## Supplementary material for "Retrospective cell lineage reconstruction in Humans using short tandem repeats": sup file4

| terid | padlock_sequence |
|---|---|
| 12780_1 | TATGAGTGTGGAGTCGTTGCGGAGAATATATCCCCTCATATTCNNNAGATCGGAAG[illegible] |
| 12780_2 | TATGAGTGTGGAGTCGTTGCGGAGAATATATCCCCTCATATTCNNNAGATCGGAAG[illegible] |
| 12780_3 | TATGAGTGTGGAGTCGTTGCGGAGAATATATCCCCTCATATTCNNNAGATCGGAAG[illegible] |
| 12780_4 | TATGAGTGTGGAGTCGTTGCGGAGAATATATCCCCTCATATTCNNNAGATCGGAAG[illegible] |
| 12780_5 | TATGAGTGTGGAGTCGTTGCGGAGAATATATCCCCTCATATTCNNNAGATCGGAAG[illegible] |
| 12780_6 | TATGAGTGTGGAGTCGTTGCGGAGAATATATCCCCTCATATTCNNNAGATCGGAAG[illegible] |
| 12780_7 | TATGAGTGTGGAGTCGTTGCGGAGAATATATCCCCTCATATTCNNNAGATCGGAAG[illegible] |
| 49555_1 | TATGAGTGTGGAGTCGTTGCGCTTTTTGGCCACATGTATGTCTNNNAGATCGGAAG[illegible] |
| 49555_2 | TATGAGTGTGGAGTCGTTGCGCTTTTTGGCCACATGTATGTCTNNNAGATCGGAAG[illegible] |
| 49555_3 | TATGAGTGTGGAGTCGTTGCGCTTTTTGGCCACATGTATGTCTNNNAGATCGGAAG[illegible] |
| 49555_4 | TATGAGTGTGGAGTCGTTGCGCTTTTTGGCCACATGTATGTCTNNNAGATCGGAAG[illegible] |
| 49555_5 | TATGAGTGTGGAGTCGTTGCGCTTTTTGGCCACATGTATGTCTNNNAGATCGGAAG[illegible] |
| 49555_6 | TATGAGTGTGGAGTCGTTGCGCTTTTTGGCCACATGTATGTCTNNNAGATCGGAAG[illegible] |
| 49555_7 | TATGAGTGTGGAGTCGTTGCGCTTTTTGGCCACATGTATGTCTNNNAGATCGGAAG[illegible] |
| 11708_1 | TATGAGTGTGGAGTCGTTGCGACACACCCACCAGATTTTCTTTNNNAGATCGGAAG[illegible] |
| 11708_2 | TATGAGTGTGGAGTCGTTGCGACACACCCACCAGATTTTCTTTNNNAGATCGGAAG[illegible] |
| 11708_3 | TATGAGTGTGGAGTCGTTGCGACACACCCACCAGATTTTCTTTNNNAGATCGGAAG[illegible] |
| 11708_4 | TATGAGTGTGGAGTCGTTGCGACACACCCACCAGATTTTCTTTNNNAGATCGGAAG[illegible] |
| 11708_5 | TATGAGTGTGGAGTCGTTGCGACACACCCACCAGATTTTCTTTNNNAGATCGGAAG[illegible] |
| 11708_6 | TATGAGTGTGGAGTCGTTGCGACACACCCACCAGATTTTCTTTNNNAGATCGGAAG[illegible] |
| 11708_7 | TATGAGTGTGGAGTCGTTGCGACACACCCACCAGATTTTCTTTNNNAGATCGGAAG[illegible] |
| 52326_1 | TATGAGTGTGGAGTCGTTGCTGAGAGACCTGGCTTGTAGACTANNNAGATCGGAA[illegible] |
| 52326_2 | TATGAGTGTGGAGTCGTTGCTGAGAGACCTGGCTTGTAGACTANNNAGATCGGAA[illegible] |
| 52326_3 | TATGAGTGTGGAGTCGTTGCTGAGAGACCTGGCTTGTAGACTANNNAGATCGGAA[illegible] |
| 52326_4 | TATGAGTGTGGAGTCGTTGCTGAGAGACCTGGCTTGTAGACTANNNAGATCGGAA[illegible] |
| 52326_5 | TATGAGTGTGGAGTCGTTGCTGAGAGACCTGGCTTGTAGACTANNNAGATCGGAA[illegible] |
| 52326_6 | TATGAGTGTGGAGTCGTTGCTGAGAGACCTGGCTTGTAGACTANNNAGATCGGAA[illegible] |
| 52326_7 | TATGAGTGTGGAGTCGTTGCTGAGAGACCTGGCTTGTAGACTANNNAGATCGGAA[illegible] |
| 33409_1 | TATGAGTGTGGAGTCGTTGCCAGCCTAGTAGGTTTTCAGCCTTNNNAGATCGGAAG[illegible] |
| 33409_2 | TATGAGTGTGGAGTCGTTGCCAGCCTAGTAGGTTTTCAGCCTTNNNAGATCGGAAG[illegible] |
| 33409_3 | TATGAGTGTGGAGTCGTTGCCAGCCTAGTAGGTTTTCAGCCTTNNNAGATCGGAAG[illegible] |
| 33409_4 | TATGAGTGTGGAGTCGTTGCCAGCCTAGTAGGTTTTCAGCCTTNNNAGATCGGAAG[illegible] |
| 33409_5 | TATGAGTGTGGAGTCGTTGCCAGCCTAGTAGGTTTTCAGCCTTNNNAGATCGGAAG[illegible] |
| 33409_6 | TATGAGTGTGGAGTCGTTGCCAGCCTAGTAGGTTTTCAGCCTTNNNAGATCGGAAG[illegible] |
| 33409_7 | TATGAGTGTGGAGTCGTTGCCAGCCTAGTAGGTTTTCAGCCTTNNNAGATCGGAAG[illegible] |
| 43136_1 | TATGAGTGTGGAGTCGTTGCCATGCCTGTAGTCTCAGCTACTTNNNAGATCGGAAG[illegible] |
| 43136_2 | TATGAGTGTGGAGTCGTTGCCATGCCTGTAGTCTCAGCTACTTNNNAGATCGGAAG[illegible] |
| 43136_3 | TATGAGTGTGGAGTCGTTGCCATGCCTGTAGTCTCAGCTACTTNNNAGATCGGAAG[illegible] |
| 43136_4 | TATGAGTGTGGAGTCGTTGCCATGCCTGTAGTCTCAGCTACTTNNNAGATCGGAAG[illegible] |
| 43136_5 | TATGAGTGTGGAGTCGTTGCCATGCCTGTAGTCTCAGCTACTTNNNAGATCGGAAG[illegible] |
| 43136_6 | TATGAGTGTGGAGTCGTTGCCATGCCTGTAGTCTCAGCTACTTNNNAGATCGGAAG[illegible] |
| 43136_7 | TATGAGTGTGGAGTCGTTGCCATGCCTGTAGTCTCAGCTACTTNNNAGATCGGAAG[illegible] |

| | |
|---|---|
| 51474_1 | TATGAGTGTGGAGTCGTTGCACAAAAATTAGCTGGGTGTGGTGNNNAGATCGGAA |
| 51474_2 | TATGAGTGTGGAGTCGTTGCACAAAAATTAGCTGGGTGTGGTGNNNAGATCGGAA |
| 51474_3 | TATGAGTGTGGAGTCGTTGCACAAAAATTAGCTGGGTGTGGTGNNNAGATCGGAA |
| 51474_4 | TATGAGTGTGGAGTCGTTGCACAAAAATTAGCTGGGTGTGGTGNNNAGATCGGAA |
| 51474_5 | TATGAGTGTGGAGTCGTTGCACAAAAATTAGCTGGGTGTGGTGNNNAGATCGGAA |
| 51474_6 | TATGAGTGTGGAGTCGTTGCACAAAAATTAGCTGGGTGTGGTGNNNAGATCGGAA |
| 51474_7 | TATGAGTGTGGAGTCGTTGCACAAAAATTAGCTGGGTGTGGTGNNNAGATCGGAA |
| 38469_1 | TATGAGTGTGGAGTCGTTGCTGATGATCCATCTTGATGCAAGGANNNAGATCGGAA |
| 38469_2 | TATGAGTGTGGAGTCGTTGCTGATGATCCATCTTGATGCAAGGANNNAGATCGGAA |
| 38469_3 | TATGAGTGTGGAGTCGTTGCTGATGATCCATCTTGATGCAAGGANNNAGATCGGAA |
| 38469_4 | TATGAGTGTGGAGTCGTTGCTGATGATCCATCTTGATGCAAGGANNNAGATCGGAA |
| 38469_5 | TATGAGTGTGGAGTCGTTGCTGATGATCCATCTTGATGCAAGGANNNAGATCGGAA |
| 38469_6 | TATGAGTGTGGAGTCGTTGCTGATGATCCATCTTGATGCAAGGANNNAGATCGGAA |
| 38469_7 | TATGAGTGTGGAGTCGTTGCTGATGATCCATCTTGATGCAAGGANNNAGATCGGAA |
| 30940_1 | TATGAGTGTGGAGTCGTTGCGCATGGCTTGTTCTCAATTTGGANNNAGATCGGAAG |
| 30940_2 | TATGAGTGTGGAGTCGTTGCGCATGGCTTGTTCTCAATTTGGANNNAGATCGGAAG |
| 30940_3 | TATGAGTGTGGAGTCGTTGCGCATGGCTTGTTCTCAATTTGGANNNAGATCGGAAG |
| 30940_4 | TATGAGTGTGGAGTCGTTGCGCATGGCTTGTTCTCAATTTGGANNNAGATCGGAAG |
| 30940_5 | TATGAGTGTGGAGTCGTTGCGCATGGCTTGTTCTCAATTTGGANNNAGATCGGAAG |
| 30940_6 | TATGAGTGTGGAGTCGTTGCGCATGGCTTGTTCTCAATTTGGANNNAGATCGGAAG |
| 30940_7 | TATGAGTGTGGAGTCGTTGCGCATGGCTTGTTCTCAATTTGGANNNAGATCGGAAG |
| 6532_1 | TATGAGTGTGGAGTCGTTGCGTAGTCCGTATCACTCGGCTTTCNNNAGATCGGAAG |
| 6532_2 | TATGAGTGTGGAGTCGTTGCGTAGTCCGTATCACTCGGCTTTCNNNAGATCGGAAG |
| 6532_3 | TATGAGTGTGGAGTCGTTGCGTAGTCCGTATCACTCGGCTTTCNNNAGATCGGAAG |
| 6532_4 | TATGAGTGTGGAGTCGTTGCGTAGTCCGTATCACTCGGCTTTCNNNAGATCGGAAG |
| 6532_5 | TATGAGTGTGGAGTCGTTGCGTAGTCCGTATCACTCGGCTTTCNNNAGATCGGAAG |
| 6532_6 | TATGAGTGTGGAGTCGTTGCGTAGTCCGTATCACTCGGCTTTCNNNAGATCGGAAG |
| 6532_7 | TATGAGTGTGGAGTCGTTGCGTAGTCCGTATCACTCGGCTTTCNNNAGATCGGAAG |
| 33963_1 | TATGAGTGTGGAGTCGTTGCTAGCTGGGATTACAGGTATGTGCNNNAGATCGGAA |
| 33963_2 | TATGAGTGTGGAGTCGTTGCTAGCTGGGATTACAGGTATGTGCNNNAGATCGGAA |
| 33963_3 | TATGAGTGTGGAGTCGTTGCTAGCTGGGATTACAGGTATGTGCNNNAGATCGGAA |
| 33963_4 | TATGAGTGTGGAGTCGTTGCTAGCTGGGATTACAGGTATGTGCNNNAGATCGGAA |
| 33963_5 | TATGAGTGTGGAGTCGTTGCTAGCTGGGATTACAGGTATGTGCNNNAGATCGGAA |
| 33963_6 | TATGAGTGTGGAGTCGTTGCTAGCTGGGATTACAGGTATGTGCNNNAGATCGGAA |
| 33963_7 | TATGAGTGTGGAGTCGTTGCTAGCTGGGATTACAGGTATGTGCNNNAGATCGGAA |
| 46008_1 | TATGAGTGTGGAGTCGTTGCGATTCTCCTGTCTTAGCCTCCTGNNNAGATCGGAAGA |
| 46008_2 | TATGAGTGTGGAGTCGTTGCGATTCTCCTGTCTTAGCCTCCTGNNNAGATCGGAAGA |
| 46008_3 | TATGAGTGTGGAGTCGTTGCGATTCTCCTGTCTTAGCCTCCTGNNNAGATCGGAAGA |
| 46008_4 | TATGAGTGTGGAGTCGTTGCGATTCTCCTGTCTTAGCCTCCTGNNNAGATCGGAAGA |
| 46008_5 | TATGAGTGTGGAGTCGTTGCGATTCTCCTGTCTTAGCCTCCTGNNNAGATCGGAAGA |
| 46008_6 | TATGAGTGTGGAGTCGTTGCGATTCTCCTGTCTTAGCCTCCTGNNNAGATCGGAAGA |
| 46008_7 | TATGAGTGTGGAGTCGTTGCGATTCTCCTGTCTTAGCCTCCTGNNNAGATCGGAAGA |
| 33176_1 | TATGAGTGTGGAGTCGTTGCTCAGTGGATTTCTGGCAATACCANNNAGATCGGAAG |

| | |
|---|---|
| 33176_2 | TATGAGTGTGGAGTCGTTGCTCAGTGGATTTCTGGCAATACCANNNAGATCGGAAG |
| 33176_3 | TATGAGTGTGGAGTCGTTGCTCAGTGGATTTCTGGCAATACCANNNAGATCGGAAG |
| 33176_4 | TATGAGTGTGGAGTCGTTGCTCAGTGGATTTCTGGCAATACCANNNAGATCGGAAG |
| 33176_5 | TATGAGTGTGGAGTCGTTGCTCAGTGGATTTCTGGCAATACCANNNAGATCGGAAG |
| 33176_6 | TATGAGTGTGGAGTCGTTGCTCAGTGGATTTCTGGCAATACCANNNAGATCGGAAG |
| 33176_7 | TATGAGTGTGGAGTCGTTGCTCAGTGGATTTCTGGCAATACCANNNAGATCGGAAG |
| 12274_1 | TATGAGTGTGGAGTCGTTGCCAAACTGAACTTGGGAACTTGCANNNAGATCGGAAG |
| 12274_2 | TATGAGTGTGGAGTCGTTGCCAAACTGAACTTGGGAACTTGCANNNAGATCGGAAG |
| 12274_3 | TATGAGTGTGGAGTCGTTGCCAAACTGAACTTGGGAACTTGCANNNAGATCGGAAG |
| 12274_4 | TATGAGTGTGGAGTCGTTGCCAAACTGAACTTGGGAACTTGCANNNAGATCGGAAG |
| 12274_5 | TATGAGTGTGGAGTCGTTGCCAAACTGAACTTGGGAACTTGCANNNAGATCGGAAG |
| 12274_6 | TATGAGTGTGGAGTCGTTGCCAAACTGAACTTGGGAACTTGCANNNAGATCGGAAG |
| 12274_7 | TATGAGTGTGGAGTCGTTGCCAAACTGAACTTGGGAACTTGCANNNAGATCGGAAG |
| 31392_1 | TATGAGTGTGGAGTCGTTGCGTGTGTGTGTGTGTGTGTTTGTANNNAGATCGGAA |
| 31392_2 | TATGAGTGTGGAGTCGTTGCGTGTGTGTGTGTGTGTGTTTGTANNNAGATCGGAA |
| 31392_3 | TATGAGTGTGGAGTCGTTGCGTGTGTGTGTGTGTGTGTTTGTANNNAGATCGGAA |
| 31392_4 | TATGAGTGTGGAGTCGTTGCGTGTGTGTGTGTGTGTGTTTGTANNNAGATCGGAA |
| 31392_5 | TATGAGTGTGGAGTCGTTGCGTGTGTGTGTGTGTGTGTTTGTANNNAGATCGGAA |
| 31392_6 | TATGAGTGTGGAGTCGTTGCGTGTGTGTGTGTGTGTGTTTGTANNNAGATCGGAA |
| 31392_7 | TATGAGTGTGGAGTCGTTGCGTGTGTGTGTGTGTGTGTTTGTANNNAGATCGGAA |
| 43114_1 | TATGAGTGTGGAGTCGTTGCTAATCCCAGCTACTCAGGAGGTTNNNAGATCGGAAG |
| 43114_2 | TATGAGTGTGGAGTCGTTGCTAATCCCAGCTACTCAGGAGGTTNNNAGATCGGAAG |
| 43114_3 | TATGAGTGTGGAGTCGTTGCTAATCCCAGCTACTCAGGAGGTTNNNAGATCGGAAG |
| 43114_4 | TATGAGTGTGGAGTCGTTGCTAATCCCAGCTACTCAGGAGGTTNNNAGATCGGAAG |
| 43114_5 | TATGAGTGTGGAGTCGTTGCTAATCCCAGCTACTCAGGAGGTTNNNAGATCGGAAG |
| 43114_6 | TATGAGTGTGGAGTCGTTGCTAATCCCAGCTACTCAGGAGGTTNNNAGATCGGAAG |
| 43114_7 | TATGAGTGTGGAGTCGTTGCTAATCCCAGCTACTCAGGAGGTTNNNAGATCGGAAG |
| 10716_1 | TATGAGTGTGGAGTCGTTGCTAGTGCCTGGTAATTGGTAGGCANNNAGATCGGAA |
| 10716_2 | TATGAGTGTGGAGTCGTTGCTAGTGCCTGGTAATTGGTAGGCANNNAGATCGGAA |
| 10716_3 | TATGAGTGTGGAGTCGTTGCTAGTGCCTGGTAATTGGTAGGCANNNAGATCGGAA |
| 10716_4 | TATGAGTGTGGAGTCGTTGCTAGTGCCTGGTAATTGGTAGGCANNNAGATCGGAA |
| 10716_5 | TATGAGTGTGGAGTCGTTGCTAGTGCCTGGTAATTGGTAGGCANNNAGATCGGAA |
| 10716_6 | TATGAGTGTGGAGTCGTTGCTAGTGCCTGGTAATTGGTAGGCANNNAGATCGGAA |
| 10716_7 | TATGAGTGTGGAGTCGTTGCTAGTGCCTGGTAATTGGTAGGCANNNAGATCGGAA |
| 6700_1 | TATGAGTGTGGAGTCGTTGCGAGATGGCAGCCAATTGCAGTTGNNNAGATCGGAA |
| 6700_2 | TATGAGTGTGGAGTCGTTGCGAGATGGCAGCCAATTGCAGTTGNNNAGATCGGAA |
| 6700_3 | TATGAGTGTGGAGTCGTTGCGAGATGGCAGCCAATTGCAGTTGNNNAGATCGGAA |
| 6700_4 | TATGAGTGTGGAGTCGTTGCGAGATGGCAGCCAATTGCAGTTGNNNAGATCGGAA |
| 6700_5 | TATGAGTGTGGAGTCGTTGCGAGATGGCAGCCAATTGCAGTTGNNNAGATCGGAA |
| 6700_6 | TATGAGTGTGGAGTCGTTGCGAGATGGCAGCCAATTGCAGTTGNNNAGATCGGAA |
| 6700_7 | TATGAGTGTGGAGTCGTTGCGAGATGGCAGCCAATTGCAGTTGNNNAGATCGGAA |
| 39690_1 | TATGAGTGTGGAGTCGTTGCTGTATGAGAACAGGATGACCAGCNNNAGATCGGAA |
| 39690_2 | TATGAGTGTGGAGTCGTTGCTGTATGAGAACAGGATGACCAGCNNNAGATCGGAA |

| | |
|---|---|
| 39690_3 | TATGAGTGTGGAGTCGTTGCTGTATGAGAACAGGATGACCAGCNNNAGATCGGAA |
| 39690_4 | TATGAGTGTGGAGTCGTTGCTGTATGAGAACAGGATGACCAGCNNNAGATCGGAA |
| 39690_5 | TATGAGTGTGGAGTCGTTGCTGTATGAGAACAGGATGACCAGCNNNAGATCGGAA |
| 39690_6 | TATGAGTGTGGAGTCGTTGCTGTATGAGAACAGGATGACCAGCNNNAGATCGGAA |
| 39690_7 | TATGAGTGTGGAGTCGTTGCTGTATGAGAACAGGATGACCAGCNNNAGATCGGAA |
| 34204_1 | TATGAGTGTGGAGTCGTTGCTGAAATGACAACCATGCTGCATTNNNAGATCGGAAG |
| 34204_2 | TATGAGTGTGGAGTCGTTGCTGAAATGACAACCATGCTGCATTNNNAGATCGGAAG |
| 34204_3 | TATGAGTGTGGAGTCGTTGCTGAAATGACAACCATGCTGCATTNNNAGATCGGAAG |
| 34204_4 | TATGAGTGTGGAGTCGTTGCTGAAATGACAACCATGCTGCATTNNNAGATCGGAAG |
| 34204_5 | TATGAGTGTGGAGTCGTTGCTGAAATGACAACCATGCTGCATTNNNAGATCGGAAG |
| 34204_6 | TATGAGTGTGGAGTCGTTGCTGAAATGACAACCATGCTGCATTNNNAGATCGGAAG |
| 34204_7 | TATGAGTGTGGAGTCGTTGCTGAAATGACAACCATGCTGCATTNNNAGATCGGAAG |
| 49177_1 | TATGAGTGTGGAGTCGTTGCTTCCTGGGTTCAAGTGATTCTCCNNNAGATCGGAAG |
| 49177_2 | TATGAGTGTGGAGTCGTTGCTTCCTGGGTTCAAGTGATTCTCCNNNAGATCGGAAG |
| 49177_3 | TATGAGTGTGGAGTCGTTGCTTCCTGGGTTCAAGTGATTCTCCNNNAGATCGGAAG |
| 49177_4 | TATGAGTGTGGAGTCGTTGCTTCCTGGGTTCAAGTGATTCTCCNNNAGATCGGAAG |
| 49177_5 | TATGAGTGTGGAGTCGTTGCTTCCTGGGTTCAAGTGATTCTCCNNNAGATCGGAAG |
| 49177_6 | TATGAGTGTGGAGTCGTTGCTTCCTGGGTTCAAGTGATTCTCCNNNAGATCGGAAG |
| 49177_7 | TATGAGTGTGGAGTCGTTGCTTCCTGGGTTCAAGTGATTCTCCNNNAGATCGGAAG |
| 32940_1 | TATGAGTGTGGAGTCGTTGCTGTGGTAGCTCACACCTGTAATCNNNAGATCGGAAG |
| 32940_2 | TATGAGTGTGGAGTCGTTGCTGTGGTAGCTCACACCTGTAATCNNNAGATCGGAAG |
| 32940_3 | TATGAGTGTGGAGTCGTTGCTGTGGTAGCTCACACCTGTAATCNNNAGATCGGAAG |
| 32940_4 | TATGAGTGTGGAGTCGTTGCTGTGGTAGCTCACACCTGTAATCNNNAGATCGGAAG |
| 32940_5 | TATGAGTGTGGAGTCGTTGCTGTGGTAGCTCACACCTGTAATCNNNAGATCGGAAG |
| 32940_6 | TATGAGTGTGGAGTCGTTGCTGTGGTAGCTCACACCTGTAATCNNNAGATCGGAAG |
| 32940_7 | TATGAGTGTGGAGTCGTTGCTGTGGTAGCTCACACCTGTAATCNNNAGATCGGAAG |
| 34567_1 | TATGAGTGTGGAGTCGTTGCGTCATGCCCTCCACGTGTTCNNNAGATCGGAAGAGC |
| 34567_2 | TATGAGTGTGGAGTCGTTGCGTCATGCCCTCCACGTGTTCNNNAGATCGGAAGAGC |
| 34567_3 | TATGAGTGTGGAGTCGTTGCGTCATGCCCTCCACGTGTTCNNNAGATCGGAAGAGC |
| 34567_4 | TATGAGTGTGGAGTCGTTGCGTCATGCCCTCCACGTGTTCNNNAGATCGGAAGAGC |
| 34567_5 | TATGAGTGTGGAGTCGTTGCGTCATGCCCTCCACGTGTTCNNNAGATCGGAAGAGC |
| 34567_6 | TATGAGTGTGGAGTCGTTGCGTCATGCCCTCCACGTGTTCNNNAGATCGGAAGAGC |
| 34567_7 | TATGAGTGTGGAGTCGTTGCGTCATGCCCTCCACGTGTTCNNNAGATCGGAAGAGC |
| 37504_1 | TATGAGTGTGGAGTCGTTGCGAGTGGCACCTCATTGTGATTTTNNNAGATCGGAAG |
| 37504_2 | TATGAGTGTGGAGTCGTTGCGAGTGGCACCTCATTGTGATTTTNNNAGATCGGAAG |
| 37504_3 | TATGAGTGTGGAGTCGTTGCGAGTGGCACCTCATTGTGATTTTNNNAGATCGGAAG |
| 37504_4 | TATGAGTGTGGAGTCGTTGCGAGTGGCACCTCATTGTGATTTTNNNAGATCGGAAG |
| 37504_5 | TATGAGTGTGGAGTCGTTGCGAGTGGCACCTCATTGTGATTTTNNNAGATCGGAAG |
| 37504_6 | TATGAGTGTGGAGTCGTTGCGAGTGGCACCTCATTGTGATTTTNNNAGATCGGAAG |
| 37504_7 | TATGAGTGTGGAGTCGTTGCGAGTGGCACCTCATTGTGATTTTNNNAGATCGGAAG |
| 35639_1 | TATGAGTGTGGAGTCGTTGCTCCTGAATACAGCACACTGATGGNNNAGATCGGAAG |
| 35639_2 | TATGAGTGTGGAGTCGTTGCTCCTGAATACAGCACACTGATGGNNNAGATCGGAAG |
| 35639_3 | TATGAGTGTGGAGTCGTTGCTCCTGAATACAGCACACTGATGGNNNAGATCGGAAG |

| | |
|---|---|
| 35639_4 | TATGAGTGTGGAGTCGTTGCTCCTGAATACAGCACACTGATGGNNNAGATCGGAAG |
| 35639_5 | TATGAGTGTGGAGTCGTTGCTCCTGAATACAGCACACTGATGGNNNAGATCGGAAG |
| 35639_6 | TATGAGTGTGGAGTCGTTGCTCCTGAATACAGCACACTGATGGNNNAGATCGGAAG |
| 35639_7 | TATGAGTGTGGAGTCGTTGCTCCTGAATACAGCACACTGATGGNNNAGATCGGAAG |
| 8403_1 | TATGAGTGTGGAGTCGTTGCCGTGTCTTAGGGATTGTCTTCCTGNNNAGATCGGAA |
| 8403_2 | TATGAGTGTGGAGTCGTTGCCGTGTCTTAGGGATTGTCTTCCTGNNNAGATCGGAA |
| 8403_3 | TATGAGTGTGGAGTCGTTGCCGTGTCTTAGGGATTGTCTTCCTGNNNAGATCGGAA |
| 8403_4 | TATGAGTGTGGAGTCGTTGCCGTGTCTTAGGGATTGTCTTCCTGNNNAGATCGGAA |
| 8403_5 | TATGAGTGTGGAGTCGTTGCCGTGTCTTAGGGATTGTCTTCCTGNNNAGATCGGAA |
| 8403_6 | TATGAGTGTGGAGTCGTTGCCGTGTCTTAGGGATTGTCTTCCTGNNNAGATCGGAA |
| 8403_7 | TATGAGTGTGGAGTCGTTGCCGTGTCTTAGGGATTGTCTTCCTGNNNAGATCGGAA |
| 38782_1 | TATGAGTGTGGAGTCGTTGCATGAAGAAGGGCTAAGACTGCTCNNNAGATCGGAA |
| 38782_2 | TATGAGTGTGGAGTCGTTGCATGAAGAAGGGCTAAGACTGCTCNNNAGATCGGAA |
| 38782_3 | TATGAGTGTGGAGTCGTTGCATGAAGAAGGGCTAAGACTGCTCNNNAGATCGGAA |
| 38782_4 | TATGAGTGTGGAGTCGTTGCATGAAGAAGGGCTAAGACTGCTCNNNAGATCGGAA |
| 38782_5 | TATGAGTGTGGAGTCGTTGCATGAAGAAGGGCTAAGACTGCTCNNNAGATCGGAA |
| 38782_6 | TATGAGTGTGGAGTCGTTGCATGAAGAAGGGCTAAGACTGCTCNNNAGATCGGAA |
| 38782_7 | TATGAGTGTGGAGTCGTTGCATGAAGAAGGGCTAAGACTGCTCNNNAGATCGGAA |
| 40807_1 | TATGAGTGTGGAGTCGTTGCAGATGAGAGATGGCATCACATGGNNNAGATCGGAA |
| 40807_2 | TATGAGTGTGGAGTCGTTGCAGATGAGAGATGGCATCACATGGNNNAGATCGGAA |
| 40807_3 | TATGAGTGTGGAGTCGTTGCAGATGAGAGATGGCATCACATGGNNNAGATCGGAA |
| 40807_4 | TATGAGTGTGGAGTCGTTGCAGATGAGAGATGGCATCACATGGNNNAGATCGGAA |
| 40807_5 | TATGAGTGTGGAGTCGTTGCAGATGAGAGATGGCATCACATGGNNNAGATCGGAA |
| 40807_6 | TATGAGTGTGGAGTCGTTGCAGATGAGAGATGGCATCACATGGNNNAGATCGGAA |
| 40807_7 | TATGAGTGTGGAGTCGTTGCAGATGAGAGATGGCATCACATGGNNNAGATCGGAA |
| 43229_1 | TATGAGTGTGGAGTCGTTGCAGGCATATGTGGTCATTTTGCACNNNAGATCGGAAG |
| 43229_2 | TATGAGTGTGGAGTCGTTGCAGGCATATGTGGTCATTTTGCACNNNAGATCGGAAG |
| 43229_3 | TATGAGTGTGGAGTCGTTGCAGGCATATGTGGTCATTTTGCACNNNAGATCGGAAG |
| 43229_4 | TATGAGTGTGGAGTCGTTGCAGGCATATGTGGTCATTTTGCACNNNAGATCGGAAG |
| 43229_5 | TATGAGTGTGGAGTCGTTGCAGGCATATGTGGTCATTTTGCACNNNAGATCGGAAG |
| 43229_6 | TATGAGTGTGGAGTCGTTGCAGGCATATGTGGTCATTTTGCACNNNAGATCGGAAG |
| 43229_7 | TATGAGTGTGGAGTCGTTGCAGGCATATGTGGTCATTTTGCACNNNAGATCGGAAG |
| 34091_1 | TATGAGTGTGGAGTCGTTGCTTTTTAGTAGAGACGGGGGCTTCNNNAGATCGGAA |
| 34091_2 | TATGAGTGTGGAGTCGTTGCTTTTTAGTAGAGACGGGGGCTTCNNNAGATCGGAA |
| 34091_3 | TATGAGTGTGGAGTCGTTGCTTTTTAGTAGAGACGGGGGCTTCNNNAGATCGGAA |
| 34091_4 | TATGAGTGTGGAGTCGTTGCTTTTTAGTAGAGACGGGGGCTTCNNNAGATCGGAA |
| 34091_5 | TATGAGTGTGGAGTCGTTGCTTTTTAGTAGAGACGGGGGCTTCNNNAGATCGGAA |
| 34091_6 | TATGAGTGTGGAGTCGTTGCTTTTTAGTAGAGACGGGGGCTTCNNNAGATCGGAA |
| 34091_7 | TATGAGTGTGGAGTCGTTGCTTTTTAGTAGAGACGGGGGCTTCNNNAGATCGGAA |
| 36968_1 | TATGAGTGTGGAGTCGTTGCAACTCGTTTCACCTGATAGTCCCNNNAGATCGGAAG |
| 36968_2 | TATGAGTGTGGAGTCGTTGCAACTCGTTTCACCTGATAGTCCCNNNAGATCGGAAG |
| 36968_3 | TATGAGTGTGGAGTCGTTGCAACTCGTTTCACCTGATAGTCCCNNNAGATCGGAAG |
| 36968_4 | TATGAGTGTGGAGTCGTTGCAACTCGTTTCACCTGATAGTCCCNNNAGATCGGAAG |

| | |
|---|---|
| 36968_5 | TATGAGTGTGGAGTCGTTGCAACTCGTTTCACCTGATAGTCCCNNNAGATCGGAAGA |
| 36968_6 | TATGAGTGTGGAGTCGTTGCAACTCGTTTCACCTGATAGTCCCNNNAGATCGGAAGA |
| 36968_7 | TATGAGTGTGGAGTCGTTGCAACTCGTTTCACCTGATAGTCCCNNNAGATCGGAAGA |
| 28964_1 | TATGAGTGTGGAGTCGTTGCACAGCAAGAAATAGCACTTTCAGNNNAGATCGGAAG |
| 28964_2 | TATGAGTGTGGAGTCGTTGCACAGCAAGAAATAGCACTTTCAGNNNAGATCGGAAG |
| 28964_3 | TATGAGTGTGGAGTCGTTGCACAGCAAGAAATAGCACTTTCAGNNNAGATCGGAAG |
| 28964_4 | TATGAGTGTGGAGTCGTTGCACAGCAAGAAATAGCACTTTCAGNNNAGATCGGAAG |
| 28964_5 | TATGAGTGTGGAGTCGTTGCACAGCAAGAAATAGCACTTTCAGNNNAGATCGGAAG |
| 28964_6 | TATGAGTGTGGAGTCGTTGCACAGCAAGAAATAGCACTTTCAGNNNAGATCGGAAG |
| 28964_7 | TATGAGTGTGGAGTCGTTGCACAGCAAGAAATAGCACTTTCAGNNNAGATCGGAAG |
| 31660_1 | TATGAGTGTGGAGTCGTTGCTAGGCTGATGTAGTTGGTTGACCNNNAGATCGGAA[illegible] |
| 31660_2 | TATGAGTGTGGAGTCGTTGCTAGGCTGATGTAGTTGGTTGACCNNNAGATCGGAA[illegible] |
| 31660_3 | TATGAGTGTGGAGTCGTTGCTAGGCTGATGTAGTTGGTTGACCNNNAGATCGGAA[illegible] |
| 31660_4 | TATGAGTGTGGAGTCGTTGCTAGGCTGATGTAGTTGGTTGACCNNNAGATCGGAA[illegible] |
| 31660_5 | TATGAGTGTGGAGTCGTTGCTAGGCTGATGTAGTTGGTTGACCNNNAGATCGGAA[illegible] |
| 31660_6 | TATGAGTGTGGAGTCGTTGCTAGGCTGATGTAGTTGGTTGACCNNNAGATCGGAA[illegible] |
| 31660_7 | TATGAGTGTGGAGTCGTTGCTAGGCTGATGTAGTTGGTTGACCNNNAGATCGGAA[illegible] |
| 36138_1 | TATGAGTGTGGAGTCGTTGCTGAAATACAGACACATGCAACGANNNAGATCGGAA[illegible] |
| 36138_2 | TATGAGTGTGGAGTCGTTGCTGAAATACAGACACATGCAACGANNNAGATCGGAA[illegible] |
| 36138_3 | TATGAGTGTGGAGTCGTTGCTGAAATACAGACACATGCAACGANNNAGATCGGAA[illegible] |
| 36138_4 | TATGAGTGTGGAGTCGTTGCTGAAATACAGACACATGCAACGANNNAGATCGGAA[illegible] |
| 36138_5 | TATGAGTGTGGAGTCGTTGCTGAAATACAGACACATGCAACGANNNAGATCGGAA[illegible] |
| 36138_6 | TATGAGTGTGGAGTCGTTGCTGAAATACAGACACATGCAACGANNNAGATCGGAA[illegible] |
| 36138_7 | TATGAGTGTGGAGTCGTTGCTGAAATACAGACACATGCAACGANNNAGATCGGAA[illegible] |
| 49380_1 | TATGAGTGTGGAGTCGTTGCCTCCTGAGAAGCTGGGATTACAGNNNAGATCGGAA[illegible] |
| 49380_2 | TATGAGTGTGGAGTCGTTGCCTCCTGAGAAGCTGGGATTACAGNNNAGATCGGAA[illegible] |
| 49380_3 | TATGAGTGTGGAGTCGTTGCCTCCTGAGAAGCTGGGATTACAGNNNAGATCGGAA[illegible] |
| 49380_4 | TATGAGTGTGGAGTCGTTGCCTCCTGAGAAGCTGGGATTACAGNNNAGATCGGAA[illegible] |
| 49380_5 | TATGAGTGTGGAGTCGTTGCCTCCTGAGAAGCTGGGATTACAGNNNAGATCGGAA[illegible] |
| 49380_6 | TATGAGTGTGGAGTCGTTGCCTCCTGAGAAGCTGGGATTACAGNNNAGATCGGAA[illegible] |
| 49380_7 | TATGAGTGTGGAGTCGTTGCCTCCTGAGAAGCTGGGATTACAGNNNAGATCGGAA[illegible] |
| 40063_1 | TATGAGTGTGGAGTCGTTGCGAAACTTCCCTAAGGTGAGTGATNNNAGATCGGAA[illegible] |
| 40063_2 | TATGAGTGTGGAGTCGTTGCGAAACTTCCCTAAGGTGAGTGATNNNAGATCGGAA[illegible] |
| 40063_3 | TATGAGTGTGGAGTCGTTGCGAAACTTCCCTAAGGTGAGTGATNNNAGATCGGAA[illegible] |
| 40063_4 | TATGAGTGTGGAGTCGTTGCGAAACTTCCCTAAGGTGAGTGATNNNAGATCGGAA[illegible] |
| 40063_5 | TATGAGTGTGGAGTCGTTGCGAAACTTCCCTAAGGTGAGTGATNNNAGATCGGAA[illegible] |
| 40063_6 | TATGAGTGTGGAGTCGTTGCGAAACTTCCCTAAGGTGAGTGATNNNAGATCGGAA[illegible] |
| 40063_7 | TATGAGTGTGGAGTCGTTGCGAAACTTCCCTAAGGTGAGTGATNNNAGATCGGAA[illegible] |
| 40785_1 | TATGAGTGTGGAGTCGTTGCGGATGAAAAAGAACCTGTAAGGGCNNNAGATCGGA |
| 40785_2 | TATGAGTGTGGAGTCGTTGCGGATGAAAAAGAACCTGTAAGGGCNNNAGATCGGA |
| 40785_3 | TATGAGTGTGGAGTCGTTGCGGATGAAAAAGAACCTGTAAGGGCNNNAGATCGGA |
| 40785_4 | TATGAGTGTGGAGTCGTTGCGGATGAAAAAGAACCTGTAAGGGCNNNAGATCGGA |
| 40785_5 | TATGAGTGTGGAGTCGTTGCGGATGAAAAAGAACCTGTAAGGGCNNNAGATCGGA |

| | |
|---|---|
| 40785_6 | TATGAGTGTGGAGTCGTTGCGGATGAAAAAGAACCTGTAAGGGCNNNAGATCGGA |
| 40785_7 | TATGAGTGTGGAGTCGTTGCGGATGAAAAAGAACCTGTAAGGGCNNNAGATCGGA |
| 33183_1 | TATGAGTGTGGAGTCGTTGCTAGTGATTCTTCCGCTGGATCTGNNNAGATCGGAAG |
| 33183_2 | TATGAGTGTGGAGTCGTTGCTAGTGATTCTTCCGCTGGATCTGNNNAGATCGGAAG |
| 33183_3 | TATGAGTGTGGAGTCGTTGCTAGTGATTCTTCCGCTGGATCTGNNNAGATCGGAAG |
| 33183_4 | TATGAGTGTGGAGTCGTTGCTAGTGATTCTTCCGCTGGATCTGNNNAGATCGGAAG |
| 33183_5 | TATGAGTGTGGAGTCGTTGCTAGTGATTCTTCCGCTGGATCTGNNNAGATCGGAAG |
| 33183_6 | TATGAGTGTGGAGTCGTTGCTAGTGATTCTTCCGCTGGATCTGNNNAGATCGGAAG |
| 33183_7 | TATGAGTGTGGAGTCGTTGCTAGTGATTCTTCCGCTGGATCTGNNNAGATCGGAAG |
| 49038_1 | TATGAGTGTGGAGTCGTTGCCACCACAGTCGCCTAATTTTTGTNNNAGATCGGAAGA |
| 49038_2 | TATGAGTGTGGAGTCGTTGCCACCACAGTCGCCTAATTTTTGTNNNAGATCGGAAGA |
| 49038_3 | TATGAGTGTGGAGTCGTTGCCACCACAGTCGCCTAATTTTTGTNNNAGATCGGAAGA |
| 49038_4 | TATGAGTGTGGAGTCGTTGCCACCACAGTCGCCTAATTTTTGTNNNAGATCGGAAGA |
| 49038_5 | TATGAGTGTGGAGTCGTTGCCACCACAGTCGCCTAATTTTTGTNNNAGATCGGAAGA |
| 49038_6 | TATGAGTGTGGAGTCGTTGCCACCACAGTCGCCTAATTTTTGTNNNAGATCGGAAGA |
| 49038_7 | TATGAGTGTGGAGTCGTTGCCACCACAGTCGCCTAATTTTTGTNNNAGATCGGAAGA |
| 30791_1 | TATGAGTGTGGAGTCGTTGCGAGAATGCAAAGGAGGGAAACACNNNAGATCGGAA |
| 30791_2 | TATGAGTGTGGAGTCGTTGCGAGAATGCAAAGGAGGGAAACACNNNAGATCGGAA |
| 30791_3 | TATGAGTGTGGAGTCGTTGCGAGAATGCAAAGGAGGGAAACACNNNAGATCGGAA |
| 30791_4 | TATGAGTGTGGAGTCGTTGCGAGAATGCAAAGGAGGGAAACACNNNAGATCGGAA |
| 30791_5 | TATGAGTGTGGAGTCGTTGCGAGAATGCAAAGGAGGGAAACACNNNAGATCGGAA |
| 30791_6 | TATGAGTGTGGAGTCGTTGCGAGAATGCAAAGGAGGGAAACACNNNAGATCGGAA |
| 30791_7 | TATGAGTGTGGAGTCGTTGCGAGAATGCAAAGGAGGGAAACACNNNAGATCGGAA |
| 33802_1 | TATGAGTGTGGAGTCGTTGCAACTGAGGTGACAGTCAATGGAGNNNAGATCGGAA |
| 33802_2 | TATGAGTGTGGAGTCGTTGCAACTGAGGTGACAGTCAATGGAGNNNAGATCGGAA |
| 33802_3 | TATGAGTGTGGAGTCGTTGCAACTGAGGTGACAGTCAATGGAGNNNAGATCGGAA |
| 33802_4 | TATGAGTGTGGAGTCGTTGCAACTGAGGTGACAGTCAATGGAGNNNAGATCGGAA |
| 33802_5 | TATGAGTGTGGAGTCGTTGCAACTGAGGTGACAGTCAATGGAGNNNAGATCGGAA |
| 33802_6 | TATGAGTGTGGAGTCGTTGCAACTGAGGTGACAGTCAATGGAGNNNAGATCGGAA |
| 33802_7 | TATGAGTGTGGAGTCGTTGCAACTGAGGTGACAGTCAATGGAGNNNAGATCGGAA |
| 43292_1 | TATGAGTGTGGAGTCGTTGCCAGGAATGGAAACCCAAACATCGNNNAGATCGGAAC |
| 43292_2 | TATGAGTGTGGAGTCGTTGCCAGGAATGGAAACCCAAACATCGNNNAGATCGGAAC |
| 43292_3 | TATGAGTGTGGAGTCGTTGCCAGGAATGGAAACCCAAACATCGNNNAGATCGGAAC |
| 43292_4 | TATGAGTGTGGAGTCGTTGCCAGGAATGGAAACCCAAACATCGNNNAGATCGGAAC |
| 43292_5 | TATGAGTGTGGAGTCGTTGCCAGGAATGGAAACCCAAACATCGNNNAGATCGGAAC |
| 43292_6 | TATGAGTGTGGAGTCGTTGCCAGGAATGGAAACCCAAACATCGNNNAGATCGGAAC |
| 43292_7 | TATGAGTGTGGAGTCGTTGCCAGGAATGGAAACCCAAACATCGNNNAGATCGGAAC |
| 45546_1 | TATGAGTGTGGAGTCGTTGCAGTGAGAATGATGTCCATGGAGGNNNAGATCGGAA |
| 45546_2 | TATGAGTGTGGAGTCGTTGCAGTGAGAATGATGTCCATGGAGGNNNAGATCGGAA |
| 45546_3 | TATGAGTGTGGAGTCGTTGCAGTGAGAATGATGTCCATGGAGGNNNAGATCGGAA |
| 45546_4 | TATGAGTGTGGAGTCGTTGCAGTGAGAATGATGTCCATGGAGGNNNAGATCGGAA |
| 45546_5 | TATGAGTGTGGAGTCGTTGCAGTGAGAATGATGTCCATGGAGGNNNAGATCGGAA |
| 45546_6 | TATGAGTGTGGAGTCGTTGCAGTGAGAATGATGTCCATGGAGGNNNAGATCGGAA |

| | |
|---|---|
| 45546_7 | TATGAGTGTGGAGTCGTTGCAGTGAGAATGATGTCCATGGAGGNNNAGATCGGAA |
| 32901_1 | TATGAGTGTGGAGTCGTTGCGCAAGACTCTGCCTCAAAAACAANNNAGATCGGAAG |
| 32901_2 | TATGAGTGTGGAGTCGTTGCGCAAGACTCTGCCTCAAAAACAANNNAGATCGGAAG |
| 32901_3 | TATGAGTGTGGAGTCGTTGCGCAAGACTCTGCCTCAAAAACAANNNAGATCGGAAG |
| 32901_4 | TATGAGTGTGGAGTCGTTGCGCAAGACTCTGCCTCAAAAACAANNNAGATCGGAAG |
| 32901_5 | TATGAGTGTGGAGTCGTTGCGCAAGACTCTGCCTCAAAAACAANNNAGATCGGAAG |
| 32901_6 | TATGAGTGTGGAGTCGTTGCGCAAGACTCTGCCTCAAAAACAANNNAGATCGGAAG |
| 32901_7 | TATGAGTGTGGAGTCGTTGCGCAAGACTCTGCCTCAAAAACAANNNAGATCGGAAG |
| 47538_1 | TATGAGTGTGGAGTCGTTGCATCTTGGCTCACTACAACCTCTGNNNAGATCGGAAG |
| 47538_2 | TATGAGTGTGGAGTCGTTGCATCTTGGCTCACTACAACCTCTGNNNAGATCGGAAG |
| 47538_3 | TATGAGTGTGGAGTCGTTGCATCTTGGCTCACTACAACCTCTGNNNAGATCGGAAG |
| 47538_4 | TATGAGTGTGGAGTCGTTGCATCTTGGCTCACTACAACCTCTGNNNAGATCGGAAG |
| 47538_5 | TATGAGTGTGGAGTCGTTGCATCTTGGCTCACTACAACCTCTGNNNAGATCGGAAG |
| 47538_6 | TATGAGTGTGGAGTCGTTGCATCTTGGCTCACTACAACCTCTGNNNAGATCGGAAG |
| 47538_7 | TATGAGTGTGGAGTCGTTGCATCTTGGCTCACTACAACCTCTGNNNAGATCGGAAG |
| 29087_1 | TATGAGTGTGGAGTCGTTGCGGTATCTAGAGACCAACTGGCAGNNNAGATCGGAA |
| 29087_2 | TATGAGTGTGGAGTCGTTGCGGTATCTAGAGACCAACTGGCAGNNNAGATCGGAA |
| 29087_3 | TATGAGTGTGGAGTCGTTGCGGTATCTAGAGACCAACTGGCAGNNNAGATCGGAA |
| 29087_4 | TATGAGTGTGGAGTCGTTGCGGTATCTAGAGACCAACTGGCAGNNNAGATCGGAA |
| 29087_5 | TATGAGTGTGGAGTCGTTGCGGTATCTAGAGACCAACTGGCAGNNNAGATCGGAA |
| 29087_6 | TATGAGTGTGGAGTCGTTGCGGTATCTAGAGACCAACTGGCAGNNNAGATCGGAA |
| 29087_7 | TATGAGTGTGGAGTCGTTGCGGTATCTAGAGACCAACTGGCAGNNNAGATCGGAA |
| 49115_1 | TATGAGTGTGGAGTCGTTGCAACCTCCTGAGTAGCTGGAACTANNNAGATCGGAAG |
| 49115_2 | TATGAGTGTGGAGTCGTTGCAACCTCCTGAGTAGCTGGAACTANNNAGATCGGAAG |
| 49115_3 | TATGAGTGTGGAGTCGTTGCAACCTCCTGAGTAGCTGGAACTANNNAGATCGGAAG |
| 49115_4 | TATGAGTGTGGAGTCGTTGCAACCTCCTGAGTAGCTGGAACTANNNAGATCGGAAG |
| 49115_5 | TATGAGTGTGGAGTCGTTGCAACCTCCTGAGTAGCTGGAACTANNNAGATCGGAAG |
| 49115_6 | TATGAGTGTGGAGTCGTTGCAACCTCCTGAGTAGCTGGAACTANNNAGATCGGAAG |
| 49115_7 | TATGAGTGTGGAGTCGTTGCAACCTCCTGAGTAGCTGGAACTANNNAGATCGGAAG |
| 51287_1 | TATGAGTGTGGAGTCGTTGCCATAGACCCTCTATTCCCAAGGCNNNAGATCGGAAG |
| 51287_2 | TATGAGTGTGGAGTCGTTGCCATAGACCCTCTATTCCCAAGGCNNNAGATCGGAAG |
| 51287_3 | TATGAGTGTGGAGTCGTTGCCATAGACCCTCTATTCCCAAGGCNNNAGATCGGAAG |
| 51287_4 | TATGAGTGTGGAGTCGTTGCCATAGACCCTCTATTCCCAAGGCNNNAGATCGGAAG |
| 51287_5 | TATGAGTGTGGAGTCGTTGCCATAGACCCTCTATTCCCAAGGCNNNAGATCGGAAG |
| 51287_6 | TATGAGTGTGGAGTCGTTGCCATAGACCCTCTATTCCCAAGGCNNNAGATCGGAAG |
| 51287_7 | TATGAGTGTGGAGTCGTTGCCATAGACCCTCTATTCCCAAGGCNNNAGATCGGAAG |
| 50214_1 | TATGAGTGTGGAGTCGTTGCCATCCACAGCACAGATCCTGAGNNNAGATCGGAAGA |
| 50214_2 | TATGAGTGTGGAGTCGTTGCCATCCACAGCACAGATCCTGAGNNNAGATCGGAAGA |
| 50214_3 | TATGAGTGTGGAGTCGTTGCCATCCACAGCACAGATCCTGAGNNNAGATCGGAAGA |
| 50214_4 | TATGAGTGTGGAGTCGTTGCCATCCACAGCACAGATCCTGAGNNNAGATCGGAAGA |
| 50214_5 | TATGAGTGTGGAGTCGTTGCCATCCACAGCACAGATCCTGAGNNNAGATCGGAAGA |
| 50214_6 | TATGAGTGTGGAGTCGTTGCCATCCACAGCACAGATCCTGAGNNNAGATCGGAAGA |
| 50214_7 | TATGAGTGTGGAGTCGTTGCCATCCACAGCACAGATCCTGAGNNNAGATCGGAAGA |

| | |
|---|---|
| 7798_1 | TATGAGTGTGGAGTCGTTGCTTCAGCGTCCTGTCCAAAGTCTANNNAGATCGGAAG |
| 7798_2 | TATGAGTGTGGAGTCGTTGCTTCAGCGTCCTGTCCAAAGTCTANNNAGATCGGAAG |
| 7798_3 | TATGAGTGTGGAGTCGTTGCTTCAGCGTCCTGTCCAAAGTCTANNNAGATCGGAAG |
| 7798_4 | TATGAGTGTGGAGTCGTTGCTTCAGCGTCCTGTCCAAAGTCTANNNAGATCGGAAG |
| 7798_5 | TATGAGTGTGGAGTCGTTGCTTCAGCGTCCTGTCCAAAGTCTANNNAGATCGGAAG |
| 7798_6 | TATGAGTGTGGAGTCGTTGCTTCAGCGTCCTGTCCAAAGTCTANNNAGATCGGAAG |
| 7798_7 | TATGAGTGTGGAGTCGTTGCTTCAGCGTCCTGTCCAAAGTCTANNNAGATCGGAAG |
| 40668_1 | TATGAGTGTGGAGTCGTTGCACATTTATGCAGCCAACAGACACNNNAGATCGGAAG |
| 40668_2 | TATGAGTGTGGAGTCGTTGCACATTTATGCAGCCAACAGACACNNNAGATCGGAAG |
| 40668_3 | TATGAGTGTGGAGTCGTTGCACATTTATGCAGCCAACAGACACNNNAGATCGGAAG |
| 40668_4 | TATGAGTGTGGAGTCGTTGCACATTTATGCAGCCAACAGACACNNNAGATCGGAAG |
| 40668_5 | TATGAGTGTGGAGTCGTTGCACATTTATGCAGCCAACAGACACNNNAGATCGGAAG |
| 40668_6 | TATGAGTGTGGAGTCGTTGCACATTTATGCAGCCAACAGACACNNNAGATCGGAAG |
| 40668_7 | TATGAGTGTGGAGTCGTTGCACATTTATGCAGCCAACAGACACNNNAGATCGGAAG |
| 52451_1 | TATGAGTGTGGAGTCGTTGCGTCACCTCTTCAGGGCTGTTTTNNNAGATCGGAAGA |
| 52451_2 | TATGAGTGTGGAGTCGTTGCGTCACCTCTTCAGGGCTGTTTTNNNAGATCGGAAGA |
| 52451_3 | TATGAGTGTGGAGTCGTTGCGTCACCTCTTCAGGGCTGTTTTNNNAGATCGGAAGA |
| 52451_4 | TATGAGTGTGGAGTCGTTGCGTCACCTCTTCAGGGCTGTTTTNNNAGATCGGAAGA |
| 52451_5 | TATGAGTGTGGAGTCGTTGCGTCACCTCTTCAGGGCTGTTTTNNNAGATCGGAAGA |
| 52451_6 | TATGAGTGTGGAGTCGTTGCGTCACCTCTTCAGGGCTGTTTTNNNAGATCGGAAGA |
| 52451_7 | TATGAGTGTGGAGTCGTTGCGTCACCTCTTCAGGGCTGTTTTNNNAGATCGGAAGA |
| 29529_1 | TATGAGTGTGGAGTCGTTGCGGCGTTGCTTTTCAAAGTGAAGANNNAGATCGGAA |
| 29529_2 | TATGAGTGTGGAGTCGTTGCGGCGTTGCTTTTCAAAGTGAAGANNNAGATCGGAA |
| 29529_3 | TATGAGTGTGGAGTCGTTGCGGCGTTGCTTTTCAAAGTGAAGANNNAGATCGGAA |
| 29529_4 | TATGAGTGTGGAGTCGTTGCGGCGTTGCTTTTCAAAGTGAAGANNNAGATCGGAA |
| 29529_5 | TATGAGTGTGGAGTCGTTGCGGCGTTGCTTTTCAAAGTGAAGANNNAGATCGGAA |
| 29529_6 | TATGAGTGTGGAGTCGTTGCGGCGTTGCTTTTCAAAGTGAAGANNNAGATCGGAA |
| 29529_7 | TATGAGTGTGGAGTCGTTGCGGCGTTGCTTTTCAAAGTGAAGANNNAGATCGGAA |
| 33068_1 | TATGAGTGTGGAGTCGTTGCTTCAAGTGATTCTCCTGCCTCAGNNNAGATCGGAAG |
| 33068_2 | TATGAGTGTGGAGTCGTTGCTTCAAGTGATTCTCCTGCCTCAGNNNAGATCGGAAG |
| 33068_3 | TATGAGTGTGGAGTCGTTGCTTCAAGTGATTCTCCTGCCTCAGNNNAGATCGGAAG |
| 33068_4 | TATGAGTGTGGAGTCGTTGCTTCAAGTGATTCTCCTGCCTCAGNNNAGATCGGAAG |
| 33068_5 | TATGAGTGTGGAGTCGTTGCTTCAAGTGATTCTCCTGCCTCAGNNNAGATCGGAAG |
| 33068_6 | TATGAGTGTGGAGTCGTTGCTTCAAGTGATTCTCCTGCCTCAGNNNAGATCGGAAG |
| 33068_7 | TATGAGTGTGGAGTCGTTGCTTCAAGTGATTCTCCTGCCTCAGNNNAGATCGGAAG |
| 34154_1 | TATGAGTGTGGAGTCGTTGCTTTTCAAATGTCCAGCCAATTGTNNNAGATCGGAAG |
| 34154_2 | TATGAGTGTGGAGTCGTTGCTTTTCAAATGTCCAGCCAATTGTNNNAGATCGGAAG |
| 34154_3 | TATGAGTGTGGAGTCGTTGCTTTTCAAATGTCCAGCCAATTGTNNNAGATCGGAAG |
| 34154_4 | TATGAGTGTGGAGTCGTTGCTTTTCAAATGTCCAGCCAATTGTNNNAGATCGGAAG |
| 34154_5 | TATGAGTGTGGAGTCGTTGCTTTTCAAATGTCCAGCCAATTGTNNNAGATCGGAAG |
| 34154_6 | TATGAGTGTGGAGTCGTTGCTTTTCAAATGTCCAGCCAATTGTNNNAGATCGGAAG |
| 34154_7 | TATGAGTGTGGAGTCGTTGCTTTTCAAATGTCCAGCCAATTGTNNNAGATCGGAAG |
| 35340_1 | TATGAGTGTGGAGTCGTTGCAGTGATTCTTCTGACTCAGCCTCNNNAGATCGGAAG |

| | |
|---|---|
| 35340_2 | TATGAGTGTGGAGTCGTTGCAGTGATTCTTCTGACTCAGCCTCNNNAGATCGGAAG |
| 35340_3 | TATGAGTGTGGAGTCGTTGCAGTGATTCTTCTGACTCAGCCTCNNNAGATCGGAAG |
| 35340_4 | TATGAGTGTGGAGTCGTTGCAGTGATTCTTCTGACTCAGCCTCNNNAGATCGGAAG |
| 35340_5 | TATGAGTGTGGAGTCGTTGCAGTGATTCTTCTGACTCAGCCTCNNNAGATCGGAAG |
| 35340_6 | TATGAGTGTGGAGTCGTTGCAGTGATTCTTCTGACTCAGCCTCNNNAGATCGGAAG |
| 35340_7 | TATGAGTGTGGAGTCGTTGCAGTGATTCTTCTGACTCAGCCTCNNNAGATCGGAAG |
| 6414_1 | TATGAGTGTGGAGTCGTTGCCCTTGGGGAGAAGTTTCAGTGTTNNNAGATCGGAA |
| 6414_2 | TATGAGTGTGGAGTCGTTGCCCTTGGGGAGAAGTTTCAGTGTTNNNAGATCGGAA |
| 6414_3 | TATGAGTGTGGAGTCGTTGCCCTTGGGGAGAAGTTTCAGTGTTNNNAGATCGGAA |
| 6414_4 | TATGAGTGTGGAGTCGTTGCCCTTGGGGAGAAGTTTCAGTGTTNNNAGATCGGAA |
| 6414_5 | TATGAGTGTGGAGTCGTTGCCCTTGGGGAGAAGTTTCAGTGTTNNNAGATCGGAA |
| 6414_6 | TATGAGTGTGGAGTCGTTGCCCTTGGGGAGAAGTTTCAGTGTTNNNAGATCGGAA |
| 6414_7 | TATGAGTGTGGAGTCGTTGCCCTTGGGGAGAAGTTTCAGTGTTNNNAGATCGGAA |
| 34021_1 | TATGAGTGTGGAGTCGTTGCGGAGTAGTGCAATGGTGTGATCTNNNAGATCGGAA |
| 34021_2 | TATGAGTGTGGAGTCGTTGCGGAGTAGTGCAATGGTGTGATCTNNNAGATCGGAA |
| 34021_3 | TATGAGTGTGGAGTCGTTGCGGAGTAGTGCAATGGTGTGATCTNNNAGATCGGAA |
| 34021_4 | TATGAGTGTGGAGTCGTTGCGGAGTAGTGCAATGGTGTGATCTNNNAGATCGGAA |
| 34021_5 | TATGAGTGTGGAGTCGTTGCGGAGTAGTGCAATGGTGTGATCTNNNAGATCGGAA |
| 34021_6 | TATGAGTGTGGAGTCGTTGCGGAGTAGTGCAATGGTGTGATCTNNNAGATCGGAA |
| 34021_7 | TATGAGTGTGGAGTCGTTGCGGAGTAGTGCAATGGTGTGATCTNNNAGATCGGAA |
| 51841_1 | TATGAGTGTGGAGTCGTTGCCTATCATCTGGCTCTTCCCACTCNNNAGATCGGAAGA |
| 51841_2 | TATGAGTGTGGAGTCGTTGCCTATCATCTGGCTCTTCCCACTCNNNAGATCGGAAGA |
| 51841_3 | TATGAGTGTGGAGTCGTTGCCTATCATCTGGCTCTTCCCACTCNNNAGATCGGAAGA |
| 51841_4 | TATGAGTGTGGAGTCGTTGCCTATCATCTGGCTCTTCCCACTCNNNAGATCGGAAGA |
| 51841_5 | TATGAGTGTGGAGTCGTTGCCTATCATCTGGCTCTTCCCACTCNNNAGATCGGAAGA |
| 51841_6 | TATGAGTGTGGAGTCGTTGCCTATCATCTGGCTCTTCCCACTCNNNAGATCGGAAGA |
| 51841_7 | TATGAGTGTGGAGTCGTTGCCTATCATCTGGCTCTTCCCACTCNNNAGATCGGAAGA |
| 34289_1 | TATGAGTGTGGAGTCGTTGCGGATTCAGCGCGTGTATGTAAAGNNNAGATCGGAA |
| 34289_2 | TATGAGTGTGGAGTCGTTGCGGATTCAGCGCGTGTATGTAAAGNNNAGATCGGAA |
| 34289_3 | TATGAGTGTGGAGTCGTTGCGGATTCAGCGCGTGTATGTAAAGNNNAGATCGGAA |
| 34289_4 | TATGAGTGTGGAGTCGTTGCGGATTCAGCGCGTGTATGTAAAGNNNAGATCGGAA |
| 34289_5 | TATGAGTGTGGAGTCGTTGCGGATTCAGCGCGTGTATGTAAAGNNNAGATCGGAA |
| 34289_6 | TATGAGTGTGGAGTCGTTGCGGATTCAGCGCGTGTATGTAAAGNNNAGATCGGAA |
| 34289_7 | TATGAGTGTGGAGTCGTTGCGGATTCAGCGCGTGTATGTAAAGNNNAGATCGGAA |
| 48627_1 | TATGAGTGTGGAGTCGTTGCCTTCCACATCCTCAGGTGTGACNNNAGATCGGAAGA |
| 48627_2 | TATGAGTGTGGAGTCGTTGCCTTCCACATCCTCAGGTGTGACNNNAGATCGGAAGA |
| 48627_3 | TATGAGTGTGGAGTCGTTGCCTTCCACATCCTCAGGTGTGACNNNAGATCGGAAGA |
| 48627_4 | TATGAGTGTGGAGTCGTTGCCTTCCACATCCTCAGGTGTGACNNNAGATCGGAAGA |
| 48627_5 | TATGAGTGTGGAGTCGTTGCCTTCCACATCCTCAGGTGTGACNNNAGATCGGAAGA |
| 48627_6 | TATGAGTGTGGAGTCGTTGCCTTCCACATCCTCAGGTGTGACNNNAGATCGGAAGA |
| 48627_7 | TATGAGTGTGGAGTCGTTGCCTTCCACATCCTCAGGTGTGACNNNAGATCGGAAGA |
| 53128_1 | TATGAGTGTGGAGTCGTTGCGCAGAAAAAAATGTGTATGTGTGTNNNAGATCGGA |
| 53128_2 | TATGAGTGTGGAGTCGTTGCGCAGAAAAAAATGTGTATGTGTGTNNNAGATCGGA |

| | |
|---|---|
| 53128_3 | TATGAGTGTGGAGTCGTTGCGCAGAAAAAAATGTGTATGTGTGTNNNAGATCGGA |
| 53128_4 | TATGAGTGTGGAGTCGTTGCGCAGAAAAAAATGTGTATGTGTGTNNNAGATCGGA |
| 53128_5 | TATGAGTGTGGAGTCGTTGCGCAGAAAAAAATGTGTATGTGTGTNNNAGATCGGA |
| 53128_6 | TATGAGTGTGGAGTCGTTGCGCAGAAAAAAATGTGTATGTGTGTNNNAGATCGGA |
| 53128_7 | TATGAGTGTGGAGTCGTTGCGCAGAAAAAAATGTGTATGTGTGTNNNAGATCGGA |
| 40790_1 | TATGAGTGTGGAGTCGTTGCTGAGATGCAGTGTTAGAAATTCTGANNNAGATCGG |
| 40790_2 | TATGAGTGTGGAGTCGTTGCTGAGATGCAGTGTTAGAAATTCTGANNNAGATCGG |
| 40790_3 | TATGAGTGTGGAGTCGTTGCTGAGATGCAGTGTTAGAAATTCTGANNNAGATCGG |
| 40790_4 | TATGAGTGTGGAGTCGTTGCTGAGATGCAGTGTTAGAAATTCTGANNNAGATCGG |
| 40790_5 | TATGAGTGTGGAGTCGTTGCTGAGATGCAGTGTTAGAAATTCTGANNNAGATCGG |
| 40790_6 | TATGAGTGTGGAGTCGTTGCTGAGATGCAGTGTTAGAAATTCTGANNNAGATCGG |
| 40790_7 | TATGAGTGTGGAGTCGTTGCTGAGATGCAGTGTTAGAAATTCTGANNNAGATCGG |
| 32467_1 | TATGAGTGTGGAGTCGTTGCGGGTGTAGTTTGGAAAATGTTTGCNNNAGATCGGA |
| 32467_2 | TATGAGTGTGGAGTCGTTGCGGGTGTAGTTTGGAAAATGTTTGCNNNAGATCGGA |
| 32467_3 | TATGAGTGTGGAGTCGTTGCGGGTGTAGTTTGGAAAATGTTTGCNNNAGATCGGA |
| 32467_4 | TATGAGTGTGGAGTCGTTGCGGGTGTAGTTTGGAAAATGTTTGCNNNAGATCGGA |
| 32467_5 | TATGAGTGTGGAGTCGTTGCGGGTGTAGTTTGGAAAATGTTTGCNNNAGATCGGA |
| 32467_6 | TATGAGTGTGGAGTCGTTGCGGGTGTAGTTTGGAAAATGTTTGCNNNAGATCGGA |
| 32467_7 | TATGAGTGTGGAGTCGTTGCGGGTGTAGTTTGGAAAATGTTTGCNNNAGATCGGA |
| 48666_1 | TATGAGTGTGGAGTCGTTGCCAAGGAACCCCAAGTATCCACGNNNAGATCGGAAG |
| 48666_2 | TATGAGTGTGGAGTCGTTGCCAAGGAACCCCAAGTATCCACGNNNAGATCGGAAG |
| 48666_3 | TATGAGTGTGGAGTCGTTGCCAAGGAACCCCAAGTATCCACGNNNAGATCGGAAG |
| 48666_4 | TATGAGTGTGGAGTCGTTGCCAAGGAACCCCAAGTATCCACGNNNAGATCGGAAG |
| 48666_5 | TATGAGTGTGGAGTCGTTGCCAAGGAACCCCAAGTATCCACGNNNAGATCGGAAG |
| 48666_6 | TATGAGTGTGGAGTCGTTGCCAAGGAACCCCAAGTATCCACGNNNAGATCGGAAG |
| 48666_7 | TATGAGTGTGGAGTCGTTGCCAAGGAACCCCAAGTATCCACGNNNAGATCGGAAG |
| 53294_1 | TATGAGTGTGGAGTCGTTGCCTGATCCAGGAGGCCGCATCNNNAGATCGGAAGAG |
| 53294_2 | TATGAGTGTGGAGTCGTTGCCTGATCCAGGAGGCCGCATCNNNAGATCGGAAGAG |
| 53294_3 | TATGAGTGTGGAGTCGTTGCCTGATCCAGGAGGCCGCATCNNNAGATCGGAAGAG |
| 53294_4 | TATGAGTGTGGAGTCGTTGCCTGATCCAGGAGGCCGCATCNNNAGATCGGAAGAG |
| 53294_5 | TATGAGTGTGGAGTCGTTGCCTGATCCAGGAGGCCGCATCNNNAGATCGGAAGAG |
| 53294_6 | TATGAGTGTGGAGTCGTTGCCTGATCCAGGAGGCCGCATCNNNAGATCGGAAGAG |
| 53294_7 | TATGAGTGTGGAGTCGTTGCCTGATCCAGGAGGCCGCATCNNNAGATCGGAAGAG |
| 33437_1 | TATGAGTGTGGAGTCGTTGCCACACACACACACACAGTATGNNNAGATCGGAAG |
| 33437_2 | TATGAGTGTGGAGTCGTTGCCACACACACACACACAGTATGNNNAGATCGGAAG |
| 33437_3 | TATGAGTGTGGAGTCGTTGCCACACACACACACACAGTATGNNNAGATCGGAAG |
| 33437_4 | TATGAGTGTGGAGTCGTTGCCACACACACACACACAGTATGNNNAGATCGGAAG |
| 33437_5 | TATGAGTGTGGAGTCGTTGCCACACACACACACACAGTATGNNNAGATCGGAAG |
| 33437_6 | TATGAGTGTGGAGTCGTTGCCACACACACACACACAGTATGNNNAGATCGGAAG |
| 33437_7 | TATGAGTGTGGAGTCGTTGCCACACACACACACACAGTATGNNNAGATCGGAAG |
| 44330_1 | TATGAGTGTGGAGTCGTTGCGATGATGTGATGCAGGTTGTACCNNNAGATCGGAA |
| 44330_2 | TATGAGTGTGGAGTCGTTGCGATGATGTGATGCAGGTTGTACCNNNAGATCGGAA |
| 44330_3 | TATGAGTGTGGAGTCGTTGCGATGATGTGATGCAGGTTGTACCNNNAGATCGGAA |

| | |
|---|---|
| 44330_4 | TATGAGTGTGGAGTCGTTGCGATGATGTGATGCAGGTTGTACCNNNAGATCGGAA |
| 44330_5 | TATGAGTGTGGAGTCGTTGCGATGATGTGATGCAGGTTGTACCNNNAGATCGGAA |
| 44330_6 | TATGAGTGTGGAGTCGTTGCGATGATGTGATGCAGGTTGTACCNNNAGATCGGAA |
| 44330_7 | TATGAGTGTGGAGTCGTTGCGATGATGTGATGCAGGTTGTACCNNNAGATCGGAA |
| 52568_1 | TATGAGTGTGGAGTCGTTGCTTGAAGGAGCAGGATTCAACCTTNNNAGATCGGAA |
| 52568_2 | TATGAGTGTGGAGTCGTTGCTTGAAGGAGCAGGATTCAACCTTNNNAGATCGGAA |
| 52568_3 | TATGAGTGTGGAGTCGTTGCTTGAAGGAGCAGGATTCAACCTTNNNAGATCGGAA |
| 52568_4 | TATGAGTGTGGAGTCGTTGCTTGAAGGAGCAGGATTCAACCTTNNNAGATCGGAA |
| 52568_5 | TATGAGTGTGGAGTCGTTGCTTGAAGGAGCAGGATTCAACCTTNNNAGATCGGAA |
| 52568_6 | TATGAGTGTGGAGTCGTTGCTTGAAGGAGCAGGATTCAACCTTNNNAGATCGGAA |
| 52568_7 | TATGAGTGTGGAGTCGTTGCTTGAAGGAGCAGGATTCAACCTTNNNAGATCGGAA |
| 42187_1 | TATGAGTGTGGAGTCGTTGCCACAACTACGCCCAGCTAATTTTNNNAGATCGGAAG |
| 42187_2 | TATGAGTGTGGAGTCGTTGCCACAACTACGCCCAGCTAATTTTNNNAGATCGGAAG |
| 42187_3 | TATGAGTGTGGAGTCGTTGCCACAACTACGCCCAGCTAATTTTNNNAGATCGGAAG |
| 42187_4 | TATGAGTGTGGAGTCGTTGCCACAACTACGCCCAGCTAATTTTNNNAGATCGGAAG |
| 42187_5 | TATGAGTGTGGAGTCGTTGCCACAACTACGCCCAGCTAATTTTNNNAGATCGGAAG |
| 42187_6 | TATGAGTGTGGAGTCGTTGCCACAACTACGCCCAGCTAATTTTNNNAGATCGGAAG |
| 42187_7 | TATGAGTGTGGAGTCGTTGCCACAACTACGCCCAGCTAATTTTNNNAGATCGGAAG |
| 13369_1 | TATGAGTGTGGAGTCGTTGCTAAAACATATTCTTGTCTCTGTCNNNAGATCGGAAG |
| 13369_2 | TATGAGTGTGGAGTCGTTGCTAAAACATATTCTTGTCTCTGTCNNNAGATCGGAAG |
| 13369_3 | TATGAGTGTGGAGTCGTTGCTAAAACATATTCTTGTCTCTGTCNNNAGATCGGAAG |
| 13369_4 | TATGAGTGTGGAGTCGTTGCTAAAACATATTCTTGTCTCTGTCNNNAGATCGGAAG |
| 13369_5 | TATGAGTGTGGAGTCGTTGCTAAAACATATTCTTGTCTCTGTCNNNAGATCGGAAG |
| 13369_6 | TATGAGTGTGGAGTCGTTGCTAAAACATATTCTTGTCTCTGTCNNNAGATCGGAAG |
| 13369_7 | TATGAGTGTGGAGTCGTTGCTAAAACATATTCTTGTCTCTGTCNNNAGATCGGAAG |
| 29310_1 | TATGAGTGTGGAGTCGTTGCGTCTCAAGCTCCCTGGAAATACANNNAGATCGGAAG |
| 29310_2 | TATGAGTGTGGAGTCGTTGCGTCTCAAGCTCCCTGGAAATACANNNAGATCGGAAG |
| 29310_3 | TATGAGTGTGGAGTCGTTGCGTCTCAAGCTCCCTGGAAATACANNNAGATCGGAAG |
| 29310_4 | TATGAGTGTGGAGTCGTTGCGTCTCAAGCTCCCTGGAAATACANNNAGATCGGAAG |
| 29310_5 | TATGAGTGTGGAGTCGTTGCGTCTCAAGCTCCCTGGAAATACANNNAGATCGGAAG |
| 29310_6 | TATGAGTGTGGAGTCGTTGCGTCTCAAGCTCCCTGGAAATACANNNAGATCGGAAG |
| 29310_7 | TATGAGTGTGGAGTCGTTGCGTCTCAAGCTCCCTGGAAATACANNNAGATCGGAAG |
| 36653_1 | TATGAGTGTGGAGTCGTTGCTCCAGGGCTTTCTTTTCAAAACCNNNAGATCGGAAG |
| 36653_2 | TATGAGTGTGGAGTCGTTGCTCCAGGGCTTTCTTTTCAAAACCNNNAGATCGGAAG |
| 36653_3 | TATGAGTGTGGAGTCGTTGCTCCAGGGCTTTCTTTTCAAAACCNNNAGATCGGAAG |
| 36653_4 | TATGAGTGTGGAGTCGTTGCTCCAGGGCTTTCTTTTCAAAACCNNNAGATCGGAAG |
| 36653_5 | TATGAGTGTGGAGTCGTTGCTCCAGGGCTTTCTTTTCAAAACCNNNAGATCGGAAG |
| 36653_6 | TATGAGTGTGGAGTCGTTGCTCCAGGGCTTTCTTTTCAAAACCNNNAGATCGGAAG |
| 36653_7 | TATGAGTGTGGAGTCGTTGCTCCAGGGCTTTCTTTTCAAAACCNNNAGATCGGAAG |
| 40756_1 | TATGAGTGTGGAGTCGTTGCAAAGTAGACTAGTGGTTGCCAGGNNNAGATCGGAA |
| 40756_2 | TATGAGTGTGGAGTCGTTGCAAAGTAGACTAGTGGTTGCCAGGNNNAGATCGGAA |
| 40756_3 | TATGAGTGTGGAGTCGTTGCAAAGTAGACTAGTGGTTGCCAGGNNNAGATCGGAA |
| 40756_4 | TATGAGTGTGGAGTCGTTGCAAAGTAGACTAGTGGTTGCCAGGNNNAGATCGGAA |

| | |
|---|---|
| 40756_5 | TATGAGTGTGGAGTCGTTGCAAAGTAGACTAGTGGTTGCCAGGNNNAGATCGGAA |
| 40756_6 | TATGAGTGTGGAGTCGTTGCAAAGTAGACTAGTGGTTGCCAGGNNNAGATCGGAA |
| 40756_7 | TATGAGTGTGGAGTCGTTGCAAAGTAGACTAGTGGTTGCCAGGNNNAGATCGGAA |
| 33716_1 | TATGAGTGTGGAGTCGTTGCATTGAGGTGGTCATGAGGAGAAGNNNAGATCGGAA |
| 33716_2 | TATGAGTGTGGAGTCGTTGCATTGAGGTGGTCATGAGGAGAAGNNNAGATCGGAA |
| 33716_3 | TATGAGTGTGGAGTCGTTGCATTGAGGTGGTCATGAGGAGAAGNNNAGATCGGAA |
| 33716_4 | TATGAGTGTGGAGTCGTTGCATTGAGGTGGTCATGAGGAGAAGNNNAGATCGGAA |
| 33716_5 | TATGAGTGTGGAGTCGTTGCATTGAGGTGGTCATGAGGAGAAGNNNAGATCGGAA |
| 33716_6 | TATGAGTGTGGAGTCGTTGCATTGAGGTGGTCATGAGGAGAAGNNNAGATCGGAA |
| 33716_7 | TATGAGTGTGGAGTCGTTGCATTGAGGTGGTCATGAGGAGAAGNNNAGATCGGAA |
| 36690_1 | TATGAGTGTGGAGTCGTTGCACTGATTCCTAGTCTTGCCAAGGNNNAGATCGGAAG |
| 36690_2 | TATGAGTGTGGAGTCGTTGCACTGATTCCTAGTCTTGCCAAGGNNNAGATCGGAAG |
| 36690_3 | TATGAGTGTGGAGTCGTTGCACTGATTCCTAGTCTTGCCAAGGNNNAGATCGGAAG |
| 36690_4 | TATGAGTGTGGAGTCGTTGCACTGATTCCTAGTCTTGCCAAGGNNNAGATCGGAAG |
| 36690_5 | TATGAGTGTGGAGTCGTTGCACTGATTCCTAGTCTTGCCAAGGNNNAGATCGGAAG |
| 36690_6 | TATGAGTGTGGAGTCGTTGCACTGATTCCTAGTCTTGCCAAGGNNNAGATCGGAAG |
| 36690_7 | TATGAGTGTGGAGTCGTTGCACTGATTCCTAGTCTTGCCAAGGNNNAGATCGGAAG |
| 30571_1 | TATGAGTGTGGAGTCGTTGCATAGATTTGCTTTTCCCAAGTTGCCNNNAGATCGGAA |
| 30571_2 | TATGAGTGTGGAGTCGTTGCATAGATTTGCTTTTCCCAAGTTGCCNNNAGATCGGAA |
| 30571_3 | TATGAGTGTGGAGTCGTTGCATAGATTTGCTTTTCCCAAGTTGCCNNNAGATCGGAA |
| 30571_4 | TATGAGTGTGGAGTCGTTGCATAGATTTGCTTTTCCCAAGTTGCCNNNAGATCGGAA |
| 30571_5 | TATGAGTGTGGAGTCGTTGCATAGATTTGCTTTTCCCAAGTTGCCNNNAGATCGGAA |
| 30571_6 | TATGAGTGTGGAGTCGTTGCATAGATTTGCTTTTCCCAAGTTGCCNNNAGATCGGAA |
| 30571_7 | TATGAGTGTGGAGTCGTTGCATAGATTTGCTTTTCCCAAGTTGCCNNNAGATCGGAA |
| 38171_1 | TATGAGTGTGGAGTCGTTGCAAGTGCTCAATGGATGTAAGCTTNNNAGATCGGAA |
| 38171_2 | TATGAGTGTGGAGTCGTTGCAAGTGCTCAATGGATGTAAGCTTNNNAGATCGGAA |
| 38171_3 | TATGAGTGTGGAGTCGTTGCAAGTGCTCAATGGATGTAAGCTTNNNAGATCGGAA |
| 38171_4 | TATGAGTGTGGAGTCGTTGCAAGTGCTCAATGGATGTAAGCTTNNNAGATCGGAA |
| 38171_5 | TATGAGTGTGGAGTCGTTGCAAGTGCTCAATGGATGTAAGCTTNNNAGATCGGAA |
| 38171_6 | TATGAGTGTGGAGTCGTTGCAAGTGCTCAATGGATGTAAGCTTNNNAGATCGGAA |
| 38171_7 | TATGAGTGTGGAGTCGTTGCAAGTGCTCAATGGATGTAAGCTTNNNAGATCGGAA |
| 30485_1 | ATGAGTGTGGAGTCGTTGCGCTACTTTTGTCTTTTAGGTACAAAATNNNAGATCGGA |
| 30485_2 | ATGAGTGTGGAGTCGTTGCGCTACTTTTGTCTTTTAGGTACAAAATNNNAGATCGGA |
| 30485_3 | ATGAGTGTGGAGTCGTTGCGCTACTTTTGTCTTTTAGGTACAAAATNNNAGATCGGA |
| 30485_4 | ATGAGTGTGGAGTCGTTGCGCTACTTTTGTCTTTTAGGTACAAAATNNNAGATCGGA |
| 30485_5 | ATGAGTGTGGAGTCGTTGCGCTACTTTTGTCTTTTAGGTACAAAATNNNAGATCGGA |
| 30485_6 | ATGAGTGTGGAGTCGTTGCGCTACTTTTGTCTTTTAGGTACAAAATNNNAGATCGGA |
| 30485_7 | ATGAGTGTGGAGTCGTTGCGCTACTTTTGTCTTTTAGGTACAAAATNNNAGATCGGA |
| 40462_1 | TATGAGTGTGGAGTCGTTGCCTTACGCCTGTAATCCCAGCANNNAGATCGGAAGAG |
| 40462_2 | TATGAGTGTGGAGTCGTTGCCTTACGCCTGTAATCCCAGCANNNAGATCGGAAGAG |
| 40462_3 | TATGAGTGTGGAGTCGTTGCCTTACGCCTGTAATCCCAGCANNNAGATCGGAAGAG |
| 40462_4 | TATGAGTGTGGAGTCGTTGCCTTACGCCTGTAATCCCAGCANNNAGATCGGAAGAG |
| 40462_5 | TATGAGTGTGGAGTCGTTGCCTTACGCCTGTAATCCCAGCANNNAGATCGGAAGAG |

| | |
|---|---|
| 40462_6 | TATGAGTGTGGAGTCGTTGCCTTACGCCTGTAATCCCAGCANNNAGATCGGAAGAG |
| 40462_7 | TATGAGTGTGGAGTCGTTGCCTTACGCCTGTAATCCCAGCANNNAGATCGGAAGAG |
| 32494_1 | TATGAGTGTGGAGTCGTTGCTTCAATCGATTCTCCTGCCTCAGNNNAGATCGGAAGA |
| 32494_2 | TATGAGTGTGGAGTCGTTGCTTCAATCGATTCTCCTGCCTCAGNNNAGATCGGAAGA |
| 32494_3 | TATGAGTGTGGAGTCGTTGCTTCAATCGATTCTCCTGCCTCAGNNNAGATCGGAAGA |
| 32494_4 | TATGAGTGTGGAGTCGTTGCTTCAATCGATTCTCCTGCCTCAGNNNAGATCGGAAGA |
| 32494_5 | TATGAGTGTGGAGTCGTTGCTTCAATCGATTCTCCTGCCTCAGNNNAGATCGGAAGA |
| 32494_6 | TATGAGTGTGGAGTCGTTGCTTCAATCGATTCTCCTGCCTCAGNNNAGATCGGAAGA |
| 32494_7 | TATGAGTGTGGAGTCGTTGCTTCAATCGATTCTCCTGCCTCAGNNNAGATCGGAAGA |
| 28931_1 | TATGAGTGTGGAGTCGTTGCGAGAGAGAGAGAGACAGACCTCCNNNAGATCGGAA |
| 28931_2 | TATGAGTGTGGAGTCGTTGCGAGAGAGAGAGAGACAGACCTCCNNNAGATCGGAA |
| 28931_3 | TATGAGTGTGGAGTCGTTGCGAGAGAGAGAGAGACAGACCTCCNNNAGATCGGAA |
| 28931_4 | TATGAGTGTGGAGTCGTTGCGAGAGAGAGAGAGACAGACCTCCNNNAGATCGGAA |
| 28931_5 | TATGAGTGTGGAGTCGTTGCGAGAGAGAGAGAGACAGACCTCCNNNAGATCGGAA |
| 28931_6 | TATGAGTGTGGAGTCGTTGCGAGAGAGAGAGAGACAGACCTCCNNNAGATCGGAA |
| 28931_7 | TATGAGTGTGGAGTCGTTGCGAGAGAGAGAGAGACAGACCTCCNNNAGATCGGAA |
| 32150_1 | TATGAGTGTGGAGTCGTTGCTGACTTTTCCTTTTGCTGTGCAANNNAGATCGGAAGA |
| 32150_2 | TATGAGTGTGGAGTCGTTGCTGACTTTTCCTTTTGCTGTGCAANNNAGATCGGAAGA |
| 32150_3 | TATGAGTGTGGAGTCGTTGCTGACTTTTCCTTTTGCTGTGCAANNNAGATCGGAAGA |
| 32150_4 | TATGAGTGTGGAGTCGTTGCTGACTTTTCCTTTTGCTGTGCAANNNAGATCGGAAGA |
| 32150_5 | TATGAGTGTGGAGTCGTTGCTGACTTTTCCTTTTGCTGTGCAANNNAGATCGGAAGA |
| 32150_6 | TATGAGTGTGGAGTCGTTGCTGACTTTTCCTTTTGCTGTGCAANNNAGATCGGAAGA |
| 32150_7 | TATGAGTGTGGAGTCGTTGCTGACTTTTCCTTTTGCTGTGCAANNNAGATCGGAAGA |
| 44763_1 | TATGAGTGTGGAGTCGTTGCAGTTGTGTGACTGGACAATCTGTNNNAGATCGGAAG |
| 44763_2 | TATGAGTGTGGAGTCGTTGCAGTTGTGTGACTGGACAATCTGTNNNAGATCGGAAG |
| 44763_3 | TATGAGTGTGGAGTCGTTGCAGTTGTGTGACTGGACAATCTGTNNNAGATCGGAAG |
| 44763_4 | TATGAGTGTGGAGTCGTTGCAGTTGTGTGACTGGACAATCTGTNNNAGATCGGAAG |
| 44763_5 | TATGAGTGTGGAGTCGTTGCAGTTGTGTGACTGGACAATCTGTNNNAGATCGGAAG |
| 44763_6 | TATGAGTGTGGAGTCGTTGCAGTTGTGTGACTGGACAATCTGTNNNAGATCGGAAG |
| 44763_7 | TATGAGTGTGGAGTCGTTGCAGTTGTGTGACTGGACAATCTGTNNNAGATCGGAAG |
| 45455_1 | TATGAGTGTGGAGTCGTTGCGAGGAAAGGGCATTCTGGGAATANNNAGATCGGAA |
| 45455_2 | TATGAGTGTGGAGTCGTTGCGAGGAAAGGGCATTCTGGGAATANNNAGATCGGAA |
| 45455_3 | TATGAGTGTGGAGTCGTTGCGAGGAAAGGGCATTCTGGGAATANNNAGATCGGAA |
| 45455_4 | TATGAGTGTGGAGTCGTTGCGAGGAAAGGGCATTCTGGGAATANNNAGATCGGAA |
| 45455_5 | TATGAGTGTGGAGTCGTTGCGAGGAAAGGGCATTCTGGGAATANNNAGATCGGAA |
| 45455_6 | TATGAGTGTGGAGTCGTTGCGAGGAAAGGGCATTCTGGGAATANNNAGATCGGAA |
| 45455_7 | TATGAGTGTGGAGTCGTTGCGAGGAAAGGGCATTCTGGGAATANNNAGATCGGAA |
| 13038_1 | TATGAGTGTGGAGTCGTTGCGGTCTGTCTTAACATTCTTATTGNNNAGATCGGAAGA |
| 13038_2 | TATGAGTGTGGAGTCGTTGCGGTCTGTCTTAACATTCTTATTGNNNAGATCGGAAGA |
| 13038_3 | TATGAGTGTGGAGTCGTTGCGGTCTGTCTTAACATTCTTATTGNNNAGATCGGAAGA |
| 13038_4 | TATGAGTGTGGAGTCGTTGCGGTCTGTCTTAACATTCTTATTGNNNAGATCGGAAGA |
| 13038_5 | TATGAGTGTGGAGTCGTTGCGGTCTGTCTTAACATTCTTATTGNNNAGATCGGAAGA |
| 13038_6 | TATGAGTGTGGAGTCGTTGCGGTCTGTCTTAACATTCTTATTGNNNAGATCGGAAGA |

| | |
|---|---|
| 13038_7 | TATGAGTGTGGAGTCGTTGCGGTCTGTCTTAACATTCTTATTGNNNAGATCGGAAG |
| 42016_1 | TATGAGTGTGGAGTCGTTGCCTGAGTAGCTGGGATTAGCAGATNNNAGATCGGAA |
| 42016_2 | TATGAGTGTGGAGTCGTTGCCTGAGTAGCTGGGATTAGCAGATNNNAGATCGGAA |
| 42016_3 | TATGAGTGTGGAGTCGTTGCCTGAGTAGCTGGGATTAGCAGATNNNAGATCGGAA |
| 42016_4 | TATGAGTGTGGAGTCGTTGCCTGAGTAGCTGGGATTAGCAGATNNNAGATCGGAA |
| 42016_5 | TATGAGTGTGGAGTCGTTGCCTGAGTAGCTGGGATTAGCAGATNNNAGATCGGAA |
| 42016_6 | TATGAGTGTGGAGTCGTTGCCTGAGTAGCTGGGATTAGCAGATNNNAGATCGGAA |
| 42016_7 | TATGAGTGTGGAGTCGTTGCCTGAGTAGCTGGGATTAGCAGATNNNAGATCGGAA |
| 38144_1 | TATGAGTGTGGAGTCGTTGCAGGGCCCTGCTTTCTCAAATTTANNNAGATCGGAAG |
| 38144_2 | TATGAGTGTGGAGTCGTTGCAGGGCCCTGCTTTCTCAAATTTANNNAGATCGGAAG |
| 38144_3 | TATGAGTGTGGAGTCGTTGCAGGGCCCTGCTTTCTCAAATTTANNNAGATCGGAAG |
| 38144_4 | TATGAGTGTGGAGTCGTTGCAGGGCCCTGCTTTCTCAAATTTANNNAGATCGGAAG |
| 38144_5 | TATGAGTGTGGAGTCGTTGCAGGGCCCTGCTTTCTCAAATTTANNNAGATCGGAAG |
| 38144_6 | TATGAGTGTGGAGTCGTTGCAGGGCCCTGCTTTCTCAAATTTANNNAGATCGGAAG |
| 38144_7 | TATGAGTGTGGAGTCGTTGCAGGGCCCTGCTTTCTCAAATTTANNNAGATCGGAAG |
| 51133_1 | TATGAGTGTGGAGTCGTTGCCCAGCACACTACAACTCCAGTATNNNAGATCGGAAG |
| 51133_2 | TATGAGTGTGGAGTCGTTGCCCAGCACACTACAACTCCAGTATNNNAGATCGGAAG |
| 51133_3 | TATGAGTGTGGAGTCGTTGCCCAGCACACTACAACTCCAGTATNNNAGATCGGAAG |
| 51133_4 | TATGAGTGTGGAGTCGTTGCCCAGCACACTACAACTCCAGTATNNNAGATCGGAAG |
| 51133_5 | TATGAGTGTGGAGTCGTTGCCCAGCACACTACAACTCCAGTATNNNAGATCGGAAG |
| 51133_6 | TATGAGTGTGGAGTCGTTGCCCAGCACACTACAACTCCAGTATNNNAGATCGGAAG |
| 51133_7 | TATGAGTGTGGAGTCGTTGCCCAGCACACTACAACTCCAGTATNNNAGATCGGAAG |
| 28975_1 | TATGAGTGTGGAGTCGTTGCTTGGCACAAAAGTAATTGCGGTTNNNAGATCGGAA |
| 28975_2 | TATGAGTGTGGAGTCGTTGCTTGGCACAAAAGTAATTGCGGTTNNNAGATCGGAA |
| 28975_3 | TATGAGTGTGGAGTCGTTGCTTGGCACAAAAGTAATTGCGGTTNNNAGATCGGAA |
| 28975_4 | TATGAGTGTGGAGTCGTTGCTTGGCACAAAAGTAATTGCGGTTNNNAGATCGGAA |
| 28975_5 | TATGAGTGTGGAGTCGTTGCTTGGCACAAAAGTAATTGCGGTTNNNAGATCGGAA |
| 28975_6 | TATGAGTGTGGAGTCGTTGCTTGGCACAAAAGTAATTGCGGTTNNNAGATCGGAA |
| 28975_7 | TATGAGTGTGGAGTCGTTGCTTGGCACAAAAGTAATTGCGGTTNNNAGATCGGAA |
| 41025_1 | TATGAGTGTGGAGTCGTTGCGAACTTGGAGTCCGATGTTCAAGNNNAGATCGGAA |
| 41025_2 | TATGAGTGTGGAGTCGTTGCGAACTTGGAGTCCGATGTTCAAGNNNAGATCGGAA |
| 41025_3 | TATGAGTGTGGAGTCGTTGCGAACTTGGAGTCCGATGTTCAAGNNNAGATCGGAA |
| 41025_4 | TATGAGTGTGGAGTCGTTGCGAACTTGGAGTCCGATGTTCAAGNNNAGATCGGAA |
| 41025_5 | TATGAGTGTGGAGTCGTTGCGAACTTGGAGTCCGATGTTCAAGNNNAGATCGGAA |
| 41025_6 | TATGAGTGTGGAGTCGTTGCGAACTTGGAGTCCGATGTTCAAGNNNAGATCGGAA |
| 41025_7 | TATGAGTGTGGAGTCGTTGCGAACTTGGAGTCCGATGTTCAAGNNNAGATCGGAA |
| 48606_1 | TATGAGTGTGGAGTCGTTGCTGAGACAGAGGAATGGCATTGAANNNAGATCGGAA |
| 48606_2 | TATGAGTGTGGAGTCGTTGCTGAGACAGAGGAATGGCATTGAANNNAGATCGGAA |
| 48606_3 | TATGAGTGTGGAGTCGTTGCTGAGACAGAGGAATGGCATTGAANNNAGATCGGAA |
| 48606_4 | TATGAGTGTGGAGTCGTTGCTGAGACAGAGGAATGGCATTGAANNNAGATCGGAA |
| 48606_5 | TATGAGTGTGGAGTCGTTGCTGAGACAGAGGAATGGCATTGAANNNAGATCGGAA |
| 48606_6 | TATGAGTGTGGAGTCGTTGCTGAGACAGAGGAATGGCATTGAANNNAGATCGGAA |
| 48606_7 | TATGAGTGTGGAGTCGTTGCTGAGACAGAGGAATGGCATTGAANNNAGATCGGAA |

| | |
|---|---|
| 38957_1 | TATGAGTGTGGAGTCGTTGCTGAGGACATGAGATTTTGGAGGGNNNAGATCGGAA |
| 38957_2 | TATGAGTGTGGAGTCGTTGCTGAGGACATGAGATTTTGGAGGGNNNAGATCGGAA |
| 38957_3 | TATGAGTGTGGAGTCGTTGCTGAGGACATGAGATTTTGGAGGGNNNAGATCGGAA |
| 38957_4 | TATGAGTGTGGAGTCGTTGCTGAGGACATGAGATTTTGGAGGGNNNAGATCGGAA |
| 38957_5 | TATGAGTGTGGAGTCGTTGCTGAGGACATGAGATTTTGGAGGGNNNAGATCGGAA |
| 38957_6 | TATGAGTGTGGAGTCGTTGCTGAGGACATGAGATTTTGGAGGGNNNAGATCGGAA |
| 38957_7 | TATGAGTGTGGAGTCGTTGCTGAGGACATGAGATTTTGGAGGGNNNAGATCGGAA |
| 45571_1 | TATGAGTGTGGAGTCGTTGCTTCAAGTGATTCTCCTGCCTCAGNNNAGATCGGAAGA |
| 45571_2 | TATGAGTGTGGAGTCGTTGCTTCAAGTGATTCTCCTGCCTCAGNNNAGATCGGAAGA |
| 45571_3 | TATGAGTGTGGAGTCGTTGCTTCAAGTGATTCTCCTGCCTCAGNNNAGATCGGAAGA |
| 45571_4 | TATGAGTGTGGAGTCGTTGCTTCAAGTGATTCTCCTGCCTCAGNNNAGATCGGAAGA |
| 45571_5 | TATGAGTGTGGAGTCGTTGCTTCAAGTGATTCTCCTGCCTCAGNNNAGATCGGAAGA |
| 45571_6 | TATGAGTGTGGAGTCGTTGCTTCAAGTGATTCTCCTGCCTCAGNNNAGATCGGAAGA |
| 45571_7 | TATGAGTGTGGAGTCGTTGCTTCAAGTGATTCTCCTGCCTCAGNNNAGATCGGAAGA |
| 41601_1 | TATGAGTGTGGAGTCGTTGCGGGTTCACGCCATTCTCCTGNNNAGATCGGAAGAGC |
| 41601_2 | TATGAGTGTGGAGTCGTTGCGGGTTCACGCCATTCTCCTGNNNAGATCGGAAGAGC |
| 41601_3 | TATGAGTGTGGAGTCGTTGCGGGTTCACGCCATTCTCCTGNNNAGATCGGAAGAGC |
| 41601_4 | TATGAGTGTGGAGTCGTTGCGGGTTCACGCCATTCTCCTGNNNAGATCGGAAGAGC |
| 41601_5 | TATGAGTGTGGAGTCGTTGCGGGTTCACGCCATTCTCCTGNNNAGATCGGAAGAGC |
| 41601_6 | TATGAGTGTGGAGTCGTTGCGGGTTCACGCCATTCTCCTGNNNAGATCGGAAGAGC |
| 41601_7 | TATGAGTGTGGAGTCGTTGCGGGTTCACGCCATTCTCCTGNNNAGATCGGAAGAGC |
| 35761_1 | TATGAGTGTGGAGTCGTTGCACAGAAAGAGGGGAAAACCTGTTNNNAGATCGGAA |
| 35761_2 | TATGAGTGTGGAGTCGTTGCACAGAAAGAGGGGAAAACCTGTTNNNAGATCGGAA |
| 35761_3 | TATGAGTGTGGAGTCGTTGCACAGAAAGAGGGGAAAACCTGTTNNNAGATCGGAA |
| 35761_4 | TATGAGTGTGGAGTCGTTGCACAGAAAGAGGGGAAAACCTGTTNNNAGATCGGAA |
| 35761_5 | TATGAGTGTGGAGTCGTTGCACAGAAAGAGGGGAAAACCTGTTNNNAGATCGGAA |
| 35761_6 | TATGAGTGTGGAGTCGTTGCACAGAAAGAGGGGAAAACCTGTTNNNAGATCGGAA |
| 35761_7 | TATGAGTGTGGAGTCGTTGCACAGAAAGAGGGGAAAACCTGTTNNNAGATCGGAA |
| 4821_1 | TATGAGTGTGGAGTCGTTGCATCCCCAGGAAGTTTTCTAGTGCNNNAGATCGGAAG |
| 4821_2 | TATGAGTGTGGAGTCGTTGCATCCCCAGGAAGTTTTCTAGTGCNNNAGATCGGAAG |
| 4821_3 | TATGAGTGTGGAGTCGTTGCATCCCCAGGAAGTTTTCTAGTGCNNNAGATCGGAAG |
| 4821_4 | TATGAGTGTGGAGTCGTTGCATCCCCAGGAAGTTTTCTAGTGCNNNAGATCGGAAG |
| 4821_5 | TATGAGTGTGGAGTCGTTGCATCCCCAGGAAGTTTTCTAGTGCNNNAGATCGGAAG |
| 4821_6 | TATGAGTGTGGAGTCGTTGCATCCCCAGGAAGTTTTCTAGTGCNNNAGATCGGAAG |
| 4821_7 | TATGAGTGTGGAGTCGTTGCATCCCCAGGAAGTTTTCTAGTGCNNNAGATCGGAAG |
| 8526_1 | TATGAGTGTGGAGTCGTTGCGCTCTGTCTTTCAGATTGGAGTGGNNNAGATCGGAA |
| 8526_2 | TATGAGTGTGGAGTCGTTGCGCTCTGTCTTTCAGATTGGAGTGGNNNAGATCGGAA |
| 8526_3 | TATGAGTGTGGAGTCGTTGCGCTCTGTCTTTCAGATTGGAGTGGNNNAGATCGGAA |
| 8526_4 | TATGAGTGTGGAGTCGTTGCGCTCTGTCTTTCAGATTGGAGTGGNNNAGATCGGAA |
| 8526_5 | TATGAGTGTGGAGTCGTTGCGCTCTGTCTTTCAGATTGGAGTGGNNNAGATCGGAA |
| 8526_6 | TATGAGTGTGGAGTCGTTGCGCTCTGTCTTTCAGATTGGAGTGGNNNAGATCGGAA |
| 8526_7 | TATGAGTGTGGAGTCGTTGCGCTCTGTCTTTCAGATTGGAGTGGNNNAGATCGGAA |
| 46090_1 | TATGAGTGTGGAGTCGTTGCTCTCCTGGGTTCAAGCAATACTCNNNAGATCGGAAGA |

| | |
|---|---|
| 46090_2 | TATGAGTGTGGAGTCGTTGCTCTCCTGGGTTCAAGCAATACTCNNNAGATCGGAAG |
| 46090_3 | TATGAGTGTGGAGTCGTTGCTCTCCTGGGTTCAAGCAATACTCNNNAGATCGGAAG |
| 46090_4 | TATGAGTGTGGAGTCGTTGCTCTCCTGGGTTCAAGCAATACTCNNNAGATCGGAAG |
| 46090_5 | TATGAGTGTGGAGTCGTTGCTCTCCTGGGTTCAAGCAATACTCNNNAGATCGGAAG |
| 46090_6 | TATGAGTGTGGAGTCGTTGCTCTCCTGGGTTCAAGCAATACTCNNNAGATCGGAAG |
| 46090_7 | TATGAGTGTGGAGTCGTTGCTCTCCTGGGTTCAAGCAATACTCNNNAGATCGGAAG |
| 39020_1 | TATGAGTGTGGAGTCGTTGCCTAGTTCAGCCACTGTGAAAAGCNNNAGATCGGAAG |
| 39020_2 | TATGAGTGTGGAGTCGTTGCCTAGTTCAGCCACTGTGAAAAGCNNNAGATCGGAAG |
| 39020_3 | TATGAGTGTGGAGTCGTTGCCTAGTTCAGCCACTGTGAAAAGCNNNAGATCGGAAG |
| 39020_4 | TATGAGTGTGGAGTCGTTGCCTAGTTCAGCCACTGTGAAAAGCNNNAGATCGGAAG |
| 39020_5 | TATGAGTGTGGAGTCGTTGCCTAGTTCAGCCACTGTGAAAAGCNNNAGATCGGAAG |
| 39020_6 | TATGAGTGTGGAGTCGTTGCCTAGTTCAGCCACTGTGAAAAGCNNNAGATCGGAAG |
| 39020_7 | TATGAGTGTGGAGTCGTTGCCTAGTTCAGCCACTGTGAAAAGCNNNAGATCGGAAG |
| 32683_1 | TATGAGTGTGGAGTCGTTGCTTGGCTCTCAGCTTGGATGTTATNNNAGATCGGAAG |
| 32683_2 | TATGAGTGTGGAGTCGTTGCTTGGCTCTCAGCTTGGATGTTATNNNAGATCGGAAG |
| 32683_3 | TATGAGTGTGGAGTCGTTGCTTGGCTCTCAGCTTGGATGTTATNNNAGATCGGAAG |
| 32683_4 | TATGAGTGTGGAGTCGTTGCTTGGCTCTCAGCTTGGATGTTATNNNAGATCGGAAG |
| 32683_5 | TATGAGTGTGGAGTCGTTGCTTGGCTCTCAGCTTGGATGTTATNNNAGATCGGAAG |
| 32683_6 | TATGAGTGTGGAGTCGTTGCTTGGCTCTCAGCTTGGATGTTATNNNAGATCGGAAG |
| 32683_7 | TATGAGTGTGGAGTCGTTGCTTGGCTCTCAGCTTGGATGTTATNNNAGATCGGAAG |
| 50124_1 | TATGAGTGTGGAGTCGTTGCCCCTCTAGAATCCCGGTGCCNNNAGATCGGAAGAGC |
| 50124_2 | TATGAGTGTGGAGTCGTTGCCCCTCTAGAATCCCGGTGCCNNNAGATCGGAAGAGC |
| 50124_3 | TATGAGTGTGGAGTCGTTGCCCCTCTAGAATCCCGGTGCCNNNAGATCGGAAGAGC |
| 50124_4 | TATGAGTGTGGAGTCGTTGCCCCTCTAGAATCCCGGTGCCNNNAGATCGGAAGAGC |
| 50124_5 | TATGAGTGTGGAGTCGTTGCCCCTCTAGAATCCCGGTGCCNNNAGATCGGAAGAGC |
| 50124_6 | TATGAGTGTGGAGTCGTTGCCCCTCTAGAATCCCGGTGCCNNNAGATCGGAAGAGC |
| 50124_7 | TATGAGTGTGGAGTCGTTGCCCCTCTAGAATCCCGGTGCCNNNAGATCGGAAGAGC |
| 30317_1 | TATGAGTGTGGAGTCGTTGCACGCAAGGAAAGGAGACTCTAGANNNAGATCGGAA |
| 30317_2 | TATGAGTGTGGAGTCGTTGCACGCAAGGAAAGGAGACTCTAGANNNAGATCGGAA |
| 30317_3 | TATGAGTGTGGAGTCGTTGCACGCAAGGAAAGGAGACTCTAGANNNAGATCGGAA |
| 30317_4 | TATGAGTGTGGAGTCGTTGCACGCAAGGAAAGGAGACTCTAGANNNAGATCGGAA |
| 30317_5 | TATGAGTGTGGAGTCGTTGCACGCAAGGAAAGGAGACTCTAGANNNAGATCGGAA |
| 30317_6 | TATGAGTGTGGAGTCGTTGCACGCAAGGAAAGGAGACTCTAGANNNAGATCGGAA |
| 30317_7 | TATGAGTGTGGAGTCGTTGCACGCAAGGAAAGGAGACTCTAGANNNAGATCGGAA |
| 9506_1 | TATGAGTGTGGAGTCGTTGCTGTGTCTGCTCCCCTTTCTTTTCNNNAGATCGGAAGA |
| 9506_2 | TATGAGTGTGGAGTCGTTGCTGTGTCTGCTCCCCTTTCTTTTCNNNAGATCGGAAGA |
| 9506_3 | TATGAGTGTGGAGTCGTTGCTGTGTCTGCTCCCCTTTCTTTTCNNNAGATCGGAAGA |
| 9506_4 | TATGAGTGTGGAGTCGTTGCTGTGTCTGCTCCCCTTTCTTTTCNNNAGATCGGAAGA |
| 9506_5 | TATGAGTGTGGAGTCGTTGCTGTGTCTGCTCCCCTTTCTTTTCNNNAGATCGGAAGA |
| 9506_6 | TATGAGTGTGGAGTCGTTGCTGTGTCTGCTCCCCTTTCTTTTCNNNAGATCGGAAGA |
| 9506_7 | TATGAGTGTGGAGTCGTTGCTGTGTCTGCTCCCCTTTCTTTTCNNNAGATCGGAAGA |
| 37584_1 | TATGAGTGTGGAGTCGTTGCGCAAGGCTTAGCATATATCCAAGTNNNAGATCGGAA |
| 37584_2 | TATGAGTGTGGAGTCGTTGCGCAAGGCTTAGCATATATCCAAGTNNNAGATCGGAA |

| | |
|---|---|
| 37584_3 | TATGAGTGTGGAGTCGTTGCGCAAGGCTTAGCATATATCCAAGTNNNAGATCGGAA |
| 37584_4 | TATGAGTGTGGAGTCGTTGCGCAAGGCTTAGCATATATCCAAGTNNNAGATCGGAA |
| 37584_5 | TATGAGTGTGGAGTCGTTGCGCAAGGCTTAGCATATATCCAAGTNNNAGATCGGAA |
| 37584_6 | TATGAGTGTGGAGTCGTTGCGCAAGGCTTAGCATATATCCAAGTNNNAGATCGGAA |
| 37584_7 | TATGAGTGTGGAGTCGTTGCGCAAGGCTTAGCATATATCCAAGTNNNAGATCGGAA |
| 53325_1 | TATGAGTGTGGAGTCGTTGCTCACACAAAAACAAGAGCAGTGTNNNAGATCGGAA |
| 53325_2 | TATGAGTGTGGAGTCGTTGCTCACACAAAAACAAGAGCAGTGTNNNAGATCGGAA |
| 53325_3 | TATGAGTGTGGAGTCGTTGCTCACACAAAAACAAGAGCAGTGTNNNAGATCGGAA |
| 53325_4 | TATGAGTGTGGAGTCGTTGCTCACACAAAAACAAGAGCAGTGTNNNAGATCGGAA |
| 53325_5 | TATGAGTGTGGAGTCGTTGCTCACACAAAAACAAGAGCAGTGTNNNAGATCGGAA |
| 53325_6 | TATGAGTGTGGAGTCGTTGCTCACACAAAAACAAGAGCAGTGTNNNAGATCGGAA |
| 53325_7 | TATGAGTGTGGAGTCGTTGCTCACACAAAAACAAGAGCAGTGTNNNAGATCGGAA |
| 33575_1 | TATGAGTGTGGAGTCGTTGCATCTCAGCTTACTGCATCCTCTGNNNAGATCGGAAGA |
| 33575_2 | TATGAGTGTGGAGTCGTTGCATCTCAGCTTACTGCATCCTCTGNNNAGATCGGAAGA |
| 33575_3 | TATGAGTGTGGAGTCGTTGCATCTCAGCTTACTGCATCCTCTGNNNAGATCGGAAGA |
| 33575_4 | TATGAGTGTGGAGTCGTTGCATCTCAGCTTACTGCATCCTCTGNNNAGATCGGAAGA |
| 33575_5 | TATGAGTGTGGAGTCGTTGCATCTCAGCTTACTGCATCCTCTGNNNAGATCGGAAGA |
| 33575_6 | TATGAGTGTGGAGTCGTTGCATCTCAGCTTACTGCATCCTCTGNNNAGATCGGAAGA |
| 33575_7 | TATGAGTGTGGAGTCGTTGCATCTCAGCTTACTGCATCCTCTGNNNAGATCGGAAGA |
| 41320_1 | TATGAGTGTGGAGTCGTTGCTCTCAGTTCATCTTTCATCCACTNNNAGATCGGAAGA |
| 41320_2 | TATGAGTGTGGAGTCGTTGCTCTCAGTTCATCTTTCATCCACTNNNAGATCGGAAGA |
| 41320_3 | TATGAGTGTGGAGTCGTTGCTCTCAGTTCATCTTTCATCCACTNNNAGATCGGAAGA |
| 41320_4 | TATGAGTGTGGAGTCGTTGCTCTCAGTTCATCTTTCATCCACTNNNAGATCGGAAGA |
| 41320_5 | TATGAGTGTGGAGTCGTTGCTCTCAGTTCATCTTTCATCCACTNNNAGATCGGAAGA |
| 41320_6 | TATGAGTGTGGAGTCGTTGCTCTCAGTTCATCTTTCATCCACTNNNAGATCGGAAGA |
| 41320_7 | TATGAGTGTGGAGTCGTTGCTCTCAGTTCATCTTTCATCCACTNNNAGATCGGAAGA |
| 50033_1 | TATGAGTGTGGAGTCGTTGCCCTGCACATTATTTGCTCCTGTTNNNAGATCGGAAGA |
| 50033_2 | TATGAGTGTGGAGTCGTTGCCCTGCACATTATTTGCTCCTGTTNNNAGATCGGAAGA |
| 50033_3 | TATGAGTGTGGAGTCGTTGCCCTGCACATTATTTGCTCCTGTTNNNAGATCGGAAGA |
| 50033_4 | TATGAGTGTGGAGTCGTTGCCCTGCACATTATTTGCTCCTGTTNNNAGATCGGAAGA |
| 50033_5 | TATGAGTGTGGAGTCGTTGCCCTGCACATTATTTGCTCCTGTTNNNAGATCGGAAGA |
| 50033_6 | TATGAGTGTGGAGTCGTTGCCCTGCACATTATTTGCTCCTGTTNNNAGATCGGAAGA |
| 50033_7 | TATGAGTGTGGAGTCGTTGCCCTGCACATTATTTGCTCCTGTTNNNAGATCGGAAGA |
| 52508_1 | TATGAGTGTGGAGTCGTTGCCCCACTGTGTACTGAAGTGATCTNNNAGATCGGAAG |
| 52508_2 | TATGAGTGTGGAGTCGTTGCCCCACTGTGTACTGAAGTGATCTNNNAGATCGGAAG |
| 52508_3 | TATGAGTGTGGAGTCGTTGCCCCACTGTGTACTGAAGTGATCTNNNAGATCGGAAG |
| 52508_4 | TATGAGTGTGGAGTCGTTGCCCCACTGTGTACTGAAGTGATCTNNNAGATCGGAAG |
| 52508_5 | TATGAGTGTGGAGTCGTTGCCCCACTGTGTACTGAAGTGATCTNNNAGATCGGAAG |
| 52508_6 | TATGAGTGTGGAGTCGTTGCCCCACTGTGTACTGAAGTGATCTNNNAGATCGGAAG |
| 52508_7 | TATGAGTGTGGAGTCGTTGCCCCACTGTGTACTGAAGTGATCTNNNAGATCGGAAG |
| 43033_1 | TATGAGTGTGGAGTCGTTGCGCATGATCTCGGCTCAATACAACNNNAGATCGGAAG |
| 43033_2 | TATGAGTGTGGAGTCGTTGCGCATGATCTCGGCTCAATACAACNNNAGATCGGAAG |
| 43033_3 | TATGAGTGTGGAGTCGTTGCGCATGATCTCGGCTCAATACAACNNNAGATCGGAAG |

| | |
|---|---|
| 43033_4 | TATGAGTGTGGAGTCGTTGCGCATGATCTCGGCTCAATACAACNNNAGATCGGAAG |
| 43033_5 | TATGAGTGTGGAGTCGTTGCGCATGATCTCGGCTCAATACAACNNNAGATCGGAAG |
| 43033_6 | TATGAGTGTGGAGTCGTTGCGCATGATCTCGGCTCAATACAACNNNAGATCGGAAG |
| 43033_7 | TATGAGTGTGGAGTCGTTGCGCATGATCTCGGCTCAATACAACNNNAGATCGGAAG |
| 29870_1 | TATGAGTGTGGAGTCGTTGCGCAGCTATCAGAGAGGGAAGACNNNAGATCGGAAG |
| 29870_2 | TATGAGTGTGGAGTCGTTGCGCAGCTATCAGAGAGGGAAGACNNNAGATCGGAAG |
| 29870_3 | TATGAGTGTGGAGTCGTTGCGCAGCTATCAGAGAGGGAAGACNNNAGATCGGAAG |
| 29870_4 | TATGAGTGTGGAGTCGTTGCGCAGCTATCAGAGAGGGAAGACNNNAGATCGGAAG |
| 29870_5 | TATGAGTGTGGAGTCGTTGCGCAGCTATCAGAGAGGGAAGACNNNAGATCGGAAG |
| 29870_6 | TATGAGTGTGGAGTCGTTGCGCAGCTATCAGAGAGGGAAGACNNNAGATCGGAAG |
| 29870_7 | TATGAGTGTGGAGTCGTTGCGCAGCTATCAGAGAGGGAAGACNNNAGATCGGAAG |
| 51331_1 | TATGAGTGTGGAGTCGTTGCGTGCACAAGAGAGAGAAAGCAAGNNNAGATCGGAA |
| 51331_2 | TATGAGTGTGGAGTCGTTGCGTGCACAAGAGAGAGAAAGCAAGNNNAGATCGGAA |
| 51331_3 | TATGAGTGTGGAGTCGTTGCGTGCACAAGAGAGAGAAAGCAAGNNNAGATCGGAA |
| 51331_4 | TATGAGTGTGGAGTCGTTGCGTGCACAAGAGAGAGAAAGCAAGNNNAGATCGGAA |
| 51331_5 | TATGAGTGTGGAGTCGTTGCGTGCACAAGAGAGAGAAAGCAAGNNNAGATCGGAA |
| 51331_6 | TATGAGTGTGGAGTCGTTGCGTGCACAAGAGAGAGAAAGCAAGNNNAGATCGGAA |
| 51331_7 | TATGAGTGTGGAGTCGTTGCGTGCACAAGAGAGAGAAAGCAAGNNNAGATCGGAA |
| 52566_1 | TATGAGTGTGGAGTCGTTGCTCCTGGCCTGGATGTTTTGATATNNNAGATCGGAAG |
| 52566_2 | TATGAGTGTGGAGTCGTTGCTCCTGGCCTGGATGTTTTGATATNNNAGATCGGAAG |
| 52566_3 | TATGAGTGTGGAGTCGTTGCTCCTGGCCTGGATGTTTTGATATNNNAGATCGGAAG |
| 52566_4 | TATGAGTGTGGAGTCGTTGCTCCTGGCCTGGATGTTTTGATATNNNAGATCGGAAG |
| 52566_5 | TATGAGTGTGGAGTCGTTGCTCCTGGCCTGGATGTTTTGATATNNNAGATCGGAAG |
| 52566_6 | TATGAGTGTGGAGTCGTTGCTCCTGGCCTGGATGTTTTGATATNNNAGATCGGAAG |
| 52566_7 | TATGAGTGTGGAGTCGTTGCTCCTGGCCTGGATGTTTTGATATNNNAGATCGGAAG |
| 30707_1 | TATGAGTGTGGAGTCGTTGCCTGCAACCTCCATCTTCTAGGTTNNNAGATCGGAAGA |
| 30707_2 | TATGAGTGTGGAGTCGTTGCCTGCAACCTCCATCTTCTAGGTTNNNAGATCGGAAGA |
| 30707_3 | TATGAGTGTGGAGTCGTTGCCTGCAACCTCCATCTTCTAGGTTNNNAGATCGGAAGA |
| 30707_4 | TATGAGTGTGGAGTCGTTGCCTGCAACCTCCATCTTCTAGGTTNNNAGATCGGAAGA |
| 30707_5 | TATGAGTGTGGAGTCGTTGCCTGCAACCTCCATCTTCTAGGTTNNNAGATCGGAAGA |
| 30707_6 | TATGAGTGTGGAGTCGTTGCCTGCAACCTCCATCTTCTAGGTTNNNAGATCGGAAGA |
| 30707_7 | TATGAGTGTGGAGTCGTTGCCTGCAACCTCCATCTTCTAGGTTNNNAGATCGGAAGA |
| 30927_1 | TATGAGTGTGGAGTCGTTGCGAGAGAGATTGGACTTGCCTGAANNNAGATCGGAA |
| 30927_2 | TATGAGTGTGGAGTCGTTGCGAGAGAGATTGGACTTGCCTGAANNNAGATCGGAA |
| 30927_3 | TATGAGTGTGGAGTCGTTGCGAGAGAGATTGGACTTGCCTGAANNNAGATCGGAA |
| 30927_4 | TATGAGTGTGGAGTCGTTGCGAGAGAGATTGGACTTGCCTGAANNNAGATCGGAA |
| 30927_5 | TATGAGTGTGGAGTCGTTGCGAGAGAGATTGGACTTGCCTGAANNNAGATCGGAA |
| 30927_6 | TATGAGTGTGGAGTCGTTGCGAGAGAGATTGGACTTGCCTGAANNNAGATCGGAA |
| 30927_7 | TATGAGTGTGGAGTCGTTGCGAGAGAGATTGGACTTGCCTGAANNNAGATCGGAA |
| 53076_1 | TATGAGTGTGGAGTCGTTGCTTTCCTACACGAACCACTTGTGANNNAGATCGGAAGA |
| 53076_2 | TATGAGTGTGGAGTCGTTGCTTTCCTACACGAACCACTTGTGANNNAGATCGGAAGA |
| 53076_3 | TATGAGTGTGGAGTCGTTGCTTTCCTACACGAACCACTTGTGANNNAGATCGGAAGA |
| 53076_4 | TATGAGTGTGGAGTCGTTGCTTTCCTACACGAACCACTTGTGANNNAGATCGGAAGA |

| | |
|---|---|
| 53076_5 | TATGAGTGTGGAGTCGTTGCTTTCCTACACGAACCACTTGTGANNNAGATCGGAAG |
| 53076_6 | TATGAGTGTGGAGTCGTTGCTTTCCTACACGAACCACTTGTGANNNAGATCGGAAG |
| 53076_7 | TATGAGTGTGGAGTCGTTGCTTTCCTACACGAACCACTTGTGANNNAGATCGGAAG |
| 46116_1 | TATGAGTGTGGAGTCGTTGCAAATGGTTCTCCTGACTCAGCTTNNNAGATCGGAAG |
| 46116_2 | TATGAGTGTGGAGTCGTTGCAAATGGTTCTCCTGACTCAGCTTNNNAGATCGGAAG |
| 46116_3 | TATGAGTGTGGAGTCGTTGCAAATGGTTCTCCTGACTCAGCTTNNNAGATCGGAAG |
| 46116_4 | TATGAGTGTGGAGTCGTTGCAAATGGTTCTCCTGACTCAGCTTNNNAGATCGGAAG |
| 46116_5 | TATGAGTGTGGAGTCGTTGCAAATGGTTCTCCTGACTCAGCTTNNNAGATCGGAAG |
| 46116_6 | TATGAGTGTGGAGTCGTTGCAAATGGTTCTCCTGACTCAGCTTNNNAGATCGGAAG |
| 46116_7 | TATGAGTGTGGAGTCGTTGCAAATGGTTCTCCTGACTCAGCTTNNNAGATCGGAAG |
| 12593_1 | TATGAGTGTGGAGTCGTTGCATTCTTCACTATGGAGCTCTTTNNNAGATCGGAAGA |
| 12593_2 | TATGAGTGTGGAGTCGTTGCATTCTTCACTATGGAGCTCTTTNNNAGATCGGAAGA |
| 12593_3 | TATGAGTGTGGAGTCGTTGCATTCTTCACTATGGAGCTCTTTNNNAGATCGGAAGA |
| 12593_4 | TATGAGTGTGGAGTCGTTGCATTCTTCACTATGGAGCTCTTTNNNAGATCGGAAGA |
| 12593_5 | TATGAGTGTGGAGTCGTTGCATTCTTCACTATGGAGCTCTTTNNNAGATCGGAAGA |
| 12593_6 | TATGAGTGTGGAGTCGTTGCATTCTTCACTATGGAGCTCTTTNNNAGATCGGAAGA |
| 12593_7 | TATGAGTGTGGAGTCGTTGCATTCTTCACTATGGAGCTCTTTNNNAGATCGGAAGA |
| 33240_1 | TATGAGTGTGGAGTCGTTGCCAATCCTCCCAGCTTACTCTCTGNNNAGATCGGAAGA |
| 33240_2 | TATGAGTGTGGAGTCGTTGCCAATCCTCCCAGCTTACTCTCTGNNNAGATCGGAAGA |
| 33240_3 | TATGAGTGTGGAGTCGTTGCCAATCCTCCCAGCTTACTCTCTGNNNAGATCGGAAGA |
| 33240_4 | TATGAGTGTGGAGTCGTTGCCAATCCTCCCAGCTTACTCTCTGNNNAGATCGGAAGA |
| 33240_5 | TATGAGTGTGGAGTCGTTGCCAATCCTCCCAGCTTACTCTCTGNNNAGATCGGAAGA |
| 33240_6 | TATGAGTGTGGAGTCGTTGCCAATCCTCCCAGCTTACTCTCTGNNNAGATCGGAAGA |
| 33240_7 | TATGAGTGTGGAGTCGTTGCCAATCCTCCCAGCTTACTCTCTGNNNAGATCGGAAGA |
| 43058_1 | TATGAGTGTGGAGTCGTTGCTATTCACAGCAACCTGGATGGAANNNAGATCGGAA |
| 43058_2 | TATGAGTGTGGAGTCGTTGCTATTCACAGCAACCTGGATGGAANNNAGATCGGAA |
| 43058_3 | TATGAGTGTGGAGTCGTTGCTATTCACAGCAACCTGGATGGAANNNAGATCGGAA |
| 43058_4 | TATGAGTGTGGAGTCGTTGCTATTCACAGCAACCTGGATGGAANNNAGATCGGAA |
| 43058_5 | TATGAGTGTGGAGTCGTTGCTATTCACAGCAACCTGGATGGAANNNAGATCGGAA |
| 43058_6 | TATGAGTGTGGAGTCGTTGCTATTCACAGCAACCTGGATGGAANNNAGATCGGAA |
| 43058_7 | TATGAGTGTGGAGTCGTTGCTATTCACAGCAACCTGGATGGAANNNAGATCGGAA |
| 47597_1 | TATGAGTGTGGAGTCGTTGCTGTACTCTTTCCTCTGCTTCCACNNNAGATCGGAAGA |
| 47597_2 | TATGAGTGTGGAGTCGTTGCTGTACTCTTTCCTCTGCTTCCACNNNAGATCGGAAGA |
| 47597_3 | TATGAGTGTGGAGTCGTTGCTGTACTCTTTCCTCTGCTTCCACNNNAGATCGGAAGA |
| 47597_4 | TATGAGTGTGGAGTCGTTGCTGTACTCTTTCCTCTGCTTCCACNNNAGATCGGAAGA |
| 47597_5 | TATGAGTGTGGAGTCGTTGCTGTACTCTTTCCTCTGCTTCCACNNNAGATCGGAAGA |
| 47597_6 | TATGAGTGTGGAGTCGTTGCTGTACTCTTTCCTCTGCTTCCACNNNAGATCGGAAGA |
| 47597_7 | TATGAGTGTGGAGTCGTTGCTGTACTCTTTCCTCTGCTTCCACNNNAGATCGGAAGA |
| 35124_1 | TATGAGTGTGGAGTCGTTGCCATTCTCCTGCCTCAGCCTCNNNAGATCGGAAGAGCA |
| 35124_2 | TATGAGTGTGGAGTCGTTGCCATTCTCCTGCCTCAGCCTCNNNAGATCGGAAGAGCA |
| 35124_3 | TATGAGTGTGGAGTCGTTGCCATTCTCCTGCCTCAGCCTCNNNAGATCGGAAGAGCA |
| 35124_4 | TATGAGTGTGGAGTCGTTGCCATTCTCCTGCCTCAGCCTCNNNAGATCGGAAGAGCA |
| 35124_5 | TATGAGTGTGGAGTCGTTGCCATTCTCCTGCCTCAGCCTCNNNAGATCGGAAGAGCA |

| | |
|---|---|
| 35124_6 | TATGAGTGTGGAGTCGTTGCCATTCTCCTGCCTCAGCCTCNNNAGATCGGAAGAGCA |
| 35124_7 | TATGAGTGTGGAGTCGTTGCCATTCTCCTGCCTCAGCCTCNNNAGATCGGAAGAGCA |
| 31932_1 | TATGAGTGTGGAGTCGTTGCAAATAATTAGCCAGGCCAGTTGGNNNAGATCGGAAC |
| 31932_2 | TATGAGTGTGGAGTCGTTGCAAATAATTAGCCAGGCCAGTTGGNNNAGATCGGAAC |
| 31932_3 | TATGAGTGTGGAGTCGTTGCAAATAATTAGCCAGGCCAGTTGGNNNAGATCGGAAC |
| 31932_4 | TATGAGTGTGGAGTCGTTGCAAATAATTAGCCAGGCCAGTTGGNNNAGATCGGAAC |
| 31932_5 | TATGAGTGTGGAGTCGTTGCAAATAATTAGCCAGGCCAGTTGGNNNAGATCGGAAC |
| 31932_6 | TATGAGTGTGGAGTCGTTGCAAATAATTAGCCAGGCCAGTTGGNNNAGATCGGAAC |
| 31932_7 | TATGAGTGTGGAGTCGTTGCAAATAATTAGCCAGGCCAGTTGGNNNAGATCGGAAC |
| 53430_1 | TATGAGTGTGGAGTCGTTGCACAGAGAGAGAGAGACTGAGACTNNNAGATCGGAA |
| 53430_2 | TATGAGTGTGGAGTCGTTGCACAGAGAGAGAGAGACTGAGACTNNNAGATCGGAA |
| 53430_3 | TATGAGTGTGGAGTCGTTGCACAGAGAGAGAGAGACTGAGACTNNNAGATCGGAA |
| 53430_4 | TATGAGTGTGGAGTCGTTGCACAGAGAGAGAGAGACTGAGACTNNNAGATCGGAA |
| 53430_5 | TATGAGTGTGGAGTCGTTGCACAGAGAGAGAGAGACTGAGACTNNNAGATCGGAA |
| 53430_6 | TATGAGTGTGGAGTCGTTGCACAGAGAGAGAGAGACTGAGACTNNNAGATCGGAA |
| 53430_7 | TATGAGTGTGGAGTCGTTGCACAGAGAGAGAGAGACTGAGACTNNNAGATCGGAA |
| 50346_1 | TATGAGTGTGGAGTCGTTGCGATCTGGGGAAGCACTGACATTNNNAGATCGGAAGA |
| 50346_2 | TATGAGTGTGGAGTCGTTGCGATCTGGGGAAGCACTGACATTNNNAGATCGGAAGA |
| 50346_3 | TATGAGTGTGGAGTCGTTGCGATCTGGGGAAGCACTGACATTNNNAGATCGGAAGA |
| 50346_4 | TATGAGTGTGGAGTCGTTGCGATCTGGGGAAGCACTGACATTNNNAGATCGGAAGA |
| 50346_5 | TATGAGTGTGGAGTCGTTGCGATCTGGGGAAGCACTGACATTNNNAGATCGGAAGA |
| 50346_6 | TATGAGTGTGGAGTCGTTGCGATCTGGGGAAGCACTGACATTNNNAGATCGGAAGA |
| 50346_7 | TATGAGTGTGGAGTCGTTGCGATCTGGGGAAGCACTGACATTNNNAGATCGGAAGA |
| 13085_1 | TATGAGTGTGGAGTCGTTGCAAAGTCTAAAGTGGTTCTTCTCNNNAGATCGGAAGA |
| 13085_2 | TATGAGTGTGGAGTCGTTGCAAAGTCTAAAGTGGTTCTTCTCNNNAGATCGGAAGA |
| 13085_3 | TATGAGTGTGGAGTCGTTGCAAAGTCTAAAGTGGTTCTTCTCNNNAGATCGGAAGA |
| 13085_4 | TATGAGTGTGGAGTCGTTGCAAAGTCTAAAGTGGTTCTTCTCNNNAGATCGGAAGA |
| 13085_5 | TATGAGTGTGGAGTCGTTGCAAAGTCTAAAGTGGTTCTTCTCNNNAGATCGGAAGA |
| 13085_6 | TATGAGTGTGGAGTCGTTGCAAAGTCTAAAGTGGTTCTTCTCNNNAGATCGGAAGA |
| 13085_7 | TATGAGTGTGGAGTCGTTGCAAAGTCTAAAGTGGTTCTTCTCNNNAGATCGGAAGA |
| 37362_1 | TATGAGTGTGGAGTCGTTGCGATGGTTTTGAGCAATAGCACGTNNNAGATCGGAAC |
| 37362_2 | TATGAGTGTGGAGTCGTTGCGATGGTTTTGAGCAATAGCACGTNNNAGATCGGAAC |
| 37362_3 | TATGAGTGTGGAGTCGTTGCGATGGTTTTGAGCAATAGCACGTNNNAGATCGGAAC |
| 37362_4 | TATGAGTGTGGAGTCGTTGCGATGGTTTTGAGCAATAGCACGTNNNAGATCGGAAC |
| 37362_5 | TATGAGTGTGGAGTCGTTGCGATGGTTTTGAGCAATAGCACGTNNNAGATCGGAAC |
| 37362_6 | TATGAGTGTGGAGTCGTTGCGATGGTTTTGAGCAATAGCACGTNNNAGATCGGAAC |
| 37362_7 | TATGAGTGTGGAGTCGTTGCGATGGTTTTGAGCAATAGCACGTNNNAGATCGGAAC |
| 30996_1 | TATGAGTGTGGAGTCGTTGCAGTCAGAGTGGTCATCAAATCCTNNNAGATCGGAAG |
| 30996_2 | TATGAGTGTGGAGTCGTTGCAGTCAGAGTGGTCATCAAATCCTNNNAGATCGGAAG |
| 30996_3 | TATGAGTGTGGAGTCGTTGCAGTCAGAGTGGTCATCAAATCCTNNNAGATCGGAAG |
| 30996_4 | TATGAGTGTGGAGTCGTTGCAGTCAGAGTGGTCATCAAATCCTNNNAGATCGGAAG |
| 30996_5 | TATGAGTGTGGAGTCGTTGCAGTCAGAGTGGTCATCAAATCCTNNNAGATCGGAAG |
| 30996_6 | TATGAGTGTGGAGTCGTTGCAGTCAGAGTGGTCATCAAATCCTNNNAGATCGGAAG |

| | |
|---|---|
| 30996_7 | TATGAGTGTGGAGTCGTTGCAGTCAGAGTGGTCATCAAATCCTNNNAGATCGGAAG |
| 30920_1 | TATGAGTGTGGAGTCGTTGCTGGACTGAACCCAGCAGAAATAANNNAGATCGGAA |
| 30920_2 | TATGAGTGTGGAGTCGTTGCTGGACTGAACCCAGCAGAAATAANNNAGATCGGAA |
| 30920_3 | TATGAGTGTGGAGTCGTTGCTGGACTGAACCCAGCAGAAATAANNNAGATCGGAA |
| 30920_4 | TATGAGTGTGGAGTCGTTGCTGGACTGAACCCAGCAGAAATAANNNAGATCGGAA |
| 30920_5 | TATGAGTGTGGAGTCGTTGCTGGACTGAACCCAGCAGAAATAANNNAGATCGGAA |
| 30920_6 | TATGAGTGTGGAGTCGTTGCTGGACTGAACCCAGCAGAAATAANNNAGATCGGAA |
| 30920_7 | TATGAGTGTGGAGTCGTTGCTGGACTGAACCCAGCAGAAATAANNNAGATCGGAA |
| 44679_1 | TATGAGTGTGGAGTCGTTGCCTTCCCGAAAAGGATGAAGCTCCNNNAGATCGGAAG |
| 44679_2 | TATGAGTGTGGAGTCGTTGCCTTCCCGAAAAGGATGAAGCTCCNNNAGATCGGAAG |
| 44679_3 | TATGAGTGTGGAGTCGTTGCCTTCCCGAAAAGGATGAAGCTCCNNNAGATCGGAAG |
| 44679_4 | TATGAGTGTGGAGTCGTTGCCTTCCCGAAAAGGATGAAGCTCCNNNAGATCGGAAG |
| 44679_5 | TATGAGTGTGGAGTCGTTGCCTTCCCGAAAAGGATGAAGCTCCNNNAGATCGGAAG |
| 44679_6 | TATGAGTGTGGAGTCGTTGCCTTCCCGAAAAGGATGAAGCTCCNNNAGATCGGAAG |
| 44679_7 | TATGAGTGTGGAGTCGTTGCCTTCCCGAAAAGGATGAAGCTCCNNNAGATCGGAAG |
| 38831_1 | TATGAGTGTGGAGTCGTTGCTGTGTGGTGGTTCATGCCTATAANNNAGATCGGAAG |
| 38831_2 | TATGAGTGTGGAGTCGTTGCTGTGTGGTGGTTCATGCCTATAANNNAGATCGGAAG |
| 38831_3 | TATGAGTGTGGAGTCGTTGCTGTGTGGTGGTTCATGCCTATAANNNAGATCGGAAG |
| 38831_4 | TATGAGTGTGGAGTCGTTGCTGTGTGGTGGTTCATGCCTATAANNNAGATCGGAAG |
| 38831_5 | TATGAGTGTGGAGTCGTTGCTGTGTGGTGGTTCATGCCTATAANNNAGATCGGAAG |
| 38831_6 | TATGAGTGTGGAGTCGTTGCTGTGTGGTGGTTCATGCCTATAANNNAGATCGGAAG |
| 38831_7 | TATGAGTGTGGAGTCGTTGCTGTGTGGTGGTTCATGCCTATAANNNAGATCGGAAG |
| 37659_1 | TATGAGTGTGGAGTCGTTGCAAAAAATTAGCTGGGCATGGTGGNNNAGATCGGAA |
| 37659_2 | TATGAGTGTGGAGTCGTTGCAAAAAATTAGCTGGGCATGGTGGNNNAGATCGGAA |
| 37659_3 | TATGAGTGTGGAGTCGTTGCAAAAAATTAGCTGGGCATGGTGGNNNAGATCGGAA |
| 37659_4 | TATGAGTGTGGAGTCGTTGCAAAAAATTAGCTGGGCATGGTGGNNNAGATCGGAA |
| 37659_5 | TATGAGTGTGGAGTCGTTGCAAAAAATTAGCTGGGCATGGTGGNNNAGATCGGAA |
| 37659_6 | TATGAGTGTGGAGTCGTTGCAAAAAATTAGCTGGGCATGGTGGNNNAGATCGGAA |
| 37659_7 | TATGAGTGTGGAGTCGTTGCAAAAAATTAGCTGGGCATGGTGGNNNAGATCGGAA |
| 48055_1 | TATGAGTGTGGAGTCGTTGCAGAGAAGGGGAGACTGAGAGAATNNNAGATCGGA |
| 48055_2 | TATGAGTGTGGAGTCGTTGCAGAGAAGGGGAGACTGAGAGAATNNNAGATCGGA |
| 48055_3 | TATGAGTGTGGAGTCGTTGCAGAGAAGGGGAGACTGAGAGAATNNNAGATCGGA |
| 48055_4 | TATGAGTGTGGAGTCGTTGCAGAGAAGGGGAGACTGAGAGAATNNNAGATCGGA |
| 48055_5 | TATGAGTGTGGAGTCGTTGCAGAGAAGGGGAGACTGAGAGAATNNNAGATCGGA |
| 48055_6 | TATGAGTGTGGAGTCGTTGCAGAGAAGGGGAGACTGAGAGAATNNNAGATCGGA |
| 48055_7 | TATGAGTGTGGAGTCGTTGCAGAGAAGGGGAGACTGAGAGAATNNNAGATCGGA |
| 37358_1 | TATGAGTGTGGAGTCGTTGCATTCTCTAGAAGCAGAGCCTGAGNNNAGATCGGAA |
| 37358_2 | TATGAGTGTGGAGTCGTTGCATTCTCTAGAAGCAGAGCCTGAGNNNAGATCGGAA |
| 37358_3 | TATGAGTGTGGAGTCGTTGCATTCTCTAGAAGCAGAGCCTGAGNNNAGATCGGAA |
| 37358_4 | TATGAGTGTGGAGTCGTTGCATTCTCTAGAAGCAGAGCCTGAGNNNAGATCGGAA |
| 37358_5 | TATGAGTGTGGAGTCGTTGCATTCTCTAGAAGCAGAGCCTGAGNNNAGATCGGAA |
| 37358_6 | TATGAGTGTGGAGTCGTTGCATTCTCTAGAAGCAGAGCCTGAGNNNAGATCGGAA |
| 37358_7 | TATGAGTGTGGAGTCGTTGCATTCTCTAGAAGCAGAGCCTGAGNNNAGATCGGAA |

| | |
|---|---|
| 34181_1 | TATGAGTGTGGAGTCGTTGCGTTTACAGCAACTGTCCAACCAANNNAGATCGGAAG |
| 34181_2 | TATGAGTGTGGAGTCGTTGCGTTTACAGCAACTGTCCAACCAANNNAGATCGGAAG |
| 34181_3 | TATGAGTGTGGAGTCGTTGCGTTTACAGCAACTGTCCAACCAANNNAGATCGGAAG |
| 34181_4 | TATGAGTGTGGAGTCGTTGCGTTTACAGCAACTGTCCAACCAANNNAGATCGGAAG |
| 34181_5 | TATGAGTGTGGAGTCGTTGCGTTTACAGCAACTGTCCAACCAANNNAGATCGGAAG |
| 34181_6 | TATGAGTGTGGAGTCGTTGCGTTTACAGCAACTGTCCAACCAANNNAGATCGGAAG |
| 34181_7 | TATGAGTGTGGAGTCGTTGCGTTTACAGCAACTGTCCAACCAANNNAGATCGGAAG |
| 52609_1 | TATGAGTGTGGAGTCGTTGCGGCGTCATATCTCATAGTGGTGTNNNAGATCGGAA |
| 52609_2 | TATGAGTGTGGAGTCGTTGCGGCGTCATATCTCATAGTGGTGTNNNAGATCGGAA |
| 52609_3 | TATGAGTGTGGAGTCGTTGCGGCGTCATATCTCATAGTGGTGTNNNAGATCGGAA |
| 52609_4 | TATGAGTGTGGAGTCGTTGCGGCGTCATATCTCATAGTGGTGTNNNAGATCGGAA |
| 52609_5 | TATGAGTGTGGAGTCGTTGCGGCGTCATATCTCATAGTGGTGTNNNAGATCGGAA |
| 52609_6 | TATGAGTGTGGAGTCGTTGCGGCGTCATATCTCATAGTGGTGTNNNAGATCGGAA |
| 52609_7 | TATGAGTGTGGAGTCGTTGCGGCGTCATATCTCATAGTGGTGTNNNAGATCGGAA |
| 8801_1 | TATGAGTGTGGAGTCGTTGCTTGTACTCCCAATACTTTGGGGGANNNAGATCGGAA |
| 8801_2 | TATGAGTGTGGAGTCGTTGCTTGTACTCCCAATACTTTGGGGGANNNAGATCGGAA |
| 8801_3 | TATGAGTGTGGAGTCGTTGCTTGTACTCCCAATACTTTGGGGGANNNAGATCGGAA |
| 8801_4 | TATGAGTGTGGAGTCGTTGCTTGTACTCCCAATACTTTGGGGGANNNAGATCGGAA |
| 8801_5 | TATGAGTGTGGAGTCGTTGCTTGTACTCCCAATACTTTGGGGGANNNAGATCGGAA |
| 8801_6 | TATGAGTGTGGAGTCGTTGCTTGTACTCCCAATACTTTGGGGGANNNAGATCGGAA |
| 8801_7 | TATGAGTGTGGAGTCGTTGCTTGTACTCCCAATACTTTGGGGGANNNAGATCGGAA |
| 42688_1 | TATGAGTGTGGAGTCGTTGCCTTTCAGTGAGTCAAGATCGCGNNNAGATCGGAAG |
| 42688_2 | TATGAGTGTGGAGTCGTTGCCTTTCAGTGAGTCAAGATCGCGNNNAGATCGGAAG |
| 42688_3 | TATGAGTGTGGAGTCGTTGCCTTTCAGTGAGTCAAGATCGCGNNNAGATCGGAAG |
| 42688_4 | TATGAGTGTGGAGTCGTTGCCTTTCAGTGAGTCAAGATCGCGNNNAGATCGGAAG |
| 42688_5 | TATGAGTGTGGAGTCGTTGCCTTTCAGTGAGTCAAGATCGCGNNNAGATCGGAAG |
| 42688_6 | TATGAGTGTGGAGTCGTTGCCTTTCAGTGAGTCAAGATCGCGNNNAGATCGGAAG |
| 42688_7 | TATGAGTGTGGAGTCGTTGCCTTTCAGTGAGTCAAGATCGCGNNNAGATCGGAAG |
| 32840_1 | TATGAGTGTGGAGTCGTTGCTGCGTAGCAAACACTTTTCACAANNNAGATCGGAAG |
| 32840_2 | TATGAGTGTGGAGTCGTTGCTGCGTAGCAAACACTTTTCACAANNNAGATCGGAAG |
| 32840_3 | TATGAGTGTGGAGTCGTTGCTGCGTAGCAAACACTTTTCACAANNNAGATCGGAAG |
| 32840_4 | TATGAGTGTGGAGTCGTTGCTGCGTAGCAAACACTTTTCACAANNNAGATCGGAAG |
| 32840_5 | TATGAGTGTGGAGTCGTTGCTGCGTAGCAAACACTTTTCACAANNNAGATCGGAAG |
| 32840_6 | TATGAGTGTGGAGTCGTTGCTGCGTAGCAAACACTTTTCACAANNNAGATCGGAAG |
| 32840_7 | TATGAGTGTGGAGTCGTTGCTGCGTAGCAAACACTTTTCACAANNNAGATCGGAAG |
| 34901_1 | TATGAGTGTGGAGTCGTTGCCTGAGTAGCTCGGATTACTGGTGNNNAGATCGGAA |
| 34901_2 | TATGAGTGTGGAGTCGTTGCCTGAGTAGCTCGGATTACTGGTGNNNAGATCGGAA |
| 34901_3 | TATGAGTGTGGAGTCGTTGCCTGAGTAGCTCGGATTACTGGTGNNNAGATCGGAA |
| 34901_4 | TATGAGTGTGGAGTCGTTGCCTGAGTAGCTCGGATTACTGGTGNNNAGATCGGAA |
| 34901_5 | TATGAGTGTGGAGTCGTTGCCTGAGTAGCTCGGATTACTGGTGNNNAGATCGGAA |
| 34901_6 | TATGAGTGTGGAGTCGTTGCCTGAGTAGCTCGGATTACTGGTGNNNAGATCGGAA |
| 34901_7 | TATGAGTGTGGAGTCGTTGCCTGAGTAGCTCGGATTACTGGTGNNNAGATCGGAA |
| 51577_1 | TATGAGTGTGGAGTCGTTGCGGAGGTCTTGGATTCTGGTTTCTNNNAGATCGGAA |

| | |
|---|---|
| 51577_2 | TATGAGTGTGGAGTCGTTGCGGAGGTCTTGGATTCTGGTTTCTNNNAGATCGGAAG |
| 51577_3 | TATGAGTGTGGAGTCGTTGCGGAGGTCTTGGATTCTGGTTTCTNNNAGATCGGAAG |
| 41402_1 | TATGAGTGTGGAGTCGTTGCAATAAATCCCAAGGCCACCATCTNNNAGATCGGAAG |
| 41402_2 | TATGAGTGTGGAGTCGTTGCAATAAATCCCAAGGCCACCATCTNNNAGATCGGAAG |
| 41402_3 | TATGAGTGTGGAGTCGTTGCAATAAATCCCAAGGCCACCATCTNNNAGATCGGAAG |
| 10127_1 | TATGAGTGTGGAGTCGTTGCCCACTCCTAACTTCGATGCTTCCNNNAGATCGGAAGA |
| 10127_2 | TATGAGTGTGGAGTCGTTGCCCACTCCTAACTTCGATGCTTCCNNNAGATCGGAAGA |
| 10127_3 | TATGAGTGTGGAGTCGTTGCCCACTCCTAACTTCGATGCTTCCNNNAGATCGGAAGA |
| 28834_1 | TATGAGTGTGGAGTCGTTGCTCCAGGGATTGAGAAGAGTTTGGNNNAGATCGGAA |
| 28834_2 | TATGAGTGTGGAGTCGTTGCTCCAGGGATTGAGAAGAGTTTGGNNNAGATCGGAA |
| 28834_3 | TATGAGTGTGGAGTCGTTGCTCCAGGGATTGAGAAGAGTTTGGNNNAGATCGGAA |
| 43431_1 | TATGAGTGTGGAGTCGTTGCTCTCCTGAGTAGCTGGGATTACANNNAGATCGGAAG |
| 43431_2 | TATGAGTGTGGAGTCGTTGCTCTCCTGAGTAGCTGGGATTACANNNAGATCGGAAG |
| 43431_3 | TATGAGTGTGGAGTCGTTGCTCTCCTGAGTAGCTGGGATTACANNNAGATCGGAAG |
| 46947_1 | TATGAGTGTGGAGTCGTTGCGAGTAGCTGGGATTACAGGCATGNNNAGATCGGAA |
| 46947_2 | TATGAGTGTGGAGTCGTTGCGAGTAGCTGGGATTACAGGCATGNNNAGATCGGAA |
| 46947_3 | TATGAGTGTGGAGTCGTTGCGAGTAGCTGGGATTACAGGCATGNNNAGATCGGAA |
| 9204_1 | TATGAGTGTGGAGTCGTTGCTTGTGCATCTGTATACCGGGTTGNNNAGATCGGAAG |
| 9204_2 | TATGAGTGTGGAGTCGTTGCTTGTGCATCTGTATACCGGGTTGNNNAGATCGGAAG |
| 9204_3 | TATGAGTGTGGAGTCGTTGCTTGTGCATCTGTATACCGGGTTGNNNAGATCGGAAG |
| 42807_1 | TATGAGTGTGGAGTCGTTGCATGCTCAGAGTAACAAGGAAGGGNNNAGATCGGAA |
| 42807_2 | TATGAGTGTGGAGTCGTTGCATGCTCAGAGTAACAAGGAAGGGNNNAGATCGGAA |
| 42807_3 | TATGAGTGTGGAGTCGTTGCATGCTCAGAGTAACAAGGAAGGGNNNAGATCGGAA |
| 13118_1 | TATGAGTGTGGAGTCGTTGCCTTTATGAAGTGTTTCACCAAANNNAGATCGGAAGA |
| 13118_2 | TATGAGTGTGGAGTCGTTGCCTTTATGAAGTGTTTCACCAAANNNAGATCGGAAGA |
| 13118_3 | TATGAGTGTGGAGTCGTTGCCTTTATGAAGTGTTTCACCAAANNNAGATCGGAAGA |
| 37926_1 | TATGAGTGTGGAGTCGTTGCAAACTCTTCTGTGTGGGAGACAGNNNAGATCGGAA |
| 37926_2 | TATGAGTGTGGAGTCGTTGCAAACTCTTCTGTGTGGGAGACAGNNNAGATCGGAA |
| 37926_3 | TATGAGTGTGGAGTCGTTGCAAACTCTTCTGTGTGGGAGACAGNNNAGATCGGAA |
| 12378_1 | TATGAGTGTGGAGTCGTTGCGGGTGAAAATCATACAGTGTGTCCNNNAGATCGGA |
| 12378_2 | TATGAGTGTGGAGTCGTTGCGGGTGAAAATCATACAGTGTGTCCNNNAGATCGGA |
| 12378_3 | TATGAGTGTGGAGTCGTTGCGGGTGAAAATCATACAGTGTGTCCNNNAGATCGGA |
| 46802_1 | TATGAGTGTGGAGTCGTTGCGCGATCTCAGCTCACAGCAAGNNNAGATCGGAAGA |
| 46802_2 | TATGAGTGTGGAGTCGTTGCGCGATCTCAGCTCACAGCAAGNNNAGATCGGAAGA |
| 46802_3 | TATGAGTGTGGAGTCGTTGCGCGATCTCAGCTCACAGCAAGNNNAGATCGGAAGA |
| 44250_1 | TATGAGTGTGGAGTCGTTGCCAGTTTGTGTCGGTATAGAGGCTNNNAGATCGGAA |
| 44250_2 | TATGAGTGTGGAGTCGTTGCCAGTTTGTGTCGGTATAGAGGCTNNNAGATCGGAA |
| 44250_3 | TATGAGTGTGGAGTCGTTGCCAGTTTGTGTCGGTATAGAGGCTNNNAGATCGGAA |
| 43109_1 | TATGAGTGTGGAGTCGTTGCGTCAAGACCAGTCTGGGTAACATNNNAGATCGGAA |
| 43109_2 | TATGAGTGTGGAGTCGTTGCGTCAAGACCAGTCTGGGTAACATNNNAGATCGGAA |
| 43109_3 | TATGAGTGTGGAGTCGTTGCGTCAAGACCAGTCTGGGTAACATNNNAGATCGGAA |
| 46888_1 | TATGAGTGTGGAGTCGTTGCTTCACGTCATTCTCCTGCCTCNNNAGATCGGAAGAGC |
| 46888_2 | TATGAGTGTGGAGTCGTTGCTTCACGTCATTCTCCTGCCTCNNNAGATCGGAAGAGC |

| | |
|---|---|
| 46888_3 | TATGAGTGTGGAGTCGTTGCTTCACGTCATTCTCCTGCCTCNNNAGATCGGAAGAGC |
| 51940_1 | TATGAGTGTGGAGTCGTTGCTCCTTCCTTCCTTTTTCCCTGAGNNNAGATCGGAAGA |
| 51940_2 | TATGAGTGTGGAGTCGTTGCTCCTTCCTTCCTTTTTCCCTGAGNNNAGATCGGAAGA |
| 51940_3 | TATGAGTGTGGAGTCGTTGCTCCTTCCTTCCTTTTTCCCTGAGNNNAGATCGGAAGA |
| 46876_1 | TATGAGTGTGGAGTCGTTGCCAGAGTGCTGGGATTATAGGCTTNNNAGATCGGAA |
| 46876_2 | TATGAGTGTGGAGTCGTTGCCAGAGTGCTGGGATTATAGGCTTNNNAGATCGGAA |
| 46876_3 | TATGAGTGTGGAGTCGTTGCCAGAGTGCTGGGATTATAGGCTTNNNAGATCGGAA |
| 37369_1 | TATGAGTGTGGAGTCGTTGCAATTTGAAACTAGCCTGGCCAACNNNAGATCGGAAG |
| 37369_2 | TATGAGTGTGGAGTCGTTGCAATTTGAAACTAGCCTGGCCAACNNNAGATCGGAAG |
| 37369_3 | TATGAGTGTGGAGTCGTTGCAATTTGAAACTAGCCTGGCCAACNNNAGATCGGAAG |
| 49413_1 | TATGAGTGTGGAGTCGTTGCCTTCCAAGATGGCCAAATAGGGANNNAGATCGGAA |
| 49413_2 | TATGAGTGTGGAGTCGTTGCCTTCCAAGATGGCCAAATAGGGANNNAGATCGGAA |
| 49413_3 | TATGAGTGTGGAGTCGTTGCCTTCCAAGATGGCCAAATAGGGANNNAGATCGGAA |
| 52801_1 | TATGAGTGTGGAGTCGTTGCGTCATCTCTGCCTGTGACCTTCNNNAGATCGGAAGA |
| 52801_2 | TATGAGTGTGGAGTCGTTGCGTCATCTCTGCCTGTGACCTTCNNNAGATCGGAAGA |
| 52801_3 | TATGAGTGTGGAGTCGTTGCGTCATCTCTGCCTGTGACCTTCNNNAGATCGGAAGA |
| 33903_1 | TATGAGTGTGGAGTCGTTGCTGCGTTCAAGTGATTCTTATGCCNNNAGATCGGAAG |
| 33903_2 | TATGAGTGTGGAGTCGTTGCTGCGTTCAAGTGATTCTTATGCCNNNAGATCGGAAG |
| 33903_3 | TATGAGTGTGGAGTCGTTGCTGCGTTCAAGTGATTCTTATGCCNNNAGATCGGAAG |
| 29925_1 | TATGAGTGTGGAGTCGTTGCTCTCCCTCCCTACACAGAATTCCNNNAGATCGGAAGA |
| 29925_2 | TATGAGTGTGGAGTCGTTGCTCTCCCTCCCTACACAGAATTCCNNNAGATCGGAAGA |
| 29925_3 | TATGAGTGTGGAGTCGTTGCTCTCCCTCCCTACACAGAATTCCNNNAGATCGGAAGA |
| 42296_1 | TATGAGTGTGGAGTCGTTGCGTTTCGGGAAATGCATCCTTAGGNNNAGATCGGAA |
| 42296_2 | TATGAGTGTGGAGTCGTTGCGTTTCGGGAAATGCATCCTTAGGNNNAGATCGGAA |
| 42296_3 | TATGAGTGTGGAGTCGTTGCGTTTCGGGAAATGCATCCTTAGGNNNAGATCGGAA |
| 42127_1 | TATGAGTGTGGAGTCGTTGCCAAAAGTGCTGGGATTACAGGTGNNNAGATCGGAA |
| 42127_2 | TATGAGTGTGGAGTCGTTGCCAAAAGTGCTGGGATTACAGGTGNNNAGATCGGAA |
| 42127_3 | TATGAGTGTGGAGTCGTTGCCAAAAGTGCTGGGATTACAGGTGNNNAGATCGGAA |
| 45221_1 | TATGAGTGTGGAGTCGTTGCCTGGATGGACAGGAAATGTAGCTNNNAGATCGGAA |
| 45221_2 | TATGAGTGTGGAGTCGTTGCCTGGATGGACAGGAAATGTAGCTNNNAGATCGGAA |
| 45221_3 | TATGAGTGTGGAGTCGTTGCCTGGATGGACAGGAAATGTAGCTNNNAGATCGGAA |
| 12709_1 | TATGAGTGTGGAGTCGTTGCAGTAAGTCAAAGACGATAAGAATNNNAGATCGGAA |
| 12709_2 | TATGAGTGTGGAGTCGTTGCAGTAAGTCAAAGACGATAAGAATNNNAGATCGGAA |
| 12709_3 | TATGAGTGTGGAGTCGTTGCAGTAAGTCAAAGACGATAAGAATNNNAGATCGGAA |
| 44995_1 | TATGAGTGTGGAGTCGTTGCTTTGGGGAAGAGAACCAAGGAAANNNAGATCGGA |
| 44995_2 | TATGAGTGTGGAGTCGTTGCTTTGGGGAAGAGAACCAAGGAAANNNAGATCGGA |
| 44995_3 | TATGAGTGTGGAGTCGTTGCTTTGGGGAAGAGAACCAAGGAAANNNAGATCGGA |
| 46988_1 | TATGAGTGTGGAGTCGTTGCGATCAAGTGATTCTCCTGCCTCANNNAGATCGGAAG |
| 46988_2 | TATGAGTGTGGAGTCGTTGCGATCAAGTGATTCTCCTGCCTCANNNAGATCGGAAG |
| 46988_3 | TATGAGTGTGGAGTCGTTGCGATCAAGTGATTCTCCTGCCTCANNNAGATCGGAAG |
| 29681_1 | TATGAGTGTGGAGTCGTTGCGAGCAAGGGAGGAGAGCTTACATNNNAGATCGGA |
| 29681_2 | TATGAGTGTGGAGTCGTTGCGAGCAAGGGAGGAGAGCTTACATNNNAGATCGGA |
| 29681_3 | TATGAGTGTGGAGTCGTTGCGAGCAAGGGAGGAGAGCTTACATNNNAGATCGGA |

| | |
|---|---|
| 33449_1 | TATGAGTGTGGAGTCGTTGCAGAAATGCTTCAGGTGGTGAGATNNNAGATCGGAA |
| 33449_2 | TATGAGTGTGGAGTCGTTGCAGAAATGCTTCAGGTGGTGAGATNNNAGATCGGAA |
| 33449_3 | TATGAGTGTGGAGTCGTTGCAGAAATGCTTCAGGTGGTGAGATNNNAGATCGGAA |
| 33530_1 | TATGAGTGTGGAGTCGTTGCCCAAACCCCTGGGATTAGATCTCNNNAGATCGGAAG |
| 33530_2 | TATGAGTGTGGAGTCGTTGCCCAAACCCCTGGGATTAGATCTCNNNAGATCGGAAG |
| 33530_3 | TATGAGTGTGGAGTCGTTGCCCAAACCCCTGGGATTAGATCTCNNNAGATCGGAAG |
| 38221_1 | TATGAGTGTGGAGTCGTTGCACCTCCCAAGATTGACCAGAAAGNNNAGATCGGAAG |
| 38221_2 | TATGAGTGTGGAGTCGTTGCACCTCCCAAGATTGACCAGAAAGNNNAGATCGGAAG |
| 38221_3 | TATGAGTGTGGAGTCGTTGCACCTCCCAAGATTGACCAGAAAGNNNAGATCGGAAG |
| 37901_1 | TATGAGTGTGGAGTCGTTGCTCCTCATACTCTTGACAGCACTGNNNAGATCGGAAG |
| 37901_2 | TATGAGTGTGGAGTCGTTGCTCCTCATACTCTTGACAGCACTGNNNAGATCGGAAG |
| 37901_3 | TATGAGTGTGGAGTCGTTGCTCCTCATACTCTTGACAGCACTGNNNAGATCGGAAG |
| 48616_1 | TATGAGTGTGGAGTCGTTGCTATACGTTGAGTTCCAAACGCCTNNNAGATCGGAAG |
| 48616_2 | TATGAGTGTGGAGTCGTTGCTATACGTTGAGTTCCAAACGCCTNNNAGATCGGAAG |
| 48616_3 | TATGAGTGTGGAGTCGTTGCTATACGTTGAGTTCCAAACGCCTNNNAGATCGGAAG |
| 47714_1 | TATGAGTGTGGAGTCGTTGCTGCCACTACACCCAGCTAATTTTNNNAGATCGGAAGA |
| 47714_2 | TATGAGTGTGGAGTCGTTGCTGCCACTACACCCAGCTAATTTTNNNAGATCGGAAGA |
| 47714_3 | TATGAGTGTGGAGTCGTTGCTGCCACTACACCCAGCTAATTTTNNNAGATCGGAAGA |
| 41834_1 | TATGAGTGTGGAGTCGTTGCGTTCAAGAGATTCTCCTGCCTCANNNAGATCGGAAG |
| 41834_2 | TATGAGTGTGGAGTCGTTGCGTTCAAGAGATTCTCCTGCCTCANNNAGATCGGAAG |
| 41834_3 | TATGAGTGTGGAGTCGTTGCGTTCAAGAGATTCTCCTGCCTCANNNAGATCGGAAG |
| 5142_1 | TATGAGTGTGGAGTCGTTGCGGGTTCTTGCCCACCTTTACTTTNNNAGATCGGAAGA |
| 5142_2 | TATGAGTGTGGAGTCGTTGCGGGTTCTTGCCCACCTTTACTTTNNNAGATCGGAAGA |
| 5142_3 | TATGAGTGTGGAGTCGTTGCGGGTTCTTGCCCACCTTTACTTTNNNAGATCGGAAGA |
| 51654_1 | TATGAGTGTGGAGTCGTTGCCATTGTGTGAATGTGTGAGCCAANNNAGATCGGAA |
| 51654_2 | TATGAGTGTGGAGTCGTTGCCATTGTGTGAATGTGTGAGCCAANNNAGATCGGAA |
| 51654_3 | TATGAGTGTGGAGTCGTTGCCATTGTGTGAATGTGTGAGCCAANNNAGATCGGAA |
| 45940_1 | TATGAGTGTGGAGTCGTTGCACACATTGGAAAGTCACTGGAGANNNAGATCGGAA |
| 45940_2 | TATGAGTGTGGAGTCGTTGCACACATTGGAAAGTCACTGGAGANNNAGATCGGAA |
| 45940_3 | TATGAGTGTGGAGTCGTTGCACACATTGGAAAGTCACTGGAGANNNAGATCGGAA |
| 31796_1 | TATGAGTGTGGAGTCGTTGCCCACCACGCCTAGCTAATTTTTTNNNAGATCGGAAGA |
| 31796_2 | TATGAGTGTGGAGTCGTTGCCCACCACGCCTAGCTAATTTTTTNNNAGATCGGAAGA |
| 31796_3 | TATGAGTGTGGAGTCGTTGCCCACCACGCCTAGCTAATTTTTTNNNAGATCGGAAGA |
| 10537_1 | TATGAGTGTGGAGTCGTTGCATGGACACCAACACAGGTGTTTTNNNAGATCGGAAG |
| 10537_2 | TATGAGTGTGGAGTCGTTGCATGGACACCAACACAGGTGTTTTNNNAGATCGGAAG |
| 10537_3 | TATGAGTGTGGAGTCGTTGCATGGACACCAACACAGGTGTTTTNNNAGATCGGAAG |
| 44727_1 | TATGAGTGTGGAGTCGTTGCCTGATACTGTGCAGAGTGGGTCNNNAGATCGGAAG |
| 44727_2 | TATGAGTGTGGAGTCGTTGCCTGATACTGTGCAGAGTGGGTCNNNAGATCGGAAG |
| 44727_3 | TATGAGTGTGGAGTCGTTGCCTGATACTGTGCAGAGTGGGTCNNNAGATCGGAAG |
| 52845_1 | TATGAGTGTGGAGTCGTTGCTCAGCCATGAGTATTATGACCCGNNNAGATCGGAAG |
| 52845_2 | TATGAGTGTGGAGTCGTTGCTCAGCCATGAGTATTATGACCCGNNNAGATCGGAAG |
| 52845_3 | TATGAGTGTGGAGTCGTTGCTCAGCCATGAGTATTATGACCCGNNNAGATCGGAAG |
| 10666_1 | TATGAGTGTGGAGTCGTTGCTTTTTGCCAGCCAATTGGACAAGNNNAGATCGGAAG |

| | |
|---|---|
| 10666_2 | TATGAGTGTGGAGTCGTTGCTTTTTGCCAGCCAATTGGACAAGNNNAGATCGGAAG |
| 10666_3 | TATGAGTGTGGAGTCGTTGCTTTTTGCCAGCCAATTGGACAAGNNNAGATCGGAAG |
| 53373_1 | TATGAGTGTGGAGTCGTTGCTCCCTCTTTCTCTCTCCTCCTTCNNNAGATCGGAAGAG |
| 53373_2 | TATGAGTGTGGAGTCGTTGCTCCCTCTTTCTCTCTCCTCCTTCNNNAGATCGGAAGAG |
| 53373_3 | TATGAGTGTGGAGTCGTTGCTCCCTCTTTCTCTCTCCTCCTTCNNNAGATCGGAAGAG |
| 52274_1 | TATGAGTGTGGAGTCGTTGCAGAAACTGTGGACAAAGAGAGCANNNAGATCGGAA |
| 52274_2 | TATGAGTGTGGAGTCGTTGCAGAAACTGTGGACAAAGAGAGCANNNAGATCGGAA |
| 52274_3 | TATGAGTGTGGAGTCGTTGCAGAAACTGTGGACAAAGAGAGCANNNAGATCGGAA |
| 39816_1 | TATGAGTGTGGAGTCGTTGCTCAGAACATCCTCTCTCTGGGAANNNAGATCGGAAG |
| 39816_2 | TATGAGTGTGGAGTCGTTGCTCAGAACATCCTCTCTCTGGGAANNNAGATCGGAAG |
| 39816_3 | TATGAGTGTGGAGTCGTTGCTCAGAACATCCTCTCTCTGGGAANNNAGATCGGAAG |
| 11090_1 | TATGAGTGTGGAGTCGTTGCAAATGTTCCCAGTTTCCCAGGTCNNNAGATCGGAAG |
| 11090_2 | TATGAGTGTGGAGTCGTTGCAAATGTTCCCAGTTTCCCAGGTCNNNAGATCGGAAG |
| 11090_3 | TATGAGTGTGGAGTCGTTGCAAATGTTCCCAGTTTCCCAGGTCNNNAGATCGGAAG |
| 30667_1 | TATGAGTGTGGAGTCGTTGCTGCCCCTTTGACCCAGTATCTNNNAGATCGGAAGAG |
| 30667_2 | TATGAGTGTGGAGTCGTTGCTGCCCCTTTGACCCAGTATCTNNNAGATCGGAAGAG |
| 30667_3 | TATGAGTGTGGAGTCGTTGCTGCCCCTTTGACCCAGTATCTNNNAGATCGGAAGAG |
| 28885_1 | TATGAGTGTGGAGTCGTTGCCGGATCAGGTCTCCATCCTTATGNNNAGATCGGAAG |
| 28885_2 | TATGAGTGTGGAGTCGTTGCCGGATCAGGTCTCCATCCTTATGNNNAGATCGGAAG |
| 28885_3 | TATGAGTGTGGAGTCGTTGCCGGATCAGGTCTCCATCCTTATGNNNAGATCGGAAG |
| 50673_1 | TATGAGTGTGGAGTCGTTGCTTGCACTGTCATCCATATGCTCTNNNAGATCGGAAGA |
| 50673_2 | TATGAGTGTGGAGTCGTTGCTTGCACTGTCATCCATATGCTCTNNNAGATCGGAAGA |
| 50673_3 | TATGAGTGTGGAGTCGTTGCTTGCACTGTCATCCATATGCTCTNNNAGATCGGAAGA |
| 43841_1 | TATGAGTGTGGAGTCGTTGCGACAGGAGAGGCTTCTTGTGAGNNNAGATCGGAAG |
| 43841_2 | TATGAGTGTGGAGTCGTTGCGACAGGAGAGGCTTCTTGTGAGNNNAGATCGGAAG |
| 43841_3 | TATGAGTGTGGAGTCGTTGCGACAGGAGAGGCTTCTTGTGAGNNNAGATCGGAAG |
| 35175_1 | TATGAGTGTGGAGTCGTTGCATCAGCTCTCTCTGGTCACTAGANNNAGATCGGAAG |
| 35175_2 | TATGAGTGTGGAGTCGTTGCATCAGCTCTCTCTGGTCACTAGANNNAGATCGGAAG |
| 35175_3 | TATGAGTGTGGAGTCGTTGCATCAGCTCTCTCTGGTCACTAGANNNAGATCGGAAG |
| 50915_1 | TATGAGTGTGGAGTCGTTGCCCAATACCAACCACATTCTCAGCNNNAGATCGGAAGA |
| 50915_2 | TATGAGTGTGGAGTCGTTGCCCAATACCAACCACATTCTCAGCNNNAGATCGGAAGA |
| 50915_3 | TATGAGTGTGGAGTCGTTGCCCAATACCAACCACATTCTCAGCNNNAGATCGGAAGA |
| 45071_1 | TATGAGTGTGGAGTCGTTGCTTAGGAGCATGCAAGAGAGCTAGNNNAGATCGGAA |
| 45071_2 | TATGAGTGTGGAGTCGTTGCTTAGGAGCATGCAAGAGAGCTAGNNNAGATCGGAA |
| 45071_3 | TATGAGTGTGGAGTCGTTGCTTAGGAGCATGCAAGAGAGCTAGNNNAGATCGGAA |
| 32179_1 | TATGAGTGTGGAGTCGTTGCTGGCAGTCCCTCAAAAATATGACNNNAGATCGGAAG |
| 32179_2 | TATGAGTGTGGAGTCGTTGCTGGCAGTCCCTCAAAAATATGACNNNAGATCGGAAG |
| 32179_3 | TATGAGTGTGGAGTCGTTGCTGGCAGTCCCTCAAAAATATGACNNNAGATCGGAAG |
| 36397_1 | TATGAGTGTGGAGTCGTTGCTTTGTTTTGTTCCTAGTCCCACANNNAGATCGGAAGA |
| 36397_2 | TATGAGTGTGGAGTCGTTGCTTTGTTTTGTTCCTAGTCCCACANNNAGATCGGAAGA |
| 36397_3 | TATGAGTGTGGAGTCGTTGCTTTGTTTTGTTCCTAGTCCCACANNNAGATCGGAAGA |
| 35946_1 | TATGAGTGTGGAGTCGTTGCTCTGTGTACATTTTTTGGCTGGGNNNAGATCGGAAG |
| 35946_2 | TATGAGTGTGGAGTCGTTGCTCTGTGTACATTTTTTGGCTGGGNNNAGATCGGAAG |

| | |
|---|---|
| 35946_3 | TATGAGTGTGGAGTCGTTGCTCTGTGTACATTTTTTGGCTGGGNNNAGATCGGAAG |
| 29442_1 | TATGAGTGTGGAGTCGTTGCGAAGATACTTACTGTAGTTCTCANNNAGATCGGAAG |
| 29442_2 | TATGAGTGTGGAGTCGTTGCGAAGATACTTACTGTAGTTCTCANNNAGATCGGAAG |
| 29442_3 | TATGAGTGTGGAGTCGTTGCGAAGATACTTACTGTAGTTCTCANNNAGATCGGAAG |
| 43152_1 | TATGAGTGTGGAGTCGTTGCCCATCCCCACTCCTTTTCAGTTANNNAGATCGGAAGA |
| 43152_2 | TATGAGTGTGGAGTCGTTGCCCATCCCCACTCCTTTTCAGTTANNNAGATCGGAAGA |
| 43152_3 | TATGAGTGTGGAGTCGTTGCCCATCCCCACTCCTTTTCAGTTANNNAGATCGGAAGA |
| 46900_1 | TATGAGTGTGGAGTCGTTGCAATCTTCCTGCCTCTTTCCATGTNNNAGATCGGAAGA |
| 46900_2 | TATGAGTGTGGAGTCGTTGCAATCTTCCTGCCTCTTTCCATGTNNNAGATCGGAAGA |
| 46900_3 | TATGAGTGTGGAGTCGTTGCAATCTTCCTGCCTCTTTCCATGTNNNAGATCGGAAGA |
| 13261_1 | TATGAGTGTGGAGTCGTTGCCTGAGAAACAGGGATATGAAAANNNAGATCGGAAG |
| 13261_2 | TATGAGTGTGGAGTCGTTGCCTGAGAAACAGGGATATGAAAANNNAGATCGGAAG |
| 13261_3 | TATGAGTGTGGAGTCGTTGCCTGAGAAACAGGGATATGAAAANNNAGATCGGAAG |
| 41105_1 | TATGAGTGTGGAGTCGTTGCAAATTTGTCCAAGCGTAGTGGTGNNNAGATCGGAA |
| 41105_2 | TATGAGTGTGGAGTCGTTGCAAATTTGTCCAAGCGTAGTGGTGNNNAGATCGGAA |
| 41105_3 | TATGAGTGTGGAGTCGTTGCAAATTTGTCCAAGCGTAGTGGTGNNNAGATCGGAA |
| 37111_1 | TATGAGTGTGGAGTCGTTGCGGAGAAATGAGCAGGAAAAAGCANNNAGATCGGA |
| 37111_2 | TATGAGTGTGGAGTCGTTGCGGAGAAATGAGCAGGAAAAAGCANNNAGATCGGA |
| 37111_3 | TATGAGTGTGGAGTCGTTGCGGAGAAATGAGCAGGAAAAAGCANNNAGATCGGA |
| 53093_1 | TATGAGTGTGGAGTCGTTGCTGTGGCAGTAAACTTGTTTCTGGNNNAGATCGGAAG |
| 53093_2 | TATGAGTGTGGAGTCGTTGCTGTGGCAGTAAACTTGTTTCTGGNNNAGATCGGAAG |
| 53093_3 | TATGAGTGTGGAGTCGTTGCTGTGGCAGTAAACTTGTTTCTGGNNNAGATCGGAAG |
| 52911_1 | TATGAGTGTGGAGTCGTTGCGTGCTTAGAGACAGAGGATGTGGNNNAGATCGGA |
| 52911_2 | TATGAGTGTGGAGTCGTTGCGTGCTTAGAGACAGAGGATGTGGNNNAGATCGGA |
| 52911_3 | TATGAGTGTGGAGTCGTTGCGTGCTTAGAGACAGAGGATGTGGNNNAGATCGGA |
| 30872_1 | TATGAGTGTGGAGTCGTTGCTGTGTGTGTGTGTGTGTGATTTCNNNAGATCGGAA |
| 30872_2 | TATGAGTGTGGAGTCGTTGCTGTGTGTGTGTGTGTGTGATTTCNNNAGATCGGAA |
| 30872_3 | TATGAGTGTGGAGTCGTTGCTGTGTGTGTGTGTGTGTGATTTCNNNAGATCGGAA |
| 33041_1 | TATGAGTGTGGAGTCGTTGCGAACTCACCTATGCTAGTGGGAGNNNAGATCGGAA |
| 33041_2 | TATGAGTGTGGAGTCGTTGCGAACTCACCTATGCTAGTGGGAGNNNAGATCGGAA |
| 33041_3 | TATGAGTGTGGAGTCGTTGCGAACTCACCTATGCTAGTGGGAGNNNAGATCGGAA |
| 30910_1 | TATGAGTGTGGAGTCGTTGCAGAGGAGAGGAGAGAGAGAGAGANNNAGATCGG |
| 30910_2 | TATGAGTGTGGAGTCGTTGCAGAGGAGAGGAGAGAGAGAGAGANNNAGATCGG |
| 30910_3 | TATGAGTGTGGAGTCGTTGCAGAGGAGAGGAGAGAGAGAGAGANNNAGATCGG |
| 30506_1 | TATGAGTGTGGAGTCGTTGCCTAAGTTCTGGGAAGCAGCTCACNNNAGATCGGAA |
| 30506_2 | TATGAGTGTGGAGTCGTTGCCTAAGTTCTGGGAAGCAGCTCACNNNAGATCGGAA |
| 30506_3 | TATGAGTGTGGAGTCGTTGCCTAAGTTCTGGGAAGCAGCTCACNNNAGATCGGAA |
| 47064_1 | TATGAGTGTGGAGTCGTTGCAGTAGCATGATCACAGCTCACGNNNAGATCGGAAG |
| 47064_2 | TATGAGTGTGGAGTCGTTGCAGTAGCATGATCACAGCTCACGNNNAGATCGGAAG |
| 47064_3 | TATGAGTGTGGAGTCGTTGCAGTAGCATGATCACAGCTCACGNNNAGATCGGAAG |
| 39801_1 | TATGAGTGTGGAGTCGTTGCTTCCGCCATTAGTAAAAGCTCCTNNNAGATCGGAAG |
| 39801_2 | TATGAGTGTGGAGTCGTTGCTTCCGCCATTAGTAAAAGCTCCTNNNAGATCGGAAG |
| 39801_3 | TATGAGTGTGGAGTCGTTGCTTCCGCCATTAGTAAAAGCTCCTNNNAGATCGGAAG |

| | |
|---|---|
| 42649_1 | TATGAGTGTGGAGTCGTTGCCACACATACGCACCATGGAATACNNNAGATCGGAAG |
| 42649_2 | TATGAGTGTGGAGTCGTTGCCACACATACGCACCATGGAATACNNNAGATCGGAAG |
| 42649_3 | TATGAGTGTGGAGTCGTTGCCACACATACGCACCATGGAATACNNNAGATCGGAAG |
| 10984_1 | TATGAGTGTGGAGTCGTTGCGCTGTCCTGTAAAATTGGTGACAANNNAGATCGGAA |
| 10984_2 | TATGAGTGTGGAGTCGTTGCGCTGTCCTGTAAAATTGGTGACAANNNAGATCGGAA |
| 10984_3 | TATGAGTGTGGAGTCGTTGCGCTGTCCTGTAAAATTGGTGACAANNNAGATCGGAA |
| 40205_1 | TATGAGTGTGGAGTCGTTGCATGGAGGCTAGGAATGGGATAGANNNAGATCGGAA |
| 40205_2 | TATGAGTGTGGAGTCGTTGCATGGAGGCTAGGAATGGGATAGANNNAGATCGGAA |
| 40205_3 | TATGAGTGTGGAGTCGTTGCATGGAGGCTAGGAATGGGATAGANNNAGATCGGAA |
| 10067_1 | TATGAGTGTGGAGTCGTTGCCCAAACGGAGAGAAGAGAAGCACNNNAGATCGGAA |
| 10067_2 | TATGAGTGTGGAGTCGTTGCCCAAACGGAGAGAAGAGAAGCACNNNAGATCGGAA |
| 10067_3 | TATGAGTGTGGAGTCGTTGCCCAAACGGAGAGAAGAGAAGCACNNNAGATCGGAA |
| 12624_1 | TATGAGTGTGGAGTCGTTGCACAGTGGTAATGTATTTTTGTGNNNAGATCGGAAGA |
| 12624_2 | TATGAGTGTGGAGTCGTTGCACAGTGGTAATGTATTTTTGTGNNNAGATCGGAAGA |
| 12624_3 | TATGAGTGTGGAGTCGTTGCACAGTGGTAATGTATTTTTGTGNNNAGATCGGAAGA |
| 41545_1 | TATGAGTGTGGAGTCGTTGCGTGTGATCTCAGTTCATGGCAACNNNAGATCGGAAG |
| 41545_2 | TATGAGTGTGGAGTCGTTGCGTGTGATCTCAGTTCATGGCAACNNNAGATCGGAAG |
| 41545_3 | TATGAGTGTGGAGTCGTTGCGTGTGATCTCAGTTCATGGCAACNNNAGATCGGAAG |
| 46058_1 | TATGAGTGTGGAGTCGTTGCGGGGTTTCACTGTTTCACTCTTGNNNAGATCGGAAGA |
| 46058_2 | TATGAGTGTGGAGTCGTTGCGGGGTTTCACTGTTTCACTCTTGNNNAGATCGGAAGA |
| 46058_3 | TATGAGTGTGGAGTCGTTGCGGGGTTTCACTGTTTCACTCTTGNNNAGATCGGAAGA |
| 43359_1 | TATGAGTGTGGAGTCGTTGCTTGGCGCTTATTCCTTATGTCCTNNNAGATCGGAAGA |
| 43359_2 | TATGAGTGTGGAGTCGTTGCTTGGCGCTTATTCCTTATGTCCTNNNAGATCGGAAGA |
| 43359_3 | TATGAGTGTGGAGTCGTTGCTTGGCGCTTATTCCTTATGTCCTNNNAGATCGGAAGA |
| 29123_1 | TATGAGTGTGGAGTCGTTGCATAGGGATCTCCGTACCATGCTANNNAGATCGGAAG |
| 29123_2 | TATGAGTGTGGAGTCGTTGCATAGGGATCTCCGTACCATGCTANNNAGATCGGAAG |
| 29123_3 | TATGAGTGTGGAGTCGTTGCATAGGGATCTCCGTACCATGCTANNNAGATCGGAAG |
| 43901_1 | TATGAGTGTGGAGTCGTTGCATGACACTTGAAGCCAGTACACTNNNAGATCGGAAG |
| 43901_2 | TATGAGTGTGGAGTCGTTGCATGACACTTGAAGCCAGTACACTNNNAGATCGGAAG |
| 43901_3 | TATGAGTGTGGAGTCGTTGCATGACACTTGAAGCCAGTACACTNNNAGATCGGAAG |
| 49683_1 | TATGAGTGTGGAGTCGTTGCTGCTGCAAACTAGTGGCATATTGNNNAGATCGGAAG |
| 49683_2 | TATGAGTGTGGAGTCGTTGCTGCTGCAAACTAGTGGCATATTGNNNAGATCGGAAG |
| 49683_3 | TATGAGTGTGGAGTCGTTGCTGCTGCAAACTAGTGGCATATTGNNNAGATCGGAAG |
| 43492_1 | TATGAGTGTGGAGTCGTTGCCAATCTTGGCTCAGTGCAACCNNNAGATCGGAAGAG |
| 43492_2 | TATGAGTGTGGAGTCGTTGCCAATCTTGGCTCAGTGCAACCNNNAGATCGGAAGAG |
| 43492_3 | TATGAGTGTGGAGTCGTTGCCAATCTTGGCTCAGTGCAACCNNNAGATCGGAAGAG |
| 35113_1 | TATGAGTGTGGAGTCGTTGCCTCCCAGGTTCATGCCATTCTCNNNAGATCGGAAGAG |
| 35113_2 | TATGAGTGTGGAGTCGTTGCCTCCCAGGTTCATGCCATTCTCNNNAGATCGGAAGAG |
| 35113_3 | TATGAGTGTGGAGTCGTTGCCTCCCAGGTTCATGCCATTCTCNNNAGATCGGAAGAG |
| 45069_1 | TATGAGTGTGGAGTCGTTGCTCAATGGAATGTACCTGGGTGAGNNNAGATCGGAAG |
| 45069_2 | TATGAGTGTGGAGTCGTTGCTCAATGGAATGTACCTGGGTGAGNNNAGATCGGAAG |
| 45069_3 | TATGAGTGTGGAGTCGTTGCTCAATGGAATGTACCTGGGTGAGNNNAGATCGGAAG |
| 31293_1 | TATGAGTGTGGAGTCGTTGCAGAAGATGGATATGACCAGTGGCNNNAGATCGGAA |

| | |
|---|---|
| 31293_2 | TATGAGTGTGGAGTCGTTGCAGAAGATGGATATGACCAGTGGCNNNAGATCGGAA |
| 31293_3 | TATGAGTGTGGAGTCGTTGCAGAAGATGGATATGACCAGTGGCNNNAGATCGGAA |
| 31106_1 | TATGAGTGTGGAGTCGTTGCGGTTCCAAGAGAAACCCTAGCAANNNAGATCGGAA |
| 31106_2 | TATGAGTGTGGAGTCGTTGCGGTTCCAAGAGAAACCCTAGCAANNNAGATCGGAA |
| 31106_3 | TATGAGTGTGGAGTCGTTGCGGTTCCAAGAGAAACCCTAGCAANNNAGATCGGAA |
| 37052_1 | TATGAGTGTGGAGTCGTTGCAAACTTAGGCTCAGAGCAGTCTGNNNAGATCGGAA |
| 37052_2 | TATGAGTGTGGAGTCGTTGCAAACTTAGGCTCAGAGCAGTCTGNNNAGATCGGAA |
| 37052_3 | TATGAGTGTGGAGTCGTTGCAAACTTAGGCTCAGAGCAGTCTGNNNAGATCGGAA |
| 33050_1 | TATGAGTGTGGAGTCGTTGCCTCCTGAGTAGCTGGGACTACAGNNNAGATCGGAA |
| 33050_2 | TATGAGTGTGGAGTCGTTGCCTCCTGAGTAGCTGGGACTACAGNNNAGATCGGAA |
| 33050_3 | TATGAGTGTGGAGTCGTTGCCTCCTGAGTAGCTGGGACTACAGNNNAGATCGGAA |
| 30455_1 | TATGAGTGTGGAGTCGTTGCGGACAAAGCACCCTGAAGTGAGANNNAGATCGGAA |
| 30455_2 | TATGAGTGTGGAGTCGTTGCGGACAAAGCACCCTGAAGTGAGANNNAGATCGGAA |
| 30455_3 | TATGAGTGTGGAGTCGTTGCGGACAAAGCACCCTGAAGTGAGANNNAGATCGGAA |
| 38832_1 | TATGAGTGTGGAGTCGTTGCACCGTAAAGACTCCACAAAACCTNNNAGATCGGAAG |
| 38832_2 | TATGAGTGTGGAGTCGTTGCACCGTAAAGACTCCACAAAACCTNNNAGATCGGAAG |
| 38832_3 | TATGAGTGTGGAGTCGTTGCACCGTAAAGACTCCACAAAACCTNNNAGATCGGAAG |
| 34517_1 | TATGAGTGTGGAGTCGTTGCCAGCTGTGAAAGAAAAGGGCTTTNNNAGATCGGAA |
| 34517_2 | TATGAGTGTGGAGTCGTTGCCAGCTGTGAAAGAAAAGGGCTTTNNNAGATCGGAA |
| 34517_3 | TATGAGTGTGGAGTCGTTGCCAGCTGTGAAAGAAAAGGGCTTTNNNAGATCGGAA |
| 38177_1 | TATGAGTGTGGAGTCGTTGCTCCTAAAGTGATGGGATTGTGGGNNNAGATCGGAA |
| 38177_2 | TATGAGTGTGGAGTCGTTGCTCCTAAAGTGATGGGATTGTGGGNNNAGATCGGAA |
| 38177_3 | TATGAGTGTGGAGTCGTTGCTCCTAAAGTGATGGGATTGTGGGNNNAGATCGGAA |
| 37577_1 | TATGAGTGTGGAGTCGTTGCGGCACAGAAACAGCTTATTGGTTNNNAGATCGGAA |
| 37577_2 | TATGAGTGTGGAGTCGTTGCGGCACAGAAACAGCTTATTGGTTNNNAGATCGGAA |
| 37577_3 | TATGAGTGTGGAGTCGTTGCGGCACAGAAACAGCTTATTGGTTNNNAGATCGGAA |
| 33323_1 | TATGAGTGTGGAGTCGTTGCAGCCTCCCAAGTAGCTAGAACTANNNAGATCGGAAG |
| 33323_2 | TATGAGTGTGGAGTCGTTGCAGCCTCCCAAGTAGCTAGAACTANNNAGATCGGAAG |
| 33323_3 | TATGAGTGTGGAGTCGTTGCAGCCTCCCAAGTAGCTAGAACTANNNAGATCGGAAG |
| 42285_1 | TATGAGTGTGGAGTCGTTGCTGCATTGGTGAGATCTCAACTCANNNAGATCGGAAG |
| 42285_2 | TATGAGTGTGGAGTCGTTGCTGCATTGGTGAGATCTCAACTCANNNAGATCGGAAG |
| 42285_3 | TATGAGTGTGGAGTCGTTGCTGCATTGGTGAGATCTCAACTCANNNAGATCGGAAG |
| 34129_1 | TATGAGTGTGGAGTCGTTGCTCCAGGAAGAATTGGTGCAAAAGNNNAGATCGGAA |
| 34129_2 | TATGAGTGTGGAGTCGTTGCTCCAGGAAGAATTGGTGCAAAAGNNNAGATCGGAA |
| 34129_3 | TATGAGTGTGGAGTCGTTGCTCCAGGAAGAATTGGTGCAAAAGNNNAGATCGGAA |
| 50486_1 | TATGAGTGTGGAGTCGTTGCGAGAGGGAGGGAAGGAGGAAGNNNAGATCGGAA |
| 50486_2 | TATGAGTGTGGAGTCGTTGCGAGAGGGAGGGAAGGAGGAAGNNNAGATCGGAA |
| 50486_3 | TATGAGTGTGGAGTCGTTGCGAGAGGGAGGGAAGGAGGAAGNNNAGATCGGAA |
| 53342_1 | TATGAGTGTGGAGTCGTTGCACAAAAATAAGCCAGGTGTGGTGNNNAGATCGGAA |
| 53342_2 | TATGAGTGTGGAGTCGTTGCACAAAAATAAGCCAGGTGTGGTGNNNAGATCGGAA |
| 53342_3 | TATGAGTGTGGAGTCGTTGCACAAAAATAAGCCAGGTGTGGTGNNNAGATCGGAA |
| 53351_1 | TATGAGTGTGGAGTCGTTGCCTTGGTCTCTCCCTCCTTCTTTCNNNAGATCGGAAGA |
| 53351_2 | TATGAGTGTGGAGTCGTTGCCTTGGTCTCTCCCTCCTTCTTTCNNNAGATCGGAAGA |

| | |
|---|---|
| 53351_3 | TATGAGTGTGGAGTCGTTGCCTTGGTCTCTCCCTCCTTCTTTCNNNAGATCGGAAGA |
| 41212_1 | TATGAGTGTGGAGTCGTTGCAAGAAATTCACCTGTCTCAGCCTNNNAGATCGGAAG |
| 41212_2 | TATGAGTGTGGAGTCGTTGCAAGAAATTCACCTGTCTCAGCCTNNNAGATCGGAAG |
| 41212_3 | TATGAGTGTGGAGTCGTTGCAAGAAATTCACCTGTCTCAGCCTNNNAGATCGGAAG |
| 32471_1 | TATGAGTGTGGAGTCGTTGCCATTCTCCTGCCTCAGCCTCNNNAGATCGGAAGAGCA |
| 32471_2 | TATGAGTGTGGAGTCGTTGCCATTCTCCTGCCTCAGCCTCNNNAGATCGGAAGAGCA |
| 32471_3 | TATGAGTGTGGAGTCGTTGCCATTCTCCTGCCTCAGCCTCNNNAGATCGGAAGAGCA |
| 30539_1 | TATGAGTGTGGAGTCGTTGCACTCTCTCTCTTGAGCGTCTTGCNNNAGATCGGAAGA |
| 30539_2 | TATGAGTGTGGAGTCGTTGCACTCTCTCTCTTGAGCGTCTTGCNNNAGATCGGAAGA |
| 30539_3 | TATGAGTGTGGAGTCGTTGCACTCTCTCTCTTGAGCGTCTTGCNNNAGATCGGAAGA |
| 31037_1 | TATGAGTGTGGAGTCGTTGCCAGTCCATTGTGAGGCCAAGATNNNAGATCGGAAG |
| 31037_2 | TATGAGTGTGGAGTCGTTGCCAGTCCATTGTGAGGCCAAGATNNNAGATCGGAAG |
| 31037_3 | TATGAGTGTGGAGTCGTTGCCAGTCCATTGTGAGGCCAAGATNNNAGATCGGAAG |
| 34573_1 | TATGAGTGTGGAGTCGTTGCCTTTGCTTCAGTAGGATCTGCACNNNAGATCGGAAG |
| 34573_2 | TATGAGTGTGGAGTCGTTGCCTTTGCTTCAGTAGGATCTGCACNNNAGATCGGAAG |
| 34573_3 | TATGAGTGTGGAGTCGTTGCCTTTGCTTCAGTAGGATCTGCACNNNAGATCGGAAG |
| 905_1 | TATGAGTGTGGAGTCGTTGCCCACTCCAGTTTCTCTGAGCATTANNNAGATCGGAAG |
| 905_2 | TATGAGTGTGGAGTCGTTGCCCACTCCAGTTTCTCTGAGCATTANNNAGATCGGAAG |
| 905_3 | TATGAGTGTGGAGTCGTTGCCCACTCCAGTTTCTCTGAGCATTANNNAGATCGGAAG |
| 32576_1 | TATGAGTGTGGAGTCGTTGCTTCAAGTGATTCTCCTGCCTCAANNNAGATCGGAAGA |
| 32576_2 | TATGAGTGTGGAGTCGTTGCTTCAAGTGATTCTCCTGCCTCAANNNAGATCGGAAGA |
| 32576_3 | TATGAGTGTGGAGTCGTTGCTTCAAGTGATTCTCCTGCCTCAANNNAGATCGGAAGA |
| 33376_1 | TATGAGTGTGGAGTCGTTGCAAATGAGGAGACTGTGCTCAGAGNNNAGATCGGAA |
| 33376_2 | TATGAGTGTGGAGTCGTTGCAAATGAGGAGACTGTGCTCAGAGNNNAGATCGGAA |
| 33376_3 | TATGAGTGTGGAGTCGTTGCAAATGAGGAGACTGTGCTCAGAGNNNAGATCGGAA |
| 46970_1 | TATGAGTGTGGAGTCGTTGCCATGATCTCAGCTCACTGCAAGNNNAGATCGGAAGA |
| 46970_2 | TATGAGTGTGGAGTCGTTGCCATGATCTCAGCTCACTGCAAGNNNAGATCGGAAGA |
| 46970_3 | TATGAGTGTGGAGTCGTTGCCATGATCTCAGCTCACTGCAAGNNNAGATCGGAAGA |
| 10708_1 | TATGAGTGTGGAGTCGTTGCCGAAGATGGGTGAAGAAGCAAGANNNAGATCGGA |
| 10708_2 | TATGAGTGTGGAGTCGTTGCCGAAGATGGGTGAAGAAGCAAGANNNAGATCGGA |
| 10708_3 | TATGAGTGTGGAGTCGTTGCCGAAGATGGGTGAAGAAGCAAGANNNAGATCGGA |
| 48120_1 | TATGAGTGTGGAGTCGTTGCCCTCCTAGGTTCAAGCGATTCTCNNNAGATCGGAAG |
| 48120_2 | TATGAGTGTGGAGTCGTTGCCCTCCTAGGTTCAAGCGATTCTCNNNAGATCGGAAG |
| 48120_3 | TATGAGTGTGGAGTCGTTGCCCTCCTAGGTTCAAGCGATTCTCNNNAGATCGGAAG |
| 37337_1 | TATGAGTGTGGAGTCGTTGCTGAATGCTCACTGTAGAGAGAGTGNNNAGATCGGA |
| 37337_2 | TATGAGTGTGGAGTCGTTGCTGAATGCTCACTGTAGAGAGAGTGNNNAGATCGGA |
| 37337_3 | TATGAGTGTGGAGTCGTTGCTGAATGCTCACTGTAGAGAGAGTGNNNAGATCGGA |
| 50987_1 | TATGAGTGTGGAGTCGTTGCCACACTCCAGGAGCGAAACANNNAGATCGGAAGAG |
| 50987_2 | TATGAGTGTGGAGTCGTTGCCACACTCCAGGAGCGAAACANNNAGATCGGAAGAG |
| 50987_3 | TATGAGTGTGGAGTCGTTGCCACACTCCAGGAGCGAAACANNNAGATCGGAAGAG |
| 47182_1 | TATGAGTGTGGAGTCGTTGCAGAGTTTGGGAGGTCACAAGTTTNNNAGATCGGAA |
| 47182_2 | TATGAGTGTGGAGTCGTTGCAGAGTTTGGGAGGTCACAAGTTTNNNAGATCGGAA |
| 47182_3 | TATGAGTGTGGAGTCGTTGCAGAGTTTGGGAGGTCACAAGTTTNNNAGATCGGAA |

| | |
|---|---|
| 39225_1 | TATGAGTGTGGAGTCGTTGCAATGGAAGAACATCTCATGCTCANNNAGATCGGAAG |
| 39225_2 | TATGAGTGTGGAGTCGTTGCAATGGAAGAACATCTCATGCTCANNNAGATCGGAAG |
| 39225_3 | TATGAGTGTGGAGTCGTTGCAATGGAAGAACATCTCATGCTCANNNAGATCGGAAG |
| 38172_1 | TATGAGTGTGGAGTCGTTGCTCTGGATTTCTGGATTCCCTTGANNNAGATCGGAAG |
| 38172_2 | TATGAGTGTGGAGTCGTTGCTCTGGATTTCTGGATTCCCTTGANNNAGATCGGAAG |
| 38172_3 | TATGAGTGTGGAGTCGTTGCTCTGGATTTCTGGATTCCCTTGANNNAGATCGGAAG |
| 33092_1 | TATGAGTGTGGAGTCGTTGCCATTCTCCTGCCTCAGCCTCNNNAGATCGGAAGAGCA |
| 33092_2 | TATGAGTGTGGAGTCGTTGCCATTCTCCTGCCTCAGCCTCNNNAGATCGGAAGAGCA |
| 33092_3 | TATGAGTGTGGAGTCGTTGCCATTCTCCTGCCTCAGCCTCNNNAGATCGGAAGAGCA |
| 36306_1 | TATGAGTGTGGAGTCGTTGCTGAGGTCAGGATTTTGAGACCAGNNNAGATCGGAA |
| 36306_2 | TATGAGTGTGGAGTCGTTGCTGAGGTCAGGATTTTGAGACCAGNNNAGATCGGAA |
| 36306_3 | TATGAGTGTGGAGTCGTTGCTGAGGTCAGGATTTTGAGACCAGNNNAGATCGGAA |
| 46299_1 | TATGAGTGTGGAGTCGTTGCTCACCCAGGTTGGAGTACAGTAGNNNAGATCGGAA |
| 46299_2 | TATGAGTGTGGAGTCGTTGCTCACCCAGGTTGGAGTACAGTAGNNNAGATCGGAA |
| 46299_3 | TATGAGTGTGGAGTCGTTGCTCACCCAGGTTGGAGTACAGTAGNNNAGATCGGAA |
| 35252_1 | TATGAGTGTGGAGTCGTTGCCCTCAATCTCTTGACCTCGTGATNNNAGATCGGAAGA |
| 35252_2 | TATGAGTGTGGAGTCGTTGCCCTCAATCTCTTGACCTCGTGATNNNAGATCGGAAGA |
| 35252_3 | TATGAGTGTGGAGTCGTTGCCCTCAATCTCTTGACCTCGTGATNNNAGATCGGAAGA |
| 34414_1 | TATGAGTGTGGAGTCGTTGCCAAGTAGCTGGGACTACAGGTGNNNAGATCGGAAG |
| 34414_2 | TATGAGTGTGGAGTCGTTGCCAAGTAGCTGGGACTACAGGTGNNNAGATCGGAAG |
| 34414_3 | TATGAGTGTGGAGTCGTTGCCAAGTAGCTGGGACTACAGGTGNNNAGATCGGAAG |
| 12584_1 | TATGAGTGTGGAGTCGTTGCCATAATGTCTTTTTCCAACAAACNNNAGATCGGAAGA |
| 12584_2 | TATGAGTGTGGAGTCGTTGCCATAATGTCTTTTTCCAACAAACNNNAGATCGGAAGA |
| 12584_3 | TATGAGTGTGGAGTCGTTGCCATAATGTCTTTTTCCAACAAACNNNAGATCGGAAGA |
| 48079_1 | TATGAGTGTGGAGTCGTTGCACAGCTATCTACACACACACGTTNNNAGATCGGAAG |
| 48079_2 | TATGAGTGTGGAGTCGTTGCACAGCTATCTACACACACACGTTNNNAGATCGGAAG |
| 48079_3 | TATGAGTGTGGAGTCGTTGCACAGCTATCTACACACACACGTTNNNAGATCGGAAG |
| 47275_1 | TATGAGTGTGGAGTCGTTGCTTTTAGTAGAGACGGGGTTTCACNNNAGATCGGAA |
| 47275_2 | TATGAGTGTGGAGTCGTTGCTTTTAGTAGAGACGGGGTTTCACNNNAGATCGGAA |
| 47275_3 | TATGAGTGTGGAGTCGTTGCTTTTAGTAGAGACGGGGTTTCACNNNAGATCGGAA |
| 44047_1 | TATGAGTGTGGAGTCGTTGCCTACAAGGGAGGAACACCTGATCNNNAGATCGGAA |
| 44047_2 | TATGAGTGTGGAGTCGTTGCCTACAAGGGAGGAACACCTGATCNNNAGATCGGAA |
| 44047_3 | TATGAGTGTGGAGTCGTTGCCTACAAGGGAGGAACACCTGATCNNNAGATCGGAA |
| 2663_1 | TATGAGTGTGGAGTCGTTGCAAAGGGCGAAATGTGTGGAGAATNNNAGATCGGA |
| 2663_2 | TATGAGTGTGGAGTCGTTGCAAAGGGCGAAATGTGTGGAGAATNNNAGATCGGA |
| 2663_3 | TATGAGTGTGGAGTCGTTGCAAAGGGCGAAATGTGTGGAGAATNNNAGATCGGA |
| 9044_1 | TATGAGTGTGGAGTCGTTGCCACAATGAAGTGAAGCAGGTCCTNNNAGATCGGAA |
| 9044_2 | TATGAGTGTGGAGTCGTTGCCACAATGAAGTGAAGCAGGTCCTNNNAGATCGGAA |
| 9044_3 | TATGAGTGTGGAGTCGTTGCCACAATGAAGTGAAGCAGGTCCTNNNAGATCGGAA |
| 32608_1 | TATGAGTGTGGAGTCGTTGCTGAAACAAAAGCAAACATCTTTTCTGANNNAGATCG |
| 32608_2 | TATGAGTGTGGAGTCGTTGCTGAAACAAAAGCAAACATCTTTTCTGANNNAGATCG |
| 32608_3 | TATGAGTGTGGAGTCGTTGCTGAAACAAAAGCAAACATCTTTTCTGANNNAGATCG |
| 32117_1 | TATGAGTGTGGAGTCGTTGCCAGGGGTTGCAATTCTAGTCTCTNNNAGATCGGAAG |

| | |
|---|---|
| 32117_2 | TATGAGTGTGGAGTCGTTGCCAGGGGTTGCAATTCTAGTCTCTNNNAGATCGGAAG |
| 32117_3 | TATGAGTGTGGAGTCGTTGCCAGGGGTTGCAATTCTAGTCTCTNNNAGATCGGAAG |
| 36040_1 | TATGAGTGTGGAGTCGTTGCATCGGGCAAGAACTAGTCAACATNNNAGATCGGAA |
| 36040_2 | TATGAGTGTGGAGTCGTTGCATCGGGCAAGAACTAGTCAACATNNNAGATCGGAA |
| 36040_3 | TATGAGTGTGGAGTCGTTGCATCGGGCAAGAACTAGTCAACATNNNAGATCGGAA |
| 30700_1 | TATGAGTGTGGAGTCGTTGCGAAAAGGCTGATTTCTGAAGGGCNNNAGATCGGAA |
| 30700_2 | TATGAGTGTGGAGTCGTTGCGAAAAGGCTGATTTCTGAAGGGCNNNAGATCGGAA |
| 30700_3 | TATGAGTGTGGAGTCGTTGCGAAAAGGCTGATTTCTGAAGGGCNNNAGATCGGAA |
| 35852_1 | TATGAGTGTGGAGTCGTTGCAAGGTCTCACTTTGTTGCCTAGGNNNAGATCGGAAG |
| 35852_2 | TATGAGTGTGGAGTCGTTGCAAGGTCTCACTTTGTTGCCTAGGNNNAGATCGGAAG |
| 35852_3 | TATGAGTGTGGAGTCGTTGCAAGGTCTCACTTTGTTGCCTAGGNNNAGATCGGAAG |
| 42395_1 | TATGAGTGTGGAGTCGTTGCTCTGTTGGTTTGTTTGGTTATGTNNNAGATCGGAAG |
| 42395_2 | TATGAGTGTGGAGTCGTTGCTCTGTTGGTTTGTTTGGTTATGTNNNAGATCGGAAG |
| 42395_3 | TATGAGTGTGGAGTCGTTGCTCTGTTGGTTTGTTTGGTTATGTNNNAGATCGGAAG |
| 30227_1 | TATGAGTGTGGAGTCGTTGCGAAAATGTCCTTGGCAGTGTCTCNNNAGATCGGAAG |
| 30227_2 | TATGAGTGTGGAGTCGTTGCGAAAATGTCCTTGGCAGTGTCTCNNNAGATCGGAAG |
| 30227_3 | TATGAGTGTGGAGTCGTTGCGAAAATGTCCTTGGCAGTGTCTCNNNAGATCGGAAG |
| 30094_1 | TATGAGTGTGGAGTCGTTGCTGATGTAAGTATCCAGCATTGCTNNNAGATCGGAAG |
| 30094_2 | TATGAGTGTGGAGTCGTTGCTGATGTAAGTATCCAGCATTGCTNNNAGATCGGAAG |
| 30094_3 | TATGAGTGTGGAGTCGTTGCTGATGTAAGTATCCAGCATTGCTNNNAGATCGGAAG |
| 33629_1 | TATGAGTGTGGAGTCGTTGCTTGGTAGATCTTCCTCCATCCCTNNNAGATCGGAAGA |
| 33629_2 | TATGAGTGTGGAGTCGTTGCTTGGTAGATCTTCCTCCATCCCTNNNAGATCGGAAGA |
| 33629_3 | TATGAGTGTGGAGTCGTTGCTTGGTAGATCTTCCTCCATCCCTNNNAGATCGGAAGA |
| 42475_1 | TATGAGTGTGGAGTCGTTGCGAAGGAAGGAAGGAAGGAAGGAANNNAGATCGG |
| 42475_2 | TATGAGTGTGGAGTCGTTGCGAAGGAAGGAAGGAAGGAAGGAANNNAGATCGG |
| 42475_3 | TATGAGTGTGGAGTCGTTGCGAAGGAAGGAAGGAAGGAAGGAANNNAGATCGG |
| 34637_1 | TATGAGTGTGGAGTCGTTGCATGTAACAAACCTGCACGTTCTGNNNAGATCGGAAG |
| 34637_2 | TATGAGTGTGGAGTCGTTGCATGTAACAAACCTGCACGTTCTGNNNAGATCGGAAG |
| 34637_3 | TATGAGTGTGGAGTCGTTGCATGTAACAAACCTGCACGTTCTGNNNAGATCGGAAG |
| 35388_1 | TATGAGTGTGGAGTCGTTGCGCTTACCGACTGAGGTCATAAGANNNAGATCGGAA |
| 35388_2 | TATGAGTGTGGAGTCGTTGCGCTTACCGACTGAGGTCATAAGANNNAGATCGGAA |
| 35388_3 | TATGAGTGTGGAGTCGTTGCGCTTACCGACTGAGGTCATAAGANNNAGATCGGAA |
| 47398_1 | TATGAGTGTGGAGTCGTTGCGGGAGAATAACTTGAGCTTGGGANNNAGATCGGA |
| 47398_2 | TATGAGTGTGGAGTCGTTGCGGGAGAATAACTTGAGCTTGGGANNNAGATCGGA |
| 47398_3 | TATGAGTGTGGAGTCGTTGCGGGAGAATAACTTGAGCTTGGGANNNAGATCGGA |
| 49072_1 | TATGAGTGTGGAGTCGTTGCCAACATACTGGCTTTCCCACATCNNNAGATCGGAAG |
| 49072_2 | TATGAGTGTGGAGTCGTTGCCAACATACTGGCTTTCCCACATCNNNAGATCGGAAG |
| 49072_3 | TATGAGTGTGGAGTCGTTGCCAACATACTGGCTTTCCCACATCNNNAGATCGGAAG |
| 39073_1 | TATGAGTGTGGAGTCGTTGCACTGTCAGCTCCCTACTTTTGAGNNNAGATCGGAAG |
| 39073_2 | TATGAGTGTGGAGTCGTTGCACTGTCAGCTCCCTACTTTTGAGNNNAGATCGGAAG |
| 39073_3 | TATGAGTGTGGAGTCGTTGCACTGTCAGCTCCCTACTTTTGAGNNNAGATCGGAAG |
| 49344_1 | TATGAGTGTGGAGTCGTTGCAGGGAGCAGCAGATTTTGTTTTCNNNAGATCGGAA |
| 49344_2 | TATGAGTGTGGAGTCGTTGCAGGGAGCAGCAGATTTTGTTTTCNNNAGATCGGAA |

| | |
|---|---|
| 49344_3 | TATGAGTGTGGAGTCGTTGCAGGGAGCAGCAGATTTTGTTTTCNNNAGATCGGAA |
| 47763_1 | TATGAGTGTGGAGTCGTTGCGGTGTGAAAAAGAACTATCGGGCNNNAGATCGGAA |
| 47763_2 | TATGAGTGTGGAGTCGTTGCGGTGTGAAAAAGAACTATCGGGCNNNAGATCGGAA |
| 47763_3 | TATGAGTGTGGAGTCGTTGCGGTGTGAAAAAGAACTATCGGGCNNNAGATCGGAA |
| 31019_1 | TATGAGTGTGGAGTCGTTGCATTGTTTTCCTGCCCTCATTTCCNNNAGATCGGAAGA |
| 31019_2 | TATGAGTGTGGAGTCGTTGCATTGTTTTCCTGCCCTCATTTCCNNNAGATCGGAAGA |
| 31019_3 | TATGAGTGTGGAGTCGTTGCATTGTTTTCCTGCCCTCATTTCCNNNAGATCGGAAGA |
| 44319_1 | TATGAGTGTGGAGTCGTTGCGAGGCATCTCTGACCACAAAGGNNNAGATCGGAAG |
| 44319_2 | TATGAGTGTGGAGTCGTTGCGAGGCATCTCTGACCACAAAGGNNNAGATCGGAAG |
| 44319_3 | TATGAGTGTGGAGTCGTTGCGAGGCATCTCTGACCACAAAGGNNNAGATCGGAAG |
| 38934_1 | TATGAGTGTGGAGTCGTTGCTTATGAAGCCAGCATCATCCTGANNNAGATCGGAAG |
| 38934_2 | TATGAGTGTGGAGTCGTTGCTTATGAAGCCAGCATCATCCTGANNNAGATCGGAAG |
| 38934_3 | TATGAGTGTGGAGTCGTTGCTTATGAAGCCAGCATCATCCTGANNNAGATCGGAAG |
| 50590_1 | TATGAGTGTGGAGTCGTTGCTGCATGCTGTTGATGTTCAGTTTNNNAGATCGGAAG |
| 50590_2 | TATGAGTGTGGAGTCGTTGCTGCATGCTGTTGATGTTCAGTTTNNNAGATCGGAAG |
| 50590_3 | TATGAGTGTGGAGTCGTTGCTGCATGCTGTTGATGTTCAGTTTNNNAGATCGGAAG |
| 245_1 | TATGAGTGTGGAGTCGTTGCTGATTTTTGTATGGGCCCAGGGTNNNAGATCGGAA |
| 245_2 | TATGAGTGTGGAGTCGTTGCTGATTTTTGTATGGGCCCAGGGTNNNAGATCGGAA |
| 245_3 | TATGAGTGTGGAGTCGTTGCTGATTTTTGTATGGGCCCAGGGTNNNAGATCGGAA |
| 42681_1 | TATGAGTGTGGAGTCGTTGCGAGAATGGTGTGAACCCAGGAGNNNAGATCGGAA |
| 42681_2 | TATGAGTGTGGAGTCGTTGCGAGAATGGTGTGAACCCAGGAGNNNAGATCGGAA |
| 42681_3 | TATGAGTGTGGAGTCGTTGCGAGAATGGTGTGAACCCAGGAGNNNAGATCGGAA |
| 29187_1 | TATGAGTGTGGAGTCGTTGCTTTTGTGCTTTTTGTCCGTGACTNNNAGATCGGAAG |
| 29187_2 | TATGAGTGTGGAGTCGTTGCTTTTGTGCTTTTTGTCCGTGACTNNNAGATCGGAAG |
| 29187_3 | TATGAGTGTGGAGTCGTTGCTTTTGTGCTTTTTGTCCGTGACTNNNAGATCGGAAG |
| 42144_1 | TATGAGTGTGGAGTCGTTGCGTTCAAGAGATTCTCCTGCCTCANNNAGATCGGAAG |
| 42144_2 | TATGAGTGTGGAGTCGTTGCGTTCAAGAGATTCTCCTGCCTCANNNAGATCGGAAG |
| 42144_3 | TATGAGTGTGGAGTCGTTGCGTTCAAGAGATTCTCCTGCCTCANNNAGATCGGAAG |
| 28987_1 | TATGAGTGTGGAGTCGTTGCGCACAAATACATTCCCCACATGTNNNAGATCGGAAG |
| 28987_2 | TATGAGTGTGGAGTCGTTGCGCACAAATACATTCCCCACATGTNNNAGATCGGAAG |
| 28987_3 | TATGAGTGTGGAGTCGTTGCGCACAAATACATTCCCCACATGTNNNAGATCGGAAG |
| 53078_1 | TATGAGTGTGGAGTCGTTGCGGGAGAGACATTAGCAATCCTGTNNNAGATCGGAA |
| 53078_2 | TATGAGTGTGGAGTCGTTGCGGGAGAGACATTAGCAATCCTGTNNNAGATCGGAA |
| 53078_3 | TATGAGTGTGGAGTCGTTGCGGGAGAGACATTAGCAATCCTGTNNNAGATCGGAA |
| 46240_1 | TATGAGTGTGGAGTCGTTGCCCACCACACCCAGTGATTTTTTTNNNAGATCGGAAGA |
| 46240_2 | TATGAGTGTGGAGTCGTTGCCCACCACACCCAGTGATTTTTTTNNNAGATCGGAAGA |
| 46240_3 | TATGAGTGTGGAGTCGTTGCCCACCACACCCAGTGATTTTTTTNNNAGATCGGAAGA |
| 34953_1 | TATGAGTGTGGAGTCGTTGCACAAAGGCAGTGGGGAATTCTTANNNAGATCGGAA |
| 34953_2 | TATGAGTGTGGAGTCGTTGCACAAAGGCAGTGGGGAATTCTTANNNAGATCGGAA |
| 34953_3 | TATGAGTGTGGAGTCGTTGCACAAAGGCAGTGGGGAATTCTTANNNAGATCGGAA |
| 50689_1 | TATGAGTGTGGAGTCGTTGCGGGAAGCCTGAAACATCCTAAGANNNAGATCGGAA |
| 50689_2 | TATGAGTGTGGAGTCGTTGCGGGAAGCCTGAAACATCCTAAGANNNAGATCGGAA |
| 50689_3 | TATGAGTGTGGAGTCGTTGCGGGAAGCCTGAAACATCCTAAGANNNAGATCGGAA |

| | |
|---|---|
| 45797_1 | TATGAGTGTGGAGTCGTTGCCCCGAGTAGCTGGTATTACAGACNNNAGATCGGAA |
| 45797_2 | TATGAGTGTGGAGTCGTTGCCCCGAGTAGCTGGTATTACAGACNNNAGATCGGAA |
| 45797_3 | TATGAGTGTGGAGTCGTTGCCCCGAGTAGCTGGTATTACAGACNNNAGATCGGAA |
| 34490_1 | TATGAGTGTGGAGTCGTTGCCTGGAGGACTCACAACTCACTACNNNAGATCGGAAG |
| 34490_2 | TATGAGTGTGGAGTCGTTGCCTGGAGGACTCACAACTCACTACNNNAGATCGGAAG |
| 34490_3 | TATGAGTGTGGAGTCGTTGCCTGGAGGACTCACAACTCACTACNNNAGATCGGAAG |
| 44154_1 | TATGAGTGTGGAGTCGTTGCCTGTCCAAACAACTCTCTCTGGANNNAGATCGGAAG |
| 44154_2 | TATGAGTGTGGAGTCGTTGCCTGTCCAAACAACTCTCTCTGGANNNAGATCGGAAG |
| 44154_3 | TATGAGTGTGGAGTCGTTGCCTGTCCAAACAACTCTCTCTGGANNNAGATCGGAAG |
| 39832_1 | TATGAGTGTGGAGTCGTTGCCTGTGGGCCCTTTTTGTGTAAATNNNAGATCGGAAG |
| 39832_2 | TATGAGTGTGGAGTCGTTGCCTGTGGGCCCTTTTTGTGTAAATNNNAGATCGGAAG |
| 39832_3 | TATGAGTGTGGAGTCGTTGCCTGTGGGCCCTTTTTGTGTAAATNNNAGATCGGAAG |
| 12860_1 | TATGAGTGTGGAGTCGTTGCGTAAGACTCTGTGACAAATCTATNNNAGATCGGAAG |
| 12860_2 | TATGAGTGTGGAGTCGTTGCGTAAGACTCTGTGACAAATCTATNNNAGATCGGAAG |
| 12860_3 | TATGAGTGTGGAGTCGTTGCGTAAGACTCTGTGACAAATCTATNNNAGATCGGAAG |
| 13170_1 | TATGAGTGTGGAGTCGTTGCTTGTTGTCCAGTTGAAAAAAGNNNAGATCGGAAGA |
| 13170_2 | TATGAGTGTGGAGTCGTTGCTTGTTGTCCAGTTGAAAAAAGNNNAGATCGGAAGA |
| 13170_3 | TATGAGTGTGGAGTCGTTGCTTGTTGTCCAGTTGAAAAAAGNNNAGATCGGAAGA |
| 28953_1 | TATGAGTGTGGAGTCGTTGCATGTGTGTCTCTGTGTGTGTACANNNAGATCGGAAG |
| 28953_2 | TATGAGTGTGGAGTCGTTGCATGTGTGTCTCTGTGTGTGTACANNNAGATCGGAAG |
| 28953_3 | TATGAGTGTGGAGTCGTTGCATGTGTGTCTCTGTGTGTGTACANNNAGATCGGAAG |
| 36218_1 | TATGAGTGTGGAGTCGTTGCGAGTACCATGAGTTCTCCAGCAANNNAGATCGGAAG |
| 36218_2 | TATGAGTGTGGAGTCGTTGCGAGTACCATGAGTTCTCCAGCAANNNAGATCGGAAG |
| 36218_3 | TATGAGTGTGGAGTCGTTGCGAGTACCATGAGTTCTCCAGCAANNNAGATCGGAAG |
| 13755_1 | TATGAGTGTGGAGTCGTTGCCTCAACAGATAGAACATTTTGAANNNAGATCGGAAG |
| 13755_2 | TATGAGTGTGGAGTCGTTGCCTCAACAGATAGAACATTTTGAANNNAGATCGGAAG |
| 13755_3 | TATGAGTGTGGAGTCGTTGCCTCAACAGATAGAACATTTTGAANNNAGATCGGAAG |
| 45352_1 | TATGAGTGTGGAGTCGTTGCGAAATCTACGTTATTGGGCCAGCNNNAGATCGGAA |
| 45352_2 | TATGAGTGTGGAGTCGTTGCGAAATCTACGTTATTGGGCCAGCNNNAGATCGGAA |
| 45352_3 | TATGAGTGTGGAGTCGTTGCGAAATCTACGTTATTGGGCCAGCNNNAGATCGGAA |
| 38324_1 | TATGAGTGTGGAGTCGTTGCGAAAGAAAGAAGGAACCTAGGGCNNNAGATCGGA |
| 38324_2 | TATGAGTGTGGAGTCGTTGCGAAAGAAAGAAGGAACCTAGGGCNNNAGATCGGA |
| 38324_3 | TATGAGTGTGGAGTCGTTGCGAAAGAAAGAAGGAACCTAGGGCNNNAGATCGGA |
| 38432_1 | TATGAGTGTGGAGTCGTTGCGAGTCCTCTAAAACAGCCAAGGANNNAGATCGGAA |
| 38432_2 | TATGAGTGTGGAGTCGTTGCGAGTCCTCTAAAACAGCCAAGGANNNAGATCGGAA |
| 38432_3 | TATGAGTGTGGAGTCGTTGCGAGTCCTCTAAAACAGCCAAGGANNNAGATCGGAA |
| 30822_1 | TATGAGTGTGGAGTCGTTGCTGGGTATTCAATGAAGTCGGAGGNNNAGATCGGAA |
| 30822_2 | TATGAGTGTGGAGTCGTTGCTGGGTATTCAATGAAGTCGGAGGNNNAGATCGGAA |
| 30822_3 | TATGAGTGTGGAGTCGTTGCTGGGTATTCAATGAAGTCGGAGGNNNAGATCGGAA |
| 48379_1 | TATGAGTGTGGAGTCGTTGCCTCTCTCTAGCTATCCCCCTCTCNNNAGATCGGAAGA |
| 48379_2 | TATGAGTGTGGAGTCGTTGCCTCTCTCTAGCTATCCCCCTCTCNNNAGATCGGAAGA |
| 48379_3 | TATGAGTGTGGAGTCGTTGCCTCTCTCTAGCTATCCCCCTCTCNNNAGATCGGAAGA |
| 43591_1 | TATGAGTGTGGAGTCGTTGCGAACCTGAGGCTGTAGGTCATGNNNAGATCGGAAG |

| | |
|---|---|
| 43591_2 | TATGAGTGTGGAGTCGTTGCGAACCTGAGGCTGTAGGTCATGNNNAGATCGGAAG |
| 43591_3 | TATGAGTGTGGAGTCGTTGCGAACCTGAGGCTGTAGGTCATGNNNAGATCGGAAG |
| 52349_1 | TATGAGTGTGGAGTCGTTGCAGCTGAGTTTAGTGGTTAGTGCANNNAGATCGGAA |
| 52349_2 | TATGAGTGTGGAGTCGTTGCAGCTGAGTTTAGTGGTTAGTGCANNNAGATCGGAA |
| 52349_3 | TATGAGTGTGGAGTCGTTGCAGCTGAGTTTAGTGGTTAGTGCANNNAGATCGGAA |
| 50523_1 | TATGAGTGTGGAGTCGTTGCTATTTTGGGGATTTCAAGGCAGCNNNAGATCGGAA |
| 50523_2 | TATGAGTGTGGAGTCGTTGCTATTTTGGGGATTTCAAGGCAGCNNNAGATCGGAA |
| 50523_3 | TATGAGTGTGGAGTCGTTGCTATTTTGGGGATTTCAAGGCAGCNNNAGATCGGAA |
| 34258_1 | TATGAGTGTGGAGTCGTTGCCCTCCTAAACTCCCCATGTTCAANNNAGATCGGAAGA |
| 34258_2 | TATGAGTGTGGAGTCGTTGCCCTCCTAAACTCCCCATGTTCAANNNAGATCGGAAGA |
| 34258_3 | TATGAGTGTGGAGTCGTTGCCCTCCTAAACTCCCCATGTTCAANNNAGATCGGAAGA |
| 50114_1 | TATGAGTGTGGAGTCGTTGCCTCCTCTTGATTTCGGGTGTCTGNNNAGATCGGAAG |
| 50114_2 | TATGAGTGTGGAGTCGTTGCCTCCTCTTGATTTCGGGTGTCTGNNNAGATCGGAAG |
| 50114_3 | TATGAGTGTGGAGTCGTTGCCTCCTCTTGATTTCGGGTGTCTGNNNAGATCGGAAG |
| 29841_1 | TATGAGTGTGGAGTCGTTGCAGAAATCACAGCAAGACTGAGGCNNNAGATCGGAA |
| 29841_2 | TATGAGTGTGGAGTCGTTGCAGAAATCACAGCAAGACTGAGGCNNNAGATCGGAA |
| 29841_3 | TATGAGTGTGGAGTCGTTGCAGAAATCACAGCAAGACTGAGGCNNNAGATCGGAA |
| 300_1 | TATGAGTGTGGAGTCGTTGCGAAGAAAGAGGCGAGGGCTAGGNNNAGATCGGAA |
| 300_2 | TATGAGTGTGGAGTCGTTGCGAAGAAAGAGGCGAGGGCTAGGNNNAGATCGGAA |
| 300_3 | TATGAGTGTGGAGTCGTTGCGAAGAAAGAGGCGAGGGCTAGGNNNAGATCGGAA |
| 12954_1 | TATGAGTGTGGAGTCGTTGCTAAAACTAGACTCAGTGAAAAGANNNAGATCGGAA |
| 12954_2 | TATGAGTGTGGAGTCGTTGCTAAAACTAGACTCAGTGAAAAGANNNAGATCGGAA |
| 12954_3 | TATGAGTGTGGAGTCGTTGCTAAAACTAGACTCAGTGAAAAGANNNAGATCGGAA |
| 32783_1 | TATGAGTGTGGAGTCGTTGCCTATGATCCTGCCACTACACTCCNNNAGATCGGAAGA |
| 32783_2 | TATGAGTGTGGAGTCGTTGCCTATGATCCTGCCACTACACTCCNNNAGATCGGAAGA |
| 32783_3 | TATGAGTGTGGAGTCGTTGCCTATGATCCTGCCACTACACTCCNNNAGATCGGAAGA |
| 3114_1 | TATGAGTGTGGAGTCGTTGCGCAGCTCTTGGCCTGAGACTNNNAGATCGGAAGAG |
| 3114_2 | TATGAGTGTGGAGTCGTTGCGCAGCTCTTGGCCTGAGACTNNNAGATCGGAAGAG |
| 3114_3 | TATGAGTGTGGAGTCGTTGCGCAGCTCTTGGCCTGAGACTNNNAGATCGGAAGAG |
| 39746_1 | TATGAGTGTGGAGTCGTTGCCCTGTAGTCCCAGCTACTTGGNNNAGATCGGAAGAG |
| 39746_2 | TATGAGTGTGGAGTCGTTGCCCTGTAGTCCCAGCTACTTGGNNNAGATCGGAAGAG |
| 39746_3 | TATGAGTGTGGAGTCGTTGCCCTGTAGTCCCAGCTACTTGGNNNAGATCGGAAGAG |
| 49908_1 | TATGAGTGTGGAGTCGTTGCTGGCTCATGATGACATTCCGTATNNNAGATCGGAAG |
| 49908_2 | TATGAGTGTGGAGTCGTTGCTGGCTCATGATGACATTCCGTATNNNAGATCGGAAG |
| 49908_3 | TATGAGTGTGGAGTCGTTGCTGGCTCATGATGACATTCCGTATNNNAGATCGGAAG |
| 42173_1 | TATGAGTGTGGAGTCGTTGCTTCAAGTGATTCTCCTGCCTCAGNNNAGATCGGAAG |
| 42173_2 | TATGAGTGTGGAGTCGTTGCTTCAAGTGATTCTCCTGCCTCAGNNNAGATCGGAAG |
| 42173_3 | TATGAGTGTGGAGTCGTTGCTTCAAGTGATTCTCCTGCCTCAGNNNAGATCGGAAG |
| 35253_1 | TATGAGTGTGGAGTCGTTGCCAGTGAAGTGGCACAATCTTGTCNNNAGATCGGAA |
| 35253_2 | TATGAGTGTGGAGTCGTTGCCAGTGAAGTGGCACAATCTTGTCNNNAGATCGGAA |
| 35253_3 | TATGAGTGTGGAGTCGTTGCCAGTGAAGTGGCACAATCTTGTCNNNAGATCGGAA |
| 48604_1 | TATGAGTGTGGAGTCGTTGCTTCAAGCGATTTTTCTGCCTCAGNNNAGATCGGAAG |
| 48604_2 | TATGAGTGTGGAGTCGTTGCTTCAAGCGATTTTTCTGCCTCAGNNNAGATCGGAAG |

| | |
|---|---|
| 48604_3 | TATGAGTGTGGAGTCGTTGCTTCAAGCGATTTTTCTGCCTCAGNNNAGATCGGAAG |
| 40059_1 | TATGAGTGTGGAGTCGTTGCTTGCAGAGCTAGTATGGTGATGGNNNAGATCGGAA |
| 40059_2 | TATGAGTGTGGAGTCGTTGCTTGCAGAGCTAGTATGGTGATGGNNNAGATCGGAA |
| 40059_3 | TATGAGTGTGGAGTCGTTGCTTGCAGAGCTAGTATGGTGATGGNNNAGATCGGAA |
| 32690_1 | TATGAGTGTGGAGTCGTTGCCCAAGAGACTAGCCCTTTTGTCANNNAGATCGGAAG |
| 32690_2 | TATGAGTGTGGAGTCGTTGCCCAAGAGACTAGCCCTTTTGTCANNNAGATCGGAAG |
| 32690_3 | TATGAGTGTGGAGTCGTTGCCCAAGAGACTAGCCCTTTTGTCANNNAGATCGGAAG |
| 48898_1 | TATGAGTGTGGAGTCGTTGCAATGAAGTGGCTTTCTCAAAGGCNNNAGATCGGAA |
| 48898_2 | TATGAGTGTGGAGTCGTTGCAATGAAGTGGCTTTCTCAAAGGCNNNAGATCGGAA |
| 48898_3 | TATGAGTGTGGAGTCGTTGCAATGAAGTGGCTTTCTCAAAGGCNNNAGATCGGAA |
| 42093_1 | TATGAGTGTGGAGTCGTTGCTTGGGTGCATGTATGTCTTCCTTNNNAGATCGGAAG |
| 42093_2 | TATGAGTGTGGAGTCGTTGCTTGGGTGCATGTATGTCTTCCTTNNNAGATCGGAAG |
| 42093_3 | TATGAGTGTGGAGTCGTTGCTTGGGTGCATGTATGTCTTCCTTNNNAGATCGGAAG |
| 41679_1 | TATGAGTGTGGAGTCGTTGCTTCAAGTGATTCTCCTGCCTCAGNNNAGATCGGAAG |
| 41679_2 | TATGAGTGTGGAGTCGTTGCTTCAAGTGATTCTCCTGCCTCAGNNNAGATCGGAAG |
| 41679_3 | TATGAGTGTGGAGTCGTTGCTTCAAGTGATTCTCCTGCCTCAGNNNAGATCGGAAG |
| 30225_1 | TATGAGTGTGGAGTCGTTGCACCACCTTGTGAGGATGAAATAGANNNAGATCGGA |
| 30225_2 | TATGAGTGTGGAGTCGTTGCACCACCTTGTGAGGATGAAATAGANNNAGATCGGA |
| 30225_3 | TATGAGTGTGGAGTCGTTGCACCACCTTGTGAGGATGAAATAGANNNAGATCGGA |
| 40055_1 | TATGAGTGTGGAGTCGTTGCAAGCACATCTTGGACTCAAAGGANNNAGATCGGAA |
| 40055_2 | TATGAGTGTGGAGTCGTTGCAAGCACATCTTGGACTCAAAGGANNNAGATCGGAA |
| 40055_3 | TATGAGTGTGGAGTCGTTGCAAGCACATCTTGGACTCAAAGGANNNAGATCGGAA |
| 32538_1 | TATGAGTGTGGAGTCGTTGCATGTTGACCTGTACCACACAAAANNNAGATCGGAAG |
| 32538_2 | TATGAGTGTGGAGTCGTTGCATGTTGACCTGTACCACACAAAANNNAGATCGGAAG |
| 32538_3 | TATGAGTGTGGAGTCGTTGCATGTTGACCTGTACCACACAAAANNNAGATCGGAAG |
| 40666_1 | TATGAGTGTGGAGTCGTTGCGCAGAAACTGGCCAAACATACATNNNAGATCGGAA |
| 40666_2 | TATGAGTGTGGAGTCGTTGCGCAGAAACTGGCCAAACATACATNNNAGATCGGAA |
| 40666_3 | TATGAGTGTGGAGTCGTTGCGCAGAAACTGGCCAAACATACATNNNAGATCGGAA |
| 35865_1 | TATGAGTGTGGAGTCGTTGCAAAAGGAAACAGGACGGTCTGATNNNAGATCGGAA |
| 35865_2 | TATGAGTGTGGAGTCGTTGCAAAAGGAAACAGGACGGTCTGATNNNAGATCGGAA |
| 35865_3 | TATGAGTGTGGAGTCGTTGCAAAAGGAAACAGGACGGTCTGATNNNAGATCGGAA |
| 41920_1 | TATGAGTGTGGAGTCGTTGCAGACACTGTCTCACTCTGTTGCNNNAGATCGGAAGA |
| 41920_2 | TATGAGTGTGGAGTCGTTGCAGACACTGTCTCACTCTGTTGCNNNAGATCGGAAGA |
| 41920_3 | TATGAGTGTGGAGTCGTTGCAGACACTGTCTCACTCTGTTGCNNNAGATCGGAAGA |
| 39048_1 | TATGAGTGTGGAGTCGTTGCAGGAAGACATTTATGTGACCAACANNNAGATCGGA |
| 39048_2 | TATGAGTGTGGAGTCGTTGCAGGAAGACATTTATGTGACCAACANNNAGATCGGA |
| 39048_3 | TATGAGTGTGGAGTCGTTGCAGGAAGACATTTATGTGACCAACANNNAGATCGGA |
| 36647_1 | TATGAGTGTGGAGTCGTTGCAAAACAATTTAGCGGAGTCCCACNNNAGATCGGAAG |
| 36647_2 | TATGAGTGTGGAGTCGTTGCAAAACAATTTAGCGGAGTCCCACNNNAGATCGGAAG |
| 36647_3 | TATGAGTGTGGAGTCGTTGCAAAACAATTTAGCGGAGTCCCACNNNAGATCGGAAG |
| 29377_1 | TATGAGTGTGGAGTCGTTGCGATTTGAGTGAGCTATGTTCGGCNNNAGATCGGAA |
| 29377_2 | TATGAGTGTGGAGTCGTTGCGATTTGAGTGAGCTATGTTCGGCNNNAGATCGGAA |
| 29377_3 | TATGAGTGTGGAGTCGTTGCGATTTGAGTGAGCTATGTTCGGCNNNAGATCGGAA |

| | |
|---|---|
| 51116_1 | TATGAGTGTGGAGTCGTTGCGCCTGTGACAGCTTCATAATGTCNNNAGATCGGAAG |
| 51116_2 | TATGAGTGTGGAGTCGTTGCGCCTGTGACAGCTTCATAATGTCNNNAGATCGGAAG |
| 51116_3 | TATGAGTGTGGAGTCGTTGCGCCTGTGACAGCTTCATAATGTCNNNAGATCGGAAG |
| 38517_1 | TATGAGTGTGGAGTCGTTGCTCACTTGAGGTCAGGAGTTCAACNNNAGATCGGAAG |
| 38517_2 | TATGAGTGTGGAGTCGTTGCTCACTTGAGGTCAGGAGTTCAACNNNAGATCGGAAG |
| 38517_3 | TATGAGTGTGGAGTCGTTGCTCACTTGAGGTCAGGAGTTCAACNNNAGATCGGAAG |
| 44085_1 | TATGAGTGTGGAGTCGTTGCTCCTACTTGTGTGGGAAATCTGGNNNAGATCGGAAG |
| 44085_2 | TATGAGTGTGGAGTCGTTGCTCCTACTTGTGTGGGAAATCTGGNNNAGATCGGAAG |
| 44085_3 | TATGAGTGTGGAGTCGTTGCTCCTACTTGTGTGGGAAATCTGGNNNAGATCGGAAG |
| 37046_1 | TATGAGTGTGGAGTCGTTGCAACAAGTTGGCTATAGTTGCAGANNNAGATCGGAA |
| 37046_2 | TATGAGTGTGGAGTCGTTGCAACAAGTTGGCTATAGTTGCAGANNNAGATCGGAA |
| 37046_3 | TATGAGTGTGGAGTCGTTGCAACAAGTTGGCTATAGTTGCAGANNNAGATCGGAA |
| 32585_1 | TATGAGTGTGGAGTCGTTGCGTTCAAGCGATTCTCTTGCCTCNNNAGATCGGAAGA |
| 32585_2 | TATGAGTGTGGAGTCGTTGCGTTCAAGCGATTCTCTTGCCTCNNNAGATCGGAAGA |
| 32585_3 | TATGAGTGTGGAGTCGTTGCGTTCAAGCGATTCTCTTGCCTCNNNAGATCGGAAGA |
| 37753_1 | TATGAGTGTGGAGTCGTTGCTGTTGCAACATGGATGAACCTTCNNNAGATCGGAAG |
| 37753_2 | TATGAGTGTGGAGTCGTTGCTGTTGCAACATGGATGAACCTTCNNNAGATCGGAAG |
| 37753_3 | TATGAGTGTGGAGTCGTTGCTGTTGCAACATGGATGAACCTTCNNNAGATCGGAAG |
| 3093_1 | TATGAGTGTGGAGTCGTTGCAGGTGCAGTCTATCTGACTTTCANNNAGATCGGAAG |
| 3093_2 | TATGAGTGTGGAGTCGTTGCAGGTGCAGTCTATCTGACTTTCANNNAGATCGGAAG |
| 3093_3 | TATGAGTGTGGAGTCGTTGCAGGTGCAGTCTATCTGACTTTCANNNAGATCGGAAG |
| 50626_1 | TATGAGTGTGGAGTCGTTGCACTAAGGCCCAATCATTCACCTTNNNAGATCGGAAGA |
| 50626_2 | TATGAGTGTGGAGTCGTTGCACTAAGGCCCAATCATTCACCTTNNNAGATCGGAAGA |
| 50626_3 | TATGAGTGTGGAGTCGTTGCACTAAGGCCCAATCATTCACCTTNNNAGATCGGAAGA |
| 32393_1 | TATGAGTGTGGAGTCGTTGCACAGCTTTTTTCCTTTGGCTCTCNNNAGATCGGAAGA |
| 32393_2 | TATGAGTGTGGAGTCGTTGCACAGCTTTTTTCCTTTGGCTCTCNNNAGATCGGAAGA |
| 32393_3 | TATGAGTGTGGAGTCGTTGCACAGCTTTTTTCCTTTGGCTCTCNNNAGATCGGAAGA |
| 35028_1 | TATGAGTGTGGAGTCGTTGCCCTCCCGAGTTCAAGAGATTCTCNNNAGATCGGAAGA |
| 35028_2 | TATGAGTGTGGAGTCGTTGCCCTCCCGAGTTCAAGAGATTCTCNNNAGATCGGAAGA |
| 35028_3 | TATGAGTGTGGAGTCGTTGCCCTCCCGAGTTCAAGAGATTCTCNNNAGATCGGAAGA |
| 43495_1 | TATGAGTGTGGAGTCGTTGCGGTCATGCCCATACACATTTGAGNNNAGATCGGAAG |
| 43495_2 | TATGAGTGTGGAGTCGTTGCGGTCATGCCCATACACATTTGAGNNNAGATCGGAAG |
| 43495_3 | TATGAGTGTGGAGTCGTTGCGGTCATGCCCATACACATTTGAGNNNAGATCGGAAG |
| 36521_1 | TATGAGTGTGGAGTCGTTGCAGATTGCAGTGAGCTGAGATCAANNNAGATCGGAA |
| 36521_2 | TATGAGTGTGGAGTCGTTGCAGATTGCAGTGAGCTGAGATCAANNNAGATCGGAA |
| 36521_3 | TATGAGTGTGGAGTCGTTGCAGATTGCAGTGAGCTGAGATCAANNNAGATCGGAA |
| 48238_1 | TATGAGTGTGGAGTCGTTGCGCTGGAATCAACGAATGTGCTTANNNAGATCGGAAG |
| 48238_2 | TATGAGTGTGGAGTCGTTGCGCTGGAATCAACGAATGTGCTTANNNAGATCGGAAG |
| 48238_3 | TATGAGTGTGGAGTCGTTGCGCTGGAATCAACGAATGTGCTTANNNAGATCGGAAG |
| 46945_1 | TATGAGTGTGGAGTCGTTGCCCACCACACCCAGCTAATTTTTTNNNAGATCGGAAGA |
| 46945_2 | TATGAGTGTGGAGTCGTTGCCCACCACACCCAGCTAATTTTTTNNNAGATCGGAAGA |
| 46945_3 | TATGAGTGTGGAGTCGTTGCCCACCACACCCAGCTAATTTTTTNNNAGATCGGAAGA |
| 46787_1 | TATGAGTGTGGAGTCGTTGCCTGAGTAGCTGTGATTACATGCGNNNAGATCGGAAG |

| | |
|---|---|
| 46787_2 | TATGAGTGTGGAGTCGTTGCCTGAGTAGCTGTGATTACATGCGNNNAGATCGGAAG |
| 46787_3 | TATGAGTGTGGAGTCGTTGCCTGAGTAGCTGTGATTACATGCGNNNAGATCGGAAG |
| 36608_1 | TATGAGTGTGGAGTCGTTGCAAGCAGGCTTTCATTTATGGCTCNNNAGATCGGAAG |
| 36608_2 | TATGAGTGTGGAGTCGTTGCAAGCAGGCTTTCATTTATGGCTCNNNAGATCGGAAG |
| 36608_3 | TATGAGTGTGGAGTCGTTGCAAGCAGGCTTTCATTTATGGCTCNNNAGATCGGAAG |
| 46513_1 | TATGAGTGTGGAGTCGTTGCCACACAGATCTCCATCTCGTAGGNNNAGATCGGAAGA |
| 46513_2 | TATGAGTGTGGAGTCGTTGCCACACAGATCTCCATCTCGTAGGNNNAGATCGGAAGA |
| 46513_3 | TATGAGTGTGGAGTCGTTGCCACACAGATCTCCATCTCGTAGGNNNAGATCGGAAGA |
| 29168_1 | TATGAGTGTGGAGTCGTTGCTGCCCTCACCACATAATTCTCTTNNNAGATCGGAAGA |
| 29168_2 | TATGAGTGTGGAGTCGTTGCTGCCCTCACCACATAATTCTCTTNNNAGATCGGAAGA |
| 29168_3 | TATGAGTGTGGAGTCGTTGCTGCCCTCACCACATAATTCTCTTNNNAGATCGGAAGA |
| 42550_1 | TATGAGTGTGGAGTCGTTGCAGAGACGAGGTTTCTCCATGTTGNNNAGATCGGAAG |
| 42550_2 | TATGAGTGTGGAGTCGTTGCAGAGACGAGGTTTCTCCATGTTGNNNAGATCGGAAG |
| 42550_3 | TATGAGTGTGGAGTCGTTGCAGAGACGAGGTTTCTCCATGTTGNNNAGATCGGAAG |
| 35934_1 | TATGAGTGTGGAGTCGTTGCTGGTAGATGACAGATTGACAGGTGNNNAGATCGGA |
| 35934_2 | TATGAGTGTGGAGTCGTTGCTGGTAGATGACAGATTGACAGGTGNNNAGATCGGA |
| 35934_3 | TATGAGTGTGGAGTCGTTGCTGGTAGATGACAGATTGACAGGTGNNNAGATCGGA |
| 29996_1 | TATGAGTGTGGAGTCGTTGCTCTATGGTAAAGTGGTTGCCTCTNNNAGATCGGAAG |
| 29996_2 | TATGAGTGTGGAGTCGTTGCTCTATGGTAAAGTGGTTGCCTCTNNNAGATCGGAAG |
| 29996_3 | TATGAGTGTGGAGTCGTTGCTCTATGGTAAAGTGGTTGCCTCTNNNAGATCGGAAG |
| 42701_1 | TATGAGTGTGGAGTCGTTGCGAGTACAGCTAATCAGGGAGGTGNNNAGATCGGAA |
| 42701_2 | TATGAGTGTGGAGTCGTTGCGAGTACAGCTAATCAGGGAGGTGNNNAGATCGGAA |
| 42701_3 | TATGAGTGTGGAGTCGTTGCGAGTACAGCTAATCAGGGAGGTGNNNAGATCGGAA |
| 8162_1 | TATGAGTGTGGAGTCGTTGCTTCTGCCATTCCATCTCCTTCTACNNNAGATCGGAAGA |
| 8162_2 | TATGAGTGTGGAGTCGTTGCTTCTGCCATTCCATCTCCTTCTACNNNAGATCGGAAGA |
| 8162_3 | TATGAGTGTGGAGTCGTTGCTTCTGCCATTCCATCTCCTTCTACNNNAGATCGGAAGA |
| 50743_1 | TATGAGTGTGGAGTCGTTGCCCCACTTTACTCTTCCCCCTTTCNNNAGATCGGAAGAG |
| 50743_2 | TATGAGTGTGGAGTCGTTGCCCCACTTTACTCTTCCCCCTTTCNNNAGATCGGAAGAG |
| 50743_3 | TATGAGTGTGGAGTCGTTGCCCCACTTTACTCTTCCCCCTTTCNNNAGATCGGAAGAG |
| 36437_1 | TATGAGTGTGGAGTCGTTGCTCCCACAAACATAAATTACTCCGTNNNAGATCGGAAG |
| 36437_2 | TATGAGTGTGGAGTCGTTGCTCCCACAAACATAAATTACTCCGTNNNAGATCGGAAG |
| 36437_3 | TATGAGTGTGGAGTCGTTGCTCCCACAAACATAAATTACTCCGTNNNAGATCGGAAG |
| 39985_1 | TATGAGTGTGGAGTCGTTGCAGAGAGGAGGAGGATGGATGATTNNNAGATCGGA |
| 39985_2 | TATGAGTGTGGAGTCGTTGCAGAGAGGAGGAGGATGGATGATTNNNAGATCGGA |
| 39985_3 | TATGAGTGTGGAGTCGTTGCAGAGAGGAGGAGGATGGATGATTNNNAGATCGGA |
| 52796_1 | TATGAGTGTGGAGTCGTTGCCTGCCTCTCTCACCACTTCATCNNNAGATCGGAAGAG |
| 52796_2 | TATGAGTGTGGAGTCGTTGCCTGCCTCTCTCACCACTTCATCNNNAGATCGGAAGAG |
| 52796_3 | TATGAGTGTGGAGTCGTTGCCTGCCTCTCTCACCACTTCATCNNNAGATCGGAAGAG |
| 40488_1 | TATGAGTGTGGAGTCGTTGCCAGCATGACACCACACCTAAAAGNNNAGATCGGAAG |
| 40488_2 | TATGAGTGTGGAGTCGTTGCCAGCATGACACCACACCTAAAAGNNNAGATCGGAAG |
| 40488_3 | TATGAGTGTGGAGTCGTTGCCAGCATGACACCACACCTAAAAGNNNAGATCGGAAG |
| 42903_1 | TATGAGTGTGGAGTCGTTGCTCCTCTTGCTGAATTGACTCCTTNNNAGATCGGAAGA |
| 42903_2 | TATGAGTGTGGAGTCGTTGCTCCTCTTGCTGAATTGACTCCTTNNNAGATCGGAAGA |

| | |
|---|---|
| 42903_3 | TATGAGTGTGGAGTCGTTGCTCCTCTTGCTGAATTGACTCCTTNNNAGATCGGAAGA |
| 45882_1 | TATGAGTGTGGAGTCGTTGCCTCACTGTACCCTCAATCTCCTGNNNAGATCGGAAGA |
| 45882_2 | TATGAGTGTGGAGTCGTTGCCTCACTGTACCCTCAATCTCCTGNNNAGATCGGAAGA |
| 45882_3 | TATGAGTGTGGAGTCGTTGCCTCACTGTACCCTCAATCTCCTGNNNAGATCGGAAGA |
| 32424_1 | TATGAGTGTGGAGTCGTTGCTGATAGAGCATCTACAGGGGTCANNNAGATCGGAA |
| 32424_2 | TATGAGTGTGGAGTCGTTGCTGATAGAGCATCTACAGGGGTCANNNAGATCGGAA |
| 32424_3 | TATGAGTGTGGAGTCGTTGCTGATAGAGCATCTACAGGGGTCANNNAGATCGGAA |
| 41357_1 | TATGAGTGTGGAGTCGTTGCCAGTGTTTCATTGTTTTGCAGCCNNNAGATCGGAAG |
| 41357_2 | TATGAGTGTGGAGTCGTTGCCAGTGTTTCATTGTTTTGCAGCCNNNAGATCGGAAG |
| 41357_3 | TATGAGTGTGGAGTCGTTGCCAGTGTTTCATTGTTTTGCAGCCNNNAGATCGGAAG |
| 45808_1 | TATGAGTGTGGAGTCGTTGCTAGTGCTCAAGTGATCCTTCTGCNNNAGATCGGAAG |
| 45808_2 | TATGAGTGTGGAGTCGTTGCTAGTGCTCAAGTGATCCTTCTGCNNNAGATCGGAAG |
| 45808_3 | TATGAGTGTGGAGTCGTTGCTAGTGCTCAAGTGATCCTTCTGCNNNAGATCGGAAG |
| 40449_1 | TATGAGTGTGGAGTCGTTGCAACTAACAAGTCAGGGCACTTTTNNNAGATCGGAAG |
| 40449_2 | TATGAGTGTGGAGTCGTTGCAACTAACAAGTCAGGGCACTTTTNNNAGATCGGAAG |
| 40449_3 | TATGAGTGTGGAGTCGTTGCAACTAACAAGTCAGGGCACTTTTNNNAGATCGGAAG |
| 37248_1 | TATGAGTGTGGAGTCGTTGCGTGAGCAGCACAGAATTCAGAATNNNAGATCGGAA |
| 37248_2 | TATGAGTGTGGAGTCGTTGCGTGAGCAGCACAGAATTCAGAATNNNAGATCGGAA |
| 37248_3 | TATGAGTGTGGAGTCGTTGCGTGAGCAGCACAGAATTCAGAATNNNAGATCGGAA |
| 39632_1 | TATGAGTGTGGAGTCGTTGCCACGCCCGGCTAATTTTTTGTANNNAGATCGGAAGA |
| 39632_2 | TATGAGTGTGGAGTCGTTGCCACGCCCGGCTAATTTTTTGTANNNAGATCGGAAGA |
| 39632_3 | TATGAGTGTGGAGTCGTTGCCACGCCCGGCTAATTTTTTGTANNNAGATCGGAAGA |
| 30316_1 | TATGAGTGTGGAGTCGTTGCCGGAACTGGAAGACAGGGAAATCNNNAGATCGGAA |
| 30316_2 | TATGAGTGTGGAGTCGTTGCCGGAACTGGAAGACAGGGAAATCNNNAGATCGGAA |
| 30316_3 | TATGAGTGTGGAGTCGTTGCCGGAACTGGAAGACAGGGAAATCNNNAGATCGGAA |
| 37627_1 | TATGAGTGTGGAGTCGTTGCATGGGATGGGGGAAAAATGATGTNNNAGATCGGA |
| 37627_2 | TATGAGTGTGGAGTCGTTGCATGGGATGGGGGAAAAATGATGTNNNAGATCGGA |
| 37627_3 | TATGAGTGTGGAGTCGTTGCATGGGATGGGGGAAAAATGATGTNNNAGATCGGA |
| 33576_1 | TATGAGTGTGGAGTCGTTGCCAGTAGCATGATCTCGGCTCACNNNAGATCGGAAGA |
| 33576_2 | TATGAGTGTGGAGTCGTTGCCAGTAGCATGATCTCGGCTCACNNNAGATCGGAAGA |
| 33576_3 | TATGAGTGTGGAGTCGTTGCCAGTAGCATGATCTCGGCTCACNNNAGATCGGAAGA |
| 41616_1 | TATGAGTGTGGAGTCGTTGCCATGATCTCGGCTCAATGCAAGNNNAGATCGGAAGA |
| 41616_2 | TATGAGTGTGGAGTCGTTGCCATGATCTCGGCTCAATGCAAGNNNAGATCGGAAGA |
| 41616_3 | TATGAGTGTGGAGTCGTTGCCATGATCTCGGCTCAATGCAAGNNNAGATCGGAAGA |
| 11107_1 | TATGAGTGTGGAGTCGTTGCGAAGCAACCACATGCATGGATTTCNNNAGATCGGAA |
| 11107_2 | TATGAGTGTGGAGTCGTTGCGAAGCAACCACATGCATGGATTTCNNNAGATCGGAA |
| 11107_3 | TATGAGTGTGGAGTCGTTGCGAAGCAACCACATGCATGGATTTCNNNAGATCGGAA |
| 50247_1 | TATGAGTGTGGAGTCGTTGCTTCTGTCTCTCCTCCCTTTCTCANNNAGATCGGAAGA |
| 50247_2 | TATGAGTGTGGAGTCGTTGCTTCTGTCTCTCCTCCCTTTCTCANNNAGATCGGAAGA |
| 50247_3 | TATGAGTGTGGAGTCGTTGCTTCTGTCTCTCCTCCCTTTCTCANNNAGATCGGAAGA |
| 38595_1 | TATGAGTGTGGAGTCGTTGCTCTGACTTTGGGGAGTTAGCAAANNNAGATCGGAA |
| 38595_2 | TATGAGTGTGGAGTCGTTGCTCTGACTTTGGGGAGTTAGCAAANNNAGATCGGAA |
| 38595_3 | TATGAGTGTGGAGTCGTTGCTCTGACTTTGGGGAGTTAGCAAANNNAGATCGGAA |

| | |
|---|---|
| 29573_1 | TATGAGTGTGGAGTCGTTGCCTTGCATCCAGGTGGTCTATCAANNNAGATCGGAAG |
| 29573_2 | TATGAGTGTGGAGTCGTTGCCTTGCATCCAGGTGGTCTATCAANNNAGATCGGAAG |
| 29573_3 | TATGAGTGTGGAGTCGTTGCCTTGCATCCAGGTGGTCTATCAANNNAGATCGGAAG |
| 32184_1 | TATGAGTGTGGAGTCGTTGCTACCAAGCTATGAAGTTGGGCATNNNAGATCGGAAG |
| 32184_2 | TATGAGTGTGGAGTCGTTGCTACCAAGCTATGAAGTTGGGCATNNNAGATCGGAAG |
| 32184_3 | TATGAGTGTGGAGTCGTTGCTACCAAGCTATGAAGTTGGGCATNNNAGATCGGAAG |
| 34728_1 | TATGAGTGTGGAGTCGTTGCCACGCCTGGGAGTTTTTGTATTTNNNAGATCGGAAG |
| 34728_2 | TATGAGTGTGGAGTCGTTGCCACGCCTGGGAGTTTTTGTATTTNNNAGATCGGAAG |
| 34728_3 | TATGAGTGTGGAGTCGTTGCCACGCCTGGGAGTTTTTGTATTTNNNAGATCGGAAG |
| 52600_1 | TATGAGTGTGGAGTCGTTGCCAGGCAGAGATATACACAGCCATNNNAGATCGGAAG |
| 52600_2 | TATGAGTGTGGAGTCGTTGCCAGGCAGAGATATACACAGCCATNNNAGATCGGAAG |
| 52600_3 | TATGAGTGTGGAGTCGTTGCCAGGCAGAGATATACACAGCCATNNNAGATCGGAAG |
| 13414_1 | TATGAGTGTGGAGTCGTTGCAAACTGATTTTTGTTGTTGTAGTNNNAGATCGGAAG |
| 13414_2 | TATGAGTGTGGAGTCGTTGCAAACTGATTTTTGTTGTTGTAGTNNNAGATCGGAAG |
| 13414_3 | TATGAGTGTGGAGTCGTTGCAAACTGATTTTTGTTGTTGTAGTNNNAGATCGGAAG |
| 38228_1 | TATGAGTGTGGAGTCGTTGCACCAAACATGATCTCACACTGTCNNNAGATCGGAAGA |
| 38228_2 | TATGAGTGTGGAGTCGTTGCACCAAACATGATCTCACACTGTCNNNAGATCGGAAGA |
| 38228_3 | TATGAGTGTGGAGTCGTTGCACCAAACATGATCTCACACTGTCNNNAGATCGGAAGA |
| 51848_1 | TATGAGTGTGGAGTCGTTGCCCCCACTTTTATCGCCAAGTTTTNNNAGATCGGAAGA |
| 51848_2 | TATGAGTGTGGAGTCGTTGCCCCCACTTTTATCGCCAAGTTTTNNNAGATCGGAAGA |
| 51848_3 | TATGAGTGTGGAGTCGTTGCCCCCACTTTTATCGCCAAGTTTTNNNAGATCGGAAGA |
| 40895_1 | TATGAGTGTGGAGTCGTTGCATCTTTTTGCCCAAGTCACAGTGNNNAGATCGGAAGA |
| 40895_2 | TATGAGTGTGGAGTCGTTGCATCTTTTTGCCCAAGTCACAGTGNNNAGATCGGAAGA |
| 40895_3 | TATGAGTGTGGAGTCGTTGCATCTTTTTGCCCAAGTCACAGTGNNNAGATCGGAAGA |
| 10647_1 | TATGAGTGTGGAGTCGTTGCATAAGATGCTGAAAGGAGGGGGNNNAGATCGGAAG |
| 10647_2 | TATGAGTGTGGAGTCGTTGCATAAGATGCTGAAAGGAGGGGGNNNAGATCGGAAG |
| 10647_3 | TATGAGTGTGGAGTCGTTGCATAAGATGCTGAAAGGAGGGGGNNNAGATCGGAAG |
| 45004_1 | TATGAGTGTGGAGTCGTTGCAGAATTAGGAAAAAGTGGCCCCANNNAGATCGGAAG |
| 45004_2 | TATGAGTGTGGAGTCGTTGCAGAATTAGGAAAAAGTGGCCCCANNNAGATCGGAAG |
| 45004_3 | TATGAGTGTGGAGTCGTTGCAGAATTAGGAAAAAGTGGCCCCANNNAGATCGGAAG |
| 47792_1 | TATGAGTGTGGAGTCGTTGCGCCCTCATTGAATCTGCATTTGANNNAGATCGGAAGA |
| 47792_2 | TATGAGTGTGGAGTCGTTGCGCCCTCATTGAATCTGCATTTGANNNAGATCGGAAGA |
| 47792_3 | TATGAGTGTGGAGTCGTTGCGCCCTCATTGAATCTGCATTTGANNNAGATCGGAAGA |
| 36078_1 | TATGAGTGTGGAGTCGTTGCGCCATAAACTACGATCATGCCACNNNAGATCGGAAGA |
| 36078_2 | TATGAGTGTGGAGTCGTTGCGCCATAAACTACGATCATGCCACNNNAGATCGGAAGA |
| 36078_3 | TATGAGTGTGGAGTCGTTGCGCCATAAACTACGATCATGCCACNNNAGATCGGAAGA |
| 31774_1 | TATGAGTGTGGAGTCGTTGCATATCCTCAGAGGCACAACACTGNNNAGATCGGAAG |
| 31774_2 | TATGAGTGTGGAGTCGTTGCATATCCTCAGAGGCACAACACTGNNNAGATCGGAAG |
| 31774_3 | TATGAGTGTGGAGTCGTTGCATATCCTCAGAGGCACAACACTGNNNAGATCGGAAG |
| 44118_1 | TATGAGTGTGGAGTCGTTGCCTGGTCTGGGCTGAGATAACTTTNNNAGATCGGAAG |
| 44118_2 | TATGAGTGTGGAGTCGTTGCCTGGTCTGGGCTGAGATAACTTTNNNAGATCGGAAG |
| 44118_3 | TATGAGTGTGGAGTCGTTGCCTGGTCTGGGCTGAGATAACTTTNNNAGATCGGAAG |
| 29607_1 | TATGAGTGTGGAGTCGTTGCTTTGGCTTCCATCAATTCCACACNNNAGATCGGAAGA |

| | |
|---|---|
| 29607_2 | TATGAGTGTGGAGTCGTTGCTTTGGCTTCCATCAATTCCACACNNNAGATCGGAAGA |
| 29607_3 | TATGAGTGTGGAGTCGTTGCTTTGGCTTCCATCAATTCCACACNNNAGATCGGAAGA |
| 32652_1 | TATGAGTGTGGAGTCGTTGCTCTCCTGGGTTCAAGCAATTCTCNNNAGATCGGAAG |
| 32652_2 | TATGAGTGTGGAGTCGTTGCTCTCCTGGGTTCAAGCAATTCTCNNNAGATCGGAAG |
| 32652_3 | TATGAGTGTGGAGTCGTTGCTCTCCTGGGTTCAAGCAATTCTCNNNAGATCGGAAG |
| 44007_1 | TATGAGTGTGGAGTCGTTGCTGGAAAACATGCTCTCTCCATCANNNAGATCGGAAG |
| 44007_2 | TATGAGTGTGGAGTCGTTGCTGGAAAACATGCTCTCTCCATCANNNAGATCGGAAG |
| 44007_3 | TATGAGTGTGGAGTCGTTGCTGGAAAACATGCTCTCTCCATCANNNAGATCGGAAG |
| 46967_1 | TATGAGTGTGGAGTCGTTGCCACCACACCTGGCCTATATTGAANNNAGATCGGAAG |
| 46967_2 | TATGAGTGTGGAGTCGTTGCCACCACACCTGGCCTATATTGAANNNAGATCGGAAG |
| 46967_3 | TATGAGTGTGGAGTCGTTGCCACCACACCTGGCCTATATTGAANNNAGATCGGAAG |
| 50407_1 | TATGAGTGTGGAGTCGTTGCCTTCCCTCATTGCAGTGAGAAAGNNNAGATCGGAAG |
| 50407_2 | TATGAGTGTGGAGTCGTTGCCTTCCCTCATTGCAGTGAGAAAGNNNAGATCGGAAG |
| 50407_3 | TATGAGTGTGGAGTCGTTGCCTTCCCTCATTGCAGTGAGAAAGNNNAGATCGGAAG |
| 36274_1 | TATGAGTGTGGAGTCGTTGCAAATGTAGGTCTATGAGCTGGGGNNNAGATCGGAA |
| 36274_2 | TATGAGTGTGGAGTCGTTGCAAATGTAGGTCTATGAGCTGGGGNNNAGATCGGAA |
| 36274_3 | TATGAGTGTGGAGTCGTTGCAAATGTAGGTCTATGAGCTGGGGNNNAGATCGGAA |
| 39096_1 | TATGAGTGTGGAGTCGTTGCGAGATTGCAGGCAAGAGTCCTATNNNAGATCGGAA |
| 39096_2 | TATGAGTGTGGAGTCGTTGCGAGATTGCAGGCAAGAGTCCTATNNNAGATCGGAA |
| 39096_3 | TATGAGTGTGGAGTCGTTGCGAGATTGCAGGCAAGAGTCCTATNNNAGATCGGAA |
| 45776_1 | TATGAGTGTGGAGTCGTTGCTGAACTCCTGAGCTCAAGTCAAGNNNAGATCGGAAG |
| 45776_2 | TATGAGTGTGGAGTCGTTGCTGAACTCCTGAGCTCAAGTCAAGNNNAGATCGGAAG |
| 45776_3 | TATGAGTGTGGAGTCGTTGCTGAACTCCTGAGCTCAAGTCAAGNNNAGATCGGAAG |
| 10462_1 | TATGAGTGTGGAGTCGTTGCTTGGCATTCTCTAGGCTTATTCANNNAGATCGGAAG |
| 10462_2 | TATGAGTGTGGAGTCGTTGCTTGGCATTCTCTAGGCTTATTCANNNAGATCGGAAG |
| 10462_3 | TATGAGTGTGGAGTCGTTGCTTGGCATTCTCTAGGCTTATTCANNNAGATCGGAAG |
| 47263_1 | TATGAGTGTGGAGTCGTTGCTCTCCCTAGCTCAAACCATCCTANNNAGATCGGAAGA |
| 47263_2 | TATGAGTGTGGAGTCGTTGCTCTCCCTAGCTCAAACCATCCTANNNAGATCGGAAGA |
| 47263_3 | TATGAGTGTGGAGTCGTTGCTCTCCCTAGCTCAAACCATCCTANNNAGATCGGAAGA |
| 45619_1 | TATGAGTGTGGAGTCGTTGCAAACTCCTGGTCTCAAGCAATCTNNNAGATCGGAAG |
| 45619_2 | TATGAGTGTGGAGTCGTTGCAAACTCCTGGTCTCAAGCAATCTNNNAGATCGGAAG |
| 45619_3 | TATGAGTGTGGAGTCGTTGCAAACTCCTGGTCTCAAGCAATCTNNNAGATCGGAAG |
| 48672_1 | TATGAGTGTGGAGTCGTTGCAACCTCCTGAGTAGCTGGGATTANNNAGATCGGAA |
| 48672_2 | TATGAGTGTGGAGTCGTTGCAACCTCCTGAGTAGCTGGGATTANNNAGATCGGAA |
| 48672_3 | TATGAGTGTGGAGTCGTTGCAACCTCCTGAGTAGCTGGGATTANNNAGATCGGAA |
| 34770_1 | TATGAGTGTGGAGTCGTTGCCCGGGTTCTAAGAGATTCTCCTGNNNAGATCGGAAG |
| 34770_2 | TATGAGTGTGGAGTCGTTGCCCGGGTTCTAAGAGATTCTCCTGNNNAGATCGGAAG |
| 34770_3 | TATGAGTGTGGAGTCGTTGCCCGGGTTCTAAGAGATTCTCCTGNNNAGATCGGAAG |
| 13712_1 | TATGAGTGTGGAGTCGTTGCAAATGTCAGCAAATGTAAAAATTNNNAGATCGGAA |
| 13712_2 | TATGAGTGTGGAGTCGTTGCAAATGTCAGCAAATGTAAAAATTNNNAGATCGGAA |
| 13712_3 | TATGAGTGTGGAGTCGTTGCAAATGTCAGCAAATGTAAAAATTNNNAGATCGGAA |
| 39707_1 | TATGAGTGTGGAGTCGTTGCACAAGAGCTGGTTTCTGAAAAGANNNAGATCGGAA |
| 39707_2 | TATGAGTGTGGAGTCGTTGCACAAGAGCTGGTTTCTGAAAAGANNNAGATCGGAA |

| | |
|---|---|
| 39707_3 | TATGAGTGTGGAGTCGTTGCACAAGAGCTGGTTTCTGAAAAGANNNAGATCGGAA |
| 32989_1 | TATGAGTGTGGAGTCGTTGCGGTCAGGCCTATAATATCCCAGCNNNAGATCGGAAG |
| 32989_2 | TATGAGTGTGGAGTCGTTGCGGTCAGGCCTATAATATCCCAGCNNNAGATCGGAAG |
| 32989_3 | TATGAGTGTGGAGTCGTTGCGGTCAGGCCTATAATATCCCAGCNNNAGATCGGAAG |
| 40086_1 | TATGAGTGTGGAGTCGTTGCGAAACAAGAGCAAACTAACCCCCNNNAGATCGGAAG |
| 40086_2 | TATGAGTGTGGAGTCGTTGCGAAACAAGAGCAAACTAACCCCCNNNAGATCGGAAG |
| 40086_3 | TATGAGTGTGGAGTCGTTGCGAAACAAGAGCAAACTAACCCCCNNNAGATCGGAAG |
| 45877_1 | TATGAGTGTGGAGTCGTTGCCACACTTACTACCTTAGGCAGCANNNAGATCGGAAG |
| 45877_2 | TATGAGTGTGGAGTCGTTGCCACACTTACTACCTTAGGCAGCANNNAGATCGGAAG |
| 45877_3 | TATGAGTGTGGAGTCGTTGCCACACTTACTACCTTAGGCAGCANNNAGATCGGAAG |
| 34748_1 | TATGAGTGTGGAGTCGTTGCTTCAAGTGATTCTCCTGCCTCAGNNNAGATCGGAAG |
| 34748_2 | TATGAGTGTGGAGTCGTTGCTTCAAGTGATTCTCCTGCCTCAGNNNAGATCGGAAG |
| 34748_3 | TATGAGTGTGGAGTCGTTGCTTCAAGTGATTCTCCTGCCTCAGNNNAGATCGGAAG |
| 46237_1 | TATGAGTGTGGAGTCGTTGCTTCACACCATTCTCCTGCCTCNNNAGATCGGAAGAGC |
| 46237_2 | TATGAGTGTGGAGTCGTTGCTTCACACCATTCTCCTGCCTCNNNAGATCGGAAGAGC |
| 46237_3 | TATGAGTGTGGAGTCGTTGCTTCACACCATTCTCCTGCCTCNNNAGATCGGAAGAGC |
| 42790_1 | TATGAGTGTGGAGTCGTTGCCCAAGTTGGAAAGCACTCTTCAGNNNAGATCGGAAG |
| 42790_2 | TATGAGTGTGGAGTCGTTGCCCAAGTTGGAAAGCACTCTTCAGNNNAGATCGGAAG |
| 42790_3 | TATGAGTGTGGAGTCGTTGCCCAAGTTGGAAAGCACTCTTCAGNNNAGATCGGAAG |
| 49075_1 | TATGAGTGTGGAGTCGTTGCCTCCTGGGTTCAAGCGATTCTCNNNAGATCGGAAGA |
| 49075_2 | TATGAGTGTGGAGTCGTTGCCTCCTGGGTTCAAGCGATTCTCNNNAGATCGGAAGA |
| 49075_3 | TATGAGTGTGGAGTCGTTGCCTCCTGGGTTCAAGCGATTCTCNNNAGATCGGAAGA |
| 2548_1 | TATGAGTGTGGAGTCGTTGCCACGCAAGCACATGAAATGTAGGNNNAGATCGGAA |
| 2548_2 | TATGAGTGTGGAGTCGTTGCCACGCAAGCACATGAAATGTAGGNNNAGATCGGAA |
| 2548_3 | TATGAGTGTGGAGTCGTTGCCACGCAAGCACATGAAATGTAGGNNNAGATCGGAA |
| 31843_1 | TATGAGTGTGGAGTCGTTGCTGCTTTGTGTATTTGGGAACTCTGNNNAGATCGGAA |
| 31843_2 | TATGAGTGTGGAGTCGTTGCTGCTTTGTGTATTTGGGAACTCTGNNNAGATCGGAA |
| 31843_3 | TATGAGTGTGGAGTCGTTGCTGCTTTGTGTATTTGGGAACTCTGNNNAGATCGGAA |
| 34820_1 | TATGAGTGTGGAGTCGTTGCAGTAGAAACAGGCTTTCACCGTGNNNAGATCGGAA |
| 34820_2 | TATGAGTGTGGAGTCGTTGCAGTAGAAACAGGCTTTCACCGTGNNNAGATCGGAA |
| 34820_3 | TATGAGTGTGGAGTCGTTGCAGTAGAAACAGGCTTTCACCGTGNNNAGATCGGAA |
| 31719_1 | TATGAGTGTGGAGTCGTTGCACAGCTGAGTTTCCTGCATTAGANNNAGATCGGAAG |
| 31719_2 | TATGAGTGTGGAGTCGTTGCACAGCTGAGTTTCCTGCATTAGANNNAGATCGGAAG |
| 31719_3 | TATGAGTGTGGAGTCGTTGCACAGCTGAGTTTCCTGCATTAGANNNAGATCGGAAG |
| 49664_1 | TATGAGTGTGGAGTCGTTGCGATTTTTCCTCGCTGCAACCTCNNNAGATCGGAAGA |
| 49664_2 | TATGAGTGTGGAGTCGTTGCGATTTTTCCTCGCTGCAACCTCNNNAGATCGGAAGA |
| 49664_3 | TATGAGTGTGGAGTCGTTGCGATTTTTCCTCGCTGCAACCTCNNNAGATCGGAAGA |
| 46175_1 | TATGAGTGTGGAGTCGTTGCCCCCCCCAAAAAGAACATAGAGTNNNAGATCGGAAG |
| 46175_2 | TATGAGTGTGGAGTCGTTGCCCCCCCCAAAAAGAACATAGAGTNNNAGATCGGAAG |
| 46175_3 | TATGAGTGTGGAGTCGTTGCCCCCCCCAAAAAGAACATAGAGTNNNAGATCGGAAG |
| 51581_1 | TATGAGTGTGGAGTCGTTGCGTACATTCACCCACATCTTTGCCNNNAGATCGGAAGA |
| 51581_2 | TATGAGTGTGGAGTCGTTGCGTACATTCACCCACATCTTTGCCNNNAGATCGGAAGA |
| 51581_3 | TATGAGTGTGGAGTCGTTGCGTACATTCACCCACATCTTTGCCNNNAGATCGGAAGA |

| | |
|---|---|
| 46372_1 | TATGAGTGTGGAGTCGTTGCCCCCAATCCTCTTCTTATCTGCTNNNAGATCGGAAGA |
| 46372_2 | TATGAGTGTGGAGTCGTTGCCCCCAATCCTCTTCTTATCTGCTNNNAGATCGGAAGA |
| 46372_3 | TATGAGTGTGGAGTCGTTGCCCCCAATCCTCTTCTTATCTGCTNNNAGATCGGAAGA |
| 40377_1 | TATGAGTGTGGAGTCGTTGCTCAATCCAAGGTATGCACTGGTTNNNAGATCGGAAG |
| 40377_2 | TATGAGTGTGGAGTCGTTGCTCAATCCAAGGTATGCACTGGTTNNNAGATCGGAAG |
| 40377_3 | TATGAGTGTGGAGTCGTTGCTCAATCCAAGGTATGCACTGGTTNNNAGATCGGAAG |
| 42246_1 | TATGAGTGTGGAGTCGTTGCGAGTGCAGTGGCGTGATCTTNNNAGATCGGAAGAG |
| 42246_2 | TATGAGTGTGGAGTCGTTGCGAGTGCAGTGGCGTGATCTTNNNAGATCGGAAGAG |
| 42246_3 | TATGAGTGTGGAGTCGTTGCGAGTGCAGTGGCGTGATCTTNNNAGATCGGAAGAG |
| 29240_1 | TATGAGTGTGGAGTCGTTGCGGGATGAGATACAGGCAATCGATNNNAGATCGGAA |
| 29240_2 | TATGAGTGTGGAGTCGTTGCGGGATGAGATACAGGCAATCGATNNNAGATCGGAA |
| 29240_3 | TATGAGTGTGGAGTCGTTGCGGGATGAGATACAGGCAATCGATNNNAGATCGGAA |
| 53245_1 | TATGAGTGTGGAGTCGTTGCTACAAGTCGCAGCCCTCTTACTTNNNAGATCGGAAG |
| 53245_2 | TATGAGTGTGGAGTCGTTGCTACAAGTCGCAGCCCTCTTACTTNNNAGATCGGAAG |
| 53245_3 | TATGAGTGTGGAGTCGTTGCTACAAGTCGCAGCCCTCTTACTTNNNAGATCGGAAG |
| 49111_1 | TATGAGTGTGGAGTCGTTGCATCTTGGCTCACTGTAACCTCTGNNNAGATCGGAAG |
| 49111_2 | TATGAGTGTGGAGTCGTTGCATCTTGGCTCACTGTAACCTCTGNNNAGATCGGAAG |
| 49111_3 | TATGAGTGTGGAGTCGTTGCATCTTGGCTCACTGTAACCTCTGNNNAGATCGGAAG |
| 35933_1 | TATGAGTGTGGAGTCGTTGCGGATCACGAGGTCAAGAGATCAANNNAGATCGGAA |
| 35933_2 | TATGAGTGTGGAGTCGTTGCGGATCACGAGGTCAAGAGATCAANNNAGATCGGAA |
| 35933_3 | TATGAGTGTGGAGTCGTTGCGGATCACGAGGTCAAGAGATCAANNNAGATCGGAA |
| 10004_1 | TATGAGTGTGGAGTCGTTGCGGCAAGAAAAATGCCCCTTTTGTNNNAGATCGGAAG |
| 10004_2 | TATGAGTGTGGAGTCGTTGCGGCAAGAAAAATGCCCCTTTTGTNNNAGATCGGAAG |
| 10004_3 | TATGAGTGTGGAGTCGTTGCGGCAAGAAAAATGCCCCTTTTGTNNNAGATCGGAAG |
| 43226_1 | TATGAGTGTGGAGTCGTTGCAGGGAGGGAGGGAAAGATAATGANNNAGATCGG |
| 43226_2 | TATGAGTGTGGAGTCGTTGCAGGGAGGGAGGGAAAGATAATGANNNAGATCGG |
| 43226_3 | TATGAGTGTGGAGTCGTTGCAGGGAGGGAGGGAAAGATAATGANNNAGATCGG |
| 46828_1 | TATGAGTGTGGAGTCGTTGCTTCCTTGACACATGGCCTCTTTANNNAGATCGGAAGA |
| 46828_2 | TATGAGTGTGGAGTCGTTGCTTCCTTGACACATGGCCTCTTTANNNAGATCGGAAGA |
| 46828_3 | TATGAGTGTGGAGTCGTTGCTTCCTTGACACATGGCCTCTTTANNNAGATCGGAAGA |
| 48535_1 | TATGAGTGTGGAGTCGTTGCCTGAAATTACAGGTGTGAGCCACNNNAGATCGGAA |
| 48535_2 | TATGAGTGTGGAGTCGTTGCCTGAAATTACAGGTGTGAGCCACNNNAGATCGGAA |
| 48535_3 | TATGAGTGTGGAGTCGTTGCCTGAAATTACAGGTGTGAGCCACNNNAGATCGGAA |
| 42959_1 | TATGAGTGTGGAGTCGTTGCTGTCTACACATTGGGAAAGGACTNNNAGATCGGAA |
| 42959_2 | TATGAGTGTGGAGTCGTTGCTGTCTACACATTGGGAAAGGACTNNNAGATCGGAA |
| 42959_3 | TATGAGTGTGGAGTCGTTGCTGTCTACACATTGGGAAAGGACTNNNAGATCGGAA |
| 30213_1 | TATGAGTGTGGAGTCGTTGCGCAGGAGGATTAACAGATGAATGGNNNAGATCGG |
| 30213_2 | TATGAGTGTGGAGTCGTTGCGCAGGAGGATTAACAGATGAATGGNNNAGATCGG |
| 30213_3 | TATGAGTGTGGAGTCGTTGCGCAGGAGGATTAACAGATGAATGGNNNAGATCGG |
| 31988_1 | TATGAGTGTGGAGTCGTTGCTTTTTGAGACAGGGTCTTGCTCTNNNAGATCGGAAG |
| 31988_2 | TATGAGTGTGGAGTCGTTGCTTTTTGAGACAGGGTCTTGCTCTNNNAGATCGGAAG |
| 31988_3 | TATGAGTGTGGAGTCGTTGCTTTTTGAGACAGGGTCTTGCTCTNNNAGATCGGAAG |
| 30099_1 | TATGAGTGTGGAGTCGTTGCGAGAGGTCTAATCCCAGAGTGCTNNNAGATCGGAA |

| | |
|---|---|
| 30099_2 | TATGAGTGTGGAGTCGTTGCGAGAGGTCTAATCCCAGAGTGCTNNNAGATCGGAA |
| 30099_3 | TATGAGTGTGGAGTCGTTGCGAGAGGTCTAATCCCAGAGTGCTNNNAGATCGGAA |
| 45672_1 | TATGAGTGTGGAGTCGTTGCCTCCTGGGTTCATGCCATTCTCNNNAGATCGGAAGAG |
| 45672_2 | TATGAGTGTGGAGTCGTTGCCTCCTGGGTTCATGCCATTCTCNNNAGATCGGAAGAG |
| 45672_3 | TATGAGTGTGGAGTCGTTGCCTCCTGGGTTCATGCCATTCTCNNNAGATCGGAAGAG |
| 38478_1 | TATGAGTGTGGAGTCGTTGCCCACCAATGTGCTTTCTGTCTTTNNNAGATCGGAAGA |
| 38478_2 | TATGAGTGTGGAGTCGTTGCCCACCAATGTGCTTTCTGTCTTTNNNAGATCGGAAGA |
| 38478_3 | TATGAGTGTGGAGTCGTTGCCCACCAATGTGCTTTCTGTCTTTNNNAGATCGGAAGA |
| 9770_1 | TATGAGTGTGGAGTCGTTGCAAGGGAGGATCACAGGCAGTTTANNNAGATCGGAA |
| 9770_2 | TATGAGTGTGGAGTCGTTGCAAGGGAGGATCACAGGCAGTTTANNNAGATCGGAA |
| 9770_3 | TATGAGTGTGGAGTCGTTGCAAGGGAGGATCACAGGCAGTTTANNNAGATCGGAA |
| 40174_1 | TATGAGTGTGGAGTCGTTGCCACAGAATGGGAGAAAATACTTGCANNNAGATCGG |
| 40174_2 | TATGAGTGTGGAGTCGTTGCCACAGAATGGGAGAAAATACTTGCANNNAGATCGG |
| 40174_3 | TATGAGTGTGGAGTCGTTGCCACAGAATGGGAGAAAATACTTGCANNNAGATCGG |
| 50173_1 | TATGAGTGTGGAGTCGTTGCGAGCCCTTTGTGCTTCAGTTTAGNNNAGATCGGAAG |
| 50173_2 | TATGAGTGTGGAGTCGTTGCGAGCCCTTTGTGCTTCAGTTTAGNNNAGATCGGAAG |
| 50173_3 | TATGAGTGTGGAGTCGTTGCGAGCCCTTTGTGCTTCAGTTTAGNNNAGATCGGAAG |
| 9989_1 | TATGAGTGTGGAGTCGTTGCCGAAGTTTCTCTCTCATCACCCANNNAGATCGGAAGA |
| 9989_2 | TATGAGTGTGGAGTCGTTGCCGAAGTTTCTCTCTCATCACCCANNNAGATCGGAAGA |
| 9989_3 | TATGAGTGTGGAGTCGTTGCCGAAGTTTCTCTCTCATCACCCANNNAGATCGGAAGA |
| 38113_1 | TATGAGTGTGGAGTCGTTGCAGCAGCTGAAGTCCTCAATCTACNNNAGATCGGAAG |
| 38113_2 | TATGAGTGTGGAGTCGTTGCAGCAGCTGAAGTCCTCAATCTACNNNAGATCGGAAG |
| 38113_3 | TATGAGTGTGGAGTCGTTGCAGCAGCTGAAGTCCTCAATCTACNNNAGATCGGAAG |
| 43936_1 | TATGAGTGTGGAGTCGTTGCAGCCATGCTTCATATCCTGGTAGNNNAGATCGGAAG |
| 43936_2 | TATGAGTGTGGAGTCGTTGCAGCCATGCTTCATATCCTGGTAGNNNAGATCGGAAG |
| 43936_3 | TATGAGTGTGGAGTCGTTGCAGCCATGCTTCATATCCTGGTAGNNNAGATCGGAAG |
| 49979_1 | TATGAGTGTGGAGTCGTTGCGAATCCTCGGGTCAGAAGAGATGNNNAGATCGGAA |
| 49979_2 | TATGAGTGTGGAGTCGTTGCGAATCCTCGGGTCAGAAGAGATGNNNAGATCGGAA |
| 49979_3 | TATGAGTGTGGAGTCGTTGCGAATCCTCGGGTCAGAAGAGATGNNNAGATCGGAA |
| 32351_1 | TATGAGTGTGGAGTCGTTGCGAGGAAACATGACAGGTGAGTCTNNNAGATCGGAA |
| 32351_2 | TATGAGTGTGGAGTCGTTGCGAGGAAACATGACAGGTGAGTCTNNNAGATCGGAA |
| 32351_3 | TATGAGTGTGGAGTCGTTGCGAGGAAACATGACAGGTGAGTCTNNNAGATCGGAA |
| 12655_1 | TATGAGTGTGGAGTCGTTGCGATTCCTAACAGACTGGTTGNNNAGATCGGAAGAG |
| 12655_2 | TATGAGTGTGGAGTCGTTGCGATTCCTAACAGACTGGTTGNNNAGATCGGAAGAG |
| 12655_3 | TATGAGTGTGGAGTCGTTGCGATTCCTAACAGACTGGTTGNNNAGATCGGAAGAG |
| 53000_1 | TATGAGTGTGGAGTCGTTGCTGTTTCTTGTCTAGCGTGAAAGCNNNAGATCGGAAG |
| 53000_2 | TATGAGTGTGGAGTCGTTGCTGTTTCTTGTCTAGCGTGAAAGCNNNAGATCGGAAG |
| 53000_3 | TATGAGTGTGGAGTCGTTGCTGTTTCTTGTCTAGCGTGAAAGCNNNAGATCGGAAG |
| 37364_1 | TATGAGTGTGGAGTCGTTGCACATTATCCTTATGGTTCCAGCANNNAGATCGGAAG |
| 37364_2 | TATGAGTGTGGAGTCGTTGCACATTATCCTTATGGTTCCAGCANNNAGATCGGAAG |
| 37364_3 | TATGAGTGTGGAGTCGTTGCACATTATCCTTATGGTTCCAGCANNNAGATCGGAAG |
| 36929_1 | TATGAGTGTGGAGTCGTTGCCACTACCCCTGAATATCACTCGGNNNAGATCGGAAG |
| 36929_2 | TATGAGTGTGGAGTCGTTGCCACTACCCCTGAATATCACTCGGNNNAGATCGGAAG |

| | |
|---|---|
| 36929_3 | TATGAGTGTGGAGTCGTTGCCACTACCCCTGAATATCACTCGGNNNAGATCGGAAG |
| 41286_1 | TATGAGTGTGGAGTCGTTGCAAACAACCAAACACCATTTCCCANNNAGATCGGAAG |
| 41286_2 | TATGAGTGTGGAGTCGTTGCAAACAACCAAACACCATTTCCCANNNAGATCGGAAG |
| 41286_3 | TATGAGTGTGGAGTCGTTGCAAACAACCAAACACCATTTCCCANNNAGATCGGAAG |
| 51623_1 | TATGAGTGTGGAGTCGTTGCTACGGTTTTGTTTGCAGGATGACNNNAGATCGGAAG |
| 51623_2 | TATGAGTGTGGAGTCGTTGCTACGGTTTTGTTTGCAGGATGACNNNAGATCGGAAG |
| 51623_3 | TATGAGTGTGGAGTCGTTGCTACGGTTTTGTTTGCAGGATGACNNNAGATCGGAAG |
| 29995_1 | TATGAGTGTGGAGTCGTTGCGTAGATGAACGAGGAGGTCTTCCNNNAGATCGGAA |
| 29995_2 | TATGAGTGTGGAGTCGTTGCGTAGATGAACGAGGAGGTCTTCCNNNAGATCGGAA |
| 29995_3 | TATGAGTGTGGAGTCGTTGCGTAGATGAACGAGGAGGTCTTCCNNNAGATCGGAA |
| 43485_1 | TATGAGTGTGGAGTCGTTGCCAATCTCAGCTCACTGCAAGTTCNNNAGATCGGAAG |
| 43485_2 | TATGAGTGTGGAGTCGTTGCCAATCTCAGCTCACTGCAAGTTCNNNAGATCGGAAG |
| 43485_3 | TATGAGTGTGGAGTCGTTGCCAATCTCAGCTCACTGCAAGTTCNNNAGATCGGAAG |
| 45015_1 | TATGAGTGTGGAGTCGTTGCCTGGAAACTGCTTTGTCGATTGTNNNAGATCGGAAG |
| 45015_2 | TATGAGTGTGGAGTCGTTGCCTGGAAACTGCTTTGTCGATTGTNNNAGATCGGAAG |
| 45015_3 | TATGAGTGTGGAGTCGTTGCCTGGAAACTGCTTTGTCGATTGTNNNAGATCGGAAG |
| 39294_1 | TATGAGTGTGGAGTCGTTGCAAAGGCATAGCTGTACATCCACANNNAGATCGGAAG |
| 39294_2 | TATGAGTGTGGAGTCGTTGCAAAGGCATAGCTGTACATCCACANNNAGATCGGAAG |
| 39294_3 | TATGAGTGTGGAGTCGTTGCAAAGGCATAGCTGTACATCCACANNNAGATCGGAAG |
| 51178_1 | TATGAGTGTGGAGTCGTTGCAAACCTGCCTCCAAACAGACATANNNAGATCGGAAG |
| 51178_2 | TATGAGTGTGGAGTCGTTGCAAACCTGCCTCCAAACAGACATANNNAGATCGGAAG |
| 51178_3 | TATGAGTGTGGAGTCGTTGCAAACCTGCCTCCAAACAGACATANNNAGATCGGAAG |
| 8495_1 | TATGAGTGTGGAGTCGTTGCTTCTATTGTGAGTGATCTCTGGTGANNNAGATCGGA |
| 8495_2 | TATGAGTGTGGAGTCGTTGCTTCTATTGTGAGTGATCTCTGGTGANNNAGATCGGA |
| 8495_3 | TATGAGTGTGGAGTCGTTGCTTCTATTGTGAGTGATCTCTGGTGANNNAGATCGGA |
| 32837_1 | TATGAGTGTGGAGTCGTTGCGTAGAGACGGACTTTCACTGTGTNNNAGATCGGAA |
| 32837_2 | TATGAGTGTGGAGTCGTTGCGTAGAGACGGACTTTCACTGTGTNNNAGATCGGAA |
| 32837_3 | TATGAGTGTGGAGTCGTTGCGTAGAGACGGACTTTCACTGTGTNNNAGATCGGAA |
| 41610_1 | TATGAGTGTGGAGTCGTTGCGCCTCCTAAGTAGCTGGGATTACNNNAGATCGGAAG |
| 41610_2 | TATGAGTGTGGAGTCGTTGCGCCTCCTAAGTAGCTGGGATTACNNNAGATCGGAAG |
| 41610_3 | TATGAGTGTGGAGTCGTTGCGCCTCCTAAGTAGCTGGGATTACNNNAGATCGGAAG |
| 51377_1 | TATGAGTGTGGAGTCGTTGCCCTTACTGAGTTCCCCCAATTTNNNAGATCGGAAGA |
| 51377_2 | TATGAGTGTGGAGTCGTTGCCCTTACTGAGTTCCCCCAATTTNNNAGATCGGAAGA |
| 51377_3 | TATGAGTGTGGAGTCGTTGCCCTTACTGAGTTCCCCCAATTTNNNAGATCGGAAGA |
| 47637_1 | TATGAGTGTGGAGTCGTTGCAGGGTCTTACCATATTGCCCATGNNNAGATCGGAAG |
| 47637_2 | TATGAGTGTGGAGTCGTTGCAGGGTCTTACCATATTGCCCATGNNNAGATCGGAAG |
| 47637_3 | TATGAGTGTGGAGTCGTTGCAGGGTCTTACCATATTGCCCATGNNNAGATCGGAAG |
| 52360_1 | TATGAGTGTGGAGTCGTTGCGGACAAATGTGGCATATCTCAGCNNNAGATCGGAA |
| 52360_2 | TATGAGTGTGGAGTCGTTGCGGACAAATGTGGCATATCTCAGCNNNAGATCGGAA |
| 52360_3 | TATGAGTGTGGAGTCGTTGCGGACAAATGTGGCATATCTCAGCNNNAGATCGGAA |
| 50126_1 | TATGAGTGTGGAGTCGTTGCTAGCTGCCTCCTCTTATCCCATANNNAGATCGGAAGA |
| 50126_2 | TATGAGTGTGGAGTCGTTGCTAGCTGCCTCCTCTTATCCCATANNNAGATCGGAAGA |
| 50126_3 | TATGAGTGTGGAGTCGTTGCTAGCTGCCTCCTCTTATCCCATANNNAGATCGGAAGA |

| | |
|---|---|
| 48384_1 | TATGAGTGTGGAGTCGTTGCTATGTAGCTGGGACTACAGGAGTNNNAGATCGGAA |
| 48384_2 | TATGAGTGTGGAGTCGTTGCTATGTAGCTGGGACTACAGGAGTNNNAGATCGGAA |
| 48384_3 | TATGAGTGTGGAGTCGTTGCTATGTAGCTGGGACTACAGGAGTNNNAGATCGGAA |
| 37571_1 | TATGAGTGTGGAGTCGTTGCTCTAGAAGCTCTAAGGGTTCAGGNNNAGATCGGAA |
| 37571_2 | TATGAGTGTGGAGTCGTTGCTCTAGAAGCTCTAAGGGTTCAGGNNNAGATCGGAA |
| 37571_3 | TATGAGTGTGGAGTCGTTGCTCTAGAAGCTCTAAGGGTTCAGGNNNAGATCGGAA |
| 34841_1 | TATGAGTGTGGAGTCGTTGCTGGTAACGGCTCAACTCTACTTCNNNAGATCGGAAG |
| 34841_2 | TATGAGTGTGGAGTCGTTGCTGGTAACGGCTCAACTCTACTTCNNNAGATCGGAAG |
| 34841_3 | TATGAGTGTGGAGTCGTTGCTGGTAACGGCTCAACTCTACTTCNNNAGATCGGAAG |
| 40789_1 | TATGAGTGTGGAGTCGTTGCACCCGAGACTGGTCAATTTACAANNNAGATCGGAAG |
| 40789_2 | TATGAGTGTGGAGTCGTTGCACCCGAGACTGGTCAATTTACAANNNAGATCGGAAG |
| 40789_3 | TATGAGTGTGGAGTCGTTGCACCCGAGACTGGTCAATTTACAANNNAGATCGGAAG |
| 43672_1 | TATGAGTGTGGAGTCGTTGCCATCCTGGAAACCCTGGTTCTTNNNAGATCGGAAGA |
| 43672_2 | TATGAGTGTGGAGTCGTTGCCATCCTGGAAACCCTGGTTCTTNNNAGATCGGAAGA |
| 43672_3 | TATGAGTGTGGAGTCGTTGCCATCCTGGAAACCCTGGTTCTTNNNAGATCGGAAGA |
| 45686_1 | TATGAGTGTGGAGTCGTTGCCTGAGTAGCTGGGATTACAGGTGNNNAGATCGGAA |
| 45686_2 | TATGAGTGTGGAGTCGTTGCCTGAGTAGCTGGGATTACAGGTGNNNAGATCGGAA |
| 45686_3 | TATGAGTGTGGAGTCGTTGCCTGAGTAGCTGGGATTACAGGTGNNNAGATCGGAA |
| 8248_1 | TATGAGTGTGGAGTCGTTGCAAAAACTGATGCTTTGGGGTGGTNNNAGATCGGAA |
| 8248_2 | TATGAGTGTGGAGTCGTTGCAAAAACTGATGCTTTGGGGTGGTNNNAGATCGGAA |
| 8248_3 | TATGAGTGTGGAGTCGTTGCAAAAACTGATGCTTTGGGGTGGTNNNAGATCGGAA |
| 38827_1 | TATGAGTGTGGAGTCGTTGCGCAACTTTGCCAGATTTGTTTACAANNNAGATCGGA |
| 38827_2 | TATGAGTGTGGAGTCGTTGCGCAACTTTGCCAGATTTGTTTACAANNNAGATCGGA |
| 38827_3 | TATGAGTGTGGAGTCGTTGCGCAACTTTGCCAGATTTGTTTACAANNNAGATCGGA |
| 44617_1 | TATGAGTGTGGAGTCGTTGCAGCCACAGAGTTGAAAAAGCAAANNNAGATCGGAA |
| 44617_2 | TATGAGTGTGGAGTCGTTGCAGCCACAGAGTTGAAAAAGCAAANNNAGATCGGAA |
| 44617_3 | TATGAGTGTGGAGTCGTTGCAGCCACAGAGTTGAAAAAGCAAANNNAGATCGGAA |
| 31007_1 | TATGAGTGTGGAGTCGTTGCTTCAAGTGATTCTCCTGCCTCAGNNNAGATCGGAAG |
| 31007_2 | TATGAGTGTGGAGTCGTTGCTTCAAGTGATTCTCCTGCCTCAGNNNAGATCGGAAG |
| 31007_3 | TATGAGTGTGGAGTCGTTGCTTCAAGTGATTCTCCTGCCTCAGNNNAGATCGGAAG |
| 40937_1 | TATGAGTGTGGAGTCGTTGCCCAGTTATTTGCCTTTCCACAGGNNNAGATCGGAAG |
| 40937_2 | TATGAGTGTGGAGTCGTTGCCCAGTTATTTGCCTTTCCACAGGNNNAGATCGGAAG |
| 40937_3 | TATGAGTGTGGAGTCGTTGCCCAGTTATTTGCCTTTCCACAGGNNNAGATCGGAAG |
| 32944_1 | TATGAGTGTGGAGTCGTTGCGGTGCACCAAAATCTCAGAAGTCNNNAGATCGGAAG |
| 32944_2 | TATGAGTGTGGAGTCGTTGCGGTGCACCAAAATCTCAGAAGTCNNNAGATCGGAAG |
| 32944_3 | TATGAGTGTGGAGTCGTTGCGGTGCACCAAAATCTCAGAAGTCNNNAGATCGGAAG |
| 35524_1 | TATGAGTGTGGAGTCGTTGCTTTATGTTTCCCAGGCTGGTCTTNNNAGATCGGAAG |
| 35524_2 | TATGAGTGTGGAGTCGTTGCTTTATGTTTCCCAGGCTGGTCTTNNNAGATCGGAAG |
| 35524_3 | TATGAGTGTGGAGTCGTTGCTTTATGTTTCCCAGGCTGGTCTTNNNAGATCGGAAG |
| 11001_1 | TATGAGTGTGGAGTCGTTGCTTCTCTACGCAGGCCCCCGTNNNAGATCGGAAGAGC |
| 11001_2 | TATGAGTGTGGAGTCGTTGCTTCTCTACGCAGGCCCCCGTNNNAGATCGGAAGAGC |
| 11001_3 | TATGAGTGTGGAGTCGTTGCTTCTCTACGCAGGCCCCCGTNNNAGATCGGAAGAGC |
| 35814_1 | TATGAGTGTGGAGTCGTTGCCCACCACATTCTTGAACACTGACNNNAGATCGGAAG |

| | |
|---|---|
| 35814_2 | TATGAGTGTGGAGTCGTTGCCCACCACATTCTTGAACACTGACNNNAGATCGGAAGA |
| 35814_3 | TATGAGTGTGGAGTCGTTGCCCACCACATTCTTGAACACTGACNNNAGATCGGAAGA |
| 31490_1 | TATGAGTGTGGAGTCGTTGCTCTGAAACCTCTTGTCCATTCCCNNNAGATCGGAAGA |
| 31490_2 | TATGAGTGTGGAGTCGTTGCTCTGAAACCTCTTGTCCATTCCCNNNAGATCGGAAGA |
| 31490_3 | TATGAGTGTGGAGTCGTTGCTCTGAAACCTCTTGTCCATTCCCNNNAGATCGGAAGA |
| 33657_1 | TATGAGTGTGGAGTCGTTGCGCTGCACCTGTTTTTGCTTTTATNNNAGATCGGAAGA |
| 33657_2 | TATGAGTGTGGAGTCGTTGCGCTGCACCTGTTTTTGCTTTTATNNNAGATCGGAAGA |
| 33657_3 | TATGAGTGTGGAGTCGTTGCGCTGCACCTGTTTTTGCTTTTATNNNAGATCGGAAGA |
| 52259_1 | TATGAGTGTGGAGTCGTTGCAGTCTTGACTGAAGTACTATGGGANNNAGATCGGAA |
| 52259_2 | TATGAGTGTGGAGTCGTTGCAGTCTTGACTGAAGTACTATGGGANNNAGATCGGAA |
| 52259_3 | TATGAGTGTGGAGTCGTTGCAGTCTTGACTGAAGTACTATGGGANNNAGATCGGAA |
| 43268_1 | TATGAGTGTGGAGTCGTTGCTTTTTTCCTCTGTATGTGTTTCTATCTNNNAGATCGGA |
| 43268_2 | TATGAGTGTGGAGTCGTTGCTTTTTTCCTCTGTATGTGTTTCTATCTNNNAGATCGGA |
| 43268_3 | TATGAGTGTGGAGTCGTTGCTTTTTTCCTCTGTATGTGTTTCTATCTNNNAGATCGGA |
| 49369_1 | TATGAGTGTGGAGTCGTTGCCTGAACTGTGGGCTATTCTGCTANNNAGATCGGAAG |
| 49369_2 | TATGAGTGTGGAGTCGTTGCCTGAACTGTGGGCTATTCTGCTANNNAGATCGGAAG |
| 49369_3 | TATGAGTGTGGAGTCGTTGCCTGAACTGTGGGCTATTCTGCTANNNAGATCGGAAG |
| 51664_1 | TATGAGTGTGGAGTCGTTGCCAGTGTATACGTGTGTGTGTTCANNNAGATCGGAAG |
| 51664_2 | TATGAGTGTGGAGTCGTTGCCAGTGTATACGTGTGTGTGTTCANNNAGATCGGAAG |
| 51664_3 | TATGAGTGTGGAGTCGTTGCCAGTGTATACGTGTGTGTGTTCANNNAGATCGGAAG |
| 45825_1 | TATGAGTGTGGAGTCGTTGCCTCAGTGATGTGAGGGGCTTTTCNNNAGATCGGAAG |
| 45825_2 | TATGAGTGTGGAGTCGTTGCCTCAGTGATGTGAGGGGCTTTTCNNNAGATCGGAAG |
| 45825_3 | TATGAGTGTGGAGTCGTTGCCTCAGTGATGTGAGGGGCTTTTCNNNAGATCGGAAG |
| 40418_1 | TATGAGTGTGGAGTCGTTGCAAACAGACAGTCCCTCAAAAGGTNNNAGATCGGAAG |
| 40418_2 | TATGAGTGTGGAGTCGTTGCAAACAGACAGTCCCTCAAAAGGTNNNAGATCGGAAG |
| 40418_3 | TATGAGTGTGGAGTCGTTGCAAACAGACAGTCCCTCAAAAGGTNNNAGATCGGAAG |
| 39763_1 | TATGAGTGTGGAGTCGTTGCACTCCAACATTTAGTGGGTGGAANNNAGATCGGAAG |
| 39763_2 | TATGAGTGTGGAGTCGTTGCACTCCAACATTTAGTGGGTGGAANNNAGATCGGAAG |
| 39763_3 | TATGAGTGTGGAGTCGTTGCACTCCAACATTTAGTGGGTGGAANNNAGATCGGAAG |
| 34329_1 | TATGAGTGTGGAGTCGTTGCGATAGTAGAGATGTCCAGCACCCNNNAGATCGGAAG |
| 34329_2 | TATGAGTGTGGAGTCGTTGCGATAGTAGAGATGTCCAGCACCCNNNAGATCGGAAG |
| 34329_3 | TATGAGTGTGGAGTCGTTGCGATAGTAGAGATGTCCAGCACCCNNNAGATCGGAAG |
| 49569_1 | TATGAGTGTGGAGTCGTTGCGCATTAGCAAACTAGACAGCTGCNNNAGATCGGAAG |
| 49569_2 | TATGAGTGTGGAGTCGTTGCGCATTAGCAAACTAGACAGCTGCNNNAGATCGGAAG |
| 49569_3 | TATGAGTGTGGAGTCGTTGCGCATTAGCAAACTAGACAGCTGCNNNAGATCGGAAG |
| 39379_1 | TATGAGTGTGGAGTCGTTGCTCAGGATAAAATGTGAAAACGGACANNNAGATCGGA |
| 39379_2 | TATGAGTGTGGAGTCGTTGCTCAGGATAAAATGTGAAAACGGACANNNAGATCGGA |
| 39379_3 | TATGAGTGTGGAGTCGTTGCTCAGGATAAAATGTGAAAACGGACANNNAGATCGGA |
| 32376_1 | TATGAGTGTGGAGTCGTTGCAATGCACCAGATCCCATCTCATTNNNAGATCGGAAGA |
| 32376_2 | TATGAGTGTGGAGTCGTTGCAATGCACCAGATCCCATCTCATTNNNAGATCGGAAGA |
| 32376_3 | TATGAGTGTGGAGTCGTTGCAATGCACCAGATCCCATCTCATTNNNAGATCGGAAGA |
| 29539_1 | TATGAGTGTGGAGTCGTTGCGTGTGTCTGTGGGGACCTCCNNNAGATCGGAAGAG |
| 29539_2 | TATGAGTGTGGAGTCGTTGCGTGTGTCTGTGGGGACCTCCNNNAGATCGGAAGAG |

| | |
|---|---|
| 29539_3 | TATGAGTGTGGAGTCGTTGCGTGTGTCTGTGGGGACCTCCNNNAGATCGGAAGAG |
| 13573_1 | TATGAGTGTGGAGTCGTTGCATCTTGCAACAGAGAGATTATCNNNAGATCGGAAGA |
| 13573_2 | TATGAGTGTGGAGTCGTTGCATCTTGCAACAGAGAGATTATCNNNAGATCGGAAGA |
| 13573_3 | TATGAGTGTGGAGTCGTTGCATCTTGCAACAGAGAGATTATCNNNAGATCGGAAGA |
| 29424_1 | TATGAGTGTGGAGTCGTTGCCAAAAGTGAGTCCCAGGTAAAATCAGNNNAGATCG |
| 29424_2 | TATGAGTGTGGAGTCGTTGCCAAAAGTGAGTCCCAGGTAAAATCAGNNNAGATCG |
| 29424_3 | TATGAGTGTGGAGTCGTTGCCAAAAGTGAGTCCCAGGTAAAATCAGNNNAGATCG |
| 3051_1 | TATGAGTGTGGAGTCGTTGCAGTACCTCAGCCCATGCCAAGNNNAGATCGGAAGAG |
| 3051_2 | TATGAGTGTGGAGTCGTTGCAGTACCTCAGCCCATGCCAAGNNNAGATCGGAAGAG |
| 3051_3 | TATGAGTGTGGAGTCGTTGCAGTACCTCAGCCCATGCCAAGNNNAGATCGGAAGAG |
| 48096_1 | TATGAGTGTGGAGTCGTTGCTTCAAGTGATTCTCCTGCCTCAGNNNAGATCGGAAG |
| 48096_2 | TATGAGTGTGGAGTCGTTGCTTCAAGTGATTCTCCTGCCTCAGNNNAGATCGGAAG |
| 48096_3 | TATGAGTGTGGAGTCGTTGCTTCAAGTGATTCTCCTGCCTCAGNNNAGATCGGAAG |
| 42232_1 | TATGAGTGTGGAGTCGTTGCCAATTGGCTCACACATTCACACANNNAGATCGGAAG |
| 42232_2 | TATGAGTGTGGAGTCGTTGCCAATTGGCTCACACATTCACACANNNAGATCGGAAG |
| 42232_3 | TATGAGTGTGGAGTCGTTGCCAATTGGCTCACACATTCACACANNNAGATCGGAAG |
| 36259_1 | TATGAGTGTGGAGTCGTTGCCAGAGGGACAAAGGAGACAAAGANNNAGATCGGA |
| 36259_2 | TATGAGTGTGGAGTCGTTGCCAGAGGGACAAAGGAGACAAAGANNNAGATCGGA |
| 36259_3 | TATGAGTGTGGAGTCGTTGCCAGAGGGACAAAGGAGACAAAGANNNAGATCGGA |
| 52837_1 | TATGAGTGTGGAGTCGTTGCATAGTACCTGGAGGAGCAGACTTNNNAGATCGGAA |
| 52837_2 | TATGAGTGTGGAGTCGTTGCATAGTACCTGGAGGAGCAGACTTNNNAGATCGGAA |
| 52837_3 | TATGAGTGTGGAGTCGTTGCATAGTACCTGGAGGAGCAGACTTNNNAGATCGGAA |
| 37983_1 | TATGAGTGTGGAGTCGTTGCAGGCATGAGAATCACTTGAACCTNNNAGATCGGAAG |
| 37983_2 | TATGAGTGTGGAGTCGTTGCAGGCATGAGAATCACTTGAACCTNNNAGATCGGAAG |
| 37983_3 | TATGAGTGTGGAGTCGTTGCAGGCATGAGAATCACTTGAACCTNNNAGATCGGAAG |
| 35069_1 | TATGAGTGTGGAGTCGTTGCTTCAAGTGATTCTCCTGCCTCAGNNNAGATCGGAAG |
| 35069_2 | TATGAGTGTGGAGTCGTTGCTTCAAGTGATTCTCCTGCCTCAGNNNAGATCGGAAG |
| 35069_3 | TATGAGTGTGGAGTCGTTGCTTCAAGTGATTCTCCTGCCTCAGNNNAGATCGGAAG |
| 46492_1 | TATGAGTGTGGAGTCGTTGCCCAAGTATCTGGGATTACAGGCANNNAGATCGGAA |
| 46492_2 | TATGAGTGTGGAGTCGTTGCCCAAGTATCTGGGATTACAGGCANNNAGATCGGAA |
| 46492_3 | TATGAGTGTGGAGTCGTTGCCCAAGTATCTGGGATTACAGGCANNNAGATCGGAA |
| 32090_1 | TATGAGTGTGGAGTCGTTGCGTCTCCCTATGTTACTCAGGCTGNNNAGATCGGAAG |
| 32090_2 | TATGAGTGTGGAGTCGTTGCGTCTCCCTATGTTACTCAGGCTGNNNAGATCGGAAG |
| 32090_3 | TATGAGTGTGGAGTCGTTGCGTCTCCCTATGTTACTCAGGCTGNNNAGATCGGAAG |
| 51697_1 | TATGAGTGTGGAGTCGTTGCCTAATAGGAACTCTCCCCCTCCANNNAGATCGGAAGA |
| 51697_2 | TATGAGTGTGGAGTCGTTGCCTAATAGGAACTCTCCCCCTCCANNNAGATCGGAAGA |
| 51697_3 | TATGAGTGTGGAGTCGTTGCCTAATAGGAACTCTCCCCCTCCANNNAGATCGGAAGA |
| 39645_1 | TATGAGTGTGGAGTCGTTGCGCCAATATCCCTGATGAACAACGNNNAGATCGGAAG |
| 39645_2 | TATGAGTGTGGAGTCGTTGCGCCAATATCCCTGATGAACAACGNNNAGATCGGAAG |
| 39645_3 | TATGAGTGTGGAGTCGTTGCGCCAATATCCCTGATGAACAACGNNNAGATCGGAAG |
| 34411_1 | TATGAGTGTGGAGTCGTTGCAGGTATGCAGACAATTGGAGTGTNNNAGATCGGAA |
| 34411_2 | TATGAGTGTGGAGTCGTTGCAGGTATGCAGACAATTGGAGTGTNNNAGATCGGAA |
| 34411_3 | TATGAGTGTGGAGTCGTTGCAGGTATGCAGACAATTGGAGTGTNNNAGATCGGAA |

| | |
|---|---|
| 39620_1 | TATGAGTGTGGAGTCGTTGCGCTGCCCAAGTATAACAGGAAAGNNNAGATCGGAA |
| 39620_2 | TATGAGTGTGGAGTCGTTGCGCTGCCCAAGTATAACAGGAAAGNNNAGATCGGAA |
| 39620_3 | TATGAGTGTGGAGTCGTTGCGCTGCCCAAGTATAACAGGAAAGNNNAGATCGGAA |
| 34525_1 | TATGAGTGTGGAGTCGTTGCCCTTCAGAAAGCTCCAATCAGGANNNAGATCGGAAG |
| 34525_2 | TATGAGTGTGGAGTCGTTGCCCTTCAGAAAGCTCCAATCAGGANNNAGATCGGAAG |
| 34525_3 | TATGAGTGTGGAGTCGTTGCCCTTCAGAAAGCTCCAATCAGGANNNAGATCGGAAG |
| 53146_1 | TATGAGTGTGGAGTCGTTGCCCCTTACATCAACCACTGCCTAANNNAGATCGGAAGA |
| 53146_2 | TATGAGTGTGGAGTCGTTGCCCCTTACATCAACCACTGCCTAANNNAGATCGGAAGA |
| 53146_3 | TATGAGTGTGGAGTCGTTGCCCCTTACATCAACCACTGCCTAANNNAGATCGGAAGA |
| 51333_1 | TATGAGTGTGGAGTCGTTGCGAAAAGAGGCCTCTGAGTTCTCANNNAGATCGGAA |
| 51333_2 | TATGAGTGTGGAGTCGTTGCGAAAAGAGGCCTCTGAGTTCTCANNNAGATCGGAA |
| 51333_3 | TATGAGTGTGGAGTCGTTGCGAAAAGAGGCCTCTGAGTTCTCANNNAGATCGGAA |
| 50528_1 | TATGAGTGTGGAGTCGTTGCCCTAGAGGGTTTGGGACAAACTGNNNAGATCGGAA |
| 50528_2 | TATGAGTGTGGAGTCGTTGCCCTAGAGGGTTTGGGACAAACTGNNNAGATCGGAA |
| 50528_3 | TATGAGTGTGGAGTCGTTGCCCTAGAGGGTTTGGGACAAACTGNNNAGATCGGAA |
| 12770_1 | TATGAGTGTGGAGTCGTTGCCAGATGAACCTGGAATTAAAAAGNNNAGATCGGAA |
| 12770_2 | TATGAGTGTGGAGTCGTTGCCAGATGAACCTGGAATTAAAAAGNNNAGATCGGAA |
| 12770_3 | TATGAGTGTGGAGTCGTTGCCAGATGAACCTGGAATTAAAAAGNNNAGATCGGAA |
| 49884_1 | TATGAGTGTGGAGTCGTTGCGCTCTTGCCCATATGTGTTCAAGNNNAGATCGGAAG |
| 49884_2 | TATGAGTGTGGAGTCGTTGCGCTCTTGCCCATATGTGTTCAAGNNNAGATCGGAAG |
| 49884_3 | TATGAGTGTGGAGTCGTTGCGCTCTTGCCCATATGTGTTCAAGNNNAGATCGGAAG |
| 8322_1 | TATGAGTGTGGAGTCGTTGCCTGTTGGGAGCTTCTGCTCTGNNNAGATCGGAAGAG |
| 8322_2 | TATGAGTGTGGAGTCGTTGCCTGTTGGGAGCTTCTGCTCTGNNNAGATCGGAAGAG |
| 8322_3 | TATGAGTGTGGAGTCGTTGCCTGTTGGGAGCTTCTGCTCTGNNNAGATCGGAAGAG |
| 52403_1 | TATGAGTGTGGAGTCGTTGCCAACTTTGCTCTTCTCCACGCNNNAGATCGGAAGAGC |
| 52403_2 | TATGAGTGTGGAGTCGTTGCCAACTTTGCTCTTCTCCACGCNNNAGATCGGAAGAGC |
| 52403_3 | TATGAGTGTGGAGTCGTTGCCAACTTTGCTCTTCTCCACGCNNNAGATCGGAAGAGC |
| 49033_1 | TATGAGTGTGGAGTCGTTGCCCAGGTTCAAGCGATTTTCCTGNNNAGATCGGAAGA |
| 49033_2 | TATGAGTGTGGAGTCGTTGCCCAGGTTCAAGCGATTTTCCTGNNNAGATCGGAAGA |
| 49033_3 | TATGAGTGTGGAGTCGTTGCCCAGGTTCAAGCGATTTTCCTGNNNAGATCGGAAGA |
| 46570_1 | TATGAGTGTGGAGTCGTTGCTATGCTGGCTTTGTCAGAGTTCTNNNAGATCGGAAG |
| 46570_2 | TATGAGTGTGGAGTCGTTGCTATGCTGGCTTTGTCAGAGTTCTNNNAGATCGGAAG |
| 46570_3 | TATGAGTGTGGAGTCGTTGCTATGCTGGCTTTGTCAGAGTTCTNNNAGATCGGAAG |
| 31757_1 | TATGAGTGTGGAGTCGTTGCCCCAGGAGATCTCAAGCCTTTAGNNNAGATCGGAAG |
| 31757_2 | TATGAGTGTGGAGTCGTTGCCCCAGGAGATCTCAAGCCTTTAGNNNAGATCGGAAG |
| 31757_3 | TATGAGTGTGGAGTCGTTGCCCCAGGAGATCTCAAGCCTTTAGNNNAGATCGGAAG |
| 37902_1 | TATGAGTGTGGAGTCGTTGCACATGGTAAAACCCCGTCTCTACNNNAGATCGGAAG |
| 37902_2 | TATGAGTGTGGAGTCGTTGCACATGGTAAAACCCCGTCTCTACNNNAGATCGGAAG |
| 37902_3 | TATGAGTGTGGAGTCGTTGCACATGGTAAAACCCCGTCTCTACNNNAGATCGGAAG |
| 29547_1 | TATGAGTGTGGAGTCGTTGCGCTGTGCTTTGGTTTAACTTCCTNNNAGATCGGAAG |
| 29547_2 | TATGAGTGTGGAGTCGTTGCGCTGTGCTTTGGTTTAACTTCCTNNNAGATCGGAAG |
| 29547_3 | TATGAGTGTGGAGTCGTTGCGCTGTGCTTTGGTTTAACTTCCTNNNAGATCGGAAG |
| 3393_1 | TATGAGTGTGGAGTCGTTGCTGGACCTGAAATATGGCTGATGANNNAGATCGGAA |

| | |
|---|---|
| 3393_2 | TATGAGTGTGGAGTCGTTGCTGGACCTGAAATATGGCTGATGANNNAGATCGGAA |
| 3393_3 | TATGAGTGTGGAGTCGTTGCTGGACCTGAAATATGGCTGATGANNNAGATCGGAA |
| 39249_1 | TATGAGTGTGGAGTCGTTGCATGACTGCATCACTGTACTCCAGNNNAGATCGGAAG |
| 39249_2 | TATGAGTGTGGAGTCGTTGCATGACTGCATCACTGTACTCCAGNNNAGATCGGAAG |
| 39249_3 | TATGAGTGTGGAGTCGTTGCATGACTGCATCACTGTACTCCAGNNNAGATCGGAAG |
| 51457_1 | TATGAGTGTGGAGTCGTTGCTAGTTTTTCCTCTAACAGGCCCCNNNAGATCGGAAGA |
| 51457_2 | TATGAGTGTGGAGTCGTTGCTAGTTTTTCCTCTAACAGGCCCCNNNAGATCGGAAGA |
| 51457_3 | TATGAGTGTGGAGTCGTTGCTAGTTTTTCCTCTAACAGGCCCCNNNAGATCGGAAGA |
| 11998_1 | TATGAGTGTGGAGTCGTTGCCTGGAGTTTCCCAAACTAAATGGCNNNAGATCGGAA |
| 11998_2 | TATGAGTGTGGAGTCGTTGCCTGGAGTTTCCCAAACTAAATGGCNNNAGATCGGAA |
| 11998_3 | TATGAGTGTGGAGTCGTTGCCTGGAGTTTCCCAAACTAAATGGCNNNAGATCGGAA |
| 41432_1 | TATGAGTGTGGAGTCGTTGCATAACAGATGTTGGCCTGGATGTNNNAGATCGGAA |
| 41432_2 | TATGAGTGTGGAGTCGTTGCATAACAGATGTTGGCCTGGATGTNNNAGATCGGAA |
| 41432_3 | TATGAGTGTGGAGTCGTTGCATAACAGATGTTGGCCTGGATGTNNNAGATCGGAA |
| 39267_1 | TATGAGTGTGGAGTCGTTGCAAAGAAAAACACTCCAGCAAACANNNAGATCGGAAG |
| 39267_2 | TATGAGTGTGGAGTCGTTGCAAAGAAAAACACTCCAGCAAACANNNAGATCGGAAG |
| 39267_3 | TATGAGTGTGGAGTCGTTGCAAAGAAAAACACTCCAGCAAACANNNAGATCGGAAG |
| 39453_1 | TATGAGTGTGGAGTCGTTGCGGCTAAGGCAGGAGAATCACTTANNNAGATCGGAA |
| 39453_2 | TATGAGTGTGGAGTCGTTGCGGCTAAGGCAGGAGAATCACTTANNNAGATCGGAA |
| 39453_3 | TATGAGTGTGGAGTCGTTGCGGCTAAGGCAGGAGAATCACTTANNNAGATCGGAA |
| 32497_1 | TATGAGTGTGGAGTCGTTGCGGTTCAAACAATTCTCCTGCCTCNNNAGATCGGAAGA |
| 32497_2 | TATGAGTGTGGAGTCGTTGCGGTTCAAACAATTCTCCTGCCTCNNNAGATCGGAAGA |
| 32497_3 | TATGAGTGTGGAGTCGTTGCGGTTCAAACAATTCTCCTGCCTCNNNAGATCGGAAGA |
| 31690_1 | TATGAGTGTGGAGTCGTTGCTGTTTGCTAATCTCCTGCTTCCTNNNAGATCGGAAGA |
| 31690_2 | TATGAGTGTGGAGTCGTTGCTGTTTGCTAATCTCCTGCTTCCTNNNAGATCGGAAGA |
| 31690_3 | TATGAGTGTGGAGTCGTTGCTGTTTGCTAATCTCCTGCTTCCTNNNAGATCGGAAGA |
| 50931_1 | TATGAGTGTGGAGTCGTTGCTCTCATCCAGGTGAAGTCACATGNNNAGATCGGAAG |
| 50931_2 | TATGAGTGTGGAGTCGTTGCTCTCATCCAGGTGAAGTCACATGNNNAGATCGGAAG |
| 50931_3 | TATGAGTGTGGAGTCGTTGCTCTCATCCAGGTGAAGTCACATGNNNAGATCGGAAG |
| 9096_1 | TATGAGTGTGGAGTCGTTGCTGGAAGTGTGGGTAAAATGGACANNNAGATCGGAA |
| 9096_2 | TATGAGTGTGGAGTCGTTGCTGGAAGTGTGGGTAAAATGGACANNNAGATCGGAA |
| 9096_3 | TATGAGTGTGGAGTCGTTGCTGGAAGTGTGGGTAAAATGGACANNNAGATCGGAA |
| 29702_1 | TATGAGTGTGGAGTCGTTGCAGAAACCAGACACACTAACATGTNNNAGATCGGAAG |
| 29702_2 | TATGAGTGTGGAGTCGTTGCAGAAACCAGACACACTAACATGTNNNAGATCGGAAG |
| 29702_3 | TATGAGTGTGGAGTCGTTGCAGAAACCAGACACACTAACATGTNNNAGATCGGAAG |
| 37903_1 | TATGAGTGTGGAGTCGTTGCGAAGTTTCACTCTTGTTGCCCAGNNNAGATCGGAAG |
| 37903_2 | TATGAGTGTGGAGTCGTTGCGAAGTTTCACTCTTGTTGCCCAGNNNAGATCGGAAG |
| 37903_3 | TATGAGTGTGGAGTCGTTGCGAAGTTTCACTCTTGTTGCCCAGNNNAGATCGGAAG |
| 32366_1 | TATGAGTGTGGAGTCGTTGCCCTTCTCTCCGTGTCTCTACTCTNNNAGATCGGAAGA |
| 32366_2 | TATGAGTGTGGAGTCGTTGCCCTTCTCTCCGTGTCTCTACTCTNNNAGATCGGAAGA |
| 32366_3 | TATGAGTGTGGAGTCGTTGCCCTTCTCTCCGTGTCTCTACTCTNNNAGATCGGAAGA |
| 45706_1 | TATGAGTGTGGAGTCGTTGCAACTCTCGACCTCAAGTGATCTGNNNAGATCGGAAG |
| 45706_2 | TATGAGTGTGGAGTCGTTGCAACTCTCGACCTCAAGTGATCTGNNNAGATCGGAAG |

| | |
|---|---|
| 45706_3 | TATGAGTGTGGAGTCGTTGCAACTCTCGACCTCAAGTGATCTGNNNAGATCGGAAG |
| 38418_1 | TATGAGTGTGGAGTCGTTGCCTAAGAATTTCAGCCAGGTGCAGNNNAGATCGGAA |
| 38418_2 | TATGAGTGTGGAGTCGTTGCCTAAGAATTTCAGCCAGGTGCAGNNNAGATCGGAA |
| 38418_3 | TATGAGTGTGGAGTCGTTGCCTAAGAATTTCAGCCAGGTGCAGNNNAGATCGGAA |
| 37587_1 | TATGAGTGTGGAGTCGTTGCACACATATAAGGGCTGATCACCCNNNAGATCGGAAG |
| 37587_2 | TATGAGTGTGGAGTCGTTGCACACATATAAGGGCTGATCACCCNNNAGATCGGAAG |
| 37587_3 | TATGAGTGTGGAGTCGTTGCACACATATAAGGGCTGATCACCCNNNAGATCGGAAG |
| 13278_1 | TATGAGTGTGGAGTCGTTGCGGACAAAAACTACAACCATATCNNNAGATCGGAAGA |
| 13278_2 | TATGAGTGTGGAGTCGTTGCGGACAAAAACTACAACCATATCNNNAGATCGGAAGA |
| 13278_3 | TATGAGTGTGGAGTCGTTGCGGACAAAAACTACAACCATATCNNNAGATCGGAAGA |
| 44372_1 | TATGAGTGTGGAGTCGTTGCAAAGATGAGATCTCGAGACCACGNNNAGATCGGAA |
| 44372_2 | TATGAGTGTGGAGTCGTTGCAAAGATGAGATCTCGAGACCACGNNNAGATCGGAA |
| 44372_3 | TATGAGTGTGGAGTCGTTGCAAAGATGAGATCTCGAGACCACGNNNAGATCGGAA |
| 34484_1 | TATGAGTGTGGAGTCGTTGCAACCTCCCACAGTTTGTACCTACNNNAGATCGGAAGA |
| 34484_2 | TATGAGTGTGGAGTCGTTGCAACCTCCCACAGTTTGTACCTACNNNAGATCGGAAGA |
| 34484_3 | TATGAGTGTGGAGTCGTTGCAACCTCCCACAGTTTGTACCTACNNNAGATCGGAAGA |
| 38207_1 | TATGAGTGTGGAGTCGTTGCAGCAAACCAGGAATAGAACTTCCNNNAGATCGGAAG |
| 38207_2 | TATGAGTGTGGAGTCGTTGCAGCAAACCAGGAATAGAACTTCCNNNAGATCGGAAG |
| 38207_3 | TATGAGTGTGGAGTCGTTGCAGCAAACCAGGAATAGAACTTCCNNNAGATCGGAAG |
| 36965_1 | TATGAGTGTGGAGTCGTTGCTTCAGGTTGGATCTTGGGTTCTTNNNAGATCGGAAG |
| 36965_2 | TATGAGTGTGGAGTCGTTGCTTCAGGTTGGATCTTGGGTTCTTNNNAGATCGGAAG |
| 36965_3 | TATGAGTGTGGAGTCGTTGCTTCAGGTTGGATCTTGGGTTCTTNNNAGATCGGAAG |
| 51869_1 | TATGAGTGTGGAGTCGTTGCCACTGTTTTCGGACCTTCCATGNNNAGATCGGAAGA |
| 51869_2 | TATGAGTGTGGAGTCGTTGCCACTGTTTTCGGACCTTCCATGNNNAGATCGGAAGA |
| 51869_3 | TATGAGTGTGGAGTCGTTGCCACTGTTTTCGGACCTTCCATGNNNAGATCGGAAGA |
| 31338_1 | TATGAGTGTGGAGTCGTTGCAGATTGAAAAGGTGGCCTAGGAGNNNAGATCGGA |
| 31338_2 | TATGAGTGTGGAGTCGTTGCAGATTGAAAAGGTGGCCTAGGAGNNNAGATCGGA |
| 31338_3 | TATGAGTGTGGAGTCGTTGCAGATTGAAAAGGTGGCCTAGGAGNNNAGATCGGA |
| 29217_1 | TATGAGTGTGGAGTCGTTGCCCTCCCGAATAGCTAGGACTACANNNAGATCGGAAG |
| 29217_2 | TATGAGTGTGGAGTCGTTGCCCTCCCGAATAGCTAGGACTACANNNAGATCGGAAG |
| 29217_3 | TATGAGTGTGGAGTCGTTGCCCTCCCGAATAGCTAGGACTACANNNAGATCGGAAG |
| 30371_1 | TATGAGTGTGGAGTCGTTGCCTGGCTTGGTTCTCAACTGTCTTNNNAGATCGGAAG |
| 30371_2 | TATGAGTGTGGAGTCGTTGCCTGGCTTGGTTCTCAACTGTCTTNNNAGATCGGAAG |
| 30371_3 | TATGAGTGTGGAGTCGTTGCCTGGCTTGGTTCTCAACTGTCTTNNNAGATCGGAAG |
| 47294_1 | TATGAGTGTGGAGTCGTTGCAACCTCCCAAGTAGCTGAGACTANNNAGATCGGAAG |
| 47294_2 | TATGAGTGTGGAGTCGTTGCAACCTCCCAAGTAGCTGAGACTANNNAGATCGGAAG |
| 47294_3 | TATGAGTGTGGAGTCGTTGCAACCTCCCAAGTAGCTGAGACTANNNAGATCGGAAG |
| 36059_1 | TATGAGTGTGGAGTCGTTGCGGAAGTCCAAGATCAAAGTGTGCNNNAGATCGGAA |
| 36059_2 | TATGAGTGTGGAGTCGTTGCGGAAGTCCAAGATCAAAGTGTGCNNNAGATCGGAA |
| 36059_3 | TATGAGTGTGGAGTCGTTGCGGAAGTCCAAGATCAAAGTGTGCNNNAGATCGGAA |
| 38302_1 | TATGAGTGTGGAGTCGTTGCTACCACACAGCTCCACATGAAATNNNAGATCGGAAG |
| 38302_2 | TATGAGTGTGGAGTCGTTGCTACCACACAGCTCCACATGAAATNNNAGATCGGAAG |
| 38302_3 | TATGAGTGTGGAGTCGTTGCTACCACACAGCTCCACATGAAATNNNAGATCGGAAG |

| | |
|---|---|
| 47916_1 | TATGAGTGTGGAGTCGTTGCCTCCTGAGTAGCTGGGATTACAGNNNAGATCGGAA |
| 47916_2 | TATGAGTGTGGAGTCGTTGCCTCCTGAGTAGCTGGGATTACAGNNNAGATCGGAA |
| 47916_3 | TATGAGTGTGGAGTCGTTGCCTCCTGAGTAGCTGGGATTACAGNNNAGATCGGAA |
| 34752_1 | TATGAGTGTGGAGTCGTTGCATGAGGCTTTCCCTAATCCCATCNNNAGATCGGAAG |
| 34752_2 | TATGAGTGTGGAGTCGTTGCATGAGGCTTTCCCTAATCCCATCNNNAGATCGGAAG |
| 34752_3 | TATGAGTGTGGAGTCGTTGCATGAGGCTTTCCCTAATCCCATCNNNAGATCGGAAG |
| 34556_1 | TATGAGTGTGGAGTCGTTGCGATGGTCCTAAGAGGAGGAGCAGNNNAGATCGGA |
| 34556_2 | TATGAGTGTGGAGTCGTTGCGATGGTCCTAAGAGGAGGAGCAGNNNAGATCGGA |
| 34556_3 | TATGAGTGTGGAGTCGTTGCGATGGTCCTAAGAGGAGGAGCAGNNNAGATCGGA |
| 36950_1 | TATGAGTGTGGAGTCGTTGCAGCCTCAACATATTTAGATGCCANNNAGATCGGAAG |
| 36950_2 | TATGAGTGTGGAGTCGTTGCAGCCTCAACATATTTAGATGCCANNNAGATCGGAAG |
| 36950_3 | TATGAGTGTGGAGTCGTTGCAGCCTCAACATATTTAGATGCCANNNAGATCGGAAG |
| 50530_1 | TATGAGTGTGGAGTCGTTGCTTTTGCCTCTTTGTGCATCAGTTNNNAGATCGGAAGA |
| 50530_2 | TATGAGTGTGGAGTCGTTGCTTTTGCCTCTTTGTGCATCAGTTNNNAGATCGGAAGA |
| 50530_3 | TATGAGTGTGGAGTCGTTGCTTTTGCCTCTTTGTGCATCAGTTNNNAGATCGGAAGA |
| 52340_1 | TATGAGTGTGGAGTCGTTGCTGTTTGGTGAGGTCATGTTTTCCNNNAGATCGGAAG |
| 52340_2 | TATGAGTGTGGAGTCGTTGCTGTTTGGTGAGGTCATGTTTTCCNNNAGATCGGAAG |
| 52340_3 | TATGAGTGTGGAGTCGTTGCTGTTTGGTGAGGTCATGTTTTCCNNNAGATCGGAAG |
| 30474_1 | TATGAGTGTGGAGTCGTTGCAGTAATGGTTTGGCATTTGGAAANNNAGATCGGAA |
| 30474_2 | TATGAGTGTGGAGTCGTTGCAGTAATGGTTTGGCATTTGGAAANNNAGATCGGAA |
| 30474_3 | TATGAGTGTGGAGTCGTTGCAGTAATGGTTTGGCATTTGGAAANNNAGATCGGAA |
| 9262_1 | TATGAGTGTGGAGTCGTTGCGGAGCACTTTACTATGGTGGTGGNNNAGATCGGAA |
| 9262_2 | TATGAGTGTGGAGTCGTTGCGGAGCACTTTACTATGGTGGTGGNNNAGATCGGAA |
| 9262_3 | TATGAGTGTGGAGTCGTTGCGGAGCACTTTACTATGGTGGTGGNNNAGATCGGAA |
| 48848_1 | TATGAGTGTGGAGTCGTTGCAGTGCTAGGATTACAGGCATGAGNNNAGATCGGAA |
| 48848_2 | TATGAGTGTGGAGTCGTTGCAGTGCTAGGATTACAGGCATGAGNNNAGATCGGAA |
| 48848_3 | TATGAGTGTGGAGTCGTTGCAGTGCTAGGATTACAGGCATGAGNNNAGATCGGAA |
| 36566_1 | TATGAGTGTGGAGTCGTTGCTAAAGGCCACCAGAACAATAGCTNNNAGATCGGAAG |
| 36566_2 | TATGAGTGTGGAGTCGTTGCTAAAGGCCACCAGAACAATAGCTNNNAGATCGGAAG |
| 36566_3 | TATGAGTGTGGAGTCGTTGCTAAAGGCCACCAGAACAATAGCTNNNAGATCGGAAG |
| 47084_1 | TATGAGTGTGGAGTCGTTGCGCCTCCTGAGTAGCTGAGATTACNNNAGATCGGAAG |
| 47084_2 | TATGAGTGTGGAGTCGTTGCGCCTCCTGAGTAGCTGAGATTACNNNAGATCGGAAG |
| 47084_3 | TATGAGTGTGGAGTCGTTGCGCCTCCTGAGTAGCTGAGATTACNNNAGATCGGAAG |
| 47348_1 | TATGAGTGTGGAGTCGTTGCCAAGTAGCTGGGACTACAGGCNNNAGATCGGAAGA |
| 47348_2 | TATGAGTGTGGAGTCGTTGCCAAGTAGCTGGGACTACAGGCNNNAGATCGGAAGA |
| 47348_3 | TATGAGTGTGGAGTCGTTGCCAAGTAGCTGGGACTACAGGCNNNAGATCGGAAGA |
| 33110_1 | TATGAGTGTGGAGTCGTTGCCATGGATGGAACTGGAGGACATTNNNAGATCGGAA |
| 33110_2 | TATGAGTGTGGAGTCGTTGCCATGGATGGAACTGGAGGACATTNNNAGATCGGAA |
| 33110_3 | TATGAGTGTGGAGTCGTTGCCATGGATGGAACTGGAGGACATTNNNAGATCGGAA |
| 12806_1 | TATGAGTGTGGAGTCGTTGCCTCCACTTCTTTATTAACAACTGNNNAGATCGGAAGA |
| 12806_2 | TATGAGTGTGGAGTCGTTGCCTCCACTTCTTTATTAACAACTGNNNAGATCGGAAGA |
| 12806_3 | TATGAGTGTGGAGTCGTTGCCTCCACTTCTTTATTAACAACTGNNNAGATCGGAAGA |
| 49760_1 | TATGAGTGTGGAGTCGTTGCTGGACAGACACTCATTGAATGGANNNAGATCGGAA |

| | |
|---|---|
| 49760_2 | TATGAGTGTGGAGTCGTTGCTGGACAGACACTCATTGAATGGANNNAGATCGGAA |
| 49760_3 | TATGAGTGTGGAGTCGTTGCTGGACAGACACTCATTGAATGGANNNAGATCGGAA |
| 46276_1 | TATGAGTGTGGAGTCGTTGCGAAATGGGGTCTCACTATGTTGCNNNAGATCGGAA |
| 46276_2 | TATGAGTGTGGAGTCGTTGCGAAATGGGGTCTCACTATGTTGCNNNAGATCGGAA |
| 46276_3 | TATGAGTGTGGAGTCGTTGCGAAATGGGGTCTCACTATGTTGCNNNAGATCGGAA |
| 48171_1 | TATGAGTGTGGAGTCGTTGCCTAGAGTGCAGTGGTGTGATGTCNNNAGATCGGAA |
| 48171_2 | TATGAGTGTGGAGTCGTTGCCTAGAGTGCAGTGGTGTGATGTCNNNAGATCGGAA |
| 48171_3 | TATGAGTGTGGAGTCGTTGCCTAGAGTGCAGTGGTGTGATGTCNNNAGATCGGAA |
| 47163_1 | TATGAGTGTGGAGTCGTTGCCAACCTCCTGAGTAGCTGGAATTNNNAGATCGGAAG |
| 47163_2 | TATGAGTGTGGAGTCGTTGCCAACCTCCTGAGTAGCTGGAATTNNNAGATCGGAAG |
| 47163_3 | TATGAGTGTGGAGTCGTTGCCAACCTCCTGAGTAGCTGGAATTNNNAGATCGGAAG |
| 28916_1 | TATGAGTGTGGAGTCGTTGCTTGCTACAAGACCCACATGTGATNNNAGATCGGAAG |
| 28916_2 | TATGAGTGTGGAGTCGTTGCTTGCTACAAGACCCACATGTGATNNNAGATCGGAAG |
| 28916_3 | TATGAGTGTGGAGTCGTTGCTTGCTACAAGACCCACATGTGATNNNAGATCGGAAG |
| 52543_1 | TATGAGTGTGGAGTCGTTGCCTTGCTTCCATCTCCTGTTCTCTNNNAGATCGGAAGA |
| 52543_2 | TATGAGTGTGGAGTCGTTGCCTTGCTTCCATCTCCTGTTCTCTNNNAGATCGGAAGA |
| 52543_3 | TATGAGTGTGGAGTCGTTGCCTTGCTTCCATCTCCTGTTCTCTNNNAGATCGGAAGA |
| 39305_1 | TATGAGTGTGGAGTCGTTGCAATTAGGCAGGAATCGGTGCTAANNNAGATCGGAA |
| 39305_2 | TATGAGTGTGGAGTCGTTGCAATTAGGCAGGAATCGGTGCTAANNNAGATCGGAA |
| 39305_3 | TATGAGTGTGGAGTCGTTGCAATTAGGCAGGAATCGGTGCTAANNNAGATCGGAA |
| 44335_1 | TATGAGTGTGGAGTCGTTGCGCAGGACACAAGTGCATTTGTNNNAGATCGGAAGA |
| 44335_2 | TATGAGTGTGGAGTCGTTGCGCAGGACACAAGTGCATTTGTNNNAGATCGGAAGA |
| 44335_3 | TATGAGTGTGGAGTCGTTGCGCAGGACACAAGTGCATTTGTNNNAGATCGGAAGA |
| 34660_1 | TATGAGTGTGGAGTCGTTGCAGAATGCTGCTATTCCTGTCCTCNNNAGATCGGAAG |
| 34660_2 | TATGAGTGTGGAGTCGTTGCAGAATGCTGCTATTCCTGTCCTCNNNAGATCGGAAG |
| 34660_3 | TATGAGTGTGGAGTCGTTGCAGAATGCTGCTATTCCTGTCCTCNNNAGATCGGAAG |
| 30021_1 | TATGAGTGTGGAGTCGTTGCACTCCACTTCTGAATTCAGTCCANNNAGATCGGAAGA |
| 30021_2 | TATGAGTGTGGAGTCGTTGCACTCCACTTCTGAATTCAGTCCANNNAGATCGGAAGA |
| 30021_3 | TATGAGTGTGGAGTCGTTGCACTCCACTTCTGAATTCAGTCCANNNAGATCGGAAGA |
| 30588_1 | TATGAGTGTGGAGTCGTTGCCATCTCACTTCTTTGGGAAACCACTCNNNAGATCGGA |
| 30588_2 | TATGAGTGTGGAGTCGTTGCCATCTCACTTCTTTGGGAAACCACTCNNNAGATCGGA |
| 30588_3 | TATGAGTGTGGAGTCGTTGCCATCTCACTTCTTTGGGAAACCACTCNNNAGATCGGA |
| 52621_1 | TATGAGTGTGGAGTCGTTGCTCAGAGGAACAGAAACACACACANNNAGATCGGAA |
| 52621_2 | TATGAGTGTGGAGTCGTTGCTCAGAGGAACAGAAACACACACANNNAGATCGGAA |
| 52621_3 | TATGAGTGTGGAGTCGTTGCTCAGAGGAACAGAAACACACACANNNAGATCGGAA |
| 34899_1 | TATGAGTGTGGAGTCGTTGCAGATGGAGTCTCTCTCTGTCACCNNNAGATCGGAAG |
| 34899_2 | TATGAGTGTGGAGTCGTTGCAGATGGAGTCTCTCTCTGTCACCNNNAGATCGGAAG |
| 34899_3 | TATGAGTGTGGAGTCGTTGCAGATGGAGTCTCTCTCTGTCACCNNNAGATCGGAAG |
| 49587_1 | TATGAGTGTGGAGTCGTTGCATTTCTCCAACTCCTTGAGAGGCNNNAGATCGGAAG |
| 49587_2 | TATGAGTGTGGAGTCGTTGCATTTCTCCAACTCCTTGAGAGGCNNNAGATCGGAAG |
| 49587_3 | TATGAGTGTGGAGTCGTTGCATTTCTCCAACTCCTTGAGAGGCNNNAGATCGGAAG |
| 39015_1 | TATGAGTGTGGAGTCGTTGCATGTCTGTGCAACTTTCAGAAGCNNNAGATCGGAAG |
| 39015_2 | TATGAGTGTGGAGTCGTTGCATGTCTGTGCAACTTTCAGAAGCNNNAGATCGGAAG |

| | |
|---|---|
| 39015_3 | TATGAGTGTGGAGTCGTTGCATGTCTGTGCAACTTTCAGAAGCNNNAGATCGGAAG |
| 5124_1 | TATGAGTGTGGAGTCGTTGCGTGTGATAGCTGCAGGAGTGAAANNNAGATCGGAA |
| 5124_2 | TATGAGTGTGGAGTCGTTGCGTGTGATAGCTGCAGGAGTGAAANNNAGATCGGAA |
| 5124_3 | TATGAGTGTGGAGTCGTTGCGTGTGATAGCTGCAGGAGTGAAANNNAGATCGGAA |
| 41038_1 | TATGAGTGTGGAGTCGTTGCTGAGACACTGTCAAAGACTGGAANNNAGATCGGAA |
| 41038_2 | TATGAGTGTGGAGTCGTTGCTGAGACACTGTCAAAGACTGGAANNNAGATCGGAA |
| 41038_3 | TATGAGTGTGGAGTCGTTGCTGAGACACTGTCAAAGACTGGAANNNAGATCGGAA |
| 29747_1 | TATGAGTGTGGAGTCGTTGCTTGTCTTCCAAATCAGCCAAACCNNNAGATCGGAAGA |
| 29747_2 | TATGAGTGTGGAGTCGTTGCTTGTCTTCCAAATCAGCCAAACCNNNAGATCGGAAGA |
| 29747_3 | TATGAGTGTGGAGTCGTTGCTTGTCTTCCAAATCAGCCAAACCNNNAGATCGGAAGA |
| 31981_1 | TATGAGTGTGGAGTCGTTGCTTGGGGGCAATCAAAATTTAGCCNNNAGATCGGAAG |
| 31981_2 | TATGAGTGTGGAGTCGTTGCTTGGGGGCAATCAAAATTTAGCCNNNAGATCGGAAG |
| 31981_3 | TATGAGTGTGGAGTCGTTGCTTGGGGGCAATCAAAATTTAGCCNNNAGATCGGAAG |
| 29128_1 | TATGAGTGTGGAGTCGTTGCTGGCTTCAAACTCTTCTACTGCANNNAGATCGGAAGA |
| 29128_2 | TATGAGTGTGGAGTCGTTGCTGGCTTCAAACTCTTCTACTGCANNNAGATCGGAAGA |
| 29128_3 | TATGAGTGTGGAGTCGTTGCTGGCTTCAAACTCTTCTACTGCANNNAGATCGGAAGA |
| 42749_1 | TATGAGTGTGGAGTCGTTGCCAATTCTTCTGTCTCAGCCTCCTNNNAGATCGGAAGA |
| 42749_2 | TATGAGTGTGGAGTCGTTGCCAATTCTTCTGTCTCAGCCTCCTNNNAGATCGGAAGA |
| 42749_3 | TATGAGTGTGGAGTCGTTGCCAATTCTTCTGTCTCAGCCTCCTNNNAGATCGGAAGA |
| 52265_1 | TATGAGTGTGGAGTCGTTGCCCTCTCCTGCTCTGTCCTCTCNNNAGATCGGAAGAGC |
| 52265_2 | TATGAGTGTGGAGTCGTTGCCCTCTCCTGCTCTGTCCTCTCNNNAGATCGGAAGAGC |
| 52265_3 | TATGAGTGTGGAGTCGTTGCCCTCTCCTGCTCTGTCCTCTCNNNAGATCGGAAGAGC |
| 40372_1 | TATGAGTGTGGAGTCGTTGCTCATTACCTCCAAAGTTAGGGCCNNNAGATCGGAAG |
| 40372_2 | TATGAGTGTGGAGTCGTTGCTCATTACCTCCAAAGTTAGGGCCNNNAGATCGGAAG |
| 40372_3 | TATGAGTGTGGAGTCGTTGCTCATTACCTCCAAAGTTAGGGCCNNNAGATCGGAAG |
| 36556_1 | TATGAGTGTGGAGTCGTTGCGTCCACATCCTACAGAATCCAGGNNNAGATCGGAAG |
| 36556_2 | TATGAGTGTGGAGTCGTTGCGTCCACATCCTACAGAATCCAGGNNNAGATCGGAAG |
| 36556_3 | TATGAGTGTGGAGTCGTTGCGTCCACATCCTACAGAATCCAGGNNNAGATCGGAAG |
| 12905_1 | TATGAGTGTGGAGTCGTTGCAACTGACCAAGATGTGTACTATNNNAGATCGGAAGA |
| 12905_2 | TATGAGTGTGGAGTCGTTGCAACTGACCAAGATGTGTACTATNNNAGATCGGAAGA |
| 12905_3 | TATGAGTGTGGAGTCGTTGCAACTGACCAAGATGTGTACTATNNNAGATCGGAAGA |
| 50883_1 | TATGAGTGTGGAGTCGTTGCTGATTGCAGACACCTCATTTTGGNNNAGATCGGAAG |
| 50883_2 | TATGAGTGTGGAGTCGTTGCTGATTGCAGACACCTCATTTTGGNNNAGATCGGAAG |
| 50883_3 | TATGAGTGTGGAGTCGTTGCTGATTGCAGACACCTCATTTTGGNNNAGATCGGAAG |
| 35342_1 | TATGAGTGTGGAGTCGTTGCCCAGTAGAGATGAGGTCTTGCTGNNNAGATCGGAA |
| 35342_2 | TATGAGTGTGGAGTCGTTGCCCAGTAGAGATGAGGTCTTGCTGNNNAGATCGGAA |
| 35342_3 | TATGAGTGTGGAGTCGTTGCCCAGTAGAGATGAGGTCTTGCTGNNNAGATCGGAA |
| 37380_1 | TATGAGTGTGGAGTCGTTGCGTGAGAACATGGCACATTAGCAGNNNAGATCGGAA |
| 37380_2 | TATGAGTGTGGAGTCGTTGCGTGAGAACATGGCACATTAGCAGNNNAGATCGGAA |
| 37380_3 | TATGAGTGTGGAGTCGTTGCGTGAGAACATGGCACATTAGCAGNNNAGATCGGAA |
| 29557_1 | TATGAGTGTGGAGTCGTTGCCTTTCCAACATGCCCACCTATTCNNNAGATCGGAAGA |
| 29557_2 | TATGAGTGTGGAGTCGTTGCCTTTCCAACATGCCCACCTATTCNNNAGATCGGAAGA |
| 29557_3 | TATGAGTGTGGAGTCGTTGCCTTTCCAACATGCCCACCTATTCNNNAGATCGGAAGA |

| | |
|---|---|
| 50290_1 | TATGAGTGTGGAGTCGTTGCCACAAACACAATAAACGCAGGGANNNAGATCGGAA |
| 50290_2 | TATGAGTGTGGAGTCGTTGCCACAAACACAATAAACGCAGGGANNNAGATCGGAA |
| 50290_3 | TATGAGTGTGGAGTCGTTGCCACAAACACAATAAACGCAGGGANNNAGATCGGAA |
| 42916_1 | TATGAGTGTGGAGTCGTTGCTTCTTTGGCTATTCAGGCTCCTTNNNAGATCGGAAG |
| 42916_2 | TATGAGTGTGGAGTCGTTGCTTCTTTGGCTATTCAGGCTCCTTNNNAGATCGGAAG |
| 42916_3 | TATGAGTGTGGAGTCGTTGCTTCTTTGGCTATTCAGGCTCCTTNNNAGATCGGAAG |
| 50776_1 | TATGAGTGTGGAGTCGTTGCGAGAACTCATTCTTGGCGTTCACNNNAGATCGGAAG |
| 50776_2 | TATGAGTGTGGAGTCGTTGCGAGAACTCATTCTTGGCGTTCACNNNAGATCGGAAG |
| 50776_3 | TATGAGTGTGGAGTCGTTGCGAGAACTCATTCTTGGCGTTCACNNNAGATCGGAAG |
| 48413_1 | TATGAGTGTGGAGTCGTTGCTCTCAAATGCCTAGACCACTTCCNNNAGATCGGAAG |
| 48413_2 | TATGAGTGTGGAGTCGTTGCTCTCAAATGCCTAGACCACTTCCNNNAGATCGGAAG |
| 48413_3 | TATGAGTGTGGAGTCGTTGCTCTCAAATGCCTAGACCACTTCCNNNAGATCGGAAG |
| 48849_1 | TATGAGTGTGGAGTCGTTGCCTCCCCTCATCCCACTAAGANNNAGATCGGAAGAG |
| 48849_2 | TATGAGTGTGGAGTCGTTGCCTCCCCTCATCCCACTAAGANNNAGATCGGAAGAG |
| 48849_3 | TATGAGTGTGGAGTCGTTGCCTCCCCTCATCCCACTAAGANNNAGATCGGAAGAG |
| 32476_1 | TATGAGTGTGGAGTCGTTGCCCTCCTGAGTAGCTGAGACTACANNNAGATCGGAAG |
| 32476_2 | TATGAGTGTGGAGTCGTTGCCCTCCTGAGTAGCTGAGACTACANNNAGATCGGAAG |
| 32476_3 | TATGAGTGTGGAGTCGTTGCCCTCCTGAGTAGCTGAGACTACANNNAGATCGGAAG |
| 31863_1 | TATGAGTGTGGAGTCGTTGCTTTGTTTTTTTGCCACAGAGCCTNNNAGATCGGAAG |
| 31863_2 | TATGAGTGTGGAGTCGTTGCTTTGTTTTTTTGCCACAGAGCCTNNNAGATCGGAAG |
| 31863_3 | TATGAGTGTGGAGTCGTTGCTTTGTTTTTTTGCCACAGAGCCTNNNAGATCGGAAG |
| 33141_1 | TATGAGTGTGGAGTCGTTGCGCGATAATGCGAGACTCCATCTCNNNAGATCGGAAG |
| 33141_2 | TATGAGTGTGGAGTCGTTGCGCGATAATGCGAGACTCCATCTCNNNAGATCGGAAG |
| 33141_3 | TATGAGTGTGGAGTCGTTGCGCGATAATGCGAGACTCCATCTCNNNAGATCGGAAG |
| 48815_1 | TATGAGTGTGGAGTCGTTGCTTCAAATGCAACCAGGCTAAACCNNNAGATCGGAAG |
| 48815_2 | TATGAGTGTGGAGTCGTTGCTTCAAATGCAACCAGGCTAAACCNNNAGATCGGAAG |
| 48815_3 | TATGAGTGTGGAGTCGTTGCTTCAAATGCAACCAGGCTAAACCNNNAGATCGGAAG |
| 38653_1 | TATGAGTGTGGAGTCGTTGCACCCCATCAGTAACCTAGAAAAGGNNNAGATCGGAA |
| 38653_2 | TATGAGTGTGGAGTCGTTGCACCCCATCAGTAACCTAGAAAAGGNNNAGATCGGAA |
| 38653_3 | TATGAGTGTGGAGTCGTTGCACCCCATCAGTAACCTAGAAAAGGNNNAGATCGGAA |
| 38529_1 | TATGAGTGTGGAGTCGTTGCAGTGCTCAAACAATGTCTAACGANNNAGATCGGAAG |
| 38529_2 | TATGAGTGTGGAGTCGTTGCAGTGCTCAAACAATGTCTAACGANNNAGATCGGAAG |
| 38529_3 | TATGAGTGTGGAGTCGTTGCAGTGCTCAAACAATGTCTAACGANNNAGATCGGAAG |
| 4688_1 | TATGAGTGTGGAGTCGTTGCTGGCTACAATGAATGCTTTTCCANNNAGATCGGAAG |
| 4688_2 | TATGAGTGTGGAGTCGTTGCTGGCTACAATGAATGCTTTTCCANNNAGATCGGAAG |
| 4688_3 | TATGAGTGTGGAGTCGTTGCTGGCTACAATGAATGCTTTTCCANNNAGATCGGAAG |
| 40932_1 | TATGAGTGTGGAGTCGTTGCCCTTGTGAATTGTGATTTACTGCCTNNNAGATCGGA |
| 40932_2 | TATGAGTGTGGAGTCGTTGCCCTTGTGAATTGTGATTTACTGCCTNNNAGATCGGA |
| 40932_3 | TATGAGTGTGGAGTCGTTGCCCTTGTGAATTGTGATTTACTGCCTNNNAGATCGGA |
| 39967_1 | TATGAGTGTGGAGTCGTTGCCCATTCCAACTTTCACTGACCAGNNNAGATCGGAAG |
| 39967_2 | TATGAGTGTGGAGTCGTTGCCCATTCCAACTTTCACTGACCAGNNNAGATCGGAAG |
| 39967_3 | TATGAGTGTGGAGTCGTTGCCCATTCCAACTTTCACTGACCAGNNNAGATCGGAAG |
| 50709_1 | TATGAGTGTGGAGTCGTTGCGTGACTTCCTTGGGAGAAGACAGNNNAGATCGGAA |

| | |
|---|---|
| 50709_2 | TATGAGTGTGGAGTCGTTGCGTGACTTCCTTGGGAGAAGACAGNNNAGATCGGAA |
| 50709_3 | TATGAGTGTGGAGTCGTTGCGTGACTTCCTTGGGAGAAGACAGNNNAGATCGGAA |
| 33677_1 | TATGAGTGTGGAGTCGTTGCGCCTCCTGAGTATACTGGGACTANNNAGATCGGAAG |
| 33677_2 | TATGAGTGTGGAGTCGTTGCGCCTCCTGAGTATACTGGGACTANNNAGATCGGAAG |
| 33677_3 | TATGAGTGTGGAGTCGTTGCGCCTCCTGAGTATACTGGGACTANNNAGATCGGAAG |
| 47425_1 | TATGAGTGTGGAGTCGTTGCATCAAAGCCAAACCCCAAATCAGNNNAGATCGGAAG |
| 47425_2 | TATGAGTGTGGAGTCGTTGCATCAAAGCCAAACCCCAAATCAGNNNAGATCGGAAG |
| 47425_3 | TATGAGTGTGGAGTCGTTGCATCAAAGCCAAACCCCAAATCAGNNNAGATCGGAAG |
| 49367_1 | TATGAGTGTGGAGTCGTTGCCTCCAAGAGTGCTGGGATTACAANNNAGATCGGAA |
| 49367_2 | TATGAGTGTGGAGTCGTTGCCTCCAAGAGTGCTGGGATTACAANNNAGATCGGAA |
| 49367_3 | TATGAGTGTGGAGTCGTTGCCTCCAAGAGTGCTGGGATTACAANNNAGATCGGAA |
| 51902_1 | TATGAGTGTGGAGTCGTTGCGATCAGCACCGTGGTCTGATGAANNNAGATCGGAA |
| 51902_2 | TATGAGTGTGGAGTCGTTGCGATCAGCACCGTGGTCTGATGAANNNAGATCGGAA |
| 51902_3 | TATGAGTGTGGAGTCGTTGCGATCAGCACCGTGGTCTGATGAANNNAGATCGGAA |
| 52792_1 | TATGAGTGTGGAGTCGTTGCCAACCCGACCTGAACTCTGACNNNAGATCGGAAGAG |
| 52792_2 | TATGAGTGTGGAGTCGTTGCCAACCCGACCTGAACTCTGACNNNAGATCGGAAGAG |
| 52792_3 | TATGAGTGTGGAGTCGTTGCCAACCCGACCTGAACTCTGACNNNAGATCGGAAGAG |
| 34095_1 | TATGAGTGTGGAGTCGTTGCATAGCTCGTGGAAAGTGGTTTCTNNNAGATCGGAA |
| 34095_2 | TATGAGTGTGGAGTCGTTGCATAGCTCGTGGAAAGTGGTTTCTNNNAGATCGGAA |
| 34095_3 | TATGAGTGTGGAGTCGTTGCATAGCTCGTGGAAAGTGGTTTCTNNNAGATCGGAA |
| 37477_1 | TATGAGTGTGGAGTCGTTGCAGCAACAGACAATCACAAAAACANNNAGATCGGAAG |
| 37477_2 | TATGAGTGTGGAGTCGTTGCAGCAACAGACAATCACAAAAACANNNAGATCGGAAG |
| 37477_3 | TATGAGTGTGGAGTCGTTGCAGCAACAGACAATCACAAAAACANNNAGATCGGAAG |
| 36517_1 | TATGAGTGTGGAGTCGTTGCATACTTTACTTCAGGCCAGCAGGNNNAGATCGGAAG |
| 36517_2 | TATGAGTGTGGAGTCGTTGCATACTTTACTTCAGGCCAGCAGGNNNAGATCGGAAG |
| 36517_3 | TATGAGTGTGGAGTCGTTGCATACTTTACTTCAGGCCAGCAGGNNNAGATCGGAAG |
| 48543_1 | TATGAGTGTGGAGTCGTTGCGGATTACATGCATGCATCACCATNNNAGATCGGAAG |
| 48543_2 | TATGAGTGTGGAGTCGTTGCGGATTACATGCATGCATCACCATNNNAGATCGGAAG |
| 48543_3 | TATGAGTGTGGAGTCGTTGCGGATTACATGCATGCATCACCATNNNAGATCGGAAG |
| 45114_1 | TATGAGTGTGGAGTCGTTGCCTATTTCCTGAGCACCTCCTCTCNNNAGATCGGAAGA |
| 45114_2 | TATGAGTGTGGAGTCGTTGCCTATTTCCTGAGCACCTCCTCTCNNNAGATCGGAAGA |
| 45114_3 | TATGAGTGTGGAGTCGTTGCCTATTTCCTGAGCACCTCCTCTCNNNAGATCGGAAGA |
| 35823_1 | TATGAGTGTGGAGTCGTTGCCCACTACTCCCGGCTAATTTTTTNNNAGATCGGAAGA |
| 35823_2 | TATGAGTGTGGAGTCGTTGCCCACTACTCCCGGCTAATTTTTTNNNAGATCGGAAGA |
| 35823_3 | TATGAGTGTGGAGTCGTTGCCCACTACTCCCGGCTAATTTTTTNNNAGATCGGAAGA |
| 43995_1 | TATGAGTGTGGAGTCGTTGCGGCTGTGATCAAAGAGGAGAGATNNNAGATCGGA |
| 43995_2 | TATGAGTGTGGAGTCGTTGCGGCTGTGATCAAAGAGGAGAGATNNNAGATCGGA |
| 43995_3 | TATGAGTGTGGAGTCGTTGCGGCTGTGATCAAAGAGGAGAGATNNNAGATCGGA |
| 9413_1 | TATGAGTGTGGAGTCGTTGCTGAGGAGACTTAGGAAGCCTACGNNNAGATCGGAA |
| 9413_2 | TATGAGTGTGGAGTCGTTGCTGAGGAGACTTAGGAAGCCTACGNNNAGATCGGAA |
| 9413_3 | TATGAGTGTGGAGTCGTTGCTGAGGAGACTTAGGAAGCCTACGNNNAGATCGGAA |
| 45536_1 | TATGAGTGTGGAGTCGTTGCTCCTGGGCTCTCTGATAGAAGGNNNAGATCGGAAG |
| 45536_2 | TATGAGTGTGGAGTCGTTGCTCCTGGGCTCTCTGATAGAAGGNNNAGATCGGAAG |

| | |
|---|---|
| 45536_3 | TATGAGTGTGGAGTCGTTGCTCCTGGGCTCTCTGATAGAAGGNNNAGATCGGAAG |
| 48903_1 | TATGAGTGTGGAGTCGTTGCCTGAGTAGCTGGGATTACAAGCANNNAGATCGGAA |
| 48903_2 | TATGAGTGTGGAGTCGTTGCCTGAGTAGCTGGGATTACAAGCANNNAGATCGGAA |
| 48903_3 | TATGAGTGTGGAGTCGTTGCCTGAGTAGCTGGGATTACAAGCANNNAGATCGGAA |
| 42331_1 | TATGAGTGTGGAGTCGTTGCTGGTTCTGTTTCCCTGAAAACCTNNNAGATCGGAAG |
| 42331_2 | TATGAGTGTGGAGTCGTTGCTGGTTCTGTTTCCCTGAAAACCTNNNAGATCGGAAG |
| 42331_3 | TATGAGTGTGGAGTCGTTGCTGGTTCTGTTTCCCTGAAAACCTNNNAGATCGGAAG |
| 37516_1 | TATGAGTGTGGAGTCGTTGCCGGAAGCAGTACAAACAAGAACANNNAGATCGGAA |
| 37516_2 | TATGAGTGTGGAGTCGTTGCCGGAAGCAGTACAAACAAGAACANNNAGATCGGAA |
| 37516_3 | TATGAGTGTGGAGTCGTTGCCGGAAGCAGTACAAACAAGAACANNNAGATCGGAA |
| 41843_1 | TATGAGTGTGGAGTCGTTGCGCTGTTGAGTTGTCATTGCTTCTNNNAGATCGGAAG |
| 41843_2 | TATGAGTGTGGAGTCGTTGCGCTGTTGAGTTGTCATTGCTTCTNNNAGATCGGAAG |
| 41843_3 | TATGAGTGTGGAGTCGTTGCGCTGTTGAGTTGTCATTGCTTCTNNNAGATCGGAAG |
| 51705_1 | TATGAGTGTGGAGTCGTTGCTTCTGGCTAATGGAATGTGGACANNNAGATCGGAA |
| 51705_2 | TATGAGTGTGGAGTCGTTGCTTCTGGCTAATGGAATGTGGACANNNAGATCGGAA |
| 51705_3 | TATGAGTGTGGAGTCGTTGCTTCTGGCTAATGGAATGTGGACANNNAGATCGGAA |
| 29067_1 | TATGAGTGTGGAGTCGTTGCTGTGTGTGTGATGGGAAAGAGAANNNAGATCGGA |
| 29067_2 | TATGAGTGTGGAGTCGTTGCTGTGTGTGTGATGGGAAAGAGAANNNAGATCGGA |
| 29067_3 | TATGAGTGTGGAGTCGTTGCTGTGTGTGTGATGGGAAAGAGAANNNAGATCGGA |
| 44318_1 | TATGAGTGTGGAGTCGTTGCGAACAGCAATGTGTTAGGAGACGNNNAGATCGGAA |
| 44318_2 | TATGAGTGTGGAGTCGTTGCGAACAGCAATGTGTTAGGAGACGNNNAGATCGGAA |
| 44318_3 | TATGAGTGTGGAGTCGTTGCGAACAGCAATGTGTTAGGAGACGNNNAGATCGGAA |
| 44449_1 | TATGAGTGTGGAGTCGTTGCGAGCTGATAGGCTGGAAGCTATTNNNAGATCGGAA |
| 44449_2 | TATGAGTGTGGAGTCGTTGCGAGCTGATAGGCTGGAAGCTATTNNNAGATCGGAA |
| 44449_3 | TATGAGTGTGGAGTCGTTGCGAGCTGATAGGCTGGAAGCTATTNNNAGATCGGAA |
| 41083_1 | TATGAGTGTGGAGTCGTTGCATACCAAGCAGTTTGTATGCAGCNNNAGATCGGAAG |
| 41083_2 | TATGAGTGTGGAGTCGTTGCATACCAAGCAGTTTGTATGCAGCNNNAGATCGGAAG |
| 41083_3 | TATGAGTGTGGAGTCGTTGCATACCAAGCAGTTTGTATGCAGCNNNAGATCGGAAG |
| 39405_1 | TATGAGTGTGGAGTCGTTGCAAAGAGGAGGGAAGATGAATGCCNNNAGATCGGA |
| 39405_2 | TATGAGTGTGGAGTCGTTGCAAAGAGGAGGGAAGATGAATGCCNNNAGATCGGA |
| 39405_3 | TATGAGTGTGGAGTCGTTGCAAAGAGGAGGGAAGATGAATGCCNNNAGATCGGA |
| 50503_1 | TATGAGTGTGGAGTCGTTGCCTTCCAGGGGTCTGTTATTAGCANNNAGATCGGAAG |
| 50503_2 | TATGAGTGTGGAGTCGTTGCCTTCCAGGGGTCTGTTATTAGCANNNAGATCGGAAG |
| 50503_3 | TATGAGTGTGGAGTCGTTGCCTTCCAGGGGTCTGTTATTAGCANNNAGATCGGAAG |
| 44905_1 | TATGAGTGTGGAGTCGTTGCAGAGAGAGGAGAGAGAGGTTTCCNNNAGATCGGA |
| 44905_2 | TATGAGTGTGGAGTCGTTGCAGAGAGAGGAGAGAGAGGTTTCCNNNAGATCGGA |
| 44905_3 | TATGAGTGTGGAGTCGTTGCAGAGAGAGGAGAGAGAGGTTTCCNNNAGATCGGA |
| 52136_1 | TATGAGTGTGGAGTCGTTGCCCTGAACTTAGTGTGCTCTGTCTNNNAGATCGGAAG |
| 52136_2 | TATGAGTGTGGAGTCGTTGCCCTGAACTTAGTGTGCTCTGTCTNNNAGATCGGAAG |
| 52136_3 | TATGAGTGTGGAGTCGTTGCCCTGAACTTAGTGTGCTCTGTCTNNNAGATCGGAAG |
| 13336_1 | TATGAGTGTGGAGTCGTTGCTTGATTGTTTTCTAAAAGGAGACNNNAGATCGGAAG |
| 13336_2 | TATGAGTGTGGAGTCGTTGCTTGATTGTTTTCTAAAAGGAGACNNNAGATCGGAAG |
| 13336_3 | TATGAGTGTGGAGTCGTTGCTTGATTGTTTTCTAAAAGGAGACNNNAGATCGGAAG |

| | |
|---|---|
| 29103_1 | TATGAGTGTGGAGTCGTTGCGAGGGAGAGAGAGAGAAACATGCNNNAGATCGGA |
| 29103_2 | TATGAGTGTGGAGTCGTTGCGAGGGAGAGAGAGAGAAACATGCNNNAGATCGGA |
| 29103_3 | TATGAGTGTGGAGTCGTTGCGAGGGAGAGAGAGAGAAACATGCNNNAGATCGGA |
| 9708_1 | TATGAGTGTGGAGTCGTTGCTTCTCCCTCTGTGATTAGCCACCNNNAGATCGGAAGA |
| 9708_2 | TATGAGTGTGGAGTCGTTGCTTCTCCCTCTGTGATTAGCCACCNNNAGATCGGAAGA |
| 9708_3 | TATGAGTGTGGAGTCGTTGCTTCTCCCTCTGTGATTAGCCACCNNNAGATCGGAAGA |
| 48018_1 | TATGAGTGTGGAGTCGTTGCGTTTCACTGTGTTAGCCAGGATGNNNAGATCGGAA |
| 48018_2 | TATGAGTGTGGAGTCGTTGCGTTTCACTGTGTTAGCCAGGATGNNNAGATCGGAA |
| 48018_3 | TATGAGTGTGGAGTCGTTGCGTTTCACTGTGTTAGCCAGGATGNNNAGATCGGAA |
| 43279_1 | TATGAGTGTGGAGTCGTTGCAATAATGGCATTTGCAGCAACCTNNNAGATCGGAAG |
| 43279_2 | TATGAGTGTGGAGTCGTTGCAATAATGGCATTTGCAGCAACCTNNNAGATCGGAAG |
| 43279_3 | TATGAGTGTGGAGTCGTTGCAATAATGGCATTTGCAGCAACCTNNNAGATCGGAAG |
| 3094_1 | TATGAGTGTGGAGTCGTTGCGCTTTTCTTCCCAGGAGCTACAGNNNAGATCGGAAG |
| 3094_2 | TATGAGTGTGGAGTCGTTGCGCTTTTCTTCCCAGGAGCTACAGNNNAGATCGGAAG |
| 3094_3 | TATGAGTGTGGAGTCGTTGCGCTTTTCTTCCCAGGAGCTACAGNNNAGATCGGAAG |
| 10416_1 | TATGAGTGTGGAGTCGTTGCTAGGGATAAGTGATCAGGTTCCTGNNNAGATCGGA |
| 10416_2 | TATGAGTGTGGAGTCGTTGCTAGGGATAAGTGATCAGGTTCCTGNNNAGATCGGA |
| 10416_3 | TATGAGTGTGGAGTCGTTGCTAGGGATAAGTGATCAGGTTCCTGNNNAGATCGGA |
| 30148_1 | TATGAGTGTGGAGTCGTTGCTGAATTCACTCCTCCTTGATGCTNNNAGATCGGAAGA |
| 30148_2 | TATGAGTGTGGAGTCGTTGCTGAATTCACTCCTCCTTGATGCTNNNAGATCGGAAGA |
| 30148_3 | TATGAGTGTGGAGTCGTTGCTGAATTCACTCCTCCTTGATGCTNNNAGATCGGAAGA |
| 47512_1 | TATGAGTGTGGAGTCGTTGCCTTGGCTCATTTCCAGAGAGTGTNNNAGATCGGAAG |
| 47512_2 | TATGAGTGTGGAGTCGTTGCCTTGGCTCATTTCCAGAGAGTGTNNNAGATCGGAAG |
| 47512_3 | TATGAGTGTGGAGTCGTTGCCTTGGCTCATTTCCAGAGAGTGTNNNAGATCGGAAG |
| 34427_1 | TATGAGTGTGGAGTCGTTGCTTAGCCTCAGTGTATGTTGTGGGNNNAGATCGGAA |
| 34427_2 | TATGAGTGTGGAGTCGTTGCTTAGCCTCAGTGTATGTTGTGGGNNNAGATCGGAA |
| 34427_3 | TATGAGTGTGGAGTCGTTGCTTAGCCTCAGTGTATGTTGTGGGNNNAGATCGGAA |
| 13023_1 | TATGAGTGTGGAGTCGTTGCGAGTATGGTGAAGAGGTTAATGNNNAGATCGGAA |
| 13023_2 | TATGAGTGTGGAGTCGTTGCGAGTATGGTGAAGAGGTTAATGNNNAGATCGGAA |
| 13023_3 | TATGAGTGTGGAGTCGTTGCGAGTATGGTGAAGAGGTTAATGNNNAGATCGGAA |
| 31016_1 | TATGAGTGTGGAGTCGTTGCACTTGTCTATCTGGTAAAGCCCGNNNAGATCGGAAG |
| 31016_2 | TATGAGTGTGGAGTCGTTGCACTTGTCTATCTGGTAAAGCCCGNNNAGATCGGAAG |
| 31016_3 | TATGAGTGTGGAGTCGTTGCACTTGTCTATCTGGTAAAGCCCGNNNAGATCGGAAG |
| 48392_1 | TATGAGTGTGGAGTCGTTGCCCATGCCCGGCTAATTTTTGTATNNNAGATCGGAAG |
| 48392_2 | TATGAGTGTGGAGTCGTTGCCCATGCCCGGCTAATTTTTGTATNNNAGATCGGAAG |
| 48392_3 | TATGAGTGTGGAGTCGTTGCCCATGCCCGGCTAATTTTTGTATNNNAGATCGGAAG |
| 48764_1 | TATGAGTGTGGAGTCGTTGCCGGATTTGTGTATTGCTTTTGGCNNNAGATCGGAAG |
| 48764_2 | TATGAGTGTGGAGTCGTTGCCGGATTTGTGTATTGCTTTTGGCNNNAGATCGGAAG |
| 48764_3 | TATGAGTGTGGAGTCGTTGCCGGATTTGTGTATTGCTTTTGGCNNNAGATCGGAAG |
| 34742_1 | TATGAGTGTGGAGTCGTTGCTTTTTGAGACAGAGTCTTGCCCTNNNAGATCGGAAG |
| 34742_2 | TATGAGTGTGGAGTCGTTGCTTTTTGAGACAGAGTCTTGCCCTNNNAGATCGGAAG |
| 34742_3 | TATGAGTGTGGAGTCGTTGCTTTTTGAGACAGAGTCTTGCCCTNNNAGATCGGAAG |
| 49114_1 | TATGAGTGTGGAGTCGTTGCGTCTTGAATTCCTGACCTCGTGANNNAGATCGGAAG |

| | |
|---|---|
| 49114_2 | TATGAGTGTGGAGTCGTTGCGTCTTGAATTCCTGACCTCGTGANNNAGATCGGAAG |
| 49114_3 | TATGAGTGTGGAGTCGTTGCGTCTTGAATTCCTGACCTCGTGANNNAGATCGGAAG |
| 45919_1 | TATGAGTGTGGAGTCGTTGCTAGGATTTACAGGTGTGAGCCACNNNAGATCGGAA |
| 45919_2 | TATGAGTGTGGAGTCGTTGCTAGGATTTACAGGTGTGAGCCACNNNAGATCGGAA |
| 45919_3 | TATGAGTGTGGAGTCGTTGCTAGGATTTACAGGTGTGAGCCACNNNAGATCGGAA |
| 39089_1 | TATGAGTGTGGAGTCGTTGCCTGTTTTTATCAAGGGGCCTGGNNNAGATCGGAAG |
| 39089_2 | TATGAGTGTGGAGTCGTTGCCTGTTTTTATCAAGGGGCCTGGNNNAGATCGGAAG |
| 39089_3 | TATGAGTGTGGAGTCGTTGCCTGTTTTTATCAAGGGGCCTGGNNNAGATCGGAAG |
| 34191_1 | TATGAGTGTGGAGTCGTTGCTGCAGACCTTTATGATGATCCACTNNNAGATCGGAA |
| 34191_2 | TATGAGTGTGGAGTCGTTGCTGCAGACCTTTATGATGATCCACTNNNAGATCGGAA |
| 34191_3 | TATGAGTGTGGAGTCGTTGCTGCAGACCTTTATGATGATCCACTNNNAGATCGGAA |
| 9314_1 | TATGAGTGTGGAGTCGTTGCCAGAGTGAAAGAGCGAGTTCTGGNNNAGATCGGA |
| 9314_2 | TATGAGTGTGGAGTCGTTGCCAGAGTGAAAGAGCGAGTTCTGGNNNAGATCGGA |
| 9314_3 | TATGAGTGTGGAGTCGTTGCCAGAGTGAAAGAGCGAGTTCTGGNNNAGATCGGA |
| 36515_1 | TATGAGTGTGGAGTCGTTGCTTTACCCTGAAAGTTCTTGGGCTNNNAGATCGGAAG |
| 36515_2 | TATGAGTGTGGAGTCGTTGCTTTACCCTGAAAGTTCTTGGGCTNNNAGATCGGAAG |
| 36515_3 | TATGAGTGTGGAGTCGTTGCTTTACCCTGAAAGTTCTTGGGCTNNNAGATCGGAAG |
| 46389_1 | TATGAGTGTGGAGTCGTTGCTTCAAGTGATTCTCCTGCCTCAGNNNAGATCGGAAG |
| 46389_2 | TATGAGTGTGGAGTCGTTGCTTCAAGTGATTCTCCTGCCTCAGNNNAGATCGGAAG |
| 46389_3 | TATGAGTGTGGAGTCGTTGCTTCAAGTGATTCTCCTGCCTCAGNNNAGATCGGAAG |
| 42041_1 | TATGAGTGTGGAGTCGTTGCTTTTCTTTCAAATACCGTGGCCANNNAGATCGGAAG |
| 42041_2 | TATGAGTGTGGAGTCGTTGCTTTTCTTTCAAATACCGTGGCCANNNAGATCGGAAG |
| 42041_3 | TATGAGTGTGGAGTCGTTGCTTTTCTTTCAAATACCGTGGCCANNNAGATCGGAAG |
| 32132_1 | TATGAGTGTGGAGTCGTTGCGGATCACTTGAGCTCAGGAGTTCNNNAGATCGGAA |
| 32132_2 | TATGAGTGTGGAGTCGTTGCGGATCACTTGAGCTCAGGAGTTCNNNAGATCGGAA |
| 32132_3 | TATGAGTGTGGAGTCGTTGCGGATCACTTGAGCTCAGGAGTTCNNNAGATCGGAA |
| 31509_1 | TATGAGTGTGGAGTCGTTGCACACACACACACACACAAATACCNNNAGATCGGAAG |
| 31509_2 | TATGAGTGTGGAGTCGTTGCACACACACACACACACAAATACCNNNAGATCGGAAG |
| 31509_3 | TATGAGTGTGGAGTCGTTGCACACACACACACACACAAATACCNNNAGATCGGAAG |
| 9275_1 | TATGAGTGTGGAGTCGTTGCAACACCTGAAAGCCAGAGAACTGNNNAGATCGGAA |
| 9275_2 | TATGAGTGTGGAGTCGTTGCAACACCTGAAAGCCAGAGAACTGNNNAGATCGGAA |
| 9275_3 | TATGAGTGTGGAGTCGTTGCAACACCTGAAAGCCAGAGAACTGNNNAGATCGGAA |
| 40838_1 | TATGAGTGTGGAGTCGTTGCAAGAGCAGACAAGACAATTGCAGNNNAGATCGGAA |
| 40838_2 | TATGAGTGTGGAGTCGTTGCAAGAGCAGACAAGACAATTGCAGNNNAGATCGGAA |
| 40838_3 | TATGAGTGTGGAGTCGTTGCAAGAGCAGACAAGACAATTGCAGNNNAGATCGGAA |
| 39203_1 | TATGAGTGTGGAGTCGTTGCTTAGCTAGTCAACCAAAGTGGGGNNNAGATCGGAA |
| 39203_2 | TATGAGTGTGGAGTCGTTGCTTAGCTAGTCAACCAAAGTGGGGNNNAGATCGGAA |
| 39203_3 | TATGAGTGTGGAGTCGTTGCTTAGCTAGTCAACCAAAGTGGGGNNNAGATCGGAA |
| 46392_1 | TATGAGTGTGGAGTCGTTGCGTTTCTTTCACCTCAGCCTCCTANNNAGATCGGAAGA |
| 46392_2 | TATGAGTGTGGAGTCGTTGCGTTTCTTTCACCTCAGCCTCCTANNNAGATCGGAAGA |
| 46392_3 | TATGAGTGTGGAGTCGTTGCGTTTCTTTCACCTCAGCCTCCTANNNAGATCGGAAGA |
| 40525_1 | TATGAGTGTGGAGTCGTTGCTTCCCTACCTAACATGATACGCANNNAGATCGGAAG |
| 40525_2 | TATGAGTGTGGAGTCGTTGCTTCCCTACCTAACATGATACGCANNNAGATCGGAAG |

| | |
|---|---|
| 40525_3 | TATGAGTGTGGAGTCGTTGCTTCCCTACCTAACATGATACGCANNNAGATCGGAAGA |
| 40463_1 | TATGAGTGTGGAGTCGTTGCACTGCCTCTGGTAATCTTTCCTGNNNAGATCGGAAGA |
| 40463_2 | TATGAGTGTGGAGTCGTTGCACTGCCTCTGGTAATCTTTCCTGNNNAGATCGGAAGA |
| 40463_3 | TATGAGTGTGGAGTCGTTGCACTGCCTCTGGTAATCTTTCCTGNNNAGATCGGAAGA |
| 44659_1 | TATGAGTGTGGAGTCGTTGCTGCTGCGGATAAGCTAGGAAATANNNAGATCGGAA |
| 44659_2 | TATGAGTGTGGAGTCGTTGCTGCTGCGGATAAGCTAGGAAATANNNAGATCGGAA |
| 44659_3 | TATGAGTGTGGAGTCGTTGCTGCTGCGGATAAGCTAGGAAATANNNAGATCGGAA |
| 30035_1 | TATGAGTGTGGAGTCGTTGCTCCTCCCACTAAATTGTGACAGGNNNAGATCGGAAG |
| 30035_2 | TATGAGTGTGGAGTCGTTGCTCCTCCCACTAAATTGTGACAGGNNNAGATCGGAAG |
| 30035_3 | TATGAGTGTGGAGTCGTTGCTCCTCCCACTAAATTGTGACAGGNNNAGATCGGAAG |
| 33966_1 | TATGAGTGTGGAGTCGTTGCTCTCCTGAGTAGCTGGGATTACANNNAGATCGGAAG |
| 33966_2 | TATGAGTGTGGAGTCGTTGCTCTCCTGAGTAGCTGGGATTACANNNAGATCGGAAG |
| 33966_3 | TATGAGTGTGGAGTCGTTGCTCTCCTGAGTAGCTGGGATTACANNNAGATCGGAAG |
| 32028_1 | TATGAGTGTGGAGTCGTTGCATGCACCTAACACTGGATATCCCNNNAGATCGGAAG |
| 32028_2 | TATGAGTGTGGAGTCGTTGCATGCACCTAACACTGGATATCCCNNNAGATCGGAAG |
| 32028_3 | TATGAGTGTGGAGTCGTTGCATGCACCTAACACTGGATATCCCNNNAGATCGGAAG |
| 165_1 | TATGAGTGTGGAGTCGTTGCCTCTGCAGCCTATGGAGCTGANNNAGATCGGAAGA |
| 165_2 | TATGAGTGTGGAGTCGTTGCCTCTGCAGCCTATGGAGCTGANNNAGATCGGAAGA |
| 165_3 | TATGAGTGTGGAGTCGTTGCCTCTGCAGCCTATGGAGCTGANNNAGATCGGAAGA |
| 42200_1 | TATGAGTGTGGAGTCGTTGCGGTAAATAGTCTCAGCTCCTCGGNNNAGATCGGAAG |
| 42200_2 | TATGAGTGTGGAGTCGTTGCGGTAAATAGTCTCAGCTCCTCGGNNNAGATCGGAAG |
| 42200_3 | TATGAGTGTGGAGTCGTTGCGGTAAATAGTCTCAGCTCCTCGGNNNAGATCGGAAG |
| 39371_1 | TATGAGTGTGGAGTCGTTGCCTCCAACTCACAGCCCATTTCTANNNAGATCGGAAGA |
| 39371_2 | TATGAGTGTGGAGTCGTTGCCTCCAACTCACAGCCCATTTCTANNNAGATCGGAAGA |
| 39371_3 | TATGAGTGTGGAGTCGTTGCCTCCAACTCACAGCCCATTTCTANNNAGATCGGAAGA |
| 35831_1 | TATGAGTGTGGAGTCGTTGCGATTCAAGCAATTCTCCTGCCTCNNNAGATCGGAAGA |
| 35831_2 | TATGAGTGTGGAGTCGTTGCGATTCAAGCAATTCTCCTGCCTCNNNAGATCGGAAGA |
| 35831_3 | TATGAGTGTGGAGTCGTTGCGATTCAAGCAATTCTCCTGCCTCNNNAGATCGGAAGA |
| 37848_1 | TATGAGTGTGGAGTCGTTGCTCACAGACTGGATAAGAAAAACCANNNAGATCGGAA |
| 37848_2 | TATGAGTGTGGAGTCGTTGCTCACAGACTGGATAAGAAAAACCANNNAGATCGGAA |
| 37848_3 | TATGAGTGTGGAGTCGTTGCTCACAGACTGGATAAGAAAAACCANNNAGATCGGAA |
| 43795_1 | TATGAGTGTGGAGTCGTTGCGAGGCAGAAGAATGGCATGAACNNNAGATCGGAA |
| 43795_2 | TATGAGTGTGGAGTCGTTGCGAGGCAGAAGAATGGCATGAACNNNAGATCGGAA |
| 43795_3 | TATGAGTGTGGAGTCGTTGCGAGGCAGAAGAATGGCATGAACNNNAGATCGGAA |
| 51473_1 | TATGAGTGTGGAGTCGTTGCGGCGAAACCCCATCTCTACTAAANNNAGATCGGAAG |
| 51473_2 | TATGAGTGTGGAGTCGTTGCGGCGAAACCCCATCTCTACTAAANNNAGATCGGAAG |
| 51473_3 | TATGAGTGTGGAGTCGTTGCGGCGAAACCCCATCTCTACTAAANNNAGATCGGAAG |
| 47590_1 | TATGAGTGTGGAGTCGTTGCGTGTGCTGGGGATTCTGTGGNNNAGATCGGAAGA |
| 47590_2 | TATGAGTGTGGAGTCGTTGCGTGTGCTGGGGATTCTGTGGNNNAGATCGGAAGA |
| 47590_3 | TATGAGTGTGGAGTCGTTGCGTGTGCTGGGGATTCTGTGGNNNAGATCGGAAGA |
| 29499_1 | TATGAGTGTGGAGTCGTTGCCATTTTGTTGCGTCTTCTTAATCNNNAGATCGGAAGA |
| 29499_2 | TATGAGTGTGGAGTCGTTGCCATTTTGTTGCGTCTTCTTAATCNNNAGATCGGAAGA |
| 29499_3 | TATGAGTGTGGAGTCGTTGCCATTTTGTTGCGTCTTCTTAATCNNNAGATCGGAAGA |

| | |
|---|---|
| 37223_1 | TATGAGTGTGGAGTCGTTGCGCCCACAAAGATGAGAAAGAACCNNNAGATCGGAA |
| 37223_2 | TATGAGTGTGGAGTCGTTGCGCCCACAAAGATGAGAAAGAACCNNNAGATCGGAA |
| 37223_3 | TATGAGTGTGGAGTCGTTGCGCCCACAAAGATGAGAAAGAACCNNNAGATCGGAA |
| 5037_1 | TATGAGTGTGGAGTCGTTGCAGAGAGAAGGAGAGAAAGGCAGTGNNNAGATCGG |
| 5037_2 | TATGAGTGTGGAGTCGTTGCAGAGAGAAGGAGAGAAAGGCAGTGNNNAGATCGG |
| 5037_3 | TATGAGTGTGGAGTCGTTGCAGAGAGAAGGAGAGAAAGGCAGTGNNNAGATCGG |
| 10039_1 | TATGAGTGTGGAGTCGTTGCTGTCCTCTTTGACCTTGAGAGCTNNNAGATCGGAAG |
| 10039_2 | TATGAGTGTGGAGTCGTTGCTGTCCTCTTTGACCTTGAGAGCTNNNAGATCGGAAG |
| 10039_3 | TATGAGTGTGGAGTCGTTGCTGTCCTCTTTGACCTTGAGAGCTNNNAGATCGGAAG |
| 52589_1 | TATGAGTGTGGAGTCGTTGCGCGTTGATTTACAGACCCATACGNNNAGATCGGAAG |
| 52589_2 | TATGAGTGTGGAGTCGTTGCGCGTTGATTTACAGACCCATACGNNNAGATCGGAAG |
| 52589_3 | TATGAGTGTGGAGTCGTTGCGCGTTGATTTACAGACCCATACGNNNAGATCGGAAG |
| 44812_1 | TATGAGTGTGGAGTCGTTGCTGTTTTTCGGATGTCAAGTGCTCNNNAGATCGGAAG |
| 44812_2 | TATGAGTGTGGAGTCGTTGCTGTTTTTCGGATGTCAAGTGCTCNNNAGATCGGAAG |
| 44812_3 | TATGAGTGTGGAGTCGTTGCTGTTTTTCGGATGTCAAGTGCTCNNNAGATCGGAAG |
| 52584_1 | TATGAGTGTGGAGTCGTTGCCCTGAGTCTGAGCCGAGAGGNNNAGATCGGAAGAG |
| 52584_2 | TATGAGTGTGGAGTCGTTGCCCTGAGTCTGAGCCGAGAGGNNNAGATCGGAAGAG |
| 52584_3 | TATGAGTGTGGAGTCGTTGCCCTGAGTCTGAGCCGAGAGGNNNAGATCGGAAGAG |
| 40030_1 | TATGAGTGTGGAGTCGTTGCCTCCCGAGTAGCTGGAATTACAGNNNAGATCGGAAG |
| 40030_2 | TATGAGTGTGGAGTCGTTGCCTCCCGAGTAGCTGGAATTACAGNNNAGATCGGAAG |
| 40030_3 | TATGAGTGTGGAGTCGTTGCCTCCCGAGTAGCTGGAATTACAGNNNAGATCGGAAG |
| 13286_1 | TATGAGTGTGGAGTCGTTGCATAACCAAAGTCACCAATACTTANNNAGATCGGAAG |
| 13286_2 | TATGAGTGTGGAGTCGTTGCATAACCAAAGTCACCAATACTTANNNAGATCGGAAG |
| 13286_3 | TATGAGTGTGGAGTCGTTGCATAACCAAAGTCACCAATACTTANNNAGATCGGAAG |
| 46298_1 | TATGAGTGTGGAGTCGTTGCAATTCTGGTATCATGCCCATGGTNNNAGATCGGAAG |
| 46298_2 | TATGAGTGTGGAGTCGTTGCAATTCTGGTATCATGCCCATGGTNNNAGATCGGAAG |
| 46298_3 | TATGAGTGTGGAGTCGTTGCAATTCTGGTATCATGCCCATGGTNNNAGATCGGAAG |
| 40273_1 | TATGAGTGTGGAGTCGTTGCGGGCTTCCAGGCTATTGGTAAATNNNAGATCGGAAG |
| 40273_2 | TATGAGTGTGGAGTCGTTGCGGGCTTCCAGGCTATTGGTAAATNNNAGATCGGAAG |
| 40273_3 | TATGAGTGTGGAGTCGTTGCGGGCTTCCAGGCTATTGGTAAATNNNAGATCGGAAG |
| 37179_1 | TATGAGTGTGGAGTCGTTGCAGGGAGTTTCAGCTAATCCACTCNNNAGATCGGAAG |
| 37179_2 | TATGAGTGTGGAGTCGTTGCAGGGAGTTTCAGCTAATCCACTCNNNAGATCGGAAG |
| 37179_3 | TATGAGTGTGGAGTCGTTGCAGGGAGTTTCAGCTAATCCACTCNNNAGATCGGAAG |
| 35068_1 | TATGAGTGTGGAGTCGTTGCGCCAAATTCCAAAGAGGTGTACANNNAGATCGGAA |
| 35068_2 | TATGAGTGTGGAGTCGTTGCGCCAAATTCCAAAGAGGTGTACANNNAGATCGGAA |
| 35068_3 | TATGAGTGTGGAGTCGTTGCGCCAAATTCCAAAGAGGTGTACANNNAGATCGGAA |
| 43917_1 | TATGAGTGTGGAGTCGTTGCCCCACCACTCCAAGGTTAGAAAANNNAGATCGGAAG |
| 43917_2 | TATGAGTGTGGAGTCGTTGCCCCACCACTCCAAGGTTAGAAAANNNAGATCGGAAG |
| 43917_3 | TATGAGTGTGGAGTCGTTGCCCCACCACTCCAAGGTTAGAAAANNNAGATCGGAAG |
| 42448_1 | TATGAGTGTGGAGTCGTTGCTTTTTTGAGACGGAGTCTTGCTCNNNAGATCGGAAG |
| 42448_2 | TATGAGTGTGGAGTCGTTGCTTTTTTGAGACGGAGTCTTGCTCNNNAGATCGGAAG |
| 42448_3 | TATGAGTGTGGAGTCGTTGCTTTTTTGAGACGGAGTCTTGCTCNNNAGATCGGAAG |
| 30938_1 | TATGAGTGTGGAGTCGTTGCTCTCTCTCTCTCTCTCTCCCTNNNAGATCGGAAGAG |

| | |
|---|---|
| 30938_2 | TATGAGTGTGGAGTCGTTGCTCTCTCTCTCTCTCTCTCTCCCTNNNAGATCGGAAGAG |
| 30938_3 | TATGAGTGTGGAGTCGTTGCTCTCTCTCTCTCTCTCTCTCCCTNNNAGATCGGAAGAG |
| 39536_1 | TATGAGTGTGGAGTCGTTGCGGATAGGTGGGAACAAGTTGACANNNAGATCGGA |
| 39536_2 | TATGAGTGTGGAGTCGTTGCGGATAGGTGGGAACAAGTTGACANNNAGATCGGA |
| 39536_3 | TATGAGTGTGGAGTCGTTGCGGATAGGTGGGAACAAGTTGACANNNAGATCGGA |
| 44017_1 | TATGAGTGTGGAGTCGTTGCCGTAACCATAGGAAGGGTACTGGNNNAGATCGGAA |
| 44017_2 | TATGAGTGTGGAGTCGTTGCCGTAACCATAGGAAGGGTACTGGNNNAGATCGGAA |
| 44017_3 | TATGAGTGTGGAGTCGTTGCCGTAACCATAGGAAGGGTACTGGNNNAGATCGGAA |
| 49196_1 | TATGAGTGTGGAGTCGTTGCAAGGCAGAGGCATGAATAGTTCANNNAGATCGGAA |
| 49196_2 | TATGAGTGTGGAGTCGTTGCAAGGCAGAGGCATGAATAGTTCANNNAGATCGGAA |
| 49196_3 | TATGAGTGTGGAGTCGTTGCAAGGCAGAGGCATGAATAGTTCANNNAGATCGGAA |
| 53390_1 | TATGAGTGTGGAGTCGTTGCACTTTCCCCCAAACAAAGTCTCTNNNAGATCGGAAGA |
| 53390_2 | TATGAGTGTGGAGTCGTTGCACTTTCCCCCAAACAAAGTCTCTNNNAGATCGGAAGA |
| 53390_3 | TATGAGTGTGGAGTCGTTGCACTTTCCCCCAAACAAAGTCTCTNNNAGATCGGAAGA |
| 37676_1 | TATGAGTGTGGAGTCGTTGCTGTGGCACTGTAACAAAAGACACNNNAGATCGGAA |
| 37676_2 | TATGAGTGTGGAGTCGTTGCTGTGGCACTGTAACAAAAGACACNNNAGATCGGAA |
| 37676_3 | TATGAGTGTGGAGTCGTTGCTGTGGCACTGTAACAAAAGACACNNNAGATCGGAA |
| 41196_1 | TATGAGTGTGGAGTCGTTGCAGAGAGAGGAGAGAGAGAGGAGANNNAGATCGG |
| 41196_2 | TATGAGTGTGGAGTCGTTGCAGAGAGAGGAGAGAGAGAGGAGANNNAGATCGG |
| 41196_3 | TATGAGTGTGGAGTCGTTGCAGAGAGAGGAGAGAGAGAGGAGANNNAGATCGG |
| 48467_1 | TATGAGTGTGGAGTCGTTGCTTGAAAGAGCCGGTCTCACAATANNNAGATCGGAA |
| 48467_2 | TATGAGTGTGGAGTCGTTGCTTGAAAGAGCCGGTCTCACAATANNNAGATCGGAA |
| 48467_3 | TATGAGTGTGGAGTCGTTGCTTGAAAGAGCCGGTCTCACAATANNNAGATCGGAA |
| 35893_1 | TATGAGTGTGGAGTCGTTGCGGTGGACTACTTGATGTCAGGAGNNNAGATCGGAA |
| 35893_2 | TATGAGTGTGGAGTCGTTGCGGTGGACTACTTGATGTCAGGAGNNNAGATCGGAA |
| 35893_3 | TATGAGTGTGGAGTCGTTGCGGTGGACTACTTGATGTCAGGAGNNNAGATCGGAA |
| 5129_1 | TATGAGTGTGGAGTCGTTGCTAGATTGTTGATAGCGTCCCTCTNNNAGATCGGAAG |
| 5129_2 | TATGAGTGTGGAGTCGTTGCTAGATTGTTGATAGCGTCCCTCTNNNAGATCGGAAG |
| 5129_3 | TATGAGTGTGGAGTCGTTGCTAGATTGTTGATAGCGTCCCTCTNNNAGATCGGAAG |
| 32355_1 | TATGAGTGTGGAGTCGTTGCTGTCTGGTTTTGAAGTGTTTTCANNNAGATCGGAAG |
| 32355_2 | TATGAGTGTGGAGTCGTTGCTGTCTGGTTTTGAAGTGTTTTCANNNAGATCGGAAG |
| 32355_3 | TATGAGTGTGGAGTCGTTGCTGTCTGGTTTTGAAGTGTTTTCANNNAGATCGGAAG |
| 43614_1 | TATGAGTGTGGAGTCGTTGCGGAGAGATGGAGGAAATGGAGACNNNAGATCGGA |
| 43614_2 | TATGAGTGTGGAGTCGTTGCGGAGAGATGGAGGAAATGGAGACNNNAGATCGGA |
| 43614_3 | TATGAGTGTGGAGTCGTTGCGGAGAGATGGAGGAAATGGAGACNNNAGATCGGA |
| 51897_1 | TATGAGTGTGGAGTCGTTGCCTCCTCGAGAATGACATAGGGAGNNNAGATCGGAA |
| 51897_2 | TATGAGTGTGGAGTCGTTGCCTCCTCGAGAATGACATAGGGAGNNNAGATCGGAA |
| 51897_3 | TATGAGTGTGGAGTCGTTGCCTCCTCGAGAATGACATAGGGAGNNNAGATCGGAA |
| 30353_1 | TATGAGTGTGGAGTCGTTGCTGGCAAATACAGCCTTCTCAGACNNNAGATCGGAAG |
| 30353_2 | TATGAGTGTGGAGTCGTTGCTGGCAAATACAGCCTTCTCAGACNNNAGATCGGAAG |
| 30353_3 | TATGAGTGTGGAGTCGTTGCTGGCAAATACAGCCTTCTCAGACNNNAGATCGGAAG |
| 33213_1 | TATGAGTGTGGAGTCGTTGCTCATTCTTCAACCTGCAAACACCNNNAGATCGGAAGA |
| 33213_2 | TATGAGTGTGGAGTCGTTGCTCATTCTTCAACCTGCAAACACCNNNAGATCGGAAGA |

| | |
|---|---|
| 33213_3 | TATGAGTGTGGAGTCGTTGCTCATTCTTCAACCTGCAAACACCNNNAGATCGGAAGA |
| 48346_1 | TATGAGTGTGGAGTCGTTGCCTGGAGTGCAATGGTGTGATCTCNNNAGATCGGAA |
| 48346_2 | TATGAGTGTGGAGTCGTTGCCTGGAGTGCAATGGTGTGATCTCNNNAGATCGGAA |
| 48346_3 | TATGAGTGTGGAGTCGTTGCCTGGAGTGCAATGGTGTGATCTCNNNAGATCGGAA |
| 31087_1 | TATGAGTGTGGAGTCGTTGCCCACATCGGACTAAAAACCACACNNNAGATCGGAAG |
| 31087_2 | TATGAGTGTGGAGTCGTTGCCCACATCGGACTAAAAACCACACNNNAGATCGGAAG |
| 31087_3 | TATGAGTGTGGAGTCGTTGCCCACATCGGACTAAAAACCACACNNNAGATCGGAAG |
| 52558_1 | TATGAGTGTGGAGTCGTTGCCACACGCGCACTTTAGCATAATANNNAGATCGGAAG |
| 52558_2 | TATGAGTGTGGAGTCGTTGCCACACGCGCACTTTAGCATAATANNNAGATCGGAAG |
| 52558_3 | TATGAGTGTGGAGTCGTTGCCACACGCGCACTTTAGCATAATANNNAGATCGGAAG |
| 32521_1 | TATGAGTGTGGAGTCGTTGCGAGCTTCAGTACAGTGTCTGGAANNNAGATCGGAA |
| 32521_2 | TATGAGTGTGGAGTCGTTGCGAGCTTCAGTACAGTGTCTGGAANNNAGATCGGAA |
| 32521_3 | TATGAGTGTGGAGTCGTTGCGAGCTTCAGTACAGTGTCTGGAANNNAGATCGGAA |
| 35772_1 | TATGAGTGTGGAGTCGTTGCGAGTTCAAGCGATTCTTTTGCCTNNNAGATCGGAAG |
| 35772_2 | TATGAGTGTGGAGTCGTTGCGAGTTCAAGCGATTCTTTTGCCTNNNAGATCGGAAG |
| 35772_3 | TATGAGTGTGGAGTCGTTGCGAGTTCAAGCGATTCTTTTGCCTNNNAGATCGGAAG |
| 34107_1 | TATGAGTGTGGAGTCGTTGCAAAAACCAGATCACCTCTATGCANNNAGATCGGAAG |
| 34107_2 | TATGAGTGTGGAGTCGTTGCAAAAACCAGATCACCTCTATGCANNNAGATCGGAAG |
| 34107_3 | TATGAGTGTGGAGTCGTTGCAAAAACCAGATCACCTCTATGCANNNAGATCGGAAG |
| 45750_1 | TATGAGTGTGGAGTCGTTGCCTGAGTATAGAGTCACCATGGGCNNNAGATCGGAA |
| 45750_2 | TATGAGTGTGGAGTCGTTGCCTGAGTATAGAGTCACCATGGGCNNNAGATCGGAA |
| 45750_3 | TATGAGTGTGGAGTCGTTGCCTGAGTATAGAGTCACCATGGGCNNNAGATCGGAA |
| 37432_1 | TATGAGTGTGGAGTCGTTGCCTGTAATCCCAGCTACTCAGGAGNNNAGATCGGAAG |
| 37432_2 | TATGAGTGTGGAGTCGTTGCCTGTAATCCCAGCTACTCAGGAGNNNAGATCGGAAG |
| 37432_3 | TATGAGTGTGGAGTCGTTGCCTGTAATCCCAGCTACTCAGGAGNNNAGATCGGAAG |
| 49810_1 | TATGAGTGTGGAGTCGTTGCTATTCCCGTGACGGAAAGTTCAANNNAGATCGGAAG |
| 49810_2 | TATGAGTGTGGAGTCGTTGCTATTCCCGTGACGGAAAGTTCAANNNAGATCGGAAG |
| 49810_3 | TATGAGTGTGGAGTCGTTGCTATTCCCGTGACGGAAAGTTCAANNNAGATCGGAAG |
| 49801_1 | TATGAGTGTGGAGTCGTTGCTAAAGCTGGGGATCCTCTTCCTANNNAGATCGGAAG |
| 49801_2 | TATGAGTGTGGAGTCGTTGCTAAAGCTGGGGATCCTCTTCCTANNNAGATCGGAAG |
| 49801_3 | TATGAGTGTGGAGTCGTTGCTAAAGCTGGGGATCCTCTTCCTANNNAGATCGGAAG |
| 30552_1 | TATGAGTGTGGAGTCGTTGCGTCATCTCCCCCATCCACAATTNNNAGATCGGAAGAG |
| 30552_2 | TATGAGTGTGGAGTCGTTGCGTCATCTCCCCCATCCACAATTNNNAGATCGGAAGAG |
| 30552_3 | TATGAGTGTGGAGTCGTTGCGTCATCTCCCCCATCCACAATTNNNAGATCGGAAGAG |
| 50710_1 | TATGAGTGTGGAGTCGTTGCCCTCTCTTTGTGTGGGCGTTAGNNNAGATCGGAAGA |
| 50710_2 | TATGAGTGTGGAGTCGTTGCCCTCTCTTTGTGTGGGCGTTAGNNNAGATCGGAAGA |
| 50710_3 | TATGAGTGTGGAGTCGTTGCCCTCTCTTTGTGTGGGCGTTAGNNNAGATCGGAAGA |
| 12627_1 | TATGAGTGTGGAGTCGTTGCGTGATTTCCAAGTGTAACAATAGNNNAGATCGGAA |
| 12627_2 | TATGAGTGTGGAGTCGTTGCGTGATTTCCAAGTGTAACAATAGNNNAGATCGGAA |
| 12627_3 | TATGAGTGTGGAGTCGTTGCGTGATTTCCAAGTGTAACAATAGNNNAGATCGGAA |
| 47521_1 | TATGAGTGTGGAGTCGTTGCCCACGCCCAGCTAATTTTTTGTANNNAGATCGGAAG |
| 47521_2 | TATGAGTGTGGAGTCGTTGCCCACGCCCAGCTAATTTTTTGTANNNAGATCGGAAG |
| 47521_3 | TATGAGTGTGGAGTCGTTGCCCACGCCCAGCTAATTTTTTGTANNNAGATCGGAAG |

| | |
|---|---|
| 35431_1 | TATGAGTGTGGAGTCGTTGCGTTCACGCTATTCTCCTGCCTNNNAGATCGGAAGAG |
| 35431_2 | TATGAGTGTGGAGTCGTTGCGTTCACGCTATTCTCCTGCCTNNNAGATCGGAAGAG |
| 35431_3 | TATGAGTGTGGAGTCGTTGCGTTCACGCTATTCTCCTGCCTNNNAGATCGGAAGAG |
| 44300_1 | TATGAGTGTGGAGTCGTTGCAGAATGAGGTTCAGTCCAAGCATNNNAGATCGGAA |
| 44300_2 | TATGAGTGTGGAGTCGTTGCAGAATGAGGTTCAGTCCAAGCATNNNAGATCGGAA |
| 44300_3 | TATGAGTGTGGAGTCGTTGCAGAATGAGGTTCAGTCCAAGCATNNNAGATCGGAA |
| 10491_1 | TATGAGTGTGGAGTCGTTGCATAGCCACTGTGCATGGAGTTTTNNNAGATCGGAAG |
| 10491_2 | TATGAGTGTGGAGTCGTTGCATAGCCACTGTGCATGGAGTTTTNNNAGATCGGAAG |
| 10491_3 | TATGAGTGTGGAGTCGTTGCATAGCCACTGTGCATGGAGTTTTNNNAGATCGGAAG |
| 45326_1 | TATGAGTGTGGAGTCGTTGCTTCCGAGGCAAAAATTGTGTGTTNNNAGATCGGAA |
| 45326_2 | TATGAGTGTGGAGTCGTTGCTTCCGAGGCAAAAATTGTGTGTTNNNAGATCGGAA |
| 45326_3 | TATGAGTGTGGAGTCGTTGCTTCCGAGGCAAAAATTGTGTGTTNNNAGATCGGAA |
| 49892_1 | TATGAGTGTGGAGTCGTTGCTTTTGAATCACTCCTGGGAAGGTNNNAGATCGGAAG |
| 49892_2 | TATGAGTGTGGAGTCGTTGCTTTTGAATCACTCCTGGGAAGGTNNNAGATCGGAAG |
| 49892_3 | TATGAGTGTGGAGTCGTTGCTTTTGAATCACTCCTGGGAAGGTNNNAGATCGGAAG |
| 37622_1 | TATGAGTGTGGAGTCGTTGCTGTTTGAGGTGATGGATATGCCANNNAGATCGGAA |
| 37622_2 | TATGAGTGTGGAGTCGTTGCTGTTTGAGGTGATGGATATGCCANNNAGATCGGAA |
| 37622_3 | TATGAGTGTGGAGTCGTTGCTGTTTGAGGTGATGGATATGCCANNNAGATCGGAA |
| 42139_1 | TATGAGTGTGGAGTCGTTGCCTGAGTAGCAGGGATTACAGGCNNNAGATCGGAAG |
| 42139_2 | TATGAGTGTGGAGTCGTTGCCTGAGTAGCAGGGATTACAGGCNNNAGATCGGAAG |
| 42139_3 | TATGAGTGTGGAGTCGTTGCCTGAGTAGCAGGGATTACAGGCNNNAGATCGGAAG |
| 2525_1 | TATGAGTGTGGAGTCGTTGCACTGACTTACTGCTGCATCAAGGNNNAGATCGGAAG |
| 2525_2 | TATGAGTGTGGAGTCGTTGCACTGACTTACTGCTGCATCAAGGNNNAGATCGGAAG |
| 2525_3 | TATGAGTGTGGAGTCGTTGCACTGACTTACTGCTGCATCAAGGNNNAGATCGGAAG |
| 38920_1 | TATGAGTGTGGAGTCGTTGCTGTCTCTTTATGTCCTTTGCCCANNNAGATCGGAAGA |
| 38920_2 | TATGAGTGTGGAGTCGTTGCTGTCTCTTTATGTCCTTTGCCCANNNAGATCGGAAGA |
| 38920_3 | TATGAGTGTGGAGTCGTTGCTGTCTCTTTATGTCCTTTGCCCANNNAGATCGGAAGA |
| 50865_1 | TATGAGTGTGGAGTCGTTGCTTTTCGTCAGTTGATTGCTGAGCNNNAGATCGGAAG |
| 50865_2 | TATGAGTGTGGAGTCGTTGCTTTTCGTCAGTTGATTGCTGAGCNNNAGATCGGAAG |
| 50865_3 | TATGAGTGTGGAGTCGTTGCTTTTCGTCAGTTGATTGCTGAGCNNNAGATCGGAAG |
| 33772_1 | TATGAGTGTGGAGTCGTTGCCATTCTCAGCAAACTATCGCCAGNNNAGATCGGAAG |
| 33772_2 | TATGAGTGTGGAGTCGTTGCCATTCTCAGCAAACTATCGCCAGNNNAGATCGGAAG |
| 33772_3 | TATGAGTGTGGAGTCGTTGCCATTCTCAGCAAACTATCGCCAGNNNAGATCGGAAG |
| 33591_1 | TATGAGTGTGGAGTCGTTGCTTCAAGTGATTCTCCTGCCTCAGNNNAGATCGGAAG |
| 33591_2 | TATGAGTGTGGAGTCGTTGCTTCAAGTGATTCTCCTGCCTCAGNNNAGATCGGAAG |
| 33591_3 | TATGAGTGTGGAGTCGTTGCTTCAAGTGATTCTCCTGCCTCAGNNNAGATCGGAAG |
| 45202_1 | TATGAGTGTGGAGTCGTTGCGTCAGTGAATACCGTGAGGGAATNNNAGATCGGAA |
| 45202_2 | TATGAGTGTGGAGTCGTTGCGTCAGTGAATACCGTGAGGGAATNNNAGATCGGAA |
| 45202_3 | TATGAGTGTGGAGTCGTTGCGTCAGTGAATACCGTGAGGGAATNNNAGATCGGAA |
| 42411_1 | TATGAGTGTGGAGTCGTTGCGAGAAGGAAGAGAGTGTTTGGGTNNNAGATCGGA |
| 42411_2 | TATGAGTGTGGAGTCGTTGCGAGAAGGAAGAGAGTGTTTGGGTNNNAGATCGGA |
| 42411_3 | TATGAGTGTGGAGTCGTTGCGAGAAGGAAGAGAGTGTTTGGGTNNNAGATCGGA |
| 50179_1 | TATGAGTGTGGAGTCGTTGCCAACCATGAGAAGATGACCCAGANNNAGATCGGAA |

| | |
|---|---|
| 50179_2 | TATGAGTGTGGAGTCGTTGCCAACCATGAGAAGATGACCCAGANNNAGATCGGAA |
| 50179_3 | TATGAGTGTGGAGTCGTTGCCAACCATGAGAAGATGACCCAGANNNAGATCGGAA |
| 52898_1 | TATGAGTGTGGAGTCGTTGCCATCTCTGCTGTCCCTGTGTCNNNAGATCGGAAGAG |
| 52898_2 | TATGAGTGTGGAGTCGTTGCCATCTCTGCTGTCCCTGTGTCNNNAGATCGGAAGAG |
| 52898_3 | TATGAGTGTGGAGTCGTTGCCATCTCTGCTGTCCCTGTGTCNNNAGATCGGAAGAG |
| 37416_1 | TATGAGTGTGGAGTCGTTGCACTCACACCTCTAAGAGTGGGATNNNAGATCGGAAG |
| 37416_2 | TATGAGTGTGGAGTCGTTGCACTCACACCTCTAAGAGTGGGATNNNAGATCGGAAG |
| 37416_3 | TATGAGTGTGGAGTCGTTGCACTCACACCTCTAAGAGTGGGATNNNAGATCGGAAG |
| 46540_1 | TATGAGTGTGGAGTCGTTGCCCTGGGTTCAAGCAATTCTCATGNNNAGATCGGAAG |
| 46540_2 | TATGAGTGTGGAGTCGTTGCCCTGGGTTCAAGCAATTCTCATGNNNAGATCGGAAG |
| 46540_3 | TATGAGTGTGGAGTCGTTGCCCTGGGTTCAAGCAATTCTCATGNNNAGATCGGAAG |
| 37510_1 | TATGAGTGTGGAGTCGTTGCGACATCCTCGCCTCATTTTGAGANNNAGATCGGAAG |
| 37510_2 | TATGAGTGTGGAGTCGTTGCGACATCCTCGCCTCATTTTGAGANNNAGATCGGAAG |
| 37510_3 | TATGAGTGTGGAGTCGTTGCGACATCCTCGCCTCATTTTGAGANNNAGATCGGAAG |
| 35109_1 | TATGAGTGTGGAGTCGTTGCGGAATTACAGGCACACACCACTANNNAGATCGGAAG |
| 35109_2 | TATGAGTGTGGAGTCGTTGCGGAATTACAGGCACACACCACTANNNAGATCGGAAG |
| 35109_3 | TATGAGTGTGGAGTCGTTGCGGAATTACAGGCACACACCACTANNNAGATCGGAAG |
| 47594_1 | TATGAGTGTGGAGTCGTTGCAGTGCTAGGATTACAGGCATGAGNNNAGATCGGAA |
| 47594_2 | TATGAGTGTGGAGTCGTTGCAGTGCTAGGATTACAGGCATGAGNNNAGATCGGAA |
| 47594_3 | TATGAGTGTGGAGTCGTTGCAGTGCTAGGATTACAGGCATGAGNNNAGATCGGAA |
| 48432_1 | TATGAGTGTGGAGTCGTTGCTACTTGGAGCTCATTCTTCCCACNNNAGATCGGAAGA |
| 48432_2 | TATGAGTGTGGAGTCGTTGCTACTTGGAGCTCATTCTTCCCACNNNAGATCGGAAGA |
| 48432_3 | TATGAGTGTGGAGTCGTTGCTACTTGGAGCTCATTCTTCCCACNNNAGATCGGAAGA |
| 41850_1 | TATGAGTGTGGAGTCGTTGCCTCCTGAGTAGCTGGGAGAGTAGNNNAGATCGGAA |
| 41850_2 | TATGAGTGTGGAGTCGTTGCCTCCTGAGTAGCTGGGAGAGTAGNNNAGATCGGAA |
| 41850_3 | TATGAGTGTGGAGTCGTTGCCTCCTGAGTAGCTGGGAGAGTAGNNNAGATCGGAA |
| 36585_1 | TATGAGTGTGGAGTCGTTGCAACTCAAAGATGGGGCCAATAGTNNNAGATCGGAA |
| 36585_2 | TATGAGTGTGGAGTCGTTGCAACTCAAAGATGGGGCCAATAGTNNNAGATCGGAA |
| 36585_3 | TATGAGTGTGGAGTCGTTGCAACTCAAAGATGGGGCCAATAGTNNNAGATCGGAA |
| 35240_1 | TATGAGTGTGGAGTCGTTGCGAGATGCTAGTAGGCAGAGACACNNNAGATCGGAA |
| 35240_2 | TATGAGTGTGGAGTCGTTGCGAGATGCTAGTAGGCAGAGACACNNNAGATCGGAA |
| 35240_3 | TATGAGTGTGGAGTCGTTGCGAGATGCTAGTAGGCAGAGACACNNNAGATCGGAA |
| 45965_1 | TATGAGTGTGGAGTCGTTGCTAGCTAGGATTACAGGCATGTGCNNNAGATCGGAA |
| 45965_2 | TATGAGTGTGGAGTCGTTGCTAGCTAGGATTACAGGCATGTGCNNNAGATCGGAA |
| 45965_3 | TATGAGTGTGGAGTCGTTGCTAGCTAGGATTACAGGCATGTGCNNNAGATCGGAA |
| 10700_1 | TATGAGTGTGGAGTCGTTGCAGGCTACAGTGATCATGCCACTANNNAGATCGGAAG |
| 10700_2 | TATGAGTGTGGAGTCGTTGCAGGCTACAGTGATCATGCCACTANNNAGATCGGAAG |
| 10700_3 | TATGAGTGTGGAGTCGTTGCAGGCTACAGTGATCATGCCACTANNNAGATCGGAAG |
| 52335_1 | TATGAGTGTGGAGTCGTTGCCCCTGTTAGCACTGCCAGTTANNNAGATCGGAAGAG |
| 52335_2 | TATGAGTGTGGAGTCGTTGCCCCTGTTAGCACTGCCAGTTANNNAGATCGGAAGAG |
| 52335_3 | TATGAGTGTGGAGTCGTTGCCCCTGTTAGCACTGCCAGTTANNNAGATCGGAAGAG |
| 40387_1 | TATGAGTGTGGAGTCGTTGCTATCAATCCCCTAGCATGACAGCNNNAGATCGGAAG |
| 40387_2 | TATGAGTGTGGAGTCGTTGCTATCAATCCCCTAGCATGACAGCNNNAGATCGGAAG |

| | |
|---|---|
| 40387_3 | TATGAGTGTGGAGTCGTTGCTATCAATCCCCTAGCATGACAGCNNNAGATCGGAAG |
| 33847_1 | TATGAGTGTGGAGTCGTTGCCACAAACAAAGCGAGGAAGGAATNNNAGATCGGAA |
| 33847_2 | TATGAGTGTGGAGTCGTTGCCACAAACAAAGCGAGGAAGGAATNNNAGATCGGAA |
| 33847_3 | TATGAGTGTGGAGTCGTTGCCACAAACAAAGCGAGGAAGGAATNNNAGATCGGAA |
| 10428_1 | TATGAGTGTGGAGTCGTTGCGACTACAAGTGGGCCCTAAAGATTNNNAGATCGGA |
| 10428_2 | TATGAGTGTGGAGTCGTTGCGACTACAAGTGGGCCCTAAAGATTNNNAGATCGGA |
| 10428_3 | TATGAGTGTGGAGTCGTTGCGACTACAAGTGGGCCCTAAAGATTNNNAGATCGGA |
| 34639_1 | TATGAGTGTGGAGTCGTTGCTTTTCTGGGAAGCCTTTTTTGGGNNNAGATCGGAAG |
| 34639_2 | TATGAGTGTGGAGTCGTTGCTTTTCTGGGAAGCCTTTTTTGGGNNNAGATCGGAAG |
| 34639_3 | TATGAGTGTGGAGTCGTTGCTTTTCTGGGAAGCCTTTTTTGGGNNNAGATCGGAAG |
| 39891_1 | TATGAGTGTGGAGTCGTTGCCACTTGTAGTCCCAGTTACTCGGNNNAGATCGGAAG |
| 39891_2 | TATGAGTGTGGAGTCGTTGCCACTTGTAGTCCCAGTTACTCGGNNNAGATCGGAAG |
| 39891_3 | TATGAGTGTGGAGTCGTTGCCACTTGTAGTCCCAGTTACTCGGNNNAGATCGGAAG |
| 36993_1 | TATGAGTGTGGAGTCGTTGCCCTTCTTGTCACTAGGCAGACTTNNNAGATCGGAAG |
| 36993_2 | TATGAGTGTGGAGTCGTTGCCCTTCTTGTCACTAGGCAGACTTNNNAGATCGGAAG |
| 36993_3 | TATGAGTGTGGAGTCGTTGCCCTTCTTGTCACTAGGCAGACTTNNNAGATCGGAAG |
| 42776_1 | TATGAGTGTGGAGTCGTTGCGTTGGGCAAGGAGGAGAGAAATANNNAGATCGGA |
| 42776_2 | TATGAGTGTGGAGTCGTTGCGTTGGGCAAGGAGGAGAGAAATANNNAGATCGGA |
| 42776_3 | TATGAGTGTGGAGTCGTTGCGTTGGGCAAGGAGGAGAGAAATANNNAGATCGGA |
| 53026_1 | TATGAGTGTGGAGTCGTTGCGAAACAACAGATACTGGGGCCTANNNAGATCGGAA |
| 53026_2 | TATGAGTGTGGAGTCGTTGCGAAACAACAGATACTGGGGCCTANNNAGATCGGAA |
| 53026_3 | TATGAGTGTGGAGTCGTTGCGAAACAACAGATACTGGGGCCTANNNAGATCGGAA |
| 40425_1 | TATGAGTGTGGAGTCGTTGCGGGAACAGATCATAGTACCTAGCANNNAGATCGGA |
| 40425_2 | TATGAGTGTGGAGTCGTTGCGGGAACAGATCATAGTACCTAGCANNNAGATCGGA |
| 40425_3 | TATGAGTGTGGAGTCGTTGCGGGAACAGATCATAGTACCTAGCANNNAGATCGGA |
| 37048_1 | TATGAGTGTGGAGTCGTTGCTTGAGTTGGAGGTAGACGTCAAGNNNAGATCGGAA |
| 37048_2 | TATGAGTGTGGAGTCGTTGCTTGAGTTGGAGGTAGACGTCAAGNNNAGATCGGAA |
| 37048_3 | TATGAGTGTGGAGTCGTTGCTTGAGTTGGAGGTAGACGTCAAGNNNAGATCGGAA |
| 43440_1 | TATGAGTGTGGAGTCGTTGCCAGCTTCCTGAGTAGTTGGGAATNNNAGATCGGAA |
| 43440_2 | TATGAGTGTGGAGTCGTTGCCAGCTTCCTGAGTAGTTGGGAATNNNAGATCGGAA |
| 43440_3 | TATGAGTGTGGAGTCGTTGCCAGCTTCCTGAGTAGTTGGGAATNNNAGATCGGAA |
| 38523_1 | TATGAGTGTGGAGTCGTTGCTGTGTATTTTACCTTTGGGCACGNNNAGATCGGAAG |
| 38523_2 | TATGAGTGTGGAGTCGTTGCTGTGTATTTTACCTTTGGGCACGNNNAGATCGGAAG |
| 38523_3 | TATGAGTGTGGAGTCGTTGCTGTGTATTTTACCTTTGGGCACGNNNAGATCGGAAG |
| 42648_1 | TATGAGTGTGGAGTCGTTGCCGCCCTCATTCTGTCTCTATGAANNNAGATCGGAAGA |
| 42648_2 | TATGAGTGTGGAGTCGTTGCCGCCCTCATTCTGTCTCTATGAANNNAGATCGGAAGA |
| 42648_3 | TATGAGTGTGGAGTCGTTGCCGCCCTCATTCTGTCTCTATGAANNNAGATCGGAAGA |
| 44067_1 | TATGAGTGTGGAGTCGTTGCCGTATTCCCTTCTACCACCTTCCNNNAGATCGGAAGA |
| 44067_2 | TATGAGTGTGGAGTCGTTGCCGTATTCCCTTCTACCACCTTCCNNNAGATCGGAAGA |
| 44067_3 | TATGAGTGTGGAGTCGTTGCCGTATTCCCTTCTACCACCTTCCNNNAGATCGGAAGA |
| 37128_1 | TATGAGTGTGGAGTCGTTGCCTAGCTATACCTGACAGTGTGGCNNNAGATCGGAAG |
| 37128_2 | TATGAGTGTGGAGTCGTTGCCTAGCTATACCTGACAGTGTGGCNNNAGATCGGAAG |
| 37128_3 | TATGAGTGTGGAGTCGTTGCCTAGCTATACCTGACAGTGTGGCNNNAGATCGGAAG |

| | |
|---|---|
| 37954_1 | TATGAGTGTGGAGTCGTTGCAGATTCTGGAAAGGAGGAACACCNNNAGATCGGAA |
| 37954_2 | TATGAGTGTGGAGTCGTTGCAGATTCTGGAAAGGAGGAACACCNNNAGATCGGAA |
| 37954_3 | TATGAGTGTGGAGTCGTTGCAGATTCTGGAAAGGAGGAACACCNNNAGATCGGAA |
| 53365_1 | TATGAGTGTGGAGTCGTTGCGTTCAAGCTATTCTCCCACCTCANNNAGATCGGAAGA |
| 53365_2 | TATGAGTGTGGAGTCGTTGCGTTCAAGCTATTCTCCCACCTCANNNAGATCGGAAGA |
| 53365_3 | TATGAGTGTGGAGTCGTTGCGTTCAAGCTATTCTCCCACCTCANNNAGATCGGAAGA |
| 40565_1 | TATGAGTGTGGAGTCGTTGCTGGCATATTGTAAGTGTGCACTCNNNAGATCGGAAG |
| 40565_2 | TATGAGTGTGGAGTCGTTGCTGGCATATTGTAAGTGTGCACTCNNNAGATCGGAAG |
| 40565_3 | TATGAGTGTGGAGTCGTTGCTGGCATATTGTAAGTGTGCACTCNNNAGATCGGAAG |
| 41733_1 | TATGAGTGTGGAGTCGTTGCCCACCACACCCAGCTAATTTTTTNNNAGATCGGAAGA |
| 41733_2 | TATGAGTGTGGAGTCGTTGCCCACCACACCCAGCTAATTTTTTNNNAGATCGGAAGA |
| 41733_3 | TATGAGTGTGGAGTCGTTGCCCACCACACCCAGCTAATTTTTTNNNAGATCGGAAGA |
| 12579_1 | TATGAGTGTGGAGTCGTTGCTATATGTATGAGAGAGAAACACGNNNAGATCGGAA |
| 12579_2 | TATGAGTGTGGAGTCGTTGCTATATGTATGAGAGAGAAACACGNNNAGATCGGAA |
| 12579_3 | TATGAGTGTGGAGTCGTTGCTATATGTATGAGAGAGAAACACGNNNAGATCGGAA |
| 30853_1 | TATGAGTGTGGAGTCGTTGCACATACAGGTACGCATGCACATANNNAGATCGGAAG |
| 30853_2 | TATGAGTGTGGAGTCGTTGCACATACAGGTACGCATGCACATANNNAGATCGGAAG |
| 30853_3 | TATGAGTGTGGAGTCGTTGCACATACAGGTACGCATGCACATANNNAGATCGGAAG |
| 38587_1 | TATGAGTGTGGAGTCGTTGCACACATGCACAGACATTTCACTGNNNAGATCGGAAG |
| 38587_2 | TATGAGTGTGGAGTCGTTGCACACATGCACAGACATTTCACTGNNNAGATCGGAAG |
| 38587_3 | TATGAGTGTGGAGTCGTTGCACACATGCACAGACATTTCACTGNNNAGATCGGAAG |
| 34194_1 | TATGAGTGTGGAGTCGTTGCCAGGTTGGTGGTTTGTGTTGATTNNNAGATCGGAA |
| 34194_2 | TATGAGTGTGGAGTCGTTGCCAGGTTGGTGGTTTGTGTTGATTNNNAGATCGGAA |
| 34194_3 | TATGAGTGTGGAGTCGTTGCCAGGTTGGTGGTTTGTGTTGATTNNNAGATCGGAA |
| 12734_1 | TATGAGTGTGGAGTCGTTGCTATTTTGGCCATCTGAAAGAANNNAGATCGGAAGA |
| 12734_2 | TATGAGTGTGGAGTCGTTGCTATTTTGGCCATCTGAAAGAANNNAGATCGGAAGA |
| 12734_3 | TATGAGTGTGGAGTCGTTGCTATTTTGGCCATCTGAAAGAANNNAGATCGGAAGA |
| 9216_1 | TATGAGTGTGGAGTCGTTGCTAGTTGAGGGTTTGAGGGTTGTGNNNAGATCGGA |
| 9216_2 | TATGAGTGTGGAGTCGTTGCTAGTTGAGGGTTTGAGGGTTGTGNNNAGATCGGA |
| 9216_3 | TATGAGTGTGGAGTCGTTGCTAGTTGAGGGTTTGAGGGTTGTGNNNAGATCGGA |
| 37287_1 | TATGAGTGTGGAGTCGTTGCTCTACTTGAGGATTGCTGTTGCTNNNAGATCGGAAG |
| 37287_2 | TATGAGTGTGGAGTCGTTGCTCTACTTGAGGATTGCTGTTGCTNNNAGATCGGAAG |
| 37287_3 | TATGAGTGTGGAGTCGTTGCTCTACTTGAGGATTGCTGTTGCTNNNAGATCGGAAG |
| 41647_1 | TATGAGTGTGGAGTCGTTGCCTGGTCTTGAACTCCTGAACTCANNNAGATCGGAAG |
| 41647_2 | TATGAGTGTGGAGTCGTTGCCTGGTCTTGAACTCCTGAACTCANNNAGATCGGAAG |
| 41647_3 | TATGAGTGTGGAGTCGTTGCCTGGTCTTGAACTCCTGAACTCANNNAGATCGGAAG |
| 47309_1 | TATGAGTGTGGAGTCGTTGCCACGCCCGGCTAATTTTTTTGTANNNAGATCGGAAG |
| 47309_2 | TATGAGTGTGGAGTCGTTGCCACGCCCGGCTAATTTTTTTGTANNNAGATCGGAAG |
| 47309_3 | TATGAGTGTGGAGTCGTTGCCACGCCCGGCTAATTTTTTTGTANNNAGATCGGAAG |
| 6125_1 | TATGAGTGTGGAGTCGTTGCTGAACCAAACAGCTGCTAGCATTNNNAGATCGGAAG |
| 6125_2 | TATGAGTGTGGAGTCGTTGCTGAACCAAACAGCTGCTAGCATTNNNAGATCGGAAG |
| 6125_3 | TATGAGTGTGGAGTCGTTGCTGAACCAAACAGCTGCTAGCATTNNNAGATCGGAAG |
| 12597_1 | TATGAGTGTGGAGTCGTTGCACCGAGATAGAAAAAAAAGTTTTNNNAGATCGGAA |

| | |
|---|---|
| 12597_2 | TATGAGTGTGGAGTCGTTGCACCGAGATAGAAAAAAAAGTTTTNNNAGATCGGAA |
| 12597_3 | TATGAGTGTGGAGTCGTTGCACCGAGATAGAAAAAAAAGTTTTNNNAGATCGGAA |
| 34245_1 | TATGAGTGTGGAGTCGTTGCCCCGACACCTCCTAATCCTAAACNNNAGATCGGAAGA |
| 34245_2 | TATGAGTGTGGAGTCGTTGCCCCGACACCTCCTAATCCTAAACNNNAGATCGGAAGA |
| 34245_3 | TATGAGTGTGGAGTCGTTGCCCCGACACCTCCTAATCCTAAACNNNAGATCGGAAGA |
| 32254_1 | TATGAGTGTGGAGTCGTTGCTCCCCAGTTTTTTTTTGCTGTGTNNNAGATCGGAAGA |
| 32254_2 | TATGAGTGTGGAGTCGTTGCTCCCCAGTTTTTTTTTGCTGTGTNNNAGATCGGAAGA |
| 32254_3 | TATGAGTGTGGAGTCGTTGCTCCCCAGTTTTTTTTTGCTGTGTNNNAGATCGGAAGA |
| 48500_1 | TATGAGTGTGGAGTCGTTGCTCCCCTTGGAATAAAACGGTGAANNNAGATCGGAAG |
| 48500_2 | TATGAGTGTGGAGTCGTTGCTCCCCTTGGAATAAAACGGTGAANNNAGATCGGAAG |
| 48500_3 | TATGAGTGTGGAGTCGTTGCTCCCCTTGGAATAAAACGGTGAANNNAGATCGGAAG |
| 36655_1 | TATGAGTGTGGAGTCGTTGCGTGCTAAGAGGATTCAATGGGGANNNAGATCGGAA |
| 36655_2 | TATGAGTGTGGAGTCGTTGCGTGCTAAGAGGATTCAATGGGGANNNAGATCGGAA |
| 36655_3 | TATGAGTGTGGAGTCGTTGCGTGCTAAGAGGATTCAATGGGGANNNAGATCGGAA |
| 41240_1 | TATGAGTGTGGAGTCGTTGCGGTGGATCATGAGGTCAAGAGATNNNAGATCGGAA |
| 41240_2 | TATGAGTGTGGAGTCGTTGCGGTGGATCATGAGGTCAAGAGATNNNAGATCGGAA |
| 41240_3 | TATGAGTGTGGAGTCGTTGCGGTGGATCATGAGGTCAAGAGATNNNAGATCGGAA |
| 32061_1 | TATGAGTGTGGAGTCGTTGCGAGTAGCTGGGACTACAGATGCNNNAGATCGGAAG |
| 32061_2 | TATGAGTGTGGAGTCGTTGCGAGTAGCTGGGACTACAGATGCNNNAGATCGGAAG |
| 32061_3 | TATGAGTGTGGAGTCGTTGCGAGTAGCTGGGACTACAGATGCNNNAGATCGGAAG |
| 34735_1 | TATGAGTGTGGAGTCGTTGCTTTTTTGAGACGGAGTCTTGCTCNNNAGATCGGAAG |
| 34735_2 | TATGAGTGTGGAGTCGTTGCTTTTTTGAGACGGAGTCTTGCTCNNNAGATCGGAAG |
| 34735_3 | TATGAGTGTGGAGTCGTTGCTTTTTTGAGACGGAGTCTTGCTCNNNAGATCGGAAG |
| 10193_1 | TATGAGTGTGGAGTCGTTGCAAAATGAAGGGGGAAGCAAGAGGNNNAGATCGGA |
| 10193_2 | TATGAGTGTGGAGTCGTTGCAAAATGAAGGGGGAAGCAAGAGGNNNAGATCGGA |
| 10193_3 | TATGAGTGTGGAGTCGTTGCAAAATGAAGGGGGAAGCAAGAGGNNNAGATCGGA |
| 37467_1 | TATGAGTGTGGAGTCGTTGCAATGAAACTGTGATAGGCTGGGANNNAGATCGGAA |
| 37467_2 | TATGAGTGTGGAGTCGTTGCAATGAAACTGTGATAGGCTGGGANNNAGATCGGAA |
| 37467_3 | TATGAGTGTGGAGTCGTTGCAATGAAACTGTGATAGGCTGGGANNNAGATCGGAA |
| 49783_1 | TATGAGTGTGGAGTCGTTGCACTGCTGCTTTTCCTTACAGTGANNNAGATCGGAAG |
| 49783_2 | TATGAGTGTGGAGTCGTTGCACTGCTGCTTTTCCTTACAGTGANNNAGATCGGAAG |
| 49783_3 | TATGAGTGTGGAGTCGTTGCACTGCTGCTTTTCCTTACAGTGANNNAGATCGGAAG |
| 48363_1 | TATGAGTGTGGAGTCGTTGCGGGTTCAAGTAATTCTCCTGCCTNNNAGATCGGAAG |
| 48363_2 | TATGAGTGTGGAGTCGTTGCGGGTTCAAGTAATTCTCCTGCCTNNNAGATCGGAAG |
| 48363_3 | TATGAGTGTGGAGTCGTTGCGGGTTCAAGTAATTCTCCTGCCTNNNAGATCGGAAG |
| 10985_1 | TATGAGTGTGGAGTCGTTGCTCGGACAGTAACCCTAAGGCTAANNNAGATCGGAAG |
| 10985_2 | TATGAGTGTGGAGTCGTTGCTCGGACAGTAACCCTAAGGCTAANNNAGATCGGAAG |
| 10985_3 | TATGAGTGTGGAGTCGTTGCTCGGACAGTAACCCTAAGGCTAANNNAGATCGGAAG |
| 31626_1 | TATGAGTGTGGAGTCGTTGCAATTGCATCACTTCCATATCGGANNNAGATCGGAAG |
| 31626_2 | TATGAGTGTGGAGTCGTTGCAATTGCATCACTTCCATATCGGANNNAGATCGGAAG |
| 31626_3 | TATGAGTGTGGAGTCGTTGCAATTGCATCACTTCCATATCGGANNNAGATCGGAAG |
| 42399_1 | TATGAGTGTGGAGTCGTTGCTTTCCCTCTTTCTGTTAGAAGCANNNAGATCGGAAGA |
| 42399_2 | TATGAGTGTGGAGTCGTTGCTTTCCCTCTTTCTGTTAGAAGCANNNAGATCGGAAGA |

| | |
|---|---|
| 42399_3 | TATGAGTGTGGAGTCGTTGCTTTCCCTCTTTCTGTTAGAAGCANNNAGATCGGAAG |
| 13485_1 | TATGAGTGTGGAGTCGTTGCCTTAAAAGTGTTGTGTTACTTTGNNNAGATCGGAAG |
| 13485_2 | TATGAGTGTGGAGTCGTTGCCTTAAAAGTGTTGTGTTACTTTGNNNAGATCGGAAG |
| 13485_3 | TATGAGTGTGGAGTCGTTGCCTTAAAAGTGTTGTGTTACTTTGNNNAGATCGGAAG |
| 46643_1 | TATGAGTGTGGAGTCGTTGCGTGTGTGTGTGTGTGTGTATTNNNAGATCGGAA |
| 46643_2 | TATGAGTGTGGAGTCGTTGCGTGTGTGTGTGTGTGTGTATTNNNAGATCGGAA |
| 46643_3 | TATGAGTGTGGAGTCGTTGCGTGTGTGTGTGTGTGTGTATTNNNAGATCGGAA |
| 51655_1 | TATGAGTGTGGAGTCGTTGCATAGGACTACAGATGCCACCATGNNNAGATCGGAA |
| 51655_2 | TATGAGTGTGGAGTCGTTGCATAGGACTACAGATGCCACCATGNNNAGATCGGAA |
| 51655_3 | TATGAGTGTGGAGTCGTTGCATAGGACTACAGATGCCACCATGNNNAGATCGGAA |
| 45163_1 | TATGAGTGTGGAGTCGTTGCGGAGAAGGAGCTAGCGTTCCNNNAGATCGGAAGA |
| 45163_2 | TATGAGTGTGGAGTCGTTGCGGAGAAGGAGCTAGCGTTCCNNNAGATCGGAAGA |
| 45163_3 | TATGAGTGTGGAGTCGTTGCGGAGAAGGAGCTAGCGTTCCNNNAGATCGGAAGA |
| 47106_1 | TATGAGTGTGGAGTCGTTGCTCTGGGTTCAAGTGATTCTCCTGNNNAGATCGGAAG |
| 47106_2 | TATGAGTGTGGAGTCGTTGCTCTGGGTTCAAGTGATTCTCCTGNNNAGATCGGAAG |
| 47106_3 | TATGAGTGTGGAGTCGTTGCTCTGGGTTCAAGTGATTCTCCTGNNNAGATCGGAAG |
| 10239_1 | TATGAGTGTGGAGTCGTTGCGGGATAGACAGAAGGACAACCCTNNNAGATCGGAA |
| 10239_2 | TATGAGTGTGGAGTCGTTGCGGGATAGACAGAAGGACAACCCTNNNAGATCGGAA |
| 10239_3 | TATGAGTGTGGAGTCGTTGCGGGATAGACAGAAGGACAACCCTNNNAGATCGGAA |
| 30495_1 | TATGAGTGTGGAGTCGTTGCTGATGTTTGTGTGTGTTTGGGTATNNNAGATCGGA |
| 30495_2 | TATGAGTGTGGAGTCGTTGCTGATGTTTGTGTGTGTTTGGGTATNNNAGATCGGA |
| 30495_3 | TATGAGTGTGGAGTCGTTGCTGATGTTTGTGTGTGTTTGGGTATNNNAGATCGGA |
| 52525_1 | TATGAGTGTGGAGTCGTTGCGTAAAGCCTTCCTCGAGCAATTCNNNAGATCGGAAG |
| 52525_2 | TATGAGTGTGGAGTCGTTGCGTAAAGCCTTCCTCGAGCAATTCNNNAGATCGGAAG |
| 52525_3 | TATGAGTGTGGAGTCGTTGCGTAAAGCCTTCCTCGAGCAATTCNNNAGATCGGAAG |
| 44773_1 | TATGAGTGTGGAGTCGTTGCACTGTTCTCAAGCTGCTACTGAANNNAGATCGGAAG |
| 44773_2 | TATGAGTGTGGAGTCGTTGCACTGTTCTCAAGCTGCTACTGAANNNAGATCGGAAG |
| 44773_3 | TATGAGTGTGGAGTCGTTGCACTGTTCTCAAGCTGCTACTGAANNNAGATCGGAAG |
| 30562_1 | ATGAGTGTGGAGTCGTTGCTACCAGCTGACTCCATTTGTTAATCTTNNNAGATCGGA |
| 30562_2 | ATGAGTGTGGAGTCGTTGCTACCAGCTGACTCCATTTGTTAATCTTNNNAGATCGGA |
| 30562_3 | ATGAGTGTGGAGTCGTTGCTACCAGCTGACTCCATTTGTTAATCTTNNNAGATCGGA |
| 29679_1 | TATGAGTGTGGAGTCGTTGCCTGCATCCATTGGCTTAGCTTTANNNAGATCGGAAG |
| 29679_2 | TATGAGTGTGGAGTCGTTGCCTGCATCCATTGGCTTAGCTTTANNNAGATCGGAAG |
| 29679_3 | TATGAGTGTGGAGTCGTTGCCTGCATCCATTGGCTTAGCTTTANNNAGATCGGAAG |
| 42085_1 | TATGAGTGTGGAGTCGTTGCCGAAGGACATGAACAGACACTTCNNNAGATCGGAA |
| 42085_2 | TATGAGTGTGGAGTCGTTGCCGAAGGACATGAACAGACACTTCNNNAGATCGGAA |
| 42085_3 | TATGAGTGTGGAGTCGTTGCCGAAGGACATGAACAGACACTTCNNNAGATCGGAA |
| 47913_1 | TATGAGTGTGGAGTCGTTGCCCATGCAAAAGAGTTAGTGGGTGNNNAGATCGGAA |
| 47913_2 | TATGAGTGTGGAGTCGTTGCCCATGCAAAAGAGTTAGTGGGTGNNNAGATCGGAA |
| 47913_3 | TATGAGTGTGGAGTCGTTGCCCATGCAAAAGAGTTAGTGGGTGNNNAGATCGGAA |
| 38686_1 | TATGAGTGTGGAGTCGTTGCGTCAAAGAAATGTCTGCACTCCCNNNAGATCGGAAG |
| 38686_2 | TATGAGTGTGGAGTCGTTGCGTCAAAGAAATGTCTGCACTCCCNNNAGATCGGAAG |
| 38686_3 | TATGAGTGTGGAGTCGTTGCGTCAAAGAAATGTCTGCACTCCCNNNAGATCGGAAG |

| | |
|---|---|
| 44015_1 | TATGAGTGTGGAGTCGTTGCCATCTACCTCTACCTCCAGGGATNNNAGATCGGAAGA |
| 44015_2 | TATGAGTGTGGAGTCGTTGCCATCTACCTCTACCTCCAGGGATNNNAGATCGGAAGA |
| 44015_3 | TATGAGTGTGGAGTCGTTGCCATCTACCTCTACCTCCAGGGATNNNAGATCGGAAGA |
| 690_1 | TATGAGTGTGGAGTCGTTGCTAATAGCATCTGTTCCCACCAGCNNNAGATCGGAAGA |
| 690_2 | TATGAGTGTGGAGTCGTTGCTAATAGCATCTGTTCCCACCAGCNNNAGATCGGAAGA |
| 690_3 | TATGAGTGTGGAGTCGTTGCTAATAGCATCTGTTCCCACCAGCNNNAGATCGGAAGA |
| 43036_1 | TATGAGTGTGGAGTCGTTGCCACATATATATGCGTGTGTGTGTGTNNNAGATCGGA |
| 43036_2 | TATGAGTGTGGAGTCGTTGCCACATATATATGCGTGTGTGTGTGTNNNAGATCGGA |
| 43036_3 | TATGAGTGTGGAGTCGTTGCCACATATATATGCGTGTGTGTGTGTNNNAGATCGGA |
| 51162_1 | TATGAGTGTGGAGTCGTTGCCCTGGGCAACAAGAATGAAACTCNNNAGATCGGAAG |
| 51162_2 | TATGAGTGTGGAGTCGTTGCCCTGGGCAACAAGAATGAAACTCNNNAGATCGGAAG |
| 51162_3 | TATGAGTGTGGAGTCGTTGCCCTGGGCAACAAGAATGAAACTCNNNAGATCGGAAG |
| 8491_1 | TATGAGTGTGGAGTCGTTGCTAGCTTACTCTGTTCTCTACTGCNNNAGATCGGAAGA |
| 8491_2 | TATGAGTGTGGAGTCGTTGCTAGCTTACTCTGTTCTCTACTGCNNNAGATCGGAAGA |
| 8491_3 | TATGAGTGTGGAGTCGTTGCTAGCTTACTCTGTTCTCTACTGCNNNAGATCGGAAGA |
| 30032_1 | TATGAGTGTGGAGTCGTTGCCCGAGACTGTGAAGGATTGTCATNNNAGATCGGAAG |
| 30032_2 | TATGAGTGTGGAGTCGTTGCCCGAGACTGTGAAGGATTGTCATNNNAGATCGGAAG |
| 30032_3 | TATGAGTGTGGAGTCGTTGCCCGAGACTGTGAAGGATTGTCATNNNAGATCGGAAG |
| 36463_1 | TATGAGTGTGGAGTCGTTGCTGCTTCTTTATGTCCATCCTTTGCNNNAGATCGGAAGA |
| 36463_2 | TATGAGTGTGGAGTCGTTGCTGCTTCTTTATGTCCATCCTTTGCNNNAGATCGGAAGA |
| 36463_3 | TATGAGTGTGGAGTCGTTGCTGCTTCTTTATGTCCATCCTTTGCNNNAGATCGGAAGA |
| 33465_1 | TATGAGTGTGGAGTCGTTGCACAGGACTGTGATCTCAATTTTTTNNNAGATCGGAAG |
| 33465_2 | TATGAGTGTGGAGTCGTTGCACAGGACTGTGATCTCAATTTTTTNNNAGATCGGAAG |
| 33465_3 | TATGAGTGTGGAGTCGTTGCACAGGACTGTGATCTCAATTTTTTNNNAGATCGGAAG |
| 42476_1 | TATGAGTGTGGAGTCGTTGCGCTCATGCCTGTAATCCCAAAACNNNAGATCGGAAGA |
| 42476_2 | TATGAGTGTGGAGTCGTTGCGCTCATGCCTGTAATCCCAAAACNNNAGATCGGAAGA |
| 42476_3 | TATGAGTGTGGAGTCGTTGCGCTCATGCCTGTAATCCCAAAACNNNAGATCGGAAGA |
| 36278_1 | TATGAGTGTGGAGTCGTTGCTCCATGTGCTATAAAGTGGGATGTNNNAGATCGGAA |
| 36278_2 | TATGAGTGTGGAGTCGTTGCTCCATGTGCTATAAAGTGGGATGTNNNAGATCGGAA |
| 36278_3 | TATGAGTGTGGAGTCGTTGCTCCATGTGCTATAAAGTGGGATGTNNNAGATCGGAA |
| 46200_1 | TATGAGTGTGGAGTCGTTGCCTTCGGAGTAGGTGGGATTACAGNNNAGATCGGAA |
| 46200_2 | TATGAGTGTGGAGTCGTTGCCTTCGGAGTAGGTGGGATTACAGNNNAGATCGGAA |
| 46200_3 | TATGAGTGTGGAGTCGTTGCCTTCGGAGTAGGTGGGATTACAGNNNAGATCGGAA |
| 45702_1 | TATGAGTGTGGAGTCGTTGCAAACAAAAGCATCCCACCAAACANNNAGATCGGAAG |
| 45702_2 | TATGAGTGTGGAGTCGTTGCAAACAAAAGCATCCCACCAAACANNNAGATCGGAAG |
| 45702_3 | TATGAGTGTGGAGTCGTTGCAAACAAAAGCATCCCACCAAACANNNAGATCGGAAG |
| 29470_1 | TATGAGTGTGGAGTCGTTGCTTTGCGGATATTAACCTCTACAGGTNNNAGATCGGAA |
| 29470_2 | TATGAGTGTGGAGTCGTTGCTTTGCGGATATTAACCTCTACAGGTNNNAGATCGGAA |
| 29470_3 | TATGAGTGTGGAGTCGTTGCTTTGCGGATATTAACCTCTACAGGTNNNAGATCGGAA |
| 49112_1 | TATGAGTGTGGAGTCGTTGCCCACCATGCCCAGCTAATTTTTANNNAGATCGGAAGA |
| 49112_2 | TATGAGTGTGGAGTCGTTGCCCACCATGCCCAGCTAATTTTTANNNAGATCGGAAGA |
| 49112_3 | TATGAGTGTGGAGTCGTTGCCCACCATGCCCAGCTAATTTTTANNNAGATCGGAAGA |
| 52519_1 | TATGAGTGTGGAGTCGTTGCGGTCATTTCAGCAGGAAAAGTCGNNNAGATCGGAAG |

| | |
|---|---|
| 52519_2 | TATGAGTGTGGAGTCGTTGCGGTCATTTCAGCAGGAAAAGTCGNNNAGATCGGAA |
| 52519_3 | TATGAGTGTGGAGTCGTTGCGGTCATTTCAGCAGGAAAAGTCGNNNAGATCGGAA |
| 40489_1 | TATGAGTGTGGAGTCGTTGCCTCCAGCATAGGTAACAGAGCAANNNAGATCGGAA |
| 40489_2 | TATGAGTGTGGAGTCGTTGCCTCCAGCATAGGTAACAGAGCAANNNAGATCGGAA |
| 40489_3 | TATGAGTGTGGAGTCGTTGCCTCCAGCATAGGTAACAGAGCAANNNAGATCGGAA |
| 39809_1 | TATGAGTGTGGAGTCGTTGCCATTCTCCTGCCTCAGCCTCNNNAGATCGGAAGAGCA |
| 39809_2 | TATGAGTGTGGAGTCGTTGCCATTCTCCTGCCTCAGCCTCNNNAGATCGGAAGAGCA |
| 39809_3 | TATGAGTGTGGAGTCGTTGCCATTCTCCTGCCTCAGCCTCNNNAGATCGGAAGAGCA |
| 39656_1 | TATGAGTGTGGAGTCGTTGCCCTGTTTTACAAGGTTGCAGGTTNNNAGATCGGAAG |
| 39656_2 | TATGAGTGTGGAGTCGTTGCCCTGTTTTACAAGGTTGCAGGTTNNNAGATCGGAAG |
| 39656_3 | TATGAGTGTGGAGTCGTTGCCCTGTTTTACAAGGTTGCAGGTTNNNAGATCGGAAG |
| 53368_1 | TATGAGTGTGGAGTCGTTGCATCAATGGGGAAGGCATAGACTTNNNAGATCGGAA |
| 53368_2 | TATGAGTGTGGAGTCGTTGCATCAATGGGGAAGGCATAGACTTNNNAGATCGGAA |
| 53368_3 | TATGAGTGTGGAGTCGTTGCATCAATGGGGAAGGCATAGACTTNNNAGATCGGAA |
| 29451_1 | TATGAGTGTGGAGTCGTTGCTCGTTTAGGTTTCTGTTGATGCTNNNAGATCGGAAG |
| 29451_2 | TATGAGTGTGGAGTCGTTGCTCGTTTAGGTTTCTGTTGATGCTNNNAGATCGGAAG |
| 29451_3 | TATGAGTGTGGAGTCGTTGCTCGTTTAGGTTTCTGTTGATGCTNNNAGATCGGAAG |
| 40067_1 | TATGAGTGTGGAGTCGTTGCGGGTGGGGAGAAATAATTGAGGTNNNAGATCGGA |
| 40067_2 | TATGAGTGTGGAGTCGTTGCGGGTGGGGAGAAATAATTGAGGTNNNAGATCGGA |
| 40067_3 | TATGAGTGTGGAGTCGTTGCGGGTGGGGAGAAATAATTGAGGTNNNAGATCGGA |
| 45099_1 | TATGAGTGTGGAGTCGTTGCGTGAGGTGCAGAGGAGAATTACANNNAGATCGGA |
| 45099_2 | TATGAGTGTGGAGTCGTTGCGTGAGGTGCAGAGGAGAATTACANNNAGATCGGA |
| 45099_3 | TATGAGTGTGGAGTCGTTGCGTGAGGTGCAGAGGAGAATTACANNNAGATCGGA |
| 10283_1 | TATGAGTGTGGAGTCGTTGCATCCACTGGGAGACACTTTGAGANNNAGATCGGAA |
| 10283_2 | TATGAGTGTGGAGTCGTTGCATCCACTGGGAGACACTTTGAGANNNAGATCGGAA |
| 10283_3 | TATGAGTGTGGAGTCGTTGCATCCACTGGGAGACACTTTGAGANNNAGATCGGAA |
| 53282_1 | TATGAGTGTGGAGTCGTTGCTGACAATAACAAGCCCTGGTTCTNNNAGATCGGAAG |
| 53282_2 | TATGAGTGTGGAGTCGTTGCTGACAATAACAAGCCCTGGTTCTNNNAGATCGGAAG |
| 53282_3 | TATGAGTGTGGAGTCGTTGCTGACAATAACAAGCCCTGGTTCTNNNAGATCGGAAG |
| 48038_1 | TATGAGTGTGGAGTCGTTGCAGAGAAGGGGAGACTGAGAGAATNNNAGATCGGA |
| 48038_2 | TATGAGTGTGGAGTCGTTGCAGAGAAGGGGAGACTGAGAGAATNNNAGATCGGA |
| 48038_3 | TATGAGTGTGGAGTCGTTGCAGAGAAGGGGAGACTGAGAGAATNNNAGATCGGA |
| 9237_1 | TATGAGTGTGGAGTCGTTGCATGAGAGCACAAGAACGTGAGTGNNNAGATCGGA |
| 9237_2 | TATGAGTGTGGAGTCGTTGCATGAGAGCACAAGAACGTGAGTGNNNAGATCGGA |
| 9237_3 | TATGAGTGTGGAGTCGTTGCATGAGAGCACAAGAACGTGAGTGNNNAGATCGGA |
| 36273_1 | TATGAGTGTGGAGTCGTTGCTGAAAAGATGTTGCCCTACTGGANNNAGATCGGAA |
| 36273_2 | TATGAGTGTGGAGTCGTTGCTGAAAAGATGTTGCCCTACTGGANNNAGATCGGAA |
| 36273_3 | TATGAGTGTGGAGTCGTTGCTGAAAAGATGTTGCCCTACTGGANNNAGATCGGAA |
| 45657_1 | TATGAGTGTGGAGTCGTTGCTTCACACCATTCTTCTGCCTCAGNNNAGATCGGAAGA |
| 45657_2 | TATGAGTGTGGAGTCGTTGCTTCACACCATTCTTCTGCCTCAGNNNAGATCGGAAGA |
| 45657_3 | TATGAGTGTGGAGTCGTTGCTTCACACCATTCTTCTGCCTCAGNNNAGATCGGAAGA |
| 13181_1 | TATGAGTGTGGAGTCGTTGCGCTGTTATTATTAAGGTAAGAGCNNNAGATCGGAA |
| 13181_2 | TATGAGTGTGGAGTCGTTGCGCTGTTATTATTAAGGTAAGAGCNNNAGATCGGAA |

| | |
|---|---|
| 13181_3 | TATGAGTGTGGAGTCGTTGCGCTGTTATTATTAAGGTAAGAGCNNNAGATCGGAA |
| 49820_1 | TATGAGTGTGGAGTCGTTGCTGGCAATGACAGAACGAGTGATANNNAGATCGGAA |
| 49820_2 | TATGAGTGTGGAGTCGTTGCTGGCAATGACAGAACGAGTGATANNNAGATCGGAA |
| 49820_3 | TATGAGTGTGGAGTCGTTGCTGGCAATGACAGAACGAGTGATANNNAGATCGGAA |
| 40325_1 | TATGAGTGTGGAGTCGTTGCAATCAGAATATCTTGGGGGTGGGNNNAGATCGGAA |
| 40325_2 | TATGAGTGTGGAGTCGTTGCAATCAGAATATCTTGGGGGTGGGNNNAGATCGGAA |
| 40325_3 | TATGAGTGTGGAGTCGTTGCAATCAGAATATCTTGGGGGTGGGNNNAGATCGGAA |
| 10346_1 | TATGAGTGTGGAGTCGTTGCAAATTCCTGCAGCTAGATGGCTCNNNAGATCGGAAG |
| 10346_2 | TATGAGTGTGGAGTCGTTGCAAATTCCTGCAGCTAGATGGCTCNNNAGATCGGAAG |
| 10346_3 | TATGAGTGTGGAGTCGTTGCAAATTCCTGCAGCTAGATGGCTCNNNAGATCGGAAG |
| 34463_1 | TATGAGTGTGGAGTCGTTGCCCACGTGATCCCATTGTTCAATTNNNAGATCGGAAG |
| 34463_2 | TATGAGTGTGGAGTCGTTGCCCACGTGATCCCATTGTTCAATTNNNAGATCGGAAG |
| 34463_3 | TATGAGTGTGGAGTCGTTGCCCACGTGATCCCATTGTTCAATTNNNAGATCGGAAG |
| 38810_1 | TATGAGTGTGGAGTCGTTGCTCTGGGTTCCAATGATTCTCCTGNNNAGATCGGAAG |
| 38810_2 | TATGAGTGTGGAGTCGTTGCTCTGGGTTCCAATGATTCTCCTGNNNAGATCGGAAG |
| 38810_3 | TATGAGTGTGGAGTCGTTGCTCTGGGTTCCAATGATTCTCCTGNNNAGATCGGAAG |
| 46764_1 | TATGAGTGTGGAGTCGTTGCCCTCCTAAGTAGCTGGGACTACANNNAGATCGGAAG |
| 46764_2 | TATGAGTGTGGAGTCGTTGCCCTCCTAAGTAGCTGGGACTACANNNAGATCGGAAG |
| 46764_3 | TATGAGTGTGGAGTCGTTGCCCTCCTAAGTAGCTGGGACTACANNNAGATCGGAAG |
| 53111_1 | TATGAGTGTGGAGTCGTTGCTGCAAAGGGATATTTAGGAGCCCNNNAGATCGGAA |
| 53111_2 | TATGAGTGTGGAGTCGTTGCTGCAAAGGGATATTTAGGAGCCCNNNAGATCGGAA |
| 53111_3 | TATGAGTGTGGAGTCGTTGCTGCAAAGGGATATTTAGGAGCCCNNNAGATCGGAA |
| 38239_1 | TATGAGTGTGGAGTCGTTGCATAAAGGACTCATTGCCTCAGCTNNNAGATCGGAAG |
| 38239_2 | TATGAGTGTGGAGTCGTTGCATAAAGGACTCATTGCCTCAGCTNNNAGATCGGAAG |
| 38239_3 | TATGAGTGTGGAGTCGTTGCATAAAGGACTCATTGCCTCAGCTNNNAGATCGGAAG |
| 30126_1 | TATGAGTGTGGAGTCGTTGCACTCAGTTTAATCCAGGCGGAATNNNAGATCGGAAG |
| 30126_2 | TATGAGTGTGGAGTCGTTGCACTCAGTTTAATCCAGGCGGAATNNNAGATCGGAAG |
| 30126_3 | TATGAGTGTGGAGTCGTTGCACTCAGTTTAATCCAGGCGGAATNNNAGATCGGAAG |
| 30162_1 | TATGAGTGTGGAGTCGTTGCGTGTGAATACGATGGATCCCAGANNNAGATCGGAA |
| 30162_2 | TATGAGTGTGGAGTCGTTGCGTGTGAATACGATGGATCCCAGANNNAGATCGGAA |
| 30162_3 | TATGAGTGTGGAGTCGTTGCGTGTGAATACGATGGATCCCAGANNNAGATCGGAA |
| 831_1 | TATGAGTGTGGAGTCGTTGCCGGTGCAAGGTGATGGGATCATNNNAGATCGGAAG |
| 831_2 | TATGAGTGTGGAGTCGTTGCCGGTGCAAGGTGATGGGATCATNNNAGATCGGAAG |
| 831_3 | TATGAGTGTGGAGTCGTTGCCGGTGCAAGGTGATGGGATCATNNNAGATCGGAAG |
| 38223_1 | TATGAGTGTGGAGTCGTTGCGACCAGACAGATTCACAGCTGAANNNAGATCGGAA |
| 38223_2 | TATGAGTGTGGAGTCGTTGCGACCAGACAGATTCACAGCTGAANNNAGATCGGAA |
| 38223_3 | TATGAGTGTGGAGTCGTTGCGACCAGACAGATTCACAGCTGAANNNAGATCGGAA |
| 12561_1 | TATGAGTGTGGAGTCGTTGCAACATGTTAGTGTTAAGGAAAATNNNAGATCGGAA |
| 12561_2 | TATGAGTGTGGAGTCGTTGCAACATGTTAGTGTTAAGGAAAATNNNAGATCGGAA |
| 12561_3 | TATGAGTGTGGAGTCGTTGCAACATGTTAGTGTTAAGGAAAATNNNAGATCGGAA |
| 31597_1 | TATGAGTGTGGAGTCGTTGCCACACACACACACACACATACTCNNNAGATCGGAAGA |
| 31597_2 | TATGAGTGTGGAGTCGTTGCCACACACACACACACACATACTCNNNAGATCGGAAGA |
| 31597_3 | TATGAGTGTGGAGTCGTTGCCACACACACACACACACATACTCNNNAGATCGGAAGA |

| | |
|---|---|
| 13269_1 | TATGAGTGTGGAGTCGTTGCAAGAAGAAGGGGGAAAATTAAGNNNAGATCGGAA |
| 13269_2 | TATGAGTGTGGAGTCGTTGCAAGAAGAAGGGGGAAAATTAAGNNNAGATCGGAA |
| 13269_3 | TATGAGTGTGGAGTCGTTGCAAGAAGAAGGGGGAAAATTAAGNNNAGATCGGAA |
| 38414_1 | TATGAGTGTGGAGTCGTTGCATCCTAGCTACTCAGAAGGCTGANNNAGATCGGAAG |
| 38414_2 | TATGAGTGTGGAGTCGTTGCATCCTAGCTACTCAGAAGGCTGANNNAGATCGGAAG |
| 38414_3 | TATGAGTGTGGAGTCGTTGCATCCTAGCTACTCAGAAGGCTGANNNAGATCGGAAG |
| 12804_1 | TATGAGTGTGGAGTCGTTGCATGAAGGTTCTAAGTAGAATTGANNNAGATCGGAA |
| 12804_2 | TATGAGTGTGGAGTCGTTGCATGAAGGTTCTAAGTAGAATTGANNNAGATCGGAA |
| 12804_3 | TATGAGTGTGGAGTCGTTGCATGAAGGTTCTAAGTAGAATTGANNNAGATCGGAA |
| 29495_1 | ATGAGTGTGGAGTCGTTGCCAGAAATGGGAGAGAAATCAAAGAAGNNNAGATCG |
| 29495_2 | ATGAGTGTGGAGTCGTTGCCAGAAATGGGAGAGAAATCAAAGAAGNNNAGATCG |
| 29495_3 | ATGAGTGTGGAGTCGTTGCCAGAAATGGGAGAGAAATCAAAGAAGNNNAGATCG |
| 49417_1 | TATGAGTGTGGAGTCGTTGCCTTATCCCTCAACACCTCTCACCNNNAGATCGGAAGA |
| 49417_2 | TATGAGTGTGGAGTCGTTGCCTTATCCCTCAACACCTCTCACCNNNAGATCGGAAGA |
| 49417_3 | TATGAGTGTGGAGTCGTTGCCTTATCCCTCAACACCTCTCACCNNNAGATCGGAAGA |
| 46557_1 | TATGAGTGTGGAGTCGTTGCCGAGTAGCTGGGATTACAGACATNNNAGATCGGAA |
| 46557_2 | TATGAGTGTGGAGTCGTTGCCGAGTAGCTGGGATTACAGACATNNNAGATCGGAA |
| 46557_3 | TATGAGTGTGGAGTCGTTGCCGAGTAGCTGGGATTACAGACATNNNAGATCGGAA |
| 29941_1 | TATGAGTGTGGAGTCGTTGCATCCTTGAGATAAGGTATGTGTGTNNNAGATCGGA |
| 29941_2 | TATGAGTGTGGAGTCGTTGCATCCTTGAGATAAGGTATGTGTGTNNNAGATCGGA |
| 29941_3 | TATGAGTGTGGAGTCGTTGCATCCTTGAGATAAGGTATGTGTGTNNNAGATCGGA |
| 45288_1 | TATGAGTGTGGAGTCGTTGCCACTTGGAAGAGGACTAACTGCTNNNAGATCGGAA |
| 45288_2 | TATGAGTGTGGAGTCGTTGCCACTTGGAAGAGGACTAACTGCTNNNAGATCGGAA |
| 45288_3 | TATGAGTGTGGAGTCGTTGCCACTTGGAAGAGGACTAACTGCTNNNAGATCGGAA |
| 6934_1 | TATGAGTGTGGAGTCGTTGCGTTCTGCTTTCAGGCTCTTTCAGNNNAGATCGGAAG |
| 6934_2 | TATGAGTGTGGAGTCGTTGCGTTCTGCTTTCAGGCTCTTTCAGNNNAGATCGGAAG |
| 6934_3 | TATGAGTGTGGAGTCGTTGCGTTCTGCTTTCAGGCTCTTTCAGNNNAGATCGGAAG |
| 47947_1 | TATGAGTGTGGAGTCGTTGCCTCGATCTCCTGACCTTGTGATCNNNAGATCGGAAG |
| 47947_2 | TATGAGTGTGGAGTCGTTGCCTCGATCTCCTGACCTTGTGATCNNNAGATCGGAAG |
| 47947_3 | TATGAGTGTGGAGTCGTTGCCTCGATCTCCTGACCTTGTGATCNNNAGATCGGAAG |
| 37894_1 | TATGAGTGTGGAGTCGTTGCTTCCCTATGGTATGTAAGCCGTGNNNAGATCGGAAG |
| 37894_2 | TATGAGTGTGGAGTCGTTGCTTCCCTATGGTATGTAAGCCGTGNNNAGATCGGAAG |
| 37894_3 | TATGAGTGTGGAGTCGTTGCTTCCCTATGGTATGTAAGCCGTGNNNAGATCGGAAG |
| 30627_1 | TATGAGTGTGGAGTCGTTGCGACAGGTGGAAGAAGACCAAGGGNNNAGATCGGA |
| 30627_2 | TATGAGTGTGGAGTCGTTGCGACAGGTGGAAGAAGACCAAGGGNNNAGATCGGA |
| 30627_3 | TATGAGTGTGGAGTCGTTGCGACAGGTGGAAGAAGACCAAGGGNNNAGATCGGA |
| 34380_1 | TATGAGTGTGGAGTCGTTGCAGTTATGGCTTTGGCTGAGAGAGNNNAGATCGGAA |
| 34380_2 | TATGAGTGTGGAGTCGTTGCAGTTATGGCTTTGGCTGAGAGAGNNNAGATCGGAA |
| 34380_3 | TATGAGTGTGGAGTCGTTGCAGTTATGGCTTTGGCTGAGAGAGNNNAGATCGGAA |
| 37193_1 | TATGAGTGTGGAGTCGTTGCTGGACACACATCGTTAGAAGACANNNAGATCGGAA |
| 37193_2 | TATGAGTGTGGAGTCGTTGCTGGACACACATCGTTAGAAGACANNNAGATCGGAA |
| 37193_3 | TATGAGTGTGGAGTCGTTGCTGGACACACATCGTTAGAAGACANNNAGATCGGAA |
| 37541_1 | TATGAGTGTGGAGTCGTTGCGTGAGCTCAAAGATGGTCATTTGANNNAGATCGGA |

| | |
|---|---|
| 37541_2 | TATGAGTGTGGAGTCGTTGCGTGAGCTCAAAGATGGTCATTTGANNNAGATCGGA |
| 37541_3 | TATGAGTGTGGAGTCGTTGCGTGAGCTCAAAGATGGTCATTTGANNNAGATCGGA |
| 36279_1 | TATGAGTGTGGAGTCGTTGCTTGTACCTTCCTCTTCAGAAGCCNNNAGATCGGAAG |
| 36279_2 | TATGAGTGTGGAGTCGTTGCTTGTACCTTCCTCTTCAGAAGCCNNNAGATCGGAAG |
| 36279_3 | TATGAGTGTGGAGTCGTTGCTTGTACCTTCCTCTTCAGAAGCCNNNAGATCGGAAG |
| 13609_1 | TATGAGTGTGGAGTCGTTGCAGACAAGGAAACTAATGATAGAANNNAGATCGGAA |
| 13609_2 | TATGAGTGTGGAGTCGTTGCAGACAAGGAAACTAATGATAGAANNNAGATCGGAA |
| 13609_3 | TATGAGTGTGGAGTCGTTGCAGACAAGGAAACTAATGATAGAANNNAGATCGGAA |
| 35851_1 | TATGAGTGTGGAGTCGTTGCCCATGATGAGACTATCAGCACCANNNAGATCGGAAG |
| 35851_2 | TATGAGTGTGGAGTCGTTGCCCATGATGAGACTATCAGCACCANNNAGATCGGAAG |
| 35851_3 | TATGAGTGTGGAGTCGTTGCCCATGATGAGACTATCAGCACCANNNAGATCGGAAG |
| 35040_1 | TATGAGTGTGGAGTCGTTGCTATCACAGATCACTGCAGTCTCCNNNAGATCGGAAG |
| 35040_2 | TATGAGTGTGGAGTCGTTGCTATCACAGATCACTGCAGTCTCCNNNAGATCGGAAG |
| 35040_3 | TATGAGTGTGGAGTCGTTGCTATCACAGATCACTGCAGTCTCCNNNAGATCGGAAG |
| 45924_1 | TATGAGTGTGGAGTCGTTGCTTCTGTTGAGAAGGTAGCCAACANNNAGATCGGAA |
| 45924_2 | TATGAGTGTGGAGTCGTTGCTTCTGTTGAGAAGGTAGCCAACANNNAGATCGGAA |
| 45924_3 | TATGAGTGTGGAGTCGTTGCTTCTGTTGAGAAGGTAGCCAACANNNAGATCGGAA |
| 13107_1 | TATGAGTGTGGAGTCGTTGCTAAGCAAATGTCATATTACCTTGNNNAGATCGGAAG |
| 13107_2 | TATGAGTGTGGAGTCGTTGCTAAGCAAATGTCATATTACCTTGNNNAGATCGGAAG |
| 13107_3 | TATGAGTGTGGAGTCGTTGCTAAGCAAATGTCATATTACCTTGNNNAGATCGGAAG |
| 43985_1 | TATGAGTGTGGAGTCGTTGCTGTAGCTAGGGATTGCCATGATCNNNAGATCGGAA |
| 43985_2 | TATGAGTGTGGAGTCGTTGCTGTAGCTAGGGATTGCCATGATCNNNAGATCGGAA |
| 43985_3 | TATGAGTGTGGAGTCGTTGCTGTAGCTAGGGATTGCCATGATCNNNAGATCGGAA |
| 43117_1 | TATGAGTGTGGAGTCGTTGCAGAACAGACATCAGCATTAGTGANNNAGATCGGAA |
| 43117_2 | TATGAGTGTGGAGTCGTTGCAGAACAGACATCAGCATTAGTGANNNAGATCGGAA |
| 43117_3 | TATGAGTGTGGAGTCGTTGCAGAACAGACATCAGCATTAGTGANNNAGATCGGAA |
| 39007_1 | TATGAGTGTGGAGTCGTTGCACGTGCCTGTAATCTCAGCTATTNNNAGATCGGAAG |
| 39007_2 | TATGAGTGTGGAGTCGTTGCACGTGCCTGTAATCTCAGCTATTNNNAGATCGGAAG |
| 39007_3 | TATGAGTGTGGAGTCGTTGCACGTGCCTGTAATCTCAGCTATTNNNAGATCGGAAG |
| 33779_1 | TATGAGTGTGGAGTCGTTGCATTTGCTTCTGTTTGGTGTGTGGNNNAGATCGGAAG |
| 33779_2 | TATGAGTGTGGAGTCGTTGCATTTGCTTCTGTTTGGTGTGTGGNNNAGATCGGAAG |
| 33779_3 | TATGAGTGTGGAGTCGTTGCATTTGCTTCTGTTTGGTGTGTGGNNNAGATCGGAAG |
| 32493_1 | TATGAGTGTGGAGTCGTTGCTCTCGATCTGACCTTGTGATCTGNNNAGATCGGAAG |
| 32493_2 | TATGAGTGTGGAGTCGTTGCTCTCGATCTGACCTTGTGATCTGNNNAGATCGGAAG |
| 32493_3 | TATGAGTGTGGAGTCGTTGCTCTCGATCTGACCTTGTGATCTGNNNAGATCGGAAG |
| 6159_1 | TATGAGTGTGGAGTCGTTGCTAGTCCTTTCAATACTGCAGCCCNNNAGATCGGAAG |
| 6159_2 | TATGAGTGTGGAGTCGTTGCTAGTCCTTTCAATACTGCAGCCCNNNAGATCGGAAG |
| 6159_3 | TATGAGTGTGGAGTCGTTGCTAGTCCTTTCAATACTGCAGCCCNNNAGATCGGAAG |
| 36784_1 | TATGAGTGTGGAGTCGTTGCATGAGCTGCTTATTCCTCAGAGGNNNAGATCGGAAG |
| 36784_2 | TATGAGTGTGGAGTCGTTGCATGAGCTGCTTATTCCTCAGAGGNNNAGATCGGAAG |
| 36784_3 | TATGAGTGTGGAGTCGTTGCATGAGCTGCTTATTCCTCAGAGGNNNAGATCGGAAG |
| 46415_1 | TATGAGTGTGGAGTCGTTGCAGTGCTGGGATTACAGACATGAGNNNAGATCGGAA |
| 46415_2 | TATGAGTGTGGAGTCGTTGCAGTGCTGGGATTACAGACATGAGNNNAGATCGGAA |

| | |
|---|---|
| 46415_3 | TATGAGTGTGGAGTCGTTGCAGTGCTGGGATTACAGACATGAGNNNAGATCGGAA |
| 34155_1 | TATGAGTGTGGAGTCGTTGCCACACCTGTAATCCCAGCACTTTNNNAGATCGGAAG |
| 34155_2 | TATGAGTGTGGAGTCGTTGCCACACCTGTAATCCCAGCACTTTNNNAGATCGGAAG |
| 34155_3 | TATGAGTGTGGAGTCGTTGCCACACCTGTAATCCCAGCACTTTNNNAGATCGGAAG |
| 38597_1 | TATGAGTGTGGAGTCGTTGCTCCTTCAGGCTGAAATGAAAGGANNNAGATCGGAA |
| 38597_2 | TATGAGTGTGGAGTCGTTGCTCCTTCAGGCTGAAATGAAAGGANNNAGATCGGAA |
| 38597_3 | TATGAGTGTGGAGTCGTTGCTCCTTCAGGCTGAAATGAAAGGANNNAGATCGGAA |
| 51117_1 | TATGAGTGTGGAGTCGTTGCGGGCTCCTCACTTCCCAGTANNNAGATCGGAAGAGC |
| 51117_2 | TATGAGTGTGGAGTCGTTGCGGGCTCCTCACTTCCCAGTANNNAGATCGGAAGAGC |
| 51117_3 | TATGAGTGTGGAGTCGTTGCGGGCTCCTCACTTCCCAGTANNNAGATCGGAAGAGC |
| 48276_1 | TATGAGTGTGGAGTCGTTGCATCTTGGCTCACTGTAACCTCTGNNNAGATCGGAAG |
| 48276_2 | TATGAGTGTGGAGTCGTTGCATCTTGGCTCACTGTAACCTCTGNNNAGATCGGAAG |
| 48276_3 | TATGAGTGTGGAGTCGTTGCATCTTGGCTCACTGTAACCTCTGNNNAGATCGGAAG |
| 45483_1 | TATGAGTGTGGAGTCGTTGCTAATTGAGACCAGCATCCCACAANNNAGATCGGAAG |
| 45483_2 | TATGAGTGTGGAGTCGTTGCTAATTGAGACCAGCATCCCACAANNNAGATCGGAAG |
| 45483_3 | TATGAGTGTGGAGTCGTTGCTAATTGAGACCAGCATCCCACAANNNAGATCGGAAG |
| 49589_1 | TATGAGTGTGGAGTCGTTGCTGTCGCCAGCTTCACATATTTCTNNNAGATCGGAAG |
| 49589_2 | TATGAGTGTGGAGTCGTTGCTGTCGCCAGCTTCACATATTTCTNNNAGATCGGAAG |
| 49589_3 | TATGAGTGTGGAGTCGTTGCTGTCGCCAGCTTCACATATTTCTNNNAGATCGGAAG |
| 45145_1 | TATGAGTGTGGAGTCGTTGCAGACCTACGACGAGAAGGGAATANNNAGATCGGAA |
| 45145_2 | TATGAGTGTGGAGTCGTTGCAGACCTACGACGAGAAGGGAATANNNAGATCGGAA |
| 45145_3 | TATGAGTGTGGAGTCGTTGCAGACCTACGACGAGAAGGGAATANNNAGATCGGAA |
| 47149_1 | TATGAGTGTGGAGTCGTTGCTTCAAGTGATTCTCCTGCCTCAGNNNAGATCGGAAG |
| 47149_2 | TATGAGTGTGGAGTCGTTGCTTCAAGTGATTCTCCTGCCTCAGNNNAGATCGGAAG |
| 47149_3 | TATGAGTGTGGAGTCGTTGCTTCAAGTGATTCTCCTGCCTCAGNNNAGATCGGAAG |
| 38327_1 | TATGAGTGTGGAGTCGTTGCTGATGTAGATCAGGTGGCGTATGNNNAGATCGGAA |
| 38327_2 | TATGAGTGTGGAGTCGTTGCTGATGTAGATCAGGTGGCGTATGNNNAGATCGGAA |
| 38327_3 | TATGAGTGTGGAGTCGTTGCTGATGTAGATCAGGTGGCGTATGNNNAGATCGGAA |
| 37373_1 | TATGAGTGTGGAGTCGTTGCAGCAGATGAGCTCAGATCATGAGNNNAGATCGGAA |
| 37373_2 | TATGAGTGTGGAGTCGTTGCAGCAGATGAGCTCAGATCATGAGNNNAGATCGGAA |
| 37373_3 | TATGAGTGTGGAGTCGTTGCAGCAGATGAGCTCAGATCATGAGNNNAGATCGGAA |
| 51045_1 | TATGAGTGTGGAGTCGTTGCGCTTTCCTTTTCACTTGGCTCTTNNNAGATCGGAAGA |
| 51045_2 | TATGAGTGTGGAGTCGTTGCGCTTTCCTTTTCACTTGGCTCTTNNNAGATCGGAAGA |
| 51045_3 | TATGAGTGTGGAGTCGTTGCGCTTTCCTTTTCACTTGGCTCTTNNNAGATCGGAAGA |
| 53171_1 | TATGAGTGTGGAGTCGTTGCATCATTGAGTCGCCTGAATCAGTNNNAGATCGGAAG |
| 53171_2 | TATGAGTGTGGAGTCGTTGCATCATTGAGTCGCCTGAATCAGTNNNAGATCGGAAG |
| 53171_3 | TATGAGTGTGGAGTCGTTGCATCATTGAGTCGCCTGAATCAGTNNNAGATCGGAAG |
| 43999_1 | TATGAGTGTGGAGTCGTTGCGAAGGAAGGAAGGAAGGAAGGAANNNAGATCGG |
| 43999_2 | TATGAGTGTGGAGTCGTTGCGAAGGAAGGAAGGAAGGAAGGAANNNAGATCGG |
| 43999_3 | TATGAGTGTGGAGTCGTTGCGAAGGAAGGAAGGAAGGAAGGAANNNAGATCGG |
| 49659_1 | TATGAGTGTGGAGTCGTTGCGTCTTGAACTCCTGACCTTGTGANNNAGATCGGAAG |
| 49659_2 | TATGAGTGTGGAGTCGTTGCGTCTTGAACTCCTGACCTTGTGANNNAGATCGGAAG |
| 49659_3 | TATGAGTGTGGAGTCGTTGCGTCTTGAACTCCTGACCTTGTGANNNAGATCGGAAG |

| | |
|---|---|
| 36470_1 | TATGAGTGTGGAGTCGTTGCAGAGTGAATTTCAAGGTGGTGGANNNAGATCGGAA |
| 36470_2 | TATGAGTGTGGAGTCGTTGCAGAGTGAATTTCAAGGTGGTGGANNNAGATCGGAA |
| 36470_3 | TATGAGTGTGGAGTCGTTGCAGAGTGAATTTCAAGGTGGTGGANNNAGATCGGAA |
| 49761_1 | TATGAGTGTGGAGTCGTTGCGAAGGCCGTGTGTGTAAAATGAANNNAGATCGGAA |
| 49761_2 | TATGAGTGTGGAGTCGTTGCGAAGGCCGTGTGTGTAAAATGAANNNAGATCGGAA |
| 49761_3 | TATGAGTGTGGAGTCGTTGCGAAGGCCGTGTGTGTAAAATGAANNNAGATCGGAA |
| 40836_1 | TATGAGTGTGGAGTCGTTGCGATCACACCACTATACTCCAGCCNNNAGATCGGAAGA |
| 40836_2 | TATGAGTGTGGAGTCGTTGCGATCACACCACTATACTCCAGCCNNNAGATCGGAAGA |
| 40836_3 | TATGAGTGTGGAGTCGTTGCGATCACACCACTATACTCCAGCCNNNAGATCGGAAGA |
| 45424_1 | TATGAGTGTGGAGTCGTTGCAAGAAAGCTTGCATGAGGTGAACNNNAGATCGGAA |
| 45424_2 | TATGAGTGTGGAGTCGTTGCAAGAAAGCTTGCATGAGGTGAACNNNAGATCGGAA |
| 45424_3 | TATGAGTGTGGAGTCGTTGCAAGAAAGCTTGCATGAGGTGAACNNNAGATCGGAA |
| 29837_1 | TATGAGTGTGGAGTCGTTGCTCACAAAGGGACCATCAAGGAATNNNAGATCGGAA |
| 29837_2 | TATGAGTGTGGAGTCGTTGCTCACAAAGGGACCATCAAGGAATNNNAGATCGGAA |
| 29837_3 | TATGAGTGTGGAGTCGTTGCTCACAAAGGGACCATCAAGGAATNNNAGATCGGAA |
| 47115_1 | TATGAGTGTGGAGTCGTTGCATGGAGTCTTGTTCTTGTCACCCNNNAGATCGGAAGA |
| 47115_2 | TATGAGTGTGGAGTCGTTGCATGGAGTCTTGTTCTTGTCACCCNNNAGATCGGAAGA |
| 47115_3 | TATGAGTGTGGAGTCGTTGCATGGAGTCTTGTTCTTGTCACCCNNNAGATCGGAAGA |
| 48106_1 | TATGAGTGTGGAGTCGTTGCACTGTTGTTGGTTTCAAGTCTGTNNNAGATCGGAAG |
| 48106_2 | TATGAGTGTGGAGTCGTTGCACTGTTGTTGGTTTCAAGTCTGTNNNAGATCGGAAG |
| 48106_3 | TATGAGTGTGGAGTCGTTGCACTGTTGTTGGTTTCAAGTCTGTNNNAGATCGGAAG |
| 46291_1 | TATGAGTGTGGAGTCGTTGCTCCCCAGTAGGTGAGATTACAGGNNNAGATCGGAA |
| 46291_2 | TATGAGTGTGGAGTCGTTGCTCCCCAGTAGGTGAGATTACAGGNNNAGATCGGAA |
| 46291_3 | TATGAGTGTGGAGTCGTTGCTCCCCAGTAGGTGAGATTACAGGNNNAGATCGGAA |
| 38735_1 | TATGAGTGTGGAGTCGTTGCCCATTTGACCCTTGGCTGAAATTNNNAGATCGGAAGA |
| 38735_2 | TATGAGTGTGGAGTCGTTGCCCATTTGACCCTTGGCTGAAATTNNNAGATCGGAAGA |
| 38735_3 | TATGAGTGTGGAGTCGTTGCCCATTTGACCCTTGGCTGAAATTNNNAGATCGGAAGA |
| 12217_1 | TATGAGTGTGGAGTCGTTGCGCTGCATGTTCATTGATGCATTGNNNAGATCGGAAG |
| 12217_2 | TATGAGTGTGGAGTCGTTGCGCTGCATGTTCATTGATGCATTGNNNAGATCGGAAG |
| 12217_3 | TATGAGTGTGGAGTCGTTGCGCTGCATGTTCATTGATGCATTGNNNAGATCGGAAG |
| 50176_1 | TATGAGTGTGGAGTCGTTGCATGATTCCATTTCCTGAGCCTGTNNNAGATCGGAAGA |
| 50176_2 | TATGAGTGTGGAGTCGTTGCATGATTCCATTTCCTGAGCCTGTNNNAGATCGGAAGA |
| 50176_3 | TATGAGTGTGGAGTCGTTGCATGATTCCATTTCCTGAGCCTGTNNNAGATCGGAAGA |
| 41618_1 | TATGAGTGTGGAGTCGTTGCTTCAAGTGATTCTCCTGCCTCAGNNNAGATCGGAAGA |
| 41618_2 | TATGAGTGTGGAGTCGTTGCTTCAAGTGATTCTCCTGCCTCAGNNNAGATCGGAAGA |
| 41618_3 | TATGAGTGTGGAGTCGTTGCTTCAAGTGATTCTCCTGCCTCAGNNNAGATCGGAAGA |
| 36327_1 | TATGAGTGTGGAGTCGTTGCCCCAGCTAACTAAGTGCACAGTANNNAGATCGGAAG |
| 36327_2 | TATGAGTGTGGAGTCGTTGCCCCAGCTAACTAAGTGCACAGTANNNAGATCGGAAG |
| 36327_3 | TATGAGTGTGGAGTCGTTGCCCCAGCTAACTAAGTGCACAGTANNNAGATCGGAAG |
| 48814_1 | TATGAGTGTGGAGTCGTTGCTAGCTGGGATTACAGGCATGTACNNNAGATCGGAA |
| 48814_2 | TATGAGTGTGGAGTCGTTGCTAGCTGGGATTACAGGCATGTACNNNAGATCGGAA |
| 48814_3 | TATGAGTGTGGAGTCGTTGCTAGCTGGGATTACAGGCATGTACNNNAGATCGGAA |
| 52248_1 | TATGAGTGTGGAGTCGTTGCACCATGTAAGAACTGCCTTTTGCNNNAGATCGGAAG |

| | |
|---|---|
| 52248_2 | TATGAGTGTGGAGTCGTTGCACCATGTAAGAACTGCCTTTTGCNNNAGATCGGAAG |
| 52248_3 | TATGAGTGTGGAGTCGTTGCACCATGTAAGAACTGCCTTTTGCNNNAGATCGGAAG |
| 13422_1 | TATGAGTGTGGAGTCGTTGCATACATGTGTAGATGGATTAGGANNNAGATCGGAA |
| 13422_2 | TATGAGTGTGGAGTCGTTGCATACATGTGTAGATGGATTAGGANNNAGATCGGAA |
| 13422_3 | TATGAGTGTGGAGTCGTTGCATACATGTGTAGATGGATTAGGANNNAGATCGGAA |
| 36593_1 | TATGAGTGTGGAGTCGTTGCCTGTAGAACTTTGCCAAGGTGTGNNNAGATCGGAA |
| 36593_2 | TATGAGTGTGGAGTCGTTGCCTGTAGAACTTTGCCAAGGTGTGNNNAGATCGGAA |
| 36593_3 | TATGAGTGTGGAGTCGTTGCCTGTAGAACTTTGCCAAGGTGTGNNNAGATCGGAA |
| 33330_1 | TATGAGTGTGGAGTCGTTGCCACAATCTTGGCTCACTGTGAACNNNAGATCGGAAG |
| 33330_2 | TATGAGTGTGGAGTCGTTGCCACAATCTTGGCTCACTGTGAACNNNAGATCGGAAG |
| 33330_3 | TATGAGTGTGGAGTCGTTGCCACAATCTTGGCTCACTGTGAACNNNAGATCGGAAG |
| 45867_1 | TATGAGTGTGGAGTCGTTGCCTCCCAGGTTCAAGCGATTCTCNNNAGATCGGAAGA |
| 45867_2 | TATGAGTGTGGAGTCGTTGCCTCCCAGGTTCAAGCGATTCTCNNNAGATCGGAAGA |
| 45867_3 | TATGAGTGTGGAGTCGTTGCCTCCCAGGTTCAAGCGATTCTCNNNAGATCGGAAGA |
| 10790_1 | TATGAGTGTGGAGTCGTTGCCTCTTCTGGATGGCCCTGTTTTCNNNAGATCGGAAG |
| 10790_2 | TATGAGTGTGGAGTCGTTGCCTCTTCTGGATGGCCCTGTTTTCNNNAGATCGGAAG |
| 10790_3 | TATGAGTGTGGAGTCGTTGCCTCTTCTGGATGGCCCTGTTTTCNNNAGATCGGAAG |
| 47473_1 | TATGAGTGTGGAGTCGTTGCCAGTGTTGTGATCCTGGCTCANNNAGATCGGAAGA |
| 47473_2 | TATGAGTGTGGAGTCGTTGCCAGTGTTGTGATCCTGGCTCANNNAGATCGGAAGA |
| 47473_3 | TATGAGTGTGGAGTCGTTGCCAGTGTTGTGATCCTGGCTCANNNAGATCGGAAGA |
| 47891_1 | TATGAGTGTGGAGTCGTTGCTGTAATTGCCTGAGGAATGGTGANNNAGATCGGAA |
| 47891_2 | TATGAGTGTGGAGTCGTTGCTGTAATTGCCTGAGGAATGGTGANNNAGATCGGAA |
| 47891_3 | TATGAGTGTGGAGTCGTTGCTGTAATTGCCTGAGGAATGGTGANNNAGATCGGAA |
| 37867_1 | TATGAGTGTGGAGTCGTTGCCGTCCTGAAAGGCATTAGTTGTCNNNAGATCGGAAG |
| 37867_2 | TATGAGTGTGGAGTCGTTGCCGTCCTGAAAGGCATTAGTTGTCNNNAGATCGGAAG |
| 37867_3 | TATGAGTGTGGAGTCGTTGCCGTCCTGAAAGGCATTAGTTGTCNNNAGATCGGAAG |
| 52752_1 | TATGAGTGTGGAGTCGTTGCCTGTGACACGTCAACTGTCTTTCNNNAGATCGGAAG |
| 52752_2 | TATGAGTGTGGAGTCGTTGCCTGTGACACGTCAACTGTCTTTCNNNAGATCGGAAG |
| 52752_3 | TATGAGTGTGGAGTCGTTGCCTGTGACACGTCAACTGTCTTTCNNNAGATCGGAAG |
| 46694_1 | TATGAGTGTGGAGTCGTTGCTTTTGTCATGTTGCTTAGGCTGGNNNAGATCGGAAG |
| 46694_2 | TATGAGTGTGGAGTCGTTGCTTTTGTCATGTTGCTTAGGCTGGNNNAGATCGGAAG |
| 46694_3 | TATGAGTGTGGAGTCGTTGCTTTTGTCATGTTGCTTAGGCTGGNNNAGATCGGAAG |
| 42626_1 | TATGAGTGTGGAGTCGTTGCGAGAGAGAGAGAGAGAGACAGGGNNNAGATCGG |
| 42626_2 | TATGAGTGTGGAGTCGTTGCGAGAGAGAGAGAGAGAGACAGGGNNNAGATCGG |
| 42626_3 | TATGAGTGTGGAGTCGTTGCGAGAGAGAGAGAGAGAGACAGGGNNNAGATCGG |
| 29160_1 | TATGAGTGTGGAGTCGTTGCGAAGAGTCTCGAAGTTCCTCCTGNNNAGATCGGAAG |
| 29160_2 | TATGAGTGTGGAGTCGTTGCGAAGAGTCTCGAAGTTCCTCCTGNNNAGATCGGAAG |
| 29160_3 | TATGAGTGTGGAGTCGTTGCGAAGAGTCTCGAAGTTCCTCCTGNNNAGATCGGAAG |
| 10779_1 | TATGAGTGTGGAGTCGTTGCATGAATAGCCCCGCATTTAGAGGNNNAGATCGGAA |
| 10779_2 | TATGAGTGTGGAGTCGTTGCATGAATAGCCCCGCATTTAGAGGNNNAGATCGGAA |
| 10779_3 | TATGAGTGTGGAGTCGTTGCATGAATAGCCCCGCATTTAGAGGNNNAGATCGGAA |
| 41799_1 | TATGAGTGTGGAGTCGTTGCCTACATTTTTCCCACAGAGCAGCNNNAGATCGGAAG |
| 41799_2 | TATGAGTGTGGAGTCGTTGCCTACATTTTTCCCACAGAGCAGCNNNAGATCGGAAG |

| | |
|---|---|
| 41799_3 | TATGAGTGTGGAGTCGTTGCCTACATTTTTCCCACAGAGCAGCNNNAGATCGGAAG |
| 39574_1 | TATGAGTGTGGAGTCGTTGCGGAGGTGGAGCTTGCAGTAAGNNNAGATCGGAAG |
| 39574_2 | TATGAGTGTGGAGTCGTTGCGGAGGTGGAGCTTGCAGTAAGNNNAGATCGGAAG |
| 39574_3 | TATGAGTGTGGAGTCGTTGCGGAGGTGGAGCTTGCAGTAAGNNNAGATCGGAAG |
| 45537_1 | TATGAGTGTGGAGTCGTTGCGATAGGGACATGATCTCACTGGGNNNAGATCGGAA |
| 45537_2 | TATGAGTGTGGAGTCGTTGCGATAGGGACATGATCTCACTGGGNNNAGATCGGAA |
| 45537_3 | TATGAGTGTGGAGTCGTTGCGATAGGGACATGATCTCACTGGGNNNAGATCGGAA |
| 37138_1 | TATGAGTGTGGAGTCGTTGCGAGTGCAATGGCGTGATCTCNNNAGATCGGAAGAG |
| 37138_2 | TATGAGTGTGGAGTCGTTGCGAGTGCAATGGCGTGATCTCNNNAGATCGGAAGAG |
| 37138_3 | TATGAGTGTGGAGTCGTTGCGAGTGCAATGGCGTGATCTCNNNAGATCGGAAGAG |
| 42791_1 | TATGAGTGTGGAGTCGTTGCGCCTAGAGGAGAATGAGGGAGTANNNAGATCGGA |
| 42791_2 | TATGAGTGTGGAGTCGTTGCGCCTAGAGGAGAATGAGGGAGTANNNAGATCGGA |
| 42791_3 | TATGAGTGTGGAGTCGTTGCGCCTAGAGGAGAATGAGGGAGTANNNAGATCGGA |
| 41734_1 | TATGAGTGTGGAGTCGTTGCGAGATGGAGTCTTGTTTGTTGGCNNNAGATCGGAA |
| 41734_2 | TATGAGTGTGGAGTCGTTGCGAGATGGAGTCTTGTTTGTTGGCNNNAGATCGGAA |
| 41734_3 | TATGAGTGTGGAGTCGTTGCGAGATGGAGTCTTGTTTGTTGGCNNNAGATCGGAA |
| 34320_1 | TATGAGTGTGGAGTCGTTGCTGAACAGGGGTGTGATATTTGGTNNNAGATCGGAA |
| 34320_2 | TATGAGTGTGGAGTCGTTGCTGAACAGGGGTGTGATATTTGGTNNNAGATCGGAA |
| 34320_3 | TATGAGTGTGGAGTCGTTGCTGAACAGGGGTGTGATATTTGGTNNNAGATCGGAA |
| 10130_1 | TATGAGTGTGGAGTCGTTGCGCTCCTTTGGCATAATGACGGTANNNAGATCGGAAG |
| 10130_2 | TATGAGTGTGGAGTCGTTGCGCTCCTTTGGCATAATGACGGTANNNAGATCGGAAG |
| 10130_3 | TATGAGTGTGGAGTCGTTGCGCTCCTTTGGCATAATGACGGTANNNAGATCGGAAG |
| 34475_1 | TATGAGTGTGGAGTCGTTGCGAGCCCAGAAATCCAAAGTCAAGNNNAGATCGGAA |
| 34475_2 | TATGAGTGTGGAGTCGTTGCGAGCCCAGAAATCCAAAGTCAAGNNNAGATCGGAA |
| 34475_3 | TATGAGTGTGGAGTCGTTGCGAGCCCAGAAATCCAAAGTCAAGNNNAGATCGGAA |
| 32778_1 | TATGAGTGTGGAGTCGTTGCCCTGTAATTCCAGCTACTTGGGANNNAGATCGGAAG |
| 32778_2 | TATGAGTGTGGAGTCGTTGCCCTGTAATTCCAGCTACTTGGGANNNAGATCGGAAG |
| 32778_3 | TATGAGTGTGGAGTCGTTGCCCTGTAATTCCAGCTACTTGGGANNNAGATCGGAAG |
| 48061_1 | TATGAGTGTGGAGTCGTTGCTGTGCAAGAAAGAGACCTGGAATNNNAGATCGGAA |
| 48061_2 | TATGAGTGTGGAGTCGTTGCTGTGCAAGAAAGAGACCTGGAATNNNAGATCGGAA |
| 48061_3 | TATGAGTGTGGAGTCGTTGCTGTGCAAGAAAGAGACCTGGAATNNNAGATCGGAA |
| 50373_1 | TATGAGTGTGGAGTCGTTGCCCAACGTGAAAGACCAAAGGAATNNNAGATCGGAA |
| 50373_2 | TATGAGTGTGGAGTCGTTGCCCAACGTGAAAGACCAAAGGAATNNNAGATCGGAA |
| 50373_3 | TATGAGTGTGGAGTCGTTGCCCAACGTGAAAGACCAAAGGAATNNNAGATCGGAA |
| 52308_1 | TATGAGTGTGGAGTCGTTGCTAAACTAAGCAAAGAGCAGCGCNNNAGATCGGAAG |
| 52308_2 | TATGAGTGTGGAGTCGTTGCTAAACTAAGCAAAGAGCAGCGCNNNAGATCGGAAG |
| 52308_3 | TATGAGTGTGGAGTCGTTGCTAAACTAAGCAAAGAGCAGCGCNNNAGATCGGAAG |
| 46973_1 | TATGAGTGTGGAGTCGTTGCGGTTCAAGCAATTATCCTGCCTCNNNAGATCGGAAG |
| 46973_2 | TATGAGTGTGGAGTCGTTGCGGTTCAAGCAATTATCCTGCCTCNNNAGATCGGAAG |
| 46973_3 | TATGAGTGTGGAGTCGTTGCGGTTCAAGCAATTATCCTGCCTCNNNAGATCGGAAG |
| 39277_1 | TATGAGTGTGGAGTCGTTGCTTTACTCTCATGTGTCTGTGGGTNNNAGATCGGAAG |
| 39277_2 | TATGAGTGTGGAGTCGTTGCTTTACTCTCATGTGTCTGTGGGTNNNAGATCGGAAG |
| 39277_3 | TATGAGTGTGGAGTCGTTGCTTTACTCTCATGTGTCTGTGGGTNNNAGATCGGAAG |

| | |
|---|---|
| 43676_1 | TATGAGTGTGGAGTCGTTGCCAAAGAAGTGCTCCCACAACAAGNNNAGATCGGAAG |
| 43676_2 | TATGAGTGTGGAGTCGTTGCCAAAGAAGTGCTCCCACAACAAGNNNAGATCGGAAG |
| 43676_3 | TATGAGTGTGGAGTCGTTGCCAAAGAAGTGCTCCCACAACAAGNNNAGATCGGAAG |
| 9975_1 | TATGAGTGTGGAGTCGTTGCTTCTCCTGGTTGAAAGGGTTTGGNNNAGATCGGAA |
| 9975_2 | TATGAGTGTGGAGTCGTTGCTTCTCCTGGTTGAAAGGGTTTGGNNNAGATCGGAA |
| 9975_3 | TATGAGTGTGGAGTCGTTGCTTCTCCTGGTTGAAAGGGTTTGGNNNAGATCGGAA |
| 50058_1 | TATGAGTGTGGAGTCGTTGCTGCCTGGAAGTGAGGACATAAAANNNAGATCGGAA |
| 50058_2 | TATGAGTGTGGAGTCGTTGCTGCCTGGAAGTGAGGACATAAAANNNAGATCGGAA |
| 50058_3 | TATGAGTGTGGAGTCGTTGCTGCCTGGAAGTGAGGACATAAAANNNAGATCGGAA |
| 34451_1 | TATGAGTGTGGAGTCGTTGCCTGTAATTTCAGATCCAGCAGCGNNNAGATCGGAAG |
| 34451_2 | TATGAGTGTGGAGTCGTTGCCTGTAATTTCAGATCCAGCAGCGNNNAGATCGGAAG |
| 34451_3 | TATGAGTGTGGAGTCGTTGCCTGTAATTTCAGATCCAGCAGCGNNNAGATCGGAAG |
| 44909_1 | TATGAGTGTGGAGTCGTTGCCTGTGAGTACTACAAGTGCTCCANNNAGATCGGAAG |
| 44909_2 | TATGAGTGTGGAGTCGTTGCCTGTGAGTACTACAAGTGCTCCANNNAGATCGGAAG |
| 44909_3 | TATGAGTGTGGAGTCGTTGCCTGTGAGTACTACAAGTGCTCCANNNAGATCGGAAG |
| 43547_1 | TATGAGTGTGGAGTCGTTGCCGCCACCACAACTAGCTAATTTTNNNAGATCGGAAG |
| 43547_2 | TATGAGTGTGGAGTCGTTGCCGCCACCACAACTAGCTAATTTTNNNAGATCGGAAG |
| 43547_3 | TATGAGTGTGGAGTCGTTGCCGCCACCACAACTAGCTAATTTTNNNAGATCGGAAG |
| 49523_1 | TATGAGTGTGGAGTCGTTGCCACGCCTGGCTAACTTTTTGTATNNNAGATCGGAAG |
| 49523_2 | TATGAGTGTGGAGTCGTTGCCACGCCTGGCTAACTTTTTGTATNNNAGATCGGAAG |
| 49523_3 | TATGAGTGTGGAGTCGTTGCCACGCCTGGCTAACTTTTTGTATNNNAGATCGGAAG |
| 30360_1 | TATGAGTGTGGAGTCGTTGCACCCCAACCTTAGTCTCTACAGTNNNAGATCGGAAGA |
| 30360_2 | TATGAGTGTGGAGTCGTTGCACCCCAACCTTAGTCTCTACAGTNNNAGATCGGAAGA |
| 30360_3 | TATGAGTGTGGAGTCGTTGCACCCCAACCTTAGTCTCTACAGTNNNAGATCGGAAGA |
| 41940_1 | TATGAGTGTGGAGTCGTTGCCGACTCCTGGATTCAAGCAATTCNNNAGATCGGAAG |
| 41940_2 | TATGAGTGTGGAGTCGTTGCCGACTCCTGGATTCAAGCAATTCNNNAGATCGGAAG |
| 41940_3 | TATGAGTGTGGAGTCGTTGCCGACTCCTGGATTCAAGCAATTCNNNAGATCGGAAG |
| 32828_1 | TATGAGTGTGGAGTCGTTGCCAGGGTCTCACTCTATTGCCAGNNNAGATCGGAAGA |
| 32828_2 | TATGAGTGTGGAGTCGTTGCCAGGGTCTCACTCTATTGCCAGNNNAGATCGGAAGA |
| 32828_3 | TATGAGTGTGGAGTCGTTGCCAGGGTCTCACTCTATTGCCAGNNNAGATCGGAAGA |
| 50031_1 | TATGAGTGTGGAGTCGTTGCGAACAAAGGAAGGAAGGAAAGGCNNNAGATCGGA |
| 50031_2 | TATGAGTGTGGAGTCGTTGCGAACAAAGGAAGGAAGGAAAGGCNNNAGATCGGA |
| 50031_3 | TATGAGTGTGGAGTCGTTGCGAACAAAGGAAGGAAGGAAAGGCNNNAGATCGGA |
| 39056_1 | TATGAGTGTGGAGTCGTTGCTGAAACTACGTGATTGCTTGCTGNNNAGATCGGAAG |
| 39056_2 | TATGAGTGTGGAGTCGTTGCTGAAACTACGTGATTGCTTGCTGNNNAGATCGGAAG |
| 39056_3 | TATGAGTGTGGAGTCGTTGCTGAAACTACGTGATTGCTTGCTGNNNAGATCGGAAG |
| 38650_1 | TATGAGTGTGGAGTCGTTGCATTACCCAAACTGCCATCATCCTNNNAGATCGGAAGA |
| 38650_2 | TATGAGTGTGGAGTCGTTGCATTACCCAAACTGCCATCATCCTNNNAGATCGGAAGA |
| 38650_3 | TATGAGTGTGGAGTCGTTGCATTACCCAAACTGCCATCATCCTNNNAGATCGGAAGA |
| 40154_1 | TATGAGTGTGGAGTCGTTGCAATACAAAATTAGCTGGGCCTGGNNNAGATCGGAA |
| 40154_2 | TATGAGTGTGGAGTCGTTGCAATACAAAATTAGCTGGGCCTGGNNNAGATCGGAA |
| 40154_3 | TATGAGTGTGGAGTCGTTGCAATACAAAATTAGCTGGGCCTGGNNNAGATCGGAA |
| 32767_1 | TATGAGTGTGGAGTCGTTGCTGAATGTTGTCCTGGATGGGAATNNNAGATCGGAA |

| | |
|---|---|
| 32767_2 | TATGAGTGTGGAGTCGTTGCTGAATGTTGTCCTGGATGGGAATNNNAGATCGGAA |
| 32767_3 | TATGAGTGTGGAGTCGTTGCTGAATGTTGTCCTGGATGGGAATNNNAGATCGGAA |
| 45605_1 | TATGAGTGTGGAGTCGTTGCTGTGTGTCTTTATAGGTAGGGGTGNNNAGATCGGA |
| 45605_2 | TATGAGTGTGGAGTCGTTGCTGTGTGTCTTTATAGGTAGGGGTGNNNAGATCGGA |
| 45605_3 | TATGAGTGTGGAGTCGTTGCTGTGTGTCTTTATAGGTAGGGGTGNNNAGATCGGA |
| 48373_1 | TATGAGTGTGGAGTCGTTGCGTCTCCTGAGTAGCTGGGACTANNNAGATCGGAAG |
| 48373_2 | TATGAGTGTGGAGTCGTTGCGTCTCCTGAGTAGCTGGGACTANNNAGATCGGAAG |
| 48373_3 | TATGAGTGTGGAGTCGTTGCGTCTCCTGAGTAGCTGGGACTANNNAGATCGGAAG |
| 47669_1 | TATGAGTGTGGAGTCGTTGCCCTGACTAGCTGGGATTATAGGCNNNAGATCGGAA |
| 47669_2 | TATGAGTGTGGAGTCGTTGCCCTGACTAGCTGGGATTATAGGCNNNAGATCGGAA |
| 47669_3 | TATGAGTGTGGAGTCGTTGCCCTGACTAGCTGGGATTATAGGCNNNAGATCGGAA |
| 50320_1 | TATGAGTGTGGAGTCGTTGCTCCAGGTACCAGGAGCAGTAATANNNAGATCGGAA |
| 50320_2 | TATGAGTGTGGAGTCGTTGCTCCAGGTACCAGGAGCAGTAATANNNAGATCGGAA |
| 50320_3 | TATGAGTGTGGAGTCGTTGCTCCAGGTACCAGGAGCAGTAATANNNAGATCGGAA |
| 38147_1 | TATGAGTGTGGAGTCGTTGCGTGTGCCTGTAACCTTAGCTACTNNNAGATCGGAAG |
| 38147_2 | TATGAGTGTGGAGTCGTTGCGTGTGCCTGTAACCTTAGCTACTNNNAGATCGGAAG |
| 38147_3 | TATGAGTGTGGAGTCGTTGCGTGTGCCTGTAACCTTAGCTACTNNNAGATCGGAAG |
| 43505_1 | TATGAGTGTGGAGTCGTTGCCCTCCTGAGTTCAAGTGATCCTCNNNAGATCGGAAG |
| 43505_2 | TATGAGTGTGGAGTCGTTGCCCTCCTGAGTTCAAGTGATCCTCNNNAGATCGGAAG |
| 43505_3 | TATGAGTGTGGAGTCGTTGCCCTCCTGAGTTCAAGTGATCCTCNNNAGATCGGAAG |
| 13167_1 | TATGAGTGTGGAGTCGTTGCGAGTTTGTCTGGTACAGTAACNNNAGATCGGAAGA |
| 13167_2 | TATGAGTGTGGAGTCGTTGCGAGTTTGTCTGGTACAGTAACNNNAGATCGGAAGA |
| 13167_3 | TATGAGTGTGGAGTCGTTGCGAGTTTGTCTGGTACAGTAACNNNAGATCGGAAGA |
| 45880_1 | TATGAGTGTGGAGTCGTTGCCTGGATTCAAGTGATTCTCGTGCNNNAGATCGGAAG |
| 45880_2 | TATGAGTGTGGAGTCGTTGCCTGGATTCAAGTGATTCTCGTGCNNNAGATCGGAAG |
| 45880_3 | TATGAGTGTGGAGTCGTTGCCTGGATTCAAGTGATTCTCGTGCNNNAGATCGGAAG |
| 13326_1 | TATGAGTGTGGAGTCGTTGCTTTTCTCTTTCAGGTTAAGCCNNNAGATCGGAAGAG |
| 13326_2 | TATGAGTGTGGAGTCGTTGCTTTTCTCTTTCAGGTTAAGCCNNNAGATCGGAAGAG |
| 13326_3 | TATGAGTGTGGAGTCGTTGCTTTTCTCTTTCAGGTTAAGCCNNNAGATCGGAAGAG |
| 35112_1 | TATGAGTGTGGAGTCGTTGCTGCAGTACAATAGCGCAATCTTGNNNAGATCGGAAG |
| 35112_2 | TATGAGTGTGGAGTCGTTGCTGCAGTACAATAGCGCAATCTTGNNNAGATCGGAAG |
| 35112_3 | TATGAGTGTGGAGTCGTTGCTGCAGTACAATAGCGCAATCTTGNNNAGATCGGAAG |
| 2579_1 | TATGAGTGTGGAGTCGTTGCTCTGTCGCAATAAGGCATCCAAANNNAGATCGGAAG |
| 2579_2 | TATGAGTGTGGAGTCGTTGCTCTGTCGCAATAAGGCATCCAAANNNAGATCGGAAG |
| 2579_3 | TATGAGTGTGGAGTCGTTGCTCTGTCGCAATAAGGCATCCAAANNNAGATCGGAAG |
| 49576_1 | TATGAGTGTGGAGTCGTTGCCCCCCATCTCTCAGAAGTCATTTNNNAGATCGGAAGA |
| 49576_2 | TATGAGTGTGGAGTCGTTGCCCCCCATCTCTCAGAAGTCATTTNNNAGATCGGAAGA |
| 49576_3 | TATGAGTGTGGAGTCGTTGCCCCCCATCTCTCAGAAGTCATTTNNNAGATCGGAAGA |
| 44755_1 | TATGAGTGTGGAGTCGTTGCCTAAGCACTTCATCTTCAGCCACNNNAGATCGGAAGA |
| 44755_2 | TATGAGTGTGGAGTCGTTGCCTAAGCACTTCATCTTCAGCCACNNNAGATCGGAAGA |
| 44755_3 | TATGAGTGTGGAGTCGTTGCCTAAGCACTTCATCTTCAGCCACNNNAGATCGGAAGA |
| 13773_1 | TATGAGTGTGGAGTCGTTGCTGATATAAAGAAAACAAGAAGCANNNAGATCGGAA |
| 13773_2 | TATGAGTGTGGAGTCGTTGCTGATATAAAGAAAACAAGAAGCANNNAGATCGGAA |

| | |
|---|---|
| 13773_3 | TATGAGTGTGGAGTCGTTGCTGATATAAAGAAAACAAGAAGCANNNAGATCGGAA |
| 40752_1 | TATGAGTGTGGAGTCGTTGCCACCCAGGAGCCTAAATTCACTANNNAGATCGGAAG |
| 40752_2 | TATGAGTGTGGAGTCGTTGCCACCCAGGAGCCTAAATTCACTANNNAGATCGGAAG |
| 40752_3 | TATGAGTGTGGAGTCGTTGCCACCCAGGAGCCTAAATTCACTANNNAGATCGGAAG |
| 42610_1 | TATGAGTGTGGAGTCGTTGCGAGAGAGAGAGAAAGAGGGAGGANNNAGATCGG |
| 42610_2 | TATGAGTGTGGAGTCGTTGCGAGAGAGAGAGAAAGAGGGAGGANNNAGATCGG |
| 42610_3 | TATGAGTGTGGAGTCGTTGCGAGAGAGAGAGAAAGAGGGAGGANNNAGATCGG |
| 13002_1 | TATGAGTGTGGAGTCGTTGCATGGAGTCATAATATTCTCAGTTNNNAGATCGGAAG |
| 13002_2 | TATGAGTGTGGAGTCGTTGCATGGAGTCATAATATTCTCAGTTNNNAGATCGGAAG |
| 13002_3 | TATGAGTGTGGAGTCGTTGCATGGAGTCATAATATTCTCAGTTNNNAGATCGGAAG |
| 38561_1 | TATGAGTGTGGAGTCGTTGCAATCTTCATTTGGCACCAGATGGNNNAGATCGGAAG |
| 38561_2 | TATGAGTGTGGAGTCGTTGCAATCTTCATTTGGCACCAGATGGNNNAGATCGGAAG |
| 38561_3 | TATGAGTGTGGAGTCGTTGCAATCTTCATTTGGCACCAGATGGNNNAGATCGGAAG |
| 52927_1 | TATGAGTGTGGAGTCGTTGCCTGTCCTCTGTGACCCTCTCTCNNNAGATCGGAAGAG |
| 52927_2 | TATGAGTGTGGAGTCGTTGCCTGTCCTCTGTGACCCTCTCTCNNNAGATCGGAAGAG |
| 52927_3 | TATGAGTGTGGAGTCGTTGCCTGTCCTCTGTGACCCTCTCTCNNNAGATCGGAAGAG |
| 37494_1 | TATGAGTGTGGAGTCGTTGCGAATTCAGGGCGAGTCCATAGACNNNAGATCGGAA |
| 37494_2 | TATGAGTGTGGAGTCGTTGCGAATTCAGGGCGAGTCCATAGACNNNAGATCGGAA |
| 37494_3 | TATGAGTGTGGAGTCGTTGCGAATTCAGGGCGAGTCCATAGACNNNAGATCGGAA |
| 38365_1 | TATGAGTGTGGAGTCGTTGCAGAGAGAGAGAGAAACCCATTGACANNNAGATCGGAA |
| 38365_2 | TATGAGTGTGGAGTCGTTGCAGAGAGAGAGAGAAACCCATTGACANNNAGATCGGAA |
| 38365_3 | TATGAGTGTGGAGTCGTTGCAGAGAGAGAGAGAAACCCATTGACANNNAGATCGGAA |
| 51183_1 | TATGAGTGTGGAGTCGTTGCTTGGAAAAGCACCTTCTTTCACCNNNAGATCGGAAG |
| 51183_2 | TATGAGTGTGGAGTCGTTGCTTGGAAAAGCACCTTCTTTCACCNNNAGATCGGAAG |
| 51183_3 | TATGAGTGTGGAGTCGTTGCTTGGAAAAGCACCTTCTTTCACCNNNAGATCGGAAG |
| 46641_1 | TATGAGTGTGGAGTCGTTGCTAGTAGAGACGGGGTTTTTCACCNNNAGATCGGAA |
| 46641_2 | TATGAGTGTGGAGTCGTTGCTAGTAGAGACGGGGTTTTTCACCNNNAGATCGGAA |
| 46641_3 | TATGAGTGTGGAGTCGTTGCTAGTAGAGACGGGGTTTTTCACCNNNAGATCGGAA |
| 36675_1 | TATGAGTGTGGAGTCGTTGCGCATCTACTTGGCTTCTAGGGAGNNNAGATCGGAA |
| 36675_2 | TATGAGTGTGGAGTCGTTGCGCATCTACTTGGCTTCTAGGGAGNNNAGATCGGAA |
| 36675_3 | TATGAGTGTGGAGTCGTTGCGCATCTACTTGGCTTCTAGGGAGNNNAGATCGGAA |
| 29505_1 | TATGAGTGTGGAGTCGTTGCCAATGAAGATGGAAATGGGAATCNNNAGATCGGAA |
| 29505_2 | TATGAGTGTGGAGTCGTTGCCAATGAAGATGGAAATGGGAATCNNNAGATCGGAA |
| 29505_3 | TATGAGTGTGGAGTCGTTGCCAATGAAGATGGAAATGGGAATCNNNAGATCGGAA |
| 33795_1 | TATGAGTGTGGAGTCGTTGCACTATCCCTGTTTGCAGATGACANNNAGATCGGAAG |
| 33795_2 | TATGAGTGTGGAGTCGTTGCACTATCCCTGTTTGCAGATGACANNNAGATCGGAAG |
| 33795_3 | TATGAGTGTGGAGTCGTTGCACTATCCCTGTTTGCAGATGACANNNAGATCGGAAG |
| 33661_1 | TATGAGTGTGGAGTCGTTGCCTGCCTCTCAAGTAGTTGGGATTNNNAGATCGGAAG |
| 33661_2 | TATGAGTGTGGAGTCGTTGCCTGCCTCTCAAGTAGTTGGGATTNNNAGATCGGAAG |
| 33661_3 | TATGAGTGTGGAGTCGTTGCCTGCCTCTCAAGTAGTTGGGATTNNNAGATCGGAAG |
| 11024_1 | TATGAGTGTGGAGTCGTTGCTTTTTCCAGTGTGGTCTGAGAGTTTNNNAGATCGGA |
| 11024_2 | TATGAGTGTGGAGTCGTTGCTTTTTCCAGTGTGGTCTGAGAGTTTNNNAGATCGGA |
| 11024_3 | TATGAGTGTGGAGTCGTTGCTTTTTCCAGTGTGGTCTGAGAGTTTNNNAGATCGGA |

| | |
|---|---|
| 41160_1 | TATGAGTGTGGAGTCGTTGCCCTGAACGAAATGCATGGGAACNNNAGATCGGAAG |
| 41160_2 | TATGAGTGTGGAGTCGTTGCCCTGAACGAAATGCATGGGAACNNNAGATCGGAAG |
| 41160_3 | TATGAGTGTGGAGTCGTTGCCCTGAACGAAATGCATGGGAACNNNAGATCGGAAG |
| 48839_1 | TATGAGTGTGGAGTCGTTGCCCAAGTAGCTGGGATTACAGGAGNNNAGATCGGAA |
| 48839_2 | TATGAGTGTGGAGTCGTTGCCCAAGTAGCTGGGATTACAGGAGNNNAGATCGGAA |
| 48839_3 | TATGAGTGTGGAGTCGTTGCCCAAGTAGCTGGGATTACAGGAGNNNAGATCGGAA |
| 37301_1 | TATGAGTGTGGAGTCGTTGCAACTGAGGTCAACAGGGAGTAAGNNNAGATCGGAA |
| 37301_2 | TATGAGTGTGGAGTCGTTGCAACTGAGGTCAACAGGGAGTAAGNNNAGATCGGAA |
| 37301_3 | TATGAGTGTGGAGTCGTTGCAACTGAGGTCAACAGGGAGTAAGNNNAGATCGGAA |
| 29787_1 | TATGAGTGTGGAGTCGTTGCCCAAAACAACTTTGCTGTGTTTTNNNAGATCGGAAG |
| 29787_2 | TATGAGTGTGGAGTCGTTGCCCAAAACAACTTTGCTGTGTTTTNNNAGATCGGAAG |
| 29787_3 | TATGAGTGTGGAGTCGTTGCCCAAAACAACTTTGCTGTGTTTTNNNAGATCGGAAG |
| 31254_1 | TATGAGTGTGGAGTCGTTGCACGTGCACACAAATAGAAACAGGNNNAGATCGGAA |
| 31254_2 | TATGAGTGTGGAGTCGTTGCACGTGCACACAAATAGAAACAGGNNNAGATCGGAA |
| 31254_3 | TATGAGTGTGGAGTCGTTGCACGTGCACACAAATAGAAACAGGNNNAGATCGGAA |
| 47448_1 | TATGAGTGTGGAGTCGTTGCTTCTCCAAAATGAGACTGGGAGGNNNAGATCGGAA |
| 47448_2 | TATGAGTGTGGAGTCGTTGCTTCTCCAAAATGAGACTGGGAGGNNNAGATCGGAA |
| 47448_3 | TATGAGTGTGGAGTCGTTGCTTCTCCAAAATGAGACTGGGAGGNNNAGATCGGAA |
| 53051_1 | TATGAGTGTGGAGTCGTTGCGTTCAGCCCATTTCTGTCTCAACNNNAGATCGGAAGA |
| 53051_2 | TATGAGTGTGGAGTCGTTGCGTTCAGCCCATTTCTGTCTCAACNNNAGATCGGAAGA |
| 53051_3 | TATGAGTGTGGAGTCGTTGCGTTCAGCCCATTTCTGTCTCAACNNNAGATCGGAAGA |
| 30747_1 | TATGAGTGTGGAGTCGTTGCATTTATAGGGGCACAGCTGGAAANNNAGATCGGAA |
| 30747_2 | TATGAGTGTGGAGTCGTTGCATTTATAGGGGCACAGCTGGAAANNNAGATCGGAA |
| 30747_3 | TATGAGTGTGGAGTCGTTGCATTTATAGGGGCACAGCTGGAAANNNAGATCGGAA |
| 44592_1 | TATGAGTGTGGAGTCGTTGCTCATCATCAAATCTGAGCTCCCCNNNAGATCGGAAGA |
| 44592_2 | TATGAGTGTGGAGTCGTTGCTCATCATCAAATCTGAGCTCCCCNNNAGATCGGAAGA |
| 44592_3 | TATGAGTGTGGAGTCGTTGCTCATCATCAAATCTGAGCTCCCCNNNAGATCGGAAGA |
| 52120_1 | TATGAGTGTGGAGTCGTTGCAGGGCAGAACAGTGTGTATAGTGNNNAGATCGGAA |
| 52120_2 | TATGAGTGTGGAGTCGTTGCAGGGCAGAACAGTGTGTATAGTGNNNAGATCGGAA |
| 52120_3 | TATGAGTGTGGAGTCGTTGCAGGGCAGAACAGTGTGTATAGTGNNNAGATCGGAA |
| 29594_1 | TATGAGTGTGGAGTCGTTGCTTTTCAAACACACACACAGAACANNNAGATCGGAAG |
| 29594_2 | TATGAGTGTGGAGTCGTTGCTTTTCAAACACACACACAGAACANNNAGATCGGAAG |
| 29594_3 | TATGAGTGTGGAGTCGTTGCTTTTCAAACACACACACAGAACANNNAGATCGGAAG |
| 34817_1 | TATGAGTGTGGAGTCGTTGCCGTCTCCCAAGTTGAAGCAATTCNNNAGATCGGAAG |
| 34817_2 | TATGAGTGTGGAGTCGTTGCCGTCTCCCAAGTTGAAGCAATTCNNNAGATCGGAAG |
| 34817_3 | TATGAGTGTGGAGTCGTTGCCGTCTCCCAAGTTGAAGCAATTCNNNAGATCGGAAG |
| 8012_1 | TATGAGTGTGGAGTCGTTGCAGCGATTTGTGGGTTCACATCACNNNAGATCGGAAG |
| 8012_2 | TATGAGTGTGGAGTCGTTGCAGCGATTTGTGGGTTCACATCACNNNAGATCGGAAG |
| 8012_3 | TATGAGTGTGGAGTCGTTGCAGCGATTTGTGGGTTCACATCACNNNAGATCGGAAG |
| 51332_1 | TATGAGTGTGGAGTCGTTGCAAGAATCTGGTCCTGCAATTCCTNNNAGATCGGAAG |
| 51332_2 | TATGAGTGTGGAGTCGTTGCAAGAATCTGGTCCTGCAATTCCTNNNAGATCGGAAG |
| 51332_3 | TATGAGTGTGGAGTCGTTGCAAGAATCTGGTCCTGCAATTCCTNNNAGATCGGAAG |
| 49873_1 | TATGAGTGTGGAGTCGTTGCTTCCCTCCAGAGCCTTCTATGTANNNAGATCGGAAGA |

| | |
|---|---|
| 49873_2 | TATGAGTGTGGAGTCGTTGCTTCCCTCCAGAGCCTTCTATGTANNNAGATCGGAAGA |
| 49873_3 | TATGAGTGTGGAGTCGTTGCTTCCCTCCAGAGCCTTCTATGTANNNAGATCGGAAGA |
| 39100_1 | TATGAGTGTGGAGTCGTTGCGAAAGAGACAGGGAAAGGGAGAANNNAGATCGGA |
| 39100_2 | TATGAGTGTGGAGTCGTTGCGAAAGAGACAGGGAAAGGGAGAANNNAGATCGGA |
| 39100_3 | TATGAGTGTGGAGTCGTTGCGAAAGAGACAGGGAAAGGGAGAANNNAGATCGGA |
| 52076_1 | TATGAGTGTGGAGTCGTTGCACAGAACCACCATAGTTCCGACNNNAGATCGGAAGA |
| 52076_2 | TATGAGTGTGGAGTCGTTGCACAGAACCACCATAGTTCCGACNNNAGATCGGAAGA |
| 52076_3 | TATGAGTGTGGAGTCGTTGCACAGAACCACCATAGTTCCGACNNNAGATCGGAAGA |
| 13511_1 | TATGAGTGTGGAGTCGTTGCGGAAAATCCCATATCTGAAGATANNNAGATCGGAAG |
| 13511_2 | TATGAGTGTGGAGTCGTTGCGGAAAATCCCATATCTGAAGATANNNAGATCGGAAG |
| 13511_3 | TATGAGTGTGGAGTCGTTGCGGAAAATCCCATATCTGAAGATANNNAGATCGGAAG |
| 41139_1 | TATGAGTGTGGAGTCGTTGCGGCAGAATGCATTGTTCCTGTAANNNAGATCGGAAG |
| 41139_2 | TATGAGTGTGGAGTCGTTGCGGCAGAATGCATTGTTCCTGTAANNNAGATCGGAAG |
| 41139_3 | TATGAGTGTGGAGTCGTTGCGGCAGAATGCATTGTTCCTGTAANNNAGATCGGAAG |
| 42978_1 | TATGAGTGTGGAGTCGTTGCCACACACACACACACACACATATNNNAGATCGGAAGA |
| 42978_2 | TATGAGTGTGGAGTCGTTGCCACACACACACACACACACATATNNNAGATCGGAAGA |
| 42978_3 | TATGAGTGTGGAGTCGTTGCCACACACACACACACACACATATNNNAGATCGGAAGA |
| 41546_1 | TATGAGTGTGGAGTCGTTGCTCAAGTGATTCTTTTGCCTCAGCNNNAGATCGGAAGA |
| 41546_2 | TATGAGTGTGGAGTCGTTGCTCAAGTGATTCTTTTGCCTCAGCNNNAGATCGGAAGA |
| 41546_3 | TATGAGTGTGGAGTCGTTGCTCAAGTGATTCTTTTGCCTCAGCNNNAGATCGGAAGA |
| 45549_1 | TATGAGTGTGGAGTCGTTGCAGGTTTGGATGGTGACAGGATTTNNNAGATCGGAA |
| 45549_2 | TATGAGTGTGGAGTCGTTGCAGGTTTGGATGGTGACAGGATTTNNNAGATCGGAA |
| 45549_3 | TATGAGTGTGGAGTCGTTGCAGGTTTGGATGGTGACAGGATTTNNNAGATCGGAA |
| 35168_1 | TATGAGTGTGGAGTCGTTGCTTAGCCAGGATGGTCTCGATTTCNNNAGATCGGAAG |
| 35168_2 | TATGAGTGTGGAGTCGTTGCTTAGCCAGGATGGTCTCGATTTCNNNAGATCGGAAG |
| 35168_3 | TATGAGTGTGGAGTCGTTGCTTAGCCAGGATGGTCTCGATTTCNNNAGATCGGAAG |
| 41302_1 | TATGAGTGTGGAGTCGTTGCAGATACAAACAAGCCCAGACAGTNNNAGATCGGAAG |
| 41302_2 | TATGAGTGTGGAGTCGTTGCAGATACAAACAAGCCCAGACAGTNNNAGATCGGAAG |
| 41302_3 | TATGAGTGTGGAGTCGTTGCAGATACAAACAAGCCCAGACAGTNNNAGATCGGAAG |
| 52447_1 | TATGAGTGTGGAGTCGTTGCGCCGAGTATCTCATCCAGTACTGNNNAGATCGGAAG |
| 52447_2 | TATGAGTGTGGAGTCGTTGCGCCGAGTATCTCATCCAGTACTGNNNAGATCGGAAG |
| 52447_3 | TATGAGTGTGGAGTCGTTGCGCCGAGTATCTCATCCAGTACTGNNNAGATCGGAAG |
| 42499_1 | TATGAGTGTGGAGTCGTTGCGTCATGGATGTACCTGAACCACANNNAGATCGGAAG |
| 42499_2 | TATGAGTGTGGAGTCGTTGCGTCATGGATGTACCTGAACCACANNNAGATCGGAAG |
| 42499_3 | TATGAGTGTGGAGTCGTTGCGTCATGGATGTACCTGAACCACANNNAGATCGGAAG |
| 37986_1 | TATGAGTGTGGAGTCGTTGCTGAGGAAAACAGACTAGACAGGCNNNAGATCGGAA |
| 37986_2 | TATGAGTGTGGAGTCGTTGCTGAGGAAAACAGACTAGACAGGCNNNAGATCGGAA |
| 37986_3 | TATGAGTGTGGAGTCGTTGCTGAGGAAAACAGACTAGACAGGCNNNAGATCGGAA |
| 47395_1 | TATGAGTGTGGAGTCGTTGCATTCCTTCTCATATGGCCTGACCNNNAGATCGGAAGA |
| 47395_2 | TATGAGTGTGGAGTCGTTGCATTCCTTCTCATATGGCCTGACCNNNAGATCGGAAGA |
| 47395_3 | TATGAGTGTGGAGTCGTTGCATTCCTTCTCATATGGCCTGACCNNNAGATCGGAAGA |
| 37592_1 | TATGAGTGTGGAGTCGTTGCAGGAAGCTAATGATGTCTGCTCCNNNAGATCGGAAG |
| 37592_2 | TATGAGTGTGGAGTCGTTGCAGGAAGCTAATGATGTCTGCTCCNNNAGATCGGAAG |

| | |
|---|---|
| 37592_3 | TATGAGTGTGGAGTCGTTGCAGGAAGCTAATGATGTCTGCTCCNNNAGATCGGAAG |
| 30919_1 | TATGAGTGTGGAGTCGTTGCGTCAGAAGGGGTTCTAAAGCCAGNNNAGATCGGAA |
| 30919_2 | TATGAGTGTGGAGTCGTTGCGTCAGAAGGGGTTCTAAAGCCAGNNNAGATCGGAA |
| 30919_3 | TATGAGTGTGGAGTCGTTGCGTCAGAAGGGGTTCTAAAGCCAGNNNAGATCGGAA |
| 36389_1 | TATGAGTGTGGAGTCGTTGCCTAACAGACTGGCTACTGTGAGGNNNAGATCGGAA |
| 36389_2 | TATGAGTGTGGAGTCGTTGCCTAACAGACTGGCTACTGTGAGGNNNAGATCGGAA |
| 36389_3 | TATGAGTGTGGAGTCGTTGCCTAACAGACTGGCTACTGTGAGGNNNAGATCGGAA |
| 43967_1 | TATGAGTGTGGAGTCGTTGCGAGAAGGAAGGAAGGAAGGAAGGNNNAGATCGG |
| 43967_2 | TATGAGTGTGGAGTCGTTGCGAGAAGGAAGGAAGGAAGGAAGGNNNAGATCGG |
| 43967_3 | TATGAGTGTGGAGTCGTTGCGAGAAGGAAGGAAGGAAGGAAGGNNNAGATCGG |
| 32758_1 | TATGAGTGTGGAGTCGTTGCAAACACAGACGTCTCCACACATANNNAGATCGGAAG |
| 32758_2 | TATGAGTGTGGAGTCGTTGCAAACACAGACGTCTCCACACATANNNAGATCGGAAG |
| 32758_3 | TATGAGTGTGGAGTCGTTGCAAACACAGACGTCTCCACACATANNNAGATCGGAAG |
| 51058_1 | TATGAGTGTGGAGTCGTTGCAGAATCCCAGAACTGAGGACAACNNNAGATCGGAA |
| 51058_2 | TATGAGTGTGGAGTCGTTGCAGAATCCCAGAACTGAGGACAACNNNAGATCGGAA |
| 51058_3 | TATGAGTGTGGAGTCGTTGCAGAATCCCAGAACTGAGGACAACNNNAGATCGGAA |
| 47256_1 | TATGAGTGTGGAGTCGTTGCCCACCACACCCAGCTAATTTTTTNNNAGATCGGAAGA |
| 47256_2 | TATGAGTGTGGAGTCGTTGCCCACCACACCCAGCTAATTTTTTNNNAGATCGGAAGA |
| 47256_3 | TATGAGTGTGGAGTCGTTGCCCACCACACCCAGCTAATTTTTTNNNAGATCGGAAGA |
| 53263_1 | TATGAGTGTGGAGTCGTTGCGCTCACTGTCACTACATCCTTGANNNAGATCGGAAG |
| 53263_2 | TATGAGTGTGGAGTCGTTGCGCTCACTGTCACTACATCCTTGANNNAGATCGGAAG |
| 53263_3 | TATGAGTGTGGAGTCGTTGCGCTCACTGTCACTACATCCTTGANNNAGATCGGAAG |
| 39867_1 | TATGAGTGTGGAGTCGTTGCAGTTACTTGAGAAGCTGAGGTGGNNNAGATCGGAA |
| 39867_2 | TATGAGTGTGGAGTCGTTGCAGTTACTTGAGAAGCTGAGGTGGNNNAGATCGGAA |
| 39867_3 | TATGAGTGTGGAGTCGTTGCAGTTACTTGAGAAGCTGAGGTGGNNNAGATCGGAA |
| 12518_1 | TATGAGTGTGGAGTCGTTGCAGAAGTCTTCTTTTGCTAATGTNNNAGATCGGAAGA |
| 12518_2 | TATGAGTGTGGAGTCGTTGCAGAAGTCTTCTTTTGCTAATGTNNNAGATCGGAAGA |
| 12518_3 | TATGAGTGTGGAGTCGTTGCAGAAGTCTTCTTTTGCTAATGTNNNAGATCGGAAGA |
| 40105_1 | TATGAGTGTGGAGTCGTTGCCTCATCAAGCTGTTTCTGTCTTTNNNAGATCGGAAGA |
| 40105_2 | TATGAGTGTGGAGTCGTTGCCTCATCAAGCTGTTTCTGTCTTTNNNAGATCGGAAGA |
| 40105_3 | TATGAGTGTGGAGTCGTTGCCTCATCAAGCTGTTTCTGTCTTTNNNAGATCGGAAGA |
| 34749_1 | TATGAGTGTGGAGTCGTTGCAATGTTGAGGTTTTGCGGAATCCNNNAGATCGGAA |
| 34749_2 | TATGAGTGTGGAGTCGTTGCAATGTTGAGGTTTTGCGGAATCCNNNAGATCGGAA |
| 34749_3 | TATGAGTGTGGAGTCGTTGCAATGTTGAGGTTTTGCGGAATCCNNNAGATCGGAA |
| 46911_1 | TATGAGTGTGGAGTCGTTGCCTGAGTAGCTGGGATTACAGGTGNNNAGATCGGAA |
| 46911_2 | TATGAGTGTGGAGTCGTTGCCTGAGTAGCTGGGATTACAGGTGNNNAGATCGGAA |
| 46911_3 | TATGAGTGTGGAGTCGTTGCCTGAGTAGCTGGGATTACAGGTGNNNAGATCGGAA |
| 10453_1 | TATGAGTGTGGAGTCGTTGCTAATTGAGGGTGCTCTCCCTTGANNNAGATCGGAAG |
| 10453_2 | TATGAGTGTGGAGTCGTTGCTAATTGAGGGTGCTCTCCCTTGANNNAGATCGGAAG |
| 10453_3 | TATGAGTGTGGAGTCGTTGCTAATTGAGGGTGCTCTCCCTTGANNNAGATCGGAAG |
| 41687_1 | TATGAGTGTGGAGTCGTTGCCCTCCCAAGTTCACACCATTCTCNNNAGATCGGAAGA |
| 41687_2 | TATGAGTGTGGAGTCGTTGCCCTCCCAAGTTCACACCATTCTCNNNAGATCGGAAGA |
| 41687_3 | TATGAGTGTGGAGTCGTTGCCCTCCCAAGTTCACACCATTCTCNNNAGATCGGAAGA |

| | |
|---|---|
| 43095_1 | TATGAGTGTGGAGTCGTTGCCTACAGTCACTCAGGATCTGGTGNNNAGATCGGAAG |
| 43095_2 | TATGAGTGTGGAGTCGTTGCCTACAGTCACTCAGGATCTGGTGNNNAGATCGGAAG |
| 43095_3 | TATGAGTGTGGAGTCGTTGCCTACAGTCACTCAGGATCTGGTGNNNAGATCGGAAG |
| 33764_1 | TATGAGTGTGGAGTCGTTGCTCATCAGAACAGATGCCTAAGGGNNNAGATCGGAA |
| 33764_2 | TATGAGTGTGGAGTCGTTGCTCATCAGAACAGATGCCTAAGGGNNNAGATCGGAA |
| 33764_3 | TATGAGTGTGGAGTCGTTGCTCATCAGAACAGATGCCTAAGGGNNNAGATCGGAA |
| 29791_1 | TATGAGTGTGGAGTCGTTGCGCCAATAAGAGGGGGCTATTNNNAGATCGGAAGAG |
| 29791_2 | TATGAGTGTGGAGTCGTTGCGCCAATAAGAGGGGGCTATTNNNAGATCGGAAGAG |
| 29791_3 | TATGAGTGTGGAGTCGTTGCGCCAATAAGAGGGGGCTATTNNNAGATCGGAAGAG |
| 41924_1 | TATGAGTGTGGAGTCGTTGCAAAGCTGGGACTTCTAATCCACCNNNAGATCGGAAG |
| 41924_2 | TATGAGTGTGGAGTCGTTGCAAAGCTGGGACTTCTAATCCACCNNNAGATCGGAAG |
| 41924_3 | TATGAGTGTGGAGTCGTTGCAAAGCTGGGACTTCTAATCCACCNNNAGATCGGAAG |
| 41262_1 | TATGAGTGTGGAGTCGTTGCCAAAAATTAGCTGGGCATGGTGANNNAGATCGGAA |
| 41262_2 | TATGAGTGTGGAGTCGTTGCCAAAAATTAGCTGGGCATGGTGANNNAGATCGGAA |
| 41262_3 | TATGAGTGTGGAGTCGTTGCCAAAAATTAGCTGGGCATGGTGANNNAGATCGGAA |
| 51550_1 | TATGAGTGTGGAGTCGTTGCCCAGAGGGGATGAAGAGGGGNNNAGATCGGAAGA |
| 51550_2 | TATGAGTGTGGAGTCGTTGCCCAGAGGGGATGAAGAGGGGNNNAGATCGGAAGA |
| 51550_3 | TATGAGTGTGGAGTCGTTGCCCAGAGGGGATGAAGAGGGGNNNAGATCGGAAGA |
| 45641_1 | TATGAGTGTGGAGTCGTTGCGTTCAAGCGATTCTCCTGCTTCNNNAGATCGGAAGA |
| 45641_2 | TATGAGTGTGGAGTCGTTGCGTTCAAGCGATTCTCCTGCTTCNNNAGATCGGAAGA |
| 45641_3 | TATGAGTGTGGAGTCGTTGCGTTCAAGCGATTCTCCTGCTTCNNNAGATCGGAAGA |
| 5769_1 | TATGAGTGTGGAGTCGTTGCCTAAGGTGTTTTGTTCCCCCTCCNNNAGATCGGAAGA |
| 5769_2 | TATGAGTGTGGAGTCGTTGCCTAAGGTGTTTTGTTCCCCCTCCNNNAGATCGGAAGA |
| 5769_3 | TATGAGTGTGGAGTCGTTGCCTAAGGTGTTTTGTTCCCCCTCCNNNAGATCGGAAGA |
| 43695_1 | TATGAGTGTGGAGTCGTTGCATGGAGGCTGCATTTTTAGAGGANNNAGATCGGAA |
| 43695_2 | TATGAGTGTGGAGTCGTTGCATGGAGGCTGCATTTTTAGAGGANNNAGATCGGAA |
| 43695_3 | TATGAGTGTGGAGTCGTTGCATGGAGGCTGCATTTTTAGAGGANNNAGATCGGAA |
| 42417_1 | TATGAGTGTGGAGTCGTTGCGGTTTTACTCTCTTCCCCTAGGCNNNAGATCGGAAGA |
| 42417_2 | TATGAGTGTGGAGTCGTTGCGGTTTTACTCTCTTCCCCTAGGCNNNAGATCGGAAGA |
| 42417_3 | TATGAGTGTGGAGTCGTTGCGGTTTTACTCTCTTCCCCTAGGCNNNAGATCGGAAGA |
| 37868_1 | TATGAGTGTGGAGTCGTTGCCAGGATGGTGCACTTCTGTAGTCNNNAGATCGGAAG |
| 37868_2 | TATGAGTGTGGAGTCGTTGCCAGGATGGTGCACTTCTGTAGTCNNNAGATCGGAAG |
| 37868_3 | TATGAGTGTGGAGTCGTTGCCAGGATGGTGCACTTCTGTAGTCNNNAGATCGGAAG |
| 53012_1 | TATGAGTGTGGAGTCGTTGCTCTTGGATTTGTAGCCTCCAGACNNNAGATCGGAAG |
| 53012_2 | TATGAGTGTGGAGTCGTTGCTCTTGGATTTGTAGCCTCCAGACNNNAGATCGGAAG |
| 53012_3 | TATGAGTGTGGAGTCGTTGCTCTTGGATTTGTAGCCTCCAGACNNNAGATCGGAAG |
| 42491_1 | TATGAGTGTGGAGTCGTTGCAAAAAGTGTGTGTGTGTGTGTGTNNNAGATCGGAA |
| 42491_2 | TATGAGTGTGGAGTCGTTGCAAAAAGTGTGTGTGTGTGTGTGTNNNAGATCGGAA |
| 42491_3 | TATGAGTGTGGAGTCGTTGCAAAAAGTGTGTGTGTGTGTGTGTNNNAGATCGGAA |
| 50195_1 | TATGAGTGTGGAGTCGTTGCCTTGCAAGAACTCCCTTCCCAGNNNAGATCGGAAGA |
| 50195_2 | TATGAGTGTGGAGTCGTTGCCTTGCAAGAACTCCCTTCCCAGNNNAGATCGGAAGA |
| 50195_3 | TATGAGTGTGGAGTCGTTGCCTTGCAAGAACTCCCTTCCCAGNNNAGATCGGAAGA |
| 34719_1 | TATGAGTGTGGAGTCGTTGCGTTCAAGTGATCCTCCTGCCTCNNNAGATCGGAAGA |

| | |
|---|---|
| 34719_2 | TATGAGTGTGGAGTCGTTGCGTTCAAGTGATCCTCCTGCCTCNNNAGATCGGAAGA |
| 34719_3 | TATGAGTGTGGAGTCGTTGCGTTCAAGTGATCCTCCTGCCTCNNNAGATCGGAAGA |
| 35921_1 | TATGAGTGTGGAGTCGTTGCAGATGACCAGTCTTGGAATTGCTNNNAGATCGGAAG |
| 35921_2 | TATGAGTGTGGAGTCGTTGCAGATGACCAGTCTTGGAATTGCTNNNAGATCGGAAG |
| 35921_3 | TATGAGTGTGGAGTCGTTGCAGATGACCAGTCTTGGAATTGCTNNNAGATCGGAAG |
| 45295_1 | TATGAGTGTGGAGTCGTTGCCAAAGAGCAGAACAGTTGTCCAGNNNAGATCGGAA |
| 45295_2 | TATGAGTGTGGAGTCGTTGCCAAAGAGCAGAACAGTTGTCCAGNNNAGATCGGAA |
| 45295_3 | TATGAGTGTGGAGTCGTTGCCAAAGAGCAGAACAGTTGTCCAGNNNAGATCGGAA |
| 8530_1 | TATGAGTGTGGAGTCGTTGCAAGTCCCTTCCCTTCCATCTATGAGNNNAGATCGGAA |
| 8530_2 | TATGAGTGTGGAGTCGTTGCAAGTCCCTTCCCTTCCATCTATGAGNNNAGATCGGAA |
| 8530_3 | TATGAGTGTGGAGTCGTTGCAAGTCCCTTCCCTTCCATCTATGAGNNNAGATCGGAA |
| 32082_1 | TATGAGTGTGGAGTCGTTGCCAGTTATTGGTTGAAGGCTGCTCNNNAGATCGGAAG |
| 32082_2 | TATGAGTGTGGAGTCGTTGCCAGTTATTGGTTGAAGGCTGCTCNNNAGATCGGAAG |
| 32082_3 | TATGAGTGTGGAGTCGTTGCCAGTTATTGGTTGAAGGCTGCTCNNNAGATCGGAAG |
| 10695_1 | TATGAGTGTGGAGTCGTTGCATCATGTAGAGCTGCTCGTTCACNNNAGATCGGAAG |
| 10695_2 | TATGAGTGTGGAGTCGTTGCATCATGTAGAGCTGCTCGTTCACNNNAGATCGGAAG |
| 10695_3 | TATGAGTGTGGAGTCGTTGCATCATGTAGAGCTGCTCGTTCACNNNAGATCGGAAG |
| 40254_1 | TATGAGTGTGGAGTCGTTGCCAGGGCCAATCAGGAAAATCTTCNNNAGATCGGAAG |
| 40254_2 | TATGAGTGTGGAGTCGTTGCCAGGGCCAATCAGGAAAATCTTCNNNAGATCGGAAG |
| 40254_3 | TATGAGTGTGGAGTCGTTGCCAGGGCCAATCAGGAAAATCTTCNNNAGATCGGAAG |
| 3163_1 | TATGAGTGTGGAGTCGTTGCTTCATTTGTTCAGCTTCTCCCCCNNNAGATCGGAAGA |
| 3163_2 | TATGAGTGTGGAGTCGTTGCTTCATTTGTTCAGCTTCTCCCCCNNNAGATCGGAAGA |
| 3163_3 | TATGAGTGTGGAGTCGTTGCTTCATTTGTTCAGCTTCTCCCCCNNNAGATCGGAAGA |
| 40867_1 | TATGAGTGTGGAGTCGTTGCGAAAATAGGCTCCCAATCGCTTCNNNAGATCGGAAG |
| 40867_2 | TATGAGTGTGGAGTCGTTGCGAAAATAGGCTCCCAATCGCTTCNNNAGATCGGAAG |
| 40867_3 | TATGAGTGTGGAGTCGTTGCGAAAATAGGCTCCCAATCGCTTCNNNAGATCGGAAG |
| 39130_1 | TATGAGTGTGGAGTCGTTGCATGGACATCAATGGAAGAGCAGTNNNAGATCGGAA |
| 39130_2 | TATGAGTGTGGAGTCGTTGCATGGACATCAATGGAAGAGCAGTNNNAGATCGGAA |
| 39130_3 | TATGAGTGTGGAGTCGTTGCATGGACATCAATGGAAGAGCAGTNNNAGATCGGAA |
| 51312_1 | TATGAGTGTGGAGTCGTTGCGAGAAAGGGGGAGTTTTATGGGTNNNAGATCGGA |
| 51312_2 | TATGAGTGTGGAGTCGTTGCGAGAAAGGGGGAGTTTTATGGGTNNNAGATCGGA |
| 51312_3 | TATGAGTGTGGAGTCGTTGCGAGAAAGGGGGAGTTTTATGGGTNNNAGATCGGA |
| 35648_1 | TATGAGTGTGGAGTCGTTGCAGACAGGTCTGGTGGTGAAAAATNNNAGATCGGAA |
| 35648_2 | TATGAGTGTGGAGTCGTTGCAGACAGGTCTGGTGGTGAAAAATNNNAGATCGGAA |
| 35648_3 | TATGAGTGTGGAGTCGTTGCAGACAGGTCTGGTGGTGAAAAATNNNAGATCGGAA |
| 33622_1 | TATGAGTGTGGAGTCGTTGCGTTACCTGCAGCAACATTTGTCTNNNAGATCGGAAG |
| 33622_2 | TATGAGTGTGGAGTCGTTGCGTTACCTGCAGCAACATTTGTCTNNNAGATCGGAAG |
| 33622_3 | TATGAGTGTGGAGTCGTTGCGTTACCTGCAGCAACATTTGTCTNNNAGATCGGAAG |
| 42011_1 | TATGAGTGTGGAGTCGTTGCAATCCTCCCACAGAAGGTTCAAANNNAGATCGGAAG |
| 42011_2 | TATGAGTGTGGAGTCGTTGCAATCCTCCCACAGAAGGTTCAAANNNAGATCGGAAG |
| 42011_3 | TATGAGTGTGGAGTCGTTGCAATCCTCCCACAGAAGGTTCAAANNNAGATCGGAAG |
| 44718_1 | TATGAGTGTGGAGTCGTTGCGACCCACCCAATCATCTCTTCTCNNNAGATCGGAAGA |
| 44718_2 | TATGAGTGTGGAGTCGTTGCGACCCACCCAATCATCTCTTCTCNNNAGATCGGAAGA |

| | |
|---|---|
| 44718_3 | TATGAGTGTGGAGTCGTTGCGACCCACCCAATCATCTCTTCTCNNNAGATCGGAAGA |
| 38732_1 | TATGAGTGTGGAGTCGTTGCAGAAGCTTCATATGCTGGTTCCANNNAGATCGGAAG |
| 38732_2 | TATGAGTGTGGAGTCGTTGCAGAAGCTTCATATGCTGGTTCCANNNAGATCGGAAG |
| 38732_3 | TATGAGTGTGGAGTCGTTGCAGAAGCTTCATATGCTGGTTCCANNNAGATCGGAAG |
| 37479_1 | TATGAGTGTGGAGTCGTTGCACCCCTCCACTTCTAGAACTGTANNNAGATCGGAAGA |
| 37479_2 | TATGAGTGTGGAGTCGTTGCACCCCTCCACTTCTAGAACTGTANNNAGATCGGAAGA |
| 37479_3 | TATGAGTGTGGAGTCGTTGCACCCCTCCACTTCTAGAACTGTANNNAGATCGGAAGA |
| 51831_1 | TATGAGTGTGGAGTCGTTGCCAGACCCAAGAACCCGAAATCTNNNAGATCGGAAGA |
| 51831_2 | TATGAGTGTGGAGTCGTTGCCAGACCCAAGAACCCGAAATCTNNNAGATCGGAAGA |
| 51831_3 | TATGAGTGTGGAGTCGTTGCCAGACCCAAGAACCCGAAATCTNNNAGATCGGAAGA |
| 48457_1 | TATGAGTGTGGAGTCGTTGCTTCAAGCGATTTTTCTGCCTCAGNNNAGATCGGAAGA |
| 48457_2 | TATGAGTGTGGAGTCGTTGCTTCAAGCGATTTTTCTGCCTCAGNNNAGATCGGAAGA |
| 48457_3 | TATGAGTGTGGAGTCGTTGCTTCAAGCGATTTTTCTGCCTCAGNNNAGATCGGAAGA |
| 50840_1 | TATGAGTGTGGAGTCGTTGCTGTCTTTTCAAATGACCTGGGGANNNAGATCGGAAG |
| 50840_2 | TATGAGTGTGGAGTCGTTGCTGTCTTTTCAAATGACCTGGGGANNNAGATCGGAAG |
| 50840_3 | TATGAGTGTGGAGTCGTTGCTGTCTTTTCAAATGACCTGGGGANNNAGATCGGAAG |
| 42594_1 | TATGAGTGTGGAGTCGTTGCCTGGGTAGAAGCTTCTGGAAGAGNNNAGATCGGAA |
| 42594_2 | TATGAGTGTGGAGTCGTTGCCTGGGTAGAAGCTTCTGGAAGAGNNNAGATCGGAA |
| 42594_3 | TATGAGTGTGGAGTCGTTGCCTGGGTAGAAGCTTCTGGAAGAGNNNAGATCGGAA |
| 30995_1 | TATGAGTGTGGAGTCGTTGCCCTCACTGAAACTCCTCTCTACCNNNAGATCGGAAGA |
| 30995_2 | TATGAGTGTGGAGTCGTTGCCCTCACTGAAACTCCTCTCTACCNNNAGATCGGAAGA |
| 30995_3 | TATGAGTGTGGAGTCGTTGCCCTCACTGAAACTCCTCTCTACCNNNAGATCGGAAGA |
| 39216_1 | TATGAGTGTGGAGTCGTTGCAACCAGAAGCCAGAAAGCAAAAGNNNAGATCGGAA |
| 39216_2 | TATGAGTGTGGAGTCGTTGCAACCAGAAGCCAGAAAGCAAAAGNNNAGATCGGAA |
| 39216_3 | TATGAGTGTGGAGTCGTTGCAACCAGAAGCCAGAAAGCAAAAGNNNAGATCGGAA |
| 28926_1 | TATGAGTGTGGAGTCGTTGCTCAGGAGTTCACAGAAAACAATGANNNAGATCGGAA |
| 28926_2 | TATGAGTGTGGAGTCGTTGCTCAGGAGTTCACAGAAAACAATGANNNAGATCGGAA |
| 28926_3 | TATGAGTGTGGAGTCGTTGCTCAGGAGTTCACAGAAAACAATGANNNAGATCGGAA |
| 40161_1 | TATGAGTGTGGAGTCGTTGCGAAGGAGAAGGAGAAGGAGAAGCNNNAGATCGGA |
| 40161_2 | TATGAGTGTGGAGTCGTTGCGAAGGAGAAGGAGAAGGAGAAGCNNNAGATCGGA |
| 40161_3 | TATGAGTGTGGAGTCGTTGCGAAGGAGAAGGAGAAGGAGAAGCNNNAGATCGGA |
| 36813_1 | TATGAGTGTGGAGTCGTTGCAGGGCAAACATAAGCGAGAAAACNNNAGATCGGAA |
| 36813_2 | TATGAGTGTGGAGTCGTTGCAGGGCAAACATAAGCGAGAAAACNNNAGATCGGAA |
| 36813_3 | TATGAGTGTGGAGTCGTTGCAGGGCAAACATAAGCGAGAAAACNNNAGATCGGAA |
| 51108_1 | TATGAGTGTGGAGTCGTTGCCCTGAGCCCTGTCTCTTATGTGNNNAGATCGGAAGA |
| 51108_2 | TATGAGTGTGGAGTCGTTGCCCTGAGCCCTGTCTCTTATGTGNNNAGATCGGAAGA |
| 51108_3 | TATGAGTGTGGAGTCGTTGCCCTGAGCCCTGTCTCTTATGTGNNNAGATCGGAAGA |
| 41279_1 | TATGAGTGTGGAGTCGTTGCCTAAAGGCACTTGTCAGGTACAGNNNAGATCGGAAG |
| 41279_2 | TATGAGTGTGGAGTCGTTGCCTAAAGGCACTTGTCAGGTACAGNNNAGATCGGAAG |
| 41279_3 | TATGAGTGTGGAGTCGTTGCCTAAAGGCACTTGTCAGGTACAGNNNAGATCGGAAG |
| 13663_1 | TATGAGTGTGGAGTCGTTGCCACTGCCTACTCTGAATAAATNNNAGATCGGAAGAG |
| 13663_2 | TATGAGTGTGGAGTCGTTGCCACTGCCTACTCTGAATAAATNNNAGATCGGAAGAG |
| 13663_3 | TATGAGTGTGGAGTCGTTGCCACTGCCTACTCTGAATAAATNNNAGATCGGAAGAG |

| | |
|---|---|
| 31025_1 | TATGAGTGTGGAGTCGTTGCCAAAGGTATCCAAAAGGCAGCAGNNNAGATCGGAA |
| 31025_2 | TATGAGTGTGGAGTCGTTGCCAAAGGTATCCAAAAGGCAGCAGNNNAGATCGGAA |
| 31025_3 | TATGAGTGTGGAGTCGTTGCCAAAGGTATCCAAAAGGCAGCAGNNNAGATCGGAA |
| 23598_1 | TATGAGTGTGGAGTCGTTGCCACAGCTGGATTAGATAACAGNNNAGATCGGAAGA |
| 23598_2 | TATGAGTGTGGAGTCGTTGCCACAGCTGGATTAGATAACAGNNNAGATCGGAAGA |
| 23598_3 | TATGAGTGTGGAGTCGTTGCCACAGCTGGATTAGATAACAGNNNAGATCGGAAGA |
| 32886_1 | TATGAGTGTGGAGTCGTTGCGACAGTGTTTGTAGGCATTCAGCNNNAGATCGGAA |
| 32886_2 | TATGAGTGTGGAGTCGTTGCGACAGTGTTTGTAGGCATTCAGCNNNAGATCGGAA |
| 32886_3 | TATGAGTGTGGAGTCGTTGCGACAGTGTTTGTAGGCATTCAGCNNNAGATCGGAA |
| 10172_1 | TATGAGTGTGGAGTCGTTGCTCAGGATGCCAAAAACCACATCTNNNAGATCGGAAG |
| 10172_2 | TATGAGTGTGGAGTCGTTGCTCAGGATGCCAAAAACCACATCTNNNAGATCGGAAG |
| 10172_3 | TATGAGTGTGGAGTCGTTGCTCAGGATGCCAAAAACCACATCTNNNAGATCGGAAG |
| 34347_1 | TATGAGTGTGGAGTCGTTGCTGTAGATGATGGGTTGATGGGTGNNNAGATCGGA |
| 34347_2 | TATGAGTGTGGAGTCGTTGCTGTAGATGATGGGTTGATGGGTGNNNAGATCGGA |
| 34347_3 | TATGAGTGTGGAGTCGTTGCTGTAGATGATGGGTTGATGGGTGNNNAGATCGGA |
| 37907_1 | TATGAGTGTGGAGTCGTTGCTGTCTATGGCAACCTGTTTGTCTNNNAGATCGGAAG |
| 37907_2 | TATGAGTGTGGAGTCGTTGCTGTCTATGGCAACCTGTTTGTCTNNNAGATCGGAAG |
| 37907_3 | TATGAGTGTGGAGTCGTTGCTGTCTATGGCAACCTGTTTGTCTNNNAGATCGGAAG |
| 50202_1 | TATGAGTGTGGAGTCGTTGCTCACTGTCAGGAAGTTTGTGGATNNNAGATCGGAA |
| 50202_2 | TATGAGTGTGGAGTCGTTGCTCACTGTCAGGAAGTTTGTGGATNNNAGATCGGAA |
| 50202_3 | TATGAGTGTGGAGTCGTTGCTCACTGTCAGGAAGTTTGTGGATNNNAGATCGGAA |
| 52949_1 | TATGAGTGTGGAGTCGTTGCTGCAGCTTTGTTGTAGCACATTTNNNAGATCGGAAG |
| 52949_2 | TATGAGTGTGGAGTCGTTGCTGCAGCTTTGTTGTAGCACATTTNNNAGATCGGAAG |
| 52949_3 | TATGAGTGTGGAGTCGTTGCTGCAGCTTTGTTGTAGCACATTTNNNAGATCGGAAG |
| 39558_1 | TATGAGTGTGGAGTCGTTGCGGAGTGACTGAGCAATGAAACAANNNAGATCGGAA |
| 39558_2 | TATGAGTGTGGAGTCGTTGCGGAGTGACTGAGCAATGAAACAANNNAGATCGGAA |
| 39558_3 | TATGAGTGTGGAGTCGTTGCGGAGTGACTGAGCAATGAAACAANNNAGATCGGAA |
| 40593_1 | TATGAGTGTGGAGTCGTTGCATCTACAAGGAACTAGGGGGACANNNAGATCGGAA |
| 40593_2 | TATGAGTGTGGAGTCGTTGCATCTACAAGGAACTAGGGGGACANNNAGATCGGAA |
| 40593_3 | TATGAGTGTGGAGTCGTTGCATCTACAAGGAACTAGGGGGACANNNAGATCGGAA |
| 53339_1 | TATGAGTGTGGAGTCGTTGCCAAGGCCCTTGATTTACCCTACTNNNAGATCGGAAG |
| 53339_2 | TATGAGTGTGGAGTCGTTGCCAAGGCCCTTGATTTACCCTACTNNNAGATCGGAAG |
| 53339_3 | TATGAGTGTGGAGTCGTTGCCAAGGCCCTTGATTTACCCTACTNNNAGATCGGAAG |
| 32547_1 | TATGAGTGTGGAGTCGTTGCTGATGTCTGTCAATTTTTGAGCTCANNNAGATCGGA |
| 32547_2 | TATGAGTGTGGAGTCGTTGCTGATGTCTGTCAATTTTTGAGCTCANNNAGATCGGA |
| 32547_3 | TATGAGTGTGGAGTCGTTGCTGATGTCTGTCAATTTTTGAGCTCANNNAGATCGGA |
| 34564_1 | TATGAGTGTGGAGTCGTTGCGGTGCAGAGAGGGAGCCAAGNNNAGATCGGAAGA |
| 34564_2 | TATGAGTGTGGAGTCGTTGCGGTGCAGAGAGGGAGCCAAGNNNAGATCGGAAGA |
| 34564_3 | TATGAGTGTGGAGTCGTTGCGGTGCAGAGAGGGAGCCAAGNNNAGATCGGAAGA |
| 31667_1 | TATGAGTGTGGAGTCGTTGCTCTTCCAGAACCTCTCCCTTCTGNNNAGATCGGAAGA |
| 31667_2 | TATGAGTGTGGAGTCGTTGCTCTTCCAGAACCTCTCCCTTCTGNNNAGATCGGAAGA |
| 31667_3 | TATGAGTGTGGAGTCGTTGCTCTTCCAGAACCTCTCCCTTCTGNNNAGATCGGAAGA |
| 30635_1 | ATGAGTGTGGAGTCGTTGCTACCTATATTCTCAAACAGTAGTAGCTNNNAGATCGG |

| | |
|---|---|
| 30635_2 | ATGAGTGTGGAGTCGTTGCTACCTATATTCTCAAACAGTAGTAGCTNNNAGATCGG |
| 30635_3 | ATGAGTGTGGAGTCGTTGCTACCTATATTCTCAAACAGTAGTAGCTNNNAGATCGG |
| 48348_1 | TATGAGTGTGGAGTCGTTGCCAGTGGTGCAAACACAGCTTGNNNAGATCGGAAGA |
| 48348_2 | TATGAGTGTGGAGTCGTTGCCAGTGGTGCAAACACAGCTTGNNNAGATCGGAAGA |
| 48348_3 | TATGAGTGTGGAGTCGTTGCCAGTGGTGCAAACACAGCTTGNNNAGATCGGAAGA |
| 10375_1 | TATGAGTGTGGAGTCGTTGCCAGTTAGGGGGAAGGAGAAGACANNNAGATCGGA |
| 10375_2 | TATGAGTGTGGAGTCGTTGCCAGTTAGGGGGAAGGAGAAGACANNNAGATCGGA |
| 10375_3 | TATGAGTGTGGAGTCGTTGCCAGTTAGGGGGAAGGAGAAGACANNNAGATCGGA |
| 48050_1 | TATGAGTGTGGAGTCGTTGCATCAGTACCATGCTGTTTTCGTTNNNAGATCGGAAG |
| 48050_2 | TATGAGTGTGGAGTCGTTGCATCAGTACCATGCTGTTTTCGTTNNNAGATCGGAAG |
| 48050_3 | TATGAGTGTGGAGTCGTTGCATCAGTACCATGCTGTTTTCGTTNNNAGATCGGAAG |
| 53109_1 | TATGAGTGTGGAGTCGTTGCTTGCTGGGTCAAATGGTAGTTCTNNNAGATCGGAA |
| 53109_2 | TATGAGTGTGGAGTCGTTGCTTGCTGGGTCAAATGGTAGTTCTNNNAGATCGGAA |
| 53109_3 | TATGAGTGTGGAGTCGTTGCTTGCTGGGTCAAATGGTAGTTCTNNNAGATCGGAA |
| 51953_1 | TATGAGTGTGGAGTCGTTGCGATGGTTTCCAGCTTCATCCATGNNNAGATCGGAAG |
| 51953_2 | TATGAGTGTGGAGTCGTTGCGATGGTTTCCAGCTTCATCCATGNNNAGATCGGAAG |
| 51953_3 | TATGAGTGTGGAGTCGTTGCGATGGTTTCCAGCTTCATCCATGNNNAGATCGGAAG |
| 49242_1 | TATGAGTGTGGAGTCGTTGCTCCTGAGTTCAAGCGATTCTCTTNNNAGATCGGAAG |
| 49242_2 | TATGAGTGTGGAGTCGTTGCTCCTGAGTTCAAGCGATTCTCTTNNNAGATCGGAAG |
| 49242_3 | TATGAGTGTGGAGTCGTTGCTCCTGAGTTCAAGCGATTCTCTTNNNAGATCGGAAG |
| 10240_1 | TATGAGTGTGGAGTCGTTGCTATTTCTCCCCAGCTTCTGACCTNNNAGATCGGAAGA |
| 10240_2 | TATGAGTGTGGAGTCGTTGCTATTTCTCCCCAGCTTCTGACCTNNNAGATCGGAAGA |
| 10240_3 | TATGAGTGTGGAGTCGTTGCTATTTCTCCCCAGCTTCTGACCTNNNAGATCGGAAGA |
| 13692_1 | TATGAGTGTGGAGTCGTTGCTCTGTGCAGATAAACAGAAATNNNAGATCGGAAGA |
| 13692_2 | TATGAGTGTGGAGTCGTTGCTCTGTGCAGATAAACAGAAATNNNAGATCGGAAGA |
| 13692_3 | TATGAGTGTGGAGTCGTTGCTCTGTGCAGATAAACAGAAATNNNAGATCGGAAGA |
| 46668_1 | TATGAGTGTGGAGTCGTTGCCAATCTCGGCTTACTGCAACTTCNNNAGATCGGAAG |
| 46668_2 | TATGAGTGTGGAGTCGTTGCCAATCTCGGCTTACTGCAACTTCNNNAGATCGGAAG |
| 46668_3 | TATGAGTGTGGAGTCGTTGCCAATCTCGGCTTACTGCAACTTCNNNAGATCGGAAG |
| 37226_1 | TATGAGTGTGGAGTCGTTGCCCACATAGTGCTCGTCTATCTCCNNNAGATCGGAAGA |
| 37226_2 | TATGAGTGTGGAGTCGTTGCCCACATAGTGCTCGTCTATCTCCNNNAGATCGGAAGA |
| 37226_3 | TATGAGTGTGGAGTCGTTGCCCACATAGTGCTCGTCTATCTCCNNNAGATCGGAAGA |
| 45179_1 | TATGAGTGTGGAGTCGTTGCCGGACAAAGAAGCCTTGTTCTTTNNNAGATCGGAAG |
| 45179_2 | TATGAGTGTGGAGTCGTTGCCGGACAAAGAAGCCTTGTTCTTTNNNAGATCGGAAG |
| 45179_3 | TATGAGTGTGGAGTCGTTGCCGGACAAAGAAGCCTTGTTCTTTNNNAGATCGGAAG |
| 52469_1 | TATGAGTGTGGAGTCGTTGCCCTCTCCCCCCTGCATTATTCNNNAGATCGGAAGAGC |
| 52469_2 | TATGAGTGTGGAGTCGTTGCCCTCTCCCCCCTGCATTATTCNNNAGATCGGAAGAGC |
| 52469_3 | TATGAGTGTGGAGTCGTTGCCCTCTCCCCCCTGCATTATTCNNNAGATCGGAAGAGC |
| 34492_1 | TATGAGTGTGGAGTCGTTGCCCCATCACTACACCTGGCTAATTNNNAGATCGGAAGA |
| 34492_2 | TATGAGTGTGGAGTCGTTGCCCCATCACTACACCTGGCTAATTNNNAGATCGGAAGA |
| 34492_3 | TATGAGTGTGGAGTCGTTGCCCCATCACTACACCTGGCTAATTNNNAGATCGGAAGA |
| 34173_1 | TATGAGTGTGGAGTCGTTGCGCCAGTTTTGGATGTCAGTGAAANNNAGATCGGAA |
| 34173_2 | TATGAGTGTGGAGTCGTTGCGCCAGTTTTGGATGTCAGTGAAANNNAGATCGGAA |

| | |
|---|---|
| 34173_3 | TATGAGTGTGGAGTCGTTGCGCCAGTTTTGGATGTCAGTGAAANNNAGATCGGAA |
| 30286_1 | TATGAGTGTGGAGTCGTTGCGCAAAACAAAACCAAACTCCACANNNAGATCGGAAG |
| 30286_2 | TATGAGTGTGGAGTCGTTGCGCAAAACAAAACCAAACTCCACANNNAGATCGGAAG |
| 30286_3 | TATGAGTGTGGAGTCGTTGCGCAAAACAAAACCAAACTCCACANNNAGATCGGAAG |
| 45695_1 | TATGAGTGTGGAGTCGTTGCCAGACAACAAGGCACAGATTAGCNNNAGATCGGAA |
| 45695_2 | TATGAGTGTGGAGTCGTTGCCAGACAACAAGGCACAGATTAGCNNNAGATCGGAA |
| 45695_3 | TATGAGTGTGGAGTCGTTGCCAGACAACAAGGCACAGATTAGCNNNAGATCGGAA |
| 36733_1 | TATGAGTGTGGAGTCGTTGCTGTCAAGAGCAAATTCTGGGGTANNNAGATCGGAA |
| 36733_2 | TATGAGTGTGGAGTCGTTGCTGTCAAGAGCAAATTCTGGGGTANNNAGATCGGAA |
| 36733_3 | TATGAGTGTGGAGTCGTTGCTGTCAAGAGCAAATTCTGGGGTANNNAGATCGGAA |
| 50435_1 | TATGAGTGTGGAGTCGTTGCCACACACCCAAAAACTTCGTGAGNNNAGATCGGAAG |
| 50435_2 | TATGAGTGTGGAGTCGTTGCCACACACCCAAAAACTTCGTGAGNNNAGATCGGAAG |
| 50435_3 | TATGAGTGTGGAGTCGTTGCCACACACCCAAAAACTTCGTGAGNNNAGATCGGAAG |
| 33886_1 | TATGAGTGTGGAGTCGTTGCCCTCAAACTTCCAGGGTCACTTANNNAGATCGGAAG |
| 33886_2 | TATGAGTGTGGAGTCGTTGCCCTCAAACTTCCAGGGTCACTTANNNAGATCGGAAG |
| 33886_3 | TATGAGTGTGGAGTCGTTGCCCTCAAACTTCCAGGGTCACTTANNNAGATCGGAAG |
| 50016_1 | TATGAGTGTGGAGTCGTTGCTCATCCCGGTTTCCACACTTTATNNNAGATCGGAAGA |
| 50016_2 | TATGAGTGTGGAGTCGTTGCTCATCCCGGTTTCCACACTTTATNNNAGATCGGAAGA |
| 50016_3 | TATGAGTGTGGAGTCGTTGCTCATCCCGGTTTCCACACTTTATNNNAGATCGGAAGA |
| 29488_1 | TATGAGTGTGGAGTCGTTGCAACAGCTTAGCCACCTCAGCAANNNAGATCGGAAGA |
| 29488_2 | TATGAGTGTGGAGTCGTTGCAACAGCTTAGCCACCTCAGCAANNNAGATCGGAAGA |
| 29488_3 | TATGAGTGTGGAGTCGTTGCAACAGCTTAGCCACCTCAGCAANNNAGATCGGAAGA |
| 42521_1 | TATGAGTGTGGAGTCGTTGCTTTCACTCTTGTCTCCCAGGCNNNAGATCGGAAGAG |
| 42521_2 | TATGAGTGTGGAGTCGTTGCTTTCACTCTTGTCTCCCAGGCNNNAGATCGGAAGAG |
| 42521_3 | TATGAGTGTGGAGTCGTTGCTTTCACTCTTGTCTCCCAGGCNNNAGATCGGAAGAG |
| 6663_1 | TATGAGTGTGGAGTCGTTGCTGGGAGGAGATGAAGTGGTGATTNNNAGATCGGA |
| 6663_2 | TATGAGTGTGGAGTCGTTGCTGGGAGGAGATGAAGTGGTGATTNNNAGATCGGA |
| 6663_3 | TATGAGTGTGGAGTCGTTGCTGGGAGGAGATGAAGTGGTGATTNNNAGATCGGA |
| 37188_1 | TATGAGTGTGGAGTCGTTGCGAGGCATGAGAATCACTTGAACCNNNAGATCGGAA |
| 37188_2 | TATGAGTGTGGAGTCGTTGCGAGGCATGAGAATCACTTGAACCNNNAGATCGGAA |
| 37188_3 | TATGAGTGTGGAGTCGTTGCGAGGCATGAGAATCACTTGAACCNNNAGATCGGAA |
| 32513_1 | TATGAGTGTGGAGTCGTTGCGAATGAGGATGAGGATGAGGACANNNAGATCGGA |
| 32513_2 | TATGAGTGTGGAGTCGTTGCGAATGAGGATGAGGATGAGGACANNNAGATCGGA |
| 32513_3 | TATGAGTGTGGAGTCGTTGCGAATGAGGATGAGGATGAGGACANNNAGATCGGA |
| 50429_1 | TATGAGTGTGGAGTCGTTGCGACTAGCTCCCAGGTATCATGTGNNNAGATCGGAAG |
| 50429_2 | TATGAGTGTGGAGTCGTTGCGACTAGCTCCCAGGTATCATGTGNNNAGATCGGAAG |
| 50429_3 | TATGAGTGTGGAGTCGTTGCGACTAGCTCCCAGGTATCATGTGNNNAGATCGGAAG |
| 52551_1 | TATGAGTGTGGAGTCGTTGCCTTGATAGGCTGCCGCATATAACNNNAGATCGGAAG |
| 52551_2 | TATGAGTGTGGAGTCGTTGCCTTGATAGGCTGCCGCATATAACNNNAGATCGGAAG |
| 52551_3 | TATGAGTGTGGAGTCGTTGCCTTGATAGGCTGCCGCATATAACNNNAGATCGGAAG |
| 32586_1 | TATGAGTGTGGAGTCGTTGCGCAAAAAAATCTCCTCCCGATCCNNNAGATCGGAAG |
| 32586_2 | TATGAGTGTGGAGTCGTTGCGCAAAAAAATCTCCTCCCGATCCNNNAGATCGGAAG |
| 32586_3 | TATGAGTGTGGAGTCGTTGCGCAAAAAAATCTCCTCCCGATCCNNNAGATCGGAAG |

| | |
|---|---|
| 37497_1 | TATGAGTGTGGAGTCGTTGCAAGATAGGCCCCTAAATTCCCACNNNAGATCGGAAG |
| 37497_2 | TATGAGTGTGGAGTCGTTGCAAGATAGGCCCCTAAATTCCCACNNNAGATCGGAAG |
| 37497_3 | TATGAGTGTGGAGTCGTTGCAAGATAGGCCCCTAAATTCCCACNNNAGATCGGAAG |
| 41691_1 | TATGAGTGTGGAGTCGTTGCCTCTGGGGTTCAGGACATTCTCNNNAGATCGGAAGA |
| 41691_2 | TATGAGTGTGGAGTCGTTGCCTCTGGGGTTCAGGACATTCTCNNNAGATCGGAAGA |
| 41691_3 | TATGAGTGTGGAGTCGTTGCCTCTGGGGTTCAGGACATTCTCNNNAGATCGGAAGA |
| 35683_1 | TATGAGTGTGGAGTCGTTGCGAGGTGATTTGGATGTTGGCATGNNNAGATCGGAA |
| 35683_2 | TATGAGTGTGGAGTCGTTGCGAGGTGATTTGGATGTTGGCATGNNNAGATCGGAA |
| 35683_3 | TATGAGTGTGGAGTCGTTGCGAGGTGATTTGGATGTTGGCATGNNNAGATCGGAA |
| 9009_1 | TATGAGTGTGGAGTCGTTGCCGGTGGAAATAGCCAGGAAATATNNNAGATCGGAA |
| 9009_2 | TATGAGTGTGGAGTCGTTGCCGGTGGAAATAGCCAGGAAATATNNNAGATCGGAA |
| 9009_3 | TATGAGTGTGGAGTCGTTGCCGGTGGAAATAGCCAGGAAATATNNNAGATCGGAA |
| 48991_1 | TATGAGTGTGGAGTCGTTGCACGATCTTAGCTCACTGCAATCTNNNAGATCGGAAG |
| 48991_2 | TATGAGTGTGGAGTCGTTGCACGATCTTAGCTCACTGCAATCTNNNAGATCGGAAG |
| 48991_3 | TATGAGTGTGGAGTCGTTGCACGATCTTAGCTCACTGCAATCTNNNAGATCGGAAG |
| 46138_1 | TATGAGTGTGGAGTCGTTGCAATGTGGGGCTTTGACCTATCATNNNAGATCGGAAG |
| 46138_2 | TATGAGTGTGGAGTCGTTGCAATGTGGGGCTTTGACCTATCATNNNAGATCGGAAG |
| 46138_3 | TATGAGTGTGGAGTCGTTGCAATGTGGGGCTTTGACCTATCATNNNAGATCGGAAG |
| 30093_1 | TATGAGTGTGGAGTCGTTGCAACTCAAGGAAACTGCAATTGCTNNNAGATCGGAAG |
| 30093_2 | TATGAGTGTGGAGTCGTTGCAACTCAAGGAAACTGCAATTGCTNNNAGATCGGAAG |
| 30093_3 | TATGAGTGTGGAGTCGTTGCAACTCAAGGAAACTGCAATTGCTNNNAGATCGGAAG |
| 38919_1 | TATGAGTGTGGAGTCGTTGCAAAAAAAGCCAGGGCAACAAAACNNNAGATCGGAA |
| 38919_2 | TATGAGTGTGGAGTCGTTGCAAAAAAAGCCAGGGCAACAAAACNNNAGATCGGAA |
| 38919_3 | TATGAGTGTGGAGTCGTTGCAAAAAAAGCCAGGGCAACAAAACNNNAGATCGGAA |
| 45663_1 | TATGAGTGTGGAGTCGTTGCCTAGAAAGTGAAGGGAACAGGCANNNAGATCGGA |
| 45663_2 | TATGAGTGTGGAGTCGTTGCCTAGAAAGTGAAGGGAACAGGCANNNAGATCGGA |
| 45663_3 | TATGAGTGTGGAGTCGTTGCCTAGAAAGTGAAGGGAACAGGCANNNAGATCGGA |
| 43709_1 | TATGAGTGTGGAGTCGTTGCCATGCGTCTGAGTGAGAAGGAAANNNAGATCGGAA |
| 43709_2 | TATGAGTGTGGAGTCGTTGCCATGCGTCTGAGTGAGAAGGAAANNNAGATCGGAA |
| 43709_3 | TATGAGTGTGGAGTCGTTGCCATGCGTCTGAGTGAGAAGGAAANNNAGATCGGAA |
| 35496_1 | TATGAGTGTGGAGTCGTTGCATCTTCCACAGGATTACCCAACANNNAGATCGGAAG |
| 35496_2 | TATGAGTGTGGAGTCGTTGCATCTTCCACAGGATTACCCAACANNNAGATCGGAAG |
| 35496_3 | TATGAGTGTGGAGTCGTTGCATCTTCCACAGGATTACCCAACANNNAGATCGGAAG |
| 39498_1 | TATGAGTGTGGAGTCGTTGCCCAATGGTGGGTATAAGAAGCCTNNNAGATCGGAA |
| 39498_2 | TATGAGTGTGGAGTCGTTGCCCAATGGTGGGTATAAGAAGCCTNNNAGATCGGAA |
| 39498_3 | TATGAGTGTGGAGTCGTTGCCCAATGGTGGGTATAAGAAGCCTNNNAGATCGGAA |
| 42223_1 | TATGAGTGTGGAGTCGTTGCGCTAGAGTGAAGTGGCATGATCTNNNAGATCGGAA |
| 42223_2 | TATGAGTGTGGAGTCGTTGCGCTAGAGTGAAGTGGCATGATCTNNNAGATCGGAA |
| 42223_3 | TATGAGTGTGGAGTCGTTGCGCTAGAGTGAAGTGGCATGATCTNNNAGATCGGAA |
| 34869_1 | TATGAGTGTGGAGTCGTTGCCTCCTGAATAGCTGGGATGACAGNNNAGATCGGAA |
| 34869_2 | TATGAGTGTGGAGTCGTTGCCTCCTGAATAGCTGGGATGACAGNNNAGATCGGAA |
| 34869_3 | TATGAGTGTGGAGTCGTTGCCTCCTGAATAGCTGGGATGACAGNNNAGATCGGAA |
| 30696_1 | TATGAGTGTGGAGTCGTTGCTTAGAAGGTTCTTTCTTCTGTGTGAGANNNAGATCG |

| | |
|---|---|
| 30696_2 | TATGAGTGTGGAGTCGTTGCTTAGAAGGTTCTTTCTTCTGTGTGAGANNNAGATCG |
| 30696_3 | TATGAGTGTGGAGTCGTTGCTTAGAAGGTTCTTTCTTCTGTGTGAGANNNAGATCG |
| 49113_1 | TATGAGTGTGGAGTCGTTGCAGCAAAACGAGAGTGATAGCTGANNNAGATCGGAA |
| 49113_2 | TATGAGTGTGGAGTCGTTGCAGCAAAACGAGAGTGATAGCTGANNNAGATCGGAA |
| 49113_3 | TATGAGTGTGGAGTCGTTGCAGCAAAACGAGAGTGATAGCTGANNNAGATCGGAA |
| 38489_1 | TATGAGTGTGGAGTCGTTGCATACCTTCCTTTCGCACCTCAATNNNAGATCGGAAGA |
| 38489_2 | TATGAGTGTGGAGTCGTTGCATACCTTCCTTTCGCACCTCAATNNNAGATCGGAAGA |
| 38489_3 | TATGAGTGTGGAGTCGTTGCATACCTTCCTTTCGCACCTCAATNNNAGATCGGAAGA |
| 13368_1 | TATGAGTGTGGAGTCGTTGCTTGAGAATACATGAGGGAGTTANNNAGATCGGAAG |
| 13368_2 | TATGAGTGTGGAGTCGTTGCTTGAGAATACATGAGGGAGTTANNNAGATCGGAAG |
| 13368_3 | TATGAGTGTGGAGTCGTTGCTTGAGAATACATGAGGGAGTTANNNAGATCGGAAG |
| 31164_1 | TATGAGTGTGGAGTCGTTGCTGAACACAGAAACTCATGTGCAGNNNAGATCGGAA |
| 31164_2 | TATGAGTGTGGAGTCGTTGCTGAACACAGAAACTCATGTGCAGNNNAGATCGGAA |
| 31164_3 | TATGAGTGTGGAGTCGTTGCTGAACACAGAAACTCATGTGCAGNNNAGATCGGAA |
| 37770_1 | TATGAGTGTGGAGTCGTTGCGGCTTAGTTTCCTCCTGTTCTCANNNAGATCGGAAGA |
| 37770_2 | TATGAGTGTGGAGTCGTTGCGGCTTAGTTTCCTCCTGTTCTCANNNAGATCGGAAGA |
| 37770_3 | TATGAGTGTGGAGTCGTTGCGGCTTAGTTTCCTCCTGTTCTCANNNAGATCGGAAGA |
| 39899_1 | TATGAGTGTGGAGTCGTTGCCACTCTATTGCTCAGGTTGGAGTNNNAGATCGGAAG |
| 39899_2 | TATGAGTGTGGAGTCGTTGCCACTCTATTGCTCAGGTTGGAGTNNNAGATCGGAAG |
| 39899_3 | TATGAGTGTGGAGTCGTTGCCACTCTATTGCTCAGGTTGGAGTNNNAGATCGGAAG |
| 49082_1 | TATGAGTGTGGAGTCGTTGCAAATGCCACTCTATGTCTGCTCANNNAGATCGGAAG |
| 49082_2 | TATGAGTGTGGAGTCGTTGCAAATGCCACTCTATGTCTGCTCANNNAGATCGGAAG |
| 49082_3 | TATGAGTGTGGAGTCGTTGCAAATGCCACTCTATGTCTGCTCANNNAGATCGGAAG |
| 33702_1 | TATGAGTGTGGAGTCGTTGCTCCCACTTTGCTACTATTTGCCTNNNAGATCGGAAGA |
| 33702_2 | TATGAGTGTGGAGTCGTTGCTCCCACTTTGCTACTATTTGCCTNNNAGATCGGAAGA |
| 33702_3 | TATGAGTGTGGAGTCGTTGCTCCCACTTTGCTACTATTTGCCTNNNAGATCGGAAGA |
| 10147_1 | TATGAGTGTGGAGTCGTTGCTCTCCATACCCATTGTTGCCATCNNNAGATCGGAAGA |
| 10147_2 | TATGAGTGTGGAGTCGTTGCTCTCCATACCCATTGTTGCCATCNNNAGATCGGAAGA |
| 10147_3 | TATGAGTGTGGAGTCGTTGCTCTCCATACCCATTGTTGCCATCNNNAGATCGGAAGA |
| 45107_1 | TATGAGTGTGGAGTCGTTGCGTGCAAGTGACTCACAGTGAATCNNNAGATCGGAA |
| 45107_2 | TATGAGTGTGGAGTCGTTGCGTGCAAGTGACTCACAGTGAATCNNNAGATCGGAA |
| 45107_3 | TATGAGTGTGGAGTCGTTGCGTGCAAGTGACTCACAGTGAATCNNNAGATCGGAA |
| 2288_1 | TATGAGTGTGGAGTCGTTGCAGGGAAGGAAATTTGACACCCACNNNAGATCGGAA |
| 2288_2 | TATGAGTGTGGAGTCGTTGCAGGGAAGGAAATTTGACACCCACNNNAGATCGGAA |
| 2288_3 | TATGAGTGTGGAGTCGTTGCAGGGAAGGAAATTTGACACCCACNNNAGATCGGAA |
| 31319_1 | TATGAGTGTGGAGTCGTTGCCTGGGAACCATGGACCATAGTAGNNNAGATCGGAA |
| 31319_2 | TATGAGTGTGGAGTCGTTGCCTGGGAACCATGGACCATAGTAGNNNAGATCGGAA |
| 31319_3 | TATGAGTGTGGAGTCGTTGCCTGGGAACCATGGACCATAGTAGNNNAGATCGGAA |
| 33332_1 | TATGAGTGTGGAGTCGTTGCCTATCTGGGCTCACTACAACCTCNNNAGATCGGAAG |
| 33332_2 | TATGAGTGTGGAGTCGTTGCCTATCTGGGCTCACTACAACCTCNNNAGATCGGAAG |
| 33332_3 | TATGAGTGTGGAGTCGTTGCCTATCTGGGCTCACTACAACCTCNNNAGATCGGAAG |
| 36500_1 | TATGAGTGTGGAGTCGTTGCCTGAGATCATGCCACTGTACTCCNNNAGATCGGAAG |
| 36500_2 | TATGAGTGTGGAGTCGTTGCCTGAGATCATGCCACTGTACTCCNNNAGATCGGAAG |

| | |
|---|---|
| 36500_3 | TATGAGTGTGGAGTCGTTGCCTGAGATCATGCCACTGTACTCCNNNAGATCGGAAG |
| 46877_1 | TATGAGTGTGGAGTCGTTGCACCATCACACCCAGCTATTCTTTNNNAGATCGGAAGA |
| 46877_2 | TATGAGTGTGGAGTCGTTGCACCATCACACCCAGCTATTCTTTNNNAGATCGGAAGA |
| 46877_3 | TATGAGTGTGGAGTCGTTGCACCATCACACCCAGCTATTCTTTNNNAGATCGGAAGA |
| 10234_1 | TATGAGTGTGGAGTCGTTGCGGAAAAGTAAAAGCGCTTGGAGTNNNAGATCGGAA |
| 10234_2 | TATGAGTGTGGAGTCGTTGCGGAAAAGTAAAAGCGCTTGGAGTNNNAGATCGGAA |
| 10234_3 | TATGAGTGTGGAGTCGTTGCGGAAAAGTAAAAGCGCTTGGAGTNNNAGATCGGAA |
| 32066_1 | TATGAGTGTGGAGTCGTTGCAGTTCTGGAGATCTGCTGTATGGNNNAGATCGGAA |
| 32066_2 | TATGAGTGTGGAGTCGTTGCAGTTCTGGAGATCTGCTGTATGGNNNAGATCGGAA |
| 32066_3 | TATGAGTGTGGAGTCGTTGCAGTTCTGGAGATCTGCTGTATGGNNNAGATCGGAA |
| 34162_1 | TATGAGTGTGGAGTCGTTGCATCTCTGCAAGCAACCTATAGGGNNNAGATCGGAAG |
| 34162_2 | TATGAGTGTGGAGTCGTTGCATCTCTGCAAGCAACCTATAGGGNNNAGATCGGAAG |
| 34162_3 | TATGAGTGTGGAGTCGTTGCATCTCTGCAAGCAACCTATAGGGNNNAGATCGGAAG |
| 46568_1 | TATGAGTGTGGAGTCGTTGCGAGTACCTGGGACTACAAGTGTGNNNAGATCGGAA |
| 46568_2 | TATGAGTGTGGAGTCGTTGCGAGTACCTGGGACTACAAGTGTGNNNAGATCGGAA |
| 46568_3 | TATGAGTGTGGAGTCGTTGCGAGTACCTGGGACTACAAGTGTGNNNAGATCGGAA |
| 50112_1 | TATGAGTGTGGAGTCGTTGCCCAACATGGTGAAATGACGTCTCNNNAGATCGGAAG |
| 50112_2 | TATGAGTGTGGAGTCGTTGCCCAACATGGTGAAATGACGTCTCNNNAGATCGGAAG |
| 50112_3 | TATGAGTGTGGAGTCGTTGCCCAACATGGTGAAATGACGTCTCNNNAGATCGGAAG |
| 44460_1 | TATGAGTGTGGAGTCGTTGCATCATTCTTGGGTGTTTCTCGGANNNAGATCGGAAG |
| 44460_2 | TATGAGTGTGGAGTCGTTGCATCATTCTTGGGTGTTTCTCGGANNNAGATCGGAAG |
| 44460_3 | TATGAGTGTGGAGTCGTTGCATCATTCTTGGGTGTTTCTCGGANNNAGATCGGAAG |
| 47408_1 | TATGAGTGTGGAGTCGTTGCTGTTGTTGTTTTTGAGACGGAGTNNNAGATCGGAA |
| 47408_2 | TATGAGTGTGGAGTCGTTGCTGTTGTTGTTTTTGAGACGGAGTNNNAGATCGGAA |
| 47408_3 | TATGAGTGTGGAGTCGTTGCTGTTGTTGTTTTTGAGACGGAGTNNNAGATCGGAA |
| 46319_1 | TATGAGTGTGGAGTCGTTGCTTATTAGAGGACTTGTCGGCACCNNNAGATCGGAAG |
| 46319_2 | TATGAGTGTGGAGTCGTTGCTTATTAGAGGACTTGTCGGCACCNNNAGATCGGAAG |
| 46319_3 | TATGAGTGTGGAGTCGTTGCTTATTAGAGGACTTGTCGGCACCNNNAGATCGGAAG |
| 34367_1 | TATGAGTGTGGAGTCGTTGCCCTACTGGGAAGTGAGGAGCCNNNAGATCGGAAGA |
| 34367_2 | TATGAGTGTGGAGTCGTTGCCCTACTGGGAAGTGAGGAGCCNNNAGATCGGAAGA |
| 34367_3 | TATGAGTGTGGAGTCGTTGCCCTACTGGGAAGTGAGGAGCCNNNAGATCGGAAGA |
| 13175_1 | TATGAGTGTGGAGTCGTTGCCTCAATGGAGAACAGGAAAGNNNAGATCGGAAGAG |
| 13175_2 | TATGAGTGTGGAGTCGTTGCCTCAATGGAGAACAGGAAAGNNNAGATCGGAAGAG |
| 13175_3 | TATGAGTGTGGAGTCGTTGCCTCAATGGAGAACAGGAAAGNNNAGATCGGAAGAG |
| 38948_1 | TATGAGTGTGGAGTCGTTGCTTGCATACTGATCCCATAGAGGCNNNAGATCGGAAG |
| 38948_2 | TATGAGTGTGGAGTCGTTGCTTGCATACTGATCCCATAGAGGCNNNAGATCGGAAG |
| 38948_3 | TATGAGTGTGGAGTCGTTGCTTGCATACTGATCCCATAGAGGCNNNAGATCGGAAG |
| 32380_1 | TATGAGTGTGGAGTCGTTGCGGCTGAGGTAGGAAGATCACTTGNNNAGATCGGAA |
| 32380_2 | TATGAGTGTGGAGTCGTTGCGGCTGAGGTAGGAAGATCACTTGNNNAGATCGGAA |
| 32380_3 | TATGAGTGTGGAGTCGTTGCGGCTGAGGTAGGAAGATCACTTGNNNAGATCGGAA |
| 46253_1 | TATGAGTGTGGAGTCGTTGCCATTCTCCTGCCTCAGCCTCNNNAGATCGGAAGAGCA |
| 46253_2 | TATGAGTGTGGAGTCGTTGCCATTCTCCTGCCTCAGCCTCNNNAGATCGGAAGAGCA |
| 46253_3 | TATGAGTGTGGAGTCGTTGCCATTCTCCTGCCTCAGCCTCNNNAGATCGGAAGAGCA |

| | |
|---|---|
| 33231_1 | TATGAGTGTGGAGTCGTTGCCTAGACTTACTGCACCCTCCATCNNNAGATCGGAAGA |
| 33231_2 | TATGAGTGTGGAGTCGTTGCCTAGACTTACTGCACCCTCCATCNNNAGATCGGAAGA |
| 33231_3 | TATGAGTGTGGAGTCGTTGCCTAGACTTACTGCACCCTCCATCNNNAGATCGGAAGA |
| 44027_1 | TATGAGTGTGGAGTCGTTGCAATGGTGTTCTGAGATGGGTGAANNNAGATCGGAA |
| 44027_2 | TATGAGTGTGGAGTCGTTGCAATGGTGTTCTGAGATGGGTGAANNNAGATCGGAA |
| 44027_3 | TATGAGTGTGGAGTCGTTGCAATGGTGTTCTGAGATGGGTGAANNNAGATCGGAA |
| 31670_1 | TATGAGTGTGGAGTCGTTGCGCACAGATAGACACAACACACACNNNAGATCGGAAG |
| 31670_2 | TATGAGTGTGGAGTCGTTGCGCACAGATAGACACAACACACACNNNAGATCGGAAG |
| 31670_3 | TATGAGTGTGGAGTCGTTGCGCACAGATAGACACAACACACACNNNAGATCGGAAG |
| 41688_1 | TATGAGTGTGGAGTCGTTGCGTTCACACCATTCTCCTACCTCANNNAGATCGGAAGA |
| 41688_2 | TATGAGTGTGGAGTCGTTGCGTTCACACCATTCTCCTACCTCANNNAGATCGGAAGA |
| 41688_3 | TATGAGTGTGGAGTCGTTGCGTTCACACCATTCTCCTACCTCANNNAGATCGGAAGA |
| 5788_1 | TATGAGTGTGGAGTCGTTGCGCCTCCCAAGCTCTAGTGATACTNNNAGATCGGAAG |
| 5788_2 | TATGAGTGTGGAGTCGTTGCGCCTCCCAAGCTCTAGTGATACTNNNAGATCGGAAG |
| 5788_3 | TATGAGTGTGGAGTCGTTGCGCCTCCCAAGCTCTAGTGATACTNNNAGATCGGAAG |
| 4234_1 | TATGAGTGTGGAGTCGTTGCTTGCACACAGTGGTACAGTAGTGNNNAGATCGGAA |
| 4234_2 | TATGAGTGTGGAGTCGTTGCTTGCACACAGTGGTACAGTAGTGNNNAGATCGGAA |
| 4234_3 | TATGAGTGTGGAGTCGTTGCTTGCACACAGTGGTACAGTAGTGNNNAGATCGGAA |
| 52228_1 | TATGAGTGTGGAGTCGTTGCTATCCTGTGTTGGGAAAGGAGTGNNNAGATCGGAA |
| 52228_2 | TATGAGTGTGGAGTCGTTGCTATCCTGTGTTGGGAAAGGAGTGNNNAGATCGGAA |
| 52228_3 | TATGAGTGTGGAGTCGTTGCTATCCTGTGTTGGGAAAGGAGTGNNNAGATCGGAA |
| 38674_1 | TATGAGTGTGGAGTCGTTGCAAAAGTCCAGAATTCTCTCCGCTNNNAGATCGGAAG |
| 38674_2 | TATGAGTGTGGAGTCGTTGCAAAAGTCCAGAATTCTCTCCGCTNNNAGATCGGAAG |
| 38674_3 | TATGAGTGTGGAGTCGTTGCAAAAGTCCAGAATTCTCTCCGCTNNNAGATCGGAAG |
| 49564_1 | TATGAGTGTGGAGTCGTTGCAGTTGGAGTCTCACTCTTGTCACNNNAGATCGGAAG |
| 49564_2 | TATGAGTGTGGAGTCGTTGCAGTTGGAGTCTCACTCTTGTCACNNNAGATCGGAAG |
| 49564_3 | TATGAGTGTGGAGTCGTTGCAGTTGGAGTCTCACTCTTGTCACNNNAGATCGGAAG |
| 48400_1 | TATGAGTGTGGAGTCGTTGCTTCTTGAGAGAAGCAGAGCACATNNNAGATCGGAA |
| 48400_2 | TATGAGTGTGGAGTCGTTGCTTCTTGAGAGAAGCAGAGCACATNNNAGATCGGAA |
| 48400_3 | TATGAGTGTGGAGTCGTTGCTTCTTGAGAGAAGCAGAGCACATNNNAGATCGGAA |
| 29155_1 | TATGAGTGTGGAGTCGTTGCCCTGTAAGTTCTCACCTAGTGCANNNAGATCGGAAG |
| 29155_2 | TATGAGTGTGGAGTCGTTGCCCTGTAAGTTCTCACCTAGTGCANNNAGATCGGAAG |
| 29155_3 | TATGAGTGTGGAGTCGTTGCCCTGTAAGTTCTCACCTAGTGCANNNAGATCGGAAG |
| 52571_1 | TATGAGTGTGGAGTCGTTGCACAAGAGAGTATGGCAGACACTGNNNAGATCGGAA |
| 52571_2 | TATGAGTGTGGAGTCGTTGCACAAGAGAGTATGGCAGACACTGNNNAGATCGGAA |
| 52571_3 | TATGAGTGTGGAGTCGTTGCACAAGAGAGTATGGCAGACACTGNNNAGATCGGAA |
| 32792_1 | TATGAGTGTGGAGTCGTTGCCAATCTTGGCTCACTGCAAACTCNNNAGATCGGAAG |
| 32792_2 | TATGAGTGTGGAGTCGTTGCCAATCTTGGCTCACTGCAAACTCNNNAGATCGGAAG |
| 32792_3 | TATGAGTGTGGAGTCGTTGCCAATCTTGGCTCACTGCAAACTCNNNAGATCGGAAG |
| 46395_1 | TATGAGTGTGGAGTCGTTGCCTCCCAGGTTCACGCTATTCTCNNNAGATCGGAAGAG |
| 46395_2 | TATGAGTGTGGAGTCGTTGCCTCCCAGGTTCACGCTATTCTCNNNAGATCGGAAGAG |
| 46395_3 | TATGAGTGTGGAGTCGTTGCCTCCCAGGTTCACGCTATTCTCNNNAGATCGGAAGAG |
| 48140_1 | TATGAGTGTGGAGTCGTTGCCTCCTAAGTTGCTGGGACTACAGNNNAGATCGGAAG |

| | |
|---|---|
| 48140_2 | TATGAGTGTGGAGTCGTTGCCTCCTAAGTTGCTGGGACTACAGNNNAGATCGGAAG |
| 48140_3 | TATGAGTGTGGAGTCGTTGCCTCCTAAGTTGCTGGGACTACAGNNNAGATCGGAAG |
| 52475_1 | TATGAGTGTGGAGTCGTTGCATTGGATAGCAATTTCTGGCTGCNNNAGATCGGAAG |
| 52475_2 | TATGAGTGTGGAGTCGTTGCATTGGATAGCAATTTCTGGCTGCNNNAGATCGGAAG |
| 52475_3 | TATGAGTGTGGAGTCGTTGCATTGGATAGCAATTTCTGGCTGCNNNAGATCGGAAG |
| 48865_1 | TATGAGTGTGGAGTCGTTGCGATTACAGGCATGAACCACCATGNNNAGATCGGAA |
| 48865_2 | TATGAGTGTGGAGTCGTTGCGATTACAGGCATGAACCACCATGNNNAGATCGGAA |
| 48865_3 | TATGAGTGTGGAGTCGTTGCGATTACAGGCATGAACCACCATGNNNAGATCGGAA |
| 13412_1 | TATGAGTGTGGAGTCGTTGCAGCTAGTTTACTGGCATTATCNNNAGATCGGAAGAG |
| 13412_2 | TATGAGTGTGGAGTCGTTGCAGCTAGTTTACTGGCATTATCNNNAGATCGGAAGAG |
| 13412_3 | TATGAGTGTGGAGTCGTTGCAGCTAGTTTACTGGCATTATCNNNAGATCGGAAGAG |
| 32309_1 | TATGAGTGTGGAGTCGTTGCATGACTTCCTTTGGGTAGACTCCNNNAGATCGGAAG |
| 32309_2 | TATGAGTGTGGAGTCGTTGCATGACTTCCTTTGGGTAGACTCCNNNAGATCGGAAG |
| 32309_3 | TATGAGTGTGGAGTCGTTGCATGACTTCCTTTGGGTAGACTCCNNNAGATCGGAAG |
| 37033_1 | TATGAGTGTGGAGTCGTTGCAGGTCACAGGATACAAGGTCAACNNNAGATCGGAA |
| 37033_2 | TATGAGTGTGGAGTCGTTGCAGGTCACAGGATACAAGGTCAACNNNAGATCGGAA |
| 37033_3 | TATGAGTGTGGAGTCGTTGCAGGTCACAGGATACAAGGTCAACNNNAGATCGGAA |
| 29388_1 | TATGAGTGTGGAGTCGTTGCAAGAACTCCTTGCTGAGACCAATNNNAGATCGGAAG |
| 29388_2 | TATGAGTGTGGAGTCGTTGCAAGAACTCCTTGCTGAGACCAATNNNAGATCGGAAG |
| 29388_3 | TATGAGTGTGGAGTCGTTGCAAGAACTCCTTGCTGAGACCAATNNNAGATCGGAAG |
| 49565_1 | TATGAGTGTGGAGTCGTTGCCAATTCTGCCTCAGTCTCCCTAGNNNAGATCGGAAG |
| 49565_2 | TATGAGTGTGGAGTCGTTGCCAATTCTGCCTCAGTCTCCCTAGNNNAGATCGGAAG |
| 49565_3 | TATGAGTGTGGAGTCGTTGCCAATTCTGCCTCAGTCTCCCTAGNNNAGATCGGAAG |
| 31248_1 | TATGAGTGTGGAGTCGTTGCCTACTGATGAAGACCAGCCTCAGNNNAGATCGGAAG |
| 31248_2 | TATGAGTGTGGAGTCGTTGCCTACTGATGAAGACCAGCCTCAGNNNAGATCGGAAG |
| 31248_3 | TATGAGTGTGGAGTCGTTGCCTACTGATGAAGACCAGCCTCAGNNNAGATCGGAAG |
| 34778_1 | TATGAGTGTGGAGTCGTTGCAAGAGTTTTTGCTCTGTCCTCCANNNAGATCGGAAG |
| 34778_2 | TATGAGTGTGGAGTCGTTGCAAGAGTTTTTGCTCTGTCCTCCANNNAGATCGGAAG |
| 34778_3 | TATGAGTGTGGAGTCGTTGCAAGAGTTTTTGCTCTGTCCTCCANNNAGATCGGAAG |
| 37639_1 | TATGAGTGTGGAGTCGTTGCCCTCTTATCTTGGCCTCCCAAATNNNAGATCGGAAGA |
| 37639_2 | TATGAGTGTGGAGTCGTTGCCCTCTTATCTTGGCCTCCCAAATNNNAGATCGGAAGA |
| 37639_3 | TATGAGTGTGGAGTCGTTGCCCTCTTATCTTGGCCTCCCAAATNNNAGATCGGAAGA |
| 44394_1 | TATGAGTGTGGAGTCGTTGCATTGCAGATGTTTCTGAAGCACCNNNAGATCGGAAG |
| 44394_2 | TATGAGTGTGGAGTCGTTGCATTGCAGATGTTTCTGAAGCACCNNNAGATCGGAAG |
| 44394_3 | TATGAGTGTGGAGTCGTTGCATTGCAGATGTTTCTGAAGCACCNNNAGATCGGAAG |
| 42979_1 | TATGAGTGTGGAGTCGTTGCCCCAGGTTCAAGCAATTCTTCTGNNNAGATCGGAAG |
| 42979_2 | TATGAGTGTGGAGTCGTTGCCCCAGGTTCAAGCAATTCTTCTGNNNAGATCGGAAG |
| 42979_3 | TATGAGTGTGGAGTCGTTGCCCCAGGTTCAAGCAATTCTTCTGNNNAGATCGGAAG |
| 13607_1 | TATGAGTGTGGAGTCGTTGCATCAGAATTAAGACTAGAGATGGNNNAGATCGGAA |
| 13607_2 | TATGAGTGTGGAGTCGTTGCATCAGAATTAAGACTAGAGATGGNNNAGATCGGAA |
| 13607_3 | TATGAGTGTGGAGTCGTTGCATCAGAATTAAGACTAGAGATGGNNNAGATCGGAA |
| 8786_1 | TATGAGTGTGGAGTCGTTGCTGTGTAGGCATGTGAATGTGCTTNNNAGATCGGAA |
| 8786_2 | TATGAGTGTGGAGTCGTTGCTGTGTAGGCATGTGAATGTGCTTNNNAGATCGGAA |

| | |
|---|---|
| 8786_3 | TATGAGTGTGGAGTCGTTGCTGTGTAGGCATGTGAATGTGCTTNNNAGATCGGAA |
| 40065_1 | TATGAGTGTGGAGTCGTTGCACACTGAAAATCTGTGGTGCTAGNNNAGATCGGAA |
| 40065_2 | TATGAGTGTGGAGTCGTTGCACACTGAAAATCTGTGGTGCTAGNNNAGATCGGAA |
| 40065_3 | TATGAGTGTGGAGTCGTTGCACACTGAAAATCTGTGGTGCTAGNNNAGATCGGAA |
| 33767_1 | TATGAGTGTGGAGTCGTTGCCGCATACACTTCTGGTGGTAGTANNNAGATCGGAAG |
| 33767_2 | TATGAGTGTGGAGTCGTTGCCGCATACACTTCTGGTGGTAGTANNNAGATCGGAAG |
| 33767_3 | TATGAGTGTGGAGTCGTTGCCGCATACACTTCTGGTGGTAGTANNNAGATCGGAAG |
| 48524_1 | TATGAGTGTGGAGTCGTTGCCTATCTCGGCTCACTGCGAGNNNAGATCGGAAGAG |
| 48524_2 | TATGAGTGTGGAGTCGTTGCCTATCTCGGCTCACTGCGAGNNNAGATCGGAAGAG |
| 48524_3 | TATGAGTGTGGAGTCGTTGCCTATCTCGGCTCACTGCGAGNNNAGATCGGAAGAG |
| 39913_1 | TATGAGTGTGGAGTCGTTGCAATAGGTCAGCCACTGTGTATCCNNNAGATCGGAAG |
| 39913_2 | TATGAGTGTGGAGTCGTTGCAATAGGTCAGCCACTGTGTATCCNNNAGATCGGAAG |
| 39913_3 | TATGAGTGTGGAGTCGTTGCAATAGGTCAGCCACTGTGTATCCNNNAGATCGGAAG |
| 12510_1 | TATGAGTGTGGAGTCGTTGCCAGAATTGGAGTCTCTGTATCNNNAGATCGGAAGA |
| 12510_2 | TATGAGTGTGGAGTCGTTGCCAGAATTGGAGTCTCTGTATCNNNAGATCGGAAGA |
| 12510_3 | TATGAGTGTGGAGTCGTTGCCAGAATTGGAGTCTCTGTATCNNNAGATCGGAAGA |
| 50808_1 | TATGAGTGTGGAGTCGTTGCGGAAGAGAAGAAAGTCTGGTCCCNNNAGATCGGA |
| 50808_2 | TATGAGTGTGGAGTCGTTGCGGAAGAGAAGAAAGTCTGGTCCCNNNAGATCGGA |
| 50808_3 | TATGAGTGTGGAGTCGTTGCGGAAGAGAAGAAAGTCTGGTCCCNNNAGATCGGA |
| 30726_1 | TATGAGTGTGGAGTCGTTGCTTATGGCAGTGAGCATAGTTTTATTTTNNNAGATCG |
| 30726_2 | TATGAGTGTGGAGTCGTTGCTTATGGCAGTGAGCATAGTTTTATTTTNNNAGATCG |
| 30726_3 | TATGAGTGTGGAGTCGTTGCTTATGGCAGTGAGCATAGTTTTATTTTNNNAGATCG |
| 38452_1 | TATGAGTGTGGAGTCGTTGCACGCCTGTAATTCCAACACTTTGNNNAGATCGGAAG |
| 38452_2 | TATGAGTGTGGAGTCGTTGCACGCCTGTAATTCCAACACTTTGNNNAGATCGGAAG |
| 38452_3 | TATGAGTGTGGAGTCGTTGCACGCCTGTAATTCCAACACTTTGNNNAGATCGGAAG |
| 45958_1 | TATGAGTGTGGAGTCGTTGCTTCTCACTTTTCATTCCCAGCCTNNNAGATCGGAAGA |
| 45958_2 | TATGAGTGTGGAGTCGTTGCTTCTCACTTTTCATTCCCAGCCTNNNAGATCGGAAGA |
| 45958_3 | TATGAGTGTGGAGTCGTTGCTTCTCACTTTTCATTCCCAGCCTNNNAGATCGGAAGA |
| 38245_1 | TATGAGTGTGGAGTCGTTGCAGTCATGCTCATCGTTGGTAAGTNNNAGATCGGAAG |
| 38245_2 | TATGAGTGTGGAGTCGTTGCAGTCATGCTCATCGTTGGTAAGTNNNAGATCGGAAG |
| 38245_3 | TATGAGTGTGGAGTCGTTGCAGTCATGCTCATCGTTGGTAAGTNNNAGATCGGAAG |
| 10614_1 | TATGAGTGTGGAGTCGTTGCTCATGGGGCAGATTTCCCTTTTTNNNAGATCGGAAG |
| 10614_2 | TATGAGTGTGGAGTCGTTGCTCATGGGGCAGATTTCCCTTTTTNNNAGATCGGAAG |
| 10614_3 | TATGAGTGTGGAGTCGTTGCTCATGGGGCAGATTTCCCTTTTTNNNAGATCGGAAG |
| 50553_1 | TATGAGTGTGGAGTCGTTGCCTCACAGGGTGTAGGAGAGAATGNNNAGATCGGA |
| 50553_2 | TATGAGTGTGGAGTCGTTGCCTCACAGGGTGTAGGAGAGAATGNNNAGATCGGA |
| 50553_3 | TATGAGTGTGGAGTCGTTGCCTCACAGGGTGTAGGAGAGAATGNNNAGATCGGA |
| 50447_1 | TATGAGTGTGGAGTCGTTGCGCCTCTTCCTTATCAGCGATACANNNAGATCGGAAG |
| 50447_2 | TATGAGTGTGGAGTCGTTGCGCCTCTTCCTTATCAGCGATACANNNAGATCGGAAG |
| 50447_3 | TATGAGTGTGGAGTCGTTGCGCCTCTTCCTTATCAGCGATACANNNAGATCGGAAG |
| 43796_1 | TATGAGTGTGGAGTCGTTGCCACTGAAGGACACCACAGATTCTNNNAGATCGGAAG |
| 43796_2 | TATGAGTGTGGAGTCGTTGCCACTGAAGGACACCACAGATTCTNNNAGATCGGAAG |
| 43796_3 | TATGAGTGTGGAGTCGTTGCCACTGAAGGACACCACAGATTCTNNNAGATCGGAAG |

| | |
|---|---|
| 41575_1 | TATGAGTGTGGAGTCGTTGCCGAGTAGCTGGGACTACAGGTNNNAGATCGGAAGA |
| 41575_2 | TATGAGTGTGGAGTCGTTGCCGAGTAGCTGGGACTACAGGTNNNAGATCGGAAGA |
| 41575_3 | TATGAGTGTGGAGTCGTTGCCGAGTAGCTGGGACTACAGGTNNNAGATCGGAAGA |
| 44374_1 | TATGAGTGTGGAGTCGTTGCTTGCCAAGAAAATAATGCTGGGGNNNAGATCGGAA |
| 44374_2 | TATGAGTGTGGAGTCGTTGCTTGCCAAGAAAATAATGCTGGGGNNNAGATCGGAA |
| 44374_3 | TATGAGTGTGGAGTCGTTGCTTGCCAAGAAAATAATGCTGGGGNNNAGATCGGAA |
| 46726_1 | TATGAGTGTGGAGTCGTTGCGGTTTCACTCCTGTCACCCAGNNNAGATCGGAAGAG |
| 46726_2 | TATGAGTGTGGAGTCGTTGCGGTTTCACTCCTGTCACCCAGNNNAGATCGGAAGAG |
| 46726_3 | TATGAGTGTGGAGTCGTTGCGGTTTCACTCCTGTCACCCAGNNNAGATCGGAAGAG |
| 17832_1 | TATGAGTGTGGAGTCGTTGCGTGTCTACTACCTAGTGTAAAGNNNAGATCGGAAGA |
| 17832_2 | TATGAGTGTGGAGTCGTTGCGTGTCTACTACCTAGTGTAAAGNNNAGATCGGAAGA |
| 17832_3 | TATGAGTGTGGAGTCGTTGCGTGTCTACTACCTAGTGTAAAGNNNAGATCGGAAGA |
| 48567_1 | TATGAGTGTGGAGTCGTTGCACCACGCCTGGCTAATTTTTGTANNNAGATCGGAAG |
| 48567_2 | TATGAGTGTGGAGTCGTTGCACCACGCCTGGCTAATTTTTGTANNNAGATCGGAAG |
| 48567_3 | TATGAGTGTGGAGTCGTTGCACCACGCCTGGCTAATTTTTGTANNNAGATCGGAAG |
| 12703_1 | TATGAGTGTGGAGTCGTTGCTTATTGGTCATCATTTCCAGTANNNAGATCGGAAGA |
| 12703_2 | TATGAGTGTGGAGTCGTTGCTTATTGGTCATCATTTCCAGTANNNAGATCGGAAGA |
| 12703_3 | TATGAGTGTGGAGTCGTTGCTTATTGGTCATCATTTCCAGTANNNAGATCGGAAGA |
| 34051_1 | TATGAGTGTGGAGTCGTTGCGAATTTTGCTCTTGCCCAGCTATNNNAGATCGGAAG |
| 34051_2 | TATGAGTGTGGAGTCGTTGCGAATTTTGCTCTTGCCCAGCTATNNNAGATCGGAAG |
| 34051_3 | TATGAGTGTGGAGTCGTTGCGAATTTTGCTCTTGCCCAGCTATNNNAGATCGGAAG |
| 50526_1 | TATGAGTGTGGAGTCGTTGCCAGAGGAAGAGAAGAGCCAAAGANNNAGATCGGA |
| 50526_2 | TATGAGTGTGGAGTCGTTGCCAGAGGAAGAGAAGAGCCAAAGANNNAGATCGGA |
| 50526_3 | TATGAGTGTGGAGTCGTTGCCAGAGGAAGAGAAGAGCCAAAGANNNAGATCGGA |
| 45584_1 | TATGAGTGTGGAGTCGTTGCGATTACAGTCACATGCCACCATGNNNAGATCGGAAG |
| 45584_2 | TATGAGTGTGGAGTCGTTGCGATTACAGTCACATGCCACCATGNNNAGATCGGAAG |
| 45584_3 | TATGAGTGTGGAGTCGTTGCGATTACAGTCACATGCCACCATGNNNAGATCGGAAG |
| 48474_1 | TATGAGTGTGGAGTCGTTGCAAATTACAGGCACCCACCACTATNNNAGATCGGAAG |
| 48474_2 | TATGAGTGTGGAGTCGTTGCAAATTACAGGCACCCACCACTATNNNAGATCGGAAG |
| 48474_3 | TATGAGTGTGGAGTCGTTGCAAATTACAGGCACCCACCACTATNNNAGATCGGAAG |
| 53090_1 | TATGAGTGTGGAGTCGTTGCGCTACTCTCTTCCTAGGGGGTAANNNAGATCGGAAG |
| 53090_2 | TATGAGTGTGGAGTCGTTGCGCTACTCTCTTCCTAGGGGGTAANNNAGATCGGAAG |
| 53090_3 | TATGAGTGTGGAGTCGTTGCGCTACTCTCTTCCTAGGGGGTAANNNAGATCGGAAG |
| 13525_1 | TATGAGTGTGGAGTCGTTGCAACTTTTACATACAGTGAGAAACNNNAGATCGGAAG |
| 13525_2 | TATGAGTGTGGAGTCGTTGCAACTTTTACATACAGTGAGAAACNNNAGATCGGAAG |
| 13525_3 | TATGAGTGTGGAGTCGTTGCAACTTTTACATACAGTGAGAAACNNNAGATCGGAAG |
| 29820_1 | TATGAGTGTGGAGTCGTTGCGAACAGGTAAGACCAAGACCCAANNNAGATCGGAA |
| 29820_2 | TATGAGTGTGGAGTCGTTGCGAACAGGTAAGACCAAGACCCAANNNAGATCGGAA |
| 29820_3 | TATGAGTGTGGAGTCGTTGCGAACAGGTAAGACCAAGACCCAANNNAGATCGGAA |
| 44220_1 | TATGAGTGTGGAGTCGTTGCGCACAGGATGGAGATGCCTATTANNNAGATCGGAA |
| 44220_2 | TATGAGTGTGGAGTCGTTGCGCACAGGATGGAGATGCCTATTANNNAGATCGGAA |
| 44220_3 | TATGAGTGTGGAGTCGTTGCGCACAGGATGGAGATGCCTATTANNNAGATCGGAA |
| 46433_1 | TATGAGTGTGGAGTCGTTGCTCAGATCACTGAGGTCTAGAGCANNNAGATCGGAA |

| | |
|---|---|
| 46433_2 | TATGAGTGTGGAGTCGTTGCTCAGATCACTGAGGTCTAGAGCANNNAGATCGGAA |
| 46433_3 | TATGAGTGTGGAGTCGTTGCTCAGATCACTGAGGTCTAGAGCANNNAGATCGGAA |
| 52187_1 | TATGAGTGTGGAGTCGTTGCGAGTGAGAATATGCGGTGTTTGGNNNAGATCGGAA |
| 52187_2 | TATGAGTGTGGAGTCGTTGCGAGTGAGAATATGCGGTGTTTGGNNNAGATCGGAA |
| 52187_3 | TATGAGTGTGGAGTCGTTGCGAGTGAGAATATGCGGTGTTTGGNNNAGATCGGAA |
| 52191_1 | TATGAGTGTGGAGTCGTTGCCAACATTTGTAGACACACCCTGGNNNAGATCGGAAG |
| 52191_2 | TATGAGTGTGGAGTCGTTGCCAACATTTGTAGACACACCCTGGNNNAGATCGGAAG |
| 52191_3 | TATGAGTGTGGAGTCGTTGCCAACATTTGTAGACACACCCTGGNNNAGATCGGAAG |
| 37634_1 | TATGAGTGTGGAGTCGTTGCTTTGAGACCAGCCTGATCAACATNNNAGATCGGAAG |
| 37634_2 | TATGAGTGTGGAGTCGTTGCTTTGAGACCAGCCTGATCAACATNNNAGATCGGAAG |
| 37634_3 | TATGAGTGTGGAGTCGTTGCTTTGAGACCAGCCTGATCAACATNNNAGATCGGAAG |
| 33560_1 | TATGAGTGTGGAGTCGTTGCGATCTCAGCTCACTGCAGTCTCNNNAGATCGGAAGA |
| 33560_2 | TATGAGTGTGGAGTCGTTGCGATCTCAGCTCACTGCAGTCTCNNNAGATCGGAAGA |
| 33560_3 | TATGAGTGTGGAGTCGTTGCGATCTCAGCTCACTGCAGTCTCNNNAGATCGGAAGA |
| 35277_1 | TATGAGTGTGGAGTCGTTGCTGTATTTTGGAGTGCATTGAGGCNNNAGATCGGAA |
| 35277_2 | TATGAGTGTGGAGTCGTTGCTGTATTTTGGAGTGCATTGAGGCNNNAGATCGGAA |
| 35277_3 | TATGAGTGTGGAGTCGTTGCTGTATTTTGGAGTGCATTGAGGCNNNAGATCGGAA |
| 10343_1 | TATGAGTGTGGAGTCGTTGCCTTACTATGTGCCAGGCGTTGTCNNNAGATCGGAAG |
| 10343_2 | TATGAGTGTGGAGTCGTTGCCTTACTATGTGCCAGGCGTTGTCNNNAGATCGGAAG |
| 10343_3 | TATGAGTGTGGAGTCGTTGCCTTACTATGTGCCAGGCGTTGTCNNNAGATCGGAAG |
| 10197_1 | TATGAGTGTGGAGTCGTTGCCCTGTGCCTGCCTTGTACTATAGNNNAGATCGGAAG |
| 10197_2 | TATGAGTGTGGAGTCGTTGCCCTGTGCCTGCCTTGTACTATAGNNNAGATCGGAAG |
| 10197_3 | TATGAGTGTGGAGTCGTTGCCCTGTGCCTGCCTTGTACTATAGNNNAGATCGGAAG |
| 38794_1 | TATGAGTGTGGAGTCGTTGCAACCTCCTGAGTAGCTGGGATTANNNAGATCGGAAG |
| 38794_2 | TATGAGTGTGGAGTCGTTGCAACCTCCTGAGTAGCTGGGATTANNNAGATCGGAAG |
| 38794_3 | TATGAGTGTGGAGTCGTTGCAACCTCCTGAGTAGCTGGGATTANNNAGATCGGAAG |
| 28848_1 | TATGAGTGTGGAGTCGTTGCAGCTCCTTTGATCTGTGTTTCGANNNAGATCGGAAG |
| 28848_2 | TATGAGTGTGGAGTCGTTGCAGCTCCTTTGATCTGTGTTTCGANNNAGATCGGAAG |
| 28848_3 | TATGAGTGTGGAGTCGTTGCAGCTCCTTTGATCTGTGTTTCGANNNAGATCGGAAG |
| 51979_1 | TATGAGTGTGGAGTCGTTGCATCAGAGGTGTTCTTGGTCTCACNNNAGATCGGAAG |
| 51979_2 | TATGAGTGTGGAGTCGTTGCATCAGAGGTGTTCTTGGTCTCACNNNAGATCGGAAG |
| 51979_3 | TATGAGTGTGGAGTCGTTGCATCAGAGGTGTTCTTGGTCTCACNNNAGATCGGAAG |
| 10098_1 | TATGAGTGTGGAGTCGTTGCCCATCCTTTTCTCCCATCTGCTCNNNAGATCGGAAGA |
| 10098_2 | TATGAGTGTGGAGTCGTTGCCCATCCTTTTCTCCCATCTGCTCNNNAGATCGGAAGA |
| 10098_3 | TATGAGTGTGGAGTCGTTGCCCATCCTTTTCTCCCATCTGCTCNNNAGATCGGAAGA |
| 47298_1 | TATGAGTGTGGAGTCGTTGCCTCACTGCAACCTCTGTCTTACANNNAGATCGGAAGA |
| 47298_2 | TATGAGTGTGGAGTCGTTGCCTCACTGCAACCTCTGTCTTACANNNAGATCGGAAGA |
| 47298_3 | TATGAGTGTGGAGTCGTTGCCTCACTGCAACCTCTGTCTTACANNNAGATCGGAAGA |
| 40309_1 | TATGAGTGTGGAGTCGTTGCTTAGAGGTGGTTTGCTATGCAGANNNAGATCGGAA |
| 40309_2 | TATGAGTGTGGAGTCGTTGCTTAGAGGTGGTTTGCTATGCAGANNNAGATCGGAA |
| 40309_3 | TATGAGTGTGGAGTCGTTGCTTAGAGGTGGTTTGCTATGCAGANNNAGATCGGAA |
| 47237_1 | TATGAGTGTGGAGTCGTTGCGGCTCAAATGCAGTAGTGTGATCNNNAGATCGGAA |
| 47237_2 | TATGAGTGTGGAGTCGTTGCGGCTCAAATGCAGTAGTGTGATCNNNAGATCGGAA |

| | |
|---|---|
| 47237_3 | TATGAGTGTGGAGTCGTTGCGGCTCAAATGCAGTAGTGTGATCNNNAGATCGGAA |
| 2118_1 | TATGAGTGTGGAGTCGTTGCTGTAAATAAAAGACAGTTGGCCAANNNAGATCGGA |
| 2118_2 | TATGAGTGTGGAGTCGTTGCTGTAAATAAAAGACAGTTGGCCAANNNAGATCGGA |
| 2118_3 | TATGAGTGTGGAGTCGTTGCTGTAAATAAAAGACAGTTGGCCAANNNAGATCGGA |
| 50291_1 | TATGAGTGTGGAGTCGTTGCTGCCATCATAGTTTCCAGAAGCANNNAGATCGGAAG |
| 50291_2 | TATGAGTGTGGAGTCGTTGCTGCCATCATAGTTTCCAGAAGCANNNAGATCGGAAG |
| 50291_3 | TATGAGTGTGGAGTCGTTGCTGCCATCATAGTTTCCAGAAGCANNNAGATCGGAAG |
| 41731_1 | TATGAGTGTGGAGTCGTTGCGAGGTGCAGTCTCTCTCTGTTGNNNAGATCGGAAG |
| 41731_2 | TATGAGTGTGGAGTCGTTGCGAGGTGCAGTCTCTCTCTGTTGNNNAGATCGGAAG |
| 41731_3 | TATGAGTGTGGAGTCGTTGCGAGGTGCAGTCTCTCTCTGTTGNNNAGATCGGAAG |
| 43783_1 | TATGAGTGTGGAGTCGTTGCTTCAACTATTTCCTGCCTTGGGANNNAGATCGGAAG |
| 43783_2 | TATGAGTGTGGAGTCGTTGCTTCAACTATTTCCTGCCTTGGGANNNAGATCGGAAG |
| 43783_3 | TATGAGTGTGGAGTCGTTGCTTCAACTATTTCCTGCCTTGGGANNNAGATCGGAAG |
| 13206_1 | TATGAGTGTGGAGTCGTTGCTTTCTGCCTTTTAGTTTTTCCNNNAGATCGGAAGAGC |
| 13206_2 | TATGAGTGTGGAGTCGTTGCTTTCTGCCTTTTAGTTTTTCCNNNAGATCGGAAGAGC |
| 13206_3 | TATGAGTGTGGAGTCGTTGCTTTCTGCCTTTTAGTTTTTCCNNNAGATCGGAAGAGC |
| 2717_1 | TATGAGTGTGGAGTCGTTGCAAGAAGGGGCAGATTTGCAAACTNNNAGATCGGAA |
| 2717_2 | TATGAGTGTGGAGTCGTTGCAAGAAGGGGCAGATTTGCAAACTNNNAGATCGGAA |
| 2717_3 | TATGAGTGTGGAGTCGTTGCAAGAAGGGGCAGATTTGCAAACTNNNAGATCGGAA |
| 32235_1 | TATGAGTGTGGAGTCGTTGCGGCTTTTCCTTCATCATTTGCCTNNNAGATCGGAAGA |
| 32235_2 | TATGAGTGTGGAGTCGTTGCGGCTTTTCCTTCATCATTTGCCTNNNAGATCGGAAGA |
| 32235_3 | TATGAGTGTGGAGTCGTTGCGGCTTTTCCTTCATCATTTGCCTNNNAGATCGGAAGA |
| 48871_1 | TATGAGTGTGGAGTCGTTGCCACCACGCCTGGCTAATTTTTTNNNAGATCGGAAGA |
| 48871_2 | TATGAGTGTGGAGTCGTTGCCACCACGCCTGGCTAATTTTTTNNNAGATCGGAAGA |
| 48871_3 | TATGAGTGTGGAGTCGTTGCCACCACGCCTGGCTAATTTTTTNNNAGATCGGAAGA |
| 32749_1 | TATGAGTGTGGAGTCGTTGCTCACACCTGTAATTCCAGCACTTNNNAGATCGGAAGA |
| 32749_2 | TATGAGTGTGGAGTCGTTGCTCACACCTGTAATTCCAGCACTTNNNAGATCGGAAGA |
| 32749_3 | TATGAGTGTGGAGTCGTTGCTCACACCTGTAATTCCAGCACTTNNNAGATCGGAAGA |
| 2339_1 | TATGAGTGTGGAGTCGTTGCTCAAGTTGTTGGGTAGGGCTTACANNNAGATCGGA |
| 2339_2 | TATGAGTGTGGAGTCGTTGCTCAAGTTGTTGGGTAGGGCTTACANNNAGATCGGA |
| 2339_3 | TATGAGTGTGGAGTCGTTGCTCAAGTTGTTGGGTAGGGCTTACANNNAGATCGGA |
| 50436_1 | TATGAGTGTGGAGTCGTTGCTGATACAATCCCAAGCCTTCTAGANNNAGATCGGAA |
| 50436_2 | TATGAGTGTGGAGTCGTTGCTGATACAATCCCAAGCCTTCTAGANNNAGATCGGAA |
| 50436_3 | TATGAGTGTGGAGTCGTTGCTGATACAATCCCAAGCCTTCTAGANNNAGATCGGAA |
| 33431_1 | TATGAGTGTGGAGTCGTTGCATCTAGGGGAAAAGCATTCCTGGNNNAGATCGGAA |
| 33431_2 | TATGAGTGTGGAGTCGTTGCATCTAGGGGAAAAGCATTCCTGGNNNAGATCGGAA |
| 33431_3 | TATGAGTGTGGAGTCGTTGCATCTAGGGGAAAAGCATTCCTGGNNNAGATCGGAA |
| 34056_1 | TATGAGTGTGGAGTCGTTGCTGAGTAGCTGAGATTACAGACGCNNNAGATCGGAA |
| 34056_2 | TATGAGTGTGGAGTCGTTGCTGAGTAGCTGAGATTACAGACGCNNNAGATCGGAA |
| 34056_3 | TATGAGTGTGGAGTCGTTGCTGAGTAGCTGAGATTACAGACGCNNNAGATCGGAA |
| 31206_1 | TATGAGTGTGGAGTCGTTGCAGTCATAGGAGCAGGAGAGATTACNNNAGATCGGA |
| 31206_2 | TATGAGTGTGGAGTCGTTGCAGTCATAGGAGCAGGAGAGATTACNNNAGATCGGA |
| 31206_3 | TATGAGTGTGGAGTCGTTGCAGTCATAGGAGCAGGAGAGATTACNNNAGATCGGA |

| | |
|---|---|
| 32451_1 | TATGAGTGTGGAGTCGTTGCGTAGTTTTCCTTTGCACAGGTCCNNNAGATCGGAAG |
| 32451_2 | TATGAGTGTGGAGTCGTTGCGTAGTTTTCCTTTGCACAGGTCCNNNAGATCGGAAG |
| 32451_3 | TATGAGTGTGGAGTCGTTGCGTAGTTTTCCTTTGCACAGGTCCNNNAGATCGGAAG |
| 43876_1 | TATGAGTGTGGAGTCGTTGCTGACAAGGGTAGGATATCTGGGTNNNAGATCGGAA |
| 43876_2 | TATGAGTGTGGAGTCGTTGCTGACAAGGGTAGGATATCTGGGTNNNAGATCGGAA |
| 43876_3 | TATGAGTGTGGAGTCGTTGCTGACAAGGGTAGGATATCTGGGTNNNAGATCGGAA |
| 37929_1 | TATGAGTGTGGAGTCGTTGCCCTTCCCAATTTTGTGTACGTGTNNNAGATCGGAAG |
| 37929_2 | TATGAGTGTGGAGTCGTTGCCCTTCCCAATTTTGTGTACGTGTNNNAGATCGGAAG |
| 37929_3 | TATGAGTGTGGAGTCGTTGCCCTTCCCAATTTTGTGTACGTGTNNNAGATCGGAAG |
| 35748_1 | TATGAGTGTGGAGTCGTTGCGAGTCAGAATCTTGCTCTGTTGCNNNAGATCGGAAG |
| 35748_2 | TATGAGTGTGGAGTCGTTGCGAGTCAGAATCTTGCTCTGTTGCNNNAGATCGGAAG |
| 35748_3 | TATGAGTGTGGAGTCGTTGCGAGTCAGAATCTTGCTCTGTTGCNNNAGATCGGAAG |
| 37071_1 | TATGAGTGTGGAGTCGTTGCAAAAACTTAGCTGAGCATGGTGGNNNAGATCGGAA |
| 37071_2 | TATGAGTGTGGAGTCGTTGCAAAAACTTAGCTGAGCATGGTGGNNNAGATCGGAA |
| 37071_3 | TATGAGTGTGGAGTCGTTGCAAAAACTTAGCTGAGCATGGTGGNNNAGATCGGAA |
| 45470_1 | TATGAGTGTGGAGTCGTTGCCTGGAACAGTTCGGTCCATTAGANNNAGATCGGAAG |
| 45470_2 | TATGAGTGTGGAGTCGTTGCCTGGAACAGTTCGGTCCATTAGANNNAGATCGGAAG |
| 45470_3 | TATGAGTGTGGAGTCGTTGCCTGGAACAGTTCGGTCCATTAGANNNAGATCGGAAG |
| 13373_1 | TATGAGTGTGGAGTCGTTGCCATTTATCTGGGATAAATGAGAANNNAGATCGGAAG |
| 13373_2 | TATGAGTGTGGAGTCGTTGCCATTTATCTGGGATAAATGAGAANNNAGATCGGAAG |
| 13373_3 | TATGAGTGTGGAGTCGTTGCCATTTATCTGGGATAAATGAGAANNNAGATCGGAAG |
| 5589_1 | TATGAGTGTGGAGTCGTTGCTGTCTGTGTTGGTTGACAATAGCANNNAGATCGGAA |
| 5589_2 | TATGAGTGTGGAGTCGTTGCTGTCTGTGTTGGTTGACAATAGCANNNAGATCGGAA |
| 5589_3 | TATGAGTGTGGAGTCGTTGCTGTCTGTGTTGGTTGACAATAGCANNNAGATCGGAA |
| 53219_1 | TATGAGTGTGGAGTCGTTGCTTTGGCCTCAAACTTCAGGTGANNNAGATCGGAAGA |
| 53219_2 | TATGAGTGTGGAGTCGTTGCTTTGGCCTCAAACTTCAGGTGANNNAGATCGGAAGA |
| 53219_3 | TATGAGTGTGGAGTCGTTGCTTTGGCCTCAAACTTCAGGTGANNNAGATCGGAAGA |
| 47805_1 | TATGAGTGTGGAGTCGTTGCGGTAACCTTGGTTTTGGCAATGANNNAGATCGGAAG |
| 47805_2 | TATGAGTGTGGAGTCGTTGCGGTAACCTTGGTTTTGGCAATGANNNAGATCGGAAG |
| 47805_3 | TATGAGTGTGGAGTCGTTGCGGTAACCTTGGTTTTGGCAATGANNNAGATCGGAAG |
| 43854_1 | TATGAGTGTGGAGTCGTTGCGGGAGTGAGTTTTTGCAATGTGANNNAGATCGGAA |
| 43854_2 | TATGAGTGTGGAGTCGTTGCGGGAGTGAGTTTTTGCAATGTGANNNAGATCGGAA |
| 43854_3 | TATGAGTGTGGAGTCGTTGCGGGAGTGAGTTTTTGCAATGTGANNNAGATCGGAA |
| 35170_1 | TATGAGTGTGGAGTCGTTGCATGAGGTTTTACCGTGTTAGCCANNNAGATCGGAAG |
| 35170_2 | TATGAGTGTGGAGTCGTTGCATGAGGTTTTACCGTGTTAGCCANNNAGATCGGAAG |
| 35170_3 | TATGAGTGTGGAGTCGTTGCATGAGGTTTTACCGTGTTAGCCANNNAGATCGGAAG |
| 51988_1 | TATGAGTGTGGAGTCGTTGCCTTTCCCTTTCTCCTCCTTGACANNNAGATCGGAAGA |
| 51988_2 | TATGAGTGTGGAGTCGTTGCCTTTCCCTTTCTCCTCCTTGACANNNAGATCGGAAGA |
| 51988_3 | TATGAGTGTGGAGTCGTTGCCTTTCCCTTTCTCCTCCTTGACANNNAGATCGGAAGA |
| 29974_1 | TATGAGTGTGGAGTCGTTGCTGTTTCTTTCAAGAGGAAATTTGTGTNNNAGATCGG |
| 29974_2 | TATGAGTGTGGAGTCGTTGCTGTTTCTTTCAAGAGGAAATTTGTGTNNNAGATCGG |
| 29974_3 | TATGAGTGTGGAGTCGTTGCTGTTTCTTTCAAGAGGAAATTTGTGTNNNAGATCGG |
| 47065_1 | TATGAGTGTGGAGTCGTTGCTTACGCGACGAACTTCAACATTCNNNAGATCGGAAG |

| | |
|---|---|
| 47065_2 | TATGAGTGTGGAGTCGTTGCTTACGCGACGAACTTCAACATTCNNNAGATCGGAAG |
| 47065_3 | TATGAGTGTGGAGTCGTTGCTTACGCGACGAACTTCAACATTCNNNAGATCGGAAG |
| 49599_1 | TATGAGTGTGGAGTCGTTGCGGAACCAGGAATCATGCCAAAAANNNAGATCGGAA |
| 49599_2 | TATGAGTGTGGAGTCGTTGCGGAACCAGGAATCATGCCAAAAANNNAGATCGGAA |
| 49599_3 | TATGAGTGTGGAGTCGTTGCGGAACCAGGAATCATGCCAAAAANNNAGATCGGAA |
| 47463_1 | TATGAGTGTGGAGTCGTTGCGAGGAAGATCAAAAGAGGGAGGGNNNAGATCGG |
| 47463_2 | TATGAGTGTGGAGTCGTTGCGAGGAAGATCAAAAGAGGGAGGGNNNAGATCGG |
| 47463_3 | TATGAGTGTGGAGTCGTTGCGAGGAAGATCAAAAGAGGGAGGGNNNAGATCGG |
| 31352_1 | TATGAGTGTGGAGTCGTTGCCCACCACTGCTAACTGGACTAATNNNAGATCGGAAG |
| 31352_2 | TATGAGTGTGGAGTCGTTGCCCACCACTGCTAACTGGACTAATNNNAGATCGGAAG |
| 31352_3 | TATGAGTGTGGAGTCGTTGCCCACCACTGCTAACTGGACTAATNNNAGATCGGAAG |
| 3200_1 | TATGAGTGTGGAGTCGTTGCAAGGGGAAAGGAATTACATGATANNNAGATCGGA |
| 3200_2 | TATGAGTGTGGAGTCGTTGCAAGGGGAAAGGAATTACATGATANNNAGATCGGA |
| 3200_3 | TATGAGTGTGGAGTCGTTGCAAGGGGAAAGGAATTACATGATANNNAGATCGGA |
| 33135_1 | TATGAGTGTGGAGTCGTTGCAAGAAGAAAGTACTCTGCCTGGGNNNAGATCGGAA |
| 33135_2 | TATGAGTGTGGAGTCGTTGCAAGAAGAAAGTACTCTGCCTGGGNNNAGATCGGAA |
| 33135_3 | TATGAGTGTGGAGTCGTTGCAAGAAGAAAGTACTCTGCCTGGGNNNAGATCGGAA |
| 34443_1 | TATGAGTGTGGAGTCGTTGCCCGGATCCCCCTAGTTTATCTTCNNNAGATCGGAAGA |
| 34443_2 | TATGAGTGTGGAGTCGTTGCCCGGATCCCCCTAGTTTATCTTCNNNAGATCGGAAGA |
| 34443_3 | TATGAGTGTGGAGTCGTTGCCCGGATCCCCCTAGTTTATCTTCNNNAGATCGGAAGA |
| 51300_1 | TATGAGTGTGGAGTCGTTGCCACTGGCAACAGGTTGATACATGNNNAGATCGGAAG |
| 51300_2 | TATGAGTGTGGAGTCGTTGCCACTGGCAACAGGTTGATACATGNNNAGATCGGAAG |
| 51300_3 | TATGAGTGTGGAGTCGTTGCCACTGGCAACAGGTTGATACATGNNNAGATCGGAAG |
| 45927_1 | TATGAGTGTGGAGTCGTTGCGCGCCCGGCTAATTTTTTGTATTNNNAGATCGGAAG |
| 45927_2 | TATGAGTGTGGAGTCGTTGCGCGCCCGGCTAATTTTTTGTATTNNNAGATCGGAAG |
| 45927_3 | TATGAGTGTGGAGTCGTTGCGCGCCCGGCTAATTTTTTGTATTNNNAGATCGGAAG |
| 34008_1 | TATGAGTGTGGAGTCGTTGCAGACAGAATCTCACTGTGTCACCNNNAGATCGGAAG |
| 34008_2 | TATGAGTGTGGAGTCGTTGCAGACAGAATCTCACTGTGTCACCNNNAGATCGGAAG |
| 34008_3 | TATGAGTGTGGAGTCGTTGCAGACAGAATCTCACTGTGTCACCNNNAGATCGGAAG |
| 37167_1 | TATGAGTGTGGAGTCGTTGCTTGTGGATTGAATAGTGTCCCCCNNNAGATCGGAAG |
| 37167_2 | TATGAGTGTGGAGTCGTTGCTTGTGGATTGAATAGTGTCCCCCNNNAGATCGGAAG |
| 37167_3 | TATGAGTGTGGAGTCGTTGCTTGTGGATTGAATAGTGTCCCCCNNNAGATCGGAAG |
| 12927_1 | TATGAGTGTGGAGTCGTTGCATTTCCCTGAAGTATAAATAAGCNNNAGATCGGAAG |
| 12927_2 | TATGAGTGTGGAGTCGTTGCATTTCCCTGAAGTATAAATAAGCNNNAGATCGGAAG |
| 12927_3 | TATGAGTGTGGAGTCGTTGCATTTCCCTGAAGTATAAATAAGCNNNAGATCGGAAG |
| 51110_1 | TATGAGTGTGGAGTCGTTGCGGTGAGCTACAGATCGGAAGTATNNNAGATCGGAA |
| 51110_2 | TATGAGTGTGGAGTCGTTGCGGTGAGCTACAGATCGGAAGTATNNNAGATCGGAA |
| 51110_3 | TATGAGTGTGGAGTCGTTGCGGTGAGCTACAGATCGGAAGTATNNNAGATCGGAA |
| 49768_1 | TATGAGTGTGGAGTCGTTGCCCTCTAAAGTACATGCCCATGGTNNNAGATCGGAAG |
| 49768_2 | TATGAGTGTGGAGTCGTTGCCCTCTAAAGTACATGCCCATGGTNNNAGATCGGAAG |
| 49768_3 | TATGAGTGTGGAGTCGTTGCCCTCTAAAGTACATGCCCATGGTNNNAGATCGGAAG |
| 35785_1 | TATGAGTGTGGAGTCGTTGCCAGAGACTCATTGTGTCACCCAGNNNAGATCGGAAG |
| 35785_2 | TATGAGTGTGGAGTCGTTGCCAGAGACTCATTGTGTCACCCAGNNNAGATCGGAAG |

| | |
|---|---|
| 35785_3 | TATGAGTGTGGAGTCGTTGCCAGAGACTCATTGTGTCACCCAGNNNAGATCGGAAG |
| 51845_1 | TATGAGTGTGGAGTCGTTGCACAATGCGGAACTGTGAGTCTATNNNAGATCGGAA |
| 51845_2 | TATGAGTGTGGAGTCGTTGCACAATGCGGAACTGTGAGTCTATNNNAGATCGGAA |
| 51845_3 | TATGAGTGTGGAGTCGTTGCACAATGCGGAACTGTGAGTCTATNNNAGATCGGAA |
| 45488_1 | TATGAGTGTGGAGTCGTTGCGCTGTCACCGAGTAGAAATTTGGNNNAGATCGGAA |
| 45488_2 | TATGAGTGTGGAGTCGTTGCGCTGTCACCGAGTAGAAATTTGGNNNAGATCGGAA |
| 45488_3 | TATGAGTGTGGAGTCGTTGCGCTGTCACCGAGTAGAAATTTGGNNNAGATCGGAA |
| 31446_1 | TATGAGTGTGGAGTCGTTGCTTCAGGCTTATCCATCTACTGCCNNNAGATCGGAAGA |
| 31446_2 | TATGAGTGTGGAGTCGTTGCTTCAGGCTTATCCATCTACTGCCNNNAGATCGGAAGA |
| 31446_3 | TATGAGTGTGGAGTCGTTGCTTCAGGCTTATCCATCTACTGCCNNNAGATCGGAAGA |
| 44703_1 | TATGAGTGTGGAGTCGTTGCCAGTGTCATGTTGAACCCCTCTANNNAGATCGGAAG |
| 44703_2 | TATGAGTGTGGAGTCGTTGCCAGTGTCATGTTGAACCCCTCTANNNAGATCGGAAG |
| 44703_3 | TATGAGTGTGGAGTCGTTGCCAGTGTCATGTTGAACCCCTCTANNNAGATCGGAAG |
| 51752_1 | TATGAGTGTGGAGTCGTTGCCTGAAGGATGGAATTGTGTGCTTNNNAGATCGGAA |
| 51752_2 | TATGAGTGTGGAGTCGTTGCCTGAAGGATGGAATTGTGTGCTTNNNAGATCGGAA |
| 51752_3 | TATGAGTGTGGAGTCGTTGCCTGAAGGATGGAATTGTGTGCTTNNNAGATCGGAA |
| 40731_1 | TATGAGTGTGGAGTCGTTGCAGGGTTTGGTCAACGTGAAAATCNNNAGATCGGAA |
| 40731_2 | TATGAGTGTGGAGTCGTTGCAGGGTTTGGTCAACGTGAAAATCNNNAGATCGGAA |
| 40731_3 | TATGAGTGTGGAGTCGTTGCAGGGTTTGGTCAACGTGAAAATCNNNAGATCGGAA |
| 51055_1 | TATGAGTGTGGAGTCGTTGCCGGAGTGGCGTATATGCTCTACNNNAGATCGGAAG |
| 51055_2 | TATGAGTGTGGAGTCGTTGCCGGAGTGGCGTATATGCTCTACNNNAGATCGGAAG |
| 51055_3 | TATGAGTGTGGAGTCGTTGCCGGAGTGGCGTATATGCTCTACNNNAGATCGGAAG |
| 51999_1 | TATGAGTGTGGAGTCGTTGCTTCTGGTAACAAAGCCTCACAGANNNAGATCGGAAG |
| 51999_2 | TATGAGTGTGGAGTCGTTGCTTCTGGTAACAAAGCCTCACAGANNNAGATCGGAAG |
| 51999_3 | TATGAGTGTGGAGTCGTTGCTTCTGGTAACAAAGCCTCACAGANNNAGATCGGAAG |
| 45715_1 | TATGAGTGTGGAGTCGTTGCTTCTTCCTGCTTCAATCTTGGGANNNAGATCGGAAGA |
| 45715_2 | TATGAGTGTGGAGTCGTTGCTTCTTCCTGCTTCAATCTTGGGANNNAGATCGGAAGA |
| 45715_3 | TATGAGTGTGGAGTCGTTGCTTCTTCCTGCTTCAATCTTGGGANNNAGATCGGAAGA |
| 29200_1 | TATGAGTGTGGAGTCGTTGCTCTCTGATTCTACATGCTCCACTNNNAGATCGGAAGA |
| 29200_2 | TATGAGTGTGGAGTCGTTGCTCTCTGATTCTACATGCTCCACTNNNAGATCGGAAGA |
| 29200_3 | TATGAGTGTGGAGTCGTTGCTCTCTGATTCTACATGCTCCACTNNNAGATCGGAAGA |
| 34310_1 | TATGAGTGTGGAGTCGTTGCCTGGCCCAAATGTATGCTTTTCANNNAGATCGGAAG |
| 34310_2 | TATGAGTGTGGAGTCGTTGCCTGGCCCAAATGTATGCTTTTCANNNAGATCGGAAG |
| 34310_3 | TATGAGTGTGGAGTCGTTGCCTGGCCCAAATGTATGCTTTTCANNNAGATCGGAAG |
| 34881_1 | TATGAGTGTGGAGTCGTTGCTTCACACCATTCTCCTGCCTCNNNAGATCGGAAGAGC |
| 34881_2 | TATGAGTGTGGAGTCGTTGCTTCACACCATTCTCCTGCCTCNNNAGATCGGAAGAGC |
| 34881_3 | TATGAGTGTGGAGTCGTTGCTTCACACCATTCTCCTGCCTCNNNAGATCGGAAGAGC |
| 47488_1 | TATGAGTGTGGAGTCGTTGCCATGTATTTGCAGAGCTTCCCATNNNAGATCGGAAG |
| 47488_2 | TATGAGTGTGGAGTCGTTGCCATGTATTTGCAGAGCTTCCCATNNNAGATCGGAAG |
| 47488_3 | TATGAGTGTGGAGTCGTTGCCATGTATTTGCAGAGCTTCCCATNNNAGATCGGAAG |
| 36676_1 | TATGAGTGTGGAGTCGTTGCCTATCAGTGTGTTGGTGTTGCTGNNNAGATCGGAAG |
| 36676_2 | TATGAGTGTGGAGTCGTTGCCTATCAGTGTGTTGGTGTTGCTGNNNAGATCGGAAG |
| 36676_3 | TATGAGTGTGGAGTCGTTGCCTATCAGTGTGTTGGTGTTGCTGNNNAGATCGGAA |

| | |
|---|---|
| 10203_1 | TATGAGTGTGGAGTCGTTGCCACCAATCCCTTTTATCTGCTGCNNNAGATCGGAAGA |
| 10203_2 | TATGAGTGTGGAGTCGTTGCCACCAATCCCTTTTATCTGCTGCNNNAGATCGGAAGA |
| 10203_3 | TATGAGTGTGGAGTCGTTGCCACCAATCCCTTTTATCTGCTGCNNNAGATCGGAAGA |
| 48642_1 | TATGAGTGTGGAGTCGTTGCCTCAGACCTCAGTTCACCCTCTNNNAGATCGGAAGAG |
| 48642_2 | TATGAGTGTGGAGTCGTTGCCTCAGACCTCAGTTCACCCTCTNNNAGATCGGAAGAG |
| 48642_3 | TATGAGTGTGGAGTCGTTGCCTCAGACCTCAGTTCACCCTCTNNNAGATCGGAAGAG |
| 48653_1 | TATGAGTGTGGAGTCGTTGCCCACCACGCCAGACTAATTTTTTNNNAGATCGGAAGA |
| 48653_2 | TATGAGTGTGGAGTCGTTGCCCACCACGCCAGACTAATTTTTTNNNAGATCGGAAGA |
| 48653_3 | TATGAGTGTGGAGTCGTTGCCCACCACGCCAGACTAATTTTTTNNNAGATCGGAAGA |
| 51961_1 | TATGAGTGTGGAGTCGTTGCAGCTGGGGTATTGGTATTTTCGTNNNAGATCGGAA |
| 51961_2 | TATGAGTGTGGAGTCGTTGCAGCTGGGGTATTGGTATTTTCGTNNNAGATCGGAA |
| 51961_3 | TATGAGTGTGGAGTCGTTGCAGCTGGGGTATTGGTATTTTCGTNNNAGATCGGAA |
| 12841_1 | TATGAGTGTGGAGTCGTTGCCTCTGGGGTAATATCTAAACTTGNNNAGATCGGAAG |
| 12841_2 | TATGAGTGTGGAGTCGTTGCCTCTGGGGTAATATCTAAACTTGNNNAGATCGGAAG |
| 12841_3 | TATGAGTGTGGAGTCGTTGCCTCTGGGGTAATATCTAAACTTGNNNAGATCGGAAG |
| 52427_1 | TATGAGTGTGGAGTCGTTGCATGTTGAAGCCCTAACTCCCAATNNNAGATCGGAAGA |
| 52427_2 | TATGAGTGTGGAGTCGTTGCATGTTGAAGCCCTAACTCCCAATNNNAGATCGGAAGA |
| 52427_3 | TATGAGTGTGGAGTCGTTGCATGTTGAAGCCCTAACTCCCAATNNNAGATCGGAAGA |
| 32827_1 | TATGAGTGTGGAGTCGTTGCCTCCCAGGTTCAAGCTATTCTCANNNAGATCGGAAGA |
| 32827_2 | TATGAGTGTGGAGTCGTTGCCTCCCAGGTTCAAGCTATTCTCANNNAGATCGGAAGA |
| 32827_3 | TATGAGTGTGGAGTCGTTGCCTCCCAGGTTCAAGCTATTCTCANNNAGATCGGAAGA |
| 39742_1 | TATGAGTGTGGAGTCGTTGCAGAAAAGAAAAATCAACTAGTGCAGANNNAGATCG |
| 39742_2 | TATGAGTGTGGAGTCGTTGCAGAAAAGAAAAATCAACTAGTGCAGANNNAGATCG |
| 39742_3 | TATGAGTGTGGAGTCGTTGCAGAAAAGAAAAATCAACTAGTGCAGANNNAGATCG |
| 38149_1 | TATGAGTGTGGAGTCGTTGCAATACAAAATTAGCTGGGCGTGGNNNAGATCGGAA |
| 38149_2 | TATGAGTGTGGAGTCGTTGCAATACAAAATTAGCTGGGCGTGGNNNAGATCGGAA |
| 38149_3 | TATGAGTGTGGAGTCGTTGCAATACAAAATTAGCTGGGCGTGGNNNAGATCGGAA |
| 13464_1 | TATGAGTGTGGAGTCGTTGCTGAAAATTAAGTGCTATGGTAAGNNNAGATCGGAA |
| 13464_2 | TATGAGTGTGGAGTCGTTGCTGAAAATTAAGTGCTATGGTAAGNNNAGATCGGAA |
| 13464_3 | TATGAGTGTGGAGTCGTTGCTGAAAATTAAGTGCTATGGTAAGNNNAGATCGGAA |
| 44722_1 | TATGAGTGTGGAGTCGTTGCAGAGGAGAGAGTCTAGACCACAGNNNAGATCGGAA |
| 44722_2 | TATGAGTGTGGAGTCGTTGCAGAGGAGAGAGTCTAGACCACAGNNNAGATCGGAA |
| 44722_3 | TATGAGTGTGGAGTCGTTGCAGAGGAGAGAGTCTAGACCACAGNNNAGATCGGAA |
| 49084_1 | TATGAGTGTGGAGTCGTTGCAAGTGATTCTTGTGCCTCAGTCTNNNAGATCGGAAGA |
| 49084_2 | TATGAGTGTGGAGTCGTTGCAAGTGATTCTTGTGCCTCAGTCTNNNAGATCGGAAGA |
| 49084_3 | TATGAGTGTGGAGTCGTTGCAAGTGATTCTTGTGCCTCAGTCTNNNAGATCGGAAGA |
| 29165_1 | TATGAGTGTGGAGTCGTTGCCATACACACACACACACACACTCNNNAGATCGGAAGA |
| 29165_2 | TATGAGTGTGGAGTCGTTGCCATACACACACACACACACACTCNNNAGATCGGAAGA |
| 29165_3 | TATGAGTGTGGAGTCGTTGCCATACACACACACACACACACTCNNNAGATCGGAAGA |
| 36303_1 | TATGAGTGTGGAGTCGTTGCAAAACAAAGAATTGAGGCCGAGGNNNAGATCGGAA |
| 36303_2 | TATGAGTGTGGAGTCGTTGCAAAACAAAGAATTGAGGCCGAGGNNNAGATCGGAA |
| 36303_3 | TATGAGTGTGGAGTCGTTGCAAAACAAAGAATTGAGGCCGAGGNNNAGATCGGAA |
| 42766_1 | TATGAGTGTGGAGTCGTTGCTGTGTGTGTGTGTGTGTTAGAGANNNAGATCGGAA |

| | |
|---|---|
| 42766_2 | TATGAGTGTGGAGTCGTTGCTGTGTGTGTGTGTGTGTTAGAGANNNAGATCGGAA |
| 42766_3 | TATGAGTGTGGAGTCGTTGCTGTGTGTGTGTGTGTGTTAGAGANNNAGATCGGAA |
| 34029_1 | TATGAGTGTGGAGTCGTTGCAATAGCTGGGATTACAGGTGTGCNNNAGATCGGAA |
| 34029_2 | TATGAGTGTGGAGTCGTTGCAATAGCTGGGATTACAGGTGTGCNNNAGATCGGAA |
| 34029_3 | TATGAGTGTGGAGTCGTTGCAATAGCTGGGATTACAGGTGTGCNNNAGATCGGAA |
| 38305_1 | TATGAGTGTGGAGTCGTTGCAAAGACAAGGGATACACTGCAGTNNNAGATCGGAA |
| 38305_2 | TATGAGTGTGGAGTCGTTGCAAAGACAAGGGATACACTGCAGTNNNAGATCGGAA |
| 38305_3 | TATGAGTGTGGAGTCGTTGCAAAGACAAGGGATACACTGCAGTNNNAGATCGGAA |
| 52958_1 | TATGAGTGTGGAGTCGTTGCTAATGGCGAATAGGGTGTCCATTNNNAGATCGGAA |
| 52958_2 | TATGAGTGTGGAGTCGTTGCTAATGGCGAATAGGGTGTCCATTNNNAGATCGGAA |
| 52958_3 | TATGAGTGTGGAGTCGTTGCTAATGGCGAATAGGGTGTCCATTNNNAGATCGGAA |
| 39614_1 | TATGAGTGTGGAGTCGTTGCAGGAAGAGGTCTACAGAACAAGTNNNAGATCGGAA |
| 39614_2 | TATGAGTGTGGAGTCGTTGCAGGAAGAGGTCTACAGAACAAGTNNNAGATCGGAA |
| 39614_3 | TATGAGTGTGGAGTCGTTGCAGGAAGAGGTCTACAGAACAAGTNNNAGATCGGAA |
| 31876_1 | TATGAGTGTGGAGTCGTTGCTCCCGCAGTCTTATTTATACCCGNNNAGATCGGAAGA |
| 31876_2 | TATGAGTGTGGAGTCGTTGCTCCCGCAGTCTTATTTATACCCGNNNAGATCGGAAGA |
| 31876_3 | TATGAGTGTGGAGTCGTTGCTCCCGCAGTCTTATTTATACCCGNNNAGATCGGAAGA |
| 50278_1 | TATGAGTGTGGAGTCGTTGCGGGATAGAATTCAATGAGGGGGGNNNAGATCGGA |
| 50278_2 | TATGAGTGTGGAGTCGTTGCGGGATAGAATTCAATGAGGGGGGNNNAGATCGGA |
| 50278_3 | TATGAGTGTGGAGTCGTTGCGGGATAGAATTCAATGAGGGGGGNNNAGATCGGA |
| 51437_1 | TATGAGTGTGGAGTCGTTGCGAGGAGAGGAGGAGGGGATCNNNAGATCGGAAG |
| 51437_2 | TATGAGTGTGGAGTCGTTGCGAGGAGAGGAGGAGGGGATCNNNAGATCGGAAG |
| 51437_3 | TATGAGTGTGGAGTCGTTGCGAGGAGAGGAGGAGGGGATCNNNAGATCGGAAG |
| 37061_1 | TATGAGTGTGGAGTCGTTGCCATCCTTCAGGTAGGTTGCCATANNNAGATCGGAAG |
| 37061_2 | TATGAGTGTGGAGTCGTTGCCATCCTTCAGGTAGGTTGCCATANNNAGATCGGAAG |
| 37061_3 | TATGAGTGTGGAGTCGTTGCCATCCTTCAGGTAGGTTGCCATANNNAGATCGGAAG |
| 36886_1 | TATGAGTGTGGAGTCGTTGCGAGAATCGCTTGAACCCAGGAAGNNNAGATCGGAA |
| 36886_2 | TATGAGTGTGGAGTCGTTGCGAGAATCGCTTGAACCCAGGAAGNNNAGATCGGAA |
| 36886_3 | TATGAGTGTGGAGTCGTTGCGAGAATCGCTTGAACCCAGGAAGNNNAGATCGGAA |
| 40212_1 | TATGAGTGTGGAGTCGTTGCAACACAAAAATGAGGTCAGGTGCNNNAGATCGGAA |
| 40212_2 | TATGAGTGTGGAGTCGTTGCAACACAAAAATGAGGTCAGGTGCNNNAGATCGGAA |
| 40212_3 | TATGAGTGTGGAGTCGTTGCAACACAAAAATGAGGTCAGGTGCNNNAGATCGGAA |
| 34244_1 | TATGAGTGTGGAGTCGTTGCCTCGAACTCCTGACCTCATGATCNNNAGATCGGAAGA |
| 34244_2 | TATGAGTGTGGAGTCGTTGCCTCGAACTCCTGACCTCATGATCNNNAGATCGGAAGA |
| 34244_3 | TATGAGTGTGGAGTCGTTGCCTCGAACTCCTGACCTCATGATCNNNAGATCGGAAGA |
| 30160_1 | TATGAGTGTGGAGTCGTTGCAAGAAGGTAAGTCAAGGATAAGGGATNNNAGATCG |
| 30160_2 | TATGAGTGTGGAGTCGTTGCAAGAAGGTAAGTCAAGGATAAGGGATNNNAGATCG |
| 30160_3 | TATGAGTGTGGAGTCGTTGCAAGAAGGTAAGTCAAGGATAAGGGATNNNAGATCG |
| 7814_1 | TATGAGTGTGGAGTCGTTGCAAGAAAGGAGGGGAAACCTTATTNNNAGATCGGAA |
| 7814_2 | TATGAGTGTGGAGTCGTTGCAAGAAAGGAGGGGAAACCTTATTNNNAGATCGGAA |
| 7814_3 | TATGAGTGTGGAGTCGTTGCAAGAAAGGAGGGGAAACCTTATTNNNAGATCGGAA |
| 30229_1 | TATGAGTGTGGAGTCGTTGCATCTGCAAACCCTCCCAGATTAGNNNAGATCGGAAG |
| 30229_2 | TATGAGTGTGGAGTCGTTGCATCTGCAAACCCTCCCAGATTAGNNNAGATCGGAAG |

| | |
|---|---|
| 30229_3 | TATGAGTGTGGAGTCGTTGCATCTGCAAACCCTCCCAGATTAGNNNAGATCGGAAG |
| 34460_1 | TATGAGTGTGGAGTCGTTGCTGTGTCTGGCTGATTGGTCAATANNNAGATCGGAA |
| 34460_2 | TATGAGTGTGGAGTCGTTGCTGTGTCTGGCTGATTGGTCAATANNNAGATCGGAA |
| 34460_3 | TATGAGTGTGGAGTCGTTGCTGTGTCTGGCTGATTGGTCAATANNNAGATCGGAA |
| 10726_1 | TATGAGTGTGGAGTCGTTGCAGGCAGCCAGAAAGAAATAACACANNNAGATCGGA |
| 10726_2 | TATGAGTGTGGAGTCGTTGCAGGCAGCCAGAAAGAAATAACACANNNAGATCGGA |
| 10726_3 | TATGAGTGTGGAGTCGTTGCAGGCAGCCAGAAAGAAATAACACANNNAGATCGGA |
| 52056_1 | TATGAGTGTGGAGTCGTTGCAAGACTCCCAGTCAAAAAGGTGTNNNAGATCGGAA |
| 52056_2 | TATGAGTGTGGAGTCGTTGCAAGACTCCCAGTCAAAAAGGTGTNNNAGATCGGAA |
| 52056_3 | TATGAGTGTGGAGTCGTTGCAAGACTCCCAGTCAAAAAGGTGTNNNAGATCGGAA |
| 30154_1 | TATGAGTGTGGAGTCGTTGCCTGTTGCAGCATCCTTGGTTATTNNNAGATCGGAAG |
| 30154_2 | TATGAGTGTGGAGTCGTTGCCTGTTGCAGCATCCTTGGTTATTNNNAGATCGGAAG |
| 30154_3 | TATGAGTGTGGAGTCGTTGCCTGTTGCAGCATCCTTGGTTATTNNNAGATCGGAAG |
| 33887_1 | TATGAGTGTGGAGTCGTTGCGGAGTGAAATGGCACGATCTTGNNNAGATCGGAAG |
| 33887_2 | TATGAGTGTGGAGTCGTTGCGGAGTGAAATGGCACGATCTTGNNNAGATCGGAAG |
| 33887_3 | TATGAGTGTGGAGTCGTTGCGGAGTGAAATGGCACGATCTTGNNNAGATCGGAAG |
| 38900_1 | TATGAGTGTGGAGTCGTTGCACCCAATAATTCATGAATCCTCCTNNNAGATCGGAAG |
| 38900_2 | TATGAGTGTGGAGTCGTTGCACCCAATAATTCATGAATCCTCCTNNNAGATCGGAAG |
| 38900_3 | TATGAGTGTGGAGTCGTTGCACCCAATAATTCATGAATCCTCCTNNNAGATCGGAAG |
| 51586_1 | TATGAGTGTGGAGTCGTTGCGGTCTTCTGTCACTCTTGCTCTTNNNAGATCGGAAGA |
| 51586_2 | TATGAGTGTGGAGTCGTTGCGGTCTTCTGTCACTCTTGCTCTTNNNAGATCGGAAGA |
| 51586_3 | TATGAGTGTGGAGTCGTTGCGGTCTTCTGTCACTCTTGCTCTTNNNAGATCGGAAGA |
| 32298_1 | TATGAGTGTGGAGTCGTTGCTACAGGGCTGAGCTACCATATCTNNNAGATCGGAAG |
| 32298_2 | TATGAGTGTGGAGTCGTTGCTACAGGGCTGAGCTACCATATCTNNNAGATCGGAAG |
| 32298_3 | TATGAGTGTGGAGTCGTTGCTACAGGGCTGAGCTACCATATCTNNNAGATCGGAAG |
| 12484_1 | TATGAGTGTGGAGTCGTTGCAAGACCTACATGTTCAGAATCNNNAGATCGGAAGAG |
| 12484_2 | TATGAGTGTGGAGTCGTTGCAAGACCTACATGTTCAGAATCNNNAGATCGGAAGAG |
| 12484_3 | TATGAGTGTGGAGTCGTTGCAAGACCTACATGTTCAGAATCNNNAGATCGGAAGAG |
| 40327_1 | TATGAGTGTGGAGTCGTTGCCTTACGCCTGTAATCCCAGCANNNAGATCGGAAGAG |
| 40327_2 | TATGAGTGTGGAGTCGTTGCCTTACGCCTGTAATCCCAGCANNNAGATCGGAAGAG |
| 40327_3 | TATGAGTGTGGAGTCGTTGCCTTACGCCTGTAATCCCAGCANNNAGATCGGAAGAG |
| 50973_1 | TATGAGTGTGGAGTCGTTGCAGCCACATGTTAGTGAAGCAGTANNNAGATCGGAA |
| 50973_2 | TATGAGTGTGGAGTCGTTGCAGCCACATGTTAGTGAAGCAGTANNNAGATCGGAA |
| 50973_3 | TATGAGTGTGGAGTCGTTGCAGCCACATGTTAGTGAAGCAGTANNNAGATCGGAA |
| 30548_1 | TATGAGTGTGGAGTCGTTGCCTGAAGATGCGCCTTACTTTGTANNNAGATCGGAAG |
| 30548_2 | TATGAGTGTGGAGTCGTTGCCTGAAGATGCGCCTTACTTTGTANNNAGATCGGAAG |
| 30548_3 | TATGAGTGTGGAGTCGTTGCCTGAAGATGCGCCTTACTTTGTANNNAGATCGGAAG |
| 30347_1 | TATGAGTGTGGAGTCGTTGCTCCACCTGAGTAAGTATTGTGAACCNNNAGATCGGA |
| 30347_2 | TATGAGTGTGGAGTCGTTGCTCCACCTGAGTAAGTATTGTGAACCNNNAGATCGGA |
| 30347_3 | TATGAGTGTGGAGTCGTTGCTCCACCTGAGTAAGTATTGTGAACCNNNAGATCGGA |
| 49534_1 | TATGAGTGTGGAGTCGTTGCCAGTTAGCTGGACAAAACTGAGCNNNAGATCGGAA |
| 49534_2 | TATGAGTGTGGAGTCGTTGCCAGTTAGCTGGACAAAACTGAGCNNNAGATCGGAA |
| 49534_3 | TATGAGTGTGGAGTCGTTGCCAGTTAGCTGGACAAAACTGAGCNNNAGATCGGAA |

| | |
|---|---|
| 31197_1 | TATGAGTGTGGAGTCGTTGCCTTTGCCATAACCTTCTTTCCCCNNNAGATCGGAAGA |
| 31197_2 | TATGAGTGTGGAGTCGTTGCCTTTGCCATAACCTTCTTTCCCCNNNAGATCGGAAGA |
| 31197_3 | TATGAGTGTGGAGTCGTTGCCTTTGCCATAACCTTCTTTCCCCNNNAGATCGGAAGA |
| 48206_1 | TATGAGTGTGGAGTCGTTGCTGAGAGAGTACTCACGGATTTGCNNNAGATCGGAA |
| 48206_2 | TATGAGTGTGGAGTCGTTGCTGAGAGAGTACTCACGGATTTGCNNNAGATCGGAA |
| 48206_3 | TATGAGTGTGGAGTCGTTGCTGAGAGAGTACTCACGGATTTGCNNNAGATCGGAA |
| 50936_1 | TATGAGTGTGGAGTCGTTGCTTGCTTCTCCTCTCCTCTACAGANNNAGATCGGAAGA |
| 50936_2 | TATGAGTGTGGAGTCGTTGCTTGCTTCTCCTCTCCTCTACAGANNNAGATCGGAAGA |
| 50936_3 | TATGAGTGTGGAGTCGTTGCTTGCTTCTCCTCTCCTCTACAGANNNAGATCGGAAGA |
| 40694_1 | TATGAGTGTGGAGTCGTTGCTGCTTAGTGAAACTTTGGTCAGGNNNAGATCGGAA |
| 40694_2 | TATGAGTGTGGAGTCGTTGCTGCTTAGTGAAACTTTGGTCAGGNNNAGATCGGAA |
| 40694_3 | TATGAGTGTGGAGTCGTTGCTGCTTAGTGAAACTTTGGTCAGGNNNAGATCGGAA |
| 49540_1 | TATGAGTGTGGAGTCGTTGCGATCTCGGCTCACTACAAGCTCNNNAGATCGGAAGA |
| 49540_2 | TATGAGTGTGGAGTCGTTGCGATCTCGGCTCACTACAAGCTCNNNAGATCGGAAGA |
| 49540_3 | TATGAGTGTGGAGTCGTTGCGATCTCGGCTCACTACAAGCTCNNNAGATCGGAAGA |
| 41950_1 | TATGAGTGTGGAGTCGTTGCCATTCTCCTGCCTCAGCCTCNNNAGATCGGAAGAGCA |
| 41950_2 | TATGAGTGTGGAGTCGTTGCCATTCTCCTGCCTCAGCCTCNNNAGATCGGAAGAGCA |
| 41950_3 | TATGAGTGTGGAGTCGTTGCCATTCTCCTGCCTCAGCCTCNNNAGATCGGAAGAGCA |
| 48714_1 | TATGAGTGTGGAGTCGTTGCTCAAGCAATTCTCCTCTCTCAGCNNNAGATCGGAAGA |
| 48714_2 | TATGAGTGTGGAGTCGTTGCTCAAGCAATTCTCCTCTCTCAGCNNNAGATCGGAAGA |
| 48714_3 | TATGAGTGTGGAGTCGTTGCTCAAGCAATTCTCCTCTCTCAGCNNNAGATCGGAAGA |
| 41007_1 | TATGAGTGTGGAGTCGTTGCCAGCATGGCTCTTCCTCAAAAAANNNAGATCGGAAG |
| 41007_2 | TATGAGTGTGGAGTCGTTGCCAGCATGGCTCTTCCTCAAAAAANNNAGATCGGAAG |
| 41007_3 | TATGAGTGTGGAGTCGTTGCCAGCATGGCTCTTCCTCAAAAAANNNAGATCGGAAG |
| 51597_1 | TATGAGTGTGGAGTCGTTGCCTAGTTCCCAGAGCTTTGCCATANNNAGATCGGAAG |
| 51597_2 | TATGAGTGTGGAGTCGTTGCCTAGTTCCCAGAGCTTTGCCATANNNAGATCGGAAG |
| 51597_3 | TATGAGTGTGGAGTCGTTGCCTAGTTCCCAGAGCTTTGCCATANNNAGATCGGAAG |
| 33643_1 | TATGAGTGTGGAGTCGTTGCGAGTAGCTGGGACTACAGGGGNNNAGATCGGAAG |
| 33643_2 | TATGAGTGTGGAGTCGTTGCGAGTAGCTGGGACTACAGGGGNNNAGATCGGAAG |
| 33643_3 | TATGAGTGTGGAGTCGTTGCGAGTAGCTGGGACTACAGGGGNNNAGATCGGAAG |
| 13564_1 | TATGAGTGTGGAGTCGTTGCGCTCCTTATGAGAATCTGATGNNNAGATCGGAAGA |
| 13564_2 | TATGAGTGTGGAGTCGTTGCGCTCCTTATGAGAATCTGATGNNNAGATCGGAAGA |
| 13564_3 | TATGAGTGTGGAGTCGTTGCGCTCCTTATGAGAATCTGATGNNNAGATCGGAAGA |
| 2078_1 | TATGAGTGTGGAGTCGTTGCTGGGTTTAGGTATATGGGCATACANNNAGATCGGA |
| 2078_2 | TATGAGTGTGGAGTCGTTGCTGGGTTTAGGTATATGGGCATACANNNAGATCGGA |
| 2078_3 | TATGAGTGTGGAGTCGTTGCTGGGTTTAGGTATATGGGCATACANNNAGATCGGA |
| 30404_1 | TATGAGTGTGGAGTCGTTGCAGAGTTTTGTTTGTTTCTAATCCNNNAGATCGGAAG |
| 30404_2 | TATGAGTGTGGAGTCGTTGCAGAGTTTTGTTTGTTTCTAATCCNNNAGATCGGAAG |
| 30404_3 | TATGAGTGTGGAGTCGTTGCAGAGTTTTGTTTGTTTCTAATCCNNNAGATCGGAAG |
| 37127_1 | TATGAGTGTGGAGTCGTTGCTACTCAGCCTCCATGACCTCTTANNNAGATCGGAAGA |
| 37127_2 | TATGAGTGTGGAGTCGTTGCTACTCAGCCTCCATGACCTCTTANNNAGATCGGAAGA |
| 37127_3 | TATGAGTGTGGAGTCGTTGCTACTCAGCCTCCATGACCTCTTANNNAGATCGGAAGA |
| 33715_1 | TATGAGTGTGGAGTCGTTGCTTGAGAGGGAGTGCTTTGAGATCNNNAGATCGGAA |

| | |
|---|---|
| 33715_2 | TATGAGTGTGGAGTCGTTGCTTGAGAGGGAGTGCTTTGAGATCNNNAGATCGGAA |
| 33715_3 | TATGAGTGTGGAGTCGTTGCTTGAGAGGGAGTGCTTTGAGATCNNNAGATCGGAA |
| 48575_1 | TATGAGTGTGGAGTCGTTGCATCATAACTCACTGCAGCCTTGANNNAGATCGGAAG |
| 48575_2 | TATGAGTGTGGAGTCGTTGCATCATAACTCACTGCAGCCTTGANNNAGATCGGAAG |
| 48575_3 | TATGAGTGTGGAGTCGTTGCATCATAACTCACTGCAGCCTTGANNNAGATCGGAAG |
| 40573_1 | TATGAGTGTGGAGTCGTTGCTAGACAGGAATGCACTTCATGGTNNNAGATCGGAA |
| 40573_2 | TATGAGTGTGGAGTCGTTGCTAGACAGGAATGCACTTCATGGTNNNAGATCGGAA |
| 40573_3 | TATGAGTGTGGAGTCGTTGCTAGACAGGAATGCACTTCATGGTNNNAGATCGGAA |
| 29196_1 | TATGAGTGTGGAGTCGTTGCGTACTCACAAACACACACATGCANNNAGATCGGAAG |
| 29196_2 | TATGAGTGTGGAGTCGTTGCGTACTCACAAACACACACATGCANNNAGATCGGAAG |
| 29196_3 | TATGAGTGTGGAGTCGTTGCGTACTCACAAACACACACATGCANNNAGATCGGAAG |
| 33207_1 | TATGAGTGTGGAGTCGTTGCAGAAAAGGACACACATTGACTGANNNAGATCGGAA |
| 33207_2 | TATGAGTGTGGAGTCGTTGCAGAAAAGGACACACATTGACTGANNNAGATCGGAA |
| 33207_3 | TATGAGTGTGGAGTCGTTGCAGAAAAGGACACACATTGACTGANNNAGATCGGAA |
| 28768_1 | TATGAGTGTGGAGTCGTTGCGTGTGTGTGTGTGTGTGTGTATGNNNAGATCGGAA |
| 28768_2 | TATGAGTGTGGAGTCGTTGCGTGTGTGTGTGTGTGTGTGTATGNNNAGATCGGAA |
| 28768_3 | TATGAGTGTGGAGTCGTTGCGTGTGTGTGTGTGTGTGTGTATGNNNAGATCGGAA |
| 43602_1 | TATGAGTGTGGAGTCGTTGCTTCAAACCACAATCATGCCTTCCNNNAGATCGGAAGA |
| 43602_2 | TATGAGTGTGGAGTCGTTGCTTCAAACCACAATCATGCCTTCCNNNAGATCGGAAGA |
| 43602_3 | TATGAGTGTGGAGTCGTTGCTTCAAACCACAATCATGCCTTCCNNNAGATCGGAAGA |
| 39124_1 | TATGAGTGTGGAGTCGTTGCTATCTTTAGCATGGCTCCCAACCNNNAGATCGGAAG |
| 39124_2 | TATGAGTGTGGAGTCGTTGCTATCTTTAGCATGGCTCCCAACCNNNAGATCGGAAG |
| 39124_3 | TATGAGTGTGGAGTCGTTGCTATCTTTAGCATGGCTCCCAACCNNNAGATCGGAAG |
| 33367_1 | TATGAGTGTGGAGTCGTTGCGGAGTGCAGTAGAGCGATCTCNNNAGATCGGAAGA |
| 33367_2 | TATGAGTGTGGAGTCGTTGCGGAGTGCAGTAGAGCGATCTCNNNAGATCGGAAGA |
| 33367_3 | TATGAGTGTGGAGTCGTTGCGGAGTGCAGTAGAGCGATCTCNNNAGATCGGAAGA |
| 30067_1 | TATGAGTGTGGAGTCGTTGCAAAAATCCCAGAGAGTGGTGTGTNNNAGATCGGAA |
| 30067_2 | TATGAGTGTGGAGTCGTTGCAAAAATCCCAGAGAGTGGTGTGTNNNAGATCGGAA |
| 30067_3 | TATGAGTGTGGAGTCGTTGCAAAAATCCCAGAGAGTGGTGTGTNNNAGATCGGAA |
| 42020_1 | TATGAGTGTGGAGTCGTTGCTTACAGAGAAGAGGGTCTCGCTANNNAGATCGGAA |
| 42020_2 | TATGAGTGTGGAGTCGTTGCTTACAGAGAAGAGGGTCTCGCTANNNAGATCGGAA |
| 42020_3 | TATGAGTGTGGAGTCGTTGCTTACAGAGAAGAGGGTCTCGCTANNNAGATCGGAA |
| 39969_1 | TATGAGTGTGGAGTCGTTGCCTCTATTGCTCAGGCTGGAGAGNNNAGATCGGAAG |
| 39969_2 | TATGAGTGTGGAGTCGTTGCCTCTATTGCTCAGGCTGGAGAGNNNAGATCGGAAG |
| 39969_3 | TATGAGTGTGGAGTCGTTGCCTCTATTGCTCAGGCTGGAGAGNNNAGATCGGAAG |
| 40484_1 | TATGAGTGTGGAGTCGTTGCGGTCCCAGTTACTCTGAAGGATGNNNAGATCGGAA |
| 40484_2 | TATGAGTGTGGAGTCGTTGCGGTCCCAGTTACTCTGAAGGATGNNNAGATCGGAA |
| 40484_3 | TATGAGTGTGGAGTCGTTGCGGTCCCAGTTACTCTGAAGGATGNNNAGATCGGAA |
| 35886_1 | TATGAGTGTGGAGTCGTTGCAAAATCTACATCCACAGGACCCCNNNAGATCGGAAG |
| 35886_2 | TATGAGTGTGGAGTCGTTGCAAAATCTACATCCACAGGACCCCNNNAGATCGGAAG |
| 35886_3 | TATGAGTGTGGAGTCGTTGCAAAATCTACATCCACAGGACCCCNNNAGATCGGAAG |
| 43762_1 | TATGAGTGTGGAGTCGTTGCGCATCCGTATTTCTGCAAAGGACNNNAGATCGGAAG |
| 43762_2 | TATGAGTGTGGAGTCGTTGCGCATCCGTATTTCTGCAAAGGACNNNAGATCGGAAG |

| | |
|---|---|
| 43762_3 | TATGAGTGTGGAGTCGTTGCGCATCCGTATTTCTGCAAAGGACNNNAGATCGGAAG |
| 24658_1 | TATGAGTGTGGAGTCGTTGCCTCCAATCTCTACTGTAATAAGANNNAGATCGGAAG |
| 24658_2 | TATGAGTGTGGAGTCGTTGCCTCCAATCTCTACTGTAATAAGANNNAGATCGGAAG |
| 24658_3 | TATGAGTGTGGAGTCGTTGCCTCCAATCTCTACTGTAATAAGANNNAGATCGGAAG |
| 41576_1 | TATGAGTGTGGAGTCGTTGCCAAGTAGCTGGGACTACAGGTGNNNAGATCGGAAG |
| 41576_2 | TATGAGTGTGGAGTCGTTGCCAAGTAGCTGGGACTACAGGTGNNNAGATCGGAAG |
| 41576_3 | TATGAGTGTGGAGTCGTTGCCAAGTAGCTGGGACTACAGGTGNNNAGATCGGAAG |
| 33596_1 | TATGAGTGTGGAGTCGTTGCTTCACATCATTCTCCTGCCTCAGNNNAGATCGGAAGA |
| 33596_2 | TATGAGTGTGGAGTCGTTGCTTCACATCATTCTCCTGCCTCAGNNNAGATCGGAAGA |
| 33596_3 | TATGAGTGTGGAGTCGTTGCTTCACATCATTCTCCTGCCTCAGNNNAGATCGGAAGA |
| 35990_1 | TATGAGTGTGGAGTCGTTGCTGATGTCGTCAAGATTCTGGTCANNNAGATCGGAAG |
| 35990_2 | TATGAGTGTGGAGTCGTTGCTGATGTCGTCAAGATTCTGGTCANNNAGATCGGAAG |
| 35990_3 | TATGAGTGTGGAGTCGTTGCTGATGTCGTCAAGATTCTGGTCANNNAGATCGGAAG |
| 2455_1 | TATGAGTGTGGAGTCGTTGCGCTCCCTTTGTGTTGCCAATTTCNNNAGATCGGAAGA |
| 2455_2 | TATGAGTGTGGAGTCGTTGCGCTCCCTTTGTGTTGCCAATTTCNNNAGATCGGAAGA |
| 2455_3 | TATGAGTGTGGAGTCGTTGCGCTCCCTTTGTGTTGCCAATTTCNNNAGATCGGAAGA |
| 35105_1 | TATGAGTGTGGAGTCGTTGCGTTCAATGGCACGATCTTGTTCANNNAGATCGGAAG |
| 35105_2 | TATGAGTGTGGAGTCGTTGCGTTCAATGGCACGATCTTGTTCANNNAGATCGGAAG |
| 35105_3 | TATGAGTGTGGAGTCGTTGCGTTCAATGGCACGATCTTGTTCANNNAGATCGGAAG |
| 49288_1 | TATGAGTGTGGAGTCGTTGCCTCCTGAGTAGCTGGGATTACAGNNNAGATCGGAAG |
| 49288_2 | TATGAGTGTGGAGTCGTTGCCTCCTGAGTAGCTGGGATTACAGNNNAGATCGGAAG |
| 49288_3 | TATGAGTGTGGAGTCGTTGCCTCCTGAGTAGCTGGGATTACAGNNNAGATCGGAAG |
| 29225_1 | TATGAGTGTGGAGTCGTTGCGAAGCTTGGAGTTTGCATGTGAANNNAGATCGGAA |
| 29225_2 | TATGAGTGTGGAGTCGTTGCGAAGCTTGGAGTTTGCATGTGAANNNAGATCGGAA |
| 29225_3 | TATGAGTGTGGAGTCGTTGCGAAGCTTGGAGTTTGCATGTGAANNNAGATCGGAA |
| 40946_1 | TATGAGTGTGGAGTCGTTGCCAATCTCAGTTCACTGCAAGCTCNNNAGATCGGAAG |
| 40946_2 | TATGAGTGTGGAGTCGTTGCCAATCTCAGTTCACTGCAAGCTCNNNAGATCGGAAG |
| 40946_3 | TATGAGTGTGGAGTCGTTGCCAATCTCAGTTCACTGCAAGCTCNNNAGATCGGAAG |
| 40814_1 | TATGAGTGTGGAGTCGTTGCCTTTTTTTGAGACAGAGGCTCGCNNNAGATCGGAAG |
| 40814_2 | TATGAGTGTGGAGTCGTTGCCTTTTTTTGAGACAGAGGCTCGCNNNAGATCGGAAG |
| 40814_3 | TATGAGTGTGGAGTCGTTGCCTTTTTTTGAGACAGAGGCTCGCNNNAGATCGGAAG |
| 13543_1 | TATGAGTGTGGAGTCGTTGCAAAAAGGTGAGCAAACTGATATNNNAGATCGGAAG |
| 13543_2 | TATGAGTGTGGAGTCGTTGCAAAAAGGTGAGCAAACTGATATNNNAGATCGGAAG |
| 13543_3 | TATGAGTGTGGAGTCGTTGCAAAAAGGTGAGCAAACTGATATNNNAGATCGGAAG |
| 8854_1 | TATGAGTGTGGAGTCGTTGCCAAGTAGGCCAATTACGTGCTGTNNNAGATCGGAAG |
| 8854_2 | TATGAGTGTGGAGTCGTTGCCAAGTAGGCCAATTACGTGCTGTNNNAGATCGGAAG |
| 8854_3 | TATGAGTGTGGAGTCGTTGCCAAGTAGGCCAATTACGTGCTGTNNNAGATCGGAAG |
| 31030_1 | TATGAGTGTGGAGTCGTTGCCTTTCCCTCTTTCCCTTTCCCTTNNNAGATCGGAAGAG |
| 31030_2 | TATGAGTGTGGAGTCGTTGCCTTTCCCTCTTTCCCTTTCCCTTNNNAGATCGGAAGAG |
| 31030_3 | TATGAGTGTGGAGTCGTTGCCTTTCCCTCTTTCCCTTTCCCTTNNNAGATCGGAAGAG |
| 51950_1 | TATGAGTGTGGAGTCGTTGCGCTGTCCTGTCATCTTGAGACTTNNNAGATCGGAAG |
| 51950_2 | TATGAGTGTGGAGTCGTTGCGCTGTCCTGTCATCTTGAGACTTNNNAGATCGGAAG |
| 51950_3 | TATGAGTGTGGAGTCGTTGCGCTGTCCTGTCATCTTGAGACTTNNNAGATCGGAAG |

| | |
|---|---|
| 37643_1 | TATGAGTGTGGAGTCGTTGCTTGCAATCCCCCTGTCTTGATAANNNAGATCGGAAG |
| 37643_2 | TATGAGTGTGGAGTCGTTGCTTGCAATCCCCCTGTCTTGATAANNNAGATCGGAAG |
| 37643_3 | TATGAGTGTGGAGTCGTTGCTTGCAATCCCCCTGTCTTGATAANNNAGATCGGAAG |
| 51565_1 | TATGAGTGTGGAGTCGTTGCGTCAGTTGATATGGGGTTGGACTNNNAGATCGGAA |
| 51565_2 | TATGAGTGTGGAGTCGTTGCGTCAGTTGATATGGGGTTGGACTNNNAGATCGGAA |
| 51565_3 | TATGAGTGTGGAGTCGTTGCGTCAGTTGATATGGGGTTGGACTNNNAGATCGGAA |
| 53358_1 | TATGAGTGTGGAGTCGTTGCCCTCCCTCCTCCTTTTCTTTTCTNNNAGATCGGAAGAG |
| 53358_2 | TATGAGTGTGGAGTCGTTGCCCTCCCTCCTCCTTTTCTTTTCTNNNAGATCGGAAGAG |
| 53358_3 | TATGAGTGTGGAGTCGTTGCCCTCCCTCCTCCTTTTCTTTTCTNNNAGATCGGAAGAG |
| 34675_1 | TATGAGTGTGGAGTCGTTGCTCTCCCTTTTGGCCCTTTATCTCNNNAGATCGGAAGA |
| 34675_2 | TATGAGTGTGGAGTCGTTGCTCTCCCTTTTGGCCCTTTATCTCNNNAGATCGGAAGA |
| 34675_3 | TATGAGTGTGGAGTCGTTGCTCTCCCTTTTGGCCCTTTATCTCNNNAGATCGGAAGA |
| 46403_1 | TATGAGTGTGGAGTCGTTGCTATAGTGGAAGGACCAGGGACTTNNNAGATCGGAA |
| 46403_2 | TATGAGTGTGGAGTCGTTGCTATAGTGGAAGGACCAGGGACTTNNNAGATCGGAA |
| 46403_3 | TATGAGTGTGGAGTCGTTGCTATAGTGGAAGGACCAGGGACTTNNNAGATCGGAA |
| 43140_1 | TATGAGTGTGGAGTCGTTGCAATCATGAGGGGAGAAAACCACANNNAGATCGGAA |
| 43140_2 | TATGAGTGTGGAGTCGTTGCAATCATGAGGGGAGAAAACCACANNNAGATCGGAA |
| 43140_3 | TATGAGTGTGGAGTCGTTGCAATCATGAGGGGAGAAAACCACANNNAGATCGGAA |
| 38950_1 | TATGAGTGTGGAGTCGTTGCCCTTTGCAGAAACATGGATGGAGNNNAGATCGGAA |
| 38950_2 | TATGAGTGTGGAGTCGTTGCCCTTTGCAGAAACATGGATGGAGNNNAGATCGGAA |
| 38950_3 | TATGAGTGTGGAGTCGTTGCCCTTTGCAGAAACATGGATGGAGNNNAGATCGGAA |
| 30925_1 | TATGAGTGTGGAGTCGTTGCCAATGATGACAGGAAGACAGCCTNNNAGATCGGAA |
| 30925_2 | TATGAGTGTGGAGTCGTTGCCAATGATGACAGGAAGACAGCCTNNNAGATCGGAA |
| 30925_3 | TATGAGTGTGGAGTCGTTGCCAATGATGACAGGAAGACAGCCTNNNAGATCGGAA |
| 32934_1 | TATGAGTGTGGAGTCGTTGCTGACATTCCTCCTTGTGAGAGTGNNNAGATCGGAAG |
| 32934_2 | TATGAGTGTGGAGTCGTTGCTGACATTCCTCCTTGTGAGAGTGNNNAGATCGGAAG |
| 32934_3 | TATGAGTGTGGAGTCGTTGCTGACATTCCTCCTTGTGAGAGTGNNNAGATCGGAAG |
| 39792_1 | TATGAGTGTGGAGTCGTTGCAGGAGCTGGTACCATTTCTTCTGNNNAGATCGGAAG |
| 39792_2 | TATGAGTGTGGAGTCGTTGCAGGAGCTGGTACCATTTCTTCTGNNNAGATCGGAAG |
| 39792_3 | TATGAGTGTGGAGTCGTTGCAGGAGCTGGTACCATTTCTTCTGNNNAGATCGGAAG |
| 48201_1 | TATGAGTGTGGAGTCGTTGCCACTCTTTCGGACGTAGCATTTCNNNAGATCGGAAG |
| 48201_2 | TATGAGTGTGGAGTCGTTGCCACTCTTTCGGACGTAGCATTTCNNNAGATCGGAAG |
| 48201_3 | TATGAGTGTGGAGTCGTTGCCACTCTTTCGGACGTAGCATTTCNNNAGATCGGAAG |
| 39447_1 | TATGAGTGTGGAGTCGTTGCTATGGCTACTTGGTGCAACAATCNNNAGATCGGAAG |
| 39447_2 | TATGAGTGTGGAGTCGTTGCTATGGCTACTTGGTGCAACAATCNNNAGATCGGAAG |
| 39447_3 | TATGAGTGTGGAGTCGTTGCTATGGCTACTTGGTGCAACAATCNNNAGATCGGAAG |
| 49855_1 | TATGAGTGTGGAGTCGTTGCCAGGCATGAGAAAGTTTCACAGGNNNAGATCGGAA |
| 49855_2 | TATGAGTGTGGAGTCGTTGCCAGGCATGAGAAAGTTTCACAGGNNNAGATCGGAA |
| 49855_3 | TATGAGTGTGGAGTCGTTGCCAGGCATGAGAAAGTTTCACAGGNNNAGATCGGAA |
| 40069_1 | TATGAGTGTGGAGTCGTTGCTGTCTACCACTCCTTCTACCACTNNNAGATCGGAAGA |
| 40069_2 | TATGAGTGTGGAGTCGTTGCTGTCTACCACTCCTTCTACCACTNNNAGATCGGAAGA |
| 40069_3 | TATGAGTGTGGAGTCGTTGCTGTCTACCACTCCTTCTACCACTNNNAGATCGGAAGA |
| 48773_1 | TATGAGTGTGGAGTCGTTGCAAAGAGGGGAGAGCTAGGTAAGTNNNAGATCGGA |

| | |
|---|---|
| 48773_2 | TATGAGTGTGGAGTCGTTGCAAAGAGGGGAGAGCTAGGTAAGTNNNAGATCGGA |
| 48773_3 | TATGAGTGTGGAGTCGTTGCAAAGAGGGGAGAGCTAGGTAAGTNNNAGATCGGA |
| 31442_1 | TATGAGTGTGGAGTCGTTGCTGAGCTCAAACTACATATGCAGTNNNAGATCGGAAG |
| 31442_2 | TATGAGTGTGGAGTCGTTGCTGAGCTCAAACTACATATGCAGTNNNAGATCGGAAG |
| 31442_3 | TATGAGTGTGGAGTCGTTGCTGAGCTCAAACTACATATGCAGTNNNAGATCGGAAG |
| 11026_1 | TATGAGTGTGGAGTCGTTGCAAGACCAAAGTGCAATATGATAGGTTGNNNAGATC |
| 11026_2 | TATGAGTGTGGAGTCGTTGCAAGACCAAAGTGCAATATGATAGGTTGNNNAGATC |
| 11026_3 | TATGAGTGTGGAGTCGTTGCAAGACCAAAGTGCAATATGATAGGTTGNNNAGATC |
| 46516_1 | TATGAGTGTGGAGTCGTTGCCTGGGTTCAAAACATGGGAAGACNNNAGATCGGAA |
| 46516_2 | TATGAGTGTGGAGTCGTTGCCTGGGTTCAAAACATGGGAAGACNNNAGATCGGAA |
| 46516_3 | TATGAGTGTGGAGTCGTTGCCTGGGTTCAAAACATGGGAAGACNNNAGATCGGAA |
| 36921_1 | TATGAGTGTGGAGTCGTTGCAACTCACTAATGTTTTGTGGCCCNNNAGATCGGAAG |
| 36921_2 | TATGAGTGTGGAGTCGTTGCAACTCACTAATGTTTTGTGGCCCNNNAGATCGGAAG |
| 36921_3 | TATGAGTGTGGAGTCGTTGCAACTCACTAATGTTTTGTGGCCCNNNAGATCGGAAG |
| 31991_1 | TATGAGTGTGGAGTCGTTGCCTGAGTCTGAAGTCCTGAGAACCNNNAGATCGGAAG |
| 31991_2 | TATGAGTGTGGAGTCGTTGCCTGAGTCTGAAGTCCTGAGAACCNNNAGATCGGAAG |
| 31991_3 | TATGAGTGTGGAGTCGTTGCCTGAGTCTGAAGTCCTGAGAACCNNNAGATCGGAAG |
| 29919_1 | TATGAGTGTGGAGTCGTTGCAAACGTTTTTAGTCATCTGGGCCNNNAGATCGGAAG |
| 29919_2 | TATGAGTGTGGAGTCGTTGCAAACGTTTTTAGTCATCTGGGCCNNNAGATCGGAAG |
| 29919_3 | TATGAGTGTGGAGTCGTTGCAAACGTTTTTAGTCATCTGGGCCNNNAGATCGGAAG |
| 51959_1 | TATGAGTGTGGAGTCGTTGCATCCTGATCAAACCATTCCTGCTNNNAGATCGGAAGA |
| 51959_2 | TATGAGTGTGGAGTCGTTGCATCCTGATCAAACCATTCCTGCTNNNAGATCGGAAGA |
| 51959_3 | TATGAGTGTGGAGTCGTTGCATCCTGATCAAACCATTCCTGCTNNNAGATCGGAAGA |
| 39722_1 | TATGAGTGTGGAGTCGTTGCAGCCTCTGGGATTGTACTTTTGTNNNAGATCGGAAG |
| 39722_2 | TATGAGTGTGGAGTCGTTGCAGCCTCTGGGATTGTACTTTTGTNNNAGATCGGAAG |
| 39722_3 | TATGAGTGTGGAGTCGTTGCAGCCTCTGGGATTGTACTTTTGTNNNAGATCGGAAG |
| 36919_1 | TATGAGTGTGGAGTCGTTGCGGGATGTAGACTAAGAAGCCATGTNNNAGATCGGA |
| 36919_2 | TATGAGTGTGGAGTCGTTGCGGGATGTAGACTAAGAAGCCATGTNNNAGATCGGA |
| 36919_3 | TATGAGTGTGGAGTCGTTGCGGGATGTAGACTAAGAAGCCATGTNNNAGATCGGA |
| 13687_1 | TATGAGTGTGGAGTCGTTGCGTAGCGTATACATTTGTTTTGGNNNAGATCGGAAGA |
| 13687_2 | TATGAGTGTGGAGTCGTTGCGTAGCGTATACATTTGTTTTGGNNNAGATCGGAAGA |
| 13687_3 | TATGAGTGTGGAGTCGTTGCGTAGCGTATACATTTGTTTTGGNNNAGATCGGAAGA |
| 32624_1 | TATGAGTGTGGAGTCGTTGCGGGGAAAAGGCTAGAAGAAATGCNNNAGATCGGA |
| 32624_2 | TATGAGTGTGGAGTCGTTGCGGGGAAAAGGCTAGAAGAAATGCNNNAGATCGGA |
| 32624_3 | TATGAGTGTGGAGTCGTTGCGGGGAAAAGGCTAGAAGAAATGCNNNAGATCGGA |
| 46265_1 | TATGAGTGTGGAGTCGTTGCAAAAGAAAAAGTAATCCGGCCGGNNNAGATCGGAA |
| 46265_2 | TATGAGTGTGGAGTCGTTGCAAAAGAAAAAGTAATCCGGCCGGNNNAGATCGGAA |
| 46265_3 | TATGAGTGTGGAGTCGTTGCAAAAGAAAAAGTAATCCGGCCGGNNNAGATCGGAA |
| 3127_1 | TATGAGTGTGGAGTCGTTGCTCTTTGTGTACGTTATGCCTTGCNNNAGATCGGAAG |
| 3127_2 | TATGAGTGTGGAGTCGTTGCTCTTTGTGTACGTTATGCCTTGCNNNAGATCGGAAG |
| 3127_3 | TATGAGTGTGGAGTCGTTGCTCTTTGTGTACGTTATGCCTTGCNNNAGATCGGAAG |
| 39252_1 | TATGAGTGTGGAGTCGTTGCGGCAGGGAAATTTTTAGGTGCATNNNAGATCGGAA |
| 39252_2 | TATGAGTGTGGAGTCGTTGCGGCAGGGAAATTTTTAGGTGCATNNNAGATCGGAA |

| | |
|---|---|
| 39252_3 | TATGAGTGTGGAGTCGTTGCGGCAGGGAAATTTTTAGGTGCATNNNAGATCGGAA |
| 43912_1 | TATGAGTGTGGAGTCGTTGCCAAGAGGGAGGTGTTGTATTGGANNNAGATCGGA |
| 43912_2 | TATGAGTGTGGAGTCGTTGCCAAGAGGGAGGTGTTGTATTGGANNNAGATCGGA |
| 43912_3 | TATGAGTGTGGAGTCGTTGCCAAGAGGGAGGTGTTGTATTGGANNNAGATCGGA |
| 41231_1 | TATGAGTGTGGAGTCGTTGCATAGCTCGTGGAAAGTGGTTTCTNNNAGATCGGAA |
| 41231_2 | TATGAGTGTGGAGTCGTTGCATAGCTCGTGGAAAGTGGTTTCTNNNAGATCGGAA |
| 41231_3 | TATGAGTGTGGAGTCGTTGCATAGCTCGTGGAAAGTGGTTTCTNNNAGATCGGAA |
| 13682_1 | TATGAGTGTGGAGTCGTTGCAATGTGGTCCTATTTAAGCCNNNAGATCGGAAGAG |
| 13682_2 | TATGAGTGTGGAGTCGTTGCAATGTGGTCCTATTTAAGCCNNNAGATCGGAAGAG |
| 13682_3 | TATGAGTGTGGAGTCGTTGCAATGTGGTCCTATTTAAGCCNNNAGATCGGAAGAG |
| 37130_1 | TATGAGTGTGGAGTCGTTGCACACACACACACATATACACACANNNAGATCGGAAG |
| 37130_2 | TATGAGTGTGGAGTCGTTGCACACACACACACATATACACACANNNAGATCGGAAG |
| 37130_3 | TATGAGTGTGGAGTCGTTGCACACACACACACATATACACACANNNAGATCGGAAG |
| 46672_1 | TATGAGTGTGGAGTCGTTGCACTCCTAAGTAGCTGGGACTACANNNAGATCGGAAG |
| 46672_2 | TATGAGTGTGGAGTCGTTGCACTCCTAAGTAGCTGGGACTACANNNAGATCGGAAG |
| 46672_3 | TATGAGTGTGGAGTCGTTGCACTCCTAAGTAGCTGGGACTACANNNAGATCGGAAG |
| 1785_1 | TATGAGTGTGGAGTCGTTGCTACTGAAATGGGCACTCTGATACATNNNAGATCGGA |
| 1785_2 | TATGAGTGTGGAGTCGTTGCTACTGAAATGGGCACTCTGATACATNNNAGATCGGA |
| 1785_3 | TATGAGTGTGGAGTCGTTGCTACTGAAATGGGCACTCTGATACATNNNAGATCGGA |
| 43697_1 | TATGAGTGTGGAGTCGTTGCTTCTCCTCTGCACACCAGAATTCNNNAGATCGGAAGA |
| 43697_2 | TATGAGTGTGGAGTCGTTGCTTCTCCTCTGCACACCAGAATTCNNNAGATCGGAAGA |
| 43697_3 | TATGAGTGTGGAGTCGTTGCTTCTCCTCTGCACACCAGAATTCNNNAGATCGGAAGA |
| 47797_1 | TATGAGTGTGGAGTCGTTGCGAAGGGCATGCCACTCACAAGNNNAGATCGGAAGA |
| 47797_2 | TATGAGTGTGGAGTCGTTGCGAAGGGCATGCCACTCACAAGNNNAGATCGGAAGA |
| 47797_3 | TATGAGTGTGGAGTCGTTGCGAAGGGCATGCCACTCACAAGNNNAGATCGGAAGA |
| 35373_1 | TATGAGTGTGGAGTCGTTGCAGGAAATTTTTTTGCCTCAGCCTNNNAGATCGGAAG |
| 35373_2 | TATGAGTGTGGAGTCGTTGCAGGAAATTTTTTTGCCTCAGCCTNNNAGATCGGAAG |
| 35373_3 | TATGAGTGTGGAGTCGTTGCAGGAAATTTTTTTGCCTCAGCCTNNNAGATCGGAAG |
| 33556_1 | TATGAGTGTGGAGTCGTTGCCGGCTCACCATAGCAGGTATATANNNAGATCGGAAG |
| 33556_2 | TATGAGTGTGGAGTCGTTGCCGGCTCACCATAGCAGGTATATANNNAGATCGGAAG |
| 33556_3 | TATGAGTGTGGAGTCGTTGCCGGCTCACCATAGCAGGTATATANNNAGATCGGAAG |
| 36170_1 | TATGAGTGTGGAGTCGTTGCCCCTGGAGAAACTCACACATGTANNNAGATCGGAAG |
| 36170_2 | TATGAGTGTGGAGTCGTTGCCCCTGGAGAAACTCACACATGTANNNAGATCGGAAG |
| 36170_3 | TATGAGTGTGGAGTCGTTGCCCCTGGAGAAACTCACACATGTANNNAGATCGGAAG |
| 29662_1 | TATGAGTGTGGAGTCGTTGCTCCAGCTATAAAGCCAGGAGATGNNNAGATCGGAA |
| 29662_2 | TATGAGTGTGGAGTCGTTGCTCCAGCTATAAAGCCAGGAGATGNNNAGATCGGAA |
| 29662_3 | TATGAGTGTGGAGTCGTTGCTCCAGCTATAAAGCCAGGAGATGNNNAGATCGGAA |
| 43221_1 | TATGAGTGTGGAGTCGTTGCAAAGAAGTGAATGTTGCAGAGTTNNNAGATCGGAA |
| 43221_2 | TATGAGTGTGGAGTCGTTGCAAAGAAGTGAATGTTGCAGAGTTNNNAGATCGGAA |
| 43221_3 | TATGAGTGTGGAGTCGTTGCAAAGAAGTGAATGTTGCAGAGTTNNNAGATCGGAA |
| 49634_1 | TATGAGTGTGGAGTCGTTGCTTCAAGTGATTCTCCTGCCTCAGNNNAGATCGGAAG |
| 49634_2 | TATGAGTGTGGAGTCGTTGCTTCAAGTGATTCTCCTGCCTCAGNNNAGATCGGAAG |
| 49634_3 | TATGAGTGTGGAGTCGTTGCTTCAAGTGATTCTCCTGCCTCAGNNNAGATCGGAAG |

| | |
|---|---|
| 37805_1 | TATGAGTGTGGAGTCGTTGCTAAAAAGGCCAAGTGCAATGGTTNNNAGATCGGAA |
| 37805_2 | TATGAGTGTGGAGTCGTTGCTAAAAAGGCCAAGTGCAATGGTTNNNAGATCGGAA |
| 37805_3 | TATGAGTGTGGAGTCGTTGCTAAAAAGGCCAAGTGCAATGGTTNNNAGATCGGAA |
| 47483_1 | TATGAGTGTGGAGTCGTTGCAGACAGTGACTAAACGGGGTTTTNNNAGATCGGAA |
| 47483_2 | TATGAGTGTGGAGTCGTTGCAGACAGTGACTAAACGGGGTTTTNNNAGATCGGAA |
| 47483_3 | TATGAGTGTGGAGTCGTTGCAGACAGTGACTAAACGGGGTTTTNNNAGATCGGAA |
| 33428_1 | TATGAGTGTGGAGTCGTTGCATAACATCTCAGCTCTTTGGCCANNNAGATCGGAAG |
| 33428_2 | TATGAGTGTGGAGTCGTTGCATAACATCTCAGCTCTTTGGCCANNNAGATCGGAAG |
| 33428_3 | TATGAGTGTGGAGTCGTTGCATAACATCTCAGCTCTTTGGCCANNNAGATCGGAAG |
| 9744_1 | TATGAGTGTGGAGTCGTTGCGGGGGAGTGTTTTAGATGGACACNNNAGATCGGA |
| 9744_2 | TATGAGTGTGGAGTCGTTGCGGGGGAGTGTTTTAGATGGACACNNNAGATCGGA |
| 9744_3 | TATGAGTGTGGAGTCGTTGCGGGGGAGTGTTTTAGATGGACACNNNAGATCGGA |
| 39846_1 | TATGAGTGTGGAGTCGTTGCCCAAGCAAACGGAAACCAAAAAGNNNAGATCGGAA |
| 39846_2 | TATGAGTGTGGAGTCGTTGCCCAAGCAAACGGAAACCAAAAAGNNNAGATCGGAA |
| 39846_3 | TATGAGTGTGGAGTCGTTGCCCAAGCAAACGGAAACCAAAAAGNNNAGATCGGAA |
| 12598_1 | TATGAGTGTGGAGTCGTTGCCTTTCCTTCTCAACAGAGTGNNNAGATCGGAAGAGC |
| 12598_2 | TATGAGTGTGGAGTCGTTGCCTTTCCTTCTCAACAGAGTGNNNAGATCGGAAGAGC |
| 12598_3 | TATGAGTGTGGAGTCGTTGCCTTTCCTTCTCAACAGAGTGNNNAGATCGGAAGAGC |
| 44172_1 | TATGAGTGTGGAGTCGTTGCGCTAAAAGTTGCTTGGGAGAGTTNNNAGATCGGAA |
| 44172_2 | TATGAGTGTGGAGTCGTTGCGCTAAAAGTTGCTTGGGAGAGTTNNNAGATCGGAA |
| 44172_3 | TATGAGTGTGGAGTCGTTGCGCTAAAAGTTGCTTGGGAGAGTTNNNAGATCGGAA |
| 42199_1 | TATGAGTGTGGAGTCGTTGCGGAGGTGGGAGAATCAATTGAGTNNNAGATCGGA |
| 42199_2 | TATGAGTGTGGAGTCGTTGCGGAGGTGGGAGAATCAATTGAGTNNNAGATCGGA |
| 42199_3 | TATGAGTGTGGAGTCGTTGCGGAGGTGGGAGAATCAATTGAGTNNNAGATCGGA |
| 47733_1 | TATGAGTGTGGAGTCGTTGCGAGTACAGGCATGCGCTACTATNNNAGATCGGAAG |
| 47733_2 | TATGAGTGTGGAGTCGTTGCGAGTACAGGCATGCGCTACTATNNNAGATCGGAAG |
| 47733_3 | TATGAGTGTGGAGTCGTTGCGAGTACAGGCATGCGCTACTATNNNAGATCGGAAG |
| 44634_1 | TATGAGTGTGGAGTCGTTGCCTTGGACCAGTACTTGGAGAGACNNNAGATCGGAA |
| 44634_2 | TATGAGTGTGGAGTCGTTGCCTTGGACCAGTACTTGGAGAGACNNNAGATCGGAA |
| 44634_3 | TATGAGTGTGGAGTCGTTGCCTTGGACCAGTACTTGGAGAGACNNNAGATCGGAA |
| 30362_1 | TATGAGTGTGGAGTCGTTGCCCCAACTTGCTGCTCTTTCTATGNNNAGATCGGAAGA |
| 30362_2 | TATGAGTGTGGAGTCGTTGCCCCAACTTGCTGCTCTTTCTATGNNNAGATCGGAAGA |
| 30362_3 | TATGAGTGTGGAGTCGTTGCCCCAACTTGCTGCTCTTTCTATGNNNAGATCGGAAGA |
| 48431_1 | TATGAGTGTGGAGTCGTTGCGCCTGTGGGTTCAAATGATTCTCNNNAGATCGGAAG |
| 48431_2 | TATGAGTGTGGAGTCGTTGCGCCTGTGGGTTCAAATGATTCTCNNNAGATCGGAAG |
| 48431_3 | TATGAGTGTGGAGTCGTTGCGCCTGTGGGTTCAAATGATTCTCNNNAGATCGGAAG |
| 50722_1 | TATGAGTGTGGAGTCGTTGCTTAGCCTGTCTTTGAGATGCCATNNNAGATCGGAAG |
| 50722_2 | TATGAGTGTGGAGTCGTTGCTTAGCCTGTCTTTGAGATGCCATNNNAGATCGGAAG |
| 50722_3 | TATGAGTGTGGAGTCGTTGCTTAGCCTGTCTTTGAGATGCCATNNNAGATCGGAAG |
| 46355_1 | TATGAGTGTGGAGTCGTTGCCATTCTCCTGCCTCAGCCTCNNNAGATCGGAAGAGCA |
| 46355_2 | TATGAGTGTGGAGTCGTTGCCATTCTCCTGCCTCAGCCTCNNNAGATCGGAAGAGCA |
| 46355_3 | TATGAGTGTGGAGTCGTTGCCATTCTCCTGCCTCAGCCTCNNNAGATCGGAAGAGCA |
| 32098_1 | TATGAGTGTGGAGTCGTTGCGTGGAAATGTAGCAAGGCTTGTTNNNAGATCGGAA |

| | |
|---|---|
| 32098_2 | TATGAGTGTGGAGTCGTTGCGTGGAAATGTAGCAAGGCTTGTTNNNAGATCGGAA |
| 32098_3 | TATGAGTGTGGAGTCGTTGCGTGGAAATGTAGCAAGGCTTGTTNNNAGATCGGAA |
| 12929_1 | TATGAGTGTGGAGTCGTTGCAACAGTCTCCAGATTATAAATCCNNNAGATCGGAAG |
| 12929_2 | TATGAGTGTGGAGTCGTTGCAACAGTCTCCAGATTATAAATCCNNNAGATCGGAAG |
| 12929_3 | TATGAGTGTGGAGTCGTTGCAACAGTCTCCAGATTATAAATCCNNNAGATCGGAAG |
| 49874_1 | TATGAGTGTGGAGTCGTTGCCACTGGAGGATCTCATTGCTCTANNNAGATCGGAAG |
| 49874_2 | TATGAGTGTGGAGTCGTTGCCACTGGAGGATCTCATTGCTCTANNNAGATCGGAAG |
| 49874_3 | TATGAGTGTGGAGTCGTTGCCACTGGAGGATCTCATTGCTCTANNNAGATCGGAAG |
| 50543_1 | TATGAGTGTGGAGTCGTTGCTCCTGTGGGTGTTGCGATTATTANNNAGATCGGAAG |
| 50543_2 | TATGAGTGTGGAGTCGTTGCTCCTGTGGGTGTTGCGATTATTANNNAGATCGGAAG |
| 50543_3 | TATGAGTGTGGAGTCGTTGCTCCTGTGGGTGTTGCGATTATTANNNAGATCGGAAG |
| 37281_1 | TATGAGTGTGGAGTCGTTGCGGAGAATGGCATGAACCTAGGAGNNNAGATCGGA |
| 37281_2 | TATGAGTGTGGAGTCGTTGCGGAGAATGGCATGAACCTAGGAGNNNAGATCGGA |
| 37281_3 | TATGAGTGTGGAGTCGTTGCGGAGAATGGCATGAACCTAGGAGNNNAGATCGGA |
| 39383_1 | TATGAGTGTGGAGTCGTTGCAAATAGCTGGGATTACAGGCACANNNAGATCGGAA |
| 39383_2 | TATGAGTGTGGAGTCGTTGCAAATAGCTGGGATTACAGGCACANNNAGATCGGAA |
| 39383_3 | TATGAGTGTGGAGTCGTTGCAAATAGCTGGGATTACAGGCACANNNAGATCGGAA |
| 11967_1 | TATGAGTGTGGAGTCGTTGCCCCTGATTTCCCTTTCTGCTCTTNNNAGATCGGAAGA |
| 11967_2 | TATGAGTGTGGAGTCGTTGCCCCTGATTTCCCTTTCTGCTCTTNNNAGATCGGAAGA |
| 11967_3 | TATGAGTGTGGAGTCGTTGCCCCTGATTTCCCTTTCTGCTCTTNNNAGATCGGAAGA |
| 42482_1 | TATGAGTGTGGAGTCGTTGCCCAAGTCAGAATGCAGGAGAAGANNNAGATCGGAA |
| 42482_2 | TATGAGTGTGGAGTCGTTGCCCAAGTCAGAATGCAGGAGAAGANNNAGATCGGAA |
| 42482_3 | TATGAGTGTGGAGTCGTTGCCCAAGTCAGAATGCAGGAGAAGANNNAGATCGGAA |
| 44361_1 | TATGAGTGTGGAGTCGTTGCGGTGCTCCTTTGCCAAATGTAAANNNAGATCGGAAG |
| 44361_2 | TATGAGTGTGGAGTCGTTGCGGTGCTCCTTTGCCAAATGTAAANNNAGATCGGAAG |
| 44361_3 | TATGAGTGTGGAGTCGTTGCGGTGCTCCTTTGCCAAATGTAAANNNAGATCGGAAG |
| 13646_1 | TATGAGTGTGGAGTCGTTGCAGAACTCTGATGTTTGTAAAATCNNNAGATCGGAAG |
| 13646_2 | TATGAGTGTGGAGTCGTTGCAGAACTCTGATGTTTGTAAAATCNNNAGATCGGAAG |
| 13646_3 | TATGAGTGTGGAGTCGTTGCAGAACTCTGATGTTTGTAAAATCNNNAGATCGGAAG |
| 13311_1 | TATGAGTGTGGAGTCGTTGCTTGCCTTTCATGAATACCAGNNNAGATCGGAAGAGC |
| 13311_2 | TATGAGTGTGGAGTCGTTGCTTGCCTTTCATGAATACCAGNNNAGATCGGAAGAGC |
| 13311_3 | TATGAGTGTGGAGTCGTTGCTTGCCTTTCATGAATACCAGNNNAGATCGGAAGAGC |
| 47221_1 | TATGAGTGTGGAGTCGTTGCATGCGTGATTTGCTTCATTGTCANNNAGATCGGAAG |
| 47221_2 | TATGAGTGTGGAGTCGTTGCATGCGTGATTTGCTTCATTGTCANNNAGATCGGAAG |
| 47221_3 | TATGAGTGTGGAGTCGTTGCATGCGTGATTTGCTTCATTGTCANNNAGATCGGAAG |
| 36799_1 | TATGAGTGTGGAGTCGTTGCCAAGTACTGGGATTCCACTGACTNNNAGATCGGAAG |
| 36799_2 | TATGAGTGTGGAGTCGTTGCCAAGTACTGGGATTCCACTGACTNNNAGATCGGAAG |
| 36799_3 | TATGAGTGTGGAGTCGTTGCCAAGTACTGGGATTCCACTGACTNNNAGATCGGAAG |
| 38235_1 | TATGAGTGTGGAGTCGTTGCCTGACTGATGCATGTGTGTTGTTNNNAGATCGGAAG |
| 38235_2 | TATGAGTGTGGAGTCGTTGCCTGACTGATGCATGTGTGTTGTTNNNAGATCGGAAG |
| 38235_3 | TATGAGTGTGGAGTCGTTGCCTGACTGATGCATGTGTGTTGTTNNNAGATCGGAAG |
| 10715_1 | TATGAGTGTGGAGTCGTTGCACTCGTATTCTCCTTCCTCTGCANNNAGATCGGAAGA |
| 10715_2 | TATGAGTGTGGAGTCGTTGCACTCGTATTCTCCTTCCTCTGCANNNAGATCGGAAGA |

| | |
|---|---|
| 10715_3 | TATGAGTGTGGAGTCGTTGCACTCGTATTCTCCTTCCTCTGCANNNAGATCGGAAGA |
| 40303_1 | TATGAGTGTGGAGTCGTTGCCACACAGCATTGATTTGTGATGCNNNAGATCGGAAG |
| 40303_2 | TATGAGTGTGGAGTCGTTGCCACACAGCATTGATTTGTGATGCNNNAGATCGGAAG |
| 40303_3 | TATGAGTGTGGAGTCGTTGCCACACAGCATTGATTTGTGATGCNNNAGATCGGAAG |
| 41692_1 | TATGAGTGTGGAGTCGTTGCCATTCTCCTGCCTCAGCCTCNNNAGATCGGAAGAGCA |
| 41692_2 | TATGAGTGTGGAGTCGTTGCCATTCTCCTGCCTCAGCCTCNNNAGATCGGAAGAGCA |
| 41692_3 | TATGAGTGTGGAGTCGTTGCCATTCTCCTGCCTCAGCCTCNNNAGATCGGAAGAGCA |
| 13128_1 | TATGAGTGTGGAGTCGTTGCCTAAAGAAAAAAAAAGTGGTAGCNNNAGATCGGAA |
| 13128_2 | TATGAGTGTGGAGTCGTTGCCTAAAGAAAAAAAAAGTGGTAGCNNNAGATCGGAA |
| 13128_3 | TATGAGTGTGGAGTCGTTGCCTAAAGAAAAAAAAAGTGGTAGCNNNAGATCGGAA |
| 38434_1 | TATGAGTGTGGAGTCGTTGCACCCGAACCATATCACCAAGTACNNNAGATCGGAAG |
| 38434_2 | TATGAGTGTGGAGTCGTTGCACCCGAACCATATCACCAAGTACNNNAGATCGGAAG |
| 38434_3 | TATGAGTGTGGAGTCGTTGCACCCGAACCATATCACCAAGTACNNNAGATCGGAAG |
| 36937_1 | TATGAGTGTGGAGTCGTTGCCCTTTCTCAAATGGAGTACGGGANNNAGATCGGAA |
| 36937_2 | TATGAGTGTGGAGTCGTTGCCCTTTCTCAAATGGAGTACGGGANNNAGATCGGAA |
| 36937_3 | TATGAGTGTGGAGTCGTTGCCCTTTCTCAAATGGAGTACGGGANNNAGATCGGAA |
| 33705_1 | TATGAGTGTGGAGTCGTTGCGTACATTCTAGAGCTCCAGGCAGNNNAGATCGGAA |
| 33705_2 | TATGAGTGTGGAGTCGTTGCGTACATTCTAGAGCTCCAGGCAGNNNAGATCGGAA |
| 33705_3 | TATGAGTGTGGAGTCGTTGCGTACATTCTAGAGCTCCAGGCAGNNNAGATCGGAA |
| 52103_1 | TATGAGTGTGGAGTCGTTGCTAACCCACAGATCCAGGAAAGTCNNNAGATCGGAAG |
| 52103_2 | TATGAGTGTGGAGTCGTTGCTAACCCACAGATCCAGGAAAGTCNNNAGATCGGAAG |
| 52103_3 | TATGAGTGTGGAGTCGTTGCTAACCCACAGATCCAGGAAAGTCNNNAGATCGGAAG |
| 45173_1 | TATGAGTGTGGAGTCGTTGCGTGACAGCTGATCAAAGAGGGTANNNAGATCGGAA |
| 45173_2 | TATGAGTGTGGAGTCGTTGCGTGACAGCTGATCAAAGAGGGTANNNAGATCGGAA |
| 45173_3 | TATGAGTGTGGAGTCGTTGCGTGACAGCTGATCAAAGAGGGTANNNAGATCGGAA |
| 49171_1 | TATGAGTGTGGAGTCGTTGCCACAATGATGGCTCACTGCAANNNAGATCGGAAGA |
| 49171_2 | TATGAGTGTGGAGTCGTTGCCACAATGATGGCTCACTGCAANNNAGATCGGAAGA |
| 49171_3 | TATGAGTGTGGAGTCGTTGCCACAATGATGGCTCACTGCAANNNAGATCGGAAGA |
| 36180_1 | TATGAGTGTGGAGTCGTTGCGAAGAAGAAGAGGAGATGAGGCCNNNAGATCGGA |
| 36180_2 | TATGAGTGTGGAGTCGTTGCGAAGAAGAAGAGGAGATGAGGCCNNNAGATCGGA |
| 36180_3 | TATGAGTGTGGAGTCGTTGCGAAGAAGAAGAGGAGATGAGGCCNNNAGATCGGA |
| 46079_1 | TATGAGTGTGGAGTCGTTGCTTCTGTGGTGCCTTCTCTATCTGNNNAGATCGGAAG |
| 46079_2 | TATGAGTGTGGAGTCGTTGCTTCTGTGGTGCCTTCTCTATCTGNNNAGATCGGAAG |
| 46079_3 | TATGAGTGTGGAGTCGTTGCTTCTGTGGTGCCTTCTCTATCTGNNNAGATCGGAAG |
| 13153_1 | TATGAGTGTGGAGTCGTTGCGACTACCACACCATCATTTAGNNNAGATCGGAAGAG |
| 13153_2 | TATGAGTGTGGAGTCGTTGCGACTACCACACCATCATTTAGNNNAGATCGGAAGAG |
| 13153_3 | TATGAGTGTGGAGTCGTTGCGACTACCACACCATCATTTAGNNNAGATCGGAAGAG |
| 53049_1 | TATGAGTGTGGAGTCGTTGCCTGAGGAAGGAGGATTGCTTGAGNNNAGATCGGA |
| 53049_2 | TATGAGTGTGGAGTCGTTGCCTGAGGAAGGAGGATTGCTTGAGNNNAGATCGGA |
| 53049_3 | TATGAGTGTGGAGTCGTTGCCTGAGGAAGGAGGATTGCTTGAGNNNAGATCGGA |
| 47602_1 | TATGAGTGTGGAGTCGTTGCGGCCTCTGAAAGTGCTAGGATTANNNAGATCGGAA |
| 47602_2 | TATGAGTGTGGAGTCGTTGCGGCCTCTGAAAGTGCTAGGATTANNNAGATCGGAA |
| 47602_3 | TATGAGTGTGGAGTCGTTGCGGCCTCTGAAAGTGCTAGGATTANNNAGATCGGAA |

| | |
|---|---|
| 4801_1 | TATGAGTGTGGAGTCGTTGCGTTTCTCCTGGTGAGTTGCCATTTNNNAGATCGGAA |
| 4801_2 | TATGAGTGTGGAGTCGTTGCGTTTCTCCTGGTGAGTTGCCATTTNNNAGATCGGAA |
| 4801_3 | TATGAGTGTGGAGTCGTTGCGTTTCTCCTGGTGAGTTGCCATTTNNNAGATCGGAA |
| 51740_1 | TATGAGTGTGGAGTCGTTGCCAGGGCCAGTTTTGTATCAAAGGNNNAGATCGGAA |
| 51740_2 | TATGAGTGTGGAGTCGTTGCCAGGGCCAGTTTTGTATCAAAGGNNNAGATCGGAA |
| 51740_3 | TATGAGTGTGGAGTCGTTGCCAGGGCCAGTTTTGTATCAAAGGNNNAGATCGGAA |
| 38952_1 | TATGAGTGTGGAGTCGTTGCAAAGGACACGAGCAGACACTTTNNNAGATCGGAAG |
| 38952_2 | TATGAGTGTGGAGTCGTTGCAAAGGACACGAGCAGACACTTTNNNAGATCGGAAG |
| 38952_3 | TATGAGTGTGGAGTCGTTGCAAAGGACACGAGCAGACACTTTNNNAGATCGGAAG |
| 29156_1 | TATGAGTGTGGAGTCGTTGCCACTGCACACAAGGCATATGAAANNNAGATCGGAAG |
| 29156_2 | TATGAGTGTGGAGTCGTTGCCACTGCACACAAGGCATATGAAANNNAGATCGGAAG |
| 29156_3 | TATGAGTGTGGAGTCGTTGCCACTGCACACAAGGCATATGAAANNNAGATCGGAAG |
| 34080_1 | TATGAGTGTGGAGTCGTTGCGTAGCTGGGACTACAGGTGAATGNNNAGATCGGAA |
| 34080_2 | TATGAGTGTGGAGTCGTTGCGTAGCTGGGACTACAGGTGAATGNNNAGATCGGAA |
| 34080_3 | TATGAGTGTGGAGTCGTTGCGTAGCTGGGACTACAGGTGAATGNNNAGATCGGAA |
| 52700_1 | TATGAGTGTGGAGTCGTTGCCCATCTTTTCCTTTTCCCACACCNNNAGATCGGAAGAG |
| 52700_2 | TATGAGTGTGGAGTCGTTGCCCATCTTTTCCTTTTCCCACACCNNNAGATCGGAAGAG |
| 52700_3 | TATGAGTGTGGAGTCGTTGCCCATCTTTTCCTTTTCCCACACCNNNAGATCGGAAGAG |
| 13432_1 | TATGAGTGTGGAGTCGTTGCCTGAAAGTCGCAAAAGTTCTNNNAGATCGGAAGAG |
| 13432_2 | TATGAGTGTGGAGTCGTTGCCTGAAAGTCGCAAAAGTTCTNNNAGATCGGAAGAG |
| 13432_3 | TATGAGTGTGGAGTCGTTGCCTGAAAGTCGCAAAAGTTCTNNNAGATCGGAAGAG |
| 40074_1 | TATGAGTGTGGAGTCGTTGCTGAGAGGTCATAACACTTGGGTTNNNAGATCGGAA |
| 40074_2 | TATGAGTGTGGAGTCGTTGCTGAGAGGTCATAACACTTGGGTTNNNAGATCGGAA |
| 40074_3 | TATGAGTGTGGAGTCGTTGCTGAGAGGTCATAACACTTGGGTTNNNAGATCGGAA |
| 36219_1 | TATGAGTGTGGAGTCGTTGCCACGCCTATAATCCCAGCTACTTNNNAGATCGGAAGA |
| 36219_2 | TATGAGTGTGGAGTCGTTGCCACGCCTATAATCCCAGCTACTTNNNAGATCGGAAGA |
| 36219_3 | TATGAGTGTGGAGTCGTTGCCACGCCTATAATCCCAGCTACTTNNNAGATCGGAAGA |
| 51201_1 | TATGAGTGTGGAGTCGTTGCTGGGTAGGAAGAGCACTCTAGAANNNAGATCGGAA |
| 51201_2 | TATGAGTGTGGAGTCGTTGCTGGGTAGGAAGAGCACTCTAGAANNNAGATCGGAA |
| 51201_3 | TATGAGTGTGGAGTCGTTGCTGGGTAGGAAGAGCACTCTAGAANNNAGATCGGAA |
| 6191_1 | TATGAGTGTGGAGTCGTTGCCATTACAGGCTTCTTCTGCTTGGAANNNAGATCGGA |
| 6191_2 | TATGAGTGTGGAGTCGTTGCCATTACAGGCTTCTTCTGCTTGGAANNNAGATCGGA |
| 6191_3 | TATGAGTGTGGAGTCGTTGCCATTACAGGCTTCTTCTGCTTGGAANNNAGATCGGA |
| 50315_1 | TATGAGTGTGGAGTCGTTGCTTATTCCTATCCTGGGACCCTGTNNNAGATCGGAAG |
| 50315_2 | TATGAGTGTGGAGTCGTTGCTTATTCCTATCCTGGGACCCTGTNNNAGATCGGAAG |
| 50315_3 | TATGAGTGTGGAGTCGTTGCTTATTCCTATCCTGGGACCCTGTNNNAGATCGGAAG |
| 13800_1 | TATGAGTGTGGAGTCGTTGCTGTTTTCTAAGTTTTTTCTCCTTNNNAGATCGGAAGA |
| 13800_2 | TATGAGTGTGGAGTCGTTGCTGTTTTCTAAGTTTTTTCTCCTTNNNAGATCGGAAGA |
| 13800_3 | TATGAGTGTGGAGTCGTTGCTGTTTTCTAAGTTTTTTCTCCTTNNNAGATCGGAAGA |
| 43773_1 | TATGAGTGTGGAGTCGTTGCATTTTTCCAGTAGAGGGCATCCCNNNAGATCGGAAG |
| 43773_2 | TATGAGTGTGGAGTCGTTGCATTTTTCCAGTAGAGGGCATCCCNNNAGATCGGAAG |
| 43773_3 | TATGAGTGTGGAGTCGTTGCATTTTTCCAGTAGAGGGCATCCCNNNAGATCGGAAG |
| 13579_1 | TATGAGTGTGGAGTCGTTGCCAAATTGAAGATCTAATAGAGGCNNNAGATCGGAA |

| | |
|---|---|
| 13579_2 | TATGAGTGTGGAGTCGTTGCCAAATTGAAGATCTAATAGAGGCNNNAGATCGGAAC |
| 13579_3 | TATGAGTGTGGAGTCGTTGCCAAATTGAAGATCTAATAGAGGCNNNAGATCGGAAC |
| 36982_1 | TATGAGTGTGGAGTCGTTGCTGGTCTGTGTTGGTTACTTAGGGNNNAGATCGGAAC |
| 36982_2 | TATGAGTGTGGAGTCGTTGCTGGTCTGTGTTGGTTACTTAGGGNNNAGATCGGAAC |
| 36982_3 | TATGAGTGTGGAGTCGTTGCTGGTCTGTGTTGGTTACTTAGGGNNNAGATCGGAAC |
| 36068_1 | TATGAGTGTGGAGTCGTTGCCCTGGATTGCAGTGGATCCTAAANNNAGATCGGAAG |
| 36068_2 | TATGAGTGTGGAGTCGTTGCCCTGGATTGCAGTGGATCCTAAANNNAGATCGGAAG |
| 36068_3 | TATGAGTGTGGAGTCGTTGCCCTGGATTGCAGTGGATCCTAAANNNAGATCGGAAG |
| 48533_1 | TATGAGTGTGGAGTCGTTGCATCTTAGCTCACTGCAACTTCCANNNAGATCGGAAGA |
| 48533_2 | TATGAGTGTGGAGTCGTTGCATCTTAGCTCACTGCAACTTCCANNNAGATCGGAAGA |
| 48533_3 | TATGAGTGTGGAGTCGTTGCATCTTAGCTCACTGCAACTTCCANNNAGATCGGAAGA |
| 33988_1 | TATGAGTGTGGAGTCGTTGCCCTACCAAAGTGCTGGGATTACANNNAGATCGGAAG |
| 33988_2 | TATGAGTGTGGAGTCGTTGCCCTACCAAAGTGCTGGGATTACANNNAGATCGGAAG |
| 33988_3 | TATGAGTGTGGAGTCGTTGCCCTACCAAAGTGCTGGGATTACANNNAGATCGGAAG |
| 39306_1 | TATGAGTGTGGAGTCGTTGCTTTCAGATCCCCTCATTGCTTCANNNAGATCGGAAGA |
| 39306_2 | TATGAGTGTGGAGTCGTTGCTTTCAGATCCCCTCATTGCTTCANNNAGATCGGAAGA |
| 39306_3 | TATGAGTGTGGAGTCGTTGCTTTCAGATCCCCTCATTGCTTCANNNAGATCGGAAGA |
| 41818_1 | TATGAGTGTGGAGTCGTTGCGTTCAAGAGATTCTCCTGCCTCANNNAGATCGGAAGA |
| 41818_2 | TATGAGTGTGGAGTCGTTGCGTTCAAGAGATTCTCCTGCCTCANNNAGATCGGAAGA |
| 41818_3 | TATGAGTGTGGAGTCGTTGCGTTCAAGAGATTCTCCTGCCTCANNNAGATCGGAAGA |
| 49848_1 | TATGAGTGTGGAGTCGTTGCGGAAGGAAAACCAAAGATGGAGCNNNAGATCGGAA |
| 49848_2 | TATGAGTGTGGAGTCGTTGCGGAAGGAAAACCAAAGATGGAGCNNNAGATCGGAA |
| 49848_3 | TATGAGTGTGGAGTCGTTGCGGAAGGAAAACCAAAGATGGAGCNNNAGATCGGAA |
| 41902_1 | TATGAGTGTGGAGTCGTTGCTATAGATCACTGCAGCCTTGAGCNNNAGATCGGAAG |
| 41902_2 | TATGAGTGTGGAGTCGTTGCTATAGATCACTGCAGCCTTGAGCNNNAGATCGGAAG |
| 41902_3 | TATGAGTGTGGAGTCGTTGCTATAGATCACTGCAGCCTTGAGCNNNAGATCGGAAG |
| 9892_1 | TATGAGTGTGGAGTCGTTGCCGTGTGTGTTTGGTGATGAAGGANNNAGATCGGAA |
| 9892_2 | TATGAGTGTGGAGTCGTTGCCGTGTGTGTTTGGTGATGAAGGANNNAGATCGGAA |
| 9892_3 | TATGAGTGTGGAGTCGTTGCCGTGTGTGTTTGGTGATGAAGGANNNAGATCGGAA |
| 40619_1 | TATGAGTGTGGAGTCGTTGCTCCCACTTGGAAATTCCCTTGAANNNAGATCGGAAGA |
| 40619_2 | TATGAGTGTGGAGTCGTTGCTCCCACTTGGAAATTCCCTTGAANNNAGATCGGAAGA |
| 40619_3 | TATGAGTGTGGAGTCGTTGCTCCCACTTGGAAATTCCCTTGAANNNAGATCGGAAGA |
| 46038_1 | TATGAGTGTGGAGTCGTTGCCTCAAGAGATTCTCCTGCCTCAGNNNAGATCGGAAGA |
| 46038_2 | TATGAGTGTGGAGTCGTTGCCTCAAGAGATTCTCCTGCCTCAGNNNAGATCGGAAGA |
| 46038_3 | TATGAGTGTGGAGTCGTTGCCTCAAGAGATTCTCCTGCCTCAGNNNAGATCGGAAGA |
| 36096_1 | TATGAGTGTGGAGTCGTTGCGTCAGTATGTCTAGACTGGGCACNNNAGATCGGAAC |
| 36096_2 | TATGAGTGTGGAGTCGTTGCGTCAGTATGTCTAGACTGGGCACNNNAGATCGGAAC |
| 36096_3 | TATGAGTGTGGAGTCGTTGCGTCAGTATGTCTAGACTGGGCACNNNAGATCGGAAC |
| 29907_1 | TATGAGTGTGGAGTCGTTGCGCCTTCATTCAAAAATCGGCAAGNNNAGATCGGAAG |
| 29907_2 | TATGAGTGTGGAGTCGTTGCGCCTTCATTCAAAAATCGGCAAGNNNAGATCGGAAG |
| 29907_3 | TATGAGTGTGGAGTCGTTGCGCCTTCATTCAAAAATCGGCAAGNNNAGATCGGAAG |
| 47738_1 | TATGAGTGTGGAGTCGTTGCCTCCTGAGTAGCTGGGACTACAGNNNAGATCGGAAC |
| 47738_2 | TATGAGTGTGGAGTCGTTGCCTCCTGAGTAGCTGGGACTACAGNNNAGATCGGAAC |

| | |
|---|---|
| 47738_3 | TATGAGTGTGGAGTCGTTGCCTCCTGAGTAGCTGGGACTACAGNNNAGATCGGAA |
| 32805_1 | TATGAGTGTGGAGTCGTTGCGCTTCCTGAGTAGCTGGGATTACNNNAGATCGGAAG |
| 32805_2 | TATGAGTGTGGAGTCGTTGCGCTTCCTGAGTAGCTGGGATTACNNNAGATCGGAAG |
| 32805_3 | TATGAGTGTGGAGTCGTTGCGCTTCCTGAGTAGCTGGGATTACNNNAGATCGGAAG |
| 33940_1 | TATGAGTGTGGAGTCGTTGCCAAGTAGCTGGGACTACAGTCGNNNAGATCGGAAG |
| 33940_2 | TATGAGTGTGGAGTCGTTGCCAAGTAGCTGGGACTACAGTCGNNNAGATCGGAAG |
| 33940_3 | TATGAGTGTGGAGTCGTTGCCAAGTAGCTGGGACTACAGTCGNNNAGATCGGAAG |
| 38610_1 | TATGAGTGTGGAGTCGTTGCAAATGATGGTGGGATAACTGGCTNNNAGATCGGAA |
| 38610_2 | TATGAGTGTGGAGTCGTTGCAAATGATGGTGGGATAACTGGCTNNNAGATCGGAA |
| 38610_3 | TATGAGTGTGGAGTCGTTGCAAATGATGGTGGGATAACTGGCTNNNAGATCGGAA |
| 32128_1 | TATGAGTGTGGAGTCGTTGCCCACTCCTGCGTATGACATTTTTNNNAGATCGGAAGA |
| 32128_2 | TATGAGTGTGGAGTCGTTGCCCACTCCTGCGTATGACATTTTTNNNAGATCGGAAGA |
| 32128_3 | TATGAGTGTGGAGTCGTTGCCCACTCCTGCGTATGACATTTTTNNNAGATCGGAAGA |
| 43473_1 | TATGAGTGTGGAGTCGTTGCCCTCCAGGGTTCAAGTGATTCTCNNNAGATCGGAAG |
| 43473_2 | TATGAGTGTGGAGTCGTTGCCCTCCAGGGTTCAAGTGATTCTCNNNAGATCGGAAG |
| 43473_3 | TATGAGTGTGGAGTCGTTGCCCTCCAGGGTTCAAGTGATTCTCNNNAGATCGGAAG |
| 46825_1 | TATGAGTGTGGAGTCGTTGCTCAAGTGATTCTCCTGCCTTAGCNNNAGATCGGAAG |
| 46825_2 | TATGAGTGTGGAGTCGTTGCTCAAGTGATTCTCCTGCCTTAGCNNNAGATCGGAAG |
| 46825_3 | TATGAGTGTGGAGTCGTTGCTCAAGTGATTCTCCTGCCTTAGCNNNAGATCGGAAG |
| 41178_1 | TATGAGTGTGGAGTCGTTGCGGGATTGCACTAAGTAGCTCGTANNNAGATCGGAA |
| 41178_2 | TATGAGTGTGGAGTCGTTGCGGGATTGCACTAAGTAGCTCGTANNNAGATCGGAA |
| 41178_3 | TATGAGTGTGGAGTCGTTGCGGGATTGCACTAAGTAGCTCGTANNNAGATCGGAA |
| 51301_1 | TATGAGTGTGGAGTCGTTGCTCTCTTCAAAGCTGTCAGACAGGNNNAGATCGGAAG |
| 51301_2 | TATGAGTGTGGAGTCGTTGCTCTCTTCAAAGCTGTCAGACAGGNNNAGATCGGAAG |
| 51301_3 | TATGAGTGTGGAGTCGTTGCTCTCTTCAAAGCTGTCAGACAGGNNNAGATCGGAAG |
| 8041_1 | TATGAGTGTGGAGTCGTTGCCGGAGTTTTCCCTCTTGTCGACCNNNAGATCGGAAG |
| 8041_2 | TATGAGTGTGGAGTCGTTGCCGGAGTTTTCCCTCTTGTCGACCNNNAGATCGGAAG |
| 8041_3 | TATGAGTGTGGAGTCGTTGCCGGAGTTTTCCCTCTTGTCGACCNNNAGATCGGAAG |
| 16_1 | TATGAGTGTGGAGTCGTTGCTTTTCTTCACTTGCTCTGCCCATNNNAGATCGGAAGA |
| 16_2 | TATGAGTGTGGAGTCGTTGCTTTTCTTCACTTGCTCTGCCCATNNNAGATCGGAAGA |
| 16_3 | TATGAGTGTGGAGTCGTTGCTTTTCTTCACTTGCTCTGCCCATNNNAGATCGGAAGA |
| 46436_1 | TATGAGTGTGGAGTCGTTGCGCCTCTCAAGTAGCTGGGAATACNNNAGATCGGAAG |
| 46436_2 | TATGAGTGTGGAGTCGTTGCGCCTCTCAAGTAGCTGGGAATACNNNAGATCGGAAG |
| 46436_3 | TATGAGTGTGGAGTCGTTGCGCCTCTCAAGTAGCTGGGAATACNNNAGATCGGAAG |
| 45092_1 | TATGAGTGTGGAGTCGTTGCAGGCAAATCCCATGAAAATGTGGNNNAGATCGGAA |
| 45092_2 | TATGAGTGTGGAGTCGTTGCAGGCAAATCCCATGAAAATGTGGNNNAGATCGGAA |
| 45092_3 | TATGAGTGTGGAGTCGTTGCAGGCAAATCCCATGAAAATGTGGNNNAGATCGGAA |
| 11730_1 | TATGAGTGTGGAGTCGTTGCATCGGGGCATAGGGATAAGAGGNNNAGATCGGAA |
| 11730_2 | TATGAGTGTGGAGTCGTTGCATCGGGGCATAGGGATAAGAGGNNNAGATCGGAA |
| 11730_3 | TATGAGTGTGGAGTCGTTGCATCGGGGCATAGGGATAAGAGGNNNAGATCGGAA |
| 52572_1 | TATGAGTGTGGAGTCGTTGCGAGAAAAGTGAGCAGGCATGTTTNNNAGATCGGAA |
| 52572_2 | TATGAGTGTGGAGTCGTTGCGAGAAAAGTGAGCAGGCATGTTTNNNAGATCGGAA |
| 52572_3 | TATGAGTGTGGAGTCGTTGCGAGAAAAGTGAGCAGGCATGTTTNNNAGATCGGAA |

| | |
|---|---|
| 52518_1 | TATGAGTGTGGAGTCGTTGCGGTCATTTCAGCAGGAAAAGTCGNNNAGATCGGAA |
| 52518_2 | TATGAGTGTGGAGTCGTTGCGGTCATTTCAGCAGGAAAAGTCGNNNAGATCGGAA |
| 52518_3 | TATGAGTGTGGAGTCGTTGCGGTCATTTCAGCAGGAAAAGTCGNNNAGATCGGAA |
| 44073_1 | TATGAGTGTGGAGTCGTTGCACCTGGTGATTGTCCCTATTGACNNNAGATCGGAAG |
| 44073_2 | TATGAGTGTGGAGTCGTTGCACCTGGTGATTGTCCCTATTGACNNNAGATCGGAAG |
| 44073_3 | TATGAGTGTGGAGTCGTTGCACCTGGTGATTGTCCCTATTGACNNNAGATCGGAAG |
| 53211_1 | TATGAGTGTGGAGTCGTTGCCTGTAAGAATCATGCAGGGGAGGNNNAGATCGGAA |
| 53211_2 | TATGAGTGTGGAGTCGTTGCCTGTAAGAATCATGCAGGGGAGGNNNAGATCGGAA |
| 53211_3 | TATGAGTGTGGAGTCGTTGCCTGTAAGAATCATGCAGGGGAGGNNNAGATCGGAA |
| 31055_1 | TATGAGTGTGGAGTCGTTGCTACAGTTGTCTCAGATTGCACCANNNAGATCGGAAG |
| 31055_2 | TATGAGTGTGGAGTCGTTGCTACAGTTGTCTCAGATTGCACCANNNAGATCGGAAG |
| 31055_3 | TATGAGTGTGGAGTCGTTGCTACAGTTGTCTCAGATTGCACCANNNAGATCGGAAG |
| 40829_1 | TATGAGTGTGGAGTCGTTGCTGTGTCCCAAATCATTGAACTGTNNNAGATCGGAAG |
| 40829_2 | TATGAGTGTGGAGTCGTTGCTGTGTCCCAAATCATTGAACTGTNNNAGATCGGAAG |
| 40829_3 | TATGAGTGTGGAGTCGTTGCTGTGTCCCAAATCATTGAACTGTNNNAGATCGGAAG |
| 35319_1 | TATGAGTGTGGAGTCGTTGCCACCTAGCCCGGCTAATTTTTTGNNNAGATCGGAAG |
| 35319_2 | TATGAGTGTGGAGTCGTTGCCACCTAGCCCGGCTAATTTTTTGNNNAGATCGGAAG |
| 35319_3 | TATGAGTGTGGAGTCGTTGCCACCTAGCCCGGCTAATTTTTTGNNNAGATCGGAAG |
| 53175_1 | TATGAGTGTGGAGTCGTTGCCAGAACAGGGATGAAACTGAGGANNNAGATCGGAA |
| 53175_2 | TATGAGTGTGGAGTCGTTGCCAGAACAGGGATGAAACTGAGGANNNAGATCGGAA |
| 53175_3 | TATGAGTGTGGAGTCGTTGCCAGAACAGGGATGAAACTGAGGANNNAGATCGGAA |
| 46603_1 | TATGAGTGTGGAGTCGTTGCGTTTCACTCTTGTTGCTCAGGCNNNAGATCGGAAGA |
| 46603_2 | TATGAGTGTGGAGTCGTTGCGTTTCACTCTTGTTGCTCAGGCNNNAGATCGGAAGA |
| 46603_3 | TATGAGTGTGGAGTCGTTGCGTTTCACTCTTGTTGCTCAGGCNNNAGATCGGAAGA |
| 52217_1 | TATGAGTGTGGAGTCGTTGCGTAACTCAGAAGGATGAGGTGGGNNNAGATCGGA |
| 52217_2 | TATGAGTGTGGAGTCGTTGCGTAACTCAGAAGGATGAGGTGGGNNNAGATCGGA |
| 52217_3 | TATGAGTGTGGAGTCGTTGCGTAACTCAGAAGGATGAGGTGGGNNNAGATCGGA |
| 29373_1 | TATGAGTGTGGAGTCGTTGCGTCTCTCTCTCTCTCTCTCTCCCNNNAGATCGGAAGAG |
| 29373_2 | TATGAGTGTGGAGTCGTTGCGTCTCTCTCTCTCTCTCTCTCCCNNNAGATCGGAAGAG |
| 29373_3 | TATGAGTGTGGAGTCGTTGCGTCTCTCTCTCTCTCTCTCTCCCNNNAGATCGGAAGAG |
| 52434_1 | TATGAGTGTGGAGTCGTTGCTCAGAGATGCATCTGCCTATGACNNNAGATCGGAAG |
| 52434_2 | TATGAGTGTGGAGTCGTTGCTCAGAGATGCATCTGCCTATGACNNNAGATCGGAAG |
| 52434_3 | TATGAGTGTGGAGTCGTTGCTCAGAGATGCATCTGCCTATGACNNNAGATCGGAAG |
| 52455_1 | TATGAGTGTGGAGTCGTTGCGCAAAAGTCGCTTTTTACAAGCCNNNAGATCGGAAG |
| 52455_2 | TATGAGTGTGGAGTCGTTGCGCAAAAGTCGCTTTTTACAAGCCNNNAGATCGGAAG |
| 52455_3 | TATGAGTGTGGAGTCGTTGCGCAAAAGTCGCTTTTTACAAGCCNNNAGATCGGAAG |
| 12848_1 | TATGAGTGTGGAGTCGTTGCGTTAACTTCAATGGGTTTTCCNNNAGATCGGAAGAG |
| 12848_2 | TATGAGTGTGGAGTCGTTGCGTTAACTTCAATGGGTTTTCCNNNAGATCGGAAGAG |
| 12848_3 | TATGAGTGTGGAGTCGTTGCGTTAACTTCAATGGGTTTTCCNNNAGATCGGAAGAG |
| 29342_1 | TATGAGTGTGGAGTCGTTGCAGCCTCTATCTGGTCTCATGACTNNNAGATCGGAAG |
| 29342_2 | TATGAGTGTGGAGTCGTTGCAGCCTCTATCTGGTCTCATGACTNNNAGATCGGAAG |
| 29342_3 | TATGAGTGTGGAGTCGTTGCAGCCTCTATCTGGTCTCATGACTNNNAGATCGGAAG |
| 47813_1 | TATGAGTGTGGAGTCGTTGCGGGTTCCTGAGTGCATGCTGNNNAGATCGGAAGAG |

| | |
|---|---|
| 47813_2 | TATGAGTGTGGAGTCGTTGCGGGTTCCTGAGTGCATGCTGNNNAGATCGGAAGAG |
| 47813_3 | TATGAGTGTGGAGTCGTTGCGGGTTCCTGAGTGCATGCTGNNNAGATCGGAAGAG |
| 47070_1 | TATGAGTGTGGAGTCGTTGCCTCAAGAAGCTGGAGTTACAGGTNNNAGATCGGAA |
| 47070_2 | TATGAGTGTGGAGTCGTTGCCTCAAGAAGCTGGAGTTACAGGTNNNAGATCGGAA |
| 47070_3 | TATGAGTGTGGAGTCGTTGCCTCAAGAAGCTGGAGTTACAGGTNNNAGATCGGAA |
| 13098_1 | TATGAGTGTGGAGTCGTTGCCATACATCTTGATTCTTTTCCATNNNAGATCGGAAGA |
| 13098_2 | TATGAGTGTGGAGTCGTTGCCATACATCTTGATTCTTTTCCATNNNAGATCGGAAGA |
| 13098_3 | TATGAGTGTGGAGTCGTTGCCATACATCTTGATTCTTTTCCATNNNAGATCGGAAGA |
| 32853_1 | TATGAGTGTGGAGTCGTTGCCCAAAGTGCTGGGATAACAGGTANNNAGATCGGAA |
| 32853_2 | TATGAGTGTGGAGTCGTTGCCCAAAGTGCTGGGATAACAGGTANNNAGATCGGAA |
| 32853_3 | TATGAGTGTGGAGTCGTTGCCCAAAGTGCTGGGATAACAGGTANNNAGATCGGAA |
| 43889_1 | TATGAGTGTGGAGTCGTTGCGAGTGCGGGGATGTGATCATAGNNNAGATCGGAA |
| 43889_2 | TATGAGTGTGGAGTCGTTGCGAGTGCGGGGATGTGATCATAGNNNAGATCGGAA |
| 43889_3 | TATGAGTGTGGAGTCGTTGCGAGTGCGGGGATGTGATCATAGNNNAGATCGGAA |
| 37336_1 | TATGAGTGTGGAGTCGTTGCTAGTACCATCTACTTGGGAGGCTNNNAGATCGGAAG |
| 37336_2 | TATGAGTGTGGAGTCGTTGCTAGTACCATCTACTTGGGAGGCTNNNAGATCGGAAG |
| 37336_3 | TATGAGTGTGGAGTCGTTGCTAGTACCATCTACTTGGGAGGCTNNNAGATCGGAAG |
| 2912_1 | TATGAGTGTGGAGTCGTTGCGTTTCTAGGAGGCAGCTGTTTGTNNNAGATCGGAA |
| 2912_2 | TATGAGTGTGGAGTCGTTGCGTTTCTAGGAGGCAGCTGTTTGTNNNAGATCGGAA |
| 2912_3 | TATGAGTGTGGAGTCGTTGCGTTTCTAGGAGGCAGCTGTTTGTNNNAGATCGGAA |
| 39599_1 | TATGAGTGTGGAGTCGTTGCAAAGGAAGTCAATCAATGTGGCANNNAGATCGGAA |
| 39599_2 | TATGAGTGTGGAGTCGTTGCAAAGGAAGTCAATCAATGTGGCANNNAGATCGGAA |
| 39599_3 | TATGAGTGTGGAGTCGTTGCAAAGGAAGTCAATCAATGTGGCANNNAGATCGGAA |
| 44626_1 | TATGAGTGTGGAGTCGTTGCTAGGAGGCAAGGATCGGGAANNNAGATCGGAAGA |
| 44626_2 | TATGAGTGTGGAGTCGTTGCTAGGAGGCAAGGATCGGGAANNNAGATCGGAAGA |
| 44626_3 | TATGAGTGTGGAGTCGTTGCTAGGAGGCAAGGATCGGGAANNNAGATCGGAAGA |
| 48872_1 | TATGAGTGTGGAGTCGTTGCATGTGCCACCATACCCAACTAATNNNAGATCGGAAG |
| 48872_2 | TATGAGTGTGGAGTCGTTGCATGTGCCACCATACCCAACTAATNNNAGATCGGAAG |
| 48872_3 | TATGAGTGTGGAGTCGTTGCATGTGCCACCATACCCAACTAATNNNAGATCGGAAG |
| 29514_1 | TATGAGTGTGGAGTCGTTGCGTTCCAAGTAAGTCAAGCTTTAANNNAGATCGGAAG |
| 29514_2 | TATGAGTGTGGAGTCGTTGCGTTCCAAGTAAGTCAAGCTTTAANNNAGATCGGAAG |
| 29514_3 | TATGAGTGTGGAGTCGTTGCGTTCCAAGTAAGTCAAGCTTTAANNNAGATCGGAAG |
| 29752_1 | TATGAGTGTGGAGTCGTTGCTTGTCGTTGTAGGTAAGAGGACANNNAGATCGGAA |
| 29752_2 | TATGAGTGTGGAGTCGTTGCTTGTCGTTGTAGGTAAGAGGACANNNAGATCGGAA |
| 29752_3 | TATGAGTGTGGAGTCGTTGCTTGTCGTTGTAGGTAAGAGGACANNNAGATCGGAA |
| 36890_1 | TATGAGTGTGGAGTCGTTGCTACCCCAAAGCCATATAAGCACANNNAGATCGGAAG |
| 36890_2 | TATGAGTGTGGAGTCGTTGCTACCCCAAAGCCATATAAGCACANNNAGATCGGAAG |
| 36890_3 | TATGAGTGTGGAGTCGTTGCTACCCCAAAGCCATATAAGCACANNNAGATCGGAAG |
| 38654_1 | TATGAGTGTGGAGTCGTTGCTTTGCCATGATTGTAAGAAGCCCNNNAGATCGGAAG |
| 38654_2 | TATGAGTGTGGAGTCGTTGCTTTGCCATGATTGTAAGAAGCCCNNNAGATCGGAAG |
| 38654_3 | TATGAGTGTGGAGTCGTTGCTTTGCCATGATTGTAAGAAGCCCNNNAGATCGGAAG |
| 29162_1 | TATGAGTGTGGAGTCGTTGCAATCTTTGCTAGCACAATGGACGNNNAGATCGGAAG |
| 29162_2 | TATGAGTGTGGAGTCGTTGCAATCTTTGCTAGCACAATGGACGNNNAGATCGGAAG |

| | |
|---|---|
| 29162_3 | TATGAGTGTGGAGTCGTTGCAATCTTTGCTAGCACAATGGACGNNNAGATCGGAAG |
| 29387_1 | TATGAGTGTGGAGTCGTTGCAGAGGTGAGAAGGGACTAAGACANNNAGATCGGA |
| 29387_2 | TATGAGTGTGGAGTCGTTGCAGAGGTGAGAAGGGACTAAGACANNNAGATCGGA |
| 29387_3 | TATGAGTGTGGAGTCGTTGCAGAGGTGAGAAGGGACTAAGACANNNAGATCGGA |
| 50521_1 | TATGAGTGTGGAGTCGTTGCCCGTGTGACGCTCAACAAATTATNNNAGATCGGAAG |
| 50521_2 | TATGAGTGTGGAGTCGTTGCCCGTGTGACGCTCAACAAATTATNNNAGATCGGAAG |
| 50521_3 | TATGAGTGTGGAGTCGTTGCCCGTGTGACGCTCAACAAATTATNNNAGATCGGAAG |
| 46955_1 | TATGAGTGTGGAGTCGTTGCCTGGAGTGCAATGGTGTGATCTTNNNAGATCGGAA |
| 46955_2 | TATGAGTGTGGAGTCGTTGCCTGGAGTGCAATGGTGTGATCTTNNNAGATCGGAA |
| 46955_3 | TATGAGTGTGGAGTCGTTGCCTGGAGTGCAATGGTGTGATCTTNNNAGATCGGAA |
| 51337_1 | TATGAGTGTGGAGTCGTTGCTCCTGTTGCTTGGTATCATGTCANNNAGATCGGAAG |
| 51337_2 | TATGAGTGTGGAGTCGTTGCTCCTGTTGCTTGGTATCATGTCANNNAGATCGGAAG |
| 51337_3 | TATGAGTGTGGAGTCGTTGCTCCTGTTGCTTGGTATCATGTCANNNAGATCGGAAG |
| 39794_1 | TATGAGTGTGGAGTCGTTGCGTGAAGAAATGCCAGGCTCTTTTNNNAGATCGGAAG |
| 39794_2 | TATGAGTGTGGAGTCGTTGCGTGAAGAAATGCCAGGCTCTTTTNNNAGATCGGAAG |
| 39794_3 | TATGAGTGTGGAGTCGTTGCGTGAAGAAATGCCAGGCTCTTTTNNNAGATCGGAAG |
| 39235_1 | TATGAGTGTGGAGTCGTTGCGCTAAGGCAGGAGAATCACTCAANNNAGATCGGAA |
| 39235_2 | TATGAGTGTGGAGTCGTTGCGCTAAGGCAGGAGAATCACTCAANNNAGATCGGAA |
| 39235_3 | TATGAGTGTGGAGTCGTTGCGCTAAGGCAGGAGAATCACTCAANNNAGATCGGAA |
| 42204_1 | TATGAGTGTGGAGTCGTTGCGGTGAGTCTGTGGTCATTTTGTGNNNAGATCGGAA |
| 42204_2 | TATGAGTGTGGAGTCGTTGCGGTGAGTCTGTGGTCATTTTGTGNNNAGATCGGAA |
| 42204_3 | TATGAGTGTGGAGTCGTTGCGGTGAGTCTGTGGTCATTTTGTGNNNAGATCGGAA |
| 6054_1 | TATGAGTGTGGAGTCGTTGCGCCACATTTATTGCTGCATTTGCNNNAGATCGGAAG |
| 6054_2 | TATGAGTGTGGAGTCGTTGCGCCACATTTATTGCTGCATTTGCNNNAGATCGGAAG |
| 6054_3 | TATGAGTGTGGAGTCGTTGCGCCACATTTATTGCTGCATTTGCNNNAGATCGGAAG |
| 36787_1 | TATGAGTGTGGAGTCGTTGCTGTGGCAATTCCTCAAAGACCTANNNAGATCGGAAG |
| 36787_2 | TATGAGTGTGGAGTCGTTGCTGTGGCAATTCCTCAAAGACCTANNNAGATCGGAAG |
| 36787_3 | TATGAGTGTGGAGTCGTTGCTGTGGCAATTCCTCAAAGACCTANNNAGATCGGAAG |
| 30235_1 | TATGAGTGTGGAGTCGTTGCAAACTGGAAAAGGCTGAGACAGANNNAGATCGGAA |
| 30235_2 | TATGAGTGTGGAGTCGTTGCAAACTGGAAAAGGCTGAGACAGANNNAGATCGGAA |
| 30235_3 | TATGAGTGTGGAGTCGTTGCAAACTGGAAAAGGCTGAGACAGANNNAGATCGGAA |
| 38884_1 | TATGAGTGTGGAGTCGTTGCAGGAAGGGGTTTGAAACAAGACANNNAGATCGGAA |
| 38884_2 | TATGAGTGTGGAGTCGTTGCAGGAAGGGGTTTGAAACAAGACANNNAGATCGGAA |
| 38884_3 | TATGAGTGTGGAGTCGTTGCAGGAAGGGGTTTGAAACAAGACANNNAGATCGGAA |
| 9574_1 | TATGAGTGTGGAGTCGTTGCATTGGTCACTGGCCTTACTATGCNNNAGATCGGAAG |
| 9574_2 | TATGAGTGTGGAGTCGTTGCATTGGTCACTGGCCTTACTATGCNNNAGATCGGAAG |
| 9574_3 | TATGAGTGTGGAGTCGTTGCATTGGTCACTGGCCTTACTATGCNNNAGATCGGAAG |
| 13190_1 | TATGAGTGTGGAGTCGTTGCACTGACTGACTTGGATCTTANNNAGATCGGAAGAG |
| 13190_2 | TATGAGTGTGGAGTCGTTGCACTGACTGACTTGGATCTTANNNAGATCGGAAGAG |
| 13190_3 | TATGAGTGTGGAGTCGTTGCACTGACTGACTTGGATCTTANNNAGATCGGAAGAG |
| 38192_1 | TATGAGTGTGGAGTCGTTGCAGTGGTTCACACCTGTAATCCCNNNAGATCGGAAGA |
| 38192_2 | TATGAGTGTGGAGTCGTTGCAGTGGTTCACACCTGTAATCCCNNNAGATCGGAAGA |
| 38192_3 | TATGAGTGTGGAGTCGTTGCAGTGGTTCACACCTGTAATCCCNNNAGATCGGAAGA |

| | |
|---|---|
| 52533_1 | TATGAGTGTGGAGTCGTTGCGGGAGCAAGTAAGACTGATCACANNNAGATCGGAA |
| 52533_2 | TATGAGTGTGGAGTCGTTGCGGGAGCAAGTAAGACTGATCACANNNAGATCGGAA |
| 52533_3 | TATGAGTGTGGAGTCGTTGCGGGAGCAAGTAAGACTGATCACANNNAGATCGGAA |
| 51945_1 | TATGAGTGTGGAGTCGTTGCCCCCTCACCTCCTGGATAGGNNNAGATCGGAAGAGC |
| 51945_2 | TATGAGTGTGGAGTCGTTGCCCCCTCACCTCCTGGATAGGNNNAGATCGGAAGAGC |
| 51945_3 | TATGAGTGTGGAGTCGTTGCCCCCTCACCTCCTGGATAGGNNNAGATCGGAAGAGC |
| 44406_1 | TATGAGTGTGGAGTCGTTGCCCCAACTATTGTCATTGTTCCAGGNNNAGATCGGAA |
| 44406_2 | TATGAGTGTGGAGTCGTTGCCCCAACTATTGTCATTGTTCCAGGNNNAGATCGGAA |
| 44406_3 | TATGAGTGTGGAGTCGTTGCCCCAACTATTGTCATTGTTCCAGGNNNAGATCGGAA |
| 33159_1 | TATGAGTGTGGAGTCGTTGCTGAACTATGATCACACCACAGCANNNAGATCGGAAG |
| 33159_2 | TATGAGTGTGGAGTCGTTGCTGAACTATGATCACACCACAGCANNNAGATCGGAAG |
| 33159_3 | TATGAGTGTGGAGTCGTTGCTGAACTATGATCACACCACAGCANNNAGATCGGAAG |
| 37114_1 | TATGAGTGTGGAGTCGTTGCAAGCACATGAATTTGTTTGGGGANNNAGATCGGAA |
| 37114_2 | TATGAGTGTGGAGTCGTTGCAAGCACATGAATTTGTTTGGGGANNNAGATCGGAA |
| 37114_3 | TATGAGTGTGGAGTCGTTGCAAGCACATGAATTTGTTTGGGGANNNAGATCGGAA |
| 50304_1 | TATGAGTGTGGAGTCGTTGCAAGCGTAATCACCATCAAAACCCNNNAGATCGGAAG |
| 50304_2 | TATGAGTGTGGAGTCGTTGCAAGCGTAATCACCATCAAAACCCNNNAGATCGGAAG |
| 50304_3 | TATGAGTGTGGAGTCGTTGCAAGCGTAATCACCATCAAAACCCNNNAGATCGGAAG |
| 43344_1 | TATGAGTGTGGAGTCGTTGCCTGAGTAGCTGGGATTATAGGCGNNNAGATCGGAA |
| 43344_2 | TATGAGTGTGGAGTCGTTGCCTGAGTAGCTGGGATTATAGGCGNNNAGATCGGAA |
| 43344_3 | TATGAGTGTGGAGTCGTTGCCTGAGTAGCTGGGATTATAGGCGNNNAGATCGGAA |
| 10868_1 | TATGAGTGTGGAGTCGTTGCAAGTTCCTGTAGCGCCGGAAAACNNNAGATCGGAA |
| 10868_2 | TATGAGTGTGGAGTCGTTGCAAGTTCCTGTAGCGCCGGAAAACNNNAGATCGGAA |
| 10868_3 | TATGAGTGTGGAGTCGTTGCAAGTTCCTGTAGCGCCGGAAAACNNNAGATCGGAA |
| 39777_1 | TATGAGTGTGGAGTCGTTGCATGCACCCAATACTAGAGCACAANNNAGATCGGAAG |
| 39777_2 | TATGAGTGTGGAGTCGTTGCATGCACCCAATACTAGAGCACAANNNAGATCGGAAG |
| 39777_3 | TATGAGTGTGGAGTCGTTGCATGCACCCAATACTAGAGCACAANNNAGATCGGAAG |
| 42633_1 | TATGAGTGTGGAGTCGTTGCAGAGAGAGACTTGTTCTGTTGCCNNNAGATCGGAA |
| 42633_2 | TATGAGTGTGGAGTCGTTGCAGAGAGAGACTTGTTCTGTTGCCNNNAGATCGGAA |
| 42633_3 | TATGAGTGTGGAGTCGTTGCAGAGAGAGACTTGTTCTGTTGCCNNNAGATCGGAA |
| 34449_1 | TATGAGTGTGGAGTCGTTGCCTCTCTCTAGCTATCCCCCTCTCNNNAGATCGGAAGA |
| 34449_2 | TATGAGTGTGGAGTCGTTGCCTCTCTCTAGCTATCCCCCTCTCNNNAGATCGGAAGA |
| 34449_3 | TATGAGTGTGGAGTCGTTGCCTCTCTCTAGCTATCCCCCTCTCNNNAGATCGGAAGA |
| 32644_1 | TATGAGTGTGGAGTCGTTGCCAGGAGAAAAGCATGCAAGTCAANNNAGATCGGAA |
| 32644_2 | TATGAGTGTGGAGTCGTTGCCAGGAGAAAAGCATGCAAGTCAANNNAGATCGGAA |
| 32644_3 | TATGAGTGTGGAGTCGTTGCCAGGAGAAAAGCATGCAAGTCAANNNAGATCGGAA |
| 3049_1 | TATGAGTGTGGAGTCGTTGCGGCTTCGGAAGTAGAAGCACTTTNNNAGATCGGAA |
| 3049_2 | TATGAGTGTGGAGTCGTTGCGGCTTCGGAAGTAGAAGCACTTTNNNAGATCGGAA |
| 3049_3 | TATGAGTGTGGAGTCGTTGCGGCTTCGGAAGTAGAAGCACTTTNNNAGATCGGAA |
| 51803_1 | TATGAGTGTGGAGTCGTTGCTCAATAGGTCCTGTCAGACAAGCNNNAGATCGGAAG |
| 51803_2 | TATGAGTGTGGAGTCGTTGCTCAATAGGTCCTGTCAGACAAGCNNNAGATCGGAAG |
| 51803_3 | TATGAGTGTGGAGTCGTTGCTCAATAGGTCCTGTCAGACAAGCNNNAGATCGGAAG |
| 36674_1 | TATGAGTGTGGAGTCGTTGCCATGTCTGTGATCCCAGGTACTTNNNAGATCGGAAG |

| | |
|---|---|
| 36674_2 | TATGAGTGTGGAGTCGTTGCCATGTCTGTGATCCCAGGTACTTNNNAGATCGGAAG |
| 36674_3 | TATGAGTGTGGAGTCGTTGCCATGTCTGTGATCCCAGGTACTTNNNAGATCGGAAG |
| 34465_1 | TATGAGTGTGGAGTCGTTGCTGACCAAAACGTTTTTATAGGGCANNNAGATCGGAA |
| 34465_2 | TATGAGTGTGGAGTCGTTGCTGACCAAAACGTTTTTATAGGGCANNNAGATCGGAA |
| 34465_3 | TATGAGTGTGGAGTCGTTGCTGACCAAAACGTTTTTATAGGGCANNNAGATCGGAA |
| 47108_1 | TATGAGTGTGGAGTCGTTGCGACGTAACCTCACCTATAGCTGGNNNAGATCGGAAG |
| 47108_2 | TATGAGTGTGGAGTCGTTGCGACGTAACCTCACCTATAGCTGGNNNAGATCGGAAG |
| 47108_3 | TATGAGTGTGGAGTCGTTGCGACGTAACCTCACCTATAGCTGGNNNAGATCGGAAG |
| 13222_1 | TATGAGTGTGGAGTCGTTGCCTATAATCCCAGCACTTTTTGNNNAGATCGGAAGAG |
| 13222_2 | TATGAGTGTGGAGTCGTTGCCTATAATCCCAGCACTTTTTGNNNAGATCGGAAGAG |
| 13222_3 | TATGAGTGTGGAGTCGTTGCCTATAATCCCAGCACTTTTTGNNNAGATCGGAAGAG |
| 9693_1 | TATGAGTGTGGAGTCGTTGCTACCCGTTTCATTTACCCAACCCNNNAGATCGGAAGA |
| 9693_2 | TATGAGTGTGGAGTCGTTGCTACCCGTTTCATTTACCCAACCCNNNAGATCGGAAGA |
| 9693_3 | TATGAGTGTGGAGTCGTTGCTACCCGTTTCATTTACCCAACCCNNNAGATCGGAAGA |
| 37589_1 | TATGAGTGTGGAGTCGTTGCGTGTCACATGCTTGTAATCCCTGNNNAGATCGGAAG |
| 37589_2 | TATGAGTGTGGAGTCGTTGCGTGTCACATGCTTGTAATCCCTGNNNAGATCGGAAG |
| 37589_3 | TATGAGTGTGGAGTCGTTGCGTGTCACATGCTTGTAATCCCTGNNNAGATCGGAAG |
| 30087_1 | TATGAGTGTGGAGTCGTTGCACCGTCTATATACAAGGCTCCAGNNNAGATCGGAAG |
| 30087_2 | TATGAGTGTGGAGTCGTTGCACCGTCTATATACAAGGCTCCAGNNNAGATCGGAAG |
| 30087_3 | TATGAGTGTGGAGTCGTTGCACCGTCTATATACAAGGCTCCAGNNNAGATCGGAAG |
| 33510_1 | TATGAGTGTGGAGTCGTTGCTTCTTTTGATTGCTCGTGGATCTNNNAGATCGGAAG |
| 33510_2 | TATGAGTGTGGAGTCGTTGCTTCTTTTGATTGCTCGTGGATCTNNNAGATCGGAAG |
| 33510_3 | TATGAGTGTGGAGTCGTTGCTTCTTTTGATTGCTCGTGGATCTNNNAGATCGGAAG |
| 31777_1 | TATGAGTGTGGAGTCGTTGCCAGGTTTGATTGGTGGTCAGAACNNNAGATCGGAA |
| 31777_2 | TATGAGTGTGGAGTCGTTGCCAGGTTTGATTGGTGGTCAGAACNNNAGATCGGAA |
| 31777_3 | TATGAGTGTGGAGTCGTTGCCAGGTTTGATTGGTGGTCAGAACNNNAGATCGGAA |
| 35238_1 | TATGAGTGTGGAGTCGTTGCCCAATTTGCTGTGGATTCACGATNNNAGATCGGAAG |
| 35238_2 | TATGAGTGTGGAGTCGTTGCCCAATTTGCTGTGGATTCACGATNNNAGATCGGAAG |
| 35238_3 | TATGAGTGTGGAGTCGTTGCCCAATTTGCTGTGGATTCACGATNNNAGATCGGAAG |
| 38739_1 | TATGAGTGTGGAGTCGTTGCTCCAACAAGGCCCTTTACCTTTANNNAGATCGGAAGA |
| 38739_2 | TATGAGTGTGGAGTCGTTGCTCCAACAAGGCCCTTTACCTTTANNNAGATCGGAAGA |
| 38739_3 | TATGAGTGTGGAGTCGTTGCTCCAACAAGGCCCTTTACCTTTANNNAGATCGGAAGA |
| 52984_1 | TATGAGTGTGGAGTCGTTGCCAGTCACACTAGCCATCACTCATNNNAGATCGGAAGA |
| 52984_2 | TATGAGTGTGGAGTCGTTGCCAGTCACACTAGCCATCACTCATNNNAGATCGGAAGA |
| 52984_3 | TATGAGTGTGGAGTCGTTGCCAGTCACACTAGCCATCACTCATNNNAGATCGGAAGA |
| 41972_1 | TATGAGTGTGGAGTCGTTGCCGAGTAGCTGGGACTACAGGTNNNAGATCGGAAGA |
| 41972_2 | TATGAGTGTGGAGTCGTTGCCGAGTAGCTGGGACTACAGGTNNNAGATCGGAAGA |
| 41972_3 | TATGAGTGTGGAGTCGTTGCCGAGTAGCTGGGACTACAGGTNNNAGATCGGAAGA |
| 10558_1 | TATGAGTGTGGAGTCGTTGCAGGAATTTTGGGGAGTGGCTTTTNNNAGATCGGAA |
| 10558_2 | TATGAGTGTGGAGTCGTTGCAGGAATTTTGGGGAGTGGCTTTTNNNAGATCGGAA |
| 10558_3 | TATGAGTGTGGAGTCGTTGCAGGAATTTTGGGGAGTGGCTTTTNNNAGATCGGAA |
| 46848_1 | TATGAGTGTGGAGTCGTTGCGGACAAGCACAGTTCCTTACTCTNNNAGATCGGAAG |
| 46848_2 | TATGAGTGTGGAGTCGTTGCGGACAAGCACAGTTCCTTACTCTNNNAGATCGGAAG |

| | |
|---|---|
| 46848_3 | TATGAGTGTGGAGTCGTTGCGGACAAGCACAGTTCCTTACTCTNNNAGATCGGAAG |
| 39807_1 | TATGAGTGTGGAGTCGTTGCGGAAAGTATGGCATAAAGGCAGGNNNAGATCGGA |
| 39807_2 | TATGAGTGTGGAGTCGTTGCGGAAAGTATGGCATAAAGGCAGGNNNAGATCGGA |
| 39807_3 | TATGAGTGTGGAGTCGTTGCGGAAAGTATGGCATAAAGGCAGGNNNAGATCGGA |
| 37609_1 | TATGAGTGTGGAGTCGTTGCCCCAGTTTTGTGCTTGAAACCTANNNAGATCGGAAG |
| 37609_2 | TATGAGTGTGGAGTCGTTGCCCCAGTTTTGTGCTTGAAACCTANNNAGATCGGAAG |
| 37609_3 | TATGAGTGTGGAGTCGTTGCCCCAGTTTTGTGCTTGAAACCTANNNAGATCGGAAG |
| 34026_1 | TATGAGTGTGGAGTCGTTGCCAGTGGTGTGATCTTGGCTCANNNAGATCGGAAGA |
| 34026_2 | TATGAGTGTGGAGTCGTTGCCAGTGGTGTGATCTTGGCTCANNNAGATCGGAAGA |
| 34026_3 | TATGAGTGTGGAGTCGTTGCCAGTGGTGTGATCTTGGCTCANNNAGATCGGAAGA |
| 50421_1 | TATGAGTGTGGAGTCGTTGCATTCAACACTAGCCTGGCTAACANNNAGATCGGAAG |
| 50421_2 | TATGAGTGTGGAGTCGTTGCATTCAACACTAGCCTGGCTAACANNNAGATCGGAAG |
| 50421_3 | TATGAGTGTGGAGTCGTTGCATTCAACACTAGCCTGGCTAACANNNAGATCGGAAG |
| 34725_1 | TATGAGTGTGGAGTCGTTGCCTTCCCAGGTTCAAGCAATTCTCNNNAGATCGGAAG |
| 34725_2 | TATGAGTGTGGAGTCGTTGCCTTCCCAGGTTCAAGCAATTCTCNNNAGATCGGAAG |
| 34725_3 | TATGAGTGTGGAGTCGTTGCCTTCCCAGGTTCAAGCAATTCTCNNNAGATCGGAAG |
| 40561_1 | TATGAGTGTGGAGTCGTTGCAGGAAAAAAAGAAAATGCCACGANNNAGATCGGAA |
| 40561_2 | TATGAGTGTGGAGTCGTTGCAGGAAAAAAAGAAAATGCCACGANNNAGATCGGAA |
| 40561_3 | TATGAGTGTGGAGTCGTTGCAGGAAAAAAAGAAAATGCCACGANNNAGATCGGAA |
| 34522_1 | TATGAGTGTGGAGTCGTTGCGACGCAGATTTGGTCTTTTCACANNNAGATCGGAAG |
| 34522_2 | TATGAGTGTGGAGTCGTTGCGACGCAGATTTGGTCTTTTCACANNNAGATCGGAAG |
| 34522_3 | TATGAGTGTGGAGTCGTTGCGACGCAGATTTGGTCTTTTCACANNNAGATCGGAAG |
| 4907_1 | TATGAGTGTGGAGTCGTTGCTCACTTGGTCCTTTCTTGTGTGTNNNAGATCGGAAG |
| 4907_2 | TATGAGTGTGGAGTCGTTGCTCACTTGGTCCTTTCTTGTGTGTNNNAGATCGGAAG |
| 4907_3 | TATGAGTGTGGAGTCGTTGCTCACTTGGTCCTTTCTTGTGTGTNNNAGATCGGAAG |
| 49921_1 | TATGAGTGTGGAGTCGTTGCTGTATTTACAGCCACTCCCCATTNNNAGATCGGAAGA |
| 49921_2 | TATGAGTGTGGAGTCGTTGCTGTATTTACAGCCACTCCCCATTNNNAGATCGGAAGA |
| 49921_3 | TATGAGTGTGGAGTCGTTGCTGTATTTACAGCCACTCCCCATTNNNAGATCGGAAGA |
| 51665_1 | TATGAGTGTGGAGTCGTTGCCTCTCCTGTATATCACAAGGCCCNNNAGATCGGAAG |
| 51665_2 | TATGAGTGTGGAGTCGTTGCCTCTCCTGTATATCACAAGGCCCNNNAGATCGGAAG |
| 51665_3 | TATGAGTGTGGAGTCGTTGCCTCTCCTGTATATCACAAGGCCCNNNAGATCGGAAG |
| 50603_1 | TATGAGTGTGGAGTCGTTGCCACTTGACAGGAGCAGAAGATCTNNNAGATCGGAA |
| 50603_2 | TATGAGTGTGGAGTCGTTGCCACTTGACAGGAGCAGAAGATCTNNNAGATCGGAA |
| 50603_3 | TATGAGTGTGGAGTCGTTGCCACTTGACAGGAGCAGAAGATCTNNNAGATCGGAA |
| 50232_1 | TATGAGTGTGGAGTCGTTGCTTCCTGGAGCTCTGTCCCTATTANNNAGATCGGAAG |
| 50232_2 | TATGAGTGTGGAGTCGTTGCTTCCTGGAGCTCTGTCCCTATTANNNAGATCGGAAG |
| 50232_3 | TATGAGTGTGGAGTCGTTGCTTCCTGGAGCTCTGTCCCTATTANNNAGATCGGAAG |
| 30603_1 | TATGAGTGTGGAGTCGTTGCTGGCATCATTCTGTCTCTCTGCTNNNAGATCGGAAGA |
| 30603_2 | TATGAGTGTGGAGTCGTTGCTGGCATCATTCTGTCTCTCTGCTNNNAGATCGGAAGA |
| 30603_3 | TATGAGTGTGGAGTCGTTGCTGGCATCATTCTGTCTCTCTGCTNNNAGATCGGAAGA |
| 39113_1 | TATGAGTGTGGAGTCGTTGCGCTTTGCAGTACAGTGCTCAAATNNNAGATCGGAAG |
| 39113_2 | TATGAGTGTGGAGTCGTTGCGCTTTGCAGTACAGTGCTCAAATNNNAGATCGGAAG |
| 39113_3 | TATGAGTGTGGAGTCGTTGCGCTTTGCAGTACAGTGCTCAAATNNNAGATCGGAAG |

| | |
|---|---|
| 36135_1 | TATGAGTGTGGAGTCGTTGCCTGGCGACAGAGTGAGACTCNNNAGATCGGAAGAG |
| 36135_2 | TATGAGTGTGGAGTCGTTGCCTGGCGACAGAGTGAGACTCNNNAGATCGGAAGAG |
| 36135_3 | TATGAGTGTGGAGTCGTTGCCTGGCGACAGAGTGAGACTCNNNAGATCGGAAGAG |
| 31104_1 | TATGAGTGTGGAGTCGTTGCAGCACTCCATGAACATCCTTGATNNNAGATCGGAAG |
| 31104_2 | TATGAGTGTGGAGTCGTTGCAGCACTCCATGAACATCCTTGATNNNAGATCGGAAG |
| 31104_3 | TATGAGTGTGGAGTCGTTGCAGCACTCCATGAACATCCTTGATNNNAGATCGGAAG |
| 44779_1 | TATGAGTGTGGAGTCGTTGCCAGGAAGGACAGAGTCATGGATGNNNAGATCGGA |
| 44779_2 | TATGAGTGTGGAGTCGTTGCCAGGAAGGACAGAGTCATGGATGNNNAGATCGGA |
| 44779_3 | TATGAGTGTGGAGTCGTTGCCAGGAAGGACAGAGTCATGGATGNNNAGATCGGA |
| 32112_1 | TATGAGTGTGGAGTCGTTGCATGGATGGAAGAGGAGGTCACTANNNAGATCGGA |
| 32112_2 | TATGAGTGTGGAGTCGTTGCATGGATGGAAGAGGAGGTCACTANNNAGATCGGA |
| 32112_3 | TATGAGTGTGGAGTCGTTGCATGGATGGAAGAGGAGGTCACTANNNAGATCGGA |
| 46579_1 | TATGAGTGTGGAGTCGTTGCGGAGCAATTTTCAGATCAGGCTCNNNAGATCGGAA |
| 46579_2 | TATGAGTGTGGAGTCGTTGCGGAGCAATTTTCAGATCAGGCTCNNNAGATCGGAA |
| 46579_3 | TATGAGTGTGGAGTCGTTGCGGAGCAATTTTCAGATCAGGCTCNNNAGATCGGAA |
| 34922_1 | TATGAGTGTGGAGTCGTTGCCCTGGGTTCAAGCAATTCTCATGNNNAGATCGGAAG |
| 34922_2 | TATGAGTGTGGAGTCGTTGCCCTGGGTTCAAGCAATTCTCATGNNNAGATCGGAAG |
| 34922_3 | TATGAGTGTGGAGTCGTTGCCCTGGGTTCAAGCAATTCTCATGNNNAGATCGGAAG |
| 38625_1 | TATGAGTGTGGAGTCGTTGCCACCTTATTCCTGCAAGAATGGCNNNAGATCGGAAG |
| 38625_2 | TATGAGTGTGGAGTCGTTGCCACCTTATTCCTGCAAGAATGGCNNNAGATCGGAAG |
| 38625_3 | TATGAGTGTGGAGTCGTTGCCACCTTATTCCTGCAAGAATGGCNNNAGATCGGAAG |
| 47612_1 | TATGAGTGTGGAGTCGTTGCGACAGTACATCCCGGAGTCTCTNNNAGATCGGAAGA |
| 47612_2 | TATGAGTGTGGAGTCGTTGCGACAGTACATCCCGGAGTCTCTNNNAGATCGGAAGA |
| 47612_3 | TATGAGTGTGGAGTCGTTGCGACAGTACATCCCGGAGTCTCTNNNAGATCGGAAGA |
| 31499_1 | TATGAGTGTGGAGTCGTTGCTCTGTTGCTATAACGGCACTAGTNNNAGATCGGAAG |
| 31499_2 | TATGAGTGTGGAGTCGTTGCTCTGTTGCTATAACGGCACTAGTNNNAGATCGGAAG |
| 31499_3 | TATGAGTGTGGAGTCGTTGCTCTGTTGCTATAACGGCACTAGTNNNAGATCGGAAG |
| 42024_1 | TATGAGTGTGGAGTCGTTGCGTGTGGAATGCATGGAAATGTCANNNAGATCGGAA |
| 42024_2 | TATGAGTGTGGAGTCGTTGCGTGTGGAATGCATGGAAATGTCANNNAGATCGGAA |
| 42024_3 | TATGAGTGTGGAGTCGTTGCGTGTGGAATGCATGGAAATGTCANNNAGATCGGAA |
| 42276_1 | TATGAGTGTGGAGTCGTTGCAGTTGGAGTCTCACTGTGTCATCNNNAGATCGGAAG |
| 42276_2 | TATGAGTGTGGAGTCGTTGCAGTTGGAGTCTCACTGTGTCATCNNNAGATCGGAAG |
| 42276_3 | TATGAGTGTGGAGTCGTTGCAGTTGGAGTCTCACTGTGTCATCNNNAGATCGGAAG |
| 46036_1 | TATGAGTGTGGAGTCGTTGCGGCCAGGGTGGTTTATTTCTAGANNNAGATCGGAA |
| 46036_2 | TATGAGTGTGGAGTCGTTGCGGCCAGGGTGGTTTATTTCTAGANNNAGATCGGAA |
| 46036_3 | TATGAGTGTGGAGTCGTTGCGGCCAGGGTGGTTTATTTCTAGANNNAGATCGGAA |
| 2247_1 | TATGAGTGTGGAGTCGTTGCCCAGGAAACCTTCTCTTCCCCTANNNAGATCGGAAGA |
| 2247_2 | TATGAGTGTGGAGTCGTTGCCCAGGAAACCTTCTCTTCCCCTANNNAGATCGGAAGA |
| 2247_3 | TATGAGTGTGGAGTCGTTGCCCAGGAAACCTTCTCTTCCCCTANNNAGATCGGAAGA |
| 38898_1 | TATGAGTGTGGAGTCGTTGCTCTGAACAAGTAGGCTTTCTCCCNNNAGATCGGAAG |
| 38898_2 | TATGAGTGTGGAGTCGTTGCTCTGAACAAGTAGGCTTTCTCCCNNNAGATCGGAAG |
| 38898_3 | TATGAGTGTGGAGTCGTTGCTCTGAACAAGTAGGCTTTCTCCCNNNAGATCGGAAG |
| 12768_1 | TATGAGTGTGGAGTCGTTGCAATTTCTTCCTAAAAGCCTACTNNNAGATCGGAAGA |

| | |
|---|---|
| 12768_2 | TATGAGTGTGGAGTCGTTGCAATTTCTTCCTAAAAGCCTACTNNNAGATCGGAAGA |
| 12768_3 | TATGAGTGTGGAGTCGTTGCAATTTCTTCCTAAAAGCCTACTNNNAGATCGGAAGA |
| 9636_1 | TATGAGTGTGGAGTCGTTGCGAAGTTTGCCTTTTCAGCTGTCCNNNAGATCGGAAG |
| 9636_2 | TATGAGTGTGGAGTCGTTGCGAAGTTTGCCTTTTCAGCTGTCCNNNAGATCGGAAG |
| 9636_3 | TATGAGTGTGGAGTCGTTGCGAAGTTTGCCTTTTCAGCTGTCCNNNAGATCGGAAG |
| 44919_1 | TATGAGTGTGGAGTCGTTGCCCACTTCACCAGTTCAGACTCATNNNAGATCGGAAG |
| 44919_2 | TATGAGTGTGGAGTCGTTGCCCACTTCACCAGTTCAGACTCATNNNAGATCGGAAG |
| 44919_3 | TATGAGTGTGGAGTCGTTGCCCACTTCACCAGTTCAGACTCATNNNAGATCGGAAG |
| 41964_1 | TATGAGTGTGGAGTCGTTGCAGCAAAGCTACTGATTTGTGTACANNNAGATCGGAA |
| 41964_2 | TATGAGTGTGGAGTCGTTGCAGCAAAGCTACTGATTTGTGTACANNNAGATCGGAA |
| 41964_3 | TATGAGTGTGGAGTCGTTGCAGCAAAGCTACTGATTTGTGTACANNNAGATCGGAA |
| 48135_1 | TATGAGTGTGGAGTCGTTGCTTCAAGTGATTCTCCCACCTCAGNNNAGATCGGAAG |
| 48135_2 | TATGAGTGTGGAGTCGTTGCTTCAAGTGATTCTCCCACCTCAGNNNAGATCGGAAG |
| 48135_3 | TATGAGTGTGGAGTCGTTGCTTCAAGTGATTCTCCCACCTCAGNNNAGATCGGAAG |
| 43650_1 | TATGAGTGTGGAGTCGTTGCGAAACCTCCAACGTGCCTCTTCNNNAGATCGGAAGA |
| 43650_2 | TATGAGTGTGGAGTCGTTGCGAAACCTCCAACGTGCCTCTTCNNNAGATCGGAAGA |
| 43650_3 | TATGAGTGTGGAGTCGTTGCGAAACCTCCAACGTGCCTCTTCNNNAGATCGGAAGA |
| 29773_1 | TATGAGTGTGGAGTCGTTGCACATGCTCCTTCTTCCCATCAATNNNAGATCGGAAGA |
| 29773_2 | TATGAGTGTGGAGTCGTTGCACATGCTCCTTCTTCCCATCAATNNNAGATCGGAAGA |
| 29773_3 | TATGAGTGTGGAGTCGTTGCACATGCTCCTTCTTCCCATCAATNNNAGATCGGAAGA |
| 13293_1 | TATGAGTGTGGAGTCGTTGCCGTGAGAATTTAGAGATACACTANNNAGATCGGAA |
| 13293_2 | TATGAGTGTGGAGTCGTTGCCGTGAGAATTTAGAGATACACTANNNAGATCGGAA |
| 13293_3 | TATGAGTGTGGAGTCGTTGCCGTGAGAATTTAGAGATACACTANNNAGATCGGAA |
| 36533_1 | TATGAGTGTGGAGTCGTTGCAAACGAAACAAAGAGATGGTTCTTNNNAGATCGGA |
| 36533_2 | TATGAGTGTGGAGTCGTTGCAAACGAAACAAAGAGATGGTTCTTNNNAGATCGGA |
| 36533_3 | TATGAGTGTGGAGTCGTTGCAAACGAAACAAAGAGATGGTTCTTNNNAGATCGGA |
| 46583_1 | TATGAGTGTGGAGTCGTTGCGTAGTGTGATCATGGCTCATTGCNNNAGATCGGAA |
| 46583_2 | TATGAGTGTGGAGTCGTTGCGTAGTGTGATCATGGCTCATTGCNNNAGATCGGAA |
| 46583_3 | TATGAGTGTGGAGTCGTTGCGTAGTGTGATCATGGCTCATTGCNNNAGATCGGAA |
| 48939_1 | TATGAGTGTGGAGTCGTTGCCTCCTGAGTAGCTGGGACTACAGNNNAGATCGGAA |
| 48939_2 | TATGAGTGTGGAGTCGTTGCCTCCTGAGTAGCTGGGACTACAGNNNAGATCGGAA |
| 48939_3 | TATGAGTGTGGAGTCGTTGCCTCCTGAGTAGCTGGGACTACAGNNNAGATCGGAA |
| 36210_1 | TATGAGTGTGGAGTCGTTGCTTCCCTGATGCCTCAAATTTTGGNNNAGATCGGAAG |
| 36210_2 | TATGAGTGTGGAGTCGTTGCTTCCCTGATGCCTCAAATTTTGGNNNAGATCGGAAG |
| 36210_3 | TATGAGTGTGGAGTCGTTGCTTCCCTGATGCCTCAAATTTTGGNNNAGATCGGAAG |
| 44515_1 | TATGAGTGTGGAGTCGTTGCATTCAACACTTATTAGCGCCTGCNNNAGATCGGAAG |
| 44515_2 | TATGAGTGTGGAGTCGTTGCATTCAACACTTATTAGCGCCTGCNNNAGATCGGAAG |
| 44515_3 | TATGAGTGTGGAGTCGTTGCATTCAACACTTATTAGCGCCTGCNNNAGATCGGAAG |
| 41978_1 | TATGAGTGTGGAGTCGTTGCCGAGTAGCTGGGATTACAGGTGNNNAGATCGGAAG |
| 41978_2 | TATGAGTGTGGAGTCGTTGCCGAGTAGCTGGGATTACAGGTGNNNAGATCGGAAG |
| 41978_3 | TATGAGTGTGGAGTCGTTGCCGAGTAGCTGGGATTACAGGTGNNNAGATCGGAAG |
| 50111_1 | TATGAGTGTGGAGTCGTTGCATTACCAATCACCAACCACCAGANNNAGATCGGAAG |
| 50111_2 | TATGAGTGTGGAGTCGTTGCATTACCAATCACCAACCACCAGANNNAGATCGGAAG |

| | |
|---|---|
| 50111_3 | TATGAGTGTGGAGTCGTTGCATTACCAATCACCAACCACCAGANNNAGATCGGAAG |
| 33509_1 | TATGAGTGTGGAGTCGTTGCGCTGTGTACAGGCTCACTACTTANNNAGATCGGAAG |
| 33509_2 | TATGAGTGTGGAGTCGTTGCGCTGTGTACAGGCTCACTACTTANNNAGATCGGAAG |
| 33509_3 | TATGAGTGTGGAGTCGTTGCGCTGTGTACAGGCTCACTACTTANNNAGATCGGAAG |
| 36343_1 | TATGAGTGTGGAGTCGTTGCAAAAGGAAAGCAAGCAAGGTCTCNNNAGATCGGAA |
| 36343_2 | TATGAGTGTGGAGTCGTTGCAAAAGGAAAGCAAGCAAGGTCTCNNNAGATCGGAA |
| 36343_3 | TATGAGTGTGGAGTCGTTGCAAAAGGAAAGCAAGCAAGGTCTCNNNAGATCGGAA |
| 2433_1 | TATGAGTGTGGAGTCGTTGCTTGCCATAAACTGCCTTTTCAGCNNNAGATCGGAAG |
| 2433_2 | TATGAGTGTGGAGTCGTTGCTTGCCATAAACTGCCTTTTCAGCNNNAGATCGGAAG |
| 2433_3 | TATGAGTGTGGAGTCGTTGCTTGCCATAAACTGCCTTTTCAGCNNNAGATCGGAAG |
| 43111_1 | TATGAGTGTGGAGTCGTTGCCAGATCACTTCAGGTCAGGAGTTNNNAGATCGGAAG |
| 43111_2 | TATGAGTGTGGAGTCGTTGCCAGATCACTTCAGGTCAGGAGTTNNNAGATCGGAAG |
| 43111_3 | TATGAGTGTGGAGTCGTTGCCAGATCACTTCAGGTCAGGAGTTNNNAGATCGGAAG |
| 49553_1 | TATGAGTGTGGAGTCGTTGCCTCGAGTGATTCTCCTACCTCAGNNNAGATCGGAAG |
| 49553_2 | TATGAGTGTGGAGTCGTTGCCTCGAGTGATTCTCCTACCTCAGNNNAGATCGGAAG |
| 49553_3 | TATGAGTGTGGAGTCGTTGCCTCGAGTGATTCTCCTACCTCAGNNNAGATCGGAAG |
| 47707_1 | TATGAGTGTGGAGTCGTTGCCCAAGTAGCTGGGATTATAGGCANNNAGATCGGAA |
| 47707_2 | TATGAGTGTGGAGTCGTTGCCCAAGTAGCTGGGATTATAGGCANNNAGATCGGAA |
| 47707_3 | TATGAGTGTGGAGTCGTTGCCCAAGTAGCTGGGATTATAGGCANNNAGATCGGAA |
| 50537_1 | TATGAGTGTGGAGTCGTTGCCACGAGCCCTTTCTTTGATGTCNNNAGATCGGAAGA |
| 50537_2 | TATGAGTGTGGAGTCGTTGCCACGAGCCCTTTCTTTGATGTCNNNAGATCGGAAGA |
| 50537_3 | TATGAGTGTGGAGTCGTTGCCACGAGCCCTTTCTTTGATGTCNNNAGATCGGAAGA |
| 10361_1 | TATGAGTGTGGAGTCGTTGCGAGAACGAAGGGAGACAGAGAGANNNAGATCGGA |
| 10361_2 | TATGAGTGTGGAGTCGTTGCGAGAACGAAGGGAGACAGAGAGANNNAGATCGGA |
| 10361_3 | TATGAGTGTGGAGTCGTTGCGAGAACGAAGGGAGACAGAGAGANNNAGATCGGA |
| 52893_1 | TATGAGTGTGGAGTCGTTGCTTTTTTTGAGACGGAGTCTTGCGNNNAGATCGGAAG |
| 52893_2 | TATGAGTGTGGAGTCGTTGCTTTTTTTGAGACGGAGTCTTGCGNNNAGATCGGAAG |
| 52893_3 | TATGAGTGTGGAGTCGTTGCTTTTTTTGAGACGGAGTCTTGCGNNNAGATCGGAAG |
| 43834_1 | TATGAGTGTGGAGTCGTTGCCTTCAGTGTGAGAGGAGTGCTAGNNNAGATCGGAA |
| 43834_2 | TATGAGTGTGGAGTCGTTGCCTTCAGTGTGAGAGGAGTGCTAGNNNAGATCGGAA |
| 43834_3 | TATGAGTGTGGAGTCGTTGCCTTCAGTGTGAGAGGAGTGCTAGNNNAGATCGGAA |
| 33043_1 | TATGAGTGTGGAGTCGTTGCAGGCTGGAGTGCAATGATATGATNNNAGATCGGAA |
| 33043_2 | TATGAGTGTGGAGTCGTTGCAGGCTGGAGTGCAATGATATGATNNNAGATCGGAA |
| 33043_3 | TATGAGTGTGGAGTCGTTGCAGGCTGGAGTGCAATGATATGATNNNAGATCGGAA |
| 40268_1 | TATGAGTGTGGAGTCGTTGCTCCATCCAACATTGTGCTTTGTTNNNAGATCGGAAGA |
| 40268_2 | TATGAGTGTGGAGTCGTTGCTCCATCCAACATTGTGCTTTGTTNNNAGATCGGAAGA |
| 40268_3 | TATGAGTGTGGAGTCGTTGCTCCATCCAACATTGTGCTTTGTTNNNAGATCGGAAGA |
| 34815_1 | TATGAGTGTGGAGTCGTTGCAGGAGGATGTGTACTGTCTCAGANNNAGATCGGAA |
| 34815_2 | TATGAGTGTGGAGTCGTTGCAGGAGGATGTGTACTGTCTCAGANNNAGATCGGAA |
| 34815_3 | TATGAGTGTGGAGTCGTTGCAGGAGGATGTGTACTGTCTCAGANNNAGATCGGAA |
| 40341_1 | TATGAGTGTGGAGTCGTTGCTGAGATGTTGGATATGGTGGCTCNNNAGATCGGAA |
| 40341_2 | TATGAGTGTGGAGTCGTTGCTGAGATGTTGGATATGGTGGCTCNNNAGATCGGAA |
| 40341_3 | TATGAGTGTGGAGTCGTTGCTGAGATGTTGGATATGGTGGCTCNNNAGATCGGAA |

| | |
|---|---|
| 49415_1 | TATGAGTGTGGAGTCGTTGCGGTGAGTACACAGGTGTTCAGTTNNNAGATCGGAA |
| 49415_2 | TATGAGTGTGGAGTCGTTGCGGTGAGTACACAGGTGTTCAGTTNNNAGATCGGAA |
| 49415_3 | TATGAGTGTGGAGTCGTTGCGGTGAGTACACAGGTGTTCAGTTNNNAGATCGGAA |
| 45896_1 | TATGAGTGTGGAGTCGTTGCTTCAAGTGATTCTCCTGCCTCAGNNNAGATCGGAAG |
| 45896_2 | TATGAGTGTGGAGTCGTTGCTTCAAGTGATTCTCCTGCCTCAGNNNAGATCGGAAG |
| 45896_3 | TATGAGTGTGGAGTCGTTGCTTCAAGTGATTCTCCTGCCTCAGNNNAGATCGGAAG |
| 51967_1 | TATGAGTGTGGAGTCGTTGCTAGTTCCTGAACAATCCCCATGGNNNAGATCGGAAG |
| 51967_2 | TATGAGTGTGGAGTCGTTGCTAGTTCCTGAACAATCCCCATGGNNNAGATCGGAAG |
| 51967_3 | TATGAGTGTGGAGTCGTTGCTAGTTCCTGAACAATCCCCATGGNNNAGATCGGAAG |
| 34670_1 | TATGAGTGTGGAGTCGTTGCGGAAGAAGAGGAAGAGTTGGAGGNNNAGATCGG |
| 34670_2 | TATGAGTGTGGAGTCGTTGCGGAAGAAGAGGAAGAGTTGGAGGNNNAGATCGG |
| 34670_3 | TATGAGTGTGGAGTCGTTGCGGAAGAAGAGGAAGAGTTGGAGGNNNAGATCGG |
| 39444_1 | TATGAGTGTGGAGTCGTTGCGTCAAACCATCACCTGACATTCCNNNAGATCGGAAG |
| 39444_2 | TATGAGTGTGGAGTCGTTGCGTCAAACCATCACCTGACATTCCNNNAGATCGGAAG |
| 39444_3 | TATGAGTGTGGAGTCGTTGCGTCAAACCATCACCTGACATTCCNNNAGATCGGAAG |
| 36294_1 | TATGAGTGTGGAGTCGTTGCATATGTGTGTGTGTGTGTGTGTGNNNAGATCGGAA |
| 36294_2 | TATGAGTGTGGAGTCGTTGCATATGTGTGTGTGTGTGTGTGTGNNNAGATCGGAA |
| 36294_3 | TATGAGTGTGGAGTCGTTGCATATGTGTGTGTGTGTGTGTGTGNNNAGATCGGAA |
| 41524_1 | TATGAGTGTGGAGTCGTTGCAATTCCAGCTACTTGGGAGGATGNNNAGATCGGAA |
| 41524_2 | TATGAGTGTGGAGTCGTTGCAATTCCAGCTACTTGGGAGGATGNNNAGATCGGAA |
| 41524_3 | TATGAGTGTGGAGTCGTTGCAATTCCAGCTACTTGGGAGGATGNNNAGATCGGAA |
| 48453_1 | TATGAGTGTGGAGTCGTTGCCTCCTGAGTAGCTGGGATTACAGNNNAGATCGGAA |
| 48453_2 | TATGAGTGTGGAGTCGTTGCCTCCTGAGTAGCTGGGATTACAGNNNAGATCGGAA |
| 48453_3 | TATGAGTGTGGAGTCGTTGCCTCCTGAGTAGCTGGGATTACAGNNNAGATCGGAA |
| 29022_1 | TATGAGTGTGGAGTCGTTGCTCCCCTCAATATATGCTTCAGGTNNNAGATCGGAAG |
| 29022_2 | TATGAGTGTGGAGTCGTTGCTCCCCTCAATATATGCTTCAGGTNNNAGATCGGAAG |
| 29022_3 | TATGAGTGTGGAGTCGTTGCTCCCCTCAATATATGCTTCAGGTNNNAGATCGGAAG |
| 51644_1 | TATGAGTGTGGAGTCGTTGCAACATGTGGGAATTATGGGAGCTNNNAGATCGGAA |
| 51644_2 | TATGAGTGTGGAGTCGTTGCAACATGTGGGAATTATGGGAGCTNNNAGATCGGAA |
| 51644_3 | TATGAGTGTGGAGTCGTTGCAACATGTGGGAATTATGGGAGCTNNNAGATCGGAA |
| 38776_1 | TATGAGTGTGGAGTCGTTGCCAAAGCAGGGTCATGGAATCATCNNNAGATCGGAA |
| 38776_2 | TATGAGTGTGGAGTCGTTGCCAAAGCAGGGTCATGGAATCATCNNNAGATCGGAA |
| 38776_3 | TATGAGTGTGGAGTCGTTGCCAAAGCAGGGTCATGGAATCATCNNNAGATCGGAA |
| 35247_1 | TATGAGTGTGGAGTCGTTGCTGTAGCTGGGACTATAGGTGCTANNNAGATCGGAA |
| 35247_2 | TATGAGTGTGGAGTCGTTGCTGTAGCTGGGACTATAGGTGCTANNNAGATCGGAA |
| 35247_3 | TATGAGTGTGGAGTCGTTGCTGTAGCTGGGACTATAGGTGCTANNNAGATCGGAA |
| 37091_1 | TATGAGTGTGGAGTCGTTGCCTACTGCACTCCAGCCTATTTGANNNAGATCGGAAG |
| 37091_2 | TATGAGTGTGGAGTCGTTGCCTACTGCACTCCAGCCTATTTGANNNAGATCGGAAG |
| 37091_3 | TATGAGTGTGGAGTCGTTGCCTACTGCACTCCAGCCTATTTGANNNAGATCGGAAG |
| 29786_1 | TATGAGTGTGGAGTCGTTGCCTCAGAGAAGGCACAGACAGTGNNNAGATCGGAAG |
| 29786_2 | TATGAGTGTGGAGTCGTTGCCTCAGAGAAGGCACAGACAGTGNNNAGATCGGAAG |
| 29786_3 | TATGAGTGTGGAGTCGTTGCCTCAGAGAAGGCACAGACAGTGNNNAGATCGGAAG |
| 45544_1 | TATGAGTGTGGAGTCGTTGCGAATGGGGACCTAGAGAGCAAAANNNAGATCGGA |

| | |
|---|---|
| 45544_2 | TATGAGTGTGGAGTCGTTGCGAATGGGGACCTAGAGAGCAAAANNNAGATCGGAA |
| 45544_3 | TATGAGTGTGGAGTCGTTGCGAATGGGGACCTAGAGAGCAAAANNNAGATCGGAA |
| 42338_1 | TATGAGTGTGGAGTCGTTGCAAAAGTTTTGCCACATGTGAGCTNNNAGATCGGAAG |
| 42338_2 | TATGAGTGTGGAGTCGTTGCAAAAGTTTTGCCACATGTGAGCTNNNAGATCGGAAG |
| 42338_3 | TATGAGTGTGGAGTCGTTGCAAAAGTTTTGCCACATGTGAGCTNNNAGATCGGAAG |
| 34830_1 | TATGAGTGTGGAGTCGTTGCCCATGCCCGGCTAATTTTTGTATNNNAGATCGGAAGA |
| 34830_2 | TATGAGTGTGGAGTCGTTGCCCATGCCCGGCTAATTTTTGTATNNNAGATCGGAAGA |
| 34830_3 | TATGAGTGTGGAGTCGTTGCCCATGCCCGGCTAATTTTTGTATNNNAGATCGGAAGA |
| 12801_1 | TATGAGTGTGGAGTCGTTGCATGACTAGAAGGTCAGATGTNNNAGATCGGAAGAG |
| 12801_2 | TATGAGTGTGGAGTCGTTGCATGACTAGAAGGTCAGATGTNNNAGATCGGAAGAG |
| 12801_3 | TATGAGTGTGGAGTCGTTGCATGACTAGAAGGTCAGATGTNNNAGATCGGAAGAG |
| 5_1 | TATGAGTGTGGAGTCGTTGCGGTTTTCACAGTACCAGAGACATNNNAGATCGGAAG |
| 5_2 | TATGAGTGTGGAGTCGTTGCGGTTTTCACAGTACCAGAGACATNNNAGATCGGAAG |
| 5_3 | TATGAGTGTGGAGTCGTTGCGGTTTTCACAGTACCAGAGACATNNNAGATCGGAAG |
| 41054_1 | TATGAGTGTGGAGTCGTTGCAAGAGTTCCATCTCCATGCAAGTNNNAGATCGGAAG |
| 41054_2 | TATGAGTGTGGAGTCGTTGCAAGAGTTCCATCTCCATGCAAGTNNNAGATCGGAAG |
| 41054_3 | TATGAGTGTGGAGTCGTTGCAAGAGTTCCATCTCCATGCAAGTNNNAGATCGGAAG |
| 35620_1 | TATGAGTGTGGAGTCGTTGCTTCACTGCACCTGGCCTATAAATNNNAGATCGGAAGA |
| 35620_2 | TATGAGTGTGGAGTCGTTGCTTCACTGCACCTGGCCTATAAATNNNAGATCGGAAGA |
| 35620_3 | TATGAGTGTGGAGTCGTTGCTTCACTGCACCTGGCCTATAAATNNNAGATCGGAAGA |
| 41174_1 | TATGAGTGTGGAGTCGTTGCGATACTCCTCACCTTCTGATGGGNNNAGATCGGAAG |
| 41174_2 | TATGAGTGTGGAGTCGTTGCGATACTCCTCACCTTCTGATGGGNNNAGATCGGAAG |
| 41174_3 | TATGAGTGTGGAGTCGTTGCGATACTCCTCACCTTCTGATGGGNNNAGATCGGAAG |
| 47447_1 | TATGAGTGTGGAGTCGTTGCAATGGTGGTGACAGGATTAGCAANNNAGATCGGAA |
| 47447_2 | TATGAGTGTGGAGTCGTTGCAATGGTGGTGACAGGATTAGCAANNNAGATCGGAA |
| 47447_3 | TATGAGTGTGGAGTCGTTGCAATGGTGGTGACAGGATTAGCAANNNAGATCGGAA |
| 7809_1 | TATGAGTGTGGAGTCGTTGCACTTCCTCTGGACCATGTTCATTNNNAGATCGGAAGA |
| 7809_2 | TATGAGTGTGGAGTCGTTGCACTTCCTCTGGACCATGTTCATTNNNAGATCGGAAGA |
| 7809_3 | TATGAGTGTGGAGTCGTTGCACTTCCTCTGGACCATGTTCATTNNNAGATCGGAAGA |
| 51118_1 | TATGAGTGTGGAGTCGTTGCCCTAGCCAACCTAGAAAAACCCTNNNAGATCGGAAGA |
| 51118_2 | TATGAGTGTGGAGTCGTTGCCCTAGCCAACCTAGAAAAACCCTNNNAGATCGGAAGA |
| 51118_3 | TATGAGTGTGGAGTCGTTGCCCTAGCCAACCTAGAAAAACCCTNNNAGATCGGAAGA |
| 42334_1 | TATGAGTGTGGAGTCGTTGCGAGAATCTCTGGAACCTGGGATGNNNAGATCGGAA |
| 42334_2 | TATGAGTGTGGAGTCGTTGCGAGAATCTCTGGAACCTGGGATGNNNAGATCGGAA |
| 42334_3 | TATGAGTGTGGAGTCGTTGCGAGAATCTCTGGAACCTGGGATGNNNAGATCGGAA |
| 37525_1 | TATGAGTGTGGAGTCGTTGCAAAGAAAGAAAAGGCCAGGTGTGNNNAGATCGGAA |
| 37525_2 | TATGAGTGTGGAGTCGTTGCAAAGAAAGAAAAGGCCAGGTGTGNNNAGATCGGAA |
| 37525_3 | TATGAGTGTGGAGTCGTTGCAAAGAAAGAAAAGGCCAGGTGTGNNNAGATCGGAA |
| 12680_1 | TATGAGTGTGGAGTCGTTGCCATCCCTGGAGAAAGTAAAGNNNAGATCGGAAGAG |
| 12680_2 | TATGAGTGTGGAGTCGTTGCCATCCCTGGAGAAAGTAAAGNNNAGATCGGAAGAG |
| 12680_3 | TATGAGTGTGGAGTCGTTGCCATCCCTGGAGAAAGTAAAGNNNAGATCGGAAGAG |
| 31549_1 | TATGAGTGTGGAGTCGTTGCGTGCCATGTTTCTTGTACAGCATNNNAGATCGGAAG |
| 31549_2 | TATGAGTGTGGAGTCGTTGCGTGCCATGTTTCTTGTACAGCATNNNAGATCGGAAG |

| | |
|---|---|
| 31549_3 | TATGAGTGTGGAGTCGTTGCGTGCCATGTTTCTTGTACAGCATNNNAGATCGGAAG |
| 42651_1 | TATGAGTGTGGAGTCGTTGCCCTTTGCAGAAACATGGATGGAGNNNAGATCGGAA |
| 42651_2 | TATGAGTGTGGAGTCGTTGCCCTTTGCAGAAACATGGATGGAGNNNAGATCGGAA |
| 42651_3 | TATGAGTGTGGAGTCGTTGCCCTTTGCAGAAACATGGATGGAGNNNAGATCGGAA |
| 47580_1 | TATGAGTGTGGAGTCGTTGCTTCAAGTGATTCTCCTGCCTCAGNNNAGATCGGAAG |
| 47580_2 | TATGAGTGTGGAGTCGTTGCTTCAAGTGATTCTCCTGCCTCAGNNNAGATCGGAAG |
| 47580_3 | TATGAGTGTGGAGTCGTTGCTTCAAGTGATTCTCCTGCCTCAGNNNAGATCGGAAG |
| 41629_1 | TATGAGTGTGGAGTCGTTGCCAGGTTCAAGCAATTCTCCTGCNNNAGATCGGAAGA |
| 41629_2 | TATGAGTGTGGAGTCGTTGCCAGGTTCAAGCAATTCTCCTGCNNNAGATCGGAAGA |
| 41629_3 | TATGAGTGTGGAGTCGTTGCCAGGTTCAAGCAATTCTCCTGCNNNAGATCGGAAGA |
| 36528_1 | TATGAGTGTGGAGTCGTTGCGCCACACAGAAGTGCTTGTTATTNNNAGATCGGAAG |
| 36528_2 | TATGAGTGTGGAGTCGTTGCGCCACACAGAAGTGCTTGTTATTNNNAGATCGGAAG |
| 36528_3 | TATGAGTGTGGAGTCGTTGCGCCACACAGAAGTGCTTGTTATTNNNAGATCGGAAG |
| 10630_1 | TATGAGTGTGGAGTCGTTGCGAGGCTACTTGCTGCTTCTGAATNNNAGATCGGAAG |
| 10630_2 | TATGAGTGTGGAGTCGTTGCGAGGCTACTTGCTGCTTCTGAATNNNAGATCGGAAG |
| 10630_3 | TATGAGTGTGGAGTCGTTGCGAGGCTACTTGCTGCTTCTGAATNNNAGATCGGAAG |
| 35288_1 | TATGAGTGTGGAGTCGTTGCACTGCAAATTCTGTCTCCTGGATNNNAGATCGGAAG |
| 35288_2 | TATGAGTGTGGAGTCGTTGCACTGCAAATTCTGTCTCCTGGATNNNAGATCGGAAG |
| 35288_3 | TATGAGTGTGGAGTCGTTGCACTGCAAATTCTGTCTCCTGGATNNNAGATCGGAAG |
| 46118_1 | TATGAGTGTGGAGTCGTTGCTTTTAGTAGAGACGGGGTTTCGCNNNAGATCGGAA |
| 46118_2 | TATGAGTGTGGAGTCGTTGCTTTTAGTAGAGACGGGGTTTCGCNNNAGATCGGAA |
| 46118_3 | TATGAGTGTGGAGTCGTTGCTTTTAGTAGAGACGGGGTTTCGCNNNAGATCGGAA |
| 33501_1 | TATGAGTGTGGAGTCGTTGCTGAGCATCCAGATGTCTCTTGTTNNNAGATCGGAAG |
| 33501_2 | TATGAGTGTGGAGTCGTTGCTGAGCATCCAGATGTCTCTTGTTNNNAGATCGGAAG |
| 33501_3 | TATGAGTGTGGAGTCGTTGCTGAGCATCCAGATGTCTCTTGTTNNNAGATCGGAAG |
| 30670_1 | TATGAGTGTGGAGTCGTTGCACCTCTCTTTCCCTCCTTCCTGNNNAGATCGGAAGAG |
| 30670_2 | TATGAGTGTGGAGTCGTTGCACCTCTCTTTCCCTCCTTCCTGNNNAGATCGGAAGAG |
| 30670_3 | TATGAGTGTGGAGTCGTTGCACCTCTCTTTCCCTCCTTCCTGNNNAGATCGGAAGAG |
| 51543_1 | TATGAGTGTGGAGTCGTTGCAGGAGGAAATTTGTAGCCTCTGGNNNAGATCGGAA |
| 51543_2 | TATGAGTGTGGAGTCGTTGCAGGAGGAAATTTGTAGCCTCTGGNNNAGATCGGAA |
| 51543_3 | TATGAGTGTGGAGTCGTTGCAGGAGGAAATTTGTAGCCTCTGGNNNAGATCGGAA |
| 31038_1 | TATGAGTGTGGAGTCGTTGCGCTCATCTCTGGTACTTCCTAGCNNNAGATCGGAAG |
| 31038_2 | TATGAGTGTGGAGTCGTTGCGCTCATCTCTGGTACTTCCTAGCNNNAGATCGGAAG |
| 31038_3 | TATGAGTGTGGAGTCGTTGCGCTCATCTCTGGTACTTCCTAGCNNNAGATCGGAAG |
| 34218_1 | TATGAGTGTGGAGTCGTTGCACCTGTAATCCCAGCACTTTAGGNNNAGATCGGAAG |
| 34218_2 | TATGAGTGTGGAGTCGTTGCACCTGTAATCCCAGCACTTTAGGNNNAGATCGGAAG |
| 34218_3 | TATGAGTGTGGAGTCGTTGCACCTGTAATCCCAGCACTTTAGGNNNAGATCGGAAG |
| 29293_1 | TATGAGTGTGGAGTCGTTGCTTGGGTCCCATTAGGCAAAATTGNNNAGATCGGAAG |
| 29293_2 | TATGAGTGTGGAGTCGTTGCTTGGGTCCCATTAGGCAAAATTGNNNAGATCGGAAG |
| 29293_3 | TATGAGTGTGGAGTCGTTGCTTGGGTCCCATTAGGCAAAATTGNNNAGATCGGAAG |
| 33861_1 | TATGAGTGTGGAGTCGTTGCCTGTAATCCCAGCTACTCAGGAGNNNAGATCGGAAG |
| 33861_2 | TATGAGTGTGGAGTCGTTGCCTGTAATCCCAGCTACTCAGGAGNNNAGATCGGAAG |
| 33861_3 | TATGAGTGTGGAGTCGTTGCCTGTAATCCCAGCTACTCAGGAGNNNAGATCGGAAG |

| | |
|---|---|
| 33150_1 | TATGAGTGTGGAGTCGTTGCAGAGAAGGCGTGTCTGTTTCCNNNAGATCGGAAGA |
| 33150_2 | TATGAGTGTGGAGTCGTTGCAGAGAAGGCGTGTCTGTTTCCNNNAGATCGGAAGA |
| 33150_3 | TATGAGTGTGGAGTCGTTGCAGAGAAGGCGTGTCTGTTTCCNNNAGATCGGAAGA |
| 43023_1 | TATGAGTGTGGAGTCGTTGCACTTCAAGTAGGGAAAGGACACCNNNAGATCGGAA |
| 43023_2 | TATGAGTGTGGAGTCGTTGCACTTCAAGTAGGGAAAGGACACCNNNAGATCGGAA |
| 43023_3 | TATGAGTGTGGAGTCGTTGCACTTCAAGTAGGGAAAGGACACCNNNAGATCGGAA |
| 52737_1 | TATGAGTGTGGAGTCGTTGCCTGGAAGAGGAAAGGTGTGGAAGNNNAGATCGGA |
| 52737_2 | TATGAGTGTGGAGTCGTTGCCTGGAAGAGGAAAGGTGTGGAAGNNNAGATCGGA |
| 52737_3 | TATGAGTGTGGAGTCGTTGCCTGGAAGAGGAAAGGTGTGGAAGNNNAGATCGGA |
| 44276_1 | TATGAGTGTGGAGTCGTTGCGGTGGGTGTCTTGGTCTTTATCTNNNAGATCGGAAG |
| 44276_2 | TATGAGTGTGGAGTCGTTGCGGTGGGTGTCTTGGTCTTTATCTNNNAGATCGGAAG |
| 44276_3 | TATGAGTGTGGAGTCGTTGCGGTGGGTGTCTTGGTCTTTATCTNNNAGATCGGAAG |
| 41198_1 | TATGAGTGTGGAGTCGTTGCGTCTCCTCCCCTCTCACCCCNNNAGATCGGAAGAGCA |
| 41198_2 | TATGAGTGTGGAGTCGTTGCGTCTCCTCCCCTCTCACCCCNNNAGATCGGAAGAGCA |
| 41198_3 | TATGAGTGTGGAGTCGTTGCGTCTCCTCCCCTCTCACCCCNNNAGATCGGAAGAGCA |
| 3164_1 | TATGAGTGTGGAGTCGTTGCAACCAGATGCTGTCATAGGCAAGNNNAGATCGGAA |
| 3164_2 | TATGAGTGTGGAGTCGTTGCAACCAGATGCTGTCATAGGCAAGNNNAGATCGGAA |
| 3164_3 | TATGAGTGTGGAGTCGTTGCAACCAGATGCTGTCATAGGCAAGNNNAGATCGGAA |
| 44256_1 | TATGAGTGTGGAGTCGTTGCTGGGTTTGGAATGTGAGACAGATNNNAGATCGGAA |
| 44256_2 | TATGAGTGTGGAGTCGTTGCTGGGTTTGGAATGTGAGACAGATNNNAGATCGGAA |
| 44256_3 | TATGAGTGTGGAGTCGTTGCTGGGTTTGGAATGTGAGACAGATNNNAGATCGGAA |
| 35722_1 | TATGAGTGTGGAGTCGTTGCGCAATTTGCCAGTCTGTGTCTTTNNNAGATCGGAAG |
| 35722_2 | TATGAGTGTGGAGTCGTTGCGCAATTTGCCAGTCTGTGTCTTTNNNAGATCGGAAG |
| 35722_3 | TATGAGTGTGGAGTCGTTGCGCAATTTGCCAGTCTGTGTCTTTNNNAGATCGGAAG |
| 35997_1 | TATGAGTGTGGAGTCGTTGCAAGCGTAATCACCATCAAAACCCNNNAGATCGGAAG |
| 35997_2 | TATGAGTGTGGAGTCGTTGCAAGCGTAATCACCATCAAAACCCNNNAGATCGGAAG |
| 35997_3 | TATGAGTGTGGAGTCGTTGCAAGCGTAATCACCATCAAAACCCNNNAGATCGGAAG |
| 32756_1 | TATGAGTGTGGAGTCGTTGCAAGACCTCCAGAAGTAAGCACTGNNNAGATCGGAA |
| 32756_2 | TATGAGTGTGGAGTCGTTGCAAGACCTCCAGAAGTAAGCACTGNNNAGATCGGAA |
| 32756_3 | TATGAGTGTGGAGTCGTTGCAAGACCTCCAGAAGTAAGCACTGNNNAGATCGGAA |
| 50532_1 | TATGAGTGTGGAGTCGTTGCGAGACCCAGGCTGAAGCTGGNNNAGATCGGAAGA |
| 50532_2 | TATGAGTGTGGAGTCGTTGCGAGACCCAGGCTGAAGCTGGNNNAGATCGGAAGA |
| 50532_3 | TATGAGTGTGGAGTCGTTGCGAGACCCAGGCTGAAGCTGGNNNAGATCGGAAGA |
| 47562_1 | TATGAGTGTGGAGTCGTTGCAAGTTCAAGTGATTCTCCTGCCTNNNAGATCGGAAG |
| 47562_2 | TATGAGTGTGGAGTCGTTGCAAGTTCAAGTGATTCTCCTGCCTNNNAGATCGGAAG |
| 47562_3 | TATGAGTGTGGAGTCGTTGCAAGTTCAAGTGATTCTCCTGCCTNNNAGATCGGAAG |
| 45319_1 | TATGAGTGTGGAGTCGTTGCAGCTGAGGGGAATGCTTTAGAAANNNAGATCGGAA |
| 45319_2 | TATGAGTGTGGAGTCGTTGCAGCTGAGGGGAATGCTTTAGAAANNNAGATCGGAA |
| 45319_3 | TATGAGTGTGGAGTCGTTGCAGCTGAGGGGAATGCTTTAGAAANNNAGATCGGAA |
| 47332_1 | TATGAGTGTGGAGTCGTTGCATGATCTTGACTCACTGCAACCTNNNAGATCGGAAG |
| 47332_2 | TATGAGTGTGGAGTCGTTGCATGATCTTGACTCACTGCAACCTNNNAGATCGGAAG |
| 47332_3 | TATGAGTGTGGAGTCGTTGCATGATCTTGACTCACTGCAACCTNNNAGATCGGAAG |
| 39104_1 | TATGAGTGTGGAGTCGTTGCCCCCACCACATAGACACAAATTGNNNAGATCGGAAG |

| | |
|---|---|
| 39104_2 | TATGAGTGTGGAGTCGTTGCCCCCACCACATAGACACAAATTGNNNAGATCGGAAG |
| 39104_3 | TATGAGTGTGGAGTCGTTGCCCCCACCACATAGACACAAATTGNNNAGATCGGAAG |
| 49000_1 | TATGAGTGTGGAGTCGTTGCTTTAGACATGAAGCGAGCTGTGNNNAGATCGGAA |
| 49000_2 | TATGAGTGTGGAGTCGTTGCTTTAGACATGAAGCGAGCTGTGNNNAGATCGGAA |
| 49000_3 | TATGAGTGTGGAGTCGTTGCTTTAGACATGAAGCGAGCTGTGNNNAGATCGGAA |
| 33630_1 | TATGAGTGTGGAGTCGTTGCGTGAATACCTGGGACCACAGGNNNAGATCGGAAGA |
| 33630_2 | TATGAGTGTGGAGTCGTTGCGTGAATACCTGGGACCACAGGNNNAGATCGGAAGA |
| 33630_3 | TATGAGTGTGGAGTCGTTGCGTGAATACCTGGGACCACAGGNNNAGATCGGAAGA |
| 39526_1 | TATGAGTGTGGAGTCGTTGCGTAATCCCAGCTACTATGGAGGCNNNAGATCGGAA |
| 39526_2 | TATGAGTGTGGAGTCGTTGCGTAATCCCAGCTACTATGGAGGCNNNAGATCGGAA |
| 39526_3 | TATGAGTGTGGAGTCGTTGCGTAATCCCAGCTACTATGGAGGCNNNAGATCGGAA |
| 35663_1 | TATGAGTGTGGAGTCGTTGCTGTTAGCCAGGATGGTCTCAATCNNNAGATCGGAAG |
| 35663_2 | TATGAGTGTGGAGTCGTTGCTGTTAGCCAGGATGGTCTCAATCNNNAGATCGGAAG |
| 35663_3 | TATGAGTGTGGAGTCGTTGCTGTTAGCCAGGATGGTCTCAATCNNNAGATCGGAAG |
| 47822_1 | TATGAGTGTGGAGTCGTTGCCTCCAGGGTTCAAGCGATTCTCNNNAGATCGGAAGA |
| 47822_2 | TATGAGTGTGGAGTCGTTGCCTCCAGGGTTCAAGCGATTCTCNNNAGATCGGAAGA |
| 47822_3 | TATGAGTGTGGAGTCGTTGCCTCCAGGGTTCAAGCGATTCTCNNNAGATCGGAAGA |
| 41841_1 | TATGAGTGTGGAGTCGTTGCCTGGTCTCAAACTTTTGGCATCANNNAGATCGGAAG |
| 41841_2 | TATGAGTGTGGAGTCGTTGCCTGGTCTCAAACTTTTGGCATCANNNAGATCGGAAG |
| 41841_3 | TATGAGTGTGGAGTCGTTGCCTGGTCTCAAACTTTTGGCATCANNNAGATCGGAAG |
| 41811_1 | TATGAGTGTGGAGTCGTTGCCATCATCTCAGCTCACTGCAACNNNAGATCGGAAGA |
| 41811_2 | TATGAGTGTGGAGTCGTTGCCATCATCTCAGCTCACTGCAACNNNAGATCGGAAGA |
| 41811_3 | TATGAGTGTGGAGTCGTTGCCATCATCTCAGCTCACTGCAACNNNAGATCGGAAGA |
| 33635_1 | TATGAGTGTGGAGTCGTTGCCTAGGCTGTAGTGCAGAAGTTCANNNAGATCGGAA |
| 33635_2 | TATGAGTGTGGAGTCGTTGCCTAGGCTGTAGTGCAGAAGTTCANNNAGATCGGAA |
| 33635_3 | TATGAGTGTGGAGTCGTTGCCTAGGCTGTAGTGCAGAAGTTCANNNAGATCGGAA |
| 50889_1 | TATGAGTGTGGAGTCGTTGCCACTGCTATGATTTTGCATGCCTNNNAGATCGGAAG |
| 50889_2 | TATGAGTGTGGAGTCGTTGCCACTGCTATGATTTTGCATGCCTNNNAGATCGGAAG |
| 50889_3 | TATGAGTGTGGAGTCGTTGCCACTGCTATGATTTTGCATGCCTNNNAGATCGGAAG |
| 50252_1 | TATGAGTGTGGAGTCGTTGCACAAGACCCTGAATGCACTAAGTNNNAGATCGGAAG |
| 50252_2 | TATGAGTGTGGAGTCGTTGCACAAGACCCTGAATGCACTAAGTNNNAGATCGGAAG |
| 50252_3 | TATGAGTGTGGAGTCGTTGCACAAGACCCTGAATGCACTAAGTNNNAGATCGGAAG |
| 52398_1 | TATGAGTGTGGAGTCGTTGCAACCTTCTACCTTCTCGTTTCCCNNNAGATCGGAAGA |
| 52398_2 | TATGAGTGTGGAGTCGTTGCAACCTTCTACCTTCTCGTTTCCCNNNAGATCGGAAGA |
| 52398_3 | TATGAGTGTGGAGTCGTTGCAACCTTCTACCTTCTCGTTTCCCNNNAGATCGGAAGA |
| 32075_1 | TATGAGTGTGGAGTCGTTGCCTCGAACTCCTGGGCTCAAGNNNAGATCGGAAGAG |
| 32075_2 | TATGAGTGTGGAGTCGTTGCCTCGAACTCCTGGGCTCAAGNNNAGATCGGAAGAG |
| 32075_3 | TATGAGTGTGGAGTCGTTGCCTCGAACTCCTGGGCTCAAGNNNAGATCGGAAGAG |
| 42641_1 | TATGAGTGTGGAGTCGTTGCTTTGGGCTGGCTCCATATTTTTGNNNAGATCGGAAG |
| 42641_2 | TATGAGTGTGGAGTCGTTGCTTTGGGCTGGCTCCATATTTTTGNNNAGATCGGAAG |
| 42641_3 | TATGAGTGTGGAGTCGTTGCTTTGGGCTGGCTCCATATTTTTGNNNAGATCGGAAG |
| 40840_1 | TATGAGTGTGGAGTCGTTGCCTGTAGTCCCAGCTACTCAGGAGNNNAGATCGGAAG |
| 40840_2 | TATGAGTGTGGAGTCGTTGCCTGTAGTCCCAGCTACTCAGGAGNNNAGATCGGAAG |

| | |
|---|---|
| 40840_3 | TATGAGTGTGGAGTCGTTGCCTGTAGTCCCAGCTACTCAGGAGNNNAGATCGGAAG |
| 39394_1 | TATGAGTGTGGAGTCGTTGCATGGTTTCGTCCTTCCCTCTTTTNNNAGATCGGAAGA |
| 39394_2 | TATGAGTGTGGAGTCGTTGCATGGTTTCGTCCTTCCCTCTTTTNNNAGATCGGAAGA |
| 39394_3 | TATGAGTGTGGAGTCGTTGCATGGTTTCGTCCTTCCCTCTTTTNNNAGATCGGAAGA |
| 49229_1 | TATGAGTGTGGAGTCGTTGCGAGTCCACCTTGGAGCATCTATTNNNAGATCGGAAG |
| 49229_2 | TATGAGTGTGGAGTCGTTGCGAGTCCACCTTGGAGCATCTATTNNNAGATCGGAAG |
| 49229_3 | TATGAGTGTGGAGTCGTTGCGAGTCCACCTTGGAGCATCTATTNNNAGATCGGAAG |
| 38323_1 | TATGAGTGTGGAGTCGTTGCCAGTGAGCTGAGATTGTACCACTNNNAGATCGGAAG |
| 38323_2 | TATGAGTGTGGAGTCGTTGCCAGTGAGCTGAGATTGTACCACTNNNAGATCGGAAG |
| 38323_3 | TATGAGTGTGGAGTCGTTGCCAGTGAGCTGAGATTGTACCACTNNNAGATCGGAAG |
| 51917_1 | TATGAGTGTGGAGTCGTTGCAACATGGTAGCTGTTTATCCCGTNNNAGATCGGAAG |
| 51917_2 | TATGAGTGTGGAGTCGTTGCAACATGGTAGCTGTTTATCCCGTNNNAGATCGGAAG |
| 51917_3 | TATGAGTGTGGAGTCGTTGCAACATGGTAGCTGTTTATCCCGTNNNAGATCGGAAG |
| 41916_1 | TATGAGTGTGGAGTCGTTGCAGACGGAGTCTCGCTCTGTCNNNAGATCGGAAGAG |
| 41916_2 | TATGAGTGTGGAGTCGTTGCAGACGGAGTCTCGCTCTGTCNNNAGATCGGAAGAG |
| 41916_3 | TATGAGTGTGGAGTCGTTGCAGACGGAGTCTCGCTCTGTCNNNAGATCGGAAGAG |
| 41118_1 | TATGAGTGTGGAGTCGTTGCAGAGATGTTAGGTGTGCTGATGGNNNAGATCGGAA |
| 41118_2 | TATGAGTGTGGAGTCGTTGCAGAGATGTTAGGTGTGCTGATGGNNNAGATCGGAA |
| 41118_3 | TATGAGTGTGGAGTCGTTGCAGAGATGTTAGGTGTGCTGATGGNNNAGATCGGAA |
| 37544_1 | TATGAGTGTGGAGTCGTTGCATAGTGCCTTGGGTTGCTCTTTANNNAGATCGGAAG |
| 37544_2 | TATGAGTGTGGAGTCGTTGCATAGTGCCTTGGGTTGCTCTTTANNNAGATCGGAAG |
| 37544_3 | TATGAGTGTGGAGTCGTTGCATAGTGCCTTGGGTTGCTCTTTANNNAGATCGGAAG |
| 36223_1 | TATGAGTGTGGAGTCGTTGCGAATCTGTGAGTCTGGCACCTTANNNAGATCGGAAG |
| 36223_2 | TATGAGTGTGGAGTCGTTGCGAATCTGTGAGTCTGGCACCTTANNNAGATCGGAAG |
| 36223_3 | TATGAGTGTGGAGTCGTTGCGAATCTGTGAGTCTGGCACCTTANNNAGATCGGAAG |
| 33453_1 | TATGAGTGTGGAGTCGTTGCAAAGGATGTCCACTTTCACCACTNNNAGATCGGAAG |
| 33453_2 | TATGAGTGTGGAGTCGTTGCAAAGGATGTCCACTTTCACCACTNNNAGATCGGAAG |
| 33453_3 | TATGAGTGTGGAGTCGTTGCAAAGGATGTCCACTTTCACCACTNNNAGATCGGAAG |
| 44192_1 | TATGAGTGTGGAGTCGTTGCCAGCCACAAACCTCAGTGAATTCNNNAGATCGGAAG |
| 44192_2 | TATGAGTGTGGAGTCGTTGCCAGCCACAAACCTCAGTGAATTCNNNAGATCGGAAG |
| 44192_3 | TATGAGTGTGGAGTCGTTGCCAGCCACAAACCTCAGTGAATTCNNNAGATCGGAAG |
| 28923_1 | TATGAGTGTGGAGTCGTTGCCCATTGCACAAGACTTGTTGCTANNNAGATCGGAAG |
| 28923_2 | TATGAGTGTGGAGTCGTTGCCCATTGCACAAGACTTGTTGCTANNNAGATCGGAAG |
| 28923_3 | TATGAGTGTGGAGTCGTTGCCCATTGCACAAGACTTGTTGCTANNNAGATCGGAAG |
| 38917_1 | TATGAGTGTGGAGTCGTTGCTGGGTCTAGTTGTTTGCCTAGTCNNNAGATCGGAAG |
| 38917_2 | TATGAGTGTGGAGTCGTTGCTGGGTCTAGTTGTTTGCCTAGTCNNNAGATCGGAAG |
| 38917_3 | TATGAGTGTGGAGTCGTTGCTGGGTCTAGTTGTTTGCCTAGTCNNNAGATCGGAAG |
| 35989_1 | TATGAGTGTGGAGTCGTTGCAAAAAAAACAGAGGGTTGGGTGCNNNAGATCGGAA |
| 35989_2 | TATGAGTGTGGAGTCGTTGCAAAAAAAACAGAGGGTTGGGTGCNNNAGATCGGAA |
| 35989_3 | TATGAGTGTGGAGTCGTTGCAAAAAAAACAGAGGGTTGGGTGCNNNAGATCGGAA |
| 50481_1 | TATGAGTGTGGAGTCGTTGCAGCAATCCACTGGGAACTCTTAGNNNAGATCGGAAG |
| 50481_2 | TATGAGTGTGGAGTCGTTGCAGCAATCCACTGGGAACTCTTAGNNNAGATCGGAAG |
| 50481_3 | TATGAGTGTGGAGTCGTTGCAGCAATCCACTGGGAACTCTTAGNNNAGATCGGAAG |

| | |
|---|---|
| 44225_1 | TATGAGTGTGGAGTCGTTGCTTTTACAACATTGGGGGCATTCCNNNAGATCGGAAG |
| 44225_2 | TATGAGTGTGGAGTCGTTGCTTTTACAACATTGGGGGCATTCCNNNAGATCGGAAG |
| 44225_3 | TATGAGTGTGGAGTCGTTGCTTTTACAACATTGGGGGCATTCCNNNAGATCGGAAG |
| 30860_1 | TATGAGTGTGGAGTCGTTGCTCTCTCTTACACACACATCCCACNNNAGATCGGAAGA |
| 30860_2 | TATGAGTGTGGAGTCGTTGCTCTCTCTTACACACACATCCCACNNNAGATCGGAAGA |
| 30860_3 | TATGAGTGTGGAGTCGTTGCTCTCTCTTACACACACATCCCACNNNAGATCGGAAGA |
| 30289_1 | TATGAGTGTGGAGTCGTTGCATTCTTGGAAACAGAGGTGAGGTNNNAGATCGGAA |
| 30289_2 | TATGAGTGTGGAGTCGTTGCATTCTTGGAAACAGAGGTGAGGTNNNAGATCGGAA |
| 30289_3 | TATGAGTGTGGAGTCGTTGCATTCTTGGAAACAGAGGTGAGGTNNNAGATCGGAA |
| 42135_1 | TATGAGTGTGGAGTCGTTGCAGACGGAATTTCACTGTTGTTGCNNNAGATCGGAAG |
| 42135_2 | TATGAGTGTGGAGTCGTTGCAGACGGAATTTCACTGTTGTTGCNNNAGATCGGAAG |
| 42135_3 | TATGAGTGTGGAGTCGTTGCAGACGGAATTTCACTGTTGTTGCNNNAGATCGGAAG |
| 11101_1 | TATGAGTGTGGAGTCGTTGCGAAGTTTACAGTCTGGGCCACTTNNNAGATCGGAAG |
| 11101_2 | TATGAGTGTGGAGTCGTTGCGAAGTTTACAGTCTGGGCCACTTNNNAGATCGGAAG |
| 11101_3 | TATGAGTGTGGAGTCGTTGCGAAGTTTACAGTCTGGGCCACTTNNNAGATCGGAAG |
| 49504_1 | TATGAGTGTGGAGTCGTTGCCCACACCCAGCTAATTTTTGTCCNNNAGATCGGAAGA |
| 49504_2 | TATGAGTGTGGAGTCGTTGCCCACACCCAGCTAATTTTTGTCCNNNAGATCGGAAGA |
| 49504_3 | TATGAGTGTGGAGTCGTTGCCCACACCCAGCTAATTTTTGTCCNNNAGATCGGAAGA |
| 49986_1 | TATGAGTGTGGAGTCGTTGCTCAGGTTTTAGTGGGTGATCCTGNNNAGATCGGAA |
| 49986_2 | TATGAGTGTGGAGTCGTTGCTCAGGTTTTAGTGGGTGATCCTGNNNAGATCGGAA |
| 49986_3 | TATGAGTGTGGAGTCGTTGCTCAGGTTTTAGTGGGTGATCCTGNNNAGATCGGAA |
| 41791_1 | TATGAGTGTGGAGTCGTTGCATCTTGGCTCACTGTTACCTCTGNNNAGATCGGAAG |
| 41791_2 | TATGAGTGTGGAGTCGTTGCATCTTGGCTCACTGTTACCTCTGNNNAGATCGGAAG |
| 41791_3 | TATGAGTGTGGAGTCGTTGCATCTTGGCTCACTGTTACCTCTGNNNAGATCGGAAG |
| 12823_1 | TATGAGTGTGGAGTCGTTGCACATTAATGTCATCACTGCANNNAGATCGGAAGAGC |
| 12823_2 | TATGAGTGTGGAGTCGTTGCACATTAATGTCATCACTGCANNNAGATCGGAAGAGC |
| 12823_3 | TATGAGTGTGGAGTCGTTGCACATTAATGTCATCACTGCANNNAGATCGGAAGAGC |
| 2221_1 | TATGAGTGTGGAGTCGTTGCTCTTGCTTGTTCCTACCTCCTGANNNAGATCGGAAGA |
| 2221_2 | TATGAGTGTGGAGTCGTTGCTCTTGCTTGTTCCTACCTCCTGANNNAGATCGGAAGA |
| 2221_3 | TATGAGTGTGGAGTCGTTGCTCTTGCTTGTTCCTACCTCCTGANNNAGATCGGAAGA |
| 47719_1 | TATGAGTGTGGAGTCGTTGCACAGAGTCTCATTTTGTCACCCANNNAGATCGGAAG |
| 47719_2 | TATGAGTGTGGAGTCGTTGCACAGAGTCTCATTTTGTCACCCANNNAGATCGGAAG |
| 47719_3 | TATGAGTGTGGAGTCGTTGCACAGAGTCTCATTTTGTCACCCANNNAGATCGGAAG |
| 52264_1 | TATGAGTGTGGAGTCGTTGCTGTCATCGCTTTCCCCCTTCNNNAGATCGGAAGAGCA |
| 52264_2 | TATGAGTGTGGAGTCGTTGCTGTCATCGCTTTCCCCCTTCNNNAGATCGGAAGAGCA |
| 52264_3 | TATGAGTGTGGAGTCGTTGCTGTCATCGCTTTCCCCCTTCNNNAGATCGGAAGAGCA |
| 2598_1 | TATGAGTGTGGAGTCGTTGCATCTATCTGCACCAGCAGTTTGGNNNAGATCGGAAG |
| 2598_2 | TATGAGTGTGGAGTCGTTGCATCTATCTGCACCAGCAGTTTGGNNNAGATCGGAAG |
| 2598_3 | TATGAGTGTGGAGTCGTTGCATCTATCTGCACCAGCAGTTTGGNNNAGATCGGAAG |
| 35265_1 | TATGAGTGTGGAGTCGTTGCGCGCTCTTCTTGACATTTCTTCANNNAGATCGGAAGA |
| 35265_2 | TATGAGTGTGGAGTCGTTGCGCGCTCTTCTTGACATTTCTTCANNNAGATCGGAAGA |
| 35265_3 | TATGAGTGTGGAGTCGTTGCGCGCTCTTCTTGACATTTCTTCANNNAGATCGGAAGA |
| 47550_1 | TATGAGTGTGGAGTCGTTGCCCTTCTAGGGCTTTAGCTCTGACNNNAGATCGGAAG |

| | |
|---|---|
| 47550_2 | TATGAGTGTGGAGTCGTTGCCCTTCTAGGGCTTTAGCTCTGACNNNAGATCGGAAG |
| 47550_3 | TATGAGTGTGGAGTCGTTGCCCTTCTAGGGCTTTAGCTCTGACNNNAGATCGGAAG |
| 43463_1 | TATGAGTGTGGAGTCGTTGCTGGCTATTCTCGATCCCTTGAAGNNNAGATCGGAAG |
| 43463_2 | TATGAGTGTGGAGTCGTTGCTGGCTATTCTCGATCCCTTGAAGNNNAGATCGGAAG |
| 43463_3 | TATGAGTGTGGAGTCGTTGCTGGCTATTCTCGATCCCTTGAAGNNNAGATCGGAAG |
| 46631_1 | TATGAGTGTGGAGTCGTTGCCACAATCTTAGCTCACTGCAACCNNNAGATCGGAAG |
| 46631_2 | TATGAGTGTGGAGTCGTTGCCACAATCTTAGCTCACTGCAACCNNNAGATCGGAAG |
| 46631_3 | TATGAGTGTGGAGTCGTTGCCACAATCTTAGCTCACTGCAACCNNNAGATCGGAAG |
| 46238_1 | TATGAGTGTGGAGTCGTTGCGATCACAAGAACCTGCCAATGTGNNNAGATCGGAA |
| 46238_2 | TATGAGTGTGGAGTCGTTGCGATCACAAGAACCTGCCAATGTGNNNAGATCGGAA |
| 46238_3 | TATGAGTGTGGAGTCGTTGCGATCACAAGAACCTGCCAATGTGNNNAGATCGGAA |
| 37657_1 | TATGAGTGTGGAGTCGTTGCTCATCTGAGGTTGGGAGTTCAAGNNNAGATCGGAA |
| 37657_2 | TATGAGTGTGGAGTCGTTGCTCATCTGAGGTTGGGAGTTCAAGNNNAGATCGGAA |
| 37657_3 | TATGAGTGTGGAGTCGTTGCTCATCTGAGGTTGGGAGTTCAAGNNNAGATCGGAA |
| 43271_1 | TATGAGTGTGGAGTCGTTGCACCCTTGATGGTCCTGCTTTTATNNNAGATCGGAAG |
| 43271_2 | TATGAGTGTGGAGTCGTTGCACCCTTGATGGTCCTGCTTTTATNNNAGATCGGAAG |
| 43271_3 | TATGAGTGTGGAGTCGTTGCACCCTTGATGGTCCTGCTTTTATNNNAGATCGGAAG |
| 49164_1 | TATGAGTGTGGAGTCGTTGCCACCACACCCAGCTAATTTTTGTNNNAGATCGGAAG |
| 49164_2 | TATGAGTGTGGAGTCGTTGCCACCACACCCAGCTAATTTTTGTNNNAGATCGGAAG |
| 49164_3 | TATGAGTGTGGAGTCGTTGCCACCACACCCAGCTAATTTTTGTNNNAGATCGGAAG |
| 52394_1 | TATGAGTGTGGAGTCGTTGCGGTTTCATTCCATTCTTCACCCCNNNAGATCGGAAGA |
| 52394_2 | TATGAGTGTGGAGTCGTTGCGGTTTCATTCCATTCTTCACCCCNNNAGATCGGAAGA |
| 52394_3 | TATGAGTGTGGAGTCGTTGCGGTTTCATTCCATTCTTCACCCCNNNAGATCGGAAGA |
| 46134_1 | TATGAGTGTGGAGTCGTTGCCCTCCCAGGTTCAAGGAATTCTCNNNAGATCGGAAG |
| 46134_2 | TATGAGTGTGGAGTCGTTGCCCTCCCAGGTTCAAGGAATTCTCNNNAGATCGGAAG |
| 46134_3 | TATGAGTGTGGAGTCGTTGCCCTCCCAGGTTCAAGGAATTCTCNNNAGATCGGAAG |
| 38155_1 | TATGAGTGTGGAGTCGTTGCATAATCCCAGCTACTCAGGAGGCNNNAGATCGGAAG |
| 38155_2 | TATGAGTGTGGAGTCGTTGCATAATCCCAGCTACTCAGGAGGCNNNAGATCGGAAG |
| 38155_3 | TATGAGTGTGGAGTCGTTGCATAATCCCAGCTACTCAGGAGGCNNNAGATCGGAAG |
| 48361_1 | TATGAGTGTGGAGTCGTTGCTAGAAGTGCAGTTTCGTTACCCTNNNAGATCGGAAG |
| 48361_2 | TATGAGTGTGGAGTCGTTGCTAGAAGTGCAGTTTCGTTACCCTNNNAGATCGGAAG |
| 48361_3 | TATGAGTGTGGAGTCGTTGCTAGAAGTGCAGTTTCGTTACCCTNNNAGATCGGAAG |
| 13020_1 | TATGAGTGTGGAGTCGTTGCTAAGCAAATGTCATATTACCTTGNNNAGATCGGAAG |
| 13020_2 | TATGAGTGTGGAGTCGTTGCTAAGCAAATGTCATATTACCTTGNNNAGATCGGAAG |
| 13020_3 | TATGAGTGTGGAGTCGTTGCTAAGCAAATGTCATATTACCTTGNNNAGATCGGAAG |
| 33091_1 | TATGAGTGTGGAGTCGTTGCCTACTGAACTTATGATCCGCCCANNNAGATCGGAAG |
| 33091_2 | TATGAGTGTGGAGTCGTTGCCTACTGAACTTATGATCCGCCCANNNAGATCGGAAG |
| 33091_3 | TATGAGTGTGGAGTCGTTGCCTACTGAACTTATGATCCGCCCANNNAGATCGGAAG |
| 28792_1 | TATGAGTGTGGAGTCGTTGCACAACCTGATTATGCAGTTCCTTGNNNAGATCGGAA |
| 28792_2 | TATGAGTGTGGAGTCGTTGCACAACCTGATTATGCAGTTCCTTGNNNAGATCGGAA |
| 28792_3 | TATGAGTGTGGAGTCGTTGCACAACCTGATTATGCAGTTCCTTGNNNAGATCGGAA |
| 47651_1 | TATGAGTGTGGAGTCGTTGCGATGGGCATCTGGAGTTCTAGAGNNNAGATCGGAA |
| 47651_2 | TATGAGTGTGGAGTCGTTGCGATGGGCATCTGGAGTTCTAGAGNNNAGATCGGAA |

| | |
|---|---|
| 47651_3 | TATGAGTGTGGAGTCGTTGCGATGGGCATCTGGAGTTCTAGAGNNNAGATCGGAA |
| 30435_1 | TATGAGTGTGGAGTCGTTGCTGCCTGATCTCAATCCTTCTGGANNNAGATCGGAAG |
| 30435_2 | TATGAGTGTGGAGTCGTTGCTGCCTGATCTCAATCCTTCTGGANNNAGATCGGAAG |
| 30435_3 | TATGAGTGTGGAGTCGTTGCTGCCTGATCTCAATCCTTCTGGANNNAGATCGGAAG |
| 30464_1 | TATGAGTGTGGAGTCGTTGCTTTCCTCAGACAGACCATACTTGANNNAGATCGGAA |
| 30464_2 | TATGAGTGTGGAGTCGTTGCTTTCCTCAGACAGACCATACTTGANNNAGATCGGAA |
| 30464_3 | TATGAGTGTGGAGTCGTTGCTTTCCTCAGACAGACCATACTTGANNNAGATCGGAA |
| 611_1 | TATGAGTGTGGAGTCGTTGCCAGCCCAGAAAAACTAGAGTGCTNNNAGATCGGAA |
| 611_2 | TATGAGTGTGGAGTCGTTGCCAGCCCAGAAAAACTAGAGTGCTNNNAGATCGGAA |
| 611_3 | TATGAGTGTGGAGTCGTTGCCAGCCCAGAAAAACTAGAGTGCTNNNAGATCGGAA |
| 50276_1 | TATGAGTGTGGAGTCGTTGCTAGCTCTCTACCGAAGGGAACTTNNNAGATCGGAAG |
| 50276_2 | TATGAGTGTGGAGTCGTTGCTAGCTCTCTACCGAAGGGAACTTNNNAGATCGGAAG |
| 50276_3 | TATGAGTGTGGAGTCGTTGCTAGCTCTCTACCGAAGGGAACTTNNNAGATCGGAAG |
| 50296_1 | TATGAGTGTGGAGTCGTTGCAGTTTGAAAGCTAAAGTGTGGGCNNNAGATCGGAA |
| 50296_2 | TATGAGTGTGGAGTCGTTGCAGTTTGAAAGCTAAAGTGTGGGCNNNAGATCGGAA |
| 50296_3 | TATGAGTGTGGAGTCGTTGCAGTTTGAAAGCTAAAGTGTGGGCNNNAGATCGGAA |
| 43810_1 | TATGAGTGTGGAGTCGTTGCAAAAAACGGGCATACCACGAATTNNNAGATCGGAA |
| 43810_2 | TATGAGTGTGGAGTCGTTGCAAAAAACGGGCATACCACGAATTNNNAGATCGGAA |
| 43810_3 | TATGAGTGTGGAGTCGTTGCAAAAAACGGGCATACCACGAATTNNNAGATCGGAA |
| 36257_1 | TATGAGTGTGGAGTCGTTGCCGCCTGTAGTCCCAGTTACTTGNNNAGATCGGAAGA |
| 36257_2 | TATGAGTGTGGAGTCGTTGCCGCCTGTAGTCCCAGTTACTTGNNNAGATCGGAAGA |
| 36257_3 | TATGAGTGTGGAGTCGTTGCCGCCTGTAGTCCCAGTTACTTGNNNAGATCGGAAGA |
| 10818_1 | TATGAGTGTGGAGTCGTTGCGAATGTGTTTTGCCAAGGCTGACNNNAGATCGGAA |
| 10818_2 | TATGAGTGTGGAGTCGTTGCGAATGTGTTTTGCCAAGGCTGACNNNAGATCGGAA |
| 10818_3 | TATGAGTGTGGAGTCGTTGCGAATGTGTTTTGCCAAGGCTGACNNNAGATCGGAA |
| 39184_1 | TATGAGTGTGGAGTCGTTGCGGACATAATGTCGAACCGGACTANNNAGATCGGAA |
| 39184_2 | TATGAGTGTGGAGTCGTTGCGGACATAATGTCGAACCGGACTANNNAGATCGGAA |
| 39184_3 | TATGAGTGTGGAGTCGTTGCGGACATAATGTCGAACCGGACTANNNAGATCGGAA |
| 40972_1 | TATGAGTGTGGAGTCGTTGCAGTGGAAAGATGGCTGATAAGCANNNAGATCGGA |
| 40972_2 | TATGAGTGTGGAGTCGTTGCAGTGGAAAGATGGCTGATAAGCANNNAGATCGGA |
| 40972_3 | TATGAGTGTGGAGTCGTTGCAGTGGAAAGATGGCTGATAAGCANNNAGATCGGA |
| 52254_1 | TATGAGTGTGGAGTCGTTGCACCCCTCTCTCACATAGAAGTGANNNAGATCGGAAG |
| 52254_2 | TATGAGTGTGGAGTCGTTGCACCCCTCTCTCACATAGAAGTGANNNAGATCGGAAG |
| 52254_3 | TATGAGTGTGGAGTCGTTGCACCCCTCTCTCACATAGAAGTGANNNAGATCGGAAG |
| 13711_1 | TATGAGTGTGGAGTCGTTGCTGGAGAAAAATGAAGGAAAAAATNNNAGATCGGA |
| 13711_2 | TATGAGTGTGGAGTCGTTGCTGGAGAAAAATGAAGGAAAAAATNNNAGATCGGA |
| 13711_3 | TATGAGTGTGGAGTCGTTGCTGGAGAAAAATGAAGGAAAAAATNNNAGATCGGA |
| 43338_1 | TATGAGTGTGGAGTCGTTGCCATGCTGGCCTACTCTCCATATCNNNAGATCGGAAGA |
| 43338_2 | TATGAGTGTGGAGTCGTTGCCATGCTGGCCTACTCTCCATATCNNNAGATCGGAAGA |
| 43338_3 | TATGAGTGTGGAGTCGTTGCCATGCTGGCCTACTCTCCATATCNNNAGATCGGAAGA |
| 40351_1 | TATGAGTGTGGAGTCGTTGCCGAGGCTCTGTCTCAAAAAACAANNNAGATCGGAAG |
| 40351_2 | TATGAGTGTGGAGTCGTTGCCGAGGCTCTGTCTCAAAAAACAANNNAGATCGGAAG |
| 40351_3 | TATGAGTGTGGAGTCGTTGCCGAGGCTCTGTCTCAAAAAACAANNNAGATCGGAAG |

| | |
|---|---|
| 42739_1 | TATGAGTGTGGAGTCGTTGCACAGTTTTGTGCTTGAATCCCAGNNNAGATCGGAAG |
| 42739_2 | TATGAGTGTGGAGTCGTTGCACAGTTTTGTGCTTGAATCCCAGNNNAGATCGGAAG |
| 42739_3 | TATGAGTGTGGAGTCGTTGCACAGTTTTGTGCTTGAATCCCAGNNNAGATCGGAAG |
| 33918_1 | TATGAGTGTGGAGTCGTTGCCAAGTGATTCTCCCAACCCAGTANNNAGATCGGAAG |
| 33918_2 | TATGAGTGTGGAGTCGTTGCCAAGTGATTCTCCCAACCCAGTANNNAGATCGGAAG |
| 33918_3 | TATGAGTGTGGAGTCGTTGCCAAGTGATTCTCCCAACCCAGTANNNAGATCGGAAG |
| 50688_1 | TATGAGTGTGGAGTCGTTGCCAGATGGGCTTGTGACTTTGAAGNNNAGATCGGAA |
| 50688_2 | TATGAGTGTGGAGTCGTTGCCAGATGGGCTTGTGACTTTGAAGNNNAGATCGGAA |
| 50688_3 | TATGAGTGTGGAGTCGTTGCCAGATGGGCTTGTGACTTTGAAGNNNAGATCGGAA |
| 35882_1 | TATGAGTGTGGAGTCGTTGCAAGAAAATTAGCTGGGCATGGTGNNNAGATCGGAA |
| 35882_2 | TATGAGTGTGGAGTCGTTGCAAGAAAATTAGCTGGGCATGGTGNNNAGATCGGAA |
| 35882_3 | TATGAGTGTGGAGTCGTTGCAAGAAAATTAGCTGGGCATGGTGNNNAGATCGGAA |
| 50813_1 | TATGAGTGTGGAGTCGTTGCATTTTTTGGCTCAATGGTGGACCNNNAGATCGGAAG |
| 50813_2 | TATGAGTGTGGAGTCGTTGCATTTTTTGGCTCAATGGTGGACCNNNAGATCGGAAG |
| 50813_3 | TATGAGTGTGGAGTCGTTGCATTTTTTGGCTCAATGGTGGACCNNNAGATCGGAAG |
| 13377_1 | TATGAGTGTGGAGTCGTTGCGAGCAATTAAAACAGATTCCTAANNNAGATCGGAAG |
| 13377_2 | TATGAGTGTGGAGTCGTTGCGAGCAATTAAAACAGATTCCTAANNNAGATCGGAAG |
| 13377_3 | TATGAGTGTGGAGTCGTTGCGAGCAATTAAAACAGATTCCTAANNNAGATCGGAAG |
| 35320_1 | TATGAGTGTGGAGTCGTTGCATTTACTGACCTGATGATCCGCCNNNAGATCGGAAG |
| 35320_2 | TATGAGTGTGGAGTCGTTGCATTTACTGACCTGATGATCCGCCNNNAGATCGGAAG |
| 35320_3 | TATGAGTGTGGAGTCGTTGCATTTACTGACCTGATGATCCGCCNNNAGATCGGAAG |
| 42068_1 | TATGAGTGTGGAGTCGTTGCAGGAGCAACTTTTACCAAATCACANNNAGATCGGAA |
| 42068_2 | TATGAGTGTGGAGTCGTTGCAGGAGCAACTTTTACCAAATCACANNNAGATCGGAA |
| 42068_3 | TATGAGTGTGGAGTCGTTGCAGGAGCAACTTTTACCAAATCACANNNAGATCGGAA |
| 50037_1 | TATGAGTGTGGAGTCGTTGCTCTTTCTGAGTGAGAACTGTGCTNNNAGATCGGAAG |
| 50037_2 | TATGAGTGTGGAGTCGTTGCTCTTTCTGAGTGAGAACTGTGCTNNNAGATCGGAAG |
| 50037_3 | TATGAGTGTGGAGTCGTTGCTCTTTCTGAGTGAGAACTGTGCTNNNAGATCGGAAG |
| 41398_1 | TATGAGTGTGGAGTCGTTGCGTGCTAGCCCACTTTTTTTCCTTNNNAGATCGGAAGA |
| 41398_2 | TATGAGTGTGGAGTCGTTGCGTGCTAGCCCACTTTTTTTCCTTNNNAGATCGGAAGA |
| 41398_3 | TATGAGTGTGGAGTCGTTGCGTGCTAGCCCACTTTTTTTCCTTNNNAGATCGGAAGA |
| 47313_1 | TATGAGTGTGGAGTCGTTGCCTCCCAAGTAGCTGGAACTACAGNNNAGATCGGAAG |
| 47313_2 | TATGAGTGTGGAGTCGTTGCCTCCCAAGTAGCTGGAACTACAGNNNAGATCGGAAG |
| 47313_3 | TATGAGTGTGGAGTCGTTGCCTCCCAAGTAGCTGGAACTACAGNNNAGATCGGAAG |
| 3071_1 | TATGAGTGTGGAGTCGTTGCCCGTCCCCTGTACCTACTACATTNNNAGATCGGAAGA |
| 3071_2 | TATGAGTGTGGAGTCGTTGCCCGTCCCCTGTACCTACTACATTNNNAGATCGGAAGA |
| 3071_3 | TATGAGTGTGGAGTCGTTGCCCGTCCCCTGTACCTACTACATTNNNAGATCGGAAGA |
| 36522_1 | TATGAGTGTGGAGTCGTTGCCAAGGCTTGTAGGATGTCACTCTNNNAGATCGGAAG |
| 36522_2 | TATGAGTGTGGAGTCGTTGCCAAGGCTTGTAGGATGTCACTCTNNNAGATCGGAAG |
| 36522_3 | TATGAGTGTGGAGTCGTTGCCAAGGCTTGTAGGATGTCACTCTNNNAGATCGGAAG |
| 43434_1 | TATGAGTGTGGAGTCGTTGCGCTCCCGGAAGCTATTTTTTGAANNNAGATCGGAAG |
| 43434_2 | TATGAGTGTGGAGTCGTTGCGCTCCCGGAAGCTATTTTTTGAANNNAGATCGGAAG |
| 43434_3 | TATGAGTGTGGAGTCGTTGCGCTCCCGGAAGCTATTTTTTGAANNNAGATCGGAAG |
| 39028_1 | TATGAGTGTGGAGTCGTTGCTGAACACGGTTCAAACAGTTAGCNNNAGATCGGAA |

| | |
|---|---|
| 39028_2 | TATGAGTGTGGAGTCGTTGCTGAACACGGTTCAAACAGTTAGCNNNAGATCGGAAG |
| 39028_3 | TATGAGTGTGGAGTCGTTGCTGAACACGGTTCAAACAGTTAGCNNNAGATCGGAAG |
| 45476_1 | TATGAGTGTGGAGTCGTTGCCCGATGACCACATGACCAAGTNNNAGATCGGAAGAG |
| 45476_2 | TATGAGTGTGGAGTCGTTGCCCGATGACCACATGACCAAGTNNNAGATCGGAAGAG |
| 45476_3 | TATGAGTGTGGAGTCGTTGCCCGATGACCACATGACCAAGTNNNAGATCGGAAGAG |
| 2786_1 | TATGAGTGTGGAGTCGTTGCAAAGGAAAGATCACAAGCAGGCTNNNAGATCGGAA |
| 2786_2 | TATGAGTGTGGAGTCGTTGCAAAGGAAAGATCACAAGCAGGCTNNNAGATCGGAA |
| 2786_3 | TATGAGTGTGGAGTCGTTGCAAAGGAAAGATCACAAGCAGGCTNNNAGATCGGAA |
| 35743_1 | TATGAGTGTGGAGTCGTTGCCAGTGCAATCATGGCTCATTGTANNNAGATCGGAAG |
| 35743_2 | TATGAGTGTGGAGTCGTTGCCAGTGCAATCATGGCTCATTGTANNNAGATCGGAAG |
| 35743_3 | TATGAGTGTGGAGTCGTTGCCAGTGCAATCATGGCTCATTGTANNNAGATCGGAAG |
| 44589_1 | TATGAGTGTGGAGTCGTTGCGAGGCAAGAGAATCGCTTGAACNNNAGATCGGAAG |
| 44589_2 | TATGAGTGTGGAGTCGTTGCGAGGCAAGAGAATCGCTTGAACNNNAGATCGGAAG |
| 44589_3 | TATGAGTGTGGAGTCGTTGCGAGGCAAGAGAATCGCTTGAACNNNAGATCGGAAG |
| 30899_1 | TATGAGTGTGGAGTCGTTGCCAGGAATAGATGGGCAATGCAAGNNNAGATCGGAA |
| 30899_2 | TATGAGTGTGGAGTCGTTGCCAGGAATAGATGGGCAATGCAAGNNNAGATCGGAA |
| 30899_3 | TATGAGTGTGGAGTCGTTGCCAGGAATAGATGGGCAATGCAAGNNNAGATCGGAA |
| 44401_1 | TATGAGTGTGGAGTCGTTGCCTGATAACCAGGGAAGCCAAAACNNNAGATCGGAA |
| 44401_2 | TATGAGTGTGGAGTCGTTGCCTGATAACCAGGGAAGCCAAAACNNNAGATCGGAA |
| 44401_3 | TATGAGTGTGGAGTCGTTGCCTGATAACCAGGGAAGCCAAAACNNNAGATCGGAA |
| 46334_1 | TATGAGTGTGGAGTCGTTGCCACCACACCCAGCTAACTTTTTTNNNAGATCGGAAGA |
| 46334_2 | TATGAGTGTGGAGTCGTTGCCACCACACCCAGCTAACTTTTTTNNNAGATCGGAAGA |
| 46334_3 | TATGAGTGTGGAGTCGTTGCCACCACACCCAGCTAACTTTTTTNNNAGATCGGAAGA |
| 29563_1 | TATGAGTGTGGAGTCGTTGCTCAGCCAAAAACATTTCCAGCATNNNAGATCGGAAGA |
| 29563_2 | TATGAGTGTGGAGTCGTTGCTCAGCCAAAAACATTTCCAGCATNNNAGATCGGAAGA |
| 29563_3 | TATGAGTGTGGAGTCGTTGCTCAGCCAAAAACATTTCCAGCATNNNAGATCGGAAGA |
| 30978_1 | TATGAGTGTGGAGTCGTTGCCACACACACACACACACACATATNNNAGATCGGAAGA |
| 30978_2 | TATGAGTGTGGAGTCGTTGCCACACACACACACACACACATATNNNAGATCGGAAGA |
| 30978_3 | TATGAGTGTGGAGTCGTTGCCACACACACACACACACACATATNNNAGATCGGAAGA |
| 36461_1 | TATGAGTGTGGAGTCGTTGCGTAGAAAAAGAGCAGGCATGGTGNNNAGATCGGAA |
| 36461_2 | TATGAGTGTGGAGTCGTTGCGTAGAAAAAGAGCAGGCATGGTGNNNAGATCGGAA |
| 36461_3 | TATGAGTGTGGAGTCGTTGCGTAGAAAAAGAGCAGGCATGGTGNNNAGATCGGAA |
| 42666_1 | TATGAGTGTGGAGTCGTTGCCTGATAGTCCCAGCTACTCAGGANNNAGATCGGAAG |
| 42666_2 | TATGAGTGTGGAGTCGTTGCCTGATAGTCCCAGCTACTCAGGANNNAGATCGGAAG |
| 42666_3 | TATGAGTGTGGAGTCGTTGCCTGATAGTCCCAGCTACTCAGGANNNAGATCGGAAG |
| 39798_1 | TATGAGTGTGGAGTCGTTGCAATGTAAGCTTGATGAGGGCAGANNNAGATCGGAA |
| 39798_2 | TATGAGTGTGGAGTCGTTGCAATGTAAGCTTGATGAGGGCAGANNNAGATCGGAA |
| 39798_3 | TATGAGTGTGGAGTCGTTGCAATGTAAGCTTGATGAGGGCAGANNNAGATCGGAA |
| 49338_1 | TATGAGTGTGGAGTCGTTGCACTGCATCTATTTCAATGATCACATNNNAGATCGGAA |
| 49338_2 | TATGAGTGTGGAGTCGTTGCACTGCATCTATTTCAATGATCACATNNNAGATCGGAA |
| 49338_3 | TATGAGTGTGGAGTCGTTGCACTGCATCTATTTCAATGATCACATNNNAGATCGGAA |
| 41583_1 | TATGAGTGTGGAGTCGTTGCCTCAAGTGATTCTCCTGCCTCAGNNNAGATCGGAAGA |
| 41583_2 | TATGAGTGTGGAGTCGTTGCCTCAAGTGATTCTCCTGCCTCAGNNNAGATCGGAAGA |

| | |
|---|---|
| 41583_3 | TATGAGTGTGGAGTCGTTGCCTCAAGTGATTCTCCTGCCTCAGNNNAGATCGGAAG |
| 10704_1 | TATGAGTGTGGAGTCGTTGCGTTAGAGAGAAGCCCCATCACCTNNNAGATCGGAAG |
| 10704_2 | TATGAGTGTGGAGTCGTTGCGTTAGAGAGAAGCCCCATCACCTNNNAGATCGGAAG |
| 10704_3 | TATGAGTGTGGAGTCGTTGCGTTAGAGAGAAGCCCCATCACCTNNNAGATCGGAAG |
| 10809_1 | TATGAGTGTGGAGTCGTTGCTATTTGTGGTTGTCGGGGTTTCCNNNAGATCGGAAG |
| 10809_2 | TATGAGTGTGGAGTCGTTGCTATTTGTGGTTGTCGGGGTTTCCNNNAGATCGGAAG |
| 10809_3 | TATGAGTGTGGAGTCGTTGCTATTTGTGGTTGTCGGGGTTTCCNNNAGATCGGAAG |
| 12946_1 | TATGAGTGTGGAGTCGTTGCTTTGAATGTTCGGTGAAGACNNNAGATCGGAAGAG |
| 12946_2 | TATGAGTGTGGAGTCGTTGCTTTGAATGTTCGGTGAAGACNNNAGATCGGAAGAG |
| 12946_3 | TATGAGTGTGGAGTCGTTGCTTTGAATGTTCGGTGAAGACNNNAGATCGGAAGAG |
| 41345_1 | TATGAGTGTGGAGTCGTTGCCTTCCACTCCACTAACCTTCCTTNNNAGATCGGAAGA |
| 41345_2 | TATGAGTGTGGAGTCGTTGCCTTCCACTCCACTAACCTTCCTTNNNAGATCGGAAGA |
| 41345_3 | TATGAGTGTGGAGTCGTTGCCTTCCACTCCACTAACCTTCCTTNNNAGATCGGAAGA |
| 40645_1 | TATGAGTGTGGAGTCGTTGCAGCTGCTTCTTTTTCCTCTCTGTNNNAGATCGGAAGA |
| 40645_2 | TATGAGTGTGGAGTCGTTGCAGCTGCTTCTTTTTCCTCTCTGTNNNAGATCGGAAGA |
| 40645_3 | TATGAGTGTGGAGTCGTTGCAGCTGCTTCTTTTTCCTCTCTGTNNNAGATCGGAAGA |
| 43541_1 | TATGAGTGTGGAGTCGTTGCTGGAGTACAGTGGCACCAATTCNNNAGATCGGAAG |
| 43541_2 | TATGAGTGTGGAGTCGTTGCTGGAGTACAGTGGCACCAATTCNNNAGATCGGAAG |
| 43541_3 | TATGAGTGTGGAGTCGTTGCTGGAGTACAGTGGCACCAATTCNNNAGATCGGAAG |
| 34457_1 | TATGAGTGTGGAGTCGTTGCGTCTCCTCCCTTCTCACCCCNNNAGATCGGAAGAGCA |
| 34457_2 | TATGAGTGTGGAGTCGTTGCGTCTCCTCCCTTCTCACCCCNNNAGATCGGAAGAGCA |
| 34457_3 | TATGAGTGTGGAGTCGTTGCGTCTCCTCCCTTCTCACCCCNNNAGATCGGAAGAGCA |
| 13096_1 | TATGAGTGTGGAGTCGTTGCAATCTCGATGAATGTGTAGATATNNNAGATCGGAA |
| 13096_2 | TATGAGTGTGGAGTCGTTGCAATCTCGATGAATGTGTAGATATNNNAGATCGGAA |
| 13096_3 | TATGAGTGTGGAGTCGTTGCAATCTCGATGAATGTGTAGATATNNNAGATCGGAA |
| 52931_1 | TATGAGTGTGGAGTCGTTGCTGACAACTGTTTAGACTTGAGGAGTNNNAGATCGG |
| 52931_2 | TATGAGTGTGGAGTCGTTGCTGACAACTGTTTAGACTTGAGGAGTNNNAGATCGG |
| 52931_3 | TATGAGTGTGGAGTCGTTGCTGACAACTGTTTAGACTTGAGGAGTNNNAGATCGG |
| 41952_1 | TATGAGTGTGGAGTCGTTGCGCTGGGACTATAGATGTACACCANNNAGATCGGAA |
| 41952_2 | TATGAGTGTGGAGTCGTTGCGCTGGGACTATAGATGTACACCANNNAGATCGGAA |
| 41952_3 | TATGAGTGTGGAGTCGTTGCGCTGGGACTATAGATGTACACCANNNAGATCGGAA |
| 43290_1 | TATGAGTGTGGAGTCGTTGCTCCATCCTCAAATTAGGCAAATTCNNNAGATCGGAA |
| 43290_2 | TATGAGTGTGGAGTCGTTGCTCCATCCTCAAATTAGGCAAATTCNNNAGATCGGAA |
| 43290_3 | TATGAGTGTGGAGTCGTTGCTCCATCCTCAAATTAGGCAAATTCNNNAGATCGGAA |
| 36444_1 | TATGAGTGTGGAGTCGTTGCCAGGCCAAAGCTTTTCTTCATCANNNAGATCGGAAG |
| 36444_2 | TATGAGTGTGGAGTCGTTGCCAGGCCAAAGCTTTTCTTCATCANNNAGATCGGAAG |
| 36444_3 | TATGAGTGTGGAGTCGTTGCCAGGCCAAAGCTTTTCTTCATCANNNAGATCGGAAG |
| 36879_1 | TATGAGTGTGGAGTCGTTGCGACAACTACCACAGACTGGGAATNNNAGATCGGAA |
| 36879_2 | TATGAGTGTGGAGTCGTTGCGACAACTACCACAGACTGGGAATNNNAGATCGGAA |
| 36879_3 | TATGAGTGTGGAGTCGTTGCGACAACTACCACAGACTGGGAATNNNAGATCGGAA |
| 49155_1 | TATGAGTGTGGAGTCGTTGCCTTCCCAGGTTCAAGCAATTCTCNNNAGATCGGAAG |
| 49155_2 | TATGAGTGTGGAGTCGTTGCCTTCCCAGGTTCAAGCAATTCTCNNNAGATCGGAAG |
| 49155_3 | TATGAGTGTGGAGTCGTTGCCTTCCCAGGTTCAAGCAATTCTCNNNAGATCGGAAG |

| | |
|---|---|
| 36696_1 | TATGAGTGTGGAGTCGTTGCCGAGACAGGACAACAAACGAAAANNNAGATCGGAA |
| 36696_2 | TATGAGTGTGGAGTCGTTGCCGAGACAGGACAACAAACGAAAANNNAGATCGGAA |
| 36696_3 | TATGAGTGTGGAGTCGTTGCCGAGACAGGACAACAAACGAAAANNNAGATCGGAA |
| 2509_1 | TATGAGTGTGGAGTCGTTGCCGGCCCATCATAGCCAAAGATAGNNNAGATCGGAAG |
| 2509_2 | TATGAGTGTGGAGTCGTTGCCGGCCCATCATAGCCAAAGATAGNNNAGATCGGAAG |
| 2509_3 | TATGAGTGTGGAGTCGTTGCCGGCCCATCATAGCCAAAGATAGNNNAGATCGGAAG |
| 36098_1 | TATGAGTGTGGAGTCGTTGCCAACATAGCAAGACCCTGTCTCTNNNAGATCGGAAG |
| 36098_2 | TATGAGTGTGGAGTCGTTGCCAACATAGCAAGACCCTGTCTCTNNNAGATCGGAAG |
| 36098_3 | TATGAGTGTGGAGTCGTTGCCAACATAGCAAGACCCTGTCTCTNNNAGATCGGAAG |
| 48682_1 | TATGAGTGTGGAGTCGTTGCCCGGCCAAAAGCACATTTCTTATNNNAGATCGGAAG |
| 48682_2 | TATGAGTGTGGAGTCGTTGCCCGGCCAAAAGCACATTTCTTATNNNAGATCGGAAG |
| 48682_3 | TATGAGTGTGGAGTCGTTGCCCGGCCAAAAGCACATTTCTTATNNNAGATCGGAAG |
| 47056_1 | TATGAGTGTGGAGTCGTTGCGGACAGTAACAATTGAAAGGTTCTGANNNAGATCG |
| 47056_2 | TATGAGTGTGGAGTCGTTGCGGACAGTAACAATTGAAAGGTTCTGANNNAGATCG |
| 47056_3 | TATGAGTGTGGAGTCGTTGCGGACAGTAACAATTGAAAGGTTCTGANNNAGATCG |
| 34339_1 | TATGAGTGTGGAGTCGTTGCCTGGAATGCAGTGGAGTGATCTCNNNAGATCGGAA |
| 34339_2 | TATGAGTGTGGAGTCGTTGCCTGGAATGCAGTGGAGTGATCTCNNNAGATCGGAA |
| 34339_3 | TATGAGTGTGGAGTCGTTGCCTGGAATGCAGTGGAGTGATCTCNNNAGATCGGAA |
| 13322_1 | TATGAGTGTGGAGTCGTTGCTTTCCAAACTTATTAAGAAGCTTNNNAGATCGGAAG |
| 13322_2 | TATGAGTGTGGAGTCGTTGCTTTCCAAACTTATTAAGAAGCTTNNNAGATCGGAAG |
| 13322_3 | TATGAGTGTGGAGTCGTTGCTTTCCAAACTTATTAAGAAGCTTNNNAGATCGGAAG |
| 42732_1 | TATGAGTGTGGAGTCGTTGCCAGCAGTTTGACAATGATGTGGTNNNAGATCGGAA |
| 42732_2 | TATGAGTGTGGAGTCGTTGCCAGCAGTTTGACAATGATGTGGTNNNAGATCGGAA |
| 42732_3 | TATGAGTGTGGAGTCGTTGCCAGCAGTTTGACAATGATGTGGTNNNAGATCGGAA |
| 32118_1 | TATGAGTGTGGAGTCGTTGCGAAAAGAAATGTAGGCTCTGGGCNNNAGATCGGAA |
| 32118_2 | TATGAGTGTGGAGTCGTTGCGAAAAGAAATGTAGGCTCTGGGCNNNAGATCGGAA |
| 32118_3 | TATGAGTGTGGAGTCGTTGCGAAAAGAAATGTAGGCTCTGGGCNNNAGATCGGAA |
| 37340_1 | TATGAGTGTGGAGTCGTTGCGAGTACATCTAGTCAGCAGAGCCNNNAGATCGGAAG |
| 37340_2 | TATGAGTGTGGAGTCGTTGCGAGTACATCTAGTCAGCAGAGCCNNNAGATCGGAAG |
| 37340_3 | TATGAGTGTGGAGTCGTTGCGAGTACATCTAGTCAGCAGAGCCNNNAGATCGGAAG |
| 13771_1 | TATGAGTGTGGAGTCGTTGCAAATGGAGAAAAGGAAACTTTAGNNNAGATCGGAA |
| 13771_2 | TATGAGTGTGGAGTCGTTGCAAATGGAGAAAAGGAAACTTTAGNNNAGATCGGAA |
| 13771_3 | TATGAGTGTGGAGTCGTTGCAAATGGAGAAAAGGAAACTTTAGNNNAGATCGGAA |
| 10266_1 | TATGAGTGTGGAGTCGTTGCTTGATCTTTGGTGCTATGAGGCCNNNAGATCGGAAG |
| 10266_2 | TATGAGTGTGGAGTCGTTGCTTGATCTTTGGTGCTATGAGGCCNNNAGATCGGAAG |
| 10266_3 | TATGAGTGTGGAGTCGTTGCTTGATCTTTGGTGCTATGAGGCCNNNAGATCGGAAG |
| 29245_1 | TATGAGTGTGGAGTCGTTGCCACCTACTACACGGCAGATACAGNNNAGATCGGAAG |
| 29245_2 | TATGAGTGTGGAGTCGTTGCCACCTACTACACGGCAGATACAGNNNAGATCGGAAG |
| 29245_3 | TATGAGTGTGGAGTCGTTGCCACCTACTACACGGCAGATACAGNNNAGATCGGAAG |
| 29408_1 | TATGAGTGTGGAGTCGTTGCATCTACAAGCAGTCACAGCACATNNNAGATCGGAAG |
| 29408_2 | TATGAGTGTGGAGTCGTTGCATCTACAAGCAGTCACAGCACATNNNAGATCGGAAG |
| 29408_3 | TATGAGTGTGGAGTCGTTGCATCTACAAGCAGTCACAGCACATNNNAGATCGGAAG |
| 53159_1 | TATGAGTGTGGAGTCGTTGCAGGCACACAATAACTGGAAGCTANNNAGATCGGAA |

| | |
|---|---|
| 53159_2 | TATGAGTGTGGAGTCGTTGCAGGCACACAATAACTGGAAGCTANNNAGATCGGAA |
| 53159_3 | TATGAGTGTGGAGTCGTTGCAGGCACACAATAACTGGAAGCTANNNAGATCGGAA |
| 42990_1 | TATGAGTGTGGAGTCGTTGCTCCCACCAACAATGTGTAAGTGTNNNAGATCGGAAG |
| 42990_2 | TATGAGTGTGGAGTCGTTGCTCCCACCAACAATGTGTAAGTGTNNNAGATCGGAAG |
| 42990_3 | TATGAGTGTGGAGTCGTTGCTCCCACCAACAATGTGTAAGTGTNNNAGATCGGAAG |
| 47383_1 | TATGAGTGTGGAGTCGTTGCTAGACATGGTGTGGTGGGTAATGNNNAGATCGGA |
| 47383_2 | TATGAGTGTGGAGTCGTTGCTAGACATGGTGTGGTGGGTAATGNNNAGATCGGA |
| 47383_3 | TATGAGTGTGGAGTCGTTGCTAGACATGGTGTGGTGGGTAATGNNNAGATCGGA |
| 43148_1 | TATGAGTGTGGAGTCGTTGCTGAAACAAATGAAAACACAACCCANNNAGATCGGAA |
| 43148_2 | TATGAGTGTGGAGTCGTTGCTGAAACAAATGAAAACACAACCCANNNAGATCGGAA |
| 43148_3 | TATGAGTGTGGAGTCGTTGCTGAAACAAATGAAAACACAACCCANNNAGATCGGAA |
| 31713_1 | TATGAGTGTGGAGTCGTTGCCTCTCTCCTCTCTCTCTGTCACANNNAGATCGGAAGA |
| 31713_2 | TATGAGTGTGGAGTCGTTGCCTCTCTCCTCTCTCTCTGTCACANNNAGATCGGAAGA |
| 31713_3 | TATGAGTGTGGAGTCGTTGCCTCTCTCCTCTCTCTCTGTCACANNNAGATCGGAAGA |
| 36731_1 | TATGAGTGTGGAGTCGTTGCATGGTATTGGCACAATTGGACACNNNAGATCGGAA |
| 36731_2 | TATGAGTGTGGAGTCGTTGCATGGTATTGGCACAATTGGACACNNNAGATCGGAA |
| 36731_3 | TATGAGTGTGGAGTCGTTGCATGGTATTGGCACAATTGGACACNNNAGATCGGAA |
| 42172_1 | TATGAGTGTGGAGTCGTTGCCTCACTGCAACCTCTCTCTTACANNNAGATCGGAAGA |
| 42172_2 | TATGAGTGTGGAGTCGTTGCCTCACTGCAACCTCTCTCTTACANNNAGATCGGAAGA |
| 42172_3 | TATGAGTGTGGAGTCGTTGCCTCACTGCAACCTCTCTCTTACANNNAGATCGGAAGA |
| 48685_1 | TATGAGTGTGGAGTCGTTGCCTCACTATGTTTCCCAGGCTCATNNNAGATCGGAAG |
| 48685_2 | TATGAGTGTGGAGTCGTTGCCTCACTATGTTTCCCAGGCTCATNNNAGATCGGAAG |
| 48685_3 | TATGAGTGTGGAGTCGTTGCCTCACTATGTTTCCCAGGCTCATNNNAGATCGGAAG |
| 40851_1 | TATGAGTGTGGAGTCGTTGCGAAGAGGAGACCACGTTAGTCAANNNAGATCGGAA |
| 40851_2 | TATGAGTGTGGAGTCGTTGCGAAGAGGAGACCACGTTAGTCAANNNAGATCGGAA |
| 40851_3 | TATGAGTGTGGAGTCGTTGCGAAGAGGAGACCACGTTAGTCAANNNAGATCGGAA |
| 36622_1 | TATGAGTGTGGAGTCGTTGCGACCCAGATCCAAACATACTGGTNNNAGATCGGAAG |
| 36622_2 | TATGAGTGTGGAGTCGTTGCGACCCAGATCCAAACATACTGGTNNNAGATCGGAAG |
| 36622_3 | TATGAGTGTGGAGTCGTTGCGACCCAGATCCAAACATACTGGTNNNAGATCGGAAG |
| 37793_1 | TATGAGTGTGGAGTCGTTGCTGTAATCACAGCACTTTGGGAGGNNNAGATCGGAA |
| 37793_2 | TATGAGTGTGGAGTCGTTGCTGTAATCACAGCACTTTGGGAGGNNNAGATCGGAA |
| 37793_3 | TATGAGTGTGGAGTCGTTGCTGTAATCACAGCACTTTGGGAGGNNNAGATCGGAA |
| 47956_1 | TATGAGTGTGGAGTCGTTGCTGCCTGGTCTGATTCCATTGTTANNNAGATCGGAAG |
| 47956_2 | TATGAGTGTGGAGTCGTTGCTGCCTGGTCTGATTCCATTGTTANNNAGATCGGAAG |
| 47956_3 | TATGAGTGTGGAGTCGTTGCTGCCTGGTCTGATTCCATTGTTANNNAGATCGGAAG |
| 29913_1 | TATGAGTGTGGAGTCGTTGCCAATCGCTAGAGGGCAAAGAAAGNNNAGATCGGAA |
| 29913_2 | TATGAGTGTGGAGTCGTTGCCAATCGCTAGAGGGCAAAGAAAGNNNAGATCGGAA |
| 29913_3 | TATGAGTGTGGAGTCGTTGCCAATCGCTAGAGGGCAAAGAAAGNNNAGATCGGAA |
| 46062_1 | TATGAGTGTGGAGTCGTTGCCAGTGGTATGATCACAGCTCACTNNNAGATCGGAAG |
| 46062_2 | TATGAGTGTGGAGTCGTTGCCAGTGGTATGATCACAGCTCACTNNNAGATCGGAAG |
| 46062_3 | TATGAGTGTGGAGTCGTTGCCAGTGGTATGATCACAGCTCACTNNNAGATCGGAAG |
| 46333_1 | TATGAGTGTGGAGTCGTTGCTAGAGTGCAATGGCTCAATCTCANNNAGATCGGAAG |
| 46333_2 | TATGAGTGTGGAGTCGTTGCTAGAGTGCAATGGCTCAATCTCANNNAGATCGGAAG |

| | |
|---|---|
| 46333_3 | TATGAGTGTGGAGTCGTTGCTAGAGTGCAATGGCTCAATCTCANNNAGATCGGAAG |
| 42787_1 | TATGAGTGTGGAGTCGTTGCCAGGCATTGTGAAGGGAACACNNNAGATCGGAAGA |
| 42787_2 | TATGAGTGTGGAGTCGTTGCCAGGCATTGTGAAGGGAACACNNNAGATCGGAAGA |
| 42787_3 | TATGAGTGTGGAGTCGTTGCCAGGCATTGTGAAGGGAACACNNNAGATCGGAAGA |
| 40272_1 | TATGAGTGTGGAGTCGTTGCCCCACCCATTACAAAATTGCCAANNNAGATCGGAAG |
| 40272_2 | TATGAGTGTGGAGTCGTTGCCCCACCCATTACAAAATTGCCAANNNAGATCGGAAG |
| 40272_3 | TATGAGTGTGGAGTCGTTGCCCCACCCATTACAAAATTGCCAANNNAGATCGGAAG |
| 49793_1 | TATGAGTGTGGAGTCGTTGCAGCACTGCAAATATGGTAGGTCANNNAGATCGGAA |
| 49793_2 | TATGAGTGTGGAGTCGTTGCAGCACTGCAAATATGGTAGGTCANNNAGATCGGAA |
| 49793_3 | TATGAGTGTGGAGTCGTTGCAGCACTGCAAATATGGTAGGTCANNNAGATCGGAA |
| 52928_1 | TATGAGTGTGGAGTCGTTGCCAGCCTCTCTCTGCAGACAGNNNAGATCGGAAGAGC |
| 52928_2 | TATGAGTGTGGAGTCGTTGCCAGCCTCTCTCTGCAGACAGNNNAGATCGGAAGAGC |
| 52928_3 | TATGAGTGTGGAGTCGTTGCCAGCCTCTCTCTGCAGACAGNNNAGATCGGAAGAGC |
| 36541_1 | TATGAGTGTGGAGTCGTTGCGAGTTTTGCTCTTGTTGCCCAGNNNAGATCGGAAGA |
| 36541_2 | TATGAGTGTGGAGTCGTTGCGAGTTTTGCTCTTGTTGCCCAGNNNAGATCGGAAGA |
| 36541_3 | TATGAGTGTGGAGTCGTTGCGAGTTTTGCTCTTGTTGCCCAGNNNAGATCGGAAGA |
| 47449_1 | TATGAGTGTGGAGTCGTTGCTTCCTGGGTTGAAGCTATTCTCCNNNAGATCGGAAG |
| 47449_2 | TATGAGTGTGGAGTCGTTGCTTCCTGGGTTGAAGCTATTCTCCNNNAGATCGGAAG |
| 47449_3 | TATGAGTGTGGAGTCGTTGCTTCCTGGGTTGAAGCTATTCTCCNNNAGATCGGAAG |
| 30542_1 | TATGAGTGTGGAGTCGTTGCTGGGGGACTGAGATGTTTTGCTTGNNNAGATCGGA |
| 30542_2 | TATGAGTGTGGAGTCGTTGCTGGGGGACTGAGATGTTTTGCTTGNNNAGATCGGA |
| 30542_3 | TATGAGTGTGGAGTCGTTGCTGGGGGACTGAGATGTTTTGCTTGNNNAGATCGGA |
| 39606_1 | TATGAGTGTGGAGTCGTTGCCGAAAGAAGGAAGGAAGGAAGGANNNAGATCGG |
| 39606_2 | TATGAGTGTGGAGTCGTTGCCGAAAGAAGGAAGGAAGGAAGGANNNAGATCGG |
| 39606_3 | TATGAGTGTGGAGTCGTTGCCGAAAGAAGGAAGGAAGGAAGGANNNAGATCGG |
| 36668_1 | TATGAGTGTGGAGTCGTTGCTGATGTATAGCAACGACCCACTTNNNAGATCGGAAG |
| 36668_2 | TATGAGTGTGGAGTCGTTGCTGATGTATAGCAACGACCCACTTNNNAGATCGGAAG |
| 36668_3 | TATGAGTGTGGAGTCGTTGCTGATGTATAGCAACGACCCACTTNNNAGATCGGAAG |
| 45399_1 | TATGAGTGTGGAGTCGTTGCGCTGCATCGAATTTAGGGAGTTCNNNAGATCGGAA |
| 45399_2 | TATGAGTGTGGAGTCGTTGCGCTGCATCGAATTTAGGGAGTTCNNNAGATCGGAA |
| 45399_3 | TATGAGTGTGGAGTCGTTGCGCTGCATCGAATTTAGGGAGTTCNNNAGATCGGAA |
| 41071_1 | TATGAGTGTGGAGTCGTTGCAAGAAGAAGAGAGAAGTTGGCGGNNNAGATCGGA |
| 41071_2 | TATGAGTGTGGAGTCGTTGCAAGAAGAAGAGAGAAGTTGGCGGNNNAGATCGGA |
| 41071_3 | TATGAGTGTGGAGTCGTTGCAAGAAGAAGAGAGAAGTTGGCGGNNNAGATCGGA |
| 36603_1 | TATGAGTGTGGAGTCGTTGCACACAATGAACTCCCTGAAAGGANNNAGATCGGAA |
| 36603_2 | TATGAGTGTGGAGTCGTTGCACACAATGAACTCCCTGAAAGGANNNAGATCGGAA |
| 36603_3 | TATGAGTGTGGAGTCGTTGCACACAATGAACTCCCTGAAAGGANNNAGATCGGAA |
| 29153_1 | TATGAGTGTGGAGTCGTTGCTGTCTCTGTATGCTTCTTTTCCTNNNAGATCGGAAGA |
| 29153_2 | TATGAGTGTGGAGTCGTTGCTGTCTCTGTATGCTTCTTTTCCTNNNAGATCGGAAGA |
| 29153_3 | TATGAGTGTGGAGTCGTTGCTGTCTCTGTATGCTTCTTTTCCTNNNAGATCGGAAGA |
| 44182_1 | TATGAGTGTGGAGTCGTTGCAAGTGGTCTCTCTCTCTCTGANNNAGATCGGAAG |
| 44182_2 | TATGAGTGTGGAGTCGTTGCAAGTGGTCTCTCTCTCTCTGANNNAGATCGGAAG |
| 44182_3 | TATGAGTGTGGAGTCGTTGCAAGTGGTCTCTCTCTCTCTGANNNAGATCGGAAG |

| | |
|---|---|
| 38218_1 | TATGAGTGTGGAGTCGTTGCCTCTCAATTGGATTGGGATGTGCNNNAGATCGGAAG |
| 38218_2 | TATGAGTGTGGAGTCGTTGCCTCTCAATTGGATTGGGATGTGCNNNAGATCGGAAG |
| 38218_3 | TATGAGTGTGGAGTCGTTGCCTCTCAATTGGATTGGGATGTGCNNNAGATCGGAAG |
| 32617_1 | TATGAGTGTGGAGTCGTTGCGTGAGGTAAGGACTGGAAAGAGGNNNAGATCGGA |
| 32617_2 | TATGAGTGTGGAGTCGTTGCGTGAGGTAAGGACTGGAAAGAGGNNNAGATCGGA |
| 32617_3 | TATGAGTGTGGAGTCGTTGCGTGAGGTAAGGACTGGAAAGAGGNNNAGATCGGA |
| 51028_1 | TATGAGTGTGGAGTCGTTGCAGCCACATTGCAGATAAAGATGCNNNAGATCGGAAG |
| 51028_2 | TATGAGTGTGGAGTCGTTGCAGCCACATTGCAGATAAAGATGCNNNAGATCGGAAG |
| 51028_3 | TATGAGTGTGGAGTCGTTGCAGCCACATTGCAGATAAAGATGCNNNAGATCGGAAG |
| 39486_1 | TATGAGTGTGGAGTCGTTGCAAACGTCAGGCATGTCTGAATTCNNNAGATCGGAAG |
| 39486_2 | TATGAGTGTGGAGTCGTTGCAAACGTCAGGCATGTCTGAATTCNNNAGATCGGAAG |
| 39486_3 | TATGAGTGTGGAGTCGTTGCAAACGTCAGGCATGTCTGAATTCNNNAGATCGGAAG |
| 38893_1 | TATGAGTGTGGAGTCGTTGCTTCTTTGCAAATGTGGATGCCTTNNNAGATCGGAAGA |
| 38893_2 | TATGAGTGTGGAGTCGTTGCTTCTTTGCAAATGTGGATGCCTTNNNAGATCGGAAGA |
| 38893_3 | TATGAGTGTGGAGTCGTTGCTTCTTTGCAAATGTGGATGCCTTNNNAGATCGGAAGA |
| 35299_1 | TATGAGTGTGGAGTCGTTGCCATTCTCCTGCCTCAGCCTCNNNAGATCGGAAGAGCA |
| 35299_2 | TATGAGTGTGGAGTCGTTGCCATTCTCCTGCCTCAGCCTCNNNAGATCGGAAGAGCA |
| 35299_3 | TATGAGTGTGGAGTCGTTGCCATTCTCCTGCCTCAGCCTCNNNAGATCGGAAGAGCA |
| 31102_1 | TATGAGTGTGGAGTCGTTGCAGAATGGATAGCTGCCTCTATGCNNNAGATCGGAAG |
| 31102_2 | TATGAGTGTGGAGTCGTTGCAGAATGGATAGCTGCCTCTATGCNNNAGATCGGAAG |
| 31102_3 | TATGAGTGTGGAGTCGTTGCAGAATGGATAGCTGCCTCTATGCNNNAGATCGGAAG |
| 50440_1 | TATGAGTGTGGAGTCGTTGCTGTAGAAGACTGAAACCTGTGCANNNAGATCGGAAG |
| 50440_2 | TATGAGTGTGGAGTCGTTGCTGTAGAAGACTGAAACCTGTGCANNNAGATCGGAAG |
| 50440_3 | TATGAGTGTGGAGTCGTTGCTGTAGAAGACTGAAACCTGTGCANNNAGATCGGAAG |
| 42047_1 | TATGAGTGTGGAGTCGTTGCATGATGGCATATGCTTCTGGTCTNNNAGATCGGAAG |
| 42047_2 | TATGAGTGTGGAGTCGTTGCATGATGGCATATGCTTCTGGTCTNNNAGATCGGAAG |
| 42047_3 | TATGAGTGTGGAGTCGTTGCATGATGGCATATGCTTCTGGTCTNNNAGATCGGAAG |
| 10169_1 | TATGAGTGTGGAGTCGTTGCCTCATTTAGGCCAGAGGCTTTGTNNNAGATCGGAAG |
| 10169_2 | TATGAGTGTGGAGTCGTTGCCTCATTTAGGCCAGAGGCTTTGTNNNAGATCGGAAG |
| 10169_3 | TATGAGTGTGGAGTCGTTGCCTCATTTAGGCCAGAGGCTTTGTNNNAGATCGGAAG |
| 33519_1 | TATGAGTGTGGAGTCGTTGCCACGTAGACCAATCGAACAGAACNNNAGATCGGAAG |
| 33519_2 | TATGAGTGTGGAGTCGTTGCCACGTAGACCAATCGAACAGAACNNNAGATCGGAAG |
| 33519_3 | TATGAGTGTGGAGTCGTTGCCACGTAGACCAATCGAACAGAACNNNAGATCGGAAG |
| 52306_1 | TATGAGTGTGGAGTCGTTGCTGACATCTCAGAATCAGGTGAGCNNNAGATCGGAAG |
| 52306_2 | TATGAGTGTGGAGTCGTTGCTGACATCTCAGAATCAGGTGAGCNNNAGATCGGAAG |
| 52306_3 | TATGAGTGTGGAGTCGTTGCTGACATCTCAGAATCAGGTGAGCNNNAGATCGGAAG |
| 29761_1 | TATGAGTGTGGAGTCGTTGCCGCTTCAGCAACTCCTGTTTATANNNAGATCGGAAGA |
| 29761_2 | TATGAGTGTGGAGTCGTTGCCGCTTCAGCAACTCCTGTTTATANNNAGATCGGAAGA |
| 29761_3 | TATGAGTGTGGAGTCGTTGCCGCTTCAGCAACTCCTGTTTATANNNAGATCGGAAGA |
| 45906_1 | TATGAGTGTGGAGTCGTTGCCTCTCAGTTCCTTCCCCTCTTTCNNNAGATCGGAAGAG |
| 45906_2 | TATGAGTGTGGAGTCGTTGCCTCTCAGTTCCTTCCCCTCTTTCNNNAGATCGGAAGAG |
| 45906_3 | TATGAGTGTGGAGTCGTTGCCTCTCAGTTCCTTCCCCTCTTTCNNNAGATCGGAAGAG |
| 46808_1 | TATGAGTGTGGAGTCGTTGCCCAGCACATATTCTGCCCAAATGNNNAGATCGGAAGA |

| | |
|---|---|
| 46808_2 | TATGAGTGTGGAGTCGTTGCCCAGCACATATTCTGCCCAAATGNNNAGATCGGAAG |
| 46808_3 | TATGAGTGTGGAGTCGTTGCCCAGCACATATTCTGCCCAAATGNNNAGATCGGAAG |
| 48454_1 | TATGAGTGTGGAGTCGTTGCTGGAGTATAGTGGCGTGATCTTGNNNAGATCGGAA |
| 48454_2 | TATGAGTGTGGAGTCGTTGCTGGAGTATAGTGGCGTGATCTTGNNNAGATCGGAA |
| 48454_3 | TATGAGTGTGGAGTCGTTGCTGGAGTATAGTGGCGTGATCTTGNNNAGATCGGAA |
| 36858_1 | TATGAGTGTGGAGTCGTTGCACCATTGATGGGCATTTAGGTTGNNNAGATCGGAAG |
| 36858_2 | TATGAGTGTGGAGTCGTTGCACCATTGATGGGCATTTAGGTTGNNNAGATCGGAAG |
| 36858_3 | TATGAGTGTGGAGTCGTTGCACCATTGATGGGCATTTAGGTTGNNNAGATCGGAAG |
| 48555_1 | TATGAGTGTGGAGTCGTTGCACCATCAGGGATTACGTTTCCAANNNAGATCGGAAG |
| 48555_2 | TATGAGTGTGGAGTCGTTGCACCATCAGGGATTACGTTTCCAANNNAGATCGGAAG |
| 48555_3 | TATGAGTGTGGAGTCGTTGCACCATCAGGGATTACGTTTCCAANNNAGATCGGAAG |
| 44399_1 | TATGAGTGTGGAGTCGTTGCCCACTTCTCAGTAAGGAATGCCTNNNAGATCGGAAG |
| 44399_2 | TATGAGTGTGGAGTCGTTGCCCACTTCTCAGTAAGGAATGCCTNNNAGATCGGAAG |
| 44399_3 | TATGAGTGTGGAGTCGTTGCCCACTTCTCAGTAAGGAATGCCTNNNAGATCGGAAG |
| 40246_1 | TATGAGTGTGGAGTCGTTGCCTCCCAGGTTCACACCATTCTCNNNAGATCGGAAGAG |
| 40246_2 | TATGAGTGTGGAGTCGTTGCCTCCCAGGTTCACACCATTCTCNNNAGATCGGAAGAG |
| 40246_3 | TATGAGTGTGGAGTCGTTGCCTCCCAGGTTCACACCATTCTCNNNAGATCGGAAGAG |
| 34122_1 | TATGAGTGTGGAGTCGTTGCGCTCGTACAGAATTCAATCCAGCNNNAGATCGGAAG |
| 34122_2 | TATGAGTGTGGAGTCGTTGCGCTCGTACAGAATTCAATCCAGCNNNAGATCGGAAG |
| 34122_3 | TATGAGTGTGGAGTCGTTGCGCTCGTACAGAATTCAATCCAGCNNNAGATCGGAAG |
| 44770_1 | TATGAGTGTGGAGTCGTTGCATGGCCCTACTCTTCCATCTCAANNNAGATCGGAAGA |
| 44770_2 | TATGAGTGTGGAGTCGTTGCATGGCCCTACTCTTCCATCTCAANNNAGATCGGAAGA |
| 44770_3 | TATGAGTGTGGAGTCGTTGCATGGCCCTACTCTTCCATCTCAANNNAGATCGGAAGA |
| 39336_1 | TATGAGTGTGGAGTCGTTGCGATCGTAGGGAGCCAAAACCTTANNNAGATCGGAA |
| 39336_2 | TATGAGTGTGGAGTCGTTGCGATCGTAGGGAGCCAAAACCTTANNNAGATCGGAA |
| 39336_3 | TATGAGTGTGGAGTCGTTGCGATCGTAGGGAGCCAAAACCTTANNNAGATCGGAA |
| 10296_1 | TATGAGTGTGGAGTCGTTGCCCAATGAGCTTCCATTCTCCCAANNNAGATCGGAAGA |
| 10296_2 | TATGAGTGTGGAGTCGTTGCCCAATGAGCTTCCATTCTCCCAANNNAGATCGGAAGA |
| 10296_3 | TATGAGTGTGGAGTCGTTGCCCAATGAGCTTCCATTCTCCCAANNNAGATCGGAAGA |
| 46125_1 | TATGAGTGTGGAGTCGTTGCCTCACTAGCAAGCTCAGCCTCNNNAGATCGGAAGAG |
| 46125_2 | TATGAGTGTGGAGTCGTTGCCTCACTAGCAAGCTCAGCCTCNNNAGATCGGAAGAG |
| 46125_3 | TATGAGTGTGGAGTCGTTGCCTCACTAGCAAGCTCAGCCTCNNNAGATCGGAAGAG |
| 45875_1 | TATGAGTGTGGAGTCGTTGCCACTTCTGAGGTTCTAGCAGGAGNNNAGATCGGAAG |
| 45875_2 | TATGAGTGTGGAGTCGTTGCCACTTCTGAGGTTCTAGCAGGAGNNNAGATCGGAAG |
| 45875_3 | TATGAGTGTGGAGTCGTTGCCACTTCTGAGGTTCTAGCAGGAGNNNAGATCGGAAG |
| 45528_1 | TATGAGTGTGGAGTCGTTGCGTGTGTATTTGGTTTGCGCTTTGNNNAGATCGGAAG |
| 45528_2 | TATGAGTGTGGAGTCGTTGCGTGTGTATTTGGTTTGCGCTTTGNNNAGATCGGAAG |
| 45528_3 | TATGAGTGTGGAGTCGTTGCGTGTGTATTTGGTTTGCGCTTTGNNNAGATCGGAAG |
| 50442_1 | TATGAGTGTGGAGTCGTTGCTTGTCAGGATATTGCATTGTGGCNNNAGATCGGAAG |
| 50442_2 | TATGAGTGTGGAGTCGTTGCTTGTCAGGATATTGCATTGTGGCNNNAGATCGGAAG |
| 50442_3 | TATGAGTGTGGAGTCGTTGCTTGTCAGGATATTGCATTGTGGCNNNAGATCGGAAG |
| 53279_1 | TATGAGTGTGGAGTCGTTGCACACACTGTAAGCCACCTACAATNNNAGATCGGAAG |
| 53279_2 | TATGAGTGTGGAGTCGTTGCACACACTGTAAGCCACCTACAATNNNAGATCGGAAG |

| | |
|---|---|
| 53279_3 | TATGAGTGTGGAGTCGTTGCACACACTGTAAGCCACCTACAATNNNAGATCGGAAG |
| 10028_1 | TATGAGTGTGGAGTCGTTGCTGGATGCCCCAACATTATGATCGNNNAGATCGGAAG |
| 10028_2 | TATGAGTGTGGAGTCGTTGCTGGATGCCCCAACATTATGATCGNNNAGATCGGAAG |
| 10028_3 | TATGAGTGTGGAGTCGTTGCTGGATGCCCCAACATTATGATCGNNNAGATCGGAAG |
| 50271_1 | TATGAGTGTGGAGTCGTTGCCTTAGGCAAGCAGCTCTTCTTTCNNNAGATCGGAAG |
| 50271_2 | TATGAGTGTGGAGTCGTTGCCTTAGGCAAGCAGCTCTTCTTTCNNNAGATCGGAAG |
| 50271_3 | TATGAGTGTGGAGTCGTTGCCTTAGGCAAGCAGCTCTTCTTTCNNNAGATCGGAAG |
| 13568_1 | TATGAGTGTGGAGTCGTTGCCTTCATCTGTATTTATAGCCACTNNNAGATCGGAAGA |
| 13568_2 | TATGAGTGTGGAGTCGTTGCCTTCATCTGTATTTATAGCCACTNNNAGATCGGAAGA |
| 13568_3 | TATGAGTGTGGAGTCGTTGCCTTCATCTGTATTTATAGCCACTNNNAGATCGGAAGA |
| 38216_1 | TATGAGTGTGGAGTCGTTGCGGGCAACTTGCTTGATGAAAGTANNNAGATCGGAA |
| 38216_2 | TATGAGTGTGGAGTCGTTGCGGGCAACTTGCTTGATGAAAGTANNNAGATCGGAA |
| 38216_3 | TATGAGTGTGGAGTCGTTGCGGGCAACTTGCTTGATGAAAGTANNNAGATCGGAA |
| 48446_1 | TATGAGTGTGGAGTCGTTGCACATAGTGTCACTCTATTGCCCGNNNAGATCGGAAG |
| 48446_2 | TATGAGTGTGGAGTCGTTGCACATAGTGTCACTCTATTGCCCGNNNAGATCGGAAG |
| 48446_3 | TATGAGTGTGGAGTCGTTGCACATAGTGTCACTCTATTGCCCGNNNAGATCGGAAG |
| 10848_1 | TATGAGTGTGGAGTCGTTGCGTGTGGAGACAGAAGGAATGGAANNNAGATCGGA |
| 10848_2 | TATGAGTGTGGAGTCGTTGCGTGTGGAGACAGAAGGAATGGAANNNAGATCGGA |
| 10848_3 | TATGAGTGTGGAGTCGTTGCGTGTGGAGACAGAAGGAATGGAANNNAGATCGGA |
| 36409_1 | TATGAGTGTGGAGTCGTTGCGGCAGGAATCATCAATAAGTGCCNNNAGATCGGAA |
| 36409_2 | TATGAGTGTGGAGTCGTTGCGGCAGGAATCATCAATAAGTGCCNNNAGATCGGAA |
| 36409_3 | TATGAGTGTGGAGTCGTTGCGGCAGGAATCATCAATAAGTGCCNNNAGATCGGAA |
| 42118_1 | TATGAGTGTGGAGTCGTTGCCCAAGTTGCTGGGATTACAAGCNNNAGATCGGAAG |
| 42118_2 | TATGAGTGTGGAGTCGTTGCCCAAGTTGCTGGGATTACAAGCNNNAGATCGGAAG |
| 42118_3 | TATGAGTGTGGAGTCGTTGCCCAAGTTGCTGGGATTACAAGCNNNAGATCGGAAG |
| 51051_1 | TATGAGTGTGGAGTCGTTGCAGTTGGCCTTGAAACAAATCTGGNNNAGATCGGAA |
| 51051_2 | TATGAGTGTGGAGTCGTTGCAGTTGGCCTTGAAACAAATCTGGNNNAGATCGGAA |
| 51051_3 | TATGAGTGTGGAGTCGTTGCAGTTGGCCTTGAAACAAATCTGGNNNAGATCGGAA |
| 48153_1 | TATGAGTGTGGAGTCGTTGCCTCGCCCGGCTAATTTTTTGTATNNNAGATCGGAAG |
| 48153_2 | TATGAGTGTGGAGTCGTTGCCTCGCCCGGCTAATTTTTTGTATNNNAGATCGGAAG |
| 48153_3 | TATGAGTGTGGAGTCGTTGCCTCGCCCGGCTAATTTTTTGTATNNNAGATCGGAAG |
| 45354_1 | TATGAGTGTGGAGTCGTTGCTCTTCATCTTACTGCCTGTCACCNNNAGATCGGAAGA |
| 45354_2 | TATGAGTGTGGAGTCGTTGCTCTTCATCTTACTGCCTGTCACCNNNAGATCGGAAGA |
| 45354_3 | TATGAGTGTGGAGTCGTTGCTCTTCATCTTACTGCCTGTCACCNNNAGATCGGAAGA |
| 41907_1 | TATGAGTGTGGAGTCGTTGCGCCTCCTAAGTAGCTGGGATTACNNNAGATCGGAAG |
| 41907_2 | TATGAGTGTGGAGTCGTTGCGCCTCCTAAGTAGCTGGGATTACNNNAGATCGGAAG |
| 41907_3 | TATGAGTGTGGAGTCGTTGCGCCTCCTAAGTAGCTGGGATTACNNNAGATCGGAAG |
| 32753_1 | TATGAGTGTGGAGTCGTTGCTCACCTTGATAGCCATTTCGGAANNNAGATCGGAAG |
| 32753_2 | TATGAGTGTGGAGTCGTTGCTCACCTTGATAGCCATTTCGGAANNNAGATCGGAAG |
| 32753_3 | TATGAGTGTGGAGTCGTTGCTCACCTTGATAGCCATTTCGGAANNNAGATCGGAAG |
| 34165_1 | TATGAGTGTGGAGTCGTTGCGTGGGTGGAGAAAGGGGAAATATNNNAGATCGGA |
| 34165_2 | TATGAGTGTGGAGTCGTTGCGTGGGTGGAGAAAGGGGAAATATNNNAGATCGGA |
| 34165_3 | TATGAGTGTGGAGTCGTTGCGTGGGTGGAGAAAGGGGAAATATNNNAGATCGGA |

| | |
|---|---|
| 34489_1 | TATGAGTGTGGAGTCGTTGCTCTTTTTACTGGAACAGTCCCCNNNAGATCGGAAGA |
| 34489_2 | TATGAGTGTGGAGTCGTTGCTCTTTTTACTGGAACAGTCCCCNNNAGATCGGAAGA |
| 34489_3 | TATGAGTGTGGAGTCGTTGCTCTTTTTACTGGAACAGTCCCCNNNAGATCGGAAGA |
| 53253_1 | TATGAGTGTGGAGTCGTTGCTTTCTAACATGCCTCTTTGCAGGNNNAGATCGGAAGA |
| 53253_2 | TATGAGTGTGGAGTCGTTGCTTTCTAACATGCCTCTTTGCAGGNNNAGATCGGAAGA |
| 53253_3 | TATGAGTGTGGAGTCGTTGCTTTCTAACATGCCTCTTTGCAGGNNNAGATCGGAAGA |
| 33247_1 | TATGAGTGTGGAGTCGTTGCCTCCTGGGTTCAAGCGATTCTCNNNAGATCGGAAGA |
| 33247_2 | TATGAGTGTGGAGTCGTTGCCTCCTGGGTTCAAGCGATTCTCNNNAGATCGGAAGA |
| 33247_3 | TATGAGTGTGGAGTCGTTGCCTCCTGGGTTCAAGCGATTCTCNNNAGATCGGAAGA |
| 38611_1 | TATGAGTGTGGAGTCGTTGCCCAGCAGCACATCAAAAAGCTTANNNAGATCGGAAG |
| 38611_2 | TATGAGTGTGGAGTCGTTGCCCAGCAGCACATCAAAAAGCTTANNNAGATCGGAAG |
| 38611_3 | TATGAGTGTGGAGTCGTTGCCCAGCAGCACATCAAAAAGCTTANNNAGATCGGAAG |
| 9939_1 | TATGAGTGTGGAGTCGTTGCTTGGGTGATGAGGGATCCAAAGANNNAGATCGGAA |
| 9939_2 | TATGAGTGTGGAGTCGTTGCTTGGGTGATGAGGGATCCAAAGANNNAGATCGGAA |
| 9939_3 | TATGAGTGTGGAGTCGTTGCTTGGGTGATGAGGGATCCAAAGANNNAGATCGGAA |
| 52388_1 | TATGAGTGTGGAGTCGTTGCACTCTAACACTGACTGCCACAAANNNAGATCGGAAGA |
| 52388_2 | TATGAGTGTGGAGTCGTTGCACTCTAACACTGACTGCCACAAANNNAGATCGGAAGA |
| 52388_3 | TATGAGTGTGGAGTCGTTGCACTCTAACACTGACTGCCACAAANNNAGATCGGAAGA |
| 44628_1 | TATGAGTGTGGAGTCGTTGCCAGCCATGACTATACTCCCCAAANNNAGATCGGAAGA |
| 44628_2 | TATGAGTGTGGAGTCGTTGCCAGCCATGACTATACTCCCCAAANNNAGATCGGAAGA |
| 44628_3 | TATGAGTGTGGAGTCGTTGCCAGCCATGACTATACTCCCCAAANNNAGATCGGAAGA |
| 38356_1 | TATGAGTGTGGAGTCGTTGCAGAAAAAATCAGGCAGATGGCACNNNAGATCGGAA |
| 38356_2 | TATGAGTGTGGAGTCGTTGCAGAAAAAATCAGGCAGATGGCACNNNAGATCGGAA |
| 38356_3 | TATGAGTGTGGAGTCGTTGCAGAAAAAATCAGGCAGATGGCACNNNAGATCGGAA |
| 46117_1 | TATGAGTGTGGAGTCGTTGCCCTGTTGCCTGGATCTCAATACTNNNAGATCGGAAGA |
| 46117_2 | TATGAGTGTGGAGTCGTTGCCCTGTTGCCTGGATCTCAATACTNNNAGATCGGAAGA |
| 46117_3 | TATGAGTGTGGAGTCGTTGCCCTGTTGCCTGGATCTCAATACTNNNAGATCGGAAGA |
| 29909_1 | TATGAGTGTGGAGTCGTTGCTCACTGTTTAACTAGAAACACACCTNNNAGATCGGAA |
| 29909_2 | TATGAGTGTGGAGTCGTTGCTCACTGTTTAACTAGAAACACACCTNNNAGATCGGAA |
| 29909_3 | TATGAGTGTGGAGTCGTTGCTCACTGTTTAACTAGAAACACACCTNNNAGATCGGAA |
| 39049_1 | TATGAGTGTGGAGTCGTTGCAGCTGTGAATTTCCTAGGTTACGANNNAGATCGGAA |
| 39049_2 | TATGAGTGTGGAGTCGTTGCAGCTGTGAATTTCCTAGGTTACGANNNAGATCGGAA |
| 39049_3 | TATGAGTGTGGAGTCGTTGCAGCTGTGAATTTCCTAGGTTACGANNNAGATCGGAA |
| 13210_1 | TATGAGTGTGGAGTCGTTGCGTTTCCTTAAAGGCAGTTACNNNAGATCGGAAGAGC |
| 13210_2 | TATGAGTGTGGAGTCGTTGCGTTTCCTTAAAGGCAGTTACNNNAGATCGGAAGAGC |
| 13210_3 | TATGAGTGTGGAGTCGTTGCGTTTCCTTAAAGGCAGTTACNNNAGATCGGAAGAGC |
| 29151_1 | TATGAGTGTGGAGTCGTTGCTGTGTGTATGAGAGAGAGAAAGCTNNNAGATCGGA |
| 29151_2 | TATGAGTGTGGAGTCGTTGCTGTGTGTATGAGAGAGAGAAAGCTNNNAGATCGGA |
| 29151_3 | TATGAGTGTGGAGTCGTTGCTGTGTGTATGAGAGAGAGAAAGCTNNNAGATCGGA |
| 39521_1 | TATGAGTGTGGAGTCGTTGCCACCTGTGCTAAAGCCAGTTATGNNNAGATCGGAAG |
| 39521_2 | TATGAGTGTGGAGTCGTTGCCACCTGTGCTAAAGCCAGTTATGNNNAGATCGGAAG |
| 39521_3 | TATGAGTGTGGAGTCGTTGCCACCTGTGCTAAAGCCAGTTATGNNNAGATCGGAAG |
| 43721_1 | TATGAGTGTGGAGTCGTTGCACTGTGGTAACGGAGGGTCTATANNNAGATCGGAA |

| | |
|---|---|
| 43721_2 | TATGAGTGTGGAGTCGTTGCACTGTGGTAACGGAGGGTCTATANNNAGATCGGAA |
| 43721_3 | TATGAGTGTGGAGTCGTTGCACTGTGGTAACGGAGGGTCTATANNNAGATCGGAA |
| 42570_1 | TATGAGTGTGGAGTCGTTGCTTTGGTATCAGCATGATGTTGGCNNNAGATCGGAAG |
| 42570_2 | TATGAGTGTGGAGTCGTTGCTTTGGTATCAGCATGATGTTGGCNNNAGATCGGAAG |
| 42570_3 | TATGAGTGTGGAGTCGTTGCTTTGGTATCAGCATGATGTTGGCNNNAGATCGGAAG |
| 51750_1 | TATGAGTGTGGAGTCGTTGCACGATCTCTGCTCATACATGCAANNNAGATCGGAAG |
| 51750_2 | TATGAGTGTGGAGTCGTTGCACGATCTCTGCTCATACATGCAANNNAGATCGGAAG |
| 51750_3 | TATGAGTGTGGAGTCGTTGCACGATCTCTGCTCATACATGCAANNNAGATCGGAAG |
| 34323_1 | TATGAGTGTGGAGTCGTTGCAACCCATAAAGCCATCCATAGCANNNAGATCGGAAG |
| 34323_2 | TATGAGTGTGGAGTCGTTGCAACCCATAAAGCCATCCATAGCANNNAGATCGGAAG |
| 34323_3 | TATGAGTGTGGAGTCGTTGCAACCCATAAAGCCATCCATAGCANNNAGATCGGAAG |
| 28990_1 | TATGAGTGTGGAGTCGTTGCGCCAGACACTTTGGTCTTTGATCNNNAGATCGGAAG |
| 28990_2 | TATGAGTGTGGAGTCGTTGCGCCAGACACTTTGGTCTTTGATCNNNAGATCGGAAG |
| 28990_3 | TATGAGTGTGGAGTCGTTGCGCCAGACACTTTGGTCTTTGATCNNNAGATCGGAAG |
| 45197_1 | TATGAGTGTGGAGTCGTTGCACTGAGAGGGACCCATCATAAGANNNAGATCGGAA |
| 45197_2 | TATGAGTGTGGAGTCGTTGCACTGAGAGGGACCCATCATAAGANNNAGATCGGAA |
| 45197_3 | TATGAGTGTGGAGTCGTTGCACTGAGAGGGACCCATCATAAGANNNAGATCGGAA |
| 52516_1 | TATGAGTGTGGAGTCGTTGCGGTCATCACCTCATCTTCTCTGGNNNAGATCGGAAG |
| 52516_2 | TATGAGTGTGGAGTCGTTGCGGTCATCACCTCATCTTCTCTGGNNNAGATCGGAAG |
| 52516_3 | TATGAGTGTGGAGTCGTTGCGGTCATCACCTCATCTTCTCTGGNNNAGATCGGAAG |
| 49867_1 | TATGAGTGTGGAGTCGTTGCGAGGTGAGGAAAACTCAGCTCTCNNNAGATCGGAA |
| 49867_2 | TATGAGTGTGGAGTCGTTGCGAGGTGAGGAAAACTCAGCTCTCNNNAGATCGGAA |
| 49867_3 | TATGAGTGTGGAGTCGTTGCGAGGTGAGGAAAACTCAGCTCTCNNNAGATCGGAA |
| 52917_1 | TATGAGTGTGGAGTCGTTGCTTCTCTGTGAAGGTGACAGGCNNNAGATCGGAAGA |
| 52917_2 | TATGAGTGTGGAGTCGTTGCTTCTCTGTGAAGGTGACAGGCNNNAGATCGGAAGA |
| 52917_3 | TATGAGTGTGGAGTCGTTGCTTCTCTGTGAAGGTGACAGGCNNNAGATCGGAAGA |
| 49705_1 | TATGAGTGTGGAGTCGTTGCAGGAAATTGACCACTGCGAATTGNNNAGATCGGAA |
| 49705_2 | TATGAGTGTGGAGTCGTTGCAGGAAATTGACCACTGCGAATTGNNNAGATCGGAA |
| 49705_3 | TATGAGTGTGGAGTCGTTGCAGGAAATTGACCACTGCGAATTGNNNAGATCGGAA |
| 36485_1 | TATGAGTGTGGAGTCGTTGCCTGAGTGAAAAAGCCCCTCCTATNNNAGATCGGAAG |
| 36485_2 | TATGAGTGTGGAGTCGTTGCCTGAGTGAAAAAGCCCCTCCTATNNNAGATCGGAAG |
| 36485_3 | TATGAGTGTGGAGTCGTTGCCTGAGTGAAAAAGCCCCTCCTATNNNAGATCGGAAG |
| 35456_1 | TATGAGTGTGGAGTCGTTGCAACTCAGCTTCTGACTCTTTGGTNNNAGATCGGAAG |
| 35456_2 | TATGAGTGTGGAGTCGTTGCAACTCAGCTTCTGACTCTTTGGTNNNAGATCGGAAG |
| 35456_3 | TATGAGTGTGGAGTCGTTGCAACTCAGCTTCTGACTCTTTGGTNNNAGATCGGAAG |
| 43448_1 | TATGAGTGTGGAGTCGTTGCCTATTTTGGCTCACTGCAACCTCNNNAGATCGGAAG |
| 43448_2 | TATGAGTGTGGAGTCGTTGCCTATTTTGGCTCACTGCAACCTCNNNAGATCGGAAG |
| 43448_3 | TATGAGTGTGGAGTCGTTGCCTATTTTGGCTCACTGCAACCTCNNNAGATCGGAAG |
| 48897_1 | TATGAGTGTGGAGTCGTTGCCCAAGTTAGAGTGCAGTAGTGGTNNNAGATCGGAA |
| 48897_2 | TATGAGTGTGGAGTCGTTGCCCAAGTTAGAGTGCAGTAGTGGTNNNAGATCGGAA |
| 48897_3 | TATGAGTGTGGAGTCGTTGCCCAAGTTAGAGTGCAGTAGTGGTNNNAGATCGGAA |
| 48111_1 | TATGAGTGTGGAGTCGTTGCTTCGAGGTGTACGTGGATAAGTCNNNAGATCGGAA |
| 48111_2 | TATGAGTGTGGAGTCGTTGCTTCGAGGTGTACGTGGATAAGTCNNNAGATCGGAA |

| | |
|---|---|
| 48111_3 | TATGAGTGTGGAGTCGTTGCTTCGAGGTGTACGTGGATAAGTCNNNAGATCGGAA |
| 32416_1 | TATGAGTGTGGAGTCGTTGCAGTTTCTGGCATTTCCCTACAGTNNNAGATCGGAAG |
| 32416_2 | TATGAGTGTGGAGTCGTTGCAGTTTCTGGCATTTCCCTACAGTNNNAGATCGGAAG |
| 32416_3 | TATGAGTGTGGAGTCGTTGCAGTTTCTGGCATTTCCCTACAGTNNNAGATCGGAAG |
| 29203_1 | TATGAGTGTGGAGTCGTTGCTCCACACACCACATTCAAGAACTNNNAGATCGGAAG |
| 29203_2 | TATGAGTGTGGAGTCGTTGCTCCACACACCACATTCAAGAACTNNNAGATCGGAAG |
| 29203_3 | TATGAGTGTGGAGTCGTTGCTCCACACACCACATTCAAGAACTNNNAGATCGGAAG |
| 40819_1 | TATGAGTGTGGAGTCGTTGCTCTTCCTCAAGAAACTGGTCCTGNNNAGATCGGAAG |
| 40819_2 | TATGAGTGTGGAGTCGTTGCTCTTCCTCAAGAAACTGGTCCTGNNNAGATCGGAAG |
| 40819_3 | TATGAGTGTGGAGTCGTTGCTCTTCCTCAAGAAACTGGTCCTGNNNAGATCGGAAG |
| 37795_1 | TATGAGTGTGGAGTCGTTGCACAAAATTATCCTGGCATGGTGGNNNAGATCGGAA |
| 37795_2 | TATGAGTGTGGAGTCGTTGCACAAAATTATCCTGGCATGGTGGNNNAGATCGGAA |
| 37795_3 | TATGAGTGTGGAGTCGTTGCACAAAATTATCCTGGCATGGTGGNNNAGATCGGAA |
| 45057_1 | TATGAGTGTGGAGTCGTTGCGTCATGTCTGAGTGTAGGCAGAANNNAGATCGGAA |
| 45057_2 | TATGAGTGTGGAGTCGTTGCGTCATGTCTGAGTGTAGGCAGAANNNAGATCGGAA |
| 45057_3 | TATGAGTGTGGAGTCGTTGCGTCATGTCTGAGTGTAGGCAGAANNNAGATCGGAA |
| 48879_1 | TATGAGTGTGGAGTCGTTGCGCCACCATGCCTAGCTAATTTTTNNNAGATCGGAAG |
| 48879_2 | TATGAGTGTGGAGTCGTTGCGCCACCATGCCTAGCTAATTTTTNNNAGATCGGAAG |
| 48879_3 | TATGAGTGTGGAGTCGTTGCGCCACCATGCCTAGCTAATTTTTNNNAGATCGGAAG |
| 41127_1 | TATGAGTGTGGAGTCGTTGCATGTACTATACCCAGTCCGTGGANNNAGATCGGAAG |
| 41127_2 | TATGAGTGTGGAGTCGTTGCATGTACTATACCCAGTCCGTGGANNNAGATCGGAAG |
| 41127_3 | TATGAGTGTGGAGTCGTTGCATGTACTATACCCAGTCCGTGGANNNAGATCGGAAG |
| 12521_1 | TATGAGTGTGGAGTCGTTGCTGACTGAGAAAGACCTTTTAATNNNAGATCGGAAGA |
| 12521_2 | TATGAGTGTGGAGTCGTTGCTGACTGAGAAAGACCTTTTAATNNNAGATCGGAAGA |
| 12521_3 | TATGAGTGTGGAGTCGTTGCTGACTGAGAAAGACCTTTTAATNNNAGATCGGAAGA |
| 32961_1 | TATGAGTGTGGAGTCGTTGCCTGAGGCAGGAGAATCACTTGAANNNAGATCGGAA |
| 32961_2 | TATGAGTGTGGAGTCGTTGCCTGAGGCAGGAGAATCACTTGAANNNAGATCGGAA |
| 32961_3 | TATGAGTGTGGAGTCGTTGCCTGAGGCAGGAGAATCACTTGAANNNAGATCGGAA |
| 42480_1 | TATGAGTGTGGAGTCGTTGCCAGGAAGACTCAGGGAGAAGATCNNNAGATCGGA |
| 42480_2 | TATGAGTGTGGAGTCGTTGCCAGGAAGACTCAGGGAGAAGATCNNNAGATCGGA |
| 42480_3 | TATGAGTGTGGAGTCGTTGCCAGGAAGACTCAGGGAGAAGATCNNNAGATCGGA |
| 52276_1 | TATGAGTGTGGAGTCGTTGCCTTCTTCCACAACCCTTCTGAGTNNNAGATCGGAAGA |
| 52276_2 | TATGAGTGTGGAGTCGTTGCCTTCTTCCACAACCCTTCTGAGTNNNAGATCGGAAGA |
| 52276_3 | TATGAGTGTGGAGTCGTTGCCTTCTTCCACAACCCTTCTGAGTNNNAGATCGGAAGA |
| 48116_1 | TATGAGTGTGGAGTCGTTGCCTGGTCCTTCCCTCTCTCTTTTGNNNAGATCGGAAGA |
| 48116_2 | TATGAGTGTGGAGTCGTTGCCTGGTCCTTCCCTCTCTCTTTTGNNNAGATCGGAAGA |
| 48116_3 | TATGAGTGTGGAGTCGTTGCCTGGTCCTTCCCTCTCTCTTTTGNNNAGATCGGAAGA |
| 45505_1 | TATGAGTGTGGAGTCGTTGCCATTGTCACGGTGATTCACATCTNNNAGATCGGAAG |
| 45505_2 | TATGAGTGTGGAGTCGTTGCCATTGTCACGGTGATTCACATCTNNNAGATCGGAAG |
| 45505_3 | TATGAGTGTGGAGTCGTTGCCATTGTCACGGTGATTCACATCTNNNAGATCGGAAG |
| 36635_1 | TATGAGTGTGGAGTCGTTGCATAATCCCAGCTACTTGGAAGGCNNNAGATCGGAAG |
| 36635_2 | TATGAGTGTGGAGTCGTTGCATAATCCCAGCTACTTGGAAGGCNNNAGATCGGAAG |
| 36635_3 | TATGAGTGTGGAGTCGTTGCATAATCCCAGCTACTTGGAAGGCNNNAGATCGGAAG |

| | |
|---|---|
| 47262_1 | TATGAGTGTGGAGTCGTTGCCTCACTCCTGGCTGTTTGATGAANNNAGATCGGAAG |
| 47262_2 | TATGAGTGTGGAGTCGTTGCCTCACTCCTGGCTGTTTGATGAANNNAGATCGGAAG |
| 47262_3 | TATGAGTGTGGAGTCGTTGCCTCACTCCTGGCTGTTTGATGAANNNAGATCGGAAG |
| 34546_1 | TATGAGTGTGGAGTCGTTGCCCATGAAACTAGCTAGGTAGGGGNNNAGATCGGAA |
| 34546_2 | TATGAGTGTGGAGTCGTTGCCCATGAAACTAGCTAGGTAGGGGNNNAGATCGGAA |
| 34546_3 | TATGAGTGTGGAGTCGTTGCCCATGAAACTAGCTAGGTAGGGGNNNAGATCGGAA |
| 34846_1 | TATGAGTGTGGAGTCGTTGCCCCCTACAGTCACAGAAACTACANNNAGATCGGAAG |
| 34846_2 | TATGAGTGTGGAGTCGTTGCCCCCTACAGTCACAGAAACTACANNNAGATCGGAAG |
| 34846_3 | TATGAGTGTGGAGTCGTTGCCCCCTACAGTCACAGAAACTACANNNAGATCGGAAG |
| 44209_1 | TATGAGTGTGGAGTCGTTGCGAGGAAATAGCTGAGAGGGCATTNNNAGATCGGA |
| 44209_2 | TATGAGTGTGGAGTCGTTGCGAGGAAATAGCTGAGAGGGCATTNNNAGATCGGA |
| 44209_3 | TATGAGTGTGGAGTCGTTGCGAGGAAATAGCTGAGAGGGCATTNNNAGATCGGA |
| 40213_1 | TATGAGTGTGGAGTCGTTGCCACAGTCACTTCTCGTTGTTGACNNNAGATCGGAAG |
| 40213_2 | TATGAGTGTGGAGTCGTTGCCACAGTCACTTCTCGTTGTTGACNNNAGATCGGAAG |
| 40213_3 | TATGAGTGTGGAGTCGTTGCCACAGTCACTTCTCGTTGTTGACNNNAGATCGGAAG |
| 10026_1 | TATGAGTGTGGAGTCGTTGCATGAGGTCTTTCCCAGTGGTCTTNNNAGATCGGAAG |
| 10026_2 | TATGAGTGTGGAGTCGTTGCATGAGGTCTTTCCCAGTGGTCTTNNNAGATCGGAAG |
| 10026_3 | TATGAGTGTGGAGTCGTTGCATGAGGTCTTTCCCAGTGGTCTTNNNAGATCGGAAG |
| 48777_1 | TATGAGTGTGGAGTCGTTGCATTTGTGACATCCTGGTGTACCCNNNAGATCGGAAG |
| 48777_2 | TATGAGTGTGGAGTCGTTGCATTTGTGACATCCTGGTGTACCCNNNAGATCGGAAG |
| 48777_3 | TATGAGTGTGGAGTCGTTGCATTTGTGACATCCTGGTGTACCCNNNAGATCGGAAG |
| 45530_1 | TATGAGTGTGGAGTCGTTGCGGGGGGAGAAGCACAAAATAATGNNNAGATCGGA |
| 45530_2 | TATGAGTGTGGAGTCGTTGCGGGGGGAGAAGCACAAAATAATGNNNAGATCGGA |
| 45530_3 | TATGAGTGTGGAGTCGTTGCGGGGGGAGAAGCACAAAATAATGNNNAGATCGGA |
| 29952_1 | TATGAGTGTGGAGTCGTTGCAACATGTGAAGACTGGATTGCTGNNNAGATCGGAA |
| 29952_2 | TATGAGTGTGGAGTCGTTGCAACATGTGAAGACTGGATTGCTGNNNAGATCGGAA |
| 29952_3 | TATGAGTGTGGAGTCGTTGCAACATGTGAAGACTGGATTGCTGNNNAGATCGGAA |
| 10683_1 | TATGAGTGTGGAGTCGTTGCATGACTTGGAGAACAGCATGTCCNNNAGATCGGAA |
| 10683_2 | TATGAGTGTGGAGTCGTTGCATGACTTGGAGAACAGCATGTCCNNNAGATCGGAA |
| 10683_3 | TATGAGTGTGGAGTCGTTGCATGACTTGGAGAACAGCATGTCCNNNAGATCGGAA |
| 2201_1 | TATGAGTGTGGAGTCGTTGCGGGTCCAAATATGTGGCATTGGANNNAGATCGGAA |
| 2201_2 | TATGAGTGTGGAGTCGTTGCGGGTCCAAATATGTGGCATTGGANNNAGATCGGAA |
| 2201_3 | TATGAGTGTGGAGTCGTTGCGGGTCCAAATATGTGGCATTGGANNNAGATCGGAA |
| 33695_1 | TATGAGTGTGGAGTCGTTGCGTATTGATGGGATGTCTTGCAGCNNNAGATCGGAA |
| 33695_2 | TATGAGTGTGGAGTCGTTGCGTATTGATGGGATGTCTTGCAGCNNNAGATCGGAA |
| 33695_3 | TATGAGTGTGGAGTCGTTGCGTATTGATGGGATGTCTTGCAGCNNNAGATCGGAA |
| 10842_1 | TATGAGTGTGGAGTCGTTGCCTAAAATAGGCTGCAGCGATCCANNNAGATCGGAA |
| 10842_2 | TATGAGTGTGGAGTCGTTGCCTAAAATAGGCTGCAGCGATCCANNNAGATCGGAA |
| 10842_3 | TATGAGTGTGGAGTCGTTGCCTAAAATAGGCTGCAGCGATCCANNNAGATCGGAA |
| 50065_1 | TATGAGTGTGGAGTCGTTGCACACACACACACACACATGAAAANNNAGATCGGAAG |
| 50065_2 | TATGAGTGTGGAGTCGTTGCACACACACACACACACATGAAAANNNAGATCGGAAG |
| 50065_3 | TATGAGTGTGGAGTCGTTGCACACACACACACACACATGAAAANNNAGATCGGAAG |
| 31432_1 | TATGAGTGTGGAGTCGTTGCATTGAGAAACAGAGCCTCTGAGGNNNAGATCGGAA |

| | |
|---|---|
| 31432_2 | TATGAGTGTGGAGTCGTTGCATTGAGAAACAGAGCCTCTGAGGNNNAGATCGGAA |
| 31432_3 | TATGAGTGTGGAGTCGTTGCATTGAGAAACAGAGCCTCTGAGGNNNAGATCGGAA |
| 45138_1 | TATGAGTGTGGAGTCGTTGCAATCACAAGCACTTTTCCTGCAGNNNAGATCGGAAG |
| 45138_2 | TATGAGTGTGGAGTCGTTGCAATCACAAGCACTTTTCCTGCAGNNNAGATCGGAAG |
| 45138_3 | TATGAGTGTGGAGTCGTTGCAATCACAAGCACTTTTCCTGCAGNNNAGATCGGAAG |
| 48579_1 | TATGAGTGTGGAGTCGTTGCTTCAAGTGATTCTCCTGCCTCAGNNNAGATCGGAAG |
| 48579_2 | TATGAGTGTGGAGTCGTTGCTTCAAGTGATTCTCCTGCCTCAGNNNAGATCGGAAG |
| 48579_3 | TATGAGTGTGGAGTCGTTGCTTCAAGTGATTCTCCTGCCTCAGNNNAGATCGGAAG |
| 36673_1 | TATGAGTGTGGAGTCGTTGCCTGGGTGACAGAGTGAGATTCTGNNNAGATCGGAA |
| 36673_2 | TATGAGTGTGGAGTCGTTGCCTGGGTGACAGAGTGAGATTCTGNNNAGATCGGAA |
| 36673_3 | TATGAGTGTGGAGTCGTTGCCTGGGTGACAGAGTGAGATTCTGNNNAGATCGGAA |
| 30081_1 | TATGAGTGTGGAGTCGTTGCTATGTCCTTTCCACACTTGCCTTNNNAGATCGGAAGA |
| 30081_2 | TATGAGTGTGGAGTCGTTGCTATGTCCTTTCCACACTTGCCTTNNNAGATCGGAAGA |
| 30081_3 | TATGAGTGTGGAGTCGTTGCTATGTCCTTTCCACACTTGCCTTNNNAGATCGGAAGA |
| 30734_1 | TATGAGTGTGGAGTCGTTGCACACAGACAAACAGGTTAGTGGTNNNAGATCGGAA |
| 30734_2 | TATGAGTGTGGAGTCGTTGCACACAGACAAACAGGTTAGTGGTNNNAGATCGGAA |
| 30734_3 | TATGAGTGTGGAGTCGTTGCACACAGACAAACAGGTTAGTGGTNNNAGATCGGAA |
| 49893_1 | TATGAGTGTGGAGTCGTTGCCTGCAGATCCCAAGATCCTTTCTNNNAGATCGGAAG |
| 49893_2 | TATGAGTGTGGAGTCGTTGCCTGCAGATCCCAAGATCCTTTCTNNNAGATCGGAAG |
| 49893_3 | TATGAGTGTGGAGTCGTTGCCTGCAGATCCCAAGATCCTTTCTNNNAGATCGGAAG |
| 49152_1 | TATGAGTGTGGAGTCGTTGCCATTCTCCTGCCTCAGCCTCNNNAGATCGGAAGAGCA |
| 49152_2 | TATGAGTGTGGAGTCGTTGCCATTCTCCTGCCTCAGCCTCNNNAGATCGGAAGAGCA |
| 49152_3 | TATGAGTGTGGAGTCGTTGCCATTCTCCTGCCTCAGCCTCNNNAGATCGGAAGAGCA |
| 49685_1 | TATGAGTGTGGAGTCGTTGCAGACGGAGTCTCTCTCTGTCACNNNAGATCGGAAGA |
| 49685_2 | TATGAGTGTGGAGTCGTTGCAGACGGAGTCTCTCTCTGTCACNNNAGATCGGAAGA |
| 49685_3 | TATGAGTGTGGAGTCGTTGCAGACGGAGTCTCTCTCTGTCACNNNAGATCGGAAGA |
| 4107_1 | TATGAGTGTGGAGTCGTTGCGGGGAAAACGTCATAGTGGCTTANNNAGATCGGAA |
| 4107_2 | TATGAGTGTGGAGTCGTTGCGGGGAAAACGTCATAGTGGCTTANNNAGATCGGAA |
| 4107_3 | TATGAGTGTGGAGTCGTTGCGGGGAAAACGTCATAGTGGCTTANNNAGATCGGAA |
| 45108_1 | TATGAGTGTGGAGTCGTTGCCCATGGACTGTACTGCATCTCAANNNAGATCGGAAG |
| 45108_2 | TATGAGTGTGGAGTCGTTGCCCATGGACTGTACTGCATCTCAANNNAGATCGGAAG |
| 45108_3 | TATGAGTGTGGAGTCGTTGCCCATGGACTGTACTGCATCTCAANNNAGATCGGAAG |
| 29292_1 | TATGAGTGTGGAGTCGTTGCTTTCTGGTCGATGATTTCTGGGTNNNAGATCGGAAG |
| 29292_2 | TATGAGTGTGGAGTCGTTGCTTTCTGGTCGATGATTTCTGGGTNNNAGATCGGAAG |
| 29292_3 | TATGAGTGTGGAGTCGTTGCTTTCTGGTCGATGATTTCTGGGTNNNAGATCGGAAG |
| 28752_1 | TATGAGTGTGGAGTCGTTGCGACAGTGGAAATGAGAGGGAACTNNNAGATCGGA |
| 28752_2 | TATGAGTGTGGAGTCGTTGCGACAGTGGAAATGAGAGGGAACTNNNAGATCGGA |
| 28752_3 | TATGAGTGTGGAGTCGTTGCGACAGTGGAAATGAGAGGGAACTNNNAGATCGGA |
| 7717_1 | TATGAGTGTGGAGTCGTTGCCAGGCTACAGTGGTGCAATCTCAANNNAGATCGGA |
| 7717_2 | TATGAGTGTGGAGTCGTTGCCAGGCTACAGTGGTGCAATCTCAANNNAGATCGGA |
| 7717_3 | TATGAGTGTGGAGTCGTTGCCAGGCTACAGTGGTGCAATCTCAANNNAGATCGGA |
| 41135_1 | TATGAGTGTGGAGTCGTTGCTGAGACCAGGAATTCAAGACCAGNNNAGATCGGAA |
| 41135_2 | TATGAGTGTGGAGTCGTTGCTGAGACCAGGAATTCAAGACCAGNNNAGATCGGAA |

| | |
|---|---|
| 41135_3 | TATGAGTGTGGAGTCGTTGCTGAGACCAGGAATTCAAGACCAGNNNAGATCGGAA |
| 49054_1 | TATGAGTGTGGAGTCGTTGCCCTTAGCCTCTTGAGTATGGGACNNNAGATCGGAAG |
| 49054_2 | TATGAGTGTGGAGTCGTTGCCCTTAGCCTCTTGAGTATGGGACNNNAGATCGGAAG |
| 49054_3 | TATGAGTGTGGAGTCGTTGCCCTTAGCCTCTTGAGTATGGGACNNNAGATCGGAAG |
| 39147_1 | TATGAGTGTGGAGTCGTTGCAAAATGGCCATACTGCACAAAGTNNNAGATCGGAAG |
| 39147_2 | TATGAGTGTGGAGTCGTTGCAAAATGGCCATACTGCACAAAGTNNNAGATCGGAAG |
| 39147_3 | TATGAGTGTGGAGTCGTTGCAAAATGGCCATACTGCACAAAGTNNNAGATCGGAAG |
| 7866_1 | TATGAGTGTGGAGTCGTTGCACGCATGAGAGTAAAGGCTCTGNNNAGATCGGAAG |
| 7866_2 | TATGAGTGTGGAGTCGTTGCACGCATGAGAGTAAAGGCTCTGNNNAGATCGGAAG |
| 7866_3 | TATGAGTGTGGAGTCGTTGCACGCATGAGAGTAAAGGCTCTGNNNAGATCGGAAG |
| 44167_1 | TATGAGTGTGGAGTCGTTGCTCCAGTATCCATTCTCCCCTTCTNNNAGATCGGAAGA |
| 44167_2 | TATGAGTGTGGAGTCGTTGCTCCAGTATCCATTCTCCCCTTCTNNNAGATCGGAAGA |
| 44167_3 | TATGAGTGTGGAGTCGTTGCTCCAGTATCCATTCTCCCCTTCTNNNAGATCGGAAGA |
| 41577_1 | TATGAGTGTGGAGTCGTTGCTTCAAGTGATCTCCTGTCTCAGCNNNAGATCGGAAG |
| 41577_2 | TATGAGTGTGGAGTCGTTGCTTCAAGTGATCTCCTGTCTCAGCNNNAGATCGGAAG |
| 41577_3 | TATGAGTGTGGAGTCGTTGCTTCAAGTGATCTCCTGTCTCAGCNNNAGATCGGAAG |
| 33020_1 | TATGAGTGTGGAGTCGTTGCCAAGTAGCTGGGATTACAGGCATNNNAGATCGGAA |
| 33020_2 | TATGAGTGTGGAGTCGTTGCCAAGTAGCTGGGATTACAGGCATNNNAGATCGGAA |
| 33020_3 | TATGAGTGTGGAGTCGTTGCCAAGTAGCTGGGATTACAGGCATNNNAGATCGGAA |
| 38875_1 | TATGAGTGTGGAGTCGTTGCTGCAAGGATAAACAAAATTGACACANNNAGATCGG |
| 38875_2 | TATGAGTGTGGAGTCGTTGCTGCAAGGATAAACAAAATTGACACANNNAGATCGG |
| 38875_3 | TATGAGTGTGGAGTCGTTGCTGCAAGGATAAACAAAATTGACACANNNAGATCGG |
| 45378_1 | TATGAGTGTGGAGTCGTTGCGCACAAGCATATTGGTGTACACANNNAGATCGGAAG |
| 45378_2 | TATGAGTGTGGAGTCGTTGCGCACAAGCATATTGGTGTACACANNNAGATCGGAAG |
| 45378_3 | TATGAGTGTGGAGTCGTTGCGCACAAGCATATTGGTGTACACANNNAGATCGGAAG |
| 42719_1 | TATGAGTGTGGAGTCGTTGCACAGTAGCTTATACCCTGCATCCNNNAGATCGGAAG |
| 42719_2 | TATGAGTGTGGAGTCGTTGCACAGTAGCTTATACCCTGCATCCNNNAGATCGGAAG |
| 42719_3 | TATGAGTGTGGAGTCGTTGCACAGTAGCTTATACCCTGCATCCNNNAGATCGGAAG |
| 49672_1 | TATGAGTGTGGAGTCGTTGCGATCTCGGCTCACTGCAACCNNNAGATCGGAAGAGC |
| 49672_2 | TATGAGTGTGGAGTCGTTGCGATCTCGGCTCACTGCAACCNNNAGATCGGAAGAGC |
| 49672_3 | TATGAGTGTGGAGTCGTTGCGATCTCGGCTCACTGCAACCNNNAGATCGGAAGAGC |
| 51458_1 | TATGAGTGTGGAGTCGTTGCTTTCTTGCACACTCTCTCCTCAANNNAGATCGGAAGA |
| 51458_2 | TATGAGTGTGGAGTCGTTGCTTTCTTGCACACTCTCTCCTCAANNNAGATCGGAAGA |
| 51458_3 | TATGAGTGTGGAGTCGTTGCTTTCTTGCACACTCTCTCCTCAANNNAGATCGGAAGA |
| 28905_1 | TATGAGTGTGGAGTCGTTGCTGGGAATGCACTGCTTACCTTNNNAGATCGGAAGAG |
| 28905_2 | TATGAGTGTGGAGTCGTTGCTGGGAATGCACTGCTTACCTTNNNAGATCGGAAGAG |
| 28905_3 | TATGAGTGTGGAGTCGTTGCTGGGAATGCACTGCTTACCTTNNNAGATCGGAAGAG |
| 12713_1 | TATGAGTGTGGAGTCGTTGCGATGGGTTCATTATCAAAGATTTNNNAGATCGGAAG |
| 12713_2 | TATGAGTGTGGAGTCGTTGCGATGGGTTCATTATCAAAGATTTNNNAGATCGGAAG |
| 12713_3 | TATGAGTGTGGAGTCGTTGCGATGGGTTCATTATCAAAGATTTNNNAGATCGGAAG |
| 43712_1 | TATGAGTGTGGAGTCGTTGCTGATTCACAGGGTTCGCTAGATTNNNAGATCGGAAG |
| 43712_2 | TATGAGTGTGGAGTCGTTGCTGATTCACAGGGTTCGCTAGATTNNNAGATCGGAAG |
| 43712_3 | TATGAGTGTGGAGTCGTTGCTGATTCACAGGGTTCGCTAGATTNNNAGATCGGAAG |

| | |
|---|---|
| 29764_1 | TATGAGTGTGGAGTCGTTGCTGAAAGAAAGGAGAGGTAACTGAANNNAGATCGG |
| 29764_2 | TATGAGTGTGGAGTCGTTGCTGAAAGAAAGGAGAGGTAACTGAANNNAGATCGG |
| 29764_3 | TATGAGTGTGGAGTCGTTGCTGAAAGAAAGGAGAGGTAACTGAANNNAGATCGG |
| 41326_1 | TATGAGTGTGGAGTCGTTGCATGCTAGAGAGGTTGTGGAGAACNNNAGATCGGAA |
| 41326_2 | TATGAGTGTGGAGTCGTTGCATGCTAGAGAGGTTGTGGAGAACNNNAGATCGGAA |
| 41326_3 | TATGAGTGTGGAGTCGTTGCATGCTAGAGAGGTTGTGGAGAACNNNAGATCGGAA |
| 45977_1 | TATGAGTGTGGAGTCGTTGCGAGTGTAGTGGCATGATCTCGATNNNAGATCGGAA |
| 45977_2 | TATGAGTGTGGAGTCGTTGCGAGTGTAGTGGCATGATCTCGATNNNAGATCGGAA |
| 45977_3 | TATGAGTGTGGAGTCGTTGCGAGTGTAGTGGCATGATCTCGATNNNAGATCGGAA |
| 53020_1 | TATGAGTGTGGAGTCGTTGCAGTATTAGGTGCCGTAGGTAACGNNNAGATCGGAA |
| 53020_2 | TATGAGTGTGGAGTCGTTGCAGTATTAGGTGCCGTAGGTAACGNNNAGATCGGAA |
| 53020_3 | TATGAGTGTGGAGTCGTTGCAGTATTAGGTGCCGTAGGTAACGNNNAGATCGGAA |
| 32108_1 | TATGAGTGTGGAGTCGTTGCAAACTCTTGACCTCAGGTGATCCNNNAGATCGGAAG |
| 32108_2 | TATGAGTGTGGAGTCGTTGCAAACTCTTGACCTCAGGTGATCCNNNAGATCGGAAG |
| 32108_3 | TATGAGTGTGGAGTCGTTGCAAACTCTTGACCTCAGGTGATCCNNNAGATCGGAAG |
| 13534_1 | TATGAGTGTGGAGTCGTTGCTGCTATCAAAGGATAAAAAATGANNNAGATCGGAA |
| 13534_2 | TATGAGTGTGGAGTCGTTGCTGCTATCAAAGGATAAAAAATGANNNAGATCGGAA |
| 13534_3 | TATGAGTGTGGAGTCGTTGCTGCTATCAAAGGATAAAAAATGANNNAGATCGGAA |
| 52523_1 | TATGAGTGTGGAGTCGTTGCTTGATGGGAAAACCAAGGGAGTCNNNAGATCGGAA |
| 52523_2 | TATGAGTGTGGAGTCGTTGCTTGATGGGAAAACCAAGGGAGTCNNNAGATCGGAA |
| 52523_3 | TATGAGTGTGGAGTCGTTGCTTGATGGGAAAACCAAGGGAGTCNNNAGATCGGAA |
| 47588_1 | TATGAGTGTGGAGTCGTTGCCTCCTGAGTAGCTGGGATTACAGNNNAGATCGGAA |
| 47588_2 | TATGAGTGTGGAGTCGTTGCCTCCTGAGTAGCTGGGATTACAGNNNAGATCGGAA |
| 47588_3 | TATGAGTGTGGAGTCGTTGCCTCCTGAGTAGCTGGGATTACAGNNNAGATCGGAA |
| 46130_1 | TATGAGTGTGGAGTCGTTGCAGATATGTGCTTGCCTGGAGAAANNNAGATCGGAA |
| 46130_2 | TATGAGTGTGGAGTCGTTGCAGATATGTGCTTGCCTGGAGAAANNNAGATCGGAA |
| 46130_3 | TATGAGTGTGGAGTCGTTGCAGATATGTGCTTGCCTGGAGAAANNNAGATCGGAA |
| 30609_1 | TATGAGTGTGGAGTCGTTGCTGCTGGCTACATCCTCTGTTGNNNAGATCGGAAGAG |
| 30609_2 | TATGAGTGTGGAGTCGTTGCTGCTGGCTACATCCTCTGTTGNNNAGATCGGAAGAG |
| 30609_3 | TATGAGTGTGGAGTCGTTGCTGCTGGCTACATCCTCTGTTGNNNAGATCGGAAGAG |
| 38010_1 | TATGAGTGTGGAGTCGTTGCTCAGAGGACAGTGAGGATCTCTTNNNAGATCGGAA |
| 38010_2 | TATGAGTGTGGAGTCGTTGCTCAGAGGACAGTGAGGATCTCTTNNNAGATCGGAA |
| 38010_3 | TATGAGTGTGGAGTCGTTGCTCAGAGGACAGTGAGGATCTCTTNNNAGATCGGAA |
| 50300_1 | TATGAGTGTGGAGTCGTTGCCTCCCTTCATCCTGCAGCTTTTCNNNAGATCGGAAGA |
| 50300_2 | TATGAGTGTGGAGTCGTTGCCTCCCTTCATCCTGCAGCTTTTCNNNAGATCGGAAGA |
| 50300_3 | TATGAGTGTGGAGTCGTTGCCTCCCTTCATCCTGCAGCTTTTCNNNAGATCGGAAGA |
| 52509_1 | TATGAGTGTGGAGTCGTTGCAGAGACATTACACTTTGTCCGCTNNNAGATCGGAAG |
| 52509_2 | TATGAGTGTGGAGTCGTTGCAGAGACATTACACTTTGTCCGCTNNNAGATCGGAAG |
| 52509_3 | TATGAGTGTGGAGTCGTTGCAGAGACATTACACTTTGTCCGCTNNNAGATCGGAAG |
| 30926_1 | TATGAGTGTGGAGTCGTTGCGCACAGATAGACACAACACACACNNNAGATCGGAAG |
| 30926_2 | TATGAGTGTGGAGTCGTTGCGCACAGATAGACACAACACACACNNNAGATCGGAAG |
| 30926_3 | TATGAGTGTGGAGTCGTTGCGCACAGATAGACACAACACACACNNNAGATCGGAAG |
| 8323_1 | TATGAGTGTGGAGTCGTTGCATGCCCACCCCAGAAATACATATNNNAGATCGGAAG |

| | |
|---|---|
| 8323_2 | TATGAGTGTGGAGTCGTTGCATGCCCACCCCAGAAATACATATNNNAGATCGGAAG |
| 8323_3 | TATGAGTGTGGAGTCGTTGCATGCCCACCCCAGAAATACATATNNNAGATCGGAAG |
| 40176_1 | TATGAGTGTGGAGTCGTTGCAGAGTACCTACTGCATGAGTCCANNNAGATCGGAAG |
| 40176_2 | TATGAGTGTGGAGTCGTTGCAGAGTACCTACTGCATGAGTCCANNNAGATCGGAAG |
| 40176_3 | TATGAGTGTGGAGTCGTTGCAGAGTACCTACTGCATGAGTCCANNNAGATCGGAAG |
| 33420_1 | TATGAGTGTGGAGTCGTTGCAGCTAGGTTTTTTCTAGATGCCCNNNAGATCGGAA |
| 33420_2 | TATGAGTGTGGAGTCGTTGCAGCTAGGTTTTTTCTAGATGCCCNNNAGATCGGAA |
| 33420_3 | TATGAGTGTGGAGTCGTTGCAGCTAGGTTTTTTCTAGATGCCCNNNAGATCGGAA |
| 42228_1 | TATGAGTGTGGAGTCGTTGCAGTTGGAGTCTCACTGTGTCATCNNNAGATCGGAAG |
| 42228_2 | TATGAGTGTGGAGTCGTTGCAGTTGGAGTCTCACTGTGTCATCNNNAGATCGGAAG |
| 42228_3 | TATGAGTGTGGAGTCGTTGCAGTTGGAGTCTCACTGTGTCATCNNNAGATCGGAAG |
| 33540_1 | TATGAGTGTGGAGTCGTTGCCCTGGGTTCAAGCAATTCTCCTANNNAGATCGGAAG |
| 33540_2 | TATGAGTGTGGAGTCGTTGCCCTGGGTTCAAGCAATTCTCCTANNNAGATCGGAAG |
| 33540_3 | TATGAGTGTGGAGTCGTTGCCCTGGGTTCAAGCAATTCTCCTANNNAGATCGGAAG |
| 52270_1 | TATGAGTGTGGAGTCGTTGCTTACACTTAGTTTCTCGGGCTGGNNNAGATCGGAAG |
| 52270_2 | TATGAGTGTGGAGTCGTTGCTTACACTTAGTTTCTCGGGCTGGNNNAGATCGGAAG |
| 52270_3 | TATGAGTGTGGAGTCGTTGCTTACACTTAGTTTCTCGGGCTGGNNNAGATCGGAAG |
| 32081_1 | TATGAGTGTGGAGTCGTTGCTGCAATCAACAGACTCAAAGTGCNNNAGATCGGAAG |
| 32081_2 | TATGAGTGTGGAGTCGTTGCTGCAATCAACAGACTCAAAGTGCNNNAGATCGGAAG |
| 32081_3 | TATGAGTGTGGAGTCGTTGCTGCAATCAACAGACTCAAAGTGCNNNAGATCGGAAG |
| 43850_1 | TATGAGTGTGGAGTCGTTGCACACATCGAATATCCAGCACAGANNNAGATCGGAAG |
| 43850_2 | TATGAGTGTGGAGTCGTTGCACACATCGAATATCCAGCACAGANNNAGATCGGAAG |
| 43850_3 | TATGAGTGTGGAGTCGTTGCACACATCGAATATCCAGCACAGANNNAGATCGGAAG |
| 13563_1 | TATGAGTGTGGAGTCGTTGCGAAATAAAGTGCACAATAAATGCNNNAGATCGGAA |
| 13563_2 | TATGAGTGTGGAGTCGTTGCGAAATAAAGTGCACAATAAATGCNNNAGATCGGAA |
| 13563_3 | TATGAGTGTGGAGTCGTTGCGAAATAAAGTGCACAATAAATGCNNNAGATCGGAA |
| 30422_1 | TATGAGTGTGGAGTCGTTGCTCCTGTGTGTGCGTAGGTGTATANNNAGATCGGAA |
| 30422_2 | TATGAGTGTGGAGTCGTTGCTCCTGTGTGTGCGTAGGTGTATANNNAGATCGGAA |
| 30422_3 | TATGAGTGTGGAGTCGTTGCTCCTGTGTGTGCGTAGGTGTATANNNAGATCGGAA |
| 13139_1 | TATGAGTGTGGAGTCGTTGCTTAATAGTTTTTCTCATCATGGCNNNAGATCGGAAGA |
| 13139_2 | TATGAGTGTGGAGTCGTTGCTTAATAGTTTTTCTCATCATGGCNNNAGATCGGAAGA |
| 13139_3 | TATGAGTGTGGAGTCGTTGCTTAATAGTTTTTCTCATCATGGCNNNAGATCGGAAGA |
| 48417_1 | TATGAGTGTGGAGTCGTTGCCATCCTGTCCTTCCAGAATCCTCNNNAGATCGGAAGA |
| 48417_2 | TATGAGTGTGGAGTCGTTGCCATCCTGTCCTTCCAGAATCCTCNNNAGATCGGAAGA |
| 48417_3 | TATGAGTGTGGAGTCGTTGCCATCCTGTCCTTCCAGAATCCTCNNNAGATCGGAAGA |
| 40631_1 | TATGAGTGTGGAGTCGTTGCTTGATTTGCAAATATTTTCCTCCCANNNAGATCGGAA |
| 40631_2 | TATGAGTGTGGAGTCGTTGCTTGATTTGCAAATATTTTCCTCCCANNNAGATCGGAA |
| 40631_3 | TATGAGTGTGGAGTCGTTGCTTGATTTGCAAATATTTTCCTCCCANNNAGATCGGAA |
| 52222_1 | TATGAGTGTGGAGTCGTTGCGTATTCTGCAACACAAGCATGGANNNAGATCGGAA |
| 52222_2 | TATGAGTGTGGAGTCGTTGCGTATTCTGCAACACAAGCATGGANNNAGATCGGAA |
| 52222_3 | TATGAGTGTGGAGTCGTTGCGTATTCTGCAACACAAGCATGGANNNAGATCGGAA |
| 10526_1 | TATGAGTGTGGAGTCGTTGCTTTTGAGTGGGGTTTCAAGGTTTNNNAGATCGGAA |
| 10526_2 | TATGAGTGTGGAGTCGTTGCTTTTGAGTGGGGTTTCAAGGTTTNNNAGATCGGAA |

| | |
|---|---|
| 10526_3 | TATGAGTGTGGAGTCGTTGCTTTTGAGTGGGGTTTCAAGGTTTNNNAGATCGGAA |
| 47915_1 | TATGAGTGTGGAGTCGTTGCCCTCCCAGGTTCAAGAGATTCTCNNNAGATCGGAAG |
| 47915_2 | TATGAGTGTGGAGTCGTTGCCCTCCCAGGTTCAAGAGATTCTCNNNAGATCGGAAG |
| 47915_3 | TATGAGTGTGGAGTCGTTGCCCTCCCAGGTTCAAGAGATTCTCNNNAGATCGGAAG |
| 32842_1 | TATGAGTGTGGAGTCGTTGCCTCCTGAGTAGCTGGGACTACAGNNNAGATCGGAA |
| 32842_2 | TATGAGTGTGGAGTCGTTGCCTCCTGAGTAGCTGGGACTACAGNNNAGATCGGAA |
| 32842_3 | TATGAGTGTGGAGTCGTTGCCTCCTGAGTAGCTGGGACTACAGNNNAGATCGGAA |
| 30136_1 | TATGAGTGTGGAGTCGTTGCCTTGACAGCCCATTCAATAAGCANNNAGATCGGAAG |
| 30136_2 | TATGAGTGTGGAGTCGTTGCCTTGACAGCCCATTCAATAAGCANNNAGATCGGAAG |
| 30136_3 | TATGAGTGTGGAGTCGTTGCCTTGACAGCCCATTCAATAAGCANNNAGATCGGAAG |
| 40727_1 | TATGAGTGTGGAGTCGTTGCCAGCCACTATGGAAAACAGTGTGNNNAGATCGGAA |
| 40727_2 | TATGAGTGTGGAGTCGTTGCCAGCCACTATGGAAAACAGTGTGNNNAGATCGGAA |
| 40727_3 | TATGAGTGTGGAGTCGTTGCCAGCCACTATGGAAAACAGTGTGNNNAGATCGGAA |
| 38503_1 | TATGAGTGTGGAGTCGTTGCTTTTTGAGACAGGGTCTTGCTCTNNNAGATCGGAAG |
| 38503_2 | TATGAGTGTGGAGTCGTTGCTTTTTGAGACAGGGTCTTGCTCTNNNAGATCGGAAG |
| 38503_3 | TATGAGTGTGGAGTCGTTGCTTTTTGAGACAGGGTCTTGCTCTNNNAGATCGGAAG |
| 47295_1 | TATGAGTGTGGAGTCGTTGCGTGGGTTTGTCAAATATGGCCTTNNNAGATCGGAA |
| 47295_2 | TATGAGTGTGGAGTCGTTGCGTGGGTTTGTCAAATATGGCCTTNNNAGATCGGAA |
| 47295_3 | TATGAGTGTGGAGTCGTTGCGTGGGTTTGTCAAATATGGCCTTNNNAGATCGGAA |
| 45900_1 | TATGAGTGTGGAGTCGTTGCCTCCTGTGTTCAAGTGATTCTGCNNNAGATCGGAAG |
| 45900_2 | TATGAGTGTGGAGTCGTTGCCTCCTGTGTTCAAGTGATTCTGCNNNAGATCGGAAG |
| 45900_3 | TATGAGTGTGGAGTCGTTGCCTCCTGTGTTCAAGTGATTCTGCNNNAGATCGGAAG |
| 49477_1 | TATGAGTGTGGAGTCGTTGCCTGTAGTGGAGTGCTGTGATCATNNNAGATCGGAA |
| 49477_2 | TATGAGTGTGGAGTCGTTGCCTGTAGTGGAGTGCTGTGATCATNNNAGATCGGAA |
| 49477_3 | TATGAGTGTGGAGTCGTTGCCTGTAGTGGAGTGCTGTGATCATNNNAGATCGGAA |
| 3018_1 | TATGAGTGTGGAGTCGTTGCTGGTACTTTACTGGGTCTGGAGGNNNAGATCGGAA |
| 3018_2 | TATGAGTGTGGAGTCGTTGCTGGTACTTTACTGGGTCTGGAGGNNNAGATCGGAA |
| 3018_3 | TATGAGTGTGGAGTCGTTGCTGGTACTTTACTGGGTCTGGAGGNNNAGATCGGAA |
| 33148_1 | TATGAGTGTGGAGTCGTTGCCCAAAAAAAAAAAAACCCTCCCGNNNAGATCGGAAG |
| 33148_2 | TATGAGTGTGGAGTCGTTGCCCAAAAAAAAAAAAACCCTCCCGNNNAGATCGGAAG |
| 33148_3 | TATGAGTGTGGAGTCGTTGCCCAAAAAAAAAAAAACCCTCCCGNNNAGATCGGAAG |
| 29671_1 | TATGAGTGTGGAGTCGTTGCAGCCATTTACAGTGGGTGACAATNNNAGATCGGAA |
| 29671_2 | TATGAGTGTGGAGTCGTTGCAGCCATTTACAGTGGGTGACAATNNNAGATCGGAA |
| 29671_3 | TATGAGTGTGGAGTCGTTGCAGCCATTTACAGTGGGTGACAATNNNAGATCGGAA |
| 32899_1 | TATGAGTGTGGAGTCGTTGCACAGAGAAGTTCGGTCTCAAACANNNAGATCGGAA |
| 32899_2 | TATGAGTGTGGAGTCGTTGCACAGAGAAGTTCGGTCTCAAACANNNAGATCGGAA |
| 32899_3 | TATGAGTGTGGAGTCGTTGCACAGAGAAGTTCGGTCTCAAACANNNAGATCGGAA |
| 39279_1 | TATGAGTGTGGAGTCGTTGCGTCCTACCTAGAGCAAGCATACCNNNAGATCGGAAG |
| 39279_2 | TATGAGTGTGGAGTCGTTGCGTCCTACCTAGAGCAAGCATACCNNNAGATCGGAAG |
| 39279_3 | TATGAGTGTGGAGTCGTTGCGTCCTACCTAGAGCAAGCATACCNNNAGATCGGAAG |
| 46561_1 | TATGAGTGTGGAGTCGTTGCTTCCAGTGATTCTCCTGCCTTACNNNAGATCGGAAGA |
| 46561_2 | TATGAGTGTGGAGTCGTTGCTTCCAGTGATTCTCCTGCCTTACNNNAGATCGGAAGA |
| 46561_3 | TATGAGTGTGGAGTCGTTGCTTCCAGTGATTCTCCTGCCTTACNNNAGATCGGAAGA |

| | |
|---|---|
| 38606_1 | TATGAGTGTGGAGTCGTTGCTTGGTTTCACTCCTGACCATTGANNNAGATCGGAAG |
| 38606_2 | TATGAGTGTGGAGTCGTTGCTTGGTTTCACTCCTGACCATTGANNNAGATCGGAAG |
| 38606_3 | TATGAGTGTGGAGTCGTTGCTTGGTTTCACTCCTGACCATTGANNNAGATCGGAAG |
| 40100_1 | TATGAGTGTGGAGTCGTTGCACACGGTGAAACCTCATCTCTACNNNAGATCGGAAG |
| 40100_2 | TATGAGTGTGGAGTCGTTGCACACGGTGAAACCTCATCTCTACNNNAGATCGGAAG |
| 40100_3 | TATGAGTGTGGAGTCGTTGCACACGGTGAAACCTCATCTCTACNNNAGATCGGAAG |
| 48803_1 | TATGAGTGTGGAGTCGTTGCGATCTCGGCTCATTGCAAACTCNNNAGATCGGAAGA |
| 48803_2 | TATGAGTGTGGAGTCGTTGCGATCTCGGCTCATTGCAAACTCNNNAGATCGGAAGA |
| 48803_3 | TATGAGTGTGGAGTCGTTGCGATCTCGGCTCATTGCAAACTCNNNAGATCGGAAGA |
| 35682_1 | TATGAGTGTGGAGTCGTTGCATCCTCACCCTTATCTGACCTCANNNAGATCGGAAGA |
| 35682_2 | TATGAGTGTGGAGTCGTTGCATCCTCACCCTTATCTGACCTCANNNAGATCGGAAGA |
| 35682_3 | TATGAGTGTGGAGTCGTTGCATCCTCACCCTTATCTGACCTCANNNAGATCGGAAGA |
| 53236_1 | TATGAGTGTGGAGTCGTTGCCTCTGGGTACTGAGAGCTCAATGNNNAGATCGGAA |
| 53236_2 | TATGAGTGTGGAGTCGTTGCCTCTGGGTACTGAGAGCTCAATGNNNAGATCGGAA |
| 53236_3 | TATGAGTGTGGAGTCGTTGCCTCTGGGTACTGAGAGCTCAATGNNNAGATCGGAA |
| 53005_1 | TATGAGTGTGGAGTCGTTGCATTCAATTACCTCCCACGACACANNNAGATCGGAAGA |
| 53005_2 | TATGAGTGTGGAGTCGTTGCATTCAATTACCTCCCACGACACANNNAGATCGGAAGA |
| 53005_3 | TATGAGTGTGGAGTCGTTGCATTCAATTACCTCCCACGACACANNNAGATCGGAAGA |
| 37260_1 | TATGAGTGTGGAGTCGTTGCACTTGCTCCTGAATGACTTTTGANNNAGATCGGAAG |
| 37260_2 | TATGAGTGTGGAGTCGTTGCACTTGCTCCTGAATGACTTTTGANNNAGATCGGAAG |
| 37260_3 | TATGAGTGTGGAGTCGTTGCACTTGCTCCTGAATGACTTTTGANNNAGATCGGAAG |
| 45333_1 | TATGAGTGTGGAGTCGTTGCCACATCTTCAGGGGCATCATAGANNNAGATCGGAAG |
| 45333_2 | TATGAGTGTGGAGTCGTTGCCACATCTTCAGGGGCATCATAGANNNAGATCGGAAG |
| 45333_3 | TATGAGTGTGGAGTCGTTGCCACATCTTCAGGGGCATCATAGANNNAGATCGGAAG |
| 44730_1 | TATGAGTGTGGAGTCGTTGCGAAAACCATGGACAGAAGAAGGCNNNAGATCGGAA |
| 44730_2 | TATGAGTGTGGAGTCGTTGCGAAAACCATGGACAGAAGAAGGCNNNAGATCGGAA |
| 44730_3 | TATGAGTGTGGAGTCGTTGCGAAAACCATGGACAGAAGAAGGCNNNAGATCGGAA |
| 42971_1 | TATGAGTGTGGAGTCGTTGCACTACTCAGCCATAAAAAGGAATGANNNAGATCGGA |
| 42971_2 | TATGAGTGTGGAGTCGTTGCACTACTCAGCCATAAAAAGGAATGANNNAGATCGGA |
| 42971_3 | TATGAGTGTGGAGTCGTTGCACTACTCAGCCATAAAAAGGAATGANNNAGATCGGA |
| 43015_1 | TATGAGTGTGGAGTCGTTGCGGACTAAAGTCAGAGCTACAGGGNNNAGATCGGAA |
| 43015_2 | TATGAGTGTGGAGTCGTTGCGGACTAAAGTCAGAGCTACAGGGNNNAGATCGGAA |
| 43015_3 | TATGAGTGTGGAGTCGTTGCGGACTAAAGTCAGAGCTACAGGGNNNAGATCGGAA |
| 44476_1 | TATGAGTGTGGAGTCGTTGCTTGGATTTGTGAATTGTGCTGCTNNNAGATCGGAAG |
| 44476_2 | TATGAGTGTGGAGTCGTTGCTTGGATTTGTGAATTGTGCTGCTNNNAGATCGGAAG |
| 44476_3 | TATGAGTGTGGAGTCGTTGCTTGGATTTGTGAATTGTGCTGCTNNNAGATCGGAAG |
| 10847_1 | TATGAGTGTGGAGTCGTTGCTGGTTGATTCTAGGCCCCACTAANNNAGATCGGAAG |
| 10847_2 | TATGAGTGTGGAGTCGTTGCTGGTTGATTCTAGGCCCCACTAANNNAGATCGGAAG |
| 10847_3 | TATGAGTGTGGAGTCGTTGCTGGTTGATTCTAGGCCCCACTAANNNAGATCGGAAG |
| 754_1 | TATGAGTGTGGAGTCGTTGCAAGAAACTTCCGGATTATGGGCANNNAGATCGGAA |
| 754_2 | TATGAGTGTGGAGTCGTTGCAAGAAACTTCCGGATTATGGGCANNNAGATCGGAA |
| 754_3 | TATGAGTGTGGAGTCGTTGCAAGAAACTTCCGGATTATGGGCANNNAGATCGGAA |
| 43107_1 | TATGAGTGTGGAGTCGTTGCGTGTAGTTTTCAAGGTCTTGGCCNNNAGATCGGAAG |

| | |
|---|---|
| 43107_2 | TATGAGTGTGGAGTCGTTGCGTGTAGTTTTCAAGGTCTTGGCCNNNAGATCGGAAG |
| 43107_3 | TATGAGTGTGGAGTCGTTGCGTGTAGTTTTCAAGGTCTTGGCCNNNAGATCGGAAG |
| 10167_1 | TATGAGTGTGGAGTCGTTGCATTTGATCTGTTGGGGGTGCATTNNNAGATCGGAAG |
| 10167_2 | TATGAGTGTGGAGTCGTTGCATTTGATCTGTTGGGGGTGCATTNNNAGATCGGAAG |
| 10167_3 | TATGAGTGTGGAGTCGTTGCATTTGATCTGTTGGGGGTGCATTNNNAGATCGGAAG |
| 52071_1 | TATGAGTGTGGAGTCGTTGCAACTTTTGTCAGCCCTCATAGCTNNNAGATCGGAAGA |
| 52071_2 | TATGAGTGTGGAGTCGTTGCAACTTTTGTCAGCCCTCATAGCTNNNAGATCGGAAGA |
| 52071_3 | TATGAGTGTGGAGTCGTTGCAACTTTTGTCAGCCCTCATAGCTNNNAGATCGGAAGA |
| 48314_1 | TATGAGTGTGGAGTCGTTGCGCTCTGGCATCTATCCTCATTCANNNAGATCGGAAGA |
| 48314_2 | TATGAGTGTGGAGTCGTTGCGCTCTGGCATCTATCCTCATTCANNNAGATCGGAAGA |
| 48314_3 | TATGAGTGTGGAGTCGTTGCGCTCTGGCATCTATCCTCATTCANNNAGATCGGAAGA |
| 38280_1 | TATGAGTGTGGAGTCGTTGCATTTGGATATGGCTCCTTTGGGANNNAGATCGGAAG |
| 38280_2 | TATGAGTGTGGAGTCGTTGCATTTGGATATGGCTCCTTTGGGANNNAGATCGGAAG |
| 38280_3 | TATGAGTGTGGAGTCGTTGCATTTGGATATGGCTCCTTTGGGANNNAGATCGGAAG |
| 43278_1 | TATGAGTGTGGAGTCGTTGCACACACACACACACTGAGAGATANNNAGATCGGAAG |
| 43278_2 | TATGAGTGTGGAGTCGTTGCACACACACACACACTGAGAGATANNNAGATCGGAAG |
| 43278_3 | TATGAGTGTGGAGTCGTTGCACACACACACACACTGAGAGATANNNAGATCGGAAG |
| 50079_1 | TATGAGTGTGGAGTCGTTGCCTTGTGGAGGTGAGGTGTGCNNNAGATCGGAAGAG |
| 50079_2 | TATGAGTGTGGAGTCGTTGCCTTGTGGAGGTGAGGTGTGCNNNAGATCGGAAGAG |
| 50079_3 | TATGAGTGTGGAGTCGTTGCCTTGTGGAGGTGAGGTGTGCNNNAGATCGGAAGAG |
| 2343_1 | TATGAGTGTGGAGTCGTTGCTGACCAAGTTCCTTATACCAGCCNNNAGATCGGAAGA |
| 2343_2 | TATGAGTGTGGAGTCGTTGCTGACCAAGTTCCTTATACCAGCCNNNAGATCGGAAGA |
| 2343_3 | TATGAGTGTGGAGTCGTTGCTGACCAAGTTCCTTATACCAGCCNNNAGATCGGAAGA |
| 48054_1 | TATGAGTGTGGAGTCGTTGCAGTAGTTGGGATTACAGGTGTGCNNNAGATCGGAA |
| 48054_2 | TATGAGTGTGGAGTCGTTGCAGTAGTTGGGATTACAGGTGTGCNNNAGATCGGAA |
| 48054_3 | TATGAGTGTGGAGTCGTTGCAGTAGTTGGGATTACAGGTGTGCNNNAGATCGGAA |
| 44915_1 | TATGAGTGTGGAGTCGTTGCATTTAGGCCTCTCAGTGTGGTTTNNNAGATCGGAAG |
| 44915_2 | TATGAGTGTGGAGTCGTTGCATTTAGGCCTCTCAGTGTGGTTTNNNAGATCGGAAG |
| 44915_3 | TATGAGTGTGGAGTCGTTGCATTTAGGCCTCTCAGTGTGGTTTNNNAGATCGGAAG |
| 34928_1 | TATGAGTGTGGAGTCGTTGCTTTAGTAGAGACGGAGTTTCGCCNNNAGATCGGAAG |
| 34928_2 | TATGAGTGTGGAGTCGTTGCTTTAGTAGAGACGGAGTTTCGCCNNNAGATCGGAAG |
| 34928_3 | TATGAGTGTGGAGTCGTTGCTTTAGTAGAGACGGAGTTTCGCCNNNAGATCGGAAG |
| 44317_1 | TATGAGTGTGGAGTCGTTGCCAGCAATTTCTTCCAGTTCTCCGNNNAGATCGGAAGA |
| 44317_2 | TATGAGTGTGGAGTCGTTGCCAGCAATTTCTTCCAGTTCTCCGNNNAGATCGGAAGA |
| 44317_3 | TATGAGTGTGGAGTCGTTGCCAGCAATTTCTTCCAGTTCTCCGNNNAGATCGGAAGA |
| 36712_1 | TATGAGTGTGGAGTCGTTGCATTTCACACCTCAAAAGTTGGCTNNNAGATCGGAAGA |
| 36712_2 | TATGAGTGTGGAGTCGTTGCATTTCACACCTCAAAAGTTGGCTNNNAGATCGGAAGA |
| 36712_3 | TATGAGTGTGGAGTCGTTGCATTTCACACCTCAAAAGTTGGCTNNNAGATCGGAAGA |
| 13629_1 | TATGAGTGTGGAGTCGTTGCCATGAGATGCTTTTGAAGATGNNNAGATCGGAAGAG |
| 13629_2 | TATGAGTGTGGAGTCGTTGCCATGAGATGCTTTTGAAGATGNNNAGATCGGAAGAG |
| 13629_3 | TATGAGTGTGGAGTCGTTGCCATGAGATGCTTTTGAAGATGNNNAGATCGGAAGAG |
| 38744_1 | TATGAGTGTGGAGTCGTTGCAAGTCTCTTGATGGTTTTTGCCANNNAGATCGGAAGA |
| 38744_2 | TATGAGTGTGGAGTCGTTGCAAGTCTCTTGATGGTTTTTGCCANNNAGATCGGAAGA |

| | |
|---|---|
| 38744_3 | TATGAGTGTGGAGTCGTTGCAAGTCTCTTGATGGTTTTTGCCANNNAGATCGGAAG |
| 39903_1 | TATGAGTGTGGAGTCGTTGCAAACTGTAGCTGTGACCGTTACTNNNAGATCGGAAG |
| 39903_2 | TATGAGTGTGGAGTCGTTGCAAACTGTAGCTGTGACCGTTACTNNNAGATCGGAAG |
| 39903_3 | TATGAGTGTGGAGTCGTTGCAAACTGTAGCTGTGACCGTTACTNNNAGATCGGAAG |
| 39659_1 | TATGAGTGTGGAGTCGTTGCAAAATTCTATGTGTTGGCCAGGCNNNAGATCGGAA |
| 39659_2 | TATGAGTGTGGAGTCGTTGCAAAATTCTATGTGTTGGCCAGGCNNNAGATCGGAA |
| 39659_3 | TATGAGTGTGGAGTCGTTGCAAAATTCTATGTGTTGGCCAGGCNNNAGATCGGAA |
| 45593_1 | TATGAGTGTGGAGTCGTTGCGTCTGGTGGGAAGTCAGAAAGATNNNAGATCGGA |
| 45593_2 | TATGAGTGTGGAGTCGTTGCGTCTGGTGGGAAGTCAGAAAGATNNNAGATCGGA |
| 45593_3 | TATGAGTGTGGAGTCGTTGCGTCTGGTGGGAAGTCAGAAAGATNNNAGATCGGA |
| 40480_1 | TATGAGTGTGGAGTCGTTGCCCAGGAAGGTGGAAGCTATACTCNNNAGATCGGAA |
| 40480_2 | TATGAGTGTGGAGTCGTTGCCCAGGAAGGTGGAAGCTATACTCNNNAGATCGGAA |
| 40480_3 | TATGAGTGTGGAGTCGTTGCCCAGGAAGGTGGAAGCTATACTCNNNAGATCGGAA |
| 31232_1 | TATGAGTGTGGAGTCGTTGCGCTGGCTGGCTTTTTTTGTTTGNNNAGATCGGAAGA |
| 31232_2 | TATGAGTGTGGAGTCGTTGCGCTGGCTGGCTTTTTTTGTTTGNNNAGATCGGAAGA |
| 31232_3 | TATGAGTGTGGAGTCGTTGCGCTGGCTGGCTTTTTTTGTTTGNNNAGATCGGAAGA |
| 36074_1 | TATGAGTGTGGAGTCGTTGCCAACTTAGGGATACCAGGAGCTGNNNAGATCGGAA |
| 36074_2 | TATGAGTGTGGAGTCGTTGCCAACTTAGGGATACCAGGAGCTGNNNAGATCGGAA |
| 36074_3 | TATGAGTGTGGAGTCGTTGCCAACTTAGGGATACCAGGAGCTGNNNAGATCGGAA |
| 40954_1 | TATGAGTGTGGAGTCGTTGCGGGAGCAGACTTCTTGGTAGAAANNNAGATCGGAA |
| 40954_2 | TATGAGTGTGGAGTCGTTGCGGGAGCAGACTTCTTGGTAGAAANNNAGATCGGAA |
| 40954_3 | TATGAGTGTGGAGTCGTTGCGGGAGCAGACTTCTTGGTAGAAANNNAGATCGGAA |
| 44570_1 | TATGAGTGTGGAGTCGTTGCCAATTATCACACCTCAGCCTCCTNNNAGATCGGAAGA |
| 44570_2 | TATGAGTGTGGAGTCGTTGCCAATTATCACACCTCAGCCTCCTNNNAGATCGGAAGA |
| 44570_3 | TATGAGTGTGGAGTCGTTGCCAATTATCACACCTCAGCCTCCTNNNAGATCGGAAGA |
| 32473_1 | TATGAGTGTGGAGTCGTTGCGTTCAAGCAATTCTCCTGCCTCNNNAGATCGGAAGA |
| 32473_2 | TATGAGTGTGGAGTCGTTGCGTTCAAGCAATTCTCCTGCCTCNNNAGATCGGAAGA |
| 32473_3 | TATGAGTGTGGAGTCGTTGCGTTCAAGCAATTCTCCTGCCTCNNNAGATCGGAAGA |
| 47664_1 | TATGAGTGTGGAGTCGTTGCGAGACAGTCTCGCTCTTGTCCNNNAGATCGGAAGAG |
| 47664_2 | TATGAGTGTGGAGTCGTTGCGAGACAGTCTCGCTCTTGTCCNNNAGATCGGAAGAG |
| 47664_3 | TATGAGTGTGGAGTCGTTGCGAGACAGTCTCGCTCTTGTCCNNNAGATCGGAAGAG |
| 43961_1 | TATGAGTGTGGAGTCGTTGCTAGTAGGTTAGCTCAGCAGGTCTNNNAGATCGGAA |
| 43961_2 | TATGAGTGTGGAGTCGTTGCTAGTAGGTTAGCTCAGCAGGTCTNNNAGATCGGAA |
| 43961_3 | TATGAGTGTGGAGTCGTTGCTAGTAGGTTAGCTCAGCAGGTCTNNNAGATCGGAA |
| 51951_1 | TATGAGTGTGGAGTCGTTGCTGACATGCTTTTCCAGGGTCATANNNAGATCGGAAG |
| 51951_2 | TATGAGTGTGGAGTCGTTGCTGACATGCTTTTCCAGGGTCATANNNAGATCGGAAG |
| 51951_3 | TATGAGTGTGGAGTCGTTGCTGACATGCTTTTCCAGGGTCATANNNAGATCGGAAG |
| 30550_1 | TATGAGTGTGGAGTCGTTGCAGCCTTCTTTTAAGCTATACCACCTNNNAGATCGGAA |
| 30550_2 | TATGAGTGTGGAGTCGTTGCAGCCTTCTTTTAAGCTATACCACCTNNNAGATCGGAA |
| 30550_3 | TATGAGTGTGGAGTCGTTGCAGCCTTCTTTTAAGCTATACCACCTNNNAGATCGGAA |
| 37224_1 | TATGAGTGTGGAGTCGTTGCCTGGAGTCTCAGCTACTCAAGAGNNNAGATCGGAA |
| 37224_2 | TATGAGTGTGGAGTCGTTGCCTGGAGTCTCAGCTACTCAAGAGNNNAGATCGGAA |
| 37224_3 | TATGAGTGTGGAGTCGTTGCCTGGAGTCTCAGCTACTCAAGAGNNNAGATCGGAA |

| | |
|---|---|
| 39008_1 | TATGAGTGTGGAGTCGTTGCGCTACAAACCATGTTTAGCCAGGNNNAGATCGGAAG |
| 39008_2 | TATGAGTGTGGAGTCGTTGCGCTACAAACCATGTTTAGCCAGGNNNAGATCGGAAG |
| 39008_3 | TATGAGTGTGGAGTCGTTGCGCTACAAACCATGTTTAGCCAGGNNNAGATCGGAAG |
| 42988_1 | TATGAGTGTGGAGTCGTTGCTCCTCACTATGCTTGATGAACTGANNNAGATCGGAA |
| 42988_2 | TATGAGTGTGGAGTCGTTGCTCCTCACTATGCTTGATGAACTGANNNAGATCGGAA |
| 42988_3 | TATGAGTGTGGAGTCGTTGCTCCTCACTATGCTTGATGAACTGANNNAGATCGGAA |
| 51205_1 | TATGAGTGTGGAGTCGTTGCATGACTGTTTTCACCATTCCCCTNNNAGATCGGAAGA |
| 51205_2 | TATGAGTGTGGAGTCGTTGCATGACTGTTTTCACCATTCCCCTNNNAGATCGGAAGA |
| 51205_3 | TATGAGTGTGGAGTCGTTGCATGACTGTTTTCACCATTCCCCTNNNAGATCGGAAGA |
| 38540_1 | TATGAGTGTGGAGTCGTTGCTGTCCCTAATCAGCTCCTTTTCCNNNAGATCGGAAGA |
| 38540_2 | TATGAGTGTGGAGTCGTTGCTGTCCCTAATCAGCTCCTTTTCCNNNAGATCGGAAGA |
| 38540_3 | TATGAGTGTGGAGTCGTTGCTGTCCCTAATCAGCTCCTTTTCCNNNAGATCGGAAGA |
| 30285_1 | TATGAGTGTGGAGTCGTTGCGTATGTCCACGAGCACAAGGATANNNAGATCGGAA |
| 30285_2 | TATGAGTGTGGAGTCGTTGCGTATGTCCACGAGCACAAGGATANNNAGATCGGAA |
| 30285_3 | TATGAGTGTGGAGTCGTTGCGTATGTCCACGAGCACAAGGATANNNAGATCGGAA |
| 48810_1 | TATGAGTGTGGAGTCGTTGCTTGAGACGGAGTCTAGCTTTGTCNNNAGATCGGAAG |
| 48810_2 | TATGAGTGTGGAGTCGTTGCTTGAGACGGAGTCTAGCTTTGTCNNNAGATCGGAAG |
| 48810_3 | TATGAGTGTGGAGTCGTTGCTTGAGACGGAGTCTAGCTTTGTCNNNAGATCGGAAG |
| 36296_1 | TATGAGTGTGGAGTCGTTGCGCAAGGGAACAAAATTGGACAGANNNAGATCGGAA |
| 36296_2 | TATGAGTGTGGAGTCGTTGCGCAAGGGAACAAAATTGGACAGANNNAGATCGGAA |
| 36296_3 | TATGAGTGTGGAGTCGTTGCGCAAGGGAACAAAATTGGACAGANNNAGATCGGAA |
| 49257_1 | TATGAGTGTGGAGTCGTTGCAGGATCACTTGAGCCAGAAGTTTNNNAGATCGGAA |
| 49257_2 | TATGAGTGTGGAGTCGTTGCAGGATCACTTGAGCCAGAAGTTTNNNAGATCGGAA |
| 49257_3 | TATGAGTGTGGAGTCGTTGCAGGATCACTTGAGCCAGAAGTTTNNNAGATCGGAA |
| 52849_1 | TATGAGTGTGGAGTCGTTGCAACACACCTGATTGTTGGAAAGGNNNAGATCGGAA |
| 52849_2 | TATGAGTGTGGAGTCGTTGCAACACACCTGATTGTTGGAAAGGNNNAGATCGGAA |
| 52849_3 | TATGAGTGTGGAGTCGTTGCAACACACCTGATTGTTGGAAAGGNNNAGATCGGAA |
| 43395_1 | TATGAGTGTGGAGTCGTTGCTAGGATTATAGGCATGCACCACCNNNAGATCGGAAG |
| 43395_2 | TATGAGTGTGGAGTCGTTGCTAGGATTATAGGCATGCACCACCNNNAGATCGGAAG |
| 43395_3 | TATGAGTGTGGAGTCGTTGCTAGGATTATAGGCATGCACCACCNNNAGATCGGAAG |
| 13569_1 | TATGAGTGTGGAGTCGTTGCAGTCTCTAGAACAGGGATTCNNNAGATCGGAAGAG |
| 13569_2 | TATGAGTGTGGAGTCGTTGCAGTCTCTAGAACAGGGATTCNNNAGATCGGAAGAG |
| 13569_3 | TATGAGTGTGGAGTCGTTGCAGTCTCTAGAACAGGGATTCNNNAGATCGGAAGAG |
| 44123_1 | TATGAGTGTGGAGTCGTTGCTTTCTCCCTGATACCCTGTGTTGNNNAGATCGGAAGA |
| 44123_2 | TATGAGTGTGGAGTCGTTGCTTTCTCCCTGATACCCTGTGTTGNNNAGATCGGAAGA |
| 44123_3 | TATGAGTGTGGAGTCGTTGCTTTCTCCCTGATACCCTGTGTTGNNNAGATCGGAAGA |
| 31243_1 | TATGAGTGTGGAGTCGTTGCTAAATAGCTCCTCTTGCCACCAGNNNAGATCGGAAG |
| 31243_2 | TATGAGTGTGGAGTCGTTGCTAAATAGCTCCTCTTGCCACCAGNNNAGATCGGAAG |
| 31243_3 | TATGAGTGTGGAGTCGTTGCTAAATAGCTCCTCTTGCCACCAGNNNAGATCGGAAG |
| 44044_1 | TATGAGTGTGGAGTCGTTGCGAGTTTTGAGAACATGGAAGGGCNNNAGATCGGAA |
| 44044_2 | TATGAGTGTGGAGTCGTTGCGAGTTTTGAGAACATGGAAGGGCNNNAGATCGGAA |
| 44044_3 | TATGAGTGTGGAGTCGTTGCGAGTTTTGAGAACATGGAAGGGCNNNAGATCGGAA |
| 13418_1 | TATGAGTGTGGAGTCGTTGCCTGTTAAGGTTTCTCTGAGGNNNAGATCGGAAGAG |

| | |
|---|---|
| 13418_2 | TATGAGTGTGGAGTCGTTGCCTGTTAAGGTTTCTCTGAGGNNNAGATCGGAAGAG |
| 13418_3 | TATGAGTGTGGAGTCGTTGCCTGTTAAGGTTTCTCTGAGGNNNAGATCGGAAGAG |
| 32019_1 | TATGAGTGTGGAGTCGTTGCCTTGGCCACACAGCTTTACAAATNNNAGATCGGAAG |
| 32019_2 | TATGAGTGTGGAGTCGTTGCCTTGGCCACACAGCTTTACAAATNNNAGATCGGAAG |
| 32019_3 | TATGAGTGTGGAGTCGTTGCCTTGGCCACACAGCTTTACAAATNNNAGATCGGAAG |
| 12738_1 | TATGAGTGTGGAGTCGTTGCAATAATCTTGTGCAGGAAGTTNNNAGATCGGAAGA |
| 12738_2 | TATGAGTGTGGAGTCGTTGCAATAATCTTGTGCAGGAAGTTNNNAGATCGGAAGA |
| 12738_3 | TATGAGTGTGGAGTCGTTGCAATAATCTTGTGCAGGAAGTTNNNAGATCGGAAGA |
| 10353_1 | TATGAGTGTGGAGTCGTTGCGCAAGGAAAACCTCAACTTGCACNNNAGATCGGAAG |
| 10353_2 | TATGAGTGTGGAGTCGTTGCGCAAGGAAAACCTCAACTTGCACNNNAGATCGGAAG |
| 10353_3 | TATGAGTGTGGAGTCGTTGCGCAAGGAAAACCTCAACTTGCACNNNAGATCGGAAG |
| 44787_1 | TATGAGTGTGGAGTCGTTGCGGAAAATGAACAGGGGGAAAGTGNNNAGATCGGA |
| 44787_2 | TATGAGTGTGGAGTCGTTGCGGAAAATGAACAGGGGGAAAGTGNNNAGATCGGA |
| 44787_3 | TATGAGTGTGGAGTCGTTGCGGAAAATGAACAGGGGGAAAGTGNNNAGATCGGA |
| 9983_1 | TATGAGTGTGGAGTCGTTGCTACATCCCCCACAAAAAGAGCACNNNAGATCGGAAG |
| 9983_2 | TATGAGTGTGGAGTCGTTGCTACATCCCCCACAAAAAGAGCACNNNAGATCGGAAG |
| 9983_3 | TATGAGTGTGGAGTCGTTGCTACATCCCCCACAAAAAGAGCACNNNAGATCGGAAG |
| 12517_1 | TATGAGTGTGGAGTCGTTGCGTTTTTTTCTGTAGGTAAACTCTNNNAGATCGGAAG |
| 12517_2 | TATGAGTGTGGAGTCGTTGCGTTTTTTTCTGTAGGTAAACTCTNNNAGATCGGAAG |
| 12517_3 | TATGAGTGTGGAGTCGTTGCGTTTTTTTCTGTAGGTAAACTCTNNNAGATCGGAAG |
| 8107_1 | TATGAGTGTGGAGTCGTTGCAACACACGCCTTCCTTGTACTTGNNNAGATCGGAAG |
| 8107_2 | TATGAGTGTGGAGTCGTTGCAACACACGCCTTCCTTGTACTTGNNNAGATCGGAAG |
| 8107_3 | TATGAGTGTGGAGTCGTTGCAACACACGCCTTCCTTGTACTTGNNNAGATCGGAAG |
| 50068_1 | TATGAGTGTGGAGTCGTTGCCTTAGATCCCGCTCCACCTCTTCNNNAGATCGGAAGA |
| 50068_2 | TATGAGTGTGGAGTCGTTGCCTTAGATCCCGCTCCACCTCTTCNNNAGATCGGAAGA |
| 50068_3 | TATGAGTGTGGAGTCGTTGCCTTAGATCCCGCTCCACCTCTTCNNNAGATCGGAAGA |
| 50070_1 | TATGAGTGTGGAGTCGTTGCGAGGATAGCTTGAGCCCAGAAATNNNAGATCGGAA |
| 50070_2 | TATGAGTGTGGAGTCGTTGCGAGGATAGCTTGAGCCCAGAAATNNNAGATCGGAA |
| 50070_3 | TATGAGTGTGGAGTCGTTGCGAGGATAGCTTGAGCCCAGAAATNNNAGATCGGAA |
| 33968_1 | TATGAGTGTGGAGTCGTTGCGAGTCTCACTCTATTGCCCAGACNNNAGATCGGAAG |
| 33968_2 | TATGAGTGTGGAGTCGTTGCGAGTCTCACTCTATTGCCCAGACNNNAGATCGGAAG |
| 33968_3 | TATGAGTGTGGAGTCGTTGCGAGTCTCACTCTATTGCCCAGACNNNAGATCGGAAG |
| 32747_1 | TATGAGTGTGGAGTCGTTGCGCGTTGAGTATGGTAGCTTTCACNNNAGATCGGAAG |
| 32747_2 | TATGAGTGTGGAGTCGTTGCGCGTTGAGTATGGTAGCTTTCACNNNAGATCGGAAG |
| 32747_3 | TATGAGTGTGGAGTCGTTGCGCGTTGAGTATGGTAGCTTTCACNNNAGATCGGAAG |
| 47189_1 | TATGAGTGTGGAGTCGTTGCGTTCAAGCGATTCTTCTGCCTCNNNAGATCGGAAGA |
| 47189_2 | TATGAGTGTGGAGTCGTTGCGTTCAAGCGATTCTTCTGCCTCNNNAGATCGGAAGA |
| 47189_3 | TATGAGTGTGGAGTCGTTGCGTTCAAGCGATTCTTCTGCCTCNNNAGATCGGAAGA |
| 36429_1 | TATGAGTGTGGAGTCGTTGCGCAACACTGTGGGCAAAAGATATNNNAGATCGGAA |
| 36429_2 | TATGAGTGTGGAGTCGTTGCGCAACACTGTGGGCAAAAGATATNNNAGATCGGAA |
| 36429_3 | TATGAGTGTGGAGTCGTTGCGCAACACTGTGGGCAAAAGATATNNNAGATCGGAA |
| 12632_1 | TATGAGTGTGGAGTCGTTGCGAAGTCACATTAATGATGAACTGNNNAGATCGGAA |
| 12632_2 | TATGAGTGTGGAGTCGTTGCGAAGTCACATTAATGATGAACTGNNNAGATCGGAA |

| | |
|---|---|
| 12632_3 | TATGAGTGTGGAGTCGTTGCGAAGTCACATTAATGATGAACTGNNNAGATCGGAA |
| 41656_1 | TATGAGTGTGGAGTCGTTGCACCCCGTCTCTACTAAAAACACCNNNAGATCGGAAGA |
| 41656_2 | TATGAGTGTGGAGTCGTTGCACCCCGTCTCTACTAAAAACACCNNNAGATCGGAAGA |
| 41656_3 | TATGAGTGTGGAGTCGTTGCACCCCGTCTCTACTAAAAACACCNNNAGATCGGAAGA |
| 33099_1 | TATGAGTGTGGAGTCGTTGCGGTGTGAGGTGGTATCTCATTGTNNNAGATCGGAA |
| 33099_2 | TATGAGTGTGGAGTCGTTGCGGTGTGAGGTGGTATCTCATTGTNNNAGATCGGAA |
| 33099_3 | TATGAGTGTGGAGTCGTTGCGGTGTGAGGTGGTATCTCATTGTNNNAGATCGGAA |
| 51008_1 | TATGAGTGTGGAGTCGTTGCTGTTCAAATGTTCATACTGCCCANNNAGATCGGAAG |
| 51008_2 | TATGAGTGTGGAGTCGTTGCTGTTCAAATGTTCATACTGCCCANNNAGATCGGAAG |
| 51008_3 | TATGAGTGTGGAGTCGTTGCTGTTCAAATGTTCATACTGCCCANNNAGATCGGAAG |
| 52610_1 | TATGAGTGTGGAGTCGTTGCCACTGTCTGGAGGAGAAACTTGANNNAGATCGGAA |
| 52610_2 | TATGAGTGTGGAGTCGTTGCCACTGTCTGGAGGAGAAACTTGANNNAGATCGGAA |
| 52610_3 | TATGAGTGTGGAGTCGTTGCCACTGTCTGGAGGAGAAACTTGANNNAGATCGGAA |
| 45664_1 | TATGAGTGTGGAGTCGTTGCAGCCTCTTGAAGTGCTGGAATTANNNAGATCGGAAG |
| 45664_2 | TATGAGTGTGGAGTCGTTGCAGCCTCTTGAAGTGCTGGAATTANNNAGATCGGAAG |
| 45664_3 | TATGAGTGTGGAGTCGTTGCAGCCTCTTGAAGTGCTGGAATTANNNAGATCGGAAG |
| 42718_1 | TATGAGTGTGGAGTCGTTGCAGGGAAGAGAACAAGAGTGTCTGNNNAGATCGGA |
| 42718_2 | TATGAGTGTGGAGTCGTTGCAGGGAAGAGAACAAGAGTGTCTGNNNAGATCGGA |
| 42718_3 | TATGAGTGTGGAGTCGTTGCAGGGAAGAGAACAAGAGTGTCTGNNNAGATCGGA |
| 34638_1 | TATGAGTGTGGAGTCGTTGCCAGAGTTTGTGAAGGGTTGCAATNNNAGATCGGAA |
| 34638_2 | TATGAGTGTGGAGTCGTTGCCAGAGTTTGTGAAGGGTTGCAATNNNAGATCGGAA |
| 34638_3 | TATGAGTGTGGAGTCGTTGCCAGAGTTTGTGAAGGGTTGCAATNNNAGATCGGAA |
| 2363_1 | TATGAGTGTGGAGTCGTTGCCCCTTTGTGTCACTTTGTAGAAGCNNNAGATCGGAA |
| 2363_2 | TATGAGTGTGGAGTCGTTGCCCCTTTGTGTCACTTTGTAGAAGCNNNAGATCGGAA |
| 2363_3 | TATGAGTGTGGAGTCGTTGCCCCTTTGTGTCACTTTGTAGAAGCNNNAGATCGGAA |
| 47876_1 | TATGAGTGTGGAGTCGTTGCGTAATCAGGGACCCAGACTGTACNNNAGATCGGAAG |
| 47876_2 | TATGAGTGTGGAGTCGTTGCGTAATCAGGGACCCAGACTGTACNNNAGATCGGAAG |
| 47876_3 | TATGAGTGTGGAGTCGTTGCGTAATCAGGGACCCAGACTGTACNNNAGATCGGAAG |
| 8119_1 | TATGAGTGTGGAGTCGTTGCGGTTCGTTGCAGCCTCGATTTNNNAGATCGGAAGAG |
| 8119_2 | TATGAGTGTGGAGTCGTTGCGGTTCGTTGCAGCCTCGATTTNNNAGATCGGAAGAG |
| 8119_3 | TATGAGTGTGGAGTCGTTGCGGTTCGTTGCAGCCTCGATTTNNNAGATCGGAAGAG |
| 44829_1 | TATGAGTGTGGAGTCGTTGCTGGGCTAGTACTGACTTAGTGGANNNAGATCGGAA |
| 44829_2 | TATGAGTGTGGAGTCGTTGCTGGGCTAGTACTGACTTAGTGGANNNAGATCGGAA |
| 44829_3 | TATGAGTGTGGAGTCGTTGCTGGGCTAGTACTGACTTAGTGGANNNAGATCGGAA |
| 35836_1 | TATGAGTGTGGAGTCGTTGCCAATTCTCCTATGTCAGCCTCCANNNAGATCGGAAGA |
| 35836_2 | TATGAGTGTGGAGTCGTTGCCAATTCTCCTATGTCAGCCTCCANNNAGATCGGAAGA |
| 35836_3 | TATGAGTGTGGAGTCGTTGCCAATTCTCCTATGTCAGCCTCCANNNAGATCGGAAGA |
| 13749_1 | TATGAGTGTGGAGTCGTTGCCAGAAGAATCTTCCTCTTTTTCNNNAGATCGGAAGAG |
| 13749_2 | TATGAGTGTGGAGTCGTTGCCAGAAGAATCTTCCTCTTTTTCNNNAGATCGGAAGAG |
| 13749_3 | TATGAGTGTGGAGTCGTTGCCAGAAGAATCTTCCTCTTTTTCNNNAGATCGGAAGAG |
| 31355_1 | TATGAGTGTGGAGTCGTTGCGCCTTCAGCTGATTAGATGATGCNNNAGATCGGAAG |
| 31355_2 | TATGAGTGTGGAGTCGTTGCGCCTTCAGCTGATTAGATGATGCNNNAGATCGGAAG |
| 31355_3 | TATGAGTGTGGAGTCGTTGCGCCTTCAGCTGATTAGATGATGCNNNAGATCGGAAG |

| | |
|---|---|
| 38338_1 | TATGAGTGTGGAGTCGTTGCATACGGTCCACATCCACAAAAGANNNAGATCGGAAG |
| 38338_2 | TATGAGTGTGGAGTCGTTGCATACGGTCCACATCCACAAAAGANNNAGATCGGAAG |
| 38338_3 | TATGAGTGTGGAGTCGTTGCATACGGTCCACATCCACAAAAGANNNAGATCGGAAG |
| 42105_1 | TATGAGTGTGGAGTCGTTGCCATCTTAGCCTCCAGAGTAGCTGNNNAGATCGGAAG |
| 42105_2 | TATGAGTGTGGAGTCGTTGCCATCTTAGCCTCCAGAGTAGCTGNNNAGATCGGAAG |
| 42105_3 | TATGAGTGTGGAGTCGTTGCCATCTTAGCCTCCAGAGTAGCTGNNNAGATCGGAAG |
| 38160_1 | TATGAGTGTGGAGTCGTTGCTCTTCACCTTCCACCATGATTGTNNNAGATCGGAAGA |
| 38160_2 | TATGAGTGTGGAGTCGTTGCTCTTCACCTTCCACCATGATTGTNNNAGATCGGAAGA |
| 38160_3 | TATGAGTGTGGAGTCGTTGCTCTTCACCTTCCACCATGATTGTNNNAGATCGGAAGA |
| 36479_1 | TATGAGTGTGGAGTCGTTGCTTTCTATGAGGCCAGGAGCATTTNNNAGATCGGAAG |
| 36479_2 | TATGAGTGTGGAGTCGTTGCTTTCTATGAGGCCAGGAGCATTTNNNAGATCGGAAG |
| 36479_3 | TATGAGTGTGGAGTCGTTGCTTTCTATGAGGCCAGGAGCATTTNNNAGATCGGAAG |
| 45654_1 | TATGAGTGTGGAGTCGTTGCCCTCCCAGGTTCAAGTGATTCTCNNNAGATCGGAAG |
| 45654_2 | TATGAGTGTGGAGTCGTTGCCCTCCCAGGTTCAAGTGATTCTCNNNAGATCGGAAG |
| 45654_3 | TATGAGTGTGGAGTCGTTGCCCTCCCAGGTTCAAGTGATTCTCNNNAGATCGGAAG |
| 30490_1 | TATGAGTGTGGAGTCGTTGCAAACTCAGGGTTGAAGAAATGGCTTANNNAGATCG |
| 30490_2 | TATGAGTGTGGAGTCGTTGCAAACTCAGGGTTGAAGAAATGGCTTANNNAGATCG |
| 30490_3 | TATGAGTGTGGAGTCGTTGCAAACTCAGGGTTGAAGAAATGGCTTANNNAGATCG |
| 48971_1 | TATGAGTGTGGAGTCGTTGCAATAGCTGAGACTACAGGTGCACNNNAGATCGGAA |
| 48971_2 | TATGAGTGTGGAGTCGTTGCAATAGCTGAGACTACAGGTGCACNNNAGATCGGAA |
| 48971_3 | TATGAGTGTGGAGTCGTTGCAATAGCTGAGACTACAGGTGCACNNNAGATCGGAA |
| 37253_1 | TATGAGTGTGGAGTCGTTGCCCTGTCTTCTGCGTACTGATCATNNNAGATCGGAAG |
| 37253_2 | TATGAGTGTGGAGTCGTTGCCCTGTCTTCTGCGTACTGATCATNNNAGATCGGAAG |
| 37253_3 | TATGAGTGTGGAGTCGTTGCCCTGTCTTCTGCGTACTGATCATNNNAGATCGGAAG |
| 35101_1 | TATGAGTGTGGAGTCGTTGCGTCACCATGCCCAGCTAATTTTTNNNAGATCGGAAG |
| 35101_2 | TATGAGTGTGGAGTCGTTGCGTCACCATGCCCAGCTAATTTTTNNNAGATCGGAAG |
| 35101_3 | TATGAGTGTGGAGTCGTTGCGTCACCATGCCCAGCTAATTTTTNNNAGATCGGAAG |
| 49334_1 | TATGAGTGTGGAGTCGTTGCTTATTGCATCTGAATTCCACGGCNNNAGATCGGAAG |
| 49334_2 | TATGAGTGTGGAGTCGTTGCTTATTGCATCTGAATTCCACGGCNNNAGATCGGAAG |
| 49334_3 | TATGAGTGTGGAGTCGTTGCTTATTGCATCTGAATTCCACGGCNNNAGATCGGAAG |
| 47903_1 | TATGAGTGTGGAGTCGTTGCACACCACTATGCCTGGGTAATTTNNNAGATCGGAAG |
| 47903_2 | TATGAGTGTGGAGTCGTTGCACACCACTATGCCTGGGTAATTTNNNAGATCGGAAG |
| 47903_3 | TATGAGTGTGGAGTCGTTGCACACCACTATGCCTGGGTAATTTNNNAGATCGGAAG |
| 41050_1 | TATGAGTGTGGAGTCGTTGCAGGAAGATCCAAACATTCCCACANNNAGATCGGAAG |
| 41050_2 | TATGAGTGTGGAGTCGTTGCAGGAAGATCCAAACATTCCCACANNNAGATCGGAAG |
| 41050_3 | TATGAGTGTGGAGTCGTTGCAGGAAGATCCAAACATTCCCACANNNAGATCGGAAG |
| 47710_1 | TATGAGTGTGGAGTCGTTGCTGTCGAAGGCTATTTGTGATGGGNNNAGATCGGAA |
| 47710_2 | TATGAGTGTGGAGTCGTTGCTGTCGAAGGCTATTTGTGATGGGNNNAGATCGGAA |
| 47710_3 | TATGAGTGTGGAGTCGTTGCTGTCGAAGGCTATTTGTGATGGGNNNAGATCGGAA |
| 42490_1 | TATGAGTGTGGAGTCGTTGCACAGGTCTAACACAGTGTCTTATGTNNNAGATCGGA |
| 42490_2 | TATGAGTGTGGAGTCGTTGCACAGGTCTAACACAGTGTCTTATGTNNNAGATCGGA |
| 42490_3 | TATGAGTGTGGAGTCGTTGCACAGGTCTAACACAGTGTCTTATGTNNNAGATCGGA |
| 50660_1 | TATGAGTGTGGAGTCGTTGCAGGTCTGTTGCATTTTCCCCTATNNNAGATCGGAAG |

| | |
|---|---|
| 50660_2 | TATGAGTGTGGAGTCGTTGCAGGTCTGTTGCATTTTCCCCTATNNNAGATCGGAAGA |
| 50660_3 | TATGAGTGTGGAGTCGTTGCAGGTCTGTTGCATTTTCCCCTATNNNAGATCGGAAGA |
| 31091_1 | TATGAGTGTGGAGTCGTTGCAGAAGAAGTGAAAGCTACCAGCANNNAGATCGGAA |
| 31091_2 | TATGAGTGTGGAGTCGTTGCAGAAGAAGTGAAAGCTACCAGCANNNAGATCGGAA |
| 31091_3 | TATGAGTGTGGAGTCGTTGCAGAAGAAGTGAAAGCTACCAGCANNNAGATCGGAA |
| 29396_1 | TATGAGTGTGGAGTCGTTGCTCAGAAAGGCTGATCACAGCAAGNNNAGATCGGAA |
| 29396_2 | TATGAGTGTGGAGTCGTTGCTCAGAAAGGCTGATCACAGCAAGNNNAGATCGGAA |
| 29396_3 | TATGAGTGTGGAGTCGTTGCTCAGAAAGGCTGATCACAGCAAGNNNAGATCGGAA |
| 28976_1 | TATGAGTGTGGAGTCGTTGCATGTTGCATGTGAGAGAAGGGATNNNAGATCGGAA |
| 28976_2 | TATGAGTGTGGAGTCGTTGCATGTTGCATGTGAGAGAAGGGATNNNAGATCGGAA |
| 28976_3 | TATGAGTGTGGAGTCGTTGCATGTTGCATGTGAGAGAAGGGATNNNAGATCGGAA |
| 35242_1 | TATGAGTGTGGAGTCGTTGCTGTGTTCTTGATTTGGTTCTCAGCNNNAGATCGGAA |
| 35242_2 | TATGAGTGTGGAGTCGTTGCTGTGTTCTTGATTTGGTTCTCAGCNNNAGATCGGAA |
| 35242_3 | TATGAGTGTGGAGTCGTTGCTGTGTTCTTGATTTGGTTCTCAGCNNNAGATCGGAA |
| 39381_1 | TATGAGTGTGGAGTCGTTGCCTAGCCCTACAATTCTGCCTCATNNNAGATCGGAAGA |
| 39381_2 | TATGAGTGTGGAGTCGTTGCCTAGCCCTACAATTCTGCCTCATNNNAGATCGGAAGA |
| 39381_3 | TATGAGTGTGGAGTCGTTGCCTAGCCCTACAATTCTGCCTCATNNNAGATCGGAAGA |
| 29414_1 | TATGAGTGTGGAGTCGTTGCCCTTTGATTGTTTTCTAAAAGGANNNAGATCGGAAG |
| 29414_2 | TATGAGTGTGGAGTCGTTGCCCTTTGATTGTTTTCTAAAAGGANNNAGATCGGAAG |
| 29414_3 | TATGAGTGTGGAGTCGTTGCCCTTTGATTGTTTTCTAAAAGGANNNAGATCGGAAG |
| 30364_1 | TATGAGTGTGGAGTCGTTGCACCTGGCTAATTTTACATGGCTNNNAGATCGGAAGA |
| 30364_2 | TATGAGTGTGGAGTCGTTGCACCTGGCTAATTTTACATGGCTNNNAGATCGGAAGA |
| 30364_3 | TATGAGTGTGGAGTCGTTGCACCTGGCTAATTTTACATGGCTNNNAGATCGGAAGA |
| 34464_1 | TATGAGTGTGGAGTCGTTGCAGAAGGTGAAGCACAACAATGTGNNNAGATCGGAA |
| 34464_2 | TATGAGTGTGGAGTCGTTGCAGAAGGTGAAGCACAACAATGTGNNNAGATCGGAA |
| 34464_3 | TATGAGTGTGGAGTCGTTGCAGAAGGTGAAGCACAACAATGTGNNNAGATCGGAA |
| 49261_1 | TATGAGTGTGGAGTCGTTGCCTATCTTGGCTCACTGCAACCTCNNNAGATCGGAAGA |
| 49261_2 | TATGAGTGTGGAGTCGTTGCCTATCTTGGCTCACTGCAACCTCNNNAGATCGGAAGA |
| 49261_3 | TATGAGTGTGGAGTCGTTGCCTATCTTGGCTCACTGCAACCTCNNNAGATCGGAAGA |
| 8189_1 | TATGAGTGTGGAGTCGTTGCAGTGTAGTGTAGTGGTGCAATCACNNNAGATCGGA |
| 8189_2 | TATGAGTGTGGAGTCGTTGCAGTGTAGTGTAGTGGTGCAATCACNNNAGATCGGA |
| 8189_3 | TATGAGTGTGGAGTCGTTGCAGTGTAGTGTAGTGGTGCAATCACNNNAGATCGGA |
| 50362_1 | TATGAGTGTGGAGTCGTTGCACACAACAGACCCATAGCTAACANNNAGATCGGAAG |
| 50362_2 | TATGAGTGTGGAGTCGTTGCACACAACAGACCCATAGCTAACANNNAGATCGGAAG |
| 50362_3 | TATGAGTGTGGAGTCGTTGCACACAACAGACCCATAGCTAACANNNAGATCGGAAG |
| 34623_1 | TATGAGTGTGGAGTCGTTGCTGAAAGGAGGAAAGGAGGAAAGGNNNAGATCGGA |
| 34623_2 | TATGAGTGTGGAGTCGTTGCTGAAAGGAGGAAAGGAGGAAAGGNNNAGATCGGA |
| 34623_3 | TATGAGTGTGGAGTCGTTGCTGAAAGGAGGAAAGGAGGAAAGGNNNAGATCGGA |
| 39678_1 | TATGAGTGTGGAGTCGTTGCGGATGGAGTACCACATAGGCATTNNNAGATCGGAA |
| 39678_2 | TATGAGTGTGGAGTCGTTGCGGATGGAGTACCACATAGGCATTNNNAGATCGGAA |
| 39678_3 | TATGAGTGTGGAGTCGTTGCGGATGGAGTACCACATAGGCATTNNNAGATCGGAA |
| 36468_1 | TATGAGTGTGGAGTCGTTGCACACAGATCATAGGGCTCAAACTNNNAGATCGGAAG |
| 36468_2 | TATGAGTGTGGAGTCGTTGCACACAGATCATAGGGCTCAAACTNNNAGATCGGAAG |

| | |
|---|---|
| 36468_3 | TATGAGTGTGGAGTCGTTGCACACAGATCATAGGGCTCAAACTNNNAGATCGGAAG |
| 43138_1 | TATGAGTGTGGAGTCGTTGCAAAACATTGGGGAAATGCTCCAGNNNAGATCGGAA |
| 43138_2 | TATGAGTGTGGAGTCGTTGCAAAACATTGGGGAAATGCTCCAGNNNAGATCGGAA |
| 43138_3 | TATGAGTGTGGAGTCGTTGCAAAACATTGGGGAAATGCTCCAGNNNAGATCGGAA |
| 50334_1 | TATGAGTGTGGAGTCGTTGCTATACACACACACACACACACACNNNAGATCGGAAG |
| 50334_2 | TATGAGTGTGGAGTCGTTGCTATACACACACACACACACACACNNNAGATCGGAAG |
| 50334_3 | TATGAGTGTGGAGTCGTTGCTATACACACACACACACACACACNNNAGATCGGAAG |
| 38791_1 | TATGAGTGTGGAGTCGTTGCTTCTCTTTACCTTCTTTTGGGCANNNAGATCGGAAGA |
| 38791_2 | TATGAGTGTGGAGTCGTTGCTTCTCTTTACCTTCTTTTGGGCANNNAGATCGGAAGA |
| 38791_3 | TATGAGTGTGGAGTCGTTGCTTCTCTTTACCTTCTTTTGGGCANNNAGATCGGAAGA |
| 37017_1 | TATGAGTGTGGAGTCGTTGCCTGAGGCAGGAGAATCACTTGAANNNAGATCGGAA |
| 37017_2 | TATGAGTGTGGAGTCGTTGCCTGAGGCAGGAGAATCACTTGAANNNAGATCGGAA |
| 37017_3 | TATGAGTGTGGAGTCGTTGCCTGAGGCAGGAGAATCACTTGAANNNAGATCGGAA |
| 44524_1 | TATGAGTGTGGAGTCGTTGCCTACACACAGGTGAGCTCTGGNNNAGATCGGAAGA |
| 44524_2 | TATGAGTGTGGAGTCGTTGCCTACACACAGGTGAGCTCTGGNNNAGATCGGAAGA |
| 44524_3 | TATGAGTGTGGAGTCGTTGCCTACACACAGGTGAGCTCTGGNNNAGATCGGAAGA |
| 44414_1 | TATGAGTGTGGAGTCGTTGCGTCCCTCTTCTTTTCTCCCAGAGNNNAGATCGGAAGA |
| 44414_2 | TATGAGTGTGGAGTCGTTGCGTCCCTCTTCTTTTCTCCCAGAGNNNAGATCGGAAGA |
| 44414_3 | TATGAGTGTGGAGTCGTTGCGTCCCTCTTCTTTTCTCCCAGAGNNNAGATCGGAAGA |
| 50918_1 | TATGAGTGTGGAGTCGTTGCCTTCTGCACCCCTGTAAAATGTGNNNAGATCGGAAG |
| 50918_2 | TATGAGTGTGGAGTCGTTGCCTTCTGCACCCCTGTAAAATGTGNNNAGATCGGAAG |
| 50918_3 | TATGAGTGTGGAGTCGTTGCCTTCTGCACCCCTGTAAAATGTGNNNAGATCGGAAG |
| 31761_1 | TATGAGTGTGGAGTCGTTGCAATCTGTATGTGTGTGTGTGTGCNNNAGATCGGAAG |
| 31761_2 | TATGAGTGTGGAGTCGTTGCAATCTGTATGTGTGTGTGTGTGCNNNAGATCGGAAG |
| 31761_3 | TATGAGTGTGGAGTCGTTGCAATCTGTATGTGTGTGTGTGTGCNNNAGATCGGAAG |
| 49637_1 | TATGAGTGTGGAGTCGTTGCATTCCTAACCCTTGGCAACTTCTNNNAGATCGGAAGA |
| 49637_2 | TATGAGTGTGGAGTCGTTGCATTCCTAACCCTTGGCAACTTCTNNNAGATCGGAAGA |
| 49637_3 | TATGAGTGTGGAGTCGTTGCATTCCTAACCCTTGGCAACTTCTNNNAGATCGGAAGA |
| 44954_1 | TATGAGTGTGGAGTCGTTGCAGGCAATTTCTTGGGCATTTACCNNNAGATCGGAAG |
| 44954_2 | TATGAGTGTGGAGTCGTTGCAGGCAATTTCTTGGGCATTTACCNNNAGATCGGAAG |
| 44954_3 | TATGAGTGTGGAGTCGTTGCAGGCAATTTCTTGGGCATTTACCNNNAGATCGGAAG |
| 37664_1 | TATGAGTGTGGAGTCGTTGCGGAGGCTGAAGCAGGAGAAAGNNNAGATCGGAAG |
| 37664_2 | TATGAGTGTGGAGTCGTTGCGGAGGCTGAAGCAGGAGAAAGNNNAGATCGGAAG |
| 37664_3 | TATGAGTGTGGAGTCGTTGCGGAGGCTGAAGCAGGAGAAAGNNNAGATCGGAAG |
| 39071_1 | TATGAGTGTGGAGTCGTTGCTGGCATTCCAACCATAAAAATGCNNNAGATCGGAAG |
| 39071_2 | TATGAGTGTGGAGTCGTTGCTGGCATTCCAACCATAAAAATGCNNNAGATCGGAAG |
| 39071_3 | TATGAGTGTGGAGTCGTTGCTGGCATTCCAACCATAAAAATGCNNNAGATCGGAAG |
| 13448_1 | TATGAGTGTGGAGTCGTTGCGAAGAGAGCTGTTGAACATGNNNAGATCGGAAGAG |
| 13448_2 | TATGAGTGTGGAGTCGTTGCGAAGAGAGCTGTTGAACATGNNNAGATCGGAAGAG |
| 13448_3 | TATGAGTGTGGAGTCGTTGCGAAGAGAGCTGTTGAACATGNNNAGATCGGAAGAG |
| 3173_1 | TATGAGTGTGGAGTCGTTGCAAAGAAATGGGGAGAGAGGTGGANNNAGATCGGA |
| 3173_2 | TATGAGTGTGGAGTCGTTGCAAAGAAATGGGGAGAGAGGTGGANNNAGATCGGA |
| 3173_3 | TATGAGTGTGGAGTCGTTGCAAAGAAATGGGGAGAGAGGTGGANNNAGATCGGA |

| | |
|---|---|
| 33033_1 | TATGAGTGTGGAGTCGTTGCCTGGAGTGCAATGGTGAGATCTCNNNAGATCGGAAC |
| 33033_2 | TATGAGTGTGGAGTCGTTGCCTGGAGTGCAATGGTGAGATCTCNNNAGATCGGAAC |
| 33033_3 | TATGAGTGTGGAGTCGTTGCCTGGAGTGCAATGGTGAGATCTCNNNAGATCGGAAC |
| 53018_1 | TATGAGTGTGGAGTCGTTGCCCTCTCCTTTCCTTTCCTTTCCTNNNAGATCGGAAGAG |
| 53018_2 | TATGAGTGTGGAGTCGTTGCCCTCTCCTTTCCTTTCCTTTCCTNNNAGATCGGAAGAG |
| 53018_3 | TATGAGTGTGGAGTCGTTGCCCTCTCCTTTCCTTTCCTTTCCTNNNAGATCGGAAGAG |
| 36462_1 | TATGAGTGTGGAGTCGTTGCCAGTCGACAGAAGCTTCAAAAGGNNNAGATCGGAAC |
| 36462_2 | TATGAGTGTGGAGTCGTTGCCAGTCGACAGAAGCTTCAAAAGGNNNAGATCGGAAC |
| 36462_3 | TATGAGTGTGGAGTCGTTGCCAGTCGACAGAAGCTTCAAAAGGNNNAGATCGGAAC |
| 47622_1 | TATGAGTGTGGAGTCGTTGCCCAAGTAGCTGGGATTACAGGTGNNNAGATCGGAAC |
| 47622_2 | TATGAGTGTGGAGTCGTTGCCCAAGTAGCTGGGATTACAGGTGNNNAGATCGGAAC |
| 47622_3 | TATGAGTGTGGAGTCGTTGCCCAAGTAGCTGGGATTACAGGTGNNNAGATCGGAAC |
| 30202_1 | TATGAGTGTGGAGTCGTTGCGTTCAATTTGTGCTTCTCGTGCANNNAGATCGGAAGA |
| 30202_2 | TATGAGTGTGGAGTCGTTGCGTTCAATTTGTGCTTCTCGTGCANNNAGATCGGAAGA |
| 30202_3 | TATGAGTGTGGAGTCGTTGCGTTCAATTTGTGCTTCTCGTGCANNNAGATCGGAAGA |
| 51771_1 | TATGAGTGTGGAGTCGTTGCCCCAAACTGGCCATAAACGAAATNNNAGATCGGAAG |
| 51771_2 | TATGAGTGTGGAGTCGTTGCCCCAAACTGGCCATAAACGAAATNNNAGATCGGAAG |
| 51771_3 | TATGAGTGTGGAGTCGTTGCCCCAAACTGGCCATAAACGAAATNNNAGATCGGAAG |
| 33427_1 | TATGAGTGTGGAGTCGTTGCCTGGAGAGAAGGCAGTACTTGAANNNAGATCGGAA |
| 33427_2 | TATGAGTGTGGAGTCGTTGCCTGGAGAGAAGGCAGTACTTGAANNNAGATCGGAA |
| 33427_3 | TATGAGTGTGGAGTCGTTGCCTGGAGAGAAGGCAGTACTTGAANNNAGATCGGAA |
| 44871_1 | TATGAGTGTGGAGTCGTTGCAGAGATGTGCAGCTGGACTAAAGNNNAGATCGGAA |
| 44871_2 | TATGAGTGTGGAGTCGTTGCAGAGATGTGCAGCTGGACTAAAGNNNAGATCGGAA |
| 44871_3 | TATGAGTGTGGAGTCGTTGCAGAGATGTGCAGCTGGACTAAAGNNNAGATCGGAA |
| 35229_1 | TATGAGTGTGGAGTCGTTGCGTGATTTGGATGATGGAGGGTCTNNNAGATCGGAA |
| 35229_2 | TATGAGTGTGGAGTCGTTGCGTGATTTGGATGATGGAGGGTCTNNNAGATCGGAA |
| 35229_3 | TATGAGTGTGGAGTCGTTGCGTGATTTGGATGATGGAGGGTCTNNNAGATCGGAA |
| 46082_1 | TATGAGTGTGGAGTCGTTGCAGAAAGGGTACTGTAATGCAGGGNNNAGATCGGAA |
| 46082_2 | TATGAGTGTGGAGTCGTTGCAGAAAGGGTACTGTAATGCAGGGNNNAGATCGGAA |
| 46082_3 | TATGAGTGTGGAGTCGTTGCAGAAAGGGTACTGTAATGCAGGGNNNAGATCGGAA |
| 36723_1 | TATGAGTGTGGAGTCGTTGCAAACTGAGGCTCGAGGAGTTTAGNNNAGATCGGAA |
| 36723_2 | TATGAGTGTGGAGTCGTTGCAAACTGAGGCTCGAGGAGTTTAGNNNAGATCGGAA |
| 36723_3 | TATGAGTGTGGAGTCGTTGCAAACTGAGGCTCGAGGAGTTTAGNNNAGATCGGAA |
| 52137_1 | TATGAGTGTGGAGTCGTTGCACAGTGACACAGTCTCATTCACANNNAGATCGGAAGA |
| 52137_2 | TATGAGTGTGGAGTCGTTGCACAGTGACACAGTCTCATTCACANNNAGATCGGAAGA |
| 52137_3 | TATGAGTGTGGAGTCGTTGCACAGTGACACAGTCTCATTCACANNNAGATCGGAAGA |
| 29030_1 | TATGAGTGTGGAGTCGTTGCAGCCTGTGATAAACTGAGTCCTGNNNAGATCGGAAG |
| 29030_2 | TATGAGTGTGGAGTCGTTGCAGCCTGTGATAAACTGAGTCCTGNNNAGATCGGAAG |
| 29030_3 | TATGAGTGTGGAGTCGTTGCAGCCTGTGATAAACTGAGTCCTGNNNAGATCGGAAG |
| 34383_1 | TATGAGTGTGGAGTCGTTGCGGGAATCTGCCCTGTGTCATCNNNAGATCGGAAGAG |
| 34383_2 | TATGAGTGTGGAGTCGTTGCGGGAATCTGCCCTGTGTCATCNNNAGATCGGAAGAG |
| 34383_3 | TATGAGTGTGGAGTCGTTGCGGGAATCTGCCCTGTGTCATCNNNAGATCGGAAGAG |
| 32981_1 | TATGAGTGTGGAGTCGTTGCGTAAGGTCTAGAGGCAACCCTTCNNNAGATCGGAAG |

| | |
|---|---|
| 32981_2 | TATGAGTGTGGAGTCGTTGCGTAAGGTCTAGAGGCAACCCTTCNNNAGATCGGAAG |
| 32981_3 | TATGAGTGTGGAGTCGTTGCGTAAGGTCTAGAGGCAACCCTTCNNNAGATCGGAAG |
| 43705_1 | TATGAGTGTGGAGTCGTTGCATGAGAGGTCATCAGGCACACNNNAGATCGGAAGA |
| 43705_2 | TATGAGTGTGGAGTCGTTGCATGAGAGGTCATCAGGCACACNNNAGATCGGAAGA |
| 43705_3 | TATGAGTGTGGAGTCGTTGCATGAGAGGTCATCAGGCACACNNNAGATCGGAAGA |
| 50025_1 | TATGAGTGTGGAGTCGTTGCCAAAAGGGGTGGGAGAGAAAGAGNNNAGATCGGA |
| 50025_2 | TATGAGTGTGGAGTCGTTGCCAAAAGGGGTGGGAGAGAAAGAGNNNAGATCGGA |
| 50025_3 | TATGAGTGTGGAGTCGTTGCCAAAAGGGGTGGGAGAGAAAGAGNNNAGATCGGA |
| 2704_1 | TATGAGTGTGGAGTCGTTGCCCCAAATTGCTCCAGATGTGGATNNNAGATCGGAAG |
| 2704_2 | TATGAGTGTGGAGTCGTTGCCCCAAATTGCTCCAGATGTGGATNNNAGATCGGAAG |
| 2704_3 | TATGAGTGTGGAGTCGTTGCCCCAAATTGCTCCAGATGTGGATNNNAGATCGGAAG |
| 45390_1 | TATGAGTGTGGAGTCGTTGCTTGTGTCCATTGCTGGATTCAACNNNAGATCGGAAG |
| 45390_2 | TATGAGTGTGGAGTCGTTGCTTGTGTCCATTGCTGGATTCAACNNNAGATCGGAAG |
| 45390_3 | TATGAGTGTGGAGTCGTTGCTTGTGTCCATTGCTGGATTCAACNNNAGATCGGAAG |
| 40942_1 | TATGAGTGTGGAGTCGTTGCTCACTGATGAACATTTCTTCCATTTNNNAGATCGGAA |
| 40942_2 | TATGAGTGTGGAGTCGTTGCTCACTGATGAACATTTCTTCCATTTNNNAGATCGGAA |
| 40942_3 | TATGAGTGTGGAGTCGTTGCTCACTGATGAACATTTCTTCCATTTNNNAGATCGGAA |
| 43429_1 | TATGAGTGTGGAGTCGTTGCAGCATGAAGCGGAGTAGCATAATNNNAGATCGGAA |
| 43429_2 | TATGAGTGTGGAGTCGTTGCAGCATGAAGCGGAGTAGCATAATNNNAGATCGGAA |
| 43429_3 | TATGAGTGTGGAGTCGTTGCAGCATGAAGCGGAGTAGCATAATNNNAGATCGGAA |
| 30535_1 | TATGAGTGTGGAGTCGTTGCCCTGTTGCACTTGATGGCTTTTTNNNAGATCGGAAG |
| 30535_2 | TATGAGTGTGGAGTCGTTGCCCTGTTGCACTTGATGGCTTTTTNNNAGATCGGAAG |
| 30535_3 | TATGAGTGTGGAGTCGTTGCCCTGTTGCACTTGATGGCTTTTTNNNAGATCGGAAG |
| 48661_1 | TATGAGTGTGGAGTCGTTGCTAGGAATGGGGATGGGAGAGTAANNNAGATCGGA |
| 48661_2 | TATGAGTGTGGAGTCGTTGCTAGGAATGGGGATGGGAGAGTAANNNAGATCGGA |
| 48661_3 | TATGAGTGTGGAGTCGTTGCTAGGAATGGGGATGGGAGAGTAANNNAGATCGGA |
| 53387_1 | TATGAGTGTGGAGTCGTTGCGTAAGACGTGACTTGGTCTTCCTNNNAGATCGGAAG |
| 53387_2 | TATGAGTGTGGAGTCGTTGCGTAAGACGTGACTTGGTCTTCCTNNNAGATCGGAAG |
| 53387_3 | TATGAGTGTGGAGTCGTTGCGTAAGACGTGACTTGGTCTTCCTNNNAGATCGGAAG |
| 49108_1 | TATGAGTGTGGAGTCGTTGCGCACCCGGCTATTTTTACTTTCTNNNAGATCGGAAGA |
| 49108_2 | TATGAGTGTGGAGTCGTTGCGCACCCGGCTATTTTTACTTTCTNNNAGATCGGAAGA |
| 49108_3 | TATGAGTGTGGAGTCGTTGCGCACCCGGCTATTTTTACTTTCTNNNAGATCGGAAGA |
| 43615_1 | TATGAGTGTGGAGTCGTTGCTCCTCACACACGACAAAACTCTCNNNAGATCGGAAGA |
| 43615_2 | TATGAGTGTGGAGTCGTTGCTCCTCACACACGACAAAACTCTCNNNAGATCGGAAGA |
| 43615_3 | TATGAGTGTGGAGTCGTTGCTCCTCACACACGACAAAACTCTCNNNAGATCGGAAGA |
| 48659_1 | TATGAGTGTGGAGTCGTTGCCTCCCACGTAGCTGAGATTACAANNNAGATCGGAAG |
| 48659_2 | TATGAGTGTGGAGTCGTTGCCTCCCACGTAGCTGAGATTACAANNNAGATCGGAAG |
| 48659_3 | TATGAGTGTGGAGTCGTTGCCTCCCACGTAGCTGAGATTACAANNNAGATCGGAAG |
| 39386_1 | TATGAGTGTGGAGTCGTTGCAGCAGCACTATTCACATCAGTCANNNAGATCGGAAG |
| 39386_2 | TATGAGTGTGGAGTCGTTGCAGCAGCACTATTCACATCAGTCANNNAGATCGGAAG |
| 39386_3 | TATGAGTGTGGAGTCGTTGCAGCAGCACTATTCACATCAGTCANNNAGATCGGAAG |
| 51587_1 | TATGAGTGTGGAGTCGTTGCATAACTGGGAAAGCTGGACAGAGNNNAGATCGGAA |
| 51587_2 | TATGAGTGTGGAGTCGTTGCATAACTGGGAAAGCTGGACAGAGNNNAGATCGGAA |

| | |
|---|---|
| 51587_3 | TATGAGTGTGGAGTCGTTGCATAACTGGGAAAGCTGGACAGAGNNNAGATCGGAA |
| 50929_1 | TATGAGTGTGGAGTCGTTGCTTTCTGACCCTAGGTGCTTTACCNNNAGATCGGAAGA |
| 50929_2 | TATGAGTGTGGAGTCGTTGCTTTCTGACCCTAGGTGCTTTACCNNNAGATCGGAAGA |
| 50929_3 | TATGAGTGTGGAGTCGTTGCTTTCTGACCCTAGGTGCTTTACCNNNAGATCGGAAGA |
| 53008_1 | TATGAGTGTGGAGTCGTTGCGAACTTCCCGTTTTGGACTTCTGNNNAGATCGGAAGA |
| 53008_2 | TATGAGTGTGGAGTCGTTGCGAACTTCCCGTTTTGGACTTCTGNNNAGATCGGAAGA |
| 53008_3 | TATGAGTGTGGAGTCGTTGCGAACTTCCCGTTTTGGACTTCTGNNNAGATCGGAAGA |
| 48825_1 | TATGAGTGTGGAGTCGTTGCGATTGGCCTCTGTCCGGAAGNNNAGATCGGAAGAG |
| 48825_2 | TATGAGTGTGGAGTCGTTGCGATTGGCCTCTGTCCGGAAGNNNAGATCGGAAGAG |
| 48825_3 | TATGAGTGTGGAGTCGTTGCGATTGGCCTCTGTCCGGAAGNNNAGATCGGAAGAG |
| 49804_1 | TATGAGTGTGGAGTCGTTGCCTCAAGTGATCCTCCCATCTCAGNNNAGATCGGAAGA |
| 49804_2 | TATGAGTGTGGAGTCGTTGCCTCAAGTGATCCTCCCATCTCAGNNNAGATCGGAAGA |
| 49804_3 | TATGAGTGTGGAGTCGTTGCCTCAAGTGATCCTCCCATCTCAGNNNAGATCGGAAGA |
| 50075_1 | TATGAGTGTGGAGTCGTTGCGAAGCGATAACACGTTGACCATCNNNAGATCGGAAG |
| 50075_2 | TATGAGTGTGGAGTCGTTGCGAAGCGATAACACGTTGACCATCNNNAGATCGGAAG |
| 50075_3 | TATGAGTGTGGAGTCGTTGCGAAGCGATAACACGTTGACCATCNNNAGATCGGAAG |
| 51721_1 | TATGAGTGTGGAGTCGTTGCATTCACGTGTTTTTATGCTGCCANNNAGATCGGAAGA |
| 51721_2 | TATGAGTGTGGAGTCGTTGCATTCACGTGTTTTTATGCTGCCANNNAGATCGGAAGA |
| 51721_3 | TATGAGTGTGGAGTCGTTGCATTCACGTGTTTTTATGCTGCCANNNAGATCGGAAGA |
| 31613_1 | TATGAGTGTGGAGTCGTTGCTGAAAATGGGACAATGGAAGGGANNNAGATCGGAA |
| 31613_2 | TATGAGTGTGGAGTCGTTGCTGAAAATGGGACAATGGAAGGGANNNAGATCGGAA |
| 31613_3 | TATGAGTGTGGAGTCGTTGCTGAAAATGGGACAATGGAAGGGANNNAGATCGGAA |
| 43308_1 | TATGAGTGTGGAGTCGTTGCCCTGGGTTCAAGCAATTCTCTTGNNNAGATCGGAAGA |
| 43308_2 | TATGAGTGTGGAGTCGTTGCCCTGGGTTCAAGCAATTCTCTTGNNNAGATCGGAAGA |
| 43308_3 | TATGAGTGTGGAGTCGTTGCCCTGGGTTCAAGCAATTCTCTTGNNNAGATCGGAAGA |
| 37220_1 | TATGAGTGTGGAGTCGTTGCTGATGCCTCCAGCTTGTTCTTATNNNAGATCGGAAGA |
| 37220_2 | TATGAGTGTGGAGTCGTTGCTGATGCCTCCAGCTTGTTCTTATNNNAGATCGGAAGA |
| 37220_3 | TATGAGTGTGGAGTCGTTGCTGATGCCTCCAGCTTGTTCTTATNNNAGATCGGAAGA |
| 10452_1 | TATGAGTGTGGAGTCGTTGCGGCTACAGCTCTGAGGTACACTANNNAGATCGGAAG |
| 10452_2 | TATGAGTGTGGAGTCGTTGCGGCTACAGCTCTGAGGTACACTANNNAGATCGGAAG |
| 10452_3 | TATGAGTGTGGAGTCGTTGCGGCTACAGCTCTGAGGTACACTANNNAGATCGGAAG |
| 50055_1 | TATGAGTGTGGAGTCGTTGCGTATGGAGAGGAGGATGAGCACNNNAGATCGGAA |
| 50055_2 | TATGAGTGTGGAGTCGTTGCGTATGGAGAGGAGGATGAGCACNNNAGATCGGAA |
| 50055_3 | TATGAGTGTGGAGTCGTTGCGTATGGAGAGGAGGATGAGCACNNNAGATCGGAA |
| 40481_1 | TATGAGTGTGGAGTCGTTGCTGAAGTTTCCCAGGCTATAGTGGNNNAGATCGGAAG |
| 40481_2 | TATGAGTGTGGAGTCGTTGCTGAAGTTTCCCAGGCTATAGTGGNNNAGATCGGAAG |
| 40481_3 | TATGAGTGTGGAGTCGTTGCTGAAGTTTCCCAGGCTATAGTGGNNNAGATCGGAAG |
| 244_1 | TATGAGTGTGGAGTCGTTGCCGGTCTGTCTCAGTCTGCTAAANNNAGATCGGAAGA |
| 244_2 | TATGAGTGTGGAGTCGTTGCCGGTCTGTCTCAGTCTGCTAAANNNAGATCGGAAGA |
| 244_3 | TATGAGTGTGGAGTCGTTGCCGGTCTGTCTCAGTCTGCTAAANNNAGATCGGAAGA |
| 32663_1 | TATGAGTGTGGAGTCGTTGCGCCTCCTTAGTAGCTGGGATTACNNNAGATCGGAAG |
| 32663_2 | TATGAGTGTGGAGTCGTTGCGCCTCCTTAGTAGCTGGGATTACNNNAGATCGGAAG |
| 32663_3 | TATGAGTGTGGAGTCGTTGCGCCTCCTTAGTAGCTGGGATTACNNNAGATCGGAAG |

| | |
|---|---|
| 37374_1 | TATGAGTGTGGAGTCGTTGCTTGAAATTGAAGTAGATGGCCGGNNNAGATCGGAA |
| 37374_2 | TATGAGTGTGGAGTCGTTGCTTGAAATTGAAGTAGATGGCCGGNNNAGATCGGAA |
| 37374_3 | TATGAGTGTGGAGTCGTTGCTTGAAATTGAAGTAGATGGCCGGNNNAGATCGGAA |
| 3091_1 | TATGAGTGTGGAGTCGTTGCTGACGCACTCACTGAGGATTTTNNNAGATCGGAAG |
| 3091_2 | TATGAGTGTGGAGTCGTTGCTGACGCACTCACTGAGGATTTTNNNAGATCGGAAG |
| 3091_3 | TATGAGTGTGGAGTCGTTGCTGACGCACTCACTGAGGATTTTNNNAGATCGGAAG |
| 33693_1 | TATGAGTGTGGAGTCGTTGCAAGCAACCTGAGAAGCTAGTTCANNNAGATCGGAA |
| 33693_2 | TATGAGTGTGGAGTCGTTGCAAGCAACCTGAGAAGCTAGTTCANNNAGATCGGAA |
| 33693_3 | TATGAGTGTGGAGTCGTTGCAAGCAACCTGAGAAGCTAGTTCANNNAGATCGGAA |
| 41191_1 | TATGAGTGTGGAGTCGTTGCATGACCCATGATGAAACCCATGANNNAGATCGGAAG |
| 41191_2 | TATGAGTGTGGAGTCGTTGCATGACCCATGATGAAACCCATGANNNAGATCGGAAG |
| 41191_3 | TATGAGTGTGGAGTCGTTGCATGACCCATGATGAAACCCATGANNNAGATCGGAAG |
| 38481_1 | TATGAGTGTGGAGTCGTTGCATCTTCTCCTGTGCCCATAATGANNNAGATCGGAAG |
| 38481_2 | TATGAGTGTGGAGTCGTTGCATCTTCTCCTGTGCCCATAATGANNNAGATCGGAAG |
| 38481_3 | TATGAGTGTGGAGTCGTTGCATCTTCTCCTGTGCCCATAATGANNNAGATCGGAAG |
| 48813_1 | TATGAGTGTGGAGTCGTTGCATCAGCTGTGATGTTGACCAAGANNNAGATCGGAA |
| 48813_2 | TATGAGTGTGGAGTCGTTGCATCAGCTGTGATGTTGACCAAGANNNAGATCGGAA |
| 48813_3 | TATGAGTGTGGAGTCGTTGCATCAGCTGTGATGTTGACCAAGANNNAGATCGGAA |
| 41350_1 | TATGAGTGTGGAGTCGTTGCTGAGACGGAGTATCACTCTGTTGNNNAGATCGGAA |
| 41350_2 | TATGAGTGTGGAGTCGTTGCTGAGACGGAGTATCACTCTGTTGNNNAGATCGGAA |
| 41350_3 | TATGAGTGTGGAGTCGTTGCTGAGACGGAGTATCACTCTGTTGNNNAGATCGGAA |
| 33147_1 | TATGAGTGTGGAGTCGTTGCGCTGTGCTCCAATAAGACTTCACNNNAGATCGGAAG |
| 33147_2 | TATGAGTGTGGAGTCGTTGCGCTGTGCTCCAATAAGACTTCACNNNAGATCGGAAG |
| 33147_3 | TATGAGTGTGGAGTCGTTGCGCTGTGCTCCAATAAGACTTCACNNNAGATCGGAAG |
| 38438_1 | TATGAGTGTGGAGTCGTTGCACATTCATGATTTTGCCACATTGCNNNAGATCGGAA |
| 38438_2 | TATGAGTGTGGAGTCGTTGCACATTCATGATTTTGCCACATTGCNNNAGATCGGAA |
| 38438_3 | TATGAGTGTGGAGTCGTTGCACATTCATGATTTTGCCACATTGCNNNAGATCGGAA |
| 52080_1 | TATGAGTGTGGAGTCGTTGCATTCTTGGGAACATAGTGGTCCCNNNAGATCGGAAG |
| 52080_2 | TATGAGTGTGGAGTCGTTGCATTCTTGGGAACATAGTGGTCCCNNNAGATCGGAAG |
| 52080_3 | TATGAGTGTGGAGTCGTTGCATTCTTGGGAACATAGTGGTCCCNNNAGATCGGAAG |
| 10850_1 | TATGAGTGTGGAGTCGTTGCTGGAGGGAACAACAAAGAAGGAANNNAGATCGGA |
| 10850_2 | TATGAGTGTGGAGTCGTTGCTGGAGGGAACAACAAAGAAGGAANNNAGATCGGA |
| 10850_3 | TATGAGTGTGGAGTCGTTGCTGGAGGGAACAACAAAGAAGGAANNNAGATCGGA |
| 29726_1 | TATGAGTGTGGAGTCGTTGCGTGAGGGTGTCTGAAACCACNNNAGATCGGAAGAG |
| 29726_2 | TATGAGTGTGGAGTCGTTGCGTGAGGGTGTCTGAAACCACNNNAGATCGGAAGAG |
| 29726_3 | TATGAGTGTGGAGTCGTTGCGTGAGGGTGTCTGAAACCACNNNAGATCGGAAGAG |
| 39993_1 | TATGAGTGTGGAGTCGTTGCTAAGTGACAGTGCCAAATACCCCNNNAGATCGGAAG |
| 39993_2 | TATGAGTGTGGAGTCGTTGCTAAGTGACAGTGCCAAATACCCCNNNAGATCGGAAG |
| 39993_3 | TATGAGTGTGGAGTCGTTGCTAAGTGACAGTGCCAAATACCCCNNNAGATCGGAAG |
| 40207_1 | TATGAGTGTGGAGTCGTTGCACTGTATTAGGCCACTCTTGCATNNNAGATCGGAAG |
| 40207_2 | TATGAGTGTGGAGTCGTTGCACTGTATTAGGCCACTCTTGCATNNNAGATCGGAAG |
| 40207_3 | TATGAGTGTGGAGTCGTTGCACTGTATTAGGCCACTCTTGCATNNNAGATCGGAAG |
| 31539_1 | TATGAGTGTGGAGTCGTTGCGTGTGTGAGACAGACTTTTGCTCNNNAGATCGGAA |

| | |
|---|---|
| 31539_2 | TATGAGTGTGGAGTCGTTGCGTGTGTGAGACAGACTTTTGCTCNNNAGATCGGAAG |
| 31539_3 | TATGAGTGTGGAGTCGTTGCGTGTGTGAGACAGACTTTTGCTCNNNAGATCGGAAG |
| 36185_1 | TATGAGTGTGGAGTCGTTGCTCATAGTACACAATGGGTTCAACANNNAGATCGGAA |
| 36185_2 | TATGAGTGTGGAGTCGTTGCTCATAGTACACAATGGGTTCAACANNNAGATCGGAA |
| 36185_3 | TATGAGTGTGGAGTCGTTGCTCATAGTACACAATGGGTTCAACANNNAGATCGGAA |
| 38164_1 | TATGAGTGTGGAGTCGTTGCGGATGACAGAGTGAGACTCCATTNNNAGATCGGAA |
| 38164_2 | TATGAGTGTGGAGTCGTTGCGGATGACAGAGTGAGACTCCATTNNNAGATCGGAA |
| 38164_3 | TATGAGTGTGGAGTCGTTGCGGATGACAGAGTGAGACTCCATTNNNAGATCGGAA |
| 48397_1 | TATGAGTGTGGAGTCGTTGCCTCCCGAGTAGCCAAGATTACAGNNNAGATCGGAAG |
| 48397_2 | TATGAGTGTGGAGTCGTTGCCTCCCGAGTAGCCAAGATTACAGNNNAGATCGGAAG |
| 48397_3 | TATGAGTGTGGAGTCGTTGCCTCCCGAGTAGCCAAGATTACAGNNNAGATCGGAAG |
| 30936_1 | TATGAGTGTGGAGTCGTTGCTGGCTGTAGATGGAAAGGAGATGNNNAGATCGGA |
| 30936_2 | TATGAGTGTGGAGTCGTTGCTGGCTGTAGATGGAAAGGAGATGNNNAGATCGGA |
| 30936_3 | TATGAGTGTGGAGTCGTTGCTGGCTGTAGATGGAAAGGAGATGNNNAGATCGGA |
| 49731_1 | TATGAGTGTGGAGTCGTTGCGCTCAAAACCCACATGAGACTTCNNNAGATCGGAAG |
| 49731_2 | TATGAGTGTGGAGTCGTTGCGCTCAAAACCCACATGAGACTTCNNNAGATCGGAAG |
| 49731_3 | TATGAGTGTGGAGTCGTTGCGCTCAAAACCCACATGAGACTTCNNNAGATCGGAAG |
| 29393_1 | TATGAGTGTGGAGTCGTTGCTGACCACTGATTTAGAACGGGAANNNAGATCGGAA |
| 29393_2 | TATGAGTGTGGAGTCGTTGCTGACCACTGATTTAGAACGGGAANNNAGATCGGAA |
| 29393_3 | TATGAGTGTGGAGTCGTTGCTGACCACTGATTTAGAACGGGAANNNAGATCGGAA |
| 44555_1 | TATGAGTGTGGAGTCGTTGCGGCTTCCACTGAAAAGTCTGTTGNNNAGATCGGAAG |
| 44555_2 | TATGAGTGTGGAGTCGTTGCGGCTTCCACTGAAAAGTCTGTTGNNNAGATCGGAAG |
| 44555_3 | TATGAGTGTGGAGTCGTTGCGGCTTCCACTGAAAAGTCTGTTGNNNAGATCGGAAG |
| 36819_1 | TATGAGTGTGGAGTCGTTGCTCTGCTAGCATCTCTGAAAAGGGNNNAGATCGGAAG |
| 36819_2 | TATGAGTGTGGAGTCGTTGCTCTGCTAGCATCTCTGAAAAGGGNNNAGATCGGAAG |
| 36819_3 | TATGAGTGTGGAGTCGTTGCTCTGCTAGCATCTCTGAAAAGGGNNNAGATCGGAAG |
| 39627_1 | TATGAGTGTGGAGTCGTTGCCATCCTGGCTAACACAGTGAAACNNNAGATCGGAAG |
| 39627_2 | TATGAGTGTGGAGTCGTTGCCATCCTGGCTAACACAGTGAAACNNNAGATCGGAAG |
| 39627_3 | TATGAGTGTGGAGTCGTTGCCATCCTGGCTAACACAGTGAAACNNNAGATCGGAAG |
| 48660_1 | TATGAGTGTGGAGTCGTTGCTTCTCCTAATGCTATCCATGCCCNNNAGATCGGAAGA |
| 48660_2 | TATGAGTGTGGAGTCGTTGCTTCTCCTAATGCTATCCATGCCCNNNAGATCGGAAGA |
| 48660_3 | TATGAGTGTGGAGTCGTTGCTTCTCCTAATGCTATCCATGCCCNNNAGATCGGAAGA |
| 33269_1 | TATGAGTGTGGAGTCGTTGCCCACCACGCACAGCTAATTTTTANNNAGATCGGAAG |
| 33269_2 | TATGAGTGTGGAGTCGTTGCCCACCACGCACAGCTAATTTTTANNNAGATCGGAAG |
| 33269_3 | TATGAGTGTGGAGTCGTTGCCCACCACGCACAGCTAATTTTTANNNAGATCGGAAG |
| 29579_1 | TATGAGTGTGGAGTCGTTGCTTGTCAGTTCCACTCTGCAACAANNNAGATCGGAAG |
| 29579_2 | TATGAGTGTGGAGTCGTTGCTTGTCAGTTCCACTCTGCAACAANNNAGATCGGAAG |
| 29579_3 | TATGAGTGTGGAGTCGTTGCTTGTCAGTTCCACTCTGCAACAANNNAGATCGGAAG |
| 52920_1 | TATGAGTGTGGAGTCGTTGCGACCCAGATCATAACGTCACTGTNNNAGATCGGAAG |
| 52920_2 | TATGAGTGTGGAGTCGTTGCGACCCAGATCATAACGTCACTGTNNNAGATCGGAAG |
| 52920_3 | TATGAGTGTGGAGTCGTTGCGACCCAGATCATAACGTCACTGTNNNAGATCGGAAG |
| 39263_1 | TATGAGTGTGGAGTCGTTGCAAGAAAGAAAGTATGCTGTGGGCNNNAGATCGGAA |
| 39263_2 | TATGAGTGTGGAGTCGTTGCAAGAAAGAAAGTATGCTGTGGGCNNNAGATCGGAA |

| | |
|---|---|
| 39263_3 | TATGAGTGTGGAGTCGTTGCAAGAAAGAAAGTATGCTGTGGGCNNNAGATCGGAA |
| 32111_1 | TATGAGTGTGGAGTCGTTGCGGAGTGCAGTGGTGTGATTTTTTNNNAGATCGGAA |
| 32111_2 | TATGAGTGTGGAGTCGTTGCGGAGTGCAGTGGTGTGATTTTTTNNNAGATCGGAA |
| 32111_3 | TATGAGTGTGGAGTCGTTGCGGAGTGCAGTGGTGTGATTTTTTNNNAGATCGGAA |
| 33929_1 | TATGAGTGTGGAGTCGTTGCGGGTTCTAGTGATTCTTGTGCCTNNNAGATCGGAAG |
| 33929_2 | TATGAGTGTGGAGTCGTTGCGGGTTCTAGTGATTCTTGTGCCTNNNAGATCGGAAG |
| 33929_3 | TATGAGTGTGGAGTCGTTGCGGGTTCTAGTGATTCTTGTGCCTNNNAGATCGGAAG |
| 36865_1 | TATGAGTGTGGAGTCGTTGCGCCTGAGTGACAAAGGGAGATAANNNAGATCGGAA |
| 36865_2 | TATGAGTGTGGAGTCGTTGCGCCTGAGTGACAAAGGGAGATAANNNAGATCGGAA |
| 36865_3 | TATGAGTGTGGAGTCGTTGCGCCTGAGTGACAAAGGGAGATAANNNAGATCGGAA |
| 44432_1 | TATGAGTGTGGAGTCGTTGCTTTATGGCACTGAGGTGGTCAATNNNAGATCGGAA |
| 44432_2 | TATGAGTGTGGAGTCGTTGCTTTATGGCACTGAGGTGGTCAATNNNAGATCGGAA |
| 44432_3 | TATGAGTGTGGAGTCGTTGCTTTATGGCACTGAGGTGGTCAATNNNAGATCGGAA |
| 49578_1 | TATGAGTGTGGAGTCGTTGCATTTCTTGCAGGCTGTGGATCTANNNAGATCGGAAG |
| 49578_2 | TATGAGTGTGGAGTCGTTGCATTTCTTGCAGGCTGTGGATCTANNNAGATCGGAAG |
| 49578_3 | TATGAGTGTGGAGTCGTTGCATTTCTTGCAGGCTGTGGATCTANNNAGATCGGAAG |
| 49220_1 | TATGAGTGTGGAGTCGTTGCGATGGAGTCTCACTCTTGTCACCNNNAGATCGGAAG |
| 49220_2 | TATGAGTGTGGAGTCGTTGCGATGGAGTCTCACTCTTGTCACCNNNAGATCGGAAG |
| 49220_3 | TATGAGTGTGGAGTCGTTGCGATGGAGTCTCACTCTTGTCACCNNNAGATCGGAAG |
| 40977_1 | TATGAGTGTGGAGTCGTTGCTTATTAGAGGACTTGTCGGCACCNNNAGATCGGAAG |
| 40977_2 | TATGAGTGTGGAGTCGTTGCTTATTAGAGGACTTGTCGGCACCNNNAGATCGGAAG |
| 40977_3 | TATGAGTGTGGAGTCGTTGCTTATTAGAGGACTTGTCGGCACCNNNAGATCGGAAG |
| 13714_1 | TATGAGTGTGGAGTCGTTGCTTATTCCGTATTACATTTTTGCANNNAGATCGGAAGA |
| 13714_2 | TATGAGTGTGGAGTCGTTGCTTATTCCGTATTACATTTTTGCANNNAGATCGGAAGA |
| 13714_3 | TATGAGTGTGGAGTCGTTGCTTATTCCGTATTACATTTTTGCANNNAGATCGGAAGA |
| 35559_1 | TATGAGTGTGGAGTCGTTGCGGCTTCCTAAAGTGCTGGGATTANNNAGATCGGAA |
| 35559_2 | TATGAGTGTGGAGTCGTTGCGGCTTCCTAAAGTGCTGGGATTANNNAGATCGGAA |
| 35559_3 | TATGAGTGTGGAGTCGTTGCGGCTTCCTAAAGTGCTGGGATTANNNAGATCGGAA |
| 49658_1 | TATGAGTGTGGAGTCGTTGCTTCAAGTGATTCTCCTGCCTCAGNNNAGATCGGAAG |
| 49658_2 | TATGAGTGTGGAGTCGTTGCTTCAAGTGATTCTCCTGCCTCAGNNNAGATCGGAAG |
| 49658_3 | TATGAGTGTGGAGTCGTTGCTTCAAGTGATTCTCCTGCCTCAGNNNAGATCGGAAG |
| 36100_1 | TATGAGTGTGGAGTCGTTGCAAAGCATTCCTCCAAGATCCACANNNAGATCGGAAG |
| 36100_2 | TATGAGTGTGGAGTCGTTGCAAAGCATTCCTCCAAGATCCACANNNAGATCGGAAG |
| 36100_3 | TATGAGTGTGGAGTCGTTGCAAAGCATTCCTCCAAGATCCACANNNAGATCGGAAG |
| 31535_1 | TATGAGTGTGGAGTCGTTGCACTTCCCAATCCAACAGTAATGANNNAGATCGGAAG |
| 31535_2 | TATGAGTGTGGAGTCGTTGCACTTCCCAATCCAACAGTAATGANNNAGATCGGAAG |
| 31535_3 | TATGAGTGTGGAGTCGTTGCACTTCCCAATCCAACAGTAATGANNNAGATCGGAAG |
| 38321_1 | TATGAGTGTGGAGTCGTTGCCTGTAGTCCCAGCTACTCAGGAGNNNAGATCGGAAG |
| 38321_2 | TATGAGTGTGGAGTCGTTGCCTGTAGTCCCAGCTACTCAGGAGNNNAGATCGGAAG |
| 38321_3 | TATGAGTGTGGAGTCGTTGCCTGTAGTCCCAGCTACTCAGGAGNNNAGATCGGAAG |
| 40140_1 | TATGAGTGTGGAGTCGTTGCTGTACTTTAGAGGCTGATGTGGGNNNAGATCGGAA |
| 40140_2 | TATGAGTGTGGAGTCGTTGCTGTACTTTAGAGGCTGATGTGGGNNNAGATCGGAA |
| 40140_3 | TATGAGTGTGGAGTCGTTGCTGTACTTTAGAGGCTGATGTGGGNNNAGATCGGAA |

| | |
|---|---|
| 9736_1 | TATGAGTGTGGAGTCGTTGCATTGTCTATCTGCCCTTCAGCACNNNAGATCGGAAGA |
| 9736_2 | TATGAGTGTGGAGTCGTTGCATTGTCTATCTGCCCTTCAGCACNNNAGATCGGAAGA |
| 9736_3 | TATGAGTGTGGAGTCGTTGCATTGTCTATCTGCCCTTCAGCACNNNAGATCGGAAGA |
| 36104_1 | TATGAGTGTGGAGTCGTTGCAAACAACCAAACACCATTTCCCANNNAGATCGGAAGA |
| 36104_2 | TATGAGTGTGGAGTCGTTGCAAACAACCAAACACCATTTCCCANNNAGATCGGAAGA |
| 36104_3 | TATGAGTGTGGAGTCGTTGCAAACAACCAAACACCATTTCCCANNNAGATCGGAAGA |
| 38568_1 | TATGAGTGTGGAGTCGTTGCGAAGGAGGAATGAGGCTGAGATCNNNAGATCGGA |
| 38568_2 | TATGAGTGTGGAGTCGTTGCGAAGGAGGAATGAGGCTGAGATCNNNAGATCGGA |
| 38568_3 | TATGAGTGTGGAGTCGTTGCGAAGGAGGAATGAGGCTGAGATCNNNAGATCGGA |
| 10474_1 | TATGAGTGTGGAGTCGTTGCTTGCCTTCTTTAGTCCTTCACCANNNAGATCGGAAGA |
| 10474_2 | TATGAGTGTGGAGTCGTTGCTTGCCTTCTTTAGTCCTTCACCANNNAGATCGGAAGA |
| 10474_3 | TATGAGTGTGGAGTCGTTGCTTGCCTTCTTTAGTCCTTCACCANNNAGATCGGAAGA |
| 49964_1 | TATGAGTGTGGAGTCGTTGCCTCACTGTTACAGACCTCTGACCNNNAGATCGGAAGA |
| 49964_2 | TATGAGTGTGGAGTCGTTGCCTCACTGTTACAGACCTCTGACCNNNAGATCGGAAGA |
| 49964_3 | TATGAGTGTGGAGTCGTTGCCTCACTGTTACAGACCTCTGACCNNNAGATCGGAAGA |
| 36758_1 | TATGAGTGTGGAGTCGTTGCACTGGACTTCTGGCCTATCACTANNNAGATCGGAAG |
| 36758_2 | TATGAGTGTGGAGTCGTTGCACTGGACTTCTGGCCTATCACTANNNAGATCGGAAG |
| 36758_3 | TATGAGTGTGGAGTCGTTGCACTGGACTTCTGGCCTATCACTANNNAGATCGGAAG |
| 40513_1 | TATGAGTGTGGAGTCGTTGCTCAGGCTAGAAAAATGAAGGCCANNNAGATCGGAA |
| 40513_2 | TATGAGTGTGGAGTCGTTGCTCAGGCTAGAAAAATGAAGGCCANNNAGATCGGAA |
| 40513_3 | TATGAGTGTGGAGTCGTTGCTCAGGCTAGAAAAATGAAGGCCANNNAGATCGGAA |
| 36963_1 | TATGAGTGTGGAGTCGTTGCTGCAGGATACAAAATCAACAAACANNNAGATCGGAA |
| 36963_2 | TATGAGTGTGGAGTCGTTGCTGCAGGATACAAAATCAACAAACANNNAGATCGGAA |
| 36963_3 | TATGAGTGTGGAGTCGTTGCTGCAGGATACAAAATCAACAAACANNNAGATCGGAA |
| 38692_1 | TATGAGTGTGGAGTCGTTGCAGTTTTGTGTTTGAAACCCAGGGNNNAGATCGGAA |
| 38692_2 | TATGAGTGTGGAGTCGTTGCAGTTTTGTGTTTGAAACCCAGGGNNNAGATCGGAA |
| 38692_3 | TATGAGTGTGGAGTCGTTGCAGTTTTGTGTTTGAAACCCAGGGNNNAGATCGGAA |
| 39286_1 | TATGAGTGTGGAGTCGTTGCGCTATGATCACACCACTGCATTCNNNAGATCGGAAG |
| 39286_2 | TATGAGTGTGGAGTCGTTGCGCTATGATCACACCACTGCATTCNNNAGATCGGAAG |
| 39286_3 | TATGAGTGTGGAGTCGTTGCGCTATGATCACACCACTGCATTCNNNAGATCGGAAG |
| 46275_1 | TATGAGTGTGGAGTCGTTGCATGATTCTTCTGCCTTAGCCTCCNNNAGATCGGAAGA |
| 46275_2 | TATGAGTGTGGAGTCGTTGCATGATTCTTCTGCCTTAGCCTCCNNNAGATCGGAAGA |
| 46275_3 | TATGAGTGTGGAGTCGTTGCATGATTCTTCTGCCTTAGCCTCCNNNAGATCGGAAGA |
| 13512_1 | TATGAGTGTGGAGTCGTTGCTTCAGATTTTTATGAGCCTTTACNNNAGATCGGAAG |
| 13512_2 | TATGAGTGTGGAGTCGTTGCTTCAGATTTTTATGAGCCTTTACNNNAGATCGGAAG |
| 13512_3 | TATGAGTGTGGAGTCGTTGCTTCAGATTTTTATGAGCCTTTACNNNAGATCGGAAG |
| 40353_1 | TATGAGTGTGGAGTCGTTGCAGAACATGGACTTTGGAACCAGANNNAGATCGGAA |
| 40353_2 | TATGAGTGTGGAGTCGTTGCAGAACATGGACTTTGGAACCAGANNNAGATCGGAA |
| 40353_3 | TATGAGTGTGGAGTCGTTGCAGAACATGGACTTTGGAACCAGANNNAGATCGGAA |
| 37978_1 | TATGAGTGTGGAGTCGTTGCGCATGAAAGACAGAAGGCACAAANNNAGATCGGAA |
| 37978_2 | TATGAGTGTGGAGTCGTTGCGCATGAAAGACAGAAGGCACAAANNNAGATCGGAA |
| 37978_3 | TATGAGTGTGGAGTCGTTGCGCATGAAAGACAGAAGGCACAAANNNAGATCGGAA |
| 10441_1 | TATGAGTGTGGAGTCGTTGCATGCTTTGAACCTAGACGTGACCNNNAGATCGGAAG |

| | |
|---|---|
| 10441_2 | TATGAGTGTGGAGTCGTTGCATGCTTTGAACCTAGACGTGACCNNNAGATCGGAAG |
| 10441_3 | TATGAGTGTGGAGTCGTTGCATGCTTTGAACCTAGACGTGACCNNNAGATCGGAAG |
| 42564_1 | TATGAGTGTGGAGTCGTTGCACTTATTGCCCTGGCTAGAAACTNNNAGATCGGAAG |
| 42564_2 | TATGAGTGTGGAGTCGTTGCACTTATTGCCCTGGCTAGAAACTNNNAGATCGGAAG |
| 42564_3 | TATGAGTGTGGAGTCGTTGCACTTATTGCCCTGGCTAGAAACTNNNAGATCGGAAG |
| 36587_1 | TATGAGTGTGGAGTCGTTGCGCAATACCGGTAACTTTGTCACCNNNAGATCGGAAG |
| 36587_2 | TATGAGTGTGGAGTCGTTGCGCAATACCGGTAACTTTGTCACCNNNAGATCGGAAG |
| 36587_3 | TATGAGTGTGGAGTCGTTGCGCAATACCGGTAACTTTGTCACCNNNAGATCGGAAG |
| 30526_1 | TATGAGTGTGGAGTCGTTGCAGCACCTTTATTCTGCCCCATGNNNAGATCGGAAGA |
| 30526_2 | TATGAGTGTGGAGTCGTTGCAGCACCTTTATTCTGCCCCATGNNNAGATCGGAAGA |
| 30526_3 | TATGAGTGTGGAGTCGTTGCAGCACCTTTATTCTGCCCCATGNNNAGATCGGAAGA |
| 35006_1 | TATGAGTGTGGAGTCGTTGCCTGAGTAGCTGGGATTATAGGCGNNNAGATCGGAA |
| 35006_2 | TATGAGTGTGGAGTCGTTGCCTGAGTAGCTGGGATTATAGGCGNNNAGATCGGAA |
| 35006_3 | TATGAGTGTGGAGTCGTTGCCTGAGTAGCTGGGATTATAGGCGNNNAGATCGGAA |
| 49847_1 | TATGAGTGTGGAGTCGTTGCTGGTATTTGCAATGATCTGTGCCNNNAGATCGGAAG |
| 49847_2 | TATGAGTGTGGAGTCGTTGCTGGTATTTGCAATGATCTGTGCCNNNAGATCGGAAG |
| 49847_3 | TATGAGTGTGGAGTCGTTGCTGGTATTTGCAATGATCTGTGCCNNNAGATCGGAAG |
| 925_1 | TATGAGTGTGGAGTCGTTGCGTAACCACACAACTCACAGATCCANNNAGATCGGAA |
| 925_2 | TATGAGTGTGGAGTCGTTGCGTAACCACACAACTCACAGATCCANNNAGATCGGAA |
| 925_3 | TATGAGTGTGGAGTCGTTGCGTAACCACACAACTCACAGATCCANNNAGATCGGAA |
| 41807_1 | TATGAGTGTGGAGTCGTTGCTCCACACCATTCTTCTGCCTCNNNAGATCGGAAGAGC |
| 41807_2 | TATGAGTGTGGAGTCGTTGCTCCACACCATTCTTCTGCCTCNNNAGATCGGAAGAGC |
| 41807_3 | TATGAGTGTGGAGTCGTTGCTCCACACCATTCTTCTGCCTCNNNAGATCGGAAGAGC |
| 45481_1 | TATGAGTGTGGAGTCGTTGCGGTGGAACAGTTTCTCACATCCTNNNAGATCGGAAG |
| 45481_2 | TATGAGTGTGGAGTCGTTGCGGTGGAACAGTTTCTCACATCCTNNNAGATCGGAAG |
| 45481_3 | TATGAGTGTGGAGTCGTTGCGGTGGAACAGTTTCTCACATCCTNNNAGATCGGAAG |
| 13585_1 | TATGAGTGTGGAGTCGTTGCTATGAGTTCTTTAGCTAAACGAANNNAGATCGGAAG |
| 13585_2 | TATGAGTGTGGAGTCGTTGCTATGAGTTCTTTAGCTAAACGAANNNAGATCGGAAG |
| 13585_3 | TATGAGTGTGGAGTCGTTGCTATGAGTTCTTTAGCTAAACGAANNNAGATCGGAAG |
| 44786_1 | TATGAGTGTGGAGTCGTTGCAGATTCTTCAATGCCACAAAGCCNNNAGATCGGAAG |
| 44786_2 | TATGAGTGTGGAGTCGTTGCAGATTCTTCAATGCCACAAAGCCNNNAGATCGGAAG |
| 44786_3 | TATGAGTGTGGAGTCGTTGCAGATTCTTCAATGCCACAAAGCCNNNAGATCGGAAG |
| 28893_1 | TATGAGTGTGGAGTCGTTGCCTCCCTCTCATACTCTCGCTTTCNNNAGATCGGAAGA |
| 28893_2 | TATGAGTGTGGAGTCGTTGCCTCCCTCTCATACTCTCGCTTTCNNNAGATCGGAAGA |
| 28893_3 | TATGAGTGTGGAGTCGTTGCCTCCCTCTCATACTCTCGCTTTCNNNAGATCGGAAGA |
| 33230_1 | TATGAGTGTGGAGTCGTTGCTCTGATTCTGGAATTGTCCCCACNNNAGATCGGAAG |
| 33230_2 | TATGAGTGTGGAGTCGTTGCTCTGATTCTGGAATTGTCCCCACNNNAGATCGGAAG |
| 33230_3 | TATGAGTGTGGAGTCGTTGCTCTGATTCTGGAATTGTCCCCACNNNAGATCGGAAG |
| 37738_1 | TATGAGTGTGGAGTCGTTGCTTGTAGGGTCTTGCTATGTTGCCNNNAGATCGGAAG |
| 37738_2 | TATGAGTGTGGAGTCGTTGCTTGTAGGGTCTTGCTATGTTGCCNNNAGATCGGAAG |
| 37738_3 | TATGAGTGTGGAGTCGTTGCTTGTAGGGTCTTGCTATGTTGCCNNNAGATCGGAAG |
| 31815_1 | TATGAGTGTGGAGTCGTTGCTTCAAGTGATTCTCCTGCCTCAGNNNAGATCGGAAG |
| 31815_2 | TATGAGTGTGGAGTCGTTGCTTCAAGTGATTCTCCTGCCTCAGNNNAGATCGGAAG |

| | |
|---|---|
| 31815_3 | TATGAGTGTGGAGTCGTTGCTTCAAGTGATTCTCCTGCCTCAGNNNAGATCGGAAG |
| 51101_1 | TATGAGTGTGGAGTCGTTGCCTACTGGGAAGTAAGGAGCCCCNNNAGATCGGAAG |
| 51101_2 | TATGAGTGTGGAGTCGTTGCCTACTGGGAAGTAAGGAGCCCCNNNAGATCGGAAG |
| 51101_3 | TATGAGTGTGGAGTCGTTGCCTACTGGGAAGTAAGGAGCCCCNNNAGATCGGAAG |
| 40401_1 | TATGAGTGTGGAGTCGTTGCCATGCCTGTAATCCCAGCTACTANNNAGATCGGAAG |
| 40401_2 | TATGAGTGTGGAGTCGTTGCCATGCCTGTAATCCCAGCTACTANNNAGATCGGAAG |
| 40401_3 | TATGAGTGTGGAGTCGTTGCCATGCCTGTAATCCCAGCTACTANNNAGATCGGAAG |
| 29256_1 | TATGAGTGTGGAGTCGTTGCGACAAGTAGAGTGGTCAGAGGTGNNNAGATCGGA |
| 29256_2 | TATGAGTGTGGAGTCGTTGCGACAAGTAGAGTGGTCAGAGGTGNNNAGATCGGA |
| 29256_3 | TATGAGTGTGGAGTCGTTGCGACAAGTAGAGTGGTCAGAGGTGNNNAGATCGGA |
| 40990_1 | TATGAGTGTGGAGTCGTTGCCCACGCCCAGCTAATTTTTGTATNNNAGATCGGAAG |
| 40990_2 | TATGAGTGTGGAGTCGTTGCCCACGCCCAGCTAATTTTTGTATNNNAGATCGGAAG |
| 40990_3 | TATGAGTGTGGAGTCGTTGCCCACGCCCAGCTAATTTTTGTATNNNAGATCGGAAG |
| 40229_1 | TATGAGTGTGGAGTCGTTGCGGAACTGGAGGCCATTATCCTAANNNAGATCGGAA |
| 40229_2 | TATGAGTGTGGAGTCGTTGCGGAACTGGAGGCCATTATCCTAANNNAGATCGGAA |
| 40229_3 | TATGAGTGTGGAGTCGTTGCGGAACTGGAGGCCATTATCCTAANNNAGATCGGAA |
| 45266_1 | TATGAGTGTGGAGTCGTTGCCAGGAAATTTGAGGAGGGTGGTANNNAGATCGGA |
| 45266_2 | TATGAGTGTGGAGTCGTTGCCAGGAAATTTGAGGAGGGTGGTANNNAGATCGGA |
| 45266_3 | TATGAGTGTGGAGTCGTTGCCAGGAAATTTGAGGAGGGTGGTANNNAGATCGGA |
| 29183_1 | TATGAGTGTGGAGTCGTTGCTCTCTCTCTCTCTCTCATGTGCANNNAGATCGGAAGA |
| 29183_2 | TATGAGTGTGGAGTCGTTGCTCTCTCTCTCTCTCTCATGTGCANNNAGATCGGAAGA |
| 29183_3 | TATGAGTGTGGAGTCGTTGCTCTCTCTCTCTCTCTCATGTGCANNNAGATCGGAAGA |
| 36266_1 | TATGAGTGTGGAGTCGTTGCCCTCTGAGGTCCATGAAAGGAATNNNAGATCGGAA |
| 36266_2 | TATGAGTGTGGAGTCGTTGCCCTCTGAGGTCCATGAAAGGAATNNNAGATCGGAA |
| 36266_3 | TATGAGTGTGGAGTCGTTGCCCTCTGAGGTCCATGAAAGGAATNNNAGATCGGAA |
| 53121_1 | TATGAGTGTGGAGTCGTTGCTGCTGGAGTCCAAGTAATGTTCANNNAGATCGGAAG |
| 53121_2 | TATGAGTGTGGAGTCGTTGCTGCTGGAGTCCAAGTAATGTTCANNNAGATCGGAAG |
| 53121_3 | TATGAGTGTGGAGTCGTTGCTGCTGGAGTCCAAGTAATGTTCANNNAGATCGGAAG |
| 42711_1 | TATGAGTGTGGAGTCGTTGCCTCCTGGGTTCAAGTGATCCTCNNNAGATCGGAAGA |
| 42711_2 | TATGAGTGTGGAGTCGTTGCCTCCTGGGTTCAAGTGATCCTCNNNAGATCGGAAGA |
| 42711_3 | TATGAGTGTGGAGTCGTTGCCTCCTGGGTTCAAGTGATCCTCNNNAGATCGGAAGA |
| 28939_1 | TATGAGTGTGGAGTCGTTGCATCTCCTTCAGGTGTATGCATGGNNNAGATCGGAAG |
| 28939_2 | TATGAGTGTGGAGTCGTTGCATCTCCTTCAGGTGTATGCATGGNNNAGATCGGAAG |
| 28939_3 | TATGAGTGTGGAGTCGTTGCATCTCCTTCAGGTGTATGCATGGNNNAGATCGGAAG |
| 36471_1 | TATGAGTGTGGAGTCGTTGCAGCTGCTACCATTCCTTCTGAAANNNAGATCGGAAG |
| 36471_2 | TATGAGTGTGGAGTCGTTGCAGCTGCTACCATTCCTTCTGAAANNNAGATCGGAAG |
| 36471_3 | TATGAGTGTGGAGTCGTTGCAGCTGCTACCATTCCTTCTGAAANNNAGATCGGAAG |
| 39470_1 | TATGAGTGTGGAGTCGTTGCTGGGAGTCGTCCTTGAAAACTAGNNNAGATCGGAA |
| 39470_2 | TATGAGTGTGGAGTCGTTGCTGGGAGTCGTCCTTGAAAACTAGNNNAGATCGGAA |
| 39470_3 | TATGAGTGTGGAGTCGTTGCTGGGAGTCGTCCTTGAAAACTAGNNNAGATCGGAA |
| 39607_1 | TATGAGTGTGGAGTCGTTGCCAGGAGTATTGCTCACATCCACTNNNAGATCGGAAG |
| 39607_2 | TATGAGTGTGGAGTCGTTGCCAGGAGTATTGCTCACATCCACTNNNAGATCGGAAG |
| 39607_3 | TATGAGTGTGGAGTCGTTGCCAGGAGTATTGCTCACATCCACTNNNAGATCGGAAG |

| | |
|---|---|
| 44089_1 | TATGAGTGTGGAGTCGTTGCCTGAATGTCTGGAATGGGACTCANNNAGATCGGAA |
| 44089_2 | TATGAGTGTGGAGTCGTTGCCTGAATGTCTGGAATGGGACTCANNNAGATCGGAA |
| 44089_3 | TATGAGTGTGGAGTCGTTGCCTGAATGTCTGGAATGGGACTCANNNAGATCGGAA |
| 38442_1 | TATGAGTGTGGAGTCGTTGCGGAGAGCTCAAGATGGAGACATCNNNAGATCGGAA |
| 38442_2 | TATGAGTGTGGAGTCGTTGCGGAGAGCTCAAGATGGAGACATCNNNAGATCGGAA |
| 38442_3 | TATGAGTGTGGAGTCGTTGCGGAGAGCTCAAGATGGAGACATCNNNAGATCGGAA |
| 41131_1 | TATGAGTGTGGAGTCGTTGCGGACTAGGATAGTGGCAGTTGAGNNNAGATCGGA |
| 41131_2 | TATGAGTGTGGAGTCGTTGCGGACTAGGATAGTGGCAGTTGAGNNNAGATCGGA |
| 41131_3 | TATGAGTGTGGAGTCGTTGCGGACTAGGATAGTGGCAGTTGAGNNNAGATCGGA |
| 49926_1 | TATGAGTGTGGAGTCGTTGCGAAGCCTGTATCGATTTTGCCTCNNNAGATCGGAAG |
| 49926_2 | TATGAGTGTGGAGTCGTTGCGAAGCCTGTATCGATTTTGCCTCNNNAGATCGGAAG |
| 49926_3 | TATGAGTGTGGAGTCGTTGCGAAGCCTGTATCGATTTTGCCTCNNNAGATCGGAAG |
| 33388_1 | TATGAGTGTGGAGTCGTTGCTATTCTGAGTGAGAAGGGAAGCCNNNAGATCGGAA |
| 33388_2 | TATGAGTGTGGAGTCGTTGCTATTCTGAGTGAGAAGGGAAGCCNNNAGATCGGAA |
| 33388_3 | TATGAGTGTGGAGTCGTTGCTATTCTGAGTGAGAAGGGAAGCCNNNAGATCGGAA |
| 39497_1 | TATGAGTGTGGAGTCGTTGCGAGTGCCCTTGTAGGAAGAAGAANNNAGATCGGAA |
| 39497_2 | TATGAGTGTGGAGTCGTTGCGAGTGCCCTTGTAGGAAGAAGAANNNAGATCGGAA |
| 39497_3 | TATGAGTGTGGAGTCGTTGCGAGTGCCCTTGTAGGAAGAAGAANNNAGATCGGAA |
| 41204_1 | TATGAGTGTGGAGTCGTTGCCATCTCCAAACCCCTCCCACNNNAGATCGGAAGAGCA |
| 41204_2 | TATGAGTGTGGAGTCGTTGCCATCTCCAAACCCCTCCCACNNNAGATCGGAAGAGCA |
| 41204_3 | TATGAGTGTGGAGTCGTTGCCATCTCCAAACCCCTCCCACNNNAGATCGGAAGAGCA |
| 44140_1 | TATGAGTGTGGAGTCGTTGCTCAGCCTTAGGTCCTGACTTCTANNNAGATCGGAAG |
| 44140_2 | TATGAGTGTGGAGTCGTTGCTCAGCCTTAGGTCCTGACTTCTANNNAGATCGGAAG |
| 44140_3 | TATGAGTGTGGAGTCGTTGCTCAGCCTTAGGTCCTGACTTCTANNNAGATCGGAAG |
| 38745_1 | TATGAGTGTGGAGTCGTTGCTGACATAATTTGCTGCTTTTCCCANNNAGATCGGAAG |
| 38745_2 | TATGAGTGTGGAGTCGTTGCTGACATAATTTGCTGCTTTTCCCANNNAGATCGGAAG |
| 38745_3 | TATGAGTGTGGAGTCGTTGCTGACATAATTTGCTGCTTTTCCCANNNAGATCGGAAG |
| 13586_1 | TATGAGTGTGGAGTCGTTGCTTGATGTAGCTGTGAAAATTAATNNNAGATCGGAA |
| 13586_2 | TATGAGTGTGGAGTCGTTGCTTGATGTAGCTGTGAAAATTAATNNNAGATCGGAA |
| 13586_3 | TATGAGTGTGGAGTCGTTGCTTGATGTAGCTGTGAAAATTAATNNNAGATCGGAA |
| 10522_1 | TATGAGTGTGGAGTCGTTGCGGACATTTGGGGAACGCATTTTTNNNAGATCGGAA |
| 10522_2 | TATGAGTGTGGAGTCGTTGCGGACATTTGGGGAACGCATTTTTNNNAGATCGGAA |
| 10522_3 | TATGAGTGTGGAGTCGTTGCGGACATTTGGGGAACGCATTTTTNNNAGATCGGAA |
| 33073_1 | TATGAGTGTGGAGTCGTTGCCTGGGATTATAGGTGTGGACCACNNNAGATCGGAA |
| 33073_2 | TATGAGTGTGGAGTCGTTGCCTGGGATTATAGGTGTGGACCACNNNAGATCGGAA |
| 33073_3 | TATGAGTGTGGAGTCGTTGCCTGGGATTATAGGTGTGGACCACNNNAGATCGGAA |
| 33707_1 | TATGAGTGTGGAGTCGTTGCAAAGATCCTCAAGTGGTAAGCGGNNNAGATCGGAA |
| 33707_2 | TATGAGTGTGGAGTCGTTGCAAAGATCCTCAAGTGGTAAGCGGNNNAGATCGGAA |
| 33707_3 | TATGAGTGTGGAGTCGTTGCAAAGATCCTCAAGTGGTAAGCGGNNNAGATCGGAA |
| 45229_1 | TATGAGTGTGGAGTCGTTGCCAGAGGTTTGAGTGTAAATGGCGNNNAGATCGGAA |
| 45229_2 | TATGAGTGTGGAGTCGTTGCCAGAGGTTTGAGTGTAAATGGCGNNNAGATCGGAA |
| 45229_3 | TATGAGTGTGGAGTCGTTGCCAGAGGTTTGAGTGTAAATGGCGNNNAGATCGGAA |
| 50550_1 | TATGAGTGTGGAGTCGTTGCGCACTCATCGGACTATTTTCAGCNNNAGATCGGAAG |

| | |
|---|---|
| 50550_2 | TATGAGTGTGGAGTCGTTGCGCACTCATCGGACTATTTTCAGCNNNAGATCGGAAG |
| 50550_3 | TATGAGTGTGGAGTCGTTGCGCACTCATCGGACTATTTTCAGCNNNAGATCGGAAG |
| 35948_1 | TATGAGTGTGGAGTCGTTGCGAAAGTGTGCACATTCTTTGGCNNNAGATCGGAAGA |
| 35948_2 | TATGAGTGTGGAGTCGTTGCGAAAGTGTGCACATTCTTTGGCNNNAGATCGGAAGA |
| 35948_3 | TATGAGTGTGGAGTCGTTGCGAAAGTGTGCACATTCTTTGGCNNNAGATCGGAAGA |
| 34593_1 | TATGAGTGTGGAGTCGTTGCGGAGTCTATGGATCTGACTTGGGNNNAGATCGGAA |
| 34593_2 | TATGAGTGTGGAGTCGTTGCGGAGTCTATGGATCTGACTTGGGNNNAGATCGGAA |
| 34593_3 | TATGAGTGTGGAGTCGTTGCGGAGTCTATGGATCTGACTTGGGNNNAGATCGGAA |
| 36446_1 | TATGAGTGTGGAGTCGTTGCAGGAACTAGAGTAAGCAGGAGGTNNNAGATCGGA |
| 36446_2 | TATGAGTGTGGAGTCGTTGCAGGAACTAGAGTAAGCAGGAGGTNNNAGATCGGA |
| 36446_3 | TATGAGTGTGGAGTCGTTGCAGGAACTAGAGTAAGCAGGAGGTNNNAGATCGGA |
| 12515_1 | TATGAGTGTGGAGTCGTTGCATTAGCCAACTACCGTTTATAAANNNAGATCGGAAG |
| 12515_2 | TATGAGTGTGGAGTCGTTGCATTAGCCAACTACCGTTTATAAANNNAGATCGGAAG |
| 12515_3 | TATGAGTGTGGAGTCGTTGCATTAGCCAACTACCGTTTATAAANNNAGATCGGAAG |
| 39589_1 | TATGAGTGTGGAGTCGTTGCAGAAACTGAATCCGAGAGGCTTTNNNAGATCGGAA |
| 39589_2 | TATGAGTGTGGAGTCGTTGCAGAAACTGAATCCGAGAGGCTTTNNNAGATCGGAA |
| 39589_3 | TATGAGTGTGGAGTCGTTGCAGAAACTGAATCCGAGAGGCTTTNNNAGATCGGAA |
| 50375_1 | TATGAGTGTGGAGTCGTTGCAAACCACTCTGCTCACCTGTTCNNNAGATCGGAAGAG |
| 50375_2 | TATGAGTGTGGAGTCGTTGCAAACCACTCTGCTCACCTGTTCNNNAGATCGGAAGAG |
| 50375_3 | TATGAGTGTGGAGTCGTTGCAAACCACTCTGCTCACCTGTTCNNNAGATCGGAAGAG |
| 40220_1 | TATGAGTGTGGAGTCGTTGCAGCACTTACCTTTTCCAGGGAATNNNAGATCGGAAG |
| 40220_2 | TATGAGTGTGGAGTCGTTGCAGCACTTACCTTTTCCAGGGAATNNNAGATCGGAAG |
| 40220_3 | TATGAGTGTGGAGTCGTTGCAGCACTTACCTTTTCCAGGGAATNNNAGATCGGAAG |
| 45592_1 | TATGAGTGTGGAGTCGTTGCTCGACTCACTATAACCTCCACCTNNNAGATCGGAAGA |
| 45592_2 | TATGAGTGTGGAGTCGTTGCTCGACTCACTATAACCTCCACCTNNNAGATCGGAAGA |
| 45592_3 | TATGAGTGTGGAGTCGTTGCTCGACTCACTATAACCTCCACCTNNNAGATCGGAAGA |
| 47315_1 | TATGAGTGTGGAGTCGTTGCAGCACCCTGCACTTTTCTTTTTTNNNAGATCGGAAGA |
| 47315_2 | TATGAGTGTGGAGTCGTTGCAGCACCCTGCACTTTTCTTTTTTNNNAGATCGGAAGA |
| 47315_3 | TATGAGTGTGGAGTCGTTGCAGCACCCTGCACTTTTCTTTTTTNNNAGATCGGAAGA |
| 38165_1 | TATGAGTGTGGAGTCGTTGCGGATCACTTGAGGTCAGGAGTTTNNNAGATCGGAA |
| 38165_2 | TATGAGTGTGGAGTCGTTGCGGATCACTTGAGGTCAGGAGTTTNNNAGATCGGAA |
| 38165_3 | TATGAGTGTGGAGTCGTTGCGGATCACTTGAGGTCAGGAGTTTNNNAGATCGGAA |
| 43544_1 | TATGAGTGTGGAGTCGTTGCGTCTCCTGGGTTCAAGTGATTCTNNNAGATCGGAAG |
| 43544_2 | TATGAGTGTGGAGTCGTTGCGTCTCCTGGGTTCAAGTGATTCTNNNAGATCGGAAG |
| 43544_3 | TATGAGTGTGGAGTCGTTGCGTCTCCTGGGTTCAAGTGATTCTNNNAGATCGGAAG |
| 30584_1 | TATGAGTGTGGAGTCGTTGCGATGATTTTTCTTTGCCTTTTTGNNNAGATCGGAAGA |
| 30584_2 | TATGAGTGTGGAGTCGTTGCGATGATTTTTCTTTGCCTTTTTGNNNAGATCGGAAGA |
| 30584_3 | TATGAGTGTGGAGTCGTTGCGATGATTTTTCTTTGCCTTTTTGNNNAGATCGGAAGA |
| 52660_1 | TATGAGTGTGGAGTCGTTGCATAAAGAGGGGATGAGCAATGGGNNNAGATCGGA |
| 52660_2 | TATGAGTGTGGAGTCGTTGCATAAAGAGGGGATGAGCAATGGGNNNAGATCGGA |
| 52660_3 | TATGAGTGTGGAGTCGTTGCATAAAGAGGGGATGAGCAATGGGNNNAGATCGGA |
| 41254_1 | TATGAGTGTGGAGTCGTTGCCCCACAGGACAAGCTGTAAACTTNNNAGATCGGAAG |
| 41254_2 | TATGAGTGTGGAGTCGTTGCCCCACAGGACAAGCTGTAAACTTNNNAGATCGGAAG |

| | |
|---|---|
| 41254_3 | TATGAGTGTGGAGTCGTTGCCCCACAGGACAAGCTGTAAACTTNNNAGATCGGAAG |
| 31849_1 | TATGAGTGTGGAGTCGTTGCGTCAGTATGTCTAGACTGGGCACNNNAGATCGGAA |
| 31849_2 | TATGAGTGTGGAGTCGTTGCGTCAGTATGTCTAGACTGGGCACNNNAGATCGGAA |
| 31849_3 | TATGAGTGTGGAGTCGTTGCGTCAGTATGTCTAGACTGGGCACNNNAGATCGGAA |
| 43530_1 | TATGAGTGTGGAGTCGTTGCACGTAGTGTGATGCTTCCAGATTNNNAGATCGGAAG |
| 43530_2 | TATGAGTGTGGAGTCGTTGCACGTAGTGTGATGCTTCCAGATTNNNAGATCGGAAG |
| 43530_3 | TATGAGTGTGGAGTCGTTGCACGTAGTGTGATGCTTCCAGATTNNNAGATCGGAAG |
| 52201_1 | TATGAGTGTGGAGTCGTTGCCATCCATCACTTCAGGTCACTCANNNAGATCGGAAG |
| 52201_2 | TATGAGTGTGGAGTCGTTGCCATCCATCACTTCAGGTCACTCANNNAGATCGGAAG |
| 52201_3 | TATGAGTGTGGAGTCGTTGCCATCCATCACTTCAGGTCACTCANNNAGATCGGAAG |
| 42325_1 | TATGAGTGTGGAGTCGTTGCCACAGAGTCACACATCCTTGAGANNNAGATCGGAAG |
| 42325_2 | TATGAGTGTGGAGTCGTTGCCACAGAGTCACACATCCTTGAGANNNAGATCGGAAG |
| 42325_3 | TATGAGTGTGGAGTCGTTGCCACAGAGTCACACATCCTTGAGANNNAGATCGGAAG |
| 47230_1 | TATGAGTGTGGAGTCGTTGCCACTCAGCCTGTCAGTTTTGTTTNNNAGATCGGAAG |
| 47230_2 | TATGAGTGTGGAGTCGTTGCCACTCAGCCTGTCAGTTTTGTTTNNNAGATCGGAAG |
| 47230_3 | TATGAGTGTGGAGTCGTTGCCACTCAGCCTGTCAGTTTTGTTTNNNAGATCGGAAG |
| 12966_1 | TATGAGTGTGGAGTCGTTGCCACTCTTCTCAAACAAAAAGTCNNNAGATCGGAAGA |
| 12966_2 | TATGAGTGTGGAGTCGTTGCCACTCTTCTCAAACAAAAAGTCNNNAGATCGGAAGA |
| 12966_3 | TATGAGTGTGGAGTCGTTGCCACTCTTCTCAAACAAAAAGTCNNNAGATCGGAAGA |
| 51931_1 | TATGAGTGTGGAGTCGTTGCGAGTGAGAATATGCGGTGTTTGGNNNAGATCGGAA |
| 51931_2 | TATGAGTGTGGAGTCGTTGCGAGTGAGAATATGCGGTGTTTGGNNNAGATCGGAA |
| 51931_3 | TATGAGTGTGGAGTCGTTGCGAGTGAGAATATGCGGTGTTTGGNNNAGATCGGAA |
| 52164_1 | TATGAGTGTGGAGTCGTTGCCCACTGTCTATGGGCCTAATTCANNNAGATCGGAAG |
| 52164_2 | TATGAGTGTGGAGTCGTTGCCCACTGTCTATGGGCCTAATTCANNNAGATCGGAAG |
| 52164_3 | TATGAGTGTGGAGTCGTTGCCCACTGTCTATGGGCCTAATTCANNNAGATCGGAAG |
| 10369_1 | TATGAGTGTGGAGTCGTTGCATCACTTCATAGTAGTGCAAGCANNNAGATCGGAAG |
| 10369_2 | TATGAGTGTGGAGTCGTTGCATCACTTCATAGTAGTGCAAGCANNNAGATCGGAAG |
| 10369_3 | TATGAGTGTGGAGTCGTTGCATCACTTCATAGTAGTGCAAGCANNNAGATCGGAAG |
| 29959_1 | TATGAGTGTGGAGTCGTTGCGTGGTGAGTTAAAGCATGGCATTNNNAGATCGGAA |
| 29959_2 | TATGAGTGTGGAGTCGTTGCGTGGTGAGTTAAAGCATGGCATTNNNAGATCGGAA |
| 29959_3 | TATGAGTGTGGAGTCGTTGCGTGGTGAGTTAAAGCATGGCATTNNNAGATCGGAA |
| 42961_1 | TATGAGTGTGGAGTCGTTGCCCCAGGAGATTGATGGTTACACANNNAGATCGGAA |
| 42961_2 | TATGAGTGTGGAGTCGTTGCCCCAGGAGATTGATGGTTACACANNNAGATCGGAA |
| 42961_3 | TATGAGTGTGGAGTCGTTGCCCCAGGAGATTGATGGTTACACANNNAGATCGGAA |
| 2111_1 | TATGAGTGTGGAGTCGTTGCTGATTATGCGTGATATGCAGGCANNNAGATCGGAA |
| 2111_2 | TATGAGTGTGGAGTCGTTGCTGATTATGCGTGATATGCAGGCANNNAGATCGGAA |
| 2111_3 | TATGAGTGTGGAGTCGTTGCTGATTATGCGTGATATGCAGGCANNNAGATCGGAA |
| 41281_1 | TATGAGTGTGGAGTCGTTGCCTCCCTCTGCTGAAAACATTTGGNNNAGATCGGAAG |
| 41281_2 | TATGAGTGTGGAGTCGTTGCCTCCCTCTGCTGAAAACATTTGGNNNAGATCGGAAG |
| 41281_3 | TATGAGTGTGGAGTCGTTGCCTCCCTCTGCTGAAAACATTTGGNNNAGATCGGAAG |
| 40882_1 | TATGAGTGTGGAGTCGTTGCGAGTTTGTAGTGAGCCAAGATCGNNNAGATCGGAA |
| 40882_2 | TATGAGTGTGGAGTCGTTGCGAGTTTGTAGTGAGCCAAGATCGNNNAGATCGGAA |
| 40882_3 | TATGAGTGTGGAGTCGTTGCGAGTTTGTAGTGAGCCAAGATCGNNNAGATCGGAA |

| | |
|---|---|
| 38834_1 | TATGAGTGTGGAGTCGTTGCGAGGTTTCAATGAGCCGAGATTGNNNAGATCGGAA |
| 38834_2 | TATGAGTGTGGAGTCGTTGCGAGGTTTCAATGAGCCGAGATTGNNNAGATCGGAA |
| 38834_3 | TATGAGTGTGGAGTCGTTGCGAGGTTTCAATGAGCCGAGATTGNNNAGATCGGAA |
| 30549_1 | TATGAGTGTGGAGTCGTTGCTCCCTTAATACCAGCTGCACTTTCNNNAGATCGGAAG |
| 30549_2 | TATGAGTGTGGAGTCGTTGCTCCCTTAATACCAGCTGCACTTTCNNNAGATCGGAAG |
| 30549_3 | TATGAGTGTGGAGTCGTTGCTCCCTTAATACCAGCTGCACTTTCNNNAGATCGGAAG |
| 48716_1 | TATGAGTGTGGAGTCGTTGCCCTCCTGAATAGCTGGGTTTACANNNAGATCGGAAG |
| 48716_2 | TATGAGTGTGGAGTCGTTGCCCTCCTGAATAGCTGGGTTTACANNNAGATCGGAAG |
| 48716_3 | TATGAGTGTGGAGTCGTTGCCCTCCTGAATAGCTGGGTTTACANNNAGATCGGAAG |
| 37976_1 | TATGAGTGTGGAGTCGTTGCCGCTAGTCTTGAACTCCTGAACTNNNAGATCGGAAG |
| 37976_2 | TATGAGTGTGGAGTCGTTGCCGCTAGTCTTGAACTCCTGAACTNNNAGATCGGAAG |
| 37976_3 | TATGAGTGTGGAGTCGTTGCCGCTAGTCTTGAACTCCTGAACTNNNAGATCGGAAG |
| 34246_1 | TATGAGTGTGGAGTCGTTGCAAGAAAGAAAGAAAGGGCCATGCNNNAGATCGGAA |
| 34246_2 | TATGAGTGTGGAGTCGTTGCAAGAAAGAAAGAAAGGGCCATGCNNNAGATCGGAA |
| 34246_3 | TATGAGTGTGGAGTCGTTGCAAGAAAGAAAGAAAGGGCCATGCNNNAGATCGGAA |
| 3138_1 | TATGAGTGTGGAGTCGTTGCCCTCCAGTGGTCTCCTTAGCTGNNNAGATCGGAAGA |
| 3138_2 | TATGAGTGTGGAGTCGTTGCCCTCCAGTGGTCTCCTTAGCTGNNNAGATCGGAAGA |
| 3138_3 | TATGAGTGTGGAGTCGTTGCCCTCCAGTGGTCTCCTTAGCTGNNNAGATCGGAAGA |
| 10590_1 | TATGAGTGTGGAGTCGTTGCGGAAGTTTTCTTTAGGGGCTGCTNNNAGATCGGAA |
| 10590_2 | TATGAGTGTGGAGTCGTTGCGGAAGTTTTCTTTAGGGGCTGCTNNNAGATCGGAA |
| 10590_3 | TATGAGTGTGGAGTCGTTGCGGAAGTTTTCTTTAGGGGCTGCTNNNAGATCGGAA |
| 53395_1 | TATGAGTGTGGAGTCGTTGCCCTTCAGAGTAGTTGGGAACACANNNAGATCGGAA |
| 53395_2 | TATGAGTGTGGAGTCGTTGCCCTTCAGAGTAGTTGGGAACACANNNAGATCGGAA |
| 53395_3 | TATGAGTGTGGAGTCGTTGCCCTTCAGAGTAGTTGGGAACACANNNAGATCGGAA |
| 29896_1 | TATGAGTGTGGAGTCGTTGCCGCTGTGTTGGTGTTGTCATTAANNNAGATCGGAAG |
| 29896_2 | TATGAGTGTGGAGTCGTTGCCGCTGTGTTGGTGTTGTCATTAANNNAGATCGGAAG |
| 29896_3 | TATGAGTGTGGAGTCGTTGCCGCTGTGTTGGTGTTGTCATTAANNNAGATCGGAAG |
| 13697_1 | TATGAGTGTGGAGTCGTTGCTACCAAATTTCAGAACACAAGAANNNAGATCGGAAG |
| 13697_2 | TATGAGTGTGGAGTCGTTGCTACCAAATTTCAGAACACAAGAANNNAGATCGGAAG |
| 13697_3 | TATGAGTGTGGAGTCGTTGCTACCAAATTTCAGAACACAAGAANNNAGATCGGAAG |
| 40787_1 | TATGAGTGTGGAGTCGTTGCTTAGAGAAGAAGCTTCCAGGCAGNNNAGATCGGAA |
| 40787_2 | TATGAGTGTGGAGTCGTTGCTTAGAGAAGAAGCTTCCAGGCAGNNNAGATCGGAA |
| 40787_3 | TATGAGTGTGGAGTCGTTGCTTAGAGAAGAAGCTTCCAGGCAGNNNAGATCGGAA |
| 45105_1 | TATGAGTGTGGAGTCGTTGCCCAAACAGTATGTGAGCTCCTCTNNNAGATCGGAAG |
| 45105_2 | TATGAGTGTGGAGTCGTTGCCCAAACAGTATGTGAGCTCCTCTNNNAGATCGGAAG |
| 45105_3 | TATGAGTGTGGAGTCGTTGCCCAAACAGTATGTGAGCTCCTCTNNNAGATCGGAAG |
| 183_1 | TATGAGTGTGGAGTCGTTGCCACCCACCATAGCTGAGGATTGAGNNNAGATCGGAA |
| 183_2 | TATGAGTGTGGAGTCGTTGCCACCCACCATAGCTGAGGATTGAGNNNAGATCGGAA |
| 183_3 | TATGAGTGTGGAGTCGTTGCCACCCACCATAGCTGAGGATTGAGNNNAGATCGGAA |
| 33316_1 | TATGAGTGTGGAGTCGTTGCGTGGCATCATCATAACTCACTGCNNNAGATCGGAAG |
| 33316_2 | TATGAGTGTGGAGTCGTTGCGTGGCATCATCATAACTCACTGCNNNAGATCGGAAG |
| 33316_3 | TATGAGTGTGGAGTCGTTGCGTGGCATCATCATAACTCACTGCNNNAGATCGGAAG |
| 11699_1 | TATGAGTGTGGAGTCGTTGCTTGTGAAGGTGATTCCCCTCCCNNNAGATCGGAAGA |

| | |
|---|---|
| 11699_2 | TATGAGTGTGGAGTCGTTGCTTGTGAAGGTGATTCCCCTCCCNNNAGATCGGAAGA |
| 11699_3 | TATGAGTGTGGAGTCGTTGCTTGTGAAGGTGATTCCCCTCCCNNNAGATCGGAAGA |
| 38331_1 | TATGAGTGTGGAGTCGTTGCTTTGACAATGGATCCTTGGAAAGNNNAGATCGGAAG |
| 38331_2 | TATGAGTGTGGAGTCGTTGCTTTGACAATGGATCCTTGGAAAGNNNAGATCGGAAG |
| 38331_3 | TATGAGTGTGGAGTCGTTGCTTTGACAATGGATCCTTGGAAAGNNNAGATCGGAAG |
| 3976_1 | TATGAGTGTGGAGTCGTTGCTACAGTGGAGCAATCTGGTTCACNNNAGATCGGAAG |
| 3976_2 | TATGAGTGTGGAGTCGTTGCTACAGTGGAGCAATCTGGTTCACNNNAGATCGGAAG |
| 3976_3 | TATGAGTGTGGAGTCGTTGCTACAGTGGAGCAATCTGGTTCACNNNAGATCGGAAG |
| 28877_1 | TATGAGTGTGGAGTCGTTGCACAGAGAGAGAGAGAGAGATGCANNNAGATCGGA |
| 28877_2 | TATGAGTGTGGAGTCGTTGCACAGAGAGAGAGAGAGAGATGCANNNAGATCGGA |
| 28877_3 | TATGAGTGTGGAGTCGTTGCACAGAGAGAGAGAGAGAGATGCANNNAGATCGGA |
| 46318_1 | TATGAGTGTGGAGTCGTTGCCAATCTTGGCTCACTGCAAACTCNNNAGATCGGAAG |
| 46318_2 | TATGAGTGTGGAGTCGTTGCCAATCTTGGCTCACTGCAAACTCNNNAGATCGGAAG |
| 46318_3 | TATGAGTGTGGAGTCGTTGCCAATCTTGGCTCACTGCAAACTCNNNAGATCGGAAG |
| 29798_1 | TATGAGTGTGGAGTCGTTGCCGGTGTATGTTAGCTTGCTTGACNNNAGATCGGAAG |
| 29798_2 | TATGAGTGTGGAGTCGTTGCCGGTGTATGTTAGCTTGCTTGACNNNAGATCGGAAG |
| 29798_3 | TATGAGTGTGGAGTCGTTGCCGGTGTATGTTAGCTTGCTTGACNNNAGATCGGAAG |
| 45365_1 | TATGAGTGTGGAGTCGTTGCGCTCCATACTGTCCCAATAAGGTNNNAGATCGGAAG |
| 45365_2 | TATGAGTGTGGAGTCGTTGCGCTCCATACTGTCCCAATAAGGTNNNAGATCGGAAG |
| 45365_3 | TATGAGTGTGGAGTCGTTGCGCTCCATACTGTCCCAATAAGGTNNNAGATCGGAAG |
| 34834_1 | TATGAGTGTGGAGTCGTTGCCTCCTGAGTAGCTGGGATTACAGNNNAGATCGGAAG |
| 34834_2 | TATGAGTGTGGAGTCGTTGCCTCCTGAGTAGCTGGGATTACAGNNNAGATCGGAAG |
| 34834_3 | TATGAGTGTGGAGTCGTTGCCTCCTGAGTAGCTGGGATTACAGNNNAGATCGGAAG |
| 50558_1 | TATGAGTGTGGAGTCGTTGCATGATGGAAGCAGAGAATCCTCCNNNAGATCGGAA |
| 50558_2 | TATGAGTGTGGAGTCGTTGCATGATGGAAGCAGAGAATCCTCCNNNAGATCGGAA |
| 50558_3 | TATGAGTGTGGAGTCGTTGCATGATGGAAGCAGAGAATCCTCCNNNAGATCGGAA |
| 37721_1 | TATGAGTGTGGAGTCGTTGCTGCAGGATACAAAACTAACACACANNNAGATCGGAA |
| 37721_2 | TATGAGTGTGGAGTCGTTGCTGCAGGATACAAAACTAACACACANNNAGATCGGAA |
| 37721_3 | TATGAGTGTGGAGTCGTTGCTGCAGGATACAAAACTAACACACANNNAGATCGGAA |
| 36848_1 | TATGAGTGTGGAGTCGTTGCTCACGAAGTCAGGAGTTCAAGACNNNAGATCGGAA |
| 36848_2 | TATGAGTGTGGAGTCGTTGCTCACGAAGTCAGGAGTTCAAGACNNNAGATCGGAA |
| 36848_3 | TATGAGTGTGGAGTCGTTGCTCACGAAGTCAGGAGTTCAAGACNNNAGATCGGAA |
| 3058_1 | TATGAGTGTGGAGTCGTTGCATGGCCCAGAAATTTCCAACTTGNNNAGATCGGAAG |
| 3058_2 | TATGAGTGTGGAGTCGTTGCATGGCCCAGAAATTTCCAACTTGNNNAGATCGGAAG |
| 3058_3 | TATGAGTGTGGAGTCGTTGCATGGCCCAGAAATTTCCAACTTGNNNAGATCGGAAG |
| 46762_1 | TATGAGTGTGGAGTCGTTGCAGTAGCTGGGATTACAGGAATGCNNNAGATCGGAA |
| 46762_2 | TATGAGTGTGGAGTCGTTGCAGTAGCTGGGATTACAGGAATGCNNNAGATCGGAA |
| 46762_3 | TATGAGTGTGGAGTCGTTGCAGTAGCTGGGATTACAGGAATGCNNNAGATCGGAA |
| 13284_1 | TATGAGTGTGGAGTCGTTGCACTTTCTTAATAAACTTGCTTGCNNNAGATCGGAAGA |
| 13284_2 | TATGAGTGTGGAGTCGTTGCACTTTCTTAATAAACTTGCTTGCNNNAGATCGGAAGA |
| 13284_3 | TATGAGTGTGGAGTCGTTGCACTTTCTTAATAAACTTGCTTGCNNNAGATCGGAAGA |
| 50644_1 | TATGAGTGTGGAGTCGTTGCCACCAATTTGGGAAAGAGGCTTGNNNAGATCGGAA |
| 50644_2 | TATGAGTGTGGAGTCGTTGCCACCAATTTGGGAAAGAGGCTTGNNNAGATCGGAA |

| | |
|---|---|
| 50644_3 | TATGAGTGTGGAGTCGTTGCCACCAATTTGGGAAAGAGGCTTGNNNAGATCGGAA |
| 44363_1 | TATGAGTGTGGAGTCGTTGCGTTTCTGAGCAGAGAGGACACAGNNNAGATCGGAA |
| 44363_2 | TATGAGTGTGGAGTCGTTGCGTTTCTGAGCAGAGAGGACACAGNNNAGATCGGAA |
| 44363_3 | TATGAGTGTGGAGTCGTTGCGTTTCTGAGCAGAGAGGACACAGNNNAGATCGGAA |
| 8088_1 | TATGAGTGTGGAGTCGTTGCGGGATTACAGGTACGGGTCAACATNNNAGATCGGA |
| 8088_2 | TATGAGTGTGGAGTCGTTGCGGGATTACAGGTACGGGTCAACATNNNAGATCGGA |
| 8088_3 | TATGAGTGTGGAGTCGTTGCGGGATTACAGGTACGGGTCAACATNNNAGATCGGA |
| 35597_1 | TATGAGTGTGGAGTCGTTGCAGTGCTGGGATTACAGATGTGAGNNNAGATCGGAA |
| 35597_2 | TATGAGTGTGGAGTCGTTGCAGTGCTGGGATTACAGATGTGAGNNNAGATCGGAA |
| 35597_3 | TATGAGTGTGGAGTCGTTGCAGTGCTGGGATTACAGATGTGAGNNNAGATCGGAA |
| 52295_1 | TATGAGTGTGGAGTCGTTGCTCCTCACACTCAGTCTTGCTTACNNNAGATCGGAAGA |
| 52295_2 | TATGAGTGTGGAGTCGTTGCTCCTCACACTCAGTCTTGCTTACNNNAGATCGGAAGA |
| 52295_3 | TATGAGTGTGGAGTCGTTGCTCCTCACACTCAGTCTTGCTTACNNNAGATCGGAAGA |
| 30890_1 | TATGAGTGTGGAGTCGTTGCAAAAACCCAAACCTGTCAACTGGNNNAGATCGGAAG |
| 30890_2 | TATGAGTGTGGAGTCGTTGCAAAAACCCAAACCTGTCAACTGGNNNAGATCGGAAG |
| 30890_3 | TATGAGTGTGGAGTCGTTGCAAAAACCCAAACCTGTCAACTGGNNNAGATCGGAAG |
| 38706_1 | TATGAGTGTGGAGTCGTTGCGGGAAACCCATCAGACTAACAGTNNNAGATCGGAA |
| 38706_2 | TATGAGTGTGGAGTCGTTGCGGGAAACCCATCAGACTAACAGTNNNAGATCGGAA |
| 38706_3 | TATGAGTGTGGAGTCGTTGCGGGAAACCCATCAGACTAACAGTNNNAGATCGGAA |
| 35827_1 | TATGAGTGTGGAGTCGTTGCAAGTGATTCTTGTTCCTCAGCCTNNNAGATCGGAAG |
| 35827_2 | TATGAGTGTGGAGTCGTTGCAAGTGATTCTTGTTCCTCAGCCTNNNAGATCGGAAG |
| 35827_3 | TATGAGTGTGGAGTCGTTGCAAGTGATTCTTGTTCCTCAGCCTNNNAGATCGGAAG |
| 32396_1 | TATGAGTGTGGAGTCGTTGCGCAAATGTATTCTTCACGGGAGGNNNAGATCGGAA |
| 32396_2 | TATGAGTGTGGAGTCGTTGCGCAAATGTATTCTTCACGGGAGGNNNAGATCGGAA |
| 32396_3 | TATGAGTGTGGAGTCGTTGCGCAAATGTATTCTTCACGGGAGGNNNAGATCGGAA |
| 40366_1 | TATGAGTGTGGAGTCGTTGCATTGAGGGGAAGTTTCAAAGGGTNNNAGATCGGA |
| 40366_2 | TATGAGTGTGGAGTCGTTGCATTGAGGGGAAGTTTCAAAGGGTNNNAGATCGGA |
| 40366_3 | TATGAGTGTGGAGTCGTTGCATTGAGGGGAAGTTTCAAAGGGTNNNAGATCGGA |
| 41956_1 | TATGAGTGTGGAGTCGTTGCGGCATGTTCATAGCTCAATGCAANNNAGATCGGAAG |
| 41956_2 | TATGAGTGTGGAGTCGTTGCGGCATGTTCATAGCTCAATGCAANNNAGATCGGAAG |
| 41956_3 | TATGAGTGTGGAGTCGTTGCGGCATGTTCATAGCTCAATGCAANNNAGATCGGAAG |
| 50263_1 | TATGAGTGTGGAGTCGTTGCCTTTGAGAGGGACCAGGAGATTCNNNAGATCGGAA |
| 50263_2 | TATGAGTGTGGAGTCGTTGCCTTTGAGAGGGACCAGGAGATTCNNNAGATCGGAA |
| 50263_3 | TATGAGTGTGGAGTCGTTGCCTTTGAGAGGGACCAGGAGATTCNNNAGATCGGAA |
| 36574_1 | TATGAGTGTGGAGTCGTTGCAGATCACCCAAGGTCAAGAGTTCNNNAGATCGGAA |
| 36574_2 | TATGAGTGTGGAGTCGTTGCAGATCACCCAAGGTCAAGAGTTCNNNAGATCGGAA |
| 36574_3 | TATGAGTGTGGAGTCGTTGCAGATCACCCAAGGTCAAGAGTTCNNNAGATCGGAA |
| 48907_1 | TATGAGTGTGGAGTCGTTGCCTCCCAAGTAGCCAGGATTACAGNNNAGATCGGAAG |
| 48907_2 | TATGAGTGTGGAGTCGTTGCCTCCCAAGTAGCCAGGATTACAGNNNAGATCGGAAG |
| 48907_3 | TATGAGTGTGGAGTCGTTGCCTCCCAAGTAGCCAGGATTACAGNNNAGATCGGAAG |
| 39181_1 | TATGAGTGTGGAGTCGTTGCAGCCAGTCATTTTGTCTCTCAGTNNNAGATCGGAAG |
| 39181_2 | TATGAGTGTGGAGTCGTTGCAGCCAGTCATTTTGTCTCTCAGTNNNAGATCGGAAG |
| 39181_3 | TATGAGTGTGGAGTCGTTGCAGCCAGTCATTTTGTCTCTCAGTNNNAGATCGGAAG |

| | |
|---|---|
| 40704_1 | TATGAGTGTGGAGTCGTTGCGAGACGGAATTTCGCTCTTGTTGNNNAGATCGGAAG |
| 40704_2 | TATGAGTGTGGAGTCGTTGCGAGACGGAATTTCGCTCTTGTTGNNNAGATCGGAAG |
| 40704_3 | TATGAGTGTGGAGTCGTTGCGAGACGGAATTTCGCTCTTGTTGNNNAGATCGGAAG |
| 45708_1 | TATGAGTGTGGAGTCGTTGCGTCTTGAACTCCTGACCTCATGANNNAGATCGGAAG |
| 45708_2 | TATGAGTGTGGAGTCGTTGCGTCTTGAACTCCTGACCTCATGANNNAGATCGGAAG |
| 45708_3 | TATGAGTGTGGAGTCGTTGCGTCTTGAACTCCTGACCTCATGANNNAGATCGGAAG |
| 47531_1 | TATGAGTGTGGAGTCGTTGCTCAATCTCCTGAGCTCAAGCAATNNNAGATCGGAAG |
| 47531_2 | TATGAGTGTGGAGTCGTTGCTCAATCTCCTGAGCTCAAGCAATNNNAGATCGGAAG |
| 47531_3 | TATGAGTGTGGAGTCGTTGCTCAATCTCCTGAGCTCAAGCAATNNNAGATCGGAAG |
| 13664_1 | TATGAGTGTGGAGTCGTTGCGGTCAACATCTTTTACATTCTGNNNAGATCGGAAGA |
| 13664_2 | TATGAGTGTGGAGTCGTTGCGGTCAACATCTTTTACATTCTGNNNAGATCGGAAGA |
| 13664_3 | TATGAGTGTGGAGTCGTTGCGGTCAACATCTTTTACATTCTGNNNAGATCGGAAGA |
| 33838_1 | TATGAGTGTGGAGTCGTTGCAAAACTTCTGCAAGGACATCTGCNNNAGATCGGAAG |
| 33838_2 | TATGAGTGTGGAGTCGTTGCAAAACTTCTGCAAGGACATCTGCNNNAGATCGGAAG |
| 33838_3 | TATGAGTGTGGAGTCGTTGCAAAACTTCTGCAAGGACATCTGCNNNAGATCGGAAG |
| 33315_1 | TATGAGTGTGGAGTCGTTGCCTCCCAGGTTCATGCCATTCTCNNNAGATCGGAAGAG |
| 33315_2 | TATGAGTGTGGAGTCGTTGCCTCCCAGGTTCATGCCATTCTCNNNAGATCGGAAGAG |
| 33315_3 | TATGAGTGTGGAGTCGTTGCCTCCCAGGTTCATGCCATTCTCNNNAGATCGGAAGAG |
| 48998_1 | TATGAGTGTGGAGTCGTTGCCCCGAGTAGCTGGGACTACAGNNNAGATCGGAAGA |
| 48998_2 | TATGAGTGTGGAGTCGTTGCCCCGAGTAGCTGGGACTACAGNNNAGATCGGAAGA |
| 48998_3 | TATGAGTGTGGAGTCGTTGCCCCGAGTAGCTGGGACTACAGNNNAGATCGGAAGA |
| 37371_1 | TATGAGTGTGGAGTCGTTGCAAGCCATAGGGTTATTTTGGGGTNNNAGATCGGAA |
| 37371_2 | TATGAGTGTGGAGTCGTTGCAAGCCATAGGGTTATTTTGGGGTNNNAGATCGGAA |
| 37371_3 | TATGAGTGTGGAGTCGTTGCAAGCCATAGGGTTATTTTGGGGTNNNAGATCGGAA |
| 41947_1 | TATGAGTGTGGAGTCGTTGCGATCTCGACTCACTGCAAGCTCNNNAGATCGGAAGA |
| 41947_2 | TATGAGTGTGGAGTCGTTGCGATCTCGACTCACTGCAAGCTCNNNAGATCGGAAGA |
| 41947_3 | TATGAGTGTGGAGTCGTTGCGATCTCGACTCACTGCAAGCTCNNNAGATCGGAAGA |
| 43311_1 | TATGAGTGTGGAGTCGTTGCCAGCCACTCTTTCTTTTGACTGGNNNAGATCGGAAGA |
| 43311_2 | TATGAGTGTGGAGTCGTTGCCAGCCACTCTTTCTTTTGACTGGNNNAGATCGGAAGA |
| 43311_3 | TATGAGTGTGGAGTCGTTGCCAGCCACTCTTTCTTTTGACTGGNNNAGATCGGAAGA |
| 40358_1 | TATGAGTGTGGAGTCGTTGCCATGGTGGTGCATGTCTGTAATCNNNAGATCGGAAG |
| 40358_2 | TATGAGTGTGGAGTCGTTGCCATGGTGGTGCATGTCTGTAATCNNNAGATCGGAAG |
| 40358_3 | TATGAGTGTGGAGTCGTTGCCATGGTGGTGCATGTCTGTAATCNNNAGATCGGAAG |
| 45942_1 | TATGAGTGTGGAGTCGTTGCGAGCAGCTGGGATTACAGACATANNNAGATCGGAA |
| 45942_2 | TATGAGTGTGGAGTCGTTGCGAGCAGCTGGGATTACAGACATANNNAGATCGGAA |
| 45942_3 | TATGAGTGTGGAGTCGTTGCGAGCAGCTGGGATTACAGACATANNNAGATCGGAA |
| 8891_1 | TATGAGTGTGGAGTCGTTGCTGGGATAGAAGTGGTTTTCTGGTNNNAGATCGGAA |
| 8891_2 | TATGAGTGTGGAGTCGTTGCTGGGATAGAAGTGGTTTTCTGGTNNNAGATCGGAA |
| 8891_3 | TATGAGTGTGGAGTCGTTGCTGGGATAGAAGTGGTTTTCTGGTNNNAGATCGGAA |
| 10641_1 | TATGAGTGTGGAGTCGTTGCCCGAGGGAATCCTGTATGTTTGGNNNAGATCGGAA |
| 10641_2 | TATGAGTGTGGAGTCGTTGCCCGAGGGAATCCTGTATGTTTGGNNNAGATCGGAA |
| 10641_3 | TATGAGTGTGGAGTCGTTGCCCGAGGGAATCCTGTATGTTTGGNNNAGATCGGAA |
| 30017_1 | TATGAGTGTGGAGTCGTTGCATCTAACTGTCCATTGGGTCCACNNNAGATCGGAAG |

| | |
|---|---|
| 30017_2 | TATGAGTGTGGAGTCGTTGCATCTAACTGTCCATTGGGTCCACNNNAGATCGGAAG |
| 30017_3 | TATGAGTGTGGAGTCGTTGCATCTAACTGTCCATTGGGTCCACNNNAGATCGGAAG |
| 35569_1 | TATGAGTGTGGAGTCGTTGCTTTCACCATGTTAGCCAGGATGANNNAGATCGGAAG |
| 35569_2 | TATGAGTGTGGAGTCGTTGCTTTCACCATGTTAGCCAGGATGANNNAGATCGGAAG |
| 35569_3 | TATGAGTGTGGAGTCGTTGCTTTCACCATGTTAGCCAGGATGANNNAGATCGGAAG |
| 38384_1 | TATGAGTGTGGAGTCGTTGCAGTCGGAATTGTGACCAGAATCANNNAGATCGGAA |
| 38384_2 | TATGAGTGTGGAGTCGTTGCAGTCGGAATTGTGACCAGAATCANNNAGATCGGAA |
| 38384_3 | TATGAGTGTGGAGTCGTTGCAGTCGGAATTGTGACCAGAATCANNNAGATCGGAA |
| 46715_1 | TATGAGTGTGGAGTCGTTGCGACAGAGTTTAGCTGTTGTTGCCNNNAGATCGGAAG |
| 46715_2 | TATGAGTGTGGAGTCGTTGCGACAGAGTTTAGCTGTTGTTGCCNNNAGATCGGAAG |
| 46715_3 | TATGAGTGTGGAGTCGTTGCGACAGAGTTTAGCTGTTGTTGCCNNNAGATCGGAAG |
| 50100_1 | TATGAGTGTGGAGTCGTTGCGGGCCAGGAACTTACACAATTTTNNNAGATCGGAAG |
| 50100_2 | TATGAGTGTGGAGTCGTTGCGGGCCAGGAACTTACACAATTTTNNNAGATCGGAAG |
| 50100_3 | TATGAGTGTGGAGTCGTTGCGGGCCAGGAACTTACACAATTTTNNNAGATCGGAAG |
| 39471_1 | TATGAGTGTGGAGTCGTTGCGAAGACAAGGGGAGGAAGAGATGNNNAGATCGGA |
| 39471_2 | TATGAGTGTGGAGTCGTTGCGAAGACAAGGGGAGGAAGAGATGNNNAGATCGGA |
| 39471_3 | TATGAGTGTGGAGTCGTTGCGAAGACAAGGGGAGGAAGAGATGNNNAGATCGGA |
| 30820_1 | TATGAGTGTGGAGTCGTTGCAAGAAAACTGGACATAGGGGGAGNNNAGATCGGA |
| 30820_2 | TATGAGTGTGGAGTCGTTGCAAGAAAACTGGACATAGGGGGAGNNNAGATCGGA |
| 30820_3 | TATGAGTGTGGAGTCGTTGCAAGAAAACTGGACATAGGGGGAGNNNAGATCGGA |
| 37628_1 | TATGAGTGTGGAGTCGTTGCTGTCAATATGTGGAGTGATTGGGNNNAGATCGGAA |
| 37628_2 | TATGAGTGTGGAGTCGTTGCTGTCAATATGTGGAGTGATTGGGNNNAGATCGGAA |
| 37628_3 | TATGAGTGTGGAGTCGTTGCTGTCAATATGTGGAGTGATTGGGNNNAGATCGGAA |
| 40287_1 | TATGAGTGTGGAGTCGTTGCAACGGAACTTCGAGTTATCTGCANNNAGATCGGAAG |
| 40287_2 | TATGAGTGTGGAGTCGTTGCAACGGAACTTCGAGTTATCTGCANNNAGATCGGAAG |
| 40287_3 | TATGAGTGTGGAGTCGTTGCAACGGAACTTCGAGTTATCTGCANNNAGATCGGAAG |
| 38091_1 | TATGAGTGTGGAGTCGTTGCTGGAAAGAGAAGTTCAAGTGGCTNNNAGATCGGAA |
| 38091_2 | TATGAGTGTGGAGTCGTTGCTGGAAAGAGAAGTTCAAGTGGCTNNNAGATCGGAA |
| 38091_3 | TATGAGTGTGGAGTCGTTGCTGGAAAGAGAAGTTCAAGTGGCTNNNAGATCGGAA |
| 34651_1 | TATGAGTGTGGAGTCGTTGCTGGAGTCCATTTGTATTCTGACANNNAGATCGGAAG |
| 34651_2 | TATGAGTGTGGAGTCGTTGCTGGAGTCCATTTGTATTCTGACANNNAGATCGGAAG |
| 34651_3 | TATGAGTGTGGAGTCGTTGCTGGAGTCCATTTGTATTCTGACANNNAGATCGGAAG |
| 51240_1 | TATGAGTGTGGAGTCGTTGCTACGGTAATTTGCCTCCAATCCANNNAGATCGGAAG |
| 51240_2 | TATGAGTGTGGAGTCGTTGCTACGGTAATTTGCCTCCAATCCANNNAGATCGGAAG |
| 51240_3 | TATGAGTGTGGAGTCGTTGCTACGGTAATTTGCCTCCAATCCANNNAGATCGGAAG |
| 48028_1 | TATGAGTGTGGAGTCGTTGCCCCATGAGCTGTAGGTCCTTTGNNNAGATCGGAAGA |
| 48028_2 | TATGAGTGTGGAGTCGTTGCCCCATGAGCTGTAGGTCCTTTGNNNAGATCGGAAGA |
| 48028_3 | TATGAGTGTGGAGTCGTTGCCCCATGAGCTGTAGGTCCTTTGNNNAGATCGGAAGA |
| 34811_1 | TATGAGTGTGGAGTCGTTGCACGCCCGGCTAATTTTTTGTATTNNNAGATCGGAAG |
| 34811_2 | TATGAGTGTGGAGTCGTTGCACGCCCGGCTAATTTTTTGTATTNNNAGATCGGAAG |
| 34811_3 | TATGAGTGTGGAGTCGTTGCACGCCCGGCTAATTTTTTGTATTNNNAGATCGGAAG |
| 32441_1 | TATGAGTGTGGAGTCGTTGCGCAGTAGTGCAATCTCAGCTCATNNNAGATCGGAAG |
| 32441_2 | TATGAGTGTGGAGTCGTTGCGCAGTAGTGCAATCTCAGCTCATNNNAGATCGGAAG |

| | |
|---|---|
| 32441_3 | TATGAGTGTGGAGTCGTTGCGCAGTAGTGCAATCTCAGCTCATNNNAGATCGGAAG |
| 32490_1 | TATGAGTGTGGAGTCGTTGCCATTCTCCTGCCTCAGCCTCNNNAGATCGGAAGAGCA |
| 32490_2 | TATGAGTGTGGAGTCGTTGCCATTCTCCTGCCTCAGCCTCNNNAGATCGGAAGAGCA |
| 32490_3 | TATGAGTGTGGAGTCGTTGCCATTCTCCTGCCTCAGCCTCNNNAGATCGGAAGAGCA |
| 12638_1 | TATGAGTGTGGAGTCGTTGCCTCTGTAGTTACATGCAAATAAANNNAGATCGGAAG |
| 12638_2 | TATGAGTGTGGAGTCGTTGCCTCTGTAGTTACATGCAAATAAANNNAGATCGGAAG |
| 12638_3 | TATGAGTGTGGAGTCGTTGCCTCTGTAGTTACATGCAAATAAANNNAGATCGGAAG |
| 50349_1 | TATGAGTGTGGAGTCGTTGCAAAGCAAAACTCTTCACAAGGCANNNAGATCGGAAG |
| 50349_2 | TATGAGTGTGGAGTCGTTGCAAAGCAAAACTCTTCACAAGGCANNNAGATCGGAAG |
| 50349_3 | TATGAGTGTGGAGTCGTTGCAAAGCAAAACTCTTCACAAGGCANNNAGATCGGAAG |
| 10699_1 | TATGAGTGTGGAGTCGTTGCCCTCTCCGAAACTGAGGAGTAGTNNNAGATCGGAAG |
| 10699_2 | TATGAGTGTGGAGTCGTTGCCCTCTCCGAAACTGAGGAGTAGTNNNAGATCGGAAG |
| 10699_3 | TATGAGTGTGGAGTCGTTGCCCTCTCCGAAACTGAGGAGTAGTNNNAGATCGGAAG |
| 53321_1 | TATGAGTGTGGAGTCGTTGCCAGTTAGCTCCTGATGGACACTTNNNAGATCGGAAG |
| 53321_2 | TATGAGTGTGGAGTCGTTGCCAGTTAGCTCCTGATGGACACTTNNNAGATCGGAAG |
| 53321_3 | TATGAGTGTGGAGTCGTTGCCAGTTAGCTCCTGATGGACACTTNNNAGATCGGAAG |
| 46984_1 | TATGAGTGTGGAGTCGTTGCTTCAAGCTATCCTCATGCCTCAGNNNAGATCGGAAGA |
| 46984_2 | TATGAGTGTGGAGTCGTTGCTTCAAGCTATCCTCATGCCTCAGNNNAGATCGGAAGA |
| 46984_3 | TATGAGTGTGGAGTCGTTGCTTCAAGCTATCCTCATGCCTCAGNNNAGATCGGAAGA |
| 49846_1 | TATGAGTGTGGAGTCGTTGCTGTCTTCTGCTGTGTGTCTTCTTNNNAGATCGGAAGA |
| 49846_2 | TATGAGTGTGGAGTCGTTGCTGTCTTCTGCTGTGTGTCTTCTTNNNAGATCGGAAGA |
| 49846_3 | TATGAGTGTGGAGTCGTTGCTGTCTTCTGCTGTGTGTCTTCTTNNNAGATCGGAAGA |
| 48688_1 | TATGAGTGTGGAGTCGTTGCCCTCTTGGGTTCATGCCATTCTCNNNAGATCGGAAGA |
| 48688_2 | TATGAGTGTGGAGTCGTTGCCCTCTTGGGTTCATGCCATTCTCNNNAGATCGGAAGA |
| 48688_3 | TATGAGTGTGGAGTCGTTGCCCTCTTGGGTTCATGCCATTCTCNNNAGATCGGAAGA |
| 9870_1 | TATGAGTGTGGAGTCGTTGCTTGGTTACTTTTGGGAGGATGGANNNAGATCGGAA |
| 9870_2 | TATGAGTGTGGAGTCGTTGCTTGGTTACTTTTGGGAGGATGGANNNAGATCGGAA |
| 9870_3 | TATGAGTGTGGAGTCGTTGCTTGGTTACTTTTGGGAGGATGGANNNAGATCGGAA |
| 2576_1 | TATGAGTGTGGAGTCGTTGCTCTGCCATTTTCTAGGCTCATGGNNNAGATCGGAAG |
| 2576_2 | TATGAGTGTGGAGTCGTTGCTCTGCCATTTTCTAGGCTCATGGNNNAGATCGGAAG |
| 2576_3 | TATGAGTGTGGAGTCGTTGCTCTGCCATTTTCTAGGCTCATGGNNNAGATCGGAAG |
| 32793_1 | TATGAGTGTGGAGTCGTTGCGGGTGATTCTCCAGCCTAAGTAGNNNAGATCGGAA |
| 32793_2 | TATGAGTGTGGAGTCGTTGCGGGTGATTCTCCAGCCTAAGTAGNNNAGATCGGAA |
| 32793_3 | TATGAGTGTGGAGTCGTTGCGGGTGATTCTCCAGCCTAAGTAGNNNAGATCGGAA |
| 43497_1 | TATGAGTGTGGAGTCGTTGCAGGTTCAAGCAATTCTCCTGTCTNNNAGATCGGAAG |
| 43497_2 | TATGAGTGTGGAGTCGTTGCAGGTTCAAGCAATTCTCCTGTCTNNNAGATCGGAAG |
| 43497_3 | TATGAGTGTGGAGTCGTTGCAGGTTCAAGCAATTCTCCTGTCTNNNAGATCGGAAG |
| 42474_1 | TATGAGTGTGGAGTCGTTGCTCTTTCCATGTGCCAGGTACTACNNNAGATCGGAAG |
| 42474_2 | TATGAGTGTGGAGTCGTTGCTCTTTCCATGTGCCAGGTACTACNNNAGATCGGAAG |
| 42474_3 | TATGAGTGTGGAGTCGTTGCTCTTTCCATGTGCCAGGTACTACNNNAGATCGGAAG |
| 41080_1 | TATGAGTGTGGAGTCGTTGCAAATGCCTGATAAGAGGAATGCANNNAGATCGGAA |
| 41080_2 | TATGAGTGTGGAGTCGTTGCAAATGCCTGATAAGAGGAATGCANNNAGATCGGAA |
| 41080_3 | TATGAGTGTGGAGTCGTTGCAAATGCCTGATAAGAGGAATGCANNNAGATCGGAA |

| | |
|---|---|
| 10086_1 | TATGAGTGTGGAGTCGTTGCGGGGATAGGATTTCCCAAGGAGTNNNAGATCGGAA |
| 10086_2 | TATGAGTGTGGAGTCGTTGCGGGGATAGGATTTCCCAAGGAGTNNNAGATCGGAA |
| 10086_3 | TATGAGTGTGGAGTCGTTGCGGGGATAGGATTTCCCAAGGAGTNNNAGATCGGAA |
| 49726_1 | TATGAGTGTGGAGTCGTTGCCTCACTACAACTTCTGCCTCCTGNNNAGATCGGAAGA |
| 49726_2 | TATGAGTGTGGAGTCGTTGCCTCACTACAACTTCTGCCTCCTGNNNAGATCGGAAGA |
| 49726_3 | TATGAGTGTGGAGTCGTTGCCTCACTACAACTTCTGCCTCCTGNNNAGATCGGAAGA |
| 37559_1 | TATGAGTGTGGAGTCGTTGCAGCAGAGGTTTGAATAGTGCTCANNNAGATCGGAA |
| 37559_2 | TATGAGTGTGGAGTCGTTGCAGCAGAGGTTTGAATAGTGCTCANNNAGATCGGAA |
| 37559_3 | TATGAGTGTGGAGTCGTTGCAGCAGAGGTTTGAATAGTGCTCANNNAGATCGGAA |
| 34333_1 | TATGAGTGTGGAGTCGTTGCATAGCACAACAGGGTGACTACAGNNNAGATCGGAA |
| 34333_2 | TATGAGTGTGGAGTCGTTGCATAGCACAACAGGGTGACTACAGNNNAGATCGGAA |
| 34333_3 | TATGAGTGTGGAGTCGTTGCATAGCACAACAGGGTGACTACAGNNNAGATCGGAA |
| 44013_1 | TATGAGTGTGGAGTCGTTGCATCCCCAGCAAGGATTCTACAAANNNAGATCGGAAG |
| 44013_2 | TATGAGTGTGGAGTCGTTGCATCCCCAGCAAGGATTCTACAAANNNAGATCGGAAG |
| 44013_3 | TATGAGTGTGGAGTCGTTGCATCCCCAGCAAGGATTCTACAAANNNAGATCGGAAG |
| 37209_1 | TATGAGTGTGGAGTCGTTGCACTCAGGAGGATAAGGCAGAAGANNNAGATCGGA |
| 37209_2 | TATGAGTGTGGAGTCGTTGCACTCAGGAGGATAAGGCAGAAGANNNAGATCGGA |
| 37209_3 | TATGAGTGTGGAGTCGTTGCACTCAGGAGGATAAGGCAGAAGANNNAGATCGGA |
| 51924_1 | TATGAGTGTGGAGTCGTTGCCCCAGAGAGATGATGTGACTGAGNNNAGATCGGAA |
| 51924_2 | TATGAGTGTGGAGTCGTTGCCCCAGAGAGATGATGTGACTGAGNNNAGATCGGAA |
| 51924_3 | TATGAGTGTGGAGTCGTTGCCCCAGAGAGATGATGTGACTGAGNNNAGATCGGAA |
| 13306_1 | TATGAGTGTGGAGTCGTTGCAATGGGAGAAATCAACAATTCTNNNAGATCGGAAG |
| 13306_2 | TATGAGTGTGGAGTCGTTGCAATGGGAGAAATCAACAATTCTNNNAGATCGGAAG |
| 13306_3 | TATGAGTGTGGAGTCGTTGCAATGGGAGAAATCAACAATTCTNNNAGATCGGAAG |
| 50313_1 | TATGAGTGTGGAGTCGTTGCAAGAATCACTTCAACCCGGGAGNNNAGATCGGAAG |
| 50313_2 | TATGAGTGTGGAGTCGTTGCAAGAATCACTTCAACCCGGGAGNNNAGATCGGAAG |
| 50313_3 | TATGAGTGTGGAGTCGTTGCAAGAATCACTTCAACCCGGGAGNNNAGATCGGAAG |
| 45048_1 | TATGAGTGTGGAGTCGTTGCTAAGGAAGGGACCATAGAGAGGGNNNAGATCGGA |
| 45048_2 | TATGAGTGTGGAGTCGTTGCTAAGGAAGGGACCATAGAGAGGGNNNAGATCGGA |
| 45048_3 | TATGAGTGTGGAGTCGTTGCTAAGGAAGGGACCATAGAGAGGGNNNAGATCGGA |
| 30325_1 | TATGAGTGTGGAGTCGTTGCGGCACCTGGTAGGTTTTAATGCANNNAGATCGGAA |
| 30325_2 | TATGAGTGTGGAGTCGTTGCGGCACCTGGTAGGTTTTAATGCANNNAGATCGGAA |
| 30325_3 | TATGAGTGTGGAGTCGTTGCGGCACCTGGTAGGTTTTAATGCANNNAGATCGGAA |
| 10610_1 | TATGAGTGTGGAGTCGTTGCCCTTCCTCTCTGTCCGCTATTTTTNNNAGATCGGAAG |
| 10610_2 | TATGAGTGTGGAGTCGTTGCCCTTCCTCTCTGTCCGCTATTTTTNNNAGATCGGAAG |
| 10610_3 | TATGAGTGTGGAGTCGTTGCCCTTCCTCTCTGTCCGCTATTTTTNNNAGATCGGAAG |
| 47434_1 | TATGAGTGTGGAGTCGTTGCCCACCACACCTGGCTAATTTTTTNNNAGATCGGAAGA |
| 47434_2 | TATGAGTGTGGAGTCGTTGCCCACCACACCTGGCTAATTTTTTNNNAGATCGGAAGA |
| 47434_3 | TATGAGTGTGGAGTCGTTGCCCACCACACCTGGCTAATTTTTTNNNAGATCGGAAGA |
| 12551_1 | TATGAGTGTGGAGTCGTTGCCAAAGACTAAGTCCACCTAAGNNNAGATCGGAAGA |
| 12551_2 | TATGAGTGTGGAGTCGTTGCCAAAGACTAAGTCCACCTAAGNNNAGATCGGAAGA |
| 12551_3 | TATGAGTGTGGAGTCGTTGCCAAAGACTAAGTCCACCTAAGNNNAGATCGGAAGA |
| 35610_1 | TATGAGTGTGGAGTCGTTGCCTCCTCAGCTTCCCAAGTAGTTGNNNAGATCGGAAG |

| | |
|---|---|
| 35610_2 | TATGAGTGTGGAGTCGTTGCCTCCTCAGCTTCCCAAGTAGTTGNNNAGATCGGAAG |
| 35610_3 | TATGAGTGTGGAGTCGTTGCCTCCTCAGCTTCCCAAGTAGTTGNNNAGATCGGAAG |
| 36037_1 | TATGAGTGTGGAGTCGTTGCAAAGGGCCACATAGATTACAGCANNNAGATCGGAA |
| 36037_2 | TATGAGTGTGGAGTCGTTGCAAAGGGCCACATAGATTACAGCANNNAGATCGGAA |
| 36037_3 | TATGAGTGTGGAGTCGTTGCAAAGGGCCACATAGATTACAGCANNNAGATCGGAA |
| 44163_1 | TATGAGTGTGGAGTCGTTGCATGCATCTAGATTTCCAAGGCCTNNNAGATCGGAAG |
| 44163_2 | TATGAGTGTGGAGTCGTTGCATGCATCTAGATTTCCAAGGCCTNNNAGATCGGAAG |
| 44163_3 | TATGAGTGTGGAGTCGTTGCATGCATCTAGATTTCCAAGGCCTNNNAGATCGGAAG |
| 29121_1 | TATGAGTGTGGAGTCGTTGCGATTGAAAAGGATGCTGGCAAGANNNAGATCGGAA |
| 29121_2 | TATGAGTGTGGAGTCGTTGCGATTGAAAAGGATGCTGGCAAGANNNAGATCGGAA |
| 29121_3 | TATGAGTGTGGAGTCGTTGCGATTGAAAAGGATGCTGGCAAGANNNAGATCGGAA |
| 44796_1 | TATGAGTGTGGAGTCGTTGCTAGGGTCAAGGAATTGTGGAGTGNNNAGATCGGAA |
| 44796_2 | TATGAGTGTGGAGTCGTTGCTAGGGTCAAGGAATTGTGGAGTGNNNAGATCGGAA |
| 44796_3 | TATGAGTGTGGAGTCGTTGCTAGGGTCAAGGAATTGTGGAGTGNNNAGATCGGAA |
| 34303_1 | TATGAGTGTGGAGTCGTTGCGAGAGTGTTGAAAGGGTCGGAGNNNAGATCGGAA |
| 34303_2 | TATGAGTGTGGAGTCGTTGCGAGAGTGTTGAAAGGGTCGGAGNNNAGATCGGAA |
| 34303_3 | TATGAGTGTGGAGTCGTTGCGAGAGTGTTGAAAGGGTCGGAGNNNAGATCGGAA |
| 13361_1 | TATGAGTGTGGAGTCGTTGCCTTGGAAGTTAACTAGCCTTTATNNNAGATCGGAAG |
| 13361_2 | TATGAGTGTGGAGTCGTTGCCTTGGAAGTTAACTAGCCTTTATNNNAGATCGGAAG |
| 13361_3 | TATGAGTGTGGAGTCGTTGCCTTGGAAGTTAACTAGCCTTTATNNNAGATCGGAAG |
| 47641_1 | TATGAGTGTGGAGTCGTTGCAATGTTGAGGTTTTGCGGAATCCNNNAGATCGGAA |
| 47641_2 | TATGAGTGTGGAGTCGTTGCAATGTTGAGGTTTTGCGGAATCCNNNAGATCGGAA |
| 47641_3 | TATGAGTGTGGAGTCGTTGCAATGTTGAGGTTTTGCGGAATCCNNNAGATCGGAA |
| 35607_1 | TATGAGTGTGGAGTCGTTGCGGCCTAGAGCTATTCAAGTTTGCNNNAGATCGGAAG |
| 35607_2 | TATGAGTGTGGAGTCGTTGCGGCCTAGAGCTATTCAAGTTTGCNNNAGATCGGAAG |
| 35607_3 | TATGAGTGTGGAGTCGTTGCGGCCTAGAGCTATTCAAGTTTGCNNNAGATCGGAAG |
| 35289_1 | TATGAGTGTGGAGTCGTTGCTTCAAGCAATTTTCCTGCCTCAGNNNAGATCGGAAG |
| 35289_2 | TATGAGTGTGGAGTCGTTGCTTCAAGCAATTTTCCTGCCTCAGNNNAGATCGGAAG |
| 35289_3 | TATGAGTGTGGAGTCGTTGCTTCAAGCAATTTTCCTGCCTCAGNNNAGATCGGAAG |
| 33706_1 | TATGAGTGTGGAGTCGTTGCGAGCAACTTCAGGTAGACTCCTNNNAGATCGGAAG |
| 33706_2 | TATGAGTGTGGAGTCGTTGCGAGCAACTTCAGGTAGACTCCTNNNAGATCGGAAG |
| 33706_3 | TATGAGTGTGGAGTCGTTGCGAGCAACTTCAGGTAGACTCCTNNNAGATCGGAAG |
| 50397_1 | TATGAGTGTGGAGTCGTTGCCAGCCTGTGAAGGGAAGGAGNNNAGATCGGAAGA |
| 50397_2 | TATGAGTGTGGAGTCGTTGCCAGCCTGTGAAGGGAAGGAGNNNAGATCGGAAGA |
| 50397_3 | TATGAGTGTGGAGTCGTTGCCAGCCTGTGAAGGGAAGGAGNNNAGATCGGAAGA |
| 30299_1 | TATGAGTGTGGAGTCGTTGCACAATCAGATCAACTTTAGCCACTNNNAGATCGGAA |
| 30299_2 | TATGAGTGTGGAGTCGTTGCACAATCAGATCAACTTTAGCCACTNNNAGATCGGAA |
| 30299_3 | TATGAGTGTGGAGTCGTTGCACAATCAGATCAACTTTAGCCACTNNNAGATCGGAA |
| 34964_1 | TATGAGTGTGGAGTCGTTGCCATGATCTCGGCTCACTGCAATNNNAGATCGGAAGA |
| 34964_2 | TATGAGTGTGGAGTCGTTGCCATGATCTCGGCTCACTGCAATNNNAGATCGGAAGA |
| 34964_3 | TATGAGTGTGGAGTCGTTGCCATGATCTCGGCTCACTGCAATNNNAGATCGGAAGA |
| 44480_1 | TATGAGTGTGGAGTCGTTGCGGCATGATCTTGGCTTATTGCAGNNNAGATCGGAAG |
| 44480_2 | TATGAGTGTGGAGTCGTTGCGGCATGATCTTGGCTTATTGCAGNNNAGATCGGAAG |

| | |
|---|---|
| 44480_3 | TATGAGTGTGGAGTCGTTGCGGCATGATCTTGGCTTATTGCAGNNNAGATCGGAA |
| 47708_1 | TATGAGTGTGGAGTCGTTGCCCGGGAAGTTTCCGAAGCTCNNNAGATCGGAAGAG |
| 47708_2 | TATGAGTGTGGAGTCGTTGCCCGGGAAGTTTCCGAAGCTCNNNAGATCGGAAGAG |
| 47708_3 | TATGAGTGTGGAGTCGTTGCCCGGGAAGTTTCCGAAGCTCNNNAGATCGGAAGAG |
| 45807_1 | TATGAGTGTGGAGTCGTTGCCCACGCCCAGCTAATTTTGTATTNNNAGATCGGAAG |
| 45807_2 | TATGAGTGTGGAGTCGTTGCCCACGCCCAGCTAATTTTGTATTNNNAGATCGGAAG |
| 45807_3 | TATGAGTGTGGAGTCGTTGCCCACGCCCAGCTAATTTTGTATTNNNAGATCGGAAG |
| 43504_1 | TATGAGTGTGGAGTCGTTGCACCTTTGATGGAAATTTGTGTTGTNNNAGATCGGAA |
| 43504_2 | TATGAGTGTGGAGTCGTTGCACCTTTGATGGAAATTTGTGTTGTNNNAGATCGGAA |
| 43504_3 | TATGAGTGTGGAGTCGTTGCACCTTTGATGGAAATTTGTGTTGTNNNAGATCGGAA |
| 45649_1 | TATGAGTGTGGAGTCGTTGCAAGCTAAGTGGGTCTCCATCAAGNNNAGATCGGAA |
| 45649_2 | TATGAGTGTGGAGTCGTTGCAAGCTAAGTGGGTCTCCATCAAGNNNAGATCGGAA |
| 45649_3 | TATGAGTGTGGAGTCGTTGCAAGCTAAGTGGGTCTCCATCAAGNNNAGATCGGAA |
| 40973_1 | TATGAGTGTGGAGTCGTTGCACTCATATGCTGAATAAATACCAAACANNNAGATCG |
| 40973_2 | TATGAGTGTGGAGTCGTTGCACTCATATGCTGAATAAATACCAAACANNNAGATCG |
| 40973_3 | TATGAGTGTGGAGTCGTTGCACTCATATGCTGAATAAATACCAAACANNNAGATCG |
| 42042_1 | TATGAGTGTGGAGTCGTTGCGAGGTCTTGCCACTGTACTCCNNNAGATCGGAAGAG |
| 42042_2 | TATGAGTGTGGAGTCGTTGCGAGGTCTTGCCACTGTACTCCNNNAGATCGGAAGAG |
| 42042_3 | TATGAGTGTGGAGTCGTTGCGAGGTCTTGCCACTGTACTCCNNNAGATCGGAAGAG |
| 48794_1 | TATGAGTGTGGAGTCGTTGCACAGCACTCCAAAAACACAAACANNNAGATCGGAAG |
| 48794_2 | TATGAGTGTGGAGTCGTTGCACAGCACTCCAAAAACACAAACANNNAGATCGGAAG |
| 48794_3 | TATGAGTGTGGAGTCGTTGCACAGCACTCCAAAAACACAAACANNNAGATCGGAAG |
| 31050_1 | TATGAGTGTGGAGTCGTTGCCCTGTGCTTTGAGGACTAGTTCTNNNAGATCGGAAG |
| 31050_2 | TATGAGTGTGGAGTCGTTGCCCTGTGCTTTGAGGACTAGTTCTNNNAGATCGGAAG |
| 31050_3 | TATGAGTGTGGAGTCGTTGCCCTGTGCTTTGAGGACTAGTTCTNNNAGATCGGAAG |
| 46010_1 | TATGAGTGTGGAGTCGTTGCGGAGTACAGTAGCACGATCTCAGNNNAGATCGGAA |
| 46010_2 | TATGAGTGTGGAGTCGTTGCGGAGTACAGTAGCACGATCTCAGNNNAGATCGGAA |
| 46010_3 | TATGAGTGTGGAGTCGTTGCGGAGTACAGTAGCACGATCTCAGNNNAGATCGGAA |
| 38772_1 | TATGAGTGTGGAGTCGTTGCAGCTCCTCTAGGTATGATGTTGANNNAGATCGGAAG |
| 38772_2 | TATGAGTGTGGAGTCGTTGCAGCTCCTCTAGGTATGATGTTGANNNAGATCGGAAG |
| 38772_3 | TATGAGTGTGGAGTCGTTGCAGCTCCTCTAGGTATGATGTTGANNNAGATCGGAAG |
| 41794_1 | TATGAGTGTGGAGTCGTTGCCCTGGGTTCAAGCAATTCTTCTGNNNAGATCGGAAG |
| 41794_2 | TATGAGTGTGGAGTCGTTGCCCTGGGTTCAAGCAATTCTTCTGNNNAGATCGGAAG |
| 41794_3 | TATGAGTGTGGAGTCGTTGCCCTGGGTTCAAGCAATTCTTCTGNNNAGATCGGAAG |
| 53092_1 | TATGAGTGTGGAGTCGTTGCCGGTAACATGAGGTGCTTGAAAGNNNAGATCGGAA |
| 53092_2 | TATGAGTGTGGAGTCGTTGCCGGTAACATGAGGTGCTTGAAAGNNNAGATCGGAA |
| 53092_3 | TATGAGTGTGGAGTCGTTGCCGGTAACATGAGGTGCTTGAAAGNNNAGATCGGAA |
| 42737_1 | TATGAGTGTGGAGTCGTTGCTGCCAGGCACTATTCTAAGTACCNNNAGATCGGAAG |
| 42737_2 | TATGAGTGTGGAGTCGTTGCTGCCAGGCACTATTCTAAGTACCNNNAGATCGGAAG |
| 42737_3 | TATGAGTGTGGAGTCGTTGCTGCCAGGCACTATTCTAAGTACCNNNAGATCGGAAG |
| 38886_1 | TATGAGTGTGGAGTCGTTGCCAAGGGCTGTAGCTTACCAAAATNNNAGATCGGAA |
| 38886_2 | TATGAGTGTGGAGTCGTTGCCAAGGGCTGTAGCTTACCAAAATNNNAGATCGGAA |
| 38886_3 | TATGAGTGTGGAGTCGTTGCCAAGGGCTGTAGCTTACCAAAATNNNAGATCGGAA |

| | |
|---|---|
| 32087_1 | TATGAGTGTGGAGTCGTTGCAGCAAAGTAATAATGAGAAGCCCTNNNAGATCGGA |
| 32087_2 | TATGAGTGTGGAGTCGTTGCAGCAAAGTAATAATGAGAAGCCCTNNNAGATCGGA |
| 32087_3 | TATGAGTGTGGAGTCGTTGCAGCAAAGTAATAATGAGAAGCCCTNNNAGATCGGA |
| 40171_1 | TATGAGTGTGGAGTCGTTGCTGTGTAGTTCAAGTAGTCACCCTNNNAGATCGGAAG |
| 40171_2 | TATGAGTGTGGAGTCGTTGCTGTGTAGTTCAAGTAGTCACCCTNNNAGATCGGAAG |
| 40171_3 | TATGAGTGTGGAGTCGTTGCTGTGTAGTTCAAGTAGTCACCCTNNNAGATCGGAAG |
| 41693_1 | TATGAGTGTGGAGTCGTTGCGGGTTCAAGTGATTCTCCTGCTNNNAGATCGGAAGA |
| 41693_2 | TATGAGTGTGGAGTCGTTGCGGGTTCAAGTGATTCTCCTGCTNNNAGATCGGAAGA |
| 41693_3 | TATGAGTGTGGAGTCGTTGCGGGTTCAAGTGATTCTCCTGCTNNNAGATCGGAAGA |
| 47874_1 | TATGAGTGTGGAGTCGTTGCCCAAAGAATGCTGGTGTTCTTGGNNNAGATCGGAA |
| 47874_2 | TATGAGTGTGGAGTCGTTGCCCAAAGAATGCTGGTGTTCTTGGNNNAGATCGGAA |
| 47874_3 | TATGAGTGTGGAGTCGTTGCCCAAAGAATGCTGGTGTTCTTGGNNNAGATCGGAA |
| 9182_1 | TATGAGTGTGGAGTCGTTGCTGCTTACCCTTTCTACATGCAACNNNAGATCGGAAGA |
| 9182_2 | TATGAGTGTGGAGTCGTTGCTGCTTACCCTTTCTACATGCAACNNNAGATCGGAAGA |
| 9182_3 | TATGAGTGTGGAGTCGTTGCTGCTTACCCTTTCTACATGCAACNNNAGATCGGAAGA |
| 49375_1 | TATGAGTGTGGAGTCGTTGCGTTAGCCAGGATGGTCTCGATCNNNAGATCGGAAG |
| 49375_2 | TATGAGTGTGGAGTCGTTGCGTTAGCCAGGATGGTCTCGATCNNNAGATCGGAAG |
| 49375_3 | TATGAGTGTGGAGTCGTTGCGTTAGCCAGGATGGTCTCGATCNNNAGATCGGAAG |
| 50154_1 | TATGAGTGTGGAGTCGTTGCCCCCAAGGTTTTTCTCTCACAGANNNAGATCGGAAG |
| 50154_2 | TATGAGTGTGGAGTCGTTGCCCCCAAGGTTTTTCTCTCACAGANNNAGATCGGAAG |
| 50154_3 | TATGAGTGTGGAGTCGTTGCCCCCAAGGTTTTTCTCTCACAGANNNAGATCGGAAG |
| 52454_1 | TATGAGTGTGGAGTCGTTGCACAGAGGGTTGGAAGTAGTTGTTNNNAGATCGGAA |
| 52454_2 | TATGAGTGTGGAGTCGTTGCACAGAGGGTTGGAAGTAGTTGTTNNNAGATCGGAA |
| 52454_3 | TATGAGTGTGGAGTCGTTGCACAGAGGGTTGGAAGTAGTTGTTNNNAGATCGGAA |
| 3023_1 | TATGAGTGTGGAGTCGTTGCGGAGCAAGTCCTTCCCTTTCATTNNNAGATCGGAAG |
| 3023_2 | TATGAGTGTGGAGTCGTTGCGGAGCAAGTCCTTCCCTTTCATTNNNAGATCGGAAG |
| 3023_3 | TATGAGTGTGGAGTCGTTGCGGAGCAAGTCCTTCCCTTTCATTNNNAGATCGGAAG |
| 31614_1 | TATGAGTGTGGAGTCGTTGCTGGAATCAGTAAATCCCCTCCTGNNNAGATCGGAAG |
| 31614_2 | TATGAGTGTGGAGTCGTTGCTGGAATCAGTAAATCCCCTCCTGNNNAGATCGGAAG |
| 31614_3 | TATGAGTGTGGAGTCGTTGCTGGAATCAGTAAATCCCCTCCTGNNNAGATCGGAAG |
| 47660_1 | TATGAGTGTGGAGTCGTTGCTTTCCTGCCTCAGATTTGCATTGNNNAGATCGGAAG |
| 47660_2 | TATGAGTGTGGAGTCGTTGCTTTCCTGCCTCAGATTTGCATTGNNNAGATCGGAAG |
| 47660_3 | TATGAGTGTGGAGTCGTTGCTTTCCTGCCTCAGATTTGCATTGNNNAGATCGGAAG |
| 31816_1 | TATGAGTGTGGAGTCGTTGCTCAAATAGGAACTCTCCATGGCCNNNAGATCGGAAG |
| 31816_2 | TATGAGTGTGGAGTCGTTGCTCAAATAGGAACTCTCCATGGCCNNNAGATCGGAAG |
| 31816_3 | TATGAGTGTGGAGTCGTTGCTCAAATAGGAACTCTCCATGGCCNNNAGATCGGAAG |
| 33259_1 | TATGAGTGTGGAGTCGTTGCCCTCCCAGGTTCAAGTGATTCTCNNNAGATCGGAAG |
| 33259_2 | TATGAGTGTGGAGTCGTTGCCCTCCCAGGTTCAAGTGATTCTCNNNAGATCGGAAG |
| 33259_3 | TATGAGTGTGGAGTCGTTGCCCTCCCAGGTTCAAGTGATTCTCNNNAGATCGGAAG |
| 44867_1 | TATGAGTGTGGAGTCGTTGCGATTGCACAAGTTAGCCATTCCANNNAGATCGGAAG |
| 44867_2 | TATGAGTGTGGAGTCGTTGCGATTGCACAAGTTAGCCATTCCANNNAGATCGGAAG |
| 44867_3 | TATGAGTGTGGAGTCGTTGCGATTGCACAAGTTAGCCATTCCANNNAGATCGGAAG |
| 36895_1 | TATGAGTGTGGAGTCGTTGCAATTGCTTGAACCCAGGAGTTTGNNNAGATCGGAA |

| | |
|---|---|
| 36895_2 | TATGAGTGTGGAGTCGTTGCAATTGCTTGAACCCAGGAGTTTGNNNAGATCGGAAG |
| 36895_3 | TATGAGTGTGGAGTCGTTGCAATTGCTTGAACCCAGGAGTTTGNNNAGATCGGAAG |
| 11031_1 | TATGAGTGTGGAGTCGTTGCTGGTCAAAATACCCCAATCCAGTAANNNAGATCGGAA |
| 11031_2 | TATGAGTGTGGAGTCGTTGCTGGTCAAAATACCCCAATCCAGTAANNNAGATCGGAA |
| 11031_3 | TATGAGTGTGGAGTCGTTGCTGGTCAAAATACCCCAATCCAGTAANNNAGATCGGAA |
| 48745_1 | TATGAGTGTGGAGTCGTTGCCTCTGGGTTTTCTCATGCCAAAANNNAGATCGGAAGA |
| 48745_2 | TATGAGTGTGGAGTCGTTGCCTCTGGGTTTTCTCATGCCAAAANNNAGATCGGAAGA |
| 48745_3 | TATGAGTGTGGAGTCGTTGCCTCTGGGTTTTCTCATGCCAAAANNNAGATCGGAAGA |
| 41042_1 | TATGAGTGTGGAGTCGTTGCAGACATAATCCGTTAGGACAGTCCNNNAGATCGGAA |
| 41042_2 | TATGAGTGTGGAGTCGTTGCAGACATAATCCGTTAGGACAGTCCNNNAGATCGGAA |
| 41042_3 | TATGAGTGTGGAGTCGTTGCAGACATAATCCGTTAGGACAGTCCNNNAGATCGGAA |
| 47824_1 | TATGAGTGTGGAGTCGTTGCTTCAGCATTGGACAGATCTTCCANNNAGATCGGAAGA |
| 47824_2 | TATGAGTGTGGAGTCGTTGCTTCAGCATTGGACAGATCTTCCANNNAGATCGGAAGA |
| 47824_3 | TATGAGTGTGGAGTCGTTGCTTCAGCATTGGACAGATCTTCCANNNAGATCGGAAGA |
| 33303_1 | TATGAGTGTGGAGTCGTTGCGACTGAGTCTCATTCTGTCACCANNNAGATCGGAAGA |
| 33303_2 | TATGAGTGTGGAGTCGTTGCGACTGAGTCTCATTCTGTCACCANNNAGATCGGAAGA |
| 33303_3 | TATGAGTGTGGAGTCGTTGCGACTGAGTCTCATTCTGTCACCANNNAGATCGGAAGA |
| 33584_1 | TATGAGTGTGGAGTCGTTGCAGTGGCATGATCTCTGCTCACNNNAGATCGGAAGAG |
| 33584_2 | TATGAGTGTGGAGTCGTTGCAGTGGCATGATCTCTGCTCACNNNAGATCGGAAGAG |
| 33584_3 | TATGAGTGTGGAGTCGTTGCAGTGGCATGATCTCTGCTCACNNNAGATCGGAAGAG |
| 40446_1 | TATGAGTGTGGAGTCGTTGCTGATAGGTGTTTGGTCTGAGTGGNNNAGATCGGAA |
| 40446_2 | TATGAGTGTGGAGTCGTTGCTGATAGGTGTTTGGTCTGAGTGGNNNAGATCGGAA |
| 40446_3 | TATGAGTGTGGAGTCGTTGCTGATAGGTGTTTGGTCTGAGTGGNNNAGATCGGAA |
| 30423_1 | TATGAGTGTGGAGTCGTTGCTGAACCATGCCTCCAACTAGATCNNNAGATCGGAAGA |
| 30423_2 | TATGAGTGTGGAGTCGTTGCTGAACCATGCCTCCAACTAGATCNNNAGATCGGAAGA |
| 30423_3 | TATGAGTGTGGAGTCGTTGCTGAACCATGCCTCCAACTAGATCNNNAGATCGGAAGA |
| 44416_1 | TATGAGTGTGGAGTCGTTGCTAGCCTAAACCTGCCCAGTAAAGNNNAGATCGGAAG |
| 44416_2 | TATGAGTGTGGAGTCGTTGCTAGCCTAAACCTGCCCAGTAAAGNNNAGATCGGAAG |
| 44416_3 | TATGAGTGTGGAGTCGTTGCTAGCCTAAACCTGCCCAGTAAAGNNNAGATCGGAAG |
| 2652_1 | TATGAGTGTGGAGTCGTTGCTCAGCTTCTCCACCATCTTGTTGNNNAGATCGGAAGA |
| 2652_2 | TATGAGTGTGGAGTCGTTGCTCAGCTTCTCCACCATCTTGTTGNNNAGATCGGAAGA |
| 2652_3 | TATGAGTGTGGAGTCGTTGCTCAGCTTCTCCACCATCTTGTTGNNNAGATCGGAAGA |
| 46926_1 | TATGAGTGTGGAGTCGTTGCTCTGGGTTCAAGTGATTCTCCTGNNNAGATCGGAAG |
| 46926_2 | TATGAGTGTGGAGTCGTTGCTCTGGGTTCAAGTGATTCTCCTGNNNAGATCGGAAG |
| 46926_3 | TATGAGTGTGGAGTCGTTGCTCTGGGTTCAAGTGATTCTCCTGNNNAGATCGGAAG |
| 30620_1 | TATGAGTGTGGAGTCGTTGCAGCAAATTAAAACACCAAGGCAGNNNAGATCGGAAG |
| 30620_2 | TATGAGTGTGGAGTCGTTGCAGCAAATTAAAACACCAAGGCAGNNNAGATCGGAAG |
| 30620_3 | TATGAGTGTGGAGTCGTTGCAGCAAATTAAAACACCAAGGCAGNNNAGATCGGAAG |
| 35458_1 | TATGAGTGTGGAGTCGTTGCGTTTCACCATGTTCCCTAAGCTGNNNAGATCGGAAGA |
| 35458_2 | TATGAGTGTGGAGTCGTTGCGTTTCACCATGTTCCCTAAGCTGNNNAGATCGGAAGA |
| 35458_3 | TATGAGTGTGGAGTCGTTGCGTTTCACCATGTTCCCTAAGCTGNNNAGATCGGAAGA |
| 48213_1 | TATGAGTGTGGAGTCGTTGCCAACCTCCATCTCCTGAGTTCAANNNAGATCGGAAGA |
| 48213_2 | TATGAGTGTGGAGTCGTTGCCAACCTCCATCTCCTGAGTTCAANNNAGATCGGAAGA |

| | |
|---|---|
| 48213_3 | TATGAGTGTGGAGTCGTTGCCAACCTCCATCTCCTGAGTTCAANNNAGATCGGAAGA |
| 29641_1 | TATGAGTGTGGAGTCGTTGCTTTTAGCTCAATGGGATGGGGAANNNAGATCGGAA |
| 29641_2 | TATGAGTGTGGAGTCGTTGCTTTTAGCTCAATGGGATGGGGAANNNAGATCGGAA |
| 29641_3 | TATGAGTGTGGAGTCGTTGCTTTTAGCTCAATGGGATGGGGAANNNAGATCGGAA |
| 43343_1 | TATGAGTGTGGAGTCGTTGCGTAGCTGGGATTACATATGCCCANNNAGATCGGAAG |
| 43343_2 | TATGAGTGTGGAGTCGTTGCGTAGCTGGGATTACATATGCCCANNNAGATCGGAAG |
| 43343_3 | TATGAGTGTGGAGTCGTTGCGTAGCTGGGATTACATATGCCCANNNAGATCGGAAG |
| 47747_1 | TATGAGTGTGGAGTCGTTGCGTGACCAGGACAGTGATCGGNNNAGATCGGAAGAG |
| 47747_2 | TATGAGTGTGGAGTCGTTGCGTGACCAGGACAGTGATCGGNNNAGATCGGAAGAG |
| 47747_3 | TATGAGTGTGGAGTCGTTGCGTGACCAGGACAGTGATCGGNNNAGATCGGAAGAG |
| 29883_1 | TATGAGTGTGGAGTCGTTGCCGTGTTTAGTGTGAGAACTGCAGNNNAGATCGGAA |
| 29883_2 | TATGAGTGTGGAGTCGTTGCCGTGTTTAGTGTGAGAACTGCAGNNNAGATCGGAA |
| 29883_3 | TATGAGTGTGGAGTCGTTGCCGTGTTTAGTGTGAGAACTGCAGNNNAGATCGGAA |
| 47193_1 | TATGAGTGTGGAGTCGTTGCATGGAGTTTCACTTTTGTTGCCCNNNAGATCGGAAG |
| 47193_2 | TATGAGTGTGGAGTCGTTGCATGGAGTTTCACTTTTGTTGCCCNNNAGATCGGAAG |
| 47193_3 | TATGAGTGTGGAGTCGTTGCATGGAGTTTCACTTTTGTTGCCCNNNAGATCGGAAG |
| 34602_1 | TATGAGTGTGGAGTCGTTGCAGAACCACTGCTCTCTTCAAAGCNNNAGATCGGAAG |
| 34602_2 | TATGAGTGTGGAGTCGTTGCAGAACCACTGCTCTCTTCAAAGCNNNAGATCGGAAG |
| 34602_3 | TATGAGTGTGGAGTCGTTGCAGAACCACTGCTCTCTTCAAAGCNNNAGATCGGAAG |
| 37882_1 | TATGAGTGTGGAGTCGTTGCAAGTGCTGGGATTACAAGTGTGANNNAGATCGGAA |
| 37882_2 | TATGAGTGTGGAGTCGTTGCAAGTGCTGGGATTACAAGTGTGANNNAGATCGGAA |
| 37882_3 | TATGAGTGTGGAGTCGTTGCAAGTGCTGGGATTACAAGTGTGANNNAGATCGGAA |
| 39682_1 | TATGAGTGTGGAGTCGTTGCCCAAAGAATGGGAGAAAAAATCTGCNNNAGATCGG |
| 39682_2 | TATGAGTGTGGAGTCGTTGCCCAAAGAATGGGAGAAAAAATCTGCNNNAGATCGG |
| 39682_3 | TATGAGTGTGGAGTCGTTGCCCAAAGAATGGGAGAAAAAATCTGCNNNAGATCGG |
| 36651_1 | TATGAGTGTGGAGTCGTTGCCTTGTTTCTTCCACAGTGCTGAGNNNAGATCGGAAG |
| 36651_2 | TATGAGTGTGGAGTCGTTGCCTTGTTTCTTCCACAGTGCTGAGNNNAGATCGGAAG |
| 36651_3 | TATGAGTGTGGAGTCGTTGCCTTGTTTCTTCCACAGTGCTGAGNNNAGATCGGAAG |
| 46230_1 | TATGAGTGTGGAGTCGTTGCTTTAGTAGAGACGGGGTTTTGCCNNNAGATCGGAA |
| 46230_2 | TATGAGTGTGGAGTCGTTGCTTTAGTAGAGACGGGGTTTTGCCNNNAGATCGGAA |
| 46230_3 | TATGAGTGTGGAGTCGTTGCTTTAGTAGAGACGGGGTTTTGCCNNNAGATCGGAA |
| 41030_1 | TATGAGTGTGGAGTCGTTGCCAGAAGGGCAAAAGTCGGAAATTNNNAGATCGGAA |
| 41030_2 | TATGAGTGTGGAGTCGTTGCCAGAAGGGCAAAAGTCGGAAATTNNNAGATCGGAA |
| 41030_3 | TATGAGTGTGGAGTCGTTGCCAGAAGGGCAAAAGTCGGAAATTNNNAGATCGGAA |
| 9064_1 | TATGAGTGTGGAGTCGTTGCTTATCAGAAGACAGAGGCAGCCTNNNAGATCGGAA |
| 9064_2 | TATGAGTGTGGAGTCGTTGCTTATCAGAAGACAGAGGCAGCCTNNNAGATCGGAA |
| 9064_3 | TATGAGTGTGGAGTCGTTGCTTATCAGAAGACAGAGGCAGCCTNNNAGATCGGAA |
| 2978_1 | TATGAGTGTGGAGTCGTTGCTTGCTTTCAACTTTGGGTGGCTNNNAGATCGGAAGA |
| 2978_2 | TATGAGTGTGGAGTCGTTGCTTGCTTTCAACTTTGGGTGGCTNNNAGATCGGAAGA |
| 2978_3 | TATGAGTGTGGAGTCGTTGCTTGCTTTCAACTTTGGGTGGCTNNNAGATCGGAAGA |
| 10644_1 | TATGAGTGTGGAGTCGTTGCGGAGCACACTTTTGATAATGCTGGNNNAGATCGGA |
| 10644_2 | TATGAGTGTGGAGTCGTTGCGGAGCACACTTTTGATAATGCTGGNNNAGATCGGA |
| 10644_3 | TATGAGTGTGGAGTCGTTGCGGAGCACACTTTTGATAATGCTGGNNNAGATCGGA |

| | |
|---|---|
| 45845_1 | TATGAGTGTGGAGTCGTTGCCCAAGTAGCTGAGATTACAGGCANNNAGATCGGAA |
| 45845_2 | TATGAGTGTGGAGTCGTTGCCCAAGTAGCTGAGATTACAGGCANNNAGATCGGAA |
| 45845_3 | TATGAGTGTGGAGTCGTTGCCCAAGTAGCTGAGATTACAGGCANNNAGATCGGAA |
| 46326_1 | TATGAGTGTGGAGTCGTTGCAGGTTGACAGTAAGCCAGACATTNNNAGATCGGAA |
| 46326_2 | TATGAGTGTGGAGTCGTTGCAGGTTGACAGTAAGCCAGACATTNNNAGATCGGAA |
| 46326_3 | TATGAGTGTGGAGTCGTTGCAGGTTGACAGTAAGCCAGACATTNNNAGATCGGAA |
| 51608_1 | TATGAGTGTGGAGTCGTTGCCAGAGACGGGTGGAACTTCAGNNNAGATCGGAAGA |
| 51608_2 | TATGAGTGTGGAGTCGTTGCCAGAGACGGGTGGAACTTCAGNNNAGATCGGAAGA |
| 51608_3 | TATGAGTGTGGAGTCGTTGCCAGAGACGGGTGGAACTTCAGNNNAGATCGGAAGA |
| 37505_1 | TATGAGTGTGGAGTCGTTGCGTTGCAGTGAGCTGAGATCATGNNNAGATCGGAAG |
| 37505_2 | TATGAGTGTGGAGTCGTTGCGTTGCAGTGAGCTGAGATCATGNNNAGATCGGAAG |
| 37505_3 | TATGAGTGTGGAGTCGTTGCGTTGCAGTGAGCTGAGATCATGNNNAGATCGGAAG |
| 45045_1 | TATGAGTGTGGAGTCGTTGCATAGTCGACAGGCTCTTTGAAGGNNNAGATCGGAA |
| 45045_2 | TATGAGTGTGGAGTCGTTGCATAGTCGACAGGCTCTTTGAAGGNNNAGATCGGAA |
| 45045_3 | TATGAGTGTGGAGTCGTTGCATAGTCGACAGGCTCTTTGAAGGNNNAGATCGGAA |
| 43860_1 | TATGAGTGTGGAGTCGTTGCTTTGAGACAGGATCTTGCTCTGTNNNAGATCGGAAG |
| 43860_2 | TATGAGTGTGGAGTCGTTGCTTTGAGACAGGATCTTGCTCTGTNNNAGATCGGAAG |
| 43860_3 | TATGAGTGTGGAGTCGTTGCTTTGAGACAGGATCTTGCTCTGTNNNAGATCGGAAG |
| 45144_1 | TATGAGTGTGGAGTCGTTGCGAAACATCTGACCTCCGCAAAAANNNAGATCGGAAG |
| 45144_2 | TATGAGTGTGGAGTCGTTGCGAAACATCTGACCTCCGCAAAAANNNAGATCGGAAG |
| 45144_3 | TATGAGTGTGGAGTCGTTGCGAAACATCTGACCTCCGCAAAAANNNAGATCGGAAG |
| 35291_1 | TATGAGTGTGGAGTCGTTGCCCTTTGGCCTTTTGAGACAAGAANNNAGATCGGAAG |
| 35291_2 | TATGAGTGTGGAGTCGTTGCCCTTTGGCCTTTTGAGACAAGAANNNAGATCGGAAG |
| 35291_3 | TATGAGTGTGGAGTCGTTGCCCTTTGGCCTTTTGAGACAAGAANNNAGATCGGAAG |
| 13764_1 | TATGAGTGTGGAGTCGTTGCCTAGAATAAAGGCTGAATGTTTANNNAGATCGGAAG |
| 13764_2 | TATGAGTGTGGAGTCGTTGCCTAGAATAAAGGCTGAATGTTTANNNAGATCGGAAG |
| 13764_3 | TATGAGTGTGGAGTCGTTGCCTAGAATAAAGGCTGAATGTTTANNNAGATCGGAAG |
| 32951_1 | TATGAGTGTGGAGTCGTTGCTCACAGGTACAGGATTGCTTCATNNNAGATCGGAAG |
| 32951_2 | TATGAGTGTGGAGTCGTTGCTCACAGGTACAGGATTGCTTCATNNNAGATCGGAAG |
| 32951_3 | TATGAGTGTGGAGTCGTTGCTCACAGGTACAGGATTGCTTCATNNNAGATCGGAAG |
| 30309_1 | TATGAGTGTGGAGTCGTTGCGCAAACTGACCCAGAGGAAGAAANNNAGATCGGAA |
| 30309_2 | TATGAGTGTGGAGTCGTTGCGCAAACTGACCCAGAGGAAGAAANNNAGATCGGAA |
| 30309_3 | TATGAGTGTGGAGTCGTTGCGCAAACTGACCCAGAGGAAGAAANNNAGATCGGAA |
| 49251_1 | TATGAGTGTGGAGTCGTTGCGTTGTTCAGGCTGGTTTCAAACTNNNAGATCGGAAG |
| 49251_2 | TATGAGTGTGGAGTCGTTGCGTTGTTCAGGCTGGTTTCAAACTNNNAGATCGGAAG |
| 49251_3 | TATGAGTGTGGAGTCGTTGCGTTGTTCAGGCTGGTTTCAAACTNNNAGATCGGAAG |
| 38800_1 | TATGAGTGTGGAGTCGTTGCTCTTTGGCTAGACTGACCAAGAANNNAGATCGGAAG |
| 38800_2 | TATGAGTGTGGAGTCGTTGCTCTTTGGCTAGACTGACCAAGAANNNAGATCGGAAG |
| 38800_3 | TATGAGTGTGGAGTCGTTGCTCTTTGGCTAGACTGACCAAGAANNNAGATCGGAAG |
| 44130_1 | TATGAGTGTGGAGTCGTTGCAGCTGTGAATGGCTTGATTGTTCNNNAGATCGGAAG |
| 44130_2 | TATGAGTGTGGAGTCGTTGCAGCTGTGAATGGCTTGATTGTTCNNNAGATCGGAAG |
| 44130_3 | TATGAGTGTGGAGTCGTTGCAGCTGTGAATGGCTTGATTGTTCNNNAGATCGGAAG |
| 48021_1 | TATGAGTGTGGAGTCGTTGCTTCAACCAATTCTCCTGCCTCAGNNNAGATCGGAAGA |

| | |
|---|---|
| 48021_2 | TATGAGTGTGGAGTCGTTGCTTCAACCAATTCTCCTGCCTCAGNNNAGATCGGAAGA |
| 48021_3 | TATGAGTGTGGAGTCGTTGCTTCAACCAATTCTCCTGCCTCAGNNNAGATCGGAAGA |
| 32923_1 | TATGAGTGTGGAGTCGTTGCATGGAACCCCAAAAGGATCTGAANNNAGATCGGAA |
| 32923_2 | TATGAGTGTGGAGTCGTTGCATGGAACCCCAAAAGGATCTGAANNNAGATCGGAA |
| 32923_3 | TATGAGTGTGGAGTCGTTGCATGGAACCCCAAAAGGATCTGAANNNAGATCGGAA |
| 10120_1 | TATGAGTGTGGAGTCGTTGCATTGAGAAGAGAGACTGGGGGAGNNNAGATCGGA |
| 10120_2 | TATGAGTGTGGAGTCGTTGCATTGAGAAGAGAGACTGGGGGAGNNNAGATCGGA |
| 10120_3 | TATGAGTGTGGAGTCGTTGCATTGAGAAGAGAGACTGGGGGAGNNNAGATCGGA |
| 36641_1 | TATGAGTGTGGAGTCGTTGCAGAAACAAACGATGGAGACTGGANNNAGATCGGA |
| 36641_2 | TATGAGTGTGGAGTCGTTGCAGAAACAAACGATGGAGACTGGANNNAGATCGGA |
| 36641_3 | TATGAGTGTGGAGTCGTTGCAGAAACAAACGATGGAGACTGGANNNAGATCGGA |
| 44380_1 | TATGAGTGTGGAGTCGTTGCATCTCTAAATCCCCTTTCCCAGCNNNAGATCGGAAGA |
| 44380_2 | TATGAGTGTGGAGTCGTTGCATCTCTAAATCCCCTTTCCCAGCNNNAGATCGGAAGA |
| 44380_3 | TATGAGTGTGGAGTCGTTGCATCTCTAAATCCCCTTTCCCAGCNNNAGATCGGAAGA |
| 2039_1 | TATGAGTGTGGAGTCGTTGCTGACTGCACATTCTAGCAATGAANNNAGATCGGAAG |
| 2039_2 | TATGAGTGTGGAGTCGTTGCTGACTGCACATTCTAGCAATGAANNNAGATCGGAAG |
| 2039_3 | TATGAGTGTGGAGTCGTTGCTGACTGCACATTCTAGCAATGAANNNAGATCGGAAG |
| 34885_1 | TATGAGTGTGGAGTCGTTGCACCACGCCTGGCTAATTTGTATTNNNAGATCGGAAG |
| 34885_2 | TATGAGTGTGGAGTCGTTGCACCACGCCTGGCTAATTTGTATTNNNAGATCGGAAG |
| 34885_3 | TATGAGTGTGGAGTCGTTGCACCACGCCTGGCTAATTTGTATTNNNAGATCGGAAG |
| 37680_1 | TATGAGTGTGGAGTCGTTGCAATGATGAGGTCGGGTGATTAGGNNNAGATCGGA |
| 37680_2 | TATGAGTGTGGAGTCGTTGCAATGATGAGGTCGGGTGATTAGGNNNAGATCGGA |
| 37680_3 | TATGAGTGTGGAGTCGTTGCAATGATGAGGTCGGGTGATTAGGNNNAGATCGGA |
| 33298_1 | TATGAGTGTGGAGTCGTTGCTTCAAGTGCTTATCCTGCCTCAGNNNAGATCGGAAG |
| 33298_2 | TATGAGTGTGGAGTCGTTGCTTCAAGTGCTTATCCTGCCTCAGNNNAGATCGGAAG |
| 33298_3 | TATGAGTGTGGAGTCGTTGCTTCAAGTGCTTATCCTGCCTCAGNNNAGATCGGAAG |
| 51912_1 | TATGAGTGTGGAGTCGTTGCCCCCTCACCTCCTGGATAGGNNNAGATCGGAAGAGC |
| 51912_2 | TATGAGTGTGGAGTCGTTGCCCCCTCACCTCCTGGATAGGNNNAGATCGGAAGAGC |
| 51912_3 | TATGAGTGTGGAGTCGTTGCCCCCTCACCTCCTGGATAGGNNNAGATCGGAAGAGC |
| 13656_1 | TATGAGTGTGGAGTCGTTGCAAGAAACATTTGGAAATAGAACANNNAGATCGGAA |
| 13656_2 | TATGAGTGTGGAGTCGTTGCAAGAAACATTTGGAAATAGAACANNNAGATCGGAA |
| 13656_3 | TATGAGTGTGGAGTCGTTGCAAGAAACATTTGGAAATAGAACANNNAGATCGGAA |
| 31178_1 | TATGAGTGTGGAGTCGTTGCAAACAGAGCCTCAAAGATGTGTGNNNAGATCGGAA |
| 31178_2 | TATGAGTGTGGAGTCGTTGCAAACAGAGCCTCAAAGATGTGTGNNNAGATCGGAA |
| 31178_3 | TATGAGTGTGGAGTCGTTGCAAACAGAGCCTCAAAGATGTGTGNNNAGATCGGAA |
| 39977_1 | TATGAGTGTGGAGTCGTTGCTGTATAGGTGAGGAAACCAAGGCNNNAGATCGGAA |
| 39977_2 | TATGAGTGTGGAGTCGTTGCTGTATAGGTGAGGAAACCAAGGCNNNAGATCGGAA |
| 39977_3 | TATGAGTGTGGAGTCGTTGCTGTATAGGTGAGGAAACCAAGGCNNNAGATCGGAA |
| 43534_1 | TATGAGTGTGGAGTCGTTGCGCACTTGTGTTGCACTACTAGAANNNAGATCGGAAG |
| 43534_2 | TATGAGTGTGGAGTCGTTGCGCACTTGTGTTGCACTACTAGAANNNAGATCGGAAG |
| 43534_3 | TATGAGTGTGGAGTCGTTGCGCACTTGTGTTGCACTACTAGAANNNAGATCGGAAG |
| 39982_1 | TATGAGTGTGGAGTCGTTGCTGTGCCTGTAATCCCAGTTACTCNNNAGATCGGAAG |
| 39982_2 | TATGAGTGTGGAGTCGTTGCTGTGCCTGTAATCCCAGTTACTCNNNAGATCGGAAG |

| | |
|---|---|
| 39982_3 | TATGAGTGTGGAGTCGTTGCTGTGCCTGTAATCCCAGTTACTCNNNAGATCGGAAG |
| 46351_1 | TATGAGTGTGGAGTCGTTGCGGCTAGGAAAGGCATTCTGAGTANNNAGATCGGAA |
| 46351_2 | TATGAGTGTGGAGTCGTTGCGGCTAGGAAAGGCATTCTGAGTANNNAGATCGGAA |
| 46351_3 | TATGAGTGTGGAGTCGTTGCGGCTAGGAAAGGCATTCTGAGTANNNAGATCGGAA |
| 37794_1 | TATGAGTGTGGAGTCGTTGCGAGGTGGAGGTGGAAGATTGTTTNNNAGATCGGA |
| 37794_2 | TATGAGTGTGGAGTCGTTGCGAGGTGGAGGTGGAAGATTGTTTNNNAGATCGGA |
| 37794_3 | TATGAGTGTGGAGTCGTTGCGAGGTGGAGGTGGAAGATTGTTTNNNAGATCGGA |
| 31044_1 | TATGAGTGTGGAGTCGTTGCAGCATCCAAGAATGTCAGGAGAGNNNAGATCGGAA |
| 31044_2 | TATGAGTGTGGAGTCGTTGCAGCATCCAAGAATGTCAGGAGAGNNNAGATCGGAA |
| 31044_3 | TATGAGTGTGGAGTCGTTGCAGCATCCAAGAATGTCAGGAGAGNNNAGATCGGAA |
| 45910_1 | TATGAGTGTGGAGTCGTTGCGAGTAGCTGGGATTATAGGTGCCNNNAGATCGGAA |
| 45910_2 | TATGAGTGTGGAGTCGTTGCGAGTAGCTGGGATTATAGGTGCCNNNAGATCGGAA |
| 45910_3 | TATGAGTGTGGAGTCGTTGCGAGTAGCTGGGATTATAGGTGCCNNNAGATCGGAA |
| 44894_1 | TATGAGTGTGGAGTCGTTGCGATAGTCCAATAGCCCTGGTAGCNNNAGATCGGAA |
| 44894_2 | TATGAGTGTGGAGTCGTTGCGATAGTCCAATAGCCCTGGTAGCNNNAGATCGGAA |
| 44894_3 | TATGAGTGTGGAGTCGTTGCGATAGTCCAATAGCCCTGGTAGCNNNAGATCGGAA |
| 34004_1 | TATGAGTGTGGAGTCGTTGCAAAAATGGACTGCCACATTCACCNNNAGATCGGAAG |
| 34004_2 | TATGAGTGTGGAGTCGTTGCAAAAATGGACTGCCACATTCACCNNNAGATCGGAAG |
| 34004_3 | TATGAGTGTGGAGTCGTTGCAAAAATGGACTGCCACATTCACCNNNAGATCGGAAG |
| 51604_1 | TATGAGTGTGGAGTCGTTGCCTTTGGAACTGTGTAACAGGCAGNNNAGATCGGAA |
| 51604_2 | TATGAGTGTGGAGTCGTTGCCTTTGGAACTGTGTAACAGGCAGNNNAGATCGGAA |
| 51604_3 | TATGAGTGTGGAGTCGTTGCCTTTGGAACTGTGTAACAGGCAGNNNAGATCGGAA |
| 48993_1 | TATGAGTGTGGAGTCGTTGCAGTCTGAGGTACATGTGCAGAACNNNAGATCGGAA |
| 48993_2 | TATGAGTGTGGAGTCGTTGCAGTCTGAGGTACATGTGCAGAACNNNAGATCGGAA |
| 48993_3 | TATGAGTGTGGAGTCGTTGCAGTCTGAGGTACATGTGCAGAACNNNAGATCGGAA |
| 31712_1 | TATGAGTGTGGAGTCGTTGCCAAAACTTTTGGGTCCATCTCCCNNNAGATCGGAAG |
| 31712_2 | TATGAGTGTGGAGTCGTTGCCAAAACTTTTGGGTCCATCTCCCNNNAGATCGGAAG |
| 31712_3 | TATGAGTGTGGAGTCGTTGCCAAAACTTTTGGGTCCATCTCCCNNNAGATCGGAAG |
| 10054_1 | TATGAGTGTGGAGTCGTTGCCGAGCTAAAATCATTTAGGGTGTTNNNAGATCGGA |
| 10054_2 | TATGAGTGTGGAGTCGTTGCCGAGCTAAAATCATTTAGGGTGTTNNNAGATCGGA |
| 10054_3 | TATGAGTGTGGAGTCGTTGCCGAGCTAAAATCATTTAGGGTGTTNNNAGATCGGA |
| 2424_1 | TATGAGTGTGGAGTCGTTGCCAGTACACACAGGCAGATGCATCNNNAGATCGGAA |
| 2424_2 | TATGAGTGTGGAGTCGTTGCCAGTACACACAGGCAGATGCATCNNNAGATCGGAA |
| 2424_3 | TATGAGTGTGGAGTCGTTGCCAGTACACACAGGCAGATGCATCNNNAGATCGGAA |
| 50085_1 | TATGAGTGTGGAGTCGTTGCAAGAGTACCTGCTTCTCTAGGGTNNNAGATCGGAAG |
| 50085_2 | TATGAGTGTGGAGTCGTTGCAAGAGTACCTGCTTCTCTAGGGTNNNAGATCGGAAG |
| 50085_3 | TATGAGTGTGGAGTCGTTGCAAGAGTACCTGCTTCTCTAGGGTNNNAGATCGGAAG |
| 42721_1 | TATGAGTGTGGAGTCGTTGCCAACATGAGACTCAGAGGTGGAANNNAGATCGGAA |
| 42721_2 | TATGAGTGTGGAGTCGTTGCCAACATGAGACTCAGAGGTGGAANNNAGATCGGAA |
| 42721_3 | TATGAGTGTGGAGTCGTTGCCAACATGAGACTCAGAGGTGGAANNNAGATCGGAA |
| 44695_1 | TATGAGTGTGGAGTCGTTGCTGTTCAGTGTCAGAGTCTCACAGNNNAGATCGGAAG |
| 44695_2 | TATGAGTGTGGAGTCGTTGCTGTTCAGTGTCAGAGTCTCACAGNNNAGATCGGAAG |
| 44695_3 | TATGAGTGTGGAGTCGTTGCTGTTCAGTGTCAGAGTCTCACAGNNNAGATCGGAAG |

| | |
|---|---|
| 47184_1 | TATGAGTGTGGAGTCGTTGCTGGTGTTCAAGTTTGTTCTCAGCNNNAGATCGGAAG |
| 47184_2 | TATGAGTGTGGAGTCGTTGCTGGTGTTCAAGTTTGTTCTCAGCNNNAGATCGGAAG |
| 47184_3 | TATGAGTGTGGAGTCGTTGCTGGTGTTCAAGTTTGTTCTCAGCNNNAGATCGGAAG |
| 41164_1 | TATGAGTGTGGAGTCGTTGCACATGGTGAAACCTCGTCTCTACNNNAGATCGGAAG |
| 41164_2 | TATGAGTGTGGAGTCGTTGCACATGGTGAAACCTCGTCTCTACNNNAGATCGGAAG |
| 41164_3 | TATGAGTGTGGAGTCGTTGCACATGGTGAAACCTCGTCTCTACNNNAGATCGGAAG |
| 39518_1 | TATGAGTGTGGAGTCGTTGCCACGTGCCTACCCGGTATTTATANNNAGATCGGAAG |
| 39518_2 | TATGAGTGTGGAGTCGTTGCCACGTGCCTACCCGGTATTTATANNNAGATCGGAAG |
| 39518_3 | TATGAGTGTGGAGTCGTTGCCACGTGCCTACCCGGTATTTATANNNAGATCGGAAG |
| 37233_1 | TATGAGTGTGGAGTCGTTGCGAGGCAGGAGAATAGCGTGAACNNNAGATCGGAA |
| 37233_2 | TATGAGTGTGGAGTCGTTGCGAGGCAGGAGAATAGCGTGAACNNNAGATCGGAA |
| 37233_3 | TATGAGTGTGGAGTCGTTGCGAGGCAGGAGAATAGCGTGAACNNNAGATCGGAA |
| 39597_1 | TATGAGTGTGGAGTCGTTGCGATTCTCCGATTCTCCAAGGTCANNNAGATCGGAAG |
| 39597_2 | TATGAGTGTGGAGTCGTTGCGATTCTCCGATTCTCCAAGGTCANNNAGATCGGAAG |
| 39597_3 | TATGAGTGTGGAGTCGTTGCGATTCTCCGATTCTCCAAGGTCANNNAGATCGGAAG |
| 31552_1 | TATGAGTGTGGAGTCGTTGCACTGCCTATTTCCCCTAACCAAGNNNAGATCGGAAGA |
| 31552_2 | TATGAGTGTGGAGTCGTTGCACTGCCTATTTCCCCTAACCAAGNNNAGATCGGAAGA |
| 31552_3 | TATGAGTGTGGAGTCGTTGCACTGCCTATTTCCCCTAACCAAGNNNAGATCGGAAGA |
| 42773_1 | TATGAGTGTGGAGTCGTTGCAAAAACCCAGCTTCTCCCATTTGNNNAGATCGGAAGA |
| 42773_2 | TATGAGTGTGGAGTCGTTGCAAAAACCCAGCTTCTCCCATTTGNNNAGATCGGAAGA |
| 42773_3 | TATGAGTGTGGAGTCGTTGCAAAAACCCAGCTTCTCCCATTTGNNNAGATCGGAAGA |
| 33078_1 | TATGAGTGTGGAGTCGTTGCGTTTCTCCAGCTTTTTGCCCTAGNNNAGATCGGAAGA |
| 33078_2 | TATGAGTGTGGAGTCGTTGCGTTTCTCCAGCTTTTTGCCCTAGNNNAGATCGGAAGA |
| 33078_3 | TATGAGTGTGGAGTCGTTGCGTTTCTCCAGCTTTTTGCCCTAGNNNAGATCGGAAGA |
| 3166_1 | TATGAGTGTGGAGTCGTTGCCACATTCCTGGCCTAGGTTTCTCNNNAGATCGGAAGA |
| 3166_2 | TATGAGTGTGGAGTCGTTGCCACATTCCTGGCCTAGGTTTCTCNNNAGATCGGAAGA |
| 3166_3 | TATGAGTGTGGAGTCGTTGCCACATTCCTGGCCTAGGTTTCTCNNNAGATCGGAAGA |
| 45990_1 | TATGAGTGTGGAGTCGTTGCCTAGGCTGGTATTGAACTCCTGGNNNAGATCGGAAG |
| 45990_2 | TATGAGTGTGGAGTCGTTGCCTAGGCTGGTATTGAACTCCTGGNNNAGATCGGAAG |
| 45990_3 | TATGAGTGTGGAGTCGTTGCCTAGGCTGGTATTGAACTCCTGGNNNAGATCGGAAG |
| 51746_1 | TATGAGTGTGGAGTCGTTGCAGGGCATAGAATCTTGGGAGTTCNNNAGATCGGAA |
| 51746_2 | TATGAGTGTGGAGTCGTTGCAGGGCATAGAATCTTGGGAGTTCNNNAGATCGGAA |
| 51746_3 | TATGAGTGTGGAGTCGTTGCAGGGCATAGAATCTTGGGAGTTCNNNAGATCGGAA |
| 5950_1 | TATGAGTGTGGAGTCGTTGCACCTTTTACACGGCTCTGTACTCNNNAGATCGGAAGA |
| 5950_2 | TATGAGTGTGGAGTCGTTGCACCTTTTACACGGCTCTGTACTCNNNAGATCGGAAGA |
| 5950_3 | TATGAGTGTGGAGTCGTTGCACCTTTTACACGGCTCTGTACTCNNNAGATCGGAAGA |
| 33728_1 | TATGAGTGTGGAGTCGTTGCGACCAACTTTCCTCGTTTTCCTGNNNAGATCGGAAGA |
| 33728_2 | TATGAGTGTGGAGTCGTTGCGACCAACTTTCCTCGTTTTCCTGNNNAGATCGGAAGA |
| 33728_3 | TATGAGTGTGGAGTCGTTGCGACCAACTTTCCTCGTTTTCCTGNNNAGATCGGAAGA |
| 39623_1 | TATGAGTGTGGAGTCGTTGCAACCAGACACATAGACCAATGGANNNAGATCGGAA |
| 39623_2 | TATGAGTGTGGAGTCGTTGCAACCAGACACATAGACCAATGGANNNAGATCGGAA |
| 39623_3 | TATGAGTGTGGAGTCGTTGCAACCAGACACATAGACCAATGGANNNAGATCGGAA |
| 48030_1 | TATGAGTGTGGAGTCGTTGCAACATATACACGCACACACACACNNNAGATCGGAAG |

| | |
|---|---|
| 48030_2 | TATGAGTGTGGAGTCGTTGCAACATATACACGCACACACACACNNNAGATCGGAAG |
| 48030_3 | TATGAGTGTGGAGTCGTTGCAACATATACACGCACACACACACNNNAGATCGGAAG |
| 3504_1 | TATGAGTGTGGAGTCGTTGCAGTTATTTCACAGGACTGTTGCTTNNNAGATCGGAA |
| 3504_2 | TATGAGTGTGGAGTCGTTGCAGTTATTTCACAGGACTGTTGCTTNNNAGATCGGAA |
| 3504_3 | TATGAGTGTGGAGTCGTTGCAGTTATTTCACAGGACTGTTGCTTNNNAGATCGGAA |
| 35211_1 | TATGAGTGTGGAGTCGTTGCTGAAAGAGCAAGAACCTGTGAGTNNNAGATCGGAA |
| 35211_2 | TATGAGTGTGGAGTCGTTGCTGAAAGAGCAAGAACCTGTGAGTNNNAGATCGGAA |
| 35211_3 | TATGAGTGTGGAGTCGTTGCTGAAAGAGCAAGAACCTGTGAGTNNNAGATCGGAA |
| 29675_1 | TATGAGTGTGGAGTCGTTGCTGAGGATGGTTCTTCACTTGGAGNNNAGATCGGAA |
| 29675_2 | TATGAGTGTGGAGTCGTTGCTGAGGATGGTTCTTCACTTGGAGNNNAGATCGGAA |
| 29675_3 | TATGAGTGTGGAGTCGTTGCTGAGGATGGTTCTTCACTTGGAGNNNAGATCGGAA |
| 34100_1 | TATGAGTGTGGAGTCGTTGCGAGATCACACCATTGCATTCCAGNNNAGATCGGAAG |
| 34100_2 | TATGAGTGTGGAGTCGTTGCGAGATCACACCATTGCATTCCAGNNNAGATCGGAAG |
| 34100_3 | TATGAGTGTGGAGTCGTTGCGAGATCACACCATTGCATTCCAGNNNAGATCGGAAG |
| 44853_1 | TATGAGTGTGGAGTCGTTGCTGTGTGATGGGACTAGGAGTTTGNNNAGATCGGAA |
| 44853_2 | TATGAGTGTGGAGTCGTTGCTGTGTGATGGGACTAGGAGTTTGNNNAGATCGGAA |
| 44853_3 | TATGAGTGTGGAGTCGTTGCTGTGTGATGGGACTAGGAGTTTGNNNAGATCGGAA |
| 39633_1 | TATGAGTGTGGAGTCGTTGCCCTGATATTCATGCTGCATAATGTNNNAGATCGGAA |
| 39633_2 | TATGAGTGTGGAGTCGTTGCCCTGATATTCATGCTGCATAATGTNNNAGATCGGAA |
| 39633_3 | TATGAGTGTGGAGTCGTTGCCCTGATATTCATGCTGCATAATGTNNNAGATCGGAA |
| 38371_1 | TATGAGTGTGGAGTCGTTGCAGGGAATCTGAACAAGGTGAGACNNNAGATCGGA |
| 38371_2 | TATGAGTGTGGAGTCGTTGCAGGGAATCTGAACAAGGTGAGACNNNAGATCGGA |
| 38371_3 | TATGAGTGTGGAGTCGTTGCAGGGAATCTGAACAAGGTGAGACNNNAGATCGGA |
| 32398_1 | TATGAGTGTGGAGTCGTTGCTCGTTATCCTGACGAAGCAGAATNNNAGATCGGAAG |
| 32398_2 | TATGAGTGTGGAGTCGTTGCTCGTTATCCTGACGAAGCAGAATNNNAGATCGGAAG |
| 32398_3 | TATGAGTGTGGAGTCGTTGCTCGTTATCCTGACGAAGCAGAATNNNAGATCGGAAG |
| 33462_1 | TATGAGTGTGGAGTCGTTGCAGGAACGCAGTGACATACAAAGANNNAGATCGGAA |
| 33462_2 | TATGAGTGTGGAGTCGTTGCAGGAACGCAGTGACATACAAAGANNNAGATCGGAA |
| 33462_3 | TATGAGTGTGGAGTCGTTGCAGGAACGCAGTGACATACAAAGANNNAGATCGGAA |
| 40651_1 | TATGAGTGTGGAGTCGTTGCCACTCCTGTAATCCCAGCACTTTNNNAGATCGGAAGA |
| 40651_2 | TATGAGTGTGGAGTCGTTGCCACTCCTGTAATCCCAGCACTTTNNNAGATCGGAAGA |
| 40651_3 | TATGAGTGTGGAGTCGTTGCCACTCCTGTAATCCCAGCACTTTNNNAGATCGGAAGA |
| 52763_1 | TATGAGTGTGGAGTCGTTGCTTTCAAGATCATCCGTTTTGGCCNNNAGATCGGAAG |
| 52763_2 | TATGAGTGTGGAGTCGTTGCTTTCAAGATCATCCGTTTTGGCCNNNAGATCGGAAG |
| 52763_3 | TATGAGTGTGGAGTCGTTGCTTTCAAGATCATCCGTTTTGGCCNNNAGATCGGAAG |
| 44490_1 | TATGAGTGTGGAGTCGTTGCCTCACCTCTACTCGCTCCATTTANNNAGATCGGAAGA |
| 44490_2 | TATGAGTGTGGAGTCGTTGCCTCACCTCTACTCGCTCCATTTANNNAGATCGGAAGA |
| 44490_3 | TATGAGTGTGGAGTCGTTGCCTCACCTCTACTCGCTCCATTTANNNAGATCGGAAGA |
| 47151_1 | TATGAGTGTGGAGTCGTTGCGGCATCCATGGACACTATGATCTNNNAGATCGGAAG |
| 47151_2 | TATGAGTGTGGAGTCGTTGCGGCATCCATGGACACTATGATCTNNNAGATCGGAAG |
| 47151_3 | TATGAGTGTGGAGTCGTTGCGGCATCCATGGACACTATGATCTNNNAGATCGGAAG |
| 35641_1 | TATGAGTGTGGAGTCGTTGCTGGTGAGTCTTGGGAATGCATAANNNAGATCGGAA |
| 35641_2 | TATGAGTGTGGAGTCGTTGCTGGTGAGTCTTGGGAATGCATAANNNAGATCGGAA |

| | |
|---|---|
| 35641_3 | TATGAGTGTGGAGTCGTTGCTGGTGAGTCTTGGGAATGCATAANNNAGATCGGAA |
| 48811_1 | TATGAGTGTGGAGTCGTTGCTCTCTCTTGTAGCTGGGACTACANNNAGATCGGAAG |
| 48811_2 | TATGAGTGTGGAGTCGTTGCTCTCTCTTGTAGCTGGGACTACANNNAGATCGGAAG |
| 48811_3 | TATGAGTGTGGAGTCGTTGCTCTCTCTTGTAGCTGGGACTACANNNAGATCGGAAG |
| 32106_1 | TATGAGTGTGGAGTCGTTGCCTCCTGAGTAGCTGGGATTACAGNNNAGATCGGAA |
| 32106_2 | TATGAGTGTGGAGTCGTTGCCTCCTGAGTAGCTGGGATTACAGNNNAGATCGGAA |
| 32106_3 | TATGAGTGTGGAGTCGTTGCCTCCTGAGTAGCTGGGATTACAGNNNAGATCGGAA |
| 43299_1 | TATGAGTGTGGAGTCGTTGCTCAAACTCCTAGGCTCAAGCAATNNNAGATCGGAAG |
| 43299_2 | TATGAGTGTGGAGTCGTTGCTCAAACTCCTAGGCTCAAGCAATNNNAGATCGGAAG |
| 43299_3 | TATGAGTGTGGAGTCGTTGCTCAAACTCCTAGGCTCAAGCAATNNNAGATCGGAAG |
| 34962_1 | TATGAGTGTGGAGTCGTTGCCACCACGCCAGGCTAATTTTTTNNNAGATCGGAAGA |
| 34962_2 | TATGAGTGTGGAGTCGTTGCCACCACGCCAGGCTAATTTTTTNNNAGATCGGAAGA |
| 34962_3 | TATGAGTGTGGAGTCGTTGCCACCACGCCAGGCTAATTTTTTNNNAGATCGGAAGA |
| 41638_1 | TATGAGTGTGGAGTCGTTGCTTTGTTGCACTGTTCTCTGAAGGNNNAGATCGGAAG |
| 41638_2 | TATGAGTGTGGAGTCGTTGCTTTGTTGCACTGTTCTCTGAAGGNNNAGATCGGAAG |
| 41638_3 | TATGAGTGTGGAGTCGTTGCTTTGTTGCACTGTTCTCTGAAGGNNNAGATCGGAAG |
| 47140_1 | TATGAGTGTGGAGTCGTTGCGATCTCCTGACCTTGTGATCCACNNNAGATCGGAAG |
| 47140_2 | TATGAGTGTGGAGTCGTTGCGATCTCCTGACCTTGTGATCCACNNNAGATCGGAAG |
| 47140_3 | TATGAGTGTGGAGTCGTTGCGATCTCCTGACCTTGTGATCCACNNNAGATCGGAAG |
| 32096_1 | TATGAGTGTGGAGTCGTTGCAACTGAATTCTGCCAACAACCAGNNNAGATCGGAAG |
| 32096_2 | TATGAGTGTGGAGTCGTTGCAACTGAATTCTGCCAACAACCAGNNNAGATCGGAAG |
| 32096_3 | TATGAGTGTGGAGTCGTTGCAACTGAATTCTGCCAACAACCAGNNNAGATCGGAAG |
| 41085_1 | TATGAGTGTGGAGTCGTTGCATGCTACTAGGTGAGGCATGATGNNNAGATCGGAA |
| 41085_2 | TATGAGTGTGGAGTCGTTGCATGCTACTAGGTGAGGCATGATGNNNAGATCGGAA |
| 41085_3 | TATGAGTGTGGAGTCGTTGCATGCTACTAGGTGAGGCATGATGNNNAGATCGGAA |
| 32484_1 | TATGAGTGTGGAGTCGTTGCAGTAGCTAGGATTACAGGCATGCNNNAGATCGGAA |
| 32484_2 | TATGAGTGTGGAGTCGTTGCAGTAGCTAGGATTACAGGCATGCNNNAGATCGGAA |
| 32484_3 | TATGAGTGTGGAGTCGTTGCAGTAGCTAGGATTACAGGCATGCNNNAGATCGGAA |
| 47981_1 | TATGAGTGTGGAGTCGTTGCCAAGTAGCTGGGATTACAGGCATNNNAGATCGGAA |
| 47981_2 | TATGAGTGTGGAGTCGTTGCCAAGTAGCTGGGATTACAGGCATNNNAGATCGGAA |
| 47981_3 | TATGAGTGTGGAGTCGTTGCCAAGTAGCTGGGATTACAGGCATNNNAGATCGGAA |
| 35330_1 | TATGAGTGTGGAGTCGTTGCACATGTGATTCCACCACTGATCANNNAGATCGGAAG |
| 35330_2 | TATGAGTGTGGAGTCGTTGCACATGTGATTCCACCACTGATCANNNAGATCGGAAG |
| 35330_3 | TATGAGTGTGGAGTCGTTGCACATGTGATTCCACCACTGATCANNNAGATCGGAAG |
| 43604_1 | TATGAGTGTGGAGTCGTTGCATTTCCTGGATCTTCCCTGACAGNNNAGATCGGAAG |
| 43604_2 | TATGAGTGTGGAGTCGTTGCATTTCCTGGATCTTCCCTGACAGNNNAGATCGGAAG |
| 43604_3 | TATGAGTGTGGAGTCGTTGCATTTCCTGGATCTTCCCTGACAGNNNAGATCGGAAG |
| 48296_1 | TATGAGTGTGGAGTCGTTGCTCACGAAGAGAGGCACACAAATANNNAGATCGGAA |
| 48296_2 | TATGAGTGTGGAGTCGTTGCTCACGAAGAGAGGCACACAAATANNNAGATCGGAA |
| 48296_3 | TATGAGTGTGGAGTCGTTGCTCACGAAGAGAGGCACACAAATANNNAGATCGGAA |
| 34715_1 | TATGAGTGTGGAGTCGTTGCCGCGCCCAGCTAATTTTTTGTATNNNAGATCGGAAG |
| 34715_2 | TATGAGTGTGGAGTCGTTGCCGCGCCCAGCTAATTTTTTGTATNNNAGATCGGAAG |
| 34715_3 | TATGAGTGTGGAGTCGTTGCCGCGCCCAGCTAATTTTTTGTATNNNAGATCGGAAG |

| | |
|---|---|
| 10844_1 | TATGAGTGTGGAGTCGTTGCATGTGTTTGGAACGTTTGAACCANNNAGATCGGAAG |
| 10844_2 | TATGAGTGTGGAGTCGTTGCATGTGTTTGGAACGTTTGAACCANNNAGATCGGAAG |
| 10844_3 | TATGAGTGTGGAGTCGTTGCATGTGTTTGGAACGTTTGAACCANNNAGATCGGAAG |
| 10729_1 | TATGAGTGTGGAGTCGTTGCGCTGGCAAAGGTAAACTACCAGANNNAGATCGGAA |
| 10729_2 | TATGAGTGTGGAGTCGTTGCGCTGGCAAAGGTAAACTACCAGANNNAGATCGGAA |
| 10729_3 | TATGAGTGTGGAGTCGTTGCGCTGGCAAAGGTAAACTACCAGANNNAGATCGGAA |
| 37699_1 | TATGAGTGTGGAGTCGTTGCTTTGGTTGAAATGCTACAGCCAGNNNAGATCGGAAG |
| 37699_2 | TATGAGTGTGGAGTCGTTGCTTTGGTTGAAATGCTACAGCCAGNNNAGATCGGAAG |
| 37699_3 | TATGAGTGTGGAGTCGTTGCTTTGGTTGAAATGCTACAGCCAGNNNAGATCGGAAG |
| 13018_1 | TATGAGTGTGGAGTCGTTGCGTATTGACATCTAATCTGCATTTNNNAGATCGGAAG |
| 13018_2 | TATGAGTGTGGAGTCGTTGCGTATTGACATCTAATCTGCATTTNNNAGATCGGAAG |
| 13018_3 | TATGAGTGTGGAGTCGTTGCGTATTGACATCTAATCTGCATTTNNNAGATCGGAAG |
| 30943_1 | TATGAGTGTGGAGTCGTTGCATCCCTGGAGTCTGCAAGTTTAGNNNAGATCGGAAG |
| 30943_2 | TATGAGTGTGGAGTCGTTGCATCCCTGGAGTCTGCAAGTTTAGNNNAGATCGGAAG |
| 30943_3 | TATGAGTGTGGAGTCGTTGCATCCCTGGAGTCTGCAAGTTTAGNNNAGATCGGAAG |
| 12731_1 | TATGAGTGTGGAGTCGTTGCGTACTGGCTCCTAACGTATTNNNAGATCGGAAGAGC |
| 12731_2 | TATGAGTGTGGAGTCGTTGCGTACTGGCTCCTAACGTATTNNNAGATCGGAAGAGC |
| 12731_3 | TATGAGTGTGGAGTCGTTGCGTACTGGCTCCTAACGTATTNNNAGATCGGAAGAGC |
| 47604_1 | TATGAGTGTGGAGTCGTTGCCTCCTGAGTAGCTGGGATTACAGNNNAGATCGGAA |
| 47604_2 | TATGAGTGTGGAGTCGTTGCCTCCTGAGTAGCTGGGATTACAGNNNAGATCGGAA |
| 47604_3 | TATGAGTGTGGAGTCGTTGCCTCCTGAGTAGCTGGGATTACAGNNNAGATCGGAA |
| 48074_1 | TATGAGTGTGGAGTCGTTGCTCACTTCCAGGTTTAGTTGTACCTNNNAGATCGGAAG |
| 48074_2 | TATGAGTGTGGAGTCGTTGCTCACTTCCAGGTTTAGTTGTACCTNNNAGATCGGAAG |
| 48074_3 | TATGAGTGTGGAGTCGTTGCTCACTTCCAGGTTTAGTTGTACCTNNNAGATCGGAAG |
| 13584_1 | TATGAGTGTGGAGTCGTTGCCTAAACGAATAAAGCAGTTACATNNNAGATCGGAAG |
| 13584_2 | TATGAGTGTGGAGTCGTTGCCTAAACGAATAAAGCAGTTACATNNNAGATCGGAAG |
| 13584_3 | TATGAGTGTGGAGTCGTTGCCTAAACGAATAAAGCAGTTACATNNNAGATCGGAAG |
| 49064_1 | TATGAGTGTGGAGTCGTTGCAGTAGCTGGGATTACAGGTATGCNNNAGATCGGAA |
| 49064_2 | TATGAGTGTGGAGTCGTTGCAGTAGCTGGGATTACAGGTATGCNNNAGATCGGAA |
| 49064_3 | TATGAGTGTGGAGTCGTTGCAGTAGCTGGGATTACAGGTATGCNNNAGATCGGAA |
| 50531_1 | TATGAGTGTGGAGTCGTTGCTCACTTTCACCGCATCTCTTAGGNNNAGATCGGAAGA |
| 50531_2 | TATGAGTGTGGAGTCGTTGCTCACTTTCACCGCATCTCTTAGGNNNAGATCGGAAGA |
| 50531_3 | TATGAGTGTGGAGTCGTTGCTCACTTTCACCGCATCTCTTAGGNNNAGATCGGAAGA |
| 31404_1 | TATGAGTGTGGAGTCGTTGCGTGTGTGTGTGTGTGTGTGATTATNNNAGATCGGA |
| 31404_2 | TATGAGTGTGGAGTCGTTGCGTGTGTGTGTGTGTGTGTGATTATNNNAGATCGGA |
| 31404_3 | TATGAGTGTGGAGTCGTTGCGTGTGTGTGTGTGTGTGTGATTATNNNAGATCGGA |
| 33827_1 | TATGAGTGTGGAGTCGTTGCAACAAGTTTGTGTCTGAGGAGGANNNAGATCGGAA |
| 33827_2 | TATGAGTGTGGAGTCGTTGCAACAAGTTTGTGTCTGAGGAGGANNNAGATCGGAA |
| 33827_3 | TATGAGTGTGGAGTCGTTGCAACAAGTTTGTGTCTGAGGAGGANNNAGATCGGAA |
| 40215_1 | TATGAGTGTGGAGTCGTTGCGTTAGCTATGATCATGCCACTGCNNNAGATCGGAAG |
| 40215_2 | TATGAGTGTGGAGTCGTTGCGTTAGCTATGATCATGCCACTGCNNNAGATCGGAAG |
| 40215_3 | TATGAGTGTGGAGTCGTTGCGTTAGCTATGATCATGCCACTGCNNNAGATCGGAAG |
| 49324_1 | TATGAGTGTGGAGTCGTTGCAAGGTCCTCGGTAATTCTTTCCCNNNAGATCGGAAG |

| | |
|---|---|
| 49324_2 | TATGAGTGTGGAGTCGTTGCAAGGTCCTCGGTAATTCTTTCCCNNNAGATCGGAAG |
| 49324_3 | TATGAGTGTGGAGTCGTTGCAAGGTCCTCGGTAATTCTTTCCCNNNAGATCGGAAG |
| 44369_1 | TATGAGTGTGGAGTCGTTGCTGGGAGACACAAAAAGAGAGGAGNNNAGATCGGA |
| 44369_2 | TATGAGTGTGGAGTCGTTGCTGGGAGACACAAAAAGAGAGGAGNNNAGATCGGA |
| 44369_3 | TATGAGTGTGGAGTCGTTGCTGGGAGACACAAAAAGAGAGGAGNNNAGATCGGA |
| 13387_1 | TATGAGTGTGGAGTCGTTGCCTTTAGGCGTTAAAACTGTAAATNNNAGATCGGAAG |
| 13387_2 | TATGAGTGTGGAGTCGTTGCCTTTAGGCGTTAAAACTGTAAATNNNAGATCGGAAG |
| 13387_3 | TATGAGTGTGGAGTCGTTGCCTTTAGGCGTTAAAACTGTAAATNNNAGATCGGAAG |
| 39070_1 | TATGAGTGTGGAGTCGTTGCTCCTGTGTTGGGTGCATATATGTNNNAGATCGGAAG |
| 39070_2 | TATGAGTGTGGAGTCGTTGCTCCTGTGTTGGGTGCATATATGTNNNAGATCGGAAG |
| 39070_3 | TATGAGTGTGGAGTCGTTGCTCCTGTGTTGGGTGCATATATGTNNNAGATCGGAAG |
| 29476_1 | TATGAGTGTGGAGTCGTTGCCAGACTCACTGGGTTCTTTCCTGNNNAGATCGGAAG |
| 29476_2 | TATGAGTGTGGAGTCGTTGCCAGACTCACTGGGTTCTTTCCTGNNNAGATCGGAAG |
| 29476_3 | TATGAGTGTGGAGTCGTTGCCAGACTCACTGGGTTCTTTCCTGNNNAGATCGGAAG |
| 10359_1 | TATGAGTGTGGAGTCGTTGCATCCCTGCCCATAACCGATATGANNNAGATCGGAAG |
| 10359_2 | TATGAGTGTGGAGTCGTTGCATCCCTGCCCATAACCGATATGANNNAGATCGGAAG |
| 10359_3 | TATGAGTGTGGAGTCGTTGCATCCCTGCCCATAACCGATATGANNNAGATCGGAAG |
| 31730_1 | TATGAGTGTGGAGTCGTTGCTGTTATATGGCTGACAGGTTGGTNNNAGATCGGAA |
| 31730_2 | TATGAGTGTGGAGTCGTTGCTGTTATATGGCTGACAGGTTGGTNNNAGATCGGAA |
| 31730_3 | TATGAGTGTGGAGTCGTTGCTGTTATATGGCTGACAGGTTGGTNNNAGATCGGAA |
| 51719_1 | TATGAGTGTGGAGTCGTTGCCTTCTCCCTCTCCTGGAAAAACANNNAGATCGGAAGA |
| 51719_2 | TATGAGTGTGGAGTCGTTGCCTTCTCCCTCTCCTGGAAAAACANNNAGATCGGAAGA |
| 51719_3 | TATGAGTGTGGAGTCGTTGCCTTCTCCCTCTCCTGGAAAAACANNNAGATCGGAAGA |
| 52287_1 | TATGAGTGTGGAGTCGTTGCGTCAGAGGACAGCGAGATTCTCNNNAGATCGGAAG |
| 52287_2 | TATGAGTGTGGAGTCGTTGCGTCAGAGGACAGCGAGATTCTCNNNAGATCGGAAG |
| 52287_3 | TATGAGTGTGGAGTCGTTGCGTCAGAGGACAGCGAGATTCTCNNNAGATCGGAAG |
| 13789_1 | TATGAGTGTGGAGTCGTTGCAAGTTATAGTCATTGGGGTAAAANNNAGATCGGAA |
| 13789_2 | TATGAGTGTGGAGTCGTTGCAAGTTATAGTCATTGGGGTAAAANNNAGATCGGAA |
| 13789_3 | TATGAGTGTGGAGTCGTTGCAAGTTATAGTCATTGGGGTAAAANNNAGATCGGAA |
| 49232_1 | TATGAGTGTGGAGTCGTTGCGCCACGCCCAACTAATTTTTTTGNNNAGATCGGAAGA |
| 49232_2 | TATGAGTGTGGAGTCGTTGCGCCACGCCCAACTAATTTTTTTGNNNAGATCGGAAGA |
| 49232_3 | TATGAGTGTGGAGTCGTTGCGCCACGCCCAACTAATTTTTTTGNNNAGATCGGAAGA |
| 51367_1 | TATGAGTGTGGAGTCGTTGCTGGATGCCTGCTCAGCTAATAATNNNAGATCGGAAG |
| 51367_2 | TATGAGTGTGGAGTCGTTGCTGGATGCCTGCTCAGCTAATAATNNNAGATCGGAAG |
| 51367_3 | TATGAGTGTGGAGTCGTTGCTGGATGCCTGCTCAGCTAATAATNNNAGATCGGAAG |
| 34266_1 | TATGAGTGTGGAGTCGTTGCTGAGTCTTTTGGTAAGCTCGTGANNNAGATCGGAAG |
| 34266_2 | TATGAGTGTGGAGTCGTTGCTGAGTCTTTTGGTAAGCTCGTGANNNAGATCGGAAG |
| 34266_3 | TATGAGTGTGGAGTCGTTGCTGAGTCTTTTGGTAAGCTCGTGANNNAGATCGGAAG |
| 42612_1 | TATGAGTGTGGAGTCGTTGCCCAGTGAAAACCACATCAAAAACANNNAGATCGGAA |
| 42612_2 | TATGAGTGTGGAGTCGTTGCCCAGTGAAAACCACATCAAAAACANNNAGATCGGAA |
| 42612_3 | TATGAGTGTGGAGTCGTTGCCCAGTGAAAACCACATCAAAAACANNNAGATCGGAA |
| 47730_1 | TATGAGTGTGGAGTCGTTGCAATCATAGCTCACTGCAACCTCANNNAGATCGGAAG |
| 47730_2 | TATGAGTGTGGAGTCGTTGCAATCATAGCTCACTGCAACCTCANNNAGATCGGAAG |

| | |
|---|---|
| 47730_3 | TATGAGTGTGGAGTCGTTGCAATCATAGCTCACTGCAACCTCANNNAGATCGGAAGA |
| 38792_1 | TATGAGTGTGGAGTCGTTGCGATCACAAGGTCAGGAGATGGAGNNNAGATCGGAA |
| 38792_2 | TATGAGTGTGGAGTCGTTGCGATCACAAGGTCAGGAGATGGAGNNNAGATCGGAA |
| 38792_3 | TATGAGTGTGGAGTCGTTGCGATCACAAGGTCAGGAGATGGAGNNNAGATCGGAA |
| 47223_1 | TATGAGTGTGGAGTCGTTGCCCTGATTCATGCCATTCTTCTGCNNNAGATCGGAAGA |
| 47223_2 | TATGAGTGTGGAGTCGTTGCCCTGATTCATGCCATTCTTCTGCNNNAGATCGGAAGA |
| 47223_3 | TATGAGTGTGGAGTCGTTGCCCTGATTCATGCCATTCTTCTGCNNNAGATCGGAAGA |
| 51041_1 | TATGAGTGTGGAGTCGTTGCAAGAAAGCTAGAAGCAAGCAAGCNNNAGATCGGAA |
| 51041_2 | TATGAGTGTGGAGTCGTTGCAAGAAAGCTAGAAGCAAGCAAGCNNNAGATCGGAA |
| 51041_3 | TATGAGTGTGGAGTCGTTGCAAGAAAGCTAGAAGCAAGCAAGCNNNAGATCGGAA |
| 31113_1 | TATGAGTGTGGAGTCGTTGCCACTGCCTAGAACTAACAGAGCANNNAGATCGGAAG |
| 31113_2 | TATGAGTGTGGAGTCGTTGCCACTGCCTAGAACTAACAGAGCANNNAGATCGGAAG |
| 31113_3 | TATGAGTGTGGAGTCGTTGCCACTGCCTAGAACTAACAGAGCANNNAGATCGGAAG |
| 40998_1 | TATGAGTGTGGAGTCGTTGCGCACCTCAAGATGCTTCAAGATTNNNAGATCGGAAG |
| 40998_2 | TATGAGTGTGGAGTCGTTGCGCACCTCAAGATGCTTCAAGATTNNNAGATCGGAAG |
| 40998_3 | TATGAGTGTGGAGTCGTTGCGCACCTCAAGATGCTTCAAGATTNNNAGATCGGAAG |
| 29371_1 | TATGAGTGTGGAGTCGTTGCGCCCCCTTTTCTCTGGAATATGANNNAGATCGGAAG |
| 29371_2 | TATGAGTGTGGAGTCGTTGCGCCCCCTTTTCTCTGGAATATGANNNAGATCGGAAG |
| 29371_3 | TATGAGTGTGGAGTCGTTGCGCCCCCTTTTCTCTGGAATATGANNNAGATCGGAAG |
| 51507_1 | TATGAGTGTGGAGTCGTTGCTGTGCTGGTTTTGTTACATGCATNNNAGATCGGAAG |
| 51507_2 | TATGAGTGTGGAGTCGTTGCTGTGCTGGTTTTGTTACATGCATNNNAGATCGGAAG |
| 51507_3 | TATGAGTGTGGAGTCGTTGCTGTGCTGGTTTTGTTACATGCATNNNAGATCGGAAG |
| 45030_1 | TATGAGTGTGGAGTCGTTGCGCACAATTGTTGAAGTGTGGAGANNNAGATCGGAA |
| 45030_2 | TATGAGTGTGGAGTCGTTGCGCACAATTGTTGAAGTGTGGAGANNNAGATCGGAA |
| 45030_3 | TATGAGTGTGGAGTCGTTGCGCACAATTGTTGAAGTGTGGAGANNNAGATCGGAA |
| 31776_1 | TATGAGTGTGGAGTCGTTGCTGTTCCTCTACATCCTTGTCAGCNNNAGATCGGAAGA |
| 31776_2 | TATGAGTGTGGAGTCGTTGCTGTTCCTCTACATCCTTGTCAGCNNNAGATCGGAAGA |
| 31776_3 | TATGAGTGTGGAGTCGTTGCTGTTCCTCTACATCCTTGTCAGCNNNAGATCGGAAGA |
| 7993_1 | TATGAGTGTGGAGTCGTTGCTTTGCGGAGATATTGAGTGAGTGNNNAGATCGGAA |
| 7993_2 | TATGAGTGTGGAGTCGTTGCTTTGCGGAGATATTGAGTGAGTGNNNAGATCGGAA |
| 7993_3 | TATGAGTGTGGAGTCGTTGCTTTGCGGAGATATTGAGTGAGTGNNNAGATCGGAA |
| 39811_1 | TATGAGTGTGGAGTCGTTGCTTTCCTCCACTGATGTCTTGAGCNNNAGATCGGAAG |
| 39811_2 | TATGAGTGTGGAGTCGTTGCTTTCCTCCACTGATGTCTTGAGCNNNAGATCGGAAG |
| 39811_3 | TATGAGTGTGGAGTCGTTGCTTTCCTCCACTGATGTCTTGAGCNNNAGATCGGAAG |
| 5355_1 | TATGAGTGTGGAGTCGTTGCATCAAAGAACACTGAGCACCTCNNNAGATCGGAAGA |
| 5355_2 | TATGAGTGTGGAGTCGTTGCATCAAAGAACACTGAGCACCTCNNNAGATCGGAAGA |
| 5355_3 | TATGAGTGTGGAGTCGTTGCATCAAAGAACACTGAGCACCTCNNNAGATCGGAAGA |
| 42536_1 | TATGAGTGTGGAGTCGTTGCGCAGTAGCATAATCACAGCTCACNNNAGATCGGAAG |
| 42536_2 | TATGAGTGTGGAGTCGTTGCGCAGTAGCATAATCACAGCTCACNNNAGATCGGAAG |
| 42536_3 | TATGAGTGTGGAGTCGTTGCGCAGTAGCATAATCACAGCTCACNNNAGATCGGAAG |
| 52221_1 | TATGAGTGTGGAGTCGTTGCGAAAACATGGAGAGCCCCCATTTNNNAGATCGGAAG |
| 52221_2 | TATGAGTGTGGAGTCGTTGCGAAAACATGGAGAGCCCCCATTTNNNAGATCGGAAG |
| 52221_3 | TATGAGTGTGGAGTCGTTGCGAAAACATGGAGAGCCCCCATTTNNNAGATCGGAAG |

| | |
|---|---|
| 35004_1 | TATGAGTGTGGAGTCGTTGCGAGTAGCTGGGATTACAGGCATGNNNAGATCGGAA |
| 35004_2 | TATGAGTGTGGAGTCGTTGCGAGTAGCTGGGATTACAGGCATGNNNAGATCGGAA |
| 35004_3 | TATGAGTGTGGAGTCGTTGCGAGTAGCTGGGATTACAGGCATGNNNAGATCGGAA |
| 48315_1 | TATGAGTGTGGAGTCGTTGCGCCAACTAAATCAGGATGATTGGGNNNAGATCGGA |
| 48315_2 | TATGAGTGTGGAGTCGTTGCGCCAACTAAATCAGGATGATTGGGNNNAGATCGGA |
| 48315_3 | TATGAGTGTGGAGTCGTTGCGCCAACTAAATCAGGATGATTGGGNNNAGATCGGA |
| 49588_1 | TATGAGTGTGGAGTCGTTGCGCCTCCTGAGTAGTTGGGATTACNNNAGATCGGAAG |
| 49588_2 | TATGAGTGTGGAGTCGTTGCGCCTCCTGAGTAGTTGGGATTACNNNAGATCGGAAG |
| 49588_3 | TATGAGTGTGGAGTCGTTGCGCCTCCTGAGTAGTTGGGATTACNNNAGATCGGAAG |
| 13704_1 | TATGAGTGTGGAGTCGTTGCTTCATTGACTTCAGGTAGATTAANNNAGATCGGAAG |
| 13704_2 | TATGAGTGTGGAGTCGTTGCTTCATTGACTTCAGGTAGATTAANNNAGATCGGAAG |
| 13704_3 | TATGAGTGTGGAGTCGTTGCTTCATTGACTTCAGGTAGATTAANNNAGATCGGAAG |
| 32400_1 | TATGAGTGTGGAGTCGTTGCATCTTTTCAGCATCCCACTCTGTNNNAGATCGGAAGA |
| 32400_2 | TATGAGTGTGGAGTCGTTGCATCTTTTCAGCATCCCACTCTGTNNNAGATCGGAAGA |
| 32400_3 | TATGAGTGTGGAGTCGTTGCATCTTTTCAGCATCCCACTCTGTNNNAGATCGGAAGA |
| 51749_1 | TATGAGTGTGGAGTCGTTGCGGAGAATCACTTGAACCTGGGAGNNNAGATCGGAA |
| 51749_2 | TATGAGTGTGGAGTCGTTGCGGAGAATCACTTGAACCTGGGAGNNNAGATCGGAA |
| 51749_3 | TATGAGTGTGGAGTCGTTGCGGAGAATCACTTGAACCTGGGAGNNNAGATCGGAA |
| 12358_1 | TATGAGTGTGGAGTCGTTGCCAACCAGACTTACATGCAGTTTCCNNNAGATCGGAAG |
| 12358_2 | TATGAGTGTGGAGTCGTTGCCAACCAGACTTACATGCAGTTTCCNNNAGATCGGAAG |
| 12358_3 | TATGAGTGTGGAGTCGTTGCCAACCAGACTTACATGCAGTTTCCNNNAGATCGGAAG |
| 5344_1 | TATGAGTGTGGAGTCGTTGCTCCCCACTCTCACGTTGTAATCTNNNAGATCGGAAGA |
| 5344_2 | TATGAGTGTGGAGTCGTTGCTCCCCACTCTCACGTTGTAATCTNNNAGATCGGAAGA |
| 5344_3 | TATGAGTGTGGAGTCGTTGCTCCCCACTCTCACGTTGTAATCTNNNAGATCGGAAGA |
| 5996_1 | TATGAGTGTGGAGTCGTTGCGGGCATTTGAAACATTTGAAGCCNNNAGATCGGAAG |
| 5996_2 | TATGAGTGTGGAGTCGTTGCGGGCATTTGAAACATTTGAAGCCNNNAGATCGGAAG |
| 5996_3 | TATGAGTGTGGAGTCGTTGCGGGCATTTGAAACATTTGAAGCCNNNAGATCGGAAG |
| 972_1 | TATGAGTGTGGAGTCGTTGCTGAAGGAGTAACCAGAAGACAGGTNNNAGATCGGA |
| 972_2 | TATGAGTGTGGAGTCGTTGCTGAAGGAGTAACCAGAAGACAGGTNNNAGATCGGA |
| 972_3 | TATGAGTGTGGAGTCGTTGCTGAAGGAGTAACCAGAAGACAGGTNNNAGATCGGA |
| 46203_1 | TATGAGTGTGGAGTCGTTGCTCCAGGAGTTTGGAGCTTTACAANNNAGATCGGAAG |
| 46203_2 | TATGAGTGTGGAGTCGTTGCTCCAGGAGTTTGGAGCTTTACAANNNAGATCGGAAG |
| 46203_3 | TATGAGTGTGGAGTCGTTGCTCCAGGAGTTTGGAGCTTTACAANNNAGATCGGAAG |
| 51254_1 | TATGAGTGTGGAGTCGTTGCGAAGACGGGTGATTTCTGCATTTNNNAGATCGGAAG |
| 51254_2 | TATGAGTGTGGAGTCGTTGCGAAGACGGGTGATTTCTGCATTTNNNAGATCGGAAG |
| 51254_3 | TATGAGTGTGGAGTCGTTGCGAAGACGGGTGATTTCTGCATTTNNNAGATCGGAAG |
| 29382_1 | TATGAGTGTGGAGTCGTTGCGTGTGAGATGATGTGTGTGTGTGNNNAGATCGGAA |
| 29382_2 | TATGAGTGTGGAGTCGTTGCGTGTGAGATGATGTGTGTGTGTGNNNAGATCGGAA |
| 29382_3 | TATGAGTGTGGAGTCGTTGCGTGTGAGATGATGTGTGTGTGTGNNNAGATCGGAA |
| 51388_1 | TATGAGTGTGGAGTCGTTGCCCTTCTTAGGCCTGACCCTAATCNNNAGATCGGAAGA |
| 51388_2 | TATGAGTGTGGAGTCGTTGCCCTTCTTAGGCCTGACCCTAATCNNNAGATCGGAAGA |
| 51388_3 | TATGAGTGTGGAGTCGTTGCCCTTCTTAGGCCTGACCCTAATCNNNAGATCGGAAGA |
| 42084_1 | TATGAGTGTGGAGTCGTTGCGCTATTGTGAACAGTGCTGCAATNNNAGATCGGAAG |

| | |
|---|---|
| 42084_2 | TATGAGTGTGGAGTCGTTGCGCTATTGTGAACAGTGCTGCAATNNNAGATCGGAAG |
| 42084_3 | TATGAGTGTGGAGTCGTTGCGCTATTGTGAACAGTGCTGCAATNNNAGATCGGAAG |
| 43664_1 | TATGAGTGTGGAGTCGTTGCAAGAGTGGAATGAAAATGCAGGCNNNAGATCGGAA |
| 43664_2 | TATGAGTGTGGAGTCGTTGCAAGAGTGGAATGAAAATGCAGGCNNNAGATCGGAA |
| 43664_3 | TATGAGTGTGGAGTCGTTGCAAGAGTGGAATGAAAATGCAGGCNNNAGATCGGAA |
| 10953_1 | TATGAGTGTGGAGTCGTTGCCTTTTGTCAGCTGCATCCTGTGNNNAGATCGGAAGA |
| 10953_2 | TATGAGTGTGGAGTCGTTGCCTTTTGTCAGCTGCATCCTGTGNNNAGATCGGAAGA |
| 10953_3 | TATGAGTGTGGAGTCGTTGCCTTTTGTCAGCTGCATCCTGTGNNNAGATCGGAAGA |
| 45088_1 | TATGAGTGTGGAGTCGTTGCAATAGAAGCACCATTCCACACCTNNNAGATCGGAAG |
| 45088_2 | TATGAGTGTGGAGTCGTTGCAATAGAAGCACCATTCCACACCTNNNAGATCGGAAG |
| 45088_3 | TATGAGTGTGGAGTCGTTGCAATAGAAGCACCATTCCACACCTNNNAGATCGGAAG |
| 37932_1 | TATGAGTGTGGAGTCGTTGCTCATCTGAAAATGGGGACAAGGANNNAGATCGGAA |
| 37932_2 | TATGAGTGTGGAGTCGTTGCTCATCTGAAAATGGGGACAAGGANNNAGATCGGAA |
| 37932_3 | TATGAGTGTGGAGTCGTTGCTCATCTGAAAATGGGGACAAGGANNNAGATCGGAA |
| 52888_1 | TATGAGTGTGGAGTCGTTGCCGAGACAAGGAGGAAGGAATTCANNNAGATCGGAA |
| 52888_2 | TATGAGTGTGGAGTCGTTGCCGAGACAAGGAGGAAGGAATTCANNNAGATCGGAA |
| 52888_3 | TATGAGTGTGGAGTCGTTGCCGAGACAAGGAGGAAGGAATTCANNNAGATCGGAA |
| 43923_1 | TATGAGTGTGGAGTCGTTGCTCTTTCTTGGGATGGAGTTTTGCNNNAGATCGGAAG |
| 43923_2 | TATGAGTGTGGAGTCGTTGCTCTTTCTTGGGATGGAGTTTTGCNNNAGATCGGAAG |
| 43923_3 | TATGAGTGTGGAGTCGTTGCTCTTTCTTGGGATGGAGTTTTGCNNNAGATCGGAAG |
| 13283_1 | TATGAGTGTGGAGTCGTTGCCACTGCCTACTCTGAATAAATNNNAGATCGGAAGAG |
| 13283_2 | TATGAGTGTGGAGTCGTTGCCACTGCCTACTCTGAATAAATNNNAGATCGGAAGAG |
| 13283_3 | TATGAGTGTGGAGTCGTTGCCACTGCCTACTCTGAATAAATNNNAGATCGGAAGAG |
| 43782_1 | TATGAGTGTGGAGTCGTTGCCTTTGCATGGCTGAAAAAAAGGTNNNAGATCGGAA |
| 43782_2 | TATGAGTGTGGAGTCGTTGCCTTTGCATGGCTGAAAAAAAGGTNNNAGATCGGAA |
| 43782_3 | TATGAGTGTGGAGTCGTTGCCTTTGCATGGCTGAAAAAAAGGTNNNAGATCGGAA |
| 46627_1 | TATGAGTGTGGAGTCGTTGCTGATCTTCTTGTCTTGGTCTCCGNNNAGATCGGAAGA |
| 46627_2 | TATGAGTGTGGAGTCGTTGCTGATCTTCTTGTCTTGGTCTCCGNNNAGATCGGAAGA |
| 46627_3 | TATGAGTGTGGAGTCGTTGCTGATCTTCTTGTCTTGGTCTCCGNNNAGATCGGAAGA |
| 49978_1 | TATGAGTGTGGAGTCGTTGCGGACACTTCTTTATCCTGGCAGANNNAGATCGGAAG |
| 49978_2 | TATGAGTGTGGAGTCGTTGCGGACACTTCTTTATCCTGGCAGANNNAGATCGGAAG |
| 49978_3 | TATGAGTGTGGAGTCGTTGCGGACACTTCTTTATCCTGGCAGANNNAGATCGGAAG |
| 41184_1 | TATGAGTGTGGAGTCGTTGCATGCAGCCAAAAGACACATGAAANNNAGATCGGAA |
| 41184_2 | TATGAGTGTGGAGTCGTTGCATGCAGCCAAAAGACACATGAAANNNAGATCGGAA |
| 41184_3 | TATGAGTGTGGAGTCGTTGCATGCAGCCAAAAGACACATGAAANNNAGATCGGAA |
| 10471_1 | TATGAGTGTGGAGTCGTTGCGTTTCCCCATACTGTTCTTGCCANNNAGATCGGAAGA |
| 10471_2 | TATGAGTGTGGAGTCGTTGCGTTTCCCCATACTGTTCTTGCCANNNAGATCGGAAGA |
| 10471_3 | TATGAGTGTGGAGTCGTTGCGTTTCCCCATACTGTTCTTGCCANNNAGATCGGAAGA |
| 51829_1 | TATGAGTGTGGAGTCGTTGCGATGTCTGCTTGCTGTCCTTTTTNNNAGATCGGAAGA |
| 51829_2 | TATGAGTGTGGAGTCGTTGCGATGTCTGCTTGCTGTCCTTTTTNNNAGATCGGAAGA |
| 51829_3 | TATGAGTGTGGAGTCGTTGCGATGTCTGCTTGCTGTCCTTTTTNNNAGATCGGAAGA |
| 32954_1 | TATGAGTGTGGAGTCGTTGCTTTTGAATACAGCACAGCCTGTGNNNAGATCGGAAG |
| 32954_2 | TATGAGTGTGGAGTCGTTGCTTTTGAATACAGCACAGCCTGTGNNNAGATCGGAAG |

| | |
|---|---|
| 32954_3 | TATGAGTGTGGAGTCGTTGCTTTTGAATACAGCACAGCCTGTGNNNAGATCGGAAG |
| 13142_1 | TATGAGTGTGGAGTCGTTGCTTCGAATTACAACTTGTAAAACTNNNAGATCGGAAG |
| 13142_2 | TATGAGTGTGGAGTCGTTGCTTCGAATTACAACTTGTAAAACTNNNAGATCGGAAG |
| 13142_3 | TATGAGTGTGGAGTCGTTGCTTCGAATTACAACTTGTAAAACTNNNAGATCGGAAG |
| 30072_1 | TATGAGTGTGGAGTCGTTGCCTCCAGGATTTGCTTTTTCCCTGNNNAGATCGGAAGA |
| 30072_2 | TATGAGTGTGGAGTCGTTGCCTCCAGGATTTGCTTTTTCCCTGNNNAGATCGGAAGA |
| 30072_3 | TATGAGTGTGGAGTCGTTGCCTCCAGGATTTGCTTTTTCCCTGNNNAGATCGGAAGA |
| 37251_1 | TATGAGTGTGGAGTCGTTGCCCCTTCTTTCCCAGCAGTAATCTNNNAGATCGGAAGA |
| 37251_2 | TATGAGTGTGGAGTCGTTGCCCCTTCTTTCCCAGCAGTAATCTNNNAGATCGGAAGA |
| 37251_3 | TATGAGTGTGGAGTCGTTGCCCCTTCTTTCCCAGCAGTAATCTNNNAGATCGGAAGA |
| 50824_1 | TATGAGTGTGGAGTCGTTGCTGCTGGTCTTCTAAGCAGATTGANNNAGATCGGAAG |
| 50824_2 | TATGAGTGTGGAGTCGTTGCTGCTGGTCTTCTAAGCAGATTGANNNAGATCGGAAG |
| 50824_3 | TATGAGTGTGGAGTCGTTGCTGCTGGTCTTCTAAGCAGATTGANNNAGATCGGAAG |
| 49210_1 | TATGAGTGTGGAGTCGTTGCCAGCAAACAAGATGGTAGGTTCANNNAGATCGGAA |
| 49210_2 | TATGAGTGTGGAGTCGTTGCCAGCAAACAAGATGGTAGGTTCANNNAGATCGGAA |
| 49210_3 | TATGAGTGTGGAGTCGTTGCCAGCAAACAAGATGGTAGGTTCANNNAGATCGGAA |
| 51514_1 | TATGAGTGTGGAGTCGTTGCATTTCTGTGGCCTGCTTGAATTCNNNAGATCGGAAG |
| 51514_2 | TATGAGTGTGGAGTCGTTGCATTTCTGTGGCCTGCTTGAATTCNNNAGATCGGAAG |
| 51514_3 | TATGAGTGTGGAGTCGTTGCATTTCTGTGGCCTGCTTGAATTCNNNAGATCGGAAG |
| 39226_1 | TATGAGTGTGGAGTCGTTGCTTATTTTCCACCTTCCACCTGCANNNAGATCGGAAGA |
| 39226_2 | TATGAGTGTGGAGTCGTTGCTTATTTTCCACCTTCCACCTGCANNNAGATCGGAAGA |
| 39226_3 | TATGAGTGTGGAGTCGTTGCTTATTTTCCACCTTCCACCTGCANNNAGATCGGAAGA |
| 31868_1 | TATGAGTGTGGAGTCGTTGCGAGAGAGAGAGAGTCTTCTTGCGNNNAGATCGGAA |
| 31868_2 | TATGAGTGTGGAGTCGTTGCGAGAGAGAGAGAGTCTTCTTGCGNNNAGATCGGAA |
| 31868_3 | TATGAGTGTGGAGTCGTTGCGAGAGAGAGAGAGTCTTCTTGCGNNNAGATCGGAA |
| 47303_1 | TATGAGTGTGGAGTCGTTGCCCTCCTAGGTTCAAGCGATTCTCNNNAGATCGGAAG |
| 47303_2 | TATGAGTGTGGAGTCGTTGCCCTCCTAGGTTCAAGCGATTCTCNNNAGATCGGAAG |
| 47303_3 | TATGAGTGTGGAGTCGTTGCCCTCCTAGGTTCAAGCGATTCTCNNNAGATCGGAAG |
| 43154_1 | TATGAGTGTGGAGTCGTTGCTTTAGTGCAGACAGGGTTTCACTNNNAGATCGGAAG |
| 43154_2 | TATGAGTGTGGAGTCGTTGCTTTAGTGCAGACAGGGTTTCACTNNNAGATCGGAAG |
| 43154_3 | TATGAGTGTGGAGTCGTTGCTTTAGTGCAGACAGGGTTTCACTNNNAGATCGGAAG |
| 47890_1 | TATGAGTGTGGAGTCGTTGCCTGGAGTACAATGGCACGATCTCNNNAGATCGGAA |
| 47890_2 | TATGAGTGTGGAGTCGTTGCCTGGAGTACAATGGCACGATCTCNNNAGATCGGAA |
| 47890_3 | TATGAGTGTGGAGTCGTTGCCTGGAGTACAATGGCACGATCTCNNNAGATCGGAA |
| 48236_1 | TATGAGTGTGGAGTCGTTGCCCTAGTAGCTGGGATTACAGGTGNNNAGATCGGAA |
| 48236_2 | TATGAGTGTGGAGTCGTTGCCCTAGTAGCTGGGATTACAGGTGNNNAGATCGGAA |
| 48236_3 | TATGAGTGTGGAGTCGTTGCCCTAGTAGCTGGGATTACAGGTGNNNAGATCGGAA |
| 42620_1 | TATGAGTGTGGAGTCGTTGCAGACAGTCATGCCATCTGTGAATNNNAGATCGGAAG |
| 42620_2 | TATGAGTGTGGAGTCGTTGCAGACAGTCATGCCATCTGTGAATNNNAGATCGGAAG |
| 42620_3 | TATGAGTGTGGAGTCGTTGCAGACAGTCATGCCATCTGTGAATNNNAGATCGGAAG |
| 29099_1 | TATGAGTGTGGAGTCGTTGCCTAATTTTGCTATCAGCCCCTGCNNNAGATCGGAAGA |
| 29099_2 | TATGAGTGTGGAGTCGTTGCCTAATTTTGCTATCAGCCCCTGCNNNAGATCGGAAGA |
| 29099_3 | TATGAGTGTGGAGTCGTTGCCTAATTTTGCTATCAGCCCCTGCNNNAGATCGGAAGA |

| | |
|---|---|
| 35468_1 | TATGAGTGTGGAGTCGTTGCATTTTATGCCTCACTAGTGCCCANNNAGATCGGAAGA |
| 35468_2 | TATGAGTGTGGAGTCGTTGCATTTTATGCCTCACTAGTGCCCANNNAGATCGGAAGA |
| 35468_3 | TATGAGTGTGGAGTCGTTGCATTTTATGCCTCACTAGTGCCCANNNAGATCGGAAGA |
| 33278_1 | TATGAGTGTGGAGTCGTTGCCATATGTTGGGCAATGGGTTCACNNNAGATCGGAAG |
| 33278_2 | TATGAGTGTGGAGTCGTTGCCATATGTTGGGCAATGGGTTCACNNNAGATCGGAAG |
| 33278_3 | TATGAGTGTGGAGTCGTTGCCATATGTTGGGCAATGGGTTCACNNNAGATCGGAAG |
| 33056_1 | TATGAGTGTGGAGTCGTTGCGATTCTCCTGCTTCAGACTCCTGNNNAGATCGGAAGA |
| 33056_2 | TATGAGTGTGGAGTCGTTGCGATTCTCCTGCTTCAGACTCCTGNNNAGATCGGAAGA |
| 33056_3 | TATGAGTGTGGAGTCGTTGCGATTCTCCTGCTTCAGACTCCTGNNNAGATCGGAAGA |
| 40991_1 | TATGAGTGTGGAGTCGTTGCGAACCCCTACAAGTCAACAACAANNNAGATCGGAAG |
| 40991_2 | TATGAGTGTGGAGTCGTTGCGAACCCCTACAAGTCAACAACAANNNAGATCGGAAG |
| 40991_3 | TATGAGTGTGGAGTCGTTGCGAACCCCTACAAGTCAACAACAANNNAGATCGGAAG |
| 51299_1 | TATGAGTGTGGAGTCGTTGCTGAGGCAGGAGAATCTCTTGAACNNNAGATCGGAAG |
| 51299_2 | TATGAGTGTGGAGTCGTTGCTGAGGCAGGAGAATCTCTTGAACNNNAGATCGGAAG |
| 51299_3 | TATGAGTGTGGAGTCGTTGCTGAGGCAGGAGAATCTCTTGAACNNNAGATCGGAAG |
| 8759_1 | TATGAGTGTGGAGTCGTTGCGTTAGGGGCTGTGGTGAGAATTTNNNAGATCGGAA |
| 8759_2 | TATGAGTGTGGAGTCGTTGCGTTAGGGGCTGTGGTGAGAATTTNNNAGATCGGAA |
| 8759_3 | TATGAGTGTGGAGTCGTTGCGTTAGGGGCTGTGGTGAGAATTTNNNAGATCGGAA |
| 40833_1 | TATGAGTGTGGAGTCGTTGCCACTGCAACCTCTACCTCCTGNNNAGATCGGAAGAGC |
| 40833_2 | TATGAGTGTGGAGTCGTTGCCACTGCAACCTCTACCTCCTGNNNAGATCGGAAGAGC |
| 40833_3 | TATGAGTGTGGAGTCGTTGCCACTGCAACCTCTACCTCCTGNNNAGATCGGAAGAGC |
| 5200_1 | TATGAGTGTGGAGTCGTTGCACAGTGAATCAGGCCAAAAGGAANNNAGATCGGAA |
| 5200_2 | TATGAGTGTGGAGTCGTTGCACAGTGAATCAGGCCAAAAGGAANNNAGATCGGAA |
| 5200_3 | TATGAGTGTGGAGTCGTTGCACAGTGAATCAGGCCAAAAGGAANNNAGATCGGAA |
| 40347_1 | TATGAGTGTGGAGTCGTTGCTAACATGGTGAAAACCCGTCTCTNNNAGATCGGAAG |
| 40347_2 | TATGAGTGTGGAGTCGTTGCTAACATGGTGAAAACCCGTCTCTNNNAGATCGGAAG |
| 40347_3 | TATGAGTGTGGAGTCGTTGCTAACATGGTGAAAACCCGTCTCTNNNAGATCGGAAG |
| 37182_1 | TATGAGTGTGGAGTCGTTGCAGAATTTTAGACAGGAAGCCCCANNNAGATCGGAAG |
| 37182_2 | TATGAGTGTGGAGTCGTTGCAGAATTTTAGACAGGAAGCCCCANNNAGATCGGAAG |
| 37182_3 | TATGAGTGTGGAGTCGTTGCAGAATTTTAGACAGGAAGCCCCANNNAGATCGGAAG |
| 38035_1 | TATGAGTGTGGAGTCGTTGCACCCAAAACAACAAGAAGGAAAANNNAGATCGGAAG |
| 38035_2 | TATGAGTGTGGAGTCGTTGCACCCAAAACAACAAGAAGGAAAANNNAGATCGGAAG |
| 38035_3 | TATGAGTGTGGAGTCGTTGCACCCAAAACAACAAGAAGGAAAANNNAGATCGGAAG |
| 3033_1 | TATGAGTGTGGAGTCGTTGCCTTCATGCTTTCCTGTGGGAGAANNNAGATCGGAAG |
| 3033_2 | TATGAGTGTGGAGTCGTTGCCTTCATGCTTTCCTGTGGGAGAANNNAGATCGGAAG |
| 3033_3 | TATGAGTGTGGAGTCGTTGCCTTCATGCTTTCCTGTGGGAGAANNNAGATCGGAAG |
| 34171_1 | TATGAGTGTGGAGTCGTTGCAAACAACAGATGCTGGAGAGGATNNNAGATCGGAA |
| 34171_2 | TATGAGTGTGGAGTCGTTGCAAACAACAGATGCTGGAGAGGATNNNAGATCGGAA |
| 34171_3 | TATGAGTGTGGAGTCGTTGCAAACAACAGATGCTGGAGAGGATNNNAGATCGGAA |
| 45079_1 | TATGAGTGTGGAGTCGTTGCAATCATTTCATTGCCGTTCTGCANNNAGATCGGAAGA |
| 45079_2 | TATGAGTGTGGAGTCGTTGCAATCATTTCATTGCCGTTCTGCANNNAGATCGGAAGA |
| 45079_3 | TATGAGTGTGGAGTCGTTGCAATCATTTCATTGCCGTTCTGCANNNAGATCGGAAGA |
| 50988_1 | TATGAGTGTGGAGTCGTTGCTGTGTTTACTGATTCCCTAGGGTNNNAGATCGGAAG |

| | |
|---|---|
| 50988_2 | TATGAGTGTGGAGTCGTTGCTGTGTTTACTGATTCCCTAGGGTNNNAGATCGGAAG |
| 50988_3 | TATGAGTGTGGAGTCGTTGCTGTGTTTACTGATTCCCTAGGGTNNNAGATCGGAAG |
| 49498_1 | TATGAGTGTGGAGTCGTTGCACATTCCCACCAACAAGTGTACANNNAGATCGGAAG |
| 49498_2 | TATGAGTGTGGAGTCGTTGCACATTCCCACCAACAAGTGTACANNNAGATCGGAAG |
| 49498_3 | TATGAGTGTGGAGTCGTTGCACATTCCCACCAACAAGTGTACANNNAGATCGGAAG |
| 13570_1 | TATGAGTGTGGAGTCGTTGCAGTCTCTAGAACAGGGATTCNNNAGATCGGAAGAG |
| 13570_2 | TATGAGTGTGGAGTCGTTGCAGTCTCTAGAACAGGGATTCNNNAGATCGGAAGAG |
| 13570_3 | TATGAGTGTGGAGTCGTTGCAGTCTCTAGAACAGGGATTCNNNAGATCGGAAGAG |
| 52204_1 | TATGAGTGTGGAGTCGTTGCCCAAAGTGCTATGATTACAGGCGNNNAGATCGGAA |
| 52204_2 | TATGAGTGTGGAGTCGTTGCCCAAAGTGCTATGATTACAGGCGNNNAGATCGGAA |
| 52204_3 | TATGAGTGTGGAGTCGTTGCCCAAAGTGCTATGATTACAGGCGNNNAGATCGGAA |
| 48537_1 | TATGAGTGTGGAGTCGTTGCCCCGGCCCAATAATTTGTTTTTTNNNAGATCGGAAG |
| 48537_2 | TATGAGTGTGGAGTCGTTGCCCCGGCCCAATAATTTGTTTTTTNNNAGATCGGAAG |
| 48537_3 | TATGAGTGTGGAGTCGTTGCCCCGGCCCAATAATTTGTTTTTTNNNAGATCGGAAG |
| 44405_1 | TATGAGTGTGGAGTCGTTGCTGATTCAACAGACTGGGCACATANNNAGATCGGAA |
| 44405_2 | TATGAGTGTGGAGTCGTTGCTGATTCAACAGACTGGGCACATANNNAGATCGGAA |
| 44405_3 | TATGAGTGTGGAGTCGTTGCTGATTCAACAGACTGGGCACATANNNAGATCGGAA |
| 49503_1 | TATGAGTGTGGAGTCGTTGCTTTGAAAATGCCAAGTGAGGGTCNNNAGATCGGAA |
| 49503_2 | TATGAGTGTGGAGTCGTTGCTTTGAAAATGCCAAGTGAGGGTCNNNAGATCGGAA |
| 49503_3 | TATGAGTGTGGAGTCGTTGCTTTGAAAATGCCAAGTGAGGGTCNNNAGATCGGAA |
| 50902_1 | TATGAGTGTGGAGTCGTTGCTGAAGTTTTCCGTAGCCTCCTACNNNAGATCGGAAG |
| 50902_2 | TATGAGTGTGGAGTCGTTGCTGAAGTTTTCCGTAGCCTCCTACNNNAGATCGGAAG |
| 50902_3 | TATGAGTGTGGAGTCGTTGCTGAAGTTTTCCGTAGCCTCCTACNNNAGATCGGAAG |
| 41300_1 | TATGAGTGTGGAGTCGTTGCAGCTGTGAATTTCCTAGGTTACGANNNAGATCGGAA |
| 41300_2 | TATGAGTGTGGAGTCGTTGCAGCTGTGAATTTCCTAGGTTACGANNNAGATCGGAA |
| 41300_3 | TATGAGTGTGGAGTCGTTGCAGCTGTGAATTTCCTAGGTTACGANNNAGATCGGAA |
| 32557_1 | TATGAGTGTGGAGTCGTTGCCAATCTTGGTTCACTGCAACCTCNNNAGATCGGAAG |
| 32557_2 | TATGAGTGTGGAGTCGTTGCCAATCTTGGTTCACTGCAACCTCNNNAGATCGGAAG |
| 32557_3 | TATGAGTGTGGAGTCGTTGCCAATCTTGGTTCACTGCAACCTCNNNAGATCGGAAG |
| 44937_1 | TATGAGTGTGGAGTCGTTGCGGAAAGGAGGGCATATGAAAAGCNNNAGATCGGA |
| 44937_2 | TATGAGTGTGGAGTCGTTGCGGAAAGGAGGGCATATGAAAAGCNNNAGATCGGA |
| 44937_3 | TATGAGTGTGGAGTCGTTGCGGAAAGGAGGGCATATGAAAAGCNNNAGATCGGA |
| 38136_1 | TATGAGTGTGGAGTCGTTGCCAGATCACGAGGTCAGGAGATCNNNAGATCGGAAG |
| 38136_2 | TATGAGTGTGGAGTCGTTGCCAGATCACGAGGTCAGGAGATCNNNAGATCGGAAG |
| 38136_3 | TATGAGTGTGGAGTCGTTGCCAGATCACGAGGTCAGGAGATCNNNAGATCGGAAG |
| 31751_1 | TATGAGTGTGGAGTCGTTGCCAACTTGAGAACCCTCTTCTGGANNNAGATCGGAAG |
| 31751_2 | TATGAGTGTGGAGTCGTTGCCAACTTGAGAACCCTCTTCTGGANNNAGATCGGAAG |
| 31751_3 | TATGAGTGTGGAGTCGTTGCCAACTTGAGAACCCTCTTCTGGANNNAGATCGGAAG |
| 43306_1 | TATGAGTGTGGAGTCGTTGCTGGAGTGTAATGGTGCAATCTCANNNAGATCGGAA |
| 43306_2 | TATGAGTGTGGAGTCGTTGCTGGAGTGTAATGGTGCAATCTCANNNAGATCGGAA |
| 43306_3 | TATGAGTGTGGAGTCGTTGCTGGAGTGTAATGGTGCAATCTCANNNAGATCGGAA |
| 40646_1 | TATGAGTGTGGAGTCGTTGCGTAGCTCATGCCTGTAATCCCAGNNNAGATCGGAAG |
| 40646_2 | TATGAGTGTGGAGTCGTTGCGTAGCTCATGCCTGTAATCCCAGNNNAGATCGGAAG |

| | |
|---|---|
| 40646_3 | TATGAGTGTGGAGTCGTTGCGTAGCTCATGCCTGTAATCCCAGNNNAGATCGGAAG |
| 41883_1 | TATGAGTGTGGAGTCGTTGCGTTCAAGAGATTCTCCTGCCTCANNNAGATCGGAAG |
| 41883_2 | TATGAGTGTGGAGTCGTTGCGTTCAAGAGATTCTCCTGCCTCANNNAGATCGGAAG |
| 41883_3 | TATGAGTGTGGAGTCGTTGCGTTCAAGAGATTCTCCTGCCTCANNNAGATCGGAAG |
| 38316_1 | TATGAGTGTGGAGTCGTTGCAGAAGAAAAAGGCTGGGTCTGAANNNAGATCGGA |
| 38316_2 | TATGAGTGTGGAGTCGTTGCAGAAGAAAAAGGCTGGGTCTGAANNNAGATCGGA |
| 38316_3 | TATGAGTGTGGAGTCGTTGCAGAAGAAAAAGGCTGGGTCTGAANNNAGATCGGA |
| 51600_1 | TATGAGTGTGGAGTCGTTGCACAGTTCTTATGCTGGGAGAAGGNNNAGATCGGAA |
| 51600_2 | TATGAGTGTGGAGTCGTTGCACAGTTCTTATGCTGGGAGAAGGNNNAGATCGGAA |
| 51600_3 | TATGAGTGTGGAGTCGTTGCACAGTTCTTATGCTGGGAGAAGGNNNAGATCGGAA |
| 45801_1 | TATGAGTGTGGAGTCGTTGCTGCTATCTCAGTTCACTGCAACTNNNAGATCGGAAG |
| 45801_2 | TATGAGTGTGGAGTCGTTGCTGCTATCTCAGTTCACTGCAACTNNNAGATCGGAAG |
| 45801_3 | TATGAGTGTGGAGTCGTTGCTGCTATCTCAGTTCACTGCAACTNNNAGATCGGAAG |
| 34491_1 | TATGAGTGTGGAGTCGTTGCCATGGTGGCTTGCACTTGTAATCNNNAGATCGGAAG |
| 34491_2 | TATGAGTGTGGAGTCGTTGCCATGGTGGCTTGCACTTGTAATCNNNAGATCGGAAG |
| 34491_3 | TATGAGTGTGGAGTCGTTGCCATGGTGGCTTGCACTTGTAATCNNNAGATCGGAAG |
| 28839_1 | TATGAGTGTGGAGTCGTTGCTTGATGCACACTAAAGTTGATTCTNNNAGATCGGAA |
| 28839_2 | TATGAGTGTGGAGTCGTTGCTTGATGCACACTAAAGTTGATTCTNNNAGATCGGAA |
| 28839_3 | TATGAGTGTGGAGTCGTTGCTTGATGCACACTAAAGTTGATTCTNNNAGATCGGAA |
| 41381_1 | TATGAGTGTGGAGTCGTTGCGCATGCCTGTAATCCTAGCTACTNNNAGATCGGAAG |
| 41381_2 | TATGAGTGTGGAGTCGTTGCGCATGCCTGTAATCCTAGCTACTNNNAGATCGGAAG |
| 41381_3 | TATGAGTGTGGAGTCGTTGCGCATGCCTGTAATCCTAGCTACTNNNAGATCGGAAG |
| 40747_1 | TATGAGTGTGGAGTCGTTGCAGAGGTTGCAGTGAGCTAAGAATNNNAGATCGGAA |
| 40747_2 | TATGAGTGTGGAGTCGTTGCAGAGGTTGCAGTGAGCTAAGAATNNNAGATCGGAA |
| 40747_3 | TATGAGTGTGGAGTCGTTGCAGAGGTTGCAGTGAGCTAAGAATNNNAGATCGGAA |
| 42590_1 | TATGAGTGTGGAGTCGTTGCGAACTCCTGACCTCAAGTGATCCNNNAGATCGGAAG |
| 42590_2 | TATGAGTGTGGAGTCGTTGCGAACTCCTGACCTCAAGTGATCCNNNAGATCGGAAG |
| 42590_3 | TATGAGTGTGGAGTCGTTGCGAACTCCTGACCTCAAGTGATCCNNNAGATCGGAAG |
| 50236_1 | TATGAGTGTGGAGTCGTTGCCAGCATGCCCCACTGAATAAAATNNNAGATCGGAAG |
| 50236_2 | TATGAGTGTGGAGTCGTTGCCAGCATGCCCCACTGAATAAAATNNNAGATCGGAAG |
| 50236_3 | TATGAGTGTGGAGTCGTTGCCAGCATGCCCCACTGAATAAAATNNNAGATCGGAAG |
| 5376_1 | TATGAGTGTGGAGTCGTTGCATCCCTCTCCTGTCTGCTTCTTTNNNAGATCGGAAGA |
| 5376_2 | TATGAGTGTGGAGTCGTTGCATCCCTCTCCTGTCTGCTTCTTTNNNAGATCGGAAGA |
| 5376_3 | TATGAGTGTGGAGTCGTTGCATCCCTCTCCTGTCTGCTTCTTTNNNAGATCGGAAGA |
| 52640_1 | TATGAGTGTGGAGTCGTTGCCTTGTGAAGACCCTGCCTACTAGNNNAGATCGGAAG |
| 52640_2 | TATGAGTGTGGAGTCGTTGCCTTGTGAAGACCCTGCCTACTAGNNNAGATCGGAAG |
| 52640_3 | TATGAGTGTGGAGTCGTTGCCTTGTGAAGACCCTGCCTACTAGNNNAGATCGGAAG |
| 34860_1 | TATGAGTGTGGAGTCGTTGCTGGTTCAAGTCATTCTCTTGCCTNNNAGATCGGAAG |
| 34860_2 | TATGAGTGTGGAGTCGTTGCTGGTTCAAGTCATTCTCTTGCCTNNNAGATCGGAAG |
| 34860_3 | TATGAGTGTGGAGTCGTTGCTGGTTCAAGTCATTCTCTTGCCTNNNAGATCGGAAG |
| 40293_1 | TATGAGTGTGGAGTCGTTGCTTATTGGACAGTGCTTGGACAGTNNNAGATCGGAA |
| 40293_2 | TATGAGTGTGGAGTCGTTGCTTATTGGACAGTGCTTGGACAGTNNNAGATCGGAA |
| 40293_3 | TATGAGTGTGGAGTCGTTGCTTATTGGACAGTGCTTGGACAGTNNNAGATCGGAA |

| | |
|---|---|
| 3086_1 | TATGAGTGTGGAGTCGTTGCCCCGTTAGGGTGTTTTTAGGCTTNNNAGATCGGAAG |
| 3086_2 | TATGAGTGTGGAGTCGTTGCCCCGTTAGGGTGTTTTTAGGCTTNNNAGATCGGAAG |
| 3086_3 | TATGAGTGTGGAGTCGTTGCCCCGTTAGGGTGTTTTTAGGCTTNNNAGATCGGAAG |
| 2734_1 | TATGAGTGTGGAGTCGTTGCGTCAAAGCATGTCGACTGAGACTNNNAGATCGGAA |
| 2734_2 | TATGAGTGTGGAGTCGTTGCGTCAAAGCATGTCGACTGAGACTNNNAGATCGGAA |
| 2734_3 | TATGAGTGTGGAGTCGTTGCGTCAAAGCATGTCGACTGAGACTNNNAGATCGGAA |
| 46321_1 | TATGAGTGTGGAGTCGTTGCCATCCCGGTTCAAGCAATTTTCTNNNAGATCGGAAG |
| 46321_2 | TATGAGTGTGGAGTCGTTGCCATCCCGGTTCAAGCAATTTTCTNNNAGATCGGAAG |
| 46321_3 | TATGAGTGTGGAGTCGTTGCCATCCCGGTTCAAGCAATTTTCTNNNAGATCGGAAG |
| 52851_1 | TATGAGTGTGGAGTCGTTGCTCCCTATCGTCTGAACAGTTGTCNNNAGATCGGAAG |
| 52851_2 | TATGAGTGTGGAGTCGTTGCTCCCTATCGTCTGAACAGTTGTCNNNAGATCGGAAG |
| 52851_3 | TATGAGTGTGGAGTCGTTGCTCCCTATCGTCTGAACAGTTGTCNNNAGATCGGAAG |
| 46782_1 | TATGAGTGTGGAGTCGTTGCCTCAAGCAATTCCCCCACTTTAGNNNAGATCGGAAGA |
| 46782_2 | TATGAGTGTGGAGTCGTTGCCTCAAGCAATTCCCCCACTTTAGNNNAGATCGGAAGA |
| 46782_3 | TATGAGTGTGGAGTCGTTGCCTCAAGCAATTCCCCCACTTTAGNNNAGATCGGAAGA |
| 40902_1 | TATGAGTGTGGAGTCGTTGCCCACTTTCTTCCGTTTGTCCTTCNNNAGATCGGAAGA |
| 40902_2 | TATGAGTGTGGAGTCGTTGCCCACTTTCTTCCGTTTGTCCTTCNNNAGATCGGAAGA |
| 40902_3 | TATGAGTGTGGAGTCGTTGCCCACTTTCTTCCGTTTGTCCTTCNNNAGATCGGAAGA |
| 34695_1 | TATGAGTGTGGAGTCGTTGCAATGGGGTCTTACCATATTGCCCNNNAGATCGGAAG |
| 34695_2 | TATGAGTGTGGAGTCGTTGCAATGGGGTCTTACCATATTGCCCNNNAGATCGGAAG |
| 34695_3 | TATGAGTGTGGAGTCGTTGCAATGGGGTCTTACCATATTGCCCNNNAGATCGGAAG |
| 39566_1 | TATGAGTGTGGAGTCGTTGCCCTTTACTGACCAACTCTCCTCCNNNAGATCGGAAGA |
| 39566_2 | TATGAGTGTGGAGTCGTTGCCCTTTACTGACCAACTCTCCTCCNNNAGATCGGAAGA |
| 39566_3 | TATGAGTGTGGAGTCGTTGCCCTTTACTGACCAACTCTCCTCCNNNAGATCGGAAGA |
| 12898_1 | TATGAGTGTGGAGTCGTTGCTTCTCCCCATATTTCACAGATNNNAGATCGGAAGAGC |
| 12898_2 | TATGAGTGTGGAGTCGTTGCTTCTCCCCATATTTCACAGATNNNAGATCGGAAGAGC |
| 12898_3 | TATGAGTGTGGAGTCGTTGCTTCTCCCCATATTTCACAGATNNNAGATCGGAAGAGC |
| 49085_1 | TATGAGTGTGGAGTCGTTGCAAGGTAGGTCAGGAAGAAATGGGNNNAGATCGGA |
| 49085_2 | TATGAGTGTGGAGTCGTTGCAAGGTAGGTCAGGAAGAAATGGGNNNAGATCGGA |
| 49085_3 | TATGAGTGTGGAGTCGTTGCAAGGTAGGTCAGGAAGAAATGGGNNNAGATCGGA |
| 44184_1 | TATGAGTGTGGAGTCGTTGCAGGTGAGTCTTATTATGTCGGCCNNNAGATCGGAA |
| 44184_2 | TATGAGTGTGGAGTCGTTGCAGGTGAGTCTTATTATGTCGGCCNNNAGATCGGAA |
| 44184_3 | TATGAGTGTGGAGTCGTTGCAGGTGAGTCTTATTATGTCGGCCNNNAGATCGGAA |
| 28858_1 | TATGAGTGTGGAGTCGTTGCAACACTGATACCTGTATGTGGGGNNNAGATCGGAA |
| 28858_2 | TATGAGTGTGGAGTCGTTGCAACACTGATACCTGTATGTGGGGNNNAGATCGGAA |
| 28858_3 | TATGAGTGTGGAGTCGTTGCAACACTGATACCTGTATGTGGGGNNNAGATCGGAA |
| 36598_1 | TATGAGTGTGGAGTCGTTGCCCAATGAAATGGCACCTCTCAAGNNNAGATCGGAAG |
| 36598_2 | TATGAGTGTGGAGTCGTTGCCCAATGAAATGGCACCTCTCAAGNNNAGATCGGAAG |
| 36598_3 | TATGAGTGTGGAGTCGTTGCCCAATGAAATGGCACCTCTCAAGNNNAGATCGGAAG |
| 32712_1 | TATGAGTGTGGAGTCGTTGCAACGAGCTAATGAATGCTGTCAANNNAGATCGGAA |
| 32712_2 | TATGAGTGTGGAGTCGTTGCAACGAGCTAATGAATGCTGTCAANNNAGATCGGAA |
| 32712_3 | TATGAGTGTGGAGTCGTTGCAACGAGCTAATGAATGCTGTCAANNNAGATCGGAA |
| 51871_1 | TATGAGTGTGGAGTCGTTGCCACTGTTTTCGGACCTTCCATGNNNAGATCGGAAGA |

| | |
|---|---|
| 51871_2 | TATGAGTGTGGAGTCGTTGCCACTGTTTTCGGACCTTCCATGNNNAGATCGGAAGA |
| 51871_3 | TATGAGTGTGGAGTCGTTGCCACTGTTTTCGGACCTTCCATGNNNAGATCGGAAGA |
| 46525_1 | TATGAGTGTGGAGTCGTTGCCCTCCCAAAGTGCTAGGATTACANNNAGATCGGAAG |
| 46525_2 | TATGAGTGTGGAGTCGTTGCCCTCCCAAAGTGCTAGGATTACANNNAGATCGGAAG |
| 46525_3 | TATGAGTGTGGAGTCGTTGCCCTCCCAAAGTGCTAGGATTACANNNAGATCGGAAG |
| 36136_1 | TATGAGTGTGGAGTCGTTGCACTCTTAGCTATTTGGGAGGCTGNNNAGATCGGAA |
| 36136_2 | TATGAGTGTGGAGTCGTTGCACTCTTAGCTATTTGGGAGGCTGNNNAGATCGGAA |
| 36136_3 | TATGAGTGTGGAGTCGTTGCACTCTTAGCTATTTGGGAGGCTGNNNAGATCGGAA |
| 30699_1 | TATGAGTGTGGAGTCGTTGCTCCAGCTTTCTTTTTCTCCCTACANNNAGATCGGAAG |
| 30699_2 | TATGAGTGTGGAGTCGTTGCTCCAGCTTTCTTTTTCTCCCTACANNNAGATCGGAAG |
| 30699_3 | TATGAGTGTGGAGTCGTTGCTCCAGCTTTCTTTTTCTCCCTACANNNAGATCGGAAG |
| 46956_1 | TATGAGTGTGGAGTCGTTGCAAACTGCCTTGCACTCTTTTTGTNNNAGATCGGAAG |
| 46956_2 | TATGAGTGTGGAGTCGTTGCAAACTGCCTTGCACTCTTTTTGTNNNAGATCGGAAG |
| 46956_3 | TATGAGTGTGGAGTCGTTGCAAACTGCCTTGCACTCTTTTTGTNNNAGATCGGAAG |
| 41291_1 | TATGAGTGTGGAGTCGTTGCGAGCTGGAGGCCATCATACTTAGNNNAGATCGGAA |
| 41291_2 | TATGAGTGTGGAGTCGTTGCGAGCTGGAGGCCATCATACTTAGNNNAGATCGGAA |
| 41291_3 | TATGAGTGTGGAGTCGTTGCGAGCTGGAGGCCATCATACTTAGNNNAGATCGGAA |
| 11041_1 | TATGAGTGTGGAGTCGTTGCGGTGGTGGGGACAGGATATTGNNNAGATCGGAAG |
| 11041_2 | TATGAGTGTGGAGTCGTTGCGGTGGTGGGGACAGGATATTGNNNAGATCGGAAG |
| 11041_3 | TATGAGTGTGGAGTCGTTGCGGTGGTGGGGACAGGATATTGNNNAGATCGGAAG |
| 41429_1 | TATGAGTGTGGAGTCGTTGCGGTAACTTCAGCCCAGGACTTTANNNAGATCGGAAG |
| 41429_2 | TATGAGTGTGGAGTCGTTGCGGTAACTTCAGCCCAGGACTTTANNNAGATCGGAAG |
| 41429_3 | TATGAGTGTGGAGTCGTTGCGGTAACTTCAGCCCAGGACTTTANNNAGATCGGAAG |
| 3010_1 | TATGAGTGTGGAGTCGTTGCTCTTGCTCAGACTGACCTTCCATNNNAGATCGGAAG |
| 3010_2 | TATGAGTGTGGAGTCGTTGCTCTTGCTCAGACTGACCTTCCATNNNAGATCGGAAG |
| 3010_3 | TATGAGTGTGGAGTCGTTGCTCTTGCTCAGACTGACCTTCCATNNNAGATCGGAAG |
| 37892_1 | TATGAGTGTGGAGTCGTTGCAACTGAGCCCTTGATTCACAGAANNNAGATCGGAAG |
| 37892_2 | TATGAGTGTGGAGTCGTTGCAACTGAGCCCTTGATTCACAGAANNNAGATCGGAAG |
| 37892_3 | TATGAGTGTGGAGTCGTTGCAACTGAGCCCTTGATTCACAGAANNNAGATCGGAAG |
| 10150_1 | TATGAGTGTGGAGTCGTTGCTAAAGATAGTCAGGCTGCTGTCCNNNAGATCGGAA |
| 10150_2 | TATGAGTGTGGAGTCGTTGCTAAAGATAGTCAGGCTGCTGTCCNNNAGATCGGAA |
| 10150_3 | TATGAGTGTGGAGTCGTTGCTAAAGATAGTCAGGCTGCTGTCCNNNAGATCGGAA |
| 31384_1 | TATGAGTGTGGAGTCGTTGCTTGGAAAGAGAAATGCACACACANNNAGATCGGAA |
| 31384_2 | TATGAGTGTGGAGTCGTTGCTTGGAAAGAGAAATGCACACACANNNAGATCGGAA |
| 31384_3 | TATGAGTGTGGAGTCGTTGCTTGGAAAGAGAAATGCACACACANNNAGATCGGAA |
| 31209_1 | TATGAGTGTGGAGTCGTTGCAAGGTGTTATAGGGGTGCAATGTNNNAGATCGGAA |
| 31209_2 | TATGAGTGTGGAGTCGTTGCAAGGTGTTATAGGGGTGCAATGTNNNAGATCGGAA |
| 31209_3 | TATGAGTGTGGAGTCGTTGCAAGGTGTTATAGGGGTGCAATGTNNNAGATCGGAA |
| 43205_1 | TATGAGTGTGGAGTCGTTGCGCATCCAGCTGCATCTATATTGCNNNAGATCGGAAG |
| 43205_2 | TATGAGTGTGGAGTCGTTGCGCATCCAGCTGCATCTATATTGCNNNAGATCGGAAG |
| 43205_3 | TATGAGTGTGGAGTCGTTGCGCATCCAGCTGCATCTATATTGCNNNAGATCGGAAG |
| 10356_1 | TATGAGTGTGGAGTCGTTGCGGGAGGGGGTGGTGATATAAGTANNNAGATCGGA |
| 10356_2 | TATGAGTGTGGAGTCGTTGCGGGAGGGGGTGGTGATATAAGTANNNAGATCGGA |

| | |
|---|---|
| 10356_3 | TATGAGTGTGGAGTCGTTGCGGGAGGGGGTGGTGATATAAGTANNNAGATCGGA |
| 53177_1 | TATGAGTGTGGAGTCGTTGCTTCAGGGCTTGATATATGCTCGTNNNAGATCGGAAG |
| 53177_2 | TATGAGTGTGGAGTCGTTGCTTCAGGGCTTGATATATGCTCGTNNNAGATCGGAAG |
| 53177_3 | TATGAGTGTGGAGTCGTTGCTTCAGGGCTTGATATATGCTCGTNNNAGATCGGAAG |
| 51267_1 | TATGAGTGTGGAGTCGTTGCCTGGACTTTGAGACTGCCATTTGNNNAGATCGGAAG |
| 51267_2 | TATGAGTGTGGAGTCGTTGCCTGGACTTTGAGACTGCCATTTGNNNAGATCGGAAG |
| 51267_3 | TATGAGTGTGGAGTCGTTGCCTGGACTTTGAGACTGCCATTTGNNNAGATCGGAAG |
| 11758_1 | TATGAGTGTGGAGTCGTTGCATCCCCCTGCATCCTTTTCTTTNNNAGATCGGAAGAG |
| 11758_2 | TATGAGTGTGGAGTCGTTGCATCCCCCTGCATCCTTTTCTTTNNNAGATCGGAAGAG |
| 11758_3 | TATGAGTGTGGAGTCGTTGCATCCCCCTGCATCCTTTTCTTTNNNAGATCGGAAGAG |
| 39568_1 | TATGAGTGTGGAGTCGTTGCTAATGGGATGTATGGAGGCATGGNNNAGATCGGA |
| 39568_2 | TATGAGTGTGGAGTCGTTGCTAATGGGATGTATGGAGGCATGGNNNAGATCGGA |
| 39568_3 | TATGAGTGTGGAGTCGTTGCTAATGGGATGTATGGAGGCATGGNNNAGATCGGA |
| 48739_1 | TATGAGTGTGGAGTCGTTGCCTCCACCTCCCGAATATCTAGGANNNAGATCGGAAG |
| 48739_2 | TATGAGTGTGGAGTCGTTGCCTCCACCTCCCGAATATCTAGGANNNAGATCGGAAG |
| 48739_3 | TATGAGTGTGGAGTCGTTGCCTCCACCTCCCGAATATCTAGGANNNAGATCGGAAG |
| 33909_1 | TATGAGTGTGGAGTCGTTGCGCCTCCCAGGTTCAAGTAATTCTNNNAGATCGGAAG |
| 33909_2 | TATGAGTGTGGAGTCGTTGCGCCTCCCAGGTTCAAGTAATTCTNNNAGATCGGAAG |
| 33909_3 | TATGAGTGTGGAGTCGTTGCGCCTCCCAGGTTCAAGTAATTCTNNNAGATCGGAAG |
| 39058_1 | TATGAGTGTGGAGTCGTTGCGTGGGCAAAGGACATAAACAGACNNNAGATCGGAA |
| 39058_2 | TATGAGTGTGGAGTCGTTGCGTGGGCAAAGGACATAAACAGACNNNAGATCGGAA |
| 39058_3 | TATGAGTGTGGAGTCGTTGCGTGGGCAAAGGACATAAACAGACNNNAGATCGGAA |
| 13073_1 | TATGAGTGTGGAGTCGTTGCCCTACAGGCTTTAAGTAGATAATNNNAGATCGGAAG |
| 13073_2 | TATGAGTGTGGAGTCGTTGCCCTACAGGCTTTAAGTAGATAATNNNAGATCGGAAG |
| 13073_3 | TATGAGTGTGGAGTCGTTGCCCTACAGGCTTTAAGTAGATAATNNNAGATCGGAAG |
| 50868_1 | TATGAGTGTGGAGTCGTTGCCACACATCAATGAGGCTCTTTGCNNNAGATCGGAAG |
| 50868_2 | TATGAGTGTGGAGTCGTTGCCACACATCAATGAGGCTCTTTGCNNNAGATCGGAAG |
| 50868_3 | TATGAGTGTGGAGTCGTTGCCACACATCAATGAGGCTCTTTGCNNNAGATCGGAAG |
| 29205_1 | TATGAGTGTGGAGTCGTTGCTGAGTACACAATGCTCCTGTTCANNNAGATCGGAAG |
| 29205_2 | TATGAGTGTGGAGTCGTTGCTGAGTACACAATGCTCCTGTTCANNNAGATCGGAAG |
| 29205_3 | TATGAGTGTGGAGTCGTTGCTGAGTACACAATGCTCCTGTTCANNNAGATCGGAAG |
| 46830_1 | TATGAGTGTGGAGTCGTTGCCCTGGGTCCATGTGATATGTGATNNNAGATCGGAA |
| 46830_2 | TATGAGTGTGGAGTCGTTGCCCTGGGTCCATGTGATATGTGATNNNAGATCGGAA |
| 46830_3 | TATGAGTGTGGAGTCGTTGCCCTGGGTCCATGTGATATGTGATNNNAGATCGGAA |
| 48349_1 | TATGAGTGTGGAGTCGTTGCCTCCTGGATTAGTGATACACCCGNNNAGATCGGAAG |
| 48349_2 | TATGAGTGTGGAGTCGTTGCCTCCTGGATTAGTGATACACCCGNNNAGATCGGAAG |
| 48349_3 | TATGAGTGTGGAGTCGTTGCCTCCTGGATTAGTGATACACCCGNNNAGATCGGAAG |
| 40170_1 | TATGAGTGTGGAGTCGTTGCTCTTTCTGTGGTTTTGTCAACCANNNAGATCGGAAG |
| 40170_2 | TATGAGTGTGGAGTCGTTGCTCTTTCTGTGGTTTTGTCAACCANNNAGATCGGAAG |
| 40170_3 | TATGAGTGTGGAGTCGTTGCTCTTTCTGTGGTTTTGTCAACCANNNAGATCGGAAG |
| 36570_1 | TATGAGTGTGGAGTCGTTGCGATTGTTTGAGTCCAGCAGTTCCNNNAGATCGGAAG |
| 36570_2 | TATGAGTGTGGAGTCGTTGCGATTGTTTGAGTCCAGCAGTTCCNNNAGATCGGAAG |
| 36570_3 | TATGAGTGTGGAGTCGTTGCGATTGTTTGAGTCCAGCAGTTCCNNNAGATCGGAAG |

| | |
|---|---|
| 49331_1 | TATGAGTGTGGAGTCGTTGCGTTCAAGCGATTCTCCTACCTCANNNAGATCGGAAG |
| 49331_2 | TATGAGTGTGGAGTCGTTGCGTTCAAGCGATTCTCCTACCTCANNNAGATCGGAAG |
| 49331_3 | TATGAGTGTGGAGTCGTTGCGTTCAAGCGATTCTCCTACCTCANNNAGATCGGAAG |
| 30056_1 | TATGAGTGTGGAGTCGTTGCCATCAATGCTGCTCTTATGGACGNNNAGATCGGAAG |
| 30056_2 | TATGAGTGTGGAGTCGTTGCCATCAATGCTGCTCTTATGGACGNNNAGATCGGAAG |
| 30056_3 | TATGAGTGTGGAGTCGTTGCCATCAATGCTGCTCTTATGGACGNNNAGATCGGAAG |
| 29560_1 | TATGAGTGTGGAGTCGTTGCCCAAATACCCCAAGGACTGTAGTNNNAGATCGGAAG |
| 29560_2 | TATGAGTGTGGAGTCGTTGCCCAAATACCCCAAGGACTGTAGTNNNAGATCGGAAG |
| 29560_3 | TATGAGTGTGGAGTCGTTGCCCAAATACCCCAAGGACTGTAGTNNNAGATCGGAAG |
| 46680_1 | TATGAGTGTGGAGTCGTTGCCAAGGTCTCACTCTGTCACTCAANNNAGATCGGAAG |
| 46680_2 | TATGAGTGTGGAGTCGTTGCCAAGGTCTCACTCTGTCACTCAANNNAGATCGGAAG |
| 46680_3 | TATGAGTGTGGAGTCGTTGCCAAGGTCTCACTCTGTCACTCAANNNAGATCGGAAG |
| 45757_1 | TATGAGTGTGGAGTCGTTGCTGACCTTCTCTGCTCAAGTGATCNNNAGATCGGAAG |
| 45757_2 | TATGAGTGTGGAGTCGTTGCTGACCTTCTCTGCTCAAGTGATCNNNAGATCGGAAG |
| 45757_3 | TATGAGTGTGGAGTCGTTGCTGACCTTCTCTGCTCAAGTGATCNNNAGATCGGAAG |
| 52321_1 | TATGAGTGTGGAGTCGTTGCGGACTAGACATTGTGTTCTGGGTNNNAGATCGGAA |
| 52321_2 | TATGAGTGTGGAGTCGTTGCGGACTAGACATTGTGTTCTGGGTNNNAGATCGGAA |
| 52321_3 | TATGAGTGTGGAGTCGTTGCGGACTAGACATTGTGTTCTGGGTNNNAGATCGGAA |
| 47204_1 | TATGAGTGTGGAGTCGTTGCTTCAAGTGATTCTGGTGCCTCAGNNNAGATCGGAAG |
| 47204_2 | TATGAGTGTGGAGTCGTTGCTTCAAGTGATTCTGGTGCCTCAGNNNAGATCGGAAG |
| 47204_3 | TATGAGTGTGGAGTCGTTGCTTCAAGTGATTCTGGTGCCTCAGNNNAGATCGGAAG |
| 13618_1 | TATGAGTGTGGAGTCGTTGCTTTATCACCATTTTACAAAGTGGNNNAGATCGGAAG |
| 13618_2 | TATGAGTGTGGAGTCGTTGCTTTATCACCATTTTACAAAGTGGNNNAGATCGGAAG |
| 13618_3 | TATGAGTGTGGAGTCGTTGCTTTATCACCATTTTACAAAGTGGNNNAGATCGGAAG |
| 40583_1 | TATGAGTGTGGAGTCGTTGCACACAAAAATTAGCTGGGCATGGNNNAGATCGGAA |
| 40583_2 | TATGAGTGTGGAGTCGTTGCACACAAAAATTAGCTGGGCATGGNNNAGATCGGAA |
| 40583_3 | TATGAGTGTGGAGTCGTTGCACACAAAAATTAGCTGGGCATGGNNNAGATCGGAA |
| 44943_1 | TATGAGTGTGGAGTCGTTGCGAGTGTGAGAGCTGAGAGGTTGNNNAGATCGGAA |
| 44943_2 | TATGAGTGTGGAGTCGTTGCGAGTGTGAGAGCTGAGAGGTTGNNNAGATCGGAA |
| 44943_3 | TATGAGTGTGGAGTCGTTGCGAGTGTGAGAGCTGAGAGGTTGNNNAGATCGGAA |
| 38612_1 | TATGAGTGTGGAGTCGTTGCACACAGAAATGCAGTATGGAGGTNNNAGATCGGAA |
| 38612_2 | TATGAGTGTGGAGTCGTTGCACACAGAAATGCAGTATGGAGGTNNNAGATCGGAA |
| 38612_3 | TATGAGTGTGGAGTCGTTGCACACAGAAATGCAGTATGGAGGTNNNAGATCGGAA |
| 36110_1 | TATGAGTGTGGAGTCGTTGCTGATCACACTACTGAACTCCAGCNNNAGATCGGAAG |
| 36110_2 | TATGAGTGTGGAGTCGTTGCTGATCACACTACTGAACTCCAGCNNNAGATCGGAAG |
| 36110_3 | TATGAGTGTGGAGTCGTTGCTGATCACACTACTGAACTCCAGCNNNAGATCGGAAG |
| 42175_1 | TATGAGTGTGGAGTCGTTGCCCACCACGCCTGTCTAATTTTTTNNNAGATCGGAAGA |
| 42175_2 | TATGAGTGTGGAGTCGTTGCCCACCACGCCTGTCTAATTTTTTNNNAGATCGGAAGA |
| 42175_3 | TATGAGTGTGGAGTCGTTGCCCACCACGCCTGTCTAATTTTTTNNNAGATCGGAAGA |
| 31124_1 | TATGAGTGTGGAGTCGTTGCTGTATTGCAAGCTTCCAAACAGANNNAGATCGGAAG |
| 31124_2 | TATGAGTGTGGAGTCGTTGCTGTATTGCAAGCTTCCAAACAGANNNAGATCGGAAG |
| 31124_3 | TATGAGTGTGGAGTCGTTGCTGTATTGCAAGCTTCCAAACAGANNNAGATCGGAAG |
| 39949_1 | TATGAGTGTGGAGTCGTTGCTATGTATTAGCTGGGCATGGTGGNNNAGATCGGAA |

| | |
|---|---|
| 39949_2 | TATGAGTGTGGAGTCGTTGCTATGTATTAGCTGGGCATGGTGGNNNAGATCGGAA |
| 39949_3 | TATGAGTGTGGAGTCGTTGCTATGTATTAGCTGGGCATGGTGGNNNAGATCGGAA |
| 45668_1 | TATGAGTGTGGAGTCGTTGCCCACCATGCCCAGCTAATTTTTANNNAGATCGGAAGA |
| 45668_2 | TATGAGTGTGGAGTCGTTGCCCACCATGCCCAGCTAATTTTTANNNAGATCGGAAGA |
| 45668_3 | TATGAGTGTGGAGTCGTTGCCCACCATGCCCAGCTAATTTTTANNNAGATCGGAAGA |
| 51262_1 | TATGAGTGTGGAGTCGTTGCAAAAAATTAGCTGGGTGTGGTGGNNNAGATCGGAA |
| 51262_2 | TATGAGTGTGGAGTCGTTGCAAAAAATTAGCTGGGTGTGGTGGNNNAGATCGGAA |
| 51262_3 | TATGAGTGTGGAGTCGTTGCAAAAAATTAGCTGGGTGTGGTGGNNNAGATCGGAA |
| 43952_1 | TATGAGTGTGGAGTCGTTGCGACACACACAGAGACAGAGAGAGNNNAGATCGGAA |
| 43952_2 | TATGAGTGTGGAGTCGTTGCGACACACACAGAGACAGAGAGAGNNNAGATCGGAA |
| 43952_3 | TATGAGTGTGGAGTCGTTGCGACACACACAGAGACAGAGAGAGNNNAGATCGGAA |
| 47083_1 | TATGAGTGTGGAGTCGTTGCTGGAATGCAGTGGTGTGATCTCNNNAGATCGGAAG |
| 47083_2 | TATGAGTGTGGAGTCGTTGCTGGAATGCAGTGGTGTGATCTCNNNAGATCGGAAG |
| 47083_3 | TATGAGTGTGGAGTCGTTGCTGGAATGCAGTGGTGTGATCTCNNNAGATCGGAAG |
| 34680_1 | TATGAGTGTGGAGTCGTTGCACAGTAATCCCCCCTTATCTCCANNNAGATCGGAAGA |
| 34680_2 | TATGAGTGTGGAGTCGTTGCACAGTAATCCCCCCTTATCTCCANNNAGATCGGAAGA |
| 34680_3 | TATGAGTGTGGAGTCGTTGCACAGTAATCCCCCCTTATCTCCANNNAGATCGGAAGA |
| 39764_1 | TATGAGTGTGGAGTCGTTGCAGTTTACATGCCACAGTTACAGTNNNAGATCGGAAG |
| 39764_2 | TATGAGTGTGGAGTCGTTGCAGTTTACATGCCACAGTTACAGTNNNAGATCGGAAG |
| 39764_3 | TATGAGTGTGGAGTCGTTGCAGTTTACATGCCACAGTTACAGTNNNAGATCGGAAG |
| 52405_1 | TATGAGTGTGGAGTCGTTGCAAGGAACTTGGAAGAAACCTGGTNNNAGATCGGAA |
| 52405_2 | TATGAGTGTGGAGTCGTTGCAAGGAACTTGGAAGAAACCTGGTNNNAGATCGGAA |
| 52405_3 | TATGAGTGTGGAGTCGTTGCAAGGAACTTGGAAGAAACCTGGTNNNAGATCGGAA |
| 38992_1 | TATGAGTGTGGAGTCGTTGCAAGATGTGGGCGTTTGCAAAATANNNAGATCGGAA |
| 38992_2 | TATGAGTGTGGAGTCGTTGCAAGATGTGGGCGTTTGCAAAATANNNAGATCGGAA |
| 38992_3 | TATGAGTGTGGAGTCGTTGCAAGATGTGGGCGTTTGCAAAATANNNAGATCGGAA |
| 42207_1 | TATGAGTGTGGAGTCGTTGCCTCCCAGGTTCAAGTGCTTCTCNNNAGATCGGAAGA |
| 42207_2 | TATGAGTGTGGAGTCGTTGCCTCCCAGGTTCAAGTGCTTCTCNNNAGATCGGAAGA |
| 42207_3 | TATGAGTGTGGAGTCGTTGCCTCCCAGGTTCAAGTGCTTCTCNNNAGATCGGAAGA |
| 47845_1 | TATGAGTGTGGAGTCGTTGCCAAGTAGCTGGGATTACAGGCATNNNAGATCGGAA |
| 47845_2 | TATGAGTGTGGAGTCGTTGCCAAGTAGCTGGGATTACAGGCATNNNAGATCGGAA |
| 47845_3 | TATGAGTGTGGAGTCGTTGCCAAGTAGCTGGGATTACAGGCATNNNAGATCGGAA |
| 41290_1 | TATGAGTGTGGAGTCGTTGCCTGTAGTCCCAGCTACTTGGAAGNNNAGATCGGAAG |
| 41290_2 | TATGAGTGTGGAGTCGTTGCCTGTAGTCCCAGCTACTTGGAAGNNNAGATCGGAAG |
| 41290_3 | TATGAGTGTGGAGTCGTTGCCTGTAGTCCCAGCTACTTGGAAGNNNAGATCGGAAG |
| 47713_1 | TATGAGTGTGGAGTCGTTGCCAGTAGCACAATCTCAGCTCACTNNNAGATCGGAAG |
| 47713_2 | TATGAGTGTGGAGTCGTTGCCAGTAGCACAATCTCAGCTCACTNNNAGATCGGAAG |
| 47713_3 | TATGAGTGTGGAGTCGTTGCCAGTAGCACAATCTCAGCTCACTNNNAGATCGGAAG |
| 44705_1 | TATGAGTGTGGAGTCGTTGCTCGGGATAAAAAAAAATGGGGGCNNNAGATCGGA |
| 44705_2 | TATGAGTGTGGAGTCGTTGCTCGGGATAAAAAAAAATGGGGGCNNNAGATCGGA |
| 44705_3 | TATGAGTGTGGAGTCGTTGCTCGGGATAAAAAAAAATGGGGGCNNNAGATCGGA |
| 46061_1 | TATGAGTGTGGAGTCGTTGCGAGGTCTCACTACATTGTCCAGGNNNAGATCGGAAG |
| 46061_2 | TATGAGTGTGGAGTCGTTGCGAGGTCTCACTACATTGTCCAGGNNNAGATCGGAAG |

| | |
|---|---|
| 46061_3 | TATGAGTGTGGAGTCGTTGCGAGGTCTCACTACATTGTCCAGGNNNAGATCGGAAG |
| 34170_1 | TATGAGTGTGGAGTCGTTGCCTAATGTGCGAGGATTGCTTGAGNNNAGATCGGAA |
| 34170_2 | TATGAGTGTGGAGTCGTTGCCTAATGTGCGAGGATTGCTTGAGNNNAGATCGGAA |
| 34170_3 | TATGAGTGTGGAGTCGTTGCCTAATGTGCGAGGATTGCTTGAGNNNAGATCGGAA |
| 48619_1 | TATGAGTGTGGAGTCGTTGCCACCCTAGAGATGCATGTTCTGTNNNAGATCGGAAG |
| 48619_2 | TATGAGTGTGGAGTCGTTGCCACCCTAGAGATGCATGTTCTGTNNNAGATCGGAAG |
| 48619_3 | TATGAGTGTGGAGTCGTTGCCACCCTAGAGATGCATGTTCTGTNNNAGATCGGAAG |
| 36833_1 | TATGAGTGTGGAGTCGTTGCGTCCTCCCAAGAAGGAAGCAATANNNAGATCGGAA |
| 36833_2 | TATGAGTGTGGAGTCGTTGCGTCCTCCCAAGAAGGAAGCAATANNNAGATCGGAA |
| 36833_3 | TATGAGTGTGGAGTCGTTGCGTCCTCCCAAGAAGGAAGCAATANNNAGATCGGAA |
| 42924_1 | TATGAGTGTGGAGTCGTTGCCAGACAGGTTCTCAGAAGACCAGNNNAGATCGGAA |
| 42924_2 | TATGAGTGTGGAGTCGTTGCCAGACAGGTTCTCAGAAGACCAGNNNAGATCGGAA |
| 42924_3 | TATGAGTGTGGAGTCGTTGCCAGACAGGTTCTCAGAAGACCAGNNNAGATCGGAA |
| 29595_1 | TATGAGTGTGGAGTCGTTGCAGAAAATGGCAGATTTCTGGCAGNNNAGATCGGAA |
| 29595_2 | TATGAGTGTGGAGTCGTTGCAGAAAATGGCAGATTTCTGGCAGNNNAGATCGGAA |
| 29595_3 | TATGAGTGTGGAGTCGTTGCAGAAAATGGCAGATTTCTGGCAGNNNAGATCGGAA |
| 34848_1 | TATGAGTGTGGAGTCGTTGCGAGAGAGAGTCTCACTGTGTTGCNNNAGATCGGAA |
| 34848_2 | TATGAGTGTGGAGTCGTTGCGAGAGAGAGTCTCACTGTGTTGCNNNAGATCGGAA |
| 34848_3 | TATGAGTGTGGAGTCGTTGCGAGAGAGAGTCTCACTGTGTTGCNNNAGATCGGAA |
| 32495_1 | TATGAGTGTGGAGTCGTTGCCTCACTGCAACCTCCACCTCNNNAGATCGGAAGAGCA |
| 32495_2 | TATGAGTGTGGAGTCGTTGCCTCACTGCAACCTCCACCTCNNNAGATCGGAAGAGCA |
| 32495_3 | TATGAGTGTGGAGTCGTTGCCTCACTGCAACCTCCACCTCNNNAGATCGGAAGAGCA |
| 50721_1 | TATGAGTGTGGAGTCGTTGCCAATCCTCTCCCGTCCTCTTCNNNAGATCGGAAGAGC |
| 50721_2 | TATGAGTGTGGAGTCGTTGCCAATCCTCTCCCGTCCTCTTCNNNAGATCGGAAGAGC |
| 50721_3 | TATGAGTGTGGAGTCGTTGCCAATCCTCTCCCGTCCTCTTCNNNAGATCGGAAGAGC |
| 42797_1 | TATGAGTGTGGAGTCGTTGCGGGATACAGCTAACTAGTGAGGTGNNNAGATCGGA |
| 42797_2 | TATGAGTGTGGAGTCGTTGCGGGATACAGCTAACTAGTGAGGTGNNNAGATCGGA |
| 42797_3 | TATGAGTGTGGAGTCGTTGCGGGATACAGCTAACTAGTGAGGTGNNNAGATCGGA |
| 46811_1 | TATGAGTGTGGAGTCGTTGCTTCAAGCGATTTTTCTGCCTCAGNNNAGATCGGAAG |
| 46811_2 | TATGAGTGTGGAGTCGTTGCTTCAAGCGATTTTTCTGCCTCAGNNNAGATCGGAAG |
| 46811_3 | TATGAGTGTGGAGTCGTTGCTTCAAGCGATTTTTCTGCCTCAGNNNAGATCGGAAG |
| 30630_1 | TATGAGTGTGGAGTCGTTGCGTCCAATGAAATTTGCCTTGCTAGGNNNAGATCGGA |
| 30630_2 | TATGAGTGTGGAGTCGTTGCGTCCAATGAAATTTGCCTTGCTAGGNNNAGATCGGA |
| 30630_3 | TATGAGTGTGGAGTCGTTGCGTCCAATGAAATTTGCCTTGCTAGGNNNAGATCGGA |
| 48884_1 | TATGAGTGTGGAGTCGTTGCCGGAGTTTTCACTCTTATTGCCCNNNAGATCGGAAG |
| 48884_2 | TATGAGTGTGGAGTCGTTGCCGGAGTTTTCACTCTTATTGCCCNNNAGATCGGAAG |
| 48884_3 | TATGAGTGTGGAGTCGTTGCCGGAGTTTTCACTCTTATTGCCCNNNAGATCGGAAG |
| 37782_1 | TATGAGTGTGGAGTCGTTGCTGGACGTCATCATTCACAAGTTGNNNAGATCGGAAG |
| 37782_2 | TATGAGTGTGGAGTCGTTGCTGGACGTCATCATTCACAAGTTGNNNAGATCGGAAG |
| 37782_3 | TATGAGTGTGGAGTCGTTGCTGGACGTCATCATTCACAAGTTGNNNAGATCGGAAG |
| 43147_1 | TATGAGTGTGGAGTCGTTGCGCAGATGACACAGAAGTGGTTTCNNNAGATCGGAA |
| 43147_2 | TATGAGTGTGGAGTCGTTGCGCAGATGACACAGAAGTGGTTTCNNNAGATCGGAA |
| 43147_3 | TATGAGTGTGGAGTCGTTGCGCAGATGACACAGAAGTGGTTTCNNNAGATCGGAA |

| | |
|---|---|
| 10543_1 | TATGAGTGTGGAGTCGTTGCTTCAAGCACAGTAGCTTTGGTTTNNNAGATCGGAAG |
| 10543_2 | TATGAGTGTGGAGTCGTTGCTTCAAGCACAGTAGCTTTGGTTTNNNAGATCGGAAG |
| 10543_3 | TATGAGTGTGGAGTCGTTGCTTCAAGCACAGTAGCTTTGGTTTNNNAGATCGGAAG |
| 10547_1 | TATGAGTGTGGAGTCGTTGCGCATGGGTGTCTGTCTCTCTTTTNNNAGATCGGAAG |
| 10547_2 | TATGAGTGTGGAGTCGTTGCGCATGGGTGTCTGTCTCTCTTTTNNNAGATCGGAAG |
| 10547_3 | TATGAGTGTGGAGTCGTTGCGCATGGGTGTCTGTCTCTCTTTTNNNAGATCGGAAG |
| 32607_1 | TATGAGTGTGGAGTCGTTGCCCAAAAAAACCCTGCATGCTACTNNNAGATCGGAAG |
| 32607_2 | TATGAGTGTGGAGTCGTTGCCCAAAAAAACCCTGCATGCTACTNNNAGATCGGAAG |
| 32607_3 | TATGAGTGTGGAGTCGTTGCCCAAAAAAACCCTGCATGCTACTNNNAGATCGGAAG |
| 30419_1 | TATGAGTGTGGAGTCGTTGCATAATGGCAATCTAGTGGCCTGGNNNAGATCGGAA |
| 30419_2 | TATGAGTGTGGAGTCGTTGCATAATGGCAATCTAGTGGCCTGGNNNAGATCGGAA |
| 30419_3 | TATGAGTGTGGAGTCGTTGCATAATGGCAATCTAGTGGCCTGGNNNAGATCGGAA |
| 3135_1 | TATGAGTGTGGAGTCGTTGCGTCAAGGCCATGGGCAACTGNNNAGATCGGAAGAG |
| 3135_2 | TATGAGTGTGGAGTCGTTGCGTCAAGGCCATGGGCAACTGNNNAGATCGGAAGAG |
| 3135_3 | TATGAGTGTGGAGTCGTTGCGTCAAGGCCATGGGCAACTGNNNAGATCGGAAGAG |
| 44677_1 | TATGAGTGTGGAGTCGTTGCTTTGGGCAAAAGTTAGATGGGTGNNNAGATCGGAA |
| 44677_2 | TATGAGTGTGGAGTCGTTGCTTTGGGCAAAAGTTAGATGGGTGNNNAGATCGGAA |
| 44677_3 | TATGAGTGTGGAGTCGTTGCTTTGGGCAAAAGTTAGATGGGTGNNNAGATCGGAA |
| 13327_1 | TATGAGTGTGGAGTCGTTGCCACGTCTGGTTAATTTTTGTATNNNAGATCGGAAGA |
| 13327_2 | TATGAGTGTGGAGTCGTTGCCACGTCTGGTTAATTTTTGTATNNNAGATCGGAAGA |
| 13327_3 | TATGAGTGTGGAGTCGTTGCCACGTCTGGTTAATTTTTGTATNNNAGATCGGAAGA |
| 46554_1 | TATGAGTGTGGAGTCGTTGCATCTCCTGGGTTCAAGCAATTCTNNNAGATCGGAAG |
| 46554_2 | TATGAGTGTGGAGTCGTTGCATCTCCTGGGTTCAAGCAATTCTNNNAGATCGGAAG |
| 46554_3 | TATGAGTGTGGAGTCGTTGCATCTCCTGGGTTCAAGCAATTCTNNNAGATCGGAAG |
| 45849_1 | TATGAGTGTGGAGTCGTTGCATTCAAGTGATTCTCCTGCCTCANNNAGATCGGAAG |
| 45849_2 | TATGAGTGTGGAGTCGTTGCATTCAAGTGATTCTCCTGCCTCANNNAGATCGGAAG |
| 45849_3 | TATGAGTGTGGAGTCGTTGCATTCAAGTGATTCTCCTGCCTCANNNAGATCGGAAG |
| 30508_1 | TATGAGTGTGGAGTCGTTGCACACACACACAGAGTTTCACTATGTNNNAGATCGGA |
| 30508_2 | TATGAGTGTGGAGTCGTTGCACACACACACAGAGTTTCACTATGTNNNAGATCGGA |
| 30508_3 | TATGAGTGTGGAGTCGTTGCACACACACACAGAGTTTCACTATGTNNNAGATCGGA |
| 40764_1 | TATGAGTGTGGAGTCGTTGCCGCAGGGACTCATACCTCTAATCNNNAGATCGGAAG |
| 40764_2 | TATGAGTGTGGAGTCGTTGCCGCAGGGACTCATACCTCTAATCNNNAGATCGGAAG |
| 40764_3 | TATGAGTGTGGAGTCGTTGCCGCAGGGACTCATACCTCTAATCNNNAGATCGGAAG |
| 35508_1 | TATGAGTGTGGAGTCGTTGCTGATCTCGATCTCCTGACCTAGTNNNAGATCGGAAG |
| 35508_2 | TATGAGTGTGGAGTCGTTGCTGATCTCGATCTCCTGACCTAGTNNNAGATCGGAAG |
| 35508_3 | TATGAGTGTGGAGTCGTTGCTGATCTCGATCTCCTGACCTAGTNNNAGATCGGAAG |
| 5327_1 | TATGAGTGTGGAGTCGTTGCAAAGAGGCCAGTCTCAAGCAAATNNNAGATCGGAA |
| 5327_2 | TATGAGTGTGGAGTCGTTGCAAAGAGGCCAGTCTCAAGCAAATNNNAGATCGGAA |
| 5327_3 | TATGAGTGTGGAGTCGTTGCAAAGAGGCCAGTCTCAAGCAAATNNNAGATCGGAA |
| 48313_1 | TATGAGTGTGGAGTCGTTGCAGATGGTGTTTTGCTCTTGTTGCNNNAGATCGGAAG |
| 48313_2 | TATGAGTGTGGAGTCGTTGCAGATGGTGTTTTGCTCTTGTTGCNNNAGATCGGAAG |
| 48313_3 | TATGAGTGTGGAGTCGTTGCAGATGGTGTTTTGCTCTTGTTGCNNNAGATCGGAAG |
| 34317_1 | TATGAGTGTGGAGTCGTTGCGCCATCATTCCTAGTTGCTGTTCNNNAGATCGGAAG |

| | |
|---|---|
| 34317_2 | TATGAGTGTGGAGTCGTTGCGCCATCATTCCTAGTTGCTGTTCNNNAGATCGGAAG[illegible] |
| 34317_3 | TATGAGTGTGGAGTCGTTGCGCCATCATTCCTAGTTGCTGTTCNNNAGATCGGAAG[illegible] |
| 31457_1 | TATGAGTGTGGAGTCGTTGCACATGATTTGGTAGGCTTTTCTCANNNAGATCGGAA[illegible] |
| 31457_2 | TATGAGTGTGGAGTCGTTGCACATGATTTGGTAGGCTTTTCTCANNNAGATCGGAA[illegible] |
| 31457_3 | TATGAGTGTGGAGTCGTTGCACATGATTTGGTAGGCTTTTCTCANNNAGATCGGAA[illegible] |
| 33306_1 | TATGAGTGTGGAGTCGTTGCCCTCCCAGGTTCAAGTGATTCTCNNNAGATCGGAAG[illegible] |
| 33306_2 | TATGAGTGTGGAGTCGTTGCCCTCCCAGGTTCAAGTGATTCTCNNNAGATCGGAAG[illegible] |
| 33306_3 | TATGAGTGTGGAGTCGTTGCCCTCCCAGGTTCAAGTGATTCTCNNNAGATCGGAAG[illegible] |
| 49268_1 | TATGAGTGTGGAGTCGTTGCCCTGGGTTCAAGCAATTCTTCTGNNNAGATCGGAAG[illegible] |
| 49268_2 | TATGAGTGTGGAGTCGTTGCCCTGGGTTCAAGCAATTCTTCTGNNNAGATCGGAAG[illegible] |
| 49268_3 | TATGAGTGTGGAGTCGTTGCCCTGGGTTCAAGCAATTCTTCTGNNNAGATCGGAAG[illegible] |
| 32614_1 | TATGAGTGTGGAGTCGTTGCCCTGAGACCCTCATTAGAAGCTGNNNAGATCGGAAG |
| 32614_2 | TATGAGTGTGGAGTCGTTGCCCTGAGACCCTCATTAGAAGCTGNNNAGATCGGAAG |
| 32614_3 | TATGAGTGTGGAGTCGTTGCCCTGAGACCCTCATTAGAAGCTGNNNAGATCGGAAG |
| 49280_1 | TATGAGTGTGGAGTCGTTGCACCACACCCAGCTAGTTTTTGTANNNAGATCGGAAG[illegible] |
| 49280_2 | TATGAGTGTGGAGTCGTTGCACCACACCCAGCTAGTTTTTGTANNNAGATCGGAAG[illegible] |
| 49280_3 | TATGAGTGTGGAGTCGTTGCACCACACCCAGCTAGTTTTTGTANNNAGATCGGAAG[illegible] |
| 48283_1 | TATGAGTGTGGAGTCGTTGCCAAGTAGCTGGGACTACAGGTGNNNAGATCGGAAG |
| 48283_2 | TATGAGTGTGGAGTCGTTGCCAAGTAGCTGGGACTACAGGTGNNNAGATCGGAAG |
| 48283_3 | TATGAGTGTGGAGTCGTTGCCAAGTAGCTGGGACTACAGGTGNNNAGATCGGAAG |
| 51005_1 | TATGAGTGTGGAGTCGTTGCAGGTGTAATTGGCTCATGGTTCTNNNAGATCGGAA[illegible] |
| 51005_2 | TATGAGTGTGGAGTCGTTGCAGGTGTAATTGGCTCATGGTTCTNNNAGATCGGAA[illegible] |
| 51005_3 | TATGAGTGTGGAGTCGTTGCAGGTGTAATTGGCTCATGGTTCTNNNAGATCGGAA[illegible] |
| 41325_1 | TATGAGTGTGGAGTCGTTGCTTTAGCAACCACCACTCTGATCANNNAGATCGGAAG[illegible] |
| 41325_2 | TATGAGTGTGGAGTCGTTGCTTTAGCAACCACCACTCTGATCANNNAGATCGGAAG[illegible] |
| 41325_3 | TATGAGTGTGGAGTCGTTGCTTTAGCAACCACCACTCTGATCANNNAGATCGGAAG[illegible] |
| 34900_1 | TATGAGTGTGGAGTCGTTGCCCAAAGTGCTGGGATTACAAGTGNNNAGATCGGAA[illegible] |
| 34900_2 | TATGAGTGTGGAGTCGTTGCCCAAAGTGCTGGGATTACAAGTGNNNAGATCGGAA[illegible] |
| 34900_3 | TATGAGTGTGGAGTCGTTGCCCAAAGTGCTGGGATTACAAGTGNNNAGATCGGAA[illegible] |
| 29684_1 | TATGAGTGTGGAGTCGTTGCAGGAAGGGGTGAAGAGGAATAGTNNNAGATCGGA |
| 29684_2 | TATGAGTGTGGAGTCGTTGCAGGAAGGGGTGAAGAGGAATAGTNNNAGATCGGA |
| 29684_3 | TATGAGTGTGGAGTCGTTGCAGGAAGGGGTGAAGAGGAATAGTNNNAGATCGGA |
| 40945_1 | TATGAGTGTGGAGTCGTTGCTGGAAAACTGCCAGAGAAAGAGANNNAGATCGGA[illegible] |
| 40945_2 | TATGAGTGTGGAGTCGTTGCTGGAAAACTGCCAGAGAAAGAGANNNAGATCGGA[illegible] |
| 40945_3 | TATGAGTGTGGAGTCGTTGCTGGAAAACTGCCAGAGAAAGAGANNNAGATCGGA[illegible] |
| 39538_1 | TATGAGTGTGGAGTCGTTGCAGGAGTCTGAGACAGGAGGATCNNNAGATCGGAA[illegible] |
| 39538_2 | TATGAGTGTGGAGTCGTTGCAGGAGTCTGAGACAGGAGGATCNNNAGATCGGAA[illegible] |
| 39538_3 | TATGAGTGTGGAGTCGTTGCAGGAGTCTGAGACAGGAGGATCNNNAGATCGGAA[illegible] |
| 53242_1 | TATGAGTGTGGAGTCGTTGCATACAGCTAAACAGGCTTCCCTCNNNAGATCGGAAG[illegible] |
| 53242_2 | TATGAGTGTGGAGTCGTTGCATACAGCTAAACAGGCTTCCCTCNNNAGATCGGAAG[illegible] |
| 53242_3 | TATGAGTGTGGAGTCGTTGCATACAGCTAAACAGGCTTCCCTCNNNAGATCGGAAG[illegible] |
| 39648_1 | TATGAGTGTGGAGTCGTTGCACTGCATGGTTTCACTTACATGTNNNAGATCGGAAG[illegible] |
| 39648_2 | TATGAGTGTGGAGTCGTTGCACTGCATGGTTTCACTTACATGTNNNAGATCGGAAG[illegible] |

| | |
|---|---|
| 39648_3 | TATGAGTGTGGAGTCGTTGCACTGCATGGTTTCACTTACATGTNNNAGATCGGAAG |
| 51866_1 | TATGAGTGTGGAGTCGTTGCCACTGTTTTCGGACCTTCCATGNNNAGATCGGAAGA |
| 51866_2 | TATGAGTGTGGAGTCGTTGCCACTGTTTTCGGACCTTCCATGNNNAGATCGGAAGA |
| 51866_3 | TATGAGTGTGGAGTCGTTGCCACTGTTTTCGGACCTTCCATGNNNAGATCGGAAGA |
| 2644_1 | TATGAGTGTGGAGTCGTTGCTCTTCTACTCCCACACCTCCTTGNNNAGATCGGAAGA |
| 2644_2 | TATGAGTGTGGAGTCGTTGCTCTTCTACTCCCACACCTCCTTGNNNAGATCGGAAGA |
| 2644_3 | TATGAGTGTGGAGTCGTTGCTCTTCTACTCCCACACCTCCTTGNNNAGATCGGAAGA |
| 49318_1 | TATGAGTGTGGAGTCGTTGCCTCGGGTTCAAGCAATTCTTCTGNNNAGATCGGAAG |
| 49318_2 | TATGAGTGTGGAGTCGTTGCCTCGGGTTCAAGCAATTCTTCTGNNNAGATCGGAAG |
| 49318_3 | TATGAGTGTGGAGTCGTTGCCTCGGGTTCAAGCAATTCTTCTGNNNAGATCGGAAG |
| 363_1 | TATGAGTGTGGAGTCGTTGCCCCACATCAAATCAAACCCCCTNNNAGATCGGAAGAG |
| 363_2 | TATGAGTGTGGAGTCGTTGCCCCACATCAAATCAAACCCCCTNNNAGATCGGAAGAG |
| 363_3 | TATGAGTGTGGAGTCGTTGCCCCACATCAAATCAAACCCCCTNNNAGATCGGAAGAG |
| 13407_1 | TATGAGTGTGGAGTCGTTGCTGCCTTGTTTTTAAGTAATACAANNNAGATCGGAAG |
| 13407_2 | TATGAGTGTGGAGTCGTTGCTGCCTTGTTTTTAAGTAATACAANNNAGATCGGAAG |
| 13407_3 | TATGAGTGTGGAGTCGTTGCTGCCTTGTTTTTAAGTAATACAANNNAGATCGGAAG |
| 41404_1 | TATGAGTGTGGAGTCGTTGCTGAAGTCGATGTGTTGACATGTCNNNAGATCGGAA |
| 41404_2 | TATGAGTGTGGAGTCGTTGCTGAAGTCGATGTGTTGACATGTCNNNAGATCGGAA |
| 41404_3 | TATGAGTGTGGAGTCGTTGCTGAAGTCGATGTGTTGACATGTCNNNAGATCGGAA |
| 34758_1 | TATGAGTGTGGAGTCGTTGCGAGTAGCTGGGATTACAGGAGTGNNNAGATCGGA |
| 34758_2 | TATGAGTGTGGAGTCGTTGCGAGTAGCTGGGATTACAGGAGTGNNNAGATCGGA |
| 34758_3 | TATGAGTGTGGAGTCGTTGCGAGTAGCTGGGATTACAGGAGTGNNNAGATCGGA |
| 51889_1 | TATGAGTGTGGAGTCGTTGCCTCTTCTCTCAAGTACACCCACCNNNAGATCGGAAGA |
| 51889_2 | TATGAGTGTGGAGTCGTTGCCTCTTCTCTCAAGTACACCCACCNNNAGATCGGAAGA |
| 51889_3 | TATGAGTGTGGAGTCGTTGCCTCTTCTCTCAAGTACACCCACCNNNAGATCGGAAGA |
| 50507_1 | TATGAGTGTGGAGTCGTTGCTTCGTTTCCATGACAACTGCATCNNNAGATCGGAAG |
| 50507_2 | TATGAGTGTGGAGTCGTTGCTTCGTTTCCATGACAACTGCATCNNNAGATCGGAAG |
| 50507_3 | TATGAGTGTGGAGTCGTTGCTTCGTTTCCATGACAACTGCATCNNNAGATCGGAAG |
| 38315_1 | TATGAGTGTGGAGTCGTTGCTCCGGAAGCTTTGGACTATGTTANNNAGATCGGAAG |
| 38315_2 | TATGAGTGTGGAGTCGTTGCTCCGGAAGCTTTGGACTATGTTANNNAGATCGGAAG |
| 38315_3 | TATGAGTGTGGAGTCGTTGCTCCGGAAGCTTTGGACTATGTTANNNAGATCGGAAG |
| 36314_1 | TATGAGTGTGGAGTCGTTGCATGAGGCATTCAGAAAAGGTCCTNNNAGATCGGAA |
| 36314_2 | TATGAGTGTGGAGTCGTTGCATGAGGCATTCAGAAAAGGTCCTNNNAGATCGGAA |
| 36314_3 | TATGAGTGTGGAGTCGTTGCATGAGGCATTCAGAAAAGGTCCTNNNAGATCGGAA |
| 33545_1 | TATGAGTGTGGAGTCGTTGCGGATACACAGTCCCACTTTGTCANNNAGATCGGAAG |
| 33545_2 | TATGAGTGTGGAGTCGTTGCGGATACACAGTCCCACTTTGTCANNNAGATCGGAAG |
| 33545_3 | TATGAGTGTGGAGTCGTTGCGGATACACAGTCCCACTTTGTCANNNAGATCGGAAG |
| 10355_1 | TATGAGTGTGGAGTCGTTGCGCAGGCTGGACAACCAAATCTTANNNAGATCGGAAG |
| 10355_2 | TATGAGTGTGGAGTCGTTGCGCAGGCTGGACAACCAAATCTTANNNAGATCGGAAG |
| 10355_3 | TATGAGTGTGGAGTCGTTGCGCAGGCTGGACAACCAAATCTTANNNAGATCGGAAG |
| 36905_1 | TATGAGTGTGGAGTCGTTGCGTCCCAGGCACTATGGTAAGTAGNNNAGATCGGAA |
| 36905_2 | TATGAGTGTGGAGTCGTTGCGTCCCAGGCACTATGGTAAGTAGNNNAGATCGGAA |
| 36905_3 | TATGAGTGTGGAGTCGTTGCGTCCCAGGCACTATGGTAAGTAGNNNAGATCGGAA |

| | |
|---|---|
| 52991_1 | TATGAGTGTGGAGTCGTTGCTTGGGGGTATGTTGATACAAGCANNNAGATCGGAA |
| 52991_2 | TATGAGTGTGGAGTCGTTGCTTGGGGGTATGTTGATACAAGCANNNAGATCGGAA |
| 52991_3 | TATGAGTGTGGAGTCGTTGCTTGGGGGTATGTTGATACAAGCANNNAGATCGGAA |
| 12815_1 | TATGAGTGTGGAGTCGTTGCCATGTTCTCTGTGTATTATTACANNNAGATCGGAAG |
| 12815_2 | TATGAGTGTGGAGTCGTTGCCATGTTCTCTGTGTATTATTACANNNAGATCGGAAG |
| 12815_3 | TATGAGTGTGGAGTCGTTGCCATGTTCTCTGTGTATTATTACANNNAGATCGGAAG |
| 32133_1 | TATGAGTGTGGAGTCGTTGCACGCCTGTAATCCCAGAACTTTNNNAGATCGGAAGA |
| 32133_2 | TATGAGTGTGGAGTCGTTGCACGCCTGTAATCCCAGAACTTTNNNAGATCGGAAGA |
| 32133_3 | TATGAGTGTGGAGTCGTTGCACGCCTGTAATCCCAGAACTTTNNNAGATCGGAAGA |
| 49139_1 | TATGAGTGTGGAGTCGTTGCCTCCCGAGTAGCTGGAATAACAGNNNAGATCGGAA |
| 49139_2 | TATGAGTGTGGAGTCGTTGCCTCCCGAGTAGCTGGAATAACAGNNNAGATCGGAA |
| 49139_3 | TATGAGTGTGGAGTCGTTGCCTCCCGAGTAGCTGGAATAACAGNNNAGATCGGAA |
| 30606_1 | TATGAGTGTGGAGTCGTTGCGCTTCCAGTGTGGATTGCACAAGNNNAGATCGGAA |
| 30606_2 | TATGAGTGTGGAGTCGTTGCGCTTCCAGTGTGGATTGCACAAGNNNAGATCGGAA |
| 30606_3 | TATGAGTGTGGAGTCGTTGCGCTTCCAGTGTGGATTGCACAAGNNNAGATCGGAA |
| 38522_1 | TATGAGTGTGGAGTCGTTGCAATAAAACTCCGGTCTCCTCCACNNNAGATCGGAAG |
| 38522_2 | TATGAGTGTGGAGTCGTTGCAATAAAACTCCGGTCTCCTCCACNNNAGATCGGAAG |
| 38522_3 | TATGAGTGTGGAGTCGTTGCAATAAAACTCCGGTCTCCTCCACNNNAGATCGGAAG |
| 43228_1 | TATGAGTGTGGAGTCGTTGCTAGGTAGGTAGGCAGGTAGATGGNNNAGATCGGA |
| 43228_2 | TATGAGTGTGGAGTCGTTGCTAGGTAGGTAGGCAGGTAGATGGNNNAGATCGGA |
| 43228_3 | TATGAGTGTGGAGTCGTTGCTAGGTAGGTAGGCAGGTAGATGGNNNAGATCGGA |
| 41864_1 | TATGAGTGTGGAGTCGTTGCGTTCACGCCATTCTCCTACCTCNNNAGATCGGAAGAG |
| 41864_2 | TATGAGTGTGGAGTCGTTGCGTTCACGCCATTCTCCTACCTCNNNAGATCGGAAGAG |
| 41864_3 | TATGAGTGTGGAGTCGTTGCGTTCACGCCATTCTCCTACCTCNNNAGATCGGAAGAG |
| 44680_1 | TATGAGTGTGGAGTCGTTGCGTGGATCGGAGGGAAGGGTGNNNAGATCGGAAGA |
| 44680_2 | TATGAGTGTGGAGTCGTTGCGTGGATCGGAGGGAAGGGTGNNNAGATCGGAAGA |
| 44680_3 | TATGAGTGTGGAGTCGTTGCGTGGATCGGAGGGAAGGGTGNNNAGATCGGAAGA |
| 49029_1 | TATGAGTGTGGAGTCGTTGCCCTTCTGGGTTTACGCCATTCTCNNNAGATCGGAAGA |
| 49029_2 | TATGAGTGTGGAGTCGTTGCCCTTCTGGGTTTACGCCATTCTCNNNAGATCGGAAGA |
| 49029_3 | TATGAGTGTGGAGTCGTTGCCCTTCTGGGTTTACGCCATTCTCNNNAGATCGGAAGA |
| 41422_1 | TATGAGTGTGGAGTCGTTGCCTCGCCCGGCTAATTTTTTGTATNNNAGATCGGAAGA |
| 41422_2 | TATGAGTGTGGAGTCGTTGCCTCGCCCGGCTAATTTTTTGTATNNNAGATCGGAAGA |
| 41422_3 | TATGAGTGTGGAGTCGTTGCCTCGCCCGGCTAATTTTTTGTATNNNAGATCGGAAGA |
| 40071_1 | TATGAGTGTGGAGTCGTTGCCCACTGAATGTTTTATCTGTTTGCCNNNAGATCGGAA |
| 40071_2 | TATGAGTGTGGAGTCGTTGCCCACTGAATGTTTTATCTGTTTGCCNNNAGATCGGAA |
| 40071_3 | TATGAGTGTGGAGTCGTTGCCCACTGAATGTTTTATCTGTTTGCCNNNAGATCGGAA |
| 10611_1 | TATGAGTGTGGAGTCGTTGCATTTCTGTGCTGCTGTAGATGCTNNNAGATCGGAAG |
| 10611_2 | TATGAGTGTGGAGTCGTTGCATTTCTGTGCTGCTGTAGATGCTNNNAGATCGGAAG |
| 10611_3 | TATGAGTGTGGAGTCGTTGCATTTCTGTGCTGCTGTAGATGCTNNNAGATCGGAAG |
| 42887_1 | TATGAGTGTGGAGTCGTTGCCCTGAACTCAATTGATTCACCCGNNNAGATCGGAAG |
| 42887_2 | TATGAGTGTGGAGTCGTTGCCCTGAACTCAATTGATTCACCCGNNNAGATCGGAAG |
| 42887_3 | TATGAGTGTGGAGTCGTTGCCCTGAACTCAATTGATTCACCCGNNNAGATCGGAAG |
| 36743_1 | TATGAGTGTGGAGTCGTTGCCTCATGCCTGTAATCCCAGCTACNNNAGATCGGAAG |

| | |
|---|---|
| 36743_2 | TATGAGTGTGGAGTCGTTGCCTCATGCCTGTAATCCCAGCTACNNNAGATCGGAAGA |
| 36743_3 | TATGAGTGTGGAGTCGTTGCCTCATGCCTGTAATCCCAGCTACNNNAGATCGGAAGA |
| 34024_1 | TATGAGTGTGGAGTCGTTGCCAATGGCATGATTTCAGCTCACTNNNAGATCGGAAG |
| 34024_2 | TATGAGTGTGGAGTCGTTGCCAATGGCATGATTTCAGCTCACTNNNAGATCGGAAG |
| 34024_3 | TATGAGTGTGGAGTCGTTGCCAATGGCATGATTTCAGCTCACTNNNAGATCGGAAG |
| 33236_1 | TATGAGTGTGGAGTCGTTGCGCCCAAAACTAGACCAAGAACACNNNAGATCGGAAG |
| 33236_2 | TATGAGTGTGGAGTCGTTGCGCCCAAAACTAGACCAAGAACACNNNAGATCGGAAG |
| 33236_3 | TATGAGTGTGGAGTCGTTGCGCCCAAAACTAGACCAAGAACACNNNAGATCGGAAG |
| 49323_1 | TATGAGTGTGGAGTCGTTGCCCCGAATACCTAGGATTACAGGCNNNAGATCGGAAG |
| 49323_2 | TATGAGTGTGGAGTCGTTGCCCCGAATACCTAGGATTACAGGCNNNAGATCGGAAG |
| 49323_3 | TATGAGTGTGGAGTCGTTGCCCCGAATACCTAGGATTACAGGCNNNAGATCGGAAG |
| 40634_1 | TATGAGTGTGGAGTCGTTGCTGGAACAGACCAACCAAAGAGAANNNAGATCGGAA |
| 40634_2 | TATGAGTGTGGAGTCGTTGCTGGAACAGACCAACCAAAGAGAANNNAGATCGGAA |
| 40634_3 | TATGAGTGTGGAGTCGTTGCTGGAACAGACCAACCAAAGAGAANNNAGATCGGAA |
| 33719_1 | TATGAGTGTGGAGTCGTTGCGGAGAATCACTTGAACCCAGGAGNNNAGATCGGAA |
| 33719_2 | TATGAGTGTGGAGTCGTTGCGGAGAATCACTTGAACCCAGGAGNNNAGATCGGAA |
| 33719_3 | TATGAGTGTGGAGTCGTTGCGGAGAATCACTTGAACCCAGGAGNNNAGATCGGAA |
| 48151_1 | TATGAGTGTGGAGTCGTTGCTTTTAGTAGAGACGGGGTTCCACNNNAGATCGGAAG |
| 48151_2 | TATGAGTGTGGAGTCGTTGCTTTTAGTAGAGACGGGGTTCCACNNNAGATCGGAAG |
| 48151_3 | TATGAGTGTGGAGTCGTTGCTTTTAGTAGAGACGGGGTTCCACNNNAGATCGGAAG |
| 43702_1 | TATGAGTGTGGAGTCGTTGCAGGAGTCTACTCTTGGGGACTACNNNAGATCGGAAG |
| 43702_2 | TATGAGTGTGGAGTCGTTGCAGGAGTCTACTCTTGGGGACTACNNNAGATCGGAAG |
| 43702_3 | TATGAGTGTGGAGTCGTTGCAGGAGTCTACTCTTGGGGACTACNNNAGATCGGAAG |
| 41682_1 | TATGAGTGTGGAGTCGTTGCGATCTCCTGACCTCGTGATCTGNNNAGATCGGAAGA |
| 41682_2 | TATGAGTGTGGAGTCGTTGCGATCTCCTGACCTCGTGATCTGNNNAGATCGGAAGA |
| 41682_3 | TATGAGTGTGGAGTCGTTGCGATCTCCTGACCTCGTGATCTGNNNAGATCGGAAGA |
| 46462_1 | TATGAGTGTGGAGTCGTTGCTTCAAGTGATTCTCCTGCCTCAGNNNAGATCGGAAGA |
| 46462_2 | TATGAGTGTGGAGTCGTTGCTTCAAGTGATTCTCCTGCCTCAGNNNAGATCGGAAGA |
| 46462_3 | TATGAGTGTGGAGTCGTTGCTTCAAGTGATTCTCCTGCCTCAGNNNAGATCGGAAGA |
| 45785_1 | TATGAGTGTGGAGTCGTTGCTAGCCCCTTTACCTTGATGCAGNNNAGATCGGAAGA |
| 45785_2 | TATGAGTGTGGAGTCGTTGCTAGCCCCTTTACCTTGATGCAGNNNAGATCGGAAGA |
| 45785_3 | TATGAGTGTGGAGTCGTTGCTAGCCCCTTTACCTTGATGCAGNNNAGATCGGAAGA |
| 41424_1 | TATGAGTGTGGAGTCGTTGCGCTCAGTAGGGCAGACTCTAATCNNNAGATCGGAAG |
| 41424_2 | TATGAGTGTGGAGTCGTTGCGCTCAGTAGGGCAGACTCTAATCNNNAGATCGGAAG |
| 41424_3 | TATGAGTGTGGAGTCGTTGCGCTCAGTAGGGCAGACTCTAATCNNNAGATCGGAAG |
| 50932_1 | TATGAGTGTGGAGTCGTTGCCCAATAAAAAAGGGGGCACCAATNNNAGATCGGAA |
| 50932_2 | TATGAGTGTGGAGTCGTTGCCCAATAAAAAAGGGGGCACCAATNNNAGATCGGAA |
| 50932_3 | TATGAGTGTGGAGTCGTTGCCCAATAAAAAAGGGGGCACCAATNNNAGATCGGAA |
| 10241_1 | TATGAGTGTGGAGTCGTTGCATTCAACAAGGAGGGGTCTGACTNNNAGATCGGAA |
| 10241_2 | TATGAGTGTGGAGTCGTTGCATTCAACAAGGAGGGGTCTGACTNNNAGATCGGAA |
| 10241_3 | TATGAGTGTGGAGTCGTTGCATTCAACAAGGAGGGGTCTGACTNNNAGATCGGAA |
| 37734_1 | TATGAGTGTGGAGTCGTTGCACCAGAGGTTGAGAACAAGTTGANNNAGATCGGAA |
| 37734_2 | TATGAGTGTGGAGTCGTTGCACCAGAGGTTGAGAACAAGTTGANNNAGATCGGAA |

| | |
|---|---|
| 37734_3 | TATGAGTGTGGAGTCGTTGCACCAGAGGTTGAGAACAAGTTGANNNAGATCGGAA |
| 35988_1 | TATGAGTGTGGAGTCGTTGCGCAGAATCATTTGAACCTGGGAGNNNAGATCGGAA |
| 35988_2 | TATGAGTGTGGAGTCGTTGCGCAGAATCATTTGAACCTGGGAGNNNAGATCGGAA |
| 35988_3 | TATGAGTGTGGAGTCGTTGCGCAGAATCATTTGAACCTGGGAGNNNAGATCGGAA |
| 40319_1 | TATGAGTGTGGAGTCGTTGCAAATGTGATTGTGGAGGGGATGANNNAGATCGGA |
| 40319_2 | TATGAGTGTGGAGTCGTTGCAAATGTGATTGTGGAGGGGATGANNNAGATCGGA |
| 40319_3 | TATGAGTGTGGAGTCGTTGCAAATGTGATTGTGGAGGGGATGANNNAGATCGGA |
| 51994_1 | TATGAGTGTGGAGTCGTTGCACTTTGCTAATCCCAAGGAGTGTNNNAGATCGGAAG |
| 51994_2 | TATGAGTGTGGAGTCGTTGCACTTTGCTAATCCCAAGGAGTGTNNNAGATCGGAAG |
| 51994_3 | TATGAGTGTGGAGTCGTTGCACTTTGCTAATCCCAAGGAGTGTNNNAGATCGGAAG |
| 47141_1 | TATGAGTGTGGAGTCGTTGCGAAAGCACATTTCCCCAACATCANNNAGATCGGAAG |
| 47141_2 | TATGAGTGTGGAGTCGTTGCGAAAGCACATTTCCCCAACATCANNNAGATCGGAAG |
| 47141_3 | TATGAGTGTGGAGTCGTTGCGAAAGCACATTTCCCCAACATCANNNAGATCGGAAG |
| 9977_1 | TATGAGTGTGGAGTCGTTGCATTCTACCCTGGATTCCCTTCCCNNNAGATCGGAAGA |
| 9977_2 | TATGAGTGTGGAGTCGTTGCATTCTACCCTGGATTCCCTTCCCNNNAGATCGGAAGA |
| 9977_3 | TATGAGTGTGGAGTCGTTGCATTCTACCCTGGATTCCCTTCCCNNNAGATCGGAAGA |
| 41771_1 | TATGAGTGTGGAGTCGTTGCAATGGCACAATCTCAGCTCACTANNNAGATCGGAAG |
| 41771_2 | TATGAGTGTGGAGTCGTTGCAATGGCACAATCTCAGCTCACTANNNAGATCGGAAG |
| 41771_3 | TATGAGTGTGGAGTCGTTGCAATGGCACAATCTCAGCTCACTANNNAGATCGGAAG |
| 30473_1 | TATGAGTGTGGAGTCGTTGCGGGGAAATTTGTTTGGGAGACTATGNNNAGATCGG |
| 30473_2 | TATGAGTGTGGAGTCGTTGCGGGGAAATTTGTTTGGGAGACTATGNNNAGATCGG |
| 30473_3 | TATGAGTGTGGAGTCGTTGCGGGGAAATTTGTTTGGGAGACTATGNNNAGATCGG |
| 12699_1 | TATGAGTGTGGAGTCGTTGCTAAAGTGCTACTCCCAACTTNNNAGATCGGAAGAGC |
| 12699_2 | TATGAGTGTGGAGTCGTTGCTAAAGTGCTACTCCCAACTTNNNAGATCGGAAGAGC |
| 12699_3 | TATGAGTGTGGAGTCGTTGCTAAAGTGCTACTCCCAACTTNNNAGATCGGAAGAGC |
| 39032_1 | TATGAGTGTGGAGTCGTTGCGTGGTGATGGATGCCTGTAGTCNNNAGATCGGAAG |
| 39032_2 | TATGAGTGTGGAGTCGTTGCGTGGTGATGGATGCCTGTAGTCNNNAGATCGGAAG |
| 39032_3 | TATGAGTGTGGAGTCGTTGCGTGGTGATGGATGCCTGTAGTCNNNAGATCGGAAG |
| 41078_1 | TATGAGTGTGGAGTCGTTGCAAAGCCAGAATGCTATGGACTGANNNAGATCGGAA |
| 41078_2 | TATGAGTGTGGAGTCGTTGCAAAGCCAGAATGCTATGGACTGANNNAGATCGGAA |
| 41078_3 | TATGAGTGTGGAGTCGTTGCAAAGCCAGAATGCTATGGACTGANNNAGATCGGAA |
| 52348_1 | TATGAGTGTGGAGTCGTTGCGAGAGATGTGTGGGCTAAGGTAGNNNAGATCGGA |
| 52348_2 | TATGAGTGTGGAGTCGTTGCGAGAGATGTGTGGGCTAAGGTAGNNNAGATCGGA |
| 52348_3 | TATGAGTGTGGAGTCGTTGCGAGAGATGTGTGGGCTAAGGTAGNNNAGATCGGA |
| 9445_1 | TATGAGTGTGGAGTCGTTGCACAGGTGAATCAGTTGGACCTGNNNAGATCGGAAG |
| 9445_2 | TATGAGTGTGGAGTCGTTGCACAGGTGAATCAGTTGGACCTGNNNAGATCGGAAG |
| 9445_3 | TATGAGTGTGGAGTCGTTGCACAGGTGAATCAGTTGGACCTGNNNAGATCGGAAG |
| 45401_1 | TATGAGTGTGGAGTCGTTGCTATGTCTGTGTGTGGGTGTGAATNNNAGATCGGAA |
| 45401_2 | TATGAGTGTGGAGTCGTTGCTATGTCTGTGTGTGGGTGTGAATNNNAGATCGGAA |
| 45401_3 | TATGAGTGTGGAGTCGTTGCTATGTCTGTGTGTGGGTGTGAATNNNAGATCGGAA |
| 47352_1 | TATGAGTGTGGAGTCGTTGCACAAATTGATTTTTGACAAGGGTGTNNNAGATCGGA |
| 47352_2 | TATGAGTGTGGAGTCGTTGCACAAATTGATTTTTGACAAGGGTGTNNNAGATCGGA |
| 47352_3 | TATGAGTGTGGAGTCGTTGCACAAATTGATTTTTGACAAGGGTGTNNNAGATCGGA |

| | |
|---|---|
| 30604_1 | TATGAGTGTGGAGTCGTTGCGGGTTTCTCTCCATGTTGGTCAGNNNAGATCGGAAG |
| 30604_2 | TATGAGTGTGGAGTCGTTGCGGGTTTCTCTCCATGTTGGTCAGNNNAGATCGGAAG |
| 30604_3 | TATGAGTGTGGAGTCGTTGCGGGTTTCTCTCCATGTTGGTCAGNNNAGATCGGAAG |
| 32888_1 | TATGAGTGTGGAGTCGTTGCGATAGTGGTGATGGTTGCACAACNNNAGATCGGAA |
| 32888_2 | TATGAGTGTGGAGTCGTTGCGATAGTGGTGATGGTTGCACAACNNNAGATCGGAA |
| 32888_3 | TATGAGTGTGGAGTCGTTGCGATAGTGGTGATGGTTGCACAACNNNAGATCGGAA |
| 43962_1 | TATGAGTGTGGAGTCGTTGCCTTCCAAGTAGCTGGGACTACAGNNNAGATCGGAA |
| 43962_2 | TATGAGTGTGGAGTCGTTGCCTTCCAAGTAGCTGGGACTACAGNNNAGATCGGAA |
| 43962_3 | TATGAGTGTGGAGTCGTTGCCTTCCAAGTAGCTGGGACTACAGNNNAGATCGGAA |
| 47674_1 | TATGAGTGTGGAGTCGTTGCGAGTCTCCCTCTATCACCCAGGNNNAGATCGGAAGA |
| 47674_2 | TATGAGTGTGGAGTCGTTGCGAGTCTCCCTCTATCACCCAGGNNNAGATCGGAAGA |
| 47674_3 | TATGAGTGTGGAGTCGTTGCGAGTCTCCCTCTATCACCCAGGNNNAGATCGGAAGA |
| 42454_1 | TATGAGTGTGGAGTCGTTGCTTGTGTGATCTCTGCACATGAGTNNNAGATCGGAAG |
| 42454_2 | TATGAGTGTGGAGTCGTTGCTTGTGTGATCTCTGCACATGAGTNNNAGATCGGAAG |
| 42454_3 | TATGAGTGTGGAGTCGTTGCTTGTGTGATCTCTGCACATGAGTNNNAGATCGGAAG |
| 30582_1 | TATGAGTGTGGAGTCGTTGCTCTTGAAAACTTCCTTCACCAGCNNNAGATCGGAAG |
| 30582_2 | TATGAGTGTGGAGTCGTTGCTCTTGAAAACTTCCTTCACCAGCNNNAGATCGGAAG |
| 30582_3 | TATGAGTGTGGAGTCGTTGCTCTTGAAAACTTCCTTCACCAGCNNNAGATCGGAAG |
| 49770_1 | TATGAGTGTGGAGTCGTTGCCCAGGGATAGAGTCACAGTGAACNNNAGATCGGAA |
| 49770_2 | TATGAGTGTGGAGTCGTTGCCCAGGGATAGAGTCACAGTGAACNNNAGATCGGAA |
| 49770_3 | TATGAGTGTGGAGTCGTTGCCCAGGGATAGAGTCACAGTGAACNNNAGATCGGAA |
| 45231_1 | TATGAGTGTGGAGTCGTTGCAAGAGAGTAAGGGGCCGTAAGAANNNAGATCGGA |
| 45231_2 | TATGAGTGTGGAGTCGTTGCAAGAGAGTAAGGGGCCGTAAGAANNNAGATCGGA |
| 45231_3 | TATGAGTGTGGAGTCGTTGCAAGAGAGTAAGGGGCCGTAAGAANNNAGATCGGA |
| 51629_1 | TATGAGTGTGGAGTCGTTGCCCCAAATCTCTATCGCTCACAGANNNAGATCGGAAG |
| 51629_2 | TATGAGTGTGGAGTCGTTGCCCCAAATCTCTATCGCTCACAGANNNAGATCGGAAG |
| 51629_3 | TATGAGTGTGGAGTCGTTGCCCCAAATCTCTATCGCTCACAGANNNAGATCGGAAG |
| 39898_1 | TATGAGTGTGGAGTCGTTGCCAACAACAAAATCAGACTTCCCCANNNAGATCGGAA |
| 39898_2 | TATGAGTGTGGAGTCGTTGCCAACAACAAAATCAGACTTCCCCANNNAGATCGGAA |
| 39898_3 | TATGAGTGTGGAGTCGTTGCCAACAACAAAATCAGACTTCCCCANNNAGATCGGAA |
| 50924_1 | TATGAGTGTGGAGTCGTTGCAGAAACACATTCGGACTCAGACANNNAGATCGGAA |
| 50924_2 | TATGAGTGTGGAGTCGTTGCAGAAACACATTCGGACTCAGACANNNAGATCGGAA |
| 50924_3 | TATGAGTGTGGAGTCGTTGCAGAAACACATTCGGACTCAGACANNNAGATCGGAA |
| 29889_1 | TATGAGTGTGGAGTCGTTGCAGGAGAAAAACAAAGCCATGCTGNNNAGATCGGAA |
| 29889_2 | TATGAGTGTGGAGTCGTTGCAGGAGAAAAACAAAGCCATGCTGNNNAGATCGGAA |
| 29889_3 | TATGAGTGTGGAGTCGTTGCAGGAGAAAAACAAAGCCATGCTGNNNAGATCGGAA |
| 31340_1 | TATGAGTGTGGAGTCGTTGCCCCCTTTTACCCCGGAGAATAAANNNAGATCGGAAG |
| 31340_2 | TATGAGTGTGGAGTCGTTGCCCCCTTTTACCCCGGAGAATAAANNNAGATCGGAAG |
| 31340_3 | TATGAGTGTGGAGTCGTTGCCCCCTTTTACCCCGGAGAATAAANNNAGATCGGAAG |
| 32078_1 | TATGAGTGTGGAGTCGTTGCCATTCTCCTGCCTCAGCCTCNNNAGATCGGAAGAGCA |
| 32078_2 | TATGAGTGTGGAGTCGTTGCCATTCTCCTGCCTCAGCCTCNNNAGATCGGAAGAGCA |
| 32078_3 | TATGAGTGTGGAGTCGTTGCCATTCTCCTGCCTCAGCCTCNNNAGATCGGAAGAGCA |
| 43906_1 | TATGAGTGTGGAGTCGTTGCCCTTCCTTGTCTCTGTCTCTTCCNNNAGATCGGAAGA |

| | |
|---|---|
| 43906_2 | TATGAGTGTGGAGTCGTTGCCCTTCCTTGTCTCTGTCTCTTCCNNNAGATCGGAAGA |
| 43906_3 | TATGAGTGTGGAGTCGTTGCCCTTCCTTGTCTCTGTCTCTTCCNNNAGATCGGAAGA |
| 39075_1 | TATGAGTGTGGAGTCGTTGCGCCTCCCAAGTAGCTAGGATTACNNNAGATCGGAAG |
| 39075_2 | TATGAGTGTGGAGTCGTTGCGCCTCCCAAGTAGCTAGGATTACNNNAGATCGGAAG |
| 39075_3 | TATGAGTGTGGAGTCGTTGCGCCTCCCAAGTAGCTAGGATTACNNNAGATCGGAAG |
| 29428_1 | TATGAGTGTGGAGTCGTTGCGAAAACACATTCTTTGCAGCNNNAGATCGGAAGAG |
| 29428_2 | TATGAGTGTGGAGTCGTTGCGAAAACACATTCTTTGCAGCNNNAGATCGGAAGAG |
| 29428_3 | TATGAGTGTGGAGTCGTTGCGAAAACACATTCTTTGCAGCNNNAGATCGGAAGAG |
| 44528_1 | TATGAGTGTGGAGTCGTTGCACACATACACACACTACTGGTCCNNNAGATCGGAAG |
| 44528_2 | TATGAGTGTGGAGTCGTTGCACACATACACACACTACTGGTCCNNNAGATCGGAAG |
| 44528_3 | TATGAGTGTGGAGTCGTTGCACACATACACACACTACTGGTCCNNNAGATCGGAAG |
| 47712_1 | TATGAGTGTGGAGTCGTTGCAAACTCACTGTTCTGTCATTGCCNNNAGATCGGAAG |
| 47712_2 | TATGAGTGTGGAGTCGTTGCAAACTCACTGTTCTGTCATTGCCNNNAGATCGGAAG |
| 47712_3 | TATGAGTGTGGAGTCGTTGCAAACTCACTGTTCTGTCATTGCCNNNAGATCGGAAG |
| 50299_1 | TATGAGTGTGGAGTCGTTGCCAAGATTATGCCCCTGTACTCCANNNAGATCGGAAG |
| 50299_2 | TATGAGTGTGGAGTCGTTGCCAAGATTATGCCCCTGTACTCCANNNAGATCGGAAG |
| 50299_3 | TATGAGTGTGGAGTCGTTGCCAAGATTATGCCCCTGTACTCCANNNAGATCGGAAG |
| 12675_1 | TATGAGTGTGGAGTCGTTGCTGGATTTCTAATTCTTTTCTGTGNNNAGATCGGAAG |
| 12675_2 | TATGAGTGTGGAGTCGTTGCTGGATTTCTAATTCTTTTCTGTGNNNAGATCGGAAG |
| 12675_3 | TATGAGTGTGGAGTCGTTGCTGGATTTCTAATTCTTTTCTGTGNNNAGATCGGAAG |
| 45025_1 | TATGAGTGTGGAGTCGTTGCGTGATGGTAGTGCAAGACTGAGANNNAGATCGGA |
| 45025_2 | TATGAGTGTGGAGTCGTTGCGTGATGGTAGTGCAAGACTGAGANNNAGATCGGA |
| 45025_3 | TATGAGTGTGGAGTCGTTGCGTGATGGTAGTGCAAGACTGAGANNNAGATCGGA |
| 52959_1 | TATGAGTGTGGAGTCGTTGCAGAGAGAGAGAGAGAGAGAGAGGGANNNAGATCGG |
| 52959_2 | TATGAGTGTGGAGTCGTTGCAGAGAGAGAGAGAGAGAGAGAGGGANNNAGATCGG |
| 52959_3 | TATGAGTGTGGAGTCGTTGCAGAGAGAGAGAGAGAGAGAGAGGGANNNAGATCGG |
| 13143_1 | TATGAGTGTGGAGTCGTTGCTAAATACATGTTTGGCATCATCNNNAGATCGGAAGA |
| 13143_2 | TATGAGTGTGGAGTCGTTGCTAAATACATGTTTGGCATCATCNNNAGATCGGAAGA |
| 13143_3 | TATGAGTGTGGAGTCGTTGCTAAATACATGTTTGGCATCATCNNNAGATCGGAAGA |
| 46234_1 | TATGAGTGTGGAGTCGTTGCAGGTTCAAGCAATTCTCCTGTCTNNNAGATCGGAAG |
| 46234_2 | TATGAGTGTGGAGTCGTTGCAGGTTCAAGCAATTCTCCTGTCTNNNAGATCGGAAG |
| 46234_3 | TATGAGTGTGGAGTCGTTGCAGGTTCAAGCAATTCTCCTGTCTNNNAGATCGGAAG |
| 41453_1 | TATGAGTGTGGAGTCGTTGCAATAAATGAAGGAAAGCCAGGCGNNNAGATCGGA |
| 41453_2 | TATGAGTGTGGAGTCGTTGCAATAAATGAAGGAAAGCCAGGCGNNNAGATCGGA |
| 41453_3 | TATGAGTGTGGAGTCGTTGCAATAAATGAAGGAAAGCCAGGCGNNNAGATCGGA |
| 37069_1 | TATGAGTGTGGAGTCGTTGCCCAAGTTCTCCCATGTTACCTGANNNAGATCGGAAG |
| 37069_2 | TATGAGTGTGGAGTCGTTGCCCAAGTTCTCCCATGTTACCTGANNNAGATCGGAAG |
| 37069_3 | TATGAGTGTGGAGTCGTTGCCCAAGTTCTCCCATGTTACCTGANNNAGATCGGAAG |
| 30439_1 | TATGAGTGTGGAGTCGTTGCGGAGTAAGGTAAAGAAGATAAGANNNAGATCGGA |
| 30439_2 | TATGAGTGTGGAGTCGTTGCGGAGTAAGGTAAAGAAGATAAGANNNAGATCGGA |
| 30439_3 | TATGAGTGTGGAGTCGTTGCGGAGTAAGGTAAAGAAGATAAGANNNAGATCGGA |
| 35269_1 | TATGAGTGTGGAGTCGTTGCCAGGCAGAGATAGTATGGACACCNNNAGATCGGAA |
| 35269_2 | TATGAGTGTGGAGTCGTTGCCAGGCAGAGATAGTATGGACACCNNNAGATCGGAA |

| | |
|---|---|
| 35269_3 | TATGAGTGTGGAGTCGTTGCCAGGCAGAGATAGTATGGACACCNNNAGATCGGAA |
| 38135_1 | TATGAGTGTGGAGTCGTTGCTCAAGTGATTCTCCTGCCTTAGCNNNAGATCGGAAG |
| 38135_2 | TATGAGTGTGGAGTCGTTGCTCAAGTGATTCTCCTGCCTTAGCNNNAGATCGGAAG |
| 38135_3 | TATGAGTGTGGAGTCGTTGCTCAAGTGATTCTCCTGCCTTAGCNNNAGATCGGAAG |
| 52603_1 | TATGAGTGTGGAGTCGTTGCTCTACGAGGGCTGATATGAGGAGNNNAGATCGGAA |
| 52603_2 | TATGAGTGTGGAGTCGTTGCTCTACGAGGGCTGATATGAGGAGNNNAGATCGGAA |
| 52603_3 | TATGAGTGTGGAGTCGTTGCTCTACGAGGGCTGATATGAGGAGNNNAGATCGGAA |
| 38252_1 | TATGAGTGTGGAGTCGTTGCGAAAAGTGGTGCAAGAACTCCAGNNNAGATCGGAA |
| 38252_2 | TATGAGTGTGGAGTCGTTGCGAAAAGTGGTGCAAGAACTCCAGNNNAGATCGGAA |
| 38252_3 | TATGAGTGTGGAGTCGTTGCGAAAAGTGGTGCAAGAACTCCAGNNNAGATCGGAA |
| 34445_1 | TATGAGTGTGGAGTCGTTGCTATACCTCTGCGAATGGAACCAGNNNAGATCGGAAG |
| 34445_2 | TATGAGTGTGGAGTCGTTGCTATACCTCTGCGAATGGAACCAGNNNAGATCGGAAG |
| 34445_3 | TATGAGTGTGGAGTCGTTGCTATACCTCTGCGAATGGAACCAGNNNAGATCGGAAG |
| 35195_1 | TATGAGTGTGGAGTCGTTGCAGAGCTAGCTTCCCTGAAAAGACNNNAGATCGGAAG |
| 35195_2 | TATGAGTGTGGAGTCGTTGCAGAGCTAGCTTCCCTGAAAAGACNNNAGATCGGAAG |
| 35195_3 | TATGAGTGTGGAGTCGTTGCAGAGCTAGCTTCCCTGAAAAGACNNNAGATCGGAAG |
| 52467_1 | TATGAGTGTGGAGTCGTTGCAATGCAGTTGGTATAAGGGAGGGNNNAGATCGGA |
| 52467_2 | TATGAGTGTGGAGTCGTTGCAATGCAGTTGGTATAAGGGAGGGNNNAGATCGGA |
| 52467_3 | TATGAGTGTGGAGTCGTTGCAATGCAGTTGGTATAAGGGAGGGNNNAGATCGGA |
| 34710_1 | TATGAGTGTGGAGTCGTTGCCACCTGGCCTCTACAAACAATTTNNNAGATCGGAAG |
| 34710_2 | TATGAGTGTGGAGTCGTTGCCACCTGGCCTCTACAAACAATTTNNNAGATCGGAAG |
| 34710_3 | TATGAGTGTGGAGTCGTTGCCACCTGGCCTCTACAAACAATTTNNNAGATCGGAAG |
| 34511_1 | TATGAGTGTGGAGTCGTTGCCAGGTTCAAGCAATTCTCCTGCNNNAGATCGGAAGA |
| 34511_2 | TATGAGTGTGGAGTCGTTGCCAGGTTCAAGCAATTCTCCTGCNNNAGATCGGAAGA |
| 34511_3 | TATGAGTGTGGAGTCGTTGCCAGGTTCAAGCAATTCTCCTGCNNNAGATCGGAAGA |
| 29823_1 | TATGAGTGTGGAGTCGTTGCACTGAAGGCCGAACAGGTAATTANNNAGATCGGAA |
| 29823_2 | TATGAGTGTGGAGTCGTTGCACTGAAGGCCGAACAGGTAATTANNNAGATCGGAA |
| 29823_3 | TATGAGTGTGGAGTCGTTGCACTGAAGGCCGAACAGGTAATTANNNAGATCGGAA |
| 35447_1 | TATGAGTGTGGAGTCGTTGCCTCCTGTGTAACTGGGACTACAGNNNAGATCGGAAG |
| 35447_2 | TATGAGTGTGGAGTCGTTGCCTCCTGTGTAACTGGGACTACAGNNNAGATCGGAAG |
| 35447_3 | TATGAGTGTGGAGTCGTTGCCTCCTGTGTAACTGGGACTACAGNNNAGATCGGAAG |
| 29177_1 | TATGAGTGTGGAGTCGTTGCGGACACATTCTATGTCAAAGCCCNNNAGATCGGAAG |
| 29177_2 | TATGAGTGTGGAGTCGTTGCGGACACATTCTATGTCAAAGCCCNNNAGATCGGAAG |
| 29177_3 | TATGAGTGTGGAGTCGTTGCGGACACATTCTATGTCAAAGCCCNNNAGATCGGAAG |
| 42794_1 | TATGAGTGTGGAGTCGTTGCAATAATGGCATTTGCAGCAACCTNNNAGATCGGAAG |
| 42794_2 | TATGAGTGTGGAGTCGTTGCAATAATGGCATTTGCAGCAACCTNNNAGATCGGAAG |
| 42794_3 | TATGAGTGTGGAGTCGTTGCAATAATGGCATTTGCAGCAACCTNNNAGATCGGAAG |
| 34575_1 | TATGAGTGTGGAGTCGTTGCGGGAAGGGAAAGGAAGGGAAGNNNAGATCGGAA |
| 34575_2 | TATGAGTGTGGAGTCGTTGCGGGAAGGGAAAGGAAGGGAAGNNNAGATCGGAA |
| 34575_3 | TATGAGTGTGGAGTCGTTGCGGGAAGGGAAAGGAAGGGAAGNNNAGATCGGAA |
| 39896_1 | TATGAGTGTGGAGTCGTTGCAGGCAAGTCACTGATAGAACTGGNNNAGATCGGAA |
| 39896_2 | TATGAGTGTGGAGTCGTTGCAGGCAAGTCACTGATAGAACTGGNNNAGATCGGAA |
| 39896_3 | TATGAGTGTGGAGTCGTTGCAGGCAAGTCACTGATAGAACTGGNNNAGATCGGAA |

| | |
|---|---|
| 40843_1 | TATGAGTGTGGAGTCGTTGCTTGTGTTGGTTTTTTCCTCCTCCNNNAGATCGGAAGA |
| 40843_2 | TATGAGTGTGGAGTCGTTGCTTGTGTTGGTTTTTTCCTCCTCCNNNAGATCGGAAGA |
| 40843_3 | TATGAGTGTGGAGTCGTTGCTTGTGTTGGTTTTTTCCTCCTCCNNNAGATCGGAAGA |
| 41128_1 | TATGAGTGTGGAGTCGTTGCTTCTGACATATGCTCCAACATGANNNAGATCGGAAG |
| 41128_2 | TATGAGTGTGGAGTCGTTGCTTCTGACATATGCTCCAACATGANNNAGATCGGAAG |
| 41128_3 | TATGAGTGTGGAGTCGTTGCTTCTGACATATGCTCCAACATGANNNAGATCGGAAG |
| 36159_1 | TATGAGTGTGGAGTCGTTGCCAGAATAGGCATCCAGGAGACTCNNNAGATCGGAA |
| 36159_2 | TATGAGTGTGGAGTCGTTGCCAGAATAGGCATCCAGGAGACTCNNNAGATCGGAA |
| 36159_3 | TATGAGTGTGGAGTCGTTGCCAGAATAGGCATCCAGGAGACTCNNNAGATCGGAA |
| 35979_1 | TATGAGTGTGGAGTCGTTGCATACATCTATGACCTCACCCCCANNNAGATCGGAAGA |
| 35979_2 | TATGAGTGTGGAGTCGTTGCATACATCTATGACCTCACCCCCANNNAGATCGGAAGA |
| 35979_3 | TATGAGTGTGGAGTCGTTGCATACATCTATGACCTCACCCCCANNNAGATCGGAAGA |
| 30047_1 | TATGAGTGTGGAGTCGTTGCTTCCAGTGGAATCTGCCCAATCNNNAGATCGGAAGA |
| 30047_2 | TATGAGTGTGGAGTCGTTGCTTCCAGTGGAATCTGCCCAATCNNNAGATCGGAAGA |
| 30047_3 | TATGAGTGTGGAGTCGTTGCTTCCAGTGGAATCTGCCCAATCNNNAGATCGGAAGA |
| 39715_1 | TATGAGTGTGGAGTCGTTGCTCAGGATTGTTTCCGAAAGGTTGNNNAGATCGGAA |
| 39715_2 | TATGAGTGTGGAGTCGTTGCTCAGGATTGTTTCCGAAAGGTTGNNNAGATCGGAA |
| 39715_3 | TATGAGTGTGGAGTCGTTGCTCAGGATTGTTTCCGAAAGGTTGNNNAGATCGGAA |
| 43812_1 | TATGAGTGTGGAGTCGTTGCCCCATATTCTCTTCCTTGCCTCANNNAGATCGGAAGA |
| 43812_2 | TATGAGTGTGGAGTCGTTGCCCCATATTCTCTTCCTTGCCTCANNNAGATCGGAAGA |
| 43812_3 | TATGAGTGTGGAGTCGTTGCCCCATATTCTCTTCCTTGCCTCANNNAGATCGGAAGA |
| 47961_1 | TATGAGTGTGGAGTCGTTGCCTGAGTAGCTAGGATTACAGGCGNNNAGATCGGAA |
| 47961_2 | TATGAGTGTGGAGTCGTTGCCTGAGTAGCTAGGATTACAGGCGNNNAGATCGGAA |
| 47961_3 | TATGAGTGTGGAGTCGTTGCCTGAGTAGCTAGGATTACAGGCGNNNAGATCGGAA |
| 50425_1 | TATGAGTGTGGAGTCGTTGCAGGGCTTCCAAGTAGATGCTTTCNNNAGATCGGAAG |
| 50425_2 | TATGAGTGTGGAGTCGTTGCAGGGCTTCCAAGTAGATGCTTTCNNNAGATCGGAAG |
| 50425_3 | TATGAGTGTGGAGTCGTTGCAGGGCTTCCAAGTAGATGCTTTCNNNAGATCGGAAG |
| 13352_1 | TATGAGTGTGGAGTCGTTGCTTCTTCTCTCCAGAACACTAGNNNAGATCGGAAGAG |
| 13352_2 | TATGAGTGTGGAGTCGTTGCTTCTTCTCTCCAGAACACTAGNNNAGATCGGAAGAG |
| 13352_3 | TATGAGTGTGGAGTCGTTGCTTCTTCTCTCCAGAACACTAGNNNAGATCGGAAGAG |
| 34503_1 | TATGAGTGTGGAGTCGTTGCCCCACTGTGCTTGACTTTTTTCANNNAGATCGGAAGA |
| 34503_2 | TATGAGTGTGGAGTCGTTGCCCCACTGTGCTTGACTTTTTTCANNNAGATCGGAAGA |
| 34503_3 | TATGAGTGTGGAGTCGTTGCCCCACTGTGCTTGACTTTTTTCANNNAGATCGGAAGA |
| 37491_1 | TATGAGTGTGGAGTCGTTGCAGAAAATCAATGAAACCGAAAGCTNNNAGATCGGA |
| 37491_2 | TATGAGTGTGGAGTCGTTGCAGAAAATCAATGAAACCGAAAGCTNNNAGATCGGA |
| 37491_3 | TATGAGTGTGGAGTCGTTGCAGAAAATCAATGAAACCGAAAGCTNNNAGATCGGA |
| 50171_1 | TATGAGTGTGGAGTCGTTGCAGGAACTCAAGAAAAAGCCATGCNNNAGATCGGAA |
| 50171_2 | TATGAGTGTGGAGTCGTTGCAGGAACTCAAGAAAAAGCCATGCNNNAGATCGGAA |
| 50171_3 | TATGAGTGTGGAGTCGTTGCAGGAACTCAAGAAAAAGCCATGCNNNAGATCGGAA |
| 37739_1 | TATGAGTGTGGAGTCGTTGCGGGCCAGAGAGACTACATTTCTTNNNAGATCGGAA |
| 37739_2 | TATGAGTGTGGAGTCGTTGCGGGCCAGAGAGACTACATTTCTTNNNAGATCGGAA |
| 37739_3 | TATGAGTGTGGAGTCGTTGCGGGCCAGAGAGACTACATTTCTTNNNAGATCGGAA |
| 42442_1 | TATGAGTGTGGAGTCGTTGCCACACCAAGAGGGATTCAGAGAGNNNAGATCGGAA |

| | |
|---|---|
| 42442_2 | TATGAGTGTGGAGTCGTTGCCACACCAAGAGGGATTCAGAGAGNNNAGATCGGAA |
| 42442_3 | TATGAGTGTGGAGTCGTTGCCACACCAAGAGGGATTCAGAGAGNNNAGATCGGAA |
| 30198_1 | TATGAGTGTGGAGTCGTTGCCGGGCTCTTTCTGAAGACTAGAANNNAGATCGGAAG |
| 30198_2 | TATGAGTGTGGAGTCGTTGCCGGGCTCTTTCTGAAGACTAGAANNNAGATCGGAAG |
| 30198_3 | TATGAGTGTGGAGTCGTTGCCGGGCTCTTTCTGAAGACTAGAANNNAGATCGGAAG |
| 40354_1 | TATGAGTGTGGAGTCGTTGCAAGAAAAAAAATTGCCGGGCATGNNNAGATCGGAA |
| 40354_2 | TATGAGTGTGGAGTCGTTGCAAGAAAAAAAATTGCCGGGCATGNNNAGATCGGAA |
| 40354_3 | TATGAGTGTGGAGTCGTTGCAAGAAAAAAAATTGCCGGGCATGNNNAGATCGGAA |
| 34101_1 | TATGAGTGTGGAGTCGTTGCATGTGCCTGTAATCCCAGCTACNNNAGATCGGAAGA |
| 34101_2 | TATGAGTGTGGAGTCGTTGCATGTGCCTGTAATCCCAGCTACNNNAGATCGGAAGA |
| 34101_3 | TATGAGTGTGGAGTCGTTGCATGTGCCTGTAATCCCAGCTACNNNAGATCGGAAGA |
| 46296_1 | TATGAGTGTGGAGTCGTTGCCAGTACTGTGATCTCGGCTCACNNNAGATCGGAAGA |
| 46296_2 | TATGAGTGTGGAGTCGTTGCCAGTACTGTGATCTCGGCTCACNNNAGATCGGAAGA |
| 46296_3 | TATGAGTGTGGAGTCGTTGCCAGTACTGTGATCTCGGCTCACNNNAGATCGGAAGA |
| 38617_1 | TATGAGTGTGGAGTCGTTGCACCCTGGGAAAAAGAGCAAACTANNNAGATCGGAA |
| 38617_2 | TATGAGTGTGGAGTCGTTGCACCCTGGGAAAAAGAGCAAACTANNNAGATCGGAA |
| 38617_3 | TATGAGTGTGGAGTCGTTGCACCCTGGGAAAAAGAGCAAACTANNNAGATCGGAA |
| 51528_1 | TATGAGTGTGGAGTCGTTGCTGGTAGTTTCTTTTGCCATGCAGNNNAGATCGGAAG |
| 51528_2 | TATGAGTGTGGAGTCGTTGCTGGTAGTTTCTTTTGCCATGCAGNNNAGATCGGAAG |
| 51528_3 | TATGAGTGTGGAGTCGTTGCTGGTAGTTTCTTTTGCCATGCAGNNNAGATCGGAAG |
| 13766_1 | TATGAGTGTGGAGTCGTTGCCACAGTAAAACATCTGTTAAAAGNNNAGATCGGAAG |
| 13766_2 | TATGAGTGTGGAGTCGTTGCCACAGTAAAACATCTGTTAAAAGNNNAGATCGGAAG |
| 13766_3 | TATGAGTGTGGAGTCGTTGCCACAGTAAAACATCTGTTAAAAGNNNAGATCGGAAG |
| 44754_1 | TATGAGTGTGGAGTCGTTGCTATTGTTCAGACCTCAGCTCCTGNNNAGATCGGAAG |
| 44754_2 | TATGAGTGTGGAGTCGTTGCTATTGTTCAGACCTCAGCTCCTGNNNAGATCGGAAG |
| 44754_3 | TATGAGTGTGGAGTCGTTGCTATTGTTCAGACCTCAGCTCCTGNNNAGATCGGAAG |
| 52309_1 | TATGAGTGTGGAGTCGTTGCTTTCTTAGGAGTGTGGGGAGTTGNNNAGATCGGAA |
| 52309_2 | TATGAGTGTGGAGTCGTTGCTTTCTTAGGAGTGTGGGGAGTTGNNNAGATCGGAA |
| 52309_3 | TATGAGTGTGGAGTCGTTGCTTTCTTAGGAGTGTGGGGAGTTGNNNAGATCGGAA |
| 40740_1 | TATGAGTGTGGAGTCGTTGCCACGTCTGTAATCCCAGGTACTCNNNAGATCGGAAG |
| 40740_2 | TATGAGTGTGGAGTCGTTGCCACGTCTGTAATCCCAGGTACTCNNNAGATCGGAAG |
| 40740_3 | TATGAGTGTGGAGTCGTTGCCACGTCTGTAATCCCAGGTACTCNNNAGATCGGAAG |
| 51025_1 | TATGAGTGTGGAGTCGTTGCTACAAAAAAATTAGCCGGGTGTGGNNNAGATCGGAA |
| 51025_2 | TATGAGTGTGGAGTCGTTGCTACAAAAAAATTAGCCGGGTGTGGNNNAGATCGGAA |
| 51025_3 | TATGAGTGTGGAGTCGTTGCTACAAAAAAATTAGCCGGGTGTGGNNNAGATCGGAA |
| 36402_1 | TATGAGTGTGGAGTCGTTGCGAGTTCAAGACTAGCCTGACCAANNNAGATCGGAAG |
| 36402_2 | TATGAGTGTGGAGTCGTTGCGAGTTCAAGACTAGCCTGACCAANNNAGATCGGAAG |
| 36402_3 | TATGAGTGTGGAGTCGTTGCGAGTTCAAGACTAGCCTGACCAANNNAGATCGGAAG |
| 39909_1 | TATGAGTGTGGAGTCGTTGCGAGAGTGAGGATTCTAGAGCAGCNNNAGATCGGAA |
| 39909_2 | TATGAGTGTGGAGTCGTTGCGAGAGTGAGGATTCTAGAGCAGCNNNAGATCGGAA |
| 39909_3 | TATGAGTGTGGAGTCGTTGCGAGAGTGAGGATTCTAGAGCAGCNNNAGATCGGAA |
| 51877_1 | TATGAGTGTGGAGTCGTTGCAAATACCACCCCTCTTTGTCCTCNNNAGATCGGAAGA |
| 51877_2 | TATGAGTGTGGAGTCGTTGCAAATACCACCCCTCTTTGTCCTCNNNAGATCGGAAGA |

| | |
|---|---|
| 51877_3 | TATGAGTGTGGAGTCGTTGCAAATACCACCCCTCTTTGTCCTCNNNAGATCGGAAGA |
| 30205_1 | TATGAGTGTGGAGTCGTTGCGCTGGAAAAAGCTTCTCAGAGTCNNNAGATCGGAA |
| 30205_2 | TATGAGTGTGGAGTCGTTGCGCTGGAAAAAGCTTCTCAGAGTCNNNAGATCGGAA |
| 30205_3 | TATGAGTGTGGAGTCGTTGCGCTGGAAAAAGCTTCTCAGAGTCNNNAGATCGGAA |
| 43527_1 | TATGAGTGTGGAGTCGTTGCCACTCTGTCACTCTCTCATCCAGNNNAGATCGGAAGA |
| 43527_2 | TATGAGTGTGGAGTCGTTGCCACTCTGTCACTCTCTCATCCAGNNNAGATCGGAAGA |
| 43527_3 | TATGAGTGTGGAGTCGTTGCCACTCTGTCACTCTCTCATCCAGNNNAGATCGGAAGA |
| 32230_1 | TATGAGTGTGGAGTCGTTGCGAAACCTTTGGCCAATCAGAGAGNNNAGATCGGAA |
| 32230_2 | TATGAGTGTGGAGTCGTTGCGAAACCTTTGGCCAATCAGAGAGNNNAGATCGGAA |
| 32230_3 | TATGAGTGTGGAGTCGTTGCGAAACCTTTGGCCAATCAGAGAGNNNAGATCGGAA |
| 45532_1 | TATGAGTGTGGAGTCGTTGCATTCTCAGCCACTGATCTTTCCCNNNAGATCGGAAGA |
| 45532_2 | TATGAGTGTGGAGTCGTTGCATTCTCAGCCACTGATCTTTCCCNNNAGATCGGAAGA |
| 45532_3 | TATGAGTGTGGAGTCGTTGCATTCTCAGCCACTGATCTTTCCCNNNAGATCGGAAGA |
| 30478_1 | TATGAGTGTGGAGTCGTTGCAGGAGAGGACTTGAGACTGGAAANNNAGATCGGA |
| 30478_2 | TATGAGTGTGGAGTCGTTGCAGGAGAGGACTTGAGACTGGAAANNNAGATCGGA |
| 30478_3 | TATGAGTGTGGAGTCGTTGCAGGAGAGGACTTGAGACTGGAAANNNAGATCGGA |
| 6148_1 | TATGAGTGTGGAGTCGTTGCTGGAAATCAAGACTGCTGTGAGCTNNNAGATCGGA |
| 6148_2 | TATGAGTGTGGAGTCGTTGCTGGAAATCAAGACTGCTGTGAGCTNNNAGATCGGA |
| 6148_3 | TATGAGTGTGGAGTCGTTGCTGGAAATCAAGACTGCTGTGAGCTNNNAGATCGGA |
| 2620_1 | TATGAGTGTGGAGTCGTTGCCTGCCTACTGTGTGCTTACTCTGNNNAGATCGGAAG |
| 2620_2 | TATGAGTGTGGAGTCGTTGCCTGCCTACTGTGTGCTTACTCTGNNNAGATCGGAAG |
| 2620_3 | TATGAGTGTGGAGTCGTTGCCTGCCTACTGTGTGCTTACTCTGNNNAGATCGGAAG |
| 43710_1 | TATGAGTGTGGAGTCGTTGCTGTCCTTCATACCCTTCCCTACTNNNAGATCGGAAGA |
| 43710_2 | TATGAGTGTGGAGTCGTTGCTGTCCTTCATACCCTTCCCTACTNNNAGATCGGAAGA |
| 43710_3 | TATGAGTGTGGAGTCGTTGCTGTCCTTCATACCCTTCCCTACTNNNAGATCGGAAGA |
| 10153_1 | TATGAGTGTGGAGTCGTTGCGAGTGTGCCTATAGCCCCAGATANNNAGATCGGAA |
| 10153_2 | TATGAGTGTGGAGTCGTTGCGAGTGTGCCTATAGCCCCAGATANNNAGATCGGAA |
| 10153_3 | TATGAGTGTGGAGTCGTTGCGAGTGTGCCTATAGCCCCAGATANNNAGATCGGAA |
| 10579_1 | TATGAGTGTGGAGTCGTTGCGACCAACAGAGTGGCATGAAGATNNNAGATCGGAA |
| 10579_2 | TATGAGTGTGGAGTCGTTGCGACCAACAGAGTGGCATGAAGATNNNAGATCGGAA |
| 10579_3 | TATGAGTGTGGAGTCGTTGCGACCAACAGAGTGGCATGAAGATNNNAGATCGGAA |
| 37455_1 | TATGAGTGTGGAGTCGTTGCCTCACACCACTCAGAATGGCTATNNNAGATCGGAAG |
| 37455_2 | TATGAGTGTGGAGTCGTTGCCTCACACCACTCAGAATGGCTATNNNAGATCGGAAG |
| 37455_3 | TATGAGTGTGGAGTCGTTGCCTCACACCACTCAGAATGGCTATNNNAGATCGGAAG |
| 30912_1 | TATGAGTGTGGAGTCGTTGCACCATTGCAGTCAGAATCTTTGCNNNAGATCGGAAG |
| 30912_2 | TATGAGTGTGGAGTCGTTGCACCATTGCAGTCAGAATCTTTGCNNNAGATCGGAAG |
| 30912_3 | TATGAGTGTGGAGTCGTTGCACCATTGCAGTCAGAATCTTTGCNNNAGATCGGAAG |
| 43340_1 | TATGAGTGTGGAGTCGTTGCCTCGATTTCCTGACCTCGTGATCNNNAGATCGGAAG |
| 43340_2 | TATGAGTGTGGAGTCGTTGCCTCGATTTCCTGACCTCGTGATCNNNAGATCGGAAG |
| 43340_3 | TATGAGTGTGGAGTCGTTGCCTCGATTTCCTGACCTCGTGATCNNNAGATCGGAAG |
| 40579_1 | TATGAGTGTGGAGTCGTTGCACTGGGAGCAACTGTAATAGCAANNNAGATCGGAA |
| 40579_2 | TATGAGTGTGGAGTCGTTGCACTGGGAGCAACTGTAATAGCAANNNAGATCGGAA |
| 40579_3 | TATGAGTGTGGAGTCGTTGCACTGGGAGCAACTGTAATAGCAANNNAGATCGGAA |

| | |
|---|---|
| 10775_1 | TATGAGTGTGGAGTCGTTGCCACACACAACAGACATAAGCTGGNNNAGATCGGAAG |
| 10775_2 | TATGAGTGTGGAGTCGTTGCCACACACAACAGACATAAGCTGGNNNAGATCGGAAG |
| 10775_3 | TATGAGTGTGGAGTCGTTGCCACACACAACAGACATAAGCTGGNNNAGATCGGAAG |
| 12084_1 | TATGAGTGTGGAGTCGTTGCGGATGGAGACTGGTTAGGAGATTNNNAGATCGGAA |
| 12084_2 | TATGAGTGTGGAGTCGTTGCGGATGGAGACTGGTTAGGAGATTNNNAGATCGGAA |
| 12084_3 | TATGAGTGTGGAGTCGTTGCGGATGGAGACTGGTTAGGAGATTNNNAGATCGGAA |
| 49511_1 | TATGAGTGTGGAGTCGTTGCACTGGACAGGAAATAGTACTAAACANNNAGATCGGA |
| 49511_2 | TATGAGTGTGGAGTCGTTGCACTGGACAGGAAATAGTACTAAACANNNAGATCGGA |
| 49511_3 | TATGAGTGTGGAGTCGTTGCACTGGACAGGAAATAGTACTAAACANNNAGATCGGA |
| 29295_1 | TATGAGTGTGGAGTCGTTGCACACTTCAGAAACTCTTAGGTGCANNNAGATCGGAA |
| 29295_2 | TATGAGTGTGGAGTCGTTGCACACTTCAGAAACTCTTAGGTGCANNNAGATCGGAA |
| 29295_3 | TATGAGTGTGGAGTCGTTGCACACTTCAGAAACTCTTAGGTGCANNNAGATCGGAA |
| 44500_1 | TATGAGTGTGGAGTCGTTGCAGAGAGGATGTGAGTGGAGGTAANNNAGATCGGA |
| 44500_2 | TATGAGTGTGGAGTCGTTGCAGAGAGGATGTGAGTGGAGGTAANNNAGATCGGA |
| 44500_3 | TATGAGTGTGGAGTCGTTGCAGAGAGGATGTGAGTGGAGGTAANNNAGATCGGA |
| 49332_1 | TATGAGTGTGGAGTCGTTGCTTTAGGTTGCCTGTTCACTCTGANNNAGATCGGAAGA |
| 49332_2 | TATGAGTGTGGAGTCGTTGCTTTAGGTTGCCTGTTCACTCTGANNNAGATCGGAAGA |
| 49332_3 | TATGAGTGTGGAGTCGTTGCTTTAGGTTGCCTGTTCACTCTGANNNAGATCGGAAGA |
| 37249_1 | TATGAGTGTGGAGTCGTTGCTGAGAAGAACGAGAGTGAAGGTGNNNAGATCGGAA |
| 37249_2 | TATGAGTGTGGAGTCGTTGCTGAGAAGAACGAGAGTGAAGGTGNNNAGATCGGAA |
| 37249_3 | TATGAGTGTGGAGTCGTTGCTGAGAAGAACGAGAGTGAAGGTGNNNAGATCGGAA |
| 12926_1 | TATGAGTGTGGAGTCGTTGCCATGGTAAAGTTACTAACACAAANNNAGATCGGAAG |
| 12926_2 | TATGAGTGTGGAGTCGTTGCCATGGTAAAGTTACTAACACAAANNNAGATCGGAAG |
| 12926_3 | TATGAGTGTGGAGTCGTTGCCATGGTAAAGTTACTAACACAAANNNAGATCGGAAG |
| 33263_1 | TATGAGTGTGGAGTCGTTGCCCCAGGTTCAAGCAATTCTTCTGNNNAGATCGGAAGA |
| 33263_2 | TATGAGTGTGGAGTCGTTGCCCCAGGTTCAAGCAATTCTTCTGNNNAGATCGGAAGA |
| 33263_3 | TATGAGTGTGGAGTCGTTGCCCCAGGTTCAAGCAATTCTTCTGNNNAGATCGGAAGA |
| 48629_1 | TATGAGTGTGGAGTCGTTGCGTTCAAGCAATTCTCTGCCTCAGNNNAGATCGGAAGA |
| 48629_2 | TATGAGTGTGGAGTCGTTGCGTTCAAGCAATTCTCTGCCTCAGNNNAGATCGGAAGA |
| 48629_3 | TATGAGTGTGGAGTCGTTGCGTTCAAGCAATTCTCTGCCTCAGNNNAGATCGGAAGA |
| 33349_1 | TATGAGTGTGGAGTCGTTGCTTCAAGTGATTCTCCTGCCTCAGNNNAGATCGGAAGA |
| 33349_2 | TATGAGTGTGGAGTCGTTGCTTCAAGTGATTCTCCTGCCTCAGNNNAGATCGGAAGA |
| 33349_3 | TATGAGTGTGGAGTCGTTGCTTCAAGTGATTCTCCTGCCTCAGNNNAGATCGGAAGA |
| 35953_1 | TATGAGTGTGGAGTCGTTGCATTTTTCCAGTAGAGGGCATCCCNNNAGATCGGAAG |
| 35953_2 | TATGAGTGTGGAGTCGTTGCATTTTTCCAGTAGAGGGCATCCCNNNAGATCGGAAG |
| 35953_3 | TATGAGTGTGGAGTCGTTGCATTTTTCCAGTAGAGGGCATCCCNNNAGATCGGAAG |
| 33721_1 | TATGAGTGTGGAGTCGTTGCTTGCAGAAAGGAAAACTATGGCCNNNAGATCGGAA |
| 33721_2 | TATGAGTGTGGAGTCGTTGCTTGCAGAAAGGAAAACTATGGCCNNNAGATCGGAA |
| 33721_3 | TATGAGTGTGGAGTCGTTGCTTGCAGAAAGGAAAACTATGGCCNNNAGATCGGAA |
| 51944_1 | TATGAGTGTGGAGTCGTTGCGGGTTTCCTCACTTCTCAGACGNNNAGATCGGAAGA |
| 51944_2 | TATGAGTGTGGAGTCGTTGCGGGTTTCCTCACTTCTCAGACGNNNAGATCGGAAGA |
| 51944_3 | TATGAGTGTGGAGTCGTTGCGGGTTTCCTCACTTCTCAGACGNNNAGATCGGAAGA |
| 31100_1 | TATGAGTGTGGAGTCGTTGCTAGAGTGAGTGCTTTCAGAAGGCNNNAGATCGGAA |

| | |
|---|---|
| 31100_2 | TATGAGTGTGGAGTCGTTGCTAGAGTGAGTGCTTTCAGAAGGCNNNAGATCGGAA |
| 31100_3 | TATGAGTGTGGAGTCGTTGCTAGAGTGAGTGCTTTCAGAAGGCNNNAGATCGGAA |
| 39091_1 | TATGAGTGTGGAGTCGTTGCTGCCAATCCTACTCAAACTACTCCNNNAGATCGGAAG |
| 39091_2 | TATGAGTGTGGAGTCGTTGCTGCCAATCCTACTCAAACTACTCCNNNAGATCGGAAG |
| 39091_3 | TATGAGTGTGGAGTCGTTGCTGCCAATCCTACTCAAACTACTCCNNNAGATCGGAAG |
| 41808_1 | TATGAGTGTGGAGTCGTTGCTTCACGCCATTCTCCTGTCTCNNNAGATCGGAAGAGC |
| 41808_2 | TATGAGTGTGGAGTCGTTGCTTCACGCCATTCTCCTGTCTCNNNAGATCGGAAGAGC |
| 41808_3 | TATGAGTGTGGAGTCGTTGCTTCACGCCATTCTCCTGTCTCNNNAGATCGGAAGAGC |
| 40502_1 | TATGAGTGTGGAGTCGTTGCATCCAAATGTCCATCAACAGCAGNNNAGATCGGAAG |
| 40502_2 | TATGAGTGTGGAGTCGTTGCATCCAAATGTCCATCAACAGCAGNNNAGATCGGAAG |
| 40502_3 | TATGAGTGTGGAGTCGTTGCATCCAAATGTCCATCAACAGCAGNNNAGATCGGAAG |
| 52869_1 | TATGAGTGTGGAGTCGTTGCTAACCGCCTATGGGGAAATTGTGNNNAGATCGGAA |
| 52869_2 | TATGAGTGTGGAGTCGTTGCTAACCGCCTATGGGGAAATTGTGNNNAGATCGGAA |
| 52869_3 | TATGAGTGTGGAGTCGTTGCTAACCGCCTATGGGGAAATTGTGNNNAGATCGGAA |
| 34510_1 | TATGAGTGTGGAGTCGTTGCTTTGTCAGATGAGTAGGTTGCGANNNAGATCGGAA |
| 34510_2 | TATGAGTGTGGAGTCGTTGCTTTGTCAGATGAGTAGGTTGCGANNNAGATCGGAA |
| 34510_3 | TATGAGTGTGGAGTCGTTGCTTTGTCAGATGAGTAGGTTGCGANNNAGATCGGAA |
| 51455_1 | TATGAGTGTGGAGTCGTTGCGTACTGCAGACATAGACTGGGAGNNNAGATCGGAA |
| 51455_2 | TATGAGTGTGGAGTCGTTGCGTACTGCAGACATAGACTGGGAGNNNAGATCGGAA |
| 51455_3 | TATGAGTGTGGAGTCGTTGCGTACTGCAGACATAGACTGGGAGNNNAGATCGGAA |
| 12803_1 | TATGAGTGTGGAGTCGTTGCCATAGACATAAAAGGACTCACTNNNAGATCGGAAGA |
| 12803_2 | TATGAGTGTGGAGTCGTTGCCATAGACATAAAAGGACTCACTNNNAGATCGGAAGA |
| 12803_3 | TATGAGTGTGGAGTCGTTGCCATAGACATAAAAGGACTCACTNNNAGATCGGAAGA |
| 53095_1 | TATGAGTGTGGAGTCGTTGCCCAGTAACCGCATTTCTAGGTCTNNNAGATCGGAAG |
| 53095_2 | TATGAGTGTGGAGTCGTTGCCCAGTAACCGCATTTCTAGGTCTNNNAGATCGGAAG |
| 53095_3 | TATGAGTGTGGAGTCGTTGCCCAGTAACCGCATTTCTAGGTCTNNNAGATCGGAAG |
| 32105_1 | TATGAGTGTGGAGTCGTTGCCTCCCAGGTTCATGCCATTCTCNNNAGATCGGAAGAG |
| 32105_2 | TATGAGTGTGGAGTCGTTGCCTCCCAGGTTCATGCCATTCTCNNNAGATCGGAAGAG |
| 32105_3 | TATGAGTGTGGAGTCGTTGCCTCCCAGGTTCATGCCATTCTCNNNAGATCGGAAGAG |
| 13030_1 | TATGAGTGTGGAGTCGTTGCCAACACTTCATAGACATCAATAANNNAGATCGGAAG |
| 13030_2 | TATGAGTGTGGAGTCGTTGCCAACACTTCATAGACATCAATAANNNAGATCGGAAG |
| 13030_3 | TATGAGTGTGGAGTCGTTGCCAACACTTCATAGACATCAATAANNNAGATCGGAAG |
| 39942_1 | TATGAGTGTGGAGTCGTTGCGAAAGGAAGGAAGGAAGGAAGGANNNAGATCGG |
| 39942_2 | TATGAGTGTGGAGTCGTTGCGAAAGGAAGGAAGGAAGGAAGGANNNAGATCGG |
| 39942_3 | TATGAGTGTGGAGTCGTTGCGAAAGGAAGGAAGGAAGGAAGGANNNAGATCGG |
| 45162_1 | TATGAGTGTGGAGTCGTTGCTCTTTCTGTTGCCTTGCTTTCAGNNNAGATCGGAAGA |
| 45162_2 | TATGAGTGTGGAGTCGTTGCTCTTTCTGTTGCCTTGCTTTCAGNNNAGATCGGAAGA |
| 45162_3 | TATGAGTGTGGAGTCGTTGCTCTTTCTGTTGCCTTGCTTTCAGNNNAGATCGGAAGA |
| 31033_1 | TATGAGTGTGGAGTCGTTGCGATCACTCTGGTTGCTGCTGATCNNNAGATCGGAAG |
| 31033_2 | TATGAGTGTGGAGTCGTTGCGATCACTCTGGTTGCTGCTGATCNNNAGATCGGAAG |
| 31033_3 | TATGAGTGTGGAGTCGTTGCGATCACTCTGGTTGCTGCTGATCNNNAGATCGGAAG |
| 32010_1 | TATGAGTGTGGAGTCGTTGCAGAGAGAGAGAGAGAGAGATTCTCNNNAGATCGG |
| 32010_2 | TATGAGTGTGGAGTCGTTGCAGAGAGAGAGAGAGAGAGATTCTCNNNAGATCGG |

| | |
|---|---|
| 32010_3 | TATGAGTGTGGAGTCGTTGCAGAGAGAGAGAGAGAGAGATTCTCNNNAGATCGGA |
| 39399_1 | TATGAGTGTGGAGTCGTTGCTGGATTGGAAGATTCAGCAGAGTNNNAGATCGGAA |
| 39399_2 | TATGAGTGTGGAGTCGTTGCTGGATTGGAAGATTCAGCAGAGTNNNAGATCGGAA |
| 39399_3 | TATGAGTGTGGAGTCGTTGCTGGATTGGAAGATTCAGCAGAGTNNNAGATCGGAA |
| 47777_1 | TATGAGTGTGGAGTCGTTGCAAGTAGCTGGGATTACAGGCACNNNAGATCGGAAG |
| 47777_2 | TATGAGTGTGGAGTCGTTGCAAGTAGCTGGGATTACAGGCACNNNAGATCGGAAG |
| 47777_3 | TATGAGTGTGGAGTCGTTGCAAGTAGCTGGGATTACAGGCACNNNAGATCGGAAG |
| 36707_1 | TATGAGTGTGGAGTCGTTGCCCAGAAAAGGCAAAACTACAGGGNNNAGATCGGAA |
| 36707_2 | TATGAGTGTGGAGTCGTTGCCCAGAAAAGGCAAAACTACAGGGNNNAGATCGGAA |
| 36707_3 | TATGAGTGTGGAGTCGTTGCCCAGAAAAGGCAAAACTACAGGGNNNAGATCGGAA |
| 36650_1 | TATGAGTGTGGAGTCGTTGCGTCTGGGAAATGGAGTTTTGGTGNNNAGATCGGAA |
| 36650_2 | TATGAGTGTGGAGTCGTTGCGTCTGGGAAATGGAGTTTTGGTGNNNAGATCGGAA |
| 36650_3 | TATGAGTGTGGAGTCGTTGCGTCTGGGAAATGGAGTTTTGGTGNNNAGATCGGAA |
| 32060_1 | TATGAGTGTGGAGTCGTTGCGTAGTCTCCTTCTGGCCCAAATTNNNAGATCGGAAGA |
| 32060_2 | TATGAGTGTGGAGTCGTTGCGTAGTCTCCTTCTGGCCCAAATTNNNAGATCGGAAGA |
| 32060_3 | TATGAGTGTGGAGTCGTTGCGTAGTCTCCTTCTGGCCCAAATTNNNAGATCGGAAGA |
| 39114_1 | TATGAGTGTGGAGTCGTTGCCAGAAGATTGAAACTGGACCCCTNNNAGATCGGAAG |
| 39114_2 | TATGAGTGTGGAGTCGTTGCCAGAAGATTGAAACTGGACCCCTNNNAGATCGGAAG |
| 39114_3 | TATGAGTGTGGAGTCGTTGCCAGAAGATTGAAACTGGACCCCTNNNAGATCGGAAG |
| 32723_1 | TATGAGTGTGGAGTCGTTGCGTTCCAGGGGGTGAAAGACTATANNNAGATCGGAA |
| 32723_2 | TATGAGTGTGGAGTCGTTGCGTTCCAGGGGGTGAAAGACTATANNNAGATCGGAA |
| 32723_3 | TATGAGTGTGGAGTCGTTGCGTTCCAGGGGGTGAAAGACTATANNNAGATCGGAA |
| 37654_1 | TATGAGTGTGGAGTCGTTGCGGTAACCTCCATATTTCCCCGTANNNAGATCGGAAGA |
| 37654_2 | TATGAGTGTGGAGTCGTTGCGGTAACCTCCATATTTCCCCGTANNNAGATCGGAAGA |
| 37654_3 | TATGAGTGTGGAGTCGTTGCGGTAACCTCCATATTTCCCCGTANNNAGATCGGAAGA |
| 35370_1 | TATGAGTGTGGAGTCGTTGCTATGCTTCTGAACCTCCCTAGTGNNNAGATCGGAAGA |
| 35370_2 | TATGAGTGTGGAGTCGTTGCTATGCTTCTGAACCTCCCTAGTGNNNAGATCGGAAGA |
| 35370_3 | TATGAGTGTGGAGTCGTTGCTATGCTTCTGAACCTCCCTAGTGNNNAGATCGGAAGA |
| 37787_1 | TATGAGTGTGGAGTCGTTGCGAAGAAAAGAAAAGGCAGGGCCNNNAGATCGGAA[illegible] |
| 37787_2 | TATGAGTGTGGAGTCGTTGCGAAGAAAAGAAAAGGCAGGGCCNNNAGATCGGAA[illegible] |
| 37787_3 | TATGAGTGTGGAGTCGTTGCGAAGAAAAGAAAAGGCAGGGCCNNNAGATCGGAA[illegible] |
| 13081_1 | TATGAGTGTGGAGTCGTTGCTATTAATGCTAAGATTTAGTGCCNNNAGATCGGAAG |
| 13081_2 | TATGAGTGTGGAGTCGTTGCTATTAATGCTAAGATTTAGTGCCNNNAGATCGGAAG |
| 13081_3 | TATGAGTGTGGAGTCGTTGCTATTAATGCTAAGATTTAGTGCCNNNAGATCGGAAG |
| 13717_1 | TATGAGTGTGGAGTCGTTGCGGTCTGTCTTAACATTCTTATTGNNNAGATCGGAAGA |
| 13717_2 | TATGAGTGTGGAGTCGTTGCGGTCTGTCTTAACATTCTTATTGNNNAGATCGGAAGA |
| 13717_3 | TATGAGTGTGGAGTCGTTGCGGTCTGTCTTAACATTCTTATTGNNNAGATCGGAAGA |
| 34307_1 | TATGAGTGTGGAGTCGTTGCGAGAAACCATGGTGTCGTCTTTGNNNAGATCGGAAG |
| 34307_2 | TATGAGTGTGGAGTCGTTGCGAGAAACCATGGTGTCGTCTTTGNNNAGATCGGAAG |
| 34307_3 | TATGAGTGTGGAGTCGTTGCGAGAAACCATGGTGTCGTCTTTGNNNAGATCGGAAG |
| 49329_1 | TATGAGTGTGGAGTCGTTGCACCAGACTCAACCTTCCAAAGAGNNNAGATCGGAAG |
| 49329_2 | TATGAGTGTGGAGTCGTTGCACCAGACTCAACCTTCCAAAGAGNNNAGATCGGAAG |
| 49329_3 | TATGAGTGTGGAGTCGTTGCACCAGACTCAACCTTCCAAAGAGNNNAGATCGGAAG |

| | |
|---|---|
| 42153_1 | TATGAGTGTGGAGTCGTTGCTTTTTTGAGACGGAGTCTTGCTCNNNAGATCGGAAG |
| 42153_2 | TATGAGTGTGGAGTCGTTGCTTTTTTGAGACGGAGTCTTGCTCNNNAGATCGGAAG |
| 42153_3 | TATGAGTGTGGAGTCGTTGCTTTTTTGAGACGGAGTCTTGCTCNNNAGATCGGAAG |
| 46699_1 | TATGAGTGTGGAGTCGTTGCCTCATTCTCTTATCCAGGCTGCANNNAGATCGGAAGA |
| 46699_2 | TATGAGTGTGGAGTCGTTGCCTCATTCTCTTATCCAGGCTGCANNNAGATCGGAAGA |
| 46699_3 | TATGAGTGTGGAGTCGTTGCCTCATTCTCTTATCCAGGCTGCANNNAGATCGGAAGA |
| 47059_1 | TATGAGTGTGGAGTCGTTGCGAGAAGAAGCTGGCAGCAATTAGNNNAGATCGGAA |
| 47059_2 | TATGAGTGTGGAGTCGTTGCGAGAAGAAGCTGGCAGCAATTAGNNNAGATCGGAA |
| 47059_3 | TATGAGTGTGGAGTCGTTGCGAGAAGAAGCTGGCAGCAATTAGNNNAGATCGGAA |
| 44333_1 | TATGAGTGTGGAGTCGTTGCGAGCTTGGTTGGGTTTTTACACANNNAGATCGGAAG |
| 44333_2 | TATGAGTGTGGAGTCGTTGCGAGCTTGGTTGGGTTTTTACACANNNAGATCGGAAG |
| 44333_3 | TATGAGTGTGGAGTCGTTGCGAGCTTGGTTGGGTTTTTACACANNNAGATCGGAAG |
| 30092_1 | TATGAGTGTGGAGTCGTTGCTGGGACAAAGAATTGGATCTGGANNNAGATCGGAA |
| 30092_2 | TATGAGTGTGGAGTCGTTGCTGGGACAAAGAATTGGATCTGGANNNAGATCGGAA |
| 30092_3 | TATGAGTGTGGAGTCGTTGCTGGGACAAAGAATTGGATCTGGANNNAGATCGGAA |
| 29586_1 | TATGAGTGTGGAGTCGTTGCTAAGCAGCATCCCTGGTCTACANNNAGATCGGAAGA |
| 29586_2 | TATGAGTGTGGAGTCGTTGCTAAGCAGCATCCCTGGTCTACANNNAGATCGGAAGA |
| 29586_3 | TATGAGTGTGGAGTCGTTGCTAAGCAGCATCCCTGGTCTACANNNAGATCGGAAGA |
| 52121_1 | TATGAGTGTGGAGTCGTTGCCCAGACCCAAACATTTTCCCCTANNNAGATCGGAAGA |
| 52121_2 | TATGAGTGTGGAGTCGTTGCCCAGACCCAAACATTTTCCCCTANNNAGATCGGAAGA |
| 52121_3 | TATGAGTGTGGAGTCGTTGCCCAGACCCAAACATTTTCCCCTANNNAGATCGGAAGA |
| 40096_1 | TATGAGTGTGGAGTCGTTGCAAAACTGGGGTATGCTTGCTTTGNNNAGATCGGAAG |
| 40096_2 | TATGAGTGTGGAGTCGTTGCAAAACTGGGGTATGCTTGCTTTGNNNAGATCGGAAG |
| 40096_3 | TATGAGTGTGGAGTCGTTGCAAAACTGGGGTATGCTTGCTTTGNNNAGATCGGAAG |
| 29612_1 | TATGAGTGTGGAGTCGTTGCACCTACAGAAATGCACGAATAGGANNNAGATCGGAA |
| 29612_2 | TATGAGTGTGGAGTCGTTGCACCTACAGAAATGCACGAATAGGANNNAGATCGGAA |
| 29612_3 | TATGAGTGTGGAGTCGTTGCACCTACAGAAATGCACGAATAGGANNNAGATCGGAA |
| 48207_1 | TATGAGTGTGGAGTCGTTGCCTCCATGACTTTCCTCTCTACGGNNNAGATCGGAAGA |
| 48207_2 | TATGAGTGTGGAGTCGTTGCCTCCATGACTTTCCTCTCTACGGNNNAGATCGGAAGA |
| 48207_3 | TATGAGTGTGGAGTCGTTGCCTCCATGACTTTCCTCTCTACGGNNNAGATCGGAAGA |
| 13110_1 | TATGAGTGTGGAGTCGTTGCCATTGGATTTCAAGTAAAACATTNNNAGATCGGAAG |
| 13110_2 | TATGAGTGTGGAGTCGTTGCCATTGGATTTCAAGTAAAACATTNNNAGATCGGAAG |
| 13110_3 | TATGAGTGTGGAGTCGTTGCCATTGGATTTCAAGTAAAACATTNNNAGATCGGAAG |
| 52448_1 | TATGAGTGTGGAGTCGTTGCTGGGAAACTGTGGGCAATTTTTTNNNAGATCGGAAG |
| 52448_2 | TATGAGTGTGGAGTCGTTGCTGGGAAACTGTGGGCAATTTTTTNNNAGATCGGAAG |
| 52448_3 | TATGAGTGTGGAGTCGTTGCTGGGAAACTGTGGGCAATTTTTTNNNAGATCGGAAG |
| 53269_1 | TATGAGTGTGGAGTCGTTGCGTTCTGGTTGCTGTTAGTTTCCCNNNAGATCGGAAGA |
| 53269_2 | TATGAGTGTGGAGTCGTTGCGTTCTGGTTGCTGTTAGTTTCCCNNNAGATCGGAAGA |
| 53269_3 | TATGAGTGTGGAGTCGTTGCGTTCTGGTTGCTGTTAGTTTCCCNNNAGATCGGAAGA |
| 51102_1 | TATGAGTGTGGAGTCGTTGCCCTACTGGGAAGTGAGGAGCCNNNAGATCGGAAGA |
| 51102_2 | TATGAGTGTGGAGTCGTTGCCCTACTGGGAAGTGAGGAGCCNNNAGATCGGAAGA |
| 51102_3 | TATGAGTGTGGAGTCGTTGCCCTACTGGGAAGTGAGGAGCCNNNAGATCGGAAGA |
| 44004_1 | TATGAGTGTGGAGTCGTTGCGAAAGTTGGAGAACCAGGAAAGCNNNAGATCGGAA |

| | |
|---|---|
| 44004_2 | TATGAGTGTGGAGTCGTTGCGAAAGTTGGAGAACCAGGAAAGCNNNAGATCGGA |
| 44004_3 | TATGAGTGTGGAGTCGTTGCGAAAGTTGGAGAACCAGGAAAGCNNNAGATCGGA |
| 45717_1 | TATGAGTGTGGAGTCGTTGCTTCAAGTGATTCTCCTGCCTCAANNNAGATCGGAAG |
| 45717_2 | TATGAGTGTGGAGTCGTTGCTTCAAGTGATTCTCCTGCCTCAANNNAGATCGGAAG |
| 45717_3 | TATGAGTGTGGAGTCGTTGCTTCAAGTGATTCTCCTGCCTCAANNNAGATCGGAAG |
| 32860_1 | TATGAGTGTGGAGTCGTTGCTGGGTTCAAGTGATTCTCCTACCNNNAGATCGGAAG |
| 32860_2 | TATGAGTGTGGAGTCGTTGCTGGGTTCAAGTGATTCTCCTACCNNNAGATCGGAAG |
| 32860_3 | TATGAGTGTGGAGTCGTTGCTGGGTTCAAGTGATTCTCCTACCNNNAGATCGGAAG |
| 41862_1 | TATGAGTGTGGAGTCGTTGCCATGATCTCAGCTCACTGCAAGNNNAGATCGGAAGA |
| 41862_2 | TATGAGTGTGGAGTCGTTGCCATGATCTCAGCTCACTGCAAGNNNAGATCGGAAGA |
| 41862_3 | TATGAGTGTGGAGTCGTTGCCATGATCTCAGCTCACTGCAAGNNNAGATCGGAAGA |
| 37839_1 | TATGAGTGTGGAGTCGTTGCGGGCTGGGACTTGAAAGATAAGANNNAGATCGGA |
| 37839_2 | TATGAGTGTGGAGTCGTTGCGGGCTGGGACTTGAAAGATAAGANNNAGATCGGA |
| 37839_3 | TATGAGTGTGGAGTCGTTGCGGGCTGGGACTTGAAAGATAAGANNNAGATCGGA |
| 51265_1 | TATGAGTGTGGAGTCGTTGCTGAGTTCAGATTCTCCTTGGGTGNNNAGATCGGAAG |
| 51265_2 | TATGAGTGTGGAGTCGTTGCTGAGTTCAGATTCTCCTTGGGTGNNNAGATCGGAAG |
| 51265_3 | TATGAGTGTGGAGTCGTTGCTGAGTTCAGATTCTCCTTGGGTGNNNAGATCGGAAG |
| 13734_1 | TATGAGTGTGGAGTCGTTGCTTGCCTTTCATGAATACCAGNNNAGATCGGAAGAGC |
| 13734_2 | TATGAGTGTGGAGTCGTTGCTTGCCTTTCATGAATACCAGNNNAGATCGGAAGAGC |
| 13734_3 | TATGAGTGTGGAGTCGTTGCTTGCCTTTCATGAATACCAGNNNAGATCGGAAGAGC |
| 33685_1 | TATGAGTGTGGAGTCGTTGCTGGAGTGCAATGGTGTGATCTCNNNAGATCGGAAG |
| 33685_2 | TATGAGTGTGGAGTCGTTGCTGGAGTGCAATGGTGTGATCTCNNNAGATCGGAAG |
| 33685_3 | TATGAGTGTGGAGTCGTTGCTGGAGTGCAATGGTGTGATCTCNNNAGATCGGAAG |
| 50462_1 | TATGAGTGTGGAGTCGTTGCCTTCTGGGAGGATTGATCTGAGGNNNAGATCGGAA |
| 50462_2 | TATGAGTGTGGAGTCGTTGCCTTCTGGGAGGATTGATCTGAGGNNNAGATCGGAA |
| 50462_3 | TATGAGTGTGGAGTCGTTGCCTTCTGGGAGGATTGATCTGAGGNNNAGATCGGAA |
| 47210_1 | TATGAGTGTGGAGTCGTTGCGATTCAAGCAATTCTCCTGCCTCNNNAGATCGGAAG |
| 47210_2 | TATGAGTGTGGAGTCGTTGCGATTCAAGCAATTCTCCTGCCTCNNNAGATCGGAAG |
| 47210_3 | TATGAGTGTGGAGTCGTTGCGATTCAAGCAATTCTCCTGCCTCNNNAGATCGGAAG |
| 12403_1 | TATGAGTGTGGAGTCGTTGCGCAAGATGTTCTCGGGCTTGATNNNAGATCGGAAG |
| 12403_2 | TATGAGTGTGGAGTCGTTGCGCAAGATGTTCTCGGGCTTGATNNNAGATCGGAAG |
| 12403_3 | TATGAGTGTGGAGTCGTTGCGCAAGATGTTCTCGGGCTTGATNNNAGATCGGAAG |
| 47247_1 | TATGAGTGTGGAGTCGTTGCGGAGTTTCGCTCTTGTTACCCAGNNNAGATCGGAAG |
| 47247_2 | TATGAGTGTGGAGTCGTTGCGGAGTTTCGCTCTTGTTACCCAGNNNAGATCGGAAG |
| 47247_3 | TATGAGTGTGGAGTCGTTGCGGAGTTTCGCTCTTGTTACCCAGNNNAGATCGGAAG |
| 50237_1 | TATGAGTGTGGAGTCGTTGCGAAGGGTCTTTCTCTGTCTCCTGNNNAGATCGGAAG |
| 50237_2 | TATGAGTGTGGAGTCGTTGCGAAGGGTCTTTCTCTGTCTCCTGNNNAGATCGGAAG |
| 50237_3 | TATGAGTGTGGAGTCGTTGCGAAGGGTCTTTCTCTGTCTCCTGNNNAGATCGGAAG |
| 42051_1 | TATGAGTGTGGAGTCGTTGCTGACCAAGTTATCAGTCTCTACCANNNAGATCGGAA |
| 42051_2 | TATGAGTGTGGAGTCGTTGCTGACCAAGTTATCAGTCTCTACCANNNAGATCGGAA |
| 42051_3 | TATGAGTGTGGAGTCGTTGCTGACCAAGTTATCAGTCTCTACCANNNAGATCGGAA |
| 29998_1 | TATGAGTGTGGAGTCGTTGCCAACAATGAAGGTGTCTGTGTCCNNNAGATCGGAA |
| 29998_2 | TATGAGTGTGGAGTCGTTGCCAACAATGAAGGTGTCTGTGTCCNNNAGATCGGAA |

| | |
|---|---|
| 29998_3 | TATGAGTGTGGAGTCGTTGCCAACAATGAAGGTGTCTGTGTCCNNNAGATCGGAAG |
| 33548_1 | TATGAGTGTGGAGTCGTTGCCCTTAGCTTCCTGAGTACTTGGGNNNAGATCGGAAG |
| 33548_2 | TATGAGTGTGGAGTCGTTGCCCTTAGCTTCCTGAGTACTTGGGNNNAGATCGGAAG |
| 33548_3 | TATGAGTGTGGAGTCGTTGCCCTTAGCTTCCTGAGTACTTGGGNNNAGATCGGAAG |
| 3125_1 | TATGAGTGTGGAGTCGTTGCATGCCACCCAACTTTTCATACGTNNNAGATCGGAAGA |
| 3125_2 | TATGAGTGTGGAGTCGTTGCATGCCACCCAACTTTTCATACGTNNNAGATCGGAAGA |
| 3125_3 | TATGAGTGTGGAGTCGTTGCATGCCACCCAACTTTTCATACGTNNNAGATCGGAAGA |
| 47587_1 | TATGAGTGTGGAGTCGTTGCAGCTTTGCAAAATTGACAAACCTNNNAGATCGGAAG |
| 47587_2 | TATGAGTGTGGAGTCGTTGCAGCTTTGCAAAATTGACAAACCTNNNAGATCGGAAG |
| 47587_3 | TATGAGTGTGGAGTCGTTGCAGCTTTGCAAAATTGACAAACCTNNNAGATCGGAAG |
| 33130_1 | TATGAGTGTGGAGTCGTTGCAACTGTGGTAGCACCTCTGTAAGNNNAGATCGGAAG |
| 33130_2 | TATGAGTGTGGAGTCGTTGCAACTGTGGTAGCACCTCTGTAAGNNNAGATCGGAAG |
| 33130_3 | TATGAGTGTGGAGTCGTTGCAACTGTGGTAGCACCTCTGTAAGNNNAGATCGGAAG |
| 33406_1 | TATGAGTGTGGAGTCGTTGCTCATTTCCAGCAGAACTCTGAGTNNNAGATCGGAAG |
| 33406_2 | TATGAGTGTGGAGTCGTTGCTCATTTCCAGCAGAACTCTGAGTNNNAGATCGGAAG |
| 33406_3 | TATGAGTGTGGAGTCGTTGCTCATTTCCAGCAGAACTCTGAGTNNNAGATCGGAAG |
| 36757_1 | TATGAGTGTGGAGTCGTTGCAACAAGATGGAGTTGGAGTTGGTNNNAGATCGGAA |
| 36757_2 | TATGAGTGTGGAGTCGTTGCAACAAGATGGAGTTGGAGTTGGTNNNAGATCGGAA |
| 36757_3 | TATGAGTGTGGAGTCGTTGCAACAAGATGGAGTTGGAGTTGGTNNNAGATCGGAA |
| 46021_1 | TATGAGTGTGGAGTCGTTGCCAATATCCGCTCACCACAACTTCNNNAGATCGGAAGA |
| 46021_2 | TATGAGTGTGGAGTCGTTGCCAATATCCGCTCACCACAACTTCNNNAGATCGGAAGA |
| 46021_3 | TATGAGTGTGGAGTCGTTGCCAATATCCGCTCACCACAACTTCNNNAGATCGGAAGA |
| 36623_1 | TATGAGTGTGGAGTCGTTGCATAGTGTGACTCAGAGATGGGGANNNAGATCGGAA |
| 36623_2 | TATGAGTGTGGAGTCGTTGCATAGTGTGACTCAGAGATGGGGANNNAGATCGGAA |
| 36623_3 | TATGAGTGTGGAGTCGTTGCATAGTGTGACTCAGAGATGGGGANNNAGATCGGAA |
| 31287_1 | TATGAGTGTGGAGTCGTTGCAACAACAAACTCTCCCAAGAAGANNNAGATCGGAAG |
| 31287_2 | TATGAGTGTGGAGTCGTTGCAACAACAAACTCTCCCAAGAAGANNNAGATCGGAAG |
| 31287_3 | TATGAGTGTGGAGTCGTTGCAACAACAAACTCTCCCAAGAAGANNNAGATCGGAAG |
| 3084_1 | TATGAGTGTGGAGTCGTTGCGTTCGGGAGCAAGAATTGGTTTGNNNAGATCGGAA |
| 3084_2 | TATGAGTGTGGAGTCGTTGCGTTCGGGAGCAAGAATTGGTTTGNNNAGATCGGAA |
| 3084_3 | TATGAGTGTGGAGTCGTTGCGTTCGGGAGCAAGAATTGGTTTGNNNAGATCGGAA |
| 37474_1 | TATGAGTGTGGAGTCGTTGCCATTACAAACAGCACTGCAGTGANNNAGATCGGAAG |
| 37474_2 | TATGAGTGTGGAGTCGTTGCCATTACAAACAGCACTGCAGTGANNNAGATCGGAAG |
| 37474_3 | TATGAGTGTGGAGTCGTTGCCATTACAAACAGCACTGCAGTGANNNAGATCGGAAG |
| 45280_1 | TATGAGTGTGGAGTCGTTGCGTTACCATCACCAACACATTGCANNNAGATCGGAAG |
| 45280_2 | TATGAGTGTGGAGTCGTTGCGTTACCATCACCAACACATTGCANNNAGATCGGAAG |
| 45280_3 | TATGAGTGTGGAGTCGTTGCGTTACCATCACCAACACATTGCANNNAGATCGGAAG |
| 41187_1 | TATGAGTGTGGAGTCGTTGCAGCTACTATGGAAACCAGTGTGGNNNAGATCGGAA |
| 41187_2 | TATGAGTGTGGAGTCGTTGCAGCTACTATGGAAACCAGTGTGGNNNAGATCGGAA |
| 41187_3 | TATGAGTGTGGAGTCGTTGCAGCTACTATGGAAACCAGTGTGGNNNAGATCGGAA |
| 50223_1 | TATGAGTGTGGAGTCGTTGCGGGTCATTATATTACAGGCGGGANNNAGATCGGAA |
| 50223_2 | TATGAGTGTGGAGTCGTTGCGGGTCATTATATTACAGGCGGGANNNAGATCGGAA |
| 50223_3 | TATGAGTGTGGAGTCGTTGCGGGTCATTATATTACAGGCGGGANNNAGATCGGAA |

| | |
|---|---|
| 32757_1 | TATGAGTGTGGAGTCGTTGCATTGAAGGCTATGGGCACAAATGNNNAGATCGGAA |
| 32757_2 | TATGAGTGTGGAGTCGTTGCATTGAAGGCTATGGGCACAAATGNNNAGATCGGAA |
| 32757_3 | TATGAGTGTGGAGTCGTTGCATTGAAGGCTATGGGCACAAATGNNNAGATCGGAA |
| 2250_1 | TATGAGTGTGGAGTCGTTGCAGAAAAGGGCTTGGTCAGTCCTANNNAGATCGGAA |
| 2250_2 | TATGAGTGTGGAGTCGTTGCAGAAAAGGGCTTGGTCAGTCCTANNNAGATCGGAA |
| 2250_3 | TATGAGTGTGGAGTCGTTGCAGAAAAGGGCTTGGTCAGTCCTANNNAGATCGGAA |
| 52240_1 | TATGAGTGTGGAGTCGTTGCTCACCACATGGCATGTATTCTGANNNAGATCGGAAG |
| 52240_2 | TATGAGTGTGGAGTCGTTGCTCACCACATGGCATGTATTCTGANNNAGATCGGAAG |
| 52240_3 | TATGAGTGTGGAGTCGTTGCTCACCACATGGCATGTATTCTGANNNAGATCGGAAG |
| 8338_1 | TATGAGTGTGGAGTCGTTGCAGTAGGTGAGGTGCAGGCACNNNAGATCGGAAGA |
| 8338_2 | TATGAGTGTGGAGTCGTTGCAGTAGGTGAGGTGCAGGCACNNNAGATCGGAAGA |
| 8338_3 | TATGAGTGTGGAGTCGTTGCAGTAGGTGAGGTGCAGGCACNNNAGATCGGAAGA |
| 44551_1 | TATGAGTGTGGAGTCGTTGCTCGACCAGTACAAGGTGTCTTGNNNAGATCGGAAG |
| 44551_2 | TATGAGTGTGGAGTCGTTGCTCGACCAGTACAAGGTGTCTTGNNNAGATCGGAAG |
| 44551_3 | TATGAGTGTGGAGTCGTTGCTCGACCAGTACAAGGTGTCTTGNNNAGATCGGAAG |
| 48491_1 | TATGAGTGTGGAGTCGTTGCCTCATTTCATCACTGTTTGGCGTNNNAGATCGGAAG |
| 48491_2 | TATGAGTGTGGAGTCGTTGCCTCATTTCATCACTGTTTGGCGTNNNAGATCGGAAG |
| 48491_3 | TATGAGTGTGGAGTCGTTGCCTCATTTCATCACTGTTTGGCGTNNNAGATCGGAAG |
| 47802_1 | TATGAGTGTGGAGTCGTTGCTGATCTTGTCTCACTGCAACCTCNNNAGATCGGAAGA |
| 47802_2 | TATGAGTGTGGAGTCGTTGCTGATCTTGTCTCACTGCAACCTCNNNAGATCGGAAGA |
| 47802_3 | TATGAGTGTGGAGTCGTTGCTGATCTTGTCTCACTGCAACCTCNNNAGATCGGAAGA |
| 29189_1 | TATGAGTGTGGAGTCGTTGCATTCCCTGAAAGCTTCTCCTGACNNNAGATCGGAAG |
| 29189_2 | TATGAGTGTGGAGTCGTTGCATTCCCTGAAAGCTTCTCCTGACNNNAGATCGGAAG |
| 29189_3 | TATGAGTGTGGAGTCGTTGCATTCCCTGAAAGCTTCTCCTGACNNNAGATCGGAAG |
| 5369_1 | TATGAGTGTGGAGTCGTTGCGTCTGTGTGTGTGTTTGTTGTTTTNNNAGATCGGAA |
| 5369_2 | TATGAGTGTGGAGTCGTTGCGTCTGTGTGTGTGTTTGTTGTTTTNNNAGATCGGAA |
| 5369_3 | TATGAGTGTGGAGTCGTTGCGTCTGTGTGTGTGTTTGTTGTTTTNNNAGATCGGAA |
| 51549_1 | TATGAGTGTGGAGTCGTTGCGCAATAGTCATCTGTGGCATGTCNNNAGATCGGAAG |
| 51549_2 | TATGAGTGTGGAGTCGTTGCGCAATAGTCATCTGTGGCATGTCNNNAGATCGGAAG |
| 51549_3 | TATGAGTGTGGAGTCGTTGCGCAATAGTCATCTGTGGCATGTCNNNAGATCGGAAG |
| 29438_1 | TATGAGTGTGGAGTCGTTGCGCAAGGAAAGAGTATTTAAAAACNNNAGATCGGAA |
| 29438_2 | TATGAGTGTGGAGTCGTTGCGCAAGGAAAGAGTATTTAAAAACNNNAGATCGGAA |
| 29438_3 | TATGAGTGTGGAGTCGTTGCGCAAGGAAAGAGTATTTAAAAACNNNAGATCGGAA |
| 42673_1 | TATGAGTGTGGAGTCGTTGCGGTTGAGAATTCCCATGATCTGCNNNAGATCGGAAG |
| 42673_2 | TATGAGTGTGGAGTCGTTGCGGTTGAGAATTCCCATGATCTGCNNNAGATCGGAAG |
| 42673_3 | TATGAGTGTGGAGTCGTTGCGGTTGAGAATTCCCATGATCTGCNNNAGATCGGAAG |
| 34116_1 | TATGAGTGTGGAGTCGTTGCGAGGAAAGAAAGGCATGCATGAANNNAGATCGGA |
| 34116_2 | TATGAGTGTGGAGTCGTTGCGAGGAAAGAAAGGCATGCATGAANNNAGATCGGA |
| 34116_3 | TATGAGTGTGGAGTCGTTGCGAGGAAAGAAAGGCATGCATGAANNNAGATCGGA |
| 52369_1 | TATGAGTGTGGAGTCGTTGCCACAGAAGGCAAACTGGATCAAGNNNAGATCGGAA |
| 52369_2 | TATGAGTGTGGAGTCGTTGCCACAGAAGGCAAACTGGATCAAGNNNAGATCGGAA |
| 52369_3 | TATGAGTGTGGAGTCGTTGCCACAGAAGGCAAACTGGATCAAGNNNAGATCGGAA |
| 50201_1 | TATGAGTGTGGAGTCGTTGCCAAGTCTTTCACCCCATTGTCACNNNAGATCGGAAGA |

| | |
|---|---|
| 50201_2 | TATGAGTGTGGAGTCGTTGCCAAGTCTTTCACCCCATTGTCACNNNAGATCGGAAGA |
| 50201_3 | TATGAGTGTGGAGTCGTTGCCAAGTCTTTCACCCCATTGTCACNNNAGATCGGAAGA |
| 10137_1 | TATGAGTGTGGAGTCGTTGCATGGGAGTTCCTTGTACTGTGCTNNNAGATCGGAAG |
| 10137_2 | TATGAGTGTGGAGTCGTTGCATGGGAGTTCCTTGTACTGTGCTNNNAGATCGGAAG |
| 10137_3 | TATGAGTGTGGAGTCGTTGCATGGGAGTTCCTTGTACTGTGCTNNNAGATCGGAAG |
| 31393_1 | TATGAGTGTGGAGTCGTTGCAAGGAAGGAGGAAGCAACAAGAANNNAGATCGGA |
| 31393_2 | TATGAGTGTGGAGTCGTTGCAAGGAAGGAGGAAGCAACAAGAANNNAGATCGGA |
| 31393_3 | TATGAGTGTGGAGTCGTTGCAAGGAAGGAGGAAGCAACAAGAANNNAGATCGGA |
| 30660_1 | TATGAGTGTGGAGTCGTTGCTAAGATCAGCTTCACTGGGCTAAANNNAGATCGGAA |
| 30660_2 | TATGAGTGTGGAGTCGTTGCTAAGATCAGCTTCACTGGGCTAAANNNAGATCGGAA |
| 30660_3 | TATGAGTGTGGAGTCGTTGCTAAGATCAGCTTCACTGGGCTAAANNNAGATCGGAA |
| 36483_1 | TATGAGTGTGGAGTCGTTGCAGACAAGGGAAGGGAAGGAAAAANNNAGATCGGA |
| 36483_2 | TATGAGTGTGGAGTCGTTGCAGACAAGGGAAGGGAAGGAAAAANNNAGATCGGA |
| 36483_3 | TATGAGTGTGGAGTCGTTGCAGACAAGGGAAGGGAAGGAAAAANNNAGATCGGA |
| 35001_1 | TATGAGTGTGGAGTCGTTGCGATCGAATGTGTCTGATGCAAGGNNNAGATCGGAA |
| 35001_2 | TATGAGTGTGGAGTCGTTGCGATCGAATGTGTCTGATGCAAGGNNNAGATCGGAA |
| 35001_3 | TATGAGTGTGGAGTCGTTGCGATCGAATGTGTCTGATGCAAGGNNNAGATCGGAA |
| 38761_1 | TATGAGTGTGGAGTCGTTGCTGTAAGTGTTTACGTGTGCAGTTNNNAGATCGGAAG |
| 38761_2 | TATGAGTGTGGAGTCGTTGCTGTAAGTGTTTACGTGTGCAGTTNNNAGATCGGAAG |
| 38761_3 | TATGAGTGTGGAGTCGTTGCTGTAAGTGTTTACGTGTGCAGTTNNNAGATCGGAAG |
| 45728_1 | TATGAGTGTGGAGTCGTTGCCTCCTGAGTAGCTGGGATTACAGNNNAGATCGGAA |
| 45728_2 | TATGAGTGTGGAGTCGTTGCCTCCTGAGTAGCTGGGATTACAGNNNAGATCGGAA |
| 45728_3 | TATGAGTGTGGAGTCGTTGCCTCCTGAGTAGCTGGGATTACAGNNNAGATCGGAA |
| 2103_1 | TATGAGTGTGGAGTCGTTGCTTGTCTTCCCTTCTCTACCAGCANNNAGATCGGAAGA |
| 2103_2 | TATGAGTGTGGAGTCGTTGCTTGTCTTCCCTTCTCTACCAGCANNNAGATCGGAAGA |
| 2103_3 | TATGAGTGTGGAGTCGTTGCTTGTCTTCCCTTCTCTACCAGCANNNAGATCGGAAGA |
| 30824_1 | TATGAGTGTGGAGTCGTTGCGGTACAGGATTGGAAACTGAGGANNNAGATCGGA |
| 30824_2 | TATGAGTGTGGAGTCGTTGCGGTACAGGATTGGAAACTGAGGANNNAGATCGGA |
| 30824_3 | TATGAGTGTGGAGTCGTTGCGGTACAGGATTGGAAACTGAGGANNNAGATCGGA |
| 52527_1 | TATGAGTGTGGAGTCGTTGCCACAGGCTGCCCTCTTATACTTANNNAGATCGGAAG |
| 52527_2 | TATGAGTGTGGAGTCGTTGCCACAGGCTGCCCTCTTATACTTANNNAGATCGGAAG |
| 52527_3 | TATGAGTGTGGAGTCGTTGCCACAGGCTGCCCTCTTATACTTANNNAGATCGGAAG |
| 48297_1 | TATGAGTGTGGAGTCGTTGCACAATTTGAGGTTGAGAGTGCAGNNNAGATCGGAA |
| 48297_2 | TATGAGTGTGGAGTCGTTGCACAATTTGAGGTTGAGAGTGCAGNNNAGATCGGAA |
| 48297_3 | TATGAGTGTGGAGTCGTTGCACAATTTGAGGTTGAGAGTGCAGNNNAGATCGGAA |
| 41059_1 | TATGAGTGTGGAGTCGTTGCTCTGTCTCTCTCCTCTCTCTCTCNNNAGATCGGAAGAG |
| 41059_2 | TATGAGTGTGGAGTCGTTGCTCTGTCTCTCTCCTCTCTCTCTCNNNAGATCGGAAGAG |
| 41059_3 | TATGAGTGTGGAGTCGTTGCTCTGTCTCTCTCCTCTCTCTCTCNNNAGATCGGAAGAG |
| 52736_1 | TATGAGTGTGGAGTCGTTGCTAACTTCCGAGCAGCTAGATCTGNNNAGATCGGAAG |
| 52736_2 | TATGAGTGTGGAGTCGTTGCTAACTTCCGAGCAGCTAGATCTGNNNAGATCGGAAG |
| 52736_3 | TATGAGTGTGGAGTCGTTGCTAACTTCCGAGCAGCTAGATCTGNNNAGATCGGAAG |
| 10017_1 | TATGAGTGTGGAGTCGTTGCTGAAGCAGCCACAACAATGAAAGNNNAGATCGGAA |
| 10017_2 | TATGAGTGTGGAGTCGTTGCTGAAGCAGCCACAACAATGAAAGNNNAGATCGGAA |

| | |
|---|---|
| 10017_3 | TATGAGTGTGGAGTCGTTGCTGAAGCAGCCACAACAATGAAAGNNNAGATCGGAA |
| 34077_1 | TATGAGTGTGGAGTCGTTGCCTTCCTGAGTAGCTGGGACTACANNNAGATCGGAAG |
| 34077_2 | TATGAGTGTGGAGTCGTTGCCTTCCTGAGTAGCTGGGACTACANNNAGATCGGAAG |
| 34077_3 | TATGAGTGTGGAGTCGTTGCCTTCCTGAGTAGCTGGGACTACANNNAGATCGGAAG |
| 43077_1 | TATGAGTGTGGAGTCGTTGCCCACACACACATCGCTAGTCATANNNAGATCGGAAG |
| 43077_2 | TATGAGTGTGGAGTCGTTGCCCACACACACATCGCTAGTCATANNNAGATCGGAAG |
| 43077_3 | TATGAGTGTGGAGTCGTTGCCCACACACACATCGCTAGTCATANNNAGATCGGAAG |
| 50370_1 | TATGAGTGTGGAGTCGTTGCACCTACTCGACTTCATCATCGTCNNNAGATCGGAAGA |
| 50370_2 | TATGAGTGTGGAGTCGTTGCACCTACTCGACTTCATCATCGTCNNNAGATCGGAAGA |
| 50370_3 | TATGAGTGTGGAGTCGTTGCACCTACTCGACTTCATCATCGTCNNNAGATCGGAAGA |
| 46448_1 | TATGAGTGTGGAGTCGTTGCTGCTTTATGGAGTTTGAGCTTGCNNNAGATCGGAAG |
| 46448_2 | TATGAGTGTGGAGTCGTTGCTGCTTTATGGAGTTTGAGCTTGCNNNAGATCGGAAG |
| 46448_3 | TATGAGTGTGGAGTCGTTGCTGCTTTATGGAGTTTGAGCTTGCNNNAGATCGGAAG |
| 10228_1 | TATGAGTGTGGAGTCGTTGCACAGGAAAACTGACATGCCACTTNNNAGATCGGAAG |
| 10228_2 | TATGAGTGTGGAGTCGTTGCACAGGAAAACTGACATGCCACTTNNNAGATCGGAAG |
| 10228_3 | TATGAGTGTGGAGTCGTTGCACAGGAAAACTGACATGCCACTTNNNAGATCGGAAG |
| 34696_1 | TATGAGTGTGGAGTCGTTGCAATGGGGTCTTACCATATTGCCCNNNAGATCGGAAG |
| 34696_2 | TATGAGTGTGGAGTCGTTGCAATGGGGTCTTACCATATTGCCCNNNAGATCGGAAG |
| 34696_3 | TATGAGTGTGGAGTCGTTGCAATGGGGTCTTACCATATTGCCCNNNAGATCGGAAG |
| 45021_1 | TATGAGTGTGGAGTCGTTGCCTGCCTAGGTAACCTGTGTATGGNNNAGATCGGAAG |
| 45021_2 | TATGAGTGTGGAGTCGTTGCCTGCCTAGGTAACCTGTGTATGGNNNAGATCGGAAG |
| 45021_3 | TATGAGTGTGGAGTCGTTGCCTGCCTAGGTAACCTGTGTATGGNNNAGATCGGAAG |
| 12674_1 | TATGAGTGTGGAGTCGTTGCATCTACTCCCACAAGATACTTNNNAGATCGGAAGAG |
| 12674_2 | TATGAGTGTGGAGTCGTTGCATCTACTCCCACAAGATACTTNNNAGATCGGAAGAG |
| 12674_3 | TATGAGTGTGGAGTCGTTGCATCTACTCCCACAAGATACTTNNNAGATCGGAAGAG |
| 30357_1 | TATGAGTGTGGAGTCGTTGCTTCAGCTTGCACGTGTTCTTTTGNNNAGATCGGAAG |
| 30357_2 | TATGAGTGTGGAGTCGTTGCTTCAGCTTGCACGTGTTCTTTTGNNNAGATCGGAAG |
| 30357_3 | TATGAGTGTGGAGTCGTTGCTTCAGCTTGCACGTGTTCTTTTGNNNAGATCGGAAG |
| 30813_1 | TATGAGTGTGGAGTCGTTGCATAGAGTATGTGTGTGTGTGCGTNNNAGATCGGAA |
| 30813_2 | TATGAGTGTGGAGTCGTTGCATAGAGTATGTGTGTGTGTGCGTNNNAGATCGGAA |
| 30813_3 | TATGAGTGTGGAGTCGTTGCATAGAGTATGTGTGTGTGTGCGTNNNAGATCGGAA |
| 40210_1 | TATGAGTGTGGAGTCGTTGCAGGTAGGAGAATTGCTTGAACCCNNNAGATCGGAA |
| 40210_2 | TATGAGTGTGGAGTCGTTGCAGGTAGGAGAATTGCTTGAACCCNNNAGATCGGAA |
| 40210_3 | TATGAGTGTGGAGTCGTTGCAGGTAGGAGAATTGCTTGAACCCNNNAGATCGGAA |
| 35770_1 | TATGAGTGTGGAGTCGTTGCAATAATAGGTGTTCCTGGCTGGGNNNAGATCGGAA |
| 35770_2 | TATGAGTGTGGAGTCGTTGCAATAATAGGTGTTCCTGGCTGGGNNNAGATCGGAA |
| 35770_3 | TATGAGTGTGGAGTCGTTGCAATAATAGGTGTTCCTGGCTGGGNNNAGATCGGAA |
| 52619_1 | TATGAGTGTGGAGTCGTTGCTTCACTGAGGTACAGCACAGATCNNNAGATCGGAAG |
| 52619_2 | TATGAGTGTGGAGTCGTTGCTTCACTGAGGTACAGCACAGATCNNNAGATCGGAAG |
| 52619_3 | TATGAGTGTGGAGTCGTTGCTTCACTGAGGTACAGCACAGATCNNNAGATCGGAAG |
| 37414_1 | TATGAGTGTGGAGTCGTTGCTTCGTGAGGTCCTAAAAGGTACGNNNAGATCGGAA |
| 37414_2 | TATGAGTGTGGAGTCGTTGCTTCGTGAGGTCCTAAAAGGTACGNNNAGATCGGAA |
| 37414_3 | TATGAGTGTGGAGTCGTTGCTTCGTGAGGTCCTAAAAGGTACGNNNAGATCGGAA |

| | |
|---|---|
| 47283_1 | TATGAGTGTGGAGTCGTTGCTATCTCGGTTCACTGCAACATCTNNNAGATCGGAAG |
| 47283_2 | TATGAGTGTGGAGTCGTTGCTATCTCGGTTCACTGCAACATCTNNNAGATCGGAAG |
| 47283_3 | TATGAGTGTGGAGTCGTTGCTATCTCGGTTCACTGCAACATCTNNNAGATCGGAAG |
| 49216_1 | TATGAGTGTGGAGTCGTTGCTTTGTAGAGACAGGGTCTTGCTNNNAGATCGGAA |
| 49216_2 | TATGAGTGTGGAGTCGTTGCTTTGTAGAGACAGGGTCTTGCTNNNAGATCGGAA |
| 49216_3 | TATGAGTGTGGAGTCGTTGCTTTGTAGAGACAGGGTCTTGCTNNNAGATCGGAA |
| 48229_1 | TATGAGTGTGGAGTCGTTGCCTGAGTAGCTGGGATTACAGGTGNNNAGATCGGAA |
| 48229_2 | TATGAGTGTGGAGTCGTTGCCTGAGTAGCTGGGATTACAGGTGNNNAGATCGGAA |
| 48229_3 | TATGAGTGTGGAGTCGTTGCCTGAGTAGCTGGGATTACAGGTGNNNAGATCGGAA |
| 31342_1 | TATGAGTGTGGAGTCGTTGCTTCTTGCCTTCTCCTTTATCCCCNNNAGATCGGAAGA |
| 31342_2 | TATGAGTGTGGAGTCGTTGCTTCTTGCCTTCTCCTTTATCCCCNNNAGATCGGAAGA |
| 31342_3 | TATGAGTGTGGAGTCGTTGCTTCTTGCCTTCTCCTTTATCCCCNNNAGATCGGAAGA |
| 35331_1 | TATGAGTGTGGAGTCGTTGCTCAAGTGATTCTCCTGTCTCAGCNNNAGATCGGAAG |
| 35331_2 | TATGAGTGTGGAGTCGTTGCTCAAGTGATTCTCCTGTCTCAGCNNNAGATCGGAAG |
| 35331_3 | TATGAGTGTGGAGTCGTTGCTCAAGTGATTCTCCTGTCTCAGCNNNAGATCGGAAG |
| 42792_1 | TATGAGTGTGGAGTCGTTGCAATAATGGCATTTGCAGCAACCTNNNAGATCGGAAG |
| 42792_2 | TATGAGTGTGGAGTCGTTGCAATAATGGCATTTGCAGCAACCTNNNAGATCGGAAG |
| 42792_3 | TATGAGTGTGGAGTCGTTGCAATAATGGCATTTGCAGCAACCTNNNAGATCGGAAG |
| 37751_1 | TATGAGTGTGGAGTCGTTGCGGCCTAGAAACATTTTGGTGCATNNNAGATCGGAA |
| 37751_2 | TATGAGTGTGGAGTCGTTGCGGCCTAGAAACATTTTGGTGCATNNNAGATCGGAA |
| 37751_3 | TATGAGTGTGGAGTCGTTGCGGCCTAGAAACATTTTGGTGCATNNNAGATCGGAA |
| 36363_1 | TATGAGTGTGGAGTCGTTGCGCTGAGGTGGAAGGATAACTTGANNNAGATCGGAA |
| 36363_2 | TATGAGTGTGGAGTCGTTGCGCTGAGGTGGAAGGATAACTTGANNNAGATCGGAA |
| 36363_3 | TATGAGTGTGGAGTCGTTGCGCTGAGGTGGAAGGATAACTTGANNNAGATCGGAA |
| 53344_1 | TATGAGTGTGGAGTCGTTGCACACACATACACACACAGACACANNNAGATCGGAAG |
| 53344_2 | TATGAGTGTGGAGTCGTTGCACACACATACACACACAGACACANNNAGATCGGAAG |
| 53344_3 | TATGAGTGTGGAGTCGTTGCACACACATACACACACAGACACANNNAGATCGGAAG |
| 47640_1 | TATGAGTGTGGAGTCGTTGCGTGACACAATCTTGGCACACTACNNNAGATCGGAAG |
| 47640_2 | TATGAGTGTGGAGTCGTTGCGTGACACAATCTTGGCACACTACNNNAGATCGGAAG |
| 47640_3 | TATGAGTGTGGAGTCGTTGCGTGACACAATCTTGGCACACTACNNNAGATCGGAAG |
| 28772_1 | TATGAGTGTGGAGTCGTTGCACATGTCTGAACAAGATCTGGCTNNNAGATCGGAAG |
| 28772_2 | TATGAGTGTGGAGTCGTTGCACATGTCTGAACAAGATCTGGCTNNNAGATCGGAAG |
| 28772_3 | TATGAGTGTGGAGTCGTTGCACATGTCTGAACAAGATCTGGCTNNNAGATCGGAAG |
| 46588_1 | TATGAGTGTGGAGTCGTTGCTGTCTCTGAGTTTCTCCCAAGTGNNNAGATCGGAAG |
| 46588_2 | TATGAGTGTGGAGTCGTTGCTGTCTCTGAGTTTCTCCCAAGTGNNNAGATCGGAAG |
| 46588_3 | TATGAGTGTGGAGTCGTTGCTGTCTCTGAGTTTCTCCCAAGTGNNNAGATCGGAAG |
| 38090_1 | TATGAGTGTGGAGTCGTTGCGAAAAAAGTACATTTGCCAGGCGNNNAGATCGGAA |
| 38090_2 | TATGAGTGTGGAGTCGTTGCGAAAAAAGTACATTTGCCAGGCGNNNAGATCGGAA |
| 38090_3 | TATGAGTGTGGAGTCGTTGCGAAAAAAGTACATTTGCCAGGCGNNNAGATCGGAA |
| 50021_1 | TATGAGTGTGGAGTCGTTGCGATGAGGAAACTGAGTCCCAGAGNNNAGATCGGAA |
| 50021_2 | TATGAGTGTGGAGTCGTTGCGATGAGGAAACTGAGTCCCAGAGNNNAGATCGGAA |
| 50021_3 | TATGAGTGTGGAGTCGTTGCGATGAGGAAACTGAGTCCCAGAGNNNAGATCGGAA |
| 50168_1 | TATGAGTGTGGAGTCGTTGCCAGCATCTTCATGGACCACTTCANNNAGATCGGAAG |

| | |
|---|---|
| 50168_2 | TATGAGTGTGGAGTCGTTGCCAGCATCTTCATGGACCACTTCANNNAGATCGGAAG |
| 50168_3 | TATGAGTGTGGAGTCGTTGCCAGCATCTTCATGGACCACTTCANNNAGATCGGAAG |
| 52498_1 | TATGAGTGTGGAGTCGTTGCTGCTTCCACCCAGTTATTAGACCNNNAGATCGGAAG |
| 52498_2 | TATGAGTGTGGAGTCGTTGCTGCTTCCACCCAGTTATTAGACCNNNAGATCGGAAG |
| 52498_3 | TATGAGTGTGGAGTCGTTGCTGCTTCCACCCAGTTATTAGACCNNNAGATCGGAAG |
| 39951_1 | TATGAGTGTGGAGTCGTTGCGAAAGGGGGAGGGATAGCATTAGNNNAGATCGGA |
| 39951_2 | TATGAGTGTGGAGTCGTTGCGAAAGGGGGAGGGATAGCATTAGNNNAGATCGGA |
| 39951_3 | TATGAGTGTGGAGTCGTTGCGAAAGGGGGAGGGATAGCATTAGNNNAGATCGGA |
| 12694_1 | TATGAGTGTGGAGTCGTTGCCAAGTCACAAGAGATACCTAGNNNAGATCGGAAGA |
| 12694_2 | TATGAGTGTGGAGTCGTTGCCAAGTCACAAGAGATACCTAGNNNAGATCGGAAGA |
| 12694_3 | TATGAGTGTGGAGTCGTTGCCAAGTCACAAGAGATACCTAGNNNAGATCGGAAGA |
| 44210_1 | TATGAGTGTGGAGTCGTTGCGAGCTGCTGGAGGACGAAGANNNAGATCGGAAGA |
| 44210_2 | TATGAGTGTGGAGTCGTTGCGAGCTGCTGGAGGACGAAGANNNAGATCGGAAGA |
| 44210_3 | TATGAGTGTGGAGTCGTTGCGAGCTGCTGGAGGACGAAGANNNAGATCGGAAGA |
| 41757_1 | TATGAGTGTGGAGTCGTTGCTTCACACCATTCTCCTGCCTCNNNAGATCGGAAGAGC |
| 41757_2 | TATGAGTGTGGAGTCGTTGCTTCACACCATTCTCCTGCCTCNNNAGATCGGAAGAGC |
| 41757_3 | TATGAGTGTGGAGTCGTTGCTTCACACCATTCTCCTGCCTCNNNAGATCGGAAGAGC |
| 52066_1 | TATGAGTGTGGAGTCGTTGCGATTGCAGTGGTACAATCTTGGCNNNAGATCGGAA |
| 52066_2 | TATGAGTGTGGAGTCGTTGCGATTGCAGTGGTACAATCTTGGCNNNAGATCGGAA |
| 52066_3 | TATGAGTGTGGAGTCGTTGCGATTGCAGTGGTACAATCTTGGCNNNAGATCGGAA |
| 51256_1 | TATGAGTGTGGAGTCGTTGCTCTACAACCTCCCCATGAAACTGNNNAGATCGGAAG |
| 51256_2 | TATGAGTGTGGAGTCGTTGCTCTACAACCTCCCCATGAAACTGNNNAGATCGGAAG |
| 51256_3 | TATGAGTGTGGAGTCGTTGCTCTACAACCTCCCCATGAAACTGNNNAGATCGGAAG |
| 31775_1 | TATGAGTGTGGAGTCGTTGCAAAGAGGGAAAGGAAGAGAGCAGNNNAGATCGGA |
| 31775_2 | TATGAGTGTGGAGTCGTTGCAAAGAGGGAAAGGAAGAGAGCAGNNNAGATCGGA |
| 31775_3 | TATGAGTGTGGAGTCGTTGCAAAGAGGGAAAGGAAGAGAGCAGNNNAGATCGGA |
| 41378_1 | TATGAGTGTGGAGTCGTTGCTTCTATTCTGGTGTCCCCTCTCTNNNAGATCGGAAGA |
| 41378_2 | TATGAGTGTGGAGTCGTTGCTTCTATTCTGGTGTCCCCTCTCTNNNAGATCGGAAGA |
| 41378_3 | TATGAGTGTGGAGTCGTTGCTTCTATTCTGGTGTCCCCTCTCTNNNAGATCGGAAGA |
| 29433_1 | TATGAGTGTGGAGTCGTTGCGAGCTGATCTTCGAATGGGGTTTNNNAGATCGGAA |
| 29433_2 | TATGAGTGTGGAGTCGTTGCGAGCTGATCTTCGAATGGGGTTTNNNAGATCGGAA |
| 29433_3 | TATGAGTGTGGAGTCGTTGCGAGCTGATCTTCGAATGGGGTTTNNNAGATCGGAA |
| 36777_1 | TATGAGTGTGGAGTCGTTGCAGGGATAGAAGCACACATACTGCNNNAGATCGGAA |
| 36777_2 | TATGAGTGTGGAGTCGTTGCAGGGATAGAAGCACACATACTGCNNNAGATCGGAA |
| 36777_3 | TATGAGTGTGGAGTCGTTGCAGGGATAGAAGCACACATACTGCNNNAGATCGGAA |
| 52704_1 | TATGAGTGTGGAGTCGTTGCGGGACCACGGAAAATTCGAATATNNNAGATCGGAA |
| 52704_2 | TATGAGTGTGGAGTCGTTGCGGGACCACGGAAAATTCGAATATNNNAGATCGGAA |
| 52704_3 | TATGAGTGTGGAGTCGTTGCGGGACCACGGAAAATTCGAATATNNNAGATCGGAA |
| 49900_1 | TATGAGTGTGGAGTCGTTGCTATCGTGTTGAGAACAGACTGCANNNAGATCGGAA |
| 49900_2 | TATGAGTGTGGAGTCGTTGCTATCGTGTTGAGAACAGACTGCANNNAGATCGGAA |
| 49900_3 | TATGAGTGTGGAGTCGTTGCTATCGTGTTGAGAACAGACTGCANNNAGATCGGAA |
| 33766_1 | TATGAGTGTGGAGTCGTTGCAAAGGCTTTGGAATCACAATGCANNNAGATCGGAA |
| 33766_2 | TATGAGTGTGGAGTCGTTGCAAAGGCTTTGGAATCACAATGCANNNAGATCGGAA |

| | |
|---|---|
| 33766_3 | TATGAGTGTGGAGTCGTTGCAAAGGCTTTGGAATCACAATGCANNNAGATCGGAA |
| 43144_1 | TATGAGTGTGGAGTCGTTGCTCCTTTGAGTGTTCGTTTTTCCTNNNAGATCGGAAGA |
| 43144_2 | TATGAGTGTGGAGTCGTTGCTCCTTTGAGTGTTCGTTTTTCCTNNNAGATCGGAAGA |
| 43144_3 | TATGAGTGTGGAGTCGTTGCTCCTTTGAGTGTTCGTTTTTCCTNNNAGATCGGAAGA |
| 36046_1 | TATGAGTGTGGAGTCGTTGCACAAGGAATACATTTGGGGTTCCNNNAGATCGGAA |
| 36046_2 | TATGAGTGTGGAGTCGTTGCACAAGGAATACATTTGGGGTTCCNNNAGATCGGAA |
| 36046_3 | TATGAGTGTGGAGTCGTTGCACAAGGAATACATTTGGGGTTCCNNNAGATCGGAA |
| 38198_1 | TATGAGTGTGGAGTCGTTGCTAGCCAGGATATACAGGGATCCCNNNAGATCGGAA |
| 38198_2 | TATGAGTGTGGAGTCGTTGCTAGCCAGGATATACAGGGATCCCNNNAGATCGGAA |
| 38198_3 | TATGAGTGTGGAGTCGTTGCTAGCCAGGATATACAGGGATCCCNNNAGATCGGAA |
| 42552_1 | TATGAGTGTGGAGTCGTTGCCTTCTGTGCCTGCCATGAAAAATNNNAGATCGGAAG |
| 42552_2 | TATGAGTGTGGAGTCGTTGCCTTCTGTGCCTGCCATGAAAAATNNNAGATCGGAAG |
| 42552_3 | TATGAGTGTGGAGTCGTTGCCTTCTGTGCCTGCCATGAAAAATNNNAGATCGGAAG |
| 33947_1 | TATGAGTGTGGAGTCGTTGCCCAAGTAGCTGGAACTACAGGTGNNNAGATCGGAA |
| 33947_2 | TATGAGTGTGGAGTCGTTGCCCAAGTAGCTGGAACTACAGGTGNNNAGATCGGAA |
| 33947_3 | TATGAGTGTGGAGTCGTTGCCCAAGTAGCTGGAACTACAGGTGNNNAGATCGGAA |
| 43686_1 | TATGAGTGTGGAGTCGTTGCTGGTCACAGAGAGAGCAAATCATNNNAGATCGGAA |
| 43686_2 | TATGAGTGTGGAGTCGTTGCTGGTCACAGAGAGAGCAAATCATNNNAGATCGGAA |
| 43686_3 | TATGAGTGTGGAGTCGTTGCTGGTCACAGAGAGAGCAAATCATNNNAGATCGGAA |
| 43749_1 | TATGAGTGTGGAGTCGTTGCATCCTTGTGCCTGCTACGTCNNNAGATCGGAAGAGC |
| 43749_2 | TATGAGTGTGGAGTCGTTGCATCCTTGTGCCTGCTACGTCNNNAGATCGGAAGAGC |
| 43749_3 | TATGAGTGTGGAGTCGTTGCATCCTTGTGCCTGCTACGTCNNNAGATCGGAAGAGC |
| 8540_1 | TATGAGTGTGGAGTCGTTGCCCGAGATTTCAATCCAACTGAATGNNNAGATCGGAA |
| 8540_2 | TATGAGTGTGGAGTCGTTGCCCGAGATTTCAATCCAACTGAATGNNNAGATCGGAA |
| 8540_3 | TATGAGTGTGGAGTCGTTGCCCGAGATTTCAATCCAACTGAATGNNNAGATCGGAA |
| 34661_1 | TATGAGTGTGGAGTCGTTGCATTGTGACTTCTTTGACCCATGGNNNAGATCGGAAG |
| 34661_2 | TATGAGTGTGGAGTCGTTGCATTGTGACTTCTTTGACCCATGGNNNAGATCGGAAG |
| 34661_3 | TATGAGTGTGGAGTCGTTGCATTGTGACTTCTTTGACCCATGGNNNAGATCGGAAG |
| 47307_1 | TATGAGTGTGGAGTCGTTGCGGTGTGATCATGGCTTATTGCAGNNNAGATCGGAA |
| 47307_2 | TATGAGTGTGGAGTCGTTGCGGTGTGATCATGGCTTATTGCAGNNNAGATCGGAA |
| 47307_3 | TATGAGTGTGGAGTCGTTGCGGTGTGATCATGGCTTATTGCAGNNNAGATCGGAA |
| 47613_1 | TATGAGTGTGGAGTCGTTGCCTCCCAGGATCAGGCAATTCTCNNNAGATCGGAAGA |
| 47613_2 | TATGAGTGTGGAGTCGTTGCCTCCCAGGATCAGGCAATTCTCNNNAGATCGGAAGA |
| 47613_3 | TATGAGTGTGGAGTCGTTGCCTCCCAGGATCAGGCAATTCTCNNNAGATCGGAAGA |
| 34153_1 | TATGAGTGTGGAGTCGTTGCGATGTGGAAGAATCGCTTGAACCNNNAGATCGGAA |
| 34153_2 | TATGAGTGTGGAGTCGTTGCGATGTGGAAGAATCGCTTGAACCNNNAGATCGGAA |
| 34153_3 | TATGAGTGTGGAGTCGTTGCGATGTGGAAGAATCGCTTGAACCNNNAGATCGGAA |
| 30171_1 | TATGAGTGTGGAGTCGTTGCATGCTAGATGACGAGTTAGTGGGNNNAGATCGGAA |
| 30171_2 | TATGAGTGTGGAGTCGTTGCATGCTAGATGACGAGTTAGTGGGNNNAGATCGGAA |
| 30171_3 | TATGAGTGTGGAGTCGTTGCATGCTAGATGACGAGTTAGTGGGNNNAGATCGGAA |
| 34562_1 | TATGAGTGTGGAGTCGTTGCGATGGTCCTAAGAGGAGGAGCAGNNNAGATCGGA |
| 34562_2 | TATGAGTGTGGAGTCGTTGCGATGGTCCTAAGAGGAGGAGCAGNNNAGATCGGA |
| 34562_3 | TATGAGTGTGGAGTCGTTGCGATGGTCCTAAGAGGAGGAGCAGNNNAGATCGGA |

| | |
|---|---|
| 53414_1 | TATGAGTGTGGAGTCGTTGCCTCAGCCTACTCAACAGGAAGTTNNNAGATCGGAAG |
| 53414_2 | TATGAGTGTGGAGTCGTTGCCTCAGCCTACTCAACAGGAAGTTNNNAGATCGGAAG |
| 53414_3 | TATGAGTGTGGAGTCGTTGCCTCAGCCTACTCAACAGGAAGTTNNNAGATCGGAAG |
| 35209_1 | TATGAGTGTGGAGTCGTTGCCATCAGCATCCTGAGTAACTGGTNNNAGATCGGAAG |
| 35209_2 | TATGAGTGTGGAGTCGTTGCCATCAGCATCCTGAGTAACTGGTNNNAGATCGGAAG |
| 35209_3 | TATGAGTGTGGAGTCGTTGCCATCAGCATCCTGAGTAACTGGTNNNAGATCGGAAG |
| 10718_1 | TATGAGTGTGGAGTCGTTGCACCAGCACCATTTATTGAAGGCANNNAGATCGGAAG |
| 10718_2 | TATGAGTGTGGAGTCGTTGCACCAGCACCATTTATTGAAGGCANNNAGATCGGAAG |
| 10718_3 | TATGAGTGTGGAGTCGTTGCACCAGCACCATTTATTGAAGGCANNNAGATCGGAAG |
| 43681_1 | TATGAGTGTGGAGTCGTTGCAGTTGGCGTTTCTATCAGGTCATNNNAGATCGGAAG |
| 43681_2 | TATGAGTGTGGAGTCGTTGCAGTTGGCGTTTCTATCAGGTCATNNNAGATCGGAAG |
| 43681_3 | TATGAGTGTGGAGTCGTTGCAGTTGGCGTTTCTATCAGGTCATNNNAGATCGGAAG |
| 9916_1 | TATGAGTGTGGAGTCGTTGCTGACCCCTTGATTTACCATCTCCTNNNAGATCGGAAG |
| 9916_2 | TATGAGTGTGGAGTCGTTGCTGACCCCTTGATTTACCATCTCCTNNNAGATCGGAAG |
| 9916_3 | TATGAGTGTGGAGTCGTTGCTGACCCCTTGATTTACCATCTCCTNNNAGATCGGAAG |
| 960_1 | TATGAGTGTGGAGTCGTTGCAGTTCTCTCTCTCCAGAATGCACNNNAGATCGGAAG |
| 960_2 | TATGAGTGTGGAGTCGTTGCAGTTCTCTCTCTCCAGAATGCACNNNAGATCGGAAG |
| 960_3 | TATGAGTGTGGAGTCGTTGCAGTTCTCTCTCTCCAGAATGCACNNNAGATCGGAAG |
| 11796_1 | TATGAGTGTGGAGTCGTTGCTGCTCTTTTCATTCCACCTCCTCNNNAGATCGGAAGA |
| 11796_2 | TATGAGTGTGGAGTCGTTGCTGCTCTTTTCATTCCACCTCCTCNNNAGATCGGAAGA |
| 11796_3 | TATGAGTGTGGAGTCGTTGCTGCTCTTTTCATTCCACCTCCTCNNNAGATCGGAAGA |
| 34458_1 | TATGAGTGTGGAGTCGTTGCACAAGAAAGTCCCACGTACTGAANNNAGATCGGAA |
| 34458_2 | TATGAGTGTGGAGTCGTTGCACAAGAAAGTCCCACGTACTGAANNNAGATCGGAA |
| 34458_3 | TATGAGTGTGGAGTCGTTGCACAAGAAAGTCCCACGTACTGAANNNAGATCGGAA |
| 35020_1 | TATGAGTGTGGAGTCGTTGCTGCCCAGCCAATTTCTTTTTCTTNNNAGATCGGAAGA |
| 35020_2 | TATGAGTGTGGAGTCGTTGCTGCCCAGCCAATTTCTTTTTCTTNNNAGATCGGAAGA |
| 35020_3 | TATGAGTGTGGAGTCGTTGCTGCCCAGCCAATTTCTTTTTCTTNNNAGATCGGAAGA |
| 33865_1 | TATGAGTGTGGAGTCGTTGCGTGCTCCACCCATATTTTCTTGGNNNAGATCGGAAG |
| 33865_2 | TATGAGTGTGGAGTCGTTGCGTGCTCCACCCATATTTTCTTGGNNNAGATCGGAAG |
| 33865_3 | TATGAGTGTGGAGTCGTTGCGTGCTCCACCCATATTTTCTTGGNNNAGATCGGAAG |
| 33358_1 | TATGAGTGTGGAGTCGTTGCCGGATTCATGCCATTCTCCTCTNNNAGATCGGAAGA |
| 33358_2 | TATGAGTGTGGAGTCGTTGCCGGATTCATGCCATTCTCCTCTNNNAGATCGGAAGA |
| 33358_3 | TATGAGTGTGGAGTCGTTGCCGGATTCATGCCATTCTCCTCTNNNAGATCGGAAGA |
| 4699_1 | TATGAGTGTGGAGTCGTTGCGGAGTGGCTCTTGCTATCTCAGGGNNNAGATCGGA |
| 4699_2 | TATGAGTGTGGAGTCGTTGCGGAGTGGCTCTTGCTATCTCAGGGNNNAGATCGGA |
| 4699_3 | TATGAGTGTGGAGTCGTTGCGGAGTGGCTCTTGCTATCTCAGGGNNNAGATCGGA |
| 39276_1 | TATGAGTGTGGAGTCGTTGCTTTTTTGAGTCAGAGTCTGGCTCNNNAGATCGGAAG |
| 39276_2 | TATGAGTGTGGAGTCGTTGCTTTTTTGAGTCAGAGTCTGGCTCNNNAGATCGGAAG |
| 39276_3 | TATGAGTGTGGAGTCGTTGCTTTTTTGAGTCAGAGTCTGGCTCNNNAGATCGGAAG |
| 34069_1 | TATGAGTGTGGAGTCGTTGCGCCTCTTGAGTAGCTGGGATTACNNNAGATCGGAA |
| 34069_2 | TATGAGTGTGGAGTCGTTGCGCCTCTTGAGTAGCTGGGATTACNNNAGATCGGAA |
| 34069_3 | TATGAGTGTGGAGTCGTTGCGCCTCTTGAGTAGCTGGGATTACNNNAGATCGGAA |
| 51903_1 | TATGAGTGTGGAGTCGTTGCTTCAGTGATCTTCCCCTCTACCTNNNAGATCGGAAGA |

| | |
|---|---|
| 51903_2 | TATGAGTGTGGAGTCGTTGCTTCAGTGATCTTCCCCTCTACCTNNNAGATCGGAAGA |
| 51903_3 | TATGAGTGTGGAGTCGTTGCTTCAGTGATCTTCCCCTCTACCTNNNAGATCGGAAGA |
| 13133_1 | TATGAGTGTGGAGTCGTTGCTAAGCAGAGAAAGACTGTCAANNNAGATCGGAAGA |
| 13133_2 | TATGAGTGTGGAGTCGTTGCTAAGCAGAGAAAGACTGTCAANNNAGATCGGAAGA |
| 13133_3 | TATGAGTGTGGAGTCGTTGCTAAGCAGAGAAAGACTGTCAANNNAGATCGGAAGA |
| 45762_1 | TATGAGTGTGGAGTCGTTGCCCTGCCAAGTTCAAGCAATTCTCNNNAGATCGGAAG |
| 45762_2 | TATGAGTGTGGAGTCGTTGCCCTGCCAAGTTCAAGCAATTCTCNNNAGATCGGAAG |
| 45762_3 | TATGAGTGTGGAGTCGTTGCCCTGCCAAGTTCAAGCAATTCTCNNNAGATCGGAAG |
| 50613_1 | TATGAGTGTGGAGTCGTTGCTAGTCCAATCTTCATACCCCCCTNNNAGATCGGAAGA |
| 50613_2 | TATGAGTGTGGAGTCGTTGCTAGTCCAATCTTCATACCCCCCTNNNAGATCGGAAGA |
| 50613_3 | TATGAGTGTGGAGTCGTTGCTAGTCCAATCTTCATACCCCCCTNNNAGATCGGAAGA |
| 35919_1 | TATGAGTGTGGAGTCGTTGCGGCTAGAGAAAGACCCCAAGTTTNNNAGATCGGAA |
| 35919_2 | TATGAGTGTGGAGTCGTTGCGGCTAGAGAAAGACCCCAAGTTTNNNAGATCGGAA |
| 35919_3 | TATGAGTGTGGAGTCGTTGCGGCTAGAGAAAGACCCCAAGTTTNNNAGATCGGAA |
| 34590_1 | TATGAGTGTGGAGTCGTTGCATATACTGTTCGGGTGATGGGTGNNNAGATCGGAA |
| 34590_2 | TATGAGTGTGGAGTCGTTGCATATACTGTTCGGGTGATGGGTGNNNAGATCGGAA |
| 34590_3 | TATGAGTGTGGAGTCGTTGCATATACTGTTCGGGTGATGGGTGNNNAGATCGGAA |
| 44941_1 | TATGAGTGTGGAGTCGTTGCAGATGACCAGGACAACTCCAAAANNNAGATCGGAA |
| 44941_2 | TATGAGTGTGGAGTCGTTGCAGATGACCAGGACAACTCCAAAANNNAGATCGGAA |
| 44941_3 | TATGAGTGTGGAGTCGTTGCAGATGACCAGGACAACTCCAAAANNNAGATCGGAA |
| 52363_1 | TATGAGTGTGGAGTCGTTGCCTCTCTGCTCAATGGCTGCTCNNNAGATCGGAAGAG |
| 52363_2 | TATGAGTGTGGAGTCGTTGCCTCTCTGCTCAATGGCTGCTCNNNAGATCGGAAGAG |
| 52363_3 | TATGAGTGTGGAGTCGTTGCCTCTCTGCTCAATGGCTGCTCNNNAGATCGGAAGAG |
| 51576_1 | TATGAGTGTGGAGTCGTTGCCTGCTCTCCCCTTCTGAATAACANNNAGATCGGAAGA |
| 51576_2 | TATGAGTGTGGAGTCGTTGCCTGCTCTCCCCTTCTGAATAACANNNAGATCGGAAGA |
| 51576_3 | TATGAGTGTGGAGTCGTTGCCTGCTCTCCCCTTCTGAATAACANNNAGATCGGAAGA |
| 52957_1 | TATGAGTGTGGAGTCGTTGCCAAGGGCAACAAGAGTGAAACTCNNNAGATCGGAA |
| 52957_2 | TATGAGTGTGGAGTCGTTGCCAAGGGCAACAAGAGTGAAACTCNNNAGATCGGAA |
| 52957_3 | TATGAGTGTGGAGTCGTTGCCAAGGGCAACAAGAGTGAAACTCNNNAGATCGGAA |
| 3969_1 | TATGAGTGTGGAGTCGTTGCTTTGAGTTTGAGGATAGCCAACTNNNAGATCGGAAG |
| 3969_2 | TATGAGTGTGGAGTCGTTGCTTTGAGTTTGAGGATAGCCAACTNNNAGATCGGAAG |
| 3969_3 | TATGAGTGTGGAGTCGTTGCTTTGAGTTTGAGGATAGCCAACTNNNAGATCGGAAG |
| 39890_1 | TATGAGTGTGGAGTCGTTGCTCCAAGATAGGAGAGCCCAAGTANNNAGATCGGAA |
| 39890_2 | TATGAGTGTGGAGTCGTTGCTCCAAGATAGGAGAGCCCAAGTANNNAGATCGGAA |
| 39890_3 | TATGAGTGTGGAGTCGTTGCTCCAAGATAGGAGAGCCCAAGTANNNAGATCGGAA |
| 36615_1 | TATGAGTGTGGAGTCGTTGCAGGTTGAGGCAGGAGAATCAATTNNNAGATCGGAA |
| 36615_2 | TATGAGTGTGGAGTCGTTGCAGGTTGAGGCAGGAGAATCAATTNNNAGATCGGAA |
| 36615_3 | TATGAGTGTGGAGTCGTTGCAGGTTGAGGCAGGAGAATCAATTNNNAGATCGGAA |
| 28999_1 | TATGAGTGTGGAGTCGTTGCGTATGGAGCCTTCACAAAATGGCNNNAGATCGGAA |
| 28999_2 | TATGAGTGTGGAGTCGTTGCGTATGGAGCCTTCACAAAATGGCNNNAGATCGGAA |
| 28999_3 | TATGAGTGTGGAGTCGTTGCGTATGGAGCCTTCACAAAATGGCNNNAGATCGGAA |
| 44648_1 | TATGAGTGTGGAGTCGTTGCCCAGAGGAGGCTAATTATGAGGCNNNAGATCGGAA |
| 44648_2 | TATGAGTGTGGAGTCGTTGCCCAGAGGAGGCTAATTATGAGGCNNNAGATCGGAA |

| | |
|---|---|
| 44648_3 | TATGAGTGTGGAGTCGTTGCCCAGAGGAGGCTAATTATGAGGCNNNAGATCGGAA |
| 47653_1 | TATGAGTGTGGAGTCGTTGCAAGTGTCTGTTGTCTAAGCCTCCNNNAGATCGGAAGA |
| 47653_2 | TATGAGTGTGGAGTCGTTGCAAGTGTCTGTTGTCTAAGCCTCCNNNAGATCGGAAGA |
| 47653_3 | TATGAGTGTGGAGTCGTTGCAAGTGTCTGTTGTCTAAGCCTCCNNNAGATCGGAAGA |
| 46566_1 | TATGAGTGTGGAGTCGTTGCTGGTTATGCTGTGTGGAGTATCCNNNAGATCGGAA |
| 46566_2 | TATGAGTGTGGAGTCGTTGCTGGTTATGCTGTGTGGAGTATCCNNNAGATCGGAA |
| 46566_3 | TATGAGTGTGGAGTCGTTGCTGGTTATGCTGTGTGGAGTATCCNNNAGATCGGAA |
| 34744_1 | TATGAGTGTGGAGTCGTTGCGTGACACAATCTTGGCACACTACNNNAGATCGGAAG |
| 34744_2 | TATGAGTGTGGAGTCGTTGCGTGACACAATCTTGGCACACTACNNNAGATCGGAAG |
| 34744_3 | TATGAGTGTGGAGTCGTTGCGTGACACAATCTTGGCACACTACNNNAGATCGGAAG |
| 12888_1 | TATGAGTGTGGAGTCGTTGCCTTGTGAAAGACAAATCTGTAAGNNNAGATCGGAA |
| 12888_2 | TATGAGTGTGGAGTCGTTGCCTTGTGAAAGACAAATCTGTAAGNNNAGATCGGAA |
| 12888_3 | TATGAGTGTGGAGTCGTTGCCTTGTGAAAGACAAATCTGTAAGNNNAGATCGGAA |
| 50818_1 | TATGAGTGTGGAGTCGTTGCTTCAAGCCAGTCTCCAAAAAAGCNNNAGATCGGAAG |
| 50818_2 | TATGAGTGTGGAGTCGTTGCTTCAAGCCAGTCTCCAAAAAAGCNNNAGATCGGAAG |
| 50818_3 | TATGAGTGTGGAGTCGTTGCTTCAAGCCAGTCTCCAAAAAAGCNNNAGATCGGAAG |
| 42152_1 | TATGAGTGTGGAGTCGTTGCAATGGGGTCTTACCATATTGCCCNNNAGATCGGAAG |
| 42152_2 | TATGAGTGTGGAGTCGTTGCAATGGGGTCTTACCATATTGCCCNNNAGATCGGAAG |
| 42152_3 | TATGAGTGTGGAGTCGTTGCAATGGGGTCTTACCATATTGCCCNNNAGATCGGAAG |
| 34324_1 | TATGAGTGTGGAGTCGTTGCCACAGCACAGAAAGAGAGAGACTNNNAGATCGGAA |
| 34324_2 | TATGAGTGTGGAGTCGTTGCCACAGCACAGAAAGAGAGAGACTNNNAGATCGGAA |
| 34324_3 | TATGAGTGTGGAGTCGTTGCCACAGCACAGAAAGAGAGAGACTNNNAGATCGGAA |
| 52070_1 | TATGAGTGTGGAGTCGTTGCCTGTTCACCAATATCCGCTGTTCNNNAGATCGGAAGA |
| 52070_2 | TATGAGTGTGGAGTCGTTGCCTGTTCACCAATATCCGCTGTTCNNNAGATCGGAAGA |
| 52070_3 | TATGAGTGTGGAGTCGTTGCCTGTTCACCAATATCCGCTGTTCNNNAGATCGGAAGA |
| 51080_1 | TATGAGTGTGGAGTCGTTGCCTACACACCACACAAACAGCCTCNNNAGATCGGAAGA |
| 51080_2 | TATGAGTGTGGAGTCGTTGCCTACACACCACACAAACAGCCTCNNNAGATCGGAAGA |
| 51080_3 | TATGAGTGTGGAGTCGTTGCCTACACACCACACAAACAGCCTCNNNAGATCGGAAGA |
| 49780_1 | TATGAGTGTGGAGTCGTTGCCCCACCCATCTAACTCCATGAATNNNAGATCGGAAGA |
| 49780_2 | TATGAGTGTGGAGTCGTTGCCCCACCCATCTAACTCCATGAATNNNAGATCGGAAGA |
| 49780_3 | TATGAGTGTGGAGTCGTTGCCCCACCCATCTAACTCCATGAATNNNAGATCGGAAGA |
| 44427_1 | TATGAGTGTGGAGTCGTTGCTTGGAGATACTACTGCATCCACGNNNAGATCGGAAG |
| 44427_2 | TATGAGTGTGGAGTCGTTGCTTGGAGATACTACTGCATCCACGNNNAGATCGGAAG |
| 44427_3 | TATGAGTGTGGAGTCGTTGCTTGGAGATACTACTGCATCCACGNNNAGATCGGAAG |
| 33938_1 | TATGAGTGTGGAGTCGTTGCCACGATCTCAGTTCACTGCAACNNNAGATCGGAAGA |
| 33938_2 | TATGAGTGTGGAGTCGTTGCCACGATCTCAGTTCACTGCAACNNNAGATCGGAAGA |
| 33938_3 | TATGAGTGTGGAGTCGTTGCCACGATCTCAGTTCACTGCAACNNNAGATCGGAAGA |
| 2674_1 | TATGAGTGTGGAGTCGTTGCATGATCCAGGAGTTGGAACCCATNNNAGATCGGAA |
| 2674_2 | TATGAGTGTGGAGTCGTTGCATGATCCAGGAGTTGGAACCCATNNNAGATCGGAA |
| 2674_3 | TATGAGTGTGGAGTCGTTGCATGATCCAGGAGTTGGAACCCATNNNAGATCGGAA |
| 33745_1 | TATGAGTGTGGAGTCGTTGCGAAGGAGGTAGTGGGAGTAGAGTNNNAGATCGGA |
| 33745_2 | TATGAGTGTGGAGTCGTTGCGAAGGAGGTAGTGGGAGTAGAGTNNNAGATCGGA |
| 33745_3 | TATGAGTGTGGAGTCGTTGCGAAGGAGGTAGTGGGAGTAGAGTNNNAGATCGGA |

| | |
|---|---|
| 47655_1 | TATGAGTGTGGAGTCGTTGCCTGGGTTCAAGCGATTCTCCTNNNAGATCGGAAGAG |
| 47655_2 | TATGAGTGTGGAGTCGTTGCCTGGGTTCAAGCGATTCTCCTNNNAGATCGGAAGAG |
| 47655_3 | TATGAGTGTGGAGTCGTTGCCTGGGTTCAAGCGATTCTCCTNNNAGATCGGAAGAG |
| 33439_1 | TATGAGTGTGGAGTCGTTGCTGCAGGATACATCTCTGAAAGAGANNNAGATCGGA |
| 33439_2 | TATGAGTGTGGAGTCGTTGCTGCAGGATACATCTCTGAAAGAGANNNAGATCGGA |
| 33439_3 | TATGAGTGTGGAGTCGTTGCTGCAGGATACATCTCTGAAAGAGANNNAGATCGGA |
| 12991_1 | TATGAGTGTGGAGTCGTTGCTTCTCTTAATAATGCTCACAATGNNNAGATCGGAAGA |
| 12991_2 | TATGAGTGTGGAGTCGTTGCTTCTCTTAATAATGCTCACAATGNNNAGATCGGAAGA |
| 12991_3 | TATGAGTGTGGAGTCGTTGCTTCTCTTAATAATGCTCACAATGNNNAGATCGGAAGA |
| 31800_1 | TATGAGTGTGGAGTCGTTGCATCGCTTAGATATGGCAGTCCTGNNNAGATCGGAAG |
| 31800_2 | TATGAGTGTGGAGTCGTTGCATCGCTTAGATATGGCAGTCCTGNNNAGATCGGAAG |
| 31800_3 | TATGAGTGTGGAGTCGTTGCATCGCTTAGATATGGCAGTCCTGNNNAGATCGGAAG |
| 37189_1 | TATGAGTGTGGAGTCGTTGCCCTTTCAAGTTTCAGTCCTTTGGGNNNAGATCGGAA |
| 37189_2 | TATGAGTGTGGAGTCGTTGCCCTTTCAAGTTTCAGTCCTTTGGGNNNAGATCGGAA |
| 37189_3 | TATGAGTGTGGAGTCGTTGCCCTTTCAAGTTTCAGTCCTTTGGGNNNAGATCGGAA |
| 45667_1 | TATGAGTGTGGAGTCGTTGCAGCCTCCAAAGTAGCTAGGACTANNNAGATCGGAA |
| 45667_2 | TATGAGTGTGGAGTCGTTGCAGCCTCCAAAGTAGCTAGGACTANNNAGATCGGAA |
| 45667_3 | TATGAGTGTGGAGTCGTTGCAGCCTCCAAAGTAGCTAGGACTANNNAGATCGGAA |
| 37873_1 | TATGAGTGTGGAGTCGTTGCGCTGAGACAGGACAATCTCTTGANNNAGATCGGAA |
| 37873_2 | TATGAGTGTGGAGTCGTTGCGCTGAGACAGGACAATCTCTTGANNNAGATCGGAA |
| 37873_3 | TATGAGTGTGGAGTCGTTGCGCTGAGACAGGACAATCTCTTGANNNAGATCGGAA |
| 11970_1 | TATGAGTGTGGAGTCGTTGCTAAAAGACACATGCACCTGCACGNNNAGATCGGAAG |
| 11970_2 | TATGAGTGTGGAGTCGTTGCTAAAAGACACATGCACCTGCACGNNNAGATCGGAAG |
| 11970_3 | TATGAGTGTGGAGTCGTTGCTAAAAGACACATGCACCTGCACGNNNAGATCGGAAG |
| 4372_1 | TATGAGTGTGGAGTCGTTGCAGCCTTGCAGAGTTAGATGACACNNNAGATCGGAA |
| 4372_2 | TATGAGTGTGGAGTCGTTGCAGCCTTGCAGAGTTAGATGACACNNNAGATCGGAA |
| 4372_3 | TATGAGTGTGGAGTCGTTGCAGCCTTGCAGAGTTAGATGACACNNNAGATCGGAA |
| 5052_1 | TATGAGTGTGGAGTCGTTGCTGGGGTTTCTATGGAAGGGCTAANNNAGATCGGAA |
| 5052_2 | TATGAGTGTGGAGTCGTTGCTGGGGTTTCTATGGAAGGGCTAANNNAGATCGGAA |
| 5052_3 | TATGAGTGTGGAGTCGTTGCTGGGGTTTCTATGGAAGGGCTAANNNAGATCGGAA |
| 32874_1 | TATGAGTGTGGAGTCGTTGCCTCCGGAATAGCTGGGACTAAAGNNNAGATCGGAA |
| 32874_2 | TATGAGTGTGGAGTCGTTGCCTCCGGAATAGCTGGGACTAAAGNNNAGATCGGAA |
| 32874_3 | TATGAGTGTGGAGTCGTTGCCTCCGGAATAGCTGGGACTAAAGNNNAGATCGGAA |
| 12884_1 | TATGAGTGTGGAGTCGTTGCTGAAGTAATAGATGAAAGGGTCNNNAGATCGGAAG |
| 12884_2 | TATGAGTGTGGAGTCGTTGCTGAAGTAATAGATGAAAGGGTCNNNAGATCGGAAG |
| 12884_3 | TATGAGTGTGGAGTCGTTGCTGAAGTAATAGATGAAAGGGTCNNNAGATCGGAAG |
| 13217_1 | TATGAGTGTGGAGTCGTTGCTTAGTGTCTGGCTTAAATTCCNNNAGATCGGAAGAG |
| 13217_2 | TATGAGTGTGGAGTCGTTGCTTAGTGTCTGGCTTAAATTCCNNNAGATCGGAAGAG |
| 13217_3 | TATGAGTGTGGAGTCGTTGCTTAGTGTCTGGCTTAAATTCCNNNAGATCGGAAGAG |
| 36393_1 | TATGAGTGTGGAGTCGTTGCAGATCCAACCTAAATCAACGCCTNNNAGATCGGAAG |
| 36393_2 | TATGAGTGTGGAGTCGTTGCAGATCCAACCTAAATCAACGCCTNNNAGATCGGAAG |
| 36393_3 | TATGAGTGTGGAGTCGTTGCAGATCCAACCTAAATCAACGCCTNNNAGATCGGAAG |
| 31227_1 | TATGAGTGTGGAGTCGTTGCAATTTGGTCTTGGGGTCTCTCTCNNNAGATCGGAAG |

| | |
|---|---|
| 31227_2 | TATGAGTGTGGAGTCGTTGCAATTTGGTCTTGGGGTCTCTCTCNNNAGATCGGAAG |
| 31227_3 | TATGAGTGTGGAGTCGTTGCAATTTGGTCTTGGGGTCTCTCTCNNNAGATCGGAAG |
| 39468_1 | TATGAGTGTGGAGTCGTTGCGTACCACTGTAGTCCAACCTGAGNNNAGATCGGAAG |
| 39468_2 | TATGAGTGTGGAGTCGTTGCGTACCACTGTAGTCCAACCTGAGNNNAGATCGGAAG |
| 39468_3 | TATGAGTGTGGAGTCGTTGCGTACCACTGTAGTCCAACCTGAGNNNAGATCGGAAG |
| 50754_1 | TATGAGTGTGGAGTCGTTGCGTTCCAAGAAATCCCACTACCCTNNNAGATCGGAAG |
| 50754_2 | TATGAGTGTGGAGTCGTTGCGTTCCAAGAAATCCCACTACCCTNNNAGATCGGAAG |
| 50754_3 | TATGAGTGTGGAGTCGTTGCGTTCCAAGAAATCCCACTACCCTNNNAGATCGGAAG |
| 43331_1 | TATGAGTGTGGAGTCGTTGCGAACCTGGGATTATAGGCACACANNNAGATCGGAA |
| 43331_2 | TATGAGTGTGGAGTCGTTGCGAACCTGGGATTATAGGCACACANNNAGATCGGAA |
| 43331_3 | TATGAGTGTGGAGTCGTTGCGAACCTGGGATTATAGGCACACANNNAGATCGGAA |
| 51939_1 | TATGAGTGTGGAGTCGTTGCGGCATGGAACAAACTGACTGAAGNNNAGATCGGAA |
| 51939_2 | TATGAGTGTGGAGTCGTTGCGGCATGGAACAAACTGACTGAAGNNNAGATCGGAA |
| 51939_3 | TATGAGTGTGGAGTCGTTGCGGCATGGAACAAACTGACTGAAGNNNAGATCGGAA |
| 47433_1 | TATGAGTGTGGAGTCGTTGCCGTCTCCCGGTTTCAAGTAATTCNNNAGATCGGAAG |
| 47433_2 | TATGAGTGTGGAGTCGTTGCCGTCTCCCGGTTTCAAGTAATTCNNNAGATCGGAAG |
| 47433_3 | TATGAGTGTGGAGTCGTTGCCGTCTCCCGGTTTCAAGTAATTCNNNAGATCGGAAG |
| 48686_1 | TATGAGTGTGGAGTCGTTGCACATTTGCAGGGAATATCTGGGTNNNAGATCGGAA |
| 48686_2 | TATGAGTGTGGAGTCGTTGCACATTTGCAGGGAATATCTGGGTNNNAGATCGGAA |
| 48686_3 | TATGAGTGTGGAGTCGTTGCACATTTGCAGGGAATATCTGGGTNNNAGATCGGAA |
| 46100_1 | TATGAGTGTGGAGTCGTTGCGTTGGTGTTGCAGTTACTAACCCNNNAGATCGGAAG |
| 46100_2 | TATGAGTGTGGAGTCGTTGCGTTGGTGTTGCAGTTACTAACCCNNNAGATCGGAAG |
| 46100_3 | TATGAGTGTGGAGTCGTTGCGTTGGTGTTGCAGTTACTAACCCNNNAGATCGGAAG |
| 45022_1 | TATGAGTGTGGAGTCGTTGCCGTTTCGCAGTGGTCAGCTCNNNAGATCGGAAGAG |
| 45022_2 | TATGAGTGTGGAGTCGTTGCCGTTTCGCAGTGGTCAGCTCNNNAGATCGGAAGAG |
| 45022_3 | TATGAGTGTGGAGTCGTTGCCGTTTCGCAGTGGTCAGCTCNNNAGATCGGAAGAG |
| 40305_1 | TATGAGTGTGGAGTCGTTGCACAATGCCCTGTTGTTAGTTGTGNNNAGATCGGAAG |
| 40305_2 | TATGAGTGTGGAGTCGTTGCACAATGCCCTGTTGTTAGTTGTGNNNAGATCGGAAG |
| 40305_3 | TATGAGTGTGGAGTCGTTGCACAATGCCCTGTTGTTAGTTGTGNNNAGATCGGAAG |
| 49815_1 | TATGAGTGTGGAGTCGTTGCCTCTTGGTGCCGTATCCTGTAATNNNAGATCGGAAG |
| 49815_2 | TATGAGTGTGGAGTCGTTGCCTCTTGGTGCCGTATCCTGTAATNNNAGATCGGAAG |
| 49815_3 | TATGAGTGTGGAGTCGTTGCCTCTTGGTGCCGTATCCTGTAATNNNAGATCGGAAG |
| 52870_1 | TATGAGTGTGGAGTCGTTGCCCCACGTGATTGACAAGTTCCTANNNAGATCGGAAG |
| 52870_2 | TATGAGTGTGGAGTCGTTGCCCCACGTGATTGACAAGTTCCTANNNAGATCGGAAG |
| 52870_3 | TATGAGTGTGGAGTCGTTGCCCCACGTGATTGACAAGTTCCTANNNAGATCGGAAG |
| 48443_1 | TATGAGTGTGGAGTCGTTGCCTCCTGAGTAGCTGGGACTACAGNNNAGATCGGAA |
| 48443_2 | TATGAGTGTGGAGTCGTTGCCTCCTGAGTAGCTGGGACTACAGNNNAGATCGGAA |
| 48443_3 | TATGAGTGTGGAGTCGTTGCCTCCTGAGTAGCTGGGACTACAGNNNAGATCGGAA |
| 40858_1 | TATGAGTGTGGAGTCGTTGCATCACCACCACCCATTTCTATGTNNNAGATCGGAAGA |
| 40858_2 | TATGAGTGTGGAGTCGTTGCATCACCACCACCCATTTCTATGTNNNAGATCGGAAGA |
| 40858_3 | TATGAGTGTGGAGTCGTTGCATCACCACCACCCATTTCTATGTNNNAGATCGGAAGA |
| 48240_1 | TATGAGTGTGGAGTCGTTGCCTCCTGAGTAGCTGGGACTACAGNNNAGATCGGAA |
| 48240_2 | TATGAGTGTGGAGTCGTTGCCTCCTGAGTAGCTGGGACTACAGNNNAGATCGGAA |

| | |
|---|---|
| 48240_3 | TATGAGTGTGGAGTCGTTGCCTCCTGAGTAGCTGGGACTACAGNNNAGATCGGAA |
| 51303_1 | TATGAGTGTGGAGTCGTTGCGCCCACTTAGATTCAAGAGGAGANNNAGATCGGAA |
| 51303_2 | TATGAGTGTGGAGTCGTTGCGCCCACTTAGATTCAAGAGGAGANNNAGATCGGAA |
| 51303_3 | TATGAGTGTGGAGTCGTTGCGCCCACTTAGATTCAAGAGGAGANNNAGATCGGAA |
| 41793_1 | TATGAGTGTGGAGTCGTTGCCTCCCAAGTAGCTGGAACTACAGNNNAGATCGGAAG |
| 41793_2 | TATGAGTGTGGAGTCGTTGCCTCCCAAGTAGCTGGAACTACAGNNNAGATCGGAAG |
| 41793_3 | TATGAGTGTGGAGTCGTTGCCTCCCAAGTAGCTGGAACTACAGNNNAGATCGGAAG |
| 33580_1 | TATGAGTGTGGAGTCGTTGCGGAGTTTCATTCTCGCTCTGTTGNNNAGATCGGAAG |
| 33580_2 | TATGAGTGTGGAGTCGTTGCGGAGTTTCATTCTCGCTCTGTTGNNNAGATCGGAAG |
| 33580_3 | TATGAGTGTGGAGTCGTTGCGGAGTTTCATTCTCGCTCTGTTGNNNAGATCGGAAG |
| 34447_1 | TATGAGTGTGGAGTCGTTGCCTCTCTCTAGCTATCCCCCTCTCNNNAGATCGGAAGA |
| 34447_2 | TATGAGTGTGGAGTCGTTGCCTCTCTCTAGCTATCCCCCTCTCNNNAGATCGGAAGA |
| 34447_3 | TATGAGTGTGGAGTCGTTGCCTCTCTCTAGCTATCCCCCTCTCNNNAGATCGGAAGA |
| 10099_1 | TATGAGTGTGGAGTCGTTGCTCATAACCTAACTGGGCATGTGCNNNAGATCGGAAG |
| 10099_2 | TATGAGTGTGGAGTCGTTGCTCATAACCTAACTGGGCATGTGCNNNAGATCGGAAG |
| 10099_3 | TATGAGTGTGGAGTCGTTGCTCATAACCTAACTGGGCATGTGCNNNAGATCGGAAG |
| 35116_1 | TATGAGTGTGGAGTCGTTGCCCTGGGTTCAAGCAATTCTTCTGNNNAGATCGGAAG |
| 35116_2 | TATGAGTGTGGAGTCGTTGCCCTGGGTTCAAGCAATTCTTCTGNNNAGATCGGAAG |
| 35116_3 | TATGAGTGTGGAGTCGTTGCCCTGGGTTCAAGCAATTCTTCTGNNNAGATCGGAAG |
| 40337_1 | TATGAGTGTGGAGTCGTTGCACCCCTTCTTTCTGAGCACTAAGNNNAGATCGGAAG |
| 40337_2 | TATGAGTGTGGAGTCGTTGCACCCCTTCTTTCTGAGCACTAAGNNNAGATCGGAAG |
| 40337_3 | TATGAGTGTGGAGTCGTTGCACCCCTTCTTTCTGAGCACTAAGNNNAGATCGGAAG |
| 36022_1 | TATGAGTGTGGAGTCGTTGCGCCTCCCTCTAACCATGAGAAAANNNAGATCGGAAG |
| 36022_2 | TATGAGTGTGGAGTCGTTGCGCCTCCCTCTAACCATGAGAAAANNNAGATCGGAAG |
| 36022_3 | TATGAGTGTGGAGTCGTTGCGCCTCCCTCTAACCATGAGAAAANNNAGATCGGAAG |
| 41109_1 | TATGAGTGTGGAGTCGTTGCAAGGTGGAGTAACTTTGTAGGGCNNNAGATCGGAA |
| 41109_2 | TATGAGTGTGGAGTCGTTGCAAGGTGGAGTAACTTTGTAGGGCNNNAGATCGGAA |
| 41109_3 | TATGAGTGTGGAGTCGTTGCAAGGTGGAGTAACTTTGTAGGGCNNNAGATCGGAA |
| 52272_1 | TATGAGTGTGGAGTCGTTGCGCTTCTCCAAAGGCTGTTTTCAANNNAGATCGGAAG |
| 52272_2 | TATGAGTGTGGAGTCGTTGCGCTTCTCCAAAGGCTGTTTTCAANNNAGATCGGAAG |
| 52272_3 | TATGAGTGTGGAGTCGTTGCGCTTCTCCAAAGGCTGTTTTCAANNNAGATCGGAAG |
| 44106_1 | TATGAGTGTGGAGTCGTTGCCTCTGAGAGTGATATGGGAGCAGNNNAGATCGGAA |
| 44106_2 | TATGAGTGTGGAGTCGTTGCCTCTGAGAGTGATATGGGAGCAGNNNAGATCGGAA |
| 44106_3 | TATGAGTGTGGAGTCGTTGCCTCTGAGAGTGATATGGGAGCAGNNNAGATCGGAA |
| 33034_1 | TATGAGTGTGGAGTCGTTGCCATGATCTCGGCTCACTGCAANNNAGATCGGAAGAG |
| 33034_2 | TATGAGTGTGGAGTCGTTGCCATGATCTCGGCTCACTGCAANNNAGATCGGAAGAG |
| 33034_3 | TATGAGTGTGGAGTCGTTGCCATGATCTCGGCTCACTGCAANNNAGATCGGAAGAG |
| 47911_1 | TATGAGTGTGGAGTCGTTGCAAACTCTTGACCTCAGGTGATCCNNNAGATCGGAAG |
| 47911_2 | TATGAGTGTGGAGTCGTTGCAAACTCTTGACCTCAGGTGATCCNNNAGATCGGAAG |
| 47911_3 | TATGAGTGTGGAGTCGTTGCAAACTCTTGACCTCAGGTGATCCNNNAGATCGGAAG |
| 42242_1 | TATGAGTGTGGAGTCGTTGCATGTCTTTTCCCTTCTCCCCTTCNNNAGATCGGAAGA |
| 42242_2 | TATGAGTGTGGAGTCGTTGCATGTCTTTTCCCTTCTCCCCTTCNNNAGATCGGAAGA |
| 42242_3 | TATGAGTGTGGAGTCGTTGCATGTCTTTTCCCTTCTCCCCTTCNNNAGATCGGAAGA |

| | |
|---|---|
| 51645_1 | TATGAGTGTGGAGTCGTTGCTTCATCCATCCATCCATCCATCCNNNAGATCGGAAGA |
| 51645_2 | TATGAGTGTGGAGTCGTTGCTTCATCCATCCATCCATCCATCCNNNAGATCGGAAGA |
| 51645_3 | TATGAGTGTGGAGTCGTTGCTTCATCCATCCATCCATCCATCCNNNAGATCGGAAGA |
| 51345_1 | TATGAGTGTGGAGTCGTTGCGCAGTATCGGGGACATTGTTTTTNNNAGATCGGAAG |
| 51345_2 | TATGAGTGTGGAGTCGTTGCGCAGTATCGGGGACATTGTTTTTNNNAGATCGGAAG |
| 51345_3 | TATGAGTGTGGAGTCGTTGCGCAGTATCGGGGACATTGTTTTTNNNAGATCGGAAG |
| 31247_1 | TATGAGTGTGGAGTCGTTGCCTCCATGTTCTGCCCTCTAAACANNNAGATCGGAAGA |
| 31247_2 | TATGAGTGTGGAGTCGTTGCCTCCATGTTCTGCCCTCTAAACANNNAGATCGGAAGA |
| 31247_3 | TATGAGTGTGGAGTCGTTGCCTCCATGTTCTGCCCTCTAAACANNNAGATCGGAAGA |
| 2148_1 | TATGAGTGTGGAGTCGTTGCGGTGTTTGGGGCATGGATATTCANNNAGATCGGAA |
| 2148_2 | TATGAGTGTGGAGTCGTTGCGGTGTTTGGGGCATGGATATTCANNNAGATCGGAA |
| 2148_3 | TATGAGTGTGGAGTCGTTGCGGTGTTTGGGGCATGGATATTCANNNAGATCGGAA |
| 32780_1 | TATGAGTGTGGAGTCGTTGCCAGGACATGTTGGAGACACAGATNNNAGATCGGAA |
| 32780_2 | TATGAGTGTGGAGTCGTTGCCAGGACATGTTGGAGACACAGATNNNAGATCGGAA |
| 32780_3 | TATGAGTGTGGAGTCGTTGCCAGGACATGTTGGAGACACAGATNNNAGATCGGAA |
| 40683_1 | TATGAGTGTGGAGTCGTTGCCTGCTTTCCCACTTGAGAGAGATNNNAGATCGGAAG |
| 40683_2 | TATGAGTGTGGAGTCGTTGCCTGCTTTCCCACTTGAGAGAGATNNNAGATCGGAAG |
| 40683_3 | TATGAGTGTGGAGTCGTTGCCTGCTTTCCCACTTGAGAGAGATNNNAGATCGGAAG |
| 32664_1 | TATGAGTGTGGAGTCGTTGCACCTCTACCTCCTGAGTTCAAGTNNNAGATCGGAAG |
| 32664_2 | TATGAGTGTGGAGTCGTTGCACCTCTACCTCCTGAGTTCAAGTNNNAGATCGGAAG |
| 32664_3 | TATGAGTGTGGAGTCGTTGCACCTCTACCTCCTGAGTTCAAGTNNNAGATCGGAAG |
| 45775_1 | TATGAGTGTGGAGTCGTTGCGGTTCAGCCTGATCTCTAGGTGNNNAGATCGGAAG |
| 45775_2 | TATGAGTGTGGAGTCGTTGCGGTTCAGCCTGATCTCTAGGTGNNNAGATCGGAAG |
| 45775_3 | TATGAGTGTGGAGTCGTTGCGGTTCAGCCTGATCTCTAGGTGNNNAGATCGGAAG |
| 34674_1 | TATGAGTGTGGAGTCGTTGCGTGTGTGTATGTGCTGTTTGTGTNNNAGATCGGAA |
| 34674_2 | TATGAGTGTGGAGTCGTTGCGTGTGTGTATGTGCTGTTTGTGTNNNAGATCGGAA |
| 34674_3 | TATGAGTGTGGAGTCGTTGCGTGTGTGTATGTGCTGTTTGTGTNNNAGATCGGAA |
| 12959_1 | TATGAGTGTGGAGTCGTTGCGTCAAATTTCTCAACTCTGTGNNNAGATCGGAAGAG |
| 12959_2 | TATGAGTGTGGAGTCGTTGCGTCAAATTTCTCAACTCTGTGNNNAGATCGGAAGAG |
| 12959_3 | TATGAGTGTGGAGTCGTTGCGTCAAATTTCTCAACTCTGTGNNNAGATCGGAAGAG |
| 2710_1 | TATGAGTGTGGAGTCGTTGCAAGCAGTTAGGGGCCTCAGATATNNNAGATCGGAA |
| 2710_2 | TATGAGTGTGGAGTCGTTGCAAGCAGTTAGGGGCCTCAGATATNNNAGATCGGAA |
| 2710_3 | TATGAGTGTGGAGTCGTTGCAAGCAGTTAGGGGCCTCAGATATNNNAGATCGGAA |
| 40138_1 | TATGAGTGTGGAGTCGTTGCATGTAATCCCAGCACTTTGAGGTNNNAGATCGGAAG |
| 40138_2 | TATGAGTGTGGAGTCGTTGCATGTAATCCCAGCACTTTGAGGTNNNAGATCGGAAG |
| 40138_3 | TATGAGTGTGGAGTCGTTGCATGTAATCCCAGCACTTTGAGGTNNNAGATCGGAAG |
| 31861_1 | TATGAGTGTGGAGTCGTTGCGCTCATGGATGGGTAGAACCAATNNNAGATCGGAA |
| 31861_2 | TATGAGTGTGGAGTCGTTGCGCTCATGGATGGGTAGAACCAATNNNAGATCGGAA |
| 31861_3 | TATGAGTGTGGAGTCGTTGCGCTCATGGATGGGTAGAACCAATNNNAGATCGGAA |
| 52962_1 | TATGAGTGTGGAGTCGTTGCCTTGTTCCCACTCCCATGATACANNNAGATCGGAAGA |
| 52962_2 | TATGAGTGTGGAGTCGTTGCCTTGTTCCCACTCCCATGATACANNNAGATCGGAAGA |
| 52962_3 | TATGAGTGTGGAGTCGTTGCCTTGTTCCCACTCCCATGATACANNNAGATCGGAAGA |
| 49946_1 | TATGAGTGTGGAGTCGTTGCCCAAATGTTCCTGGGGAAGACCNNNAGATCGGAAG |

| | |
|---|---|
| 49946_2 | TATGAGTGTGGAGTCGTTGCCCAAATGTTCCTGGGGAAGACCNNNAGATCGGAAG |
| 49946_3 | TATGAGTGTGGAGTCGTTGCCCAAATGTTCCTGGGGAAGACCNNNAGATCGGAAG |
| 40936_1 | TATGAGTGTGGAGTCGTTGCTGCCAAAGTCTCTGAAATGTGTCNNNAGATCGGAAG |
| 40936_2 | TATGAGTGTGGAGTCGTTGCTGCCAAAGTCTCTGAAATGTGTCNNNAGATCGGAAG |
| 40936_3 | TATGAGTGTGGAGTCGTTGCTGCCAAAGTCTCTGAAATGTGTCNNNAGATCGGAAG |
| 48233_1 | TATGAGTGTGGAGTCGTTGCCTCCCAGGTTCACACCATTCTCNNNAGATCGGAAGAG |
| 48233_2 | TATGAGTGTGGAGTCGTTGCCTCCCAGGTTCACACCATTCTCNNNAGATCGGAAGAG |
| 48233_3 | TATGAGTGTGGAGTCGTTGCCTCCCAGGTTCACACCATTCTCNNNAGATCGGAAGAG |
| 13552_1 | TATGAGTGTGGAGTCGTTGCATTATGCATCGTCCATAACTTTNNNAGATCGGAAGA |
| 13552_2 | TATGAGTGTGGAGTCGTTGCATTATGCATCGTCCATAACTTTNNNAGATCGGAAGA |
| 13552_3 | TATGAGTGTGGAGTCGTTGCATTATGCATCGTCCATAACTTTNNNAGATCGGAAGA |
| 39718_1 | TATGAGTGTGGAGTCGTTGCCTGCTGGAGACATCTCTTCTGTTNNNAGATCGGAAG |
| 39718_2 | TATGAGTGTGGAGTCGTTGCCTGCTGGAGACATCTCTTCTGTTNNNAGATCGGAAG |
| 39718_3 | TATGAGTGTGGAGTCGTTGCCTGCTGGAGACATCTCTTCTGTTNNNAGATCGGAAG |
| 31532_1 | TATGAGTGTGGAGTCGTTGCTCTGAGGTGTAGGAAAGAAAGCCNNNAGATCGGAA |
| 31532_2 | TATGAGTGTGGAGTCGTTGCTCTGAGGTGTAGGAAAGAAAGCCNNNAGATCGGAA |
| 31532_3 | TATGAGTGTGGAGTCGTTGCTCTGAGGTGTAGGAAAGAAAGCCNNNAGATCGGAA |
| 35745_1 | TATGAGTGTGGAGTCGTTGCGAGACGGAGTCTTGTTCTGTCACNNNAGATCGGAA |
| 35745_2 | TATGAGTGTGGAGTCGTTGCGAGACGGAGTCTTGTTCTGTCACNNNAGATCGGAA |
| 35745_3 | TATGAGTGTGGAGTCGTTGCGAGACGGAGTCTTGTTCTGTCACNNNAGATCGGAA |
| 37376_1 | TATGAGTGTGGAGTCGTTGCGGGGGATTTCAGATCTCTGTACCNNNAGATCGGAA |
| 37376_2 | TATGAGTGTGGAGTCGTTGCGGGGGATTTCAGATCTCTGTACCNNNAGATCGGAA |
| 37376_3 | TATGAGTGTGGAGTCGTTGCGGGGGATTTCAGATCTCTGTACCNNNAGATCGGAA |
| 35150_1 | TATGAGTGTGGAGTCGTTGCCCGGGTTCACTTTGATTCTGTTGNNNAGATCGGAAG |
| 35150_2 | TATGAGTGTGGAGTCGTTGCCCGGGTTCACTTTGATTCTGTTGNNNAGATCGGAAG |
| 35150_3 | TATGAGTGTGGAGTCGTTGCCCGGGTTCACTTTGATTCTGTTGNNNAGATCGGAAG |
| 45100_1 | TATGAGTGTGGAGTCGTTGCAAATCCAAACCCCCTGAATAGCTNNNAGATCGGAAG |
| 45100_2 | TATGAGTGTGGAGTCGTTGCAAATCCAAACCCCCTGAATAGCTNNNAGATCGGAAG |
| 45100_3 | TATGAGTGTGGAGTCGTTGCAAATCCAAACCCCCTGAATAGCTNNNAGATCGGAAG |
| 41969_1 | TATGAGTGTGGAGTCGTTGCAGAGTAGCTGGGACTACAGGCNNNAGATCGGAAGA |
| 41969_2 | TATGAGTGTGGAGTCGTTGCAGAGTAGCTGGGACTACAGGCNNNAGATCGGAAGA |
| 41969_3 | TATGAGTGTGGAGTCGTTGCAGAGTAGCTGGGACTACAGGCNNNAGATCGGAAGA |
| 32438_1 | TATGAGTGTGGAGTCGTTGCGTGATCTCGGTTCATTGCAACCNNNAGATCGGAAGA |
| 32438_2 | TATGAGTGTGGAGTCGTTGCGTGATCTCGGTTCATTGCAACCNNNAGATCGGAAGA |
| 32438_3 | TATGAGTGTGGAGTCGTTGCGTGATCTCGGTTCATTGCAACCNNNAGATCGGAAGA |
| 10723_1 | TATGAGTGTGGAGTCGTTGCGCTGTTGCCATCTGGTTGATCTANNNAGATCGGAAG |
| 10723_2 | TATGAGTGTGGAGTCGTTGCGCTGTTGCCATCTGGTTGATCTANNNAGATCGGAAG |
| 10723_3 | TATGAGTGTGGAGTCGTTGCGCTGTTGCCATCTGGTTGATCTANNNAGATCGGAAG |
| 50133_1 | TATGAGTGTGGAGTCGTTGCACCACCAAACCTTCCTATGTACCNNNAGATCGGAAGA |
| 50133_2 | TATGAGTGTGGAGTCGTTGCACCACCAAACCTTCCTATGTACCNNNAGATCGGAAGA |
| 50133_3 | TATGAGTGTGGAGTCGTTGCACCACCAAACCTTCCTATGTACCNNNAGATCGGAAGA |
| 50000_1 | TATGAGTGTGGAGTCGTTGCCATTTCTCCAGTTGATCACGGTCNNNAGATCGGAAG |
| 50000_2 | TATGAGTGTGGAGTCGTTGCCATTTCTCCAGTTGATCACGGTCNNNAGATCGGAAG |

| | |
|---|---|
| 50000_3 | TATGAGTGTGGAGTCGTTGCCATTTCTCCAGTTGATCACGGTCNNNAGATCGGAAG |
| 38378_1 | TATGAGTGTGGAGTCGTTGCTGGGTGATAGAGTGAGACTCCATNNNAGATCGGAA |
| 38378_2 | TATGAGTGTGGAGTCGTTGCTGGGTGATAGAGTGAGACTCCATNNNAGATCGGAA |
| 38378_3 | TATGAGTGTGGAGTCGTTGCTGGGTGATAGAGTGAGACTCCATNNNAGATCGGAA |
| 40771_1 | TATGAGTGTGGAGTCGTTGCTGTAAGCATCAATGTCAATCGGGNNNAGATCGGAA |
| 40771_2 | TATGAGTGTGGAGTCGTTGCTGTAAGCATCAATGTCAATCGGGNNNAGATCGGAA |
| 40771_3 | TATGAGTGTGGAGTCGTTGCTGTAAGCATCAATGTCAATCGGGNNNAGATCGGAA |
| 28983_1 | TATGAGTGTGGAGTCGTTGCTGGGAGATTGTGTCTTCATGTCCNNNAGATCGGAAG |
| 28983_2 | TATGAGTGTGGAGTCGTTGCTGGGAGATTGTGTCTTCATGTCCNNNAGATCGGAAG |
| 28983_3 | TATGAGTGTGGAGTCGTTGCTGGGAGATTGTGTCTTCATGTCCNNNAGATCGGAAG |
| 30971_1 | TATGAGTGTGGAGTCGTTGCTTTTTTGACTCCTTCCAATGCCCNNNAGATCGGAAGA |
| 30971_2 | TATGAGTGTGGAGTCGTTGCTTTTTTGACTCCTTCCAATGCCCNNNAGATCGGAAGA |
| 30971_3 | TATGAGTGTGGAGTCGTTGCTTTTTTGACTCCTTCCAATGCCCNNNAGATCGGAAGA |
| 51466_1 | TATGAGTGTGGAGTCGTTGCGGGAGATACTCAGTTCTGCACATNNNAGATCGGAA |
| 51466_2 | TATGAGTGTGGAGTCGTTGCGGGAGATACTCAGTTCTGCACATNNNAGATCGGAA |
| 51466_3 | TATGAGTGTGGAGTCGTTGCGGGAGATACTCAGTTCTGCACATNNNAGATCGGAA |
| 45245_1 | TATGAGTGTGGAGTCGTTGCGAAGACCTCCAGTATATCGCAGGNNNAGATCGGAA |
| 45245_2 | TATGAGTGTGGAGTCGTTGCGAAGACCTCCAGTATATCGCAGGNNNAGATCGGAA |
| 45245_3 | TATGAGTGTGGAGTCGTTGCGAAGACCTCCAGTATATCGCAGGNNNAGATCGGAA |
| 51440_1 | TATGAGTGTGGAGTCGTTGCACATGTGTTACACTGTGAGCTCTNNNAGATCGGAAG |
| 51440_2 | TATGAGTGTGGAGTCGTTGCACATGTGTTACACTGTGAGCTCTNNNAGATCGGAAG |
| 51440_3 | TATGAGTGTGGAGTCGTTGCACATGTGTTACACTGTGAGCTCTNNNAGATCGGAAG |
| 36323_1 | TATGAGTGTGGAGTCGTTGCATGACTTCGTATAGATCCTGCCTNNNAGATCGGAAG |
| 36323_2 | TATGAGTGTGGAGTCGTTGCATGACTTCGTATAGATCCTGCCTNNNAGATCGGAAG |
| 36323_3 | TATGAGTGTGGAGTCGTTGCATGACTTCGTATAGATCCTGCCTNNNAGATCGGAAG |
| 37192_1 | TATGAGTGTGGAGTCGTTGCCATGGCTTCAAACAGATCGCATTNNNAGATCGGAAG |
| 37192_2 | TATGAGTGTGGAGTCGTTGCCATGGCTTCAAACAGATCGCATTNNNAGATCGGAAG |
| 37192_3 | TATGAGTGTGGAGTCGTTGCCATGGCTTCAAACAGATCGCATTNNNAGATCGGAAG |
| 34708_1 | TATGAGTGTGGAGTCGTTGCGCGCCCGGCCTATTTATTTTATTNNNAGATCGGAAG |
| 34708_2 | TATGAGTGTGGAGTCGTTGCGCGCCCGGCCTATTTATTTTATTNNNAGATCGGAAG |
| 34708_3 | TATGAGTGTGGAGTCGTTGCGCGCCCGGCCTATTTATTTTATTNNNAGATCGGAAG |
| 50777_1 | TATGAGTGTGGAGTCGTTGCAGGAGAGCTCATTACTTTTCGGGNNNAGATCGGAA |
| 50777_2 | TATGAGTGTGGAGTCGTTGCAGGAGAGCTCATTACTTTTCGGGNNNAGATCGGAA |
| 50777_3 | TATGAGTGTGGAGTCGTTGCAGGAGAGCTCATTACTTTTCGGGNNNAGATCGGAA |
| 42579_1 | TATGAGTGTGGAGTCGTTGCCTGGCTGATCTCGAATTCCTGAANNNAGATCGGAAG |
| 42579_2 | TATGAGTGTGGAGTCGTTGCCTGGCTGATCTCGAATTCCTGAANNNAGATCGGAAG |
| 42579_3 | TATGAGTGTGGAGTCGTTGCCTGGCTGATCTCGAATTCCTGAANNNAGATCGGAAG |
| 47704_1 | TATGAGTGTGGAGTCGTTGCGGAGTGGAGTGGCATGATCATAGNNNAGATCGGA |
| 47704_2 | TATGAGTGTGGAGTCGTTGCGGAGTGGAGTGGCATGATCATAGNNNAGATCGGA |
| 47704_3 | TATGAGTGTGGAGTCGTTGCGGAGTGGAGTGGCATGATCATAGNNNAGATCGGA |
| 38175_1 | TATGAGTGTGGAGTCGTTGCGACGAAGCTGGGCAAAATAATGANNNAGATCGGAA |
| 38175_2 | TATGAGTGTGGAGTCGTTGCGACGAAGCTGGGCAAAATAATGANNNAGATCGGAA |
| 38175_3 | TATGAGTGTGGAGTCGTTGCGACGAAGCTGGGCAAAATAATGANNNAGATCGGAA |

| | |
|---|---|
| 36826_1 | TATGAGTGTGGAGTCGTTGCTGGCTCTTATAATCCCCTTCTGCNNNAGATCGGAAGA |
| 36826_2 | TATGAGTGTGGAGTCGTTGCTGGCTCTTATAATCCCCTTCTGCNNNAGATCGGAAGA |
| 36826_3 | TATGAGTGTGGAGTCGTTGCTGGCTCTTATAATCCCCTTCTGCNNNAGATCGGAAGA |
| 51789_1 | TATGAGTGTGGAGTCGTTGCCTACACAATCTAGCCCCCTCATCNNNAGATCGGAAGA |
| 51789_2 | TATGAGTGTGGAGTCGTTGCCTACACAATCTAGCCCCCTCATCNNNAGATCGGAAGA |
| 51789_3 | TATGAGTGTGGAGTCGTTGCCTACACAATCTAGCCCCCTCATCNNNAGATCGGAAGA |
| 44846_1 | TATGAGTGTGGAGTCGTTGCAGGAAGCCTTTAGTCATCGGTTTNNNAGATCGGAAG |
| 44846_2 | TATGAGTGTGGAGTCGTTGCAGGAAGCCTTTAGTCATCGGTTTNNNAGATCGGAAG |
| 44846_3 | TATGAGTGTGGAGTCGTTGCAGGAAGCCTTTAGTCATCGGTTTNNNAGATCGGAAG |
| 10545_1 | TATGAGTGTGGAGTCGTTGCACCTCTCACACGGTTTCTGTTTTNNNAGATCGGAAGA |
| 10545_2 | TATGAGTGTGGAGTCGTTGCACCTCTCACACGGTTTCTGTTTTNNNAGATCGGAAGA |
| 10545_3 | TATGAGTGTGGAGTCGTTGCACCTCTCACACGGTTTCTGTTTTNNNAGATCGGAAGA |
| 46416_1 | TATGAGTGTGGAGTCGTTGCTCCCTGCTGGTTCTCAAAGTTTANNNAGATCGGAAG |
| 46416_2 | TATGAGTGTGGAGTCGTTGCTCCCTGCTGGTTCTCAAAGTTTANNNAGATCGGAAG |
| 46416_3 | TATGAGTGTGGAGTCGTTGCTCCCTGCTGGTTCTCAAAGTTTANNNAGATCGGAAG |
| 39067_1 | TATGAGTGTGGAGTCGTTGCAATGTCCATTGTTGAAGCCACACNNNAGATCGGAAG |
| 39067_2 | TATGAGTGTGGAGTCGTTGCAATGTCCATTGTTGAAGCCACACNNNAGATCGGAAG |
| 39067_3 | TATGAGTGTGGAGTCGTTGCAATGTCCATTGTTGAAGCCACACNNNAGATCGGAAG |
| 29112_1 | TATGAGTGTGGAGTCGTTGCTCTGAGGAGCATTGAAACCTACCNNNAGATCGGAAG |
| 29112_2 | TATGAGTGTGGAGTCGTTGCTCTGAGGAGCATTGAAACCTACCNNNAGATCGGAAG |
| 29112_3 | TATGAGTGTGGAGTCGTTGCTCTGAGGAGCATTGAAACCTACCNNNAGATCGGAAG |
| 37534_1 | TATGAGTGTGGAGTCGTTGCAGTTGTCTTCAATGGGAGACATCANNNAGATCGGAA |
| 37534_2 | TATGAGTGTGGAGTCGTTGCAGTTGTCTTCAATGGGAGACATCANNNAGATCGGAA |
| 37534_3 | TATGAGTGTGGAGTCGTTGCAGTTGTCTTCAATGGGAGACATCANNNAGATCGGAA |
| 12989_1 | TATGAGTGTGGAGTCGTTGCCTCAGGTGAAGGACAAAAAGNNNAGATCGGAAGA |
| 12989_2 | TATGAGTGTGGAGTCGTTGCCTCAGGTGAAGGACAAAAAGNNNAGATCGGAAGA |
| 12989_3 | TATGAGTGTGGAGTCGTTGCCTCAGGTGAAGGACAAAAAGNNNAGATCGGAAGA |
| 41110_1 | TATGAGTGTGGAGTCGTTGCGTTCAAAAGATTCTTGTGCCCCANNNAGATCGGAAG |
| 41110_2 | TATGAGTGTGGAGTCGTTGCGTTCAAAAGATTCTTGTGCCCCANNNAGATCGGAAG |
| 41110_3 | TATGAGTGTGGAGTCGTTGCGTTCAAAAGATTCTTGTGCCCCANNNAGATCGGAAG |
| 30413_1 | TATGAGTGTGGAGTCGTTGCAGAAACACATACACAAGGATGCACTNNNAGATCGG |
| 30413_2 | TATGAGTGTGGAGTCGTTGCAGAAACACATACACAAGGATGCACTNNNAGATCGG |
| 30413_3 | TATGAGTGTGGAGTCGTTGCAGAAACACATACACAAGGATGCACTNNNAGATCGG |
| 43755_1 | TATGAGTGTGGAGTCGTTGCCTCGGAGAAGTAGAGCGCACNNNAGATCGGAAGAG |
| 43755_2 | TATGAGTGTGGAGTCGTTGCCTCGGAGAAGTAGAGCGCACNNNAGATCGGAAGAG |
| 43755_3 | TATGAGTGTGGAGTCGTTGCCTCGGAGAAGTAGAGCGCACNNNAGATCGGAAGAG |
| 35421_1 | TATGAGTGTGGAGTCGTTGCGCCTGTAGCTAGAACAATAGGCANNNAGATCGGAA |
| 35421_2 | TATGAGTGTGGAGTCGTTGCGCCTGTAGCTAGAACAATAGGCANNNAGATCGGAA |
| 35421_3 | TATGAGTGTGGAGTCGTTGCGCCTGTAGCTAGAACAATAGGCANNNAGATCGGAA |
| 2951_1 | TATGAGTGTGGAGTCGTTGCCACACACACCTTGGGGTATTCTTNNNAGATCGGAAG |
| 2951_2 | TATGAGTGTGGAGTCGTTGCCACACACACCTTGGGGTATTCTTNNNAGATCGGAAG |
| 2951_3 | TATGAGTGTGGAGTCGTTGCCACACACACCTTGGGGTATTCTTNNNAGATCGGAAG |
| 2211_1 | TATGAGTGTGGAGTCGTTGCACACCAGAAAGGCTTCAAGATGANNNAGATCGGAA |

| | |
|---|---|
| 2211_2 | TATGAGTGTGGAGTCGTTGCACACCAGAAAGGCTTCAAGATGANNNAGATCGGAA |
| 2211_3 | TATGAGTGTGGAGTCGTTGCACACCAGAAAGGCTTCAAGATGANNNAGATCGGAA |
| 8499_1 | TATGAGTGTGGAGTCGTTGCGCCATTATGTGTAGGGACCATTCTNNNAGATCGGAA |
| 8499_2 | TATGAGTGTGGAGTCGTTGCGCCATTATGTGTAGGGACCATTCTNNNAGATCGGAA |
| 8499_3 | TATGAGTGTGGAGTCGTTGCGCCATTATGTGTAGGGACCATTCTNNNAGATCGGAA |
| 48496_1 | TATGAGTGTGGAGTCGTTGCAATTGAGCCCAACTGACAAACTGNNNAGATCGGAAG |
| 48496_2 | TATGAGTGTGGAGTCGTTGCAATTGAGCCCAACTGACAAACTGNNNAGATCGGAAG |
| 48496_3 | TATGAGTGTGGAGTCGTTGCAATTGAGCCCAACTGACAAACTGNNNAGATCGGAAG |
| 32512_1 | TATGAGTGTGGAGTCGTTGCACCATCCAGTGAACAAAAGACCTNNNAGATCGGAAG |
| 32512_2 | TATGAGTGTGGAGTCGTTGCACCATCCAGTGAACAAAAGACCTNNNAGATCGGAAG |
| 32512_3 | TATGAGTGTGGAGTCGTTGCACCATCCAGTGAACAAAAGACCTNNNAGATCGGAAG |
| 29785_1 | TATGAGTGTGGAGTCGTTGCGCGTTCAGCATCTACATTCACTANNNAGATCGGAAG |
| 29785_2 | TATGAGTGTGGAGTCGTTGCGCGTTCAGCATCTACATTCACTANNNAGATCGGAAG |
| 29785_3 | TATGAGTGTGGAGTCGTTGCGCGTTCAGCATCTACATTCACTANNNAGATCGGAAG |
| 2179_1 | TATGAGTGTGGAGTCGTTGCATGGCTCTGGTGCATCTCTTAGANNNAGATCGGAAG |
| 2179_2 | TATGAGTGTGGAGTCGTTGCATGGCTCTGGTGCATCTCTTAGANNNAGATCGGAAG |
| 2179_3 | TATGAGTGTGGAGTCGTTGCATGGCTCTGGTGCATCTCTTAGANNNAGATCGGAAG |
| 49487_1 | TATGAGTGTGGAGTCGTTGCGCCTCAAAATACCTCTCCATCCTNNNAGATCGGAAGA |
| 49487_2 | TATGAGTGTGGAGTCGTTGCGCCTCAAAATACCTCTCCATCCTNNNAGATCGGAAGA |
| 49487_3 | TATGAGTGTGGAGTCGTTGCGCCTCAAAATACCTCTCCATCCTNNNAGATCGGAAGA |
| 45462_1 | TATGAGTGTGGAGTCGTTGCCAGATTTACCTTTCAGTTCCGCCNNNAGATCGGAAG |
| 45462_2 | TATGAGTGTGGAGTCGTTGCCAGATTTACCTTTCAGTTCCGCCNNNAGATCGGAAG |
| 45462_3 | TATGAGTGTGGAGTCGTTGCCAGATTTACCTTTCAGTTCCGCCNNNAGATCGGAAG |
| 51527_1 | TATGAGTGTGGAGTCGTTGCTTCTTCAATAGGCTGATGGTGCANNNAGATCGGAAG |
| 51527_2 | TATGAGTGTGGAGTCGTTGCTTCTTCAATAGGCTGATGGTGCANNNAGATCGGAAG |
| 51527_3 | TATGAGTGTGGAGTCGTTGCTTCTTCAATAGGCTGATGGTGCANNNAGATCGGAAG |
| 47900_1 | TATGAGTGTGGAGTCGTTGCACACCACTACACCTGGCTAATTTNNNAGATCGGAAG |
| 47900_2 | TATGAGTGTGGAGTCGTTGCACACCACTACACCTGGCTAATTTNNNAGATCGGAAG |
| 47900_3 | TATGAGTGTGGAGTCGTTGCACACCACTACACCTGGCTAATTTNNNAGATCGGAAG |
| 45406_1 | TATGAGTGTGGAGTCGTTGCACCAGCTACAAATGACAGTTCCANNNAGATCGGAAG |
| 45406_2 | TATGAGTGTGGAGTCGTTGCACCAGCTACAAATGACAGTTCCANNNAGATCGGAAG |
| 45406_3 | TATGAGTGTGGAGTCGTTGCACCAGCTACAAATGACAGTTCCANNNAGATCGGAAG |
| 12834_1 | TATGAGTGTGGAGTCGTTGCACCTTCTATTTTAATCTCTCTCCNNNAGATCGGAAGA |
| 12834_2 | TATGAGTGTGGAGTCGTTGCACCTTCTATTTTAATCTCTCTCCNNNAGATCGGAAGA |
| 12834_3 | TATGAGTGTGGAGTCGTTGCACCTTCTATTTTAATCTCTCTCCNNNAGATCGGAAGA |
| 38729_1 | TATGAGTGTGGAGTCGTTGCGGAGGAATCACATTACCGGACTTNNNAGATCGGAA |
| 38729_2 | TATGAGTGTGGAGTCGTTGCGGAGGAATCACATTACCGGACTTNNNAGATCGGAA |
| 38729_3 | TATGAGTGTGGAGTCGTTGCGGAGGAATCACATTACCGGACTTNNNAGATCGGAA |
| 40211_1 | TATGAGTGTGGAGTCGTTGCTGGGACAAACTTATGTCTCCTGANNNAGATCGGAAG |
| 40211_2 | TATGAGTGTGGAGTCGTTGCTGGGACAAACTTATGTCTCCTGANNNAGATCGGAAG |
| 40211_3 | TATGAGTGTGGAGTCGTTGCTGGGACAAACTTATGTCTCCTGANNNAGATCGGAAG |
| 30284_1 | TATGAGTGTGGAGTCGTTGCAGCTAAGGTAAGTGCCTGTCAATNNNAGATCGGAA |
| 30284_2 | TATGAGTGTGGAGTCGTTGCAGCTAAGGTAAGTGCCTGTCAATNNNAGATCGGAA |

| | |
|---|---|
| 30284_3 | TATGAGTGTGGAGTCGTTGCAGCTAAGGTAAGTGCCTGTCAATNNNAGATCGGAA |
| 34306_1 | TATGAGTGTGGAGTCGTTGCAACAAAAGGCATGAAAACCCAANNNAGATCGGAA |
| 34306_2 | TATGAGTGTGGAGTCGTTGCAACAAAAGGCATGAAAACCCAANNNAGATCGGAA |
| 34306_3 | TATGAGTGTGGAGTCGTTGCAACAAAAGGCATGAAAACCCAANNNAGATCGGAA |
| 38225_1 | TATGAGTGTGGAGTCGTTGCTGCTAAGCGAAATAAGCCAGATGNNNAGATCGGAA |
| 38225_2 | TATGAGTGTGGAGTCGTTGCTGCTAAGCGAAATAAGCCAGATGNNNAGATCGGAA |
| 38225_3 | TATGAGTGTGGAGTCGTTGCTGCTAAGCGAAATAAGCCAGATGNNNAGATCGGAA |
| 47994_1 | TATGAGTGTGGAGTCGTTGCTCCATTCAGCCAGGCTATTTCTTNNNAGATCGGAAG |
| 47994_2 | TATGAGTGTGGAGTCGTTGCTCCATTCAGCCAGGCTATTTCTTNNNAGATCGGAAG |
| 47994_3 | TATGAGTGTGGAGTCGTTGCTCCATTCAGCCAGGCTATTTCTTNNNAGATCGGAAG |
| 29927_1 | TATGAGTGTGGAGTCGTTGCACAATTAGCCAGGTGACAGAGAGNNNAGATCGGAA |
| 29927_2 | TATGAGTGTGGAGTCGTTGCACAATTAGCCAGGTGACAGAGAGNNNAGATCGGAA |
| 29927_3 | TATGAGTGTGGAGTCGTTGCACAATTAGCCAGGTGACAGAGAGNNNAGATCGGAA |
| 42351_1 | TATGAGTGTGGAGTCGTTGCACTTGGGATATTGAGGCAGGAAANNNAGATCGGAA |
| 42351_2 | TATGAGTGTGGAGTCGTTGCACTTGGGATATTGAGGCAGGAAANNNAGATCGGAA |
| 42351_3 | TATGAGTGTGGAGTCGTTGCACTTGGGATATTGAGGCAGGAAANNNAGATCGGAA |
| 36380_1 | TATGAGTGTGGAGTCGTTGCTGGCAGTAAAGTGGGTTCTGAATNNNAGATCGGAA |
| 36380_2 | TATGAGTGTGGAGTCGTTGCTGGCAGTAAAGTGGGTTCTGAATNNNAGATCGGAA |
| 36380_3 | TATGAGTGTGGAGTCGTTGCTGGCAGTAAAGTGGGTTCTGAATNNNAGATCGGAA |
| 48312_1 | TATGAGTGTGGAGTCGTTGCGGCCGCCTGTATTTTCTTTTTCTNNNAGATCGGAAGA |
| 48312_2 | TATGAGTGTGGAGTCGTTGCGGCCGCCTGTATTTTCTTTTTCTNNNAGATCGGAAGA |
| 48312_3 | TATGAGTGTGGAGTCGTTGCGGCCGCCTGTATTTTCTTTTTCTNNNAGATCGGAAGA |
| 38569_1 | TATGAGTGTGGAGTCGTTGCAGCTTGGCATAACCTCAATACCANNNAGATCGGAAG |
| 38569_2 | TATGAGTGTGGAGTCGTTGCAGCTTGGCATAACCTCAATACCANNNAGATCGGAAG |
| 38569_3 | TATGAGTGTGGAGTCGTTGCAGCTTGGCATAACCTCAATACCANNNAGATCGGAAG |
| 49575_1 | TATGAGTGTGGAGTCGTTGCTCCTGTTACTTTTCTGTGGAACTNNNAGATCGGAAG |
| 49575_2 | TATGAGTGTGGAGTCGTTGCTCCTGTTACTTTTCTGTGGAACTNNNAGATCGGAAG |
| 49575_3 | TATGAGTGTGGAGTCGTTGCTCCTGTTACTTTTCTGTGGAACTNNNAGATCGGAAG |
| 29362_1 | TATGAGTGTGGAGTCGTTGCAATGTAGCAGAGCATGTGACAGANNNAGATCGGAA |
| 29362_2 | TATGAGTGTGGAGTCGTTGCAATGTAGCAGAGCATGTGACAGANNNAGATCGGAA |
| 29362_3 | TATGAGTGTGGAGTCGTTGCAATGTAGCAGAGCATGTGACAGANNNAGATCGGAA |
| 34301_1 | TATGAGTGTGGAGTCGTTGCAAACTGTGGGGTGTTGAGAGAATNNNAGATCGGA |
| 34301_2 | TATGAGTGTGGAGTCGTTGCAAACTGTGGGGTGTTGAGAGAATNNNAGATCGGA |
| 34301_3 | TATGAGTGTGGAGTCGTTGCAAACTGTGGGGTGTTGAGAGAATNNNAGATCGGA |
| 31065_1 | TATGAGTGTGGAGTCGTTGCCCTGTGAGCTCTATTTTTCTCTGCNNNAGATCGGAAG |
| 31065_2 | TATGAGTGTGGAGTCGTTGCCCTGTGAGCTCTATTTTTCTCTGCNNNAGATCGGAAG |
| 31065_3 | TATGAGTGTGGAGTCGTTGCCCTGTGAGCTCTATTTTTCTCTGCNNNAGATCGGAAG |
| 41144_1 | TATGAGTGTGGAGTCGTTGCTGTAGTGGCACCCATCTGTAATCNNNAGATCGGAAG |
| 41144_2 | TATGAGTGTGGAGTCGTTGCTGTAGTGGCACCCATCTGTAATCNNNAGATCGGAAG |
| 41144_3 | TATGAGTGTGGAGTCGTTGCTGTAGTGGCACCCATCTGTAATCNNNAGATCGGAAG |
| 49825_1 | TATGAGTGTGGAGTCGTTGCGTGTCAAACTGCTAAGACTGCTGNNNAGATCGGAA |
| 49825_2 | TATGAGTGTGGAGTCGTTGCGTGTCAAACTGCTAAGACTGCTGNNNAGATCGGAA |
| 49825_3 | TATGAGTGTGGAGTCGTTGCGTGTCAAACTGCTAAGACTGCTGNNNAGATCGGAA |

| | |
|---|---|
| 31635_1 | TATGAGTGTGGAGTCGTTGCAGGGCCTCATGTTTTATCTGGTTNNNAGATCGGAAG |
| 31635_2 | TATGAGTGTGGAGTCGTTGCAGGGCCTCATGTTTTATCTGGTTNNNAGATCGGAAG |
| 31635_3 | TATGAGTGTGGAGTCGTTGCAGGGCCTCATGTTTTATCTGGTTNNNAGATCGGAAG |
| 44897_1 | TATGAGTGTGGAGTCGTTGCGGAAGAGCAGAGATGGGATTCTGNNNAGATCGGA |
| 44897_2 | TATGAGTGTGGAGTCGTTGCGGAAGAGCAGAGATGGGATTCTGNNNAGATCGGA |
| 44897_3 | TATGAGTGTGGAGTCGTTGCGGAAGAGCAGAGATGGGATTCTGNNNAGATCGGA |
| 9908_1 | TATGAGTGTGGAGTCGTTGCACTGCTCTCTCAAAGCTCAGAGANNNAGATCGGAAG |
| 9908_2 | TATGAGTGTGGAGTCGTTGCACTGCTCTCTCAAAGCTCAGAGANNNAGATCGGAAG |
| 9908_3 | TATGAGTGTGGAGTCGTTGCACTGCTCTCTCAAAGCTCAGAGANNNAGATCGGAAG |
| 47899_1 | TATGAGTGTGGAGTCGTTGCCCTCCTGGGTTCAGATGATTCTCNNNAGATCGGAAG |
| 47899_2 | TATGAGTGTGGAGTCGTTGCCCTCCTGGGTTCAGATGATTCTCNNNAGATCGGAAG |
| 47899_3 | TATGAGTGTGGAGTCGTTGCCCTCCTGGGTTCAGATGATTCTCNNNAGATCGGAAG |
| 50908_1 | TATGAGTGTGGAGTCGTTGCTGCTCCAGGGAAAAATATGTGGANNNAGATCGGAA |
| 50908_2 | TATGAGTGTGGAGTCGTTGCTGCTCCAGGGAAAAATATGTGGANNNAGATCGGAA |
| 50908_3 | TATGAGTGTGGAGTCGTTGCTGCTCCAGGGAAAAATATGTGGANNNAGATCGGAA |
| 53218_1 | TATGAGTGTGGAGTCGTTGCTTTGGCCTCAAACTTCAGGTGANNNAGATCGGAAGA |
| 53218_2 | TATGAGTGTGGAGTCGTTGCTTTGGCCTCAAACTTCAGGTGANNNAGATCGGAAGA |
| 53218_3 | TATGAGTGTGGAGTCGTTGCTTTGGCCTCAAACTTCAGGTGANNNAGATCGGAAGA |
| 37401_1 | TATGAGTGTGGAGTCGTTGCGAGACAGGGCTTATCATTCCCAANNNAGATCGGAAG |
| 37401_2 | TATGAGTGTGGAGTCGTTGCGAGACAGGGCTTATCATTCCCAANNNAGATCGGAAG |
| 37401_3 | TATGAGTGTGGAGTCGTTGCGAGACAGGGCTTATCATTCCCAANNNAGATCGGAAG |
| 38230_1 | TATGAGTGTGGAGTCGTTGCACCACTGCAAATTCTGTTTGACCNNNAGATCGGAAG |
| 38230_2 | TATGAGTGTGGAGTCGTTGCACCACTGCAAATTCTGTTTGACCNNNAGATCGGAAG |
| 38230_3 | TATGAGTGTGGAGTCGTTGCACCACTGCAAATTCTGTTTGACCNNNAGATCGGAAG |
| 49458_1 | TATGAGTGTGGAGTCGTTGCAGGAGATATTACCACTGACCCCANNNAGATCGGAAG |
| 49458_2 | TATGAGTGTGGAGTCGTTGCAGGAGATATTACCACTGACCCCANNNAGATCGGAAG |
| 49458_3 | TATGAGTGTGGAGTCGTTGCAGGAGATATTACCACTGACCCCANNNAGATCGGAAG |
| 36830_1 | TATGAGTGTGGAGTCGTTGCGTTAGGGGTGTTTCTTGGTGTTGNNNAGATCGGAA |
| 36830_2 | TATGAGTGTGGAGTCGTTGCGTTAGGGGTGTTTCTTGGTGTTGNNNAGATCGGAA |
| 36830_3 | TATGAGTGTGGAGTCGTTGCGTTAGGGGTGTTTCTTGGTGTTGNNNAGATCGGAA |
| 41659_1 | TATGAGTGTGGAGTCGTTGCTCTCCCAAGTAGCTGAGACTACANNNAGATCGGAAG |
| 41659_2 | TATGAGTGTGGAGTCGTTGCTCTCCCAAGTAGCTGAGACTACANNNAGATCGGAAG |
| 41659_3 | TATGAGTGTGGAGTCGTTGCTCTCCCAAGTAGCTGAGACTACANNNAGATCGGAAG |
| 52529_1 | TATGAGTGTGGAGTCGTTGCCCGGGCATCATCCAAATCTTTTTNNNAGATCGGAAG |
| 52529_2 | TATGAGTGTGGAGTCGTTGCCCGGGCATCATCCAAATCTTTTTNNNAGATCGGAAG |
| 52529_3 | TATGAGTGTGGAGTCGTTGCCCGGGCATCATCCAAATCTTTTTNNNAGATCGGAAG |
| 37749_1 | TATGAGTGTGGAGTCGTTGCGAGAATTGCTTGAACCTGGGAAGNNNAGATCGGAA |
| 37749_2 | TATGAGTGTGGAGTCGTTGCGAGAATTGCTTGAACCTGGGAAGNNNAGATCGGAA |
| 37749_3 | TATGAGTGTGGAGTCGTTGCGAGAATTGCTTGAACCTGGGAAGNNNAGATCGGAA |
| 49312_1 | TATGAGTGTGGAGTCGTTGCTGCCTCTTGGTTTGTTGGTGATANNNAGATCGGAAG |
| 49312_2 | TATGAGTGTGGAGTCGTTGCTGCCTCTTGGTTTGTTGGTGATANNNAGATCGGAAG |
| 49312_3 | TATGAGTGTGGAGTCGTTGCTGCCTCTTGGTTTGTTGGTGATANNNAGATCGGAAG |
| 31966_1 | TATGAGTGTGGAGTCGTTGCAGACAGAGAAAAGAAAGCCCACANNNAGATCGGAA |

| | |
|---|---|
| 31966_2 | TATGAGTGTGGAGTCGTTGCAGACAGAGAAAAGAAAGCCCACANNNAGATCGGAA |
| 31966_3 | TATGAGTGTGGAGTCGTTGCAGACAGAGAAAAGAAAGCCCACANNNAGATCGGAA |
| 46777_1 | TATGAGTGTGGAGTCGTTGCCTCCTGAGTAGCTGGGATCACAGNNNAGATCGGAA |
| 46777_2 | TATGAGTGTGGAGTCGTTGCCTCCTGAGTAGCTGGGATCACAGNNNAGATCGGAA |
| 46777_3 | TATGAGTGTGGAGTCGTTGCCTCCTGAGTAGCTGGGATCACAGNNNAGATCGGAA |
| 51241_1 | TATGAGTGTGGAGTCGTTGCACAAAGAGCAAGTCAGAGAGAGGNNNAGATCGGA |
| 51241_2 | TATGAGTGTGGAGTCGTTGCACAAAGAGCAAGTCAGAGAGAGGNNNAGATCGGA |
| 51241_3 | TATGAGTGTGGAGTCGTTGCACAAAGAGCAAGTCAGAGAGAGGNNNAGATCGGA |
| 29559_1 | TATGAGTGTGGAGTCGTTGCCCACCCCAGTTTGTTGAAAATGTNNNAGATCGGAAG |
| 29559_2 | TATGAGTGTGGAGTCGTTGCCCACCCCAGTTTGTTGAAAATGTNNNAGATCGGAAG |
| 29559_3 | TATGAGTGTGGAGTCGTTGCCCACCCCAGTTTGTTGAAAATGTNNNAGATCGGAAG |
| 46609_1 | TATGAGTGTGGAGTCGTTGCGTTCAATCGATTCTCCTCCCTCANNNAGATCGGAAGA |
| 46609_2 | TATGAGTGTGGAGTCGTTGCGTTCAATCGATTCTCCTCCCTCANNNAGATCGGAAGA |
| 46609_3 | TATGAGTGTGGAGTCGTTGCGTTCAATCGATTCTCCTCCCTCANNNAGATCGGAAGA |
| 51561_1 | TATGAGTGTGGAGTCGTTGCTGACTCTGAGCTTGCAAATCTCTNNNAGATCGGAAG |
| 51561_2 | TATGAGTGTGGAGTCGTTGCTGACTCTGAGCTTGCAAATCTCTNNNAGATCGGAAG |
| 51561_3 | TATGAGTGTGGAGTCGTTGCTGACTCTGAGCTTGCAAATCTCTNNNAGATCGGAAG |
| 44029_1 | TATGAGTGTGGAGTCGTTGCCATAAAAGGATGCCAGCTGTGTTNNNAGATCGGAA |
| 44029_2 | TATGAGTGTGGAGTCGTTGCCATAAAAGGATGCCAGCTGTGTTNNNAGATCGGAA |
| 44029_3 | TATGAGTGTGGAGTCGTTGCCATAAAAGGATGCCAGCTGTGTTNNNAGATCGGAA |
| 43049_1 | TATGAGTGTGGAGTCGTTGCCAAGTCCAGGAGTGAAGACATGANNNAGATCGGAA |
| 43049_2 | TATGAGTGTGGAGTCGTTGCCAAGTCCAGGAGTGAAGACATGANNNAGATCGGAA |
| 43049_3 | TATGAGTGTGGAGTCGTTGCCAAGTCCAGGAGTGAAGACATGANNNAGATCGGAA |
| 52685_1 | TATGAGTGTGGAGTCGTTGCCCAGGCTGCCATAGTATGTTACTNNNAGATCGGAAG |
| 52685_2 | TATGAGTGTGGAGTCGTTGCCCAGGCTGCCATAGTATGTTACTNNNAGATCGGAAG |
| 52685_3 | TATGAGTGTGGAGTCGTTGCCCAGGCTGCCATAGTATGTTACTNNNAGATCGGAAG |
| 470_1 | TATGAGTGTGGAGTCGTTGCAAGTAGCTGTCTCCACCCCTTTNNNAGATCGGAAGA |
| 470_2 | TATGAGTGTGGAGTCGTTGCAAGTAGCTGTCTCCACCCCTTTNNNAGATCGGAAGA |
| 470_3 | TATGAGTGTGGAGTCGTTGCAAGTAGCTGTCTCCACCCCTTTNNNAGATCGGAAGA |
| 41441_1 | TATGAGTGTGGAGTCGTTGCGTGAGCCCAGATCACTACACTACNNNAGATCGGAAG |
| 41441_2 | TATGAGTGTGGAGTCGTTGCGTGAGCCCAGATCACTACACTACNNNAGATCGGAAG |
| 41441_3 | TATGAGTGTGGAGTCGTTGCGTGAGCCCAGATCACTACACTACNNNAGATCGGAAG |
| 52779_1 | TATGAGTGTGGAGTCGTTGCTTGTGGATGAAAAACCCCCTCTCNNNAGATCGGAAG |
| 52779_2 | TATGAGTGTGGAGTCGTTGCTTGTGGATGAAAAACCCCCTCTCNNNAGATCGGAAG |
| 52779_3 | TATGAGTGTGGAGTCGTTGCTTGTGGATGAAAAACCCCCTCTCNNNAGATCGGAAG |
| 35293_1 | TATGAGTGTGGAGTCGTTGCTGGTCTCAAAATCCTGAGCTCAANNNAGATCGGAAG |
| 35293_2 | TATGAGTGTGGAGTCGTTGCTGGTCTCAAAATCCTGAGCTCAANNNAGATCGGAAG |
| 35293_3 | TATGAGTGTGGAGTCGTTGCTGGTCTCAAAATCCTGAGCTCAANNNAGATCGGAAG |
| 44338_1 | TATGAGTGTGGAGTCGTTGCCCAACAGGATGTTTGTGTTAGCCNNNAGATCGGAAG |
| 44338_2 | TATGAGTGTGGAGTCGTTGCCCAACAGGATGTTTGTGTTAGCCNNNAGATCGGAAG |
| 44338_3 | TATGAGTGTGGAGTCGTTGCCCAACAGGATGTTTGTGTTAGCCNNNAGATCGGAAG |
| 36410_1 | TATGAGTGTGGAGTCGTTGCATTTGGTTGAAAGGTCAGTGCAGNNNAGATCGGAA |
| 36410_2 | TATGAGTGTGGAGTCGTTGCATTTGGTTGAAAGGTCAGTGCAGNNNAGATCGGAA |

| | |
|---|---|
| 36410_3 | TATGAGTGTGGAGTCGTTGCATTTGGTTGAAAGGTCAGTGCAGNNNAGATCGGAA |
| 42603_1 | TATGAGTGTGGAGTCGTTGCTTGGTTGAGAATTATGTCCCCCCNNNAGATCGGAAG |
| 42603_2 | TATGAGTGTGGAGTCGTTGCTTGGTTGAGAATTATGTCCCCCCNNNAGATCGGAAG |
| 42603_3 | TATGAGTGTGGAGTCGTTGCTTGGTTGAGAATTATGTCCCCCCNNNAGATCGGAAG |
| 41745_1 | TATGAGTGTGGAGTCGTTGCCTCAGCCTCCCGAGTAGTTGNNNAGATCGGAAGAG |
| 41745_2 | TATGAGTGTGGAGTCGTTGCCTCAGCCTCCCGAGTAGTTGNNNAGATCGGAAGAG |
| 41745_3 | TATGAGTGTGGAGTCGTTGCCTCAGCCTCCCGAGTAGTTGNNNAGATCGGAAGAG |
| 46273_1 | TATGAGTGTGGAGTCGTTGCCCTTCTCAGAGGTGGAGATTTCCNNNAGATCGGAAG |
| 46273_2 | TATGAGTGTGGAGTCGTTGCCCTTCTCAGAGGTGGAGATTTCCNNNAGATCGGAAG |
| 46273_3 | TATGAGTGTGGAGTCGTTGCCCTTCTCAGAGGTGGAGATTTCCNNNAGATCGGAAG |
| 40743_1 | TATGAGTGTGGAGTCGTTGCGAGCTTCAGCCAAGTATCTCCATNNNAGATCGGAAG |
| 40743_2 | TATGAGTGTGGAGTCGTTGCGAGCTTCAGCCAAGTATCTCCATNNNAGATCGGAAG |
| 40743_3 | TATGAGTGTGGAGTCGTTGCGAGCTTCAGCCAAGTATCTCCATNNNAGATCGGAAG |
| 35762_1 | TATGAGTGTGGAGTCGTTGCAATCTCGGCTCACTGCAATCTCNNNAGATCGGAAGA |
| 35762_2 | TATGAGTGTGGAGTCGTTGCAATCTCGGCTCACTGCAATCTCNNNAGATCGGAAGA |
| 35762_3 | TATGAGTGTGGAGTCGTTGCAATCTCGGCTCACTGCAATCTCNNNAGATCGGAAGA |
| 36057_1 | TATGAGTGTGGAGTCGTTGCTCTCATCAGAACACATGGACTCGNNNAGATCGGAAG |
| 36057_2 | TATGAGTGTGGAGTCGTTGCTCTCATCAGAACACATGGACTCGNNNAGATCGGAAG |
| 36057_3 | TATGAGTGTGGAGTCGTTGCTCTCATCAGAACACATGGACTCGNNNAGATCGGAAG |
| 32926_1 | TATGAGTGTGGAGTCGTTGCGCCTGTAGTCCCAGCTATTTGGNNNAGATCGGAAGA |
| 32926_2 | TATGAGTGTGGAGTCGTTGCGCCTGTAGTCCCAGCTATTTGGNNNAGATCGGAAGA |
| 32926_3 | TATGAGTGTGGAGTCGTTGCGCCTGTAGTCCCAGCTATTTGGNNNAGATCGGAAGA |
| 41392_1 | TATGAGTGTGGAGTCGTTGCGCCTGTAATCCCAGTACTTTGGANNNAGATCGGAAG |
| 41392_2 | TATGAGTGTGGAGTCGTTGCGCCTGTAATCCCAGTACTTTGGANNNAGATCGGAAG |
| 41392_3 | TATGAGTGTGGAGTCGTTGCGCCTGTAATCCCAGTACTTTGGANNNAGATCGGAAG |
| 35540_1 | TATGAGTGTGGAGTCGTTGCTTCTGATTCACATGGACTTCCCCNNNAGATCGGAAGA |
| 35540_2 | TATGAGTGTGGAGTCGTTGCTTCTGATTCACATGGACTTCCCCNNNAGATCGGAAGA |
| 35540_3 | TATGAGTGTGGAGTCGTTGCTTCTGATTCACATGGACTTCCCCNNNAGATCGGAAGA |
| 48940_1 | TATGAGTGTGGAGTCGTTGCGGCTTCACATTTCTTCCTCTTGGNNNAGATCGGAAGA |
| 48940_2 | TATGAGTGTGGAGTCGTTGCGGCTTCACATTTCTTCCTCTTGGNNNAGATCGGAAGA |
| 48940_3 | TATGAGTGTGGAGTCGTTGCGGCTTCACATTTCTTCCTCTTGGNNNAGATCGGAAGA |
| 30252_1 | TATGAGTGTGGAGTCGTTGCATACTCCCAACCCCCAGGAANNNAGATCGGAAGAGC |
| 30252_2 | TATGAGTGTGGAGTCGTTGCATACTCCCAACCCCCAGGAANNNAGATCGGAAGAGC |
| 30252_3 | TATGAGTGTGGAGTCGTTGCATACTCCCAACCCCCAGGAANNNAGATCGGAAGAGC |
| 46022_1 | TATGAGTGTGGAGTCGTTGCCCACAGTGCCCAGCTTATTTTTTNNNAGATCGGAAGA |
| 46022_2 | TATGAGTGTGGAGTCGTTGCCCACAGTGCCCAGCTTATTTTTTNNNAGATCGGAAGA |
| 46022_3 | TATGAGTGTGGAGTCGTTGCCCACAGTGCCCAGCTTATTTTTTNNNAGATCGGAAGA |
| 45374_1 | TATGAGTGTGGAGTCGTTGCGTGACCTCTACCCTGCAAACATANNNAGATCGGAAG |
| 45374_2 | TATGAGTGTGGAGTCGTTGCGTGACCTCTACCCTGCAAACATANNNAGATCGGAAG |
| 45374_3 | TATGAGTGTGGAGTCGTTGCGTGACCTCTACCCTGCAAACATANNNAGATCGGAAG |
| 40367_1 | TATGAGTGTGGAGTCGTTGCCTGCTTACAGAGATGGGGGTAAGNNNAGATCGGAA |
| 40367_2 | TATGAGTGTGGAGTCGTTGCCTGCTTACAGAGATGGGGGTAAGNNNAGATCGGAA |
| 40367_3 | TATGAGTGTGGAGTCGTTGCCTGCTTACAGAGATGGGGGTAAGNNNAGATCGGAA |

| | |
|---|---|
| 35361_1 | TATGAGTGTGGAGTCGTTGCGATCTCAGCTCACTGCAACATCTNNNAGATCGGAAGA |
| 35361_2 | TATGAGTGTGGAGTCGTTGCGATCTCAGCTCACTGCAACATCTNNNAGATCGGAAGA |
| 35361_3 | TATGAGTGTGGAGTCGTTGCGATCTCAGCTCACTGCAACATCTNNNAGATCGGAAGA |
| 45068_1 | TATGAGTGTGGAGTCGTTGCAGGAGGTCATTAGGGAAATGCAGNNNAGATCGGAA |
| 45068_2 | TATGAGTGTGGAGTCGTTGCAGGAGGTCATTAGGGAAATGCAGNNNAGATCGGAA |
| 45068_3 | TATGAGTGTGGAGTCGTTGCAGGAGGTCATTAGGGAAATGCAGNNNAGATCGGAA |
| 29875_1 | TATGAGTGTGGAGTCGTTGCGCCTTCAGATAAAATTCCCGAAGCNNNAGATCGGAA[illegible] |
| 29875_2 | TATGAGTGTGGAGTCGTTGCGCCTTCAGATAAAATTCCCGAAGCNNNAGATCGGAA[illegible] |
| 29875_3 | TATGAGTGTGGAGTCGTTGCGCCTTCAGATAAAATTCCCGAAGCNNNAGATCGGAA[illegible] |
| 3145_1 | TATGAGTGTGGAGTCGTTGCCCACCCATGCCTTCTTCTCTTTTNNNAGATCGGAAGAG |
| 3145_2 | TATGAGTGTGGAGTCGTTGCCCACCCATGCCTTCTTCTCTTTTNNNAGATCGGAAGAG |
| 3145_3 | TATGAGTGTGGAGTCGTTGCCCACCCATGCCTTCTTCTCTTTTNNNAGATCGGAAGAG |
| 29175_1 | TATGAGTGTGGAGTCGTTGCTTGAGCTCTTCATCACCCATGAANNNAGATCGGAAGA |
| 29175_2 | TATGAGTGTGGAGTCGTTGCTTGAGCTCTTCATCACCCATGAANNNAGATCGGAAGA |
| 29175_3 | TATGAGTGTGGAGTCGTTGCTTGAGCTCTTCATCACCCATGAANNNAGATCGGAAGA |
| 12641_1 | TATGAGTGTGGAGTCGTTGCGGAAAGAAAAACTGGAATATAGTNNNAGATCGGAA |
| 12641_2 | TATGAGTGTGGAGTCGTTGCGGAAAGAAAAACTGGAATATAGTNNNAGATCGGAA |
| 12641_3 | TATGAGTGTGGAGTCGTTGCGGAAAGAAAAACTGGAATATAGTNNNAGATCGGAA |
| 10658_1 | TATGAGTGTGGAGTCGTTGCATTCCCTCAGTTCAACACAGCAGNNNAGATCGGAAGA |
| 10658_2 | TATGAGTGTGGAGTCGTTGCATTCCCTCAGTTCAACACAGCAGNNNAGATCGGAAGA |
| 10658_3 | TATGAGTGTGGAGTCGTTGCATTCCCTCAGTTCAACACAGCAGNNNAGATCGGAAGA |
| 47940_1 | TATGAGTGTGGAGTCGTTGCGAGACGGAATCTAGCTCTGTCACNNNAGATCGGAAG |
| 47940_2 | TATGAGTGTGGAGTCGTTGCGAGACGGAATCTAGCTCTGTCACNNNAGATCGGAAG |
| 47940_3 | TATGAGTGTGGAGTCGTTGCGAGACGGAATCTAGCTCTGTCACNNNAGATCGGAAG |
| 40184_1 | TATGAGTGTGGAGTCGTTGCGCCTGCAACTTATCCTCAAATGGNNNAGATCGGAAGA |
| 40184_2 | TATGAGTGTGGAGTCGTTGCGCCTGCAACTTATCCTCAAATGGNNNAGATCGGAAGA |
| 40184_3 | TATGAGTGTGGAGTCGTTGCGCCTGCAACTTATCCTCAAATGGNNNAGATCGGAAGA |
| 39397_1 | TATGAGTGTGGAGTCGTTGCGCCTGTAATCCCAGCACTTTGNNNAGATCGGAAGAG |
| 39397_2 | TATGAGTGTGGAGTCGTTGCGCCTGTAATCCCAGCACTTTGNNNAGATCGGAAGAG |
| 39397_3 | TATGAGTGTGGAGTCGTTGCGCCTGTAATCCCAGCACTTTGNNNAGATCGGAAGAG |
| 52479_1 | TATGAGTGTGGAGTCGTTGCTGCCTCTTTGCAGCATCTGTATANNNAGATCGGAAGA |
| 52479_2 | TATGAGTGTGGAGTCGTTGCTGCCTCTTTGCAGCATCTGTATANNNAGATCGGAAGA |
| 52479_3 | TATGAGTGTGGAGTCGTTGCTGCCTCTTTGCAGCATCTGTATANNNAGATCGGAAGA |
| 52456_1 | TATGAGTGTGGAGTCGTTGCTGGTACACTGCAACTGATACCAANNNAGATCGGAAG |
| 52456_2 | TATGAGTGTGGAGTCGTTGCTGGTACACTGCAACTGATACCAANNNAGATCGGAAG |
| 52456_3 | TATGAGTGTGGAGTCGTTGCTGGTACACTGCAACTGATACCAANNNAGATCGGAAG |
| 13499_1 | TATGAGTGTGGAGTCGTTGCTTCTTGCTTCTGTAAGAGATGNNNAGATCGGAAGAG |
| 13499_2 | TATGAGTGTGGAGTCGTTGCTTCTTGCTTCTGTAAGAGATGNNNAGATCGGAAGAG |
| 13499_3 | TATGAGTGTGGAGTCGTTGCTTCTTGCTTCTGTAAGAGATGNNNAGATCGGAAGAG |
| 29020_1 | TATGAGTGTGGAGTCGTTGCCTGCAAGACAAATGAGGTCAAGGNNNAGATCGGAA |
| 29020_2 | TATGAGTGTGGAGTCGTTGCCTGCAAGACAAATGAGGTCAAGGNNNAGATCGGAA |
| 29020_3 | TATGAGTGTGGAGTCGTTGCCTGCAAGACAAATGAGGTCAAGGNNNAGATCGGAA |
| 40702_1 | TATGAGTGTGGAGTCGTTGCAGCTGCCTTATACAGTAGTTGTGANNNAGATCGGAA |

| | |
|---|---|
| 40702_2 | TATGAGTGTGGAGTCGTTGCAGCTGCCTTATACAGTAGTTGTGANNNAGATCGGAA |
| 40702_3 | TATGAGTGTGGAGTCGTTGCAGCTGCCTTATACAGTAGTTGTGANNNAGATCGGAA |
| 36956_1 | TATGAGTGTGGAGTCGTTGCTGGCTTCTCCTTTATGTGAGTCCNNNAGATCGGAAG |
| 36956_2 | TATGAGTGTGGAGTCGTTGCTGGCTTCTCCTTTATGTGAGTCCNNNAGATCGGAAG |
| 36956_3 | TATGAGTGTGGAGTCGTTGCTGGCTTCTCCTTTATGTGAGTCCNNNAGATCGGAAG |
| 36030_1 | TATGAGTGTGGAGTCGTTGCTTTTCTTTCAAATACCGTGGCCANNNAGATCGGAAG |
| 36030_2 | TATGAGTGTGGAGTCGTTGCTTTTCTTTCAAATACCGTGGCCANNNAGATCGGAAG |
| 36030_3 | TATGAGTGTGGAGTCGTTGCTTTTCTTTCAAATACCGTGGCCANNNAGATCGGAAG |
| 37662_1 | TATGAGTGTGGAGTCGTTGCGAGATGAGGTTTTGCCATGTTGTNNNAGATCGGAA |
| 37662_2 | TATGAGTGTGGAGTCGTTGCGAGATGAGGTTTTGCCATGTTGTNNNAGATCGGAA |
| 37662_3 | TATGAGTGTGGAGTCGTTGCGAGATGAGGTTTTGCCATGTTGTNNNAGATCGGAA |
| 45279_1 | TATGAGTGTGGAGTCGTTGCACTCTAATGGGTACATGCTTGCANNNAGATCGGAAG |
| 45279_2 | TATGAGTGTGGAGTCGTTGCACTCTAATGGGTACATGCTTGCANNNAGATCGGAAG |
| 45279_3 | TATGAGTGTGGAGTCGTTGCACTCTAATGGGTACATGCTTGCANNNAGATCGGAAG |
| 34515_1 | TATGAGTGTGGAGTCGTTGCCACTGTCTCTGGGCCTAATTCAANNNAGATCGGAAG |
| 34515_2 | TATGAGTGTGGAGTCGTTGCCACTGTCTCTGGGCCTAATTCAANNNAGATCGGAAG |
| 34515_3 | TATGAGTGTGGAGTCGTTGCCACTGTCTCTGGGCCTAATTCAANNNAGATCGGAAG |
| 52500_1 | TATGAGTGTGGAGTCGTTGCACAAACAGGAGACTAATGGCTGANNNAGATCGGAA |
| 52500_2 | TATGAGTGTGGAGTCGTTGCACAAACAGGAGACTAATGGCTGANNNAGATCGGAA |
| 52500_3 | TATGAGTGTGGAGTCGTTGCACAAACAGGAGACTAATGGCTGANNNAGATCGGAA |
| 38456_1 | TATGAGTGTGGAGTCGTTGCTAGCTACTCAGGATCACTTGAGCNNNAGATCGGAAG |
| 38456_2 | TATGAGTGTGGAGTCGTTGCTAGCTACTCAGGATCACTTGAGCNNNAGATCGGAAG |
| 38456_3 | TATGAGTGTGGAGTCGTTGCTAGCTACTCAGGATCACTTGAGCNNNAGATCGGAAG |
| 31303_1 | TATGAGTGTGGAGTCGTTGCAGTTGGGGTAGGGAGTAGAGAAANNNAGATCGGA |
| 31303_2 | TATGAGTGTGGAGTCGTTGCAGTTGGGGTAGGGAGTAGAGAAANNNAGATCGGA |
| 31303_3 | TATGAGTGTGGAGTCGTTGCAGTTGGGGTAGGGAGTAGAGAAANNNAGATCGGA |
| 52758_1 | TATGAGTGTGGAGTCGTTGCAACAAGTAACAACAGGGGTGGAANNNAGATCGGAA |
| 52758_2 | TATGAGTGTGGAGTCGTTGCAACAAGTAACAACAGGGGTGGAANNNAGATCGGAA |
| 52758_3 | TATGAGTGTGGAGTCGTTGCAACAAGTAACAACAGGGGTGGAANNNAGATCGGAA |
| 39442_1 | TATGAGTGTGGAGTCGTTGCCTTGCAGATAGTTGGCTTTGCTTNNNAGATCGGAAG |
| 39442_2 | TATGAGTGTGGAGTCGTTGCCTTGCAGATAGTTGGCTTTGCTTNNNAGATCGGAAG |
| 39442_3 | TATGAGTGTGGAGTCGTTGCCTTGCAGATAGTTGGCTTTGCTTNNNAGATCGGAAG |
| 44126_1 | TATGAGTGTGGAGTCGTTGCATGTGGACCTGAAAAGAGCTCAANNNAGATCGGAA |
| 44126_2 | TATGAGTGTGGAGTCGTTGCATGTGGACCTGAAAAGAGCTCAANNNAGATCGGAA |
| 44126_3 | TATGAGTGTGGAGTCGTTGCATGTGGACCTGAAAAGAGCTCAANNNAGATCGGAA |
| 43845_1 | TATGAGTGTGGAGTCGTTGCCTGCTGAGTAGAACTGACAGGAGNNNAGATCGGAA |
| 43845_2 | TATGAGTGTGGAGTCGTTGCCTGCTGAGTAGAACTGACAGGAGNNNAGATCGGAA |
| 43845_3 | TATGAGTGTGGAGTCGTTGCCTGCTGAGTAGAACTGACAGGAGNNNAGATCGGAA |
| 47095_1 | TATGAGTGTGGAGTCGTTGCATTCAAGTGATCCTCCTGCTTGANNNAGATCGGAAG |
| 47095_2 | TATGAGTGTGGAGTCGTTGCATTCAAGTGATCCTCCTGCTTGANNNAGATCGGAAG |
| 47095_3 | TATGAGTGTGGAGTCGTTGCATTCAAGTGATCCTCCTGCTTGANNNAGATCGGAAG |
| 12607_1 | TATGAGTGTGGAGTCGTTGCTATCTTCAAAACAACCTGATAGANNNAGATCGGAAG |
| 12607_2 | TATGAGTGTGGAGTCGTTGCTATCTTCAAAACAACCTGATAGANNNAGATCGGAAG |

| | |
|---|---|
| 12607_3 | TATGAGTGTGGAGTCGTTGCTATCTTCAAAACAACCTGATAGANNNAGATCGGAAG |
| 47758_1 | TATGAGTGTGGAGTCGTTGCCCTGGTCAAGAACAAAGAAGCAGNNNAGATCGGAA |
| 47758_2 | TATGAGTGTGGAGTCGTTGCCCTGGTCAAGAACAAAGAAGCAGNNNAGATCGGAA |
| 47758_3 | TATGAGTGTGGAGTCGTTGCCCTGGTCAAGAACAAAGAAGCAGNNNAGATCGGAA |
| 33956_1 | TATGAGTGTGGAGTCGTTGCGGTTCAAGTGATTCTCCTGCTTCNNNAGATCGGAAG |
| 33956_2 | TATGAGTGTGGAGTCGTTGCGGTTCAAGTGATTCTCCTGCTTCNNNAGATCGGAAG |
| 33956_3 | TATGAGTGTGGAGTCGTTGCGGTTCAAGTGATTCTCCTGCTTCNNNAGATCGGAAG |
| 45372_1 | TATGAGTGTGGAGTCGTTGCGTCTGTAAAAGAGTGCTTGAGGCNNNAGATCGGAA |
| 45372_2 | TATGAGTGTGGAGTCGTTGCGTCTGTAAAAGAGTGCTTGAGGCNNNAGATCGGAA |
| 45372_3 | TATGAGTGTGGAGTCGTTGCGTCTGTAAAAGAGTGCTTGAGGCNNNAGATCGGAA |
| 30315_1 | TATGAGTGTGGAGTCGTTGCAAGTCAGCTACAGTTGTTGCAGTNNNAGATCGGAAG |
| 30315_2 | TATGAGTGTGGAGTCGTTGCAAGTCAGCTACAGTTGTTGCAGTNNNAGATCGGAAG |
| 30315_3 | TATGAGTGTGGAGTCGTTGCAAGTCAGCTACAGTTGTTGCAGTNNNAGATCGGAAG |
| 46135_1 | TATGAGTGTGGAGTCGTTGCTTTGGTAGAGATGGGGTTTCTCCNNNAGATCGGAA |
| 46135_2 | TATGAGTGTGGAGTCGTTGCTTTGGTAGAGATGGGGTTTCTCCNNNAGATCGGAA |
| 46135_3 | TATGAGTGTGGAGTCGTTGCTTTGGTAGAGATGGGGTTTCTCCNNNAGATCGGAA |
| 53231_1 | TATGAGTGTGGAGTCGTTGCCCCACAAACGACAAACAAACAAANNNAGATCGGAAG |
| 53231_2 | TATGAGTGTGGAGTCGTTGCCCCACAAACGACAAACAAACAAANNNAGATCGGAAG |
| 53231_3 | TATGAGTGTGGAGTCGTTGCCCCACAAACGACAAACAAACAAANNNAGATCGGAAG |
| 41649_1 | TATGAGTGTGGAGTCGTTGCACCATGCCCACCTAATGTTTGTANNNAGATCGGAAG |
| 41649_2 | TATGAGTGTGGAGTCGTTGCACCATGCCCACCTAATGTTTGTANNNAGATCGGAAG |
| 41649_3 | TATGAGTGTGGAGTCGTTGCACCATGCCCACCTAATGTTTGTANNNAGATCGGAAG |
| 35850_1 | TATGAGTGTGGAGTCGTTGCCCTCTGGAGTATCTGGGACTACANNNAGATCGGAAG |
| 35850_2 | TATGAGTGTGGAGTCGTTGCCCTCTGGAGTATCTGGGACTACANNNAGATCGGAAG |
| 35850_3 | TATGAGTGTGGAGTCGTTGCCCTCTGGAGTATCTGGGACTACANNNAGATCGGAAG |
| 51092_1 | TATGAGTGTGGAGTCGTTGCTGTGGAGAAATCTGTGCATCTCANNNAGATCGGAAG |
| 51092_2 | TATGAGTGTGGAGTCGTTGCTGTGGAGAAATCTGTGCATCTCANNNAGATCGGAAG |
| 51092_3 | TATGAGTGTGGAGTCGTTGCTGTGGAGAAATCTGTGCATCTCANNNAGATCGGAAG |
| 3119_1 | TATGAGTGTGGAGTCGTTGCCTTCCACACTCTGTCCCCATCTGNNNAGATCGGAAGA |
| 3119_2 | TATGAGTGTGGAGTCGTTGCCTTCCACACTCTGTCCCCATCTGNNNAGATCGGAAGA |
| 3119_3 | TATGAGTGTGGAGTCGTTGCCTTCCACACTCTGTCCCCATCTGNNNAGATCGGAAGA |
| 39622_1 | TATGAGTGTGGAGTCGTTGCTCAAAAGCTAGCAGAAGACAAGANNNAGATCGGAA |
| 39622_2 | TATGAGTGTGGAGTCGTTGCTCAAAAGCTAGCAGAAGACAAGANNNAGATCGGAA |
| 39622_3 | TATGAGTGTGGAGTCGTTGCTCAAAAGCTAGCAGAAGACAAGANNNAGATCGGAA |
| 47592_1 | TATGAGTGTGGAGTCGTTGCGTTAGCCAGGATGGTCTCAATCTNNNAGATCGGAAG |
| 47592_2 | TATGAGTGTGGAGTCGTTGCGTTAGCCAGGATGGTCTCAATCTNNNAGATCGGAAG |
| 47592_3 | TATGAGTGTGGAGTCGTTGCGTTAGCCAGGATGGTCTCAATCTNNNAGATCGGAAG |
| 10198_1 | TATGAGTGTGGAGTCGTTGCTTTTCCCAGTGGGTTTGGATCAGNNNAGATCGGAAG |
| 10198_2 | TATGAGTGTGGAGTCGTTGCTTTTCCCAGTGGGTTTGGATCAGNNNAGATCGGAAG |
| 10198_3 | TATGAGTGTGGAGTCGTTGCTTTTCCCAGTGGGTTTGGATCAGNNNAGATCGGAAG |
| 30025_1 | TATGAGTGTGGAGTCGTTGCCACAGCATTTGTTCCAGAGTCAGNNNAGATCGGAAG |
| 30025_2 | TATGAGTGTGGAGTCGTTGCCACAGCATTTGTTCCAGAGTCAGNNNAGATCGGAAG |
| 30025_3 | TATGAGTGTGGAGTCGTTGCCACAGCATTTGTTCCAGAGTCAGNNNAGATCGGAAG |

| | |
|---|---|
| 10752_1 | TATGAGTGTGGAGTCGTTGCTGGAAATTCTGGAGTTGAAAAGGTNNNAGATCGGA |
| 10752_2 | TATGAGTGTGGAGTCGTTGCTGGAAATTCTGGAGTTGAAAAGGTNNNAGATCGGA |
| 10752_3 | TATGAGTGTGGAGTCGTTGCTGGAAATTCTGGAGTTGAAAAGGTNNNAGATCGGA |
| 34474_1 | TATGAGTGTGGAGTCGTTGCACCCCCCCAAAAAAAGAAAACTGNNNAGATCGGAAG |
| 34474_2 | TATGAGTGTGGAGTCGTTGCACCCCCCCAAAAAAAGAAAACTGNNNAGATCGGAAG |
| 34474_3 | TATGAGTGTGGAGTCGTTGCACCCCCCCAAAAAAAGAAAACTGNNNAGATCGGAAG |
| 39389_1 | TATGAGTGTGGAGTCGTTGCAATGGTACAAACCCTAAGTGCCTNNNAGATCGGAAG |
| 39389_2 | TATGAGTGTGGAGTCGTTGCAATGGTACAAACCCTAAGTGCCTNNNAGATCGGAAG |
| 39389_3 | TATGAGTGTGGAGTCGTTGCAATGGTACAAACCCTAAGTGCCTNNNAGATCGGAAG |
| 51479_1 | TATGAGTGTGGAGTCGTTGCTGCTTCCTTTCATGTACCAGTCANNNAGATCGGAAGA |
| 51479_2 | TATGAGTGTGGAGTCGTTGCTGCTTCCTTTCATGTACCAGTCANNNAGATCGGAAGA |
| 51479_3 | TATGAGTGTGGAGTCGTTGCTGCTTCCTTTCATGTACCAGTCANNNAGATCGGAAGA |
| 37205_1 | TATGAGTGTGGAGTCGTTGCAGCATTCCTTTGAGTCCCAAGATNNNAGATCGGAAG |
| 37205_2 | TATGAGTGTGGAGTCGTTGCAGCATTCCTTTGAGTCCCAAGATNNNAGATCGGAAG |
| 37205_3 | TATGAGTGTGGAGTCGTTGCAGCATTCCTTTGAGTCCCAAGATNNNAGATCGGAAG |
| 38465_1 | TATGAGTGTGGAGTCGTTGCTGATATGGTTGTTGGATACAGGTNNNAGATCGGAA |
| 38465_2 | TATGAGTGTGGAGTCGTTGCTGATATGGTTGTTGGATACAGGTNNNAGATCGGAA |
| 38465_3 | TATGAGTGTGGAGTCGTTGCTGATATGGTTGTTGGATACAGGTNNNAGATCGGAA |
| 45888_1 | TATGAGTGTGGAGTCGTTGCCTCAAGTGATCCGCCCATCTTAGNNNAGATCGGAAG |
| 45888_2 | TATGAGTGTGGAGTCGTTGCCTCAAGTGATCCGCCCATCTTAGNNNAGATCGGAAG |
| 45888_3 | TATGAGTGTGGAGTCGTTGCCTCAAGTGATCCGCCCATCTTAGNNNAGATCGGAAG |
| 36029_1 | TATGAGTGTGGAGTCGTTGCTAATCCCAGCTACTCAGGAGGTTNNNAGATCGGAAG |
| 36029_2 | TATGAGTGTGGAGTCGTTGCTAATCCCAGCTACTCAGGAGGTTNNNAGATCGGAAG |
| 36029_3 | TATGAGTGTGGAGTCGTTGCTAATCCCAGCTACTCAGGAGGTTNNNAGATCGGAAG |
| 29898_1 | TATGAGTGTGGAGTCGTTGCCTTCTAGGCCCGTGCAGATATTNNNAGATCGGAAGA |
| 29898_2 | TATGAGTGTGGAGTCGTTGCCTTCTAGGCCCGTGCAGATATTNNNAGATCGGAAGA |
| 29898_3 | TATGAGTGTGGAGTCGTTGCCTTCTAGGCCCGTGCAGATATTNNNAGATCGGAAGA |
| 41650_1 | TATGAGTGTGGAGTCGTTGCTGGGTTCAAGCAATTCTCTGTCTNNNAGATCGGAAG |
| 41650_2 | TATGAGTGTGGAGTCGTTGCTGGGTTCAAGCAATTCTCTGTCTNNNAGATCGGAAG |
| 41650_3 | TATGAGTGTGGAGTCGTTGCTGGGTTCAAGCAATTCTCTGTCTNNNAGATCGGAAG |
| 31351_1 | TATGAGTGTGGAGTCGTTGCGAGAGAGGGAGGGTGTAAATGTGNNNAGATCGGA |
| 31351_2 | TATGAGTGTGGAGTCGTTGCGAGAGAGGGAGGGTGTAAATGTGNNNAGATCGGA |
| 31351_3 | TATGAGTGTGGAGTCGTTGCGAGAGAGGGAGGGTGTAAATGTGNNNAGATCGGA |
| 32846_1 | TATGAGTGTGGAGTCGTTGCTTCAAGTGATTCTCCTGCCTCAGNNNAGATCGGAAG |
| 32846_2 | TATGAGTGTGGAGTCGTTGCTTCAAGTGATTCTCCTGCCTCAGNNNAGATCGGAAG |
| 32846_3 | TATGAGTGTGGAGTCGTTGCTTCAAGTGATTCTCCTGCCTCAGNNNAGATCGGAAG |
| 36321_1 | TATGAGTGTGGAGTCGTTGCGATGGCCCCACACCAAAAAATTANNNAGATCGGAAG |
| 36321_2 | TATGAGTGTGGAGTCGTTGCGATGGCCCCACACCAAAAAATTANNNAGATCGGAAG |
| 36321_3 | TATGAGTGTGGAGTCGTTGCGATGGCCCCACACCAAAAAATTANNNAGATCGGAAG |
| 46679_1 | TATGAGTGTGGAGTCGTTGCCCAAGTATCTGGGATTACAGGCANNNAGATCGGAA |
| 46679_2 | TATGAGTGTGGAGTCGTTGCCCAAGTATCTGGGATTACAGGCANNNAGATCGGAA |
| 46679_3 | TATGAGTGTGGAGTCGTTGCCCAAGTATCTGGGATTACAGGCANNNAGATCGGAA |
| 52319_1 | TATGAGTGTGGAGTCGTTGCCTTCACTTCTGACTCTTCCCCAGNNNAGATCGGAAGA |

| | |
|---|---|
| 52319_2 | TATGAGTGTGGAGTCGTTGCCTTCACTTCTGACTCTTCCCCAGNNNAGATCGGAAGA |
| 52319_3 | TATGAGTGTGGAGTCGTTGCCTTCACTTCTGACTCTTCCCCAGNNNAGATCGGAAGA |
| 36439_1 | TATGAGTGTGGAGTCGTTGCGAGATTGCACCACTGTACTCCAGNNNAGATCGGAAG |
| 36439_2 | TATGAGTGTGGAGTCGTTGCGAGATTGCACCACTGTACTCCAGNNNAGATCGGAAG |
| 36439_3 | TATGAGTGTGGAGTCGTTGCGAGATTGCACCACTGTACTCCAGNNNAGATCGGAAG |
| 43742_1 | TATGAGTGTGGAGTCGTTGCAGCCTTCTAGGTGAGCAATGAAANNNAGATCGGAA |
| 43742_2 | TATGAGTGTGGAGTCGTTGCAGCCTTCTAGGTGAGCAATGAAANNNAGATCGGAA |
| 43742_3 | TATGAGTGTGGAGTCGTTGCAGCCTTCTAGGTGAGCAATGAAANNNAGATCGGAA |
| 38953_1 | TATGAGTGTGGAGTCGTTGCTGATCACATGGGGGTAGAGTGTANNNAGATCGGAA |
| 38953_2 | TATGAGTGTGGAGTCGTTGCTGATCACATGGGGGTAGAGTGTANNNAGATCGGAA |
| 38953_3 | TATGAGTGTGGAGTCGTTGCTGATCACATGGGGGTAGAGTGTANNNAGATCGGAA |
| 40709_1 | TATGAGTGTGGAGTCGTTGCATAGAACGCAATAGAGAGGCCAGNNNAGATCGGAA |
| 40709_2 | TATGAGTGTGGAGTCGTTGCATAGAACGCAATAGAGAGGCCAGNNNAGATCGGAA |
| 40709_3 | TATGAGTGTGGAGTCGTTGCATAGAACGCAATAGAGAGGCCAGNNNAGATCGGAA |
| 47317_1 | TATGAGTGTGGAGTCGTTGCGATCAGGATGATGCTGGTCTCATNNNAGATCGGAA |
| 47317_2 | TATGAGTGTGGAGTCGTTGCGATCAGGATGATGCTGGTCTCATNNNAGATCGGAA |
| 47317_3 | TATGAGTGTGGAGTCGTTGCGATCAGGATGATGCTGGTCTCATNNNAGATCGGAA |
| 42839_1 | TATGAGTGTGGAGTCGTTGCTCCTAGGAACAGAATTGCCAGTCNNNAGATCGGAAG |
| 42839_2 | TATGAGTGTGGAGTCGTTGCTCCTAGGAACAGAATTGCCAGTCNNNAGATCGGAAG |
| 42839_3 | TATGAGTGTGGAGTCGTTGCTCCTAGGAACAGAATTGCCAGTCNNNAGATCGGAAG |
| 51064_1 | TATGAGTGTGGAGTCGTTGCAACCACAATCAATCAATCCCAGCNNNAGATCGGAAG |
| 51064_2 | TATGAGTGTGGAGTCGTTGCAACCACAATCAATCAATCCCAGCNNNAGATCGGAAG |
| 51064_3 | TATGAGTGTGGAGTCGTTGCAACCACAATCAATCAATCCCAGCNNNAGATCGGAAG |
| 41445_1 | TATGAGTGTGGAGTCGTTGCATCGTGAAATCTAGGTGGTAGGTNNNAGATCGGAA |
| 41445_2 | TATGAGTGTGGAGTCGTTGCATCGTGAAATCTAGGTGGTAGGTNNNAGATCGGAA |
| 41445_3 | TATGAGTGTGGAGTCGTTGCATCGTGAAATCTAGGTGGTAGGTNNNAGATCGGAA |
| 36577_1 | TATGAGTGTGGAGTCGTTGCTTGAATCCTTTGTGCAGCTGAAANNNAGATCGGAAG |
| 36577_2 | TATGAGTGTGGAGTCGTTGCTTGAATCCTTTGTGCAGCTGAAANNNAGATCGGAAG |
| 36577_3 | TATGAGTGTGGAGTCGTTGCTTGAATCCTTTGTGCAGCTGAAANNNAGATCGGAAG |
| 35950_1 | TATGAGTGTGGAGTCGTTGCTTGTGTACATTTTTTGGCTGGGCNNNAGATCGGAAG |
| 35950_2 | TATGAGTGTGGAGTCGTTGCTTGTGTACATTTTTTGGCTGGGCNNNAGATCGGAAG |
| 35950_3 | TATGAGTGTGGAGTCGTTGCTTGTGTACATTTTTTGGCTGGGCNNNAGATCGGAAG |
| 43252_1 | TATGAGTGTGGAGTCGTTGCCCCGGATTCTAGTGATTCTCCTGNNNAGATCGGAAG |
| 43252_2 | TATGAGTGTGGAGTCGTTGCCCCGGATTCTAGTGATTCTCCTGNNNAGATCGGAAG |
| 43252_3 | TATGAGTGTGGAGTCGTTGCCCCGGATTCTAGTGATTCTCCTGNNNAGATCGGAAG |
| 42669_1 | TATGAGTGTGGAGTCGTTGCTCACAAAACACACACACAGACACNNNAGATCGGAAG |
| 42669_2 | TATGAGTGTGGAGTCGTTGCTCACAAAACACACACACAGACACNNNAGATCGGAAG |
| 42669_3 | TATGAGTGTGGAGTCGTTGCTCACAAAACACACACACAGACACNNNAGATCGGAAG |
| 33108_1 | TATGAGTGTGGAGTCGTTGCCTGAGGCAGGAGAATCACTTGANNNAGATCGGAAG |
| 33108_2 | TATGAGTGTGGAGTCGTTGCCTGAGGCAGGAGAATCACTTGANNNAGATCGGAAG |
| 33108_3 | TATGAGTGTGGAGTCGTTGCCTGAGGCAGGAGAATCACTTGANNNAGATCGGAAG |
| 31350_1 | TATGAGTGTGGAGTCGTTGCTGTGTTTCTACCTATTGTGTGTGANNNAGATCGGAA |
| 31350_2 | TATGAGTGTGGAGTCGTTGCTGTGTTTCTACCTATTGTGTGTGANNNAGATCGGAA |

| | |
|---|---|
| 31350_3 | TATGAGTGTGGAGTCGTTGCTGTGTTTCTACCTATTGTGTGTGANNNAGATCGGAA[illegible] |
| 12476_1 | TATGAGTGTGGAGTCGTTGCTTTTCAAATCTTCTACAGTTCTGNNNAGATCGGAAGA |
| 12476_2 | TATGAGTGTGGAGTCGTTGCTTTTCAAATCTTCTACAGTTCTGNNNAGATCGGAAGA |
| 12476_3 | TATGAGTGTGGAGTCGTTGCTTTTCAAATCTTCTACAGTTCTGNNNAGATCGGAAGA |
| 10145_1 | TATGAGTGTGGAGTCGTTGCACAGCAGAGGGCTCTTTGAATTCNNNAGATCGGAAG |
| 10145_2 | TATGAGTGTGGAGTCGTTGCACAGCAGAGGGCTCTTTGAATTCNNNAGATCGGAAG |
| 10145_3 | TATGAGTGTGGAGTCGTTGCACAGCAGAGGGCTCTTTGAATTCNNNAGATCGGAAG |
| 48109_1 | TATGAGTGTGGAGTCGTTGCACATCTTGAATGCTTTGCTGCTTNNNAGATCGGAAG[illegible] |
| 48109_2 | TATGAGTGTGGAGTCGTTGCACATCTTGAATGCTTTGCTGCTTNNNAGATCGGAAG[illegible] |
| 48109_3 | TATGAGTGTGGAGTCGTTGCACATCTTGAATGCTTTGCTGCTTNNNAGATCGGAAG[illegible] |
| 33302_1 | TATGAGTGTGGAGTCGTTGCCATTCTCCTGCCTCAGCCTCNNNAGATCGGAAGAGCA |
| 33302_2 | TATGAGTGTGGAGTCGTTGCCATTCTCCTGCCTCAGCCTCNNNAGATCGGAAGAGCA |
| 33302_3 | TATGAGTGTGGAGTCGTTGCCATTCTCCTGCCTCAGCCTCNNNAGATCGGAAGAGCA |
| 10331_1 | TATGAGTGTGGAGTCGTTGCGGGGCTGGCAAATCCAAATTATGNNNAGATCGGAA[illegible] |
| 10331_2 | TATGAGTGTGGAGTCGTTGCGGGGCTGGCAAATCCAAATTATGNNNAGATCGGAA[illegible] |
| 10331_3 | TATGAGTGTGGAGTCGTTGCGGGGCTGGCAAATCCAAATTATGNNNAGATCGGAA[illegible] |
| 43717_1 | TATGAGTGTGGAGTCGTTGCTTCTCTGCAGAGCATATGGGAAGNNNAGATCGGAA[illegible] |
| 43717_2 | TATGAGTGTGGAGTCGTTGCTTCTCTGCAGAGCATATGGGAAGNNNAGATCGGAA[illegible] |
| 43717_3 | TATGAGTGTGGAGTCGTTGCTTCTCTGCAGAGCATATGGGAAGNNNAGATCGGAA[illegible] |
| 37387_1 | TATGAGTGTGGAGTCGTTGCAAGAAAAAGTTAGCCAGACATGGNNNAGATCGGAA |
| 37387_2 | TATGAGTGTGGAGTCGTTGCAAGAAAAAGTTAGCCAGACATGGNNNAGATCGGAA |
| 37387_3 | TATGAGTGTGGAGTCGTTGCAAGAAAAAGTTAGCCAGACATGGNNNAGATCGGAA |
| 37004_1 | TATGAGTGTGGAGTCGTTGCATGGAAAATAGTGTGGCGCTTTCNNNAGATCGGAA[illegible] |
| 37004_2 | TATGAGTGTGGAGTCGTTGCATGGAAAATAGTGTGGCGCTTTCNNNAGATCGGAA[illegible] |
| 37004_3 | TATGAGTGTGGAGTCGTTGCATGGAAAATAGTGTGGCGCTTTCNNNAGATCGGAA[illegible] |
| 47197_1 | TATGAGTGTGGAGTCGTTGCGAGAAAGAGTCTTGCCCTGTCTCNNNAGATCGGAAG |
| 47197_2 | TATGAGTGTGGAGTCGTTGCGAGAAAGAGTCTTGCCCTGTCTCNNNAGATCGGAAG |
| 47197_3 | TATGAGTGTGGAGTCGTTGCGAGAAAGAGTCTTGCCCTGTCTCNNNAGATCGGAAG |
| 46983_1 | TATGAGTGTGGAGTCGTTGCTCTCAGTTCACTGCAACCTCTACNNNAGATCGGAAGA |
| 46983_2 | TATGAGTGTGGAGTCGTTGCTCTCAGTTCACTGCAACCTCTACNNNAGATCGGAAGA |
| 46983_3 | TATGAGTGTGGAGTCGTTGCTCTCAGTTCACTGCAACCTCTACNNNAGATCGGAAGA |
| 45885_1 | TATGAGTGTGGAGTCGTTGCGAGTCTTGCTCTGTCGCCTAGNNNAGATCGGAAGAG |
| 45885_2 | TATGAGTGTGGAGTCGTTGCGAGTCTTGCTCTGTCGCCTAGNNNAGATCGGAAGAG |
| 45885_3 | TATGAGTGTGGAGTCGTTGCGAGTCTTGCTCTGTCGCCTAGNNNAGATCGGAAGAG |
| 40278_1 | TATGAGTGTGGAGTCGTTGCTGCCTAGGAAGGGAAATTCTGTTNNNAGATCGGAA[illegible] |
| 40278_2 | TATGAGTGTGGAGTCGTTGCTGCCTAGGAAGGGAAATTCTGTTNNNAGATCGGAA[illegible] |
| 40278_3 | TATGAGTGTGGAGTCGTTGCTGCCTAGGAAGGGAAATTCTGTTNNNAGATCGGAA[illegible] |
| 40291_1 | TATGAGTGTGGAGTCGTTGCTCAACCAAATACTAGCCAACCGANNNAGATCGGAAG[illegible] |
| 40291_2 | TATGAGTGTGGAGTCGTTGCTCAACCAAATACTAGCCAACCGANNNAGATCGGAAG[illegible] |
| 40291_3 | TATGAGTGTGGAGTCGTTGCTCAACCAAATACTAGCCAACCGANNNAGATCGGAAG[illegible] |
| 32092_1 | TATGAGTGTGGAGTCGTTGCCCAATCAATTTGCCAGTCTGTGTNNNAGATCGGAAG[illegible] |
| 32092_2 | TATGAGTGTGGAGTCGTTGCCCAATCAATTTGCCAGTCTGTGTNNNAGATCGGAAG[illegible] |
| 32092_3 | TATGAGTGTGGAGTCGTTGCCCAATCAATTTGCCAGTCTGTGTNNNAGATCGGAAG[illegible] |

| | |
|---|---|
| 51329_1 | TATGAGTGTGGAGTCGTTGCATTCAAGGCTTCACTGGGATCAGNNNAGATCGGAAG |
| 51329_2 | TATGAGTGTGGAGTCGTTGCATTCAAGGCTTCACTGGGATCAGNNNAGATCGGAAG |
| 51329_3 | TATGAGTGTGGAGTCGTTGCATTCAAGGCTTCACTGGGATCAGNNNAGATCGGAAG |
| 31013_1 | TATGAGTGTGGAGTCGTTGCAGGAAAGGGGAAAGGAAAAAGGANNNAGATCGGA |
| 31013_2 | TATGAGTGTGGAGTCGTTGCAGGAAAGGGGAAAGGAAAAAGGANNNAGATCGGA |
| 31013_3 | TATGAGTGTGGAGTCGTTGCAGGAAAGGGGAAAGGAAAAAGGANNNAGATCGGA |
| 33574_1 | TATGAGTGTGGAGTCGTTGCGTGTGGAGTCATTTCTAGGGTGTNNNAGATCGGAA |
| 33574_2 | TATGAGTGTGGAGTCGTTGCGTGTGGAGTCATTTCTAGGGTGTNNNAGATCGGAA |
| 33574_3 | TATGAGTGTGGAGTCGTTGCGTGTGGAGTCATTTCTAGGGTGTNNNAGATCGGAA |
| 31566_1 | TATGAGTGTGGAGTCGTTGCTGATGGGTTAGTTTTGCATCAGTNNNAGATCGGAAG |
| 31566_2 | TATGAGTGTGGAGTCGTTGCTGATGGGTTAGTTTTGCATCAGTNNNAGATCGGAAG |
| 31566_3 | TATGAGTGTGGAGTCGTTGCTGATGGGTTAGTTTTGCATCAGTNNNAGATCGGAAG |
| 52253_1 | TATGAGTGTGGAGTCGTTGCCAGTCCCATCAACCACCCAAGNNNAGATCGGAAGAG |
| 52253_2 | TATGAGTGTGGAGTCGTTGCCAGTCCCATCAACCACCCAAGNNNAGATCGGAAGAG |
| 52253_3 | TATGAGTGTGGAGTCGTTGCCAGTCCCATCAACCACCCAAGNNNAGATCGGAAGAG |
| 33914_1 | TATGAGTGTGGAGTCGTTGCAAATGATTTGCCTGCCTTGTCTCNNNAGATCGGAAGA |
| 33914_2 | TATGAGTGTGGAGTCGTTGCAAATGATTTGCCTGCCTTGTCTCNNNAGATCGGAAGA |
| 33914_3 | TATGAGTGTGGAGTCGTTGCAAATGATTTGCCTGCCTTGTCTCNNNAGATCGGAAGA |
| 13135_1 | TATGAGTGTGGAGTCGTTGCCGCCAGCGAATAAATAATAAATANNNAGATCGGAAG |
| 13135_2 | TATGAGTGTGGAGTCGTTGCCGCCAGCGAATAAATAATAAATANNNAGATCGGAAG |
| 13135_3 | TATGAGTGTGGAGTCGTTGCCGCCAGCGAATAAATAATAAATANNNAGATCGGAAG |
| 38816_1 | TATGAGTGTGGAGTCGTTGCAACAAACAAACAAACGAACAGCCNNNAGATCGGAAG |
| 38816_2 | TATGAGTGTGGAGTCGTTGCAACAAACAAACAAACGAACAGCCNNNAGATCGGAAG |
| 38816_3 | TATGAGTGTGGAGTCGTTGCAACAAACAAACAAACGAACAGCCNNNAGATCGGAAG |
| 36683_1 | TATGAGTGTGGAGTCGTTGCCACGCCTGTAATCCCAACATTTTNNNAGATCGGAAGA |
| 36683_2 | TATGAGTGTGGAGTCGTTGCCACGCCTGTAATCCCAACATTTTNNNAGATCGGAAGA |
| 36683_3 | TATGAGTGTGGAGTCGTTGCCACGCCTGTAATCCCAACATTTTNNNAGATCGGAAGA |
| 53317_1 | TATGAGTGTGGAGTCGTTGCCACAGAGTCTCTCATGTGCTACANNNAGATCGGAAGA |
| 53317_2 | TATGAGTGTGGAGTCGTTGCCACAGAGTCTCTCATGTGCTACANNNAGATCGGAAGA |
| 53317_3 | TATGAGTGTGGAGTCGTTGCCACAGAGTCTCTCATGTGCTACANNNAGATCGGAAGA |
| 48294_1 | TATGAGTGTGGAGTCGTTGCTTGACTTCCCCGTTCAAGAGATCNNNAGATCGGAAGA |
| 48294_2 | TATGAGTGTGGAGTCGTTGCTTGACTTCCCCGTTCAAGAGATCNNNAGATCGGAAGA |
| 48294_3 | TATGAGTGTGGAGTCGTTGCTTGACTTCCCCGTTCAAGAGATCNNNAGATCGGAAGA |
| 46465_1 | TATGAGTGTGGAGTCGTTGCGATTCAAGCGATTCTCCTGCTTCNNNAGATCGGAAGA |
| 46465_2 | TATGAGTGTGGAGTCGTTGCGATTCAAGCGATTCTCCTGCTTCNNNAGATCGGAAGA |
| 46465_3 | TATGAGTGTGGAGTCGTTGCGATTCAAGCGATTCTCCTGCTTCNNNAGATCGGAAGA |
| 41887_1 | TATGAGTGTGGAGTCGTTGCAGACTGCACATTCTCTTTTTGGCNNNAGATCGGAAGA |
| 41887_2 | TATGAGTGTGGAGTCGTTGCAGACTGCACATTCTCTTTTTGGCNNNAGATCGGAAGA |
| 41887_3 | TATGAGTGTGGAGTCGTTGCAGACTGCACATTCTCTTTTTGGCNNNAGATCGGAAGA |
| 31684_1 | TATGAGTGTGGAGTCGTTGCACACAACCTTGGAGGAAGAAAGANNNAGATCGGAA |
| 31684_2 | TATGAGTGTGGAGTCGTTGCACACAACCTTGGAGGAAGAAAGANNNAGATCGGAA |
| 31684_3 | TATGAGTGTGGAGTCGTTGCACACAACCTTGGAGGAAGAAAGANNNAGATCGGAA |
| 49838_1 | TATGAGTGTGGAGTCGTTGCACCAGACTGCCTCTATTTTCAGGNNNAGATCGGAAGA |

| | |
|---|---|
| 49838_2 | TATGAGTGTGGAGTCGTTGCACCAGACTGCCTCTATTTTCAGGNNNAGATCGGAAG |
| 49838_3 | TATGAGTGTGGAGTCGTTGCACCAGACTGCCTCTATTTTCAGGNNNAGATCGGAAG |
| 42543_1 | TATGAGTGTGGAGTCGTTGCCAAGCACCAGAGAAGCAGAAATCNNNAGATCGGAA |
| 42543_2 | TATGAGTGTGGAGTCGTTGCCAAGCACCAGAGAAGCAGAAATCNNNAGATCGGAA |
| 42543_3 | TATGAGTGTGGAGTCGTTGCCAAGCACCAGAGAAGCAGAAATCNNNAGATCGGAA |
| 39910_1 | TATGAGTGTGGAGTCGTTGCCCTGCATGTCCCTCTAGTTTTCANNNAGATCGGAAGA |
| 39910_2 | TATGAGTGTGGAGTCGTTGCCCTGCATGTCCCTCTAGTTTTCANNNAGATCGGAAGA |
| 39910_3 | TATGAGTGTGGAGTCGTTGCCCTGCATGTCCCTCTAGTTTTCANNNAGATCGGAAGA |
| 5283_1 | TATGAGTGTGGAGTCGTTGCGCACCAGTAGAGACTGGGTTTCACNNNAGATCGGA |
| 5283_2 | TATGAGTGTGGAGTCGTTGCGCACCAGTAGAGACTGGGTTTCACNNNAGATCGGA |
| 5283_3 | TATGAGTGTGGAGTCGTTGCGCACCAGTAGAGACTGGGTTTCACNNNAGATCGGA |
| 31231_1 | TATGAGTGTGGAGTCGTTGCTTACCAGTAACCAGAAGGCAGTCNNNAGATCGGAA |
| 31231_2 | TATGAGTGTGGAGTCGTTGCTTACCAGTAACCAGAAGGCAGTCNNNAGATCGGAA |
| 31231_3 | TATGAGTGTGGAGTCGTTGCTTACCAGTAACCAGAAGGCAGTCNNNAGATCGGAA |
| 45175_1 | TATGAGTGTGGAGTCGTTGCGATTTGAGCGACATCTTCTCTGCNNNAGATCGGAAG |
| 45175_2 | TATGAGTGTGGAGTCGTTGCGATTTGAGCGACATCTTCTCTGCNNNAGATCGGAAG |
| 45175_3 | TATGAGTGTGGAGTCGTTGCGATTTGAGCGACATCTTCTCTGCNNNAGATCGGAAG |
| 47897_1 | TATGAGTGTGGAGTCGTTGCTGTCTTACTTGCTCTGATCTGGCNNNAGATCGGAAG |
| 47897_2 | TATGAGTGTGGAGTCGTTGCTGTCTTACTTGCTCTGATCTGGCNNNAGATCGGAAG |
| 47897_3 | TATGAGTGTGGAGTCGTTGCTGTCTTACTTGCTCTGATCTGGCNNNAGATCGGAAG |
| 31674_1 | TATGAGTGTGGAGTCGTTGCGACTCAGGAAGAGCCTTTCTGAANNNAGATCGGAA |
| 31674_2 | TATGAGTGTGGAGTCGTTGCGACTCAGGAAGAGCCTTTCTGAANNNAGATCGGAA |
| 31674_3 | TATGAGTGTGGAGTCGTTGCGACTCAGGAAGAGCCTTTCTGAANNNAGATCGGAA |
| 10793_1 | TATGAGTGTGGAGTCGTTGCCACAGATGAGATTGAGGCAGGAGNNNAGATCGGA |
| 10793_2 | TATGAGTGTGGAGTCGTTGCCACAGATGAGATTGAGGCAGGAGNNNAGATCGGA |
| 10793_3 | TATGAGTGTGGAGTCGTTGCCACAGATGAGATTGAGGCAGGAGNNNAGATCGGA |
| 51431_1 | TATGAGTGTGGAGTCGTTGCTAGTGTGGGAGGGACTTAGAGAGNNNAGATCGGA |
| 51431_2 | TATGAGTGTGGAGTCGTTGCTAGTGTGGGAGGGACTTAGAGAGNNNAGATCGGA |
| 51431_3 | TATGAGTGTGGAGTCGTTGCTAGTGTGGGAGGGACTTAGAGAGNNNAGATCGGA |
| 33446_1 | TATGAGTGTGGAGTCGTTGCAGATGTGGATGATGATGGTGATGNNNAGATCGGA |
| 33446_2 | TATGAGTGTGGAGTCGTTGCAGATGTGGATGATGATGGTGATGNNNAGATCGGA |
| 33446_3 | TATGAGTGTGGAGTCGTTGCAGATGTGGATGATGATGGTGATGNNNAGATCGGA |
| 13613_1 | TATGAGTGTGGAGTCGTTGCCCCATATGCTATCCCTTATATTTNNNAGATCGGAAGA |
| 13613_2 | TATGAGTGTGGAGTCGTTGCCCCATATGCTATCCCTTATATTTNNNAGATCGGAAGA |
| 13613_3 | TATGAGTGTGGAGTCGTTGCCCCATATGCTATCCCTTATATTTNNNAGATCGGAAGA |
| 45637_1 | TATGAGTGTGGAGTCGTTGCTTCAAACGATTCTCCTGCCTCAGNNNAGATCGGAAG |
| 45637_2 | TATGAGTGTGGAGTCGTTGCTTCAAACGATTCTCCTGCCTCAGNNNAGATCGGAAG |
| 45637_3 | TATGAGTGTGGAGTCGTTGCTTCAAACGATTCTCCTGCCTCAGNNNAGATCGGAAG |
| 42083_1 | TATGAGTGTGGAGTCGTTGCTCTTCAAGCTAGCTCAGTGTCTCNNNAGATCGGAAG |
| 42083_2 | TATGAGTGTGGAGTCGTTGCTCTTCAAGCTAGCTCAGTGTCTCNNNAGATCGGAAG |
| 42083_3 | TATGAGTGTGGAGTCGTTGCTCTTCAAGCTAGCTCAGTGTCTCNNNAGATCGGAAG |
| 49308_1 | TATGAGTGTGGAGTCGTTGCCAAAGAGGTATTACGCGTCTGGTNNNAGATCGGAA |
| 49308_2 | TATGAGTGTGGAGTCGTTGCCAAAGAGGTATTACGCGTCTGGTNNNAGATCGGAA |

| | |
|---|---|
| 49308_3 | TATGAGTGTGGAGTCGTTGCCAAAGAGGTATTACGCGTCTGGTNNNAGATCGGAA |
| 46826_1 | TATGAGTGTGGAGTCGTTGCCTCCTGAGTAGCTGGGATTACAGNNNAGATCGGAA |
| 46826_2 | TATGAGTGTGGAGTCGTTGCCTCCTGAGTAGCTGGGATTACAGNNNAGATCGGAA |
| 46826_3 | TATGAGTGTGGAGTCGTTGCCTCCTGAGTAGCTGGGATTACAGNNNAGATCGGAA |
| 47037_1 | TATGAGTGTGGAGTCGTTGCAAGAAAGGAAAAGCTCCCTGTCTNNNAGATCGGAA |
| 47037_2 | TATGAGTGTGGAGTCGTTGCAAGAAAGGAAAAGCTCCCTGTCTNNNAGATCGGAA |
| 47037_3 | TATGAGTGTGGAGTCGTTGCAAGAAAGGAAAAGCTCCCTGTCTNNNAGATCGGAA |
| 44283_1 | TATGAGTGTGGAGTCGTTGCATCAGGTTTGAGTTTGCTTGCTCNNNAGATCGGAAG |
| 44283_2 | TATGAGTGTGGAGTCGTTGCATCAGGTTTGAGTTTGCTTGCTCNNNAGATCGGAAG |
| 44283_3 | TATGAGTGTGGAGTCGTTGCATCAGGTTTGAGTTTGCTTGCTCNNNAGATCGGAAG |
| 48832_1 | TATGAGTGTGGAGTCGTTGCCCTCCCAAGAAGCTGTACTACAGNNNAGATCGGAAG |
| 48832_2 | TATGAGTGTGGAGTCGTTGCCCTCCCAAGAAGCTGTACTACAGNNNAGATCGGAAG |
| 48832_3 | TATGAGTGTGGAGTCGTTGCCCTCCCAAGAAGCTGTACTACAGNNNAGATCGGAAG |
| 42407_1 | TATGAGTGTGGAGTCGTTGCGATAGAGTTTTGCTCCTGTTGCCNNNAGATCGGAAG |
| 42407_2 | TATGAGTGTGGAGTCGTTGCGATAGAGTTTTGCTCCTGTTGCCNNNAGATCGGAAG |
| 42407_3 | TATGAGTGTGGAGTCGTTGCGATAGAGTTTTGCTCCTGTTGCCNNNAGATCGGAAG |
| 45003_1 | TATGAGTGTGGAGTCGTTGCAGGGGGACTGCAAATAGTGTATGNNNAGATCGGAA |
| 45003_2 | TATGAGTGTGGAGTCGTTGCAGGGGGACTGCAAATAGTGTATGNNNAGATCGGAA |
| 45003_3 | TATGAGTGTGGAGTCGTTGCAGGGGGACTGCAAATAGTGTATGNNNAGATCGGAA |
| 33697_1 | TATGAGTGTGGAGTCGTTGCCTGTTCAGAGAGGCCAAGAAGAGNNNAGATCGGAA |
| 33697_2 | TATGAGTGTGGAGTCGTTGCCTGTTCAGAGAGGCCAAGAAGAGNNNAGATCGGAA |
| 33697_3 | TATGAGTGTGGAGTCGTTGCCTGTTCAGAGAGGCCAAGAAGAGNNNAGATCGGAA |
| 29515_1 | TATGAGTGTGGAGTCGTTGCCTGGGCCCACCTAAATTCTCNNNAGATCGGAAGAGC |
| 29515_2 | TATGAGTGTGGAGTCGTTGCCTGGGCCCACCTAAATTCTCNNNAGATCGGAAGAGC |
| 29515_3 | TATGAGTGTGGAGTCGTTGCCTGGGCCCACCTAAATTCTCNNNAGATCGGAAGAGC |
| 50077_1 | TATGAGTGTGGAGTCGTTGCGCGATTGTGATTGTGCTGCTATANNNAGATCGGAA |
| 50077_2 | TATGAGTGTGGAGTCGTTGCGCGATTGTGATTGTGCTGCTATANNNAGATCGGAA |
| 50077_3 | TATGAGTGTGGAGTCGTTGCGCGATTGTGATTGTGCTGCTATANNNAGATCGGAA |
| 50368_1 | TATGAGTGTGGAGTCGTTGCAAGGAACTAGAGAGGCAAGAACANNNAGATCGGA |
| 50368_2 | TATGAGTGTGGAGTCGTTGCAAGGAACTAGAGAGGCAAGAACANNNAGATCGGA |
| 50368_3 | TATGAGTGTGGAGTCGTTGCAAGGAACTAGAGAGGCAAGAACANNNAGATCGGA |
| 48731_1 | TATGAGTGTGGAGTCGTTGCTTTCTGCAACAATGATGGGAAGGNNNAGATCGGAA |
| 48731_2 | TATGAGTGTGGAGTCGTTGCTTTCTGCAACAATGATGGGAAGGNNNAGATCGGAA |
| 48731_3 | TATGAGTGTGGAGTCGTTGCTTTCTGCAACAATGATGGGAAGGNNNAGATCGGAA |
| 32626_1 | TATGAGTGTGGAGTCGTTGCTTACACCATGATTGTGAGGCCTTNNNAGATCGGAAG |
| 32626_2 | TATGAGTGTGGAGTCGTTGCTTACACCATGATTGTGAGGCCTTNNNAGATCGGAAG |
| 32626_3 | TATGAGTGTGGAGTCGTTGCTTACACCATGATTGTGAGGCCTTNNNAGATCGGAAG |
| 40541_1 | TATGAGTGTGGAGTCGTTGCAACACTGCTCAAAAGAAAGAAGTNNNAGATCGGAA |
| 40541_2 | TATGAGTGTGGAGTCGTTGCAACACTGCTCAAAAGAAAGAAGTNNNAGATCGGAA |
| 40541_3 | TATGAGTGTGGAGTCGTTGCAACACTGCTCAAAAGAAAGAAGTNNNAGATCGGAA |
| 42877_1 | TATGAGTGTGGAGTCGTTGCTGTGATATATGTGTGTGTGTGCANNNAGATCGGAA |
| 42877_2 | TATGAGTGTGGAGTCGTTGCTGTGATATATGTGTGTGTGTGCANNNAGATCGGAA |
| 42877_3 | TATGAGTGTGGAGTCGTTGCTGTGATATATGTGTGTGTGTGCANNNAGATCGGAA |

| | |
|---|---|
| 41200_1 | TATGAGTGTGGAGTCGTTGCGAGAAAGGGTGAAGGCAAACATCNNNAGATCGGA |
| 41200_2 | TATGAGTGTGGAGTCGTTGCGAGAAAGGGTGAAGGCAAACATCNNNAGATCGGA |
| 41200_3 | TATGAGTGTGGAGTCGTTGCGAGAAAGGGTGAAGGCAAACATCNNNAGATCGGA |
| 30644_1 | TATGAGTGTGGAGTCGTTGCCTGGGATGGCTTTCTCTGCTTCTTNNNAGATCGGAA |
| 30644_2 | TATGAGTGTGGAGTCGTTGCCTGGGATGGCTTTCTCTGCTTCTTNNNAGATCGGAA |
| 30644_3 | TATGAGTGTGGAGTCGTTGCCTGGGATGGCTTTCTCTGCTTCTTNNNAGATCGGAA |
| 44980_1 | TATGAGTGTGGAGTCGTTGCAGAGAGAGACAGACAAGGACAGANNNAGATCGGA |
| 44980_2 | TATGAGTGTGGAGTCGTTGCAGAGAGAGACAGACAAGGACAGANNNAGATCGGA |
| 44980_3 | TATGAGTGTGGAGTCGTTGCAGAGAGAGACAGACAAGGACAGANNNAGATCGGA |
| 42439_1 | TATGAGTGTGGAGTCGTTGCGGGAGAGAGAGAGGAAGAGAGAGNNNAGATCGG |
| 42439_2 | TATGAGTGTGGAGTCGTTGCGGGAGAGAGAGAGGAAGAGAGAGNNNAGATCGG |
| 42439_3 | TATGAGTGTGGAGTCGTTGCGGGAGAGAGAGAGGAAGAGAGAGNNNAGATCGG |
| 39679_1 | TATGAGTGTGGAGTCGTTGCCAACATCATTCATTTACCGGCCANNNAGATCGGAAG |
| 39679_2 | TATGAGTGTGGAGTCGTTGCCAACATCATTCATTTACCGGCCANNNAGATCGGAAG |
| 39679_3 | TATGAGTGTGGAGTCGTTGCCAACATCATTCATTTACCGGCCANNNAGATCGGAAG |
| 40202_1 | TATGAGTGTGGAGTCGTTGCTCCCATTCAAACTCTCCAACACTNNNAGATCGGAAGA |
| 40202_2 | TATGAGTGTGGAGTCGTTGCTCCCATTCAAACTCTCCAACACTNNNAGATCGGAAGA |
| 40202_3 | TATGAGTGTGGAGTCGTTGCTCCCATTCAAACTCTCCAACACTNNNAGATCGGAAGA |
| 43736_1 | TATGAGTGTGGAGTCGTTGCTGCAGAATTCAAAGGAGGACTGANNNAGATCGGAA |
| 43736_2 | TATGAGTGTGGAGTCGTTGCTGCAGAATTCAAAGGAGGACTGANNNAGATCGGAA |
| 43736_3 | TATGAGTGTGGAGTCGTTGCTGCAGAATTCAAAGGAGGACTGANNNAGATCGGAA |
| 42964_1 | TATGAGTGTGGAGTCGTTGCGCTTCCCAAGTAGCTAGGACTACNNNAGATCGGAAG |
| 42964_2 | TATGAGTGTGGAGTCGTTGCGCTTCCCAAGTAGCTAGGACTACNNNAGATCGGAAG |
| 42964_3 | TATGAGTGTGGAGTCGTTGCGCTTCCCAAGTAGCTAGGACTACNNNAGATCGGAAG |
| 46840_1 | TATGAGTGTGGAGTCGTTGCTTTCACTTTTGTTGCCTAGGCTGNNNAGATCGGAAG |
| 46840_2 | TATGAGTGTGGAGTCGTTGCTTTCACTTTTGTTGCCTAGGCTGNNNAGATCGGAAG |
| 46840_3 | TATGAGTGTGGAGTCGTTGCTTTCACTTTTGTTGCCTAGGCTGNNNAGATCGGAAG |
| 49728_1 | TATGAGTGTGGAGTCGTTGCTAACAGAACCGTGTTAGCTGTCANNNAGATCGGAAG |
| 49728_2 | TATGAGTGTGGAGTCGTTGCTAACAGAACCGTGTTAGCTGTCANNNAGATCGGAAG |
| 49728_3 | TATGAGTGTGGAGTCGTTGCTAACAGAACCGTGTTAGCTGTCANNNAGATCGGAAG |
| 32974_1 | TATGAGTGTGGAGTCGTTGCAAAAATTGAGGAGGAGGGTCTCCNNNAGATCGGAA |
| 32974_2 | TATGAGTGTGGAGTCGTTGCAAAAATTGAGGAGGAGGGTCTCCNNNAGATCGGAA |
| 32974_3 | TATGAGTGTGGAGTCGTTGCAAAAATTGAGGAGGAGGGTCTCCNNNAGATCGGAA |
| 36613_1 | TATGAGTGTGGAGTCGTTGCTAAAGGCCACTAGGAAACAGTCCNNNAGATCGGAA |
| 36613_2 | TATGAGTGTGGAGTCGTTGCTAAAGGCCACTAGGAAACAGTCCNNNAGATCGGAA |
| 36613_3 | TATGAGTGTGGAGTCGTTGCTAAAGGCCACTAGGAAACAGTCCNNNAGATCGGAA |
| 50045_1 | TATGAGTGTGGAGTCGTTGCATTCCCACACAATTGAAACGTCCNNNAGATCGGAAG |
| 50045_2 | TATGAGTGTGGAGTCGTTGCATTCCCACACAATTGAAACGTCCNNNAGATCGGAAG |
| 50045_3 | TATGAGTGTGGAGTCGTTGCATTCCCACACAATTGAAACGTCCNNNAGATCGGAAG |
| 41259_1 | TATGAGTGTGGAGTCGTTGCGGAAAATCTGGCGAAATGCAAACNNNAGATCGGAA |
| 41259_2 | TATGAGTGTGGAGTCGTTGCGGAAAATCTGGCGAAATGCAAACNNNAGATCGGAA |
| 41259_3 | TATGAGTGTGGAGTCGTTGCGGAAAATCTGGCGAAATGCAAACNNNAGATCGGAA |
| 32154_1 | TATGAGTGTGGAGTCGTTGCGAGAATCACTTGAACCAGGGAGTNNNAGATCGGAA |

| | |
|---|---|
| 32154_2 | TATGAGTGTGGAGTCGTTGCGAGAATCACTTGAACCAGGGAGTNNNAGATCGGAA |
| 32154_3 | TATGAGTGTGGAGTCGTTGCGAGAATCACTTGAACCAGGGAGTNNNAGATCGGAA |
| 49522_1 | TATGAGTGTGGAGTCGTTGCCTCGCCCGGCTAATTTTTTGTATNNNAGATCGGAAG |
| 49522_2 | TATGAGTGTGGAGTCGTTGCCTCGCCCGGCTAATTTTTTGTATNNNAGATCGGAAG |
| 49522_3 | TATGAGTGTGGAGTCGTTGCCTCGCCCGGCTAATTTTTTGTATNNNAGATCGGAAG |
| 38320_1 | TATGAGTGTGGAGTCGTTGCTGTGGAATCAAAAACAAACCCGANNNAGATCGGAA |
| 38320_2 | TATGAGTGTGGAGTCGTTGCTGTGGAATCAAAAACAAACCCGANNNAGATCGGAA |
| 38320_3 | TATGAGTGTGGAGTCGTTGCTGTGGAATCAAAAACAAACCCGANNNAGATCGGAA |
| 45445_1 | TATGAGTGTGGAGTCGTTGCAGGGGGTGTGCAAATCATAATCANNNAGATCGGAA |
| 45445_2 | TATGAGTGTGGAGTCGTTGCAGGGGGTGTGCAAATCATAATCANNNAGATCGGAA |
| 45445_3 | TATGAGTGTGGAGTCGTTGCAGGGGGTGTGCAAATCATAATCANNNAGATCGGAA |
| 32024_1 | TATGAGTGTGGAGTCGTTGCGAGACAAAGGCCTTGCTCTATTGNNNAGATCGGAA |
| 32024_2 | TATGAGTGTGGAGTCGTTGCGAGACAAAGGCCTTGCTCTATTGNNNAGATCGGAA |
| 32024_3 | TATGAGTGTGGAGTCGTTGCGAGACAAAGGCCTTGCTCTATTGNNNAGATCGGAA |
| 46481_1 | TATGAGTGTGGAGTCGTTGCAACACAGGCTACAGAGGACTTGANNNAGATCGGAA |
| 46481_2 | TATGAGTGTGGAGTCGTTGCAACACAGGCTACAGAGGACTTGANNNAGATCGGAA |
| 46481_3 | TATGAGTGTGGAGTCGTTGCAACACAGGCTACAGAGGACTTGANNNAGATCGGAA |
| 31276_1 | TATGAGTGTGGAGTCGTTGCAGGAAATGAGGCTAAGCAATCCANNNAGATCGGAA |
| 31276_2 | TATGAGTGTGGAGTCGTTGCAGGAAATGAGGCTAAGCAATCCANNNAGATCGGAA |
| 31276_3 | TATGAGTGTGGAGTCGTTGCAGGAAATGAGGCTAAGCAATCCANNNAGATCGGAA |
| 52814_1 | TATGAGTGTGGAGTCGTTGCCATACCCCCTTTCTGCTCCTGNNNAGATCGGAAGAGC |
| 52814_2 | TATGAGTGTGGAGTCGTTGCCATACCCCCTTTCTGCTCCTGNNNAGATCGGAAGAGC |
| 52814_3 | TATGAGTGTGGAGTCGTTGCCATACCCCCTTTCTGCTCCTGNNNAGATCGGAAGAGC |
| 34893_1 | TATGAGTGTGGAGTCGTTGCGTCTCCCGAGTAGCTGGAATTACNNNAGATCGGAAG |
| 34893_2 | TATGAGTGTGGAGTCGTTGCGTCTCCCGAGTAGCTGGAATTACNNNAGATCGGAAG |
| 34893_3 | TATGAGTGTGGAGTCGTTGCGTCTCCCGAGTAGCTGGAATTACNNNAGATCGGAAG |
| 28762_1 | TATGAGTGTGGAGTCGTTGCCCAAGTAGCTGGGATTACAGGTGNNNAGATCGGAA |
| 28762_2 | TATGAGTGTGGAGTCGTTGCCCAAGTAGCTGGGATTACAGGTGNNNAGATCGGAA |
| 28762_3 | TATGAGTGTGGAGTCGTTGCCCAAGTAGCTGGGATTACAGGTGNNNAGATCGGAA |
| 7772_1 | TATGAGTGTGGAGTCGTTGCCGGGGAGGAAAATATGGGTGTTGNNNAGATCGGA |
| 7772_2 | TATGAGTGTGGAGTCGTTGCCGGGGAGGAAAATATGGGTGTTGNNNAGATCGGA |
| 7772_3 | TATGAGTGTGGAGTCGTTGCCGGGGAGGAAAATATGGGTGTTGNNNAGATCGGA |
| 40395_1 | TATGAGTGTGGAGTCGTTGCGGACTCCACACTATCAACACCTTNNNAGATCGGAAG |
| 40395_2 | TATGAGTGTGGAGTCGTTGCGGACTCCACACTATCAACACCTTNNNAGATCGGAAG |
| 40395_3 | TATGAGTGTGGAGTCGTTGCGGACTCCACACTATCAACACCTTNNNAGATCGGAAG |
| 40979_1 | TATGAGTGTGGAGTCGTTGCGTCTCTTCACACTGCGTTTCTTGNNNAGATCGGAAGA |
| 40979_2 | TATGAGTGTGGAGTCGTTGCGTCTCTTCACACTGCGTTTCTTGNNNAGATCGGAAGA |
| 40979_3 | TATGAGTGTGGAGTCGTTGCGTCTCTTCACACTGCGTTTCTTGNNNAGATCGGAAGA |
| 33683_1 | TATGAGTGTGGAGTCGTTGCCCAAAGTGCTGGGATTACAAGTGNNNAGATCGGAA |
| 33683_2 | TATGAGTGTGGAGTCGTTGCCCAAAGTGCTGGGATTACAAGTGNNNAGATCGGAA |
| 33683_3 | TATGAGTGTGGAGTCGTTGCCCAAAGTGCTGGGATTACAAGTGNNNAGATCGGAA |
| 3029_1 | TATGAGTGTGGAGTCGTTGCGCTCAGGATTCCAAGTGCTTTCANNNAGATCGGAAG |
| 3029_2 | TATGAGTGTGGAGTCGTTGCGCTCAGGATTCCAAGTGCTTTCANNNAGATCGGAAG |

| | |
|---|---|
| 3029_3 | TATGAGTGTGGAGTCGTTGCGCTCAGGATTCCAAGTGCTTTCANNNAGATCGGAAG |
| 46502_1 | TATGAGTGTGGAGTCGTTGCCTCCTGAGTAGCTGGGACTACTNNNAGATCGGAAGA |
| 46502_2 | TATGAGTGTGGAGTCGTTGCCTCCTGAGTAGCTGGGACTACTNNNAGATCGGAAGA |
| 46502_3 | TATGAGTGTGGAGTCGTTGCCTCCTGAGTAGCTGGGACTACTNNNAGATCGGAAGA |
| 5809_1 | TATGAGTGTGGAGTCGTTGCTGAACGGAATAGCCTGTACTTCCNNNAGATCGGAAG |
| 5809_2 | TATGAGTGTGGAGTCGTTGCTGAACGGAATAGCCTGTACTTCCNNNAGATCGGAAG |
| 5809_3 | TATGAGTGTGGAGTCGTTGCTGAACGGAATAGCCTGTACTTCCNNNAGATCGGAAG |
| 42512_1 | TATGAGTGTGGAGTCGTTGCTGTATTTTCCCTTCTCCCCTTCCNNNAGATCGGAAGAG |
| 42512_2 | TATGAGTGTGGAGTCGTTGCTGTATTTTCCCTTCTCCCCTTCCNNNAGATCGGAAGAG |
| 42512_3 | TATGAGTGTGGAGTCGTTGCTGTATTTTCCCTTCTCCCCTTCCNNNAGATCGGAAGAG |
| 34878_1 | TATGAGTGTGGAGTCGTTGCCAGTGGTGCAGTCCTAGCTCNNNAGATCGGAAGAG |
| 34878_2 | TATGAGTGTGGAGTCGTTGCCAGTGGTGCAGTCCTAGCTCNNNAGATCGGAAGAG |
| 34878_3 | TATGAGTGTGGAGTCGTTGCCAGTGGTGCAGTCCTAGCTCNNNAGATCGGAAGAG |
| 50478_1 | TATGAGTGTGGAGTCGTTGCCTGCTTTACAAGAGACAGCACACNNNAGATCGGAAG |
| 50478_2 | TATGAGTGTGGAGTCGTTGCCTGCTTTACAAGAGACAGCACACNNNAGATCGGAAG |
| 50478_3 | TATGAGTGTGGAGTCGTTGCCTGCTTTACAAGAGACAGCACACNNNAGATCGGAAG |
| 36596_1 | TATGAGTGTGGAGTCGTTGCAGGAGCTATTACAACTGACACCANNNAGATCGGAAG |
| 36596_2 | TATGAGTGTGGAGTCGTTGCAGGAGCTATTACAACTGACACCANNNAGATCGGAAG |
| 36596_3 | TATGAGTGTGGAGTCGTTGCAGGAGCTATTACAACTGACACCANNNAGATCGGAAG |
| 29479_1 | TATGAGTGTGGAGTCGTTGCTTGTACTGCATTTCTGCCAGTGANNNAGATCGGAAG |
| 29479_2 | TATGAGTGTGGAGTCGTTGCTTGTACTGCATTTCTGCCAGTGANNNAGATCGGAAG |
| 29479_3 | TATGAGTGTGGAGTCGTTGCTTGTACTGCATTTCTGCCAGTGANNNAGATCGGAAG |
| 30825_1 | TATGAGTGTGGAGTCGTTGCTCTCTCTCTCTCTCTGTGTGTGTNNNAGATCGGAAGA |
| 30825_2 | TATGAGTGTGGAGTCGTTGCTCTCTCTCTCTCTCTGTGTGTGTNNNAGATCGGAAGA |
| 30825_3 | TATGAGTGTGGAGTCGTTGCTCTCTCTCTCTCTCTGTGTGTGTNNNAGATCGGAAGA |
| 12659_1 | TATGAGTGTGGAGTCGTTGCAGCTATGAATACAGAGGTATATGNNNAGATCGGAA |
| 12659_2 | TATGAGTGTGGAGTCGTTGCAGCTATGAATACAGAGGTATATGNNNAGATCGGAA |
| 12659_3 | TATGAGTGTGGAGTCGTTGCAGCTATGAATACAGAGGTATATGNNNAGATCGGAA |
| 45835_1 | TATGAGTGTGGAGTCGTTGCCGAGTAGCTGGGATTACAGGTGNNNAGATCGGAAG |
| 45835_2 | TATGAGTGTGGAGTCGTTGCCGAGTAGCTGGGATTACAGGTGNNNAGATCGGAAG |
| 45835_3 | TATGAGTGTGGAGTCGTTGCCGAGTAGCTGGGATTACAGGTGNNNAGATCGGAAG |
| 40860_1 | TATGAGTGTGGAGTCGTTGCAGATCTGGGAGTCTTTGAGGAGANNNAGATCGGAA |
| 40860_2 | TATGAGTGTGGAGTCGTTGCAGATCTGGGAGTCTTTGAGGAGANNNAGATCGGAA |
| 40860_3 | TATGAGTGTGGAGTCGTTGCAGATCTGGGAGTCTTTGAGGAGANNNAGATCGGAA |
| 42481_1 | TATGAGTGTGGAGTCGTTGCTTATGAGAACAGCATGGGGGAAANNNAGATCGGAA |
| 42481_2 | TATGAGTGTGGAGTCGTTGCTTATGAGAACAGCATGGGGGAAANNNAGATCGGAA |
| 42481_3 | TATGAGTGTGGAGTCGTTGCTTATGAGAACAGCATGGGGGAAANNNAGATCGGAA |
| 42569_1 | TATGAGTGTGGAGTCGTTGCTCCTCTTGTTGAATTGACCCCTTNNNAGATCGGAAGA |
| 42569_2 | TATGAGTGTGGAGTCGTTGCTCCTCTTGTTGAATTGACCCCTTNNNAGATCGGAAGA |
| 42569_3 | TATGAGTGTGGAGTCGTTGCTCCTCTTGTTGAATTGACCCCTTNNNAGATCGGAAGA |
| 30509_1 | TATGAGTGTGGAGTCGTTGCGAAGAGCCTGATACTTGGGTTGGNNNAGATCGGAA |
| 30509_2 | TATGAGTGTGGAGTCGTTGCGAAGAGCCTGATACTTGGGTTGGNNNAGATCGGAA |
| 30509_3 | TATGAGTGTGGAGTCGTTGCGAAGAGCCTGATACTTGGGTTGGNNNAGATCGGAA |

| | |
|---|---|
| 52310_1 | TATGAGTGTGGAGTCGTTGCGACTCCATTGACTCCTGAGATCCNNNAGATCGGAAG |
| 52310_2 | TATGAGTGTGGAGTCGTTGCGACTCCATTGACTCCTGAGATCCNNNAGATCGGAAG |
| 52310_3 | TATGAGTGTGGAGTCGTTGCGACTCCATTGACTCCTGAGATCCNNNAGATCGGAAG |
| 51787_1 | TATGAGTGTGGAGTCGTTGCCTCCTACACACAGCTGCTAGATTNNNAGATCGGAAG |
| 51787_2 | TATGAGTGTGGAGTCGTTGCCTCCTACACACAGCTGCTAGATTNNNAGATCGGAAG |
| 51787_3 | TATGAGTGTGGAGTCGTTGCCTCCTACACACAGCTGCTAGATTNNNAGATCGGAAG |
| 41087_1 | TATGAGTGTGGAGTCGTTGCGACTTCGATCCTAGATATGGCCANNNAGATCGGAAG |
| 41087_2 | TATGAGTGTGGAGTCGTTGCGACTTCGATCCTAGATATGGCCANNNAGATCGGAAG |
| 41087_3 | TATGAGTGTGGAGTCGTTGCGACTTCGATCCTAGATATGGCCANNNAGATCGGAAG |
| 42950_1 | TATGAGTGTGGAGTCGTTGCAAAATTAGCTGGATATGGTGGCGNNNAGATCGGAA |
| 42950_2 | TATGAGTGTGGAGTCGTTGCAAAATTAGCTGGATATGGTGGCGNNNAGATCGGAA |
| 42950_3 | TATGAGTGTGGAGTCGTTGCAAAATTAGCTGGATATGGTGGCGNNNAGATCGGAA |
| 41498_1 | TATGAGTGTGGAGTCGTTGCTTCGAATTTGTTTGCTCTTGCCTNNNAGATCGGAAGA |
| 41498_2 | TATGAGTGTGGAGTCGTTGCTTCGAATTTGTTTGCTCTTGCCTNNNAGATCGGAAGA |
| 41498_3 | TATGAGTGTGGAGTCGTTGCTTCGAATTTGTTTGCTCTTGCCTNNNAGATCGGAAGA |
| 45430_1 | TATGAGTGTGGAGTCGTTGCTGTGTGTGTGTGTGTGGTTTTTTNNNAGATCGGAAG |
| 45430_2 | TATGAGTGTGGAGTCGTTGCTGTGTGTGTGTGTGTGGTTTTTTNNNAGATCGGAAG |
| 45430_3 | TATGAGTGTGGAGTCGTTGCTGTGTGTGTGTGTGTGGTTTTTTNNNAGATCGGAAG |
| 34895_1 | TATGAGTGTGGAGTCGTTGCGATTTTGAGACGGAGTCTTGCTCNNNAGATCGGAAG |
| 34895_2 | TATGAGTGTGGAGTCGTTGCGATTTTGAGACGGAGTCTTGCTCNNNAGATCGGAAG |
| 34895_3 | TATGAGTGTGGAGTCGTTGCGATTTTGAGACGGAGTCTTGCTCNNNAGATCGGAAG |
| 44489_1 | TATGAGTGTGGAGTCGTTGCTATGTCTCAGGCCTATGGTAGCTNNNAGATCGGAAG |
| 44489_2 | TATGAGTGTGGAGTCGTTGCTATGTCTCAGGCCTATGGTAGCTNNNAGATCGGAAG |
| 44489_3 | TATGAGTGTGGAGTCGTTGCTATGTCTCAGGCCTATGGTAGCTNNNAGATCGGAAG |
| 35769_1 | TATGAGTGTGGAGTCGTTGCTACTTGAGGCTATACCCAGGTGANNNAGATCGGAAG |
| 35769_2 | TATGAGTGTGGAGTCGTTGCTACTTGAGGCTATACCCAGGTGANNNAGATCGGAAG |
| 35769_3 | TATGAGTGTGGAGTCGTTGCTACTTGAGGCTATACCCAGGTGANNNAGATCGGAAG |
| 47388_1 | TATGAGTGTGGAGTCGTTGCTGTGCGGGAGAAAGGAGTTTTATNNNAGATCGGAA |
| 47388_2 | TATGAGTGTGGAGTCGTTGCTGTGCGGGAGAAAGGAGTTTTATNNNAGATCGGAA |
| 47388_3 | TATGAGTGTGGAGTCGTTGCTGTGCGGGAGAAAGGAGTTTTATNNNAGATCGGAA |
| 34544_1 | TATGAGTGTGGAGTCGTTGCCAAGCTCTGACCCCACAGTCNNNAGATCGGAAGAGC |
| 34544_2 | TATGAGTGTGGAGTCGTTGCCAAGCTCTGACCCCACAGTCNNNAGATCGGAAGAGC |
| 34544_3 | TATGAGTGTGGAGTCGTTGCCAAGCTCTGACCCCACAGTCNNNAGATCGGAAGAGC |
| 13619_1 | TATGAGTGTGGAGTCGTTGCGATGCCATAAGAACAAAGATACNNNAGATCGGAAG |
| 13619_2 | TATGAGTGTGGAGTCGTTGCGATGCCATAAGAACAAAGATACNNNAGATCGGAAG |
| 13619_3 | TATGAGTGTGGAGTCGTTGCGATGCCATAAGAACAAAGATACNNNAGATCGGAAG |
| 50594_1 | TATGAGTGTGGAGTCGTTGCCCCAAATCAACCACCACTCTAGANNNAGATCGGAAGA |
| 50594_2 | TATGAGTGTGGAGTCGTTGCCCCAAATCAACCACCACTCTAGANNNAGATCGGAAGA |
| 50594_3 | TATGAGTGTGGAGTCGTTGCCCCAAATCAACCACCACTCTAGANNNAGATCGGAAGA |
| 36031_1 | TATGAGTGTGGAGTCGTTGCGTTCAGCCCATTTCTGTCTCAACNNNAGATCGGAAGA |
| 36031_2 | TATGAGTGTGGAGTCGTTGCGTTCAGCCCATTTCTGTCTCAACNNNAGATCGGAAGA |
| 36031_3 | TATGAGTGTGGAGTCGTTGCGTTCAGCCCATTTCTGTCTCAACNNNAGATCGGAAGA |
| 36817_1 | TATGAGTGTGGAGTCGTTGCATGCATCAGAGCTCTGTCCATAGNNNAGATCGGAAG |

| | |
|---|---|
| 36817_2 | TATGAGTGTGGAGTCGTTGCATGCATCAGAGCTCTGTCCATAGNNNAGATCGGAAG |
| 36817_3 | TATGAGTGTGGAGTCGTTGCATGCATCAGAGCTCTGTCCATAGNNNAGATCGGAAG |
| 40027_1 | TATGAGTGTGGAGTCGTTGCAAGCAAGGTAATGGGCTAAGAGTNNNAGATCGGAA |
| 40027_2 | TATGAGTGTGGAGTCGTTGCAAGCAAGGTAATGGGCTAAGAGTNNNAGATCGGAA |
| 40027_3 | TATGAGTGTGGAGTCGTTGCAAGCAAGGTAATGGGCTAAGAGTNNNAGATCGGAA |
| 45342_1 | TATGAGTGTGGAGTCGTTGCTTGTGTAAAAGGGCACTCAGAGANNNAGATCGGAA |
| 45342_2 | TATGAGTGTGGAGTCGTTGCTTGTGTAAAAGGGCACTCAGAGANNNAGATCGGAA |
| 45342_3 | TATGAGTGTGGAGTCGTTGCTTGTGTAAAAGGGCACTCAGAGANNNAGATCGGAA |
| 31859_1 | TATGAGTGTGGAGTCGTTGCAAACTGCAAACAACCTCAGAACCNNNAGATCGGAAG |
| 31859_2 | TATGAGTGTGGAGTCGTTGCAAACTGCAAACAACCTCAGAACCNNNAGATCGGAAG |
| 31859_3 | TATGAGTGTGGAGTCGTTGCAAACTGCAAACAACCTCAGAACCNNNAGATCGGAAG |
| 36697_1 | TATGAGTGTGGAGTCGTTGCGGCACCTATAATCCCAGCTACTCNNNAGATCGGAAG |
| 36697_2 | TATGAGTGTGGAGTCGTTGCGGCACCTATAATCCCAGCTACTCNNNAGATCGGAAG |
| 36697_3 | TATGAGTGTGGAGTCGTTGCGGCACCTATAATCCCAGCTACTCNNNAGATCGGAAG |
| 47371_1 | TATGAGTGTGGAGTCGTTGCATGCAATCTCAGCTTACTGCAACNNNAGATCGGAAG |
| 47371_2 | TATGAGTGTGGAGTCGTTGCATGCAATCTCAGCTTACTGCAACNNNAGATCGGAAG |
| 47371_3 | TATGAGTGTGGAGTCGTTGCATGCAATCTCAGCTTACTGCAACNNNAGATCGGAAG |
| 40147_1 | TATGAGTGTGGAGTCGTTGCAGGGTTTCTTACTGCCTCTAGGANNNAGATCGGAAG |
| 40147_2 | TATGAGTGTGGAGTCGTTGCAGGGTTTCTTACTGCCTCTAGGANNNAGATCGGAAG |
| 40147_3 | TATGAGTGTGGAGTCGTTGCAGGGTTTCTTACTGCCTCTAGGANNNAGATCGGAAG |
| 40494_1 | TATGAGTGTGGAGTCGTTGCGGTTCAAGTGATTTTCCTGCCTCNNNAGATCGGAAG |
| 40494_2 | TATGAGTGTGGAGTCGTTGCGGTTCAAGTGATTTTCCTGCCTCNNNAGATCGGAAG |
| 40494_3 | TATGAGTGTGGAGTCGTTGCGGTTCAAGTGATTTTCCTGCCTCNNNAGATCGGAAG |
| 40414_1 | TATGAGTGTGGAGTCGTTGCATATTACTGTCACCGACTGTCCCNNNAGATCGGAAG |
| 40414_2 | TATGAGTGTGGAGTCGTTGCATATTACTGTCACCGACTGTCCCNNNAGATCGGAAG |
| 40414_3 | TATGAGTGTGGAGTCGTTGCATATTACTGTCACCGACTGTCCCNNNAGATCGGAAG |
| 39311_1 | TATGAGTGTGGAGTCGTTGCAACTGCATATTGGGTACCGTGTANNNAGATCGGAAG |
| 39311_2 | TATGAGTGTGGAGTCGTTGCAACTGCATATTGGGTACCGTGTANNNAGATCGGAAG |
| 39311_3 | TATGAGTGTGGAGTCGTTGCAACTGCATATTGGGTACCGTGTANNNAGATCGGAAG |
| 37772_1 | TATGAGTGTGGAGTCGTTGCCTGAGAGGTGAAGTTTGCAGTGANNNAGATCGGAA |
| 37772_2 | TATGAGTGTGGAGTCGTTGCCTGAGAGGTGAAGTTTGCAGTGANNNAGATCGGAA |
| 37772_3 | TATGAGTGTGGAGTCGTTGCCTGAGAGGTGAAGTTTGCAGTGANNNAGATCGGAA |
| 10021_1 | TATGAGTGTGGAGTCGTTGCGGTCAAGAGCATCACTGAGCTTTNNNAGATCGGAAG |
| 10021_2 | TATGAGTGTGGAGTCGTTGCGGTCAAGAGCATCACTGAGCTTTNNNAGATCGGAAG |
| 10021_3 | TATGAGTGTGGAGTCGTTGCGGTCAAGAGCATCACTGAGCTTTNNNAGATCGGAAG |
| 44698_1 | TATGAGTGTGGAGTCGTTGCGAAAAAGAGAGCACACACTCCACNNNAGATCGGAAG |
| 44698_2 | TATGAGTGTGGAGTCGTTGCGAAAAAGAGAGCACACACTCCACNNNAGATCGGAAG |
| 44698_3 | TATGAGTGTGGAGTCGTTGCGAAAAAGAGAGCACACACTCCACNNNAGATCGGAAG |
| 34139_1 | TATGAGTGTGGAGTCGTTGCAAGCCTGAAGTGTGGAAACAAACNNNAGATCGGAA |
| 34139_2 | TATGAGTGTGGAGTCGTTGCAAGCCTGAAGTGTGGAAACAAACNNNAGATCGGAA |
| 34139_3 | TATGAGTGTGGAGTCGTTGCAAGCCTGAAGTGTGGAAACAAACNNNAGATCGGAA |
| 37703_1 | TATGAGTGTGGAGTCGTTGCCACACCTAGAGGAACTAGAAGAGCNNNAGATCGGA |
| 37703_2 | TATGAGTGTGGAGTCGTTGCCACACCTAGAGGAACTAGAAGAGCNNNAGATCGGA |

| | |
|---|---|
| 37703_3 | TATGAGTGTGGAGTCGTTGCCACACCTAGAGGAACTAGAAGAGCNNNAGATCGGA |
| 42159_1 | TATGAGTGTGGAGTCGTTGCCTCCCGAGTAGCTGGAATTACAGNNNAGATCGGAAG |
| 42159_2 | TATGAGTGTGGAGTCGTTGCCTCCCGAGTAGCTGGAATTACAGNNNAGATCGGAAG |
| 42159_3 | TATGAGTGTGGAGTCGTTGCCTCCCGAGTAGCTGGAATTACAGNNNAGATCGGAAG |
| 45001_1 | TATGAGTGTGGAGTCGTTGCAGTTTCCGATGAGCCTGTATGTTNNNAGATCGGAAG |
| 45001_2 | TATGAGTGTGGAGTCGTTGCAGTTTCCGATGAGCCTGTATGTTNNNAGATCGGAAG |
| 45001_3 | TATGAGTGTGGAGTCGTTGCAGTTTCCGATGAGCCTGTATGTTNNNAGATCGGAAG |
| 49528_1 | TATGAGTGTGGAGTCGTTGCGGATCTCGGCTCACTGCAAGNNNAGATCGGAAGAG |
| 49528_2 | TATGAGTGTGGAGTCGTTGCGGATCTCGGCTCACTGCAAGNNNAGATCGGAAGAG |
| 49528_3 | TATGAGTGTGGAGTCGTTGCGGATCTCGGCTCACTGCAAGNNNAGATCGGAAGAG |
| 52436_1 | TATGAGTGTGGAGTCGTTGCCTTACAGATTCCACTGCCAATGCNNNAGATCGGAAG |
| 52436_2 | TATGAGTGTGGAGTCGTTGCCTTACAGATTCCACTGCCAATGCNNNAGATCGGAAG |
| 52436_3 | TATGAGTGTGGAGTCGTTGCCTTACAGATTCCACTGCCAATGCNNNAGATCGGAAG |
| 37858_1 | TATGAGTGTGGAGTCGTTGCTCTGATCATACACCTTGTGGCAANNNAGATCGGAAG |
| 37858_2 | TATGAGTGTGGAGTCGTTGCTCTGATCATACACCTTGTGGCAANNNAGATCGGAAG |
| 37858_3 | TATGAGTGTGGAGTCGTTGCTCTGATCATACACCTTGTGGCAANNNAGATCGGAAG |
| 32006_1 | TATGAGTGTGGAGTCGTTGCGAGAGGGAACCAACTTGTACTGTNNNAGATCGGAA |
| 32006_2 | TATGAGTGTGGAGTCGTTGCGAGAGGGAACCAACTTGTACTGTNNNAGATCGGAA |
| 32006_3 | TATGAGTGTGGAGTCGTTGCGAGAGGGAACCAACTTGTACTGTNNNAGATCGGAA |
| 38509_1 | TATGAGTGTGGAGTCGTTGCTTGACAGGACGAAATCAGGACTTNNNAGATCGGAA |
| 38509_2 | TATGAGTGTGGAGTCGTTGCTTGACAGGACGAAATCAGGACTTNNNAGATCGGAA |
| 38509_3 | TATGAGTGTGGAGTCGTTGCTTGACAGGACGAAATCAGGACTTNNNAGATCGGAA |
| 48983_1 | TATGAGTGTGGAGTCGTTGCCGGCATCAACATTGAAATCCAGANNNAGATCGGAAG |
| 48983_2 | TATGAGTGTGGAGTCGTTGCCGGCATCAACATTGAAATCCAGANNNAGATCGGAAG |
| 48983_3 | TATGAGTGTGGAGTCGTTGCCGGCATCAACATTGAAATCCAGANNNAGATCGGAAG |
| 52953_1 | TATGAGTGTGGAGTCGTTGCGGAGTTGCAAGAAGGGAAACATCNNNAGATCGGAA |
| 52953_2 | TATGAGTGTGGAGTCGTTGCGGAGTTGCAAGAAGGGAAACATCNNNAGATCGGAA |
| 52953_3 | TATGAGTGTGGAGTCGTTGCGGAGTTGCAAGAAGGGAAACATCNNNAGATCGGAA |
| 41407_1 | TATGAGTGTGGAGTCGTTGCAGGTAGGGGATTATGTGTGTAAGCNNNAGATCGGA |
| 41407_2 | TATGAGTGTGGAGTCGTTGCAGGTAGGGGATTATGTGTGTAAGCNNNAGATCGGA |
| 41407_3 | TATGAGTGTGGAGTCGTTGCAGGTAGGGGATTATGTGTGTAAGCNNNAGATCGGA |
| 30386_1 | TATGAGTGTGGAGTCGTTGCCCGGTATGTAATTTAACATTCAANNNAGATCGGAAG |
| 30386_2 | TATGAGTGTGGAGTCGTTGCCCGGTATGTAATTTAACATTCAANNNAGATCGGAAG |
| 30386_3 | TATGAGTGTGGAGTCGTTGCCCGGTATGTAATTTAACATTCAANNNAGATCGGAAG |
| 42833_1 | TATGAGTGTGGAGTCGTTGCGGTAAATAGTCTCAGCTCCTCGGNNNAGATCGGAAG |
| 42833_2 | TATGAGTGTGGAGTCGTTGCGGTAAATAGTCTCAGCTCCTCGGNNNAGATCGGAAG |
| 42833_3 | TATGAGTGTGGAGTCGTTGCGGTAAATAGTCTCAGCTCCTCGGNNNAGATCGGAAG |
| 48727_1 | TATGAGTGTGGAGTCGTTGCCAATCTTGGCTCACTGCAATCTCNNNAGATCGGAAG |
| 48727_2 | TATGAGTGTGGAGTCGTTGCCAATCTTGGCTCACTGCAATCTCNNNAGATCGGAAG |
| 48727_3 | TATGAGTGTGGAGTCGTTGCCAATCTTGGCTCACTGCAATCTCNNNAGATCGGAAG |
| 46329_1 | TATGAGTGTGGAGTCGTTGCAAGTTCAAGTGATTCTCCTGCCTNNNAGATCGGAAG |
| 46329_2 | TATGAGTGTGGAGTCGTTGCAAGTTCAAGTGATTCTCCTGCCTNNNAGATCGGAAG |
| 46329_3 | TATGAGTGTGGAGTCGTTGCAAGTTCAAGTGATTCTCCTGCCTNNNAGATCGGAAG |

| | |
|---|---|
| 34939_1 | TATGAGTGTGGAGTCGTTGCTGTTCTTCTCATTCAGTCTGGGGNNNAGATCGGAAG |
| 34939_2 | TATGAGTGTGGAGTCGTTGCTGTTCTTCTCATTCAGTCTGGGGNNNAGATCGGAAG |
| 34939_3 | TATGAGTGTGGAGTCGTTGCTGTTCTTCTCATTCAGTCTGGGGNNNAGATCGGAAG |
| 44393_1 | TATGAGTGTGGAGTCGTTGCAAGTGGCCCCAACTTATCTTTCTNNNAGATCGGAAG |
| 44393_2 | TATGAGTGTGGAGTCGTTGCAAGTGGCCCCAACTTATCTTTCTNNNAGATCGGAAG |
| 44393_3 | TATGAGTGTGGAGTCGTTGCAAGTGGCCCCAACTTATCTTTCTNNNAGATCGGAAG |
| 38887_1 | TATGAGTGTGGAGTCGTTGCACCAGATGTACAAAGAAGGGCTGNNNAGATCGGAA |
| 38887_2 | TATGAGTGTGGAGTCGTTGCACCAGATGTACAAAGAAGGGCTGNNNAGATCGGAA |
| 38887_3 | TATGAGTGTGGAGTCGTTGCACCAGATGTACAAAGAAGGGCTGNNNAGATCGGAA |
| 44238_1 | TATGAGTGTGGAGTCGTTGCCTAGACTTATGTGCTGGGACCAGNNNAGATCGGAA |
| 44238_2 | TATGAGTGTGGAGTCGTTGCCTAGACTTATGTGCTGGGACCAGNNNAGATCGGAA |
| 44238_3 | TATGAGTGTGGAGTCGTTGCCTAGACTTATGTGCTGGGACCAGNNNAGATCGGAA |
| 8815_1 | TATGAGTGTGGAGTCGTTGCCTTTGGAAGGGTAGTGTGGCATTNNNAGATCGGAA |
| 8815_2 | TATGAGTGTGGAGTCGTTGCCTTTGGAAGGGTAGTGTGGCATTNNNAGATCGGAA |
| 8815_3 | TATGAGTGTGGAGTCGTTGCCTTTGGAAGGGTAGTGTGGCATTNNNAGATCGGAA |
| 2796_1 | TATGAGTGTGGAGTCGTTGCTTGGCATGATTGTTATGCTTGCTNNNAGATCGGAAG |
| 2796_2 | TATGAGTGTGGAGTCGTTGCTTGGCATGATTGTTATGCTTGCTNNNAGATCGGAAG |
| 2796_3 | TATGAGTGTGGAGTCGTTGCTTGGCATGATTGTTATGCTTGCTNNNAGATCGGAAG |
| 5912_1 | TATGAGTGTGGAGTCGTTGCATCACCGTGCCATGAATCAATCANNNAGATCGGAAG |
| 5912_2 | TATGAGTGTGGAGTCGTTGCATCACCGTGCCATGAATCAATCANNNAGATCGGAAG |
| 5912_3 | TATGAGTGTGGAGTCGTTGCATCACCGTGCCATGAATCAATCANNNAGATCGGAAG |
| 29359_1 | TATGAGTGTGGAGTCGTTGCTTTGTATGTCCCTTCCTCCACAGNNNAGATCGGAAGA |
| 29359_2 | TATGAGTGTGGAGTCGTTGCTTTGTATGTCCCTTCCTCCACAGNNNAGATCGGAAGA |
| 29359_3 | TATGAGTGTGGAGTCGTTGCTTTGTATGTCCCTTCCTCCACAGNNNAGATCGGAAGA |
| 46733_1 | TATGAGTGTGGAGTCGTTGCGGTTCAAGTGATTTTCCTGCCTCNNNAGATCGGAAG |
| 46733_2 | TATGAGTGTGGAGTCGTTGCGGTTCAAGTGATTTTCCTGCCTCNNNAGATCGGAAG |
| 46733_3 | TATGAGTGTGGAGTCGTTGCGGTTCAAGTGATTTTCCTGCCTCNNNAGATCGGAAG |
| 13051_1 | TATGAGTGTGGAGTCGTTGCCTAATCCAGAACCACTTTGTANNNAGATCGGAAGAG |
| 13051_2 | TATGAGTGTGGAGTCGTTGCCTAATCCAGAACCACTTTGTANNNAGATCGGAAGAG |
| 13051_3 | TATGAGTGTGGAGTCGTTGCCTAATCCAGAACCACTTTGTANNNAGATCGGAAGAG |
| 45222_1 | TATGAGTGTGGAGTCGTTGCCTGGTCACCATCGCAAACATCNNNAGATCGGAAGAG |
| 45222_2 | TATGAGTGTGGAGTCGTTGCCTGGTCACCATCGCAAACATCNNNAGATCGGAAGAG |
| 45222_3 | TATGAGTGTGGAGTCGTTGCCTGGTCACCATCGCAAACATCNNNAGATCGGAAGAG |
| 52142_1 | TATGAGTGTGGAGTCGTTGCCAAAGGTGGTGAATTCGGTTTCTNNNAGATCGGAA |
| 52142_2 | TATGAGTGTGGAGTCGTTGCCAAAGGTGGTGAATTCGGTTTCTNNNAGATCGGAA |
| 52142_3 | TATGAGTGTGGAGTCGTTGCCAAAGGTGGTGAATTCGGTTTCTNNNAGATCGGAA |
| 44344_1 | TATGAGTGTGGAGTCGTTGCCAGCCTTGGGGTATCTTACTCTCNNNAGATCGGAAG |
| 44344_2 | TATGAGTGTGGAGTCGTTGCCAGCCTTGGGGTATCTTACTCTCNNNAGATCGGAAG |
| 44344_3 | TATGAGTGTGGAGTCGTTGCCAGCCTTGGGGTATCTTACTCTCNNNAGATCGGAAG |
| 32414_1 | TATGAGTGTGGAGTCGTTGCAAAAAAACAAACCTAGCCAGGCANNNAGATCGGAA |
| 32414_2 | TATGAGTGTGGAGTCGTTGCAAAAAAACAAACCTAGCCAGGCANNNAGATCGGAA |
| 32414_3 | TATGAGTGTGGAGTCGTTGCAAAAAAACAAACCTAGCCAGGCANNNAGATCGGAA |
| 39146_1 | TATGAGTGTGGAGTCGTTGCGGTGCAAGGGACATGAATAGACANNNAGATCGGA |

| | |
|---|---|
| 39146_2 | TATGAGTGTGGAGTCGTTGCGGTGCAAGGGACATGAATAGACANNNAGATCGGAA |
| 39146_3 | TATGAGTGTGGAGTCGTTGCGGTGCAAGGGACATGAATAGACANNNAGATCGGAA |
| 50158_1 | TATGAGTGTGGAGTCGTTGCCAAGCCTAGGAATCTTGGTGCTANNNAGATCGGAAG |
| 50158_2 | TATGAGTGTGGAGTCGTTGCCAAGCCTAGGAATCTTGGTGCTANNNAGATCGGAAG |
| 50158_3 | TATGAGTGTGGAGTCGTTGCCAAGCCTAGGAATCTTGGTGCTANNNAGATCGGAAG |
| 919_1 | TATGAGTGTGGAGTCGTTGCGGGTGGTTTTAGGGCTCCATCTNNNAGATCGGAAGA |
| 919_2 | TATGAGTGTGGAGTCGTTGCGGGTGGTTTTAGGGCTCCATCTNNNAGATCGGAAGA |
| 919_3 | TATGAGTGTGGAGTCGTTGCGGGTGGTTTTAGGGCTCCATCTNNNAGATCGGAAGA |
| 46660_1 | TATGAGTGTGGAGTCGTTGCTTCAAGTGATTCTCCTGCCTCAGNNNAGATCGGAAGA |
| 46660_2 | TATGAGTGTGGAGTCGTTGCTTCAAGTGATTCTCCTGCCTCAGNNNAGATCGGAAGA |
| 46660_3 | TATGAGTGTGGAGTCGTTGCTTCAAGTGATTCTCCTGCCTCAGNNNAGATCGGAAGA |
| 38428_1 | TATGAGTGTGGAGTCGTTGCTAAGTCAGAATCCTTGTGGGTGGNNNAGATCGGAA |
| 38428_2 | TATGAGTGTGGAGTCGTTGCTAAGTCAGAATCCTTGTGGGTGGNNNAGATCGGAA |
| 38428_3 | TATGAGTGTGGAGTCGTTGCTAAGTCAGAATCCTTGTGGGTGGNNNAGATCGGAA |
| 48945_1 | TATGAGTGTGGAGTCGTTGCCTTCCCAGGTTCAAGCAATTCTCNNNAGATCGGAAGA |
| 48945_2 | TATGAGTGTGGAGTCGTTGCCTTCCCAGGTTCAAGCAATTCTCNNNAGATCGGAAGA |
| 48945_3 | TATGAGTGTGGAGTCGTTGCCTTCCCAGGTTCAAGCAATTCTCNNNAGATCGGAAGA |
| 36646_1 | TATGAGTGTGGAGTCGTTGCTGGAATCCTGTAAAACGGTCAGTNNNAGATCGGAAG |
| 36646_2 | TATGAGTGTGGAGTCGTTGCTGGAATCCTGTAAAACGGTCAGTNNNAGATCGGAAG |
| 36646_3 | TATGAGTGTGGAGTCGTTGCTGGAATCCTGTAAAACGGTCAGTNNNAGATCGGAAG |
| 48086_1 | TATGAGTGTGGAGTCGTTGCCTAAATGGGATTACAAGTGCGCCNNNAGATCGGAAG |
| 48086_2 | TATGAGTGTGGAGTCGTTGCCTAAATGGGATTACAAGTGCGCCNNNAGATCGGAAG |
| 48086_3 | TATGAGTGTGGAGTCGTTGCCTAAATGGGATTACAAGTGCGCCNNNAGATCGGAAG |
| 39194_1 | TATGAGTGTGGAGTCGTTGCATGCACTCATAGCACTGTTCAGANNNAGATCGGAAG |
| 39194_2 | TATGAGTGTGGAGTCGTTGCATGCACTCATAGCACTGTTCAGANNNAGATCGGAAG |
| 39194_3 | TATGAGTGTGGAGTCGTTGCATGCACTCATAGCACTGTTCAGANNNAGATCGGAAG |
| 46844_1 | TATGAGTGTGGAGTCGTTGCTCCATTTTTACTGTGCATACACGTNNNAGATCGGAAG |
| 46844_2 | TATGAGTGTGGAGTCGTTGCTCCATTTTTACTGTGCATACACGTNNNAGATCGGAAG |
| 46844_3 | TATGAGTGTGGAGTCGTTGCTCCATTTTTACTGTGCATACACGTNNNAGATCGGAAG |
| 37555_1 | TATGAGTGTGGAGTCGTTGCCATGAGCAGCTGAGCAACAATAGNNNAGATCGGAA |
| 37555_2 | TATGAGTGTGGAGTCGTTGCCATGAGCAGCTGAGCAACAATAGNNNAGATCGGAA |
| 37555_3 | TATGAGTGTGGAGTCGTTGCCATGAGCAGCTGAGCAACAATAGNNNAGATCGGAA |
| 52981_1 | TATGAGTGTGGAGTCGTTGCCCCCCACTAGTGACCATGTATACNNNAGATCGGAAGA |
| 52981_2 | TATGAGTGTGGAGTCGTTGCCCCCCACTAGTGACCATGTATACNNNAGATCGGAAGA |
| 52981_3 | TATGAGTGTGGAGTCGTTGCCCCCCACTAGTGACCATGTATACNNNAGATCGGAAGA |
| 53316_1 | TATGAGTGTGGAGTCGTTGCGGTGTGTGTATCTGTGTGTGTTTNNNAGATCGGAA |
| 53316_2 | TATGAGTGTGGAGTCGTTGCGGTGTGTGTATCTGTGTGTGTTTNNNAGATCGGAA |
| 53316_3 | TATGAGTGTGGAGTCGTTGCGGTGTGTGTATCTGTGTGTGTTTNNNAGATCGGAA |
| 30952_1 | TATGAGTGTGGAGTCGTTGCCACACACACACAGACATACACACNNNAGATCGGAAGA |
| 30952_2 | TATGAGTGTGGAGTCGTTGCCACACACACACAGACATACACACNNNAGATCGGAAGA |
| 30952_3 | TATGAGTGTGGAGTCGTTGCCACACACACACAGACATACACACNNNAGATCGGAAGA |
| 52797_1 | TATGAGTGTGGAGTCGTTGCGAGTGGGAGTGGAGGCGAAGNNNAGATCGGAAGA |
| 52797_2 | TATGAGTGTGGAGTCGTTGCGAGTGGGAGTGGAGGCGAAGNNNAGATCGGAAGA |

| | |
|---|---|
| 52797_3 | TATGAGTGTGGAGTCGTTGCGAGTGGGAGTGGAGGCGAAGNNNAGATCGGAAG |
| 12324_1 | TATGAGTGTGGAGTCGTTGCGCTTCACAATCCTAGGCAGTTATNNNAGATCGGAAG |
| 12324_2 | TATGAGTGTGGAGTCGTTGCGCTTCACAATCCTAGGCAGTTATNNNAGATCGGAAG |
| 12324_3 | TATGAGTGTGGAGTCGTTGCGCTTCACAATCCTAGGCAGTTATNNNAGATCGGAAG |
| 41946_1 | TATGAGTGTGGAGTCGTTGCATCTCCTGGGTTCAAGCAATTCTNNNAGATCGGAAG |
| 41946_2 | TATGAGTGTGGAGTCGTTGCATCTCCTGGGTTCAAGCAATTCTNNNAGATCGGAAG |
| 41946_3 | TATGAGTGTGGAGTCGTTGCATCTCCTGGGTTCAAGCAATTCTNNNAGATCGGAAG |
| 36560_1 | TATGAGTGTGGAGTCGTTGCTTGAGCCCAAGAGTTCAAGATCANNNAGATCGGAA |
| 36560_2 | TATGAGTGTGGAGTCGTTGCTTGAGCCCAAGAGTTCAAGATCANNNAGATCGGAA |
| 36560_3 | TATGAGTGTGGAGTCGTTGCTTGAGCCCAAGAGTTCAAGATCANNNAGATCGGAA |
| 40746_1 | TATGAGTGTGGAGTCGTTGCAGCTTTGAAGTAATCGAGATGCTNNNAGATCGGAA |
| 40746_2 | TATGAGTGTGGAGTCGTTGCAGCTTTGAAGTAATCGAGATGCTNNNAGATCGGAA |
| 40746_3 | TATGAGTGTGGAGTCGTTGCAGCTTTGAAGTAATCGAGATGCTNNNAGATCGGAA |
| 42522_1 | TATGAGTGTGGAGTCGTTGCATATGTGTGTGTGTGTGTGTGTGNNNAGATCGGAA |
| 42522_2 | TATGAGTGTGGAGTCGTTGCATATGTGTGTGTGTGTGTGTGTGNNNAGATCGGAA |
| 42522_3 | TATGAGTGTGGAGTCGTTGCATATGTGTGTGTGTGTGTGTGTGNNNAGATCGGAA |
| 31651_1 | TATGAGTGTGGAGTCGTTGCTATAGACGTGTGTGTGTGTGTGTNNNAGATCGGAA |
| 31651_2 | TATGAGTGTGGAGTCGTTGCTATAGACGTGTGTGTGTGTGTGTNNNAGATCGGAA |
| 31651_3 | TATGAGTGTGGAGTCGTTGCTATAGACGTGTGTGTGTGTGTGTNNNAGATCGGAA |
| 51568_1 | TATGAGTGTGGAGTCGTTGCAGAGCCGAGAATGTTCCAAGATTNNNAGATCGGAA |
| 51568_2 | TATGAGTGTGGAGTCGTTGCAGAGCCGAGAATGTTCCAAGATTNNNAGATCGGAA |
| 51568_3 | TATGAGTGTGGAGTCGTTGCAGAGCCGAGAATGTTCCAAGATTNNNAGATCGGAA |
| 2752_1 | TATGAGTGTGGAGTCGTTGCACAACAGATTTGACTGAATACCCTNNNAGATCGGAA |
| 2752_2 | TATGAGTGTGGAGTCGTTGCACAACAGATTTGACTGAATACCCTNNNAGATCGGAA |
| 2752_3 | TATGAGTGTGGAGTCGTTGCACAACAGATTTGACTGAATACCCTNNNAGATCGGAA |
| 43592_1 | TATGAGTGTGGAGTCGTTGCATCACGGCTGAAAACAAGAGAGANNNAGATCGGAA |
| 43592_2 | TATGAGTGTGGAGTCGTTGCATCACGGCTGAAAACAAGAGAGANNNAGATCGGAA |
| 43592_3 | TATGAGTGTGGAGTCGTTGCATCACGGCTGAAAACAAGAGAGANNNAGATCGGAA |
| 29596_1 | TATGAGTGTGGAGTCGTTGCAGATAGGAACAAAATCCCTGGCANNNAGATCGGAA |
| 29596_2 | TATGAGTGTGGAGTCGTTGCAGATAGGAACAAAATCCCTGGCANNNAGATCGGAA |
| 29596_3 | TATGAGTGTGGAGTCGTTGCAGATAGGAACAAAATCCCTGGCANNNAGATCGGAA |
| 41860_1 | TATGAGTGTGGAGTCGTTGCGATGGAGTCTCGCTGTCACCNNNAGATCGGAAGAG |
| 41860_2 | TATGAGTGTGGAGTCGTTGCGATGGAGTCTCGCTGTCACCNNNAGATCGGAAGAG |
| 41860_3 | TATGAGTGTGGAGTCGTTGCGATGGAGTCTCGCTGTCACCNNNAGATCGGAAGAG |
| 52149_1 | TATGAGTGTGGAGTCGTTGCCCACTCCAACACGTGCTCTCNNNAGATCGGAAGAGC |
| 52149_2 | TATGAGTGTGGAGTCGTTGCCCACTCCAACACGTGCTCTCNNNAGATCGGAAGAGC |
| 52149_3 | TATGAGTGTGGAGTCGTTGCCCACTCCAACACGTGCTCTCNNNAGATCGGAAGAGC |
| 3096_1 | TATGAGTGTGGAGTCGTTGCGGGATGTCAGGACACATGTCTTCNNNAGATCGGAA |
| 3096_2 | TATGAGTGTGGAGTCGTTGCGGGATGTCAGGACACATGTCTTCNNNAGATCGGAA |
| 3096_3 | TATGAGTGTGGAGTCGTTGCGGGATGTCAGGACACATGTCTTCNNNAGATCGGAA |
| 47363_1 | TATGAGTGTGGAGTCGTTGCGGGGAAATGCCACCTTCAGGNNNAGATCGGAAGAG |
| 47363_2 | TATGAGTGTGGAGTCGTTGCGGGGAAATGCCACCTTCAGGNNNAGATCGGAAGAG |
| 47363_3 | TATGAGTGTGGAGTCGTTGCGGGGAAATGCCACCTTCAGGNNNAGATCGGAAGAG |

| | |
|---|---|
| 5538_1 | TATGAGTGTGGAGTCGTTGCGTGCAGGTAACTCTTCATCCGTTNNNAGATCGGAAG |
| 5538_2 | TATGAGTGTGGAGTCGTTGCGTGCAGGTAACTCTTCATCCGTTNNNAGATCGGAAG |
| 5538_3 | TATGAGTGTGGAGTCGTTGCGTGCAGGTAACTCTTCATCCGTTNNNAGATCGGAAG |
| 32582_1 | TATGAGTGTGGAGTCGTTGCCTCAAGTAGCTGGGATCACAGTTNNNAGATCGGAAG |
| 32582_2 | TATGAGTGTGGAGTCGTTGCCTCAAGTAGCTGGGATCACAGTTNNNAGATCGGAAG |
| 32582_3 | TATGAGTGTGGAGTCGTTGCCTCAAGTAGCTGGGATCACAGTTNNNAGATCGGAAG |
| 32032_1 | TATGAGTGTGGAGTCGTTGCTCAATTGGCAGAATAAATGCTCAGTNNNAGATCGGA |
| 32032_2 | TATGAGTGTGGAGTCGTTGCTCAATTGGCAGAATAAATGCTCAGTNNNAGATCGGA |
| 32032_3 | TATGAGTGTGGAGTCGTTGCTCAATTGGCAGAATAAATGCTCAGTNNNAGATCGGA |
| 40093_1 | TATGAGTGTGGAGTCGTTGCAGAGGAGGAATGTGTCCACTCTANNNAGATCGGAA |
| 40093_2 | TATGAGTGTGGAGTCGTTGCAGAGGAGGAATGTGTCCACTCTANNNAGATCGGAA |
| 40093_3 | TATGAGTGTGGAGTCGTTGCAGAGGAGGAATGTGTCCACTCTANNNAGATCGGAA |
| 28841_1 | TATGAGTGTGGAGTCGTTGCGGTGTGCAACTTCTCTTCTCCTANNNAGATCGGAAGA |
| 28841_2 | TATGAGTGTGGAGTCGTTGCGGTGTGCAACTTCTCTTCTCCTANNNAGATCGGAAGA |
| 28841_3 | TATGAGTGTGGAGTCGTTGCGGTGTGCAACTTCTCTTCTCCTANNNAGATCGGAAGA |
| 39585_1 | TATGAGTGTGGAGTCGTTGCTCCATTACCACATTCAAGGTCATNNNAGATCGGAAGA |
| 39585_2 | TATGAGTGTGGAGTCGTTGCTCCATTACCACATTCAAGGTCATNNNAGATCGGAAGA |
| 39585_3 | TATGAGTGTGGAGTCGTTGCTCCATTACCACATTCAAGGTCATNNNAGATCGGAAGA |
| 35488_1 | TATGAGTGTGGAGTCGTTGCAGTGCTGGGATTACAGACATGAGNNNAGATCGGAA |
| 35488_2 | TATGAGTGTGGAGTCGTTGCAGTGCTGGGATTACAGACATGAGNNNAGATCGGAA |
| 35488_3 | TATGAGTGTGGAGTCGTTGCAGTGCTGGGATTACAGACATGAGNNNAGATCGGAA |
| 13535_1 | TATGAGTGTGGAGTCGTTGCCTCCTTTAAAATAATGGTTCACANNNAGATCGGAAGA |
| 13535_2 | TATGAGTGTGGAGTCGTTGCCTCCTTTAAAATAATGGTTCACANNNAGATCGGAAGA |
| 13535_3 | TATGAGTGTGGAGTCGTTGCCTCCTTTAAAATAATGGTTCACANNNAGATCGGAAGA |
| 53277_1 | TATGAGTGTGGAGTCGTTGCTTTTCACCTGTCTCACACTCCATNNNAGATCGGAAGA |
| 53277_2 | TATGAGTGTGGAGTCGTTGCTTTTCACCTGTCTCACACTCCATNNNAGATCGGAAGA |
| 53277_3 | TATGAGTGTGGAGTCGTTGCTTTTCACCTGTCTCACACTCCATNNNAGATCGGAAGA |
| 30105_1 | TATGAGTGTGGAGTCGTTGCATAATGGGCAGTGAGCACTAACANNNAGATCGGAA |
| 30105_2 | TATGAGTGTGGAGTCGTTGCATAATGGGCAGTGAGCACTAACANNNAGATCGGAA |
| 30105_3 | TATGAGTGTGGAGTCGTTGCATAATGGGCAGTGAGCACTAACANNNAGATCGGAA |
| 12673_1 | TATGAGTGTGGAGTCGTTGCAAGGTCAGATAAGGAAAATGAAANNNAGATCGGAA |
| 12673_2 | TATGAGTGTGGAGTCGTTGCAAGGTCAGATAAGGAAAATGAAANNNAGATCGGAA |
| 12673_3 | TATGAGTGTGGAGTCGTTGCAAGGTCAGATAAGGAAAATGAAANNNAGATCGGAA |
| 36687_1 | TATGAGTGTGGAGTCGTTGCAAGAAGAAAGAAAATGCGCTGGGNNNAGATCGGAA |
| 36687_2 | TATGAGTGTGGAGTCGTTGCAAGAAGAAAGAAAATGCGCTGGGNNNAGATCGGAA |
| 36687_3 | TATGAGTGTGGAGTCGTTGCAAGAAGAAAGAAAATGCGCTGGGNNNAGATCGGAA |
| 37517_1 | TATGAGTGTGGAGTCGTTGCCTGTAGTCCCAGCTACTTGAGAGNNNAGATCGGAAG |
| 37517_2 | TATGAGTGTGGAGTCGTTGCCTGTAGTCCCAGCTACTTGAGAGNNNAGATCGGAAG |
| 37517_3 | TATGAGTGTGGAGTCGTTGCCTGTAGTCCCAGCTACTTGAGAGNNNAGATCGGAAG |
| 53256_1 | TATGAGTGTGGAGTCGTTGCAAGAATGAAAGCATTGGGACAGGNNNAGATCGGAA |
| 53256_2 | TATGAGTGTGGAGTCGTTGCAAGAATGAAAGCATTGGGACAGGNNNAGATCGGAA |
| 53256_3 | TATGAGTGTGGAGTCGTTGCAAGAATGAAAGCATTGGGACAGGNNNAGATCGGAA |
| 38064_1 | TATGAGTGTGGAGTCGTTGCGCTTAGGTGATCACTTGAGTCCANNNAGATCGGAAG |

| | |
|---|---|
| 38064_2 | TATGAGTGTGGAGTCGTTGCGCTTAGGTGATCACTTGAGTCCANNNAGATCGGAAG |
| 38064_3 | TATGAGTGTGGAGTCGTTGCGCTTAGGTGATCACTTGAGTCCANNNAGATCGGAAG |
| 41856_1 | TATGAGTGTGGAGTCGTTGCACTCCCGAGTAGCTGGAACTATANNNAGATCGGAAG |
| 41856_2 | TATGAGTGTGGAGTCGTTGCACTCCCGAGTAGCTGGAACTATANNNAGATCGGAAG |
| 41856_3 | TATGAGTGTGGAGTCGTTGCACTCCCGAGTAGCTGGAACTATANNNAGATCGGAAG |
| 52484_1 | TATGAGTGTGGAGTCGTTGCGCAAGACGTTCAGTCCTACAGAANNNAGATCGGAAG |
| 52484_2 | TATGAGTGTGGAGTCGTTGCGCAAGACGTTCAGTCCTACAGAANNNAGATCGGAAG |
| 52484_3 | TATGAGTGTGGAGTCGTTGCGCAAGACGTTCAGTCCTACAGAANNNAGATCGGAAG |
| 36184_1 | TATGAGTGTGGAGTCGTTGCAATTTCCTAGCACTGCCCAGTAANNNAGATCGGAAGA |
| 36184_2 | TATGAGTGTGGAGTCGTTGCAATTTCCTAGCACTGCCCAGTAANNNAGATCGGAAGA |
| 36184_3 | TATGAGTGTGGAGTCGTTGCAATTTCCTAGCACTGCCCAGTAANNNAGATCGGAAGA |
| 31711_1 | TATGAGTGTGGAGTCGTTGCCCATCTCCCACTCAGAGTCTTTCNNNAGATCGGAAGA |
| 31711_2 | TATGAGTGTGGAGTCGTTGCCCATCTCCCACTCAGAGTCTTTCNNNAGATCGGAAGA |
| 31711_3 | TATGAGTGTGGAGTCGTTGCCCATCTCCCACTCAGAGTCTTTCNNNAGATCGGAAGA |
| 53240_1 | TATGAGTGTGGAGTCGTTGCAAATAGTCCCTCTTGTCTGCAGGNNNAGATCGGAAG |
| 53240_2 | TATGAGTGTGGAGTCGTTGCAAATAGTCCCTCTTGTCTGCAGGNNNAGATCGGAAG |
| 53240_3 | TATGAGTGTGGAGTCGTTGCAAATAGTCCCTCTTGTCTGCAGGNNNAGATCGGAAG |
| 53038_1 | TATGAGTGTGGAGTCGTTGCGCCTCATATTGTCCTGCTTTTCCNNNAGATCGGAAGA |
| 53038_2 | TATGAGTGTGGAGTCGTTGCGCCTCATATTGTCCTGCTTTTCCNNNAGATCGGAAGA |
| 53038_3 | TATGAGTGTGGAGTCGTTGCGCCTCATATTGTCCTGCTTTTCCNNNAGATCGGAAGA |
| 43217_1 | TATGAGTGTGGAGTCGTTGCGACTGTACTGCCGTGATCTCAGNNNAGATCGGAAGA |
| 43217_2 | TATGAGTGTGGAGTCGTTGCGACTGTACTGCCGTGATCTCAGNNNAGATCGGAAGA |
| 43217_3 | TATGAGTGTGGAGTCGTTGCGACTGTACTGCCGTGATCTCAGNNNAGATCGGAAGA |
| 50901_1 | TATGAGTGTGGAGTCGTTGCTTACCAGTTCACTTGTCCAGAGGNNNAGATCGGAAG |
| 50901_2 | TATGAGTGTGGAGTCGTTGCTTACCAGTTCACTTGTCCAGAGGNNNAGATCGGAAG |
| 50901_3 | TATGAGTGTGGAGTCGTTGCTTACCAGTTCACTTGTCCAGAGGNNNAGATCGGAAG |
| 34030_1 | TATGAGTGTGGAGTCGTTGCGTTTCACCATGTTAGCCAGGATGNNNAGATCGGAAG |
| 34030_2 | TATGAGTGTGGAGTCGTTGCGTTTCACCATGTTAGCCAGGATGNNNAGATCGGAAG |
| 34030_3 | TATGAGTGTGGAGTCGTTGCGTTTCACCATGTTAGCCAGGATGNNNAGATCGGAAG |
| 41531_1 | TATGAGTGTGGAGTCGTTGCACCAAATGATACCAAAACATGCTGNNNAGATCGGAA |
| 41531_2 | TATGAGTGTGGAGTCGTTGCACCAAATGATACCAAAACATGCTGNNNAGATCGGAA |
| 41531_3 | TATGAGTGTGGAGTCGTTGCACCAAATGATACCAAAACATGCTGNNNAGATCGGAA |
| 40409_1 | TATGAGTGTGGAGTCGTTGCTCACGTCTGTAATCCCAGCATTTNNNAGATCGGAAGA |
| 40409_2 | TATGAGTGTGGAGTCGTTGCTCACGTCTGTAATCCCAGCATTTNNNAGATCGGAAGA |
| 40409_3 | TATGAGTGTGGAGTCGTTGCTCACGTCTGTAATCCCAGCATTTNNNAGATCGGAAGA |
| 2637_1 | TATGAGTGTGGAGTCGTTGCAGGAATACGAAGTTGACGGTGTGNNNAGATCGGAA |
| 2637_2 | TATGAGTGTGGAGTCGTTGCAGGAATACGAAGTTGACGGTGTGNNNAGATCGGAA |
| 2637_3 | TATGAGTGTGGAGTCGTTGCAGGAATACGAAGTTGACGGTGTGNNNAGATCGGAA |
| 39641_1 | TATGAGTGTGGAGTCGTTGCCTGTCATTTGCACCAACATGGATNNNAGATCGGAAG |
| 39641_2 | TATGAGTGTGGAGTCGTTGCCTGTCATTTGCACCAACATGGATNNNAGATCGGAAG |
| 39641_3 | TATGAGTGTGGAGTCGTTGCCTGTCATTTGCACCAACATGGATNNNAGATCGGAAG |
| 42571_1 | TATGAGTGTGGAGTCGTTGCTTCAGTTCAGCTCTGACTTTGGTNNNAGATCGGAAG |
| 42571_2 | TATGAGTGTGGAGTCGTTGCTTCAGTTCAGCTCTGACTTTGGTNNNAGATCGGAAG |

| | |
|---|---|
| 42571_3 | TATGAGTGTGGAGTCGTTGCTTCAGTTCAGCTCTGACTTTGGTNNNAGATCGGAAG |
| 37672_1 | TATGAGTGTGGAGTCGTTGCTGTGTGTATAAGCATGTCTGTGCNNNAGATCGGAAG |
| 37672_2 | TATGAGTGTGGAGTCGTTGCTGTGTGTATAAGCATGTCTGTGCNNNAGATCGGAAG |
| 37672_3 | TATGAGTGTGGAGTCGTTGCTGTGTGTATAAGCATGTCTGTGCNNNAGATCGGAAG |
| 38367_1 | TATGAGTGTGGAGTCGTTGCCGTGACAGAGTGAGACTTCATCTNNNAGATCGGAAG |
| 38367_2 | TATGAGTGTGGAGTCGTTGCCGTGACAGAGTGAGACTTCATCTNNNAGATCGGAAG |
| 38367_3 | TATGAGTGTGGAGTCGTTGCCGTGACAGAGTGAGACTTCATCTNNNAGATCGGAAG |
| 36995_1 | TATGAGTGTGGAGTCGTTGCCTGAATGCTCGAGGAGACTGATTNNNAGATCGGAA |
| 36995_2 | TATGAGTGTGGAGTCGTTGCCTGAATGCTCGAGGAGACTGATTNNNAGATCGGAA |
| 36995_3 | TATGAGTGTGGAGTCGTTGCCTGAATGCTCGAGGAGACTGATTNNNAGATCGGAA |
| 32360_1 | TATGAGTGTGGAGTCGTTGCTCTCAGTGGAAGTTGGGATTTCCNNNAGATCGGAAG |
| 32360_2 | TATGAGTGTGGAGTCGTTGCTCTCAGTGGAAGTTGGGATTTCCNNNAGATCGGAAG |
| 32360_3 | TATGAGTGTGGAGTCGTTGCTCTCAGTGGAAGTTGGGATTTCCNNNAGATCGGAAG |
| 46820_1 | TATGAGTGTGGAGTCGTTGCCATTTTCCTGCCTCAGCCTTCTNNNAGATCGGAAGAG |
| 46820_2 | TATGAGTGTGGAGTCGTTGCCATTTTCCTGCCTCAGCCTTCTNNNAGATCGGAAGAG |
| 46820_3 | TATGAGTGTGGAGTCGTTGCCATTTTCCTGCCTCAGCCTTCTNNNAGATCGGAAGAG |
| 38550_1 | TATGAGTGTGGAGTCGTTGCTGCATGGATTTGTTTCTGGGTTCNNNAGATCGGAAG |
| 38550_2 | TATGAGTGTGGAGTCGTTGCTGCATGGATTTGTTTCTGGGTTCNNNAGATCGGAAG |
| 38550_3 | TATGAGTGTGGAGTCGTTGCTGCATGGATTTGTTTCTGGGTTCNNNAGATCGGAAG |
| 30781_1 | TATGAGTGTGGAGTCGTTGCGGACCATGGGCAGATAGAAAGAANNNAGATCGGA |
| 30781_2 | TATGAGTGTGGAGTCGTTGCGGACCATGGGCAGATAGAAAGAANNNAGATCGGA |
| 30781_3 | TATGAGTGTGGAGTCGTTGCGGACCATGGGCAGATAGAAAGAANNNAGATCGGA |
| 46841_1 | TATGAGTGTGGAGTCGTTGCGCAGGGATAGCATGTTGTCAAATNNNAGATCGGAA |
| 46841_2 | TATGAGTGTGGAGTCGTTGCGCAGGGATAGCATGTTGTCAAATNNNAGATCGGAA |
| 46841_3 | TATGAGTGTGGAGTCGTTGCGCAGGGATAGCATGTTGTCAAATNNNAGATCGGAA |
| 32347_1 | TATGAGTGTGGAGTCGTTGCTGTTGTTGTTGTTGAGACAGAGTNNNAGATCGGAA |
| 32347_2 | TATGAGTGTGGAGTCGTTGCTGTTGTTGTTGTTGAGACAGAGTNNNAGATCGGAA |
| 32347_3 | TATGAGTGTGGAGTCGTTGCTGTTGTTGTTGTTGAGACAGAGTNNNAGATCGGAA |
| 49866_1 | TATGAGTGTGGAGTCGTTGCTCTTGCTGTGTGAATGTTATGGCNNNAGATCGGAAG |
| 49866_2 | TATGAGTGTGGAGTCGTTGCTCTTGCTGTGTGAATGTTATGGCNNNAGATCGGAAG |
| 49866_3 | TATGAGTGTGGAGTCGTTGCTCTTGCTGTGTGAATGTTATGGCNNNAGATCGGAAG |
| 48656_1 | TATGAGTGTGGAGTCGTTGCTTTCCTGCACAACACCTTGTTTTNNNAGATCGGAAGA |
| 48656_2 | TATGAGTGTGGAGTCGTTGCTTTCCTGCACAACACCTTGTTTTNNNAGATCGGAAGA |
| 48656_3 | TATGAGTGTGGAGTCGTTGCTTTCCTGCACAACACCTTGTTTTNNNAGATCGGAAGA |
| 41694_1 | TATGAGTGTGGAGTCGTTGCCTCCTGAGTAGCTGGGACTACAGNNNAGATCGGAA |
| 41694_2 | TATGAGTGTGGAGTCGTTGCCTCCTGAGTAGCTGGGACTACAGNNNAGATCGGAA |
| 41694_3 | TATGAGTGTGGAGTCGTTGCCTCCTGAGTAGCTGGGACTACAGNNNAGATCGGAA |
| 36852_1 | TATGAGTGTGGAGTCGTTGCAAACTTTGATTCTGTGCGTCCACNNNAGATCGGAAG |
| 36852_2 | TATGAGTGTGGAGTCGTTGCAAACTTTGATTCTGTGCGTCCACNNNAGATCGGAAG |
| 36852_3 | TATGAGTGTGGAGTCGTTGCAAACTTTGATTCTGTGCGTCCACNNNAGATCGGAAG |
| 43828_1 | TATGAGTGTGGAGTCGTTGCCCTTCCTTGAGAGTGGTTTCAGTNNNAGATCGGAAG |
| 43828_2 | TATGAGTGTGGAGTCGTTGCCCTTCCTTGAGAGTGGTTTCAGTNNNAGATCGGAAG |
| 43828_3 | TATGAGTGTGGAGTCGTTGCCCTTCCTTGAGAGTGGTTTCAGTNNNAGATCGGAAG |

| | |
|---|---|
| 29530_1 | TATGAGTGTGGAGTCGTTGCTAAGGAAGCTGTTGCTGGTTTGTNNNAGATCGGAA |
| 29530_2 | TATGAGTGTGGAGTCGTTGCTAAGGAAGCTGTTGCTGGTTTGTNNNAGATCGGAA |
| 29530_3 | TATGAGTGTGGAGTCGTTGCTAAGGAAGCTGTTGCTGGTTTGTNNNAGATCGGAA |
| 30477_1 | TATGAGTGTGGAGTCGTTGCGAAGGGGAGGGGGTTTAGATTGANNNAGATCGGA |
| 30477_2 | TATGAGTGTGGAGTCGTTGCGAAGGGGAGGGGGTTTAGATTGANNNAGATCGGA |
| 30477_3 | TATGAGTGTGGAGTCGTTGCGAAGGGGAGGGGGTTTAGATTGANNNAGATCGGA |
| 45287_1 | TATGAGTGTGGAGTCGTTGCAGGACAGCAGTAGAAATCACACTNNNAGATCGGAA |
| 45287_2 | TATGAGTGTGGAGTCGTTGCAGGACAGCAGTAGAAATCACACTNNNAGATCGGAA |
| 45287_3 | TATGAGTGTGGAGTCGTTGCAGGACAGCAGTAGAAATCACACTNNNAGATCGGAA |
| 52361_1 | TATGAGTGTGGAGTCGTTGCACTTCTCAAATGCTACCCCAGTTNNNAGATCGGAAG |
| 52361_2 | TATGAGTGTGGAGTCGTTGCACTTCTCAAATGCTACCCCAGTTNNNAGATCGGAAG |
| 52361_3 | TATGAGTGTGGAGTCGTTGCACTTCTCAAATGCTACCCCAGTTNNNAGATCGGAAG |
| 33582_1 | TATGAGTGTGGAGTCGTTGCGTTCAAGAGATTCTCCTGCCTCANNNAGATCGGAAG |
| 33582_2 | TATGAGTGTGGAGTCGTTGCGTTCAAGAGATTCTCCTGCCTCANNNAGATCGGAAG |
| 33582_3 | TATGAGTGTGGAGTCGTTGCGTTCAAGAGATTCTCCTGCCTCANNNAGATCGGAAG |
| 43496_1 | TATGAGTGTGGAGTCGTTGCAGCCTCTCGAGTAGTTGGTACTANNNAGATCGGAAG |
| 43496_2 | TATGAGTGTGGAGTCGTTGCAGCCTCTCGAGTAGTTGGTACTANNNAGATCGGAAG |
| 43496_3 | TATGAGTGTGGAGTCGTTGCAGCCTCTCGAGTAGTTGGTACTANNNAGATCGGAAG |
| 41775_1 | TATGAGTGTGGAGTCGTTGCCTCTCAAGTAGCTGGGACTACAGNNNAGATCGGAA |
| 41775_2 | TATGAGTGTGGAGTCGTTGCCTCTCAAGTAGCTGGGACTACAGNNNAGATCGGAA |
| 41775_3 | TATGAGTGTGGAGTCGTTGCCTCTCAAGTAGCTGGGACTACAGNNNAGATCGGAA |
| 12746_1 | TATGAGTGTGGAGTCGTTGCGTTCTAACCTAATCACCTATCTTNNNAGATCGGAAGA |
| 12746_2 | TATGAGTGTGGAGTCGTTGCGTTCTAACCTAATCACCTATCTTNNNAGATCGGAAGA |
| 12746_3 | TATGAGTGTGGAGTCGTTGCGTTCTAACCTAATCACCTATCTTNNNAGATCGGAAGA |
| 50785_1 | TATGAGTGTGGAGTCGTTGCATAAAAGATGCAGCTGTTGTGGCNNNAGATCGGAA |
| 50785_2 | TATGAGTGTGGAGTCGTTGCATAAAAGATGCAGCTGTTGTGGCNNNAGATCGGAA |
| 50785_3 | TATGAGTGTGGAGTCGTTGCATAAAAGATGCAGCTGTTGTGGCNNNAGATCGGAA |
| 43637_1 | TATGAGTGTGGAGTCGTTGCATCATTTCTCCCCACTCTGCATCNNNAGATCGGAAGA |
| 43637_2 | TATGAGTGTGGAGTCGTTGCATCATTTCTCCCCACTCTGCATCNNNAGATCGGAAGA |
| 43637_3 | TATGAGTGTGGAGTCGTTGCATCATTTCTCCCCACTCTGCATCNNNAGATCGGAAGA |
| 35516_1 | TATGAGTGTGGAGTCGTTGCAGAGTTGGGATTACAGGCATGAGNNNAGATCGGAA |
| 35516_2 | TATGAGTGTGGAGTCGTTGCAGAGTTGGGATTACAGGCATGAGNNNAGATCGGAA |
| 35516_3 | TATGAGTGTGGAGTCGTTGCAGAGTTGGGATTACAGGCATGAGNNNAGATCGGAA |
| 43679_1 | TATGAGTGTGGAGTCGTTGCCCTCGTCCTCCTCCATCCTAGNNNAGATCGGAAGAGC |
| 43679_2 | TATGAGTGTGGAGTCGTTGCCCTCGTCCTCCTCCATCCTAGNNNAGATCGGAAGAGC |
| 43679_3 | TATGAGTGTGGAGTCGTTGCCCTCGTCCTCCTCCATCCTAGNNNAGATCGGAAGAGC |
| 38057_1 | TATGAGTGTGGAGTCGTTGCGGGGAAAAGAAGGATGAGACCAGNNNAGATCGGA |
| 38057_2 | TATGAGTGTGGAGTCGTTGCGGGGAAAAGAAGGATGAGACCAGNNNAGATCGGA |
| 38057_3 | TATGAGTGTGGAGTCGTTGCGGGGAAAAGAAGGATGAGACCAGNNNAGATCGGA |
| 45738_1 | TATGAGTGTGGAGTCGTTGCCACCACGCCCAGCTAATTTTTTNNNAGATCGGAAGA |
| 45738_2 | TATGAGTGTGGAGTCGTTGCCACCACGCCCAGCTAATTTTTTNNNAGATCGGAAGA |
| 45738_3 | TATGAGTGTGGAGTCGTTGCCACCACGCCCAGCTAATTTTTTNNNAGATCGGAAGA |
| 50052_1 | TATGAGTGTGGAGTCGTTGCAGAGTCAACCTGCCTTGAGAAGNNNAGATCGGAAG |

| | |
|---|---|
| 50052_2 | TATGAGTGTGGAGTCGTTGCAGAGTCAACCTGCCTTGAGAAGNNNAGATCGGAAG |
| 50052_3 | TATGAGTGTGGAGTCGTTGCAGAGTCAACCTGCCTTGAGAAGNNNAGATCGGAAG |
| 30554_1 | TATGAGTGTGGAGTCGTTGCTTGAGTGAGGAGGGGAGATCAAGNNNAGATCGGA |
| 30554_2 | TATGAGTGTGGAGTCGTTGCTTGAGTGAGGAGGGGAGATCAAGNNNAGATCGGA |
| 30554_3 | TATGAGTGTGGAGTCGTTGCTTGAGTGAGGAGGGGAGATCAAGNNNAGATCGGA |
| 45215_1 | TATGAGTGTGGAGTCGTTGCTCCCTGTCCAGAGAAGACTAGACNNNAGATCGGAAG |
| 45215_2 | TATGAGTGTGGAGTCGTTGCTCCCTGTCCAGAGAAGACTAGACNNNAGATCGGAAG |
| 45215_3 | TATGAGTGTGGAGTCGTTGCTCCCTGTCCAGAGAAGACTAGACNNNAGATCGGAAG |
| 44178_1 | TATGAGTGTGGAGTCGTTGCCCAGGACTAGGAGGTGGGAGNNNAGATCGGAAGA |
| 44178_2 | TATGAGTGTGGAGTCGTTGCCCAGGACTAGGAGGTGGGAGNNNAGATCGGAAGA |
| 44178_3 | TATGAGTGTGGAGTCGTTGCCCAGGACTAGGAGGTGGGAGNNNAGATCGGAAGA |
| 33125_1 | TATGAGTGTGGAGTCGTTGCTCTGGTATTCCCTAGCTCTGTCANNNAGATCGGAAG |
| 33125_2 | TATGAGTGTGGAGTCGTTGCTCTGGTATTCCCTAGCTCTGTCANNNAGATCGGAAG |
| 33125_3 | TATGAGTGTGGAGTCGTTGCTCTGGTATTCCCTAGCTCTGTCANNNAGATCGGAAG |
| 46702_1 | TATGAGTGTGGAGTCGTTGCAAGTACAGTAGGGCCATCATCACNNNAGATCGGAA |
| 46702_2 | TATGAGTGTGGAGTCGTTGCAAGTACAGTAGGGCCATCATCACNNNAGATCGGAA |
| 46702_3 | TATGAGTGTGGAGTCGTTGCAAGTACAGTAGGGCCATCATCACNNNAGATCGGAA |
| 29009_1 | TATGAGTGTGGAGTCGTTGCATGGAAGCTGAGAAGTTCCATGANNNAGATCGGAA |
| 29009_2 | TATGAGTGTGGAGTCGTTGCATGGAAGCTGAGAAGTTCCATGANNNAGATCGGAA |
| 29009_3 | TATGAGTGTGGAGTCGTTGCATGGAAGCTGAGAAGTTCCATGANNNAGATCGGAA |
| 38121_1 | TATGAGTGTGGAGTCGTTGCCTGAGGTAGGAGGATTGCTTGAGNNNAGATCGGA |
| 38121_2 | TATGAGTGTGGAGTCGTTGCCTGAGGTAGGAGGATTGCTTGAGNNNAGATCGGA |
| 38121_3 | TATGAGTGTGGAGTCGTTGCCTGAGGTAGGAGGATTGCTTGAGNNNAGATCGGA |
| 43856_1 | TATGAGTGTGGAGTCGTTGCCATCCTCTTAGCCACACAAGACTNNNAGATCGGAAG |
| 43856_2 | TATGAGTGTGGAGTCGTTGCCATCCTCTTAGCCACACAAGACTNNNAGATCGGAAG |
| 43856_3 | TATGAGTGTGGAGTCGTTGCCATCCTCTTAGCCACACAAGACTNNNAGATCGGAAG |
| 12671_1 | TATGAGTGTGGAGTCGTTGCATGTTTTAGTTCAACAGAATACCNNNAGATCGGAAG |
| 12671_2 | TATGAGTGTGGAGTCGTTGCATGTTTTAGTTCAACAGAATACCNNNAGATCGGAAG |
| 12671_3 | TATGAGTGTGGAGTCGTTGCATGTTTTAGTTCAACAGAATACCNNNAGATCGGAAG |
| 5026_1 | TATGAGTGTGGAGTCGTTGCTACCCCAATCTCCAAAGTTTGCCNNNAGATCGGAAG |
| 5026_2 | TATGAGTGTGGAGTCGTTGCTACCCCAATCTCCAAAGTTTGCCNNNAGATCGGAAG |
| 5026_3 | TATGAGTGTGGAGTCGTTGCTACCCCAATCTCCAAAGTTTGCCNNNAGATCGGAAG |
| 13769_1 | TATGAGTGTGGAGTCGTTGCAACTTTTGTCTTTCTTATGAACCNNNAGATCGGAAGA |
| 13769_2 | TATGAGTGTGGAGTCGTTGCAACTTTTGTCTTTCTTATGAACCNNNAGATCGGAAGA |
| 13769_3 | TATGAGTGTGGAGTCGTTGCAACTTTTGTCTTTCTTATGAACCNNNAGATCGGAAGA |
| 44314_1 | TATGAGTGTGGAGTCGTTGCCCAATGTTTGTCATGGGTGTGAANNNAGATCGGAA |
| 44314_2 | TATGAGTGTGGAGTCGTTGCCCAATGTTTGTCATGGGTGTGAANNNAGATCGGAA |
| 44314_3 | TATGAGTGTGGAGTCGTTGCCCAATGTTTGTCATGGGTGTGAANNNAGATCGGAA |
| 50463_1 | TATGAGTGTGGAGTCGTTGCAAAGCCTTCGACGTCATGAGTAANNNAGATCGGAA |
| 50463_2 | TATGAGTGTGGAGTCGTTGCAAAGCCTTCGACGTCATGAGTAANNNAGATCGGAA |
| 50463_3 | TATGAGTGTGGAGTCGTTGCAAAGCCTTCGACGTCATGAGTAANNNAGATCGGAA |
| 39410_1 | TATGAGTGTGGAGTCGTTGCACTGTCATGGAAAAATGCTCGTGNNNAGATCGGAA |
| 39410_2 | TATGAGTGTGGAGTCGTTGCACTGTCATGGAAAAATGCTCGTGNNNAGATCGGAA |

| | |
|---|---|
| 39410_3 | TATGAGTGTGGAGTCGTTGCACTGTCATGGAAAAATGCTCGTGNNNAGATCGGAA |
| 33395_1 | TATGAGTGTGGAGTCGTTGCGAAGAAGACCTGACAATGCCTTGNNNAGATCGGAA |
| 33395_2 | TATGAGTGTGGAGTCGTTGCGAAGAAGACCTGACAATGCCTTGNNNAGATCGGAA |
| 33395_3 | TATGAGTGTGGAGTCGTTGCGAAGAAGACCTGACAATGCCTTGNNNAGATCGGAA |
| 38712_1 | TATGAGTGTGGAGTCGTTGCGCAATCCTCCTGCCTCATTTTTANNNAGATCGGAAGA |
| 38712_2 | TATGAGTGTGGAGTCGTTGCGCAATCCTCCTGCCTCATTTTTANNNAGATCGGAAGA |
| 38712_3 | TATGAGTGTGGAGTCGTTGCGCAATCCTCCTGCCTCATTTTTANNNAGATCGGAAGA |
| 52872_1 | TATGAGTGTGGAGTCGTTGCAGGGCCTCTGGTTTGTTGTATTANNNAGATCGGAAG |
| 52872_2 | TATGAGTGTGGAGTCGTTGCAGGGCCTCTGGTTTGTTGTATTANNNAGATCGGAAG |
| 52872_3 | TATGAGTGTGGAGTCGTTGCAGGGCCTCTGGTTTGTTGTATTANNNAGATCGGAAG |
| 53154_1 | TATGAGTGTGGAGTCGTTGCTTTCCATCACTGGGAGGAGAAAGNNNAGATCGGAA |
| 53154_2 | TATGAGTGTGGAGTCGTTGCTTTCCATCACTGGGAGGAGAAAGNNNAGATCGGAA |
| 53154_3 | TATGAGTGTGGAGTCGTTGCTTTCCATCACTGGGAGGAGAAAGNNNAGATCGGAA |
| 32574_1 | TATGAGTGTGGAGTCGTTGCATCCTATCAACTCAGCCTCCTCTNNNAGATCGGAAGA |
| 32574_2 | TATGAGTGTGGAGTCGTTGCATCCTATCAACTCAGCCTCCTCTNNNAGATCGGAAGA |
| 32574_3 | TATGAGTGTGGAGTCGTTGCATCCTATCAACTCAGCCTCCTCTNNNAGATCGGAAGA |
| 51311_1 | TATGAGTGTGGAGTCGTTGCGAGTGGGAAGATGATATTGCCCTNNNAGATCGGAA |
| 51311_2 | TATGAGTGTGGAGTCGTTGCGAGTGGGAAGATGATATTGCCCTNNNAGATCGGAA |
| 51311_3 | TATGAGTGTGGAGTCGTTGCGAGTGGGAAGATGATATTGCCCTNNNAGATCGGAA |
| 50175_1 | TATGAGTGTGGAGTCGTTGCTTTATCCATCCACCCATCCATCCNNNAGATCGGAAGA |
| 50175_2 | TATGAGTGTGGAGTCGTTGCTTTATCCATCCACCCATCCATCCNNNAGATCGGAAGA |
| 50175_3 | TATGAGTGTGGAGTCGTTGCTTTATCCATCCACCCATCCATCCNNNAGATCGGAAGA |
| 11907_1 | TATGAGTGTGGAGTCGTTGCACTTACCTGCTACCTGGAGCGTTNNNAGATCGGAAGA |
| 11907_2 | TATGAGTGTGGAGTCGTTGCACTTACCTGCTACCTGGAGCGTTNNNAGATCGGAAGA |
| 11907_3 | TATGAGTGTGGAGTCGTTGCACTTACCTGCTACCTGGAGCGTTNNNAGATCGGAAGA |
| 46796_1 | TATGAGTGTGGAGTCGTTGCCTCCTAGGCTCAAGTGATTCTCCNNNAGATCGGAAGA |
| 46796_2 | TATGAGTGTGGAGTCGTTGCCTCCTAGGCTCAAGTGATTCTCCNNNAGATCGGAAGA |
| 46796_3 | TATGAGTGTGGAGTCGTTGCCTCCTAGGCTCAAGTGATTCTCCNNNAGATCGGAAGA |
| 36058_1 | TATGAGTGTGGAGTCGTTGCTTGGAGACAGAGAGACACAGAGANNNAGATCGGAA |
| 36058_2 | TATGAGTGTGGAGTCGTTGCTTGGAGACAGAGAGACACAGAGANNNAGATCGGAA |
| 36058_3 | TATGAGTGTGGAGTCGTTGCTTGGAGACAGAGAGACACAGAGANNNAGATCGGAA |
| 33016_1 | TATGAGTGTGGAGTCGTTGCCGCACGGCCATTTTTTTGTATTTNNNAGATCGGAAGA |
| 33016_2 | TATGAGTGTGGAGTCGTTGCCGCACGGCCATTTTTTTGTATTTNNNAGATCGGAAGA |
| 33016_3 | TATGAGTGTGGAGTCGTTGCCGCACGGCCATTTTTTTGTATTTNNNAGATCGGAAGA |
| 43075_1 | TATGAGTGTGGAGTCGTTGCGAAGCAGAAAAACTTGCCTTCCTNNNAGATCGGAAG |
| 43075_2 | TATGAGTGTGGAGTCGTTGCGAAGCAGAAAAACTTGCCTTCCTNNNAGATCGGAAG |
| 43075_3 | TATGAGTGTGGAGTCGTTGCGAAGCAGAAAAACTTGCCTTCCTNNNAGATCGGAAG |
| 45680_1 | TATGAGTGTGGAGTCGTTGCCGAGTAGCTGGGATTACAGGCNNNAGATCGGAAGA |
| 45680_2 | TATGAGTGTGGAGTCGTTGCCGAGTAGCTGGGATTACAGGCNNNAGATCGGAAGA |
| 45680_3 | TATGAGTGTGGAGTCGTTGCCGAGTAGCTGGGATTACAGGCNNNAGATCGGAAGA |
| 35091_1 | TATGAGTGTGGAGTCGTTGCCAAAGTGCTAGGATTACAGGGGTNNNAGATCGGAA |
| 35091_2 | TATGAGTGTGGAGTCGTTGCCAAAGTGCTAGGATTACAGGGGTNNNAGATCGGAA |
| 35091_3 | TATGAGTGTGGAGTCGTTGCCAAAGTGCTAGGATTACAGGGGTNNNAGATCGGAA |

| | |
|---|---|
| 39116_1 | TATGAGTGTGGAGTCGTTGCATGCTAGAGAGGTTGTGGAGAACNNNAGATCGGA |
| 39116_2 | TATGAGTGTGGAGTCGTTGCATGCTAGAGAGGTTGTGGAGAACNNNAGATCGGA |
| 39116_3 | TATGAGTGTGGAGTCGTTGCATGCTAGAGAGGTTGTGGAGAACNNNAGATCGGA |
| 30557_1 | TATGAGTGTGGAGTCGTTGCTCATCACCTGGGTCAATGCCTGNNNAGATCGGAAGA |
| 30557_2 | TATGAGTGTGGAGTCGTTGCTCATCACCTGGGTCAATGCCTGNNNAGATCGGAAGA |
| 30557_3 | TATGAGTGTGGAGTCGTTGCTCATCACCTGGGTCAATGCCTGNNNAGATCGGAAGA |
| 49617_1 | TATGAGTGTGGAGTCGTTGCAATCTGAGGGCAGGAACTATGTCNNNAGATCGGAA |
| 49617_2 | TATGAGTGTGGAGTCGTTGCAATCTGAGGGCAGGAACTATGTCNNNAGATCGGAA |
| 49617_3 | TATGAGTGTGGAGTCGTTGCAATCTGAGGGCAGGAACTATGTCNNNAGATCGGAA |
| 32491_1 | TATGAGTGTGGAGTCGTTGCTGGTGGCTTATCCCTGTAATTCCNNNAGATCGGAAG |
| 32491_2 | TATGAGTGTGGAGTCGTTGCTGGTGGCTTATCCCTGTAATTCCNNNAGATCGGAAG |
| 32491_3 | TATGAGTGTGGAGTCGTTGCTGGTGGCTTATCCCTGTAATTCCNNNAGATCGGAAG |
| 50166_1 | TATGAGTGTGGAGTCGTTGCCCCAGATAACTGCTGCTGACATANNNAGATCGGAAG |
| 50166_2 | TATGAGTGTGGAGTCGTTGCCCCAGATAACTGCTGCTGACATANNNAGATCGGAAG |
| 50166_3 | TATGAGTGTGGAGTCGTTGCCCCAGATAACTGCTGCTGACATANNNAGATCGGAAG |
| 34234_1 | TATGAGTGTGGAGTCGTTGCTGGTGCCATTCCATTCTACAAGTNNNAGATCGGAAG |
| 34234_2 | TATGAGTGTGGAGTCGTTGCTGGTGCCATTCCATTCTACAAGTNNNAGATCGGAAG |
| 34234_3 | TATGAGTGTGGAGTCGTTGCTGGTGCCATTCCATTCTACAAGTNNNAGATCGGAAG |
| 52908_1 | TATGAGTGTGGAGTCGTTGCAGAAAAGCTATGGGAGAAAGCCTNNNAGATCGGAA |
| 52908_2 | TATGAGTGTGGAGTCGTTGCAGAAAAGCTATGGGAGAAAGCCTNNNAGATCGGAA |
| 52908_3 | TATGAGTGTGGAGTCGTTGCAGAAAAGCTATGGGAGAAAGCCTNNNAGATCGGAA |
| 50881_1 | TATGAGTGTGGAGTCGTTGCCTTTGAGTTTGTGAGCTTCCCTGNNNAGATCGGAAG |
| 50881_2 | TATGAGTGTGGAGTCGTTGCCTTTGAGTTTGTGAGCTTCCCTGNNNAGATCGGAAG |
| 50881_3 | TATGAGTGTGGAGTCGTTGCCTTTGAGTTTGTGAGCTTCCCTGNNNAGATCGGAAG |
| 50376_1 | TATGAGTGTGGAGTCGTTGCGCTTATAGACCTCCTCCTTCCTGNNNAGATCGGAAGA |
| 50376_2 | TATGAGTGTGGAGTCGTTGCGCTTATAGACCTCCTCCTTCCTGNNNAGATCGGAAGA |
| 50376_3 | TATGAGTGTGGAGTCGTTGCGCTTATAGACCTCCTCCTTCCTGNNNAGATCGGAAGA |
| 45811_1 | TATGAGTGTGGAGTCGTTGCACTATCTGTGCTCATGGAACGTGNNNAGATCGGAAG |
| 45811_2 | TATGAGTGTGGAGTCGTTGCACTATCTGTGCTCATGGAACGTGNNNAGATCGGAAG |
| 45811_3 | TATGAGTGTGGAGTCGTTGCACTATCTGTGCTCATGGAACGTGNNNAGATCGGAAG |
| 42630_1 | TATGAGTGTGGAGTCGTTGCGAGAGAGCGAGAGAGAGAGAGAANNNAGATCGG |
| 42630_2 | TATGAGTGTGGAGTCGTTGCGAGAGAGCGAGAGAGAGAGAGAANNNAGATCGG |
| 42630_3 | TATGAGTGTGGAGTCGTTGCGAGAGAGCGAGAGAGAGAGAGAANNNAGATCGG |
| 43450_1 | TATGAGTGTGGAGTCGTTGCCTTCCGAGTAGCTGGGATTACAGNNNAGATCGGAA |
| 43450_2 | TATGAGTGTGGAGTCGTTGCCTTCCGAGTAGCTGGGATTACAGNNNAGATCGGAA |
| 43450_3 | TATGAGTGTGGAGTCGTTGCCTTCCGAGTAGCTGGGATTACAGNNNAGATCGGAA |
| 29452_1 | TATGAGTGTGGAGTCGTTGCAAAAAATGTAATAAACAGAACCANNNAGATCGGAA |
| 29452_2 | TATGAGTGTGGAGTCGTTGCAAAAAATGTAATAAACAGAACCANNNAGATCGGAA |
| 29452_3 | TATGAGTGTGGAGTCGTTGCAAAAAATGTAATAAACAGAACCANNNAGATCGGAA |
| 46812_1 | TATGAGTGTGGAGTCGTTGCCTACTCTCTGACCCCCAGTGTNNNAGATCGGAAGAG |
| 46812_2 | TATGAGTGTGGAGTCGTTGCCTACTCTCTGACCCCCAGTGTNNNAGATCGGAAGAG |
| 46812_3 | TATGAGTGTGGAGTCGTTGCCTACTCTCTGACCCCCAGTGTNNNAGATCGGAAGAG |
| 32324_1 | TATGAGTGTGGAGTCGTTGCATTTCACACCTTCTGCTCTTTGCNNNAGATCGGAAGA |

| | |
|---|---|
| 32324_2 | TATGAGTGTGGAGTCGTTGCATTTCACACCTTCTGCTCTTTGCNNNAGATCGGAAGA |
| 32324_3 | TATGAGTGTGGAGTCGTTGCATTTCACACCTTCTGCTCTTTGCNNNAGATCGGAAGA |
| 45085_1 | TATGAGTGTGGAGTCGTTGCTTCTGTCATTTGGAGGTCCCTTTNNNAGATCGGAAG |
| 45085_2 | TATGAGTGTGGAGTCGTTGCTTCTGTCATTTGGAGGTCCCTTTNNNAGATCGGAAG |
| 45085_3 | TATGAGTGTGGAGTCGTTGCTTCTGTCATTTGGAGGTCCCTTTNNNAGATCGGAAG |
| 52643_1 | TATGAGTGTGGAGTCGTTGCAAAGACGACTGCATAGGAGTCTCNNNAGATCGGAA |
| 52643_2 | TATGAGTGTGGAGTCGTTGCAAAGACGACTGCATAGGAGTCTCNNNAGATCGGAA |
| 52643_3 | TATGAGTGTGGAGTCGTTGCAAAGACGACTGCATAGGAGTCTCNNNAGATCGGAA |
| 41645_1 | TATGAGTGTGGAGTCGTTGCAGAGTAGCTGGGACTATAGGCATNNNAGATCGGAA |
| 41645_2 | TATGAGTGTGGAGTCGTTGCAGAGTAGCTGGGACTATAGGCATNNNAGATCGGAA |
| 41645_3 | TATGAGTGTGGAGTCGTTGCAGAGTAGCTGGGACTATAGGCATNNNAGATCGGAA |
| 32536_1 | TATGAGTGTGGAGTCGTTGCGCTACAGAATCCTCTCCCCAATCNNNAGATCGGAAG |
| 32536_2 | TATGAGTGTGGAGTCGTTGCGCTACAGAATCCTCTCCCCAATCNNNAGATCGGAAG |
| 32536_3 | TATGAGTGTGGAGTCGTTGCGCTACAGAATCCTCTCCCCAATCNNNAGATCGGAAG |
| 47770_1 | TATGAGTGTGGAGTCGTTGCGACGCCCGGCTAATTTTTTGTATNNNAGATCGGAAG |
| 47770_2 | TATGAGTGTGGAGTCGTTGCGACGCCCGGCTAATTTTTTGTATNNNAGATCGGAAG |
| 47770_3 | TATGAGTGTGGAGTCGTTGCGACGCCCGGCTAATTTTTTGTATNNNAGATCGGAAG |
| 45538_1 | TATGAGTGTGGAGTCGTTGCGGAGACAGAGAAACAGGAGAAGGNNNAGATCGGA |
| 45538_2 | TATGAGTGTGGAGTCGTTGCGGAGACAGAGAAACAGGAGAAGGNNNAGATCGGA |
| 45538_3 | TATGAGTGTGGAGTCGTTGCGGAGACAGAGAAACAGGAGAAGGNNNAGATCGGA |
| 42337_1 | TATGAGTGTGGAGTCGTTGCACATTCAAGTAGTCCAGGTAGGCNNNAGATCGGAA |
| 42337_2 | TATGAGTGTGGAGTCGTTGCACATTCAAGTAGTCCAGGTAGGCNNNAGATCGGAA |
| 42337_3 | TATGAGTGTGGAGTCGTTGCACATTCAAGTAGTCCAGGTAGGCNNNAGATCGGAA |
| 30057_1 | TATGAGTGTGGAGTCGTTGCGACTCCCAGACTAACTCTTCACANNNAGATCGGAAG |
| 30057_2 | TATGAGTGTGGAGTCGTTGCGACTCCCAGACTAACTCTTCACANNNAGATCGGAAG |
| 30057_3 | TATGAGTGTGGAGTCGTTGCGACTCCCAGACTAACTCTTCACANNNAGATCGGAAG |
| 12935_1 | TATGAGTGTGGAGTCGTTGCGGAAGAAATTAATGTTATGCTCANNNAGATCGGAA |
| 12935_2 | TATGAGTGTGGAGTCGTTGCGGAAGAAATTAATGTTATGCTCANNNAGATCGGAA |
| 12935_3 | TATGAGTGTGGAGTCGTTGCGGAAGAAATTAATGTTATGCTCANNNAGATCGGAA |
| 31503_1 | TATGAGTGTGGAGTCGTTGCACCCCTATTTTCCTCTCTCACCTNNNAGATCGGAAGA |
| 31503_2 | TATGAGTGTGGAGTCGTTGCACCCCTATTTTCCTCTCTCACCTNNNAGATCGGAAGA |
| 31503_3 | TATGAGTGTGGAGTCGTTGCACCCCTATTTTCCTCTCTCACCTNNNAGATCGGAAGA |
| 35710_1 | TATGAGTGTGGAGTCGTTGCGATCTTGGCTCACTGAAAGCTCTNNNAGATCGGAAG |
| 35710_2 | TATGAGTGTGGAGTCGTTGCGATCTTGGCTCACTGAAAGCTCTNNNAGATCGGAAG |
| 35710_3 | TATGAGTGTGGAGTCGTTGCGATCTTGGCTCACTGAAAGCTCTNNNAGATCGGAAG |
| 45749_1 | TATGAGTGTGGAGTCGTTGCAAACTTGACTGCAGGCTAAATCGNNNAGATCGGAA |
| 45749_2 | TATGAGTGTGGAGTCGTTGCAAACTTGACTGCAGGCTAAATCGNNNAGATCGGAA |
| 45749_3 | TATGAGTGTGGAGTCGTTGCAAACTTGACTGCAGGCTAAATCGNNNAGATCGGAA |
| 45398_1 | TATGAGTGTGGAGTCGTTGCCTGCACAGCACTATCTAGCTCTTNNNAGATCGGAAG |
| 45398_2 | TATGAGTGTGGAGTCGTTGCCTGCACAGCACTATCTAGCTCTTNNNAGATCGGAAG |
| 45398_3 | TATGAGTGTGGAGTCGTTGCCTGCACAGCACTATCTAGCTCTTNNNAGATCGGAAG |
| 31720_1 | TATGAGTGTGGAGTCGTTGCCTGTTGCTCTCTGGAGACTTGACNNNAGATCGGAAG |
| 31720_2 | TATGAGTGTGGAGTCGTTGCCTGTTGCTCTCTGGAGACTTGACNNNAGATCGGAAG |

| | |
|---|---|
| 31720_3 | TATGAGTGTGGAGTCGTTGCCTGTTGCTCTCTGGAGACTTGACNNNAGATCGGAAG |
| 42840_1 | TATGAGTGTGGAGTCGTTGCTGATGCAATGACATTTGGATTCTGTNNNAGATCGGA |
| 42840_2 | TATGAGTGTGGAGTCGTTGCTGATGCAATGACATTTGGATTCTGTNNNAGATCGGA |
| 42840_3 | TATGAGTGTGGAGTCGTTGCTGATGCAATGACATTTGGATTCTGTNNNAGATCGGA |
| 41176_1 | TATGAGTGTGGAGTCGTTGCACTGCCACCAGAAAGAGAAATCTNNNAGATCGGAA |
| 41176_2 | TATGAGTGTGGAGTCGTTGCACTGCCACCAGAAAGAGAAATCTNNNAGATCGGAA |
| 41176_3 | TATGAGTGTGGAGTCGTTGCACTGCCACCAGAAAGAGAAATCTNNNAGATCGGAA |
| 29161_1 | TATGAGTGTGGAGTCGTTGCCTCCTCCAGTCAAGCTTTGAGATNNNAGATCGGAAG |
| 29161_2 | TATGAGTGTGGAGTCGTTGCCTCCTCCAGTCAAGCTTTGAGATNNNAGATCGGAAG |
| 29161_3 | TATGAGTGTGGAGTCGTTGCCTCCTCCAGTCAAGCTTTGAGATNNNAGATCGGAAG |
| 38408_1 | TATGAGTGTGGAGTCGTTGCCACCTGTAATTCCAGCACTTTGGNNNAGATCGGAAG |
| 38408_2 | TATGAGTGTGGAGTCGTTGCCACCTGTAATTCCAGCACTTTGGNNNAGATCGGAAG |
| 38408_3 | TATGAGTGTGGAGTCGTTGCCACCTGTAATTCCAGCACTTTGGNNNAGATCGGAAG |
| 48545_1 | TATGAGTGTGGAGTCGTTGCCTTTTCCTGGGGTCTGGAATACANNNAGATCGGAAG |
| 48545_2 | TATGAGTGTGGAGTCGTTGCCTTTTCCTGGGGTCTGGAATACANNNAGATCGGAAG |
| 48545_3 | TATGAGTGTGGAGTCGTTGCCTTTTCCTGGGGTCTGGAATACANNNAGATCGGAAG |
| 38476_1 | TATGAGTGTGGAGTCGTTGCTGGAAAACGGTATGGAGGTTTCTNNNAGATCGGAA |
| 38476_2 | TATGAGTGTGGAGTCGTTGCTGGAAAACGGTATGGAGGTTTCTNNNAGATCGGAA |
| 38476_3 | TATGAGTGTGGAGTCGTTGCTGGAAAACGGTATGGAGGTTTCTNNNAGATCGGAA |
| 46786_1 | TATGAGTGTGGAGTCGTTGCCGGGTTCATGCCATTCTCCTNNNAGATCGGAAGAGC |
| 46786_2 | TATGAGTGTGGAGTCGTTGCCGGGTTCATGCCATTCTCCTNNNAGATCGGAAGAGC |
| 46786_3 | TATGAGTGTGGAGTCGTTGCCGGGTTCATGCCATTCTCCTNNNAGATCGGAAGAGC |
| 47810_1 | TATGAGTGTGGAGTCGTTGCAGATGGAATTTCGCTCTTGTTGCNNNAGATCGGAAG |
| 47810_2 | TATGAGTGTGGAGTCGTTGCAGATGGAATTTCGCTCTTGTTGCNNNAGATCGGAAG |
| 47810_3 | TATGAGTGTGGAGTCGTTGCAGATGGAATTTCGCTCTTGTTGCNNNAGATCGGAAG |
| 42841_1 | TATGAGTGTGGAGTCGTTGCCAGTCAGGGGACCATTGTGATAANNNAGATCGGAA |
| 42841_2 | TATGAGTGTGGAGTCGTTGCCAGTCAGGGGACCATTGTGATAANNNAGATCGGAA |
| 42841_3 | TATGAGTGTGGAGTCGTTGCCAGTCAGGGGACCATTGTGATAANNNAGATCGGAA |
| 45855_1 | TATGAGTGTGGAGTCGTTGCGAGTGCAGTGACACGATCTCAGNNNAGATCGGAAG |
| 45855_2 | TATGAGTGTGGAGTCGTTGCGAGTGCAGTGACACGATCTCAGNNNAGATCGGAAG |
| 45855_3 | TATGAGTGTGGAGTCGTTGCGAGTGCAGTGACACGATCTCAGNNNAGATCGGAAG |
| 50898_1 | TATGAGTGTGGAGTCGTTGCAGCCTGGAAGAAAGCTAAGACTCNNNAGATCGGAA |
| 50898_2 | TATGAGTGTGGAGTCGTTGCAGCCTGGAAGAAAGCTAAGACTCNNNAGATCGGAA |
| 50898_3 | TATGAGTGTGGAGTCGTTGCAGCCTGGAAGAAAGCTAAGACTCNNNAGATCGGAA |
| 37282_1 | TATGAGTGTGGAGTCGTTGCCTCCCCTTCACCTTCACCAATAANNNAGATCGGAAGA |
| 37282_2 | TATGAGTGTGGAGTCGTTGCCTCCCCTTCACCTTCACCAATAANNNAGATCGGAAGA |
| 37282_3 | TATGAGTGTGGAGTCGTTGCCTCCCCTTCACCTTCACCAATAANNNAGATCGGAAGA |
| 39168_1 | TATGAGTGTGGAGTCGTTGCATTCAAGCGATTCTCCTGCCTTANNNAGATCGGAAG |
| 39168_2 | TATGAGTGTGGAGTCGTTGCATTCAAGCGATTCTCCTGCCTTANNNAGATCGGAAG |
| 39168_3 | TATGAGTGTGGAGTCGTTGCATTCAAGCGATTCTCCTGCCTTANNNAGATCGGAAG |
| 35333_1 | TATGAGTGTGGAGTCGTTGCAACCTCTGCCTCCTTAGTTCAAGNNNAGATCGGAAG |
| 35333_2 | TATGAGTGTGGAGTCGTTGCAACCTCTGCCTCCTTAGTTCAAGNNNAGATCGGAAG |
| 35333_3 | TATGAGTGTGGAGTCGTTGCAACCTCTGCCTCCTTAGTTCAAGNNNAGATCGGAAG |

| | |
|---|---|
| 48003_1 | TATGAGTGTGGAGTCGTTGCCAAAAAGAAAGCCGTGAAGGAGTNNNAGATCGGAA |
| 48003_2 | TATGAGTGTGGAGTCGTTGCCAAAAAGAAAGCCGTGAAGGAGTNNNAGATCGGAA |
| 48003_3 | TATGAGTGTGGAGTCGTTGCCAAAAAGAAAGCCGTGAAGGAGTNNNAGATCGGAA |
| 37036_1 | TATGAGTGTGGAGTCGTTGCTTAGCTAATGGAGTCTAGCAGCANNNAGATCGGAAG |
| 37036_2 | TATGAGTGTGGAGTCGTTGCTTAGCTAATGGAGTCTAGCAGCANNNAGATCGGAAG |
| 37036_3 | TATGAGTGTGGAGTCGTTGCTTAGCTAATGGAGTCTAGCAGCANNNAGATCGGAAG |
| 51712_1 | TATGAGTGTGGAGTCGTTGCACTTATTTGACGACTCTGCCTCCNNNAGATCGGAAGA |
| 51712_2 | TATGAGTGTGGAGTCGTTGCACTTATTTGACGACTCTGCCTCCNNNAGATCGGAAGA |
| 51712_3 | TATGAGTGTGGAGTCGTTGCACTTATTTGACGACTCTGCCTCCNNNAGATCGGAAGA |
| 28869_1 | TATGAGTGTGGAGTCGTTGCGCAGTGATGAGATCGTGTTCAAGNNNAGATCGGAA |
| 28869_2 | TATGAGTGTGGAGTCGTTGCGCAGTGATGAGATCGTGTTCAAGNNNAGATCGGAA |
| 28869_3 | TATGAGTGTGGAGTCGTTGCGCAGTGATGAGATCGTGTTCAAGNNNAGATCGGAA |
| 36369_1 | TATGAGTGTGGAGTCGTTGCAGAGTGATTATTTCTGCATCAGGTNNNAGATCGGAA |
| 36369_2 | TATGAGTGTGGAGTCGTTGCAGAGTGATTATTTCTGCATCAGGTNNNAGATCGGAA |
| 36369_3 | TATGAGTGTGGAGTCGTTGCAGAGTGATTATTTCTGCATCAGGTNNNAGATCGGAA |
| 48198_1 | TATGAGTGTGGAGTCGTTGCCTCCTGAGTAGCTGGGATTACAGNNNAGATCGGAAG |
| 48198_2 | TATGAGTGTGGAGTCGTTGCCTCCTGAGTAGCTGGGATTACAGNNNAGATCGGAAG |
| 48198_3 | TATGAGTGTGGAGTCGTTGCCTCCTGAGTAGCTGGGATTACAGNNNAGATCGGAAG |
| 46026_1 | TATGAGTGTGGAGTCGTTGCCTGGAGTACAGTGGAGCAATCTCNNNAGATCGGAA |
| 46026_2 | TATGAGTGTGGAGTCGTTGCCTGGAGTACAGTGGAGCAATCTCNNNAGATCGGAA |
| 46026_3 | TATGAGTGTGGAGTCGTTGCCTGGAGTACAGTGGAGCAATCTCNNNAGATCGGAA |
| 35811_1 | TATGAGTGTGGAGTCGTTGCCACAATCTCGGTTCACTGCAAGNNNAGATCGGAAGA |
| 35811_2 | TATGAGTGTGGAGTCGTTGCCACAATCTCGGTTCACTGCAAGNNNAGATCGGAAGA |
| 35811_3 | TATGAGTGTGGAGTCGTTGCCACAATCTCGGTTCACTGCAAGNNNAGATCGGAAGA |
| 40933_1 | TATGAGTGTGGAGTCGTTGCTTCCAAAAGCCTCCTCTAACTGANNNAGATCGGAAGA |
| 40933_2 | TATGAGTGTGGAGTCGTTGCTTCCAAAAGCCTCCTCTAACTGANNNAGATCGGAAGA |
| 40933_3 | TATGAGTGTGGAGTCGTTGCTTCCAAAAGCCTCCTCTAACTGANNNAGATCGGAAGA |
| 12996_1 | TATGAGTGTGGAGTCGTTGCGAATGTAGAACGATTTCAGAATGNNNAGATCGGAA |
| 12996_2 | TATGAGTGTGGAGTCGTTGCGAATGTAGAACGATTTCAGAATGNNNAGATCGGAA |
| 12996_3 | TATGAGTGTGGAGTCGTTGCGAATGTAGAACGATTTCAGAATGNNNAGATCGGAA |
| 3043_1 | TATGAGTGTGGAGTCGTTGCCCTCTGCCTCACAGTTCAATTCTNNNAGATCGGAAGA |
| 3043_2 | TATGAGTGTGGAGTCGTTGCCCTCTGCCTCACAGTTCAATTCTNNNAGATCGGAAGA |
| 3043_3 | TATGAGTGTGGAGTCGTTGCCCTCTGCCTCACAGTTCAATTCTNNNAGATCGGAAGA |
| 35861_1 | TATGAGTGTGGAGTCGTTGCCTGTCATCTCAGCACTTTGGGAGNNNAGATCGGAAG |
| 35861_2 | TATGAGTGTGGAGTCGTTGCCTGTCATCTCAGCACTTTGGGAGNNNAGATCGGAAG |
| 35861_3 | TATGAGTGTGGAGTCGTTGCCTGTCATCTCAGCACTTTGGGAGNNNAGATCGGAAG |
| 35003_1 | TATGAGTGTGGAGTCGTTGCGATTACAGGCATGCACCATCATGNNNAGATCGGAAG |
| 35003_2 | TATGAGTGTGGAGTCGTTGCGATTACAGGCATGCACCATCATGNNNAGATCGGAAG |
| 35003_3 | TATGAGTGTGGAGTCGTTGCGATTACAGGCATGCACCATCATGNNNAGATCGGAAG |
| 35349_1 | TATGAGTGTGGAGTCGTTGCCATTGCTAAGAACCAGCTACAGCNNNAGATCGGAAG |
| 35349_2 | TATGAGTGTGGAGTCGTTGCCATTGCTAAGAACCAGCTACAGCNNNAGATCGGAAG |
| 35349_3 | TATGAGTGTGGAGTCGTTGCCATTGCTAAGAACCAGCTACAGCNNNAGATCGGAAG |
| 35866_1 | TATGAGTGTGGAGTCGTTGCCAGTGGTATGATCTCGGCTCACNNNAGATCGGAAGA |

| | |
|---|---|
| 35866_2 | TATGAGTGTGGAGTCGTTGCCAGTGGTATGATCTCGGCTCACNNNAGATCGGAAGA |
| 35866_3 | TATGAGTGTGGAGTCGTTGCCAGTGGTATGATCTCGGCTCACNNNAGATCGGAAGA |
| 39026_1 | TATGAGTGTGGAGTCGTTGCACTTGGCTGCTTTTCTTCTACTGNNNAGATCGGAAGA |
| 39026_2 | TATGAGTGTGGAGTCGTTGCACTTGGCTGCTTTTCTTCTACTGNNNAGATCGGAAGA |
| 39026_3 | TATGAGTGTGGAGTCGTTGCACTTGGCTGCTTTTCTTCTACTGNNNAGATCGGAAGA |
| 38531_1 | TATGAGTGTGGAGTCGTTGCACTGTGAAACCCCGTCTCTACTANNNAGATCGGAAGA |
| 38531_2 | TATGAGTGTGGAGTCGTTGCACTGTGAAACCCCGTCTCTACTANNNAGATCGGAAGA |
| 38531_3 | TATGAGTGTGGAGTCGTTGCACTGTGAAACCCCGTCTCTACTANNNAGATCGGAAGA |
| 52632_1 | TATGAGTGTGGAGTCGTTGCATAGTGGGTGACAGTATCCAAGCNNNAGATCGGAA |
| 52632_2 | TATGAGTGTGGAGTCGTTGCATAGTGGGTGACAGTATCCAAGCNNNAGATCGGAA |
| 52632_3 | TATGAGTGTGGAGTCGTTGCATAGTGGGTGACAGTATCCAAGCNNNAGATCGGAA |
| 44275_1 | TATGAGTGTGGAGTCGTTGCTTTAGCAGCTGAAGGGAGAACATNNNAGATCGGAA |
| 44275_2 | TATGAGTGTGGAGTCGTTGCTTTAGCAGCTGAAGGGAGAACATNNNAGATCGGAA |
| 44275_3 | TATGAGTGTGGAGTCGTTGCTTTAGCAGCTGAAGGGAGAACATNNNAGATCGGAA |
| 43766_1 | TATGAGTGTGGAGTCGTTGCAATTCTCCAGGAATGAGCACTGTNNNAGATCGGAAG |
| 43766_2 | TATGAGTGTGGAGTCGTTGCAATTCTCCAGGAATGAGCACTGTNNNAGATCGGAAG |
| 43766_3 | TATGAGTGTGGAGTCGTTGCAATTCTCCAGGAATGAGCACTGTNNNAGATCGGAAG |
| 39236_1 | TATGAGTGTGGAGTCGTTGCCCTGCCTCCAATCAATCACAAACNNNAGATCGGAAGA |
| 39236_2 | TATGAGTGTGGAGTCGTTGCCCTGCCTCCAATCAATCACAAACNNNAGATCGGAAGA |
| 39236_3 | TATGAGTGTGGAGTCGTTGCCCTGCCTCCAATCAATCACAAACNNNAGATCGGAAGA |
| 31936_1 | TATGAGTGTGGAGTCGTTGCAATTCTGAAGACTCCCAGACAGGNNNAGATCGGAAG |
| 31936_2 | TATGAGTGTGGAGTCGTTGCAATTCTGAAGACTCCCAGACAGGNNNAGATCGGAAG |
| 31936_3 | TATGAGTGTGGAGTCGTTGCAATTCTGAAGACTCCCAGACAGGNNNAGATCGGAAG |
| 50482_1 | TATGAGTGTGGAGTCGTTGCTCTTGGTGAGATCCATTGCTGAANNNAGATCGGAAG |
| 50482_2 | TATGAGTGTGGAGTCGTTGCTCTTGGTGAGATCCATTGCTGAANNNAGATCGGAAG |
| 50482_3 | TATGAGTGTGGAGTCGTTGCTCTTGGTGAGATCCATTGCTGAANNNAGATCGGAAG |
| 1585_1 | TATGAGTGTGGAGTCGTTGCTGGCAGCAACTCTGATTTTGGTANNNAGATCGGAAG |
| 1585_2 | TATGAGTGTGGAGTCGTTGCTGGCAGCAACTCTGATTTTGGTANNNAGATCGGAAG |
| 1585_3 | TATGAGTGTGGAGTCGTTGCTGGCAGCAACTCTGATTTTGGTANNNAGATCGGAAG |
| 36844_1 | TATGAGTGTGGAGTCGTTGCTCTCTCAGGATGGGGCATTAGTANNNAGATCGGAA |
| 36844_2 | TATGAGTGTGGAGTCGTTGCTCTCTCAGGATGGGGCATTAGTANNNAGATCGGAA |
| 36844_3 | TATGAGTGTGGAGTCGTTGCTCTCTCAGGATGGGGCATTAGTANNNAGATCGGAA |
| 41896_1 | TATGAGTGTGGAGTCGTTGCCAGGTTTTGGAGCATTTCGGATTNNNAGATCGGAA |
| 41896_2 | TATGAGTGTGGAGTCGTTGCCAGGTTTTGGAGCATTTCGGATTNNNAGATCGGAA |
| 41896_3 | TATGAGTGTGGAGTCGTTGCCAGGTTTTGGAGCATTTCGGATTNNNAGATCGGAA |
| 31081_1 | TATGAGTGTGGAGTCGTTGCGGACCCATAATTCCTCCAGATGGNNNAGATCGGAAG |
| 31081_2 | TATGAGTGTGGAGTCGTTGCGGACCCATAATTCCTCCAGATGGNNNAGATCGGAAG |
| 31081_3 | TATGAGTGTGGAGTCGTTGCGGACCCATAATTCCTCCAGATGGNNNAGATCGGAAG |
| 44173_1 | TATGAGTGTGGAGTCGTTGCCCCCATCATTCTAGCACAGAACANNNAGATCGGAAGA |
| 44173_2 | TATGAGTGTGGAGTCGTTGCCCCCATCATTCTAGCACAGAACANNNAGATCGGAAGA |
| 44173_3 | TATGAGTGTGGAGTCGTTGCCCCCATCATTCTAGCACAGAACANNNAGATCGGAAGA |
| 44306_1 | TATGAGTGTGGAGTCGTTGCGTCGAATGCTGGCCTTTAGTGNNNAGATCGGAAGA |
| 44306_2 | TATGAGTGTGGAGTCGTTGCGTCGAATGCTGGCCTTTAGTGNNNAGATCGGAAGA |

| | |
|---|---|
| 44306_3 | TATGAGTGTGGAGTCGTTGCGTCGAATGCTGGCCTTTAGTGNNNAGATCGGAAGA |
| 39400_1 | TATGAGTGTGGAGTCGTTGCAATTGTTTTGTTCGAGGAGGCTGNNNAGATCGGAA |
| 39400_2 | TATGAGTGTGGAGTCGTTGCAATTGTTTTGTTCGAGGAGGCTGNNNAGATCGGAA |
| 39400_3 | TATGAGTGTGGAGTCGTTGCAATTGTTTTGTTCGAGGAGGCTGNNNAGATCGGAA |
| 52879_1 | TATGAGTGTGGAGTCGTTGCCCAACATGAGTTCAAACCAAGGTNNNAGATCGGAAG |
| 52879_2 | TATGAGTGTGGAGTCGTTGCCCAACATGAGTTCAAACCAAGGTNNNAGATCGGAAG |
| 52879_3 | TATGAGTGTGGAGTCGTTGCCCAACATGAGTTCAAACCAAGGTNNNAGATCGGAAG |
| 40318_1 | TATGAGTGTGGAGTCGTTGCAAACAAAAAAGCCAGGCATACCTNNNAGATCGGAAG |
| 40318_2 | TATGAGTGTGGAGTCGTTGCAAACAAAAAAGCCAGGCATACCTNNNAGATCGGAAG |
| 40318_3 | TATGAGTGTGGAGTCGTTGCAAACAAAAAAGCCAGGCATACCTNNNAGATCGGAAG |
| 40622_1 | TATGAGTGTGGAGTCGTTGCCAAAAGCCAGTACTGTGCTTGAGNNNAGATCGGAA |
| 40622_2 | TATGAGTGTGGAGTCGTTGCCAAAAGCCAGTACTGTGCTTGAGNNNAGATCGGAA |
| 40622_3 | TATGAGTGTGGAGTCGTTGCCAAAAGCCAGTACTGTGCTTGAGNNNAGATCGGAA |
| 42294_1 | TATGAGTGTGGAGTCGTTGCAGACTGTAGGAAACGTGAACACANNNAGATCGGAA |
| 42294_2 | TATGAGTGTGGAGTCGTTGCAGACTGTAGGAAACGTGAACACANNNAGATCGGAA |
| 42294_3 | TATGAGTGTGGAGTCGTTGCAGACTGTAGGAAACGTGAACACANNNAGATCGGAA |
| 52099_1 | TATGAGTGTGGAGTCGTTGCACTTTCCAGACAGGGTTAGGTTCNNNAGATCGGAAG |
| 52099_2 | TATGAGTGTGGAGTCGTTGCACTTTCCAGACAGGGTTAGGTTCNNNAGATCGGAAG |
| 52099_3 | TATGAGTGTGGAGTCGTTGCACTTTCCAGACAGGGTTAGGTTCNNNAGATCGGAAG |
| 39702_1 | TATGAGTGTGGAGTCGTTGCTGTCAGCTCTCCATTTTGTCTCANNNAGATCGGAAGA |
| 39702_2 | TATGAGTGTGGAGTCGTTGCTGTCAGCTCTCCATTTTGTCTCANNNAGATCGGAAGA |
| 39702_3 | TATGAGTGTGGAGTCGTTGCTGTCAGCTCTCCATTTTGTCTCANNNAGATCGGAAGA |
| 49192_1 | TATGAGTGTGGAGTCGTTGCGAGACGGAGTCTTGCTCTGTCNNNAGATCGGAAGA |
| 49192_2 | TATGAGTGTGGAGTCGTTGCGAGACGGAGTCTTGCTCTGTCNNNAGATCGGAAGA |
| 49192_3 | TATGAGTGTGGAGTCGTTGCGAGACGGAGTCTTGCTCTGTCNNNAGATCGGAAGA |
| 12687_1 | TATGAGTGTGGAGTCGTTGCCAGTTGCTTTGTACTTAACATCNNNAGATCGGAAGA |
| 12687_2 | TATGAGTGTGGAGTCGTTGCCAGTTGCTTTGTACTTAACATCNNNAGATCGGAAGA |
| 12687_3 | TATGAGTGTGGAGTCGTTGCCAGTTGCTTTGTACTTAACATCNNNAGATCGGAAGA |
| 41355_1 | TATGAGTGTGGAGTCGTTGCCATTGTTGAGGTCCTGAGTCCTANNNAGATCGGAAG |
| 41355_2 | TATGAGTGTGGAGTCGTTGCCATTGTTGAGGTCCTGAGTCCTANNNAGATCGGAAG |
| 41355_3 | TATGAGTGTGGAGTCGTTGCCATTGTTGAGGTCCTGAGTCCTANNNAGATCGGAAG |
| 44733_1 | TATGAGTGTGGAGTCGTTGCCTGATACTGTGCAGAGTGGGTCNNNAGATCGGAAG |
| 44733_2 | TATGAGTGTGGAGTCGTTGCCTGATACTGTGCAGAGTGGGTCNNNAGATCGGAAG |
| 44733_3 | TATGAGTGTGGAGTCGTTGCCTGATACTGTGCAGAGTGGGTCNNNAGATCGGAAG |
| 32701_1 | TATGAGTGTGGAGTCGTTGCCCACCTCCTGAGTTCAAGCTATTNNNAGATCGGAAG |
| 32701_2 | TATGAGTGTGGAGTCGTTGCCCACCTCCTGAGTTCAAGCTATTNNNAGATCGGAAG |
| 32701_3 | TATGAGTGTGGAGTCGTTGCCCACCTCCTGAGTTCAAGCTATTNNNAGATCGGAAG |
| 38251_1 | TATGAGTGTGGAGTCGTTGCCACTGGCAACCATGACAAAGAAANNNAGATCGGAA |
| 38251_2 | TATGAGTGTGGAGTCGTTGCCACTGGCAACCATGACAAAGAAANNNAGATCGGAA |
| 38251_3 | TATGAGTGTGGAGTCGTTGCCACTGGCAACCATGACAAAGAAANNNAGATCGGAA |
| 33104_1 | TATGAGTGTGGAGTCGTTGCCAAGCACTTGGAGGGAAGAAAAGNNNAGATCGGA |
| 33104_2 | TATGAGTGTGGAGTCGTTGCCAAGCACTTGGAGGGAAGAAAAGNNNAGATCGGA |
| 33104_3 | TATGAGTGTGGAGTCGTTGCCAAGCACTTGGAGGGAAGAAAAGNNNAGATCGGA |

| | |
|---|---|
| 31215_1 | TATGAGTGTGGAGTCGTTGCCATGTTGATGTTCTGGCTGACAGNNNAGATCGGAAG |
| 31215_2 | TATGAGTGTGGAGTCGTTGCCATGTTGATGTTCTGGCTGACAGNNNAGATCGGAAG |
| 31215_3 | TATGAGTGTGGAGTCGTTGCCATGTTGATGTTCTGGCTGACAGNNNAGATCGGAAG |
| 36902_1 | TATGAGTGTGGAGTCGTTGCGCTAGGAGAACCCAGGAAATTCANNNAGATCGGAA |
| 36902_2 | TATGAGTGTGGAGTCGTTGCGCTAGGAGAACCCAGGAAATTCANNNAGATCGGAA |
| 36902_3 | TATGAGTGTGGAGTCGTTGCGCTAGGAGAACCCAGGAAATTCANNNAGATCGGAA |
| 13363_1 | TATGAGTGTGGAGTCGTTGCTCCATTGTTTCTAAGGTCTTCNNNAGATCGGAAGAG |
| 13363_2 | TATGAGTGTGGAGTCGTTGCTCCATTGTTTCTAAGGTCTTCNNNAGATCGGAAGAG |
| 13363_3 | TATGAGTGTGGAGTCGTTGCTCCATTGTTTCTAAGGTCTTCNNNAGATCGGAAGAG |
| 46939_1 | TATGAGTGTGGAGTCGTTGCGACAGAGTTTCACTTTGTTGCCCNNNAGATCGGAAG |
| 46939_2 | TATGAGTGTGGAGTCGTTGCGACAGAGTTTCACTTTGTTGCCCNNNAGATCGGAAG |
| 46939_3 | TATGAGTGTGGAGTCGTTGCGACAGAGTTTCACTTTGTTGCCCNNNAGATCGGAAG |
| 29268_1 | TATGAGTGTGGAGTCGTTGCCATCCCAAAAGCCACATCTTCAGNNNAGATCGGAAG |
| 29268_2 | TATGAGTGTGGAGTCGTTGCCATCCCAAAAGCCACATCTTCAGNNNAGATCGGAAG |
| 29268_3 | TATGAGTGTGGAGTCGTTGCCATCCCAAAAGCCACATCTTCAGNNNAGATCGGAAG |
| 31782_1 | TATGAGTGTGGAGTCGTTGCCACCTGACCTCTGCAGACTTTATNNNAGATCGGAAG |
| 31782_2 | TATGAGTGTGGAGTCGTTGCCACCTGACCTCTGCAGACTTTATNNNAGATCGGAAG |
| 31782_3 | TATGAGTGTGGAGTCGTTGCCACCTGACCTCTGCAGACTTTATNNNAGATCGGAAG |
| 32707_1 | TATGAGTGTGGAGTCGTTGCGTTCAAGCGATTCTCTTGCCTCNNNAGATCGGAAGA |
| 32707_2 | TATGAGTGTGGAGTCGTTGCGTTCAAGCGATTCTCTTGCCTCNNNAGATCGGAAGA |
| 32707_3 | TATGAGTGTGGAGTCGTTGCGTTCAAGCGATTCTCTTGCCTCNNNAGATCGGAAGA |
| 12855_1 | TATGAGTGTGGAGTCGTTGCGGTGAAGATTTTTGGATTCTTTNNNAGATCGGAAGA |
| 12855_2 | TATGAGTGTGGAGTCGTTGCGGTGAAGATTTTTGGATTCTTTNNNAGATCGGAAGA |
| 12855_3 | TATGAGTGTGGAGTCGTTGCGGTGAAGATTTTTGGATTCTTTNNNAGATCGGAAGA |
| 35391_1 | TATGAGTGTGGAGTCGTTGCGGGCTCAAATGATTCTCCTGTCTNNNAGATCGGAAG |
| 35391_2 | TATGAGTGTGGAGTCGTTGCGGGCTCAAATGATTCTCCTGTCTNNNAGATCGGAAG |
| 35391_3 | TATGAGTGTGGAGTCGTTGCGGGCTCAAATGATTCTCCTGTCTNNNAGATCGGAAG |
| 6139_1 | TATGAGTGTGGAGTCGTTGCTCTGGAGCGGTTTATCACAAGTGNNNAGATCGGAAG |
| 6139_2 | TATGAGTGTGGAGTCGTTGCTCTGGAGCGGTTTATCACAAGTGNNNAGATCGGAAG |
| 6139_3 | TATGAGTGTGGAGTCGTTGCTCTGGAGCGGTTTATCACAAGTGNNNAGATCGGAAG |
| 31655_1 | TATGAGTGTGGAGTCGTTGCTTTCGCAGCCAGACCAGAAAGNNNAGATCGGAAGA |
| 31655_2 | TATGAGTGTGGAGTCGTTGCTTTCGCAGCCAGACCAGAAAGNNNAGATCGGAAGA |
| 31655_3 | TATGAGTGTGGAGTCGTTGCTTTCGCAGCCAGACCAGAAAGNNNAGATCGGAAGA |
| 10166_1 | TATGAGTGTGGAGTCGTTGCCTCTCACCCACTCCCTATTCCTTNNNAGATCGGAAGAG |
| 10166_2 | TATGAGTGTGGAGTCGTTGCCTCTCACCCACTCCCTATTCCTTNNNAGATCGGAAGAG |
| 10166_3 | TATGAGTGTGGAGTCGTTGCCTCTCACCCACTCCCTATTCCTTNNNAGATCGGAAGAG |
| 12556_1 | TATGAGTGTGGAGTCGTTGCAAAGAAGTAACCCAAACGTAANNNAGATCGGAAGA |
| 12556_2 | TATGAGTGTGGAGTCGTTGCAAAGAAGTAACCCAAACGTAANNNAGATCGGAAGA |
| 12556_3 | TATGAGTGTGGAGTCGTTGCAAAGAAGTAACCCAAACGTAANNNAGATCGGAAGA |
| 30345_1 | TATGAGTGTGGAGTCGTTGCGCCCTCAACTGAAGCAATCTTGANNNAGATCGGAAG |
| 30345_2 | TATGAGTGTGGAGTCGTTGCGCCCTCAACTGAAGCAATCTTGANNNAGATCGGAAG |
| 30345_3 | TATGAGTGTGGAGTCGTTGCGCCCTCAACTGAAGCAATCTTGANNNAGATCGGAAG |
| 51687_1 | TATGAGTGTGGAGTCGTTGCCAGTTTTATTGCCGTGTAGGACANNNAGATCGGAAG |

| | |
|---|---|
| 51687_2 | TATGAGTGTGGAGTCGTTGCCAGTTTTATTGCCGTGTAGGACANNNAGATCGGAAG |
| 51687_3 | TATGAGTGTGGAGTCGTTGCCAGTTTTATTGCCGTGTAGGACANNNAGATCGGAAG |
| 38590_1 | TATGAGTGTGGAGTCGTTGCTGAGGAAGCTGTGTGAGAATTGANNNAGATCGGAA |
| 38590_2 | TATGAGTGTGGAGTCGTTGCTGAGGAAGCTGTGTGAGAATTGANNNAGATCGGAA |
| 38590_3 | TATGAGTGTGGAGTCGTTGCTGAGGAAGCTGTGTGAGAATTGANNNAGATCGGAA |
| 42827_1 | TATGAGTGTGGAGTCGTTGCCATGGATGGAACTGGAGGTCATTNNNAGATCGGAA |
| 42827_2 | TATGAGTGTGGAGTCGTTGCCATGGATGGAACTGGAGGTCATTNNNAGATCGGAA |
| 42827_3 | TATGAGTGTGGAGTCGTTGCCATGGATGGAACTGGAGGTCATTNNNAGATCGGAA |
| 51079_1 | TATGAGTGTGGAGTCGTTGCGAAGCACCATGCCAGCAATCNNNAGATCGGAAGAG |
| 51079_2 | TATGAGTGTGGAGTCGTTGCGAAGCACCATGCCAGCAATCNNNAGATCGGAAGAG |
| 51079_3 | TATGAGTGTGGAGTCGTTGCGAAGCACCATGCCAGCAATCNNNAGATCGGAAGAG |
| 10252_1 | TATGAGTGTGGAGTCGTTGCACTCTTTCCTCTATATGCACGTGGNNNAGATCGGAAG |
| 10252_2 | TATGAGTGTGGAGTCGTTGCACTCTTTCCTCTATATGCACGTGGNNNAGATCGGAAG |
| 10252_3 | TATGAGTGTGGAGTCGTTGCACTCTTTCCTCTATATGCACGTGGNNNAGATCGGAAG |
| 38983_1 | TATGAGTGTGGAGTCGTTGCATTTTGTCCCTCATCTCAGTGGGNNNAGATCGGAAG |
| 38983_2 | TATGAGTGTGGAGTCGTTGCATTTTGTCCCTCATCTCAGTGGGNNNAGATCGGAAG |
| 38983_3 | TATGAGTGTGGAGTCGTTGCATTTTGTCCCTCATCTCAGTGGGNNNAGATCGGAAG |
| 10221_1 | TATGAGTGTGGAGTCGTTGCCCCTCCCACGTGACATCTAGTAANNNAGATCGGAAG |
| 10221_2 | TATGAGTGTGGAGTCGTTGCCCCTCCCACGTGACATCTAGTAANNNAGATCGGAAG |
| 10221_3 | TATGAGTGTGGAGTCGTTGCCCCTCCCACGTGACATCTAGTAANNNAGATCGGAAG |
| 33941_1 | TATGAGTGTGGAGTCGTTGCCTCTGGAGTAGCTGGGACTACAGNNNAGATCGGAA |
| 33941_2 | TATGAGTGTGGAGTCGTTGCCTCTGGAGTAGCTGGGACTACAGNNNAGATCGGAA |
| 33941_3 | TATGAGTGTGGAGTCGTTGCCTCTGGAGTAGCTGGGACTACAGNNNAGATCGGAA |
| 12947_1 | TATGAGTGTGGAGTCGTTGCTGGAAGAATAGATACAAGCATAANNNAGATCGGAA |
| 12947_2 | TATGAGTGTGGAGTCGTTGCTGGAAGAATAGATACAAGCATAANNNAGATCGGAA |
| 12947_3 | TATGAGTGTGGAGTCGTTGCTGGAAGAATAGATACAAGCATAANNNAGATCGGAA |
| 30652_1 | TATGAGTGTGGAGTCGTTGCGAAAAGATAAAATGAGAGATGGANNNAGATCGGA |
| 30652_2 | TATGAGTGTGGAGTCGTTGCGAAAAGATAAAATGAGAGATGGANNNAGATCGGA |
| 30652_3 | TATGAGTGTGGAGTCGTTGCGAAAAGATAAAATGAGAGATGGANNNAGATCGGA |
| 38084_1 | TATGAGTGTGGAGTCGTTGCACTTCTGTTGATGTTCAGTCTCCTNNNAGATCGGAAG |
| 38084_2 | TATGAGTGTGGAGTCGTTGCACTTCTGTTGATGTTCAGTCTCCTNNNAGATCGGAAG |
| 38084_3 | TATGAGTGTGGAGTCGTTGCACTTCTGTTGATGTTCAGTCTCCTNNNAGATCGGAAG |
| 5526_1 | TATGAGTGTGGAGTCGTTGCCAGTGTGGGAAAGCATTCAGTTTNNNAGATCGGAA |
| 5526_2 | TATGAGTGTGGAGTCGTTGCCAGTGTGGGAAAGCATTCAGTTTNNNAGATCGGAA |
| 5526_3 | TATGAGTGTGGAGTCGTTGCCAGTGTGGGAAAGCATTCAGTTTNNNAGATCGGAA |
| 48318_1 | TATGAGTGTGGAGTCGTTGCAACTCCGGAGCTCAAGTTATCTGNNNAGATCGGAAG |
| 48318_2 | TATGAGTGTGGAGTCGTTGCAACTCCGGAGCTCAAGTTATCTGNNNAGATCGGAAG |
| 48318_3 | TATGAGTGTGGAGTCGTTGCAACTCCGGAGCTCAAGTTATCTGNNNAGATCGGAAG |
| 9531_1 | TATGAGTGTGGAGTCGTTGCGAAGGTTGAGAAGTCCCATGCTCNNNAGATCGGAA |
| 9531_2 | TATGAGTGTGGAGTCGTTGCGAAGGTTGAGAAGTCCCATGCTCNNNAGATCGGAA |
| 9531_3 | TATGAGTGTGGAGTCGTTGCGAAGGTTGAGAAGTCCCATGCTCNNNAGATCGGAA |
| 8210_1 | TATGAGTGTGGAGTCGTTGCGTTGGAGGGCATACTTGTTGAANNNAGATCGGAAG |
| 8210_2 | TATGAGTGTGGAGTCGTTGCGTTGGAGGGCATACTTGTTGAANNNAGATCGGAAG |

| | |
|---|---|
| 8210_3 | TATGAGTGTGGAGTCGTTGCGTTGGAGGGCATACTTGTTGAANNNAGATCGGAAG |
| 48369_1 | TATGAGTGTGGAGTCGTTGCTTCAAGTGATTCTCCTGCCTCAGNNNAGATCGGAAG |
| 48369_2 | TATGAGTGTGGAGTCGTTGCTTCAAGTGATTCTCCTGCCTCAGNNNAGATCGGAAG |
| 48369_3 | TATGAGTGTGGAGTCGTTGCTTCAAGTGATTCTCCTGCCTCAGNNNAGATCGGAAG |
| 41800_1 | TATGAGTGTGGAGTCGTTGCAAGTGATCTCAAAGTGCTGGGATNNNAGATCGGAA |
| 41800_2 | TATGAGTGTGGAGTCGTTGCAAGTGATCTCAAAGTGCTGGGATNNNAGATCGGAA |
| 41800_3 | TATGAGTGTGGAGTCGTTGCAAGTGATCTCAAAGTGCTGGGATNNNAGATCGGAA |
| 34743_1 | TATGAGTGTGGAGTCGTTGCGAGACGAAGTCTTGCTCTGTCANNNAGATCGGAAG |
| 34743_2 | TATGAGTGTGGAGTCGTTGCGAGACGAAGTCTTGCTCTGTCANNNAGATCGGAAG |
| 34743_3 | TATGAGTGTGGAGTCGTTGCGAGACGAAGTCTTGCTCTGTCANNNAGATCGGAAG |
| 33165_1 | TATGAGTGTGGAGTCGTTGCACTACATGGAAATGGAACGACCTNNNAGATCGGAA |
| 33165_2 | TATGAGTGTGGAGTCGTTGCACTACATGGAAATGGAACGACCTNNNAGATCGGAA |
| 33165_3 | TATGAGTGTGGAGTCGTTGCACTACATGGAAATGGAACGACCTNNNAGATCGGAA |
| 44487_1 | TATGAGTGTGGAGTCGTTGCCAAGAGTATGACGGGGCTGTACNNNAGATCGGAAG |
| 44487_2 | TATGAGTGTGGAGTCGTTGCCAAGAGTATGACGGGGCTGTACNNNAGATCGGAAG |
| 44487_3 | TATGAGTGTGGAGTCGTTGCCAAGAGTATGACGGGGCTGTACNNNAGATCGGAAG |
| 44701_1 | TATGAGTGTGGAGTCGTTGCATGGGTCAGGCTGTAAAAGATGTNNNAGATCGGAA |
| 44701_2 | TATGAGTGTGGAGTCGTTGCATGGGTCAGGCTGTAAAAGATGTNNNAGATCGGAA |
| 44701_3 | TATGAGTGTGGAGTCGTTGCATGGGTCAGGCTGTAAAAGATGTNNNAGATCGGAA |
| 33739_1 | TATGAGTGTGGAGTCGTTGCCTCTTCTGTGGCAAACCCTAGAGNNNAGATCGGAAG |
| 33739_2 | TATGAGTGTGGAGTCGTTGCCTCTTCTGTGGCAAACCCTAGAGNNNAGATCGGAAG |
| 33739_3 | TATGAGTGTGGAGTCGTTGCCTCTTCTGTGGCAAACCCTAGAGNNNAGATCGGAAG |
| 40135_1 | TATGAGTGTGGAGTCGTTGCTGGCATATACCTGTAGTTCCAGCNNNAGATCGGAAG |
| 40135_2 | TATGAGTGTGGAGTCGTTGCTGGCATATACCTGTAGTTCCAGCNNNAGATCGGAAG |
| 40135_3 | TATGAGTGTGGAGTCGTTGCTGGCATATACCTGTAGTTCCAGCNNNAGATCGGAAG |
| 47168_1 | TATGAGTGTGGAGTCGTTGCTTCAAGTGATTCTCCTGCCTCAGNNNAGATCGGAAG |
| 47168_2 | TATGAGTGTGGAGTCGTTGCTTCAAGTGATTCTCCTGCCTCAGNNNAGATCGGAAG |
| 47168_3 | TATGAGTGTGGAGTCGTTGCTTCAAGTGATTCTCCTGCCTCAGNNNAGATCGGAAG |
| 9763_1 | TATGAGTGTGGAGTCGTTGCGTAAGCCCTGAAACCAGCAAAACNNNAGATCGGAA |
| 9763_2 | TATGAGTGTGGAGTCGTTGCGTAAGCCCTGAAACCAGCAAAACNNNAGATCGGAA |
| 9763_3 | TATGAGTGTGGAGTCGTTGCGTAAGCCCTGAAACCAGCAAAACNNNAGATCGGAA |
| 48057_1 | TATGAGTGTGGAGTCGTTGCCAAGCTGGAGTGCAATGGTAAAANNNAGATCGGAA |
| 48057_2 | TATGAGTGTGGAGTCGTTGCCAAGCTGGAGTGCAATGGTAAAANNNAGATCGGAA |
| 48057_3 | TATGAGTGTGGAGTCGTTGCCAAGCTGGAGTGCAATGGTAAAANNNAGATCGGAA |
| 49129_1 | TATGAGTGTGGAGTCGTTGCTTTTTGAGACAGATTCTTGCCCCNNNAGATCGGAAG |
| 49129_2 | TATGAGTGTGGAGTCGTTGCTTTTTGAGACAGATTCTTGCCCCNNNAGATCGGAAG |
| 49129_3 | TATGAGTGTGGAGTCGTTGCTTTTTGAGACAGATTCTTGCCCCNNNAGATCGGAAG |
| 48550_1 | TATGAGTGTGGAGTCGTTGCCCATCAGAAGCTCCACTATGGTCNNNAGATCGGAAG |
| 48550_2 | TATGAGTGTGGAGTCGTTGCCCATCAGAAGCTCCACTATGGTCNNNAGATCGGAAG |
| 48550_3 | TATGAGTGTGGAGTCGTTGCCCATCAGAAGCTCCACTATGGTCNNNAGATCGGAAG |
| 30935_1 | TATGAGTGTGGAGTCGTTGCATCTTTTTTTCTGTGGAGTGCGGNNNAGATCGGAAG |
| 30935_2 | TATGAGTGTGGAGTCGTTGCATCTTTTTTTCTGTGGAGTGCGGNNNAGATCGGAAG |
| 30935_3 | TATGAGTGTGGAGTCGTTGCATCTTTTTTTCTGTGGAGTGCGGNNNAGATCGGAAG |

| | |
|---|---|
| 45647_1 | TATGAGTGTGGAGTCGTTGCCTCCCAAGTAGCTGGCATTACAGNNNAGATCGGAAG |
| 45647_2 | TATGAGTGTGGAGTCGTTGCCTCCCAAGTAGCTGGCATTACAGNNNAGATCGGAAG |
| 45647_3 | TATGAGTGTGGAGTCGTTGCCTCCCAAGTAGCTGGCATTACAGNNNAGATCGGAAG |
| 52990_1 | TATGAGTGTGGAGTCGTTGCAACTCCTGACCTCAAGTGATCTGNNNAGATCGGAAG |
| 52990_2 | TATGAGTGTGGAGTCGTTGCAACTCCTGACCTCAAGTGATCTGNNNAGATCGGAAG |
| 52990_3 | TATGAGTGTGGAGTCGTTGCAACTCCTGACCTCAAGTGATCTGNNNAGATCGGAAG |
| 45027_1 | TATGAGTGTGGAGTCGTTGCGAACATCTCTGCAGCTTTCACAGNNNAGATCGGAAG |
| 45027_2 | TATGAGTGTGGAGTCGTTGCGAACATCTCTGCAGCTTTCACAGNNNAGATCGGAAG |
| 45027_3 | TATGAGTGTGGAGTCGTTGCGAACATCTCTGCAGCTTTCACAGNNNAGATCGGAAG |
| 31139_1 | TATGAGTGTGGAGTCGTTGCATGGATCTCAACATCATACGTCANNNAGATCGGAAG |
| 31139_2 | TATGAGTGTGGAGTCGTTGCATGGATCTCAACATCATACGTCANNNAGATCGGAAG |
| 31139_3 | TATGAGTGTGGAGTCGTTGCATGGATCTCAACATCATACGTCANNNAGATCGGAAG |
| 50755_1 | TATGAGTGTGGAGTCGTTGCCTACTAAAGACCCACCTACCTGCNNNAGATCGGAAG |
| 50755_2 | TATGAGTGTGGAGTCGTTGCCTACTAAAGACCCACCTACCTGCNNNAGATCGGAAG |
| 50755_3 | TATGAGTGTGGAGTCGTTGCCTACTAAAGACCCACCTACCTGCNNNAGATCGGAAG |
| 42542_1 | TATGAGTGTGGAGTCGTTGCTGTGTGTGTGTGTGTGTGTACTANNNAGATCGGAA |
| 42542_2 | TATGAGTGTGGAGTCGTTGCTGTGTGTGTGTGTGTGTGTACTANNNAGATCGGAA |
| 42542_3 | TATGAGTGTGGAGTCGTTGCTGTGTGTGTGTGTGTGTGTACTANNNAGATCGGAA |
| 45856_1 | TATGAGTGTGGAGTCGTTGCTGACAGTATCTCACTCTTTCGCCNNNAGATCGGAAG |
| 45856_2 | TATGAGTGTGGAGTCGTTGCTGACAGTATCTCACTCTTTCGCCNNNAGATCGGAAG |
| 45856_3 | TATGAGTGTGGAGTCGTTGCTGACAGTATCTCACTCTTTCGCCNNNAGATCGGAAG |
| 46407_1 | TATGAGTGTGGAGTCGTTGCCATTCTCCTGCCTCAGCCTCNNNAGATCGGAAGAGCA |
| 46407_2 | TATGAGTGTGGAGTCGTTGCCATTCTCCTGCCTCAGCCTCNNNAGATCGGAAGAGCA |
| 46407_3 | TATGAGTGTGGAGTCGTTGCCATTCTCCTGCCTCAGCCTCNNNAGATCGGAAGAGCA |
| 41327_1 | TATGAGTGTGGAGTCGTTGCTGGCTTTGGGATAATGCTTGAACNNNAGATCGGAA |
| 41327_2 | TATGAGTGTGGAGTCGTTGCTGGCTTTGGGATAATGCTTGAACNNNAGATCGGAA |
| 41327_3 | TATGAGTGTGGAGTCGTTGCTGGCTTTGGGATAATGCTTGAACNNNAGATCGGAA |
| 36276_1 | TATGAGTGTGGAGTCGTTGCAGGTACTTGCTGTATGAGGCATCNNNAGATCGGAA |
| 36276_2 | TATGAGTGTGGAGTCGTTGCAGGTACTTGCTGTATGAGGCATCNNNAGATCGGAA |
| 36276_3 | TATGAGTGTGGAGTCGTTGCAGGTACTTGCTGTATGAGGCATCNNNAGATCGGAA |
| 33765_1 | TATGAGTGTGGAGTCGTTGCTTTTACTCAGAGGGGGGACTACTNNNAGATCGGAA |
| 33765_2 | TATGAGTGTGGAGTCGTTGCTTTTACTCAGAGGGGGGACTACTNNNAGATCGGAA |
| 33765_3 | TATGAGTGTGGAGTCGTTGCTTTTACTCAGAGGGGGGACTACTNNNAGATCGGAA |
| 29552_1 | TATGAGTGTGGAGTCGTTGCTTTTACCTGCAATTGGGGCATTTNNNAGATCGGAAG |
| 29552_2 | TATGAGTGTGGAGTCGTTGCTTTTACCTGCAATTGGGGCATTTNNNAGATCGGAAG |
| 29552_3 | TATGAGTGTGGAGTCGTTGCTTTTACCTGCAATTGGGGCATTTNNNAGATCGGAAG |
| 36459_1 | TATGAGTGTGGAGTCGTTGCACAAAAATTATCCAGGCGTGGTGNNNAGATCGGAA |
| 36459_2 | TATGAGTGTGGAGTCGTTGCACAAAAATTATCCAGGCGTGGTGNNNAGATCGGAA |
| 36459_3 | TATGAGTGTGGAGTCGTTGCACAAAAATTATCCAGGCGTGGTGNNNAGATCGGAA |
| 52906_1 | TATGAGTGTGGAGTCGTTGCGTGCTTAGAGACAGAGGATGTGGNNNAGATCGGA |
| 52906_2 | TATGAGTGTGGAGTCGTTGCGTGCTTAGAGACAGAGGATGTGGNNNAGATCGGA |
| 52906_3 | TATGAGTGTGGAGTCGTTGCGTGCTTAGAGACAGAGGATGTGGNNNAGATCGGA |
| 50360_1 | TATGAGTGTGGAGTCGTTGCGAGTGTGAGTCTAGGGAGTGAGANNNAGATCGGA |

| | |
|---|---|
| 50360_2 | TATGAGTGTGGAGTCGTTGCGAGTGTGAGTCTAGGGAGTGAGANNNAGATCGGA |
| 50360_3 | TATGAGTGTGGAGTCGTTGCGAGTGTGAGTCTAGGGAGTGAGANNNAGATCGGA |
| 29777_1 | TATGAGTGTGGAGTCGTTGCTAAGATTGCCAGTTTCCCAGGACNNNAGATCGGAAG |
| 29777_2 | TATGAGTGTGGAGTCGTTGCTAAGATTGCCAGTTTCCCAGGACNNNAGATCGGAAG |
| 29777_3 | TATGAGTGTGGAGTCGTTGCTAAGATTGCCAGTTTCCCAGGACNNNAGATCGGAAG |
| 33820_1 | TATGAGTGTGGAGTCGTTGCGGAGGCTCTCTTGGAAGAAATCANNNAGATCGGAA |
| 33820_2 | TATGAGTGTGGAGTCGTTGCGGAGGCTCTCTTGGAAGAAATCANNNAGATCGGAA |
| 33820_3 | TATGAGTGTGGAGTCGTTGCGGAGGCTCTCTTGGAAGAAATCANNNAGATCGGAA |
| 2290_1 | TATGAGTGTGGAGTCGTTGCCTGGGCTACTAAAACCTTCCCACNNNAGATCGGAAG |
| 2290_2 | TATGAGTGTGGAGTCGTTGCCTGGGCTACTAAAACCTTCCCACNNNAGATCGGAAG |
| 2290_3 | TATGAGTGTGGAGTCGTTGCCTGGGCTACTAAAACCTTCCCACNNNAGATCGGAAG |
| 30100_1 | TATGAGTGTGGAGTCGTTGCGAGGACAGTGGTGAGTCAGTTTTNNNAGATCGGAA |
| 30100_2 | TATGAGTGTGGAGTCGTTGCGAGGACAGTGGTGAGTCAGTTTTNNNAGATCGGAA |
| 30100_3 | TATGAGTGTGGAGTCGTTGCGAGGACAGTGGTGAGTCAGTTTTNNNAGATCGGAA |
| 53228_1 | TATGAGTGTGGAGTCGTTGCGTGAACCCACAATTCCGGAATAGNNNAGATCGGAA |
| 53228_2 | TATGAGTGTGGAGTCGTTGCGTGAACCCACAATTCCGGAATAGNNNAGATCGGAA |
| 53228_3 | TATGAGTGTGGAGTCGTTGCGTGAACCCACAATTCCGGAATAGNNNAGATCGGAA |
| 36083_1 | TATGAGTGTGGAGTCGTTGCAAGGACACTAGTGTAGCCATAGCNNNAGATCGGAA |
| 36083_2 | TATGAGTGTGGAGTCGTTGCAAGGACACTAGTGTAGCCATAGCNNNAGATCGGAA |
| 36083_3 | TATGAGTGTGGAGTCGTTGCAAGGACACTAGTGTAGCCATAGCNNNAGATCGGAA |
| 51336_1 | TATGAGTGTGGAGTCGTTGCCTCCCACCCCTTTTCCACTTTCNNNAGATCGGAAGAG |
| 51336_2 | TATGAGTGTGGAGTCGTTGCCTCCCACCCCTTTTCCACTTTCNNNAGATCGGAAGAG |
| 51336_3 | TATGAGTGTGGAGTCGTTGCCTCCCACCCCTTTTCCACTTTCNNNAGATCGGAAGAG |
| 46994_1 | TATGAGTGTGGAGTCGTTGCCCTCCTGGGTTCAAGCAATTCTCNNNAGATCGGAAG |
| 46994_2 | TATGAGTGTGGAGTCGTTGCCCTCCTGGGTTCAAGCAATTCTCNNNAGATCGGAAG |
| 46994_3 | TATGAGTGTGGAGTCGTTGCCCTCCTGGGTTCAAGCAATTCTCNNNAGATCGGAAG |
| 12807_1 | TATGAGTGTGGAGTCGTTGCCTTCTTTTCCACTTTGCTCTNNNAGATCGGAAGAGCA |
| 12807_2 | TATGAGTGTGGAGTCGTTGCCTTCTTTTCCACTTTGCTCTNNNAGATCGGAAGAGCA |
| 12807_3 | TATGAGTGTGGAGTCGTTGCCTTCTTTTCCACTTTGCTCTNNNAGATCGGAAGAGCA |
| 38204_1 | TATGAGTGTGGAGTCGTTGCAGCCGCAGAATTCAGAATTTGACNNNAGATCGGAA |
| 38204_2 | TATGAGTGTGGAGTCGTTGCAGCCGCAGAATTCAGAATTTGACNNNAGATCGGAA |
| 38204_3 | TATGAGTGTGGAGTCGTTGCAGCCGCAGAATTCAGAATTTGACNNNAGATCGGAA |
| 45989_1 | TATGAGTGTGGAGTCGTTGCGCACAATCTCAGCTCACTGTAACNNNAGATCGGAAG |
| 45989_2 | TATGAGTGTGGAGTCGTTGCGCACAATCTCAGCTCACTGTAACNNNAGATCGGAAG |
| 45989_3 | TATGAGTGTGGAGTCGTTGCGCACAATCTCAGCTCACTGTAACNNNAGATCGGAAG |
| 31149_1 | TATGAGTGTGGAGTCGTTGCATCATTACAGCATAATGGGGCCANNNAGATCGGAA |
| 31149_2 | TATGAGTGTGGAGTCGTTGCATCATTACAGCATAATGGGGCCANNNAGATCGGAA |
| 31149_3 | TATGAGTGTGGAGTCGTTGCATCATTACAGCATAATGGGGCCANNNAGATCGGAA |
| 36928_1 | TATGAGTGTGGAGTCGTTGCCTACTTGGGAGGCTGAGACATGNNNAGATCGGAAG |
| 36928_2 | TATGAGTGTGGAGTCGTTGCCTACTTGGGAGGCTGAGACATGNNNAGATCGGAAG |
| 36928_3 | TATGAGTGTGGAGTCGTTGCCTACTTGGGAGGCTGAGACATGNNNAGATCGGAAG |
| 42547_1 | TATGAGTGTGGAGTCGTTGCCTGAAATGCTGAGTTGCTTGGAANNNAGATCGGAA |
| 42547_2 | TATGAGTGTGGAGTCGTTGCCTGAAATGCTGAGTTGCTTGGAANNNAGATCGGAA |

| | |
|---|---|
| 42547_3 | TATGAGTGTGGAGTCGTTGCCTGAAATGCTGAGTTGCTTGGAANNNAGATCGGAA |
| 50740_1 | TATGAGTGTGGAGTCGTTGCCCTTCTTTTCCCTACCTCCTTCCNNNAGATCGGAAGAG |
| 50740_2 | TATGAGTGTGGAGTCGTTGCCCTTCTTTTCCCTACCTCCTTCCNNNAGATCGGAAGAG |
| 50740_3 | TATGAGTGTGGAGTCGTTGCCCTTCTTTTCCCTACCTCCTTCCNNNAGATCGGAAGAG |
| 50731_1 | TATGAGTGTGGAGTCGTTGCCTCATACCAGTCCCGGTTGGTANNNAGATCGGAAGA |
| 50731_2 | TATGAGTGTGGAGTCGTTGCCTCATACCAGTCCCGGTTGGTANNNAGATCGGAAGA |
| 50731_3 | TATGAGTGTGGAGTCGTTGCCTCATACCAGTCCCGGTTGGTANNNAGATCGGAAGA |
| 40673_1 | TATGAGTGTGGAGTCGTTGCATCACTTGAGTCTGCAAGGTCAANNNAGATCGGAAG |
| 40673_2 | TATGAGTGTGGAGTCGTTGCATCACTTGAGTCTGCAAGGTCAANNNAGATCGGAAG |
| 40673_3 | TATGAGTGTGGAGTCGTTGCATCACTTGAGTCTGCAAGGTCAANNNAGATCGGAAG |
| 51596_1 | TATGAGTGTGGAGTCGTTGCCAGAACACCATCCCTGGACCNNNAGATCGGAAGAGC |
| 51596_2 | TATGAGTGTGGAGTCGTTGCCAGAACACCATCCCTGGACCNNNAGATCGGAAGAGC |
| 51596_3 | TATGAGTGTGGAGTCGTTGCCAGAACACCATCCCTGGACCNNNAGATCGGAAGAGC |
| 32164_1 | TATGAGTGTGGAGTCGTTGCGAGACGGAGTCTTGCTCTGTCNNNAGATCGGAAGA |
| 32164_2 | TATGAGTGTGGAGTCGTTGCGAGACGGAGTCTTGCTCTGTCNNNAGATCGGAAGA |
| 32164_3 | TATGAGTGTGGAGTCGTTGCGAGACGGAGTCTTGCTCTGTCNNNAGATCGGAAGA |
| 13338_1 | TATGAGTGTGGAGTCGTTGCTTCTCTAACTCTGAATCATTCATNNNAGATCGGAAGA |
| 13338_2 | TATGAGTGTGGAGTCGTTGCTTCTCTAACTCTGAATCATTCATNNNAGATCGGAAGA |
| 13338_3 | TATGAGTGTGGAGTCGTTGCTTCTCTAACTCTGAATCATTCATNNNAGATCGGAAGA |
| 29309_1 | TATGAGTGTGGAGTCGTTGCTGGATTCCAGGGCCTATATGTTGNNNAGATCGGAAG |
| 29309_2 | TATGAGTGTGGAGTCGTTGCTGGATTCCAGGGCCTATATGTTGNNNAGATCGGAAG |
| 29309_3 | TATGAGTGTGGAGTCGTTGCTGGATTCCAGGGCCTATATGTTGNNNAGATCGGAAG |
| 45609_1 | TATGAGTGTGGAGTCGTTGCTCAGCCTCCTGAATAGCTAGGATNNNAGATCGGAAG |
| 45609_2 | TATGAGTGTGGAGTCGTTGCTCAGCCTCCTGAATAGCTAGGATNNNAGATCGGAAG |
| 45609_3 | TATGAGTGTGGAGTCGTTGCTCAGCCTCCTGAATAGCTAGGATNNNAGATCGGAAG |
| 39176_1 | TATGAGTGTGGAGTCGTTGCGATGGATGGAACTGGAGGTCATTNNNAGATCGGAA |
| 39176_2 | TATGAGTGTGGAGTCGTTGCGATGGATGGAACTGGAGGTCATTNNNAGATCGGAA |
| 39176_3 | TATGAGTGTGGAGTCGTTGCGATGGATGGAACTGGAGGTCATTNNNAGATCGGAA |
| 36983_1 | TATGAGTGTGGAGTCGTTGCTTATCCTGAAACCATCCCAACCTNNNAGATCGGAAGA |
| 36983_2 | TATGAGTGTGGAGTCGTTGCTTATCCTGAAACCATCCCAACCTNNNAGATCGGAAGA |
| 36983_3 | TATGAGTGTGGAGTCGTTGCTTATCCTGAAACCATCCCAACCTNNNAGATCGGAAGA |
| 37750_1 | TATGAGTGTGGAGTCGTTGCGATTGGTTTTTGTGTGCTGACCTNNNAGATCGGAAG |
| 37750_2 | TATGAGTGTGGAGTCGTTGCGATTGGTTTTTGTGTGCTGACCTNNNAGATCGGAAG |
| 37750_3 | TATGAGTGTGGAGTCGTTGCGATTGGTTTTTGTGTGCTGACCTNNNAGATCGGAAG |
| 33220_1 | TATGAGTGTGGAGTCGTTGCTATACCTCTGCGAATGGAACCAGNNNAGATCGGAAG |
| 33220_2 | TATGAGTGTGGAGTCGTTGCTATACCTCTGCGAATGGAACCAGNNNAGATCGGAAG |
| 33220_3 | TATGAGTGTGGAGTCGTTGCTATACCTCTGCGAATGGAACCAGNNNAGATCGGAAG |
| 49590_1 | TATGAGTGTGGAGTCGTTGCCTCCTGAGTAGCTGGGACTACAGNNNAGATCGGAA |
| 49590_2 | TATGAGTGTGGAGTCGTTGCCTCCTGAGTAGCTGGGACTACAGNNNAGATCGGAA |
| 49590_3 | TATGAGTGTGGAGTCGTTGCCTCCTGAGTAGCTGGGACTACAGNNNAGATCGGAA |
| 31430_1 | TATGAGTGTGGAGTCGTTGCAGAACATTCTATCGGAGATGGGCNNNAGATCGGAA |
| 31430_2 | TATGAGTGTGGAGTCGTTGCAGAACATTCTATCGGAGATGGGCNNNAGATCGGAA |
| 31430_3 | TATGAGTGTGGAGTCGTTGCAGAACATTCTATCGGAGATGGGCNNNAGATCGGAA |

| | |
|---|---|
| 44503_1 | TATGAGTGTGGAGTCGTTGCACTATCTCAGCTCACTGCAAGCNNNAGATCGGAAGA |
| 44503_2 | TATGAGTGTGGAGTCGTTGCACTATCTCAGCTCACTGCAAGCNNNAGATCGGAAGA |
| 44503_3 | TATGAGTGTGGAGTCGTTGCACTATCTCAGCTCACTGCAAGCNNNAGATCGGAAGA |
| 30947_1 | TATGAGTGTGGAGTCGTTGCAAAGCAGGAGAAGGGACAAGAATNNNAGATCGGA |
| 30947_2 | TATGAGTGTGGAGTCGTTGCAAAGCAGGAGAAGGGACAAGAATNNNAGATCGGA |
| 30947_3 | TATGAGTGTGGAGTCGTTGCAAAGCAGGAGAAGGGACAAGAATNNNAGATCGGA |
| 35023_1 | TATGAGTGTGGAGTCGTTGCCTCCCAGGTTCAAGTGATTCTCANNNAGATCGGAAG |
| 35023_2 | TATGAGTGTGGAGTCGTTGCCTCCCAGGTTCAAGTGATTCTCANNNAGATCGGAAG |
| 35023_3 | TATGAGTGTGGAGTCGTTGCCTCCCAGGTTCAAGTGATTCTCANNNAGATCGGAAG |
| 35766_1 | TATGAGTGTGGAGTCGTTGCTCCCTTTATTTCTCAGACCAGCCNNNAGATCGGAAGA |
| 35766_2 | TATGAGTGTGGAGTCGTTGCTCCCTTTATTTCTCAGACCAGCCNNNAGATCGGAAGA |
| 35766_3 | TATGAGTGTGGAGTCGTTGCTCCCTTTATTTCTCAGACCAGCCNNNAGATCGGAAGA |
| 52554_1 | TATGAGTGTGGAGTCGTTGCTCCAGCTCAAACTCCTCTTCTTCNNNAGATCGGAAGA |
| 52554_2 | TATGAGTGTGGAGTCGTTGCTCCAGCTCAAACTCCTCTTCTTCNNNAGATCGGAAGA |
| 52554_3 | TATGAGTGTGGAGTCGTTGCTCCAGCTCAAACTCCTCTTCTTCNNNAGATCGGAAGA |
| 40498_1 | TATGAGTGTGGAGTCGTTGCTTTTTTCTGGAGACGGGGTTTTGNNNAGATCGGAA |
| 40498_2 | TATGAGTGTGGAGTCGTTGCTTTTTTCTGGAGACGGGGTTTTGNNNAGATCGGAA |
| 40498_3 | TATGAGTGTGGAGTCGTTGCTTTTTTCTGGAGACGGGGTTTTGNNNAGATCGGAA |
| 52381_1 | TATGAGTGTGGAGTCGTTGCATTGGGACAGTACTAGCAAAGGGNNNAGATCGGAA |
| 52381_2 | TATGAGTGTGGAGTCGTTGCATTGGGACAGTACTAGCAAAGGGNNNAGATCGGAA |
| 52381_3 | TATGAGTGTGGAGTCGTTGCATTGGGACAGTACTAGCAAAGGGNNNAGATCGGAA |
| 36727_1 | TATGAGTGTGGAGTCGTTGCGAAATAACCAGACACCACCCTCTNNNAGATCGGAAG |
| 36727_2 | TATGAGTGTGGAGTCGTTGCGAAATAACCAGACACCACCCTCTNNNAGATCGGAAG |
| 36727_3 | TATGAGTGTGGAGTCGTTGCGAAATAACCAGACACCACCCTCTNNNAGATCGGAAG |
| 50287_1 | TATGAGTGTGGAGTCGTTGCCGACCTCTCTTTTTCTTTCCCCTNNNAGATCGGAAGA |
| 50287_2 | TATGAGTGTGGAGTCGTTGCCGACCTCTCTTTTTCTTTCCCCTNNNAGATCGGAAGA |
| 50287_3 | TATGAGTGTGGAGTCGTTGCCGACCTCTCTTTTTCTTTCCCCTNNNAGATCGGAAGA |
| 50985_1 | TATGAGTGTGGAGTCGTTGCCTCTTCTTGTGGAAATTCTGCCCNNNAGATCGGAAG |
| 50985_2 | TATGAGTGTGGAGTCGTTGCCTCTTCTTGTGGAAATTCTGCCCNNNAGATCGGAAG |
| 50985_3 | TATGAGTGTGGAGTCGTTGCCTCTTCTTGTGGAAATTCTGCCCNNNAGATCGGAAG |
| 41835_1 | TATGAGTGTGGAGTCGTTGCGTTCAAGCGATTCTCCTGCTTCNNNAGATCGGAAGA |
| 41835_2 | TATGAGTGTGGAGTCGTTGCGTTCAAGCGATTCTCCTGCTTCNNNAGATCGGAAGA |
| 41835_3 | TATGAGTGTGGAGTCGTTGCGTTCAAGCGATTCTCCTGCTTCNNNAGATCGGAAGA |
| 42937_1 | TATGAGTGTGGAGTCGTTGCGCGTTTCTGGTCCTAGTTCTTCTNNNAGATCGGAAG |
| 42937_2 | TATGAGTGTGGAGTCGTTGCGCGTTTCTGGTCCTAGTTCTTCTNNNAGATCGGAAG |
| 42937_3 | TATGAGTGTGGAGTCGTTGCGCGTTTCTGGTCCTAGTTCTTCTNNNAGATCGGAAG |
| 39111_1 | TATGAGTGTGGAGTCGTTGCACATGACATATCACGCGAGATCANNNAGATCGGAAG |
| 39111_2 | TATGAGTGTGGAGTCGTTGCACATGACATATCACGCGAGATCANNNAGATCGGAAG |
| 39111_3 | TATGAGTGTGGAGTCGTTGCACATGACATATCACGCGAGATCANNNAGATCGGAAG |
| 46210_1 | TATGAGTGTGGAGTCGTTGCGGCATTGAAAATGCAAAGCACAANNNAGATCGGAA |
| 46210_2 | TATGAGTGTGGAGTCGTTGCGGCATTGAAAATGCAAAGCACAANNNAGATCGGAA |
| 46210_3 | TATGAGTGTGGAGTCGTTGCGGCATTGAAAATGCAAAGCACAANNNAGATCGGAA |
| 45178_1 | TATGAGTGTGGAGTCGTTGCGTAAGAAGAGCCATCCACCCTAGNNNAGATCGGAA |

| | |
|---|---|
| 45178_2 | TATGAGTGTGGAGTCGTTGCGTAAGAAGAGCCATCCACCCTAGNNNAGATCGGAAG |
| 45178_3 | TATGAGTGTGGAGTCGTTGCGTAAGAAGAGCCATCCACCCTAGNNNAGATCGGAAG |
| 41207_1 | TATGAGTGTGGAGTCGTTGCCATGAGGGGAAGGTGAGGTTTGNNNAGATCGGAA |
| 41207_2 | TATGAGTGTGGAGTCGTTGCCATGAGGGGAAGGTGAGGTTTGNNNAGATCGGAA |
| 41207_3 | TATGAGTGTGGAGTCGTTGCCATGAGGGGAAGGTGAGGTTTGNNNAGATCGGAA |
| 29990_1 | TATGAGTGTGGAGTCGTTGCGCCTCATAAGAGACTCCCTGTCNNNAGATCGGAAGA |
| 29990_2 | TATGAGTGTGGAGTCGTTGCGCCTCATAAGAGACTCCCTGTCNNNAGATCGGAAGA |
| 29990_3 | TATGAGTGTGGAGTCGTTGCGCCTCATAAGAGACTCCCTGTCNNNAGATCGGAAGA |
| 51326_1 | TATGAGTGTGGAGTCGTTGCGCATGGCACATGTATGCATATGTNNNAGATCGGAAG |
| 51326_2 | TATGAGTGTGGAGTCGTTGCGCATGGCACATGTATGCATATGTNNNAGATCGGAAG |
| 51326_3 | TATGAGTGTGGAGTCGTTGCGCATGGCACATGTATGCATATGTNNNAGATCGGAAG |
| 48452_1 | TATGAGTGTGGAGTCGTTGCCCCGAGTAGCTGGGATTACAATTNNNAGATCGGAAG |
| 48452_2 | TATGAGTGTGGAGTCGTTGCCCCGAGTAGCTGGGATTACAATTNNNAGATCGGAAG |
| 48452_3 | TATGAGTGTGGAGTCGTTGCCCCGAGTAGCTGGGATTACAATTNNNAGATCGGAAG |
| 38099_1 | TATGAGTGTGGAGTCGTTGCGAGAAAATGATGGCCAGACACAGNNNAGATCGGAA |
| 38099_2 | TATGAGTGTGGAGTCGTTGCGAGAAAATGATGGCCAGACACAGNNNAGATCGGAA |
| 38099_3 | TATGAGTGTGGAGTCGTTGCGAGAAAATGATGGCCAGACACAGNNNAGATCGGAA |
| 32458_1 | TATGAGTGTGGAGTCGTTGCGGTGAAACCCCATCTCTACCAAANNNAGATCGGAAG |
| 32458_2 | TATGAGTGTGGAGTCGTTGCGGTGAAACCCCATCTCTACCAAANNNAGATCGGAAG |
| 32458_3 | TATGAGTGTGGAGTCGTTGCGGTGAAACCCCATCTCTACCAAANNNAGATCGGAAG |
| 30622_1 | TATGAGTGTGGAGTCGTTGCAGTAAAGGGAGATCAAGCTCCTGTNNNAGATCGGA |
| 30622_2 | TATGAGTGTGGAGTCGTTGCAGTAAAGGGAGATCAAGCTCCTGTNNNAGATCGGA |
| 30622_3 | TATGAGTGTGGAGTCGTTGCAGTAAAGGGAGATCAAGCTCCTGTNNNAGATCGGA |
| 43142_1 | TATGAGTGTGGAGTCGTTGCGCAGAAGCCATTACACTTCAGTTNNNAGATCGGAAG |
| 43142_2 | TATGAGTGTGGAGTCGTTGCGCAGAAGCCATTACACTTCAGTTNNNAGATCGGAAG |
| 43142_3 | TATGAGTGTGGAGTCGTTGCGCAGAAGCCATTACACTTCAGTTNNNAGATCGGAAG |
| 2496_1 | TATGAGTGTGGAGTCGTTGCGTCAACAGAGCAGGCAGATAGAGNNNAGATCGGAA |
| 2496_2 | TATGAGTGTGGAGTCGTTGCGTCAACAGAGCAGGCAGATAGAGNNNAGATCGGAA |
| 2496_3 | TATGAGTGTGGAGTCGTTGCGTCAACAGAGCAGGCAGATAGAGNNNAGATCGGAA |
| 37649_1 | TATGAGTGTGGAGTCGTTGCAGAAAGCTGTACTGATACGTGCTNNNAGATCGGAA |
| 37649_2 | TATGAGTGTGGAGTCGTTGCAGAAAGCTGTACTGATACGTGCTNNNAGATCGGAA |
| 37649_3 | TATGAGTGTGGAGTCGTTGCAGAAAGCTGTACTGATACGTGCTNNNAGATCGGAA |
| 750_1 | TATGAGTGTGGAGTCGTTGCACTCTCATTGCTGGCCTTGAAGANNNAGATCGGAAG |
| 750_2 | TATGAGTGTGGAGTCGTTGCACTCTCATTGCTGGCCTTGAAGANNNAGATCGGAAG |
| 750_3 | TATGAGTGTGGAGTCGTTGCACTCTCATTGCTGGCCTTGAAGANNNAGATCGGAAG |
| 32515_1 | TATGAGTGTGGAGTCGTTGCATTCTCGTGTATGGCTTGAGGAANNNAGATCGGAA |
| 32515_2 | TATGAGTGTGGAGTCGTTGCATTCTCGTGTATGGCTTGAGGAANNNAGATCGGAA |
| 32515_3 | TATGAGTGTGGAGTCGTTGCATTCTCGTGTATGGCTTGAGGAANNNAGATCGGAA |
| 42507_1 | TATGAGTGTGGAGTCGTTGCGGAGGTGGGAGAATCAATTGAGTNNNAGATCGGA |
| 42507_2 | TATGAGTGTGGAGTCGTTGCGGAGGTGGGAGAATCAATTGAGTNNNAGATCGGA |
| 42507_3 | TATGAGTGTGGAGTCGTTGCGGAGGTGGGAGAATCAATTGAGTNNNAGATCGGA |
| 2497_1 | TATGAGTGTGGAGTCGTTGCTGGCTTGTGAGACAGAAAAGGAGNNNAGATCGGA |
| 2497_2 | TATGAGTGTGGAGTCGTTGCTGGCTTGTGAGACAGAAAAGGAGNNNAGATCGGA |

| | |
|---|---|
| 2497_3 | TATGAGTGTGGAGTCGTTGCTGGCTTGTGAGACAGAAAAGGAGNNNAGATCGGA |
| 52127_1 | TATGAGTGTGGAGTCGTTGCGCAGTCTGCTTGCTGTTTCTTANNNAGATCGGAAG |
| 52127_2 | TATGAGTGTGGAGTCGTTGCGCAGTCTGCTTGCTGTTTCTTANNNAGATCGGAAG |
| 52127_3 | TATGAGTGTGGAGTCGTTGCGCAGTCTGCTTGCTGTTTCTTANNNAGATCGGAAG |
| 31846_1 | TATGAGTGTGGAGTCGTTGCGGATGGATACTTGCTGCTTTTCCNNNAGATCGGAAG |
| 31846_2 | TATGAGTGTGGAGTCGTTGCGGATGGATACTTGCTGCTTTTCCNNNAGATCGGAAG |
| 31846_3 | TATGAGTGTGGAGTCGTTGCGGATGGATACTTGCTGCTTTTCCNNNAGATCGGAAG |
| 39448_1 | TATGAGTGTGGAGTCGTTGCTCTGCCACATAAAAACTGCATGCNNNAGATCGGAAG |
| 39448_2 | TATGAGTGTGGAGTCGTTGCTCTGCCACATAAAAACTGCATGCNNNAGATCGGAAG |
| 39448_3 | TATGAGTGTGGAGTCGTTGCTCTGCCACATAAAAACTGCATGCNNNAGATCGGAAG |
| 33487_1 | TATGAGTGTGGAGTCGTTGCGTGAGTCTGAGGGATGACAAGTTNNNAGATCGGAA |
| 33487_2 | TATGAGTGTGGAGTCGTTGCGTGAGTCTGAGGGATGACAAGTTNNNAGATCGGAA |
| 33487_3 | TATGAGTGTGGAGTCGTTGCGTGAGTCTGAGGGATGACAAGTTNNNAGATCGGAA |
| 51673_1 | TATGAGTGTGGAGTCGTTGCTCTGCCTCTTTCTGTTTCCAAGANNNAGATCGGAAGA |
| 51673_2 | TATGAGTGTGGAGTCGTTGCTCTGCCTCTTTCTGTTTCCAAGANNNAGATCGGAAGA |
| 51673_3 | TATGAGTGTGGAGTCGTTGCTCTGCCTCTTTCTGTTTCCAAGANNNAGATCGGAAGA |
| 50991_1 | TATGAGTGTGGAGTCGTTGCGTGATCATGAGCCTACACCAAGANNNAGATCGGAA |
| 50991_2 | TATGAGTGTGGAGTCGTTGCGTGATCATGAGCCTACACCAAGANNNAGATCGGAA |
| 50991_3 | TATGAGTGTGGAGTCGTTGCGTGATCATGAGCCTACACCAAGANNNAGATCGGAA |
| 31638_1 | TATGAGTGTGGAGTCGTTGCTTAGCACCAGGGACCTACTTAGANNNAGATCGGAAG |
| 31638_2 | TATGAGTGTGGAGTCGTTGCTTAGCACCAGGGACCTACTTAGANNNAGATCGGAAG |
| 31638_3 | TATGAGTGTGGAGTCGTTGCTTAGCACCAGGGACCTACTTAGANNNAGATCGGAAG |
| 3498_1 | TATGAGTGTGGAGTCGTTGCCCTCCCAGATTCTCTCCCACCTTNNNAGATCGGAAGA |
| 3498_2 | TATGAGTGTGGAGTCGTTGCCCTCCCAGATTCTCTCCCACCTTNNNAGATCGGAAGA |
| 3498_3 | TATGAGTGTGGAGTCGTTGCCCTCCCAGATTCTCTCCCACCTTNNNAGATCGGAAGA |
| 29619_1 | TATGAGTGTGGAGTCGTTGCACATGCACCAAAAATTTCAAGGTNNNAGATCGGAAG |
| 29619_2 | TATGAGTGTGGAGTCGTTGCACATGCACCAAAAATTTCAAGGTNNNAGATCGGAAG |
| 29619_3 | TATGAGTGTGGAGTCGTTGCACATGCACCAAAAATTTCAAGGTNNNAGATCGGAAG |
| 13648_1 | TATGAGTGTGGAGTCGTTGCGCTAGGATCTTTAGAGAAGTTTANNNAGATCGGAA |
| 13648_2 | TATGAGTGTGGAGTCGTTGCGCTAGGATCTTTAGAGAAGTTTANNNAGATCGGAA |
| 13648_3 | TATGAGTGTGGAGTCGTTGCGCTAGGATCTTTAGAGAAGTTTANNNAGATCGGAA |
| 46425_1 | TATGAGTGTGGAGTCGTTGCCTCTCGGATTCACGCCATTCTCNNNAGATCGGAAGA |
| 46425_2 | TATGAGTGTGGAGTCGTTGCCTCTCGGATTCACGCCATTCTCNNNAGATCGGAAGA |
| 46425_3 | TATGAGTGTGGAGTCGTTGCCTCTCGGATTCACGCCATTCTCNNNAGATCGGAAGA |
| 12531_1 | TATGAGTGTGGAGTCGTTGCGTACAGAAGACTCAAAAACTTCNNNAGATCGGAAG |
| 12531_2 | TATGAGTGTGGAGTCGTTGCGTACAGAAGACTCAAAAACTTCNNNAGATCGGAAG |
| 12531_3 | TATGAGTGTGGAGTCGTTGCGTACAGAAGACTCAAAAACTTCNNNAGATCGGAAG |
| 43931_1 | TATGAGTGTGGAGTCGTTGCAAAGGATGAATGGAAGCAGGGATNNNAGATCGGA |
| 43931_2 | TATGAGTGTGGAGTCGTTGCAAAGGATGAATGGAAGCAGGGATNNNAGATCGGA |
| 43931_3 | TATGAGTGTGGAGTCGTTGCAAAGGATGAATGGAAGCAGGGATNNNAGATCGGA |
| 49390_1 | TATGAGTGTGGAGTCGTTGCGCACAAATGAACACCTCAGACTCNNNAGATCGGAAG |
| 49390_2 | TATGAGTGTGGAGTCGTTGCGCACAAATGAACACCTCAGACTCNNNAGATCGGAAG |
| 49390_3 | TATGAGTGTGGAGTCGTTGCGCACAAATGAACACCTCAGACTCNNNAGATCGGAAG |

| | |
|---|---|
| 2519_1 | TATGAGTGTGGAGTCGTTGCTTACCACTCTGTCTCTCTTCCCGNNNAGATCGGAAGA |
| 2519_2 | TATGAGTGTGGAGTCGTTGCTTACCACTCTGTCTCTCTTCCCGNNNAGATCGGAAGA |
| 2519_3 | TATGAGTGTGGAGTCGTTGCTTACCACTCTGTCTCTCTTCCCGNNNAGATCGGAAGA |
| 29398_1 | TATGAGTGTGGAGTCGTTGCTTGCTTGTCAGATGCCTATGACANNNAGATCGGAAG |
| 29398_2 | TATGAGTGTGGAGTCGTTGCTTGCTTGTCAGATGCCTATGACANNNAGATCGGAAG |
| 29398_3 | TATGAGTGTGGAGTCGTTGCTTGCTTGTCAGATGCCTATGACANNNAGATCGGAAG |
| 36337_1 | TATGAGTGTGGAGTCGTTGCACATTGCGCTAAGTGAAAGAAGCNNNAGATCGGAA |
| 36337_2 | TATGAGTGTGGAGTCGTTGCACATTGCGCTAAGTGAAAGAAGCNNNAGATCGGAA |
| 36337_3 | TATGAGTGTGGAGTCGTTGCACATTGCGCTAAGTGAAAGAAGCNNNAGATCGGAA |
| 12546_1 | TATGAGTGTGGAGTCGTTGCCAATGGACTACTTGGAATTTTTANNNAGATCGGAAG |
| 12546_2 | TATGAGTGTGGAGTCGTTGCCAATGGACTACTTGGAATTTTTANNNAGATCGGAAG |
| 12546_3 | TATGAGTGTGGAGTCGTTGCCAATGGACTACTTGGAATTTTTANNNAGATCGGAAG |
| 46871_1 | TATGAGTGTGGAGTCGTTGCGTAGCTGGGATTACAGGCGTATGNNNAGATCGGAA |
| 46871_2 | TATGAGTGTGGAGTCGTTGCGTAGCTGGGATTACAGGCGTATGNNNAGATCGGAA |
| 46871_3 | TATGAGTGTGGAGTCGTTGCGTAGCTGGGATTACAGGCGTATGNNNAGATCGGAA |
| 10194_1 | TATGAGTGTGGAGTCGTTGCATAGTCCAAAGATGAGCTGCAGGNNNAGATCGGAA |
| 10194_2 | TATGAGTGTGGAGTCGTTGCATAGTCCAAAGATGAGCTGCAGGNNNAGATCGGAA |
| 10194_3 | TATGAGTGTGGAGTCGTTGCATAGTCCAAAGATGAGCTGCAGGNNNAGATCGGAA |
| 52774_1 | TATGAGTGTGGAGTCGTTGCCTCTCCAACTGCCCAGAGACNNNAGATCGGAAGAGC |
| 52774_2 | TATGAGTGTGGAGTCGTTGCCTCTCCAACTGCCCAGAGACNNNAGATCGGAAGAGC |
| 52774_3 | TATGAGTGTGGAGTCGTTGCCTCTCCAACTGCCCAGAGACNNNAGATCGGAAGAGC |
| 35339_1 | TATGAGTGTGGAGTCGTTGCCTCCTGAGTACCTGGGACTACAGNNNAGATCGGAAG |
| 35339_2 | TATGAGTGTGGAGTCGTTGCCTCCTGAGTACCTGGGACTACAGNNNAGATCGGAAG |
| 35339_3 | TATGAGTGTGGAGTCGTTGCCTCCTGAGTACCTGGGACTACAGNNNAGATCGGAAG |
| 45411_1 | TATGAGTGTGGAGTCGTTGCACAAAGTTCTGACCAAAACACCGNNNAGATCGGAAG |
| 45411_2 | TATGAGTGTGGAGTCGTTGCACAAAGTTCTGACCAAAACACCGNNNAGATCGGAAG |
| 45411_3 | TATGAGTGTGGAGTCGTTGCACAAAGTTCTGACCAAAACACCGNNNAGATCGGAAG |
| 37080_1 | TATGAGTGTGGAGTCGTTGCAGCAATGAGGCAGTATGCAAAATNNNAGATCGGAA |
| 37080_2 | TATGAGTGTGGAGTCGTTGCAGCAATGAGGCAGTATGCAAAATNNNAGATCGGAA |
| 37080_3 | TATGAGTGTGGAGTCGTTGCAGCAATGAGGCAGTATGCAAAATNNNAGATCGGAA |
| 36656_1 | TATGAGTGTGGAGTCGTTGCAGTTCAGGAAAATGTGATGCTTCANNNAGATCGGA |
| 36656_2 | TATGAGTGTGGAGTCGTTGCAGTTCAGGAAAATGTGATGCTTCANNNAGATCGGA |
| 36656_3 | TATGAGTGTGGAGTCGTTGCAGTTCAGGAAAATGTGATGCTTCANNNAGATCGGA |
| 2441_1 | TATGAGTGTGGAGTCGTTGCCTTCCTCCCTCACCATCTCTCTCNNNAGATCGGAAGAG |
| 2441_2 | TATGAGTGTGGAGTCGTTGCCTTCCTCCCTCACCATCTCTCTCNNNAGATCGGAAGAG |
| 2441_3 | TATGAGTGTGGAGTCGTTGCCTTCCTCCCTCACCATCTCTCTCNNNAGATCGGAAGAG |
| 47919_1 | TATGAGTGTGGAGTCGTTGCTGAGTAGTTGGGATTACAGGCACNNNAGATCGGAA |
| 47919_2 | TATGAGTGTGGAGTCGTTGCTGAGTAGTTGGGATTACAGGCACNNNAGATCGGAA |
| 47919_3 | TATGAGTGTGGAGTCGTTGCTGAGTAGTTGGGATTACAGGCACNNNAGATCGGAA |
| 47767_1 | TATGAGTGTGGAGTCGTTGCCACGCCCAGCTAAGTTTTTGTATNNNAGATCGGAAG |
| 47767_2 | TATGAGTGTGGAGTCGTTGCCACGCCCAGCTAAGTTTTTGTATNNNAGATCGGAAG |
| 47767_3 | TATGAGTGTGGAGTCGTTGCCACGCCCAGCTAAGTTTTTGTATNNNAGATCGGAAG |
| 52334_1 | TATGAGTGTGGAGTCGTTGCGGACCTTCATCGAGTCACTGTAGNNNAGATCGGAAG |

| | |
|---|---|
| 52334_2 | TATGAGTGTGGAGTCGTTGCGGACCTTCATCGAGTCACTGTAGNNNAGATCGGAAG |
| 52334_3 | TATGAGTGTGGAGTCGTTGCGGACCTTCATCGAGTCACTGTAGNNNAGATCGGAAG |
| 34396_1 | TATGAGTGTGGAGTCGTTGCGACTTGGAACTTTGACTCTTGGCNNNAGATCGGAAG |
| 34396_2 | TATGAGTGTGGAGTCGTTGCGACTTGGAACTTTGACTCTTGGCNNNAGATCGGAAG |
| 34396_3 | TATGAGTGTGGAGTCGTTGCGACTTGGAACTTTGACTCTTGGCNNNAGATCGGAAG |
| 50081_1 | TATGAGTGTGGAGTCGTTGCTATCCTTAGGAAGCACAGTCAGGNNNAGATCGGAA |
| 50081_2 | TATGAGTGTGGAGTCGTTGCTATCCTTAGGAAGCACAGTCAGGNNNAGATCGGAA |
| 50081_3 | TATGAGTGTGGAGTCGTTGCTATCCTTAGGAAGCACAGTCAGGNNNAGATCGGAA |
| 32511_1 | TATGAGTGTGGAGTCGTTGCGAAATGGCCCCACAATCTAACCNNNAGATCGGAAGA |
| 32511_2 | TATGAGTGTGGAGTCGTTGCGAAATGGCCCCACAATCTAACCNNNAGATCGGAAGA |
| 32511_3 | TATGAGTGTGGAGTCGTTGCGAAATGGCCCCACAATCTAACCNNNAGATCGGAAGA |
| 42440_1 | TATGAGTGTGGAGTCGTTGCGGCTGTGATCAAAGAGGAGAGATNNNAGATCGGA |
| 42440_2 | TATGAGTGTGGAGTCGTTGCGGCTGTGATCAAAGAGGAGAGATNNNAGATCGGA |
| 42440_3 | TATGAGTGTGGAGTCGTTGCGGCTGTGATCAAAGAGGAGAGATNNNAGATCGGA |
| 34319_1 | TATGAGTGTGGAGTCGTTGCGCTCCTCACTTCTCAGACGGNNNAGATCGGAAGAGC |
| 34319_2 | TATGAGTGTGGAGTCGTTGCGCTCCTCACTTCTCAGACGGNNNAGATCGGAAGAGC |
| 34319_3 | TATGAGTGTGGAGTCGTTGCGCTCCTCACTTCTCAGACGGNNNAGATCGGAAGAGC |
| 45091_1 | TATGAGTGTGGAGTCGTTGCTCAGTCCCTCACCAAACTGAATCNNNAGATCGGAAG |
| 45091_2 | TATGAGTGTGGAGTCGTTGCTCAGTCCCTCACCAAACTGAATCNNNAGATCGGAAG |
| 45091_3 | TATGAGTGTGGAGTCGTTGCTCAGTCCCTCACCAAACTGAATCNNNAGATCGGAAG |
| 32176_1 | TATGAGTGTGGAGTCGTTGCCTGAAGCGGGTGGATAATTTGAGNNNAGATCGGAA |
| 32176_2 | TATGAGTGTGGAGTCGTTGCCTGAAGCGGGTGGATAATTTGAGNNNAGATCGGAA |
| 32176_3 | TATGAGTGTGGAGTCGTTGCCTGAAGCGGGTGGATAATTTGAGNNNAGATCGGAA |
| 36795_1 | TATGAGTGTGGAGTCGTTGCAAGAAAGCTTCATTTGAGCAGGTNNNAGATCGGAA |
| 36795_2 | TATGAGTGTGGAGTCGTTGCAAGAAAGCTTCATTTGAGCAGGTNNNAGATCGGAA |
| 36795_3 | TATGAGTGTGGAGTCGTTGCAAGAAAGCTTCATTTGAGCAGGTNNNAGATCGGAA |
| 51062_1 | TATGAGTGTGGAGTCGTTGCGTAGCCTTGGAGTTCCCTAGAACNNNAGATCGGAAG |
| 51062_2 | TATGAGTGTGGAGTCGTTGCGTAGCCTTGGAGTTCCCTAGAACNNNAGATCGGAAG |
| 51062_3 | TATGAGTGTGGAGTCGTTGCGTAGCCTTGGAGTTCCCTAGAACNNNAGATCGGAAG |
| 50394_1 | TATGAGTGTGGAGTCGTTGCCAGCCTGTGAAGGGAAGGAGNNNAGATCGGAAGA |
| 50394_2 | TATGAGTGTGGAGTCGTTGCCAGCCTGTGAAGGGAAGGAGNNNAGATCGGAAGA |
| 50394_3 | TATGAGTGTGGAGTCGTTGCCAGCCTGTGAAGGGAAGGAGNNNAGATCGGAAGA |
| 42555_1 | TATGAGTGTGGAGTCGTTGCCACCACCGCAATACTATGCAATCNNNAGATCGGAAG |
| 42555_2 | TATGAGTGTGGAGTCGTTGCCACCACCGCAATACTATGCAATCNNNAGATCGGAAG |
| 42555_3 | TATGAGTGTGGAGTCGTTGCCACCACCGCAATACTATGCAATCNNNAGATCGGAAG |
| 35630_1 | TATGAGTGTGGAGTCGTTGCGTTCATGCCATTCTCCTACCTCANNNAGATCGGAAGA |
| 35630_2 | TATGAGTGTGGAGTCGTTGCGTTCATGCCATTCTCCTACCTCANNNAGATCGGAAGA |
| 35630_3 | TATGAGTGTGGAGTCGTTGCGTTCATGCCATTCTCCTACCTCANNNAGATCGGAAGA |
| 35913_1 | TATGAGTGTGGAGTCGTTGCTGCCATCATAGTTTCCAGAAGCANNNAGATCGGAAG |
| 35913_2 | TATGAGTGTGGAGTCGTTGCTGCCATCATAGTTTCCAGAAGCANNNAGATCGGAAG |
| 35913_3 | TATGAGTGTGGAGTCGTTGCTGCCATCATAGTTTCCAGAAGCANNNAGATCGGAAG |
| 49200_1 | TATGAGTGTGGAGTCGTTGCTATTTCTGTGGGATCAGTGGCAANNNAGATCGGAA |
| 49200_2 | TATGAGTGTGGAGTCGTTGCTATTTCTGTGGGATCAGTGGCAANNNAGATCGGAA |

| | |
|---|---|
| 49200_3 | TATGAGTGTGGAGTCGTTGCTATTTCTGTGGGATCAGTGGCAANNNAGATCGGAA |
| 34810_1 | TATGAGTGTGGAGTCGTTGCTCCTCTGCTTCCTGAGTTCAATCNNNAGATCGGAAGA |
| 34810_2 | TATGAGTGTGGAGTCGTTGCTCCTCTGCTTCCTGAGTTCAATCNNNAGATCGGAAGA |
| 34810_3 | TATGAGTGTGGAGTCGTTGCTCCTCTGCTTCCTGAGTTCAATCNNNAGATCGGAAGA |
| 35194_1 | TATGAGTGTGGAGTCGTTGCAGATGGGGTTTCATTCTTGTTGCNNNAGATCGGAAG |
| 35194_2 | TATGAGTGTGGAGTCGTTGCAGATGGGGTTTCATTCTTGTTGCNNNAGATCGGAAG |
| 35194_3 | TATGAGTGTGGAGTCGTTGCAGATGGGGTTTCATTCTTGTTGCNNNAGATCGGAAG |
| 40106_1 | TATGAGTGTGGAGTCGTTGCTCTTACCTCCGAATGATTGCACTNNNAGATCGGAAGA |
| 40106_2 | TATGAGTGTGGAGTCGTTGCTCTTACCTCCGAATGATTGCACTNNNAGATCGGAAGA |
| 40106_3 | TATGAGTGTGGAGTCGTTGCTCTTACCTCCGAATGATTGCACTNNNAGATCGGAAGA |
| 39915_1 | TATGAGTGTGGAGTCGTTGCGAGGTTGCAGTGAGCTGAGATTNNNAGATCGGAAG |
| 39915_2 | TATGAGTGTGGAGTCGTTGCGAGGTTGCAGTGAGCTGAGATTNNNAGATCGGAAG |
| 39915_3 | TATGAGTGTGGAGTCGTTGCGAGGTTGCAGTGAGCTGAGATTNNNAGATCGGAAG |
| 43874_1 | TATGAGTGTGGAGTCGTTGCAATCATCCCCAAGTCATGCTTCTNNNAGATCGGAAGA |
| 43874_2 | TATGAGTGTGGAGTCGTTGCAATCATCCCCAAGTCATGCTTCTNNNAGATCGGAAGA |
| 43874_3 | TATGAGTGTGGAGTCGTTGCAATCATCCCCAAGTCATGCTTCTNNNAGATCGGAAGA |
| 50301_1 | TATGAGTGTGGAGTCGTTGCGTCATGAGATGTAGGGACCAGCNNNAGATCGGAAG |
| 50301_2 | TATGAGTGTGGAGTCGTTGCGTCATGAGATGTAGGGACCAGCNNNAGATCGGAAG |
| 50301_3 | TATGAGTGTGGAGTCGTTGCGTCATGAGATGTAGGGACCAGCNNNAGATCGGAAG |
| 49719_1 | TATGAGTGTGGAGTCGTTGCTTCAAGTGATTCTCCTGCCTCAGNNNAGATCGGAAGA |
| 49719_2 | TATGAGTGTGGAGTCGTTGCTTCAAGTGATTCTCCTGCCTCAGNNNAGATCGGAAGA |
| 49719_3 | TATGAGTGTGGAGTCGTTGCTTCAAGTGATTCTCCTGCCTCAGNNNAGATCGGAAGA |
| 36701_1 | TATGAGTGTGGAGTCGTTGCTGAGAGGCAATAGTGACAATGCTNNNAGATCGGAA |
| 36701_2 | TATGAGTGTGGAGTCGTTGCTGAGAGGCAATAGTGACAATGCTNNNAGATCGGAA |
| 36701_3 | TATGAGTGTGGAGTCGTTGCTGAGAGGCAATAGTGACAATGCTNNNAGATCGGAA |
| 34495_1 | TATGAGTGTGGAGTCGTTGCGAGAGGTCCTATGGGAAGGGATCNNNAGATCGGAA |
| 34495_2 | TATGAGTGTGGAGTCGTTGCGAGAGGTCCTATGGGAAGGGATCNNNAGATCGGAA |
| 34495_3 | TATGAGTGTGGAGTCGTTGCGAGAGGTCCTATGGGAAGGGATCNNNAGATCGGAA |
| 49621_1 | TATGAGTGTGGAGTCGTTGCGCCACCACACCAAGCTAATTTTTNNNAGATCGGAAGA |
| 49621_2 | TATGAGTGTGGAGTCGTTGCGCCACCACACCAAGCTAATTTTTNNNAGATCGGAAGA |
| 49621_3 | TATGAGTGTGGAGTCGTTGCGCCACCACACCAAGCTAATTTTTNNNAGATCGGAAGA |
| 34071_1 | TATGAGTGTGGAGTCGTTGCGAGACGGAGTCTTGCTCTGTCNNNAGATCGGAAGA |
| 34071_2 | TATGAGTGTGGAGTCGTTGCGAGACGGAGTCTTGCTCTGTCNNNAGATCGGAAGA |
| 34071_3 | TATGAGTGTGGAGTCGTTGCGAGACGGAGTCTTGCTCTGTCNNNAGATCGGAAGA |
| 50905_1 | TATGAGTGTGGAGTCGTTGCCCAGTTCAATGATGGTCACATGCNNNAGATCGGAAG |
| 50905_2 | TATGAGTGTGGAGTCGTTGCCCAGTTCAATGATGGTCACATGCNNNAGATCGGAAG |
| 50905_3 | TATGAGTGTGGAGTCGTTGCCCAGTTCAATGATGGTCACATGCNNNAGATCGGAAG |
| 36308_1 | TATGAGTGTGGAGTCGTTGCGAGGCTGATCGGAAAGTATCACTNNNAGATCGGAA |
| 36308_2 | TATGAGTGTGGAGTCGTTGCGAGGCTGATCGGAAAGTATCACTNNNAGATCGGAA |
| 36308_3 | TATGAGTGTGGAGTCGTTGCGAGGCTGATCGGAAAGTATCACTNNNAGATCGGAA |
| 52541_1 | TATGAGTGTGGAGTCGTTGCTGTTCCTGCCAGTTTACACCTTANNNAGATCGGAAGA |
| 52541_2 | TATGAGTGTGGAGTCGTTGCTGTTCCTGCCAGTTTACACCTTANNNAGATCGGAAGA |
| 52541_3 | TATGAGTGTGGAGTCGTTGCTGTTCCTGCCAGTTTACACCTTANNNAGATCGGAAGA |

| | |
|---|---|
| 42554_1 | TATGAGTGTGGAGTCGTTGCCACCATGGAATAGTACTCAGCCANNNAGATCGGAAG |
| 42554_2 | TATGAGTGTGGAGTCGTTGCCACCATGGAATAGTACTCAGCCANNNAGATCGGAAG |
| 42554_3 | TATGAGTGTGGAGTCGTTGCCACCATGGAATAGTACTCAGCCANNNAGATCGGAAG |
| 48786_1 | TATGAGTGTGGAGTCGTTGCTTCAAGCGATTCTCCTGCCTCNNNAGATCGGAAGAG |
| 48786_2 | TATGAGTGTGGAGTCGTTGCTTCAAGCGATTCTCCTGCCTCNNNAGATCGGAAGAG |
| 48786_3 | TATGAGTGTGGAGTCGTTGCTTCAAGCGATTCTCCTGCCTCNNNAGATCGGAAGAG |
| 40247_1 | TATGAGTGTGGAGTCGTTGCATTCATTTGCACTTGGGAGGTTGNNNAGATCGGAA |
| 40247_2 | TATGAGTGTGGAGTCGTTGCATTCATTTGCACTTGGGAGGTTGNNNAGATCGGAA |
| 40247_3 | TATGAGTGTGGAGTCGTTGCATTCATTTGCACTTGGGAGGTTGNNNAGATCGGAA |
| 51167_1 | TATGAGTGTGGAGTCGTTGCCCTTCCATTTGAGGAGAGGAGACNNNAGATCGGAA |
| 51167_2 | TATGAGTGTGGAGTCGTTGCCCTTCCATTTGAGGAGAGGAGACNNNAGATCGGAA |
| 51167_3 | TATGAGTGTGGAGTCGTTGCCCTTCCATTTGAGGAGAGGAGACNNNAGATCGGAA |
| 31770_1 | TATGAGTGTGGAGTCGTTGCCACACAGAGAGAGATGAGGGAACNNNAGATCGGA |
| 31770_2 | TATGAGTGTGGAGTCGTTGCCACACAGAGAGAGATGAGGGAACNNNAGATCGGA |
| 31770_3 | TATGAGTGTGGAGTCGTTGCCACACAGAGAGAGATGAGGGAACNNNAGATCGGA |
| 31947_1 | TATGAGTGTGGAGTCGTTGCTCTGAAAAAAAATGGCTGTGAAGCNNNAGATCGGA |
| 31947_2 | TATGAGTGTGGAGTCGTTGCTCTGAAAAAAAATGGCTGTGAAGCNNNAGATCGGA |
| 31947_3 | TATGAGTGTGGAGTCGTTGCTCTGAAAAAAAATGGCTGTGAAGCNNNAGATCGGA |
| 9940_1 | TATGAGTGTGGAGTCGTTGCAGTATGAACACAGCGATTTCCCNNNAGATCGGAAG |
| 9940_2 | TATGAGTGTGGAGTCGTTGCAGTATGAACACAGCGATTTCCCNNNAGATCGGAAG |
| 9940_3 | TATGAGTGTGGAGTCGTTGCAGTATGAACACAGCGATTTCCCNNNAGATCGGAAG |
| 53134_1 | TATGAGTGTGGAGTCGTTGCGAAAATTCTGTCGCAACGGAGACNNNAGATCGGAA |
| 53134_2 | TATGAGTGTGGAGTCGTTGCGAAAATTCTGTCGCAACGGAGACNNNAGATCGGAA |
| 53134_3 | TATGAGTGTGGAGTCGTTGCGAAAATTCTGTCGCAACGGAGACNNNAGATCGGAA |
| 32552_1 | TATGAGTGTGGAGTCGTTGCGGGTTCACGCCATGATAAGAATGNNNAGATCGGAA |
| 32552_2 | TATGAGTGTGGAGTCGTTGCGGGTTCACGCCATGATAAGAATGNNNAGATCGGAA |
| 32552_3 | TATGAGTGTGGAGTCGTTGCGGGTTCACGCCATGATAAGAATGNNNAGATCGGAA |
| 10066_1 | TATGAGTGTGGAGTCGTTGCCTCCTGTTTGAGTGCTCCTGAAGNNNAGATCGGAAG |
| 10066_2 | TATGAGTGTGGAGTCGTTGCCTCCTGTTTGAGTGCTCCTGAAGNNNAGATCGGAAG |
| 10066_3 | TATGAGTGTGGAGTCGTTGCCTCCTGTTTGAGTGCTCCTGAAGNNNAGATCGGAAG |
| 37650_1 | TATGAGTGTGGAGTCGTTGCCAAACTCATAGGGTCCAGACGATNNNAGATCGGAA |
| 37650_2 | TATGAGTGTGGAGTCGTTGCCAAACTCATAGGGTCCAGACGATNNNAGATCGGAA |
| 37650_3 | TATGAGTGTGGAGTCGTTGCCAAACTCATAGGGTCCAGACGATNNNAGATCGGAA |
| 47377_1 | TATGAGTGTGGAGTCGTTGCGGTGATTCCTCCAGTTTTGTTCCNNNAGATCGGAAG |
| 47377_2 | TATGAGTGTGGAGTCGTTGCGGTGATTCCTCCAGTTTTGTTCCNNNAGATCGGAAG |
| 47377_3 | TATGAGTGTGGAGTCGTTGCGGTGATTCCTCCAGTTTTGTTCCNNNAGATCGGAAG |
| 49648_1 | TATGAGTGTGGAGTCGTTGCTCAAGCGATTCTCCTATCTCAGCNNNAGATCGGAAG |
| 49648_2 | TATGAGTGTGGAGTCGTTGCTCAAGCGATTCTCCTATCTCAGCNNNAGATCGGAAG |
| 49648_3 | TATGAGTGTGGAGTCGTTGCTCAAGCGATTCTCCTATCTCAGCNNNAGATCGGAAG |
| 30497_1 | TATGAGTGTGGAGTCGTTGCTGTTGACTTTAAAAGAGCCAAGGTNNNAGATCGGA |
| 30497_2 | TATGAGTGTGGAGTCGTTGCTGTTGACTTTAAAAGAGCCAAGGTNNNAGATCGGA |
| 30497_3 | TATGAGTGTGGAGTCGTTGCTGTTGACTTTAAAAGAGCCAAGGTNNNAGATCGGA |
| 41266_1 | TATGAGTGTGGAGTCGTTGCCATATGGGGAAGATCCAACAGCTNNNAGATCGGAA |

| | |
|---|---|
| 41266_2 | TATGAGTGTGGAGTCGTTGCCATATGGGGAAGATCCAACAGCTNNNAGATCGGAA |
| 41266_3 | TATGAGTGTGGAGTCGTTGCCATATGGGGAAGATCCAACAGCTNNNAGATCGGAA |
| 44199_1 | TATGAGTGTGGAGTCGTTGCACATGGAAAAATTACCCGCAAGGNNNAGATCGGAA |
| 44199_2 | TATGAGTGTGGAGTCGTTGCACATGGAAAAATTACCCGCAAGGNNNAGATCGGAA |
| 44199_3 | TATGAGTGTGGAGTCGTTGCACATGGAAAAATTACCCGCAAGGNNNAGATCGGAA |
| 31750_1 | TATGAGTGTGGAGTCGTTGCCCAAAGTGGAAGGTTTGTGATGGNNNAGATCGGAA |
| 31750_2 | TATGAGTGTGGAGTCGTTGCCCAAAGTGGAAGGTTTGTGATGGNNNAGATCGGAA |
| 31750_3 | TATGAGTGTGGAGTCGTTGCCCAAAGTGGAAGGTTTGTGATGGNNNAGATCGGAA |
| 33875_1 | TATGAGTGTGGAGTCGTTGCTCCCTTTCCCATAAGAGGAGATGNNNAGATCGGAAG |
| 33875_2 | TATGAGTGTGGAGTCGTTGCTCCCTTTCCCATAAGAGGAGATGNNNAGATCGGAAG |
| 33875_3 | TATGAGTGTGGAGTCGTTGCTCCCTTTCCCATAAGAGGAGATGNNNAGATCGGAAG |
| 11812_1 | TATGAGTGTGGAGTCGTTGCCTGCCCTGTAGACTTATGACCCNNNAGATCGGAAGA |
| 11812_2 | TATGAGTGTGGAGTCGTTGCCTGCCCTGTAGACTTATGACCCNNNAGATCGGAAGA |
| 11812_3 | TATGAGTGTGGAGTCGTTGCCTGCCCTGTAGACTTATGACCCNNNAGATCGGAAGA |
| 29307_1 | TATGAGTGTGGAGTCGTTGCCCCTTATCCAGTCCCTTGTTGATNNNAGATCGGAAGA |
| 29307_2 | TATGAGTGTGGAGTCGTTGCCCCTTATCCAGTCCCTTGTTGATNNNAGATCGGAAGA |
| 29307_3 | TATGAGTGTGGAGTCGTTGCCCCTTATCCAGTCCCTTGTTGATNNNAGATCGGAAGA |
| 51408_1 | TATGAGTGTGGAGTCGTTGCGAAGGGAGAGAGAGAAAGGGAGANNNAGATCGG |
| 51408_2 | TATGAGTGTGGAGTCGTTGCGAAGGGAGAGAGAGAAAGGGAGANNNAGATCGG |
| 51408_3 | TATGAGTGTGGAGTCGTTGCGAAGGGAGAGAGAGAAAGGGAGANNNAGATCGG |
| 46151_1 | TATGAGTGTGGAGTCGTTGCTGGAGTGTAGTGGTGTGATCATGNNNAGATCGGAA |
| 46151_2 | TATGAGTGTGGAGTCGTTGCTGGAGTGTAGTGGTGTGATCATGNNNAGATCGGAA |
| 46151_3 | TATGAGTGTGGAGTCGTTGCTGGAGTGTAGTGGTGTGATCATGNNNAGATCGGAA |
| 34984_1 | TATGAGTGTGGAGTCGTTGCCAGTAGTACGATCTTGGCTCACTNNNAGATCGGAAG |
| 34984_2 | TATGAGTGTGGAGTCGTTGCCAGTAGTACGATCTTGGCTCACTNNNAGATCGGAAG |
| 34984_3 | TATGAGTGTGGAGTCGTTGCCAGTAGTACGATCTTGGCTCACTNNNAGATCGGAAG |
| 35911_1 | TATGAGTGTGGAGTCGTTGCAAATGTCTGCCTTCTCATCTGGTNNNAGATCGGAAG |
| 35911_2 | TATGAGTGTGGAGTCGTTGCAAATGTCTGCCTTCTCATCTGGTNNNAGATCGGAAG |
| 35911_3 | TATGAGTGTGGAGTCGTTGCAAATGTCTGCCTTCTCATCTGGTNNNAGATCGGAAG |
| 34859_1 | TATGAGTGTGGAGTCGTTGCCCTCCTAGGTTCACATGCTTCTCNNNAGATCGGAAGA |
| 34859_2 | TATGAGTGTGGAGTCGTTGCCCTCCTAGGTTCACATGCTTCTCNNNAGATCGGAAGA |
| 34859_3 | TATGAGTGTGGAGTCGTTGCCCTCCTAGGTTCACATGCTTCTCNNNAGATCGGAAGA |
| 30783_1 | TATGAGTGTGGAGTCGTTGCTCATATTCTGCTGCTGTCTCTCCNNNAGATCGGAAGA |
| 30783_2 | TATGAGTGTGGAGTCGTTGCTCATATTCTGCTGCTGTCTCTCCNNNAGATCGGAAGA |
| 30783_3 | TATGAGTGTGGAGTCGTTGCTCATATTCTGCTGCTGTCTCTCCNNNAGATCGGAAGA |
| 32388_1 | TATGAGTGTGGAGTCGTTGCTTCAAGCAATTTTCCTGCCTCAGNNNAGATCGGAAG |
| 32388_2 | TATGAGTGTGGAGTCGTTGCTTCAAGCAATTTTCCTGCCTCAGNNNAGATCGGAAG |
| 32388_3 | TATGAGTGTGGAGTCGTTGCTTCAAGCAATTTTCCTGCCTCAGNNNAGATCGGAAG |
| 48172_1 | TATGAGTGTGGAGTCGTTGCGATCTCCTGACCTCATGATCCACNNNAGATCGGAAG |
| 48172_2 | TATGAGTGTGGAGTCGTTGCGATCTCCTGACCTCATGATCCACNNNAGATCGGAAG |
| 48172_3 | TATGAGTGTGGAGTCGTTGCGATCTCCTGACCTCATGATCCACNNNAGATCGGAAG |
| 50662_1 | TATGAGTGTGGAGTCGTTGCACACAACCTGCATTCTTGAACAGNNNAGATCGGAAG |
| 50662_2 | TATGAGTGTGGAGTCGTTGCACACAACCTGCATTCTTGAACAGNNNAGATCGGAAG |

| | |
|---|---|
| 50662_3 | TATGAGTGTGGAGTCGTTGCACACAACCTGCATTCTTGAACAGNNNAGATCGGAAG |
| 13531_1 | TATGAGTGTGGAGTCGTTGCAAAGCTAACACTTAGTGAATCTANNNAGATCGGAAG |
| 13531_2 | TATGAGTGTGGAGTCGTTGCAAAGCTAACACTTAGTGAATCTANNNAGATCGGAAG |
| 13531_3 | TATGAGTGTGGAGTCGTTGCAAAGCTAACACTTAGTGAATCTANNNAGATCGGAAG |
| 7971_1 | TATGAGTGTGGAGTCGTTGCCAACAGGGCCACTGCCTTTGANNNAGATCGGAAGAG |
| 7971_2 | TATGAGTGTGGAGTCGTTGCCAACAGGGCCACTGCCTTTGANNNAGATCGGAAGAG |
| 7971_3 | TATGAGTGTGGAGTCGTTGCCAACAGGGCCACTGCCTTTGANNNAGATCGGAAGAG |
| 48689_1 | TATGAGTGTGGAGTCGTTGCCTCAGTGATTGAAAGGCTTGGACNNNAGATCGGAA |
| 48689_2 | TATGAGTGTGGAGTCGTTGCCTCAGTGATTGAAAGGCTTGGACNNNAGATCGGAA |
| 48689_3 | TATGAGTGTGGAGTCGTTGCCTCAGTGATTGAAAGGCTTGGACNNNAGATCGGAA |
| 45796_1 | TATGAGTGTGGAGTCGTTGCCCCACTCCTGGATTCAAGTGATTNNNAGATCGGAAG |
| 45796_2 | TATGAGTGTGGAGTCGTTGCCCCACTCCTGGATTCAAGTGATTNNNAGATCGGAAG |
| 45796_3 | TATGAGTGTGGAGTCGTTGCCCCACTCCTGGATTCAAGTGATTNNNAGATCGGAAG |
| 35259_1 | TATGAGTGTGGAGTCGTTGCATACAATCTCTGCCACCCATGTTNNNAGATCGGAAGA |
| 35259_2 | TATGAGTGTGGAGTCGTTGCATACAATCTCTGCCACCCATGTTNNNAGATCGGAAGA |
| 35259_3 | TATGAGTGTGGAGTCGTTGCATACAATCTCTGCCACCCATGTTNNNAGATCGGAAGA |
| 47110_1 | TATGAGTGTGGAGTCGTTGCCTTCCAAAGTGCTGGGATTACAGNNNAGATCGGAAG |
| 47110_2 | TATGAGTGTGGAGTCGTTGCCTTCCAAAGTGCTGGGATTACAGNNNAGATCGGAAG |
| 47110_3 | TATGAGTGTGGAGTCGTTGCCTTCCAAAGTGCTGGGATTACAGNNNAGATCGGAAG |
| 34756_1 | TATGAGTGTGGAGTCGTTGCCGCCTGATCCAAACGATAATTGTNNNAGATCGGAAG |
| 34756_2 | TATGAGTGTGGAGTCGTTGCCGCCTGATCCAAACGATAATTGTNNNAGATCGGAAG |
| 34756_3 | TATGAGTGTGGAGTCGTTGCCGCCTGATCCAAACGATAATTGTNNNAGATCGGAAG |
| 45955_1 | TATGAGTGTGGAGTCGTTGCCTCCCAGGTTCACACCATTCTCNNNAGATCGGAAGAG |
| 45955_2 | TATGAGTGTGGAGTCGTTGCCTCCCAGGTTCACACCATTCTCNNNAGATCGGAAGAG |
| 45955_3 | TATGAGTGTGGAGTCGTTGCCTCCCAGGTTCACACCATTCTCNNNAGATCGGAAGAG |
| 43656_1 | TATGAGTGTGGAGTCGTTGCAAGGAAAGATTTGGGGTTCCAGANNNAGATCGGAA |
| 43656_2 | TATGAGTGTGGAGTCGTTGCAAGGAAAGATTTGGGGTTCCAGANNNAGATCGGAA |
| 43656_3 | TATGAGTGTGGAGTCGTTGCAAGGAAAGATTTGGGGTTCCAGANNNAGATCGGAA |
| 45805_1 | TATGAGTGTGGAGTCGTTGCCATTCTCCTGCCTCAGCCTCNNNAGATCGGAAGAGCA |
| 45805_2 | TATGAGTGTGGAGTCGTTGCCATTCTCCTGCCTCAGCCTCNNNAGATCGGAAGAGCA |
| 45805_3 | TATGAGTGTGGAGTCGTTGCCATTCTCCTGCCTCAGCCTCNNNAGATCGGAAGAGCA |
| 34372_1 | TATGAGTGTGGAGTCGTTGCGCCTTGTCTTTTGAACTTCTCCCNNNAGATCGGAAGA |
| 34372_2 | TATGAGTGTGGAGTCGTTGCGCCTTGTCTTTTGAACTTCTCCCNNNAGATCGGAAGA |
| 34372_3 | TATGAGTGTGGAGTCGTTGCGCCTTGTCTTTTGAACTTCTCCCNNNAGATCGGAAGA |
| 37921_1 | TATGAGTGTGGAGTCGTTGCCCTTCTATTTTCTGTCAGCTGGCNNNAGATCGGAAGA |
| 37921_2 | TATGAGTGTGGAGTCGTTGCCCTTCTATTTTCTGTCAGCTGGCNNNAGATCGGAAGA |
| 37921_3 | TATGAGTGTGGAGTCGTTGCCCTTCTATTTTCTGTCAGCTGGCNNNAGATCGGAAGA |
| 38850_1 | TATGAGTGTGGAGTCGTTGCTGCATGAGTCTTCCAAAGGAGAANNNAGATCGGAA |
| 38850_2 | TATGAGTGTGGAGTCGTTGCTGCATGAGTCTTCCAAAGGAGAANNNAGATCGGAA |
| 38850_3 | TATGAGTGTGGAGTCGTTGCTGCATGAGTCTTCCAAAGGAGAANNNAGATCGGAA |
| 42194_1 | TATGAGTGTGGAGTCGTTGCCTTTTTGTGTGTGCAGGCATTTCNNNAGATCGGAAG |
| 42194_2 | TATGAGTGTGGAGTCGTTGCCTTTTTGTGTGTGCAGGCATTTCNNNAGATCGGAAG |
| 42194_3 | TATGAGTGTGGAGTCGTTGCCTTTTTGTGTGTGCAGGCATTTCNNNAGATCGGAAG |

| | |
|---|---|
| 9729_1 | TATGAGTGTGGAGTCGTTGCAGAGAGAGAGGCAGAGATGTCACNNNAGATCGGA |
| 9729_2 | TATGAGTGTGGAGTCGTTGCAGAGAGAGAGGCAGAGATGTCACNNNAGATCGGA |
| 9729_3 | TATGAGTGTGGAGTCGTTGCAGAGAGAGAGGCAGAGATGTCACNNNAGATCGGA |
| 5387_1 | TATGAGTGTGGAGTCGTTGCTGTGGATGGTAAGAATTGAGGCTNNNAGATCGGAA |
| 5387_2 | TATGAGTGTGGAGTCGTTGCTGTGGATGGTAAGAATTGAGGCTNNNAGATCGGAA |
| 5387_3 | TATGAGTGTGGAGTCGTTGCTGTGGATGGTAAGAATTGAGGCTNNNAGATCGGAA |
| 45890_1 | TATGAGTGTGGAGTCGTTGCTTGGATTAGGGAGGGGTTTGAAGNNNAGATCGGA |
| 45890_2 | TATGAGTGTGGAGTCGTTGCTTGGATTAGGGAGGGGTTTGAAGNNNAGATCGGA |
| 45890_3 | TATGAGTGTGGAGTCGTTGCTTGGATTAGGGAGGGGTTTGAAGNNNAGATCGGA |
| 38262_1 | TATGAGTGTGGAGTCGTTGCAGGTGTTTCTGTCCTTAGAGTGCNNNAGATCGGAAG |
| 38262_2 | TATGAGTGTGGAGTCGTTGCAGGTGTTTCTGTCCTTAGAGTGCNNNAGATCGGAAG |
| 38262_3 | TATGAGTGTGGAGTCGTTGCAGGTGTTTCTGTCCTTAGAGTGCNNNAGATCGGAAG |
| 42523_1 | TATGAGTGTGGAGTCGTTGCCCTCCTAGGTTCAAGCGATTCTCNNNAGATCGGAAG |
| 42523_2 | TATGAGTGTGGAGTCGTTGCCCTCCTAGGTTCAAGCGATTCTCNNNAGATCGGAAG |
| 42523_3 | TATGAGTGTGGAGTCGTTGCCCTCCTAGGTTCAAGCGATTCTCNNNAGATCGGAAG |
| 49882_1 | TATGAGTGTGGAGTCGTTGCTCATGGGGAAATCACTGCTTGTANNNAGATCGGAA |
| 49882_2 | TATGAGTGTGGAGTCGTTGCTCATGGGGAAATCACTGCTTGTANNNAGATCGGAA |
| 49882_3 | TATGAGTGTGGAGTCGTTGCTCATGGGGAAATCACTGCTTGTANNNAGATCGGAA |
| 29751_1 | TATGAGTGTGGAGTCGTTGCCCCTTCTTCGAAGGTAACGCTAANNNAGATCGGAAG |
| 29751_2 | TATGAGTGTGGAGTCGTTGCCCCTTCTTCGAAGGTAACGCTAANNNAGATCGGAAG |
| 29751_3 | TATGAGTGTGGAGTCGTTGCCCCTTCTTCGAAGGTAACGCTAANNNAGATCGGAAG |
| 39671_1 | TATGAGTGTGGAGTCGTTGCGGTCTCCTCTATACTCCATACCCANNNAGATCGGAAG |
| 39671_2 | TATGAGTGTGGAGTCGTTGCGGTCTCCTCTATACTCCATACCCANNNAGATCGGAAG |
| 39671_3 | TATGAGTGTGGAGTCGTTGCGGTCTCCTCTATACTCCATACCCANNNAGATCGGAAG |
| 32822_1 | TATGAGTGTGGAGTCGTTGCCTCATTGCAACCTCTACCTCCTGNNNAGATCGGAAGA |
| 32822_2 | TATGAGTGTGGAGTCGTTGCCTCATTGCAACCTCTACCTCCTGNNNAGATCGGAAGA |
| 32822_3 | TATGAGTGTGGAGTCGTTGCCTCATTGCAACCTCTACCTCCTGNNNAGATCGGAAGA |
| 39506_1 | TATGAGTGTGGAGTCGTTGCAGCAACCTATGTGACAGTGTCTCNNNAGATCGGAAG |
| 39506_2 | TATGAGTGTGGAGTCGTTGCAGCAACCTATGTGACAGTGTCTCNNNAGATCGGAAG |
| 39506_3 | TATGAGTGTGGAGTCGTTGCAGCAACCTATGTGACAGTGTCTCNNNAGATCGGAAG |
| 45246_1 | TATGAGTGTGGAGTCGTTGCTCCAAGGGGGAAAAAAGAAAACTGNNNAGATCGGA |
| 45246_2 | TATGAGTGTGGAGTCGTTGCTCCAAGGGGGAAAAAAGAAAACTGNNNAGATCGGA |
| 45246_3 | TATGAGTGTGGAGTCGTTGCTCCAAGGGGGAAAAAAGAAAACTGNNNAGATCGGA |
| 13295_1 | TATGAGTGTGGAGTCGTTGCGAGTTCATAGATAGATGCAAAACNNNAGATCGGAA |
| 13295_2 | TATGAGTGTGGAGTCGTTGCGAGTTCATAGATAGATGCAAAACNNNAGATCGGAA |
| 13295_3 | TATGAGTGTGGAGTCGTTGCGAGTTCATAGATAGATGCAAAACNNNAGATCGGAA |
| 45979_1 | TATGAGTGTGGAGTCGTTGCCCTACCGGGTTCAAGCAATTCTTNNNAGATCGGAAG |
| 45979_2 | TATGAGTGTGGAGTCGTTGCCCTACCGGGTTCAAGCAATTCTTNNNAGATCGGAAG |
| 45979_3 | TATGAGTGTGGAGTCGTTGCCCTACCGGGTTCAAGCAATTCTTNNNAGATCGGAAG |
| 46361_1 | TATGAGTGTGGAGTCGTTGCATCTTGGCTCACTGCAACCTCNNNAGATCGGAAGAG |
| 46361_2 | TATGAGTGTGGAGTCGTTGCATCTTGGCTCACTGCAACCTCNNNAGATCGGAAGAG |
| 46361_3 | TATGAGTGTGGAGTCGTTGCATCTTGGCTCACTGCAACCTCNNNAGATCGGAAGAG |
| 2654_1 | TATGAGTGTGGAGTCGTTGCACTCGCCTTTGCATGACATTTTGNNNAGATCGGAAG |

| | |
|---|---|
| 2654_2 | TATGAGTGTGGAGTCGTTGCACTCGCCTTTGCATGACATTTTGNNNAGATCGGAAG |
| 2654_3 | TATGAGTGTGGAGTCGTTGCACTCGCCTTTGCATGACATTTTGNNNAGATCGGAAG |
| 46046_1 | TATGAGTGTGGAGTCGTTGCGTTTCACTCTTGTTGCCCAGGNNNAGATCGGAAGAG |
| 46046_2 | TATGAGTGTGGAGTCGTTGCGTTTCACTCTTGTTGCCCAGGNNNAGATCGGAAGAG |
| 46046_3 | TATGAGTGTGGAGTCGTTGCGTTTCACTCTTGTTGCCCAGGNNNAGATCGGAAGAG |
| 37475_1 | TATGAGTGTGGAGTCGTTGCATCCTAGCTACTTGAGAGGCTGANNNAGATCGGAA |
| 37475_2 | TATGAGTGTGGAGTCGTTGCATCCTAGCTACTTGAGAGGCTGANNNAGATCGGAA |
| 37475_3 | TATGAGTGTGGAGTCGTTGCATCCTAGCTACTTGAGAGGCTGANNNAGATCGGAA |
| 36267_1 | TATGAGTGTGGAGTCGTTGCGCATACAGTTGGCACTTCTGAAGNNNAGATCGGAA |
| 36267_2 | TATGAGTGTGGAGTCGTTGCGCATACAGTTGGCACTTCTGAAGNNNAGATCGGAA |
| 36267_3 | TATGAGTGTGGAGTCGTTGCGCATACAGTTGGCACTTCTGAAGNNNAGATCGGAA |
| 47203_1 | TATGAGTGTGGAGTCGTTGCGCCTCCCAAGTAACTGGTACTACNNNAGATCGGAAG |
| 47203_2 | TATGAGTGTGGAGTCGTTGCGCCTCCCAAGTAACTGGTACTACNNNAGATCGGAAG |
| 47203_3 | TATGAGTGTGGAGTCGTTGCGCCTCCCAAGTAACTGGTACTACNNNAGATCGGAAG |
| 51016_1 | TATGAGTGTGGAGTCGTTGCCCTAGTGGAGCTGTGAGAAGAAGNNNAGATCGGA |
| 51016_2 | TATGAGTGTGGAGTCGTTGCCCTAGTGGAGCTGTGAGAAGAAGNNNAGATCGGA |
| 51016_3 | TATGAGTGTGGAGTCGTTGCCCTAGTGGAGCTGTGAGAAGAAGNNNAGATCGGA |
| 33196_1 | TATGAGTGTGGAGTCGTTGCTGTCCTTGTCCTTGAAATGTCGANNNAGATCGGAAG |
| 33196_2 | TATGAGTGTGGAGTCGTTGCTGTCCTTGTCCTTGAAATGTCGANNNAGATCGGAAG |
| 33196_3 | TATGAGTGTGGAGTCGTTGCTGTCCTTGTCCTTGAAATGTCGANNNAGATCGGAAG |
| 35479_1 | TATGAGTGTGGAGTCGTTGCACATTCATTTTGCTGGTGGATGGNNNAGATCGGAA |
| 35479_2 | TATGAGTGTGGAGTCGTTGCACATTCATTTTGCTGGTGGATGGNNNAGATCGGAA |
| 35479_3 | TATGAGTGTGGAGTCGTTGCACATTCATTTTGCTGGTGGATGGNNNAGATCGGAA |
| 38524_1 | TATGAGTGTGGAGTCGTTGCAATACAAAATTAGCCAGGCGTGGNNNAGATCGGAA |
| 38524_2 | TATGAGTGTGGAGTCGTTGCAATACAAAATTAGCCAGGCGTGGNNNAGATCGGAA |
| 38524_3 | TATGAGTGTGGAGTCGTTGCAATACAAAATTAGCCAGGCGTGGNNNAGATCGGAA |
| 29844_1 | TATGAGTGTGGAGTCGTTGCGGTGAACTAAAGCAGGGTGAACANNNAGATCGGA |
| 29844_2 | TATGAGTGTGGAGTCGTTGCGGTGAACTAAAGCAGGGTGAACANNNAGATCGGA |
| 29844_3 | TATGAGTGTGGAGTCGTTGCGGTGAACTAAAGCAGGGTGAACANNNAGATCGGA |
| 2236_1 | TATGAGTGTGGAGTCGTTGCCACTGCAAACCCCCAGGTAAATANNNAGATCGGAAG |
| 2236_2 | TATGAGTGTGGAGTCGTTGCCACTGCAAACCCCCAGGTAAATANNNAGATCGGAAG |
| 2236_3 | TATGAGTGTGGAGTCGTTGCCACTGCAAACCCCCAGGTAAATANNNAGATCGGAAG |
| 33336_1 | TATGAGTGTGGAGTCGTTGCCTCCTGAGTAGCTGGGATTACAGNNNAGATCGGAA |
| 33336_2 | TATGAGTGTGGAGTCGTTGCCTCCTGAGTAGCTGGGATTACAGNNNAGATCGGAA |
| 33336_3 | TATGAGTGTGGAGTCGTTGCCTCCTGAGTAGCTGGGATTACAGNNNAGATCGGAA |
| 29654_1 | TATGAGTGTGGAGTCGTTGCGCAACTAACGAAACTGCCCATACNNNAGATCGGAAG |
| 29654_2 | TATGAGTGTGGAGTCGTTGCGCAACTAACGAAACTGCCCATACNNNAGATCGGAAG |
| 29654_3 | TATGAGTGTGGAGTCGTTGCGCAACTAACGAAACTGCCCATACNNNAGATCGGAAG |
| 8516_1 | TATGAGTGTGGAGTCGTTGCAAGTTGAAGCGCGGCACGAATCNNNAGATCGGAAG |
| 8516_2 | TATGAGTGTGGAGTCGTTGCAAGTTGAAGCGCGGCACGAATCNNNAGATCGGAAG |
| 8516_3 | TATGAGTGTGGAGTCGTTGCAAGTTGAAGCGCGGCACGAATCNNNAGATCGGAAG |
| 45970_1 | TATGAGTGTGGAGTCGTTGCCTCCTGAGTAGCTGGGACTACAGNNNAGATCGGAA |
| 45970_2 | TATGAGTGTGGAGTCGTTGCCTCCTGAGTAGCTGGGACTACAGNNNAGATCGGAA |

| | |
|---|---|
| 45970_3 | TATGAGTGTGGAGTCGTTGCCTCCTGAGTAGCTGGGACTACAGNNNAGATCGGAA |
| 48888_1 | TATGAGTGTGGAGTCGTTGCGTATGTTGCCCAGACTGACCTTANNNAGATCGGAAG |
| 48888_2 | TATGAGTGTGGAGTCGTTGCGTATGTTGCCCAGACTGACCTTANNNAGATCGGAAG |
| 48888_3 | TATGAGTGTGGAGTCGTTGCGTATGTTGCCCAGACTGACCTTANNNAGATCGGAAG |
| 41742_1 | TATGAGTGTGGAGTCGTTGCATCTCCCAGGTTCAAGCAATTCTNNNAGATCGGAAG |
| 41742_2 | TATGAGTGTGGAGTCGTTGCATCTCCCAGGTTCAAGCAATTCTNNNAGATCGGAAG |
| 41742_3 | TATGAGTGTGGAGTCGTTGCATCTCCCAGGTTCAAGCAATTCTNNNAGATCGGAAG |
| 29150_1 | TATGAGTGTGGAGTCGTTGCTGCTGATAACAAGGCAGAAGATGNNNAGATCGGAA |
| 29150_2 | TATGAGTGTGGAGTCGTTGCTGCTGATAACAAGGCAGAAGATGNNNAGATCGGAA |
| 29150_3 | TATGAGTGTGGAGTCGTTGCTGCTGATAACAAGGCAGAAGATGNNNAGATCGGAA |
| 48571_1 | TATGAGTGTGGAGTCGTTGCCTCCCAAGTTCAAGCGTTTCTCNNNAGATCGGAAGA |
| 48571_2 | TATGAGTGTGGAGTCGTTGCCTCCCAAGTTCAAGCGTTTCTCNNNAGATCGGAAGA |
| 48571_3 | TATGAGTGTGGAGTCGTTGCCTCCCAAGTTCAAGCGTTTCTCNNNAGATCGGAAGA |
| 39922_1 | TATGAGTGTGGAGTCGTTGCCAGGAATGTGACCTTTGGCTCTANNNAGATCGGAAG |
| 39922_2 | TATGAGTGTGGAGTCGTTGCCAGGAATGTGACCTTTGGCTCTANNNAGATCGGAAG |
| 39922_3 | TATGAGTGTGGAGTCGTTGCCAGGAATGTGACCTTTGGCTCTANNNAGATCGGAAG |
| 40113_1 | TATGAGTGTGGAGTCGTTGCTGGGAGATTAGGGATTCTCTGGANNNAGATCGGAA |
| 40113_2 | TATGAGTGTGGAGTCGTTGCTGGGAGATTAGGGATTCTCTGGANNNAGATCGGAA |
| 40113_3 | TATGAGTGTGGAGTCGTTGCTGGGAGATTAGGGATTCTCTGGANNNAGATCGGAA |
| 30005_1 | TATGAGTGTGGAGTCGTTGCAGACCCCAGTAATAGGATTCAGGANNNAGATCGGA |
| 30005_2 | TATGAGTGTGGAGTCGTTGCAGACCCCAGTAATAGGATTCAGGANNNAGATCGGA |
| 30005_3 | TATGAGTGTGGAGTCGTTGCAGACCCCAGTAATAGGATTCAGGANNNAGATCGGA |
| 41831_1 | TATGAGTGTGGAGTCGTTGCAAGTGATTCTCCTGCTTCAACCTNNNAGATCGGAAG |
| 41831_2 | TATGAGTGTGGAGTCGTTGCAAGTGATTCTCCTGCTTCAACCTNNNAGATCGGAAG |
| 41831_3 | TATGAGTGTGGAGTCGTTGCAAGTGATTCTCCTGCTTCAACCTNNNAGATCGGAAG |
| 30836_1 | TATGAGTGTGGAGTCGTTGCTCCAGGCACTAATCACCTCATTTNNNAGATCGGAAGA |
| 30836_2 | TATGAGTGTGGAGTCGTTGCTCCAGGCACTAATCACCTCATTTNNNAGATCGGAAGA |
| 30836_3 | TATGAGTGTGGAGTCGTTGCTCCAGGCACTAATCACCTCATTTNNNAGATCGGAAGA |
| 48916_1 | TATGAGTGTGGAGTCGTTGCCTCCTGAGTAGCTGGGACTACAGNNNAGATCGGAA |
| 48916_2 | TATGAGTGTGGAGTCGTTGCCTCCTGAGTAGCTGGGACTACAGNNNAGATCGGAA |
| 48916_3 | TATGAGTGTGGAGTCGTTGCCTCCTGAGTAGCTGGGACTACAGNNNAGATCGGAA |
| 42238_1 | TATGAGTGTGGAGTCGTTGCTGCATCTGGCCAGTTTATAGTGANNNAGATCGGAAG |
| 42238_2 | TATGAGTGTGGAGTCGTTGCTGCATCTGGCCAGTTTATAGTGANNNAGATCGGAAG |
| 42238_3 | TATGAGTGTGGAGTCGTTGCTGCATCTGGCCAGTTTATAGTGANNNAGATCGGAAG |
| 49537_1 | TATGAGTGTGGAGTCGTTGCTGTGATCGACTTGATGCAATGTGNNNAGATCGGAA |
| 49537_2 | TATGAGTGTGGAGTCGTTGCTGTGATCGACTTGATGCAATGTGNNNAGATCGGAA |
| 49537_3 | TATGAGTGTGGAGTCGTTGCTGTGATCGACTTGATGCAATGTGNNNAGATCGGAA |
| 30226_1 | TATGAGTGTGGAGTCGTTGCGTCCATGGTGGGTTTGAAGATTGNNNAGATCGGAA |
| 30226_2 | TATGAGTGTGGAGTCGTTGCGTCCATGGTGGGTTTGAAGATTGNNNAGATCGGAA |
| 30226_3 | TATGAGTGTGGAGTCGTTGCGTCCATGGTGGGTTTGAAGATTGNNNAGATCGGAA |
| 49189_1 | TATGAGTGTGGAGTCGTTGCCAGGTTCACGCCATTCTCCTNNNAGATCGGAAGAGC |
| 49189_2 | TATGAGTGTGGAGTCGTTGCCAGGTTCACGCCATTCTCCTNNNAGATCGGAAGAGC |
| 49189_3 | TATGAGTGTGGAGTCGTTGCCAGGTTCACGCCATTCTCCTNNNAGATCGGAAGAGC |

| | |
|---|---|
| 52856_1 | TATGAGTGTGGAGTCGTTGCTTTTTAGGTCTGTACAGCACCCANNNAGATCGGAAG |
| 52856_2 | TATGAGTGTGGAGTCGTTGCTTTTTAGGTCTGTACAGCACCCANNNAGATCGGAAG |
| 52856_3 | TATGAGTGTGGAGTCGTTGCTTTTTAGGTCTGTACAGCACCCANNNAGATCGGAAG |
| 31095_1 | TATGAGTGTGGAGTCGTTGCGGGAGGCCTCTTCAAGAATAAGTNNNAGATCGGAA |
| 31095_2 | TATGAGTGTGGAGTCGTTGCGGGAGGCCTCTTCAAGAATAAGTNNNAGATCGGAA |
| 31095_3 | TATGAGTGTGGAGTCGTTGCGGGAGGCCTCTTCAAGAATAAGTNNNAGATCGGAA |
| 42919_1 | TATGAGTGTGGAGTCGTTGCCCACTTGTTGATTGATGGGCATTNNNAGATCGGAAG |
| 42919_2 | TATGAGTGTGGAGTCGTTGCCCACTTGTTGATTGATGGGCATTNNNAGATCGGAAG |
| 42919_3 | TATGAGTGTGGAGTCGTTGCCCACTTGTTGATTGATGGGCATTNNNAGATCGGAAG |
| 38304_1 | TATGAGTGTGGAGTCGTTGCACCCGAGTAGCAAACACAAAATGNNNAGATCGGAA |
| 38304_2 | TATGAGTGTGGAGTCGTTGCACCCGAGTAGCAAACACAAAATGNNNAGATCGGAA |
| 38304_3 | TATGAGTGTGGAGTCGTTGCACCCGAGTAGCAAACACAAAATGNNNAGATCGGAA |
| 32593_1 | TATGAGTGTGGAGTCGTTGCTCGGTCTCAAATACCTAAGGCTGNNNAGATCGGAAG |
| 32593_2 | TATGAGTGTGGAGTCGTTGCTCGGTCTCAAATACCTAAGGCTGNNNAGATCGGAAG |
| 32593_3 | TATGAGTGTGGAGTCGTTGCTCGGTCTCAAATACCTAAGGCTGNNNAGATCGGAAG |
| 43765_1 | TATGAGTGTGGAGTCGTTGCTACACAGGAGGCTTCAAAAGGAANNNAGATCGGAA |
| 43765_2 | TATGAGTGTGGAGTCGTTGCTACACAGGAGGCTTCAAAAGGAANNNAGATCGGAA |
| 43765_3 | TATGAGTGTGGAGTCGTTGCTACACAGGAGGCTTCAAAAGGAANNNAGATCGGAA |
| 31610_1 | TATGAGTGTGGAGTCGTTGCGACCCTGTGAAAAACCCTAAAGCNNNAGATCGGAAG |
| 31610_2 | TATGAGTGTGGAGTCGTTGCGACCCTGTGAAAAACCCTAAAGCNNNAGATCGGAAG |
| 31610_3 | TATGAGTGTGGAGTCGTTGCGACCCTGTGAAAAACCCTAAAGCNNNAGATCGGAAG |
| 41552_1 | TATGAGTGTGGAGTCGTTGCCCAAAGTGCTGGGATTATAAGCGNNNAGATCGGAA |
| 41552_2 | TATGAGTGTGGAGTCGTTGCCCAAAGTGCTGGGATTATAAGCGNNNAGATCGGAA |
| 41552_3 | TATGAGTGTGGAGTCGTTGCCCAAAGTGCTGGGATTATAAGCGNNNAGATCGGAA |
| 44655_1 | TATGAGTGTGGAGTCGTTGCAAAGGGGACATATAAGGAGGTGGNNNAGATCGGA |
| 44655_2 | TATGAGTGTGGAGTCGTTGCAAAGGGGACATATAAGGAGGTGGNNNAGATCGGA |
| 44655_3 | TATGAGTGTGGAGTCGTTGCAAAGGGGACATATAAGGAGGTGGNNNAGATCGGA |
| 50823_1 | TATGAGTGTGGAGTCGTTGCAACAAAAGGGAGAAGTAGAGGGCNNNAGATCGGA |
| 50823_2 | TATGAGTGTGGAGTCGTTGCAACAAAAGGGAGAAGTAGAGGGCNNNAGATCGGA |
| 50823_3 | TATGAGTGTGGAGTCGTTGCAACAAAAGGGAGAAGTAGAGGGCNNNAGATCGGA |
| 29546_1 | TATGAGTGTGGAGTCGTTGCCCTGAGCTGTACTTTGTCAAGGTNNNAGATCGGAAG |
| 29546_2 | TATGAGTGTGGAGTCGTTGCCCTGAGCTGTACTTTGTCAAGGTNNNAGATCGGAAG |
| 29546_3 | TATGAGTGTGGAGTCGTTGCCCTGAGCTGTACTTTGTCAAGGTNNNAGATCGGAAG |
| 42912_1 | TATGAGTGTGGAGTCGTTGCACACTATGTACTAGGCTCTGTGCNNNAGATCGGAAG |
| 42912_2 | TATGAGTGTGGAGTCGTTGCACACTATGTACTAGGCTCTGTGCNNNAGATCGGAAG |
| 42912_3 | TATGAGTGTGGAGTCGTTGCACACTATGTACTAGGCTCTGTGCNNNAGATCGGAAG |
| 40899_1 | TATGAGTGTGGAGTCGTTGCTAGTGGGAAAACAACCTTCCTCANNNAGATCGGAAG |
| 40899_2 | TATGAGTGTGGAGTCGTTGCTAGTGGGAAAACAACCTTCCTCANNNAGATCGGAAG |
| 40899_3 | TATGAGTGTGGAGTCGTTGCTAGTGGGAAAACAACCTTCCTCANNNAGATCGGAAG |
| 49373_1 | TATGAGTGTGGAGTCGTTGCCAAAGTGCTGGGATTATAGGCCTNNNAGATCGGAA |
| 49373_2 | TATGAGTGTGGAGTCGTTGCCAAAGTGCTGGGATTATAGGCCTNNNAGATCGGAA |
| 49373_3 | TATGAGTGTGGAGTCGTTGCCAAAGTGCTGGGATTATAGGCCTNNNAGATCGGAA |
| 47830_1 | TATGAGTGTGGAGTCGTTGCGGTTTCACTGTGTTAGCCAGAACNNNAGATCGGAAG |

| | |
|---|---|
| 47830_2 | TATGAGTGTGGAGTCGTTGCGGTTTCACTGTGTTAGCCAGAACNNNAGATCGGAAG |
| 47830_3 | TATGAGTGTGGAGTCGTTGCGGTTTCACTGTGTTAGCCAGAACNNNAGATCGGAAG |
| 52545_1 | TATGAGTGTGGAGTCGTTGCCCCAGGAATGAAGATTGGGGTATNNNAGATCGGAA |
| 52545_2 | TATGAGTGTGGAGTCGTTGCCCCAGGAATGAAGATTGGGGTATNNNAGATCGGAA |
| 52545_3 | TATGAGTGTGGAGTCGTTGCCCCAGGAATGAAGATTGGGGTATNNNAGATCGGAA |
| 2333_1 | TATGAGTGTGGAGTCGTTGCCCAAATACTCCCGACCCTTCCTNNNAGATCGGAAGAG |
| 2333_2 | TATGAGTGTGGAGTCGTTGCCCAAATACTCCCGACCCTTCCTNNNAGATCGGAAGAG |
| 2333_3 | TATGAGTGTGGAGTCGTTGCCCAAATACTCCCGACCCTTCCTNNNAGATCGGAAGAG |
| 42036_1 | TATGAGTGTGGAGTCGTTGCAAGAAACCTTCCCTTTCACCTCCNNNAGATCGGAAGA |
| 42036_2 | TATGAGTGTGGAGTCGTTGCAAGAAACCTTCCCTTTCACCTCCNNNAGATCGGAAGA |
| 42036_3 | TATGAGTGTGGAGTCGTTGCAAGAAACCTTCCCTTTCACCTCCNNNAGATCGGAAGA |
| 38487_1 | TATGAGTGTGGAGTCGTTGCGAGGTTTTTTGCTGTTGCTGTTGNNNAGATCGGAAG |
| 38487_2 | TATGAGTGTGGAGTCGTTGCGAGGTTTTTTGCTGTTGCTGTTGNNNAGATCGGAAG |
| 38487_3 | TATGAGTGTGGAGTCGTTGCGAGGTTTTTTGCTGTTGCTGTTGNNNAGATCGGAAG |
| 42332_1 | TATGAGTGTGGAGTCGTTGCAACTCCTGAACTAGTGGAGGAAANNNAGATCGGAA |
| 42332_2 | TATGAGTGTGGAGTCGTTGCAACTCCTGAACTAGTGGAGGAAANNNAGATCGGAA |
| 42332_3 | TATGAGTGTGGAGTCGTTGCAACTCCTGAACTAGTGGAGGAAANNNAGATCGGAA |
| 37996_1 | TATGAGTGTGGAGTCGTTGCACCGGACTGAAGAAAAGCAGATANNNAGATCGGAA |
| 37996_2 | TATGAGTGTGGAGTCGTTGCACCGGACTGAAGAAAAGCAGATANNNAGATCGGAA |
| 37996_3 | TATGAGTGTGGAGTCGTTGCACCGGACTGAAGAAAAGCAGATANNNAGATCGGAA |
| 31036_1 | TATGAGTGTGGAGTCGTTGCCAGAGAGCTGATTCGTAGTTCCANNNAGATCGGAAG |
| 31036_2 | TATGAGTGTGGAGTCGTTGCCAGAGAGCTGATTCGTAGTTCCANNNAGATCGGAAG |
| 31036_3 | TATGAGTGTGGAGTCGTTGCCAGAGAGCTGATTCGTAGTTCCANNNAGATCGGAAG |
| 41787_1 | TATGAGTGTGGAGTCGTTGCCTTCCGAGTAGCTGGGATTACAGNNNAGATCGGAAG |
| 41787_2 | TATGAGTGTGGAGTCGTTGCCTTCCGAGTAGCTGGGATTACAGNNNAGATCGGAAG |
| 41787_3 | TATGAGTGTGGAGTCGTTGCCTTCCGAGTAGCTGGGATTACAGNNNAGATCGGAAG |
| 34268_1 | TATGAGTGTGGAGTCGTTGCCCTACTGGGAAGTGAGGAGCNNNAGATCGGAAGAG |
| 34268_2 | TATGAGTGTGGAGTCGTTGCCCTACTGGGAAGTGAGGAGCNNNAGATCGGAAGAG |
| 34268_3 | TATGAGTGTGGAGTCGTTGCCCTACTGGGAAGTGAGGAGCNNNAGATCGGAAGAG |
| 32115_1 | TATGAGTGTGGAGTCGTTGCACACATTTTAGGGAGGCATGAGANNNAGATCGGAA |
| 32115_2 | TATGAGTGTGGAGTCGTTGCACACATTTTAGGGAGGCATGAGANNNAGATCGGAA |
| 32115_3 | TATGAGTGTGGAGTCGTTGCACACATTTTAGGGAGGCATGAGANNNAGATCGGAA |
| 43238_1 | TATGAGTGTGGAGTCGTTGCCTTCCGAGTAACTGGGACTACAGNNNAGATCGGAAG |
| 43238_2 | TATGAGTGTGGAGTCGTTGCCTTCCGAGTAACTGGGACTACAGNNNAGATCGGAAG |
| 43238_3 | TATGAGTGTGGAGTCGTTGCCTTCCGAGTAACTGGGACTACAGNNNAGATCGGAAG |
| 51140_1 | TATGAGTGTGGAGTCGTTGCGAGAGAGAGAGAGAGCAAACCAGNNNAGATCGGA |
| 51140_2 | TATGAGTGTGGAGTCGTTGCGAGAGAGAGAGAGAGCAAACCAGNNNAGATCGGA |
| 51140_3 | TATGAGTGTGGAGTCGTTGCGAGAGAGAGAGAGAGCAAACCAGNNNAGATCGGA |
| 42189_1 | TATGAGTGTGGAGTCGTTGCCACCAGGCCCAGCTAATTTTTTGNNNAGATCGGAAG |
| 42189_2 | TATGAGTGTGGAGTCGTTGCCACCAGGCCCAGCTAATTTTTTGNNNAGATCGGAAG |
| 42189_3 | TATGAGTGTGGAGTCGTTGCCACCAGGCCCAGCTAATTTTTTGNNNAGATCGGAAG |
| 48875_1 | TATGAGTGTGGAGTCGTTGCAGTATCTGGGACTATAGGCTCGTNNNAGATCGGAA |
| 48875_2 | TATGAGTGTGGAGTCGTTGCAGTATCTGGGACTATAGGCTCGTNNNAGATCGGAA |

| | |
|---|---|
| 48875_3 | TATGAGTGTGGAGTCGTTGCAGTATCTGGGACTATAGGCTCGTNNNAGATCGGAAG |
| 32694_1 | TATGAGTGTGGAGTCGTTGCCTCCTGAGTAGCTGGGACTACAGNNNAGATCGGAAG |
| 32694_2 | TATGAGTGTGGAGTCGTTGCCTCCTGAGTAGCTGGGACTACAGNNNAGATCGGAAG |
| 32694_3 | TATGAGTGTGGAGTCGTTGCCTCCTGAGTAGCTGGGACTACAGNNNAGATCGGAAG |
| 39490_1 | TATGAGTGTGGAGTCGTTGCGGGATGATTCCGAAATTCACGTTNNNAGATCGGAAG |
| 39490_2 | TATGAGTGTGGAGTCGTTGCGGGATGATTCCGAAATTCACGTTNNNAGATCGGAAG |
| 39490_3 | TATGAGTGTGGAGTCGTTGCGGGATGATTCCGAAATTCACGTTNNNAGATCGGAAG |
| 43598_1 | TATGAGTGTGGAGTCGTTGCACAAATTTGCATGTTGAGGGAGGNNNAGATCGGAA |
| 43598_2 | TATGAGTGTGGAGTCGTTGCACAAATTTGCATGTTGAGGGAGGNNNAGATCGGAA |
| 43598_3 | TATGAGTGTGGAGTCGTTGCACAAATTTGCATGTTGAGGGAGGNNNAGATCGGAA |
| 29511_1 | TATGAGTGTGGAGTCGTTGCCTCCAGCCCGGAAAATAGCATNNNAGATCGGAAGAG |
| 29511_2 | TATGAGTGTGGAGTCGTTGCCTCCAGCCCGGAAAATAGCATNNNAGATCGGAAGAG |
| 29511_3 | TATGAGTGTGGAGTCGTTGCCTCCAGCCCGGAAAATAGCATNNNAGATCGGAAGAG |
| 50335_1 | TATGAGTGTGGAGTCGTTGCGAAAGTGTCTTCGCATAAGCCCNNNAGATCGGAAGA |
| 50335_2 | TATGAGTGTGGAGTCGTTGCGAAAGTGTCTTCGCATAAGCCCNNNAGATCGGAAGA |
| 50335_3 | TATGAGTGTGGAGTCGTTGCGAAAGTGTCTTCGCATAAGCCCNNNAGATCGGAAGA |
| 40036_1 | TATGAGTGTGGAGTCGTTGCGGATCACAAGGTCAGAAGATCGANNNAGATCGGAA |
| 40036_2 | TATGAGTGTGGAGTCGTTGCGGATCACAAGGTCAGAAGATCGANNNAGATCGGAA |
| 40036_3 | TATGAGTGTGGAGTCGTTGCGGATCACAAGGTCAGAAGATCGANNNAGATCGGAA |
| 2225_1 | TATGAGTGTGGAGTCGTTGCGCCCTGGAAATAGAGGTAGATGGNNNAGATCGGAA |
| 2225_2 | TATGAGTGTGGAGTCGTTGCGCCCTGGAAATAGAGGTAGATGGNNNAGATCGGAA |
| 2225_3 | TATGAGTGTGGAGTCGTTGCGCCCTGGAAATAGAGGTAGATGGNNNAGATCGGAA |
| 38209_1 | TATGAGTGTGGAGTCGTTGCAAAACACAGAGACTGGCACATTGNNNAGATCGGAAG |
| 38209_2 | TATGAGTGTGGAGTCGTTGCAAAACACAGAGACTGGCACATTGNNNAGATCGGAAG |
| 38209_3 | TATGAGTGTGGAGTCGTTGCAAAACACAGAGACTGGCACATTGNNNAGATCGGAAG |
| 42572_1 | TATGAGTGTGGAGTCGTTGCGAGTTTTGCTCTTGTTGCCCAGNNNAGATCGGAAGA |
| 42572_2 | TATGAGTGTGGAGTCGTTGCGAGTTTTGCTCTTGTTGCCCAGNNNAGATCGGAAGA |
| 42572_3 | TATGAGTGTGGAGTCGTTGCGAGTTTTGCTCTTGTTGCCCAGNNNAGATCGGAAGA |
| 43209_1 | TATGAGTGTGGAGTCGTTGCCGCCTCCTGGATTCAAATGATTCNNNAGATCGGAAGA |
| 43209_2 | TATGAGTGTGGAGTCGTTGCCGCCTCCTGGATTCAAATGATTCNNNAGATCGGAAGA |
| 43209_3 | TATGAGTGTGGAGTCGTTGCCGCCTCCTGGATTCAAATGATTCNNNAGATCGGAAGA |
| 35972_1 | TATGAGTGTGGAGTCGTTGCACGTATGTTTTTGCAGCACTGTTNNNAGATCGGAAGA |
| 35972_2 | TATGAGTGTGGAGTCGTTGCACGTATGTTTTTGCAGCACTGTTNNNAGATCGGAAGA |
| 35972_3 | TATGAGTGTGGAGTCGTTGCACGTATGTTTTTGCAGCACTGTTNNNAGATCGGAAGA |
| 36627_1 | TATGAGTGTGGAGTCGTTGCGTCCAGAACATGAGAAACCAGGANNNAGATCGGAAG |
| 36627_2 | TATGAGTGTGGAGTCGTTGCGTCCAGAACATGAGAAACCAGGANNNAGATCGGAAG |
| 36627_3 | TATGAGTGTGGAGTCGTTGCGTCCAGAACATGAGAAACCAGGANNNAGATCGGAAG |
| 12545_1 | TATGAGTGTGGAGTCGTTGCAGTTTACAAATGCCTACAAGTTANNNAGATCGGAAG |
| 12545_2 | TATGAGTGTGGAGTCGTTGCAGTTTACAAATGCCTACAAGTTANNNAGATCGGAAG |
| 12545_3 | TATGAGTGTGGAGTCGTTGCAGTTTACAAATGCCTACAAGTTANNNAGATCGGAAG |
| 33352_1 | TATGAGTGTGGAGTCGTTGCCTGGGTTCAAGTGATTCATCTGCNNNAGATCGGAAG |
| 33352_2 | TATGAGTGTGGAGTCGTTGCCTGGGTTCAAGTGATTCATCTGCNNNAGATCGGAAG |
| 33352_3 | TATGAGTGTGGAGTCGTTGCCTGGGTTCAAGTGATTCATCTGCNNNAGATCGGAAG |

| | |
|---|---|
| 44961_1 | TATGAGTGTGGAGTCGTTGCTTTTCTCAGGCTCTAGCAGGGTANNNAGATCGGAAG |
| 44961_2 | TATGAGTGTGGAGTCGTTGCTTTTCTCAGGCTCTAGCAGGGTANNNAGATCGGAAG |
| 44961_3 | TATGAGTGTGGAGTCGTTGCTTTTCTCAGGCTCTAGCAGGGTANNNAGATCGGAAG |
| 47176_1 | TATGAGTGTGGAGTCGTTGCACTCCTGACATCAAGTGATCCACNNNAGATCGGAAG |
| 47176_2 | TATGAGTGTGGAGTCGTTGCACTCCTGACATCAAGTGATCCACNNNAGATCGGAAG |
| 47176_3 | TATGAGTGTGGAGTCGTTGCACTCCTGACATCAAGTGATCCACNNNAGATCGGAAG |
| 48034_1 | TATGAGTGTGGAGTCGTTGCATGAAAAGGACAAGGGTACTCCGNNNAGATCGGAA |
| 48034_2 | TATGAGTGTGGAGTCGTTGCATGAAAAGGACAAGGGTACTCCGNNNAGATCGGAA |
| 48034_3 | TATGAGTGTGGAGTCGTTGCATGAAAAGGACAAGGGTACTCCGNNNAGATCGGAA |
| 43981_1 | TATGAGTGTGGAGTCGTTGCGAAAGGAAGGAAGGAAGGAAGGANNNAGATCGG |
| 43981_2 | TATGAGTGTGGAGTCGTTGCGAAAGGAAGGAAGGAAGGAAGGANNNAGATCGG |
| 43981_3 | TATGAGTGTGGAGTCGTTGCGAAAGGAAGGAAGGAAGGAAGGANNNAGATCGG |
| 49766_1 | TATGAGTGTGGAGTCGTTGCCTTGCTGCTGTCAGAGAGAACTANNNAGATCGGAA |
| 49766_2 | TATGAGTGTGGAGTCGTTGCCTTGCTGCTGTCAGAGAGAACTANNNAGATCGGAA |
| 49766_3 | TATGAGTGTGGAGTCGTTGCCTTGCTGCTGTCAGAGAGAACTANNNAGATCGGAA |
| 39522_1 | TATGAGTGTGGAGTCGTTGCGTTTCAGAAAAGAAAGCCCCAGGNNNAGATCGGAA |
| 39522_2 | TATGAGTGTGGAGTCGTTGCGTTTCAGAAAAGAAAGCCCCAGGNNNAGATCGGAA |
| 39522_3 | TATGAGTGTGGAGTCGTTGCGTTTCAGAAAAGAAAGCCCCAGGNNNAGATCGGAA |
| 33489_1 | TATGAGTGTGGAGTCGTTGCAAAGAGAAAGGGGGATAAGGCAGNNNAGATCGGA |
| 33489_2 | TATGAGTGTGGAGTCGTTGCAAAGAGAAAGGGGGATAAGGCAGNNNAGATCGGA |
| 33489_3 | TATGAGTGTGGAGTCGTTGCAAAGAGAAAGGGGGATAAGGCAGNNNAGATCGGA |
| 11631_1 | TATGAGTGTGGAGTCGTTGCGCATGTGTTGGTGTTAGTGTCTGTNNNAGATCGGA |
| 11631_2 | TATGAGTGTGGAGTCGTTGCGCATGTGTTGGTGTTAGTGTCTGTNNNAGATCGGA |
| 11631_3 | TATGAGTGTGGAGTCGTTGCGCATGTGTTGGTGTTAGTGTCTGTNNNAGATCGGA |
| 13484_1 | TATGAGTGTGGAGTCGTTGCGATTCAAGTGCTGTTTCAACNNNAGATCGGAAGAG |
| 13484_2 | TATGAGTGTGGAGTCGTTGCGATTCAAGTGCTGTTTCAACNNNAGATCGGAAGAG |
| 13484_3 | TATGAGTGTGGAGTCGTTGCGATTCAAGTGCTGTTTCAACNNNAGATCGGAAGAG |
| 44033_1 | TATGAGTGTGGAGTCGTTGCGTCCCCCTGTGTTTAGGCTAAATNNNAGATCGGAAG |
| 44033_2 | TATGAGTGTGGAGTCGTTGCGTCCCCCTGTGTTTAGGCTAAATNNNAGATCGGAAG |
| 44033_3 | TATGAGTGTGGAGTCGTTGCGTCCCCCTGTGTTTAGGCTAAATNNNAGATCGGAAG |
| 52793_1 | TATGAGTGTGGAGTCGTTGCGAGGAGGACTCTACAGGGACTCNNNAGATCGGAAG |
| 52793_2 | TATGAGTGTGGAGTCGTTGCGAGGAGGACTCTACAGGGACTCNNNAGATCGGAAG |
| 52793_3 | TATGAGTGTGGAGTCGTTGCGAGGAGGACTCTACAGGGACTCNNNAGATCGGAAG |
| 42715_1 | TATGAGTGTGGAGTCGTTGCACTTGAATAGACACTTCTCCACAGNNNAGATCGGAA |
| 42715_2 | TATGAGTGTGGAGTCGTTGCACTTGAATAGACACTTCTCCACAGNNNAGATCGGAA |
| 42715_3 | TATGAGTGTGGAGTCGTTGCACTTGAATAGACACTTCTCCACAGNNNAGATCGGAA |
| 682_1 | TATGAGTGTGGAGTCGTTGCACTGGAAAAATATTGTGACCCCCANNNAGATCGGAA |
| 682_2 | TATGAGTGTGGAGTCGTTGCACTGGAAAAATATTGTGACCCCCANNNAGATCGGAA |
| 682_3 | TATGAGTGTGGAGTCGTTGCACTGGAAAAATATTGTGACCCCCANNNAGATCGGAA |
| 42077_1 | TATGAGTGTGGAGTCGTTGCCACCTCACATTACACGACTCTGANNNAGATCGGAAG |
| 42077_2 | TATGAGTGTGGAGTCGTTGCCACCTCACATTACACGACTCTGANNNAGATCGGAAG |
| 42077_3 | TATGAGTGTGGAGTCGTTGCCACCTCACATTACACGACTCTGANNNAGATCGGAAG |
| 29394_1 | TATGAGTGTGGAGTCGTTGCAAATTAATCCTGGTTTTCAAGAGNNNAGATCGGAAG |

| | |
|---|---|
| 29394_2 | TATGAGTGTGGAGTCGTTGCAAATTAATCCTGGTTTTCAAGAGNNNAGATCGGAAG |
| 29394_3 | TATGAGTGTGGAGTCGTTGCAAATTAATCCTGGTTTTCAAGAGNNNAGATCGGAAG |
| 45251_1 | TATGAGTGTGGAGTCGTTGCGAGTCTCTTCTCACGACCCTTTTNNNAGATCGGAAGA |
| 45251_2 | TATGAGTGTGGAGTCGTTGCGAGTCTCTTCTCACGACCCTTTTNNNAGATCGGAAGA |
| 45251_3 | TATGAGTGTGGAGTCGTTGCGAGTCTCTTCTCACGACCCTTTTNNNAGATCGGAAGA |
| 42650_1 | TATGAGTGTGGAGTCGTTGCCAGCTACTTGGGAGGTACAAGAGNNNAGATCGGAA |
| 42650_2 | TATGAGTGTGGAGTCGTTGCCAGCTACTTGGGAGGTACAAGAGNNNAGATCGGAA |
| 42650_3 | TATGAGTGTGGAGTCGTTGCCAGCTACTTGGGAGGTACAAGAGNNNAGATCGGAA |
| 33770_1 | TATGAGTGTGGAGTCGTTGCTCATGAAACAACAGATGCTGGTGNNNAGATCGGAA |
| 33770_2 | TATGAGTGTGGAGTCGTTGCTCATGAAACAACAGATGCTGGTGNNNAGATCGGAA |
| 33770_3 | TATGAGTGTGGAGTCGTTGCTCATGAAACAACAGATGCTGGTGNNNAGATCGGAA |
| 51350_1 | TATGAGTGTGGAGTCGTTGCGGTGGTTGGCGAAGAAAAAAAACNNNAGATCGGA |
| 51350_2 | TATGAGTGTGGAGTCGTTGCGGTGGTTGGCGAAGAAAAAAAACNNNAGATCGGA |
| 51350_3 | TATGAGTGTGGAGTCGTTGCGGTGGTTGGCGAAGAAAAAAAACNNNAGATCGGA |
| 37468_1 | TATGAGTGTGGAGTCGTTGCAGCCTCTGTTTGTGTTCTTCAGTNNNAGATCGGAAG |
| 37468_2 | TATGAGTGTGGAGTCGTTGCAGCCTCTGTTTGTGTTCTTCAGTNNNAGATCGGAAG |
| 37468_3 | TATGAGTGTGGAGTCGTTGCAGCCTCTGTTTGTGTTCTTCAGTNNNAGATCGGAAG |
| 41435_1 | TATGAGTGTGGAGTCGTTGCGTAGTAACATGGATGGTGCTGGANNNAGATCGGAA |
| 41435_2 | TATGAGTGTGGAGTCGTTGCGTAGTAACATGGATGGTGCTGGANNNAGATCGGAA |
| 41435_3 | TATGAGTGTGGAGTCGTTGCGTAGTAACATGGATGGTGCTGGANNNAGATCGGAA |
| 32505_1 | TATGAGTGTGGAGTCGTTGCTAGCTGGGATTATAGGCACATGCNNNAGATCGGAA |
| 32505_2 | TATGAGTGTGGAGTCGTTGCTAGCTGGGATTATAGGCACATGCNNNAGATCGGAA |
| 32505_3 | TATGAGTGTGGAGTCGTTGCTAGCTGGGATTATAGGCACATGCNNNAGATCGGAA |
| 33864_1 | TATGAGTGTGGAGTCGTTGCGTAGGGGCACATAGGTGATAAGGNNNAGATCGGA |
| 33864_2 | TATGAGTGTGGAGTCGTTGCGTAGGGGCACATAGGTGATAAGGNNNAGATCGGA |
| 33864_3 | TATGAGTGTGGAGTCGTTGCGTAGGGGCACATAGGTGATAAGGNNNAGATCGGA |
| 35892_1 | TATGAGTGTGGAGTCGTTGCTCATGTCTGTAATCCCAGCATGTNNNAGATCGGAAG |
| 35892_2 | TATGAGTGTGGAGTCGTTGCTCATGTCTGTAATCCCAGCATGTNNNAGATCGGAAG |
| 35892_3 | TATGAGTGTGGAGTCGTTGCTCATGTCTGTAATCCCAGCATGTNNNAGATCGGAAG |
| 38473_1 | TATGAGTGTGGAGTCGTTGCGTTTTGTCTTTCTGCAGTGTCCANNNAGATCGGAAG |
| 38473_2 | TATGAGTGTGGAGTCGTTGCGTTTTGTCTTTCTGCAGTGTCCANNNAGATCGGAAG |
| 38473_3 | TATGAGTGTGGAGTCGTTGCGTTTTGTCTTTCTGCAGTGTCCANNNAGATCGGAAG |
| 33819_1 | TATGAGTGTGGAGTCGTTGCAAGTGGCATGGTGGTTATGTTTGNNNAGATCGGAA |
| 33819_2 | TATGAGTGTGGAGTCGTTGCAAGTGGCATGGTGGTTATGTTTGNNNAGATCGGAA |
| 33819_3 | TATGAGTGTGGAGTCGTTGCAAGTGGCATGGTGGTTATGTTTGNNNAGATCGGAA |
| 10495_1 | TATGAGTGTGGAGTCGTTGCCAAACACACGCACTCACTCATTTNNNAGATCGGAAGA |
| 10495_2 | TATGAGTGTGGAGTCGTTGCCAAACACACGCACTCACTCATTTNNNAGATCGGAAGA |
| 10495_3 | TATGAGTGTGGAGTCGTTGCCAAACACACGCACTCACTCATTTNNNAGATCGGAAGA |
| 47356_1 | TATGAGTGTGGAGTCGTTGCAGGCTGGGACAGTCAAGTATTTTNNNAGATCGGAA |
| 47356_2 | TATGAGTGTGGAGTCGTTGCAGGCTGGGACAGTCAAGTATTTTNNNAGATCGGAA |
| 47356_3 | TATGAGTGTGGAGTCGTTGCAGGCTGGGACAGTCAAGTATTTTNNNAGATCGGAA |
| 42932_1 | TATGAGTGTGGAGTCGTTGCTGGTGGCTCATTCCTGTAATCCNNNAGATCGGAAGA |
| 42932_2 | TATGAGTGTGGAGTCGTTGCTGGTGGCTCATTCCTGTAATCCNNNAGATCGGAAGA |

| | |
|---|---|
| 42932_3 | TATGAGTGTGGAGTCGTTGCTGGTGGCTCATTCCTGTAATCCNNNAGATCGGAAGA |
| 32605_1 | TATGAGTGTGGAGTCGTTGCAAAGAGACGGATGGCATTTTTCCNNNAGATCGGAA |
| 32605_2 | TATGAGTGTGGAGTCGTTGCAAAGAGACGGATGGCATTTTTCCNNNAGATCGGAA |
| 32605_3 | TATGAGTGTGGAGTCGTTGCAAAGAGACGGATGGCATTTTTCCNNNAGATCGGAA |
| 52803_1 | TATGAGTGTGGAGTCGTTGCCGGATCAGACCCCTCTCCAGNNNAGATCGGAAGAGC |
| 52803_2 | TATGAGTGTGGAGTCGTTGCCGGATCAGACCCCTCTCCAGNNNAGATCGGAAGAGC |
| 52803_3 | TATGAGTGTGGAGTCGTTGCCGGATCAGACCCCTCTCCAGNNNAGATCGGAAGAGC |
| 43362_1 | TATGAGTGTGGAGTCGTTGCGATCTCAGCTCACTGCAACCANNNAGATCGGAAGAG |
| 43362_2 | TATGAGTGTGGAGTCGTTGCGATCTCAGCTCACTGCAACCANNNAGATCGGAAGAG |
| 43362_3 | TATGAGTGTGGAGTCGTTGCGATCTCAGCTCACTGCAACCANNNAGATCGGAAGAG |
| 42170_1 | TATGAGTGTGGAGTCGTTGCCTCAAGTGATCCTCCTGTCTCAGNNNAGATCGGAAG |
| 42170_2 | TATGAGTGTGGAGTCGTTGCCTCAAGTGATCCTCCTGTCTCAGNNNAGATCGGAAG |
| 42170_3 | TATGAGTGTGGAGTCGTTGCCTCAAGTGATCCTCCTGTCTCAGNNNAGATCGGAAG |
| 39981_1 | TATGAGTGTGGAGTCGTTGCCTTGTGTGGAACAACTGCAGATGNNNAGATCGGAA |
| 39981_2 | TATGAGTGTGGAGTCGTTGCCTTGTGTGGAACAACTGCAGATGNNNAGATCGGAA |
| 39981_3 | TATGAGTGTGGAGTCGTTGCCTTGTGTGGAACAACTGCAGATGNNNAGATCGGAA |
| 38993_1 | TATGAGTGTGGAGTCGTTGCCAAGTGTTAGGATCCTGGCATCTNNNAGATCGGAAG |
| 38993_2 | TATGAGTGTGGAGTCGTTGCCAAGTGTTAGGATCCTGGCATCTNNNAGATCGGAAG |
| 38993_3 | TATGAGTGTGGAGTCGTTGCCAAGTGTTAGGATCCTGGCATCTNNNAGATCGGAAG |
| 49275_1 | TATGAGTGTGGAGTCGTTGCCCACGCCCAGCTAATTTTTGTATNNNAGATCGGAAG |
| 49275_2 | TATGAGTGTGGAGTCGTTGCCCACGCCCAGCTAATTTTTGTATNNNAGATCGGAAG |
| 49275_3 | TATGAGTGTGGAGTCGTTGCCCACGCCCAGCTAATTTTTGTATNNNAGATCGGAAG |
| 48752_1 | TATGAGTGTGGAGTCGTTGCATTTTTATCGGGTCAGGTGCCATNNNAGATCGGAAG |
| 48752_2 | TATGAGTGTGGAGTCGTTGCATTTTTATCGGGTCAGGTGCCATNNNAGATCGGAAG |
| 48752_3 | TATGAGTGTGGAGTCGTTGCATTTTTATCGGGTCAGGTGCCATNNNAGATCGGAAG |
| 52860_1 | TATGAGTGTGGAGTCGTTGCATCACCTTCCTCCTCCTTCTAGGNNNAGATCGGAAGA |
| 52860_2 | TATGAGTGTGGAGTCGTTGCATCACCTTCCTCCTCCTTCTAGGNNNAGATCGGAAGA |
| 52860_3 | TATGAGTGTGGAGTCGTTGCATCACCTTCCTCCTCCTTCTAGGNNNAGATCGGAAGA |
| 44341_1 | TATGAGTGTGGAGTCGTTGCCTTCCTCAGTACACTCCCTTTGGNNNAGATCGGAAGA |
| 44341_2 | TATGAGTGTGGAGTCGTTGCCTTCCTCAGTACACTCCCTTTGGNNNAGATCGGAAGA |
| 44341_3 | TATGAGTGTGGAGTCGTTGCCTTCCTCAGTACACTCCCTTTGGNNNAGATCGGAAGA |
| 39545_1 | TATGAGTGTGGAGTCGTTGCAGAAGCTAACTGAAGAGGCACAANNNAGATCGGAA |
| 39545_2 | TATGAGTGTGGAGTCGTTGCAGAAGCTAACTGAAGAGGCACAANNNAGATCGGAA |
| 39545_3 | TATGAGTGTGGAGTCGTTGCAGAAGCTAACTGAAGAGGCACAANNNAGATCGGAA |
| 41674_1 | TATGAGTGTGGAGTCGTTGCCCTGAGTAGCTAGGATTACAGGCNNNAGATCGGAA |
| 41674_2 | TATGAGTGTGGAGTCGTTGCCCTGAGTAGCTAGGATTACAGGCNNNAGATCGGAA |
| 41674_3 | TATGAGTGTGGAGTCGTTGCCCTGAGTAGCTAGGATTACAGGCNNNAGATCGGAA |
| 48461_1 | TATGAGTGTGGAGTCGTTGCTTCAAGCAATTCTCCCACCTCAGNNNAGATCGGAAGA |
| 48461_2 | TATGAGTGTGGAGTCGTTGCTTCAAGCAATTCTCCCACCTCAGNNNAGATCGGAAGA |
| 48461_3 | TATGAGTGTGGAGTCGTTGCTTCAAGCAATTCTCCCACCTCAGNNNAGATCGGAAGA |
| 13675_1 | TATGAGTGTGGAGTCGTTGCCACACACTAGGAAGATACCTNNNAGATCGGAAGAG |
| 13675_2 | TATGAGTGTGGAGTCGTTGCCACACACTAGGAAGATACCTNNNAGATCGGAAGAG |
| 13675_3 | TATGAGTGTGGAGTCGTTGCCACACACTAGGAAGATACCTNNNAGATCGGAAGAG |

| | |
|---|---|
| 32969_1 | TATGAGTGTGGAGTCGTTGCACTGTTTGTTTTTGAGCTAGGGCNNNAGATCGGAAG |
| 32969_2 | TATGAGTGTGGAGTCGTTGCACTGTTTGTTTTTGAGCTAGGGCNNNAGATCGGAAG |
| 32969_3 | TATGAGTGTGGAGTCGTTGCACTGTTTGTTTTTGAGCTAGGGCNNNAGATCGGAAG |
| 30821_1 | TATGAGTGTGGAGTCGTTGCCTCATGTCTCAGCTAAGGAAGGGNNNAGATCGGAA |
| 30821_2 | TATGAGTGTGGAGTCGTTGCCTCATGTCTCAGCTAAGGAAGGGNNNAGATCGGAA |
| 30821_3 | TATGAGTGTGGAGTCGTTGCCTCATGTCTCAGCTAAGGAAGGGNNNAGATCGGAA |
| 51801_1 | TATGAGTGTGGAGTCGTTGCCCAGAGCGGAGTGAAATCTTAGANNNAGATCGGAA |
| 51801_2 | TATGAGTGTGGAGTCGTTGCCCAGAGCGGAGTGAAATCTTAGANNNAGATCGGAA |
| 51801_3 | TATGAGTGTGGAGTCGTTGCCCAGAGCGGAGTGAAATCTTAGANNNAGATCGGAA |
| 36670_1 | TATGAGTGTGGAGTCGTTGCAGTCAAAGAGACCAGTGGATCACNNNAGATCGGAA |
| 36670_2 | TATGAGTGTGGAGTCGTTGCAGTCAAAGAGACCAGTGGATCACNNNAGATCGGAA |
| 36670_3 | TATGAGTGTGGAGTCGTTGCAGTCAAAGAGACCAGTGGATCACNNNAGATCGGAA |
| 46684_1 | TATGAGTGTGGAGTCGTTGCGCCTTCTGAGTAGCTAGGATCACNNNAGATCGGAAG |
| 46684_2 | TATGAGTGTGGAGTCGTTGCGCCTTCTGAGTAGCTAGGATCACNNNAGATCGGAAG |
| 46684_3 | TATGAGTGTGGAGTCGTTGCGCCTTCTGAGTAGCTAGGATCACNNNAGATCGGAAG |
| 42855_1 | TATGAGTGTGGAGTCGTTGCAGACGCCCGGCTAATTTTTTTTTNNNAGATCGGAAG |
| 42855_2 | TATGAGTGTGGAGTCGTTGCAGACGCCCGGCTAATTTTTTTTTNNNAGATCGGAAG |
| 42855_3 | TATGAGTGTGGAGTCGTTGCAGACGCCCGGCTAATTTTTTTTTNNNAGATCGGAAG |
| 45299_1 | TATGAGTGTGGAGTCGTTGCAGGGATTTGCCATTCTCTTCCATNNNAGATCGGAAG |
| 45299_2 | TATGAGTGTGGAGTCGTTGCAGGGATTTGCCATTCTCTTCCATNNNAGATCGGAAG |
| 45299_3 | TATGAGTGTGGAGTCGTTGCAGGGATTTGCCATTCTCTTCCATNNNAGATCGGAAG |
| 51139_1 | TATGAGTGTGGAGTCGTTGCCCACCATCCCACTTGTTCAAATCNNNAGATCGGAAGA |
| 51139_2 | TATGAGTGTGGAGTCGTTGCCCACCATCCCACTTGTTCAAATCNNNAGATCGGAAGA |
| 51139_3 | TATGAGTGTGGAGTCGTTGCCCACCATCCCACTTGTTCAAATCNNNAGATCGGAAGA |
| 33889_1 | TATGAGTGTGGAGTCGTTGCCCTCATATCTCAGCCTCTCAAGGNNNAGATCGGAAG |
| 33889_2 | TATGAGTGTGGAGTCGTTGCCCTCATATCTCAGCCTCTCAAGGNNNAGATCGGAAG |
| 33889_3 | TATGAGTGTGGAGTCGTTGCCCTCATATCTCAGCCTCTCAAGGNNNAGATCGGAAG |
| 50010_1 | TATGAGTGTGGAGTCGTTGCCTGTGCCTCATTTCCTAGCAGTNNNAGATCGGAAGA |
| 50010_2 | TATGAGTGTGGAGTCGTTGCCTGTGCCTCATTTCCTAGCAGTNNNAGATCGGAAGA |
| 50010_3 | TATGAGTGTGGAGTCGTTGCCTGTGCCTCATTTCCTAGCAGTNNNAGATCGGAAGA |
| 43157_1 | TATGAGTGTGGAGTCGTTGCTGTTCTTCTCATTCAGTCTGGGGNNNAGATCGGAAG |
| 43157_2 | TATGAGTGTGGAGTCGTTGCTGTTCTTCTCATTCAGTCTGGGGNNNAGATCGGAAG |
| 43157_3 | TATGAGTGTGGAGTCGTTGCTGTTCTTCTCATTCAGTCTGGGGNNNAGATCGGAAG |
| 37000_1 | TATGAGTGTGGAGTCGTTGCTATCTCGTGACTTGGATACCAGCNNNAGATCGGAAG |
| 37000_2 | TATGAGTGTGGAGTCGTTGCTATCTCGTGACTTGGATACCAGCNNNAGATCGGAAG |
| 37000_3 | TATGAGTGTGGAGTCGTTGCTATCTCGTGACTTGGATACCAGCNNNAGATCGGAAG |
| 13723_1 | TATGAGTGTGGAGTCGTTGCAAGATAGTCTCCACTTTTTAATGNNNAGATCGGAAG |
| 13723_2 | TATGAGTGTGGAGTCGTTGCAAGATAGTCTCCACTTTTTAATGNNNAGATCGGAAG |
| 13723_3 | TATGAGTGTGGAGTCGTTGCAAGATAGTCTCCACTTTTTAATGNNNAGATCGGAAG |
| 13001_1 | TATGAGTGTGGAGTCGTTGCATCCTTTATGTACTAGAAAAGGTNNNAGATCGGAAG |
| 13001_2 | TATGAGTGTGGAGTCGTTGCATCCTTTATGTACTAGAAAAGGTNNNAGATCGGAAG |
| 13001_3 | TATGAGTGTGGAGTCGTTGCATCCTTTATGTACTAGAAAAGGTNNNAGATCGGAAG |
| 35937_1 | TATGAGTGTGGAGTCGTTGCCACCATGGAATACTACTCAGCCANNNAGATCGGAAG |

| | |
|---|---|
| 35937_2 | TATGAGTGTGGAGTCGTTGCCACCATGGAATACTACTCAGCCANNNAGATCGGAAG |
| 35937_3 | TATGAGTGTGGAGTCGTTGCCACCATGGAATACTACTCAGCCANNNAGATCGGAAG |
| 28894_1 | TATGAGTGTGGAGTCGTTGCCCAGTGACTATTTTTGTGGCTGGNNNAGATCGGAAG |
| 28894_2 | TATGAGTGTGGAGTCGTTGCCCAGTGACTATTTTTGTGGCTGGNNNAGATCGGAAG |
| 28894_3 | TATGAGTGTGGAGTCGTTGCCCAGTGACTATTTTTGTGGCTGGNNNAGATCGGAAG |
| 40397_1 | TATGAGTGTGGAGTCGTTGCAATACAGCTGACCCTTGAACCATNNNAGATCGGAAG |
| 40397_2 | TATGAGTGTGGAGTCGTTGCAATACAGCTGACCCTTGAACCATNNNAGATCGGAAG |
| 40397_3 | TATGAGTGTGGAGTCGTTGCAATACAGCTGACCCTTGAACCATNNNAGATCGGAAG |
| 36780_1 | TATGAGTGTGGAGTCGTTGCAGGATTGGTAAGAAGTTTCAGCANNNAGATCGGAA |
| 36780_2 | TATGAGTGTGGAGTCGTTGCAGGATTGGTAAGAAGTTTCAGCANNNAGATCGGAA |
| 36780_3 | TATGAGTGTGGAGTCGTTGCAGGATTGGTAAGAAGTTTCAGCANNNAGATCGGAA |
| 51385_1 | TATGAGTGTGGAGTCGTTGCTGATTGATCAGCTAAGGTAGGGCNNNAGATCGGAA |
| 51385_2 | TATGAGTGTGGAGTCGTTGCTGATTGATCAGCTAAGGTAGGGCNNNAGATCGGAA |
| 51385_3 | TATGAGTGTGGAGTCGTTGCTGATTGATCAGCTAAGGTAGGGCNNNAGATCGGAA |
| 38343_1 | TATGAGTGTGGAGTCGTTGCGCTTTTGGCAGTGACATACTTGTNNNAGATCGGAAG |
| 38343_2 | TATGAGTGTGGAGTCGTTGCGCTTTTGGCAGTGACATACTTGTNNNAGATCGGAAG |
| 38343_3 | TATGAGTGTGGAGTCGTTGCGCTTTTGGCAGTGACATACTTGTNNNAGATCGGAAG |
| 34284_1 | TATGAGTGTGGAGTCGTTGCATGCCTGTAATCTCAGCTACTCGNNNAGATCGGAAG |
| 34284_2 | TATGAGTGTGGAGTCGTTGCATGCCTGTAATCTCAGCTACTCGNNNAGATCGGAAG |
| 34284_3 | TATGAGTGTGGAGTCGTTGCATGCCTGTAATCTCAGCTACTCGNNNAGATCGGAAG |
| 48802_1 | TATGAGTGTGGAGTCGTTGCGTCATTTCTGGGATTTTGGTGCANNNAGATCGGAAG |
| 48802_2 | TATGAGTGTGGAGTCGTTGCGTCATTTCTGGGATTTTGGTGCANNNAGATCGGAAG |
| 48802_3 | TATGAGTGTGGAGTCGTTGCGTCATTTCTGGGATTTTGGTGCANNNAGATCGGAAG |
| 40638_1 | TATGAGTGTGGAGTCGTTGCACATTCACTGAAAAGGACAACGGNNNAGATCGGAA |
| 40638_2 | TATGAGTGTGGAGTCGTTGCACATTCACTGAAAAGGACAACGGNNNAGATCGGAA |
| 40638_3 | TATGAGTGTGGAGTCGTTGCACATTCACTGAAAAGGACAACGGNNNAGATCGGAA |
| 35856_1 | TATGAGTGTGGAGTCGTTGCGCTCCTGGCCTCTTTTCTTTTTTNNNAGATCGGAAGA |
| 35856_2 | TATGAGTGTGGAGTCGTTGCGCTCCTGGCCTCTTTTCTTTTTTNNNAGATCGGAAGA |
| 35856_3 | TATGAGTGTGGAGTCGTTGCGCTCCTGGCCTCTTTTCTTTTTTNNNAGATCGGAAGA |
| 53408_1 | TATGAGTGTGGAGTCGTTGCTCCTTAGCTAATAATCAGGGAAATGTNNNAGATCGG |
| 53408_2 | TATGAGTGTGGAGTCGTTGCTCCTTAGCTAATAATCAGGGAAATGTNNNAGATCGG |
| 53408_3 | TATGAGTGTGGAGTCGTTGCTCCTTAGCTAATAATCAGGGAAATGTNNNAGATCGG |
| 29058_1 | TATGAGTGTGGAGTCGTTGCTCTTTTTCAATTTGGGTGCTTCTNNNAGATCGGAAGA |
| 29058_2 | TATGAGTGTGGAGTCGTTGCTCTTTTTCAATTTGGGTGCTTCTNNNAGATCGGAAGA |
| 29058_3 | TATGAGTGTGGAGTCGTTGCTCTTTTTCAATTTGGGTGCTTCTNNNAGATCGGAAGA |
| 31219_1 | TATGAGTGTGGAGTCGTTGCCCAAATTAGCCAGAAACCACCAGNNNAGATCGGAAG |
| 31219_2 | TATGAGTGTGGAGTCGTTGCCCAAATTAGCCAGAAACCACCAGNNNAGATCGGAAG |
| 31219_3 | TATGAGTGTGGAGTCGTTGCCCAAATTAGCCAGAAACCACCAGNNNAGATCGGAAG |
| 13751_1 | TATGAGTGTGGAGTCGTTGCAAAGAATGTTTGTATACAAAGCTNNNAGATCGGAA |
| 13751_2 | TATGAGTGTGGAGTCGTTGCAAAGAATGTTTGTATACAAAGCTNNNAGATCGGAA |
| 13751_3 | TATGAGTGTGGAGTCGTTGCAAAGAATGTTTGTATACAAAGCTNNNAGATCGGAA |
| 2234_1 | TATGAGTGTGGAGTCGTTGCGGAGGTGGGAAGAGGGAGAAAATANNNAGATCG |
| 2234_2 | TATGAGTGTGGAGTCGTTGCGGAGGTGGGAAGAGGGAGAAAATANNNAGATCG |

| | |
|---|---|
| 2234_3 | TATGAGTGTGGAGTCGTTGCGGAGGTGGGAAGAGGGAGAAAATANNNAGATCGG |
| 33920_1 | TATGAGTGTGGAGTCGTTGCGTTCACTCCATTCTCCTGGCTCNNNAGATCGGAAGAG |
| 33920_2 | TATGAGTGTGGAGTCGTTGCGTTCACTCCATTCTCCTGGCTCNNNAGATCGGAAGAG |
| 33920_3 | TATGAGTGTGGAGTCGTTGCGTTCACTCCATTCTCCTGGCTCNNNAGATCGGAAGAG |
| 46368_1 | TATGAGTGTGGAGTCGTTGCGAGTTTTGCTCTTGTTGCCCAGNNNAGATCGGAAGA |
| 46368_2 | TATGAGTGTGGAGTCGTTGCGAGTTTTGCTCTTGTTGCCCAGNNNAGATCGGAAGA |
| 46368_3 | TATGAGTGTGGAGTCGTTGCGAGTTTTGCTCTTGTTGCCCAGNNNAGATCGGAAGA |
| 45831_1 | TATGAGTGTGGAGTCGTTGCCTTGAACTCCTGACCTCGTGATCNNNAGATCGGAAGA |
| 45831_2 | TATGAGTGTGGAGTCGTTGCCTTGAACTCCTGACCTCGTGATCNNNAGATCGGAAGA |
| 45831_3 | TATGAGTGTGGAGTCGTTGCCTTGAACTCCTGACCTCGTGATCNNNAGATCGGAAGA |
| 45725_1 | TATGAGTGTGGAGTCGTTGCTTGGGTTCAAGCGAGTCTACTGNNNAGATCGGAAGA |
| 45725_2 | TATGAGTGTGGAGTCGTTGCTTGGGTTCAAGCGAGTCTACTGNNNAGATCGGAAGA |
| 45725_3 | TATGAGTGTGGAGTCGTTGCTTGGGTTCAAGCGAGTCTACTGNNNAGATCGGAAGA |
| 47476_1 | TATGAGTGTGGAGTCGTTGCCAGGCACAATCATGGAACACTACNNNAGATCGGAAG |
| 47476_2 | TATGAGTGTGGAGTCGTTGCCAGGCACAATCATGGAACACTACNNNAGATCGGAAG |
| 47476_3 | TATGAGTGTGGAGTCGTTGCCAGGCACAATCATGGAACACTACNNNAGATCGGAAG |
| 51498_1 | TATGAGTGTGGAGTCGTTGCATACTTGGCAGTCACTTACGAGGNNNAGATCGGAAG |
| 51498_2 | TATGAGTGTGGAGTCGTTGCATACTTGGCAGTCACTTACGAGGNNNAGATCGGAAG |
| 51498_3 | TATGAGTGTGGAGTCGTTGCATACTTGGCAGTCACTTACGAGGNNNAGATCGGAAG |
| 44533_1 | TATGAGTGTGGAGTCGTTGCCTTGTACATCCTCACTTCCCCAANNNAGATCGGAAGA |
| 44533_2 | TATGAGTGTGGAGTCGTTGCCTTGTACATCCTCACTTCCCCAANNNAGATCGGAAGA |
| 44533_3 | TATGAGTGTGGAGTCGTTGCCTTGTACATCCTCACTTCCCCAANNNAGATCGGAAGA |
| 34337_1 | TATGAGTGTGGAGTCGTTGCTTGGCTGTTGTTGGTGTATAGGANNNAGATCGGAA |
| 34337_2 | TATGAGTGTGGAGTCGTTGCTTGGCTGTTGTTGGTGTATAGGANNNAGATCGGAA |
| 34337_3 | TATGAGTGTGGAGTCGTTGCTTGGCTGTTGTTGGTGTATAGGANNNAGATCGGAA |
| 38000_1 | TATGAGTGTGGAGTCGTTGCGCAGGGTGAACACATTTAGCAATNNNAGATCGGAA |
| 38000_2 | TATGAGTGTGGAGTCGTTGCGCAGGGTGAACACATTTAGCAATNNNAGATCGGAA |
| 38000_3 | TATGAGTGTGGAGTCGTTGCGCAGGGTGAACACATTTAGCAATNNNAGATCGGAA |
| 33227_1 | TATGAGTGTGGAGTCGTTGCAATAGCTGGGATTACAGACACCCNNNAGATCGGAAG |
| 33227_2 | TATGAGTGTGGAGTCGTTGCAATAGCTGGGATTACAGACACCCNNNAGATCGGAAG |
| 33227_3 | TATGAGTGTGGAGTCGTTGCAATAGCTGGGATTACAGACACCCNNNAGATCGGAAG |
| 13770_1 | TATGAGTGTGGAGTCGTTGCCATTATACCAAAAAGATACACGANNNAGATCGGAAG |
| 13770_2 | TATGAGTGTGGAGTCGTTGCCATTATACCAAAAAGATACACGANNNAGATCGGAAG |
| 13770_3 | TATGAGTGTGGAGTCGTTGCCATTATACCAAAAAGATACACGANNNAGATCGGAAG |
| 46053_1 | TATGAGTGTGGAGTCGTTGCCTCTCGAGTAGCTGGGACTACAGNNNAGATCGGAA |
| 46053_2 | TATGAGTGTGGAGTCGTTGCCTCTCGAGTAGCTGGGACTACAGNNNAGATCGGAA |
| 46053_3 | TATGAGTGTGGAGTCGTTGCCTCTCGAGTAGCTGGGACTACAGNNNAGATCGGAA |
| 30790_1 | TATGAGTGTGGAGTCGTTGCACACACACATCTGCCTACGTTATNNNAGATCGGAAGA |
| 30790_2 | TATGAGTGTGGAGTCGTTGCACACACACATCTGCCTACGTTATNNNAGATCGGAAGA |
| 30790_3 | TATGAGTGTGGAGTCGTTGCACACACACATCTGCCTACGTTATNNNAGATCGGAAGA |
| 10813_1 | TATGAGTGTGGAGTCGTTGCGCAGAAGACCAAAAATGGATGCCNNNAGATCGGAA |
| 10813_2 | TATGAGTGTGGAGTCGTTGCGCAGAAGACCAAAAATGGATGCCNNNAGATCGGAA |
| 10813_3 | TATGAGTGTGGAGTCGTTGCGCAGAAGACCAAAAATGGATGCCNNNAGATCGGAA |

| | |
|---|---|
| 10827_1 | TATGAGTGTGGAGTCGTTGCAGAGGCTGACTTTGCAGAGACTANNNAGATCGGAA |
| 10827_2 | TATGAGTGTGGAGTCGTTGCAGAGGCTGACTTTGCAGAGACTANNNAGATCGGAA |
| 10827_3 | TATGAGTGTGGAGTCGTTGCAGAGGCTGACTTTGCAGAGACTANNNAGATCGGAA |
| 10385_1 | TATGAGTGTGGAGTCGTTGCACAAAAATCACCTGGAGAGCTGTTNNNAGATCGGA |
| 10385_2 | TATGAGTGTGGAGTCGTTGCACAAAAATCACCTGGAGAGCTGTTNNNAGATCGGA |
| 10385_3 | TATGAGTGTGGAGTCGTTGCACAAAAATCACCTGGAGAGCTGTTNNNAGATCGGA |
| 10349_1 | TATGAGTGTGGAGTCGTTGCTAGGGCACCACCAAATGAATAGCNNNAGATCGGAA |
| 10349_2 | TATGAGTGTGGAGTCGTTGCTAGGGCACCACCAAATGAATAGCNNNAGATCGGAA |
| 10349_3 | TATGAGTGTGGAGTCGTTGCTAGGGCACCACCAAATGAATAGCNNNAGATCGGAA |
| 43696_1 | TATGAGTGTGGAGTCGTTGCAGGCAGCATGTTCATTATAGGGANNNAGATCGGAA |
| 43696_2 | TATGAGTGTGGAGTCGTTGCAGGCAGCATGTTCATTATAGGGANNNAGATCGGAA |
| 43696_3 | TATGAGTGTGGAGTCGTTGCAGGCAGCATGTTCATTATAGGGANNNAGATCGGAA |
| 44675_1 | TATGAGTGTGGAGTCGTTGCCTCATCAGAAACGTCCTCTGTCANNNAGATCGGAAG |
| 44675_2 | TATGAGTGTGGAGTCGTTGCCTCATCAGAAACGTCCTCTGTCANNNAGATCGGAAG |
| 44675_3 | TATGAGTGTGGAGTCGTTGCCTCATCAGAAACGTCCTCTGTCANNNAGATCGGAAG |
| 47978_1 | TATGAGTGTGGAGTCGTTGCGCTATGCTGGGTCTTTTGTTGTTNNNAGATCGGAAG |
| 47978_2 | TATGAGTGTGGAGTCGTTGCGCTATGCTGGGTCTTTTGTTGTTNNNAGATCGGAAG |
| 47978_3 | TATGAGTGTGGAGTCGTTGCGCTATGCTGGGTCTTTTGTTGTTNNNAGATCGGAAG |
| 35905_1 | TATGAGTGTGGAGTCGTTGCCTGAGGCAGGAGAATCACTTGAANNNAGATCGGAA |
| 35905_2 | TATGAGTGTGGAGTCGTTGCCTGAGGCAGGAGAATCACTTGAANNNAGATCGGAA |
| 35905_3 | TATGAGTGTGGAGTCGTTGCCTGAGGCAGGAGAATCACTTGAANNNAGATCGGAA |
| 43552_1 | TATGAGTGTGGAGTCGTTGCCTGGTCTTGAACTCCTGATCTCANNNAGATCGGAAG |
| 43552_2 | TATGAGTGTGGAGTCGTTGCCTGGTCTTGAACTCCTGATCTCANNNAGATCGGAAG |
| 43552_3 | TATGAGTGTGGAGTCGTTGCCTGGTCTTGAACTCCTGATCTCANNNAGATCGGAAG |
| 43151_1 | TATGAGTGTGGAGTCGTTGCAATACTCGACCAGGTACATTGGCNNNAGATCGGAAG |
| 43151_2 | TATGAGTGTGGAGTCGTTGCAATACTCGACCAGGTACATTGGCNNNAGATCGGAAG |
| 43151_3 | TATGAGTGTGGAGTCGTTGCAATACTCGACCAGGTACATTGGCNNNAGATCGGAAG |
| 48435_1 | TATGAGTGTGGAGTCGTTGCGATCTCAGCTCACTGCAACCANNNAGATCGGAAGAG |
| 48435_2 | TATGAGTGTGGAGTCGTTGCGATCTCAGCTCACTGCAACCANNNAGATCGGAAGAG |
| 48435_3 | TATGAGTGTGGAGTCGTTGCGATCTCAGCTCACTGCAACCANNNAGATCGGAAGAG |
| 52042_1 | TATGAGTGTGGAGTCGTTGCCAGTCCCATCAACCACCCAAGNNNAGATCGGAAGAG |
| 52042_2 | TATGAGTGTGGAGTCGTTGCCAGTCCCATCAACCACCCAAGNNNAGATCGGAAGAG |
| 52042_3 | TATGAGTGTGGAGTCGTTGCCAGTCCCATCAACCACCCAAGNNNAGATCGGAAGAG |
| 43442_1 | TATGAGTGTGGAGTCGTTGCCCACCACACCTGGATAATTTTCGNNNAGATCGGAAG |
| 43442_2 | TATGAGTGTGGAGTCGTTGCCCACCACACCTGGATAATTTTCGNNNAGATCGGAAG |
| 43442_3 | TATGAGTGTGGAGTCGTTGCCCACCACACCTGGATAATTTTCGNNNAGATCGGAAG |
| 37507_1 | TATGAGTGTGGAGTCGTTGCGGAACTCCTGGGATTACAACAGTNNNAGATCGGAA |
| 37507_2 | TATGAGTGTGGAGTCGTTGCGGAACTCCTGGGATTACAACAGTNNNAGATCGGAA |
| 37507_3 | TATGAGTGTGGAGTCGTTGCGGAACTCCTGGGATTACAACAGTNNNAGATCGGAA |
| 48709_1 | TATGAGTGTGGAGTCGTTGCGTTCAAGCGATTCTTCTGCCTCNNNAGATCGGAAGA |
| 48709_2 | TATGAGTGTGGAGTCGTTGCGTTCAAGCGATTCTTCTGCCTCNNNAGATCGGAAGA |
| 48709_3 | TATGAGTGTGGAGTCGTTGCGTTCAAGCGATTCTTCTGCCTCNNNAGATCGGAAGA |
| 13245_1 | TATGAGTGTGGAGTCGTTGCTTTGAGATATTACATGTTCAAGCNNNAGATCGGAAG |

| | |
|---|---|
| 13245_2 | TATGAGTGTGGAGTCGTTGCTTTGAGATATTACATGTTCAAGCNNNAGATCGGAAG |
| 13245_3 | TATGAGTGTGGAGTCGTTGCTTTGAGATATTACATGTTCAAGCNNNAGATCGGAAG |
| 33964_1 | TATGAGTGTGGAGTCGTTGCCTGAAGTGCAATGGCGAAATCTCNNNAGATCGGAA |
| 33964_2 | TATGAGTGTGGAGTCGTTGCCTGAAGTGCAATGGCGAAATCTCNNNAGATCGGAA |
| 33964_3 | TATGAGTGTGGAGTCGTTGCCTGAAGTGCAATGGCGAAATCTCNNNAGATCGGAA |
| 29981_1 | TATGAGTGTGGAGTCGTTGCGGTAGTGAGAACTGCCAAAAACCNNNAGATCGGAA |
| 29981_2 | TATGAGTGTGGAGTCGTTGCGGTAGTGAGAACTGCCAAAAACCNNNAGATCGGAA |
| 29981_3 | TATGAGTGTGGAGTCGTTGCGGTAGTGAGAACTGCCAAAAACCNNNAGATCGGAA |
| 41440_1 | TATGAGTGTGGAGTCGTTGCGGATGGCATGGAAAACACAAGTTNNNAGATCGGAA |
| 41440_2 | TATGAGTGTGGAGTCGTTGCGGATGGCATGGAAAACACAAGTTNNNAGATCGGAA |
| 41440_3 | TATGAGTGTGGAGTCGTTGCGGATGGCATGGAAAACACAAGTTNNNAGATCGGAA |
| 7451_1 | TATGAGTGTGGAGTCGTTGCTTGATCCCTTTTGGACCAGTTCGNNNAGATCGGAAG |
| 7451_2 | TATGAGTGTGGAGTCGTTGCTTGATCCCTTTTGGACCAGTTCGNNNAGATCGGAAG |
| 7451_3 | TATGAGTGTGGAGTCGTTGCTTGATCCCTTTTGGACCAGTTCGNNNAGATCGGAAG |
| 37875_1 | TATGAGTGTGGAGTCGTTGCGCTAACTGACAGAACATGAGGGANNNAGATCGGAA |
| 37875_2 | TATGAGTGTGGAGTCGTTGCGCTAACTGACAGAACATGAGGGANNNAGATCGGAA |
| 37875_3 | TATGAGTGTGGAGTCGTTGCGCTAACTGACAGAACATGAGGGANNNAGATCGGAA |
| 28889_1 | TATGAGTGTGGAGTCGTTGCCCTTCTTAGGGCCATTCCAAGATNNNAGATCGGAAG |
| 28889_2 | TATGAGTGTGGAGTCGTTGCCCTTCTTAGGGCCATTCCAAGATNNNAGATCGGAAG |
| 28889_3 | TATGAGTGTGGAGTCGTTGCCCTTCTTAGGGCCATTCCAAGATNNNAGATCGGAAG |
| 46419_1 | TATGAGTGTGGAGTCGTTGCCAAGTAGCTGGGACTACAGGCNNNAGATCGGAAGA |
| 46419_2 | TATGAGTGTGGAGTCGTTGCCAAGTAGCTGGGACTACAGGCNNNAGATCGGAAGA |
| 46419_3 | TATGAGTGTGGAGTCGTTGCCAAGTAGCTGGGACTACAGGCNNNAGATCGGAAGA |
| 35059_1 | TATGAGTGTGGAGTCGTTGCAATGGCGAGATCACACCTCATTNNNAGATCGGAAGA |
| 35059_2 | TATGAGTGTGGAGTCGTTGCAATGGCGAGATCACACCTCATTNNNAGATCGGAAGA |
| 35059_3 | TATGAGTGTGGAGTCGTTGCAATGGCGAGATCACACCTCATTNNNAGATCGGAAGA |
| 52998_1 | TATGAGTGTGGAGTCGTTGCTGGTGTCTCTTCTACTTCCCGTANNNAGATCGGAAGA |
| 52998_2 | TATGAGTGTGGAGTCGTTGCTGGTGTCTCTTCTACTTCCCGTANNNAGATCGGAAGA |
| 52998_3 | TATGAGTGTGGAGTCGTTGCTGGTGTCTCTTCTACTTCCCGTANNNAGATCGGAAGA |
| 8350_1 | TATGAGTGTGGAGTCGTTGCCCCCATAAAAGGAGTTAGGGCANNNAGATCGGAAG |
| 8350_2 | TATGAGTGTGGAGTCGTTGCCCCCATAAAAGGAGTTAGGGCANNNAGATCGGAAG |
| 8350_3 | TATGAGTGTGGAGTCGTTGCCCCCATAAAAGGAGTTAGGGCANNNAGATCGGAAG |
| 29048_1 | TATGAGTGTGGAGTCGTTGCGCCTCAAACACCTTGGAAAATCANNNAGATCGGAAG |
| 29048_2 | TATGAGTGTGGAGTCGTTGCGCCTCAAACACCTTGGAAAATCANNNAGATCGGAAG |
| 29048_3 | TATGAGTGTGGAGTCGTTGCGCCTCAAACACCTTGGAAAATCANNNAGATCGGAAG |
| 43409_1 | TATGAGTGTGGAGTCGTTGCAGATGAAGTCTTTCTCTGTCGCCNNNAGATCGGAAG |
| 43409_2 | TATGAGTGTGGAGTCGTTGCAGATGAAGTCTTTCTCTGTCGCCNNNAGATCGGAAG |
| 43409_3 | TATGAGTGTGGAGTCGTTGCAGATGAAGTCTTTCTCTGTCGCCNNNAGATCGGAAG |
| 45578_1 | TATGAGTGTGGAGTCGTTGCTAGCTGGGATTATAGGTGTGTGCNNNAGATCGGAA |
| 45578_2 | TATGAGTGTGGAGTCGTTGCTAGCTGGGATTATAGGTGTGTGCNNNAGATCGGAA |
| 45578_3 | TATGAGTGTGGAGTCGTTGCTAGCTGGGATTATAGGTGTGTGCNNNAGATCGGAA |
| 38589_1 | TATGAGTGTGGAGTCGTTGCTCTGTAATTCCAGCTGAGGAAGGNNNAGATCGGAA |
| 38589_2 | TATGAGTGTGGAGTCGTTGCTCTGTAATTCCAGCTGAGGAAGGNNNAGATCGGAA |

| | |
|---|---|
| 38589_3 | TATGAGTGTGGAGTCGTTGCTCTGTAATTCCAGCTGAGGAAGGNNNAGATCGGAA |
| 7956_1 | TATGAGTGTGGAGTCGTTGCCTTTCTGTCCCTACTTCCTGGCTNNNAGATCGGAAGA |
| 7956_2 | TATGAGTGTGGAGTCGTTGCCTTTCTGTCCCTACTTCCTGGCTNNNAGATCGGAAGA |
| 7956_3 | TATGAGTGTGGAGTCGTTGCCTTTCTGTCCCTACTTCCTGGCTNNNAGATCGGAAGA |
| 32999_1 | TATGAGTGTGGAGTCGTTGCCCACAAAAGACAGCTTTACAGGGNNNAGATCGGAA |
| 32999_2 | TATGAGTGTGGAGTCGTTGCCCACAAAAGACAGCTTTACAGGGNNNAGATCGGAA |
| 32999_3 | TATGAGTGTGGAGTCGTTGCCCACAAAAGACAGCTTTACAGGGNNNAGATCGGAA |
| 42578_1 | TATGAGTGTGGAGTCGTTGCGATCACGAGGTCAGGAGTTCAAGNNNAGATCGGAA |
| 42578_2 | TATGAGTGTGGAGTCGTTGCGATCACGAGGTCAGGAGTTCAAGNNNAGATCGGAA |
| 42578_3 | TATGAGTGTGGAGTCGTTGCGATCACGAGGTCAGGAGTTCAAGNNNAGATCGGAA |
| 41578_1 | TATGAGTGTGGAGTCGTTGCCTCCTGACCTCAAGTGATACACCNNNAGATCGGAAG |
| 41578_2 | TATGAGTGTGGAGTCGTTGCCTCCTGACCTCAAGTGATACACCNNNAGATCGGAAG |
| 41578_3 | TATGAGTGTGGAGTCGTTGCCTCCTGACCTCAAGTGATACACCNNNAGATCGGAAG |
| 38649_1 | TATGAGTGTGGAGTCGTTGCGCTGATCACCTGAAGTTAGGAGTNNNAGATCGGAA |
| 38649_2 | TATGAGTGTGGAGTCGTTGCGCTGATCACCTGAAGTTAGGAGTNNNAGATCGGAA |
| 38649_3 | TATGAGTGTGGAGTCGTTGCGCTGATCACCTGAAGTTAGGAGTNNNAGATCGGAA |
| 35107_1 | TATGAGTGTGGAGTCGTTGCCCACCTAAGCTCAAGTGATCCTTNNNAGATCGGAAG |
| 35107_2 | TATGAGTGTGGAGTCGTTGCCCACCTAAGCTCAAGTGATCCTTNNNAGATCGGAAG |
| 35107_3 | TATGAGTGTGGAGTCGTTGCCCACCTAAGCTCAAGTGATCCTTNNNAGATCGGAAG |
| 51977_1 | TATGAGTGTGGAGTCGTTGCACAAAAATTAGCCGGATGTGGTGNNNAGATCGGAA |
| 51977_2 | TATGAGTGTGGAGTCGTTGCACAAAAATTAGCCGGATGTGGTGNNNAGATCGGAA |
| 51977_3 | TATGAGTGTGGAGTCGTTGCACAAAAATTAGCCGGATGTGGTGNNNAGATCGGAA |
| 12853_1 | TATGAGTGTGGAGTCGTTGCATAGGATGCAGATATTGTGATAANNNAGATCGGAA |
| 12853_2 | TATGAGTGTGGAGTCGTTGCATAGGATGCAGATATTGTGATAANNNAGATCGGAA |
| 12853_3 | TATGAGTGTGGAGTCGTTGCATAGGATGCAGATATTGTGATAANNNAGATCGGAA |
| 41579_1 | TATGAGTGTGGAGTCGTTGCGAGTTTTGCTCTTGTTGCCCAGNNNAGATCGGAAGA |
| 41579_2 | TATGAGTGTGGAGTCGTTGCGAGTTTTGCTCTTGTTGCCCAGNNNAGATCGGAAGA |
| 41579_3 | TATGAGTGTGGAGTCGTTGCGAGTTTTGCTCTTGTTGCCCAGNNNAGATCGGAAGA |
| 39677_1 | TATGAGTGTGGAGTCGTTGCGAAGGAAGGAAGGAAGGAAGGAGNNNAGATCGG |
| 39677_2 | TATGAGTGTGGAGTCGTTGCGAAGGAAGGAAGGAAGGAAGGAGNNNAGATCGG |
| 39677_3 | TATGAGTGTGGAGTCGTTGCGAAGGAAGGAAGGAAGGAAGGAGNNNAGATCGG |
| 48713_1 | TATGAGTGTGGAGTCGTTGCACACCGTGACTTGTCTTACACTTNNNAGATCGGAAG |
| 48713_2 | TATGAGTGTGGAGTCGTTGCACACCGTGACTTGTCTTACACTTNNNAGATCGGAAG |
| 48713_3 | TATGAGTGTGGAGTCGTTGCACACCGTGACTTGTCTTACACTTNNNAGATCGGAAG |
| 53162_1 | TATGAGTGTGGAGTCGTTGCAGATGGCAGTAAAGAGCTGGAAANNNAGATCGGA |
| 53162_2 | TATGAGTGTGGAGTCGTTGCAGATGGCAGTAAAGAGCTGGAAANNNAGATCGGA |
| 53162_3 | TATGAGTGTGGAGTCGTTGCAGATGGCAGTAAAGAGCTGGAAANNNAGATCGGA |
| 36240_1 | TATGAGTGTGGAGTCGTTGCTCCCCAGTTTCTTCAGAACAACANNNAGATCGGAAG |
| 36240_2 | TATGAGTGTGGAGTCGTTGCTCCCCAGTTTCTTCAGAACAACANNNAGATCGGAAG |
| 36240_3 | TATGAGTGTGGAGTCGTTGCTCCCCAGTTTCTTCAGAACAACANNNAGATCGGAAG |
| 28816_1 | TATGAGTGTGGAGTCGTTGCATCCACTCATTGAATGACGGGAANNNAGATCGGAAG |
| 28816_2 | TATGAGTGTGGAGTCGTTGCATCCACTCATTGAATGACGGGAANNNAGATCGGAAG |
| 28816_3 | TATGAGTGTGGAGTCGTTGCATCCACTCATTGAATGACGGGAANNNAGATCGGAAG |

| | |
|---|---|
| 36026_1 | TATGAGTGTGGAGTCGTTGCCAGTGAGCCAAGATGACCAATGNNNAGATCGGAAG |
| 36026_2 | TATGAGTGTGGAGTCGTTGCCAGTGAGCCAAGATGACCAATGNNNAGATCGGAAG |
| 36026_3 | TATGAGTGTGGAGTCGTTGCCAGTGAGCCAAGATGACCAATGNNNAGATCGGAAG |
| 31593_1 | TATGAGTGTGGAGTCGTTGCATTGTAATCTCAGCAGTCCCCACNNNAGATCGGAAG |
| 31593_2 | TATGAGTGTGGAGTCGTTGCATTGTAATCTCAGCAGTCCCCACNNNAGATCGGAAG |
| 31593_3 | TATGAGTGTGGAGTCGTTGCATTGTAATCTCAGCAGTCCCCACNNNAGATCGGAAG |
| 49243_1 | TATGAGTGTGGAGTCGTTGCGCAGCATGCTGAGTTGTTAGAAANNNAGATCGGAA |
| 49243_2 | TATGAGTGTGGAGTCGTTGCGCAGCATGCTGAGTTGTTAGAAANNNAGATCGGAA |
| 49243_3 | TATGAGTGTGGAGTCGTTGCGCAGCATGCTGAGTTGTTAGAAANNNAGATCGGAA |
| 49382_1 | TATGAGTGTGGAGTCGTTGCCAAGTAGCTGGGATTACAGACGTNNNAGATCGGAA |
| 49382_2 | TATGAGTGTGGAGTCGTTGCCAAGTAGCTGGGATTACAGACGTNNNAGATCGGAA |
| 49382_3 | TATGAGTGTGGAGTCGTTGCCAAGTAGCTGGGATTACAGACGTNNNAGATCGGAA |
| 10653_1 | TATGAGTGTGGAGTCGTTGCTACATTCATAGTGAGGACCAGCGNNNAGATCGGAAG |
| 10653_2 | TATGAGTGTGGAGTCGTTGCTACATTCATAGTGAGGACCAGCGNNNAGATCGGAAG |
| 10653_3 | TATGAGTGTGGAGTCGTTGCTACATTCATAGTGAGGACCAGCGNNNAGATCGGAAG |
| 45386_1 | TATGAGTGTGGAGTCGTTGCAAGGTTCCAGTCACCTAAGAAGGNNNAGATCGGAA |
| 45386_2 | TATGAGTGTGGAGTCGTTGCAAGGTTCCAGTCACCTAAGAAGGNNNAGATCGGAA |
| 45386_3 | TATGAGTGTGGAGTCGTTGCAAGGTTCCAGTCACCTAAGAAGGNNNAGATCGGAA |
| 29301_1 | TATGAGTGTGGAGTCGTTGCGTCTGCTGTTCCTCAATTCTCCTNNNAGATCGGAAGA |
| 29301_2 | TATGAGTGTGGAGTCGTTGCGTCTGCTGTTCCTCAATTCTCCTNNNAGATCGGAAGA |
| 29301_3 | TATGAGTGTGGAGTCGTTGCGTCTGCTGTTCCTCAATTCTCCTNNNAGATCGGAAGA |
| 46033_1 | TATGAGTGTGGAGTCGTTGCTGGCCAGAAATAATGTTTACCAGCNNNAGATCGGAA |
| 46033_2 | TATGAGTGTGGAGTCGTTGCTGGCCAGAAATAATGTTTACCAGCNNNAGATCGGAA |
| 46033_3 | TATGAGTGTGGAGTCGTTGCTGGCCAGAAATAATGTTTACCAGCNNNAGATCGGAA |
| 46803_1 | TATGAGTGTGGAGTCGTTGCCATTCTCCTGCCTCAGCCTCNNNAGATCGGAAGAGCA |
| 46803_2 | TATGAGTGTGGAGTCGTTGCCATTCTCCTGCCTCAGCCTCNNNAGATCGGAAGAGCA |
| 46803_3 | TATGAGTGTGGAGTCGTTGCCATTCTCCTGCCTCAGCCTCNNNAGATCGGAAGAGCA |
| 47598_1 | TATGAGTGTGGAGTCGTTGCGTTGTGGTCTAGGTGGAGTTCATNNNAGATCGGAA |
| 47598_2 | TATGAGTGTGGAGTCGTTGCGTTGTGGTCTAGGTGGAGTTCATNNNAGATCGGAA |
| 47598_3 | TATGAGTGTGGAGTCGTTGCGTTGTGGTCTAGGTGGAGTTCATNNNAGATCGGAA |
| 37243_1 | TATGAGTGTGGAGTCGTTGCAATGGCAACTGTAATCCCAGCTANNNAGATCGGAAG |
| 37243_2 | TATGAGTGTGGAGTCGTTGCAATGGCAACTGTAATCCCAGCTANNNAGATCGGAAG |
| 37243_3 | TATGAGTGTGGAGTCGTTGCAATGGCAACTGTAATCCCAGCTANNNAGATCGGAAG |
| 9879_1 | TATGAGTGTGGAGTCGTTGCTTTCATGTGTTTCCTGCAAGGGANNNAGATCGGAAG |
| 9879_2 | TATGAGTGTGGAGTCGTTGCTTTCATGTGTTTCCTGCAAGGGANNNAGATCGGAAG |
| 9879_3 | TATGAGTGTGGAGTCGTTGCTTTCATGTGTTTCCTGCAAGGGANNNAGATCGGAAG |
| 43441_1 | TATGAGTGTGGAGTCGTTGCCTCCTGAGTAGCTGGGATTACAGNNNAGATCGGAA |
| 43441_2 | TATGAGTGTGGAGTCGTTGCCTCCTGAGTAGCTGGGATTACAGNNNAGATCGGAA |
| 43441_3 | TATGAGTGTGGAGTCGTTGCCTCCTGAGTAGCTGGGATTACAGNNNAGATCGGAA |
| 35426_1 | TATGAGTGTGGAGTCGTTGCACCTCCCAGATTCAAGCAATTCTNNNAGATCGGAAG |
| 35426_2 | TATGAGTGTGGAGTCGTTGCACCTCCCAGATTCAAGCAATTCTNNNAGATCGGAAG |
| 35426_3 | TATGAGTGTGGAGTCGTTGCACCTCCCAGATTCAAGCAATTCTNNNAGATCGGAAG |
| 30428_1 | TATGAGTGTGGAGTCGTTGCCTGAGACCTGACGTTTGGAAGAGNNNAGATCGGAA |

| | |
|---|---|
| 30428_2 | TATGAGTGTGGAGTCGTTGCCTGAGACCTGACGTTTGGAAGAGNNNAGATCGGAA |
| 30428_3 | TATGAGTGTGGAGTCGTTGCCTGAGACCTGACGTTTGGAAGAGNNNAGATCGGAA |
| 36754_1 | TATGAGTGTGGAGTCGTTGCGCTCAGCAAGTAACCCCAAAAATNNNAGATCGGAAG |
| 36754_2 | TATGAGTGTGGAGTCGTTGCGCTCAGCAAGTAACCCCAAAAATNNNAGATCGGAAG |
| 36754_3 | TATGAGTGTGGAGTCGTTGCGCTCAGCAAGTAACCCCAAAAATNNNAGATCGGAAG |
| 52007_1 | TATGAGTGTGGAGTCGTTGCGCTGCTGGGTGGAGACACTCNNNAGATCGGAAGAG |
| 52007_2 | TATGAGTGTGGAGTCGTTGCGCTGCTGGGTGGAGACACTCNNNAGATCGGAAGAG |
| 52007_3 | TATGAGTGTGGAGTCGTTGCGCTGCTGGGTGGAGACACTCNNNAGATCGGAAGAG |
| 43811_1 | TATGAGTGTGGAGTCGTTGCGTGAGAGAGACAGCTTTCTTCCANNNAGATCGGAA |
| 43811_2 | TATGAGTGTGGAGTCGTTGCGTGAGAGAGACAGCTTTCTTCCANNNAGATCGGAA |
| 43811_3 | TATGAGTGTGGAGTCGTTGCGTGAGAGAGACAGCTTTCTTCCANNNAGATCGGAA |
| 40715_1 | TATGAGTGTGGAGTCGTTGCTGGGTGTTTGAGGTGAGAAAATGNNNAGATCGGA |
| 40715_2 | TATGAGTGTGGAGTCGTTGCTGGGTGTTTGAGGTGAGAAAATGNNNAGATCGGA |
| 40715_3 | TATGAGTGTGGAGTCGTTGCTGGGTGTTTGAGGTGAGAAAATGNNNAGATCGGA |
| 52942_1 | TATGAGTGTGGAGTCGTTGCGATGCGTCCACTGATGAGTTCANNNAGATCGGAAG |
| 52942_2 | TATGAGTGTGGAGTCGTTGCGATGCGTCCACTGATGAGTTCANNNAGATCGGAAG |
| 52942_3 | TATGAGTGTGGAGTCGTTGCGATGCGTCCACTGATGAGTTCANNNAGATCGGAAG |
| 2990_1 | TATGAGTGTGGAGTCGTTGCCCAGAACAGACCACTATCCCAANNNAGATCGGAAGA |
| 2990_2 | TATGAGTGTGGAGTCGTTGCCCAGAACAGACCACTATCCCAANNNAGATCGGAAGA |
| 2990_3 | TATGAGTGTGGAGTCGTTGCCCAGAACAGACCACTATCCCAANNNAGATCGGAAGA |
| 32895_1 | TATGAGTGTGGAGTCGTTGCACTCCTATTCTGCTAGCACGTTTNNNAGATCGGAAG |
| 32895_2 | TATGAGTGTGGAGTCGTTGCACTCCTATTCTGCTAGCACGTTTNNNAGATCGGAAG |
| 32895_3 | TATGAGTGTGGAGTCGTTGCACTCCTATTCTGCTAGCACGTTTNNNAGATCGGAAG |
| 4811_1 | TATGAGTGTGGAGTCGTTGCGAGCCACAGCCTATTTCCATCTCNNNAGATCGGAAG |
| 4811_2 | TATGAGTGTGGAGTCGTTGCGAGCCACAGCCTATTTCCATCTCNNNAGATCGGAAG |
| 4811_3 | TATGAGTGTGGAGTCGTTGCGAGCCACAGCCTATTTCCATCTCNNNAGATCGGAAG |
| 40310_1 | TATGAGTGTGGAGTCGTTGCAAGAAAGTCTCCAGGAAGTGTGGNNNAGATCGGAA |
| 40310_2 | TATGAGTGTGGAGTCGTTGCAAGAAAGTCTCCAGGAAGTGTGGNNNAGATCGGAA |
| 40310_3 | TATGAGTGTGGAGTCGTTGCAAGAAAGTCTCCAGGAAGTGTGGNNNAGATCGGAA |
| 46157_1 | TATGAGTGTGGAGTCGTTGCGCCACCACATCCAGCTAATTTTTNNNAGATCGGAAG |
| 46157_2 | TATGAGTGTGGAGTCGTTGCGCCACCACATCCAGCTAATTTTTNNNAGATCGGAAG |
| 46157_3 | TATGAGTGTGGAGTCGTTGCGCCACCACATCCAGCTAATTTTTNNNAGATCGGAAG |
| 29888_1 | TATGAGTGTGGAGTCGTTGCTGTCATGTTGCAGAGAAAACGACNNNAGATCGGAA |
| 29888_2 | TATGAGTGTGGAGTCGTTGCTGTCATGTTGCAGAGAAAACGACNNNAGATCGGAA |
| 29888_3 | TATGAGTGTGGAGTCGTTGCTGTCATGTTGCAGAGAAAACGACNNNAGATCGGAA |
| 3146_1 | TATGAGTGTGGAGTCGTTGCTCACCATTTGTCTTTGCAGGGAANNNAGATCGGAAG |
| 3146_2 | TATGAGTGTGGAGTCGTTGCTCACCATTTGTCTTTGCAGGGAANNNAGATCGGAAG |
| 3146_3 | TATGAGTGTGGAGTCGTTGCTCACCATTTGTCTTTGCAGGGAANNNAGATCGGAAG |
| 50587_1 | TATGAGTGTGGAGTCGTTGCAAGGTAAGTGAGCTGGGTAATGGNNNAGATCGGA |
| 50587_2 | TATGAGTGTGGAGTCGTTGCAAGGTAAGTGAGCTGGGTAATGGNNNAGATCGGA |
| 50587_3 | TATGAGTGTGGAGTCGTTGCAAGGTAAGTGAGCTGGGTAATGGNNNAGATCGGA |
| 36235_1 | TATGAGTGTGGAGTCGTTGCACCTATGATGGGTCTGGATGTACNNNAGATCGGAA |
| 36235_2 | TATGAGTGTGGAGTCGTTGCACCTATGATGGGTCTGGATGTACNNNAGATCGGAA |

| | |
|---|---|
| 36235_3 | TATGAGTGTGGAGTCGTTGCACCTATGATGGGTCTGGATGTACNNNAGATCGGAA |
| 52702_1 | TATGAGTGTGGAGTCGTTGCCTTGAGTAAGGAATGCCAGGACTNNNAGATCGGAA |
| 52702_2 | TATGAGTGTGGAGTCGTTGCCTTGAGTAAGGAATGCCAGGACTNNNAGATCGGAA |
| 52702_3 | TATGAGTGTGGAGTCGTTGCCTTGAGTAAGGAATGCCAGGACTNNNAGATCGGAA |
| 31427_1 | TATGAGTGTGGAGTCGTTGCAACCATTTCATCTGCTTTGAGGCNNNAGATCGGAAG |
| 31427_2 | TATGAGTGTGGAGTCGTTGCAACCATTTCATCTGCTTTGAGGCNNNAGATCGGAAG |
| 31427_3 | TATGAGTGTGGAGTCGTTGCAACCATTTCATCTGCTTTGAGGCNNNAGATCGGAAG |
| 2993_1 | TATGAGTGTGGAGTCGTTGCAGCAGTGGCCAGAGGAGATGAANNNAGATCGGAA |
| 2993_2 | TATGAGTGTGGAGTCGTTGCAGCAGTGGCCAGAGGAGATGAANNNAGATCGGAA |
| 2993_3 | TATGAGTGTGGAGTCGTTGCAGCAGTGGCCAGAGGAGATGAANNNAGATCGGAA |
| 51260_1 | TATGAGTGTGGAGTCGTTGCCTAAAGAGAGGAAGAAGGGGCTGNNNAGATCGGA |
| 51260_2 | TATGAGTGTGGAGTCGTTGCCTAAAGAGAGGAAGAAGGGGCTGNNNAGATCGGA |
| 51260_3 | TATGAGTGTGGAGTCGTTGCCTAAAGAGAGGAAGAAGGGGCTGNNNAGATCGGA |
| 52978_1 | TATGAGTGTGGAGTCGTTGCAGTAGCTGGGACTATAGGTGGTTNNNAGATCGGAA |
| 52978_2 | TATGAGTGTGGAGTCGTTGCAGTAGCTGGGACTATAGGTGGTTNNNAGATCGGAA |
| 52978_3 | TATGAGTGTGGAGTCGTTGCAGTAGCTGGGACTATAGGTGGTTNNNAGATCGGAA |
| 47068_1 | TATGAGTGTGGAGTCGTTGCTTTCACTATGTTGCCCAGGCTAGNNNAGATCGGAAG |
| 47068_2 | TATGAGTGTGGAGTCGTTGCTTTCACTATGTTGCCCAGGCTAGNNNAGATCGGAAG |
| 47068_3 | TATGAGTGTGGAGTCGTTGCTTTCACTATGTTGCCCAGGCTAGNNNAGATCGGAAG |
| 37975_1 | TATGAGTGTGGAGTCGTTGCAGGAGCTAGGAATGGGAGAAAAANNNAGATCGGA |
| 37975_2 | TATGAGTGTGGAGTCGTTGCAGGAGCTAGGAATGGGAGAAAAANNNAGATCGGA |
| 37975_3 | TATGAGTGTGGAGTCGTTGCAGGAGCTAGGAATGGGAGAAAAANNNAGATCGGA |
| 52281_1 | TATGAGTGTGGAGTCGTTGCCTGATCCAGGAGGCCGCATCNNNAGATCGGAAGAG |
| 52281_2 | TATGAGTGTGGAGTCGTTGCCTGATCCAGGAGGCCGCATCNNNAGATCGGAAGAG |
| 52281_3 | TATGAGTGTGGAGTCGTTGCCTGATCCAGGAGGCCGCATCNNNAGATCGGAAGAG |
| 29985_1 | TATGAGTGTGGAGTCGTTGCCACTCTGAGCATTGACCTGAAACNNNAGATCGGAAG |
| 29985_2 | TATGAGTGTGGAGTCGTTGCCACTCTGAGCATTGACCTGAAACNNNAGATCGGAAG |
| 29985_3 | TATGAGTGTGGAGTCGTTGCCACTCTGAGCATTGACCTGAAACNNNAGATCGGAAG |
| 45290_1 | TATGAGTGTGGAGTCGTTGCCTCACTTGATCAACTGCGAGAACNNNAGATCGGAAG |
| 45290_2 | TATGAGTGTGGAGTCGTTGCCTCACTTGATCAACTGCGAGAACNNNAGATCGGAAG |
| 45290_3 | TATGAGTGTGGAGTCGTTGCCTCACTTGATCAACTGCGAGAACNNNAGATCGGAAG |
| 35273_1 | TATGAGTGTGGAGTCGTTGCCCACACTGGATTGCATTGGTATCNNNAGATCGGAAG |
| 35273_2 | TATGAGTGTGGAGTCGTTGCCCACACTGGATTGCATTGGTATCNNNAGATCGGAAG |
| 35273_3 | TATGAGTGTGGAGTCGTTGCCCACACTGGATTGCATTGGTATCNNNAGATCGGAAG |
| 31704_1 | TATGAGTGTGGAGTCGTTGCATGACAATACCTGCATCTCTGCANNNAGATCGGAAG |
| 31704_2 | TATGAGTGTGGAGTCGTTGCATGACAATACCTGCATCTCTGCANNNAGATCGGAAG |
| 31704_3 | TATGAGTGTGGAGTCGTTGCATGACAATACCTGCATCTCTGCANNNAGATCGGAAG |
| 34395_1 | TATGAGTGTGGAGTCGTTGCGAGTTCACACAATCCCCCATTCNNNAGATCGGAAGA |
| 34395_2 | TATGAGTGTGGAGTCGTTGCGAGTTCACACAATCCCCCATTCNNNAGATCGGAAGA |
| 34395_3 | TATGAGTGTGGAGTCGTTGCGAGTTCACACAATCCCCCATTCNNNAGATCGGAAGA |
| 43098_1 | TATGAGTGTGGAGTCGTTGCGCCTCCTTAGTAGCTGGGATTACNNNAGATCGGAAG |
| 43098_2 | TATGAGTGTGGAGTCGTTGCGCCTCCTTAGTAGCTGGGATTACNNNAGATCGGAAG |
| 43098_3 | TATGAGTGTGGAGTCGTTGCGCCTCCTTAGTAGCTGGGATTACNNNAGATCGGAAG |

| | |
|---|---|
| 51784_1 | TATGAGTGTGGAGTCGTTGCACAGCTAGAAGCACCTTCCTAACNNNAGATCGGAAG |
| 51784_2 | TATGAGTGTGGAGTCGTTGCACAGCTAGAAGCACCTTCCTAACNNNAGATCGGAAG |
| 51784_3 | TATGAGTGTGGAGTCGTTGCACAGCTAGAAGCACCTTCCTAACNNNAGATCGGAAG |
| 45960_1 | TATGAGTGTGGAGTCGTTGCCACCACGCCTGGCTAATTTTTTNNNAGATCGGAAGA |
| 45960_2 | TATGAGTGTGGAGTCGTTGCCACCACGCCTGGCTAATTTTTTNNNAGATCGGAAGA |
| 45960_3 | TATGAGTGTGGAGTCGTTGCCACCACGCCTGGCTAATTTTTTNNNAGATCGGAAGA |
| 43220_1 | TATGAGTGTGGAGTCGTTGCCAATCTCAGCTCATTGCAACCTCNNNAGATCGGAAGA |
| 43220_2 | TATGAGTGTGGAGTCGTTGCCAATCTCAGCTCATTGCAACCTCNNNAGATCGGAAGA |
| 43220_3 | TATGAGTGTGGAGTCGTTGCCAATCTCAGCTCATTGCAACCTCNNNAGATCGGAAGA |
| 51185_1 | TATGAGTGTGGAGTCGTTGCAACCACAGGCAGTTGTTTCTTTCNNNAGATCGGAAG |
| 51185_2 | TATGAGTGTGGAGTCGTTGCAACCACAGGCAGTTGTTTCTTTCNNNAGATCGGAAG |
| 51185_3 | TATGAGTGTGGAGTCGTTGCAACCACAGGCAGTTGTTTCTTTCNNNAGATCGGAAG |
| 32227_1 | TATGAGTGTGGAGTCGTTGCTGGAGGATACATTTGCTGTTGGANNNAGATCGGAA |
| 32227_2 | TATGAGTGTGGAGTCGTTGCTGGAGGATACATTTGCTGTTGGANNNAGATCGGAA |
| 32227_3 | TATGAGTGTGGAGTCGTTGCTGGAGGATACATTTGCTGTTGGANNNAGATCGGAA |
| 48352_1 | TATGAGTGTGGAGTCGTTGCAACCTCTGACTCCCAACTTCAAGNNNAGATCGGAAG |
| 48352_2 | TATGAGTGTGGAGTCGTTGCAACCTCTGACTCCCAACTTCAAGNNNAGATCGGAAG |
| 48352_3 | TATGAGTGTGGAGTCGTTGCAACCTCTGACTCCCAACTTCAAGNNNAGATCGGAAG |
| 9325_1 | TATGAGTGTGGAGTCGTTGCTGGAGACTGAGAGAGAGAGATGGNNNAGATCGGA |
| 9325_2 | TATGAGTGTGGAGTCGTTGCTGGAGACTGAGAGAGAGAGATGGNNNAGATCGGA |
| 9325_3 | TATGAGTGTGGAGTCGTTGCTGGAGACTGAGAGAGAGAGATGGNNNAGATCGGA |
| 38846_1 | TATGAGTGTGGAGTCGTTGCATGATGTTAGCTCAGGTGGGAAGNNNAGATCGGAA |
| 38846_2 | TATGAGTGTGGAGTCGTTGCATGATGTTAGCTCAGGTGGGAAGNNNAGATCGGAA |
| 38846_3 | TATGAGTGTGGAGTCGTTGCATGATGTTAGCTCAGGTGGGAAGNNNAGATCGGAA |
| 47079_1 | TATGAGTGTGGAGTCGTTGCCAGGCTGGAGTGTAATGGTATGANNNAGATCGGAA |
| 47079_2 | TATGAGTGTGGAGTCGTTGCCAGGCTGGAGTGTAATGGTATGANNNAGATCGGAA |
| 47079_3 | TATGAGTGTGGAGTCGTTGCCAGGCTGGAGTGTAATGGTATGANNNAGATCGGAA |
| 50268_1 | TATGAGTGTGGAGTCGTTGCCCTTTGGTAAGCCTCCTTTGTCTNNNAGATCGGAAGA |
| 50268_2 | TATGAGTGTGGAGTCGTTGCCCTTTGGTAAGCCTCCTTTGTCTNNNAGATCGGAAGA |
| 50268_3 | TATGAGTGTGGAGTCGTTGCCCTTTGGTAAGCCTCCTTTGTCTNNNAGATCGGAAGA |
| 53007_1 | TATGAGTGTGGAGTCGTTGCGTGTGTGGACATGTGTGTATGTGNNNAGATCGGAA |
| 53007_2 | TATGAGTGTGGAGTCGTTGCGTGTGTGGACATGTGTGTATGTGNNNAGATCGGAA |
| 53007_3 | TATGAGTGTGGAGTCGTTGCGTGTGTGGACATGTGTGTATGTGNNNAGATCGGAA |
| 37038_1 | TATGAGTGTGGAGTCGTTGCAGTTGTTTTGTTAGAAGAGGGAGANNNAGATCGGA |
| 37038_2 | TATGAGTGTGGAGTCGTTGCAGTTGTTTTGTTAGAAGAGGGAGANNNAGATCGGA |
| 37038_3 | TATGAGTGTGGAGTCGTTGCAGTTGTTTTGTTAGAAGAGGGAGANNNAGATCGGA |
| 37089_1 | TATGAGTGTGGAGTCGTTGCACCAAAATCTATGGGCTACAGCANNNAGATCGGAAG |
| 37089_2 | TATGAGTGTGGAGTCGTTGCACCAAAATCTATGGGCTACAGCANNNAGATCGGAAG |
| 37089_3 | TATGAGTGTGGAGTCGTTGCACCAAAATCTATGGGCTACAGCANNNAGATCGGAAG |
| 35008_1 | TATGAGTGTGGAGTCGTTGCAATTCTTGTGCCTCCCGAGTAGNNNAGATCGGAAGA |
| 35008_2 | TATGAGTGTGGAGTCGTTGCAATTCTTGTGCCTCCCGAGTAGNNNAGATCGGAAGA |
| 35008_3 | TATGAGTGTGGAGTCGTTGCAATTCTTGTGCCTCCCGAGTAGNNNAGATCGGAAGA |
| 51204_1 | TATGAGTGTGGAGTCGTTGCAGTGTTCACTGCTCAGATGACAGNNNAGATCGGAAG |

| | |
|---|---|
| 51204_2 | TATGAGTGTGGAGTCGTTGCAGTGTTCACTGCTCAGATGACAGNNNAGATCGGAAG |
| 51204_3 | TATGAGTGTGGAGTCGTTGCAGTGTTCACTGCTCAGATGACAGNNNAGATCGGAAG |
| 30487_1 | TATGAGTGTGGAGTCGTTGCTTTATTACCGCCAGCCTGACGCNNNAGATCGGAAGA |
| 30487_2 | TATGAGTGTGGAGTCGTTGCTTTATTACCGCCAGCCTGACGCNNNAGATCGGAAGA |
| 30487_3 | TATGAGTGTGGAGTCGTTGCTTTATTACCGCCAGCCTGACGCNNNAGATCGGAAGA |
| 28947_1 | TATGAGTGTGGAGTCGTTGCATTTATGCACACACACACACACTNNNAGATCGGAAGA |
| 28947_2 | TATGAGTGTGGAGTCGTTGCATTTATGCACACACACACACACTNNNAGATCGGAAGA |
| 28947_3 | TATGAGTGTGGAGTCGTTGCATTTATGCACACACACACACACTNNNAGATCGGAAGA |
| 47992_1 | TATGAGTGTGGAGTCGTTGCCTGAGTAGCTGGGATTATAGGCGNNNAGATCGGAA |
| 47992_2 | TATGAGTGTGGAGTCGTTGCCTGAGTAGCTGGGATTATAGGCGNNNAGATCGGAA |
| 47992_3 | TATGAGTGTGGAGTCGTTGCCTGAGTAGCTGGGATTATAGGCGNNNAGATCGGAA |
| 12138_1 | TATGAGTGTGGAGTCGTTGCCTTTTTGCATCTCCTGACCACAGNNNAGATCGGAAGA |
| 12138_2 | TATGAGTGTGGAGTCGTTGCCTTTTTGCATCTCCTGACCACAGNNNAGATCGGAAGA |
| 12138_3 | TATGAGTGTGGAGTCGTTGCCTTTTTGCATCTCCTGACCACAGNNNAGATCGGAAGA |
| 45510_1 | TATGAGTGTGGAGTCGTTGCCATCAAGGATCACTCCCTCCACNNNAGATCGGAAGAG |
| 45510_2 | TATGAGTGTGGAGTCGTTGCCATCAAGGATCACTCCCTCCACNNNAGATCGGAAGAG |
| 45510_3 | TATGAGTGTGGAGTCGTTGCCATCAAGGATCACTCCCTCCACNNNAGATCGGAAGAG |
| 35153_1 | TATGAGTGTGGAGTCGTTGCGACACAGGGTCTCACTCTGTTGNNNAGATCGGAAGA |
| 35153_2 | TATGAGTGTGGAGTCGTTGCGACACAGGGTCTCACTCTGTTGNNNAGATCGGAAGA |
| 35153_3 | TATGAGTGTGGAGTCGTTGCGACACAGGGTCTCACTCTGTTGNNNAGATCGGAAGA |
| 30754_1 | TATGAGTGTGGAGTCGTTGCAGAGAGAGAGGGAGATGTTTGGANNNAGATCGGA |
| 30754_2 | TATGAGTGTGGAGTCGTTGCAGAGAGAGAGGGAGATGTTTGGANNNAGATCGGA |
| 30754_3 | TATGAGTGTGGAGTCGTTGCAGAGAGAGAGGGAGATGTTTGGANNNAGATCGGA |
| 45274_1 | TATGAGTGTGGAGTCGTTGCGAGATTTGCTCTTGTTGCCCAGNNNAGATCGGAAGA |
| 45274_2 | TATGAGTGTGGAGTCGTTGCGAGATTTGCTCTTGTTGCCCAGNNNAGATCGGAAGA |
| 45274_3 | TATGAGTGTGGAGTCGTTGCGAGATTTGCTCTTGTTGCCCAGNNNAGATCGGAAGA |
| 12874_1 | TATGAGTGTGGAGTCGTTGCGTTTTGTTGCTGACACAAATNNNAGATCGGAAGAGC |
| 12874_2 | TATGAGTGTGGAGTCGTTGCGTTTTGTTGCTGACACAAATNNNAGATCGGAAGAGC |
| 12874_3 | TATGAGTGTGGAGTCGTTGCGTTTTGTTGCTGACACAAATNNNAGATCGGAAGAGC |
| 10663_1 | TATGAGTGTGGAGTCGTTGCTGCTATTGGCTTTCAAGCAACAGNNNAGATCGGAAG |
| 10663_2 | TATGAGTGTGGAGTCGTTGCTGCTATTGGCTTTCAAGCAACAGNNNAGATCGGAAG |
| 10663_3 | TATGAGTGTGGAGTCGTTGCTGCTATTGGCTTTCAAGCAACAGNNNAGATCGGAAG |
| 45210_1 | TATGAGTGTGGAGTCGTTGCAAGCTTTTTGATGATGCAAGGCTNNNAGATCGGAAG |
| 45210_2 | TATGAGTGTGGAGTCGTTGCAAGCTTTTTGATGATGCAAGGCTNNNAGATCGGAAG |
| 45210_3 | TATGAGTGTGGAGTCGTTGCAAGCTTTTTGATGATGCAAGGCTNNNAGATCGGAAG |
| 47119_1 | TATGAGTGTGGAGTCGTTGCTTGGCTTCTTCACTTACCTACCCNNNAGATCGGAAGA |
| 47119_2 | TATGAGTGTGGAGTCGTTGCTTGGCTTCTTCACTTACCTACCCNNNAGATCGGAAGA |
| 47119_3 | TATGAGTGTGGAGTCGTTGCTTGGCTTCTTCACTTACCTACCCNNNAGATCGGAAGA |
| 52838_1 | TATGAGTGTGGAGTCGTTGCTCTGCTCTGCTAGTCTTTCCACNNNAGATCGGAAGAG |
| 52838_2 | TATGAGTGTGGAGTCGTTGCTCTGCTCTGCTAGTCTTTCCACNNNAGATCGGAAGAG |
| 52838_3 | TATGAGTGTGGAGTCGTTGCTCTGCTCTGCTAGTCTTTCCACNNNAGATCGGAAGAG |
| 35398_1 | TATGAGTGTGGAGTCGTTGCGATTCAAGCGATTCTCCTGCCTCNNNAGATCGGAAGA |
| 35398_2 | TATGAGTGTGGAGTCGTTGCGATTCAAGCGATTCTCCTGCCTCNNNAGATCGGAAGA |

| | |
|---|---|
| 35398_3 | TATGAGTGTGGAGTCGTTGCGATTCAAGCGATTCTCCTGCCTCNNNAGATCGGAAG |
| 36945_1 | TATGAGTGTGGAGTCGTTGCTTTTCTACTTCTGTGAGCCGGAANNNAGATCGGAAG |
| 36945_2 | TATGAGTGTGGAGTCGTTGCTTTTCTACTTCTGTGAGCCGGAANNNAGATCGGAAG |
| 36945_3 | TATGAGTGTGGAGTCGTTGCTTTTCTACTTCTGTGAGCCGGAANNNAGATCGGAAG |
| 34358_1 | TATGAGTGTGGAGTCGTTGCACCAGCATGGCACATGTATACATNNNAGATCGGAAG |
| 34358_2 | TATGAGTGTGGAGTCGTTGCACCAGCATGGCACATGTATACATNNNAGATCGGAAG |
| 34358_3 | TATGAGTGTGGAGTCGTTGCACCAGCATGGCACATGTATACATNNNAGATCGGAAG |
| 30168_1 | TATGAGTGTGGAGTCGTTGCACCGAGTAAGAGAGAAAGGTCAGNNNAGATCGGA |
| 30168_2 | TATGAGTGTGGAGTCGTTGCACCGAGTAAGAGAGAAAGGTCAGNNNAGATCGGA |
| 30168_3 | TATGAGTGTGGAGTCGTTGCACCGAGTAAGAGAGAAAGGTCAGNNNAGATCGGA |
| 36899_1 | TATGAGTGTGGAGTCGTTGCTGCACAAAAAACCCACAGAATGGNNNAGATCGGAA |
| 36899_2 | TATGAGTGTGGAGTCGTTGCTGCACAAAAAACCCACAGAATGGNNNAGATCGGAA |
| 36899_3 | TATGAGTGTGGAGTCGTTGCTGCACAAAAAACCCACAGAATGGNNNAGATCGGAA |
| 39407_1 | TATGAGTGTGGAGTCGTTGCCACTGGGGGGTTTCATACACTAGNNNAGATCGGAA |
| 39407_2 | TATGAGTGTGGAGTCGTTGCCACTGGGGGGTTTCATACACTAGNNNAGATCGGAA |
| 39407_3 | TATGAGTGTGGAGTCGTTGCCACTGGGGGGTTTCATACACTAGNNNAGATCGGAA |
| 38882_1 | TATGAGTGTGGAGTCGTTGCCTGAGGAAGGAGAATGGTGTGAANNNAGATCGGA |
| 38882_2 | TATGAGTGTGGAGTCGTTGCCTGAGGAAGGAGAATGGTGTGAANNNAGATCGGA |
| 38882_3 | TATGAGTGTGGAGTCGTTGCCTGAGGAAGGAGAATGGTGTGAANNNAGATCGGA |
| 43852_1 | TATGAGTGTGGAGTCGTTGCTCGCTAGGTGACTCAGAGGTAGNNNAGATCGGAAG |
| 43852_2 | TATGAGTGTGGAGTCGTTGCTCGCTAGGTGACTCAGAGGTAGNNNAGATCGGAAG |
| 43852_3 | TATGAGTGTGGAGTCGTTGCTCGCTAGGTGACTCAGAGGTAGNNNAGATCGGAAG |
| 12925_1 | TATGAGTGTGGAGTCGTTGCTTCTCACCTTGTCATTAGAAAANNNAGATCGGAAGA |
| 12925_2 | TATGAGTGTGGAGTCGTTGCTTCTCACCTTGTCATTAGAAAANNNAGATCGGAAGA |
| 12925_3 | TATGAGTGTGGAGTCGTTGCTTCTCACCTTGTCATTAGAAAANNNAGATCGGAAGA |
| 30109_1 | TATGAGTGTGGAGTCGTTGCAGCCTTTGTATGTACATGAAATCCCNNNAGATCGGA |
| 30109_2 | TATGAGTGTGGAGTCGTTGCAGCCTTTGTATGTACATGAAATCCCNNNAGATCGGA |
| 30109_3 | TATGAGTGTGGAGTCGTTGCAGCCTTTGTATGTACATGAAATCCCNNNAGATCGGA |
| 47632_1 | TATGAGTGTGGAGTCGTTGCAAGGTCTCACTTTGTTGCCTAGGNNNAGATCGGAAG |
| 47632_2 | TATGAGTGTGGAGTCGTTGCAAGGTCTCACTTTGTTGCCTAGGNNNAGATCGGAAG |
| 47632_3 | TATGAGTGTGGAGTCGTTGCAAGGTCTCACTTTGTTGCCTAGGNNNAGATCGGAAG |
| 37204_1 | TATGAGTGTGGAGTCGTTGCGTCATATGGTGTTTCCCCTTCCTNNNAGATCGGAAGA |
| 37204_2 | TATGAGTGTGGAGTCGTTGCGTCATATGGTGTTTCCCCTTCCTNNNAGATCGGAAGA |
| 37204_3 | TATGAGTGTGGAGTCGTTGCGTCATATGGTGTTTCCCCTTCCTNNNAGATCGGAAGA |
| 2545_1 | TATGAGTGTGGAGTCGTTGCTCAAGTTTCTGCCAGATTGGAGGNNNAGATCGGAA |
| 2545_2 | TATGAGTGTGGAGTCGTTGCTCAAGTTTCTGCCAGATTGGAGGNNNAGATCGGAA |
| 2545_3 | TATGAGTGTGGAGTCGTTGCTCAAGTTTCTGCCAGATTGGAGGNNNAGATCGGAA |
| 52329_1 | TATGAGTGTGGAGTCGTTGCAGCCATCCTATCTGACTGGGTNNNAGATCGGAAGAG |
| 52329_2 | TATGAGTGTGGAGTCGTTGCAGCCATCCTATCTGACTGGGTNNNAGATCGGAAGAG |
| 52329_3 | TATGAGTGTGGAGTCGTTGCAGCCATCCTATCTGACTGGGTNNNAGATCGGAAGAG |
| 47589_1 | TATGAGTGTGGAGTCGTTGCTCCAGCTTGTCTCAGTTCCAATTNNNAGATCGGAAGA |
| 47589_2 | TATGAGTGTGGAGTCGTTGCTCCAGCTTGTCTCAGTTCCAATTNNNAGATCGGAAGA |
| 47589_3 | TATGAGTGTGGAGTCGTTGCTCCAGCTTGTCTCAGTTCCAATTNNNAGATCGGAAGA |

| | |
|---|---|
| 31864_1 | TATGAGTGTGGAGTCGTTGCCTGGAAGTTATGGTATGGCCCTTNNNAGATCGGAAG |
| 31864_2 | TATGAGTGTGGAGTCGTTGCCTGGAAGTTATGGTATGGCCCTTNNNAGATCGGAAG |
| 31864_3 | TATGAGTGTGGAGTCGTTGCCTGGAAGTTATGGTATGGCCCTTNNNAGATCGGAAG |
| 37896_1 | TATGAGTGTGGAGTCGTTGCGTCCAAAGTTAGCAGAAAGATGGANNNAGATCGGA |
| 37896_2 | TATGAGTGTGGAGTCGTTGCGTCCAAAGTTAGCAGAAAGATGGANNNAGATCGGA |
| 37896_3 | TATGAGTGTGGAGTCGTTGCGTCCAAAGTTAGCAGAAAGATGGANNNAGATCGGA |
| 48016_1 | TATGAGTGTGGAGTCGTTGCCGTCCCTCCTCTCTACCCTCNNNAGATCGGAAGAGCA |
| 48016_2 | TATGAGTGTGGAGTCGTTGCCGTCCCTCCTCTCTACCCTCNNNAGATCGGAAGAGCA |
| 48016_3 | TATGAGTGTGGAGTCGTTGCCGTCCCTCCTCTCTACCCTCNNNAGATCGGAAGAGCA |
| 35652_1 | TATGAGTGTGGAGTCGTTGCGATTTTTCTGCCTCAGCTTCCTGNNNAGATCGGAAGA |
| 35652_2 | TATGAGTGTGGAGTCGTTGCGATTTTTCTGCCTCAGCTTCCTGNNNAGATCGGAAGA |
| 35652_3 | TATGAGTGTGGAGTCGTTGCGATTTTTCTGCCTCAGCTTCCTGNNNAGATCGGAAGA |
| 48718_1 | TATGAGTGTGGAGTCGTTGCCAATCGCCTCTCACTGCAAGNNNAGATCGGAAGAGC |
| 48718_2 | TATGAGTGTGGAGTCGTTGCCAATCGCCTCTCACTGCAAGNNNAGATCGGAAGAGC |
| 48718_3 | TATGAGTGTGGAGTCGTTGCCAATCGCCTCTCACTGCAAGNNNAGATCGGAAGAGC |
| 10057_1 | TATGAGTGTGGAGTCGTTGCCATCCAACCCCTGTCTGTTTTCANNNAGATCGGAAGA |
| 10057_2 | TATGAGTGTGGAGTCGTTGCCATCCAACCCCTGTCTGTTTTCANNNAGATCGGAAGA |
| 10057_3 | TATGAGTGTGGAGTCGTTGCCATCCAACCCCTGTCTGTTTTCANNNAGATCGGAAGA |
| 37333_1 | TATGAGTGTGGAGTCGTTGCCAGATCATGTGGTCAGGAGTTCANNNAGATCGGAA |
| 37333_2 | TATGAGTGTGGAGTCGTTGCCAGATCATGTGGTCAGGAGTTCANNNAGATCGGAA |
| 37333_3 | TATGAGTGTGGAGTCGTTGCCAGATCATGTGGTCAGGAGTTCANNNAGATCGGAA |
| 31032_1 | TATGAGTGTGGAGTCGTTGCTCTCTGGAGAAGGATTCCCATCTNNNAGATCGGAAG |
| 31032_2 | TATGAGTGTGGAGTCGTTGCTCTCTGGAGAAGGATTCCCATCTNNNAGATCGGAAG |
| 31032_3 | TATGAGTGTGGAGTCGTTGCTCTCTGGAGAAGGATTCCCATCTNNNAGATCGGAAG |
| 30875_1 | TATGAGTGTGGAGTCGTTGCCAGACGACGGAGGATCTTATTCANNNAGATCGGAAG |
| 30875_2 | TATGAGTGTGGAGTCGTTGCCAGACGACGGAGGATCTTATTCANNNAGATCGGAAG |
| 30875_3 | TATGAGTGTGGAGTCGTTGCCAGACGACGGAGGATCTTATTCANNNAGATCGGAAG |
| 47823_1 | TATGAGTGTGGAGTCGTTGCCAACATCTGAAGCCATCCAGGANNNAGATCGGAAGA |
| 47823_2 | TATGAGTGTGGAGTCGTTGCCAACATCTGAAGCCATCCAGGANNNAGATCGGAAGA |
| 47823_3 | TATGAGTGTGGAGTCGTTGCCAACATCTGAAGCCATCCAGGANNNAGATCGGAAGA |
| 52252_1 | TATGAGTGTGGAGTCGTTGCGGTGCTTCCACTCTTCATTGAAANNNAGATCGGAAG |
| 52252_2 | TATGAGTGTGGAGTCGTTGCGGTGCTTCCACTCTTCATTGAAANNNAGATCGGAAG |
| 52252_3 | TATGAGTGTGGAGTCGTTGCGGTGCTTCCACTCTTCATTGAAANNNAGATCGGAAG |
| 45772_1 | TATGAGTGTGGAGTCGTTGCGCCAGGTCCTTTATCTTCTCCAANNNAGATCGGAAGA |
| 45772_2 | TATGAGTGTGGAGTCGTTGCGCCAGGTCCTTTATCTTCTCCAANNNAGATCGGAAGA |
| 45772_3 | TATGAGTGTGGAGTCGTTGCGCCAGGTCCTTTATCTTCTCCAANNNAGATCGGAAGA |
| 9444_1 | TATGAGTGTGGAGTCGTTGCACGGATGTGGAACTTAGGTGAGANNNAGATCGGAA |
| 9444_2 | TATGAGTGTGGAGTCGTTGCACGGATGTGGAACTTAGGTGAGANNNAGATCGGAA |
| 9444_3 | TATGAGTGTGGAGTCGTTGCACGGATGTGGAACTTAGGTGAGANNNAGATCGGAA |
| 42784_1 | TATGAGTGTGGAGTCGTTGCCCACCATGAGATACTGCTACACANNNAGATCGGAAG |
| 42784_2 | TATGAGTGTGGAGTCGTTGCCCACCATGAGATACTGCTACACANNNAGATCGGAAG |
| 42784_3 | TATGAGTGTGGAGTCGTTGCCCACCATGAGATACTGCTACACANNNAGATCGGAAG |
| 10728_1 | TATGAGTGTGGAGTCGTTGCGTGCAAACACATGCATTCTGGAANNNAGATCGGAAG |

| | |
|---|---|
| 10728_2 | TATGAGTGTGGAGTCGTTGCGTGCAAACACATGCATTCTGGAANNNAGATCGGAAG |
| 10728_3 | TATGAGTGTGGAGTCGTTGCGTGCAAACACATGCATTCTGGAANNNAGATCGGAAG |
| 38635_1 | TATGAGTGTGGAGTCGTTGCATGCCTGTAATCTCAGCTACTCGNNNAGATCGGAAGA |
| 38635_2 | TATGAGTGTGGAGTCGTTGCATGCCTGTAATCTCAGCTACTCGNNNAGATCGGAAGA |
| 38635_3 | TATGAGTGTGGAGTCGTTGCATGCCTGTAATCTCAGCTACTCGNNNAGATCGGAAGA |
| 38635_4 | TATGAGTGTGGAGTCGTTGCATGCCTGTAATCTCAGCTACTCGNNNAGATCGGAAGA |
| 38635_5 | TATGAGTGTGGAGTCGTTGCATGCCTGTAATCTCAGCTACTCGNNNAGATCGGAAGA |
| 38635_6 | TATGAGTGTGGAGTCGTTGCATGCCTGTAATCTCAGCTACTCGNNNAGATCGGAAGA |
| 50280_1 | TATGAGTGTGGAGTCGTTGCGAAACTCCTCGAGTGTGGGAGNNNAGATCGGAAGA |
| 50280_2 | TATGAGTGTGGAGTCGTTGCGAAACTCCTCGAGTGTGGGAGNNNAGATCGGAAGA |
| 50280_3 | TATGAGTGTGGAGTCGTTGCGAAACTCCTCGAGTGTGGGAGNNNAGATCGGAAGA |
| 50280_4 | TATGAGTGTGGAGTCGTTGCGAAACTCCTCGAGTGTGGGAGNNNAGATCGGAAGA |
| 50280_5 | TATGAGTGTGGAGTCGTTGCGAAACTCCTCGAGTGTGGGAGNNNAGATCGGAAGA |
| 50280_6 | TATGAGTGTGGAGTCGTTGCGAAACTCCTCGAGTGTGGGAGNNNAGATCGGAAGA |
| 31788_1 | TATGAGTGTGGAGTCGTTGCGGAGTGAGCATGTACTCTGATGTNNNAGATCGGAA |
| 31788_2 | TATGAGTGTGGAGTCGTTGCGGAGTGAGCATGTACTCTGATGTNNNAGATCGGAA |
| 31788_3 | TATGAGTGTGGAGTCGTTGCGGAGTGAGCATGTACTCTGATGTNNNAGATCGGAA |
| 31788_4 | TATGAGTGTGGAGTCGTTGCGGAGTGAGCATGTACTCTGATGTNNNAGATCGGAA |
| 31788_5 | TATGAGTGTGGAGTCGTTGCGGAGTGAGCATGTACTCTGATGTNNNAGATCGGAA |
| 31788_6 | TATGAGTGTGGAGTCGTTGCGGAGTGAGCATGTACTCTGATGTNNNAGATCGGAA |
| 42614_1 | TATGAGTGTGGAGTCGTTGCAGAGAGAGAAGAGAGATGGAGACANNNAGATCGG |
| 42614_2 | TATGAGTGTGGAGTCGTTGCAGAGAGAGAAGAGAGATGGAGACANNNAGATCGG |
| 42614_3 | TATGAGTGTGGAGTCGTTGCAGAGAGAGAAGAGAGATGGAGACANNNAGATCGG |
| 42614_4 | TATGAGTGTGGAGTCGTTGCAGAGAGAGAAGAGAGATGGAGACANNNAGATCGG |
| 42614_5 | TATGAGTGTGGAGTCGTTGCAGAGAGAGAAGAGAGATGGAGACANNNAGATCGG |
| 42614_6 | TATGAGTGTGGAGTCGTTGCAGAGAGAGAAGAGAGATGGAGACANNNAGATCGG |
| 40257_1 | TATGAGTGTGGAGTCGTTGCCAGATAACTGGCAAGCCATATGCNNNAGATCGGAAG |
| 40257_2 | TATGAGTGTGGAGTCGTTGCCAGATAACTGGCAAGCCATATGCNNNAGATCGGAAG |
| 40257_3 | TATGAGTGTGGAGTCGTTGCCAGATAACTGGCAAGCCATATGCNNNAGATCGGAAG |
| 40257_4 | TATGAGTGTGGAGTCGTTGCCAGATAACTGGCAAGCCATATGCNNNAGATCGGAAG |
| 40257_5 | TATGAGTGTGGAGTCGTTGCCAGATAACTGGCAAGCCATATGCNNNAGATCGGAAG |
| 40257_6 | TATGAGTGTGGAGTCGTTGCCAGATAACTGGCAAGCCATATGCNNNAGATCGGAAG |
| 51556_1 | TATGAGTGTGGAGTCGTTGCGGTTTTTGGGAGTGGGTTGAAAGNNNAGATCGGAA |
| 51556_2 | TATGAGTGTGGAGTCGTTGCGGTTTTTGGGAGTGGGTTGAAAGNNNAGATCGGAA |
| 51556_3 | TATGAGTGTGGAGTCGTTGCGGTTTTTGGGAGTGGGTTGAAAGNNNAGATCGGAA |
| 51556_4 | TATGAGTGTGGAGTCGTTGCGGTTTTTGGGAGTGGGTTGAAAGNNNAGATCGGAA |
| 51556_5 | TATGAGTGTGGAGTCGTTGCGGTTTTTGGGAGTGGGTTGAAAGNNNAGATCGGAA |
| 51556_6 | TATGAGTGTGGAGTCGTTGCGGTTTTTGGGAGTGGGTTGAAAGNNNAGATCGGAA |
| 2186_1 | TATGAGTGTGGAGTCGTTGCGAAAACATGTGTGTGGTCAGGGANNNAGATCGGAA |
| 2186_2 | TATGAGTGTGGAGTCGTTGCGAAAACATGTGTGTGGTCAGGGANNNAGATCGGAA |
| 2186_3 | TATGAGTGTGGAGTCGTTGCGAAAACATGTGTGTGGTCAGGGANNNAGATCGGAA |
| 2186_4 | TATGAGTGTGGAGTCGTTGCGAAAACATGTGTGTGGTCAGGGANNNAGATCGGAA |
| 2186_5 | TATGAGTGTGGAGTCGTTGCGAAAACATGTGTGTGGTCAGGGANNNAGATCGGAA |

| | |
|---|---|
| 2186_6 | TATGAGTGTGGAGTCGTTGCGAAAACATGTGTGTGGTCAGGGANNNAGATCGGAA |
| 51797_1 | TATGAGTGTGGAGTCGTTGCACCAGTGATCTTGAGCACTTTCTNNNAGATCGGAAGA |
| 51797_2 | TATGAGTGTGGAGTCGTTGCACCAGTGATCTTGAGCACTTTCTNNNAGATCGGAAGA |
| 51797_3 | TATGAGTGTGGAGTCGTTGCACCAGTGATCTTGAGCACTTTCTNNNAGATCGGAAGA |
| 51797_4 | TATGAGTGTGGAGTCGTTGCACCAGTGATCTTGAGCACTTTCTNNNAGATCGGAAGA |
| 51797_5 | TATGAGTGTGGAGTCGTTGCACCAGTGATCTTGAGCACTTTCTNNNAGATCGGAAGA |
| 51797_6 | TATGAGTGTGGAGTCGTTGCACCAGTGATCTTGAGCACTTTCTNNNAGATCGGAAGA |
| 32034_1 | TATGAGTGTGGAGTCGTTGCAGTTCAAGACCACCCTGGATAACNNNAGATCGGAAG |
| 32034_2 | TATGAGTGTGGAGTCGTTGCAGTTCAAGACCACCCTGGATAACNNNAGATCGGAAG |
| 32034_3 | TATGAGTGTGGAGTCGTTGCAGTTCAAGACCACCCTGGATAACNNNAGATCGGAAG |
| 32034_4 | TATGAGTGTGGAGTCGTTGCAGTTCAAGACCACCCTGGATAACNNNAGATCGGAAG |
| 32034_5 | TATGAGTGTGGAGTCGTTGCAGTTCAAGACCACCCTGGATAACNNNAGATCGGAAG |
| 32034_6 | TATGAGTGTGGAGTCGTTGCAGTTCAAGACCACCCTGGATAACNNNAGATCGGAAG |
| 48572_1 | TATGAGTGTGGAGTCGTTGCGGACATACGGTCTCATCTCTGTCNNNAGATCGGAAGA |
| 48572_2 | TATGAGTGTGGAGTCGTTGCGGACATACGGTCTCATCTCTGTCNNNAGATCGGAAGA |
| 48572_3 | TATGAGTGTGGAGTCGTTGCGGACATACGGTCTCATCTCTGTCNNNAGATCGGAAGA |
| 48572_4 | TATGAGTGTGGAGTCGTTGCGGACATACGGTCTCATCTCTGTCNNNAGATCGGAAGA |
| 48572_5 | TATGAGTGTGGAGTCGTTGCGGACATACGGTCTCATCTCTGTCNNNAGATCGGAAGA |
| 48572_6 | TATGAGTGTGGAGTCGTTGCGGACATACGGTCTCATCTCTGTCNNNAGATCGGAAGA |
| 52013_1 | TATGAGTGTGGAGTCGTTGCTTCCTGCATCATGTGAAGAAGGANNNAGATCGGAAG |
| 52013_2 | TATGAGTGTGGAGTCGTTGCTTCCTGCATCATGTGAAGAAGGANNNAGATCGGAAG |
| 52013_3 | TATGAGTGTGGAGTCGTTGCTTCCTGCATCATGTGAAGAAGGANNNAGATCGGAAG |
| 52013_4 | TATGAGTGTGGAGTCGTTGCTTCCTGCATCATGTGAAGAAGGANNNAGATCGGAAG |
| 52013_5 | TATGAGTGTGGAGTCGTTGCTTCCTGCATCATGTGAAGAAGGANNNAGATCGGAAG |
| 52013_6 | TATGAGTGTGGAGTCGTTGCTTCCTGCATCATGTGAAGAAGGANNNAGATCGGAAG |
| 44938_1 | TATGAGTGTGGAGTCGTTGCCCTTCTTGTGGACAGAAAAGCTGNNNAGATCGGAAG |
| 44938_2 | TATGAGTGTGGAGTCGTTGCCCTTCTTGTGGACAGAAAAGCTGNNNAGATCGGAAG |
| 44938_3 | TATGAGTGTGGAGTCGTTGCCCTTCTTGTGGACAGAAAAGCTGNNNAGATCGGAAG |
| 44938_4 | TATGAGTGTGGAGTCGTTGCCCTTCTTGTGGACAGAAAAGCTGNNNAGATCGGAAG |
| 44938_5 | TATGAGTGTGGAGTCGTTGCCCTTCTTGTGGACAGAAAAGCTGNNNAGATCGGAAG |
| 44938_6 | TATGAGTGTGGAGTCGTTGCCCTTCTTGTGGACAGAAAAGCTGNNNAGATCGGAAG |
| 35983_1 | TATGAGTGTGGAGTCGTTGCGTCATGAGATGTAGGGACCAGCNNNAGATCGGAAG |
| 35983_2 | TATGAGTGTGGAGTCGTTGCGTCATGAGATGTAGGGACCAGCNNNAGATCGGAAG |
| 35983_3 | TATGAGTGTGGAGTCGTTGCGTCATGAGATGTAGGGACCAGCNNNAGATCGGAAG |
| 35983_4 | TATGAGTGTGGAGTCGTTGCGTCATGAGATGTAGGGACCAGCNNNAGATCGGAAG |
| 35983_5 | TATGAGTGTGGAGTCGTTGCGTCATGAGATGTAGGGACCAGCNNNAGATCGGAAG |
| 35983_6 | TATGAGTGTGGAGTCGTTGCGTCATGAGATGTAGGGACCAGCNNNAGATCGGAAG |
| 13093_1 | TATGAGTGTGGAGTCGTTGCATTCCTGTATTTCTTTAGTTGTCNNNAGATCGGAAGA |
| 13093_2 | TATGAGTGTGGAGTCGTTGCATTCCTGTATTTCTTTAGTTGTCNNNAGATCGGAAGA |
| 13093_3 | TATGAGTGTGGAGTCGTTGCATTCCTGTATTTCTTTAGTTGTCNNNAGATCGGAAGA |
| 13093_4 | TATGAGTGTGGAGTCGTTGCATTCCTGTATTTCTTTAGTTGTCNNNAGATCGGAAGA |
| 13093_5 | TATGAGTGTGGAGTCGTTGCATTCCTGTATTTCTTTAGTTGTCNNNAGATCGGAAGA |
| 13093_6 | TATGAGTGTGGAGTCGTTGCATTCCTGTATTTCTTTAGTTGTCNNNAGATCGGAAGA |

| | |
|---|---|
| 2641_1 | TATGAGTGTGGAGTCGTTGCATTCAGATTTGTGTTGGTGGCCTNNNAGATCGGAAG |
| 2641_2 | TATGAGTGTGGAGTCGTTGCATTCAGATTTGTGTTGGTGGCCTNNNAGATCGGAAG |
| 2641_3 | TATGAGTGTGGAGTCGTTGCATTCAGATTTGTGTTGGTGGCCTNNNAGATCGGAAG |
| 2641_4 | TATGAGTGTGGAGTCGTTGCATTCAGATTTGTGTTGGTGGCCTNNNAGATCGGAAG |
| 2641_5 | TATGAGTGTGGAGTCGTTGCATTCAGATTTGTGTTGGTGGCCTNNNAGATCGGAAG |
| 2641_6 | TATGAGTGTGGAGTCGTTGCATTCAGATTTGTGTTGGTGGCCTNNNAGATCGGAAG |
| 37300_1 | TATGAGTGTGGAGTCGTTGCGATTGGTGTACCTGAAAGTGACGNNNAGATCGGAA |
| 37300_2 | TATGAGTGTGGAGTCGTTGCGATTGGTGTACCTGAAAGTGACGNNNAGATCGGAA |
| 37300_3 | TATGAGTGTGGAGTCGTTGCGATTGGTGTACCTGAAAGTGACGNNNAGATCGGAA |
| 37300_4 | TATGAGTGTGGAGTCGTTGCGATTGGTGTACCTGAAAGTGACGNNNAGATCGGAA |
| 37300_5 | TATGAGTGTGGAGTCGTTGCGATTGGTGTACCTGAAAGTGACGNNNAGATCGGAA |
| 37300_6 | TATGAGTGTGGAGTCGTTGCGATTGGTGTACCTGAAAGTGACGNNNAGATCGGAA |
| 39042_1 | TATGAGTGTGGAGTCGTTGCAGTACTGAAAACTGGCTCTGTCANNNAGATCGGAAG |
| 39042_2 | TATGAGTGTGGAGTCGTTGCAGTACTGAAAACTGGCTCTGTCANNNAGATCGGAAG |
| 39042_3 | TATGAGTGTGGAGTCGTTGCAGTACTGAAAACTGGCTCTGTCANNNAGATCGGAAG |
| 39042_4 | TATGAGTGTGGAGTCGTTGCAGTACTGAAAACTGGCTCTGTCANNNAGATCGGAAG |
| 39042_5 | TATGAGTGTGGAGTCGTTGCAGTACTGAAAACTGGCTCTGTCANNNAGATCGGAAG |
| 39042_6 | TATGAGTGTGGAGTCGTTGCAGTACTGAAAACTGGCTCTGTCANNNAGATCGGAAG |
| 39734_1 | TATGAGTGTGGAGTCGTTGCTTGAATTTCAACTTCACCACTTACTNNNAGATCGGAA |
| 39734_2 | TATGAGTGTGGAGTCGTTGCTTGAATTTCAACTTCACCACTTACTNNNAGATCGGAA |
| 39734_3 | TATGAGTGTGGAGTCGTTGCTTGAATTTCAACTTCACCACTTACTNNNAGATCGGAA |
| 39734_4 | TATGAGTGTGGAGTCGTTGCTTGAATTTCAACTTCACCACTTACTNNNAGATCGGAA |
| 39734_5 | TATGAGTGTGGAGTCGTTGCTTGAATTTCAACTTCACCACTTACTNNNAGATCGGAA |
| 39734_6 | TATGAGTGTGGAGTCGTTGCTTGAATTTCAACTTCACCACTTACTNNNAGATCGGAA |
| 33957_1 | TATGAGTGTGGAGTCGTTGCGAGATCTTGACTCGCCACAACTTNNNAGATCGGAAG |
| 33957_2 | TATGAGTGTGGAGTCGTTGCGAGATCTTGACTCGCCACAACTTNNNAGATCGGAAG |
| 33957_3 | TATGAGTGTGGAGTCGTTGCGAGATCTTGACTCGCCACAACTTNNNAGATCGGAAG |
| 33957_4 | TATGAGTGTGGAGTCGTTGCGAGATCTTGACTCGCCACAACTTNNNAGATCGGAAG |
| 33957_5 | TATGAGTGTGGAGTCGTTGCGAGATCTTGACTCGCCACAACTTNNNAGATCGGAAG |
| 33957_6 | TATGAGTGTGGAGTCGTTGCGAGATCTTGACTCGCCACAACTTNNNAGATCGGAAG |
| 34010_1 | TATGAGTGTGGAGTCGTTGCGAGTGCAATGGCGAGATCACANNNAGATCGGAAGA |
| 34010_2 | TATGAGTGTGGAGTCGTTGCGAGTGCAATGGCGAGATCACANNNAGATCGGAAGA |
| 34010_3 | TATGAGTGTGGAGTCGTTGCGAGTGCAATGGCGAGATCACANNNAGATCGGAAGA |
| 34010_4 | TATGAGTGTGGAGTCGTTGCGAGTGCAATGGCGAGATCACANNNAGATCGGAAGA |
| 34010_5 | TATGAGTGTGGAGTCGTTGCGAGTGCAATGGCGAGATCACANNNAGATCGGAAGA |
| 34010_6 | TATGAGTGTGGAGTCGTTGCGAGTGCAATGGCGAGATCACANNNAGATCGGAAGA |
| 48858_1 | TATGAGTGTGGAGTCGTTGCCTAGAGTACAGTGGCCCAATCACNNNAGATCGGAAG |
| 48858_2 | TATGAGTGTGGAGTCGTTGCCTAGAGTACAGTGGCCCAATCACNNNAGATCGGAAG |
| 48858_3 | TATGAGTGTGGAGTCGTTGCCTAGAGTACAGTGGCCCAATCACNNNAGATCGGAAG |
| 48858_4 | TATGAGTGTGGAGTCGTTGCCTAGAGTACAGTGGCCCAATCACNNNAGATCGGAAG |
| 48858_5 | TATGAGTGTGGAGTCGTTGCCTAGAGTACAGTGGCCCAATCACNNNAGATCGGAAG |
| 48858_6 | TATGAGTGTGGAGTCGTTGCCTAGAGTACAGTGGCCCAATCACNNNAGATCGGAAG |
| 33523_1 | TATGAGTGTGGAGTCGTTGCTTCACACCATGACAAAAACTGCANNNAGATCGGAAG |

| | |
|---|---|
| 33523_2 | TATGAGTGTGGAGTCGTTGCTTCACACCATGACAAAAACTGCANNNAGATCGGAAG |
| 33523_3 | TATGAGTGTGGAGTCGTTGCTTCACACCATGACAAAAACTGCANNNAGATCGGAAG |
| 33523_4 | TATGAGTGTGGAGTCGTTGCTTCACACCATGACAAAAACTGCANNNAGATCGGAAG |
| 33523_5 | TATGAGTGTGGAGTCGTTGCTTCACACCATGACAAAAACTGCANNNAGATCGGAAG |
| 33523_6 | TATGAGTGTGGAGTCGTTGCTTCACACCATGACAAAAACTGCANNNAGATCGGAAG |
| 46345_1 | TATGAGTGTGGAGTCGTTGCCTCTGTCTCCCAAGTTCAAGTGANNNAGATCGGAAG |
| 46345_2 | TATGAGTGTGGAGTCGTTGCCTCTGTCTCCCAAGTTCAAGTGANNNAGATCGGAAG |
| 46345_3 | TATGAGTGTGGAGTCGTTGCCTCTGTCTCCCAAGTTCAAGTGANNNAGATCGGAAG |
| 46345_4 | TATGAGTGTGGAGTCGTTGCCTCTGTCTCCCAAGTTCAAGTGANNNAGATCGGAAG |
| 46345_5 | TATGAGTGTGGAGTCGTTGCCTCTGTCTCCCAAGTTCAAGTGANNNAGATCGGAAG |
| 46345_6 | TATGAGTGTGGAGTCGTTGCCTCTGTCTCCCAAGTTCAAGTGANNNAGATCGGAAG |
| 5570_1 | TATGAGTGTGGAGTCGTTGCAACAAAGGTAGGTGCCAGAATGGNNNAGATCGGAA |
| 5570_2 | TATGAGTGTGGAGTCGTTGCAACAAAGGTAGGTGCCAGAATGGNNNAGATCGGAA |
| 5570_3 | TATGAGTGTGGAGTCGTTGCAACAAAGGTAGGTGCCAGAATGGNNNAGATCGGAA |
| 5570_4 | TATGAGTGTGGAGTCGTTGCAACAAAGGTAGGTGCCAGAATGGNNNAGATCGGAA |
| 5570_5 | TATGAGTGTGGAGTCGTTGCAACAAAGGTAGGTGCCAGAATGGNNNAGATCGGAA |
| 5570_6 | TATGAGTGTGGAGTCGTTGCAACAAAGGTAGGTGCCAGAATGGNNNAGATCGGAA |
| 46895_1 | TATGAGTGTGGAGTCGTTGCACAGGTGTAAACCATTGCACTTGNNNAGATCGGAAG |
| 46895_2 | TATGAGTGTGGAGTCGTTGCACAGGTGTAAACCATTGCACTTGNNNAGATCGGAAG |
| 46895_3 | TATGAGTGTGGAGTCGTTGCACAGGTGTAAACCATTGCACTTGNNNAGATCGGAAG |
| 46895_4 | TATGAGTGTGGAGTCGTTGCACAGGTGTAAACCATTGCACTTGNNNAGATCGGAAG |
| 46895_5 | TATGAGTGTGGAGTCGTTGCACAGGTGTAAACCATTGCACTTGNNNAGATCGGAAG |
| 46895_6 | TATGAGTGTGGAGTCGTTGCACAGGTGTAAACCATTGCACTTGNNNAGATCGGAAG |
| 42420_1 | TATGAGTGTGGAGTCGTTGCGTCCCTTTGACAATTCCAAGTGGNNNAGATCGGAAG |
| 42420_2 | TATGAGTGTGGAGTCGTTGCGTCCCTTTGACAATTCCAAGTGGNNNAGATCGGAAG |
| 42420_3 | TATGAGTGTGGAGTCGTTGCGTCCCTTTGACAATTCCAAGTGGNNNAGATCGGAAG |
| 42420_4 | TATGAGTGTGGAGTCGTTGCGTCCCTTTGACAATTCCAAGTGGNNNAGATCGGAAG |
| 42420_5 | TATGAGTGTGGAGTCGTTGCGTCCCTTTGACAATTCCAAGTGGNNNAGATCGGAAG |
| 42420_6 | TATGAGTGTGGAGTCGTTGCGTCCCTTTGACAATTCCAAGTGGNNNAGATCGGAAG |
| 9682_1 | TATGAGTGTGGAGTCGTTGCACGGTAGTGCTCTACTACATCCCNNNAGATCGGAAG |
| 9682_2 | TATGAGTGTGGAGTCGTTGCACGGTAGTGCTCTACTACATCCCNNNAGATCGGAAG |
| 9682_3 | TATGAGTGTGGAGTCGTTGCACGGTAGTGCTCTACTACATCCCNNNAGATCGGAAG |
| 9682_4 | TATGAGTGTGGAGTCGTTGCACGGTAGTGCTCTACTACATCCCNNNAGATCGGAAG |
| 9682_5 | TATGAGTGTGGAGTCGTTGCACGGTAGTGCTCTACTACATCCCNNNAGATCGGAAG |
| 9682_6 | TATGAGTGTGGAGTCGTTGCACGGTAGTGCTCTACTACATCCCNNNAGATCGGAAG |
| 33353_1 | TATGAGTGTGGAGTCGTTGCGAGGTAGGGTCTCAATCTGTCACNNNAGATCGGAA |
| 33353_2 | TATGAGTGTGGAGTCGTTGCGAGGTAGGGTCTCAATCTGTCACNNNAGATCGGAA |
| 33353_3 | TATGAGTGTGGAGTCGTTGCGAGGTAGGGTCTCAATCTGTCACNNNAGATCGGAA |
| 33353_4 | TATGAGTGTGGAGTCGTTGCGAGGTAGGGTCTCAATCTGTCACNNNAGATCGGAA |
| 33353_5 | TATGAGTGTGGAGTCGTTGCGAGGTAGGGTCTCAATCTGTCACNNNAGATCGGAA |
| 33353_6 | TATGAGTGTGGAGTCGTTGCGAGGTAGGGTCTCAATCTGTCACNNNAGATCGGAA |
| 2917_1 | TATGAGTGTGGAGTCGTTGCCTTCCTGTCTTTGGCACCTCATTNNNAGATCGGAAGA |
| 2917_2 | TATGAGTGTGGAGTCGTTGCCTTCCTGTCTTTGGCACCTCATTNNNAGATCGGAAGA |

| | |
|---|---|
| 2917_3 | TATGAGTGTGGAGTCGTTGCCTTCCTGTCTTTGGCACCTCATTNNNAGATCGGAAGA |
| 2917_4 | TATGAGTGTGGAGTCGTTGCCTTCCTGTCTTTGGCACCTCATTNNNAGATCGGAAGA |
| 2917_5 | TATGAGTGTGGAGTCGTTGCCTTCCTGTCTTTGGCACCTCATTNNNAGATCGGAAGA |
| 2917_6 | TATGAGTGTGGAGTCGTTGCCTTCCTGTCTTTGGCACCTCATTNNNAGATCGGAAGA |
| 8858_1 | TATGAGTGTGGAGTCGTTGCTACTGTGGAAGAGATATGCCTGTNNNAGATCGGAAC |
| 8858_2 | TATGAGTGTGGAGTCGTTGCTACTGTGGAAGAGATATGCCTGTNNNAGATCGGAAC |
| 8858_3 | TATGAGTGTGGAGTCGTTGCTACTGTGGAAGAGATATGCCTGTNNNAGATCGGAAC |
| 8858_4 | TATGAGTGTGGAGTCGTTGCTACTGTGGAAGAGATATGCCTGTNNNAGATCGGAAC |
| 8858_5 | TATGAGTGTGGAGTCGTTGCTACTGTGGAAGAGATATGCCTGTNNNAGATCGGAAC |
| 8858_6 | TATGAGTGTGGAGTCGTTGCTACTGTGGAAGAGATATGCCTGTNNNAGATCGGAAC |
| 30643_1 | TATGAGTGTGGAGTCGTTGCCCCTGGAGCCCGTGTTTTATTTNNNAGATCGGAAGAC |
| 30643_2 | TATGAGTGTGGAGTCGTTGCCCCTGGAGCCCGTGTTTTATTTNNNAGATCGGAAGAC |
| 30643_3 | TATGAGTGTGGAGTCGTTGCCCCTGGAGCCCGTGTTTTATTTNNNAGATCGGAAGAC |
| 30643_4 | TATGAGTGTGGAGTCGTTGCCCCTGGAGCCCGTGTTTTATTTNNNAGATCGGAAGAC |
| 30643_5 | TATGAGTGTGGAGTCGTTGCCCCTGGAGCCCGTGTTTTATTTNNNAGATCGGAAGAC |
| 30643_6 | TATGAGTGTGGAGTCGTTGCCCCTGGAGCCCGTGTTTTATTTNNNAGATCGGAAGAC |
| 32338_1 | TATGAGTGTGGAGTCGTTGCAGAGTTTGAGCCAGAGTCATGTTNNNAGATCGGAAC |
| 32338_2 | TATGAGTGTGGAGTCGTTGCAGAGTTTGAGCCAGAGTCATGTTNNNAGATCGGAAC |
| 32338_3 | TATGAGTGTGGAGTCGTTGCAGAGTTTGAGCCAGAGTCATGTTNNNAGATCGGAAC |
| 32338_4 | TATGAGTGTGGAGTCGTTGCAGAGTTTGAGCCAGAGTCATGTTNNNAGATCGGAAC |
| 32338_5 | TATGAGTGTGGAGTCGTTGCAGAGTTTGAGCCAGAGTCATGTTNNNAGATCGGAAC |
| 32338_6 | TATGAGTGTGGAGTCGTTGCAGAGTTTGAGCCAGAGTCATGTTNNNAGATCGGAAC |
| 10163_1 | TATGAGTGTGGAGTCGTTGCGGCTCGTACACATGGAACTCTTTNNNAGATCGGAAG |
| 10163_2 | TATGAGTGTGGAGTCGTTGCGGCTCGTACACATGGAACTCTTTNNNAGATCGGAAG |
| 10163_3 | TATGAGTGTGGAGTCGTTGCGGCTCGTACACATGGAACTCTTTNNNAGATCGGAAG |
| 10163_4 | TATGAGTGTGGAGTCGTTGCGGCTCGTACACATGGAACTCTTTNNNAGATCGGAAG |
| 10163_5 | TATGAGTGTGGAGTCGTTGCGGCTCGTACACATGGAACTCTTTNNNAGATCGGAAG |
| 10163_6 | TATGAGTGTGGAGTCGTTGCGGCTCGTACACATGGAACTCTTTNNNAGATCGGAAG |
| 50652_1 | TATGAGTGTGGAGTCGTTGCATCAGGGGAAGAAAAGGAGCAAANNNAGATCGGAA |
| 50652_2 | TATGAGTGTGGAGTCGTTGCATCAGGGGAAGAAAAGGAGCAAANNNAGATCGGAA |
| 50652_3 | TATGAGTGTGGAGTCGTTGCATCAGGGGAAGAAAAGGAGCAAANNNAGATCGGAA |
| 50652_4 | TATGAGTGTGGAGTCGTTGCATCAGGGGAAGAAAAGGAGCAAANNNAGATCGGAA |
| 50652_5 | TATGAGTGTGGAGTCGTTGCATCAGGGGAAGAAAAGGAGCAAANNNAGATCGGAA |
| 50652_6 | TATGAGTGTGGAGTCGTTGCATCAGGGGAAGAAAAGGAGCAAANNNAGATCGGAA |
| 82_1 | TATGAGTGTGGAGTCGTTGCCATAACCTCAATTGCCAGCCATANNNAGATCGGAAGA |
| 82_2 | TATGAGTGTGGAGTCGTTGCCATAACCTCAATTGCCAGCCATANNNAGATCGGAAGA |
| 82_3 | TATGAGTGTGGAGTCGTTGCCATAACCTCAATTGCCAGCCATANNNAGATCGGAAGA |
| 82_4 | TATGAGTGTGGAGTCGTTGCCATAACCTCAATTGCCAGCCATANNNAGATCGGAAGA |
| 82_5 | TATGAGTGTGGAGTCGTTGCCATAACCTCAATTGCCAGCCATANNNAGATCGGAAGA |
| 82_6 | TATGAGTGTGGAGTCGTTGCCATAACCTCAATTGCCAGCCATANNNAGATCGGAAGA |
| 35402_1 | TATGAGTGTGGAGTCGTTGCTTCACGCCATTCTCCTGACTCNNNAGATCGGAAGAGC |
| 35402_2 | TATGAGTGTGGAGTCGTTGCTTCACGCCATTCTCCTGACTCNNNAGATCGGAAGAGC |
| 35402_3 | TATGAGTGTGGAGTCGTTGCTTCACGCCATTCTCCTGACTCNNNAGATCGGAAGAGC |

| | |
|---|---|
| 35402_4 | TATGAGTGTGGAGTCGTTGCTTCACGCCATTCTCCTGACTCNNNAGATCGGAAGAGC |
| 35402_5 | TATGAGTGTGGAGTCGTTGCTTCACGCCATTCTCCTGACTCNNNAGATCGGAAGAGC |
| 35402_6 | TATGAGTGTGGAGTCGTTGCTTCACGCCATTCTCCTGACTCNNNAGATCGGAAGAGC |
| 35596_1 | TATGAGTGTGGAGTCGTTGCCAATCTCGGCTCACTGCAAACNNNAGATCGGAAGAG |
| 35596_2 | TATGAGTGTGGAGTCGTTGCCAATCTCGGCTCACTGCAAACNNNAGATCGGAAGAG |
| 35596_3 | TATGAGTGTGGAGTCGTTGCCAATCTCGGCTCACTGCAAACNNNAGATCGGAAGAG |
| 35596_4 | TATGAGTGTGGAGTCGTTGCCAATCTCGGCTCACTGCAAACNNNAGATCGGAAGAG |
| 35596_5 | TATGAGTGTGGAGTCGTTGCCAATCTCGGCTCACTGCAAACNNNAGATCGGAAGAG |
| 35596_6 | TATGAGTGTGGAGTCGTTGCCAATCTCGGCTCACTGCAAACNNNAGATCGGAAGAG |
| 39610_1 | TATGAGTGTGGAGTCGTTGCGAGGTAGCAGTGAGGAGAGATTGNNNAGATCGGA |
| 39610_2 | TATGAGTGTGGAGTCGTTGCGAGGTAGCAGTGAGGAGAGATTGNNNAGATCGGA |
| 39610_3 | TATGAGTGTGGAGTCGTTGCGAGGTAGCAGTGAGGAGAGATTGNNNAGATCGGA |
| 39610_4 | TATGAGTGTGGAGTCGTTGCGAGGTAGCAGTGAGGAGAGATTGNNNAGATCGGA |
| 39610_5 | TATGAGTGTGGAGTCGTTGCGAGGTAGCAGTGAGGAGAGATTGNNNAGATCGGA |
| 39610_6 | TATGAGTGTGGAGTCGTTGCGAGGTAGCAGTGAGGAGAGATTGNNNAGATCGGA |
| 49465_1 | TATGAGTGTGGAGTCGTTGCTCCTACGCAGAACACACATACAANNNAGATCGGAAG |
| 49465_2 | TATGAGTGTGGAGTCGTTGCTCCTACGCAGAACACACATACAANNNAGATCGGAAG |
| 49465_3 | TATGAGTGTGGAGTCGTTGCTCCTACGCAGAACACACATACAANNNAGATCGGAAG |
| 49465_4 | TATGAGTGTGGAGTCGTTGCTCCTACGCAGAACACACATACAANNNAGATCGGAAG |
| 49465_5 | TATGAGTGTGGAGTCGTTGCTCCTACGCAGAACACACATACAANNNAGATCGGAAG |
| 49465_6 | TATGAGTGTGGAGTCGTTGCTCCTACGCAGAACACACATACAANNNAGATCGGAAG |
| 40475_1 | TATGAGTGTGGAGTCGTTGCGATAAGGTTCTGCTTGAGGGTCANNNAGATCGGAA |
| 40475_2 | TATGAGTGTGGAGTCGTTGCGATAAGGTTCTGCTTGAGGGTCANNNAGATCGGAA |
| 40475_3 | TATGAGTGTGGAGTCGTTGCGATAAGGTTCTGCTTGAGGGTCANNNAGATCGGAA |
| 40475_4 | TATGAGTGTGGAGTCGTTGCGATAAGGTTCTGCTTGAGGGTCANNNAGATCGGAA |
| 40475_5 | TATGAGTGTGGAGTCGTTGCGATAAGGTTCTGCTTGAGGGTCANNNAGATCGGAA |
| 40475_6 | TATGAGTGTGGAGTCGTTGCGATAAGGTTCTGCTTGAGGGTCANNNAGATCGGAA |
| 41481_1 | TATGAGTGTGGAGTCGTTGCAGTACATCAACGTGGAAGGAACANNNAGATCGGAA |
| 41481_2 | TATGAGTGTGGAGTCGTTGCAGTACATCAACGTGGAAGGAACANNNAGATCGGAA |
| 41481_3 | TATGAGTGTGGAGTCGTTGCAGTACATCAACGTGGAAGGAACANNNAGATCGGAA |
| 41481_4 | TATGAGTGTGGAGTCGTTGCAGTACATCAACGTGGAAGGAACANNNAGATCGGAA |
| 41481_5 | TATGAGTGTGGAGTCGTTGCAGTACATCAACGTGGAAGGAACANNNAGATCGGAA |
| 41481_6 | TATGAGTGTGGAGTCGTTGCAGTACATCAACGTGGAAGGAACANNNAGATCGGAA |
| 42658_1 | TATGAGTGTGGAGTCGTTGCACCTGTAGTCCTAGCTACTCAGGNNNAGATCGGAAG |
| 42658_2 | TATGAGTGTGGAGTCGTTGCACCTGTAGTCCTAGCTACTCAGGNNNAGATCGGAAG |
| 42658_3 | TATGAGTGTGGAGTCGTTGCACCTGTAGTCCTAGCTACTCAGGNNNAGATCGGAAG |
| 42658_4 | TATGAGTGTGGAGTCGTTGCACCTGTAGTCCTAGCTACTCAGGNNNAGATCGGAAG |
| 42658_5 | TATGAGTGTGGAGTCGTTGCACCTGTAGTCCTAGCTACTCAGGNNNAGATCGGAAG |
| 42658_6 | TATGAGTGTGGAGTCGTTGCACCTGTAGTCCTAGCTACTCAGGNNNAGATCGGAAG |
| 28767_1 | TATGAGTGTGGAGTCGTTGCGAGACAGGAGGATCACTTGAGTCNNNAGATCGGAA |
| 28767_2 | TATGAGTGTGGAGTCGTTGCGAGACAGGAGGATCACTTGAGTCNNNAGATCGGAA |
| 28767_3 | TATGAGTGTGGAGTCGTTGCGAGACAGGAGGATCACTTGAGTCNNNAGATCGGAA |
| 28767_4 | TATGAGTGTGGAGTCGTTGCGAGACAGGAGGATCACTTGAGTCNNNAGATCGGAA |

| | |
|---|---|
| 28767_5 | TATGAGTGTGGAGTCGTTGCGAGACAGGAGGATCACTTGAGTCNNNAGATCGGAA |
| 28767_6 | TATGAGTGTGGAGTCGTTGCGAGACAGGAGGATCACTTGAGTCNNNAGATCGGAA |
| 10156_1 | TATGAGTGTGGAGTCGTTGCCAGTGACTTGGATTGCAGTGAGANNNAGATCGGAA |
| 10156_2 | TATGAGTGTGGAGTCGTTGCCAGTGACTTGGATTGCAGTGAGANNNAGATCGGAA |
| 10156_3 | TATGAGTGTGGAGTCGTTGCCAGTGACTTGGATTGCAGTGAGANNNAGATCGGAA |
| 10156_4 | TATGAGTGTGGAGTCGTTGCCAGTGACTTGGATTGCAGTGAGANNNAGATCGGAA |
| 10156_5 | TATGAGTGTGGAGTCGTTGCCAGTGACTTGGATTGCAGTGAGANNNAGATCGGAA |
| 10156_6 | TATGAGTGTGGAGTCGTTGCCAGTGACTTGGATTGCAGTGAGANNNAGATCGGAA |
| 7852_1 | TATGAGTGTGGAGTCGTTGCACTGCAAGCTTTGCCAAGCGANNNAGATCGGAAGA |
| 7852_2 | TATGAGTGTGGAGTCGTTGCACTGCAAGCTTTGCCAAGCGANNNAGATCGGAAGA |
| 7852_3 | TATGAGTGTGGAGTCGTTGCACTGCAAGCTTTGCCAAGCGANNNAGATCGGAAGA |
| 7852_4 | TATGAGTGTGGAGTCGTTGCACTGCAAGCTTTGCCAAGCGANNNAGATCGGAAGA |
| 7852_5 | TATGAGTGTGGAGTCGTTGCACTGCAAGCTTTGCCAAGCGANNNAGATCGGAAGA |
| 7852_6 | TATGAGTGTGGAGTCGTTGCACTGCAAGCTTTGCCAAGCGANNNAGATCGGAAGA |
| 48145_1 | TATGAGTGTGGAGTCGTTGCTTCAGTAGAGATGGGGTTTCAGCNNNAGATCGGAA |
| 48145_2 | TATGAGTGTGGAGTCGTTGCTTCAGTAGAGATGGGGTTTCAGCNNNAGATCGGAA |
| 48145_3 | TATGAGTGTGGAGTCGTTGCTTCAGTAGAGATGGGGTTTCAGCNNNAGATCGGAA |
| 48145_4 | TATGAGTGTGGAGTCGTTGCTTCAGTAGAGATGGGGTTTCAGCNNNAGATCGGAA |
| 48145_5 | TATGAGTGTGGAGTCGTTGCTTCAGTAGAGATGGGGTTTCAGCNNNAGATCGGAA |
| 48145_6 | TATGAGTGTGGAGTCGTTGCTTCAGTAGAGATGGGGTTTCAGCNNNAGATCGGAA |
| 33612_1 | TATGAGTGTGGAGTCGTTGCTCAAACAATTTTCCGACCTCAGCNNNAGATCGGAAG |
| 33612_2 | TATGAGTGTGGAGTCGTTGCTCAAACAATTTTCCGACCTCAGCNNNAGATCGGAAG |
| 33612_3 | TATGAGTGTGGAGTCGTTGCTCAAACAATTTTCCGACCTCAGCNNNAGATCGGAAG |
| 33612_4 | TATGAGTGTGGAGTCGTTGCTCAAACAATTTTCCGACCTCAGCNNNAGATCGGAAG |
| 33612_5 | TATGAGTGTGGAGTCGTTGCTCAAACAATTTTCCGACCTCAGCNNNAGATCGGAAG |
| 33612_6 | TATGAGTGTGGAGTCGTTGCTCAAACAATTTTCCGACCTCAGCNNNAGATCGGAAG |
| 42745_1 | TATGAGTGTGGAGTCGTTGCTCATGAAAGTGACACAGGATTGTNNNAGATCGGAA |
| 42745_2 | TATGAGTGTGGAGTCGTTGCTCATGAAAGTGACACAGGATTGTNNNAGATCGGAA |
| 42745_3 | TATGAGTGTGGAGTCGTTGCTCATGAAAGTGACACAGGATTGTNNNAGATCGGAA |
| 42745_4 | TATGAGTGTGGAGTCGTTGCTCATGAAAGTGACACAGGATTGTNNNAGATCGGAA |
| 42745_5 | TATGAGTGTGGAGTCGTTGCTCATGAAAGTGACACAGGATTGTNNNAGATCGGAA |
| 42745_6 | TATGAGTGTGGAGTCGTTGCTCATGAAAGTGACACAGGATTGTNNNAGATCGGAA |
| 41044_1 | TATGAGTGTGGAGTCGTTGCAGGCCCATATATTCAGGCTTGAGNNNAGATCGGAA |
| 41044_2 | TATGAGTGTGGAGTCGTTGCAGGCCCATATATTCAGGCTTGAGNNNAGATCGGAA |
| 41044_3 | TATGAGTGTGGAGTCGTTGCAGGCCCATATATTCAGGCTTGAGNNNAGATCGGAA |
| 41044_4 | TATGAGTGTGGAGTCGTTGCAGGCCCATATATTCAGGCTTGAGNNNAGATCGGAA |
| 41044_5 | TATGAGTGTGGAGTCGTTGCAGGCCCATATATTCAGGCTTGAGNNNAGATCGGAA |
| 41044_6 | TATGAGTGTGGAGTCGTTGCAGGCCCATATATTCAGGCTTGAGNNNAGATCGGAA |
| 47656_1 | TATGAGTGTGGAGTCGTTGCTTTAGTAGAGACGGAGTTTCGCCNNNAGATCGGAA |
| 47656_2 | TATGAGTGTGGAGTCGTTGCTTTAGTAGAGACGGAGTTTCGCCNNNAGATCGGAA |
| 47656_3 | TATGAGTGTGGAGTCGTTGCTTTAGTAGAGACGGAGTTTCGCCNNNAGATCGGAA |
| 47656_4 | TATGAGTGTGGAGTCGTTGCTTTAGTAGAGACGGAGTTTCGCCNNNAGATCGGAA |
| 47656_5 | TATGAGTGTGGAGTCGTTGCTTTAGTAGAGACGGAGTTTCGCCNNNAGATCGGAA |

| | |
|---|---|
| 47656_6 | TATGAGTGTGGAGTCGTTGCTTTAGTAGAGACGGAGTTTCGCCNNNAGATCGGAA |
| 47626_1 | TATGAGTGTGGAGTCGTTGCGCTCACTACAGCCACATTTTGTTNNNAGATCGGAAG |
| 47626_2 | TATGAGTGTGGAGTCGTTGCGCTCACTACAGCCACATTTTGTTNNNAGATCGGAAG |
| 47626_3 | TATGAGTGTGGAGTCGTTGCGCTCACTACAGCCACATTTTGTTNNNAGATCGGAAG |
| 47626_4 | TATGAGTGTGGAGTCGTTGCGCTCACTACAGCCACATTTTGTTNNNAGATCGGAAG |
| 47626_5 | TATGAGTGTGGAGTCGTTGCGCTCACTACAGCCACATTTTGTTNNNAGATCGGAAG |
| 47626_6 | TATGAGTGTGGAGTCGTTGCGCTCACTACAGCCACATTTTGTTNNNAGATCGGAAG |
| 37236_1 | TATGAGTGTGGAGTCGTTGCAAAGTTCAGGCCAATATCCCTGTNNNAGATCGGAAG |
| 37236_2 | TATGAGTGTGGAGTCGTTGCAAAGTTCAGGCCAATATCCCTGTNNNAGATCGGAAG |
| 37236_3 | TATGAGTGTGGAGTCGTTGCAAAGTTCAGGCCAATATCCCTGTNNNAGATCGGAAG |
| 37236_4 | TATGAGTGTGGAGTCGTTGCAAAGTTCAGGCCAATATCCCTGTNNNAGATCGGAAG |
| 37236_5 | TATGAGTGTGGAGTCGTTGCAAAGTTCAGGCCAATATCCCTGTNNNAGATCGGAAG |
| 37236_6 | TATGAGTGTGGAGTCGTTGCAAAGTTCAGGCCAATATCCCTGTNNNAGATCGGAAG |
| 35279_1 | TATGAGTGTGGAGTCGTTGCTTGAACTGTTTGTGGCCAATTGTNNNAGATCGGAAG |
| 35279_2 | TATGAGTGTGGAGTCGTTGCTTGAACTGTTTGTGGCCAATTGTNNNAGATCGGAAG |
| 35279_3 | TATGAGTGTGGAGTCGTTGCTTGAACTGTTTGTGGCCAATTGTNNNAGATCGGAAG |
| 35279_4 | TATGAGTGTGGAGTCGTTGCTTGAACTGTTTGTGGCCAATTGTNNNAGATCGGAAG |
| 35279_5 | TATGAGTGTGGAGTCGTTGCTTGAACTGTTTGTGGCCAATTGTNNNAGATCGGAAG |
| 35279_6 | TATGAGTGTGGAGTCGTTGCTTGAACTGTTTGTGGCCAATTGTNNNAGATCGGAAG |
| 45379_1 | TATGAGTGTGGAGTCGTTGCTCTTCTGAAAGAACGTGGAGACTNNNAGATCGGAA |
| 45379_2 | TATGAGTGTGGAGTCGTTGCTCTTCTGAAAGAACGTGGAGACTNNNAGATCGGAA |
| 45379_3 | TATGAGTGTGGAGTCGTTGCTCTTCTGAAAGAACGTGGAGACTNNNAGATCGGAA |
| 45379_4 | TATGAGTGTGGAGTCGTTGCTCTTCTGAAAGAACGTGGAGACTNNNAGATCGGAA |
| 45379_5 | TATGAGTGTGGAGTCGTTGCTCTTCTGAAAGAACGTGGAGACTNNNAGATCGGAA |
| 45379_6 | TATGAGTGTGGAGTCGTTGCTCTTCTGAAAGAACGTGGAGACTNNNAGATCGGAA |
| 47393_1 | TATGAGTGTGGAGTCGTTGCAGTGCTAGGATTACAGGCATGAGNNNAGATCGGAA |
| 47393_2 | TATGAGTGTGGAGTCGTTGCAGTGCTAGGATTACAGGCATGAGNNNAGATCGGAA |
| 47393_3 | TATGAGTGTGGAGTCGTTGCAGTGCTAGGATTACAGGCATGAGNNNAGATCGGAA |
| 47393_4 | TATGAGTGTGGAGTCGTTGCAGTGCTAGGATTACAGGCATGAGNNNAGATCGGAA |
| 47393_5 | TATGAGTGTGGAGTCGTTGCAGTGCTAGGATTACAGGCATGAGNNNAGATCGGAA |
| 47393_6 | TATGAGTGTGGAGTCGTTGCAGTGCTAGGATTACAGGCATGAGNNNAGATCGGAA |
| 37244_1 | TATGAGTGTGGAGTCGTTGCCTTGCAGTGAGCCGAGATCANNNAGATCGGAAGAG |
| 37244_2 | TATGAGTGTGGAGTCGTTGCCTTGCAGTGAGCCGAGATCANNNAGATCGGAAGAG |
| 37244_3 | TATGAGTGTGGAGTCGTTGCCTTGCAGTGAGCCGAGATCANNNAGATCGGAAGAG |
| 37244_4 | TATGAGTGTGGAGTCGTTGCCTTGCAGTGAGCCGAGATCANNNAGATCGGAAGAG |
| 37244_5 | TATGAGTGTGGAGTCGTTGCCTTGCAGTGAGCCGAGATCANNNAGATCGGAAGAG |
| 37244_6 | TATGAGTGTGGAGTCGTTGCCTTGCAGTGAGCCGAGATCANNNAGATCGGAAGAG |
| 42694_1 | TATGAGTGTGGAGTCGTTGCCACACACGCAGAAAACTATAGGCNNNAGATCGGAA |
| 42694_2 | TATGAGTGTGGAGTCGTTGCCACACACGCAGAAAACTATAGGCNNNAGATCGGAA |
| 42694_3 | TATGAGTGTGGAGTCGTTGCCACACACGCAGAAAACTATAGGCNNNAGATCGGAA |
| 42694_4 | TATGAGTGTGGAGTCGTTGCCACACACGCAGAAAACTATAGGCNNNAGATCGGAA |
| 42694_5 | TATGAGTGTGGAGTCGTTGCCACACACGCAGAAAACTATAGGCNNNAGATCGGAA |
| 42694_6 | TATGAGTGTGGAGTCGTTGCCACACACGCAGAAAACTATAGGCNNNAGATCGGAA |

| | |
|---|---|
| 47827_1 | TATGAGTGTGGAGTCGTTGCCTGTCACCTCCTTAGTGTCCTTGNNNAGATCGGAAGA |
| 47827_2 | TATGAGTGTGGAGTCGTTGCCTGTCACCTCCTTAGTGTCCTTGNNNAGATCGGAAGA |
| 47827_3 | TATGAGTGTGGAGTCGTTGCCTGTCACCTCCTTAGTGTCCTTGNNNAGATCGGAAGA |
| 47827_4 | TATGAGTGTGGAGTCGTTGCCTGTCACCTCCTTAGTGTCCTTGNNNAGATCGGAAGA |
| 47827_5 | TATGAGTGTGGAGTCGTTGCCTGTCACCTCCTTAGTGTCCTTGNNNAGATCGGAAGA |
| 47827_6 | TATGAGTGTGGAGTCGTTGCCTGTCACCTCCTTAGTGTCCTTGNNNAGATCGGAAGA |
| 38807_1 | TATGAGTGTGGAGTCGTTGCCAATAGCCCTGAAAGAAGCAATGTNNNAGATCGGAA |
| 38807_2 | TATGAGTGTGGAGTCGTTGCCAATAGCCCTGAAAGAAGCAATGTNNNAGATCGGAA |
| 38807_3 | TATGAGTGTGGAGTCGTTGCCAATAGCCCTGAAAGAAGCAATGTNNNAGATCGGAA |
| 38807_4 | TATGAGTGTGGAGTCGTTGCCAATAGCCCTGAAAGAAGCAATGTNNNAGATCGGAA |
| 38807_5 | TATGAGTGTGGAGTCGTTGCCAATAGCCCTGAAAGAAGCAATGTNNNAGATCGGAA |
| 38807_6 | TATGAGTGTGGAGTCGTTGCCAATAGCCCTGAAAGAAGCAATGTNNNAGATCGGAA |
| 35709_1 | TATGAGTGTGGAGTCGTTGCACTTGAGAAGCTGGGATTACAGGNNNAGATCGGAA |
| 35709_2 | TATGAGTGTGGAGTCGTTGCACTTGAGAAGCTGGGATTACAGGNNNAGATCGGAA |
| 35709_3 | TATGAGTGTGGAGTCGTTGCACTTGAGAAGCTGGGATTACAGGNNNAGATCGGAA |
| 35709_4 | TATGAGTGTGGAGTCGTTGCACTTGAGAAGCTGGGATTACAGGNNNAGATCGGAA |
| 35709_5 | TATGAGTGTGGAGTCGTTGCACTTGAGAAGCTGGGATTACAGGNNNAGATCGGAA |
| 35709_6 | TATGAGTGTGGAGTCGTTGCACTTGAGAAGCTGGGATTACAGGNNNAGATCGGAA |
| 43820_1 | TATGAGTGTGGAGTCGTTGCGGCTTTTTTTTTTCCGGACTGCNNNAGATCGGAAGA |
| 43820_2 | TATGAGTGTGGAGTCGTTGCGGCTTTTTTTTTTCCGGACTGCNNNAGATCGGAAGA |
| 43820_3 | TATGAGTGTGGAGTCGTTGCGGCTTTTTTTTTTCCGGACTGCNNNAGATCGGAAGA |
| 43820_4 | TATGAGTGTGGAGTCGTTGCGGCTTTTTTTTTTCCGGACTGCNNNAGATCGGAAGA |
| 43820_5 | TATGAGTGTGGAGTCGTTGCGGCTTTTTTTTTTCCGGACTGCNNNAGATCGGAAGA |
| 43820_6 | TATGAGTGTGGAGTCGTTGCGGCTTTTTTTTTTCCGGACTGCNNNAGATCGGAAGA |
| 38148_1 | TATGAGTGTGGAGTCGTTGCGCTCAGACAGGAGAATCACTTGANNNAGATCGGAAC |
| 38148_2 | TATGAGTGTGGAGTCGTTGCGCTCAGACAGGAGAATCACTTGANNNAGATCGGAAC |
| 38148_3 | TATGAGTGTGGAGTCGTTGCGCTCAGACAGGAGAATCACTTGANNNAGATCGGAAC |
| 38148_4 | TATGAGTGTGGAGTCGTTGCGCTCAGACAGGAGAATCACTTGANNNAGATCGGAAC |
| 38148_5 | TATGAGTGTGGAGTCGTTGCGCTCAGACAGGAGAATCACTTGANNNAGATCGGAAC |
| 38148_6 | TATGAGTGTGGAGTCGTTGCGCTCAGACAGGAGAATCACTTGANNNAGATCGGAAC |
| 10828_1 | TATGAGTGTGGAGTCGTTGCGAGACTGTGTGTGTCTGTGCATANNNAGATCGGAAC |
| 10828_2 | TATGAGTGTGGAGTCGTTGCGAGACTGTGTGTGTCTGTGCATANNNAGATCGGAAC |
| 10828_3 | TATGAGTGTGGAGTCGTTGCGAGACTGTGTGTGTCTGTGCATANNNAGATCGGAAC |
| 10828_4 | TATGAGTGTGGAGTCGTTGCGAGACTGTGTGTGTCTGTGCATANNNAGATCGGAAC |
| 10828_5 | TATGAGTGTGGAGTCGTTGCGAGACTGTGTGTGTCTGTGCATANNNAGATCGGAAC |
| 10828_6 | TATGAGTGTGGAGTCGTTGCGAGACTGTGTGTGTCTGTGCATANNNAGATCGGAAC |
| 44711_1 | TATGAGTGTGGAGTCGTTGCCTGTCCCTTTTTCCTGAGTAGCTNNNAGATCGGAAGA |
| 44711_2 | TATGAGTGTGGAGTCGTTGCCTGTCCCTTTTTCCTGAGTAGCTNNNAGATCGGAAGA |
| 44711_3 | TATGAGTGTGGAGTCGTTGCCTGTCCCTTTTTCCTGAGTAGCTNNNAGATCGGAAGA |
| 44711_4 | TATGAGTGTGGAGTCGTTGCCTGTCCCTTTTTCCTGAGTAGCTNNNAGATCGGAAGA |
| 44711_5 | TATGAGTGTGGAGTCGTTGCCTGTCCCTTTTTCCTGAGTAGCTNNNAGATCGGAAGA |
| 44711_6 | TATGAGTGTGGAGTCGTTGCCTGTCCCTTTTTCCTGAGTAGCTNNNAGATCGGAAGA |
| 41657_1 | TATGAGTGTGGAGTCGTTGCCAAGTGATTCTCTTGCTTCAGCCNNNAGATCGGAAGA |

| | |
|---|---|
| 41657_2 | TATGAGTGTGGAGTCGTTGCCAAGTGATTCTCTTGCTTCAGCCNNNAGATCGGAAG |
| 41657_3 | TATGAGTGTGGAGTCGTTGCCAAGTGATTCTCTTGCTTCAGCCNNNAGATCGGAAG |
| 41657_4 | TATGAGTGTGGAGTCGTTGCCAAGTGATTCTCTTGCTTCAGCCNNNAGATCGGAAG |
| 41657_5 | TATGAGTGTGGAGTCGTTGCCAAGTGATTCTCTTGCTTCAGCCNNNAGATCGGAAG |
| 41657_6 | TATGAGTGTGGAGTCGTTGCCAAGTGATTCTCTTGCTTCAGCCNNNAGATCGGAAG |
| 43384_1 | TATGAGTGTGGAGTCGTTGCCCAAGAGCAGTGATCCTCCTGNNNAGATCGGAAGA |
| 43384_2 | TATGAGTGTGGAGTCGTTGCCCAAGAGCAGTGATCCTCCTGNNNAGATCGGAAGA |
| 43384_3 | TATGAGTGTGGAGTCGTTGCCCAAGAGCAGTGATCCTCCTGNNNAGATCGGAAGA |
| 43384_4 | TATGAGTGTGGAGTCGTTGCCCAAGAGCAGTGATCCTCCTGNNNAGATCGGAAGA |
| 43384_5 | TATGAGTGTGGAGTCGTTGCCCAAGAGCAGTGATCCTCCTGNNNAGATCGGAAGA |
| 43384_6 | TATGAGTGTGGAGTCGTTGCCCAAGAGCAGTGATCCTCCTGNNNAGATCGGAAGA |
| 28787_1 | TATGAGTGTGGAGTCGTTGCTATACACACACACACACACACACNNNAGATCGGAAG |
| 28787_2 | TATGAGTGTGGAGTCGTTGCTATACACACACACACACACACACNNNAGATCGGAAG |
| 28787_3 | TATGAGTGTGGAGTCGTTGCTATACACACACACACACACACACNNNAGATCGGAAG |
| 28787_4 | TATGAGTGTGGAGTCGTTGCTATACACACACACACACACACACNNNAGATCGGAAG |
| 28787_5 | TATGAGTGTGGAGTCGTTGCTATACACACACACACACACACACNNNAGATCGGAAG |
| 28787_6 | TATGAGTGTGGAGTCGTTGCTATACACACACACACACACACACNNNAGATCGGAAG |
| 12821_1 | TATGAGTGTGGAGTCGTTGCCCTAGGAGAGAATGGGTTTGNNNAGATCGGAAGA |
| 12821_2 | TATGAGTGTGGAGTCGTTGCCCTAGGAGAGAATGGGTTTGNNNAGATCGGAAGA |
| 12821_3 | TATGAGTGTGGAGTCGTTGCCCTAGGAGAGAATGGGTTTGNNNAGATCGGAAGA |
| 12821_4 | TATGAGTGTGGAGTCGTTGCCCTAGGAGAGAATGGGTTTGNNNAGATCGGAAGA |
| 12821_5 | TATGAGTGTGGAGTCGTTGCCCTAGGAGAGAATGGGTTTGNNNAGATCGGAAGA |
| 12821_6 | TATGAGTGTGGAGTCGTTGCCCTAGGAGAGAATGGGTTTGNNNAGATCGGAAGA |
| 39489_1 | TATGAGTGTGGAGTCGTTGCGATACCAGAGTACCAGAGATGCGNNNAGATCGGAA |
| 39489_2 | TATGAGTGTGGAGTCGTTGCGATACCAGAGTACCAGAGATGCGNNNAGATCGGAA |
| 39489_3 | TATGAGTGTGGAGTCGTTGCGATACCAGAGTACCAGAGATGCGNNNAGATCGGAA |
| 39489_4 | TATGAGTGTGGAGTCGTTGCGATACCAGAGTACCAGAGATGCGNNNAGATCGGAA |
| 39489_5 | TATGAGTGTGGAGTCGTTGCGATACCAGAGTACCAGAGATGCGNNNAGATCGGAA |
| 39489_6 | TATGAGTGTGGAGTCGTTGCGATACCAGAGTACCAGAGATGCGNNNAGATCGGAA |
| 38828_1 | TATGAGTGTGGAGTCGTTGCGGGTGGGAGGAAGTAAGCATAAANNNAGATCGGA |
| 38828_2 | TATGAGTGTGGAGTCGTTGCGGGTGGGAGGAAGTAAGCATAAANNNAGATCGGA |
| 38828_3 | TATGAGTGTGGAGTCGTTGCGGGTGGGAGGAAGTAAGCATAAANNNAGATCGGA |
| 38828_4 | TATGAGTGTGGAGTCGTTGCGGGTGGGAGGAAGTAAGCATAAANNNAGATCGGA |
| 38828_5 | TATGAGTGTGGAGTCGTTGCGGGTGGGAGGAAGTAAGCATAAANNNAGATCGGA |
| 38828_6 | TATGAGTGTGGAGTCGTTGCGGGTGGGAGGAAGTAAGCATAAANNNAGATCGGA |
| 10632_1 | TATGAGTGTGGAGTCGTTGCGAGGTCCAACTGAACAGCTCAATNNNAGATCGGAA |
| 10632_2 | TATGAGTGTGGAGTCGTTGCGAGGTCCAACTGAACAGCTCAATNNNAGATCGGAA |
| 10632_3 | TATGAGTGTGGAGTCGTTGCGAGGTCCAACTGAACAGCTCAATNNNAGATCGGAA |
| 10632_4 | TATGAGTGTGGAGTCGTTGCGAGGTCCAACTGAACAGCTCAATNNNAGATCGGAA |
| 10632_5 | TATGAGTGTGGAGTCGTTGCGAGGTCCAACTGAACAGCTCAATNNNAGATCGGAA |
| 10632_6 | TATGAGTGTGGAGTCGTTGCGAGGTCCAACTGAACAGCTCAATNNNAGATCGGAA |
| 2878_1 | TATGAGTGTGGAGTCGTTGCTAATCTGCTGAGTGGAGGGCATANNNAGATCGGAA |
| 2878_2 | TATGAGTGTGGAGTCGTTGCTAATCTGCTGAGTGGAGGGCATANNNAGATCGGAA |

| | |
|---|---|
| 2878_3 | TATGAGTGTGGAGTCGTTGCTAATCTGCTGAGTGGAGGGCATANNNAGATCGGAA |
| 2878_4 | TATGAGTGTGGAGTCGTTGCTAATCTGCTGAGTGGAGGGCATANNNAGATCGGAA |
| 2878_5 | TATGAGTGTGGAGTCGTTGCTAATCTGCTGAGTGGAGGGCATANNNAGATCGGAA |
| 2878_6 | TATGAGTGTGGAGTCGTTGCTAATCTGCTGAGTGGAGGGCATANNNAGATCGGAA |
| 47643_1 | TATGAGTGTGGAGTCGTTGCGGTTCCTGCAGCATATACCACTANNNAGATCGGAAG |
| 47643_2 | TATGAGTGTGGAGTCGTTGCGGTTCCTGCAGCATATACCACTANNNAGATCGGAAG |
| 47643_3 | TATGAGTGTGGAGTCGTTGCGGTTCCTGCAGCATATACCACTANNNAGATCGGAAG |
| 47643_4 | TATGAGTGTGGAGTCGTTGCGGTTCCTGCAGCATATACCACTANNNAGATCGGAAG |
| 47643_5 | TATGAGTGTGGAGTCGTTGCGGTTCCTGCAGCATATACCACTANNNAGATCGGAAG |
| 47643_6 | TATGAGTGTGGAGTCGTTGCGGTTCCTGCAGCATATACCACTANNNAGATCGGAAG |
| 33744_1 | TATGAGTGTGGAGTCGTTGCAAAAGAAAAGAGAAACCCACGCGNNNAGATCGGAA |
| 33744_2 | TATGAGTGTGGAGTCGTTGCAAAAGAAAAGAGAAACCCACGCGNNNAGATCGGAA |
| 33744_3 | TATGAGTGTGGAGTCGTTGCAAAAGAAAAGAGAAACCCACGCGNNNAGATCGGAA |
| 33744_4 | TATGAGTGTGGAGTCGTTGCAAAAGAAAAGAGAAACCCACGCGNNNAGATCGGAA |
| 33744_5 | TATGAGTGTGGAGTCGTTGCAAAAGAAAAGAGAAACCCACGCGNNNAGATCGGAA |
| 33744_6 | TATGAGTGTGGAGTCGTTGCAAAAGAAAAGAGAAACCCACGCGNNNAGATCGGAA |
| 44678_1 | TATGAGTGTGGAGTCGTTGCACTTTGAAGTCCAGTTGAGTCCANNNAGATCGGAAG |
| 44678_2 | TATGAGTGTGGAGTCGTTGCACTTTGAAGTCCAGTTGAGTCCANNNAGATCGGAAG |
| 44678_3 | TATGAGTGTGGAGTCGTTGCACTTTGAAGTCCAGTTGAGTCCANNNAGATCGGAAG |
| 44678_4 | TATGAGTGTGGAGTCGTTGCACTTTGAAGTCCAGTTGAGTCCANNNAGATCGGAAG |
| 44678_5 | TATGAGTGTGGAGTCGTTGCACTTTGAAGTCCAGTTGAGTCCANNNAGATCGGAAG |
| 44678_6 | TATGAGTGTGGAGTCGTTGCACTTTGAAGTCCAGTTGAGTCCANNNAGATCGGAAG |
| 40075_1 | TATGAGTGTGGAGTCGTTGCCAATGGGATTTTTGGGGCAGTAANNNAGATCGGAA |
| 40075_2 | TATGAGTGTGGAGTCGTTGCCAATGGGATTTTTGGGGCAGTAANNNAGATCGGAA |
| 40075_3 | TATGAGTGTGGAGTCGTTGCCAATGGGATTTTTGGGGCAGTAANNNAGATCGGAA |
| 40075_4 | TATGAGTGTGGAGTCGTTGCCAATGGGATTTTTGGGGCAGTAANNNAGATCGGAA |
| 40075_5 | TATGAGTGTGGAGTCGTTGCCAATGGGATTTTTGGGGCAGTAANNNAGATCGGAA |
| 40075_6 | TATGAGTGTGGAGTCGTTGCCAATGGGATTTTTGGGGCAGTAANNNAGATCGGAA |
| 51769_1 | TATGAGTGTGGAGTCGTTGCTCACCAATCTCAGTGCTATTGCTNNNAGATCGGAAG |
| 51769_2 | TATGAGTGTGGAGTCGTTGCTCACCAATCTCAGTGCTATTGCTNNNAGATCGGAAG |
| 51769_3 | TATGAGTGTGGAGTCGTTGCTCACCAATCTCAGTGCTATTGCTNNNAGATCGGAAG |
| 51769_4 | TATGAGTGTGGAGTCGTTGCTCACCAATCTCAGTGCTATTGCTNNNAGATCGGAAG |
| 51769_5 | TATGAGTGTGGAGTCGTTGCTCACCAATCTCAGTGCTATTGCTNNNAGATCGGAAG |
| 51769_6 | TATGAGTGTGGAGTCGTTGCTCACCAATCTCAGTGCTATTGCTNNNAGATCGGAAG |
| 45236_1 | TATGAGTGTGGAGTCGTTGCGGAGATGGGGTAGGTGAGGAAANNNAGATCGGA |
| 45236_2 | TATGAGTGTGGAGTCGTTGCGGAGATGGGGTAGGTGAGGAAANNNAGATCGGA |
| 45236_3 | TATGAGTGTGGAGTCGTTGCGGAGATGGGGTAGGTGAGGAAANNNAGATCGGA |
| 45236_4 | TATGAGTGTGGAGTCGTTGCGGAGATGGGGTAGGTGAGGAAANNNAGATCGGA |
| 45236_5 | TATGAGTGTGGAGTCGTTGCGGAGATGGGGTAGGTGAGGAAANNNAGATCGGA |
| 45236_6 | TATGAGTGTGGAGTCGTTGCGGAGATGGGGTAGGTGAGGAAANNNAGATCGGA |
| 52166_1 | TATGAGTGTGGAGTCGTTGCTTTGCTGAGCATTTCTCTCTCCTNNNAGATCGGAAGA |
| 52166_2 | TATGAGTGTGGAGTCGTTGCTTTGCTGAGCATTTCTCTCTCCTNNNAGATCGGAAGA |
| 52166_3 | TATGAGTGTGGAGTCGTTGCTTTGCTGAGCATTTCTCTCTCCTNNNAGATCGGAAGA |

| | |
|---|---|
| 52166_4 | TATGAGTGTGGAGTCGTTGCTTTGCTGAGCATTTCTCTCTCCTNNNAGATCGGAAGA |
| 52166_5 | TATGAGTGTGGAGTCGTTGCTTTGCTGAGCATTTCTCTCTCCTNNNAGATCGGAAGA |
| 52166_6 | TATGAGTGTGGAGTCGTTGCTTTGCTGAGCATTTCTCTCTCCTNNNAGATCGGAAGA |
| 35676_1 | TATGAGTGTGGAGTCGTTGCCTTGATCTCCTGACCTCGTGATCNNNAGATCGGAAGA |
| 35676_2 | TATGAGTGTGGAGTCGTTGCCTTGATCTCCTGACCTCGTGATCNNNAGATCGGAAGA |
| 35676_3 | TATGAGTGTGGAGTCGTTGCCTTGATCTCCTGACCTCGTGATCNNNAGATCGGAAGA |
| 35676_4 | TATGAGTGTGGAGTCGTTGCCTTGATCTCCTGACCTCGTGATCNNNAGATCGGAAGA |
| 35676_5 | TATGAGTGTGGAGTCGTTGCCTTGATCTCCTGACCTCGTGATCNNNAGATCGGAAGA |
| 35676_6 | TATGAGTGTGGAGTCGTTGCCTTGATCTCCTGACCTCGTGATCNNNAGATCGGAAGA |
| 35977_1 | TATGAGTGTGGAGTCGTTGCTGAGGGAGTAGATTCGCTTGAACNNNAGATCGGAA |
| 35977_2 | TATGAGTGTGGAGTCGTTGCTGAGGGAGTAGATTCGCTTGAACNNNAGATCGGAA |
| 35977_3 | TATGAGTGTGGAGTCGTTGCTGAGGGAGTAGATTCGCTTGAACNNNAGATCGGAA |
| 35977_4 | TATGAGTGTGGAGTCGTTGCTGAGGGAGTAGATTCGCTTGAACNNNAGATCGGAA |
| 35977_5 | TATGAGTGTGGAGTCGTTGCTGAGGGAGTAGATTCGCTTGAACNNNAGATCGGAA |
| 35977_6 | TATGAGTGTGGAGTCGTTGCTGAGGGAGTAGATTCGCTTGAACNNNAGATCGGAA |
| 50847_1 | TATGAGTGTGGAGTCGTTGCAAGAAAGAGATCTGGGAGTCGTGNNNAGATCGGAA |
| 50847_2 | TATGAGTGTGGAGTCGTTGCAAGAAAGAGATCTGGGAGTCGTGNNNAGATCGGAA |
| 50847_3 | TATGAGTGTGGAGTCGTTGCAAGAAAGAGATCTGGGAGTCGTGNNNAGATCGGAA |
| 50847_4 | TATGAGTGTGGAGTCGTTGCAAGAAAGAGATCTGGGAGTCGTGNNNAGATCGGAA |
| 50847_5 | TATGAGTGTGGAGTCGTTGCAAGAAAGAGATCTGGGAGTCGTGNNNAGATCGGAA |
| 50847_6 | TATGAGTGTGGAGTCGTTGCAAGAAAGAGATCTGGGAGTCGTGNNNAGATCGGAA |
| 45139_1 | TATGAGTGTGGAGTCGTTGCTGTGGAACAGAACAAGCAGAATGNNNAGATCGGAA |
| 45139_2 | TATGAGTGTGGAGTCGTTGCTGTGGAACAGAACAAGCAGAATGNNNAGATCGGAA |
| 45139_3 | TATGAGTGTGGAGTCGTTGCTGTGGAACAGAACAAGCAGAATGNNNAGATCGGAA |
| 45139_4 | TATGAGTGTGGAGTCGTTGCTGTGGAACAGAACAAGCAGAATGNNNAGATCGGAA |
| 45139_5 | TATGAGTGTGGAGTCGTTGCTGTGGAACAGAACAAGCAGAATGNNNAGATCGGAA |
| 45139_6 | TATGAGTGTGGAGTCGTTGCTGTGGAACAGAACAAGCAGAATGNNNAGATCGGAA |
| 35551_1 | TATGAGTGTGGAGTCGTTGCTGGGAATAATGACCTGCAGAACANNNAGATCGGAA |
| 35551_2 | TATGAGTGTGGAGTCGTTGCTGGGAATAATGACCTGCAGAACANNNAGATCGGAA |
| 35551_3 | TATGAGTGTGGAGTCGTTGCTGGGAATAATGACCTGCAGAACANNNAGATCGGAA |
| 35551_4 | TATGAGTGTGGAGTCGTTGCTGGGAATAATGACCTGCAGAACANNNAGATCGGAA |
| 35551_5 | TATGAGTGTGGAGTCGTTGCTGGGAATAATGACCTGCAGAACANNNAGATCGGAA |
| 35551_6 | TATGAGTGTGGAGTCGTTGCTGGGAATAATGACCTGCAGAACANNNAGATCGGAA |
| 36618_1 | TATGAGTGTGGAGTCGTTGCAAGTTCAAGAACAAGGAAAGCCANNNAGATCGGAA |
| 36618_2 | TATGAGTGTGGAGTCGTTGCAAGTTCAAGAACAAGGAAAGCCANNNAGATCGGAA |
| 36618_3 | TATGAGTGTGGAGTCGTTGCAAGTTCAAGAACAAGGAAAGCCANNNAGATCGGAA |
| 36618_4 | TATGAGTGTGGAGTCGTTGCAAGTTCAAGAACAAGGAAAGCCANNNAGATCGGAA |
| 36618_5 | TATGAGTGTGGAGTCGTTGCAAGTTCAAGAACAAGGAAAGCCANNNAGATCGGAA |
| 36618_6 | TATGAGTGTGGAGTCGTTGCAAGTTCAAGAACAAGGAAAGCCANNNAGATCGGAA |
| 36236_1 | TATGAGTGTGGAGTCGTTGCACTGACAAAGTGAAAAGGAAGCANNNAGATCGGAA |
| 36236_2 | TATGAGTGTGGAGTCGTTGCACTGACAAAGTGAAAAGGAAGCANNNAGATCGGAA |
| 36236_3 | TATGAGTGTGGAGTCGTTGCACTGACAAAGTGAAAAGGAAGCANNNAGATCGGAA |
| 36236_4 | TATGAGTGTGGAGTCGTTGCACTGACAAAGTGAAAAGGAAGCANNNAGATCGGAA |

| | |
|---|---|
| 36236_5 | TATGAGTGTGGAGTCGTTGCACTGACAAAGTGAAAAGGAAGCANNNAGATCGGAA |
| 36236_6 | TATGAGTGTGGAGTCGTTGCACTGACAAAGTGAAAAGGAAGCANNNAGATCGGAA |
| 49604_1 | TATGAGTGTGGAGTCGTTGCTACTATCTCAGCTCACTGCAACCNNNAGATCGGAAGA |
| 49604_2 | TATGAGTGTGGAGTCGTTGCTACTATCTCAGCTCACTGCAACCNNNAGATCGGAAGA |
| 49604_3 | TATGAGTGTGGAGTCGTTGCTACTATCTCAGCTCACTGCAACCNNNAGATCGGAAGA |
| 49604_4 | TATGAGTGTGGAGTCGTTGCTACTATCTCAGCTCACTGCAACCNNNAGATCGGAAGA |
| 49604_5 | TATGAGTGTGGAGTCGTTGCTACTATCTCAGCTCACTGCAACCNNNAGATCGGAAGA |
| 49604_6 | TATGAGTGTGGAGTCGTTGCTACTATCTCAGCTCACTGCAACCNNNAGATCGGAAGA |
| 7537_1 | TATGAGTGTGGAGTCGTTGCTCAACCCCCACTCTTGAGAACATGNNNAGATCGGAAG |
| 7537_2 | TATGAGTGTGGAGTCGTTGCTCAACCCCCACTCTTGAGAACATGNNNAGATCGGAAG |
| 7537_3 | TATGAGTGTGGAGTCGTTGCTCAACCCCCACTCTTGAGAACATGNNNAGATCGGAAG |
| 7537_4 | TATGAGTGTGGAGTCGTTGCTCAACCCCCACTCTTGAGAACATGNNNAGATCGGAAG |
| 7537_5 | TATGAGTGTGGAGTCGTTGCTCAACCCCCACTCTTGAGAACATGNNNAGATCGGAAG |
| 7537_6 | TATGAGTGTGGAGTCGTTGCTCAACCCCCACTCTTGAGAACATGNNNAGATCGGAAG |
| 33255_1 | TATGAGTGTGGAGTCGTTGCATCTTGGCTCAATGCAACCTCTANNNAGATCGGAAGA |
| 33255_2 | TATGAGTGTGGAGTCGTTGCATCTTGGCTCAATGCAACCTCTANNNAGATCGGAAGA |
| 33255_3 | TATGAGTGTGGAGTCGTTGCATCTTGGCTCAATGCAACCTCTANNNAGATCGGAAGA |
| 33255_4 | TATGAGTGTGGAGTCGTTGCATCTTGGCTCAATGCAACCTCTANNNAGATCGGAAGA |
| 33255_5 | TATGAGTGTGGAGTCGTTGCATCTTGGCTCAATGCAACCTCTANNNAGATCGGAAGA |
| 33255_6 | TATGAGTGTGGAGTCGTTGCATCTTGGCTCAATGCAACCTCTANNNAGATCGGAAGA |
| 5437_1 | TATGAGTGTGGAGTCGTTGCTTGAGACATTACCACCACCTTGGNNNAGATCGGAAGA |
| 5437_2 | TATGAGTGTGGAGTCGTTGCTTGAGACATTACCACCACCTTGGNNNAGATCGGAAGA |
| 5437_3 | TATGAGTGTGGAGTCGTTGCTTGAGACATTACCACCACCTTGGNNNAGATCGGAAGA |
| 5437_4 | TATGAGTGTGGAGTCGTTGCTTGAGACATTACCACCACCTTGGNNNAGATCGGAAGA |
| 5437_5 | TATGAGTGTGGAGTCGTTGCTTGAGACATTACCACCACCTTGGNNNAGATCGGAAGA |
| 5437_6 | TATGAGTGTGGAGTCGTTGCTTGAGACATTACCACCACCTTGGNNNAGATCGGAAGA |
| 48601_1 | TATGAGTGTGGAGTCGTTGCTTGTTGGTAGTGGGATTTGGAGTNNNAGATCGGAA |
| 48601_2 | TATGAGTGTGGAGTCGTTGCTTGTTGGTAGTGGGATTTGGAGTNNNAGATCGGAA |
| 48601_3 | TATGAGTGTGGAGTCGTTGCTTGTTGGTAGTGGGATTTGGAGTNNNAGATCGGAA |
| 48601_4 | TATGAGTGTGGAGTCGTTGCTTGTTGGTAGTGGGATTTGGAGTNNNAGATCGGAA |
| 48601_5 | TATGAGTGTGGAGTCGTTGCTTGTTGGTAGTGGGATTTGGAGTNNNAGATCGGAA |
| 48601_6 | TATGAGTGTGGAGTCGTTGCTTGTTGGTAGTGGGATTTGGAGTNNNAGATCGGAA |
| 52677_1 | TATGAGTGTGGAGTCGTTGCCCCCTCCTTTCCCTTCAACTTTANNNAGATCGGAAGAG |
| 52677_2 | TATGAGTGTGGAGTCGTTGCCCCCTCCTTTCCCTTCAACTTTANNNAGATCGGAAGAG |
| 52677_3 | TATGAGTGTGGAGTCGTTGCCCCCTCCTTTCCCTTCAACTTTANNNAGATCGGAAGAG |
| 52677_4 | TATGAGTGTGGAGTCGTTGCCCCCTCCTTTCCCTTCAACTTTANNNAGATCGGAAGAG |
| 52677_5 | TATGAGTGTGGAGTCGTTGCCCCCTCCTTTCCCTTCAACTTTANNNAGATCGGAAGAG |
| 52677_6 | TATGAGTGTGGAGTCGTTGCCCCCTCCTTTCCCTTCAACTTTANNNAGATCGGAAGAG |
| 38328_1 | TATGAGTGTGGAGTCGTTGCCTACTCTGCAAATGTGATGGCTGNNNAGATCGGAAG |
| 38328_2 | TATGAGTGTGGAGTCGTTGCCTACTCTGCAAATGTGATGGCTGNNNAGATCGGAAG |
| 38328_3 | TATGAGTGTGGAGTCGTTGCCTACTCTGCAAATGTGATGGCTGNNNAGATCGGAAG |
| 38328_4 | TATGAGTGTGGAGTCGTTGCCTACTCTGCAAATGTGATGGCTGNNNAGATCGGAAG |
| 38328_5 | TATGAGTGTGGAGTCGTTGCCTACTCTGCAAATGTGATGGCTGNNNAGATCGGAAG |

| | |
|---|---|
| 38328_6 | TATGAGTGTGGAGTCGTTGCCTACTCTGCAAATGTGATGGCTGNNNAGATCGGAAG |
| 41954_1 | TATGAGTGTGGAGTCGTTGCGTAGCTGGGACTACAGGCATGNNNAGATCGGAAGA |
| 41954_2 | TATGAGTGTGGAGTCGTTGCGTAGCTGGGACTACAGGCATGNNNAGATCGGAAGA |
| 41954_3 | TATGAGTGTGGAGTCGTTGCGTAGCTGGGACTACAGGCATGNNNAGATCGGAAGA |
| 41954_4 | TATGAGTGTGGAGTCGTTGCGTAGCTGGGACTACAGGCATGNNNAGATCGGAAGA |
| 41954_5 | TATGAGTGTGGAGTCGTTGCGTAGCTGGGACTACAGGCATGNNNAGATCGGAAGA |
| 41954_6 | TATGAGTGTGGAGTCGTTGCGTAGCTGGGACTACAGGCATGNNNAGATCGGAAGA |
| 50125_1 | TATGAGTGTGGAGTCGTTGCAACAAGCCAAGTAATTTGAGCCCNNNAGATCGGAAG |
| 50125_2 | TATGAGTGTGGAGTCGTTGCAACAAGCCAAGTAATTTGAGCCCNNNAGATCGGAAG |
| 50125_3 | TATGAGTGTGGAGTCGTTGCAACAAGCCAAGTAATTTGAGCCCNNNAGATCGGAAG |
| 50125_4 | TATGAGTGTGGAGTCGTTGCAACAAGCCAAGTAATTTGAGCCCNNNAGATCGGAAG |
| 50125_5 | TATGAGTGTGGAGTCGTTGCAACAAGCCAAGTAATTTGAGCCCNNNAGATCGGAAG |
| 50125_6 | TATGAGTGTGGAGTCGTTGCAACAAGCCAAGTAATTTGAGCCCNNNAGATCGGAAG |
| 36981_1 | TATGAGTGTGGAGTCGTTGCGAGAATCGCTTGAACCAGGGAGNNNAGATCGGAAG |
| 36981_2 | TATGAGTGTGGAGTCGTTGCGAGAATCGCTTGAACCAGGGAGNNNAGATCGGAAG |
| 36981_3 | TATGAGTGTGGAGTCGTTGCGAGAATCGCTTGAACCAGGGAGNNNAGATCGGAAG |
| 36981_4 | TATGAGTGTGGAGTCGTTGCGAGAATCGCTTGAACCAGGGAGNNNAGATCGGAAG |
| 36981_5 | TATGAGTGTGGAGTCGTTGCGAGAATCGCTTGAACCAGGGAGNNNAGATCGGAAG |
| 36981_6 | TATGAGTGTGGAGTCGTTGCGAGAATCGCTTGAACCAGGGAGNNNAGATCGGAAG |
| 51701_1 | TATGAGTGTGGAGTCGTTGCCCTCTTGAAGCACGTAACCAATGNNNAGATCGGAAG |
| 51701_2 | TATGAGTGTGGAGTCGTTGCCCTCTTGAAGCACGTAACCAATGNNNAGATCGGAAG |
| 51701_3 | TATGAGTGTGGAGTCGTTGCCCTCTTGAAGCACGTAACCAATGNNNAGATCGGAAG |
| 51701_4 | TATGAGTGTGGAGTCGTTGCCCTCTTGAAGCACGTAACCAATGNNNAGATCGGAAG |
| 51701_5 | TATGAGTGTGGAGTCGTTGCCCTCTTGAAGCACGTAACCAATGNNNAGATCGGAAG |
| 51701_6 | TATGAGTGTGGAGTCGTTGCCCTCTTGAAGCACGTAACCAATGNNNAGATCGGAAG |
| 52482_1 | TATGAGTGTGGAGTCGTTGCCACCCCCTGTACGACTGAAGNNNAGATCGGAAGAGC |
| 52482_2 | TATGAGTGTGGAGTCGTTGCCACCCCCTGTACGACTGAAGNNNAGATCGGAAGAGC |
| 52482_3 | TATGAGTGTGGAGTCGTTGCCACCCCCTGTACGACTGAAGNNNAGATCGGAAGAGC |
| 52482_4 | TATGAGTGTGGAGTCGTTGCCACCCCCTGTACGACTGAAGNNNAGATCGGAAGAGC |
| 52482_5 | TATGAGTGTGGAGTCGTTGCCACCCCCTGTACGACTGAAGNNNAGATCGGAAGAGC |
| 52482_6 | TATGAGTGTGGAGTCGTTGCCACCCCCTGTACGACTGAAGNNNAGATCGGAAGAGC |
| 40485_1 | TATGAGTGTGGAGTCGTTGCCCATGTCGGGCACTTTACCTATANNNAGATCGGAAG/ |
| 40485_2 | TATGAGTGTGGAGTCGTTGCCCATGTCGGGCACTTTACCTATANNNAGATCGGAAG/ |
| 40485_3 | TATGAGTGTGGAGTCGTTGCCCATGTCGGGCACTTTACCTATANNNAGATCGGAAG/ |
| 40485_4 | TATGAGTGTGGAGTCGTTGCCCATGTCGGGCACTTTACCTATANNNAGATCGGAAG/ |
| 40485_5 | TATGAGTGTGGAGTCGTTGCCCATGTCGGGCACTTTACCTATANNNAGATCGGAAG/ |
| 40485_6 | TATGAGTGTGGAGTCGTTGCCCATGTCGGGCACTTTACCTATANNNAGATCGGAAG/ |
| 35185_1 | TATGAGTGTGGAGTCGTTGCTTTTGGCTCACTGTAACCTCAGANNNAGATCGGAAG/ |
| 35185_2 | TATGAGTGTGGAGTCGTTGCTTTTGGCTCACTGTAACCTCAGANNNAGATCGGAAG/ |
| 35185_3 | TATGAGTGTGGAGTCGTTGCTTTTGGCTCACTGTAACCTCAGANNNAGATCGGAAG/ |
| 35185_4 | TATGAGTGTGGAGTCGTTGCTTTTGGCTCACTGTAACCTCAGANNNAGATCGGAAG/ |
| 35185_5 | TATGAGTGTGGAGTCGTTGCTTTTGGCTCACTGTAACCTCAGANNNAGATCGGAAG/ |
| 35185_6 | TATGAGTGTGGAGTCGTTGCTTTTGGCTCACTGTAACCTCAGANNNAGATCGGAAG/ |

| | |
|---|---|
| 35285_1 | TATGAGTGTGGAGTCGTTGCTTCAAGCTATTCTCCTGCCTCTGNNNAGATCGGAAGA |
| 35285_2 | TATGAGTGTGGAGTCGTTGCTTCAAGCTATTCTCCTGCCTCTGNNNAGATCGGAAGA |
| 35285_3 | TATGAGTGTGGAGTCGTTGCTTCAAGCTATTCTCCTGCCTCTGNNNAGATCGGAAGA |
| 35285_4 | TATGAGTGTGGAGTCGTTGCTTCAAGCTATTCTCCTGCCTCTGNNNAGATCGGAAGA |
| 35285_5 | TATGAGTGTGGAGTCGTTGCTTCAAGCTATTCTCCTGCCTCTGNNNAGATCGGAAGA |
| 35285_6 | TATGAGTGTGGAGTCGTTGCTTCAAGCTATTCTCCTGCCTCTGNNNAGATCGGAAGA |
| 12582_1 | TATGAGTGTGGAGTCGTTGCCTTCTTCTTCATCACTTTCCTTNNNAGATCGGAAGAG |
| 12582_2 | TATGAGTGTGGAGTCGTTGCCTTCTTCTTCATCACTTTCCTTNNNAGATCGGAAGAG |
| 12582_3 | TATGAGTGTGGAGTCGTTGCCTTCTTCTTCATCACTTTCCTTNNNAGATCGGAAGAG |
| 12582_4 | TATGAGTGTGGAGTCGTTGCCTTCTTCTTCATCACTTTCCTTNNNAGATCGGAAGAG |
| 12582_5 | TATGAGTGTGGAGTCGTTGCCTTCTTCTTCATCACTTTCCTTNNNAGATCGGAAGAG |
| 12582_6 | TATGAGTGTGGAGTCGTTGCCTTCTTCTTCATCACTTTCCTTNNNAGATCGGAAGAG |
| 35119_1 | TATGAGTGTGGAGTCGTTGCGTTTCACCATGTTGGCCAAGATGNNNAGATCGGAAG |
| 35119_2 | TATGAGTGTGGAGTCGTTGCGTTTCACCATGTTGGCCAAGATGNNNAGATCGGAAG |
| 35119_3 | TATGAGTGTGGAGTCGTTGCGTTTCACCATGTTGGCCAAGATGNNNAGATCGGAAG |
| 35119_4 | TATGAGTGTGGAGTCGTTGCGTTTCACCATGTTGGCCAAGATGNNNAGATCGGAAG |
| 35119_5 | TATGAGTGTGGAGTCGTTGCGTTTCACCATGTTGGCCAAGATGNNNAGATCGGAAG |
| 35119_6 | TATGAGTGTGGAGTCGTTGCGTTTCACCATGTTGGCCAAGATGNNNAGATCGGAAG |
| 42097_1 | TATGAGTGTGGAGTCGTTGCTGGCTGCAGTGACACAGATAATANNNAGATCGGAA |
| 42097_2 | TATGAGTGTGGAGTCGTTGCTGGCTGCAGTGACACAGATAATANNNAGATCGGAA |
| 42097_3 | TATGAGTGTGGAGTCGTTGCTGGCTGCAGTGACACAGATAATANNNAGATCGGAA |
| 42097_4 | TATGAGTGTGGAGTCGTTGCTGGCTGCAGTGACACAGATAATANNNAGATCGGAA |
| 42097_5 | TATGAGTGTGGAGTCGTTGCTGGCTGCAGTGACACAGATAATANNNAGATCGGAA |
| 42097_6 | TATGAGTGTGGAGTCGTTGCTGGCTGCAGTGACACAGATAATANNNAGATCGGAA |
| 45285_1 | TATGAGTGTGGAGTCGTTGCTACACTCAAGACACAGTCATGCANNNAGATCGGAAG |
| 45285_2 | TATGAGTGTGGAGTCGTTGCTACACTCAAGACACAGTCATGCANNNAGATCGGAAG |
| 45285_3 | TATGAGTGTGGAGTCGTTGCTACACTCAAGACACAGTCATGCANNNAGATCGGAAG |
| 45285_4 | TATGAGTGTGGAGTCGTTGCTACACTCAAGACACAGTCATGCANNNAGATCGGAAG |
| 45285_5 | TATGAGTGTGGAGTCGTTGCTACACTCAAGACACAGTCATGCANNNAGATCGGAAG |
| 45285_6 | TATGAGTGTGGAGTCGTTGCTACACTCAAGACACAGTCATGCANNNAGATCGGAAG |
| 7277_1 | TATGAGTGTGGAGTCGTTGCAAGAAAGGGGCGAGACCCGGTNNNAGATCGGAAGA |
| 7277_2 | TATGAGTGTGGAGTCGTTGCAAGAAAGGGGCGAGACCCGGTNNNAGATCGGAAGA |
| 7277_3 | TATGAGTGTGGAGTCGTTGCAAGAAAGGGGCGAGACCCGGTNNNAGATCGGAAGA |
| 7277_4 | TATGAGTGTGGAGTCGTTGCAAGAAAGGGGCGAGACCCGGTNNNAGATCGGAAGA |
| 7277_5 | TATGAGTGTGGAGTCGTTGCAAGAAAGGGGCGAGACCCGGTNNNAGATCGGAAGA |
| 7277_6 | TATGAGTGTGGAGTCGTTGCAAGAAAGGGGCGAGACCCGGTNNNAGATCGGAAGA |
| 45434_1 | TATGAGTGTGGAGTCGTTGCTGCAGTTGTCCAGTGGATTCTAGNNNAGATCGGAAG |
| 45434_2 | TATGAGTGTGGAGTCGTTGCTGCAGTTGTCCAGTGGATTCTAGNNNAGATCGGAAG |
| 45434_3 | TATGAGTGTGGAGTCGTTGCTGCAGTTGTCCAGTGGATTCTAGNNNAGATCGGAAG |
| 45434_4 | TATGAGTGTGGAGTCGTTGCTGCAGTTGTCCAGTGGATTCTAGNNNAGATCGGAAG |
| 45434_5 | TATGAGTGTGGAGTCGTTGCTGCAGTTGTCCAGTGGATTCTAGNNNAGATCGGAAG |
| 45434_6 | TATGAGTGTGGAGTCGTTGCTGCAGTTGTCCAGTGGATTCTAGNNNAGATCGGAAG |
| 39460_1 | TATGAGTGTGGAGTCGTTGCATGCCCACTTTCACCACTTCTATNNNAGATCGGAAGA |

| | |
|---|---|
| 39460_2 | TATGAGTGTGGAGTCGTTGCATGCCCACTTTCACCACTTCTATNNNAGATCGGAAGA |
| 39460_3 | TATGAGTGTGGAGTCGTTGCATGCCCACTTTCACCACTTCTATNNNAGATCGGAAGA |
| 39460_4 | TATGAGTGTGGAGTCGTTGCATGCCCACTTTCACCACTTCTATNNNAGATCGGAAGA |
| 39460_5 | TATGAGTGTGGAGTCGTTGCATGCCCACTTTCACCACTTCTATNNNAGATCGGAAGA |
| 39460_6 | TATGAGTGTGGAGTCGTTGCATGCCCACTTTCACCACTTCTATNNNAGATCGGAAGA |
| 33840_1 | TATGAGTGTGGAGTCGTTGCAGAGGTTGCGATGAGTCAAGATTNNNAGATCGGAA |
| 33840_2 | TATGAGTGTGGAGTCGTTGCAGAGGTTGCGATGAGTCAAGATTNNNAGATCGGAA |
| 33840_3 | TATGAGTGTGGAGTCGTTGCAGAGGTTGCGATGAGTCAAGATTNNNAGATCGGAA |
| 33840_4 | TATGAGTGTGGAGTCGTTGCAGAGGTTGCGATGAGTCAAGATTNNNAGATCGGAA |
| 33840_5 | TATGAGTGTGGAGTCGTTGCAGAGGTTGCGATGAGTCAAGATTNNNAGATCGGAA |
| 33840_6 | TATGAGTGTGGAGTCGTTGCAGAGGTTGCGATGAGTCAAGATTNNNAGATCGGAA |
| 37606_1 | TATGAGTGTGGAGTCGTTGCATGGCGGGCATCTATAATCTCAGNNNAGATCGGAAG |
| 37606_2 | TATGAGTGTGGAGTCGTTGCATGGCGGGCATCTATAATCTCAGNNNAGATCGGAAG |
| 37606_3 | TATGAGTGTGGAGTCGTTGCATGGCGGGCATCTATAATCTCAGNNNAGATCGGAAG |
| 37606_4 | TATGAGTGTGGAGTCGTTGCATGGCGGGCATCTATAATCTCAGNNNAGATCGGAAG |
| 37606_5 | TATGAGTGTGGAGTCGTTGCATGGCGGGCATCTATAATCTCAGNNNAGATCGGAAG |
| 37606_6 | TATGAGTGTGGAGTCGTTGCATGGCGGGCATCTATAATCTCAGNNNAGATCGGAAG |
| 30844_1 | TATGAGTGTGGAGTCGTTGCCAAATCAAGGCAAGGAGTTGGAGNNNAGATCGGAA |
| 30844_2 | TATGAGTGTGGAGTCGTTGCCAAATCAAGGCAAGGAGTTGGAGNNNAGATCGGAA |
| 30844_3 | TATGAGTGTGGAGTCGTTGCCAAATCAAGGCAAGGAGTTGGAGNNNAGATCGGAA |
| 30844_4 | TATGAGTGTGGAGTCGTTGCCAAATCAAGGCAAGGAGTTGGAGNNNAGATCGGAA |
| 30844_5 | TATGAGTGTGGAGTCGTTGCCAAATCAAGGCAAGGAGTTGGAGNNNAGATCGGAA |
| 30844_6 | TATGAGTGTGGAGTCGTTGCCAAATCAAGGCAAGGAGTTGGAGNNNAGATCGGAA |
| 13713_1 | TATGAGTGTGGAGTCGTTGCTTATTCCGTATTACATTTTTGCANNNAGATCGGAAGA |
| 13713_2 | TATGAGTGTGGAGTCGTTGCTTATTCCGTATTACATTTTTGCANNNAGATCGGAAGA |
| 13713_3 | TATGAGTGTGGAGTCGTTGCTTATTCCGTATTACATTTTTGCANNNAGATCGGAAGA |
| 13713_4 | TATGAGTGTGGAGTCGTTGCTTATTCCGTATTACATTTTTGCANNNAGATCGGAAGA |
| 13713_5 | TATGAGTGTGGAGTCGTTGCTTATTCCGTATTACATTTTTGCANNNAGATCGGAAGA |
| 13713_6 | TATGAGTGTGGAGTCGTTGCTTATTCCGTATTACATTTTTGCANNNAGATCGGAAGA |
| 40112_1 | TATGAGTGTGGAGTCGTTGCCCCCATCTCACTGACTGAATCTCNNNAGATCGGAAGA |
| 40112_2 | TATGAGTGTGGAGTCGTTGCCCCCATCTCACTGACTGAATCTCNNNAGATCGGAAGA |
| 40112_3 | TATGAGTGTGGAGTCGTTGCCCCCATCTCACTGACTGAATCTCNNNAGATCGGAAGA |
| 40112_4 | TATGAGTGTGGAGTCGTTGCCCCCATCTCACTGACTGAATCTCNNNAGATCGGAAGA |
| 40112_5 | TATGAGTGTGGAGTCGTTGCCCCCATCTCACTGACTGAATCTCNNNAGATCGGAAGA |
| 40112_6 | TATGAGTGTGGAGTCGTTGCCCCCATCTCACTGACTGAATCTCNNNAGATCGGAAGA |
| 41006_1 | TATGAGTGTGGAGTCGTTGCTTCACCTACACTGACAGCATGAANNNAGATCGGAAG |
| 41006_2 | TATGAGTGTGGAGTCGTTGCTTCACCTACACTGACAGCATGAANNNAGATCGGAAG |
| 41006_3 | TATGAGTGTGGAGTCGTTGCTTCACCTACACTGACAGCATGAANNNAGATCGGAAG |
| 41006_4 | TATGAGTGTGGAGTCGTTGCTTCACCTACACTGACAGCATGAANNNAGATCGGAAG |
| 41006_5 | TATGAGTGTGGAGTCGTTGCTTCACCTACACTGACAGCATGAANNNAGATCGGAAG |
| 41006_6 | TATGAGTGTGGAGTCGTTGCTTCACCTACACTGACAGCATGAANNNAGATCGGAAG |
| 32228_1 | TATGAGTGTGGAGTCGTTGCAGTTACAGCACCTAAAACCCCAANNNAGATCGGAAG |
| 32228_2 | TATGAGTGTGGAGTCGTTGCAGTTACAGCACCTAAAACCCCAANNNAGATCGGAAG |

| | |
|---|---|
| 32228_3 | TATGAGTGTGGAGTCGTTGCAGTTACAGCACCTAAAACCCCAANNNAGATCGGAAG |
| 32228_4 | TATGAGTGTGGAGTCGTTGCAGTTACAGCACCTAAAACCCCAANNNAGATCGGAAG |
| 32228_5 | TATGAGTGTGGAGTCGTTGCAGTTACAGCACCTAAAACCCCAANNNAGATCGGAAG |
| 32228_6 | TATGAGTGTGGAGTCGTTGCAGTTACAGCACCTAAAACCCCAANNNAGATCGGAAG |
| 36152_1 | TATGAGTGTGGAGTCGTTGCTATTCCATACACACCTGCTCACANNNAGATCGGAAGA |
| 36152_2 | TATGAGTGTGGAGTCGTTGCTATTCCATACACACCTGCTCACANNNAGATCGGAAGA |
| 36152_3 | TATGAGTGTGGAGTCGTTGCTATTCCATACACACCTGCTCACANNNAGATCGGAAGA |
| 36152_4 | TATGAGTGTGGAGTCGTTGCTATTCCATACACACCTGCTCACANNNAGATCGGAAGA |
| 36152_5 | TATGAGTGTGGAGTCGTTGCTATTCCATACACACCTGCTCACANNNAGATCGGAAGA |
| 36152_6 | TATGAGTGTGGAGTCGTTGCTATTCCATACACACCTGCTCACANNNAGATCGGAAGA |
| 49263_1 | TATGAGTGTGGAGTCGTTGCCAATCTCAGCTCACTGCAATCTGNNNAGATCGGAAG |
| 49263_2 | TATGAGTGTGGAGTCGTTGCCAATCTCAGCTCACTGCAATCTGNNNAGATCGGAAG |
| 49263_3 | TATGAGTGTGGAGTCGTTGCCAATCTCAGCTCACTGCAATCTGNNNAGATCGGAAG |
| 49263_4 | TATGAGTGTGGAGTCGTTGCCAATCTCAGCTCACTGCAATCTGNNNAGATCGGAAG |
| 49263_5 | TATGAGTGTGGAGTCGTTGCCAATCTCAGCTCACTGCAATCTGNNNAGATCGGAAG |
| 49263_6 | TATGAGTGTGGAGTCGTTGCCAATCTCAGCTCACTGCAATCTGNNNAGATCGGAAG |
| 53428_1 | TATGAGTGTGGAGTCGTTGCAAAAAGAGGGGAGAGAAAGTCCCNNNAGATCGGA |
| 53428_2 | TATGAGTGTGGAGTCGTTGCAAAAAGAGGGGAGAGAAAGTCCCNNNAGATCGGA |
| 53428_3 | TATGAGTGTGGAGTCGTTGCAAAAAGAGGGGAGAGAAAGTCCCNNNAGATCGGA |
| 53428_4 | TATGAGTGTGGAGTCGTTGCAAAAAGAGGGGAGAGAAAGTCCCNNNAGATCGGA |
| 53428_5 | TATGAGTGTGGAGTCGTTGCAAAAAGAGGGGAGAGAAAGTCCCNNNAGATCGGA |
| 53428_6 | TATGAGTGTGGAGTCGTTGCAAAAAGAGGGGAGAGAAAGTCCCNNNAGATCGGA |
| 28942_1 | TATGAGTGTGGAGTCGTTGCTCACCCTCTCCAAGCAGAAATTTNNNAGATCGGAAG |
| 28942_2 | TATGAGTGTGGAGTCGTTGCTCACCCTCTCCAAGCAGAAATTTNNNAGATCGGAAG |
| 28942_3 | TATGAGTGTGGAGTCGTTGCTCACCCTCTCCAAGCAGAAATTTNNNAGATCGGAAG |
| 28942_4 | TATGAGTGTGGAGTCGTTGCTCACCCTCTCCAAGCAGAAATTTNNNAGATCGGAAG |
| 28942_5 | TATGAGTGTGGAGTCGTTGCTCACCCTCTCCAAGCAGAAATTTNNNAGATCGGAAG |
| 28942_6 | TATGAGTGTGGAGTCGTTGCTCACCCTCTCCAAGCAGAAATTTNNNAGATCGGAAG |
| 40217_1 | TATGAGTGTGGAGTCGTTGCGCTTTCTTGCAATCCTGAAAGGANNNAGATCGGAAG |
| 40217_2 | TATGAGTGTGGAGTCGTTGCGCTTTCTTGCAATCCTGAAAGGANNNAGATCGGAAG |
| 40217_3 | TATGAGTGTGGAGTCGTTGCGCTTTCTTGCAATCCTGAAAGGANNNAGATCGGAAG |
| 40217_4 | TATGAGTGTGGAGTCGTTGCGCTTTCTTGCAATCCTGAAAGGANNNAGATCGGAAG |
| 40217_5 | TATGAGTGTGGAGTCGTTGCGCTTTCTTGCAATCCTGAAAGGANNNAGATCGGAAG |
| 40217_6 | TATGAGTGTGGAGTCGTTGCGCTTTCTTGCAATCCTGAAAGGANNNAGATCGGAAG |
| 12901_1 | TATGAGTGTGGAGTCGTTGCAGCACAGGATCCGAAATTAANNNAGATCGGAAGAG |
| 12901_2 | TATGAGTGTGGAGTCGTTGCAGCACAGGATCCGAAATTAANNNAGATCGGAAGAG |
| 12901_3 | TATGAGTGTGGAGTCGTTGCAGCACAGGATCCGAAATTAANNNAGATCGGAAGAG |
| 12901_4 | TATGAGTGTGGAGTCGTTGCAGCACAGGATCCGAAATTAANNNAGATCGGAAGAG |
| 12901_5 | TATGAGTGTGGAGTCGTTGCAGCACAGGATCCGAAATTAANNNAGATCGGAAGAG |
| 12901_6 | TATGAGTGTGGAGTCGTTGCAGCACAGGATCCGAAATTAANNNAGATCGGAAGAG |
| 49057_1 | TATGAGTGTGGAGTCGTTGCAGAGCAGGAAAGAGAAGGCTTACNNNAGATCGGA |
| 49057_2 | TATGAGTGTGGAGTCGTTGCAGAGCAGGAAAGAGAAGGCTTACNNNAGATCGGA |
| 49057_3 | TATGAGTGTGGAGTCGTTGCAGAGCAGGAAAGAGAAGGCTTACNNNAGATCGGA |

| | |
|---|---|
| 49057_4 | TATGAGTGTGGAGTCGTTGCAGAGCAGGAAAGAGAAGGCTTACNNNAGATCGGAA |
| 49057_5 | TATGAGTGTGGAGTCGTTGCAGAGCAGGAAAGAGAAGGCTTACNNNAGATCGGAA |
| 49057_6 | TATGAGTGTGGAGTCGTTGCAGAGCAGGAAAGAGAAGGCTTACNNNAGATCGGAA |
| 43310_1 | TATGAGTGTGGAGTCGTTGCCCACCACACCTGGCTATTTTTTTNNNAGATCGGAAGA |
| 43310_2 | TATGAGTGTGGAGTCGTTGCCCACCACACCTGGCTATTTTTTTNNNAGATCGGAAGA |
| 43310_3 | TATGAGTGTGGAGTCGTTGCCCACCACACCTGGCTATTTTTTTNNNAGATCGGAAGA |
| 43310_4 | TATGAGTGTGGAGTCGTTGCCCACCACACCTGGCTATTTTTTTNNNAGATCGGAAGA |
| 43310_5 | TATGAGTGTGGAGTCGTTGCCCACCACACCTGGCTATTTTTTTNNNAGATCGGAAGA |
| 43310_6 | TATGAGTGTGGAGTCGTTGCCCACCACACCTGGCTATTTTTTTNNNAGATCGGAAGA |
| 40012_1 | TATGAGTGTGGAGTCGTTGCCCATGTAGCTGGGATTACAGGTGNNNAGATCGGAAC |
| 40012_2 | TATGAGTGTGGAGTCGTTGCCCATGTAGCTGGGATTACAGGTGNNNAGATCGGAAC |
| 40012_3 | TATGAGTGTGGAGTCGTTGCCCATGTAGCTGGGATTACAGGTGNNNAGATCGGAAC |
| 40012_4 | TATGAGTGTGGAGTCGTTGCCCATGTAGCTGGGATTACAGGTGNNNAGATCGGAAC |
| 40012_5 | TATGAGTGTGGAGTCGTTGCCCATGTAGCTGGGATTACAGGTGNNNAGATCGGAAC |
| 40012_6 | TATGAGTGTGGAGTCGTTGCCCATGTAGCTGGGATTACAGGTGNNNAGATCGGAAC |
| 49552_1 | TATGAGTGTGGAGTCGTTGCAAGTCATGGGAAGTGTGAGAAGGNNNAGATCGGAA |
| 49552_2 | TATGAGTGTGGAGTCGTTGCAAGTCATGGGAAGTGTGAGAAGGNNNAGATCGGAA |
| 49552_3 | TATGAGTGTGGAGTCGTTGCAAGTCATGGGAAGTGTGAGAAGGNNNAGATCGGAA |
| 49552_4 | TATGAGTGTGGAGTCGTTGCAAGTCATGGGAAGTGTGAGAAGGNNNAGATCGGAA |
| 49552_5 | TATGAGTGTGGAGTCGTTGCAAGTCATGGGAAGTGTGAGAAGGNNNAGATCGGAA |
| 49552_6 | TATGAGTGTGGAGTCGTTGCAAGTCATGGGAAGTGTGAGAAGGNNNAGATCGGAA |
| 13324_1 | TATGAGTGTGGAGTCGTTGCCTTAACCTGAATTTGATTAAAACANNNAGATCGGAAC |
| 13324_2 | TATGAGTGTGGAGTCGTTGCCTTAACCTGAATTTGATTAAAACANNNAGATCGGAAC |
| 13324_3 | TATGAGTGTGGAGTCGTTGCCTTAACCTGAATTTGATTAAAACANNNAGATCGGAAC |
| 13324_4 | TATGAGTGTGGAGTCGTTGCCTTAACCTGAATTTGATTAAAACANNNAGATCGGAAC |
| 13324_5 | TATGAGTGTGGAGTCGTTGCCTTAACCTGAATTTGATTAAAACANNNAGATCGGAAC |
| 13324_6 | TATGAGTGTGGAGTCGTTGCCTTAACCTGAATTTGATTAAAACANNNAGATCGGAAC |
| 30859_1 | TATGAGTGTGGAGTCGTTGCTTCTTCTATCAAAGTGGTGGCCTNNNAGATCGGAAGA |
| 30859_2 | TATGAGTGTGGAGTCGTTGCTTCTTCTATCAAAGTGGTGGCCTNNNAGATCGGAAGA |
| 30859_3 | TATGAGTGTGGAGTCGTTGCTTCTTCTATCAAAGTGGTGGCCTNNNAGATCGGAAGA |
| 30859_4 | TATGAGTGTGGAGTCGTTGCTTCTTCTATCAAAGTGGTGGCCTNNNAGATCGGAAGA |
| 30859_5 | TATGAGTGTGGAGTCGTTGCTTCTTCTATCAAAGTGGTGGCCTNNNAGATCGGAAGA |
| 30859_6 | TATGAGTGTGGAGTCGTTGCTTCTTCTATCAAAGTGGTGGCCTNNNAGATCGGAAGA |
| 45291_1 | TATGAGTGTGGAGTCGTTGCGAATAATCTGGCACAACCGGAAGNNNAGATCGGAAC |
| 45291_2 | TATGAGTGTGGAGTCGTTGCGAATAATCTGGCACAACCGGAAGNNNAGATCGGAAC |
| 45291_3 | TATGAGTGTGGAGTCGTTGCGAATAATCTGGCACAACCGGAAGNNNAGATCGGAAC |
| 45291_4 | TATGAGTGTGGAGTCGTTGCGAATAATCTGGCACAACCGGAAGNNNAGATCGGAAC |
| 45291_5 | TATGAGTGTGGAGTCGTTGCGAATAATCTGGCACAACCGGAAGNNNAGATCGGAAC |
| 45291_6 | TATGAGTGTGGAGTCGTTGCGAATAATCTGGCACAACCGGAAGNNNAGATCGGAAC |
| 41838_1 | TATGAGTGTGGAGTCGTTGCCTCCTGAGTAGCTGGGATTACAGNNNAGATCGGAAC |
| 41838_2 | TATGAGTGTGGAGTCGTTGCCTCCTGAGTAGCTGGGATTACAGNNNAGATCGGAAC |
| 41838_3 | TATGAGTGTGGAGTCGTTGCCTCCTGAGTAGCTGGGATTACAGNNNAGATCGGAAC |
| 41838_4 | TATGAGTGTGGAGTCGTTGCCTCCTGAGTAGCTGGGATTACAGNNNAGATCGGAAC |

41838_5 TATGAGTGTGGAGTCGTTGCCTCCTGAGTAGCTGGGATTACAGNNNAGATCGGAA
41838_6 TATGAGTGTGGAGTCGTTGCCTCCTGAGTAGCTGGGATTACAGNNNAGATCGGAA
35055_1 TATGAGTGTGGAGTCGTTGCTGTGCTGTTTCCCTCTATGAGTCNNNAGATCGGAAG
35055_2 TATGAGTGTGGAGTCGTTGCTGTGCTGTTTCCCTCTATGAGTCNNNAGATCGGAAG
35055_3 TATGAGTGTGGAGTCGTTGCTGTGCTGTTTCCCTCTATGAGTCNNNAGATCGGAAG
35055_4 TATGAGTGTGGAGTCGTTGCTGTGCTGTTTCCCTCTATGAGTCNNNAGATCGGAAG
35055_5 TATGAGTGTGGAGTCGTTGCTGTGCTGTTTCCCTCTATGAGTCNNNAGATCGGAAG
35055_6 TATGAGTGTGGAGTCGTTGCTGTGCTGTTTCCCTCTATGAGTCNNNAGATCGGAAG
38056_1 TATGAGTGTGGAGTCGTTGCCCTGGCTCTGGACACATTTTTTTNNNAGATCGGAAG
38056_2 TATGAGTGTGGAGTCGTTGCCCTGGCTCTGGACACATTTTTTTNNNAGATCGGAAG
38056_3 TATGAGTGTGGAGTCGTTGCCCTGGCTCTGGACACATTTTTTTNNNAGATCGGAAG
38056_4 TATGAGTGTGGAGTCGTTGCCCTGGCTCTGGACACATTTTTTTNNNAGATCGGAAG
38056_5 TATGAGTGTGGAGTCGTTGCCCTGGCTCTGGACACATTTTTTTNNNAGATCGGAAG
38056_6 TATGAGTGTGGAGTCGTTGCCCTGGCTCTGGACACATTTTTTTNNNAGATCGGAAG
45413_1 TATGAGTGTGGAGTCGTTGCACAAAGTTCTGACCAAAACACCGNNNAGATCGGAAG
45413_2 TATGAGTGTGGAGTCGTTGCACAAAGTTCTGACCAAAACACCGNNNAGATCGGAAG
45413_3 TATGAGTGTGGAGTCGTTGCACAAAGTTCTGACCAAAACACCGNNNAGATCGGAAG
45413_4 TATGAGTGTGGAGTCGTTGCACAAAGTTCTGACCAAAACACCGNNNAGATCGGAAG
45413_5 TATGAGTGTGGAGTCGTTGCACAAAGTTCTGACCAAAACACCGNNNAGATCGGAAG
45413_6 TATGAGTGTGGAGTCGTTGCACAAAGTTCTGACCAAAACACCGNNNAGATCGGAAG
41268_1 TATGAGTGTGGAGTCGTTGCGTATTGATGGGATGTCTTGCAGCNNNAGATCGGAA
41268_2 TATGAGTGTGGAGTCGTTGCGTATTGATGGGATGTCTTGCAGCNNNAGATCGGAA
41268_3 TATGAGTGTGGAGTCGTTGCGTATTGATGGGATGTCTTGCAGCNNNAGATCGGAA
41268_4 TATGAGTGTGGAGTCGTTGCGTATTGATGGGATGTCTTGCAGCNNNAGATCGGAA
41268_5 TATGAGTGTGGAGTCGTTGCGTATTGATGGGATGTCTTGCAGCNNNAGATCGGAA
41268_6 TATGAGTGTGGAGTCGTTGCGTATTGATGGGATGTCTTGCAGCNNNAGATCGGAA
50544_1 TATGAGTGTGGAGTCGTTGCTCTGGGAATTACATCATTGGGCTNNNAGATCGGAAG
50544_2 TATGAGTGTGGAGTCGTTGCTCTGGGAATTACATCATTGGGCTNNNAGATCGGAAG
50544_3 TATGAGTGTGGAGTCGTTGCTCTGGGAATTACATCATTGGGCTNNNAGATCGGAAG
50544_4 TATGAGTGTGGAGTCGTTGCTCTGGGAATTACATCATTGGGCTNNNAGATCGGAAG
50544_5 TATGAGTGTGGAGTCGTTGCTCTGGGAATTACATCATTGGGCTNNNAGATCGGAAG
50544_6 TATGAGTGTGGAGTCGTTGCTCTGGGAATTACATCATTGGGCTNNNAGATCGGAAG
28979_1 TATGAGTGTGGAGTCGTTGCACTGAGAATGGAAACTGTAAGAGANNNAGATCGGA
28979_2 TATGAGTGTGGAGTCGTTGCACTGAGAATGGAAACTGTAAGAGANNNAGATCGGA
28979_3 TATGAGTGTGGAGTCGTTGCACTGAGAATGGAAACTGTAAGAGANNNAGATCGGA
28979_4 TATGAGTGTGGAGTCGTTGCACTGAGAATGGAAACTGTAAGAGANNNAGATCGGA
28979_5 TATGAGTGTGGAGTCGTTGCACTGAGAATGGAAACTGTAAGAGANNNAGATCGGA
28979_6 TATGAGTGTGGAGTCGTTGCACTGAGAATGGAAACTGTAAGAGANNNAGATCGGA
45806_1 TATGAGTGTGGAGTCGTTGCAACTCCTGACCTCAAGTGATCTGNNNAGATCGGAAG
45806_2 TATGAGTGTGGAGTCGTTGCAACTCCTGACCTCAAGTGATCTGNNNAGATCGGAAG
45806_3 TATGAGTGTGGAGTCGTTGCAACTCCTGACCTCAAGTGATCTGNNNAGATCGGAAG
45806_4 TATGAGTGTGGAGTCGTTGCAACTCCTGACCTCAAGTGATCTGNNNAGATCGGAAG
45806_5 TATGAGTGTGGAGTCGTTGCAACTCCTGACCTCAAGTGATCTGNNNAGATCGGAAG

| | |
|---|---|
| 45806_6 | TATGAGTGTGGAGTCGTTGCAACTCCTGACCTCAAGTGATCTGNNNAGATCGGAAG |
| 49147_1 | TATGAGTGTGGAGTCGTTGCCCAGGTTCAAGTGATTCTCCTCTNNNAGATCGGAAG |
| 49147_2 | TATGAGTGTGGAGTCGTTGCCCAGGTTCAAGTGATTCTCCTCTNNNAGATCGGAAG |
| 49147_3 | TATGAGTGTGGAGTCGTTGCCCAGGTTCAAGTGATTCTCCTCTNNNAGATCGGAAG |
| 49147_4 | TATGAGTGTGGAGTCGTTGCCCAGGTTCAAGTGATTCTCCTCTNNNAGATCGGAAG |
| 49147_5 | TATGAGTGTGGAGTCGTTGCCCAGGTTCAAGTGATTCTCCTCTNNNAGATCGGAAG |
| 49147_6 | TATGAGTGTGGAGTCGTTGCCCAGGTTCAAGTGATTCTCCTCTNNNAGATCGGAAG |
| 47325_1 | TATGAGTGTGGAGTCGTTGCGCCATTCTGACTAGTGTGACATGNNNAGATCGGAAG |
| 47325_2 | TATGAGTGTGGAGTCGTTGCGCCATTCTGACTAGTGTGACATGNNNAGATCGGAAG |
| 47325_3 | TATGAGTGTGGAGTCGTTGCGCCATTCTGACTAGTGTGACATGNNNAGATCGGAAG |
| 47325_4 | TATGAGTGTGGAGTCGTTGCGCCATTCTGACTAGTGTGACATGNNNAGATCGGAAG |
| 47325_5 | TATGAGTGTGGAGTCGTTGCGCCATTCTGACTAGTGTGACATGNNNAGATCGGAAG |
| 47325_6 | TATGAGTGTGGAGTCGTTGCGCCATTCTGACTAGTGTGACATGNNNAGATCGGAAG |
| 44221_1 | TATGAGTGTGGAGTCGTTGCCCACCCAACACTGTCCTTGTACNNNAGATCGGAAGA |
| 44221_2 | TATGAGTGTGGAGTCGTTGCCCACCCAACACTGTCCTTGTACNNNAGATCGGAAGA |
| 44221_3 | TATGAGTGTGGAGTCGTTGCCCACCCAACACTGTCCTTGTACNNNAGATCGGAAGA |
| 44221_4 | TATGAGTGTGGAGTCGTTGCCCACCCAACACTGTCCTTGTACNNNAGATCGGAAGA |
| 44221_5 | TATGAGTGTGGAGTCGTTGCCCACCCAACACTGTCCTTGTACNNNAGATCGGAAGA |
| 44221_6 | TATGAGTGTGGAGTCGTTGCCCACCCAACACTGTCCTTGTACNNNAGATCGGAAGA |
| 36447_1 | TATGAGTGTGGAGTCGTTGCACTAGGCAATTGGGAAGTGTAGGNNNAGATCGGAA |
| 36447_2 | TATGAGTGTGGAGTCGTTGCACTAGGCAATTGGGAAGTGTAGGNNNAGATCGGAA |
| 36447_3 | TATGAGTGTGGAGTCGTTGCACTAGGCAATTGGGAAGTGTAGGNNNAGATCGGAA |
| 36447_4 | TATGAGTGTGGAGTCGTTGCACTAGGCAATTGGGAAGTGTAGGNNNAGATCGGAA |
| 36447_5 | TATGAGTGTGGAGTCGTTGCACTAGGCAATTGGGAAGTGTAGGNNNAGATCGGAA |
| 36447_6 | TATGAGTGTGGAGTCGTTGCACTAGGCAATTGGGAAGTGTAGGNNNAGATCGGAA |
| 38188_1 | TATGAGTGTGGAGTCGTTGCTGATACTAGCTGTGGGTCTGTCANNNAGATCGGAA |
| 38188_2 | TATGAGTGTGGAGTCGTTGCTGATACTAGCTGTGGGTCTGTCANNNAGATCGGAA |
| 38188_3 | TATGAGTGTGGAGTCGTTGCTGATACTAGCTGTGGGTCTGTCANNNAGATCGGAA |
| 38188_4 | TATGAGTGTGGAGTCGTTGCTGATACTAGCTGTGGGTCTGTCANNNAGATCGGAA |
| 38188_5 | TATGAGTGTGGAGTCGTTGCTGATACTAGCTGTGGGTCTGTCANNNAGATCGGAA |
| 38188_6 | TATGAGTGTGGAGTCGTTGCTGATACTAGCTGTGGGTCTGTCANNNAGATCGGAA |
| 47340_1 | TATGAGTGTGGAGTCGTTGCGTTTTGCTGTGGCTATTTGAGGANNNAGATCGGAA |
| 47340_2 | TATGAGTGTGGAGTCGTTGCGTTTTGCTGTGGCTATTTGAGGANNNAGATCGGAA |
| 47340_3 | TATGAGTGTGGAGTCGTTGCGTTTTGCTGTGGCTATTTGAGGANNNAGATCGGAA |
| 47340_4 | TATGAGTGTGGAGTCGTTGCGTTTTGCTGTGGCTATTTGAGGANNNAGATCGGAA |
| 47340_5 | TATGAGTGTGGAGTCGTTGCGTTTTGCTGTGGCTATTTGAGGANNNAGATCGGAA |
| 47340_6 | TATGAGTGTGGAGTCGTTGCGTTTTGCTGTGGCTATTTGAGGANNNAGATCGGAA |
| 38757_1 | TATGAGTGTGGAGTCGTTGCGTCACCATAGCTCACTGTAACCTNNNAGATCGGAAG |
| 38757_2 | TATGAGTGTGGAGTCGTTGCGTCACCATAGCTCACTGTAACCTNNNAGATCGGAAG |
| 38757_3 | TATGAGTGTGGAGTCGTTGCGTCACCATAGCTCACTGTAACCTNNNAGATCGGAAG |
| 38757_4 | TATGAGTGTGGAGTCGTTGCGTCACCATAGCTCACTGTAACCTNNNAGATCGGAAG |
| 38757_5 | TATGAGTGTGGAGTCGTTGCGTCACCATAGCTCACTGTAACCTNNNAGATCGGAAG |
| 38757_6 | TATGAGTGTGGAGTCGTTGCGTCACCATAGCTCACTGTAACCTNNNAGATCGGAAG |

| | |
|---|---|
| 49278_1 | TATGAGTGTGGAGTCGTTGCCCACACCCGGCTAATTTTTGTACNNNAGATCGGAAGA |
| 49278_2 | TATGAGTGTGGAGTCGTTGCCCACACCCGGCTAATTTTTGTACNNNAGATCGGAAGA |
| 49278_3 | TATGAGTGTGGAGTCGTTGCCCACACCCGGCTAATTTTTGTACNNNAGATCGGAAGA |
| 49278_4 | TATGAGTGTGGAGTCGTTGCCCACACCCGGCTAATTTTTGTACNNNAGATCGGAAGA |
| 49278_5 | TATGAGTGTGGAGTCGTTGCCCACACCCGGCTAATTTTTGTACNNNAGATCGGAAGA |
| 49278_6 | TATGAGTGTGGAGTCGTTGCCCACACCCGGCTAATTTTTGTACNNNAGATCGGAAGA |
| 12784_1 | TATGAGTGTGGAGTCGTTGCTCATTGTTGTAGGACTCCAGNNNAGATCGGAAGAGC |
| 12784_2 | TATGAGTGTGGAGTCGTTGCTCATTGTTGTAGGACTCCAGNNNAGATCGGAAGAGC |
| 12784_3 | TATGAGTGTGGAGTCGTTGCTCATTGTTGTAGGACTCCAGNNNAGATCGGAAGAGC |
| 12784_4 | TATGAGTGTGGAGTCGTTGCTCATTGTTGTAGGACTCCAGNNNAGATCGGAAGAGC |
| 12784_5 | TATGAGTGTGGAGTCGTTGCTCATTGTTGTAGGACTCCAGNNNAGATCGGAAGAGC |
| 12784_6 | TATGAGTGTGGAGTCGTTGCTCATTGTTGTAGGACTCCAGNNNAGATCGGAAGAGC |
| 46051_1 | TATGAGTGTGGAGTCGTTGCCAATCTCCTGACCTCATGATCCGNNNAGATCGGAAGA |
| 46051_2 | TATGAGTGTGGAGTCGTTGCCAATCTCCTGACCTCATGATCCGNNNAGATCGGAAGA |
| 46051_3 | TATGAGTGTGGAGTCGTTGCCAATCTCCTGACCTCATGATCCGNNNAGATCGGAAGA |
| 46051_4 | TATGAGTGTGGAGTCGTTGCCAATCTCCTGACCTCATGATCCGNNNAGATCGGAAGA |
| 46051_5 | TATGAGTGTGGAGTCGTTGCCAATCTCCTGACCTCATGATCCGNNNAGATCGGAAGA |
| 46051_6 | TATGAGTGTGGAGTCGTTGCCAATCTCCTGACCTCATGATCCGNNNAGATCGGAAGA |
| 36783_1 | TATGAGTGTGGAGTCGTTGCGACCATTTTTGCTGCATTGAAGCNNNAGATCGGAAG |
| 36783_2 | TATGAGTGTGGAGTCGTTGCGACCATTTTTGCTGCATTGAAGCNNNAGATCGGAAG |
| 36783_3 | TATGAGTGTGGAGTCGTTGCGACCATTTTTGCTGCATTGAAGCNNNAGATCGGAAG |
| 36783_4 | TATGAGTGTGGAGTCGTTGCGACCATTTTTGCTGCATTGAAGCNNNAGATCGGAAG |
| 36783_5 | TATGAGTGTGGAGTCGTTGCGACCATTTTTGCTGCATTGAAGCNNNAGATCGGAAG |
| 36783_6 | TATGAGTGTGGAGTCGTTGCGACCATTTTTGCTGCATTGAAGCNNNAGATCGGAAG |
| 52923_1 | TATGAGTGTGGAGTCGTTGCGAGCCTGACAAGTCTTATGGGAANNNAGATCGGAAC |
| 52923_2 | TATGAGTGTGGAGTCGTTGCGAGCCTGACAAGTCTTATGGGAANNNAGATCGGAAC |
| 52923_3 | TATGAGTGTGGAGTCGTTGCGAGCCTGACAAGTCTTATGGGAANNNAGATCGGAAC |
| 52923_4 | TATGAGTGTGGAGTCGTTGCGAGCCTGACAAGTCTTATGGGAANNNAGATCGGAAC |
| 52923_5 | TATGAGTGTGGAGTCGTTGCGAGCCTGACAAGTCTTATGGGAANNNAGATCGGAAC |
| 52923_6 | TATGAGTGTGGAGTCGTTGCGAGCCTGACAAGTCTTATGGGAANNNAGATCGGAAC |
| 48409_1 | TATGAGTGTGGAGTCGTTGCTGGGAATTGCCTTGATAATCCCTNNNAGATCGGAAG |
| 48409_2 | TATGAGTGTGGAGTCGTTGCTGGGAATTGCCTTGATAATCCCTNNNAGATCGGAAG |
| 48409_3 | TATGAGTGTGGAGTCGTTGCTGGGAATTGCCTTGATAATCCCTNNNAGATCGGAAG |
| 48409_4 | TATGAGTGTGGAGTCGTTGCTGGGAATTGCCTTGATAATCCCTNNNAGATCGGAAG |
| 48409_5 | TATGAGTGTGGAGTCGTTGCTGGGAATTGCCTTGATAATCCCTNNNAGATCGGAAG |
| 48409_6 | TATGAGTGTGGAGTCGTTGCTGGGAATTGCCTTGATAATCCCTNNNAGATCGGAAG |
| 30264_1 | TATGAGTGTGGAGTCGTTGCTACAAGGCTGTCTGTCATGAAGCNNNAGATCGGAAG |
| 30264_2 | TATGAGTGTGGAGTCGTTGCTACAAGGCTGTCTGTCATGAAGCNNNAGATCGGAAG |
| 30264_3 | TATGAGTGTGGAGTCGTTGCTACAAGGCTGTCTGTCATGAAGCNNNAGATCGGAAG |
| 30264_4 | TATGAGTGTGGAGTCGTTGCTACAAGGCTGTCTGTCATGAAGCNNNAGATCGGAAG |
| 30264_5 | TATGAGTGTGGAGTCGTTGCTACAAGGCTGTCTGTCATGAAGCNNNAGATCGGAAG |
| 30264_6 | TATGAGTGTGGAGTCGTTGCTACAAGGCTGTCTGTCATGAAGCNNNAGATCGGAAG |
| 46352_1 | TATGAGTGTGGAGTCGTTGCAAGTAGCTGGGATTACAGGCACNNNAGATCGGAAG |

| | |
|---|---|
| 46352_2 | TATGAGTGTGGAGTCGTTGCAAGTAGCTGGGATTACAGGCACNNNAGATCGGAAG |
| 46352_3 | TATGAGTGTGGAGTCGTTGCAAGTAGCTGGGATTACAGGCACNNNAGATCGGAAG |
| 46352_4 | TATGAGTGTGGAGTCGTTGCAAGTAGCTGGGATTACAGGCACNNNAGATCGGAAG |
| 46352_5 | TATGAGTGTGGAGTCGTTGCAAGTAGCTGGGATTACAGGCACNNNAGATCGGAAG |
| 46352_6 | TATGAGTGTGGAGTCGTTGCAAGTAGCTGGGATTACAGGCACNNNAGATCGGAAG |
| 48771_1 | TATGAGTGTGGAGTCGTTGCGACCAACTGCTGTGTAATGTGACNNNAGATCGGAAG |
| 48771_2 | TATGAGTGTGGAGTCGTTGCGACCAACTGCTGTGTAATGTGACNNNAGATCGGAAG |
| 48771_3 | TATGAGTGTGGAGTCGTTGCGACCAACTGCTGTGTAATGTGACNNNAGATCGGAAG |
| 48771_4 | TATGAGTGTGGAGTCGTTGCGACCAACTGCTGTGTAATGTGACNNNAGATCGGAAG |
| 48771_5 | TATGAGTGTGGAGTCGTTGCGACCAACTGCTGTGTAATGTGACNNNAGATCGGAAG |
| 48771_6 | TATGAGTGTGGAGTCGTTGCGACCAACTGCTGTGTAATGTGACNNNAGATCGGAAG |
| 40505_1 | TATGAGTGTGGAGTCGTTGCATACCTTCCACTGTAACCTTGCANNNAGATCGGAAGA |
| 40505_2 | TATGAGTGTGGAGTCGTTGCATACCTTCCACTGTAACCTTGCANNNAGATCGGAAGA |
| 40505_3 | TATGAGTGTGGAGTCGTTGCATACCTTCCACTGTAACCTTGCANNNAGATCGGAAGA |
| 40505_4 | TATGAGTGTGGAGTCGTTGCATACCTTCCACTGTAACCTTGCANNNAGATCGGAAGA |
| 40505_5 | TATGAGTGTGGAGTCGTTGCATACCTTCCACTGTAACCTTGCANNNAGATCGGAAGA |
| 40505_6 | TATGAGTGTGGAGTCGTTGCATACCTTCCACTGTAACCTTGCANNNAGATCGGAAGA |
| 10350_1 | TATGAGTGTGGAGTCGTTGCGACCCAAGGTTTGGAAAAACAGCNNNAGATCGGAAG |
| 10350_2 | TATGAGTGTGGAGTCGTTGCGACCCAAGGTTTGGAAAAACAGCNNNAGATCGGAAG |
| 10350_3 | TATGAGTGTGGAGTCGTTGCGACCCAAGGTTTGGAAAAACAGCNNNAGATCGGAAG |
| 10350_4 | TATGAGTGTGGAGTCGTTGCGACCCAAGGTTTGGAAAAACAGCNNNAGATCGGAAG |
| 10350_5 | TATGAGTGTGGAGTCGTTGCGACCCAAGGTTTGGAAAAACAGCNNNAGATCGGAAG |
| 10350_6 | TATGAGTGTGGAGTCGTTGCGACCCAAGGTTTGGAAAAACAGCNNNAGATCGGAAG |
| 29854_1 | TATGAGTGTGGAGTCGTTGCATCTTCACCTTTAGCTGGCAGACNNNAGATCGGAAGA |
| 29854_2 | TATGAGTGTGGAGTCGTTGCATCTTCACCTTTAGCTGGCAGACNNNAGATCGGAAGA |
| 29854_3 | TATGAGTGTGGAGTCGTTGCATCTTCACCTTTAGCTGGCAGACNNNAGATCGGAAGA |
| 29854_4 | TATGAGTGTGGAGTCGTTGCATCTTCACCTTTAGCTGGCAGACNNNAGATCGGAAGA |
| 29854_5 | TATGAGTGTGGAGTCGTTGCATCTTCACCTTTAGCTGGCAGACNNNAGATCGGAAGA |
| 29854_6 | TATGAGTGTGGAGTCGTTGCATCTTCACCTTTAGCTGGCAGACNNNAGATCGGAAGA |
| 37219_1 | TATGAGTGTGGAGTCGTTGCCATTCGTTGCCAGGTCTCTTTAGNNNAGATCGGAAGA |
| 37219_2 | TATGAGTGTGGAGTCGTTGCCATTCGTTGCCAGGTCTCTTTAGNNNAGATCGGAAGA |
| 37219_3 | TATGAGTGTGGAGTCGTTGCCATTCGTTGCCAGGTCTCTTTAGNNNAGATCGGAAGA |
| 37219_4 | TATGAGTGTGGAGTCGTTGCCATTCGTTGCCAGGTCTCTTTAGNNNAGATCGGAAGA |
| 37219_5 | TATGAGTGTGGAGTCGTTGCCATTCGTTGCCAGGTCTCTTTAGNNNAGATCGGAAGA |
| 37219_6 | TATGAGTGTGGAGTCGTTGCCATTCGTTGCCAGGTCTCTTTAGNNNAGATCGGAAGA |
| 31737_1 | TATGAGTGTGGAGTCGTTGCGTGTGTGTGTGATGCTTGCTTATNNNAGATCGGAAG |
| 31737_2 | TATGAGTGTGGAGTCGTTGCGTGTGTGTGTGATGCTTGCTTATNNNAGATCGGAAG |
| 31737_3 | TATGAGTGTGGAGTCGTTGCGTGTGTGTGTGATGCTTGCTTATNNNAGATCGGAAG |
| 31737_4 | TATGAGTGTGGAGTCGTTGCGTGTGTGTGTGATGCTTGCTTATNNNAGATCGGAAG |
| 31737_5 | TATGAGTGTGGAGTCGTTGCGTGTGTGTGTGATGCTTGCTTATNNNAGATCGGAAG |
| 31737_6 | TATGAGTGTGGAGTCGTTGCGTGTGTGTGTGATGCTTGCTTATNNNAGATCGGAAG |
| 28912_1 | TATGAGTGTGGAGTCGTTGCAGATGGCACAAACAACTGAACTTNNNAGATCGGAAG |
| 28912_2 | TATGAGTGTGGAGTCGTTGCAGATGGCACAAACAACTGAACTTNNNAGATCGGAAG |

| | |
|---|---|
| 28912_3 | TATGAGTGTGGAGTCGTTGCAGATGGCACAAACAACTGAACTTNNNAGATCGGAAG |
| 28912_4 | TATGAGTGTGGAGTCGTTGCAGATGGCACAAACAACTGAACTTNNNAGATCGGAAG |
| 28912_5 | TATGAGTGTGGAGTCGTTGCAGATGGCACAAACAACTGAACTTNNNAGATCGGAAG |
| 28912_6 | TATGAGTGTGGAGTCGTTGCAGATGGCACAAACAACTGAACTTNNNAGATCGGAAG |
| 37040_1 | TATGAGTGTGGAGTCGTTGCTGTGGTCAACAAGCATACAGTGANNNAGATCGGAA |
| 37040_2 | TATGAGTGTGGAGTCGTTGCTGTGGTCAACAAGCATACAGTGANNNAGATCGGAA |
| 37040_3 | TATGAGTGTGGAGTCGTTGCTGTGGTCAACAAGCATACAGTGANNNAGATCGGAA |
| 37040_4 | TATGAGTGTGGAGTCGTTGCTGTGGTCAACAAGCATACAGTGANNNAGATCGGAA |
| 37040_5 | TATGAGTGTGGAGTCGTTGCTGTGGTCAACAAGCATACAGTGANNNAGATCGGAA |
| 37040_6 | TATGAGTGTGGAGTCGTTGCTGTGGTCAACAAGCATACAGTGANNNAGATCGGAA |
| 40684_1 | TATGAGTGTGGAGTCGTTGCACCTGGACTGTTCTGGATTGTTTNNNAGATCGGAAG |
| 40684_2 | TATGAGTGTGGAGTCGTTGCACCTGGACTGTTCTGGATTGTTTNNNAGATCGGAAG |
| 40684_3 | TATGAGTGTGGAGTCGTTGCACCTGGACTGTTCTGGATTGTTTNNNAGATCGGAAG |
| 40684_4 | TATGAGTGTGGAGTCGTTGCACCTGGACTGTTCTGGATTGTTTNNNAGATCGGAAG |
| 40684_5 | TATGAGTGTGGAGTCGTTGCACCTGGACTGTTCTGGATTGTTTNNNAGATCGGAAG |
| 40684_6 | TATGAGTGTGGAGTCGTTGCACCTGGACTGTTCTGGATTGTTTNNNAGATCGGAAG |
| 46095_1 | TATGAGTGTGGAGTCGTTGCGAGTGAAATGGCGTGATCTCAACNNNAGATCGGAA |
| 46095_2 | TATGAGTGTGGAGTCGTTGCGAGTGAAATGGCGTGATCTCAACNNNAGATCGGAA |
| 46095_3 | TATGAGTGTGGAGTCGTTGCGAGTGAAATGGCGTGATCTCAACNNNAGATCGGAA |
| 46095_4 | TATGAGTGTGGAGTCGTTGCGAGTGAAATGGCGTGATCTCAACNNNAGATCGGAA |
| 46095_5 | TATGAGTGTGGAGTCGTTGCGAGTGAAATGGCGTGATCTCAACNNNAGATCGGAA |
| 46095_6 | TATGAGTGTGGAGTCGTTGCGAGTGAAATGGCGTGATCTCAACNNNAGATCGGAA |
| 12741_1 | TATGAGTGTGGAGTCGTTGCTTACTCCATGCAAAATCTCCNNNAGATCGGAAGAGCA |
| 12741_2 | TATGAGTGTGGAGTCGTTGCTTACTCCATGCAAAATCTCCNNNAGATCGGAAGAGCA |
| 12741_3 | TATGAGTGTGGAGTCGTTGCTTACTCCATGCAAAATCTCCNNNAGATCGGAAGAGCA |
| 12741_4 | TATGAGTGTGGAGTCGTTGCTTACTCCATGCAAAATCTCCNNNAGATCGGAAGAGCA |
| 12741_5 | TATGAGTGTGGAGTCGTTGCTTACTCCATGCAAAATCTCCNNNAGATCGGAAGAGCA |
| 12741_6 | TATGAGTGTGGAGTCGTTGCTTACTCCATGCAAAATCTCCNNNAGATCGGAAGAGCA |
| 30678_1 | TATGAGTGTGGAGTCGTTGCGAAGAGGTGAAGATGGAGTTCGANNNAGATCGGA |
| 30678_2 | TATGAGTGTGGAGTCGTTGCGAAGAGGTGAAGATGGAGTTCGANNNAGATCGGA |
| 30678_3 | TATGAGTGTGGAGTCGTTGCGAAGAGGTGAAGATGGAGTTCGANNNAGATCGGA |
| 30678_4 | TATGAGTGTGGAGTCGTTGCGAAGAGGTGAAGATGGAGTTCGANNNAGATCGGA |
| 30678_5 | TATGAGTGTGGAGTCGTTGCGAAGAGGTGAAGATGGAGTTCGANNNAGATCGGA |
| 30678_6 | TATGAGTGTGGAGTCGTTGCGAAGAGGTGAAGATGGAGTTCGANNNAGATCGGA |
| 51454_1 | TATGAGTGTGGAGTCGTTGCATAGCGCCTTTCTCTTTACTGGGNNNAGATCGGAAG |
| 51454_2 | TATGAGTGTGGAGTCGTTGCATAGCGCCTTTCTCTTTACTGGGNNNAGATCGGAAG |
| 51454_3 | TATGAGTGTGGAGTCGTTGCATAGCGCCTTTCTCTTTACTGGGNNNAGATCGGAAG |
| 51454_4 | TATGAGTGTGGAGTCGTTGCATAGCGCCTTTCTCTTTACTGGGNNNAGATCGGAAG |
| 51454_5 | TATGAGTGTGGAGTCGTTGCATAGCGCCTTTCTCTTTACTGGGNNNAGATCGGAAG |
| 51454_6 | TATGAGTGTGGAGTCGTTGCATAGCGCCTTTCTCTTTACTGGGNNNAGATCGGAAG |
| 51643_1 | TATGAGTGTGGAGTCGTTGCCCCTCTACCCCTTCATCCATCTANNNAGATCGGAAGA |
| 51643_2 | TATGAGTGTGGAGTCGTTGCCCCTCTACCCCTTCATCCATCTANNNAGATCGGAAGA |
| 51643_3 | TATGAGTGTGGAGTCGTTGCCCCTCTACCCCTTCATCCATCTANNNAGATCGGAAGA |

| | |
|---|---|
| 51643_4 | TATGAGTGTGGAGTCGTTGCCCCTCTACCCCTTCATCCATCTANNNAGATCGGAAGA |
| 51643_5 | TATGAGTGTGGAGTCGTTGCCCCTCTACCCCTTCATCCATCTANNNAGATCGGAAGA |
| 51643_6 | TATGAGTGTGGAGTCGTTGCCCCTCTACCCCTTCATCCATCTANNNAGATCGGAAGA |
| 38047_1 | TATGAGTGTGGAGTCGTTGCGGATCCCTTGAGTTCAGGAGTTCNNNAGATCGGAA |
| 38047_2 | TATGAGTGTGGAGTCGTTGCGGATCCCTTGAGTTCAGGAGTTCNNNAGATCGGAA |
| 38047_3 | TATGAGTGTGGAGTCGTTGCGGATCCCTTGAGTTCAGGAGTTCNNNAGATCGGAA |
| 38047_4 | TATGAGTGTGGAGTCGTTGCGGATCCCTTGAGTTCAGGAGTTCNNNAGATCGGAA |
| 38047_5 | TATGAGTGTGGAGTCGTTGCGGATCCCTTGAGTTCAGGAGTTCNNNAGATCGGAA |
| 38047_6 | TATGAGTGTGGAGTCGTTGCGGATCCCTTGAGTTCAGGAGTTCNNNAGATCGGAA |
| 46581_1 | TATGAGTGTGGAGTCGTTGCGCTTCTTTTCCAAAGCCATGACTNNNAGATCGGAAG |
| 46581_2 | TATGAGTGTGGAGTCGTTGCGCTTCTTTTCCAAAGCCATGACTNNNAGATCGGAAG |
| 46581_3 | TATGAGTGTGGAGTCGTTGCGCTTCTTTTCCAAAGCCATGACTNNNAGATCGGAAG |
| 46581_4 | TATGAGTGTGGAGTCGTTGCGCTTCTTTTCCAAAGCCATGACTNNNAGATCGGAAG |
| 46581_5 | TATGAGTGTGGAGTCGTTGCGCTTCTTTTCCAAAGCCATGACTNNNAGATCGGAAG |
| 46581_6 | TATGAGTGTGGAGTCGTTGCGCTTCTTTTCCAAAGCCATGACTNNNAGATCGGAAG |
| 36752_1 | TATGAGTGTGGAGTCGTTGCCCCCTCTAAAAATAATGGCCAGCNNNAGATCGGAAG |
| 36752_2 | TATGAGTGTGGAGTCGTTGCCCCCTCTAAAAATAATGGCCAGCNNNAGATCGGAAG |
| 36752_3 | TATGAGTGTGGAGTCGTTGCCCCCTCTAAAAATAATGGCCAGCNNNAGATCGGAAG |
| 36752_4 | TATGAGTGTGGAGTCGTTGCCCCCTCTAAAAATAATGGCCAGCNNNAGATCGGAAG |
| 36752_5 | TATGAGTGTGGAGTCGTTGCCCCCTCTAAAAATAATGGCCAGCNNNAGATCGGAAG |
| 36752_6 | TATGAGTGTGGAGTCGTTGCCCCCTCTAAAAATAATGGCCAGCNNNAGATCGGAAG |
| 45665_1 | TATGAGTGTGGAGTCGTTGCTTTCACTCTTGTTACCCAGGCTGNNNAGATCGGAAG |
| 45665_2 | TATGAGTGTGGAGTCGTTGCTTTCACTCTTGTTACCCAGGCTGNNNAGATCGGAAG |
| 45665_3 | TATGAGTGTGGAGTCGTTGCTTTCACTCTTGTTACCCAGGCTGNNNAGATCGGAAG |
| 45665_4 | TATGAGTGTGGAGTCGTTGCTTTCACTCTTGTTACCCAGGCTGNNNAGATCGGAAG |
| 45665_5 | TATGAGTGTGGAGTCGTTGCTTTCACTCTTGTTACCCAGGCTGNNNAGATCGGAAG |
| 45665_6 | TATGAGTGTGGAGTCGTTGCTTTCACTCTTGTTACCCAGGCTGNNNAGATCGGAAG |
| 31611_1 | TATGAGTGTGGAGTCGTTGCCACACACATTCACACACACACATNNNAGATCGGAAG |
| 31611_2 | TATGAGTGTGGAGTCGTTGCCACACACATTCACACACACACATNNNAGATCGGAAG |
| 31611_3 | TATGAGTGTGGAGTCGTTGCCACACACATTCACACACACACATNNNAGATCGGAAG |
| 31611_4 | TATGAGTGTGGAGTCGTTGCCACACACATTCACACACACACATNNNAGATCGGAAG |
| 31611_5 | TATGAGTGTGGAGTCGTTGCCACACACATTCACACACACACATNNNAGATCGGAAG |
| 31611_6 | TATGAGTGTGGAGTCGTTGCCACACACATTCACACACACACATNNNAGATCGGAAG |
| 40554_1 | TATGAGTGTGGAGTCGTTGCGCCACACTGGAAGAAGAATTGTCNNNAGATCGGAA |
| 40554_2 | TATGAGTGTGGAGTCGTTGCGCCACACTGGAAGAAGAATTGTCNNNAGATCGGAA |
| 40554_3 | TATGAGTGTGGAGTCGTTGCGCCACACTGGAAGAAGAATTGTCNNNAGATCGGAA |
| 40554_4 | TATGAGTGTGGAGTCGTTGCGCCACACTGGAAGAAGAATTGTCNNNAGATCGGAA |
| 40554_5 | TATGAGTGTGGAGTCGTTGCGCCACACTGGAAGAAGAATTGTCNNNAGATCGGAA |
| 40554_6 | TATGAGTGTGGAGTCGTTGCGCCACACTGGAAGAAGAATTGTCNNNAGATCGGAA |
| 35341_1 | TATGAGTGTGGAGTCGTTGCAGTAGAGACAGATTTCGCCATGTNNNAGATCGGAA |
| 35341_2 | TATGAGTGTGGAGTCGTTGCAGTAGAGACAGATTTCGCCATGTNNNAGATCGGAA |
| 35341_3 | TATGAGTGTGGAGTCGTTGCAGTAGAGACAGATTTCGCCATGTNNNAGATCGGAA |
| 35341_4 | TATGAGTGTGGAGTCGTTGCAGTAGAGACAGATTTCGCCATGTNNNAGATCGGAA |

| | |
|---|---|
| 35341_5 | TATGAGTGTGGAGTCGTTGCAGTAGAGACAGATTTCGCCATGTNNNAGATCGGAAG |
| 35341_6 | TATGAGTGTGGAGTCGTTGCAGTAGAGACAGATTTCGCCATGTNNNAGATCGGAAG |
| 43600_1 | TATGAGTGTGGAGTCGTTGCACCTGAAGACTTCACTGCTGAATNNNAGATCGGAAG |
| 43600_2 | TATGAGTGTGGAGTCGTTGCACCTGAAGACTTCACTGCTGAATNNNAGATCGGAAG |
| 43600_3 | TATGAGTGTGGAGTCGTTGCACCTGAAGACTTCACTGCTGAATNNNAGATCGGAAG |
| 43600_4 | TATGAGTGTGGAGTCGTTGCACCTGAAGACTTCACTGCTGAATNNNAGATCGGAAG |
| 43600_5 | TATGAGTGTGGAGTCGTTGCACCTGAAGACTTCACTGCTGAATNNNAGATCGGAAG |
| 43600_6 | TATGAGTGTGGAGTCGTTGCACCTGAAGACTTCACTGCTGAATNNNAGATCGGAAG |
| 12961_1 | TATGAGTGTGGAGTCGTTGCCTAAATTCCATCATAAGTCGAAANNNAGATCGGAAGA |
| 12961_2 | TATGAGTGTGGAGTCGTTGCCTAAATTCCATCATAAGTCGAAANNNAGATCGGAAGA |
| 12961_3 | TATGAGTGTGGAGTCGTTGCCTAAATTCCATCATAAGTCGAAANNNAGATCGGAAGA |
| 12961_4 | TATGAGTGTGGAGTCGTTGCCTAAATTCCATCATAAGTCGAAANNNAGATCGGAAGA |
| 12961_5 | TATGAGTGTGGAGTCGTTGCCTAAATTCCATCATAAGTCGAAANNNAGATCGGAAGA |
| 12961_6 | TATGAGTGTGGAGTCGTTGCCTAAATTCCATCATAAGTCGAAANNNAGATCGGAAGA |
| 51481_1 | TATGAGTGTGGAGTCGTTGCGATTGATGAGTCTGCCTTTCACGNNNAGATCGGAAG |
| 51481_2 | TATGAGTGTGGAGTCGTTGCGATTGATGAGTCTGCCTTTCACGNNNAGATCGGAAG |
| 51481_3 | TATGAGTGTGGAGTCGTTGCGATTGATGAGTCTGCCTTTCACGNNNAGATCGGAAG |
| 51481_4 | TATGAGTGTGGAGTCGTTGCGATTGATGAGTCTGCCTTTCACGNNNAGATCGGAAG |
| 51481_5 | TATGAGTGTGGAGTCGTTGCGATTGATGAGTCTGCCTTTCACGNNNAGATCGGAAG |
| 51481_6 | TATGAGTGTGGAGTCGTTGCGATTGATGAGTCTGCCTTTCACGNNNAGATCGGAAG |
| 35182_1 | TATGAGTGTGGAGTCGTTGCCACCACGCCCAGCTAATTTTATNNNAGATCGGAAGA |
| 35182_2 | TATGAGTGTGGAGTCGTTGCCACCACGCCCAGCTAATTTTATNNNAGATCGGAAGA |
| 35182_3 | TATGAGTGTGGAGTCGTTGCCACCACGCCCAGCTAATTTTATNNNAGATCGGAAGA |
| 35182_4 | TATGAGTGTGGAGTCGTTGCCACCACGCCCAGCTAATTTTATNNNAGATCGGAAGA |
| 35182_5 | TATGAGTGTGGAGTCGTTGCCACCACGCCCAGCTAATTTTATNNNAGATCGGAAGA |
| 35182_6 | TATGAGTGTGGAGTCGTTGCCACCACGCCCAGCTAATTTTATNNNAGATCGGAAGA |
| 9281_1 | TATGAGTGTGGAGTCGTTGCTGGCATGTTTGCAACAACTGATGNNNAGATCGGAAG |
| 9281_2 | TATGAGTGTGGAGTCGTTGCTGGCATGTTTGCAACAACTGATGNNNAGATCGGAAG |
| 9281_3 | TATGAGTGTGGAGTCGTTGCTGGCATGTTTGCAACAACTGATGNNNAGATCGGAAG |
| 9281_4 | TATGAGTGTGGAGTCGTTGCTGGCATGTTTGCAACAACTGATGNNNAGATCGGAAG |
| 9281_5 | TATGAGTGTGGAGTCGTTGCTGGCATGTTTGCAACAACTGATGNNNAGATCGGAAG |
| 9281_6 | TATGAGTGTGGAGTCGTTGCTGGCATGTTTGCAACAACTGATGNNNAGATCGGAAG |
| 39755_1 | TATGAGTGTGGAGTCGTTGCTGGAAACCAGTGGCTCATCATAANNNAGATCGGAAG |
| 39755_2 | TATGAGTGTGGAGTCGTTGCTGGAAACCAGTGGCTCATCATAANNNAGATCGGAAG |
| 39755_3 | TATGAGTGTGGAGTCGTTGCTGGAAACCAGTGGCTCATCATAANNNAGATCGGAAG |
| 39755_4 | TATGAGTGTGGAGTCGTTGCTGGAAACCAGTGGCTCATCATAANNNAGATCGGAAG |
| 39755_5 | TATGAGTGTGGAGTCGTTGCTGGAAACCAGTGGCTCATCATAANNNAGATCGGAAG |
| 39755_6 | TATGAGTGTGGAGTCGTTGCTGGAAACCAGTGGCTCATCATAANNNAGATCGGAAG |
| 39756_1 | TATGAGTGTGGAGTCGTTGCCACTGAGTCAAGATCACGTCACTNNNAGATCGGAAG |
| 39756_2 | TATGAGTGTGGAGTCGTTGCCACTGAGTCAAGATCACGTCACTNNNAGATCGGAAG |
| 39756_3 | TATGAGTGTGGAGTCGTTGCCACTGAGTCAAGATCACGTCACTNNNAGATCGGAAG |
| 39756_4 | TATGAGTGTGGAGTCGTTGCCACTGAGTCAAGATCACGTCACTNNNAGATCGGAAG |
| 39756_5 | TATGAGTGTGGAGTCGTTGCCACTGAGTCAAGATCACGTCACTNNNAGATCGGAAG |

| | |
|---|---|
| 39756_6 | TATGAGTGTGGAGTCGTTGCCACTGAGTCAAGATCACGTCACTNNNAGATCGGAAG |
| 47355_1 | TATGAGTGTGGAGTCGTTGCTGCCCTGAAAACCACAAATTCTCNNNAGATCGGAAG |
| 47355_2 | TATGAGTGTGGAGTCGTTGCTGCCCTGAAAACCACAAATTCTCNNNAGATCGGAAG |
| 47355_3 | TATGAGTGTGGAGTCGTTGCTGCCCTGAAAACCACAAATTCTCNNNAGATCGGAAG |
| 47355_4 | TATGAGTGTGGAGTCGTTGCTGCCCTGAAAACCACAAATTCTCNNNAGATCGGAAG |
| 47355_5 | TATGAGTGTGGAGTCGTTGCTGCCCTGAAAACCACAAATTCTCNNNAGATCGGAAG |
| 47355_6 | TATGAGTGTGGAGTCGTTGCTGCCCTGAAAACCACAAATTCTCNNNAGATCGGAAG |
| 34373_1 | TATGAGTGTGGAGTCGTTGCGCCTGTAATCCTAGCTACTCTGGNNNAGATCGGAAG |
| 34373_2 | TATGAGTGTGGAGTCGTTGCGCCTGTAATCCTAGCTACTCTGGNNNAGATCGGAAG |
| 34373_3 | TATGAGTGTGGAGTCGTTGCGCCTGTAATCCTAGCTACTCTGGNNNAGATCGGAAG |
| 34373_4 | TATGAGTGTGGAGTCGTTGCGCCTGTAATCCTAGCTACTCTGGNNNAGATCGGAAG |
| 34373_5 | TATGAGTGTGGAGTCGTTGCGCCTGTAATCCTAGCTACTCTGGNNNAGATCGGAAG |
| 34373_6 | TATGAGTGTGGAGTCGTTGCGCCTGTAATCCTAGCTACTCTGGNNNAGATCGGAAG |
| 10846_1 | TATGAGTGTGGAGTCGTTGCATTGGCCCCTGAAGGTTCACTAANNNAGATCGGAAG |
| 10846_2 | TATGAGTGTGGAGTCGTTGCATTGGCCCCTGAAGGTTCACTAANNNAGATCGGAAG |
| 10846_3 | TATGAGTGTGGAGTCGTTGCATTGGCCCCTGAAGGTTCACTAANNNAGATCGGAAG |
| 10846_4 | TATGAGTGTGGAGTCGTTGCATTGGCCCCTGAAGGTTCACTAANNNAGATCGGAAG |
| 10846_5 | TATGAGTGTGGAGTCGTTGCATTGGCCCCTGAAGGTTCACTAANNNAGATCGGAAG |
| 10846_6 | TATGAGTGTGGAGTCGTTGCATTGGCCCCTGAAGGTTCACTAANNNAGATCGGAAG |
| 37010_1 | TATGAGTGTGGAGTCGTTGCCACACATGCACATACACATACACANNNAGATCGGAA |
| 37010_2 | TATGAGTGTGGAGTCGTTGCCACACATGCACATACACATACACANNNAGATCGGAA |
| 37010_3 | TATGAGTGTGGAGTCGTTGCCACACATGCACATACACATACACANNNAGATCGGAA |
| 37010_4 | TATGAGTGTGGAGTCGTTGCCACACATGCACATACACATACACANNNAGATCGGAA |
| 37010_5 | TATGAGTGTGGAGTCGTTGCCACACATGCACATACACATACACANNNAGATCGGAA |
| 37010_6 | TATGAGTGTGGAGTCGTTGCCACACATGCACATACACATACACANNNAGATCGGAA |
| 35280_1 | TATGAGTGTGGAGTCGTTGCTGTAATTCCCTGGTTCAAGCGATNNNAGATCGGAAG |
| 35280_2 | TATGAGTGTGGAGTCGTTGCTGTAATTCCCTGGTTCAAGCGATNNNAGATCGGAAG |
| 35280_3 | TATGAGTGTGGAGTCGTTGCTGTAATTCCCTGGTTCAAGCGATNNNAGATCGGAAG |
| 35280_4 | TATGAGTGTGGAGTCGTTGCTGTAATTCCCTGGTTCAAGCGATNNNAGATCGGAAG |
| 35280_5 | TATGAGTGTGGAGTCGTTGCTGTAATTCCCTGGTTCAAGCGATNNNAGATCGGAAG |
| 35280_6 | TATGAGTGTGGAGTCGTTGCTGTAATTCCCTGGTTCAAGCGATNNNAGATCGGAAG |
| 40129_1 | TATGAGTGTGGAGTCGTTGCGAGTGAATGAAAAGAAGAGGGGANNNAGATCGGA |
| 40129_2 | TATGAGTGTGGAGTCGTTGCGAGTGAATGAAAAGAAGAGGGGANNNAGATCGGA |
| 40129_3 | TATGAGTGTGGAGTCGTTGCGAGTGAATGAAAAGAAGAGGGGANNNAGATCGGA |
| 40129_4 | TATGAGTGTGGAGTCGTTGCGAGTGAATGAAAAGAAGAGGGGANNNAGATCGGA |
| 40129_5 | TATGAGTGTGGAGTCGTTGCGAGTGAATGAAAAGAAGAGGGGANNNAGATCGGA |
| 40129_6 | TATGAGTGTGGAGTCGTTGCGAGTGAATGAAAAGAAGAGGGGANNNAGATCGGA |
| 49100_1 | TATGAGTGTGGAGTCGTTGCACCCAGTCTTCAATTGATGAGCTNNNAGATCGGAAG |
| 49100_2 | TATGAGTGTGGAGTCGTTGCACCCAGTCTTCAATTGATGAGCTNNNAGATCGGAAG |
| 49100_3 | TATGAGTGTGGAGTCGTTGCACCCAGTCTTCAATTGATGAGCTNNNAGATCGGAAG |
| 49100_4 | TATGAGTGTGGAGTCGTTGCACCCAGTCTTCAATTGATGAGCTNNNAGATCGGAAG |
| 49100_5 | TATGAGTGTGGAGTCGTTGCACCCAGTCTTCAATTGATGAGCTNNNAGATCGGAAG |
| 49100_6 | TATGAGTGTGGAGTCGTTGCACCCAGTCTTCAATTGATGAGCTNNNAGATCGGAAG |

| | |
|---|---|
| 47816_1 | TATGAGTGTGGAGTCGTTGCCTCTGCTCTGCTAGTCTTTCCACNNNAGATCGGAAGA |
| 47816_2 | TATGAGTGTGGAGTCGTTGCCTCTGCTCTGCTAGTCTTTCCACNNNAGATCGGAAGA |
| 47816_3 | TATGAGTGTGGAGTCGTTGCCTCTGCTCTGCTAGTCTTTCCACNNNAGATCGGAAGA |
| 47816_4 | TATGAGTGTGGAGTCGTTGCCTCTGCTCTGCTAGTCTTTCCACNNNAGATCGGAAGA |
| 47816_5 | TATGAGTGTGGAGTCGTTGCCTCTGCTCTGCTAGTCTTTCCACNNNAGATCGGAAGA |
| 47816_6 | TATGAGTGTGGAGTCGTTGCCTCTGCTCTGCTAGTCTTTCCACNNNAGATCGGAAGA |
| 32774_1 | TATGAGTGTGGAGTCGTTGCCATGATCATATCACTGCCCTCCANNNAGATCGGAAGA |
| 32774_2 | TATGAGTGTGGAGTCGTTGCCATGATCATATCACTGCCCTCCANNNAGATCGGAAGA |
| 32774_3 | TATGAGTGTGGAGTCGTTGCCATGATCATATCACTGCCCTCCANNNAGATCGGAAGA |
| 32774_4 | TATGAGTGTGGAGTCGTTGCCATGATCATATCACTGCCCTCCANNNAGATCGGAAGA |
| 32774_5 | TATGAGTGTGGAGTCGTTGCCATGATCATATCACTGCCCTCCANNNAGATCGGAAGA |
| 32774_6 | TATGAGTGTGGAGTCGTTGCCATGATCATATCACTGCCCTCCANNNAGATCGGAAGA |
| 53115_1 | TATGAGTGTGGAGTCGTTGCGGTCAGCACCAAGGTAGAACTTANNNAGATCGGAA |
| 53115_2 | TATGAGTGTGGAGTCGTTGCGGTCAGCACCAAGGTAGAACTTANNNAGATCGGAA |
| 53115_3 | TATGAGTGTGGAGTCGTTGCGGTCAGCACCAAGGTAGAACTTANNNAGATCGGAA |
| 53115_4 | TATGAGTGTGGAGTCGTTGCGGTCAGCACCAAGGTAGAACTTANNNAGATCGGAA |
| 53115_5 | TATGAGTGTGGAGTCGTTGCGGTCAGCACCAAGGTAGAACTTANNNAGATCGGAA |
| 53115_6 | TATGAGTGTGGAGTCGTTGCGGTCAGCACCAAGGTAGAACTTANNNAGATCGGAA |
| 48380_1 | TATGAGTGTGGAGTCGTTGCTTTGAGACAGGGTCTTGTTCTGTNNNAGATCGGAAG |
| 48380_2 | TATGAGTGTGGAGTCGTTGCTTTGAGACAGGGTCTTGTTCTGTNNNAGATCGGAAG |
| 48380_3 | TATGAGTGTGGAGTCGTTGCTTTGAGACAGGGTCTTGTTCTGTNNNAGATCGGAAG |
| 48380_4 | TATGAGTGTGGAGTCGTTGCTTTGAGACAGGGTCTTGTTCTGTNNNAGATCGGAAG |
| 48380_5 | TATGAGTGTGGAGTCGTTGCTTTGAGACAGGGTCTTGTTCTGTNNNAGATCGGAAG |
| 48380_6 | TATGAGTGTGGAGTCGTTGCTTTGAGACAGGGTCTTGTTCTGTNNNAGATCGGAAG |
| 30398_1 | TATGAGTGTGGAGTCGTTGCGCCAGAAGTCCACAATCAGTTTCNNNAGATCGGAAG |
| 30398_2 | TATGAGTGTGGAGTCGTTGCGCCAGAAGTCCACAATCAGTTTCNNNAGATCGGAAG |
| 30398_3 | TATGAGTGTGGAGTCGTTGCGCCAGAAGTCCACAATCAGTTTCNNNAGATCGGAAG |
| 30398_4 | TATGAGTGTGGAGTCGTTGCGCCAGAAGTCCACAATCAGTTTCNNNAGATCGGAAG |
| 30398_5 | TATGAGTGTGGAGTCGTTGCGCCAGAAGTCCACAATCAGTTTCNNNAGATCGGAAG |
| 30398_6 | TATGAGTGTGGAGTCGTTGCGCCAGAAGTCCACAATCAGTTTCNNNAGATCGGAAG |
| 32771_1 | TATGAGTGTGGAGTCGTTGCTGAAGTCTTATAGTGGCTTCCCTNNNAGATCGGAAG |
| 32771_2 | TATGAGTGTGGAGTCGTTGCTGAAGTCTTATAGTGGCTTCCCTNNNAGATCGGAAG |
| 32771_3 | TATGAGTGTGGAGTCGTTGCTGAAGTCTTATAGTGGCTTCCCTNNNAGATCGGAAG |
| 32771_4 | TATGAGTGTGGAGTCGTTGCTGAAGTCTTATAGTGGCTTCCCTNNNAGATCGGAAG |
| 32771_5 | TATGAGTGTGGAGTCGTTGCTGAAGTCTTATAGTGGCTTCCCTNNNAGATCGGAAG |
| 32771_6 | TATGAGTGTGGAGTCGTTGCTGAAGTCTTATAGTGGCTTCCCTNNNAGATCGGAAG |
| 39462_1 | TATGAGTGTGGAGTCGTTGCCCAATGACCAAAGCTGGAACAATNNNAGATCGGAAG |
| 39462_2 | TATGAGTGTGGAGTCGTTGCCCAATGACCAAAGCTGGAACAATNNNAGATCGGAAG |
| 39462_3 | TATGAGTGTGGAGTCGTTGCCCAATGACCAAAGCTGGAACAATNNNAGATCGGAAG |
| 39462_4 | TATGAGTGTGGAGTCGTTGCCCAATGACCAAAGCTGGAACAATNNNAGATCGGAAG |
| 39462_5 | TATGAGTGTGGAGTCGTTGCCCAATGACCAAAGCTGGAACAATNNNAGATCGGAAG |
| 39462_6 | TATGAGTGTGGAGTCGTTGCCCAATGACCAAAGCTGGAACAATNNNAGATCGGAAG |
| 31177_1 | TATGAGTGTGGAGTCGTTGCGTGTGTCTGTGTGGTAAATGAGCNNNAGATCGGAA |

| | |
|---|---|
| 31177_2 | TATGAGTGTGGAGTCGTTGCGTGTGTCTGTGTGGTAAATGAGCNNNAGATCGGAA |
| 31177_3 | TATGAGTGTGGAGTCGTTGCGTGTGTCTGTGTGGTAAATGAGCNNNAGATCGGAA |
| 31177_4 | TATGAGTGTGGAGTCGTTGCGTGTGTCTGTGTGGTAAATGAGCNNNAGATCGGAA |
| 31177_5 | TATGAGTGTGGAGTCGTTGCGTGTGTCTGTGTGGTAAATGAGCNNNAGATCGGAA |
| 31177_6 | TATGAGTGTGGAGTCGTTGCGTGTGTCTGTGTGGTAAATGAGCNNNAGATCGGAA |
| 50267_1 | TATGAGTGTGGAGTCGTTGCATAGACAAGAGGCACACAGAGACNNNAGATCGGAA |
| 50267_2 | TATGAGTGTGGAGTCGTTGCATAGACAAGAGGCACACAGAGACNNNAGATCGGAA |
| 50267_3 | TATGAGTGTGGAGTCGTTGCATAGACAAGAGGCACACAGAGACNNNAGATCGGAA |
| 50267_4 | TATGAGTGTGGAGTCGTTGCATAGACAAGAGGCACACAGAGACNNNAGATCGGAA |
| 50267_5 | TATGAGTGTGGAGTCGTTGCATAGACAAGAGGCACACAGAGACNNNAGATCGGAA |
| 50267_6 | TATGAGTGTGGAGTCGTTGCATAGACAAGAGGCACACAGAGACNNNAGATCGGAA |
| 45261_1 | TATGAGTGTGGAGTCGTTGCAGGGAGCTGGTAGACAGAACTATNNNAGATCGGAA |
| 45261_2 | TATGAGTGTGGAGTCGTTGCAGGGAGCTGGTAGACAGAACTATNNNAGATCGGAA |
| 45261_3 | TATGAGTGTGGAGTCGTTGCAGGGAGCTGGTAGACAGAACTATNNNAGATCGGAA |
| 45261_4 | TATGAGTGTGGAGTCGTTGCAGGGAGCTGGTAGACAGAACTATNNNAGATCGGAA |
| 45261_5 | TATGAGTGTGGAGTCGTTGCAGGGAGCTGGTAGACAGAACTATNNNAGATCGGAA |
| 45261_6 | TATGAGTGTGGAGTCGTTGCAGGGAGCTGGTAGACAGAACTATNNNAGATCGGAA |
| 37271_1 | TATGAGTGTGGAGTCGTTGCTGTCTCTTTTCCTATCACTGGCCNNNAGATCGGAAGA |
| 37271_2 | TATGAGTGTGGAGTCGTTGCTGTCTCTTTTCCTATCACTGGCCNNNAGATCGGAAGA |
| 37271_3 | TATGAGTGTGGAGTCGTTGCTGTCTCTTTTCCTATCACTGGCCNNNAGATCGGAAGA |
| 37271_4 | TATGAGTGTGGAGTCGTTGCTGTCTCTTTTCCTATCACTGGCCNNNAGATCGGAAGA |
| 37271_5 | TATGAGTGTGGAGTCGTTGCTGTCTCTTTTCCTATCACTGGCCNNNAGATCGGAAGA |
| 37271_6 | TATGAGTGTGGAGTCGTTGCTGTCTCTTTTCCTATCACTGGCCNNNAGATCGGAAGA |
| 35528_1 | TATGAGTGTGGAGTCGTTGCCAATTGATTCTTCTGCCTCAGCCNNNAGATCGGAAGA |
| 35528_2 | TATGAGTGTGGAGTCGTTGCCAATTGATTCTTCTGCCTCAGCCNNNAGATCGGAAGA |
| 35528_3 | TATGAGTGTGGAGTCGTTGCCAATTGATTCTTCTGCCTCAGCCNNNAGATCGGAAGA |
| 35528_4 | TATGAGTGTGGAGTCGTTGCCAATTGATTCTTCTGCCTCAGCCNNNAGATCGGAAGA |
| 35528_5 | TATGAGTGTGGAGTCGTTGCCAATTGATTCTTCTGCCTCAGCCNNNAGATCGGAAGA |
| 35528_6 | TATGAGTGTGGAGTCGTTGCCAATTGATTCTTCTGCCTCAGCCNNNAGATCGGAAGA |
| 43128_1 | TATGAGTGTGGAGTCGTTGCACAAAAGGGGAAAGGGAAGTTCTNNNAGATCGGAA |
| 43128_2 | TATGAGTGTGGAGTCGTTGCACAAAAGGGGAAAGGGAAGTTCTNNNAGATCGGAA |
| 43128_3 | TATGAGTGTGGAGTCGTTGCACAAAAGGGGAAAGGGAAGTTCTNNNAGATCGGAA |
| 43128_4 | TATGAGTGTGGAGTCGTTGCACAAAAGGGGAAAGGGAAGTTCTNNNAGATCGGAA |
| 43128_5 | TATGAGTGTGGAGTCGTTGCACAAAAGGGGAAAGGGAAGTTCTNNNAGATCGGAA |
| 43128_6 | TATGAGTGTGGAGTCGTTGCACAAAAGGGGAAAGGGAAGTTCTNNNAGATCGGAA |
| 39437_1 | TATGAGTGTGGAGTCGTTGCTCTAGTTCTATTGCCCAGGTTGGNNNAGATCGGAAG |
| 39437_2 | TATGAGTGTGGAGTCGTTGCTCTAGTTCTATTGCCCAGGTTGGNNNAGATCGGAAG |
| 39437_3 | TATGAGTGTGGAGTCGTTGCTCTAGTTCTATTGCCCAGGTTGGNNNAGATCGGAAG |
| 39437_4 | TATGAGTGTGGAGTCGTTGCTCTAGTTCTATTGCCCAGGTTGGNNNAGATCGGAAG |
| 39437_5 | TATGAGTGTGGAGTCGTTGCTCTAGTTCTATTGCCCAGGTTGGNNNAGATCGGAAG |
| 39437_6 | TATGAGTGTGGAGTCGTTGCTCTAGTTCTATTGCCCAGGTTGGNNNAGATCGGAAG |
| 41606_1 | TATGAGTGTGGAGTCGTTGCCTCCTGAGTAGCTGGGATTACAGNNNAGATCGGAAG |
| 41606_2 | TATGAGTGTGGAGTCGTTGCCTCCTGAGTAGCTGGGATTACAGNNNAGATCGGAAG |

| | |
|---|---|
| 41606_3 | TATGAGTGTGGAGTCGTTGCCTCCTGAGTAGCTGGGATTACAGNNNAGATCGGAA |
| 41606_4 | TATGAGTGTGGAGTCGTTGCCTCCTGAGTAGCTGGGATTACAGNNNAGATCGGAA |
| 41606_5 | TATGAGTGTGGAGTCGTTGCCTCCTGAGTAGCTGGGATTACAGNNNAGATCGGAA |
| 41606_6 | TATGAGTGTGGAGTCGTTGCCTCCTGAGTAGCTGGGATTACAGNNNAGATCGGAA |
| 12745_1 | TATGAGTGTGGAGTCGTTGCCTGCAGGAGTAACTAAGATTATANNNAGATCGGAA |
| 12745_2 | TATGAGTGTGGAGTCGTTGCCTGCAGGAGTAACTAAGATTATANNNAGATCGGAA |
| 12745_3 | TATGAGTGTGGAGTCGTTGCCTGCAGGAGTAACTAAGATTATANNNAGATCGGAA |
| 12745_4 | TATGAGTGTGGAGTCGTTGCCTGCAGGAGTAACTAAGATTATANNNAGATCGGAA |
| 12745_5 | TATGAGTGTGGAGTCGTTGCCTGCAGGAGTAACTAAGATTATANNNAGATCGGAA |
| 12745_6 | TATGAGTGTGGAGTCGTTGCCTGCAGGAGTAACTAAGATTATANNNAGATCGGAA |
| 41447_1 | TATGAGTGTGGAGTCGTTGCTCTTTGAGTTATAGAAGACGGCGTNNNAGATCGGA |
| 41447_2 | TATGAGTGTGGAGTCGTTGCTCTTTGAGTTATAGAAGACGGCGTNNNAGATCGGA |
| 41447_3 | TATGAGTGTGGAGTCGTTGCTCTTTGAGTTATAGAAGACGGCGTNNNAGATCGGA |
| 41447_4 | TATGAGTGTGGAGTCGTTGCTCTTTGAGTTATAGAAGACGGCGTNNNAGATCGGA |
| 41447_5 | TATGAGTGTGGAGTCGTTGCTCTTTGAGTTATAGAAGACGGCGTNNNAGATCGGA |
| 41447_6 | TATGAGTGTGGAGTCGTTGCTCTTTGAGTTATAGAAGACGGCGTNNNAGATCGGA |
| 2551_1 | TATGAGTGTGGAGTCGTTGCAGGTCTAGCATAAATGTTTTCTCAGGANNNAGATCG |
| 2551_2 | TATGAGTGTGGAGTCGTTGCAGGTCTAGCATAAATGTTTTCTCAGGANNNAGATCG |
| 2551_3 | TATGAGTGTGGAGTCGTTGCAGGTCTAGCATAAATGTTTTCTCAGGANNNAGATCG |
| 2551_4 | TATGAGTGTGGAGTCGTTGCAGGTCTAGCATAAATGTTTTCTCAGGANNNAGATCG |
| 2551_5 | TATGAGTGTGGAGTCGTTGCAGGTCTAGCATAAATGTTTTCTCAGGANNNAGATCG |
| 2551_6 | TATGAGTGTGGAGTCGTTGCAGGTCTAGCATAAATGTTTTCTCAGGANNNAGATCG |
| 35788_1 | TATGAGTGTGGAGTCGTTGCCTGGTATGTTCCTGTTTCCCTCANNNAGATCGGAAGA |
| 35788_2 | TATGAGTGTGGAGTCGTTGCCTGGTATGTTCCTGTTTCCCTCANNNAGATCGGAAGA |
| 35788_3 | TATGAGTGTGGAGTCGTTGCCTGGTATGTTCCTGTTTCCCTCANNNAGATCGGAAGA |
| 35788_4 | TATGAGTGTGGAGTCGTTGCCTGGTATGTTCCTGTTTCCCTCANNNAGATCGGAAGA |
| 35788_5 | TATGAGTGTGGAGTCGTTGCCTGGTATGTTCCTGTTTCCCTCANNNAGATCGGAAGA |
| 35788_6 | TATGAGTGTGGAGTCGTTGCCTGGTATGTTCCTGTTTCCCTCANNNAGATCGGAAGA |
| 40330_1 | TATGAGTGTGGAGTCGTTGCAATTGCCACATCTTCCACAATGGNNNAGATCGGAAG |
| 40330_2 | TATGAGTGTGGAGTCGTTGCAATTGCCACATCTTCCACAATGGNNNAGATCGGAAG |
| 40330_3 | TATGAGTGTGGAGTCGTTGCAATTGCCACATCTTCCACAATGGNNNAGATCGGAAG |
| 40330_4 | TATGAGTGTGGAGTCGTTGCAATTGCCACATCTTCCACAATGGNNNAGATCGGAAG |
| 40330_5 | TATGAGTGTGGAGTCGTTGCAATTGCCACATCTTCCACAATGGNNNAGATCGGAAG |
| 40330_6 | TATGAGTGTGGAGTCGTTGCAATTGCCACATCTTCCACAATGGNNNAGATCGGAAG |
| 38578_1 | TATGAGTGTGGAGTCGTTGCAAGGGGATACTGGTGCCAAAATTNNNAGATCGGAA |
| 38578_2 | TATGAGTGTGGAGTCGTTGCAAGGGGATACTGGTGCCAAAATTNNNAGATCGGAA |
| 38578_3 | TATGAGTGTGGAGTCGTTGCAAGGGGATACTGGTGCCAAAATTNNNAGATCGGAA |
| 38578_4 | TATGAGTGTGGAGTCGTTGCAAGGGGATACTGGTGCCAAAATTNNNAGATCGGAA |
| 38578_5 | TATGAGTGTGGAGTCGTTGCAAGGGGATACTGGTGCCAAAATTNNNAGATCGGAA |
| 38578_6 | TATGAGTGTGGAGTCGTTGCAAGGGGATACTGGTGCCAAAATTNNNAGATCGGAA |
| 10414_1 | TATGAGTGTGGAGTCGTTGCTATGCGGGATGTGTGTGTATGTGNNNAGATCGGAA |
| 10414_2 | TATGAGTGTGGAGTCGTTGCTATGCGGGATGTGTGTGTATGTGNNNAGATCGGAA |
| 10414_3 | TATGAGTGTGGAGTCGTTGCTATGCGGGATGTGTGTGTATGTGNNNAGATCGGAA |

| | |
|---|---|
| 10414_4 | TATGAGTGTGGAGTCGTTGCTATGCGGGATGTGTGTGTATGTGNNNAGATCGGAA |
| 10414_5 | TATGAGTGTGGAGTCGTTGCTATGCGGGATGTGTGTGTATGTGNNNAGATCGGAA |
| 10414_6 | TATGAGTGTGGAGTCGTTGCTATGCGGGATGTGTGTGTATGTGNNNAGATCGGAA |
| 31554_1 | TATGAGTGTGGAGTCGTTGCACATCAAAATTCGTGGTCCCTCTNNNAGATCGGAAG |
| 31554_2 | TATGAGTGTGGAGTCGTTGCACATCAAAATTCGTGGTCCCTCTNNNAGATCGGAAG |
| 31554_3 | TATGAGTGTGGAGTCGTTGCACATCAAAATTCGTGGTCCCTCTNNNAGATCGGAAG |
| 31554_4 | TATGAGTGTGGAGTCGTTGCACATCAAAATTCGTGGTCCCTCTNNNAGATCGGAAG |
| 31554_5 | TATGAGTGTGGAGTCGTTGCACATCAAAATTCGTGGTCCCTCTNNNAGATCGGAAG |
| 31554_6 | TATGAGTGTGGAGTCGTTGCACATCAAAATTCGTGGTCCCTCTNNNAGATCGGAAG |
| 29109_1 | TATGAGTGTGGAGTCGTTGCTGAGACTGCAAAAACCATAGTGCNNNAGATCGGAA |
| 29109_2 | TATGAGTGTGGAGTCGTTGCTGAGACTGCAAAAACCATAGTGCNNNAGATCGGAA |
| 29109_3 | TATGAGTGTGGAGTCGTTGCTGAGACTGCAAAAACCATAGTGCNNNAGATCGGAA |
| 29109_4 | TATGAGTGTGGAGTCGTTGCTGAGACTGCAAAAACCATAGTGCNNNAGATCGGAA |
| 29109_5 | TATGAGTGTGGAGTCGTTGCTGAGACTGCAAAAACCATAGTGCNNNAGATCGGAA |
| 29109_6 | TATGAGTGTGGAGTCGTTGCTGAGACTGCAAAAACCATAGTGCNNNAGATCGGAA |
| 13481_1 | TATGAGTGTGGAGTCGTTGCCATGAATGTAATCTTAGCTTTCANNNAGATCGGAAG |
| 13481_2 | TATGAGTGTGGAGTCGTTGCCATGAATGTAATCTTAGCTTTCANNNAGATCGGAAG |
| 13481_3 | TATGAGTGTGGAGTCGTTGCCATGAATGTAATCTTAGCTTTCANNNAGATCGGAAG |
| 13481_4 | TATGAGTGTGGAGTCGTTGCCATGAATGTAATCTTAGCTTTCANNNAGATCGGAAG |
| 13481_5 | TATGAGTGTGGAGTCGTTGCCATGAATGTAATCTTAGCTTTCANNNAGATCGGAAG |
| 13481_6 | TATGAGTGTGGAGTCGTTGCCATGAATGTAATCTTAGCTTTCANNNAGATCGGAAG |
| 47889_1 | TATGAGTGTGGAGTCGTTGCAACTTCCACTTTCCAGGCTCAGNNNAGATCGGAAGA |
| 47889_2 | TATGAGTGTGGAGTCGTTGCAACTTCCACTTTCCAGGCTCAGNNNAGATCGGAAGA |
| 47889_3 | TATGAGTGTGGAGTCGTTGCAACTTCCACTTTCCAGGCTCAGNNNAGATCGGAAGA |
| 47889_4 | TATGAGTGTGGAGTCGTTGCAACTTCCACTTTCCAGGCTCAGNNNAGATCGGAAGA |
| 47889_5 | TATGAGTGTGGAGTCGTTGCAACTTCCACTTTCCAGGCTCAGNNNAGATCGGAAGA |
| 47889_6 | TATGAGTGTGGAGTCGTTGCAACTTCCACTTTCCAGGCTCAGNNNAGATCGGAAGA |
| 47881_1 | TATGAGTGTGGAGTCGTTGCCACCACACCTGGCTAATTTTTGTNNNAGATCGGAAG |
| 47881_2 | TATGAGTGTGGAGTCGTTGCCACCACACCTGGCTAATTTTTGTNNNAGATCGGAAG |
| 47881_3 | TATGAGTGTGGAGTCGTTGCCACCACACCTGGCTAATTTTTGTNNNAGATCGGAAG |
| 47881_4 | TATGAGTGTGGAGTCGTTGCCACCACACCTGGCTAATTTTTGTNNNAGATCGGAAG |
| 47881_5 | TATGAGTGTGGAGTCGTTGCCACCACACCTGGCTAATTTTTGTNNNAGATCGGAAG |
| 47881_6 | TATGAGTGTGGAGTCGTTGCCACCACACCTGGCTAATTTTTGTNNNAGATCGGAAG |
| 29721_1 | TATGAGTGTGGAGTCGTTGCTCACAGATTCCATTAACTAGCGGTNNNAGATCGGAA |
| 29721_2 | TATGAGTGTGGAGTCGTTGCTCACAGATTCCATTAACTAGCGGTNNNAGATCGGAA |
| 29721_3 | TATGAGTGTGGAGTCGTTGCTCACAGATTCCATTAACTAGCGGTNNNAGATCGGAA |
| 29721_4 | TATGAGTGTGGAGTCGTTGCTCACAGATTCCATTAACTAGCGGTNNNAGATCGGAA |
| 29721_5 | TATGAGTGTGGAGTCGTTGCTCACAGATTCCATTAACTAGCGGTNNNAGATCGGAA |
| 29721_6 | TATGAGTGTGGAGTCGTTGCTCACAGATTCCATTAACTAGCGGTNNNAGATCGGAA |
| 53170_1 | TATGAGTGTGGAGTCGTTGCCCCATTTTGAAGGTGGAGTTGTGNNNAGATCGGAA |
| 53170_2 | TATGAGTGTGGAGTCGTTGCCCCATTTTGAAGGTGGAGTTGTGNNNAGATCGGAA |
| 53170_3 | TATGAGTGTGGAGTCGTTGCCCCATTTTGAAGGTGGAGTTGTGNNNAGATCGGAA |
| 53170_4 | TATGAGTGTGGAGTCGTTGCCCCATTTTGAAGGTGGAGTTGTGNNNAGATCGGAA |

| | |
|---|---|
| 53170_5 | TATGAGTGTGGAGTCGTTGCCCCATTTTGAAGGTGGAGTTGTGNNNAGATCGGAAC |
| 53170_6 | TATGAGTGTGGAGTCGTTGCCCCATTTTGAAGGTGGAGTTGTGNNNAGATCGGAAC |
| 2835_1 | TATGAGTGTGGAGTCGTTGCCTCTTGCCCATTCTTCCAACAGTNNNAGATCGGAAGA |
| 2835_2 | TATGAGTGTGGAGTCGTTGCCTCTTGCCCATTCTTCCAACAGTNNNAGATCGGAAGA |
| 2835_3 | TATGAGTGTGGAGTCGTTGCCTCTTGCCCATTCTTCCAACAGTNNNAGATCGGAAGA |
| 2835_4 | TATGAGTGTGGAGTCGTTGCCTCTTGCCCATTCTTCCAACAGTNNNAGATCGGAAGA |
| 2835_5 | TATGAGTGTGGAGTCGTTGCCTCTTGCCCATTCTTCCAACAGTNNNAGATCGGAAGA |
| 2835_6 | TATGAGTGTGGAGTCGTTGCCTCTTGCCCATTCTTCCAACAGTNNNAGATCGGAAGA |
| 12981_1 | TATGAGTGTGGAGTCGTTGCAAAAGTCCAGTTTGTATGTTTTGNNNAGATCGGAAG |
| 12981_2 | TATGAGTGTGGAGTCGTTGCAAAAGTCCAGTTTGTATGTTTTGNNNAGATCGGAAG |
| 12981_3 | TATGAGTGTGGAGTCGTTGCAAAAGTCCAGTTTGTATGTTTTGNNNAGATCGGAAG |
| 12981_4 | TATGAGTGTGGAGTCGTTGCAAAAGTCCAGTTTGTATGTTTTGNNNAGATCGGAAG |
| 12981_5 | TATGAGTGTGGAGTCGTTGCAAAAGTCCAGTTTGTATGTTTTGNNNAGATCGGAAG |
| 12981_6 | TATGAGTGTGGAGTCGTTGCAAAAGTCCAGTTTGTATGTTTTGNNNAGATCGGAAG |
| 30271_1 | TATGAGTGTGGAGTCGTTGCCACATTGACCAAGCTCCTGAAAGNNNAGATCGGAAG |
| 30271_2 | TATGAGTGTGGAGTCGTTGCCACATTGACCAAGCTCCTGAAAGNNNAGATCGGAAG |
| 30271_3 | TATGAGTGTGGAGTCGTTGCCACATTGACCAAGCTCCTGAAAGNNNAGATCGGAAG |
| 30271_4 | TATGAGTGTGGAGTCGTTGCCACATTGACCAAGCTCCTGAAAGNNNAGATCGGAAG |
| 30271_5 | TATGAGTGTGGAGTCGTTGCCACATTGACCAAGCTCCTGAAAGNNNAGATCGGAAG |
| 30271_6 | TATGAGTGTGGAGTCGTTGCCACATTGACCAAGCTCCTGAAAGNNNAGATCGGAAG |
| 28865_1 | TATGAGTGTGGAGTCGTTGCTGACACTTGTTTGAAGCTTTGCTNNNAGATCGGAAGA |
| 28865_2 | TATGAGTGTGGAGTCGTTGCTGACACTTGTTTGAAGCTTTGCTNNNAGATCGGAAGA |
| 28865_3 | TATGAGTGTGGAGTCGTTGCTGACACTTGTTTGAAGCTTTGCTNNNAGATCGGAAGA |
| 28865_4 | TATGAGTGTGGAGTCGTTGCTGACACTTGTTTGAAGCTTTGCTNNNAGATCGGAAGA |
| 28865_5 | TATGAGTGTGGAGTCGTTGCTGACACTTGTTTGAAGCTTTGCTNNNAGATCGGAAGA |
| 28865_6 | TATGAGTGTGGAGTCGTTGCTGACACTTGTTTGAAGCTTTGCTNNNAGATCGGAAGA |
| 32848_1 | TATGAGTGTGGAGTCGTTGCCAGAGTCTCATTCTGTCTCCCGNNNAGATCGGAAGAC |
| 32848_2 | TATGAGTGTGGAGTCGTTGCCAGAGTCTCATTCTGTCTCCCGNNNAGATCGGAAGAC |
| 32848_3 | TATGAGTGTGGAGTCGTTGCCAGAGTCTCATTCTGTCTCCCGNNNAGATCGGAAGAC |
| 32848_4 | TATGAGTGTGGAGTCGTTGCCAGAGTCTCATTCTGTCTCCCGNNNAGATCGGAAGAC |
| 32848_5 | TATGAGTGTGGAGTCGTTGCCAGAGTCTCATTCTGTCTCCCGNNNAGATCGGAAGAC |
| 32848_6 | TATGAGTGTGGAGTCGTTGCCAGAGTCTCATTCTGTCTCCCGNNNAGATCGGAAGAC |
| 51348_1 | TATGAGTGTGGAGTCGTTGCCACTGCACATGGTCAAATTTGGTNNNAGATCGGAAG |
| 51348_2 | TATGAGTGTGGAGTCGTTGCCACTGCACATGGTCAAATTTGGTNNNAGATCGGAAG |
| 51348_3 | TATGAGTGTGGAGTCGTTGCCACTGCACATGGTCAAATTTGGTNNNAGATCGGAAG |
| 51348_4 | TATGAGTGTGGAGTCGTTGCCACTGCACATGGTCAAATTTGGTNNNAGATCGGAAG |
| 51348_5 | TATGAGTGTGGAGTCGTTGCCACTGCACATGGTCAAATTTGGTNNNAGATCGGAAG |
| 51348_6 | TATGAGTGTGGAGTCGTTGCCACTGCACATGGTCAAATTTGGTNNNAGATCGGAAG |
| 42986_1 | TATGAGTGTGGAGTCGTTGCTAGAGTACCGAAGCCACTTATGCNNNAGATCGGAAG |
| 42986_2 | TATGAGTGTGGAGTCGTTGCTAGAGTACCGAAGCCACTTATGCNNNAGATCGGAAG |
| 42986_3 | TATGAGTGTGGAGTCGTTGCTAGAGTACCGAAGCCACTTATGCNNNAGATCGGAAG |
| 42986_4 | TATGAGTGTGGAGTCGTTGCTAGAGTACCGAAGCCACTTATGCNNNAGATCGGAAG |
| 42986_5 | TATGAGTGTGGAGTCGTTGCTAGAGTACCGAAGCCACTTATGCNNNAGATCGGAAG |

| | |
|---|---|
| 42986_6 | TATGAGTGTGGAGTCGTTGCTAGAGTACCGAAGCCACTTATGCNNNAGATCGGAAG |
| 46600_1 | TATGAGTGTGGAGTCGTTGCCACCATGCCCAGCTAATTTTTGTNNNAGATCGGAAGA |
| 46600_2 | TATGAGTGTGGAGTCGTTGCCACCATGCCCAGCTAATTTTTGTNNNAGATCGGAAGA |
| 46600_3 | TATGAGTGTGGAGTCGTTGCCACCATGCCCAGCTAATTTTTGTNNNAGATCGGAAGA |
| 46600_4 | TATGAGTGTGGAGTCGTTGCCACCATGCCCAGCTAATTTTTGTNNNAGATCGGAAGA |
| 46600_5 | TATGAGTGTGGAGTCGTTGCCACCATGCCCAGCTAATTTTTGTNNNAGATCGGAAGA |
| 46600_6 | TATGAGTGTGGAGTCGTTGCCACCATGCCCAGCTAATTTTTGTNNNAGATCGGAAGA |
| 45817_1 | TATGAGTGTGGAGTCGTTGCAAATGGACGGCTGGCAATTTTTANNNAGATCGGAA[illegible] |
| 45817_2 | TATGAGTGTGGAGTCGTTGCAAATGGACGGCTGGCAATTTTTANNNAGATCGGAA[illegible] |
| 45817_3 | TATGAGTGTGGAGTCGTTGCAAATGGACGGCTGGCAATTTTTANNNAGATCGGAA[illegible] |
| 45817_4 | TATGAGTGTGGAGTCGTTGCAAATGGACGGCTGGCAATTTTTANNNAGATCGGAA[illegible] |
| 45817_5 | TATGAGTGTGGAGTCGTTGCAAATGGACGGCTGGCAATTTTTANNNAGATCGGAA[illegible] |
| 45817_6 | TATGAGTGTGGAGTCGTTGCAAATGGACGGCTGGCAATTTTTANNNAGATCGGAA[illegible] |
| 43318_1 | TATGAGTGTGGAGTCGTTGCAAAATACTCCTGGGGCTTCCTTTNNNAGATCGGAAGA |
| 43318_2 | TATGAGTGTGGAGTCGTTGCAAAATACTCCTGGGGCTTCCTTTNNNAGATCGGAAGA |
| 43318_3 | TATGAGTGTGGAGTCGTTGCAAAATACTCCTGGGGCTTCCTTTNNNAGATCGGAAGA |
| 43318_4 | TATGAGTGTGGAGTCGTTGCAAAATACTCCTGGGGCTTCCTTTNNNAGATCGGAAGA |
| 43318_5 | TATGAGTGTGGAGTCGTTGCAAAATACTCCTGGGGCTTCCTTTNNNAGATCGGAAGA |
| 43318_6 | TATGAGTGTGGAGTCGTTGCAAAATACTCCTGGGGCTTCCTTTNNNAGATCGGAAGA |
| 36552_1 | TATGAGTGTGGAGTCGTTGCATGAAACAACTCCCCATTCCTGTNNNAGATCGGAAGA |
| 36552_2 | TATGAGTGTGGAGTCGTTGCATGAAACAACTCCCCATTCCTGTNNNAGATCGGAAGA |
| 36552_3 | TATGAGTGTGGAGTCGTTGCATGAAACAACTCCCCATTCCTGTNNNAGATCGGAAGA |
| 36552_4 | TATGAGTGTGGAGTCGTTGCATGAAACAACTCCCCATTCCTGTNNNAGATCGGAAGA |
| 36552_5 | TATGAGTGTGGAGTCGTTGCATGAAACAACTCCCCATTCCTGTNNNAGATCGGAAGA |
| 36552_6 | TATGAGTGTGGAGTCGTTGCATGAAACAACTCCCCATTCCTGTNNNAGATCGGAAGA |
| 44043_1 | TATGAGTGTGGAGTCGTTGCTTCCTACATCCAAGAATGTGCCANNNAGATCGGAAGA |
| 44043_2 | TATGAGTGTGGAGTCGTTGCTTCCTACATCCAAGAATGTGCCANNNAGATCGGAAGA |
| 44043_3 | TATGAGTGTGGAGTCGTTGCTTCCTACATCCAAGAATGTGCCANNNAGATCGGAAGA |
| 44043_4 | TATGAGTGTGGAGTCGTTGCTTCCTACATCCAAGAATGTGCCANNNAGATCGGAAGA |
| 44043_5 | TATGAGTGTGGAGTCGTTGCTTCCTACATCCAAGAATGTGCCANNNAGATCGGAAGA |
| 44043_6 | TATGAGTGTGGAGTCGTTGCTTCCTACATCCAAGAATGTGCCANNNAGATCGGAAGA |
| 42386_1 | TATGAGTGTGGAGTCGTTGCAGTGATTCTACTGCCTCAACCTCNNNAGATCGGAAGA |
| 42386_2 | TATGAGTGTGGAGTCGTTGCAGTGATTCTACTGCCTCAACCTCNNNAGATCGGAAGA |
| 42386_3 | TATGAGTGTGGAGTCGTTGCAGTGATTCTACTGCCTCAACCTCNNNAGATCGGAAGA |
| 42386_4 | TATGAGTGTGGAGTCGTTGCAGTGATTCTACTGCCTCAACCTCNNNAGATCGGAAGA |
| 42386_5 | TATGAGTGTGGAGTCGTTGCAGTGATTCTACTGCCTCAACCTCNNNAGATCGGAAGA |
| 42386_6 | TATGAGTGTGGAGTCGTTGCAGTGATTCTACTGCCTCAACCTCNNNAGATCGGAAGA |
| 51974_1 | TATGAGTGTGGAGTCGTTGCAAAAAGACCCTGGAGTCCTGGATNNNAGATCGGAA[illegible] |
| 51974_2 | TATGAGTGTGGAGTCGTTGCAAAAAGACCCTGGAGTCCTGGATNNNAGATCGGAA[illegible] |
| 51974_3 | TATGAGTGTGGAGTCGTTGCAAAAAGACCCTGGAGTCCTGGATNNNAGATCGGAA[illegible] |
| 51974_4 | TATGAGTGTGGAGTCGTTGCAAAAAGACCCTGGAGTCCTGGATNNNAGATCGGAA[illegible] |
| 51974_5 | TATGAGTGTGGAGTCGTTGCAAAAAGACCCTGGAGTCCTGGATNNNAGATCGGAA[illegible] |
| 51974_6 | TATGAGTGTGGAGTCGTTGCAAAAAGACCCTGGAGTCCTGGATNNNAGATCGGAA[illegible] |

| | |
|---|---|
| 32226_1 | TATGAGTGTGGAGTCGTTGCGTCTCCTCCCCTCTCACCCCNNNAGATCGGAAGAGCA |
| 32226_2 | TATGAGTGTGGAGTCGTTGCGTCTCCTCCCCTCTCACCCCNNNAGATCGGAAGAGCA |
| 32226_3 | TATGAGTGTGGAGTCGTTGCGTCTCCTCCCCTCTCACCCCNNNAGATCGGAAGAGCA |
| 32226_4 | TATGAGTGTGGAGTCGTTGCGTCTCCTCCCCTCTCACCCCNNNAGATCGGAAGAGCA |
| 32226_5 | TATGAGTGTGGAGTCGTTGCGTCTCCTCCCCTCTCACCCCNNNAGATCGGAAGAGCA |
| 32226_6 | TATGAGTGTGGAGTCGTTGCGTCTCCTCCCCTCTCACCCCNNNAGATCGGAAGAGCA |
| 42754_1 | TATGAGTGTGGAGTCGTTGCGCAATTATGGAGGCTGACAAGTCNNNAGATCGGAA |
| 42754_2 | TATGAGTGTGGAGTCGTTGCGCAATTATGGAGGCTGACAAGTCNNNAGATCGGAA |
| 42754_3 | TATGAGTGTGGAGTCGTTGCGCAATTATGGAGGCTGACAAGTCNNNAGATCGGAA |
| 42754_4 | TATGAGTGTGGAGTCGTTGCGCAATTATGGAGGCTGACAAGTCNNNAGATCGGAA |
| 42754_5 | TATGAGTGTGGAGTCGTTGCGCAATTATGGAGGCTGACAAGTCNNNAGATCGGAA |
| 42754_6 | TATGAGTGTGGAGTCGTTGCGCAATTATGGAGGCTGACAAGTCNNNAGATCGGAA |
| 35281_1 | TATGAGTGTGGAGTCGTTGCCATTCTCCTGCCTCAGCCTCNNNAGATCGGAAGAGCA |
| 35281_2 | TATGAGTGTGGAGTCGTTGCCATTCTCCTGCCTCAGCCTCNNNAGATCGGAAGAGCA |
| 35281_3 | TATGAGTGTGGAGTCGTTGCCATTCTCCTGCCTCAGCCTCNNNAGATCGGAAGAGCA |
| 35281_4 | TATGAGTGTGGAGTCGTTGCCATTCTCCTGCCTCAGCCTCNNNAGATCGGAAGAGCA |
| 35281_5 | TATGAGTGTGGAGTCGTTGCCATTCTCCTGCCTCAGCCTCNNNAGATCGGAAGAGCA |
| 35281_6 | TATGAGTGTGGAGTCGTTGCCATTCTCCTGCCTCAGCCTCNNNAGATCGGAAGAGCA |
| 12721_1 | TATGAGTGTGGAGTCGTTGCAAAATGGAAAACGTAAACACAANNNAGATCGGAAG |
| 12721_2 | TATGAGTGTGGAGTCGTTGCAAAATGGAAAACGTAAACACAANNNAGATCGGAAG |
| 12721_3 | TATGAGTGTGGAGTCGTTGCAAAATGGAAAACGTAAACACAANNNAGATCGGAAG |
| 12721_4 | TATGAGTGTGGAGTCGTTGCAAAATGGAAAACGTAAACACAANNNAGATCGGAAG |
| 12721_5 | TATGAGTGTGGAGTCGTTGCAAAATGGAAAACGTAAACACAANNNAGATCGGAAG |
| 12721_6 | TATGAGTGTGGAGTCGTTGCAAAATGGAAAACGTAAACACAANNNAGATCGGAAG |
| 45286_1 | TATGAGTGTGGAGTCGTTGCTCTGGTGAAATTGAAGAACCCCANNNAGATCGGAAG |
| 45286_2 | TATGAGTGTGGAGTCGTTGCTCTGGTGAAATTGAAGAACCCCANNNAGATCGGAAG |
| 45286_3 | TATGAGTGTGGAGTCGTTGCTCTGGTGAAATTGAAGAACCCCANNNAGATCGGAAG |
| 45286_4 | TATGAGTGTGGAGTCGTTGCTCTGGTGAAATTGAAGAACCCCANNNAGATCGGAAG |
| 45286_5 | TATGAGTGTGGAGTCGTTGCTCTGGTGAAATTGAAGAACCCCANNNAGATCGGAAG |
| 45286_6 | TATGAGTGTGGAGTCGTTGCTCTGGTGAAATTGAAGAACCCCANNNAGATCGGAAG |
| 41623_1 | TATGAGTGTGGAGTCGTTGCTCCCAGGTTCAAGTGATTCTCTGNNNAGATCGGAAG |
| 41623_2 | TATGAGTGTGGAGTCGTTGCTCCCAGGTTCAAGTGATTCTCTGNNNAGATCGGAAG |
| 41623_3 | TATGAGTGTGGAGTCGTTGCTCCCAGGTTCAAGTGATTCTCTGNNNAGATCGGAAG |
| 41623_4 | TATGAGTGTGGAGTCGTTGCTCCCAGGTTCAAGTGATTCTCTGNNNAGATCGGAAG |
| 41623_5 | TATGAGTGTGGAGTCGTTGCTCCCAGGTTCAAGTGATTCTCTGNNNAGATCGGAAG |
| 41623_6 | TATGAGTGTGGAGTCGTTGCTCCCAGGTTCAAGTGATTCTCTGNNNAGATCGGAAG |
| 53399_1 | TATGAGTGTGGAGTCGTTGCCCTGGGAACAAATGGGAAGATTGNNNAGATCGGAA |
| 53399_2 | TATGAGTGTGGAGTCGTTGCCCTGGGAACAAATGGGAAGATTGNNNAGATCGGAA |
| 53399_3 | TATGAGTGTGGAGTCGTTGCCCTGGGAACAAATGGGAAGATTGNNNAGATCGGAA |
| 53399_4 | TATGAGTGTGGAGTCGTTGCCCTGGGAACAAATGGGAAGATTGNNNAGATCGGAA |
| 53399_5 | TATGAGTGTGGAGTCGTTGCCCTGGGAACAAATGGGAAGATTGNNNAGATCGGAA |
| 53399_6 | TATGAGTGTGGAGTCGTTGCCCTGGGAACAAATGGGAAGATTGNNNAGATCGGAA |
| 38577_1 | TATGAGTGTGGAGTCGTTGCGAGGCAATGTTCTGAGAGCTTTCNNNAGATCGGAA |

| | |
|---|---|
| 38577_2 | TATGAGTGTGGAGTCGTTGCGAGGCAATGTTCTGAGAGCTTTCNNNAGATCGGAA |
| 38577_3 | TATGAGTGTGGAGTCGTTGCGAGGCAATGTTCTGAGAGCTTTCNNNAGATCGGAA |
| 38577_4 | TATGAGTGTGGAGTCGTTGCGAGGCAATGTTCTGAGAGCTTTCNNNAGATCGGAA |
| 38577_5 | TATGAGTGTGGAGTCGTTGCGAGGCAATGTTCTGAGAGCTTTCNNNAGATCGGAA |
| 38577_6 | TATGAGTGTGGAGTCGTTGCGAGGCAATGTTCTGAGAGCTTTCNNNAGATCGGAA |
| 41949_1 | TATGAGTGTGGAGTCGTTGCTTATTTTGTCTCACCCTCCCGAGNNNAGATCGGAAGA |
| 41949_2 | TATGAGTGTGGAGTCGTTGCTTATTTTGTCTCACCCTCCCGAGNNNAGATCGGAAGA |
| 41949_3 | TATGAGTGTGGAGTCGTTGCTTATTTTGTCTCACCCTCCCGAGNNNAGATCGGAAGA |
| 41949_4 | TATGAGTGTGGAGTCGTTGCTTATTTTGTCTCACCCTCCCGAGNNNAGATCGGAAGA |
| 41949_5 | TATGAGTGTGGAGTCGTTGCTTATTTTGTCTCACCCTCCCGAGNNNAGATCGGAAGA |
| 41949_6 | TATGAGTGTGGAGTCGTTGCTTATTTTGTCTCACCCTCCCGAGNNNAGATCGGAAGA |
| 41428_1 | TATGAGTGTGGAGTCGTTGCGCTTCTGATCACTGCTGTTTGAANNNAGATCGGAAG |
| 41428_2 | TATGAGTGTGGAGTCGTTGCGCTTCTGATCACTGCTGTTTGAANNNAGATCGGAAG |
| 41428_3 | TATGAGTGTGGAGTCGTTGCGCTTCTGATCACTGCTGTTTGAANNNAGATCGGAAG |
| 41428_4 | TATGAGTGTGGAGTCGTTGCGCTTCTGATCACTGCTGTTTGAANNNAGATCGGAAG |
| 41428_5 | TATGAGTGTGGAGTCGTTGCGCTTCTGATCACTGCTGTTTGAANNNAGATCGGAAG |
| 41428_6 | TATGAGTGTGGAGTCGTTGCGCTTCTGATCACTGCTGTTTGAANNNAGATCGGAAG |
| 33636_1 | TATGAGTGTGGAGTCGTTGCCAGACTGCAGTGGTGCTATCTCNNNAGATCGGAAGA |
| 33636_2 | TATGAGTGTGGAGTCGTTGCCAGACTGCAGTGGTGCTATCTCNNNAGATCGGAAGA |
| 33636_3 | TATGAGTGTGGAGTCGTTGCCAGACTGCAGTGGTGCTATCTCNNNAGATCGGAAGA |
| 33636_4 | TATGAGTGTGGAGTCGTTGCCAGACTGCAGTGGTGCTATCTCNNNAGATCGGAAGA |
| 33636_5 | TATGAGTGTGGAGTCGTTGCCAGACTGCAGTGGTGCTATCTCNNNAGATCGGAAGA |
| 33636_6 | TATGAGTGTGGAGTCGTTGCCAGACTGCAGTGGTGCTATCTCNNNAGATCGGAAGA |
| 51453_1 | TATGAGTGTGGAGTCGTTGCGGGATGGTTTTGGGATGAAACTGNNNAGATCGGAA |
| 51453_2 | TATGAGTGTGGAGTCGTTGCGGGATGGTTTTGGGATGAAACTGNNNAGATCGGAA |
| 51453_3 | TATGAGTGTGGAGTCGTTGCGGGATGGTTTTGGGATGAAACTGNNNAGATCGGAA |
| 51453_4 | TATGAGTGTGGAGTCGTTGCGGGATGGTTTTGGGATGAAACTGNNNAGATCGGAA |
| 51453_5 | TATGAGTGTGGAGTCGTTGCGGGATGGTTTTGGGATGAAACTGNNNAGATCGGAA |
| 51453_6 | TATGAGTGTGGAGTCGTTGCGGGATGGTTTTGGGATGAAACTGNNNAGATCGGAA |
| 34102_1 | TATGAGTGTGGAGTCGTTGCCCAGGTGATGGATACCCCAAATANNNAGATCGGAA |
| 34102_2 | TATGAGTGTGGAGTCGTTGCCCAGGTGATGGATACCCCAAATANNNAGATCGGAA |
| 34102_3 | TATGAGTGTGGAGTCGTTGCCCAGGTGATGGATACCCCAAATANNNAGATCGGAA |
| 34102_4 | TATGAGTGTGGAGTCGTTGCCCAGGTGATGGATACCCCAAATANNNAGATCGGAA |
| 34102_5 | TATGAGTGTGGAGTCGTTGCCCAGGTGATGGATACCCCAAATANNNAGATCGGAA |
| 34102_6 | TATGAGTGTGGAGTCGTTGCCCAGGTGATGGATACCCCAAATANNNAGATCGGAA |
| 31881_1 | TATGAGTGTGGAGTCGTTGCTGTTGTGAGTCGTGCTATCAAGANNNAGATCGGAA |
| 31881_2 | TATGAGTGTGGAGTCGTTGCTGTTGTGAGTCGTGCTATCAAGANNNAGATCGGAA |
| 31881_3 | TATGAGTGTGGAGTCGTTGCTGTTGTGAGTCGTGCTATCAAGANNNAGATCGGAA |
| 31881_4 | TATGAGTGTGGAGTCGTTGCTGTTGTGAGTCGTGCTATCAAGANNNAGATCGGAA |
| 31881_5 | TATGAGTGTGGAGTCGTTGCTGTTGTGAGTCGTGCTATCAAGANNNAGATCGGAA |
| 31881_6 | TATGAGTGTGGAGTCGTTGCTGTTGTGAGTCGTGCTATCAAGANNNAGATCGGAA |
| 48411_1 | TATGAGTGTGGAGTCGTTGCTTGCCACGTTCGTGCTGATGNNNAGATCGGAAGAG |
| 48411_2 | TATGAGTGTGGAGTCGTTGCTTGCCACGTTCGTGCTGATGNNNAGATCGGAAGAG |

| | |
|---|---|
| 48411_3 | TATGAGTGTGGAGTCGTTGCTTGCCACGTTCGTGCTGATGNNNAGATCGGAAGAG |
| 48411_4 | TATGAGTGTGGAGTCGTTGCTTGCCACGTTCGTGCTGATGNNNAGATCGGAAGAG |
| 48411_5 | TATGAGTGTGGAGTCGTTGCTTGCCACGTTCGTGCTGATGNNNAGATCGGAAGAG |
| 48411_6 | TATGAGTGTGGAGTCGTTGCTTGCCACGTTCGTGCTGATGNNNAGATCGGAAGAG |
| 50143_1 | TATGAGTGTGGAGTCGTTGCTTGGCACTCCTTTTTCCAATTCCNNNAGATCGGAAGA |
| 50143_2 | TATGAGTGTGGAGTCGTTGCTTGGCACTCCTTTTTCCAATTCCNNNAGATCGGAAGA |
| 50143_3 | TATGAGTGTGGAGTCGTTGCTTGGCACTCCTTTTTCCAATTCCNNNAGATCGGAAGA |
| 50143_4 | TATGAGTGTGGAGTCGTTGCTTGGCACTCCTTTTTCCAATTCCNNNAGATCGGAAGA |
| 50143_5 | TATGAGTGTGGAGTCGTTGCTTGGCACTCCTTTTTCCAATTCCNNNAGATCGGAAGA |
| 50143_6 | TATGAGTGTGGAGTCGTTGCTTGGCACTCCTTTTTCCAATTCCNNNAGATCGGAAGA |
| 48502_1 | TATGAGTGTGGAGTCGTTGCGGTTCAAACGATTCTCTTGCCTCNNNAGATCGGAAG |
| 48502_2 | TATGAGTGTGGAGTCGTTGCGGTTCAAACGATTCTCTTGCCTCNNNAGATCGGAAG |
| 48502_3 | TATGAGTGTGGAGTCGTTGCGGTTCAAACGATTCTCTTGCCTCNNNAGATCGGAAG |
| 48502_4 | TATGAGTGTGGAGTCGTTGCGGTTCAAACGATTCTCTTGCCTCNNNAGATCGGAAG |
| 48502_5 | TATGAGTGTGGAGTCGTTGCGGTTCAAACGATTCTCTTGCCTCNNNAGATCGGAAG |
| 48502_6 | TATGAGTGTGGAGTCGTTGCGGTTCAAACGATTCTCTTGCCTCNNNAGATCGGAAG |
| 45952_1 | TATGAGTGTGGAGTCGTTGCCAAGTAGCTGGGACTACAGGCNNNAGATCGGAAGA |
| 45952_2 | TATGAGTGTGGAGTCGTTGCCAAGTAGCTGGGACTACAGGCNNNAGATCGGAAGA |
| 45952_3 | TATGAGTGTGGAGTCGTTGCCAAGTAGCTGGGACTACAGGCNNNAGATCGGAAGA |
| 45952_4 | TATGAGTGTGGAGTCGTTGCCAAGTAGCTGGGACTACAGGCNNNAGATCGGAAGA |
| 45952_5 | TATGAGTGTGGAGTCGTTGCCAAGTAGCTGGGACTACAGGCNNNAGATCGGAAGA |
| 45952_6 | TATGAGTGTGGAGTCGTTGCCAAGTAGCTGGGACTACAGGCNNNAGATCGGAAGA |
| 31143_1 | TATGAGTGTGGAGTCGTTGCGTGGTTCTGACAGCTGTTTCTTCNNNAGATCGGAAG |
| 31143_2 | TATGAGTGTGGAGTCGTTGCGTGGTTCTGACAGCTGTTTCTTCNNNAGATCGGAAG |
| 31143_3 | TATGAGTGTGGAGTCGTTGCGTGGTTCTGACAGCTGTTTCTTCNNNAGATCGGAAG |
| 31143_4 | TATGAGTGTGGAGTCGTTGCGTGGTTCTGACAGCTGTTTCTTCNNNAGATCGGAAG |
| 31143_5 | TATGAGTGTGGAGTCGTTGCGTGGTTCTGACAGCTGTTTCTTCNNNAGATCGGAAG |
| 31143_6 | TATGAGTGTGGAGTCGTTGCGTGGTTCTGACAGCTGTTTCTTCNNNAGATCGGAAG |
| 4650_1 | TATGAGTGTGGAGTCGTTGCTCCAGCATACAATGTGTTTCACCANNNAGATCGGAA |
| 4650_2 | TATGAGTGTGGAGTCGTTGCTCCAGCATACAATGTGTTTCACCANNNAGATCGGAA |
| 4650_3 | TATGAGTGTGGAGTCGTTGCTCCAGCATACAATGTGTTTCACCANNNAGATCGGAA |
| 4650_4 | TATGAGTGTGGAGTCGTTGCTCCAGCATACAATGTGTTTCACCANNNAGATCGGAA |
| 4650_5 | TATGAGTGTGGAGTCGTTGCTCCAGCATACAATGTGTTTCACCANNNAGATCGGAA |
| 4650_6 | TATGAGTGTGGAGTCGTTGCTCCAGCATACAATGTGTTTCACCANNNAGATCGGAA |
| 51733_1 | TATGAGTGTGGAGTCGTTGCTTTTTTTTGAGACGGAGTCTGGCNNNAGATCGGAAG |
| 51733_2 | TATGAGTGTGGAGTCGTTGCTTTTTTTTGAGACGGAGTCTGGCNNNAGATCGGAAG |
| 51733_3 | TATGAGTGTGGAGTCGTTGCTTTTTTTTGAGACGGAGTCTGGCNNNAGATCGGAAG |
| 51733_4 | TATGAGTGTGGAGTCGTTGCTTTTTTTTGAGACGGAGTCTGGCNNNAGATCGGAAG |
| 51733_5 | TATGAGTGTGGAGTCGTTGCTTTTTTTTGAGACGGAGTCTGGCNNNAGATCGGAAG |
| 51733_6 | TATGAGTGTGGAGTCGTTGCTTTTTTTTGAGACGGAGTCTGGCNNNAGATCGGAAG |
| 12495_1 | TATGAGTGTGGAGTCGTTGCGAGGAGGACTATAAAAGAACAAANNNAGATCGGAA |
| 12495_2 | TATGAGTGTGGAGTCGTTGCGAGGAGGACTATAAAAGAACAAANNNAGATCGGAA |
| 12495_3 | TATGAGTGTGGAGTCGTTGCGAGGAGGACTATAAAAGAACAAANNNAGATCGGAA |

| | |
|---|---|
| 12495_4 | TATGAGTGTGGAGTCGTTGCGAGGAGGACTATAAAAGAACAAANNNAGATCGGAA |
| 12495_5 | TATGAGTGTGGAGTCGTTGCGAGGAGGACTATAAAAGAACAAANNNAGATCGGAA |
| 12495_6 | TATGAGTGTGGAGTCGTTGCGAGGAGGACTATAAAAGAACAAANNNAGATCGGAA |
| 37542_1 | TATGAGTGTGGAGTCGTTGCTCTAGAGAGAAGTGACCCCTCAGNNNAGATCGGAAG |
| 37542_2 | TATGAGTGTGGAGTCGTTGCTCTAGAGAGAAGTGACCCCTCAGNNNAGATCGGAAG |
| 37542_3 | TATGAGTGTGGAGTCGTTGCTCTAGAGAGAAGTGACCCCTCAGNNNAGATCGGAAG |
| 37542_4 | TATGAGTGTGGAGTCGTTGCTCTAGAGAGAAGTGACCCCTCAGNNNAGATCGGAAG |
| 37542_5 | TATGAGTGTGGAGTCGTTGCTCTAGAGAGAAGTGACCCCTCAGNNNAGATCGGAAG |
| 37542_6 | TATGAGTGTGGAGTCGTTGCTCTAGAGAGAAGTGACCCCTCAGNNNAGATCGGAAG |
| 45191_1 | TATGAGTGTGGAGTCGTTGCATGGGATTGTGGGGATATGGATGNNNAGATCGGAA |
| 45191_2 | TATGAGTGTGGAGTCGTTGCATGGGATTGTGGGGATATGGATGNNNAGATCGGAA |
| 45191_3 | TATGAGTGTGGAGTCGTTGCATGGGATTGTGGGGATATGGATGNNNAGATCGGAA |
| 45191_4 | TATGAGTGTGGAGTCGTTGCATGGGATTGTGGGGATATGGATGNNNAGATCGGAA |
| 45191_5 | TATGAGTGTGGAGTCGTTGCATGGGATTGTGGGGATATGGATGNNNAGATCGGAA |
| 45191_6 | TATGAGTGTGGAGTCGTTGCATGGGATTGTGGGGATATGGATGNNNAGATCGGAA |
| 40165_1 | TATGAGTGTGGAGTCGTTGCGTGACACAGCAAGACAATGTCTCNNNAGATCGGAAG |
| 40165_2 | TATGAGTGTGGAGTCGTTGCGTGACACAGCAAGACAATGTCTCNNNAGATCGGAAG |
| 40165_3 | TATGAGTGTGGAGTCGTTGCGTGACACAGCAAGACAATGTCTCNNNAGATCGGAAG |
| 40165_4 | TATGAGTGTGGAGTCGTTGCGTGACACAGCAAGACAATGTCTCNNNAGATCGGAAG |
| 40165_5 | TATGAGTGTGGAGTCGTTGCGTGACACAGCAAGACAATGTCTCNNNAGATCGGAAG |
| 40165_6 | TATGAGTGTGGAGTCGTTGCGTGACACAGCAAGACAATGTCTCNNNAGATCGGAAG |
| 42366_1 | TATGAGTGTGGAGTCGTTGCCATTCTCCTGCCTCAGCCTCNNNAGATCGGAAGAGCA |
| 42366_2 | TATGAGTGTGGAGTCGTTGCCATTCTCCTGCCTCAGCCTCNNNAGATCGGAAGAGCA |
| 42366_3 | TATGAGTGTGGAGTCGTTGCCATTCTCCTGCCTCAGCCTCNNNAGATCGGAAGAGCA |
| 42366_4 | TATGAGTGTGGAGTCGTTGCCATTCTCCTGCCTCAGCCTCNNNAGATCGGAAGAGCA |
| 42366_5 | TATGAGTGTGGAGTCGTTGCCATTCTCCTGCCTCAGCCTCNNNAGATCGGAAGAGCA |
| 42366_6 | TATGAGTGTGGAGTCGTTGCCATTCTCCTGCCTCAGCCTCNNNAGATCGGAAGAGCA |
| 12990_1 | TATGAGTGTGGAGTCGTTGCGAAACTTCATAATTAGCACAGATNNNAGATCGGAAG |
| 12990_2 | TATGAGTGTGGAGTCGTTGCGAAACTTCATAATTAGCACAGATNNNAGATCGGAAG |
| 12990_3 | TATGAGTGTGGAGTCGTTGCGAAACTTCATAATTAGCACAGATNNNAGATCGGAAG |
| 12990_4 | TATGAGTGTGGAGTCGTTGCGAAACTTCATAATTAGCACAGATNNNAGATCGGAAG |
| 12990_5 | TATGAGTGTGGAGTCGTTGCGAAACTTCATAATTAGCACAGATNNNAGATCGGAAG |
| 12990_6 | TATGAGTGTGGAGTCGTTGCGAAACTTCATAATTAGCACAGATNNNAGATCGGAAG |
| 43022_1 | TATGAGTGTGGAGTCGTTGCTTTCACTTGATAGTCAGGGCTGTNNNAGATCGGAAG |
| 43022_2 | TATGAGTGTGGAGTCGTTGCTTTCACTTGATAGTCAGGGCTGTNNNAGATCGGAAG |
| 43022_3 | TATGAGTGTGGAGTCGTTGCTTTCACTTGATAGTCAGGGCTGTNNNAGATCGGAAG |
| 43022_4 | TATGAGTGTGGAGTCGTTGCTTTCACTTGATAGTCAGGGCTGTNNNAGATCGGAAG |
| 43022_5 | TATGAGTGTGGAGTCGTTGCTTTCACTTGATAGTCAGGGCTGTNNNAGATCGGAAG |
| 43022_6 | TATGAGTGTGGAGTCGTTGCTTTCACTTGATAGTCAGGGCTGTNNNAGATCGGAAG |
| 28874_1 | TATGAGTGTGGAGTCGTTGCGTTACTGCAGCCTTGAACTCTTGNNNAGATCGGAAGA |
| 28874_2 | TATGAGTGTGGAGTCGTTGCGTTACTGCAGCCTTGAACTCTTGNNNAGATCGGAAGA |
| 28874_3 | TATGAGTGTGGAGTCGTTGCGTTACTGCAGCCTTGAACTCTTGNNNAGATCGGAAGA |
| 28874_4 | TATGAGTGTGGAGTCGTTGCGTTACTGCAGCCTTGAACTCTTGNNNAGATCGGAAGA |

| | |
|---|---|
| 28874_5 | TATGAGTGTGGAGTCGTTGCGTTACTGCAGCCTTGAACTCTTGNNNAGATCGGAAG |
| 28874_6 | TATGAGTGTGGAGTCGTTGCGTTACTGCAGCCTTGAACTCTTGNNNAGATCGGAAG |
| 36695_1 | TATGAGTGTGGAGTCGTTGCCTACCCCCAGAGTGCATAGAATCNNNAGATCGGAAG |
| 36695_2 | TATGAGTGTGGAGTCGTTGCCTACCCCCAGAGTGCATAGAATCNNNAGATCGGAAG |
| 36695_3 | TATGAGTGTGGAGTCGTTGCCTACCCCCAGAGTGCATAGAATCNNNAGATCGGAAG |
| 36695_4 | TATGAGTGTGGAGTCGTTGCCTACCCCCAGAGTGCATAGAATCNNNAGATCGGAAG |
| 36695_5 | TATGAGTGTGGAGTCGTTGCCTACCCCCAGAGTGCATAGAATCNNNAGATCGGAAG |
| 36695_6 | TATGAGTGTGGAGTCGTTGCCTACCCCCAGAGTGCATAGAATCNNNAGATCGGAAG |
| 31443_1 | TATGAGTGTGGAGTCGTTGCCATTCTCACTAAGCCTGACCTGTNNNAGATCGGAAG |
| 31443_2 | TATGAGTGTGGAGTCGTTGCCATTCTCACTAAGCCTGACCTGTNNNAGATCGGAAG |
| 31443_3 | TATGAGTGTGGAGTCGTTGCCATTCTCACTAAGCCTGACCTGTNNNAGATCGGAAG |
| 31443_4 | TATGAGTGTGGAGTCGTTGCCATTCTCACTAAGCCTGACCTGTNNNAGATCGGAAG |
| 31443_5 | TATGAGTGTGGAGTCGTTGCCATTCTCACTAAGCCTGACCTGTNNNAGATCGGAAG |
| 31443_6 | TATGAGTGTGGAGTCGTTGCCATTCTCACTAAGCCTGACCTGTNNNAGATCGGAAG |
| 52968_1 | TATGAGTGTGGAGTCGTTGCAGCAGACGTTCTTCTTCCCTTTCNNNAGATCGGAAGA |
| 52968_2 | TATGAGTGTGGAGTCGTTGCAGCAGACGTTCTTCTTCCCTTTCNNNAGATCGGAAGA |
| 52968_3 | TATGAGTGTGGAGTCGTTGCAGCAGACGTTCTTCTTCCCTTTCNNNAGATCGGAAGA |
| 52968_4 | TATGAGTGTGGAGTCGTTGCAGCAGACGTTCTTCTTCCCTTTCNNNAGATCGGAAGA |
| 52968_5 | TATGAGTGTGGAGTCGTTGCAGCAGACGTTCTTCTTCCCTTTCNNNAGATCGGAAGA |
| 52968_6 | TATGAGTGTGGAGTCGTTGCAGCAGACGTTCTTCTTCCCTTTCNNNAGATCGGAAGA |
| 39505_1 | TATGAGTGTGGAGTCGTTGCAAAAGAAAAAAATGGCCGTGGGANNNAGATCGGAA |
| 39505_2 | TATGAGTGTGGAGTCGTTGCAAAAGAAAAAAATGGCCGTGGGANNNAGATCGGAA |
| 39505_3 | TATGAGTGTGGAGTCGTTGCAAAAGAAAAAAATGGCCGTGGGANNNAGATCGGAA |
| 39505_4 | TATGAGTGTGGAGTCGTTGCAAAAGAAAAAAATGGCCGTGGGANNNAGATCGGAA |
| 39505_5 | TATGAGTGTGGAGTCGTTGCAAAAGAAAAAAATGGCCGTGGGANNNAGATCGGAA |
| 39505_6 | TATGAGTGTGGAGTCGTTGCAAAAGAAAAAAATGGCCGTGGGANNNAGATCGGAA |
| 39505_7 | TATGAGTGTGGAGTCGTTGCAAAAGAAAAAAATGGCCGTGGGANNNAGATCGGAA |
| 39505_8 | TATGAGTGTGGAGTCGTTGCAAAAGAAAAAAATGGCCGTGGGANNNAGATCGGAA |
| 39505_9 | TATGAGTGTGGAGTCGTTGCAAAAGAAAAAAATGGCCGTGGGANNNAGATCGGAA |
| 41427_1 | TATGAGTGTGGAGTCGTTGCATTGAGGACTGCTGATACAAGCANNNAGATCGGAA |
| 41427_2 | TATGAGTGTGGAGTCGTTGCATTGAGGACTGCTGATACAAGCANNNAGATCGGAA |
| 41427_3 | TATGAGTGTGGAGTCGTTGCATTGAGGACTGCTGATACAAGCANNNAGATCGGAA |
| 41427_4 | TATGAGTGTGGAGTCGTTGCATTGAGGACTGCTGATACAAGCANNNAGATCGGAA |
| 41427_5 | TATGAGTGTGGAGTCGTTGCATTGAGGACTGCTGATACAAGCANNNAGATCGGAA |
| 41427_6 | TATGAGTGTGGAGTCGTTGCATTGAGGACTGCTGATACAAGCANNNAGATCGGAA |
| 41427_7 | TATGAGTGTGGAGTCGTTGCATTGAGGACTGCTGATACAAGCANNNAGATCGGAA |
| 41427_8 | TATGAGTGTGGAGTCGTTGCATTGAGGACTGCTGATACAAGCANNNAGATCGGAA |
| 41427_9 | TATGAGTGTGGAGTCGTTGCATTGAGGACTGCTGATACAAGCANNNAGATCGGAA |
| 39888_1 | TATGAGTGTGGAGTCGTTGCAAGGATATGAGGACTTGGTGCTGNNNAGATCGGAA |
| 39888_2 | TATGAGTGTGGAGTCGTTGCAAGGATATGAGGACTTGGTGCTGNNNAGATCGGAA |
| 39888_3 | TATGAGTGTGGAGTCGTTGCAAGGATATGAGGACTTGGTGCTGNNNAGATCGGAA |
| 39888_4 | TATGAGTGTGGAGTCGTTGCAAGGATATGAGGACTTGGTGCTGNNNAGATCGGAA |
| 39888_5 | TATGAGTGTGGAGTCGTTGCAAGGATATGAGGACTTGGTGCTGNNNAGATCGGAA |

| | |
|---|---|
| 39888_6 | TATGAGTGTGGAGTCGTTGCAAGGATATGAGGACTTGGTGCTGNNNAGATCGGAA |
| 39888_7 | TATGAGTGTGGAGTCGTTGCAAGGATATGAGGACTTGGTGCTGNNNAGATCGGAA |
| 39888_8 | TATGAGTGTGGAGTCGTTGCAAGGATATGAGGACTTGGTGCTGNNNAGATCGGAA |
| 39888_9 | TATGAGTGTGGAGTCGTTGCAAGGATATGAGGACTTGGTGCTGNNNAGATCGGAA |
| 35574_1 | TATGAGTGTGGAGTCGTTGCGCTCATGTCATCTCCCCTCTTAGNNNAGATCGGAAGA |
| 35574_2 | TATGAGTGTGGAGTCGTTGCGCTCATGTCATCTCCCCTCTTAGNNNAGATCGGAAGA |
| 35574_3 | TATGAGTGTGGAGTCGTTGCGCTCATGTCATCTCCCCTCTTAGNNNAGATCGGAAGA |
| 35574_4 | TATGAGTGTGGAGTCGTTGCGCTCATGTCATCTCCCCTCTTAGNNNAGATCGGAAGA |
| 35574_5 | TATGAGTGTGGAGTCGTTGCGCTCATGTCATCTCCCCTCTTAGNNNAGATCGGAAGA |
| 35574_6 | TATGAGTGTGGAGTCGTTGCGCTCATGTCATCTCCCCTCTTAGNNNAGATCGGAAGA |
| 35574_7 | TATGAGTGTGGAGTCGTTGCGCTCATGTCATCTCCCCTCTTAGNNNAGATCGGAAGA |
| 35574_8 | TATGAGTGTGGAGTCGTTGCGCTCATGTCATCTCCCCTCTTAGNNNAGATCGGAAGA |
| 35574_9 | TATGAGTGTGGAGTCGTTGCGCTCATGTCATCTCCCCTCTTAGNNNAGATCGGAAGA |
| 32637_1 | TATGAGTGTGGAGTCGTTGCGTTCAAGCGATTCTCCTGCTTCNNNAGATCGGAAGA |
| 32637_2 | TATGAGTGTGGAGTCGTTGCGTTCAAGCGATTCTCCTGCTTCNNNAGATCGGAAGA |
| 32637_3 | TATGAGTGTGGAGTCGTTGCGTTCAAGCGATTCTCCTGCTTCNNNAGATCGGAAGA |
| 32637_4 | TATGAGTGTGGAGTCGTTGCGTTCAAGCGATTCTCCTGCTTCNNNAGATCGGAAGA |
| 32637_5 | TATGAGTGTGGAGTCGTTGCGTTCAAGCGATTCTCCTGCTTCNNNAGATCGGAAGA |
| 32637_6 | TATGAGTGTGGAGTCGTTGCGTTCAAGCGATTCTCCTGCTTCNNNAGATCGGAAGA |
| 32637_7 | TATGAGTGTGGAGTCGTTGCGTTCAAGCGATTCTCCTGCTTCNNNAGATCGGAAGA |
| 32637_8 | TATGAGTGTGGAGTCGTTGCGTTCAAGCGATTCTCCTGCTTCNNNAGATCGGAAGA |
| 32637_9 | TATGAGTGTGGAGTCGTTGCGTTCAAGCGATTCTCCTGCTTCNNNAGATCGGAAGA |
| 2587_1 | TATGAGTGTGGAGTCGTTGCAGCCAAGGTTCTTGAAAGAGTGGNNNAGATCGGAA |
| 2587_2 | TATGAGTGTGGAGTCGTTGCAGCCAAGGTTCTTGAAAGAGTGGNNNAGATCGGAA |
| 2587_3 | TATGAGTGTGGAGTCGTTGCAGCCAAGGTTCTTGAAAGAGTGGNNNAGATCGGAA |
| 2587_4 | TATGAGTGTGGAGTCGTTGCAGCCAAGGTTCTTGAAAGAGTGGNNNAGATCGGAA |
| 2587_5 | TATGAGTGTGGAGTCGTTGCAGCCAAGGTTCTTGAAAGAGTGGNNNAGATCGGAA |
| 2587_6 | TATGAGTGTGGAGTCGTTGCAGCCAAGGTTCTTGAAAGAGTGGNNNAGATCGGAA |
| 2587_7 | TATGAGTGTGGAGTCGTTGCAGCCAAGGTTCTTGAAAGAGTGGNNNAGATCGGAA |
| 2587_8 | TATGAGTGTGGAGTCGTTGCAGCCAAGGTTCTTGAAAGAGTGGNNNAGATCGGAA |
| 2587_9 | TATGAGTGTGGAGTCGTTGCAGCCAAGGTTCTTGAAAGAGTGGNNNAGATCGGAA |
| 2190_1 | TATGAGTGTGGAGTCGTTGCCTTGCCCATGAGATGAAATGGGANNNAGATCGGAA |
| 2190_2 | TATGAGTGTGGAGTCGTTGCCTTGCCCATGAGATGAAATGGGANNNAGATCGGAA |
| 2190_3 | TATGAGTGTGGAGTCGTTGCCTTGCCCATGAGATGAAATGGGANNNAGATCGGAA |
| 2190_4 | TATGAGTGTGGAGTCGTTGCCTTGCCCATGAGATGAAATGGGANNNAGATCGGAA |
| 2190_5 | TATGAGTGTGGAGTCGTTGCCTTGCCCATGAGATGAAATGGGANNNAGATCGGAA |
| 2190_6 | TATGAGTGTGGAGTCGTTGCCTTGCCCATGAGATGAAATGGGANNNAGATCGGAA |
| 2190_7 | TATGAGTGTGGAGTCGTTGCCTTGCCCATGAGATGAAATGGGANNNAGATCGGAA |
| 2190_8 | TATGAGTGTGGAGTCGTTGCCTTGCCCATGAGATGAAATGGGANNNAGATCGGAA |
| 2190_9 | TATGAGTGTGGAGTCGTTGCCTTGCCCATGAGATGAAATGGGANNNAGATCGGAA |
| 30127_1 | ATGAGTGTGGAGTCGTTGCAACCACTATTGCAAATCCTATTACAANNNAGATCGGAA |
| 30127_2 | ATGAGTGTGGAGTCGTTGCAACCACTATTGCAAATCCTATTACAANNNAGATCGGAA |
| 30127_3 | ATGAGTGTGGAGTCGTTGCAACCACTATTGCAAATCCTATTACAANNNAGATCGGAA |

| | |
|---|---|
| 30127_4 | ATGAGTGTGGAGTCGTTGCAACCACTATTGCAAATCCTATTACAANNNAGATCGGAA |
| 30127_5 | ATGAGTGTGGAGTCGTTGCAACCACTATTGCAAATCCTATTACAANNNAGATCGGAA |
| 30127_6 | ATGAGTGTGGAGTCGTTGCAACCACTATTGCAAATCCTATTACAANNNAGATCGGAA |
| 30127_7 | ATGAGTGTGGAGTCGTTGCAACCACTATTGCAAATCCTATTACAANNNAGATCGGAA |
| 30127_8 | ATGAGTGTGGAGTCGTTGCAACCACTATTGCAAATCCTATTACAANNNAGATCGGAA |
| 30127_9 | ATGAGTGTGGAGTCGTTGCAACCACTATTGCAAATCCTATTACAANNNAGATCGGAA |
| 43508_1 | TATGAGTGTGGAGTCGTTGCGAGTAACTGGGATTAGAGGTGCCNNNAGATCGGAA |
| 43508_2 | TATGAGTGTGGAGTCGTTGCGAGTAACTGGGATTAGAGGTGCCNNNAGATCGGAA |
| 43508_3 | TATGAGTGTGGAGTCGTTGCGAGTAACTGGGATTAGAGGTGCCNNNAGATCGGAA |
| 43508_4 | TATGAGTGTGGAGTCGTTGCGAGTAACTGGGATTAGAGGTGCCNNNAGATCGGAA |
| 43508_5 | TATGAGTGTGGAGTCGTTGCGAGTAACTGGGATTAGAGGTGCCNNNAGATCGGAA |
| 43508_6 | TATGAGTGTGGAGTCGTTGCGAGTAACTGGGATTAGAGGTGCCNNNAGATCGGAA |
| 43508_7 | TATGAGTGTGGAGTCGTTGCGAGTAACTGGGATTAGAGGTGCCNNNAGATCGGAA |
| 43508_8 | TATGAGTGTGGAGTCGTTGCGAGTAACTGGGATTAGAGGTGCCNNNAGATCGGAA |
| 43508_9 | TATGAGTGTGGAGTCGTTGCGAGTAACTGGGATTAGAGGTGCCNNNAGATCGGAA |
| 47718_1 | TATGAGTGTGGAGTCGTTGCAGTGTTGATTCTGGCATCTTCCTNNNAGATCGGAAGA |
| 47718_2 | TATGAGTGTGGAGTCGTTGCAGTGTTGATTCTGGCATCTTCCTNNNAGATCGGAAGA |
| 47718_3 | TATGAGTGTGGAGTCGTTGCAGTGTTGATTCTGGCATCTTCCTNNNAGATCGGAAGA |
| 47718_4 | TATGAGTGTGGAGTCGTTGCAGTGTTGATTCTGGCATCTTCCTNNNAGATCGGAAGA |
| 47718_5 | TATGAGTGTGGAGTCGTTGCAGTGTTGATTCTGGCATCTTCCTNNNAGATCGGAAGA |
| 47718_6 | TATGAGTGTGGAGTCGTTGCAGTGTTGATTCTGGCATCTTCCTNNNAGATCGGAAGA |
| 47718_7 | TATGAGTGTGGAGTCGTTGCAGTGTTGATTCTGGCATCTTCCTNNNAGATCGGAAGA |
| 47718_8 | TATGAGTGTGGAGTCGTTGCAGTGTTGATTCTGGCATCTTCCTNNNAGATCGGAAGA |
| 47718_9 | TATGAGTGTGGAGTCGTTGCAGTGTTGATTCTGGCATCTTCCTNNNAGATCGGAAGA |
| 36729_1 | TATGAGTGTGGAGTCGTTGCTCTAAAAGCAGCACTGGGTTTTGNNNAGATCGGAAG |
| 36729_2 | TATGAGTGTGGAGTCGTTGCTCTAAAAGCAGCACTGGGTTTTGNNNAGATCGGAAG |
| 36729_3 | TATGAGTGTGGAGTCGTTGCTCTAAAAGCAGCACTGGGTTTTGNNNAGATCGGAAG |
| 36729_4 | TATGAGTGTGGAGTCGTTGCTCTAAAAGCAGCACTGGGTTTTGNNNAGATCGGAAG |
| 36729_5 | TATGAGTGTGGAGTCGTTGCTCTAAAAGCAGCACTGGGTTTTGNNNAGATCGGAAG |
| 36729_6 | TATGAGTGTGGAGTCGTTGCTCTAAAAGCAGCACTGGGTTTTGNNNAGATCGGAAG |
| 36729_7 | TATGAGTGTGGAGTCGTTGCTCTAAAAGCAGCACTGGGTTTTGNNNAGATCGGAAG |
| 36729_8 | TATGAGTGTGGAGTCGTTGCTCTAAAAGCAGCACTGGGTTTTGNNNAGATCGGAAG |
| 36729_9 | TATGAGTGTGGAGTCGTTGCTCTAAAAGCAGCACTGGGTTTTGNNNAGATCGGAAG |
| 40930_1 | TATGAGTGTGGAGTCGTTGCACGAAAAACTGCTGAATCACACTNNNAGATCGGAAG |
| 40930_2 | TATGAGTGTGGAGTCGTTGCACGAAAAACTGCTGAATCACACTNNNAGATCGGAAG |
| 40930_3 | TATGAGTGTGGAGTCGTTGCACGAAAAACTGCTGAATCACACTNNNAGATCGGAAG |
| 40930_4 | TATGAGTGTGGAGTCGTTGCACGAAAAACTGCTGAATCACACTNNNAGATCGGAAG |
| 40930_5 | TATGAGTGTGGAGTCGTTGCACGAAAAACTGCTGAATCACACTNNNAGATCGGAAG |
| 40930_6 | TATGAGTGTGGAGTCGTTGCACGAAAAACTGCTGAATCACACTNNNAGATCGGAAG |
| 40930_7 | TATGAGTGTGGAGTCGTTGCACGAAAAACTGCTGAATCACACTNNNAGATCGGAAG |
| 40930_8 | TATGAGTGTGGAGTCGTTGCACGAAAAACTGCTGAATCACACTNNNAGATCGGAAG |
| 40930_9 | TATGAGTGTGGAGTCGTTGCACGAAAAACTGCTGAATCACACTNNNAGATCGGAAG |
| 34673_1 | TATGAGTGTGGAGTCGTTGCGCATGGCTCTTAGCAATCTTAGCNNNAGATCGGAAG |

| | |
|---|---|
| 34673_2 | TATGAGTGTGGAGTCGTTGCGCATGGCTCTTAGCAATCTTAGCNNNAGATCGGAAG |
| 34673_3 | TATGAGTGTGGAGTCGTTGCGCATGGCTCTTAGCAATCTTAGCNNNAGATCGGAAG |
| 34673_4 | TATGAGTGTGGAGTCGTTGCGCATGGCTCTTAGCAATCTTAGCNNNAGATCGGAAG |
| 34673_5 | TATGAGTGTGGAGTCGTTGCGCATGGCTCTTAGCAATCTTAGCNNNAGATCGGAAG |
| 34673_6 | TATGAGTGTGGAGTCGTTGCGCATGGCTCTTAGCAATCTTAGCNNNAGATCGGAAG |
| 34673_7 | TATGAGTGTGGAGTCGTTGCGCATGGCTCTTAGCAATCTTAGCNNNAGATCGGAAG |
| 34673_8 | TATGAGTGTGGAGTCGTTGCGCATGGCTCTTAGCAATCTTAGCNNNAGATCGGAAG |
| 34673_9 | TATGAGTGTGGAGTCGTTGCGCATGGCTCTTAGCAATCTTAGCNNNAGATCGGAAG |
| 34858_1 | TATGAGTGTGGAGTCGTTGCCCTCCTGAGTAACTGGGATTACANNNAGATCGGAAG |
| 34858_2 | TATGAGTGTGGAGTCGTTGCCCTCCTGAGTAACTGGGATTACANNNAGATCGGAAG |
| 34858_3 | TATGAGTGTGGAGTCGTTGCCCTCCTGAGTAACTGGGATTACANNNAGATCGGAAG |
| 34858_4 | TATGAGTGTGGAGTCGTTGCCCTCCTGAGTAACTGGGATTACANNNAGATCGGAAG |
| 34858_5 | TATGAGTGTGGAGTCGTTGCCCTCCTGAGTAACTGGGATTACANNNAGATCGGAAG |
| 34858_6 | TATGAGTGTGGAGTCGTTGCCCTCCTGAGTAACTGGGATTACANNNAGATCGGAAG |
| 34858_7 | TATGAGTGTGGAGTCGTTGCCCTCCTGAGTAACTGGGATTACANNNAGATCGGAAG |
| 34858_8 | TATGAGTGTGGAGTCGTTGCCCTCCTGAGTAACTGGGATTACANNNAGATCGGAAG |
| 34858_9 | TATGAGTGTGGAGTCGTTGCCCTCCTGAGTAACTGGGATTACANNNAGATCGGAAG |
| 9491_1 | TATGAGTGTGGAGTCGTTGCACACACACTGTTCAGGGTGAAGNNNAGATCGGAAG |
| 9491_2 | TATGAGTGTGGAGTCGTTGCACACACACTGTTCAGGGTGAAGNNNAGATCGGAAG |
| 9491_3 | TATGAGTGTGGAGTCGTTGCACACACACTGTTCAGGGTGAAGNNNAGATCGGAAG |
| 9491_4 | TATGAGTGTGGAGTCGTTGCACACACACTGTTCAGGGTGAAGNNNAGATCGGAAG |
| 9491_5 | TATGAGTGTGGAGTCGTTGCACACACACTGTTCAGGGTGAAGNNNAGATCGGAAG |
| 9491_6 | TATGAGTGTGGAGTCGTTGCACACACACTGTTCAGGGTGAAGNNNAGATCGGAAG |
| 9491_7 | TATGAGTGTGGAGTCGTTGCACACACACTGTTCAGGGTGAAGNNNAGATCGGAAG |
| 9491_8 | TATGAGTGTGGAGTCGTTGCACACACACTGTTCAGGGTGAAGNNNAGATCGGAAG |
| 9491_9 | TATGAGTGTGGAGTCGTTGCACACACACTGTTCAGGGTGAAGNNNAGATCGGAAG |
| 35264_1 | TATGAGTGTGGAGTCGTTGCAATTTCAAGGCAATGCTGGAGTGNNNAGATCGGAA |
| 35264_2 | TATGAGTGTGGAGTCGTTGCAATTTCAAGGCAATGCTGGAGTGNNNAGATCGGAA |
| 35264_3 | TATGAGTGTGGAGTCGTTGCAATTTCAAGGCAATGCTGGAGTGNNNAGATCGGAA |
| 35264_4 | TATGAGTGTGGAGTCGTTGCAATTTCAAGGCAATGCTGGAGTGNNNAGATCGGAA |
| 35264_5 | TATGAGTGTGGAGTCGTTGCAATTTCAAGGCAATGCTGGAGTGNNNAGATCGGAA |
| 35264_6 | TATGAGTGTGGAGTCGTTGCAATTTCAAGGCAATGCTGGAGTGNNNAGATCGGAA |
| 35264_7 | TATGAGTGTGGAGTCGTTGCAATTTCAAGGCAATGCTGGAGTGNNNAGATCGGAA |
| 35264_8 | TATGAGTGTGGAGTCGTTGCAATTTCAAGGCAATGCTGGAGTGNNNAGATCGGAA |
| 35264_9 | TATGAGTGTGGAGTCGTTGCAATTTCAAGGCAATGCTGGAGTGNNNAGATCGGAA |
| 29485_1 | ATGAGTGTGGAGTCGTTGCCAAACTAGAAAAGATAAAAACACACCANNNAGATCGG |
| 29485_2 | ATGAGTGTGGAGTCGTTGCCAAACTAGAAAAGATAAAAACACACCANNNAGATCGG |
| 29485_3 | ATGAGTGTGGAGTCGTTGCCAAACTAGAAAAGATAAAAACACACCANNNAGATCGG |
| 29485_4 | ATGAGTGTGGAGTCGTTGCCAAACTAGAAAAGATAAAAACACACCANNNAGATCGG |
| 29485_5 | ATGAGTGTGGAGTCGTTGCCAAACTAGAAAAGATAAAAACACACCANNNAGATCGG |
| 29485_6 | ATGAGTGTGGAGTCGTTGCCAAACTAGAAAAGATAAAAACACACCANNNAGATCGG |
| 29485_7 | ATGAGTGTGGAGTCGTTGCCAAACTAGAAAAGATAAAAACACACCANNNAGATCGG |
| 29485_8 | ATGAGTGTGGAGTCGTTGCCAAACTAGAAAAGATAAAAACACACCANNNAGATCGG |

| | |
|---|---|
| 29485_9 | ATGAGTGTGGAGTCGTTGCCAAACTAGAAAAGATAAAAACACACCANNNAGATCGG |
| 38678_1 | TATGAGTGTGGAGTCGTTGCAACCTGAACGTATTTCCAGCTCTNNNAGATCGGAAG |
| 38678_2 | TATGAGTGTGGAGTCGTTGCAACCTGAACGTATTTCCAGCTCTNNNAGATCGGAAG |
| 38678_3 | TATGAGTGTGGAGTCGTTGCAACCTGAACGTATTTCCAGCTCTNNNAGATCGGAAG |
| 38678_4 | TATGAGTGTGGAGTCGTTGCAACCTGAACGTATTTCCAGCTCTNNNAGATCGGAAG |
| 38678_5 | TATGAGTGTGGAGTCGTTGCAACCTGAACGTATTTCCAGCTCTNNNAGATCGGAAG |
| 38678_6 | TATGAGTGTGGAGTCGTTGCAACCTGAACGTATTTCCAGCTCTNNNAGATCGGAAG |
| 38678_7 | TATGAGTGTGGAGTCGTTGCAACCTGAACGTATTTCCAGCTCTNNNAGATCGGAAG |
| 38678_8 | TATGAGTGTGGAGTCGTTGCAACCTGAACGTATTTCCAGCTCTNNNAGATCGGAAG |
| 38678_9 | TATGAGTGTGGAGTCGTTGCAACCTGAACGTATTTCCAGCTCTNNNAGATCGGAAG |
| 47985_1 | TATGAGTGTGGAGTCGTTGCCACCTTGCCTGGCTAATTTCTTGNNNAGATCGGAAG |
| 47985_2 | TATGAGTGTGGAGTCGTTGCCACCTTGCCTGGCTAATTTCTTGNNNAGATCGGAAG |
| 47985_3 | TATGAGTGTGGAGTCGTTGCCACCTTGCCTGGCTAATTTCTTGNNNAGATCGGAAG |
| 47985_4 | TATGAGTGTGGAGTCGTTGCCACCTTGCCTGGCTAATTTCTTGNNNAGATCGGAAG |
| 47985_5 | TATGAGTGTGGAGTCGTTGCCACCTTGCCTGGCTAATTTCTTGNNNAGATCGGAAG |
| 47985_6 | TATGAGTGTGGAGTCGTTGCCACCTTGCCTGGCTAATTTCTTGNNNAGATCGGAAG |
| 47985_7 | TATGAGTGTGGAGTCGTTGCCACCTTGCCTGGCTAATTTCTTGNNNAGATCGGAAG |
| 47985_8 | TATGAGTGTGGAGTCGTTGCCACCTTGCCTGGCTAATTTCTTGNNNAGATCGGAAG |
| 47985_9 | TATGAGTGTGGAGTCGTTGCCACCTTGCCTGGCTAATTTCTTGNNNAGATCGGAAG |
| 32482_1 | TATGAGTGTGGAGTCGTTGCAAATTTTGCTCTATTGCCCAGGCNNNAGATCGGAAG |
| 32482_2 | TATGAGTGTGGAGTCGTTGCAAATTTTGCTCTATTGCCCAGGCNNNAGATCGGAAG |
| 32482_3 | TATGAGTGTGGAGTCGTTGCAAATTTTGCTCTATTGCCCAGGCNNNAGATCGGAAG |
| 32482_4 | TATGAGTGTGGAGTCGTTGCAAATTTTGCTCTATTGCCCAGGCNNNAGATCGGAAG |
| 32482_5 | TATGAGTGTGGAGTCGTTGCAAATTTTGCTCTATTGCCCAGGCNNNAGATCGGAAG |
| 32482_6 | TATGAGTGTGGAGTCGTTGCAAATTTTGCTCTATTGCCCAGGCNNNAGATCGGAAG |
| 32482_7 | TATGAGTGTGGAGTCGTTGCAAATTTTGCTCTATTGCCCAGGCNNNAGATCGGAAG |
| 32482_8 | TATGAGTGTGGAGTCGTTGCAAATTTTGCTCTATTGCCCAGGCNNNAGATCGGAAG |
| 32482_9 | TATGAGTGTGGAGTCGTTGCAAATTTTGCTCTATTGCCCAGGCNNNAGATCGGAAG |
| 49355_1 | TATGAGTGTGGAGTCGTTGCTCCAGGTTAGGAAAGGATTCAGCNNNAGATCGGAA |
| 49355_2 | TATGAGTGTGGAGTCGTTGCTCCAGGTTAGGAAAGGATTCAGCNNNAGATCGGAA |
| 49355_3 | TATGAGTGTGGAGTCGTTGCTCCAGGTTAGGAAAGGATTCAGCNNNAGATCGGAA |
| 49355_4 | TATGAGTGTGGAGTCGTTGCTCCAGGTTAGGAAAGGATTCAGCNNNAGATCGGAA |
| 49355_5 | TATGAGTGTGGAGTCGTTGCTCCAGGTTAGGAAAGGATTCAGCNNNAGATCGGAA |
| 49355_6 | TATGAGTGTGGAGTCGTTGCTCCAGGTTAGGAAAGGATTCAGCNNNAGATCGGAA |
| 49355_7 | TATGAGTGTGGAGTCGTTGCTCCAGGTTAGGAAAGGATTCAGCNNNAGATCGGAA |
| 49355_8 | TATGAGTGTGGAGTCGTTGCTCCAGGTTAGGAAAGGATTCAGCNNNAGATCGGAA |
| 49355_9 | TATGAGTGTGGAGTCGTTGCTCCAGGTTAGGAAAGGATTCAGCNNNAGATCGGAA |
| 40181_1 | TATGAGTGTGGAGTCGTTGCTTGAGGCAGGAGAATCACTTGAANNNAGATCGGAA |
| 40181_2 | TATGAGTGTGGAGTCGTTGCTTGAGGCAGGAGAATCACTTGAANNNAGATCGGAA |
| 40181_3 | TATGAGTGTGGAGTCGTTGCTTGAGGCAGGAGAATCACTTGAANNNAGATCGGAA |
| 40181_4 | TATGAGTGTGGAGTCGTTGCTTGAGGCAGGAGAATCACTTGAANNNAGATCGGAA |
| 40181_5 | TATGAGTGTGGAGTCGTTGCTTGAGGCAGGAGAATCACTTGAANNNAGATCGGAA |
| 40181_6 | TATGAGTGTGGAGTCGTTGCTTGAGGCAGGAGAATCACTTGAANNNAGATCGGAA |

| | |
|---|---|
| 40181_7 | TATGAGTGTGGAGTCGTTGCTTGAGGCAGGAGAATCACTTGAANNNAGATCGGAA |
| 40181_8 | TATGAGTGTGGAGTCGTTGCTTGAGGCAGGAGAATCACTTGAANNNAGATCGGAA |
| 40181_9 | TATGAGTGTGGAGTCGTTGCTTGAGGCAGGAGAATCACTTGAANNNAGATCGGAA |
| 35060_1 | TATGAGTGTGGAGTCGTTGCCTATCTCGACTCACTGCAAGCTCNNNAGATCGGAAG |
| 35060_2 | TATGAGTGTGGAGTCGTTGCCTATCTCGACTCACTGCAAGCTCNNNAGATCGGAAG |
| 35060_3 | TATGAGTGTGGAGTCGTTGCCTATCTCGACTCACTGCAAGCTCNNNAGATCGGAAG |
| 35060_4 | TATGAGTGTGGAGTCGTTGCCTATCTCGACTCACTGCAAGCTCNNNAGATCGGAAG |
| 35060_5 | TATGAGTGTGGAGTCGTTGCCTATCTCGACTCACTGCAAGCTCNNNAGATCGGAAG |
| 35060_6 | TATGAGTGTGGAGTCGTTGCCTATCTCGACTCACTGCAAGCTCNNNAGATCGGAAG |
| 35060_7 | TATGAGTGTGGAGTCGTTGCCTATCTCGACTCACTGCAAGCTCNNNAGATCGGAAG |
| 35060_8 | TATGAGTGTGGAGTCGTTGCCTATCTCGACTCACTGCAAGCTCNNNAGATCGGAAG |
| 35060_9 | TATGAGTGTGGAGTCGTTGCCTATCTCGACTCACTGCAAGCTCNNNAGATCGGAAG |
| 30279_1 | TATGAGTGTGGAGTCGTTGCGGGTGCTACTGCTGAGATTTTGNNNAGATCGGAA |
| 30279_2 | TATGAGTGTGGAGTCGTTGCGGGTGCTACTGCTGAGATTTTGNNNAGATCGGAA |
| 30279_3 | TATGAGTGTGGAGTCGTTGCGGGTGCTACTGCTGAGATTTTGNNNAGATCGGAA |
| 30279_4 | TATGAGTGTGGAGTCGTTGCGGGTGCTACTGCTGAGATTTTGNNNAGATCGGAA |
| 30279_5 | TATGAGTGTGGAGTCGTTGCGGGTGCTACTGCTGAGATTTTGNNNAGATCGGAA |
| 30279_6 | TATGAGTGTGGAGTCGTTGCGGGTGCTACTGCTGAGATTTTGNNNAGATCGGAA |
| 30279_7 | TATGAGTGTGGAGTCGTTGCGGGTGCTACTGCTGAGATTTTGNNNAGATCGGAA |
| 30279_8 | TATGAGTGTGGAGTCGTTGCGGGTGCTACTGCTGAGATTTTGNNNAGATCGGAA |
| 30279_9 | TATGAGTGTGGAGTCGTTGCGGGTGCTACTGCTGAGATTTTGNNNAGATCGGAA |
| 29120_1 | TATGAGTGTGGAGTCGTTGCACCAAAGCTAAGAAGACAGCAGANNNAGATCGGAA |
| 29120_2 | TATGAGTGTGGAGTCGTTGCACCAAAGCTAAGAAGACAGCAGANNNAGATCGGAA |
| 29120_3 | TATGAGTGTGGAGTCGTTGCACCAAAGCTAAGAAGACAGCAGANNNAGATCGGAA |
| 29120_4 | TATGAGTGTGGAGTCGTTGCACCAAAGCTAAGAAGACAGCAGANNNAGATCGGAA |
| 29120_5 | TATGAGTGTGGAGTCGTTGCACCAAAGCTAAGAAGACAGCAGANNNAGATCGGAA |
| 29120_6 | TATGAGTGTGGAGTCGTTGCACCAAAGCTAAGAAGACAGCAGANNNAGATCGGAA |
| 29120_7 | TATGAGTGTGGAGTCGTTGCACCAAAGCTAAGAAGACAGCAGANNNAGATCGGAA |
| 29120_8 | TATGAGTGTGGAGTCGTTGCACCAAAGCTAAGAAGACAGCAGANNNAGATCGGAA |
| 29120_9 | TATGAGTGTGGAGTCGTTGCACCAAAGCTAAGAAGACAGCAGANNNAGATCGGAA |
| 45986_1 | TATGAGTGTGGAGTCGTTGCACCTGCTGGATTCAAGTGATTCTNNNAGATCGGAAG |
| 45986_2 | TATGAGTGTGGAGTCGTTGCACCTGCTGGATTCAAGTGATTCTNNNAGATCGGAAG |
| 45986_3 | TATGAGTGTGGAGTCGTTGCACCTGCTGGATTCAAGTGATTCTNNNAGATCGGAAG |
| 45986_4 | TATGAGTGTGGAGTCGTTGCACCTGCTGGATTCAAGTGATTCTNNNAGATCGGAAG |
| 45986_5 | TATGAGTGTGGAGTCGTTGCACCTGCTGGATTCAAGTGATTCTNNNAGATCGGAAG |
| 45986_6 | TATGAGTGTGGAGTCGTTGCACCTGCTGGATTCAAGTGATTCTNNNAGATCGGAAG |
| 45986_7 | TATGAGTGTGGAGTCGTTGCACCTGCTGGATTCAAGTGATTCTNNNAGATCGGAAG |
| 45986_8 | TATGAGTGTGGAGTCGTTGCACCTGCTGGATTCAAGTGATTCTNNNAGATCGGAAG |
| 45986_9 | TATGAGTGTGGAGTCGTTGCACCTGCTGGATTCAAGTGATTCTNNNAGATCGGAAG |
| 52379_1 | TATGAGTGTGGAGTCGTTGCAGTCAGCTACATCAATGGCAAGANNNAGATCGGAA |
| 52379_2 | TATGAGTGTGGAGTCGTTGCAGTCAGCTACATCAATGGCAAGANNNAGATCGGAA |
| 52379_3 | TATGAGTGTGGAGTCGTTGCAGTCAGCTACATCAATGGCAAGANNNAGATCGGAA |
| 52379_4 | TATGAGTGTGGAGTCGTTGCAGTCAGCTACATCAATGGCAAGANNNAGATCGGAA |

| | |
|---|---|
| 52379_5 | TATGAGTGTGGAGTCGTTGCAGTCAGCTACATCAATGGCAAGANNNAGATCGGAA |
| 52379_6 | TATGAGTGTGGAGTCGTTGCAGTCAGCTACATCAATGGCAAGANNNAGATCGGAA |
| 52379_7 | TATGAGTGTGGAGTCGTTGCAGTCAGCTACATCAATGGCAAGANNNAGATCGGAA |
| 52379_8 | TATGAGTGTGGAGTCGTTGCAGTCAGCTACATCAATGGCAAGANNNAGATCGGAA |
| 52379_9 | TATGAGTGTGGAGTCGTTGCAGTCAGCTACATCAATGGCAAGANNNAGATCGGAA |
| 29355_1 | TATGAGTGTGGAGTCGTTGCATGAAGTTTACAGTCCCAGGAGCNNNAGATCGGAA |
| 29355_2 | TATGAGTGTGGAGTCGTTGCATGAAGTTTACAGTCCCAGGAGCNNNAGATCGGAA |
| 29355_3 | TATGAGTGTGGAGTCGTTGCATGAAGTTTACAGTCCCAGGAGCNNNAGATCGGAA |
| 29355_4 | TATGAGTGTGGAGTCGTTGCATGAAGTTTACAGTCCCAGGAGCNNNAGATCGGAA |
| 29355_5 | TATGAGTGTGGAGTCGTTGCATGAAGTTTACAGTCCCAGGAGCNNNAGATCGGAA |
| 29355_6 | TATGAGTGTGGAGTCGTTGCATGAAGTTTACAGTCCCAGGAGCNNNAGATCGGAA |
| 29355_7 | TATGAGTGTGGAGTCGTTGCATGAAGTTTACAGTCCCAGGAGCNNNAGATCGGAA |
| 29355_8 | TATGAGTGTGGAGTCGTTGCATGAAGTTTACAGTCCCAGGAGCNNNAGATCGGAA |
| 29355_9 | TATGAGTGTGGAGTCGTTGCATGAAGTTTACAGTCCCAGGAGCNNNAGATCGGAA |
| 29644_1 | TATGAGTGTGGAGTCGTTGCCAATAGGGTGGTGGGTGATAGAGNNNAGATCGGA |
| 29644_2 | TATGAGTGTGGAGTCGTTGCCAATAGGGTGGTGGGTGATAGAGNNNAGATCGGA |
| 29644_3 | TATGAGTGTGGAGTCGTTGCCAATAGGGTGGTGGGTGATAGAGNNNAGATCGGA |
| 29644_4 | TATGAGTGTGGAGTCGTTGCCAATAGGGTGGTGGGTGATAGAGNNNAGATCGGA |
| 29644_5 | TATGAGTGTGGAGTCGTTGCCAATAGGGTGGTGGGTGATAGAGNNNAGATCGGA |
| 29644_6 | TATGAGTGTGGAGTCGTTGCCAATAGGGTGGTGGGTGATAGAGNNNAGATCGGA |
| 29644_7 | TATGAGTGTGGAGTCGTTGCCAATAGGGTGGTGGGTGATAGAGNNNAGATCGGA |
| 29644_8 | TATGAGTGTGGAGTCGTTGCCAATAGGGTGGTGGGTGATAGAGNNNAGATCGGA |
| 29644_9 | TATGAGTGTGGAGTCGTTGCCAATAGGGTGGTGGGTGATAGAGNNNAGATCGGA |
| 43016_1 | TATGAGTGTGGAGTCGTTGCTGTGTGTGTGTTGTGTATGTGTGNNNAGATCGGAA |
| 43016_2 | TATGAGTGTGGAGTCGTTGCTGTGTGTGTGTTGTGTATGTGTGNNNAGATCGGAA |
| 43016_3 | TATGAGTGTGGAGTCGTTGCTGTGTGTGTGTTGTGTATGTGTGNNNAGATCGGAA |
| 43016_4 | TATGAGTGTGGAGTCGTTGCTGTGTGTGTGTTGTGTATGTGTGNNNAGATCGGAA |
| 43016_5 | TATGAGTGTGGAGTCGTTGCTGTGTGTGTGTTGTGTATGTGTGNNNAGATCGGAA |
| 43016_6 | TATGAGTGTGGAGTCGTTGCTGTGTGTGTGTTGTGTATGTGTGNNNAGATCGGAA |
| 43016_7 | TATGAGTGTGGAGTCGTTGCTGTGTGTGTGTTGTGTATGTGTGNNNAGATCGGAA |
| 43016_8 | TATGAGTGTGGAGTCGTTGCTGTGTGTGTGTTGTGTATGTGTGNNNAGATCGGAA |
| 43016_9 | TATGAGTGTGGAGTCGTTGCTGTGTGTGTGTTGTGTATGTGTGNNNAGATCGGAA |
| 33660_1 | TATGAGTGTGGAGTCGTTGCATAGTGGCATGATCTTGGCTCATNNNAGATCGGAAG |
| 33660_2 | TATGAGTGTGGAGTCGTTGCATAGTGGCATGATCTTGGCTCATNNNAGATCGGAAG |
| 33660_3 | TATGAGTGTGGAGTCGTTGCATAGTGGCATGATCTTGGCTCATNNNAGATCGGAAG |
| 33660_4 | TATGAGTGTGGAGTCGTTGCATAGTGGCATGATCTTGGCTCATNNNAGATCGGAAG |
| 33660_5 | TATGAGTGTGGAGTCGTTGCATAGTGGCATGATCTTGGCTCATNNNAGATCGGAAG |
| 33660_6 | TATGAGTGTGGAGTCGTTGCATAGTGGCATGATCTTGGCTCATNNNAGATCGGAAG |
| 33660_7 | TATGAGTGTGGAGTCGTTGCATAGTGGCATGATCTTGGCTCATNNNAGATCGGAAG |
| 33660_8 | TATGAGTGTGGAGTCGTTGCATAGTGGCATGATCTTGGCTCATNNNAGATCGGAAG |
| 33660_9 | TATGAGTGTGGAGTCGTTGCATAGTGGCATGATCTTGGCTCATNNNAGATCGGAAG |
| 10461_1 | TATGAGTGTGGAGTCGTTGCAGGCCATGCTAGCCAGAGTATAANNNAGATCGGAA |
| 10461_2 | TATGAGTGTGGAGTCGTTGCAGGCCATGCTAGCCAGAGTATAANNNAGATCGGAA |

| | |
|---|---|
| 10461_3 | TATGAGTGTGGAGTCGTTGCAGGCCATGCTAGCCAGAGTATAANNNAGATCGGAA |
| 10461_4 | TATGAGTGTGGAGTCGTTGCAGGCCATGCTAGCCAGAGTATAANNNAGATCGGAA |
| 10461_5 | TATGAGTGTGGAGTCGTTGCAGGCCATGCTAGCCAGAGTATAANNNAGATCGGAA |
| 10461_6 | TATGAGTGTGGAGTCGTTGCAGGCCATGCTAGCCAGAGTATAANNNAGATCGGAA |
| 10461_7 | TATGAGTGTGGAGTCGTTGCAGGCCATGCTAGCCAGAGTATAANNNAGATCGGAA |
| 10461_8 | TATGAGTGTGGAGTCGTTGCAGGCCATGCTAGCCAGAGTATAANNNAGATCGGAA |
| 10461_9 | TATGAGTGTGGAGTCGTTGCAGGCCATGCTAGCCAGAGTATAANNNAGATCGGAA |
| 36436_1 | TATGAGTGTGGAGTCGTTGCTGTTGTGAAGTGCTTCCCTTTTCNNNAGATCGGAAG |
| 36436_2 | TATGAGTGTGGAGTCGTTGCTGTTGTGAAGTGCTTCCCTTTTCNNNAGATCGGAAG |
| 36436_3 | TATGAGTGTGGAGTCGTTGCTGTTGTGAAGTGCTTCCCTTTTCNNNAGATCGGAAG |
| 36436_4 | TATGAGTGTGGAGTCGTTGCTGTTGTGAAGTGCTTCCCTTTTCNNNAGATCGGAAG |
| 36436_5 | TATGAGTGTGGAGTCGTTGCTGTTGTGAAGTGCTTCCCTTTTCNNNAGATCGGAAG |
| 36436_6 | TATGAGTGTGGAGTCGTTGCTGTTGTGAAGTGCTTCCCTTTTCNNNAGATCGGAAG |
| 36436_7 | TATGAGTGTGGAGTCGTTGCTGTTGTGAAGTGCTTCCCTTTTCNNNAGATCGGAAG |
| 36436_8 | TATGAGTGTGGAGTCGTTGCTGTTGTGAAGTGCTTCCCTTTTCNNNAGATCGGAAG |
| 36436_9 | TATGAGTGTGGAGTCGTTGCTGTTGTGAAGTGCTTCCCTTTTCNNNAGATCGGAAG |
| 34262_1 | TATGAGTGTGGAGTCGTTGCCTCATCTTGGTGATCTTGGGGAANNNAGATCGGAAG |
| 34262_2 | TATGAGTGTGGAGTCGTTGCCTCATCTTGGTGATCTTGGGGAANNNAGATCGGAAG |
| 34262_3 | TATGAGTGTGGAGTCGTTGCCTCATCTTGGTGATCTTGGGGAANNNAGATCGGAAG |
| 34262_4 | TATGAGTGTGGAGTCGTTGCCTCATCTTGGTGATCTTGGGGAANNNAGATCGGAAG |
| 34262_5 | TATGAGTGTGGAGTCGTTGCCTCATCTTGGTGATCTTGGGGAANNNAGATCGGAAG |
| 34262_6 | TATGAGTGTGGAGTCGTTGCCTCATCTTGGTGATCTTGGGGAANNNAGATCGGAAG |
| 34262_7 | TATGAGTGTGGAGTCGTTGCCTCATCTTGGTGATCTTGGGGAANNNAGATCGGAAG |
| 34262_8 | TATGAGTGTGGAGTCGTTGCCTCATCTTGGTGATCTTGGGGAANNNAGATCGGAAG |
| 34262_9 | TATGAGTGTGGAGTCGTTGCCTCATCTTGGTGATCTTGGGGAANNNAGATCGGAAG |
| 50946_1 | TATGAGTGTGGAGTCGTTGCAATCCTCTTCTTTCACCCCAACANNNAGATCGGAAGA |
| 50946_2 | TATGAGTGTGGAGTCGTTGCAATCCTCTTCTTTCACCCCAACANNNAGATCGGAAGA |
| 50946_3 | TATGAGTGTGGAGTCGTTGCAATCCTCTTCTTTCACCCCAACANNNAGATCGGAAGA |
| 50946_4 | TATGAGTGTGGAGTCGTTGCAATCCTCTTCTTTCACCCCAACANNNAGATCGGAAGA |
| 50946_5 | TATGAGTGTGGAGTCGTTGCAATCCTCTTCTTTCACCCCAACANNNAGATCGGAAGA |
| 50946_6 | TATGAGTGTGGAGTCGTTGCAATCCTCTTCTTTCACCCCAACANNNAGATCGGAAGA |
| 50946_7 | TATGAGTGTGGAGTCGTTGCAATCCTCTTCTTTCACCCCAACANNNAGATCGGAAGA |
| 50946_8 | TATGAGTGTGGAGTCGTTGCAATCCTCTTCTTTCACCCCAACANNNAGATCGGAAGA |
| 50946_9 | TATGAGTGTGGAGTCGTTGCAATCCTCTTCTTTCACCCCAACANNNAGATCGGAAGA |
| 48692_1 | TATGAGTGTGGAGTCGTTGCCAAATCCATGTTCTTGCTCGTGTNNNAGATCGGAAG |
| 48692_2 | TATGAGTGTGGAGTCGTTGCCAAATCCATGTTCTTGCTCGTGTNNNAGATCGGAAG |
| 48692_3 | TATGAGTGTGGAGTCGTTGCCAAATCCATGTTCTTGCTCGTGTNNNAGATCGGAAG |
| 48692_4 | TATGAGTGTGGAGTCGTTGCCAAATCCATGTTCTTGCTCGTGTNNNAGATCGGAAG |
| 48692_5 | TATGAGTGTGGAGTCGTTGCCAAATCCATGTTCTTGCTCGTGTNNNAGATCGGAAG |
| 48692_6 | TATGAGTGTGGAGTCGTTGCCAAATCCATGTTCTTGCTCGTGTNNNAGATCGGAAG |
| 48692_7 | TATGAGTGTGGAGTCGTTGCCAAATCCATGTTCTTGCTCGTGTNNNAGATCGGAAG |
| 48692_8 | TATGAGTGTGGAGTCGTTGCCAAATCCATGTTCTTGCTCGTGTNNNAGATCGGAAG |
| 48692_9 | TATGAGTGTGGAGTCGTTGCCAAATCCATGTTCTTGCTCGTGTNNNAGATCGGAAG |

| | |
|---|---|
| 9121_1 | TATGAGTGTGGAGTCGTTGCAAGTCACCACAAAGACCTTGTCANNNAGATCGGAAG |
| 9121_2 | TATGAGTGTGGAGTCGTTGCAAGTCACCACAAAGACCTTGTCANNNAGATCGGAAG |
| 9121_3 | TATGAGTGTGGAGTCGTTGCAAGTCACCACAAAGACCTTGTCANNNAGATCGGAAG |
| 9121_4 | TATGAGTGTGGAGTCGTTGCAAGTCACCACAAAGACCTTGTCANNNAGATCGGAAG |
| 9121_5 | TATGAGTGTGGAGTCGTTGCAAGTCACCACAAAGACCTTGTCANNNAGATCGGAAG |
| 9121_6 | TATGAGTGTGGAGTCGTTGCAAGTCACCACAAAGACCTTGTCANNNAGATCGGAAG |
| 9121_7 | TATGAGTGTGGAGTCGTTGCAAGTCACCACAAAGACCTTGTCANNNAGATCGGAAG |
| 9121_8 | TATGAGTGTGGAGTCGTTGCAAGTCACCACAAAGACCTTGTCANNNAGATCGGAAG |
| 9121_9 | TATGAGTGTGGAGTCGTTGCAAGTCACCACAAAGACCTTGTCANNNAGATCGGAAG |
| 28755_1 | TATGAGTGTGGAGTCGTTGCACCAACCTTCCTCACACACTATGNNNAGATCGGAAGA |
| 28755_2 | TATGAGTGTGGAGTCGTTGCACCAACCTTCCTCACACACTATGNNNAGATCGGAAGA |
| 28755_3 | TATGAGTGTGGAGTCGTTGCACCAACCTTCCTCACACACTATGNNNAGATCGGAAGA |
| 28755_4 | TATGAGTGTGGAGTCGTTGCACCAACCTTCCTCACACACTATGNNNAGATCGGAAGA |
| 28755_5 | TATGAGTGTGGAGTCGTTGCACCAACCTTCCTCACACACTATGNNNAGATCGGAAGA |
| 28755_6 | TATGAGTGTGGAGTCGTTGCACCAACCTTCCTCACACACTATGNNNAGATCGGAAGA |
| 28755_7 | TATGAGTGTGGAGTCGTTGCACCAACCTTCCTCACACACTATGNNNAGATCGGAAGA |
| 28755_8 | TATGAGTGTGGAGTCGTTGCACCAACCTTCCTCACACACTATGNNNAGATCGGAAGA |
| 28755_9 | TATGAGTGTGGAGTCGTTGCACCAACCTTCCTCACACACTATGNNNAGATCGGAAGA |
| 52258_1 | TATGAGTGTGGAGTCGTTGCATCACGATAGAATCCTGCCTTCCNNNAGATCGGAAGA |
| 52258_2 | TATGAGTGTGGAGTCGTTGCATCACGATAGAATCCTGCCTTCCNNNAGATCGGAAGA |
| 52258_3 | TATGAGTGTGGAGTCGTTGCATCACGATAGAATCCTGCCTTCCNNNAGATCGGAAGA |
| 52258_4 | TATGAGTGTGGAGTCGTTGCATCACGATAGAATCCTGCCTTCCNNNAGATCGGAAGA |
| 52258_5 | TATGAGTGTGGAGTCGTTGCATCACGATAGAATCCTGCCTTCCNNNAGATCGGAAGA |
| 52258_6 | TATGAGTGTGGAGTCGTTGCATCACGATAGAATCCTGCCTTCCNNNAGATCGGAAGA |
| 52258_7 | TATGAGTGTGGAGTCGTTGCATCACGATAGAATCCTGCCTTCCNNNAGATCGGAAGA |
| 52258_8 | TATGAGTGTGGAGTCGTTGCATCACGATAGAATCCTGCCTTCCNNNAGATCGGAAGA |
| 52258_9 | TATGAGTGTGGAGTCGTTGCATCACGATAGAATCCTGCCTTCCNNNAGATCGGAAGA |
| 10768_1 | TATGAGTGTGGAGTCGTTGCTTGCAGTTTCTCTGGAGAACCATNNNAGATCGGAAG |
| 10768_2 | TATGAGTGTGGAGTCGTTGCTTGCAGTTTCTCTGGAGAACCATNNNAGATCGGAAG |
| 10768_3 | TATGAGTGTGGAGTCGTTGCTTGCAGTTTCTCTGGAGAACCATNNNAGATCGGAAG |
| 10768_4 | TATGAGTGTGGAGTCGTTGCTTGCAGTTTCTCTGGAGAACCATNNNAGATCGGAAG |
| 10768_5 | TATGAGTGTGGAGTCGTTGCTTGCAGTTTCTCTGGAGAACCATNNNAGATCGGAAG |
| 10768_6 | TATGAGTGTGGAGTCGTTGCTTGCAGTTTCTCTGGAGAACCATNNNAGATCGGAAG |
| 10768_7 | TATGAGTGTGGAGTCGTTGCTTGCAGTTTCTCTGGAGAACCATNNNAGATCGGAAG |
| 10768_8 | TATGAGTGTGGAGTCGTTGCTTGCAGTTTCTCTGGAGAACCATNNNAGATCGGAAG |
| 10768_9 | TATGAGTGTGGAGTCGTTGCTTGCAGTTTCTCTGGAGAACCATNNNAGATCGGAAG |
| 45721_1 | TATGAGTGTGGAGTCGTTGCGTGCAGTCTCAGCTCACTTCAANNNAGATCGGAAGA |
| 45721_2 | TATGAGTGTGGAGTCGTTGCGTGCAGTCTCAGCTCACTTCAANNNAGATCGGAAGA |
| 45721_3 | TATGAGTGTGGAGTCGTTGCGTGCAGTCTCAGCTCACTTCAANNNAGATCGGAAGA |
| 45721_4 | TATGAGTGTGGAGTCGTTGCGTGCAGTCTCAGCTCACTTCAANNNAGATCGGAAGA |
| 45721_5 | TATGAGTGTGGAGTCGTTGCGTGCAGTCTCAGCTCACTTCAANNNAGATCGGAAGA |
| 45721_6 | TATGAGTGTGGAGTCGTTGCGTGCAGTCTCAGCTCACTTCAANNNAGATCGGAAGA |
| 45721_7 | TATGAGTGTGGAGTCGTTGCGTGCAGTCTCAGCTCACTTCAANNNAGATCGGAAGA |

| | |
|---|---|
| 45721_8 | TATGAGTGTGGAGTCGTTGCGTGCAGTCTCAGCTCACTTCAANNNAGATCGGAAGA |
| 45721_9 | TATGAGTGTGGAGTCGTTGCGTGCAGTCTCAGCTCACTTCAANNNAGATCGGAAGA |
| 51245_1 | TATGAGTGTGGAGTCGTTGCTCTCAAACCTCCAAAGAACAGCTNNNAGATCGGAAG |
| 51245_2 | TATGAGTGTGGAGTCGTTGCTCTCAAACCTCCAAAGAACAGCTNNNAGATCGGAAG |
| 51245_3 | TATGAGTGTGGAGTCGTTGCTCTCAAACCTCCAAAGAACAGCTNNNAGATCGGAAG |
| 51245_4 | TATGAGTGTGGAGTCGTTGCTCTCAAACCTCCAAAGAACAGCTNNNAGATCGGAAG |
| 51245_5 | TATGAGTGTGGAGTCGTTGCTCTCAAACCTCCAAAGAACAGCTNNNAGATCGGAAG |
| 51245_6 | TATGAGTGTGGAGTCGTTGCTCTCAAACCTCCAAAGAACAGCTNNNAGATCGGAAG |
| 51245_7 | TATGAGTGTGGAGTCGTTGCTCTCAAACCTCCAAAGAACAGCTNNNAGATCGGAAG |
| 51245_8 | TATGAGTGTGGAGTCGTTGCTCTCAAACCTCCAAAGAACAGCTNNNAGATCGGAAG |
| 51245_9 | TATGAGTGTGGAGTCGTTGCTCTCAAACCTCCAAAGAACAGCTNNNAGATCGGAAG |
| 40491_1 | TATGAGTGTGGAGTCGTTGCCTTTGCACAGTCTCCCTCTGATNNNAGATCGGAAGA |
| 40491_2 | TATGAGTGTGGAGTCGTTGCCTTTGCACAGTCTCCCTCTGATNNNAGATCGGAAGA |
| 40491_3 | TATGAGTGTGGAGTCGTTGCCTTTGCACAGTCTCCCTCTGATNNNAGATCGGAAGA |
| 40491_4 | TATGAGTGTGGAGTCGTTGCCTTTGCACAGTCTCCCTCTGATNNNAGATCGGAAGA |
| 40491_5 | TATGAGTGTGGAGTCGTTGCCTTTGCACAGTCTCCCTCTGATNNNAGATCGGAAGA |
| 40491_6 | TATGAGTGTGGAGTCGTTGCCTTTGCACAGTCTCCCTCTGATNNNAGATCGGAAGA |
| 40491_7 | TATGAGTGTGGAGTCGTTGCCTTTGCACAGTCTCCCTCTGATNNNAGATCGGAAGA |
| 40491_8 | TATGAGTGTGGAGTCGTTGCCTTTGCACAGTCTCCCTCTGATNNNAGATCGGAAGA |
| 40491_9 | TATGAGTGTGGAGTCGTTGCCTTTGCACAGTCTCCCTCTGATNNNAGATCGGAAGA |
| 33692_1 | TATGAGTGTGGAGTCGTTGCCAAAAATTAGCTGGGCATGGTGANNNAGATCGGAA |
| 33692_2 | TATGAGTGTGGAGTCGTTGCCAAAAATTAGCTGGGCATGGTGANNNAGATCGGAA |
| 33692_3 | TATGAGTGTGGAGTCGTTGCCAAAAATTAGCTGGGCATGGTGANNNAGATCGGAA |
| 33692_4 | TATGAGTGTGGAGTCGTTGCCAAAAATTAGCTGGGCATGGTGANNNAGATCGGAA |
| 33692_5 | TATGAGTGTGGAGTCGTTGCCAAAAATTAGCTGGGCATGGTGANNNAGATCGGAA |
| 33692_6 | TATGAGTGTGGAGTCGTTGCCAAAAATTAGCTGGGCATGGTGANNNAGATCGGAA |
| 33692_7 | TATGAGTGTGGAGTCGTTGCCAAAAATTAGCTGGGCATGGTGANNNAGATCGGAA |
| 33692_8 | TATGAGTGTGGAGTCGTTGCCAAAAATTAGCTGGGCATGGTGANNNAGATCGGAA |
| 33692_9 | TATGAGTGTGGAGTCGTTGCCAAAAATTAGCTGGGCATGGTGANNNAGATCGGAA |
| 40957_1 | TATGAGTGTGGAGTCGTTGCGTCTGGCATCCCATCAAAAATTANNNAGATCGGAAG |
| 40957_2 | TATGAGTGTGGAGTCGTTGCGTCTGGCATCCCATCAAAAATTANNNAGATCGGAAG |
| 40957_3 | TATGAGTGTGGAGTCGTTGCGTCTGGCATCCCATCAAAAATTANNNAGATCGGAAG |
| 40957_4 | TATGAGTGTGGAGTCGTTGCGTCTGGCATCCCATCAAAAATTANNNAGATCGGAAG |
| 40957_5 | TATGAGTGTGGAGTCGTTGCGTCTGGCATCCCATCAAAAATTANNNAGATCGGAAG |
| 40957_6 | TATGAGTGTGGAGTCGTTGCGTCTGGCATCCCATCAAAAATTANNNAGATCGGAAG |
| 40957_7 | TATGAGTGTGGAGTCGTTGCGTCTGGCATCCCATCAAAAATTANNNAGATCGGAAG |
| 40957_8 | TATGAGTGTGGAGTCGTTGCGTCTGGCATCCCATCAAAAATTANNNAGATCGGAAG |
| 40957_9 | TATGAGTGTGGAGTCGTTGCGTCTGGCATCCCATCAAAAATTANNNAGATCGGAAG |
| 38116_1 | TATGAGTGTGGAGTCGTTGCCACAGACTGGGAGAAAACATTGTNNNAGATCGGAA |
| 38116_2 | TATGAGTGTGGAGTCGTTGCCACAGACTGGGAGAAAACATTGTNNNAGATCGGAA |
| 38116_3 | TATGAGTGTGGAGTCGTTGCCACAGACTGGGAGAAAACATTGTNNNAGATCGGAA |
| 38116_4 | TATGAGTGTGGAGTCGTTGCCACAGACTGGGAGAAAACATTGTNNNAGATCGGAA |
| 38116_5 | TATGAGTGTGGAGTCGTTGCCACAGACTGGGAGAAAACATTGTNNNAGATCGGAA |

| | |
|---|---|
| 38116_6 | TATGAGTGTGGAGTCGTTGCCACAGACTGGGAGAAAACATTGTNNNAGATCGGAA |
| 38116_7 | TATGAGTGTGGAGTCGTTGCCACAGACTGGGAGAAAACATTGTNNNAGATCGGAA |
| 38116_8 | TATGAGTGTGGAGTCGTTGCCACAGACTGGGAGAAAACATTGTNNNAGATCGGAA |
| 38116_9 | TATGAGTGTGGAGTCGTTGCCACAGACTGGGAGAAAACATTGTNNNAGATCGGAA |
| 45560_1 | TATGAGTGTGGAGTCGTTGCAACATGTTAGTTGCTGCAGCTTTNNNAGATCGGAAG |
| 45560_2 | TATGAGTGTGGAGTCGTTGCAACATGTTAGTTGCTGCAGCTTTNNNAGATCGGAAG |
| 45560_3 | TATGAGTGTGGAGTCGTTGCAACATGTTAGTTGCTGCAGCTTTNNNAGATCGGAAG |
| 45560_4 | TATGAGTGTGGAGTCGTTGCAACATGTTAGTTGCTGCAGCTTTNNNAGATCGGAAG |
| 45560_5 | TATGAGTGTGGAGTCGTTGCAACATGTTAGTTGCTGCAGCTTTNNNAGATCGGAAG |
| 45560_6 | TATGAGTGTGGAGTCGTTGCAACATGTTAGTTGCTGCAGCTTTNNNAGATCGGAAG |
| 45560_7 | TATGAGTGTGGAGTCGTTGCAACATGTTAGTTGCTGCAGCTTTNNNAGATCGGAAG |
| 45560_8 | TATGAGTGTGGAGTCGTTGCAACATGTTAGTTGCTGCAGCTTTNNNAGATCGGAAG |
| 45560_9 | TATGAGTGTGGAGTCGTTGCAACATGTTAGTTGCTGCAGCTTTNNNAGATCGGAAG |
| 40831_1 | TATGAGTGTGGAGTCGTTGCAGCTGCATGGTTTTGTAATACCANNNAGATCGGAAG |
| 40831_2 | TATGAGTGTGGAGTCGTTGCAGCTGCATGGTTTTGTAATACCANNNAGATCGGAAG |
| 40831_3 | TATGAGTGTGGAGTCGTTGCAGCTGCATGGTTTTGTAATACCANNNAGATCGGAAG |
| 40831_4 | TATGAGTGTGGAGTCGTTGCAGCTGCATGGTTTTGTAATACCANNNAGATCGGAAG |
| 40831_5 | TATGAGTGTGGAGTCGTTGCAGCTGCATGGTTTTGTAATACCANNNAGATCGGAAG |
| 40831_6 | TATGAGTGTGGAGTCGTTGCAGCTGCATGGTTTTGTAATACCANNNAGATCGGAAG |
| 40831_7 | TATGAGTGTGGAGTCGTTGCAGCTGCATGGTTTTGTAATACCANNNAGATCGGAAG |
| 40831_8 | TATGAGTGTGGAGTCGTTGCAGCTGCATGGTTTTGTAATACCANNNAGATCGGAAG |
| 40831_9 | TATGAGTGTGGAGTCGTTGCAGCTGCATGGTTTTGTAATACCANNNAGATCGGAAG |
| 37293_1 | TATGAGTGTGGAGTCGTTGCAAAGAGGCATGATCCATGAAAGANNNAGATCGGAA |
| 37293_2 | TATGAGTGTGGAGTCGTTGCAAAGAGGCATGATCCATGAAAGANNNAGATCGGAA |
| 37293_3 | TATGAGTGTGGAGTCGTTGCAAAGAGGCATGATCCATGAAAGANNNAGATCGGAA |
| 37293_4 | TATGAGTGTGGAGTCGTTGCAAAGAGGCATGATCCATGAAAGANNNAGATCGGAA |
| 37293_5 | TATGAGTGTGGAGTCGTTGCAAAGAGGCATGATCCATGAAAGANNNAGATCGGAA |
| 37293_6 | TATGAGTGTGGAGTCGTTGCAAAGAGGCATGATCCATGAAAGANNNAGATCGGAA |
| 37293_7 | TATGAGTGTGGAGTCGTTGCAAAGAGGCATGATCCATGAAAGANNNAGATCGGAA |
| 37293_8 | TATGAGTGTGGAGTCGTTGCAAAGAGGCATGATCCATGAAAGANNNAGATCGGAA |
| 37293_9 | TATGAGTGTGGAGTCGTTGCAAAGAGGCATGATCCATGAAAGANNNAGATCGGAA |
| 41318_1 | TATGAGTGTGGAGTCGTTGCATCATAGATTTTGCCAGCACGTCNNNAGATCGGAAG |
| 41318_2 | TATGAGTGTGGAGTCGTTGCATCATAGATTTTGCCAGCACGTCNNNAGATCGGAAG |
| 41318_3 | TATGAGTGTGGAGTCGTTGCATCATAGATTTTGCCAGCACGTCNNNAGATCGGAAG |
| 41318_4 | TATGAGTGTGGAGTCGTTGCATCATAGATTTTGCCAGCACGTCNNNAGATCGGAAG |
| 41318_5 | TATGAGTGTGGAGTCGTTGCATCATAGATTTTGCCAGCACGTCNNNAGATCGGAAG |
| 41318_6 | TATGAGTGTGGAGTCGTTGCATCATAGATTTTGCCAGCACGTCNNNAGATCGGAAG |
| 41318_7 | TATGAGTGTGGAGTCGTTGCATCATAGATTTTGCCAGCACGTCNNNAGATCGGAAG |
| 41318_8 | TATGAGTGTGGAGTCGTTGCATCATAGATTTTGCCAGCACGTCNNNAGATCGGAAG |
| 41318_9 | TATGAGTGTGGAGTCGTTGCATCATAGATTTTGCCAGCACGTCNNNAGATCGGAAG |
| 46445_1 | TATGAGTGTGGAGTCGTTGCCCAAAGTGCTGGGATTACAAGTGNNNAGATCGGAA |
| 46445_2 | TATGAGTGTGGAGTCGTTGCCCAAAGTGCTGGGATTACAAGTGNNNAGATCGGAA |
| 46445_3 | TATGAGTGTGGAGTCGTTGCCCAAAGTGCTGGGATTACAAGTGNNNAGATCGGAA |

| | |
|---|---|
| 46445_4 | TATGAGTGTGGAGTCGTTGCCCAAAGTGCTGGGATTACAAGTGNNNAGATCGGAA |
| 46445_5 | TATGAGTGTGGAGTCGTTGCCCAAAGTGCTGGGATTACAAGTGNNNAGATCGGAA |
| 46445_6 | TATGAGTGTGGAGTCGTTGCCCAAAGTGCTGGGATTACAAGTGNNNAGATCGGAA |
| 46445_7 | TATGAGTGTGGAGTCGTTGCCCAAAGTGCTGGGATTACAAGTGNNNAGATCGGAA |
| 46445_8 | TATGAGTGTGGAGTCGTTGCCCAAAGTGCTGGGATTACAAGTGNNNAGATCGGAA |
| 46445_9 | TATGAGTGTGGAGTCGTTGCCCAAAGTGCTGGGATTACAAGTGNNNAGATCGGAA |
| 48697_1 | TATGAGTGTGGAGTCGTTGCTTTTTGGTAGAGACGGGGTTTCANNNAGATCGGAA |
| 48697_2 | TATGAGTGTGGAGTCGTTGCTTTTTGGTAGAGACGGGGTTTCANNNAGATCGGAA |
| 48697_3 | TATGAGTGTGGAGTCGTTGCTTTTTGGTAGAGACGGGGTTTCANNNAGATCGGAA |
| 48697_4 | TATGAGTGTGGAGTCGTTGCTTTTTGGTAGAGACGGGGTTTCANNNAGATCGGAA |
| 48697_5 | TATGAGTGTGGAGTCGTTGCTTTTTGGTAGAGACGGGGTTTCANNNAGATCGGAA |
| 48697_6 | TATGAGTGTGGAGTCGTTGCTTTTTGGTAGAGACGGGGTTTCANNNAGATCGGAA |
| 48697_7 | TATGAGTGTGGAGTCGTTGCTTTTTGGTAGAGACGGGGTTTCANNNAGATCGGAA |
| 48697_8 | TATGAGTGTGGAGTCGTTGCTTTTTGGTAGAGACGGGGTTTCANNNAGATCGGAA |
| 48697_9 | TATGAGTGTGGAGTCGTTGCTTTTTGGTAGAGACGGGGTTTCANNNAGATCGGAA |
| 42939_1 | TATGAGTGTGGAGTCGTTGCATGCCCATCAATCAACAAGTAGANNNAGATCGGAAG |
| 42939_2 | TATGAGTGTGGAGTCGTTGCATGCCCATCAATCAACAAGTAGANNNAGATCGGAAG |
| 42939_3 | TATGAGTGTGGAGTCGTTGCATGCCCATCAATCAACAAGTAGANNNAGATCGGAAG |
| 42939_4 | TATGAGTGTGGAGTCGTTGCATGCCCATCAATCAACAAGTAGANNNAGATCGGAAG |
| 1367_1 | TATGAGTGTGGAGTCGTTGCCAGAACACCATGTGATGTAGGCANNNAGATCGGAA |
| 1367_2 | TATGAGTGTGGAGTCGTTGCCAGAACACCATGTGATGTAGGCANNNAGATCGGAA |
| 1367_3 | TATGAGTGTGGAGTCGTTGCCAGAACACCATGTGATGTAGGCANNNAGATCGGAA |
| 1367_4 | TATGAGTGTGGAGTCGTTGCCAGAACACCATGTGATGTAGGCANNNAGATCGGAA |
| 12736_1 | TATGAGTGTGGAGTCGTTGCAACAACAAAAAAATTCAACTCTGNNNAGATCGGAAG |
| 12736_2 | TATGAGTGTGGAGTCGTTGCAACAACAAAAAAATTCAACTCTGNNNAGATCGGAAG |
| 12736_3 | TATGAGTGTGGAGTCGTTGCAACAACAAAAAAATTCAACTCTGNNNAGATCGGAAG |
| 12736_4 | TATGAGTGTGGAGTCGTTGCAACAACAAAAAAATTCAACTCTGNNNAGATCGGAAG |
| 45948_1 | TATGAGTGTGGAGTCGTTGCGGACAGTGAGTAGCCTTTGTCTTNNNAGATCGGAAG |
| 45948_2 | TATGAGTGTGGAGTCGTTGCGGACAGTGAGTAGCCTTTGTCTTNNNAGATCGGAAG |
| 45948_3 | TATGAGTGTGGAGTCGTTGCGGACAGTGAGTAGCCTTTGTCTTNNNAGATCGGAAG |
| 45948_4 | TATGAGTGTGGAGTCGTTGCGGACAGTGAGTAGCCTTTGTCTTNNNAGATCGGAAG |
| 43025_1 | TATGAGTGTGGAGTCGTTGCCCCTGAGGAACATAGATGCAGAANNNAGATCGGAA |
| 43025_2 | TATGAGTGTGGAGTCGTTGCCCCTGAGGAACATAGATGCAGAANNNAGATCGGAA |
| 43025_3 | TATGAGTGTGGAGTCGTTGCCCCTGAGGAACATAGATGCAGAANNNAGATCGGAA |
| 43025_4 | TATGAGTGTGGAGTCGTTGCCCCTGAGGAACATAGATGCAGAANNNAGATCGGAA |
| 31648_1 | TATGAGTGTGGAGTCGTTGCACCACACTACACACAACCCTTAGNNNAGATCGGAAGA |
| 31648_2 | TATGAGTGTGGAGTCGTTGCACCACACTACACACAACCCTTAGNNNAGATCGGAAGA |
| 31648_3 | TATGAGTGTGGAGTCGTTGCACCACACTACACACAACCCTTAGNNNAGATCGGAAGA |
| 31648_4 | TATGAGTGTGGAGTCGTTGCACCACACTACACACAACCCTTAGNNNAGATCGGAAGA |
| 41276_1 | TATGAGTGTGGAGTCGTTGCCTGAAGCGGGTGGATAATTTGAGNNNAGATCGGAA |
| 41276_2 | TATGAGTGTGGAGTCGTTGCCTGAAGCGGGTGGATAATTTGAGNNNAGATCGGAA |
| 41276_3 | TATGAGTGTGGAGTCGTTGCCTGAAGCGGGTGGATAATTTGAGNNNAGATCGGAA |
| 41276_4 | TATGAGTGTGGAGTCGTTGCCTGAAGCGGGTGGATAATTTGAGNNNAGATCGGAA |

| | |
|---|---|
| 36954_1 | TATGAGTGTGGAGTCGTTGCTGTCACACAGAAGACACAAAGGANNNAGATCGGAA |
| 36954_2 | TATGAGTGTGGAGTCGTTGCTGTCACACAGAAGACACAAAGGANNNAGATCGGAA |
| 36954_3 | TATGAGTGTGGAGTCGTTGCTGTCACACAGAAGACACAAAGGANNNAGATCGGAA |
| 36954_4 | TATGAGTGTGGAGTCGTTGCTGTCACACAGAAGACACAAAGGANNNAGATCGGAA |
| 38298_1 | TATGAGTGTGGAGTCGTTGCAAATCACCTCTAAGATCCAGGGCNNNAGATCGGAAG |
| 38298_2 | TATGAGTGTGGAGTCGTTGCAAATCACCTCTAAGATCCAGGGCNNNAGATCGGAAG |
| 38298_3 | TATGAGTGTGGAGTCGTTGCAAATCACCTCTAAGATCCAGGGCNNNAGATCGGAAG |
| 38298_4 | TATGAGTGTGGAGTCGTTGCAAATCACCTCTAAGATCCAGGGCNNNAGATCGGAAG |
| 35754_1 | TATGAGTGTGGAGTCGTTGCCAAGTAGCTGGGACTACAGGCNNNAGATCGGAAGA |
| 35754_2 | TATGAGTGTGGAGTCGTTGCCAAGTAGCTGGGACTACAGGCNNNAGATCGGAAGA |
| 35754_3 | TATGAGTGTGGAGTCGTTGCCAAGTAGCTGGGACTACAGGCNNNAGATCGGAAGA |
| 35754_4 | TATGAGTGTGGAGTCGTTGCCAAGTAGCTGGGACTACAGGCNNNAGATCGGAAGA |
| 36412_1 | TATGAGTGTGGAGTCGTTGCCGCCTATAGTCCCAGCTACTTGNNNAGATCGGAAGA |
| 36412_2 | TATGAGTGTGGAGTCGTTGCCGCCTATAGTCCCAGCTACTTGNNNAGATCGGAAGA |
| 36412_3 | TATGAGTGTGGAGTCGTTGCCGCCTATAGTCCCAGCTACTTGNNNAGATCGGAAGA |
| 36412_4 | TATGAGTGTGGAGTCGTTGCCGCCTATAGTCCCAGCTACTTGNNNAGATCGGAAGA |
| 51732_1 | TATGAGTGTGGAGTCGTTGCCAATACTATCCTCCAGGTGCCTCNNNAGATCGGAAG |
| 51732_2 | TATGAGTGTGGAGTCGTTGCCAATACTATCCTCCAGGTGCCTCNNNAGATCGGAAG |
| 51732_3 | TATGAGTGTGGAGTCGTTGCCAATACTATCCTCCAGGTGCCTCNNNAGATCGGAAG |
| 51732_4 | TATGAGTGTGGAGTCGTTGCCAATACTATCCTCCAGGTGCCTCNNNAGATCGGAAG |
| 46207_1 | TATGAGTGTGGAGTCGTTGCAATTCTTGTGTCTCAGCCTCCCNNNAGATCGGAAGA |
| 46207_2 | TATGAGTGTGGAGTCGTTGCAATTCTTGTGTCTCAGCCTCCCNNNAGATCGGAAGA |
| 46207_3 | TATGAGTGTGGAGTCGTTGCAATTCTTGTGTCTCAGCCTCCCNNNAGATCGGAAGA |
| 46207_4 | TATGAGTGTGGAGTCGTTGCAATTCTTGTGTCTCAGCCTCCCNNNAGATCGGAAGA |
| 30175_1 | TATGAGTGTGGAGTCGTTGCACATACCATGCAGATGTTAGGCANNNAGATCGGAA |
| 30175_2 | TATGAGTGTGGAGTCGTTGCACATACCATGCAGATGTTAGGCANNNAGATCGGAA |
| 30175_3 | TATGAGTGTGGAGTCGTTGCACATACCATGCAGATGTTAGGCANNNAGATCGGAA |
| 30175_4 | TATGAGTGTGGAGTCGTTGCACATACCATGCAGATGTTAGGCANNNAGATCGGAA |
| 28924_1 | TATGAGTGTGGAGTCGTTGCATCATCACCATCAACAGTCACCANNNAGATCGGAAG |
| 28924_2 | TATGAGTGTGGAGTCGTTGCATCATCACCATCAACAGTCACCANNNAGATCGGAAG |
| 28924_3 | TATGAGTGTGGAGTCGTTGCATCATCACCATCAACAGTCACCANNNAGATCGGAAG |
| 28924_4 | TATGAGTGTGGAGTCGTTGCATCATCACCATCAACAGTCACCANNNAGATCGGAAG |
| 52380_1 | TATGAGTGTGGAGTCGTTGCGGTCTCTAAATCTGGTCACAGCANNNAGATCGGAAG |
| 52380_2 | TATGAGTGTGGAGTCGTTGCGGTCTCTAAATCTGGTCACAGCANNNAGATCGGAAG |
| 52380_3 | TATGAGTGTGGAGTCGTTGCGGTCTCTAAATCTGGTCACAGCANNNAGATCGGAAG |
| 52380_4 | TATGAGTGTGGAGTCGTTGCGGTCTCTAAATCTGGTCACAGCANNNAGATCGGAAG |
| 36458_1 | TATGAGTGTGGAGTCGTTGCCTAATCCCCAGAACCTGTGAACANNNAGATCGGAAG |
| 36458_2 | TATGAGTGTGGAGTCGTTGCCTAATCCCCAGAACCTGTGAACANNNAGATCGGAAG |
| 36458_3 | TATGAGTGTGGAGTCGTTGCCTAATCCCCAGAACCTGTGAACANNNAGATCGGAAG |
| 36458_4 | TATGAGTGTGGAGTCGTTGCCTAATCCCCAGAACCTGTGAACANNNAGATCGGAAG |
| 12490_1 | TATGAGTGTGGAGTCGTTGCGGGCAAAAAAAGTTTATTGAAAANNNAGATCGGAA |
| 12490_2 | TATGAGTGTGGAGTCGTTGCGGGCAAAAAAAGTTTATTGAAAANNNAGATCGGAA |
| 12490_3 | TATGAGTGTGGAGTCGTTGCGGGCAAAAAAAGTTTATTGAAAANNNAGATCGGAA |

| | |
|---|---|
| 12490_4 | TATGAGTGTGGAGTCGTTGCGGGCAAAAAAAGTTTATTGAAAANNNAGATCGGAA |
| 4450_1 | TATGAGTGTGGAGTCGTTGCATCTAATTGCCTCCTGTGCTCCANNNAGATCGGAAG |
| 4450_2 | TATGAGTGTGGAGTCGTTGCATCTAATTGCCTCCTGTGCTCCANNNAGATCGGAAG |
| 4450_3 | TATGAGTGTGGAGTCGTTGCATCTAATTGCCTCCTGTGCTCCANNNAGATCGGAAG |
| 4450_4 | TATGAGTGTGGAGTCGTTGCATCTAATTGCCTCCTGTGCTCCANNNAGATCGGAAG |
| 46635_1 | TATGAGTGTGGAGTCGTTGCTTCATGCCATTCTCCTGTGTCAGNNNAGATCGGAAG |
| 46635_2 | TATGAGTGTGGAGTCGTTGCTTCATGCCATTCTCCTGTGTCAGNNNAGATCGGAAG |
| 46635_3 | TATGAGTGTGGAGTCGTTGCTTCATGCCATTCTCCTGTGTCAGNNNAGATCGGAAG |
| 46635_4 | TATGAGTGTGGAGTCGTTGCTTCATGCCATTCTCCTGTGTCAGNNNAGATCGGAAG |
| 41469_1 | TATGAGTGTGGAGTCGTTGCAAACAAATCAAGCTGAAGGCACANNNAGATCGGAA |
| 41469_2 | TATGAGTGTGGAGTCGTTGCAAACAAATCAAGCTGAAGGCACANNNAGATCGGAA |
| 41469_3 | TATGAGTGTGGAGTCGTTGCAAACAAATCAAGCTGAAGGCACANNNAGATCGGAA |
| 41469_4 | TATGAGTGTGGAGTCGTTGCAAACAAATCAAGCTGAAGGCACANNNAGATCGGAA |
| 47477_1 | TATGAGTGTGGAGTCGTTGCGCCTCCTGAGTAGCTGGTATTACNNNAGATCGGAAG |
| 47477_2 | TATGAGTGTGGAGTCGTTGCGCCTCCTGAGTAGCTGGTATTACNNNAGATCGGAAG |
| 47477_3 | TATGAGTGTGGAGTCGTTGCGCCTCCTGAGTAGCTGGTATTACNNNAGATCGGAAG |
| 47477_4 | TATGAGTGTGGAGTCGTTGCGCCTCCTGAGTAGCTGGTATTACNNNAGATCGGAAG |
| 33445_1 | TATGAGTGTGGAGTCGTTGCTTGGAAATGCACCAATCTCCAACNNNAGATCGGAAG |
| 33445_2 | TATGAGTGTGGAGTCGTTGCTTGGAAATGCACCAATCTCCAACNNNAGATCGGAAG |
| 33445_3 | TATGAGTGTGGAGTCGTTGCTTGGAAATGCACCAATCTCCAACNNNAGATCGGAAG |
| 33445_4 | TATGAGTGTGGAGTCGTTGCTTGGAAATGCACCAATCTCCAACNNNAGATCGGAAG |
| 50672_1 | TATGAGTGTGGAGTCGTTGCAGAGCAGGTTATACTGAGGGCTANNNAGATCGGAA |
| 50672_2 | TATGAGTGTGGAGTCGTTGCAGAGCAGGTTATACTGAGGGCTANNNAGATCGGAA |
| 50672_3 | TATGAGTGTGGAGTCGTTGCAGAGCAGGTTATACTGAGGGCTANNNAGATCGGAA |
| 50672_4 | TATGAGTGTGGAGTCGTTGCAGAGCAGGTTATACTGAGGGCTANNNAGATCGGAA |
| 40062_1 | TATGAGTGTGGAGTCGTTGCCAATCTTGGCTCACTGCAACCNNNAGATCGGAAGAG |
| 40062_2 | TATGAGTGTGGAGTCGTTGCCAATCTTGGCTCACTGCAACCNNNAGATCGGAAGAG |
| 40062_3 | TATGAGTGTGGAGTCGTTGCCAATCTTGGCTCACTGCAACCNNNAGATCGGAAGAG |
| 40062_4 | TATGAGTGTGGAGTCGTTGCCAATCTTGGCTCACTGCAACCNNNAGATCGGAAGAG |
| 33484_1 | TATGAGTGTGGAGTCGTTGCAGTTGAAGAATGGGATGCCTCTTNNNAGATCGGAA |
| 33484_2 | TATGAGTGTGGAGTCGTTGCAGTTGAAGAATGGGATGCCTCTTNNNAGATCGGAA |
| 33484_3 | TATGAGTGTGGAGTCGTTGCAGTTGAAGAATGGGATGCCTCTTNNNAGATCGGAA |
| 33484_4 | TATGAGTGTGGAGTCGTTGCAGTTGAAGAATGGGATGCCTCTTNNNAGATCGGAA |
| 36514_1 | TATGAGTGTGGAGTCGTTGCTCACTTGAATACACAAGCCCTCANNNAGATCGGAAG |
| 36514_2 | TATGAGTGTGGAGTCGTTGCTCACTTGAATACACAAGCCCTCANNNAGATCGGAAG |
| 36514_3 | TATGAGTGTGGAGTCGTTGCTCACTTGAATACACAAGCCCTCANNNAGATCGGAAG |
| 36514_4 | TATGAGTGTGGAGTCGTTGCTCACTTGAATACACAAGCCCTCANNNAGATCGGAAG |
| 51276_1 | TATGAGTGTGGAGTCGTTGCTGGGTGCCATGTGTAGAAAGTATNNNAGATCGGAA |
| 51276_2 | TATGAGTGTGGAGTCGTTGCTGGGTGCCATGTGTAGAAAGTATNNNAGATCGGAA |
| 51276_3 | TATGAGTGTGGAGTCGTTGCTGGGTGCCATGTGTAGAAAGTATNNNAGATCGGAA |
| 51276_4 | TATGAGTGTGGAGTCGTTGCTGGGTGCCATGTGTAGAAAGTATNNNAGATCGGAA |
| 13070_1 | TATGAGTGTGGAGTCGTTGCAGACAGTGAAAAAAGATGAAAATNNNAGATCGGAA |
| 13070_2 | TATGAGTGTGGAGTCGTTGCAGACAGTGAAAAAAGATGAAAATNNNAGATCGGAA |

| | |
|---|---|
| 13070_3 | TATGAGTGTGGAGTCGTTGCAGACAGTGAAAAAAGATGAAAATNNNAGATCGGAA |
| 13070_4 | TATGAGTGTGGAGTCGTTGCAGACAGTGAAAAAAGATGAAAATNNNAGATCGGAA |
| 7658_1 | TATGAGTGTGGAGTCGTTGCAAGGCACTGGAGTTTCTGAGNNNAGATCGGAAGAG |
| 7658_2 | TATGAGTGTGGAGTCGTTGCAAGGCACTGGAGTTTCTGAGNNNAGATCGGAAGAG |
| 7658_3 | TATGAGTGTGGAGTCGTTGCAAGGCACTGGAGTTTCTGAGNNNAGATCGGAAGAG |
| 7658_4 | TATGAGTGTGGAGTCGTTGCAAGGCACTGGAGTTTCTGAGNNNAGATCGGAAGAG |
| 49675_1 | TATGAGTGTGGAGTCGTTGCACATGTAATATGCCCCCCCATCNNNAGATCGGAAGA |
| 49675_2 | TATGAGTGTGGAGTCGTTGCACATGTAATATGCCCCCCCATCNNNAGATCGGAAGA |
| 49675_3 | TATGAGTGTGGAGTCGTTGCACATGTAATATGCCCCCCCATCNNNAGATCGGAAGA |
| 49675_4 | TATGAGTGTGGAGTCGTTGCACATGTAATATGCCCCCCCATCNNNAGATCGGAAGA |
| 47865_1 | TATGAGTGTGGAGTCGTTGCCTCACTATAGCCTTGACCTGGACNNNAGATCGGAAG |
| 47865_2 | TATGAGTGTGGAGTCGTTGCCTCACTATAGCCTTGACCTGGACNNNAGATCGGAAG |
| 47865_3 | TATGAGTGTGGAGTCGTTGCCTCACTATAGCCTTGACCTGGACNNNAGATCGGAAG |
| 47865_4 | TATGAGTGTGGAGTCGTTGCCTCACTATAGCCTTGACCTGGACNNNAGATCGGAAG |
| 44832_1 | TATGAGTGTGGAGTCGTTGCCAGAAGTCCTGGGGAATGAGAGNNNAGATCGGAA |
| 44832_2 | TATGAGTGTGGAGTCGTTGCCAGAAGTCCTGGGGAATGAGAGNNNAGATCGGAA |
| 44832_3 | TATGAGTGTGGAGTCGTTGCCAGAAGTCCTGGGGAATGAGAGNNNAGATCGGAA |
| 44832_4 | TATGAGTGTGGAGTCGTTGCCAGAAGTCCTGGGGAATGAGAGNNNAGATCGGAA |
| 35099_1 | TATGAGTGTGGAGTCGTTGCCTCCCTGGTTCAAGTGATCCTCNNNAGATCGGAAGA |
| 35099_2 | TATGAGTGTGGAGTCGTTGCCTCCCTGGTTCAAGTGATCCTCNNNAGATCGGAAGA |
| 35099_3 | TATGAGTGTGGAGTCGTTGCCTCCCTGGTTCAAGTGATCCTCNNNAGATCGGAAGA |
| 35099_4 | TATGAGTGTGGAGTCGTTGCCTCCCTGGTTCAAGTGATCCTCNNNAGATCGGAAGA |
| 41214_1 | TATGAGTGTGGAGTCGTTGCCAGTGGTATGATCTCGGCTCACNNNAGATCGGAAGA |
| 41214_2 | TATGAGTGTGGAGTCGTTGCCAGTGGTATGATCTCGGCTCACNNNAGATCGGAAGA |
| 41214_3 | TATGAGTGTGGAGTCGTTGCCAGTGGTATGATCTCGGCTCACNNNAGATCGGAAGA |
| 41214_4 | TATGAGTGTGGAGTCGTTGCCAGTGGTATGATCTCGGCTCACNNNAGATCGGAAGA |
| 33945_1 | TATGAGTGTGGAGTCGTTGCTCTCCAAAAGTGCTGGGATTACANNNAGATCGGAAG |
| 33945_2 | TATGAGTGTGGAGTCGTTGCTCTCCAAAAGTGCTGGGATTACANNNAGATCGGAAG |
| 33945_3 | TATGAGTGTGGAGTCGTTGCTCTCCAAAAGTGCTGGGATTACANNNAGATCGGAAG |
| 33945_4 | TATGAGTGTGGAGTCGTTGCTCTCCAAAAGTGCTGGGATTACANNNAGATCGGAAG |
| 48304_1 | TATGAGTGTGGAGTCGTTGCGGTTCTCCTTTGGCTCTATCTAANNNAGATCGGAAG |
| 48304_2 | TATGAGTGTGGAGTCGTTGCGGTTCTCCTTTGGCTCTATCTAANNNAGATCGGAAG |
| 48304_3 | TATGAGTGTGGAGTCGTTGCGGTTCTCCTTTGGCTCTATCTAANNNAGATCGGAAG |
| 48304_4 | TATGAGTGTGGAGTCGTTGCGGTTCTCCTTTGGCTCTATCTAANNNAGATCGGAAG |
| 50831_1 | TATGAGTGTGGAGTCGTTGCGACTGGTCACGTGAAAAGGAACNNNAGATCGGAAG |
| 50831_2 | TATGAGTGTGGAGTCGTTGCGACTGGTCACGTGAAAAGGAACNNNAGATCGGAAG |
| 50831_3 | TATGAGTGTGGAGTCGTTGCGACTGGTCACGTGAAAAGGAACNNNAGATCGGAAG |
| 50831_4 | TATGAGTGTGGAGTCGTTGCGACTGGTCACGTGAAAAGGAACNNNAGATCGGAAG |
| 34017_1 | TATGAGTGTGGAGTCGTTGCGATCTTGGCTTATTGCAACCTCCNNNAGATCGGAAG |
| 34017_2 | TATGAGTGTGGAGTCGTTGCGATCTTGGCTTATTGCAACCTCCNNNAGATCGGAAG |
| 34017_3 | TATGAGTGTGGAGTCGTTGCGATCTTGGCTTATTGCAACCTCCNNNAGATCGGAAG |
| 34017_4 | TATGAGTGTGGAGTCGTTGCGATCTTGGCTTATTGCAACCTCCNNNAGATCGGAAG |
| 3088_1 | TATGAGTGTGGAGTCGTTGCCAGCCCAAAACCCTAATTTCAGCNNNAGATCGGAAG |

| | |
|---|---|
| 3088_2 | TATGAGTGTGGAGTCGTTGCCAGCCCAAAACCCTAATTTCAGCNNNAGATCGGAAG |
| 3088_3 | TATGAGTGTGGAGTCGTTGCCAGCCCAAAACCCTAATTTCAGCNNNAGATCGGAAG |
| 3088_4 | TATGAGTGTGGAGTCGTTGCCAGCCCAAAACCCTAATTTCAGCNNNAGATCGGAAG |
| 34915_1 | TATGAGTGTGGAGTCGTTGCGCCATCACACCCAGCTAATTTTTNNNAGATCGGAAGA |
| 34915_2 | TATGAGTGTGGAGTCGTTGCGCCATCACACCCAGCTAATTTTTNNNAGATCGGAAGA |
| 34915_3 | TATGAGTGTGGAGTCGTTGCGCCATCACACCCAGCTAATTTTTNNNAGATCGGAAGA |
| 34915_4 | TATGAGTGTGGAGTCGTTGCGCCATCACACCCAGCTAATTTTTNNNAGATCGGAAGA |
| 41439_1 | TATGAGTGTGGAGTCGTTGCCTAGGGGAGGGATAGCATTAGGANNNAGATCGGA |
| 41439_2 | TATGAGTGTGGAGTCGTTGCCTAGGGGAGGGATAGCATTAGGANNNAGATCGGA |
| 41439_3 | TATGAGTGTGGAGTCGTTGCCTAGGGGAGGGATAGCATTAGGANNNAGATCGGA |
| 41439_4 | TATGAGTGTGGAGTCGTTGCCTAGGGGAGGGATAGCATTAGGANNNAGATCGGA |
| 30951_1 | TATGAGTGTGGAGTCGTTGCAGGGATGACTCTCCAACACAATCNNNAGATCGGAAG |
| 30951_2 | TATGAGTGTGGAGTCGTTGCAGGGATGACTCTCCAACACAATCNNNAGATCGGAAG |
| 30951_3 | TATGAGTGTGGAGTCGTTGCAGGGATGACTCTCCAACACAATCNNNAGATCGGAAG |
| 30951_4 | TATGAGTGTGGAGTCGTTGCAGGGATGACTCTCCAACACAATCNNNAGATCGGAAG |
| 29687_1 | TATGAGTGTGGAGTCGTTGCGAGCTGATCGTGAAAGGTGAGTNNNAGATCGGAA |
| 29687_2 | TATGAGTGTGGAGTCGTTGCGAGCTGATCGTGAAAGGTGAGTNNNAGATCGGAA |
| 29687_3 | TATGAGTGTGGAGTCGTTGCGAGCTGATCGTGAAAGGTGAGTNNNAGATCGGAA |
| 29687_4 | TATGAGTGTGGAGTCGTTGCGAGCTGATCGTGAAAGGTGAGTNNNAGATCGGAA |
| 37724_1 | TATGAGTGTGGAGTCGTTGCAGCAACAGAAGACCTTTCAGTCANNNAGATCGGAAG |
| 37724_2 | TATGAGTGTGGAGTCGTTGCAGCAACAGAAGACCTTTCAGTCANNNAGATCGGAAG |
| 37724_3 | TATGAGTGTGGAGTCGTTGCAGCAACAGAAGACCTTTCAGTCANNNAGATCGGAAG |
| 37724_4 | TATGAGTGTGGAGTCGTTGCAGCAACAGAAGACCTTTCAGTCANNNAGATCGGAAG |
| 2062_1 | TATGAGTGTGGAGTCGTTGCATCCACCAAATTTCCACCACACANNNAGATCGGAAGA |
| 2062_2 | TATGAGTGTGGAGTCGTTGCATCCACCAAATTTCCACCACACANNNAGATCGGAAGA |
| 2062_3 | TATGAGTGTGGAGTCGTTGCATCCACCAAATTTCCACCACACANNNAGATCGGAAGA |
| 2062_4 | TATGAGTGTGGAGTCGTTGCATCCACCAAATTTCCACCACACANNNAGATCGGAAGA |
| 31616_1 | TATGAGTGTGGAGTCGTTGCCTAGCAGGAGACTCAGACAGTTCNNNAGATCGGAA |
| 31616_2 | TATGAGTGTGGAGTCGTTGCCTAGCAGGAGACTCAGACAGTTCNNNAGATCGGAA |
| 31616_3 | TATGAGTGTGGAGTCGTTGCCTAGCAGGAGACTCAGACAGTTCNNNAGATCGGAA |
| 31616_4 | TATGAGTGTGGAGTCGTTGCCTAGCAGGAGACTCAGACAGTTCNNNAGATCGGAA |
| 12896_1 | TATGAGTGTGGAGTCGTTGCCACATATCATGGTAGTGACGNNNAGATCGGAAGAG |
| 12896_2 | TATGAGTGTGGAGTCGTTGCCACATATCATGGTAGTGACGNNNAGATCGGAAGAG |
| 12896_3 | TATGAGTGTGGAGTCGTTGCCACATATCATGGTAGTGACGNNNAGATCGGAAGAG |
| 12896_4 | TATGAGTGTGGAGTCGTTGCCACATATCATGGTAGTGACGNNNAGATCGGAAGAG |
| 32050_1 | TATGAGTGTGGAGTCGTTGCTCTATTAGGCCAGCATCATCCTGNNNAGATCGGAAG |
| 32050_2 | TATGAGTGTGGAGTCGTTGCTCTATTAGGCCAGCATCATCCTGNNNAGATCGGAAG |
| 32050_3 | TATGAGTGTGGAGTCGTTGCTCTATTAGGCCAGCATCATCCTGNNNAGATCGGAAG |
| 32050_4 | TATGAGTGTGGAGTCGTTGCTCTATTAGGCCAGCATCATCCTGNNNAGATCGGAAG |
| 41389_1 | TATGAGTGTGGAGTCGTTGCGCCATCACACCAGGCTAATTTTTNNNAGATCGGAAG |
| 41389_2 | TATGAGTGTGGAGTCGTTGCGCCATCACACCAGGCTAATTTTTNNNAGATCGGAAG |
| 41389_3 | TATGAGTGTGGAGTCGTTGCGCCATCACACCAGGCTAATTTTTNNNAGATCGGAAG |
| 41389_4 | TATGAGTGTGGAGTCGTTGCGCCATCACACCAGGCTAATTTTTNNNAGATCGGAAG |

| | |
|---|---|
| 37836_1 | TATGAGTGTGGAGTCGTTGCGGGACAAAAAGGGTGTGATCTTGNNNAGATCGGAA |
| 37836_2 | TATGAGTGTGGAGTCGTTGCGGGACAAAAAGGGTGTGATCTTGNNNAGATCGGAA |
| 37836_3 | TATGAGTGTGGAGTCGTTGCGGGACAAAAAGGGTGTGATCTTGNNNAGATCGGAA |
| 37836_4 | TATGAGTGTGGAGTCGTTGCGGGACAAAAAGGGTGTGATCTTGNNNAGATCGGAA |
| 41405_1 | TATGAGTGTGGAGTCGTTGCTGTTCACCAACAAAAAGTGAGCTNNNAGATCGGAAG |
| 41405_2 | TATGAGTGTGGAGTCGTTGCTGTTCACCAACAAAAAGTGAGCTNNNAGATCGGAAG |
| 41405_3 | TATGAGTGTGGAGTCGTTGCTGTTCACCAACAAAAAGTGAGCTNNNAGATCGGAAG |
| 41405_4 | TATGAGTGTGGAGTCGTTGCTGTTCACCAACAAAAAGTGAGCTNNNAGATCGGAAG |
| 41397_1 | TATGAGTGTGGAGTCGTTGCACTGTAATGTCATAGAACAGGGGTNNNAGATCGGA |
| 41397_2 | TATGAGTGTGGAGTCGTTGCACTGTAATGTCATAGAACAGGGGTNNNAGATCGGA |
| 41397_3 | TATGAGTGTGGAGTCGTTGCACTGTAATGTCATAGAACAGGGGTNNNAGATCGGA |
| 41397_4 | TATGAGTGTGGAGTCGTTGCACTGTAATGTCATAGAACAGGGGTNNNAGATCGGA |
| 49400_1 | TATGAGTGTGGAGTCGTTGCACCACGCCTTGCTAATTTTTTGTNNNAGATCGGAAGA |
| 49400_2 | TATGAGTGTGGAGTCGTTGCACCACGCCTTGCTAATTTTTTGTNNNAGATCGGAAGA |
| 49400_3 | TATGAGTGTGGAGTCGTTGCACCACGCCTTGCTAATTTTTTGTNNNAGATCGGAAGA |
| 49400_4 | TATGAGTGTGGAGTCGTTGCACCACGCCTTGCTAATTTTTTGTNNNAGATCGGAAGA |
| 12371_1 | TATGAGTGTGGAGTCGTTGCAGCCAATGCTAAAGATGCCTCAANNNAGATCGGAAG |
| 12371_2 | TATGAGTGTGGAGTCGTTGCAGCCAATGCTAAAGATGCCTCAANNNAGATCGGAAG |
| 12371_3 | TATGAGTGTGGAGTCGTTGCAGCCAATGCTAAAGATGCCTCAANNNAGATCGGAAG |
| 12371_4 | TATGAGTGTGGAGTCGTTGCAGCCAATGCTAAAGATGCCTCAANNNAGATCGGAAG |
| 35262_1 | TATGAGTGTGGAGTCGTTGCGGGTCCACTTTGATTCTGTTGTGNNNAGATCGGAAG |
| 35262_2 | TATGAGTGTGGAGTCGTTGCGGGTCCACTTTGATTCTGTTGTGNNNAGATCGGAAG |
| 35262_3 | TATGAGTGTGGAGTCGTTGCGGGTCCACTTTGATTCTGTTGTGNNNAGATCGGAAG |
| 35262_4 | TATGAGTGTGGAGTCGTTGCGGGTCCACTTTGATTCTGTTGTGNNNAGATCGGAAG |
| 31625_1 | TATGAGTGTGGAGTCGTTGCATCATGGCCGGTAGGGTCAGNNNAGATCGGAAGAG |
| 31625_2 | TATGAGTGTGGAGTCGTTGCATCATGGCCGGTAGGGTCAGNNNAGATCGGAAGAG |
| 31625_3 | TATGAGTGTGGAGTCGTTGCATCATGGCCGGTAGGGTCAGNNNAGATCGGAAGAG |
| 31625_4 | TATGAGTGTGGAGTCGTTGCATCATGGCCGGTAGGGTCAGNNNAGATCGGAAGAG |
| 36027_1 | TATGAGTGTGGAGTCGTTGCGCCCATGTTCTCAGAACCTTCTANNNAGATCGGAAGA |
| 36027_2 | TATGAGTGTGGAGTCGTTGCGCCCATGTTCTCAGAACCTTCTANNNAGATCGGAAGA |
| 36027_3 | TATGAGTGTGGAGTCGTTGCGCCCATGTTCTCAGAACCTTCTANNNAGATCGGAAGA |
| 36027_4 | TATGAGTGTGGAGTCGTTGCGCCCATGTTCTCAGAACCTTCTANNNAGATCGGAAGA |
| 34217_1 | TATGAGTGTGGAGTCGTTGCATTTGGCCATATATGCAAGGGTTNNNAGATCGGAAG |
| 34217_2 | TATGAGTGTGGAGTCGTTGCATTTGGCCATATATGCAAGGGTTNNNAGATCGGAAG |
| 34217_3 | TATGAGTGTGGAGTCGTTGCATTTGGCCATATATGCAAGGGTTNNNAGATCGGAAG |
| 34217_4 | TATGAGTGTGGAGTCGTTGCATTTGGCCATATATGCAAGGGTTNNNAGATCGGAAG |
| 36741_1 | TATGAGTGTGGAGTCGTTGCGTGGATCATTTGAGGTCAGGAGTNNNAGATCGGAA |
| 36741_2 | TATGAGTGTGGAGTCGTTGCGTGGATCATTTGAGGTCAGGAGTNNNAGATCGGAA |
| 36741_3 | TATGAGTGTGGAGTCGTTGCGTGGATCATTTGAGGTCAGGAGTNNNAGATCGGAA |
| 36741_4 | TATGAGTGTGGAGTCGTTGCGTGGATCATTTGAGGTCAGGAGTNNNAGATCGGAA |
| 43375_1 | TATGAGTGTGGAGTCGTTGCTAGCACAATCTCAGCTTACTGCANNNAGATCGGAAG |
| 43375_2 | TATGAGTGTGGAGTCGTTGCTAGCACAATCTCAGCTTACTGCANNNAGATCGGAAG |
| 43375_3 | TATGAGTGTGGAGTCGTTGCTAGCACAATCTCAGCTTACTGCANNNAGATCGGAAG |

| | |
|---|---|
| 43375_4 | TATGAGTGTGGAGTCGTTGCTAGCACAATCTCAGCTTACTGCANNNAGATCGGAAG |
| 29716_1 | TATGAGTGTGGAGTCGTTGCAGCTGGGAGAATAAGCTTACCACNNNAGATCGGAA |
| 29716_2 | TATGAGTGTGGAGTCGTTGCAGCTGGGAGAATAAGCTTACCACNNNAGATCGGAA |
| 29716_3 | TATGAGTGTGGAGTCGTTGCAGCTGGGAGAATAAGCTTACCACNNNAGATCGGAA |
| 29716_4 | TATGAGTGTGGAGTCGTTGCAGCTGGGAGAATAAGCTTACCACNNNAGATCGGAA |
| 48501_1 | TATGAGTGTGGAGTCGTTGCTCTGTTGTTTGTCTGTTCAGGGANNNAGATCGGAAG |
| 48501_2 | TATGAGTGTGGAGTCGTTGCTCTGTTGTTTGTCTGTTCAGGGANNNAGATCGGAAG |
| 48501_3 | TATGAGTGTGGAGTCGTTGCTCTGTTGTTTGTCTGTTCAGGGANNNAGATCGGAAG |
| 48501_4 | TATGAGTGTGGAGTCGTTGCTCTGTTGTTTGTCTGTTCAGGGANNNAGATCGGAAG |
| 37974_1 | TATGAGTGTGGAGTCGTTGCCACCAATAGTCACAGCACTTTGGNNNAGATCGGAAG |
| 37974_2 | TATGAGTGTGGAGTCGTTGCCACCAATAGTCACAGCACTTTGGNNNAGATCGGAAG |
| 37974_3 | TATGAGTGTGGAGTCGTTGCCACCAATAGTCACAGCACTTTGGNNNAGATCGGAAG |
| 37974_4 | TATGAGTGTGGAGTCGTTGCCACCAATAGTCACAGCACTTTGGNNNAGATCGGAAG |
| 44114_1 | TATGAGTGTGGAGTCGTTGCCCTCTGGGTATTGGCCATCTGNNNAGATCGGAAGAG |
| 44114_2 | TATGAGTGTGGAGTCGTTGCCCTCTGGGTATTGGCCATCTGNNNAGATCGGAAGAG |
| 44114_3 | TATGAGTGTGGAGTCGTTGCCCTCTGGGTATTGGCCATCTGNNNAGATCGGAAGAG |
| 44114_4 | TATGAGTGTGGAGTCGTTGCCCTCTGGGTATTGGCCATCTGNNNAGATCGGAAGAG |
| 42008_1 | TATGAGTGTGGAGTCGTTGCGAGTGCAGGGTGTGATCTCAGNNNAGATCGGAAGA |
| 42008_2 | TATGAGTGTGGAGTCGTTGCGAGTGCAGGGTGTGATCTCAGNNNAGATCGGAAGA |
| 42008_3 | TATGAGTGTGGAGTCGTTGCGAGTGCAGGGTGTGATCTCAGNNNAGATCGGAAGA |
| 42008_4 | TATGAGTGTGGAGTCGTTGCGAGTGCAGGGTGTGATCTCAGNNNAGATCGGAAGA |
| 52757_1 | TATGAGTGTGGAGTCGTTGCGCTTGTTCTCATTCAGGCAAACTNNNAGATCGGAAG |
| 52757_2 | TATGAGTGTGGAGTCGTTGCGCTTGTTCTCATTCAGGCAAACTNNNAGATCGGAAG |
| 52757_3 | TATGAGTGTGGAGTCGTTGCGCTTGTTCTCATTCAGGCAAACTNNNAGATCGGAAG |
| 52757_4 | TATGAGTGTGGAGTCGTTGCGCTTGTTCTCATTCAGGCAAACTNNNAGATCGGAAG |
| 10760_1 | TATGAGTGTGGAGTCGTTGCCACATCCTGTTGCCATTGTACCTNNNAGATCGGAAGA |
| 10760_2 | TATGAGTGTGGAGTCGTTGCCACATCCTGTTGCCATTGTACCTNNNAGATCGGAAGA |
| 10760_3 | TATGAGTGTGGAGTCGTTGCCACATCCTGTTGCCATTGTACCTNNNAGATCGGAAGA |
| 10760_4 | TATGAGTGTGGAGTCGTTGCCACATCCTGTTGCCATTGTACCTNNNAGATCGGAAGA |
| 29095_1 | TATGAGTGTGGAGTCGTTGCCCCACAGACTCCACAAATAGGAANNNAGATCGGAAG |
| 29095_2 | TATGAGTGTGGAGTCGTTGCCCCACAGACTCCACAAATAGGAANNNAGATCGGAAG |
| 29095_3 | TATGAGTGTGGAGTCGTTGCCCCACAGACTCCACAAATAGGAANNNAGATCGGAAG |
| 29095_4 | TATGAGTGTGGAGTCGTTGCCCCACAGACTCCACAAATAGGAANNNAGATCGGAAG |
| 46683_1 | TATGAGTGTGGAGTCGTTGCGAGAAAGGGGGCCATAACAAAACNNNAGATCGGAA |
| 46683_2 | TATGAGTGTGGAGTCGTTGCGAGAAAGGGGGCCATAACAAAACNNNAGATCGGAA |
| 46683_3 | TATGAGTGTGGAGTCGTTGCGAGAAAGGGGGCCATAACAAAACNNNAGATCGGAA |
| 46683_4 | TATGAGTGTGGAGTCGTTGCGAGAAAGGGGGCCATAACAAAACNNNAGATCGGAA |
| 10805_1 | TATGAGTGTGGAGTCGTTGCAAACAGAAATGAAAAACCCTGGGCNNNAGATCGGA |
| 10805_2 | TATGAGTGTGGAGTCGTTGCAAACAGAAATGAAAAACCCTGGGCNNNAGATCGGA |
| 10805_3 | TATGAGTGTGGAGTCGTTGCAAACAGAAATGAAAAACCCTGGGCNNNAGATCGGA |
| 10805_4 | TATGAGTGTGGAGTCGTTGCAAACAGAAATGAAAAACCCTGGGCNNNAGATCGGA |
| 32427_1 | TATGAGTGTGGAGTCGTTGCAAAGTGCTGGGATTACAGGCTTNNNAGATCGGAAG |
| 32427_2 | TATGAGTGTGGAGTCGTTGCAAAGTGCTGGGATTACAGGCTTNNNAGATCGGAAG |

| | |
|---|---|
| 32427_3 | TATGAGTGTGGAGTCGTTGCAAAGTGCTGGGATTACAGGCTTNNNAGATCGGAAG |
| 32427_4 | TATGAGTGTGGAGTCGTTGCAAAGTGCTGGGATTACAGGCTTNNNAGATCGGAAG |
| 11896_1 | TATGAGTGTGGAGTCGTTGCACCTCTGGCTAGCCCTTGAAATCNNNAGATCGGAAG |
| 11896_2 | TATGAGTGTGGAGTCGTTGCACCTCTGGCTAGCCCTTGAAATCNNNAGATCGGAAG |
| 11896_3 | TATGAGTGTGGAGTCGTTGCACCTCTGGCTAGCCCTTGAAATCNNNAGATCGGAAG |
| 11896_4 | TATGAGTGTGGAGTCGTTGCACCTCTGGCTAGCCCTTGAAATCNNNAGATCGGAAG |
| 33930_1 | TATGAGTGTGGAGTCGTTGCCTAAAGTGCAGTGGTGCGATTTCNNNAGATCGGAA |
| 33930_2 | TATGAGTGTGGAGTCGTTGCCTAAAGTGCAGTGGTGCGATTTCNNNAGATCGGAA |
| 33930_3 | TATGAGTGTGGAGTCGTTGCCTAAAGTGCAGTGGTGCGATTTCNNNAGATCGGAA |
| 33930_4 | TATGAGTGTGGAGTCGTTGCCTAAAGTGCAGTGGTGCGATTTCNNNAGATCGGAA |
| 41289_1 | TATGAGTGTGGAGTCGTTGCCAGCCAAGAAAGTACACATGCAANNNAGATCGGAA |
| 41289_2 | TATGAGTGTGGAGTCGTTGCCAGCCAAGAAAGTACACATGCAANNNAGATCGGAA |
| 41289_3 | TATGAGTGTGGAGTCGTTGCCAGCCAAGAAAGTACACATGCAANNNAGATCGGAA |
| 41289_4 | TATGAGTGTGGAGTCGTTGCCAGCCAAGAAAGTACACATGCAANNNAGATCGGAA |
| 35702_1 | TATGAGTGTGGAGTCGTTGCGCTCAATTGATCCTCCCACCTTANNNAGATCGGAAGA |
| 35702_2 | TATGAGTGTGGAGTCGTTGCGCTCAATTGATCCTCCCACCTTANNNAGATCGGAAGA |
| 35702_3 | TATGAGTGTGGAGTCGTTGCGCTCAATTGATCCTCCCACCTTANNNAGATCGGAAGA |
| 35702_4 | TATGAGTGTGGAGTCGTTGCGCTCAATTGATCCTCCCACCTTANNNAGATCGGAAGA |
| 39676_1 | TATGAGTGTGGAGTCGTTGCCAATTGCACCGGGAATTAGGAAGNNNAGATCGGAA |
| 39676_2 | TATGAGTGTGGAGTCGTTGCCAATTGCACCGGGAATTAGGAAGNNNAGATCGGAA |
| 39676_3 | TATGAGTGTGGAGTCGTTGCCAATTGCACCGGGAATTAGGAAGNNNAGATCGGAA |
| 39676_4 | TATGAGTGTGGAGTCGTTGCCAATTGCACCGGGAATTAGGAAGNNNAGATCGGAA |
| 46639_1 | TATGAGTGTGGAGTCGTTGCGATCTTGGCTTACTGCAACCTCTNNNAGATCGGAAG |
| 46639_2 | TATGAGTGTGGAGTCGTTGCGATCTTGGCTTACTGCAACCTCTNNNAGATCGGAAG |
| 46639_3 | TATGAGTGTGGAGTCGTTGCGATCTTGGCTTACTGCAACCTCTNNNAGATCGGAAG |
| 46639_4 | TATGAGTGTGGAGTCGTTGCGATCTTGGCTTACTGCAACCTCTNNNAGATCGGAAG |
| 48133_1 | TATGAGTGTGGAGTCGTTGCCCACTCTGGGTTGTCTGTTTACTNNNAGATCGGAAG |
| 48133_2 | TATGAGTGTGGAGTCGTTGCCCACTCTGGGTTGTCTGTTTACTNNNAGATCGGAAG |
| 48133_3 | TATGAGTGTGGAGTCGTTGCCCACTCTGGGTTGTCTGTTTACTNNNAGATCGGAAG |
| 48133_4 | TATGAGTGTGGAGTCGTTGCCCACTCTGGGTTGTCTGTTTACTNNNAGATCGGAAG |
| 37967_1 | TATGAGTGTGGAGTCGTTGCCTAGGATTACAGGTGTGAGCCTCNNNAGATCGGAA |
| 37967_2 | TATGAGTGTGGAGTCGTTGCCTAGGATTACAGGTGTGAGCCTCNNNAGATCGGAA |
| 37967_3 | TATGAGTGTGGAGTCGTTGCCTAGGATTACAGGTGTGAGCCTCNNNAGATCGGAA |
| 37967_4 | TATGAGTGTGGAGTCGTTGCCTAGGATTACAGGTGTGAGCCTCNNNAGATCGGAA |
| 49017_1 | TATGAGTGTGGAGTCGTTGCTCTAGATGTCAAGTGGAGAGGGTNNNAGATCGGAA |
| 49017_2 | TATGAGTGTGGAGTCGTTGCTCTAGATGTCAAGTGGAGAGGGTNNNAGATCGGAA |
| 49017_3 | TATGAGTGTGGAGTCGTTGCTCTAGATGTCAAGTGGAGAGGGTNNNAGATCGGAA |
| 49017_4 | TATGAGTGTGGAGTCGTTGCTCTAGATGTCAAGTGGAGAGGGTNNNAGATCGGAA |
| 31288_1 | TATGAGTGTGGAGTCGTTGCAAAAAGCAGGAAGTGGTGGTAGANNNAGATCGGA |
| 31288_2 | TATGAGTGTGGAGTCGTTGCAAAAAGCAGGAAGTGGTGGTAGANNNAGATCGGA |
| 31288_3 | TATGAGTGTGGAGTCGTTGCAAAAAGCAGGAAGTGGTGGTAGANNNAGATCGGA |
| 31288_4 | TATGAGTGTGGAGTCGTTGCAAAAAGCAGGAAGTGGTGGTAGANNNAGATCGGA |
| 9154_1 | TATGAGTGTGGAGTCGTTGCCTCCGAACCAATAGCATCAGCATNNNAGATCGGAAG |

| | |
|---|---|
| 9154_2 | TATGAGTGTGGAGTCGTTGCCTCCGAACCAATAGCATCAGCATNNNAGATCGGAAG |
| 9154_3 | TATGAGTGTGGAGTCGTTGCCTCCGAACCAATAGCATCAGCATNNNAGATCGGAAG |
| 9154_4 | TATGAGTGTGGAGTCGTTGCCTCCGAACCAATAGCATCAGCATNNNAGATCGGAAG |
| 28756_1 | TATGAGTGTGGAGTCGTTGCCCTATAGTCCCAGCTACTCAGGANNNAGATCGGAAG |
| 28756_2 | TATGAGTGTGGAGTCGTTGCCCTATAGTCCCAGCTACTCAGGANNNAGATCGGAAG |
| 28756_3 | TATGAGTGTGGAGTCGTTGCCCTATAGTCCCAGCTACTCAGGANNNAGATCGGAAG |
| 28756_4 | TATGAGTGTGGAGTCGTTGCCCTATAGTCCCAGCTACTCAGGANNNAGATCGGAAG |
| 39378_1 | TATGAGTGTGGAGTCGTTGCTCGTCCAGGGCATTTATGATGATNNNAGATCGGAAG |
| 39378_2 | TATGAGTGTGGAGTCGTTGCTCGTCCAGGGCATTTATGATGATNNNAGATCGGAAG |
| 39378_3 | TATGAGTGTGGAGTCGTTGCTCGTCCAGGGCATTTATGATGATNNNAGATCGGAAG |
| 39378_4 | TATGAGTGTGGAGTCGTTGCTCGTCCAGGGCATTTATGATGATNNNAGATCGGAAG |
| 45851_1 | TATGAGTGTGGAGTCGTTGCCCTCCCAGGTTCAAGTGATTCTCNNNAGATCGGAAG |
| 45851_2 | TATGAGTGTGGAGTCGTTGCCCTCCCAGGTTCAAGTGATTCTCNNNAGATCGGAAG |
| 45851_3 | TATGAGTGTGGAGTCGTTGCCCTCCCAGGTTCAAGTGATTCTCNNNAGATCGGAAG |
| 45851_4 | TATGAGTGTGGAGTCGTTGCCCTCCCAGGTTCAAGTGATTCTCNNNAGATCGGAAG |
| 3509_1 | TATGAGTGTGGAGTCGTTGCTGATGGGAAGCTCAACTAGCATGGNNNAGATCGGA |
| 3509_2 | TATGAGTGTGGAGTCGTTGCTGATGGGAAGCTCAACTAGCATGGNNNAGATCGGA |
| 3509_3 | TATGAGTGTGGAGTCGTTGCTGATGGGAAGCTCAACTAGCATGGNNNAGATCGGA |
| 3509_4 | TATGAGTGTGGAGTCGTTGCTGATGGGAAGCTCAACTAGCATGGNNNAGATCGGA |
| 49909_1 | TATGAGTGTGGAGTCGTTGCGACCACTCTTTAGGGGTATGCAANNNAGATCGGAAG |
| 49909_2 | TATGAGTGTGGAGTCGTTGCGACCACTCTTTAGGGGTATGCAANNNAGATCGGAAG |
| 49909_3 | TATGAGTGTGGAGTCGTTGCGACCACTCTTTAGGGGTATGCAANNNAGATCGGAAG |
| 49909_4 | TATGAGTGTGGAGTCGTTGCGACCACTCTTTAGGGGTATGCAANNNAGATCGGAAG |
| 35996_1 | TATGAGTGTGGAGTCGTTGCAGATCAAAGCATGCCACATTCAGNNNAGATCGGAAG |
| 35996_2 | TATGAGTGTGGAGTCGTTGCAGATCAAAGCATGCCACATTCAGNNNAGATCGGAAG |
| 35996_3 | TATGAGTGTGGAGTCGTTGCAGATCAAAGCATGCCACATTCAGNNNAGATCGGAAG |
| 35996_4 | TATGAGTGTGGAGTCGTTGCAGATCAAAGCATGCCACATTCAGNNNAGATCGGAAG |
| 10535_1 | TATGAGTGTGGAGTCGTTGCACTGCTCTTTCTCACAACTCTTTNNNAGATCGGAAGA |
| 10535_2 | TATGAGTGTGGAGTCGTTGCACTGCTCTTTCTCACAACTCTTTNNNAGATCGGAAGA |
| 10535_3 | TATGAGTGTGGAGTCGTTGCACTGCTCTTTCTCACAACTCTTTNNNAGATCGGAAGA |
| 10535_4 | TATGAGTGTGGAGTCGTTGCACTGCTCTTTCTCACAACTCTTTNNNAGATCGGAAGA |
| 38567_1 | TATGAGTGTGGAGTCGTTGCAATGGTAACAACTGCAGGATCCANNNAGATCGGAA |
| 38567_2 | TATGAGTGTGGAGTCGTTGCAATGGTAACAACTGCAGGATCCANNNAGATCGGAA |
| 38567_3 | TATGAGTGTGGAGTCGTTGCAATGGTAACAACTGCAGGATCCANNNAGATCGGAA |
| 38567_4 | TATGAGTGTGGAGTCGTTGCAATGGTAACAACTGCAGGATCCANNNAGATCGGAA |
| 37982_1 | TATGAGTGTGGAGTCGTTGCACAAGAATCCCAGCTAGCTTGTTNNNAGATCGGAAG |
| 37982_2 | TATGAGTGTGGAGTCGTTGCACAAGAATCCCAGCTAGCTTGTTNNNAGATCGGAAG |
| 37982_3 | TATGAGTGTGGAGTCGTTGCACAAGAATCCCAGCTAGCTTGTTNNNAGATCGGAAG |
| 37982_4 | TATGAGTGTGGAGTCGTTGCACAAGAATCCCAGCTAGCTTGTTNNNAGATCGGAAG |
| 37707_1 | TATGAGTGTGGAGTCGTTGCACATTGAGAGCCTTCTAGGTTGGNNNAGATCGGAA |
| 37707_2 | TATGAGTGTGGAGTCGTTGCACATTGAGAGCCTTCTAGGTTGGNNNAGATCGGAA |
| 37707_3 | TATGAGTGTGGAGTCGTTGCACATTGAGAGCCTTCTAGGTTGGNNNAGATCGGAA |
| 37707_4 | TATGAGTGTGGAGTCGTTGCACATTGAGAGCCTTCTAGGTTGGNNNAGATCGGAA |

| | |
|---|---|
| 482_1 | TATGAGTGTGGAGTCGTTGCCCCTGCAATCCAAAAACAGGAAAGNNNAGATCGGAA |
| 482_2 | TATGAGTGTGGAGTCGTTGCCCCTGCAATCCAAAAACAGGAAAGNNNAGATCGGAA |
| 482_3 | TATGAGTGTGGAGTCGTTGCCCCTGCAATCCAAAAACAGGAAAGNNNAGATCGGAA |
| 482_4 | TATGAGTGTGGAGTCGTTGCCCCTGCAATCCAAAAACAGGAAAGNNNAGATCGGAA |
| 53037_1 | TATGAGTGTGGAGTCGTTGCAACAGAAGTCAGATGTCCTCAGGNNNAGATCGGAA |
| 53037_2 | TATGAGTGTGGAGTCGTTGCAACAGAAGTCAGATGTCCTCAGGNNNAGATCGGAA |
| 53037_3 | TATGAGTGTGGAGTCGTTGCAACAGAAGTCAGATGTCCTCAGGNNNAGATCGGAA |
| 53037_4 | TATGAGTGTGGAGTCGTTGCAACAGAAGTCAGATGTCCTCAGGNNNAGATCGGAA |
| 9603_1 | TATGAGTGTGGAGTCGTTGCAAGGAGCATAATGCCTAACAGGCNNNAGATCGGAA |
| 9603_2 | TATGAGTGTGGAGTCGTTGCAAGGAGCATAATGCCTAACAGGCNNNAGATCGGAA |
| 9603_3 | TATGAGTGTGGAGTCGTTGCAAGGAGCATAATGCCTAACAGGCNNNAGATCGGAA |
| 9603_4 | TATGAGTGTGGAGTCGTTGCAAGGAGCATAATGCCTAACAGGCNNNAGATCGGAA |
| 45074_1 | TATGAGTGTGGAGTCGTTGCACGTTACAAAAATCAAGCTGCCANNNAGATCGGAAG |
| 45074_2 | TATGAGTGTGGAGTCGTTGCACGTTACAAAAATCAAGCTGCCANNNAGATCGGAAG |
| 45074_3 | TATGAGTGTGGAGTCGTTGCACGTTACAAAAATCAAGCTGCCANNNAGATCGGAAG |
| 45074_4 | TATGAGTGTGGAGTCGTTGCACGTTACAAAAATCAAGCTGCCANNNAGATCGGAAG |
| 37470_1 | TATGAGTGTGGAGTCGTTGCGATACATAGGGCAAGGTATGGGGNNNAGATCGGAA |
| 37470_2 | TATGAGTGTGGAGTCGTTGCGATACATAGGGCAAGGTATGGGGNNNAGATCGGAA |
| 37470_3 | TATGAGTGTGGAGTCGTTGCGATACATAGGGCAAGGTATGGGGNNNAGATCGGAA |
| 37470_4 | TATGAGTGTGGAGTCGTTGCGATACATAGGGCAAGGTATGGGGNNNAGATCGGAA |
| 41813_1 | TATGAGTGTGGAGTCGTTGCGCCTCCTGAATAGCTGGGAATACNNNAGATCGGAAG |
| 41813_2 | TATGAGTGTGGAGTCGTTGCGCCTCCTGAATAGCTGGGAATACNNNAGATCGGAAG |
| 41813_3 | TATGAGTGTGGAGTCGTTGCGCCTCCTGAATAGCTGGGAATACNNNAGATCGGAAG |
| 41813_4 | TATGAGTGTGGAGTCGTTGCGCCTCCTGAATAGCTGGGAATACNNNAGATCGGAAG |
| 50309_1 | TATGAGTGTGGAGTCGTTGCGCTTTTATGGGGTTTGCAGGAATNNNAGATCGGAA |
| 50309_2 | TATGAGTGTGGAGTCGTTGCGCTTTTATGGGGTTTGCAGGAATNNNAGATCGGAA |
| 50309_3 | TATGAGTGTGGAGTCGTTGCGCTTTTATGGGGTTTGCAGGAATNNNAGATCGGAA |
| 50309_4 | TATGAGTGTGGAGTCGTTGCGCTTTTATGGGGTTTGCAGGAATNNNAGATCGGAA |
| 37013_1 | TATGAGTGTGGAGTCGTTGCATGCACATGAAGAACTCTCCTGTNNNAGATCGGAAG |
| 37013_2 | TATGAGTGTGGAGTCGTTGCATGCACATGAAGAACTCTCCTGTNNNAGATCGGAAG |
| 37013_3 | TATGAGTGTGGAGTCGTTGCATGCACATGAAGAACTCTCCTGTNNNAGATCGGAAG |
| 37013_4 | TATGAGTGTGGAGTCGTTGCATGCACATGAAGAACTCTCCTGTNNNAGATCGGAAG |
| 52692_1 | TATGAGTGTGGAGTCGTTGCCTCCTTGTTGGTCTGAGTTTCCCNNNAGATCGGAAGA |
| 52692_2 | TATGAGTGTGGAGTCGTTGCCTCCTTGTTGGTCTGAGTTTCCCNNNAGATCGGAAGA |
| 52692_3 | TATGAGTGTGGAGTCGTTGCCTCCTTGTTGGTCTGAGTTTCCCNNNAGATCGGAAGA |
| 52692_4 | TATGAGTGTGGAGTCGTTGCCTCCTTGTTGGTCTGAGTTTCCCNNNAGATCGGAAGA |
| 31454_1 | TATGAGTGTGGAGTCGTTGCAAGCATGTCAATCTTACCCCCAANNNAGATCGGAAGA |
| 31454_2 | TATGAGTGTGGAGTCGTTGCAAGCATGTCAATCTTACCCCCAANNNAGATCGGAAGA |
| 31454_3 | TATGAGTGTGGAGTCGTTGCAAGCATGTCAATCTTACCCCCAANNNAGATCGGAAGA |
| 31454_4 | TATGAGTGTGGAGTCGTTGCAAGCATGTCAATCTTACCCCCAANNNAGATCGGAAGA |
| 35845_1 | TATGAGTGTGGAGTCGTTGCACGATCTCAGCTCACTGCAAGNNNAGATCGGAAGAG |
| 35845_2 | TATGAGTGTGGAGTCGTTGCACGATCTCAGCTCACTGCAAGNNNAGATCGGAAGAG |
| 35845_3 | TATGAGTGTGGAGTCGTTGCACGATCTCAGCTCACTGCAAGNNNAGATCGGAAGAG |

| | |
|---|---|
| 35845_4 | TATGAGTGTGGAGTCGTTGCACGATCTCAGCTCACTGCAAGNNNAGATCGGAAGAG |
| 48215_1 | TATGAGTGTGGAGTCGTTGCCCAAGTAGCTGGGATTATAGGCANNNAGATCGGAA |
| 48215_2 | TATGAGTGTGGAGTCGTTGCCCAAGTAGCTGGGATTATAGGCANNNAGATCGGAA |
| 48215_3 | TATGAGTGTGGAGTCGTTGCCCAAGTAGCTGGGATTATAGGCANNNAGATCGGAA |
| 48215_4 | TATGAGTGTGGAGTCGTTGCCCAAGTAGCTGGGATTATAGGCANNNAGATCGGAA |
| 43468_1 | TATGAGTGTGGAGTCGTTGCGTCACCCAGGCTAGAGTACAATGNNNAGATCGGAA |
| 43468_2 | TATGAGTGTGGAGTCGTTGCGTCACCCAGGCTAGAGTACAATGNNNAGATCGGAA |
| 43468_3 | TATGAGTGTGGAGTCGTTGCGTCACCCAGGCTAGAGTACAATGNNNAGATCGGAA |
| 43468_4 | TATGAGTGTGGAGTCGTTGCGTCACCCAGGCTAGAGTACAATGNNNAGATCGGAA |
| 35534_1 | TATGAGTGTGGAGTCGTTGCTCACTCAAGCTGGAGTACAGTTCNNNAGATCGGAAG |
| 35534_2 | TATGAGTGTGGAGTCGTTGCTCACTCAAGCTGGAGTACAGTTCNNNAGATCGGAAG |
| 35534_3 | TATGAGTGTGGAGTCGTTGCTCACTCAAGCTGGAGTACAGTTCNNNAGATCGGAAG |
| 35534_4 | TATGAGTGTGGAGTCGTTGCTCACTCAAGCTGGAGTACAGTTCNNNAGATCGGAAG |
| 41369_1 | TATGAGTGTGGAGTCGTTGCCAGGAGAATGGCGTGAAAAGAAANNNAGATCGGA |
| 41369_2 | TATGAGTGTGGAGTCGTTGCCAGGAGAATGGCGTGAAAAGAAANNNAGATCGGA |
| 41369_3 | TATGAGTGTGGAGTCGTTGCCAGGAGAATGGCGTGAAAAGAAANNNAGATCGGA |
| 41369_4 | TATGAGTGTGGAGTCGTTGCCAGGAGAATGGCGTGAAAAGAAANNNAGATCGGA |
| 52356_1 | TATGAGTGTGGAGTCGTTGCGTTCCCAGCACCACCATTTTGNNNAGATCGGAAGAG |
| 52356_2 | TATGAGTGTGGAGTCGTTGCGTTCCCAGCACCACCATTTTGNNNAGATCGGAAGAG |
| 52356_3 | TATGAGTGTGGAGTCGTTGCGTTCCCAGCACCACCATTTTGNNNAGATCGGAAGAG |
| 52356_4 | TATGAGTGTGGAGTCGTTGCGTTCCCAGCACCACCATTTTGNNNAGATCGGAAGAG |
| 48378_1 | TATGAGTGTGGAGTCGTTGCGGCTGCATACTATTCCACTCTGTNNNAGATCGGAAG |
| 48378_2 | TATGAGTGTGGAGTCGTTGCGGCTGCATACTATTCCACTCTGTNNNAGATCGGAAG |
| 48378_3 | TATGAGTGTGGAGTCGTTGCGGCTGCATACTATTCCACTCTGTNNNAGATCGGAAG |
| 48378_4 | TATGAGTGTGGAGTCGTTGCGGCTGCATACTATTCCACTCTGTNNNAGATCGGAAG |
| 34882_1 | TATGAGTGTGGAGTCGTTGCCCAGGTTCAAGCGATTTTCCTGNNNAGATCGGAAGA |
| 34882_2 | TATGAGTGTGGAGTCGTTGCCCAGGTTCAAGCGATTTTCCTGNNNAGATCGGAAGA |
| 34882_3 | TATGAGTGTGGAGTCGTTGCCCAGGTTCAAGCGATTTTCCTGNNNAGATCGGAAGA |
| 34882_4 | TATGAGTGTGGAGTCGTTGCCCAGGTTCAAGCGATTTTCCTGNNNAGATCGGAAGA |
| 44747_1 | TATGAGTGTGGAGTCGTTGCGATAGAGCATCAGTGGGGAGTCNNNAGATCGGAA |
| 44747_2 | TATGAGTGTGGAGTCGTTGCGATAGAGCATCAGTGGGGAGTCNNNAGATCGGAA |
| 44747_3 | TATGAGTGTGGAGTCGTTGCGATAGAGCATCAGTGGGGAGTCNNNAGATCGGAA |
| 44747_4 | TATGAGTGTGGAGTCGTTGCGATAGAGCATCAGTGGGGAGTCNNNAGATCGGAA |
| 48066_1 | TATGAGTGTGGAGTCGTTGCAGCATGAAAGGCTGTTGAACTTTNNNAGATCGGAA |
| 48066_2 | TATGAGTGTGGAGTCGTTGCAGCATGAAAGGCTGTTGAACTTTNNNAGATCGGAA |
| 48066_3 | TATGAGTGTGGAGTCGTTGCAGCATGAAAGGCTGTTGAACTTTNNNAGATCGGAA |
| 48066_4 | TATGAGTGTGGAGTCGTTGCAGCATGAAAGGCTGTTGAACTTTNNNAGATCGGAA |
| 7919_1 | TATGAGTGTGGAGTCGTTGCAGTGGGTCATGGGAGCATCTAGTANNNAGATCGGA |
| 7919_2 | TATGAGTGTGGAGTCGTTGCAGTGGGTCATGGGAGCATCTAGTANNNAGATCGGA |
| 7919_3 | TATGAGTGTGGAGTCGTTGCAGTGGGTCATGGGAGCATCTAGTANNNAGATCGGA |
| 7919_4 | TATGAGTGTGGAGTCGTTGCAGTGGGTCATGGGAGCATCTAGTANNNAGATCGGA |
| 37824_1 | TATGAGTGTGGAGTCGTTGCTCCTTCTTCCTCTTTGCCTTCTGNNNAGATCGGAAGA |
| 37824_2 | TATGAGTGTGGAGTCGTTGCTCCTTCTTCCTCTTTGCCTTCTGNNNAGATCGGAAGA |

| | |
|---|---|
| 37824_3 | TATGAGTGTGGAGTCGTTGCTCCTTCTTCCTCTTTGCCTTCTGNNNAGATCGGAAGA |
| 37824_4 | TATGAGTGTGGAGTCGTTGCTCCTTCTTCCTCTTTGCCTTCTGNNNAGATCGGAAGA |
| 49437_1 | TATGAGTGTGGAGTCGTTGCTACTTCCAAAGGGCACAGTTTTGNNNAGATCGGAAG |
| 49437_2 | TATGAGTGTGGAGTCGTTGCTACTTCCAAAGGGCACAGTTTTGNNNAGATCGGAAG |
| 49437_3 | TATGAGTGTGGAGTCGTTGCTACTTCCAAAGGGCACAGTTTTGNNNAGATCGGAAG |
| 49437_4 | TATGAGTGTGGAGTCGTTGCTACTTCCAAAGGGCACAGTTTTGNNNAGATCGGAAG |
| 9978_1 | TATGAGTGTGGAGTCGTTGCTTGAACAGGGGATCCTTACACCTNNNAGATCGGAAG |
| 9978_2 | TATGAGTGTGGAGTCGTTGCTTGAACAGGGGATCCTTACACCTNNNAGATCGGAAG |
| 9978_3 | TATGAGTGTGGAGTCGTTGCTTGAACAGGGGATCCTTACACCTNNNAGATCGGAAG |
| 9978_4 | TATGAGTGTGGAGTCGTTGCTTGAACAGGGGATCCTTACACCTNNNAGATCGGAAG |
| 30137_1 | TATGAGTGTGGAGTCGTTGCACTCTGGGTCTTATCTCTCCGTANNNAGATCGGAAG |
| 30137_2 | TATGAGTGTGGAGTCGTTGCACTCTGGGTCTTATCTCTCCGTANNNAGATCGGAAG |
| 30137_3 | TATGAGTGTGGAGTCGTTGCACTCTGGGTCTTATCTCTCCGTANNNAGATCGGAAG |
| 30137_4 | TATGAGTGTGGAGTCGTTGCACTCTGGGTCTTATCTCTCCGTANNNAGATCGGAAG |
| 49953_1 | TATGAGTGTGGAGTCGTTGCGCATCAAATGGATTTCACGAGGTNNNAGATCGGAA |
| 49953_2 | TATGAGTGTGGAGTCGTTGCGCATCAAATGGATTTCACGAGGTNNNAGATCGGAA |
| 49953_3 | TATGAGTGTGGAGTCGTTGCGCATCAAATGGATTTCACGAGGTNNNAGATCGGAA |
| 49953_4 | TATGAGTGTGGAGTCGTTGCGCATCAAATGGATTTCACGAGGTNNNAGATCGGAA |
| 53419_1 | TATGAGTGTGGAGTCGTTGCAAATGAATTGTGTTGCCTGGACCNNNAGATCGGAAG |
| 53419_2 | TATGAGTGTGGAGTCGTTGCAAATGAATTGTGTTGCCTGGACCNNNAGATCGGAAG |
| 53419_3 | TATGAGTGTGGAGTCGTTGCAAATGAATTGTGTTGCCTGGACCNNNAGATCGGAAG |
| 53419_4 | TATGAGTGTGGAGTCGTTGCAAATGAATTGTGTTGCCTGGACCNNNAGATCGGAAG |
| 8156_1 | TATGAGTGTGGAGTCGTTGCTGTTCCACTCTCTTCTGGTCTTTNNNAGATCGGAAGA |
| 8156_2 | TATGAGTGTGGAGTCGTTGCTGTTCCACTCTCTTCTGGTCTTTNNNAGATCGGAAGA |
| 8156_3 | TATGAGTGTGGAGTCGTTGCTGTTCCACTCTCTTCTGGTCTTTNNNAGATCGGAAGA |
| 8156_4 | TATGAGTGTGGAGTCGTTGCTGTTCCACTCTCTTCTGGTCTTTNNNAGATCGGAAGA |
| 42589_1 | TATGAGTGTGGAGTCGTTGCGAGTTTTGCTCTTGTTGCTCAGGNNNAGATCGGAAG |
| 42589_2 | TATGAGTGTGGAGTCGTTGCGAGTTTTGCTCTTGTTGCTCAGGNNNAGATCGGAAG |
| 42589_3 | TATGAGTGTGGAGTCGTTGCGAGTTTTGCTCTTGTTGCTCAGGNNNAGATCGGAAG |
| 42589_4 | TATGAGTGTGGAGTCGTTGCGAGTTTTGCTCTTGTTGCTCAGGNNNAGATCGGAAG |
| 12710_1 | TATGAGTGTGGAGTCGTTGCTTTAATGATCTTCTGGTCATTTGNNNAGATCGGAAG |
| 12710_2 | TATGAGTGTGGAGTCGTTGCTTTAATGATCTTCTGGTCATTTGNNNAGATCGGAAG |
| 12710_3 | TATGAGTGTGGAGTCGTTGCTTTAATGATCTTCTGGTCATTTGNNNAGATCGGAAG |
| 12710_4 | TATGAGTGTGGAGTCGTTGCTTTAATGATCTTCTGGTCATTTGNNNAGATCGGAAG |
| 46040_1 | TATGAGTGTGGAGTCGTTGCAGCCTCCTGAGTAGCTAGGATTANNNAGATCGGAA |
| 46040_2 | TATGAGTGTGGAGTCGTTGCAGCCTCCTGAGTAGCTAGGATTANNNAGATCGGAA |
| 46040_3 | TATGAGTGTGGAGTCGTTGCAGCCTCCTGAGTAGCTAGGATTANNNAGATCGGAA |
| 46040_4 | TATGAGTGTGGAGTCGTTGCAGCCTCCTGAGTAGCTAGGATTANNNAGATCGGAA |
| 33711_1 | TATGAGTGTGGAGTCGTTGCAACAGATACTGGTTTTCTAGACCANNNAGATCGGAA |
| 33711_2 | TATGAGTGTGGAGTCGTTGCAACAGATACTGGTTTTCTAGACCANNNAGATCGGAA |
| 33711_3 | TATGAGTGTGGAGTCGTTGCAACAGATACTGGTTTTCTAGACCANNNAGATCGGAA |
| 33711_4 | TATGAGTGTGGAGTCGTTGCAACAGATACTGGTTTTCTAGACCANNNAGATCGGAA |
| 45951_1 | TATGAGTGTGGAGTCGTTGCGAGTCAAGGACTGTGTGGAGAATNNNAGATCGGA |

| | |
|---|---|
| 45951_2 | TATGAGTGTGGAGTCGTTGCGAGTCAAGGACTGTGTGGAGAATNNNAGATCGGAA |
| 45951_3 | TATGAGTGTGGAGTCGTTGCGAGTCAAGGACTGTGTGGAGAATNNNAGATCGGAA |
| 45951_4 | TATGAGTGTGGAGTCGTTGCGAGTCAAGGACTGTGTGGAGAATNNNAGATCGGAA |
| 50393_1 | TATGAGTGTGGAGTCGTTGCCAGCCTGTGAAGGGAAGGAGNNNAGATCGGAAGA |
| 50393_2 | TATGAGTGTGGAGTCGTTGCCAGCCTGTGAAGGGAAGGAGNNNAGATCGGAAGA |
| 50393_3 | TATGAGTGTGGAGTCGTTGCCAGCCTGTGAAGGGAAGGAGNNNAGATCGGAAGA |
| 50393_4 | TATGAGTGTGGAGTCGTTGCCAGCCTGTGAAGGGAAGGAGNNNAGATCGGAAGA |
| 38646_1 | TATGAGTGTGGAGTCGTTGCGCCAGGCTCACAGAAATAGAGAANNNAGATCGGAA |
| 38646_2 | TATGAGTGTGGAGTCGTTGCGCCAGGCTCACAGAAATAGAGAANNNAGATCGGAA |
| 38646_3 | TATGAGTGTGGAGTCGTTGCGCCAGGCTCACAGAAATAGAGAANNNAGATCGGAA |
| 38646_4 | TATGAGTGTGGAGTCGTTGCGCCAGGCTCACAGAAATAGAGAANNNAGATCGGAA |
| 16390_1 | TATGAGTGTGGAGTCGTTGCAGTATCTGTTATATTTCTGTGCTNNNAGATCGGAAG |
| 16390_2 | TATGAGTGTGGAGTCGTTGCAGTATCTGTTATATTTCTGTGCTNNNAGATCGGAAG |
| 16390_3 | TATGAGTGTGGAGTCGTTGCAGTATCTGTTATATTTCTGTGCTNNNAGATCGGAAG |
| 16390_4 | TATGAGTGTGGAGTCGTTGCAGTATCTGTTATATTTCTGTGCTNNNAGATCGGAAG |
| 34389_1 | TATGAGTGTGGAGTCGTTGCCTGAGTAGCTGGGATTACAGGTGNNNAGATCGGAA |
| 34389_2 | TATGAGTGTGGAGTCGTTGCCTGAGTAGCTGGGATTACAGGTGNNNAGATCGGAA |
| 34389_3 | TATGAGTGTGGAGTCGTTGCCTGAGTAGCTGGGATTACAGGTGNNNAGATCGGAA |
| 34389_4 | TATGAGTGTGGAGTCGTTGCCTGAGTAGCTGGGATTACAGGTGNNNAGATCGGAA |
| 52396_1 | TATGAGTGTGGAGTCGTTGCTGATGTGTTTTTCTGAGTGTGGCNNNAGATCGGAAG |
| 52396_2 | TATGAGTGTGGAGTCGTTGCTGATGTGTTTTTCTGAGTGTGGCNNNAGATCGGAAG |
| 52396_3 | TATGAGTGTGGAGTCGTTGCTGATGTGTTTTTCTGAGTGTGGCNNNAGATCGGAAG |
| 52396_4 | TATGAGTGTGGAGTCGTTGCTGATGTGTTTTTCTGAGTGTGGCNNNAGATCGGAAG |
| 36111_1 | TATGAGTGTGGAGTCGTTGCCACAGGAGGAGTATGAAGGAACCNNNAGATCGGAA |
| 36111_2 | TATGAGTGTGGAGTCGTTGCCACAGGAGGAGTATGAAGGAACCNNNAGATCGGAA |
| 36111_3 | TATGAGTGTGGAGTCGTTGCCACAGGAGGAGTATGAAGGAACCNNNAGATCGGAA |
| 36111_4 | TATGAGTGTGGAGTCGTTGCCACAGGAGGAGTATGAAGGAACCNNNAGATCGGAA |
| 50062_1 | TATGAGTGTGGAGTCGTTGCACCTTTCTCTTGCTGTCCTTGAANNNAGATCGGAAGA |
| 50062_2 | TATGAGTGTGGAGTCGTTGCACCTTTCTCTTGCTGTCCTTGAANNNAGATCGGAAGA |
| 50062_3 | TATGAGTGTGGAGTCGTTGCACCTTTCTCTTGCTGTCCTTGAANNNAGATCGGAAGA |
| 50062_4 | TATGAGTGTGGAGTCGTTGCACCTTTCTCTTGCTGTCCTTGAANNNAGATCGGAAGA |
| 49269_1 | TATGAGTGTGGAGTCGTTGCAGGTGGAGTTTTGTTCTTGTTGCNNNAGATCGGAAG |
| 49269_2 | TATGAGTGTGGAGTCGTTGCAGGTGGAGTTTTGTTCTTGTTGCNNNAGATCGGAAG |
| 49269_3 | TATGAGTGTGGAGTCGTTGCAGGTGGAGTTTTGTTCTTGTTGCNNNAGATCGGAAG |
| 49269_4 | TATGAGTGTGGAGTCGTTGCAGGTGGAGTTTTGTTCTTGTTGCNNNAGATCGGAAG |
| 49440_1 | TATGAGTGTGGAGTCGTTGCTTGTCCAGGTCTCCAATGAACATNNNAGATCGGAAG |
| 49440_2 | TATGAGTGTGGAGTCGTTGCTTGTCCAGGTCTCCAATGAACATNNNAGATCGGAAG |
| 49440_3 | TATGAGTGTGGAGTCGTTGCTTGTCCAGGTCTCCAATGAACATNNNAGATCGGAAG |
| 49440_4 | TATGAGTGTGGAGTCGTTGCTTGTCCAGGTCTCCAATGAACATNNNAGATCGGAAG |
| 49230_1 | TATGAGTGTGGAGTCGTTGCTAGCTGGGATTATAGACGCACACNNNAGATCGGAA |
| 49230_2 | TATGAGTGTGGAGTCGTTGCTAGCTGGGATTATAGACGCACACNNNAGATCGGAA |
| 49230_3 | TATGAGTGTGGAGTCGTTGCTAGCTGGGATTATAGACGCACACNNNAGATCGGAA |
| 49230_4 | TATGAGTGTGGAGTCGTTGCTAGCTGGGATTATAGACGCACACNNNAGATCGGAA |

| | |
|---|---|
| 40385_1 | TATGAGTGTGGAGTCGTTGCGCTTCTAGTCCCAGCTACTCAGANNNAGATCGGAAG |
| 40385_2 | TATGAGTGTGGAGTCGTTGCGCTTCTAGTCCCAGCTACTCAGANNNAGATCGGAAG |
| 40385_3 | TATGAGTGTGGAGTCGTTGCGCTTCTAGTCCCAGCTACTCAGANNNAGATCGGAAG |
| 40385_4 | TATGAGTGTGGAGTCGTTGCGCTTCTAGTCCCAGCTACTCAGANNNAGATCGGAAG |
| 32920_1 | TATGAGTGTGGAGTCGTTGCGAGTCAGGAAAACTGCTTCAACCNNNAGATCGGAAG |
| 32920_2 | TATGAGTGTGGAGTCGTTGCGAGTCAGGAAAACTGCTTCAACCNNNAGATCGGAAG |
| 32920_3 | TATGAGTGTGGAGTCGTTGCGAGTCAGGAAAACTGCTTCAACCNNNAGATCGGAAG |
| 32920_4 | TATGAGTGTGGAGTCGTTGCGAGTCAGGAAAACTGCTTCAACCNNNAGATCGGAAG |
| 37063_1 | TATGAGTGTGGAGTCGTTGCAGCCAGCTAAGTAATTGTGGGAANNNAGATCGGAA |
| 37063_2 | TATGAGTGTGGAGTCGTTGCAGCCAGCTAAGTAATTGTGGGAANNNAGATCGGAA |
| 37063_3 | TATGAGTGTGGAGTCGTTGCAGCCAGCTAAGTAATTGTGGGAANNNAGATCGGAA |
| 37063_4 | TATGAGTGTGGAGTCGTTGCAGCCAGCTAAGTAATTGTGGGAANNNAGATCGGAA |
| 33280_1 | TATGAGTGTGGAGTCGTTGCGAGTGCAGTGGTACAATTATGGCNNNAGATCGGAA |
| 33280_2 | TATGAGTGTGGAGTCGTTGCGAGTGCAGTGGTACAATTATGGCNNNAGATCGGAA |
| 33280_3 | TATGAGTGTGGAGTCGTTGCGAGTGCAGTGGTACAATTATGGCNNNAGATCGGAA |
| 33280_4 | TATGAGTGTGGAGTCGTTGCGAGTGCAGTGGTACAATTATGGCNNNAGATCGGAA |
| 10864_1 | TATGAGTGTGGAGTCGTTGCGCCAAACCATCTGGAAAAAGCCNNNAGATCGGAAG |
| 10864_2 | TATGAGTGTGGAGTCGTTGCGCCAAACCATCTGGAAAAAGCCNNNAGATCGGAAG |
| 10864_3 | TATGAGTGTGGAGTCGTTGCGCCAAACCATCTGGAAAAAGCCNNNAGATCGGAAG |
| 10864_4 | TATGAGTGTGGAGTCGTTGCGCCAAACCATCTGGAAAAAGCCNNNAGATCGGAAG |
| 32399_1 | TATGAGTGTGGAGTCGTTGCTGCTCAGTTGTTAGTGGCTTATGNNNAGATCGGAAG |
| 32399_2 | TATGAGTGTGGAGTCGTTGCTGCTCAGTTGTTAGTGGCTTATGNNNAGATCGGAAG |
| 32399_3 | TATGAGTGTGGAGTCGTTGCTGCTCAGTTGTTAGTGGCTTATGNNNAGATCGGAAG |
| 32399_4 | TATGAGTGTGGAGTCGTTGCTGCTCAGTTGTTAGTGGCTTATGNNNAGATCGGAAG |
| 34437_1 | TATGAGTGTGGAGTCGTTGCGAAGTAGAGACTGCGCATGTAGANNNAGATCGGAA |
| 34437_2 | TATGAGTGTGGAGTCGTTGCGAAGTAGAGACTGCGCATGTAGANNNAGATCGGAA |
| 34437_3 | TATGAGTGTGGAGTCGTTGCGAAGTAGAGACTGCGCATGTAGANNNAGATCGGAA |
| 34437_4 | TATGAGTGTGGAGTCGTTGCGAAGTAGAGACTGCGCATGTAGANNNAGATCGGAA |
| 47789_1 | TATGAGTGTGGAGTCGTTGCTCTTTGCTCACCATCCTCAAACTNNNAGATCGGAAGA |
| 47789_2 | TATGAGTGTGGAGTCGTTGCTCTTTGCTCACCATCCTCAAACTNNNAGATCGGAAGA |
| 47789_3 | TATGAGTGTGGAGTCGTTGCTCTTTGCTCACCATCCTCAAACTNNNAGATCGGAAGA |
| 47789_4 | TATGAGTGTGGAGTCGTTGCTCTTTGCTCACCATCCTCAAACTNNNAGATCGGAAGA |
| 38363_1 | TATGAGTGTGGAGTCGTTGCAATTTCAGGCCAACCAATATGCCNNNAGATCGGAAG |
| 38363_2 | TATGAGTGTGGAGTCGTTGCAATTTCAGGCCAACCAATATGCCNNNAGATCGGAAG |
| 38363_3 | TATGAGTGTGGAGTCGTTGCAATTTCAGGCCAACCAATATGCCNNNAGATCGGAAG |
| 38363_4 | TATGAGTGTGGAGTCGTTGCAATTTCAGGCCAACCAATATGCCNNNAGATCGGAAG |
| 13163_1 | TATGAGTGTGGAGTCGTTGCTATTTCCACTAAACCTAGTCTTTNNNAGATCGGAAGA |
| 13163_2 | TATGAGTGTGGAGTCGTTGCTATTTCCACTAAACCTAGTCTTTNNNAGATCGGAAGA |
| 13163_3 | TATGAGTGTGGAGTCGTTGCTATTTCCACTAAACCTAGTCTTTNNNAGATCGGAAGA |
| 13163_4 | TATGAGTGTGGAGTCGTTGCTATTTCCACTAAACCTAGTCTTTNNNAGATCGGAAGA |
| 47305_1 | TATGAGTGTGGAGTCGTTGCTTTTGGACAAGTCTAAGGCTCCANNNAGATCGGAAG |
| 47305_2 | TATGAGTGTGGAGTCGTTGCTTTTGGACAAGTCTAAGGCTCCANNNAGATCGGAAG |
| 47305_3 | TATGAGTGTGGAGTCGTTGCTTTTGGACAAGTCTAAGGCTCCANNNAGATCGGAAG |

| | |
|---|---|
| 47305_4 | TATGAGTGTGGAGTCGTTGCTTTTGGACAAGTCTAAGGCTCCANNNAGATCGGAAG |
| 34053_1 | TATGAGTGTGGAGTCGTTGCCTCCTGAGTAGCTGGGATTACAGNNNAGATCGGAA |
| 34053_2 | TATGAGTGTGGAGTCGTTGCCTCCTGAGTAGCTGGGATTACAGNNNAGATCGGAA |
| 34053_3 | TATGAGTGTGGAGTCGTTGCCTCCTGAGTAGCTGGGATTACAGNNNAGATCGGAA |
| 34053_4 | TATGAGTGTGGAGTCGTTGCCTCCTGAGTAGCTGGGATTACAGNNNAGATCGGAA |
| 47545_1 | TATGAGTGTGGAGTCGTTGCGTCCTCAGGAGCTGGGTATTTAGNNNAGATCGGAA |
| 47545_2 | TATGAGTGTGGAGTCGTTGCGTCCTCAGGAGCTGGGTATTTAGNNNAGATCGGAA |
| 47545_3 | TATGAGTGTGGAGTCGTTGCGTCCTCAGGAGCTGGGTATTTAGNNNAGATCGGAA |
| 47545_4 | TATGAGTGTGGAGTCGTTGCGTCCTCAGGAGCTGGGTATTTAGNNNAGATCGGAA |
| 30661_1 | TATGAGTGTGGAGTCGTTGCTAACCAAGTGGTGGTACTGGGAANNNAGATCGGAA |
| 30661_2 | TATGAGTGTGGAGTCGTTGCTAACCAAGTGGTGGTACTGGGAANNNAGATCGGAA |
| 30661_3 | TATGAGTGTGGAGTCGTTGCTAACCAAGTGGTGGTACTGGGAANNNAGATCGGAA |
| 30661_4 | TATGAGTGTGGAGTCGTTGCTAACCAAGTGGTGGTACTGGGAANNNAGATCGGAA |
| 52846_1 | TATGAGTGTGGAGTCGTTGCGAGATTGGAGAAGTTGGTCCCTGNNNAGATCGGAA |
| 52846_2 | TATGAGTGTGGAGTCGTTGCGAGATTGGAGAAGTTGGTCCCTGNNNAGATCGGAA |
| 52846_3 | TATGAGTGTGGAGTCGTTGCGAGATTGGAGAAGTTGGTCCCTGNNNAGATCGGAA |
| 52846_4 | TATGAGTGTGGAGTCGTTGCGAGATTGGAGAAGTTGGTCCCTGNNNAGATCGGAA |
| 44464_1 | TATGAGTGTGGAGTCGTTGCATATGGGAGAGGGGATAGCCAGNNNAGATCGGAA |
| 44464_2 | TATGAGTGTGGAGTCGTTGCATATGGGAGAGGGGATAGCCAGNNNAGATCGGAA |
| 44464_3 | TATGAGTGTGGAGTCGTTGCATATGGGAGAGGGGATAGCCAGNNNAGATCGGAA |
| 44464_4 | TATGAGTGTGGAGTCGTTGCATATGGGAGAGGGGATAGCCAGNNNAGATCGGAA |
| 8778_1 | TATGAGTGTGGAGTCGTTGCCCTGGAGCCTGTGAATGTTTCTTNNNAGATCGGAAG |
| 8778_2 | TATGAGTGTGGAGTCGTTGCCCTGGAGCCTGTGAATGTTTCTTNNNAGATCGGAAG |
| 8778_3 | TATGAGTGTGGAGTCGTTGCCCTGGAGCCTGTGAATGTTTCTTNNNAGATCGGAAG |
| 8778_4 | TATGAGTGTGGAGTCGTTGCCCTGGAGCCTGTGAATGTTTCTTNNNAGATCGGAAG |
| 31818_1 | TATGAGTGTGGAGTCGTTGCCAGTGGGGTAATTATGGCTCACTNNNAGATCGGAA |
| 31818_2 | TATGAGTGTGGAGTCGTTGCCAGTGGGGTAATTATGGCTCACTNNNAGATCGGAA |
| 31818_3 | TATGAGTGTGGAGTCGTTGCCAGTGGGGTAATTATGGCTCACTNNNAGATCGGAA |
| 31818_4 | TATGAGTGTGGAGTCGTTGCCAGTGGGGTAATTATGGCTCACTNNNAGATCGGAA |
| 52510_1 | TATGAGTGTGGAGTCGTTGCAAAGAGCTTTCCTCCAGCCTTCNNNAGATCGGAAGA |
| 52510_2 | TATGAGTGTGGAGTCGTTGCAAAGAGCTTTCCTCCAGCCTTCNNNAGATCGGAAGA |
| 52510_3 | TATGAGTGTGGAGTCGTTGCAAAGAGCTTTCCTCCAGCCTTCNNNAGATCGGAAGA |
| 52510_4 | TATGAGTGTGGAGTCGTTGCAAAGAGCTTTCCTCCAGCCTTCNNNAGATCGGAAGA |
| 41893_1 | TATGAGTGTGGAGTCGTTGCAACTTCCACTCCAAGGGTTCAAGNNNAGATCGGAAG |
| 41893_2 | TATGAGTGTGGAGTCGTTGCAACTTCCACTCCAAGGGTTCAAGNNNAGATCGGAAG |
| 41893_3 | TATGAGTGTGGAGTCGTTGCAACTTCCACTCCAAGGGTTCAAGNNNAGATCGGAAG |
| 41893_4 | TATGAGTGTGGAGTCGTTGCAACTTCCACTCCAAGGGTTCAAGNNNAGATCGGAAG |
| 52341_1 | TATGAGTGTGGAGTCGTTGCGCATCTGCTGTATATGTCGATGANNNAGATCGGAAG |
| 52341_2 | TATGAGTGTGGAGTCGTTGCGCATCTGCTGTATATGTCGATGANNNAGATCGGAAG |
| 52341_3 | TATGAGTGTGGAGTCGTTGCGCATCTGCTGTATATGTCGATGANNNAGATCGGAAG |
| 52341_4 | TATGAGTGTGGAGTCGTTGCGCATCTGCTGTATATGTCGATGANNNAGATCGGAAG |
| 36942_1 | TATGAGTGTGGAGTCGTTGCTAGATTAGGGACACTCAACCTGTNNNAGATCGGAAG |
| 36942_2 | TATGAGTGTGGAGTCGTTGCTAGATTAGGGACACTCAACCTGTNNNAGATCGGAAG |

| | |
|---|---|
| 36942_3 | TATGAGTGTGGAGTCGTTGCTAGATTAGGGACACTCAACCTGTNNNAGATCGGAAG |
| 36942_4 | TATGAGTGTGGAGTCGTTGCTAGATTAGGGACACTCAACCTGTNNNAGATCGGAAG |
| 30709_1 | TATGAGTGTGGAGTCGTTGCTCCCTGTTTTACTGACTGGTTGGNNNAGATCGGAAG |
| 30709_2 | TATGAGTGTGGAGTCGTTGCTCCCTGTTTTACTGACTGGTTGGNNNAGATCGGAAG |
| 30709_3 | TATGAGTGTGGAGTCGTTGCTCCCTGTTTTACTGACTGGTTGGNNNAGATCGGAAG |
| 30709_4 | TATGAGTGTGGAGTCGTTGCTCCCTGTTTTACTGACTGGTTGGNNNAGATCGGAAG |
| 49542_1 | TATGAGTGTGGAGTCGTTGCGCACACCCGGCTAATTTTTGTATNNNAGATCGGAAG |
| 49542_2 | TATGAGTGTGGAGTCGTTGCGCACACCCGGCTAATTTTTGTATNNNAGATCGGAAG |
| 49542_3 | TATGAGTGTGGAGTCGTTGCGCACACCCGGCTAATTTTTGTATNNNAGATCGGAAG |
| 49542_4 | TATGAGTGTGGAGTCGTTGCGCACACCCGGCTAATTTTTGTATNNNAGATCGGAAG |
| 13631_1 | TATGAGTGTGGAGTCGTTGCTTGTACTAAAGACCTGACAACNNNAGATCGGAAGAG |
| 13631_2 | TATGAGTGTGGAGTCGTTGCTTGTACTAAAGACCTGACAACNNNAGATCGGAAGAG |
| 13631_3 | TATGAGTGTGGAGTCGTTGCTTGTACTAAAGACCTGACAACNNNAGATCGGAAGAG |
| 13631_4 | TATGAGTGTGGAGTCGTTGCTTGTACTAAAGACCTGACAACNNNAGATCGGAAGAG |
| 40517_1 | TATGAGTGTGGAGTCGTTGCAGTAAGCCAAGTAGGTGAGGAAANNNAGATCGGA |
| 40517_2 | TATGAGTGTGGAGTCGTTGCAGTAAGCCAAGTAGGTGAGGAAANNNAGATCGGA |
| 40517_3 | TATGAGTGTGGAGTCGTTGCAGTAAGCCAAGTAGGTGAGGAAANNNAGATCGGA |
| 40517_4 | TATGAGTGTGGAGTCGTTGCAGTAAGCCAAGTAGGTGAGGAAANNNAGATCGGA |
| 36990_1 | TATGAGTGTGGAGTCGTTGCGCATGAGAATCACTTGAACCCAGNNNAGATCGGAAG |
| 36990_2 | TATGAGTGTGGAGTCGTTGCGCATGAGAATCACTTGAACCCAGNNNAGATCGGAAG |
| 36990_3 | TATGAGTGTGGAGTCGTTGCGCATGAGAATCACTTGAACCCAGNNNAGATCGGAAG |
| 36990_4 | TATGAGTGTGGAGTCGTTGCGCATGAGAATCACTTGAACCCAGNNNAGATCGGAAG |
| 2666_1 | TATGAGTGTGGAGTCGTTGCGGTGGTTTCAGAGCTGACTGAATNNNAGATCGGAA |
| 2666_2 | TATGAGTGTGGAGTCGTTGCGGTGGTTTCAGAGCTGACTGAATNNNAGATCGGAA |
| 2666_3 | TATGAGTGTGGAGTCGTTGCGGTGGTTTCAGAGCTGACTGAATNNNAGATCGGAA |
| 2666_4 | TATGAGTGTGGAGTCGTTGCGGTGGTTTCAGAGCTGACTGAATNNNAGATCGGAA |
| 33296_1 | TATGAGTGTGGAGTCGTTGCTAGGACAATAGTGGAGGGAAGGTNNNAGATCGGA |
| 33296_2 | TATGAGTGTGGAGTCGTTGCTAGGACAATAGTGGAGGGAAGGTNNNAGATCGGA |
| 33296_3 | TATGAGTGTGGAGTCGTTGCTAGGACAATAGTGGAGGGAAGGTNNNAGATCGGA |
| 33296_4 | TATGAGTGTGGAGTCGTTGCTAGGACAATAGTGGAGGGAAGGTNNNAGATCGGA |
| 46716_1 | TATGAGTGTGGAGTCGTTGCTAGGTTCAAGTGATTCTCCTGCCNNNAGATCGGAAG |
| 46716_2 | TATGAGTGTGGAGTCGTTGCTAGGTTCAAGTGATTCTCCTGCCNNNAGATCGGAAG |
| 46716_3 | TATGAGTGTGGAGTCGTTGCTAGGTTCAAGTGATTCTCCTGCCNNNAGATCGGAAG |
| 46716_4 | TATGAGTGTGGAGTCGTTGCTAGGTTCAAGTGATTCTCCTGCCNNNAGATCGGAAG |
| 31226_1 | TATGAGTGTGGAGTCGTTGCGCCAAATCACTTGAAAGCCTTCTNNNAGATCGGAAG |
| 31226_2 | TATGAGTGTGGAGTCGTTGCGCCAAATCACTTGAAAGCCTTCTNNNAGATCGGAAG |
| 31226_3 | TATGAGTGTGGAGTCGTTGCGCCAAATCACTTGAAAGCCTTCTNNNAGATCGGAAG |
| 31226_4 | TATGAGTGTGGAGTCGTTGCGCCAAATCACTTGAAAGCCTTCTNNNAGATCGGAAG |
| 47268_1 | TATGAGTGTGGAGTCGTTGCCAATGGCACAATCTTGGCTCACNNNAGATCGGAAGA |
| 47268_2 | TATGAGTGTGGAGTCGTTGCCAATGGCACAATCTTGGCTCACNNNAGATCGGAAGA |
| 47268_3 | TATGAGTGTGGAGTCGTTGCCAATGGCACAATCTTGGCTCACNNNAGATCGGAAGA |
| 47268_4 | TATGAGTGTGGAGTCGTTGCCAATGGCACAATCTTGGCTCACNNNAGATCGGAAGA |
| 36372_1 | TATGAGTGTGGAGTCGTTGCCTTGAATGTAGAGCCCAGGTCTTNNNAGATCGGAAG |

| | |
|---|---|
| 36372_2 | TATGAGTGTGGAGTCGTTGCCTTGAATGTAGAGCCCAGGTCTTNNNAGATCGGAAG |
| 36372_3 | TATGAGTGTGGAGTCGTTGCCTTGAATGTAGAGCCCAGGTCTTNNNAGATCGGAAG |
| 36372_4 | TATGAGTGTGGAGTCGTTGCCTTGAATGTAGAGCCCAGGTCTTNNNAGATCGGAAG |
| 49436_1 | TATGAGTGTGGAGTCGTTGCCCAAAGTGCTGGAATTACAGGTGNNNAGATCGGAA |
| 49436_2 | TATGAGTGTGGAGTCGTTGCCCAAAGTGCTGGAATTACAGGTGNNNAGATCGGAA |
| 49436_3 | TATGAGTGTGGAGTCGTTGCCCAAAGTGCTGGAATTACAGGTGNNNAGATCGGAA |
| 49436_4 | TATGAGTGTGGAGTCGTTGCCCAAAGTGCTGGAATTACAGGTGNNNAGATCGGAA |
| 31689_1 | TATGAGTGTGGAGTCGTTGCGTTGGCAGTGACCTCCATAAGTANNNAGATCGGAAG |
| 31689_2 | TATGAGTGTGGAGTCGTTGCGTTGGCAGTGACCTCCATAAGTANNNAGATCGGAAG |
| 31689_3 | TATGAGTGTGGAGTCGTTGCGTTGGCAGTGACCTCCATAAGTANNNAGATCGGAAG |
| 31689_4 | TATGAGTGTGGAGTCGTTGCGTTGGCAGTGACCTCCATAAGTANNNAGATCGGAAG |
| 35689_1 | TATGAGTGTGGAGTCGTTGCGCGATGGTATGCAATCTCAGTTCNNNAGATCGGAAG |
| 35689_2 | TATGAGTGTGGAGTCGTTGCGCGATGGTATGCAATCTCAGTTCNNNAGATCGGAAG |
| 35689_3 | TATGAGTGTGGAGTCGTTGCGCGATGGTATGCAATCTCAGTTCNNNAGATCGGAAG |
| 35689_4 | TATGAGTGTGGAGTCGTTGCGCGATGGTATGCAATCTCAGTTCNNNAGATCGGAAG |
| 49915_1 | TATGAGTGTGGAGTCGTTGCAAATGAGGAACTTGAGGTCCCAANNNAGATCGGAA |
| 49915_2 | TATGAGTGTGGAGTCGTTGCAAATGAGGAACTTGAGGTCCCAANNNAGATCGGAA |
| 49915_3 | TATGAGTGTGGAGTCGTTGCAAATGAGGAACTTGAGGTCCCAANNNAGATCGGAA |
| 49915_4 | TATGAGTGTGGAGTCGTTGCAAATGAGGAACTTGAGGTCCCAANNNAGATCGGAA |
| 48232_1 | TATGAGTGTGGAGTCGTTGCCGTGAGTTTTTGTCATGAAGGGANNNAGATCGGAA |
| 48232_2 | TATGAGTGTGGAGTCGTTGCCGTGAGTTTTTGTCATGAAGGGANNNAGATCGGAA |
| 48232_3 | TATGAGTGTGGAGTCGTTGCCGTGAGTTTTTGTCATGAAGGGANNNAGATCGGAA |
| 48232_4 | TATGAGTGTGGAGTCGTTGCCGTGAGTTTTTGTCATGAAGGGANNNAGATCGGAA |
| 31460_1 | TATGAGTGTGGAGTCGTTGCTGATACAGTATTTCTGAGCCCTCANNNAGATCGGAA |
| 31460_2 | TATGAGTGTGGAGTCGTTGCTGATACAGTATTTCTGAGCCCTCANNNAGATCGGAA |
| 31460_3 | TATGAGTGTGGAGTCGTTGCTGATACAGTATTTCTGAGCCCTCANNNAGATCGGAA |
| 31460_4 | TATGAGTGTGGAGTCGTTGCTGATACAGTATTTCTGAGCCCTCANNNAGATCGGAA |
| 45361_1 | TATGAGTGTGGAGTCGTTGCAGAGGCACTTAGGTCAATTCTGTNNNAGATCGGAAG |
| 45361_2 | TATGAGTGTGGAGTCGTTGCAGAGGCACTTAGGTCAATTCTGTNNNAGATCGGAAG |
| 45361_3 | TATGAGTGTGGAGTCGTTGCAGAGGCACTTAGGTCAATTCTGTNNNAGATCGGAAG |
| 45361_4 | TATGAGTGTGGAGTCGTTGCAGAGGCACTTAGGTCAATTCTGTNNNAGATCGGAAG |
| 36160_1 | TATGAGTGTGGAGTCGTTGCAAACCTTGGGGGAAATACAGAGGNNNAGATCGGA |
| 36160_2 | TATGAGTGTGGAGTCGTTGCAAACCTTGGGGGAAATACAGAGGNNNAGATCGGA |
| 36160_3 | TATGAGTGTGGAGTCGTTGCAAACCTTGGGGGAAATACAGAGGNNNAGATCGGA |
| 36160_4 | TATGAGTGTGGAGTCGTTGCAAACCTTGGGGGAAATACAGAGGNNNAGATCGGA |
| 42511_1 | TATGAGTGTGGAGTCGTTGCCTCCCAGGTTCAAGTGCTTCTCNNNAGATCGGAAGA |
| 42511_2 | TATGAGTGTGGAGTCGTTGCCTCCCAGGTTCAAGTGCTTCTCNNNAGATCGGAAGA |
| 42511_3 | TATGAGTGTGGAGTCGTTGCCTCCCAGGTTCAAGTGCTTCTCNNNAGATCGGAAGA |
| 42511_4 | TATGAGTGTGGAGTCGTTGCCTCCCAGGTTCAAGTGCTTCTCNNNAGATCGGAAGA |
| 34959_1 | TATGAGTGTGGAGTCGTTGCGGAGAGATGGCTCTGATTTTTGCNNNAGATCGGAA |
| 34959_2 | TATGAGTGTGGAGTCGTTGCGGAGAGATGGCTCTGATTTTTGCNNNAGATCGGAA |
| 34959_3 | TATGAGTGTGGAGTCGTTGCGGAGAGATGGCTCTGATTTTTGCNNNAGATCGGAA |
| 34959_4 | TATGAGTGTGGAGTCGTTGCGGAGAGATGGCTCTGATTTTTGCNNNAGATCGGAA |

| | |
|---|---|
| 38903_1 | TATGAGTGTGGAGTCGTTGCACCCCCCAAAAATCACACTAGTTNNNAGATCGGAAG |
| 38903_2 | TATGAGTGTGGAGTCGTTGCACCCCCCAAAAATCACACTAGTTNNNAGATCGGAAG |
| 38903_3 | TATGAGTGTGGAGTCGTTGCACCCCCCAAAAATCACACTAGTTNNNAGATCGGAAG |
| 38903_4 | TATGAGTGTGGAGTCGTTGCACCCCCCAAAAATCACACTAGTTNNNAGATCGGAAG |
| 50428_1 | TATGAGTGTGGAGTCGTTGCCTTTCTCCAGCTCCGTGTCATTNNNAGATCGGAAGAG |
| 50428_2 | TATGAGTGTGGAGTCGTTGCCTTTCTCCAGCTCCGTGTCATTNNNAGATCGGAAGAG |
| 50428_3 | TATGAGTGTGGAGTCGTTGCCTTTCTCCAGCTCCGTGTCATTNNNAGATCGGAAGAG |
| 50428_4 | TATGAGTGTGGAGTCGTTGCCTTTCTCCAGCTCCGTGTCATTNNNAGATCGGAAGAG |
| 38750_1 | TATGAGTGTGGAGTCGTTGCTCACAGCACTATTCACAGTAGCANNNAGATCGGAAG |
| 38750_2 | TATGAGTGTGGAGTCGTTGCTCACAGCACTATTCACAGTAGCANNNAGATCGGAAG |
| 38750_3 | TATGAGTGTGGAGTCGTTGCTCACAGCACTATTCACAGTAGCANNNAGATCGGAAG |
| 38750_4 | TATGAGTGTGGAGTCGTTGCTCACAGCACTATTCACAGTAGCANNNAGATCGGAAG |
| 29076_1 | TATGAGTGTGGAGTCGTTGCCATGTGTGTGTGTGTGTGTGTATNNNAGATCGGAA |
| 29076_2 | TATGAGTGTGGAGTCGTTGCCATGTGTGTGTGTGTGTGTGTATNNNAGATCGGAA |
| 29076_3 | TATGAGTGTGGAGTCGTTGCCATGTGTGTGTGTGTGTGTGTATNNNAGATCGGAA |
| 29076_4 | TATGAGTGTGGAGTCGTTGCCATGTGTGTGTGTGTGTGTGTATNNNAGATCGGAA |
| 12648_1 | TATGAGTGTGGAGTCGTTGCATCTATTAGTACACGAGTAACTCNNNAGATCGGAAG |
| 12648_2 | TATGAGTGTGGAGTCGTTGCATCTATTAGTACACGAGTAACTCNNNAGATCGGAAG |
| 12648_3 | TATGAGTGTGGAGTCGTTGCATCTATTAGTACACGAGTAACTCNNNAGATCGGAAG |
| 12648_4 | TATGAGTGTGGAGTCGTTGCATCTATTAGTACACGAGTAACTCNNNAGATCGGAAG |
| 40983_1 | TATGAGTGTGGAGTCGTTGCCCTTGCCATTGAAGAGCTTAGTGNNNAGATCGGAAG |
| 40983_2 | TATGAGTGTGGAGTCGTTGCCCTTGCCATTGAAGAGCTTAGTGNNNAGATCGGAAG |
| 40983_3 | TATGAGTGTGGAGTCGTTGCCCTTGCCATTGAAGAGCTTAGTGNNNAGATCGGAAG |
| 40983_4 | TATGAGTGTGGAGTCGTTGCCCTTGCCATTGAAGAGCTTAGTGNNNAGATCGGAAG |
| 33178_1 | TATGAGTGTGGAGTCGTTGCCCATAGTGAATGTTCCCTGCAAGNNNAGATCGGAAG |
| 33178_2 | TATGAGTGTGGAGTCGTTGCCCATAGTGAATGTTCCCTGCAAGNNNAGATCGGAAG |
| 33178_3 | TATGAGTGTGGAGTCGTTGCCCATAGTGAATGTTCCCTGCAAGNNNAGATCGGAAG |
| 33178_4 | TATGAGTGTGGAGTCGTTGCCCATAGTGAATGTTCCCTGCAAGNNNAGATCGGAAG |
| 51344_1 | TATGAGTGTGGAGTCGTTGCCAAGTCTCTTCCACTATCCACCCNNNAGATCGGAAGA |
| 51344_2 | TATGAGTGTGGAGTCGTTGCCAAGTCTCTTCCACTATCCACCCNNNAGATCGGAAGA |
| 51344_3 | TATGAGTGTGGAGTCGTTGCCAAGTCTCTTCCACTATCCACCCNNNAGATCGGAAGA |
| 51344_4 | TATGAGTGTGGAGTCGTTGCCAAGTCTCTTCCACTATCCACCCNNNAGATCGGAAGA |
| 40701_1 | TATGAGTGTGGAGTCGTTGCCACCAAGCCCATGAGTACATAGANNNAGATCGGAAG |
| 40701_2 | TATGAGTGTGGAGTCGTTGCCACCAAGCCCATGAGTACATAGANNNAGATCGGAAG |
| 40701_3 | TATGAGTGTGGAGTCGTTGCCACCAAGCCCATGAGTACATAGANNNAGATCGGAAG |
| 40701_4 | TATGAGTGTGGAGTCGTTGCCACCAAGCCCATGAGTACATAGANNNAGATCGGAAG |
| 38558_1 | TATGAGTGTGGAGTCGTTGCCAGGAGAATCGCTGGAACCTGNNNAGATCGGAAGA |
| 38558_2 | TATGAGTGTGGAGTCGTTGCCAGGAGAATCGCTGGAACCTGNNNAGATCGGAAGA |
| 38558_3 | TATGAGTGTGGAGTCGTTGCCAGGAGAATCGCTGGAACCTGNNNAGATCGGAAGA |
| 38558_4 | TATGAGTGTGGAGTCGTTGCCAGGAGAATCGCTGGAACCTGNNNAGATCGGAAGA |
| 33590_1 | TATGAGTGTGGAGTCGTTGCCATGATCTTGGCTCACTGCAACNNNAGATCGGAAGA |
| 33590_2 | TATGAGTGTGGAGTCGTTGCCATGATCTTGGCTCACTGCAACNNNAGATCGGAAGA |
| 33590_3 | TATGAGTGTGGAGTCGTTGCCATGATCTTGGCTCACTGCAACNNNAGATCGGAAGA |

| | |
|---|---|
| 33590_4 | TATGAGTGTGGAGTCGTTGCCATGATCTTGGCTCACTGCAACNNNAGATCGGAAGA |
| 48658_1 | TATGAGTGTGGAGTCGTTGCTGCTTTCAAGGTTTATTCTGGCANNNAGATCGGAAG |
| 48658_2 | TATGAGTGTGGAGTCGTTGCTGCTTTCAAGGTTTATTCTGGCANNNAGATCGGAAG |
| 48658_3 | TATGAGTGTGGAGTCGTTGCTGCTTTCAAGGTTTATTCTGGCANNNAGATCGGAAG |
| 48658_4 | TATGAGTGTGGAGTCGTTGCTGCTTTCAAGGTTTATTCTGGCANNNAGATCGGAAG |
| 51925_1 | TATGAGTGTGGAGTCGTTGCCTGAATGGATCCCTTCTTGACCANNNAGATCGGAAG |
| 51925_2 | TATGAGTGTGGAGTCGTTGCCTGAATGGATCCCTTCTTGACCANNNAGATCGGAAG |
| 51925_3 | TATGAGTGTGGAGTCGTTGCCTGAATGGATCCCTTCTTGACCANNNAGATCGGAAG |
| 51925_4 | TATGAGTGTGGAGTCGTTGCCTGAATGGATCCCTTCTTGACCANNNAGATCGGAAG |
| 39142_1 | TATGAGTGTGGAGTCGTTGCCGATCTCAGCTACTGCAACTTCTNNNAGATCGGAAG |
| 39142_2 | TATGAGTGTGGAGTCGTTGCCGATCTCAGCTACTGCAACTTCTNNNAGATCGGAAG |
| 39142_3 | TATGAGTGTGGAGTCGTTGCCGATCTCAGCTACTGCAACTTCTNNNAGATCGGAAG |
| 39142_4 | TATGAGTGTGGAGTCGTTGCCGATCTCAGCTACTGCAACTTCTNNNAGATCGGAAG |
| 36094_1 | TATGAGTGTGGAGTCGTTGCCTCCCTCTGCTGAAAACATTTGGNNNAGATCGGAAG |
| 36094_2 | TATGAGTGTGGAGTCGTTGCCTCCCTCTGCTGAAAACATTTGGNNNAGATCGGAAG |
| 36094_3 | TATGAGTGTGGAGTCGTTGCCTCCCTCTGCTGAAAACATTTGGNNNAGATCGGAAG |
| 36094_4 | TATGAGTGTGGAGTCGTTGCCTCCCTCTGCTGAAAACATTTGGNNNAGATCGGAAG |
| 43747_1 | TATGAGTGTGGAGTCGTTGCTGCTCATCAGAACTACCATCTCGNNNAGATCGGAAG |
| 43747_2 | TATGAGTGTGGAGTCGTTGCTGCTCATCAGAACTACCATCTCGNNNAGATCGGAAG |
| 43747_3 | TATGAGTGTGGAGTCGTTGCTGCTCATCAGAACTACCATCTCGNNNAGATCGGAAG |
| 43747_4 | TATGAGTGTGGAGTCGTTGCTGCTCATCAGAACTACCATCTCGNNNAGATCGGAAG |
| 36751_1 | TATGAGTGTGGAGTCGTTGCGAGAGAGCCCATGTACCCTTTACNNNAGATCGGAAG |
| 36751_2 | TATGAGTGTGGAGTCGTTGCGAGAGAGCCCATGTACCCTTTACNNNAGATCGGAAG |
| 36751_3 | TATGAGTGTGGAGTCGTTGCGAGAGAGCCCATGTACCCTTTACNNNAGATCGGAAG |
| 36751_4 | TATGAGTGTGGAGTCGTTGCGAGAGAGCCCATGTACCCTTTACNNNAGATCGGAAG |
| 29004_1 | TATGAGTGTGGAGTCGTTGCCACACACATACACACACACACTCNNNAGATCGGAAG |
| 29004_2 | TATGAGTGTGGAGTCGTTGCCACACACATACACACACACACTCNNNAGATCGGAAG |
| 29004_3 | TATGAGTGTGGAGTCGTTGCCACACACATACACACACACACTCNNNAGATCGGAAG |
| 29004_4 | TATGAGTGTGGAGTCGTTGCCACACACATACACACACACACTCNNNAGATCGGAAG |
| 36823_1 | TATGAGTGTGGAGTCGTTGCGTAGGGCAGAAAAACCTATGCACNNNAGATCGGAA |
| 36823_2 | TATGAGTGTGGAGTCGTTGCGTAGGGCAGAAAAACCTATGCACNNNAGATCGGAA |
| 36823_3 | TATGAGTGTGGAGTCGTTGCGTAGGGCAGAAAAACCTATGCACNNNAGATCGGAA |
| 36823_4 | TATGAGTGTGGAGTCGTTGCGTAGGGCAGAAAAACCTATGCACNNNAGATCGGAA |
| 34947_1 | TATGAGTGTGGAGTCGTTGCATCTCCCAGGTTTGAGCAATTCTNNNAGATCGGAAG |
| 34947_2 | TATGAGTGTGGAGTCGTTGCATCTCCCAGGTTTGAGCAATTCTNNNAGATCGGAAG |
| 34947_3 | TATGAGTGTGGAGTCGTTGCATCTCCCAGGTTTGAGCAATTCTNNNAGATCGGAAG |
| 34947_4 | TATGAGTGTGGAGTCGTTGCATCTCCCAGGTTTGAGCAATTCTNNNAGATCGGAAG |
| 29744_1 | TATGAGTGTGGAGTCGTTGCTTGTTCCCTTAGGTCTTGGGCTANNNAGATCGGAAG |
| 29744_2 | TATGAGTGTGGAGTCGTTGCTTGTTCCCTTAGGTCTTGGGCTANNNAGATCGGAAG |
| 29744_3 | TATGAGTGTGGAGTCGTTGCTTGTTCCCTTAGGTCTTGGGCTANNNAGATCGGAAG |
| 29744_4 | TATGAGTGTGGAGTCGTTGCTTGTTCCCTTAGGTCTTGGGCTANNNAGATCGGAAG |
| 46661_1 | TATGAGTGTGGAGTCGTTGCTTCAAGTGATTCTCCTGCCTCAGNNNAGATCGGAAG |
| 46661_2 | TATGAGTGTGGAGTCGTTGCTTCAAGTGATTCTCCTGCCTCAGNNNAGATCGGAAG |

| | |
|---|---|
| 46661_3 | TATGAGTGTGGAGTCGTTGCTTCAAGTGATTCTCCTGCCTCAGNNNAGATCGGAAG |
| 46661_4 | TATGAGTGTGGAGTCGTTGCTTCAAGTGATTCTCCTGCCTCAGNNNAGATCGGAAG |
| 41324_1 | TATGAGTGTGGAGTCGTTGCCCATAGTGAAAGTTCCCTGCAAGNNNAGATCGGAAG |
| 41324_2 | TATGAGTGTGGAGTCGTTGCCCATAGTGAAAGTTCCCTGCAAGNNNAGATCGGAAG |
| 41324_3 | TATGAGTGTGGAGTCGTTGCCCATAGTGAAAGTTCCCTGCAAGNNNAGATCGGAAG |
| 41324_4 | TATGAGTGTGGAGTCGTTGCCCATAGTGAAAGTTCCCTGCAAGNNNAGATCGGAAG |
| 37673_1 | TATGAGTGTGGAGTCGTTGCAATATTGACTGCCTTTGGGGAGGNNNAGATCGGAA |
| 37673_2 | TATGAGTGTGGAGTCGTTGCAATATTGACTGCCTTTGGGGAGGNNNAGATCGGAA |
| 37673_3 | TATGAGTGTGGAGTCGTTGCAATATTGACTGCCTTTGGGGAGGNNNAGATCGGAA |
| 37673_4 | TATGAGTGTGGAGTCGTTGCAATATTGACTGCCTTTGGGGAGGNNNAGATCGGAA |
| 39391_1 | TATGAGTGTGGAGTCGTTGCGAGAATCGTTTGAACTCAGGAGCNNNAGATCGGAA |
| 39391_2 | TATGAGTGTGGAGTCGTTGCGAGAATCGTTTGAACTCAGGAGCNNNAGATCGGAA |
| 39391_3 | TATGAGTGTGGAGTCGTTGCGAGAATCGTTTGAACTCAGGAGCNNNAGATCGGAA |
| 39391_4 | TATGAGTGTGGAGTCGTTGCGAGAATCGTTTGAACTCAGGAGCNNNAGATCGGAA |
| 33368_1 | TATGAGTGTGGAGTCGTTGCCATTCTCCTGCCTCAGCCTCNNNAGATCGGAAGAGCA |
| 33368_2 | TATGAGTGTGGAGTCGTTGCCATTCTCCTGCCTCAGCCTCNNNAGATCGGAAGAGCA |
| 33368_3 | TATGAGTGTGGAGTCGTTGCCATTCTCCTGCCTCAGCCTCNNNAGATCGGAAGAGCA |
| 33368_4 | TATGAGTGTGGAGTCGTTGCCATTCTCCTGCCTCAGCCTCNNNAGATCGGAAGAGCA |
| 30879_1 | TATGAGTGTGGAGTCGTTGCGTGTGTGTGTGTGTTGGGATAAGNNNAGATCGGAA |
| 30879_2 | TATGAGTGTGGAGTCGTTGCGTGTGTGTGTGTGTTGGGATAAGNNNAGATCGGAA |
| 30879_3 | TATGAGTGTGGAGTCGTTGCGTGTGTGTGTGTGTTGGGATAAGNNNAGATCGGAA |
| 30879_4 | TATGAGTGTGGAGTCGTTGCGTGTGTGTGTGTGTTGGGATAAGNNNAGATCGGAA |
| 36326_1 | TATGAGTGTGGAGTCGTTGCAAAAATTTGTCAGGTGTGGTGGCNNNAGATCGGAA |
| 36326_2 | TATGAGTGTGGAGTCGTTGCAAAAATTTGTCAGGTGTGGTGGCNNNAGATCGGAA |
| 36326_3 | TATGAGTGTGGAGTCGTTGCAAAAATTTGTCAGGTGTGGTGGCNNNAGATCGGAA |
| 36326_4 | TATGAGTGTGGAGTCGTTGCAAAAATTTGTCAGGTGTGGTGGCNNNAGATCGGAA |
| 6445_1 | TATGAGTGTGGAGTCGTTGCTCACATCTTCACCTTTCAGGGTTNNNAGATCGGAAGA |
| 6445_2 | TATGAGTGTGGAGTCGTTGCTCACATCTTCACCTTTCAGGGTTNNNAGATCGGAAGA |
| 6445_3 | TATGAGTGTGGAGTCGTTGCTCACATCTTCACCTTTCAGGGTTNNNAGATCGGAAGA |
| 6445_4 | TATGAGTGTGGAGTCGTTGCTCACATCTTCACCTTTCAGGGTTNNNAGATCGGAAGA |
| 49525_1 | TATGAGTGTGGAGTCGTTGCAACAAAGAACATGGCTTCAAAGCNNNAGATCGGAA |
| 49525_2 | TATGAGTGTGGAGTCGTTGCAACAAAGAACATGGCTTCAAAGCNNNAGATCGGAA |
| 49525_3 | TATGAGTGTGGAGTCGTTGCAACAAAGAACATGGCTTCAAAGCNNNAGATCGGAA |
| 49525_4 | TATGAGTGTGGAGTCGTTGCAACAAAGAACATGGCTTCAAAGCNNNAGATCGGAA |
| 34455_1 | TATGAGTGTGGAGTCGTTGCCCCATCTCCTCCCCTCTCACNNNAGATCGGAAGAGCA |
| 34455_2 | TATGAGTGTGGAGTCGTTGCCCCATCTCCTCCCCTCTCACNNNAGATCGGAAGAGCA |
| 34455_3 | TATGAGTGTGGAGTCGTTGCCCCATCTCCTCCCCTCTCACNNNAGATCGGAAGAGCA |
| 34455_4 | TATGAGTGTGGAGTCGTTGCCCCATCTCCTCCCCTCTCACNNNAGATCGGAAGAGCA |
| 12679_1 | TATGAGTGTGGAGTCGTTGCCTGATACTTCTTGAAATACACACNNNAGATCGGAAG |
| 12679_2 | TATGAGTGTGGAGTCGTTGCCTGATACTTCTTGAAATACACACNNNAGATCGGAAG |
| 12679_3 | TATGAGTGTGGAGTCGTTGCCTGATACTTCTTGAAATACACACNNNAGATCGGAAG |
| 12679_4 | TATGAGTGTGGAGTCGTTGCCTGATACTTCTTGAAATACACACNNNAGATCGGAAG |
| 10005_1 | TATGAGTGTGGAGTCGTTGCTTCTGTGCTCAAGCAATCAGGTTNNNAGATCGGAAG |

| | |
|---|---|
| 10005_2 | TATGAGTGTGGAGTCGTTGCTTCTGTGCTCAAGCAATCAGGTTNNNAGATCGGAAG |
| 10005_3 | TATGAGTGTGGAGTCGTTGCTTCTGTGCTCAAGCAATCAGGTTNNNAGATCGGAAG |
| 10005_4 | TATGAGTGTGGAGTCGTTGCTTCTGTGCTCAAGCAATCAGGTTNNNAGATCGGAAG |
| 31027_1 | TATGAGTGTGGAGTCGTTGCGATGTTCTCAGTTTTGTCCTGGCNNNAGATCGGAAG |
| 31027_2 | TATGAGTGTGGAGTCGTTGCGATGTTCTCAGTTTTGTCCTGGCNNNAGATCGGAAG |
| 31027_3 | TATGAGTGTGGAGTCGTTGCGATGTTCTCAGTTTTGTCCTGGCNNNAGATCGGAAG |
| 31027_4 | TATGAGTGTGGAGTCGTTGCGATGTTCTCAGTTTTGTCCTGGCNNNAGATCGGAAG |
| 35224_1 | TATGAGTGTGGAGTCGTTGCATGTAGGGCCATCAATGTTCTGTNNNAGATCGGAAG |
| 35224_2 | TATGAGTGTGGAGTCGTTGCATGTAGGGCCATCAATGTTCTGTNNNAGATCGGAAG |
| 35224_3 | TATGAGTGTGGAGTCGTTGCATGTAGGGCCATCAATGTTCTGTNNNAGATCGGAAG |
| 35224_4 | TATGAGTGTGGAGTCGTTGCATGTAGGGCCATCAATGTTCTGTNNNAGATCGGAAG |
| 48333_1 | TATGAGTGTGGAGTCGTTGCGGCTTGCAAGTCTGAAATGATGTNNNAGATCGGAA |
| 48333_2 | TATGAGTGTGGAGTCGTTGCGGCTTGCAAGTCTGAAATGATGTNNNAGATCGGAA |
| 48333_3 | TATGAGTGTGGAGTCGTTGCGGCTTGCAAGTCTGAAATGATGTNNNAGATCGGAA |
| 48333_4 | TATGAGTGTGGAGTCGTTGCGGCTTGCAAGTCTGAAATGATGTNNNAGATCGGAA |
| 4948_1 | TATGAGTGTGGAGTCGTTGCCTGAACTTCCACAAGGAAGCACTNNNAGATCGGAAG |
| 4948_2 | TATGAGTGTGGAGTCGTTGCCTGAACTTCCACAAGGAAGCACTNNNAGATCGGAAG |
| 4948_3 | TATGAGTGTGGAGTCGTTGCCTGAACTTCCACAAGGAAGCACTNNNAGATCGGAAG |
| 4948_4 | TATGAGTGTGGAGTCGTTGCCTGAACTTCCACAAGGAAGCACTNNNAGATCGGAAG |
| 48938_1 | TATGAGTGTGGAGTCGTTGCTAGCCAGGGCAGTAGTATACAGANNNAGATCGGAA |
| 48938_2 | TATGAGTGTGGAGTCGTTGCTAGCCAGGGCAGTAGTATACAGANNNAGATCGGAA |
| 48938_3 | TATGAGTGTGGAGTCGTTGCTAGCCAGGGCAGTAGTATACAGANNNAGATCGGAA |
| 48938_4 | TATGAGTGTGGAGTCGTTGCTAGCCAGGGCAGTAGTATACAGANNNAGATCGGAA |
| 48538_1 | TATGAGTGTGGAGTCGTTGCCACGCCCAGCTAATTTTTGTACTNNNAGATCGGAAG |
| 48538_2 | TATGAGTGTGGAGTCGTTGCCACGCCCAGCTAATTTTTGTACTNNNAGATCGGAAG |
| 48538_3 | TATGAGTGTGGAGTCGTTGCCACGCCCAGCTAATTTTTGTACTNNNAGATCGGAAG |
| 48538_4 | TATGAGTGTGGAGTCGTTGCCACGCCCAGCTAATTTTTGTACTNNNAGATCGGAAG |
| 40258_1 | TATGAGTGTGGAGTCGTTGCTCAGAGAAGTGAAAATGAAAACCACANNNAGATCG |
| 40258_2 | TATGAGTGTGGAGTCGTTGCTCAGAGAAGTGAAAATGAAAACCACANNNAGATCG |
| 40258_3 | TATGAGTGTGGAGTCGTTGCTCAGAGAAGTGAAAATGAAAACCACANNNAGATCG |
| 40258_4 | TATGAGTGTGGAGTCGTTGCTCAGAGAAGTGAAAATGAAAACCACANNNAGATCG |
| 43522_1 | TATGAGTGTGGAGTCGTTGCGATCTTGGCTCACTGCAATCTCTNNNAGATCGGAAG |
| 43522_2 | TATGAGTGTGGAGTCGTTGCGATCTTGGCTCACTGCAATCTCTNNNAGATCGGAAG |
| 43522_3 | TATGAGTGTGGAGTCGTTGCGATCTTGGCTCACTGCAATCTCTNNNAGATCGGAAG |
| 43522_4 | TATGAGTGTGGAGTCGTTGCGATCTTGGCTCACTGCAATCTCTNNNAGATCGGAAG |
| 3140_1 | TATGAGTGTGGAGTCGTTGCGCGTTAGTGACAAGAGGGACATCNNNAGATCGGAA |
| 3140_2 | TATGAGTGTGGAGTCGTTGCGCGTTAGTGACAAGAGGGACATCNNNAGATCGGAA |
| 3140_3 | TATGAGTGTGGAGTCGTTGCGCGTTAGTGACAAGAGGGACATCNNNAGATCGGAA |
| 3140_4 | TATGAGTGTGGAGTCGTTGCGCGTTAGTGACAAGAGGGACATCNNNAGATCGGAA |
| 4110_1 | TATGAGTGTGGAGTCGTTGCTCAGGTCCCCATCCCAAACTAAANNNAGATCGGAAG |
| 4110_2 | TATGAGTGTGGAGTCGTTGCTCAGGTCCCCATCCCAAACTAAANNNAGATCGGAAG |
| 4110_3 | TATGAGTGTGGAGTCGTTGCTCAGGTCCCCATCCCAAACTAAANNNAGATCGGAAG |
| 4110_4 | TATGAGTGTGGAGTCGTTGCTCAGGTCCCCATCCCAAACTAAANNNAGATCGGAAG |

| | |
|---|---|
| 29584_1 | TATGAGTGTGGAGTCGTTGCTCTCTCATGTGCTGAGGTCGTTANNNAGATCGGAAG |
| 29584_2 | TATGAGTGTGGAGTCGTTGCTCTCTCATGTGCTGAGGTCGTTANNNAGATCGGAAG |
| 29584_3 | TATGAGTGTGGAGTCGTTGCTCTCTCATGTGCTGAGGTCGTTANNNAGATCGGAAG |
| 29584_4 | TATGAGTGTGGAGTCGTTGCTCTCTCATGTGCTGAGGTCGTTANNNAGATCGGAAG |
| 40649_1 | TATGAGTGTGGAGTCGTTGCGAGGGCCAGTAACATTATCAGCTNNNAGATCGGAA |
| 40649_2 | TATGAGTGTGGAGTCGTTGCGAGGGCCAGTAACATTATCAGCTNNNAGATCGGAA |
| 40649_3 | TATGAGTGTGGAGTCGTTGCGAGGGCCAGTAACATTATCAGCTNNNAGATCGGAA |
| 40649_4 | TATGAGTGTGGAGTCGTTGCGAGGGCCAGTAACATTATCAGCTNNNAGATCGGAA |
| 3123_1 | TATGAGTGTGGAGTCGTTGCCATCGGTTACTACAAACTGGCCANNNAGATCGGAAG |
| 3123_2 | TATGAGTGTGGAGTCGTTGCCATCGGTTACTACAAACTGGCCANNNAGATCGGAAG |
| 3123_3 | TATGAGTGTGGAGTCGTTGCCATCGGTTACTACAAACTGGCCANNNAGATCGGAAG |
| 3123_4 | TATGAGTGTGGAGTCGTTGCCATCGGTTACTACAAACTGGCCANNNAGATCGGAAG |
| 38185_1 | TATGAGTGTGGAGTCGTTGCGCATGAAACAATGCTTTTGAGGTNNNAGATCGGAA |
| 38185_2 | TATGAGTGTGGAGTCGTTGCGCATGAAACAATGCTTTTGAGGTNNNAGATCGGAA |
| 38185_3 | TATGAGTGTGGAGTCGTTGCGCATGAAACAATGCTTTTGAGGTNNNAGATCGGAA |
| 38185_4 | TATGAGTGTGGAGTCGTTGCGCATGAAACAATGCTTTTGAGGTNNNAGATCGGAA |
| 28910_1 | TATGAGTGTGGAGTCGTTGCACTGGGGGAAGAATTACATCCACNNNAGATCGGAA |
| 28910_2 | TATGAGTGTGGAGTCGTTGCACTGGGGGAAGAATTACATCCACNNNAGATCGGAA |
| 28910_3 | TATGAGTGTGGAGTCGTTGCACTGGGGGAAGAATTACATCCACNNNAGATCGGAA |
| 28910_4 | TATGAGTGTGGAGTCGTTGCACTGGGGGAAGAATTACATCCACNNNAGATCGGAA |
| 45508_1 | TATGAGTGTGGAGTCGTTGCATCTGAGAGTCCCATCCCTGAGNNNAGATCGGAAGA |
| 45508_2 | TATGAGTGTGGAGTCGTTGCATCTGAGAGTCCCATCCCTGAGNNNAGATCGGAAGA |
| 45508_3 | TATGAGTGTGGAGTCGTTGCATCTGAGAGTCCCATCCCTGAGNNNAGATCGGAAGA |
| 45508_4 | TATGAGTGTGGAGTCGTTGCATCTGAGAGTCCCATCCCTGAGNNNAGATCGGAAGA |
| 47511_1 | TATGAGTGTGGAGTCGTTGCTCAGTCTCTCGAGTAGGTAGGACNNNAGATCGGAA |
| 47511_2 | TATGAGTGTGGAGTCGTTGCTCAGTCTCTCGAGTAGGTAGGACNNNAGATCGGAA |
| 47511_3 | TATGAGTGTGGAGTCGTTGCTCAGTCTCTCGAGTAGGTAGGACNNNAGATCGGAA |
| 47511_4 | TATGAGTGTGGAGTCGTTGCTCAGTCTCTCGAGTAGGTAGGACNNNAGATCGGAA |
| 32834_1 | TATGAGTGTGGAGTCGTTGCAGATGGAGTCTCGCTTTGTCACNNNAGATCGGAAGA |
| 32834_2 | TATGAGTGTGGAGTCGTTGCAGATGGAGTCTCGCTTTGTCACNNNAGATCGGAAGA |
| 32834_3 | TATGAGTGTGGAGTCGTTGCAGATGGAGTCTCGCTTTGTCACNNNAGATCGGAAGA |
| 32834_4 | TATGAGTGTGGAGTCGTTGCAGATGGAGTCTCGCTTTGTCACNNNAGATCGGAAGA |
| 44586_1 | TATGAGTGTGGAGTCGTTGCCACACACACACATACACACACAANNNAGATCGGAAG |
| 44586_2 | TATGAGTGTGGAGTCGTTGCCACACACACACATACACACACAANNNAGATCGGAAG |
| 44586_3 | TATGAGTGTGGAGTCGTTGCCACACACACACATACACACACAANNNAGATCGGAAG |
| 44586_4 | TATGAGTGTGGAGTCGTTGCCACACACACACATACACACACAANNNAGATCGGAAG |
| 44771_1 | TATGAGTGTGGAGTCGTTGCCTCTCCATCATACCTGGGATTGGNNNAGATCGGAAG |
| 44771_2 | TATGAGTGTGGAGTCGTTGCCTCTCCATCATACCTGGGATTGGNNNAGATCGGAAG |
| 44771_3 | TATGAGTGTGGAGTCGTTGCCTCTCCATCATACCTGGGATTGGNNNAGATCGGAAG |
| 44771_4 | TATGAGTGTGGAGTCGTTGCCTCTCCATCATACCTGGGATTGGNNNAGATCGGAAG |
| 35433_1 | TATGAGTGTGGAGTCGTTGCCTCCTGAGTAGCTGGGACTACAGNNNAGATCGGAA |
| 35433_2 | TATGAGTGTGGAGTCGTTGCCTCCTGAGTAGCTGGGACTACAGNNNAGATCGGAA |
| 35433_3 | TATGAGTGTGGAGTCGTTGCCTCCTGAGTAGCTGGGACTACAGNNNAGATCGGAA |

| | |
|---|---|
| 35433_4 | TATGAGTGTGGAGTCGTTGCCTCCTGAGTAGCTGGGACTACAGNNNAGATCGGAAG |
| 44437_1 | TATGAGTGTGGAGTCGTTGCGGTTACAAAGTAAACACAGGCCCNNNAGATCGGAAG |
| 44437_2 | TATGAGTGTGGAGTCGTTGCGGTTACAAAGTAAACACAGGCCCNNNAGATCGGAAG |
| 44437_3 | TATGAGTGTGGAGTCGTTGCGGTTACAAAGTAAACACAGGCCCNNNAGATCGGAAG |
| 44437_4 | TATGAGTGTGGAGTCGTTGCGGTTACAAAGTAAACACAGGCCCNNNAGATCGGAAG |
| 35840_1 | TATGAGTGTGGAGTCGTTGCAGGTTCAAAAGATTCTCCTGCCTNNNAGATCGGAAGA |
| 35840_2 | TATGAGTGTGGAGTCGTTGCAGGTTCAAAAGATTCTCCTGCCTNNNAGATCGGAAGA |
| 35840_3 | TATGAGTGTGGAGTCGTTGCAGGTTCAAAAGATTCTCCTGCCTNNNAGATCGGAAGA |
| 35840_4 | TATGAGTGTGGAGTCGTTGCAGGTTCAAAAGATTCTCCTGCCTNNNAGATCGGAAGA |
| 38134_1 | TATGAGTGTGGAGTCGTTGCTCGCAATCTTTCATGTTGTGACCNNNAGATCGGAAGA |
| 38134_2 | TATGAGTGTGGAGTCGTTGCTCGCAATCTTTCATGTTGTGACCNNNAGATCGGAAGA |
| 38134_3 | TATGAGTGTGGAGTCGTTGCTCGCAATCTTTCATGTTGTGACCNNNAGATCGGAAGA |
| 38134_4 | TATGAGTGTGGAGTCGTTGCTCGCAATCTTTCATGTTGTGACCNNNAGATCGGAAGA |
| 31066_1 | TATGAGTGTGGAGTCGTTGCATGTCTGAACTTGTCTGGAAGCANNNAGATCGGAAG |
| 31066_2 | TATGAGTGTGGAGTCGTTGCATGTCTGAACTTGTCTGGAAGCANNNAGATCGGAAG |
| 31066_3 | TATGAGTGTGGAGTCGTTGCATGTCTGAACTTGTCTGGAAGCANNNAGATCGGAAG |
| 31066_4 | TATGAGTGTGGAGTCGTTGCATGTCTGAACTTGTCTGGAAGCANNNAGATCGGAAG |
| 50794_1 | TATGAGTGTGGAGTCGTTGCTTCCACCGACTTTGCTAATCTCCNNNAGATCGGAAGA |
| 50794_2 | TATGAGTGTGGAGTCGTTGCTTCCACCGACTTTGCTAATCTCCNNNAGATCGGAAGA |
| 50794_3 | TATGAGTGTGGAGTCGTTGCTTCCACCGACTTTGCTAATCTCCNNNAGATCGGAAGA |
| 50794_4 | TATGAGTGTGGAGTCGTTGCTTCCACCGACTTTGCTAATCTCCNNNAGATCGGAAGA |
| 30218_1 | TATGAGTGTGGAGTCGTTGCTGAGTAGAAAGGTTCTCATGCCANNNAGATCGGAAG |
| 30218_2 | TATGAGTGTGGAGTCGTTGCTGAGTAGAAAGGTTCTCATGCCANNNAGATCGGAAG |
| 30218_3 | TATGAGTGTGGAGTCGTTGCTGAGTAGAAAGGTTCTCATGCCANNNAGATCGGAAG |
| 30218_4 | TATGAGTGTGGAGTCGTTGCTGAGTAGAAAGGTTCTCATGCCANNNAGATCGGAAG |
| 53199_1 | TATGAGTGTGGAGTCGTTGCTTCGTTGGCCCCTAACTAACTACNNNAGATCGGAAGA |
| 53199_2 | TATGAGTGTGGAGTCGTTGCTTCGTTGGCCCCTAACTAACTACNNNAGATCGGAAGA |
| 53199_3 | TATGAGTGTGGAGTCGTTGCTTCGTTGGCCCCTAACTAACTACNNNAGATCGGAAGA |
| 53199_4 | TATGAGTGTGGAGTCGTTGCTTCGTTGGCCCCTAACTAACTACNNNAGATCGGAAGA |
| 39663_1 | TATGAGTGTGGAGTCGTTGCAGAATGGAGATTCCTTGCACACTNNNAGATCGGAAG |
| 39663_2 | TATGAGTGTGGAGTCGTTGCAGAATGGAGATTCCTTGCACACTNNNAGATCGGAAG |
| 39663_3 | TATGAGTGTGGAGTCGTTGCAGAATGGAGATTCCTTGCACACTNNNAGATCGGAAG |
| 39663_4 | TATGAGTGTGGAGTCGTTGCAGAATGGAGATTCCTTGCACACTNNNAGATCGGAAG |
| 46831_1 | TATGAGTGTGGAGTCGTTGCGCCTCCTGAGTAGCTGAGATTACNNNAGATCGGAAG |
| 46831_2 | TATGAGTGTGGAGTCGTTGCGCCTCCTGAGTAGCTGAGATTACNNNAGATCGGAAG |
| 46831_3 | TATGAGTGTGGAGTCGTTGCGCCTCCTGAGTAGCTGAGATTACNNNAGATCGGAAG |
| 46831_4 | TATGAGTGTGGAGTCGTTGCGCCTCCTGAGTAGCTGAGATTACNNNAGATCGGAAG |
| 44152_1 | TATGAGTGTGGAGTCGTTGCGCTACTTGTTGCACTGGTAAGTGNNNAGATCGGAAG |
| 44152_2 | TATGAGTGTGGAGTCGTTGCGCTACTTGTTGCACTGGTAAGTGNNNAGATCGGAAG |
| 44152_3 | TATGAGTGTGGAGTCGTTGCGCTACTTGTTGCACTGGTAAGTGNNNAGATCGGAAG |
| 44152_4 | TATGAGTGTGGAGTCGTTGCGCTACTTGTTGCACTGGTAAGTGNNNAGATCGGAAG |
| 29972_1 | TATGAGTGTGGAGTCGTTGCAGACGGAGTCTCGCTCTGTCNNNAGATCGGAAGAG |
| 29972_2 | TATGAGTGTGGAGTCGTTGCAGACGGAGTCTCGCTCTGTCNNNAGATCGGAAGAG |

| | |
|---|---|
| 29972_3 | TATGAGTGTGGAGTCGTTGCAGACGGAGTCTCGCTCTGTCNNNAGATCGGAAGAG |
| 29972_4 | TATGAGTGTGGAGTCGTTGCAGACGGAGTCTCGCTCTGTCNNNAGATCGGAAGAG |
| 30984_1 | TATGAGTGTGGAGTCGTTGCCAGCATGTCACCCTCCATATTCTNNNAGATCGGAAGA |
| 30984_2 | TATGAGTGTGGAGTCGTTGCCAGCATGTCACCCTCCATATTCTNNNAGATCGGAAGA |
| 30984_3 | TATGAGTGTGGAGTCGTTGCCAGCATGTCACCCTCCATATTCTNNNAGATCGGAAGA |
| 30984_4 | TATGAGTGTGGAGTCGTTGCCAGCATGTCACCCTCCATATTCTNNNAGATCGGAAGA |
| 35521_1 | TATGAGTGTGGAGTCGTTGCCTCCTGAGCAGCTGATACTACAGNNNAGATCGGAAG |
| 35521_2 | TATGAGTGTGGAGTCGTTGCCTCCTGAGCAGCTGATACTACAGNNNAGATCGGAAG |
| 35521_3 | TATGAGTGTGGAGTCGTTGCCTCCTGAGCAGCTGATACTACAGNNNAGATCGGAAG |
| 35521_4 | TATGAGTGTGGAGTCGTTGCCTCCTGAGCAGCTGATACTACAGNNNAGATCGGAAG |
| 28965_1 | TATGAGTGTGGAGTCGTTGCAAAGGAGAAGTAGGCGAGAAGTGNNNAGATCGGA |
| 28965_2 | TATGAGTGTGGAGTCGTTGCAAAGGAGAAGTAGGCGAGAAGTGNNNAGATCGGA |
| 28965_3 | TATGAGTGTGGAGTCGTTGCAAAGGAGAAGTAGGCGAGAAGTGNNNAGATCGGA |
| 28965_4 | TATGAGTGTGGAGTCGTTGCAAAGGAGAAGTAGGCGAGAAGTGNNNAGATCGGA |
| 35903_1 | TATGAGTGTGGAGTCGTTGCAGCAACTTCAGGGAAGCATTTTCNNNAGATCGGAAG |
| 35903_2 | TATGAGTGTGGAGTCGTTGCAGCAACTTCAGGGAAGCATTTTCNNNAGATCGGAAG |
| 35903_3 | TATGAGTGTGGAGTCGTTGCAGCAACTTCAGGGAAGCATTTTCNNNAGATCGGAAG |
| 35903_4 | TATGAGTGTGGAGTCGTTGCAGCAACTTCAGGGAAGCATTTTCNNNAGATCGGAAG |
| 38451_1 | TATGAGTGTGGAGTCGTTGCATGGTGAAGTCCCGTCTCTACTANNNAGATCGGAAG |
| 38451_2 | TATGAGTGTGGAGTCGTTGCATGGTGAAGTCCCGTCTCTACTANNNAGATCGGAAG |
| 38451_3 | TATGAGTGTGGAGTCGTTGCATGGTGAAGTCCCGTCTCTACTANNNAGATCGGAAG |
| 38451_4 | TATGAGTGTGGAGTCGTTGCATGGTGAAGTCCCGTCTCTACTANNNAGATCGGAAG |
| 33273_1 | TATGAGTGTGGAGTCGTTGCCAATCTCGGCTCACTGCAAACNNNAGATCGGAAGAG |
| 33273_2 | TATGAGTGTGGAGTCGTTGCCAATCTCGGCTCACTGCAAACNNNAGATCGGAAGAG |
| 33273_3 | TATGAGTGTGGAGTCGTTGCCAATCTCGGCTCACTGCAAACNNNAGATCGGAAGAG |
| 33273_4 | TATGAGTGTGGAGTCGTTGCCAATCTCGGCTCACTGCAAACNNNAGATCGGAAGAG |
| 31977_1 | TATGAGTGTGGAGTCGTTGCGCTTCAGGCTCTCTAATCAACGANNNAGATCGGAAG |
| 31977_2 | TATGAGTGTGGAGTCGTTGCGCTTCAGGCTCTCTAATCAACGANNNAGATCGGAAG |
| 31977_3 | TATGAGTGTGGAGTCGTTGCGCTTCAGGCTCTCTAATCAACGANNNAGATCGGAAG |
| 31977_4 | TATGAGTGTGGAGTCGTTGCGCTTCAGGCTCTCTAATCAACGANNNAGATCGGAAG |
| 48100_1 | TATGAGTGTGGAGTCGTTGCGTTCAATGGCACGATCTTGGTTCNNNAGATCGGAAG |
| 48100_2 | TATGAGTGTGGAGTCGTTGCGTTCAATGGCACGATCTTGGTTCNNNAGATCGGAAG |
| 48100_3 | TATGAGTGTGGAGTCGTTGCGTTCAATGGCACGATCTTGGTTCNNNAGATCGGAAG |
| 48100_4 | TATGAGTGTGGAGTCGTTGCGTTCAATGGCACGATCTTGGTTCNNNAGATCGGAAG |
| 51863_1 | TATGAGTGTGGAGTCGTTGCGGACCATGGCCCCATCTATAATTNNNAGATCGGAAG |
| 51863_2 | TATGAGTGTGGAGTCGTTGCGGACCATGGCCCCATCTATAATTNNNAGATCGGAAG |
| 51863_3 | TATGAGTGTGGAGTCGTTGCGGACCATGGCCCCATCTATAATTNNNAGATCGGAAG |
| 51863_4 | TATGAGTGTGGAGTCGTTGCGGACCATGGCCCCATCTATAATTNNNAGATCGGAAG |
| 41179_1 | TATGAGTGTGGAGTCGTTGCATGTGAGAAAGGACTGAACAGACNNNAGATCGGAA |
| 41179_2 | TATGAGTGTGGAGTCGTTGCATGTGAGAAAGGACTGAACAGACNNNAGATCGGAA |
| 41179_3 | TATGAGTGTGGAGTCGTTGCATGTGAGAAAGGACTGAACAGACNNNAGATCGGAA |
| 41179_4 | TATGAGTGTGGAGTCGTTGCATGTGAGAAAGGACTGAACAGACNNNAGATCGGAA |
| 35073_1 | TATGAGTGTGGAGTCGTTGCTTTTTTCTTGTCTCCTGCTAGCTNNNAGATCGGAAGA |

| | |
|---|---|
| 35073_2 | TATGAGTGTGGAGTCGTTGCTTTTTTCTTGTCTCCTGCTAGCTNNNAGATCGGAAGA |
| 35073_3 | TATGAGTGTGGAGTCGTTGCTTTTTTCTTGTCTCCTGCTAGCTNNNAGATCGGAAGA |
| 35073_4 | TATGAGTGTGGAGTCGTTGCTTTTTTCTTGTCTCCTGCTAGCTNNNAGATCGGAAGA |
| 38535_1 | TATGAGTGTGGAGTCGTTGCTTATGCCTGTAATCCCAGCACTTNNNAGATCGGAAGA |
| 38535_2 | TATGAGTGTGGAGTCGTTGCTTATGCCTGTAATCCCAGCACTTNNNAGATCGGAAGA |
| 38535_3 | TATGAGTGTGGAGTCGTTGCTTATGCCTGTAATCCCAGCACTTNNNAGATCGGAAGA |
| 38535_4 | TATGAGTGTGGAGTCGTTGCTTATGCCTGTAATCCCAGCACTTNNNAGATCGGAAGA |
| 51294_1 | TATGAGTGTGGAGTCGTTGCCTCAGTTATTTGGGGGAGGGAAANNNAGATCGGAA |
| 51294_2 | TATGAGTGTGGAGTCGTTGCCTCAGTTATTTGGGGGAGGGAAANNNAGATCGGAA |
| 51294_3 | TATGAGTGTGGAGTCGTTGCCTCAGTTATTTGGGGGAGGGAAANNNAGATCGGAA |
| 51294_4 | TATGAGTGTGGAGTCGTTGCCTCAGTTATTTGGGGGAGGGAAANNNAGATCGGAA |
| 36567_1 | TATGAGTGTGGAGTCGTTGCCTGAGGCAGGAAAATCGTTTGAANNNAGATCGGAA |
| 36567_2 | TATGAGTGTGGAGTCGTTGCCTGAGGCAGGAAAATCGTTTGAANNNAGATCGGAA |
| 36567_3 | TATGAGTGTGGAGTCGTTGCCTGAGGCAGGAAAATCGTTTGAANNNAGATCGGAA |
| 36567_4 | TATGAGTGTGGAGTCGTTGCCTGAGGCAGGAAAATCGTTTGAANNNAGATCGGAA |
| 50027_1 | TATGAGTGTGGAGTCGTTGCCACACACACACACATACACACTCNNNAGATCGGAAGA |
| 50027_2 | TATGAGTGTGGAGTCGTTGCCACACACACACACATACACACTCNNNAGATCGGAAGA |
| 50027_3 | TATGAGTGTGGAGTCGTTGCCACACACACACACATACACACTCNNNAGATCGGAAGA |
| 50027_4 | TATGAGTGTGGAGTCGTTGCCACACACACACACATACACACTCNNNAGATCGGAAGA |
| 41603_1 | TATGAGTGTGGAGTCGTTGCGAGCCTCCCAAAGTGATGAGATTNNNAGATCGGAAG |
| 41603_2 | TATGAGTGTGGAGTCGTTGCGAGCCTCCCAAAGTGATGAGATTNNNAGATCGGAAG |
| 41603_3 | TATGAGTGTGGAGTCGTTGCGAGCCTCCCAAAGTGATGAGATTNNNAGATCGGAAG |
| 41603_4 | TATGAGTGTGGAGTCGTTGCGAGCCTCCCAAAGTGATGAGATTNNNAGATCGGAAG |
| 28822_1 | TATGAGTGTGGAGTCGTTGCATCCACTCATTAGTTGATGGGCANNNAGATCGGAAG |
| 28822_2 | TATGAGTGTGGAGTCGTTGCATCCACTCATTAGTTGATGGGCANNNAGATCGGAAG |
| 28822_3 | TATGAGTGTGGAGTCGTTGCATCCACTCATTAGTTGATGGGCANNNAGATCGGAAG |
| 28822_4 | TATGAGTGTGGAGTCGTTGCATCCACTCATTAGTTGATGGGCANNNAGATCGGAAG |
| 53268_1 | TATGAGTGTGGAGTCGTTGCGAGAGTTGATCTTCTCCCCTCAGNNNAGATCGGAAG |
| 53268_2 | TATGAGTGTGGAGTCGTTGCGAGAGTTGATCTTCTCCCCTCAGNNNAGATCGGAAG |
| 53268_3 | TATGAGTGTGGAGTCGTTGCGAGAGTTGATCTTCTCCCCTCAGNNNAGATCGGAAG |
| 53268_4 | TATGAGTGTGGAGTCGTTGCGAGAGTTGATCTTCTCCCCTCAGNNNAGATCGGAAG |
| 13236_1 | TATGAGTGTGGAGTCGTTGCTTTGAGAGATTTCCTTTTTCAAANNNAGATCGGAAGA |
| 13236_2 | TATGAGTGTGGAGTCGTTGCTTTGAGAGATTTCCTTTTTCAAANNNAGATCGGAAGA |
| 13236_3 | TATGAGTGTGGAGTCGTTGCTTTGAGAGATTTCCTTTTTCAAANNNAGATCGGAAGA |
| 13236_4 | TATGAGTGTGGAGTCGTTGCTTTGAGAGATTTCCTTTTTCAAANNNAGATCGGAAGA |
| 30907_1 | TATGAGTGTGGAGTCGTTGCTGTGTTTTGGACTAAAGGCAGTGNNNAGATCGGAA |
| 30907_2 | TATGAGTGTGGAGTCGTTGCTGTGTTTTGGACTAAAGGCAGTGNNNAGATCGGAA |
| 30907_3 | TATGAGTGTGGAGTCGTTGCTGTGTTTTGGACTAAAGGCAGTGNNNAGATCGGAA |
| 30907_4 | TATGAGTGTGGAGTCGTTGCTGTGTTTTGGACTAAAGGCAGTGNNNAGATCGGAA |
| 46685_1 | TATGAGTGTGGAGTCGTTGCTCCATTTCTAAGGGCTGCTGTAGNNNAGATCGGAAG |
| 46685_2 | TATGAGTGTGGAGTCGTTGCTCCATTTCTAAGGGCTGCTGTAGNNNAGATCGGAAG |
| 46685_3 | TATGAGTGTGGAGTCGTTGCTCCATTTCTAAGGGCTGCTGTAGNNNAGATCGGAAG |
| 46685_4 | TATGAGTGTGGAGTCGTTGCTCCATTTCTAAGGGCTGCTGTAGNNNAGATCGGAAG |

| | |
|---|---|
| 35510_1 | TATGAGTGTGGAGTCGTTGCCACTATTGTTGCCCAGGTTTGAGNNNAGATCGGAAG |
| 35510_2 | TATGAGTGTGGAGTCGTTGCCACTATTGTTGCCCAGGTTTGAGNNNAGATCGGAAG |
| 35510_3 | TATGAGTGTGGAGTCGTTGCCACTATTGTTGCCCAGGTTTGAGNNNAGATCGGAAG |
| 35510_4 | TATGAGTGTGGAGTCGTTGCCACTATTGTTGCCCAGGTTTGAGNNNAGATCGGAAG |
| 34971_1 | TATGAGTGTGGAGTCGTTGCGGGCTCTGTGTTTCCTTAGGTAANNNAGATCGGAAG |
| 34971_2 | TATGAGTGTGGAGTCGTTGCGGGCTCTGTGTTTCCTTAGGTAANNNAGATCGGAAG |
| 34971_3 | TATGAGTGTGGAGTCGTTGCGGGCTCTGTGTTTCCTTAGGTAANNNAGATCGGAAG |
| 34971_4 | TATGAGTGTGGAGTCGTTGCGGGCTCTGTGTTTCCTTAGGTAANNNAGATCGGAAG |
| 3189_1 | TATGAGTGTGGAGTCGTTGCGGTGCAGTTATGCCTCAGATTCANNNAGATCGGAAG |
| 3189_2 | TATGAGTGTGGAGTCGTTGCGGTGCAGTTATGCCTCAGATTCANNNAGATCGGAAG |
| 3189_3 | TATGAGTGTGGAGTCGTTGCGGTGCAGTTATGCCTCAGATTCANNNAGATCGGAAG |
| 3189_4 | TATGAGTGTGGAGTCGTTGCGGTGCAGTTATGCCTCAGATTCANNNAGATCGGAAG |
| 43146_1 | TATGAGTGTGGAGTCGTTGCTTGTCCAGAGTCCAAACCAGAAANNNAGATCGGAAG |
| 43146_2 | TATGAGTGTGGAGTCGTTGCTTGTCCAGAGTCCAAACCAGAAANNNAGATCGGAAG |
| 43146_3 | TATGAGTGTGGAGTCGTTGCTTGTCCAGAGTCCAAACCAGAAANNNAGATCGGAAG |
| 43146_4 | TATGAGTGTGGAGTCGTTGCTTGTCCAGAGTCCAAACCAGAAANNNAGATCGGAAG |
| 51032_1 | TATGAGTGTGGAGTCGTTGCCGGGTGAGGGATAAAAGACTACANNNAGATCGGAA |
| 51032_2 | TATGAGTGTGGAGTCGTTGCCGGGTGAGGGATAAAAGACTACANNNAGATCGGAA |
| 51032_3 | TATGAGTGTGGAGTCGTTGCCGGGTGAGGGATAAAAGACTACANNNAGATCGGAA |
| 51032_4 | TATGAGTGTGGAGTCGTTGCCGGGTGAGGGATAAAAGACTACANNNAGATCGGAA |
| 38043_1 | TATGAGTGTGGAGTCGTTGCATCAGAACTTGACGGAACTTCAGNNNAGATCGGAAG |
| 38043_2 | TATGAGTGTGGAGTCGTTGCATCAGAACTTGACGGAACTTCAGNNNAGATCGGAAG |
| 38043_3 | TATGAGTGTGGAGTCGTTGCATCAGAACTTGACGGAACTTCAGNNNAGATCGGAAG |
| 38043_4 | TATGAGTGTGGAGTCGTTGCATCAGAACTTGACGGAACTTCAGNNNAGATCGGAAG |
| 49456_1 | TATGAGTGTGGAGTCGTTGCAGTGATAGTAGAGGGTTGAGGGANNNAGATCGGA |
| 49456_2 | TATGAGTGTGGAGTCGTTGCAGTGATAGTAGAGGGTTGAGGGANNNAGATCGGA |
| 49456_3 | TATGAGTGTGGAGTCGTTGCAGTGATAGTAGAGGGTTGAGGGANNNAGATCGGA |
| 49456_4 | TATGAGTGTGGAGTCGTTGCAGTGATAGTAGAGGGTTGAGGGANNNAGATCGGA |
| 48626_1 | TATGAGTGTGGAGTCGTTGCTTCTGACCTCATAATCTGTCCGCNNNAGATCGGAAGA |
| 48626_2 | TATGAGTGTGGAGTCGTTGCTTCTGACCTCATAATCTGTCCGCNNNAGATCGGAAGA |
| 48626_3 | TATGAGTGTGGAGTCGTTGCTTCTGACCTCATAATCTGTCCGCNNNAGATCGGAAGA |
| 48626_4 | TATGAGTGTGGAGTCGTTGCTTCTGACCTCATAATCTGTCCGCNNNAGATCGGAAGA |
| 33058_1 | TATGAGTGTGGAGTCGTTGCCCTCCAGTGTTCACTCCATTCTCNNNAGATCGGAAGA |
| 33058_2 | TATGAGTGTGGAGTCGTTGCCCTCCAGTGTTCACTCCATTCTCNNNAGATCGGAAGA |
| 33058_3 | TATGAGTGTGGAGTCGTTGCCCTCCAGTGTTCACTCCATTCTCNNNAGATCGGAAGA |
| 33058_4 | TATGAGTGTGGAGTCGTTGCCCTCCAGTGTTCACTCCATTCTCNNNAGATCGGAAGA |
| 37705_1 | TATGAGTGTGGAGTCGTTGCAATGAGGGCATATTGTCCAGAGCNNNAGATCGGAA |
| 37705_2 | TATGAGTGTGGAGTCGTTGCAATGAGGGCATATTGTCCAGAGCNNNAGATCGGAA |
| 37705_3 | TATGAGTGTGGAGTCGTTGCAATGAGGGCATATTGTCCAGAGCNNNAGATCGGAA |
| 37705_4 | TATGAGTGTGGAGTCGTTGCAATGAGGGCATATTGTCCAGAGCNNNAGATCGGAA |
| 46384_1 | TATGAGTGTGGAGTCGTTGCGCTTCATCAGTACCATCCAGGAANNNAGATCGGAAG |
| 46384_2 | TATGAGTGTGGAGTCGTTGCGCTTCATCAGTACCATCCAGGAANNNAGATCGGAAG |
| 46384_3 | TATGAGTGTGGAGTCGTTGCGCTTCATCAGTACCATCCAGGAANNNAGATCGGAAG |

| | |
|---|---|
| 46384_4 | TATGAGTGTGGAGTCGTTGCGCTTCATCAGTACCATCCAGGAANNNAGATCGGAAG |
| 39246_1 | TATGAGTGTGGAGTCGTTGCCAAGGCTATTTGGTGGCTTACAGNNNAGATCGGAAG |
| 39246_2 | TATGAGTGTGGAGTCGTTGCCAAGGCTATTTGGTGGCTTACAGNNNAGATCGGAAG |
| 39246_3 | TATGAGTGTGGAGTCGTTGCCAAGGCTATTTGGTGGCTTACAGNNNAGATCGGAAG |
| 39246_4 | TATGAGTGTGGAGTCGTTGCCAAGGCTATTTGGTGGCTTACAGNNNAGATCGGAAG |
| 38741_1 | TATGAGTGTGGAGTCGTTGCGTGGGAGGATCACTTGAGATCAGNNNAGATCGGAA |
| 38741_2 | TATGAGTGTGGAGTCGTTGCGTGGGAGGATCACTTGAGATCAGNNNAGATCGGAA |
| 38741_3 | TATGAGTGTGGAGTCGTTGCGTGGGAGGATCACTTGAGATCAGNNNAGATCGGAA |
| 38741_4 | TATGAGTGTGGAGTCGTTGCGTGGGAGGATCACTTGAGATCAGNNNAGATCGGAA |
| 40120_1 | TATGAGTGTGGAGTCGTTGCCATGGATGCAGCTGGAGTTTATTNNNAGATCGGAAG |
| 40120_2 | TATGAGTGTGGAGTCGTTGCCATGGATGCAGCTGGAGTTTATTNNNAGATCGGAAG |
| 40120_3 | TATGAGTGTGGAGTCGTTGCCATGGATGCAGCTGGAGTTTATTNNNAGATCGGAAG |
| 40120_4 | TATGAGTGTGGAGTCGTTGCCATGGATGCAGCTGGAGTTTATTNNNAGATCGGAAG |
| 48770_1 | TATGAGTGTGGAGTCGTTGCTTTCACCATATTGGCCAGGCTNNNAGATCGGAAGAG |
| 48770_2 | TATGAGTGTGGAGTCGTTGCTTTCACCATATTGGCCAGGCTNNNAGATCGGAAGAG |
| 48770_3 | TATGAGTGTGGAGTCGTTGCTTTCACCATATTGGCCAGGCTNNNAGATCGGAAGAG |
| 48770_4 | TATGAGTGTGGAGTCGTTGCTTTCACCATATTGGCCAGGCTNNNAGATCGGAAGAG |
| 51816_1 | TATGAGTGTGGAGTCGTTGCATGTCAAGCAACCATCAGGAGATNNNAGATCGGAAG |
| 51816_2 | TATGAGTGTGGAGTCGTTGCATGTCAAGCAACCATCAGGAGATNNNAGATCGGAAG |
| 51816_3 | TATGAGTGTGGAGTCGTTGCATGTCAAGCAACCATCAGGAGATNNNAGATCGGAAG |
| 51816_4 | TATGAGTGTGGAGTCGTTGCATGTCAAGCAACCATCAGGAGATNNNAGATCGGAAG |
| 29050_1 | TATGAGTGTGGAGTCGTTGCCCCCTTCCATTTATCTGGGAGTCNNNAGATCGGAAGA |
| 29050_2 | TATGAGTGTGGAGTCGTTGCCCCCTTCCATTTATCTGGGAGTCNNNAGATCGGAAGA |
| 29050_3 | TATGAGTGTGGAGTCGTTGCCCCCTTCCATTTATCTGGGAGTCNNNAGATCGGAAGA |
| 29050_4 | TATGAGTGTGGAGTCGTTGCCCCCTTCCATTTATCTGGGAGTCNNNAGATCGGAAGA |
| 48924_1 | TATGAGTGTGGAGTCGTTGCGCACACACATCTTCAAGAAGCTCNNNAGATCGGAAGA |
| 48924_2 | TATGAGTGTGGAGTCGTTGCGCACACACATCTTCAAGAAGCTCNNNAGATCGGAAGA |
| 48924_3 | TATGAGTGTGGAGTCGTTGCGCACACACATCTTCAAGAAGCTCNNNAGATCGGAAGA |
| 48924_4 | TATGAGTGTGGAGTCGTTGCGCACACACATCTTCAAGAAGCTCNNNAGATCGGAAGA |
| 40648_1 | TATGAGTGTGGAGTCGTTGCTCTTTAGTCCCAGCTACTCAGGANNNAGATCGGAAGA |
| 40648_2 | TATGAGTGTGGAGTCGTTGCTCTTTAGTCCCAGCTACTCAGGANNNAGATCGGAAGA |
| 40648_3 | TATGAGTGTGGAGTCGTTGCTCTTTAGTCCCAGCTACTCAGGANNNAGATCGGAAGA |
| 40648_4 | TATGAGTGTGGAGTCGTTGCTCTTTAGTCCCAGCTACTCAGGANNNAGATCGGAAGA |
| 10744_1 | TATGAGTGTGGAGTCGTTGCAATAAAGCCTAGATATTGCCCTGTNNNAGATCGGAAG |
| 10744_2 | TATGAGTGTGGAGTCGTTGCAATAAAGCCTAGATATTGCCCTGTNNNAGATCGGAAG |
| 10744_3 | TATGAGTGTGGAGTCGTTGCAATAAAGCCTAGATATTGCCCTGTNNNAGATCGGAAG |
| 10744_4 | TATGAGTGTGGAGTCGTTGCAATAAAGCCTAGATATTGCCCTGTNNNAGATCGGAAG |
| 51469_1 | TATGAGTGTGGAGTCGTTGCATTAGGGCCTGGAATTTTGTCCTNNNAGATCGGAAG |
| 51469_2 | TATGAGTGTGGAGTCGTTGCATTAGGGCCTGGAATTTTGTCCTNNNAGATCGGAAG |
| 51469_3 | TATGAGTGTGGAGTCGTTGCATTAGGGCCTGGAATTTTGTCCTNNNAGATCGGAAG |
| 51469_4 | TATGAGTGTGGAGTCGTTGCATTAGGGCCTGGAATTTTGTCCTNNNAGATCGGAAG |
| 31605_1 | TATGAGTGTGGAGTCGTTGCCTCTCACTTGTGCCTTCCTAAGTNNNAGATCGGAAGA |
| 31605_2 | TATGAGTGTGGAGTCGTTGCCTCTCACTTGTGCCTTCCTAAGTNNNAGATCGGAAGA |

| | |
|---|---|
| 31605_3 | TATGAGTGTGGAGTCGTTGCCTCTCACTTGTGCCTTCCTAAGTNNNAGATCGGAAGA |
| 31605_4 | TATGAGTGTGGAGTCGTTGCCTCTCACTTGTGCCTTCCTAAGTNNNAGATCGGAAGA |
| 5970_1 | TATGAGTGTGGAGTCGTTGCCATTTTAGACACATGACTTCCGCNNNAGATCGGAAG |
| 5970_2 | TATGAGTGTGGAGTCGTTGCCATTTTAGACACATGACTTCCGCNNNAGATCGGAAG |
| 5970_3 | TATGAGTGTGGAGTCGTTGCCATTTTAGACACATGACTTCCGCNNNAGATCGGAAG |
| 5970_4 | TATGAGTGTGGAGTCGTTGCCATTTTAGACACATGACTTCCGCNNNAGATCGGAAG |
| 33638_1 | TATGAGTGTGGAGTCGTTGCTTAGGGTACATGTGCACATTGTGNNNAGATCGGAA |
| 33638_2 | TATGAGTGTGGAGTCGTTGCTTAGGGTACATGTGCACATTGTGNNNAGATCGGAA |
| 33638_3 | TATGAGTGTGGAGTCGTTGCTTAGGGTACATGTGCACATTGTGNNNAGATCGGAA |
| 33638_4 | TATGAGTGTGGAGTCGTTGCTTAGGGTACATGTGCACATTGTGNNNAGATCGGAA |
| 30438_1 | TATGAGTGTGGAGTCGTTGCTAGAGAGAAGAGAGAGGGGGAAANNNAGATCGG |
| 30438_2 | TATGAGTGTGGAGTCGTTGCTAGAGAGAAGAGAGAGGGGGAAANNNAGATCGG |
| 30438_3 | TATGAGTGTGGAGTCGTTGCTAGAGAGAAGAGAGAGGGGGAAANNNAGATCGG |
| 30438_4 | TATGAGTGTGGAGTCGTTGCTAGAGAGAAGAGAGAGGGGGAAANNNAGATCGG |
| 46194_1 | TATGAGTGTGGAGTCGTTGCACAAGTCTCATTCTGTCGCCTAGNNNAGATCGGAAG |
| 46194_2 | TATGAGTGTGGAGTCGTTGCACAAGTCTCATTCTGTCGCCTAGNNNAGATCGGAAG |
| 46194_3 | TATGAGTGTGGAGTCGTTGCACAAGTCTCATTCTGTCGCCTAGNNNAGATCGGAAG |
| 46194_4 | TATGAGTGTGGAGTCGTTGCACAAGTCTCATTCTGTCGCCTAGNNNAGATCGGAAG |
| 51915_1 | TATGAGTGTGGAGTCGTTGCAGACAAAACTTCCAGAGGAACGANNNAGATCGGAA |
| 51915_2 | TATGAGTGTGGAGTCGTTGCAGACAAAACTTCCAGAGGAACGANNNAGATCGGAA |
| 51915_3 | TATGAGTGTGGAGTCGTTGCAGACAAAACTTCCAGAGGAACGANNNAGATCGGAA |
| 51915_4 | TATGAGTGTGGAGTCGTTGCAGACAAAACTTCCAGAGGAACGANNNAGATCGGAA |
| 33791_1 | TATGAGTGTGGAGTCGTTGCTGGGATCTCAACATCCTTCCAAGNNNAGATCGGAAG |
| 33791_2 | TATGAGTGTGGAGTCGTTGCTGGGATCTCAACATCCTTCCAAGNNNAGATCGGAAG |
| 33791_3 | TATGAGTGTGGAGTCGTTGCTGGGATCTCAACATCCTTCCAAGNNNAGATCGGAAG |
| 33791_4 | TATGAGTGTGGAGTCGTTGCTGGGATCTCAACATCCTTCCAAGNNNAGATCGGAAG |
| 2515_1 | TATGAGTGTGGAGTCGTTGCATGGGAAACTAGGGGAGGCTTAGNNNAGATCGGA |
| 2515_2 | TATGAGTGTGGAGTCGTTGCATGGGAAACTAGGGGAGGCTTAGNNNAGATCGGA |
| 2515_3 | TATGAGTGTGGAGTCGTTGCATGGGAAACTAGGGGAGGCTTAGNNNAGATCGGA |
| 2515_4 | TATGAGTGTGGAGTCGTTGCATGGGAAACTAGGGGAGGCTTAGNNNAGATCGGA |
| 52197_1 | TATGAGTGTGGAGTCGTTGCTGTCTGACATGTAGTAAGTGCCANNNAGATCGGAA |
| 52197_2 | TATGAGTGTGGAGTCGTTGCTGTCTGACATGTAGTAAGTGCCANNNAGATCGGAA |
| 52197_3 | TATGAGTGTGGAGTCGTTGCTGTCTGACATGTAGTAAGTGCCANNNAGATCGGAA |
| 52197_4 | TATGAGTGTGGAGTCGTTGCTGTCTGACATGTAGTAAGTGCCANNNAGATCGGAA |
| 43669_1 | TATGAGTGTGGAGTCGTTGCCAGGTTTAGAAGGGGTGTCTGGNNNAGATCGGAA |
| 43669_2 | TATGAGTGTGGAGTCGTTGCCAGGTTTAGAAGGGGTGTCTGGNNNAGATCGGAA |
| 43669_3 | TATGAGTGTGGAGTCGTTGCCAGGTTTAGAAGGGGTGTCTGGNNNAGATCGGAA |
| 43669_4 | TATGAGTGTGGAGTCGTTGCCAGGTTTAGAAGGGGTGTCTGGNNNAGATCGGAA |
| 48891_1 | TATGAGTGTGGAGTCGTTGCGAAAGAAGGGGAAAGGAGGGAAANNNAGATCGG |
| 48891_2 | TATGAGTGTGGAGTCGTTGCGAAAGAAGGGGAAAGGAGGGAAANNNAGATCGG |
| 48891_3 | TATGAGTGTGGAGTCGTTGCGAAAGAAGGGGAAAGGAGGGAAANNNAGATCGG |
| 48891_4 | TATGAGTGTGGAGTCGTTGCGAAAGAAGGGGAAAGGAGGGAAANNNAGATCGG |
| 34390_1 | TATGAGTGTGGAGTCGTTGCCTTACCACAAAGCCCTTTTGGAGNNNAGATCGGAAG |

| | |
|---|---|
| 34390_2 | TATGAGTGTGGAGTCGTTGCCTTACCACAAAGCCCTTTTGGAGNNNAGATCGGAAG |
| 34390_3 | TATGAGTGTGGAGTCGTTGCCTTACCACAAAGCCCTTTTGGAGNNNAGATCGGAAG |
| 34390_4 | TATGAGTGTGGAGTCGTTGCCTTACCACAAAGCCCTTTTGGAGNNNAGATCGGAAG |
| 42506_1 | TATGAGTGTGGAGTCGTTGCGGGGTGCTGGAACATGGTAAATANNNAGATCGGAA |
| 42506_2 | TATGAGTGTGGAGTCGTTGCGGGGTGCTGGAACATGGTAAATANNNAGATCGGAA |
| 42506_3 | TATGAGTGTGGAGTCGTTGCGGGGTGCTGGAACATGGTAAATANNNAGATCGGAA |
| 42506_4 | TATGAGTGTGGAGTCGTTGCGGGGTGCTGGAACATGGTAAATANNNAGATCGGAA |
| 30193_1 | TATGAGTGTGGAGTCGTTGCTTCAGTGATGAAGCCCAGATCAANNNAGATCGGAA |
| 30193_2 | TATGAGTGTGGAGTCGTTGCTTCAGTGATGAAGCCCAGATCAANNNAGATCGGAA |
| 30193_3 | TATGAGTGTGGAGTCGTTGCTTCAGTGATGAAGCCCAGATCAANNNAGATCGGAA |
| 30193_4 | TATGAGTGTGGAGTCGTTGCTTCAGTGATGAAGCCCAGATCAANNNAGATCGGAA |
| 36226_1 | TATGAGTGTGGAGTCGTTGCGCACTTACAAGCTGTGTGTTCTTNNNAGATCGGAAG |
| 36226_2 | TATGAGTGTGGAGTCGTTGCGCACTTACAAGCTGTGTGTTCTTNNNAGATCGGAAG |
| 36226_3 | TATGAGTGTGGAGTCGTTGCGCACTTACAAGCTGTGTGTTCTTNNNAGATCGGAAG |
| 36226_4 | TATGAGTGTGGAGTCGTTGCGCACTTACAAGCTGTGTGTTCTTNNNAGATCGGAAG |
| 49193_1 | TATGAGTGTGGAGTCGTTGCTCAAACTTCTGGCCTCATGCTATNNNAGATCGGAAG |
| 49193_2 | TATGAGTGTGGAGTCGTTGCTCAAACTTCTGGCCTCATGCTATNNNAGATCGGAAG |
| 49193_3 | TATGAGTGTGGAGTCGTTGCTCAAACTTCTGGCCTCATGCTATNNNAGATCGGAAG |
| 49193_4 | TATGAGTGTGGAGTCGTTGCTCAAACTTCTGGCCTCATGCTATNNNAGATCGGAAG |
| 30496_1 | TATGAGTGTGGAGTCGTTGCGACCAGGTTAGTCTCCAACTAGGGNNNAGATCGGA |
| 30496_2 | TATGAGTGTGGAGTCGTTGCGACCAGGTTAGTCTCCAACTAGGGNNNAGATCGGA |
| 30496_3 | TATGAGTGTGGAGTCGTTGCGACCAGGTTAGTCTCCAACTAGGGNNNAGATCGGA |
| 30496_4 | TATGAGTGTGGAGTCGTTGCGACCAGGTTAGTCTCCAACTAGGGNNNAGATCGGA |
| 47524_1 | TATGAGTGTGGAGTCGTTGCCCCGAGTAGCTGGAACTACAAATNNNAGATCGGAA |
| 47524_2 | TATGAGTGTGGAGTCGTTGCCCCGAGTAGCTGGAACTACAAATNNNAGATCGGAA |
| 47524_3 | TATGAGTGTGGAGTCGTTGCCCCGAGTAGCTGGAACTACAAATNNNAGATCGGAA |
| 47524_4 | TATGAGTGTGGAGTCGTTGCCCCGAGTAGCTGGAACTACAAATNNNAGATCGGAA |
| 30151_1 | TATGAGTGTGGAGTCGTTGCTCTCAAGACTGTCAAGACCTTCCNNNAGATCGGAAG |
| 30151_2 | TATGAGTGTGGAGTCGTTGCTCTCAAGACTGTCAAGACCTTCCNNNAGATCGGAAG |
| 30151_3 | TATGAGTGTGGAGTCGTTGCTCTCAAGACTGTCAAGACCTTCCNNNAGATCGGAAG |
| 30151_4 | TATGAGTGTGGAGTCGTTGCTCTCAAGACTGTCAAGACCTTCCNNNAGATCGGAAG |
| 52885_1 | TATGAGTGTGGAGTCGTTGCCCCTTGGACTTCTCTGAGGAATTNNNAGATCGGAAG |
| 52885_2 | TATGAGTGTGGAGTCGTTGCCCCTTGGACTTCTCTGAGGAATTNNNAGATCGGAAG |
| 52885_3 | TATGAGTGTGGAGTCGTTGCCCCTTGGACTTCTCTGAGGAATTNNNAGATCGGAAG |
| 52885_4 | TATGAGTGTGGAGTCGTTGCCCCTTGGACTTCTCTGAGGAATTNNNAGATCGGAAG |
| 35076_1 | TATGAGTGTGGAGTCGTTGCCAAGCTGTTTTCGTTACTGTGGTNNNAGATCGGAAG |
| 35076_2 | TATGAGTGTGGAGTCGTTGCCAAGCTGTTTTCGTTACTGTGGTNNNAGATCGGAAG |
| 35076_3 | TATGAGTGTGGAGTCGTTGCCAAGCTGTTTTCGTTACTGTGGTNNNAGATCGGAAG |
| 35076_4 | TATGAGTGTGGAGTCGTTGCCAAGCTGTTTTCGTTACTGTGGTNNNAGATCGGAAG |
| 44424_1 | TATGAGTGTGGAGTCGTTGCATGGGCATGAACCATGAGAAGTANNNAGATCGGAA |
| 44424_2 | TATGAGTGTGGAGTCGTTGCATGGGCATGAACCATGAGAAGTANNNAGATCGGAA |
| 44424_3 | TATGAGTGTGGAGTCGTTGCATGGGCATGAACCATGAGAAGTANNNAGATCGGAA |
| 44424_4 | TATGAGTGTGGAGTCGTTGCATGGGCATGAACCATGAGAAGTANNNAGATCGGAA |

| | |
|---|---|
| 33818_1 | TATGAGTGTGGAGTCGTTGCGCCAATTGCAAGCCAAGAAAATANNNAGATCGGAAG |
| 33818_2 | TATGAGTGTGGAGTCGTTGCGCCAATTGCAAGCCAAGAAAATANNNAGATCGGAAG |
| 33818_3 | TATGAGTGTGGAGTCGTTGCGCCAATTGCAAGCCAAGAAAATANNNAGATCGGAAG |
| 33818_4 | TATGAGTGTGGAGTCGTTGCGCCAATTGCAAGCCAAGAAAATANNNAGATCGGAAG |
| 51217_1 | TATGAGTGTGGAGTCGTTGCAAATGCTTATAGTGGTGCATGGCNNNAGATCGGAA |
| 51217_2 | TATGAGTGTGGAGTCGTTGCAAATGCTTATAGTGGTGCATGGCNNNAGATCGGAA |
| 51217_3 | TATGAGTGTGGAGTCGTTGCAAATGCTTATAGTGGTGCATGGCNNNAGATCGGAA |
| 51217_4 | TATGAGTGTGGAGTCGTTGCAAATGCTTATAGTGGTGCATGGCNNNAGATCGGAA |
| 38562_1 | TATGAGTGTGGAGTCGTTGCAGACTTAGCCACTTCTTTACGGCNNNAGATCGGAAG |
| 38562_2 | TATGAGTGTGGAGTCGTTGCAGACTTAGCCACTTCTTTACGGCNNNAGATCGGAAG |
| 38562_3 | TATGAGTGTGGAGTCGTTGCAGACTTAGCCACTTCTTTACGGCNNNAGATCGGAAG |
| 38562_4 | TATGAGTGTGGAGTCGTTGCAGACTTAGCCACTTCTTTACGGCNNNAGATCGGAAG |
| 53124_1 | TATGAGTGTGGAGTCGTTGCAGAGCTAAAAGCAGAACTCTCCTNNNAGATCGGAAG |
| 53124_2 | TATGAGTGTGGAGTCGTTGCAGAGCTAAAAGCAGAACTCTCCTNNNAGATCGGAAG |
| 53124_3 | TATGAGTGTGGAGTCGTTGCAGAGCTAAAAGCAGAACTCTCCTNNNAGATCGGAAG |
| 53124_4 | TATGAGTGTGGAGTCGTTGCAGAGCTAAAAGCAGAACTCTCCTNNNAGATCGGAAG |
| 38176_1 | TATGAGTGTGGAGTCGTTGCTTCTTCTGTTGTCCTTATCGGGTNNNAGATCGGAAG |
| 38176_2 | TATGAGTGTGGAGTCGTTGCTTCTTCTGTTGTCCTTATCGGGTNNNAGATCGGAAG |
| 38176_3 | TATGAGTGTGGAGTCGTTGCTTCTTCTGTTGTCCTTATCGGGTNNNAGATCGGAAG |
| 38176_4 | TATGAGTGTGGAGTCGTTGCTTCTTCTGTTGTCCTTATCGGGTNNNAGATCGGAAG |
| 32579_1 | TATGAGTGTGGAGTCGTTGCTCTCCTGAGTAGCTGGGATTACANNNAGATCGGAAG |
| 32579_2 | TATGAGTGTGGAGTCGTTGCTCTCCTGAGTAGCTGGGATTACANNNAGATCGGAAG |
| 32579_3 | TATGAGTGTGGAGTCGTTGCTCTCCTGAGTAGCTGGGATTACANNNAGATCGGAAG |
| 32579_4 | TATGAGTGTGGAGTCGTTGCTCTCCTGAGTAGCTGGGATTACANNNAGATCGGAAG |
| 42245_1 | TATGAGTGTGGAGTCGTTGCCCAAGGGAATGGATGCTAAAGATGNNNAGATCGGA |
| 42245_2 | TATGAGTGTGGAGTCGTTGCCCAAGGGAATGGATGCTAAAGATGNNNAGATCGGA |
| 42245_3 | TATGAGTGTGGAGTCGTTGCCCAAGGGAATGGATGCTAAAGATGNNNAGATCGGA |
| 42245_4 | TATGAGTGTGGAGTCGTTGCCCAAGGGAATGGATGCTAAAGATGNNNAGATCGGA |
| 32264_1 | TATGAGTGTGGAGTCGTTGCCTGGCTGAAGTCTATGGTCCTTGNNNAGATCGGAAG |
| 32264_2 | TATGAGTGTGGAGTCGTTGCCTGGCTGAAGTCTATGGTCCTTGNNNAGATCGGAAG |
| 32264_3 | TATGAGTGTGGAGTCGTTGCCTGGCTGAAGTCTATGGTCCTTGNNNAGATCGGAAG |
| 32264_4 | TATGAGTGTGGAGTCGTTGCCTGGCTGAAGTCTATGGTCCTTGNNNAGATCGGAAG |
| 37462_1 | TATGAGTGTGGAGTCGTTGCCTAATGGTCTACAAGGCCCTCAANNNAGATCGGAAG |
| 37462_2 | TATGAGTGTGGAGTCGTTGCCTAATGGTCTACAAGGCCCTCAANNNAGATCGGAAG |
| 37462_3 | TATGAGTGTGGAGTCGTTGCCTAATGGTCTACAAGGCCCTCAANNNAGATCGGAAG |
| 37462_4 | TATGAGTGTGGAGTCGTTGCCTAATGGTCTACAAGGCCCTCAANNNAGATCGGAAG |
| 32933_1 | TATGAGTGTGGAGTCGTTGCCAACACACACACACACACACATANNNAGATCGGAAG |
| 32933_2 | TATGAGTGTGGAGTCGTTGCCAACACACACACACACACACATANNNAGATCGGAAG |
| 32933_3 | TATGAGTGTGGAGTCGTTGCCAACACACACACACACACACATANNNAGATCGGAAG |
| 32933_4 | TATGAGTGTGGAGTCGTTGCCAACACACACACACACACACATANNNAGATCGGAAG |
| 40850_1 | TATGAGTGTGGAGTCGTTGCCTGAGACAGGAGAATCGCTTGAANNNAGATCGGAA |
| 40850_2 | TATGAGTGTGGAGTCGTTGCCTGAGACAGGAGAATCGCTTGAANNNAGATCGGAA |
| 40850_3 | TATGAGTGTGGAGTCGTTGCCTGAGACAGGAGAATCGCTTGAANNNAGATCGGAA |

| | |
|---|---|
| 40850_4 | TATGAGTGTGGAGTCGTTGCCTGAGACAGGAGAATCGCTTGAANNNAGATCGGAA |
| 37876_1 | TATGAGTGTGGAGTCGTTGCCTCCTCCTATCCACCATCCCTAANNNAGATCGGAAGA |
| 37876_2 | TATGAGTGTGGAGTCGTTGCCTCCTCCTATCCACCATCCCTAANNNAGATCGGAAGA |
| 37876_3 | TATGAGTGTGGAGTCGTTGCCTCCTCCTATCCACCATCCCTAANNNAGATCGGAAGA |
| 37876_4 | TATGAGTGTGGAGTCGTTGCCTCCTCCTATCCACCATCCCTAANNNAGATCGGAAGA |
| 32069_1 | TATGAGTGTGGAGTCGTTGCCACGCCTGTAATCTCAGCAATTNNNAGATCGGAAG |
| 32069_2 | TATGAGTGTGGAGTCGTTGCCACGCCTGTAATCTCAGCAATTNNNAGATCGGAAG |
| 32069_3 | TATGAGTGTGGAGTCGTTGCCACGCCTGTAATCTCAGCAATTNNNAGATCGGAAG |
| 32069_4 | TATGAGTGTGGAGTCGTTGCCACGCCTGTAATCTCAGCAATTNNNAGATCGGAAG |
| 36174_1 | TATGAGTGTGGAGTCGTTGCAGACCCCATCACGAAATTTGAGTNNNAGATCGGAAG |
| 36174_2 | TATGAGTGTGGAGTCGTTGCAGACCCCATCACGAAATTTGAGTNNNAGATCGGAAG |
| 36174_3 | TATGAGTGTGGAGTCGTTGCAGACCCCATCACGAAATTTGAGTNNNAGATCGGAAG |
| 36174_4 | TATGAGTGTGGAGTCGTTGCAGACCCCATCACGAAATTTGAGTNNNAGATCGGAAG |
| 44891_1 | TATGAGTGTGGAGTCGTTGCCCCATGTCCATTTCCTGATCTCANNNAGATCGGAAGA |
| 44891_2 | TATGAGTGTGGAGTCGTTGCCCCATGTCCATTTCCTGATCTCANNNAGATCGGAAGA |
| 44891_3 | TATGAGTGTGGAGTCGTTGCCCCATGTCCATTTCCTGATCTCANNNAGATCGGAAGA |
| 44891_4 | TATGAGTGTGGAGTCGTTGCCCCATGTCCATTTCCTGATCTCANNNAGATCGGAAGA |
| 51921_1 | TATGAGTGTGGAGTCGTTGCCTCTTTACCTACCGGCCGTAGNNNAGATCGGAAGAG |
| 51921_2 | TATGAGTGTGGAGTCGTTGCCTCTTTACCTACCGGCCGTAGNNNAGATCGGAAGAG |
| 51921_3 | TATGAGTGTGGAGTCGTTGCCTCTTTACCTACCGGCCGTAGNNNAGATCGGAAGAG |
| 51921_4 | TATGAGTGTGGAGTCGTTGCCTCTTTACCTACCGGCCGTAGNNNAGATCGGAAGAG |
| 9230_1 | TATGAGTGTGGAGTCGTTGCGACAGAGACAGAGAGCAAAGGTGNNNAGATCGGA |
| 9230_2 | TATGAGTGTGGAGTCGTTGCGACAGAGACAGAGAGCAAAGGTGNNNAGATCGGA |
| 9230_3 | TATGAGTGTGGAGTCGTTGCGACAGAGACAGAGAGCAAAGGTGNNNAGATCGGA |
| 9230_4 | TATGAGTGTGGAGTCGTTGCGACAGAGACAGAGAGCAAAGGTGNNNAGATCGGA |
| 51674_1 | TATGAGTGTGGAGTCGTTGCTTTAGTAGAGACGGGGGTTTCACNNNAGATCGGAA |
| 51674_2 | TATGAGTGTGGAGTCGTTGCTTTAGTAGAGACGGGGGTTTCACNNNAGATCGGAA |
| 51674_3 | TATGAGTGTGGAGTCGTTGCTTTAGTAGAGACGGGGGTTTCACNNNAGATCGGAA |
| 51674_4 | TATGAGTGTGGAGTCGTTGCTTTAGTAGAGACGGGGGTTTCACNNNAGATCGGAA |
| 46402_1 | TATGAGTGTGGAGTCGTTGCCTCCCAAGTAGCTGAGACTACAGNNNAGATCGGAAG |
| 46402_2 | TATGAGTGTGGAGTCGTTGCCTCCCAAGTAGCTGAGACTACAGNNNAGATCGGAAG |
| 46402_3 | TATGAGTGTGGAGTCGTTGCCTCCCAAGTAGCTGAGACTACAGNNNAGATCGGAAG |
| 46402_4 | TATGAGTGTGGAGTCGTTGCCTCCCAAGTAGCTGAGACTACAGNNNAGATCGGAAG |
| 40919_1 | TATGAGTGTGGAGTCGTTGCTATGGGACTAGGAGGAAGACAGGNNNAGATCGGA |
| 40919_2 | TATGAGTGTGGAGTCGTTGCTATGGGACTAGGAGGAAGACAGGNNNAGATCGGA |
| 40919_3 | TATGAGTGTGGAGTCGTTGCTATGGGACTAGGAGGAAGACAGGNNNAGATCGGA |
| 40919_4 | TATGAGTGTGGAGTCGTTGCTATGGGACTAGGAGGAAGACAGGNNNAGATCGGA |
| 30108_1 | TATGAGTGTGGAGTCGTTGCTGTCATTATTGAGTGAGATGAGTCTNNNAGATCGGA |
| 30108_2 | TATGAGTGTGGAGTCGTTGCTGTCATTATTGAGTGAGATGAGTCTNNNAGATCGGA |
| 30108_3 | TATGAGTGTGGAGTCGTTGCTGTCATTATTGAGTGAGATGAGTCTNNNAGATCGGA |
| 30108_4 | TATGAGTGTGGAGTCGTTGCTGTCATTATTGAGTGAGATGAGTCTNNNAGATCGGA |
| 9114_1 | TATGAGTGTGGAGTCGTTGCACCATCAGGACACCCTTGTAGGNNNAGATCGGAAGA |
| 9114_2 | TATGAGTGTGGAGTCGTTGCACCATCAGGACACCCTTGTAGGNNNAGATCGGAAGA |

| | |
|---|---|
| 9114_3 | TATGAGTGTGGAGTCGTTGCACCATCAGGACACCCTTGTAGGNNNAGATCGGAAGA |
| 9114_4 | TATGAGTGTGGAGTCGTTGCACCATCAGGACACCCTTGTAGGNNNAGATCGGAAGA |
| 31360_1 | TATGAGTGTGGAGTCGTTGCCCCACTCCAAAAACATTGCTGAANNNAGATCGGAAG |
| 31360_2 | TATGAGTGTGGAGTCGTTGCCCCACTCCAAAAACATTGCTGAANNNAGATCGGAAG |
| 31360_3 | TATGAGTGTGGAGTCGTTGCCCCACTCCAAAAACATTGCTGAANNNAGATCGGAAG |
| 31360_4 | TATGAGTGTGGAGTCGTTGCCCCACTCCAAAAACATTGCTGAANNNAGATCGGAAG |
| 49118_1 | TATGAGTGTGGAGTCGTTGCGAACTTCTGACCTCAGGTGATCCNNNAGATCGGAAG |
| 49118_2 | TATGAGTGTGGAGTCGTTGCGAACTTCTGACCTCAGGTGATCCNNNAGATCGGAAG |
| 49118_3 | TATGAGTGTGGAGTCGTTGCGAACTTCTGACCTCAGGTGATCCNNNAGATCGGAAG |
| 49118_4 | TATGAGTGTGGAGTCGTTGCGAACTTCTGACCTCAGGTGATCCNNNAGATCGGAAG |
| 37278_1 | TATGAGTGTGGAGTCGTTGCTGTAGCCATTTCTGGAGATTTTGCNNNAGATCGGAA |
| 37278_2 | TATGAGTGTGGAGTCGTTGCTGTAGCCATTTCTGGAGATTTTGCNNNAGATCGGAA |
| 37278_3 | TATGAGTGTGGAGTCGTTGCTGTAGCCATTTCTGGAGATTTTGCNNNAGATCGGAA |
| 37278_4 | TATGAGTGTGGAGTCGTTGCTGTAGCCATTTCTGGAGATTTTGCNNNAGATCGGAA |
| 38710_1 | TATGAGTGTGGAGTCGTTGCAGAAGAAAAAAAACCCTGCACGTNNNAGATCGGAA |
| 38710_2 | TATGAGTGTGGAGTCGTTGCAGAAGAAAAAAAACCCTGCACGTNNNAGATCGGAA |
| 38710_3 | TATGAGTGTGGAGTCGTTGCAGAAGAAAAAAAACCCTGCACGTNNNAGATCGGAA |
| 38710_4 | TATGAGTGTGGAGTCGTTGCAGAAGAAAAAAAACCCTGCACGTNNNAGATCGGAA |
| 37421_1 | TATGAGTGTGGAGTCGTTGCAATTGCTGGGTCATATGGTGTCTNNNAGATCGGAAG |
| 37421_2 | TATGAGTGTGGAGTCGTTGCAATTGCTGGGTCATATGGTGTCTNNNAGATCGGAAG |
| 37421_3 | TATGAGTGTGGAGTCGTTGCAATTGCTGGGTCATATGGTGTCTNNNAGATCGGAAG |
| 37421_4 | TATGAGTGTGGAGTCGTTGCAATTGCTGGGTCATATGGTGTCTNNNAGATCGGAAG |
| 52648_1 | TATGAGTGTGGAGTCGTTGCCTGAAGTTTGCGTGTAAGACTGGNNNAGATCGGAA |
| 52648_2 | TATGAGTGTGGAGTCGTTGCCTGAAGTTTGCGTGTAAGACTGGNNNAGATCGGAA |
| 52648_3 | TATGAGTGTGGAGTCGTTGCCTGAAGTTTGCGTGTAAGACTGGNNNAGATCGGAA |
| 52648_4 | TATGAGTGTGGAGTCGTTGCCTGAAGTTTGCGTGTAAGACTGGNNNAGATCGGAA |
| 29243_1 | TATGAGTGTGGAGTCGTTGCAAAATCTGGGGGAGGGAGAGATANNNAGATCGGA |
| 29243_2 | TATGAGTGTGGAGTCGTTGCAAAATCTGGGGGAGGGAGAGATANNNAGATCGGA |
| 29243_3 | TATGAGTGTGGAGTCGTTGCAAAATCTGGGGGAGGGAGAGATANNNAGATCGGA |
| 29243_4 | TATGAGTGTGGAGTCGTTGCAAAATCTGGGGGAGGGAGAGATANNNAGATCGGA |
| 44410_1 | TATGAGTGTGGAGTCGTTGCGCAAGGGCTCTCATGAAGAAAAANNNAGATCGGAA |
| 44410_2 | TATGAGTGTGGAGTCGTTGCGCAAGGGCTCTCATGAAGAAAAANNNAGATCGGAA |
| 44410_3 | TATGAGTGTGGAGTCGTTGCGCAAGGGCTCTCATGAAGAAAAANNNAGATCGGAA |
| 44410_4 | TATGAGTGTGGAGTCGTTGCGCAAGGGCTCTCATGAAGAAAAANNNAGATCGGAA |
| 45380_1 | TATGAGTGTGGAGTCGTTGCAGATAGGGTCTCACTCTGTCACCNNNAGATCGGAAG |
| 45380_2 | TATGAGTGTGGAGTCGTTGCAGATAGGGTCTCACTCTGTCACCNNNAGATCGGAAG |
| 45380_3 | TATGAGTGTGGAGTCGTTGCAGATAGGGTCTCACTCTGTCACCNNNAGATCGGAAG |
| 45380_4 | TATGAGTGTGGAGTCGTTGCAGATAGGGTCTCACTCTGTCACCNNNAGATCGGAAG |
| 29412_1 | TATGAGTGTGGAGTCGTTGCGGGAATGTTGTGGGAAAAGGGATNNNAGATCGGA |
| 29412_2 | TATGAGTGTGGAGTCGTTGCGGGAATGTTGTGGGAAAAGGGATNNNAGATCGGA |
| 29412_3 | TATGAGTGTGGAGTCGTTGCGGGAATGTTGTGGGAAAAGGGATNNNAGATCGGA |
| 29412_4 | TATGAGTGTGGAGTCGTTGCGGGAATGTTGTGGGAAAAGGGATNNNAGATCGGA |
| 52646_1 | TATGAGTGTGGAGTCGTTGCGTTTCAAGGAGCAGAGGAGAGAANNNAGATCGGA |

| | |
|---|---|
| 52646_2 | TATGAGTGTGGAGTCGTTGCGTTTCAAGGAGCAGAGGAGAGAANNNAGATCGGA |
| 52646_3 | TATGAGTGTGGAGTCGTTGCGTTTCAAGGAGCAGAGGAGAGAANNNAGATCGGA |
| 52646_4 | TATGAGTGTGGAGTCGTTGCGTTTCAAGGAGCAGAGGAGAGAANNNAGATCGGA |
| 38726_1 | TATGAGTGTGGAGTCGTTGCGCGCAAAACTGGATGGATAATGANNNAGATCGGAA |
| 38726_2 | TATGAGTGTGGAGTCGTTGCGCGCAAAACTGGATGGATAATGANNNAGATCGGAA |
| 38726_3 | TATGAGTGTGGAGTCGTTGCGCGCAAAACTGGATGGATAATGANNNAGATCGGAA |
| 38726_4 | TATGAGTGTGGAGTCGTTGCGCGCAAAACTGGATGGATAATGANNNAGATCGGAA |
| 33698_1 | TATGAGTGTGGAGTCGTTGCGACACCTAGCTGGCCACATTATANNNAGATCGGAAG |
| 33698_2 | TATGAGTGTGGAGTCGTTGCGACACCTAGCTGGCCACATTATANNNAGATCGGAAG |
| 33698_3 | TATGAGTGTGGAGTCGTTGCGACACCTAGCTGGCCACATTATANNNAGATCGGAAG |
| 33698_4 | TATGAGTGTGGAGTCGTTGCGACACCTAGCTGGCCACATTATANNNAGATCGGAAG |
| 43819_1 | TATGAGTGTGGAGTCGTTGCATGCAGCCAAAAGACACATGAAANNNAGATCGGAA |
| 43819_2 | TATGAGTGTGGAGTCGTTGCATGCAGCCAAAAGACACATGAAANNNAGATCGGAA |
| 43819_3 | TATGAGTGTGGAGTCGTTGCATGCAGCCAAAAGACACATGAAANNNAGATCGGAA |
| 43819_4 | TATGAGTGTGGAGTCGTTGCATGCAGCCAAAAGACACATGAAANNNAGATCGGAA |
| 39758_1 | TATGAGTGTGGAGTCGTTGCGATCAAGAGGTCAGGAGATCGTGNNNAGATCGGAA |
| 39758_2 | TATGAGTGTGGAGTCGTTGCGATCAAGAGGTCAGGAGATCGTGNNNAGATCGGAA |
| 39758_3 | TATGAGTGTGGAGTCGTTGCGATCAAGAGGTCAGGAGATCGTGNNNAGATCGGAA |
| 39758_4 | TATGAGTGTGGAGTCGTTGCGATCAAGAGGTCAGGAGATCGTGNNNAGATCGGAA |
| 44526_1 | TATGAGTGTGGAGTCGTTGCGATCTCGGCTCACTGAAACCTCNNNAGATCGGAAGA |
| 44526_2 | TATGAGTGTGGAGTCGTTGCGATCTCGGCTCACTGAAACCTCNNNAGATCGGAAGA |
| 44526_3 | TATGAGTGTGGAGTCGTTGCGATCTCGGCTCACTGAAACCTCNNNAGATCGGAAGA |
| 44526_4 | TATGAGTGTGGAGTCGTTGCGATCTCGGCTCACTGAAACCTCNNNAGATCGGAAGA |
| 40125_1 | TATGAGTGTGGAGTCGTTGCCATAGAGAGATGAACAGGTGCCANNNAGATCGGAA |
| 40125_2 | TATGAGTGTGGAGTCGTTGCCATAGAGAGATGAACAGGTGCCANNNAGATCGGAA |
| 40125_3 | TATGAGTGTGGAGTCGTTGCCATAGAGAGATGAACAGGTGCCANNNAGATCGGAA |
| 40125_4 | TATGAGTGTGGAGTCGTTGCCATAGAGAGATGAACAGGTGCCANNNAGATCGGAA |
| 38737_1 | TATGAGTGTGGAGTCGTTGCAGGTGTTATGTAGCAAGACCCTGNNNAGATCGGAA |
| 38737_2 | TATGAGTGTGGAGTCGTTGCAGGTGTTATGTAGCAAGACCCTGNNNAGATCGGAA |
| 38737_3 | TATGAGTGTGGAGTCGTTGCAGGTGTTATGTAGCAAGACCCTGNNNAGATCGGAA |
| 38737_4 | TATGAGTGTGGAGTCGTTGCAGGTGTTATGTAGCAAGACCCTGNNNAGATCGGAA |
| 40342_1 | TATGAGTGTGGAGTCGTTGCAGGCAAAGGATATGAATTGGCAANNNAGATCGGAA |
| 40342_2 | TATGAGTGTGGAGTCGTTGCAGGCAAAGGATATGAATTGGCAANNNAGATCGGAA |
| 40342_3 | TATGAGTGTGGAGTCGTTGCAGGCAAAGGATATGAATTGGCAANNNAGATCGGAA |
| 40342_4 | TATGAGTGTGGAGTCGTTGCAGGCAAAGGATATGAATTGGCAANNNAGATCGGAA |
| 13532_1 | TATGAGTGTGGAGTCGTTGCGCAACCAAAAGATGTATAAAGTTNNNAGATCGGAA |
| 13532_2 | TATGAGTGTGGAGTCGTTGCGCAACCAAAAGATGTATAAAGTTNNNAGATCGGAA |
| 13532_3 | TATGAGTGTGGAGTCGTTGCGCAACCAAAAGATGTATAAAGTTNNNAGATCGGAA |
| 13532_4 | TATGAGTGTGGAGTCGTTGCGCAACCAAAAGATGTATAAAGTTNNNAGATCGGAA |
| 13482_1 | TATGAGTGTGGAGTCGTTGCCACATGAGCTTTTGGTAAATGNNNAGATCGGAAGA |
| 13482_2 | TATGAGTGTGGAGTCGTTGCCACATGAGCTTTTGGTAAATGNNNAGATCGGAAGA |
| 13482_3 | TATGAGTGTGGAGTCGTTGCCACATGAGCTTTTGGTAAATGNNNAGATCGGAAGA |
| 13482_4 | TATGAGTGTGGAGTCGTTGCCACATGAGCTTTTGGTAAATGNNNAGATCGGAAGA |

| | |
|---|---|
| 47839_1 | TATGAGTGTGGAGTCGTTGCCGTTTCCTGGATTCAAGCGATTCNNNAGATCGGAAG |
| 47839_2 | TATGAGTGTGGAGTCGTTGCCGTTTCCTGGATTCAAGCGATTCNNNAGATCGGAAG |
| 47839_3 | TATGAGTGTGGAGTCGTTGCCGTTTCCTGGATTCAAGCGATTCNNNAGATCGGAAG |
| 47839_4 | TATGAGTGTGGAGTCGTTGCCGTTTCCTGGATTCAAGCGATTCNNNAGATCGGAAG |
| 32246_1 | TATGAGTGTGGAGTCGTTGCCTGAGCCAGCCAATCAGGAAAGNNNAGATCGGAAG |
| 32246_2 | TATGAGTGTGGAGTCGTTGCCTGAGCCAGCCAATCAGGAAAGNNNAGATCGGAAG |
| 32246_3 | TATGAGTGTGGAGTCGTTGCCTGAGCCAGCCAATCAGGAAAGNNNAGATCGGAAG |
| 32246_4 | TATGAGTGTGGAGTCGTTGCCTGAGCCAGCCAATCAGGAAAGNNNAGATCGGAAG |
| 30172_1 | TATGAGTGTGGAGTCGTTGCTCTTCCAAGTTTTATGTGAGTTTTCTNNNAGATCGGA |
| 30172_2 | TATGAGTGTGGAGTCGTTGCTCTTCCAAGTTTTATGTGAGTTTTCTNNNAGATCGGA |
| 30172_3 | TATGAGTGTGGAGTCGTTGCTCTTCCAAGTTTTATGTGAGTTTTCTNNNAGATCGGA |
| 30172_4 | TATGAGTGTGGAGTCGTTGCTCTTCCAAGTTTTATGTGAGTTTTCTNNNAGATCGGA |
| 31117_1 | TATGAGTGTGGAGTCGTTGCTTGAGACAGAGTGCAATCTCAGCNNNAGATCGGAA |
| 31117_2 | TATGAGTGTGGAGTCGTTGCTTGAGACAGAGTGCAATCTCAGCNNNAGATCGGAA |
| 31117_3 | TATGAGTGTGGAGTCGTTGCTTGAGACAGAGTGCAATCTCAGCNNNAGATCGGAA |
| 31117_4 | TATGAGTGTGGAGTCGTTGCTTGAGACAGAGTGCAATCTCAGCNNNAGATCGGAA |
| 13305_1 | TATGAGTGTGGAGTCGTTGCCTTGGGATTCAGAATTTTGAGNNNAGATCGGAAGA |
| 13305_2 | TATGAGTGTGGAGTCGTTGCCTTGGGATTCAGAATTTTGAGNNNAGATCGGAAGA |
| 13305_3 | TATGAGTGTGGAGTCGTTGCCTTGGGATTCAGAATTTTGAGNNNAGATCGGAAGA |
| 13305_4 | TATGAGTGTGGAGTCGTTGCCTTGGGATTCAGAATTTTGAGNNNAGATCGGAAGA |
| 37139_1 | TATGAGTGTGGAGTCGTTGCGGGTGGATTTCGCATTGTTGTATNNNAGATCGGAA |
| 37139_2 | TATGAGTGTGGAGTCGTTGCGGGTGGATTTCGCATTGTTGTATNNNAGATCGGAA |
| 37139_3 | TATGAGTGTGGAGTCGTTGCGGGTGGATTTCGCATTGTTGTATNNNAGATCGGAA |
| 37139_4 | TATGAGTGTGGAGTCGTTGCGGGTGGATTTCGCATTGTTGTATNNNAGATCGGAA |
| 34569_1 | TATGAGTGTGGAGTCGTTGCGCCACATCATTTTTAGCCCTAGGNNNAGATCGGAAG |
| 34569_2 | TATGAGTGTGGAGTCGTTGCGCCACATCATTTTTAGCCCTAGGNNNAGATCGGAAG |
| 34569_3 | TATGAGTGTGGAGTCGTTGCGCCACATCATTTTTAGCCCTAGGNNNAGATCGGAAG |
| 34569_4 | TATGAGTGTGGAGTCGTTGCGCCACATCATTTTTAGCCCTAGGNNNAGATCGGAAG |
| 30818_1 | TATGAGTGTGGAGTCGTTGCTCTTGTAAGGCAGTGTAGATGCANNNAGATCGGAA |
| 30818_2 | TATGAGTGTGGAGTCGTTGCTCTTGTAAGGCAGTGTAGATGCANNNAGATCGGAA |
| 30818_3 | TATGAGTGTGGAGTCGTTGCTCTTGTAAGGCAGTGTAGATGCANNNAGATCGGAA |
| 30818_4 | TATGAGTGTGGAGTCGTTGCTCTTGTAAGGCAGTGTAGATGCANNNAGATCGGAA |
| 29138_1 | TATGAGTGTGGAGTCGTTGCGTGGCCTTCAAATGCATTAGGAGNNNAGATCGGAA |
| 29138_2 | TATGAGTGTGGAGTCGTTGCGTGGCCTTCAAATGCATTAGGAGNNNAGATCGGAA |
| 29138_3 | TATGAGTGTGGAGTCGTTGCGTGGCCTTCAAATGCATTAGGAGNNNAGATCGGAA |
| 29138_4 | TATGAGTGTGGAGTCGTTGCGTGGCCTTCAAATGCATTAGGAGNNNAGATCGGAA |
| 39420_1 | TATGAGTGTGGAGTCGTTGCCTCTGATCACCCCTGGAGAATTCNNNAGATCGGAAG |
| 39420_2 | TATGAGTGTGGAGTCGTTGCCTCTGATCACCCCTGGAGAATTCNNNAGATCGGAAG |
| 39420_3 | TATGAGTGTGGAGTCGTTGCCTCTGATCACCCCTGGAGAATTCNNNAGATCGGAAG |
| 39420_4 | TATGAGTGTGGAGTCGTTGCCTCTGATCACCCCTGGAGAATTCNNNAGATCGGAAG |
| 39527_1 | TATGAGTGTGGAGTCGTTGCCCACACCTATAATCCCAGCACTTNNNAGATCGGAAGA |
| 39527_2 | TATGAGTGTGGAGTCGTTGCCCACACCTATAATCCCAGCACTTNNNAGATCGGAAGA |
| 39527_3 | TATGAGTGTGGAGTCGTTGCCCACACCTATAATCCCAGCACTTNNNAGATCGGAAGA |

| | |
|---|---|
| 39527_4 | TATGAGTGTGGAGTCGTTGCCCACACCTATAATCCCAGCACTTNNNAGATCGGAAGA |
| 44198_1 | TATGAGTGTGGAGTCGTTGCCATGTGTTTGCATGGTCTCTCAGNNNAGATCGGAAG |
| 44198_2 | TATGAGTGTGGAGTCGTTGCCATGTGTTTGCATGGTCTCTCAGNNNAGATCGGAAG |
| 44198_3 | TATGAGTGTGGAGTCGTTGCCATGTGTTTGCATGGTCTCTCAGNNNAGATCGGAAG |
| 44198_4 | TATGAGTGTGGAGTCGTTGCCATGTGTTTGCATGGTCTCTCAGNNNAGATCGGAAG |
| 41390_1 | TATGAGTGTGGAGTCGTTGCCATGGCTCCAGGTCTCAATTTTCNNNAGATCGGAAGA |
| 41390_2 | TATGAGTGTGGAGTCGTTGCCATGGCTCCAGGTCTCAATTTTCNNNAGATCGGAAGA |
| 41390_3 | TATGAGTGTGGAGTCGTTGCCATGGCTCCAGGTCTCAATTTTCNNNAGATCGGAAGA |
| 41390_4 | TATGAGTGTGGAGTCGTTGCCATGGCTCCAGGTCTCAATTTTCNNNAGATCGGAAGA |
| 42301_1 | TATGAGTGTGGAGTCGTTGCGCAGCCCAATTTGTATTCCAGAGNNNAGATCGGAAG |
| 42301_2 | TATGAGTGTGGAGTCGTTGCGCAGCCCAATTTGTATTCCAGAGNNNAGATCGGAAG |
| 42301_3 | TATGAGTGTGGAGTCGTTGCGCAGCCCAATTTGTATTCCAGAGNNNAGATCGGAAG |
| 42301_4 | TATGAGTGTGGAGTCGTTGCGCAGCCCAATTTGTATTCCAGAGNNNAGATCGGAAG |
| 2824_1 | TATGAGTGTGGAGTCGTTGCCCTGTCCTTGAGCAAGTCAAAGTNNNAGATCGGAAG |
| 2824_2 | TATGAGTGTGGAGTCGTTGCCCTGTCCTTGAGCAAGTCAAAGTNNNAGATCGGAAG |
| 2824_3 | TATGAGTGTGGAGTCGTTGCCCTGTCCTTGAGCAAGTCAAAGTNNNAGATCGGAAG |
| 2824_4 | TATGAGTGTGGAGTCGTTGCCCTGTCCTTGAGCAAGTCAAAGTNNNAGATCGGAAG |
| 34041_1 | TATGAGTGTGGAGTCGTTGCCAAGTAGCCTAGTAGCTGGGATTNNNAGATCGGAAG |
| 34041_2 | TATGAGTGTGGAGTCGTTGCCAAGTAGCCTAGTAGCTGGGATTNNNAGATCGGAAG |
| 34041_3 | TATGAGTGTGGAGTCGTTGCCAAGTAGCCTAGTAGCTGGGATTNNNAGATCGGAAG |
| 34041_4 | TATGAGTGTGGAGTCGTTGCCAAGTAGCCTAGTAGCTGGGATTNNNAGATCGGAAG |
| 40959_1 | TATGAGTGTGGAGTCGTTGCTTGAATTTCAGGTTGAAGGTCCTNNNAGATCGGAAG |
| 40959_2 | TATGAGTGTGGAGTCGTTGCTTGAATTTCAGGTTGAAGGTCCTNNNAGATCGGAAG |
| 40959_3 | TATGAGTGTGGAGTCGTTGCTTGAATTTCAGGTTGAAGGTCCTNNNAGATCGGAAG |
| 40959_4 | TATGAGTGTGGAGTCGTTGCTTGAATTTCAGGTTGAAGGTCCTNNNAGATCGGAAG |
| 42663_1 | TATGAGTGTGGAGTCGTTGCGCTTGTAGATGAAACTGCCAAGTNNNAGATCGGAAG |
| 42663_2 | TATGAGTGTGGAGTCGTTGCGCTTGTAGATGAAACTGCCAAGTNNNAGATCGGAAG |
| 42663_3 | TATGAGTGTGGAGTCGTTGCGCTTGTAGATGAAACTGCCAAGTNNNAGATCGGAAG |
| 42663_4 | TATGAGTGTGGAGTCGTTGCGCTTGTAGATGAAACTGCCAAGTNNNAGATCGGAAG |
| 30016_1 | TATGAGTGTGGAGTCGTTGCAAATTGGGGCATTATCATTGACANNNAGATCGGAAG |
| 30016_2 | TATGAGTGTGGAGTCGTTGCAAATTGGGGCATTATCATTGACANNNAGATCGGAAG |
| 30016_3 | TATGAGTGTGGAGTCGTTGCAAATTGGGGCATTATCATTGACANNNAGATCGGAAG |
| 30016_4 | TATGAGTGTGGAGTCGTTGCAAATTGGGGCATTATCATTGACANNNAGATCGGAAG |
| 33228_1 | TATGAGTGTGGAGTCGTTGCAGTATTCCCACCTAGTCAGAGCTNNNAGATCGGAAGA |
| 33228_2 | TATGAGTGTGGAGTCGTTGCAGTATTCCCACCTAGTCAGAGCTNNNAGATCGGAAGA |
| 33228_3 | TATGAGTGTGGAGTCGTTGCAGTATTCCCACCTAGTCAGAGCTNNNAGATCGGAAGA |
| 33228_4 | TATGAGTGTGGAGTCGTTGCAGTATTCCCACCTAGTCAGAGCTNNNAGATCGGAAGA |
| 32095_1 | TATGAGTGTGGAGTCGTTGCTGACAAAAAAGCCCCATTTGAACTNNNAGATCGGAAG |
| 32095_2 | TATGAGTGTGGAGTCGTTGCTGACAAAAAAGCCCCATTTGAACTNNNAGATCGGAAG |
| 32095_3 | TATGAGTGTGGAGTCGTTGCTGACAAAAAAGCCCCATTTGAACTNNNAGATCGGAAG |
| 32095_4 | TATGAGTGTGGAGTCGTTGCTGACAAAAAAGCCCCATTTGAACTNNNAGATCGGAAG |
| 44967_1 | TATGAGTGTGGAGTCGTTGCAAAAACAAACTGGGAAAGGGAGCNNNAGATCGGAAA |
| 44967_2 | TATGAGTGTGGAGTCGTTGCAAAAACAAACTGGGAAAGGGAGCNNNAGATCGGAAA |

| | |
|---|---|
| 44967_3 | TATGAGTGTGGAGTCGTTGCAAAAACAAACTGGGAAAGGGAGCNNNAGATCGGA |
| 44967_4 | TATGAGTGTGGAGTCGTTGCAAAAACAAACTGGGAAAGGGAGCNNNAGATCGGA |
| 51575_1 | TATGAGTGTGGAGTCGTTGCCCCCAGCTCACTAGTATTTTGGTNNNAGATCGGAAG |
| 51575_2 | TATGAGTGTGGAGTCGTTGCCCCCAGCTCACTAGTATTTTGGTNNNAGATCGGAAG |
| 51575_3 | TATGAGTGTGGAGTCGTTGCCCCCAGCTCACTAGTATTTTGGTNNNAGATCGGAAG |
| 51575_4 | TATGAGTGTGGAGTCGTTGCCCCCAGCTCACTAGTATTTTGGTNNNAGATCGGAAG |
| 42693_1 | TATGAGTGTGGAGTCGTTGCCTCTCTCTCTTTCTCCCTCTCCANNNAGATCGGAAGAG |
| 42693_2 | TATGAGTGTGGAGTCGTTGCCTCTCTCTCTTTCTCCCTCTCCANNNAGATCGGAAGAG |
| 42693_3 | TATGAGTGTGGAGTCGTTGCCTCTCTCTCTTTCTCCCTCTCCANNNAGATCGGAAGAG |
| 42693_4 | TATGAGTGTGGAGTCGTTGCCTCTCTCTCTTTCTCCCTCTCCANNNAGATCGGAAGAG |
| 52036_1 | TATGAGTGTGGAGTCGTTGCATGGGAGTAGATTCCTGGTTTGGNNNAGATCGGAA |
| 52036_2 | TATGAGTGTGGAGTCGTTGCATGGGAGTAGATTCCTGGTTTGGNNNAGATCGGAA |
| 52036_3 | TATGAGTGTGGAGTCGTTGCATGGGAGTAGATTCCTGGTTTGGNNNAGATCGGAA |
| 52036_4 | TATGAGTGTGGAGTCGTTGCATGGGAGTAGATTCCTGGTTTGGNNNAGATCGGAA |
| 42640_1 | TATGAGTGTGGAGTCGTTGCCAGATGGACTGGAGGTAGAGAGANNNAGATCGGA |
| 42640_2 | TATGAGTGTGGAGTCGTTGCCAGATGGACTGGAGGTAGAGAGANNNAGATCGGA |
| 42640_3 | TATGAGTGTGGAGTCGTTGCCAGATGGACTGGAGGTAGAGAGANNNAGATCGGA |
| 42640_4 | TATGAGTGTGGAGTCGTTGCCAGATGGACTGGAGGTAGAGAGANNNAGATCGGA |
| 43822_1 | TATGAGTGTGGAGTCGTTGCTGAATCATACCTCGTTTCGGGTTNNNAGATCGGAAG |
| 43822_2 | TATGAGTGTGGAGTCGTTGCTGAATCATACCTCGTTTCGGGTTNNNAGATCGGAAG |
| 43822_3 | TATGAGTGTGGAGTCGTTGCTGAATCATACCTCGTTTCGGGTTNNNAGATCGGAAG |
| 43822_4 | TATGAGTGTGGAGTCGTTGCTGAATCATACCTCGTTTCGGGTTNNNAGATCGGAAG |
| 37030_1 | TATGAGTGTGGAGTCGTTGCCATGGACATGGGTCAAAACTGTGNNNAGATCGGAA |
| 37030_2 | TATGAGTGTGGAGTCGTTGCCATGGACATGGGTCAAAACTGTGNNNAGATCGGAA |
| 37030_3 | TATGAGTGTGGAGTCGTTGCCATGGACATGGGTCAAAACTGTGNNNAGATCGGAA |
| 37030_4 | TATGAGTGTGGAGTCGTTGCCATGGACATGGGTCAAAACTGTGNNNAGATCGGAA |
| 40925_1 | TATGAGTGTGGAGTCGTTGCAGAAGAAAAGAATCCTGCCACCTNNNAGATCGGAAG |
| 40925_2 | TATGAGTGTGGAGTCGTTGCAGAAGAAAAGAATCCTGCCACCTNNNAGATCGGAAG |
| 40925_3 | TATGAGTGTGGAGTCGTTGCAGAAGAAAAGAATCCTGCCACCTNNNAGATCGGAAG |
| 40925_4 | TATGAGTGTGGAGTCGTTGCAGAAGAAAAGAATCCTGCCACCTNNNAGATCGGAAG |
| 28791_1 | TATGAGTGTGGAGTCGTTGCAGAAGGTAGCACACTTCCAACATNNNAGATCGGAAG |
| 28791_2 | TATGAGTGTGGAGTCGTTGCAGAAGGTAGCACACTTCCAACATNNNAGATCGGAAG |
| 28791_3 | TATGAGTGTGGAGTCGTTGCAGAAGGTAGCACACTTCCAACATNNNAGATCGGAAG |
| 28791_4 | TATGAGTGTGGAGTCGTTGCAGAAGGTAGCACACTTCCAACATNNNAGATCGGAAG |
| 12864_1 | TATGAGTGTGGAGTCGTTGCACTGTCTAGAGCTTTTTGATTCNNNAGATCGGAAGA |
| 12864_2 | TATGAGTGTGGAGTCGTTGCACTGTCTAGAGCTTTTTGATTCNNNAGATCGGAAGA |
| 12864_3 | TATGAGTGTGGAGTCGTTGCACTGTCTAGAGCTTTTTGATTCNNNAGATCGGAAGA |
| 12864_4 | TATGAGTGTGGAGTCGTTGCACTGTCTAGAGCTTTTTGATTCNNNAGATCGGAAGA |
| 44545_1 | TATGAGTGTGGAGTCGTTGCACACACACACACACACACAATTTNNNAGATCGGAAG |
| 44545_2 | TATGAGTGTGGAGTCGTTGCACACACACACACACACACAATTTNNNAGATCGGAAG |
| 44545_3 | TATGAGTGTGGAGTCGTTGCACACACACACACACACACAATTTNNNAGATCGGAAG |
| 44545_4 | TATGAGTGTGGAGTCGTTGCACACACACACACACACACAATTTNNNAGATCGGAAG |
| 50692_1 | TATGAGTGTGGAGTCGTTGCGTCTTGAATTTGCTCTTGGCCTTNNNAGATCGGAAG |

| | |
|---|---|
| 50692_2 | TATGAGTGTGGAGTCGTTGCGTCTTGAATTTGCTCTTGGCCTTNNNAGATCGGAAG |
| 50692_3 | TATGAGTGTGGAGTCGTTGCGTCTTGAATTTGCTCTTGGCCTTNNNAGATCGGAAG |
| 50692_4 | TATGAGTGTGGAGTCGTTGCGTCTTGAATTTGCTCTTGGCCTTNNNAGATCGGAAG |
| 47431_1 | TATGAGTGTGGAGTCGTTGCTCTGTGTCGGGAAAGCAGAATAANNNAGATCGGAA |
| 47431_2 | TATGAGTGTGGAGTCGTTGCTCTGTGTCGGGAAAGCAGAATAANNNAGATCGGAA |
| 47431_3 | TATGAGTGTGGAGTCGTTGCTCTGTGTCGGGAAAGCAGAATAANNNAGATCGGAA |
| 47431_4 | TATGAGTGTGGAGTCGTTGCTCTGTGTCGGGAAAGCAGAATAANNNAGATCGGAA |
| 40714_1 | TATGAGTGTGGAGTCGTTGCAGAGAGAGAAGGAAGGAGGGAAANNNAGATCGG |
| 40714_2 | TATGAGTGTGGAGTCGTTGCAGAGAGAGAAGGAAGGAGGGAAANNNAGATCGG |
| 40714_3 | TATGAGTGTGGAGTCGTTGCAGAGAGAGAAGGAAGGAGGGAAANNNAGATCGG |
| 40714_4 | TATGAGTGTGGAGTCGTTGCAGAGAGAGAAGGAAGGAGGGAAANNNAGATCGG |
| 10055_1 | TATGAGTGTGGAGTCGTTGCTTCAATGGATTCTGGTTGGGGACNNNAGATCGGAA |
| 10055_2 | TATGAGTGTGGAGTCGTTGCTTCAATGGATTCTGGTTGGGGACNNNAGATCGGAA |
| 10055_3 | TATGAGTGTGGAGTCGTTGCTTCAATGGATTCTGGTTGGGGACNNNAGATCGGAA |
| 10055_4 | TATGAGTGTGGAGTCGTTGCTTCAATGGATTCTGGTTGGGGACNNNAGATCGGAA |
| 38082_1 | TATGAGTGTGGAGTCGTTGCAATGAGGGCAACTTGTTGTTCTGNNNAGATCGGAA |
| 38082_2 | TATGAGTGTGGAGTCGTTGCAATGAGGGCAACTTGTTGTTCTGNNNAGATCGGAA |
| 38082_3 | TATGAGTGTGGAGTCGTTGCAATGAGGGCAACTTGTTGTTCTGNNNAGATCGGAA |
| 38082_4 | TATGAGTGTGGAGTCGTTGCAATGAGGGCAACTTGTTGTTCTGNNNAGATCGGAA |
| 45366_1 | TATGAGTGTGGAGTCGTTGCCACACTGTCTTCCACAATGGTTGNNNAGATCGGAAG |
| 45366_2 | TATGAGTGTGGAGTCGTTGCCACACTGTCTTCCACAATGGTTGNNNAGATCGGAAG |
| 45366_3 | TATGAGTGTGGAGTCGTTGCCACACTGTCTTCCACAATGGTTGNNNAGATCGGAAG |
| 45366_4 | TATGAGTGTGGAGTCGTTGCCACACTGTCTTCCACAATGGTTGNNNAGATCGGAAG |
| 43314_1 | TATGAGTGTGGAGTCGTTGCCATTCTCCTGCCTCAGCCTCNNNAGATCGGAAGAGCA |
| 43314_2 | TATGAGTGTGGAGTCGTTGCCATTCTCCTGCCTCAGCCTCNNNAGATCGGAAGAGCA |
| 43314_3 | TATGAGTGTGGAGTCGTTGCCATTCTCCTGCCTCAGCCTCNNNAGATCGGAAGAGCA |
| 43314_4 | TATGAGTGTGGAGTCGTTGCCATTCTCCTGCCTCAGCCTCNNNAGATCGGAAGAGCA |
| 10041_1 | TATGAGTGTGGAGTCGTTGCAAAGAAGAGGGTACCTGAGCCTTNNNAGATCGGAA |
| 10041_2 | TATGAGTGTGGAGTCGTTGCAAAGAAGAGGGTACCTGAGCCTTNNNAGATCGGAA |
| 10041_3 | TATGAGTGTGGAGTCGTTGCAAAGAAGAGGGTACCTGAGCCTTNNNAGATCGGAA |
| 10041_4 | TATGAGTGTGGAGTCGTTGCAAAGAAGAGGGTACCTGAGCCTTNNNAGATCGGAA |
| 13707_1 | TATGAGTGTGGAGTCGTTGCCAGACTCTTACCAGCATATATAGNNNAGATCGGAAG |
| 13707_2 | TATGAGTGTGGAGTCGTTGCCAGACTCTTACCAGCATATATAGNNNAGATCGGAAG |
| 13707_3 | TATGAGTGTGGAGTCGTTGCCAGACTCTTACCAGCATATATAGNNNAGATCGGAAG |
| 13707_4 | TATGAGTGTGGAGTCGTTGCCAGACTCTTACCAGCATATATAGNNNAGATCGGAAG |
| 41106_1 | TATGAGTGTGGAGTCGTTGCAGTGCCTAACAACAAAGTGTTCANNNAGATCGGAAG |
| 41106_2 | TATGAGTGTGGAGTCGTTGCAGTGCCTAACAACAAAGTGTTCANNNAGATCGGAAG |
| 41106_3 | TATGAGTGTGGAGTCGTTGCAGTGCCTAACAACAAAGTGTTCANNNAGATCGGAAG |
| 41106_4 | TATGAGTGTGGAGTCGTTGCAGTGCCTAACAACAAAGTGTTCANNNAGATCGGAAG |
| 51579_1 | TATGAGTGTGGAGTCGTTGCCTCATGCCTATAATCCCAGCACTNNNAGATCGGAAGA |
| 51579_2 | TATGAGTGTGGAGTCGTTGCCTCATGCCTATAATCCCAGCACTNNNAGATCGGAAGA |
| 51579_3 | TATGAGTGTGGAGTCGTTGCCTCATGCCTATAATCCCAGCACTNNNAGATCGGAAGA |
| 51579_4 | TATGAGTGTGGAGTCGTTGCCTCATGCCTATAATCCCAGCACTNNNAGATCGGAAGA |

| | |
|---|---|
| 13034_1 | TATGAGTGTGGAGTCGTTGCCATCATCTGCAAAAACATCCNNNAGATCGGAAGAGC |
| 13034_2 | TATGAGTGTGGAGTCGTTGCCATCATCTGCAAAAACATCCNNNAGATCGGAAGAGC |
| 13034_3 | TATGAGTGTGGAGTCGTTGCCATCATCTGCAAAAACATCCNNNAGATCGGAAGAGC |
| 13034_4 | TATGAGTGTGGAGTCGTTGCCATCATCTGCAAAAACATCCNNNAGATCGGAAGAGC |
| 41513_1 | TATGAGTGTGGAGTCGTTGCGGGCACTTGTAATCTCAGCTACTNNNAGATCGGAAG |
| 41513_2 | TATGAGTGTGGAGTCGTTGCGGGCACTTGTAATCTCAGCTACTNNNAGATCGGAAG |
| 41513_3 | TATGAGTGTGGAGTCGTTGCGGGCACTTGTAATCTCAGCTACTNNNAGATCGGAAG |
| 41513_4 | TATGAGTGTGGAGTCGTTGCGGGCACTTGTAATCTCAGCTACTNNNAGATCGGAAG |
| 41521_1 | TATGAGTGTGGAGTCGTTGCATCTTGCCAAAGCTGTCCTAGATNNNAGATCGGAAG |
| 41521_2 | TATGAGTGTGGAGTCGTTGCATCTTGCCAAAGCTGTCCTAGATNNNAGATCGGAAG |
| 41521_3 | TATGAGTGTGGAGTCGTTGCATCTTGCCAAAGCTGTCCTAGATNNNAGATCGGAAG |
| 41521_4 | TATGAGTGTGGAGTCGTTGCATCTTGCCAAAGCTGTCCTAGATNNNAGATCGGAAG |
| 9796_1 | TATGAGTGTGGAGTCGTTGCTGTAATCTCAGCCCCAGCATCTANNNAGATCGGAAG |
| 9796_2 | TATGAGTGTGGAGTCGTTGCTGTAATCTCAGCCCCAGCATCTANNNAGATCGGAAG |
| 9796_3 | TATGAGTGTGGAGTCGTTGCTGTAATCTCAGCCCCAGCATCTANNNAGATCGGAAG |
| 9796_4 | TATGAGTGTGGAGTCGTTGCTGTAATCTCAGCCCCAGCATCTANNNAGATCGGAAG |
| 37218_1 | TATGAGTGTGGAGTCGTTGCACAAACAGACATGCTACAAGTCTNNNAGATCGGAAG |
| 37218_2 | TATGAGTGTGGAGTCGTTGCACAAACAGACATGCTACAAGTCTNNNAGATCGGAAG |
| 37218_3 | TATGAGTGTGGAGTCGTTGCACAAACAGACATGCTACAAGTCTNNNAGATCGGAAG |
| 37218_4 | TATGAGTGTGGAGTCGTTGCACAAACAGACATGCTACAAGTCTNNNAGATCGGAAG |
| 43156_1 | TATGAGTGTGGAGTCGTTGCCGTGCCTGGCTAATTTGTGTATTNNNAGATCGGAAG |
| 43156_2 | TATGAGTGTGGAGTCGTTGCCGTGCCTGGCTAATTTGTGTATTNNNAGATCGGAAG |
| 43156_3 | TATGAGTGTGGAGTCGTTGCCGTGCCTGGCTAATTTGTGTATTNNNAGATCGGAAG |
| 43156_4 | TATGAGTGTGGAGTCGTTGCCGTGCCTGGCTAATTTGTGTATTNNNAGATCGGAAG |
| 37327_1 | TATGAGTGTGGAGTCGTTGCCTGGGTGACAGAGCAAGATCTTANNNAGATCGGAA |
| 37327_2 | TATGAGTGTGGAGTCGTTGCCTGGGTGACAGAGCAAGATCTTANNNAGATCGGAA |
| 37327_3 | TATGAGTGTGGAGTCGTTGCCTGGGTGACAGAGCAAGATCTTANNNAGATCGGAA |
| 37327_4 | TATGAGTGTGGAGTCGTTGCCTGGGTGACAGAGCAAGATCTTANNNAGATCGGAA |
| 33001_1 | TATGAGTGTGGAGTCGTTGCTTGCCTAGTCTGGAGTGTAATGGNNNAGATCGGAA |
| 33001_2 | TATGAGTGTGGAGTCGTTGCTTGCCTAGTCTGGAGTGTAATGGNNNAGATCGGAA |
| 33001_3 | TATGAGTGTGGAGTCGTTGCTTGCCTAGTCTGGAGTGTAATGGNNNAGATCGGAA |
| 33001_4 | TATGAGTGTGGAGTCGTTGCTTGCCTAGTCTGGAGTGTAATGGNNNAGATCGGAA |
| 43570_1 | TATGAGTGTGGAGTCGTTGCTTCAAGTGATTCTCCTGCCTCAGNNNAGATCGGAAG |
| 43570_2 | TATGAGTGTGGAGTCGTTGCTTCAAGTGATTCTCCTGCCTCAGNNNAGATCGGAAG |
| 43570_3 | TATGAGTGTGGAGTCGTTGCTTCAAGTGATTCTCCTGCCTCAGNNNAGATCGGAAG |
| 43570_4 | TATGAGTGTGGAGTCGTTGCTTCAAGTGATTCTCCTGCCTCAGNNNAGATCGGAAG |
| 31151_1 | TATGAGTGTGGAGTCGTTGCCTGAAACTTGGCTGTCCTCAAAGNNNAGATCGGAAG |
| 31151_2 | TATGAGTGTGGAGTCGTTGCCTGAAACTTGGCTGTCCTCAAAGNNNAGATCGGAAG |
| 31151_3 | TATGAGTGTGGAGTCGTTGCCTGAAACTTGGCTGTCCTCAAAGNNNAGATCGGAAG |
| 31151_4 | TATGAGTGTGGAGTCGTTGCCTGAAACTTGGCTGTCCTCAAAGNNNAGATCGGAAG |
| 34943_1 | TATGAGTGTGGAGTCGTTGCCTCCCAAGTAGCTGGAACTACAGNNNAGATCGGAAG |
| 34943_2 | TATGAGTGTGGAGTCGTTGCCTCCCAAGTAGCTGGAACTACAGNNNAGATCGGAAG |
| 34943_3 | TATGAGTGTGGAGTCGTTGCCTCCCAAGTAGCTGGAACTACAGNNNAGATCGGAAG |

| | |
|---|---|
| 34943_4 | TATGAGTGTGGAGTCGTTGCCTCCCAAGTAGCTGGAACTACAGNNNAGATCGGAAG |
| 48185_1 | TATGAGTGTGGAGTCGTTGCCTCTGGTACTGCTCACAGGTCNNNAGATCGGAAGAG |
| 48185_2 | TATGAGTGTGGAGTCGTTGCCTCTGGTACTGCTCACAGGTCNNNAGATCGGAAGAG |
| 48185_3 | TATGAGTGTGGAGTCGTTGCCTCTGGTACTGCTCACAGGTCNNNAGATCGGAAGAG |
| 48185_4 | TATGAGTGTGGAGTCGTTGCCTCTGGTACTGCTCACAGGTCNNNAGATCGGAAGAG |
| 49359_1 | TATGAGTGTGGAGTCGTTGCGGTCTGAATGCATTCCACACAATNNNAGATCGGAAG |
| 49359_2 | TATGAGTGTGGAGTCGTTGCGGTCTGAATGCATTCCACACAATNNNAGATCGGAAG |
| 49359_3 | TATGAGTGTGGAGTCGTTGCGGTCTGAATGCATTCCACACAATNNNAGATCGGAAG |
| 49359_4 | TATGAGTGTGGAGTCGTTGCGGTCTGAATGCATTCCACACAATNNNAGATCGGAAG |
| 46541_1 | TATGAGTGTGGAGTCGTTGCCCTCCTGAGTAGCTGAGACTACANNNAGATCGGAAG |
| 46541_2 | TATGAGTGTGGAGTCGTTGCCCTCCTGAGTAGCTGAGACTACANNNAGATCGGAAG |
| 46541_3 | TATGAGTGTGGAGTCGTTGCCCTCCTGAGTAGCTGAGACTACANNNAGATCGGAAG |
| 46541_4 | TATGAGTGTGGAGTCGTTGCCCTCCTGAGTAGCTGAGACTACANNNAGATCGGAAG |
| 51879_1 | TATGAGTGTGGAGTCGTTGCTGTGGTGTTTGGTTTTCTGTTCCNNNAGATCGGAAG |
| 51879_2 | TATGAGTGTGGAGTCGTTGCTGTGGTGTTTGGTTTTCTGTTCCNNNAGATCGGAAG |
| 51879_3 | TATGAGTGTGGAGTCGTTGCTGTGGTGTTTGGTTTTCTGTTCCNNNAGATCGGAAG |
| 51879_4 | TATGAGTGTGGAGTCGTTGCTGTGGTGTTTGGTTTTCTGTTCCNNNAGATCGGAAG |
| 44908_1 | TATGAGTGTGGAGTCGTTGCGCCTGTTTTCTAAAGAGCTTCCCNNNAGATCGGAAG |
| 44908_2 | TATGAGTGTGGAGTCGTTGCGCCTGTTTTCTAAAGAGCTTCCCNNNAGATCGGAAG |
| 44908_3 | TATGAGTGTGGAGTCGTTGCGCCTGTTTTCTAAAGAGCTTCCCNNNAGATCGGAAG |
| 44908_4 | TATGAGTGTGGAGTCGTTGCGCCTGTTTTCTAAAGAGCTTCCCNNNAGATCGGAAG |
| 3171_1 | TATGAGTGTGGAGTCGTTGCCTGCACACAGAAACATAAAGCCANNNAGATCGGAAG |
| 3171_2 | TATGAGTGTGGAGTCGTTGCCTGCACACAGAAACATAAAGCCANNNAGATCGGAAG |
| 3171_3 | TATGAGTGTGGAGTCGTTGCCTGCACACAGAAACATAAAGCCANNNAGATCGGAAG |
| 3171_4 | TATGAGTGTGGAGTCGTTGCCTGCACACAGAAACATAAAGCCANNNAGATCGGAAG |
| 33594_1 | TATGAGTGTGGAGTCGTTGCGGTTCAAGTGATTCTTCTGCCTCNNNAGATCGGAAG |
| 33594_2 | TATGAGTGTGGAGTCGTTGCGGTTCAAGTGATTCTTCTGCCTCNNNAGATCGGAAG |
| 33594_3 | TATGAGTGTGGAGTCGTTGCGGTTCAAGTGATTCTTCTGCCTCNNNAGATCGGAAG |
| 33594_4 | TATGAGTGTGGAGTCGTTGCGGTTCAAGTGATTCTTCTGCCTCNNNAGATCGGAAG |
| 46328_1 | TATGAGTGTGGAGTCGTTGCGCACCAACACATCAGGGTAATTTNNNAGATCGGAAG |
| 46328_2 | TATGAGTGTGGAGTCGTTGCGCACCAACACATCAGGGTAATTTNNNAGATCGGAAG |
| 46328_3 | TATGAGTGTGGAGTCGTTGCGCACCAACACATCAGGGTAATTTNNNAGATCGGAAG |
| 46328_4 | TATGAGTGTGGAGTCGTTGCGCACCAACACATCAGGGTAATTTNNNAGATCGGAAG |
| 33890_1 | TATGAGTGTGGAGTCGTTGCGCCTCCCAAGTAGCTGGATTATANNNAGATCGGAAG |
| 33890_2 | TATGAGTGTGGAGTCGTTGCGCCTCCCAAGTAGCTGGATTATANNNAGATCGGAAG |
| 33890_3 | TATGAGTGTGGAGTCGTTGCGCCTCCCAAGTAGCTGGATTATANNNAGATCGGAAG |
| 33890_4 | TATGAGTGTGGAGTCGTTGCGCCTCCCAAGTAGCTGGATTATANNNAGATCGGAAG |
| 38839_1 | TATGAGTGTGGAGTCGTTGCGTGCTTGAGAGAAGGGAACTACANNNAGATCGGAA |
| 38839_2 | TATGAGTGTGGAGTCGTTGCGTGCTTGAGAGAAGGGAACTACANNNAGATCGGAA |
| 38839_3 | TATGAGTGTGGAGTCGTTGCGTGCTTGAGAGAAGGGAACTACANNNAGATCGGAA |
| 38839_4 | TATGAGTGTGGAGTCGTTGCGTGCTTGAGAGAAGGGAACTACANNNAGATCGGAA |
| 32102_1 | TATGAGTGTGGAGTCGTTGCTCTCCTGGAGCTTTTTCTGTCTGNNNAGATCGGAAG |
| 32102_2 | TATGAGTGTGGAGTCGTTGCTCTCCTGGAGCTTTTTCTGTCTGNNNAGATCGGAAG |

| | |
|---|---|
| 32102_3 | TATGAGTGTGGAGTCGTTGCTCTCCTGGAGCTTTTTCTGTCTGNNNAGATCGGAAGA |
| 32102_4 | TATGAGTGTGGAGTCGTTGCTCTCCTGGAGCTTTTTCTGTCTGNNNAGATCGGAAGA |
| 52138_1 | TATGAGTGTGGAGTCGTTGCTGCTCTTTAGGAAGGCACAATCTNNNAGATCGGAAG |
| 52138_2 | TATGAGTGTGGAGTCGTTGCTGCTCTTTAGGAAGGCACAATCTNNNAGATCGGAAG |
| 52138_3 | TATGAGTGTGGAGTCGTTGCTGCTCTTTAGGAAGGCACAATCTNNNAGATCGGAAG |
| 52138_4 | TATGAGTGTGGAGTCGTTGCTGCTCTTTAGGAAGGCACAATCTNNNAGATCGGAAG |
| 29980_1 | TATGAGTGTGGAGTCGTTGCTGGCGGTGTCTAAGATGACTTGNNNAGATCGGAAGA |
| 29980_2 | TATGAGTGTGGAGTCGTTGCTGGCGGTGTCTAAGATGACTTGNNNAGATCGGAAGA |
| 29980_3 | TATGAGTGTGGAGTCGTTGCTGGCGGTGTCTAAGATGACTTGNNNAGATCGGAAGA |
| 29980_4 | TATGAGTGTGGAGTCGTTGCTGGCGGTGTCTAAGATGACTTGNNNAGATCGGAAGA |
| 2503_1 | TATGAGTGTGGAGTCGTTGCGAGGCTGAGAAGTTCTGATGGAGNNNAGATCGGAA |
| 2503_2 | TATGAGTGTGGAGTCGTTGCGAGGCTGAGAAGTTCTGATGGAGNNNAGATCGGAA |
| 2503_3 | TATGAGTGTGGAGTCGTTGCGAGGCTGAGAAGTTCTGATGGAGNNNAGATCGGAA |
| 2503_4 | TATGAGTGTGGAGTCGTTGCGAGGCTGAGAAGTTCTGATGGAGNNNAGATCGGAA |
| 13392_1 | TATGAGTGTGGAGTCGTTGCGGATACCTATAATTGGATATTTCCNNNAGATCGGAA[illegible] |
| 13392_2 | TATGAGTGTGGAGTCGTTGCGGATACCTATAATTGGATATTTCCNNNAGATCGGAA[illegible] |
| 13392_3 | TATGAGTGTGGAGTCGTTGCGGATACCTATAATTGGATATTTCCNNNAGATCGGAA[illegible] |
| 13392_4 | TATGAGTGTGGAGTCGTTGCGGATACCTATAATTGGATATTTCCNNNAGATCGGAA[illegible] |
| 33807_1 | TATGAGTGTGGAGTCGTTGCGAGGCAGGAGAATCGCTAGAACNNNAGATCGGAAG |
| 33807_2 | TATGAGTGTGGAGTCGTTGCGAGGCAGGAGAATCGCTAGAACNNNAGATCGGAAG |
| 33807_3 | TATGAGTGTGGAGTCGTTGCGAGGCAGGAGAATCGCTAGAACNNNAGATCGGAAG |
| 33807_4 | TATGAGTGTGGAGTCGTTGCGAGGCAGGAGAATCGCTAGAACNNNAGATCGGAAG |
| 37572_1 | TATGAGTGTGGAGTCGTTGCGAGCAGCAGATCTTTGAGGGATANNNAGATCGGAA |
| 37572_2 | TATGAGTGTGGAGTCGTTGCGAGCAGCAGATCTTTGAGGGATANNNAGATCGGAA |
| 37572_3 | TATGAGTGTGGAGTCGTTGCGAGCAGCAGATCTTTGAGGGATANNNAGATCGGAA |
| 37572_4 | TATGAGTGTGGAGTCGTTGCGAGCAGCAGATCTTTGAGGGATANNNAGATCGGAA |
| 50821_1 | TATGAGTGTGGAGTCGTTGCCGAAACCCCGTCTCTACAAAAAANNNAGATCGGAAGA |
| 50821_2 | TATGAGTGTGGAGTCGTTGCCGAAACCCCGTCTCTACAAAAAANNNAGATCGGAAGA |
| 50821_3 | TATGAGTGTGGAGTCGTTGCCGAAACCCCGTCTCTACAAAAAANNNAGATCGGAAGA |
| 50821_4 | TATGAGTGTGGAGTCGTTGCCGAAACCCCGTCTCTACAAAAAANNNAGATCGGAAGA |
| 48451_1 | TATGAGTGTGGAGTCGTTGCACCTAGTAGCTGGGATTACAGGTNNNAGATCGGAA[illegible] |
| 48451_2 | TATGAGTGTGGAGTCGTTGCACCTAGTAGCTGGGATTACAGGTNNNAGATCGGAA[illegible] |
| 48451_3 | TATGAGTGTGGAGTCGTTGCACCTAGTAGCTGGGATTACAGGTNNNAGATCGGAA[illegible] |
| 48451_4 | TATGAGTGTGGAGTCGTTGCACCTAGTAGCTGGGATTACAGGTNNNAGATCGGAA[illegible] |
| 31980_1 | TATGAGTGTGGAGTCGTTGCCACTCATTCCAACTACTCAGCCANNNAGATCGGAAGA |
| 31980_2 | TATGAGTGTGGAGTCGTTGCCACTCATTCCAACTACTCAGCCANNNAGATCGGAAGA |
| 31980_3 | TATGAGTGTGGAGTCGTTGCCACTCATTCCAACTACTCAGCCANNNAGATCGGAAGA |
| 31980_4 | TATGAGTGTGGAGTCGTTGCCACTCATTCCAACTACTCAGCCANNNAGATCGGAAGA |
| 47980_1 | TATGAGTGTGGAGTCGTTGCTGTGACAGGTAGAACAGAGAGGANNNAGATCGGAA |
| 47980_2 | TATGAGTGTGGAGTCGTTGCTGTGACAGGTAGAACAGAGAGGANNNAGATCGGAA |
| 47980_3 | TATGAGTGTGGAGTCGTTGCTGTGACAGGTAGAACAGAGAGGANNNAGATCGGAA |
| 47980_4 | TATGAGTGTGGAGTCGTTGCTGTGACAGGTAGAACAGAGAGGANNNAGATCGGAA |
| 28818_1 | TATGAGTGTGGAGTCGTTGCACTTGGATGAAGCTGGAGACAATNNNAGATCGGAA[illegible] |

| | |
|---|---|
| 28818_2 | TATGAGTGTGGAGTCGTTGCACTTGGATGAAGCTGGAGACAATNNNAGATCGGAA |
| 28818_3 | TATGAGTGTGGAGTCGTTGCACTTGGATGAAGCTGGAGACAATNNNAGATCGGAA |
| 28818_4 | TATGAGTGTGGAGTCGTTGCACTTGGATGAAGCTGGAGACAATNNNAGATCGGAA |
| 49333_1 | TATGAGTGTGGAGTCGTTGCGGAAGAGTTGGGGTTAGTTGAGTNNNAGATCGGA |
| 49333_2 | TATGAGTGTGGAGTCGTTGCGGAAGAGTTGGGGTTAGTTGAGTNNNAGATCGGA |
| 49333_3 | TATGAGTGTGGAGTCGTTGCGGAAGAGTTGGGGTTAGTTGAGTNNNAGATCGGA |
| 49333_4 | TATGAGTGTGGAGTCGTTGCGGAAGAGTTGGGGTTAGTTGAGTNNNAGATCGGA |
| 37626_1 | TATGAGTGTGGAGTCGTTGCATGCCTATGGTTCCTATGGTTCCNNNAGATCGGAAG |
| 37626_2 | TATGAGTGTGGAGTCGTTGCATGCCTATGGTTCCTATGGTTCCNNNAGATCGGAAG |
| 37626_3 | TATGAGTGTGGAGTCGTTGCATGCCTATGGTTCCTATGGTTCCNNNAGATCGGAAG |
| 37626_4 | TATGAGTGTGGAGTCGTTGCATGCCTATGGTTCCTATGGTTCCNNNAGATCGGAAG |
| 50495_1 | TATGAGTGTGGAGTCGTTGCACCACCAAGTACCACAGGTAAAGNNNAGATCGGAAG |
| 50495_2 | TATGAGTGTGGAGTCGTTGCACCACCAAGTACCACAGGTAAAGNNNAGATCGGAAG |
| 50495_3 | TATGAGTGTGGAGTCGTTGCACCACCAAGTACCACAGGTAAAGNNNAGATCGGAAG |
| 50495_4 | TATGAGTGTGGAGTCGTTGCACCACCAAGTACCACAGGTAAAGNNNAGATCGGAAG |
| 51029_1 | TATGAGTGTGGAGTCGTTGCCCCCTACTAGGAAGTGAGGACCNNNAGATCGGAAG |
| 51029_2 | TATGAGTGTGGAGTCGTTGCCCCCTACTAGGAAGTGAGGACCNNNAGATCGGAAG |
| 51029_3 | TATGAGTGTGGAGTCGTTGCCCCCTACTAGGAAGTGAGGACCNNNAGATCGGAAG |
| 51029_4 | TATGAGTGTGGAGTCGTTGCCCCCTACTAGGAAGTGAGGACCNNNAGATCGGAAG |
| 35227_1 | TATGAGTGTGGAGTCGTTGCGGATGAAGTCTCCCTCTGTCACNNNAGATCGGAAGA |
| 35227_2 | TATGAGTGTGGAGTCGTTGCGGATGAAGTCTCCCTCTGTCACNNNAGATCGGAAGA |
| 35227_3 | TATGAGTGTGGAGTCGTTGCGGATGAAGTCTCCCTCTGTCACNNNAGATCGGAAGA |
| 35227_4 | TATGAGTGTGGAGTCGTTGCGGATGAAGTCTCCCTCTGTCACNNNAGATCGGAAGA |
| 32640_1 | TATGAGTGTGGAGTCGTTGCCTCCAGAGTGCCAGGACTACAGNNNAGATCGGAAG |
| 32640_2 | TATGAGTGTGGAGTCGTTGCCTCCAGAGTGCCAGGACTACAGNNNAGATCGGAAG |
| 32640_3 | TATGAGTGTGGAGTCGTTGCCTCCAGAGTGCCAGGACTACAGNNNAGATCGGAAG |
| 32640_4 | TATGAGTGTGGAGTCGTTGCCTCCAGAGTGCCAGGACTACAGNNNAGATCGGAAG |
| 41905_1 | TATGAGTGTGGAGTCGTTGCCATTCTCCTGCCTCAGCCTCNNNAGATCGGAAGAGCA |
| 41905_2 | TATGAGTGTGGAGTCGTTGCCATTCTCCTGCCTCAGCCTCNNNAGATCGGAAGAGCA |
| 41905_3 | TATGAGTGTGGAGTCGTTGCCATTCTCCTGCCTCAGCCTCNNNAGATCGGAAGAGCA |
| 41905_4 | TATGAGTGTGGAGTCGTTGCCATTCTCCTGCCTCAGCCTCNNNAGATCGGAAGAGCA |
| 47244_1 | TATGAGTGTGGAGTCGTTGCCGCAATCTCAGCTTACTGCAAGNNNAGATCGGAAGA |
| 47244_2 | TATGAGTGTGGAGTCGTTGCCGCAATCTCAGCTTACTGCAAGNNNAGATCGGAAGA |
| 47244_3 | TATGAGTGTGGAGTCGTTGCCGCAATCTCAGCTTACTGCAAGNNNAGATCGGAAGA |
| 47244_4 | TATGAGTGTGGAGTCGTTGCCGCAATCTCAGCTTACTGCAAGNNNAGATCGGAAGA |
| 50524_1 | TATGAGTGTGGAGTCGTTGCGATGAGGGGACAAGAGAGTCTTCNNNAGATCGGAA |
| 50524_2 | TATGAGTGTGGAGTCGTTGCGATGAGGGGACAAGAGAGTCTTCNNNAGATCGGAA |
| 50524_3 | TATGAGTGTGGAGTCGTTGCGATGAGGGGACAAGAGAGTCTTCNNNAGATCGGAA |
| 50524_4 | TATGAGTGTGGAGTCGTTGCGATGAGGGGACAAGAGAGTCTTCNNNAGATCGGAA |
| 10209_1 | TATGAGTGTGGAGTCGTTGCAAAGGACACGCTGAAGTCCTAACNNNAGATCGGAA |
| 10209_2 | TATGAGTGTGGAGTCGTTGCAAAGGACACGCTGAAGTCCTAACNNNAGATCGGAA |
| 10209_3 | TATGAGTGTGGAGTCGTTGCAAAGGACACGCTGAAGTCCTAACNNNAGATCGGAA |
| 10209_4 | TATGAGTGTGGAGTCGTTGCAAAGGACACGCTGAAGTCCTAACNNNAGATCGGAA |

| | |
|---|---|
| 44951_1 | TATGAGTGTGGAGTCGTTGCCCCAGCTCCAAACCATTTCTAGANNNAGATCGGAAG |
| 44951_2 | TATGAGTGTGGAGTCGTTGCCCCAGCTCCAAACCATTTCTAGANNNAGATCGGAAG |
| 44951_3 | TATGAGTGTGGAGTCGTTGCCCCAGCTCCAAACCATTTCTAGANNNAGATCGGAAG |
| 44951_4 | TATGAGTGTGGAGTCGTTGCCCCAGCTCCAAACCATTTCTAGANNNAGATCGGAAG |
| 32317_1 | TATGAGTGTGGAGTCGTTGCGGTAGCTGTGTTGTTCTTATGCCNNNAGATCGGAAG |
| 32317_2 | TATGAGTGTGGAGTCGTTGCGGTAGCTGTGTTGTTCTTATGCCNNNAGATCGGAAG |
| 32317_3 | TATGAGTGTGGAGTCGTTGCGGTAGCTGTGTTGTTCTTATGCCNNNAGATCGGAAG |
| 32317_4 | TATGAGTGTGGAGTCGTTGCGGTAGCTGTGTTGTTCTTATGCCNNNAGATCGGAAG |
| 13478_1 | TATGAGTGTGGAGTCGTTGCGTTGATACTGAAGAATTGATGGNNNAGATCGGAAG |
| 13478_2 | TATGAGTGTGGAGTCGTTGCGTTGATACTGAAGAATTGATGGNNNAGATCGGAAG |
| 13478_3 | TATGAGTGTGGAGTCGTTGCGTTGATACTGAAGAATTGATGGNNNAGATCGGAAG |
| 13478_4 | TATGAGTGTGGAGTCGTTGCGTTGATACTGAAGAATTGATGGNNNAGATCGGAAG |
| 46817_1 | TATGAGTGTGGAGTCGTTGCATGCCAGACAGAAGTGCTGATTANNNAGATCGGAA |
| 46817_2 | TATGAGTGTGGAGTCGTTGCATGCCAGACAGAAGTGCTGATTANNNAGATCGGAA |
| 46817_3 | TATGAGTGTGGAGTCGTTGCATGCCAGACAGAAGTGCTGATTANNNAGATCGGAA |
| 46817_4 | TATGAGTGTGGAGTCGTTGCATGCCAGACAGAAGTGCTGATTANNNAGATCGGAA |
| 51823_1 | TATGAGTGTGGAGTCGTTGCGAGTGGATAAGTGCTCCTACACCNNNAGATCGGAA |
| 51823_2 | TATGAGTGTGGAGTCGTTGCGAGTGGATAAGTGCTCCTACACCNNNAGATCGGAA |
| 51823_3 | TATGAGTGTGGAGTCGTTGCGAGTGGATAAGTGCTCCTACACCNNNAGATCGGAA |
| 51823_4 | TATGAGTGTGGAGTCGTTGCGAGTGGATAAGTGCTCCTACACCNNNAGATCGGAA |
| 42832_1 | TATGAGTGTGGAGTCGTTGCAAACAAGGTCTTGCTCTGTTGTCNNNAGATCGGAAG |
| 42832_2 | TATGAGTGTGGAGTCGTTGCAAACAAGGTCTTGCTCTGTTGTCNNNAGATCGGAAG |
| 42832_3 | TATGAGTGTGGAGTCGTTGCAAACAAGGTCTTGCTCTGTTGTCNNNAGATCGGAAG |
| 42832_4 | TATGAGTGTGGAGTCGTTGCAAACAAGGTCTTGCTCTGTTGTCNNNAGATCGGAAG |
| 51131_1 | TATGAGTGTGGAGTCGTTGCAAGTTGGAGGGTTTGGAAGATGTNNNAGATCGGA |
| 51131_2 | TATGAGTGTGGAGTCGTTGCAAGTTGGAGGGTTTGGAAGATGTNNNAGATCGGA |
| 51131_3 | TATGAGTGTGGAGTCGTTGCAAGTTGGAGGGTTTGGAAGATGTNNNAGATCGGA |
| 51131_4 | TATGAGTGTGGAGTCGTTGCAAGTTGGAGGGTTTGGAAGATGTNNNAGATCGGA |
| 9831_1 | TATGAGTGTGGAGTCGTTGCGCCAGCTTTGAGAACTTCCTTGANNNAGATCGGAAG |
| 9831_2 | TATGAGTGTGGAGTCGTTGCGCCAGCTTTGAGAACTTCCTTGANNNAGATCGGAAG |
| 9831_3 | TATGAGTGTGGAGTCGTTGCGCCAGCTTTGAGAACTTCCTTGANNNAGATCGGAAG |
| 9831_4 | TATGAGTGTGGAGTCGTTGCGCCAGCTTTGAGAACTTCCTTGANNNAGATCGGAAG |
| 37466_1 | TATGAGTGTGGAGTCGTTGCAAGCACAAGCACAATGAGGTAACNNNAGATCGGAA |
| 37466_2 | TATGAGTGTGGAGTCGTTGCAAGCACAAGCACAATGAGGTAACNNNAGATCGGAA |
| 37466_3 | TATGAGTGTGGAGTCGTTGCAAGCACAAGCACAATGAGGTAACNNNAGATCGGAA |
| 37466_4 | TATGAGTGTGGAGTCGTTGCAAGCACAAGCACAATGAGGTAACNNNAGATCGGAA |
| 29526_1 | TATGAGTGTGGAGTCGTTGCGCTGCTAAACATCGACTCCATGANNNAGATCGGAAG |
| 29526_2 | TATGAGTGTGGAGTCGTTGCGCTGCTAAACATCGACTCCATGANNNAGATCGGAAG |
| 29526_3 | TATGAGTGTGGAGTCGTTGCGCTGCTAAACATCGACTCCATGANNNAGATCGGAAG |
| 29526_4 | TATGAGTGTGGAGTCGTTGCGCTGCTAAACATCGACTCCATGANNNAGATCGGAAG |
| 40034_1 | TATGAGTGTGGAGTCGTTGCCAGAGCAAGGCTGTCTCAAAAAANNNAGATCGGAA |
| 40034_2 | TATGAGTGTGGAGTCGTTGCCAGAGCAAGGCTGTCTCAAAAAANNNAGATCGGAA |
| 40034_3 | TATGAGTGTGGAGTCGTTGCCAGAGCAAGGCTGTCTCAAAAAANNNAGATCGGAA |

| | |
|---|---|
| 40034_4 | TATGAGTGTGGAGTCGTTGCCAGAGCAAGGCTGTCTCAAAAAANNNAGATCGGAA |
| 41476_1 | TATGAGTGTGGAGTCGTTGCTTTCTTCTCACAGTGATGGGACCNNNAGATCGGAAG |
| 41476_2 | TATGAGTGTGGAGTCGTTGCTTTCTTCTCACAGTGATGGGACCNNNAGATCGGAAG |
| 41476_3 | TATGAGTGTGGAGTCGTTGCTTTCTTCTCACAGTGATGGGACCNNNAGATCGGAAG |
| 41476_4 | TATGAGTGTGGAGTCGTTGCTTTCTTCTCACAGTGATGGGACCNNNAGATCGGAAG |
| 2194_1 | TATGAGTGTGGAGTCGTTGCTTTTGGTTGATGAGCAGCTCCTTNNNAGATCGGAAG |
| 2194_2 | TATGAGTGTGGAGTCGTTGCTTTTGGTTGATGAGCAGCTCCTTNNNAGATCGGAAG |
| 2194_3 | TATGAGTGTGGAGTCGTTGCTTTTGGTTGATGAGCAGCTCCTTNNNAGATCGGAAG |
| 2194_4 | TATGAGTGTGGAGTCGTTGCTTTTGGTTGATGAGCAGCTCCTTNNNAGATCGGAAG |
| 37097_1 | TATGAGTGTGGAGTCGTTGCGGCCTTAGGATCTGTGTAAATGCNNNAGATCGGAA |
| 37097_2 | TATGAGTGTGGAGTCGTTGCGGCCTTAGGATCTGTGTAAATGCNNNAGATCGGAA |
| 37097_3 | TATGAGTGTGGAGTCGTTGCGGCCTTAGGATCTGTGTAAATGCNNNAGATCGGAA |
| 37097_4 | TATGAGTGTGGAGTCGTTGCGGCCTTAGGATCTGTGTAAATGCNNNAGATCGGAA |
| 49886_1 | TATGAGTGTGGAGTCGTTGCTTTTGTAGAAGTTCTGGGGAGCANNNAGATCGGAA |
| 49886_2 | TATGAGTGTGGAGTCGTTGCTTTTGTAGAAGTTCTGGGGAGCANNNAGATCGGAA |
| 49886_3 | TATGAGTGTGGAGTCGTTGCTTTTGTAGAAGTTCTGGGGAGCANNNAGATCGGAA |
| 49886_4 | TATGAGTGTGGAGTCGTTGCTTTTGTAGAAGTTCTGGGGAGCANNNAGATCGGAA |
| 37020_1 | TATGAGTGTGGAGTCGTTGCGAATAGCTCCAGTCTACAGCTCCNNNAGATCGGAAG |
| 37020_2 | TATGAGTGTGGAGTCGTTGCGAATAGCTCCAGTCTACAGCTCCNNNAGATCGGAAG |
| 37020_3 | TATGAGTGTGGAGTCGTTGCGAATAGCTCCAGTCTACAGCTCCNNNAGATCGGAAG |
| 37020_4 | TATGAGTGTGGAGTCGTTGCGAATAGCTCCAGTCTACAGCTCCNNNAGATCGGAAG |
| 51932_1 | TATGAGTGTGGAGTCGTTGCCCATTGCTGCTGGTACATGAAAANNNAGATCGGAAG |
| 51932_2 | TATGAGTGTGGAGTCGTTGCCCATTGCTGCTGGTACATGAAAANNNAGATCGGAAG |
| 51932_3 | TATGAGTGTGGAGTCGTTGCCCATTGCTGCTGGTACATGAAAANNNAGATCGGAAG |
| 51932_4 | TATGAGTGTGGAGTCGTTGCCCATTGCTGCTGGTACATGAAAANNNAGATCGGAAG |
| 30963_1 | TATGAGTGTGGAGTCGTTGCACCTTTCCTCTCCCCTAGAATCANNNAGATCGGAAGA |
| 30963_2 | TATGAGTGTGGAGTCGTTGCACCTTTCCTCTCCCCTAGAATCANNNAGATCGGAAGA |
| 30963_3 | TATGAGTGTGGAGTCGTTGCACCTTTCCTCTCCCCTAGAATCANNNAGATCGGAAGA |
| 30963_4 | TATGAGTGTGGAGTCGTTGCACCTTTCCTCTCCCCTAGAATCANNNAGATCGGAAGA |
| 43885_1 | TATGAGTGTGGAGTCGTTGCAGAATCAACCCCATGTCCACTTCNNNAGATCGGAAGA |
| 43885_2 | TATGAGTGTGGAGTCGTTGCAGAATCAACCCCATGTCCACTTCNNNAGATCGGAAGA |
| 43885_3 | TATGAGTGTGGAGTCGTTGCAGAATCAACCCCATGTCCACTTCNNNAGATCGGAAGA |
| 43885_4 | TATGAGTGTGGAGTCGTTGCAGAATCAACCCCATGTCCACTTCNNNAGATCGGAAGA |
| 41744_1 | TATGAGTGTGGAGTCGTTGCGTTCAGGGATTCGACTTCTTCCTNNNAGATCGGAAG |
| 41744_2 | TATGAGTGTGGAGTCGTTGCGTTCAGGGATTCGACTTCTTCCTNNNAGATCGGAAG |
| 41744_3 | TATGAGTGTGGAGTCGTTGCGTTCAGGGATTCGACTTCTTCCTNNNAGATCGGAAG |
| 41744_4 | TATGAGTGTGGAGTCGTTGCGTTCAGGGATTCGACTTCTTCCTNNNAGATCGGAAG |
| 47379_1 | TATGAGTGTGGAGTCGTTGCGATCTCAGCTCACTGCAACCTCNNNAGATCGGAAGA |
| 47379_2 | TATGAGTGTGGAGTCGTTGCGATCTCAGCTCACTGCAACCTCNNNAGATCGGAAGA |
| 47379_3 | TATGAGTGTGGAGTCGTTGCGATCTCAGCTCACTGCAACCTCNNNAGATCGGAAGA |
| 47379_4 | TATGAGTGTGGAGTCGTTGCGATCTCAGCTCACTGCAACCTCNNNAGATCGGAAGA |
| 43748_1 | TATGAGTGTGGAGTCGTTGCGTCACACAGAAAACCATGGTCAGNNNAGATCGGAA |
| 43748_2 | TATGAGTGTGGAGTCGTTGCGTCACACAGAAAACCATGGTCAGNNNAGATCGGAA |

| | |
|---|---|
| 43748_3 | TATGAGTGTGGAGTCGTTGCGTCACACAGAAAACCATGGTCAGNNNAGATCGGAA |
| 43748_4 | TATGAGTGTGGAGTCGTTGCGTCACACAGAAAACCATGGTCAGNNNAGATCGGAA |
| 49891_1 | TATGAGTGTGGAGTCGTTGCTGTAAACACACACACACACACACNNNAGATCGGAAG |
| 49891_2 | TATGAGTGTGGAGTCGTTGCTGTAAACACACACACACACACACNNNAGATCGGAAG |
| 49891_3 | TATGAGTGTGGAGTCGTTGCTGTAAACACACACACACACACACNNNAGATCGGAAG |
| 49891_4 | TATGAGTGTGGAGTCGTTGCTGTAAACACACACACACACACACNNNAGATCGGAAG |
| 34701_1 | TATGAGTGTGGAGTCGTTGCAGGGTCTTACCATATTGCCCATGNNNAGATCGGAAG |
| 34701_2 | TATGAGTGTGGAGTCGTTGCAGGGTCTTACCATATTGCCCATGNNNAGATCGGAAG |
| 34701_3 | TATGAGTGTGGAGTCGTTGCAGGGTCTTACCATATTGCCCATGNNNAGATCGGAAG |
| 34701_4 | TATGAGTGTGGAGTCGTTGCAGGGTCTTACCATATTGCCCATGNNNAGATCGGAAG |
| 46649_1 | TATGAGTGTGGAGTCGTTGCGCCACCACACCTGCTAATTATTTNNNAGATCGGAAGA |
| 46649_2 | TATGAGTGTGGAGTCGTTGCGCCACCACACCTGCTAATTATTTNNNAGATCGGAAGA |
| 46649_3 | TATGAGTGTGGAGTCGTTGCGCCACCACACCTGCTAATTATTTNNNAGATCGGAAGA |
| 46649_4 | TATGAGTGTGGAGTCGTTGCGCCACCACACCTGCTAATTATTTNNNAGATCGGAAGA |
| 51598_1 | TATGAGTGTGGAGTCGTTGCTCCAAAATGAAGTCCATCCCCTTNNNAGATCGGAAG |
| 51598_2 | TATGAGTGTGGAGTCGTTGCTCCAAAATGAAGTCCATCCCCTTNNNAGATCGGAAG |
| 51598_3 | TATGAGTGTGGAGTCGTTGCTCCAAAATGAAGTCCATCCCCTTNNNAGATCGGAAG |
| 51598_4 | TATGAGTGTGGAGTCGTTGCTCCAAAATGAAGTCCATCCCCTTNNNAGATCGGAAG |
| 48341_1 | TATGAGTGTGGAGTCGTTGCTTCTCAGGAGGTTCAGTCTCAGANNNAGATCGGAAG |
| 48341_2 | TATGAGTGTGGAGTCGTTGCTTCTCAGGAGGTTCAGTCTCAGANNNAGATCGGAAG |
| 48341_3 | TATGAGTGTGGAGTCGTTGCTTCTCAGGAGGTTCAGTCTCAGANNNAGATCGGAAG |
| 48341_4 | TATGAGTGTGGAGTCGTTGCTTCTCAGGAGGTTCAGTCTCAGANNNAGATCGGAAG |
| 31411_1 | TATGAGTGTGGAGTCGTTGCGACAAGGTCTCACTCTGTCACCNNNAGATCGGAAGA |
| 31411_2 | TATGAGTGTGGAGTCGTTGCGACAAGGTCTCACTCTGTCACCNNNAGATCGGAAGA |
| 31411_3 | TATGAGTGTGGAGTCGTTGCGACAAGGTCTCACTCTGTCACCNNNAGATCGGAAGA |
| 31411_4 | TATGAGTGTGGAGTCGTTGCGACAAGGTCTCACTCTGTCACCNNNAGATCGGAAGA |
| 861_1 | TATGAGTGTGGAGTCGTTGCGTTGGTGGACACTCGGGTTATTTNNNAGATCGGAA |
| 861_2 | TATGAGTGTGGAGTCGTTGCGTTGGTGGACACTCGGGTTATTTNNNAGATCGGAA |
| 861_3 | TATGAGTGTGGAGTCGTTGCGTTGGTGGACACTCGGGTTATTTNNNAGATCGGAA |
| 861_4 | TATGAGTGTGGAGTCGTTGCGTTGGTGGACACTCGGGTTATTTNNNAGATCGGAA |
| 29570_1 | TATGAGTGTGGAGTCGTTGCATATTCCTGGCTAGCTACACAGCNNNAGATCGGAAG |
| 29570_2 | TATGAGTGTGGAGTCGTTGCATATTCCTGGCTAGCTACACAGCNNNAGATCGGAAG |
| 29570_3 | TATGAGTGTGGAGTCGTTGCATATTCCTGGCTAGCTACACAGCNNNAGATCGGAAG |
| 29570_4 | TATGAGTGTGGAGTCGTTGCATATTCCTGGCTAGCTACACAGCNNNAGATCGGAAG |
| 31929_1 | TATGAGTGTGGAGTCGTTGCAAGGCAGTCAGATGGGAGAAATTNNNAGATCGGAA |
| 31929_2 | TATGAGTGTGGAGTCGTTGCAAGGCAGTCAGATGGGAGAAATTNNNAGATCGGAA |
| 31929_3 | TATGAGTGTGGAGTCGTTGCAAGGCAGTCAGATGGGAGAAATTNNNAGATCGGAA |
| 31929_4 | TATGAGTGTGGAGTCGTTGCAAGGCAGTCAGATGGGAGAAATTNNNAGATCGGAA |
| 44002_1 | TATGAGTGTGGAGTCGTTGCTTCCAGCCTATATCCTTGTCACANNNAGATCGGAAGA |
| 44002_2 | TATGAGTGTGGAGTCGTTGCTTCCAGCCTATATCCTTGTCACANNNAGATCGGAAGA |
| 44002_3 | TATGAGTGTGGAGTCGTTGCTTCCAGCCTATATCCTTGTCACANNNAGATCGGAAGA |
| 44002_4 | TATGAGTGTGGAGTCGTTGCTTCCAGCCTATATCCTTGTCACANNNAGATCGGAAGA |
| 44794_1 | TATGAGTGTGGAGTCGTTGCTGCATGGGGAAAGAATCACAGATNNNAGATCGGAA |

| | |
|---|---|
| 44794_2 | TATGAGTGTGGAGTCGTTGCTGCATGGGGAAAGAATCACAGATNNNAGATCGGAA |
| 44794_3 | TATGAGTGTGGAGTCGTTGCTGCATGGGGAAAGAATCACAGATNNNAGATCGGAA |
| 44794_4 | TATGAGTGTGGAGTCGTTGCTGCATGGGGAAAGAATCACAGATNNNAGATCGGAA |
| 8772_1 | TATGAGTGTGGAGTCGTTGCGGTGTGTGTGGGTGTGTTTAGTTNNNAGATCGGAA |
| 8772_2 | TATGAGTGTGGAGTCGTTGCGGTGTGTGTGGGTGTGTTTAGTTNNNAGATCGGAA |
| 8772_3 | TATGAGTGTGGAGTCGTTGCGGTGTGTGTGGGTGTGTTTAGTTNNNAGATCGGAA |
| 8772_4 | TATGAGTGTGGAGTCGTTGCGGTGTGTGTGGGTGTGTTTAGTTNNNAGATCGGAA |
| 48305_1 | TATGAGTGTGGAGTCGTTGCCTGGGTAGCTGGGATTACAAGTGNNNAGATCGGAA |
| 48305_2 | TATGAGTGTGGAGTCGTTGCCTGGGTAGCTGGGATTACAAGTGNNNAGATCGGAA |
| 48305_3 | TATGAGTGTGGAGTCGTTGCCTGGGTAGCTGGGATTACAAGTGNNNAGATCGGAA |
| 48305_4 | TATGAGTGTGGAGTCGTTGCCTGGGTAGCTGGGATTACAAGTGNNNAGATCGGAA |
| 44525_1 | TATGAGTGTGGAGTCGTTGCCCAACCCTTTGAATTGGCGGNNNAGATCGGAAGAG |
| 44525_2 | TATGAGTGTGGAGTCGTTGCCCAACCCTTTGAATTGGCGGNNNAGATCGGAAGAG |
| 44525_3 | TATGAGTGTGGAGTCGTTGCCCAACCCTTTGAATTGGCGGNNNAGATCGGAAGAG |
| 44525_4 | TATGAGTGTGGAGTCGTTGCCCAACCCTTTGAATTGGCGGNNNAGATCGGAAGAG |
| 40047_1 | TATGAGTGTGGAGTCGTTGCAATTCAATAATTAGCCGGGCGTGNNNAGATCGGAA |
| 40047_2 | TATGAGTGTGGAGTCGTTGCAATTCAATAATTAGCCGGGCGTGNNNAGATCGGAA |
| 40047_3 | TATGAGTGTGGAGTCGTTGCAATTCAATAATTAGCCGGGCGTGNNNAGATCGGAA |
| 40047_4 | TATGAGTGTGGAGTCGTTGCAATTCAATAATTAGCCGGGCGTGNNNAGATCGGAA |
| 38865_1 | TATGAGTGTGGAGTCGTTGCATCTCAGTGCTGACTCCTAGGATNNNAGATCGGAAG |
| 38865_2 | TATGAGTGTGGAGTCGTTGCATCTCAGTGCTGACTCCTAGGATNNNAGATCGGAAG |
| 38865_3 | TATGAGTGTGGAGTCGTTGCATCTCAGTGCTGACTCCTAGGATNNNAGATCGGAAG |
| 38865_4 | TATGAGTGTGGAGTCGTTGCATCTCAGTGCTGACTCCTAGGATNNNAGATCGGAAG |
| 10465_1 | TATGAGTGTGGAGTCGTTGCCTGCTGAACCTGATGCATAGTCANNNAGATCGGAAG |
| 10465_2 | TATGAGTGTGGAGTCGTTGCCTGCTGAACCTGATGCATAGTCANNNAGATCGGAAG |
| 10465_3 | TATGAGTGTGGAGTCGTTGCCTGCTGAACCTGATGCATAGTCANNNAGATCGGAAG |
| 10465_4 | TATGAGTGTGGAGTCGTTGCCTGCTGAACCTGATGCATAGTCANNNAGATCGGAAG |
| 3007_1 | TATGAGTGTGGAGTCGTTGCCTAAAATGGATCCACCAGCCCACNNNAGATCGGAAG |
| 3007_2 | TATGAGTGTGGAGTCGTTGCCTAAAATGGATCCACCAGCCCACNNNAGATCGGAAG |
| 3007_3 | TATGAGTGTGGAGTCGTTGCCTAAAATGGATCCACCAGCCCACNNNAGATCGGAAG |
| 3007_4 | TATGAGTGTGGAGTCGTTGCCTAAAATGGATCCACCAGCCCACNNNAGATCGGAAG |
| 38533_1 | TATGAGTGTGGAGTCGTTGCTGGCCATTCTCTAAGGTTCAATCTNNNAGATCGGAA |
| 38533_2 | TATGAGTGTGGAGTCGTTGCTGGCCATTCTCTAAGGTTCAATCTNNNAGATCGGAA |
| 38533_3 | TATGAGTGTGGAGTCGTTGCTGGCCATTCTCTAAGGTTCAATCTNNNAGATCGGAA |
| 38533_4 | TATGAGTGTGGAGTCGTTGCTGGCCATTCTCTAAGGTTCAATCTNNNAGATCGGAA |
| 49938_1 | TATGAGTGTGGAGTCGTTGCGCATCCATTGGTGTCTTTGTGAANNNAGATCGGAAG |
| 49938_2 | TATGAGTGTGGAGTCGTTGCGCATCCATTGGTGTCTTTGTGAANNNAGATCGGAAG |
| 49938_3 | TATGAGTGTGGAGTCGTTGCGCATCCATTGGTGTCTTTGTGAANNNAGATCGGAAG |
| 49938_4 | TATGAGTGTGGAGTCGTTGCGCATCCATTGGTGTCTTTGTGAANNNAGATCGGAAG |
| 28790_1 | TATGAGTGTGGAGTCGTTGCTGGAGAGAGAGAGAGAGAGAGAANNNAGATCGG |
| 28790_2 | TATGAGTGTGGAGTCGTTGCTGGAGAGAGAGAGAGAGAGAGAANNNAGATCGG |
| 28790_3 | TATGAGTGTGGAGTCGTTGCTGGAGAGAGAGAGAGAGAGAGAANNNAGATCGG |
| 28790_4 | TATGAGTGTGGAGTCGTTGCTGGAGAGAGAGAGAGAGAGAGAANNNAGATCGG |

| | |
|---|---|
| 2698_1 | TATGAGTGTGGAGTCGTTGCTGAAGAGACGAGGAAGCTGAGATNNNAGATCGGA |
| 2698_2 | TATGAGTGTGGAGTCGTTGCTGAAGAGACGAGGAAGCTGAGATNNNAGATCGGA |
| 2698_3 | TATGAGTGTGGAGTCGTTGCTGAAGAGACGAGGAAGCTGAGATNNNAGATCGGA |
| 2698_4 | TATGAGTGTGGAGTCGTTGCTGAAGAGACGAGGAAGCTGAGATNNNAGATCGGA |
| 38078_1 | TATGAGTGTGGAGTCGTTGCAAGGATTTGGGTAGCTAAGAGGCNNNAGATCGGA |
| 38078_2 | TATGAGTGTGGAGTCGTTGCAAGGATTTGGGTAGCTAAGAGGCNNNAGATCGGA |
| 38078_3 | TATGAGTGTGGAGTCGTTGCAAGGATTTGGGTAGCTAAGAGGCNNNAGATCGGA |
| 38078_4 | TATGAGTGTGGAGTCGTTGCAAGGATTTGGGTAGCTAAGAGGCNNNAGATCGGA |
| 46618_1 | TATGAGTGTGGAGTCGTTGCGCATGATCTTGTTCTCAGCTCACNNNAGATCGGAAG |
| 46618_2 | TATGAGTGTGGAGTCGTTGCGCATGATCTTGTTCTCAGCTCACNNNAGATCGGAAG |
| 46618_3 | TATGAGTGTGGAGTCGTTGCGCATGATCTTGTTCTCAGCTCACNNNAGATCGGAAG |
| 46618_4 | TATGAGTGTGGAGTCGTTGCGCATGATCTTGTTCTCAGCTCACNNNAGATCGGAAG |
| 38937_1 | TATGAGTGTGGAGTCGTTGCGTAATCCTAGCTACTTGGGAGGCNNNAGATCGGAA |
| 38937_2 | TATGAGTGTGGAGTCGTTGCGTAATCCTAGCTACTTGGGAGGCNNNAGATCGGAA |
| 38937_3 | TATGAGTGTGGAGTCGTTGCGTAATCCTAGCTACTTGGGAGGCNNNAGATCGGAA |
| 38937_4 | TATGAGTGTGGAGTCGTTGCGTAATCCTAGCTACTTGGGAGGCNNNAGATCGGAA |
| 43524_1 | TATGAGTGTGGAGTCGTTGCCAAACTCGGCTCACTGCAACNNNAGATCGGAAGAGC |
| 43524_2 | TATGAGTGTGGAGTCGTTGCCAAACTCGGCTCACTGCAACNNNAGATCGGAAGAGC |
| 43524_3 | TATGAGTGTGGAGTCGTTGCCAAACTCGGCTCACTGCAACNNNAGATCGGAAGAGC |
| 43524_4 | TATGAGTGTGGAGTCGTTGCCAAACTCGGCTCACTGCAACNNNAGATCGGAAGAGC |
| 34826_1 | TATGAGTGTGGAGTCGTTGCTGGTTTTGAACTCCTGACCTCAANNNAGATCGGAAG |
| 34826_2 | TATGAGTGTGGAGTCGTTGCTGGTTTTGAACTCCTGACCTCAANNNAGATCGGAAG |
| 34826_3 | TATGAGTGTGGAGTCGTTGCTGGTTTTGAACTCCTGACCTCAANNNAGATCGGAAG |
| 34826_4 | TATGAGTGTGGAGTCGTTGCTGGTTTTGAACTCCTGACCTCAANNNAGATCGGAAG |
| 51819_1 | TATGAGTGTGGAGTCGTTGCTACCTACTTGCCTGCCTTTCATTNNNAGATCGGAAGA |
| 51819_2 | TATGAGTGTGGAGTCGTTGCTACCTACTTGCCTGCCTTTCATTNNNAGATCGGAAGA |
| 51819_3 | TATGAGTGTGGAGTCGTTGCTACCTACTTGCCTGCCTTTCATTNNNAGATCGGAAGA |
| 51819_4 | TATGAGTGTGGAGTCGTTGCTACCTACTTGCCTGCCTTTCATTNNNAGATCGGAAGA |
| 47977_1 | TATGAGTGTGGAGTCGTTGCTGGTGGAGTATGGGAAATACAGCNNNAGATCGGA |
| 47977_2 | TATGAGTGTGGAGTCGTTGCTGGTGGAGTATGGGAAATACAGCNNNAGATCGGA |
| 47977_3 | TATGAGTGTGGAGTCGTTGCTGGTGGAGTATGGGAAATACAGCNNNAGATCGGA |
| 47977_4 | TATGAGTGTGGAGTCGTTGCTGGTGGAGTATGGGAAATACAGCNNNAGATCGGA |
| 46083_1 | TATGAGTGTGGAGTCGTTGCCTCCCAGGTTCAAGTGATTCTCANNNAGATCGGAAG |
| 46083_2 | TATGAGTGTGGAGTCGTTGCCTCCCAGGTTCAAGTGATTCTCANNNAGATCGGAAG |
| 46083_3 | TATGAGTGTGGAGTCGTTGCCTCCCAGGTTCAAGTGATTCTCANNNAGATCGGAAG |
| 46083_4 | TATGAGTGTGGAGTCGTTGCCTCCCAGGTTCAAGTGATTCTCANNNAGATCGGAAG |
| 29395_1 | TATGAGTGTGGAGTCGTTGCTCAGAAAGGCTGATCACAGCAAGNNNAGATCGGAA |
| 29395_2 | TATGAGTGTGGAGTCGTTGCTCAGAAAGGCTGATCACAGCAAGNNNAGATCGGAA |
| 29395_3 | TATGAGTGTGGAGTCGTTGCTCAGAAAGGCTGATCACAGCAAGNNNAGATCGGAA |
| 29395_4 | TATGAGTGTGGAGTCGTTGCTCAGAAAGGCTGATCACAGCAAGNNNAGATCGGAA |
| 47962_1 | TATGAGTGTGGAGTCGTTGCGTGTAAAGAAGTCAGTGCCAAGGNNNAGATCGGAA |
| 47962_2 | TATGAGTGTGGAGTCGTTGCGTGTAAAGAAGTCAGTGCCAAGGNNNAGATCGGAA |
| 47962_3 | TATGAGTGTGGAGTCGTTGCGTGTAAAGAAGTCAGTGCCAAGGNNNAGATCGGAA |

| | |
|---|---|
| 47962_4 | TATGAGTGTGGAGTCGTTGCGTGTAAAGAAGTCAGTGCCAAGGNNNAGATCGGAA |
| 47759_1 | TATGAGTGTGGAGTCGTTGCGTTCAAGAGATTCTCCTGCCTCANNNAGATCGGAAG |
| 47759_2 | TATGAGTGTGGAGTCGTTGCGTTCAAGAGATTCTCCTGCCTCANNNAGATCGGAAG |
| 47759_3 | TATGAGTGTGGAGTCGTTGCGTTCAAGAGATTCTCCTGCCTCANNNAGATCGGAAG |
| 47759_4 | TATGAGTGTGGAGTCGTTGCGTTCAAGAGATTCTCCTGCCTCANNNAGATCGGAAG |
| 12879_1 | TATGAGTGTGGAGTCGTTGCAAGTTACTTCCATCCACTTTCNNNAGATCGGAAGAGC |
| 12879_2 | TATGAGTGTGGAGTCGTTGCAAGTTACTTCCATCCACTTTCNNNAGATCGGAAGAGC |
| 12879_3 | TATGAGTGTGGAGTCGTTGCAAGTTACTTCCATCCACTTTCNNNAGATCGGAAGAGC |
| 12879_4 | TATGAGTGTGGAGTCGTTGCAAGTTACTTCCATCCACTTTCNNNAGATCGGAAGAGC |
| 49454_1 | TATGAGTGTGGAGTCGTTGCCCCAAATCCCACTAGTGCTAAGTNNNAGATCGGAAG |
| 49454_2 | TATGAGTGTGGAGTCGTTGCCCCAAATCCCACTAGTGCTAAGTNNNAGATCGGAAG |
| 49454_3 | TATGAGTGTGGAGTCGTTGCCCCAAATCCCACTAGTGCTAAGTNNNAGATCGGAAG |
| 49454_4 | TATGAGTGTGGAGTCGTTGCCCCAAATCCCACTAGTGCTAAGTNNNAGATCGGAAG |
| 35031_1 | TATGAGTGTGGAGTCGTTGCCTGATCTCAGCTCACTGCAAGCNNNAGATCGGAAGA |
| 35031_2 | TATGAGTGTGGAGTCGTTGCCTGATCTCAGCTCACTGCAAGCNNNAGATCGGAAGA |
| 35031_3 | TATGAGTGTGGAGTCGTTGCCTGATCTCAGCTCACTGCAAGCNNNAGATCGGAAGA |
| 35031_4 | TATGAGTGTGGAGTCGTTGCCTGATCTCAGCTCACTGCAAGCNNNAGATCGGAAGA |
| 29895_1 | TATGAGTGTGGAGTCGTTGCCACTAATTTCCAACCCAGGCTTGNNNAGATCGGAAG |
| 29895_2 | TATGAGTGTGGAGTCGTTGCCACTAATTTCCAACCCAGGCTTGNNNAGATCGGAAG |
| 29895_3 | TATGAGTGTGGAGTCGTTGCCACTAATTTCCAACCCAGGCTTGNNNAGATCGGAAG |
| 29895_4 | TATGAGTGTGGAGTCGTTGCCACTAATTTCCAACCCAGGCTTGNNNAGATCGGAAG |
| 29812_1 | TATGAGTGTGGAGTCGTTGCAATTGCTTCCTCAGAGGTCATGTNNNAGATCGGAAG |
| 29812_2 | TATGAGTGTGGAGTCGTTGCAATTGCTTCCTCAGAGGTCATGTNNNAGATCGGAAG |
| 29812_3 | TATGAGTGTGGAGTCGTTGCAATTGCTTCCTCAGAGGTCATGTNNNAGATCGGAAG |
| 29812_4 | TATGAGTGTGGAGTCGTTGCAATTGCTTCCTCAGAGGTCATGTNNNAGATCGGAAG |
| 37688_1 | TATGAGTGTGGAGTCGTTGCGTTCATATGCTCGAATTCCACCCNNNAGATCGGAAG |
| 37688_2 | TATGAGTGTGGAGTCGTTGCGTTCATATGCTCGAATTCCACCCNNNAGATCGGAAG |
| 37688_3 | TATGAGTGTGGAGTCGTTGCGTTCATATGCTCGAATTCCACCCNNNAGATCGGAAG |
| 37688_4 | TATGAGTGTGGAGTCGTTGCGTTCATATGCTCGAATTCCACCCNNNAGATCGGAAG |
| 38297_1 | TATGAGTGTGGAGTCGTTGCGATGACAGCTGCTGTTTGCTAAANNNAGATCGGAA |
| 38297_2 | TATGAGTGTGGAGTCGTTGCGATGACAGCTGCTGTTTGCTAAANNNAGATCGGAA |
| 38297_3 | TATGAGTGTGGAGTCGTTGCGATGACAGCTGCTGTTTGCTAAANNNAGATCGGAA |
| 38297_4 | TATGAGTGTGGAGTCGTTGCGATGACAGCTGCTGTTTGCTAAANNNAGATCGGAA |
| 4287_1 | TATGAGTGTGGAGTCGTTGCGGAGACCATGTCTATGCAAGAGAAANNNAGATCGG |
| 4287_2 | TATGAGTGTGGAGTCGTTGCGGAGACCATGTCTATGCAAGAGAAANNNAGATCGG |
| 4287_3 | TATGAGTGTGGAGTCGTTGCGGAGACCATGTCTATGCAAGAGAAANNNAGATCGG |
| 4287_4 | TATGAGTGTGGAGTCGTTGCGGAGACCATGTCTATGCAAGAGAAANNNAGATCGG |
| 48076_1 | TATGAGTGTGGAGTCGTTGCTGAGCCTTTGAAATCAAACGAGTNNNAGATCGGAA |
| 48076_2 | TATGAGTGTGGAGTCGTTGCTGAGCCTTTGAAATCAAACGAGTNNNAGATCGGAA |
| 48076_3 | TATGAGTGTGGAGTCGTTGCTGAGCCTTTGAAATCAAACGAGTNNNAGATCGGAA |
| 48076_4 | TATGAGTGTGGAGTCGTTGCTGAGCCTTTGAAATCAAACGAGTNNNAGATCGGAA |
| 40029_1 | TATGAGTGTGGAGTCGTTGCGGAGGCTGAAGCAGGAAAATTTCNNNAGATCGGAA |
| 40029_2 | TATGAGTGTGGAGTCGTTGCGGAGGCTGAAGCAGGAAAATTTCNNNAGATCGGAA |

| | |
|---|---|
| 40029_3 | TATGAGTGTGGAGTCGTTGCGGAGGCTGAAGCAGGAAAATTTCNNNAGATCGGAA |
| 40029_4 | TATGAGTGTGGAGTCGTTGCGGAGGCTGAAGCAGGAAAATTTCNNNAGATCGGAA |
| 38146_1 | TATGAGTGTGGAGTCGTTGCTATCCAGTGGCTCTTCATGTCACNNNAGATCGGAAG |
| 38146_2 | TATGAGTGTGGAGTCGTTGCTATCCAGTGGCTCTTCATGTCACNNNAGATCGGAAG |
| 38146_3 | TATGAGTGTGGAGTCGTTGCTATCCAGTGGCTCTTCATGTCACNNNAGATCGGAAG |
| 38146_4 | TATGAGTGTGGAGTCGTTGCTATCCAGTGGCTCTTCATGTCACNNNAGATCGGAAG |
| 49297_1 | TATGAGTGTGGAGTCGTTGCGTCCAGGAAGATGGTAATAGGGCNNNAGATCGGAA |
| 49297_2 | TATGAGTGTGGAGTCGTTGCGTCCAGGAAGATGGTAATAGGGCNNNAGATCGGAA |
| 49297_3 | TATGAGTGTGGAGTCGTTGCGTCCAGGAAGATGGTAATAGGGCNNNAGATCGGAA |
| 49297_4 | TATGAGTGTGGAGTCGTTGCGTCCAGGAAGATGGTAATAGGGCNNNAGATCGGAA |
| 38962_1 | TATGAGTGTGGAGTCGTTGCCCTGGAATGTCTGCAAAAACGATNNNAGATCGGAA |
| 38962_2 | TATGAGTGTGGAGTCGTTGCCCTGGAATGTCTGCAAAAACGATNNNAGATCGGAA |
| 38962_3 | TATGAGTGTGGAGTCGTTGCCCTGGAATGTCTGCAAAAACGATNNNAGATCGGAA |
| 38962_4 | TATGAGTGTGGAGTCGTTGCCCTGGAATGTCTGCAAAAACGATNNNAGATCGGAA |
| 12683_1 | TATGAGTGTGGAGTCGTTGCCTTCATGAAATCAGCATGAGANNNAGATCGGAAGA |
| 12683_2 | TATGAGTGTGGAGTCGTTGCCTTCATGAAATCAGCATGAGANNNAGATCGGAAGA |
| 12683_3 | TATGAGTGTGGAGTCGTTGCCTTCATGAAATCAGCATGAGANNNAGATCGGAAGA |
| 12683_4 | TATGAGTGTGGAGTCGTTGCCTTCATGAAATCAGCATGAGANNNAGATCGGAAGA |
| 43507_1 | TATGAGTGTGGAGTCGTTGCGGGCTCCTACTTCCTGATTTCAANNNAGATCGGAAG |
| 43507_2 | TATGAGTGTGGAGTCGTTGCGGGCTCCTACTTCCTGATTTCAANNNAGATCGGAAG |
| 43507_3 | TATGAGTGTGGAGTCGTTGCGGGCTCCTACTTCCTGATTTCAANNNAGATCGGAAG |
| 43507_4 | TATGAGTGTGGAGTCGTTGCGGGCTCCTACTTCCTGATTTCAANNNAGATCGGAAG |
| 52173_1 | TATGAGTGTGGAGTCGTTGCCTGAGGCAGGAGAATCACTTGAANNNAGATCGGAA |
| 52173_2 | TATGAGTGTGGAGTCGTTGCCTGAGGCAGGAGAATCACTTGAANNNAGATCGGAA |
| 52173_3 | TATGAGTGTGGAGTCGTTGCCTGAGGCAGGAGAATCACTTGAANNNAGATCGGAA |
| 52173_4 | TATGAGTGTGGAGTCGTTGCCTGAGGCAGGAGAATCACTTGAANNNAGATCGGAA |
| 48820_1 | TATGAGTGTGGAGTCGTTGCACCACACCCAGCCTAAAATTCTTNNNAGATCGGAAGA |
| 48820_2 | TATGAGTGTGGAGTCGTTGCACCACACCCAGCCTAAAATTCTTNNNAGATCGGAAGA |
| 48820_3 | TATGAGTGTGGAGTCGTTGCACCACACCCAGCCTAAAATTCTTNNNAGATCGGAAGA |
| 48820_4 | TATGAGTGTGGAGTCGTTGCACCACACCCAGCCTAAAATTCTTNNNAGATCGGAAGA |
| 39398_1 | TATGAGTGTGGAGTCGTTGCACTCCATACACCCCAACAATGAANNNAGATCGGAAG |
| 39398_2 | TATGAGTGTGGAGTCGTTGCACTCCATACACCCCAACAATGAANNNAGATCGGAAG |
| 39398_3 | TATGAGTGTGGAGTCGTTGCACTCCATACACCCCAACAATGAANNNAGATCGGAAG |
| 39398_4 | TATGAGTGTGGAGTCGTTGCACTCCATACACCCCAACAATGAANNNAGATCGGAAG |
| 45963_1 | TATGAGTGTGGAGTCGTTGCGAGTCTCGCTCTGTTTCCCAAGNNNAGATCGGAAGA |
| 45963_2 | TATGAGTGTGGAGTCGTTGCGAGTCTCGCTCTGTTTCCCAAGNNNAGATCGGAAGA |
| 45963_3 | TATGAGTGTGGAGTCGTTGCGAGTCTCGCTCTGTTTCCCAAGNNNAGATCGGAAGA |
| 45963_4 | TATGAGTGTGGAGTCGTTGCGAGTCTCGCTCTGTTTCCCAAGNNNAGATCGGAAGA |
| 29263_1 | TATGAGTGTGGAGTCGTTGCCCTGGCACCACTTTGATACATTGNNNAGATCGGAAG |
| 29263_2 | TATGAGTGTGGAGTCGTTGCCCTGGCACCACTTTGATACATTGNNNAGATCGGAAG |
| 29263_3 | TATGAGTGTGGAGTCGTTGCCCTGGCACCACTTTGATACATTGNNNAGATCGGAAG |
| 29263_4 | TATGAGTGTGGAGTCGTTGCCCTGGCACCACTTTGATACATTGNNNAGATCGGAAG |
| 38909_1 | TATGAGTGTGGAGTCGTTGCATGTCCTATGCAAAATGGCACACNNNAGATCGGAAG |

| | |
|---|---|
| 38909_2 | TATGAGTGTGGAGTCGTTGCATGTCCTATGCAAAATGGCACACNNNAGATCGGAAG |
| 38909_3 | TATGAGTGTGGAGTCGTTGCATGTCCTATGCAAAATGGCACACNNNAGATCGGAAG |
| 38909_4 | TATGAGTGTGGAGTCGTTGCATGTCCTATGCAAAATGGCACACNNNAGATCGGAAG |
| 30897_1 | TATGAGTGTGGAGTCGTTGCAATCCCAGAGAGATTCGGAGAGANNNAGATCGGAA |
| 30897_2 | TATGAGTGTGGAGTCGTTGCAATCCCAGAGAGATTCGGAGAGANNNAGATCGGAA |
| 30897_3 | TATGAGTGTGGAGTCGTTGCAATCCCAGAGAGATTCGGAGAGANNNAGATCGGAA |
| 30897_4 | TATGAGTGTGGAGTCGTTGCAATCCCAGAGAGATTCGGAGAGANNNAGATCGGAA |
| 10046_1 | TATGAGTGTGGAGTCGTTGCCAGGGGCCAGATAGAGGAAGATANNNAGATCGGAA |
| 10046_2 | TATGAGTGTGGAGTCGTTGCCAGGGGCCAGATAGAGGAAGATANNNAGATCGGAA |
| 10046_3 | TATGAGTGTGGAGTCGTTGCCAGGGGCCAGATAGAGGAAGATANNNAGATCGGAA |
| 10046_4 | TATGAGTGTGGAGTCGTTGCCAGGGGCCAGATAGAGGAAGATANNNAGATCGGAA |
| 33823_1 | TATGAGTGTGGAGTCGTTGCATTTACAACTCTGCAACATGGGGNNNAGATCGGAAG |
| 33823_2 | TATGAGTGTGGAGTCGTTGCATTTACAACTCTGCAACATGGGGNNNAGATCGGAAG |
| 33823_3 | TATGAGTGTGGAGTCGTTGCATTTACAACTCTGCAACATGGGGNNNAGATCGGAAG |
| 33823_4 | TATGAGTGTGGAGTCGTTGCATTTACAACTCTGCAACATGGGGNNNAGATCGGAAG |
| 51764_1 | TATGAGTGTGGAGTCGTTGCTGAGGCAGGAGAATCACTTGAAANNNAGATCGGAA |
| 51764_2 | TATGAGTGTGGAGTCGTTGCTGAGGCAGGAGAATCACTTGAAANNNAGATCGGAA |
| 51764_3 | TATGAGTGTGGAGTCGTTGCTGAGGCAGGAGAATCACTTGAAANNNAGATCGGAA |
| 51764_4 | TATGAGTGTGGAGTCGTTGCTGAGGCAGGAGAATCACTTGAAANNNAGATCGGAA |
| 47150_1 | TATGAGTGTGGAGTCGTTGCCTCGCTGTAGCCTCTACCTCTTNNNAGATCGGAAGAG |
| 47150_2 | TATGAGTGTGGAGTCGTTGCCTCGCTGTAGCCTCTACCTCTTNNNAGATCGGAAGAG |
| 47150_3 | TATGAGTGTGGAGTCGTTGCCTCGCTGTAGCCTCTACCTCTTNNNAGATCGGAAGAG |
| 47150_4 | TATGAGTGTGGAGTCGTTGCCTCGCTGTAGCCTCTACCTCTTNNNAGATCGGAAGAG |
| 48649_1 | TATGAGTGTGGAGTCGTTGCTGTGTTCCAGGATGCGTACTACNNNAGATCGGAAGA |
| 48649_2 | TATGAGTGTGGAGTCGTTGCTGTGTTCCAGGATGCGTACTACNNNAGATCGGAAGA |
| 48649_3 | TATGAGTGTGGAGTCGTTGCTGTGTTCCAGGATGCGTACTACNNNAGATCGGAAGA |
| 48649_4 | TATGAGTGTGGAGTCGTTGCTGTGTTCCAGGATGCGTACTACNNNAGATCGGAAGA |
| 38565_1 | TATGAGTGTGGAGTCGTTGCAACTCTTCTGGACATAGGCCTTGNNNAGATCGGAAG |
| 38565_2 | TATGAGTGTGGAGTCGTTGCAACTCTTCTGGACATAGGCCTTGNNNAGATCGGAAG |
| 38565_3 | TATGAGTGTGGAGTCGTTGCAACTCTTCTGGACATAGGCCTTGNNNAGATCGGAAG |
| 38565_4 | TATGAGTGTGGAGTCGTTGCAACTCTTCTGGACATAGGCCTTGNNNAGATCGGAAG |
| 50585_1 | TATGAGTGTGGAGTCGTTGCATATTCACAGGCCAGACTCAGTGNNNAGATCGGAAG |
| 50585_2 | TATGAGTGTGGAGTCGTTGCATATTCACAGGCCAGACTCAGTGNNNAGATCGGAAG |
| 50585_3 | TATGAGTGTGGAGTCGTTGCATATTCACAGGCCAGACTCAGTGNNNAGATCGGAAG |
| 50585_4 | TATGAGTGTGGAGTCGTTGCATATTCACAGGCCAGACTCAGTGNNNAGATCGGAAG |
| 32955_1 | TATGAGTGTGGAGTCGTTGCCACGCCCAGCTACTTTTTGTATTNNNAGATCGGAAGA |
| 32955_2 | TATGAGTGTGGAGTCGTTGCCACGCCCAGCTACTTTTTGTATTNNNAGATCGGAAGA |
| 32955_3 | TATGAGTGTGGAGTCGTTGCCACGCCCAGCTACTTTTTGTATTNNNAGATCGGAAGA |
| 32955_4 | TATGAGTGTGGAGTCGTTGCCACGCCCAGCTACTTTTTGTATTNNNAGATCGGAAGA |
| 43891_1 | TATGAGTGTGGAGTCGTTGCAGACGGAGTCTCACTCTGTCGNNNAGATCGGAAGAG |
| 43891_2 | TATGAGTGTGGAGTCGTTGCAGACGGAGTCTCACTCTGTCGNNNAGATCGGAAGAG |
| 43891_3 | TATGAGTGTGGAGTCGTTGCAGACGGAGTCTCACTCTGTCGNNNAGATCGGAAGAG |
| 43891_4 | TATGAGTGTGGAGTCGTTGCAGACGGAGTCTCACTCTGTCGNNNAGATCGGAAGAG |

| | |
|---|---|
| 51844_1 | TATGAGTGTGGAGTCGTTGCGGTCCATTGTATGAGGGTGAAGTNNNAGATCGGAA |
| 51844_2 | TATGAGTGTGGAGTCGTTGCGGTCCATTGTATGAGGGTGAAGTNNNAGATCGGAA |
| 51844_3 | TATGAGTGTGGAGTCGTTGCGGTCCATTGTATGAGGGTGAAGTNNNAGATCGGAA |
| 51844_4 | TATGAGTGTGGAGTCGTTGCGGTCCATTGTATGAGGGTGAAGTNNNAGATCGGAA |
| 12868_1 | TATGAGTGTGGAGTCGTTGCCACAAAATCAACCTTGTCAGNNNAGATCGGAAGAGC |
| 12868_2 | TATGAGTGTGGAGTCGTTGCCACAAAATCAACCTTGTCAGNNNAGATCGGAAGAGC |
| 12868_3 | TATGAGTGTGGAGTCGTTGCCACAAAATCAACCTTGTCAGNNNAGATCGGAAGAGC |
| 12868_4 | TATGAGTGTGGAGTCGTTGCCACAAAATCAACCTTGTCAGNNNAGATCGGAAGAGC |
| 29358_1 | TATGAGTGTGGAGTCGTTGCACCATTTGGTGAGATGAGTCTTGTNNNAGATCGGA |
| 29358_2 | TATGAGTGTGGAGTCGTTGCACCATTTGGTGAGATGAGTCTTGTNNNAGATCGGA |
| 29358_3 | TATGAGTGTGGAGTCGTTGCACCATTTGGTGAGATGAGTCTTGTNNNAGATCGGA |
| 29358_4 | TATGAGTGTGGAGTCGTTGCACCATTTGGTGAGATGAGTCTTGTNNNAGATCGGA |
| 47481_1 | TATGAGTGTGGAGTCGTTGCCTGGAGTGCAATGGTGTGATCTCNNNAGATCGGAA |
| 47481_2 | TATGAGTGTGGAGTCGTTGCCTGGAGTGCAATGGTGTGATCTCNNNAGATCGGAA |
| 47481_3 | TATGAGTGTGGAGTCGTTGCCTGGAGTGCAATGGTGTGATCTCNNNAGATCGGAA |
| 47481_4 | TATGAGTGTGGAGTCGTTGCCTGGAGTGCAATGGTGTGATCTCNNNAGATCGGAA |
| 45321_1 | TATGAGTGTGGAGTCGTTGCTCCCTTTACTTGGCCGTATTTCANNNAGATCGGAAGA |
| 45321_2 | TATGAGTGTGGAGTCGTTGCTCCCTTTACTTGGCCGTATTTCANNNAGATCGGAAGA |
| 45321_3 | TATGAGTGTGGAGTCGTTGCTCCCTTTACTTGGCCGTATTTCANNNAGATCGGAAGA |
| 45321_4 | TATGAGTGTGGAGTCGTTGCTCCCTTTACTTGGCCGTATTTCANNNAGATCGGAAGA |
| 42373_1 | TATGAGTGTGGAGTCGTTGCTATTGACCAGATGCCATGGTTCANNNAGATCGGAAG |
| 42373_2 | TATGAGTGTGGAGTCGTTGCTATTGACCAGATGCCATGGTTCANNNAGATCGGAAG |
| 42373_3 | TATGAGTGTGGAGTCGTTGCTATTGACCAGATGCCATGGTTCANNNAGATCGGAAG |
| 42373_4 | TATGAGTGTGGAGTCGTTGCTATTGACCAGATGCCATGGTTCANNNAGATCGGAAG |
| 12839_1 | TATGAGTGTGGAGTCGTTGCGACTGTTTCAAGGAGCTACTNNNAGATCGGAAGAG |
| 12839_2 | TATGAGTGTGGAGTCGTTGCGACTGTTTCAAGGAGCTACTNNNAGATCGGAAGAG |
| 12839_3 | TATGAGTGTGGAGTCGTTGCGACTGTTTCAAGGAGCTACTNNNAGATCGGAAGAG |
| 12839_4 | TATGAGTGTGGAGTCGTTGCGACTGTTTCAAGGAGCTACTNNNAGATCGGAAGAG |
| 8069_1 | TATGAGTGTGGAGTCGTTGCGGAAGCCACATGAAGAAGGTGAANNNAGATCGGA |
| 8069_2 | TATGAGTGTGGAGTCGTTGCGGAAGCCACATGAAGAAGGTGAANNNAGATCGGA |
| 8069_3 | TATGAGTGTGGAGTCGTTGCGGAAGCCACATGAAGAAGGTGAANNNAGATCGGA |
| 8069_4 | TATGAGTGTGGAGTCGTTGCGGAAGCCACATGAAGAAGGTGAANNNAGATCGGA |
| 41499_1 | TATGAGTGTGGAGTCGTTGCTTGAACCCAGGAGACATAGGTTGNNNAGATCGGAA |
| 41499_2 | TATGAGTGTGGAGTCGTTGCTTGAACCCAGGAGACATAGGTTGNNNAGATCGGAA |
| 41499_3 | TATGAGTGTGGAGTCGTTGCTTGAACCCAGGAGACATAGGTTGNNNAGATCGGAA |
| 41499_4 | TATGAGTGTGGAGTCGTTGCTTGAACCCAGGAGACATAGGTTGNNNAGATCGGAA |
| 37761_1 | TATGAGTGTGGAGTCGTTGCTCAAGAGATCTGGAGTAGAGCCTNNNAGATCGGAA |
| 37761_2 | TATGAGTGTGGAGTCGTTGCTCAAGAGATCTGGAGTAGAGCCTNNNAGATCGGAA |
| 37761_3 | TATGAGTGTGGAGTCGTTGCTCAAGAGATCTGGAGTAGAGCCTNNNAGATCGGAA |
| 37761_4 | TATGAGTGTGGAGTCGTTGCTCAAGAGATCTGGAGTAGAGCCTNNNAGATCGGAA |
| 46977_1 | TATGAGTGTGGAGTCGTTGCGTTCACACCATTCTCCTACCTCANNNAGATCGGAAGA |
| 46977_2 | TATGAGTGTGGAGTCGTTGCGTTCACACCATTCTCCTACCTCANNNAGATCGGAAGA |
| 46977_3 | TATGAGTGTGGAGTCGTTGCGTTCACACCATTCTCCTACCTCANNNAGATCGGAAGA |

| | |
|---|---|
| 46977_4 | TATGAGTGTGGAGTCGTTGCGTTCACACCATTCTCCTACCTCANNNAGATCGGAAGA |
| 37804_1 | TATGAGTGTGGAGTCGTTGCAGTAAGCATGAGATGTGTCAGGGNNNAGATCGGAA |
| 37804_2 | TATGAGTGTGGAGTCGTTGCAGTAAGCATGAGATGTGTCAGGGNNNAGATCGGAA |
| 37804_3 | TATGAGTGTGGAGTCGTTGCAGTAAGCATGAGATGTGTCAGGGNNNAGATCGGAA |
| 37804_4 | TATGAGTGTGGAGTCGTTGCAGTAAGCATGAGATGTGTCAGGGNNNAGATCGGAA |
| 37920_1 | TATGAGTGTGGAGTCGTTGCCACTCCTGTAATCCCAGCACTTTNNNAGATCGGAAGA |
| 37920_2 | TATGAGTGTGGAGTCGTTGCCACTCCTGTAATCCCAGCACTTTNNNAGATCGGAAGA |
| 37920_3 | TATGAGTGTGGAGTCGTTGCCACTCCTGTAATCCCAGCACTTTNNNAGATCGGAAGA |
| 37920_4 | TATGAGTGTGGAGTCGTTGCCACTCCTGTAATCCCAGCACTTTNNNAGATCGGAAGA |
| 35147_1 | TATGAGTGTGGAGTCGTTGCATTGACTGGCAGTGAAGGTATGTNNNAGATCGGAA |
| 35147_2 | TATGAGTGTGGAGTCGTTGCATTGACTGGCAGTGAAGGTATGTNNNAGATCGGAA |
| 35147_3 | TATGAGTGTGGAGTCGTTGCATTGACTGGCAGTGAAGGTATGTNNNAGATCGGAA |
| 35147_4 | TATGAGTGTGGAGTCGTTGCATTGACTGGCAGTGAAGGTATGTNNNAGATCGGAA |
| 37593_1 | TATGAGTGTGGAGTCGTTGCAGAAGAGCAAAAAGCAAGAGCACNNNAGATCGGAA |
| 37593_2 | TATGAGTGTGGAGTCGTTGCAGAAGAGCAAAAAGCAAGAGCACNNNAGATCGGAA |
| 37593_3 | TATGAGTGTGGAGTCGTTGCAGAAGAGCAAAAAGCAAGAGCACNNNAGATCGGAA |
| 37593_4 | TATGAGTGTGGAGTCGTTGCAGAAGAGCAAAAAGCAAGAGCACNNNAGATCGGAA |
| 35249_1 | TATGAGTGTGGAGTCGTTGCGACACAGGATTTCACCACATTGGNNNAGATCGGAAG |
| 35249_2 | TATGAGTGTGGAGTCGTTGCGACACAGGATTTCACCACATTGGNNNAGATCGGAAG |
| 35249_3 | TATGAGTGTGGAGTCGTTGCGACACAGGATTTCACCACATTGGNNNAGATCGGAAG |
| 35249_4 | TATGAGTGTGGAGTCGTTGCGACACAGGATTTCACCACATTGGNNNAGATCGGAAG |
| 29446_1 | TATGAGTGTGGAGTCGTTGCAAACCATCCTATCCTGGGACTCCNNNAGATCGGAAG |
| 29446_2 | TATGAGTGTGGAGTCGTTGCAAACCATCCTATCCTGGGACTCCNNNAGATCGGAAG |
| 29446_3 | TATGAGTGTGGAGTCGTTGCAAACCATCCTATCCTGGGACTCCNNNAGATCGGAAG |
| 29446_4 | TATGAGTGTGGAGTCGTTGCAAACCATCCTATCCTGGGACTCCNNNAGATCGGAAG |
| 29831_1 | TATGAGTGTGGAGTCGTTGCGTGTGAGTGGAGTAACGGTGAANNNAGATCGGAA |
| 29831_2 | TATGAGTGTGGAGTCGTTGCGTGTGAGTGGAGTAACGGTGAANNNAGATCGGAA |
| 29831_3 | TATGAGTGTGGAGTCGTTGCGTGTGAGTGGAGTAACGGTGAANNNAGATCGGAA |
| 29831_4 | TATGAGTGTGGAGTCGTTGCGTGTGAGTGGAGTAACGGTGAANNNAGATCGGAA |
| 32008_1 | TATGAGTGTGGAGTCGTTGCACACCTCCAACCTGTCTCAATTTNNNAGATCGGAAGA |
| 32008_2 | TATGAGTGTGGAGTCGTTGCACACCTCCAACCTGTCTCAATTTNNNAGATCGGAAGA |
| 32008_3 | TATGAGTGTGGAGTCGTTGCACACCTCCAACCTGTCTCAATTTNNNAGATCGGAAGA |
| 32008_4 | TATGAGTGTGGAGTCGTTGCACACCTCCAACCTGTCTCAATTTNNNAGATCGGAAGA |
| 4568_1 | TATGAGTGTGGAGTCGTTGCCGCCCTTGGCTAAGTAGTTGGTTNNNAGATCGGAAG |
| 4568_2 | TATGAGTGTGGAGTCGTTGCCGCCCTTGGCTAAGTAGTTGGTTNNNAGATCGGAAG |
| 4568_3 | TATGAGTGTGGAGTCGTTGCCGCCCTTGGCTAAGTAGTTGGTTNNNAGATCGGAAG |
| 4568_4 | TATGAGTGTGGAGTCGTTGCCGCCCTTGGCTAAGTAGTTGGTTNNNAGATCGGAAG |
| 43179_1 | TATGAGTGTGGAGTCGTTGCGGTTCAAGCAATTATCCTGCCTCNNNAGATCGGAAG |
| 43179_2 | TATGAGTGTGGAGTCGTTGCGGTTCAAGCAATTATCCTGCCTCNNNAGATCGGAAG |
| 43179_3 | TATGAGTGTGGAGTCGTTGCGGTTCAAGCAATTATCCTGCCTCNNNAGATCGGAAG |
| 43179_4 | TATGAGTGTGGAGTCGTTGCGGTTCAAGCAATTATCCTGCCTCNNNAGATCGGAAG |
| 43030_1 | TATGAGTGTGGAGTCGTTGCAGAGAGAGAGAGAGAGTATGTCCCNNNAGATCGG |
| 43030_2 | TATGAGTGTGGAGTCGTTGCAGAGAGAGAGAGAGAGTATGTCCCNNNAGATCGG |

| | |
|---|---|
| 43030_3 | TATGAGTGTGGAGTCGTTGCAGAGAGAGAGAGAGAGTATGTCCCNNNAGATCGG |
| 43030_4 | TATGAGTGTGGAGTCGTTGCAGAGAGAGAGAGAGAGTATGTCCCNNNAGATCGG |
| 42388_1 | TATGAGTGTGGAGTCGTTGCTGTAGGTCTCTGCCTTTCTGATGNNNAGATCGGAAG |
| 42388_2 | TATGAGTGTGGAGTCGTTGCTGTAGGTCTCTGCCTTTCTGATGNNNAGATCGGAAG |
| 42388_3 | TATGAGTGTGGAGTCGTTGCTGTAGGTCTCTGCCTTTCTGATGNNNAGATCGGAAG |
| 42388_4 | TATGAGTGTGGAGTCGTTGCTGTAGGTCTCTGCCTTTCTGATGNNNAGATCGGAAG |
| 45652_1 | TATGAGTGTGGAGTCGTTGCTGATCTTGGCTCACTGTAACCTCNNNAGATCGGAAG |
| 45652_2 | TATGAGTGTGGAGTCGTTGCTGATCTTGGCTCACTGTAACCTCNNNAGATCGGAAG |
| 45652_3 | TATGAGTGTGGAGTCGTTGCTGATCTTGGCTCACTGTAACCTCNNNAGATCGGAAG |
| 45652_4 | TATGAGTGTGGAGTCGTTGCTGATCTTGGCTCACTGTAACCTCNNNAGATCGGAAG |
| 3149_1 | TATGAGTGTGGAGTCGTTGCGCTGCCAAGTCTCTGTGAATACANNNAGATCGGAAG |
| 3149_2 | TATGAGTGTGGAGTCGTTGCGCTGCCAAGTCTCTGTGAATACANNNAGATCGGAAG |
| 3149_3 | TATGAGTGTGGAGTCGTTGCGCTGCCAAGTCTCTGTGAATACANNNAGATCGGAAG |
| 3149_4 | TATGAGTGTGGAGTCGTTGCGCTGCCAAGTCTCTGTGAATACANNNAGATCGGAAG |
| 43099_1 | TATGAGTGTGGAGTCGTTGCAGGGATAGAGTGGTGGAGAATCANNNAGATCGGA |
| 43099_2 | TATGAGTGTGGAGTCGTTGCAGGGATAGAGTGGTGGAGAATCANNNAGATCGGA |
| 43099_3 | TATGAGTGTGGAGTCGTTGCAGGGATAGAGTGGTGGAGAATCANNNAGATCGGA |
| 43099_4 | TATGAGTGTGGAGTCGTTGCAGGGATAGAGTGGTGGAGAATCANNNAGATCGGA |
| 44845_1 | TATGAGTGTGGAGTCGTTGCGATCTCAGCTCACTGTAACCTCCNNNAGATCGGAAG |
| 44845_2 | TATGAGTGTGGAGTCGTTGCGATCTCAGCTCACTGTAACCTCCNNNAGATCGGAAG |
| 44845_3 | TATGAGTGTGGAGTCGTTGCGATCTCAGCTCACTGTAACCTCCNNNAGATCGGAAG |
| 44845_4 | TATGAGTGTGGAGTCGTTGCGATCTCAGCTCACTGTAACCTCCNNNAGATCGGAAG |
| 48426_1 | TATGAGTGTGGAGTCGTTGCCCATGTCCCGCTAGTTTTTGTTCNNNAGATCGGAAG |
| 48426_2 | TATGAGTGTGGAGTCGTTGCCCATGTCCCGCTAGTTTTTGTTCNNNAGATCGGAAG |
| 48426_3 | TATGAGTGTGGAGTCGTTGCCCATGTCCCGCTAGTTTTTGTTCNNNAGATCGGAAG |
| 48426_4 | TATGAGTGTGGAGTCGTTGCCCATGTCCCGCTAGTTTTTGTTCNNNAGATCGGAAG |
| 39030_1 | TATGAGTGTGGAGTCGTTGCTGTTCAGATTTCTGTGACCACCANNNAGATCGGAAG |
| 39030_2 | TATGAGTGTGGAGTCGTTGCTGTTCAGATTTCTGTGACCACCANNNAGATCGGAAG |
| 39030_3 | TATGAGTGTGGAGTCGTTGCTGTTCAGATTTCTGTGACCACCANNNAGATCGGAAG |
| 39030_4 | TATGAGTGTGGAGTCGTTGCTGTTCAGATTTCTGTGACCACCANNNAGATCGGAAG |
| 36867_1 | TATGAGTGTGGAGTCGTTGCAAAGAGTGGTTGAGACAGGATCANNNAGATCGGA |
| 36867_2 | TATGAGTGTGGAGTCGTTGCAAAGAGTGGTTGAGACAGGATCANNNAGATCGGA |
| 36867_3 | TATGAGTGTGGAGTCGTTGCAAAGAGTGGTTGAGACAGGATCANNNAGATCGGA |
| 36867_4 | TATGAGTGTGGAGTCGTTGCAAAGAGTGGTTGAGACAGGATCANNNAGATCGGA |
| 29260_1 | TATGAGTGTGGAGTCGTTGCCTCACCTTGGCTCTTAGACACAANNNAGATCGGAAG |
| 29260_2 | TATGAGTGTGGAGTCGTTGCCTCACCTTGGCTCTTAGACACAANNNAGATCGGAAG |
| 29260_3 | TATGAGTGTGGAGTCGTTGCCTCACCTTGGCTCTTAGACACAANNNAGATCGGAAG |
| 29260_4 | TATGAGTGTGGAGTCGTTGCCTCACCTTGGCTCTTAGACACAANNNAGATCGGAAG |
| 30441_1 | TATGAGTGTGGAGTCGTTGCATGATGAGGCTGTAACTGAGTGANNNAGATCGGAA |
| 30441_2 | TATGAGTGTGGAGTCGTTGCATGATGAGGCTGTAACTGAGTGANNNAGATCGGAA |
| 30441_3 | TATGAGTGTGGAGTCGTTGCATGATGAGGCTGTAACTGAGTGANNNAGATCGGAA |
| 30441_4 | TATGAGTGTGGAGTCGTTGCATGATGAGGCTGTAACTGAGTGANNNAGATCGGAA |
| 33410_1 | TATGAGTGTGGAGTCGTTGCGTGGCTCACACCTGTAATCCTAANNNAGATCGGAAG |

| | |
|---|---|
| 33410_2 | TATGAGTGTGGAGTCGTTGCGTGGCTCACACCTGTAATCCTAANNNAGATCGGAAGA |
| 33410_3 | TATGAGTGTGGAGTCGTTGCGTGGCTCACACCTGTAATCCTAANNNAGATCGGAAGA |
| 33410_4 | TATGAGTGTGGAGTCGTTGCGTGGCTCACACCTGTAATCCTAANNNAGATCGGAAGA |
| 32965_1 | TATGAGTGTGGAGTCGTTGCGCCAAATAAAGCCAAAACTGCTGNNNAGATCGGAAG |
| 32965_2 | TATGAGTGTGGAGTCGTTGCGCCAAATAAAGCCAAAACTGCTGNNNAGATCGGAAG |
| 32965_3 | TATGAGTGTGGAGTCGTTGCGCCAAATAAAGCCAAAACTGCTGNNNAGATCGGAAG |
| 32965_4 | TATGAGTGTGGAGTCGTTGCGCCAAATAAAGCCAAAACTGCTGNNNAGATCGGAAG |
| 44816_1 | TATGAGTGTGGAGTCGTTGCGCTAGGAAAAGAGCGAGGAGATTNNNAGATCGGAA |
| 44816_2 | TATGAGTGTGGAGTCGTTGCGCTAGGAAAAGAGCGAGGAGATTNNNAGATCGGAA |
| 44816_3 | TATGAGTGTGGAGTCGTTGCGCTAGGAAAAGAGCGAGGAGATTNNNAGATCGGAA |
| 44816_4 | TATGAGTGTGGAGTCGTTGCGCTAGGAAAAGAGCGAGGAGATTNNNAGATCGGAA |
| 44359_1 | TATGAGTGTGGAGTCGTTGCGAAAGAACACAATCACTGGCTCCNNNAGATCGGAAG |
| 44359_2 | TATGAGTGTGGAGTCGTTGCGAAAGAACACAATCACTGGCTCCNNNAGATCGGAAG |
| 44359_3 | TATGAGTGTGGAGTCGTTGCGAAAGAACACAATCACTGGCTCCNNNAGATCGGAAG |
| 44359_4 | TATGAGTGTGGAGTCGTTGCGAAAGAACACAATCACTGGCTCCNNNAGATCGGAAG |
| 52124_1 | TATGAGTGTGGAGTCGTTGCTGATCTCAAAACCCTGTCTCCTGNNNAGATCGGAAGA |
| 52124_2 | TATGAGTGTGGAGTCGTTGCTGATCTCAAAACCCTGTCTCCTGNNNAGATCGGAAGA |
| 52124_3 | TATGAGTGTGGAGTCGTTGCTGATCTCAAAACCCTGTCTCCTGNNNAGATCGGAAGA |
| 52124_4 | TATGAGTGTGGAGTCGTTGCTGATCTCAAAACCCTGTCTCCTGNNNAGATCGGAAGA |
| 47129_1 | TATGAGTGTGGAGTCGTTGCCCACGCCCAGCTAATTTTTTGTANNNAGATCGGAAGA |
| 47129_2 | TATGAGTGTGGAGTCGTTGCCCACGCCCAGCTAATTTTTTGTANNNAGATCGGAAGA |
| 47129_3 | TATGAGTGTGGAGTCGTTGCCCACGCCCAGCTAATTTTTTGTANNNAGATCGGAAGA |
| 47129_4 | TATGAGTGTGGAGTCGTTGCCCACGCCCAGCTAATTTTTTGTANNNAGATCGGAAGA |
| 46145_1 | TATGAGTGTGGAGTCGTTGCCCTTCCATCTCCTTGGCCAACNNNAGATCGGAAGAGC |
| 46145_2 | TATGAGTGTGGAGTCGTTGCCCTTCCATCTCCTTGGCCAACNNNAGATCGGAAGAGC |
| 46145_3 | TATGAGTGTGGAGTCGTTGCCCTTCCATCTCCTTGGCCAACNNNAGATCGGAAGAGC |
| 46145_4 | TATGAGTGTGGAGTCGTTGCCCTTCCATCTCCTTGGCCAACNNNAGATCGGAAGAGC |
| 46722_1 | TATGAGTGTGGAGTCGTTGCGGCAAAACCTTGTCTTCAAACCTNNNAGATCGGAAGA |
| 46722_2 | TATGAGTGTGGAGTCGTTGCGGCAAAACCTTGTCTTCAAACCTNNNAGATCGGAAGA |
| 46722_3 | TATGAGTGTGGAGTCGTTGCGGCAAAACCTTGTCTTCAAACCTNNNAGATCGGAAGA |
| 46722_4 | TATGAGTGTGGAGTCGTTGCGGCAAAACCTTGTCTTCAAACCTNNNAGATCGGAAGA |
| 53116_1 | TATGAGTGTGGAGTCGTTGCTTCCCCTCCCTGGTTAGAACTATNNNAGATCGGAAGA |
| 53116_2 | TATGAGTGTGGAGTCGTTGCTTCCCCTCCCTGGTTAGAACTATNNNAGATCGGAAGA |
| 53116_3 | TATGAGTGTGGAGTCGTTGCTTCCCCTCCCTGGTTAGAACTATNNNAGATCGGAAGA |
| 53116_4 | TATGAGTGTGGAGTCGTTGCTTCCCCTCCCTGGTTAGAACTATNNNAGATCGGAAGA |
| 44840_1 | TATGAGTGTGGAGTCGTTGCCTCTGAAAGTCCCTCAGCCTTCNNNAGATCGGAAGAG |
| 44840_2 | TATGAGTGTGGAGTCGTTGCCTCTGAAAGTCCCTCAGCCTTCNNNAGATCGGAAGAG |
| 44840_3 | TATGAGTGTGGAGTCGTTGCCTCTGAAAGTCCCTCAGCCTTCNNNAGATCGGAAGAG |
| 44840_4 | TATGAGTGTGGAGTCGTTGCCTCTGAAAGTCCCTCAGCCTTCNNNAGATCGGAAGAG |
| 35646_1 | TATGAGTGTGGAGTCGTTGCATCATCTCAGCTCAACACAACCTNNNAGATCGGAAGA |
| 35646_2 | TATGAGTGTGGAGTCGTTGCATCATCTCAGCTCAACACAACCTNNNAGATCGGAAGA |
| 35646_3 | TATGAGTGTGGAGTCGTTGCATCATCTCAGCTCAACACAACCTNNNAGATCGGAAGA |
| 35646_4 | TATGAGTGTGGAGTCGTTGCATCATCTCAGCTCAACACAACCTNNNAGATCGGAAGA |

| | |
|---|---|
| 38105_1 | TATGAGTGTGGAGTCGTTGCTACTGGTCTAGTCTGCTTTGCTGNNNAGATCGGAAG |
| 38105_2 | TATGAGTGTGGAGTCGTTGCTACTGGTCTAGTCTGCTTTGCTGNNNAGATCGGAAG |
| 38105_3 | TATGAGTGTGGAGTCGTTGCTACTGGTCTAGTCTGCTTTGCTGNNNAGATCGGAAG |
| 38105_4 | TATGAGTGTGGAGTCGTTGCTACTGGTCTAGTCTGCTTTGCTGNNNAGATCGGAAG |
| 50444_1 | TATGAGTGTGGAGTCGTTGCGACAGCACTCACAGCCTTCTCNNNAGATCGGAAGAG |
| 50444_2 | TATGAGTGTGGAGTCGTTGCGACAGCACTCACAGCCTTCTCNNNAGATCGGAAGAG |
| 50444_3 | TATGAGTGTGGAGTCGTTGCGACAGCACTCACAGCCTTCTCNNNAGATCGGAAGAG |
| 50444_4 | TATGAGTGTGGAGTCGTTGCGACAGCACTCACAGCCTTCTCNNNAGATCGGAAGAG |
| 47161_1 | TATGAGTGTGGAGTCGTTGCCTCCTGAGTAGCTGGAACAACAGNNNAGATCGGAA |
| 47161_2 | TATGAGTGTGGAGTCGTTGCCTCCTGAGTAGCTGGAACAACAGNNNAGATCGGAA |
| 47161_3 | TATGAGTGTGGAGTCGTTGCCTCCTGAGTAGCTGGAACAACAGNNNAGATCGGAA |
| 47161_4 | TATGAGTGTGGAGTCGTTGCCTCCTGAGTAGCTGGAACAACAGNNNAGATCGGAA |
| 49929_1 | TATGAGTGTGGAGTCGTTGCCCAGCCTTGCCAAATAATGTCTCNNNAGATCGGAAG |
| 49929_2 | TATGAGTGTGGAGTCGTTGCCCAGCCTTGCCAAATAATGTCTCNNNAGATCGGAAG |
| 49929_3 | TATGAGTGTGGAGTCGTTGCCCAGCCTTGCCAAATAATGTCTCNNNAGATCGGAAG |
| 49929_4 | TATGAGTGTGGAGTCGTTGCCCAGCCTTGCCAAATAATGTCTCNNNAGATCGGAAG |
| 47400_1 | TATGAGTGTGGAGTCGTTGCCAAAGTGCTGGAATTACAGGCATNNNAGATCGGAA |
| 47400_2 | TATGAGTGTGGAGTCGTTGCCAAAGTGCTGGAATTACAGGCATNNNAGATCGGAA |
| 47400_3 | TATGAGTGTGGAGTCGTTGCCAAAGTGCTGGAATTACAGGCATNNNAGATCGGAA |
| 47400_4 | TATGAGTGTGGAGTCGTTGCCAAAGTGCTGGAATTACAGGCATNNNAGATCGGAA |
| 12849_1 | TATGAGTGTGGAGTCGTTGCCTGTTTCAGGTTTCTTATTGGNNNAGATCGGAAGAG |
| 12849_2 | TATGAGTGTGGAGTCGTTGCCTGTTTCAGGTTTCTTATTGGNNNAGATCGGAAGAG |
| 12849_3 | TATGAGTGTGGAGTCGTTGCCTGTTTCAGGTTTCTTATTGGNNNAGATCGGAAGAG |
| 12849_4 | TATGAGTGTGGAGTCGTTGCCTGTTTCAGGTTTCTTATTGGNNNAGATCGGAAGAG |
| 30656_1 | TATGAGTGTGGAGTCGTTGCCTCAGGAAAGCAAGCTACACCANNNAGATCGGAAG |
| 30656_2 | TATGAGTGTGGAGTCGTTGCCTCAGGAAAGCAAGCTACACCANNNAGATCGGAAG |
| 30656_3 | TATGAGTGTGGAGTCGTTGCCTCAGGAAAGCAAGCTACACCANNNAGATCGGAAG |
| 30656_4 | TATGAGTGTGGAGTCGTTGCCTCAGGAAAGCAAGCTACACCANNNAGATCGGAAG |
| 30367_1 | TATGAGTGTGGAGTCGTTGCGGCTCTCATGTACCCCAGAAATTNNNAGATCGGAAG |
| 30367_2 | TATGAGTGTGGAGTCGTTGCGGCTCTCATGTACCCCAGAAATTNNNAGATCGGAAG |
| 30367_3 | TATGAGTGTGGAGTCGTTGCGGCTCTCATGTACCCCAGAAATTNNNAGATCGGAAG |
| 30367_4 | TATGAGTGTGGAGTCGTTGCGGCTCTCATGTACCCCAGAAATTNNNAGATCGGAAG |
| 34014_1 | TATGAGTGTGGAGTCGTTGCGAGTAGCTGGGACTTCAGGCNNNAGATCGGAAGAG |
| 34014_2 | TATGAGTGTGGAGTCGTTGCGAGTAGCTGGGACTTCAGGCNNNAGATCGGAAGAG |
| 34014_3 | TATGAGTGTGGAGTCGTTGCGAGTAGCTGGGACTTCAGGCNNNAGATCGGAAGAG |
| 34014_4 | TATGAGTGTGGAGTCGTTGCGAGTAGCTGGGACTTCAGGCNNNAGATCGGAAGAG |
| 43365_1 | TATGAGTGTGGAGTCGTTGCAAGCTAAGATCCACTGGCTTGAANNNAGATCGGAA |
| 43365_2 | TATGAGTGTGGAGTCGTTGCAAGCTAAGATCCACTGGCTTGAANNNAGATCGGAA |
| 43365_3 | TATGAGTGTGGAGTCGTTGCAAGCTAAGATCCACTGGCTTGAANNNAGATCGGAA |
| 43365_4 | TATGAGTGTGGAGTCGTTGCAAGCTAAGATCCACTGGCTTGAANNNAGATCGGAA |
| 13545_1 | TATGAGTGTGGAGTCGTTGCATTTATCTCATAAGTGGAAATGCNNNAGATCGGAAG |
| 13545_2 | TATGAGTGTGGAGTCGTTGCATTTATCTCATAAGTGGAAATGCNNNAGATCGGAAG |
| 13545_3 | TATGAGTGTGGAGTCGTTGCATTTATCTCATAAGTGGAAATGCNNNAGATCGGAAG |

| | |
|---|---|
| 13545_4 | TATGAGTGTGGAGTCGTTGCATTTATCTCATAAGTGGAAATGCNNNAGATCGGAAG |
| 51327_1 | TATGAGTGTGGAGTCGTTGCAGAGAGAAAGAGAGAGCATGTGCNNNAGATCGGA |
| 51327_2 | TATGAGTGTGGAGTCGTTGCAGAGAGAAAGAGAGAGCATGTGCNNNAGATCGGA |
| 51327_3 | TATGAGTGTGGAGTCGTTGCAGAGAGAAAGAGAGAGCATGTGCNNNAGATCGGA |
| 51327_4 | TATGAGTGTGGAGTCGTTGCAGAGAGAAAGAGAGAGCATGTGCNNNAGATCGGA |
| 34766_1 | TATGAGTGTGGAGTCGTTGCAGGAGATGATGACACAGACACACNNNAGATCGGAA |
| 34766_2 | TATGAGTGTGGAGTCGTTGCAGGAGATGATGACACAGACACACNNNAGATCGGAA |
| 34766_3 | TATGAGTGTGGAGTCGTTGCAGGAGATGATGACACAGACACACNNNAGATCGGAA |
| 34766_4 | TATGAGTGTGGAGTCGTTGCAGGAGATGATGACACAGACACACNNNAGATCGGAA |
| 53017_1 | TATGAGTGTGGAGTCGTTGCATTCATGAAACACAGAGGGAGGGNNNAGATCGGAA |
| 53017_2 | TATGAGTGTGGAGTCGTTGCATTCATGAAACACAGAGGGAGGGNNNAGATCGGAA |
| 53017_3 | TATGAGTGTGGAGTCGTTGCATTCATGAAACACAGAGGGAGGGNNNAGATCGGAA |
| 53017_4 | TATGAGTGTGGAGTCGTTGCATTCATGAAACACAGAGGGAGGGNNNAGATCGGAA |
| 42218_1 | TATGAGTGTGGAGTCGTTGCCTCCAGCTTCAAAGGATTCCTCTNNNAGATCGGAAGA |
| 42218_2 | TATGAGTGTGGAGTCGTTGCCTCCAGCTTCAAAGGATTCCTCTNNNAGATCGGAAGA |
| 42218_3 | TATGAGTGTGGAGTCGTTGCCTCCAGCTTCAAAGGATTCCTCTNNNAGATCGGAAGA |
| 42218_4 | TATGAGTGTGGAGTCGTTGCCTCCAGCTTCAAAGGATTCCTCTNNNAGATCGGAAGA |
| 45158_1 | TATGAGTGTGGAGTCGTTGCCTCCTCTCTTCAGCTCTCCTCTNNNAGATCGGAAGAG |
| 45158_2 | TATGAGTGTGGAGTCGTTGCCTCCTCTCTTCAGCTCTCCTCTNNNAGATCGGAAGAG |
| 45158_3 | TATGAGTGTGGAGTCGTTGCCTCCTCTCTTCAGCTCTCCTCTNNNAGATCGGAAGAG |
| 45158_4 | TATGAGTGTGGAGTCGTTGCCTCCTCTCTTCAGCTCTCCTCTNNNAGATCGGAAGAG |
| 32865_1 | TATGAGTGTGGAGTCGTTGCGATAGAGCCTCATTCTGTCACCCNNNAGATCGGAAG |
| 32865_2 | TATGAGTGTGGAGTCGTTGCGATAGAGCCTCATTCTGTCACCCNNNAGATCGGAAG |
| 32865_3 | TATGAGTGTGGAGTCGTTGCGATAGAGCCTCATTCTGTCACCCNNNAGATCGGAAG |
| 32865_4 | TATGAGTGTGGAGTCGTTGCGATAGAGCCTCATTCTGTCACCCNNNAGATCGGAAG |
| 49876_1 | TATGAGTGTGGAGTCGTTGCGCTGGCACTTTACAAATTCGTCTNNNAGATCGGAAG |
| 49876_2 | TATGAGTGTGGAGTCGTTGCGCTGGCACTTTACAAATTCGTCTNNNAGATCGGAAG |
| 49876_3 | TATGAGTGTGGAGTCGTTGCGCTGGCACTTTACAAATTCGTCTNNNAGATCGGAAG |
| 49876_4 | TATGAGTGTGGAGTCGTTGCGCTGGCACTTTACAAATTCGTCTNNNAGATCGGAAG |
| 46963_1 | TATGAGTGTGGAGTCGTTGCCACCACACCTGGCTCATTTTTTGNNNAGATCGGAAGA |
| 46963_2 | TATGAGTGTGGAGTCGTTGCCACCACACCTGGCTCATTTTTTGNNNAGATCGGAAGA |
| 46963_3 | TATGAGTGTGGAGTCGTTGCCACCACACCTGGCTCATTTTTTGNNNAGATCGGAAGA |
| 46963_4 | TATGAGTGTGGAGTCGTTGCCACCACACCTGGCTCATTTTTTGNNNAGATCGGAAGA |
| 36017_1 | TATGAGTGTGGAGTCGTTGCATCATTGAGTCGCCTGAATCAGTNNNAGATCGGAAG |
| 36017_2 | TATGAGTGTGGAGTCGTTGCATCATTGAGTCGCCTGAATCAGTNNNAGATCGGAAG |
| 36017_3 | TATGAGTGTGGAGTCGTTGCATCATTGAGTCGCCTGAATCAGTNNNAGATCGGAAG |
| 36017_4 | TATGAGTGTGGAGTCGTTGCATCATTGAGTCGCCTGAATCAGTNNNAGATCGGAAG |
| 46127_1 | TATGAGTGTGGAGTCGTTGCCACCACACACCCAGCTAATTTTTNNNAGATCGGAAGA |
| 46127_2 | TATGAGTGTGGAGTCGTTGCCACCACACACCCAGCTAATTTTTNNNAGATCGGAAGA |
| 46127_3 | TATGAGTGTGGAGTCGTTGCCACCACACACCCAGCTAATTTTTNNNAGATCGGAAGA |
| 46127_4 | TATGAGTGTGGAGTCGTTGCCACCACACACCCAGCTAATTTTTNNNAGATCGGAAGA |
| 42995_1 | TATGAGTGTGGAGTCGTTGCTTGGTTCTGAGTCTGTGGAGAACNNNAGATCGGAA |
| 42995_2 | TATGAGTGTGGAGTCGTTGCTTGGTTCTGAGTCTGTGGAGAACNNNAGATCGGAA |

| | |
|---|---|
| 42995_3 | TATGAGTGTGGAGTCGTTGCTTGGTTCTGAGTCTGTGGAGAACNNNAGATCGGAA |
| 42995_4 | TATGAGTGTGGAGTCGTTGCTTGGTTCTGAGTCTGTGGAGAACNNNAGATCGGAA |
| 13489_1 | TATGAGTGTGGAGTCGTTGCGCTTGATTTTGAATACTGATTTTNNNAGATCGGAAG |
| 13489_2 | TATGAGTGTGGAGTCGTTGCGCTTGATTTTGAATACTGATTTTNNNAGATCGGAAG |
| 13489_3 | TATGAGTGTGGAGTCGTTGCGCTTGATTTTGAATACTGATTTTNNNAGATCGGAAG |
| 13489_4 | TATGAGTGTGGAGTCGTTGCGCTTGATTTTGAATACTGATTTTNNNAGATCGGAAG |
| 47170_1 | TATGAGTGTGGAGTCGTTGCCTGTGCCTGGCCTTTATTTTTCANNNAGATCGGAAGA |
| 47170_2 | TATGAGTGTGGAGTCGTTGCCTGTGCCTGGCCTTTATTTTTCANNNAGATCGGAAGA |
| 47170_3 | TATGAGTGTGGAGTCGTTGCCTGTGCCTGGCCTTTATTTTTCANNNAGATCGGAAGA |
| 47170_4 | TATGAGTGTGGAGTCGTTGCCTGTGCCTGGCCTTTATTTTTCANNNAGATCGGAAGA |
| 30710_1 | TATGAGTGTGGAGTCGTTGCGTGTGTGTCTGATCTCCTTGAACTCNNNAGATCGGA |
| 30710_2 | TATGAGTGTGGAGTCGTTGCGTGTGTGTCTGATCTCCTTGAACTCNNNAGATCGGA |
| 30710_3 | TATGAGTGTGGAGTCGTTGCGTGTGTGTCTGATCTCCTTGAACTCNNNAGATCGGA |
| 30710_4 | TATGAGTGTGGAGTCGTTGCGTGTGTGTCTGATCTCCTTGAACTCNNNAGATCGGA |
| 13509_1 | TATGAGTGTGGAGTCGTTGCGAGTATCTAGTAATTTGGAAGGTNNNAGATCGGAA |
| 13509_2 | TATGAGTGTGGAGTCGTTGCGAGTATCTAGTAATTTGGAAGGTNNNAGATCGGAA |
| 13509_3 | TATGAGTGTGGAGTCGTTGCGAGTATCTAGTAATTTGGAAGGTNNNAGATCGGAA |
| 13509_4 | TATGAGTGTGGAGTCGTTGCGAGTATCTAGTAATTTGGAAGGTNNNAGATCGGAA |
| 49182_1 | TATGAGTGTGGAGTCGTTGCCCTCCCAAGTAGCTCGGATTATGNNNAGATCGGAAG |
| 49182_2 | TATGAGTGTGGAGTCGTTGCCCTCCCAAGTAGCTCGGATTATGNNNAGATCGGAAG |
| 49182_3 | TATGAGTGTGGAGTCGTTGCCCTCCCAAGTAGCTCGGATTATGNNNAGATCGGAAG |
| 49182_4 | TATGAGTGTGGAGTCGTTGCCCTCCCAAGTAGCTCGGATTATGNNNAGATCGGAAG |
| 35248_1 | TATGAGTGTGGAGTCGTTGCCTGAAGACTGGTCCTGGATATGGNNNAGATCGGAA |
| 35248_2 | TATGAGTGTGGAGTCGTTGCCTGAAGACTGGTCCTGGATATGGNNNAGATCGGAA |
| 35248_3 | TATGAGTGTGGAGTCGTTGCCTGAAGACTGGTCCTGGATATGGNNNAGATCGGAA |
| 35248_4 | TATGAGTGTGGAGTCGTTGCCTGAAGACTGGTCCTGGATATGGNNNAGATCGGAA |
| 13248_1 | TATGAGTGTGGAGTCGTTGCCTCTACATTCAAAACCTATTGTTNNNAGATCGGAAGA |
| 13248_2 | TATGAGTGTGGAGTCGTTGCCTCTACATTCAAAACCTATTGTTNNNAGATCGGAAGA |
| 13248_3 | TATGAGTGTGGAGTCGTTGCCTCTACATTCAAAACCTATTGTTNNNAGATCGGAAGA |
| 13248_4 | TATGAGTGTGGAGTCGTTGCCTCTACATTCAAAACCTATTGTTNNNAGATCGGAAGA |
| 34371_1 | TATGAGTGTGGAGTCGTTGCGACACCACCCCTGCTAATTTTTGNNNAGATCGGAAG |
| 34371_2 | TATGAGTGTGGAGTCGTTGCGACACCACCCCTGCTAATTTTTGNNNAGATCGGAAG |
| 34371_3 | TATGAGTGTGGAGTCGTTGCGACACCACCCCTGCTAATTTTTGNNNAGATCGGAAG |
| 34371_4 | TATGAGTGTGGAGTCGTTGCGACACCACCCCTGCTAATTTTTGNNNAGATCGGAAG |
| 47974_1 | TATGAGTGTGGAGTCGTTGCGAAATATGACATTCCTGCCACGGNNNAGATCGGAAG |
| 47974_2 | TATGAGTGTGGAGTCGTTGCGAAATATGACATTCCTGCCACGGNNNAGATCGGAAG |
| 47974_3 | TATGAGTGTGGAGTCGTTGCGAAATATGACATTCCTGCCACGGNNNAGATCGGAAG |
| 47974_4 | TATGAGTGTGGAGTCGTTGCGAAATATGACATTCCTGCCACGGNNNAGATCGGAAG |
| 29719_1 | TATGAGTGTGGAGTCGTTGCTTTGTATCCCCATTGCTGGTGTTNNNAGATCGGAAG |
| 29719_2 | TATGAGTGTGGAGTCGTTGCTTTGTATCCCCATTGCTGGTGTTNNNAGATCGGAAG |
| 29719_3 | TATGAGTGTGGAGTCGTTGCTTTGTATCCCCATTGCTGGTGTTNNNAGATCGGAAG |
| 29719_4 | TATGAGTGTGGAGTCGTTGCTTTGTATCCCCATTGCTGGTGTTNNNAGATCGGAAG |
| 48002_1 | TATGAGTGTGGAGTCGTTGCGTCCCTTGAACTACTGGGTGTAGNNNAGATCGGAA |

| | |
|---|---|
| 48002_2 | TATGAGTGTGGAGTCGTTGCGTCCCTTGAACTACTGGGTGTAGNNNAGATCGGAAG |
| 48002_3 | TATGAGTGTGGAGTCGTTGCGTCCCTTGAACTACTGGGTGTAGNNNAGATCGGAAG |
| 48002_4 | TATGAGTGTGGAGTCGTTGCGTCCCTTGAACTACTGGGTGTAGNNNAGATCGGAAG |
| 40594_1 | TATGAGTGTGGAGTCGTTGCACTGACATGCATTGATGTTTCAGTNNNAGATCGGAA |
| 40594_2 | TATGAGTGTGGAGTCGTTGCACTGACATGCATTGATGTTTCAGTNNNAGATCGGAA |
| 40594_3 | TATGAGTGTGGAGTCGTTGCACTGACATGCATTGATGTTTCAGTNNNAGATCGGAA |
| 40594_4 | TATGAGTGTGGAGTCGTTGCACTGACATGCATTGATGTTTCAGTNNNAGATCGGAA |
| 32937_1 | TATGAGTGTGGAGTCGTTGCAAAAGCCCAGACATTCAACTCACNNNAGATCGGAAG |
| 32937_2 | TATGAGTGTGGAGTCGTTGCAAAAGCCCAGACATTCAACTCACNNNAGATCGGAAG |
| 32937_3 | TATGAGTGTGGAGTCGTTGCAAAAGCCCAGACATTCAACTCACNNNAGATCGGAAG |
| 32937_4 | TATGAGTGTGGAGTCGTTGCAAAAGCCCAGACATTCAACTCACNNNAGATCGGAAG |
| 44968_1 | TATGAGTGTGGAGTCGTTGCGTAAAATGAGGGGGCTGGAGTAANNNAGATCGGA |
| 44968_2 | TATGAGTGTGGAGTCGTTGCGTAAAATGAGGGGGCTGGAGTAANNNAGATCGGA |
| 44968_3 | TATGAGTGTGGAGTCGTTGCGTAAAATGAGGGGGCTGGAGTAANNNAGATCGGA |
| 44968_4 | TATGAGTGTGGAGTCGTTGCGTAAAATGAGGGGGCTGGAGTAANNNAGATCGGA |
| 4426_1 | TATGAGTGTGGAGTCGTTGCCCACTCATCCATCAACAGTCACTNNNAGATCGGAAGA |
| 4426_2 | TATGAGTGTGGAGTCGTTGCCCACTCATCCATCAACAGTCACTNNNAGATCGGAAGA |
| 4426_3 | TATGAGTGTGGAGTCGTTGCCCACTCATCCATCAACAGTCACTNNNAGATCGGAAGA |
| 4426_4 | TATGAGTGTGGAGTCGTTGCCCACTCATCCATCAACAGTCACTNNNAGATCGGAAGA |
| 45165_1 | TATGAGTGTGGAGTCGTTGCCATGGATTAGGGTCTCAGACTGGNNNAGATCGGAA |
| 45165_2 | TATGAGTGTGGAGTCGTTGCCATGGATTAGGGTCTCAGACTGGNNNAGATCGGAA |
| 45165_3 | TATGAGTGTGGAGTCGTTGCCATGGATTAGGGTCTCAGACTGGNNNAGATCGGAA |
| 45165_4 | TATGAGTGTGGAGTCGTTGCCATGGATTAGGGTCTCAGACTGGNNNAGATCGGAA |
| 31573_1 | TATGAGTGTGGAGTCGTTGCGCACTCACTGAATTCCACACAAANNNAGATCGGAAG |
| 31573_2 | TATGAGTGTGGAGTCGTTGCGCACTCACTGAATTCCACACAAANNNAGATCGGAAG |
| 31573_3 | TATGAGTGTGGAGTCGTTGCGCACTCACTGAATTCCACACAAANNNAGATCGGAAG |
| 31573_4 | TATGAGTGTGGAGTCGTTGCGCACTCACTGAATTCCACACAAANNNAGATCGGAAG |
| 41518_1 | TATGAGTGTGGAGTCGTTGCTCACTGCTAATGACTCCTCACACNNNAGATCGGAAGA |
| 41518_2 | TATGAGTGTGGAGTCGTTGCTCACTGCTAATGACTCCTCACACNNNAGATCGGAAGA |
| 41518_3 | TATGAGTGTGGAGTCGTTGCTCACTGCTAATGACTCCTCACACNNNAGATCGGAAGA |
| 41518_4 | TATGAGTGTGGAGTCGTTGCTCACTGCTAATGACTCCTCACACNNNAGATCGGAAGA |
| 49934_1 | TATGAGTGTGGAGTCGTTGCGTCATCCAAACCAACGTTGTCATNNNAGATCGGAAG |
| 49934_2 | TATGAGTGTGGAGTCGTTGCGTCATCCAAACCAACGTTGTCATNNNAGATCGGAAG |
| 49934_3 | TATGAGTGTGGAGTCGTTGCGTCATCCAAACCAACGTTGTCATNNNAGATCGGAAG |
| 49934_4 | TATGAGTGTGGAGTCGTTGCGTCATCCAAACCAACGTTGTCATNNNAGATCGGAAG |
| 53074_1 | TATGAGTGTGGAGTCGTTGCAAAGCAACCTATCAGTGGCTCATNNNAGATCGGAAG |
| 53074_2 | TATGAGTGTGGAGTCGTTGCAAAGCAACCTATCAGTGGCTCATNNNAGATCGGAAG |
| 53074_3 | TATGAGTGTGGAGTCGTTGCAAAGCAACCTATCAGTGGCTCATNNNAGATCGGAAG |
| 53074_4 | TATGAGTGTGGAGTCGTTGCAAAGCAACCTATCAGTGGCTCATNNNAGATCGGAAG |
| 49021_1 | TATGAGTGTGGAGTCGTTGCCCACCACGCCCAGATAATTTTTTNNNAGATCGGAAGA |
| 49021_2 | TATGAGTGTGGAGTCGTTGCCCACCACGCCCAGATAATTTTTTNNNAGATCGGAAGA |
| 49021_3 | TATGAGTGTGGAGTCGTTGCCCACCACGCCCAGATAATTTTTTNNNAGATCGGAAGA |
| 49021_4 | TATGAGTGTGGAGTCGTTGCCCACCACGCCCAGATAATTTTTTNNNAGATCGGAAGA |

| | |
|---|---|
| 35440_1 | TATGAGTGTGGAGTCGTTGCCTCAAACTCCTGACCTCAAGTGANNNAGATCGGAAG |
| 35440_2 | TATGAGTGTGGAGTCGTTGCCTCAAACTCCTGACCTCAAGTGANNNAGATCGGAAG |
| 35440_3 | TATGAGTGTGGAGTCGTTGCCTCAAACTCCTGACCTCAAGTGANNNAGATCGGAAG |
| 35440_4 | TATGAGTGTGGAGTCGTTGCCTCAAACTCCTGACCTCAAGTGANNNAGATCGGAAG |
| 47399_1 | TATGAGTGTGGAGTCGTTGCCATCCTGGGCCTCTGAATATTCANNNAGATCGGAAG |
| 47399_2 | TATGAGTGTGGAGTCGTTGCCATCCTGGGCCTCTGAATATTCANNNAGATCGGAAG |
| 47399_3 | TATGAGTGTGGAGTCGTTGCCATCCTGGGCCTCTGAATATTCANNNAGATCGGAAG |
| 47399_4 | TATGAGTGTGGAGTCGTTGCCATCCTGGGCCTCTGAATATTCANNNAGATCGGAAG |
| 44254_1 | TATGAGTGTGGAGTCGTTGCTCTTGCCTTTATGACCACTGACANNNAGATCGGAAG |
| 44254_2 | TATGAGTGTGGAGTCGTTGCTCTTGCCTTTATGACCACTGACANNNAGATCGGAAG |
| 44254_3 | TATGAGTGTGGAGTCGTTGCTCTTGCCTTTATGACCACTGACANNNAGATCGGAAG |
| 44254_4 | TATGAGTGTGGAGTCGTTGCTCTTGCCTTTATGACCACTGACANNNAGATCGGAAG |
| 45239_1 | TATGAGTGTGGAGTCGTTGCCTTGGAGAGCTCTAATGGGTGACNNNAGATCGGAA |
| 45239_2 | TATGAGTGTGGAGTCGTTGCCTTGGAGAGCTCTAATGGGTGACNNNAGATCGGAA |
| 45239_3 | TATGAGTGTGGAGTCGTTGCCTTGGAGAGCTCTAATGGGTGACNNNAGATCGGAA |
| 45239_4 | TATGAGTGTGGAGTCGTTGCCTTGGAGAGCTCTAATGGGTGACNNNAGATCGGAA |
| 32516_1 | TATGAGTGTGGAGTCGTTGCAGAGCGAAACGAACATCAACAATNNNAGATCGGAA |
| 32516_2 | TATGAGTGTGGAGTCGTTGCAGAGCGAAACGAACATCAACAATNNNAGATCGGAA |
| 32516_3 | TATGAGTGTGGAGTCGTTGCAGAGCGAAACGAACATCAACAATNNNAGATCGGAA |
| 32516_4 | TATGAGTGTGGAGTCGTTGCAGAGCGAAACGAACATCAACAATNNNAGATCGGAA |
| 41394_1 | TATGAGTGTGGAGTCGTTGCCCTGTAGTCCCAGCTACTCGGNNNAGATCGGAAGAG |
| 41394_2 | TATGAGTGTGGAGTCGTTGCCCTGTAGTCCCAGCTACTCGGNNNAGATCGGAAGAG |
| 41394_3 | TATGAGTGTGGAGTCGTTGCCCTGTAGTCCCAGCTACTCGGNNNAGATCGGAAGAG |
| 41394_4 | TATGAGTGTGGAGTCGTTGCCCTGTAGTCCCAGCTACTCGGNNNAGATCGGAAGAG |
| 47437_1 | TATGAGTGTGGAGTCGTTGCAGTGAGACAGACAAGGCTATTGGNNNAGATCGGAA |
| 47437_2 | TATGAGTGTGGAGTCGTTGCAGTGAGACAGACAAGGCTATTGGNNNAGATCGGAA |
| 47437_3 | TATGAGTGTGGAGTCGTTGCAGTGAGACAGACAAGGCTATTGGNNNAGATCGGAA |
| 47437_4 | TATGAGTGTGGAGTCGTTGCAGTGAGACAGACAAGGCTATTGGNNNAGATCGGAA |
| 29982_1 | TATGAGTGTGGAGTCGTTGCCATTCTCATCTACCGAGGCGTAGNNNAGATCGGAAG |
| 29982_2 | TATGAGTGTGGAGTCGTTGCCATTCTCATCTACCGAGGCGTAGNNNAGATCGGAAG |
| 29982_3 | TATGAGTGTGGAGTCGTTGCCATTCTCATCTACCGAGGCGTAGNNNAGATCGGAAG |
| 29982_4 | TATGAGTGTGGAGTCGTTGCCATTCTCATCTACCGAGGCGTAGNNNAGATCGGAAG |
| 9961_1 | TATGAGTGTGGAGTCGTTGCCCAGGCTGAGAGTACAGTGACTANNNAGATCGGAA |
| 9961_2 | TATGAGTGTGGAGTCGTTGCCCAGGCTGAGAGTACAGTGACTANNNAGATCGGAA |
| 9961_3 | TATGAGTGTGGAGTCGTTGCCCAGGCTGAGAGTACAGTGACTANNNAGATCGGAA |
| 9961_4 | TATGAGTGTGGAGTCGTTGCCCAGGCTGAGAGTACAGTGACTANNNAGATCGGAA |
| 48573_1 | TATGAGTGTGGAGTCGTTGCACTACAGGTATGTGCCACCATACNNNAGATCGGAAG |
| 48573_2 | TATGAGTGTGGAGTCGTTGCACTACAGGTATGTGCCACCATACNNNAGATCGGAAG |
| 48573_3 | TATGAGTGTGGAGTCGTTGCACTACAGGTATGTGCCACCATACNNNAGATCGGAAG |
| 48573_4 | TATGAGTGTGGAGTCGTTGCACTACAGGTATGTGCCACCATACNNNAGATCGGAAG |
| 53400_1 | TATGAGTGTGGAGTCGTTGCGAGAGGAGAGACAGAGACAGAGANNNAGATCGGA |
| 53400_2 | TATGAGTGTGGAGTCGTTGCGAGAGGAGAGACAGAGACAGAGANNNAGATCGGA |
| 53400_3 | TATGAGTGTGGAGTCGTTGCGAGAGGAGAGACAGAGACAGAGANNNAGATCGGA |

| | |
|---|---|
| 53400_4 | TATGAGTGTGGAGTCGTTGCGAGAGGAGAGACAGAGACAGAGANNNAGATCGGA |
| 46260_1 | TATGAGTGTGGAGTCGTTGCCAAGTAGCTGGGATTACAGGCATNNNAGATCGGAA |
| 46260_2 | TATGAGTGTGGAGTCGTTGCCAAGTAGCTGGGATTACAGGCATNNNAGATCGGAA |
| 46260_3 | TATGAGTGTGGAGTCGTTGCCAAGTAGCTGGGATTACAGGCATNNNAGATCGGAA |
| 46260_4 | TATGAGTGTGGAGTCGTTGCCAAGTAGCTGGGATTACAGGCATNNNAGATCGGAA |
| 36382_1 | TATGAGTGTGGAGTCGTTGCTTCTTGGGACACAATAAGCCCTTNNNAGATCGGAAG |
| 36382_2 | TATGAGTGTGGAGTCGTTGCTTCTTGGGACACAATAAGCCCTTNNNAGATCGGAAG |
| 36382_3 | TATGAGTGTGGAGTCGTTGCTTCTTGGGACACAATAAGCCCTTNNNAGATCGGAAG |
| 36382_4 | TATGAGTGTGGAGTCGTTGCTTCTTGGGACACAATAAGCCCTTNNNAGATCGGAAG |
| 52905_1 | TATGAGTGTGGAGTCGTTGCTCAGGGTATGTTTGACTCAAGGGNNNAGATCGGAA |
| 52905_2 | TATGAGTGTGGAGTCGTTGCTCAGGGTATGTTTGACTCAAGGGNNNAGATCGGAA |
| 52905_3 | TATGAGTGTGGAGTCGTTGCTCAGGGTATGTTTGACTCAAGGGNNNAGATCGGAA |
| 52905_4 | TATGAGTGTGGAGTCGTTGCTCAGGGTATGTTTGACTCAAGGGNNNAGATCGGAA |
| 41708_1 | TATGAGTGTGGAGTCGTTGCGGTTCAAGCGATTCCATGTTTCANNNAGATCGGAAG |
| 41708_2 | TATGAGTGTGGAGTCGTTGCGGTTCAAGCGATTCCATGTTTCANNNAGATCGGAAG |
| 41708_3 | TATGAGTGTGGAGTCGTTGCGGTTCAAGCGATTCCATGTTTCANNNAGATCGGAAG |
| 41708_4 | TATGAGTGTGGAGTCGTTGCGGTTCAAGCGATTCCATGTTTCANNNAGATCGGAAG |
| 48675_1 | TATGAGTGTGGAGTCGTTGCTTGGAAAGACAGGGTTTATGTGANNNAGATCGGAA |
| 48675_2 | TATGAGTGTGGAGTCGTTGCTTGGAAAGACAGGGTTTATGTGANNNAGATCGGAA |
| 48675_3 | TATGAGTGTGGAGTCGTTGCTTGGAAAGACAGGGTTTATGTGANNNAGATCGGAA |
| 48675_4 | TATGAGTGTGGAGTCGTTGCTTGGAAAGACAGGGTTTATGTGANNNAGATCGGAA |
| 49560_1 | TATGAGTGTGGAGTCGTTGCAACAGTTCCCTTTTCTCCACACCNNNAGATCGGAAGA |
| 49560_2 | TATGAGTGTGGAGTCGTTGCAACAGTTCCCTTTTCTCCACACCNNNAGATCGGAAGA |
| 49560_3 | TATGAGTGTGGAGTCGTTGCAACAGTTCCCTTTTCTCCACACCNNNAGATCGGAAGA |
| 49560_4 | TATGAGTGTGGAGTCGTTGCAACAGTTCCCTTTTCTCCACACCNNNAGATCGGAAGA |
| 51601_1 | TATGAGTGTGGAGTCGTTGCCACGTGATCCTAGAATCTGCTGTNNNAGATCGGAAG |
| 51601_2 | TATGAGTGTGGAGTCGTTGCCACGTGATCCTAGAATCTGCTGTNNNAGATCGGAAG |
| 51601_3 | TATGAGTGTGGAGTCGTTGCCACGTGATCCTAGAATCTGCTGTNNNAGATCGGAAG |
| 51601_4 | TATGAGTGTGGAGTCGTTGCCACGTGATCCTAGAATCTGCTGTNNNAGATCGGAAG |
| 31900_1 | TATGAGTGTGGAGTCGTTGCCTGTCTGGACCTGGAGTTTCTTTNNNAGATCGGAAG |
| 31900_2 | TATGAGTGTGGAGTCGTTGCCTGTCTGGACCTGGAGTTTCTTTNNNAGATCGGAAG |
| 31900_3 | TATGAGTGTGGAGTCGTTGCCTGTCTGGACCTGGAGTTTCTTTNNNAGATCGGAAG |
| 31900_4 | TATGAGTGTGGAGTCGTTGCCTGTCTGGACCTGGAGTTTCTTTNNNAGATCGGAAG |
| 51324_1 | TATGAGTGTGGAGTCGTTGCTGAAGCCCAGACTCTCATTTTCANNNAGATCGGAAG |
| 51324_2 | TATGAGTGTGGAGTCGTTGCTGAAGCCCAGACTCTCATTTTCANNNAGATCGGAAG |
| 51324_3 | TATGAGTGTGGAGTCGTTGCTGAAGCCCAGACTCTCATTTTCANNNAGATCGGAAG |
| 51324_4 | TATGAGTGTGGAGTCGTTGCTGAAGCCCAGACTCTCATTTTCANNNAGATCGGAAG |
| 49781_1 | TATGAGTGTGGAGTCGTTGCTTTTTTTGCGATGGAGTCTTGCTNNNAGATCGGAAG |
| 49781_2 | TATGAGTGTGGAGTCGTTGCTTTTTTTGCGATGGAGTCTTGCTNNNAGATCGGAAG |
| 49781_3 | TATGAGTGTGGAGTCGTTGCTTTTTTTGCGATGGAGTCTTGCTNNNAGATCGGAAG |
| 49781_4 | TATGAGTGTGGAGTCGTTGCTTTTTTTGCGATGGAGTCTTGCTNNNAGATCGGAAG |
| 45601_1 | TATGAGTGTGGAGTCGTTGCGGCTGAGAATGGGGTTTTCTTTGNNNAGATCGGAA |
| 45601_2 | TATGAGTGTGGAGTCGTTGCGGCTGAGAATGGGGTTTTCTTTGNNNAGATCGGAA |

| | |
|---|---|
| 45601_3 | TATGAGTGTGGAGTCGTTGCGGCTGAGAATGGGGTTTTCTTTGNNNAGATCGGAA |
| 45601_4 | TATGAGTGTGGAGTCGTTGCGGCTGAGAATGGGGTTTTCTTTGNNNAGATCGGAA |
| 43055_1 | TATGAGTGTGGAGTCGTTGCGTAAAAAACTCTTGAACGGGGCANNNAGATCGGAA |
| 43055_2 | TATGAGTGTGGAGTCGTTGCGTAAAAAACTCTTGAACGGGGCANNNAGATCGGAA |
| 43055_3 | TATGAGTGTGGAGTCGTTGCGTAAAAAACTCTTGAACGGGGCANNNAGATCGGAA |
| 43055_4 | TATGAGTGTGGAGTCGTTGCGTAAAAAACTCTTGAACGGGGCANNNAGATCGGAA |
| 29274_1 | TATGAGTGTGGAGTCGTTGCATCTCAATCTGTGTGCCTGTCATNNNAGATCGGAAG |
| 29274_2 | TATGAGTGTGGAGTCGTTGCATCTCAATCTGTGTGCCTGTCATNNNAGATCGGAAG |
| 29274_3 | TATGAGTGTGGAGTCGTTGCATCTCAATCTGTGTGCCTGTCATNNNAGATCGGAAG |
| 29274_4 | TATGAGTGTGGAGTCGTTGCATCTCAATCTGTGTGCCTGTCATNNNAGATCGGAAG |
| 28881_1 | TATGAGTGTGGAGTCGTTGCTTCTCTCTCTCACACACACACACNNNAGATCGGAAGA |
| 28881_2 | TATGAGTGTGGAGTCGTTGCTTCTCTCTCTCACACACACACACNNNAGATCGGAAGA |
| 28881_3 | TATGAGTGTGGAGTCGTTGCTTCTCTCTCTCACACACACACACNNNAGATCGGAAGA |
| 28881_4 | TATGAGTGTGGAGTCGTTGCTTCTCTCTCTCACACACACACACNNNAGATCGGAAGA |
| 44795_1 | TATGAGTGTGGAGTCGTTGCACCAGCAGACATTCTTACTCTGGNNNAGATCGGAAG |
| 44795_2 | TATGAGTGTGGAGTCGTTGCACCAGCAGACATTCTTACTCTGGNNNAGATCGGAAG |
| 44795_3 | TATGAGTGTGGAGTCGTTGCACCAGCAGACATTCTTACTCTGGNNNAGATCGGAAG |
| 44795_4 | TATGAGTGTGGAGTCGTTGCACCAGCAGACATTCTTACTCTGGNNNAGATCGGAAG |
| 50449_1 | TATGAGTGTGGAGTCGTTGCCTTCCGAAGGCTCTGAGGATATGNNNAGATCGGAA |
| 50449_2 | TATGAGTGTGGAGTCGTTGCCTTCCGAAGGCTCTGAGGATATGNNNAGATCGGAA |
| 50449_3 | TATGAGTGTGGAGTCGTTGCCTTCCGAAGGCTCTGAGGATATGNNNAGATCGGAA |
| 50449_4 | TATGAGTGTGGAGTCGTTGCCTTCCGAAGGCTCTGAGGATATGNNNAGATCGGAA |
| 47089_1 | TATGAGTGTGGAGTCGTTGCGGATCACTTGAGGCTAGGAGTTCNNNAGATCGGAA |
| 47089_2 | TATGAGTGTGGAGTCGTTGCGGATCACTTGAGGCTAGGAGTTCNNNAGATCGGAA |
| 47089_3 | TATGAGTGTGGAGTCGTTGCGGATCACTTGAGGCTAGGAGTTCNNNAGATCGGAA |
| 47089_4 | TATGAGTGTGGAGTCGTTGCGGATCACTTGAGGCTAGGAGTTCNNNAGATCGGAA |
| 38243_1 | TATGAGTGTGGAGTCGTTGCACGGTTTTTCACATGTCCAACTCNNNAGATCGGAAGA |
| 38243_2 | TATGAGTGTGGAGTCGTTGCACGGTTTTTCACATGTCCAACTCNNNAGATCGGAAGA |
| 38243_3 | TATGAGTGTGGAGTCGTTGCACGGTTTTTCACATGTCCAACTCNNNAGATCGGAAGA |
| 38243_4 | TATGAGTGTGGAGTCGTTGCACGGTTTTTCACATGTCCAACTCNNNAGATCGGAAGA |
| 13477_1 | TATGAGTGTGGAGTCGTTGCGTTGATACTGAAGAATTGATGGNNNAGATCGGAAG |
| 13477_2 | TATGAGTGTGGAGTCGTTGCGTTGATACTGAAGAATTGATGGNNNAGATCGGAAG |
| 13477_3 | TATGAGTGTGGAGTCGTTGCGTTGATACTGAAGAATTGATGGNNNAGATCGGAAG |
| 13477_4 | TATGAGTGTGGAGTCGTTGCGTTGATACTGAAGAATTGATGGNNNAGATCGGAAG |
| 32148_1 | TATGAGTGTGGAGTCGTTGCCTCACTGCAAGCTCCACCTCNNNAGATCGGAAGAGC |
| 32148_2 | TATGAGTGTGGAGTCGTTGCCTCACTGCAAGCTCCACCTCNNNAGATCGGAAGAGC |
| 32148_3 | TATGAGTGTGGAGTCGTTGCCTCACTGCAAGCTCCACCTCNNNAGATCGGAAGAGC |
| 32148_4 | TATGAGTGTGGAGTCGTTGCCTCACTGCAAGCTCCACCTCNNNAGATCGGAAGAGC |
| 37389_1 | TATGAGTGTGGAGTCGTTGCGAGGATTGTTTGAGCACAGGAAGNNNAGATCGGAA |
| 37389_2 | TATGAGTGTGGAGTCGTTGCGAGGATTGTTTGAGCACAGGAAGNNNAGATCGGAA |
| 37389_3 | TATGAGTGTGGAGTCGTTGCGAGGATTGTTTGAGCACAGGAAGNNNAGATCGGAA |
| 37389_4 | TATGAGTGTGGAGTCGTTGCGAGGATTGTTTGAGCACAGGAAGNNNAGATCGGAA |
| 52734_1 | TATGAGTGTGGAGTCGTTGCGCCTATGACTCACCACACTGTTANNNAGATCGGAAG |

| | |
|---|---|
| 52734_2 | TATGAGTGTGGAGTCGTTGCGCCTATGACTCACCACACTGTTANNNAGATCGGAAG |
| 52734_3 | TATGAGTGTGGAGTCGTTGCGCCTATGACTCACCACACTGTTANNNAGATCGGAAG |
| 52734_4 | TATGAGTGTGGAGTCGTTGCGCCTATGACTCACCACACTGTTANNNAGATCGGAAG |
| 40439_1 | TATGAGTGTGGAGTCGTTGCATCTTCTGTGCTGAAAAGGCAAGNNNAGATCGGAA |
| 40439_2 | TATGAGTGTGGAGTCGTTGCATCTTCTGTGCTGAAAAGGCAAGNNNAGATCGGAA |
| 40439_3 | TATGAGTGTGGAGTCGTTGCATCTTCTGTGCTGAAAAGGCAAGNNNAGATCGGAA |
| 40439_4 | TATGAGTGTGGAGTCGTTGCATCTTCTGTGCTGAAAAGGCAAGNNNAGATCGGAA |
| 29107_1 | TATGAGTGTGGAGTCGTTGCTTTGAGCCATAGGGAAGTAGAGCNNNAGATCGGAA |
| 29107_2 | TATGAGTGTGGAGTCGTTGCTTTGAGCCATAGGGAAGTAGAGCNNNAGATCGGAA |
| 29107_3 | TATGAGTGTGGAGTCGTTGCTTTGAGCCATAGGGAAGTAGAGCNNNAGATCGGAA |
| 29107_4 | TATGAGTGTGGAGTCGTTGCTTTGAGCCATAGGGAAGTAGAGCNNNAGATCGGAA |
| 29186_1 | TATGAGTGTGGAGTCGTTGCGTCCCATGATGACAGAGTCTTGTNNNAGATCGGAAG |
| 29186_2 | TATGAGTGTGGAGTCGTTGCGTCCCATGATGACAGAGTCTTGTNNNAGATCGGAAG |
| 29186_3 | TATGAGTGTGGAGTCGTTGCGTCCCATGATGACAGAGTCTTGTNNNAGATCGGAAG |
| 29186_4 | TATGAGTGTGGAGTCGTTGCGTCCCATGATGACAGAGTCTTGTNNNAGATCGGAAG |
| 43861_1 | TATGAGTGTGGAGTCGTTGCGACAGAGAGGGGATGAAAAGCAGNNNAGATCGGA |
| 43861_2 | TATGAGTGTGGAGTCGTTGCGACAGAGAGGGGATGAAAAGCAGNNNAGATCGGA |
| 43861_3 | TATGAGTGTGGAGTCGTTGCGACAGAGAGGGGATGAAAAGCAGNNNAGATCGGA |
| 43861_4 | TATGAGTGTGGAGTCGTTGCGACAGAGAGGGGATGAAAAGCAGNNNAGATCGGA |
| 6130_1 | TATGAGTGTGGAGTCGTTGCGTCATCAGTCATTTCCTCCAGTTTNNNAGATCGGAAG |
| 6130_2 | TATGAGTGTGGAGTCGTTGCGTCATCAGTCATTTCCTCCAGTTTNNNAGATCGGAAG |
| 6130_3 | TATGAGTGTGGAGTCGTTGCGTCATCAGTCATTTCCTCCAGTTTNNNAGATCGGAAG |
| 6130_4 | TATGAGTGTGGAGTCGTTGCGTCATCAGTCATTTCCTCCAGTTTNNNAGATCGGAAG |
| 29624_1 | TATGAGTGTGGAGTCGTTGCCTGGTTTGCAATGGTTAAGGTGANNNAGATCGGAA |
| 29624_2 | TATGAGTGTGGAGTCGTTGCCTGGTTTGCAATGGTTAAGGTGANNNAGATCGGAA |
| 29624_3 | TATGAGTGTGGAGTCGTTGCCTGGTTTGCAATGGTTAAGGTGANNNAGATCGGAA |
| 29624_4 | TATGAGTGTGGAGTCGTTGCCTGGTTTGCAATGGTTAAGGTGANNNAGATCGGAA |
| 29450_1 | TATGAGTGTGGAGTCGTTGCCCTTTGGTATTGCCCTATAATAANNNAGATCGGAAG |
| 29450_2 | TATGAGTGTGGAGTCGTTGCCCTTTGGTATTGCCCTATAATAANNNAGATCGGAAG |
| 29450_3 | TATGAGTGTGGAGTCGTTGCCCTTTGGTATTGCCCTATAATAANNNAGATCGGAAG |
| 29450_4 | TATGAGTGTGGAGTCGTTGCCCTTTGGTATTGCCCTATAATAANNNAGATCGGAAG |
| 46754_1 | TATGAGTGTGGAGTCGTTGCGATTACAGGCACCCGCTATCATANNNAGATCGGAAG |
| 46754_2 | TATGAGTGTGGAGTCGTTGCGATTACAGGCACCCGCTATCATANNNAGATCGGAAG |
| 46754_3 | TATGAGTGTGGAGTCGTTGCGATTACAGGCACCCGCTATCATANNNAGATCGGAAG |
| 46754_4 | TATGAGTGTGGAGTCGTTGCGATTACAGGCACCCGCTATCATANNNAGATCGGAAG |
| 52631_1 | TATGAGTGTGGAGTCGTTGCTCTCATTCTCTAGTGACCCCCTTNNNAGATCGGAAGA |
| 52631_2 | TATGAGTGTGGAGTCGTTGCTCTCATTCTCTAGTGACCCCCTTNNNAGATCGGAAGA |
| 52631_3 | TATGAGTGTGGAGTCGTTGCTCTCATTCTCTAGTGACCCCCTTNNNAGATCGGAAGA |
| 52631_4 | TATGAGTGTGGAGTCGTTGCTCTCATTCTCTAGTGACCCCCTTNNNAGATCGGAAGA |
| 49501_1 | TATGAGTGTGGAGTCGTTGCTTGCTGCTGTAACAAATAGCCACNNNAGATCGGAAG |
| 49501_2 | TATGAGTGTGGAGTCGTTGCTTGCTGCTGTAACAAATAGCCACNNNAGATCGGAAG |
| 49501_3 | TATGAGTGTGGAGTCGTTGCTTGCTGCTGTAACAAATAGCCACNNNAGATCGGAAG |
| 49501_4 | TATGAGTGTGGAGTCGTTGCTTGCTGCTGTAACAAATAGCCACNNNAGATCGGAAG |

| | |
|---|---|
| 53290_1 | TATGAGTGTGGAGTCGTTGCCTGATCCAGGAGGCCGCATCNNNAGATCGGAAGAG |
| 53290_2 | TATGAGTGTGGAGTCGTTGCCTGATCCAGGAGGCCGCATCNNNAGATCGGAAGAG |
| 53290_3 | TATGAGTGTGGAGTCGTTGCCTGATCCAGGAGGCCGCATCNNNAGATCGGAAGAG |
| 53290_4 | TATGAGTGTGGAGTCGTTGCCTGATCCAGGAGGCCGCATCNNNAGATCGGAAGAG |
| 36451_1 | TATGAGTGTGGAGTCGTTGCGGCTTTTACGCGAAAGGTAAAGTNNNAGATCGGAA |
| 36451_2 | TATGAGTGTGGAGTCGTTGCGGCTTTTACGCGAAAGGTAAAGTNNNAGATCGGAA |
| 36451_3 | TATGAGTGTGGAGTCGTTGCGGCTTTTACGCGAAAGGTAAAGTNNNAGATCGGAA |
| 36451_4 | TATGAGTGTGGAGTCGTTGCGGCTTTTACGCGAAAGGTAAAGTNNNAGATCGGAA |
| 46411_1 | TATGAGTGTGGAGTCGTTGCCACAATCTCAGCTCACTGCAACNNNAGATCGGAAGA |
| 46411_2 | TATGAGTGTGGAGTCGTTGCCACAATCTCAGCTCACTGCAACNNNAGATCGGAAGA |
| 46411_3 | TATGAGTGTGGAGTCGTTGCCACAATCTCAGCTCACTGCAACNNNAGATCGGAAGA |
| 46411_4 | TATGAGTGTGGAGTCGTTGCCACAATCTCAGCTCACTGCAACNNNAGATCGGAAGA |
| 36325_1 | TATGAGTGTGGAGTCGTTGCCCACACACACACACAGAAAAACTNNNAGATCGGAAG |
| 36325_2 | TATGAGTGTGGAGTCGTTGCCCACACACACACACAGAAAAACTNNNAGATCGGAAG |
| 36325_3 | TATGAGTGTGGAGTCGTTGCCCACACACACACACAGAAAAACTNNNAGATCGGAAG |
| 36325_4 | TATGAGTGTGGAGTCGTTGCCCACACACACACACAGAAAAACTNNNAGATCGGAAG |
| 49410_1 | TATGAGTGTGGAGTCGTTGCCAGAAAGCCGTTGCGTCTAGATNNNAGATCGGAAG |
| 49410_2 | TATGAGTGTGGAGTCGTTGCCAGAAAGCCGTTGCGTCTAGATNNNAGATCGGAAG |
| 49410_3 | TATGAGTGTGGAGTCGTTGCCAGAAAGCCGTTGCGTCTAGATNNNAGATCGGAAG |
| 49410_4 | TATGAGTGTGGAGTCGTTGCCAGAAAGCCGTTGCGTCTAGATNNNAGATCGGAAG |
| 41142_1 | TATGAGTGTGGAGTCGTTGCTGAATGTAACTGATGCCACAGGANNNAGATCGGAA |
| 41142_2 | TATGAGTGTGGAGTCGTTGCTGAATGTAACTGATGCCACAGGANNNAGATCGGAA |
| 41142_3 | TATGAGTGTGGAGTCGTTGCTGAATGTAACTGATGCCACAGGANNNAGATCGGAA |
| 41142_4 | TATGAGTGTGGAGTCGTTGCTGAATGTAACTGATGCCACAGGANNNAGATCGGAA |
| 43824_1 | TATGAGTGTGGAGTCGTTGCTCAACGACATCACACTGATCACTNNNAGATCGGAAG |
| 43824_2 | TATGAGTGTGGAGTCGTTGCTCAACGACATCACACTGATCACTNNNAGATCGGAAG |
| 43824_3 | TATGAGTGTGGAGTCGTTGCTCAACGACATCACACTGATCACTNNNAGATCGGAAG |
| 43824_4 | TATGAGTGTGGAGTCGTTGCTCAACGACATCACACTGATCACTNNNAGATCGGAAG |
| 9747_1 | TATGAGTGTGGAGTCGTTGCCCAGCCCAAGAAATACAACACACNNNAGATCGGAAG |
| 9747_2 | TATGAGTGTGGAGTCGTTGCCCAGCCCAAGAAATACAACACACNNNAGATCGGAAG |
| 9747_3 | TATGAGTGTGGAGTCGTTGCCCAGCCCAAGAAATACAACACACNNNAGATCGGAAG |
| 9747_4 | TATGAGTGTGGAGTCGTTGCCCAGCCCAAGAAATACAACACACNNNAGATCGGAAG |
| 33140_1 | TATGAGTGTGGAGTCGTTGCTGACTTTGCCTGTAACCTCTACCNNNAGATCGGAAGA |
| 33140_2 | TATGAGTGTGGAGTCGTTGCTGACTTTGCCTGTAACCTCTACCNNNAGATCGGAAGA |
| 33140_3 | TATGAGTGTGGAGTCGTTGCTGACTTTGCCTGTAACCTCTACCNNNAGATCGGAAGA |
| 33140_4 | TATGAGTGTGGAGTCGTTGCTGACTTTGCCTGTAACCTCTACCNNNAGATCGGAAGA |
| 52145_1 | TATGAGTGTGGAGTCGTTGCGGGCTCCTCACTTCCCAGTANNNAGATCGGAAGAGC |
| 52145_2 | TATGAGTGTGGAGTCGTTGCGGGCTCCTCACTTCCCAGTANNNAGATCGGAAGAGC |
| 52145_3 | TATGAGTGTGGAGTCGTTGCGGGCTCCTCACTTCCCAGTANNNAGATCGGAAGAGC |
| 52145_4 | TATGAGTGTGGAGTCGTTGCGGGCTCCTCACTTCCCAGTANNNAGATCGGAAGAGC |
| 6297_1 | TATGAGTGTGGAGTCGTTGCCCCCAGGCTTGAGAAGACTAAAANNNAGATCGGAA |
| 6297_2 | TATGAGTGTGGAGTCGTTGCCCCCAGGCTTGAGAAGACTAAAANNNAGATCGGAA |
| 6297_3 | TATGAGTGTGGAGTCGTTGCCCCCAGGCTTGAGAAGACTAAAANNNAGATCGGAA |

| | |
|---|---|
| 6297_4 | TATGAGTGTGGAGTCGTTGCCCCCAGGCTTGAGAAGACTAAAANNNAGATCGGAAG |
| 42866_1 | TATGAGTGTGGAGTCGTTGCCATTCTCCTGCCTCAGCCTCNNNAGATCGGAAGAGCA |
| 42866_2 | TATGAGTGTGGAGTCGTTGCCATTCTCCTGCCTCAGCCTCNNNAGATCGGAAGAGCA |
| 42866_3 | TATGAGTGTGGAGTCGTTGCCATTCTCCTGCCTCAGCCTCNNNAGATCGGAAGAGCA |
| 42866_4 | TATGAGTGTGGAGTCGTTGCCATTCTCCTGCCTCAGCCTCNNNAGATCGGAAGAGCA |
| 40461_1 | TATGAGTGTGGAGTCGTTGCGCAACTACTCTTGACCACTGTCTNNNAGATCGGAAGA |
| 40461_2 | TATGAGTGTGGAGTCGTTGCGCAACTACTCTTGACCACTGTCTNNNAGATCGGAAGA |
| 40461_3 | TATGAGTGTGGAGTCGTTGCGCAACTACTCTTGACCACTGTCTNNNAGATCGGAAGA |
| 40461_4 | TATGAGTGTGGAGTCGTTGCGCAACTACTCTTGACCACTGTCTNNNAGATCGGAAGA |
| 48115_1 | TATGAGTGTGGAGTCGTTGCGTTAGGTGGCAAGTGGGTCAGNNNAGATCGGAAGA |
| 48115_2 | TATGAGTGTGGAGTCGTTGCGTTAGGTGGCAAGTGGGTCAGNNNAGATCGGAAGA |
| 48115_3 | TATGAGTGTGGAGTCGTTGCGTTAGGTGGCAAGTGGGTCAGNNNAGATCGGAAGA |
| 48115_4 | TATGAGTGTGGAGTCGTTGCGTTAGGTGGCAAGTGGGTCAGNNNAGATCGGAAGA |
| 33177_1 | TATGAGTGTGGAGTCGTTGCGCCCATTTTCTGTGGAGGAATTCNNNAGATCGGAAG |
| 33177_2 | TATGAGTGTGGAGTCGTTGCGCCCATTTTCTGTGGAGGAATTCNNNAGATCGGAAG |
| 33177_3 | TATGAGTGTGGAGTCGTTGCGCCCATTTTCTGTGGAGGAATTCNNNAGATCGGAAG |
| 33177_4 | TATGAGTGTGGAGTCGTTGCGCCCATTTTCTGTGGAGGAATTCNNNAGATCGGAAG |
| 40233_1 | TATGAGTGTGGAGTCGTTGCGCCTTTGAGCTCGTTGATTCTTTNNNAGATCGGAAGA |
| 40233_2 | TATGAGTGTGGAGTCGTTGCGCCTTTGAGCTCGTTGATTCTTTNNNAGATCGGAAGA |
| 40233_3 | TATGAGTGTGGAGTCGTTGCGCCTTTGAGCTCGTTGATTCTTTNNNAGATCGGAAGA |
| 40233_4 | TATGAGTGTGGAGTCGTTGCGCCTTTGAGCTCGTTGATTCTTTNNNAGATCGGAAGA |
| 39530_1 | TATGAGTGTGGAGTCGTTGCCCTACTGAAGAGTGTCCTTGGACNNNAGATCGGAAG |
| 39530_2 | TATGAGTGTGGAGTCGTTGCCCTACTGAAGAGTGTCCTTGGACNNNAGATCGGAAG |
| 39530_3 | TATGAGTGTGGAGTCGTTGCCCTACTGAAGAGTGTCCTTGGACNNNAGATCGGAAG |
| 39530_4 | TATGAGTGTGGAGTCGTTGCCCTACTGAAGAGTGTCCTTGGACNNNAGATCGGAAG |
| 10829_1 | TATGAGTGTGGAGTCGTTGCTAGTGCTTTTGCTGCCCAACATANNNAGATCGGAAGA |
| 10829_2 | TATGAGTGTGGAGTCGTTGCTAGTGCTTTTGCTGCCCAACATANNNAGATCGGAAGA |
| 10829_3 | TATGAGTGTGGAGTCGTTGCTAGTGCTTTTGCTGCCCAACATANNNAGATCGGAAGA |
| 10829_4 | TATGAGTGTGGAGTCGTTGCTAGTGCTTTTGCTGCCCAACATANNNAGATCGGAAGA |
| 33094_1 | TATGAGTGTGGAGTCGTTGCAGCCAACACAACACACTTTGAAANNNAGATCGGAAG |
| 33094_2 | TATGAGTGTGGAGTCGTTGCAGCCAACACAACACACTTTGAAANNNAGATCGGAAG |
| 33094_3 | TATGAGTGTGGAGTCGTTGCAGCCAACACAACACACTTTGAAANNNAGATCGGAAG |
| 33094_4 | TATGAGTGTGGAGTCGTTGCAGCCAACACAACACACTTTGAAANNNAGATCGGAAG |
| 40331_1 | TATGAGTGTGGAGTCGTTGCTGGTGGGTTTCCTACATCATGTTNNNAGATCGGAAG |
| 40331_2 | TATGAGTGTGGAGTCGTTGCTGGTGGGTTTCCTACATCATGTTNNNAGATCGGAAG |
| 40331_3 | TATGAGTGTGGAGTCGTTGCTGGTGGGTTTCCTACATCATGTTNNNAGATCGGAAG |
| 40331_4 | TATGAGTGTGGAGTCGTTGCTGGTGGGTTTCCTACATCATGTTNNNAGATCGGAAG |
| 40133_1 | TATGAGTGTGGAGTCGTTGCGAAAAGCCAACTTATGGGATGGGNNNAGATCGGAA |
| 40133_2 | TATGAGTGTGGAGTCGTTGCGAAAAGCCAACTTATGGGATGGGNNNAGATCGGAA |
| 40133_3 | TATGAGTGTGGAGTCGTTGCGAAAAGCCAACTTATGGGATGGGNNNAGATCGGAA |
| 40133_4 | TATGAGTGTGGAGTCGTTGCGAAAAGCCAACTTATGGGATGGGNNNAGATCGGAA |
| 37674_1 | TATGAGTGTGGAGTCGTTGCGGACATACCAAGCCAACCAAAAANNNAGATCGGAAG |
| 37674_2 | TATGAGTGTGGAGTCGTTGCGGACATACCAAGCCAACCAAAAANNNAGATCGGAAG |

| | |
|---|---|
| 37674_3 | TATGAGTGTGGAGTCGTTGCGGACATACCAAGCCAACCAAAAANNNAGATCGGAAG |
| 37674_4 | TATGAGTGTGGAGTCGTTGCGGACATACCAAGCCAACCAAAAANNNAGATCGGAAG |
| 47169_1 | TATGAGTGTGGAGTCGTTGCAGACAGAGTCTCACTGTGTTTCCNNNAGATCGGAAG |
| 47169_2 | TATGAGTGTGGAGTCGTTGCAGACAGAGTCTCACTGTGTTTCCNNNAGATCGGAAG |
| 47169_3 | TATGAGTGTGGAGTCGTTGCAGACAGAGTCTCACTGTGTTTCCNNNAGATCGGAAG |
| 47169_4 | TATGAGTGTGGAGTCGTTGCAGACAGAGTCTCACTGTGTTTCCNNNAGATCGGAAG |
| 32391_1 | TATGAGTGTGGAGTCGTTGCCATTCTCCTGCCTCAGCCTCNNNAGATCGGAAGAGCA |
| 32391_2 | TATGAGTGTGGAGTCGTTGCCATTCTCCTGCCTCAGCCTCNNNAGATCGGAAGAGCA |
| 32391_3 | TATGAGTGTGGAGTCGTTGCCATTCTCCTGCCTCAGCCTCNNNAGATCGGAAGAGCA |
| 32391_4 | TATGAGTGTGGAGTCGTTGCCATTCTCCTGCCTCAGCCTCNNNAGATCGGAAGAGCA |
| 42665_1 | TATGAGTGTGGAGTCGTTGCTTTTGAGACAGAGTCACGTTCTGNNNAGATCGGAAG |
| 42665_2 | TATGAGTGTGGAGTCGTTGCTTTTGAGACAGAGTCACGTTCTGNNNAGATCGGAAG |
| 42665_3 | TATGAGTGTGGAGTCGTTGCTTTTGAGACAGAGTCACGTTCTGNNNAGATCGGAAG |
| 42665_4 | TATGAGTGTGGAGTCGTTGCTTTTGAGACAGAGTCACGTTCTGNNNAGATCGGAAG |
| 30319_1 | TATGAGTGTGGAGTCGTTGCCATCAACAACCTCATGTCCTCCTNNNAGATCGGAAGA |
| 30319_2 | TATGAGTGTGGAGTCGTTGCCATCAACAACCTCATGTCCTCCTNNNAGATCGGAAGA |
| 30319_3 | TATGAGTGTGGAGTCGTTGCCATCAACAACCTCATGTCCTCCTNNNAGATCGGAAGA |
| 30319_4 | TATGAGTGTGGAGTCGTTGCCATCAACAACCTCATGTCCTCCTNNNAGATCGGAAGA |
| 50895_1 | TATGAGTGTGGAGTCGTTGCTGGCTGTGTGATAATGAGGAAGTNNNAGATCGGAA |
| 50895_2 | TATGAGTGTGGAGTCGTTGCTGGCTGTGTGATAATGAGGAAGTNNNAGATCGGAA |
| 50895_3 | TATGAGTGTGGAGTCGTTGCTGGCTGTGTGATAATGAGGAAGTNNNAGATCGGAA |
| 50895_4 | TATGAGTGTGGAGTCGTTGCTGGCTGTGTGATAATGAGGAAGTNNNAGATCGGAA |
| 44580_1 | TATGAGTGTGGAGTCGTTGCCAGTGGCACAATCTTGGTTCATTNNNAGATCGGAAG |
| 44580_2 | TATGAGTGTGGAGTCGTTGCCAGTGGCACAATCTTGGTTCATTNNNAGATCGGAAG |
| 44580_3 | TATGAGTGTGGAGTCGTTGCCAGTGGCACAATCTTGGTTCATTNNNAGATCGGAAG |
| 44580_4 | TATGAGTGTGGAGTCGTTGCCAGTGGCACAATCTTGGTTCATTNNNAGATCGGAAG |
| 42826_1 | TATGAGTGTGGAGTCGTTGCCACCTTCAAAGTTTCCTCATGCCNNNAGATCGGAAGA |
| 42826_2 | TATGAGTGTGGAGTCGTTGCCACCTTCAAAGTTTCCTCATGCCNNNAGATCGGAAGA |
| 42826_3 | TATGAGTGTGGAGTCGTTGCCACCTTCAAAGTTTCCTCATGCCNNNAGATCGGAAGA |
| 42826_4 | TATGAGTGTGGAGTCGTTGCCACCTTCAAAGTTTCCTCATGCCNNNAGATCGGAAGA |
| 35888_1 | TATGAGTGTGGAGTCGTTGCGAGAGAGAGAGAGAGAGACCTCANNNAGATCGGAA |
| 35888_2 | TATGAGTGTGGAGTCGTTGCGAGAGAGAGAGAGAGAGACCTCANNNAGATCGGAA |
| 35888_3 | TATGAGTGTGGAGTCGTTGCGAGAGAGAGAGAGAGAGACCTCANNNAGATCGGAA |
| 35888_4 | TATGAGTGTGGAGTCGTTGCGAGAGAGAGAGAGAGAGACCTCANNNAGATCGGAA |
| 37405_1 | TATGAGTGTGGAGTCGTTGCTGGTGCACCAAATGGAAAAAATTNNNAGATCGGAAG |
| 37405_2 | TATGAGTGTGGAGTCGTTGCTGGTGCACCAAATGGAAAAAATTNNNAGATCGGAAG |
| 37405_3 | TATGAGTGTGGAGTCGTTGCTGGTGCACCAAATGGAAAAAATTNNNAGATCGGAAG |
| 37405_4 | TATGAGTGTGGAGTCGTTGCTGGTGCACCAAATGGAAAAAATTNNNAGATCGGAAG |
| 35176_1 | TATGAGTGTGGAGTCGTTGCCAAGTAGCTGGGACTACAGGCNNNAGATCGGAAGA |
| 35176_2 | TATGAGTGTGGAGTCGTTGCCAAGTAGCTGGGACTACAGGCNNNAGATCGGAAGA |
| 35176_3 | TATGAGTGTGGAGTCGTTGCCAAGTAGCTGGGACTACAGGCNNNAGATCGGAAGA |
| 35176_4 | TATGAGTGTGGAGTCGTTGCCAAGTAGCTGGGACTACAGGCNNNAGATCGGAAGA |
| 52408_1 | TATGAGTGTGGAGTCGTTGCGCCAGTGTTCTCTCAGTAAAGGANNNAGATCGGAAG |

| | |
|---|---|
| 52408_2 | TATGAGTGTGGAGTCGTTGCGCCAGTGTTCTCTCAGTAAAGGANNNAGATCGGAAG |
| 52408_3 | TATGAGTGTGGAGTCGTTGCGCCAGTGTTCTCTCAGTAAAGGANNNAGATCGGAAG |
| 52408_4 | TATGAGTGTGGAGTCGTTGCGCCAGTGTTCTCTCAGTAAAGGANNNAGATCGGAAG |
| 51486_1 | TATGAGTGTGGAGTCGTTGCTGGGGATGAGGATGTCAAAATGTNNNAGATCGGAA |
| 51486_2 | TATGAGTGTGGAGTCGTTGCTGGGGATGAGGATGTCAAAATGTNNNAGATCGGAA |
| 51486_3 | TATGAGTGTGGAGTCGTTGCTGGGGATGAGGATGTCAAAATGTNNNAGATCGGAA |
| 51486_4 | TATGAGTGTGGAGTCGTTGCTGGGGATGAGGATGTCAAAATGTNNNAGATCGGAA |
| 38988_1 | TATGAGTGTGGAGTCGTTGCGGTTTCAGAAAGCCGAGATCAACNNNAGATCGGAAC |
| 38988_2 | TATGAGTGTGGAGTCGTTGCGGTTTCAGAAAGCCGAGATCAACNNNAGATCGGAAC |
| 38988_3 | TATGAGTGTGGAGTCGTTGCGGTTTCAGAAAGCCGAGATCAACNNNAGATCGGAAC |
| 38988_4 | TATGAGTGTGGAGTCGTTGCGGTTTCAGAAAGCCGAGATCAACNNNAGATCGGAAC |
| 28757_1 | TATGAGTGTGGAGTCGTTGCAGGACAGTTCATCCATGTTGCTANNNAGATCGGAAG |
| 28757_2 | TATGAGTGTGGAGTCGTTGCAGGACAGTTCATCCATGTTGCTANNNAGATCGGAAG |
| 28757_3 | TATGAGTGTGGAGTCGTTGCAGGACAGTTCATCCATGTTGCTANNNAGATCGGAAG |
| 28757_4 | TATGAGTGTGGAGTCGTTGCAGGACAGTTCATCCATGTTGCTANNNAGATCGGAAG |
| 13157_1 | TATGAGTGTGGAGTCGTTGCCAACAAGATTCTGAAGAACATGNNNAGATCGGAAGA |
| 13157_2 | TATGAGTGTGGAGTCGTTGCCAACAAGATTCTGAAGAACATGNNNAGATCGGAAGA |
| 13157_3 | TATGAGTGTGGAGTCGTTGCCAACAAGATTCTGAAGAACATGNNNAGATCGGAAGA |
| 13157_4 | TATGAGTGTGGAGTCGTTGCCAACAAGATTCTGAAGAACATGNNNAGATCGGAAGA |
| 32751_1 | TATGAGTGTGGAGTCGTTGCGCAACTCAACAAAAAGACAACCCNNNAGATCGGAAG |
| 32751_2 | TATGAGTGTGGAGTCGTTGCGCAACTCAACAAAAAGACAACCCNNNAGATCGGAAG |
| 32751_3 | TATGAGTGTGGAGTCGTTGCGCAACTCAACAAAAAGACAACCCNNNAGATCGGAAG |
| 32751_4 | TATGAGTGTGGAGTCGTTGCGCAACTCAACAAAAAGACAACCCNNNAGATCGGAAG |
| 52834_1 | TATGAGTGTGGAGTCGTTGCCACCCACCTGTGAACTCCTTGNNNAGATCGGAAGAGC |
| 52834_2 | TATGAGTGTGGAGTCGTTGCCACCCACCTGTGAACTCCTTGNNNAGATCGGAAGAGC |
| 52834_3 | TATGAGTGTGGAGTCGTTGCCACCCACCTGTGAACTCCTTGNNNAGATCGGAAGAGC |
| 52834_4 | TATGAGTGTGGAGTCGTTGCCACCCACCTGTGAACTCCTTGNNNAGATCGGAAGAGC |
| 33876_1 | TATGAGTGTGGAGTCGTTGCCATTGTCATACCTACCTGCCTCANNNAGATCGGAAGA |
| 33876_2 | TATGAGTGTGGAGTCGTTGCCATTGTCATACCTACCTGCCTCANNNAGATCGGAAGA |
| 33876_3 | TATGAGTGTGGAGTCGTTGCCATTGTCATACCTACCTGCCTCANNNAGATCGGAAGA |
| 33876_4 | TATGAGTGTGGAGTCGTTGCCATTGTCATACCTACCTGCCTCANNNAGATCGGAAGA |
| 31180_1 | TATGAGTGTGGAGTCGTTGCTCCAAAAGCCTGAGGATTCTGAANNNAGATCGGAAG |
| 31180_2 | TATGAGTGTGGAGTCGTTGCTCCAAAAGCCTGAGGATTCTGAANNNAGATCGGAAG |
| 31180_3 | TATGAGTGTGGAGTCGTTGCTCCAAAAGCCTGAGGATTCTGAANNNAGATCGGAAG |
| 31180_4 | TATGAGTGTGGAGTCGTTGCTCCAAAAGCCTGAGGATTCTGAANNNAGATCGGAAG |
| 36820_1 | TATGAGTGTGGAGTCGTTGCCGCCTGGCCTGAAACTAATATTTNNNAGATCGGAAGA |
| 36820_2 | TATGAGTGTGGAGTCGTTGCCGCCTGGCCTGAAACTAATATTTNNNAGATCGGAAGA |
| 36820_3 | TATGAGTGTGGAGTCGTTGCCGCCTGGCCTGAAACTAATATTTNNNAGATCGGAAGA |
| 36820_4 | TATGAGTGTGGAGTCGTTGCCGCCTGGCCTGAAACTAATATTTNNNAGATCGGAAGA |
| 31585_1 | TATGAGTGTGGAGTCGTTGCAGACCTTTTCACACTCCTCACTCNNNAGATCGGAAGAC |
| 31585_2 | TATGAGTGTGGAGTCGTTGCAGACCTTTTCACACTCCTCACTCNNNAGATCGGAAGAC |
| 31585_3 | TATGAGTGTGGAGTCGTTGCAGACCTTTTCACACTCCTCACTCNNNAGATCGGAAGAC |
| 31585_4 | TATGAGTGTGGAGTCGTTGCAGACCTTTTCACACTCCTCACTCNNNAGATCGGAAGAC |

| | |
|---|---|
| 33674_1 | TATGAGTGTGGAGTCGTTGCTTCAAGTGATTCTCCTGCCTCAGNNNAGATCGGAAG |
| 33674_2 | TATGAGTGTGGAGTCGTTGCTTCAAGTGATTCTCCTGCCTCAGNNNAGATCGGAAG |
| 33674_3 | TATGAGTGTGGAGTCGTTGCTTCAAGTGATTCTCCTGCCTCAGNNNAGATCGGAAG |
| 33674_4 | TATGAGTGTGGAGTCGTTGCTTCAAGTGATTCTCCTGCCTCAGNNNAGATCGGAAG |
| 33905_1 | TATGAGTGTGGAGTCGTTGCTTCAAGCTATTCTCCTGCCTCAGNNNAGATCGGAAGA |
| 33905_2 | TATGAGTGTGGAGTCGTTGCTTCAAGCTATTCTCCTGCCTCAGNNNAGATCGGAAGA |
| 33905_3 | TATGAGTGTGGAGTCGTTGCTTCAAGCTATTCTCCTGCCTCAGNNNAGATCGGAAGA |
| 33905_4 | TATGAGTGTGGAGTCGTTGCTTCAAGCTATTCTCCTGCCTCAGNNNAGATCGGAAGA |
| 52074_1 | TATGAGTGTGGAGTCGTTGCCTGTGGACGCAGGCATCTCTNNNAGATCGGAAGAG |
| 52074_2 | TATGAGTGTGGAGTCGTTGCCTGTGGACGCAGGCATCTCTNNNAGATCGGAAGAG |
| 52074_3 | TATGAGTGTGGAGTCGTTGCCTGTGGACGCAGGCATCTCTNNNAGATCGGAAGAG |
| 52074_4 | TATGAGTGTGGAGTCGTTGCCTGTGGACGCAGGCATCTCTNNNAGATCGGAAGAG |
| 38694_1 | TATGAGTGTGGAGTCGTTGCAGGATGAACAAAGATATGCCATGTNNNAGATCGGA |
| 38694_2 | TATGAGTGTGGAGTCGTTGCAGGATGAACAAAGATATGCCATGTNNNAGATCGGA |
| 38694_3 | TATGAGTGTGGAGTCGTTGCAGGATGAACAAAGATATGCCATGTNNNAGATCGGA |
| 38694_4 | TATGAGTGTGGAGTCGTTGCAGGATGAACAAAGATATGCCATGTNNNAGATCGGA |
| 35362_1 | TATGAGTGTGGAGTCGTTGCCAGGTTCAAGCAATTCTCCTGCNNNAGATCGGAAGA |
| 35362_2 | TATGAGTGTGGAGTCGTTGCCAGGTTCAAGCAATTCTCCTGCNNNAGATCGGAAGA |
| 35362_3 | TATGAGTGTGGAGTCGTTGCCAGGTTCAAGCAATTCTCCTGCNNNAGATCGGAAGA |
| 35362_4 | TATGAGTGTGGAGTCGTTGCCAGGTTCAAGCAATTCTCCTGCNNNAGATCGGAAGA |
| 30220_1 | TATGAGTGTGGAGTCGTTGCGGTCCCCGAGATTGTATACACAGNNNAGATCGGAAG |
| 30220_2 | TATGAGTGTGGAGTCGTTGCGGTCCCCGAGATTGTATACACAGNNNAGATCGGAAG |
| 30220_3 | TATGAGTGTGGAGTCGTTGCGGTCCCCGAGATTGTATACACAGNNNAGATCGGAAG |
| 30220_4 | TATGAGTGTGGAGTCGTTGCGGTCCCCGAGATTGTATACACAGNNNAGATCGGAAG |
| 44382_1 | TATGAGTGTGGAGTCGTTGCCTGCTATTGAGTGAGGGCCATTANNNAGATCGGAA |
| 44382_2 | TATGAGTGTGGAGTCGTTGCCTGCTATTGAGTGAGGGCCATTANNNAGATCGGAA |
| 44382_3 | TATGAGTGTGGAGTCGTTGCCTGCTATTGAGTGAGGGCCATTANNNAGATCGGAA |
| 44382_4 | TATGAGTGTGGAGTCGTTGCCTGCTATTGAGTGAGGGCCATTANNNAGATCGGAA |
| 29740_1 | TATGAGTGTGGAGTCGTTGCAGACCCTTGTAGAAGACTCAGGTNNNAGATCGGAA |
| 29740_2 | TATGAGTGTGGAGTCGTTGCAGACCCTTGTAGAAGACTCAGGTNNNAGATCGGAA |
| 29740_3 | TATGAGTGTGGAGTCGTTGCAGACCCTTGTAGAAGACTCAGGTNNNAGATCGGAA |
| 29740_4 | TATGAGTGTGGAGTCGTTGCAGACCCTTGTAGAAGACTCAGGTNNNAGATCGGAA |
| 32703_1 | TATGAGTGTGGAGTCGTTGCCCAAAGTGCTGGGATTACAAGTGNNNAGATCGGAA |
| 32703_2 | TATGAGTGTGGAGTCGTTGCCCAAAGTGCTGGGATTACAAGTGNNNAGATCGGAA |
| 32703_3 | TATGAGTGTGGAGTCGTTGCCCAAAGTGCTGGGATTACAAGTGNNNAGATCGGAA |
| 32703_4 | TATGAGTGTGGAGTCGTTGCCCAAAGTGCTGGGATTACAAGTGNNNAGATCGGAA |
| 51343_1 | TATGAGTGTGGAGTCGTTGCGGGCATTCCTCATTTGGTTCAAANNNAGATCGGAAG |
| 51343_2 | TATGAGTGTGGAGTCGTTGCGGGCATTCCTCATTTGGTTCAAANNNAGATCGGAAG |
| 51343_3 | TATGAGTGTGGAGTCGTTGCGGGCATTCCTCATTTGGTTCAAANNNAGATCGGAAG |
| 51343_4 | TATGAGTGTGGAGTCGTTGCGGGCATTCCTCATTTGGTTCAAANNNAGATCGGAAG |
| 46608_1 | TATGAGTGTGGAGTCGTTGCCTCCCAGGTTCAAGCAATTCTTCNNNAGATCGGAAG |
| 46608_2 | TATGAGTGTGGAGTCGTTGCCTCCCAGGTTCAAGCAATTCTTCNNNAGATCGGAAG |
| 46608_3 | TATGAGTGTGGAGTCGTTGCCTCCCAGGTTCAAGCAATTCTTCNNNAGATCGGAAG |

| | |
|---|---|
| 46608_4 | TATGAGTGTGGAGTCGTTGCCTCCCAGGTTCAAGCAATTCTTCNNNAGATCGGAAGA |
| 47044_1 | TATGAGTGTGGAGTCGTTGCTTTTTTGAGACGGAGTCTTGCTCNNNAGATCGGAAG |
| 47044_2 | TATGAGTGTGGAGTCGTTGCTTTTTTGAGACGGAGTCTTGCTCNNNAGATCGGAAG |
| 47044_3 | TATGAGTGTGGAGTCGTTGCTTTTTTGAGACGGAGTCTTGCTCNNNAGATCGGAAG |
| 47044_4 | TATGAGTGTGGAGTCGTTGCTTTTTTGAGACGGAGTCTTGCTCNNNAGATCGGAAG |
| 36452_1 | TATGAGTGTGGAGTCGTTGCCACCATGGAATACTACTCAGCCANNNAGATCGGAAG |
| 36452_2 | TATGAGTGTGGAGTCGTTGCCACCATGGAATACTACTCAGCCANNNAGATCGGAAG |
| 36452_3 | TATGAGTGTGGAGTCGTTGCCACCATGGAATACTACTCAGCCANNNAGATCGGAAG |
| 36452_4 | TATGAGTGTGGAGTCGTTGCCACCATGGAATACTACTCAGCCANNNAGATCGGAAG |
| 52203_1 | TATGAGTGTGGAGTCGTTGCCAAACTTGGCATGAGTCAAGGAGNNNAGATCGGAA |
| 52203_2 | TATGAGTGTGGAGTCGTTGCCAAACTTGGCATGAGTCAAGGAGNNNAGATCGGAA |
| 52203_3 | TATGAGTGTGGAGTCGTTGCCAAACTTGGCATGAGTCAAGGAGNNNAGATCGGAA |
| 52203_4 | TATGAGTGTGGAGTCGTTGCCAAACTTGGCATGAGTCAAGGAGNNNAGATCGGAA |
| 48060_1 | TATGAGTGTGGAGTCGTTGCCTTGGCTCCCCTTCTACACTATCNNNAGATCGGAAGA |
| 48060_2 | TATGAGTGTGGAGTCGTTGCCTTGGCTCCCCTTCTACACTATCNNNAGATCGGAAGA |
| 48060_3 | TATGAGTGTGGAGTCGTTGCCTTGGCTCCCCTTCTACACTATCNNNAGATCGGAAGA |
| 48060_4 | TATGAGTGTGGAGTCGTTGCCTTGGCTCCCCTTCTACACTATCNNNAGATCGGAAGA |
| 48393_1 | TATGAGTGTGGAGTCGTTGCACATGAGAGATGGAATTGCTGTCTNNNAGATCGGA |
| 48393_2 | TATGAGTGTGGAGTCGTTGCACATGAGAGATGGAATTGCTGTCTNNNAGATCGGA |
| 48393_3 | TATGAGTGTGGAGTCGTTGCACATGAGAGATGGAATTGCTGTCTNNNAGATCGGA |
| 48393_4 | TATGAGTGTGGAGTCGTTGCACATGAGAGATGGAATTGCTGTCTNNNAGATCGGA |
| 52280_1 | TATGAGTGTGGAGTCGTTGCCTAGGTACCGGTTCTGAGCATAGNNNAGATCGGAA |
| 52280_2 | TATGAGTGTGGAGTCGTTGCCTAGGTACCGGTTCTGAGCATAGNNNAGATCGGAA |
| 52280_3 | TATGAGTGTGGAGTCGTTGCCTAGGTACCGGTTCTGAGCATAGNNNAGATCGGAA |
| 52280_4 | TATGAGTGTGGAGTCGTTGCCTAGGTACCGGTTCTGAGCATAGNNNAGATCGGAA |
| 921_1 | TATGAGTGTGGAGTCGTTGCGCTAGGGGGAATCCTAGGGTAGANNNAGATCGGA |
| 921_2 | TATGAGTGTGGAGTCGTTGCGCTAGGGGGAATCCTAGGGTAGANNNAGATCGGA |
| 921_3 | TATGAGTGTGGAGTCGTTGCGCTAGGGGGAATCCTAGGGTAGANNNAGATCGGA |
| 921_4 | TATGAGTGTGGAGTCGTTGCGCTAGGGGGAATCCTAGGGTAGANNNAGATCGGA |
| 34002_1 | TATGAGTGTGGAGTCGTTGCGAGTAGCTGGGATTACAGGCATGNNNAGATCGGAA |
| 34002_2 | TATGAGTGTGGAGTCGTTGCGAGTAGCTGGGATTACAGGCATGNNNAGATCGGAA |
| 34002_3 | TATGAGTGTGGAGTCGTTGCGAGTAGCTGGGATTACAGGCATGNNNAGATCGGAA |
| 34002_4 | TATGAGTGTGGAGTCGTTGCGAGTAGCTGGGATTACAGGCATGNNNAGATCGGAA |
| 34911_1 | TATGAGTGTGGAGTCGTTGCAGTAGATGGGATTACAGGCACACNNNAGATCGGAA |
| 34911_2 | TATGAGTGTGGAGTCGTTGCAGTAGATGGGATTACAGGCACACNNNAGATCGGAA |
| 34911_3 | TATGAGTGTGGAGTCGTTGCAGTAGATGGGATTACAGGCACACNNNAGATCGGAA |
| 34911_4 | TATGAGTGTGGAGTCGTTGCAGTAGATGGGATTACAGGCACACNNNAGATCGGAA |
| 10660_1 | TATGAGTGTGGAGTCGTTGCGGCTTTCCCTGATCTGAAGACAGNNNAGATCGGAAG |
| 10660_2 | TATGAGTGTGGAGTCGTTGCGGCTTTCCCTGATCTGAAGACAGNNNAGATCGGAAG |
| 10660_3 | TATGAGTGTGGAGTCGTTGCGGCTTTCCCTGATCTGAAGACAGNNNAGATCGGAAG |
| 10660_4 | TATGAGTGTGGAGTCGTTGCGGCTTTCCCTGATCTGAAGACAGNNNAGATCGGAAG |
| 41149_1 | TATGAGTGTGGAGTCGTTGCCACACACACACACACACATACATNNNAGATCGGAAG |
| 41149_2 | TATGAGTGTGGAGTCGTTGCCACACACACACACACACATACATNNNAGATCGGAAG |

| | |
|---|---|
| 41149_3 | TATGAGTGTGGAGTCGTTGCCACACACACACACACACATACATNNNAGATCGGAAG |
| 41149_4 | TATGAGTGTGGAGTCGTTGCCACACACACACACACACATACATNNNAGATCGGAAG |
| 36317_1 | TATGAGTGTGGAGTCGTTGCCACGCTCTGCTAATTTGAGATGTNNNAGATCGGAAG |
| 36317_2 | TATGAGTGTGGAGTCGTTGCCACGCTCTGCTAATTTGAGATGTNNNAGATCGGAAG |
| 36317_3 | TATGAGTGTGGAGTCGTTGCCACGCTCTGCTAATTTGAGATGTNNNAGATCGGAAG |
| 36317_4 | TATGAGTGTGGAGTCGTTGCCACGCTCTGCTAATTTGAGATGTNNNAGATCGGAAG |
| 30129_1 | TATGAGTGTGGAGTCGTTGCCACCGGGATACACAAGTCTAACTNNNAGATCGGAAG |
| 30129_2 | TATGAGTGTGGAGTCGTTGCCACCGGGATACACAAGTCTAACTNNNAGATCGGAAG |
| 30129_3 | TATGAGTGTGGAGTCGTTGCCACCGGGATACACAAGTCTAACTNNNAGATCGGAAG |
| 30129_4 | TATGAGTGTGGAGTCGTTGCCACCGGGATACACAAGTCTAACTNNNAGATCGGAAG |
| 33652_1 | TATGAGTGTGGAGTCGTTGCAACATATTTATTTGGGGCTGGGCNNNAGATCGGAA |
| 33652_2 | TATGAGTGTGGAGTCGTTGCAACATATTTATTTGGGGCTGGGCNNNAGATCGGAA |
| 33652_3 | TATGAGTGTGGAGTCGTTGCAACATATTTATTTGGGGCTGGGCNNNAGATCGGAA |
| 33652_4 | TATGAGTGTGGAGTCGTTGCAACATATTTATTTGGGGCTGGGCNNNAGATCGGAA |
| 35453_1 | TATGAGTGTGGAGTCGTTGCGACCTCTCATTCTCTGAAGCCAANNNAGATCGGAAG |
| 35453_2 | TATGAGTGTGGAGTCGTTGCGACCTCTCATTCTCTGAAGCCAANNNAGATCGGAAG |
| 35453_3 | TATGAGTGTGGAGTCGTTGCGACCTCTCATTCTCTGAAGCCAANNNAGATCGGAAG |
| 35453_4 | TATGAGTGTGGAGTCGTTGCGACCTCTCATTCTCTGAAGCCAANNNAGATCGGAAG |
| 10188_1 | TATGAGTGTGGAGTCGTTGCCTGTGGATAAGAGCTCACTGCTGNNNAGATCGGAA |
| 10188_2 | TATGAGTGTGGAGTCGTTGCCTGTGGATAAGAGCTCACTGCTGNNNAGATCGGAA |
| 10188_3 | TATGAGTGTGGAGTCGTTGCCTGTGGATAAGAGCTCACTGCTGNNNAGATCGGAA |
| 10188_4 | TATGAGTGTGGAGTCGTTGCCTGTGGATAAGAGCTCACTGCTGNNNAGATCGGAA |
| 36175_1 | TATGAGTGTGGAGTCGTTGCAGTGGATTACAAGGTCAAAGGCTNNNAGATCGGAA |
| 36175_2 | TATGAGTGTGGAGTCGTTGCAGTGGATTACAAGGTCAAAGGCTNNNAGATCGGAA |
| 36175_3 | TATGAGTGTGGAGTCGTTGCAGTGGATTACAAGGTCAAAGGCTNNNAGATCGGAA |
| 36175_4 | TATGAGTGTGGAGTCGTTGCAGTGGATTACAAGGTCAAAGGCTNNNAGATCGGAA |
| 45311_1 | TATGAGTGTGGAGTCGTTGCAGTTGGAGGATTGGCCTTTAGAGNNNAGATCGGAA |
| 45311_2 | TATGAGTGTGGAGTCGTTGCAGTTGGAGGATTGGCCTTTAGAGNNNAGATCGGAA |
| 45311_3 | TATGAGTGTGGAGTCGTTGCAGTTGGAGGATTGGCCTTTAGAGNNNAGATCGGAA |
| 45311_4 | TATGAGTGTGGAGTCGTTGCAGTTGGAGGATTGGCCTTTAGAGNNNAGATCGGAA |
| 31388_1 | TATGAGTGTGGAGTCGTTGCTAAGGCCCACTGTACTAAGGAGANNNAGATCGGAA |
| 31388_2 | TATGAGTGTGGAGTCGTTGCTAAGGCCCACTGTACTAAGGAGANNNAGATCGGAA |
| 31388_3 | TATGAGTGTGGAGTCGTTGCTAAGGCCCACTGTACTAAGGAGANNNAGATCGGAA |
| 31388_4 | TATGAGTGTGGAGTCGTTGCTAAGGCCCACTGTACTAAGGAGANNNAGATCGGAA |
| 50997_1 | TATGAGTGTGGAGTCGTTGCCAAGGCTGTTTATTTCACCTGGGNNNAGATCGGAAG |
| 50997_2 | TATGAGTGTGGAGTCGTTGCCAAGGCTGTTTATTTCACCTGGGNNNAGATCGGAAG |
| 50997_3 | TATGAGTGTGGAGTCGTTGCCAAGGCTGTTTATTTCACCTGGGNNNAGATCGGAAG |
| 50997_4 | TATGAGTGTGGAGTCGTTGCCAAGGCTGTTTATTTCACCTGGGNNNAGATCGGAAG |
| 30053_1 | TATGAGTGTGGAGTCGTTGCTGGAAAAGTGGCAGGAGGAATGNNNAGATCGGAA |
| 30053_2 | TATGAGTGTGGAGTCGTTGCTGGAAAAGTGGCAGGAGGAATGNNNAGATCGGAA |
| 30053_3 | TATGAGTGTGGAGTCGTTGCTGGAAAAGTGGCAGGAGGAATGNNNAGATCGGAA |
| 30053_4 | TATGAGTGTGGAGTCGTTGCTGGAAAAGTGGCAGGAGGAATGNNNAGATCGGAA |
| 36241_1 | TATGAGTGTGGAGTCGTTGCAAAAGAATGAGAGCCTCCTCTGGNNNAGATCGGAA |

| | |
|---|---|
| 36241_2 | TATGAGTGTGGAGTCGTTGCAAAAGAATGAGAGCCTCCTCTGGNNNAGATCGGAA |
| 36241_3 | TATGAGTGTGGAGTCGTTGCAAAAGAATGAGAGCCTCCTCTGGNNNAGATCGGAA |
| 36241_4 | TATGAGTGTGGAGTCGTTGCAAAAGAATGAGAGCCTCCTCTGGNNNAGATCGGAA |
| 37350_1 | TATGAGTGTGGAGTCGTTGCCACGTGAAGGCCTTTGATTCTTNNNAGATCGGAAG |
| 37350_2 | TATGAGTGTGGAGTCGTTGCCACGTGAAGGCCTTTGATTCTTNNNAGATCGGAAG |
| 37350_3 | TATGAGTGTGGAGTCGTTGCCACGTGAAGGCCTTTGATTCTTNNNAGATCGGAAG |
| 37350_4 | TATGAGTGTGGAGTCGTTGCCACGTGAAGGCCTTTGATTCTTNNNAGATCGGAAG |
| 49679_1 | TATGAGTGTGGAGTCGTTGCCCCCCTGAGTAGCTGGGATTATANNNAGATCGGAAG |
| 49679_2 | TATGAGTGTGGAGTCGTTGCCCCCCTGAGTAGCTGGGATTATANNNAGATCGGAAG |
| 49679_3 | TATGAGTGTGGAGTCGTTGCCCCCCTGAGTAGCTGGGATTATANNNAGATCGGAAG |
| 49679_4 | TATGAGTGTGGAGTCGTTGCCCCCCTGAGTAGCTGGGATTATANNNAGATCGGAAG |
| 52661_1 | TATGAGTGTGGAGTCGTTGCGGAAGTGACTTTGGTGCTAGGTANNNAGATCGGAA |
| 52661_2 | TATGAGTGTGGAGTCGTTGCGGAAGTGACTTTGGTGCTAGGTANNNAGATCGGAA |
| 52661_3 | TATGAGTGTGGAGTCGTTGCGGAAGTGACTTTGGTGCTAGGTANNNAGATCGGAA |
| 52661_4 | TATGAGTGTGGAGTCGTTGCGGAAGTGACTTTGGTGCTAGGTANNNAGATCGGAA |
| 38738_1 | TATGAGTGTGGAGTCGTTGCGCTGAGCCAAGATAATTGCTTGTNNNAGATCGGAAG |
| 38738_2 | TATGAGTGTGGAGTCGTTGCGCTGAGCCAAGATAATTGCTTGTNNNAGATCGGAAG |
| 38738_3 | TATGAGTGTGGAGTCGTTGCGCTGAGCCAAGATAATTGCTTGTNNNAGATCGGAAG |
| 38738_4 | TATGAGTGTGGAGTCGTTGCGCTGAGCCAAGATAATTGCTTGTNNNAGATCGGAAG |
| 30572_1 | TATGAGTGTGGAGTCGTTGCAGTGGGGGCTACAGACAACACNNNAGATCGGAAGA |
| 30572_2 | TATGAGTGTGGAGTCGTTGCAGTGGGGGCTACAGACAACACNNNAGATCGGAAGA |
| 30572_3 | TATGAGTGTGGAGTCGTTGCAGTGGGGGCTACAGACAACACNNNAGATCGGAAGA |
| 30572_4 | TATGAGTGTGGAGTCGTTGCAGTGGGGGCTACAGACAACACNNNAGATCGGAAGA |
| 52296_1 | TATGAGTGTGGAGTCGTTGCGTATCTCCTCACCTCATCTGCAGNNNAGATCGGAAGA |
| 52296_2 | TATGAGTGTGGAGTCGTTGCGTATCTCCTCACCTCATCTGCAGNNNAGATCGGAAGA |
| 52296_3 | TATGAGTGTGGAGTCGTTGCGTATCTCCTCACCTCATCTGCAGNNNAGATCGGAAGA |
| 52296_4 | TATGAGTGTGGAGTCGTTGCGTATCTCCTCACCTCATCTGCAGNNNAGATCGGAAGA |
| 33841_1 | TATGAGTGTGGAGTCGTTGCTTTTGCCTCAAAGCAGGAATCTGNNNAGATCGGAAG |
| 33841_2 | TATGAGTGTGGAGTCGTTGCTTTTGCCTCAAAGCAGGAATCTGNNNAGATCGGAAG |
| 33841_3 | TATGAGTGTGGAGTCGTTGCTTTTGCCTCAAAGCAGGAATCTGNNNAGATCGGAAG |
| 33841_4 | TATGAGTGTGGAGTCGTTGCTTTTGCCTCAAAGCAGGAATCTGNNNAGATCGGAAG |
| 32572_1 | TATGAGTGTGGAGTCGTTGCGAGAAGCTAGGACTACAGGTGTGNNNAGATCGGAA |
| 32572_2 | TATGAGTGTGGAGTCGTTGCGAGAAGCTAGGACTACAGGTGTGNNNAGATCGGAA |
| 32572_3 | TATGAGTGTGGAGTCGTTGCGAGAAGCTAGGACTACAGGTGTGNNNAGATCGGAA |
| 32572_4 | TATGAGTGTGGAGTCGTTGCGAGAAGCTAGGACTACAGGTGTGNNNAGATCGGAA |
| 12810_1 | TATGAGTGTGGAGTCGTTGCGGAAGACAGTGTATTCCAAANNNAGATCGGAAGAG |
| 12810_2 | TATGAGTGTGGAGTCGTTGCGGAAGACAGTGTATTCCAAANNNAGATCGGAAGAG |
| 12810_3 | TATGAGTGTGGAGTCGTTGCGGAAGACAGTGTATTCCAAANNNAGATCGGAAGAG |
| 12810_4 | TATGAGTGTGGAGTCGTTGCGGAAGACAGTGTATTCCAAANNNAGATCGGAAGAG |
| 34066_1 | TATGAGTGTGGAGTCGTTGCGAGACGGAGTCTTGCTCTGTCNNNAGATCGGAAGA |
| 34066_2 | TATGAGTGTGGAGTCGTTGCGAGACGGAGTCTTGCTCTGTCNNNAGATCGGAAGA |
| 34066_3 | TATGAGTGTGGAGTCGTTGCGAGACGGAGTCTTGCTCTGTCNNNAGATCGGAAGA |
| 34066_4 | TATGAGTGTGGAGTCGTTGCGAGACGGAGTCTTGCTCTGTCNNNAGATCGGAAGA |

| | |
|---|---|
| 34316_1 | TATGAGTGTGGAGTCGTTGCTGTAGCCGGCCTCATGTTTATTANNNAGATCGGAAG |
| 34316_2 | TATGAGTGTGGAGTCGTTGCTGTAGCCGGCCTCATGTTTATTANNNAGATCGGAAG |
| 34316_3 | TATGAGTGTGGAGTCGTTGCTGTAGCCGGCCTCATGTTTATTANNNAGATCGGAAG |
| 34316_4 | TATGAGTGTGGAGTCGTTGCTGTAGCCGGCCTCATGTTTATTANNNAGATCGGAAG |
| 43987_1 | TATGAGTGTGGAGTCGTTGCAGATGGTGCATGTCTGTAATCCCNNNAGATCGGAAG |
| 43987_2 | TATGAGTGTGGAGTCGTTGCAGATGGTGCATGTCTGTAATCCCNNNAGATCGGAAG |
| 43987_3 | TATGAGTGTGGAGTCGTTGCAGATGGTGCATGTCTGTAATCCCNNNAGATCGGAAG |
| 43987_4 | TATGAGTGTGGAGTCGTTGCAGATGGTGCATGTCTGTAATCCCNNNAGATCGGAAG |
| 32244_1 | TATGAGTGTGGAGTCGTTGCGCAGTCTGTCCATTCTCAGATCTNNNAGATCGGAAG |
| 32244_2 | TATGAGTGTGGAGTCGTTGCGCAGTCTGTCCATTCTCAGATCTNNNAGATCGGAAG |
| 32244_3 | TATGAGTGTGGAGTCGTTGCGCAGTCTGTCCATTCTCAGATCTNNNAGATCGGAAG |
| 32244_4 | TATGAGTGTGGAGTCGTTGCGCAGTCTGTCCATTCTCAGATCTNNNAGATCGGAAG |
| 52605_1 | TATGAGTGTGGAGTCGTTGCATTGCTTTGAGAATGCAAGCACANNNAGATCGGAA |
| 52605_2 | TATGAGTGTGGAGTCGTTGCATTGCTTTGAGAATGCAAGCACANNNAGATCGGAA |
| 52605_3 | TATGAGTGTGGAGTCGTTGCATTGCTTTGAGAATGCAAGCACANNNAGATCGGAA |
| 52605_4 | TATGAGTGTGGAGTCGTTGCATTGCTTTGAGAATGCAAGCACANNNAGATCGGAA |
| 47426_1 | TATGAGTGTGGAGTCGTTGCCCCGGATTCAAGTGATTCTCCTANNNAGATCGGAAG |
| 47426_2 | TATGAGTGTGGAGTCGTTGCCCCGGATTCAAGTGATTCTCCTANNNAGATCGGAAG |
| 47426_3 | TATGAGTGTGGAGTCGTTGCCCCGGATTCAAGTGATTCTCCTANNNAGATCGGAAG |
| 47426_4 | TATGAGTGTGGAGTCGTTGCCCCGGATTCAAGTGATTCTCCTANNNAGATCGGAAG |
| 44716_1 | TATGAGTGTGGAGTCGTTGCCATTGGAGTTCCCGAGAAGGTAANNNAGATCGGAA |
| 44716_2 | TATGAGTGTGGAGTCGTTGCCATTGGAGTTCCCGAGAAGGTAANNNAGATCGGAA |
| 44716_3 | TATGAGTGTGGAGTCGTTGCCATTGGAGTTCCCGAGAAGGTAANNNAGATCGGAA |
| 44716_4 | TATGAGTGTGGAGTCGTTGCCATTGGAGTTCCCGAGAAGGTAANNNAGATCGGAA |
| 52139_1 | TATGAGTGTGGAGTCGTTGCCTGGAGTTCTGGATGAGCGTGNNNAGATCGGAAGA |
| 52139_2 | TATGAGTGTGGAGTCGTTGCCTGGAGTTCTGGATGAGCGTGNNNAGATCGGAAGA |
| 52139_3 | TATGAGTGTGGAGTCGTTGCCTGGAGTTCTGGATGAGCGTGNNNAGATCGGAAGA |
| 52139_4 | TATGAGTGTGGAGTCGTTGCCTGGAGTTCTGGATGAGCGTGNNNAGATCGGAAGA |
| 35800_1 | TATGAGTGTGGAGTCGTTGCTGCAGTTGTACACCCTTGTATCANNNAGATCGGAAG |
| 35800_2 | TATGAGTGTGGAGTCGTTGCTGCAGTTGTACACCCTTGTATCANNNAGATCGGAAG |
| 35800_3 | TATGAGTGTGGAGTCGTTGCTGCAGTTGTACACCCTTGTATCANNNAGATCGGAAG |
| 35800_4 | TATGAGTGTGGAGTCGTTGCTGCAGTTGTACACCCTTGTATCANNNAGATCGGAAG |
| 45981_1 | TATGAGTGTGGAGTCGTTGCCACCATGCCCAGCTAATTTTTGTNNNAGATCGGAAG |
| 45981_2 | TATGAGTGTGGAGTCGTTGCCACCATGCCCAGCTAATTTTTGTNNNAGATCGGAAG |
| 45981_3 | TATGAGTGTGGAGTCGTTGCCACCATGCCCAGCTAATTTTTGTNNNAGATCGGAAG |
| 45981_4 | TATGAGTGTGGAGTCGTTGCCACCATGCCCAGCTAATTTTTGTNNNAGATCGGAAG |
| 46336_1 | TATGAGTGTGGAGTCGTTGCCTTAGCCTCCTGAGTAGCTAGGANNNAGATCGGAAG |
| 46336_2 | TATGAGTGTGGAGTCGTTGCCTTAGCCTCCTGAGTAGCTAGGANNNAGATCGGAAG |
| 46336_3 | TATGAGTGTGGAGTCGTTGCCTTAGCCTCCTGAGTAGCTAGGANNNAGATCGGAAG |
| 46336_4 | TATGAGTGTGGAGTCGTTGCCTTAGCCTCCTGAGTAGCTAGGANNNAGATCGGAAG |
| 28866_1 | TATGAGTGTGGAGTCGTTGCAGTGCCACAATCTCAGCTCTCNNNAGATCGGAAGAG |
| 28866_2 | TATGAGTGTGGAGTCGTTGCAGTGCCACAATCTCAGCTCTCNNNAGATCGGAAGAG |
| 28866_3 | TATGAGTGTGGAGTCGTTGCAGTGCCACAATCTCAGCTCTCNNNAGATCGGAAGAG |

| | |
|---|---|
| 28866_4 | TATGAGTGTGGAGTCGTTGCAGTGCCACAATCTCAGCTCTCNNNAGATCGGAAGAG |
| 37924_1 | TATGAGTGTGGAGTCGTTGCGCTATGCCTGTAATCCTAGCACTNNNAGATCGGAAG |
| 37924_2 | TATGAGTGTGGAGTCGTTGCGCTATGCCTGTAATCCTAGCACTNNNAGATCGGAAG |
| 37924_3 | TATGAGTGTGGAGTCGTTGCGCTATGCCTGTAATCCTAGCACTNNNAGATCGGAAG |
| 37924_4 | TATGAGTGTGGAGTCGTTGCGCTATGCCTGTAATCCTAGCACTNNNAGATCGGAAG |
| 33844_1 | TATGAGTGTGGAGTCGTTGCTGCTTACTTATGCCTTCTACTTGCNNNAGATCGGAAG |
| 33844_2 | TATGAGTGTGGAGTCGTTGCTGCTTACTTATGCCTTCTACTTGCNNNAGATCGGAAG |
| 33844_3 | TATGAGTGTGGAGTCGTTGCTGCTTACTTATGCCTTCTACTTGCNNNAGATCGGAAG |
| 33844_4 | TATGAGTGTGGAGTCGTTGCTGCTTACTTATGCCTTCTACTTGCNNNAGATCGGAAG |
| 44822_1 | TATGAGTGTGGAGTCGTTGCCTACTACCCTGCCTGATTTGTGTNNNAGATCGGAAG |
| 44822_2 | TATGAGTGTGGAGTCGTTGCCTACTACCCTGCCTGATTTGTGTNNNAGATCGGAAG |
| 44822_3 | TATGAGTGTGGAGTCGTTGCCTACTACCCTGCCTGATTTGTGTNNNAGATCGGAAG |
| 44822_4 | TATGAGTGTGGAGTCGTTGCCTACTACCCTGCCTGATTTGTGTNNNAGATCGGAAG |
| 35470_1 | TATGAGTGTGGAGTCGTTGCATAAGCCACAGCATCTGGACTACNNNAGATCGGAAG |
| 35470_2 | TATGAGTGTGGAGTCGTTGCATAAGCCACAGCATCTGGACTACNNNAGATCGGAAG |
| 35470_3 | TATGAGTGTGGAGTCGTTGCATAAGCCACAGCATCTGGACTACNNNAGATCGGAAG |
| 35470_4 | TATGAGTGTGGAGTCGTTGCATAAGCCACAGCATCTGGACTACNNNAGATCGGAAG |
| 10418_1 | TATGAGTGTGGAGTCGTTGCTTCACCAGTTGGTTTCTCCACTGNNNAGATCGGAAG |
| 10418_2 | TATGAGTGTGGAGTCGTTGCTTCACCAGTTGGTTTCTCCACTGNNNAGATCGGAAG |
| 10418_3 | TATGAGTGTGGAGTCGTTGCTTCACCAGTTGGTTTCTCCACTGNNNAGATCGGAAG |
| 10418_4 | TATGAGTGTGGAGTCGTTGCTTCACCAGTTGGTTTCTCCACTGNNNAGATCGGAAG |
| 52826_1 | TATGAGTGTGGAGTCGTTGCGCTCGCTATCACATGGATCCTCNNNAGATCGGAAGA |
| 52826_2 | TATGAGTGTGGAGTCGTTGCGCTCGCTATCACATGGATCCTCNNNAGATCGGAAGA |
| 52826_3 | TATGAGTGTGGAGTCGTTGCGCTCGCTATCACATGGATCCTCNNNAGATCGGAAGA |
| 52826_4 | TATGAGTGTGGAGTCGTTGCGCTCGCTATCACATGGATCCTCNNNAGATCGGAAGA |
| 35863_1 | TATGAGTGTGGAGTCGTTGCAAAGAAAGGTGTTGTCTCTGCTGNNNAGATCGGAA |
| 35863_2 | TATGAGTGTGGAGTCGTTGCAAAGAAAGGTGTTGTCTCTGCTGNNNAGATCGGAA |
| 35863_3 | TATGAGTGTGGAGTCGTTGCAAAGAAAGGTGTTGTCTCTGCTGNNNAGATCGGAA |
| 35863_4 | TATGAGTGTGGAGTCGTTGCAAAGAAAGGTGTTGTCTCTGCTGNNNAGATCGGAA |
| 45461_1 | TATGAGTGTGGAGTCGTTGCAGGGAGCAGCTAAATAGGTTGTTNNNAGATCGGAA |
| 45461_2 | TATGAGTGTGGAGTCGTTGCAGGGAGCAGCTAAATAGGTTGTTNNNAGATCGGAA |
| 45461_3 | TATGAGTGTGGAGTCGTTGCAGGGAGCAGCTAAATAGGTTGTTNNNAGATCGGAA |
| 45461_4 | TATGAGTGTGGAGTCGTTGCAGGGAGCAGCTAAATAGGTTGTTNNNAGATCGGAA |
| 34812_1 | TATGAGTGTGGAGTCGTTGCGGTGTGATCTCAACACATTGCAANNNAGATCGGAAG |
| 34812_2 | TATGAGTGTGGAGTCGTTGCGGTGTGATCTCAACACATTGCAANNNAGATCGGAAG |
| 34812_3 | TATGAGTGTGGAGTCGTTGCGGTGTGATCTCAACACATTGCAANNNAGATCGGAAG |
| 34812_4 | TATGAGTGTGGAGTCGTTGCGGTGTGATCTCAACACATTGCAANNNAGATCGGAAG |
| 38095_1 | TATGAGTGTGGAGTCGTTGCAAGGGGGAAATGTGTATGGGTACNNNAGATCGGA |
| 38095_2 | TATGAGTGTGGAGTCGTTGCAAGGGGGAAATGTGTATGGGTACNNNAGATCGGA |
| 38095_3 | TATGAGTGTGGAGTCGTTGCAAGGGGGAAATGTGTATGGGTACNNNAGATCGGA |
| 38095_4 | TATGAGTGTGGAGTCGTTGCAAGGGGGAAATGTGTATGGGTACNNNAGATCGGA |
| 10940_1 | TATGAGTGTGGAGTCGTTGCGGACAATAGCTGGCTGGGGAAGTNNNAGATCGGA |
| 10940_2 | TATGAGTGTGGAGTCGTTGCGGACAATAGCTGGCTGGGGAAGTNNNAGATCGGA |

| | |
|---|---|
| 10940_3 | TATGAGTGTGGAGTCGTTGCGGACAATAGCTGGCTGGGGAAGTNNNAGATCGGAA |
| 10940_4 | TATGAGTGTGGAGTCGTTGCGGACAATAGCTGGCTGGGGAAGTNNNAGATCGGAA |
| 47490_1 | TATGAGTGTGGAGTCGTTGCTCCTGTGTTCAAGCTATTCTCGTNNNAGATCGGAAG |
| 47490_2 | TATGAGTGTGGAGTCGTTGCTCCTGTGTTCAAGCTATTCTCGTNNNAGATCGGAAG |
| 47490_3 | TATGAGTGTGGAGTCGTTGCTCCTGTGTTCAAGCTATTCTCGTNNNAGATCGGAAG |
| 47490_4 | TATGAGTGTGGAGTCGTTGCTCCTGTGTTCAAGCTATTCTCGTNNNAGATCGGAAG |
| 51002_1 | TATGAGTGTGGAGTCGTTGCTGATCAATCAAGCGGCATTTACCNNNAGATCGGAAG |
| 51002_2 | TATGAGTGTGGAGTCGTTGCTGATCAATCAAGCGGCATTTACCNNNAGATCGGAAG |
| 51002_3 | TATGAGTGTGGAGTCGTTGCTGATCAATCAAGCGGCATTTACCNNNAGATCGGAAG |
| 51002_4 | TATGAGTGTGGAGTCGTTGCTGATCAATCAAGCGGCATTTACCNNNAGATCGGAAG |
| 40417_1 | TATGAGTGTGGAGTCGTTGCATGAGGCAGCAGAATCACTTGAANNNAGATCGGAA |
| 40417_2 | TATGAGTGTGGAGTCGTTGCATGAGGCAGCAGAATCACTTGAANNNAGATCGGAA |
| 40417_3 | TATGAGTGTGGAGTCGTTGCATGAGGCAGCAGAATCACTTGAANNNAGATCGGAA |
| 40417_4 | TATGAGTGTGGAGTCGTTGCATGAGGCAGCAGAATCACTTGAANNNAGATCGGAA |
| 36866_1 | TATGAGTGTGGAGTCGTTGCGTGTCATCAAGGACCCATCTCTTNNNAGATCGGAAG |
| 36866_2 | TATGAGTGTGGAGTCGTTGCGTGTCATCAAGGACCCATCTCTTNNNAGATCGGAAG |
| 36866_3 | TATGAGTGTGGAGTCGTTGCGTGTCATCAAGGACCCATCTCTTNNNAGATCGGAAG |
| 36866_4 | TATGAGTGTGGAGTCGTTGCGTGTCATCAAGGACCCATCTCTTNNNAGATCGGAAG |
| 13434_1 | TATGAGTGTGGAGTCGTTGCGGTACATATATACTGTGGGATACNNNAGATCGGAA |
| 13434_2 | TATGAGTGTGGAGTCGTTGCGGTACATATATACTGTGGGATACNNNAGATCGGAA |
| 13434_3 | TATGAGTGTGGAGTCGTTGCGGTACATATATACTGTGGGATACNNNAGATCGGAA |
| 13434_4 | TATGAGTGTGGAGTCGTTGCGGTACATATATACTGTGGGATACNNNAGATCGGAA |
| 32682_1 | TATGAGTGTGGAGTCGTTGCCAGTGTCTTGCTCTGTCAACCANNNAGATCGGAAGA |
| 32682_2 | TATGAGTGTGGAGTCGTTGCCAGTGTCTTGCTCTGTCAACCANNNAGATCGGAAGA |
| 32682_3 | TATGAGTGTGGAGTCGTTGCCAGTGTCTTGCTCTGTCAACCANNNAGATCGGAAGA |
| 32682_4 | TATGAGTGTGGAGTCGTTGCCAGTGTCTTGCTCTGTCAACCANNNAGATCGGAAGA |
| 12752_1 | TATGAGTGTGGAGTCGTTGCAAACTGCTTAATGATGGATTTAGNNNAGATCGGAA |
| 12752_2 | TATGAGTGTGGAGTCGTTGCAAACTGCTTAATGATGGATTTAGNNNAGATCGGAA |
| 12752_3 | TATGAGTGTGGAGTCGTTGCAAACTGCTTAATGATGGATTTAGNNNAGATCGGAA |
| 12752_4 | TATGAGTGTGGAGTCGTTGCAAACTGCTTAATGATGGATTTAGNNNAGATCGGAA |
| 51382_1 | TATGAGTGTGGAGTCGTTGCGCTGAGTCTCCATCTATTCCCTGNNNAGATCGGAAG |
| 51382_2 | TATGAGTGTGGAGTCGTTGCGCTGAGTCTCCATCTATTCCCTGNNNAGATCGGAAG |
| 51382_3 | TATGAGTGTGGAGTCGTTGCGCTGAGTCTCCATCTATTCCCTGNNNAGATCGGAAG |
| 51382_4 | TATGAGTGTGGAGTCGTTGCGCTGAGTCTCCATCTATTCCCTGNNNAGATCGGAAG |
| 30450_1 | TATGAGTGTGGAGTCGTTGCGGATCAGAGTGTTGTTCTGAGGCNNNAGATCGGAA |
| 30450_2 | TATGAGTGTGGAGTCGTTGCGGATCAGAGTGTTGTTCTGAGGCNNNAGATCGGAA |
| 30450_3 | TATGAGTGTGGAGTCGTTGCGGATCAGAGTGTTGTTCTGAGGCNNNAGATCGGAA |
| 30450_4 | TATGAGTGTGGAGTCGTTGCGGATCAGAGTGTTGTTCTGAGGCNNNAGATCGGAA |
| 6720_1 | TATGAGTGTGGAGTCGTTGCTTTTGACAGCCTCCTTCTGCTGNNNAGATCGGAAGA |
| 6720_2 | TATGAGTGTGGAGTCGTTGCTTTTGACAGCCTCCTTCTGCTGNNNAGATCGGAAGA |
| 6720_3 | TATGAGTGTGGAGTCGTTGCTTTTGACAGCCTCCTTCTGCTGNNNAGATCGGAAGA |
| 6720_4 | TATGAGTGTGGAGTCGTTGCTTTTGACAGCCTCCTTCTGCTGNNNAGATCGGAAGA |
| 36487_1 | TATGAGTGTGGAGTCGTTGCTGTAACAGAACTCAAGGTTTCCANNNAGATCGGAAG |

| | |
|---|---|
| 36487_2 | TATGAGTGTGGAGTCGTTGCTGTAACAGAACTCAAGGTTTCCANNNAGATCGGAAG |
| 36487_3 | TATGAGTGTGGAGTCGTTGCTGTAACAGAACTCAAGGTTTCCANNNAGATCGGAAG |
| 36487_4 | TATGAGTGTGGAGTCGTTGCTGTAACAGAACTCAAGGTTTCCANNNAGATCGGAAG |
| 40222_1 | TATGAGTGTGGAGTCGTTGCCTCTGAGGCAGCATAATGAGACTNNNAGATCGGAA |
| 40222_2 | TATGAGTGTGGAGTCGTTGCCTCTGAGGCAGCATAATGAGACTNNNAGATCGGAA |
| 40222_3 | TATGAGTGTGGAGTCGTTGCCTCTGAGGCAGCATAATGAGACTNNNAGATCGGAA |
| 40222_4 | TATGAGTGTGGAGTCGTTGCCTCTGAGGCAGCATAATGAGACTNNNAGATCGGAA |
| 30664_1 | TATGAGTGTGGAGTCGTTGCTGTGTGTCTCCATCCGAGACATTNNNAGATCGGAAG |
| 30664_2 | TATGAGTGTGGAGTCGTTGCTGTGTGTCTCCATCCGAGACATTNNNAGATCGGAAG |
| 30664_3 | TATGAGTGTGGAGTCGTTGCTGTGTGTCTCCATCCGAGACATTNNNAGATCGGAAG |
| 30664_4 | TATGAGTGTGGAGTCGTTGCTGTGTGTCTCCATCCGAGACATTNNNAGATCGGAAG |
| 30449_1 | TATGAGTGTGGAGTCGTTGCCTCACCAGAGTAGTCCTGGCTTNNNAGATCGGAAGA |
| 30449_2 | TATGAGTGTGGAGTCGTTGCCTCACCAGAGTAGTCCTGGCTTNNNAGATCGGAAGA |
| 30449_3 | TATGAGTGTGGAGTCGTTGCCTCACCAGAGTAGTCCTGGCTTNNNAGATCGGAAGA |
| 30449_4 | TATGAGTGTGGAGTCGTTGCCTCACCAGAGTAGTCCTGGCTTNNNAGATCGGAAGA |
| 29824_1 | TATGAGTGTGGAGTCGTTGCGGATGGATGGTTGTGGGATGATTNNNAGATCGGA |
| 29824_2 | TATGAGTGTGGAGTCGTTGCGGATGGATGGTTGTGGGATGATTNNNAGATCGGA |
| 29824_3 | TATGAGTGTGGAGTCGTTGCGGATGGATGGTTGTGGGATGATTNNNAGATCGGA |
| 29824_4 | TATGAGTGTGGAGTCGTTGCGGATGGATGGTTGTGGGATGATTNNNAGATCGGA |
| 35868_1 | TATGAGTGTGGAGTCGTTGCTTACAGAGAAGAGGGTCTCGCTANNNAGATCGGAA |
| 35868_2 | TATGAGTGTGGAGTCGTTGCTTACAGAGAAGAGGGTCTCGCTANNNAGATCGGAA |
| 35868_3 | TATGAGTGTGGAGTCGTTGCTTACAGAGAAGAGGGTCTCGCTANNNAGATCGGAA |
| 35868_4 | TATGAGTGTGGAGTCGTTGCTTACAGAGAAGAGGGTCTCGCTANNNAGATCGGAA |
| 49870_1 | TATGAGTGTGGAGTCGTTGCATAATCAGGTGGGTTTTCAGCCTNNNAGATCGGAAG |
| 49870_2 | TATGAGTGTGGAGTCGTTGCATAATCAGGTGGGTTTTCAGCCTNNNAGATCGGAAG |
| 49870_3 | TATGAGTGTGGAGTCGTTGCATAATCAGGTGGGTTTTCAGCCTNNNAGATCGGAAG |
| 49870_4 | TATGAGTGTGGAGTCGTTGCATAATCAGGTGGGTTTTCAGCCTNNNAGATCGGAAG |
| 10432_1 | TATGAGTGTGGAGTCGTTGCGGTAGCACTTGCTGTAGGGAATCNNNAGATCGGAA |
| 10432_2 | TATGAGTGTGGAGTCGTTGCGGTAGCACTTGCTGTAGGGAATCNNNAGATCGGAA |
| 10432_3 | TATGAGTGTGGAGTCGTTGCGGTAGCACTTGCTGTAGGGAATCNNNAGATCGGAA |
| 10432_4 | TATGAGTGTGGAGTCGTTGCGGTAGCACTTGCTGTAGGGAATCNNNAGATCGGAA |
| 45454_1 | TATGAGTGTGGAGTCGTTGCGCCTCCCAAATAGCTGAGACTACNNNAGATCGGAAG |
| 45454_2 | TATGAGTGTGGAGTCGTTGCGCCTCCCAAATAGCTGAGACTACNNNAGATCGGAAG |
| 45454_3 | TATGAGTGTGGAGTCGTTGCGCCTCCCAAATAGCTGAGACTACNNNAGATCGGAAG |
| 45454_4 | TATGAGTGTGGAGTCGTTGCGCCTCCCAAATAGCTGAGACTACNNNAGATCGGAAG |
| 32636_1 | TATGAGTGTGGAGTCGTTGCCTCCTGAGTAGCTGGGATTACGNNNAGATCGGAAG |
| 32636_2 | TATGAGTGTGGAGTCGTTGCCTCCTGAGTAGCTGGGATTACGNNNAGATCGGAAG |
| 32636_3 | TATGAGTGTGGAGTCGTTGCCTCCTGAGTAGCTGGGATTACGNNNAGATCGGAAG |
| 32636_4 | TATGAGTGTGGAGTCGTTGCCTCCTGAGTAGCTGGGATTACGNNNAGATCGGAAG |
| 51813_1 | TATGAGTGTGGAGTCGTTGCTTCTGTTTTGGCACTTGCTTTCTNNNAGATCGGAAGA |
| 51813_2 | TATGAGTGTGGAGTCGTTGCTTCTGTTTTGGCACTTGCTTTCTNNNAGATCGGAAGA |
| 51813_3 | TATGAGTGTGGAGTCGTTGCTTCTGTTTTGGCACTTGCTTTCTNNNAGATCGGAAGA |
| 51813_4 | TATGAGTGTGGAGTCGTTGCTTCTGTTTTGGCACTTGCTTTCTNNNAGATCGGAAGA |

| | |
|---|---|
| 49958_1 | TATGAGTGTGGAGTCGTTGCGCTTTGACTGCATCACAATCCATNNNAGATCGGAAG |
| 49958_2 | TATGAGTGTGGAGTCGTTGCGCTTTGACTGCATCACAATCCATNNNAGATCGGAAG |
| 49958_3 | TATGAGTGTGGAGTCGTTGCGCTTTGACTGCATCACAATCCATNNNAGATCGGAAG |
| 49958_4 | TATGAGTGTGGAGTCGTTGCGCTTTGACTGCATCACAATCCATNNNAGATCGGAAG |
| 41927_1 | TATGAGTGTGGAGTCGTTGCCTCCCAGGTTCATGCCATTCTCNNNAGATCGGAAGA |
| 41927_2 | TATGAGTGTGGAGTCGTTGCCTCCCAGGTTCATGCCATTCTCNNNAGATCGGAAGA |
| 41927_3 | TATGAGTGTGGAGTCGTTGCCTCCCAGGTTCATGCCATTCTCNNNAGATCGGAAGA |
| 41927_4 | TATGAGTGTGGAGTCGTTGCCTCCCAGGTTCATGCCATTCTCNNNAGATCGGAAGA |
| 37675_1 | TATGAGTGTGGAGTCGTTGCGGCCAAACAAATATGTCCAGAGANNNAGATCGGAA |
| 37675_2 | TATGAGTGTGGAGTCGTTGCGGCCAAACAAATATGTCCAGAGANNNAGATCGGAA |
| 37675_3 | TATGAGTGTGGAGTCGTTGCGGCCAAACAAATATGTCCAGAGANNNAGATCGGAA |
| 37675_4 | TATGAGTGTGGAGTCGTTGCGGCCAAACAAATATGTCCAGAGANNNAGATCGGAA |
| 34542_1 | TATGAGTGTGGAGTCGTTGCAATGCACAAATGTTACCCTTGCANNNAGATCGGAAG |
| 34542_2 | TATGAGTGTGGAGTCGTTGCAATGCACAAATGTTACCCTTGCANNNAGATCGGAAG |
| 34542_3 | TATGAGTGTGGAGTCGTTGCAATGCACAAATGTTACCCTTGCANNNAGATCGGAAG |
| 34542_4 | TATGAGTGTGGAGTCGTTGCAATGCACAAATGTTACCCTTGCANNNAGATCGGAAG |
| 49644_1 | TATGAGTGTGGAGTCGTTGCTTTCAAGCGATTCTCCTGCCTTANNNAGATCGGAAGA |
| 49644_2 | TATGAGTGTGGAGTCGTTGCTTTCAAGCGATTCTCCTGCCTTANNNAGATCGGAAGA |
| 49644_3 | TATGAGTGTGGAGTCGTTGCTTTCAAGCGATTCTCCTGCCTTANNNAGATCGGAAGA |
| 49644_4 | TATGAGTGTGGAGTCGTTGCTTTCAAGCGATTCTCCTGCCTTANNNAGATCGGAAGA |
| 38956_1 | TATGAGTGTGGAGTCGTTGCCCGGCCAAACTAAGCTTCATAAGNNNAGATCGGAAG |
| 38956_2 | TATGAGTGTGGAGTCGTTGCCCGGCCAAACTAAGCTTCATAAGNNNAGATCGGAAG |
| 38956_3 | TATGAGTGTGGAGTCGTTGCCCGGCCAAACTAAGCTTCATAAGNNNAGATCGGAAG |
| 38956_4 | TATGAGTGTGGAGTCGTTGCCCGGCCAAACTAAGCTTCATAAGNNNAGATCGGAAG |
| 51414_1 | TATGAGTGTGGAGTCGTTGCCTCCCAAATCACCACCAAAGAACNNNAGATCGGAAG |
| 51414_2 | TATGAGTGTGGAGTCGTTGCCTCCCAAATCACCACCAAAGAACNNNAGATCGGAAG |
| 51414_3 | TATGAGTGTGGAGTCGTTGCCTCCCAAATCACCACCAAAGAACNNNAGATCGGAAG |
| 51414_4 | TATGAGTGTGGAGTCGTTGCCTCCCAAATCACCACCAAAGAACNNNAGATCGGAAG |
| 39149_1 | TATGAGTGTGGAGTCGTTGCGCGACAGACTGAGACTCCTTCNNNAGATCGGAAGA |
| 39149_2 | TATGAGTGTGGAGTCGTTGCGCGACAGACTGAGACTCCTTCNNNAGATCGGAAGA |
| 39149_3 | TATGAGTGTGGAGTCGTTGCGCGACAGACTGAGACTCCTTCNNNAGATCGGAAGA |
| 39149_4 | TATGAGTGTGGAGTCGTTGCGCGACAGACTGAGACTCCTTCNNNAGATCGGAAGA |
| 39186_1 | TATGAGTGTGGAGTCGTTGCGGTGATGATTGCACAACTGTGAANNNAGATCGGAA |
| 39186_2 | TATGAGTGTGGAGTCGTTGCGGTGATGATTGCACAACTGTGAANNNAGATCGGAA |
| 39186_3 | TATGAGTGTGGAGTCGTTGCGGTGATGATTGCACAACTGTGAANNNAGATCGGAA |
| 39186_4 | TATGAGTGTGGAGTCGTTGCGGTGATGATTGCACAACTGTGAANNNAGATCGGAA |
| 46309_1 | TATGAGTGTGGAGTCGTTGCAGACTGAGTCTCACTTTGTCACCNNNAGATCGGAAG |
| 46309_2 | TATGAGTGTGGAGTCGTTGCAGACTGAGTCTCACTTTGTCACCNNNAGATCGGAAG |
| 46309_3 | TATGAGTGTGGAGTCGTTGCAGACTGAGTCTCACTTTGTCACCNNNAGATCGGAAG |
| 46309_4 | TATGAGTGTGGAGTCGTTGCAGACTGAGTCTCACTTTGTCACCNNNAGATCGGAAG |
| 32745_1 | TATGAGTGTGGAGTCGTTGCGAAATCCATGCACAGGCTTCTTTNNNAGATCGGAAG |
| 32745_2 | TATGAGTGTGGAGTCGTTGCGAAATCCATGCACAGGCTTCTTTNNNAGATCGGAAG |
| 32745_3 | TATGAGTGTGGAGTCGTTGCGAAATCCATGCACAGGCTTCTTTNNNAGATCGGAAG |

| | |
|---|---|
| 32745_4 | TATGAGTGTGGAGTCGTTGCGAAATCCATGCACAGGCTTCTTTNNNAGATCGGAAG |
| 33214_1 | TATGAGTGTGGAGTCGTTGCGCATTGTAGTGCATGCCTGTAATNNNAGATCGGAAG |
| 33214_2 | TATGAGTGTGGAGTCGTTGCGCATTGTAGTGCATGCCTGTAATNNNAGATCGGAAG |
| 33214_3 | TATGAGTGTGGAGTCGTTGCGCATTGTAGTGCATGCCTGTAATNNNAGATCGGAAG |
| 33214_4 | TATGAGTGTGGAGTCGTTGCGCATTGTAGTGCATGCCTGTAATNNNAGATCGGAAG |
| 31122_1 | TATGAGTGTGGAGTCGTTGCAGAGAGACTAGAAGCCAGAGTCANNNAGATCGGAA |
| 31122_2 | TATGAGTGTGGAGTCGTTGCAGAGAGACTAGAAGCCAGAGTCANNNAGATCGGAA |
| 31122_3 | TATGAGTGTGGAGTCGTTGCAGAGAGACTAGAAGCCAGAGTCANNNAGATCGGAA |
| 31122_4 | TATGAGTGTGGAGTCGTTGCAGAGAGACTAGAAGCCAGAGTCANNNAGATCGGAA |
| 50691_1 | TATGAGTGTGGAGTCGTTGCGAGTGGCCCTTGCATAGAATAGTNNNAGATCGGAA |
| 50691_2 | TATGAGTGTGGAGTCGTTGCGAGTGGCCCTTGCATAGAATAGTNNNAGATCGGAA |
| 50691_3 | TATGAGTGTGGAGTCGTTGCGAGTGGCCCTTGCATAGAATAGTNNNAGATCGGAA |
| 50691_4 | TATGAGTGTGGAGTCGTTGCGAGTGGCCCTTGCATAGAATAGTNNNAGATCGGAA |
| 12150_1 | TATGAGTGTGGAGTCGTTGCTCCCCTTTTCCTTGGCATTTCTCNNNAGATCGGAAGA |
| 12150_2 | TATGAGTGTGGAGTCGTTGCTCCCCTTTTCCTTGGCATTTCTCNNNAGATCGGAAGA |
| 12150_3 | TATGAGTGTGGAGTCGTTGCTCCCCTTTTCCTTGGCATTTCTCNNNAGATCGGAAGA |
| 12150_4 | TATGAGTGTGGAGTCGTTGCTCCCCTTTTCCTTGGCATTTCTCNNNAGATCGGAAGA |
| 10949_1 | TATGAGTGTGGAGTCGTTGCGGTCCAGCCCCAATCCTTTGAGNNNAGATCGGAAGA |
| 10949_2 | TATGAGTGTGGAGTCGTTGCGGTCCAGCCCCAATCCTTTGAGNNNAGATCGGAAGA |
| 10949_3 | TATGAGTGTGGAGTCGTTGCGGTCCAGCCCCAATCCTTTGAGNNNAGATCGGAAGA |
| 10949_4 | TATGAGTGTGGAGTCGTTGCGGTCCAGCCCCAATCCTTTGAGNNNAGATCGGAAGA |
| 34642_1 | TATGAGTGTGGAGTCGTTGCCCTATGTTTTCCTGCACTTGGTGNNNAGATCGGAAG |
| 34642_2 | TATGAGTGTGGAGTCGTTGCCCTATGTTTTCCTGCACTTGGTGNNNAGATCGGAAG |
| 34642_3 | TATGAGTGTGGAGTCGTTGCCCTATGTTTTCCTGCACTTGGTGNNNAGATCGGAAG |
| 34642_4 | TATGAGTGTGGAGTCGTTGCCCTATGTTTTCCTGCACTTGGTGNNNAGATCGGAAG |
| 50832_1 | TATGAGTGTGGAGTCGTTGCTCACCACTCCCTATTTGGTCTTCNNNAGATCGGAAGA |
| 50832_2 | TATGAGTGTGGAGTCGTTGCTCACCACTCCCTATTTGGTCTTCNNNAGATCGGAAGA |
| 50832_3 | TATGAGTGTGGAGTCGTTGCTCACCACTCCCTATTTGGTCTTCNNNAGATCGGAAGA |
| 50832_4 | TATGAGTGTGGAGTCGTTGCTCACCACTCCCTATTTGGTCTTCNNNAGATCGGAAGA |
| 32693_1 | TATGAGTGTGGAGTCGTTGCCCTCTTATCTCTCACAGCTTCCCNNNAGATCGGAAGA |
| 32693_2 | TATGAGTGTGGAGTCGTTGCCCTCTTATCTCTCACAGCTTCCCNNNAGATCGGAAGA |
| 32693_3 | TATGAGTGTGGAGTCGTTGCCCTCTTATCTCTCACAGCTTCCCNNNAGATCGGAAGA |
| 32693_4 | TATGAGTGTGGAGTCGTTGCCCTCTTATCTCTCACAGCTTCCCNNNAGATCGGAAGA |
| 42128_1 | TATGAGTGTGGAGTCGTTGCCTCCTGGGTTCACACCATTCTCNNNAGATCGGAAGAG |
| 42128_2 | TATGAGTGTGGAGTCGTTGCCTCCTGGGTTCACACCATTCTCNNNAGATCGGAAGAG |
| 42128_3 | TATGAGTGTGGAGTCGTTGCCTCCTGGGTTCACACCATTCTCNNNAGATCGGAAGAG |
| 42128_4 | TATGAGTGTGGAGTCGTTGCCTCCTGGGTTCACACCATTCTCNNNAGATCGGAAGAG |
| 46171_1 | TATGAGTGTGGAGTCGTTGCACAAAGGCAGTGGGGAATTCTTANNNAGATCGGAA |
| 46171_2 | TATGAGTGTGGAGTCGTTGCACAAAGGCAGTGGGGAATTCTTANNNAGATCGGAA |
| 46171_3 | TATGAGTGTGGAGTCGTTGCACAAAGGCAGTGGGGAATTCTTANNNAGATCGGAA |
| 46171_4 | TATGAGTGTGGAGTCGTTGCACAAAGGCAGTGGGGAATTCTTANNNAGATCGGAA |
| 45591_1 | TATGAGTGTGGAGTCGTTGCCTCCCTGAGTAGCTTGGACTACANNNAGATCGGAAG |
| 45591_2 | TATGAGTGTGGAGTCGTTGCCTCCCTGAGTAGCTTGGACTACANNNAGATCGGAAG |

| | |
|---|---|
| 45591_3 | TATGAGTGTGGAGTCGTTGCCTCCCTGAGTAGCTTGGACTACANNNAGATCGGAAG |
| 45591_4 | TATGAGTGTGGAGTCGTTGCCTCCCTGAGTAGCTTGGACTACANNNAGATCGGAAG |
| 29249_1 | TATGAGTGTGGAGTCGTTGCAAGGCTGTGCAATAGTCAAAGATNNNAGATCGGAA |
| 29249_2 | TATGAGTGTGGAGTCGTTGCAAGGCTGTGCAATAGTCAAAGATNNNAGATCGGAA |
| 29249_3 | TATGAGTGTGGAGTCGTTGCAAGGCTGTGCAATAGTCAAAGATNNNAGATCGGAA |
| 29249_4 | TATGAGTGTGGAGTCGTTGCAAGGCTGTGCAATAGTCAAAGATNNNAGATCGGAA |
| 37136_1 | TATGAGTGTGGAGTCGTTGCAAGTCAAGAAGATTGTTGGGGGTNNNAGATCGGA |
| 37136_2 | TATGAGTGTGGAGTCGTTGCAAGTCAAGAAGATTGTTGGGGGTNNNAGATCGGA |
| 37136_3 | TATGAGTGTGGAGTCGTTGCAAGTCAAGAAGATTGTTGGGGGTNNNAGATCGGA |
| 37136_4 | TATGAGTGTGGAGTCGTTGCAAGTCAAGAAGATTGTTGGGGGTNNNAGATCGGA |
| 50500_1 | TATGAGTGTGGAGTCGTTGCGGGGTTGGGAGAATGTATACCTGNNNAGATCGGA |
| 50500_2 | TATGAGTGTGGAGTCGTTGCGGGGTTGGGAGAATGTATACCTGNNNAGATCGGA |
| 50500_3 | TATGAGTGTGGAGTCGTTGCGGGGTTGGGAGAATGTATACCTGNNNAGATCGGA |
| 50500_4 | TATGAGTGTGGAGTCGTTGCGGGGTTGGGAGAATGTATACCTGNNNAGATCGGA |
| 45360_1 | TATGAGTGTGGAGTCGTTGCTCGGGTCATGTTCTTGGAATTCTNNNAGATCGGAAG |
| 45360_2 | TATGAGTGTGGAGTCGTTGCTCGGGTCATGTTCTTGGAATTCTNNNAGATCGGAAG |
| 45360_3 | TATGAGTGTGGAGTCGTTGCTCGGGTCATGTTCTTGGAATTCTNNNAGATCGGAAG |
| 45360_4 | TATGAGTGTGGAGTCGTTGCTCGGGTCATGTTCTTGGAATTCTNNNAGATCGGAAG |
| 34267_1 | TATGAGTGTGGAGTCGTTGCGGTCCCAGATTTGAATCTAGCCANNNAGATCGGAAG |
| 34267_2 | TATGAGTGTGGAGTCGTTGCGGTCCCAGATTTGAATCTAGCCANNNAGATCGGAAG |
| 34267_3 | TATGAGTGTGGAGTCGTTGCGGTCCCAGATTTGAATCTAGCCANNNAGATCGGAAG |
| 34267_4 | TATGAGTGTGGAGTCGTTGCGGTCCCAGATTTGAATCTAGCCANNNAGATCGGAAG |
| 52474_1 | TATGAGTGTGGAGTCGTTGCGTGGCTGTATAGATGGCTCATGANNNAGATCGGAA |
| 52474_2 | TATGAGTGTGGAGTCGTTGCGTGGCTGTATAGATGGCTCATGANNNAGATCGGAA |
| 52474_3 | TATGAGTGTGGAGTCGTTGCGTGGCTGTATAGATGGCTCATGANNNAGATCGGAA |
| 52474_4 | TATGAGTGTGGAGTCGTTGCGTGGCTGTATAGATGGCTCATGANNNAGATCGGAA |
| 50146_1 | TATGAGTGTGGAGTCGTTGCGTTCCGCTTCAGGAGAAATTCTGNNNAGATCGGAAG |
| 50146_2 | TATGAGTGTGGAGTCGTTGCGTTCCGCTTCAGGAGAAATTCTGNNNAGATCGGAAG |
| 50146_3 | TATGAGTGTGGAGTCGTTGCGTTCCGCTTCAGGAGAAATTCTGNNNAGATCGGAAG |
| 50146_4 | TATGAGTGTGGAGTCGTTGCGTTCCGCTTCAGGAGAAATTCTGNNNAGATCGGAAG |
| 35149_1 | TATGAGTGTGGAGTCGTTGCGAGTCTTGTTCTTTTGCCTAGGCNNNAGATCGGAAG |
| 35149_2 | TATGAGTGTGGAGTCGTTGCGAGTCTTGTTCTTTTGCCTAGGCNNNAGATCGGAAG |
| 35149_3 | TATGAGTGTGGAGTCGTTGCGAGTCTTGTTCTTTTGCCTAGGCNNNAGATCGGAAG |
| 35149_4 | TATGAGTGTGGAGTCGTTGCGAGTCTTGTTCTTTTGCCTAGGCNNNAGATCGGAAG |
| 45396_1 | TATGAGTGTGGAGTCGTTGCCAATAAACACCTGGGTGAGAGGANNNAGATCGGAA |
| 45396_2 | TATGAGTGTGGAGTCGTTGCCAATAAACACCTGGGTGAGAGGANNNAGATCGGAA |
| 45396_3 | TATGAGTGTGGAGTCGTTGCCAATAAACACCTGGGTGAGAGGANNNAGATCGGAA |
| 45396_4 | TATGAGTGTGGAGTCGTTGCCAATAAACACCTGGGTGAGAGGANNNAGATCGGAA |
| 50885_1 | TATGAGTGTGGAGTCGTTGCCCACTCCACACAAAGAAACTTGGNNNAGATCGGAAG |
| 50885_2 | TATGAGTGTGGAGTCGTTGCCCACTCCACACAAAGAAACTTGGNNNAGATCGGAAG |
| 50885_3 | TATGAGTGTGGAGTCGTTGCCCACTCCACACAAAGAAACTTGGNNNAGATCGGAAG |
| 50885_4 | TATGAGTGTGGAGTCGTTGCCCACTCCACACAAAGAAACTTGGNNNAGATCGGAAG |
| 43893_1 | TATGAGTGTGGAGTCGTTGCTCTCTCTCTCTCTGTACAGCTNNNAGATCGGAAGA |

| | |
|---|---|
| 43893_2 | TATGAGTGTGGAGTCGTTGCTCTCTCTCTCTCTCTGTACAGCTNNNAGATCGGAAGA |
| 43893_3 | TATGAGTGTGGAGTCGTTGCTCTCTCTCTCTCTCTGTACAGCTNNNAGATCGGAAGA |
| 43893_4 | TATGAGTGTGGAGTCGTTGCTCTCTCTCTCTCTCTGTACAGCTNNNAGATCGGAAGA |
| 51236_1 | TATGAGTGTGGAGTCGTTGCATTTGAGGTCAGGAGTTCGAGACNNNAGATCGGAA |
| 51236_2 | TATGAGTGTGGAGTCGTTGCATTTGAGGTCAGGAGTTCGAGACNNNAGATCGGAA |
| 51236_3 | TATGAGTGTGGAGTCGTTGCATTTGAGGTCAGGAGTTCGAGACNNNAGATCGGAA |
| 51236_4 | TATGAGTGTGGAGTCGTTGCATTTGAGGTCAGGAGTTCGAGACNNNAGATCGGAA |
| 43398_1 | TATGAGTGTGGAGTCGTTGCTGCTGCCTGGACATAAACTTAGTNNNAGATCGGAAG |
| 43398_2 | TATGAGTGTGGAGTCGTTGCTGCTGCCTGGACATAAACTTAGTNNNAGATCGGAAG |
| 43398_3 | TATGAGTGTGGAGTCGTTGCTGCTGCCTGGACATAAACTTAGTNNNAGATCGGAAG |
| 43398_4 | TATGAGTGTGGAGTCGTTGCTGCTGCCTGGACATAAACTTAGTNNNAGATCGGAAG |
| 49865_1 | TATGAGTGTGGAGTCGTTGCTGAAAGAAGCCAGGGAAAAAACANNNAGATCGGAA |
| 49865_2 | TATGAGTGTGGAGTCGTTGCTGAAAGAAGCCAGGGAAAAAACANNNAGATCGGAA |
| 49865_3 | TATGAGTGTGGAGTCGTTGCTGAAAGAAGCCAGGGAAAAAACANNNAGATCGGAA |
| 49865_4 | TATGAGTGTGGAGTCGTTGCTGAAAGAAGCCAGGGAAAAAACANNNAGATCGGAA |
| 40493_1 | TATGAGTGTGGAGTCGTTGCATTTTCATGCCCAAAGTCTTGCCNNNAGATCGGAAG |
| 40493_2 | TATGAGTGTGGAGTCGTTGCATTTTCATGCCCAAAGTCTTGCCNNNAGATCGGAAG |
| 40493_3 | TATGAGTGTGGAGTCGTTGCATTTTCATGCCCAAAGTCTTGCCNNNAGATCGGAAG |
| 40493_4 | TATGAGTGTGGAGTCGTTGCATTTTCATGCCCAAAGTCTTGCCNNNAGATCGGAAG |
| 30827_1 | TATGAGTGTGGAGTCGTTGCGGATGCTTCTGAATTTCCAGCTCNNNAGATCGGAAG |
| 30827_2 | TATGAGTGTGGAGTCGTTGCGGATGCTTCTGAATTTCCAGCTCNNNAGATCGGAAG |
| 30827_3 | TATGAGTGTGGAGTCGTTGCGGATGCTTCTGAATTTCCAGCTCNNNAGATCGGAAG |
| 30827_4 | TATGAGTGTGGAGTCGTTGCGGATGCTTCTGAATTTCCAGCTCNNNAGATCGGAAG |
| 36843_1 | TATGAGTGTGGAGTCGTTGCGTCTTGTGTGGCATGAGGTTTTNNNAGATCGGAAG |
| 36843_2 | TATGAGTGTGGAGTCGTTGCGTCTTGTGTGGCATGAGGTTTTNNNAGATCGGAAG |
| 36843_3 | TATGAGTGTGGAGTCGTTGCGTCTTGTGTGGCATGAGGTTTTNNNAGATCGGAAG |
| 36843_4 | TATGAGTGTGGAGTCGTTGCGTCTTGTGTGGCATGAGGTTTTNNNAGATCGGAAG |
| 30784_1 | TATGAGTGTGGAGTCGTTGCCCCCAGGTCCTAAAAAGTTGAGANNNAGATCGGAAG |
| 30784_2 | TATGAGTGTGGAGTCGTTGCCCCCAGGTCCTAAAAAGTTGAGANNNAGATCGGAAG |
| 30784_3 | TATGAGTGTGGAGTCGTTGCCCCCAGGTCCTAAAAAGTTGAGANNNAGATCGGAAG |
| 30784_4 | TATGAGTGTGGAGTCGTTGCCCCCAGGTCCTAAAAAGTTGAGANNNAGATCGGAAG |
| 41529_1 | TATGAGTGTGGAGTCGTTGCTCCAAACATAAAGAAGGGAGGCANNNAGATCGGAA |
| 41529_2 | TATGAGTGTGGAGTCGTTGCTCCAAACATAAAGAAGGGAGGCANNNAGATCGGAA |
| 41529_3 | TATGAGTGTGGAGTCGTTGCTCCAAACATAAAGAAGGGAGGCANNNAGATCGGAA |
| 41529_4 | TATGAGTGTGGAGTCGTTGCTCCAAACATAAAGAAGGGAGGCANNNAGATCGGAA |
| 51929_1 | TATGAGTGTGGAGTCGTTGCTCAGTTGACATTGGACATTGTGCNNNAGATCGGAAG |
| 51929_2 | TATGAGTGTGGAGTCGTTGCTCAGTTGACATTGGACATTGTGCNNNAGATCGGAAG |
| 51929_3 | TATGAGTGTGGAGTCGTTGCTCAGTTGACATTGGACATTGTGCNNNAGATCGGAAG |
| 51929_4 | TATGAGTGTGGAGTCGTTGCTCAGTTGACATTGGACATTGTGCNNNAGATCGGAAG |
| 40857_1 | TATGAGTGTGGAGTCGTTGCTGTTGTTGTTTTCGTAAGTGCCANNNAGATCGGAAG |
| 40857_2 | TATGAGTGTGGAGTCGTTGCTGTTGTTGTTTTCGTAAGTGCCANNNAGATCGGAAG |
| 40857_3 | TATGAGTGTGGAGTCGTTGCTGTTGTTGTTTTCGTAAGTGCCANNNAGATCGGAAG |
| 40857_4 | TATGAGTGTGGAGTCGTTGCTGTTGTTGTTTTCGTAAGTGCCANNNAGATCGGAAG |

| | |
|---|---|
| 50153_1 | TATGAGTGTGGAGTCGTTGCCCAGCCCTCAGACTTCCATATTTNNNAGATCGGAAGA |
| 50153_2 | TATGAGTGTGGAGTCGTTGCCCAGCCCTCAGACTTCCATATTTNNNAGATCGGAAGA |
| 50153_3 | TATGAGTGTGGAGTCGTTGCCCAGCCCTCAGACTTCCATATTTNNNAGATCGGAAGA |
| 50153_4 | TATGAGTGTGGAGTCGTTGCCCAGCCCTCAGACTTCCATATTTNNNAGATCGGAAGA |
| 51434_1 | TATGAGTGTGGAGTCGTTGCCGAGTGAGGAAGAAGGAGGAAGNNNAGATCGGAA |
| 51434_2 | TATGAGTGTGGAGTCGTTGCCGAGTGAGGAAGAAGGAGGAAGNNNAGATCGGAA |
| 51434_3 | TATGAGTGTGGAGTCGTTGCCGAGTGAGGAAGAAGGAGGAAGNNNAGATCGGAA |
| 51434_4 | TATGAGTGTGGAGTCGTTGCCGAGTGAGGAAGAAGGAGGAAGNNNAGATCGGAA |
| 29781_1 | TATGAGTGTGGAGTCGTTGCGGTGAAGGAGGCGATGGAATTNNNAGATCGGAAG |
| 29781_2 | TATGAGTGTGGAGTCGTTGCGGTGAAGGAGGCGATGGAATTNNNAGATCGGAAG |
| 29781_3 | TATGAGTGTGGAGTCGTTGCGGTGAAGGAGGCGATGGAATTNNNAGATCGGAAG |
| 29781_4 | TATGAGTGTGGAGTCGTTGCGGTGAAGGAGGCGATGGAATTNNNAGATCGGAAG |
| 7580_1 | TATGAGTGTGGAGTCGTTGCGGGCTATCAGAGAAAATGTGCTTTCNNNAGATCGG |
| 7580_2 | TATGAGTGTGGAGTCGTTGCGGGCTATCAGAGAAAATGTGCTTTCNNNAGATCGG |
| 7580_3 | TATGAGTGTGGAGTCGTTGCGGGCTATCAGAGAAAATGTGCTTTCNNNAGATCGG |
| 7580_4 | TATGAGTGTGGAGTCGTTGCGGGCTATCAGAGAAAATGTGCTTTCNNNAGATCGG |
| 49734_1 | TATGAGTGTGGAGTCGTTGCAGAGAGGAGAGGCACCTTCTAATNNNAGATCGGAA |
| 49734_2 | TATGAGTGTGGAGTCGTTGCAGAGAGGAGAGGCACCTTCTAATNNNAGATCGGAA |
| 49734_3 | TATGAGTGTGGAGTCGTTGCAGAGAGGAGAGGCACCTTCTAATNNNAGATCGGAA |
| 49734_4 | TATGAGTGTGGAGTCGTTGCAGAGAGGAGAGGCACCTTCTAATNNNAGATCGGAA |
| 52028_1 | TATGAGTGTGGAGTCGTTGCAGATATGGGTCCAGTTTCAAGGGNNNAGATCGGAA |
| 52028_2 | TATGAGTGTGGAGTCGTTGCAGATATGGGTCCAGTTTCAAGGGNNNAGATCGGAA |
| 52028_3 | TATGAGTGTGGAGTCGTTGCAGATATGGGTCCAGTTTCAAGGGNNNAGATCGGAA |
| 52028_4 | TATGAGTGTGGAGTCGTTGCAGATATGGGTCCAGTTTCAAGGGNNNAGATCGGAA |
| 51386_1 | TATGAGTGTGGAGTCGTTGCAGAAGTTCTCACCCCAGGATCTANNNAGATCGGAAG |
| 51386_2 | TATGAGTGTGGAGTCGTTGCAGAAGTTCTCACCCCAGGATCTANNNAGATCGGAAG |
| 51386_3 | TATGAGTGTGGAGTCGTTGCAGAAGTTCTCACCCCAGGATCTANNNAGATCGGAAG |
| 51386_4 | TATGAGTGTGGAGTCGTTGCAGAAGTTCTCACCCCAGGATCTANNNAGATCGGAAG |
| 29259_1 | TATGAGTGTGGAGTCGTTGCTGCTGTCCAACTGATTTACTCCTNNNAGATCGGAAGA |
| 29259_2 | TATGAGTGTGGAGTCGTTGCTGCTGTCCAACTGATTTACTCCTNNNAGATCGGAAGA |
| 29259_3 | TATGAGTGTGGAGTCGTTGCTGCTGTCCAACTGATTTACTCCTNNNAGATCGGAAGA |
| 29259_4 | TATGAGTGTGGAGTCGTTGCTGCTGTCCAACTGATTTACTCCTNNNAGATCGGAAGA |
| 29940_1 | TATGAGTGTGGAGTCGTTGCACCATGTCTGTCTACACACTAGCNNNAGATCGGAAG |
| 29940_2 | TATGAGTGTGGAGTCGTTGCACCATGTCTGTCTACACACTAGCNNNAGATCGGAAG |
| 29940_3 | TATGAGTGTGGAGTCGTTGCACCATGTCTGTCTACACACTAGCNNNAGATCGGAAG |
| 29940_4 | TATGAGTGTGGAGTCGTTGCACCATGTCTGTCTACACACTAGCNNNAGATCGGAAG |
| 48922_1 | TATGAGTGTGGAGTCGTTGCTCACATCCCCATTTTACAGCTGANNNAGATCGGAAGA |
| 48922_2 | TATGAGTGTGGAGTCGTTGCTCACATCCCCATTTTACAGCTGANNNAGATCGGAAGA |
| 48922_3 | TATGAGTGTGGAGTCGTTGCTCACATCCCCATTTTACAGCTGANNNAGATCGGAAGA |
| 48922_4 | TATGAGTGTGGAGTCGTTGCTCACATCCCCATTTTACAGCTGANNNAGATCGGAAGA |
| 13617_1 | TATGAGTGTGGAGTCGTTGCCCTAGTTGTCACCTTTGTATAATNNNAGATCGGAAG |
| 13617_2 | TATGAGTGTGGAGTCGTTGCCCTAGTTGTCACCTTTGTATAATNNNAGATCGGAAG |
| 13617_3 | TATGAGTGTGGAGTCGTTGCCCTAGTTGTCACCTTTGTATAATNNNAGATCGGAAG |

| | |
|---|---|
| 13617_4 | TATGAGTGTGGAGTCGTTGCCCTAGTTGTCACCTTTGTATAATNNNAGATCGGAAG |
| 32684_1 | TATGAGTGTGGAGTCGTTGCTGCAATCTCGGCTCACTGAAANNNAGATCGGAAGAG |
| 32684_2 | TATGAGTGTGGAGTCGTTGCTGCAATCTCGGCTCACTGAAANNNAGATCGGAAGAG |
| 32684_3 | TATGAGTGTGGAGTCGTTGCTGCAATCTCGGCTCACTGAAANNNAGATCGGAAGAG |
| 32684_4 | TATGAGTGTGGAGTCGTTGCTGCAATCTCGGCTCACTGAAANNNAGATCGGAAGAG |
| 52613_1 | TATGAGTGTGGAGTCGTTGCATAATGTCTTTCTGGCCTCTCCCNNNAGATCGGAAGA |
| 52613_2 | TATGAGTGTGGAGTCGTTGCATAATGTCTTTCTGGCCTCTCCCNNNAGATCGGAAGA |
| 52613_3 | TATGAGTGTGGAGTCGTTGCATAATGTCTTTCTGGCCTCTCCCNNNAGATCGGAAGA |
| 52613_4 | TATGAGTGTGGAGTCGTTGCATAATGTCTTTCTGGCCTCTCCCNNNAGATCGGAAGA |
| 45718_1 | TATGAGTGTGGAGTCGTTGCTTCAAGTGATTCTCCTGCCTCAGNNNAGATCGGAAG |
| 45718_2 | TATGAGTGTGGAGTCGTTGCTTCAAGTGATTCTCCTGCCTCAGNNNAGATCGGAAG |
| 45718_3 | TATGAGTGTGGAGTCGTTGCTTCAAGTGATTCTCCTGCCTCAGNNNAGATCGGAAG |
| 45718_4 | TATGAGTGTGGAGTCGTTGCTTCAAGTGATTCTCCTGCCTCAGNNNAGATCGGAAG |
| 43619_1 | TATGAGTGTGGAGTCGTTGCTAGACAGGATCTTCTCAGGAGCANNNAGATCGGAA |
| 43619_2 | TATGAGTGTGGAGTCGTTGCTAGACAGGATCTTCTCAGGAGCANNNAGATCGGAA |
| 43619_3 | TATGAGTGTGGAGTCGTTGCTAGACAGGATCTTCTCAGGAGCANNNAGATCGGAA |
| 43619_4 | TATGAGTGTGGAGTCGTTGCTAGACAGGATCTTCTCAGGAGCANNNAGATCGGAA |
| 51876_1 | TATGAGTGTGGAGTCGTTGCTCCCTGGTACTCATGATCTCCTTNNNAGATCGGAAGA |
| 51876_2 | TATGAGTGTGGAGTCGTTGCTCCCTGGTACTCATGATCTCCTTNNNAGATCGGAAGA |
| 51876_3 | TATGAGTGTGGAGTCGTTGCTCCCTGGTACTCATGATCTCCTTNNNAGATCGGAAGA |
| 51876_4 | TATGAGTGTGGAGTCGTTGCTCCCTGGTACTCATGATCTCCTTNNNAGATCGGAAGA |
| 49919_1 | TATGAGTGTGGAGTCGTTGCTTCCAATTTCCATCCATCAGGGTNNNAGATCGGAAG |
| 49919_2 | TATGAGTGTGGAGTCGTTGCTTCCAATTTCCATCCATCAGGGTNNNAGATCGGAAG |
| 49919_3 | TATGAGTGTGGAGTCGTTGCTTCCAATTTCCATCCATCAGGGTNNNAGATCGGAAG |
| 49919_4 | TATGAGTGTGGAGTCGTTGCTTCCAATTTCCATCCATCAGGGTNNNAGATCGGAAG |
| 42875_1 | TATGAGTGTGGAGTCGTTGCAATGTCATTTGCAGCAACATGGANNNAGATCGGAA |
| 42875_2 | TATGAGTGTGGAGTCGTTGCAATGTCATTTGCAGCAACATGGANNNAGATCGGAA |
| 42875_3 | TATGAGTGTGGAGTCGTTGCAATGTCATTTGCAGCAACATGGANNNAGATCGGAA |
| 42875_4 | TATGAGTGTGGAGTCGTTGCAATGTCATTTGCAGCAACATGGANNNAGATCGGAA |
| 36162_1 | TATGAGTGTGGAGTCGTTGCCTGGGAAATGATTGAGTGACTGCNNNAGATCGGAA |
| 36162_2 | TATGAGTGTGGAGTCGTTGCCTGGGAAATGATTGAGTGACTGCNNNAGATCGGAA |
| 36162_3 | TATGAGTGTGGAGTCGTTGCCTGGGAAATGATTGAGTGACTGCNNNAGATCGGAA |
| 36162_4 | TATGAGTGTGGAGTCGTTGCCTGGGAAATGATTGAGTGACTGCNNNAGATCGGAA |
| 37960_1 | TATGAGTGTGGAGTCGTTGCCTGAATCATGCCACGCTTTCTAGNNNAGATCGGAAG |
| 37960_2 | TATGAGTGTGGAGTCGTTGCCTGAATCATGCCACGCTTTCTAGNNNAGATCGGAAG |
| 37960_3 | TATGAGTGTGGAGTCGTTGCCTGAATCATGCCACGCTTTCTAGNNNAGATCGGAAG |
| 37960_4 | TATGAGTGTGGAGTCGTTGCCTGAATCATGCCACGCTTTCTAGNNNAGATCGGAAG |
| 40501_1 | TATGAGTGTGGAGTCGTTGCCTACTTAGGAAGCTGAGGTGGGANNNAGATCGGAA |
| 40501_2 | TATGAGTGTGGAGTCGTTGCCTACTTAGGAAGCTGAGGTGGGANNNAGATCGGAA |
| 40501_3 | TATGAGTGTGGAGTCGTTGCCTACTTAGGAAGCTGAGGTGGGANNNAGATCGGAA |
| 40501_4 | TATGAGTGTGGAGTCGTTGCCTACTTAGGAAGCTGAGGTGGGANNNAGATCGGAA |
| 47304_1 | TATGAGTGTGGAGTCGTTGCGTTCAAGTACTTCTCCTGCCTCANNNAGATCGGAAGA |
| 47304_2 | TATGAGTGTGGAGTCGTTGCGTTCAAGTACTTCTCCTGCCTCANNNAGATCGGAAGA |

| | |
|---|---|
| 47304_3 | TATGAGTGTGGAGTCGTTGCGTTCAAGTACTTCTCCTGCCTCANNNAGATCGGAAGA |
| 47304_4 | TATGAGTGTGGAGTCGTTGCGTTCAAGTACTTCTCCTGCCTCANNNAGATCGGAAGA |
| 12544_1 | TATGAGTGTGGAGTCGTTGCCCTTACATGCTTTATCAACTAAGNNNAGATCGGAAGA |
| 12544_2 | TATGAGTGTGGAGTCGTTGCCCTTACATGCTTTATCAACTAAGNNNAGATCGGAAGA |
| 12544_3 | TATGAGTGTGGAGTCGTTGCCCTTACATGCTTTATCAACTAAGNNNAGATCGGAAGA |
| 12544_4 | TATGAGTGTGGAGTCGTTGCCCTTACATGCTTTATCAACTAAGNNNAGATCGGAAGA |
| 32994_1 | TATGAGTGTGGAGTCGTTGCCTCAAGTGATCCTCCCATCTCAGNNNAGATCGGAAGA |
| 32994_2 | TATGAGTGTGGAGTCGTTGCCTCAAGTGATCCTCCCATCTCAGNNNAGATCGGAAGA |
| 32994_3 | TATGAGTGTGGAGTCGTTGCCTCAAGTGATCCTCCCATCTCAGNNNAGATCGGAAGA |
| 32994_4 | TATGAGTGTGGAGTCGTTGCCTCAAGTGATCCTCCCATCTCAGNNNAGATCGGAAGA |
| 38699_1 | TATGAGTGTGGAGTCGTTGCTATGCCAGTTGAAAGACACCAGANNNAGATCGGAAG |
| 38699_2 | TATGAGTGTGGAGTCGTTGCTATGCCAGTTGAAAGACACCAGANNNAGATCGGAAG |
| 38699_3 | TATGAGTGTGGAGTCGTTGCTATGCCAGTTGAAAGACACCAGANNNAGATCGGAAG |
| 38699_4 | TATGAGTGTGGAGTCGTTGCTATGCCAGTTGAAAGACACCAGANNNAGATCGGAAG |
| 37944_1 | TATGAGTGTGGAGTCGTTGCTATAGTTCTGGAGGCTGGGAAGTNNNAGATCGGAA |
| 37944_2 | TATGAGTGTGGAGTCGTTGCTATAGTTCTGGAGGCTGGGAAGTNNNAGATCGGAA |
| 37944_3 | TATGAGTGTGGAGTCGTTGCTATAGTTCTGGAGGCTGGGAAGTNNNAGATCGGAA |
| 37944_4 | TATGAGTGTGGAGTCGTTGCTATAGTTCTGGAGGCTGGGAAGTNNNAGATCGGAA |
| 12512_1 | TATGAGTGTGGAGTCGTTGCTTACTTGTTTGTCCAAAGTTATTNNNAGATCGGAAGA |
| 12512_2 | TATGAGTGTGGAGTCGTTGCTTACTTGTTTGTCCAAAGTTATTNNNAGATCGGAAGA |
| 12512_3 | TATGAGTGTGGAGTCGTTGCTTACTTGTTTGTCCAAAGTTATTNNNAGATCGGAAGA |
| 12512_4 | TATGAGTGTGGAGTCGTTGCTTACTTGTTTGTCCAAAGTTATTNNNAGATCGGAAGA |
| 36375_1 | TATGAGTGTGGAGTCGTTGCAGGCACGAGAATCTCTTCAACTTNNNAGATCGGAAG |
| 36375_2 | TATGAGTGTGGAGTCGTTGCAGGCACGAGAATCTCTTCAACTTNNNAGATCGGAAG |
| 36375_3 | TATGAGTGTGGAGTCGTTGCAGGCACGAGAATCTCTTCAACTTNNNAGATCGGAAG |
| 36375_4 | TATGAGTGTGGAGTCGTTGCAGGCACGAGAATCTCTTCAACTTNNNAGATCGGAAG |
| 41584_1 | TATGAGTGTGGAGTCGTTGCATTCTCCTACCTCAACCTCCTGANNNAGATCGGAAGA |
| 41584_2 | TATGAGTGTGGAGTCGTTGCATTCTCCTACCTCAACCTCCTGANNNAGATCGGAAGA |
| 41584_3 | TATGAGTGTGGAGTCGTTGCATTCTCCTACCTCAACCTCCTGANNNAGATCGGAAGA |
| 41584_4 | TATGAGTGTGGAGTCGTTGCATTCTCCTACCTCAACCTCCTGANNNAGATCGGAAGA |
| 36836_1 | TATGAGTGTGGAGTCGTTGCGGCAAGAGAGAATTACCACCTGANNNAGATCGGAA |
| 36836_2 | TATGAGTGTGGAGTCGTTGCGGCAAGAGAGAATTACCACCTGANNNAGATCGGAA |
| 36836_3 | TATGAGTGTGGAGTCGTTGCGGCAAGAGAGAATTACCACCTGANNNAGATCGGAA |
| 36836_4 | TATGAGTGTGGAGTCGTTGCGGCAAGAGAGAATTACCACCTGANNNAGATCGGAA |
| 38899_1 | TATGAGTGTGGAGTCGTTGCACTGTTCGTGGGACTGTAAATTANNNAGATCGGAAG |
| 38899_2 | TATGAGTGTGGAGTCGTTGCACTGTTCGTGGGACTGTAAATTANNNAGATCGGAAG |
| 38899_3 | TATGAGTGTGGAGTCGTTGCACTGTTCGTGGGACTGTAAATTANNNAGATCGGAAG |
| 38899_4 | TATGAGTGTGGAGTCGTTGCACTGTTCGTGGGACTGTAAATTANNNAGATCGGAAG |
| 47537_1 | TATGAGTGTGGAGTCGTTGCGAACTCCCGAAGTCAAGTGATCTNNNAGATCGGAAG |
| 47537_2 | TATGAGTGTGGAGTCGTTGCGAACTCCCGAAGTCAAGTGATCTNNNAGATCGGAAG |
| 47537_3 | TATGAGTGTGGAGTCGTTGCGAACTCCCGAAGTCAAGTGATCTNNNAGATCGGAAG |
| 47537_4 | TATGAGTGTGGAGTCGTTGCGAACTCCCGAAGTCAAGTGATCTNNNAGATCGGAAG |
| 52904_1 | TATGAGTGTGGAGTCGTTGCAGAAAAGCTATGGGAGAAAGCCTNNNAGATCGGAA |

| | |
|---|---|
| 52904_2 | TATGAGTGTGGAGTCGTTGCAGAAAAGCTATGGGAGAAAGCCTNNNAGATCGGAA |
| 52904_3 | TATGAGTGTGGAGTCGTTGCAGAAAAGCTATGGGAGAAAGCCTNNNAGATCGGAA |
| 52904_4 | TATGAGTGTGGAGTCGTTGCAGAAAAGCTATGGGAGAAAGCCTNNNAGATCGGAA |
| 29826_1 | TATGAGTGTGGAGTCGTTGCCATTCAGGCACAGTCAAAAGGTTNNNAGATCGGAAG |
| 29826_2 | TATGAGTGTGGAGTCGTTGCCATTCAGGCACAGTCAAAAGGTTNNNAGATCGGAAG |
| 29826_3 | TATGAGTGTGGAGTCGTTGCCATTCAGGCACAGTCAAAAGGTTNNNAGATCGGAAG |
| 29826_4 | TATGAGTGTGGAGTCGTTGCCATTCAGGCACAGTCAAAAGGTTNNNAGATCGGAAG |
| 12994_1 | TATGAGTGTGGAGTCGTTGCGATAGGACTTGAACAGAAGATTNNNAGATCGGAAG |
| 12994_2 | TATGAGTGTGGAGTCGTTGCGATAGGACTTGAACAGAAGATTNNNAGATCGGAAG |
| 12994_3 | TATGAGTGTGGAGTCGTTGCGATAGGACTTGAACAGAAGATTNNNAGATCGGAAG |
| 12994_4 | TATGAGTGTGGAGTCGTTGCGATAGGACTTGAACAGAAGATTNNNAGATCGGAAG |
| 33492_1 | TATGAGTGTGGAGTCGTTGCGCTCTGTGGGACTAGTTTAGGAGNNNAGATCGGAA |
| 33492_2 | TATGAGTGTGGAGTCGTTGCGCTCTGTGGGACTAGTTTAGGAGNNNAGATCGGAA |
| 33492_3 | TATGAGTGTGGAGTCGTTGCGCTCTGTGGGACTAGTTTAGGAGNNNAGATCGGAA |
| 33492_4 | TATGAGTGTGGAGTCGTTGCGCTCTGTGGGACTAGTTTAGGAGNNNAGATCGGAA |
| 40326_1 | TATGAGTGTGGAGTCGTTGCACTCTCACACACTGTTGGAGAAANNNAGATCGGAAG |
| 40326_2 | TATGAGTGTGGAGTCGTTGCACTCTCACACACTGTTGGAGAAANNNAGATCGGAAG |
| 40326_3 | TATGAGTGTGGAGTCGTTGCACTCTCACACACTGTTGGAGAAANNNAGATCGGAAG |
| 40326_4 | TATGAGTGTGGAGTCGTTGCACTCTCACACACTGTTGGAGAAANNNAGATCGGAAG |
| 35681_1 | TATGAGTGTGGAGTCGTTGCTCCTGAGTAGCTGGGATTACAGTNNNAGATCGGAA |
| 35681_2 | TATGAGTGTGGAGTCGTTGCTCCTGAGTAGCTGGGATTACAGTNNNAGATCGGAA |
| 35681_3 | TATGAGTGTGGAGTCGTTGCTCCTGAGTAGCTGGGATTACAGTNNNAGATCGGAA |
| 35681_4 | TATGAGTGTGGAGTCGTTGCTCCTGAGTAGCTGGGATTACAGTNNNAGATCGGAA |
| 41460_1 | TATGAGTGTGGAGTCGTTGCCCAGCAGGACAGCAATTACATTTNNNAGATCGGAAG |
| 41460_2 | TATGAGTGTGGAGTCGTTGCCCAGCAGGACAGCAATTACATTTNNNAGATCGGAAG |
| 41460_3 | TATGAGTGTGGAGTCGTTGCCCAGCAGGACAGCAATTACATTTNNNAGATCGGAAG |
| 41460_4 | TATGAGTGTGGAGTCGTTGCCCAGCAGGACAGCAATTACATTTNNNAGATCGGAAG |
| 30267_1 | TATGAGTGTGGAGTCGTTGCTGGAGACCAGACAAATCGGATACNNNAGATCGGAA |
| 30267_2 | TATGAGTGTGGAGTCGTTGCTGGAGACCAGACAAATCGGATACNNNAGATCGGAA |
| 30267_3 | TATGAGTGTGGAGTCGTTGCTGGAGACCAGACAAATCGGATACNNNAGATCGGAA |
| 30267_4 | TATGAGTGTGGAGTCGTTGCTGGAGACCAGACAAATCGGATACNNNAGATCGGAA |
| 37582_1 | TATGAGTGTGGAGTCGTTGCATTACTCCAACTCCAAGAGGTGCNNNAGATCGGAAG |
| 37582_2 | TATGAGTGTGGAGTCGTTGCATTACTCCAACTCCAAGAGGTGCNNNAGATCGGAAG |
| 37582_3 | TATGAGTGTGGAGTCGTTGCATTACTCCAACTCCAAGAGGTGCNNNAGATCGGAAG |
| 37582_4 | TATGAGTGTGGAGTCGTTGCATTACTCCAACTCCAAGAGGTGCNNNAGATCGGAAG |
| 49526_1 | TATGAGTGTGGAGTCGTTGCCACTCTGTTGTGGACCAGTATGTNNNAGATCGGAAG |
| 49526_2 | TATGAGTGTGGAGTCGTTGCCACTCTGTTGTGGACCAGTATGTNNNAGATCGGAAG |
| 49526_3 | TATGAGTGTGGAGTCGTTGCCACTCTGTTGTGGACCAGTATGTNNNAGATCGGAAG |
| 49526_4 | TATGAGTGTGGAGTCGTTGCCACTCTGTTGTGGACCAGTATGTNNNAGATCGGAAG |
| 30767_1 | TATGAGTGTGGAGTCGTTGCTCATTGAGATGGATTGCTGGTGANNNAGATCGGAA |
| 30767_2 | TATGAGTGTGGAGTCGTTGCTCATTGAGATGGATTGCTGGTGANNNAGATCGGAA |
| 30767_3 | TATGAGTGTGGAGTCGTTGCTCATTGAGATGGATTGCTGGTGANNNAGATCGGAA |
| 30767_4 | TATGAGTGTGGAGTCGTTGCTCATTGAGATGGATTGCTGGTGANNNAGATCGGAA |

| | |
|---|---|
| 47577_1 | TATGAGTGTGGAGTCGTTGCTGGAGTGCAATGACACAATCTTGNNNAGATCGGAA |
| 47577_2 | TATGAGTGTGGAGTCGTTGCTGGAGTGCAATGACACAATCTTGNNNAGATCGGAA |
| 47577_3 | TATGAGTGTGGAGTCGTTGCTGGAGTGCAATGACACAATCTTGNNNAGATCGGAA |
| 47577_4 | TATGAGTGTGGAGTCGTTGCTGGAGTGCAATGACACAATCTTGNNNAGATCGGAA |
| 48382_1 | TATGAGTGTGGAGTCGTTGCAGGTTCAAGCAATTATCCCGTCTNNNAGATCGGAAG |
| 48382_2 | TATGAGTGTGGAGTCGTTGCAGGTTCAAGCAATTATCCCGTCTNNNAGATCGGAAG |
| 48382_3 | TATGAGTGTGGAGTCGTTGCAGGTTCAAGCAATTATCCCGTCTNNNAGATCGGAAG |
| 48382_4 | TATGAGTGTGGAGTCGTTGCAGGTTCAAGCAATTATCCCGTCTNNNAGATCGGAAG |
| 48143_1 | TATGAGTGTGGAGTCGTTGCGGTTCAAGCAATTCTCCTGTCTCNNNAGATCGGAAG |
| 48143_2 | TATGAGTGTGGAGTCGTTGCGGTTCAAGCAATTCTCCTGTCTCNNNAGATCGGAAG |
| 48143_3 | TATGAGTGTGGAGTCGTTGCGGTTCAAGCAATTCTCCTGTCTCNNNAGATCGGAAG |
| 48143_4 | TATGAGTGTGGAGTCGTTGCGGTTCAAGCAATTCTCCTGTCTCNNNAGATCGGAAG |
| 32127_1 | TATGAGTGTGGAGTCGTTGCACGATCTCGGCTTACTGCAAGNNNAGATCGGAAGA |
| 32127_2 | TATGAGTGTGGAGTCGTTGCACGATCTCGGCTTACTGCAAGNNNAGATCGGAAGA |
| 32127_3 | TATGAGTGTGGAGTCGTTGCACGATCTCGGCTTACTGCAAGNNNAGATCGGAAGA |
| 32127_4 | TATGAGTGTGGAGTCGTTGCACGATCTCGGCTTACTGCAAGNNNAGATCGGAAGA |
| 45089_1 | TATGAGTGTGGAGTCGTTGCTGTAGCATATAGGAAGCAATGCCTNNNAGATCGGA |
| 45089_2 | TATGAGTGTGGAGTCGTTGCTGTAGCATATAGGAAGCAATGCCTNNNAGATCGGA |
| 45089_3 | TATGAGTGTGGAGTCGTTGCTGTAGCATATAGGAAGCAATGCCTNNNAGATCGGA |
| 45089_4 | TATGAGTGTGGAGTCGTTGCTGTAGCATATAGGAAGCAATGCCTNNNAGATCGGA |
| 41415_1 | TATGAGTGTGGAGTCGTTGCTCAGTCATTTGGTAAGCAGACCANNNAGATCGGAAG |
| 41415_2 | TATGAGTGTGGAGTCGTTGCTCAGTCATTTGGTAAGCAGACCANNNAGATCGGAAG |
| 41415_3 | TATGAGTGTGGAGTCGTTGCTCAGTCATTTGGTAAGCAGACCANNNAGATCGGAAG |
| 41415_4 | TATGAGTGTGGAGTCGTTGCTCAGTCATTTGGTAAGCAGACCANNNAGATCGGAAG |
| 46449_1 | TATGAGTGTGGAGTCGTTGCGAAAGCCAGCAAGACAGAACAATNNNAGATCGGAA |
| 46449_2 | TATGAGTGTGGAGTCGTTGCGAAAGCCAGCAAGACAGAACAATNNNAGATCGGAA |
| 46449_3 | TATGAGTGTGGAGTCGTTGCGAAAGCCAGCAAGACAGAACAATNNNAGATCGGAA |
| 46449_4 | TATGAGTGTGGAGTCGTTGCGAAAGCCAGCAAGACAGAACAATNNNAGATCGGAA |
| 32991_1 | TATGAGTGTGGAGTCGTTGCTTCACACCATTCTCCTGCCTCNNNAGATCGGAAGAGC |
| 32991_2 | TATGAGTGTGGAGTCGTTGCTTCACACCATTCTCCTGCCTCNNNAGATCGGAAGAGC |
| 32991_3 | TATGAGTGTGGAGTCGTTGCTTCACACCATTCTCCTGCCTCNNNAGATCGGAAGAGC |
| 32991_4 | TATGAGTGTGGAGTCGTTGCTTCACACCATTCTCCTGCCTCNNNAGATCGGAAGAGC |
| 51252_1 | TATGAGTGTGGAGTCGTTGCGGAGGGATGTAAGTTCATCACCANNNAGATCGGAA |
| 51252_2 | TATGAGTGTGGAGTCGTTGCGGAGGGATGTAAGTTCATCACCANNNAGATCGGAA |
| 51252_3 | TATGAGTGTGGAGTCGTTGCGGAGGGATGTAAGTTCATCACCANNNAGATCGGAA |
| 51252_4 | TATGAGTGTGGAGTCGTTGCGGAGGGATGTAAGTTCATCACCANNNAGATCGGAA |
| 47292_1 | TATGAGTGTGGAGTCGTTGCGACAGAATCTCACTCTATCGCCCNNNAGATCGGAAG |
| 47292_2 | TATGAGTGTGGAGTCGTTGCGACAGAATCTCACTCTATCGCCCNNNAGATCGGAAG |
| 47292_3 | TATGAGTGTGGAGTCGTTGCGACAGAATCTCACTCTATCGCCCNNNAGATCGGAAG |
| 47292_4 | TATGAGTGTGGAGTCGTTGCGACAGAATCTCACTCTATCGCCCNNNAGATCGGAAG |
| 31426_1 | TATGAGTGTGGAGTCGTTGCCCAGGACTCAGATTTTTTTTCCCANNNAGATCGGAAG |
| 31426_2 | TATGAGTGTGGAGTCGTTGCCCAGGACTCAGATTTTTTTTCCCANNNAGATCGGAAG |
| 31426_3 | TATGAGTGTGGAGTCGTTGCCCAGGACTCAGATTTTTTTTCCCANNNAGATCGGAAG |

| | |
|---|---|
| 31426_4 | TATGAGTGTGGAGTCGTTGCCCAGGACTCAGATTTTTTTCCCANNNAGATCGGAAG |
| 42676_1 | TATGAGTGTGGAGTCGTTGCTTGGCTTGGGTCAAATAGCTCTANNNAGATCGGAAG |
| 42676_2 | TATGAGTGTGGAGTCGTTGCTTGGCTTGGGTCAAATAGCTCTANNNAGATCGGAAG |
| 42676_3 | TATGAGTGTGGAGTCGTTGCTTGGCTTGGGTCAAATAGCTCTANNNAGATCGGAAG |
| 42676_4 | TATGAGTGTGGAGTCGTTGCTTGGCTTGGGTCAAATAGCTCTANNNAGATCGGAAG |
| 42591_1 | TATGAGTGTGGAGTCGTTGCACCACCATCTCTCTCTAGTTCCANNNAGATCGGAAGA |
| 42591_2 | TATGAGTGTGGAGTCGTTGCACCACCATCTCTCTCTAGTTCCANNNAGATCGGAAGA |
| 42591_3 | TATGAGTGTGGAGTCGTTGCACCACCATCTCTCTCTAGTTCCANNNAGATCGGAAGA |
| 42591_4 | TATGAGTGTGGAGTCGTTGCACCACCATCTCTCTCTAGTTCCANNNAGATCGGAAGA |
| 31261_1 | TATGAGTGTGGAGTCGTTGCCCTATCCCACTTTGGACACTTGANNNAGATCGGAAGA |
| 31261_2 | TATGAGTGTGGAGTCGTTGCCCTATCCCACTTTGGACACTTGANNNAGATCGGAAGA |
| 31261_3 | TATGAGTGTGGAGTCGTTGCCCTATCCCACTTTGGACACTTGANNNAGATCGGAAGA |
| 31261_4 | TATGAGTGTGGAGTCGTTGCCCTATCCCACTTTGGACACTTGANNNAGATCGGAAGA |
| 32975_1 | TATGAGTGTGGAGTCGTTGCGAAAATCACCACTTTGCGACCTTNNNAGATCGGAAGA |
| 32975_2 | TATGAGTGTGGAGTCGTTGCGAAAATCACCACTTTGCGACCTTNNNAGATCGGAAGA |
| 32975_3 | TATGAGTGTGGAGTCGTTGCGAAAATCACCACTTTGCGACCTTNNNAGATCGGAAGA |
| 32975_4 | TATGAGTGTGGAGTCGTTGCGAAAATCACCACTTTGCGACCTTNNNAGATCGGAAGA |
| 13436_1 | TATGAGTGTGGAGTCGTTGCATAGATATCTGAAGTGTGTGAAGNNNAGATCGGAA |
| 13436_2 | TATGAGTGTGGAGTCGTTGCATAGATATCTGAAGTGTGTGAAGNNNAGATCGGAA |
| 13436_3 | TATGAGTGTGGAGTCGTTGCATAGATATCTGAAGTGTGTGAAGNNNAGATCGGAA |
| 13436_4 | TATGAGTGTGGAGTCGTTGCATAGATATCTGAAGTGTGTGAAGNNNAGATCGGAA |
| 51678_1 | TATGAGTGTGGAGTCGTTGCGAAACCACCACTACCATTTGCATNNNAGATCGGAAGA |
| 51678_2 | TATGAGTGTGGAGTCGTTGCGAAACCACCACTACCATTTGCATNNNAGATCGGAAGA |
| 51678_3 | TATGAGTGTGGAGTCGTTGCGAAACCACCACTACCATTTGCATNNNAGATCGGAAGA |
| 51678_4 | TATGAGTGTGGAGTCGTTGCGAAACCACCACTACCATTTGCATNNNAGATCGGAAGA |
| 52301_1 | TATGAGTGTGGAGTCGTTGCCCAAGCCAACAGTAATCCCGAGNNNAGATCGGAAGA |
| 52301_2 | TATGAGTGTGGAGTCGTTGCCCAAGCCAACAGTAATCCCGAGNNNAGATCGGAAGA |
| 52301_3 | TATGAGTGTGGAGTCGTTGCCCAAGCCAACAGTAATCCCGAGNNNAGATCGGAAGA |
| 52301_4 | TATGAGTGTGGAGTCGTTGCCCAAGCCAACAGTAATCCCGAGNNNAGATCGGAAGA |
| 41215_1 | TATGAGTGTGGAGTCGTTGCCAAGCACTTGGAGGGAAGAAAAGNNNAGATCGGAA |
| 41215_2 | TATGAGTGTGGAGTCGTTGCCAAGCACTTGGAGGGAAGAAAAGNNNAGATCGGAA |
| 41215_3 | TATGAGTGTGGAGTCGTTGCCAAGCACTTGGAGGGAAGAAAAGNNNAGATCGGAA |
| 41215_4 | TATGAGTGTGGAGTCGTTGCCAAGCACTTGGAGGGAAGAAAAGNNNAGATCGGAA |
| 50535_1 | TATGAGTGTGGAGTCGTTGCGCCTAGAAACACTTCTTCCCTGANNNAGATCGGAAGA |
| 50535_2 | TATGAGTGTGGAGTCGTTGCGCCTAGAAACACTTCTTCCCTGANNNAGATCGGAAGA |
| 50535_3 | TATGAGTGTGGAGTCGTTGCGCCTAGAAACACTTCTTCCCTGANNNAGATCGGAAGA |
| 50535_4 | TATGAGTGTGGAGTCGTTGCGCCTAGAAACACTTCTTCCCTGANNNAGATCGGAAGA |
| 51609_1 | TATGAGTGTGGAGTCGTTGCCCAAGATCTGTCCATTCTCCTGGNNNAGATCGGAAGA |
| 51609_2 | TATGAGTGTGGAGTCGTTGCCCAAGATCTGTCCATTCTCCTGGNNNAGATCGGAAGA |
| 51609_3 | TATGAGTGTGGAGTCGTTGCCCAAGATCTGTCCATTCTCCTGGNNNAGATCGGAAGA |
| 51609_4 | TATGAGTGTGGAGTCGTTGCCCAAGATCTGTCCATTCTCCTGGNNNAGATCGGAAGA |
| 12781_1 | TATGAGTGTGGAGTCGTTGCCATATCTTCAAACATGTCATCAANNNAGATCGGAAGA |
| 12781_2 | TATGAGTGTGGAGTCGTTGCCATATCTTCAAACATGTCATCAANNNAGATCGGAAGA |

| | |
|---|---|
| 12781_3 | TATGAGTGTGGAGTCGTTGCCATATCTTCAAACATGTCATCAANNNAGATCGGAAG |
| 12781_4 | TATGAGTGTGGAGTCGTTGCCATATCTTCAAACATGTCATCAANNNAGATCGGAAG |
| 36262_1 | TATGAGTGTGGAGTCGTTGCGCACATGCCTGTAATCCTAGCTANNNAGATCGGAAG |
| 36262_2 | TATGAGTGTGGAGTCGTTGCGCACATGCCTGTAATCCTAGCTANNNAGATCGGAAG |
| 36262_3 | TATGAGTGTGGAGTCGTTGCGCACATGCCTGTAATCCTAGCTANNNAGATCGGAAG |
| 36262_4 | TATGAGTGTGGAGTCGTTGCGCACATGCCTGTAATCCTAGCTANNNAGATCGGAAG |
| 51442_1 | TATGAGTGTGGAGTCGTTGCTTTGCAACTATGTCTGGCATTCCNNNAGATCGGAAG |
| 51442_2 | TATGAGTGTGGAGTCGTTGCTTTGCAACTATGTCTGGCATTCCNNNAGATCGGAAG |
| 51442_3 | TATGAGTGTGGAGTCGTTGCTTTGCAACTATGTCTGGCATTCCNNNAGATCGGAAG |
| 51442_4 | TATGAGTGTGGAGTCGTTGCTTTGCAACTATGTCTGGCATTCCNNNAGATCGGAAG |
| 10387_1 | TATGAGTGTGGAGTCGTTGCGTGGGAGACTTTAGCATGCTCTTNNNAGATCGGAA |
| 10387_2 | TATGAGTGTGGAGTCGTTGCGTGGGAGACTTTAGCATGCTCTTNNNAGATCGGAA |
| 10387_3 | TATGAGTGTGGAGTCGTTGCGTGGGAGACTTTAGCATGCTCTTNNNAGATCGGAA |
| 10387_4 | TATGAGTGTGGAGTCGTTGCGTGGGAGACTTTAGCATGCTCTTNNNAGATCGGAA |
| 44285_1 | TATGAGTGTGGAGTCGTTGCTTTCCTAGTCCTCCCCATTCCTCNNNAGATCGGAAGA |
| 44285_2 | TATGAGTGTGGAGTCGTTGCTTTCCTAGTCCTCCCCATTCCTCNNNAGATCGGAAGA |
| 44285_3 | TATGAGTGTGGAGTCGTTGCTTTCCTAGTCCTCCCCATTCCTCNNNAGATCGGAAGA |
| 44285_4 | TATGAGTGTGGAGTCGTTGCTTTCCTAGTCCTCCCCATTCCTCNNNAGATCGGAAGA |
| 41287_1 | TATGAGTGTGGAGTCGTTGCCTGTTCAGAGAGGCCAAGAAGAGNNNAGATCGGAA |
| 41287_2 | TATGAGTGTGGAGTCGTTGCCTGTTCAGAGAGGCCAAGAAGAGNNNAGATCGGAA |
| 41287_3 | TATGAGTGTGGAGTCGTTGCCTGTTCAGAGAGGCCAAGAAGAGNNNAGATCGGAA |
| 41287_4 | TATGAGTGTGGAGTCGTTGCCTGTTCAGAGAGGCCAAGAAGAGNNNAGATCGGAA |
| 9236_1 | TATGAGTGTGGAGTCGTTGCGAGAGAGCGACAGAGATTGAGTGNNNAGATCGGA |
| 9236_2 | TATGAGTGTGGAGTCGTTGCGAGAGAGCGACAGAGATTGAGTGNNNAGATCGGA |
| 9236_3 | TATGAGTGTGGAGTCGTTGCGAGAGAGCGACAGAGATTGAGTGNNNAGATCGGA |
| 9236_4 | TATGAGTGTGGAGTCGTTGCGAGAGAGCGACAGAGATTGAGTGNNNAGATCGGA |
| 36714_1 | TATGAGTGTGGAGTCGTTGCTCCTGAGATGAATACGTTTCCCGNNNAGATCGGAAG |
| 36714_2 | TATGAGTGTGGAGTCGTTGCTCCTGAGATGAATACGTTTCCCGNNNAGATCGGAAG |
| 36714_3 | TATGAGTGTGGAGTCGTTGCTCCTGAGATGAATACGTTTCCCGNNNAGATCGGAAG |
| 36714_4 | TATGAGTGTGGAGTCGTTGCTCCTGAGATGAATACGTTTCCCGNNNAGATCGGAAG |
| 34765_1 | TATGAGTGTGGAGTCGTTGCCCGTGGTTCATAAGCAAATCCTCNNNAGATCGGAAG |
| 34765_2 | TATGAGTGTGGAGTCGTTGCCCGTGGTTCATAAGCAAATCCTCNNNAGATCGGAAG |
| 34765_3 | TATGAGTGTGGAGTCGTTGCCCGTGGTTCATAAGCAAATCCTCNNNAGATCGGAAG |
| 34765_4 | TATGAGTGTGGAGTCGTTGCCCGTGGTTCATAAGCAAATCCTCNNNAGATCGGAAG |
| 40134_1 | TATGAGTGTGGAGTCGTTGCATAAAACAAGAGGACCTGTGCCANNNAGATCGGAA |
| 40134_2 | TATGAGTGTGGAGTCGTTGCATAAAACAAGAGGACCTGTGCCANNNAGATCGGAA |
| 40134_3 | TATGAGTGTGGAGTCGTTGCATAAAACAAGAGGACCTGTGCCANNNAGATCGGAA |
| 40134_4 | TATGAGTGTGGAGTCGTTGCATAAAACAAGAGGACCTGTGCCANNNAGATCGGAA |
| 50546_1 | TATGAGTGTGGAGTCGTTGCCTCTGGAGCTCTAGAATCTGTGCNNNAGATCGGAAG |
| 50546_2 | TATGAGTGTGGAGTCGTTGCCTCTGGAGCTCTAGAATCTGTGCNNNAGATCGGAAG |
| 50546_3 | TATGAGTGTGGAGTCGTTGCCTCTGGAGCTCTAGAATCTGTGCNNNAGATCGGAAG |
| 50546_4 | TATGAGTGTGGAGTCGTTGCCTCTGGAGCTCTAGAATCTGTGCNNNAGATCGGAAG |
| 12978_1 | TATGAGTGTGGAGTCGTTGCAAATCTCAGCCGTTTAAAACTTNNNAGATCGGAAGA |

| | |
|---|---|
| 12978_2 | TATGAGTGTGGAGTCGTTGCAAATCTCAGCCGTTTAAAACTTNNNAGATCGGAAGA |
| 12978_3 | TATGAGTGTGGAGTCGTTGCAAATCTCAGCCGTTTAAAACTTNNNAGATCGGAAGA |
| 12978_4 | TATGAGTGTGGAGTCGTTGCAAATCTCAGCCGTTTAAAACTTNNNAGATCGGAAGA |
| 44056_1 | TATGAGTGTGGAGTCGTTGCAAAGGGCTTGGAGATCAGAAAGTNNNAGATCGGAA |
| 44056_2 | TATGAGTGTGGAGTCGTTGCAAAGGGCTTGGAGATCAGAAAGTNNNAGATCGGAA |
| 44056_3 | TATGAGTGTGGAGTCGTTGCAAAGGGCTTGGAGATCAGAAAGTNNNAGATCGGAA |
| 44056_4 | TATGAGTGTGGAGTCGTTGCAAAGGGCTTGGAGATCAGAAAGTNNNAGATCGGAA |
| 44041_1 | TATGAGTGTGGAGTCGTTGCATCTGCAAAGCTCTACAAGGTCANNNAGATCGGAAG |
| 44041_2 | TATGAGTGTGGAGTCGTTGCATCTGCAAAGCTCTACAAGGTCANNNAGATCGGAAG |
| 44041_3 | TATGAGTGTGGAGTCGTTGCATCTGCAAAGCTCTACAAGGTCANNNAGATCGGAAG |
| 44041_4 | TATGAGTGTGGAGTCGTTGCATCTGCAAAGCTCTACAAGGTCANNNAGATCGGAAG |
| 38435_1 | TATGAGTGTGGAGTCGTTGCTTGGAGTAAGCTCACTACAACCTNNNAGATCGGAAG |
| 38435_2 | TATGAGTGTGGAGTCGTTGCTTGGAGTAAGCTCACTACAACCTNNNAGATCGGAAG |
| 38435_3 | TATGAGTGTGGAGTCGTTGCTTGGAGTAAGCTCACTACAACCTNNNAGATCGGAAG |
| 38435_4 | TATGAGTGTGGAGTCGTTGCTTGGAGTAAGCTCACTACAACCTNNNAGATCGGAAG |
| 39857_1 | TATGAGTGTGGAGTCGTTGCGCAAACATGGACAAATGGGATCANNNAGATCGGAA |
| 39857_2 | TATGAGTGTGGAGTCGTTGCGCAAACATGGACAAATGGGATCANNNAGATCGGAA |
| 39857_3 | TATGAGTGTGGAGTCGTTGCGCAAACATGGACAAATGGGATCANNNAGATCGGAA |
| 39857_4 | TATGAGTGTGGAGTCGTTGCGCAAACATGGACAAATGGGATCANNNAGATCGGAA |
| 35372_1 | TATGAGTGTGGAGTCGTTGCAAGTGCTGAGATTACAGGCATGANNNAGATCGGAA |
| 35372_2 | TATGAGTGTGGAGTCGTTGCAAGTGCTGAGATTACAGGCATGANNNAGATCGGAA |
| 35372_3 | TATGAGTGTGGAGTCGTTGCAAGTGCTGAGATTACAGGCATGANNNAGATCGGAA |
| 35372_4 | TATGAGTGTGGAGTCGTTGCAAGTGCTGAGATTACAGGCATGANNNAGATCGGAA |
| 53119_1 | TATGAGTGTGGAGTCGTTGCGCATGCTTATTTTGCTGACATCCNNNAGATCGGAAG |
| 53119_2 | TATGAGTGTGGAGTCGTTGCGCATGCTTATTTTGCTGACATCCNNNAGATCGGAAG |
| 53119_3 | TATGAGTGTGGAGTCGTTGCGCATGCTTATTTTGCTGACATCCNNNAGATCGGAAG |
| 53119_4 | TATGAGTGTGGAGTCGTTGCGCATGCTTATTTTGCTGACATCCNNNAGATCGGAAG |
| 40574_1 | TATGAGTGTGGAGTCGTTGCAGGAGCAACTGGACTATGTGATCNNNAGATCGGAA |
| 40574_2 | TATGAGTGTGGAGTCGTTGCAGGAGCAACTGGACTATGTGATCNNNAGATCGGAA |
| 40574_3 | TATGAGTGTGGAGTCGTTGCAGGAGCAACTGGACTATGTGATCNNNAGATCGGAA |
| 40574_4 | TATGAGTGTGGAGTCGTTGCAGGAGCAACTGGACTATGTGATCNNNAGATCGGAA |
| 39575_1 | TATGAGTGTGGAGTCGTTGCATACACTGTCCCAAGAATGACCCNNNAGATCGGAAG |
| 39575_2 | TATGAGTGTGGAGTCGTTGCATACACTGTCCCAAGAATGACCCNNNAGATCGGAAG |
| 39575_3 | TATGAGTGTGGAGTCGTTGCATACACTGTCCCAAGAATGACCCNNNAGATCGGAAG |
| 39575_4 | TATGAGTGTGGAGTCGTTGCATACACTGTCCCAAGAATGACCCNNNAGATCGGAAG |
| 49627_1 | TATGAGTGTGGAGTCGTTGCAAATTACTTTGGGCAGTATGGCCNNNAGATCGGAA |
| 49627_2 | TATGAGTGTGGAGTCGTTGCAAATTACTTTGGGCAGTATGGCCNNNAGATCGGAA |
| 49627_3 | TATGAGTGTGGAGTCGTTGCAAATTACTTTGGGCAGTATGGCCNNNAGATCGGAA |
| 49627_4 | TATGAGTGTGGAGTCGTTGCAAATTACTTTGGGCAGTATGGCCNNNAGATCGGAA |
| 42509_1 | TATGAGTGTGGAGTCGTTGCGGTGAGTCTGTGGTCATTTTGTGNNNAGATCGGAA |
| 42509_2 | TATGAGTGTGGAGTCGTTGCGGTGAGTCTGTGGTCATTTTGTGNNNAGATCGGAA |
| 42509_3 | TATGAGTGTGGAGTCGTTGCGGTGAGTCTGTGGTCATTTTGTGNNNAGATCGGAA |
| 42509_4 | TATGAGTGTGGAGTCGTTGCGGTGAGTCTGTGGTCATTTTGTGNNNAGATCGGAA |

| | |
|---|---|
| 37958_1 | TATGAGTGTGGAGTCGTTGCAAAAGAAAAGAAAAGGCTGGGCANNNAGATCGGA |
| 37958_2 | TATGAGTGTGGAGTCGTTGCAAAAGAAAAGAAAAGGCTGGGCANNNAGATCGGA |
| 37958_3 | TATGAGTGTGGAGTCGTTGCAAAAGAAAAGAAAAGGCTGGGCANNNAGATCGGA |
| 37958_4 | TATGAGTGTGGAGTCGTTGCAAAAGAAAAGAAAAGGCTGGGCANNNAGATCGGA |
| 44640_1 | TATGAGTGTGGAGTCGTTGCCTCCACTGCAGTGAACAAATCAGNNNAGATCGGAAG |
| 44640_2 | TATGAGTGTGGAGTCGTTGCCTCCACTGCAGTGAACAAATCAGNNNAGATCGGAAG |
| 44640_3 | TATGAGTGTGGAGTCGTTGCCTCCACTGCAGTGAACAAATCAGNNNAGATCGGAAG |
| 44640_4 | TATGAGTGTGGAGTCGTTGCCTCCACTGCAGTGAACAAATCAGNNNAGATCGGAAG |
| 35257_1 | TATGAGTGTGGAGTCGTTGCATACAATCTCTGCCACCCATGTTNNNAGATCGGAAGA |
| 35257_2 | TATGAGTGTGGAGTCGTTGCATACAATCTCTGCCACCCATGTTNNNAGATCGGAAGA |
| 35257_3 | TATGAGTGTGGAGTCGTTGCATACAATCTCTGCCACCCATGTTNNNAGATCGGAAGA |
| 35257_4 | TATGAGTGTGGAGTCGTTGCATACAATCTCTGCCACCCATGTTNNNAGATCGGAAGA |
| 45703_1 | TATGAGTGTGGAGTCGTTGCCTCAAGCAATTCTCATGCCTCAGNNNAGATCGGAAG |
| 45703_2 | TATGAGTGTGGAGTCGTTGCCTCAAGCAATTCTCATGCCTCAGNNNAGATCGGAAG |
| 45703_3 | TATGAGTGTGGAGTCGTTGCCTCAAGCAATTCTCATGCCTCAGNNNAGATCGGAAG |
| 45703_4 | TATGAGTGTGGAGTCGTTGCCTCAAGCAATTCTCATGCCTCAGNNNAGATCGGAAG |
| 36118_1 | TATGAGTGTGGAGTCGTTGCCACGTCTATGTAGCTCCAGAACANNNAGATCGGAAG |
| 36118_2 | TATGAGTGTGGAGTCGTTGCCACGTCTATGTAGCTCCAGAACANNNAGATCGGAAG |
| 36118_3 | TATGAGTGTGGAGTCGTTGCCACGTCTATGTAGCTCCAGAACANNNAGATCGGAAG |
| 36118_4 | TATGAGTGTGGAGTCGTTGCCACGTCTATGTAGCTCCAGAACANNNAGATCGGAAG |
| 52790_1 | TATGAGTGTGGAGTCGTTGCGCAGCTTCAAAGGGAGCAATTTTNNNAGATCGGAA |
| 52790_2 | TATGAGTGTGGAGTCGTTGCGCAGCTTCAAAGGGAGCAATTTTNNNAGATCGGAA |
| 52790_3 | TATGAGTGTGGAGTCGTTGCGCAGCTTCAAAGGGAGCAATTTTNNNAGATCGGAA |
| 52790_4 | TATGAGTGTGGAGTCGTTGCGCAGCTTCAAAGGGAGCAATTTTNNNAGATCGGAA |
| 40410_1 | TATGAGTGTGGAGTCGTTGCCAATTCTCCTATCTCAGCCTCCCNNNAGATCGGAAGA |
| 40410_2 | TATGAGTGTGGAGTCGTTGCCAATTCTCCTATCTCAGCCTCCCNNNAGATCGGAAGA |
| 40410_3 | TATGAGTGTGGAGTCGTTGCCAATTCTCCTATCTCAGCCTCCCNNNAGATCGGAAGA |
| 40410_4 | TATGAGTGTGGAGTCGTTGCCAATTCTCCTATCTCAGCCTCCCNNNAGATCGGAAGA |
| 40963_1 | TATGAGTGTGGAGTCGTTGCTGCTTTGTGATACCTTGTTCATGGNNNAGATCGGAA |
| 40963_2 | TATGAGTGTGGAGTCGTTGCTGCTTTGTGATACCTTGTTCATGGNNNAGATCGGAA |
| 40963_3 | TATGAGTGTGGAGTCGTTGCTGCTTTGTGATACCTTGTTCATGGNNNAGATCGGAA |
| 40963_4 | TATGAGTGTGGAGTCGTTGCTGCTTTGTGATACCTTGTTCATGGNNNAGATCGGAA |
| 38929_1 | TATGAGTGTGGAGTCGTTGCATGTGGAAGTATGAACTTGGCCTNNNAGATCGGAA |
| 38929_2 | TATGAGTGTGGAGTCGTTGCATGTGGAAGTATGAACTTGGCCTNNNAGATCGGAA |
| 38929_3 | TATGAGTGTGGAGTCGTTGCATGTGGAAGTATGAACTTGGCCTNNNAGATCGGAA |
| 38929_4 | TATGAGTGTGGAGTCGTTGCATGTGGAAGTATGAACTTGGCCTNNNAGATCGGAA |
| 51361_1 | TATGAGTGTGGAGTCGTTGCGGTGGTGAAACAACCCATTGTAGNNNAGATCGGAA |
| 51361_2 | TATGAGTGTGGAGTCGTTGCGGTGGTGAAACAACCCATTGTAGNNNAGATCGGAA |
| 51361_3 | TATGAGTGTGGAGTCGTTGCGGTGGTGAAACAACCCATTGTAGNNNAGATCGGAA |
| 51361_4 | TATGAGTGTGGAGTCGTTGCGGTGGTGAAACAACCCATTGTAGNNNAGATCGGAA |
| 45167_1 | TATGAGTGTGGAGTCGTTGCCGGGTCCTCTCTAGCGTTCCNNNAGATCGGAAGAGC |
| 45167_2 | TATGAGTGTGGAGTCGTTGCCGGGTCCTCTCTAGCGTTCCNNNAGATCGGAAGAGC |
| 45167_3 | TATGAGTGTGGAGTCGTTGCCGGGTCCTCTCTAGCGTTCCNNNAGATCGGAAGAGC |

| | |
|---|---|
| 45167_4 | TATGAGTGTGGAGTCGTTGCCGGGTCCTCTCTAGCGTTCCNNNAGATCGGAAGAGC |
| 46545_1 | TATGAGTGTGGAGTCGTTGCTTCAAGTGATTCTCCTGCCTCAGNNNAGATCGGAAG |
| 46545_2 | TATGAGTGTGGAGTCGTTGCTTCAAGTGATTCTCCTGCCTCAGNNNAGATCGGAAG |
| 46545_3 | TATGAGTGTGGAGTCGTTGCTTCAAGTGATTCTCCTGCCTCAGNNNAGATCGGAAG |
| 46545_4 | TATGAGTGTGGAGTCGTTGCTTCAAGTGATTCTCCTGCCTCAGNNNAGATCGGAAG |
| 3037_1 | TATGAGTGTGGAGTCGTTGCCGCATCTCTAAGGAGGAGTACCANNNAGATCGGAA |
| 3037_2 | TATGAGTGTGGAGTCGTTGCCGCATCTCTAAGGAGGAGTACCANNNAGATCGGAA |
| 3037_3 | TATGAGTGTGGAGTCGTTGCCGCATCTCTAAGGAGGAGTACCANNNAGATCGGAA |
| 3037_4 | TATGAGTGTGGAGTCGTTGCCGCATCTCTAAGGAGGAGTACCANNNAGATCGGAA |
| 33248_1 | TATGAGTGTGGAGTCGTTGCGAGATGGGGTCTTTCTCTGTCACNNNAGATCGGAAG |
| 33248_2 | TATGAGTGTGGAGTCGTTGCGAGATGGGGTCTTTCTCTGTCACNNNAGATCGGAAG |
| 33248_3 | TATGAGTGTGGAGTCGTTGCGAGATGGGGTCTTTCTCTGTCACNNNAGATCGGAAG |
| 33248_4 | TATGAGTGTGGAGTCGTTGCGAGATGGGGTCTTTCTCTGTCACNNNAGATCGGAAG |
| 36524_1 | TATGAGTGTGGAGTCGTTGCTGACATTCTCTAGGTACCTGCAGNNNAGATCGGAAG |
| 36524_2 | TATGAGTGTGGAGTCGTTGCTGACATTCTCTAGGTACCTGCAGNNNAGATCGGAAG |
| 36524_3 | TATGAGTGTGGAGTCGTTGCTGACATTCTCTAGGTACCTGCAGNNNAGATCGGAAG |
| 36524_4 | TATGAGTGTGGAGTCGTTGCTGACATTCTCTAGGTACCTGCAGNNNAGATCGGAAG |
| 49505_1 | TATGAGTGTGGAGTCGTTGCGAGACGGAGTCTTGCTCTGTCNNNAGATCGGAAGA |
| 49505_2 | TATGAGTGTGGAGTCGTTGCGAGACGGAGTCTTGCTCTGTCNNNAGATCGGAAGA |
| 49505_3 | TATGAGTGTGGAGTCGTTGCGAGACGGAGTCTTGCTCTGTCNNNAGATCGGAAGA |
| 49505_4 | TATGAGTGTGGAGTCGTTGCGAGACGGAGTCTTGCTCTGTCNNNAGATCGGAAGA |
| 35694_1 | TATGAGTGTGGAGTCGTTGCGGAGTCTTACTCTATTGCCCAGGNNNAGATCGGAAG |
| 35694_2 | TATGAGTGTGGAGTCGTTGCGGAGTCTTACTCTATTGCCCAGGNNNAGATCGGAAG |
| 35694_3 | TATGAGTGTGGAGTCGTTGCGGAGTCTTACTCTATTGCCCAGGNNNAGATCGGAAG |
| 35694_4 | TATGAGTGTGGAGTCGTTGCGGAGTCTTACTCTATTGCCCAGGNNNAGATCGGAAG |
| 48212_1 | TATGAGTGTGGAGTCGTTGCGTTCAAGCGATTCTCATGCCTCNNNAGATCGGAAGA |
| 48212_2 | TATGAGTGTGGAGTCGTTGCGTTCAAGCGATTCTCATGCCTCNNNAGATCGGAAGA |
| 48212_3 | TATGAGTGTGGAGTCGTTGCGTTCAAGCGATTCTCATGCCTCNNNAGATCGGAAGA |
| 48212_4 | TATGAGTGTGGAGTCGTTGCGTTCAAGCGATTCTCATGCCTCNNNAGATCGGAAGA |
| 36850_1 | TATGAGTGTGGAGTCGTTGCGTTGTAGTGAGCAGAGATTGTGCNNNAGATCGGAA |
| 36850_2 | TATGAGTGTGGAGTCGTTGCGTTGTAGTGAGCAGAGATTGTGCNNNAGATCGGAA |
| 36850_3 | TATGAGTGTGGAGTCGTTGCGTTGTAGTGAGCAGAGATTGTGCNNNAGATCGGAA |
| 36850_4 | TATGAGTGTGGAGTCGTTGCGTTGTAGTGAGCAGAGATTGTGCNNNAGATCGGAA |
| 50115_1 | TATGAGTGTGGAGTCGTTGCGCGTGCCAAGATGAATGCTAAATNNNAGATCGGAA |
| 50115_2 | TATGAGTGTGGAGTCGTTGCGCGTGCCAAGATGAATGCTAAATNNNAGATCGGAA |
| 50115_3 | TATGAGTGTGGAGTCGTTGCGCGTGCCAAGATGAATGCTAAATNNNAGATCGGAA |
| 50115_4 | TATGAGTGTGGAGTCGTTGCGCGTGCCAAGATGAATGCTAAATNNNAGATCGGAA |
| 44765_1 | TATGAGTGTGGAGTCGTTGCCTTGGCTAGTTCCTCAAAGTCCANNNAGATCGGAAG |
| 44765_2 | TATGAGTGTGGAGTCGTTGCCTTGGCTAGTTCCTCAAAGTCCANNNAGATCGGAAG |
| 44765_3 | TATGAGTGTGGAGTCGTTGCCTTGGCTAGTTCCTCAAAGTCCANNNAGATCGGAAG |
| 44765_4 | TATGAGTGTGGAGTCGTTGCCTTGGCTAGTTCCTCAAAGTCCANNNAGATCGGAAG |
| 43674_1 | TATGAGTGTGGAGTCGTTGCTCCAGGTTTAGGAGGGGTCTCNNNAGATCGGAAGA |
| 43674_2 | TATGAGTGTGGAGTCGTTGCTCCAGGTTTAGGAGGGGTCTCNNNAGATCGGAAGA |

| | |
|---|---|
| 43674_3 | TATGAGTGTGGAGTCGTTGCTCCAGGTTTAGGAGGGGTCTCNNNAGATCGGAAGA |
| 43674_4 | TATGAGTGTGGAGTCGTTGCTCCAGGTTTAGGAGGGGTCTCNNNAGATCGGAAGA |
| 35599_1 | TATGAGTGTGGAGTCGTTGCCAAGGGTATACAGAAGGTCCCACNNNAGATCGGAA |
| 35599_2 | TATGAGTGTGGAGTCGTTGCCAAGGGTATACAGAAGGTCCCACNNNAGATCGGAA |
| 35599_3 | TATGAGTGTGGAGTCGTTGCCAAGGGTATACAGAAGGTCCCACNNNAGATCGGAA |
| 35599_4 | TATGAGTGTGGAGTCGTTGCCAAGGGTATACAGAAGGTCCCACNNNAGATCGGAA |
| 30519_1 | TATGAGTGTGGAGTCGTTGCGTCTAAGTGAGATCGTGTGGGATTCNNNAGATCGG |
| 30519_2 | TATGAGTGTGGAGTCGTTGCGTCTAAGTGAGATCGTGTGGGATTCNNNAGATCGG |
| 30519_3 | TATGAGTGTGGAGTCGTTGCGTCTAAGTGAGATCGTGTGGGATTCNNNAGATCGG |
| 30519_4 | TATGAGTGTGGAGTCGTTGCGTCTAAGTGAGATCGTGTGGGATTCNNNAGATCGG |
| 42562_1 | TATGAGTGTGGAGTCGTTGCTGTGTGTGTTGATGTATGTGTGTGNNNAGATCGGA |
| 42562_2 | TATGAGTGTGGAGTCGTTGCTGTGTGTGTTGATGTATGTGTGTGNNNAGATCGGA |
| 42562_3 | TATGAGTGTGGAGTCGTTGCTGTGTGTGTTGATGTATGTGTGTGNNNAGATCGGA |
| 42562_4 | TATGAGTGTGGAGTCGTTGCTGTGTGTGTTGATGTATGTGTGTGNNNAGATCGGA |
| 52623_1 | TATGAGTGTGGAGTCGTTGCGGAACTTGGCAAGGAAGAGAAACNNNAGATCGGA |
| 52623_2 | TATGAGTGTGGAGTCGTTGCGGAACTTGGCAAGGAAGAGAAACNNNAGATCGGA |
| 52623_3 | TATGAGTGTGGAGTCGTTGCGGAACTTGGCAAGGAAGAGAAACNNNAGATCGGA |
| 52623_4 | TATGAGTGTGGAGTCGTTGCGGAACTTGGCAAGGAAGAGAAACNNNAGATCGGA |
| 39618_1 | TATGAGTGTGGAGTCGTTGCCTGAGGTACTGGGTTCATCTCACNNNAGATCGGAAG |
| 39618_2 | TATGAGTGTGGAGTCGTTGCCTGAGGTACTGGGTTCATCTCACNNNAGATCGGAAG |
| 39618_3 | TATGAGTGTGGAGTCGTTGCCTGAGGTACTGGGTTCATCTCACNNNAGATCGGAAG |
| 39618_4 | TATGAGTGTGGAGTCGTTGCCTGAGGTACTGGGTTCATCTCACNNNAGATCGGAAG |
| 48758_1 | TATGAGTGTGGAGTCGTTGCCTGGATTCTGATTCCCACTCCAANNNAGATCGGAAG |
| 48758_2 | TATGAGTGTGGAGTCGTTGCCTGGATTCTGATTCCCACTCCAANNNAGATCGGAAG |
| 48758_3 | TATGAGTGTGGAGTCGTTGCCTGGATTCTGATTCCCACTCCAANNNAGATCGGAAG |
| 48758_4 | TATGAGTGTGGAGTCGTTGCCTGGATTCTGATTCCCACTCCAANNNAGATCGGAAG |
| 30663_1 | TATGAGTGTGGAGTCGTTGCGTCCTAGGTTTCTGGCATAGAGCTNNNAGATCGGAA |
| 30663_2 | TATGAGTGTGGAGTCGTTGCGTCCTAGGTTTCTGGCATAGAGCTNNNAGATCGGAA |
| 30663_3 | TATGAGTGTGGAGTCGTTGCGTCCTAGGTTTCTGGCATAGAGCTNNNAGATCGGAA |
| 30663_4 | TATGAGTGTGGAGTCGTTGCGTCCTAGGTTTCTGGCATAGAGCTNNNAGATCGGAA |
| 13194_1 | TATGAGTGTGGAGTCGTTGCCTATCTCTTAGACTTGTTCTGAANNNAGATCGGAAG |
| 13194_2 | TATGAGTGTGGAGTCGTTGCCTATCTCTTAGACTTGTTCTGAANNNAGATCGGAAG |
| 13194_3 | TATGAGTGTGGAGTCGTTGCCTATCTCTTAGACTTGTTCTGAANNNAGATCGGAAG |
| 13194_4 | TATGAGTGTGGAGTCGTTGCCTATCTCTTAGACTTGTTCTGAANNNAGATCGGAAG |
| 44062_1 | TATGAGTGTGGAGTCGTTGCGCTAGGCGATCCTTGGTCTATTTNNNAGATCGGAAG |
| 44062_2 | TATGAGTGTGGAGTCGTTGCGCTAGGCGATCCTTGGTCTATTTNNNAGATCGGAAG |
| 44062_3 | TATGAGTGTGGAGTCGTTGCGCTAGGCGATCCTTGGTCTATTTNNNAGATCGGAAG |
| 44062_4 | TATGAGTGTGGAGTCGTTGCGCTAGGCGATCCTTGGTCTATTTNNNAGATCGGAAG |
| 42329_1 | TATGAGTGTGGAGTCGTTGCATATGGTGTCTTGCTCTGTCACCNNNAGATCGGAAG |
| 42329_2 | TATGAGTGTGGAGTCGTTGCATATGGTGTCTTGCTCTGTCACCNNNAGATCGGAAG |
| 42329_3 | TATGAGTGTGGAGTCGTTGCATATGGTGTCTTGCTCTGTCACCNNNAGATCGGAAG |
| 42329_4 | TATGAGTGTGGAGTCGTTGCATATGGTGTCTTGCTCTGTCACCNNNAGATCGGAAG |
| 9269_1 | TATGAGTGTGGAGTCGTTGCCAAGGCTGAGAAACCCTAACCTGNNNAGATCGGAA |

| | |
|---|---|
| 9269_2 | TATGAGTGTGGAGTCGTTGCCAAGGCTGAGAAACCCTAACCTGNNNAGATCGGAAG |
| 9269_3 | TATGAGTGTGGAGTCGTTGCCAAGGCTGAGAAACCCTAACCTGNNNAGATCGGAAG |
| 9269_4 | TATGAGTGTGGAGTCGTTGCCAAGGCTGAGAAACCCTAACCTGNNNAGATCGGAAG |
| 28934_1 | TATGAGTGTGGAGTCGTTGCTTCGGACTCTTGGATTTGTAGCANNNAGATCGGAAG |
| 28934_2 | TATGAGTGTGGAGTCGTTGCTTCGGACTCTTGGATTTGTAGCANNNAGATCGGAAG |
| 28934_3 | TATGAGTGTGGAGTCGTTGCTTCGGACTCTTGGATTTGTAGCANNNAGATCGGAAG |
| 28934_4 | TATGAGTGTGGAGTCGTTGCTTCGGACTCTTGGATTTGTAGCANNNAGATCGGAAG |
| 31192_1 | TATGAGTGTGGAGTCGTTGCCCTCGTTTCTCTCCATTTTGCTCNNNAGATCGGAAGA |
| 31192_2 | TATGAGTGTGGAGTCGTTGCCCTCGTTTCTCTCCATTTTGCTCNNNAGATCGGAAGA |
| 31192_3 | TATGAGTGTGGAGTCGTTGCCCTCGTTTCTCTCCATTTTGCTCNNNAGATCGGAAGA |
| 31192_4 | TATGAGTGTGGAGTCGTTGCCCTCGTTTCTCTCCATTTTGCTCNNNAGATCGGAAGA |
| 35318_1 | TATGAGTGTGGAGTCGTTGCGACCGGAGTGTAGTTGTCTCAATNNNAGATCGGAA |
| 35318_2 | TATGAGTGTGGAGTCGTTGCGACCGGAGTGTAGTTGTCTCAATNNNAGATCGGAA |
| 35318_3 | TATGAGTGTGGAGTCGTTGCGACCGGAGTGTAGTTGTCTCAATNNNAGATCGGAA |
| 35318_4 | TATGAGTGTGGAGTCGTTGCGACCGGAGTGTAGTTGTCTCAATNNNAGATCGGAA |
| 48147_1 | TATGAGTGTGGAGTCGTTGCCCCAGGTCAAAAATGCGAATCTTNNNAGATCGGAAG |
| 48147_2 | TATGAGTGTGGAGTCGTTGCCCCAGGTCAAAAATGCGAATCTTNNNAGATCGGAAG |
| 48147_3 | TATGAGTGTGGAGTCGTTGCCCCAGGTCAAAAATGCGAATCTTNNNAGATCGGAAG |
| 48147_4 | TATGAGTGTGGAGTCGTTGCCCCAGGTCAAAAATGCGAATCTTNNNAGATCGGAAG |
| 34550_1 | TATGAGTGTGGAGTCGTTGCGAGACACACACACACAGAAACACNNNAGATCGGAAG |
| 34550_2 | TATGAGTGTGGAGTCGTTGCGAGACACACACACACAGAAACACNNNAGATCGGAAG |
| 34550_3 | TATGAGTGTGGAGTCGTTGCGAGACACACACACACAGAAACACNNNAGATCGGAAG |
| 34550_4 | TATGAGTGTGGAGTCGTTGCGAGACACACACACACAGAAACACNNNAGATCGGAAG |
| 50004_1 | TATGAGTGTGGAGTCGTTGCCAGAGTGTGTGTGACATTGTGTGNNNAGATCGGAA |
| 50004_2 | TATGAGTGTGGAGTCGTTGCCAGAGTGTGTGTGACATTGTGTGNNNAGATCGGAA |
| 50004_3 | TATGAGTGTGGAGTCGTTGCCAGAGTGTGTGTGACATTGTGTGNNNAGATCGGAA |
| 50004_4 | TATGAGTGTGGAGTCGTTGCCAGAGTGTGTGTGACATTGTGTGNNNAGATCGGAA |
| 41754_1 | TATGAGTGTGGAGTCGTTGCTGTGTCTGGTGGATTCAAGTGATNNNAGATCGGAA |
| 41754_2 | TATGAGTGTGGAGTCGTTGCTGTGTCTGGTGGATTCAAGTGATNNNAGATCGGAA |
| 41754_3 | TATGAGTGTGGAGTCGTTGCTGTGTCTGGTGGATTCAAGTGATNNNAGATCGGAA |
| 41754_4 | TATGAGTGTGGAGTCGTTGCTGTGTCTGGTGGATTCAAGTGATNNNAGATCGGAA |
| 8754_1 | TATGAGTGTGGAGTCGTTGCGCTCAGAGGTACCAAAAACCTTTNNNAGATCGGAAG |
| 8754_2 | TATGAGTGTGGAGTCGTTGCGCTCAGAGGTACCAAAAACCTTTNNNAGATCGGAAG |
| 8754_3 | TATGAGTGTGGAGTCGTTGCGCTCAGAGGTACCAAAAACCTTTNNNAGATCGGAAG |
| 8754_4 | TATGAGTGTGGAGTCGTTGCGCTCAGAGGTACCAAAAACCTTTNNNAGATCGGAAG |
| 40282_1 | TATGAGTGTGGAGTCGTTGCCTGATACTTCAGGGTGTCACCAANNNAGATCGGAAG |
| 40282_2 | TATGAGTGTGGAGTCGTTGCCTGATACTTCAGGGTGTCACCAANNNAGATCGGAAG |
| 40282_3 | TATGAGTGTGGAGTCGTTGCCTGATACTTCAGGGTGTCACCAANNNAGATCGGAAG |
| 40282_4 | TATGAGTGTGGAGTCGTTGCCTGATACTTCAGGGTGTCACCAANNNAGATCGGAAG |
| 36455_1 | TATGAGTGTGGAGTCGTTGCTCGATTAGTTGAGCCTGTAGTCCNNNAGATCGGAAG |
| 36455_2 | TATGAGTGTGGAGTCGTTGCTCGATTAGTTGAGCCTGTAGTCCNNNAGATCGGAAG |
| 36455_3 | TATGAGTGTGGAGTCGTTGCTCGATTAGTTGAGCCTGTAGTCCNNNAGATCGGAAG |
| 36455_4 | TATGAGTGTGGAGTCGTTGCTCGATTAGTTGAGCCTGTAGTCCNNNAGATCGGAAG |

| | |
|---|---|
| 50909_1 | TATGAGTGTGGAGTCGTTGCATTTGCCACCTTTTTGATGGTCANNNAGATCGGAAG |
| 50909_2 | TATGAGTGTGGAGTCGTTGCATTTGCCACCTTTTTGATGGTCANNNAGATCGGAAG |
| 50909_3 | TATGAGTGTGGAGTCGTTGCATTTGCCACCTTTTTGATGGTCANNNAGATCGGAAG |
| 50909_4 | TATGAGTGTGGAGTCGTTGCATTTGCCACCTTTTTGATGGTCANNNAGATCGGAAG |
| 48968_1 | TATGAGTGTGGAGTCGTTGCGGAGACCTGGAACTCTATCACCTNNNAGATCGGAAG |
| 48968_2 | TATGAGTGTGGAGTCGTTGCGGAGACCTGGAACTCTATCACCTNNNAGATCGGAAG |
| 48968_3 | TATGAGTGTGGAGTCGTTGCGGAGACCTGGAACTCTATCACCTNNNAGATCGGAAG |
| 48968_4 | TATGAGTGTGGAGTCGTTGCGGAGACCTGGAACTCTATCACCTNNNAGATCGGAAG |
| 52912_1 | TATGAGTGTGGAGTCGTTGCCCCCACTCAACTCAAGTCTCTTTNNNAGATCGGAAGA |
| 52912_2 | TATGAGTGTGGAGTCGTTGCCCCCACTCAACTCAAGTCTCTTTNNNAGATCGGAAGA |
| 52912_3 | TATGAGTGTGGAGTCGTTGCCCCCACTCAACTCAAGTCTCTTTNNNAGATCGGAAGA |
| 52912_4 | TATGAGTGTGGAGTCGTTGCCCCCACTCAACTCAAGTCTCTTTNNNAGATCGGAAGA |
| 40918_1 | TATGAGTGTGGAGTCGTTGCCTGGATTTGAGCATTCATTTGGCNNNAGATCGGAAG |
| 40918_2 | TATGAGTGTGGAGTCGTTGCCTGGATTTGAGCATTCATTTGGCNNNAGATCGGAAG |
| 40918_3 | TATGAGTGTGGAGTCGTTGCCTGGATTTGAGCATTCATTTGGCNNNAGATCGGAAG |
| 40918_4 | TATGAGTGTGGAGTCGTTGCCTGGATTTGAGCATTCATTTGGCNNNAGATCGGAAG |
| 49551_1 | TATGAGTGTGGAGTCGTTGCTGCCCAGTTATGCTTACCTGATTNNNAGATCGGAAG |
| 49551_2 | TATGAGTGTGGAGTCGTTGCTGCCCAGTTATGCTTACCTGATTNNNAGATCGGAAG |
| 49551_3 | TATGAGTGTGGAGTCGTTGCTGCCCAGTTATGCTTACCTGATTNNNAGATCGGAAG |
| 49551_4 | TATGAGTGTGGAGTCGTTGCTGCCCAGTTATGCTTACCTGATTNNNAGATCGGAAG |
| 48480_1 | TATGAGTGTGGAGTCGTTGCCCTCCTGAGTAGCTGAGACTACANNNAGATCGGAAG |
| 48480_2 | TATGAGTGTGGAGTCGTTGCCCTCCTGAGTAGCTGAGACTACANNNAGATCGGAAG |
| 48480_3 | TATGAGTGTGGAGTCGTTGCCCTCCTGAGTAGCTGAGACTACANNNAGATCGGAAG |
| 48480_4 | TATGAGTGTGGAGTCGTTGCCCTCCTGAGTAGCTGAGACTACANNNAGATCGGAAG |
| 41633_1 | TATGAGTGTGGAGTCGTTGCTATCTTGGCTCACTGCAACCTCNNNAGATCGGAAGA |
| 41633_2 | TATGAGTGTGGAGTCGTTGCTATCTTGGCTCACTGCAACCTCNNNAGATCGGAAGA |
| 41633_3 | TATGAGTGTGGAGTCGTTGCTATCTTGGCTCACTGCAACCTCNNNAGATCGGAAGA |
| 41633_4 | TATGAGTGTGGAGTCGTTGCTATCTTGGCTCACTGCAACCTCNNNAGATCGGAAGA |
| 29498_1 | TATGAGTGTGGAGTCGTTGCCTCACTGCTCTTGGTATGGCATCTNNNAGATCGGAA |
| 29498_2 | TATGAGTGTGGAGTCGTTGCCTCACTGCTCTTGGTATGGCATCTNNNAGATCGGAA |
| 29498_3 | TATGAGTGTGGAGTCGTTGCCTCACTGCTCTTGGTATGGCATCTNNNAGATCGGAA |
| 29498_4 | TATGAGTGTGGAGTCGTTGCCTCACTGCTCTTGGTATGGCATCTNNNAGATCGGAA |
| 2790_1 | TATGAGTGTGGAGTCGTTGCCCATATTTTCTGGCCAGGATGCTNNNAGATCGGAAG |
| 2790_2 | TATGAGTGTGGAGTCGTTGCCCATATTTTCTGGCCAGGATGCTNNNAGATCGGAAG |
| 2790_3 | TATGAGTGTGGAGTCGTTGCCCATATTTTCTGGCCAGGATGCTNNNAGATCGGAAG |
| 2790_4 | TATGAGTGTGGAGTCGTTGCCCATATTTTCTGGCCAGGATGCTNNNAGATCGGAAG |
| 38596_1 | TATGAGTGTGGAGTCGTTGCAAAGACAACTCACAGAATGGGGANNNAGATCGGAA |
| 38596_2 | TATGAGTGTGGAGTCGTTGCAAAGACAACTCACAGAATGGGGANNNAGATCGGAA |
| 38596_3 | TATGAGTGTGGAGTCGTTGCAAAGACAACTCACAGAATGGGGANNNAGATCGGAA |
| 38596_4 | TATGAGTGTGGAGTCGTTGCAAAGACAACTCACAGAATGGGGANNNAGATCGGAA |
| 49087_1 | TATGAGTGTGGAGTCGTTGCCCTTGAGACAGTAGCTCCAACATNNNAGATCGGAAG |
| 49087_2 | TATGAGTGTGGAGTCGTTGCCCTTGAGACAGTAGCTCCAACATNNNAGATCGGAAG |
| 49087_3 | TATGAGTGTGGAGTCGTTGCCCTTGAGACAGTAGCTCCAACATNNNAGATCGGAAG |

| | |
|---|---|
| 49087_4 | TATGAGTGTGGAGTCGTTGCCCTTGAGACAGTAGCTCCAACATNNNAGATCGGAAG |
| 35332_1 | TATGAGTGTGGAGTCGTTGCGCCTCCTGAGTAGCTGGTATTACNNNAGATCGGAAG |
| 35332_2 | TATGAGTGTGGAGTCGTTGCGCCTCCTGAGTAGCTGGTATTACNNNAGATCGGAAG |
| 35332_3 | TATGAGTGTGGAGTCGTTGCGCCTCCTGAGTAGCTGGTATTACNNNAGATCGGAAG |
| 35332_4 | TATGAGTGTGGAGTCGTTGCGCCTCCTGAGTAGCTGGTATTACNNNAGATCGGAAG |
| 39159_1 | TATGAGTGTGGAGTCGTTGCTCATCGGATCCTTGGTGATTACTNNNAGATCGGAAG |
| 39159_2 | TATGAGTGTGGAGTCGTTGCTCATCGGATCCTTGGTGATTACTNNNAGATCGGAAG |
| 39159_3 | TATGAGTGTGGAGTCGTTGCTCATCGGATCCTTGGTGATTACTNNNAGATCGGAAG |
| 39159_4 | TATGAGTGTGGAGTCGTTGCTCATCGGATCCTTGGTGATTACTNNNAGATCGGAAG |
| 43348_1 | TATGAGTGTGGAGTCGTTGCTGAGAGTAAAAAGGCCAGAGCATNNNAGATCGGAA |
| 43348_2 | TATGAGTGTGGAGTCGTTGCTGAGAGTAAAAAGGCCAGAGCATNNNAGATCGGAA |
| 43348_3 | TATGAGTGTGGAGTCGTTGCTGAGAGTAAAAAGGCCAGAGCATNNNAGATCGGAA |
| 43348_4 | TATGAGTGTGGAGTCGTTGCTGAGAGTAAAAAGGCCAGAGCATNNNAGATCGGAA |
| 45323_1 | TATGAGTGTGGAGTCGTTGCTAACTGTCCGGAGCTTCTCTTTCNNNAGATCGGAAG |
| 45323_2 | TATGAGTGTGGAGTCGTTGCTAACTGTCCGGAGCTTCTCTTTCNNNAGATCGGAAG |
| 45323_3 | TATGAGTGTGGAGTCGTTGCTAACTGTCCGGAGCTTCTCTTTCNNNAGATCGGAAG |
| 45323_4 | TATGAGTGTGGAGTCGTTGCTAACTGTCCGGAGCTTCTCTTTCNNNAGATCGGAAG |
| 50568_1 | TATGAGTGTGGAGTCGTTGCAAAACAAGCTCTGCAACAACTGANNNAGATCGGAAG |
| 50568_2 | TATGAGTGTGGAGTCGTTGCAAAACAAGCTCTGCAACAACTGANNNAGATCGGAAG |
| 50568_3 | TATGAGTGTGGAGTCGTTGCAAAACAAGCTCTGCAACAACTGANNNAGATCGGAAG |
| 50568_4 | TATGAGTGTGGAGTCGTTGCAAAACAAGCTCTGCAACAACTGANNNAGATCGGAAG |
| 48854_1 | TATGAGTGTGGAGTCGTTGCTGGGGATTCACCATGTTGACTAGNNNAGATCGGAA |
| 48854_2 | TATGAGTGTGGAGTCGTTGCTGGGGATTCACCATGTTGACTAGNNNAGATCGGAA |
| 48854_3 | TATGAGTGTGGAGTCGTTGCTGGGGATTCACCATGTTGACTAGNNNAGATCGGAA |
| 48854_4 | TATGAGTGTGGAGTCGTTGCTGGGGATTCACCATGTTGACTAGNNNAGATCGGAA |
| 46620_1 | TATGAGTGTGGAGTCGTTGCGGATAGACGCTAACACCACAGAANNNAGATCGGAA |
| 46620_2 | TATGAGTGTGGAGTCGTTGCGGATAGACGCTAACACCACAGAANNNAGATCGGAA |
| 46620_3 | TATGAGTGTGGAGTCGTTGCGGATAGACGCTAACACCACAGAANNNAGATCGGAA |
| 46620_4 | TATGAGTGTGGAGTCGTTGCGGATAGACGCTAACACCACAGAANNNAGATCGGAA |
| 37632_1 | TATGAGTGTGGAGTCGTTGCTCCCTGTAGCAGCCATTCTTTTANNNAGATCGGAAG |
| 37632_2 | TATGAGTGTGGAGTCGTTGCTCCCTGTAGCAGCCATTCTTTTANNNAGATCGGAAG |
| 37632_3 | TATGAGTGTGGAGTCGTTGCTCCCTGTAGCAGCCATTCTTTTANNNAGATCGGAAG |
| 37632_4 | TATGAGTGTGGAGTCGTTGCTCCCTGTAGCAGCCATTCTTTTANNNAGATCGGAAG |
| 11807_1 | TATGAGTGTGGAGTCGTTGCTGATGTTTGCCTTCCCTGAGACTNNNAGATCGGAAG |
| 11807_2 | TATGAGTGTGGAGTCGTTGCTGATGTTTGCCTTCCCTGAGACTNNNAGATCGGAAG |
| 11807_3 | TATGAGTGTGGAGTCGTTGCTGATGTTTGCCTTCCCTGAGACTNNNAGATCGGAAG |
| 11807_4 | TATGAGTGTGGAGTCGTTGCTGATGTTTGCCTTCCCTGAGACTNNNAGATCGGAAG |
| 2259_1 | TATGAGTGTGGAGTCGTTGCGAGTGGATGTGCGTGCATATGTANNNAGATCGGAA |
| 2259_2 | TATGAGTGTGGAGTCGTTGCGAGTGGATGTGCGTGCATATGTANNNAGATCGGAA |
| 2259_3 | TATGAGTGTGGAGTCGTTGCGAGTGGATGTGCGTGCATATGTANNNAGATCGGAA |
| 2259_4 | TATGAGTGTGGAGTCGTTGCGAGTGGATGTGCGTGCATATGTANNNAGATCGGAA |
| 37417_1 | TATGAGTGTGGAGTCGTTGCAATCATGGAAGGTCAGAGTGTGTNNNAGATCGGAA |
| 37417_2 | TATGAGTGTGGAGTCGTTGCAATCATGGAAGGTCAGAGTGTGTNNNAGATCGGAA |

| | |
|---|---|
| 37417_3 | TATGAGTGTGGAGTCGTTGCAATCATGGAAGGTCAGAGTGTGTNNNAGATCGGAA |
| 37417_4 | TATGAGTGTGGAGTCGTTGCAATCATGGAAGGTCAGAGTGTGTNNNAGATCGGAA |
| 40569_1 | TATGAGTGTGGAGTCGTTGCCTGTCTTTTCTCCATCCTCCCTCNNNAGATCGGAAGAG |
| 40569_2 | TATGAGTGTGGAGTCGTTGCCTGTCTTTTCTCCATCCTCCCTCNNNAGATCGGAAGAG |
| 40569_3 | TATGAGTGTGGAGTCGTTGCCTGTCTTTTCTCCATCCTCCCTCNNNAGATCGGAAGAG |
| 40569_4 | TATGAGTGTGGAGTCGTTGCCTGTCTTTTCTCCATCCTCCCTCNNNAGATCGGAAGAG |
| 9166_1 | TATGAGTGTGGAGTCGTTGCTTTCTCCTGTTCTTGCTCTGTCGNNNAGATCGGAAGA |
| 9166_2 | TATGAGTGTGGAGTCGTTGCTTTCTCCTGTTCTTGCTCTGTCGNNNAGATCGGAAGA |
| 9166_3 | TATGAGTGTGGAGTCGTTGCTTTCTCCTGTTCTTGCTCTGTCGNNNAGATCGGAAGA |
| 9166_4 | TATGAGTGTGGAGTCGTTGCTTTCTCCTGTTCTTGCTCTGTCGNNNAGATCGGAAGA |
| 36938_1 | TATGAGTGTGGAGTCGTTGCAGGTGAGCAGATTACTTGAGGTGNNNAGATCGGAA |
| 36938_2 | TATGAGTGTGGAGTCGTTGCAGGTGAGCAGATTACTTGAGGTGNNNAGATCGGAA |
| 36938_3 | TATGAGTGTGGAGTCGTTGCAGGTGAGCAGATTACTTGAGGTGNNNAGATCGGAA |
| 36938_4 | TATGAGTGTGGAGTCGTTGCAGGTGAGCAGATTACTTGAGGTGNNNAGATCGGAA |
| 42531_1 | TATGAGTGTGGAGTCGTTGCAATGGCACAATCTCAGCTCACTANNNAGATCGGAAG |
| 42531_2 | TATGAGTGTGGAGTCGTTGCAATGGCACAATCTCAGCTCACTANNNAGATCGGAAG |
| 42531_3 | TATGAGTGTGGAGTCGTTGCAATGGCACAATCTCAGCTCACTANNNAGATCGGAAG |
| 42531_4 | TATGAGTGTGGAGTCGTTGCAATGGCACAATCTCAGCTCACTANNNAGATCGGAAG |
| 4673_1 | TATGAGTGTGGAGTCGTTGCGGGCCTCAAAGACAGTTTCAAGANNNAGATCGGAA |
| 4673_2 | TATGAGTGTGGAGTCGTTGCGGGCCTCAAAGACAGTTTCAAGANNNAGATCGGAA |
| 4673_3 | TATGAGTGTGGAGTCGTTGCGGGCCTCAAAGACAGTTTCAAGANNNAGATCGGAA |
| 4673_4 | TATGAGTGTGGAGTCGTTGCGGGCCTCAAAGACAGTTTCAAGANNNAGATCGGAA |
| 7180_1 | TATGAGTGTGGAGTCGTTGCCTCGGACCAGGAGCCAGAGGANNNAGATCGGAAGA |
| 7180_2 | TATGAGTGTGGAGTCGTTGCCTCGGACCAGGAGCCAGAGGANNNAGATCGGAAGA |
| 7180_3 | TATGAGTGTGGAGTCGTTGCCTCGGACCAGGAGCCAGAGGANNNAGATCGGAAGA |
| 7180_4 | TATGAGTGTGGAGTCGTTGCCTCGGACCAGGAGCCAGAGGANNNAGATCGGAAGA |
| 41941_1 | TATGAGTGTGGAGTCGTTGCAGATAGGGTCTTGCTATGTTGCCNNNAGATCGGAAG |
| 41941_2 | TATGAGTGTGGAGTCGTTGCAGATAGGGTCTTGCTATGTTGCCNNNAGATCGGAAG |
| 41941_3 | TATGAGTGTGGAGTCGTTGCAGATAGGGTCTTGCTATGTTGCCNNNAGATCGGAAG |
| 41941_4 | TATGAGTGTGGAGTCGTTGCAGATAGGGTCTTGCTATGTTGCCNNNAGATCGGAAG |
| 13132_1 | TATGAGTGTGGAGTCGTTGCGCCAAGCTATTAAGTTTGTAAATNNNAGATCGGAAG |
| 13132_2 | TATGAGTGTGGAGTCGTTGCGCCAAGCTATTAAGTTTGTAAATNNNAGATCGGAAG |
| 13132_3 | TATGAGTGTGGAGTCGTTGCGCCAAGCTATTAAGTTTGTAAATNNNAGATCGGAAG |
| 13132_4 | TATGAGTGTGGAGTCGTTGCGCCAAGCTATTAAGTTTGTAAATNNNAGATCGGAAG |
| 41828_1 | TATGAGTGTGGAGTCGTTGCCCTAGTTCAGCCTCTCAAGTAGCNNNAGATCGGAAG |
| 41828_2 | TATGAGTGTGGAGTCGTTGCCCTAGTTCAGCCTCTCAAGTAGCNNNAGATCGGAAG |
| 41828_3 | TATGAGTGTGGAGTCGTTGCCCTAGTTCAGCCTCTCAAGTAGCNNNAGATCGGAAG |
| 41828_4 | TATGAGTGTGGAGTCGTTGCCCTAGTTCAGCCTCTCAAGTAGCNNNAGATCGGAAG |
| 40891_1 | TATGAGTGTGGAGTCGTTGCGCAAGGCTCCATCTCAAAAGAAANNNAGATCGGAAG |
| 40891_2 | TATGAGTGTGGAGTCGTTGCGCAAGGCTCCATCTCAAAAGAAANNNAGATCGGAAG |
| 40891_3 | TATGAGTGTGGAGTCGTTGCGCAAGGCTCCATCTCAAAAGAAANNNAGATCGGAAG |
| 40891_4 | TATGAGTGTGGAGTCGTTGCGCAAGGCTCCATCTCAAAAGAAANNNAGATCGGAAG |
| 52020_1 | TATGAGTGTGGAGTCGTTGCCAGTTTGACTTCAGACTGCTGTGNNNAGATCGGAAG |

| | |
|---|---|
| 52020_2 | TATGAGTGTGGAGTCGTTGCCAGTTTGACTTCAGACTGCTGTGNNNAGATCGGAAG |
| 52020_3 | TATGAGTGTGGAGTCGTTGCCAGTTTGACTTCAGACTGCTGTGNNNAGATCGGAAG |
| 52020_4 | TATGAGTGTGGAGTCGTTGCCAGTTTGACTTCAGACTGCTGTGNNNAGATCGGAAG |
| 38630_1 | TATGAGTGTGGAGTCGTTGCTTCCATGCTCATGAGTTGGAAGANNNAGATCGGAA |
| 38630_2 | TATGAGTGTGGAGTCGTTGCTTCCATGCTCATGAGTTGGAAGANNNAGATCGGAA |
| 38630_3 | TATGAGTGTGGAGTCGTTGCTTCCATGCTCATGAGTTGGAAGANNNAGATCGGAA |
| 38630_4 | TATGAGTGTGGAGTCGTTGCTTCCATGCTCATGAGTTGGAAGANNNAGATCGGAA |
| 29456_1 | TATGAGTGTGGAGTCGTTGCTTAGCTTGCAAGTCAATTGTGCANNNAGATCGGAAG |
| 29456_2 | TATGAGTGTGGAGTCGTTGCTTAGCTTGCAAGTCAATTGTGCANNNAGATCGGAAG |
| 29456_3 | TATGAGTGTGGAGTCGTTGCTTAGCTTGCAAGTCAATTGTGCANNNAGATCGGAAG |
| 29456_4 | TATGAGTGTGGAGTCGTTGCTTAGCTTGCAAGTCAATTGTGCANNNAGATCGGAAG |
| 46287_1 | TATGAGTGTGGAGTCGTTGCAGGTTCAAGCAATTCTCCTGTCTNNNAGATCGGAAG |
| 46287_2 | TATGAGTGTGGAGTCGTTGCAGGTTCAAGCAATTCTCCTGTCTNNNAGATCGGAAG |
| 46287_3 | TATGAGTGTGGAGTCGTTGCAGGTTCAAGCAATTCTCCTGTCTNNNAGATCGGAAG |
| 46287_4 | TATGAGTGTGGAGTCGTTGCAGGTTCAAGCAATTCTCCTGTCTNNNAGATCGGAAG |
| 35317_1 | TATGAGTGTGGAGTCGTTGCGGGAAGAGAGGAGGAATAGGAGANNNAGATCGG |
| 35317_2 | TATGAGTGTGGAGTCGTTGCGGGAAGAGAGGAGGAATAGGAGANNNAGATCGG |
| 35317_3 | TATGAGTGTGGAGTCGTTGCGGGAAGAGAGGAGGAATAGGAGANNNAGATCGG |
| 35317_4 | TATGAGTGTGGAGTCGTTGCGGGAAGAGAGGAGGAATAGGAGANNNAGATCGG |
| 33907_1 | TATGAGTGTGGAGTCGTTGCAATCTCGACTCACTTCAAGCTCCNNNAGATCGGAAG |
| 33907_2 | TATGAGTGTGGAGTCGTTGCAATCTCGACTCACTTCAAGCTCCNNNAGATCGGAAG |
| 33907_3 | TATGAGTGTGGAGTCGTTGCAATCTCGACTCACTTCAAGCTCCNNNAGATCGGAAG |
| 33907_4 | TATGAGTGTGGAGTCGTTGCAATCTCGACTCACTTCAAGCTCCNNNAGATCGGAAG |
| 10867_1 | TATGAGTGTGGAGTCGTTGCCCCTCGAATGTTCTCCATTGCTTNNNAGATCGGAAGA |
| 10867_2 | TATGAGTGTGGAGTCGTTGCCCCTCGAATGTTCTCCATTGCTTNNNAGATCGGAAGA |
| 10867_3 | TATGAGTGTGGAGTCGTTGCCCCTCGAATGTTCTCCATTGCTTNNNAGATCGGAAGA |
| 10867_4 | TATGAGTGTGGAGTCGTTGCCCCTCGAATGTTCTCCATTGCTTNNNAGATCGGAAGA |
| 33722_1 | TATGAGTGTGGAGTCGTTGCTCTGCCTCTCTCCTAGCATCATANNNAGATCGGAAGA |
| 33722_2 | TATGAGTGTGGAGTCGTTGCTCTGCCTCTCTCCTAGCATCATANNNAGATCGGAAGA |
| 33722_3 | TATGAGTGTGGAGTCGTTGCTCTGCCTCTCTCCTAGCATCATANNNAGATCGGAAGA |
| 33722_4 | TATGAGTGTGGAGTCGTTGCTCTGCCTCTCTCCTAGCATCATANNNAGATCGGAAGA |
| 37053_1 | TATGAGTGTGGAGTCGTTGCAGGTCAGTAGTAGTTCGAGACCANNNAGATCGGAA |
| 37053_2 | TATGAGTGTGGAGTCGTTGCAGGTCAGTAGTAGTTCGAGACCANNNAGATCGGAA |
| 37053_3 | TATGAGTGTGGAGTCGTTGCAGGTCAGTAGTAGTTCGAGACCANNNAGATCGGAA |
| 37053_4 | TATGAGTGTGGAGTCGTTGCAGGTCAGTAGTAGTTCGAGACCANNNAGATCGGAA |
| 34618_1 | TATGAGTGTGGAGTCGTTGCATGTGGAAAGTTCTGGAAGCTCTNNNAGATCGGAA |
| 34618_2 | TATGAGTGTGGAGTCGTTGCATGTGGAAAGTTCTGGAAGCTCTNNNAGATCGGAA |
| 34618_3 | TATGAGTGTGGAGTCGTTGCATGTGGAAAGTTCTGGAAGCTCTNNNAGATCGGAA |
| 34618_4 | TATGAGTGTGGAGTCGTTGCATGTGGAAAGTTCTGGAAGCTCTNNNAGATCGGAA |
| 2347_1 | TATGAGTGTGGAGTCGTTGCACACTTCTTCCACAGCATCATCCNNNAGATCGGAAGA |
| 2347_2 | TATGAGTGTGGAGTCGTTGCACACTTCTTCCACAGCATCATCCNNNAGATCGGAAGA |
| 2347_3 | TATGAGTGTGGAGTCGTTGCACACTTCTTCCACAGCATCATCCNNNAGATCGGAAGA |
| 2347_4 | TATGAGTGTGGAGTCGTTGCACACTTCTTCCACAGCATCATCCNNNAGATCGGAAGA |

| | |
|---|---|
| 38857_1 | TATGAGTGTGGAGTCGTTGCAAAACAGCACACCTACAACCATCNNNAGATCGGAAG |
| 38857_2 | TATGAGTGTGGAGTCGTTGCAAAACAGCACACCTACAACCATCNNNAGATCGGAAG |
| 38857_3 | TATGAGTGTGGAGTCGTTGCAAAACAGCACACCTACAACCATCNNNAGATCGGAAG |
| 38857_4 | TATGAGTGTGGAGTCGTTGCAAAACAGCACACCTACAACCATCNNNAGATCGGAAG |
| 40761_1 | TATGAGTGTGGAGTCGTTGCACCAGAGGTACAAATAAGAGCTGGNNNAGATCGGA |
| 40761_2 | TATGAGTGTGGAGTCGTTGCACCAGAGGTACAAATAAGAGCTGGNNNAGATCGGA |
| 40761_3 | TATGAGTGTGGAGTCGTTGCACCAGAGGTACAAATAAGAGCTGGNNNAGATCGGA |
| 40761_4 | TATGAGTGTGGAGTCGTTGCACCAGAGGTACAAATAAGAGCTGGNNNAGATCGGA |
| 42251_1 | TATGAGTGTGGAGTCGTTGCGTTTCACAATATTTGCTCGGGCTNNNAGATCGGAAG |
| 42251_2 | TATGAGTGTGGAGTCGTTGCGTTTCACAATATTTGCTCGGGCTNNNAGATCGGAAG |
| 42251_3 | TATGAGTGTGGAGTCGTTGCGTTTCACAATATTTGCTCGGGCTNNNAGATCGGAAG |
| 42251_4 | TATGAGTGTGGAGTCGTTGCGTTTCACAATATTTGCTCGGGCTNNNAGATCGGAAG |
| 53234_1 | TATGAGTGTGGAGTCGTTGCAGAGATCAAGTGTACGGTTGACCNNNAGATCGGAA |
| 53234_2 | TATGAGTGTGGAGTCGTTGCAGAGATCAAGTGTACGGTTGACCNNNAGATCGGAA |
| 53234_3 | TATGAGTGTGGAGTCGTTGCAGAGATCAAGTGTACGGTTGACCNNNAGATCGGAA |
| 53234_4 | TATGAGTGTGGAGTCGTTGCAGAGATCAAGTGTACGGTTGACCNNNAGATCGGAA |
| 12943_1 | TATGAGTGTGGAGTCGTTGCTTTTCTTACTCCATTCTTGCAANNNAGATCGGAAGAG |
| 12943_2 | TATGAGTGTGGAGTCGTTGCTTTTCTTACTCCATTCTTGCAANNNAGATCGGAAGAG |
| 12943_3 | TATGAGTGTGGAGTCGTTGCTTTTCTTACTCCATTCTTGCAANNNAGATCGGAAGAG |
| 12943_4 | TATGAGTGTGGAGTCGTTGCTTTTCTTACTCCATTCTTGCAANNNAGATCGGAAGAG |
| 35007_1 | TATGAGTGTGGAGTCGTTGCCAAGTCTTGCTCTGTTACCCAAGNNNAGATCGGAAG |
| 35007_2 | TATGAGTGTGGAGTCGTTGCCAAGTCTTGCTCTGTTACCCAAGNNNAGATCGGAAG |
| 35007_3 | TATGAGTGTGGAGTCGTTGCCAAGTCTTGCTCTGTTACCCAAGNNNAGATCGGAAG |
| 35007_4 | TATGAGTGTGGAGTCGTTGCCAAGTCTTGCTCTGTTACCCAAGNNNAGATCGGAAG |
| 35771_1 | TATGAGTGTGGAGTCGTTGCGCCTTCTGAGTAGCTAGGATGACNNNAGATCGGAA |
| 35771_2 | TATGAGTGTGGAGTCGTTGCGCCTTCTGAGTAGCTAGGATGACNNNAGATCGGAA |
| 35771_3 | TATGAGTGTGGAGTCGTTGCGCCTTCTGAGTAGCTAGGATGACNNNAGATCGGAA |
| 35771_4 | TATGAGTGTGGAGTCGTTGCGCCTTCTGAGTAGCTAGGATGACNNNAGATCGGAA |
| 38018_1 | TATGAGTGTGGAGTCGTTGCAGCTACTCAGGGAGTTGAGGTNNNAGATCGGAAGA |
| 38018_2 | TATGAGTGTGGAGTCGTTGCAGCTACTCAGGGAGTTGAGGTNNNAGATCGGAAGA |
| 38018_3 | TATGAGTGTGGAGTCGTTGCAGCTACTCAGGGAGTTGAGGTNNNAGATCGGAAGA |
| 38018_4 | TATGAGTGTGGAGTCGTTGCAGCTACTCAGGGAGTTGAGGTNNNAGATCGGAAGA |
| 32007_1 | TATGAGTGTGGAGTCGTTGCCAACAGATATCAGCTGGACTCCANNNAGATCGGAAG |
| 32007_2 | TATGAGTGTGGAGTCGTTGCCAACAGATATCAGCTGGACTCCANNNAGATCGGAAG |
| 32007_3 | TATGAGTGTGGAGTCGTTGCCAACAGATATCAGCTGGACTCCANNNAGATCGGAAG |
| 32007_4 | TATGAGTGTGGAGTCGTTGCCAACAGATATCAGCTGGACTCCANNNAGATCGGAAG |
| 41061_1 | TATGAGTGTGGAGTCGTTGCCACCCACCCTATTCCCAACTTATNNNAGATCGGAAGA |
| 41061_2 | TATGAGTGTGGAGTCGTTGCCACCCACCCTATTCCCAACTTATNNNAGATCGGAAGA |
| 41061_3 | TATGAGTGTGGAGTCGTTGCCACCCACCCTATTCCCAACTTATNNNAGATCGGAAGA |
| 41061_4 | TATGAGTGTGGAGTCGTTGCCACCCACCCTATTCCCAACTTATNNNAGATCGGAAGA |
| 34914_1 | TATGAGTGTGGAGTCGTTGCCGGGGTTCTAAGAGATTCTCCTGNNNAGATCGGAA |
| 34914_2 | TATGAGTGTGGAGTCGTTGCCGGGGTTCTAAGAGATTCTCCTGNNNAGATCGGAA |
| 34914_3 | TATGAGTGTGGAGTCGTTGCCGGGGTTCTAAGAGATTCTCCTGNNNAGATCGGAA |

| | |
|---|---|
| 34914_4 | TATGAGTGTGGAGTCGTTGCCGGGGTTCTAAGAGATTCTCCTGNNNAGATCGGAAG |
| 35628_1 | TATGAGTGTGGAGTCGTTGCATCAGCAAATTCTCTTCGGTCCTNNNAGATCGGAAGA |
| 35628_2 | TATGAGTGTGGAGTCGTTGCATCAGCAAATTCTCTTCGGTCCTNNNAGATCGGAAGA |
| 35628_3 | TATGAGTGTGGAGTCGTTGCATCAGCAAATTCTCTTCGGTCCTNNNAGATCGGAAGA |
| 35628_4 | TATGAGTGTGGAGTCGTTGCATCAGCAAATTCTCTTCGGTCCTNNNAGATCGGAAGA |
| 51484_1 | TATGAGTGTGGAGTCGTTGCTGTACCCAACAGCTCATTGAGAANNNAGATCGGAAG |
| 51484_2 | TATGAGTGTGGAGTCGTTGCTGTACCCAACAGCTCATTGAGAANNNAGATCGGAAG |
| 51484_3 | TATGAGTGTGGAGTCGTTGCTGTACCCAACAGCTCATTGAGAANNNAGATCGGAAG |
| 51484_4 | TATGAGTGTGGAGTCGTTGCTGTACCCAACAGCTCATTGAGAANNNAGATCGGAAG |
| 47013_1 | TATGAGTGTGGAGTCGTTGCCAAGTAGCTGGGACTACAGTCTCNNNAGATCGGAAG |
| 47013_2 | TATGAGTGTGGAGTCGTTGCCAAGTAGCTGGGACTACAGTCTCNNNAGATCGGAAG |
| 47013_3 | TATGAGTGTGGAGTCGTTGCCAAGTAGCTGGGACTACAGTCTCNNNAGATCGGAAG |
| 47013_4 | TATGAGTGTGGAGTCGTTGCCAAGTAGCTGGGACTACAGTCTCNNNAGATCGGAAG |
| 8295_1 | TATGAGTGTGGAGTCGTTGCGTGGGTTCAAGCACTTCTCAANNNAGATCGGAAGAG |
| 8295_2 | TATGAGTGTGGAGTCGTTGCGTGGGTTCAAGCACTTCTCAANNNAGATCGGAAGAG |
| 8295_3 | TATGAGTGTGGAGTCGTTGCGTGGGTTCAAGCACTTCTCAANNNAGATCGGAAGAG |
| 8295_4 | TATGAGTGTGGAGTCGTTGCGTGGGTTCAAGCACTTCTCAANNNAGATCGGAAGAG |
| 50028_1 | TATGAGTGTGGAGTCGTTGCGCCTTCCAAGTTCAAGCAATTCTNNNAGATCGGAAGA |
| 50028_2 | TATGAGTGTGGAGTCGTTGCGCCTTCCAAGTTCAAGCAATTCTNNNAGATCGGAAGA |
| 50028_3 | TATGAGTGTGGAGTCGTTGCGCCTTCCAAGTTCAAGCAATTCTNNNAGATCGGAAGA |
| 50028_4 | TATGAGTGTGGAGTCGTTGCGCCTTCCAAGTTCAAGCAATTCTNNNAGATCGGAAGA |
| 12119_1 | TATGAGTGTGGAGTCGTTGCCTGGAGCTCATTACTGGTACAACNNNAGATCGGAAG |
| 12119_2 | TATGAGTGTGGAGTCGTTGCCTGGAGCTCATTACTGGTACAACNNNAGATCGGAAG |
| 12119_3 | TATGAGTGTGGAGTCGTTGCCTGGAGCTCATTACTGGTACAACNNNAGATCGGAAG |
| 12119_4 | TATGAGTGTGGAGTCGTTGCCTGGAGCTCATTACTGGTACAACNNNAGATCGGAAG |
| 50389_1 | TATGAGTGTGGAGTCGTTGCCAGCCTGTGAAGGGAAGGAGNNNAGATCGGAAGAG |
| 50389_2 | TATGAGTGTGGAGTCGTTGCCAGCCTGTGAAGGGAAGGAGNNNAGATCGGAAGAG |
| 50389_3 | TATGAGTGTGGAGTCGTTGCCAGCCTGTGAAGGGAAGGAGNNNAGATCGGAAGAG |
| 50389_4 | TATGAGTGTGGAGTCGTTGCCAGCCTGTGAAGGGAAGGAGNNNAGATCGGAAGAG |
| 12488_1 | TATGAGTGTGGAGTCGTTGCAATCTGGTCTTCATCACCTANNNAGATCGGAAGAGCA |
| 12488_2 | TATGAGTGTGGAGTCGTTGCAATCTGGTCTTCATCACCTANNNAGATCGGAAGAGCA |
| 12488_3 | TATGAGTGTGGAGTCGTTGCAATCTGGTCTTCATCACCTANNNAGATCGGAAGAGCA |
| 12488_4 | TATGAGTGTGGAGTCGTTGCAATCTGGTCTTCATCACCTANNNAGATCGGAAGAGCA |
| 50728_1 | TATGAGTGTGGAGTCGTTGCTCAGTCTCAAACACAGTAGCTCCNNNAGATCGGAAGA |
| 50728_2 | TATGAGTGTGGAGTCGTTGCTCAGTCTCAAACACAGTAGCTCCNNNAGATCGGAAGA |
| 50728_3 | TATGAGTGTGGAGTCGTTGCTCAGTCTCAAACACAGTAGCTCCNNNAGATCGGAAGA |
| 50728_4 | TATGAGTGTGGAGTCGTTGCTCAGTCTCAAACACAGTAGCTCCNNNAGATCGGAAGA |
| 40079_1 | TATGAGTGTGGAGTCGTTGCGAATTGGACTAGGGAAACTGTGCNNNAGATCGGAA |
| 40079_2 | TATGAGTGTGGAGTCGTTGCGAATTGGACTAGGGAAACTGTGCNNNAGATCGGAA |
| 40079_3 | TATGAGTGTGGAGTCGTTGCGAATTGGACTAGGGAAACTGTGCNNNAGATCGGAA |
| 40079_4 | TATGAGTGTGGAGTCGTTGCGAATTGGACTAGGGAAACTGTGCNNNAGATCGGAA |
| 39813_1 | TATGAGTGTGGAGTCGTTGCTGATACGTTTGTCCACATGGAGANNNAGATCGGAAG |
| 39813_2 | TATGAGTGTGGAGTCGTTGCTGATACGTTTGTCCACATGGAGANNNAGATCGGAAG |

| | |
|---|---|
| 39813_3 | TATGAGTGTGGAGTCGTTGCTGATACGTTTGTCCACATGGAGANNNAGATCGGAAG |
| 39813_4 | TATGAGTGTGGAGTCGTTGCTGATACGTTTGTCCACATGGAGANNNAGATCGGAAG |
| 50056_1 | TATGAGTGTGGAGTCGTTGCTCCTGTTGGTTCTGTTTCTCTGGNNNAGATCGGAAG |
| 50056_2 | TATGAGTGTGGAGTCGTTGCTCCTGTTGGTTCTGTTTCTCTGGNNNAGATCGGAAG |
| 50056_3 | TATGAGTGTGGAGTCGTTGCTCCTGTTGGTTCTGTTTCTCTGGNNNAGATCGGAAG |
| 50056_4 | TATGAGTGTGGAGTCGTTGCTCCTGTTGGTTCTGTTTCTCTGGNNNAGATCGGAAG |
| 33906_1 | TATGAGTGTGGAGTCGTTGCTGATTCTTATGCCTCAACCTCCCNNNAGATCGGAAGA |
| 33906_2 | TATGAGTGTGGAGTCGTTGCTGATTCTTATGCCTCAACCTCCCNNNAGATCGGAAGA |
| 33906_3 | TATGAGTGTGGAGTCGTTGCTGATTCTTATGCCTCAACCTCCCNNNAGATCGGAAGA |
| 33906_4 | TATGAGTGTGGAGTCGTTGCTGATTCTTATGCCTCAACCTCCCNNNAGATCGGAAGA |
| 47951_1 | TATGAGTGTGGAGTCGTTGCCACCTCTATTCCAACACCCTCAANNNAGATCGGAAGA |
| 47951_2 | TATGAGTGTGGAGTCGTTGCCACCTCTATTCCAACACCCTCAANNNAGATCGGAAGA |
| 47951_3 | TATGAGTGTGGAGTCGTTGCCACCTCTATTCCAACACCCTCAANNNAGATCGGAAGA |
| 47951_4 | TATGAGTGTGGAGTCGTTGCCACCTCTATTCCAACACCCTCAANNNAGATCGGAAGA |
| 33164_1 | TATGAGTGTGGAGTCGTTGCTTCCTATACACCAACAACAGGCANNNAGATCGGAAG |
| 33164_2 | TATGAGTGTGGAGTCGTTGCTTCCTATACACCAACAACAGGCANNNAGATCGGAAG |
| 33164_3 | TATGAGTGTGGAGTCGTTGCTTCCTATACACCAACAACAGGCANNNAGATCGGAAG |
| 33164_4 | TATGAGTGTGGAGTCGTTGCTTCCTATACACCAACAACAGGCANNNAGATCGGAAG |
| 38676_1 | TATGAGTGTGGAGTCGTTGCTGCACCTGTAATCCCAACTACTCNNNAGATCGGAAGA |
| 38676_2 | TATGAGTGTGGAGTCGTTGCTGCACCTGTAATCCCAACTACTCNNNAGATCGGAAGA |
| 38676_3 | TATGAGTGTGGAGTCGTTGCTGCACCTGTAATCCCAACTACTCNNNAGATCGGAAGA |
| 38676_4 | TATGAGTGTGGAGTCGTTGCTGCACCTGTAATCCCAACTACTCNNNAGATCGGAAGA |
| 45954_1 | TATGAGTGTGGAGTCGTTGCTGTAAAGGGCTTAGCACATTCCTNNNAGATCGGAAG |
| 45954_2 | TATGAGTGTGGAGTCGTTGCTGTAAAGGGCTTAGCACATTCCTNNNAGATCGGAAG |
| 45954_3 | TATGAGTGTGGAGTCGTTGCTGTAAAGGGCTTAGCACATTCCTNNNAGATCGGAAG |
| 45954_4 | TATGAGTGTGGAGTCGTTGCTGTAAAGGGCTTAGCACATTCCTNNNAGATCGGAAG |
| 31910_1 | TATGAGTGTGGAGTCGTTGCGAGCCGGTATATAAGAGCTGCTTNNNAGATCGGAA |
| 31910_2 | TATGAGTGTGGAGTCGTTGCGAGCCGGTATATAAGAGCTGCTTNNNAGATCGGAA |
| 31910_3 | TATGAGTGTGGAGTCGTTGCGAGCCGGTATATAAGAGCTGCTTNNNAGATCGGAA |
| 31910_4 | TATGAGTGTGGAGTCGTTGCGAGCCGGTATATAAGAGCTGCTTNNNAGATCGGAA |
| 45142_1 | TATGAGTGTGGAGTCGTTGCGACAAAGTGAGTCACCATTTGGGNNNAGATCGGAA |
| 45142_2 | TATGAGTGTGGAGTCGTTGCGACAAAGTGAGTCACCATTTGGGNNNAGATCGGAA |
| 45142_3 | TATGAGTGTGGAGTCGTTGCGACAAAGTGAGTCACCATTTGGGNNNAGATCGGAA |
| 45142_4 | TATGAGTGTGGAGTCGTTGCGACAAAGTGAGTCACCATTTGGGNNNAGATCGGAA |
| 53014_1 | TATGAGTGTGGAGTCGTTGCGTTGCTGAGGTCTCACTCCCNNNAGATCGGAAGAGC |
| 53014_2 | TATGAGTGTGGAGTCGTTGCGTTGCTGAGGTCTCACTCCCNNNAGATCGGAAGAGC |
| 53014_3 | TATGAGTGTGGAGTCGTTGCGTTGCTGAGGTCTCACTCCCNNNAGATCGGAAGAGC |
| 53014_4 | TATGAGTGTGGAGTCGTTGCGTTGCTGAGGTCTCACTCCCNNNAGATCGGAAGAGC |
| 29893_1 | TATGAGTGTGGAGTCGTTGCGAGCATGGTGGTGTTTTGTTTTGNNNAGATCGGAA |
| 29893_2 | TATGAGTGTGGAGTCGTTGCGAGCATGGTGGTGTTTTGTTTTGNNNAGATCGGAA |
| 29893_3 | TATGAGTGTGGAGTCGTTGCGAGCATGGTGGTGTTTTGTTTTGNNNAGATCGGAA |
| 29893_4 | TATGAGTGTGGAGTCGTTGCGAGCATGGTGGTGTTTTGTTTTGNNNAGATCGGAA |
| 52550_1 | TATGAGTGTGGAGTCGTTGCAACACAGCATAGTGCCTAGTACCNNNAGATCGGAAG |

| | |
|---|---|
| 52550_2 | TATGAGTGTGGAGTCGTTGCAACACAGCATAGTGCCTAGTACCNNNAGATCGGAAG |
| 52550_3 | TATGAGTGTGGAGTCGTTGCAACACAGCATAGTGCCTAGTACCNNNAGATCGGAAG |
| 52550_4 | TATGAGTGTGGAGTCGTTGCAACACAGCATAGTGCCTAGTACCNNNAGATCGGAAG |
| 48052_1 | TATGAGTGTGGAGTCGTTGCCTTCCTCTCTAGCTGCCTTCAATNNNAGATCGGAAGA |
| 48052_2 | TATGAGTGTGGAGTCGTTGCCTTCCTCTCTAGCTGCCTTCAATNNNAGATCGGAAGA |
| 48052_3 | TATGAGTGTGGAGTCGTTGCCTTCCTCTCTAGCTGCCTTCAATNNNAGATCGGAAGA |
| 48052_4 | TATGAGTGTGGAGTCGTTGCCTTCCTCTCTAGCTGCCTTCAATNNNAGATCGGAAGA |
| 40762_1 | TATGAGTGTGGAGTCGTTGCGCAGCAAAGGCACTTTATAAGCCNNNAGATCGGAAG |
| 40762_2 | TATGAGTGTGGAGTCGTTGCGCAGCAAAGGCACTTTATAAGCCNNNAGATCGGAAG |
| 40762_3 | TATGAGTGTGGAGTCGTTGCGCAGCAAAGGCACTTTATAAGCCNNNAGATCGGAAG |
| 40762_4 | TATGAGTGTGGAGTCGTTGCGCAGCAAAGGCACTTTATAAGCCNNNAGATCGGAAG |
| 1896_1 | TATGAGTGTGGAGTCGTTGCCAAGAAAGGGAGATGGGGCTTTTNNNAGATCGGAA |
| 1896_2 | TATGAGTGTGGAGTCGTTGCCAAGAAAGGGAGATGGGGCTTTTNNNAGATCGGAA |
| 1896_3 | TATGAGTGTGGAGTCGTTGCCAAGAAAGGGAGATGGGGCTTTTNNNAGATCGGAA |
| 1896_4 | TATGAGTGTGGAGTCGTTGCCAAGAAAGGGAGATGGGGCTTTTNNNAGATCGGAA |
| 42328_1 | TATGAGTGTGGAGTCGTTGCTTCGAGCAATTCTCCTGTCTCAGNNNAGATCGGAAG |
| 42328_2 | TATGAGTGTGGAGTCGTTGCTTCGAGCAATTCTCCTGTCTCAGNNNAGATCGGAAG |
| 42328_3 | TATGAGTGTGGAGTCGTTGCTTCGAGCAATTCTCCTGTCTCAGNNNAGATCGGAAG |
| 42328_4 | TATGAGTGTGGAGTCGTTGCTTCGAGCAATTCTCCTGTCTCAGNNNAGATCGGAAG |
| 31909_1 | TATGAGTGTGGAGTCGTTGCCCTCCGATATAACCCACCCAAATNNNAGATCGGAAGA |
| 31909_2 | TATGAGTGTGGAGTCGTTGCCCTCCGATATAACCCACCCAAATNNNAGATCGGAAGA |
| 31909_3 | TATGAGTGTGGAGTCGTTGCCCTCCGATATAACCCACCCAAATNNNAGATCGGAAGA |
| 31909_4 | TATGAGTGTGGAGTCGTTGCCCTCCGATATAACCCACCCAAATNNNAGATCGGAAGA |
| 43932_1 | TATGAGTGTGGAGTCGTTGCGATCTCAGCTCACAGCAAACTCTNNNAGATCGGAAG |
| 43932_2 | TATGAGTGTGGAGTCGTTGCGATCTCAGCTCACAGCAAACTCTNNNAGATCGGAAG |
| 43932_3 | TATGAGTGTGGAGTCGTTGCGATCTCAGCTCACAGCAAACTCTNNNAGATCGGAAG |
| 43932_4 | TATGAGTGTGGAGTCGTTGCGATCTCAGCTCACAGCAAACTCTNNNAGATCGGAAG |
| 38211_1 | TATGAGTGTGGAGTCGTTGCCATCCTTGTCTTGTGCTGGTTTTNNNAGATCGGAAGA |
| 38211_2 | TATGAGTGTGGAGTCGTTGCCATCCTTGTCTTGTGCTGGTTTTNNNAGATCGGAAGA |
| 38211_3 | TATGAGTGTGGAGTCGTTGCCATCCTTGTCTTGTGCTGGTTTTNNNAGATCGGAAGA |
| 38211_4 | TATGAGTGTGGAGTCGTTGCCATCCTTGTCTTGTGCTGGTTTTNNNAGATCGGAAGA |
| 29554_1 | TATGAGTGTGGAGTCGTTGCCAGGGTTTTAAATTCCTGCAGCANNNAGATCGGAAG |
| 29554_2 | TATGAGTGTGGAGTCGTTGCCAGGGTTTTAAATTCCTGCAGCANNNAGATCGGAAG |
| 29554_3 | TATGAGTGTGGAGTCGTTGCCAGGGTTTTAAATTCCTGCAGCANNNAGATCGGAAG |
| 29554_4 | TATGAGTGTGGAGTCGTTGCCAGGGTTTTAAATTCCTGCAGCANNNAGATCGGAAG |
| 47523_1 | TATGAGTGTGGAGTCGTTGCGAAGGGTCTTATCCTCTCTGCAGNNNAGATCGGAAG |
| 47523_2 | TATGAGTGTGGAGTCGTTGCGAAGGGTCTTATCCTCTCTGCAGNNNAGATCGGAAG |
| 47523_3 | TATGAGTGTGGAGTCGTTGCGAAGGGTCTTATCCTCTCTGCAGNNNAGATCGGAAG |
| 47523_4 | TATGAGTGTGGAGTCGTTGCGAAGGGTCTTATCCTCTCTGCAGNNNAGATCGGAAG |
| 36399_1 | TATGAGTGTGGAGTCGTTGCAAGAAAGGCAACCCAAAGAATGGNNNAGATCGGAA |
| 36399_2 | TATGAGTGTGGAGTCGTTGCAAGAAAGGCAACCCAAAGAATGGNNNAGATCGGAA |
| 36399_3 | TATGAGTGTGGAGTCGTTGCAAGAAAGGCAACCCAAAGAATGGNNNAGATCGGAA |
| 36399_4 | TATGAGTGTGGAGTCGTTGCAAGAAAGGCAACCCAAAGAATGGNNNAGATCGGAA |

| | |
|---|---|
| 31168_1 | TATGAGTGTGGAGTCGTTGCCCTCTTCTGCTTTCACACACTTGNNNAGATCGGAAGA |
| 31168_2 | TATGAGTGTGGAGTCGTTGCCCTCTTCTGCTTTCACACACTTGNNNAGATCGGAAGA |
| 31168_3 | TATGAGTGTGGAGTCGTTGCCCTCTTCTGCTTTCACACACTTGNNNAGATCGGAAGA |
| 31168_4 | TATGAGTGTGGAGTCGTTGCCCTCTTCTGCTTTCACACACTTGNNNAGATCGGAAGA |
| 32802_1 | TATGAGTGTGGAGTCGTTGCGAAACAGAGTCTCACGCTGTTGNNNAGATCGGAAG |
| 32802_2 | TATGAGTGTGGAGTCGTTGCGAAACAGAGTCTCACGCTGTTGNNNAGATCGGAAG |
| 32802_3 | TATGAGTGTGGAGTCGTTGCGAAACAGAGTCTCACGCTGTTGNNNAGATCGGAAG |
| 32802_4 | TATGAGTGTGGAGTCGTTGCGAAACAGAGTCTCACGCTGTTGNNNAGATCGGAAG |
| 40085_1 | TATGAGTGTGGAGTCGTTGCAAAGAAAATGAAACCTCAGCCCCNNNAGATCGGAAG |
| 40085_2 | TATGAGTGTGGAGTCGTTGCAAAGAAAATGAAACCTCAGCCCCNNNAGATCGGAAG |
| 40085_3 | TATGAGTGTGGAGTCGTTGCAAAGAAAATGAAACCTCAGCCCCNNNAGATCGGAAG |
| 40085_4 | TATGAGTGTGGAGTCGTTGCAAAGAAAATGAAACCTCAGCCCCNNNAGATCGGAAG |
| 52739_1 | TATGAGTGTGGAGTCGTTGCATCCAATGGCGACTCGACAGNNNAGATCGGAAGAG |
| 52739_2 | TATGAGTGTGGAGTCGTTGCATCCAATGGCGACTCGACAGNNNAGATCGGAAGAG |
| 52739_3 | TATGAGTGTGGAGTCGTTGCATCCAATGGCGACTCGACAGNNNAGATCGGAAGAG |
| 52739_4 | TATGAGTGTGGAGTCGTTGCATCCAATGGCGACTCGACAGNNNAGATCGGAAGAG |
| 52795_1 | TATGAGTGTGGAGTCGTTGCCATTGGCATCCCCCCTAGAAGNNNAGATCGGAAGAG |
| 52795_2 | TATGAGTGTGGAGTCGTTGCCATTGGCATCCCCCCTAGAAGNNNAGATCGGAAGAG |
| 52795_3 | TATGAGTGTGGAGTCGTTGCCATTGGCATCCCCCCTAGAAGNNNAGATCGGAAGAG |
| 52795_4 | TATGAGTGTGGAGTCGTTGCCATTGGCATCCCCCCTAGAAGNNNAGATCGGAAGAG |
| 39936_1 | TATGAGTGTGGAGTCGTTGCGCGGATCATGAGATCAGGAGATTNNNAGATCGGAA |
| 39936_2 | TATGAGTGTGGAGTCGTTGCGCGGATCATGAGATCAGGAGATTNNNAGATCGGAA |
| 39936_3 | TATGAGTGTGGAGTCGTTGCGCGGATCATGAGATCAGGAGATTNNNAGATCGGAA |
| 39936_4 | TATGAGTGTGGAGTCGTTGCGCGGATCATGAGATCAGGAGATTNNNAGATCGGAA |
| 45499_1 | TATGAGTGTGGAGTCGTTGCTCTAACACTCGGAGTTGCTTACCNNNAGATCGGAAG |
| 45499_2 | TATGAGTGTGGAGTCGTTGCTCTAACACTCGGAGTTGCTTACCNNNAGATCGGAAG |
| 45499_3 | TATGAGTGTGGAGTCGTTGCTCTAACACTCGGAGTTGCTTACCNNNAGATCGGAAG |
| 45499_4 | TATGAGTGTGGAGTCGTTGCTCTAACACTCGGAGTTGCTTACCNNNAGATCGGAAG |
| 2283_1 | TATGAGTGTGGAGTCGTTGCTGAAGTGGGCAAAGAGAAGGAACNNNAGATCGGA |
| 2283_2 | TATGAGTGTGGAGTCGTTGCTGAAGTGGGCAAAGAGAAGGAACNNNAGATCGGA |
| 2283_3 | TATGAGTGTGGAGTCGTTGCTGAAGTGGGCAAAGAGAAGGAACNNNAGATCGGA |
| 2283_4 | TATGAGTGTGGAGTCGTTGCTGAAGTGGGCAAAGAGAAGGAACNNNAGATCGGA |
| 38682_1 | TATGAGTGTGGAGTCGTTGCGCAAAGGACATGAACAGACACTTNNNAGATCGGAA |
| 38682_2 | TATGAGTGTGGAGTCGTTGCGCAAAGGACATGAACAGACACTTNNNAGATCGGAA |
| 38682_3 | TATGAGTGTGGAGTCGTTGCGCAAAGGACATGAACAGACACTTNNNAGATCGGAA |
| 38682_4 | TATGAGTGTGGAGTCGTTGCGCAAAGGACATGAACAGACACTTNNNAGATCGGAA |
| 49790_1 | TATGAGTGTGGAGTCGTTGCTGCTACTTGGTGCCAATTTTGAGNNNAGATCGGAAG |
| 49790_2 | TATGAGTGTGGAGTCGTTGCTGCTACTTGGTGCCAATTTTGAGNNNAGATCGGAAG |
| 49790_3 | TATGAGTGTGGAGTCGTTGCTGCTACTTGGTGCCAATTTTGAGNNNAGATCGGAAG |
| 49790_4 | TATGAGTGTGGAGTCGTTGCTGCTACTTGGTGCCAATTTTGAGNNNAGATCGGAAG |
| 39847_1 | TATGAGTGTGGAGTCGTTGCTTCAAGATCCTTGGTGCAGAGAANNNAGATCGGAA |
| 39847_2 | TATGAGTGTGGAGTCGTTGCTTCAAGATCCTTGGTGCAGAGAANNNAGATCGGAA |
| 39847_3 | TATGAGTGTGGAGTCGTTGCTTCAAGATCCTTGGTGCAGAGAANNNAGATCGGAA |

| | |
|---|---|
| 39847_4 | TATGAGTGTGGAGTCGTTGCTTCAAGATCCTTGGTGCAGAGAANNNAGATCGGAA[illegible] |
| 41644_1 | TATGAGTGTGGAGTCGTTGCGATGGTGTCTCACTCTGTTACCCNNNAGATCGGAAG[illegible] |
| 41644_2 | TATGAGTGTGGAGTCGTTGCGATGGTGTCTCACTCTGTTACCCNNNAGATCGGAAG[illegible] |
| 41644_3 | TATGAGTGTGGAGTCGTTGCGATGGTGTCTCACTCTGTTACCCNNNAGATCGGAAG[illegible] |
| 41644_4 | TATGAGTGTGGAGTCGTTGCGATGGTGTCTCACTCTGTTACCCNNNAGATCGGAAG[illegible] |
| 12492_1 | TATGAGTGTGGAGTCGTTGCGAGGAACATAACAAACTTTACATNNNAGATCGGAAG |
| 12492_2 | TATGAGTGTGGAGTCGTTGCGAGGAACATAACAAACTTTACATNNNAGATCGGAAG |
| 12492_3 | TATGAGTGTGGAGTCGTTGCGAGGAACATAACAAACTTTACATNNNAGATCGGAAG |
| 12492_4 | TATGAGTGTGGAGTCGTTGCGAGGAACATAACAAACTTTACATNNNAGATCGGAAG |
| 46616_1 | TATGAGTGTGGAGTCGTTGCTAGTTGGGACTATAGGCATGTGCNNNAGATCGGAA[illegible] |
| 46616_2 | TATGAGTGTGGAGTCGTTGCTAGTTGGGACTATAGGCATGTGCNNNAGATCGGAA[illegible] |
| 46616_3 | TATGAGTGTGGAGTCGTTGCTAGTTGGGACTATAGGCATGTGCNNNAGATCGGAA[illegible] |
| 46616_4 | TATGAGTGTGGAGTCGTTGCTAGTTGGGACTATAGGCATGTGCNNNAGATCGGAA[illegible] |
| 36293_1 | TATGAGTGTGGAGTCGTTGCCACCACCACCAACAAAAACCTAGNNNAGATCGGAAG[illegible] |
| 36293_2 | TATGAGTGTGGAGTCGTTGCCACCACCACCAACAAAAACCTAGNNNAGATCGGAAG[illegible] |
| 36293_3 | TATGAGTGTGGAGTCGTTGCCACCACCACCAACAAAAACCTAGNNNAGATCGGAAG[illegible] |
| 36293_4 | TATGAGTGTGGAGTCGTTGCCACCACCACCAACAAAAACCTAGNNNAGATCGGAAG[illegible] |
| 37351_1 | TATGAGTGTGGAGTCGTTGCAGCCAAAACGTGGAAACTAATGTNNNAGATCGGAA[illegible] |
| 37351_2 | TATGAGTGTGGAGTCGTTGCAGCCAAAACGTGGAAACTAATGTNNNAGATCGGAA[illegible] |
| 37351_3 | TATGAGTGTGGAGTCGTTGCAGCCAAAACGTGGAAACTAATGTNNNAGATCGGAA[illegible] |
| 37351_4 | TATGAGTGTGGAGTCGTTGCAGCCAAAACGTGGAAACTAATGTNNNAGATCGGAA[illegible] |
| 7978_1 | TATGAGTGTGGAGTCGTTGCACGACAGAGTTTCACTTTCTTGNNNAGATCGGAAGA[illegible] |
| 7978_2 | TATGAGTGTGGAGTCGTTGCACGACAGAGTTTCACTTTCTTGNNNAGATCGGAAGA[illegible] |
| 7978_3 | TATGAGTGTGGAGTCGTTGCACGACAGAGTTTCACTTTCTTGNNNAGATCGGAAGA[illegible] |
| 7978_4 | TATGAGTGTGGAGTCGTTGCACGACAGAGTTTCACTTTCTTGNNNAGATCGGAAGA[illegible] |
| 42753_1 | TATGAGTGTGGAGTCGTTGCTAATGGTATGCGTCCCAGTTAGGNNNAGATCGGAA[illegible] |
| 42753_2 | TATGAGTGTGGAGTCGTTGCTAATGGTATGCGTCCCAGTTAGGNNNAGATCGGAA[illegible] |
| 42753_3 | TATGAGTGTGGAGTCGTTGCTAATGGTATGCGTCCCAGTTAGGNNNAGATCGGAA[illegible] |
| 42753_4 | TATGAGTGTGGAGTCGTTGCTAATGGTATGCGTCCCAGTTAGGNNNAGATCGGAA[illegible] |
| 50156_1 | TATGAGTGTGGAGTCGTTGCTGTTTTTACTCACTTCCTGCCCTNNNAGATCGGAAGA[illegible] |
| 50156_2 | TATGAGTGTGGAGTCGTTGCTGTTTTTACTCACTTCCTGCCCTNNNAGATCGGAAGA[illegible] |
| 50156_3 | TATGAGTGTGGAGTCGTTGCTGTTTTTACTCACTTCCTGCCCTNNNAGATCGGAAGA[illegible] |
| 50156_4 | TATGAGTGTGGAGTCGTTGCTGTTTTTACTCACTTCCTGCCCTNNNAGATCGGAAGA[illegible] |
| 43649_1 | TATGAGTGTGGAGTCGTTGCGACTTTGGCTCACAACGTTCTCNNNAGATCGGAAGA[illegible] |
| 43649_2 | TATGAGTGTGGAGTCGTTGCGACTTTGGCTCACAACGTTCTCNNNAGATCGGAAGA[illegible] |
| 43649_3 | TATGAGTGTGGAGTCGTTGCGACTTTGGCTCACAACGTTCTCNNNAGATCGGAAGA[illegible] |
| 43649_4 | TATGAGTGTGGAGTCGTTGCGACTTTGGCTCACAACGTTCTCNNNAGATCGGAAGA[illegible] |
| 46126_1 | TATGAGTGTGGAGTCGTTGCATAAAGTCTTGCTCTGTCACCCANNNAGATCGGAAG[illegible] |
| 46126_2 | TATGAGTGTGGAGTCGTTGCATAAAGTCTTGCTCTGTCACCCANNNAGATCGGAAG[illegible] |
| 46126_3 | TATGAGTGTGGAGTCGTTGCATAAAGTCTTGCTCTGTCACCCANNNAGATCGGAAG[illegible] |
| 46126_4 | TATGAGTGTGGAGTCGTTGCATAAAGTCTTGCTCTGTCACCCANNNAGATCGGAAG[illegible] |
| 42836_1 | TATGAGTGTGGAGTCGTTGCCGTAACGTGGATGAGAAGTCTGANNNAGATCGGAA |
| 42836_2 | TATGAGTGTGGAGTCGTTGCCGTAACGTGGATGAGAAGTCTGANNNAGATCGGAA |

| | |
|---|---|
| 42836_3 | TATGAGTGTGGAGTCGTTGCCGTAACGTGGATGAGAAGTCTGANNNAGATCGGAA |
| 42836_4 | TATGAGTGTGGAGTCGTTGCCGTAACGTGGATGAGAAGTCTGANNNAGATCGGAA |
| 40400_1 | TATGAGTGTGGAGTCGTTGCCAACATGTCATGGCTCTTTCAGCNNNAGATCGGAAG |
| 40400_2 | TATGAGTGTGGAGTCGTTGCCAACATGTCATGGCTCTTTCAGCNNNAGATCGGAAG |
| 40400_3 | TATGAGTGTGGAGTCGTTGCCAACATGTCATGGCTCTTTCAGCNNNAGATCGGAAG |
| 40400_4 | TATGAGTGTGGAGTCGTTGCCAACATGTCATGGCTCTTTCAGCNNNAGATCGGAAG |
| 9968_1 | TATGAGTGTGGAGTCGTTGCAAAGGGGTAACCCAGCAAAGAAGNNNAGATCGGAA |
| 9968_2 | TATGAGTGTGGAGTCGTTGCAAAGGGGTAACCCAGCAAAGAAGNNNAGATCGGAA |
| 9968_3 | TATGAGTGTGGAGTCGTTGCAAAGGGGTAACCCAGCAAAGAAGNNNAGATCGGAA |
| 9968_4 | TATGAGTGTGGAGTCGTTGCAAAGGGGTAACCCAGCAAAGAAGNNNAGATCGGAA |
| 34197_1 | TATGAGTGTGGAGTCGTTGCACCACTGATCCCACAGAAATACANNNAGATCGGAAG |
| 34197_2 | TATGAGTGTGGAGTCGTTGCACCACTGATCCCACAGAAATACANNNAGATCGGAAG |
| 34197_3 | TATGAGTGTGGAGTCGTTGCACCACTGATCCCACAGAAATACANNNAGATCGGAAG |
| 34197_4 | TATGAGTGTGGAGTCGTTGCACCACTGATCCCACAGAAATACANNNAGATCGGAAG |
| 41124_1 | TATGAGTGTGGAGTCGTTGCATTTTTCTGTGCAATCACCCACANNNAGATCGGAAG |
| 41124_2 | TATGAGTGTGGAGTCGTTGCATTTTTCTGTGCAATCACCCACANNNAGATCGGAAG |
| 41124_3 | TATGAGTGTGGAGTCGTTGCATTTTTCTGTGCAATCACCCACANNNAGATCGGAAG |
| 41124_4 | TATGAGTGTGGAGTCGTTGCATTTTTCTGTGCAATCACCCACANNNAGATCGGAAG |
| 37437_1 | TATGAGTGTGGAGTCGTTGCATAGTTGAGCCCTCTGTTGTACCNNNAGATCGGAAG |
| 37437_2 | TATGAGTGTGGAGTCGTTGCATAGTTGAGCCCTCTGTTGTACCNNNAGATCGGAAG |
| 37437_3 | TATGAGTGTGGAGTCGTTGCATAGTTGAGCCCTCTGTTGTACCNNNAGATCGGAAG |
| 37437_4 | TATGAGTGTGGAGTCGTTGCATAGTTGAGCCCTCTGTTGTACCNNNAGATCGGAAG |
| 34874_1 | TATGAGTGTGGAGTCGTTGCGATCTCGTCTCACTGCAACAGAGNNNAGATCGGAAG |
| 34874_2 | TATGAGTGTGGAGTCGTTGCGATCTCGTCTCACTGCAACAGAGNNNAGATCGGAAG |
| 34874_3 | TATGAGTGTGGAGTCGTTGCGATCTCGTCTCACTGCAACAGAGNNNAGATCGGAAG |
| 34874_4 | TATGAGTGTGGAGTCGTTGCGATCTCGTCTCACTGCAACAGAGNNNAGATCGGAAG |
| 30446_1 | TATGAGTGTGGAGTCGTTGCGGAGAAATTGAGATAGAGTAGTTNNNAGATCGGAA |
| 30446_2 | TATGAGTGTGGAGTCGTTGCGGAGAAATTGAGATAGAGTAGTTNNNAGATCGGAA |
| 30446_3 | TATGAGTGTGGAGTCGTTGCGGAGAAATTGAGATAGAGTAGTTNNNAGATCGGAA |
| 30446_4 | TATGAGTGTGGAGTCGTTGCGGAGAAATTGAGATAGAGTAGTTNNNAGATCGGAA |
| 39131_1 | TATGAGTGTGGAGTCGTTGCCTGTGGTTGACAATGATTTGGGGNNNAGATCGGAA |
| 39131_2 | TATGAGTGTGGAGTCGTTGCCTGTGGTTGACAATGATTTGGGGNNNAGATCGGAA |
| 39131_3 | TATGAGTGTGGAGTCGTTGCCTGTGGTTGACAATGATTTGGGGNNNAGATCGGAA |
| 39131_4 | TATGAGTGTGGAGTCGTTGCCTGTGGTTGACAATGATTTGGGGNNNAGATCGGAA |
| 43694_1 | TATGAGTGTGGAGTCGTTGCCCTTTTCAGGGATGAGGTCTTGGNNNAGATCGGAA |
| 43694_2 | TATGAGTGTGGAGTCGTTGCCCTTTTCAGGGATGAGGTCTTGGNNNAGATCGGAA |
| 43694_3 | TATGAGTGTGGAGTCGTTGCCCTTTTCAGGGATGAGGTCTTGGNNNAGATCGGAA |
| 43694_4 | TATGAGTGTGGAGTCGTTGCCCTTTTCAGGGATGAGGTCTTGGNNNAGATCGGAA |
| 43266_1 | TATGAGTGTGGAGTCGTTGCGAGTAGCTGGGACTACAGATGTGNNNAGATCGGAA |
| 43266_2 | TATGAGTGTGGAGTCGTTGCGAGTAGCTGGGACTACAGATGTGNNNAGATCGGAA |
| 43266_3 | TATGAGTGTGGAGTCGTTGCGAGTAGCTGGGACTACAGATGTGNNNAGATCGGAA |
| 43266_4 | TATGAGTGTGGAGTCGTTGCGAGTAGCTGGGACTACAGATGTGNNNAGATCGGAA |
| 41171_1 | TATGAGTGTGGAGTCGTTGCTAGGAACTTGTCAGCCACTCAAANNNAGATCGGAAG |

| | |
|---|---|
| 41171_2 | TATGAGTGTGGAGTCGTTGCTAGGAACTTGTCAGCCACTCAAANNNAGATCGGAAG |
| 41171_3 | TATGAGTGTGGAGTCGTTGCTAGGAACTTGTCAGCCACTCAAANNNAGATCGGAAG |
| 41171_4 | TATGAGTGTGGAGTCGTTGCTAGGAACTTGTCAGCCACTCAAANNNAGATCGGAAG |
| 43715_1 | TATGAGTGTGGAGTCGTTGCTGTGCGTTTATACATGTGTGTGTNNNAGATCGGAAG |
| 43715_2 | TATGAGTGTGGAGTCGTTGCTGTGCGTTTATACATGTGTGTGTNNNAGATCGGAAG |
| 43715_3 | TATGAGTGTGGAGTCGTTGCTGTGCGTTTATACATGTGTGTGTNNNAGATCGGAAG |
| 43715_4 | TATGAGTGTGGAGTCGTTGCTGTGCGTTTATACATGTGTGTGTNNNAGATCGGAAG |
| 32910_1 | TATGAGTGTGGAGTCGTTGCAGTTGTGGCAATTTGAACCCATTNNNAGATCGGAAG |
| 32910_2 | TATGAGTGTGGAGTCGTTGCAGTTGTGGCAATTTGAACCCATTNNNAGATCGGAAG |
| 32910_3 | TATGAGTGTGGAGTCGTTGCAGTTGTGGCAATTTGAACCCATTNNNAGATCGGAAG |
| 32910_4 | TATGAGTGTGGAGTCGTTGCAGTTGTGGCAATTTGAACCCATTNNNAGATCGGAAG |
| 12046_1 | TATGAGTGTGGAGTCGTTGCATGTCTGAGGGCACAGGAGCCNNNAGATCGGAAGA |
| 12046_2 | TATGAGTGTGGAGTCGTTGCATGTCTGAGGGCACAGGAGCCNNNAGATCGGAAGA |
| 12046_3 | TATGAGTGTGGAGTCGTTGCATGTCTGAGGGCACAGGAGCCNNNAGATCGGAAGA |
| 12046_4 | TATGAGTGTGGAGTCGTTGCATGTCTGAGGGCACAGGAGCCNNNAGATCGGAAGA |
| 37231_1 | TATGAGTGTGGAGTCGTTGCTTGTTTCTCCCTCTCCTTGTTGANNNAGATCGGAAGA |
| 37231_2 | TATGAGTGTGGAGTCGTTGCTTGTTTCTCCCTCTCCTTGTTGANNNAGATCGGAAGA |
| 37231_3 | TATGAGTGTGGAGTCGTTGCTTGTTTCTCCCTCTCCTTGTTGANNNAGATCGGAAGA |
| 37231_4 | TATGAGTGTGGAGTCGTTGCTTGTTTCTCCCTCTCCTTGTTGANNNAGATCGGAAGA |
| 39109_1 | TATGAGTGTGGAGTCGTTGCAGATACTGACTAACCGACCTCCTNNNAGATCGGAAG |
| 39109_2 | TATGAGTGTGGAGTCGTTGCAGATACTGACTAACCGACCTCCTNNNAGATCGGAAG |
| 39109_3 | TATGAGTGTGGAGTCGTTGCAGATACTGACTAACCGACCTCCTNNNAGATCGGAAG |
| 39109_4 | TATGAGTGTGGAGTCGTTGCAGATACTGACTAACCGACCTCCTNNNAGATCGGAAG |
| 44900_1 | TATGAGTGTGGAGTCGTTGCGAACAGAGTGACTAAAACGTGCCNNNAGATCGGAA |
| 44900_2 | TATGAGTGTGGAGTCGTTGCGAACAGAGTGACTAAAACGTGCCNNNAGATCGGAA |
| 44900_3 | TATGAGTGTGGAGTCGTTGCGAACAGAGTGACTAAAACGTGCCNNNAGATCGGAA |
| 44900_4 | TATGAGTGTGGAGTCGTTGCGAACAGAGTGACTAAAACGTGCCNNNAGATCGGAA |
| 37779_1 | TATGAGTGTGGAGTCGTTGCGGGTTGAGACATTGCACCAACNNNAGATCGGAAGA |
| 37779_2 | TATGAGTGTGGAGTCGTTGCGGGTTGAGACATTGCACCAACNNNAGATCGGAAGA |
| 37779_3 | TATGAGTGTGGAGTCGTTGCGGGTTGAGACATTGCACCAACNNNAGATCGGAAGA |
| 37779_4 | TATGAGTGTGGAGTCGTTGCGGGTTGAGACATTGCACCAACNNNAGATCGGAAGA |
| 38500_1 | TATGAGTGTGGAGTCGTTGCAAGTTCAGCATCCTGAGAATCCCNNNAGATCGGAAG |
| 38500_2 | TATGAGTGTGGAGTCGTTGCAAGTTCAGCATCCTGAGAATCCCNNNAGATCGGAAG |
| 38500_3 | TATGAGTGTGGAGTCGTTGCAAGTTCAGCATCCTGAGAATCCCNNNAGATCGGAAG |
| 38500_4 | TATGAGTGTGGAGTCGTTGCAAGTTCAGCATCCTGAGAATCCCNNNAGATCGGAAG |
| 41431_1 | TATGAGTGTGGAGTCGTTGCGGCATGGGCTACTGACTGAATATNNNAGATCGGAA |
| 41431_2 | TATGAGTGTGGAGTCGTTGCGGCATGGGCTACTGACTGAATATNNNAGATCGGAA |
| 41431_3 | TATGAGTGTGGAGTCGTTGCGGCATGGGCTACTGACTGAATATNNNAGATCGGAA |
| 41431_4 | TATGAGTGTGGAGTCGTTGCGGCATGGGCTACTGACTGAATATNNNAGATCGGAA |
| 43054_1 | TATGAGTGTGGAGTCGTTGCCTCTAAGCTAAGGACAGCAGACCNNNAGATCGGAAG |
| 43054_2 | TATGAGTGTGGAGTCGTTGCCTCTAAGCTAAGGACAGCAGACCNNNAGATCGGAAG |
| 43054_3 | TATGAGTGTGGAGTCGTTGCCTCTAAGCTAAGGACAGCAGACCNNNAGATCGGAAG |
| 43054_4 | TATGAGTGTGGAGTCGTTGCCTCTAAGCTAAGGACAGCAGACCNNNAGATCGGAAG |

| | |
|---|---|
| 36310_1 | TATGAGTGTGGAGTCGTTGCAGAAGAAGGCGAAGAAGGAGAAGNNNAGATCGGA |
| 36310_2 | TATGAGTGTGGAGTCGTTGCAGAAGAAGGCGAAGAAGGAGAAGNNNAGATCGGA |
| 36310_3 | TATGAGTGTGGAGTCGTTGCAGAAGAAGGCGAAGAAGGAGAAGNNNAGATCGGA |
| 36310_4 | TATGAGTGTGGAGTCGTTGCAGAAGAAGGCGAAGAAGGAGAAGNNNAGATCGGA |
| 38665_1 | TATGAGTGTGGAGTCGTTGCTGTAAACTACAATGCCCACACACNNNAGATCGGAAG |
| 38665_2 | TATGAGTGTGGAGTCGTTGCTGTAAACTACAATGCCCACACACNNNAGATCGGAAG |
| 38665_3 | TATGAGTGTGGAGTCGTTGCTGTAAACTACAATGCCCACACACNNNAGATCGGAAG |
| 38665_4 | TATGAGTGTGGAGTCGTTGCTGTAAACTACAATGCCCACACACNNNAGATCGGAAG |
| 45974_1 | TATGAGTGTGGAGTCGTTGCGTCTCACTCTATCACCCAGGTTGNNNAGATCGGAAG |
| 45974_2 | TATGAGTGTGGAGTCGTTGCGTCTCACTCTATCACCCAGGTTGNNNAGATCGGAAG |
| 45974_3 | TATGAGTGTGGAGTCGTTGCGTCTCACTCTATCACCCAGGTTGNNNAGATCGGAAG |
| 45974_4 | TATGAGTGTGGAGTCGTTGCGTCTCACTCTATCACCCAGGTTGNNNAGATCGGAAG |
| 31934_1 | TATGAGTGTGGAGTCGTTGCTCCTTGAAACATGGAGATGGAGGNNNAGATCGGAA |
| 31934_2 | TATGAGTGTGGAGTCGTTGCTCCTTGAAACATGGAGATGGAGGNNNAGATCGGAA |
| 31934_3 | TATGAGTGTGGAGTCGTTGCTCCTTGAAACATGGAGATGGAGGNNNAGATCGGAA |
| 31934_4 | TATGAGTGTGGAGTCGTTGCTCCTTGAAACATGGAGATGGAGGNNNAGATCGGAA |
| 45967_1 | TATGAGTGTGGAGTCGTTGCGATTCAAGCGATTCTCCTGCTTCNNNAGATCGGAAG |
| 45967_2 | TATGAGTGTGGAGTCGTTGCGATTCAAGCGATTCTCCTGCTTCNNNAGATCGGAAG |
| 45967_3 | TATGAGTGTGGAGTCGTTGCGATTCAAGCGATTCTCCTGCTTCNNNAGATCGGAAG |
| 45967_4 | TATGAGTGTGGAGTCGTTGCGATTCAAGCGATTCTCCTGCTTCNNNAGATCGGAAG |
| 38659_1 | TATGAGTGTGGAGTCGTTGCCCACTGACTCTTCATCCACCTGNNNAGATCGGAAGAG |
| 38659_2 | TATGAGTGTGGAGTCGTTGCCCACTGACTCTTCATCCACCTGNNNAGATCGGAAGAG |
| 38659_3 | TATGAGTGTGGAGTCGTTGCCCACTGACTCTTCATCCACCTGNNNAGATCGGAAGAG |
| 38659_4 | TATGAGTGTGGAGTCGTTGCCCACTGACTCTTCATCCACCTGNNNAGATCGGAAGAG |
| 49135_1 | TATGAGTGTGGAGTCGTTGCGCCTCTTGAGTAGCTGGGATTACNNNAGATCGGAAG |
| 49135_2 | TATGAGTGTGGAGTCGTTGCGCCTCTTGAGTAGCTGGGATTACNNNAGATCGGAAG |
| 49135_3 | TATGAGTGTGGAGTCGTTGCGCCTCTTGAGTAGCTGGGATTACNNNAGATCGGAAG |
| 49135_4 | TATGAGTGTGGAGTCGTTGCGCCTCTTGAGTAGCTGGGATTACNNNAGATCGGAAG |
| 45281_1 | TATGAGTGTGGAGTCGTTGCTTATCCTTCCAATGTGGGCAGAANNNAGATCGGAAG |
| 45281_2 | TATGAGTGTGGAGTCGTTGCTTATCCTTCCAATGTGGGCAGAANNNAGATCGGAAG |
| 45281_3 | TATGAGTGTGGAGTCGTTGCTTATCCTTCCAATGTGGGCAGAANNNAGATCGGAAG |
| 45281_4 | TATGAGTGTGGAGTCGTTGCTTATCCTTCCAATGTGGGCAGAANNNAGATCGGAAG |
| 41475_1 | TATGAGTGTGGAGTCGTTGCCCTGAAAATGTGGAAGCGACTTCNNNAGATCGGAA |
| 41475_2 | TATGAGTGTGGAGTCGTTGCCCTGAAAATGTGGAAGCGACTTCNNNAGATCGGAA |
| 41475_3 | TATGAGTGTGGAGTCGTTGCCCTGAAAATGTGGAAGCGACTTCNNNAGATCGGAA |
| 41475_4 | TATGAGTGTGGAGTCGTTGCCCTGAAAATGTGGAAGCGACTTCNNNAGATCGGAA |
| 47359_1 | TATGAGTGTGGAGTCGTTGCTAGTAGAGATGGGGTTTCTCCGTNNNAGATCGGAA |
| 47359_2 | TATGAGTGTGGAGTCGTTGCTAGTAGAGATGGGGTTTCTCCGTNNNAGATCGGAA |
| 47359_3 | TATGAGTGTGGAGTCGTTGCTAGTAGAGATGGGGTTTCTCCGTNNNAGATCGGAA |
| 47359_4 | TATGAGTGTGGAGTCGTTGCTAGTAGAGATGGGGTTTCTCCGTNNNAGATCGGAA |
| 10053_1 | TATGAGTGTGGAGTCGTTGCCACAAGACAAGAAACTGAGGCCTNNNAGATCGGAA |
| 10053_2 | TATGAGTGTGGAGTCGTTGCCACAAGACAAGAAACTGAGGCCTNNNAGATCGGAA |
| 10053_3 | TATGAGTGTGGAGTCGTTGCCACAAGACAAGAAACTGAGGCCTNNNAGATCGGAA |

| | |
|---|---|
| 10053_4 | TATGAGTGTGGAGTCGTTGCCACAAGACAAGAAACTGAGGCCTNNNAGATCGGAA |
| 45475_1 | TATGAGTGTGGAGTCGTTGCATATGGCAGGAAAATGGGAACCTNNNAGATCGGAA |
| 45475_2 | TATGAGTGTGGAGTCGTTGCATATGGCAGGAAAATGGGAACCTNNNAGATCGGAA |
| 45475_3 | TATGAGTGTGGAGTCGTTGCATATGGCAGGAAAATGGGAACCTNNNAGATCGGAA |
| 45475_4 | TATGAGTGTGGAGTCGTTGCATATGGCAGGAAAATGGGAACCTNNNAGATCGGAA |
| 36763_1 | TATGAGTGTGGAGTCGTTGCATGTCTCTCTGAGGTGAAGCTCTNNNAGATCGGAAG |
| 36763_2 | TATGAGTGTGGAGTCGTTGCATGTCTCTCTGAGGTGAAGCTCTNNNAGATCGGAAG |
| 36763_3 | TATGAGTGTGGAGTCGTTGCATGTCTCTCTGAGGTGAAGCTCTNNNAGATCGGAAG |
| 36763_4 | TATGAGTGTGGAGTCGTTGCATGTCTCTCTGAGGTGAAGCTCTNNNAGATCGGAAG |
| 29682_1 | TATGAGTGTGGAGTCGTTGCTACCAGTATGTGTTCAACCAGCANNNAGATCGGAAG |
| 29682_2 | TATGAGTGTGGAGTCGTTGCTACCAGTATGTGTTCAACCAGCANNNAGATCGGAAG |
| 29682_3 | TATGAGTGTGGAGTCGTTGCTACCAGTATGTGTTCAACCAGCANNNAGATCGGAAG |
| 29682_4 | TATGAGTGTGGAGTCGTTGCTACCAGTATGTGTTCAACCAGCANNNAGATCGGAAG |
| 38604_1 | TATGAGTGTGGAGTCGTTGCCTGAGGCAGGAGAATCACTTGAANNNAGATCGGAA |
| 38604_2 | TATGAGTGTGGAGTCGTTGCCTGAGGCAGGAGAATCACTTGAANNNAGATCGGAA |
| 38604_3 | TATGAGTGTGGAGTCGTTGCCTGAGGCAGGAGAATCACTTGAANNNAGATCGGAA |
| 38604_4 | TATGAGTGTGGAGTCGTTGCCTGAGGCAGGAGAATCACTTGAANNNAGATCGGAA |
| 52822_1 | TATGAGTGTGGAGTCGTTGCAAAGATTTCTCAAGTGGGAGCCANNNAGATCGGAA |
| 52822_2 | TATGAGTGTGGAGTCGTTGCAAAGATTTCTCAAGTGGGAGCCANNNAGATCGGAA |
| 52822_3 | TATGAGTGTGGAGTCGTTGCAAAGATTTCTCAAGTGGGAGCCANNNAGATCGGAA |
| 52822_4 | TATGAGTGTGGAGTCGTTGCAAAGATTTCTCAAGTGGGAGCCANNNAGATCGGAA |
| 31550_1 | TATGAGTGTGGAGTCGTTGCCCATGATTCCAACCCTCTACCATNNNAGATCGGAAGA |
| 31550_2 | TATGAGTGTGGAGTCGTTGCCCATGATTCCAACCCTCTACCATNNNAGATCGGAAGA |
| 31550_3 | TATGAGTGTGGAGTCGTTGCCCATGATTCCAACCCTCTACCATNNNAGATCGGAAGA |
| 31550_4 | TATGAGTGTGGAGTCGTTGCCCATGATTCCAACCCTCTACCATNNNAGATCGGAAGA |
| 43067_1 | TATGAGTGTGGAGTCGTTGCAACATGAATCAACCTGGAGGACANNNAGATCGGAA |
| 43067_2 | TATGAGTGTGGAGTCGTTGCAACATGAATCAACCTGGAGGACANNNAGATCGGAA |
| 43067_3 | TATGAGTGTGGAGTCGTTGCAACATGAATCAACCTGGAGGACANNNAGATCGGAA |
| 43067_4 | TATGAGTGTGGAGTCGTTGCAACATGAATCAACCTGGAGGACANNNAGATCGGAA |
| 31959_1 | TATGAGTGTGGAGTCGTTGCAGCTGCATACAAATTGACTCTCCNNNAGATCGGAAG |
| 31959_2 | TATGAGTGTGGAGTCGTTGCAGCTGCATACAAATTGACTCTCCNNNAGATCGGAAG |
| 31959_3 | TATGAGTGTGGAGTCGTTGCAGCTGCATACAAATTGACTCTCCNNNAGATCGGAAG |
| 31959_4 | TATGAGTGTGGAGTCGTTGCAGCTGCATACAAATTGACTCTCCNNNAGATCGGAAG |
| 50088_1 | TATGAGTGTGGAGTCGTTGCCACTTCTGAGCTCCTGACCTAGNNNAGATCGGAAGA |
| 50088_2 | TATGAGTGTGGAGTCGTTGCCACTTCTGAGCTCCTGACCTAGNNNAGATCGGAAGA |
| 50088_3 | TATGAGTGTGGAGTCGTTGCCACTTCTGAGCTCCTGACCTAGNNNAGATCGGAAGA |
| 50088_4 | TATGAGTGTGGAGTCGTTGCCACTTCTGAGCTCCTGACCTAGNNNAGATCGGAAGA |
| 37339_1 | TATGAGTGTGGAGTCGTTGCAGATACAAGACTACGCCCTTATCCNNNAGATCGGAA |
| 37339_2 | TATGAGTGTGGAGTCGTTGCAGATACAAGACTACGCCCTTATCCNNNAGATCGGAA |
| 37339_3 | TATGAGTGTGGAGTCGTTGCAGATACAAGACTACGCCCTTATCCNNNAGATCGGAA |
| 37339_4 | TATGAGTGTGGAGTCGTTGCAGATACAAGACTACGCCCTTATCCNNNAGATCGGAA |
| 42806_1 | TATGAGTGTGGAGTCGTTGCGGCTCTGAGGAAAGGCTAAGTAGNNNAGATCGGAA |
| 42806_2 | TATGAGTGTGGAGTCGTTGCGGCTCTGAGGAAAGGCTAAGTAGNNNAGATCGGAA |

| | |
|---|---|
| 42806_3 | TATGAGTGTGGAGTCGTTGCGGCTCTGAGGAAAGGCTAAGTAGNNNAGATCGGAA |
| 42806_4 | TATGAGTGTGGAGTCGTTGCGGCTCTGAGGAAAGGCTAAGTAGNNNAGATCGGAA |
| 35723_1 | TATGAGTGTGGAGTCGTTGCTCTGGGCTCAAGTGATTTGTTCTNNNAGATCGGAAG |
| 35723_2 | TATGAGTGTGGAGTCGTTGCTCTGGGCTCAAGTGATTTGTTCTNNNAGATCGGAAG |
| 35723_3 | TATGAGTGTGGAGTCGTTGCTCTGGGCTCAAGTGATTTGTTCTNNNAGATCGGAAG |
| 35723_4 | TATGAGTGTGGAGTCGTTGCTCTGGGCTCAAGTGATTTGTTCTNNNAGATCGGAAG |
| 13783_1 | TATGAGTGTGGAGTCGTTGCCTCTAGCATCTGTAGTTGATGNNNAGATCGGAAGAG |
| 13783_2 | TATGAGTGTGGAGTCGTTGCCTCTAGCATCTGTAGTTGATGNNNAGATCGGAAGAG |
| 13783_3 | TATGAGTGTGGAGTCGTTGCCTCTAGCATCTGTAGTTGATGNNNAGATCGGAAGAG |
| 13783_4 | TATGAGTGTGGAGTCGTTGCCTCTAGCATCTGTAGTTGATGNNNAGATCGGAAGAG |
| 2787_1 | TATGAGTGTGGAGTCGTTGCGCAATAAGAAGTCACAGCAGGCTNNNAGATCGGAA |
| 2787_2 | TATGAGTGTGGAGTCGTTGCGCAATAAGAAGTCACAGCAGGCTNNNAGATCGGAA |
| 2787_3 | TATGAGTGTGGAGTCGTTGCGCAATAAGAAGTCACAGCAGGCTNNNAGATCGGAA |
| 2787_4 | TATGAGTGTGGAGTCGTTGCGCAATAAGAAGTCACAGCAGGCTNNNAGATCGGAA |
| 35684_1 | TATGAGTGTGGAGTCGTTGCGCAGGCTCATGATTCACAGAAAANNNAGATCGGAAG |
| 35684_2 | TATGAGTGTGGAGTCGTTGCGCAGGCTCATGATTCACAGAAAANNNAGATCGGAAG |
| 35684_3 | TATGAGTGTGGAGTCGTTGCGCAGGCTCATGATTCACAGAAAANNNAGATCGGAAG |
| 35684_4 | TATGAGTGTGGAGTCGTTGCGCAGGCTCATGATTCACAGAAAANNNAGATCGGAAG |
| 52588_1 | TATGAGTGTGGAGTCGTTGCAATCCTTGCTGCTGTGATTGGAGNNNAGATCGGAAG |
| 52588_2 | TATGAGTGTGGAGTCGTTGCAATCCTTGCTGCTGTGATTGGAGNNNAGATCGGAAG |
| 52588_3 | TATGAGTGTGGAGTCGTTGCAATCCTTGCTGCTGTGATTGGAGNNNAGATCGGAAG |
| 52588_4 | TATGAGTGTGGAGTCGTTGCAATCCTTGCTGCTGTGATTGGAGNNNAGATCGGAAG |
| 31893_1 | TATGAGTGTGGAGTCGTTGCGCCATAAACTACGATCATGCCACNNNAGATCGGAAG |
| 31893_2 | TATGAGTGTGGAGTCGTTGCGCCATAAACTACGATCATGCCACNNNAGATCGGAAG |
| 31893_3 | TATGAGTGTGGAGTCGTTGCGCCATAAACTACGATCATGCCACNNNAGATCGGAAG |
| 31893_4 | TATGAGTGTGGAGTCGTTGCGCCATAAACTACGATCATGCCACNNNAGATCGGAAG |
| 31418_1 | TATGAGTGTGGAGTCGTTGCTGTGTGTGACTATCAATGCAGGANNNAGATCGGAA |
| 31418_2 | TATGAGTGTGGAGTCGTTGCTGTGTGTGACTATCAATGCAGGANNNAGATCGGAA |
| 31418_3 | TATGAGTGTGGAGTCGTTGCTGTGTGTGACTATCAATGCAGGANNNAGATCGGAA |
| 31418_4 | TATGAGTGTGGAGTCGTTGCTGTGTGTGACTATCAATGCAGGANNNAGATCGGAA |
| 7796_1 | TATGAGTGTGGAGTCGTTGCGGAATGTGGAAGGCTATTGGCNNNAGATCGGAAG |
| 7796_2 | TATGAGTGTGGAGTCGTTGCGGAATGTGGAAGGCTATTGGCNNNAGATCGGAAG |
| 7796_3 | TATGAGTGTGGAGTCGTTGCGGAATGTGGAAGGCTATTGGCNNNAGATCGGAAG |
| 7796_4 | TATGAGTGTGGAGTCGTTGCGGAATGTGGAAGGCTATTGGCNNNAGATCGGAAG |
| 43378_1 | TATGAGTGTGGAGTCGTTGCAACAATCACCATCAGTTCCTCCANNNAGATCGGAAG |
| 43378_2 | TATGAGTGTGGAGTCGTTGCAACAATCACCATCAGTTCCTCCANNNAGATCGGAAG |
| 43378_3 | TATGAGTGTGGAGTCGTTGCAACAATCACCATCAGTTCCTCCANNNAGATCGGAAG |
| 43378_4 | TATGAGTGTGGAGTCGTTGCAACAATCACCATCAGTTCCTCCANNNAGATCGGAAG |
| 12785_1 | TATGAGTGTGGAGTCGTTGCCATTTTGTCTAACTTTGGAGTGNNNAGATCGGAAGA |
| 12785_2 | TATGAGTGTGGAGTCGTTGCCATTTTGTCTAACTTTGGAGTGNNNAGATCGGAAGA |
| 12785_3 | TATGAGTGTGGAGTCGTTGCCATTTTGTCTAACTTTGGAGTGNNNAGATCGGAAGA |
| 12785_4 | TATGAGTGTGGAGTCGTTGCCATTTTGTCTAACTTTGGAGTGNNNAGATCGGAAGA |
| 48755_1 | TATGAGTGTGGAGTCGTTGCTCTGGTTCAGTCTTGAGAGGTTGNNNAGATCGGAA |

| | |
|---|---|
| 48755_2 | TATGAGTGTGGAGTCGTTGCTCTGGTTCAGTCTTGAGAGGTTGNNNAGATCGGAA |
| 48755_3 | TATGAGTGTGGAGTCGTTGCTCTGGTTCAGTCTTGAGAGGTTGNNNAGATCGGAA |
| 48755_4 | TATGAGTGTGGAGTCGTTGCTCTGGTTCAGTCTTGAGAGGTTGNNNAGATCGGAA |
| 47922_1 | TATGAGTGTGGAGTCGTTGCGGGGGTCTAGTTATGTTTCCCAGNNNAGATCGGAA |
| 47922_2 | TATGAGTGTGGAGTCGTTGCGGGGGTCTAGTTATGTTTCCCAGNNNAGATCGGAA |
| 47922_3 | TATGAGTGTGGAGTCGTTGCGGGGGTCTAGTTATGTTTCCCAGNNNAGATCGGAA |
| 47922_4 | TATGAGTGTGGAGTCGTTGCGGGGGTCTAGTTATGTTTCCCAGNNNAGATCGGAA |
| 30328_1 | TATGAGTGTGGAGTCGTTGCGGTTGATGGAGCGGTACTACAAGNNNAGATCGGAA |
| 30328_2 | TATGAGTGTGGAGTCGTTGCGGTTGATGGAGCGGTACTACAAGNNNAGATCGGAA |
| 30328_3 | TATGAGTGTGGAGTCGTTGCGGTTGATGGAGCGGTACTACAAGNNNAGATCGGAA |
| 30328_4 | TATGAGTGTGGAGTCGTTGCGGTTGATGGAGCGGTACTACAAGNNNAGATCGGAA |
| 2106_1 | TATGAGTGTGGAGTCGTTGCCAACAGCTACCTGGACAAGGTGNNNAGATCGGAAG |
| 2106_2 | TATGAGTGTGGAGTCGTTGCCAACAGCTACCTGGACAAGGTGNNNAGATCGGAAG |
| 2106_3 | TATGAGTGTGGAGTCGTTGCCAACAGCTACCTGGACAAGGTGNNNAGATCGGAAG |
| 2106_4 | TATGAGTGTGGAGTCGTTGCCAACAGCTACCTGGACAAGGTGNNNAGATCGGAAG |
| 49613_1 | TATGAGTGTGGAGTCGTTGCGCCTCCCAAGTAGCTAGGATTACNNNAGATCGGAAG |
| 49613_2 | TATGAGTGTGGAGTCGTTGCGCCTCCCAAGTAGCTAGGATTACNNNAGATCGGAAG |
| 49613_3 | TATGAGTGTGGAGTCGTTGCGCCTCCCAAGTAGCTAGGATTACNNNAGATCGGAAG |
| 49613_4 | TATGAGTGTGGAGTCGTTGCGCCTCCCAAGTAGCTAGGATTACNNNAGATCGGAAG |
| 47783_1 | TATGAGTGTGGAGTCGTTGCAGTGCTGGGAATACAGACATGAGNNNAGATCGGAA |
| 47783_2 | TATGAGTGTGGAGTCGTTGCAGTGCTGGGAATACAGACATGAGNNNAGATCGGAA |
| 47783_3 | TATGAGTGTGGAGTCGTTGCAGTGCTGGGAATACAGACATGAGNNNAGATCGGAA |
| 47783_4 | TATGAGTGTGGAGTCGTTGCAGTGCTGGGAATACAGACATGAGNNNAGATCGGAA |
| 327_1 | TATGAGTGTGGAGTCGTTGCCCCAGAATGTGTGAGCAGTTCATNNNAGATCGGAA |
| 327_2 | TATGAGTGTGGAGTCGTTGCCCCAGAATGTGTGAGCAGTTCATNNNAGATCGGAA |
| 327_3 | TATGAGTGTGGAGTCGTTGCCCCAGAATGTGTGAGCAGTTCATNNNAGATCGGAA |
| 327_4 | TATGAGTGTGGAGTCGTTGCCCCAGAATGTGTGAGCAGTTCATNNNAGATCGGAA |
| 52787_1 | TATGAGTGTGGAGTCGTTGCTCAGAACTTACACAGCTCCTCACNNNAGATCGGAAG |
| 52787_2 | TATGAGTGTGGAGTCGTTGCTCAGAACTTACACAGCTCCTCACNNNAGATCGGAAG |
| 52787_3 | TATGAGTGTGGAGTCGTTGCTCAGAACTTACACAGCTCCTCACNNNAGATCGGAAG |
| 52787_4 | TATGAGTGTGGAGTCGTTGCTCAGAACTTACACAGCTCCTCACNNNAGATCGGAAG |
| 50117_1 | TATGAGTGTGGAGTCGTTGCCACGAGCCCCTTCACTTCAGNNNAGATCGGAAGAGC |
| 50117_2 | TATGAGTGTGGAGTCGTTGCCACGAGCCCCTTCACTTCAGNNNAGATCGGAAGAGC |
| 50117_3 | TATGAGTGTGGAGTCGTTGCCACGAGCCCCTTCACTTCAGNNNAGATCGGAAGAGC |
| 50117_4 | TATGAGTGTGGAGTCGTTGCCACGAGCCCCTTCACTTCAGNNNAGATCGGAAGAGC |
| 39625_1 | TATGAGTGTGGAGTCGTTGCACAGAAAAGGGGTAGCCTAGAGANNNAGATCGGA |
| 39625_2 | TATGAGTGTGGAGTCGTTGCACAGAAAAGGGGTAGCCTAGAGANNNAGATCGGA |
| 39625_3 | TATGAGTGTGGAGTCGTTGCACAGAAAAGGGGTAGCCTAGAGANNNAGATCGGA |
| 39625_4 | TATGAGTGTGGAGTCGTTGCACAGAAAAGGGGTAGCCTAGAGANNNAGATCGGA |
| 37666_1 | TATGAGTGTGGAGTCGTTGCAAATGCATGGGATTAGACACCCTNNNAGATCGGAA |
| 37666_2 | TATGAGTGTGGAGTCGTTGCAAATGCATGGGATTAGACACCCTNNNAGATCGGAA |
| 37666_3 | TATGAGTGTGGAGTCGTTGCAAATGCATGGGATTAGACACCCTNNNAGATCGGAA |
| 37666_4 | TATGAGTGTGGAGTCGTTGCAAATGCATGGGATTAGACACCCTNNNAGATCGGAA |

| | |
|---|---|
| 13117_1 | TATGAGTGTGGAGTCGTTGCAATAATTTCCCTATTAGCCAGAANNNAGATCGGAAG |
| 13117_2 | TATGAGTGTGGAGTCGTTGCAATAATTTCCCTATTAGCCAGAANNNAGATCGGAAG |
| 13117_3 | TATGAGTGTGGAGTCGTTGCAATAATTTCCCTATTAGCCAGAANNNAGATCGGAAG |
| 13117_4 | TATGAGTGTGGAGTCGTTGCAATAATTTCCCTATTAGCCAGAANNNAGATCGGAAG |
| 10339_1 | TATGAGTGTGGAGTCGTTGCTCCTGTGACTTCCTCAGACAGAGNNNAGATCGGAAG |
| 10339_2 | TATGAGTGTGGAGTCGTTGCTCCTGTGACTTCCTCAGACAGAGNNNAGATCGGAAG |
| 10339_3 | TATGAGTGTGGAGTCGTTGCTCCTGTGACTTCCTCAGACAGAGNNNAGATCGGAAG |
| 10339_4 | TATGAGTGTGGAGTCGTTGCTCCTGTGACTTCCTCAGACAGAGNNNAGATCGGAAG |
| 41076_1 | TATGAGTGTGGAGTCGTTGCTACAAAAATTAGCTGGGCATGGCNNNAGATCGGAA |
| 41076_2 | TATGAGTGTGGAGTCGTTGCTACAAAAATTAGCTGGGCATGGCNNNAGATCGGAA |
| 41076_3 | TATGAGTGTGGAGTCGTTGCTACAAAAATTAGCTGGGCATGGCNNNAGATCGGAA |
| 41076_4 | TATGAGTGTGGAGTCGTTGCTACAAAAATTAGCTGGGCATGGCNNNAGATCGGAA |
| 45119_1 | TATGAGTGTGGAGTCGTTGCAAAACTTGGCAGGATTAGACCCTNNNAGATCGGAAG |
| 45119_2 | TATGAGTGTGGAGTCGTTGCAAAACTTGGCAGGATTAGACCCTNNNAGATCGGAAG |
| 45119_3 | TATGAGTGTGGAGTCGTTGCAAAACTTGGCAGGATTAGACCCTNNNAGATCGGAAG |
| 45119_4 | TATGAGTGTGGAGTCGTTGCAAAACTTGGCAGGATTAGACCCTNNNAGATCGGAAG |
| 46254_1 | TATGAGTGTGGAGTCGTTGCCTTCCAAAGTGCTGGGATTACAGNNNAGATCGGAAG |
| 46254_2 | TATGAGTGTGGAGTCGTTGCCTTCCAAAGTGCTGGGATTACAGNNNAGATCGGAAG |
| 46254_3 | TATGAGTGTGGAGTCGTTGCCTTCCAAAGTGCTGGGATTACAGNNNAGATCGGAAG |
| 46254_4 | TATGAGTGTGGAGTCGTTGCCTTCCAAAGTGCTGGGATTACAGNNNAGATCGGAAG |
| 37785_1 | TATGAGTGTGGAGTCGTTGCAGGCTATCTGGAAACACAGGAAGNNNAGATCGGAA |
| 37785_2 | TATGAGTGTGGAGTCGTTGCAGGCTATCTGGAAACACAGGAAGNNNAGATCGGAA |
| 37785_3 | TATGAGTGTGGAGTCGTTGCAGGCTATCTGGAAACACAGGAAGNNNAGATCGGAA |
| 37785_4 | TATGAGTGTGGAGTCGTTGCAGGCTATCTGGAAACACAGGAAGNNNAGATCGGAA |
| 41457_1 | TATGAGTGTGGAGTCGTTGCGGCACCTGTATCTCAACTATTTGGNNNAGATCGGAA |
| 41457_2 | TATGAGTGTGGAGTCGTTGCGGCACCTGTATCTCAACTATTTGGNNNAGATCGGAA |
| 41457_3 | TATGAGTGTGGAGTCGTTGCGGCACCTGTATCTCAACTATTTGGNNNAGATCGGAA |
| 41457_4 | TATGAGTGTGGAGTCGTTGCGGCACCTGTATCTCAACTATTTGGNNNAGATCGGAA |
| 46235_1 | TATGAGTGTGGAGTCGTTGCATTCCAAGGAGTGTAGGGAGACANNNAGATCGGAA |
| 46235_2 | TATGAGTGTGGAGTCGTTGCATTCCAAGGAGTGTAGGGAGACANNNAGATCGGAA |
| 46235_3 | TATGAGTGTGGAGTCGTTGCATTCCAAGGAGTGTAGGGAGACANNNAGATCGGAA |
| 46235_4 | TATGAGTGTGGAGTCGTTGCATTCCAAGGAGTGTAGGGAGACANNNAGATCGGAA |
| 48951_1 | TATGAGTGTGGAGTCGTTGCGGCTTCACATTTCTTCCTCTTGGNNNAGATCGGAAG |
| 48951_2 | TATGAGTGTGGAGTCGTTGCGGCTTCACATTTCTTCCTCTTGGNNNAGATCGGAAG |
| 48951_3 | TATGAGTGTGGAGTCGTTGCGGCTTCACATTTCTTCCTCTTGGNNNAGATCGGAAG |
| 48951_4 | TATGAGTGTGGAGTCGTTGCGGCTTCACATTTCTTCCTCTTGGNNNAGATCGGAAG |
| 2569_1 | TATGAGTGTGGAGTCGTTGCAAAAGTCTATGGTGTTGGCTGGGNNNAGATCGGAA |
| 2569_2 | TATGAGTGTGGAGTCGTTGCAAAAGTCTATGGTGTTGGCTGGGNNNAGATCGGAA |
| 2569_3 | TATGAGTGTGGAGTCGTTGCAAAAGTCTATGGTGTTGGCTGGGNNNAGATCGGAA |
| 2569_4 | TATGAGTGTGGAGTCGTTGCAAAAGTCTATGGTGTTGGCTGGGNNNAGATCGGAA |
| 32142_1 | TATGAGTGTGGAGTCGTTGCTGATCTTGGCTCACTACAACCTCNNNAGATCGGAAG |
| 32142_2 | TATGAGTGTGGAGTCGTTGCTGATCTTGGCTCACTACAACCTCNNNAGATCGGAAG |
| 32142_3 | TATGAGTGTGGAGTCGTTGCTGATCTTGGCTCACTACAACCTCNNNAGATCGGAAG |

| | |
|---|---|
| 32142_4 | TATGAGTGTGGAGTCGTTGCTGATCTTGGCTCACTACAACCTCNNNAGATCGGAAG |
| 49611_1 | TATGAGTGTGGAGTCGTTGCCATGCCCGGCTAAGTTTTGTATTNNNAGATCGGAAG |
| 49611_2 | TATGAGTGTGGAGTCGTTGCCATGCCCGGCTAAGTTTTGTATTNNNAGATCGGAAG |
| 49611_3 | TATGAGTGTGGAGTCGTTGCCATGCCCGGCTAAGTTTTGTATTNNNAGATCGGAAG |
| 49611_4 | TATGAGTGTGGAGTCGTTGCCATGCCCGGCTAAGTTTTGTATTNNNAGATCGGAAG |
| 33432_1 | TATGAGTGTGGAGTCGTTGCCCAACCAGATACCTAACCCAACANNNAGATCGGAAG |
| 33432_2 | TATGAGTGTGGAGTCGTTGCCCAACCAGATACCTAACCCAACANNNAGATCGGAAG |
| 33432_3 | TATGAGTGTGGAGTCGTTGCCCAACCAGATACCTAACCCAACANNNAGATCGGAAG |
| 33432_4 | TATGAGTGTGGAGTCGTTGCCCAACCAGATACCTAACCCAACANNNAGATCGGAAG |
| 39364_1 | TATGAGTGTGGAGTCGTTGCTTCTGGGTTCAAGAAATCCTCCCNNNAGATCGGAAG |
| 39364_2 | TATGAGTGTGGAGTCGTTGCTTCTGGGTTCAAGAAATCCTCCCNNNAGATCGGAAG |
| 39364_3 | TATGAGTGTGGAGTCGTTGCTTCTGGGTTCAAGAAATCCTCCCNNNAGATCGGAAG |
| 39364_4 | TATGAGTGTGGAGTCGTTGCTTCTGGGTTCAAGAAATCCTCCCNNNAGATCGGAAG |
| 50279_1 | TATGAGTGTGGAGTCGTTGCGGTCCAAGGAATAGTCCAAGAGGNNNAGATCGGAA |
| 50279_2 | TATGAGTGTGGAGTCGTTGCGGTCCAAGGAATAGTCCAAGAGGNNNAGATCGGAA |
| 50279_3 | TATGAGTGTGGAGTCGTTGCGGTCCAAGGAATAGTCCAAGAGGNNNAGATCGGAA |
| 50279_4 | TATGAGTGTGGAGTCGTTGCGGTCCAAGGAATAGTCCAAGAGGNNNAGATCGGAA |
| 30224_1 | TATGAGTGTGGAGTCGTTGCCTCTGAGGAGAGGCCTACAAATGNNNAGATCGGAA |
| 30224_2 | TATGAGTGTGGAGTCGTTGCCTCTGAGGAGAGGCCTACAAATGNNNAGATCGGAA |
| 30224_3 | TATGAGTGTGGAGTCGTTGCCTCTGAGGAGAGGCCTACAAATGNNNAGATCGGAA |
| 30224_4 | TATGAGTGTGGAGTCGTTGCCTCTGAGGAGAGGCCTACAAATGNNNAGATCGGAA |
| 12059_1 | TATGAGTGTGGAGTCGTTGCACCTGTCAAAACTCATCAAACACTNNNAGATCGGAAG |
| 12059_2 | TATGAGTGTGGAGTCGTTGCACCTGTCAAAACTCATCAAACACTNNNAGATCGGAAG |
| 12059_3 | TATGAGTGTGGAGTCGTTGCACCTGTCAAAACTCATCAAACACTNNNAGATCGGAAG |
| 12059_4 | TATGAGTGTGGAGTCGTTGCACCTGTCAAAACTCATCAAACACTNNNAGATCGGAAG |
| 43804_1 | TATGAGTGTGGAGTCGTTGCATGATTCCACTACTGCACTCCAGNNNAGATCGGAAG |
| 43804_2 | TATGAGTGTGGAGTCGTTGCATGATTCCACTACTGCACTCCAGNNNAGATCGGAAG |
| 43804_3 | TATGAGTGTGGAGTCGTTGCATGATTCCACTACTGCACTCCAGNNNAGATCGGAAG |
| 43804_4 | TATGAGTGTGGAGTCGTTGCATGATTCCACTACTGCACTCCAGNNNAGATCGGAAG |
| 50005_1 | TATGAGTGTGGAGTCGTTGCTGTTCCAGCATTCCCATCAAGAANNNAGATCGGAAG |
| 50005_2 | TATGAGTGTGGAGTCGTTGCTGTTCCAGCATTCCCATCAAGAANNNAGATCGGAAG |
| 50005_3 | TATGAGTGTGGAGTCGTTGCTGTTCCAGCATTCCCATCAAGAANNNAGATCGGAAG |
| 50005_4 | TATGAGTGTGGAGTCGTTGCTGTTCCAGCATTCCCATCAAGAANNNAGATCGGAAG |
| 37568_1 | TATGAGTGTGGAGTCGTTGCGGGACATGGAGCTTGCAGTAAGNNNAGATCGGAAG |
| 37568_2 | TATGAGTGTGGAGTCGTTGCGGGACATGGAGCTTGCAGTAAGNNNAGATCGGAAG |
| 37568_3 | TATGAGTGTGGAGTCGTTGCGGGACATGGAGCTTGCAGTAAGNNNAGATCGGAAG |
| 37568_4 | TATGAGTGTGGAGTCGTTGCGGGACATGGAGCTTGCAGTAAGNNNAGATCGGAAG |
| 47639_1 | TATGAGTGTGGAGTCGTTGCGAGACGAAGTCTTGCTCTGTCANNNAGATCGGAAG |
| 47639_2 | TATGAGTGTGGAGTCGTTGCGAGACGAAGTCTTGCTCTGTCANNNAGATCGGAAG |
| 47639_3 | TATGAGTGTGGAGTCGTTGCGAGACGAAGTCTTGCTCTGTCANNNAGATCGGAAG |
| 47639_4 | TATGAGTGTGGAGTCGTTGCGAGACGAAGTCTTGCTCTGTCANNNAGATCGGAAG |
| 33399_1 | TATGAGTGTGGAGTCGTTGCCGGCTTGATATTTGCTGCTTACANNNAGATCGGAAG |
| 33399_2 | TATGAGTGTGGAGTCGTTGCCGGCTTGATATTTGCTGCTTACANNNAGATCGGAAG |

| | |
|---|---|
| 33399_3 | TATGAGTGTGGAGTCGTTGCCGGCTTGATATTTGCTGCTTACANNNAGATCGGAAG |
| 33399_4 | TATGAGTGTGGAGTCGTTGCCGGCTTGATATTTGCTGCTTACANNNAGATCGGAAG |
| 35968_1 | TATGAGTGTGGAGTCGTTGCCTGGGTAACAGAGTGAGACACTGNNNAGATCGGAA |
| 35968_2 | TATGAGTGTGGAGTCGTTGCCTGGGTAACAGAGTGAGACACTGNNNAGATCGGAA |
| 35968_3 | TATGAGTGTGGAGTCGTTGCCTGGGTAACAGAGTGAGACACTGNNNAGATCGGAA |
| 35968_4 | TATGAGTGTGGAGTCGTTGCCTGGGTAACAGAGTGAGACACTGNNNAGATCGGAA |
| 12589_1 | TATGAGTGTGGAGTCGTTGCTTTTAAATTCCAAGTCATTCCTCNNNAGATCGGAAGA |
| 12589_2 | TATGAGTGTGGAGTCGTTGCTTTTAAATTCCAAGTCATTCCTCNNNAGATCGGAAGA |
| 12589_3 | TATGAGTGTGGAGTCGTTGCTTTTAAATTCCAAGTCATTCCTCNNNAGATCGGAAGA |
| 12589_4 | TATGAGTGTGGAGTCGTTGCTTTTAAATTCCAAGTCATTCCTCNNNAGATCGGAAGA |
| 43004_1 | TATGAGTGTGGAGTCGTTGCAGAGGTCATGACAGTCAACGAATNNNAGATCGGAA |
| 43004_2 | TATGAGTGTGGAGTCGTTGCAGAGGTCATGACAGTCAACGAATNNNAGATCGGAA |
| 43004_3 | TATGAGTGTGGAGTCGTTGCAGAGGTCATGACAGTCAACGAATNNNAGATCGGAA |
| 43004_4 | TATGAGTGTGGAGTCGTTGCAGAGGTCATGACAGTCAACGAATNNNAGATCGGAA |
| 51707_1 | TATGAGTGTGGAGTCGTTGCGGTTTCGAACACCAGTTGAGATGNNNAGATCGGAA |
| 51707_2 | TATGAGTGTGGAGTCGTTGCGGTTTCGAACACCAGTTGAGATGNNNAGATCGGAA |
| 51707_3 | TATGAGTGTGGAGTCGTTGCGGTTTCGAACACCAGTTGAGATGNNNAGATCGGAA |
| 51707_4 | TATGAGTGTGGAGTCGTTGCGGTTTCGAACACCAGTTGAGATGNNNAGATCGGAA |
| 44177_1 | TATGAGTGTGGAGTCGTTGCACAGATGTGCTGGGAACTTTACANNNAGATCGGAA |
| 44177_2 | TATGAGTGTGGAGTCGTTGCACAGATGTGCTGGGAACTTTACANNNAGATCGGAA |
| 44177_3 | TATGAGTGTGGAGTCGTTGCACAGATGTGCTGGGAACTTTACANNNAGATCGGAA |
| 44177_4 | TATGAGTGTGGAGTCGTTGCACAGATGTGCTGGGAACTTTACANNNAGATCGGAA |
| 31561_1 | TATGAGTGTGGAGTCGTTGCCCCATAACCCAGAGTCAACTACANNNAGATCGGAAG |
| 31561_2 | TATGAGTGTGGAGTCGTTGCCCCATAACCCAGAGTCAACTACANNNAGATCGGAAG |
| 31561_3 | TATGAGTGTGGAGTCGTTGCCCCATAACCCAGAGTCAACTACANNNAGATCGGAAG |
| 31561_4 | TATGAGTGTGGAGTCGTTGCCCCATAACCCAGAGTCAACTACANNNAGATCGGAAG |
| 35155_1 | TATGAGTGTGGAGTCGTTGCTGGCCAGAAATAATGTTTACCAGCNNNAGATCGGAA |
| 35155_2 | TATGAGTGTGGAGTCGTTGCTGGCCAGAAATAATGTTTACCAGCNNNAGATCGGAA |
| 35155_3 | TATGAGTGTGGAGTCGTTGCTGGCCAGAAATAATGTTTACCAGCNNNAGATCGGAA |
| 35155_4 | TATGAGTGTGGAGTCGTTGCTGGCCAGAAATAATGTTTACCAGCNNNAGATCGGAA |
| 34421_1 | TATGAGTGTGGAGTCGTTGCCAGGAAGAACTGGAACTGGTCTGNNNAGATCGGAA |
| 34421_2 | TATGAGTGTGGAGTCGTTGCCAGGAAGAACTGGAACTGGTCTGNNNAGATCGGAA |
| 34421_3 | TATGAGTGTGGAGTCGTTGCCAGGAAGAACTGGAACTGGTCTGNNNAGATCGGAA |
| 34421_4 | TATGAGTGTGGAGTCGTTGCCAGGAAGAACTGGAACTGGTCTGNNNAGATCGGAA |
| 35570_1 | TATGAGTGTGGAGTCGTTGCCCTCAAACCATCCTTCCACCTAANNNAGATCGGAAGA |
| 35570_2 | TATGAGTGTGGAGTCGTTGCCCTCAAACCATCCTTCCACCTAANNNAGATCGGAAGA |
| 35570_3 | TATGAGTGTGGAGTCGTTGCCCTCAAACCATCCTTCCACCTAANNNAGATCGGAAGA |
| 35570_4 | TATGAGTGTGGAGTCGTTGCCCTCAAACCATCCTTCCACCTAANNNAGATCGGAAGA |
| 29461_1 | TATGAGTGTGGAGTCGTTGCAATTGTTGGCATTTTTTTAACAANNNAGATCGGAAG |
| 29461_2 | TATGAGTGTGGAGTCGTTGCAATTGTTGGCATTTTTTTAACAANNNAGATCGGAAG |
| 29461_3 | TATGAGTGTGGAGTCGTTGCAATTGTTGGCATTTTTTTAACAANNNAGATCGGAAG |
| 29461_4 | TATGAGTGTGGAGTCGTTGCAATTGTTGGCATTTTTTTAACAANNNAGATCGGAAG |
| 9677_1 | TATGAGTGTGGAGTCGTTGCCACACTCATTTTCCCTCTCTCCCNNNAGATCGGAAGA |

| | |
|---|---|
| 9677_2 | TATGAGTGTGGAGTCGTTGCCACACTCATTTTCCCTCTCTCCCNNNAGATCGGAAGAG |
| 9677_3 | TATGAGTGTGGAGTCGTTGCCACACTCATTTTCCCTCTCTCCCNNNAGATCGGAAGAG |
| 9677_4 | TATGAGTGTGGAGTCGTTGCCACACTCATTTTCCCTCTCTCCCNNNAGATCGGAAGAG |
| 50860_1 | TATGAGTGTGGAGTCGTTGCTCCACTTCCCTACATGTAGTCCTNNNAGATCGGAAGA |
| 50860_2 | TATGAGTGTGGAGTCGTTGCTCCACTTCCCTACATGTAGTCCTNNNAGATCGGAAGA |
| 50860_3 | TATGAGTGTGGAGTCGTTGCTCCACTTCCCTACATGTAGTCCTNNNAGATCGGAAGA |
| 50860_4 | TATGAGTGTGGAGTCGTTGCTCCACTTCCCTACATGTAGTCCTNNNAGATCGGAAGA |
| 51867_1 | TATGAGTGTGGAGTCGTTGCCACTGTTTTCGGACCTTCCATGNNNAGATCGGAAGA[illegible] |
| 51867_2 | TATGAGTGTGGAGTCGTTGCCACTGTTTTCGGACCTTCCATGNNNAGATCGGAAGA[illegible] |
| 51867_3 | TATGAGTGTGGAGTCGTTGCCACTGTTTTCGGACCTTCCATGNNNAGATCGGAAGA[illegible] |
| 51867_4 | TATGAGTGTGGAGTCGTTGCCACTGTTTTCGGACCTTCCATGNNNAGATCGGAAGA[illegible] |
| 41913_1 | TATGAGTGTGGAGTCGTTGCTTCAAGCAATTCTCCTGCCTCAGNNNAGATCGGAAGA |
| 41913_2 | TATGAGTGTGGAGTCGTTGCTTCAAGCAATTCTCCTGCCTCAGNNNAGATCGGAAGA |
| 41913_3 | TATGAGTGTGGAGTCGTTGCTTCAAGCAATTCTCCTGCCTCAGNNNAGATCGGAAGA |
| 41913_4 | TATGAGTGTGGAGTCGTTGCTTCAAGCAATTCTCCTGCCTCAGNNNAGATCGGAAGA |
| 49125_1 | TATGAGTGTGGAGTCGTTGCATTCTGCCTTAGCCTCCCAAGTANNNAGATCGGAAGA |
| 49125_2 | TATGAGTGTGGAGTCGTTGCATTCTGCCTTAGCCTCCCAAGTANNNAGATCGGAAGA |
| 49125_3 | TATGAGTGTGGAGTCGTTGCATTCTGCCTTAGCCTCCCAAGTANNNAGATCGGAAGA |
| 49125_4 | TATGAGTGTGGAGTCGTTGCATTCTGCCTTAGCCTCCCAAGTANNNAGATCGGAAGA |
| 42256_1 | TATGAGTGTGGAGTCGTTGCACTGTGCTGGAGATTCTCAAACTNNNAGATCGGAAG |
| 42256_2 | TATGAGTGTGGAGTCGTTGCACTGTGCTGGAGATTCTCAAACTNNNAGATCGGAAG |
| 42256_3 | TATGAGTGTGGAGTCGTTGCACTGTGCTGGAGATTCTCAAACTNNNAGATCGGAAG |
| 42256_4 | TATGAGTGTGGAGTCGTTGCACTGTGCTGGAGATTCTCAAACTNNNAGATCGGAAG |
| 50727_1 | TATGAGTGTGGAGTCGTTGCATTTCCTCCAGTGGCAGCTATACNNNAGATCGGAAGA |
| 50727_2 | TATGAGTGTGGAGTCGTTGCATTTCCTCCAGTGGCAGCTATACNNNAGATCGGAAGA |
| 50727_3 | TATGAGTGTGGAGTCGTTGCATTTCCTCCAGTGGCAGCTATACNNNAGATCGGAAGA |
| 50727_4 | TATGAGTGTGGAGTCGTTGCATTTCCTCCAGTGGCAGCTATACNNNAGATCGGAAGA |
| 39634_1 | TATGAGTGTGGAGTCGTTGCAAGACCTACATCACCATCTCTGCNNNAGATCGGAAGA |
| 39634_2 | TATGAGTGTGGAGTCGTTGCAAGACCTACATCACCATCTCTGCNNNAGATCGGAAGA |
| 39634_3 | TATGAGTGTGGAGTCGTTGCAAGACCTACATCACCATCTCTGCNNNAGATCGGAAGA |
| 39634_4 | TATGAGTGTGGAGTCGTTGCAAGACCTACATCACCATCTCTGCNNNAGATCGGAAGA |
| 30545_1 | TATGAGTGTGGAGTCGTTGCAGTTCCAGAAAACAGCAGAACTCAGNNNAGATCGGA |
| 30545_2 | TATGAGTGTGGAGTCGTTGCAGTTCCAGAAAACAGCAGAACTCAGNNNAGATCGGA |
| 30545_3 | TATGAGTGTGGAGTCGTTGCAGTTCCAGAAAACAGCAGAACTCAGNNNAGATCGGA |
| 30545_4 | TATGAGTGTGGAGTCGTTGCAGTTCCAGAAAACAGCAGAACTCAGNNNAGATCGGA |
| 49184_1 | TATGAGTGTGGAGTCGTTGCCTAGTAGCTGGGATTACAGGTGCNNNAGATCGGAA[illegible] |
| 49184_2 | TATGAGTGTGGAGTCGTTGCCTAGTAGCTGGGATTACAGGTGCNNNAGATCGGAA[illegible] |
| 49184_3 | TATGAGTGTGGAGTCGTTGCCTAGTAGCTGGGATTACAGGTGCNNNAGATCGGAA[illegible] |
| 49184_4 | TATGAGTGTGGAGTCGTTGCCTAGTAGCTGGGATTACAGGTGCNNNAGATCGGAA[illegible] |
| 42625_1 | TATGAGTGTGGAGTCGTTGCGCAAAATGTCAACAGTGGTCAGANNNAGATCGGAA[illegible] |
| 42625_2 | TATGAGTGTGGAGTCGTTGCGCAAAATGTCAACAGTGGTCAGANNNAGATCGGAA[illegible] |
| 42625_3 | TATGAGTGTGGAGTCGTTGCGCAAAATGTCAACAGTGGTCAGANNNAGATCGGAA[illegible] |
| 42625_4 | TATGAGTGTGGAGTCGTTGCGCAAAATGTCAACAGTGGTCAGANNNAGATCGGAA[illegible] |

| | |
|---|---|
| 46610_1 | TATGAGTGTGGAGTCGTTGCCTGAGCTCAAGCGATCCTTCTGNNNAGATCGGAAGA |
| 46610_2 | TATGAGTGTGGAGTCGTTGCCTGAGCTCAAGCGATCCTTCTGNNNAGATCGGAAGA |
| 46610_3 | TATGAGTGTGGAGTCGTTGCCTGAGCTCAAGCGATCCTTCTGNNNAGATCGGAAGA |
| 46610_4 | TATGAGTGTGGAGTCGTTGCCTGAGCTCAAGCGATCCTTCTGNNNAGATCGGAAGA |
| 39160_1 | TATGAGTGTGGAGTCGTTGCTAGTCACTTCCTCTTCTAGCACTNNNAGATCGGAAGA |
| 39160_2 | TATGAGTGTGGAGTCGTTGCTAGTCACTTCCTCTTCTAGCACTNNNAGATCGGAAGA |
| 39160_3 | TATGAGTGTGGAGTCGTTGCTAGTCACTTCCTCTTCTAGCACTNNNAGATCGGAAGA |
| 39160_4 | TATGAGTGTGGAGTCGTTGCTAGTCACTTCCTCTTCTAGCACTNNNAGATCGGAAGA |
| 8334_1 | TATGAGTGTGGAGTCGTTGCGAGTGGAGACTGGGTTTCATTATGNNNAGATCGGA |
| 8334_2 | TATGAGTGTGGAGTCGTTGCGAGTGGAGACTGGGTTTCATTATGNNNAGATCGGA |
| 8334_3 | TATGAGTGTGGAGTCGTTGCGAGTGGAGACTGGGTTTCATTATGNNNAGATCGGA |
| 8334_4 | TATGAGTGTGGAGTCGTTGCGAGTGGAGACTGGGTTTCATTATGNNNAGATCGGA |
| 37615_1 | TATGAGTGTGGAGTCGTTGCAGGCATGAACAAACCAAACTGTTNNNAGATCGGAA |
| 37615_2 | TATGAGTGTGGAGTCGTTGCAGGCATGAACAAACCAAACTGTTNNNAGATCGGAA |
| 37615_3 | TATGAGTGTGGAGTCGTTGCAGGCATGAACAAACCAAACTGTTNNNAGATCGGAA |
| 37615_4 | TATGAGTGTGGAGTCGTTGCAGGCATGAACAAACCAAACTGTTNNNAGATCGGAA |
| 13275_1 | TATGAGTGTGGAGTCGTTGCTCATCTCACTTAATAAGACATGTNNNAGATCGGAAG |
| 13275_2 | TATGAGTGTGGAGTCGTTGCTCATCTCACTTAATAAGACATGTNNNAGATCGGAAG |
| 13275_3 | TATGAGTGTGGAGTCGTTGCTCATCTCACTTAATAAGACATGTNNNAGATCGGAAG |
| 13275_4 | TATGAGTGTGGAGTCGTTGCTCATCTCACTTAATAAGACATGTNNNAGATCGGAAG |
| 29688_1 | TATGAGTGTGGAGTCGTTGCAAAAGAAGGACTGGAAGGAAGCANNNAGATCGGA |
| 29688_2 | TATGAGTGTGGAGTCGTTGCAAAAGAAGGACTGGAAGGAAGCANNNAGATCGGA |
| 29688_3 | TATGAGTGTGGAGTCGTTGCAAAAGAAGGACTGGAAGGAAGCANNNAGATCGGA |
| 29688_4 | TATGAGTGTGGAGTCGTTGCAAAAGAAGGACTGGAAGGAAGCANNNAGATCGGA |
| 53273_1 | TATGAGTGTGGAGTCGTTGCCTGAGGTCAGGAGTTTGAGACCNNNAGATCGGAAG |
| 53273_2 | TATGAGTGTGGAGTCGTTGCCTGAGGTCAGGAGTTTGAGACCNNNAGATCGGAAG |
| 53273_3 | TATGAGTGTGGAGTCGTTGCCTGAGGTCAGGAGTTTGAGACCNNNAGATCGGAAG |
| 53273_4 | TATGAGTGTGGAGTCGTTGCCTGAGGTCAGGAGTTTGAGACCNNNAGATCGGAAG |
| 46217_1 | TATGAGTGTGGAGTCGTTGCTCCCAGGTTCAAATGATTCTCGTNNNAGATCGGAAG |
| 46217_2 | TATGAGTGTGGAGTCGTTGCTCCCAGGTTCAAATGATTCTCGTNNNAGATCGGAAG |
| 46217_3 | TATGAGTGTGGAGTCGTTGCTCCCAGGTTCAAATGATTCTCGTNNNAGATCGGAAG |
| 46217_4 | TATGAGTGTGGAGTCGTTGCTCCCAGGTTCAAATGATTCTCGTNNNAGATCGGAAG |
| 46745_1 | TATGAGTGTGGAGTCGTTGCGGCACTCTGAGCATTTTCATTGTNNNAGATCGGAAG |
| 46745_2 | TATGAGTGTGGAGTCGTTGCGGCACTCTGAGCATTTTCATTGTNNNAGATCGGAAG |
| 46745_3 | TATGAGTGTGGAGTCGTTGCGGCACTCTGAGCATTTTCATTGTNNNAGATCGGAAG |
| 46745_4 | TATGAGTGTGGAGTCGTTGCGGCACTCTGAGCATTTTCATTGTNNNAGATCGGAAG |
| 42655_1 | TATGAGTGTGGAGTCGTTGCCCTAGCTACTTGGGAGGTTGATGNNNAGATCGGAA |
| 42655_2 | TATGAGTGTGGAGTCGTTGCCCTAGCTACTTGGGAGGTTGATGNNNAGATCGGAA |
| 42655_3 | TATGAGTGTGGAGTCGTTGCCCTAGCTACTTGGGAGGTTGATGNNNAGATCGGAA |
| 42655_4 | TATGAGTGTGGAGTCGTTGCCCTAGCTACTTGGGAGGTTGATGNNNAGATCGGAA |
| 36417_1 | TATGAGTGTGGAGTCGTTGCGGGAAAAGCATAGTATCTGGGCTNNNAGATCGGAA |
| 36417_2 | TATGAGTGTGGAGTCGTTGCGGGAAAAGCATAGTATCTGGGCTNNNAGATCGGAA |
| 36417_3 | TATGAGTGTGGAGTCGTTGCGGGAAAAGCATAGTATCTGGGCTNNNAGATCGGAA |

| | |
|---|---|
| 36417_4 | TATGAGTGTGGAGTCGTTGCGGGAAAAGCATAGTATCTGGGCTNNNAGATCGGAA |
| 31772_1 | TATGAGTGTGGAGTCGTTGCGGTGGATCACTTGAGATCAGGAGNNNAGATCGGAA |
| 31772_2 | TATGAGTGTGGAGTCGTTGCGGTGGATCACTTGAGATCAGGAGNNNAGATCGGAA |
| 31772_3 | TATGAGTGTGGAGTCGTTGCGGTGGATCACTTGAGATCAGGAGNNNAGATCGGAA |
| 31772_4 | TATGAGTGTGGAGTCGTTGCGGTGGATCACTTGAGATCAGGAGNNNAGATCGGAA |
| 35724_1 | TATGAGTGTGGAGTCGTTGCCAGCCTCCTCAATACCTGACATTNNNAGATCGGAAGA |
| 35724_2 | TATGAGTGTGGAGTCGTTGCCAGCCTCCTCAATACCTGACATTNNNAGATCGGAAGA |
| 35724_3 | TATGAGTGTGGAGTCGTTGCCAGCCTCCTCAATACCTGACATTNNNAGATCGGAAGA |
| 35724_4 | TATGAGTGTGGAGTCGTTGCCAGCCTCCTCAATACCTGACATTNNNAGATCGGAAGA |
| 38607_1 | TATGAGTGTGGAGTCGTTGCTGATGTGCAACGGACTAATTTCCNNNAGATCGGAAG |
| 38607_2 | TATGAGTGTGGAGTCGTTGCTGATGTGCAACGGACTAATTTCCNNNAGATCGGAAG |
| 38607_3 | TATGAGTGTGGAGTCGTTGCTGATGTGCAACGGACTAATTTCCNNNAGATCGGAAG |
| 38607_4 | TATGAGTGTGGAGTCGTTGCTGATGTGCAACGGACTAATTTCCNNNAGATCGGAAG |
| 31456_1 | TATGAGTGTGGAGTCGTTGCTCAGGGAAATGCAAACCAAATCCNNNAGATCGGAAG |
| 31456_2 | TATGAGTGTGGAGTCGTTGCTCAGGGAAATGCAAACCAAATCCNNNAGATCGGAAG |
| 31456_3 | TATGAGTGTGGAGTCGTTGCTCAGGGAAATGCAAACCAAATCCNNNAGATCGGAAG |
| 31456_4 | TATGAGTGTGGAGTCGTTGCTCAGGGAAATGCAAACCAAATCCNNNAGATCGGAAG |
| 34530_1 | TATGAGTGTGGAGTCGTTGCGCAAGTCAGACTCTGAGACAGAANNNAGATCGGAA |
| 34530_2 | TATGAGTGTGGAGTCGTTGCGCAAGTCAGACTCTGAGACAGAANNNAGATCGGAA |
| 34530_3 | TATGAGTGTGGAGTCGTTGCGCAAGTCAGACTCTGAGACAGAANNNAGATCGGAA |
| 34530_4 | TATGAGTGTGGAGTCGTTGCGCAAGTCAGACTCTGAGACAGAANNNAGATCGGAA |
| 49076_1 | TATGAGTGTGGAGTCGTTGCATCTCAGCTCACTGTAACCTCTGNNNAGATCGGAAGA |
| 49076_2 | TATGAGTGTGGAGTCGTTGCATCTCAGCTCACTGTAACCTCTGNNNAGATCGGAAGA |
| 49076_3 | TATGAGTGTGGAGTCGTTGCATCTCAGCTCACTGTAACCTCTGNNNAGATCGGAAGA |
| 49076_4 | TATGAGTGTGGAGTCGTTGCATCTCAGCTCACTGTAACCTCTGNNNAGATCGGAAGA |
| 39587_1 | TATGAGTGTGGAGTCGTTGCTCCTGACTGACTGTTTTCTAGCCNNNAGATCGGAAGA |
| 39587_2 | TATGAGTGTGGAGTCGTTGCTCCTGACTGACTGTTTTCTAGCCNNNAGATCGGAAGA |
| 39587_3 | TATGAGTGTGGAGTCGTTGCTCCTGACTGACTGTTTTCTAGCCNNNAGATCGGAAGA |
| 39587_4 | TATGAGTGTGGAGTCGTTGCTCCTGACTGACTGTTTTCTAGCCNNNAGATCGGAAGA |
| 43224_1 | TATGAGTGTGGAGTCGTTGCCCCGAATAGCTAGCAATACAGGTNNNAGATCGGAAG |
| 43224_2 | TATGAGTGTGGAGTCGTTGCCCCGAATAGCTAGCAATACAGGTNNNAGATCGGAAG |
| 43224_3 | TATGAGTGTGGAGTCGTTGCCCCGAATAGCTAGCAATACAGGTNNNAGATCGGAAG |
| 43224_4 | TATGAGTGTGGAGTCGTTGCCCCGAATAGCTAGCAATACAGGTNNNAGATCGGAAG |
| 35414_1 | TATGAGTGTGGAGTCGTTGCGGTATGATCTCAGCTCACTCCAGNNNAGATCGGAAG |
| 35414_2 | TATGAGTGTGGAGTCGTTGCGGTATGATCTCAGCTCACTCCAGNNNAGATCGGAAG |
| 35414_3 | TATGAGTGTGGAGTCGTTGCGGTATGATCTCAGCTCACTCCAGNNNAGATCGGAAG |
| 35414_4 | TATGAGTGTGGAGTCGTTGCGGTATGATCTCAGCTCACTCCAGNNNAGATCGGAAG |
| 52823_1 | TATGAGTGTGGAGTCGTTGCCAGCAACAGCATGAAGAAGAACANNNAGATCGGAA |
| 52823_2 | TATGAGTGTGGAGTCGTTGCCAGCAACAGCATGAAGAAGAACANNNAGATCGGAA |
| 52823_3 | TATGAGTGTGGAGTCGTTGCCAGCAACAGCATGAAGAAGAACANNNAGATCGGAA |
| 52823_4 | TATGAGTGTGGAGTCGTTGCCAGCAACAGCATGAAGAAGAACANNNAGATCGGAA |
| 51590_1 | TATGAGTGTGGAGTCGTTGCCGACTCGAGCTTTAGCCCAGNNNAGATCGGAAGAG |
| 51590_2 | TATGAGTGTGGAGTCGTTGCCGACTCGAGCTTTAGCCCAGNNNAGATCGGAAGAG |

| | |
|---|---|
| 51590_3 | TATGAGTGTGGAGTCGTTGCCGACTCGAGCTTTAGCCCAGNNNAGATCGGAAGAGC |
| 51590_4 | TATGAGTGTGGAGTCGTTGCCGACTCGAGCTTTAGCCCAGNNNAGATCGGAAGAGC |
| 44581_1 | TATGAGTGTGGAGTCGTTGCAACTCCGTAAGAGAACCTTCCACNNNAGATCGGAAGA |
| 44581_2 | TATGAGTGTGGAGTCGTTGCAACTCCGTAAGAGAACCTTCCACNNNAGATCGGAAGA |
| 44581_3 | TATGAGTGTGGAGTCGTTGCAACTCCGTAAGAGAACCTTCCACNNNAGATCGGAAGA |
| 44581_4 | TATGAGTGTGGAGTCGTTGCAACTCCGTAAGAGAACCTTCCACNNNAGATCGGAAGA |
| 46749_1 | TATGAGTGTGGAGTCGTTGCCTCCTGAGTAGCTGGAACTACAGNNNAGATCGGAAG |
| 46749_2 | TATGAGTGTGGAGTCGTTGCCTCCTGAGTAGCTGGAACTACAGNNNAGATCGGAAG |
| 46749_3 | TATGAGTGTGGAGTCGTTGCCTCCTGAGTAGCTGGAACTACAGNNNAGATCGGAAG |
| 46749_4 | TATGAGTGTGGAGTCGTTGCCTCCTGAGTAGCTGGAACTACAGNNNAGATCGGAAG |
| 44094_1 | TATGAGTGTGGAGTCGTTGCAAATGCAAGAGAGGCTACAAGGANNNAGATCGGAA |
| 44094_2 | TATGAGTGTGGAGTCGTTGCAAATGCAAGAGAGGCTACAAGGANNNAGATCGGAA |
| 44094_3 | TATGAGTGTGGAGTCGTTGCAAATGCAAGAGAGGCTACAAGGANNNAGATCGGAA |
| 44094_4 | TATGAGTGTGGAGTCGTTGCAAATGCAAGAGAGGCTACAAGGANNNAGATCGGAA |
| 51614_1 | TATGAGTGTGGAGTCGTTGCGCCAAAGCAAGGTTGAGATGAATNNNAGATCGGAA |
| 51614_2 | TATGAGTGTGGAGTCGTTGCGCCAAAGCAAGGTTGAGATGAATNNNAGATCGGAA |
| 51614_3 | TATGAGTGTGGAGTCGTTGCGCCAAAGCAAGGTTGAGATGAATNNNAGATCGGAA |
| 51614_4 | TATGAGTGTGGAGTCGTTGCGCCAAAGCAAGGTTGAGATGAATNNNAGATCGGAA |
| 32173_1 | TATGAGTGTGGAGTCGTTGCGATATGGAGTCTCCCACTGTCACNNNAGATCGGAAG |
| 32173_2 | TATGAGTGTGGAGTCGTTGCGATATGGAGTCTCCCACTGTCACNNNAGATCGGAAG |
| 32173_3 | TATGAGTGTGGAGTCGTTGCGATATGGAGTCTCCCACTGTCACNNNAGATCGGAAG |
| 32173_4 | TATGAGTGTGGAGTCGTTGCGATATGGAGTCTCCCACTGTCACNNNAGATCGGAAG |
| 10118_1 | TATGAGTGTGGAGTCGTTGCCTCCAAATCCTAGCACCATTCATNNNAGATCGGAAGA |
| 10118_2 | TATGAGTGTGGAGTCGTTGCCTCCAAATCCTAGCACCATTCATNNNAGATCGGAAGA |
| 10118_3 | TATGAGTGTGGAGTCGTTGCCTCCAAATCCTAGCACCATTCATNNNAGATCGGAAGA |
| 10118_4 | TATGAGTGTGGAGTCGTTGCCTCCAAATCCTAGCACCATTCATNNNAGATCGGAAGA |
| 36609_1 | TATGAGTGTGGAGTCGTTGCTTTGGGGAAAAACTGTGCTGTTTNNNAGATCGGAAG |
| 36609_2 | TATGAGTGTGGAGTCGTTGCTTTGGGGAAAAACTGTGCTGTTTNNNAGATCGGAAG |
| 36609_3 | TATGAGTGTGGAGTCGTTGCTTTGGGGAAAAACTGTGCTGTTTNNNAGATCGGAAG |
| 36609_4 | TATGAGTGTGGAGTCGTTGCTTTGGGGAAAAACTGTGCTGTTTNNNAGATCGGAAG |
| 29565_1 | TATGAGTGTGGAGTCGTTGCTTAGCCTCATCACCTTCATCTGGNNNAGATCGGAAGA |
| 29565_2 | TATGAGTGTGGAGTCGTTGCTTAGCCTCATCACCTTCATCTGGNNNAGATCGGAAGA |
| 29565_3 | TATGAGTGTGGAGTCGTTGCTTAGCCTCATCACCTTCATCTGGNNNAGATCGGAAGA |
| 29565_4 | TATGAGTGTGGAGTCGTTGCTTAGCCTCATCACCTTCATCTGGNNNAGATCGGAAGA |
| 33293_1 | TATGAGTGTGGAGTCGTTGCTTCAAGTGATTCTCCTGCCTCAGNNNAGATCGGAAGA |
| 33293_2 | TATGAGTGTGGAGTCGTTGCTTCAAGTGATTCTCCTGCCTCAGNNNAGATCGGAAGA |
| 33293_3 | TATGAGTGTGGAGTCGTTGCTTCAAGTGATTCTCCTGCCTCAGNNNAGATCGGAAGA |
| 33293_4 | TATGAGTGTGGAGTCGTTGCTTCAAGTGATTCTCCTGCCTCAGNNNAGATCGGAAGA |
| 52866_1 | TATGAGTGTGGAGTCGTTGCCTTTGGCCACAGAGCTGGAGNNNAGATCGGAAGAG |
| 52866_2 | TATGAGTGTGGAGTCGTTGCCTTTGGCCACAGAGCTGGAGNNNAGATCGGAAGAG |
| 52866_3 | TATGAGTGTGGAGTCGTTGCCTTTGGCCACAGAGCTGGAGNNNAGATCGGAAGAG |
| 52866_4 | TATGAGTGTGGAGTCGTTGCCTTTGGCCACAGAGCTGGAGNNNAGATCGGAAGAG |
| 33027_1 | TATGAGTGTGGAGTCGTTGCCAAGTAGCTGGGACTACAGGCNNNAGATCGGAAGA |

| | |
|---|---|
| 33027_2 | TATGAGTGTGGAGTCGTTGCCAAGTAGCTGGGACTACAGGCNNNAGATCGGAAGA |
| 33027_3 | TATGAGTGTGGAGTCGTTGCCAAGTAGCTGGGACTACAGGCNNNAGATCGGAAGA |
| 33027_4 | TATGAGTGTGGAGTCGTTGCCAAGTAGCTGGGACTACAGGCNNNAGATCGGAAGA |
| 13059_1 | TATGAGTGTGGAGTCGTTGCATCTATAGAACTCTGAACCACTANNNAGATCGGAAG |
| 13059_2 | TATGAGTGTGGAGTCGTTGCATCTATAGAACTCTGAACCACTANNNAGATCGGAAG |
| 13059_3 | TATGAGTGTGGAGTCGTTGCATCTATAGAACTCTGAACCACTANNNAGATCGGAAG |
| 13059_4 | TATGAGTGTGGAGTCGTTGCATCTATAGAACTCTGAACCACTANNNAGATCGGAAG |
| 39965_1 | TATGAGTGTGGAGTCGTTGCGAGAGAATAGAGAATGGGGGCTGNNNAGATCGGA |
| 39965_2 | TATGAGTGTGGAGTCGTTGCGAGAGAATAGAGAATGGGGGCTGNNNAGATCGGA |
| 39965_3 | TATGAGTGTGGAGTCGTTGCGAGAGAATAGAGAATGGGGGCTGNNNAGATCGGA |
| 39965_4 | TATGAGTGTGGAGTCGTTGCGAGAGAATAGAGAATGGGGGCTGNNNAGATCGGA |
| 37811_1 | TATGAGTGTGGAGTCGTTGCTAACCTGGCAAACATGGTGAAACNNNAGATCGGAA |
| 37811_2 | TATGAGTGTGGAGTCGTTGCTAACCTGGCAAACATGGTGAAACNNNAGATCGGAA |
| 37811_3 | TATGAGTGTGGAGTCGTTGCTAACCTGGCAAACATGGTGAAACNNNAGATCGGAA |
| 37811_4 | TATGAGTGTGGAGTCGTTGCTAACCTGGCAAACATGGTGAAACNNNAGATCGGAA |
| 42973_1 | TATGAGTGTGGAGTCGTTGCACTATTTGCCCATTTTGACCACGNNNAGATCGGAAG |
| 42973_2 | TATGAGTGTGGAGTCGTTGCACTATTTGCCCATTTTGACCACGNNNAGATCGGAAG |
| 42973_3 | TATGAGTGTGGAGTCGTTGCACTATTTGCCCATTTTGACCACGNNNAGATCGGAAG |
| 42973_4 | TATGAGTGTGGAGTCGTTGCACTATTTGCCCATTTTGACCACGNNNAGATCGGAAG |
| 30378_1 | TATGAGTGTGGAGTCGTTGCCCGGTATGTAATTTAACATTCAANNNAGATCGGAAG |
| 30378_2 | TATGAGTGTGGAGTCGTTGCCCGGTATGTAATTTAACATTCAANNNAGATCGGAAG |
| 30378_3 | TATGAGTGTGGAGTCGTTGCCCGGTATGTAATTTAACATTCAANNNAGATCGGAAG |
| 30378_4 | TATGAGTGTGGAGTCGTTGCCCGGTATGTAATTTAACATTCAANNNAGATCGGAAG |
| 51691_1 | TATGAGTGTGGAGTCGTTGCTTCCAAGCAAATCCCAGATCTCANNNAGATCGGAAG |
| 51691_2 | TATGAGTGTGGAGTCGTTGCTTCCAAGCAAATCCCAGATCTCANNNAGATCGGAAG |
| 51691_3 | TATGAGTGTGGAGTCGTTGCTTCCAAGCAAATCCCAGATCTCANNNAGATCGGAAG |
| 51691_4 | TATGAGTGTGGAGTCGTTGCTTCCAAGCAAATCCCAGATCTCANNNAGATCGGAAG |
| 33121_1 | TATGAGTGTGGAGTCGTTGCAGGATTCCATTGCTCAGAAACCTNNNAGATCGGAAG |
| 33121_2 | TATGAGTGTGGAGTCGTTGCAGGATTCCATTGCTCAGAAACCTNNNAGATCGGAAG |
| 33121_3 | TATGAGTGTGGAGTCGTTGCAGGATTCCATTGCTCAGAAACCTNNNAGATCGGAAG |
| 33121_4 | TATGAGTGTGGAGTCGTTGCAGGATTCCATTGCTCAGAAACCTNNNAGATCGGAAG |
| 37685_1 | TATGAGTGTGGAGTCGTTGCAAAAAACCACACACACACACACANNNAGATCGGAAG |
| 37685_2 | TATGAGTGTGGAGTCGTTGCAAAAAACCACACACACACACACANNNAGATCGGAAG |
| 37685_3 | TATGAGTGTGGAGTCGTTGCAAAAAACCACACACACACACACANNNAGATCGGAAG |
| 37685_4 | TATGAGTGTGGAGTCGTTGCAAAAAACCACACACACACACACANNNAGATCGGAAG |
| 48260_1 | TATGAGTGTGGAGTCGTTGCCACGCCCGGCTAATTTTTTGTATNNNAGATCGGAAG |
| 48260_2 | TATGAGTGTGGAGTCGTTGCCACGCCCGGCTAATTTTTTGTATNNNAGATCGGAAG |
| 48260_3 | TATGAGTGTGGAGTCGTTGCCACGCCCGGCTAATTTTTTGTATNNNAGATCGGAAG |
| 48260_4 | TATGAGTGTGGAGTCGTTGCCACGCCCGGCTAATTTTTTGTATNNNAGATCGGAAG |
| 3038_1 | TATGAGTGTGGAGTCGTTGCGCTTGTTCTCGGCATTATGATGCNNNAGATCGGAAG |
| 3038_2 | TATGAGTGTGGAGTCGTTGCGCTTGTTCTCGGCATTATGATGCNNNAGATCGGAAG |
| 3038_3 | TATGAGTGTGGAGTCGTTGCGCTTGTTCTCGGCATTATGATGCNNNAGATCGGAAG |
| 3038_4 | TATGAGTGTGGAGTCGTTGCGCTTGTTCTCGGCATTATGATGCNNNAGATCGGAAG |

| | |
|---|---|
| 49706_1 | TATGAGTGTGGAGTCGTTGCAATTCCAGGGCAACTGAGTTACTNNNAGATCGGAAG |
| 49706_2 | TATGAGTGTGGAGTCGTTGCAATTCCAGGGCAACTGAGTTACTNNNAGATCGGAAG |
| 49706_3 | TATGAGTGTGGAGTCGTTGCAATTCCAGGGCAACTGAGTTACTNNNAGATCGGAAG |
| 49706_4 | TATGAGTGTGGAGTCGTTGCAATTCCAGGGCAACTGAGTTACTNNNAGATCGGAAG |
| 47234_1 | TATGAGTGTGGAGTCGTTGCCCTCCTGAGTAGCTGGGACTATANNNAGATCGGAAG |
| 47234_2 | TATGAGTGTGGAGTCGTTGCCCTCCTGAGTAGCTGGGACTATANNNAGATCGGAAG |
| 47234_3 | TATGAGTGTGGAGTCGTTGCCCTCCTGAGTAGCTGGGACTATANNNAGATCGGAAG |
| 47234_4 | TATGAGTGTGGAGTCGTTGCCCTCCTGAGTAGCTGGGACTATANNNAGATCGGAAG |
| 52083_1 | TATGAGTGTGGAGTCGTTGCTACTAAAATGGATGGACGGCTCANNNAGATCGGAA |
| 52083_2 | TATGAGTGTGGAGTCGTTGCTACTAAAATGGATGGACGGCTCANNNAGATCGGAA |
| 52083_3 | TATGAGTGTGGAGTCGTTGCTACTAAAATGGATGGACGGCTCANNNAGATCGGAA |
| 52083_4 | TATGAGTGTGGAGTCGTTGCTACTAAAATGGATGGACGGCTCANNNAGATCGGAA |
| 40004_1 | TATGAGTGTGGAGTCGTTGCCCTCATGCTTCATCTCATGGACTNNNAGATCGGAAGA |
| 40004_2 | TATGAGTGTGGAGTCGTTGCCCTCATGCTTCATCTCATGGACTNNNAGATCGGAAGA |
| 40004_3 | TATGAGTGTGGAGTCGTTGCCCTCATGCTTCATCTCATGGACTNNNAGATCGGAAGA |
| 40004_4 | TATGAGTGTGGAGTCGTTGCCCTCATGCTTCATCTCATGGACTNNNAGATCGGAAGA |
| 32431_1 | TATGAGTGTGGAGTCGTTGCATTCAGGGGCAGTCAGTTGATAGNNNAGATCGGAA |
| 32431_2 | TATGAGTGTGGAGTCGTTGCATTCAGGGGCAGTCAGTTGATAGNNNAGATCGGAA |
| 32431_3 | TATGAGTGTGGAGTCGTTGCATTCAGGGGCAGTCAGTTGATAGNNNAGATCGGAA |
| 32431_4 | TATGAGTGTGGAGTCGTTGCATTCAGGGGCAGTCAGTTGATAGNNNAGATCGGAA |
| 44031_1 | TATGAGTGTGGAGTCGTTGCTCCTTTGCCTTGTCATTTGAAGGNNNAGATCGGAAG |
| 44031_2 | TATGAGTGTGGAGTCGTTGCTCCTTTGCCTTGTCATTTGAAGGNNNAGATCGGAAG |
| 44031_3 | TATGAGTGTGGAGTCGTTGCTCCTTTGCCTTGTCATTTGAAGGNNNAGATCGGAAG |
| 44031_4 | TATGAGTGTGGAGTCGTTGCTCCTTTGCCTTGTCATTTGAAGGNNNAGATCGGAAG |
| 35163_1 | TATGAGTGTGGAGTCGTTGCGTCTCCCGGATTCAAGTGATTCTNNNAGATCGGAAG |
| 35163_2 | TATGAGTGTGGAGTCGTTGCGTCTCCCGGATTCAAGTGATTCTNNNAGATCGGAAG |
| 35163_3 | TATGAGTGTGGAGTCGTTGCGTCTCCCGGATTCAAGTGATTCTNNNAGATCGGAAG |
| 35163_4 | TATGAGTGTGGAGTCGTTGCGTCTCCCGGATTCAAGTGATTCTNNNAGATCGGAAG |
| 10201_1 | TATGAGTGTGGAGTCGTTGCTGGAGGGAGCAGAGGTGTTTATCNNNAGATCGGAA |
| 10201_2 | TATGAGTGTGGAGTCGTTGCTGGAGGGAGCAGAGGTGTTTATCNNNAGATCGGAA |
| 10201_3 | TATGAGTGTGGAGTCGTTGCTGGAGGGAGCAGAGGTGTTTATCNNNAGATCGGAA |
| 10201_4 | TATGAGTGTGGAGTCGTTGCTGGAGGGAGCAGAGGTGTTTATCNNNAGATCGGAA |
| 47811_1 | TATGAGTGTGGAGTCGTTGCGAGTGCAGTGGCTTAGTGCAAGNNNAGATCGGAAG |
| 47811_2 | TATGAGTGTGGAGTCGTTGCGAGTGCAGTGGCTTAGTGCAAGNNNAGATCGGAAG |
| 47811_3 | TATGAGTGTGGAGTCGTTGCGAGTGCAGTGGCTTAGTGCAAGNNNAGATCGGAAG |
| 47811_4 | TATGAGTGTGGAGTCGTTGCGAGTGCAGTGGCTTAGTGCAAGNNNAGATCGGAAG |
| 52674_1 | TATGAGTGTGGAGTCGTTGCAGCCATGTGATATCCATTGAGCTNNNAGATCGGAAG |
| 52674_2 | TATGAGTGTGGAGTCGTTGCAGCCATGTGATATCCATTGAGCTNNNAGATCGGAAG |
| 52674_3 | TATGAGTGTGGAGTCGTTGCAGCCATGTGATATCCATTGAGCTNNNAGATCGGAAG |
| 52674_4 | TATGAGTGTGGAGTCGTTGCAGCCATGTGATATCCATTGAGCTNNNAGATCGGAAG |
| 30102_1 | TATGAGTGTGGAGTCGTTGCCACCAGTCCATCCTCTGTATCAGNNNAGATCGGAAG |
| 30102_2 | TATGAGTGTGGAGTCGTTGCCACCAGTCCATCCTCTGTATCAGNNNAGATCGGAAG |
| 30102_3 | TATGAGTGTGGAGTCGTTGCCACCAGTCCATCCTCTGTATCAGNNNAGATCGGAAG |

| | |
|---|---|
| 30102_4 | TATGAGTGTGGAGTCGTTGCCACCAGTCCATCCTCTGTATCAGNNNAGATCGGAAGA |
| 30992_1 | TATGAGTGTGGAGTCGTTGCGGAATGAATGAGATTGGGGGTGANNNAGATCGGAA |
| 30992_2 | TATGAGTGTGGAGTCGTTGCGGAATGAATGAGATTGGGGGTGANNNAGATCGGAA |
| 30992_3 | TATGAGTGTGGAGTCGTTGCGGAATGAATGAGATTGGGGGTGANNNAGATCGGAA |
| 30992_4 | TATGAGTGTGGAGTCGTTGCGGAATGAATGAGATTGGGGGTGANNNAGATCGGAA |
| 39494_1 | TATGAGTGTGGAGTCGTTGCCTGTGTACATGCCAGCATTTGTTNNNAGATCGGAAGA |
| 39494_2 | TATGAGTGTGGAGTCGTTGCCTGTGTACATGCCAGCATTTGTTNNNAGATCGGAAGA |
| 39494_3 | TATGAGTGTGGAGTCGTTGCCTGTGTACATGCCAGCATTTGTTNNNAGATCGGAAGA |
| 39494_4 | TATGAGTGTGGAGTCGTTGCCTGTGTACATGCCAGCATTTGTTNNNAGATCGGAAGA |
| 40227_1 | TATGAGTGTGGAGTCGTTGCGAGAATTGCTTGAACCTGGGAGANNNAGATCGGAA |
| 40227_2 | TATGAGTGTGGAGTCGTTGCGAGAATTGCTTGAACCTGGGAGANNNAGATCGGAA |
| 40227_3 | TATGAGTGTGGAGTCGTTGCGAGAATTGCTTGAACCTGGGAGANNNAGATCGGAA |
| 40227_4 | TATGAGTGTGGAGTCGTTGCGAGAATTGCTTGAACCTGGGAGANNNAGATCGGAA |
| 42419_1 | TATGAGTGTGGAGTCGTTGCCCTCATATATCGCTAGTCTGGCANNNAGATCGGAAGA |
| 42419_2 | TATGAGTGTGGAGTCGTTGCCCTCATATATCGCTAGTCTGGCANNNAGATCGGAAGA |
| 42419_3 | TATGAGTGTGGAGTCGTTGCCCTCATATATCGCTAGTCTGGCANNNAGATCGGAAGA |
| 42419_4 | TATGAGTGTGGAGTCGTTGCCCTCATATATCGCTAGTCTGGCANNNAGATCGGAAGA |
| 33182_1 | TATGAGTGTGGAGTCGTTGCTCACCCAACATTCTTTGATTGAGGNNNAGATCGGAAC |
| 33182_2 | TATGAGTGTGGAGTCGTTGCTCACCCAACATTCTTTGATTGAGGNNNAGATCGGAAC |
| 33182_3 | TATGAGTGTGGAGTCGTTGCTCACCCAACATTCTTTGATTGAGGNNNAGATCGGAAC |
| 33182_4 | TATGAGTGTGGAGTCGTTGCTCACCCAACATTCTTTGATTGAGGNNNAGATCGGAAC |
| 45182_1 | TATGAGTGTGGAGTCGTTGCACTTTTTCTGAGGGAGGCAAAGANNNAGATCGGAAC |
| 45182_2 | TATGAGTGTGGAGTCGTTGCACTTTTTCTGAGGGAGGCAAAGANNNAGATCGGAAC |
| 45182_3 | TATGAGTGTGGAGTCGTTGCACTTTTTCTGAGGGAGGCAAAGANNNAGATCGGAAC |
| 45182_4 | TATGAGTGTGGAGTCGTTGCACTTTTTCTGAGGGAGGCAAAGANNNAGATCGGAAC |
| 44670_1 | TATGAGTGTGGAGTCGTTGCTTGGTGGCCTTTTCCTGAGCNNNAGATCGGAAGAGC |
| 44670_2 | TATGAGTGTGGAGTCGTTGCTTGGTGGCCTTTTCCTGAGCNNNAGATCGGAAGAGC |
| 44670_3 | TATGAGTGTGGAGTCGTTGCTTGGTGGCCTTTTCCTGAGCNNNAGATCGGAAGAGC |
| 44670_4 | TATGAGTGTGGAGTCGTTGCTTGGTGGCCTTTTCCTGAGCNNNAGATCGGAAGAGC |
| 10662_1 | TATGAGTGTGGAGTCGTTGCCCCAGAATCACTGGTCCTAACAGNNNAGATCGGAAG |
| 10662_2 | TATGAGTGTGGAGTCGTTGCCCCAGAATCACTGGTCCTAACAGNNNAGATCGGAAG |
| 10662_3 | TATGAGTGTGGAGTCGTTGCCCCAGAATCACTGGTCCTAACAGNNNAGATCGGAAG |
| 10662_4 | TATGAGTGTGGAGTCGTTGCCCCAGAATCACTGGTCCTAACAGNNNAGATCGGAAG |
| 38152_1 | TATGAGTGTGGAGTCGTTGCGAATAAATAAATGGGTGGCCGGGNNNAGATCGGAA |
| 38152_2 | TATGAGTGTGGAGTCGTTGCGAATAAATAAATGGGTGGCCGGGNNNAGATCGGAA |
| 38152_3 | TATGAGTGTGGAGTCGTTGCGAATAAATAAATGGGTGGCCGGGNNNAGATCGGAA |
| 38152_4 | TATGAGTGTGGAGTCGTTGCGAATAAATAAATGGGTGGCCGGGNNNAGATCGGAA |
| 49715_1 | TATGAGTGTGGAGTCGTTGCGAGACAGGGATTTGCCATGTTGNNNAGATCGGAAG |
| 49715_2 | TATGAGTGTGGAGTCGTTGCGAGACAGGGATTTGCCATGTTGNNNAGATCGGAAG |
| 49715_3 | TATGAGTGTGGAGTCGTTGCGAGACAGGGATTTGCCATGTTGNNNAGATCGGAAG |
| 49715_4 | TATGAGTGTGGAGTCGTTGCGAGACAGGGATTTGCCATGTTGNNNAGATCGGAAG |
| 31769_1 | TATGAGTGTGGAGTCGTTGCGATGAGGAGTGGAGTCAGGTATGNNNAGATCGGAA |
| 31769_2 | TATGAGTGTGGAGTCGTTGCGATGAGGAGTGGAGTCAGGTATGNNNAGATCGGAA |

| | |
|---|---|
| 31769_3 | TATGAGTGTGGAGTCGTTGCGATGAGGAGTGGAGTCAGGTATGNNNAGATCGGA |
| 31769_4 | TATGAGTGTGGAGTCGTTGCGATGAGGAGTGGAGTCAGGTATGNNNAGATCGGA |
| 33658_1 | TATGAGTGTGGAGTCGTTGCCCATTTCTCCTGACTCAGTCTCCNNNAGATCGGAAGA |
| 33658_2 | TATGAGTGTGGAGTCGTTGCCCATTTCTCCTGACTCAGTCTCCNNNAGATCGGAAGA |
| 33658_3 | TATGAGTGTGGAGTCGTTGCCCATTTCTCCTGACTCAGTCTCCNNNAGATCGGAAGA |
| 33658_4 | TATGAGTGTGGAGTCGTTGCCCATTTCTCCTGACTCAGTCTCCNNNAGATCGGAAGA |
| 37535_1 | TATGAGTGTGGAGTCGTTGCTTCTTAGTGTACAACCTTTGCCANNNAGATCGGAAGA |
| 37535_2 | TATGAGTGTGGAGTCGTTGCTTCTTAGTGTACAACCTTTGCCANNNAGATCGGAAGA |
| 37535_3 | TATGAGTGTGGAGTCGTTGCTTCTTAGTGTACAACCTTTGCCANNNAGATCGGAAGA |
| 37535_4 | TATGAGTGTGGAGTCGTTGCTTCTTAGTGTACAACCTTTGCCANNNAGATCGGAAGA |
| 51258_1 | TATGAGTGTGGAGTCGTTGCTCTAGGGCTCATGCTCTTCATTCNNNAGATCGGAAGA |
| 51258_2 | TATGAGTGTGGAGTCGTTGCTCTAGGGCTCATGCTCTTCATTCNNNAGATCGGAAGA |
| 51258_3 | TATGAGTGTGGAGTCGTTGCTCTAGGGCTCATGCTCTTCATTCNNNAGATCGGAAGA |
| 51258_4 | TATGAGTGTGGAGTCGTTGCTCTAGGGCTCATGCTCTTCATTCNNNAGATCGGAAGA |
| 39209_1 | TATGAGTGTGGAGTCGTTGCAAGCCAATCAAGAGAAACAGCACNNNAGATCGGAA |
| 39209_2 | TATGAGTGTGGAGTCGTTGCAAGCCAATCAAGAGAAACAGCACNNNAGATCGGAA |
| 39209_3 | TATGAGTGTGGAGTCGTTGCAAGCCAATCAAGAGAAACAGCACNNNAGATCGGAA |
| 39209_4 | TATGAGTGTGGAGTCGTTGCAAGCCAATCAAGAGAAACAGCACNNNAGATCGGAA |
| 43599_1 | TATGAGTGTGGAGTCGTTGCCCTACTGGGAAGTGAGGAGCCNNNAGATCGGAAGA |
| 43599_2 | TATGAGTGTGGAGTCGTTGCCCTACTGGGAAGTGAGGAGCCNNNAGATCGGAAGA |
| 43599_3 | TATGAGTGTGGAGTCGTTGCCCTACTGGGAAGTGAGGAGCCNNNAGATCGGAAGA |
| 43599_4 | TATGAGTGTGGAGTCGTTGCCCTACTGGGAAGTGAGGAGCCNNNAGATCGGAAGA |
| 51043_1 | TATGAGTGTGGAGTCGTTGCGCCTTCAAAAGACTCTTGGACATNNNAGATCGGAAG |
| 51043_2 | TATGAGTGTGGAGTCGTTGCGCCTTCAAAAGACTCTTGGACATNNNAGATCGGAAG |
| 51043_3 | TATGAGTGTGGAGTCGTTGCGCCTTCAAAAGACTCTTGGACATNNNAGATCGGAAG |
| 51043_4 | TATGAGTGTGGAGTCGTTGCGCCTTCAAAAGACTCTTGGACATNNNAGATCGGAAG |
| 36033_1 | TATGAGTGTGGAGTCGTTGCTCTAACAAGTAATGGAAGGCGGGNNNAGATCGGAA |
| 36033_2 | TATGAGTGTGGAGTCGTTGCTCTAACAAGTAATGGAAGGCGGGNNNAGATCGGAA |
| 36033_3 | TATGAGTGTGGAGTCGTTGCTCTAACAAGTAATGGAAGGCGGGNNNAGATCGGAA |
| 36033_4 | TATGAGTGTGGAGTCGTTGCTCTAACAAGTAATGGAAGGCGGGNNNAGATCGGAA |
| 31673_1 | TATGAGTGTGGAGTCGTTGCATGATTGTTTGGGCTAACAAGGCNNNAGATCGGAA |
| 31673_2 | TATGAGTGTGGAGTCGTTGCATGATTGTTTGGGCTAACAAGGCNNNAGATCGGAA |
| 31673_3 | TATGAGTGTGGAGTCGTTGCATGATTGTTTGGGCTAACAAGGCNNNAGATCGGAA |
| 31673_4 | TATGAGTGTGGAGTCGTTGCATGATTGTTTGGGCTAACAAGGCNNNAGATCGGAA |
| 45581_1 | TATGAGTGTGGAGTCGTTGCCGGTTGGTGGTGAATCAATCAAANNNAGATCGGAA |
| 45581_2 | TATGAGTGTGGAGTCGTTGCCGGTTGGTGGTGAATCAATCAAANNNAGATCGGAA |
| 45581_3 | TATGAGTGTGGAGTCGTTGCCGGTTGGTGGTGAATCAATCAAANNNAGATCGGAA |
| 45581_4 | TATGAGTGTGGAGTCGTTGCCGGTTGGTGGTGAATCAATCAAANNNAGATCGGAA |
| 36320_1 | TATGAGTGTGGAGTCGTTGCTCCTACATTGGGAAGAACAAGCTNNNAGATCGGAAG |
| 36320_2 | TATGAGTGTGGAGTCGTTGCTCCTACATTGGGAAGAACAAGCTNNNAGATCGGAAG |
| 36320_3 | TATGAGTGTGGAGTCGTTGCTCCTACATTGGGAAGAACAAGCTNNNAGATCGGAAG |
| 36320_4 | TATGAGTGTGGAGTCGTTGCTCCTACATTGGGAAGAACAAGCTNNNAGATCGGAAG |
| 48279_1 | TATGAGTGTGGAGTCGTTGCCCTCCTAAGTAGCTGGGACTACANNNAGATCGGAAG |

| | |
|---|---|
| 48279_2 | TATGAGTGTGGAGTCGTTGCCCTCCTAAGTAGCTGGGACTACANNNAGATCGGAAG |
| 48279_3 | TATGAGTGTGGAGTCGTTGCCCTCCTAAGTAGCTGGGACTACANNNAGATCGGAAG |
| 48279_4 | TATGAGTGTGGAGTCGTTGCCCTCCTAAGTAGCTGGGACTACANNNAGATCGGAAG |
| 52367_1 | TATGAGTGTGGAGTCGTTGCATTCCTACCAGACCCTTAGCCATNNNAGATCGGAAGA |
| 52367_2 | TATGAGTGTGGAGTCGTTGCATTCCTACCAGACCCTTAGCCATNNNAGATCGGAAGA |
| 52367_3 | TATGAGTGTGGAGTCGTTGCATTCCTACCAGACCCTTAGCCATNNNAGATCGGAAGA |
| 52367_4 | TATGAGTGTGGAGTCGTTGCATTCCTACCAGACCCTTAGCCATNNNAGATCGGAAGA |
| 30953_1 | TATGAGTGTGGAGTCGTTGCAGGGACTGTGAGAAAGTAGGAGANNNAGATCGGAA |
| 30953_2 | TATGAGTGTGGAGTCGTTGCAGGGACTGTGAGAAAGTAGGAGANNNAGATCGGAA |
| 30953_3 | TATGAGTGTGGAGTCGTTGCAGGGACTGTGAGAAAGTAGGAGANNNAGATCGGAA |
| 30953_4 | TATGAGTGTGGAGTCGTTGCAGGGACTGTGAGAAAGTAGGAGANNNAGATCGGAA |
| 31256_1 | TATGAGTGTGGAGTCGTTGCACACACACACACACAGTCTAAGTNNNAGATCGGAAGA |
| 31256_2 | TATGAGTGTGGAGTCGTTGCACACACACACACACAGTCTAAGTNNNAGATCGGAAGA |
| 31256_3 | TATGAGTGTGGAGTCGTTGCACACACACACACACAGTCTAAGTNNNAGATCGGAAGA |
| 31256_4 | TATGAGTGTGGAGTCGTTGCACACACACACACACAGTCTAAGTNNNAGATCGGAAGA |
| 31374_1 | TATGAGTGTGGAGTCGTTGCCCTAAGATGAAAGAGACAGGGCANNNAGATCGGAA |
| 31374_2 | TATGAGTGTGGAGTCGTTGCCCTAAGATGAAAGAGACAGGGCANNNAGATCGGAA |
| 31374_3 | TATGAGTGTGGAGTCGTTGCCCTAAGATGAAAGAGACAGGGCANNNAGATCGGAA |
| 31374_4 | TATGAGTGTGGAGTCGTTGCCCTAAGATGAAAGAGACAGGGCANNNAGATCGGAA |
| 39280_1 | TATGAGTGTGGAGTCGTTGCCAAACTCCTGGGATCAAACAACCNNNAGATCGGAAG |
| 39280_2 | TATGAGTGTGGAGTCGTTGCCAAACTCCTGGGATCAAACAACCNNNAGATCGGAAG |
| 39280_3 | TATGAGTGTGGAGTCGTTGCCAAACTCCTGGGATCAAACAACCNNNAGATCGGAAG |
| 39280_4 | TATGAGTGTGGAGTCGTTGCCAAACTCCTGGGATCAAACAACCNNNAGATCGGAAG |
| 29233_1 | TATGAGTGTGGAGTCGTTGCGAATGGAACAGAAATCCCCAAGCNNNAGATCGGAA |
| 29233_2 | TATGAGTGTGGAGTCGTTGCGAATGGAACAGAAATCCCCAAGCNNNAGATCGGAA |
| 29233_3 | TATGAGTGTGGAGTCGTTGCGAATGGAACAGAAATCCCCAAGCNNNAGATCGGAA |
| 29233_4 | TATGAGTGTGGAGTCGTTGCGAATGGAACAGAAATCCCCAAGCNNNAGATCGGAA |
| 39287_1 | TATGAGTGTGGAGTCGTTGCTATGATGACAGTGTGCCTCTCAGNNNAGATCGGAAG |
| 39287_2 | TATGAGTGTGGAGTCGTTGCTATGATGACAGTGTGCCTCTCAGNNNAGATCGGAAG |
| 39287_3 | TATGAGTGTGGAGTCGTTGCTATGATGACAGTGTGCCTCTCAGNNNAGATCGGAAG |
| 39287_4 | TATGAGTGTGGAGTCGTTGCTATGATGACAGTGTGCCTCTCAGNNNAGATCGGAAG |
| 42533_1 | TATGAGTGTGGAGTCGTTGCATAGTTGGAGACTTCACCACACCNNNAGATCGGAAG |
| 42533_2 | TATGAGTGTGGAGTCGTTGCATAGTTGGAGACTTCACCACACCNNNAGATCGGAAG |
| 42533_3 | TATGAGTGTGGAGTCGTTGCATAGTTGGAGACTTCACCACACCNNNAGATCGGAAG |
| 42533_4 | TATGAGTGTGGAGTCGTTGCATAGTTGGAGACTTCACCACACCNNNAGATCGGAAG |
| 35080_1 | TATGAGTGTGGAGTCGTTGCTAGTAGCTGGGATTACAGGCATGNNNAGATCGGAA |
| 35080_2 | TATGAGTGTGGAGTCGTTGCTAGTAGCTGGGATTACAGGCATGNNNAGATCGGAA |
| 35080_3 | TATGAGTGTGGAGTCGTTGCTAGTAGCTGGGATTACAGGCATGNNNAGATCGGAA |
| 35080_4 | TATGAGTGTGGAGTCGTTGCTAGTAGCTGGGATTACAGGCATGNNNAGATCGGAA |
| 7891_1 | TATGAGTGTGGAGTCGTTGCCCTGTAGCCACCAGAATTGTTGANNNAGATCGGAAG |
| 7891_2 | TATGAGTGTGGAGTCGTTGCCCTGTAGCCACCAGAATTGTTGANNNAGATCGGAAG |
| 7891_3 | TATGAGTGTGGAGTCGTTGCCCTGTAGCCACCAGAATTGTTGANNNAGATCGGAAG |
| 7891_4 | TATGAGTGTGGAGTCGTTGCCCTGTAGCCACCAGAATTGTTGANNNAGATCGGAAG |

| | |
|---|---|
| 48219_1 | TATGAGTGTGGAGTCGTTGCCACTGAAACTTCCACCTCCTAGGNNNAGATCGGAAG |
| 48219_2 | TATGAGTGTGGAGTCGTTGCCACTGAAACTTCCACCTCCTAGGNNNAGATCGGAAG |
| 48219_3 | TATGAGTGTGGAGTCGTTGCCACTGAAACTTCCACCTCCTAGGNNNAGATCGGAAG |
| 48219_4 | TATGAGTGTGGAGTCGTTGCCACTGAAACTTCCACCTCCTAGGNNNAGATCGGAAG |
| 30846_1 | TATGAGTGTGGAGTCGTTGCCTGACACTTCTAGCGGTACACAANNNAGATCGGAAG |
| 30846_2 | TATGAGTGTGGAGTCGTTGCCTGACACTTCTAGCGGTACACAANNNAGATCGGAAG |
| 30846_3 | TATGAGTGTGGAGTCGTTGCCTGACACTTCTAGCGGTACACAANNNAGATCGGAAG |
| 30846_4 | TATGAGTGTGGAGTCGTTGCCTGACACTTCTAGCGGTACACAANNNAGATCGGAAG |
| 37775_1 | TATGAGTGTGGAGTCGTTGCAAAAAAGGAGGCTTATTGAGGCCNNNAGATCGGAA |
| 37775_2 | TATGAGTGTGGAGTCGTTGCAAAAAAGGAGGCTTATTGAGGCCNNNAGATCGGAA |
| 37775_3 | TATGAGTGTGGAGTCGTTGCAAAAAAGGAGGCTTATTGAGGCCNNNAGATCGGAA |
| 37775_4 | TATGAGTGTGGAGTCGTTGCAAAAAAGGAGGCTTATTGAGGCCNNNAGATCGGAA |
| 48155_1 | TATGAGTGTGGAGTCGTTGCCTCCTGAGTAGCTGGGACTACAGNNNAGATCGGAA |
| 48155_2 | TATGAGTGTGGAGTCGTTGCCTCCTGAGTAGCTGGGACTACAGNNNAGATCGGAA |
| 48155_3 | TATGAGTGTGGAGTCGTTGCCTCCTGAGTAGCTGGGACTACAGNNNAGATCGGAA |
| 48155_4 | TATGAGTGTGGAGTCGTTGCCTCCTGAGTAGCTGGGACTACAGNNNAGATCGGAA |
| 51956_1 | TATGAGTGTGGAGTCGTTGCCTATCCTCACAGGCTCTGTTTGTNNNAGATCGGAAG |
| 51956_2 | TATGAGTGTGGAGTCGTTGCCTATCCTCACAGGCTCTGTTTGTNNNAGATCGGAAG |
| 51956_3 | TATGAGTGTGGAGTCGTTGCCTATCCTCACAGGCTCTGTTTGTNNNAGATCGGAAG |
| 51956_4 | TATGAGTGTGGAGTCGTTGCCTATCCTCACAGGCTCTGTTTGTNNNAGATCGGAAG |
| 53113_1 | TATGAGTGTGGAGTCGTTGCTCCTCTTACTGATCTTTGGTTTGCNNNAGATCGGAAG |
| 53113_2 | TATGAGTGTGGAGTCGTTGCTCCTCTTACTGATCTTTGGTTTGCNNNAGATCGGAAG |
| 53113_3 | TATGAGTGTGGAGTCGTTGCTCCTCTTACTGATCTTTGGTTTGCNNNAGATCGGAAG |
| 53113_4 | TATGAGTGTGGAGTCGTTGCTCCTCTTACTGATCTTTGGTTTGCNNNAGATCGGAAG |
| 52225_1 | TATGAGTGTGGAGTCGTTGCTTCATTCCTTCTCTGCATGTTGCNNNAGATCGGAAGA |
| 52225_2 | TATGAGTGTGGAGTCGTTGCTTCATTCCTTCTCTGCATGTTGCNNNAGATCGGAAGA |
| 52225_3 | TATGAGTGTGGAGTCGTTGCTTCATTCCTTCTCTGCATGTTGCNNNAGATCGGAAGA |
| 52225_4 | TATGAGTGTGGAGTCGTTGCTTCATTCCTTCTCTGCATGTTGCNNNAGATCGGAAGA |
| 31093_1 | TATGAGTGTGGAGTCGTTGCAGAGAGAGAGAGAAAGAGGGAGACANNNAGATCGG |
| 31093_2 | TATGAGTGTGGAGTCGTTGCAGAGAGAGAGAGAAAGAGGGAGACANNNAGATCGG |
| 31093_3 | TATGAGTGTGGAGTCGTTGCAGAGAGAGAGAGAAAGAGGGAGACANNNAGATCGG |
| 31093_4 | TATGAGTGTGGAGTCGTTGCAGAGAGAGAGAGAAAGAGGGAGACANNNAGATCGG |
| 52748_1 | TATGAGTGTGGAGTCGTTGCCAACCCCATTTGGTATCAACGTGNNNAGATCGGAAG |
| 52748_2 | TATGAGTGTGGAGTCGTTGCCAACCCCATTTGGTATCAACGTGNNNAGATCGGAAG |
| 52748_3 | TATGAGTGTGGAGTCGTTGCCAACCCCATTTGGTATCAACGTGNNNAGATCGGAAG |
| 52748_4 | TATGAGTGTGGAGTCGTTGCCAACCCCATTTGGTATCAACGTGNNNAGATCGGAAG |
| 40415_1 | TATGAGTGTGGAGTCGTTGCCTTGCCTTGACCATCATTCTTCCNNNAGATCGGAAGA |
| 40415_2 | TATGAGTGTGGAGTCGTTGCCTTGCCTTGACCATCATTCTTCCNNNAGATCGGAAGA |
| 40415_3 | TATGAGTGTGGAGTCGTTGCCTTGCCTTGACCATCATTCTTCCNNNAGATCGGAAGA |
| 40415_4 | TATGAGTGTGGAGTCGTTGCCTTGCCTTGACCATCATTCTTCCNNNAGATCGGAAGA |
| 40993_1 | TATGAGTGTGGAGTCGTTGCGTTGCAGTACTTCCACTACCTGANNNAGATCGGAAG |
| 40993_2 | TATGAGTGTGGAGTCGTTGCGTTGCAGTACTTCCACTACCTGANNNAGATCGGAAG |
| 40993_3 | TATGAGTGTGGAGTCGTTGCGTTGCAGTACTTCCACTACCTGANNNAGATCGGAAG |

| | |
|---|---|
| 40993_4 | TATGAGTGTGGAGTCGTTGCGTTGCAGTACTTCCACTACCTGANNNAGATCGGAAG |
| 42305_1 | TATGAGTGTGGAGTCGTTGCGAAAGAAGGGGACTGGTGAAGAGNNNAGATCGGA |
| 42305_2 | TATGAGTGTGGAGTCGTTGCGAAAGAAGGGGACTGGTGAAGAGNNNAGATCGGA |
| 42305_3 | TATGAGTGTGGAGTCGTTGCGAAAGAAGGGGACTGGTGAAGAGNNNAGATCGGA |
| 42305_4 | TATGAGTGTGGAGTCGTTGCGAAAGAAGGGGACTGGTGAAGAGNNNAGATCGGA |
| 29320_1 | TATGAGTGTGGAGTCGTTGCAAGGGGTTGAGGGTCTCCTATAANNNAGATCGGAA |
| 29320_2 | TATGAGTGTGGAGTCGTTGCAAGGGGTTGAGGGTCTCCTATAANNNAGATCGGAA |
| 29320_3 | TATGAGTGTGGAGTCGTTGCAAGGGGTTGAGGGTCTCCTATAANNNAGATCGGAA |
| 29320_4 | TATGAGTGTGGAGTCGTTGCAAGGGGTTGAGGGTCTCCTATAANNNAGATCGGAA |
| 49630_1 | TATGAGTGTGGAGTCGTTGCAAAATGAGATTGGTTCTGCTGCCNNNAGATCGGAAG |
| 49630_2 | TATGAGTGTGGAGTCGTTGCAAAATGAGATTGGTTCTGCTGCCNNNAGATCGGAAG |
| 49630_3 | TATGAGTGTGGAGTCGTTGCAAAATGAGATTGGTTCTGCTGCCNNNAGATCGGAAG |
| 49630_4 | TATGAGTGTGGAGTCGTTGCAAAATGAGATTGGTTCTGCTGCCNNNAGATCGGAAG |
| 29291_1 | TATGAGTGTGGAGTCGTTGCTGTGTGTGACCAATCCCCAAATANNNAGATCGGAAG |
| 29291_2 | TATGAGTGTGGAGTCGTTGCTGTGTGTGACCAATCCCCAAATANNNAGATCGGAAG |
| 29291_3 | TATGAGTGTGGAGTCGTTGCTGTGTGTGACCAATCCCCAAATANNNAGATCGGAAG |
| 29291_4 | TATGAGTGTGGAGTCGTTGCTGTGTGTGACCAATCCCCAAATANNNAGATCGGAAG |
| 39423_1 | TATGAGTGTGGAGTCGTTGCAATGTTCTGAAGCAGGTTCATGCNNNAGATCGGAAG |
| 39423_2 | TATGAGTGTGGAGTCGTTGCAATGTTCTGAAGCAGGTTCATGCNNNAGATCGGAAG |
| 39423_3 | TATGAGTGTGGAGTCGTTGCAATGTTCTGAAGCAGGTTCATGCNNNAGATCGGAAG |
| 39423_4 | TATGAGTGTGGAGTCGTTGCAATGTTCTGAAGCAGGTTCATGCNNNAGATCGGAAG |
| 50774_1 | TATGAGTGTGGAGTCGTTGCCTCTCAGGGAAGGTGTTGGATTCNNNAGATCGGAA |
| 50774_2 | TATGAGTGTGGAGTCGTTGCCTCTCAGGGAAGGTGTTGGATTCNNNAGATCGGAA |
| 50774_3 | TATGAGTGTGGAGTCGTTGCCTCTCAGGGAAGGTGTTGGATTCNNNAGATCGGAA |
| 50774_4 | TATGAGTGTGGAGTCGTTGCCTCTCAGGGAAGGTGTTGGATTCNNNAGATCGGAA |
| 42282_1 | TATGAGTGTGGAGTCGTTGCACACTGATGTCTTCCAATGGGTTNNNAGATCGGAAG |
| 42282_2 | TATGAGTGTGGAGTCGTTGCACACTGATGTCTTCCAATGGGTTNNNAGATCGGAAG |
| 42282_3 | TATGAGTGTGGAGTCGTTGCACACTGATGTCTTCCAATGGGTTNNNAGATCGGAAG |
| 42282_4 | TATGAGTGTGGAGTCGTTGCACACTGATGTCTTCCAATGGGTTNNNAGATCGGAAG |
| 42497_1 | TATGAGTGTGGAGTCGTTGCGAGAGAGGAGAGAGAGAGAGACCNNNAGATCGGA |
| 42497_2 | TATGAGTGTGGAGTCGTTGCGAGAGAGGAGAGAGAGAGAGACCNNNAGATCGGA |
| 42497_3 | TATGAGTGTGGAGTCGTTGCGAGAGAGGAGAGAGAGAGAGACCNNNAGATCGGA |
| 42497_4 | TATGAGTGTGGAGTCGTTGCGAGAGAGGAGAGAGAGAGAGACCNNNAGATCGGA |
| 49121_1 | TATGAGTGTGGAGTCGTTGCTCATTCCTGCCTCAGTTTAGCTTNNNAGATCGGAAGA |
| 49121_2 | TATGAGTGTGGAGTCGTTGCTCATTCCTGCCTCAGTTTAGCTTNNNAGATCGGAAGA |
| 49121_3 | TATGAGTGTGGAGTCGTTGCTCATTCCTGCCTCAGTTTAGCTTNNNAGATCGGAAGA |
| 49121_4 | TATGAGTGTGGAGTCGTTGCTCATTCCTGCCTCAGTTTAGCTTNNNAGATCGGAAGA |
| 30241_1 | TATGAGTGTGGAGTCGTTGCCCTCTAGAATGCACGCTTTTCTGNNNAGATCGGAAG |
| 30241_2 | TATGAGTGTGGAGTCGTTGCCCTCTAGAATGCACGCTTTTCTGNNNAGATCGGAAG |
| 30241_3 | TATGAGTGTGGAGTCGTTGCCCTCTAGAATGCACGCTTTTCTGNNNAGATCGGAAG |
| 30241_4 | TATGAGTGTGGAGTCGTTGCCCTCTAGAATGCACGCTTTTCTGNNNAGATCGGAAG |
| 32418_1 | TATGAGTGTGGAGTCGTTGCTCTTGGAGTACAGTGGTGCAATCNNNAGATCGGAA |
| 32418_2 | TATGAGTGTGGAGTCGTTGCTCTTGGAGTACAGTGGTGCAATCNNNAGATCGGAA |

| | |
|---|---|
| 32418_3 | TATGAGTGTGGAGTCGTTGCTCTTGGAGTACAGTGGTGCAATCNNNAGATCGGAA |
| 32418_4 | TATGAGTGTGGAGTCGTTGCTCTTGGAGTACAGTGGTGCAATCNNNAGATCGGAA |
| 35923_1 | TATGAGTGTGGAGTCGTTGCTTCTCTCTGGATTTTCAGGAGGCNNNAGATCGGAAG |
| 35923_2 | TATGAGTGTGGAGTCGTTGCTTCTCTCTGGATTTTCAGGAGGCNNNAGATCGGAAG |
| 35923_3 | TATGAGTGTGGAGTCGTTGCTTCTCTCTGGATTTTCAGGAGGCNNNAGATCGGAAG |
| 35923_4 | TATGAGTGTGGAGTCGTTGCTTCTCTCTGGATTTTCAGGAGGCNNNAGATCGGAAG |
| 42802_1 | TATGAGTGTGGAGTCGTTGCAGAAGATTCGGCACTTGTAGGAGNNNAGATCGGAA |
| 42802_2 | TATGAGTGTGGAGTCGTTGCAGAAGATTCGGCACTTGTAGGAGNNNAGATCGGAA |
| 42802_3 | TATGAGTGTGGAGTCGTTGCAGAAGATTCGGCACTTGTAGGAGNNNAGATCGGAA |
| 42802_4 | TATGAGTGTGGAGTCGTTGCAGAAGATTCGGCACTTGTAGGAGNNNAGATCGGAA |
| 35377_1 | TATGAGTGTGGAGTCGTTGCTGGTTTCGAACTTCTGACCTCAANNNAGATCGGAAG |
| 35377_2 | TATGAGTGTGGAGTCGTTGCTGGTTTCGAACTTCTGACCTCAANNNAGATCGGAAG |
| 35377_3 | TATGAGTGTGGAGTCGTTGCTGGTTTCGAACTTCTGACCTCAANNNAGATCGGAAG |
| 35377_4 | TATGAGTGTGGAGTCGTTGCTGGTTTCGAACTTCTGACCTCAANNNAGATCGGAAG |
| 52890_1 | TATGAGTGTGGAGTCGTTGCGTCACTTTATTTCATTCGGGCCCNNNAGATCGGAAG |
| 52890_2 | TATGAGTGTGGAGTCGTTGCGTCACTTTATTTCATTCGGGCCCNNNAGATCGGAAG |
| 52890_3 | TATGAGTGTGGAGTCGTTGCGTCACTTTATTTCATTCGGGCCCNNNAGATCGGAAG |
| 52890_4 | TATGAGTGTGGAGTCGTTGCGTCACTTTATTTCATTCGGGCCCNNNAGATCGGAAG |
| 44560_1 | TATGAGTGTGGAGTCGTTGCCACTTAGAGAGAATGTGGGGCATNNNAGATCGGAA |
| 44560_2 | TATGAGTGTGGAGTCGTTGCCACTTAGAGAGAATGTGGGGCATNNNAGATCGGAA |
| 44560_3 | TATGAGTGTGGAGTCGTTGCCACTTAGAGAGAATGTGGGGCATNNNAGATCGGAA |
| 44560_4 | TATGAGTGTGGAGTCGTTGCCACTTAGAGAGAATGTGGGGCATNNNAGATCGGAA |
| 50826_1 | TATGAGTGTGGAGTCGTTGCATTGAGGTATGAGTGACACGGATNNNAGATCGGAA |
| 50826_2 | TATGAGTGTGGAGTCGTTGCATTGAGGTATGAGTGACACGGATNNNAGATCGGAA |
| 50826_3 | TATGAGTGTGGAGTCGTTGCATTGAGGTATGAGTGACACGGATNNNAGATCGGAA |
| 50826_4 | TATGAGTGTGGAGTCGTTGCATTGAGGTATGAGTGACACGGATNNNAGATCGGAA |
| 37238_1 | TATGAGTGTGGAGTCGTTGCAGGCCAATAAAATCCTGCTTGATGNNNAGATCGGAA |
| 37238_2 | TATGAGTGTGGAGTCGTTGCAGGCCAATAAAATCCTGCTTGATGNNNAGATCGGAA |
| 37238_3 | TATGAGTGTGGAGTCGTTGCAGGCCAATAAAATCCTGCTTGATGNNNAGATCGGAA |
| 37238_4 | TATGAGTGTGGAGTCGTTGCAGGCCAATAAAATCCTGCTTGATGNNNAGATCGGAA |
| 45444_1 | TATGAGTGTGGAGTCGTTGCTGGAGGACAGAAAACATGGCATANNNAGATCGGAA |
| 45444_2 | TATGAGTGTGGAGTCGTTGCTGGAGGACAGAAAACATGGCATANNNAGATCGGAA |
| 45444_3 | TATGAGTGTGGAGTCGTTGCTGGAGGACAGAAAACATGGCATANNNAGATCGGAA |
| 45444_4 | TATGAGTGTGGAGTCGTTGCTGGAGGACAGAAAACATGGCATANNNAGATCGGAA |
| 43641_1 | TATGAGTGTGGAGTCGTTGCTCAGAAAACATCAGGAGGGGAAGNNNAGATCGGA |
| 43641_2 | TATGAGTGTGGAGTCGTTGCTCAGAAAACATCAGGAGGGGAAGNNNAGATCGGA |
| 43641_3 | TATGAGTGTGGAGTCGTTGCTCAGAAAACATCAGGAGGGGAAGNNNAGATCGGA |
| 43641_4 | TATGAGTGTGGAGTCGTTGCTCAGAAAACATCAGGAGGGGAAGNNNAGATCGGA |
| 30681_1 | TATGAGTGTGGAGTCGTTGCCCCGGTTGTAAATTGTTATAGTGGNNNAGATCGGA |
| 30681_2 | TATGAGTGTGGAGTCGTTGCCCCGGTTGTAAATTGTTATAGTGGNNNAGATCGGA |
| 30681_3 | TATGAGTGTGGAGTCGTTGCCCCGGTTGTAAATTGTTATAGTGGNNNAGATCGGA |
| 30681_4 | TATGAGTGTGGAGTCGTTGCCCCGGTTGTAAATTGTTATAGTGGNNNAGATCGGA |
| 38598_1 | TATGAGTGTGGAGTCGTTGCAAGAAAAGGTGGGAAAGAGCTGANNNAGATCGGA |

| | |
|---|---|
| 38598_2 | TATGAGTGTGGAGTCGTTGCAAGAAAAGGTGGGAAAGAGCTGANNNAGATCGGA |
| 38598_3 | TATGAGTGTGGAGTCGTTGCAAGAAAAGGTGGGAAAGAGCTGANNNAGATCGGA |
| 38598_4 | TATGAGTGTGGAGTCGTTGCAAGAAAAGGTGGGAAAGAGCTGANNNAGATCGGA |
| 44621_1 | TATGAGTGTGGAGTCGTTGCTTCCGAAATGGTGAGTAACTGGTNNNAGATCGGAA |
| 44621_2 | TATGAGTGTGGAGTCGTTGCTTCCGAAATGGTGAGTAACTGGTNNNAGATCGGAA |
| 44621_3 | TATGAGTGTGGAGTCGTTGCTTCCGAAATGGTGAGTAACTGGTNNNAGATCGGAA |
| 44621_4 | TATGAGTGTGGAGTCGTTGCTTCCGAAATGGTGAGTAACTGGTNNNAGATCGGAA |
| 34967_1 | TATGAGTGTGGAGTCGTTGCTTCAAGCCATTCTCCTGTCTCAGNNNAGATCGGAAG |
| 34967_2 | TATGAGTGTGGAGTCGTTGCTTCAAGCCATTCTCCTGTCTCAGNNNAGATCGGAAG |
| 34967_3 | TATGAGTGTGGAGTCGTTGCTTCAAGCCATTCTCCTGTCTCAGNNNAGATCGGAAG |
| 34967_4 | TATGAGTGTGGAGTCGTTGCTTCAAGCCATTCTCCTGTCTCAGNNNAGATCGGAAG |
| 9956_1 | TATGAGTGTGGAGTCGTTGCTCGTGACCATGTGTTAGGTAGGTNNNAGATCGGAA |
| 9956_2 | TATGAGTGTGGAGTCGTTGCTCGTGACCATGTGTTAGGTAGGTNNNAGATCGGAA |
| 9956_3 | TATGAGTGTGGAGTCGTTGCTCGTGACCATGTGTTAGGTAGGTNNNAGATCGGAA |
| 9956_4 | TATGAGTGTGGAGTCGTTGCTCGTGACCATGTGTTAGGTAGGTNNNAGATCGGAA |
| 51143_1 | TATGAGTGTGGAGTCGTTGCTGCCACTTGGATCAATGACAAAANNNAGATCGGAAG |
| 51143_2 | TATGAGTGTGGAGTCGTTGCTGCCACTTGGATCAATGACAAAANNNAGATCGGAAG |
| 51143_3 | TATGAGTGTGGAGTCGTTGCTGCCACTTGGATCAATGACAAAANNNAGATCGGAAG |
| 51143_4 | TATGAGTGTGGAGTCGTTGCTGCCACTTGGATCAATGACAAAANNNAGATCGGAAG |
| 45774_1 | TATGAGTGTGGAGTCGTTGCCCATGCCCGGCTAATTTTTTGTANNNAGATCGGAAG |
| 45774_2 | TATGAGTGTGGAGTCGTTGCCCATGCCCGGCTAATTTTTTGTANNNAGATCGGAAG |
| 45774_3 | TATGAGTGTGGAGTCGTTGCCCATGCCCGGCTAATTTTTTGTANNNAGATCGGAAG |
| 45774_4 | TATGAGTGTGGAGTCGTTGCCCATGCCCGGCTAATTTTTTGTANNNAGATCGGAAG |
| 49403_1 | TATGAGTGTGGAGTCGTTGCAACATGACCTCCTAGGAATGAGCNNNAGATCGGAA |
| 49403_2 | TATGAGTGTGGAGTCGTTGCAACATGACCTCCTAGGAATGAGCNNNAGATCGGAA |
| 49403_3 | TATGAGTGTGGAGTCGTTGCAACATGACCTCCTAGGAATGAGCNNNAGATCGGAA |
| 49403_4 | TATGAGTGTGGAGTCGTTGCAACATGACCTCCTAGGAATGAGCNNNAGATCGGAA |
| 10140_1 | TATGAGTGTGGAGTCGTTGCAACACATTCACACAGTCAGAGCANNNAGATCGGAAG |
| 10140_2 | TATGAGTGTGGAGTCGTTGCAACACATTCACACAGTCAGAGCANNNAGATCGGAAG |
| 10140_3 | TATGAGTGTGGAGTCGTTGCAACACATTCACACAGTCAGAGCANNNAGATCGGAAG |
| 10140_4 | TATGAGTGTGGAGTCGTTGCAACACATTCACACAGTCAGAGCANNNAGATCGGAAG |
| 2591_1 | TATGAGTGTGGAGTCGTTGCAGAGATGATAGTGGGTCCTGTGGNNNAGATCGGAA |
| 2591_2 | TATGAGTGTGGAGTCGTTGCAGAGATGATAGTGGGTCCTGTGGNNNAGATCGGAA |
| 2591_3 | TATGAGTGTGGAGTCGTTGCAGAGATGATAGTGGGTCCTGTGGNNNAGATCGGAA |
| 2591_4 | TATGAGTGTGGAGTCGTTGCAGAGATGATAGTGGGTCCTGTGGNNNAGATCGGAA |
| 47376_1 | TATGAGTGTGGAGTCGTTGCCCTCCTGGTTCACTCCATTCTCNNNAGATCGGAAGAG |
| 47376_2 | TATGAGTGTGGAGTCGTTGCCCTCCTGGTTCACTCCATTCTCNNNAGATCGGAAGAG |
| 47376_3 | TATGAGTGTGGAGTCGTTGCCCTCCTGGTTCACTCCATTCTCNNNAGATCGGAAGAG |
| 47376_4 | TATGAGTGTGGAGTCGTTGCCCTCCTGGTTCACTCCATTCTCNNNAGATCGGAAGAG |
| 32323_1 | TATGAGTGTGGAGTCGTTGCTGTGTGTCTTCCGTGCATATGTANNNAGATCGGAAG |
| 32323_2 | TATGAGTGTGGAGTCGTTGCTGTGTGTCTTCCGTGCATATGTANNNAGATCGGAAG |
| 32323_3 | TATGAGTGTGGAGTCGTTGCTGTGTGTCTTCCGTGCATATGTANNNAGATCGGAAG |
| 32323_4 | TATGAGTGTGGAGTCGTTGCTGTGTGTCTTCCGTGCATATGTANNNAGATCGGAAG |

| | |
|---|---|
| 37263_1 | TATGAGTGTGGAGTCGTTGCTAGGAGGAACTGAAGGCTGTTTCNNNAGATCGGAA |
| 37263_2 | TATGAGTGTGGAGTCGTTGCTAGGAGGAACTGAAGGCTGTTTCNNNAGATCGGAA |
| 37263_3 | TATGAGTGTGGAGTCGTTGCTAGGAGGAACTGAAGGCTGTTTCNNNAGATCGGAA |
| 37263_4 | TATGAGTGTGGAGTCGTTGCTAGGAGGAACTGAAGGCTGTTTCNNNAGATCGGAA |
| 41293_1 | TATGAGTGTGGAGTCGTTGCGTTCTGTGTGGGAAATGCATGAANNNAGATCGGAA |
| 41293_2 | TATGAGTGTGGAGTCGTTGCGTTCTGTGTGGGAAATGCATGAANNNAGATCGGAA |
| 41293_3 | TATGAGTGTGGAGTCGTTGCGTTCTGTGTGGGAAATGCATGAANNNAGATCGGAA |
| 41293_4 | TATGAGTGTGGAGTCGTTGCGTTCTGTGTGGGAAATGCATGAANNNAGATCGGAA |
| 45557_1 | TATGAGTGTGGAGTCGTTGCGAAGGAAGGAAGCGGGGAAGNNNAGATCGGAAG |
| 45557_2 | TATGAGTGTGGAGTCGTTGCGAAGGAAGGAAGCGGGGAAGNNNAGATCGGAAG |
| 45557_3 | TATGAGTGTGGAGTCGTTGCGAAGGAAGGAAGCGGGGAAGNNNAGATCGGAAG |
| 45557_4 | TATGAGTGTGGAGTCGTTGCGAAGGAAGGAAGCGGGGAAGNNNAGATCGGAAG |
| 7749_1 | TATGAGTGTGGAGTCGTTGCGGGTGCACTGCAGTAGTGTAATNNNAGATCGGAAG |
| 7749_2 | TATGAGTGTGGAGTCGTTGCGGGTGCACTGCAGTAGTGTAATNNNAGATCGGAAG |
| 7749_3 | TATGAGTGTGGAGTCGTTGCGGGTGCACTGCAGTAGTGTAATNNNAGATCGGAAG |
| 7749_4 | TATGAGTGTGGAGTCGTTGCGGGTGCACTGCAGTAGTGTAATNNNAGATCGGAAG |
| 29278_1 | TATGAGTGTGGAGTCGTTGCTTGTCTGAGTGTCCCAGAATCTGNNNAGATCGGAAG |
| 29278_2 | TATGAGTGTGGAGTCGTTGCTTGTCTGAGTGTCCCAGAATCTGNNNAGATCGGAAG |
| 29278_3 | TATGAGTGTGGAGTCGTTGCTTGTCTGAGTGTCCCAGAATCTGNNNAGATCGGAAG |
| 29278_4 | TATGAGTGTGGAGTCGTTGCTTGTCTGAGTGTCCCAGAATCTGNNNAGATCGGAAG |
| 49681_1 | TATGAGTGTGGAGTCGTTGCGTAGTAGCTGGGATTACAGGAGCNNNAGATCGGAA |
| 49681_2 | TATGAGTGTGGAGTCGTTGCGTAGTAGCTGGGATTACAGGAGCNNNAGATCGGAA |
| 49681_3 | TATGAGTGTGGAGTCGTTGCGTAGTAGCTGGGATTACAGGAGCNNNAGATCGGAA |
| 49681_4 | TATGAGTGTGGAGTCGTTGCGTAGTAGCTGGGATTACAGGAGCNNNAGATCGGAA |
| 8921_1 | TATGAGTGTGGAGTCGTTGCTCTCCATTCTCCACTTTCCTCGTNNNAGATCGGAAGA |
| 8921_2 | TATGAGTGTGGAGTCGTTGCTCTCCATTCTCCACTTTCCTCGTNNNAGATCGGAAGA |
| 8921_3 | TATGAGTGTGGAGTCGTTGCTCTCCATTCTCCACTTTCCTCGTNNNAGATCGGAAGA |
| 8921_4 | TATGAGTGTGGAGTCGTTGCTCTCCATTCTCCACTTTCCTCGTNNNAGATCGGAAGA |
| 45383_1 | TATGAGTGTGGAGTCGTTGCGGAGAGCCAACTGTACACAACTANNNAGATCGGAA |
| 45383_2 | TATGAGTGTGGAGTCGTTGCGGAGAGCCAACTGTACACAACTANNNAGATCGGAA |
| 45383_3 | TATGAGTGTGGAGTCGTTGCGGAGAGCCAACTGTACACAACTANNNAGATCGGAA |
| 45383_4 | TATGAGTGTGGAGTCGTTGCGGAGAGCCAACTGTACACAACTANNNAGATCGGAA |
| 33979_1 | TATGAGTGTGGAGTCGTTGCGAGACGGAGTCTTGCTTTGTCGNNNAGATCGGAAG |
| 33979_2 | TATGAGTGTGGAGTCGTTGCGAGACGGAGTCTTGCTTTGTCGNNNAGATCGGAAG |
| 33979_3 | TATGAGTGTGGAGTCGTTGCGAGACGGAGTCTTGCTTTGTCGNNNAGATCGGAAG |
| 33979_4 | TATGAGTGTGGAGTCGTTGCGAGACGGAGTCTTGCTTTGTCGNNNAGATCGGAAG |
| 51671_1 | TATGAGTGTGGAGTCGTTGCGCCCTGAGTCATGAAACCATTTTNNNAGATCGGAAG |
| 51671_2 | TATGAGTGTGGAGTCGTTGCGCCCTGAGTCATGAAACCATTTTNNNAGATCGGAAG |
| 51671_3 | TATGAGTGTGGAGTCGTTGCGCCCTGAGTCATGAAACCATTTTNNNAGATCGGAAG |
| 51671_4 | TATGAGTGTGGAGTCGTTGCGCCCTGAGTCATGAAACCATTTTNNNAGATCGGAAG |
| 48647_1 | TATGAGTGTGGAGTCGTTGCTATATCCTGACTCACCCTCCACCNNNAGATCGGAAGA |
| 48647_2 | TATGAGTGTGGAGTCGTTGCTATATCCTGACTCACCCTCCACCNNNAGATCGGAAGA |
| 48647_3 | TATGAGTGTGGAGTCGTTGCTATATCCTGACTCACCCTCCACCNNNAGATCGGAAGA |

| | |
|---|---|
| 48647_4 | TATGAGTGTGGAGTCGTTGCTATATCCTGACTCACCCTCCACCNNNAGATCGGAAGA |
| 39719_1 | TATGAGTGTGGAGTCGTTGCTGGCACATTGAATTACCTACACTNNNAGATCGGAAG |
| 39719_2 | TATGAGTGTGGAGTCGTTGCTGGCACATTGAATTACCTACACTNNNAGATCGGAAG |
| 39719_3 | TATGAGTGTGGAGTCGTTGCTGGCACATTGAATTACCTACACTNNNAGATCGGAAG |
| 39719_4 | TATGAGTGTGGAGTCGTTGCTGGCACATTGAATTACCTACACTNNNAGATCGGAAG |
| 33459_1 | TATGAGTGTGGAGTCGTTGCAAAGAGAGGCTTTGTTTCTGTGCNNNAGATCGGAA |
| 33459_2 | TATGAGTGTGGAGTCGTTGCAAAGAGAGGCTTTGTTTCTGTGCNNNAGATCGGAA |
| 33459_3 | TATGAGTGTGGAGTCGTTGCAAAGAGAGGCTTTGTTTCTGTGCNNNAGATCGGAA |
| 33459_4 | TATGAGTGTGGAGTCGTTGCAAAGAGAGGCTTTGTTTCTGTGCNNNAGATCGGAA |
| 44502_1 | TATGAGTGTGGAGTCGTTGCTTCTGTGAAGGCTGTCATGAGTANNNAGATCGGAA |
| 44502_2 | TATGAGTGTGGAGTCGTTGCTTCTGTGAAGGCTGTCATGAGTANNNAGATCGGAA |
| 44502_3 | TATGAGTGTGGAGTCGTTGCTTCTGTGAAGGCTGTCATGAGTANNNAGATCGGAA |
| 44502_4 | TATGAGTGTGGAGTCGTTGCTTCTGTGAAGGCTGTCATGAGTANNNAGATCGGAA |
| 39765_1 | TATGAGTGTGGAGTCGTTGCCTGTAGTCCCAGCTACTCAGGAGNNNAGATCGGAAG |
| 39765_2 | TATGAGTGTGGAGTCGTTGCCTGTAGTCCCAGCTACTCAGGAGNNNAGATCGGAAG |
| 39765_3 | TATGAGTGTGGAGTCGTTGCCTGTAGTCCCAGCTACTCAGGAGNNNAGATCGGAAG |
| 39765_4 | TATGAGTGTGGAGTCGTTGCCTGTAGTCCCAGCTACTCAGGAGNNNAGATCGGAAG |
| 48589_1 | TATGAGTGTGGAGTCGTTGCAAGCCCTTATCTTCCCTTACTGCNNNAGATCGGAAGA |
| 48589_2 | TATGAGTGTGGAGTCGTTGCAAGCCCTTATCTTCCCTTACTGCNNNAGATCGGAAGA |
| 48589_3 | TATGAGTGTGGAGTCGTTGCAAGCCCTTATCTTCCCTTACTGCNNNAGATCGGAAGA |
| 48589_4 | TATGAGTGTGGAGTCGTTGCAAGCCCTTATCTTCCCTTACTGCNNNAGATCGGAAGA |
| 31609_1 | TATGAGTGTGGAGTCGTTGCACACCTGTGCACAATAACATCCTNNNAGATCGGAAG |
| 31609_2 | TATGAGTGTGGAGTCGTTGCACACCTGTGCACAATAACATCCTNNNAGATCGGAAG |
| 31609_3 | TATGAGTGTGGAGTCGTTGCACACCTGTGCACAATAACATCCTNNNAGATCGGAAG |
| 31609_4 | TATGAGTGTGGAGTCGTTGCACACCTGTGCACAATAACATCCTNNNAGATCGGAAG |
| 45832_1 | TATGAGTGTGGAGTCGTTGCGAGTAGCTGGGATTACAGGCATGNNNAGATCGGAA |
| 45832_2 | TATGAGTGTGGAGTCGTTGCGAGTAGCTGGGATTACAGGCATGNNNAGATCGGAA |
| 45832_3 | TATGAGTGTGGAGTCGTTGCGAGTAGCTGGGATTACAGGCATGNNNAGATCGGAA |
| 45832_4 | TATGAGTGTGGAGTCGTTGCGAGTAGCTGGGATTACAGGCATGNNNAGATCGGAA |
| 45304_1 | TATGAGTGTGGAGTCGTTGCTTTGTGCCAGGCATTTTATACCCNNNAGATCGGAAG |
| 45304_2 | TATGAGTGTGGAGTCGTTGCTTTGTGCCAGGCATTTTATACCCNNNAGATCGGAAG |
| 45304_3 | TATGAGTGTGGAGTCGTTGCTTTGTGCCAGGCATTTTATACCCNNNAGATCGGAAG |
| 45304_4 | TATGAGTGTGGAGTCGTTGCTTTGTGCCAGGCATTTTATACCCNNNAGATCGGAAG |
| 45983_1 | TATGAGTGTGGAGTCGTTGCGAGTAGCTGGGATTACAGGCATGNNNAGATCGGAA |
| 45983_2 | TATGAGTGTGGAGTCGTTGCGAGTAGCTGGGATTACAGGCATGNNNAGATCGGAA |
| 45983_3 | TATGAGTGTGGAGTCGTTGCGAGTAGCTGGGATTACAGGCATGNNNAGATCGGAA |
| 45983_4 | TATGAGTGTGGAGTCGTTGCGAGTAGCTGGGATTACAGGCATGNNNAGATCGGAA |
| 10061_1 | TATGAGTGTGGAGTCGTTGCAGTAGCCCCATTATCTTGGGCTTNNNAGATCGGAAG |
| 10061_2 | TATGAGTGTGGAGTCGTTGCAGTAGCCCCATTATCTTGGGCTTNNNAGATCGGAAG |
| 10061_3 | TATGAGTGTGGAGTCGTTGCAGTAGCCCCATTATCTTGGGCTTNNNAGATCGGAAG |
| 10061_4 | TATGAGTGTGGAGTCGTTGCAGTAGCCCCATTATCTTGGGCTTNNNAGATCGGAAG |
| 46555_1 | TATGAGTGTGGAGTCGTTGCAGCCTCCTGAGTATCTGGGATTANNNAGATCGGAAG |
| 46555_2 | TATGAGTGTGGAGTCGTTGCAGCCTCCTGAGTATCTGGGATTANNNAGATCGGAAG |

| | |
|---|---|
| 46555_3 | TATGAGTGTGGAGTCGTTGCAGCCTCCTGAGTATCTGGGATTANNNAGATCGGAAG |
| 46555_4 | TATGAGTGTGGAGTCGTTGCAGCCTCCTGAGTATCTGGGATTANNNAGATCGGAAG |
| 29540_1 | TATGAGTGTGGAGTCGTTGCTGGGGGTTGATTTGCTTTTGATGNNNAGATCGGAA |
| 29540_2 | TATGAGTGTGGAGTCGTTGCTGGGGGTTGATTTGCTTTTGATGNNNAGATCGGAA |
| 29540_3 | TATGAGTGTGGAGTCGTTGCTGGGGGTTGATTTGCTTTTGATGNNNAGATCGGAA |
| 29540_4 | TATGAGTGTGGAGTCGTTGCTGGGGGTTGATTTGCTTTTGATGNNNAGATCGGAA |
| 47450_1 | TATGAGTGTGGAGTCGTTGCCACGCTTGTAATCTCAGCACTTTNNNAGATCGGAAGA |
| 47450_2 | TATGAGTGTGGAGTCGTTGCCACGCTTGTAATCTCAGCACTTTNNNAGATCGGAAGA |
| 47450_3 | TATGAGTGTGGAGTCGTTGCCACGCTTGTAATCTCAGCACTTTNNNAGATCGGAAGA |
| 47450_4 | TATGAGTGTGGAGTCGTTGCCACGCTTGTAATCTCAGCACTTTNNNAGATCGGAAGA |
| 47450_5 | TATGAGTGTGGAGTCGTTGCCACGCTTGTAATCTCAGCACTTTNNNAGATCGGAAGA |
| 47450_6 | TATGAGTGTGGAGTCGTTGCCACGCTTGTAATCTCAGCACTTTNNNAGATCGGAAGA |
| 47450_7 | TATGAGTGTGGAGTCGTTGCCACGCTTGTAATCTCAGCACTTTNNNAGATCGGAAGA |
| 47450_8 | TATGAGTGTGGAGTCGTTGCCACGCTTGTAATCTCAGCACTTTNNNAGATCGGAAGA |
| 47450_9 | TATGAGTGTGGAGTCGTTGCCACGCTTGTAATCTCAGCACTTTNNNAGATCGGAAGA |
| 47450_10 | TATGAGTGTGGAGTCGTTGCCACGCTTGTAATCTCAGCACTTTNNNAGATCGGAAGA |
| 34237_1 | TATGAGTGTGGAGTCGTTGCGCAAGTGAGGTGATAGGCGATATNNNAGATCGGAA |
| 34237_2 | TATGAGTGTGGAGTCGTTGCGCAAGTGAGGTGATAGGCGATATNNNAGATCGGAA |
| 34237_3 | TATGAGTGTGGAGTCGTTGCGCAAGTGAGGTGATAGGCGATATNNNAGATCGGAA |
| 34237_4 | TATGAGTGTGGAGTCGTTGCGCAAGTGAGGTGATAGGCGATATNNNAGATCGGAA |
| 34237_5 | TATGAGTGTGGAGTCGTTGCGCAAGTGAGGTGATAGGCGATATNNNAGATCGGAA |
| 34237_6 | TATGAGTGTGGAGTCGTTGCGCAAGTGAGGTGATAGGCGATATNNNAGATCGGAA |
| 34237_7 | TATGAGTGTGGAGTCGTTGCGCAAGTGAGGTGATAGGCGATATNNNAGATCGGAA |
| 34237_8 | TATGAGTGTGGAGTCGTTGCGCAAGTGAGGTGATAGGCGATATNNNAGATCGGAA |
| 34237_9 | TATGAGTGTGGAGTCGTTGCGCAAGTGAGGTGATAGGCGATATNNNAGATCGGAA |
| 34237_10 | TATGAGTGTGGAGTCGTTGCGCAAGTGAGGTGATAGGCGATATNNNAGATCGGAA |
| 47035_1 | TATGAGTGTGGAGTCGTTGCTCTCCTCATCTCATGTCACTTGCNNNAGATCGGAAGA |
| 47035_2 | TATGAGTGTGGAGTCGTTGCTCTCCTCATCTCATGTCACTTGCNNNAGATCGGAAGA |
| 47035_3 | TATGAGTGTGGAGTCGTTGCTCTCCTCATCTCATGTCACTTGCNNNAGATCGGAAGA |
| 47035_4 | TATGAGTGTGGAGTCGTTGCTCTCCTCATCTCATGTCACTTGCNNNAGATCGGAAGA |
| 47035_5 | TATGAGTGTGGAGTCGTTGCTCTCCTCATCTCATGTCACTTGCNNNAGATCGGAAGA |
| 47035_6 | TATGAGTGTGGAGTCGTTGCTCTCCTCATCTCATGTCACTTGCNNNAGATCGGAAGA |
| 47035_7 | TATGAGTGTGGAGTCGTTGCTCTCCTCATCTCATGTCACTTGCNNNAGATCGGAAGA |
| 47035_8 | TATGAGTGTGGAGTCGTTGCTCTCCTCATCTCATGTCACTTGCNNNAGATCGGAAGA |
| 47035_9 | TATGAGTGTGGAGTCGTTGCTCTCCTCATCTCATGTCACTTGCNNNAGATCGGAAGA |
| 47035_10 | TATGAGTGTGGAGTCGTTGCTCTCCTCATCTCATGTCACTTGCNNNAGATCGGAAGA |
| 41075_1 | TATGAGTGTGGAGTCGTTGCGCGACAGAACAAGACTCAGTCTANNNAGATCGGAA |
| 41075_2 | TATGAGTGTGGAGTCGTTGCGCGACAGAACAAGACTCAGTCTANNNAGATCGGAA |
| 41075_3 | TATGAGTGTGGAGTCGTTGCGCGACAGAACAAGACTCAGTCTANNNAGATCGGAA |
| 41075_4 | TATGAGTGTGGAGTCGTTGCGCGACAGAACAAGACTCAGTCTANNNAGATCGGAA |
| 41075_5 | TATGAGTGTGGAGTCGTTGCGCGACAGAACAAGACTCAGTCTANNNAGATCGGAA |
| 41075_6 | TATGAGTGTGGAGTCGTTGCGCGACAGAACAAGACTCAGTCTANNNAGATCGGAA |
| 41075_7 | TATGAGTGTGGAGTCGTTGCGCGACAGAACAAGACTCAGTCTANNNAGATCGGAA |

| | |
|---|---|
| 41075_8 | TATGAGTGTGGAGTCGTTGCGCGACAGAACAAGACTCAGTCTANNNAGATCGGAAG |
| 41075_9 | TATGAGTGTGGAGTCGTTGCGCGACAGAACAAGACTCAGTCTANNNAGATCGGAAG |
| 41075_10 | TATGAGTGTGGAGTCGTTGCGCGACAGAACAAGACTCAGTCTANNNAGATCGGAAG |
| 33291_1 | TATGAGTGTGGAGTCGTTGCTGTAATGGCATGATCTCAGCTCANNNAGATCGGAAG |
| 33291_2 | TATGAGTGTGGAGTCGTTGCTGTAATGGCATGATCTCAGCTCANNNAGATCGGAAG |
| 33291_3 | TATGAGTGTGGAGTCGTTGCTGTAATGGCATGATCTCAGCTCANNNAGATCGGAAG |
| 33291_4 | TATGAGTGTGGAGTCGTTGCTGTAATGGCATGATCTCAGCTCANNNAGATCGGAAG |
| 33291_5 | TATGAGTGTGGAGTCGTTGCTGTAATGGCATGATCTCAGCTCANNNAGATCGGAAG |
| 33291_6 | TATGAGTGTGGAGTCGTTGCTGTAATGGCATGATCTCAGCTCANNNAGATCGGAAG |
| 33291_7 | TATGAGTGTGGAGTCGTTGCTGTAATGGCATGATCTCAGCTCANNNAGATCGGAAG |
| 33291_8 | TATGAGTGTGGAGTCGTTGCTGTAATGGCATGATCTCAGCTCANNNAGATCGGAAG |
| 33291_9 | TATGAGTGTGGAGTCGTTGCTGTAATGGCATGATCTCAGCTCANNNAGATCGGAAG |
| 33291_10 | TATGAGTGTGGAGTCGTTGCTGTAATGGCATGATCTCAGCTCANNNAGATCGGAAG |
| 35753_1 | TATGAGTGTGGAGTCGTTGCGTACTAGGGTTACAGGTGTGAGCNNNAGATCGGAA |
| 35753_2 | TATGAGTGTGGAGTCGTTGCGTACTAGGGTTACAGGTGTGAGCNNNAGATCGGAA |
| 35753_3 | TATGAGTGTGGAGTCGTTGCGTACTAGGGTTACAGGTGTGAGCNNNAGATCGGAA |
| 35753_4 | TATGAGTGTGGAGTCGTTGCGTACTAGGGTTACAGGTGTGAGCNNNAGATCGGAA |
| 35753_5 | TATGAGTGTGGAGTCGTTGCGTACTAGGGTTACAGGTGTGAGCNNNAGATCGGAA |
| 35753_6 | TATGAGTGTGGAGTCGTTGCGTACTAGGGTTACAGGTGTGAGCNNNAGATCGGAA |
| 35753_7 | TATGAGTGTGGAGTCGTTGCGTACTAGGGTTACAGGTGTGAGCNNNAGATCGGAA |
| 35753_8 | TATGAGTGTGGAGTCGTTGCGTACTAGGGTTACAGGTGTGAGCNNNAGATCGGAA |
| 35753_9 | TATGAGTGTGGAGTCGTTGCGTACTAGGGTTACAGGTGTGAGCNNNAGATCGGAA |
| 35753_10 | TATGAGTGTGGAGTCGTTGCGTACTAGGGTTACAGGTGTGAGCNNNAGATCGGAA |
| 41812_1 | TATGAGTGTGGAGTCGTTGCGATTTCAGCTCACTGCAACCTTTNNNAGATCGGAAGA |
| 41812_2 | TATGAGTGTGGAGTCGTTGCGATTTCAGCTCACTGCAACCTTTNNNAGATCGGAAGA |
| 41812_3 | TATGAGTGTGGAGTCGTTGCGATTTCAGCTCACTGCAACCTTTNNNAGATCGGAAGA |
| 41812_4 | TATGAGTGTGGAGTCGTTGCGATTTCAGCTCACTGCAACCTTTNNNAGATCGGAAGA |
| 41812_5 | TATGAGTGTGGAGTCGTTGCGATTTCAGCTCACTGCAACCTTTNNNAGATCGGAAGA |
| 41812_6 | TATGAGTGTGGAGTCGTTGCGATTTCAGCTCACTGCAACCTTTNNNAGATCGGAAGA |
| 41812_7 | TATGAGTGTGGAGTCGTTGCGATTTCAGCTCACTGCAACCTTTNNNAGATCGGAAGA |
| 41812_8 | TATGAGTGTGGAGTCGTTGCGATTTCAGCTCACTGCAACCTTTNNNAGATCGGAAGA |
| 41812_9 | TATGAGTGTGGAGTCGTTGCGATTTCAGCTCACTGCAACCTTTNNNAGATCGGAAGA |
| 41812_10 | TATGAGTGTGGAGTCGTTGCGATTTCAGCTCACTGCAACCTTTNNNAGATCGGAAGA |
| 30841_1 | TATGAGTGTGGAGTCGTTGCTCAAGCACAGGAATGCATTGAAGNNNAGATCGGAAG |
| 30841_2 | TATGAGTGTGGAGTCGTTGCTCAAGCACAGGAATGCATTGAAGNNNAGATCGGAAG |
| 30841_3 | TATGAGTGTGGAGTCGTTGCTCAAGCACAGGAATGCATTGAAGNNNAGATCGGAAG |
| 30841_4 | TATGAGTGTGGAGTCGTTGCTCAAGCACAGGAATGCATTGAAGNNNAGATCGGAAG |
| 30841_5 | TATGAGTGTGGAGTCGTTGCTCAAGCACAGGAATGCATTGAAGNNNAGATCGGAAG |
| 30841_6 | TATGAGTGTGGAGTCGTTGCTCAAGCACAGGAATGCATTGAAGNNNAGATCGGAAG |
| 30841_7 | TATGAGTGTGGAGTCGTTGCTCAAGCACAGGAATGCATTGAAGNNNAGATCGGAAG |
| 30841_8 | TATGAGTGTGGAGTCGTTGCTCAAGCACAGGAATGCATTGAAGNNNAGATCGGAAG |
| 30841_9 | TATGAGTGTGGAGTCGTTGCTCAAGCACAGGAATGCATTGAAGNNNAGATCGGAAG |
| 30841_10 | TATGAGTGTGGAGTCGTTGCTCAAGCACAGGAATGCATTGAAGNNNAGATCGGAAG |

| | |
|---|---|
| 42983_1 | TATGAGTGTGGAGTCGTTGCCACACACACAAGGCTACAAAGAGNNNAGATCGGAAG |
| 42983_2 | TATGAGTGTGGAGTCGTTGCCACACACACAAGGCTACAAAGAGNNNAGATCGGAAG |
| 42983_3 | TATGAGTGTGGAGTCGTTGCCACACACACAAGGCTACAAAGAGNNNAGATCGGAAG |
| 42983_4 | TATGAGTGTGGAGTCGTTGCCACACACACAAGGCTACAAAGAGNNNAGATCGGAAG |
| 42983_5 | TATGAGTGTGGAGTCGTTGCCACACACACAAGGCTACAAAGAGNNNAGATCGGAAG |
| 42983_6 | TATGAGTGTGGAGTCGTTGCCACACACACAAGGCTACAAAGAGNNNAGATCGGAAG |
| 42983_7 | TATGAGTGTGGAGTCGTTGCCACACACACAAGGCTACAAAGAGNNNAGATCGGAAG |
| 42983_8 | TATGAGTGTGGAGTCGTTGCCACACACACAAGGCTACAAAGAGNNNAGATCGGAAG |
| 42983_9 | TATGAGTGTGGAGTCGTTGCCACACACACAAGGCTACAAAGAGNNNAGATCGGAAG |
| 42983_10 | TATGAGTGTGGAGTCGTTGCCACACACACAAGGCTACAAAGAGNNNAGATCGGAAG |
| 9932_1 | TATGAGTGTGGAGTCGTTGCACCAGTTCTAGCGGGAACTAAGANNNAGATCGGAA[illegible] |
| 9932_2 | TATGAGTGTGGAGTCGTTGCACCAGTTCTAGCGGGAACTAAGANNNAGATCGGAA[illegible] |
| 9932_3 | TATGAGTGTGGAGTCGTTGCACCAGTTCTAGCGGGAACTAAGANNNAGATCGGAA[illegible] |
| 9932_4 | TATGAGTGTGGAGTCGTTGCACCAGTTCTAGCGGGAACTAAGANNNAGATCGGAA[illegible] |
| 9932_5 | TATGAGTGTGGAGTCGTTGCACCAGTTCTAGCGGGAACTAAGANNNAGATCGGAA[illegible] |
| 9932_6 | TATGAGTGTGGAGTCGTTGCACCAGTTCTAGCGGGAACTAAGANNNAGATCGGAA[illegible] |
| 9932_7 | TATGAGTGTGGAGTCGTTGCACCAGTTCTAGCGGGAACTAAGANNNAGATCGGAA[illegible] |
| 9932_8 | TATGAGTGTGGAGTCGTTGCACCAGTTCTAGCGGGAACTAAGANNNAGATCGGAA[illegible] |
| 9932_9 | TATGAGTGTGGAGTCGTTGCACCAGTTCTAGCGGGAACTAAGANNNAGATCGGAA[illegible] |
| 9932_10 | TATGAGTGTGGAGTCGTTGCACCAGTTCTAGCGGGAACTAAGANNNAGATCGGAA[illegible] |
| 32465_1 | TATGAGTGTGGAGTCGTTGCAAAGGGTTCGCTGTAGATTCAGTNNNAGATCGGAA[illegible] |
| 32465_2 | TATGAGTGTGGAGTCGTTGCAAAGGGTTCGCTGTAGATTCAGTNNNAGATCGGAA[illegible] |
| 32465_3 | TATGAGTGTGGAGTCGTTGCAAAGGGTTCGCTGTAGATTCAGTNNNAGATCGGAA[illegible] |
| 32465_4 | TATGAGTGTGGAGTCGTTGCAAAGGGTTCGCTGTAGATTCAGTNNNAGATCGGAA[illegible] |
| 32465_5 | TATGAGTGTGGAGTCGTTGCAAAGGGTTCGCTGTAGATTCAGTNNNAGATCGGAA[illegible] |
| 32465_6 | TATGAGTGTGGAGTCGTTGCAAAGGGTTCGCTGTAGATTCAGTNNNAGATCGGAA[illegible] |
| 32465_7 | TATGAGTGTGGAGTCGTTGCAAAGGGTTCGCTGTAGATTCAGTNNNAGATCGGAA[illegible] |
| 32465_8 | TATGAGTGTGGAGTCGTTGCAAAGGGTTCGCTGTAGATTCAGTNNNAGATCGGAA[illegible] |
| 32465_9 | TATGAGTGTGGAGTCGTTGCAAAGGGTTCGCTGTAGATTCAGTNNNAGATCGGAA[illegible] |
| 32465_10 | TATGAGTGTGGAGTCGTTGCAAAGGGTTCGCTGTAGATTCAGTNNNAGATCGGAA[illegible] |
| 45278_1 | TATGAGTGTGGAGTCGTTGCTTCCAATCAGAAGATGCCAAGGANNNAGATCGGAAG |
| 45278_2 | TATGAGTGTGGAGTCGTTGCTTCCAATCAGAAGATGCCAAGGANNNAGATCGGAAG |
| 45278_3 | TATGAGTGTGGAGTCGTTGCTTCCAATCAGAAGATGCCAAGGANNNAGATCGGAAG |
| 45278_4 | TATGAGTGTGGAGTCGTTGCTTCCAATCAGAAGATGCCAAGGANNNAGATCGGAAG |
| 45278_5 | TATGAGTGTGGAGTCGTTGCTTCCAATCAGAAGATGCCAAGGANNNAGATCGGAAG |
| 45278_6 | TATGAGTGTGGAGTCGTTGCTTCCAATCAGAAGATGCCAAGGANNNAGATCGGAAG |
| 45278_7 | TATGAGTGTGGAGTCGTTGCTTCCAATCAGAAGATGCCAAGGANNNAGATCGGAAG |
| 45278_8 | TATGAGTGTGGAGTCGTTGCTTCCAATCAGAAGATGCCAAGGANNNAGATCGGAAG |
| 45278_9 | TATGAGTGTGGAGTCGTTGCTTCCAATCAGAAGATGCCAAGGANNNAGATCGGAAG |
| 45278_10 | TATGAGTGTGGAGTCGTTGCTTCCAATCAGAAGATGCCAAGGANNNAGATCGGAAG |
| 48303_1 | TATGAGTGTGGAGTCGTTGCGAGTCTCGCTCTATTGCCAGGNNNAGATCGGAAGAG |
| 48303_2 | TATGAGTGTGGAGTCGTTGCGAGTCTCGCTCTATTGCCAGGNNNAGATCGGAAGAG |
| 48303_3 | TATGAGTGTGGAGTCGTTGCGAGTCTCGCTCTATTGCCAGGNNNAGATCGGAAGAG |

| | |
|---|---|
| 48303_4 | TATGAGTGTGGAGTCGTTGCGAGTCTCGCTCTATTGCCAGGNNNAGATCGGAAGAG |
| 48303_5 | TATGAGTGTGGAGTCGTTGCGAGTCTCGCTCTATTGCCAGGNNNAGATCGGAAGAG |
| 48303_6 | TATGAGTGTGGAGTCGTTGCGAGTCTCGCTCTATTGCCAGGNNNAGATCGGAAGAG |
| 48303_7 | TATGAGTGTGGAGTCGTTGCGAGTCTCGCTCTATTGCCAGGNNNAGATCGGAAGAG |
| 48303_8 | TATGAGTGTGGAGTCGTTGCGAGTCTCGCTCTATTGCCAGGNNNAGATCGGAAGAG |
| 48303_9 | TATGAGTGTGGAGTCGTTGCGAGTCTCGCTCTATTGCCAGGNNNAGATCGGAAGAG |
| 48303_10 | TATGAGTGTGGAGTCGTTGCGAGTCTCGCTCTATTGCCAGGNNNAGATCGGAAGAG |
| 39802_1 | TATGAGTGTGGAGTCGTTGCACAGAAACTCACCTGTTATGTGTNNNAGATCGGAAG |
| 39802_2 | TATGAGTGTGGAGTCGTTGCACAGAAACTCACCTGTTATGTGTNNNAGATCGGAAG |
| 39802_3 | TATGAGTGTGGAGTCGTTGCACAGAAACTCACCTGTTATGTGTNNNAGATCGGAAG |
| 39802_4 | TATGAGTGTGGAGTCGTTGCACAGAAACTCACCTGTTATGTGTNNNAGATCGGAAG |
| 39802_5 | TATGAGTGTGGAGTCGTTGCACAGAAACTCACCTGTTATGTGTNNNAGATCGGAAG |
| 39802_6 | TATGAGTGTGGAGTCGTTGCACAGAAACTCACCTGTTATGTGTNNNAGATCGGAAG |
| 39802_7 | TATGAGTGTGGAGTCGTTGCACAGAAACTCACCTGTTATGTGTNNNAGATCGGAAG |
| 39802_8 | TATGAGTGTGGAGTCGTTGCACAGAAACTCACCTGTTATGTGTNNNAGATCGGAAG |
| 39802_9 | TATGAGTGTGGAGTCGTTGCACAGAAACTCACCTGTTATGTGTNNNAGATCGGAAG |
| 39802_10 | TATGAGTGTGGAGTCGTTGCACAGAAACTCACCTGTTATGTGTNNNAGATCGGAAG |
| 34477_1 | TATGAGTGTGGAGTCGTTGCTTGTGATCAATCTAGCCCAGCAANNNAGATCGGAAG |
| 34477_2 | TATGAGTGTGGAGTCGTTGCTTGTGATCAATCTAGCCCAGCAANNNAGATCGGAAG |
| 34477_3 | TATGAGTGTGGAGTCGTTGCTTGTGATCAATCTAGCCCAGCAANNNAGATCGGAAG |
| 34477_4 | TATGAGTGTGGAGTCGTTGCTTGTGATCAATCTAGCCCAGCAANNNAGATCGGAAG |
| 34477_5 | TATGAGTGTGGAGTCGTTGCTTGTGATCAATCTAGCCCAGCAANNNAGATCGGAAG |
| 34477_6 | TATGAGTGTGGAGTCGTTGCTTGTGATCAATCTAGCCCAGCAANNNAGATCGGAAG |
| 34477_7 | TATGAGTGTGGAGTCGTTGCTTGTGATCAATCTAGCCCAGCAANNNAGATCGGAAG |
| 34477_8 | TATGAGTGTGGAGTCGTTGCTTGTGATCAATCTAGCCCAGCAANNNAGATCGGAAG |
| 34477_9 | TATGAGTGTGGAGTCGTTGCTTGTGATCAATCTAGCCCAGCAANNNAGATCGGAAG |
| 34477_10 | TATGAGTGTGGAGTCGTTGCTTGTGATCAATCTAGCCCAGCAANNNAGATCGGAAG |
| 31955_1 | TATGAGTGTGGAGTCGTTGCGAAGGAAGGAAGGAAAGGAAGGANNNAGATCGGA |
| 31955_2 | TATGAGTGTGGAGTCGTTGCGAAGGAAGGAAGGAAAGGAAGGANNNAGATCGGA |
| 31955_3 | TATGAGTGTGGAGTCGTTGCGAAGGAAGGAAGGAAAGGAAGGANNNAGATCGGA |
| 31955_4 | TATGAGTGTGGAGTCGTTGCGAAGGAAGGAAGGAAAGGAAGGANNNAGATCGGA |
| 31955_5 | TATGAGTGTGGAGTCGTTGCGAAGGAAGGAAGGAAAGGAAGGANNNAGATCGGA |
| 31955_6 | TATGAGTGTGGAGTCGTTGCGAAGGAAGGAAGGAAAGGAAGGANNNAGATCGGA |
| 31955_7 | TATGAGTGTGGAGTCGTTGCGAAGGAAGGAAGGAAAGGAAGGANNNAGATCGGA |
| 31955_8 | TATGAGTGTGGAGTCGTTGCGAAGGAAGGAAGGAAAGGAAGGANNNAGATCGGA |
| 31955_9 | TATGAGTGTGGAGTCGTTGCGAAGGAAGGAAGGAAAGGAAGGANNNAGATCGGA |
| 31955_10 | TATGAGTGTGGAGTCGTTGCGAAGGAAGGAAGGAAAGGAAGGANNNAGATCGGA |
| 41073_1 | TATGAGTGTGGAGTCGTTGCTTCACTATATTGCCCAGGCTTGTNNNAGATCGGAAGA |
| 41073_2 | TATGAGTGTGGAGTCGTTGCTTCACTATATTGCCCAGGCTTGTNNNAGATCGGAAGA |
| 41073_3 | TATGAGTGTGGAGTCGTTGCTTCACTATATTGCCCAGGCTTGTNNNAGATCGGAAGA |
| 41073_4 | TATGAGTGTGGAGTCGTTGCTTCACTATATTGCCCAGGCTTGTNNNAGATCGGAAGA |
| 41073_5 | TATGAGTGTGGAGTCGTTGCTTCACTATATTGCCCAGGCTTGTNNNAGATCGGAAGA |
| 41073_6 | TATGAGTGTGGAGTCGTTGCTTCACTATATTGCCCAGGCTTGTNNNAGATCGGAAGA |

| | |
|---|---|
| 41073_7 | TATGAGTGTGGAGTCGTTGCTTCACTATATTGCCCAGGCTTGTNNNAGATCGGAAGA |
| 41073_8 | TATGAGTGTGGAGTCGTTGCTTCACTATATTGCCCAGGCTTGTNNNAGATCGGAAGA |
| 41073_9 | TATGAGTGTGGAGTCGTTGCTTCACTATATTGCCCAGGCTTGTNNNAGATCGGAAGA |
| 41073_10 | TATGAGTGTGGAGTCGTTGCTTCACTATATTGCCCAGGCTTGTNNNAGATCGGAAGA |
| 42265_1 | TATGAGTGTGGAGTCGTTGCCCTCAGCAGTGTACATTTGCTTCNNNAGATCGGAAGA |
| 42265_2 | TATGAGTGTGGAGTCGTTGCCCTCAGCAGTGTACATTTGCTTCNNNAGATCGGAAGA |
| 42265_3 | TATGAGTGTGGAGTCGTTGCCCTCAGCAGTGTACATTTGCTTCNNNAGATCGGAAGA |
| 42265_4 | TATGAGTGTGGAGTCGTTGCCCTCAGCAGTGTACATTTGCTTCNNNAGATCGGAAGA |
| 42265_5 | TATGAGTGTGGAGTCGTTGCCCTCAGCAGTGTACATTTGCTTCNNNAGATCGGAAGA |
| 42265_6 | TATGAGTGTGGAGTCGTTGCCCTCAGCAGTGTACATTTGCTTCNNNAGATCGGAAGA |
| 42265_7 | TATGAGTGTGGAGTCGTTGCCCTCAGCAGTGTACATTTGCTTCNNNAGATCGGAAGA |
| 42265_8 | TATGAGTGTGGAGTCGTTGCCCTCAGCAGTGTACATTTGCTTCNNNAGATCGGAAGA |
| 42265_9 | TATGAGTGTGGAGTCGTTGCCCTCAGCAGTGTACATTTGCTTCNNNAGATCGGAAGA |
| 42265_10 | TATGAGTGTGGAGTCGTTGCCCTCAGCAGTGTACATTTGCTTCNNNAGATCGGAAGA |
| 51555_1 | TATGAGTGTGGAGTCGTTGCGGTTTTTGGGAGTGGGTTGAAAGNNNAGATCGGAA |
| 51555_2 | TATGAGTGTGGAGTCGTTGCGGTTTTTGGGAGTGGGTTGAAAGNNNAGATCGGAA |
| 51555_3 | TATGAGTGTGGAGTCGTTGCGGTTTTTGGGAGTGGGTTGAAAGNNNAGATCGGAA |
| 51555_4 | TATGAGTGTGGAGTCGTTGCGGTTTTTGGGAGTGGGTTGAAAGNNNAGATCGGAA |
| 51555_5 | TATGAGTGTGGAGTCGTTGCGGTTTTTGGGAGTGGGTTGAAAGNNNAGATCGGAA |
| 51555_6 | TATGAGTGTGGAGTCGTTGCGGTTTTTGGGAGTGGGTTGAAAGNNNAGATCGGAA |
| 51555_7 | TATGAGTGTGGAGTCGTTGCGGTTTTTGGGAGTGGGTTGAAAGNNNAGATCGGAA |
| 51555_8 | TATGAGTGTGGAGTCGTTGCGGTTTTTGGGAGTGGGTTGAAAGNNNAGATCGGAA |
| 51555_9 | TATGAGTGTGGAGTCGTTGCGGTTTTTGGGAGTGGGTTGAAAGNNNAGATCGGAA |
| 51555_10 | TATGAGTGTGGAGTCGTTGCGGTTTTTGGGAGTGGGTTGAAAGNNNAGATCGGAA |
| 35941_1 | TATGAGTGTGGAGTCGTTGCTCCCGCAGTCTTATTTATACCCGNNNAGATCGGAAGA |
| 35941_2 | TATGAGTGTGGAGTCGTTGCTCCCGCAGTCTTATTTATACCCGNNNAGATCGGAAGA |
| 35941_3 | TATGAGTGTGGAGTCGTTGCTCCCGCAGTCTTATTTATACCCGNNNAGATCGGAAGA |
| 35941_4 | TATGAGTGTGGAGTCGTTGCTCCCGCAGTCTTATTTATACCCGNNNAGATCGGAAGA |
| 35941_5 | TATGAGTGTGGAGTCGTTGCTCCCGCAGTCTTATTTATACCCGNNNAGATCGGAAGA |
| 35941_6 | TATGAGTGTGGAGTCGTTGCTCCCGCAGTCTTATTTATACCCGNNNAGATCGGAAGA |
| 35941_7 | TATGAGTGTGGAGTCGTTGCTCCCGCAGTCTTATTTATACCCGNNNAGATCGGAAGA |
| 35941_8 | TATGAGTGTGGAGTCGTTGCTCCCGCAGTCTTATTTATACCCGNNNAGATCGGAAGA |
| 35941_9 | TATGAGTGTGGAGTCGTTGCTCCCGCAGTCTTATTTATACCCGNNNAGATCGGAAGA |
| 35941_10 | TATGAGTGTGGAGTCGTTGCTCCCGCAGTCTTATTTATACCCGNNNAGATCGGAAGA |
| 47875_1 | TATGAGTGTGGAGTCGTTGCCAGCCTCCCAGACATCATAACTTNNNAGATCGGAAGA |
| 47875_2 | TATGAGTGTGGAGTCGTTGCCAGCCTCCCAGACATCATAACTTNNNAGATCGGAAGA |
| 47875_3 | TATGAGTGTGGAGTCGTTGCCAGCCTCCCAGACATCATAACTTNNNAGATCGGAAGA |
| 47875_4 | TATGAGTGTGGAGTCGTTGCCAGCCTCCCAGACATCATAACTTNNNAGATCGGAAGA |
| 47875_5 | TATGAGTGTGGAGTCGTTGCCAGCCTCCCAGACATCATAACTTNNNAGATCGGAAGA |
| 47875_6 | TATGAGTGTGGAGTCGTTGCCAGCCTCCCAGACATCATAACTTNNNAGATCGGAAGA |
| 47875_7 | TATGAGTGTGGAGTCGTTGCCAGCCTCCCAGACATCATAACTTNNNAGATCGGAAGA |
| 47875_8 | TATGAGTGTGGAGTCGTTGCCAGCCTCCCAGACATCATAACTTNNNAGATCGGAAGA |
| 47875_9 | TATGAGTGTGGAGTCGTTGCCAGCCTCCCAGACATCATAACTTNNNAGATCGGAAGA |

| | |
|---|---|
| 47875_10 | TATGAGTGTGGAGTCGTTGCCAGCCTCCCAGACATCATAACTTNNNAGATCGGAAGA |
| 13378_1 | TATGAGTGTGGAGTCGTTGCGAAAGTCTGAGAACGGATTCNNNAGATCGGAAGAG |
| 13378_2 | TATGAGTGTGGAGTCGTTGCGAAAGTCTGAGAACGGATTCNNNAGATCGGAAGAG |
| 13378_3 | TATGAGTGTGGAGTCGTTGCGAAAGTCTGAGAACGGATTCNNNAGATCGGAAGAG |
| 13378_4 | TATGAGTGTGGAGTCGTTGCGAAAGTCTGAGAACGGATTCNNNAGATCGGAAGAG |
| 13378_5 | TATGAGTGTGGAGTCGTTGCGAAAGTCTGAGAACGGATTCNNNAGATCGGAAGAG |
| 13378_6 | TATGAGTGTGGAGTCGTTGCGAAAGTCTGAGAACGGATTCNNNAGATCGGAAGAG |
| 13378_7 | TATGAGTGTGGAGTCGTTGCGAAAGTCTGAGAACGGATTCNNNAGATCGGAAGAG |
| 13378_8 | TATGAGTGTGGAGTCGTTGCGAAAGTCTGAGAACGGATTCNNNAGATCGGAAGAG |
| 13378_9 | TATGAGTGTGGAGTCGTTGCGAAAGTCTGAGAACGGATTCNNNAGATCGGAAGAG |
| 13378_10 | TATGAGTGTGGAGTCGTTGCGAAAGTCTGAGAACGGATTCNNNAGATCGGAAGAG |
| 31996_1 | TATGAGTGTGGAGTCGTTGCGCTCCATCCTTATGACCTCACTTNNNAGATCGGAAGA |
| 31996_2 | TATGAGTGTGGAGTCGTTGCGCTCCATCCTTATGACCTCACTTNNNAGATCGGAAGA |
| 31996_3 | TATGAGTGTGGAGTCGTTGCGCTCCATCCTTATGACCTCACTTNNNAGATCGGAAGA |
| 31996_4 | TATGAGTGTGGAGTCGTTGCGCTCCATCCTTATGACCTCACTTNNNAGATCGGAAGA |
| 31996_5 | TATGAGTGTGGAGTCGTTGCGCTCCATCCTTATGACCTCACTTNNNAGATCGGAAGA |
| 31996_6 | TATGAGTGTGGAGTCGTTGCGCTCCATCCTTATGACCTCACTTNNNAGATCGGAAGA |
| 31996_7 | TATGAGTGTGGAGTCGTTGCGCTCCATCCTTATGACCTCACTTNNNAGATCGGAAGA |
| 31996_8 | TATGAGTGTGGAGTCGTTGCGCTCCATCCTTATGACCTCACTTNNNAGATCGGAAGA |
| 31996_9 | TATGAGTGTGGAGTCGTTGCGCTCCATCCTTATGACCTCACTTNNNAGATCGGAAGA |
| 31996_10 | TATGAGTGTGGAGTCGTTGCGCTCCATCCTTATGACCTCACTTNNNAGATCGGAAGA |
| 32120_1 | TATGAGTGTGGAGTCGTTGCCCCATAAGTGACCCAGATGGAAANNNAGATCGGAAG |
| 32120_2 | TATGAGTGTGGAGTCGTTGCCCCATAAGTGACCCAGATGGAAANNNAGATCGGAAG |
| 32120_3 | TATGAGTGTGGAGTCGTTGCCCCATAAGTGACCCAGATGGAAANNNAGATCGGAAG |
| 32120_4 | TATGAGTGTGGAGTCGTTGCCCCATAAGTGACCCAGATGGAAANNNAGATCGGAAG |
| 32120_5 | TATGAGTGTGGAGTCGTTGCCCCATAAGTGACCCAGATGGAAANNNAGATCGGAAG |
| 32120_6 | TATGAGTGTGGAGTCGTTGCCCCATAAGTGACCCAGATGGAAANNNAGATCGGAAG |
| 32120_7 | TATGAGTGTGGAGTCGTTGCCCCATAAGTGACCCAGATGGAAANNNAGATCGGAAG |
| 32120_8 | TATGAGTGTGGAGTCGTTGCCCCATAAGTGACCCAGATGGAAANNNAGATCGGAAG |
| 32120_9 | TATGAGTGTGGAGTCGTTGCCCCATAAGTGACCCAGATGGAAANNNAGATCGGAAG |
| 32120_10 | TATGAGTGTGGAGTCGTTGCCCCATAAGTGACCCAGATGGAAANNNAGATCGGAAG |
| 34999_1 | TATGAGTGTGGAGTCGTTGCGCCTACAGAGTAGCTGGGATTACNNNAGATCGGAAG |
| 34999_2 | TATGAGTGTGGAGTCGTTGCGCCTACAGAGTAGCTGGGATTACNNNAGATCGGAAG |
| 34999_3 | TATGAGTGTGGAGTCGTTGCGCCTACAGAGTAGCTGGGATTACNNNAGATCGGAAG |
| 34999_4 | TATGAGTGTGGAGTCGTTGCGCCTACAGAGTAGCTGGGATTACNNNAGATCGGAAG |
| 34999_5 | TATGAGTGTGGAGTCGTTGCGCCTACAGAGTAGCTGGGATTACNNNAGATCGGAAG |
| 34999_6 | TATGAGTGTGGAGTCGTTGCGCCTACAGAGTAGCTGGGATTACNNNAGATCGGAAG |
| 34999_7 | TATGAGTGTGGAGTCGTTGCGCCTACAGAGTAGCTGGGATTACNNNAGATCGGAAG |
| 34999_8 | TATGAGTGTGGAGTCGTTGCGCCTACAGAGTAGCTGGGATTACNNNAGATCGGAAG |
| 34999_9 | TATGAGTGTGGAGTCGTTGCGCCTACAGAGTAGCTGGGATTACNNNAGATCGGAAG |
| 34999_10 | TATGAGTGTGGAGTCGTTGCGCCTACAGAGTAGCTGGGATTACNNNAGATCGGAAG |
| 35908_1 | TATGAGTGTGGAGTCGTTGCCACACATATCCACAGCAGCAATGNNNAGATCGGAAG |
| 35908_2 | TATGAGTGTGGAGTCGTTGCCACACATATCCACAGCAGCAATGNNNAGATCGGAAG |

| | |
|---|---|
| 35908_3 | TATGAGTGTGGAGTCGTTGCCACACATATCCACAGCAGCAATGNNNAGATCGGAAG |
| 35908_4 | TATGAGTGTGGAGTCGTTGCCACACATATCCACAGCAGCAATGNNNAGATCGGAAG |
| 35908_5 | TATGAGTGTGGAGTCGTTGCCACACATATCCACAGCAGCAATGNNNAGATCGGAAG |
| 35908_6 | TATGAGTGTGGAGTCGTTGCCACACATATCCACAGCAGCAATGNNNAGATCGGAAG |
| 35908_7 | TATGAGTGTGGAGTCGTTGCCACACATATCCACAGCAGCAATGNNNAGATCGGAAG |
| 35908_8 | TATGAGTGTGGAGTCGTTGCCACACATATCCACAGCAGCAATGNNNAGATCGGAAG |
| 35908_9 | TATGAGTGTGGAGTCGTTGCCACACATATCCACAGCAGCAATGNNNAGATCGGAAG |
| 35908_10 | TATGAGTGTGGAGTCGTTGCCACACATATCCACAGCAGCAATGNNNAGATCGGAAG |
| 12550_1 | TATGAGTGTGGAGTCGTTGCGTGAATGAAATAGAAAAAATCGTTNNNAGATCGGA |
| 12550_2 | TATGAGTGTGGAGTCGTTGCGTGAATGAAATAGAAAAAATCGTTNNNAGATCGGA |
| 12550_3 | TATGAGTGTGGAGTCGTTGCGTGAATGAAATAGAAAAAATCGTTNNNAGATCGGA |
| 12550_4 | TATGAGTGTGGAGTCGTTGCGTGAATGAAATAGAAAAAATCGTTNNNAGATCGGA |
| 12550_5 | TATGAGTGTGGAGTCGTTGCGTGAATGAAATAGAAAAAATCGTTNNNAGATCGGA |
| 12550_6 | TATGAGTGTGGAGTCGTTGCGTGAATGAAATAGAAAAAATCGTTNNNAGATCGGA |
| 12550_7 | TATGAGTGTGGAGTCGTTGCGTGAATGAAATAGAAAAAATCGTTNNNAGATCGGA |
| 12550_8 | TATGAGTGTGGAGTCGTTGCGTGAATGAAATAGAAAAAATCGTTNNNAGATCGGA |
| 12550_9 | TATGAGTGTGGAGTCGTTGCGTGAATGAAATAGAAAAAATCGTTNNNAGATCGGA |
| 12550_10 | TATGAGTGTGGAGTCGTTGCGTGAATGAAATAGAAAAAATCGTTNNNAGATCGGA |
| 52453_1 | TATGAGTGTGGAGTCGTTGCTCCACCCCTAACTTGACCTCTATNNNAGATCGGAAGA |
| 52453_2 | TATGAGTGTGGAGTCGTTGCTCCACCCCTAACTTGACCTCTATNNNAGATCGGAAGA |
| 52453_3 | TATGAGTGTGGAGTCGTTGCTCCACCCCTAACTTGACCTCTATNNNAGATCGGAAGA |
| 52453_4 | TATGAGTGTGGAGTCGTTGCTCCACCCCTAACTTGACCTCTATNNNAGATCGGAAGA |
| 52453_5 | TATGAGTGTGGAGTCGTTGCTCCACCCCTAACTTGACCTCTATNNNAGATCGGAAGA |
| 52453_6 | TATGAGTGTGGAGTCGTTGCTCCACCCCTAACTTGACCTCTATNNNAGATCGGAAGA |
| 52453_7 | TATGAGTGTGGAGTCGTTGCTCCACCCCTAACTTGACCTCTATNNNAGATCGGAAGA |
| 52453_8 | TATGAGTGTGGAGTCGTTGCTCCACCCCTAACTTGACCTCTATNNNAGATCGGAAGA |
| 52453_9 | TATGAGTGTGGAGTCGTTGCTCCACCCCTAACTTGACCTCTATNNNAGATCGGAAGA |
| 52453_10 | TATGAGTGTGGAGTCGTTGCTCCACCCCTAACTTGACCTCTATNNNAGATCGGAAGA |
| 45671_1 | TATGAGTGTGGAGTCGTTGCCCTCCTGGTTCAAGAGATTCTCCNNNAGATCGGAAG |
| 45671_2 | TATGAGTGTGGAGTCGTTGCCCTCCTGGTTCAAGAGATTCTCCNNNAGATCGGAAG |
| 45671_3 | TATGAGTGTGGAGTCGTTGCCCTCCTGGTTCAAGAGATTCTCCNNNAGATCGGAAG |
| 45671_4 | TATGAGTGTGGAGTCGTTGCCCTCCTGGTTCAAGAGATTCTCCNNNAGATCGGAAG |
| 45671_5 | TATGAGTGTGGAGTCGTTGCCCTCCTGGTTCAAGAGATTCTCCNNNAGATCGGAAG |
| 45671_6 | TATGAGTGTGGAGTCGTTGCCCTCCTGGTTCAAGAGATTCTCCNNNAGATCGGAAG |
| 45671_7 | TATGAGTGTGGAGTCGTTGCCCTCCTGGTTCAAGAGATTCTCCNNNAGATCGGAAG |
| 45671_8 | TATGAGTGTGGAGTCGTTGCCCTCCTGGTTCAAGAGATTCTCCNNNAGATCGGAAG |
| 45671_9 | TATGAGTGTGGAGTCGTTGCCCTCCTGGTTCAAGAGATTCTCCNNNAGATCGGAAG |
| 45671_10 | TATGAGTGTGGAGTCGTTGCCCTCCTGGTTCAAGAGATTCTCCNNNAGATCGGAAG |
| 13701_1 | TATGAGTGTGGAGTCGTTGCACTCATGTCCCTATTTATGATTTNNNAGATCGGAAGA |
| 13701_2 | TATGAGTGTGGAGTCGTTGCACTCATGTCCCTATTTATGATTTNNNAGATCGGAAGA |
| 13701_3 | TATGAGTGTGGAGTCGTTGCACTCATGTCCCTATTTATGATTTNNNAGATCGGAAGA |
| 13701_4 | TATGAGTGTGGAGTCGTTGCACTCATGTCCCTATTTATGATTTNNNAGATCGGAAGA |
| 13701_5 | TATGAGTGTGGAGTCGTTGCACTCATGTCCCTATTTATGATTTNNNAGATCGGAAGA |

| | |
|---|---|
| 13701_6 | TATGAGTGTGGAGTCGTTGCACTCATGTCCCTATTTATGATTTNNNAGATCGGAAGA |
| 13701_7 | TATGAGTGTGGAGTCGTTGCACTCATGTCCCTATTTATGATTTNNNAGATCGGAAGA |
| 13701_8 | TATGAGTGTGGAGTCGTTGCACTCATGTCCCTATTTATGATTTNNNAGATCGGAAGA |
| 13701_9 | TATGAGTGTGGAGTCGTTGCACTCATGTCCCTATTTATGATTTNNNAGATCGGAAGA |
| 13701_10 | TATGAGTGTGGAGTCGTTGCACTCATGTCCCTATTTATGATTTNNNAGATCGGAAGA |
| 52892_1 | TATGAGTGTGGAGTCGTTGCCACCACTCAGAGTCCCCAAAGNNNAGATCGGAAGAG |
| 52892_2 | TATGAGTGTGGAGTCGTTGCCACCACTCAGAGTCCCCAAAGNNNAGATCGGAAGAG |
| 52892_3 | TATGAGTGTGGAGTCGTTGCCACCACTCAGAGTCCCCAAAGNNNAGATCGGAAGAG |
| 52892_4 | TATGAGTGTGGAGTCGTTGCCACCACTCAGAGTCCCCAAAGNNNAGATCGGAAGAG |
| 52892_5 | TATGAGTGTGGAGTCGTTGCCACCACTCAGAGTCCCCAAAGNNNAGATCGGAAGAG |
| 52892_6 | TATGAGTGTGGAGTCGTTGCCACCACTCAGAGTCCCCAAAGNNNAGATCGGAAGAG |
| 52892_7 | TATGAGTGTGGAGTCGTTGCCACCACTCAGAGTCCCCAAAGNNNAGATCGGAAGAG |
| 52892_8 | TATGAGTGTGGAGTCGTTGCCACCACTCAGAGTCCCCAAAGNNNAGATCGGAAGAG |
| 52892_9 | TATGAGTGTGGAGTCGTTGCCACCACTCAGAGTCCCCAAAGNNNAGATCGGAAGAG |
| 52892_10 | TATGAGTGTGGAGTCGTTGCCACCACTCAGAGTCCCCAAAGNNNAGATCGGAAGAG |
| 38720_1 | TATGAGTGTGGAGTCGTTGCGGCTAGCCATATGCAGAAAACTGNNNAGATCGGAA[illegible] |
| 38720_2 | TATGAGTGTGGAGTCGTTGCGGCTAGCCATATGCAGAAAACTGNNNAGATCGGAA[illegible] |
| 38720_3 | TATGAGTGTGGAGTCGTTGCGGCTAGCCATATGCAGAAAACTGNNNAGATCGGAA[illegible] |
| 38720_4 | TATGAGTGTGGAGTCGTTGCGGCTAGCCATATGCAGAAAACTGNNNAGATCGGAA[illegible] |
| 38720_5 | TATGAGTGTGGAGTCGTTGCGGCTAGCCATATGCAGAAAACTGNNNAGATCGGAA[illegible] |
| 38720_6 | TATGAGTGTGGAGTCGTTGCGGCTAGCCATATGCAGAAAACTGNNNAGATCGGAA[illegible] |
| 38720_7 | TATGAGTGTGGAGTCGTTGCGGCTAGCCATATGCAGAAAACTGNNNAGATCGGAA[illegible] |
| 38720_8 | TATGAGTGTGGAGTCGTTGCGGCTAGCCATATGCAGAAAACTGNNNAGATCGGAA[illegible] |
| 38720_9 | TATGAGTGTGGAGTCGTTGCGGCTAGCCATATGCAGAAAACTGNNNAGATCGGAA[illegible] |
| 38720_10 | TATGAGTGTGGAGTCGTTGCGGCTAGCCATATGCAGAAAACTGNNNAGATCGGAA[illegible] |
| 31746_1 | TATGAGTGTGGAGTCGTTGCCAACACTGCAGTCATGGTCTTGNNNAGATCGGAAGA |
| 31746_2 | TATGAGTGTGGAGTCGTTGCCAACACTGCAGTCATGGTCTTGNNNAGATCGGAAGA |
| 31746_3 | TATGAGTGTGGAGTCGTTGCCAACACTGCAGTCATGGTCTTGNNNAGATCGGAAGA |
| 31746_4 | TATGAGTGTGGAGTCGTTGCCAACACTGCAGTCATGGTCTTGNNNAGATCGGAAGA |
| 31746_5 | TATGAGTGTGGAGTCGTTGCCAACACTGCAGTCATGGTCTTGNNNAGATCGGAAGA |
| 31746_6 | TATGAGTGTGGAGTCGTTGCCAACACTGCAGTCATGGTCTTGNNNAGATCGGAAGA |
| 31746_7 | TATGAGTGTGGAGTCGTTGCCAACACTGCAGTCATGGTCTTGNNNAGATCGGAAGA |
| 31746_8 | TATGAGTGTGGAGTCGTTGCCAACACTGCAGTCATGGTCTTGNNNAGATCGGAAGA |
| 31746_9 | TATGAGTGTGGAGTCGTTGCCAACACTGCAGTCATGGTCTTGNNNAGATCGGAAGA |
| 31746_10 | TATGAGTGTGGAGTCGTTGCCAACACTGCAGTCATGGTCTTGNNNAGATCGGAAGA |
| 29917_1 | TATGAGTGTGGAGTCGTTGCCTGATGCAGAATGGGACTGGTTANNNAGATCGGAA[illegible] |
| 29917_2 | TATGAGTGTGGAGTCGTTGCCTGATGCAGAATGGGACTGGTTANNNAGATCGGAA[illegible] |
| 29917_3 | TATGAGTGTGGAGTCGTTGCCTGATGCAGAATGGGACTGGTTANNNAGATCGGAA[illegible] |
| 29917_4 | TATGAGTGTGGAGTCGTTGCCTGATGCAGAATGGGACTGGTTANNNAGATCGGAA[illegible] |
| 29917_5 | TATGAGTGTGGAGTCGTTGCCTGATGCAGAATGGGACTGGTTANNNAGATCGGAA[illegible] |
| 29917_6 | TATGAGTGTGGAGTCGTTGCCTGATGCAGAATGGGACTGGTTANNNAGATCGGAA[illegible] |
| 29917_7 | TATGAGTGTGGAGTCGTTGCCTGATGCAGAATGGGACTGGTTANNNAGATCGGAA[illegible] |
| 29917_8 | TATGAGTGTGGAGTCGTTGCCTGATGCAGAATGGGACTGGTTANNNAGATCGGAA[illegible] |

| | |
|---|---|
| 29917_9 | TATGAGTGTGGAGTCGTTGCCTGATGCAGAATGGGACTGGTTANNNAGATCGGAA |
| 29917_10 | TATGAGTGTGGAGTCGTTGCCTGATGCAGAATGGGACTGGTTANNNAGATCGGAA |
| 44879_1 | TATGAGTGTGGAGTCGTTGCAGGACCATTTAGATAACAGCCCCNNNAGATCGGAAG |
| 44879_2 | TATGAGTGTGGAGTCGTTGCAGGACCATTTAGATAACAGCCCCNNNAGATCGGAAG |
| 31967_1 | TATGAGTGTGGAGTCGTTGCGCCTAGGTGCTCATATGGACTTANNNAGATCGGAAG |
| 31967_2 | TATGAGTGTGGAGTCGTTGCGCCTAGGTGCTCATATGGACTTANNNAGATCGGAAG |
| 33199_1 | TATGAGTGTGGAGTCGTTGCGCTCAGGTGAAGTGAATGAGAGANNNAGATCGGA |
| 33199_2 | TATGAGTGTGGAGTCGTTGCGCTCAGGTGAAGTGAATGAGAGANNNAGATCGGA |
| 51206_1 | TATGAGTGTGGAGTCGTTGCGTGGAGAGGAGGAGGACAAAGNNNAGATCGGAA |
| 51206_2 | TATGAGTGTGGAGTCGTTGCGTGGAGAGGAGGAGGACAAAGNNNAGATCGGAA |
| 12625_1 | TATGAGTGTGGAGTCGTTGCCTAGGATGCCAACAAAGAAANNNAGATCGGAAGAG |
| 12625_2 | TATGAGTGTGGAGTCGTTGCCTAGGATGCCAACAAAGAAANNNAGATCGGAAGAG |
| 41028_1 | TATGAGTGTGGAGTCGTTGCATTGAAAGCAGGGACTGAACAGANNNAGATCGGAA |
| 41028_2 | TATGAGTGTGGAGTCGTTGCATTGAAAGCAGGGACTGAACAGANNNAGATCGGAA |
| 36295_1 | TATGAGTGTGGAGTCGTTGCAAGAAAATTAGCTGAGCGTGGTGNNNAGATCGGAA |
| 36295_2 | TATGAGTGTGGAGTCGTTGCAAGAAAATTAGCTGAGCGTGGTGNNNAGATCGGAA |
| 46129_1 | TATGAGTGTGGAGTCGTTGCTTGGTATTAGGGTGATACTGGCANNNAGATCGGAA |
| 46129_2 | TATGAGTGTGGAGTCGTTGCTTGGTATTAGGGTGATACTGGCANNNAGATCGGAA |
| 41010_1 | TATGAGTGTGGAGTCGTTGCGTGAAACCCCCGTCTCTACTAAANNNAGATCGGAAG |
| 41010_2 | TATGAGTGTGGAGTCGTTGCGTGAAACCCCCGTCTCTACTAAANNNAGATCGGAAG |
| 41295_1 | TATGAGTGTGGAGTCGTTGCCAGGAATGGCTCTCAGTTTCCTNNNAGATCGGAAGA |
| 41295_2 | TATGAGTGTGGAGTCGTTGCCAGGAATGGCTCTCAGTTTCCTNNNAGATCGGAAGA |
| 29601_1 | TATGAGTGTGGAGTCGTTGCAAGGCTTGTTACTTTTTCTTTAAATGTNNNAGATCGG |
| 29601_2 | TATGAGTGTGGAGTCGTTGCAAGGCTTGTTACTTTTTCTTTAAATGTNNNAGATCGG |
| 29872_1 | TATGAGTGTGGAGTCGTTGCGCTGATGACTCCACTAGGACTTCNNNAGATCGGAAG |
| 29872_2 | TATGAGTGTGGAGTCGTTGCGCTGATGACTCCACTAGGACTTCNNNAGATCGGAAG |
| 50419_1 | TATGAGTGTGGAGTCGTTGCTTGCCTGTGAATGTAAGAGGTGTNNNAGATCGGAA |
| 50419_2 | TATGAGTGTGGAGTCGTTGCTTGCCTGTGAATGTAAGAGGTGTNNNAGATCGGAA |
| 12902_1 | TATGAGTGTGGAGTCGTTGCTGGTTTTCTTACATAAACATGAGNNNAGATCGGAAG |
| 12902_2 | TATGAGTGTGGAGTCGTTGCTGGTTTTCTTACATAAACATGAGNNNAGATCGGAAG |
| 46921_1 | TATGAGTGTGGAGTCGTTGCTTCAAGTGATTCTCCTGCCTCAGNNNAGATCGGAAG |
| 46921_2 | TATGAGTGTGGAGTCGTTGCTTCAAGTGATTCTCCTGCCTCAGNNNAGATCGGAAG |
| 2251_1 | TATGAGTGTGGAGTCGTTGCAGAGTGGAGGCTTGAAGTTCCTANNNAGATCGGAA |
| 2251_2 | TATGAGTGTGGAGTCGTTGCAGAGTGGAGGCTTGAAGTTCCTANNNAGATCGGAA |
| 31097_1 | TATGAGTGTGGAGTCGTTGCAGCAAACAATGTTGAGGCTCAAANNNAGATCGGAA |
| 31097_2 | TATGAGTGTGGAGTCGTTGCAGCAAACAATGTTGAGGCTCAAANNNAGATCGGAA |
| 36876_1 | TATGAGTGTGGAGTCGTTGCGTTTCATGCTTTGACCAGGAGTCNNNAGATCGGAAG |
| 36876_2 | TATGAGTGTGGAGTCGTTGCGTTTCATGCTTTGACCAGGAGTCNNNAGATCGGAAG |
| 2480_1 | TATGAGTGTGGAGTCGTTGCTTCCTTGTGCTGGAAAGTACCAGNNNAGATCGGAAG |
| 2480_2 | TATGAGTGTGGAGTCGTTGCTTCCTTGTGCTGGAAAGTACCAGNNNAGATCGGAAG |
| 44847_1 | TATGAGTGTGGAGTCGTTGCATGCAAATCACCTCAGCTAAGGANNNAGATCGGAAG |
| 44847_2 | TATGAGTGTGGAGTCGTTGCATGCAAATCACCTCAGCTAAGGANNNAGATCGGAAG |
| 37837_1 | TATGAGTGTGGAGTCGTTGCTCCCTGTCTAGTTGTTTTACCCANNNAGATCGGAAGA |

| | |
|---|---|
| 37837_2 | TATGAGTGTGGAGTCGTTGCTCCCTGTCTAGTTGTTTACCCANNNAGATCGGAAGA |
| 37723_1 | TATGAGTGTGGAGTCGTTGCAAAAACCTCACTGGATGGGATCANNNAGATCGGAAG |
| 37723_2 | TATGAGTGTGGAGTCGTTGCAAAAACCTCACTGGATGGGATCANNNAGATCGGAAG |
| 52151_1 | TATGAGTGTGGAGTCGTTGCTCCATCCTACTGCTTTGGATCACNNNAGATCGGAAGA |
| 52151_2 | TATGAGTGTGGAGTCGTTGCTCCATCCTACTGCTTTGGATCACNNNAGATCGGAAGA |
| 32599_1 | TATGAGTGTGGAGTCGTTGCGACCCTGAGTCTGGAGAAATTGTNNNAGATCGGAA |
| 32599_2 | TATGAGTGTGGAGTCGTTGCGACCCTGAGTCTGGAGAAATTGTNNNAGATCGGAA |
| 37915_1 | TATGAGTGTGGAGTCGTTGCTTCCCTGTGTTTCCATAGTGTGTNNNAGATCGGAAGA |
| 37915_2 | TATGAGTGTGGAGTCGTTGCTTCCCTGTGTTTCCATAGTGTGTNNNAGATCGGAAGA |
| 45790_1 | TATGAGTGTGGAGTCGTTGCTTGTTGAGATGGCTTTCTCCCTTNNNAGATCGGAAGA |
| 45790_2 | TATGAGTGTGGAGTCGTTGCTTGTTGAGATGGCTTTCTCCCTTNNNAGATCGGAAGA |
| 29757_1 | TATGAGTGTGGAGTCGTTGCAAGTGATCGGAAATGACACTGGANNNAGATCGGAA |
| 29757_2 | TATGAGTGTGGAGTCGTTGCAAGTGATCGGAAATGACACTGGANNNAGATCGGAA |
| 39477_1 | TATGAGTGTGGAGTCGTTGCAAGGTTTCTTGCAGCATCATTGTNNNAGATCGGAAG |
| 39477_2 | TATGAGTGTGGAGTCGTTGCAAGGTTTCTTGCAGCATCATTGTNNNAGATCGGAAG |
| 30090_1 | TATGAGTGTGGAGTCGTTGCTTGTAGAAACAGAGTATGTCTGAGTNNNAGATCGGA |
| 30090_2 | TATGAGTGTGGAGTCGTTGCTTGTAGAAACAGAGTATGTCTGAGTNNNAGATCGGA |
| 29811_1 | TATGAGTGTGGAGTCGTTGCTTAAGCTTGGAGCAGAGATGGAGNNNAGATCGGAA |
| 29811_2 | TATGAGTGTGGAGTCGTTGCTTAAGCTTGGAGCAGAGATGGAGNNNAGATCGGAA |
| 29025_1 | TATGAGTGTGGAGTCGTTGCACACACAAACTTACATGGACAGGNNNAGATCGGAAG |
| 29025_2 | TATGAGTGTGGAGTCGTTGCACACACAAACTTACATGGACAGGNNNAGATCGGAAG |
| 44150_1 | TATGAGTGTGGAGTCGTTGCTTCTCTAACAGTGGTGCTTTCCANNNAGATCGGAAGA |
| 44150_2 | TATGAGTGTGGAGTCGTTGCTTCTCTAACAGTGGTGCTTTCCANNNAGATCGGAAGA |
| 41511_1 | TATGAGTGTGGAGTCGTTGCAGACTGGCCTTACAAGAAATGGTNNNAGATCGGAA |
| 41511_2 | TATGAGTGTGGAGTCGTTGCAGACTGGCCTTACAAGAAATGGTNNNAGATCGGAA |
| 13090_1 | TATGAGTGTGGAGTCGTTGCATGAGTGATTCCAACTATGTTGNNNAGATCGGAAGA |
| 13090_2 | TATGAGTGTGGAGTCGTTGCATGAGTGATTCCAACTATGTTGNNNAGATCGGAAGA |
| 49595_1 | TATGAGTGTGGAGTCGTTGCAATGTTTCGCTCTTGTTTCCCAGNNNAGATCGGAAGA |
| 49595_2 | TATGAGTGTGGAGTCGTTGCAATGTTTCGCTCTTGTTTCCCAGNNNAGATCGGAAGA |
| 29443_1 | TATGAGTGTGGAGTCGTTGCATGTGGGAGCAGACAGTGAAAAANNNAGATCGGAA |
| 29443_2 | TATGAGTGTGGAGTCGTTGCATGTGGGAGCAGACAGTGAAAAANNNAGATCGGAA |
| 45509_1 | TATGAGTGTGGAGTCGTTGCCCCTGCTCTCTCTCTCTCTTTNNNAGATCGGAAGAG |
| 45509_2 | TATGAGTGTGGAGTCGTTGCCCCTGCTCTCTCTCTCTCTTTNNNAGATCGGAAGAG |
| 34205_1 | TATGAGTGTGGAGTCGTTGCGAGGATCACAAGGTCAGGAGATGNNNAGATCGGAA |
| 34205_2 | TATGAGTGTGGAGTCGTTGCGAGGATCACAAGGTCAGGAGATGNNNAGATCGGAA |
| 46611_1 | TATGAGTGTGGAGTCGTTGCGAATTTCACTCTTGTTGCCCAGGNNNAGATCGGAAG |
| 46611_2 | TATGAGTGTGGAGTCGTTGCGAATTTCACTCTTGTTGCCCAGGNNNAGATCGGAAG |
| 42502_1 | TATGAGTGTGGAGTCGTTGCGACCATTTGGAACTGAAAACGGTNNNAGATCGGAA |
| 42502_2 | TATGAGTGTGGAGTCGTTGCGACCATTTGGAACTGAAAACGGTNNNAGATCGGAA |
| 37222_1 | TATGAGTGTGGAGTCGTTGCGCCCTCACAAAATAGTACTGGGANNNAGATCGGAAG |
| 37222_2 | TATGAGTGTGGAGTCGTTGCGCCCTCACAAAATAGTACTGGGANNNAGATCGGAAG |
| 53418_1 | TATGAGTGTGGAGTCGTTGCGAGGGAGAAAGGAAGGAAGGAAGNNNAGATCGGA |
| 53418_2 | TATGAGTGTGGAGTCGTTGCGAGGGAGAAAGGAAGGAAGGAAGNNNAGATCGGA |

| | |
|---|---|
| 50641_1 | TATGAGTGTGGAGTCGTTGCTCAGAGGCCTCATCTTCAGAAACNNNAGATCGGAAG |
| 50641_2 | TATGAGTGTGGAGTCGTTGCTCAGAGGCCTCATCTTCAGAAACNNNAGATCGGAAG |
| 37459_1 | TATGAGTGTGGAGTCGTTGCATGGAGCTCTTTCCTCTGTTAGCNNNAGATCGGAAG |
| 37459_2 | TATGAGTGTGGAGTCGTTGCATGGAGCTCTTTCCTCTGTTAGCNNNAGATCGGAAG |
| 10619_1 | TATGAGTGTGGAGTCGTTGCAGGCTCATGGATTTCCTGGCNNNAGATCGGAAGAG |
| 10619_2 | TATGAGTGTGGAGTCGTTGCAGGCTCATGGATTTCCTGGCNNNAGATCGGAAGAG |
| 49422_1 | TATGAGTGTGGAGTCGTTGCTTCAAGCTATTCTCCTGCCTCAGNNNAGATCGGAAGA |
| 49422_2 | TATGAGTGTGGAGTCGTTGCTTCAAGCTATTCTCCTGCCTCAGNNNAGATCGGAAGA |
| 52106_1 | TATGAGTGTGGAGTCGTTGCTTACCGTGCAAGAGACTTTTCCANNNAGATCGGAAG |
| 52106_2 | TATGAGTGTGGAGTCGTTGCTTACCGTGCAAGAGACTTTTCCANNNAGATCGGAAG |
| 31982_1 | TATGAGTGTGGAGTCGTTGCATGAGCGTCAAAAGTACTCCTGTNNNAGATCGGAAG |
| 31982_2 | TATGAGTGTGGAGTCGTTGCATGAGCGTCAAAAGTACTCCTGTNNNAGATCGGAAG |
| 48084_1 | TATGAGTGTGGAGTCGTTGCGTCTTGAACTCCTGACCTCATGANNNAGATCGGAAG |
| 48084_2 | TATGAGTGTGGAGTCGTTGCGTCTTGAACTCCTGACCTCATGANNNAGATCGGAAG |
| 50516_1 | TATGAGTGTGGAGTCGTTGCGCATAATCCTGCTGTGACTTCTGNNNAGATCGGAAG |
| 50516_2 | TATGAGTGTGGAGTCGTTGCGCATAATCCTGCTGTGACTTCTGNNNAGATCGGAAG |
| 45318_1 | TATGAGTGTGGAGTCGTTGCCCTTGACTGAAAGGATTAGGGCTNNNAGATCGGAA |
| 45318_2 | TATGAGTGTGGAGTCGTTGCCCTTGACTGAAAGGATTAGGGCTNNNAGATCGGAA |
| 38949_1 | TATGAGTGTGGAGTCGTTGCCTAGGCCAGTGTTCTCATTGAGTNNNAGATCGGAAG |
| 38949_2 | TATGAGTGTGGAGTCGTTGCCTAGGCCAGTGTTCTCATTGAGTNNNAGATCGGAAG |
| 42031_1 | TATGAGTGTGGAGTCGTTGCGGTCTCTCTTAGCCATCACAGAGNNNAGATCGGAAG |
| 42031_2 | TATGAGTGTGGAGTCGTTGCGGTCTCTCTTAGCCATCACAGAGNNNAGATCGGAAG |
| 31367_1 | TATGAGTGTGGAGTCGTTGCTATTGTGTGGGGTGCATACAAGANNNAGATCGGAA |
| 31367_2 | TATGAGTGTGGAGTCGTTGCTATTGTGTGGGGTGCATACAAGANNNAGATCGGAA |
| 40868_1 | TATGAGTGTGGAGTCGTTGCCTATTGTGAACAGTGCTGCAACANNNAGATCGGAAG |
| 40868_2 | TATGAGTGTGGAGTCGTTGCCTATTGTGAACAGTGCTGCAACANNNAGATCGGAAG |
| 38055_1 | TATGAGTGTGGAGTCGTTGCTGTGGGGTTCTGACATAGGTAGANNNAGATCGGAA |
| 38055_2 | TATGAGTGTGGAGTCGTTGCTGTGGGGTTCTGACATAGGTAGANNNAGATCGGAA |
| 31266_1 | TATGAGTGTGGAGTCGTTGCATTTTCTCGTTGCCTTGCTTCAANNNAGATCGGAAGA |
| 31266_2 | TATGAGTGTGGAGTCGTTGCATTTTCTCGTTGCCTTGCTTCAANNNAGATCGGAAGA |
| 39318_1 | TATGAGTGTGGAGTCGTTGCGGAGTATGTAGTTGCTGAGGGAGNNNAGATCGGA |
| 39318_2 | TATGAGTGTGGAGTCGTTGCGGAGTATGTAGTTGCTGAGGGAGNNNAGATCGGA |
| 46106_1 | TATGAGTGTGGAGTCGTTGCAGCCTCCCAAGTAGCTAGGATTANNNAGATCGGAAG |
| 46106_2 | TATGAGTGTGGAGTCGTTGCAGCCTCCCAAGTAGCTAGGATTANNNAGATCGGAAG |
| 42891_1 | TATGAGTGTGGAGTCGTTGCTGTGTGTGTGTGTTTTGCCTTAGNNNAGATCGGAAG |
| 42891_2 | TATGAGTGTGGAGTCGTTGCTGTGTGTGTGTGTTTTGCCTTAGNNNAGATCGGAAG |
| 44562_1 | TATGAGTGTGGAGTCGTTGCGTTTAGTGGTGGTGATGTTGGTGNNNAGATCGGAA |
| 44562_2 | TATGAGTGTGGAGTCGTTGCGTTTAGTGGTGGTGATGTTGGTGNNNAGATCGGAA |
| 51735_1 | TATGAGTGTGGAGTCGTTGCTCCTGCTATGGAATGATGAAGCANNNAGATCGGAA |
| 51735_2 | TATGAGTGTGGAGTCGTTGCTCCTGCTATGGAATGATGAAGCANNNAGATCGGAA |
| 13774_1 | TATGAGTGTGGAGTCGTTGCATAAGGAAGTAACTCAAAGGAATNNNAGATCGGAA |
| 13774_2 | TATGAGTGTGGAGTCGTTGCATAAGGAAGTAACTCAAAGGAATNNNAGATCGGAA |
| 46213_1 | TATGAGTGTGGAGTCGTTGCCTCCTCCATCCTCCTCTTCAGGNNNAGATCGGAAGAG |

| | |
|---|---|
| 46213_2 | TATGAGTGTGGAGTCGTTGCCTCCTCCATCCTCCTCTTCAGGNNNAGATCGGAAGAG |
| 33679_1 | TATGAGTGTGGAGTCGTTGCAGATGGAGTCTCTGTTCCCCANNNAGATCGGAAGAG |
| 33679_2 | TATGAGTGTGGAGTCGTTGCAGATGGAGTCTCTGTTCCCCANNNAGATCGGAAGAG |
| 49142_1 | TATGAGTGTGGAGTCGTTGCCCCTAAGAGCTGGGATTATAGGCNNNAGATCGGAA |
| 49142_2 | TATGAGTGTGGAGTCGTTGCCCCTAAGAGCTGGGATTATAGGCNNNAGATCGGAA |
| 30442_1 | TATGAGTGTGGAGTCGTTGCAAATACCGCAGTGAACACAGGACTGNNNAGATCGG |
| 30442_2 | TATGAGTGTGGAGTCGTTGCAAATACCGCAGTGAACACAGGACTGNNNAGATCGG |
| 37446_1 | TATGAGTGTGGAGTCGTTGCATTTCCTGCCCCAGATCTGAAATNNNAGATCGGAAG |
| 37446_2 | TATGAGTGTGGAGTCGTTGCATTTCCTGCCCCAGATCTGAAATNNNAGATCGGAAG |
| 46231_1 | TATGAGTGTGGAGTCGTTGCCTGGAGTGCAATGGTGAGATCTTNNNAGATCGGAA |
| 46231_2 | TATGAGTGTGGAGTCGTTGCCTGGAGTGCAATGGTGAGATCTTNNNAGATCGGAA |
| 43840_1 | TATGAGTGTGGAGTCGTTGCTTGGCCATTTTCGAGTTGAACAANNNAGATCGGAAG |
| 43840_2 | TATGAGTGTGGAGTCGTTGCTTGGCCATTTTCGAGTTGAACAANNNAGATCGGAAG |
| 49353_1 | TATGAGTGTGGAGTCGTTGCAAGTACTGGCATTACAGGCATGANNNAGATCGGAA |
| 49353_2 | TATGAGTGTGGAGTCGTTGCAAGTACTGGCATTACAGGCATGANNNAGATCGGAA |
| 2559_1 | TATGAGTGTGGAGTCGTTGCGATAAAAACAAGTTTGAGGCCGGGNNNAGATCGGA |
| 2559_2 | TATGAGTGTGGAGTCGTTGCGATAAAAACAAGTTTGAGGCCGGGNNNAGATCGGA |
| 46771_1 | TATGAGTGTGGAGTCGTTGCCAAGGCTTTGACTGGAAGGGTATNNNAGATCGGAA |
| 46771_2 | TATGAGTGTGGAGTCGTTGCCAAGGCTTTGACTGGAAGGGTATNNNAGATCGGAA |
| 2999_1 | TATGAGTGTGGAGTCGTTGCGTGCGCTTGGATCTGATAAACGAGGNNNAGATCGG |
| 2999_2 | TATGAGTGTGGAGTCGTTGCGTGCGCTTGGATCTGATAAACGAGGNNNAGATCGG |
| 46388_1 | TATGAGTGTGGAGTCGTTGCAGAAGGGATCTCGCTATGTTTCCNNNAGATCGGAAG |
| 46388_2 | TATGAGTGTGGAGTCGTTGCAGAAGGGATCTCGCTATGTTTCCNNNAGATCGGAAG |
| 49507_1 | TATGAGTGTGGAGTCGTTGCTAAGAGACAGAGCTCCACTGACTNNNAGATCGGAAG |
| 49507_2 | TATGAGTGTGGAGTCGTTGCTAAGAGACAGAGCTCCACTGACTNNNAGATCGGAAG |
| 47723_1 | TATGAGTGTGGAGTCGTTGCAACTTGATCTTGGAGTCGTGGAANNNAGATCGGAA |
| 47723_2 | TATGAGTGTGGAGTCGTTGCAACTTGATCTTGGAGTCGTGGAANNNAGATCGGAA |
| 31086_1 | TATGAGTGTGGAGTCGTTGCTTAGGTTTGGCCATGCAACATTTNNNAGATCGGAAG |
| 31086_2 | TATGAGTGTGGAGTCGTTGCTTAGGTTTGGCCATGCAACATTTNNNAGATCGGAAG |
| 37142_1 | TATGAGTGTGGAGTCGTTGCTGGTGTATGAGCTAGATGTGGTGNNNAGATCGGAA |
| 37142_2 | TATGAGTGTGGAGTCGTTGCTGGTGTATGAGCTAGATGTGGTGNNNAGATCGGAA |
| 29224_1 | TATGAGTGTGGAGTCGTTGCAGGGTGCAAGGGGTCTCTATATANNNAGATCGGAA |
| 29224_2 | TATGAGTGTGGAGTCGTTGCAGGGTGCAAGGGGTCTCTATATANNNAGATCGGAA |
| 2338_1 | TATGAGTGTGGAGTCGTTGCCCTTCATCCTTGCATTTTTGCCCNNNAGATCGGAAGA |
| 2338_2 | TATGAGTGTGGAGTCGTTGCCCTTCATCCTTGCATTTTTGCCCNNNAGATCGGAAGA |
| 44935_1 | TATGAGTGTGGAGTCGTTGCATTGAAAATCACAGGGCAGTTGGNNNAGATCGGAA |
| 44935_2 | TATGAGTGTGGAGTCGTTGCATTGAAAATCACAGGGCAGTTGGNNNAGATCGGAA |
| 29346_1 | TATGAGTGTGGAGTCGTTGCATCTCGGCTTACTAATGCAACCTNNNAGATCGGAAG |
| 29346_2 | TATGAGTGTGGAGTCGTTGCATCTCGGCTTACTAATGCAACCTNNNAGATCGGAAG |
| 43910_1 | TATGAGTGTGGAGTCGTTGCCAATTATCACACCTCAGCCTCCTNNNAGATCGGAAGA |
| 43910_2 | TATGAGTGTGGAGTCGTTGCCAATTATCACACCTCAGCCTCCTNNNAGATCGGAAGA |
| 33061_1 | TATGAGTGTGGAGTCGTTGCCAGTGTTGCAATCTCGGTTCACNNNAGATCGGAAGA |
| 33061_2 | TATGAGTGTGGAGTCGTTGCCAGTGTTGCAATCTCGGTTCACNNNAGATCGGAAGA |

| | |
|---|---|
| 7442_1 | TATGAGTGTGGAGTCGTTGCCTCAGTAGTCTTCCACGTCATCANNNAGATCGGAAGA |
| 7442_2 | TATGAGTGTGGAGTCGTTGCCTCAGTAGTCTTCCACGTCATCANNNAGATCGGAAGA |
| 32590_1 | TATGAGTGTGGAGTCGTTGCCAAGTGATCTGGCCTCCTTACTTNNNAGATCGGAAGA |
| 32590_2 | TATGAGTGTGGAGTCGTTGCCAAGTGATCTGGCCTCCTTACTTNNNAGATCGGAAGA |
| 38202_1 | TATGAGTGTGGAGTCGTTGCAGAAGGGAAAATGTGGCTATTTCTNNNAGATCGGAA |
| 38202_2 | TATGAGTGTGGAGTCGTTGCAGAAGGGAAAATGTGGCTATTTCTNNNAGATCGGAA |
| 43892_1 | TATGAGTGTGGAGTCGTTGCGAAACCAGCTCCTAGCAAACATGNNNAGATCGGAAG |
| 43892_2 | TATGAGTGTGGAGTCGTTGCGAAACCAGCTCCTAGCAAACATGNNNAGATCGGAAG |
| 45201_1 | TATGAGTGTGGAGTCGTTGCGGGAGGGATACATGGGACTTTTTNNNAGATCGGAA |
| 45201_2 | TATGAGTGTGGAGTCGTTGCGGGAGGGATACATGGGACTTTTTNNNAGATCGGAA |
| 34741_1 | TATGAGTGTGGAGTCGTTGCATTCAAGTGATTCTCCTGCCTCANNNAGATCGGAAGA |
| 34741_2 | TATGAGTGTGGAGTCGTTGCATTCAAGTGATTCTCCTGCCTCANNNAGATCGGAAGA |
| 44440_1 | TATGAGTGTGGAGTCGTTGCCCACCATCCACCTACTAAGGAAGNNNAGATCGGAAGA |
| 44440_2 | TATGAGTGTGGAGTCGTTGCCCACCATCCACCTACTAAGGAAGNNNAGATCGGAAGA |
| 47055_1 | TATGAGTGTGGAGTCGTTGCTGCAACTACTAGAAAGCTAGGCANNNAGATCGGAAG |
| 47055_2 | TATGAGTGTGGAGTCGTTGCTGCAACTACTAGAAAGCTAGGCANNNAGATCGGAAG |
| 39855_1 | TATGAGTGTGGAGTCGTTGCATTCAAGTGATTCTCCTGCCTCANNNAGATCGGAAGA |
| 39855_2 | TATGAGTGTGGAGTCGTTGCATTCAAGTGATTCTCCTGCCTCANNNAGATCGGAAGA |
| 46214_1 | TATGAGTGTGGAGTCGTTGCGAAGCTAATCAAGTCTTCGCGACNNNAGATCGGAAG |
| 46214_2 | TATGAGTGTGGAGTCGTTGCGAAGCTAATCAAGTCTTCGCGACNNNAGATCGGAAG |
| 30881_1 | TATGAGTGTGGAGTCGTTGCCAGAAGCTGGAAAAGGCAAAGAANNNAGATCGGAA |
| 30881_2 | TATGAGTGTGGAGTCGTTGCCAGAAGCTGGAAAAGGCAAAGAANNNAGATCGGAA |
| 34393_1 | TATGAGTGTGGAGTCGTTGCGCAAAGAAGAAGCGGGAGCTNNNAGATCGGAAGAG |
| 34393_2 | TATGAGTGTGGAGTCGTTGCGCAAAGAAGAAGCGGGAGCTNNNAGATCGGAAGAG |
| 35271_1 | TATGAGTGTGGAGTCGTTGCCTCCTGAGTAGCTGGGACTACAGNNNAGATCGGAAG |
| 35271_2 | TATGAGTGTGGAGTCGTTGCCTCCTGAGTAGCTGGGACTACAGNNNAGATCGGAAG |
| 39986_1 | TATGAGTGTGGAGTCGTTGCACTCTTGACCCAAATTCACCAGTNNNAGATCGGAAGA |
| 39986_2 | TATGAGTGTGGAGTCGTTGCACTCTTGACCCAAATTCACCAGTNNNAGATCGGAAGA |
| 44573_1 | TATGAGTGTGGAGTCGTTGCATTCTCCATCACAGGACCCAAAANNNAGATCGGAAGA |
| 44573_2 | TATGAGTGTGGAGTCGTTGCATTCTCCATCACAGGACCCAAAANNNAGATCGGAAGA |
| 35118_1 | TATGAGTGTGGAGTCGTTGCCCACGCCCAGCTAATTTTTTGTANNNAGATCGGAAGA |
| 35118_2 | TATGAGTGTGGAGTCGTTGCCCACGCCCAGCTAATTTTTTGTANNNAGATCGGAAGA |
| 38815_1 | TATGAGTGTGGAGTCGTTGCATGTTTGACCAAAAAGCAACACANNNAGATCGGAAG |
| 38815_2 | TATGAGTGTGGAGTCGTTGCATGTTTGACCAAAAAGCAACACANNNAGATCGGAAG |
| 47224_1 | TATGAGTGTGGAGTCGTTGCTTGTTTGTTCCTGCAGCAAAAGANNNAGATCGGAAG |
| 47224_2 | TATGAGTGTGGAGTCGTTGCTTGTTTGTTCCTGCAGCAAAAGANNNAGATCGGAAG |
| 40522_1 | TATGAGTGTGGAGTCGTTGCTGTCAACCCATTTCCCCAACTATNNNAGATCGGAAGA |
| 40522_2 | TATGAGTGTGGAGTCGTTGCTGTCAACCCATTTCCCCAACTATNNNAGATCGGAAGA |
| 5595_1 | TATGAGTGTGGAGTCGTTGCTAAGAGTTGCCGAGAAAATGCCANNNAGATCGGAAG |
| 5595_2 | TATGAGTGTGGAGTCGTTGCTAAGAGTTGCCGAGAAAATGCCANNNAGATCGGAAG |
| 44407_1 | TATGAGTGTGGAGTCGTTGCCTTTTCTTTTCATGGTGGCTGGTNNNAGATCGGAAGA |
| 44407_2 | TATGAGTGTGGAGTCGTTGCCTTTTCTTTTCATGGTGGCTGGTNNNAGATCGGAAGA |
| 46599_1 | TATGAGTGTGGAGTCGTTGCGCCTCCTGAGTAGCTGAGATTACNNNAGATCGGAAG |

| | |
|---|---|
| 46599_2 | TATGAGTGTGGAGTCGTTGCGCCTCCTGAGTAGCTGAGATTACNNNAGATCGGAAG |
| 44700_1 | TATGAGTGTGGAGTCGTTGCCAGAGACTGGCGCAGAATCGNNNAGATCGGAAGAG |
| 44700_2 | TATGAGTGTGGAGTCGTTGCCAGAGACTGGCGCAGAATCGNNNAGATCGGAAGAG |
| 47351_1 | TATGAGTGTGGAGTCGTTGCTGAGAAACTGTGATTGGCTCAGTNNNAGATCGGAA |
| 47351_2 | TATGAGTGTGGAGTCGTTGCTGAGAAACTGTGATTGGCTCAGTNNNAGATCGGAA |
| 39543_1 | TATGAGTGTGGAGTCGTTGCTGATCTCGGCTCACTGCAACNNNAGATCGGAAGAGC |
| 39543_2 | TATGAGTGTGGAGTCGTTGCTGATCTCGGCTCACTGCAACNNNAGATCGGAAGAGC |
| 38267_1 | TATGAGTGTGGAGTCGTTGCAAGGCCAAGGACAAAGAGAGAATNNNAGATCGGA |
| 38267_2 | TATGAGTGTGGAGTCGTTGCAAGGCCAAGGACAAAGAGAGAATNNNAGATCGGA |
| 49496_1 | TATGAGTGTGGAGTCGTTGCCTCCATCTCCTGACCTTGTGATCNNNAGATCGGAAGA |
| 49496_2 | TATGAGTGTGGAGTCGTTGCCTCCATCTCCTGACCTTGTGATCNNNAGATCGGAAGA |
| 47800_1 | TATGAGTGTGGAGTCGTTGCAGACAGAGTCTCCCTGTGTCTCNNNAGATCGGAAGA |
| 47800_2 | TATGAGTGTGGAGTCGTTGCAGACAGAGTCTCCCTGTGTCTCNNNAGATCGGAAGA |
| 39542_1 | TATGAGTGTGGAGTCGTTGCCTGTAGTCCCAGCTACTCAAGAGNNNAGATCGGAAG |
| 39542_2 | TATGAGTGTGGAGTCGTTGCCTGTAGTCCCAGCTACTCAAGAGNNNAGATCGGAAG |
| 38691_1 | TATGAGTGTGGAGTCGTTGCATAAAAGGCCTTGTCCAACAGGANNNAGATCGGAA |
| 38691_2 | TATGAGTGTGGAGTCGTTGCATAAAAGGCCTTGTCCAACAGGANNNAGATCGGAA |
| 52041_1 | TATGAGTGTGGAGTCGTTGCCACACTCAAATGCACACATGTCANNNAGATCGGAAG |
| 52041_2 | TATGAGTGTGGAGTCGTTGCCACACTCAAATGCACACATGTCANNNAGATCGGAAG |
| 29805_1 | TATGAGTGTGGAGTCGTTGCTTATGGCACTTTCCCTTTTCTGCNNNAGATCGGAAGA |
| 29805_2 | TATGAGTGTGGAGTCGTTGCTTATGGCACTTTCCCTTTTCTGCNNNAGATCGGAAGA |
| 12596_1 | TATGAGTGTGGAGTCGTTGCGAACTAGATTTACTGGAAGACTTNNNAGATCGGAAG |
| 12596_2 | TATGAGTGTGGAGTCGTTGCGAACTAGATTTACTGGAAGACTTNNNAGATCGGAAG |
| 48122_1 | TATGAGTGTGGAGTCGTTGCCTCTCGAGTAGCTGGGATTACAGNNNAGATCGGAA |
| 48122_2 | TATGAGTGTGGAGTCGTTGCCTCTCGAGTAGCTGGGATTACAGNNNAGATCGGAA |
| 3102_1 | TATGAGTGTGGAGTCGTTGCAATGGTGGGAGAAAAGGCTTTGTNNNAGATCGGAA |
| 3102_2 | TATGAGTGTGGAGTCGTTGCAATGGTGGGAGAAAAGGCTTTGTNNNAGATCGGAA |
| 52224_1 | TATGAGTGTGGAGTCGTTGCATATGTGTGTGTGTGTGTGTGTGNNNAGATCGGAA |
| 52224_2 | TATGAGTGTGGAGTCGTTGCATATGTGTGTGTGTGTGTGTGTGNNNAGATCGGAA |
| 42275_1 | TATGAGTGTGGAGTCGTTGCGAGAGAGAGAGAGAGAGAGAGGTNNNAGATCGG |
| 42275_2 | TATGAGTGTGGAGTCGTTGCGAGAGAGAGAGAGAGAGAGAGGTNNNAGATCGG |
| 43758_1 | TATGAGTGTGGAGTCGTTGCTAGCTGCTATTAGGTTGGTGCAANNNAGATCGGAA |
| 43758_2 | TATGAGTGTGGAGTCGTTGCTAGCTGCTATTAGGTTGGTGCAANNNAGATCGGAA |
| 41458_1 | TATGAGTGTGGAGTCGTTGCCACCCCAAAAAATCCAATGTAGCTNNNAGATCGGAA |
| 41458_2 | TATGAGTGTGGAGTCGTTGCCACCCCAAAAAATCCAATGTAGCTNNNAGATCGGAA |
| 32706_1 | TATGAGTGTGGAGTCGTTGCCTTGAACTCCTGACCTCGTGATCNNNAGATCGGAAG |
| 32706_2 | TATGAGTGTGGAGTCGTTGCCTTGAACTCCTGACCTCGTGATCNNNAGATCGGAAG |
| 50623_1 | TATGAGTGTGGAGTCGTTGCATGCTATAGTGGCAAGAAGGCTTNNNAGATCGGAA |
| 50623_2 | TATGAGTGTGGAGTCGTTGCATGCTATAGTGGCAAGAAGGCTTNNNAGATCGGAA |
| 45912_1 | TATGAGTGTGGAGTCGTTGCTCATAGTTCACTGCAACCTCGAANNNAGATCGGAAG |
| 45912_2 | TATGAGTGTGGAGTCGTTGCTCATAGTTCACTGCAACCTCGAANNNAGATCGGAAG |
| 44478_1 | TATGAGTGTGGAGTCGTTGCGCTGGTGGCAAGTAAATCAGAAGNNNAGATCGGAA |
| 44478_2 | TATGAGTGTGGAGTCGTTGCGCTGGTGGCAAGTAAATCAGAAGNNNAGATCGGAA |

| | |
|---|---|
| 39201_1 | TATGAGTGTGGAGTCGTTGCAAACAACCAAACTACACAAGCCTNNNAGATCGGAAG |
| 39201_2 | TATGAGTGTGGAGTCGTTGCAAACAACCAAACTACACAAGCCTNNNAGATCGGAAG |
| 11815_1 | TATGAGTGTGGAGTCGTTGCCCCAGCCATTGTCCCCGTCGNNNAGATCGGAAGAGC |
| 11815_2 | TATGAGTGTGGAGTCGTTGCCCCAGCCATTGTCCCCGTCGNNNAGATCGGAAGAGC |
| 89_1 | TATGAGTGTGGAGTCGTTGCTCTCTCAGCTCTCGGAAGGACCNNNAGATCGGAAGA |
| 89_2 | TATGAGTGTGGAGTCGTTGCTCTCTCAGCTCTCGGAAGGACCNNNAGATCGGAAGA |
| 43735_1 | TATGAGTGTGGAGTCGTTGCGAGGAGGAGAGAGGAGGGAAGNNNAGATCGGAA |
| 43735_2 | TATGAGTGTGGAGTCGTTGCGAGGAGGAGAGAGGAGGGAAGNNNAGATCGGAA |
| 9462_1 | TATGAGTGTGGAGTCGTTGCATTTAGGAGGCACATGCACAGAGNNNAGATCGGAA |
| 9462_2 | TATGAGTGTGGAGTCGTTGCATTTAGGAGGCACATGCACAGAGNNNAGATCGGAA |
| 52211_1 | TATGAGTGTGGAGTCGTTGCGGAGATTAGAAAGTGTGTGTGTGCNNNAGATCGG |
| 52211_2 | TATGAGTGTGGAGTCGTTGCGGAGATTAGAAAGTGTGTGTGTGCNNNAGATCGG |
| 48942_1 | TATGAGTGTGGAGTCGTTGCTAATCTCGGCTCAATGCAACCTCNNNAGATCGGAAG |
| 48942_2 | TATGAGTGTGGAGTCGTTGCTAATCTCGGCTCAATGCAACCTCNNNAGATCGGAAG |
| 47154_1 | TATGAGTGTGGAGTCGTTGCAAGCAACTTTGAAACTGGGGAACNNNAGATCGGAA |
| 47154_2 | TATGAGTGTGGAGTCGTTGCAAGCAACTTTGAAACTGGGGAACNNNAGATCGGAA |
| 53156_1 | TATGAGTGTGGAGTCGTTGCGGGAGGAAGCCAGAGGGAAGNNNAGATCGGAAG |
| 53156_2 | TATGAGTGTGGAGTCGTTGCGGGAGGAAGCCAGAGGGAAGNNNAGATCGGAAG |
| 10694_1 | TATGAGTGTGGAGTCGTTGCTAGCTTCTTTCAGCTTCCTCCCTNNNAGATCGGAAGA |
| 10694_2 | TATGAGTGTGGAGTCGTTGCTAGCTTCTTTCAGCTTCCTCCCTNNNAGATCGGAAGA |
| 49614_1 | TATGAGTGTGGAGTCGTTGCCTGAAGGGTCTGGGTCATTTGTANNNAGATCGGAA |
| 49614_2 | TATGAGTGTGGAGTCGTTGCCTGAAGGGTCTGGGTCATTTGTANNNAGATCGGAA |
| 53213_1 | TATGAGTGTGGAGTCGTTGCACTGTACCTCAGATCTCACCTGANNNAGATCGGAAG |
| 53213_2 | TATGAGTGTGGAGTCGTTGCACTGTACCTCAGATCTCACCTGANNNAGATCGGAAG |
| 29431_1 | TATGAGTGTGGAGTCGTTGCGGTCCCCAAATGGAGCAGTACTANNNAGATCGGAA |
| 29431_2 | TATGAGTGTGGAGTCGTTGCGGTCCCCAAATGGAGCAGTACTANNNAGATCGGAA |
| 37109_1 | TATGAGTGTGGAGTCGTTGCTAGAGAGAGACATCGGTGTGAGANNNAGATCGGA |
| 37109_2 | TATGAGTGTGGAGTCGTTGCTAGAGAGAGACATCGGTGTGAGANNNAGATCGGA |
| 43162_1 | TATGAGTGTGGAGTCGTTGCGTTTTTCTGGGAATGGCTGGAAANNNAGATCGGAA |
| 43162_2 | TATGAGTGTGGAGTCGTTGCGTTTTTCTGGGAATGGCTGGAAANNNAGATCGGAA |
| 9425_1 | TATGAGTGTGGAGTCGTTGCGCTCTTCTCCAGACCCTCTCTAGNNNAGATCGGAAGA |
| 9425_2 | TATGAGTGTGGAGTCGTTGCGCTCTTCTCCAGACCCTCTCTAGNNNAGATCGGAAGA |
| 5356_1 | TATGAGTGTGGAGTCGTTGCTCAAAATGTGGGGGATGAGACAANNNAGATCGGA |
| 5356_2 | TATGAGTGTGGAGTCGTTGCTCAAAATGTGGGGGATGAGACAANNNAGATCGGA |
| 44035_1 | TATGAGTGTGGAGTCGTTGCGCTCAAAACAAGGGAACCATGAGNNNAGATCGGAA |
| 44035_2 | TATGAGTGTGGAGTCGTTGCGCTCAAAACAAGGGAACCATGAGNNNAGATCGGAA |
| 32947_1 | TATGAGTGTGGAGTCGTTGCGAAAGAAAGAAAGCAGGAAGGCANNNAGATCGGA |
| 32947_2 | TATGAGTGTGGAGTCGTTGCGAAAGAAAGAAAGCAGGAAGGCANNNAGATCGGA |
| 46110_1 | TATGAGTGTGGAGTCGTTGCCTCAAGCAATCCTCCTGTCTCAGNNNAGATCGGAAG |
| 46110_2 | TATGAGTGTGGAGTCGTTGCCTCAAGCAATCCTCCTGTCTCAGNNNAGATCGGAAG |
| 33972_1 | TATGAGTGTGGAGTCGTTGCATCCCAAGTAACTGGAACCACAGNNNAGATCGGAAG |
| 33972_2 | TATGAGTGTGGAGTCGTTGCATCCCAAGTAACTGGAACCACAGNNNAGATCGGAAG |
| 8194_1 | TATGAGTGTGGAGTCGTTGCGCACCCAGCCTCAATATTGTTTCNNNAGATCGGAAG |

| | |
|---|---|
| 8194_2 | TATGAGTGTGGAGTCGTTGCGCACCCAGCCTCAATATTGTTTCNNNAGATCGGAAG |
| 50720_1 | TATGAGTGTGGAGTCGTTGCGTGCAACCTTCCTTGTCTCTCTANNNAGATCGGAAGA |
| 50720_2 | TATGAGTGTGGAGTCGTTGCGTGCAACCTTCCTTGTCTCTCTANNNAGATCGGAAGA |
| 51122_1 | TATGAGTGTGGAGTCGTTGCTTGTGGGGTGAATGATGGAAGATNNNAGATCGGA |
| 51122_2 | TATGAGTGTGGAGTCGTTGCTTGTGGGGTGAATGATGGAAGATNNNAGATCGGA |
| 8909_1 | TATGAGTGTGGAGTCGTTGCGGAAGGCAAAAACGTGAAATCGTNNNAGATCGGAA |
| 8909_2 | TATGAGTGTGGAGTCGTTGCGGAAGGCAAAAACGTGAAATCGTNNNAGATCGGAA |
| 32327_1 | TATGAGTGTGGAGTCGTTGCCTTAGAAGAAGGTATCGGGCCTCNNNAGATCGGAA |
| 32327_2 | TATGAGTGTGGAGTCGTTGCCTTAGAAGAAGGTATCGGGCCTCNNNAGATCGGAA |
| 46528_1 | TATGAGTGTGGAGTCGTTGCGTTGCAGTGCAGTGGTATGATCNNNAGATCGGAAG |
| 46528_2 | TATGAGTGTGGAGTCGTTGCGTTGCAGTGCAGTGGTATGATCNNNAGATCGGAAG |
| 43914_1 | TATGAGTGTGGAGTCGTTGCTGCAAAACCAACACCTTTTCCTTNNNAGATCGGAAGA |
| 43914_2 | TATGAGTGTGGAGTCGTTGCTGCAAAACCAACACCTTTTCCTTNNNAGATCGGAAGA |
| 32881_1 | TATGAGTGTGGAGTCGTTGCGTTTCACCTTGTTAGCCAGGATGNNNAGATCGGAAG |
| 32881_2 | TATGAGTGTGGAGTCGTTGCGTTTCACCTTGTTAGCCAGGATGNNNAGATCGGAAG |
| 34682_1 | TATGAGTGTGGAGTCGTTGCAAGGGTAAGCCAGGAAGAACAAANNNAGATCGGA |
| 34682_2 | TATGAGTGTGGAGTCGTTGCAAGGGTAAGCCAGGAAGAACAAANNNAGATCGGA |
| 47179_1 | TATGAGTGTGGAGTCGTTGCCCTCCCAGGTTCAAGAGATTCTCNNNAGATCGGAAG |
| 47179_2 | TATGAGTGTGGAGTCGTTGCCCTCCCAGGTTCAAGAGATTCTCNNNAGATCGGAAG |
| 49805_1 | TATGAGTGTGGAGTCGTTGCGGCTTCCCCAATTCTTTTTCTCCNNNAGATCGGAAGA |
| 49805_2 | TATGAGTGTGGAGTCGTTGCGGCTTCCCCAATTCTTTTTCTCCNNNAGATCGGAAGA |
| 50926_1 | TATGAGTGTGGAGTCGTTGCCTAAGGGCTGACCTGGTCTTGNNNAGATCGGAAGA |
| 50926_2 | TATGAGTGTGGAGTCGTTGCCTAAGGGCTGACCTGGTCTTGNNNAGATCGGAAGA |
| 47284_1 | TATGAGTGTGGAGTCGTTGCTCAGACTCCTGAGTAGCTGGAATNNNAGATCGGAA |
| 47284_2 | TATGAGTGTGGAGTCGTTGCTCAGACTCCTGAGTAGCTGGAATNNNAGATCGGAA |
| 52324_1 | TATGAGTGTGGAGTCGTTGCCATGAACACTCAATGCTCCTTGTNNNAGATCGGAAG |
| 52324_2 | TATGAGTGTGGAGTCGTTGCCATGAACACTCAATGCTCCTTGTNNNAGATCGGAAG |
| 29375_1 | TATGAGTGTGGAGTCGTTGCCTGATGTCTGTGTTTCAGGCATGNNNAGATCGGAAG |
| 29375_2 | TATGAGTGTGGAGTCGTTGCCTGATGTCTGTGTTTCAGGCATGNNNAGATCGGAAG |
| 40560_1 | TATGAGTGTGGAGTCGTTGCAATACACACACACACACACACACNNNAGATCGGAAG |
| 40560_2 | TATGAGTGTGGAGTCGTTGCAATACACACACACACACACACACNNNAGATCGGAAG |
| 50321_1 | TATGAGTGTGGAGTCGTTGCTCAACTTGTAGCGTAATTTTCCTGTNNNAGATCGGA |
| 50321_2 | TATGAGTGTGGAGTCGTTGCTCAACTTGTAGCGTAATTTTCCTGTNNNAGATCGGA |
| 11288_1 | TATGAGTGTGGAGTCGTTGCCTGAGAGATGGAGAGAGAAGCCCNNNAGATCGGA |
| 11288_2 | TATGAGTGTGGAGTCGTTGCCTGAGAGATGGAGAGAGAAGCCCNNNAGATCGGA |
| 49615_1 | TATGAGTGTGGAGTCGTTGCCCCGGCCAAGATGTTTACATTTTNNNAGATCGGAAG |
| 49615_2 | TATGAGTGTGGAGTCGTTGCCCCGGCCAAGATGTTTACATTTTNNNAGATCGGAAG |
| 48069_1 | TATGAGTGTGGAGTCGTTGCGAGATCTCGGATCTTGGCTCACNNNAGATCGGAAGA |
| 48069_2 | TATGAGTGTGGAGTCGTTGCGAGATCTCGGATCTTGGCTCACNNNAGATCGGAAGA |
| 40274_1 | TATGAGTGTGGAGTCGTTGCAGATTCCTAAAGCCCTCGATGTCNNNAGATCGGAAG |
| 40274_2 | TATGAGTGTGGAGTCGTTGCAGATTCCTAAAGCCCTCGATGTCNNNAGATCGGAAG |
| 49185_1 | TATGAGTGTGGAGTCGTTGCCAGCACTGATAACTAGGCAGCTANNNAGATCGGAA |
| 49185_2 | TATGAGTGTGGAGTCGTTGCCAGCACTGATAACTAGGCAGCTANNNAGATCGGAA |

| | |
|---|---|
| 33723_1 | TATGAGTGTGGAGTCGTTGCGATGAGGCTATGCTGAGGTCTGNNNAGATCGGAAG |
| 33723_2 | TATGAGTGTGGAGTCGTTGCGATGAGGCTATGCTGAGGTCTGNNNAGATCGGAAG |
| 50830_1 | TATGAGTGTGGAGTCGTTGCTTTGAGACGGAGTTTTGCTGTTGNNNAGATCGGAA |
| 50830_2 | TATGAGTGTGGAGTCGTTGCTTTGAGACGGAGTTTTGCTGTTGNNNAGATCGGAA |
| 42348_1 | TATGAGTGTGGAGTCGTTGCGGAAAGGCATTTCACATTCATGGANNNAGATCGGA |
| 42348_2 | TATGAGTGTGGAGTCGTTGCGGAAAGGCATTTCACATTCATGGANNNAGATCGGA |
| 49009_1 | TATGAGTGTGGAGTCGTTGCCCACGCCCAGCTAATTTTTGTATNNNAGATCGGAAG |
| 49009_2 | TATGAGTGTGGAGTCGTTGCCCACGCCCAGCTAATTTTTGTATNNNAGATCGGAAG |
| 39275_1 | TATGAGTGTGGAGTCGTTGCAGCTCCACAGCCTAAAGAAATCANNNAGATCGGAAG |
| 39275_2 | TATGAGTGTGGAGTCGTTGCAGCTCCACAGCCTAAAGAAATCANNNAGATCGGAAG |
| 34552_1 | TATGAGTGTGGAGTCGTTGCGAGCCAAATTCTTCCCAAGAGTCNNNAGATCGGAAG |
| 34552_2 | TATGAGTGTGGAGTCGTTGCGAGCCAAATTCTTCCCAAGAGTCNNNAGATCGGAAG |
| 53297_1 | TATGAGTGTGGAGTCGTTGCTGAACTTGTCACCTGAGATGGAGNNNAGATCGGAA |
| 53297_2 | TATGAGTGTGGAGTCGTTGCTGAACTTGTCACCTGAGATGGAGNNNAGATCGGAA |
| 45714_1 | TATGAGTGTGGAGTCGTTGCAGTGCTGAGATTATAGGCGTGAGNNNAGATCGGAA |
| 45714_2 | TATGAGTGTGGAGTCGTTGCAGTGCTGAGATTATAGGCGTGAGNNNAGATCGGAA |
| 34003_1 | TATGAGTGTGGAGTCGTTGCTGGAGTGCAGTGGTGTCATATCNNNAGATCGGAAG |
| 34003_2 | TATGAGTGTGGAGTCGTTGCTGGAGTGCAGTGGTGTCATATCNNNAGATCGGAAG |
| 28830_1 | TATGAGTGTGGAGTCGTTGCTTTTTTTTTTTTGCGACGGGGTCNNNAGATCGGAAG |
| 28830_2 | TATGAGTGTGGAGTCGTTGCTTTTTTTTTTTTGCGACGGGGTCNNNAGATCGGAAG |
| 51144_1 | TATGAGTGTGGAGTCGTTGCGGATGGGTGATTTCTGCATTTCCNNNAGATCGGAAG |
| 51144_2 | TATGAGTGTGGAGTCGTTGCGGATGGGTGATTTCTGCATTTCCNNNAGATCGGAAG |
| 47975_1 | TATGAGTGTGGAGTCGTTGCCTTTGTGGATGTCCAGGTCCTAANNNAGATCGGAAG |
| 47975_2 | TATGAGTGTGGAGTCGTTGCCTTTGTGGATGTCCAGGTCCTAANNNAGATCGGAAG |
| 13557_1 | TATGAGTGTGGAGTCGTTGCGGCAGCAATGGAAAACTAATNNNAGATCGGAAGAG |
| 13557_2 | TATGAGTGTGGAGTCGTTGCGGCAGCAATGGAAAACTAATNNNAGATCGGAAGAG |
| 43871_1 | TATGAGTGTGGAGTCGTTGCCAAAGCCTCCTCTGTCTCATCATNNNAGATCGGAAGA |
| 43871_2 | TATGAGTGTGGAGTCGTTGCCAAAGCCTCCTCTGTCTCATCATNNNAGATCGGAAGA |
| 46059_1 | TATGAGTGTGGAGTCGTTGCAATCTTGGCTCATTGCAACTTCCNNNAGATCGGAAG |
| 46059_2 | TATGAGTGTGGAGTCGTTGCAATCTTGGCTCATTGCAACTTCCNNNAGATCGGAAG |
| 29527_1 | TATGAGTGTGGAGTCGTTGCATGAGTCAGGCTGCCACCAATANNNAGATCGGAAG |
| 29527_2 | TATGAGTGTGGAGTCGTTGCATGAGTCAGGCTGCCACCAATANNNAGATCGGAAG |
| 49352_1 | TATGAGTGTGGAGTCGTTGCTTCCAGCGATTCTTGAACCTCAGNNNAGATCGGAAG |
| 49352_2 | TATGAGTGTGGAGTCGTTGCTTCCAGCGATTCTTGAACCTCAGNNNAGATCGGAAG |
| 42250_1 | TATGAGTGTGGAGTCGTTGCACCAAATGGAATGTGTGTGTGTGNNNAGATCGGAA |
| 42250_2 | TATGAGTGTGGAGTCGTTGCACCAAATGGAATGTGTGTGTGTGNNNAGATCGGAA |
| 45517_1 | TATGAGTGTGGAGTCGTTGCTTTGGCCCAGAGTTGACATAAGTNNNAGATCGGAA |
| 45517_2 | TATGAGTGTGGAGTCGTTGCTTTGGCCCAGAGTTGACATAAGTNNNAGATCGGAA |
| 51652_1 | TATGAGTGTGGAGTCGTTGCAAACCAGGAAAGTTTGACCTCCTNNNAGATCGGAAG |
| 51652_2 | TATGAGTGTGGAGTCGTTGCAAACCAGGAAAGTTTGACCTCCTNNNAGATCGGAAG |
| 40951_1 | TATGAGTGTGGAGTCGTTGCACAACAAAAAGAAAGTTGGGGGGNNNAGATCGGA |
| 40951_2 | TATGAGTGTGGAGTCGTTGCACAACAAAAAGAAAGTTGGGGGGNNNAGATCGGA |
| 31871_1 | TATGAGTGTGGAGTCGTTGCCACACATATCCACAGCAGCAATGNNNAGATCGGAAG |

| | |
|---|---|
| 31871_2 | TATGAGTGTGGAGTCGTTGCCACACATATCCACAGCAGCAATGNNNAGATCGGAAG |
| 48200_1 | TATGAGTGTGGAGTCGTTGCTAGGGCTCTAAAAGGCTCTCTCTNNNAGATCGGAAG |
| 48200_2 | TATGAGTGTGGAGTCGTTGCTAGGGCTCTAAAAGGCTCTCTCTNNNAGATCGGAAG |
| 10426_1 | TATGAGTGTGGAGTCGTTGCACAGGGGAGATACATTCCTGGAGNNNAGATCGGAA |
| 10426_2 | TATGAGTGTGGAGTCGTTGCACAGGGGAGATACATTCCTGGAGNNNAGATCGGAA |
| 51594_1 | TATGAGTGTGGAGTCGTTGCCTGAGCAGCATCACCAGCAGNNNAGATCGGAAGAG |
| 51594_2 | TATGAGTGTGGAGTCGTTGCCTGAGCAGCATCACCAGCAGNNNAGATCGGAAGAG |
| 32662_1 | TATGAGTGTGGAGTCGTTGCCCAAAGTAGCTGGGATTACAGGTNNNAGATCGGAA |
| 32662_2 | TATGAGTGTGGAGTCGTTGCCCAAAGTAGCTGGGATTACAGGTNNNAGATCGGAA |
| 52980_1 | TATGAGTGTGGAGTCGTTGCGGTGTGATGTCTGTGTCTAGGTTNNNAGATCGGAA |
| 52980_2 | TATGAGTGTGGAGTCGTTGCGGTGTGATGTCTGTGTCTAGGTTNNNAGATCGGAA |
| 46626_1 | TATGAGTGTGGAGTCGTTGCAATCTCGGCTCACTGTAAGCTCNNNAGATCGGAAGA |
| 46626_2 | TATGAGTGTGGAGTCGTTGCAATCTCGGCTCACTGTAAGCTCNNNAGATCGGAAGA |
| 45582_1 | TATGAGTGTGGAGTCGTTGCGGTTCAAGCAATTCTCCTGTCTCNNNAGATCGGAAG |
| 45582_2 | TATGAGTGTGGAGTCGTTGCGGTTCAAGCAATTCTCCTGTCTCNNNAGATCGGAAG |
| 32795_1 | TATGAGTGTGGAGTCGTTGCTTCAAGCTATTCTCCTGCCTCAGNNNAGATCGGAAGA |
| 32795_2 | TATGAGTGTGGAGTCGTTGCTTCAAGCTATTCTCCTGCCTCAGNNNAGATCGGAAGA |
| 48964_1 | TATGAGTGTGGAGTCGTTGCGTCTTGAACTCCTGACCTTGTGANNNAGATCGGAAG |
| 48964_2 | TATGAGTGTGGAGTCGTTGCGTCTTGAACTCCTGACCTTGTGANNNAGATCGGAAG |
| 52602_1 | TATGAGTGTGGAGTCGTTGCTCCACAGAGTTCTCTTCTCGTTGNNNAGATCGGAAG |
| 52602_2 | TATGAGTGTGGAGTCGTTGCTCCACAGAGTTCTCTTCTCGTTGNNNAGATCGGAAG |
| 39088_1 | TATGAGTGTGGAGTCGTTGCATGAGAAGGTGCTCAACATCACTNNNAGATCGGAAG |
| 39088_2 | TATGAGTGTGGAGTCGTTGCATGAGAAGGTGCTCAACATCACTNNNAGATCGGAAG |
| 47945_1 | TATGAGTGTGGAGTCGTTGCTTTTTAGTGGAGACGGGGTTTCANNNAGATCGGAA |
| 47945_2 | TATGAGTGTGGAGTCGTTGCTTTTTAGTGGAGACGGGGTTTCANNNAGATCGGAA |
| 36881_1 | TATGAGTGTGGAGTCGTTGCGAGGCAGGAGAATGGTGTGAANNNAGATCGGAAG |
| 36881_2 | TATGAGTGTGGAGTCGTTGCGAGGCAGGAGAATGGTGTGAANNNAGATCGGAAG |
| 13134_1 | TATGAGTGTGGAGTCGTTGCCATACATACTTGACTTCTCATCTNNNAGATCGGAAGA |
| 13134_2 | TATGAGTGTGGAGTCGTTGCCATACATACTTGACTTCTCATCTNNNAGATCGGAAGA |
| 39476_1 | TATGAGTGTGGAGTCGTTGCATGATGGCAGGGAGAGTGAAAATNNNAGATCGGA |
| 39476_2 | TATGAGTGTGGAGTCGTTGCATGATGGCAGGGAGAGTGAAAATNNNAGATCGGA |
| 47687_1 | TATGAGTGTGGAGTCGTTGCGAAGGATGTAGGGGCAGGAGTNNNAGATCGGAAG |
| 47687_2 | TATGAGTGTGGAGTCGTTGCGAAGGATGTAGGGGCAGGAGTNNNAGATCGGAAG |
| 39895_1 | TATGAGTGTGGAGTCGTTGCTATACAGCAAATCCCAAGAGCGTNNNAGATCGGAAG |
| 39895_2 | TATGAGTGTGGAGTCGTTGCTATACAGCAAATCCCAAGAGCGTNNNAGATCGGAAG |
| 31477_1 | TATGAGTGTGGAGTCGTTGCGTGTGTGTGTGTGTGTGTGTTTANNNAGATCGGAA |
| 31477_2 | TATGAGTGTGGAGTCGTTGCGTGTGTGTGTGTGTGTGTGTTTANNNAGATCGGAA |
| 30686_1 | TATGAGTGTGGAGTCGTTGCATTATGCTTTCATGCCTCCTCCCNNNAGATCGGAAGA |
| 30686_2 | TATGAGTGTGGAGTCGTTGCATTATGCTTTCATGCCTCCTCCCNNNAGATCGGAAGA |
| 42671_1 | TATGAGTGTGGAGTCGTTGCGAGGCAGGAAAATCACTTGAACCNNNAGATCGGAA |
| 42671_2 | TATGAGTGTGGAGTCGTTGCGAGGCAGGAAAATCACTTGAACCNNNAGATCGGAA |
| 37514_1 | TATGAGTGTGGAGTCGTTGCACCAAAATTAGTTTCAGCCCAATNNNAGATCGGAAG |
| 37514_2 | TATGAGTGTGGAGTCGTTGCACCAAAATTAGTTTCAGCCCAATNNNAGATCGGAAG |

| | |
|---|---|
| 45484_1 | TATGAGTGTGGAGTCGTTGCCTCACCTTTGTCCTAGAACAGCTNNNAGATCGGAAGA |
| 45484_2 | TATGAGTGTGGAGTCGTTGCCTCACCTTTGTCCTAGAACAGCTNNNAGATCGGAAGA |
| 52009_1 | TATGAGTGTGGAGTCGTTGCCCCCATTTTATCCCAAGCCTTTCNNNAGATCGGAAGA |
| 52009_2 | TATGAGTGTGGAGTCGTTGCCCCCATTTTATCCCAAGCCTTTCNNNAGATCGGAAGA |
| 31343_1 | TATGAGTGTGGAGTCGTTGCCAGAGATGCTGCTAAACACCCTANNNAGATCGGAAG |
| 31343_2 | TATGAGTGTGGAGTCGTTGCCAGAGATGCTGCTAAACACCCTANNNAGATCGGAAG |
| 10027_1 | TATGAGTGTGGAGTCGTTGCACCACCACCACTACCATGACTTANNNAGATCGGAAGA |
| 10027_2 | TATGAGTGTGGAGTCGTTGCACCACCACCACTACCATGACTTANNNAGATCGGAAGA |
| 50083_1 | TATGAGTGTGGAGTCGTTGCCCTGGCAGTGACACTGATTAGATNNNAGATCGGAAG |
| 50083_2 | TATGAGTGTGGAGTCGTTGCCCTGGCAGTGACACTGATTAGATNNNAGATCGGAAG |
| 40349_1 | TATGAGTGTGGAGTCGTTGCAAGAAGAAGAAGGAGGAGGAGGANNNAGATCGG |
| 40349_2 | TATGAGTGTGGAGTCGTTGCAAGAAGAAGAAGGAGGAGGAGGANNNAGATCGG |
| 36062_1 | TATGAGTGTGGAGTCGTTGCGCTGGTTCCAATCTGACTGAAACNNNAGATCGGAAG |
| 36062_2 | TATGAGTGTGGAGTCGTTGCGCTGGTTCCAATCTGACTGAAACNNNAGATCGGAAG |
| 49828_1 | TATGAGTGTGGAGTCGTTGCATGCATTCTGGCCTCCTAATCTTNNNAGATCGGAAGA |
| 49828_2 | TATGAGTGTGGAGTCGTTGCATGCATTCTGGCCTCCTAATCTTNNNAGATCGGAAGA |
| 32218_1 | TATGAGTGTGGAGTCGTTGCTGTGTTAGACGTTCTAGAGCCTGNNNAGATCGGAAG |
| 32218_2 | TATGAGTGTGGAGTCGTTGCTGTGTTAGACGTTCTAGAGCCTGNNNAGATCGGAAG |
| 3064_1 | TATGAGTGTGGAGTCGTTGCATTACCTGTTCCAGCACCATGTCNNNAGATCGGAAGA |
| 3064_2 | TATGAGTGTGGAGTCGTTGCATTACCTGTTCCAGCACCATGTCNNNAGATCGGAAGA |
| 43447_1 | TATGAGTGTGGAGTCGTTGCGGGTCACATAAACCATTTTAGAGGGNNNAGATCGG |
| 43447_2 | TATGAGTGTGGAGTCGTTGCGGGTCACATAAACCATTTTAGAGGGNNNAGATCGG |
| 52279_1 | TATGAGTGTGGAGTCGTTGCCATGCCCTTTCACTACTGTACCTNNNAGATCGGAAGA |
| 52279_2 | TATGAGTGTGGAGTCGTTGCCATGCCCTTTCACTACTGTACCTNNNAGATCGGAAGA |
| 34247_1 | TATGAGTGTGGAGTCGTTGCCACACAAGCCATATGAAGTGTCCNNNAGATCGGAAG |
| 34247_2 | TATGAGTGTGGAGTCGTTGCCACACAAGCCATATGAAGTGTCCNNNAGATCGGAAG |
| 38159_1 | TATGAGTGTGGAGTCGTTGCCATGCCTGTAGTCCCAGCTATTTNNNAGATCGGAAGA |
| 38159_2 | TATGAGTGTGGAGTCGTTGCCATGCCTGTAGTCCCAGCTATTTNNNAGATCGGAAGA |
| 37411_1 | TATGAGTGTGGAGTCGTTGCATGGGAAGAAGCCACCATCTAGANNNAGATCGGAA |
| 37411_2 | TATGAGTGTGGAGTCGTTGCATGGGAAGAAGCCACCATCTAGANNNAGATCGGAA |
| 37268_1 | TATGAGTGTGGAGTCGTTGCGCTTTCAGTTAGCTACAGTCCAANNNAGATCGGAAG |
| 37268_2 | TATGAGTGTGGAGTCGTTGCGCTTTCAGTTAGCTACAGTCCAANNNAGATCGGAAG |
| 51490_1 | TATGAGTGTGGAGTCGTTGCGAGACCTGTATGACTCAGCAGAGNNNAGATCGGAA |
| 51490_2 | TATGAGTGTGGAGTCGTTGCGAGACCTGTATGACTCAGCAGAGNNNAGATCGGAA |
| 38357_1 | TATGAGTGTGGAGTCGTTGCGAGGCTGAGACATGAGAATCACTNNNAGATCGGAA |
| 38357_2 | TATGAGTGTGGAGTCGTTGCGAGGCTGAGACATGAGAATCACTNNNAGATCGGAA |
| 46931_1 | TATGAGTGTGGAGTCGTTGCCGATAAGCTGGGACTGGATTACANNNAGATCGGAA |
| 46931_2 | TATGAGTGTGGAGTCGTTGCCGATAAGCTGGGACTGGATTACANNNAGATCGGAA |
| 40662_1 | TATGAGTGTGGAGTCGTTGCTGAAGTTGTTTCTGCTGATGTGTNNNAGATCGGAAG |
| 40662_2 | TATGAGTGTGGAGTCGTTGCTGAAGTTGTTTCTGCTGATGTGTNNNAGATCGGAAG |
| 34300_1 | TATGAGTGTGGAGTCGTTGCCTGTTCCCCCACACCATCTCNNNAGATCGGAAGAGC |
| 34300_2 | TATGAGTGTGGAGTCGTTGCCTGTTCCCCCACACCATCTCNNNAGATCGGAAGAGC |
| 2528_1 | TATGAGTGTGGAGTCGTTGCCAAGAGTAGGAGGGAGAGGAAGGNNNAGATCGG |

| | |
|---|---|
| 2528_2 | TATGAGTGTGGAGTCGTTGCCAAGAGTAGGAGGGAGAGGAAGGNNNAGATCGG |
| 32478_1 | TATGAGTGTGGAGTCGTTGCAACCTCTGCTTCCTGTGATTCTCNNNAGATCGGAAGA |
| 32478_2 | TATGAGTGTGGAGTCGTTGCAACCTCTGCTTCCTGTGATTCTCNNNAGATCGGAAGA |
| 45852_1 | TATGAGTGTGGAGTCGTTGCAAGGACACTGTAAACAGCAAAGCNNNAGATCGGAA |
| 45852_2 | TATGAGTGTGGAGTCGTTGCAAGGACACTGTAAACAGCAAAGCNNNAGATCGGAA |
| 4364_1 | TATGAGTGTGGAGTCGTTGCCTCACCGTTCGCAGTACTGAATANNNAGATCGGAAG |
| 4364_2 | TATGAGTGTGGAGTCGTTGCCTCACCGTTCGCAGTACTGAATANNNAGATCGGAAG |
| 51093_1 | TATGAGTGTGGAGTCGTTGCCCCCATACTGTTCTTGTGGTAGTNNNAGATCGGAAG |
| 51093_2 | TATGAGTGTGGAGTCGTTGCCCCCATACTGTTCTTGTGGTAGTNNNAGATCGGAAG |
| 38285_1 | TATGAGTGTGGAGTCGTTGCTCCTACAGCTATGTTGGTGTCTGNNNAGATCGGAAG |
| 38285_2 | TATGAGTGTGGAGTCGTTGCTCCTACAGCTATGTTGGTGTCTGNNNAGATCGGAAG |
| 39611_1 | TATGAGTGTGGAGTCGTTGCCCACCATGCCAGGCTAATTTTTANNNAGATCGGAAG |
| 39611_2 | TATGAGTGTGGAGTCGTTGCCCACCATGCCAGGCTAATTTTTANNNAGATCGGAAG |
| 40374_1 | TATGAGTGTGGAGTCGTTGCTGAGTTCTGTGTGTCCTGGATACNNNAGATCGGAAG |
| 40374_2 | TATGAGTGTGGAGTCGTTGCTGAGTTCTGTGTGTCCTGGATACNNNAGATCGGAAG |
| 32765_1 | TATGAGTGTGGAGTCGTTGCTGTGCTGGCATAAAGAGGCATATNNNAGATCGGAA |
| 32765_2 | TATGAGTGTGGAGTCGTTGCTGTGCTGGCATAAAGAGGCATATNNNAGATCGGAA |
| 50465_1 | TATGAGTGTGGAGTCGTTGCCAAGTTCTCTCCCAAGGTGAGTCNNNAGATCGGAAG |
| 50465_2 | TATGAGTGTGGAGTCGTTGCCAAGTTCTCTCCCAAGGTGAGTCNNNAGATCGGAAG |
| 35034_1 | TATGAGTGTGGAGTCGTTGCTCTCAGTACATTGACCAAGCTGGNNNAGATCGGAAG |
| 35034_2 | TATGAGTGTGGAGTCGTTGCTCTCAGTACATTGACCAAGCTGGNNNAGATCGGAAG |
| 45568_1 | TATGAGTGTGGAGTCGTTGCGTGAGAGGGGGATTTTACCATGTNNNAGATCGGAA |
| 45568_2 | TATGAGTGTGGAGTCGTTGCGTGAGAGGGGGATTTTACCATGTNNNAGATCGGAA |
| 49490_1 | TATGAGTGTGGAGTCGTTGCACCCAATCTTCCTTGCTGATTCTNNNAGATCGGAAGA |
| 49490_2 | TATGAGTGTGGAGTCGTTGCACCCAATCTTCCTTGCTGATTCTNNNAGATCGGAAGA |
| 34126_1 | TATGAGTGTGGAGTCGTTGCCAACCCAAGATTTCCTCAAAACCANNNAGATCGGAA |
| 34126_2 | TATGAGTGTGGAGTCGTTGCCAACCCAAGATTTCCTCAAAACCANNNAGATCGGAA |
| 39564_1 | TATGAGTGTGGAGTCGTTGCGTCTGTAAATGACCCCCTTAGCANNNAGATCGGAAG |
| 39564_2 | TATGAGTGTGGAGTCGTTGCGTCTGTAAATGACCCCCTTAGCANNNAGATCGGAAG |
| 30565_1 | TATGAGTGTGGAGTCGTTGCGACCAAGATGCTAGACCTGCTAGANNNAGATCGGA |
| 30565_2 | TATGAGTGTGGAGTCGTTGCGACCAAGATGCTAGACCTGCTAGANNNAGATCGGA |
| 40263_1 | TATGAGTGTGGAGTCGTTGCTATGAGACGTGGCAAAGTAGGTCNNNAGATCGGAA |
| 40263_2 | TATGAGTGTGGAGTCGTTGCTATGAGACGTGGCAAAGTAGGTCNNNAGATCGGAA |
| 35396_1 | TATGAGTGTGGAGTCGTTGCGATTTCGGCTCACTGCAAGCNNNAGATCGGAAGAG |
| 35396_2 | TATGAGTGTGGAGTCGTTGCGATTTCGGCTCACTGCAAGCNNNAGATCGGAAGAG |
| 51805_1 | TATGAGTGTGGAGTCGTTGCAGGTCACTAATCAGTCGGGATTCNNNAGATCGGAA |
| 51805_2 | TATGAGTGTGGAGTCGTTGCAGGTCACTAATCAGTCGGGATTCNNNAGATCGGAA |
| 52645_1 | TATGAGTGTGGAGTCGTTGCTGCCAGTGGATGTCTCTAAATCCNNNAGATCGGAAG |
| 52645_2 | TATGAGTGTGGAGTCGTTGCTGCCAGTGGATGTCTCTAAATCCNNNAGATCGGAAG |
| 48159_1 | TATGAGTGTGGAGTCGTTGCAATCTCGGCTAACCACAACCTCNNNAGATCGGAAGA |
| 48159_2 | TATGAGTGTGGAGTCGTTGCAATCTCGGCTAACCACAACCTCNNNAGATCGGAAGA |
| 3036_1 | TATGAGTGTGGAGTCGTTGCGCTGTACCTCATGAGTTGGTTGTNNNAGATCGGAAG |
| 3036_2 | TATGAGTGTGGAGTCGTTGCGCTGTACCTCATGAGTTGGTTGTNNNAGATCGGAAG |

| | |
|---|---|
| 2099_1 | TATGAGTGTGGAGTCGTTGCTAGGCAAAATGATGCGGTAAGCANNNAGATCGGAA |
| 2099_2 | TATGAGTGTGGAGTCGTTGCTAGGCAAAATGATGCGGTAAGCANNNAGATCGGAA |
| 32548_1 | TATGAGTGTGGAGTCGTTGCAAAAGACATAGAGTGGCTGAGGGNNNAGATCGGA |
| 32548_2 | TATGAGTGTGGAGTCGTTGCAAAAGACATAGAGTGGCTGAGGGNNNAGATCGGA |
| 44203_1 | TATGAGTGTGGAGTCGTTGCTGGAGGAGGGGAGAAACAGTATANNNAGATCGGA |
| 44203_2 | TATGAGTGTGGAGTCGTTGCTGGAGGAGGGGAGAAACAGTATANNNAGATCGGA |
| 35758_1 | TATGAGTGTGGAGTCGTTGCTCCATTCTGCTAGCCTAAGTCTTNNNAGATCGGAAGA |
| 35758_2 | TATGAGTGTGGAGTCGTTGCTCCATTCTGCTAGCCTAAGTCTTNNNAGATCGGAAGA |
| 53054_1 | TATGAGTGTGGAGTCGTTGCTCTTTTTGGGAGACCACAGTTCANNNAGATCGGAAG |
| 53054_2 | TATGAGTGTGGAGTCGTTGCTCTTTTTGGGAGACCACAGTTCANNNAGATCGGAAG |
| 49069_1 | TATGAGTGTGGAGTCGTTGCGTGTTCAAGCAATTCTTCTGCCTNNNAGATCGGAAG |
| 49069_2 | TATGAGTGTGGAGTCGTTGCGTGTTCAAGCAATTCTTCTGCCTNNNAGATCGGAAG |
| 36092_1 | TATGAGTGTGGAGTCGTTGCAGAGTTGCAGTCACTCGAATCTTNNNAGATCGGAAG |
| 36092_2 | TATGAGTGTGGAGTCGTTGCAGAGTTGCAGTCACTCGAATCTTNNNAGATCGGAAG |
| 2296_1 | TATGAGTGTGGAGTCGTTGCCCAACAGACAACTTGAACAGCACNNNAGATCGGAAG |
| 2296_2 | TATGAGTGTGGAGTCGTTGCCCAACAGACAACTTGAACAGCACNNNAGATCGGAAG |
| 34269_1 | TATGAGTGTGGAGTCGTTGCAACACTAAGTGAGCACAGAACCANNNAGATCGGAA |
| 34269_2 | TATGAGTGTGGAGTCGTTGCAACACTAAGTGAGCACAGAACCANNNAGATCGGAA |
| 46632_1 | TATGAGTGTGGAGTCGTTGCATAGAATCTCACTCTGTTGCCCANNNAGATCGGAAG |
| 46632_2 | TATGAGTGTGGAGTCGTTGCATAGAATCTCACTCTGTTGCCCANNNAGATCGGAAG |
| 10044_1 | TATGAGTGTGGAGTCGTTGCGCAGTAGGGCCCATTAGTCTAGANNNAGATCGGAA |
| 10044_2 | TATGAGTGTGGAGTCGTTGCGCAGTAGGGCCCATTAGTCTAGANNNAGATCGGAA |
| 44506_1 | TATGAGTGTGGAGTCGTTGCGGCAAGCAAGGAGAATGCAATAANNNAGATCGGAA |
| 44506_2 | TATGAGTGTGGAGTCGTTGCGGCAAGCAAGGAGAATGCAATAANNNAGATCGGAA |
| 37246_1 | TATGAGTGTGGAGTCGTTGCTGAAAGAAAAGGATGTCAATGAGCANNNAGATCGG |
| 37246_2 | TATGAGTGTGGAGTCGTTGCTGAAAGAAAAGGATGTCAATGAGCANNNAGATCGG |
| 32300_1 | TATGAGTGTGGAGTCGTTGCTATAAACGCCCTCATCTTGCCATNNNAGATCGGAAGA |
| 32300_2 | TATGAGTGTGGAGTCGTTGCTATAAACGCCCTCATCTTGCCATNNNAGATCGGAAGA |
| 41230_1 | TATGAGTGTGGAGTCGTTGCCATTTCTGAAACTGAGAGGGGGANNNAGATCGGAA |
| 41230_2 | TATGAGTGTGGAGTCGTTGCCATTTCTGAAACTGAGAGGGGGANNNAGATCGGAA |
| 42099_1 | TATGAGTGTGGAGTCGTTGCAGCTACTATGGAAACCAGTGTGGNNNAGATCGGAA |
| 42099_2 | TATGAGTGTGGAGTCGTTGCAGCTACTATGGAAACCAGTGTGGNNNAGATCGGAA |
| 36607_1 | TATGAGTGTGGAGTCGTTGCGAAGAAACGAAACCTGGATCAGGNNNAGATCGGAA |
| 36607_2 | TATGAGTGTGGAGTCGTTGCGAAGAAACGAAACCTGGATCAGGNNNAGATCGGAA |
| 10242_1 | TATGAGTGTGGAGTCGTTGCACATGCTATCAGGGTTACTGACTNNNAGATCGGAAG |
| 10242_2 | TATGAGTGTGGAGTCGTTGCACATGCTATCAGGGTTACTGACTNNNAGATCGGAAG |
| 38627_1 | TATGAGTGTGGAGTCGTTGCAGTCAAAGGAGGTTGTTCTCTGGNNNAGATCGGAA |
| 38627_2 | TATGAGTGTGGAGTCGTTGCAGTCAAAGGAGGTTGTTCTCTGGNNNAGATCGGAA |
| 53302_1 | TATGAGTGTGGAGTCGTTGCGGAGGCCACTATCCTAAGTGAAGNNNAGATCGGAA |
| 53302_2 | TATGAGTGTGGAGTCGTTGCGGAGGCCACTATCCTAAGTGAAGNNNAGATCGGAA |
| 32435_1 | TATGAGTGTGGAGTCGTTGCAACTCCTGGGCTCAAAAGATTCTNNNAGATCGGAAG |
| 32435_2 | TATGAGTGTGGAGTCGTTGCAACTCCTGGGCTCAAAAGATTCTNNNAGATCGGAAG |
| 38110_1 | TATGAGTGTGGAGTCGTTGCTAACAAGCAAACCTCAACCCCAANNNAGATCGGAAG |

| | |
|---|---|
| 38110_2 | TATGAGTGTGGAGTCGTTGCTAACAAGCAAACCTCAACCCCAANNNAGATCGGAAG |
| 51277_1 | TATGAGTGTGGAGTCGTTGCTATAGCGCTTACTATGTGCCAGGNNNAGATCGGAAG |
| 51277_2 | TATGAGTGTGGAGTCGTTGCTATAGCGCTTACTATGTGCCAGGNNNAGATCGGAAG |
| 51298_1 | TATGAGTGTGGAGTCGTTGCCGCCAAAATCTCACAAATCACCANNNAGATCGGAAG |
| 51298_2 | TATGAGTGTGGAGTCGTTGCCGCCAAAATCTCACAAATCACCANNNAGATCGGAAG |
| 51832_1 | TATGAGTGTGGAGTCGTTGCCTTATGCCTTTGCTTCCTCATGGNNNAGATCGGAAG |
| 51832_2 | TATGAGTGTGGAGTCGTTGCCTTATGCCTTTGCTTCCTCATGGNNNAGATCGGAAG |
| 46719_1 | TATGAGTGTGGAGTCGTTGCTACAGTGGTGCAATATCAGCTCANNNAGATCGGAAG |
| 46719_2 | TATGAGTGTGGAGTCGTTGCTACAGTGGTGCAATATCAGCTCANNNAGATCGGAAG |
| 12693_1 | TATGAGTGTGGAGTCGTTGCGAAGTCACTCCTTCTACATTATANNNAGATCGGAAG |
| 12693_2 | TATGAGTGTGGAGTCGTTGCGAAGTCACTCCTTCTACATTATANNNAGATCGGAAG |
| 46861_1 | TATGAGTGTGGAGTCGTTGCAGCCTCCTGGATTCAAGTGATTCNNNAGATCGGAAG |
| 46861_2 | TATGAGTGTGGAGTCGTTGCAGCCTCCTGGATTCAAGTGATTCNNNAGATCGGAAG |
| 39135_1 | TATGAGTGTGGAGTCGTTGCTCCCTGATTATAGCCATTGGAGCNNNAGATCGGAAG |
| 39135_2 | TATGAGTGTGGAGTCGTTGCTCCCTGATTATAGCCATTGGAGCNNNAGATCGGAAG |
| 46048_1 | TATGAGTGTGGAGTCGTTGCTTCAGGTGATTCTTCTGCCTCAGNNNAGATCGGAAG |
| 46048_2 | TATGAGTGTGGAGTCGTTGCTTCAGGTGATTCTTCTGCCTCAGNNNAGATCGGAAG |
| 46338_1 | TATGAGTGTGGAGTCGTTGCTATTGACGTAAGCAGCTCCATGTNNNAGATCGGAAG |
| 46338_2 | TATGAGTGTGGAGTCGTTGCTATTGACGTAAGCAGCTCCATGTNNNAGATCGGAAG |
| 32750_1 | TATGAGTGTGGAGTCGTTGCCAGAGCGAGTTTCCATCTCAAAANNNAGATCGGAAG |
| 32750_2 | TATGAGTGTGGAGTCGTTGCCAGAGCGAGTTTCCATCTCAAAANNNAGATCGGAAG |
| 32776_1 | TATGAGTGTGGAGTCGTTGCATTCCATCTCCAACCTGACCAATNNNAGATCGGAAGA |
| 32776_2 | TATGAGTGTGGAGTCGTTGCATTCCATCTCCAACCTGACCAATNNNAGATCGGAAGA |
| 30632_1 | TATGAGTGTGGAGTCGTTGCATGAGCCAGCAAACTGGTTAGTAACNNNAGATCGG |
| 30632_2 | TATGAGTGTGGAGTCGTTGCATGAGCCAGCAAACTGGTTAGTAACNNNAGATCGG |
| 44311_1 | TATGAGTGTGGAGTCGTTGCTCCGAAGAAGCAGATCTTTGTCANNNAGATCGGAAG |
| 44311_2 | TATGAGTGTGGAGTCGTTGCTCCGAAGAAGCAGATCTTTGTCANNNAGATCGGAAG |
| 31578_1 | TATGAGTGTGGAGTCGTTGCGTGACTCCACAACAAACTGGATGNNNAGATCGGAA |
| 31578_2 | TATGAGTGTGGAGTCGTTGCGTGACTCCACAACAAACTGGATGNNNAGATCGGAA |
| 37211_1 | TATGAGTGTGGAGTCGTTGCTTCCCAACTTCCATGTCACGATANNNAGATCGGAAG |
| 37211_2 | TATGAGTGTGGAGTCGTTGCTTCCCAACTTCCATGTCACGATANNNAGATCGGAAG |
| 49298_1 | TATGAGTGTGGAGTCGTTGCAGGGATTCTCACTCTGTCACCTANNNAGATCGGAAG |
| 49298_2 | TATGAGTGTGGAGTCGTTGCAGGGATTCTCACTCTGTCACCTANNNAGATCGGAAG |
| 41593_1 | TATGAGTGTGGAGTCGTTGCGATCTCAGCTCACTGCAACCTNNNAGATCGGAAGAG |
| 41593_2 | TATGAGTGTGGAGTCGTTGCGATCTCAGCTCACTGCAACCTNNNAGATCGGAAGAG |
| 48162_1 | TATGAGTGTGGAGTCGTTGCCTCACGGGTTCAAGCAATTCTCNNNAGATCGGAAGA |
| 48162_2 | TATGAGTGTGGAGTCGTTGCCTCACGGGTTCAAGCAATTCTCNNNAGATCGGAAGA |
| 47009_1 | TATGAGTGTGGAGTCGTTGCCTCAAGCAATTCTCCTGCCTCAGNNNAGATCGGAAG |
| 47009_2 | TATGAGTGTGGAGTCGTTGCCTCAAGCAATTCTCCTGCCTCAGNNNAGATCGGAAG |
| 8534_1 | TATGAGTGTGGAGTCGTTGCACTTACCTGTTTGGATGCCTAGCNNNAGATCGGAAG |
| 8534_2 | TATGAGTGTGGAGTCGTTGCACTTACCTGTTTGGATGCCTAGCNNNAGATCGGAAG |
| 49914_1 | TATGAGTGTGGAGTCGTTGCCCCTTTTCGCCTTTTTTCTTTGCNNNAGATCGGAAGA |
| 49914_2 | TATGAGTGTGGAGTCGTTGCCCCTTTTCGCCTTTTTTCTTTGCNNNAGATCGGAAGA |

| | |
|---|---|
| 53386_1 | TATGAGTGTGGAGTCGTTGCCCTTAGCCATGAGGAACTGTGATNNNAGATCGGAAG |
| 53386_2 | TATGAGTGTGGAGTCGTTGCCCTTAGCCATGAGGAACTGTGATNNNAGATCGGAAG |
| 48166_1 | TATGAGTGTGGAGTCGTTGCATGGGGTTTCACTATGTTGGCTANNNAGATCGGAAG |
| 48166_2 | TATGAGTGTGGAGTCGTTGCATGGGGTTTCACTATGTTGGCTANNNAGATCGGAAG |
| 44752_1 | TATGAGTGTGGAGTCGTTGCCCTCCAGGGTTCAAGTGATTCTNNNAGATCGGAAGA |
| 44752_2 | TATGAGTGTGGAGTCGTTGCCCTCCAGGGTTCAAGTGATTCTNNNAGATCGGAAGA |
| 32445_1 | TATGAGTGTGGAGTCGTTGCCCTCCTGGATTCAAGCGATTCTCNNNAGATCGGAAGA |
| 32445_2 | TATGAGTGTGGAGTCGTTGCCCTCCTGGATTCAAGCGATTCTCNNNAGATCGGAAGA |
| 42409_1 | TATGAGTGTGGAGTCGTTGCTTTGTCATGAACCATCTGTTGCCNNNAGATCGGAAGA |
| 42409_2 | TATGAGTGTGGAGTCGTTGCTTTGTCATGAACCATCTGTTGCCNNNAGATCGGAAGA |
| 45298_1 | TATGAGTGTGGAGTCGTTGCGCCATTGCCCTAGCTTATGAATGNNNAGATCGGAAG |
| 45298_2 | TATGAGTGTGGAGTCGTTGCGCCATTGCCCTAGCTTATGAATGNNNAGATCGGAAG |
| 44216_1 | TATGAGTGTGGAGTCGTTGCTAACCAGTAAGTTCTGCTGCCTTNNNAGATCGGAAGA |
| 44216_2 | TATGAGTGTGGAGTCGTTGCTAACCAGTAAGTTCTGCTGCCTTNNNAGATCGGAAGA |
| 30823_1 | TATGAGTGTGGAGTCGTTGCAGACAAGGAGAAGAGGAGGAGAANNNAGATCGGA |
| 30823_2 | TATGAGTGTGGAGTCGTTGCAGACAAGGAGAAGAGGAGGAGAANNNAGATCGGA |
| 35179_1 | TATGAGTGTGGAGTCGTTGCCTCTGAGGTGGCATCTAGCTCTNNNAGATCGGAAGA |
| 35179_2 | TATGAGTGTGGAGTCGTTGCCTCTGAGGTGGCATCTAGCTCTNNNAGATCGGAAGA |
| 2470_1 | TATGAGTGTGGAGTCGTTGCTGTGTGCGTGTGTATGAAAGAAGNNNAGATCGGAA |
| 2470_2 | TATGAGTGTGGAGTCGTTGCTGTGTGCGTGTGTATGAAAGAAGNNNAGATCGGAA |
| 13798_1 | TATGAGTGTGGAGTCGTTGCCCACACTATCTCTTTATCCATAANNNAGATCGGAAGA |
| 13798_2 | TATGAGTGTGGAGTCGTTGCCCACACTATCTCTTTATCCATAANNNAGATCGGAAGA |
| 40403_1 | TATGAGTGTGGAGTCGTTGCGCAGAGGTAAAGAAGCACGAAAGNNNAGATCGGA |
| 40403_2 | TATGAGTGTGGAGTCGTTGCGCAGAGGTAAAGAAGCACGAAAGNNNAGATCGGA |
| 29535_1 | TATGAGTGTGGAGTCGTTGCCCGAGAGCAGACACCAGCAGNNNAGATCGGAAGAG |
| 29535_2 | TATGAGTGTGGAGTCGTTGCCCGAGAGCAGACACCAGCAGNNNAGATCGGAAGAG |
| 33617_1 | TATGAGTGTGGAGTCGTTGCCTCCCGAGTAGCTGAGACTACANNNAGATCGGAAGA |
| 33617_2 | TATGAGTGTGGAGTCGTTGCCTCCCGAGTAGCTGAGACTACANNNAGATCGGAAGA |
| 35228_1 | TATGAGTGTGGAGTCGTTGCCATTTTGGAGGTTGCCTGTTCATNNNAGATCGGAAG |
| 35228_2 | TATGAGTGTGGAGTCGTTGCCATTTTGGAGGTTGCCTGTTCATNNNAGATCGGAAG |
| 48246_1 | TATGAGTGTGGAGTCGTTGCCGTATCTGGCCTTTGCATACTTTNNNAGATCGGAAGA |
| 48246_2 | TATGAGTGTGGAGTCGTTGCCGTATCTGGCCTTTGCATACTTTNNNAGATCGGAAGA |
| 48466_1 | TATGAGTGTGGAGTCGTTGCACAATCTTGGCTCAGTGTAACCTNNNAGATCGGAAGA |
| 48466_2 | TATGAGTGTGGAGTCGTTGCACAATCTTGGCTCAGTGTAACCTNNNAGATCGGAAGA |
| 31533_1 | TATGAGTGTGGAGTCGTTGCATCTGTTTCCCCATTCTTTTGCANNNAGATCGGAAGA |
| 31533_2 | TATGAGTGTGGAGTCGTTGCATCTGTTTCCCCATTCTTTTGCANNNAGATCGGAAGA |
| 40588_1 | TATGAGTGTGGAGTCGTTGCTCCTAGTACAACCAATCCACTCTNNNAGATCGGAAGA |
| 40588_2 | TATGAGTGTGGAGTCGTTGCTCCTAGTACAACCAATCCACTCTNNNAGATCGGAAGA |
| 39751_1 | TATGAGTGTGGAGTCGTTGCACAAGGAAAACTACAAAATGCACTGNNNAGATCGGA |
| 39751_2 | TATGAGTGTGGAGTCGTTGCACAAGGAAAACTACAAAATGCACTGNNNAGATCGGA |
| 42113_1 | TATGAGTGTGGAGTCGTTGCGACTCCTGGGTTCAAGTGATTCTNNNAGATCGGAAG |
| 42113_2 | TATGAGTGTGGAGTCGTTGCGACTCCTGGGTTCAAGTGATTCTNNNAGATCGGAAG |
| 31042_1 | TATGAGTGTGGAGTCGTTGCTTGCCTCTCTGAATATACCACCGNNNAGATCGGAAGA |

| | |
|---|---|
| 31042_2 | TATGAGTGTGGAGTCGTTGCTTGCCTCTCTGAATATACCACCGNNNAGATCGGAAGA |
| 40892_1 | TATGAGTGTGGAGTCGTTGCGAGTTGGTGTGAGAGTGAGCATANNNAGATCGGAA |
| 40892_2 | TATGAGTGTGGAGTCGTTGCGAGTTGGTGTGAGAGTGAGCATANNNAGATCGGAA |
| 32943_1 | TATGAGTGTGGAGTCGTTGCCCATCCAAAGGTCATCAACATCAANNNAGATCGGAA |
| 32943_2 | TATGAGTGTGGAGTCGTTGCCCATCCAAAGGTCATCAACATCAANNNAGATCGGAA |
| 53406_1 | TATGAGTGTGGAGTCGTTGCGTACAGTCCTACCTAGGGGAACANNNAGATCGGAA |
| 53406_2 | TATGAGTGTGGAGTCGTTGCGTACAGTCCTACCTAGGGGAACANNNAGATCGGAA |
| 41681_1 | TATGAGTGTGGAGTCGTTGCTCTTGAACTCCTGACCTCAAGTGNNNAGATCGGAAG |
| 41681_2 | TATGAGTGTGGAGTCGTTGCTCTTGAACTCCTGACCTCAAGTGNNNAGATCGGAAG |
| 3099_1 | TATGAGTGTGGAGTCGTTGCAGGGAGTGACAGTAGGCTCTTTGNNNAGATCGGAA |
| 3099_2 | TATGAGTGTGGAGTCGTTGCAGGGAGTGACAGTAGGCTCTTTGNNNAGATCGGAA |
| 29329_1 | TATGAGTGTGGAGTCGTTGCATTGGGGTTTTTGGACTTAGGGTNNNAGATCGGAA |
| 29329_2 | TATGAGTGTGGAGTCGTTGCATTGGGGTTTTTGGACTTAGGGTNNNAGATCGGAA |
| 12795_1 | TATGAGTGTGGAGTCGTTGCATATTAGTGACGAACATCTTACANNNAGATCGGAAG |
| 12795_2 | TATGAGTGTGGAGTCGTTGCATATTAGTGACGAACATCTTACANNNAGATCGGAAG |
| 12600_1 | TATGAGTGTGGAGTCGTTGCGAAATTCTTTAATAGCAGCTCCNNNAGATCGGAAGA |
| 12600_2 | TATGAGTGTGGAGTCGTTGCGAAATTCTTTAATAGCAGCTCCNNNAGATCGGAAGA |
| 37447_1 | TATGAGTGTGGAGTCGTTGCCGTCATGCCAAAAAGAAACCCTANNNAGATCGGAAG |
| 37447_2 | TATGAGTGTGGAGTCGTTGCCGTCATGCCAAAAAGAAACCCTANNNAGATCGGAAG |
| 50422_1 | TATGAGTGTGGAGTCGTTGCAAAATGCAAATTTCCCTGGCAGANNNAGATCGGAAG |
| 50422_2 | TATGAGTGTGGAGTCGTTGCAAAATGCAAATTTCCCTGGCAGANNNAGATCGGAAG |
| 37363_1 | TATGAGTGTGGAGTCGTTGCGGTGCATAGGGTGTTGTCTTAGTNNNAGATCGGAA |
| 37363_2 | TATGAGTGTGGAGTCGTTGCGGTGCATAGGGTGTTGTCTTAGTNNNAGATCGGAA |
| 32158_1 | TATGAGTGTGGAGTCGTTGCGATTTTCCAGACTGGCTACAAGCNNNAGATCGGAAG |
| 32158_2 | TATGAGTGTGGAGTCGTTGCGATTTTCCAGACTGGCTACAAGCNNNAGATCGGAAG |
| 1023_1 | TATGAGTGTGGAGTCGTTGCCTGTAGATGTTGAGCAGTGTTCCTNNNAGATCGGAA |
| 1023_2 | TATGAGTGTGGAGTCGTTGCCTGTAGATGTTGAGCAGTGTTCCTNNNAGATCGGAA |
| 36411_1 | TATGAGTGTGGAGTCGTTGCGTGCTTTTCCTCATTGGTTCCTCNNNAGATCGGAAGA |
| 36411_2 | TATGAGTGTGGAGTCGTTGCGTGCTTTTCCTCATTGGTTCCTCNNNAGATCGGAAGA |
| 31098_1 | TATGAGTGTGGAGTCGTTGCTGGTAATCCAGAGAATTTGCCCANNNAGATCGGAAG |
| 31098_2 | TATGAGTGTGGAGTCGTTGCTGGTAATCCAGAGAATTTGCCCANNNAGATCGGAAG |
| 44115_1 | TATGAGTGTGGAGTCGTTGCCGTGTCCCCCAGTATAGATGTTTNNNAGATCGGAAG |
| 44115_2 | TATGAGTGTGGAGTCGTTGCCGTGTCCCCCAGTATAGATGTTTNNNAGATCGGAAG |
| 37652_1 | TATGAGTGTGGAGTCGTTGCAACCAATCCCCTTTTTGTCTCCTNNNAGATCGGAAGA |
| 37652_2 | TATGAGTGTGGAGTCGTTGCAACCAATCCCCTTTTTGTCTCCTNNNAGATCGGAAGA |
| 29140_1 | TATGAGTGTGGAGTCGTTGCTGGATCCAGTGATAAGCTACCGANNNAGATCGGAA |
| 29140_2 | TATGAGTGTGGAGTCGTTGCTGGATCCAGTGATAAGCTACCGANNNAGATCGGAA |
| 32724_1 | TATGAGTGTGGAGTCGTTGCGTTTGAAACCTCCCCCCAAATTTNNNAGATCGGAAGA |
| 32724_2 | TATGAGTGTGGAGTCGTTGCGTTTGAAACCTCCCCCCAAATTTNNNAGATCGGAAGA |
| 51219_1 | TATGAGTGTGGAGTCGTTGCGGTAAGAATCCACCTTTTGCCTCNNNAGATCGGAAG |
| 51219_2 | TATGAGTGTGGAGTCGTTGCGGTAAGAATCCACCTTTTGCCTCNNNAGATCGGAAG |
| 41965_1 | TATGAGTGTGGAGTCGTTGCTGGTGGTTGTTTGTATTTCCGTGNNNAGATCGGAAG |
| 41965_2 | TATGAGTGTGGAGTCGTTGCTGGTGGTTGTTTGTATTTCCGTGNNNAGATCGGAAG |

| | |
|---|---|
| 36198_1 | TATGAGTGTGGAGTCGTTGCATAACAGACCACGTGTAAGCTGTNNNAGATCGGAAG |
| 36198_2 | TATGAGTGTGGAGTCGTTGCATAACAGACCACGTGTAAGCTGTNNNAGATCGGAAG |
| 42935_1 | TATGAGTGTGGAGTCGTTGCTTCTGGACCTGCTATCCTTTTCCNNNAGATCGGAAGA |
| 42935_2 | TATGAGTGTGGAGTCGTTGCTTCTGGACCTGCTATCCTTTTCCNNNAGATCGGAAGA |
| 37343_1 | TATGAGTGTGGAGTCGTTGCGGGCAAAACACCTGTATAGACATNNNAGATCGGAAG |
| 37343_2 | TATGAGTGTGGAGTCGTTGCGGGCAAAACACCTGTATAGACATNNNAGATCGGAAG |
| 52404_1 | TATGAGTGTGGAGTCGTTGCTTCCTTCTCCCAGCACCTCGNNNAGATCGGAAGAGCA |
| 52404_2 | TATGAGTGTGGAGTCGTTGCTTCCTTCTCCCAGCACCTCGNNNAGATCGGAAGAGCA |
| 6899_1 | TATGAGTGTGGAGTCGTTGCAATGAGGCTCAAAGCAGGTCACANNNAGATCGGAA |
| 6899_2 | TATGAGTGTGGAGTCGTTGCAATGAGGCTCAAAGCAGGTCACANNNAGATCGGAA |
| 43488_1 | TATGAGTGTGGAGTCGTTGCCAGTATCTGGGATCACAGGTGCNNNAGATCGGAAG |
| 43488_2 | TATGAGTGTGGAGTCGTTGCCAGTATCTGGGATCACAGGTGCNNNAGATCGGAAG |
| 41108_1 | TATGAGTGTGGAGTCGTTGCAGGAAGGAAGCTGTACCTCTACTNNNAGATCGGAAG |
| 41108_2 | TATGAGTGTGGAGTCGTTGCAGGAAGGAAGCTGTACCTCTACTNNNAGATCGGAAG |
| 10298_1 | TATGAGTGTGGAGTCGTTGCGTTGGCAGAGATTAGCCCTGAAANNNAGATCGGAA |
| 10298_2 | TATGAGTGTGGAGTCGTTGCGTTGGCAGAGATTAGCCCTGAAANNNAGATCGGAA |
| 51232_1 | TATGAGTGTGGAGTCGTTGCCATTCCAATTAGAGCAAGCCAGGNNNAGATCGGAAG |
| 51232_2 | TATGAGTGTGGAGTCGTTGCCATTCCAATTAGAGCAAGCCAGGNNNAGATCGGAAG |
| 31781_1 | TATGAGTGTGGAGTCGTTGCACCCCATTATGTTGAAAACCCCANNNAGATCGGAAG |
| 31781_2 | TATGAGTGTGGAGTCGTTGCACCCCATTATGTTGAAAACCCCANNNAGATCGGAAG |
| 42044_1 | TATGAGTGTGGAGTCGTTGCCATATGATCGCTACCACAACAGCNNNAGATCGGAAG |
| 42044_2 | TATGAGTGTGGAGTCGTTGCCATATGATCGCTACCACAACAGCNNNAGATCGGAAG |
| 47631_1 | TATGAGTGTGGAGTCGTTGCCTCCTGAGTAGCTGGGATTACAGNNNAGATCGGAAG |
| 47631_2 | TATGAGTGTGGAGTCGTTGCCTCCTGAGTAGCTGGGATTACAGNNNAGATCGGAAG |
| 39567_1 | TATGAGTGTGGAGTCGTTGCAGTTTCAGGTTTTGGAACATTTCANNNAGATCGGAA |
| 39567_2 | TATGAGTGTGGAGTCGTTGCAGTTTCAGGTTTTGGAACATTTCANNNAGATCGGAA |
| 48019_1 | TATGAGTGTGGAGTCGTTGCCCACGCCCAGCTAATTTTTGTATNNNAGATCGGAAGA |
| 48019_2 | TATGAGTGTGGAGTCGTTGCCCACGCCCAGCTAATTTTTGTATNNNAGATCGGAAGA |
| 35561_1 | TATGAGTGTGGAGTCGTTGCCCTCCTGGTTACATGCCATTCTCNNNAGATCGGAAGA |
| 35561_2 | TATGAGTGTGGAGTCGTTGCCCTCCTGGTTACATGCCATTCTCNNNAGATCGGAAGA |
| 44262_1 | TATGAGTGTGGAGTCGTTGCATTTGCTGATCCAAACTACCCCANNNAGATCGGAAGA |
| 44262_2 | TATGAGTGTGGAGTCGTTGCATTTGCTGATCCAAACTACCCCANNNAGATCGGAAGA |
| 49070_1 | TATGAGTGTGGAGTCGTTGCGCAAAAGGAATTTTCCCTTCCCTNNNAGATCGGAAGA |
| 49070_2 | TATGAGTGTGGAGTCGTTGCGCAAAAGGAATTTTCCCTTCCCTNNNAGATCGGAAGA |
| 34496_1 | TATGAGTGTGGAGTCGTTGCTCTTCCCACTGCTCCTTCTTTTTNNNAGATCGGAAGAG |
| 34496_2 | TATGAGTGTGGAGTCGTTGCTCTTCCCACTGCTCCTTCTTTTTNNNAGATCGGAAGAG |
| 13208_1 | TATGAGTGTGGAGTCGTTGCAGAAAGAACAAAACTCCATCTNNNAGATCGGAAGAG |
| 13208_2 | TATGAGTGTGGAGTCGTTGCAGAAAGAACAAAACTCCATCTNNNAGATCGGAAGAG |
| 29926_1 | TATGAGTGTGGAGTCGTTGCTCTCCCTCCCTACACAGAATTCCNNNAGATCGGAAGA |
| 29926_2 | TATGAGTGTGGAGTCGTTGCTCTCCCTCCCTACACAGAATTCCNNNAGATCGGAAGA |
| 39137_1 | TATGAGTGTGGAGTCGTTGCCTCCCAGGTTCAAGCCATTCTCNNNAGATCGGAAGAG |
| 39137_2 | TATGAGTGTGGAGTCGTTGCCTCCCAGGTTCAAGCCATTCTCNNNAGATCGGAAGAG |
| 30533_1 | TATGAGTGTGGAGTCGTTGCTACATAGAGTTGATGGATAGAGCAAANNNAGATCG |

| | |
|---|---|
| 30533_2 | TATGAGTGTGGAGTCGTTGCTACATAGAGTTGATGGATAGAGCAAANNNAGATCGG |
| 30749_1 | TATGAGTGTGGAGTCGTTGCGTCCTGTGGATTTTGACATGCATNNNAGATCGGAAG |
| 30749_2 | TATGAGTGTGGAGTCGTTGCGTCCTGTGGATTTTGACATGCATNNNAGATCGGAAG |
| 44034_1 | TATGAGTGTGGAGTCGTTGCCACATGACGGGGTGCATATTTACNNNAGATCGGAAG |
| 44034_2 | TATGAGTGTGGAGTCGTTGCCACATGACGGGGTGCATATTTACNNNAGATCGGAAG |
| 32988_1 | TATGAGTGTGGAGTCGTTGCAGAGATGTTAGGTGTGCTGATGGNNNAGATCGGAA |
| 32988_2 | TATGAGTGTGGAGTCGTTGCAGAGATGTTAGGTGTGCTGATGGNNNAGATCGGAA |
| 46020_1 | TATGAGTGTGGAGTCGTTGCACTCAAGCGATTATCTTGCCTCANNNAGATCGGAAGA |
| 46020_2 | TATGAGTGTGGAGTCGTTGCACTCAAGCGATTATCTTGCCTCANNNAGATCGGAAGA |
| 43484_1 | TATGAGTGTGGAGTCGTTGCAAGAGGAAGGAACTTCGGTATCGNNNAGATCGGAA |
| 43484_2 | TATGAGTGTGGAGTCGTTGCAAGAGGAAGGAACTTCGGTATCGNNNAGATCGGAA |
| 9681_1 | TATGAGTGTGGAGTCGTTGCTTGTGACCTAGCATTTGCATCCCNNNAGATCGGAAGA |
| 9681_2 | TATGAGTGTGGAGTCGTTGCTTGTGACCTAGCATTTGCATCCCNNNAGATCGGAAGA |
| 52788_1 | TATGAGTGTGGAGTCGTTGCTCAATGAGGTTGTTCTCCAGGTGNNNAGATCGGAAG |
| 52788_2 | TATGAGTGTGGAGTCGTTGCTCAATGAGGTTGTTCTCCAGGTGNNNAGATCGGAAG |
| 12302_1 | TATGAGTGTGGAGTCGTTGCACGGACCAGGGATTAGGAAAAACNNNAGATCGGAA |
| 12302_2 | TATGAGTGTGGAGTCGTTGCACGGACCAGGGATTAGGAAAAACNNNAGATCGGAA |
| 36640_1 | TATGAGTGTGGAGTCGTTGCCCCCTGATGGCCTACTTTTCATANNNAGATCGGAAGA |
| 36640_2 | TATGAGTGTGGAGTCGTTGCCCCCTGATGGCCTACTTTTCATANNNAGATCGGAAGA |
| 13444_1 | TATGAGTGTGGAGTCGTTGCATTTCCCTACATGATGTACATTTNNNAGATCGGAAGA |
| 13444_2 | TATGAGTGTGGAGTCGTTGCATTTCCCTACATGATGTACATTTNNNAGATCGGAAGA |
| 52655_1 | TATGAGTGTGGAGTCGTTGCGCATTCCTAGCCCTTGAAGTAGTNNNAGATCGGAAG |
| 52655_2 | TATGAGTGTGGAGTCGTTGCGCATTCCTAGCCCTTGAAGTAGTNNNAGATCGGAAG |
| 44514_1 | TATGAGTGTGGAGTCGTTGCACATTGTGTCTGGAACTTTGTCCNNNAGATCGGAAGA |
| 44514_2 | TATGAGTGTGGAGTCGTTGCACATTGTGTCTGGAACTTTGTCCNNNAGATCGGAAGA |
| 50281_1 | TATGAGTGTGGAGTCGTTGCAAATAGACTGTGGTGATGGTGGCNNNAGATCGGAA |
| 50281_2 | TATGAGTGTGGAGTCGTTGCAAATAGACTGTGGTGATGGTGGCNNNAGATCGGAA |
| 31452_1 | TATGAGTGTGGAGTCGTTGCCTTCACAGTAGACTGCAGCAAAGNNNAGATCGGAAG |
| 31452_2 | TATGAGTGTGGAGTCGTTGCCTTCACAGTAGACTGCAGCAAAGNNNAGATCGGAAG |
| 46514_1 | TATGAGTGTGGAGTCGTTGCGAGGTAGAGTCTCACTCCTCACTNNNAGATCGGAAG |
| 46514_2 | TATGAGTGTGGAGTCGTTGCGAGGTAGAGTCTCACTCCTCACTNNNAGATCGGAAG |
| 50761_1 | TATGAGTGTGGAGTCGTTGCCAAACTGTTCTGCTTGTCACCTTNNNAGATCGGAAGA |
| 50761_2 | TATGAGTGTGGAGTCGTTGCCAAACTGTTCTGCTTGTCACCTTNNNAGATCGGAAGA |
| 52045_1 | TATGAGTGTGGAGTCGTTGCGTTTCAGGAAAACAAGCTCAGGGNNNAGATCGGAAG |
| 52045_2 | TATGAGTGTGGAGTCGTTGCGTTTCAGGAAAACAAGCTCAGGGNNNAGATCGGAAG |
| 9133_1 | TATGAGTGTGGAGTCGTTGCCCAGGAGTTCCACATCTAGGGATNNNAGATCGGAAG |
| 9133_2 | TATGAGTGTGGAGTCGTTGCCCAGGAGTTCCACATCTAGGGATNNNAGATCGGAAG |
| 33755_1 | TATGAGTGTGGAGTCGTTGCATTAGGTCAGGAGATGGAGACCANNNAGATCGGAA |
| 33755_2 | TATGAGTGTGGAGTCGTTGCATTAGGTCAGGAGATGGAGACCANNNAGATCGGAA |
| 32862_1 | TATGAGTGTGGAGTCGTTGCAATCTCAGCTCACTGCAACCTCNNNAGATCGGAAGAG |
| 32862_2 | TATGAGTGTGGAGTCGTTGCAATCTCAGCTCACTGCAACCTCNNNAGATCGGAAGAG |
| 40532_1 | TATGAGTGTGGAGTCGTTGCTAAACTCTTTCCCCTTGACCCAGNNNAGATCGGAAGA |
| 40532_2 | TATGAGTGTGGAGTCGTTGCTAAACTCTTTCCCCTTGACCCAGNNNAGATCGGAAGA |

| | |
|---|---|
| 2462_1 | TATGAGTGTGGAGTCGTTGCGTGGCCCAGATGAGTTTAGTGTCNNNAGATCGGAA |
| 2462_2 | TATGAGTGTGGAGTCGTTGCGTGGCCCAGATGAGTTTAGTGTCNNNAGATCGGAA |
| 47218_1 | TATGAGTGTGGAGTCGTTGCCCACCATACCCGGCTAATTTTGNNNAGATCGGAAG |
| 47218_2 | TATGAGTGTGGAGTCGTTGCCCACCATACCCGGCTAATTTTGNNNAGATCGGAAG |
| 46878_1 | TATGAGTGTGGAGTCGTTGCAGAGGTTTCTCCATGTTGGTCAGNNNAGATCGGAAG |
| 46878_2 | TATGAGTGTGGAGTCGTTGCAGAGGTTTCTCCATGTTGGTCAGNNNAGATCGGAAG |
| 12502_1 | TATGAGTGTGGAGTCGTTGCTTATTTTTCCTTCCTTTTCTTGCNNNAGATCGGAAGAG |
| 12502_2 | TATGAGTGTGGAGTCGTTGCTTATTTTTCCTTCCTTTTCTTGCNNNAGATCGGAAGAG |
| 46236_1 | TATGAGTGTGGAGTCGTTGCCAGTCTGGTAATCACTGTCACCTNNNAGATCGGAAG |
| 46236_2 | TATGAGTGTGGAGTCGTTGCCAGTCTGGTAATCACTGTCACCTNNNAGATCGGAAG |
| 50627_1 | TATGAGTGTGGAGTCGTTGCTTGCACCCACTTTTCATGAACTCNNNAGATCGGAAGA |
| 50627_2 | TATGAGTGTGGAGTCGTTGCTTGCACCCACTTTTCATGAACTCNNNAGATCGGAAGA |
| 43922_1 | TATGAGTGTGGAGTCGTTGCTCTTTCTTGGGATGGAGTTTTGCNNNAGATCGGAAG |
| 43922_2 | TATGAGTGTGGAGTCGTTGCTCTTTCTTGGGATGGAGTTTTGCNNNAGATCGGAAG |
| 36803_1 | TATGAGTGTGGAGTCGTTGCTGTGGACACTGGTTGTTACAGATNNNAGATCGGAA |
| 36803_2 | TATGAGTGTGGAGTCGTTGCTGTGGACACTGGTTGTTACAGATNNNAGATCGGAA |
| 40626_1 | TATGAGTGTGGAGTCGTTGCAACATGAAGAGGCAACTGAGTCTNNNAGATCGGAA |
| 40626_2 | TATGAGTGTGGAGTCGTTGCAACATGAAGAGGCAACTGAGTCTNNNAGATCGGAA |
| 48578_1 | TATGAGTGTGGAGTCGTTGCGAGTAGCTGGGATTATAGGTGCCNNNAGATCGGAA |
| 48578_2 | TATGAGTGTGGAGTCGTTGCGAGTAGCTGGGATTATAGGTGCCNNNAGATCGGAA |
| 51530_1 | TATGAGTGTGGAGTCGTTGCATAATTTTCTGTCCCCCATCCCCNNNAGATCGGAAGA |
| 51530_2 | TATGAGTGTGGAGTCGTTGCATAATTTTCTGTCCCCCATCCCCNNNAGATCGGAAGA |
| 35165_1 | TATGAGTGTGGAGTCGTTGCCGCCTCTCTTTCTCCATTGGAAANNNAGATCGGAAGA |
| 35165_2 | TATGAGTGTGGAGTCGTTGCCGCCTCTCTTTCTCCATTGGAAANNNAGATCGGAAGA |
| 29971_1 | TATGAGTGTGGAGTCGTTGCTCGATCCATTCAACAACTGGGTANNNAGATCGGAAG |
| 29971_2 | TATGAGTGTGGAGTCGTTGCTCGATCCATTCAACAACTGGGTANNNAGATCGGAAG |
| 38080_1 | TATGAGTGTGGAGTCGTTGCATGGGACTCTTAGGACTTTGCAGNNNAGATCGGAAG |
| 38080_2 | TATGAGTGTGGAGTCGTTGCATGGGACTCTTAGGACTTTGCAGNNNAGATCGGAAG |
| 45493_1 | TATGAGTGTGGAGTCGTTGCCCAATGGGAAGGGAGAACACTTCNNNAGATCGGAA |
| 45493_2 | TATGAGTGTGGAGTCGTTGCCCAATGGGAAGGGAGAACACTTCNNNAGATCGGAA |
| 52464_1 | TATGAGTGTGGAGTCGTTGCTCACAGATAAGGAAGTTGAGCCCNNNAGATCGGAA |
| 52464_2 | TATGAGTGTGGAGTCGTTGCTCACAGATAAGGAAGTTGAGCCCNNNAGATCGGAA |
| 41253_1 | TATGAGTGTGGAGTCGTTGCTCCCACATTGGAAAAGCAGTACTNNNAGATCGGAAG |
| 41253_2 | TATGAGTGTGGAGTCGTTGCTCCCACATTGGAAAAGCAGTACTNNNAGATCGGAAG |
| 38614_1 | TATGAGTGTGGAGTCGTTGCCAACTGTAATCCCAGCACTTTGGNNNAGATCGGAAG |
| 38614_2 | TATGAGTGTGGAGTCGTTGCCAACTGTAATCCCAGCACTTTGGNNNAGATCGGAAG |
| 32464_1 | TATGAGTGTGGAGTCGTTGCCTGAAGTGAGTGGATCTCCTGAGNNNAGATCGGAA |
| 32464_2 | TATGAGTGTGGAGTCGTTGCCTGAAGTGAGTGGATCTCCTGAGNNNAGATCGGAA |
| 29484_1 | TATGAGTGTGGAGTCGTTGCGACACGCGGGCATTGATGACNNNAGATCGGAAGAG |
| 29484_2 | TATGAGTGTGGAGTCGTTGCGACACGCGGGCATTGATGACNNNAGATCGGAAGAG |
| 52891_1 | TATGAGTGTGGAGTCGTTGCCAAAGAGGAGAAGATGCCCAGAGNNNAGATCGGA |
| 52891_2 | TATGAGTGTGGAGTCGTTGCCAAAGAGGAGAAGATGCCCAGAGNNNAGATCGGA |
| 52176_1 | TATGAGTGTGGAGTCGTTGCCTCAAGTGATTCTCCTGCCTCAGNNNAGATCGGAAG |

| | |
|---|---|
| 52176_2 | TATGAGTGTGGAGTCGTTGCCTCAAGTGATTCTCCTGCCTCAGNNNAGATCGGAAG |
| 35957_1 | TATGAGTGTGGAGTCGTTGCGAACAGAGAGGTGAAGGACCATTNNNAGATCGGAA |
| 35957_2 | TATGAGTGTGGAGTCGTTGCGAACAGAGAGGTGAAGGACCATTNNNAGATCGGAA |
| 39861_1 | TATGAGTGTGGAGTCGTTGCAAATGTATAGAAGCACCAGGCCANNNAGATCGGAA |
| 39861_2 | TATGAGTGTGGAGTCGTTGCAAATGTATAGAAGCACCAGGCCANNNAGATCGGAA |
| 31354_1 | TATGAGTGTGGAGTCGTTGCGCAGAACTGGTGACACATTTACCNNNAGATCGGAAG |
| 31354_2 | TATGAGTGTGGAGTCGTTGCGCAGAACTGGTGACACATTTACCNNNAGATCGGAAG |
| 47286_1 | TATGAGTGTGGAGTCGTTGCGAAACAGGGTCTCACTCTGTTGCNNNAGATCGGAAG |
| 47286_2 | TATGAGTGTGGAGTCGTTGCGAAACAGGGTCTCACTCTGTTGCNNNAGATCGGAAG |
| 2467_1 | TATGAGTGTGGAGTCGTTGCGAACTGGGGAGTGGTGATGCAAGNNNAGATCGGA |
| 2467_2 | TATGAGTGTGGAGTCGTTGCGAACTGGGGAGTGGTGATGCAAGNNNAGATCGGA |
| 49376_1 | TATGAGTGTGGAGTCGTTGCCTGGGTTCAAGTGATTCTCCTGANNNAGATCGGAAG |
| 49376_2 | TATGAGTGTGGAGTCGTTGCCTGGGTTCAAGTGATTCTCCTGANNNAGATCGGAAG |
| 4463_1 | TATGAGTGTGGAGTCGTTGCCAAGGCCACAGCATTTTTGGAAGNNNAGATCGGAA |
| 4463_2 | TATGAGTGTGGAGTCGTTGCCAAGGCCACAGCATTTTTGGAAGNNNAGATCGGAA |
| 49947_1 | TATGAGTGTGGAGTCGTTGCCAAGTAGCTGGGACTACTGGAATNNNAGATCGGAA |
| 49947_2 | TATGAGTGTGGAGTCGTTGCCAAGTAGCTGGGACTACTGGAATNNNAGATCGGAA |
| 33600_1 | TATGAGTGTGGAGTCGTTGCGTTCTGGGTTCATGCCATTCTCNNNAGATCGGAAGA |
| 33600_2 | TATGAGTGTGGAGTCGTTGCGTTCTGGGTTCATGCCATTCTCNNNAGATCGGAAGA |
| 32734_1 | TATGAGTGTGGAGTCGTTGCTTTCCCTTCCCTTGAATCTAGGCNNNAGATCGGAAGA |
| 32734_2 | TATGAGTGTGGAGTCGTTGCTTTCCCTTCCCTTGAATCTAGGCNNNAGATCGGAAGA |
| 31912_1 | TATGAGTGTGGAGTCGTTGCTTTCTCCAGCCTGGTCTTTTACANNNAGATCGGAAGA |
| 31912_2 | TATGAGTGTGGAGTCGTTGCTTTCTCCAGCCTGGTCTTTTACANNNAGATCGGAAGA |
| 51916_1 | TATGAGTGTGGAGTCGTTGCGGGCTCCTCACTTCCCAGTANNNAGATCGGAAGAGC |
| 51916_2 | TATGAGTGTGGAGTCGTTGCGGGCTCCTCACTTCCCAGTANNNAGATCGGAAGAGC |
| 49854_1 | TATGAGTGTGGAGTCGTTGCAGTTGTTGCCTTCACACCAAATTNNNAGATCGGAAG |
| 49854_2 | TATGAGTGTGGAGTCGTTGCAGTTGTTGCCTTCACACCAAATTNNNAGATCGGAAG |
| 49217_1 | TATGAGTGTGGAGTCGTTGCTTCAAGTGATTCTCCTGCCTCAGNNNAGATCGGAAG |
| 49217_2 | TATGAGTGTGGAGTCGTTGCTTCAAGTGATTCTCCTGCCTCAGNNNAGATCGGAAG |
| 34007_1 | TATGAGTGTGGAGTCGTTGCTTTAGTAGAGACGGAGTTTCGCCNNNAGATCGGAA |
| 34007_2 | TATGAGTGTGGAGTCGTTGCTTTAGTAGAGACGGAGTTTCGCCNNNAGATCGGAA |
| 10720_1 | TATGAGTGTGGAGTCGTTGCAAAAGGGAGCAGGAGACAGGTAANNNAGATCGGA |
| 10720_2 | TATGAGTGTGGAGTCGTTGCAAAAGGGAGCAGGAGACAGGTAANNNAGATCGGA |
| 32258_1 | TATGAGTGTGGAGTCGTTGCGGAGCAATTTCTTACCTCCCTCANNNAGATCGGAAG |
| 32258_2 | TATGAGTGTGGAGTCGTTGCGGAGCAATTTCTTACCTCCCTCANNNAGATCGGAAG |
| 33103_1 | TATGAGTGTGGAGTCGTTGCGTTCAAGCGATTCTCCTACCTCANNNAGATCGGAAG |
| 33103_2 | TATGAGTGTGGAGTCGTTGCGTTCAAGCGATTCTCCTACCTCANNNAGATCGGAAG |
| 39261_1 | TATGAGTGTGGAGTCGTTGCTTGGCTAGTGCTGAAGAGGTTAGNNNAGATCGGAA |
| 39261_2 | TATGAGTGTGGAGTCGTTGCTTGGCTAGTGCTGAAGAGGTTAGNNNAGATCGGAA |
| 48469_1 | TATGAGTGTGGAGTCGTTGCGTTCAAGCAATTCTCTGCCTCAGNNNAGATCGGAAG |
| 48469_2 | TATGAGTGTGGAGTCGTTGCGTTCAAGCAATTCTCTGCCTCAGNNNAGATCGGAAG |
| 12838_1 | TATGAGTGTGGAGTCGTTGCGTATTCTTGGTTTCTGATGAAATNNNAGATCGGAAG |
| 12838_2 | TATGAGTGTGGAGTCGTTGCGTATTCTTGGTTTCTGATGAAATNNNAGATCGGAAG |

| | |
|---|---|
| 30166_1 | TATGAGTGTGGAGTCGTTGCAAGGCCATATGAGAATCCACGTTNNNAGATCGGAAG |
| 30166_2 | TATGAGTGTGGAGTCGTTGCAAGGCCATATGAGAATCCACGTTNNNAGATCGGAAG |
| 35258_1 | TATGAGTGTGGAGTCGTTGCATACAATCTCTGCCACCCATGTTNNNAGATCGGAAGA |
| 35258_2 | TATGAGTGTGGAGTCGTTGCATACAATCTCTGCCACCCATGTTNNNAGATCGGAAGA |
| 31391_1 | TATGAGTGTGGAGTCGTTGCATCACCACTTGTACTCTTCTGGGNNNAGATCGGAAG |
| 31391_2 | TATGAGTGTGGAGTCGTTGCATCACCACTTGTACTCTTCTGGGNNNAGATCGGAAG |
| 31270_1 | TATGAGTGTGGAGTCGTTGCCCTCCACCCATCTCTCTTTGAAANNNAGATCGGAAGA |
| 31270_2 | TATGAGTGTGGAGTCGTTGCCCTCCACCCATCTCTCTTTGAAANNNAGATCGGAAGA |
| 42646_1 | TATGAGTGTGGAGTCGTTGCCCTCTCAGGTTCAAGTGATCCTCNNNAGATCGGAAG |
| 42646_2 | TATGAGTGTGGAGTCGTTGCCCTCTCAGGTTCAAGTGATCCTCNNNAGATCGGAAG |
| 39810_1 | TATGAGTGTGGAGTCGTTGCAAGTATTTGCTAATAGCTTGGCAANNNAGATCGGAA |
| 39810_2 | TATGAGTGTGGAGTCGTTGCAAGTATTTGCTAATAGCTTGGCAANNNAGATCGGAA |
| 53144_1 | TATGAGTGTGGAGTCGTTGCTAGCTAACTTGGACCTCAGAGGANNNAGATCGGAA |
| 53144_2 | TATGAGTGTGGAGTCGTTGCTAGCTAACTTGGACCTCAGAGGANNNAGATCGGAA |
| 32529_1 | TATGAGTGTGGAGTCGTTGCTCCCTCTAGCCTCTAGAACTTCANNNAGATCGGAAGA |
| 32529_2 | TATGAGTGTGGAGTCGTTGCTCCCTCTAGCCTCTAGAACTTCANNNAGATCGGAAGA |
| 38872_1 | TATGAGTGTGGAGTCGTTGCTGCAAAAACTGTGGGAAAAGTGTNNNAGATCGGAA |
| 38872_2 | TATGAGTGTGGAGTCGTTGCTGCAAAAACTGTGGGAAAAGTGTNNNAGATCGGAA |
| 48590_1 | TATGAGTGTGGAGTCGTTGCGAGTTTCGATCTTGTTTCCCAGGNNNAGATCGGAAG |
| 48590_2 | TATGAGTGTGGAGTCGTTGCGAGTTTCGATCTTGTTTCCCAGGNNNAGATCGGAAG |
| 46470_1 | TATGAGTGTGGAGTCGTTGCCATCAGCTCACCACAACCTCTNNNAGATCGGAAGAG |
| 46470_2 | TATGAGTGTGGAGTCGTTGCCATCAGCTCACCACAACCTCTNNNAGATCGGAAGAG |
| 34331_1 | TATGAGTGTGGAGTCGTTGCCTGGCCAAAGGAGAATGTTCATCNNNAGATCGGAA |
| 34331_2 | TATGAGTGTGGAGTCGTTGCCTGGCCAAAGGAGAATGTTCATCNNNAGATCGGAA |
| 2244_1 | TATGAGTGTGGAGTCGTTGCGGGGGTGAACTCGCCATTTTATANNNAGATCGGAA |
| 2244_2 | TATGAGTGTGGAGTCGTTGCGGGGGTGAACTCGCCATTTTATANNNAGATCGGAA |
| 34013_1 | TATGAGTGTGGAGTCGTTGCCTCCTATTCTCAAGCCATCCTCCNNNAGATCGGAAGA |
| 34013_2 | TATGAGTGTGGAGTCGTTGCCTCCTATTCTCAAGCCATCCTCCNNNAGATCGGAAGA |
| 35947_1 | TATGAGTGTGGAGTCGTTGCTTGTGTACATTTTTTGGCTGGGCNNNAGATCGGAAG |
| 35947_2 | TATGAGTGTGGAGTCGTTGCTTGTGTACATTTTTTGGCTGGGCNNNAGATCGGAAG |
| 32796_1 | TATGAGTGTGGAGTCGTTGCGGCAAAATCTTAGCTCACTGCAANNNAGATCGGAAG |
| 32796_2 | TATGAGTGTGGAGTCGTTGCGGCAAAATCTTAGCTCACTGCAANNNAGATCGGAAG |
| 50152_1 | TATGAGTGTGGAGTCGTTGCTTCCATTCCCTCAATTCCAGGAGNNNAGATCGGAAG |
| 50152_2 | TATGAGTGTGGAGTCGTTGCTTCCATTCCCTCAATTCCAGGAGNNNAGATCGGAAG |
| 47156_1 | TATGAGTGTGGAGTCGTTGCGAGTGCAGTGGTGTGATCTCAGNNNAGATCGGAAG |
| 47156_2 | TATGAGTGTGGAGTCGTTGCGAGTGCAGTGGTGTGATCTCAGNNNAGATCGGAAG |
| 38764_1 | TATGAGTGTGGAGTCGTTGCAGAATCACTTGAACTCAGGAGGCNNNAGATCGGAA |
| 38764_2 | TATGAGTGTGGAGTCGTTGCAGAATCACTTGAACTCAGGAGGCNNNAGATCGGAA |
| 42414_1 | TATGAGTGTGGAGTCGTTGCTCTCTCTTGTCAGTCTCTACCCANNNAGATCGGAAGA |
| 42414_2 | TATGAGTGTGGAGTCGTTGCTCTCTCTTGTCAGTCTCTACCCANNNAGATCGGAAGA |
| 42545_1 | TATGAGTGTGGAGTCGTTGCATGATAATCGTCCACAAGGCTCANNNAGATCGGAAG |
| 42545_2 | TATGAGTGTGGAGTCGTTGCATGATAATCGTCCACAAGGCTCANNNAGATCGGAAG |
| 10754_1 | TATGAGTGTGGAGTCGTTGCGTTATGGAATTCAGCTGGGCTCTNNNAGATCGGAA |

| | |
|---|---|
| 10754_2 | TATGAGTGTGGAGTCGTTGCGTTATGGAATTCAGCTGGGCTCTNNNAGATCGGAAG |
| 46248_1 | TATGAGTGTGGAGTCGTTGCCTCACAAGTGAATGTACCCTGGANNNAGATCGGAAG |
| 46248_2 | TATGAGTGTGGAGTCGTTGCCTCACAAGTGAATGTACCCTGGANNNAGATCGGAAG |
| 49875_1 | TATGAGTGTGGAGTCGTTGCTTAGCACTTGCCATCTTGGTTTCNNNAGATCGGAAGA |
| 49875_2 | TATGAGTGTGGAGTCGTTGCTTAGCACTTGCCATCTTGGTTTCNNNAGATCGGAAGA |
| 41558_1 | TATGAGTGTGGAGTCGTTGCTGGATTATAGATGTGAGCCACCGNNNAGATCGGAAG |
| 41558_2 | TATGAGTGTGGAGTCGTTGCTGGATTATAGATGTGAGCCACCGNNNAGATCGGAAG |
| 37799_1 | TATGAGTGTGGAGTCGTTGCATGGATTGGAAGACAACACGGTANNNAGATCGGAA |
| 37799_2 | TATGAGTGTGGAGTCGTTGCATGGATTGGAAGACAACACGGTANNNAGATCGGAA |
| 29932_1 | TATGAGTGTGGAGTCGTTGCCTCACAGTGGATATGGAGATGGGNNNAGATCGGAA |
| 29932_2 | TATGAGTGTGGAGTCGTTGCCTCACAGTGGATATGGAGATGGGNNNAGATCGGAA |
| 42656_1 | TATGAGTGTGGAGTCGTTGCACTGAGGTAGGAGGATCACTTGANNNAGATCGGAA |
| 42656_2 | TATGAGTGTGGAGTCGTTGCACTGAGGTAGGAGGATCACTTGANNNAGATCGGAA |
| 3154_1 | TATGAGTGTGGAGTCGTTGCAGTCAATACTTGGGTGGTACCCANNNAGATCGGAAG |
| 3154_2 | TATGAGTGTGGAGTCGTTGCAGTCAATACTTGGGTGGTACCCANNNAGATCGGAAG |
| 31826_1 | TATGAGTGTGGAGTCGTTGCGTGCACAACCACACATACATAGGNNNAGATCGGAAG |
| 31826_2 | TATGAGTGTGGAGTCGTTGCGTGCACAACCACACATACATAGGNNNAGATCGGAAG |
| 40283_1 | TATGAGTGTGGAGTCGTTGCCCTGCTCTTGAACTACTCTAGTAANNNAGATCGGAAG |
| 40283_2 | TATGAGTGTGGAGTCGTTGCCCTGCTCTTGAACTACTCTAGTAANNNAGATCGGAAG |
| 45827_1 | TATGAGTGTGGAGTCGTTGCTGCCAAACACAACCTGATGATTCNNNAGATCGGAAGA |
| 45827_2 | TATGAGTGTGGAGTCGTTGCTGCCAAACACAACCTGATGATTCNNNAGATCGGAAGA |
| 1639_1 | TATGAGTGTGGAGTCGTTGCCCTGGAGTGCCTAGCATTCTTNNNAGATCGGAAGAG |
| 1639_2 | TATGAGTGTGGAGTCGTTGCCCTGGAGTGCCTAGCATTCTTNNNAGATCGGAAGAG |
| 43839_1 | TATGAGTGTGGAGTCGTTGCCCAGTGTCCGTGGATAAAAGGTANNNAGATCGGAAG |
| 43839_2 | TATGAGTGTGGAGTCGTTGCCCAGTGTCCGTGGATAAAAGGTANNNAGATCGGAAG |
| 35518_1 | TATGAGTGTGGAGTCGTTGCCTGAGTAGGTGGGATTACAGGTGNNNAGATCGGAA |
| 35518_2 | TATGAGTGTGGAGTCGTTGCCTGAGTAGGTGGGATTACAGGTGNNNAGATCGGAA |
| 32410_1 | TATGAGTGTGGAGTCGTTGCCCATCTCACACCAATCAGAATGGNNNAGATCGGAAGA |
| 32410_2 | TATGAGTGTGGAGTCGTTGCCCATCTCACACCAATCAGAATGGNNNAGATCGGAAGA |
| 47143_1 | TATGAGTGTGGAGTCGTTGCCCCAGGTATGAAACTCAGTACCCNNNAGATCGGAAG |
| 47143_2 | TATGAGTGTGGAGTCGTTGCCCCAGGTATGAAACTCAGTACCCNNNAGATCGGAAG |
| 37817_1 | TATGAGTGTGGAGTCGTTGCGGGTGAACATGAACATTGGGAAGNNNAGATCGGAA |
| 37817_2 | TATGAGTGTGGAGTCGTTGCGGGTGAACATGAACATTGGGAAGNNNAGATCGGAA |
| 32797_1 | TATGAGTGTGGAGTCGTTGCGATCTCGGCTCAGTATAAGCTCCNNNAGATCGGAAG |
| 32797_2 | TATGAGTGTGGAGTCGTTGCGATCTCGGCTCAGTATAAGCTCCNNNAGATCGGAAG |
| 35787_1 | TATGAGTGTGGAGTCGTTGCGATCTCAGCTCATCACAACCTCCNNNAGATCGGAAGA |
| 35787_2 | TATGAGTGTGGAGTCGTTGCGATCTCAGCTCATCACAACCTCCNNNAGATCGGAAGA |
| 45843_1 | TATGAGTGTGGAGTCGTTGCCCAAGTAGCTGAGATTACAGGCANNNAGATCGGAAG |
| 45843_2 | TATGAGTGTGGAGTCGTTGCCCAAGTAGCTGAGATTACAGGCANNNAGATCGGAAG |
| 29384_1 | TATGAGTGTGGAGTCGTTGCATCATAGAGGCACAGAGGAGGAANNNAGATCGGAA |
| 29384_2 | TATGAGTGTGGAGTCGTTGCATCATAGAGGCACAGAGGAGGAANNNAGATCGGAA |
| 29012_1 | TATGAGTGTGGAGTCGTTGCTGACAACTCCAAAATCTAGGGGTNNNAGATCGGAAG |
| 29012_2 | TATGAGTGTGGAGTCGTTGCTGACAACTCCAAAATCTAGGGGTNNNAGATCGGAAG |

| | |
|---|---|
| 49221_1 | TATGAGTGTGGAGTCGTTGCGTACAAGTGATTCTCCTGCCTCANNNAGATCGGAAG |
| 49221_2 | TATGAGTGTGGAGTCGTTGCGTACAAGTGATTCTCCTGCCTCANNNAGATCGGAAG |
| 51392_1 | TATGAGTGTGGAGTCGTTGCGGTTTCCTGGACAGTGTGTAGAANNNAGATCGGAA |
| 51392_2 | TATGAGTGTGGAGTCGTTGCGGTTTCCTGGACAGTGTGTAGAANNNAGATCGGAA |
| 44332_1 | TATGAGTGTGGAGTCGTTGCGAGCAGATGGCAACTTTGTCTCNNNAGATCGGAAGA |
| 44332_2 | TATGAGTGTGGAGTCGTTGCGAGCAGATGGCAACTTTGTCTCNNNAGATCGGAAGA |
| 35657_1 | TATGAGTGTGGAGTCGTTGCCTTCTGGGTCAAGCTCTTCTTCTNNNAGATCGGAAGA |
| 35657_2 | TATGAGTGTGGAGTCGTTGCCTTCTGGGTCAAGCTCTTCTTCTNNNAGATCGGAAGA |
| 36824_1 | TATGAGTGTGGAGTCGTTGCTTCAGAAGTGATTCTCAGCCACANNNAGATCGGAAG |
| 36824_2 | TATGAGTGTGGAGTCGTTGCTTCAGAAGTGATTCTCAGCCACANNNAGATCGGAAG |
| 38463_1 | TATGAGTGTGGAGTCGTTGCCAGAGCAAGACTCCATCTCGAAANNNAGATCGGAAG |
| 38463_2 | TATGAGTGTGGAGTCGTTGCCAGAGCAAGACTCCATCTCGAAANNNAGATCGGAAG |
| 32815_1 | TATGAGTGTGGAGTCGTTGCTTCAAGTGATTCTCCTGCCTCAGNNNAGATCGGAAG |
| 32815_2 | TATGAGTGTGGAGTCGTTGCTTCAAGTGATTCTCCTGCCTCAGNNNAGATCGGAAG |
| 13696_1 | TATGAGTGTGGAGTCGTTGCTACCAAATTTCAGAACACAAGAANNNAGATCGGAAG |
| 13696_2 | TATGAGTGTGGAGTCGTTGCTACCAAATTTCAGAACACAAGAANNNAGATCGGAAG |
| 29762_1 | TATGAGTGTGGAGTCGTTGCTGTCTGCTTACTTGGTTGACACTNNNAGATCGGAAG |
| 29762_2 | TATGAGTGTGGAGTCGTTGCTGTCTGCTTACTTGGTTGACACTNNNAGATCGGAAG |
| 37275_1 | TATGAGTGTGGAGTCGTTGCTCCCAGCTCCTAACTCAGTCTAANNNAGATCGGAAGA |
| 37275_2 | TATGAGTGTGGAGTCGTTGCTCCCAGCTCCTAACTCAGTCTAANNNAGATCGGAAGA |
| 43941_1 | TATGAGTGTGGAGTCGTTGCAATTGAGGTCACAGGGCAAACTANNNAGATCGGAA |
| 43941_2 | TATGAGTGTGGAGTCGTTGCAATTGAGGTCACAGGGCAAACTANNNAGATCGGAA |
| 43397_1 | TATGAGTGTGGAGTCGTTGCCTGAGTTCAAGCAATTCTCCCACNNNAGATCGGAAG |
| 43397_2 | TATGAGTGTGGAGTCGTTGCCTGAGTTCAAGCAATTCTCCCACNNNAGATCGGAAG |
| 29948_1 | TATGAGTGTGGAGTCGTTGCGCAATGGCTATTGATACCTCTAACCNNNAGATCGGA |
| 29948_2 | TATGAGTGTGGAGTCGTTGCGCAATGGCTATTGATACCTCTAACCNNNAGATCGGA |
| 41100_1 | TATGAGTGTGGAGTCGTTGCGACTAACTCCCCACCACTCTCCNNNAGATCGGAAGAG |
| 41100_2 | TATGAGTGTGGAGTCGTTGCGACTAACTCCCCACCACTCTCCNNNAGATCGGAAGAG |
| 45250_1 | TATGAGTGTGGAGTCGTTGCTGCTGAAATGGAGTTTGAGGAGANNNAGATCGGAA |
| 45250_2 | TATGAGTGTGGAGTCGTTGCTGCTGAAATGGAGTTTGAGGAGANNNAGATCGGAA |
| 53212_1 | TATGAGTGTGGAGTCGTTGCATCTACCACTCAGGAAGGTGAGANNNAGATCGGAA |
| 53212_2 | TATGAGTGTGGAGTCGTTGCATCTACCACTCAGGAAGGTGAGANNNAGATCGGAA |
| 43141_1 | TATGAGTGTGGAGTCGTTGCACTCATACAGGGTGCAGAGAAAANNNAGATCGGAA |
| 43141_2 | TATGAGTGTGGAGTCGTTGCACTCATACAGGGTGCAGAGAAAANNNAGATCGGAA |
| 50084_1 | TATGAGTGTGGAGTCGTTGCCCGAAACCTGGAAAAGTAGGGTANNNAGATCGGAA |
| 50084_2 | TATGAGTGTGGAGTCGTTGCCCGAAACCTGGAAAAGTAGGGTANNNAGATCGGAA |
| 44851_1 | TATGAGTGTGGAGTCGTTGCTAGCAAGAAGAGCAGAGTTGTGTNNNAGATCGGAA |
| 44851_2 | TATGAGTGTGGAGTCGTTGCTAGCAAGAAGAGCAGAGTTGTGTNNNAGATCGGAA |
| 30131_1 | TATGAGTGTGGAGTCGTTGCTCTGATGAAGCCCCTGATATGTGNNNAGATCGGAAG |
| 30131_2 | TATGAGTGTGGAGTCGTTGCTCTGATGAAGCCCCTGATATGTGNNNAGATCGGAAG |
| 50675_1 | TATGAGTGTGGAGTCGTTGCTGTAAGAGAAACGGGAAGGTCACNNNAGATCGGAA |
| 50675_2 | TATGAGTGTGGAGTCGTTGCTGTAAGAGAAACGGGAAGGTCACNNNAGATCGGAA |
| 41911_1 | TATGAGTGTGGAGTCGTTGCCTCCTGGGTTCTCATGACTCTTCNNNAGATCGGAAGA |

| | |
|---|---|
| 41911_2 | TATGAGTGTGGAGTCGTTGCCTCCTGGGTTCTCATGACTCTTCNNNAGATCGGAAG |
| 43007_1 | TATGAGTGTGGAGTCGTTGCACACACACATATATACACACACTCGNNNAGATCGGA |
| 43007_2 | TATGAGTGTGGAGTCGTTGCACACACACATATATACACACACTCGNNNAGATCGGA |
| 33618_1 | TATGAGTGTGGAGTCGTTGCACTGGGTTCAAGTGATTCTCCTGNNNAGATCGGAAG |
| 33618_2 | TATGAGTGTGGAGTCGTTGCACTGGGTTCAAGTGATTCTCCTGNNNAGATCGGAAG |
| 51097_1 | TATGAGTGTGGAGTCGTTGCCAGCCACTGAGTGTTTACTAGGTNNNAGATCGGAAG |
| 51097_2 | TATGAGTGTGGAGTCGTTGCCAGCCACTGAGTGTTTACTAGGTNNNAGATCGGAAG |
| 30403_1 | TATGAGTGTGGAGTCGTTGCACTCTGGTGCATTCTTCAGTATGTNNNAGATCGGAA |
| 30403_2 | TATGAGTGTGGAGTCGTTGCACTCTGGTGCATTCTTCAGTATGTNNNAGATCGGAA |
| 41772_1 | TATGAGTGTGGAGTCGTTGCGATCTTGGCTCACTGCAAGCTCNNNAGATCGGAAGA |
| 41772_2 | TATGAGTGTGGAGTCGTTGCGATCTTGGCTCACTGCAAGCTCNNNAGATCGGAAGA |
| 50277_1 | TATGAGTGTGGAGTCGTTGCCAGGTTTTTCAGGTCTGCCATTCNNNAGATCGGAAG |
| 50277_2 | TATGAGTGTGGAGTCGTTGCCAGGTTTTTCAGGTCTGCCATTCNNNAGATCGGAAG |
| 33668_1 | TATGAGTGTGGAGTCGTTGCCTCCTGAGTAGCTGGGATTACAGNNNAGATCGGAA |
| 33668_2 | TATGAGTGTGGAGTCGTTGCCTCCTGAGTAGCTGGGATTACAGNNNAGATCGGAA |
| 29380_1 | TATGAGTGTGGAGTCGTTGCAATCTTTACAGACCACTGGCAGTNNNAGATCGGAAG |
| 29380_2 | TATGAGTGTGGAGTCGTTGCAATCTTTACAGACCACTGGCAGTNNNAGATCGGAAG |
| 466_1 | TATGAGTGTGGAGTCGTTGCAGTGTTTCAGACTTCGGCTCTGTNNNAGATCGGAAG |
| 466_2 | TATGAGTGTGGAGTCGTTGCAGTGTTTCAGACTTCGGCTCTGTNNNAGATCGGAAG |
| 44509_1 | TATGAGTGTGGAGTCGTTGCATATTAGGGGGTCATGGGTTTGGNNNAGATCGGAA |
| 44509_2 | TATGAGTGTGGAGTCGTTGCATATTAGGGGGTCATGGGTTTGGNNNAGATCGGAA |
| 32340_1 | TATGAGTGTGGAGTCGTTGCGGCAGGACTGACAATAAATCAGCNNNAGATCGGAA |
| 32340_2 | TATGAGTGTGGAGTCGTTGCGGCAGGACTGACAATAAATCAGCNNNAGATCGGAA |
| 31723_1 | TATGAGTGTGGAGTCGTTGCACTCTCCTCACCCCACTACAATANNNAGATCGGAAGA |
| 31723_2 | TATGAGTGTGGAGTCGTTGCACTCTCCTCACCCCACTACAATANNNAGATCGGAAGA |
| 3083_1 | TATGAGTGTGGAGTCGTTGCGGTCGCTAAGGTCAGGTTGTATCNNNAGATCGGAA |
| 3083_2 | TATGAGTGTGGAGTCGTTGCGGTCGCTAAGGTCAGGTTGTATCNNNAGATCGGAA |
| 48097_1 | TATGAGTGTGGAGTCGTTGCGCCTCCTGAGTAGTTGGGACTANNNAGATCGGAAG |
| 48097_2 | TATGAGTGTGGAGTCGTTGCGCCTCCTGAGTAGTTGGGACTANNNAGATCGGAAG |
| 48595_1 | TATGAGTGTGGAGTCGTTGCCACGCCCGGCTAATTTTTGTATTNNNAGATCGGAAG |
| 48595_2 | TATGAGTGTGGAGTCGTTGCCACGCCCGGCTAATTTTTGTATTNNNAGATCGGAAG |
| 8113_1 | TATGAGTGTGGAGTCGTTGCACTTGTTTCTGGACTTTGCCTTGNNNAGATCGGAAG |
| 8113_2 | TATGAGTGTGGAGTCGTTGCACTTGTTTCTGGACTTTGCCTTGNNNAGATCGGAAG |
| 965_1 | TATGAGTGTGGAGTCGTTGCGTCAGGAGTTCTCCATATTGGTCNNNAGATCGGAAG |
| 965_2 | TATGAGTGTGGAGTCGTTGCGTCAGGAGTTCTCCATATTGGTCNNNAGATCGGAAG |
| 50209_1 | TATGAGTGTGGAGTCGTTGCCCCAGAGGATTTCTTGACACTCANNNAGATCGGAAG |
| 50209_2 | TATGAGTGTGGAGTCGTTGCCCCAGAGGATTTCTTGACACTCANNNAGATCGGAAG |
| 51221_1 | TATGAGTGTGGAGTCGTTGCATCTGAAGACCTCCAGTATCCCTNNNAGATCGGAAG |
| 51221_2 | TATGAGTGTGGAGTCGTTGCATCTGAAGACCTCCAGTATCCCTNNNAGATCGGAAG |
| 32507_1 | TATGAGTGTGGAGTCGTTGCAAACCTCACGTCACAGCAAAAATNNNAGATCGGAAG |
| 32507_2 | TATGAGTGTGGAGTCGTTGCAAACCTCACGTCACAGCAAAAATNNNAGATCGGAAG |
| 50803_1 | TATGAGTGTGGAGTCGTTGCCAGAAGTGTTCAGACGCCTAAGANNNAGATCGGAA |
| 50803_2 | TATGAGTGTGGAGTCGTTGCCAGAAGTGTTCAGACGCCTAAGANNNAGATCGGAA |

| | |
|---|---|
| 42212_1 | TATGAGTGTGGAGTCGTTGCGACAACCTGATCTAACACAGCCTNNNAGATCGGAAG |
| 42212_2 | TATGAGTGTGGAGTCGTTGCGACAACCTGATCTAACACAGCCTNNNAGATCGGAAG |
| 50007_1 | TATGAGTGTGGAGTCGTTGCTCTCTTGTTCCTGATGACTGTGGNNNAGATCGGAAG |
| 50007_2 | TATGAGTGTGGAGTCGTTGCTCTCTTGTTCCTGATGACTGTGGNNNAGATCGGAAG |
| 50744_1 | TATGAGTGTGGAGTCGTTGCAACGATCTAGTGCAAGGAAAAGCNNNAGATCGGAA |
| 50744_2 | TATGAGTGTGGAGTCGTTGCAACGATCTAGTGCAAGGAAAAGCNNNAGATCGGAA |
| 35474_1 | TATGAGTGTGGAGTCGTTGCGCCTCTTGGGTATCTGTGACTACNNNAGATCGGAAG |
| 35474_2 | TATGAGTGTGGAGTCGTTGCGCCTCTTGGGTATCTGTGACTACNNNAGATCGGAAG |
| 47029_1 | TATGAGTGTGGAGTCGTTGCTACTCTATCACCCAGGTTGGAGTNNNAGATCGGAAG |
| 47029_2 | TATGAGTGTGGAGTCGTTGCTACTCTATCACCCAGGTTGGAGTNNNAGATCGGAAG |
| 32294_1 | TATGAGTGTGGAGTCGTTGCCAGCTCACTTGGGAGATCTGAAANNNAGATCGGAAG |
| 32294_2 | TATGAGTGTGGAGTCGTTGCCAGCTCACTTGGGAGATCTGAAANNNAGATCGGAAG |
| 35755_1 | TATGAGTGTGGAGTCGTTGCATGACAGACATTGAGGAAAGCCANNNAGATCGGAA |
| 35755_2 | TATGAGTGTGGAGTCGTTGCATGACAGACATTGAGGAAAGCCANNNAGATCGGAA |
| 51033_1 | TATGAGTGTGGAGTCGTTGCAGTTTCATGCAAGCCTCTTCCTANNNAGATCGGAAG |
| 51033_2 | TATGAGTGTGGAGTCGTTGCAGTTTCATGCAAGCCTCTTCCTANNNAGATCGGAAG |
| 40970_1 | TATGAGTGTGGAGTCGTTGCGAAAGAAAAATGGACTGGGGTGGNNNAGATCGGA |
| 40970_2 | TATGAGTGTGGAGTCGTTGCGAAAGAAAAATGGACTGGGGTGGNNNAGATCGGA |
| 43005_1 | TATGAGTGTGGAGTCGTTGCGTGTAACAGAGCGAGACCCTATCNNNAGATCGGAAG |
| 43005_2 | TATGAGTGTGGAGTCGTTGCGTGTAACAGAGCGAGACCCTATCNNNAGATCGGAAG |
| 4716_1 | TATGAGTGTGGAGTCGTTGCTTCTTGTCTGTTGACCTGTTGATAGANNNAGATCGG |
| 4716_2 | TATGAGTGTGGAGTCGTTGCTTCTTGTCTGTTGACCTGTTGATAGANNNAGATCGG |
| 37845_1 | TATGAGTGTGGAGTCGTTGCGTTGCAGTGAGTTGAGAACACACNNNAGATCGGAA |
| 37845_2 | TATGAGTGTGGAGTCGTTGCGTTGCAGTGAGTTGAGAACACACNNNAGATCGGAA |
| 35828_1 | TATGAGTGTGGAGTCGTTGCTCCATACTGTTTTCCATAGAGGCTNNNAGATCGGAAG |
| 35828_2 | TATGAGTGTGGAGTCGTTGCTCCATACTGTTTTCCATAGAGGCTNNNAGATCGGAAG |
| 51248_1 | TATGAGTGTGGAGTCGTTGCGGAGGGGGAGAAGCTAGGTATATNNNAGATCGGA |
| 51248_2 | TATGAGTGTGGAGTCGTTGCGGAGGGGGAGAAGCTAGGTATATNNNAGATCGGA |
| 36652_1 | TATGAGTGTGGAGTCGTTGCGGCAGTGAGTCTCTAGCATGAATNNNAGATCGGAAG |
| 36652_2 | TATGAGTGTGGAGTCGTTGCGGCAGTGAGTCTCTAGCATGAATNNNAGATCGGAAG |
| 29587_1 | TATGAGTGTGGAGTCGTTGCGGGAATTCATCAGCTTCGTCAAGNNNAGATCGGAAG |
| 29587_2 | TATGAGTGTGGAGTCGTTGCGGGAATTCATCAGCTTCGTCAAGNNNAGATCGGAAG |
| 39513_1 | TATGAGTGTGGAGTCGTTGCCATTTGACATGTGGCTGGTGTAANNNAGATCGGAAG |
| 39513_2 | TATGAGTGTGGAGTCGTTGCCATTTGACATGTGGCTGGTGTAANNNAGATCGGAAG |
| 50387_1 | TATGAGTGTGGAGTCGTTGCCAGCCTGTGAAGGGAAGGAGNNNAGATCGGAAGA |
| 50387_2 | TATGAGTGTGGAGTCGTTGCCAGCCTGTGAAGGGAAGGAGNNNAGATCGGAAGA |
| 37273_1 | TATGAGTGTGGAGTCGTTGCCTTGGTGGTGCATGTTTGTAGTCNNNAGATCGGAAG |
| 37273_2 | TATGAGTGTGGAGTCGTTGCCTTGGTGGTGCATGTTTGTAGTCNNNAGATCGGAAG |
| 31012_1 | TATGAGTGTGGAGTCGTTGCTCTATCCAGTGGCTTGAGGAGTANNNAGATCGGAAG |
| 31012_2 | TATGAGTGTGGAGTCGTTGCTCTATCCAGTGGCTTGAGGAGTANNNAGATCGGAAG |
| 44046_1 | TATGAGTGTGGAGTCGTTGCCAACAGGGCAACATAGAGAGACTNNNAGATCGGAA |
| 44046_2 | TATGAGTGTGGAGTCGTTGCCAACAGGGCAACATAGAGAGACTNNNAGATCGGAA |
| 40571_1 | TATGAGTGTGGAGTCGTTGCCTTCACAGTGGTTAGGAGGACTGNNNAGATCGGAA |

| | |
|---|---|
| 40571_2 | TATGAGTGTGGAGTCGTTGCCTTCACAGTGGTTAGGAGGACTGNNNAGATCGGAA |
| 23524_1 | TATGAGTGTGGAGTCGTTGCATGTGTTTCTTTGACAGGTAANNNAGATCGGAAGAG |
| 23524_2 | TATGAGTGTGGAGTCGTTGCATGTGTTTCTTTGACAGGTAANNNAGATCGGAAGAG |
| 37816_1 | TATGAGTGTGGAGTCGTTGCCCCATGATACTGAGACCATCACCNNNAGATCGGAAG |
| 37816_2 | TATGAGTGTGGAGTCGTTGCCCCATGATACTGAGACCATCACCNNNAGATCGGAAG |
| 36629_1 | TATGAGTGTGGAGTCGTTGCAAACAAACAACTTGGCCTGCAATNNNAGATCGGAAG |
| 36629_2 | TATGAGTGTGGAGTCGTTGCAAACAAACAACTTGGCCTGCAATNNNAGATCGGAAG |
| 10904_1 | TATGAGTGTGGAGTCGTTGCAAGTCTGCCACAAGGACAAAGAGNNNAGATCGGAA |
| 10904_2 | TATGAGTGTGGAGTCGTTGCAAGTCTGCCACAAGGACAAAGAGNNNAGATCGGAA |
| 49307_1 | TATGAGTGTGGAGTCGTTGCCTCAGCTCCATCCCTTACCAAAGNNNAGATCGGAAGA |
| 49307_2 | TATGAGTGTGGAGTCGTTGCCTCAGCTCCATCCCTTACCAAAGNNNAGATCGGAAGA |
| 43802_1 | TATGAGTGTGGAGTCGTTGCTGGATTCCAATGCTCTCTGTTAGTNNNAGATCGGAA |
| 43802_2 | TATGAGTGTGGAGTCGTTGCTGGATTCCAATGCTCTCTGTTAGTNNNAGATCGGAA |
| 36419_1 | TATGAGTGTGGAGTCGTTGCTGAGATCTCAATTGCCAGCTCATNNNAGATCGGAAG |
| 36419_2 | TATGAGTGTGGAGTCGTTGCTGAGATCTCAATTGCCAGCTCATNNNAGATCGGAAG |
| 9068_1 | TATGAGTGTGGAGTCGTTGCGCACCACATAATGACATTCCCCTNNNAGATCGGAAGA |
| 9068_2 | TATGAGTGTGGAGTCGTTGCGCACCACATAATGACATTCCCCTNNNAGATCGGAAGA |
| 47848_1 | TATGAGTGTGGAGTCGTTGCGGTTCAAGTGATTCTCCTGCCTANNNAGATCGGAAG |
| 47848_2 | TATGAGTGTGGAGTCGTTGCGGTTCAAGTGATTCTCCTGCCTANNNAGATCGGAAG |
| 46295_1 | TATGAGTGTGGAGTCGTTGCAGTGGTGCAATCATGGCTTACTANNNAGATCGGAAG |
| 46295_2 | TATGAGTGTGGAGTCGTTGCAGTGGTGCAATCATGGCTTACTANNNAGATCGGAAG |
| 8256_1 | TATGAGTGTGGAGTCGTTGCGAGTTTCATTCACTCTGTCGCCAANNNAGATCGGAA |
| 8256_2 | TATGAGTGTGGAGTCGTTGCGAGTTTCATTCACTCTGTCGCCAANNNAGATCGGAA |
| 28891_1 | TATGAGTGTGGAGTCGTTGCATGTCACTCTGGTAGTTGTCACCNNNAGATCGGAAG |
| 28891_2 | TATGAGTGTGGAGTCGTTGCATGTCACTCTGGTAGTTGTCACCNNNAGATCGGAAG |
| 50601_1 | TATGAGTGTGGAGTCGTTGCGGGTAACTAAGGCCTAGTGACAGNNNAGATCGGAA |
| 50601_2 | TATGAGTGTGGAGTCGTTGCGGGTAACTAAGGCCTAGTGACAGNNNAGATCGGAA |
| 46592_1 | TATGAGTGTGGAGTCGTTGCGCTGTCCAAGTAGCTGAGACTACNNNAGATCGGAAG |
| 46592_2 | TATGAGTGTGGAGTCGTTGCGCTGTCCAAGTAGCTGAGACTACNNNAGATCGGAAG |
| 12635_1 | TATGAGTGTGGAGTCGTTGCATTATGAATAAATGAGGAAGGGCNNNAGATCGGAA |
| 12635_2 | TATGAGTGTGGAGTCGTTGCATTATGAATAAATGAGGAAGGGCNNNAGATCGGAA |
| 44738_1 | TATGAGTGTGGAGTCGTTGCTCCTCCGAATCATTTCCTTCCTCNNNAGATCGGAAGA |
| 44738_2 | TATGAGTGTGGAGTCGTTGCTCCTCCGAATCATTTCCTTCCTCNNNAGATCGGAAGA |
| 2820_1 | TATGAGTGTGGAGTCGTTGCTCTCTCCCACTCAGTCTCTTTCTNNNAGATCGGAAGA |
| 2820_2 | TATGAGTGTGGAGTCGTTGCTCTCTCCCACTCAGTCTCTTTCTNNNAGATCGGAAGA |
| 48582_1 | TATGAGTGTGGAGTCGTTGCAATCTCAGTTCACTGCAACCTCTNNNAGATCGGAAGA |
| 48582_2 | TATGAGTGTGGAGTCGTTGCAATCTCAGTTCACTGCAACCTCTNNNAGATCGGAAGA |
| 41574_1 | TATGAGTGTGGAGTCGTTGCCTGTGTTCAAGTGATTCTCCTGCNNNAGATCGGAAG |
| 41574_2 | TATGAGTGTGGAGTCGTTGCCTGTGTTCAAGTGATTCTCCTGCNNNAGATCGGAAG |
| 10134_1 | TATGAGTGTGGAGTCGTTGCACCCTGTGCATTTGAGAACACTTNNNAGATCGGAAG |
| 10134_2 | TATGAGTGTGGAGTCGTTGCACCCTGTGCATTTGAGAACACTTNNNAGATCGGAAG |
| 13504_1 | TATGAGTGTGGAGTCGTTGCCTTATGGTCAAATAACGACTTGNNNAGATCGGAAGA |
| 13504_2 | TATGAGTGTGGAGTCGTTGCCTTATGGTCAAATAACGACTTGNNNAGATCGGAAGA |

| | |
|---|---|
| 36616_1 | TATGAGTGTGGAGTCGTTGCTATCTTGTGAGAGCAGAGGGAGANNNAGATCGGAA |
| 36616_2 | TATGAGTGTGGAGTCGTTGCTATCTTGTGAGAGCAGAGGGAGANNNAGATCGGAA |
| 31621_1 | TATGAGTGTGGAGTCGTTGCCCCCGATACAATCATAGCTGGAGNNNAGATCGGAAG |
| 31621_2 | TATGAGTGTGGAGTCGTTGCCCCCGATACAATCATAGCTGGAGNNNAGATCGGAAG |
| 42637_1 | TATGAGTGTGGAGTCGTTGCACTGAATATGCTTTGGGGAGGTTNNNAGATCGGAA |
| 42637_2 | TATGAGTGTGGAGTCGTTGCACTGAATATGCTTTGGGGAGGTTNNNAGATCGGAA |
| 37338_1 | TATGAGTGTGGAGTCGTTGCCAGGACCAGATGTATTCACAGCTNNNAGATCGGAAG |
| 37338_2 | TATGAGTGTGGAGTCGTTGCCAGGACCAGATGTATTCACAGCTNNNAGATCGGAAG |
| 31852_1 | TATGAGTGTGGAGTCGTTGCAAATTCAAGCCAGCTGCAGAAATNNNAGATCGGAAG |
| 31852_2 | TATGAGTGTGGAGTCGTTGCAAATTCAAGCCAGCTGCAGAAATNNNAGATCGGAAG |
| 13667_1 | TATGAGTGTGGAGTCGTTGCTTTTTTCTTTTTCTTTGAGACTGNNNAGATCGGAAGA |
| 13667_2 | TATGAGTGTGGAGTCGTTGCTTTTTTCTTTTTCTTTGAGACTGNNNAGATCGGAAGA |
| 44030_1 | TATGAGTGTGGAGTCGTTGCTTCACTGGAATCCTCAAGGTAGCNNNAGATCGGAAG |
| 44030_2 | TATGAGTGTGGAGTCGTTGCTTCACTGGAATCCTCAAGGTAGCNNNAGATCGGAAG |
| 41898_1 | TATGAGTGTGGAGTCGTTGCGAGTAGCTGAGACTACAGGCATGNNNAGATCGGAA |
| 41898_2 | TATGAGTGTGGAGTCGTTGCGAGTAGCTGAGACTACAGGCATGNNNAGATCGGAA |
| 35478_1 | TATGAGTGTGGAGTCGTTGCCTCCTTTTGCTGTAGGGCACTAGNNNAGATCGGAAG |
| 35478_2 | TATGAGTGTGGAGTCGTTGCCTCCTTTTGCTGTAGGGCACTAGNNNAGATCGGAAG |
| 31745_1 | TATGAGTGTGGAGTCGTTGCGCAGATGGCTTCCTTGAAACATTNNNAGATCGGAAG |
| 31745_2 | TATGAGTGTGGAGTCGTTGCGCAGATGGCTTCCTTGAAACATTNNNAGATCGGAAG |
| 35924_1 | TATGAGTGTGGAGTCGTTGCAATGAGATTTCTGAGCAGCTGGANNNAGATCGGAA |
| 35924_2 | TATGAGTGTGGAGTCGTTGCAATGAGATTTCTGAGCAGCTGGANNNAGATCGGAA |
| 43191_1 | TATGAGTGTGGAGTCGTTGCTCAGTTCAAGCAATTCTTCAGCCNNNAGATCGGAAGA |
| 43191_2 | TATGAGTGTGGAGTCGTTGCTCAGTTCAAGCAATTCTTCAGCCNNNAGATCGGAAGA |
| 47559_1 | TATGAGTGTGGAGTCGTTGCCCTCCAGAGTAGCTAGGACTACANNNAGATCGGAAG |
| 47559_2 | TATGAGTGTGGAGTCGTTGCCCTCCAGAGTAGCTAGGACTACANNNAGATCGGAAG |
| 47760_1 | TATGAGTGTGGAGTCGTTGCTTGAAGTGATTCTCCCGTCTCAGNNNAGATCGGAAG |
| 47760_2 | TATGAGTGTGGAGTCGTTGCTTGAAGTGATTCTCCCGTCTCAGNNNAGATCGGAAG |
| 50964_1 | TATGAGTGTGGAGTCGTTGCAGGAAGACCAGAAAGAACAAGCANNNAGATCGGAA |
| 50964_2 | TATGAGTGTGGAGTCGTTGCAGGAAGACCAGAAAGAACAAGCANNNAGATCGGAA |
| 48046_1 | TATGAGTGTGGAGTCGTTGCTGTGCAAGAAAGAGACCTGGAATNNNAGATCGGAA |
| 48046_2 | TATGAGTGTGGAGTCGTTGCTGTGCAAGAAAGAGACCTGGAATNNNAGATCGGAA |
| 46729_1 | TATGAGTGTGGAGTCGTTGCGTCTCCAGGGTTCAAGTGATTCTNNNAGATCGGAAG |
| 46729_2 | TATGAGTGTGGAGTCGTTGCGTCTCCAGGGTTCAAGTGATTCTNNNAGATCGGAAG |
| 32219_1 | TATGAGTGTGGAGTCGTTGCCCACTGTAACCACTACCTGTCTTNNNAGATCGGAAGA |
| 32219_2 | TATGAGTGTGGAGTCGTTGCCCACTGTAACCACTACCTGTCTTNNNAGATCGGAAGA |
| 53415_1 | TATGAGTGTGGAGTCGTTGCAGAGAGACACAGGTAGAGACACANNNAGATCGGAA |
| 53415_2 | TATGAGTGTGGAGTCGTTGCAGAGAGACACAGGTAGAGACACANNNAGATCGGAA |
| 2624_1 | TATGAGTGTGGAGTCGTTGCGGGGCACAAGGAAATGTTTTCTGNNNAGATCGGAA |
| 2624_2 | TATGAGTGTGGAGTCGTTGCGGGGCACAAGGAAATGTTTTCTGNNNAGATCGGAA |
| 49889_1 | TATGAGTGTGGAGTCGTTGCAGCAAACCTACACAAAATGAACGANNNAGATCGGAA |
| 49889_2 | TATGAGTGTGGAGTCGTTGCAGCAAACCTACACAAAATGAACGANNNAGATCGGAA |
| 48631_1 | TATGAGTGTGGAGTCGTTGCCTAGTCCAGGCCATTTCACTCTTNNNAGATCGGAAGA |

| | |
|---|---|
| 48631_2 | TATGAGTGTGGAGTCGTTGCCTAGTCCAGGCCATTTCACTCTTNNNAGATCGGAAGA |
| 36460_1 | TATGAGTGTGGAGTCGTTGCAATATTACGCATCCTGTCCTGGCNNNAGATCGGAAG |
| 36460_2 | TATGAGTGTGGAGTCGTTGCAATATTACGCATCCTGTCCTGGCNNNAGATCGGAAG |
| 29549_1 | TATGAGTGTGGAGTCGTTGCCTTCTCGAAGTCCTGGAACACCNNNAGATCGGAAGA |
| 29549_2 | TATGAGTGTGGAGTCGTTGCCTTCTCGAAGTCCTGGAACACCNNNAGATCGGAAGA |
| 47884_1 | TATGAGTGTGGAGTCGTTGCGCCTCCCAAGTAGCTAGGATTACNNNAGATCGGAAG |
| 47884_2 | TATGAGTGTGGAGTCGTTGCGCCTCCCAAGTAGCTAGGATTACNNNAGATCGGAAG |
| 38127_1 | TATGAGTGTGGAGTCGTTGCAGGGATGGATTATTCATGCCTCCNNNAGATCGGAAG |
| 38127_2 | TATGAGTGTGGAGTCGTTGCAGGGATGGATTATTCATGCCTCCNNNAGATCGGAAG |
| 37153_1 | TATGAGTGTGGAGTCGTTGCTAAAGGGAGGAATGGTTTGCGTANNNAGATCGGAA |
| 37153_2 | TATGAGTGTGGAGTCGTTGCTAAAGGGAGGAATGGTTTGCGTANNNAGATCGGAA |
| 29664_1 | TATGAGTGTGGAGTCGTTGCGTAGGAAATGCAACATCTCAGGCNNNAGATCGGAA |
| 29664_2 | TATGAGTGTGGAGTCGTTGCGTAGGAAATGCAACATCTCAGGCNNNAGATCGGAA |
| 13466_1 | TATGAGTGTGGAGTCGTTGCTTACATATCATCTCGTCCAATAANNNAGATCGGAAGA |
| 13466_2 | TATGAGTGTGGAGTCGTTGCTTACATATCATCTCGTCCAATAANNNAGATCGGAAGA |
| 739_1 | TATGAGTGTGGAGTCGTTGCCTGGGAATCAGTGTGATGATTGGNNNAGATCGGAA |
| 739_2 | TATGAGTGTGGAGTCGTTGCCTGGGAATCAGTGTGATGATTGGNNNAGATCGGAA |
| 38447_1 | TATGAGTGTGGAGTCGTTGCGATGTTTCCAAGGCTTCGTCTTTNNNAGATCGGAAG |
| 38447_2 | TATGAGTGTGGAGTCGTTGCGATGTTTCCAAGGCTTCGTCTTTNNNAGATCGGAAG |
| 46222_1 | TATGAGTGTGGAGTCGTTGCTCAAGCAATTCTCCTACCTCAGCNNNAGATCGGAAGA |
| 46222_2 | TATGAGTGTGGAGTCGTTGCTCAAGCAATTCTCCTACCTCAGCNNNAGATCGGAAGA |
| 42427_1 | TATGAGTGTGGAGTCGTTGCAGCCAAGAGTGCTAGAGGAAAAANNNAGATCGGAA |
| 42427_2 | TATGAGTGTGGAGTCGTTGCAGCCAAGAGTGCTAGAGGAAAAANNNAGATCGGAA |
| 48973_1 | TATGAGTGTGGAGTCGTTGCATCCAATTTGCCAGTCTGAGTCTNNNAGATCGGAAG |
| 48973_2 | TATGAGTGTGGAGTCGTTGCATCCAATTTGCCAGTCTGAGTCTNNNAGATCGGAAG |
| 52599_1 | TATGAGTGTGGAGTCGTTGCCAGGCAGAGATATACACAGCCATNNNAGATCGGAA |
| 52599_2 | TATGAGTGTGGAGTCGTTGCCAGGCAGAGATATACACAGCCATNNNAGATCGGAA |
| 47469_1 | TATGAGTGTGGAGTCGTTGCCCAGGGTGAGGCTTTTTCTTTAGNNNAGATCGGAAG |
| 47469_2 | TATGAGTGTGGAGTCGTTGCCCAGGGTGAGGCTTTTTCTTTAGNNNAGATCGGAAG |
| 38197_1 | TATGAGTGTGGAGTCGTTGCGCCGCTTACCTACAACTATCTGANNNAGATCGGAAG |
| 38197_2 | TATGAGTGTGGAGTCGTTGCGCCGCTTACCTACAACTATCTGANNNAGATCGGAAG |
| 49094_1 | TATGAGTGTGGAGTCGTTGCCACCAGGCCCAGCTAATTTTTTTNNNAGATCGGAAGA |
| 49094_2 | TATGAGTGTGGAGTCGTTGCCACCAGGCCCAGCTAATTTTTTTNNNAGATCGGAAGA |
| 42253_1 | TATGAGTGTGGAGTCGTTGCACTGTGCTGGAGATTCTCAAACTNNNAGATCGGAAG |
| 42253_2 | TATGAGTGTGGAGTCGTTGCACTGTGCTGGAGATTCTCAAACTNNNAGATCGGAAG |
| 39174_1 | TATGAGTGTGGAGTCGTTGCCATATCCAGCCTCTACATGTCCCNNNAGATCGGAAGA |
| 39174_2 | TATGAGTGTGGAGTCGTTGCCATATCCAGCCTCTACATGTCCCNNNAGATCGGAAGA |
| 33371_1 | TATGAGTGTGGAGTCGTTGCGGAGGTCTCACTTTGTTGCCTAGNNNAGATCGGAAG |
| 33371_2 | TATGAGTGTGGAGTCGTTGCGGAGGTCTCACTTTGTTGCCTAGNNNAGATCGGAAG |
| 31348_1 | TATGAGTGTGGAGTCGTTGCCTCTGTGCCCTCTAGTTGAAGAGNNNAGATCGGAAG |
| 31348_2 | TATGAGTGTGGAGTCGTTGCCTCTGTGCCCTCTAGTTGAAGAGNNNAGATCGGAAG |
| 45133_1 | TATGAGTGTGGAGTCGTTGCGTTGGTGTAGAGGACTGGGTATCNNNAGATCGGAA |
| 45133_2 | TATGAGTGTGGAGTCGTTGCGTTGGTGTAGAGGACTGGGTATCNNNAGATCGGAA |

| | |
|---|---|
| 40621_1 | TATGAGTGTGGAGTCGTTGCTAAGGCATGAAAAGGTGGGTTGANNNAGATCGGA |
| 40621_2 | TATGAGTGTGGAGTCGTTGCTAAGGCATGAAAAGGTGGGTTGANNNAGATCGGA |
| 364_1 | TATGAGTGTGGAGTCGTTGCATAGCCAGTTGGAAAGGGTGGAAANNNAGATCGG |
| 364_2 | TATGAGTGTGGAGTCGTTGCATAGCCAGTTGGAAAGGGTGGAAANNNAGATCGG |
| 44397_1 | TATGAGTGTGGAGTCGTTGCGTTATTGGGTAGAGGTGGTCCTGNNNAGATCGGAA |
| 44397_2 | TATGAGTGTGGAGTCGTTGCGTTATTGGGTAGAGGTGGTCCTGNNNAGATCGGAA |
| 47607_1 | TATGAGTGTGGAGTCGTTGCCTGGAGTGCAATGGTGTGATCTCNNNAGATCGGAA |
| 47607_2 | TATGAGTGTGGAGTCGTTGCCTGGAGTGCAATGGTGTGATCTCNNNAGATCGGAA |
| 43461_1 | TATGAGTGTGGAGTCGTTGCGCACAATCTCAGTTCACTGCAATNNNAGATCGGAAG |
| 43461_2 | TATGAGTGTGGAGTCGTTGCGCACAATCTCAGTTCACTGCAATNNNAGATCGGAAG |
| 40039_1 | TATGAGTGTGGAGTCGTTGCCAAGAAGTAGAGGCTGCAGTGAGNNNAGATCGGA |
| 40039_2 | TATGAGTGTGGAGTCGTTGCCAAGAAGTAGAGGCTGCAGTGAGNNNAGATCGGA |
| 31514_1 | TATGAGTGTGGAGTCGTTGCACTGACAGAGAGGTGTTGAAGTCNNNAGATCGGAA |
| 31514_2 | TATGAGTGTGGAGTCGTTGCACTGACAGAGAGGTGTTGAAGTCNNNAGATCGGAA |
| 52771_1 | TATGAGTGTGGAGTCGTTGCTCCCTTAGAGAACTACCTAGGGCNNNAGATCGGAAG |
| 52771_2 | TATGAGTGTGGAGTCGTTGCTCCCTTAGAGAACTACCTAGGGCNNNAGATCGGAAG |
| 31623_1 | TATGAGTGTGGAGTCGTTGCTTGCCTACCCCAACAGTACTTACNNNAGATCGGAAGA |
| 31623_2 | TATGAGTGTGGAGTCGTTGCTTGCCTACCCCAACAGTACTTACNNNAGATCGGAAGA |
| 51804_1 | TATGAGTGTGGAGTCGTTGCGGGCTCCTCACTTCCCAGTAGNNNAGATCGGAAGAG |
| 51804_2 | TATGAGTGTGGAGTCGTTGCGGGCTCCTCACTTCCCAGTAGNNNAGATCGGAAGAG |
| 31872_1 | TATGAGTGTGGAGTCGTTGCGGCTAGAGAAAGACCCCAAGTTTNNNAGATCGGAA |
| 31872_2 | TATGAGTGTGGAGTCGTTGCGGCTAGAGAAAGACCCCAAGTTTNNNAGATCGGAA |
| 39918_1 | TATGAGTGTGGAGTCGTTGCGAGGAAGGAAGGAAGGAAGGAAGNNNAGATCGG |
| 39918_2 | TATGAGTGTGGAGTCGTTGCGAGGAAGGAAGGAAGGAAGGAAGNNNAGATCGG |
| 37424_1 | TATGAGTGTGGAGTCGTTGCACTTCAGCATCACATAGACCACANNNAGATCGGAAG |
| 37424_2 | TATGAGTGTGGAGTCGTTGCACTTCAGCATCACATAGACCACANNNAGATCGGAAG |
| 32314_1 | TATGAGTGTGGAGTCGTTGCCAACAAGAATGAAACTCCGTCTCANNNAGATCGGAA |
| 32314_2 | TATGAGTGTGGAGTCGTTGCCAACAAGAATGAAACTCCGTCTCANNNAGATCGGAA |
| 45704_1 | TATGAGTGTGGAGTCGTTGCCACTATCTTGGCTCACTGCAAGNNNAGATCGGAAGA |
| 45704_2 | TATGAGTGTGGAGTCGTTGCCACTATCTTGGCTCACTGCAAGNNNAGATCGGAAGA |
| 40500_1 | TATGAGTGTGGAGTCGTTGCGTTGTTAGTCAAAAGGGGGCTTCNNNAGATCGGAA |
| 40500_2 | TATGAGTGTGGAGTCGTTGCGTTGTTAGTCAAAAGGGGGCTTCNNNAGATCGGAA |
| 49277_1 | TATGAGTGTGGAGTCGTTGCCTCACATAAGCACATGAACACCCNNNAGATCGGAAG |
| 49277_2 | TATGAGTGTGGAGTCGTTGCCTCACATAAGCACATGAACACCCNNNAGATCGGAAG |
| 30187_1 | TATGAGTGTGGAGTCGTTGCGCAATCATCTAAAGTGCAGCCAANNNAGATCGGAAG |
| 30187_2 | TATGAGTGTGGAGTCGTTGCGCAATCATCTAAAGTGCAGCCAANNNAGATCGGAAG |
| 13693_1 | TATGAGTGTGGAGTCGTTGCAGTAAATAGACAGCCTTTCTGNNNAGATCGGAAGAG |
| 13693_2 | TATGAGTGTGGAGTCGTTGCAGTAAATAGACAGCCTTTCTGNNNAGATCGGAAGAG |
| 47336_1 | TATGAGTGTGGAGTCGTTGCAGACGGAGTTTTGTTCTTGTTGCNNNAGATCGGAAG |
| 47336_2 | TATGAGTGTGGAGTCGTTGCAGACGGAGTTTTGTTCTTGTTGCNNNAGATCGGAAG |
| 50583_1 | TATGAGTGTGGAGTCGTTGCCTGCAATCTCTAAGCAAGCCAAANNNAGATCGGAAG |
| 50583_2 | TATGAGTGTGGAGTCGTTGCCTGCAATCTCTAAGCAAGCCAAANNNAGATCGGAAG |
| 44171_1 | TATGAGTGTGGAGTCGTTGCAAGGGGCAAGAATGAAGAAGGATNNNAGATCGGA |

| | |
|---|---|
| 44171_2 | TATGAGTGTGGAGTCGTTGCAAGGGGCAAGAATGAAGAAGGATNNNAGATCGGA |
| 47455_1 | TATGAGTGTGGAGTCGTTGCCAAAGGGCTGGGATTATAGGCATNNNAGATCGGAA |
| 47455_2 | TATGAGTGTGGAGTCGTTGCCAAAGGGCTGGGATTATAGGCATNNNAGATCGGAA |
| 36931_1 | TATGAGTGTGGAGTCGTTGCCTGTGTGTGTCCATGAGGGTATANNNAGATCGGAA |
| 36931_2 | TATGAGTGTGGAGTCGTTGCCTGTGTGTGTCCATGAGGGTATANNNAGATCGGAA |
| 45953_1 | TATGAGTGTGGAGTCGTTGCAGATGGAGTTTCGTTCTTGTTGCNNNAGATCGGAAG |
| 45953_2 | TATGAGTGTGGAGTCGTTGCAGATGGAGTTTCGTTCTTGTTGCNNNAGATCGGAAG |
| 33882_1 | TATGAGTGTGGAGTCGTTGCCTCCTGAGTAGCTGGGATTACAGNNNAGATCGGAAG |
| 33882_2 | TATGAGTGTGGAGTCGTTGCCTCCTGAGTAGCTGGGATTACAGNNNAGATCGGAAG |
| 11085_1 | TATGAGTGTGGAGTCGTTGCTTTTAGAGACAGAGCTTTGCCAGNNNAGATCGGAAG |
| 11085_2 | TATGAGTGTGGAGTCGTTGCTTTTAGAGACAGAGCTTTGCCAGNNNAGATCGGAAG |
| 38200_1 | TATGAGTGTGGAGTCGTTGCTGGAAGGTCTGACATGATGATCANNNAGATCGGAA |
| 38200_2 | TATGAGTGTGGAGTCGTTGCTGGAAGGTCTGACATGATGATCANNNAGATCGGAA |
| 46466_1 | TATGAGTGTGGAGTCGTTGCTAGGACAATAGTGGAGGGAAGGTNNNAGATCGGA |
| 46466_2 | TATGAGTGTGGAGTCGTTGCTAGGACAATAGTGGAGGGAAGGTNNNAGATCGGA |
| 48347_1 | TATGAGTGTGGAGTCGTTGCTTTTTTGAGACGGAGTCTTGCTCNNNAGATCGGAAG |
| 48347_2 | TATGAGTGTGGAGTCGTTGCTTTTTTGAGACGGAGTCTTGCTCNNNAGATCGGAAG |
| 37579_1 | TATGAGTGTGGAGTCGTTGCGGAAGATATCTGGGTGATGGGTCNNNAGATCGGAA |
| 37579_2 | TATGAGTGTGGAGTCGTTGCGGAAGATATCTGGGTGATGGGTCNNNAGATCGGAA |
| 30444_1 | TATGAGTGTGGAGTCGTTGCCTTCACTCAGCCTAGAAATGCAAANNNAGATCGGAA |
| 30444_2 | TATGAGTGTGGAGTCGTTGCCTTCACTCAGCCTAGAAATGCAAANNNAGATCGGAA |
| 29832_1 | TATGAGTGTGGAGTCGTTGCAGATCTGAGTACATGAGACCCCTNNNAGATCGGAAG |
| 29832_2 | TATGAGTGTGGAGTCGTTGCAGATCTGAGTACATGAGACCCCTNNNAGATCGGAAG |
| 38299_1 | TATGAGTGTGGAGTCGTTGCCTATCTTGTAGGAAGCCAGGGTCNNNAGATCGGAAG |
| 38299_2 | TATGAGTGTGGAGTCGTTGCCTATCTTGTAGGAAGCCAGGGTCNNNAGATCGGAAG |
| 30307_1 | TATGAGTGTGGAGTCGTTGCCCCGATTTCGCTACCTCAATGAANNNAGATCGGAAG |
| 30307_2 | TATGAGTGTGGAGTCGTTGCCCCGATTTCGCTACCTCAATGAANNNAGATCGGAAG |
| 34354_1 | TATGAGTGTGGAGTCGTTGCCCCCCATGATTCAATTACCTCCANNNAGATCGGAAGA |
| 34354_2 | TATGAGTGTGGAGTCGTTGCCCCCCATGATTCAATTACCTCCANNNAGATCGGAAGA |
| 34259_1 | TATGAGTGTGGAGTCGTTGCCCATCAGGACGGGATACTTGGNNNAGATCGGAAGA |
| 34259_2 | TATGAGTGTGGAGTCGTTGCCCATCAGGACGGGATACTTGGNNNAGATCGGAAGA |
| 44753_1 | TATGAGTGTGGAGTCGTTGCAGACAAGGCCAGTGTATATGCAANNNAGATCGGAA |
| 44753_2 | TATGAGTGTGGAGTCGTTGCAGACAAGGCCAGTGTATATGCAANNNAGATCGGAA |
| 37741_1 | TATGAGTGTGGAGTCGTTGCATGTGCAGACCCTGAAAAATGTGNNNAGATCGGAA |
| 37741_2 | TATGAGTGTGGAGTCGTTGCATGTGCAGACCCTGAAAAATGTGNNNAGATCGGAA |
| 53186_1 | TATGAGTGTGGAGTCGTTGCTCTCTAAGGAATCCTGGCTGAGANNNAGATCGGAAG |
| 53186_2 | TATGAGTGTGGAGTCGTTGCTCTCTAAGGAATCCTGGCTGAGANNNAGATCGGAAG |
| 13604_1 | TATGAGTGTGGAGTCGTTGCGAACTACATGAGATGTAAGAATGNNNAGATCGGAA |
| 13604_2 | TATGAGTGTGGAGTCGTTGCGAACTACATGAGATGTAAGAATGNNNAGATCGGAA |
| 53024_1 | TATGAGTGTGGAGTCGTTGCCAGCAGACCTTGGAGAAATCTCTNNNAGATCGGAAG |
| 53024_2 | TATGAGTGTGGAGTCGTTGCCAGCAGACCTTGGAGAAATCTCTNNNAGATCGGAAG |
| 52896_1 | TATGAGTGTGGAGTCGTTGCTCCCCTATCAACCTCCAAGTAGTNNNAGATCGGAAGA |
| 52896_2 | TATGAGTGTGGAGTCGTTGCTCCCCTATCAACCTCCAAGTAGTNNNAGATCGGAAGA |

| | |
|---|---|
| 12812_1 | TATGAGTGTGGAGTCGTTGCATAAAATGAGATTCCAGCCAGNNNAGATCGGAAGA |
| 12812_2 | TATGAGTGTGGAGTCGTTGCATAAAATGAGATTCCAGCCAGNNNAGATCGGAAGA |
| 50172_1 | TATGAGTGTGGAGTCGTTGCTACCAGAAACCTTCTCACCCAAGNNNAGATCGGAAG |
| 50172_2 | TATGAGTGTGGAGTCGTTGCTACCAGAAACCTTCTCACCCAAGNNNAGATCGGAAG |
| 43096_1 | TATGAGTGTGGAGTCGTTGCATGCACAACCAGGCACTATTCTANNNAGATCGGAAG |
| 43096_2 | TATGAGTGTGGAGTCGTTGCATGCACAACCAGGCACTATTCTANNNAGATCGGAAG |
| 46371_1 | TATGAGTGTGGAGTCGTTGCGATCTCACCTTGCTGCAACCTCNNNAGATCGGAAGA |
| 46371_2 | TATGAGTGTGGAGTCGTTGCGATCTCACCTTGCTGCAACCTCNNNAGATCGGAAGA |
| 34620_1 | TATGAGTGTGGAGTCGTTGCATGATCCCTGTTCTTTTGTGGGANNNAGATCGGAAG |
| 34620_2 | TATGAGTGTGGAGTCGTTGCATGATCCCTGTTCTTTTGTGGGANNNAGATCGGAAG |
| 37323_1 | TATGAGTGTGGAGTCGTTGCACCAATTCCTAGATAATCAAGGCANNNAGATCGGAA |
| 37323_2 | TATGAGTGTGGAGTCGTTGCACCAATTCCTAGATAATCAAGGCANNNAGATCGGAA |
| 42951_1 | TATGAGTGTGGAGTCGTTGCACACCAGGACTCAAGTGATTCTCNNNAGATCGGAAG |
| 42951_2 | TATGAGTGTGGAGTCGTTGCACACCAGGACTCAAGTGATTCTCNNNAGATCGGAAG |
| 43084_1 | TATGAGTGTGGAGTCGTTGCACAGTGCTACAGTGAACATGAGANNNAGATCGGAA |
| 43084_2 | TATGAGTGTGGAGTCGTTGCACAGTGCTACAGTGAACATGAGANNNAGATCGGAA |
| 33137_1 | TATGAGTGTGGAGTCGTTGCATGAAAGGACATAGAGGCTGGTCNNNAGATCGGAA |
| 33137_2 | TATGAGTGTGGAGTCGTTGCATGAAAGGACATAGAGGCTGGTCNNNAGATCGGAA |
| 29021_1 | TATGAGTGTGGAGTCGTTGCGCAATTTCTTCCATCGAGGTTTTGNNNAGATCGGAA |
| 29021_2 | TATGAGTGTGGAGTCGTTGCGCAATTTCTTCCATCGAGGTTTTGNNNAGATCGGAA |
| 35964_1 | TATGAGTGTGGAGTCGTTGCCATCCTGGCTAACATGGTGAAACNNNAGATCGGAAG |
| 35964_2 | TATGAGTGTGGAGTCGTTGCCATCCTGGCTAACATGGTGAAACNNNAGATCGGAAG |
| 47301_1 | TATGAGTGTGGAGTCGTTGCAGGTCAGAGATTGCTTCATGTGANNNAGATCGGAA |
| 47301_2 | TATGAGTGTGGAGTCGTTGCAGGTCAGAGATTGCTTCATGTGANNNAGATCGGAA |
| 12555_1 | TATGAGTGTGGAGTCGTTGCTACATTCAAACTCAATATCCATGNNNAGATCGGAAG |
| 12555_2 | TATGAGTGTGGAGTCGTTGCTACATTCAAACTCAATATCCATGNNNAGATCGGAAG |
| 30294_1 | TATGAGTGTGGAGTCGTTGCCGCTTCAGTAGAGGAAGGATCTGNNNAGATCGGAA |
| 30294_2 | TATGAGTGTGGAGTCGTTGCCGCTTCAGTAGAGGAAGGATCTGNNNAGATCGGAA |
| 34609_1 | TATGAGTGTGGAGTCGTTGCCTCAGTCCAGGAGGGAGGTGNNNAGATCGGAAGA |
| 34609_2 | TATGAGTGTGGAGTCGTTGCCTCAGTCCAGGAGGGAGGTGNNNAGATCGGAAGA |
| 37764_1 | TATGAGTGTGGAGTCGTTGCACAGCCCACCTAGAAGAACTAGANNNAGATCGGAA |
| 37764_2 | TATGAGTGTGGAGTCGTTGCACAGCCCACCTAGAAGAACTAGANNNAGATCGGAA |
| 45572_1 | TATGAGTGTGGAGTCGTTGCGATCTCAGCTCACTGCAACCTCNNNAGATCGGAAGA |
| 45572_2 | TATGAGTGTGGAGTCGTTGCGATCTCAGCTCACTGCAACCTCNNNAGATCGGAAGA |
| 10265_1 | TATGAGTGTGGAGTCGTTGCGACCAGTTAGAAGGCAAGAGAGCNNNAGATCGGAA |
| 10265_2 | TATGAGTGTGGAGTCGTTGCGACCAGTTAGAAGGCAAGAGAGCNNNAGATCGGAA |
| 50361_1 | TATGAGTGTGGAGTCGTTGCTGTAGTCACTGGTTGATCTTTGANNNAGATCGGAAG |
| 50361_2 | TATGAGTGTGGAGTCGTTGCTGTAGTCACTGGTTGATCTTTGANNNAGATCGGAAG |
| 47453_1 | TATGAGTGTGGAGTCGTTGCAGAGAACATTTGGACCCCATTGTNNNAGATCGGAAG |
| 47453_2 | TATGAGTGTGGAGTCGTTGCAGAGAACATTTGGACCCCATTGTNNNAGATCGGAAG |
| 33797_1 | TATGAGTGTGGAGTCGTTGCTTCGTCGACAGTATCTTCCACTGNNNAGATCGGAAG |
| 33797_2 | TATGAGTGTGGAGTCGTTGCTTCGTCGACAGTATCTTCCACTGNNNAGATCGGAAG |
| 31914_1 | TATGAGTGTGGAGTCGTTGCCCACCAAAAGGCAACTAGAACTGNNNAGATCGGAA |

| | |
|---|---|
| 31914_2 | TATGAGTGTGGAGTCGTTGCCCACCAAAAGGCAACTAGAACTGNNNAGATCGGAAG |
| 31405_1 | TATGAGTGTGGAGTCGTTGCCCTTTACCTGCAATAACCTCCCTNNNAGATCGGAAGA |
| 31405_2 | TATGAGTGTGGAGTCGTTGCCCTTTACCTGCAATAACCTCCCTNNNAGATCGGAAGA |
| 10505_1 | TATGAGTGTGGAGTCGTTGCAAGTATTGCTCTTCAGCAAGGGTNNNAGATCGGAAG |
| 10505_2 | TATGAGTGTGGAGTCGTTGCAAGTATTGCTCTTCAGCAAGGGTNNNAGATCGGAAG |
| 9985_1 | TATGAGTGTGGAGTCGTTGCATCAAGCAAACAAAAAGGCCGTCNNNAGATCGGAAG |
| 9985_2 | TATGAGTGTGGAGTCGTTGCATCAAGCAAACAAAAAGGCCGTCNNNAGATCGGAAG |
| 32369_1 | TATGAGTGTGGAGTCGTTGCTCAACGAATCCAGGAGTTGACTTNNNAGATCGGAAG |
| 32369_2 | TATGAGTGTGGAGTCGTTGCTCAACGAATCCAGGAGTTGACTTNNNAGATCGGAAG |
| 39876_1 | TATGAGTGTGGAGTCGTTGCTGTGATGTCATGCAAACCTTTCCNNNAGATCGGAAGA |
| 39876_2 | TATGAGTGTGGAGTCGTTGCTGTGATGTCATGCAAACCTTTCCNNNAGATCGGAAGA |
| 3074_1 | TATGAGTGTGGAGTCGTTGCAGGCCGCATGATTGTATTTTCCTNNNAGATCGGAAGA |
| 3074_2 | TATGAGTGTGGAGTCGTTGCAGGCCGCATGATTGTATTTTCCTNNNAGATCGGAAGA |
| 42725_1 | TATGAGTGTGGAGTCGTTGCAGTTTGCCTCAGTACCCATAAGGNNNAGATCGGAAG |
| 42725_2 | TATGAGTGTGGAGTCGTTGCAGTTTGCCTCAGTACCCATAAGGNNNAGATCGGAAG |
| 40188_1 | TATGAGTGTGGAGTCGTTGCTGAAGCCAGAATATTTGTCTTGGANNNAGATCGGAA |
| 40188_2 | TATGAGTGTGGAGTCGTTGCTGAAGCCAGAATATTTGTCTTGGANNNAGATCGGAA |
| 10319_1 | TATGAGTGTGGAGTCGTTGCAGTGGTGCTTTATTTTTCAGCTCTNNNAGATCGGAAG |
| 10319_2 | TATGAGTGTGGAGTCGTTGCAGTGGTGCTTTATTTTTCAGCTCTNNNAGATCGGAAG |
| 41374_1 | TATGAGTGTGGAGTCGTTGCCCCAACTCAGTGTAACCCATTCTNNNAGATCGGAAGA |
| 41374_2 | TATGAGTGTGGAGTCGTTGCCCCAACTCAGTGTAACCCATTCTNNNAGATCGGAAGA |
| 46548_1 | TATGAGTGTGGAGTCGTTGCCACCACACTCGGCTAATTTTGTTNNNAGATCGGAAGA |
| 46548_2 | TATGAGTGTGGAGTCGTTGCCACCACACTCGGCTAATTTTGTTNNNAGATCGGAAGA |
| 36825_1 | TATGAGTGTGGAGTCGTTGCCCATTCTAGCCTAGCTTCTGGAGNNNAGATCGGAAGA |
| 36825_2 | TATGAGTGTGGAGTCGTTGCCCATTCTAGCCTAGCTTCTGGAGNNNAGATCGGAAGA |
| 30702_1 | TATGAGTGTGGAGTCGTTGCTTCAGTAGGAACGGTACCACTTCNNNAGATCGGAAG |
| 30702_2 | TATGAGTGTGGAGTCGTTGCTTCAGTAGGAACGGTACCACTTCNNNAGATCGGAAG |
| 32656_1 | TATGAGTGTGGAGTCGTTGCCACCCAGCCTCAATGGAAATTTTNNNAGATCGGAAGA |
| 32656_2 | TATGAGTGTGGAGTCGTTGCCACCCAGCCTCAATGGAAATTTTNNNAGATCGGAAGA |
| 45879_1 | TATGAGTGTGGAGTCGTTGCTAGCACAGACGATCCCATACATGNNNAGATCGGAAG |
| 45879_2 | TATGAGTGTGGAGTCGTTGCTAGCACAGACGATCCCATACATGNNNAGATCGGAAG |
| 35295_1 | TATGAGTGTGGAGTCGTTGCAGAAGGAACTTGCCTTGAGTCTTNNNAGATCGGAAG |
| 35295_2 | TATGAGTGTGGAGTCGTTGCAGAAGGAACTTGCCTTGAGTCTTNNNAGATCGGAAG |
| 44585_1 | TATGAGTGTGGAGTCGTTGCACACACACCCACACACACATATANNNAGATCGGAAGA |
| 44585_2 | TATGAGTGTGGAGTCGTTGCACACACACCCACACACACATATANNNAGATCGGAAGA |
| 49759_1 | TATGAGTGTGGAGTCGTTGCACCCTTCCAGATTCTGAGCAAATNNNAGATCGGAAGA |
| 49759_2 | TATGAGTGTGGAGTCGTTGCACCCTTCCAGATTCTGAGCAAATNNNAGATCGGAAGA |
| 52126_1 | TATGAGTGTGGAGTCGTTGCTTGGAATACTGCAGAGCTCAGTTNNNAGATCGGAAG |
| 52126_2 | TATGAGTGTGGAGTCGTTGCTTGGAATACTGCAGAGCTCAGTTNNNAGATCGGAAG |
| 45554_1 | TATGAGTGTGGAGTCGTTGCAGATGAACCAATCTGGGTGTCTCNNNAGATCGGAAG |
| 45554_2 | TATGAGTGTGGAGTCGTTGCAGATGAACCAATCTGGGTGTCTCNNNAGATCGGAAG |
| 43859_1 | TATGAGTGTGGAGTCGTTGCGGTGGGGGGTAGATAGTTCTTTCNNNAGATCGGAA |
| 43859_2 | TATGAGTGTGGAGTCGTTGCGGTGGGGGGTAGATAGTTCTTTCNNNAGATCGGAA |

| | |
|---|---|
| 34818_1 | TATGAGTGTGGAGTCGTTGCCGTCTCCCAAGTTGAAGCAATTCNNNAGATCGGAAG |
| 34818_2 | TATGAGTGTGGAGTCGTTGCCGTCTCCCAAGTTGAAGCAATTCNNNAGATCGGAAG |
| 42132_1 | TATGAGTGTGGAGTCGTTGCCATGTAGCTGGGATTACAAGTGCNNNAGATCGGAA |
| 42132_2 | TATGAGTGTGGAGTCGTTGCCATGTAGCTGGGATTACAAGTGCNNNAGATCGGAA |
| 41047_1 | TATGAGTGTGGAGTCGTTGCCACCTATATTCCCAGCTACTCGGNNNAGATCGGAAGA |
| 41047_2 | TATGAGTGTGGAGTCGTTGCCACCTATATTCCCAGCTACTCGGNNNAGATCGGAAGA |
| 46013_1 | TATGAGTGTGGAGTCGTTGCCAGGTTCACGCCATTCTCCTNNNAGATCGGAAGAGC |
| 46013_2 | TATGAGTGTGGAGTCGTTGCCAGGTTCACGCCATTCTCCTNNNAGATCGGAAGAGC |
| 30197_1 | TATGAGTGTGGAGTCGTTGCTCAAGGCAGTTTGTGACAATCTGNNNAGATCGGAAG |
| 30197_2 | TATGAGTGTGGAGTCGTTGCTCAAGGCAGTTTGTGACAATCTGNNNAGATCGGAAG |
| 10029_1 | TATGAGTGTGGAGTCGTTGCTGTCCAGGATGCCTTCAATGTTCNNNAGATCGGAAG |
| 10029_2 | TATGAGTGTGGAGTCGTTGCTGTCCAGGATGCCTTCAATGTTCNNNAGATCGGAAG |
| 49748_1 | TATGAGTGTGGAGTCGTTGCAATCGTCCTCCACTTCATCAACANNNAGATCGGAAGA |
| 49748_2 | TATGAGTGTGGAGTCGTTGCAATCGTCCTCCACTTCATCAACANNNAGATCGGAAGA |
| 10168_1 | TATGAGTGTGGAGTCGTTGCGCTCTGCCCTAAGAACTTCCATTNNNAGATCGGAAGA |
| 10168_2 | TATGAGTGTGGAGTCGTTGCGCTCTGCCCTAAGAACTTCCATTNNNAGATCGGAAGA |
| 41781_1 | TATGAGTGTGGAGTCGTTGCCTCAGCCTCCAGATTAGCTAGGANNNAGATCGGAAG |
| 41781_2 | TATGAGTGTGGAGTCGTTGCCTCAGCCTCCAGATTAGCTAGGANNNAGATCGGAAG |
| 9235_1 | TATGAGTGTGGAGTCGTTGCGCATGGTTTCTGCCTTCATAGTGNNNAGATCGGAAG |
| 9235_2 | TATGAGTGTGGAGTCGTTGCGCATGGTTTCTGCCTTCATAGTGNNNAGATCGGAAG |
| 39865_1 | TATGAGTGTGGAGTCGTTGCATGATTCCTGACCTTCCTTAGCCNNNAGATCGGAAGA |
| 39865_2 | TATGAGTGTGGAGTCGTTGCATGATTCCTGACCTTCCTTAGCCNNNAGATCGGAAGA |
| 33464_1 | TATGAGTGTGGAGTCGTTGCCAACAGTGAGTTTGCCACATCAANNNAGATCGGAAG |
| 33464_2 | TATGAGTGTGGAGTCGTTGCCAACAGTGAGTTTGCCACATCAANNNAGATCGGAAG |
| 40528_1 | TATGAGTGTGGAGTCGTTGCCACAAAGAAATGAAACACGCAGCNNNAGATCGGAA |
| 40528_2 | TATGAGTGTGGAGTCGTTGCCACAAAGAAATGAAACACGCAGCNNNAGATCGGAA |
| 45276_1 | TATGAGTGTGGAGTCGTTGCGAGTCAAGCTTTACCTACCCCAANNNAGATCGGAAG |
| 45276_2 | TATGAGTGTGGAGTCGTTGCGAGTCAAGCTTTACCTACCCCAANNNAGATCGGAAG |
| 49916_1 | TATGAGTGTGGAGTCGTTGCAACTTCAATCCCCTCCTCTCCAANNNAGATCGGAAGA |
| 49916_2 | TATGAGTGTGGAGTCGTTGCAACTTCAATCCCCTCCTCTCCAANNNAGATCGGAAGA |
| 45758_1 | TATGAGTGTGGAGTCGTTGCGTTGACGCCATTCTCTTGCTTCNNNAGATCGGAAGA |
| 45758_2 | TATGAGTGTGGAGTCGTTGCGTTGACGCCATTCTCTTGCTTCNNNAGATCGGAAGA |
| 31683_1 | TATGAGTGTGGAGTCGTTGCCACAGACAGTGGAAATGCCAAAANNNAGATCGGAA |
| 31683_2 | TATGAGTGTGGAGTCGTTGCCACAGACAGTGGAAATGCCAAAANNNAGATCGGAA |
| 32616_1 | TATGAGTGTGGAGTCGTTGCTCACAAGGAGATACACCTCACACNNNAGATCGGAAG |
| 32616_2 | TATGAGTGTGGAGTCGTTGCTCACAAGGAGATACACCTCACACNNNAGATCGGAAG |
| 42940_1 | TATGAGTGTGGAGTCGTTGCGGTGATTCTCCTAACTCAGCCTCNNNAGATCGGAAG |
| 42940_2 | TATGAGTGTGGAGTCGTTGCGGTGATTCTCCTAACTCAGCCTCNNNAGATCGGAAG |
| 46584_1 | TATGAGTGTGGAGTCGTTGCGGGTTCAAGTGATTCTCCTGTCTNNNAGATCGGAAG |
| 46584_2 | TATGAGTGTGGAGTCGTTGCGGGTTCAAGTGATTCTCCTGTCTNNNAGATCGGAAG |
| 11299_1 | TATGAGTGTGGAGTCGTTGCCCGCATGTAAGATGTGTCTTTCTCNNNAGATCGGAA |
| 11299_2 | TATGAGTGTGGAGTCGTTGCCCGCATGTAAGATGTGTCTTTCTCNNNAGATCGGAA |
| 42456_1 | TATGAGTGTGGAGTCGTTGCCAGGGTGATGCTGACCTCATAATNNNAGATCGGAA |

| | |
|---|---|
| 42456_2 | TATGAGTGTGGAGTCGTTGCCAGGGTGATGCTGACCTCATAATNNNAGATCGGAAG |
| 36940_1 | TATGAGTGTGGAGTCGTTGCGCAGGCATTCAAACAAAAACTTGTNNNAGATCGGAA |
| 36940_2 | TATGAGTGTGGAGTCGTTGCGCAGGCATTCAAACAAAAACTTGTNNNAGATCGGAA |
| 39143_1 | TATGAGTGTGGAGTCGTTGCTCCTGAACAGACCAATAACAACTNNNAGATCGGAAG |
| 39143_2 | TATGAGTGTGGAGTCGTTGCTCCTGAACAGACCAATAACAACTNNNAGATCGGAAG |
| 47102_1 | TATGAGTGTGGAGTCGTTGCGCTTCTCAAAGTGCTGGGATTACNNNAGATCGGAAG |
| 47102_2 | TATGAGTGTGGAGTCGTTGCGCTTCTCAAAGTGCTGGGATTACNNNAGATCGGAAG |
| 50239_1 | TATGAGTGTGGAGTCGTTGCTGAGGGGAGAGAGAAATCAGAGTNNNAGATCGGA |
| 50239_2 | TATGAGTGTGGAGTCGTTGCTGAGGGGAGAGAGAAATCAGAGTNNNAGATCGGA |
| 48001_1 | TATGAGTGTGGAGTCGTTGCACTCCCTCTATCCATGAGTTCAANNNAGATCGGAAGA |
| 48001_2 | TATGAGTGTGGAGTCGTTGCACTCCCTCTATCCATGAGTTCAANNNAGATCGGAAGA |
| 41869_1 | TATGAGTGTGGAGTCGTTGCCATGATCTCAGCTCACTGCAACNNNAGATCGGAAGA |
| 41869_2 | TATGAGTGTGGAGTCGTTGCCATGATCTCAGCTCACTGCAACNNNAGATCGGAAGA |
| 48674_1 | TATGAGTGTGGAGTCGTTGCATGCAAGGTTCCGAATGAAACTGNNNAGATCGGAA |
| 48674_2 | TATGAGTGTGGAGTCGTTGCATGCAAGGTTCCGAATGAAACTGNNNAGATCGGAA |
| 52157_1 | TATGAGTGTGGAGTCGTTGCCTCCTGAGTAGCTGGGATTACAGNNNAGATCGGAA |
| 52157_2 | TATGAGTGTGGAGTCGTTGCCTCCTGAGTAGCTGGGATTACAGNNNAGATCGGAA |
| 48254_1 | TATGAGTGTGGAGTCGTTGCCTTGACCCAGGCTCTGATGGNNNAGATCGGAAGAG |
| 48254_2 | TATGAGTGTGGAGTCGTTGCCTTGACCCAGGCTCTGATGGNNNAGATCGGAAGAG |
| 37949_1 | TATGAGTGTGGAGTCGTTGCAATTAGCTTAGCATGGTTGTGGGNNNAGATCGGAA |
| 37949_2 | TATGAGTGTGGAGTCGTTGCAATTAGCTTAGCATGGTTGTGGGNNNAGATCGGAA |
| 39517_1 | TATGAGTGTGGAGTCGTTGCCTAGAAACCCCAAGGATTCTGCTNNNAGATCGGAAG |
| 39517_2 | TATGAGTGTGGAGTCGTTGCCTAGAAACCCCAAGGATTCTGCTNNNAGATCGGAAG |
| 29743_1 | TGAGTGTGGAGTCGTTGCTTTGCTCTAATTGAAGGAAAAAACAAANNNAGATCGGA |
| 29743_2 | TGAGTGTGGAGTCGTTGCTTTGCTCTAATTGAAGGAAAAAACAAANNNAGATCGGA |
| 31060_1 | TATGAGTGTGGAGTCGTTGCATGAATTTTGGAACGTGCCCTTTNNNAGATCGGAAG |
| 31060_2 | TATGAGTGTGGAGTCGTTGCATGAATTTTGGAACGTGCCCTTTNNNAGATCGGAAG |
| 53223_1 | TATGAGTGTGGAGTCGTTGCTTTGGCCTCAAACTTCAGGTGANNNAGATCGGAAGA |
| 53223_2 | TATGAGTGTGGAGTCGTTGCTTTGGCCTCAAACTTCAGGTGANNNAGATCGGAAGA |
| 31847_1 | TATGAGTGTGGAGTCGTTGCTCCACAAAAGCAATTGAAAACTCTNNNAGATCGGAA |
| 31847_2 | TATGAGTGTGGAGTCGTTGCTCCACAAAAGCAATTGAAAACTCTNNNAGATCGGAA |
| 48808_1 | TATGAGTGTGGAGTCGTTGCGAGTAGCTGGGATTATAGGCACCNNNAGATCGGAA |
| 48808_2 | TATGAGTGTGGAGTCGTTGCGAGTAGCTGGGATTATAGGCACCNNNAGATCGGAA |
| 52347_1 | TATGAGTGTGGAGTCGTTGCTTTCTGGCATAAGTGAGGTTGGANNNAGATCGGAA |
| 52347_2 | TATGAGTGTGGAGTCGTTGCTTTCTGGCATAAGTGAGGTTGGANNNAGATCGGAA |
| 37221_1 | TATGAGTGTGGAGTCGTTGCATGTTTGGGGTGATGAATATGCTNNNAGATCGGAA |
| 37221_2 | TATGAGTGTGGAGTCGTTGCATGTTTGGGGTGATGAATATGCTNNNAGATCGGAA |
| 40146_1 | TATGAGTGTGGAGTCGTTGCTCCACTCAAGGATACCACTTTGCNNNAGATCGGAAG |
| 40146_2 | TATGAGTGTGGAGTCGTTGCTCCACTCAAGGATACCACTTTGCNNNAGATCGGAAG |
| 41530_1 | TATGAGTGTGGAGTCGTTGCACATGTGCAGGTTTGTTACATGGNNNAGATCGGAA |
| 41530_2 | TATGAGTGTGGAGTCGTTGCACATGTGCAGGTTTGTTACATGGNNNAGATCGGAA |
| 49651_1 | TATGAGTGTGGAGTCGTTGCGGAGTACAGTGGCATGATCATGANNNAGATCGGAA |
| 49651_2 | TATGAGTGTGGAGTCGTTGCGGAGTACAGTGGCATGATCATGANNNAGATCGGAA |

| | |
|---|---|
| 42063_1 | TATGAGTGTGGAGTCGTTGCTGTATACCCCCAATCAGGCAAAANNNAGATCGGAAG |
| 42063_2 | TATGAGTGTGGAGTCGTTGCTGTATACCCCCAATCAGGCAAAANNNAGATCGGAAG |
| 30488_1 | TATGAGTGTGGAGTCGTTGCGCCTCTCCATCATTGCTCTTTATTNNNAGATCGGAAG |
| 30488_2 | TATGAGTGTGGAGTCGTTGCGCCTCTCCATCATTGCTCTTTATTNNNAGATCGGAAG |
| 33522_1 | TATGAGTGTGGAGTCGTTGCCCCACCTGGAAAGACAGAATAGTNNNAGATCGGAA |
| 33522_2 | TATGAGTGTGGAGTCGTTGCCCCACCTGGAAAGACAGAATAGTNNNAGATCGGAA |
| 50453_1 | TATGAGTGTGGAGTCGTTGCTGCAGAGTCTCATTTCTCTACCGNNNAGATCGGAAG |
| 50453_2 | TATGAGTGTGGAGTCGTTGCTGCAGAGTCTCATTTCTCTACCGNNNAGATCGGAAG |
| 32803_1 | TATGAGTGTGGAGTCGTTGCTTCAAGCAATTTTCCTGCCTCAGNNNAGATCGGAAG |
| 32803_2 | TATGAGTGTGGAGTCGTTGCTTCAAGCAATTTTCCTGCCTCAGNNNAGATCGGAAG |
| 39105_1 | TATGAGTGTGGAGTCGTTGCAGTGCTGATAGCCAAGACAATGANNNAGATCGGAA |
| 39105_2 | TATGAGTGTGGAGTCGTTGCAGTGCTGATAGCCAAGACAATGANNNAGATCGGAA |
| 50043_1 | TATGAGTGTGGAGTCGTTGCAGGAAGGAAGGAAGGAGAAAGAANNNAGATCGG |
| 50043_2 | TATGAGTGTGGAGTCGTTGCAGGAAGGAAGGAAGGAGAAAGAANNNAGATCGG |
| 36499_1 | TATGAGTGTGGAGTCGTTGCAAAGAAGCCAGTTGCAAAGTACCNNNAGATCGGAA |
| 36499_2 | TATGAGTGTGGAGTCGTTGCAAAGAAGCCAGTTGCAAAGTACCNNNAGATCGGAA |
| 29367_1 | TATGAGTGTGGAGTCGTTGCAACCATTTCTCAGCTGAACACTGNNNAGATCGGAAG |
| 29367_2 | TATGAGTGTGGAGTCGTTGCAACCATTTCTCAGCTGAACACTGNNNAGATCGGAAG |
| 53088_1 | TATGAGTGTGGAGTCGTTGCTTCACCAATTTGACAAGGATGGCNNNAGATCGGAAG |
| 53088_2 | TATGAGTGTGGAGTCGTTGCTTCACCAATTTGACAAGGATGGCNNNAGATCGGAAG |
| 36735_1 | TATGAGTGTGGAGTCGTTGCGGGGACAGCAGCTATATTCACTTNNNAGATCGGAA |
| 36735_2 | TATGAGTGTGGAGTCGTTGCGGGGACAGCAGCTATATTCACTTNNNAGATCGGAA |
| 44785_1 | TATGAGTGTGGAGTCGTTGCCGTGAGGCTAGCACTCATCCNNNAGATCGGAAGAG |
| 44785_2 | TATGAGTGTGGAGTCGTTGCCGTGAGGCTAGCACTCATCCNNNAGATCGGAAGAG |
| 46432_1 | TATGAGTGTGGAGTCGTTGCCCTTCTGGGTTCACACCATTCTCNNNAGATCGGAAGA |
| 46432_2 | TATGAGTGTGGAGTCGTTGCCCTTCTGGGTTCACACCATTCTCNNNAGATCGGAAGA |
| 50418_1 | TATGAGTGTGGAGTCGTTGCTGAGATCTCTTCGGAGGACAAAGNNNAGATCGGAA |
| 50418_2 | TATGAGTGTGGAGTCGTTGCTGAGATCTCTTCGGAGGACAAAGNNNAGATCGGAA |
| 45889_1 | TATGAGTGTGGAGTCGTTGCGAGTAGCTGGGACTACAGGTGTCNNNAGATCGGAA |
| 45889_2 | TATGAGTGTGGAGTCGTTGCGAGTAGCTGGGACTACAGGTGTCNNNAGATCGGAA |
| 3501_1 | TATGAGTGTGGAGTCGTTGCTTCTGTTTTCTTCTGGGTTGGAANNNAGATCGGAAG |
| 3501_2 | TATGAGTGTGGAGTCGTTGCTTCTGTTTTCTTCTGGGTTGGAANNNAGATCGGAAG |
| 29638_1 | TATGAGTGTGGAGTCGTTGCAACCAAAACCTGCAACAAATCAANNNAGATCGGAAG |
| 29638_2 | TATGAGTGTGGAGTCGTTGCAACCAAAACCTGCAACAAATCAANNNAGATCGGAAG |
| 51253_1 | TATGAGTGTGGAGTCGTTGCGAGAGGAGAGGAGAGGAGAGAAANNNAGATCGG |
| 51253_2 | TATGAGTGTGGAGTCGTTGCGAGAGGAGAGGAGAGGAGAGAAANNNAGATCGG |
| 36324_1 | TATGAGTGTGGAGTCGTTGCTTAGAATCACCAATACCTGCCCCNNNAGATCGGAAG |
| 36324_2 | TATGAGTGTGGAGTCGTTGCTTAGAATCACCAATACCTGCCCCNNNAGATCGGAAG |
| 41904_1 | TATGAGTGTGGAGTCGTTGCAGTAGAGATGGGGTTTTGCTCTGNNNAGATCGGAA |
| 41904_2 | TATGAGTGTGGAGTCGTTGCAGTAGAGATGGGGTTTTGCTCTGNNNAGATCGGAA |
| 29508_1 | TATGAGTGTGGAGTCGTTGCAGGAGCTCACTATCGGTAGTNNNAGATCGGAAGAG |
| 29508_2 | TATGAGTGTGGAGTCGTTGCAGGAGCTCACTATCGGTAGTNNNAGATCGGAAGAG |
| 35416_1 | TATGAGTGTGGAGTCGTTGCATCATTGGGTGGAGGAAGAAACANNNAGATCGGA |

| | |
|---|---|
| 35416_2 | TATGAGTGTGGAGTCGTTGCATCATTGGGTGGAGGAAGAAACANNNAGATCGGAA |
| 29955_1 | TATGAGTGTGGAGTCGTTGCCCCAAACAGACTGTACCATCTGTNNNAGATCGGAAG |
| 29955_2 | TATGAGTGTGGAGTCGTTGCCCCAAACAGACTGTACCATCTGTNNNAGATCGGAAG |
| 52424_1 | TATGAGTGTGGAGTCGTTGCTAGTATGCAGCTCTCTGACTTGCNNNAGATCGGAAG |
| 52424_2 | TATGAGTGTGGAGTCGTTGCTAGTATGCAGCTCTCTGACTTGCNNNAGATCGGAAG |
| 34286_1 | TATGAGTGTGGAGTCGTTGCGGGTGCACCAAAATCTCAGAAATNNNAGATCGGAA |
| 34286_2 | TATGAGTGTGGAGTCGTTGCGGGTGCACCAAAATCTCAGAAATNNNAGATCGGAA |
| 29674_1 | TATGAGTGTGGAGTCGTTGCGCCCAAGACCCGAAGTTCTCNNNAGATCGGAAGAG |
| 29674_2 | TATGAGTGTGGAGTCGTTGCGCCCAAGACCCGAAGTTCTCNNNAGATCGGAAGAG |
| 2036_1 | TATGAGTGTGGAGTCGTTGCTTATGTGCTCCCCTTCCTGAGAANNNAGATCGGAAG |
| 2036_2 | TATGAGTGTGGAGTCGTTGCTTATGTGCTCCCCTTCCTGAGAANNNAGATCGGAAG |
| 50185_1 | TATGAGTGTGGAGTCGTTGCTGGGGTCTGAAGCTATTGTAACANNNAGATCGGAA |
| 50185_2 | TATGAGTGTGGAGTCGTTGCTGGGGTCTGAAGCTATTGTAACANNNAGATCGGAA |
| 52389_1 | TATGAGTGTGGAGTCGTTGCGGAGATTCAGACCGTACCTGAAGNNNAGATCGGAA |
| 52389_2 | TATGAGTGTGGAGTCGTTGCGGAGATTCAGACCGTACCTGAAGNNNAGATCGGAA |
| 48205_1 | TATGAGTGTGGAGTCGTTGCCGCCCAGCTAATTTTTTTGCATTNNNAGATCGGAAGA |
| 48205_2 | TATGAGTGTGGAGTCGTTGCCGCCCAGCTAATTTTTTTGCATTNNNAGATCGGAAGA |
| 47166_1 | TATGAGTGTGGAGTCGTTGCCTCCTGAGTAGCTGGGATTACAGNNNAGATCGGAA |
| 47166_2 | TATGAGTGTGGAGTCGTTGCCTCCTGAGTAGCTGGGATTACAGNNNAGATCGGAA |
| 35867_1 | TATGAGTGTGGAGTCGTTGCGAAGAAAAGCACCGTGTGGGNNNAGATCGGAAGA |
| 35867_2 | TATGAGTGTGGAGTCGTTGCGAAGAAAAGCACCGTGTGGGNNNAGATCGGAAGA |
| 12792_1 | TATGAGTGTGGAGTCGTTGCCTTGAGATGACAAGATGTTCTNNNAGATCGGAAGA |
| 12792_2 | TATGAGTGTGGAGTCGTTGCCTTGAGATGACAAGATGTTCTNNNAGATCGGAAGA |
| 13716_1 | TATGAGTGTGGAGTCGTTGCGTATACTCAGGGACCATGTCNNNAGATCGGAAGAG |
| 13716_2 | TATGAGTGTGGAGTCGTTGCGTATACTCAGGGACCATGTCNNNAGATCGGAAGAG |
| 40689_1 | TATGAGTGTGGAGTCGTTGCCACACACACACACACACATGTATNNNAGATCGGAAG |
| 40689_2 | TATGAGTGTGGAGTCGTTGCCACACACACACACACACATGTATNNNAGATCGGAAG |
| 32686_1 | TATGAGTGTGGAGTCGTTGCCAAGTAGCTGGGACTACAGGCNNNAGATCGGAAGA |
| 32686_2 | TATGAGTGTGGAGTCGTTGCCAAGTAGCTGGGACTACAGGCNNNAGATCGGAAGA |
| 33457_1 | TATGAGTGTGGAGTCGTTGCGCTTGTGTTCCAACAAATCAGGTNNNAGATCGGAAG |
| 33457_2 | TATGAGTGTGGAGTCGTTGCGCTTGTGTTCCAACAAATCAGGTNNNAGATCGGAAG |
| 42765_1 | TATGAGTGTGGAGTCGTTGCGTGAAGTCTGAATAAGGCGTGTGNNNAGATCGGAA |
| 42765_2 | TATGAGTGTGGAGTCGTTGCGTGAAGTCTGAATAAGGCGTGTGNNNAGATCGGAA |
| 49811_1 | TATGAGTGTGGAGTCGTTGCCCACTTCTGTTCTCCTCCATGATNNNAGATCGGAAGA |
| 49811_2 | TATGAGTGTGGAGTCGTTGCCCACTTCTGTTCTCCTCCATGATNNNAGATCGGAAGA |
| 43382_1 | TATGAGTGTGGAGTCGTTGCTTTTCTTTGAGACAGGCTGGAGTNNNAGATCGGAAG |
| 43382_2 | TATGAGTGTGGAGTCGTTGCTTTTCTTTGAGACAGGCTGGAGTNNNAGATCGGAAG |
| 46681_1 | TATGAGTGTGGAGTCGTTGCCCTCCTGGGTTCAAGCTATTCTCNNNAGATCGGAAG |
| 46681_2 | TATGAGTGTGGAGTCGTTGCCCTCCTGGGTTCAAGCTATTCTCNNNAGATCGGAAG |
| 30086_1 | TATGAGTGTGGAGTCGTTGCTGAGAGAGTTCAGGAGAGTAGCANNNAGATCGGAA |
| 30086_2 | TATGAGTGTGGAGTCGTTGCTGAGAGAGTTCAGGAGAGTAGCANNNAGATCGGAA |
| 34137_1 | TATGAGTGTGGAGTCGTTGCTGTCATTTGTGTTCATGGACGTGNNNAGATCGGAAG |
| 34137_2 | TATGAGTGTGGAGTCGTTGCTGTCATTTGTGTTCATGGACGTGNNNAGATCGGAAG |

| | |
|---|---|
| 10200_1 | TATGAGTGTGGAGTCGTTGCGATTCAACTGTGAGGTCTGGGTCNNNAGATCGGAA |
| 10200_2 | TATGAGTGTGGAGTCGTTGCGATTCAACTGTGAGGTCTGGGTCNNNAGATCGGAA |
| 8199_1 | TATGAGTGTGGAGTCGTTGCCTCTCCCCTCATGGTTACGAATCNNNAGATCGGAAGA |
| 8199_2 | TATGAGTGTGGAGTCGTTGCCTCTCCCCTCATGGTTACGAATCNNNAGATCGGAAGA |
| 40399_1 | TATGAGTGTGGAGTCGTTGCCCCAAACACTATCAGGCACAATGNNNAGATCGGAAG |
| 40399_2 | TATGAGTGTGGAGTCGTTGCCCCAAACACTATCAGGCACAATGNNNAGATCGGAAG |
| 41451_1 | TATGAGTGTGGAGTCGTTGCCTGAGGCAGAAGAATGGTGTGAANNNAGATCGGAA |
| 41451_2 | TATGAGTGTGGAGTCGTTGCCTGAGGCAGAAGAATGGTGTGAANNNAGATCGGAA |
| 8480_1 | TATGAGTGTGGAGTCGTTGCAAACACAATGCCACAGAAGCACNNNAGATCGGAAGA |
| 8480_2 | TATGAGTGTGGAGTCGTTGCAAACACAATGCCACAGAAGCACNNNAGATCGGAAGA |
| 43446_1 | TATGAGTGTGGAGTCGTTGCACCACATCCAGTTATTTCCCTCCNNNAGATCGGAAGA |
| 43446_2 | TATGAGTGTGGAGTCGTTGCACCACATCCAGTTATTTCCCTCCNNNAGATCGGAAGA |
| 52637_1 | TATGAGTGTGGAGTCGTTGCTCAGGGACTGCATGTTACAGAAANNNAGATCGGAA |
| 52637_2 | TATGAGTGTGGAGTCGTTGCTCAGGGACTGCATGTTACAGAAANNNAGATCGGAA |
| 43262_1 | TATGAGTGTGGAGTCGTTGCTCAAGTGATTCTCCTGTCTCAGCNNNAGATCGGAAGA |
| 43262_2 | TATGAGTGTGGAGTCGTTGCTCAAGTGATTCTCCTGTCTCAGCNNNAGATCGGAAGA |
| 42575_1 | TATGAGTGTGGAGTCGTTGCCCTGTAGTCCTAGCTACTTGGGANNNAGATCGGAAG |
| 42575_2 | TATGAGTGTGGAGTCGTTGCCCTGTAGTCCTAGCTACTTGGGANNNAGATCGGAAG |
| 30847_1 | TATGAGTGTGGAGTCGTTGCTGAAGGTTTACAGTACACCAGGGNNNAGATCGGAA |
| 30847_2 | TATGAGTGTGGAGTCGTTGCTGAAGGTTTACAGTACACCAGGGNNNAGATCGGAA |
| 33554_1 | TATGAGTGTGGAGTCGTTGCGGAGATGTGGCTTTGCTATGTTGNNNAGATCGGAA |
| 33554_2 | TATGAGTGTGGAGTCGTTGCGGAGATGTGGCTTTGCTATGTTGNNNAGATCGGAA |
| 39234_1 | TATGAGTGTGGAGTCGTTGCTAGGTTACACGGACATGGTGAAGNNNAGATCGGAA |
| 39234_2 | TATGAGTGTGGAGTCGTTGCTAGGTTACACGGACATGGTGAAGNNNAGATCGGAA |
| 41229_1 | TATGAGTGTGGAGTCGTTGCGGATCTTGCTATGTTTTCCAGGCNNNAGATCGGAAG |
| 41229_2 | TATGAGTGTGGAGTCGTTGCGGATCTTGCTATGTTTTCCAGGCNNNAGATCGGAAG |
| 46317_1 | TATGAGTGTGGAGTCGTTGCCCTGAGTAGCTGGGATTATAGGCNNNAGATCGGAA |
| 46317_2 | TATGAGTGTGGAGTCGTTGCCCTGAGTAGCTGGGATTATAGGCNNNAGATCGGAA |
| 42392_1 | TATGAGTGTGGAGTCGTTGCCACACACACACACACACATACATNNNAGATCGGAAGA |
| 42392_2 | TATGAGTGTGGAGTCGTTGCCACACACACACACACACATACATNNNAGATCGGAAGA |
| 51135_1 | TATGAGTGTGGAGTCGTTGCTTCTTTTGCAGTGAAAAGCACCANNNAGATCGGAAG |
| 51135_2 | TATGAGTGTGGAGTCGTTGCTTCTTTTGCAGTGAAAAGCACCANNNAGATCGGAAG |
| 42741_1 | TATGAGTGTGGAGTCGTTGCACCCTTCAGCCATGAAAAAAAGGNNNAGATCGGAAG |
| 42741_2 | TATGAGTGTGGAGTCGTTGCACCCTTCAGCCATGAAAAAAAGGNNNAGATCGGAAG |
| 45002_1 | TATGAGTGTGGAGTCGTTGCGGGTGAGAACCTATGCTCTTGAANNNAGATCGGAA |
| 45002_2 | TATGAGTGTGGAGTCGTTGCGGGTGAGAACCTATGCTCTTGAANNNAGATCGGAA |
| 38322_1 | TATGAGTGTGGAGTCGTTGCGTGTGGAACAAGTTGGCTAAGACNNNAGATCGGAA |
| 38322_2 | TATGAGTGTGGAGTCGTTGCGTGTGGAACAAGTTGGCTAAGACNNNAGATCGGAA |
| 37808_1 | TATGAGTGTGGAGTCGTTGCCCCCATCTCAAAAAAAAATGCCCNNNAGATCGGAAGA |
| 37808_2 | TATGAGTGTGGAGTCGTTGCCCCCATCTCAAAAAAAAATGCCCNNNAGATCGGAAGA |
| 50227_1 | TATGAGTGTGGAGTCGTTGCTTTACGCTTTCACACACCCAATCNNNAGATCGGAAGA |
| 50227_2 | TATGAGTGTGGAGTCGTTGCTTTACGCTTTCACACACCCAATCNNNAGATCGGAAGA |
| 52821_1 | TATGAGTGTGGAGTCGTTGCCCTAGCAAGAGCCAGCTTGTTNNNAGATCGGAAGAG |

| | |
|---|---|
| 52821_2 | TATGAGTGTGGAGTCGTTGCCCTAGCAAGAGCCAGCTTGTTNNNAGATCGGAAGAG |
| 143_1 | TATGAGTGTGGAGTCGTTGCCAATCAATTCTGCCACCTCAGTTTNNNAGATCGGAAG |
| 143_2 | TATGAGTGTGGAGTCGTTGCCAATCAATTCTGCCACCTCAGTTTNNNAGATCGGAAG |
| 33042_1 | TATGAGTGTGGAGTCGTTGCTTGCAAATGGAGAGCCTAGACTTNNNAGATCGGAA |
| 33042_2 | TATGAGTGTGGAGTCGTTGCTTGCAAATGGAGAGCCTAGACTTNNNAGATCGGAA |
| 29531_1 | TATGAGTGTGGAGTCGTTGCGGTTCAGCAGGATGTCCTTGTGNNNAGATCGGAAG |
| 29531_2 | TATGAGTGTGGAGTCGTTGCGGTTCAGCAGGATGTCCTTGTGNNNAGATCGGAAG |
| 44519_1 | TATGAGTGTGGAGTCGTTGCCAATGGCACATCTCGGCTCANNNAGATCGGAAGAG |
| 44519_2 | TATGAGTGTGGAGTCGTTGCCAATGGCACATCTCGGCTCANNNAGATCGGAAGAG |
| 46165_1 | TATGAGTGTGGAGTCGTTGCCACCACACCCGGCTAATTTTTNNNAGATCGGAAGAG |
| 46165_2 | TATGAGTGTGGAGTCGTTGCCACCACACCCGGCTAATTTTTNNNAGATCGGAAGAG |
| 30779_1 | TATGAGTGTGGAGTCGTTGCAGTCTGGTCTGACAATGCAGTAGNNNAGATCGGAA |
| 30779_2 | TATGAGTGTGGAGTCGTTGCAGTCTGGTCTGACAATGCAGTAGNNNAGATCGGAA |
| 13621_1 | TATGAGTGTGGAGTCGTTGCCTAATAGATTACCTTTATCGCCTNNNAGATCGGAAGA |
| 13621_2 | TATGAGTGTGGAGTCGTTGCCTAATAGATTACCTTTATCGCCTNNNAGATCGGAAGA |
| 10070_1 | TATGAGTGTGGAGTCGTTGCGCATGGGCTTGGGGCTCAGTNNNAGATCGGAAGAG |
| 10070_2 | TATGAGTGTGGAGTCGTTGCGCATGGGCTTGGGGCTCAGTNNNAGATCGGAAGAG |
| 30258_1 | TATGAGTGTGGAGTCGTTGCCCCTTTGGTCTTGATATTTCTATTGGANNNAGATCGG |
| 30258_2 | TATGAGTGTGGAGTCGTTGCCCCTTTGGTCTTGATATTTCTATTGGANNNAGATCGG |
| 37689_1 | TATGAGTGTGGAGTCGTTGCACACAACCATAAGCCTGAATTCANNNAGATCGGAAG |
| 37689_2 | TATGAGTGTGGAGTCGTTGCACACAACCATAAGCCTGAATTCANNNAGATCGGAAG |
| 32262_1 | TATGAGTGTGGAGTCGTTGCCGCCAATACGTAAAAGAAGCCATNNNAGATCGGAAG |
| 32262_2 | TATGAGTGTGGAGTCGTTGCCGCCAATACGTAAAAGAAGCCATNNNAGATCGGAAG |
| 29142_1 | TATGAGTGTGGAGTCGTTGCCAGCTCTGGATCCCTAGTCATTCNNNAGATCGGAAG |
| 29142_2 | TATGAGTGTGGAGTCGTTGCCAGCTCTGGATCCCTAGTCATTCNNNAGATCGGAAG |
| 39537_1 | TATGAGTGTGGAGTCGTTGCAGGTTGCAGTGAACTGAGATCATNNNAGATCGGAA |
| 39537_2 | TATGAGTGTGGAGTCGTTGCAGGTTGCAGTGAACTGAGATCATNNNAGATCGGAA |
| 40535_1 | TATGAGTGTGGAGTCGTTGCCAGCCTGGCCAATATGGTAAAACNNNAGATCGGAAG |
| 40535_2 | TATGAGTGTGGAGTCGTTGCCAGCCTGGCCAATATGGTAAAACNNNAGATCGGAAG |
| 32764_1 | TATGAGTGTGGAGTCGTTGCGCTCAAATGTTCCTGATGGCTTTNNNAGATCGGAAG |
| 32764_2 | TATGAGTGTGGAGTCGTTGCGCTCAAATGTTCCTGATGGCTTTNNNAGATCGGAAG |
| 30268_1 | TATGAGTGTGGAGTCGTTGCGTGCCAGAACTGCAAGTAAGTTTNNNAGATCGGAA |
| 30268_2 | TATGAGTGTGGAGTCGTTGCGTGCCAGAACTGCAAGTAAGTTTNNNAGATCGGAA |
| 34501_1 | TATGAGTGTGGAGTCGTTGCAACTCATTGTTCTCACTCGAGCTNNNAGATCGGAAGA |
| 34501_2 | TATGAGTGTGGAGTCGTTGCAACTCATTGTTCTCACTCGAGCTNNNAGATCGGAAGA |
| 11327_1 | TATGAGTGTGGAGTCGTTGCTGCGATGTGTGTTCCAATGACNNNAGATCGGAAGA |
| 11327_2 | TATGAGTGTGGAGTCGTTGCTGCGATGTGTGTTCCAATGACNNNAGATCGGAAGA |
| 35057_1 | TATGAGTGTGGAGTCGTTGCAAATGGAGGAGTTGGACTTTGCTNNNAGATCGGAA |
| 35057_2 | TATGAGTGTGGAGTCGTTGCAAATGGAGGAGTTGGACTTTGCTNNNAGATCGGAA |
| 35070_1 | TATGAGTGTGGAGTCGTTGCCCTCCTGAGTAGCTGGGACTATANNNAGATCGGAAG |
| 35070_2 | TATGAGTGTGGAGTCGTTGCCCTCCTGAGTAGCTGGGACTATANNNAGATCGGAAG |
| 53265_1 | TATGAGTGTGGAGTCGTTGCCTACCGCCGTCTTGGAGATGNNNAGATCGGAAGAG |
| 53265_2 | TATGAGTGTGGAGTCGTTGCCTACCGCCGTCTTGGAGATGNNNAGATCGGAAGAG |

| | |
|---|---|
| 45521_1 | TATGAGTGTGGAGTCGTTGCACTGAGTAGAGTAGATGGGTCCANNNAGATCGGAA |
| 45521_2 | TATGAGTGTGGAGTCGTTGCACTGAGTAGAGTAGATGGGTCCANNNAGATCGGAA |
| 35738_1 | TATGAGTGTGGAGTCGTTGCCTAGGTTCAAGCAATTCTCCTGCNNNAGATCGGAAG |
| 35738_2 | TATGAGTGTGGAGTCGTTGCCTAGGTTCAAGCAATTCTCCTGCNNNAGATCGGAAG |
| 53139_1 | TATGAGTGTGGAGTCGTTGCCCAGGCCAGTGTATTAGCAGTTANNNAGATCGGAA |
| 53139_2 | TATGAGTGTGGAGTCGTTGCCCAGGCCAGTGTATTAGCAGTTANNNAGATCGGAA |
| 44635_1 | TATGAGTGTGGAGTCGTTGCGGAGGACAGGTAATCCAATTGGTNNNAGATCGGAA |
| 44635_2 | TATGAGTGTGGAGTCGTTGCGGAGGACAGGTAATCCAATTGGTNNNAGATCGGAA |
| 38117_1 | TATGAGTGTGGAGTCGTTGCTATAGTACCAGGGACCAAGAGCTNNNAGATCGGAA |
| 38117_2 | TATGAGTGTGGAGTCGTTGCTATAGTACCAGGGACCAAGAGCTNNNAGATCGGAA |
| 6144_1 | TATGAGTGTGGAGTCGTTGCTTGAGCAGCAGAAGTGACAGATCNNNAGATCGGAA |
| 6144_2 | TATGAGTGTGGAGTCGTTGCTTGAGCAGCAGAAGTGACAGATCNNNAGATCGGAA |
| 29881_1 | TATGAGTGTGGAGTCGTTGCTCTAACCCATAAGTCACTGGCAANNNAGATCGGAAG |
| 29881_2 | TATGAGTGTGGAGTCGTTGCTCTAACCCATAAGTCACTGGCAANNNAGATCGGAAG |
| 5566_1 | TATGAGTGTGGAGTCGTTGCCACCACATACACACACACCTTCANNNAGATCGGAAGA |
| 5566_2 | TATGAGTGTGGAGTCGTTGCCACCACATACACACACACCTTCANNNAGATCGGAAGA |
| 51653_1 | TATGAGTGTGGAGTCGTTGCTGGCTGCTCATCAAGAAACATTCNNNAGATCGGAAG |
| 51653_2 | TATGAGTGTGGAGTCGTTGCTGGCTGCTCATCAAGAAACATTCNNNAGATCGGAAG |
| 34328_1 | TATGAGTGTGGAGTCGTTGCTGGTTGACATTGGACCGTATCATNNNAGATCGGAAG |
| 34328_2 | TATGAGTGTGGAGTCGTTGCTGGTTGACATTGGACCGTATCATNNNAGATCGGAAG |
| 40479_1 | TATGAGTGTGGAGTCGTTGCAAAAAGAAAAAAAGGTGGGGGGGNNNAGATCGG |
| 40479_2 | TATGAGTGTGGAGTCGTTGCAAAAAGAAAAAAAGGTGGGGGGGNNNAGATCGG |
| 36002_1 | TATGAGTGTGGAGTCGTTGCCAGGTCTCTACCCCATTTCTGAGNNNAGATCGGAAG |
| 36002_2 | TATGAGTGTGGAGTCGTTGCCAGGTCTCTACCCCATTTCTGAGNNNAGATCGGAAG |
| 38123_1 | TATGAGTGTGGAGTCGTTGCACATGCCAAGAGTGAAAAATCTGANNNAGATCGGA |
| 38123_2 | TATGAGTGTGGAGTCGTTGCACATGCCAAGAGTGAAAAATCTGANNNAGATCGGA |
| 43105_1 | TATGAGTGTGGAGTCGTTGCCTGTAGTCCCAGCTACTCAGGAGNNNAGATCGGAAG |
| 43105_2 | TATGAGTGTGGAGTCGTTGCCTGTAGTCCCAGCTACTCAGGAGNNNAGATCGGAAG |
| 48684_1 | TATGAGTGTGGAGTCGTTGCTTTGCAGAAGTGAATCTGGGGTANNNAGATCGGAA |
| 48684_2 | TATGAGTGTGGAGTCGTTGCTTTGCAGAAGTGAATCTGGGGTANNNAGATCGGAA |
| 33282_1 | TATGAGTGTGGAGTCGTTGCCAATCTTGGCTCACTGCAATCTCNNNAGATCGGAAG |
| 33282_2 | TATGAGTGTGGAGTCGTTGCCAATCTTGGCTCACTGCAATCTCNNNAGATCGGAAG |
| 30843_1 | TATGAGTGTGGAGTCGTTGCCACACACACTACTCTACCCTTCCNNNAGATCGGAAGA |
| 30843_2 | TATGAGTGTGGAGTCGTTGCCACACACACTACTCTACCCTTCCNNNAGATCGGAAGA |
| 33980_1 | TATGAGTGTGGAGTCGTTGCTCTCGAGTAGCTGGAACTACAGANNNAGATCGGAA |
| 33980_2 | TATGAGTGTGGAGTCGTTGCTCTCGAGTAGCTGGAACTACAGANNNAGATCGGAA |
| 36874_1 | TATGAGTGTGGAGTCGTTGCTGCTGATGGAAGTGGTGGAAATANNNAGATCGGA |
| 36874_2 | TATGAGTGTGGAGTCGTTGCTGCTGATGGAAGTGGTGGAAATANNNAGATCGGA |
| 34942_1 | TATGAGTGTGGAGTCGTTGCGATCTCTGCTCAGTACAGCCTACNNNAGATCGGAAG |
| 34942_2 | TATGAGTGTGGAGTCGTTGCGATCTCTGCTCAGTACAGCCTACNNNAGATCGGAAG |
| 50164_1 | TATGAGTGTGGAGTCGTTGCAGTACAAGGCAGCTTTCTACTCCNNNAGATCGGAAG |
| 50164_2 | TATGAGTGTGGAGTCGTTGCAGTACAAGGCAGCTTTCTACTCCNNNAGATCGGAAG |
| 33563_1 | TATGAGTGTGGAGTCGTTGCTCCTAAGTAGCTGGGATCACAGANNNAGATCGGAA |

| | |
|---|---|
| 33563_2 | TATGAGTGTGGAGTCGTTGCTCCTAAGTAGCTGGGATCACAGANNNAGATCGGAA |
| 45034_1 | TATGAGTGTGGAGTCGTTGCCTAGCTCCTGGGATGTCTTCATCNNNAGATCGGAAG |
| 45034_2 | TATGAGTGTGGAGTCGTTGCCTAGCTCCTGGGATGTCTTCATCNNNAGATCGGAAG |
| 33357_1 | TATGAGTGTGGAGTCGTTGCGAGACGGAGTCTTACTCTGTCGNNNAGATCGGAAG |
| 33357_2 | TATGAGTGTGGAGTCGTTGCGAGACGGAGTCTTACTCTGTCGNNNAGATCGGAAG |
| 31169_1 | TATGAGTGTGGAGTCGTTGCTGTAGAGCCACACTTGAGAGATGNNNAGATCGGAA |
| 31169_2 | TATGAGTGTGGAGTCGTTGCTGTAGAGCCACACTTGAGAGATGNNNAGATCGGAA |
| 3042_1 | TATGAGTGTGGAGTCGTTGCGGTAGTAGCTCTTGAAGCACCAANNNAGATCGGAA |
| 3042_2 | TATGAGTGTGGAGTCGTTGCGGTAGTAGCTCTTGAAGCACCAANNNAGATCGGAA |
| 10276_1 | TATGAGTGTGGAGTCGTTGCAGAGAACAGCCCATCTGAGTGTANNNAGATCGGAA |
| 10276_2 | TATGAGTGTGGAGTCGTTGCAGAGAACAGCCCATCTGAGTGTANNNAGATCGGAA |
| 46357_1 | TATGAGTGTGGAGTCGTTGCACCACCACACTCGGCTAATTTNNNAGATCGGAAGAG |
| 46357_2 | TATGAGTGTGGAGTCGTTGCACCACCACACTCGGCTAATTTNNNAGATCGGAAGAG |
| 35302_1 | TATGAGTGTGGAGTCGTTGCGAGCCATCTTGGAGTCCCTTTTANNNAGATCGGAAG |
| 35302_2 | TATGAGTGTGGAGTCGTTGCGAGCCATCTTGGAGTCCCTTTTANNNAGATCGGAAG |
| 43947_1 | TATGAGTGTGGAGTCGTTGCGTGCTCCAGTTTTGGATGCAAATNNNAGATCGGAAG |
| 43947_2 | TATGAGTGTGGAGTCGTTGCGTGCTCCAGTTTTGGATGCAAATNNNAGATCGGAAG |
| 48732_1 | TATGAGTGTGGAGTCGTTGCAATGGGGTTTTTACATGTTGGCCNNNAGATCGGAA |
| 48732_2 | TATGAGTGTGGAGTCGTTGCAATGGGGTTTTTACATGTTGGCCNNNAGATCGGAA |
| 50135_1 | TATGAGTGTGGAGTCGTTGCAAAATACCATCCTGCCTCACCTTNNNAGATCGGAAGA |
| 50135_2 | TATGAGTGTGGAGTCGTTGCAAAATACCATCCTGCCTCACCTTNNNAGATCGGAAGA |
| 2005_1 | TATGAGTGTGGAGTCGTTGCGGCTGATGAGAGCGTCTTACAAANNNAGATCGGAA |
| 2005_2 | TATGAGTGTGGAGTCGTTGCGGCTGATGAGAGCGTCTTACAAANNNAGATCGGAA |
| 49213_1 | TATGAGTGTGGAGTCGTTGCAGCCTTTACTTCCTGAGCTCAAGNNNAGATCGGAAG |
| 49213_2 | TATGAGTGTGGAGTCGTTGCAGCCTTTACTTCCTGAGCTCAAGNNNAGATCGGAAG |
| 45026_1 | TATGAGTGTGGAGTCGTTGCAGACAGAACAAGGCAGCTCATAANNNAGATCGGAA |
| 45026_2 | TATGAGTGTGGAGTCGTTGCAGACAGAACAAGGCAGCTCATAANNNAGATCGGAA |
| 51213_1 | TATGAGTGTGGAGTCGTTGCGAGTCTCTCCATCCAACGCAATANNNAGATCGGAAG |
| 51213_2 | TATGAGTGTGGAGTCGTTGCGAGTCTCTCCATCCAACGCAATANNNAGATCGGAAG |
| 32587_1 | TATGAGTGTGGAGTCGTTGCAAAAGCAAATGTTTGACTGGGCANNNAGATCGGAA |
| 32587_2 | TATGAGTGTGGAGTCGTTGCAAAAGCAAATGTTTGACTGGGCANNNAGATCGGAA |
| 42478_1 | TATGAGTGTGGAGTCGTTGCATGAAGAACAATTGCCTCAGTGCNNNAGATCGGAA |
| 42478_2 | TATGAGTGTGGAGTCGTTGCATGAAGAACAATTGCCTCAGTGCNNNAGATCGGAA |
| 48308_1 | TATGAGTGTGGAGTCGTTGCATTCCACACAGATGGGGATATGGNNNAGATCGGAA |
| 48308_2 | TATGAGTGTGGAGTCGTTGCATTCCACACAGATGGGGATATGGNNNAGATCGGAA |
| 33516_1 | TATGAGTGTGGAGTCGTTGCGAGAAGACACTGACATTCGGAGGNNNAGATCGGAA |
| 33516_2 | TATGAGTGTGGAGTCGTTGCGAGAAGACACTGACATTCGGAGGNNNAGATCGGAA |
| 50380_1 | TATGAGTGTGGAGTCGTTGCGCTAAGTAATTCCAGGAGCACCTNNNAGATCGGAAG |
| 50380_2 | TATGAGTGTGGAGTCGTTGCGCTAAGTAATTCCAGGAGCACCTNNNAGATCGGAAG |
| 29882_1 | TATGAGTGTGGAGTCGTTGCAATAGACAACTGAGGACACTTCANNNAGATCGGAA |
| 29882_2 | TATGAGTGTGGAGTCGTTGCAATAGACAACTGAGGACACTTCANNNAGATCGGAA |
| 13314_1 | TATGAGTGTGGAGTCGTTGCCATCATTTCTGGAGTCTCAGNNNAGATCGGAAGAGC |
| 13314_2 | TATGAGTGTGGAGTCGTTGCCATCATTTCTGGAGTCTCAGNNNAGATCGGAAGAGC |

9695_1 TATGAGTGTGGAGTCGTTGCTGTGGTTGGTAGGAAGAGGTACCNNNAGATCGGAA
9695_2 TATGAGTGTGGAGTCGTTGCTGTGGTTGGTAGGAAGAGGTACCNNNAGATCGGAA
9824_1 TATGAGTGTGGAGTCGTTGCAAGAACAGGCTGTCAACCAAGACNNNAGATCGGAA
9824_2 TATGAGTGTGGAGTCGTTGCAAGAACAGGCTGTCAACCAAGACNNNAGATCGGAA
34641_1 TATGAGTGTGGAGTCGTTGCGAACCATCTCTCTCAGGACGCNNNAGATCGGAAGAG
34641_2 TATGAGTGTGGAGTCGTTGCGAACCATCTCTCTCAGGACGCNNNAGATCGGAAGAG
53145_1 TATGAGTGTGGAGTCGTTGCGAAAAGCTGCTGAGTTCCAAAGTNNNAGATCGGAA
53145_2 TATGAGTGTGGAGTCGTTGCGAAAAGCTGCTGAGTTCCAAAGTNNNAGATCGGAA
49443_1 TATGAGTGTGGAGTCGTTGCGCCTCCTGAGTAGCTGAGATTACNNNAGATCGGAAG
49443_2 TATGAGTGTGGAGTCGTTGCGCCTCCTGAGTAGCTGAGATTACNNNAGATCGGAAG
38477_1 TATGAGTGTGGAGTCGTTGCATGGAGCACTACACAGCCATAAANNNAGATCGGAAG
38477_2 TATGAGTGTGGAGTCGTTGCATGGAGCACTACACAGCCATAAANNNAGATCGGAAG
44889_1 TATGAGTGTGGAGTCGTTGCAAATCATTTCCCCTCCTGGATCCNNNAGATCGGAAGA
44889_2 TATGAGTGTGGAGTCGTTGCAAATCATTTCCCCTCCTGGATCCNNNAGATCGGAAGA
40192_1 TATGAGTGTGGAGTCGTTGCGGATCACTTGAGGTCAGGAGTTCNNNAGATCGGAA
40192_2 TATGAGTGTGGAGTCGTTGCGGATCACTTGAGGTCAGGAGTTCNNNAGATCGGAA
33577_1 TATGAGTGTGGAGTCGTTGCGTAGCTAGGATTACAGGCATGCANNNAGATCGGAA
33577_2 TATGAGTGTGGAGTCGTTGCGTAGCTAGGATTACAGGCATGCANNNAGATCGGAA
33608_1 TATGAGTGTGGAGTCGTTGCAACACCGACCATCTTTTCATGTGNNNAGATCGGAAGA
33608_2 TATGAGTGTGGAGTCGTTGCAACACCGACCATCTTTTCATGTGNNNAGATCGGAAGA
31386_1 TATGAGTGTGGAGTCGTTGCGGAATGTGTTTGTGCGTATGGAANNNAGATCGGAA
31386_2 TATGAGTGTGGAGTCGTTGCGGAATGTGTTTGTGCGTATGGAANNNAGATCGGAA
29862_1 TATGAGTGTGGAGTCGTTGCCCCCTTGATTCTTCCTTCCCACCNNNAGATCGGAAGA
29862_2 TATGAGTGTGGAGTCGTTGCCCCCTTGATTCTTCCTTCCCACCNNNAGATCGGAAGA
51182_1 TATGAGTGTGGAGTCGTTGCATGGGTCCACCAAAATCTCAGAANNNAGATCGGAAG
51182_2 TATGAGTGTGGAGTCGTTGCATGGGTCCACCAAAATCTCAGAANNNAGATCGGAAG
53225_1 TATGAGTGTGGAGTCGTTGCTTTGGCCTCAAACTTCAGGTGANNNAGATCGGAAGA
53225_2 TATGAGTGTGGAGTCGTTGCTTTGGCCTCAAACTTCAGGTGANNNAGATCGGAAGA
51231_1 TATGAGTGTGGAGTCGTTGCTTATGTTGGTCAGGCTGATCTCGNNNAGATCGGAAG
51231_2 TATGAGTGTGGAGTCGTTGCTTATGTTGGTCAGGCTGATCTCGNNNAGATCGGAAG
42853_1 TATGAGTGTGGAGTCGTTGCCTCCCAGGTTCAAGTGATTCTCANNNAGATCGGAAGA
42853_2 TATGAGTGTGGAGTCGTTGCCTCCCAGGTTCAAGTGATTCTCANNNAGATCGGAAGA
49981_1 TATGAGTGTGGAGTCGTTGCCTCAGACAGACCCCAAATCCTCNNNAGATCGGAAGA
49981_2 TATGAGTGTGGAGTCGTTGCCTCAGACAGACCCCAAATCCTCNNNAGATCGGAAGA
42221_1 TATGAGTGTGGAGTCGTTGCTGCATAACCGGTGGAAGAAACTANNNAGATCGGAA
42221_2 TATGAGTGTGGAGTCGTTGCTGCATAACCGGTGGAAGAAACTANNNAGATCGGAA
51034_1 TATGAGTGTGGAGTCGTTGCTTTCCATGTGCAGATCAGATGGANNNAGATCGGAAG
51034_2 TATGAGTGTGGAGTCGTTGCTTTCCATGTGCAGATCAGATGGANNNAGATCGGAAG
30523_1 TATGAGTGTGGAGTCGTTGCACACGGACAACAGATTCAGCTGANNNAGATCGGAA
30523_2 TATGAGTGTGGAGTCGTTGCACACGGACAACAGATTCAGCTGANNNAGATCGGAA
39503_1 TATGAGTGTGGAGTCGTTGCACCTTCCTGGGATTAGGCTTTTTNNNAGATCGGAAGA
39503_2 TATGAGTGTGGAGTCGTTGCACCTTCCTGGGATTAGGCTTTTTNNNAGATCGGAAGA
42308_1 TATGAGTGTGGAGTCGTTGCGAGCCTCTTTCTCACCACATACANNNAGATCGGAAGA

| | |
|---|---|
| 42308_2 | TATGAGTGTGGAGTCGTTGCGAGCCTCTTTCTCACCACATACANNNAGATCGGAAGA |
| 30443_1 | TATGAGTGTGGAGTCGTTGCCCCTTATTAAAGTGAGTGGCCCCNNNAGATCGGAAG |
| 30443_2 | TATGAGTGTGGAGTCGTTGCCCCTTATTAAAGTGAGTGGCCCCNNNAGATCGGAAG |
| 44009_1 | TATGAGTGTGGAGTCGTTGCACCACCAAAACATGCATTTCTAGGNNNAGATCGGAA |
| 44009_2 | TATGAGTGTGGAGTCGTTGCACCACCAAAACATGCATTTCTAGGNNNAGATCGGAA |
| 34523_1 | TATGAGTGTGGAGTCGTTGCAAGTAAGTTTCTGAGTGGCTCCTNNNAGATCGGAAG |
| 34523_2 | TATGAGTGTGGAGTCGTTGCAAGTAAGTTTCTGAGTGGCTCCTNNNAGATCGGAAG |
| 30133_1 | TATGAGTGTGGAGTCGTTGCGGATGGCAGTGATGAAGAGAACTNNNAGATCGGAA |
| 30133_2 | TATGAGTGTGGAGTCGTTGCGGATGGCAGTGATGAAGAGAACTNNNAGATCGGAA |
| 39583_1 | TATGAGTGTGGAGTCGTTGCAAAAGTTGTCAGATCACTTGCCANNNAGATCGGAAG |
| 39583_2 | TATGAGTGTGGAGTCGTTGCAAAAGTTGTCAGATCACTTGCCANNNAGATCGGAAG |
| 38088_1 | TATGAGTGTGGAGTCGTTGCGGGATTTTGTTTGGGATTGTGCTNNNAGATCGGAA |
| 38088_2 | TATGAGTGTGGAGTCGTTGCGGGATTTTGTTTGGGATTGTGCTNNNAGATCGGAA |
| 45737_1 | TATGAGTGTGGAGTCGTTGCGCCTCCTGAATAGCTGGGAATACNNNAGATCGGAAG |
| 45737_2 | TATGAGTGTGGAGTCGTTGCGCCTCCTGAATAGCTGGGAATACNNNAGATCGGAAG |
| 32539_1 | TATGAGTGTGGAGTCGTTGCGAGACTTCAGGGATCATGGTTGTNNNAGATCGGAA |
| 32539_2 | TATGAGTGTGGAGTCGTTGCGAGACTTCAGGGATCATGGTTGTNNNAGATCGGAA |
| 33416_1 | TATGAGTGTGGAGTCGTTGCAGTATGGCATCTCCTCATCTCCTNNNAGATCGGAAGA |
| 33416_2 | TATGAGTGTGGAGTCGTTGCAGTATGGCATCTCCTCATCTCCTNNNAGATCGGAAGA |
| 34093_1 | TATGAGTGTGGAGTCGTTGCCCAAAGCAAGCAGATCTCTTCAGNNNAGATCGGAAG |
| 34093_2 | TATGAGTGTGGAGTCGTTGCCCAAAGCAAGCAGATCTCTTCAGNNNAGATCGGAAG |
| 46339_1 | TATGAGTGTGGAGTCGTTGCTGTATTTCTGTGGCATCAGTGGANNNAGATCGGAAG |
| 46339_2 | TATGAGTGTGGAGTCGTTGCTGTATTTCTGTGGCATCAGTGGANNNAGATCGGAAG |
| 4050_1 | TATGAGTGTGGAGTCGTTGCTGAACTACTTGTGCGTGTTAGGTNNNAGATCGGAAG |
| 4050_2 | TATGAGTGTGGAGTCGTTGCTGAACTACTTGTGCGTGTTAGGTNNNAGATCGGAAG |
| 6425_1 | TATGAGTGTGGAGTCGTTGCTACACATATGCTCCCATCCAGGCNNNAGATCGGAAGA |
| 6425_2 | TATGAGTGTGGAGTCGTTGCTACACATATGCTCCCATCCAGGCNNNAGATCGGAAGA |
| 42850_1 | TATGAGTGTGGAGTCGTTGCCTCTCCTTTTCCTCATCTGCCTTNNNAGATCGGAAGAG |
| 42850_2 | TATGAGTGTGGAGTCGTTGCCTCTCCTTTTCCTCATCTGCCTTNNNAGATCGGAAGAG |
| 44186_1 | TATGAGTGTGGAGTCGTTGCTGGTGTTGGAGAAGGAGATAAACANNNAGATCGGA |
| 44186_2 | TATGAGTGTGGAGTCGTTGCTGGTGTTGGAGAAGGAGATAAACANNNAGATCGGA |
| 44005_1 | TATGAGTGTGGAGTCGTTGCCGAGAGAGAGAGAGAGAGAGAGANNNAGATCGGA |
| 44005_2 | TATGAGTGTGGAGTCGTTGCCGAGAGAGAGAGAGAGAGAGAGANNNAGATCGGA |
| 32302_1 | TATGAGTGTGGAGTCGTTGCGGAGGACAGGTAATCCAATTGGTNNNAGATCGGAA |
| 32302_2 | TATGAGTGTGGAGTCGTTGCGGAGGACAGGTAATCCAATTGGTNNNAGATCGGAA |
| 47563_1 | TATGAGTGTGGAGTCGTTGCTGATGTTCCAGAACTGCATCAGANNNAGATCGGAAG |
| 47563_2 | TATGAGTGTGGAGTCGTTGCTGATGTTCCAGAACTGCATCAGANNNAGATCGGAAG |
| 44737_1 | TATGAGTGTGGAGTCGTTGCTGTTGAAGAGAAATGAGCATCGCNNNAGATCGGAA |
| 44737_2 | TATGAGTGTGGAGTCGTTGCTGTTGAAGAGAAATGAGCATCGCNNNAGATCGGAA |
| 12649_1 | TATGAGTGTGGAGTCGTTGCAACAGGTCATAGTTTGAACCNNNAGATCGGAAGAG |
| 12649_2 | TATGAGTGTGGAGTCGTTGCAACAGGTCATAGTTTGAACCNNNAGATCGGAAGAG |
| 35122_1 | TATGAGTGTGGAGTCGTTGCTACAAGTCTGCAAGAACCACAGTNNNAGATCGGAAG |
| 35122_2 | TATGAGTGTGGAGTCGTTGCTACAAGTCTGCAAGAACCACAGTNNNAGATCGGAAG |

| | |
|---|---|
| 40046_1 | TATGAGTGTGGAGTCGTTGCGGATGGGCTCCATATTGTGATCANNNAGATCGGAA |
| 40046_2 | TATGAGTGTGGAGTCGTTGCGGATGGGCTCCATATTGTGATCANNNAGATCGGAA |
| 51039_1 | TATGAGTGTGGAGTCGTTGCTTCACTTTCTTGTTTCAGCTGGCNNNAGATCGGAAG |
| 51039_2 | TATGAGTGTGGAGTCGTTGCTTCACTTTCTTGTTTCAGCTGGCNNNAGATCGGAAG |
| 10618_1 | TATGAGTGTGGAGTCGTTGCGGCAAGTTTTACCTACAGCTCTNNNAGATCGGAAGA |
| 10618_2 | TATGAGTGTGGAGTCGTTGCGGCAAGTTTTACCTACAGCTCTNNNAGATCGGAAGA |
| 31569_1 | TATGAGTGTGGAGTCGTTGCTTCACAGGGCTAAAGGTTATGGGNNNAGATCGGAA |
| 31569_2 | TATGAGTGTGGAGTCGTTGCTTCACAGGGCTAAAGGTTATGGGNNNAGATCGGAA |
| 37154_1 | TATGAGTGTGGAGTCGTTGCTCCAGACACCTCACTCTTCTAAGNNNAGATCGGAAG |
| 37154_2 | TATGAGTGTGGAGTCGTTGCTCCAGACACCTCACTCTTCTAAGNNNAGATCGGAAG |
| 52402_1 | TATGAGTGTGGAGTCGTTGCGTTTTAGGCAGCCCCAAAGTGNNNAGATCGGAAGA |
| 52402_2 | TATGAGTGTGGAGTCGTTGCGTTTTAGGCAGCCCCAAAGTGNNNAGATCGGAAGA |
| 43938_1 | TATGAGTGTGGAGTCGTTGCGAGAAGGAAGAGAGTGTTTGGGTNNNAGATCGGA |
| 43938_2 | TATGAGTGTGGAGTCGTTGCGAGAAGGAAGAGAGTGTTTGGGTNNNAGATCGGA |
| 41572_1 | TATGAGTGTGGAGTCGTTGCAAACTCCTAACCTCATGATCCGCNNNAGATCGGAAG |
| 41572_2 | TATGAGTGTGGAGTCGTTGCAAACTCCTAACCTCATGATCCGCNNNAGATCGGAAG |
| 12539_1 | TATGAGTGTGGAGTCGTTGCCTTTTCCTTGTACCCAACAANNNAGATCGGAAGAGC |
| 12539_2 | TATGAGTGTGGAGTCGTTGCCTTTTCCTTGTACCCAACAANNNAGATCGGAAGAGC |
| 32919_1 | TATGAGTGTGGAGTCGTTGCTCAGAACAAATGCATCCAGGAGANNNAGATCGGAA |
| 32919_2 | TATGAGTGTGGAGTCGTTGCTCAGAACAAATGCATCCAGGAGANNNAGATCGGAA |
| 34671_1 | TATGAGTGTGGAGTCGTTGCGCAGCAGGAAGCCTTATAACATTNNNAGATCGGAA |
| 34671_2 | TATGAGTGTGGAGTCGTTGCGCAGCAGGAAGCCTTATAACATTNNNAGATCGGAA |
| 33637_1 | TATGAGTGTGGAGTCGTTGCCGGGTTCATGCCATTCTCCTNNNAGATCGGAAGAGC |
| 33637_2 | TATGAGTGTGGAGTCGTTGCCGGGTTCATGCCATTCTCCTNNNAGATCGGAAGAGC |
| 33848_1 | TATGAGTGTGGAGTCGTTGCAGACTGAGGCAGGAGAATTGTTTNNNAGATCGGAA |
| 33848_2 | TATGAGTGTGGAGTCGTTGCAGACTGAGGCAGGAGAATTGTTTNNNAGATCGGAA |
| 37354_1 | TATGAGTGTGGAGTCGTTGCATGCTTTGAGAGGGATCTTCAGTNNNAGATCGGAA |
| 37354_2 | TATGAGTGTGGAGTCGTTGCATGCTTTGAGAGGGATCTTCAGTNNNAGATCGGAA |
| 43063_1 | TATGAGTGTGGAGTCGTTGCTGGAGTCAATTGAAAGAGGTCCCNNNAGATCGGAA |
| 43063_2 | TATGAGTGTGGAGTCGTTGCTGGAGTCAATTGAAAGAGGTCCCNNNAGATCGGAA |
| 38421_1 | TATGAGTGTGGAGTCGTTGCGCCTTTCCTCTAGTATCTGGAGCNNNAGATCGGAAG |
| 38421_2 | TATGAGTGTGGAGTCGTTGCGCCTTTCCTCTAGTATCTGGAGCNNNAGATCGGAAG |
| 39860_1 | TATGAGTGTGGAGTCGTTGCCTTATCCAGCCCAGTGGTGATTNNNAGATCGGAAGA |
| 39860_2 | TATGAGTGTGGAGTCGTTGCCTTATCCAGCCCAGTGGTGATTNNNAGATCGGAAGA |
| 40887_1 | TATGAGTGTGGAGTCGTTGCCTGACTCATGCTAAATGGCCATGNNNAGATCGGAAG |
| 40887_2 | TATGAGTGTGGAGTCGTTGCCTGACTCATGCTAAATGGCCATGNNNAGATCGGAAG |
| 47330_1 | TATGAGTGTGGAGTCGTTGCCCGAGTAGCTGGAATTATAGGCANNNAGATCGGAA |
| 47330_2 | TATGAGTGTGGAGTCGTTGCCCGAGTAGCTGGAATTATAGGCANNNAGATCGGAA |
| 29159_1 | TATGAGTGTGGAGTCGTTGCGAAGAGTCTCGAAGTTCCTCCTGNNNAGATCGGAAG |
| 29159_2 | TATGAGTGTGGAGTCGTTGCGAAGAGTCTCGAAGTTCCTCCTGNNNAGATCGGAAG |
| 45067_1 | TATGAGTGTGGAGTCGTTGCCTCAAATTGTGTGCCTGTGACAANNNAGATCGGAAG |
| 45067_2 | TATGAGTGTGGAGTCGTTGCCTCAAATTGTGTGCCTGTGACAANNNAGATCGGAAG |
| 44571_1 | TATGAGTGTGGAGTCGTTGCCAGATCTGGGGAACCTCTGTTGNNNAGATCGGAAG |

| | |
|---|---|
| 44571_2 | TATGAGTGTGGAGTCGTTGCCAGATCTGGGGAACCTCTGTTGNNNAGATCGGAAGA |
| 39392_1 | TATGAGTGTGGAGTCGTTGCAAAGACGGACTGACAGGGTTTTANNNAGATCGGAA |
| 39392_2 | TATGAGTGTGGAGTCGTTGCAAAGACGGACTGACAGGGTTTTANNNAGATCGGAA |
| 34794_1 | TATGAGTGTGGAGTCGTTGCCAGTGGTTCAATCTCGGCTCANNNAGATCGGAAGAG |
| 34794_2 | TATGAGTGTGGAGTCGTTGCCAGTGGTTCAATCTCGGCTCANNNAGATCGGAAGAG |
| 48910_1 | TATGAGTGTGGAGTCGTTGCGTGCCTAACCTGAAATGCCGANNNAGATCGGAAGAG |
| 48910_2 | TATGAGTGTGGAGTCGTTGCGTGCCTAACCTGAAATGCCGANNNAGATCGGAAGAG |
| 39866_1 | TATGAGTGTGGAGTCGTTGCTTGCTGCCTATCTCTCTTCTGACNNNAGATCGGAAGA |
| 39866_2 | TATGAGTGTGGAGTCGTTGCTTGCTGCCTATCTCTCTTCTGACNNNAGATCGGAAGA |
| 49680_1 | TATGAGTGTGGAGTCGTTGCAGTCTCCCTCTGAAGTACAGTGTNNNAGATCGGAAG |
| 49680_2 | TATGAGTGTGGAGTCGTTGCAGTCTCCCTCTGAAGTACAGTGTNNNAGATCGGAAG |
| 44748_1 | TATGAGTGTGGAGTCGTTGCACTGTAGGGACAACCATGTTGTTNNNAGATCGGAAG |
| 44748_2 | TATGAGTGTGGAGTCGTTGCACTGTAGGGACAACCATGTTGTTNNNAGATCGGAAG |
| 9299_1 | TATGAGTGTGGAGTCGTTGCTTTGGATTACAGCCAGGTGTTGGNNNAGATCGGAA |
| 9299_2 | TATGAGTGTGGAGTCGTTGCTTTGGATTACAGCCAGGTGTTGGNNNAGATCGGAA |
| 29568_1 | TATGAGTGTGGAGTCGTTGCAATCAGAAAGGAGTAGAGGACGCNNNAGATCGGAA |
| 29568_2 | TATGAGTGTGGAGTCGTTGCAATCAGAAAGGAGTAGAGGACGCNNNAGATCGGAA |
| 31531_1 | TATGAGTGTGGAGTCGTTGCCCCCCATATATATCCAGATAGCCANNNAGATCGGAAG |
| 31531_2 | TATGAGTGTGGAGTCGTTGCCCCCCATATATATCCAGATAGCCANNNAGATCGGAAG |
| 10528_1 | TATGAGTGTGGAGTCGTTGCGTGTGGCCAATGCATCAGGATTTNNNAGATCGGAAG |
| 10528_2 | TATGAGTGTGGAGTCGTTGCGTGTGGCCAATGCATCAGGATTTNNNAGATCGGAAG |
| 46522_1 | TATGAGTGTGGAGTCGTTGCCTCACTGCAATCTCCACCTTCTNNNAGATCGGAAGAG |
| 46522_2 | TATGAGTGTGGAGTCGTTGCCTCACTGCAATCTCCACCTTCTNNNAGATCGGAAGAG |
| 52535_1 | TATGAGTGTGGAGTCGTTGCTAGTTAGGGCACTTTGAATCCCCNNNAGATCGGAAG |
| 52535_2 | TATGAGTGTGGAGTCGTTGCTAGTTAGGGCACTTTGAATCCCCNNNAGATCGGAAG |
| 13514_1 | TATGAGTGTGGAGTCGTTGCTAACTTGACAAAAAATGTAAGCTNNNAGATCGGAAG |
| 13514_2 | TATGAGTGTGGAGTCGTTGCTAACTTGACAAAAAATGTAAGCTNNNAGATCGGAAG |
| 12496_1 | TATGAGTGTGGAGTCGTTGCTTCTCTATGTTCACCTAACTCTANNNAGATCGGAAGA |
| 12496_2 | TATGAGTGTGGAGTCGTTGCTTCTCTATGTTCACCTAACTCTANNNAGATCGGAAGA |
| 2008_1 | TATGAGTGTGGAGTCGTTGCGGCAGGAGAGACAAGACAAGAAANNNAGATCGGA |
| 2008_2 | TATGAGTGTGGAGTCGTTGCGGCAGGAGAGACAAGACAAGAAANNNAGATCGGA |
| 46556_1 | TATGAGTGTGGAGTCGTTGCATGGAGTTTTGTCCTTTTGCCTGNNNAGATCGGAAG |
| 46556_2 | TATGAGTGTGGAGTCGTTGCATGGAGTTTTGTCCTTTTGCCTGNNNAGATCGGAAG |
| 43232_1 | TATGAGTGTGGAGTCGTTGCAGGAAGGAAGGAACAGAGAGAGANNNAGATCGGA |
| 43232_2 | TATGAGTGTGGAGTCGTTGCAGGAAGGAAGGAACAGAGAGAGANNNAGATCGGA |
| 45492_1 | TATGAGTGTGGAGTCGTTGCCCAATGGGAAGGGAGAACACTTCNNNAGATCGGAA |
| 45492_2 | TATGAGTGTGGAGTCGTTGCCCAATGGGAAGGGAGAACACTTCNNNAGATCGGAA |
| 50381_1 | TATGAGTGTGGAGTCGTTGCGGATGTGAGAAGGTCTTCGAAGANNNAGATCGGAA |
| 50381_2 | TATGAGTGTGGAGTCGTTGCGGATGTGAGAAGGTCTTCGAAGANNNAGATCGGAA |
| 38031_1 | TATGAGTGTGGAGTCGTTGCCAAATACACACGGGAACCCAATTNNNAGATCGGAAG |
| 38031_2 | TATGAGTGTGGAGTCGTTGCCAAATACACACGGGAACCCAATTNNNAGATCGGAAG |
| 33468_1 | TATGAGTGTGGAGTCGTTGCATTCCTGTGTTTCATCCCAGACCNNNAGATCGGAAGA |
| 33468_2 | TATGAGTGTGGAGTCGTTGCATTCCTGTGTTTCATCCCAGACCNNNAGATCGGAAGA |

| | |
|---|---|
| 32641_1 | TATGAGTGTGGAGTCGTTGCCTGGTCTCAAACTCCTGACTTCANNNAGATCGGAAGA |
| 32641_2 | TATGAGTGTGGAGTCGTTGCCTGGTCTCAAACTCCTGACTTCANNNAGATCGGAAGA |
| 41620_1 | TATGAGTGTGGAGTCGTTGCACGAGAACAAAGGTCTCTGGTTTNNNAGATCGGAAG |
| 41620_2 | TATGAGTGTGGAGTCGTTGCACGAGAACAAAGGTCTCTGGTTTNNNAGATCGGAAG |
| 53034_1 | TATGAGTGTGGAGTCGTTGCTGAAAACAACGAGGTCTCAGAGTNNNAGATCGGAA |
| 53034_2 | TATGAGTGTGGAGTCGTTGCTGAAAACAACGAGGTCTCAGAGTNNNAGATCGGAA |
| 52171_1 | TATGAGTGTGGAGTCGTTGCAGCTTCATCTATGTCCCTGCAAANNNAGATCGGAAGA |
| 52171_2 | TATGAGTGTGGAGTCGTTGCAGCTTCATCTATGTCCCTGCAAANNNAGATCGGAAGA |
| 40370_1 | TATGAGTGTGGAGTCGTTGCAGACAGACAAGAAGGAGAGAGGANNNAGATCGGA |
| 40370_2 | TATGAGTGTGGAGTCGTTGCAGACAGACAAGAAGGAGAGAGGANNNAGATCGGA |
| 46094_1 | TATGAGTGTGGAGTCGTTGCTTGTTTTTAGACGGAGTCTCGCTNNNAGATCGGAAG |
| 46094_2 | TATGAGTGTGGAGTCGTTGCTTGTTTTTAGACGGAGTCTCGCTNNNAGATCGGAAG |
| 52536_1 | TATGAGTGTGGAGTCGTTGCACCAGAGTTAGGAGCTTTTTCCTNNNAGATCGGAAG |
| 52536_2 | TATGAGTGTGGAGTCGTTGCACCAGAGTTAGGAGCTTTTTCCTNNNAGATCGGAAG |
| 45086_1 | TATGAGTGTGGAGTCGTTGCTCTTGGTGGAAGGCTTTTTACCANNNAGATCGGAAG |
| 45086_2 | TATGAGTGTGGAGTCGTTGCTCTTGGTGGAAGGCTTTTTACCANNNAGATCGGAAG |
| 41598_1 | TATGAGTGTGGAGTCGTTGCCATTCTCCTGCCTCAGCCTCNNNAGATCGGAAGAGCA |
| 41598_2 | TATGAGTGTGGAGTCGTTGCCATTCTCCTGCCTCAGCCTCNNNAGATCGGAAGAGCA |
| 42033_1 | TATGAGTGTGGAGTCGTTGCTTCAAGTGATTCTCCTGCCTCAGNNNAGATCGGAAGA |
| 42033_2 | TATGAGTGTGGAGTCGTTGCTTCAAGTGATTCTCCTGCCTCAGNNNAGATCGGAAGA |
| 41403_1 | TATGAGTGTGGAGTCGTTGCGGGGTTGCAATCCTAGTCTATGANNNAGATCGGAAG |
| 41403_2 | TATGAGTGTGGAGTCGTTGCGGGGTTGCAATCCTAGTCTATGANNNAGATCGGAAG |
| 33955_1 | TATGAGTGTGGAGTCGTTGCCTTCCCAAGTTCAAGCGATTCTCNNNAGATCGGAAGA |
| 33955_2 | TATGAGTGTGGAGTCGTTGCCTTCCCAAGTTCAAGCGATTCTCNNNAGATCGGAAGA |
| 9507_1 | TATGAGTGTGGAGTCGTTGCCTCTCTGGGGAACTTCGGGTTTCNNNAGATCGGAAG |
| 9507_2 | TATGAGTGTGGAGTCGTTGCCTCTCTGGGGAACTTCGGGTTTCNNNAGATCGGAAG |
| 29614_1 | TATGAGTGTGGAGTCGTTGCTCAAGGCTCTCTGATGGTTCTAACNNNAGATCGGAA |
| 29614_2 | TATGAGTGTGGAGTCGTTGCTCAAGGCTCTCTGATGGTTCTAACNNNAGATCGGAA |
| 45962_1 | TATGAGTGTGGAGTCGTTGCTAGCTCATTGCAGCCTTGAACTANNNAGATCGGAAGA |
| 45962_2 | TATGAGTGTGGAGTCGTTGCTAGCTCATTGCAGCCTTGAACTANNNAGATCGGAAGA |
| 44741_1 | TATGAGTGTGGAGTCGTTGCTCGGGTGCTAGTTCTACCTCTTANNNAGATCGGAAGA |
| 44741_2 | TATGAGTGTGGAGTCGTTGCTCGGGTGCTAGTTCTACCTCTTANNNAGATCGGAAGA |
| 10671_1 | TATGAGTGTGGAGTCGTTGCATAGCATCCAAGGAGTGGTCCTCNNNAGATCGGAAG |
| 10671_2 | TATGAGTGTGGAGTCGTTGCATAGCATCCAAGGAGTGGTCCTCNNNAGATCGGAAG |
| 48482_1 | TATGAGTGTGGAGTCGTTGCAAAGCGGTGGTATTACAGGCATANNNAGATCGGAA |
| 48482_2 | TATGAGTGTGGAGTCGTTGCAAAGCGGTGGTATTACAGGCATANNNAGATCGGAA |
| 43377_1 | TATGAGTGTGGAGTCGTTGCCTGAGTAGCTGGGATTACAGGTGNNNAGATCGGAA |
| 43377_2 | TATGAGTGTGGAGTCGTTGCCTGAGTAGCTGGGATTACAGGTGNNNAGATCGGAA |
| 48932_1 | TATGAGTGTGGAGTCGTTGCGTCACCAAACTTCCTGTGGATCANNNAGATCGGAAGA |
| 48932_2 | TATGAGTGTGGAGTCGTTGCGTCACCAAACTTCCTGTGGATCANNNAGATCGGAAGA |
| 52697_1 | TATGAGTGTGGAGTCGTTGCCATCTGGCAAGGGAGATAGCATANNNAGATCGGAA |
| 52697_2 | TATGAGTGTGGAGTCGTTGCCATCTGGCAAGGGAGATAGCATANNNAGATCGGAA |
| 37465_1 | TATGAGTGTGGAGTCGTTGCTCAGGAGAATCACTTGAACCCAGNNNAGATCGGAAG |

| | |
|---|---|
| 37465_2 | TATGAGTGTGGAGTCGTTGCTCAGGAGAATCACTTGAACCCAGNNNAGATCGGAAG |
| 36704_1 | TATGAGTGTGGAGTCGTTGCTGTGTAGAATTCCCCCATTGAGGNNNAGATCGGAAG |
| 36704_2 | TATGAGTGTGGAGTCGTTGCTGTGTAGAATTCCCCCATTGAGGNNNAGATCGGAAG |
| 46158_1 | TATGAGTGTGGAGTCGTTGCTTGGTCCTAAGTCATGGTTCTGGNNNAGATCGGAAG |
| 46158_2 | TATGAGTGTGGAGTCGTTGCTTGGTCCTAAGTCATGGTTCTGGNNNAGATCGGAAG |
| 35013_1 | TATGAGTGTGGAGTCGTTGCAATAGCTGGGATTACAGACACCCNNNAGATCGGAAG |
| 35013_2 | TATGAGTGTGGAGTCGTTGCAATAGCTGGGATTACAGACACCCNNNAGATCGGAAG |
| 49798_1 | TATGAGTGTGGAGTCGTTGCGGCTGGCTATTTCTTACCAAAGCNNNAGATCGGAAG |
| 49798_2 | TATGAGTGTGGAGTCGTTGCGGCTGGCTATTTCTTACCAAAGCNNNAGATCGGAAG |
| 47546_1 | TATGAGTGTGGAGTCGTTGCCTTGGTGTCAGTTGGTGTCAGATNNNAGATCGGAAG |
| 47546_2 | TATGAGTGTGGAGTCGTTGCCTTGGTGTCAGTTGGTGTCAGATNNNAGATCGGAAG |
| 43219_1 | TATGAGTGTGGAGTCGTTGCCTTCCCCTTCCACCTTGATTGTANNNAGATCGGAAGAG |
| 43219_2 | TATGAGTGTGGAGTCGTTGCCTTCCCCTTCCACCTTGATTGTANNNAGATCGGAAGAG |
| 47595_1 | TATGAGTGTGGAGTCGTTGCTTATGGACACAAAAGCTAAGGCCNNNAGATCGGAAG |
| 47595_2 | TATGAGTGTGGAGTCGTTGCTTATGGACACAAAAGCTAAGGCCNNNAGATCGGAAG |
| 43100_1 | TATGAGTGTGGAGTCGTTGCCTCCTGGGTTCACACCATTCTCNNNAGATCGGAAGAG |
| 43100_2 | TATGAGTGTGGAGTCGTTGCCTCCTGGGTTCACACCATTCTCNNNAGATCGGAAGAG |
| 39800_1 | TATGAGTGTGGAGTCGTTGCAAGTGGGAACTGGAAATGGGATTNNNAGATCGGAA |
| 39800_2 | TATGAGTGTGGAGTCGTTGCAAGTGGGAACTGGAAATGGGATTNNNAGATCGGAA |
| 49104_1 | TATGAGTGTGGAGTCGTTGCCCCAGTTTTGTCAGGTGTAACACNNNAGATCGGAAG |
| 49104_2 | TATGAGTGTGGAGTCGTTGCCCCAGTTTTGTCAGGTGTAACACNNNAGATCGGAAG |
| 32344_1 | TATGAGTGTGGAGTCGTTGCGGGGCAATCTCTAAGGGTTACATNNNAGATCGGAAG |
| 32344_2 | TATGAGTGTGGAGTCGTTGCGGGGCAATCTCTAAGGGTTACATNNNAGATCGGAAG |
| 10267_1 | TATGAGTGTGGAGTCGTTGCCAGTTGCAGAGGAATCCACTACCNNNAGATCGGAAG |
| 10267_2 | TATGAGTGTGGAGTCGTTGCCAGTTGCAGAGGAATCCACTACCNNNAGATCGGAAG |
| 34208_1 | TATGAGTGTGGAGTCGTTGCCCCACCACATTGGAAATCATGATNNNAGATCGGAAGA |
| 34208_2 | TATGAGTGTGGAGTCGTTGCCCCACCACATTGGAAATCATGATNNNAGATCGGAAGA |
| 32859_1 | TATGAGTGTGGAGTCGTTGCCACAGGTTTTCACCATGTTCTCCNNNAGATCGGAAGA |
| 32859_2 | TATGAGTGTGGAGTCGTTGCCACAGGTTTTCACCATGTTCTCCNNNAGATCGGAAGA |
| 10819_1 | TATGAGTGTGGAGTCGTTGCCCCAGTACTCTTTCACAGCTGACNNNAGATCGGAAGA |
| 10819_2 | TATGAGTGTGGAGTCGTTGCCCCAGTACTCTTTCACAGCTGACNNNAGATCGGAAGA |
| 30832_1 | TATGAGTGTGGAGTCGTTGCTCAAGTGAAGAGCAAGACTCCAANNNAGATCGGAAG |
| 30832_2 | TATGAGTGTGGAGTCGTTGCTCAAGTGAAGAGCAAGACTCCAANNNAGATCGGAAG |
| 13126_1 | TATGAGTGTGGAGTCGTTGCTTTAAACCAACAATTAAGACAGANNNAGATCGGAAG |
| 13126_2 | TATGAGTGTGGAGTCGTTGCTTTAAACCAACAATTAAGACAGANNNAGATCGGAAG |
| 39563_1 | TATGAGTGTGGAGTCGTTGCAGATCAATGAATTCAGGAGCTAGTNNNAGATCGGAA |
| 39563_2 | TATGAGTGTGGAGTCGTTGCAGATCAATGAATTCAGGAGCTAGTNNNAGATCGGAA |
| 38668_1 | TATGAGTGTGGAGTCGTTGCTACTCTCTACTGCCCAAACCAACNNNAGATCGGAAGA |
| 38668_2 | TATGAGTGTGGAGTCGTTGCTACTCTCTACTGCCCAAACCAACNNNAGATCGGAAGA |
| 39126_1 | TATGAGTGTGGAGTCGTTGCTTCCCTTTCTCAAACCCTTGACANNNAGATCGGAAGAG |
| 39126_2 | TATGAGTGTGGAGTCGTTGCTTCCCTTTCTCAAACCCTTGACANNNAGATCGGAAGAG |
| 28796_1 | TATGAGTGTGGAGTCGTTGCTGATGGAATACTACTCAGCCATAANNNAGATCGGAA |
| 28796_2 | TATGAGTGTGGAGTCGTTGCTGATGGAATACTACTCAGCCATAANNNAGATCGGAA |

| | |
|---|---|
| 29884_1 | TATGAGTGTGGAGTCGTTGCGTCAACGTCACTACAGACCCTACNNNAGATCGGAAG |
| 29884_2 | TATGAGTGTGGAGTCGTTGCGTCAACGTCACTACAGACCCTACNNNAGATCGGAAG |
| 52096_1 | TATGAGTGTGGAGTCGTTGCCCACACTCCCATAGCATTTTGTTNNNAGATCGGAAGA |
| 52096_2 | TATGAGTGTGGAGTCGTTGCCCACACTCCCATAGCATTTTGTTNNNAGATCGGAAGA |
| 142_1 | TATGAGTGTGGAGTCGTTGCCACAAACTAAGTGGCTCCAACAANNNAGATCGGAAG |
| 142_2 | TATGAGTGTGGAGTCGTTGCCACAAACTAAGTGGCTCCAACAANNNAGATCGGAAG |
| 51288_1 | TATGAGTGTGGAGTCGTTGCATGCTTCTCCCATGGATCTTTGTNNNAGATCGGAAG |
| 51288_2 | TATGAGTGTGGAGTCGTTGCATGCTTCTCCCATGGATCTTTGTNNNAGATCGGAAG |
| 30312_1 | TATGAGTGTGGAGTCGTTGCGCCTGGTGATTTTCTTCTCCGAGNNNAGATCGGAAG |
| 30312_2 | TATGAGTGTGGAGTCGTTGCGCCTGGTGATTTTCTTCTCCGAGNNNAGATCGGAAG |
| 40433_1 | TATGAGTGTGGAGTCGTTGCACTGAAAAGTTTCCATGAACACAGTNNNAGATCGGA |
| 40433_2 | TATGAGTGTGGAGTCGTTGCACTGAAAAGTTTCCATGAACACAGTNNNAGATCGGA |
| 53322_1 | TATGAGTGTGGAGTCGTTGCATTTCTTGCCTAATTGCTCTGGCNNNAGATCGGAAG |
| 53322_2 | TATGAGTGTGGAGTCGTTGCATTTCTTGCCTAATTGCTCTGGCNNNAGATCGGAAG |
| 42631_1 | TATGAGTGTGGAGTCGTTGCTTGTAGAGACCAGGTTTTGCCATNNNAGATCGGAAG |
| 42631_2 | TATGAGTGTGGAGTCGTTGCTTGTAGAGACCAGGTTTTGCCATNNNAGATCGGAAG |
| 45937_1 | TATGAGTGTGGAGTCGTTGCTTCAAGCAATTCTCCTGCCTCAGNNNAGATCGGAAG |
| 45937_2 | TATGAGTGTGGAGTCGTTGCTTCAAGCAATTCTCCTGCCTCAGNNNAGATCGGAAG |
| 38910_1 | TATGAGTGTGGAGTCGTTGCGAATGAGTTCACGTCCTTTGCAGNNNAGATCGGAAG |
| 38910_2 | TATGAGTGTGGAGTCGTTGCGAATGAGTTCACGTCCTTTGCAGNNNAGATCGGAAG |
| 53337_1 | TATGAGTGTGGAGTCGTTGCACAAAATCAGAAATGACAAAGGGGANNNAGATCGG |
| 53337_2 | TATGAGTGTGGAGTCGTTGCACAAAATCAGAAATGACAAAGGGGANNNAGATCGG |
| 46807_1 | TATGAGTGTGGAGTCGTTGCCACCACACCCAGCTAATTTTTGTNNNAGATCGGAAG |
| 46807_2 | TATGAGTGTGGAGTCGTTGCCACCACACCCAGCTAATTTTTGTNNNAGATCGGAAG |
| 51163_1 | TATGAGTGTGGAGTCGTTGCTAGGGAGAACTCAAGCTGGAAACNNNAGATCGGAA |
| 51163_2 | TATGAGTGTGGAGTCGTTGCTAGGGAGAACTCAAGCTGGAAACNNNAGATCGGAA |
| 50753_1 | TATGAGTGTGGAGTCGTTGCTTGCCAGGTAGGTTGCCTAATTANNNAGATCGGAAG |
| 50753_2 | TATGAGTGTGGAGTCGTTGCTTGCCAGGTAGGTTGCCTAATTANNNAGATCGGAAG |
| 30882_1 | TATGAGTGTGGAGTCGTTGCATGATACAAGCGGAGTTCCAGAGNNNAGATCGGAA |
| 30882_2 | TATGAGTGTGGAGTCGTTGCATGATACAAGCGGAGTTCCAGAGNNNAGATCGGAA |
| 39998_1 | TATGAGTGTGGAGTCGTTGCAGAAGAATTTTTGTGGCCTGCTGNNNAGATCGGAA |
| 39998_2 | TATGAGTGTGGAGTCGTTGCAGAAGAATTTTTGTGGCCTGCTGNNNAGATCGGAA |
| 38771_1 | TATGAGTGTGGAGTCGTTGCTAGAGAAAGGAACTCATGCACTTNNNAGATCGGAA |
| 38771_2 | TATGAGTGTGGAGTCGTTGCTAGAGAAAGGAACTCATGCACTTNNNAGATCGGAA |
| 47814_1 | TATGAGTGTGGAGTCGTTGCGATCTCCTGACCTCATGATCCACNNNAGATCGGAAG |
| 47814_2 | TATGAGTGTGGAGTCGTTGCGATCTCCTGACCTCATGATCCACNNNAGATCGGAAG |
| 44155_1 | TATGAGTGTGGAGTCGTTGCAGCCAGACATAATTCCCTCTTCCNNNAGATCGGAAGA |
| 44155_2 | TATGAGTGTGGAGTCGTTGCAGCCAGACATAATTCCCTCTTCCNNNAGATCGGAAGA |
| 43487_1 | TATGAGTGTGGAGTCGTTGCTGCACCCTCAACATATATCCAGGNNNAGATCGGAAG |
| 43487_2 | TATGAGTGTGGAGTCGTTGCTGCACCCTCAACATATATCCAGGNNNAGATCGGAAG |
| 49925_1 | TATGAGTGTGGAGTCGTTGCCACCAAGACAGCTGAGAAGATCTNNNAGATCGGAA |
| 49925_2 | TATGAGTGTGGAGTCGTTGCCACCAAGACAGCTGAGAAGATCTNNNAGATCGGAA |
| 45249_1 | TATGAGTGTGGAGTCGTTGCTCATGTCACTTCACTGCAACCTANNNAGATCGGAAGA |

| | |
|---|---|
| 45249_2 | TATGAGTGTGGAGTCGTTGCTCATGTCACTTCACTGCAACCTANNNAGATCGGAAGA |
| 40589_1 | TATGAGTGTGGAGTCGTTGCTGCAGTCTTGAATTTCTCTGAGTGNNNAGATCGGAA |
| 40589_2 | TATGAGTGTGGAGTCGTTGCTGCAGTCTTGAATTTCTCTGAGTGNNNAGATCGGAA |
| 36158_1 | TATGAGTGTGGAGTCGTTGCAAAAGTGAAAACCTGGTTGGGTGNNNAGATCGGAA |
| 36158_2 | TATGAGTGTGGAGTCGTTGCAAAAGTGAAAACCTGGTTGGGTGNNNAGATCGGAA |
| 40590_1 | TATGAGTGTGGAGTCGTTGCCAGGAAATGTCTTCAAGCACCAGNNNAGATCGGAAG |
| 40590_2 | TATGAGTGTGGAGTCGTTGCCAGGAAATGTCTTCAAGCACCAGNNNAGATCGGAAG |
| 50260_1 | TATGAGTGTGGAGTCGTTGCGGAGAGATCGACGAGGGTGGNNNAGATCGGAAGA |
| 50260_2 | TATGAGTGTGGAGTCGTTGCGGAGAGATCGACGAGGGTGGNNNAGATCGGAAGA |
| 46261_1 | TATGAGTGTGGAGTCGTTGCCAGGTTCAAGCAATTCTCCTGCNNNAGATCGGAAGA |
| 46261_2 | TATGAGTGTGGAGTCGTTGCCAGGTTCAAGCAATTCTCCTGCNNNAGATCGGAAGA |
| 50273_1 | TATGAGTGTGGAGTCGTTGCCATACCTACACCCAAGGTTGTCANNNAGATCGGAAG |
| 50273_2 | TATGAGTGTGGAGTCGTTGCCATACCTACACCCAAGGTTGTCANNNAGATCGGAAG |
| 44375_1 | TATGAGTGTGGAGTCGTTGCGGATCTGATGAGGTCTGAAGAGCNNNAGATCGGAA |
| 44375_2 | TATGAGTGTGGAGTCGTTGCGGATCTGATGAGGTCTGAAGAGCNNNAGATCGGAA |
| 36546_1 | TATGAGTGTGGAGTCGTTGCACTTGAAGGAATTAGGAGCCCAGNNNAGATCGGAA |
| 36546_2 | TATGAGTGTGGAGTCGTTGCACTTGAAGGAATTAGGAGCCCAGNNNAGATCGGAA |
| 51378_1 | TATGAGTGTGGAGTCGTTGCACACCAACGTCTATGATACAGGTNNNAGATCGGAAG |
| 51378_2 | TATGAGTGTGGAGTCGTTGCACACCAACGTCTATGATACAGGTNNNAGATCGGAAG |
| 45096_1 | TATGAGTGTGGAGTCGTTGCAGATGTGAAGGTTTCAGGTCACANNNAGATCGGAA |
| 45096_2 | TATGAGTGTGGAGTCGTTGCAGATGTGAAGGTTTCAGGTCACANNNAGATCGGAA |
| 47112_1 | TATGAGTGTGGAGTCGTTGCTTCGATTCTCCCACTTCATCCTCNNNAGATCGGAAGA |
| 47112_2 | TATGAGTGTGGAGTCGTTGCTTCGATTCTCCCACTTCATCCTCNNNAGATCGGAAGA |
| 50396_1 | TATGAGTGTGGAGTCGTTGCCAGCCTGTGAAGGGAAGGAGNNNAGATCGGAAGA |
| 50396_2 | TATGAGTGTGGAGTCGTTGCCAGCCTGTGAAGGGAAGGAGNNNAGATCGGAAGA |
| 39202_1 | TATGAGTGTGGAGTCGTTGCTGCAGCCTAGATAATGCGTAACCNNNAGATCGGAAG |
| 39202_2 | TATGAGTGTGGAGTCGTTGCTGCAGCCTAGATAATGCGTAACCNNNAGATCGGAAG |
| 32730_1 | TATGAGTGTGGAGTCGTTGCAATGCTTTTACACTGTTGGTGGGNNNAGATCGGAAG |
| 32730_2 | TATGAGTGTGGAGTCGTTGCAATGCTTTTACACTGTTGGTGGGNNNAGATCGGAAG |
| 34175_1 | TATGAGTGTGGAGTCGTTGCCCAAAGTAGGTGGATCACAAGGTNNNAGATCGGAA |
| 34175_2 | TATGAGTGTGGAGTCGTTGCCCAAAGTAGGTGGATCACAAGGTNNNAGATCGGAA |
| 41210_1 | TATGAGTGTGGAGTCGTTGCGGTAGTGGAGTGTTGACCAATCANNNAGATCGGAA |
| 41210_2 | TATGAGTGTGGAGTCGTTGCGGTAGTGGAGTGTTGACCAATCANNNAGATCGGAA |
| 9290_1 | TATGAGTGTGGAGTCGTTGCCACTGTTCACAGCCAGAACATCANNNAGATCGGAAG |
| 9290_2 | TATGAGTGTGGAGTCGTTGCCACTGTTCACAGCCAGAACATCANNNAGATCGGAAG |
| 29392_1 | TATGAGTGTGGAGTCGTTGCTTTTTTTTTTTTTTGAGAGGGAGNNNAGATCGGAAG |
| 29392_2 | TATGAGTGTGGAGTCGTTGCTTTTTTTTTTTTTTGAGAGGGAGNNNAGATCGGAAG |
| 10392_1 | TATGAGTGTGGAGTCGTTGCGTGTGCCTTGTACCAAGTGATGTNNNAGATCGGAAG |
| 10392_2 | TATGAGTGTGGAGTCGTTGCGTGTGCCTTGTACCAAGTGATGTNNNAGATCGGAAG |
| 51919_1 | TATGAGTGTGGAGTCGTTGCGATCGTCGCCTATCCTGCTTGNNNAGATCGGAAGAG |
| 51919_2 | TATGAGTGTGGAGTCGTTGCGATCGTCGCCTATCCTGCTTGNNNAGATCGGAAGAG |
| 10635_1 | TATGAGTGTGGAGTCGTTGCTACAGCCAGACTTCACACGATTGNNNAGATCGGAAG |
| 10635_2 | TATGAGTGTGGAGTCGTTGCTACAGCCAGACTTCACACGATTGNNNAGATCGGAAG |

| | |
|---|---|
| 53044_1 | TATGAGTGTGGAGTCGTTGCTCTGTGTACATTTTTTGGCTGGGNNNAGATCGGAAG |
| 53044_2 | TATGAGTGTGGAGTCGTTGCTCTGTGTACATTTTTTGGCTGGGNNNAGATCGGAAG |
| 34132_1 | TATGAGTGTGGAGTCGTTGCGACTTGCTTTGGCCAACAAAATGNNNAGATCGGAAG |
| 34132_2 | TATGAGTGTGGAGTCGTTGCGACTTGCTTTGGCCAACAAAATGNNNAGATCGGAAG |
| 12697_1 | TATGAGTGTGGAGTCGTTGCTGAATACTACCAAACATTCTTTGNNNAGATCGGAAG |
| 12697_2 | TATGAGTGTGGAGTCGTTGCTGAATACTACCAAACATTCTTTGNNNAGATCGGAAG |
| 36789_1 | TATGAGTGTGGAGTCGTTGCTACACCATGGGATACTATGCAGCNNNAGATCGGAAG |
| 36789_2 | TATGAGTGTGGAGTCGTTGCTACACCATGGGATACTATGCAGCNNNAGATCGGAAG |
| 10942_1 | TATGAGTGTGGAGTCGTTGCCGGCCACTACAAGCTGCTTTTNNNAGATCGGAAGAG |
| 10942_2 | TATGAGTGTGGAGTCGTTGCCGGCCACTACAAGCTGCTTTTNNNAGATCGGAAGAG |
| 44201_1 | TATGAGTGTGGAGTCGTTGCGTGTAGAGCTAGAGGGAGCATTCNNNAGATCGGAA |
| 44201_2 | TATGAGTGTGGAGTCGTTGCGTGTAGAGCTAGAGGGAGCATTCNNNAGATCGGAA |
| 46220_1 | TATGAGTGTGGAGTCGTTGCCTCAATCTCCTGACCTCGTGATCNNNAGATCGGAAGA |
| 46220_2 | TATGAGTGTGGAGTCGTTGCCTCAATCTCCTGACCTCGTGATCNNNAGATCGGAAGA |
| 42401_1 | TATGAGTGTGGAGTCGTTGCTCAAGCAATTCTCCTGCCTTAGTNNNAGATCGGAAG |
| 42401_2 | TATGAGTGTGGAGTCGTTGCTCAAGCAATTCTCCTGCCTTAGTNNNAGATCGGAAG |
| 48901_1 | TATGAGTGTGGAGTCGTTGCCTAGTAGCTGGGATTACAGGCACNNNAGATCGGAA |
| 48901_2 | TATGAGTGTGGAGTCGTTGCCTAGTAGCTGGGATTACAGGCACNNNAGATCGGAA |
| 10713_1 | TATGAGTGTGGAGTCGTTGCATGCTCACTTCCTCCCATCATCANNNAGATCGGAAGA |
| 10713_2 | TATGAGTGTGGAGTCGTTGCATGCTCACTTCCTCCCATCATCANNNAGATCGGAAGA |
| 47705_1 | TATGAGTGTGGAGTCGTTGCGTCCACACCCGGCTAATTTTTATNNNAGATCGGAAGA |
| 47705_2 | TATGAGTGTGGAGTCGTTGCGTCCACACCCGGCTAATTTTTATNNNAGATCGGAAGA |
| 34838_1 | TATGAGTGTGGAGTCGTTGCGCGCCCAGCCAAATTTTTTTTANNNAGATCGGAAGA |
| 34838_2 | TATGAGTGTGGAGTCGTTGCGCGCCCAGCCAAATTTTTTTTANNNAGATCGGAAGA |
| 32194_1 | TATGAGTGTGGAGTCGTTGCATCTCCCAGGTTCAAGCAATTCTNNNAGATCGGAAG |
| 32194_2 | TATGAGTGTGGAGTCGTTGCATCTCCCAGGTTCAAGCAATTCTNNNAGATCGGAAG |
| 47276_1 | TATGAGTGTGGAGTCGTTGCCAAGTAGCTGGGACTATAGGCACNNNAGATCGGAA |
| 47276_2 | TATGAGTGTGGAGTCGTTGCCAAGTAGCTGGGACTATAGGCACNNNAGATCGGAA |
| 53100_1 | TATGAGTGTGGAGTCGTTGCCTCATCTTTTGGGTGGACACAACNNNAGATCGGAAG |
| 53100_2 | TATGAGTGTGGAGTCGTTGCCTCATCTTTTGGGTGGACACAACNNNAGATCGGAAG |
| 41380_1 | TATGAGTGTGGAGTCGTTGCAGTGCTGAAAAGGAATGTTATTGCANNNAGATCGG |
| 41380_2 | TATGAGTGTGGAGTCGTTGCAGTGCTGAAAAGGAATGTTATTGCANNNAGATCGG |
| 29846_1 | TATGAGTGTGGAGTCGTTGCGGTGTTTCATTTGAGTTCCAGGGNNNAGATCGGAA |
| 29846_2 | TATGAGTGTGGAGTCGTTGCGGTGTTTCATTTGAGTTCCAGGGNNNAGATCGGAA |
| 48670_1 | TATGAGTGTGGAGTCGTTGCGGTTCAAGTGATTTTCCTGCCTCNNNAGATCGGAAG |
| 48670_2 | TATGAGTGTGGAGTCGTTGCGGTTCAAGTGATTTTCCTGCCTCNNNAGATCGGAAG |
| 33400_1 | TATGAGTGTGGAGTCGTTGCAACATATACACACAGACACACGTNNNAGATCGGAAG |
| 33400_2 | TATGAGTGTGGAGTCGTTGCAACATATACACACAGACACACGTNNNAGATCGGAAG |
| 32229_1 | TATGAGTGTGGAGTCGTTGCTGAGAAACAAGGACAGGACACAANNNAGATCGGAA |
| 32229_2 | TATGAGTGTGGAGTCGTTGCTGAGAAACAAGGACAGGACACAANNNAGATCGGAA |
| 35425_1 | TATGAGTGTGGAGTCGTTGCCTCACTGCAACCTCAAATTCCTGNNNAGATCGGAAGA |
| 35425_2 | TATGAGTGTGGAGTCGTTGCCTCACTGCAACCTCAAATTCCTGNNNAGATCGGAAGA |
| 30528_1 | TATGAGTGTGGAGTCGTTGCTATTTGTTATCTGTCTCTAGCTCCAGNNNAGATCGGA |

30528_2 TATGAGTGTGGAGTCGTTGCTATTTGTTATCTGTCTCTAGCTCCAGNNNAGATCGGA
33616_1 TATGAGTGTGGAGTCGTTGCCATTCTCCTGCCTCAGCCTCNNNAGATCGGAAGAGCA
33616_2 TATGAGTGTGGAGTCGTTGCCATTCTCCTGCCTCAGCCTCNNNAGATCGGAAGAGCA
44158_1 TATGAGTGTGGAGTCGTTGCTGCACCTGGCCAGTTATATCATTNNNAGATCGGAAG
44158_2 TATGAGTGTGGAGTCGTTGCTGCACCTGGCCAGTTATATCATTNNNAGATCGGAAG
43953_1 TATGAGTGTGGAGTCGTTGCAAACATTTGAAGACGAGCAGCATNNNAGATCGGAA
43953_2 TATGAGTGTGGAGTCGTTGCAAACATTTGAAGACGAGCAGCATNNNAGATCGGAA
41181_1 TATGAGTGTGGAGTCGTTGCAATACAGCACACTGATGGGTCTTNNNAGATCGGAAG
41181_2 TATGAGTGTGGAGTCGTTGCAATACAGCACACTGATGGGTCTTNNNAGATCGGAAG
35568_1 TATGAGTGTGGAGTCGTTGCTAATCCTCTTGTTTCAGCCTCCCNNNAGATCGGAAGA
35568_2 TATGAGTGTGGAGTCGTTGCTAATCCTCTTGTTTCAGCCTCCCNNNAGATCGGAAGA
37742_1 TATGAGTGTGGAGTCGTTGCCCATGGGCTTTGGAAGTTTGAAANNNAGATCGGAA
37742_2 TATGAGTGTGGAGTCGTTGCCCATGGGCTTTGGAAGTTTGAAANNNAGATCGGAA
8768_1 TATGAGTGTGGAGTCGTTGCGCAGCTCTCTACAGTCTCTTTCCNNNAGATCGGAAGA
8768_2 TATGAGTGTGGAGTCGTTGCGCAGCTCTCTACAGTCTCTTTCCNNNAGATCGGAAGA
49738_1 TATGAGTGTGGAGTCGTTGCCCAGGTTCAAGCGATTGTCTTGNNNAGATCGGAAGA
49738_2 TATGAGTGTGGAGTCGTTGCCCAGGTTCAAGCGATTGTCTTGNNNAGATCGGAAGA
12945_1 TATGAGTGTGGAGTCGTTGCTATAACTGTGAAGCTTAATTCAGNNNAGATCGGAAG
12945_2 TATGAGTGTGGAGTCGTTGCTATAACTGTGAAGCTTAATTCAGNNNAGATCGGAAG
50139_1 TATGAGTGTGGAGTCGTTGCATGATACCACCCACTCCCTAAGANNNAGATCGGAAG
50139_2 TATGAGTGTGGAGTCGTTGCATGATACCACCCACTCCCTAAGANNNAGATCGGAAG
34221_1 TATGAGTGTGGAGTCGTTGCGGGAGAAGAGAAGAGAGAACACANNNAGATCGGA
34221_2 TATGAGTGTGGAGTCGTTGCGGGAGAAGAGAAGAGAGAACACANNNAGATCGGA
43425_1 TATGAGTGTGGAGTCGTTGCGCCCGGCTAATTTTTTTGTACTTNNNAGATCGGAAG
43425_2 TATGAGTGTGGAGTCGTTGCGCCCGGCTAATTTTTTTGTACTTNNNAGATCGGAAG
32498_1 TATGAGTGTGGAGTCGTTGCGTAGTAACAGGCCTTGGCTTTTCNNNAGATCGGAAG
32498_2 TATGAGTGTGGAGTCGTTGCGTAGTAACAGGCCTTGGCTTTTCNNNAGATCGGAAG
32001_1 TATGAGTGTGGAGTCGTTGCATACTGGTTTGTGTCTCACTGCTNNNAGATCGGAAG
32001_2 TATGAGTGTGGAGTCGTTGCATACTGGTTTGTGTCTCACTGCTNNNAGATCGGAAG
53348_1 TATGAGTGTGGAGTCGTTGCGCTATTGCTGCACTGTCTTGCNNNAGATCGGAAGAG
53348_2 TATGAGTGTGGAGTCGTTGCGCTATTGCTGCACTGTCTTGCNNNAGATCGGAAGAG
31887_1 TATGAGTGTGGAGTCGTTGCAAAGGGCCACATAGATTACAGCANNNAGATCGGAA
31887_2 TATGAGTGTGGAGTCGTTGCAAAGGGCCACATAGATTACAGCANNNAGATCGGAA
46225_1 TATGAGTGTGGAGTCGTTGCCAGAGTAGCTGGGATTACAAGCANNNAGATCGGAA
46225_2 TATGAGTGTGGAGTCGTTGCCAGAGTAGCTGGGATTACAAGCANNNAGATCGGAA
33488_1 TATGAGTGTGGAGTCGTTGCCAGAGTAGCTACAAGGCAAGACTNNNAGATCGGAA
33488_2 TATGAGTGTGGAGTCGTTGCCAGAGTAGCTACAAGGCAAGACTNNNAGATCGGAA
37428_1 TATGAGTGTGGAGTCGTTGCCTGAGACAGGAGAATCGCTTGAANNNAGATCGGAA
37428_2 TATGAGTGTGGAGTCGTTGCCTGAGACAGGAGAATCGCTTGAANNNAGATCGGAA
47681_1 TATGAGTGTGGAGTCGTTGCCCTGGAACTCTTGGAACAATCCTNNNAGATCGGAAG
47681_2 TATGAGTGTGGAGTCGTTGCCCTGGAACTCTTGGAACAATCCTNNNAGATCGGAAG
39440_1 TATGAGTGTGGAGTCGTTGCAAAGTCTTGAAAAAAGCCTGGGANNNAGATCGGAA
39440_2 TATGAGTGTGGAGTCGTTGCAAAGTCTTGAAAAAAGCCTGGGANNNAGATCGGAA

| | |
|---|---|
| 44352_1 | TATGAGTGTGGAGTCGTTGCTTCCTGTGATGGTTCTGAGAAGGNNNAGATCGGAA |
| 44352_2 | TATGAGTGTGGAGTCGTTGCTTCCTGTGATGGTTCTGAGAAGGNNNAGATCGGAA |
| 31928_1 | TATGAGTGTGGAGTCGTTGCAGGACTACACATCACCAAACCAANNNAGATCGGAAG |
| 31928_2 | TATGAGTGTGGAGTCGTTGCAGGACTACACATCACCAAACCAANNNAGATCGGAAG |
| 38138_1 | TATGAGTGTGGAGTCGTTGCATAAAAAAATAGGCCGGGCACAGNNNAGATCGGAA |
| 38138_2 | TATGAGTGTGGAGTCGTTGCATAAAAAAATAGGCCGGGCACAGNNNAGATCGGAA |
| 46605_1 | TATGAGTGTGGAGTCGTTGCTCTACTCCTCCAGCAATCTGAGANNNAGATCGGAAG |
| 46605_2 | TATGAGTGTGGAGTCGTTGCTCTACTCCTCCAGCAATCTGAGANNNAGATCGGAAG |
| 40066_1 | TATGAGTGTGGAGTCGTTGCGGATCACAAGGTCAGGAGATCAANNNAGATCGGAA |
| 40066_2 | TATGAGTGTGGAGTCGTTGCGGATCACAAGGTCAGGAGATCAANNNAGATCGGAA |
| 49086_1 | TATGAGTGTGGAGTCGTTGCCCACACTGGGCTGTTTTGTATTTNNNAGATCGGAAG |
| 49086_2 | TATGAGTGTGGAGTCGTTGCCCACACTGGGCTGTTTTGTATTTNNNAGATCGGAAG |
| 30088_1 | TATGAGTGTGGAGTCGTTGCGGTGATATCTCTGGTCCACCTTGNNNAGATCGGAAG |
| 30088_2 | TATGAGTGTGGAGTCGTTGCGGTGATATCTCTGGTCCACCTTGNNNAGATCGGAAG |
| 46572_1 | TATGAGTGTGGAGTCGTTGCCTCCCCCGTGTTGATTTAGTAGANNNAGATCGGAAG |
| 46572_2 | TATGAGTGTGGAGTCGTTGCCTCCCCCGTGTTGATTTAGTAGANNNAGATCGGAAG |
| 34521_1 | TATGAGTGTGGAGTCGTTGCTAAGTCTCACGAGATCTGAGGGTNNNAGATCGGAA |
| 34521_2 | TATGAGTGTGGAGTCGTTGCTAAGTCTCACGAGATCTGAGGGTNNNAGATCGGAA |
| 32469_1 | TATGAGTGTGGAGTCGTTGCCACACAGAGAAATCAGGTCCTCANNNAGATCGGAAG |
| 32469_2 | TATGAGTGTGGAGTCGTTGCCACACAGAGAAATCAGGTCCTCANNNAGATCGGAAG |
| 34485_1 | TATGAGTGTGGAGTCGTTGCCTCCTTCCAACCCAGACTACATGNNNAGATCGGAAGA |
| 34485_2 | TATGAGTGTGGAGTCGTTGCCTCCTTCCAACCCAGACTACATGNNNAGATCGGAAGA |
| 33285_1 | TATGAGTGTGGAGTCGTTGCGATGGAGTCTCACTCTGTCATCCNNNAGATCGGAAG |
| 33285_2 | TATGAGTGTGGAGTCGTTGCGATGGAGTCTCACTCTGTCATCCNNNAGATCGGAAG |
| 32161_1 | TATGAGTGTGGAGTCGTTGCCTCCTGAGTAGCTGGGATTACAGNNNAGATCGGAA |
| 32161_2 | TATGAGTGTGGAGTCGTTGCCTCCTGAGTAGCTGGGATTACAGNNNAGATCGGAA |
| 3128_1 | TATGAGTGTGGAGTCGTTGCTGGTGACCCTCTTCCTCTTACAANNNAGATCGGAAGA |
| 3128_2 | TATGAGTGTGGAGTCGTTGCTGGTGACCCTCTTCCTCTTACAANNNAGATCGGAAGA |
| 29673_1 | TATGAGTGTGGAGTCGTTGCGCCCAAGACCCGAAGTTCTCNNNAGATCGGAAGAGC |
| 29673_2 | TATGAGTGTGGAGTCGTTGCGCCCAAGACCCGAAGTTCTCNNNAGATCGGAAGAGC |
| 31099_1 | TATGAGTGTGGAGTCGTTGCCACCATTTCACATTCTCACCACCNNNAGATCGGAAGA |
| 31099_2 | TATGAGTGTGGAGTCGTTGCCACCATTTCACATTCTCACCACCNNNAGATCGGAAGA |
| 30300_1 | TATGAGTGTGGAGTCGTTGCGATTGGTCCATTTCATGCCTCTGNNNAGATCGGAAG |
| 30300_2 | TATGAGTGTGGAGTCGTTGCGATTGGTCCATTTCATGCCTCTGNNNAGATCGGAAG |
| 44744_1 | TATGAGTGTGGAGTCGTTGCGAGAGGATTGGGACAGAGAAAGGNNNAGATCGGA |
| 44744_2 | TATGAGTGTGGAGTCGTTGCGAGAGGATTGGGACAGAGAAAGGNNNAGATCGGA |
| 42279_1 | TATGAGTGTGGAGTCGTTGCACATTCCACTTACTGCAACCTCTNNNAGATCGGAAGA |
| 42279_2 | TATGAGTGTGGAGTCGTTGCACATTCCACTTACTGCAACCTCTNNNAGATCGGAAGA |
| 43778_1 | TATGAGTGTGGAGTCGTTGCTCCAGTAAAGTTGACAATGCACANNNAGATCGGAAG |
| 43778_2 | TATGAGTGTGGAGTCGTTGCTCCAGTAAAGTTGACAATGCACANNNAGATCGGAAG |
| 48677_1 | TATGAGTGTGGAGTCGTTGCACCACACCCGGCTAATTTTTGTANNNAGATCGGAAG |
| 48677_2 | TATGAGTGTGGAGTCGTTGCACCACACCCGGCTAATTTTTGTANNNAGATCGGAAG |
| 184_1 | TATGAGTGTGGAGTCGTTGCCTCGGGAAAGAACTGTAAAAACCCCNNNAGATCGGA |

| | |
|---|---|
| 184_2 | TATGAGTGTGGAGTCGTTGCCTCGGGAAAGAACTGTAAAAACCCCNNNAGATCGG |
| 38896_1 | TATGAGTGTGGAGTCGTTGCCTTGTGCTGACATTTCCTGTACTNNNAGATCGGAAG |
| 38896_2 | TATGAGTGTGGAGTCGTTGCCTTGTGCTGACATTTCCTGTACTNNNAGATCGGAAG |
| 52499_1 | TATGAGTGTGGAGTCGTTGCGAAAAAGAAATAGCCCTGGACCANNNAGATCGGAA |
| 52499_2 | TATGAGTGTGGAGTCGTTGCGAAAAAGAAATAGCCCTGGACCANNNAGATCGGAA |
| 31459_1 | TATGAGTGTGGAGTCGTTGCTCAAAATAACCAAGGGGCTCAGANNNAGATCGGAA |
| 31459_2 | TATGAGTGTGGAGTCGTTGCTCAAAATAACCAAGGGGCTCAGANNNAGATCGGAA |
| 37636_1 | TATGAGTGTGGAGTCGTTGCGGCATGATCTCTGCTCACTGTAANNNAGATCGGAAG |
| 37636_2 | TATGAGTGTGGAGTCGTTGCGGCATGATCTCTGCTCACTGTAANNNAGATCGGAAG |
| 40102_1 | TATGAGTGTGGAGTCGTTGCGACTCAGTTAGACACAAGATCCANNNAGATCGGAAG |
| 40102_2 | TATGAGTGTGGAGTCGTTGCGACTCAGTTAGACACAAGATCCANNNAGATCGGAAG |
| 41175_1 | TATGAGTGTGGAGTCGTTGCTGCATGGACCCATTTCTAGATGTNNNAGATCGGAAG |
| 41175_2 | TATGAGTGTGGAGTCGTTGCTGCATGGACCCATTTCTAGATGTNNNAGATCGGAAG |
| 35222_1 | TATGAGTGTGGAGTCGTTGCACCCGGCCTGTTATATGTTGTAANNNAGATCGGAAG |
| 35222_2 | TATGAGTGTGGAGTCGTTGCACCCGGCCTGTTATATGTTGTAANNNAGATCGGAAG |
| 33044_1 | TATGAGTGTGGAGTCGTTGCTCAAGGAATTGAATCCAGCCTCANNNAGATCGGAAG |
| 33044_2 | TATGAGTGTGGAGTCGTTGCTCAAGGAATTGAATCCAGCCTCANNNAGATCGGAAG |
| 32966_1 | TATGAGTGTGGAGTCGTTGCTGGCTTTCCTGCTGAATTCTACANNNAGATCGGAAG |
| 32966_2 | TATGAGTGTGGAGTCGTTGCTGGCTTTCCTGCTGAATTCTACANNNAGATCGGAAG |
| 37731_1 | TATGAGTGTGGAGTCGTTGCTAAGCAGTTGCAGAAGAGCTGTANNNAGATCGGAA |
| 37731_2 | TATGAGTGTGGAGTCGTTGCTAAGCAGTTGCAGAAGAGCTGTANNNAGATCGGAA |
| 45073_1 | TATGAGTGTGGAGTCGTTGCCCTGCAGATACCGGATGGAGNNNAGATCGGAAGAG |
| 45073_2 | TATGAGTGTGGAGTCGTTGCCCTGCAGATACCGGATGGAGNNNAGATCGGAAGAG |
| 45661_1 | TATGAGTGTGGAGTCGTTGCGGCTGAAGTACAATGGCATGATCNNNAGATCGGAA |
| 45661_2 | TATGAGTGTGGAGTCGTTGCGGCTGAAGTACAATGGCATGATCNNNAGATCGGAA |
| 45653_1 | TATGAGTGTGGAGTCGTTGCCCCACCACAATGTCCAACTAATTNNNAGATCGGAAG |
| 45653_2 | TATGAGTGTGGAGTCGTTGCCCCACCACAATGTCCAACTAATTNNNAGATCGGAAG |
| 32035_1 | TATGAGTGTGGAGTCGTTGCCAGTATACACGATGGATCCAGCANNNAGATCGGAA |
| 32035_2 | TATGAGTGTGGAGTCGTTGCCAGTATACACGATGGATCCAGCANNNAGATCGGAA |
| 35017_1 | TATGAGTGTGGAGTCGTTGCCGAGTAGCTGGGATTACAGACATNNNAGATCGGAA |
| 35017_2 | TATGAGTGTGGAGTCGTTGCCGAGTAGCTGGGATTACAGACATNNNAGATCGGAA |
| 40316_1 | TATGAGTGTGGAGTCGTTGCCTTACCTCAGGGCCTTTACACTTNNNAGATCGGAAG |
| 40316_2 | TATGAGTGTGGAGTCGTTGCCTTACCTCAGGGCCTTTACACTTNNNAGATCGGAAG |
| 39608_1 | TATGAGTGTGGAGTCGTTGCCCCAGGTGATTCTAATGTAGCCANNNAGATCGGAAG |
| 39608_2 | TATGAGTGTGGAGTCGTTGCCCCAGGTGATTCTAATGTAGCCANNNAGATCGGAAG |
| 48990_1 | TATGAGTGTGGAGTCGTTGCGCCTCCCTAACTGCTGGAATTATNNNAGATCGGAAG |
| 48990_2 | TATGAGTGTGGAGTCGTTGCGCCTCCCTAACTGCTGGAATTATNNNAGATCGGAAG |
| 38416_1 | TATGAGTGTGGAGTCGTTGCACCCTTTGTAGCCTGAAATTTGANNNAGATCGGAAG |
| 38416_2 | TATGAGTGTGGAGTCGTTGCACCCTTTGTAGCCTGAAATTTGANNNAGATCGGAAG |
| 40346_1 | TATGAGTGTGGAGTCGTTGCCTGGATGGAACTGGAGAGCATTANNNAGATCGGAA |
| 40346_2 | TATGAGTGTGGAGTCGTTGCCTGGATGGAACTGGAGAGCATTANNNAGATCGGAA |
| 44878_1 | TATGAGTGTGGAGTCGTTGCTAAGCCAGTCCAGAGCACTAAAGNNNAGATCGGAA |
| 44878_2 | TATGAGTGTGGAGTCGTTGCTAAGCCAGTCCAGAGCACTAAAGNNNAGATCGGAA |

| | |
|---|---|
| 10091_1 | TATGAGTGTGGAGTCGTTGCTACCTCTTCTCTTGGCACTCACANNNAGATCGGAAGA |
| 10091_2 | TATGAGTGTGGAGTCGTTGCTACCTCTTCTCTTGGCACTCACANNNAGATCGGAAGA |
| 52015_1 | TATGAGTGTGGAGTCGTTGCTTTCTTCCACGAATCCTTGCAACNNNAGATCGGAAGA |
| 52015_2 | TATGAGTGTGGAGTCGTTGCTTTCTTCCACGAATCCTTGCAACNNNAGATCGGAAGA |
| 49927_1 | TATGAGTGTGGAGTCGTTGCCATGTATGTGCAGCCTACAATGGNNNAGATCGGAAG |
| 49927_2 | TATGAGTGTGGAGTCGTTGCCATGTATGTGCAGCCTACAATGGNNNAGATCGGAAG |
| 52102_1 | TATGAGTGTGGAGTCGTTGCAGGTGATTCAGACCAGAGAGAGANNNAGATCGGAA |
| 52102_2 | TATGAGTGTGGAGTCGTTGCAGGTGATTCAGACCAGAGAGAGANNNAGATCGGAA |
| 9360_1 | TATGAGTGTGGAGTCGTTGCCTCCTTGGGCCAATCAAGTTAGGNNNAGATCGGAAG |
| 9360_2 | TATGAGTGTGGAGTCGTTGCCTCCTTGGGCCAATCAAGTTAGGNNNAGATCGGAAG |
| 34894_1 | TATGAGTGTGGAGTCGTTGCGCCACCACATCAGGCTAATTTTTNNNAGATCGGAAG |
| 34894_2 | TATGAGTGTGGAGTCGTTGCGCCACCACATCAGGCTAATTTTTNNNAGATCGGAAG |
| 29191_1 | TATGAGTGTGGAGTCGTTGCGCGAGAGAAAAAAATGTGCCTGANNNAGATCGGAA |
| 29191_2 | TATGAGTGTGGAGTCGTTGCGCGAGAGAAAAAAATGTGCCTGANNNAGATCGGAA |
| 9886_1 | TATGAGTGTGGAGTCGTTGCTCTCTCTCTTCAGTTGAGCAGGANNNAGATCGGAAG |
| 9886_2 | TATGAGTGTGGAGTCGTTGCTCTCTCTCTTCAGTTGAGCAGGANNNAGATCGGAAG |
| 39069_1 | TATGAGTGTGGAGTCGTTGCTGACTCCCATCCTATACAACCCTNNNAGATCGGAAGA |
| 39069_2 | TATGAGTGTGGAGTCGTTGCTGACTCCCATCCTATACAACCCTNNNAGATCGGAAGA |
| 32202_1 | TATGAGTGTGGAGTCGTTGCCATTCTCCTGCCTCAGCCTCNNNAGATCGGAAGAGCA |
| 32202_2 | TATGAGTGTGGAGTCGTTGCCATTCTCCTGCCTCAGCCTCNNNAGATCGGAAGAGCA |
| 47208_1 | TATGAGTGTGGAGTCGTTGCCTAAGTGCTGGGATTACAGGTGTNNNAGATCGGAA |
| 47208_2 | TATGAGTGTGGAGTCGTTGCCTAAGTGCTGGGATTACAGGTGTNNNAGATCGGAA |
| 45972_1 | TATGAGTGTGGAGTCGTTGCGGTACAAAAGTGACAACTGGTGGNNNAGATCGGAA |
| 45972_2 | TATGAGTGTGGAGTCGTTGCGGTACAAAAGTGACAACTGGTGGNNNAGATCGGAA |
| 3079_1 | TATGAGTGTGGAGTCGTTGCGGATACGTTCACCAGAAGGAAGGNNNAGATCGGAA |
| 3079_2 | TATGAGTGTGGAGTCGTTGCGGATACGTTCACCAGAAGGAAGGNNNAGATCGGAA |
| 41264_1 | TATGAGTGTGGAGTCGTTGCAACGAGCTAATGAATGCTGTCAANNNAGATCGGAA |
| 41264_2 | TATGAGTGTGGAGTCGTTGCAACGAGCTAATGAATGCTGTCAANNNAGATCGGAA |
| 42040_1 | TATGAGTGTGGAGTCGTTGCTCTCATCAGAACACATGGACTCGNNNAGATCGGAAG |
| 42040_2 | TATGAGTGTGGAGTCGTTGCTCTCATCAGAACACATGGACTCGNNNAGATCGGAAG |
| 52158_1 | TATGAGTGTGGAGTCGTTGCAAAAAATTAGCTGGGCATGGTGGNNNAGATCGGAA |
| 52158_2 | TATGAGTGTGGAGTCGTTGCAAAAAATTAGCTGGGCATGGTGGNNNAGATCGGAA |
| 51497_1 | TATGAGTGTGGAGTCGTTGCACTTCTGTGAAGTCGAGTTAGGGNNNAGATCGGAA |
| 51497_2 | TATGAGTGTGGAGTCGTTGCACTTCTGTGAAGTCGAGTTAGGGNNNAGATCGGAA |
| 41639_1 | TATGAGTGTGGAGTCGTTGCGGTTCAAGTGATTCTTCTGCCTCNNNAGATCGGAAG |
| 41639_2 | TATGAGTGTGGAGTCGTTGCGGTTCAAGTGATTCTTCTGCCTCNNNAGATCGGAAG |
| 37377_1 | TATGAGTGTGGAGTCGTTGCGAGGCAGGAGAATTGTTTGAACCNNNAGATCGGAA |
| 37377_2 | TATGAGTGTGGAGTCGTTGCGAGGCAGGAGAATTGTTTGAACCNNNAGATCGGAA |
| 44953_1 | TATGAGTGTGGAGTCGTTGCGAAATGGGTGAAGAGGGGGAAATNNNAGATCGGA |
| 44953_2 | TATGAGTGTGGAGTCGTTGCGAAATGGGTGAAGAGGGGGAAATNNNAGATCGGA |
| 46459_1 | TATGAGTGTGGAGTCGTTGCCATGTAGCTGGGATTACGGTTGTNNNAGATCGGAA |
| 46459_2 | TATGAGTGTGGAGTCGTTGCCATGTAGCTGGGATTACGGTTGTNNNAGATCGGAA |
| 13458_1 | TATGAGTGTGGAGTCGTTGCGATTTCTGAGGCACATAGTCNNNAGATCGGAAGAG |

| | |
|---|---|
| 13458_2 | TATGAGTGTGGAGTCGTTGCGATTTCTGAGGCACATAGTCNNNAGATCGGAAGAG |
| 45516_1 | TATGAGTGTGGAGTCGTTGCCCACACAGAGGCTAGGAGAAGNNNAGATCGGAAGA |
| 45516_2 | TATGAGTGTGGAGTCGTTGCCCACACAGAGGCTAGGAGAAGNNNAGATCGGAAGA |
| 28955_1 | TATGAGTGTGGAGTCGTTGCGCCTCTCTACCCTTCCTTGAATCNNNAGATCGGAAGA |
| 28955_2 | TATGAGTGTGGAGTCGTTGCGCCTCTCTACCCTTCCTTGAATCNNNAGATCGGAAGA |
| 8762_1 | TATGAGTGTGGAGTCGTTGCCAGCATGACAAGAGCTGATTGTTNNNAGATCGGAAG |
| 8762_2 | TATGAGTGTGGAGTCGTTGCCAGCATGACAAGAGCTGATTGTTNNNAGATCGGAAG |
| 37235_1 | TATGAGTGTGGAGTCGTTGCGACCTAGTTCTAACCCAGCAGACNNNAGATCGGAAG |
| 37235_2 | TATGAGTGTGGAGTCGTTGCGACCTAGTTCTAACCCAGCAGACNNNAGATCGGAAG |
| 38624_1 | TATGAGTGTGGAGTCGTTGCACAGAAATCTGGAGAACAGGCATNNNAGATCGGAA |
| 38624_2 | TATGAGTGTGGAGTCGTTGCACAGAAATCTGGAGAACAGGCATNNNAGATCGGAA |
| 34298_1 | TATGAGTGTGGAGTCGTTGCGCCCAACAGCTCATTGAGAACNNNAGATCGGAAGAG |
| 34298_2 | TATGAGTGTGGAGTCGTTGCGCCCAACAGCTCATTGAGAACNNNAGATCGGAAGAG |
| 48785_1 | TATGAGTGTGGAGTCGTTGCAAAGCATGTCAACATTCCCAGTGNNNAGATCGGAAG |
| 48785_2 | TATGAGTGTGGAGTCGTTGCAAAGCATGTCAACATTCCCAGTGNNNAGATCGGAAG |
| 44651_1 | TATGAGTGTGGAGTCGTTGCAGGATACCTAGAGGGGGAATTGGNNNAGATCGGA |
| 44651_2 | TATGAGTGTGGAGTCGTTGCAGGATACCTAGAGGGGGAATTGGNNNAGATCGGA |
| 10550_1 | TATGAGTGTGGAGTCGTTGCCGGGCAGTTTCTAGAACAAGTTTNNNAGATCGGAAG |
| 10550_2 | TATGAGTGTGGAGTCGTTGCCGGGCAGTTTCTAGAACAAGTTTNNNAGATCGGAAG |
| 49131_1 | TATGAGTGTGGAGTCGTTGCGATGGAGTACAGAGGCACAATCANNNAGATCGGAA |
| 49131_2 | TATGAGTGTGGAGTCGTTGCGATGGAGTACAGAGGCACAATCANNNAGATCGGAA |
| 40540_1 | TATGAGTGTGGAGTCGTTGCAACACATCTAGAGGCTGCAAAGANNNAGATCGGAA |
| 40540_2 | TATGAGTGTGGAGTCGTTGCAACACATCTAGAGGCTGCAAAGANNNAGATCGGAA |
| 45745_1 | TATGAGTGTGGAGTCGTTGCTTCAAGTGATTCTCCTGCCTCAGNNNAGATCGGAAG |
| 45745_2 | TATGAGTGTGGAGTCGTTGCTTCAAGTGATTCTCCTGCCTCAGNNNAGATCGGAAG |
| 43293_1 | TATGAGTGTGGAGTCGTTGCGAAACCCCCTAAGTACCTGTCACNNNAGATCGGAAG |
| 43293_2 | TATGAGTGTGGAGTCGTTGCGAAACCCCCTAAGTACCTGTCACNNNAGATCGGAAG |
| 3110_1 | TATGAGTGTGGAGTCGTTGCCTGAGCCTCTTTCTCATCCCAAGNNNAGATCGGAAGA |
| 3110_2 | TATGAGTGTGGAGTCGTTGCCTGAGCCTCTTTCTCATCCCAAGNNNAGATCGGAAGA |
| 40080_1 | TATGAGTGTGGAGTCGTTGCTTGCAAATCTACCCTTGTCTCCANNNAGATCGGAAGA |
| 40080_2 | TATGAGTGTGGAGTCGTTGCTTGCAAATCTACCCTTGTCTCCANNNAGATCGGAAGA |
| 41048_1 | TATGAGTGTGGAGTCGTTGCTTCAACCTATGCCTGCAGTTACTNNNAGATCGGAAG |
| 41048_2 | TATGAGTGTGGAGTCGTTGCTTCAACCTATGCCTGCAGTTACTNNNAGATCGGAAG |
| 36171_1 | TATGAGTGTGGAGTCGTTGCAGAATTGCTTGAATCCAGGAGGTNNNAGATCGGAA |
| 36171_2 | TATGAGTGTGGAGTCGTTGCAGAATTGCTTGAATCCAGGAGGTNNNAGATCGGAA |
| 5308_1 | TATGAGTGTGGAGTCGTTGCTAGGAGGGAAGGGTTCAAACTCCNNNAGATCGGAA |
| 5308_2 | TATGAGTGTGGAGTCGTTGCTAGGAGGGAAGGGTTCAAACTCCNNNAGATCGGAA |
| 48008_1 | TATGAGTGTGGAGTCGTTGCCTGGAGTGTAGTGGCATGATCTCNNNAGATCGGAA |
| 48008_2 | TATGAGTGTGGAGTCGTTGCCTGGAGTGTAGTGGCATGATCTCNNNAGATCGGAA |
| 34206_1 | TATGAGTGTGGAGTCGTTGCTCAGTGGCATGTAGTCCAAGAAANNNAGATCGGAA |
| 34206_2 | TATGAGTGTGGAGTCGTTGCTCAGTGGCATGTAGTCCAAGAAANNNAGATCGGAA |
| 41033_1 | TATGAGTGTGGAGTCGTTGCAAGGTCAACAGGAGAAAATCCCANNNAGATCGGAA |
| 41033_2 | TATGAGTGTGGAGTCGTTGCAAGGTCAACAGGAGAAAATCCCANNNAGATCGGAA |

| | |
|---|---|
| 10446_1 | TATGAGTGTGGAGTCGTTGCACTCCTGACAAATACACTGCCACNNNAGATCGGAAG |
| 10446_2 | TATGAGTGTGGAGTCGTTGCACTCCTGACAAATACACTGCCACNNNAGATCGGAAG |
| 42426_1 | TATGAGTGTGGAGTCGTTGCGACACACACAGAGACAGAGAGAGNNNAGATCGGAA |
| 42426_2 | TATGAGTGTGGAGTCGTTGCGACACACACAGAGACAGAGAGAGNNNAGATCGGAA |
| 42355_1 | TATGAGTGTGGAGTCGTTGCACTTGGGATATTGAGGCAGGAAANNNAGATCGGAA |
| 42355_2 | TATGAGTGTGGAGTCGTTGCACTTGGGATATTGAGGCAGGAAANNNAGATCGGAA |
| 53349_1 | TATGAGTGTGGAGTCGTTGCAGAGAGAGAGAGAGAGGGAGAGANNNAGATCGG |
| 53349_2 | TATGAGTGTGGAGTCGTTGCAGAGAGAGAGAGAGAGGGAGAGANNNAGATCGG |
| 35297_1 | TATGAGTGTGGAGTCGTTGCCACCGCCTGCTAATTTTTGTCTTNNNAGATCGGAAGA |
| 35297_2 | TATGAGTGTGGAGTCGTTGCCACCGCCTGCTAATTTTTGTCTTNNNAGATCGGAAGA |
| 46562_1 | TATGAGTGTGGAGTCGTTGCCTGAGTAGCTGGGATTACAGGTGNNNAGATCGGAA |
| 46562_2 | TATGAGTGTGGAGTCGTTGCCTGAGTAGCTGGGATTACAGGTGNNNAGATCGGAA |
| 31153_1 | TATGAGTGTGGAGTCGTTGCTGCCATTGTCACTACAGGACTACNNNAGATCGGAAG |
| 31153_2 | TATGAGTGTGGAGTCGTTGCTGCCATTGTCACTACAGGACTACNNNAGATCGGAAG |
| 13251_1 | TATGAGTGTGGAGTCGTTGCAAAATTTATCAACTAAGCACGTANNNAGATCGGAAG |
| 13251_2 | TATGAGTGTGGAGTCGTTGCAAAATTTATCAACTAAGCACGTANNNAGATCGGAAG |
| 52212_1 | TATGAGTGTGGAGTCGTTGCGGAGATTAGAAAGTGTGTGTGTGCNNNAGATCGGA |
| 52212_2 | TATGAGTGTGGAGTCGTTGCGGAGATTAGAAAGTGTGTGTGTGCNNNAGATCGGA |
| 35498_1 | TATGAGTGTGGAGTCGTTGCTGCAACAGGAGAGAGAAGATGACNNNAGATCGGAA |
| 35498_2 | TATGAGTGTGGAGTCGTTGCTGCAACAGGAGAGAGAAGATGACNNNAGATCGGAA |
| 33718_1 | TATGAGTGTGGAGTCGTTGCGTGCCTGTTGGAGAAAGAAGAACNNNAGATCGGAA |
| 33718_2 | TATGAGTGTGGAGTCGTTGCGTGCCTGTTGGAGAAAGAAGAACNNNAGATCGGAA |
| 42146_1 | TATGAGTGTGGAGTCGTTGCGAATTCAAGCGATTCTCCTGCCNNNAGATCGGAAGA |
| 42146_2 | TATGAGTGTGGAGTCGTTGCGAATTCAAGCGATTCTCCTGCCNNNAGATCGGAAGA |
| 41488_1 | TATGAGTGTGGAGTCGTTGCTGCTAATGATACGAGGAGATGAACANNNAGATCGG |
| 41488_2 | TATGAGTGTGGAGTCGTTGCTGCTAATGATACGAGGAGATGAACANNNAGATCGG |
| 40344_1 | TATGAGTGTGGAGTCGTTGCGAAAAAGATAGAGGGGCAGGGAANNNAGATCGGA |
| 40344_2 | TATGAGTGTGGAGTCGTTGCGAAAAAGATAGAGGGGCAGGGAANNNAGATCGGA |
| 33503_1 | TATGAGTGTGGAGTCGTTGCGATAACTTGGGGCTGTTTACTGCNNNAGATCGGAA |
| 33503_2 | TATGAGTGTGGAGTCGTTGCGATAACTTGGGGCTGTTTACTGCNNNAGATCGGAA |
| 43383_1 | TATGAGTGTGGAGTCGTTGCCTGAGTAGCTGGGATTACAGGTGNNNAGATCGGAA |
| 43383_2 | TATGAGTGTGGAGTCGTTGCCTGAGTAGCTGGGATTACAGGTGNNNAGATCGGAA |
| 35984_1 | TATGAGTGTGGAGTCGTTGCAAAATCTGAAAGTCACGGAAGGGNNNAGATCGGAA |
| 35984_2 | TATGAGTGTGGAGTCGTTGCAAAATCTGAAAGTCACGGAAGGGNNNAGATCGGAA |
| 11949_1 | TATGAGTGTGGAGTCGTTGCTCTTCTTTGAGGTGGTAAGGCCNNNAGATCGGAAGA |
| 11949_2 | TATGAGTGTGGAGTCGTTGCTCTTCTTTGAGGTGGTAAGGCCNNNAGATCGGAAGA |
| 50770_1 | TATGAGTGTGGAGTCGTTGCTAAGGTCCTCAATTTCCCTCTGCNNNAGATCGGAAGA |
| 50770_2 | TATGAGTGTGGAGTCGTTGCTAAGGTCCTCAATTTCCCTCTGCNNNAGATCGGAAGA |
| 8520_1 | TATGAGTGTGGAGTCGTTGCTGTAATTGACCTAGATGCCTGTCAACNNNAGATCGG |
| 8520_2 | TATGAGTGTGGAGTCGTTGCTGTAATTGACCTAGATGCCTGTCAACNNNAGATCGG |
| 28753_1 | TATGAGTGTGGAGTCGTTGCCATCAAGCAATTCTCTGCCTCAGNNNAGATCGGAAG |
| 28753_2 | TATGAGTGTGGAGTCGTTGCCATCAAGCAATTCTCTGCCTCAGNNNAGATCGGAAG |
| 42585_1 | TATGAGTGTGGAGTCGTTGCATTTGGCATACACATCTCTGCAGNNNAGATCGGAAG |

| | |
|---|---|
| 42585_2 | TATGAGTGTGGAGTCGTTGCATTTGGCATACACATCTCTGCAGNNNAGATCGGAAG |
| 32714_1 | TATGAGTGTGGAGTCGTTGCTAAGCAGTTTCCTCTCTTGCAGANNNAGATCGGAAG |
| 32714_2 | TATGAGTGTGGAGTCGTTGCTAAGCAGTTTCCTCTCTTGCAGANNNAGATCGGAAG |
| 13768_1 | TATGAGTGTGGAGTCGTTGCCATGTATGGATTGGTGGATCNNNAGATCGGAAGAG |
| 13768_2 | TATGAGTGTGGAGTCGTTGCCATGTATGGATTGGTGGATCNNNAGATCGGAAGAG |
| 44000_1 | TATGAGTGTGGAGTCGTTGCGAAATCTGCCCCCATGATCCAATNNNAGATCGGAAG |
| 44000_2 | TATGAGTGTGGAGTCGTTGCGAAATCTGCCCCCATGATCCAATNNNAGATCGGAAG |
| 37532_1 | TATGAGTGTGGAGTCGTTGCCTGGGCCACTTGCTTATACATTTNNNAGATCGGAAG |
| 37532_2 | TATGAGTGTGGAGTCGTTGCCTGGGCCACTTGCTTATACATTTNNNAGATCGGAAG |
| 33471_1 | TATGAGTGTGGAGTCGTTGCGCAAAGCAAATCCAGCAGCATATNNNAGATCGGAAG |
| 33471_2 | TATGAGTGTGGAGTCGTTGCGCAAAGCAAATCCAGCAGCATATNNNAGATCGGAAG |
| 46018_1 | TATGAGTGTGGAGTCGTTGCCTCCTGAGTAGGTGGGACTACAGNNNAGATCGGAA |
| 46018_2 | TATGAGTGTGGAGTCGTTGCCTCCTGAGTAGGTGGGACTACAGNNNAGATCGGAA |
| 50715_1 | TATGAGTGTGGAGTCGTTGCTACCTTAGTGGGTAGGGACCAAANNNAGATCGGAA |
| 50715_2 | TATGAGTGTGGAGTCGTTGCTACCTTAGTGGGTAGGGACCAAANNNAGATCGGAA |
| 40966_1 | TATGAGTGTGGAGTCGTTGCTTGTCAGCCAGAAGTCTTCTTGANNNAGATCGGAAG |
| 40966_2 | TATGAGTGTGGAGTCGTTGCTTGTCAGCCAGAAGTCTTCTTGANNNAGATCGGAAG |
| 33343_1 | TATGAGTGTGGAGTCGTTGCGGGTACCTACTGATCGTTTGTGTNNNAGATCGGAAG |
| 33343_2 | TATGAGTGTGGAGTCGTTGCGGGTACCTACTGATCGTTTGTGTNNNAGATCGGAAG |
| 50476_1 | TATGAGTGTGGAGTCGTTGCACTCAAAAAAAGGAGATGCCTGCNNNAGATCGGAA |
| 50476_2 | TATGAGTGTGGAGTCGTTGCACTCAAAAAAAGGAGATGCCTGCNNNAGATCGGAA |
| 53140_1 | TATGAGTGTGGAGTCGTTGCGCCACACCCCTAGATCATAGAAGNNNAGATCGGAAG |
| 53140_2 | TATGAGTGTGGAGTCGTTGCGCCACACCCCTAGATCATAGAAGNNNAGATCGGAAG |
| 35058_1 | TATGAGTGTGGAGTCGTTGCTTGAGTTTATCAGGAGGGGAAGCNNNAGATCGGAA |
| 35058_2 | TATGAGTGTGGAGTCGTTGCTTGAGTTTATCAGGAGGGGAAGCNNNAGATCGGAA |
| 52975_1 | TATGAGTGTGGAGTCGTTGCAGCAATATATGGCAGTGTTGGCTNNNAGATCGGAA |
| 52975_2 | TATGAGTGTGGAGTCGTTGCAGCAATATATGGCAGTGTTGGCTNNNAGATCGGAA |
| 33746_1 | TATGAGTGTGGAGTCGTTGCATAAAATTCCAAGCCCCACAACGNNNAGATCGGAAG |
| 33746_2 | TATGAGTGTGGAGTCGTTGCATAAAATTCCAAGCCCCACAACGNNNAGATCGGAAG |
| 50937_1 | TATGAGTGTGGAGTCGTTGCCTAAGGGCTGACCTGGTCTTGNNNAGATCGGAAGA |
| 50937_2 | TATGAGTGTGGAGTCGTTGCCTAAGGGCTGACCTGGTCTTGNNNAGATCGGAAGA |
| 43585_1 | TATGAGTGTGGAGTCGTTGCGTCTTGAACTCCTGGCCTAAAGANNNAGATCGGAAG |
| 43585_2 | TATGAGTGTGGAGTCGTTGCGTCTTGAACTCCTGGCCTAAAGANNNAGATCGGAAG |
| 30115_1 | TATGAGTGTGGAGTCGTTGCCTGCAATGATGACAGCCTGTTAGNNNAGATCGGAA |
| 30115_2 | TATGAGTGTGGAGTCGTTGCCTGCAATGATGACAGCCTGTTAGNNNAGATCGGAA |
| 40416_1 | TATGAGTGTGGAGTCGTTGCACCAGATTTCAGAGAAAGCAGCTNNNAGATCGGAA |
| 40416_2 | TATGAGTGTGGAGTCGTTGCACCAGATTTCAGAGAAAGCAGCTNNNAGATCGGAA |
| 34012_1 | TATGAGTGTGGAGTCGTTGCTCTGGTGATTTTGTGGTGTTCACNNNAGATCGGAAG |
| 34012_2 | TATGAGTGTGGAGTCGTTGCTCTGGTGATTTTGTGGTGTTCACNNNAGATCGGAAG |
| 52457_1 | TATGAGTGTGGAGTCGTTGCGCCCTACCTCCTTCTTTCAACTTNNNAGATCGGAAGA |
| 52457_2 | TATGAGTGTGGAGTCGTTGCGCCCTACCTCCTTCTTTCAACTTNNNAGATCGGAAGA |
| 50589_1 | TATGAGTGTGGAGTCGTTGCGCTTTTCTTTGCTCTTCTCCTGGNNNAGATCGGAAGA |
| 50589_2 | TATGAGTGTGGAGTCGTTGCGCTTTTCTTTGCTCTTCTCCTGGNNNAGATCGGAAGA |

| | |
|---|---|
| 41910_1 | TATGAGTGTGGAGTCGTTGCCCGGGTGAGATCACATCTTACTANNNAGATCGGAAG |
| 41910_2 | TATGAGTGTGGAGTCGTTGCCCGGGTGAGATCACATCTTACTANNNAGATCGGAAG |
| 30867_1 | TATGAGTGTGGAGTCGTTGCAGCTACAGATTTTTGGACCACTCNNNAGATCGGAAG |
| 30867_2 | TATGAGTGTGGAGTCGTTGCAGCTACAGATTTTTGGACCACTCNNNAGATCGGAAG |
| 49598_1 | TATGAGTGTGGAGTCGTTGCACGCCCGGCTAATTTTTTGTATTNNNAGATCGGAAG |
| 49598_2 | TATGAGTGTGGAGTCGTTGCACGCCCGGCTAATTTTTTGTATTNNNAGATCGGAAG |
| 51773_1 | TATGAGTGTGGAGTCGTTGCAGCTTGGCTGCCTCTTCAAGNNNAGATCGGAAGAG |
| 51773_2 | TATGAGTGTGGAGTCGTTGCAGCTTGGCTGCCTCTTCAAGNNNAGATCGGAAGAG |
| 45586_1 | TATGAGTGTGGAGTCGTTGCGAAGAGAACCATCGACCTGACTTNNNAGATCGGAA |
| 45586_2 | TATGAGTGTGGAGTCGTTGCGAAGAGAACCATCGACCTGACTTNNNAGATCGGAA |
| 30705_1 | TATGAGTGTGGAGTCGTTGCTGATGCAGTTGACAGATGCTGAANNNAGATCGGAA |
| 30705_2 | TATGAGTGTGGAGTCGTTGCTGATGCAGTTGACAGATGCTGAANNNAGATCGGAA |
| 10732_1 | TATGAGTGTGGAGTCGTTGCTGACCAGGAGAGGAGAGTTAGTGNNNAGATCGGA |
| 10732_2 | TATGAGTGTGGAGTCGTTGCTGACCAGGAGAGGAGAGTTAGTGNNNAGATCGGA |
| 45730_1 | TATGAGTGTGGAGTCGTTGCGTTCACACTATTCTCCTGCCTCANNNAGATCGGAAGA |
| 45730_2 | TATGAGTGTGGAGTCGTTGCGTTCACACTATTCTCCTGCCTCANNNAGATCGGAAGA |
| 52783_1 | TATGAGTGTGGAGTCGTTGCAGAACTCTGATTGGTGACTTCCCNNNAGATCGGAAG |
| 52783_2 | TATGAGTGTGGAGTCGTTGCAGAACTCTGATTGGTGACTTCCCNNNAGATCGGAAG |
| 52494_1 | TATGAGTGTGGAGTCGTTGCCCTGTAATCCTGTAATCCGAGCANNNAGATCGGAAG |
| 52494_2 | TATGAGTGTGGAGTCGTTGCCCTGTAATCCTGTAATCCGAGCANNNAGATCGGAAG |
| 45103_1 | TATGAGTGTGGAGTCGTTGCAAAGAGTATGGAGCAGCAATCCANNNAGATCGGAA |
| 45103_2 | TATGAGTGTGGAGTCGTTGCAAAGAGTATGGAGCAGCAATCCANNNAGATCGGAA |
| 29753_1 | TATGAGTGTGGAGTCGTTGCAGCTAATATTAAGATGGGTAACCTGTNNNAGATCGG |
| 29753_2 | TATGAGTGTGGAGTCGTTGCAGCTAATATTAAGATGGGTAACCTGTNNNAGATCGG |
| 36520_1 | TATGAGTGTGGAGTCGTTGCTGATTCAAGGTAAACCGTCCACTNNNAGATCGGAAG |
| 36520_2 | TATGAGTGTGGAGTCGTTGCTGATTCAAGGTAAACCGTCCACTNNNAGATCGGAAG |
| 33558_1 | TATGAGTGTGGAGTCGTTGCCTGAGTAGTTGGGATTGTAGGCANNNAGATCGGAA |
| 33558_2 | TATGAGTGTGGAGTCGTTGCCTGAGTAGTTGGGATTGTAGGCANNNAGATCGGAA |
| 34611_1 | TATGAGTGTGGAGTCGTTGCCCTACTGGGAAGTGAGGAGCCNNNAGATCGGAAGA |
| 34611_2 | TATGAGTGTGGAGTCGTTGCCCTACTGGGAAGTGAGGAGCCNNNAGATCGGAAGA |
| 44391_1 | TATGAGTGTGGAGTCGTTGCTTTCACACGCTGTAATCTGGAGANNNAGATCGGAAG |
| 44391_2 | TATGAGTGTGGAGTCGTTGCTTTCACACGCTGTAATCTGGAGANNNAGATCGGAAG |
| 40483_1 | TATGAGTGTGGAGTCGTTGCCTGTGATCACACCACTGCATTACNNNAGATCGGAAG |
| 40483_2 | TATGAGTGTGGAGTCGTTGCCTGTGATCACACCACTGCATTACNNNAGATCGGAAG |
| 34055_1 | TATGAGTGTGGAGTCGTTGCCTCAAGTGATTCTCGTGCCTCAGNNNAGATCGGAAG |
| 34055_2 | TATGAGTGTGGAGTCGTTGCCTCAAGTGATTCTCGTGCCTCAGNNNAGATCGGAAG |
| 52897_1 | TATGAGTGTGGAGTCGTTGCGGAAGGTAAGAGACTCAAGGAGCNNNAGATCGGA |
| 52897_2 | TATGAGTGTGGAGTCGTTGCGGAAGGTAAGAGACTCAAGGAGCNNNAGATCGGA |
| 32237_1 | TATGAGTGTGGAGTCGTTGCTCCCCAGATTCCTCTCTAGAGTCNNNAGATCGGAAGA |
| 32237_2 | TATGAGTGTGGAGTCGTTGCTCCCCAGATTCCTCTCTAGAGTCNNNAGATCGGAAGA |
| 48862_1 | TATGAGTGTGGAGTCGTTGCGGCTTCTCCATGTTGCTCATTTTNNNAGATCGGAAGA |
| 48862_2 | TATGAGTGTGGAGTCGTTGCGGCTTCTCCATGTTGCTCATTTTNNNAGATCGGAAGA |
| 34732_1 | TATGAGTGTGGAGTCGTTGCCCACCAAGCTTGAATCCAGAAAGNNNAGATCGGAAG |

| | |
|---|---|
| 34732_2 | TATGAGTGTGGAGTCGTTGCCCACCAAGCTTGAATCCAGAAAGNNNAGATCGGAAG |
| 43385_1 | TATGAGTGTGGAGTCGTTGCCAGTGGTGTGATCTCAGCTCANNNAGATCGGAAGAG |
| 43385_2 | TATGAGTGTGGAGTCGTTGCCAGTGGTGTGATCTCAGCTCANNNAGATCGGAAGAG |
| 45193_1 | TATGAGTGTGGAGTCGTTGCGTGAGATTCAGAGGGATGGAGTCNNNAGATCGGAA |
| 45193_2 | TATGAGTGTGGAGTCGTTGCGTGAGATTCAGAGGGATGGAGTCNNNAGATCGGAA |
| 34065_1 | TATGAGTGTGGAGTCGTTGCTTAGGGTACATGTGCACATTGTGNNNAGATCGGAAG |
| 34065_2 | TATGAGTGTGGAGTCGTTGCTTAGGGTACATGTGCACATTGTGNNNAGATCGGAAG |
| 39266_1 | TATGAGTGTGGAGTCGTTGCAAGCAAGGTTTCTCACTGTCTGANNNAGATCGGAAG |
| 39266_2 | TATGAGTGTGGAGTCGTTGCAAGCAAGGTTTCTCACTGTCTGANNNAGATCGGAAG |
| 32589_1 | TATGAGTGTGGAGTCGTTGCGCTGGAGAGGATGTGGAGAAATANNNAGATCGGAA |
| 32589_2 | TATGAGTGTGGAGTCGTTGCGCTGGAGAGGATGTGGAGAAATANNNAGATCGGAA |
| 30293_1 | TATGAGTGTGGAGTCGTTGCAGACGGAGTCTCGCTCTGTCNNNAGATCGGAAGAG |
| 30293_2 | TATGAGTGTGGAGTCGTTGCAGACGGAGTCTCGCTCTGTCNNNAGATCGGAAGAG |
| 30759_1 | TATGAGTGTGGAGTCGTTGCCTGGCAGAAGAGGATTCAGACATNNNAGATCGGAA |
| 30759_2 | TATGAGTGTGGAGTCGTTGCCTGGCAGAAGAGGATTCAGACATNNNAGATCGGAA |
| 32361_1 | TATGAGTGTGGAGTCGTTGCCACACAGACAAACACACACACATNNNAGATCGGAAGA |
| 32361_2 | TATGAGTGTGGAGTCGTTGCCACACAGACAAACACACACACATNNNAGATCGGAAGA |
| 48515_1 | TATGAGTGTGGAGTCGTTGCCCTGGGTTCAAGCAATTCTCTTGNNNAGATCGGAAGA |
| 48515_2 | TATGAGTGTGGAGTCGTTGCCCTGGGTTCAAGCAATTCTCTTGNNNAGATCGGAAGA |
| 49234_1 | TATGAGTGTGGAGTCGTTGCCCAGAGTAGCTGAGATTACAGGCNNNAGATCGGAA |
| 49234_2 | TATGAGTGTGGAGTCGTTGCCCAGAGTAGCTGAGATTACAGGCNNNAGATCGGAA |
| 47238_1 | TATGAGTGTGGAGTCGTTGCTTCAAGTGATTCTCCTGCCTCAGNNNAGATCGGAAGA |
| 47238_2 | TATGAGTGTGGAGTCGTTGCTTCAAGTGATTCTCCTGCCTCAGNNNAGATCGGAAGA |
| 35324_1 | TATGAGTGTGGAGTCGTTGCCTCCTGAGTAGCTGGGACTACAGNNNAGATCGGAAG |
| 35324_2 | TATGAGTGTGGAGTCGTTGCCTCCTGAGTAGCTGGGACTACAGNNNAGATCGGAAG |
| 37596_1 | TATGAGTGTGGAGTCGTTGCAACTATCTGGAAGCTTCAAGGCANNNAGATCGGAAG |
| 37596_2 | TATGAGTGTGGAGTCGTTGCAACTATCTGGAAGCTTCAAGGCANNNAGATCGGAAG |
| 50600_1 | TATGAGTGTGGAGTCGTTGCATCCAAGAGCAGAAGCAGAATCANNNAGATCGGAA |
| 50600_2 | TATGAGTGTGGAGTCGTTGCATCCAAGAGCAGAAGCAGAATCANNNAGATCGGAA |
| 38325_1 | TATGAGTGTGGAGTCGTTGCCTGTAATCCCAGCTACTCAGGAGNNNAGATCGGAAG |
| 38325_2 | TATGAGTGTGGAGTCGTTGCCTGTAATCCCAGCTACTCAGGAGNNNAGATCGGAAG |
| 29441_1 | TATGAGTGTGGAGTCGTTGCACCCAATCAAAAATGGGCTCTTGNNNAGATCGGAAG |
| 29441_2 | TATGAGTGTGGAGTCGTTGCACCCAATCAAAAATGGGCTCTTGNNNAGATCGGAAG |
| 31952_1 | TATGAGTGTGGAGTCGTTGCAGGGACTACCAATCTGCATTGTCNNNAGATCGGAAG |
| 31952_2 | TATGAGTGTGGAGTCGTTGCAGGGACTACCAATCTGCATTGTCNNNAGATCGGAAG |
| 31927_1 | TATGAGTGTGGAGTCGTTGCCAGAAACTGTCCTTGGAGAAACANNNAGATCGGAAG |
| 31927_2 | TATGAGTGTGGAGTCGTTGCCAGAAACTGTCCTTGGAGAAACANNNAGATCGGAAG |
| 29516_1 | TATGAGTGTGGAGTCGTTGCGGAAACTATCCTAGCATAATGCNNNAGATCGGAAGA |
| 29516_2 | TATGAGTGTGGAGTCGTTGCGGAAACTATCCTAGCATAATGCNNNAGATCGGAAGA |
| 51846_1 | TATGAGTGTGGAGTCGTTGCTTAGACATCCAGCTGCCTTCTTTNNNAGATCGGAAGA |
| 51846_2 | TATGAGTGTGGAGTCGTTGCTTAGACATCCAGCTGCCTTCTTTNNNAGATCGGAAGA |
| 48850_1 | TATGAGTGTGGAGTCGTTGCCCGATTTCTAAGAAGGCCCCTTCNNNAGATCGGAAGA |
| 48850_2 | TATGAGTGTGGAGTCGTTGCCCGATTTCTAAGAAGGCCCCTTCNNNAGATCGGAAGA |

| | |
|---|---|
| 29703_1 | TATGAGTGTGGAGTCGTTGCATATACTAAAGCCATGTGGAGGCNNNAGATCGGAA |
| 29703_2 | TATGAGTGTGGAGTCGTTGCATATACTAAAGCCATGTGGAGGCNNNAGATCGGAA |
| 51852_1 | TATGAGTGTGGAGTCGTTGCACTCATACATAGCAAGCACCCTCNNNAGATCGGAAG |
| 51852_2 | TATGAGTGTGGAGTCGTTGCACTCATACATAGCAAGCACCCTCNNNAGATCGGAAG |
| 51233_1 | TATGAGTGTGGAGTCGTTGCTATTTTGATGAGCATGCGGGATGNNNAGATCGGAA |
| 51233_2 | TATGAGTGTGGAGTCGTTGCTATTTTGATGAGCATGCGGGATGNNNAGATCGGAA |
| 45063_1 | TATGAGTGTGGAGTCGTTGCTATTGCAGCTGTAGGTTCAGGTTNNNAGATCGGAAG |
| 45063_2 | TATGAGTGTGGAGTCGTTGCTATTGCAGCTGTAGGTTCAGGTTNNNAGATCGGAAG |
| 48597_1 | TATGAGTGTGGAGTCGTTGCAACTTAGCTGTATGTGGTGGTGTNNNAGATCGGAA |
| 48597_2 | TATGAGTGTGGAGTCGTTGCAACTTAGCTGTATGTGGTGGTGTNNNAGATCGGAA |
| 32913_1 | TATGAGTGTGGAGTCGTTGCGAAAGACTGAAAGCAGGGACCTTNNNAGATCGGAA |
| 32913_2 | TATGAGTGTGGAGTCGTTGCGAAAGACTGAAAGCAGGGACCTTNNNAGATCGGAA |
| 31258_1 | TATGAGTGTGGAGTCGTTGCCCCCAATGAATGATGAACCACTGNNNAGATCGGAAG |
| 31258_2 | TATGAGTGTGGAGTCGTTGCCCCCAATGAATGATGAACCACTGNNNAGATCGGAAG |
| 51310_1 | TATGAGTGTGGAGTCGTTGCCCTAACCTAGCTTGAGTGGGTTTNNNAGATCGGAAG |
| 51310_2 | TATGAGTGTGGAGTCGTTGCCCTAACCTAGCTTGAGTGGGTTTNNNAGATCGGAAG |
| 40114_1 | TATGAGTGTGGAGTCGTTGCTGACCCCACCGAAATACAAACTANNNAGATCGGAAG |
| 40114_2 | TATGAGTGTGGAGTCGTTGCTGACCCCACCGAAATACAAACTANNNAGATCGGAAG |
| 2152_1 | TATGAGTGTGGAGTCGTTGCTGGCTGTGTTCGTTTCTCTTTCANNNAGATCGGAAG |
| 2152_2 | TATGAGTGTGGAGTCGTTGCTGGCTGTGTTCGTTTCTCTTTCANNNAGATCGGAAG |
| 33703_1 | TATGAGTGTGGAGTCGTTGCAAAGCACAAAAGAGAAAAGGGGGNNNAGATCGGA |
| 33703_2 | TATGAGTGTGGAGTCGTTGCAAAGCACAAAAGAGAAAAGGGGGNNNAGATCGGA |
| 45751_1 | TATGAGTGTGGAGTCGTTGCGGGGATCTGCAGTAGTCACAGNNNAGATCGGAAGA |
| 45751_2 | TATGAGTGTGGAGTCGTTGCGGGGATCTGCAGTAGTCACAGNNNAGATCGGAAGA |
| 48956_1 | TATGAGTGTGGAGTCGTTGCAGGGATATCTTGGTTGCTTCCAANNNAGATCGGAAG |
| 48956_2 | TATGAGTGTGGAGTCGTTGCAGGGATATCTTGGTTGCTTCCAANNNAGATCGGAAG |
| 34998_1 | TATGAGTGTGGAGTCGTTGCCTCAAGCAATTCTCCTGCCTCAGNNNAGATCGGAAG |
| 34998_2 | TATGAGTGTGGAGTCGTTGCCTCAAGCAATTCTCCTGCCTCAGNNNAGATCGGAAG |
| 42009_1 | TATGAGTGTGGAGTCGTTGCCCGAGTAGCTGGGATTACAAGTGNNNAGATCGGAA |
| 42009_2 | TATGAGTGTGGAGTCGTTGCCCGAGTAGCTGGGATTACAAGTGNNNAGATCGGAA |
| 51647_1 | TATGAGTGTGGAGTCGTTGCCACTGCACTTCTGAATGTTCCTGNNNAGATCGGAAG |
| 51647_2 | TATGAGTGTGGAGTCGTTGCCACTGCACTTCTGAATGTTCCTGNNNAGATCGGAAG |
| 32499_1 | TATGAGTGTGGAGTCGTTGCCCTCCATAGTAGCTGAGACCTCANNNAGATCGGAAG |
| 32499_2 | TATGAGTGTGGAGTCGTTGCCCTCCATAGTAGCTGAGACCTCANNNAGATCGGAAG |
| 35550_1 | TATGAGTGTGGAGTCGTTGCCCTCCCAGGTTCATGACATTCTCNNNAGATCGGAAGA |
| 35550_2 | TATGAGTGTGGAGTCGTTGCCCTCCCAGGTTCATGACATTCTCNNNAGATCGGAAGA |
| 39187_1 | TATGAGTGTGGAGTCGTTGCGCGGGTGGAAATATAGCTTTCNNNAGATCGGAAGA |
| 39187_2 | TATGAGTGTGGAGTCGTTGCGCGGGTGGAAATATAGCTTTCNNNAGATCGGAAGA |
| 46980_1 | TATGAGTGTGGAGTCGTTGCTTCAAGCGATTCTTCTCCCTCAGNNNAGATCGGAAGA |
| 46980_2 | TATGAGTGTGGAGTCGTTGCTTCAAGCGATTCTTCTCCCTCAGNNNAGATCGGAAGA |
| 46870_1 | TATGAGTGTGGAGTCGTTGCCCAGTAGTTGGGATTACAGGTGCNNNAGATCGGAA |
| 46870_2 | TATGAGTGTGGAGTCGTTGCCCAGTAGTTGGGATTACAGGTGCNNNAGATCGGAA |
| 40296_1 | TATGAGTGTGGAGTCGTTGCGACTTGTTGAATGGCTTTGACCANNNAGATCGGAAG |

| | |
|---|---|
| 40296_2 | TATGAGTGTGGAGTCGTTGCGACTTGTTGAATGGCTTTGACCANNNAGATCGGAAG |
| 30955_1 | TATGAGTGTGGAGTCGTTGCTGACAGACCTGACACTTAGCTTCNNNAGATCGGAAG |
| 30955_2 | TATGAGTGTGGAGTCGTTGCTGACAGACCTGACACTTAGCTTCNNNAGATCGGAAG |
| 42163_1 | TATGAGTGTGGAGTCGTTGCCTCAAGTGATCCTTCCACCTCAGNNNAGATCGGAAG |
| 42163_2 | TATGAGTGTGGAGTCGTTGCCTCAAGTGATCCTTCCACCTCAGNNNAGATCGGAAG |
| 40300_1 | TATGAGTGTGGAGTCGTTGCTGTTGTTTTATTTTCTGTTGCGTTNNNAGATCGGAAG |
| 40300_2 | TATGAGTGTGGAGTCGTTGCTGTTGTTTTATTTTCTGTTGCGTTNNNAGATCGGAAG |
| 34426_1 | TATGAGTGTGGAGTCGTTGCGCATGAGGGAAGCTTCTAGACTTNNNAGATCGGAA |
| 34426_2 | TATGAGTGTGGAGTCGTTGCGCATGAGGGAAGCTTCTAGACTTNNNAGATCGGAA |
| 32924_1 | TATGAGTGTGGAGTCGTTGCTAGTCCTAGCTACTTGGAAGGCTNNNAGATCGGAAG |
| 32924_2 | TATGAGTGTGGAGTCGTTGCTAGTCCTAGCTACTTGGAAGGCTNNNAGATCGGAAG |
| 44918_1 | TATGAGTGTGGAGTCGTTGCGTGGTCTGTCCAATCAGTTACCANNNAGATCGGAAG |
| 44918_2 | TATGAGTGTGGAGTCGTTGCGTGGTCTGTCCAATCAGTTACCANNNAGATCGGAAG |
| 35051_1 | TATGAGTGTGGAGTCGTTGCTTGCTTTGGTTATTCAGGGTGTTNNNAGATCGGAAG |
| 35051_2 | TATGAGTGTGGAGTCGTTGCTTGCTTTGGTTATTCAGGGTGTTNNNAGATCGGAAG |
| 39796_1 | TATGAGTGTGGAGTCGTTGCCTTGCAGTGAGCCGAGATTGTNNNAGATCGGAAGA |
| 39796_2 | TATGAGTGTGGAGTCGTTGCCTTGCAGTGAGCCGAGATTGTNNNAGATCGGAAGA |
| 2479_1 | TATGAGTGTGGAGTCGTTGCTAGGGTTCTTGTCAGGAGACCAGNNNAGATCGGAA |
| 2479_2 | TATGAGTGTGGAGTCGTTGCTAGGGTTCTTGTCAGGAGACCAGNNNAGATCGGAA |
| 49202_1 | TATGAGTGTGGAGTCGTTGCCTAGTCCAGCCCAAAGGATGTAANNNAGATCGGAAG |
| 49202_2 | TATGAGTGTGGAGTCGTTGCCTAGTCCAGCCCAAAGGATGTAANNNAGATCGGAAG |
| 43339_1 | TATGAGTGTGGAGTCGTTGCCAGTGACATGATCTCGGTTCACTNNNAGATCGGAAG |
| 43339_2 | TATGAGTGTGGAGTCGTTGCCAGTGACATGATCTCGGTTCACTNNNAGATCGGAAG |
| 52727_1 | TATGAGTGTGGAGTCGTTGCCCTTAGTCCCCAGAAGGTAGAGANNNAGATCGGAA |
| 52727_2 | TATGAGTGTGGAGTCGTTGCCCTTAGTCCCCAGAAGGTAGAGANNNAGATCGGAA |
| 49520_1 | TATGAGTGTGGAGTCGTTGCCTGATCAGCTTCCTTGGGTTTTGNNNAGATCGGAAG |
| 49520_2 | TATGAGTGTGGAGTCGTTGCCTGATCAGCTTCCTTGGGTTTTGNNNAGATCGGAAG |
| 37296_1 | TATGAGTGTGGAGTCGTTGCTGTGATCGTGTTGTCAGAAAGTTNNNAGATCGGAA |
| 37296_2 | TATGAGTGTGGAGTCGTTGCTGTGATCGTGTTGTCAGAAAGTTNNNAGATCGGAA |
| 53142_1 | TATGAGTGTGGAGTCGTTGCCTCAGCTACCAGGACTAGGTTTGNNNAGATCGGAAG |
| 53142_2 | TATGAGTGTGGAGTCGTTGCCTCAGCTACCAGGACTAGGTTTGNNNAGATCGGAAG |
| 42758_1 | TATGAGTGTGGAGTCGTTGCTTTTTGAGACAGGGTCTTGCTCTNNNAGATCGGAAG |
| 42758_2 | TATGAGTGTGGAGTCGTTGCTTTTTGAGACAGGGTCTTGCTCTNNNAGATCGGAAG |
| 13394_1 | TATGAGTGTGGAGTCGTTGCATGTGTTTATTTCTATCTATCTATCTNNNAGATCGGA |
| 13394_2 | TATGAGTGTGGAGTCGTTGCATGTGTTTATTTCTATCTATCTATCTNNNAGATCGGA |
| 49226_1 | TATGAGTGTGGAGTCGTTGCGCCACCATGCCTAGCTAATTTTTNNNAGATCGGAAG |
| 49226_2 | TATGAGTGTGGAGTCGTTGCGCCACCATGCCTAGCTAATTTTTNNNAGATCGGAAG |
| 31223_1 | TATGAGTGTGGAGTCGTTGCATCTGGTGTTACAACTTCCCCTCNNNAGATCGGAAG |
| 31223_2 | TATGAGTGTGGAGTCGTTGCATCTGGTGTTACAACTTCCCCTCNNNAGATCGGAAG |
| 38636_1 | TATGAGTGTGGAGTCGTTGCAATATAAAAACACCTGGGCCAGGNNNAGATCGGAA |
| 38636_2 | TATGAGTGTGGAGTCGTTGCAATATAAAAACACCTGGGCCAGGNNNAGATCGGAA |
| 30342_1 | TATGAGTGTGGAGTCGTTGCCTTCAGAGCCTACCTAGGACCTCNNNAGATCGGAAG |
| 30342_2 | TATGAGTGTGGAGTCGTTGCCTTCAGAGCCTACCTAGGACCTCNNNAGATCGGAAG |

| | |
|---|---|
| 34325_1 | TATGAGTGTGGAGTCGTTGCCTTTCTGGTAGCCTTTCCCTGTANNNAGATCGGAAGA |
| 34325_2 | TATGAGTGTGGAGTCGTTGCCTTTCTGGTAGCCTTTCCCTGTANNNAGATCGGAAGA |
| 40378_1 | TATGAGTGTGGAGTCGTTGCATATGGTTAGACTTCGTGTCCCCNNNAGATCGGAAG |
| 40378_2 | TATGAGTGTGGAGTCGTTGCATATGGTTAGACTTCGTGTCCCCNNNAGATCGGAAG |
| 51107_1 | TATGAGTGTGGAGTCGTTGCGTAGTCGTAGCTACTCAGGAAGCNNNAGATCGGAA |
| 51107_2 | TATGAGTGTGGAGTCGTTGCGTAGTCGTAGCTACTCAGGAAGCNNNAGATCGGAA |
| 45780_1 | TATGAGTGTGGAGTCGTTGCCCTCCTGAGTAACTGGGACTACANNNAGATCGGAAG |
| 45780_2 | TATGAGTGTGGAGTCGTTGCCCTCCTGAGTAACTGGGACTACANNNAGATCGGAAG |
| 32573_1 | TATGAGTGTGGAGTCGTTGCGGGTTCAAGCAATTCTCCTACCTNNNAGATCGGAAG |
| 32573_2 | TATGAGTGTGGAGTCGTTGCGGGTTCAAGCAATTCTCCTACCTNNNAGATCGGAAG |
| 47391_1 | TATGAGTGTGGAGTCGTTGCACCATGCGGGAGCTAATTTTTTTNNNAGATCGGAAG |
| 47391_2 | TATGAGTGTGGAGTCGTTGCACCATGCGGGAGCTAATTTTTTTNNNAGATCGGAAG |
| 34308_1 | TATGAGTGTGGAGTCGTTGCACCAACTAGTTAGTTACTTGCGCNNNAGATCGGAAG |
| 34308_2 | TATGAGTGTGGAGTCGTTGCACCAACTAGTTAGTTACTTGCGCNNNAGATCGGAAG |
| 50224_1 | TATGAGTGTGGAGTCGTTGCAGAGAGTGGGTTCAAAAGACTCCNNNAGATCGGAA |
| 50224_2 | TATGAGTGTGGAGTCGTTGCAGAGAGTGGGTTCAAAAGACTCCNNNAGATCGGAA |
| 29942_1 | TATGAGTGTGGAGTCGTTGCTAGCACCTCACTTCCAGAAAGCNNNAGATCGGAAGA |
| 29942_2 | TATGAGTGTGGAGTCGTTGCTAGCACCTCACTTCCAGAAAGCNNNAGATCGGAAGA |
| 43190_1 | TATGAGTGTGGAGTCGTTGCATAACAGTGGCCAAGGGACATAGNNNAGATCGGAA |
| 43190_2 | TATGAGTGTGGAGTCGTTGCATAACAGTGGCCAAGGGACATAGNNNAGATCGGAA |
| 46414_1 | TATGAGTGTGGAGTCGTTGCCAAAGTGCTGGGATTATAGGCGTNNNAGATCGGAA |
| 46414_2 | TATGAGTGTGGAGTCGTTGCCAAAGTGCTGGGATTATAGGCGTNNNAGATCGGAA |
| 39854_1 | TATGAGTGTGGAGTCGTTGCGACCGATGGAACAGAGTAGAGAGNNNAGATCGGA |
| 39854_2 | TATGAGTGTGGAGTCGTTGCGACCGATGGAACAGAGTAGAGAGNNNAGATCGGA |
| 38843_1 | TATGAGTGTGGAGTCGTTGCAACACTTTTTTGGGTGCAAGGAANNNAGATCGGAA |
| 38843_2 | TATGAGTGTGGAGTCGTTGCAACACTTTTTTGGGTGCAAGGAANNNAGATCGGAA |
| 32534_1 | TATGAGTGTGGAGTCGTTGCGGAGTCTGAGGCACAATAATTGCNNNAGATCGGAA |
| 32534_2 | TATGAGTGTGGAGTCGTTGCGGAGTCTGAGGCACAATAATTGCNNNAGATCGGAA |
| 38226_1 | TATGAGTGTGGAGTCGTTGCACTTCACACTCACCAACTTAGGTNNNAGATCGGAAGA |
| 38226_2 | TATGAGTGTGGAGTCGTTGCACTTCACACTCACCAACTTAGGTNNNAGATCGGAAGA |
| 27610_1 | TATGAGTGTGGAGTCGTTGCTTTTTTTTTTTTTGAGACACAGTNNNAGATCGGAAGA |
| 27610_2 | TATGAGTGTGGAGTCGTTGCTTTTTTTTTTTTTGAGACACAGTNNNAGATCGGAAGA |
| 43998_1 | TATGAGTGTGGAGTCGTTGCAGAGGGAGAGAGAGAGAGAGAGANNNAGATCGG |
| 43998_2 | TATGAGTGTGGAGTCGTTGCAGAGGGAGAGAGAGAGAGAGAGANNNAGATCGG |
| 12717_1 | TATGAGTGTGGAGTCGTTGCTTGTATTGTTTCTAAGACAAAGGNNNAGATCGGAAG |
| 12717_2 | TATGAGTGTGGAGTCGTTGCTTGTATTGTTTCTAAGACAAAGGNNNAGATCGGAAG |
| 45594_1 | TATGAGTGTGGAGTCGTTGCCTCAAGTGATCCTCCTGCTTCAGNNNAGATCGGAAG |
| 45594_2 | TATGAGTGTGGAGTCGTTGCCTCAAGTGATCCTCCTGCTTCAGNNNAGATCGGAAG |
| 49455_1 | TATGAGTGTGGAGTCGTTGCGCAATCTGCATTCACTCCCAAGNNNAGATCGGAAGA |
| 49455_2 | TATGAGTGTGGAGTCGTTGCGCAATCTGCATTCACTCCCAAGNNNAGATCGGAAGA |
| 39869_1 | TATGAGTGTGGAGTCGTTGCGAGGTCAGGAGATTGAGACCATCNNNAGATCGGAA |
| 39869_2 | TATGAGTGTGGAGTCGTTGCGAGGTCAGGAGATTGAGACCATCNNNAGATCGGAA |
| 53251_1 | TATGAGTGTGGAGTCGTTGCATTACCTGTTGCAAGCTTGAGTGNNNAGATCGGAAG |

| | |
|---|---|
| 53251_2 | TATGAGTGTGGAGTCGTTGCATTACCTGTTGCAAGCTTGAGTGNNNAGATCGGAAG |
| 31045_1 | TATGAGTGTGGAGTCGTTGCTGGGTTCCTCTTCTTCTTTGCTTNNNAGATCGGAAGA |
| 31045_2 | TATGAGTGTGGAGTCGTTGCTGGGTTCCTCTTCTTCTTTGCTTNNNAGATCGGAAGA |
| 45016_1 | TATGAGTGTGGAGTCGTTGCGGTGAATGGCATAACTTATGGCCNNNAGATCGGAA |
| 45016_2 | TATGAGTGTGGAGTCGTTGCGGTGAATGGCATAACTTATGGCCNNNAGATCGGAA |
| 12692_1 | TATGAGTGTGGAGTCGTTGCTCCTTTGAATGTTACTTAACCTANNNAGATCGGAAG |
| 12692_2 | TATGAGTGTGGAGTCGTTGCTCCTTTGAATGTTACTTAACCTANNNAGATCGGAAG |
| 35255_1 | TATGAGTGTGGAGTCGTTGCCAGGTTCAAGCAATTCTCCTGCNNNAGATCGGAAGA |
| 35255_2 | TATGAGTGTGGAGTCGTTGCCAGGTTCAAGCAATTCTCCTGCNNNAGATCGGAAGA |
| 33932_1 | TATGAGTGTGGAGTCGTTGCGTGGTACAATGTCAGCTTACTGCNNNAGATCGGAAG |
| 33932_2 | TATGAGTGTGGAGTCGTTGCGTGGTACAATGTCAGCTTACTGCNNNAGATCGGAAG |
| 45154_1 | TATGAGTGTGGAGTCGTTGCTAAGGGAGCACATGATCAGAGTGNNNAGATCGGAA |
| 45154_2 | TATGAGTGTGGAGTCGTTGCTAAGGGAGCACATGATCAGAGTGNNNAGATCGGAA |
| 42220_1 | TATGAGTGTGGAGTCGTTGCGATTGGGCAAAAAACCAAAAGCCNNNAGATCGGAA |
| 42220_2 | TATGAGTGTGGAGTCGTTGCGATTGGGCAAAAAACCAAAAGCCNNNAGATCGGAA |
| 29944_1 | TATGAGTGTGGAGTCGTTGCGAAGGCTCACTCTCCTTGTTACTNNNAGATCGGAAG |
| 29944_2 | TATGAGTGTGGAGTCGTTGCGAAGGCTCACTCTCCTTGTTACTNNNAGATCGGAAG |
| 44036_1 | TATGAGTGTGGAGTCGTTGCAGGATTTTAGGTGGCTACAGAGTNNNAGATCGGAA |
| 44036_2 | TATGAGTGTGGAGTCGTTGCAGGATTTTAGGTGGCTACAGAGTNNNAGATCGGAA |
| 12478_1 | TATGAGTGTGGAGTCGTTGCATACATGTTCAATAAGGACAATGNNNAGATCGGAAG |
| 12478_2 | TATGAGTGTGGAGTCGTTGCATACATGTTCAATAAGGACAATGNNNAGATCGGAAG |
| 31373_1 | TATGAGTGTGGAGTCGTTGCTCAAAAAGTATGCTGCGTGAGTGNNNAGATCGGAA |
| 31373_2 | TATGAGTGTGGAGTCGTTGCTCAAAAAGTATGCTGCGTGAGTGNNNAGATCGGAA |
| 50210_1 | TATGAGTGTGGAGTCGTTGCTGAGATGTACTGCTGTGTGAAGTNNNAGATCGGAA |
| 50210_2 | TATGAGTGTGGAGTCGTTGCTGAGATGTACTGCTGTGTGAAGTNNNAGATCGGAA |
| 12541_1 | TATGAGTGTGGAGTCGTTGCATAGTTGAGTGTAACAATCTCATNNNAGATCGGAAG |
| 12541_2 | TATGAGTGTGGAGTCGTTGCATAGTTGAGTGTAACAATCTCATNNNAGATCGGAAG |
| 50087_1 | TATGAGTGTGGAGTCGTTGCACTATGCTAAAAAGTCCCCTCCCNNNAGATCGGAAG |
| 50087_2 | TATGAGTGTGGAGTCGTTGCACTATGCTAAAAAGTCCCCTCCCNNNAGATCGGAAG |
| 31691_1 | TATGAGTGTGGAGTCGTTGCATGTGTGTGGTAGGTACATTGCTNNNAGATCGGAA |
| 31691_2 | TATGAGTGTGGAGTCGTTGCATGTGTGTGGTAGGTACATTGCTNNNAGATCGGAA |
| 43847_1 | TATGAGTGTGGAGTCGTTGCCTTCCAATCAAAGGAGCCAACTGNNNAGATCGGAAG |
| 43847_2 | TATGAGTGTGGAGTCGTTGCCTTCCAATCAAAGGAGCCAACTGNNNAGATCGGAAG |
| 45980_1 | TATGAGTGTGGAGTCGTTGCGTAGCTGGGATTACAGGTGCATNNNAGATCGGAAG |
| 45980_2 | TATGAGTGTGGAGTCGTTGCGTAGCTGGGATTACAGGTGCATNNNAGATCGGAAG |
| 49966_1 | TATGAGTGTGGAGTCGTTGCACTCTGGCTACTCTTCTGCTAGANNNAGATCGGAAG |
| 49966_2 | TATGAGTGTGGAGTCGTTGCACTCTGGCTACTCTTCTGCTAGANNNAGATCGGAAG |
| 42949_1 | TATGAGTGTGGAGTCGTTGCGTGTGTGTGATGTCTTTGTCTGGNNNAGATCGGAA |
| 42949_2 | TATGAGTGTGGAGTCGTTGCGTGTGTGTGATGTCTTTGTCTGGNNNAGATCGGAA |
| 51837_1 | TATGAGTGTGGAGTCGTTGCCGTCACCACACTCAGCTAATTTTNNNAGATCGGAAGA |
| 51837_2 | TATGAGTGTGGAGTCGTTGCCGTCACCACACTCAGCTAATTTTNNNAGATCGGAAGA |
| 49912_1 | TATGAGTGTGGAGTCGTTGCGCATTTTTCCCATGACCACTACCNNNAGATCGGAAGA |
| 49912_2 | TATGAGTGTGGAGTCGTTGCGCATTTTTCCCATGACCACTACCNNNAGATCGGAAGA |

| | |
|---|---|
| 46954_1 | TATGAGTGTGGAGTCGTTGCACGCCCGGCTAATTTTTCTTGTANNNAGATCGGAAG |
| 46954_2 | TATGAGTGTGGAGTCGTTGCACGCCCGGCTAATTTTTCTTGTANNNAGATCGGAAG |
| 30190_1 | TATGAGTGTGGAGTCGTTGCACAGGGATGCACTACCATTACCNNNAGATCGGAAGA |
| 30190_2 | TATGAGTGTGGAGTCGTTGCACAGGGATGCACTACCATTACCNNNAGATCGGAAGA |
| 30641_1 | TATGAGTGTGGAGTCGTTGCCAGCATGACAAGAGCTGATTGTTNNNAGATCGGAA |
| 30641_2 | TATGAGTGTGGAGTCGTTGCCAGCATGACAAGAGCTGATTGTTNNNAGATCGGAA |
| 41446_1 | TATGAGTGTGGAGTCGTTGCGACAGCCTTTCATTTTCAGAGGTNNNAGATCGGAAG |
| 41446_2 | TATGAGTGTGGAGTCGTTGCGACAGCCTTTCATTTTCAGAGGTNNNAGATCGGAAG |
| 41370_1 | TATGAGTGTGGAGTCGTTGCAGCTTACAATCCCTGGAACTTGTNNNAGATCGGAAG |
| 41370_2 | TATGAGTGTGGAGTCGTTGCAGCTTACAATCCCTGGAACTTGTNNNAGATCGGAAG |
| 39078_1 | TATGAGTGTGGAGTCGTTGCCCTTACTCCTGCAAGAATAGCCANNNAGATCGGAAG |
| 39078_2 | TATGAGTGTGGAGTCGTTGCCCTTACTCCTGCAAGAATAGCCANNNAGATCGGAAG |
| 40514_1 | TATGAGTGTGGAGTCGTTGCCTGTAGTTCCAGCTACTCAGGAGNNNAGATCGGAAG |
| 40514_2 | TATGAGTGTGGAGTCGTTGCCTGTAGTTCCAGCTACTCAGGAGNNNAGATCGGAAG |
| 31031_1 | TATGAGTGTGGAGTCGTTGCTACGAGTGAACACCCATTCTCAGNNNAGATCGGAAG |
| 31031_2 | TATGAGTGTGGAGTCGTTGCTACGAGTGAACACCCATTCTCAGNNNAGATCGGAAG |
| 30753_1 | TATGAGTGTGGAGTCGTTGCTTCCCCTCCAAAGATAGTGGATGNNNAGATCGGAAG |
| 30753_2 | TATGAGTGTGGAGTCGTTGCTTCCCCTCCAAAGATAGTGGATGNNNAGATCGGAAG |
| 33166_1 | TATGAGTGTGGAGTCGTTGCATGGCATGTCATTTTTGTCACCTNNNAGATCGGAAG |
| 33166_2 | TATGAGTGTGGAGTCGTTGCATGGCATGTCATTTTTGTCACCTNNNAGATCGGAAG |
| 34558_1 | TATGAGTGTGGAGTCGTTGCGATGGTCCTAAGAGGAGGAGCAGNNNAGATCGGA |
| 34558_2 | TATGAGTGTGGAGTCGTTGCGATGGTCCTAAGAGGAGGAGCAGNNNAGATCGGA |
| 40392_1 | TATGAGTGTGGAGTCGTTGCGATGCTGAAGGTGCTGATAAGAANNNAGATCGGAA |
| 40392_2 | TATGAGTGTGGAGTCGTTGCGATGCTGAAGGTGCTGATAAGAANNNAGATCGGAA |
| 53327_1 | TATGAGTGTGGAGTCGTTGCTCGTAGATGAGATGCTGAATGGGNNNAGATCGGAA |
| 53327_2 | TATGAGTGTGGAGTCGTTGCTCGTAGATGAGATGCTGAATGGGNNNAGATCGGAA |
| 51966_1 | TATGAGTGTGGAGTCGTTGCCTTCCTGGGTTCAAGCGATTCTCNNNAGATCGGAAG |
| 51966_2 | TATGAGTGTGGAGTCGTTGCCTTCCTGGGTTCAAGCGATTCTCNNNAGATCGGAAG |
| 32709_1 | TATGAGTGTGGAGTCGTTGCCCTCCTAGGTTCAAGCGATTCTCNNNAGATCGGAAG |
| 32709_2 | TATGAGTGTGGAGTCGTTGCCCTCCTAGGTTCAAGCGATTCTCNNNAGATCGGAAG |
| 12819_1 | TATGAGTGTGGAGTCGTTGCTAATACAAACAAACAGCAAGAAANNNAGATCGGAA |
| 12819_2 | TATGAGTGTGGAGTCGTTGCTAATACAAACAAACAGCAAGAAANNNAGATCGGAA |
| 36800_1 | TATGAGTGTGGAGTCGTTGCGATTTTGAAGCACCTATGGACGCNNNAGATCGGAA |
| 36800_2 | TATGAGTGTGGAGTCGTTGCGATTTTGAAGCACCTATGGACGCNNNAGATCGGAA |
| 36660_1 | TATGAGTGTGGAGTCGTTGCGAGGACAGAAAACAGACACAACGNNNAGATCGGAA |
| 36660_2 | TATGAGTGTGGAGTCGTTGCGAGGACAGAAAACAGACACAACGNNNAGATCGGAA |
| 49011_1 | TATGAGTGTGGAGTCGTTGCGCTCCTGGCCTCTTTTCTTTTTTNNNAGATCGGAAGA |
| 49011_2 | TATGAGTGTGGAGTCGTTGCGCTCCTGGCCTCTTTTCTTTTTTNNNAGATCGGAAGA |
| 37486_1 | TATGAGTGTGGAGTCGTTGCGCATTATGACGTTGTGTGACCATNNNAGATCGGAAG |
| 37486_2 | TATGAGTGTGGAGTCGTTGCGCATTATGACGTTGTGTGACCATNNNAGATCGGAAG |
| 52057_1 | TATGAGTGTGGAGTCGTTGCGCTCAGACTACTGTCCACATCATNNNAGATCGGAAG |
| 52057_2 | TATGAGTGTGGAGTCGTTGCGCTCAGACTACTGTCCACATCATNNNAGATCGGAAG |
| 41166_1 | TATGAGTGTGGAGTCGTTGCGTGTCACAATGAAACACTGCAGANNNAGATCGGAA |

| | |
|---|---|
| 41166_2 | TATGAGTGTGGAGTCGTTGCGTGTCACAATGAAACACTGCAGANNNAGATCGGAA |
| 50983_1 | TATGAGTGTGGAGTCGTTGCCTGCAGCCTCAAATTCTTGGATTNNNAGATCGGAAG |
| 50983_2 | TATGAGTGTGGAGTCGTTGCCTGCAGCCTCAAATTCTTGGATTNNNAGATCGGAAG |
| 50563_1 | TATGAGTGTGGAGTCGTTGCAGCACAGATGAAGACAGGAAACANNNAGATCGGAA |
| 50563_2 | TATGAGTGTGGAGTCGTTGCAGCACAGATGAAGACAGGAAACANNNAGATCGGAA |
| 46224_1 | TATGAGTGTGGAGTCGTTGCCTGCCTGAGTAGCTGGGATTACNNNAGATCGGAAG |
| 46224_2 | TATGAGTGTGGAGTCGTTGCCTGCCTGAGTAGCTGGGATTACNNNAGATCGGAAG |
| 47846_1 | TATGAGTGTGGAGTCGTTGCCGACATCACCTTCGAGCATATCANNNAGATCGGAAG |
| 47846_2 | TATGAGTGTGGAGTCGTTGCCGACATCACCTTCGAGCATATCANNNAGATCGGAAG |
| 13062_1 | TATGAGTGTGGAGTCGTTGCAATCATTTGAAGATATTCACCATNNNAGATCGGAAG |
| 13062_2 | TATGAGTGTGGAGTCGTTGCAATCATTTGAAGATATTCACCATNNNAGATCGGAAG |
| 48065_1 | TATGAGTGTGGAGTCGTTGCTGGCTATGTGGGCTCTTTTTTTGNNNAGATCGGAAG |
| 48065_2 | TATGAGTGTGGAGTCGTTGCTGGCTATGTGGGCTCTTTTTTTGNNNAGATCGGAAG |
| 29760_1 | TATGAGTGTGGAGTCGTTGCATCTTCAGCTTTCTGGTACAGCANNNAGATCGGAAG |
| 29760_2 | TATGAGTGTGGAGTCGTTGCATCTTCAGCTTTCTGGTACAGCANNNAGATCGGAAG |
| 38576_1 | TATGAGTGTGGAGTCGTTGCAAACCGTTCAACTTACAAAAGCANNNAGATCGGAAG |
| 38576_2 | TATGAGTGTGGAGTCGTTGCAAACCGTTCAACTTACAAAAGCANNNAGATCGGAAG |
| 51962_1 | TATGAGTGTGGAGTCGTTGCCTACTGTCACTCTTCCCGTGTCNNNAGATCGGAAGAG |
| 51962_2 | TATGAGTGTGGAGTCGTTGCCTACTGTCACTCTTCCCGTGTCNNNAGATCGGAAGAG |
| 52597_1 | TATGAGTGTGGAGTCGTTGCGAGGATACAGTCGTGCACTACATNNNAGATCGGAA |
| 52597_2 | TATGAGTGTGGAGTCGTTGCGAGGATACAGTCGTGCACTACATNNNAGATCGGAA |
| 49291_1 | TATGAGTGTGGAGTCGTTGCCTCCTGGGTTCAAGCGATTCTCNNNAGATCGGAAGA |
| 49291_2 | TATGAGTGTGGAGTCGTTGCCTCCTGGGTTCAAGCGATTCTCNNNAGATCGGAAGA |
| 38020_1 | TATGAGTGTGGAGTCGTTGCCTGCAAGAAACACACTTCACCTANNNAGATCGGAAG |
| 38020_2 | TATGAGTGTGGAGTCGTTGCCTGCAAGAAACACACTTCACCTANNNAGATCGGAAG |
| 41425_1 | TATGAGTGTGGAGTCGTTGCATGGACATTTGTTCAGCTGTGTGNNNAGATCGGAA |
| 41425_2 | TATGAGTGTGGAGTCGTTGCATGGACATTTGTTCAGCTGTGTGNNNAGATCGGAA |
| 41631_1 | TATGAGTGTGGAGTCGTTGCGCTTCTAGATATCCAATTGTTCTGGNNNAGATCGGA |
| 41631_2 | TATGAGTGTGGAGTCGTTGCGCTTCTAGATATCCAATTGTTCTGGNNNAGATCGGA |
| 40570_1 | TATGAGTGTGGAGTCGTTGCAAAGAAGGGCAGGTCACTTACATNNNAGATCGGAA |
| 40570_2 | TATGAGTGTGGAGTCGTTGCAAAGAAGGGCAGGTCACTTACATNNNAGATCGGAA |
| 46015_1 | TATGAGTGTGGAGTCGTTGCATGTGTAACCAATGAACATGGCCNNNAGATCGGAA |
| 46015_2 | TATGAGTGTGGAGTCGTTGCATGTGTAACCAATGAACATGGCCNNNAGATCGGAA |
| 51099_1 | TATGAGTGTGGAGTCGTTGCTAGCCAGACCCTCATCTCTACAANNNAGATCGGAAG |
| 51099_2 | TATGAGTGTGGAGTCGTTGCTAGCCAGACCCTCATCTCTACAANNNAGATCGGAAG |
| 51150_1 | TATGAGTGTGGAGTCGTTGCAGTCTACAGATTCTGAGCAGCTGNNNAGATCGGAA |
| 51150_2 | TATGAGTGTGGAGTCGTTGCAGTCTACAGATTCTGAGCAGCTGNNNAGATCGGAA |
| 8859_1 | TATGAGTGTGGAGTCGTTGCCAACTGAGAGCAATAACCCCTGTNNNAGATCGGAAG |
| 8859_2 | TATGAGTGTGGAGTCGTTGCCAACTGAGAGCAATAACCCCTGTNNNAGATCGGAAG |
| 42074_1 | TATGAGTGTGGAGTCGTTGCCTGTAATCCCAGCTACTCAGGAGNNNAGATCGGAAG |
| 42074_2 | TATGAGTGTGGAGTCGTTGCCTGTAATCCCAGCTACTCAGGAGNNNAGATCGGAAG |
| 37375_1 | TATGAGTGTGGAGTCGTTGCGACCTTTCTTCCTACAGGGGTTTNNNAGATCGGAAG |
| 37375_2 | TATGAGTGTGGAGTCGTTGCGACCTTTCTTCCTACAGGGGTTTNNNAGATCGGAAG |

| | |
|---|---|
| 4777_1 | TATGAGTGTGGAGTCGTTGCTCGGTCAGGCTACACATTTTCCANNNAGATCGGAAG |
| 4777_2 | TATGAGTGTGGAGTCGTTGCTCGGTCAGGCTACACATTTTCCANNNAGATCGGAAG |
| 51973_1 | TATGAGTGTGGAGTCGTTGCCTCAACATTTAGGCTGCCTTCTGNNNAGATCGGAAG |
| 51973_2 | TATGAGTGTGGAGTCGTTGCCTCAACATTTAGGCTGCCTTCTGNNNAGATCGGAAG |
| 49510_1 | TATGAGTGTGGAGTCGTTGCTCTCCTTCACCCTTCAAAGTGTCNNNAGATCGGAAGA |
| 49510_2 | TATGAGTGTGGAGTCGTTGCTCTCCTTCACCCTTCAAAGTGTCNNNAGATCGGAAGA |
| 51115_1 | TATGAGTGTGGAGTCGTTGCTGGGCCAATTCAGTGTATGGTTANNNAGATCGGAA |
| 51115_2 | TATGAGTGTGGAGTCGTTGCTGGGCCAATTCAGTGTATGGTTANNNAGATCGGAA |
| 47734_1 | TATGAGTGTGGAGTCGTTGCCCAGGTTCAAGCAATTCTCGTGNNNAGATCGGAAGA |
| 47734_2 | TATGAGTGTGGAGTCGTTGCCCAGGTTCAAGCAATTCTCGTGNNNAGATCGGAAGA |
| 51946_1 | TATGAGTGTGGAGTCGTTGCGAAAACGTGGACTTGAATGACGTNNNAGATCGGAA |
| 51946_2 | TATGAGTGTGGAGTCGTTGCGAAAACGTGGACTTGAATGACGTNNNAGATCGGAA |
| 35296_1 | TATGAGTGTGGAGTCGTTGCTGGAGTAGCTGGGATTACAGGTANNNAGATCGGAA |
| 35296_2 | TATGAGTGTGGAGTCGTTGCTGGAGTAGCTGGGATTACAGGTANNNAGATCGGAA |
| 29158_1 | TATGAGTGTGGAGTCGTTGCTTCCCTTTCCTCCTTTCCAGATGNNNAGATCGGAAGA |
| 29158_2 | TATGAGTGTGGAGTCGTTGCTTCCCTTTCCTCCTTTCCAGATGNNNAGATCGGAAGA |
| 30335_1 | TATGAGTGTGGAGTCGTTGCAAAAGCACACGGAGTACTTGCATNNNAGATCGGAA |
| 30335_2 | TATGAGTGTGGAGTCGTTGCAAAAGCACACGGAGTACTTGCATNNNAGATCGGAA |
| 37383_1 | TATGAGTGTGGAGTCGTTGCATGATGGCATACACCTGTAGTCCNNNAGATCGGAAG |
| 37383_2 | TATGAGTGTGGAGTCGTTGCATGATGGCATACACCTGTAGTCCNNNAGATCGGAAG |
| 47073_1 | TATGAGTGTGGAGTCGTTGCCGAGTAGCTGAGATTACAGGCATNNNAGATCGGAA |
| 47073_2 | TATGAGTGTGGAGTCGTTGCCGAGTAGCTGAGATTACAGGCATNNNAGATCGGAA |
| 47720_1 | TATGAGTGTGGAGTCGTTGCCTTTTTTTGAGACAGGGCCTCAGNNNAGATCGGAAG |
| 47720_2 | TATGAGTGTGGAGTCGTTGCCTTTTTTTGAGACAGGGCCTCAGNNNAGATCGGAAG |
| 42756_1 | TATGAGTGTGGAGTCGTTGCGTAGCCTCCAGAACTGTGAGAAANNNAGATCGGAA |
| 42756_2 | TATGAGTGTGGAGTCGTTGCGTAGCCTCCAGAACTGTGAGAAANNNAGATCGGAA |
| 33996_1 | TATGAGTGTGGAGTCGTTGCCCCTGTTTCAAGCAATTCTCCTGNNNAGATCGGAAGA |
| 33996_2 | TATGAGTGTGGAGTCGTTGCCCCTGTTTCAAGCAATTCTCCTGNNNAGATCGGAAGA |
| 35445_1 | TATGAGTGTGGAGTCGTTGCTGAAAGAACCTCTAAGCCTGTCCNNNAGATCGGAAG |
| 35445_2 | TATGAGTGTGGAGTCGTTGCTGAAAGAACCTCTAAGCCTGTCCNNNAGATCGGAAG |
| 32085_1 | TATGAGTGTGGAGTCGTTGCAGACCCTTATCTCTCACCGTACANNNAGATCGGAAGA |
| 32085_2 | TATGAGTGTGGAGTCGTTGCAGACCCTTATCTCTCACCGTACANNNAGATCGGAAGA |
| 40901_1 | TATGAGTGTGGAGTCGTTGCGTTTGTGCATGCTCTACAGTCTGNNNAGATCGGAAG |
| 40901_2 | TATGAGTGTGGAGTCGTTGCGTTTGTGCATGCTCTACAGTCTGNNNAGATCGGAAG |
| 50177_1 | TATGAGTGTGGAGTCGTTGCGAGAGTGAGAGGACAGGACTCAGNNNAGATCGGA |
| 50177_2 | TATGAGTGTGGAGTCGTTGCGAGAGTGAGAGGACAGGACTCAGNNNAGATCGGA |
| 13642_1 | TATGAGTGTGGAGTCGTTGCTCTTTAATAGAAAACCTTTGGACNNNAGATCGGAAG |
| 13642_2 | TATGAGTGTGGAGTCGTTGCTCTTTAATAGAAAACCTTTGGACNNNAGATCGGAAG |
| 51943_1 | TATGAGTGTGGAGTCGTTGCCTCATTTTTTGGGGAGTCGCATTNNNAGATCGGAAG |
| 51943_2 | TATGAGTGTGGAGTCGTTGCCTCATTTTTTGGGGAGTCGCATTNNNAGATCGGAAG |
| 10278_1 | TATGAGTGTGGAGTCGTTGCAATATCTGTGGCAGCAACACTGANNNAGATCGGAA |
| 10278_2 | TATGAGTGTGGAGTCGTTGCAATATCTGTGGCAGCAACACTGANNNAGATCGGAA |
| 51878_1 | TATGAGTGTGGAGTCGTTGCACTGCATTTGAATTTCTAGGCCTNNNAGATCGGAAG |

| | |
|---|---|
| 51878_2 | TATGAGTGTGGAGTCGTTGCACTGCATTTGAATTTCTAGGCCTNNNAGATCGGAAG |
| 41121_1 | TATGAGTGTGGAGTCGTTGCAGTGCCAAGTGGGACATTTCTAGNNNAGATCGGAA |
| 41121_2 | TATGAGTGTGGAGTCGTTGCAGTGCCAAGTGGGACATTTCTAGNNNAGATCGGAA |
| 41823_1 | TATGAGTGTGGAGTCGTTGCGTCCAGCCAACTCTCTGTAGTTTNNNAGATCGGAAG |
| 41823_2 | TATGAGTGTGGAGTCGTTGCGTCCAGCCAACTCTCTGTAGTTTNNNAGATCGGAAG |
| 49122_1 | TATGAGTGTGGAGTCGTTGCCCACCACACCCAGCTAATTTTTTNNNAGATCGGAAGA |
| 49122_2 | TATGAGTGTGGAGTCGTTGCCCACCACACCCAGCTAATTTTTTNNNAGATCGGAAGA |
| 52561_1 | TATGAGTGTGGAGTCGTTGCGGCCAGTACTTCACAAGCAAAATNNNAGATCGGAAG |
| 52561_2 | TATGAGTGTGGAGTCGTTGCGGCCAGTACTTCACAAGCAAAATNNNAGATCGGAAG |
| 43342_1 | TATGAGTGTGGAGTCGTTGCAGGATTGTTTTGGCTATTTGGGCNNNAGATCGGAA |
| 43342_2 | TATGAGTGTGGAGTCGTTGCAGGATTGTTTTGGCTATTTGGGCNNNAGATCGGAA |
| 52633_1 | TATGAGTGTGGAGTCGTTGCGTGCCGGGTCGAAAGACAGANNNAGATCGGAAGA |
| 52633_2 | TATGAGTGTGGAGTCGTTGCGTGCCGGGTCGAAAGACAGANNNAGATCGGAAGA |
| 43175_1 | TATGAGTGTGGAGTCGTTGCCATTCTCCTGCCTCAGCCTCNNNAGATCGGAAGAGCA |
| 43175_2 | TATGAGTGTGGAGTCGTTGCCATTCTCCTGCCTCAGCCTCNNNAGATCGGAAGAGCA |
| 51009_1 | TATGAGTGTGGAGTCGTTGCTTTGTATAGGGCAAGCACAGCTANNNAGATCGGAA |
| 51009_2 | TATGAGTGTGGAGTCGTTGCTTTGTATAGGGCAAGCACAGCTANNNAGATCGGAA |
| 42467_1 | TATGAGTGTGGAGTCGTTGCCTATCACTGTGCTGGGTCAGACNNNAGATCGGAAGA |
| 42467_2 | TATGAGTGTGGAGTCGTTGCCTATCACTGTGCTGGGTCAGACNNNAGATCGGAAGA |
| 13060_1 | TATGAGTGTGGAGTCGTTGCTTTTGCTACAGGAAAACAGAAANNNAGATCGGAAG |
| 13060_2 | TATGAGTGTGGAGTCGTTGCTTTTGCTACAGGAAAACAGAAANNNAGATCGGAAG |
| 12696_1 | TATGAGTGTGGAGTCGTTGCCTTCACTCCTACCACTGAAANNNAGATCGGAAGAGC |
| 12696_2 | TATGAGTGTGGAGTCGTTGCCTTCACTCCTACCACTGAAANNNAGATCGGAAGAGC |
| 45918_1 | TATGAGTGTGGAGTCGTTGCTTCAAGTGATTCTCCTGCCTCAGNNNAGATCGGAAG |
| 45918_2 | TATGAGTGTGGAGTCGTTGCTTCAAGTGATTCTCCTGCCTCAGNNNAGATCGGAAG |
| 50718_1 | TATGAGTGTGGAGTCGTTGCCTTTGTGTTCTGGGCATGGTTAGNNNAGATCGGAAG |
| 50718_2 | TATGAGTGTGGAGTCGTTGCCTTTGTGTTCTGGGCATGGTTAGNNNAGATCGGAAG |
| 43066_1 | TATGAGTGTGGAGTCGTTGCCATGACCACGTGGGATTTATTCGNNNAGATCGGAAG |
| 43066_2 | TATGAGTGTGGAGTCGTTGCCATGACCACGTGGGATTTATTCGNNNAGATCGGAAG |
| 50685_1 | TATGAGTGTGGAGTCGTTGCATCCTCAAGGTTCCAGTGCTTCNNNAGATCGGAAGA |
| 50685_2 | TATGAGTGTGGAGTCGTTGCATCCTCAAGGTTCCAGTGCTTCNNNAGATCGGAAGA |
| 2561_1 | TATGAGTGTGGAGTCGTTGCTGCCCAAACTTATCAATCCTCCCNNNAGATCGGAAGA |
| 2561_2 | TATGAGTGTGGAGTCGTTGCTGCCCAAACTTATCAATCCTCCCNNNAGATCGGAAGA |
| 1996_1 | TATGAGTGTGGAGTCGTTGCTCACCATCACCTAGGTTGGAGTGNNNAGATCGGAAG |
| 1996_2 | TATGAGTGTGGAGTCGTTGCTCACCATCACCTAGGTTGGAGTGNNNAGATCGGAAG |
| 36575_1 | TATGAGTGTGGAGTCGTTGCCATAAATTAGCCAGGTGTGGTGGNNNAGATCGGAA |
| 36575_2 | TATGAGTGTGGAGTCGTTGCCATAAATTAGCCAGGTGTGGTGGNNNAGATCGGAA |
| 41982_1 | TATGAGTGTGGAGTCGTTGCTATCATTCTCCTGCATCAGCCTCNNNAGATCGGAAGA |
| 41982_2 | TATGAGTGTGGAGTCGTTGCTATCATTCTCCTGCATCAGCCTCNNNAGATCGGAAGA |
| 33532_1 | TATGAGTGTGGAGTCGTTGCGACAGATTCTCACTCTCACCCAGNNNAGATCGGAAG |
| 33532_2 | TATGAGTGTGGAGTCGTTGCGACAGATTCTCACTCTCACCCAGNNNAGATCGGAAG |
| 51112_1 | TATGAGTGTGGAGTCGTTGCGGGGTTGGTAAGCTTAGGGATTTNNNAGATCGGAA |
| 51112_2 | TATGAGTGTGGAGTCGTTGCGGGGTTGGTAAGCTTAGGGATTTNNNAGATCGGAA |

| | |
|---|---|
| 50812_1 | TATGAGTGTGGAGTCGTTGCACTCTGTCTGAGATCAAGTCAGCNNNAGATCGGAAG |
| 50812_2 | TATGAGTGTGGAGTCGTTGCACTCTGTCTGAGATCAAGTCAGCNNNAGATCGGAAG |
| 51330_1 | TATGAGTGTGGAGTCGTTGCGGGAAACCCTGTTCAGACTACATNNNAGATCGGAAG |
| 51330_2 | TATGAGTGTGGAGTCGTTGCGGGAAACCCTGTTCAGACTACATNNNAGATCGGAAG |
| 2945_1 | TATGAGTGTGGAGTCGTTGCATTGGCTTGGAAAATAAGTGCTTNNNAGATCGGAA |
| 2945_2 | TATGAGTGTGGAGTCGTTGCATTGGCTTGGAAAATAAGTGCTTNNNAGATCGGAA |
| 53031_1 | TATGAGTGTGGAGTCGTTGCCATGGTTTGGAGTTTGAGGGAAANNNAGATCGGAA |
| 53031_2 | TATGAGTGTGGAGTCGTTGCCATGGTTTGGAGTTTGAGGGAAANNNAGATCGGAA |
| 32824_1 | TATGAGTGTGGAGTCGTTGCGTGCAATCTCGGTTCAGTGTAACNNNAGATCGGAAG |
| 32824_2 | TATGAGTGTGGAGTCGTTGCGTGCAATCTCGGTTCAGTGTAACNNNAGATCGGAAG |
| 37262_1 | TATGAGTGTGGAGTCGTTGCAGTGCTTATCAGACTGGTGGAAGNNNAGATCGGAA |
| 37262_2 | TATGAGTGTGGAGTCGTTGCAGTGCTTATCAGACTGGTGGAAGNNNAGATCGGAA |
| 12775_1 | TATGAGTGTGGAGTCGTTGCTCTTGAATTTCAAAAGGATTCTCNNNAGATCGGAAG |
| 12775_2 | TATGAGTGTGGAGTCGTTGCTCTTGAATTTCAAAAGGATTCTCNNNAGATCGGAAG |
| 51690_1 | TATGAGTGTGGAGTCGTTGCCACATTCAGTGGGTCTAGAGCTCNNNAGATCGGAAG |
| 51690_2 | TATGAGTGTGGAGTCGTTGCCACATTCAGTGGGTCTAGAGCTCNNNAGATCGGAAG |
| 33057_1 | TATGAGTGTGGAGTCGTTGCCAAAATTTTGGCTCACTGCAACCNNNAGATCGGAAG |
| 33057_2 | TATGAGTGTGGAGTCGTTGCCAAAATTTTGGCTCACTGCAACCNNNAGATCGGAAG |
| 35680_1 | TATGAGTGTGGAGTCGTTGCCCACACCCGGCTAATTTTTTGTANNNAGATCGGAAG |
| 35680_2 | TATGAGTGTGGAGTCGTTGCCCACACCCGGCTAATTTTTTGTANNNAGATCGGAAG |
| 29866_1 | TATGAGTGTGGAGTCGTTGCCCCACAGGCATTAAAACTGAACCNNNAGATCGGAAG |
| 29866_2 | TATGAGTGTGGAGTCGTTGCCCCACAGGCATTAAAACTGAACCNNNAGATCGGAAG |
| 5294_1 | TATGAGTGTGGAGTCGTTGCGGATCTGGGCGCTATGAAAAGTGTNNNAGATCGGA |
| 5294_2 | TATGAGTGTGGAGTCGTTGCGGATCTGGGCGCTATGAAAAGTGTNNNAGATCGGA |
| 49842_1 | TATGAGTGTGGAGTCGTTGCCTATGTGACTGATGGAGAGGCAGNNNAGATCGGAA |
| 49842_2 | TATGAGTGTGGAGTCGTTGCCTATGTGACTGATGGAGAGGCAGNNNAGATCGGAA |
| 45562_1 | TATGAGTGTGGAGTCGTTGCGCATGTTTCTTCTTAGACTGGCCNNNAGATCGGAAG |
| 45562_2 | TATGAGTGTGGAGTCGTTGCGCATGTTTCTTCTTAGACTGGCCNNNAGATCGGAAG |
| 12645_1 | TATGAGTGTGGAGTCGTTGCCACACCTAGTAAGAAAATTTTGTNNNAGATCGGAAG |
| 12645_2 | TATGAGTGTGGAGTCGTTGCCACACCTAGTAAGAAAATTTTGTNNNAGATCGGAAG |
| 48868_1 | TATGAGTGTGGAGTCGTTGCCTCAGCATTCACATTCACTGCACNNNAGATCGGAAG |
| 48868_2 | TATGAGTGTGGAGTCGTTGCCTCAGCATTCACATTCACTGCACNNNAGATCGGAAG |
| 51249_1 | TATGAGTGTGGAGTCGTTGCAGCCAGTTATCCCACCATCATTTNNNAGATCGGAAG |
| 51249_2 | TATGAGTGTGGAGTCGTTGCAGCCAGTTATCCCACCATCATTTNNNAGATCGGAAG |
| 13309_1 | TATGAGTGTGGAGTCGTTGCGTAACAGATGACAAGATCAAAGNNNAGATCGGAAG |
| 13309_2 | TATGAGTGTGGAGTCGTTGCGTAACAGATGACAAGATCAAAGNNNAGATCGGAAG |
| 41652_1 | TATGAGTGTGGAGTCGTTGCTCAGGGATTCAACTTCTTCCTGGNNNAGATCGGAAG |
| 41652_2 | TATGAGTGTGGAGTCGTTGCTCAGGGATTCAACTTCTTCCTGGNNNAGATCGGAAG |
| 46070_1 | TATGAGTGTGGAGTCGTTGCCTCCGGAGTAGCTGGGACTACNNNAGATCGGAAGA |
| 46070_2 | TATGAGTGTGGAGTCGTTGCCTCCGGAGTAGCTGGGACTACNNNAGATCGGAAGA |
| 30139_1 | TATGAGTGTGGAGTCGTTGCTTTTCAACTGATTATGTCTGTGTGANNNAGATCGGA |
| 30139_2 | TATGAGTGTGGAGTCGTTGCTTTTCAACTGATTATGTCTGTGTGANNNAGATCGGA |
| 38818_1 | TATGAGTGTGGAGTCGTTGCCTGTAGTCCCAGCTACTCAGGAGNNNAGATCGGAAG |

| | |
|---|---|
| 38818_2 | TATGAGTGTGGAGTCGTTGCCTGTAGTCCCAGCTACTCAGGAGNNNAGATCGGAAG |
| 38025_1 | TATGAGTGTGGAGTCGTTGCCGTATTGTCTTCCACGAAACTGGNNNAGATCGGAAG |
| 38025_2 | TATGAGTGTGGAGTCGTTGCCGTATTGTCTTCCACGAAACTGGNNNAGATCGGAAG |
| 43406_1 | TATGAGTGTGGAGTCGTTGCTTCAAGCTATTCTCCTGCCTCAGNNNAGATCGGAAGA |
| 43406_2 | TATGAGTGTGGAGTCGTTGCTTCAAGCTATTCTCCTGCCTCAGNNNAGATCGGAAGA |
| 2707_1 | TATGAGTGTGGAGTCGTTGCGGGCAACCAGGTAGATCCCATATNNNAGATCGGAA |
| 2707_2 | TATGAGTGTGGAGTCGTTGCGGGCAACCAGGTAGATCCCATATNNNAGATCGGAA |
| 50923_1 | TATGAGTGTGGAGTCGTTGCCCACACCAGGCCTTATTACTCTTNNNAGATCGGAAGA |
| 50923_2 | TATGAGTGTGGAGTCGTTGCCCACACCAGGCCTTATTACTCTTNNNAGATCGGAAGA |
| 53310_1 | TATGAGTGTGGAGTCGTTGCCACAGCAACCTGGATGTAATTGGNNNAGATCGGAA |
| 53310_2 | TATGAGTGTGGAGTCGTTGCCACAGCAACCTGGATGTAATTGGNNNAGATCGGAA |
| 41746_1 | TATGAGTGTGGAGTCGTTGCTGGTCTTGAACTTCTGAGCTCAANNNAGATCGGAAG |
| 41746_2 | TATGAGTGTGGAGTCGTTGCTGGTCTTGAACTTCTGAGCTCAANNNAGATCGGAAG |
| 34200_1 | TATGAGTGTGGAGTCGTTGCTGATGCAACATGGATGAACCTTGNNNAGATCGGAA |
| 34200_2 | TATGAGTGTGGAGTCGTTGCTGATGCAACATGGATGAACCTTGNNNAGATCGGAA |
| 42014_1 | TATGAGTGTGGAGTCGTTGCTTTTAGTAGGGACGGGGTTTCACNNNAGATCGGAA |
| 42014_2 | TATGAGTGTGGAGTCGTTGCTTTTAGTAGGGACGGGGTTTCACNNNAGATCGGAA |
| 30104_1 | TATGAGTGTGGAGTCGTTGCCATGTCCGTTTCATCACAGAACCNNNAGATCGGAAG |
| 30104_2 | TATGAGTGTGGAGTCGTTGCCATGTCCGTTTCATCACAGAACCNNNAGATCGGAAG |
| 50792_1 | TATGAGTGTGGAGTCGTTGCCAGCTCACAGAAATGCTTGTCTCNNNAGATCGGAAG |
| 50792_2 | TATGAGTGTGGAGTCGTTGCCAGCTCACAGAAATGCTTGTCTCNNNAGATCGGAAG |
| 30044_1 | TATGAGTGTGGAGTCGTTGCGCCAAGTGAGTAATTTACAGCTGCNNNAGATCGGAA |
| 30044_2 | TATGAGTGTGGAGTCGTTGCGCCAAGTGAGTAATTTACAGCTGCNNNAGATCGGAA |
| 42137_1 | TATGAGTGTGGAGTCGTTGCGTTAGGAGGAACGAGAGACTTGGNNNAGATCGGA |
| 42137_2 | TATGAGTGTGGAGTCGTTGCGTTAGGAGGAACGAGAGACTTGGNNNAGATCGGA |
| 40426_1 | TATGAGTGTGGAGTCGTTGCTCACGCTCCTCAAAGGTTTCTTANNNAGATCGGAAG |
| 40426_2 | TATGAGTGTGGAGTCGTTGCTCACGCTCCTCAAAGGTTTCTTANNNAGATCGGAAG |
| 31190_1 | TATGAGTGTGGAGTCGTTGCCCTTTCCAATTTGGGTGTCTTCCNNNAGATCGGAAGA |
| 31190_2 | TATGAGTGTGGAGTCGTTGCCCTTTCCAATTTGGGTGTCTTCCNNNAGATCGGAAGA |
| 48154_1 | TATGAGTGTGGAGTCGTTGCACATCCTCCAGTTCAATTCAGCTNNNAGATCGGAAGA |
| 48154_2 | TATGAGTGTGGAGTCGTTGCACATCCTCCAGTTCAATTCAGCTNNNAGATCGGAAGA |
| 36794_1 | TATGAGTGTGGAGTCGTTGCGTAAGCAATAGGGGGCTAGGATCNNNAGATCGGAA |
| 36794_2 | TATGAGTGTGGAGTCGTTGCGTAAGCAATAGGGGGCTAGGATCNNNAGATCGGAA |
| 44808_1 | TATGAGTGTGGAGTCGTTGCTATTTCTGGACACTAGCCTCGTGNNNAGATCGGAAG |
| 44808_2 | TATGAGTGTGGAGTCGTTGCTATTTCTGGACACTAGCCTCGTGNNNAGATCGGAAG |
| 34060_1 | TATGAGTGTGGAGTCGTTGCTCTTGAGTACCTGGGACTACAGANNNAGATCGGAA |
| 34060_2 | TATGAGTGTGGAGTCGTTGCTCTTGAGTACCTGGGACTACAGANNNAGATCGGAA |
| 43047_1 | TATGAGTGTGGAGTCGTTGCATAAGATTAGCCGAGCGTAGTGGNNNAGATCGGAA |
| 43047_2 | TATGAGTGTGGAGTCGTTGCATAAGATTAGCCGAGCGTAGTGGNNNAGATCGGAA |
| 43558_1 | TATGAGTGTGGAGTCGTTGCCTAGTAGCTGGGATTACAGGCACNNNAGATCGGAA |
| 43558_2 | TATGAGTGTGGAGTCGTTGCCTAGTAGCTGGGATTACAGGCACNNNAGATCGGAA |
| 46700_1 | TATGAGTGTGGAGTCGTTGCGATGGAGTACAACTCTGTCACCCNNNAGATCGGAAG |
| 46700_2 | TATGAGTGTGGAGTCGTTGCGATGGAGTACAACTCTGTCACCCNNNAGATCGGAAG |

| | |
|---|---|
| 32643_1 | TATGAGTGTGGAGTCGTTGCCATTCTCCTGCCTCAGCCTCNNNAGATCGGAAGAGCA |
| 32643_2 | TATGAGTGTGGAGTCGTTGCCATTCTCCTGCCTCAGCCTCNNNAGATCGGAAGAGCA |
| 39173_1 | TATGAGTGTGGAGTCGTTGCGAAAGACAAAAGACAAGCTGCCTNNNAGATCGGAA |
| 39173_2 | TATGAGTGTGGAGTCGTTGCGAAAGACAAAAGACAAGCTGCCTNNNAGATCGGAA |
| 11113_1 | TATGAGTGTGGAGTCGTTGCTCAGCCTAGGAGTTCATGGTTGCNNNAGATCGGAAG |
| 11113_2 | TATGAGTGTGGAGTCGTTGCTCAGCCTAGGAGTTCATGGTTGCNNNAGATCGGAAG |
| 31813_1 | TATGAGTGTGGAGTCGTTGCACACAGCCATAAAAAGGGATGAANNNAGATCGGAA |
| 31813_2 | TATGAGTGTGGAGTCGTTGCACACAGCCATAAAAAGGGATGAANNNAGATCGGAA |
| 31517_1 | TATGAGTGTGGAGTCGTTGCATATGTGTGTGTGTGTGTGTGTGNNNAGATCGGAA |
| 31517_2 | TATGAGTGTGGAGTCGTTGCATATGTGTGTGTGTGTGTGTGTGNNNAGATCGGAA |
| 32129_1 | TATGAGTGTGGAGTCGTTGCCAACAGAAAAATCCGTGACCCAANNNAGATCGGAAG |
| 32129_2 | TATGAGTGTGGAGTCGTTGCCAACAGAAAAATCCGTGACCCAANNNAGATCGGAAG |
| 48636_1 | TATGAGTGTGGAGTCGTTGCTGGAGTGCAATAGTGTGATCTCGNNNAGATCGGAA |
| 48636_2 | TATGAGTGTGGAGTCGTTGCTGGAGTGCAATAGTGTGATCTCGNNNAGATCGGAA |
| 49850_1 | TATGAGTGTGGAGTCGTTGCTCAGGCATCCCTTTGTCACATTANNNAGATCGGAAGA |
| 49850_2 | TATGAGTGTGGAGTCGTTGCTCAGGCATCCCTTTGTCACATTANNNAGATCGGAAGA |
| 49739_1 | TATGAGTGTGGAGTCGTTGCTGATCTCAGTTCACTGCAACCTCNNNAGATCGGAAGA |
| 49739_2 | TATGAGTGTGGAGTCGTTGCTGATCTCAGTTCACTGCAACCTCNNNAGATCGGAAGA |
| 42984_1 | TATGAGTGTGGAGTCGTTGCACACACACACACAAAATATAGGCTNNNAGATCGGAA |
| 42984_2 | TATGAGTGTGGAGTCGTTGCACACACACACACAAAATATAGGCTNNNAGATCGGAA |
| 39478_1 | TATGAGTGTGGAGTCGTTGCTCAGAAGGTTCTTGGGCATGTTANNNAGATCGGAAG |
| 39478_2 | TATGAGTGTGGAGTCGTTGCTCAGAAGGTTCTTGGGCATGTTANNNAGATCGGAAG |
| 30838_1 | TATGAGTGTGGAGTCGTTGCTGCAATCAGAGGCTCTTCTTTCTNNNAGATCGGAAGA |
| 30838_2 | TATGAGTGTGGAGTCGTTGCTGCAATCAGAGGCTCTTCTTTCTNNNAGATCGGAAGA |
| 30735_1 | TATGAGTGTGGAGTCGTTGCCCTCCCGAGTTCAAGAGATTCTTNNNAGATCGGAAGA |
| 30735_2 | TATGAGTGTGGAGTCGTTGCCCTCCCGAGTTCAAGAGATTCTTNNNAGATCGGAAGA |
| 13522_1 | TATGAGTGTGGAGTCGTTGCAAATGGAGAAAAGGAAACTTTAGNNNAGATCGGAA |
| 13522_2 | TATGAGTGTGGAGTCGTTGCAAATGGAGAAAAGGAAACTTTAGNNNAGATCGGAA |
| 30906_1 | TATGAGTGTGGAGTCGTTGCGAGACAAAGAGGGAGTCAAGGAGNNNAGATCGGA |
| 30906_2 | TATGAGTGTGGAGTCGTTGCGAGACAAAGAGGGAGTCAAGGAGNNNAGATCGGA |
| 3183_1 | TATGAGTGTGGAGTCGTTGCTGAAGCTTGTTGTGACTTCTGCTNNNAGATCGGAAG |
| 3183_2 | TATGAGTGTGGAGTCGTTGCTGAAGCTTGTTGTGACTTCTGCTNNNAGATCGGAAG |
| 32986_1 | TATGAGTGTGGAGTCGTTGCCACACCTGTAATCCCAGCACTTNNNAGATCGGAAGA |
| 32986_2 | TATGAGTGTGGAGTCGTTGCCACACCTGTAATCCCAGCACTTNNNAGATCGGAAGA |
| 3954_1 | TATGAGTGTGGAGTCGTTGCTCCGAGTGTTCCAGTGTCTTTTTNNNAGATCGGAAGA |
| 3954_2 | TATGAGTGTGGAGTCGTTGCTCCGAGTGTTCCAGTGTCTTTTTNNNAGATCGGAAGA |
| 49764_1 | TATGAGTGTGGAGTCGTTGCAAGCTGGAAGATGAAGCTCAAGANNNAGATCGGAA |
| 49764_2 | TATGAGTGTGGAGTCGTTGCAAGCTGGAAGATGAAGCTCAAGANNNAGATCGGAA |
| 51091_1 | TATGAGTGTGGAGTCGTTGCGAGGCTGAGGCAGAATAATCACTNNNAGATCGGAA |
| 51091_2 | TATGAGTGTGGAGTCGTTGCGAGGCTGAGGCAGAATAATCACTNNNAGATCGGAA |
| 4245_1 | TATGAGTGTGGAGTCGTTGCATTGATTCTCCTTCCTCTGCCTTCNNNAGATCGGAAGA |
| 4245_2 | TATGAGTGTGGAGTCGTTGCATTGATTCTCCTTCCTCTGCCTTCNNNAGATCGGAAGA |
| 32645_1 | TATGAGTGTGGAGTCGTTGCTTCAAGTGATTCTCCTGCCTCAGNNNAGATCGGAAGA |

| | |
|---|---|
| 32645_2 | TATGAGTGTGGAGTCGTTGCTTCAAGTGATTCTCCTGCCTCAGNNNAGATCGGAAG |
| 36967_1 | TATGAGTGTGGAGTCGTTGCATGCCTAGATGGTGTCTACACACNNNAGATCGGAAG |
| 36967_2 | TATGAGTGTGGAGTCGTTGCATGCCTAGATGGTGTCTACACACNNNAGATCGGAAG |
| 35052_1 | TATGAGTGTGGAGTCGTTGCCAGAGAGAGGGCATCTTTGTCTTNNNAGATCGGAA |
| 35052_2 | TATGAGTGTGGAGTCGTTGCCAGAGAGAGGGCATCTTTGTCTTNNNAGATCGGAA |
| 50707_1 | TATGAGTGTGGAGTCGTTGCCTGTTTGGGGATGTTTCCAGTCNNNAGATCGGAAGA |
| 50707_2 | TATGAGTGTGGAGTCGTTGCCTGTTTGGGGATGTTTCCAGTCNNNAGATCGGAAGA |
| 44916_1 | TATGAGTGTGGAGTCGTTGCGGGAGAAGAGACAGGGTAGAGTANNNAGATCGGA |
| 44916_2 | TATGAGTGTGGAGTCGTTGCGGGAGAAGAGACAGGGTAGAGTANNNAGATCGGA |
| 34566_1 | TATGAGTGTGGAGTCGTTGCCAGATACAGGGGATGAGGCAAAGNNNAGATCGGA |
| 34566_2 | TATGAGTGTGGAGTCGTTGCCAGATACAGGGGATGAGGCAAAGNNNAGATCGGA |
| 46420_1 | TATGAGTGTGGAGTCGTTGCACCACGCCCAGCTAATTTTTTTTNNNAGATCGGAAGA |
| 46420_2 | TATGAGTGTGGAGTCGTTGCACCACGCCCAGCTAATTTTTTTTNNNAGATCGGAAGA |
| 36861_1 | TATGAGTGTGGAGTCGTTGCAAACACTATGCCTCTCAGTGGAGNNNAGATCGGAAG |
| 36861_2 | TATGAGTGTGGAGTCGTTGCAAACACTATGCCTCTCAGTGGAGNNNAGATCGGAAG |
| 44096_1 | TATGAGTGTGGAGTCGTTGCTCAGAAGGGAAGAGCTCTTGTTCNNNAGATCGGAA |
| 44096_2 | TATGAGTGTGGAGTCGTTGCTCAGAAGGGAAGAGCTCTTGTTCNNNAGATCGGAA |
| 31811_1 | TATGAGTGTGGAGTCGTTGCTTTGAGACAAGGTCTTGCTCTGTNNNAGATCGGAAG |
| 31811_2 | TATGAGTGTGGAGTCGTTGCTTTGAGACAAGGTCTTGCTCTGTNNNAGATCGGAAG |
| 34238_1 | TATGAGTGTGGAGTCGTTGCTAAATTGTGTTGGGCAGTATGGCNNNAGATCGGAA |
| 34238_2 | TATGAGTGTGGAGTCGTTGCTAAATTGTGTTGGGCAGTATGGCNNNAGATCGGAA |
| 36486_1 | TATGAGTGTGGAGTCGTTGCACACACACACACACACACAAAAANNNAGATCGGAAG |
| 36486_2 | TATGAGTGTGGAGTCGTTGCACACACACACACACACACAAAAANNNAGATCGGAAG |
| 43596_1 | TATGAGTGTGGAGTCGTTGCTAGTAACCTGATGATTGTCGCCCNNNAGATCGGAAG |
| 43596_2 | TATGAGTGTGGAGTCGTTGCTAGTAACCTGATGATTGTCGCCCNNNAGATCGGAAG |
| 46493_1 | TATGAGTGTGGAGTCGTTGCGGGTTGAAGTAATTCTCCTGCCTNNNAGATCGGAAG |
| 46493_2 | TATGAGTGTGGAGTCGTTGCGGGTTGAAGTAATTCTCCTGCCTNNNAGATCGGAAG |
| 12828_1 | TATGAGTGTGGAGTCGTTGCAGTTTATAATCTGGAATCTCCTCNNNAGATCGGAAG |
| 12828_2 | TATGAGTGTGGAGTCGTTGCAGTTTATAATCTGGAATCTCCTCNNNAGATCGGAAG |
| 31870_1 | TATGAGTGTGGAGTCGTTGCCAACGCATCTTCACTGATGGACNNNAGATCGGAAGA |
| 31870_2 | TATGAGTGTGGAGTCGTTGCCAACGCATCTTCACTGATGGACNNNAGATCGGAAGA |
| 35962_1 | TATGAGTGTGGAGTCGTTGCTACAAAAAATTAGCCAGGCGTGGNNNAGATCGGAA |
| 35962_2 | TATGAGTGTGGAGTCGTTGCTACAAAAAATTAGCCAGGCGTGGNNNAGATCGGAA |
| 39528_1 | TATGAGTGTGGAGTCGTTGCGGGATCAATGCAACAAGAAGAGCNNNAGATCGGAA |
| 39528_2 | TATGAGTGTGGAGTCGTTGCGGGATCAATGCAACAAGAAGAGCNNNAGATCGGAA |
| 29633_1 | TATGAGTGTGGAGTCGTTGCGGAAAACGGTGGCAGGTTAAAAANNNAGATCGGA |
| 29633_2 | TATGAGTGTGGAGTCGTTGCGGAAAACGGTGGCAGGTTAAAAANNNAGATCGGA |
| 5559_1 | TATGAGTGTGGAGTCGTTGCTACTTGGTGGCTAATTCAGCAGGNNNAGATCGGAA |
| 5559_2 | TATGAGTGTGGAGTCGTTGCTACTTGGTGGCTAATTCAGCAGGNNNAGATCGGAA |
| 30383_1 | TATGAGTGTGGAGTCGTTGCCCGGTATGTAATTTAACATTCAANNNAGATCGGAAG |
| 30383_2 | TATGAGTGTGGAGTCGTTGCCCGGTATGTAATTTAACATTCAANNNAGATCGGAAG |
| 48885_1 | TATGAGTGTGGAGTCGTTGCCTCCTGAGTAGCTGGGATTACAGNNNAGATCGGAA |
| 48885_2 | TATGAGTGTGGAGTCGTTGCCTCCTGAGTAGCTGGGATTACAGNNNAGATCGGAA |

| | |
|---|---|
| 10834_1 | TATGAGTGTGGAGTCGTTGCTGATAGTTCTTTGCCCCTCCAGANNNAGATCGGAAG |
| 10834_2 | TATGAGTGTGGAGTCGTTGCTGATAGTTCTTTGCCCCTCCAGANNNAGATCGGAAG |
| 36093_1 | TATGAGTGTGGAGTCGTTGCTGAAAAGCCTCACATAATGCGTCNNNAGATCGGAAG |
| 36093_2 | TATGAGTGTGGAGTCGTTGCTGAAAAGCCTCACATAATGCGTCNNNAGATCGGAAG |
| 40955_1 | TATGAGTGTGGAGTCGTTGCAAGACTCTCTAAGCAAACATACCTNNNAGATCGGAA |
| 40955_2 | TATGAGTGTGGAGTCGTTGCAAGACTCTCTAAGCAAACATACCTNNNAGATCGGAA |
| 41255_1 | TATGAGTGTGGAGTCGTTGCCTGGGCAATATAGCAAGACTCCANNNAGATCGGAA |
| 41255_2 | TATGAGTGTGGAGTCGTTGCCTGGGCAATATAGCAAGACTCCANNNAGATCGGAA |
| 42458_1 | TATGAGTGTGGAGTCGTTGCCACACCGTGCTTTATGATCATCCNNNAGATCGGAAG |
| 42458_2 | TATGAGTGTGGAGTCGTTGCCACACCGTGCTTTATGATCATCCNNNAGATCGGAAG |
| 33528_1 | TATGAGTGTGGAGTCGTTGCATATACTTTGGGACAAGGCCAGCNNNAGATCGGAA |
| 33528_2 | TATGAGTGTGGAGTCGTTGCATATACTTTGGGACAAGGCCAGCNNNAGATCGGAA |
| 51626_1 | TATGAGTGTGGAGTCGTTGCGGAGAGACACAGAAAGACCACATNNNAGATCGGAA |
| 51626_2 | TATGAGTGTGGAGTCGTTGCGGAGAGACACAGAAAGACCACATNNNAGATCGGAA |
| 51436_1 | TATGAGTGTGGAGTCGTTGCGGAAGAACGGAGCCACCTTCNNNAGATCGGAAGAG |
| 51436_2 | TATGAGTGTGGAGTCGTTGCGGAAGAACGGAGCCACCTTCNNNAGATCGGAAGAG |
| 40363_1 | TATGAGTGTGGAGTCGTTGCATCCTGACTAACACAGTGAACCCNNNAGATCGGAAG |
| 40363_2 | TATGAGTGTGGAGTCGTTGCATCCTGACTAACACAGTGAACCCNNNAGATCGGAAG |
| 35411_1 | TATGAGTGTGGAGTCGTTGCCATTACCCAAGAACCTGCAACAGNNNAGATCGGAAG |
| 35411_2 | TATGAGTGTGGAGTCGTTGCCATTACCCAAGAACCTGCAACAGNNNAGATCGGAAG |
| 13732_1 | TATGAGTGTGGAGTCGTTGCTAATATCTACTTTTGCTCAGCTTNNNAGATCGGAAGA |
| 13732_2 | TATGAGTGTGGAGTCGTTGCTAATATCTACTTTTGCTCAGCTTNNNAGATCGGAAGA |
| 51909_1 | TATGAGTGTGGAGTCGTTGCGGCTCCTCACTTCCCAGAAGNNNAGATCGGAAGAGC |
| 51909_2 | TATGAGTGTGGAGTCGTTGCGGCTCCTCACTTCCCAGAAGNNNAGATCGGAAGAGC |
| 30807_1 | TATGAGTGTGGAGTCGTTGCCTCTGTCTCTCTCTCTCTCTCCCNNNAGATCGGAAGAG |
| 30807_2 | TATGAGTGTGGAGTCGTTGCCTCTGTCTCTCTCTCTCTCTCCCNNNAGATCGGAAGAG |
| 40874_1 | TATGAGTGTGGAGTCGTTGCCCACGACCGGCTAATTTTTGTATNNNAGATCGGAAG |
| 40874_2 | TATGAGTGTGGAGTCGTTGCCCACGACCGGCTAATTTTTGTATNNNAGATCGGAAG |
| 35236_1 | TATGAGTGTGGAGTCGTTGCGTTCAGTCAGCCTTCCAATTCACNNNAGATCGGAAG |
| 35236_2 | TATGAGTGTGGAGTCGTTGCGTTCAGTCAGCCTTCCAATTCACNNNAGATCGGAAG |
| 42406_1 | TATGAGTGTGGAGTCGTTGCACACAAACACACACACACACATTNNNAGATCGGAAG |
| 42406_2 | TATGAGTGTGGAGTCGTTGCACACAAACACACACACACACATTNNNAGATCGGAAG |
| 10216_1 | TATGAGTGTGGAGTCGTTGCAGGACACCCTAGCAAATAGCTCANNNAGATCGGAAG |
| 10216_2 | TATGAGTGTGGAGTCGTTGCAGGACACCCTAGCAAATAGCTCANNNAGATCGGAAG |
| 34290_1 | TATGAGTGTGGAGTCGTTGCTCTCTCTCTCTCTCTTTCTGCCANNNAGATCGGAAGA |
| 34290_2 | TATGAGTGTGGAGTCGTTGCTCTCTCTCTCTCTCTTTCTGCCANNNAGATCGGAAGA |
| 36255_1 | TATGAGTGTGGAGTCGTTGCGGAGAATGGCATGAACCAGTGAGNNNAGATCGGA |
| 36255_2 | TATGAGTGTGGAGTCGTTGCGGAGAATGGCATGAACCAGTGAGNNNAGATCGGA |
| 39356_1 | TATGAGTGTGGAGTCGTTGCATGTGTTTAGTACCCTTGGCTGGNNNAGATCGGAAG |
| 39356_2 | TATGAGTGTGGAGTCGTTGCATGTGTTTAGTACCCTTGGCTGGNNNAGATCGGAAG |
| 53372_1 | TATGAGTGTGGAGTCGTTGCAGCCCATGTTGAGAGCTGTATACNNNAGATCGGAA |
| 53372_2 | TATGAGTGTGGAGTCGTTGCAGCCCATGTTGAGAGCTGTATACNNNAGATCGGAA |
| 51247_1 | TATGAGTGTGGAGTCGTTGCCATTGAAGTCCTGTTGGTCCCTANNNAGATCGGAAG |

| | |
|---|---|
| 51247_2 | TATGAGTGTGGAGTCGTTGCCATTGAAGTCCTGTTGGTCCCTANNNAGATCGGAAG |
| 33168_1 | TATGAGTGTGGAGTCGTTGCAAACTGTTATCACCACTGACCCANNNAGATCGGAAG |
| 33168_2 | TATGAGTGTGGAGTCGTTGCAAACTGTTATCACCACTGACCCANNNAGATCGGAAG |
| 52198_1 | TATGAGTGTGGAGTCGTTGCATAGCCCGGTTGTCTCTCATTTTNNNAGATCGGAAG |
| 52198_2 | TATGAGTGTGGAGTCGTTGCATAGCCCGGTTGTCTCTCATTTTNNNAGATCGGAAG |
| 46965_1 | TATGAGTGTGGAGTCGTTGCAACCAATGGGGAACTAAGACTCCNNNAGATCGGAAG |
| 46965_2 | TATGAGTGTGGAGTCGTTGCAACCAATGGGGAACTAAGACTCCNNNAGATCGGAAG |
| 33732_1 | TATGAGTGTGGAGTCGTTGCGCTGTCACTTTCTACCCCTGAAANNNAGATCGGAAG |
| 33732_2 | TATGAGTGTGGAGTCGTTGCGCTGTCACTTTCTACCCCTGAAANNNAGATCGGAAG |
| 50464_1 | TATGAGTGTGGAGTCGTTGCACACGCTACAACATGGATGAAACNNNAGATCGGAAG |
| 50464_2 | TATGAGTGTGGAGTCGTTGCACACGCTACAACATGGATGAAACNNNAGATCGGAAG |
| 13388_1 | TATGAGTGTGGAGTCGTTGCCTGTTTGCATCCCAACATACNNNAGATCGGAAGAGC |
| 13388_2 | TATGAGTGTGGAGTCGTTGCCTGTTTGCATCCCAACATACNNNAGATCGGAAGAGC |
| 36088_1 | TATGAGTGTGGAGTCGTTGCATCTGAACTTTTGGGGGTCAACTNNNAGATCGGAAG |
| 36088_2 | TATGAGTGTGGAGTCGTTGCATCTGAACTTTTGGGGGTCAACTNNNAGATCGGAAG |
| 36544_1 | TATGAGTGTGGAGTCGTTGCATAAAGACAGGATCCAGCTCTGCNNNAGATCGGAAG |
| 36544_2 | TATGAGTGTGGAGTCGTTGCATAAAGACAGGATCCAGCTCTGCNNNAGATCGGAAG |
| 40448_1 | TATGAGTGTGGAGTCGTTGCCCAGTCAGTAGTGCAAGTCAGAANNNAGATCGGAA |
| 40448_2 | TATGAGTGTGGAGTCGTTGCCCAGTCAGTAGTGCAAGTCAGAANNNAGATCGGAA |
| 50238_1 | TATGAGTGTGGAGTCGTTGCATGGAACTGCAGGATCCAAAGATNNNAGATCGGAA |
| 50238_2 | TATGAGTGTGGAGTCGTTGCATGGAACTGCAGGATCCAAAGATNNNAGATCGGAA |
| 35117_1 | TATGAGTGTGGAGTCGTTGCCCTCTCAGGTTCAAGCGATTCTCNNNAGATCGGAAG |
| 35117_2 | TATGAGTGTGGAGTCGTTGCCCTCTCAGGTTCAAGCGATTCTCNNNAGATCGGAAG |
| 8183_1 | TATGAGTGTGGAGTCGTTGCATTCGGTTTTGCTGTTTGCTCANNNAGATCGGAAGA |
| 8183_2 | TATGAGTGTGGAGTCGTTGCATTCGGTTTTGCTGTTTGCTCANNNAGATCGGAAGA |
| 43826_1 | TATGAGTGTGGAGTCGTTGCTAGGAAGACGCATCTCACAAGAGNNNAGATCGGAA |
| 43826_2 | TATGAGTGTGGAGTCGTTGCTAGGAAGACGCATCTCACAAGAGNNNAGATCGGAA |
| 46582_1 | TATGAGTGTGGAGTCGTTGCCTCACTATATTGCCCAGTCTGGTNNNAGATCGGAAG |
| 46582_2 | TATGAGTGTGGAGTCGTTGCCTCACTATATTGCCCAGTCTGGTNNNAGATCGGAAG |
| 12742_1 | TATGAGTGTGGAGTCGTTGCCATTAACAGGATTCAGACTAAAANNNAGATCGGAAG |
| 12742_2 | TATGAGTGTGGAGTCGTTGCCATTAACAGGATTCAGACTAAAANNNAGATCGGAAG |
| 44252_1 | TATGAGTGTGGAGTCGTTGCGATTGTGCACTGTGTGAAACTGANNNAGATCGGAA |
| 44252_2 | TATGAGTGTGGAGTCGTTGCGATTGTGCACTGTGTGAAACTGANNNAGATCGGAA |
| 35815_1 | TATGAGTGTGGAGTCGTTGCAAACAAGAGCTTTGCATTCAGGCNNNAGATCGGAAG |
| 35815_2 | TATGAGTGTGGAGTCGTTGCAAACAAGAGCTTTGCATTCAGGCNNNAGATCGGAAG |
| 31743_1 | TATGAGTGTGGAGTCGTTGCTCACTCAACAGTATTCTTGGGCANNNAGATCGGAAG |
| 31743_2 | TATGAGTGTGGAGTCGTTGCTCACTCAACAGTATTCTTGGGCANNNAGATCGGAAG |
| 35239_1 | TATGAGTGTGGAGTCGTTGCCCCTTTAGTTGATGACCTGTCCANNNAGATCGGAAG |
| 35239_2 | TATGAGTGTGGAGTCGTTGCCCCTTTAGTTGATGACCTGTCCANNNAGATCGGAAG |
| 12984_1 | TATGAGTGTGGAGTCGTTGCTCTGGAGAAATAGAAGAAAAGATNNNAGATCGGAA |
| 12984_2 | TATGAGTGTGGAGTCGTTGCTCTGGAGAAATAGAAGAAAAGATNNNAGATCGGAA |
| 40476_1 | TATGAGTGTGGAGTCGTTGCACAGGGATGTAAGCTCCTTGAAGNNNAGATCGGAA |
| 40476_2 | TATGAGTGTGGAGTCGTTGCACAGGGATGTAAGCTCCTTGAAGNNNAGATCGGAA |

| | |
|---|---|
| 48292_1 | TATGAGTGTGGAGTCGTTGCGATTTTCTACAGGTGCCGAATGGNNNAGATCGGAA |
| 48292_2 | TATGAGTGTGGAGTCGTTGCGATTTTCTACAGGTGCCGAATGGNNNAGATCGGAA |
| 32575_1 | TATGAGTGTGGAGTCGTTGCCAGGTTCAAGCAATTCTCCTGCNNNAGATCGGAAGA |
| 32575_2 | TATGAGTGTGGAGTCGTTGCCAGGTTCAAGCAATTCTCCTGCNNNAGATCGGAAGA |
| 50943_1 | TATGAGTGTGGAGTCGTTGCTATAAGGACTGCTGGGTGTTTGCNNNAGATCGGAA |
| 50943_2 | TATGAGTGTGGAGTCGTTGCTATAAGGACTGCTGGGTGTTTGCNNNAGATCGGAA |
| 5388_1 | TATGAGTGTGGAGTCGTTGCAGCATGAGGAATAGCAGAATGGCNNNAGATCGGAA |
| 5388_2 | TATGAGTGTGGAGTCGTTGCAGCATGAGGAATAGCAGAATGGCNNNAGATCGGAA |
| 52365_1 | TATGAGTGTGGAGTCGTTGCGAAACTTAGCTTTGTCCGCCTTCNNNAGATCGGAAG |
| 52365_2 | TATGAGTGTGGAGTCGTTGCGAAACTTAGCTTTGTCCGCCTTCNNNAGATCGGAAG |
| 31362_1 | TATGAGTGTGGAGTCGTTGCGAGTCAAAGTTCTCCTGTTCCCTNNNAGATCGGAAG |
| 31362_2 | TATGAGTGTGGAGTCGTTGCGAGTCAAAGTTCTCCTGTTCCCTNNNAGATCGGAAG |
| 8191_1 | TATGAGTGTGGAGTCGTTGCACACACAGACACAATTCATGCCTNNNAGATCGGAAG |
| 8191_2 | TATGAGTGTGGAGTCGTTGCACACACAGACACAATTCATGCCTNNNAGATCGGAAG |
| 41560_1 | TATGAGTGTGGAGTCGTTGCCATGATCTTGGCTCACTGCAACNNNAGATCGGAAGA |
| 41560_2 | TATGAGTGTGGAGTCGTTGCCATGATCTTGGCTCACTGCAACNNNAGATCGGAAGA |
| 45256_1 | TATGAGTGTGGAGTCGTTGCGCAGCTATCTCTTTCCATTGCTGNNNAGATCGGAAG |
| 45256_2 | TATGAGTGTGGAGTCGTTGCGCAGCTATCTCTTTCCATTGCTGNNNAGATCGGAAG |
| 40465_1 | TATGAGTGTGGAGTCGTTGCATGATTCATATTCCCAGGGTGGCNNNAGATCGGAAG |
| 40465_2 | TATGAGTGTGGAGTCGTTGCATGATTCATATTCCCAGGGTGGCNNNAGATCGGAAG |
| 40189_1 | TATGAGTGTGGAGTCGTTGCTCCCATAAGATTATAAGGGAGCTGCNNNAGATCGGA |
| 40189_2 | TATGAGTGTGGAGTCGTTGCTCCCATAAGATTATAAGGGAGCTGCNNNAGATCGGA |
| 50775_1 | TATGAGTGTGGAGTCGTTGCCCAGAACCTGCCTAGATGTCTCNNNAGATCGGAAGA |
| 50775_2 | TATGAGTGTGGAGTCGTTGCCCAGAACCTGCCTAGATGTCTCNNNAGATCGGAAGA |
| 48004_1 | TATGAGTGTGGAGTCGTTGCGGTTCATGCCATTCTCCTGACTNNNAGATCGGAAGA |
| 48004_2 | TATGAGTGTGGAGTCGTTGCGGTTCATGCCATTCTCCTGACTNNNAGATCGGAAGA |
| 7913_1 | TATGAGTGTGGAGTCGTTGCAGAGCCTCACTCTGTTACCTAGAGNNNAGATCGGAA |
| 7913_2 | TATGAGTGTGGAGTCGTTGCAGAGCCTCACTCTGTTACCTAGAGNNNAGATCGGAA |
| 34909_1 | TATGAGTGTGGAGTCGTTGCCAAGTAGCTGGGATTACAAGTGCNNNAGATCGGAA |
| 34909_2 | TATGAGTGTGGAGTCGTTGCCAAGTAGCTGGGATTACAAGTGCNNNAGATCGGAA |
| 37395_1 | TATGAGTGTGGAGTCGTTGCATGCCTGTAATCCCAGCACTTTNNNAGATCGGAAGA |
| 37395_2 | TATGAGTGTGGAGTCGTTGCATGCCTGTAATCCCAGCACTTTNNNAGATCGGAAGA |
| 30657_1 | TATGAGTGTGGAGTCGTTGCCAGATAGGTGTGTGCTTATACTTCTNNNAGATCGGA |
| 30657_2 | TATGAGTGTGGAGTCGTTGCCAGATAGGTGTGTGCTTATACTTCTNNNAGATCGGA |
| 50790_1 | TATGAGTGTGGAGTCGTTGCCTAAAAAGGCCATTCGCAAGCTCNNNAGATCGGAAG |
| 50790_2 | TATGAGTGTGGAGTCGTTGCCTAAAAAGGCCATTCGCAAGCTCNNNAGATCGGAAG |
| 41216_1 | TATGAGTGTGGAGTCGTTGCTACATGACCCAAATGTCGTCCTTNNNAGATCGGAAG |
| 41216_2 | TATGAGTGTGGAGTCGTTGCTACATGACCCAAATGTCGTCCTTNNNAGATCGGAAG |
| 52433_1 | TATGAGTGTGGAGTCGTTGCCAACTACCTCATTGTTGGGGAGANNNAGATCGGAAG |
| 52433_2 | TATGAGTGTGGAGTCGTTGCCAACTACCTCATTGTTGGGGAGANNNAGATCGGAAG |
| 45382_1 | TATGAGTGTGGAGTCGTTGCTTTGGGGTCTGGGTGATGAATAGNNNAGATCGGAA |
| 45382_2 | TATGAGTGTGGAGTCGTTGCTTTGGGGTCTGGGTGATGAATAGNNNAGATCGGAA |
| 50441_1 | TATGAGTGTGGAGTCGTTGCCCGCTTGTTCACTCATACTTCTTGNNNAGATCGGAAG |

| | |
|---|---|
| 50441_2 | TATGAGTGTGGAGTCGTTGCCCGCTTGTTCACTCATACTTCTTGNNNAGATCGGAAG |
| 49045_1 | TATGAGTGTGGAGTCGTTGCTGGAGTACAGTGACACGATCTTGNNNAGATCGGAA |
| 49045_2 | TATGAGTGTGGAGTCGTTGCTGGAGTACAGTGACACGATCTTGNNNAGATCGGAA |
| 29964_1 | TATGAGTGTGGAGTCGTTGCCCACAAGCCAAGCGAATCATTTANNNAGATCGGAAG |
| 29964_2 | TATGAGTGTGGAGTCGTTGCCCACAAGCCAAGCGAATCATTTANNNAGATCGGAAG |
| 48310_1 | TATGAGTGTGGAGTCGTTGCGAGCCTAATCTGATGTTTGCAGGNNNAGATCGGAA |
| 48310_2 | TATGAGTGTGGAGTCGTTGCGAGCCTAATCTGATGTTTGCAGGNNNAGATCGGAA |
| 10450_1 | TATGAGTGTGGAGTCGTTGCGTTGCTGAAATCCATGAGGGCTANNNAGATCGGAA |
| 10450_2 | TATGAGTGTGGAGTCGTTGCGTTGCTGAAATCCATGAGGGCTANNNAGATCGGAA |
| 13660_1 | TATGAGTGTGGAGTCGTTGCCAATAAAACTTTGCTGCCACNNNAGATCGGAAGAGC |
| 13660_2 | TATGAGTGTGGAGTCGTTGCCAATAAAACTTTGCTGCCACNNNAGATCGGAAGAGC |
| 49699_1 | TATGAGTGTGGAGTCGTTGCAACACCAGTGAGACAGAACTGTTNNNAGATCGGAA |
| 49699_2 | TATGAGTGTGGAGTCGTTGCAACACCAGTGAGACAGAACTGTTNNNAGATCGGAA |
| 36920_1 | TATGAGTGTGGAGTCGTTGCTTGACCTCTTAGGCTCAAGCAATNNNAGATCGGAAG |
| 36920_2 | TATGAGTGTGGAGTCGTTGCTTGACCTCTTAGGCTCAAGCAATNNNAGATCGGAAG |
| 47567_1 | TATGAGTGTGGAGTCGTTGCGTTCAAGCGATTCTCCTACCTCANNNAGATCGGAAG |
| 47567_2 | TATGAGTGTGGAGTCGTTGCGTTCAAGCGATTCTCCTACCTCANNNAGATCGGAAG |
| 42945_1 | TATGAGTGTGGAGTCGTTGCTTCAGAAGGAATGGTACCAGCTCNNNAGATCGGAA |
| 42945_2 | TATGAGTGTGGAGTCGTTGCTTCAGAAGGAATGGTACCAGCTCNNNAGATCGGAA |
| 35812_1 | TATGAGTGTGGAGTCGTTGCCAGAACACAGAATGGAATGCCTGNNNAGATCGGAA |
| 35812_2 | TATGAGTGTGGAGTCGTTGCCAGAACACAGAATGGAATGCCTGNNNAGATCGGAA |
| 48191_1 | TATGAGTGTGGAGTCGTTGCCCACCACTCCTGGCTAATTTTTGNNNAGATCGGAAGA |
| 48191_2 | TATGAGTGTGGAGTCGTTGCCCACCACTCCTGGCTAATTTTTGNNNAGATCGGAAGA |
| 44309_1 | TATGAGTGTGGAGTCGTTGCAGTTGTTCACACCACCTCCTAAANNNAGATCGGAAG |
| 44309_2 | TATGAGTGTGGAGTCGTTGCAGTTGTTCACACCACCTCCTAAANNNAGATCGGAAG |
| 35902_1 | TATGAGTGTGGAGTCGTTGCATCATGAGGTCAGGAGATCGAGANNNAGATCGGAA |
| 35902_2 | TATGAGTGTGGAGTCGTTGCATCATGAGGTCAGGAGATCGAGANNNAGATCGGAA |
| 53035_1 | TATGAGTGTGGAGTCGTTGCATGAGATGTTCAGGTCACGTTGANNNAGATCGGAA |
| 53035_2 | TATGAGTGTGGAGTCGTTGCATGAGATGTTCAGGTCACGTTGANNNAGATCGGAA |
| 5094_1 | TATGAGTGTGGAGTCGTTGCAGTGGTTTGAATGGTTGTTCCCANNNAGATCGGAA |
| 5094_2 | TATGAGTGTGGAGTCGTTGCAGTGGTTTGAATGGTTGTTCCCANNNAGATCGGAA |
| 46612_1 | TATGAGTGTGGAGTCGTTGCGGAAGTGACTCTGAGCTGGTTTANNNAGATCGGAA |
| 46612_2 | TATGAGTGTGGAGTCGTTGCGGAAGTGACTCTGAGCTGGTTTANNNAGATCGGAA |
| 46136_1 | TATGAGTGTGGAGTCGTTGCTAGCTGGGATTACAGGTATGTGCNNNAGATCGGAA |
| 46136_2 | TATGAGTGTGGAGTCGTTGCTAGCTGGGATTACAGGTATGTGCNNNAGATCGGAA |
| 12837_1 | TATGAGTGTGGAGTCGTTGCAATATTTGTAAGTTCGAAATGGGNNNAGATCGGAA |
| 12837_2 | TATGAGTGTGGAGTCGTTGCAATATTTGTAAGTTCGAAATGGGNNNAGATCGGAA |
| 2985_1 | TATGAGTGTGGAGTCGTTGCAGGGGAATCAGGGCCTGCAAGNNNAGATCGGAAG |
| 2985_2 | TATGAGTGTGGAGTCGTTGCAGGGGAATCAGGGCCTGCAAGNNNAGATCGGAAG |
| 41917_1 | TATGAGTGTGGAGTCGTTGCTTTTTAGTGGAGACGGGGTTTCANNNAGATCGGAA |
| 41917_2 | TATGAGTGTGGAGTCGTTGCTTTTTAGTGGAGACGGGGTTTCANNNAGATCGGAA |
| 29027_1 | TATGAGTGTGGAGTCGTTGCCCCGGCCCACTCTGTATATTTTANNNAGATCGGAAGA |
| 29027_2 | TATGAGTGTGGAGTCGTTGCCCCGGCCCACTCTGTATATTTTANNNAGATCGGAAGA |

| | |
|---|---|
| 52617_1 | TATGAGTGTGGAGTCGTTGCACTTTCAGCCTCTCAAGTTCCAANNNAGATCGGAAGA |
| 52617_2 | TATGAGTGTGGAGTCGTTGCACTTTCAGCCTCTCAAGTTCCAANNNAGATCGGAAGA |
| 32199_1 | TATGAGTGTGGAGTCGTTGCGAGTAGCTGGGATTACAGGCATGNNNAGATCGGAA |
| 32199_2 | TATGAGTGTGGAGTCGTTGCGAGTAGCTGGGATTACAGGCATGNNNAGATCGGAA |
| 38345_1 | TATGAGTGTGGAGTCGTTGCAAGAGATGTCAGTCTTCTGGAGCNNNAGATCGGAA |
| 38345_2 | TATGAGTGTGGAGTCGTTGCAAGAGATGTCAGTCTTCTGGAGCNNNAGATCGGAA |
| 29734_1 | TATGAGTGTGGAGTCGTTGCAGGAGAAGTCCCTCAGTGATGTANNNAGATCGGAA |
| 29734_2 | TATGAGTGTGGAGTCGTTGCAGGAGAAGTCCCTCAGTGATGTANNNAGATCGGAA |
| 53122_1 | TATGAGTGTGGAGTCGTTGCTGGTTTTAGGGTTCAACAGGAGTNNNAGATCGGAA |
| 53122_2 | TATGAGTGTGGAGTCGTTGCTGGTTTTAGGGTTCAACAGGAGTNNNAGATCGGAA |
| 13691_1 | TATGAGTGTGGAGTCGTTGCATACCTGTTTCTTTGGGACATNNNAGATCGGAAGAG |
| 13691_2 | TATGAGTGTGGAGTCGTTGCATACCTGTTTCTTTGGGACATNNNAGATCGGAAGAG |
| 48853_1 | TATGAGTGTGGAGTCGTTGCCAACATCCTTTTTGCCTCTCAGGNNNAGATCGGAAGA |
| 48853_2 | TATGAGTGTGGAGTCGTTGCCAACATCCTTTTTGCCTCTCAGGNNNAGATCGGAAGA |
| 43540_1 | TATGAGTGTGGAGTCGTTGCCATTCTCCTGCCTCAGCCTCNNNAGATCGGAAGAGCA |
| 43540_2 | TATGAGTGTGGAGTCGTTGCCATTCTCCTGCCTCAGCCTCNNNAGATCGGAAGAGCA |
| 47023_1 | TATGAGTGTGGAGTCGTTGCCCGCCATGAACTCTTTCTACAACNNNAGATCGGAAGA |
| 47023_2 | TATGAGTGTGGAGTCGTTGCCCGCCATGAACTCTTTCTACAACNNNAGATCGGAAGA |
| 30840_1 | TATGAGTGTGGAGTCGTTGCAGCTGACACAGAGCACTCAATAANNNAGATCGGAAG |
| 30840_2 | TATGAGTGTGGAGTCGTTGCAGCTGACACAGAGCACTCAATAANNNAGATCGGAAG |
| 50001_1 | TATGAGTGTGGAGTCGTTGCAATACCGCGTGCAATGTATTCACNNNAGATCGGAAG |
| 50001_2 | TATGAGTGTGGAGTCGTTGCAATACCGCGTGCAATGTATTCACNNNAGATCGGAAG |
| 51283_1 | TATGAGTGTGGAGTCGTTGCCCTTTGAGTAGGCAAGAAGGGATNNNAGATCGGAA |
| 51283_2 | TATGAGTGTGGAGTCGTTGCCCTTTGAGTAGGCAAGAAGGGATNNNAGATCGGAA |
| 33152_1 | TATGAGTGTGGAGTCGTTGCCACAAAAGCTGGAGAGTGACTTGNNNAGATCGGAA |
| 33152_2 | TATGAGTGTGGAGTCGTTGCCACAAAAGCTGGAGAGTGACTTGNNNAGATCGGAA |
| 44858_1 | TATGAGTGTGGAGTCGTTGCCTGTGCTGTGCCTAATGAAACAANNNAGATCGGAAG |
| 44858_2 | TATGAGTGTGGAGTCGTTGCCTGTGCTGTGCCTAATGAAACAANNNAGATCGGAAG |
| 47435_1 | TATGAGTGTGGAGTCGTTGCGCACCTTGCTTGGTTACAGAATGNNNAGATCGGAAG |
| 47435_2 | TATGAGTGTGGAGTCGTTGCGCACCTTGCTTGGTTACAGAATGNNNAGATCGGAAG |
| 50878_1 | TATGAGTGTGGAGTCGTTGCGGGAGAGATGGAGGTAAGCTTAGNNNAGATCGGA |
| 50878_2 | TATGAGTGTGGAGTCGTTGCGGGAGAGATGGAGGTAAGCTTAGNNNAGATCGGA |
| 29091_1 | TATGAGTGTGGAGTCGTTGCTAAGGCAAATGACGAGATTGTGCNNNAGATCGGAA |
| 29091_2 | TATGAGTGTGGAGTCGTTGCTAAGGCAAATGACGAGATTGTGCNNNAGATCGGAA |
| 35585_1 | TATGAGTGTGGAGTCGTTGCTTCAAGTGATTCTCCTGCCTCAGNNNAGATCGGAAG |
| 35585_2 | TATGAGTGTGGAGTCGTTGCTTCAAGTGATTCTCCTGCCTCAGNNNAGATCGGAAG |
| 42260_1 | TATGAGTGTGGAGTCGTTGCTGTGACAGATAAGAAGCACCTCCNNNAGATCGGAAG |
| 42260_2 | TATGAGTGTGGAGTCGTTGCTGTGACAGATAAGAAGCACCTCCNNNAGATCGGAAG |
| 40108_1 | TATGAGTGTGGAGTCGTTGCAGTGATGGTTGGTCAACTCTATGANNNAGATCGGA |
| 40108_2 | TATGAGTGTGGAGTCGTTGCAGTGATGGTTGGTCAACTCTATGANNNAGATCGGA |
| 44389_1 | TATGAGTGTGGAGTCGTTGCTGAATCTCTTCTTCATGGTGGGGNNNAGATCGGAAG |
| 44389_2 | TATGAGTGTGGAGTCGTTGCTGAATCTCTTCTTCATGGTGGGGNNNAGATCGGAAG |
| 31290_1 | TATGAGTGTGGAGTCGTTGCAGGGGGAGAATTAGGAGAACAGANNNAGATCGGA |

| | |
|---|---|
| 31290_2 | TATGAGTGTGGAGTCGTTGCAGGGGGAGAATTAGGAGAACAGANNNAGATCGGA |
| 38756_1 | TATGAGTGTGGAGTCGTTGCCATGCCTGTTATCCCAGCTAGTTNNNAGATCGGAAG |
| 38756_2 | TATGAGTGTGGAGTCGTTGCCATGCCTGTTATCCCAGCTAGTTNNNAGATCGGAAG |
| 52460_1 | TATGAGTGTGGAGTCGTTGCACCATTGCCAGTCATTTCAACAGNNNAGATCGGAAG |
| 52460_2 | TATGAGTGTGGAGTCGTTGCACCATTGCCAGTCATTTCAACAGNNNAGATCGGAAG |
| 51677_1 | TATGAGTGTGGAGTCGTTGCCCAGTGGATGTCTGTTTAGTGGANNNAGATCGGAA |
| 51677_2 | TATGAGTGTGGAGTCGTTGCCCAGTGGATGTCTGTTTAGTGGANNNAGATCGGAA |
| 10204_1 | TATGAGTGTGGAGTCGTTGCCCCACTCCAGTTACCATGTTAGCNNNAGATCGGAAG |
| 10204_2 | TATGAGTGTGGAGTCGTTGCCCCACTCCAGTTACCATGTTAGCNNNAGATCGGAAG |
| 36163_1 | TATGAGTGTGGAGTCGTTGCTATCCAACATGGTGAAACCCCATNNNAGATCGGAAG |
| 36163_2 | TATGAGTGTGGAGTCGTTGCTATCCAACATGGTGAAACCCCATNNNAGATCGGAAG |
| 39572_1 | TATGAGTGTGGAGTCGTTGCGTGATTTTGGCTCACTGCAACCNNNAGATCGGAAGA |
| 39572_2 | TATGAGTGTGGAGTCGTTGCGTGATTTTGGCTCACTGCAACCNNNAGATCGGAAGA |
| 44850_1 | TATGAGTGTGGAGTCGTTGCACTTGCACAGTAAATGGTGATGGNNNAGATCGGAA |
| 44850_2 | TATGAGTGTGGAGTCGTTGCACTTGCACAGTAAATGGTGATGGNNNAGATCGGAA |
| 44343_1 | TATGAGTGTGGAGTCGTTGCAACATCAGTTTCGAAGCACTTGGNNNAGATCGGAAG |
| 44343_2 | TATGAGTGTGGAGTCGTTGCAACATCAGTTTCGAAGCACTTGGNNNAGATCGGAAG |
| 36392_1 | TATGAGTGTGGAGTCGTTGCACAATCGTAGGTCACTGAGGAACNNNAGATCGGAA |
| 36392_2 | TATGAGTGTGGAGTCGTTGCACAATCGTAGGTCACTGAGGAACNNNAGATCGGAA |
| 38404_1 | TATGAGTGTGGAGTCGTTGCGGAAGATGCCATGGATAGGAGAGNNNAGATCGGA |
| 38404_2 | TATGAGTGTGGAGTCGTTGCGGAAGATGCCATGGATAGGAGAGNNNAGATCGGA |
| 49459_1 | TATGAGTGTGGAGTCGTTGCTCACTGTAACCTCAAACTCCTGGNNNAGATCGGAAG |
| 49459_2 | TATGAGTGTGGAGTCGTTGCTCACTGTAACCTCAAACTCCTGGNNNAGATCGGAAG |
| 12776_1 | TATGAGTGTGGAGTCGTTGCTATAGAAGTTGAAACAAGAAGGANNNAGATCGGAA |
| 12776_2 | TATGAGTGTGGAGTCGTTGCTATAGAAGTTGAAACAAGAAGGANNNAGATCGGAA |
| 47883_1 | TATGAGTGTGGAGTCGTTGCTGCTGGGATTACAAGTATGAGCCNNNAGATCGGAA |
| 47883_2 | TATGAGTGTGGAGTCGTTGCTGCTGGGATTACAAGTATGAGCCNNNAGATCGGAA |
| 50593_1 | TATGAGTGTGGAGTCGTTGCGAACAGGGAAAGGCACTAGAGAANNNAGATCGGA |
| 50593_2 | TATGAGTGTGGAGTCGTTGCGAACAGGGAAAGGCACTAGAGAANNNAGATCGGA |
| 49795_1 | TATGAGTGTGGAGTCGTTGCACTCAGAGACCTTGTTACAATCCANNNAGATCGGAA |
| 49795_2 | TATGAGTGTGGAGTCGTTGCACTCAGAGACCTTGTTACAATCCANNNAGATCGGAA |
| 53422_1 | TATGAGTGTGGAGTCGTTGCAGCTAATTGGGTGATGTTTGCACNNNAGATCGGAA |
| 53422_2 | TATGAGTGTGGAGTCGTTGCAGCTAATTGGGTGATGTTTGCACNNNAGATCGGAA |
| 48360_1 | TATGAGTGTGGAGTCGTTGCCAGTGGTGCAGTCCTAGCTCNNNAGATCGGAAGAG |
| 48360_2 | TATGAGTGTGGAGTCGTTGCCAGTGGTGCAGTCCTAGCTCNNNAGATCGGAAGAG |
| 42674_1 | TATGAGTGTGGAGTCGTTGCAATTATGGAGGCTGAGATGTCCCNNNAGATCGGAA |
| 42674_2 | TATGAGTGTGGAGTCGTTGCAATTATGGAGGCTGAGATGTCCCNNNAGATCGGAA |
| 38133_1 | TATGAGTGTGGAGTCGTTGCTGAGATAGGCATTGAGCTTGAGCNNNAGATCGGAA |
| 38133_2 | TATGAGTGTGGAGTCGTTGCTGAGATAGGCATTGAGCTTGAGCNNNAGATCGGAA |
| 46025_1 | TATGAGTGTGGAGTCGTTGCGAGCTCATCATGCCCCTATTTGGNNNAGATCGGAAG |
| 46025_2 | TATGAGTGTGGAGTCGTTGCGAGCTCATCATGCCCCTATTTGGNNNAGATCGGAAG |
| 36435_1 | TATGAGTGTGGAGTCGTTGCTTCAACTTCCATAGTCTCAGGCCNNNAGATCGGAAG |
| 36435_2 | TATGAGTGTGGAGTCGTTGCTTCAACTTCCATAGTCTCAGGCCNNNAGATCGGAAG |

| | |
|---|---|
| 29313_1 | TATGAGTGTGGAGTCGTTGCTGGGTGAGAATCTGCACAGTTTANNNAGATCGGAA |
| 29313_2 | TATGAGTGTGGAGTCGTTGCTGGGTGAGAATCTGCACAGTTTANNNAGATCGGAA |
| 46167_1 | TATGAGTGTGGAGTCGTTGCCCTCCCACGTTCAAATGATTCTCNNNAGATCGGAAGA |
| 46167_2 | TATGAGTGTGGAGTCGTTGCCCTCCCACGTTCAAATGATTCTCNNNAGATCGGAAGA |
| 34369_1 | TATGAGTGTGGAGTCGTTGCCACTGGAGCTCTCTGGTAATCAGNNNAGATCGGAAG |
| 34369_2 | TATGAGTGTGGAGTCGTTGCCACTGGAGCTCTCTGGTAATCAGNNNAGATCGGAAG |
| 30354_1 | TATGAGTGTGGAGTCGTTGCCGATGTGCCAGAATAGGCATTCNNNAGATCGGAAG |
| 30354_2 | TATGAGTGTGGAGTCGTTGCCGATGTGCCAGAATAGGCATTCNNNAGATCGGAAG |
| 42266_1 | TATGAGTGTGGAGTCGTTGCGAACTCCTGGCTTCAAGCAATCNNNAGATCGGAAGA |
| 42266_2 | TATGAGTGTGGAGTCGTTGCGAACTCCTGGCTTCAAGCAATCNNNAGATCGGAAGA |
| 9208_1 | TATGAGTGTGGAGTCGTTGCACATGATTCTTTCTGTGCCAGCANNNAGATCGGAAG |
| 9208_2 | TATGAGTGTGGAGTCGTTGCACATGATTCTTTCTGTGCCAGCANNNAGATCGGAAG |
| 49089_1 | TATGAGTGTGGAGTCGTTGCGAAAAGCAGTTCGTAGAGGGAGANNNAGATCGGA |
| 49089_2 | TATGAGTGTGGAGTCGTTGCGAAAAGCAGTTCGTAGAGGGAGANNNAGATCGGA |
| 51972_1 | TATGAGTGTGGAGTCGTTGCCTCCTCCTCACTCACTTGACTTGNNNAGATCGGAAGA |
| 51972_2 | TATGAGTGTGGAGTCGTTGCCTCCTCCTCACTCACTTGACTTGNNNAGATCGGAAGA |
| 2690_1 | TATGAGTGTGGAGTCGTTGCAAGTCTCCTGAAGGCTGGTTCATNNNAGATCGGAAG |
| 2690_2 | TATGAGTGTGGAGTCGTTGCAAGTCTCCTGAAGGCTGGTTCATNNNAGATCGGAAG |
| 47414_1 | TATGAGTGTGGAGTCGTTGCGGAGTGCAGTAGTATGATCTCGGNNNAGATCGGAA |
| 47414_2 | TATGAGTGTGGAGTCGTTGCGGAGTGCAGTAGTATGATCTCGGNNNAGATCGGAA |
| 50022_1 | TATGAGTGTGGAGTCGTTGCCCATCAAGGACCTAGGCTTTCTTNNNAGATCGGAAG |
| 50022_2 | TATGAGTGTGGAGTCGTTGCCCATCAAGGACCTAGGCTTTCTTNNNAGATCGGAAG |
| 52848_1 | TATGAGTGTGGAGTCGTTGCTGATTGTTGGAAAGGTCTTCCCANNNAGATCGGAAG |
| 52848_2 | TATGAGTGTGGAGTCGTTGCTGATTGTTGGAAAGGTCTTCCCANNNAGATCGGAAG |
| 42072_1 | TATGAGTGTGGAGTCGTTGCAGGAGACAATGGTTGCAGTGAGNNNAGATCGGAA |
| 42072_2 | TATGAGTGTGGAGTCGTTGCAGGAGACAATGGTTGCAGTGAGNNNAGATCGGAA |
| 36254_1 | TATGAGTGTGGAGTCGTTGCATGAAGGCGATCCAGTCTAAAGGNNNAGATCGGAA |
| 36254_2 | TATGAGTGTGGAGTCGTTGCATGAAGGCGATCCAGTCTAAAGGNNNAGATCGGAA |
| 29617_1 | TATGAGTGTGGAGTCGTTGCAAACAGCACCAAAAAGACCTCAGNNNAGATCGGAA |
| 29617_2 | TATGAGTGTGGAGTCGTTGCAAACAGCACCAAAAAGACCTCAGNNNAGATCGGAA |
| 13702_1 | TATGAGTGTGGAGTCGTTGCCTCAGGAAAGTTAAGGTATCAANNNAGATCGGAAG |
| 13702_2 | TATGAGTGTGGAGTCGTTGCCTCAGGAAAGTTAAGGTATCAANNNAGATCGGAAG |
| 10776_1 | TATGAGTGTGGAGTCGTTGCACCTTCTGACCTATTTTGTCCTGGNNNAGATCGGAAG |
| 10776_2 | TATGAGTGTGGAGTCGTTGCACCTTCTGACCTATTTTGTCCTGGNNNAGATCGGAAG |
| 50572_1 | TATGAGTGTGGAGTCGTTGCGGATAATGAGTTGATTGCTGGGCNNNAGATCGGAA |
| 50572_2 | TATGAGTGTGGAGTCGTTGCGGATAATGAGTTGATTGCTGGGCNNNAGATCGGAA |
| 34861_1 | TATGAGTGTGGAGTCGTTGCCTCCTGAGTAGCTGGGATTACAGNNNAGATCGGAA |
| 34861_2 | TATGAGTGTGGAGTCGTTGCCTCCTGAGTAGCTGGGATTACAGNNNAGATCGGAA |
| 44170_1 | TATGAGTGTGGAGTCGTTGCGCCAATATTGACCACCACTCATGNNNAGATCGGAAG |
| 44170_2 | TATGAGTGTGGAGTCGTTGCGCCAATATTGACCACCACTCATGNNNAGATCGGAAG |
| 45742_1 | TATGAGTGTGGAGTCGTTGCCACTACCCCCGGCTAATTTTTTGNNNAGATCGGAAGA |
| 45742_2 | TATGAGTGTGGAGTCGTTGCCACTACCCCCGGCTAATTTTTTGNNNAGATCGGAAGA |
| 43223_1 | TATGAGTGTGGAGTCGTTGCGGACTTCCAGGACTAACAGTGAANNNAGATCGGAA |

| | |
|---|---|
| 43223_2 | TATGAGTGTGGAGTCGTTGCGGACTTCCAGGACTAACAGTGAANNNAGATCGGAA |
| 13323_1 | TATGAGTGTGGAGTCGTTGCCTAATGCTACAAAGGTTACTCTTNNNAGATCGGAAG |
| 13323_2 | TATGAGTGTGGAGTCGTTGCCTAATGCTACAAAGGTTACTCTTNNNAGATCGGAAG |
| 39204_1 | TATGAGTGTGGAGTCGTTGCGTTCCCAGTGACTCAGTTCCTAGNNNAGATCGGAAG |
| 39204_2 | TATGAGTGTGGAGTCGTTGCGTTCCCAGTGACTCAGTTCCTAGNNNAGATCGGAAG |
| 41648_1 | TATGAGTGTGGAGTCGTTGCATTTATGTGCGTTGCAGAGTGACNNNAGATCGGAAG |
| 41648_2 | TATGAGTGTGGAGTCGTTGCATTTATGTGCGTTGCAGAGTGACNNNAGATCGGAAG |
| 43803_1 | TATGAGTGTGGAGTCGTTGCTGCTTCTGATGGAAATTCTGGGANNNAGATCGGAAG |
| 43803_2 | TATGAGTGTGGAGTCGTTGCTGCTTCTGATGGAAATTCTGGGANNNAGATCGGAAG |
| 49295_1 | TATGAGTGTGGAGTCGTTGCGAGAGCAGTGGTGCAATCTCAGNNNAGATCGGAAG |
| 49295_2 | TATGAGTGTGGAGTCGTTGCGAGAGCAGTGGTGCAATCTCAGNNNAGATCGGAAG |
| 40682_1 | TATGAGTGTGGAGTCGTTGCTTCCAAACTACTGAAGAGGCTCCNNNAGATCGGAAG |
| 40682_2 | TATGAGTGTGGAGTCGTTGCTTCCAAACTACTGAAGAGGCTCCNNNAGATCGGAAG |
| 38257_1 | TATGAGTGTGGAGTCGTTGCAGGTCCCTTTTATGTTCCCTTGTNNNAGATCGGAAGA |
| 38257_2 | TATGAGTGTGGAGTCGTTGCAGGTCCCTTTTATGTTCCCTTGTNNNAGATCGGAAGA |
| 48556_1 | TATGAGTGTGGAGTCGTTGCAGTAGCTGGGATTATAGGCATGCNNNAGATCGGAA |
| 48556_2 | TATGAGTGTGGAGTCGTTGCAGTAGCTGGGATTATAGGCATGCNNNAGATCGGAA |
| 53082_1 | TATGAGTGTGGAGTCGTTGCGGATCATGAGGTCGGAAGATCAANNNAGATCGGAA |
| 53082_2 | TATGAGTGTGGAGTCGTTGCGGATCATGAGGTCGGAAGATCAANNNAGATCGGAA |
| 53355_1 | TATGAGTGTGGAGTCGTTGCTCTCAACGTGGGATTTGAAGAGTNNNAGATCGGAA |
| 53355_2 | TATGAGTGTGGAGTCGTTGCTCTCAACGTGGGATTTGAAGAGTNNNAGATCGGAA |
| 30387_1 | TATGAGTGTGGAGTCGTTGCAAATCAAGGACCACGGTCTTTCCNNNAGATCGGAAG |
| 30387_2 | TATGAGTGTGGAGTCGTTGCAAATCAAGGACCACGGTCTTTCCNNNAGATCGGAAG |
| 33307_1 | TATGAGTGTGGAGTCGTTGCCATTCTCCTGCCTCAGCCTCNNNAGATCGGAAGAGCA |
| 33307_2 | TATGAGTGTGGAGTCGTTGCCATTCTCCTGCCTCAGCCTCNNNAGATCGGAAGAGCA |
| 46056_1 | TATGAGTGTGGAGTCGTTGCGATCTCGGCTCACTGCAACTTCNNNAGATCGGAAGA |
| 46056_2 | TATGAGTGTGGAGTCGTTGCGATCTCGGCTCACTGCAACTTCNNNAGATCGGAAGA |
| 48278_1 | TATGAGTGTGGAGTCGTTGCCAGTGGCACAATGATATTGGCTCNNNAGATCGGAAG |
| 48278_2 | TATGAGTGTGGAGTCGTTGCCAGTGGCACAATGATATTGGCTCNNNAGATCGGAAG |
| 44362_1 | TATGAGTGTGGAGTCGTTGCGAAAAGACAGGCTTGGCTTGAATNNNAGATCGGAA |
| 44362_2 | TATGAGTGTGGAGTCGTTGCGAAAAGACAGGCTTGGCTTGAATNNNAGATCGGAA |
| 41382_1 | TATGAGTGTGGAGTCGTTGCAATGCAAGTGTTCTAAGCCCCTANNNAGATCGGAAG |
| 41382_2 | TATGAGTGTGGAGTCGTTGCAATGCAAGTGTTCTAAGCCCCTANNNAGATCGGAAG |
| 44088_1 | TATGAGTGTGGAGTCGTTGCGACTGGAATGGGAGGATGTGAATNNNAGATCGGA |
| 44088_2 | TATGAGTGTGGAGTCGTTGCGACTGGAATGGGAGGATGTGAATNNNAGATCGGA |
| 29699_1 | TATGAGTGTGGAGTCGTTGCTCCAGATAACAGTGAAAGCACCANNNAGATCGGAAG |
| 29699_2 | TATGAGTGTGGAGTCGTTGCTCCAGATAACAGTGAAAGCACCANNNAGATCGGAAG |
| 49361_1 | TATGAGTGTGGAGTCGTTGCAAAATGCTGGGATTATACGCGTGNNNAGATCGGAA |
| 49361_2 | TATGAGTGTGGAGTCGTTGCAAAATGCTGGGATTATACGCGTGNNNAGATCGGAA |
| 51758_1 | TATGAGTGTGGAGTCGTTGCCTGTGAGTGAGAGTATGCAGTGTNNNAGATCGGAA |
| 51758_2 | TATGAGTGTGGAGTCGTTGCCTGTGAGTGAGAGTATGCAGTGTNNNAGATCGGAA |
| 43548_1 | TATGAGTGTGGAGTCGTTGCGAGTCTCGCTCTGTCACCAAGNNNAGATCGGAAGAG |
| 43548_2 | TATGAGTGTGGAGTCGTTGCGAGTCTCGCTCTGTCACCAAGNNNAGATCGGAAGAG |

| | |
|---|---|
| 33634_1 | TATGAGTGTGGAGTCGTTGCATCTTGGCTCACTACAACCTCTGNNNAGATCGGAAG |
| 33634_2 | TATGAGTGTGGAGTCGTTGCATCTTGGCTCACTACAACCTCTGNNNAGATCGGAAG |
| 41437_1 | TATGAGTGTGGAGTCGTTGCTCCGGTCAAAGTCTACCTAAACANNNAGATCGGAAG |
| 41437_2 | TATGAGTGTGGAGTCGTTGCTCCGGTCAAAGTCTACCTAAACANNNAGATCGGAAG |
| 31575_1 | TATGAGTGTGGAGTCGTTGCACTAGAACAGTTGGGCACATCTGNNNAGATCGGAA |
| 31575_2 | TATGAGTGTGGAGTCGTTGCACTAGAACAGTTGGGCACATCTGNNNAGATCGGAA |
| 38039_1 | TATGAGTGTGGAGTCGTTGCGGCTCATCACAACAGACTCCTAANNNAGATCGGAAG |
| 38039_2 | TATGAGTGTGGAGTCGTTGCGGCTCATCACAACAGACTCCTAANNNAGATCGGAAG |
| 44175_1 | TATGAGTGTGGAGTCGTTGCATCATCTCATGCCTCAGTGATCCNNNAGATCGGAAG |
| 44175_2 | TATGAGTGTGGAGTCGTTGCATCATCTCATGCCTCAGTGATCCNNNAGATCGGAAG |
| 31230_1 | TATGAGTGTGGAGTCGTTGCTTACCAGTAACCAGAAGGCAGTCNNNAGATCGGAA |
| 31230_2 | TATGAGTGTGGAGTCGTTGCTTACCAGTAACCAGAAGGCAGTCNNNAGATCGGAA |
| 45887_1 | TATGAGTGTGGAGTCGTTGCCTCCTGAGTAGCTGGGACTACAGNNNAGATCGGAA |
| 45887_2 | TATGAGTGTGGAGTCGTTGCCTCCTGAGTAGCTGGGACTACAGNNNAGATCGGAA |
| 50256_1 | TATGAGTGTGGAGTCGTTGCAGTGCCTAAGTCCTCAGTCTTTGNNNAGATCGGAAG |
| 50256_2 | TATGAGTGTGGAGTCGTTGCAGTGCCTAAGTCCTCAGTCTTTGNNNAGATCGGAAG |
| 12695_1 | TATGAGTGTGGAGTCGTTGCCTCGAATTTAAGGTATGGTATTGNNNAGATCGGAA |
| 12695_2 | TATGAGTGTGGAGTCGTTGCCTCGAATTTAAGGTATGGTATTGNNNAGATCGGAA |
| 29154_1 | TATGAGTGTGGAGTCGTTGCATCAATTCACTGAGACCACAGCANNNAGATCGGAAG |
| 29154_2 | TATGAGTGTGGAGTCGTTGCATCAATTCACTGAGACCACAGCANNNAGATCGGAAG |
| 34853_1 | TATGAGTGTGGAGTCGTTGCCACCACGCCTGGCTAATTTTTANNNAGATCGGAAG |
| 34853_2 | TATGAGTGTGGAGTCGTTGCCACCACGCCTGGCTAATTTTTANNNAGATCGGAAG |
| 32870_1 | TATGAGTGTGGAGTCGTTGCACAGCTCCTGGATTTCTTGATCTNNNAGATCGGAAG |
| 32870_2 | TATGAGTGTGGAGTCGTTGCACAGCTCCTGGATTTCTTGATCTNNNAGATCGGAAG |
| 36010_1 | TATGAGTGTGGAGTCGTTGCCAGATCACTTCAGGTCAGGAGTTNNNAGATCGGAA |
| 36010_2 | TATGAGTGTGGAGTCGTTGCCAGATCACTTCAGGTCAGGAGTTNNNAGATCGGAA |
| 13036_1 | TATGAGTGTGGAGTCGTTGCGAAGTAAGTGCATCCACTTGNNNAGATCGGAAGAG |
| 13036_2 | TATGAGTGTGGAGTCGTTGCGAAGTAAGTGCATCCACTTGNNNAGATCGGAAGAG |
| 52458_1 | TATGAGTGTGGAGTCGTTGCGCTAGAGATGCTGACACCATGTANNNAGATCGGAA |
| 52458_2 | TATGAGTGTGGAGTCGTTGCGCTAGAGATGCTGACACCATGTANNNAGATCGGAA |
| 35695_1 | TATGAGTGTGGAGTCGTTGCCCACCACATTCTTGAACACTGACNNNAGATCGGAAG |
| 35695_2 | TATGAGTGTGGAGTCGTTGCCCACCACATTCTTGAACACTGACNNNAGATCGGAAG |
| 33006_1 | TATGAGTGTGGAGTCGTTGCACAATCTTCTCAACTCAGCCTCCNNNAGATCGGAAGA |
| 33006_2 | TATGAGTGTGGAGTCGTTGCACAATCTTCTCAACTCAGCCTCCNNNAGATCGGAAGA |
| 49427_1 | TATGAGTGTGGAGTCGTTGCCCGAGTAGCTGGGATTACAAGAGNNNAGATCGGAA |
| 49427_2 | TATGAGTGTGGAGTCGTTGCCCGAGTAGCTGGGATTACAAGAGNNNAGATCGGAA |
| 40566_1 | TATGAGTGTGGAGTCGTTGCGCCTGTAGTCCCAGCTACTTGNNNAGATCGGAAGAG |
| 40566_2 | TATGAGTGTGGAGTCGTTGCGCCTGTAGTCCCAGCTACTTGNNNAGATCGGAAGAG |
| 38942_1 | TATGAGTGTGGAGTCGTTGCTATTCCCAAACTCCCAAACTTGCNNNAGATCGGAAGA |
| 38942_2 | TATGAGTGTGGAGTCGTTGCTATTCCCAAACTCCCAAACTTGCNNNAGATCGGAAGA |
| 45482_1 | TATGAGTGTGGAGTCGTTGCAGTCTCATCGTTATGTGTGGCTGNNNAGATCGGAAG |
| 45482_2 | TATGAGTGTGGAGTCGTTGCAGTCTCATCGTTATGTGTGGCTGNNNAGATCGGAAG |
| 44944_1 | TATGAGTGTGGAGTCGTTGCCTAAGCCCTCTCACATCTGGAGNNNAGATCGGAAGA |

| | |
|---|---|
| 44944_2 | TATGAGTGTGGAGTCGTTGCCTAAGCCCTCTCACATCTGGAGNNNAGATCGGAAGA |
| 44849_1 | TATGAGTGTGGAGTCGTTGCCCTGGGGAGAAGACAAACAGTAANNNAGATCGGAA |
| 44849_2 | TATGAGTGTGGAGTCGTTGCCCTGGGGAGAAGACAAACAGTAANNNAGATCGGAA |
| 53291_1 | TATGAGTGTGGAGTCGTTGCCTGATCCAGGAGGCCGCATCNNNAGATCGGAAGAG |
| 53291_2 | TATGAGTGTGGAGTCGTTGCCTGATCCAGGAGGCCGCATCNNNAGATCGGAAGAG |
| 51658_1 | TATGAGTGTGGAGTCGTTGCTCTTGGTGTGTTTTCTGTGGAGANNNAGATCGGAAG |
| 51658_2 | TATGAGTGTGGAGTCGTTGCTCTTGGTGTGTTTTCTGTGGAGANNNAGATCGGAAG |
| 47362_1 | TATGAGTGTGGAGTCGTTGCAAGCCCAAAGCAGTAGAAAGGATNNNAGATCGGAA |
| 47362_2 | TATGAGTGTGGAGTCGTTGCAAGCCCAAAGCAGTAGAAAGGATNNNAGATCGGAA |
| 33369_1 | TATGAGTGTGGAGTCGTTGCTTCAAGTGATTCTCCTGCCTCAGNNNAGATCGGAAG |
| 33369_2 | TATGAGTGTGGAGTCGTTGCTTCAAGTGATTCTCCTGCCTCAGNNNAGATCGGAAG |
| 52963_1 | TATGAGTGTGGAGTCGTTGCTTATCCACCTATGACCTGGAAGCNNNAGATCGGAAG |
| 52963_2 | TATGAGTGTGGAGTCGTTGCTTATCCACCTATGACCTGGAAGCNNNAGATCGGAAG |
| 42182_1 | TATGAGTGTGGAGTCGTTGCCGGGATCTCAGCTCACTGTAAGNNNAGATCGGAAG |
| 42182_2 | TATGAGTGTGGAGTCGTTGCCGGGATCTCAGCTCACTGTAAGNNNAGATCGGAAG |
| 40315_1 | TATGAGTGTGGAGTCGTTGCTCCTGAGCTAGTTTTGCAAAATTGNNNAGATCGGAA |
| 40315_2 | TATGAGTGTGGAGTCGTTGCTCCTGAGCTAGTTTTGCAAAATTGNNNAGATCGGAA |
| 50995_1 | TATGAGTGTGGAGTCGTTGCTCTGGGTATTGTCTTTGCTTGTGNNNAGATCGGAAG |
| 50995_2 | TATGAGTGTGGAGTCGTTGCTCTGGGTATTGTCTTTGCTTGTGNNNAGATCGGAAG |
| 30215_1 | TATGAGTGTGGAGTCGTTGCTAAGCAGACATCTTTGAGCACCTNNNAGATCGGAAG |
| 30215_2 | TATGAGTGTGGAGTCGTTGCTAAGCAGACATCTTTGAGCACCTNNNAGATCGGAAG |
| 47618_1 | TATGAGTGTGGAGTCGTTGCGAGACAGGGTTTCAACATGTTGGNNNAGATCGGAA |
| 47618_2 | TATGAGTGTGGAGTCGTTGCGAGACAGGGTTTCAACATGTTGGNNNAGATCGGAA |
| 9509_1 | TATGAGTGTGGAGTCGTTGCCTCTCTAGCTTCCGCTTCCTTTCNNNAGATCGGAAGA |
| 9509_2 | TATGAGTGTGGAGTCGTTGCCTCTCTAGCTTCCGCTTCCTTTCNNNAGATCGGAAGA |
| 12976_1 | TATGAGTGTGGAGTCGTTGCATGAAAGAAACACAAAAACCTTANNNAGATCGGAAG |
| 12976_2 | TATGAGTGTGGAGTCGTTGCATGAAAGAAACACAAAAACCTTANNNAGATCGGAAG |
| 13381_1 | TATGAGTGTGGAGTCGTTGCATATATAGTGGAGAGACTCAGAANNNAGATCGGAA |
| 13381_2 | TATGAGTGTGGAGTCGTTGCATATATAGTGGAGAGACTCAGAANNNAGATCGGAA |
| 36644_1 | TATGAGTGTGGAGTCGTTGCAGACTGAACATTGTGACAGAACTNNNAGATCGGAAG |
| 36644_2 | TATGAGTGTGGAGTCGTTGCAGACTGAACATTGTGACAGAACTNNNAGATCGGAAG |
| 49326_1 | TATGAGTGTGGAGTCGTTGCTCCTGTGCCTGACTTATTTCACTNNNAGATCGGAAGA |
| 49326_2 | TATGAGTGTGGAGTCGTTGCTCCTGTGCCTGACTTATTTCACTNNNAGATCGGAAGA |
| 36146_1 | TATGAGTGTGGAGTCGTTGCAAACTGGACTGCACCAAAAACAANNNAGATCGGAAG |
| 36146_2 | TATGAGTGTGGAGTCGTTGCAAACTGGACTGCACCAAAAACAANNNAGATCGGAAG |
| 46630_1 | TATGAGTGTGGAGTCGTTGCGGACGAGAACCTTTCAACACTTGNNNAGATCGGAAG |
| 46630_2 | TATGAGTGTGGAGTCGTTGCGGACGAGAACCTTTCAACACTTGNNNAGATCGGAAG |
| 40665_1 | TATGAGTGTGGAGTCGTTGCCTCAGGAAAGACGACCCATTCTANNNAGATCGGAAG |
| 40665_2 | TATGAGTGTGGAGTCGTTGCCTCAGGAAAGACGACCCATTCTANNNAGATCGGAAG |
| 34391_1 | TATGAGTGTGGAGTCGTTGCCACCCTCCCCATTTCCTCTTGNNNAGATCGGAAGAGC |
| 34391_2 | TATGAGTGTGGAGTCGTTGCCACCCTCCCCATTTCCTCTTGNNNAGATCGGAAGAGC |
| 48985_1 | TATGAGTGTGGAGTCGTTGCCTTCATTCCCACATCACAGGAGANNNAGATCGGAAG |
| 48985_2 | TATGAGTGTGGAGTCGTTGCCTTCATTCCCACATCACAGGAGANNNAGATCGGAAG |

| | |
|---|---|
| 48835_1 | TATGAGTGTGGAGTCGTTGCGCCAACACTTTTGGGCTTATCTTNNNAGATCGGAAG |
| 48835_2 | TATGAGTGTGGAGTCGTTGCGCCAACACTTTTGGGCTTATCTTNNNAGATCGGAAG |
| 34679_1 | TATGAGTGTGGAGTCGTTGCTGGTAACTCATGCCTGTAATCCCNNNAGATCGGAAG |
| 34679_2 | TATGAGTGTGGAGTCGTTGCTGGTAACTCATGCCTGTAATCCCNNNAGATCGGAAG |
| 40784_1 | TATGAGTGTGGAGTCGTTGCGAGTGGAGGAACACCAGATTCTTNNNAGATCGGAA |
| 40784_2 | TATGAGTGTGGAGTCGTTGCGAGTGGAGGAACACCAGATTCTTNNNAGATCGGAA |
| 51767_1 | TATGAGTGTGGAGTCGTTGCAGGTTGTGTCCATCTTCACTTGTNNNAGATCGGAAG |
| 51767_2 | TATGAGTGTGGAGTCGTTGCAGGTTGTGTCCATCTTCACTTGTNNNAGATCGGAAG |
| 52302_1 | TATGAGTGTGGAGTCGTTGCCAACAGTCTGGGATCTCTGATGTNNNAGATCGGAAG |
| 52302_2 | TATGAGTGTGGAGTCGTTGCCAACAGTCTGGGATCTCTGATGTNNNAGATCGGAAG |
| 50670_1 | TATGAGTGTGGAGTCGTTGCAAGCCATTGTTTTCAGACCATGGNNNAGATCGGAAG |
| 50670_2 | TATGAGTGTGGAGTCGTTGCAAGCCATTGTTTTCAGACCATGGNNNAGATCGGAAG |
| 1936_1 | TATGAGTGTGGAGTCGTTGCGTTGTGTGTGTCTGCAAGGTTTTNNNAGATCGGAA |
| 1936_2 | TATGAGTGTGGAGTCGTTGCGTTGTGTGTGTCTGCAAGGTTTTNNNAGATCGGAA |
| 45899_1 | TATGAGTGTGGAGTCGTTGCCTCCTGAGTAGCTGGGACTACAGNNNAGATCGGAA |
| 45899_2 | TATGAGTGTGGAGTCGTTGCCTCCTGAGTAGCTGGGACTACAGNNNAGATCGGAA |
| 40518_1 | TATGAGTGTGGAGTCGTTGCAGCTTAGTAAGGTTGTAGGATGCANNNAGATCGGA |
| 40518_2 | TATGAGTGTGGAGTCGTTGCAGCTTAGTAAGGTTGTAGGATGCANNNAGATCGGA |
| 34401_1 | TATGAGTGTGGAGTCGTTGCGTGTACCCAACAGCTCATTGAGANNNAGATCGGAAG |
| 34401_2 | TATGAGTGTGGAGTCGTTGCGTGTACCCAACAGCTCATTGAGANNNAGATCGGAAG |
| 45358_1 | TATGAGTGTGGAGTCGTTGCATCTCCCTCTCTTCTTCACATGCNNNAGATCGGAAGA |
| 45358_2 | TATGAGTGTGGAGTCGTTGCATCTCCCTCTCTTCTTCACATGCNNNAGATCGGAAGA |
| 36851_1 | TATGAGTGTGGAGTCGTTGCTGAGACAGTTTCGCTATTGTTGCNNNAGATCGGAAG |
| 36851_2 | TATGAGTGTGGAGTCGTTGCTGAGACAGTTTCGCTATTGTTGCNNNAGATCGGAAG |
| 7881_1 | TATGAGTGTGGAGTCGTTGCGTGGGGGTGGTTGTGTGTTTTNNNAGATCGGAAGA |
| 7881_2 | TATGAGTGTGGAGTCGTTGCGTGGGGGTGGTTGTGTGTTTTNNNAGATCGGAAGA |
| 34082_1 | TATGAGTGTGGAGTCGTTGCAGATGGAGCTTTGCTTTTGTCACNNNAGATCGGAAG |
| 34082_2 | TATGAGTGTGGAGTCGTTGCAGATGGAGCTTTGCTTTTGTCACNNNAGATCGGAAG |
| 38947_1 | TATGAGTGTGGAGTCGTTGCGCATTTGCCCAATTTTTCATGGGNNNAGATCGGAAG |
| 38947_2 | TATGAGTGTGGAGTCGTTGCGCATTTGCCCAATTTTTCATGGGNNNAGATCGGAAG |
| 39035_1 | TATGAGTGTGGAGTCGTTGCTGTGTTCTGTCTACTACAATGGGTNNNAGATCGGAA |
| 39035_2 | TATGAGTGTGGAGTCGTTGCTGTGTTCTGTCTACTACAATGGGTNNNAGATCGGAA |
| 47145_1 | TATGAGTGTGGAGTCGTTGCTGGACATATTGTGTTTGTGGTGCNNNAGATCGGAA |
| 47145_2 | TATGAGTGTGGAGTCGTTGCTGGACATATTGTGTTTGTGGTGCNNNAGATCGGAA |
| 40308_1 | TATGAGTGTGGAGTCGTTGCGCCTAGCACATAGTAGGAGTTTGANNNAGATCGGA |
| 40308_2 | TATGAGTGTGGAGTCGTTGCGCCTAGCACATAGTAGGAGTTTGANNNAGATCGGA |
| 52784_1 | TATGAGTGTGGAGTCGTTGCTGGAATGTCCCTTAGAGATGTGCNNNAGATCGGAA |
| 52784_2 | TATGAGTGTGGAGTCGTTGCTGGAATGTCCCTTAGAGATGTGCNNNAGATCGGAA |
| 51395_1 | TATGAGTGTGGAGTCGTTGCCAGCCAAGGTTTCAATTACAGCANNNAGATCGGAAG |
| 51395_2 | TATGAGTGTGGAGTCGTTGCCAGCCAAGGTTTCAATTACAGCANNNAGATCGGAAG |
| 51145_1 | TATGAGTGTGGAGTCGTTGCAAAGGAGGCCAGATCACTTCTTTNNNAGATCGGAAG |
| 51145_2 | TATGAGTGTGGAGTCGTTGCAAAGGAGGCCAGATCACTTCTTTNNNAGATCGGAAG |
| 41347_1 | TATGAGTGTGGAGTCGTTGCGCAGATAGCACTAAGAAGCGTTGNNNAGATCGGAA |

| | |
|---|---|
| 41347_2 | TATGAGTGTGGAGTCGTTGCGCAGATAGCACTAAGAAGCGTTGNNNAGATCGGAA |
| 51261_1 | TATGAGTGTGGAGTCGTTGCGCTAGAAGTTGCACATTGTTCCANNNAGATCGGAAG |
| 51261_2 | TATGAGTGTGGAGTCGTTGCGCTAGAAGTTGCACATTGTTCCANNNAGATCGGAAG |
| 49053_1 | TATGAGTGTGGAGTCGTTGCCCTTAGCCTCTTGAGTATGGGACNNNAGATCGGAAG |
| 49053_2 | TATGAGTGTGGAGTCGTTGCCCTTAGCCTCTTGAGTATGGGACNNNAGATCGGAAG |
| 3191_1 | TATGAGTGTGGAGTCGTTGCGGTTGGAAAGCCATAGGATTCCANNNAGATCGGAA |
| 3191_2 | TATGAGTGTGGAGTCGTTGCGGTTGGAAAGCCATAGGATTCCANNNAGATCGGAA |
| 44516_1 | TATGAGTGTGGAGTCGTTGCGATCCCCCTACGTTGTCTACTTGNNNAGATCGGAAG |
| 44516_2 | TATGAGTGTGGAGTCGTTGCGATCCCCCTACGTTGTCTACTTGNNNAGATCGGAAG |
| 42755_1 | TATGAGTGTGGAGTCGTTGCTTCCGAGTGTGTGCTCATGTATGNNNAGATCGGAAG |
| 42755_2 | TATGAGTGTGGAGTCGTTGCTTCCGAGTGTGTGCTCATGTATGNNNAGATCGGAAG |
| 41203_1 | TATGAGTGTGGAGTCGTTGCCTTTCTCTTTCTGCTCCCCCTACNNNAGATCGGAAGA |
| 41203_2 | TATGAGTGTGGAGTCGTTGCCTTTCTCTTTCTGCTCCCCCTACNNNAGATCGGAAGA |
| 52955_1 | TATGAGTGTGGAGTCGTTGCCACCTTATCCTACACCTCCCATGNNNAGATCGGAAGA |
| 52955_2 | TATGAGTGTGGAGTCGTTGCCACCTTATCCTACACCTCCCATGNNNAGATCGGAAGA |
| 41974_1 | TATGAGTGTGGAGTCGTTGCGAGTAGCTGGGATTACAGGCATGNNNAGATCGGAA |
| 41974_2 | TATGAGTGTGGAGTCGTTGCGAGTAGCTGGGATTACAGGCATGNNNAGATCGGAA |
| 10600_1 | TATGAGTGTGGAGTCGTTGCGTTGCTGTCTCTGTCCATGACAANNNAGATCGGAAG |
| 10600_2 | TATGAGTGTGGAGTCGTTGCGTTGCTGTCTCTGTCCATGACAANNNAGATCGGAAG |
| 31512_1 | TATGAGTGTGGAGTCGTTGCCAGGGTCTCATCAAGAGCATTCTNNNAGATCGGAAG |
| 31512_2 | TATGAGTGTGGAGTCGTTGCCAGGGTCTCATCAAGAGCATTCTNNNAGATCGGAAG |
| 13358_1 | TATGAGTGTGGAGTCGTTGCCATGTAACAGTGAAGTCAGAGNNNAGATCGGAAGA |
| 13358_2 | TATGAGTGTGGAGTCGTTGCCATGTAACAGTGAAGTCAGAGNNNAGATCGGAAGA |
| 29783_1 | ATGAGTGTGGAGTCGTTGCTCCGAGTTACATATTTCTCATAGTACANNNAGATCGGA |
| 29783_2 | ATGAGTGTGGAGTCGTTGCTCCGAGTTACATATTTCTCATAGTACANNNAGATCGGA |
| 31584_1 | TATGAGTGTGGAGTCGTTGCCTCACGTGAGGGATGATCAAGTCNNNAGATCGGAA |
| 31584_2 | TATGAGTGTGGAGTCGTTGCCTCACGTGAGGGATGATCAAGTCNNNAGATCGGAA |
| 51445_1 | TATGAGTGTGGAGTCGTTGCTTACAAATGCCTCCAGGAATGGTNNNAGATCGGAAG |
| 51445_2 | TATGAGTGTGGAGTCGTTGCTTACAAATGCCTCCAGGAATGGTNNNAGATCGGAAG |
| 43934_1 | TATGAGTGTGGAGTCGTTGCCACATACACACACACACATCAGTNNNAGATCGGAAG |
| 43934_2 | TATGAGTGTGGAGTCGTTGCCACATACACACACACACATCAGTNNNAGATCGGAAG |
| 45692_1 | TATGAGTGTGGAGTCGTTGCCCTCCCAGGTTCAAGTGATTCTCNNNAGATCGGAAG |
| 45692_2 | TATGAGTGTGGAGTCGTTGCCCTCCCAGGTTCAAGTGATTCTCNNNAGATCGGAAG |
| 48410_1 | TATGAGTGTGGAGTCGTTGCGGGACTTTCACTTTGGAAATGGCNNNAGATCGGAA |
| 48410_2 | TATGAGTGTGGAGTCGTTGCGGGACTTTCACTTTGGAAATGGCNNNAGATCGGAA |
| 42360_1 | TATGAGTGTGGAGTCGTTGCCTCAGCTCTAGCACTGGGAATCNNNAGATCGGAAGA |
| 42360_2 | TATGAGTGTGGAGTCGTTGCCTCAGCTCTAGCACTGGGAATCNNNAGATCGGAAGA |
| 39372_1 | TATGAGTGTGGAGTCGTTGCCAGCATATAAGCCAGGAGCAGTANNNAGATCGGAA |
| 39372_2 | TATGAGTGTGGAGTCGTTGCCAGCATATAAGCCAGGAGCAGTANNNAGATCGGAA |
| 35565_1 | TATGAGTGTGGAGTCGTTGCCAATCTCAGCTCACTGCAACTTCNNNAGATCGGAAGA |
| 35565_2 | TATGAGTGTGGAGTCGTTGCCAATCTCAGCTCACTGCAACTTCNNNAGATCGGAAGA |
| 10811_1 | TATGAGTGTGGAGTCGTTGCCCCTCCTGGCTATTATCCTCAGGNNNAGATCGGAAG |
| 10811_2 | TATGAGTGTGGAGTCGTTGCCCCTCCTGGCTATTATCCTCAGGNNNAGATCGGAAG |

| | |
|---|---|
| 30077_1 | TATGAGTGTGGAGTCGTTGCATCACCAGAACAAAATGTGCAGGNNNAGATCGGAA |
| 30077_2 | TATGAGTGTGGAGTCGTTGCATCACCAGAACAAAATGTGCAGGNNNAGATCGGAA |
| 51799_1 | TATGAGTGTGGAGTCGTTGCATGTGACCAGCTTTTCCTCTTCTNNNAGATCGGAAGA |
| 51799_2 | TATGAGTGTGGAGTCGTTGCATGTGACCAGCTTTTCCTCTTCTNNNAGATCGGAAGA |
| 10689_1 | TATGAGTGTGGAGTCGTTGCAGCAGATCAGTGCATTCTTTCCCNNNAGATCGGAAG |
| 10689_2 | TATGAGTGTGGAGTCGTTGCAGCAGATCAGTGCATTCTTTCCCNNNAGATCGGAAG |
| 31395_1 | TATGAGTGTGGAGTCGTTGCGTATTTTCCTCCCCTTTATGCGCNNNAGATCGGAAGA |
| 31395_2 | TATGAGTGTGGAGTCGTTGCGTATTTTCCTCCCCTTTATGCGCNNNAGATCGGAAGA |
| 29715_1 | TATGAGTGTGGAGTCGTTGCGCATTCGACTGTTGAGGTTGAACNNNAGATCGGAA |
| 29715_2 | TATGAGTGTGGAGTCGTTGCGCATTCGACTGTTGAGGTTGAACNNNAGATCGGAA |
| 13166_1 | TATGAGTGTGGAGTCGTTGCGCTGTTTAGATTTATATTGCTGTNNNAGATCGGAAG |
| 13166_2 | TATGAGTGTGGAGTCGTTGCGCTGTTTAGATTTATATTGCTGTNNNAGATCGGAAG |
| 43683_1 | TATGAGTGTGGAGTCGTTGCGTGGTGTGTGTGTATCTATGGGTNNNAGATCGGAA |
| 43683_2 | TATGAGTGTGGAGTCGTTGCGTGGTGTGTGTGTATCTATGGGTNNNAGATCGGAA |
| 49446_1 | TATGAGTGTGGAGTCGTTGCAAAAACAATGGTGAAAGTGGGCANNNAGATCGGA |
| 49446_2 | TATGAGTGTGGAGTCGTTGCAAAAACAATGGTGAAAGTGGGCANNNAGATCGGA |
| 46282_1 | TATGAGTGTGGAGTCGTTGCGAGTAGCTGGGATTACAGGTGTGNNNAGATCGGA |
| 46282_2 | TATGAGTGTGGAGTCGTTGCGAGTAGCTGGGATTACAGGTGTGNNNAGATCGGA |
| 36367_1 | TATGAGTGTGGAGTCGTTGCACCCTTTGGCTCAATATTGTACTNNNAGATCGGAAG |
| 36367_2 | TATGAGTGTGGAGTCGTTGCACCCTTTGGCTCAATATTGTACTNNNAGATCGGAAG |
| 35854_1 | TATGAGTGTGGAGTCGTTGCTTCAAGTGATCCTCCTGCTTCAGNNNAGATCGGAAG |
| 35854_2 | TATGAGTGTGGAGTCGTTGCTTCAAGTGATCCTCCTGCTTCAGNNNAGATCGGAAG |
| 38681_1 | TATGAGTGTGGAGTCGTTGCGCTGTAACAAAAACTTCCCCAGTNNNAGATCGGAAG |
| 38681_2 | TATGAGTGTGGAGTCGTTGCGCTGTAACAAAAACTTCCCCAGTNNNAGATCGGAAG |
| 30091_1 | TATGAGTGTGGAGTCGTTGCTTGTGAAGGAGTGAGCAGTGCNNNAGATCGGAAG |
| 30091_2 | TATGAGTGTGGAGTCGTTGCTTGTGAAGGAGTGAGCAGTGCNNNAGATCGGAAG |
| 34667_1 | TATGAGTGTGGAGTCGTTGCTCTTCACACTACAGCTGGAGAACNNNAGATCGGAAG |
| 34667_2 | TATGAGTGTGGAGTCGTTGCTCTTCACACTACAGCTGGAGAACNNNAGATCGGAAG |
| 49605_1 | TATGAGTGTGGAGTCGTTGCCCTCTCCCTATGCCTGAGAAATCNNNAGATCGGAAG |
| 49605_2 | TATGAGTGTGGAGTCGTTGCCCTCTCCCTATGCCTGAGAAATCNNNAGATCGGAAG |
| 13754_1 | TATGAGTGTGGAGTCGTTGCCCACTATTTTCAATTTATCTGTCTNNNAGATCGGAAG |
| 13754_2 | TATGAGTGTGGAGTCGTTGCCCACTATTTTCAATTTATCTGTCTNNNAGATCGGAAG |
| 45752_1 | TATGAGTGTGGAGTCGTTGCCTAGTAGCTGGGATTACAGGCGNNNAGATCGGAAG |
| 45752_2 | TATGAGTGTGGAGTCGTTGCCTAGTAGCTGGGATTACAGGCGNNNAGATCGGAAG |
| 47504_1 | TATGAGTGTGGAGTCGTTGCAAACTCCTGACCTCAAGTGATCCNNNAGATCGGAAG |
| 47504_2 | TATGAGTGTGGAGTCGTTGCAAACTCCTGACCTCAAGTGATCCNNNAGATCGGAAG |
| 28938_1 | TATGAGTGTGGAGTCGTTGCAGGGTCTTCAAACATATTCCCTGTNNNAGATCGGAA |
| 28938_2 | TATGAGTGTGGAGTCGTTGCAGGGTCTTCAAACATATTCCCTGTNNNAGATCGGAA |
| 44357_1 | TATGAGTGTGGAGTCGTTGCCGTCTGAACCCCAAGAATGGAGNNNAGATCGGAAG |
| 44357_2 | TATGAGTGTGGAGTCGTTGCCGTCTGAACCCCAAGAATGGAGNNNAGATCGGAAG |
| 51420_1 | TATGAGTGTGGAGTCGTTGCCAGCCTGGGTGATAGAATGAGATNNNAGATCGGAA |
| 51420_2 | TATGAGTGTGGAGTCGTTGCCAGCCTGGGTGATAGAATGAGATNNNAGATCGGAA |
| 52705_1 | TATGAGTGTGGAGTCGTTGCCTTGCCTCTCTCTGCCGTTCNNNAGATCGGAAGAGCA |

| | |
|---|---|
| 52705_2 | TATGAGTGTGGAGTCGTTGCCTTGCCTCTCTCTGCCGTTCNNNAGATCGGAAGAGCA |
| 41822_1 | TATGAGTGTGGAGTCGTTGCTTCTGCCTTAGCTCCAATTGTCTNNNAGATCGGAAGA |
| 41822_2 | TATGAGTGTGGAGTCGTTGCTTCTGCCTTAGCTCCAATTGTCTNNNAGATCGGAAGA |
| 35692_1 | TATGAGTGTGGAGTCGTTGCCTTCTGGGTTCAGGCCATACTCNNNAGATCGGAAGA |
| 35692_2 | TATGAGTGTGGAGTCGTTGCCTTCTGGGTTCAGGCCATACTCNNNAGATCGGAAGA |
| 12999_1 | TATGAGTGTGGAGTCGTTGCAAGACATTATTTTGGGTGAATAGNNNAGATCGGAA |
| 12999_2 | TATGAGTGTGGAGTCGTTGCAAGACATTATTTTGGGTGAATAGNNNAGATCGGAA |
| 12656_1 | TATGAGTGTGGAGTCGTTGCATTCAGTCTTCTGTTCCCTATNNNAGATCGGAAGAGC |
| 12656_2 | TATGAGTGTGGAGTCGTTGCATTCAGTCTTCTGTTCCCTATNNNAGATCGGAAGAGC |
| 33997_1 | TATGAGTGTGGAGTCGTTGCCCACTTCAGTCTTCCAAGTAGCTNNNAGATCGGAAGA |
| 33997_2 | TATGAGTGTGGAGTCGTTGCCCACTTCAGTCTTCCAAGTAGCTNNNAGATCGGAAGA |
| 32284_1 | TATGAGTGTGGAGTCGTTGCCTCAGCTTGCAGGGAGGTGTNNNAGATCGGAAGAG |
| 32284_2 | TATGAGTGTGGAGTCGTTGCCTCAGCTTGCAGGGAGGTGTNNNAGATCGGAAGAG |
| 12986_1 | TATGAGTGTGGAGTCGTTGCAATTATTAGCATTGATCTGAGTGNNNAGATCGGAAG |
| 12986_2 | TATGAGTGTGGAGTCGTTGCAATTATTAGCATTGATCTGAGTGNNNAGATCGGAAG |
| 12893_1 | TATGAGTGTGGAGTCGTTGCCTCAATAGTTACCAGCCCATNNNAGATCGGAAGAGCA |
| 12893_2 | TATGAGTGTGGAGTCGTTGCCTCAATAGTTACCAGCCCATNNNAGATCGGAAGAGCA |
| 41573_1 | TATGAGTGTGGAGTCGTTGCGGGTTTGAGTGATTCTCCTGTCTNNNAGATCGGAAG |
| 41573_2 | TATGAGTGTGGAGTCGTTGCGGGTTTGAGTGATTCTCCTGTCTNNNAGATCGGAAG |
| 5393_1 | TATGAGTGTGGAGTCGTTGCATTGCCTCTGTTCTTGCCTTTGANNNAGATCGGAAGA |
| 5393_2 | TATGAGTGTGGAGTCGTTGCATTGCCTCTGTTCTTGCCTTTGANNNAGATCGGAAGA |
| 42259_1 | TATGAGTGTGGAGTCGTTGCATAGCTGGGATTACAGGCATGAGNNNAGATCGGAA |
| 42259_2 | TATGAGTGTGGAGTCGTTGCATAGCTGGGATTACAGGCATGAGNNNAGATCGGAA |
| 40285_1 | TATGAGTGTGGAGTCGTTGCAATGTTGGCTCCATCAAGTCAACNNNAGATCGGAAG |
| 40285_2 | TATGAGTGTGGAGTCGTTGCAATGTTGGCTCCATCAAGTCAACNNNAGATCGGAAG |
| 48906_1 | TATGAGTGTGGAGTCGTTGCGCCTTGGATATTCAGGCTCCTTTNNNAGATCGGAAGA |
| 48906_2 | TATGAGTGTGGAGTCGTTGCGCCTTGGATATTCAGGCTCCTTTNNNAGATCGGAAGA |
| 38284_1 | TATGAGTGTGGAGTCGTTGCGCCAAACCCACGATGATCTAGATNNNAGATCGGAAG |
| 38284_2 | TATGAGTGTGGAGTCGTTGCGCCAAACCCACGATGATCTAGATNNNAGATCGGAAG |
| 43050_1 | TATGAGTGTGGAGTCGTTGCCACATGCACACACACACACATNNNAGATCGGAAGAG |
| 43050_2 | TATGAGTGTGGAGTCGTTGCCACATGCACACACACACACATNNNAGATCGGAAGAG |
| 48424_1 | TATGAGTGTGGAGTCGTTGCTGGAGTGGCATGATCATGATTCANNNAGATCGGAA |
| 48424_2 | TATGAGTGTGGAGTCGTTGCTGGAGTGGCATGATCATGATTCANNNAGATCGGAA |
| 50372_1 | TATGAGTGTGGAGTCGTTGCCCAAAGGAAACCCAACCTCAATCNNNAGATCGGAAG |
| 50372_2 | TATGAGTGTGGAGTCGTTGCCCAAAGGAAACCCAACCTCAATCNNNAGATCGGAAG |
| 48121_1 | TATGAGTGTGGAGTCGTTGCCCAGGAACTTGTGTGGACTCAGNNNAGATCGGAAG |
| 48121_2 | TATGAGTGTGGAGTCGTTGCCCAGGAACTTGTGTGGACTCAGNNNAGATCGGAAG |
| 47667_1 | TATGAGTGTGGAGTCGTTGCTTCAAGCTATTCTCCTGCCTCAGNNNAGATCGGAAGA |
| 47667_2 | TATGAGTGTGGAGTCGTTGCTTCAAGCTATTCTCCTGCCTCAGNNNAGATCGGAAGA |
| 47209_1 | TATGAGTGTGGAGTCGTTGCTCATAGCTTACTGCAGCCTTGAANNNAGATCGGAAG |
| 47209_2 | TATGAGTGTGGAGTCGTTGCTCATAGCTTACTGCAGCCTTGAANNNAGATCGGAAG |
| 2141_1 | TATGAGTGTGGAGTCGTTGCGCTTTGAGCACAGTGAGTTCTCANNNAGATCGGAAG |
| 2141_2 | TATGAGTGTGGAGTCGTTGCGCTTTGAGCACAGTGAGTTCTCANNNAGATCGGAAG |

| | |
|---|---|
| 50994_1 | TATGAGTGTGGAGTCGTTGCACACACACACACACACACAAAATNNNAGATCGGAAG |
| 50994_2 | TATGAGTGTGGAGTCGTTGCACACACACACACACACACAAAATNNNAGATCGGAAG |
| 30967_1 | TATGAGTGTGGAGTCGTTGCCCATGGCCGGCTAATTTTTGTATNNNAGATCGGAAG |
| 30967_2 | TATGAGTGTGGAGTCGTTGCCCATGGCCGGCTAATTTTTGTATNNNAGATCGGAAG |
| 46080_1 | TATGAGTGTGGAGTCGTTGCCTGAGTAGCTGGGATTACAGGTGNNNAGATCGGAA |
| 46080_2 | TATGAGTGTGGAGTCGTTGCCTGAGTAGCTGGGATTACAGGTGNNNAGATCGGAA |
| 42657_1 | TATGAGTGTGGAGTCGTTGCACAAGGAGGAACTGGTACCATTCNNNAGATCGGAA |
| 42657_2 | TATGAGTGTGGAGTCGTTGCACAAGGAGGAACTGGTACCATTCNNNAGATCGGAA |
| 38253_1 | TATGAGTGTGGAGTCGTTGCAGGCTAGGAAGAAACTGCATCAANNNAGATCGGAA |
| 38253_2 | TATGAGTGTGGAGTCGTTGCAGGCTAGGAAGAAACTGCATCAANNNAGATCGGAA |
| 46985_1 | TATGAGTGTGGAGTCGTTGCCCCCACCTGGCTAATTTTTGTTTNNNAGATCGGAAGA |
| 46985_2 | TATGAGTGTGGAGTCGTTGCCCCCACCTGGCTAATTTTTGTTTNNNAGATCGGAAGA |
| 41070_1 | TATGAGTGTGGAGTCGTTGCTTGAATGATACTGAGGGCCTGTCNNNAGATCGGAA |
| 41070_2 | TATGAGTGTGGAGTCGTTGCTTGAATGATACTGAGGGCCTGTCNNNAGATCGGAA |
| 43039_1 | TATGAGTGTGGAGTCGTTGCTGCTCAAAGTGATTGACATCCCANNNAGATCGGAAG |
| 43039_2 | TATGAGTGTGGAGTCGTTGCTGCTCAAAGTGATTGACATCCCANNNAGATCGGAAG |
| 37291_1 | TATGAGTGTGGAGTCGTTGCATACATAACATCCAGCGTGGTGANNNAGATCGGAA |
| 37291_2 | TATGAGTGTGGAGTCGTTGCATACATAACATCCAGCGTGGTGANNNAGATCGGAA |
| 32062_1 | TATGAGTGTGGAGTCGTTGCTAGGTTGGCTAATCGAGGTTTCTNNNAGATCGGAA |
| 32062_2 | TATGAGTGTGGAGTCGTTGCTAGGTTGGCTAATCGAGGTTTCTNNNAGATCGGAA |
| 50350_1 | TATGAGTGTGGAGTCGTTGCGATTACAGGTGTAAGCAGCAGTGNNNAGATCGGAA |
| 50350_2 | TATGAGTGTGGAGTCGTTGCGATTACAGGTGTAAGCAGCAGTGNNNAGATCGGAA |
| 29886_1 | TATGAGTGTGGAGTCGTTGCAGCAAAACAAATAGAAAAGCCGTNNNAGATCGGAA |
| 29886_2 | TATGAGTGTGGAGTCGTTGCAGCAAAACAAATAGAAAAGCCGTNNNAGATCGGAA |
| 43222_1 | TATGAGTGTGGAGTCGTTGCAGAGTAGCTGGGACTACAGGAGNNNAGATCGGAA |
| 43222_2 | TATGAGTGTGGAGTCGTTGCAGAGTAGCTGGGACTACAGGAGNNNAGATCGGAA |
| 10332_1 | TATGAGTGTGGAGTCGTTGCGGATTTATGCATGCAGTGTTGGGNNNAGATCGGAA |
| 10332_2 | TATGAGTGTGGAGTCGTTGCGGATTTATGCATGCAGTGTTGGGNNNAGATCGGAA |
| 43427_1 | TATGAGTGTGGAGTCGTTGCTAAGCTTTTTGATGTGGTGCTGGNNNAGATCGGAA |
| 43427_2 | TATGAGTGTGGAGTCGTTGCTAAGCTTTTTGATGTGGTGCTGGNNNAGATCGGAA |
| 30046_1 | TATGAGTGTGGAGTCGTTGCCCCACAAAGGCAGAGAGATCTCNNNAGATCGGAAG |
| 30046_2 | TATGAGTGTGGAGTCGTTGCCCCACAAAGGCAGAGAGATCTCNNNAGATCGGAAG |
| 44799_1 | TATGAGTGTGGAGTCGTTGCACCAGAGCAACTCCATCTTGAATNNNAGATCGGAAG |
| 44799_2 | TATGAGTGTGGAGTCGTTGCACCAGAGCAACTCCATCTTGAATNNNAGATCGGAAG |
| 33536_1 | TATGAGTGTGGAGTCGTTGCCAGCTGGGATAACAGGTGCATNNNAGATCGGAAGA |
| 33536_2 | TATGAGTGTGGAGTCGTTGCCAGCTGGGATAACAGGTGCATNNNAGATCGGAAGA |
| 36730_1 | TATGAGTGTGGAGTCGTTGCCTGTCTCCTCAGGCTTCTTTCTTNNNAGATCGGAAGA |
| 36730_2 | TATGAGTGTGGAGTCGTTGCCTGTCTCCTCAGGCTTCTTTCTTNNNAGATCGGAAGA |
| 50477_1 | TATGAGTGTGGAGTCGTTGCAGACATTCCTTCTTTCCCTTCCCNNNAGATCGGAAGA |
| 50477_2 | TATGAGTGTGGAGTCGTTGCAGACATTCCTTCTTTCCCTTCCCNNNAGATCGGAAGA |
| 52620_1 | TATGAGTGTGGAGTCGTTGCTCCCCCAATGATTCAATGCCTTANNNAGATCGGAAGA |
| 52620_2 | TATGAGTGTGGAGTCGTTGCTCCCCCAATGATTCAATGCCTTANNNAGATCGGAAGA |
| 41143_1 | TATGAGTGTGGAGTCGTTGCTCTATCTTTCTCCAAAGCCAGCANNNAGATCGGAAGA |

| | |
|---|---|
| 41143_2 | TATGAGTGTGGAGTCGTTGCTCTATCTTTCTCCAAAGCCAGCANNNAGATCGGAAG |
| 39167_1 | TATGAGTGTGGAGTCGTTGCAGTAGAAGGAAGAAGTGAGGGAACNNNAGATCGG |
| 39167_2 | TATGAGTGTGGAGTCGTTGCAGTAGAAGGAAGAAGTGAGGGAACNNNAGATCGG |
| 49266_1 | TATGAGTGTGGAGTCGTTGCTACGGGAAGGCTTGAATATCCAGNNNAGATCGGAA |
| 49266_2 | TATGAGTGTGGAGTCGTTGCTACGGGAAGGCTTGAATATCCAGNNNAGATCGGAA |
| 42091_1 | TATGAGTGTGGAGTCGTTGCTTCAATATTAGGTTGGCTGAATGCNNNAGATCGGAA |
| 42091_2 | TATGAGTGTGGAGTCGTTGCTTCAATATTAGGTTGGCTGAATGCNNNAGATCGGAA |
| 41011_1 | TATGAGTGTGGAGTCGTTGCACTTCTGGGTTCAAGTGATCCTCNNNAGATCGGAAG |
| 41011_2 | TATGAGTGTGGAGTCGTTGCACTTCTGGGTTCAAGTGATCCTCNNNAGATCGGAAG |
| 12654_1 | TATGAGTGTGGAGTCGTTGCCCAAATACCAAAAACGTTGAGNNNAGATCGGAAGA |
| 12654_2 | TATGAGTGTGGAGTCGTTGCCCAAATACCAAAAACGTTGAGNNNAGATCGGAAGA |
| 46792_1 | TATGAGTGTGGAGTCGTTGCTTGGTATCAGGATGATGCTGACCNNNAGATCGGAA |
| 46792_2 | TATGAGTGTGGAGTCGTTGCTTGGTATCAGGATGATGCTGACCNNNAGATCGGAA |
| 50130_1 | TATGAGTGTGGAGTCGTTGCGCACTTCTGAACTCCTGACCTAGNNNAGATCGGAAG |
| 50130_2 | TATGAGTGTGGAGTCGTTGCGCACTTCTGAACTCCTGACCTAGNNNAGATCGGAAG |
| 43045_1 | TATGAGTGTGGAGTCGTTGCGAGAAAACCAAATCCAGCCGTACNNNAGATCGGAAG |
| 43045_2 | TATGAGTGTGGAGTCGTTGCGAGAAAACCAAATCCAGCCGTACNNNAGATCGGAAG |
| 44825_1 | TATGAGTGTGGAGTCGTTGCCAGAACAGAGACTGACATCTCCCNNNAGATCGGAAG |
| 44825_2 | TATGAGTGTGGAGTCGTTGCCAGAACAGAGACTGACATCTCCCNNNAGATCGGAAG |
| 36048_1 | TATGAGTGTGGAGTCGTTGCACAACATGGTCCTCAGTTAGTCGNNNAGATCGGAAG |
| 36048_2 | TATGAGTGTGGAGTCGTTGCACAACATGGTCCTCAGTTAGTCGNNNAGATCGGAAG |
| 47965_1 | TATGAGTGTGGAGTCGTTGCACCAGGCCCAGCTAATTTTTGTANNNAGATCGGAAG |
| 47965_2 | TATGAGTGTGGAGTCGTTGCACCAGGCCCAGCTAATTTTTGTANNNAGATCGGAAG |
| 33862_1 | TATGAGTGTGGAGTCGTTGCTTTGGGAGAAAGTAAGGGACGAGNNNAGATCGGA |
| 33862_2 | TATGAGTGTGGAGTCGTTGCTTTGGGAGAAAGTAAGGGACGAGNNNAGATCGGA |
| 44761_1 | TATGAGTGTGGAGTCGTTGCCCGTGAGAGAAATATGTGTGCACNNNAGATCGGAA |
| 44761_2 | TATGAGTGTGGAGTCGTTGCCCGTGAGAGAAATATGTGTGCACNNNAGATCGGAA |
| 38224_1 | TATGAGTGTGGAGTCGTTGCGAGGTCAGGAGTTCAAGACGAGNNNAGATCGGAAG |
| 38224_2 | TATGAGTGTGGAGTCGTTGCGAGGTCAGGAGTTCAAGACGAGNNNAGATCGGAAG |
| 47568_1 | TATGAGTGTGGAGTCGTTGCGGACTCAAGTGATTCTTCTGCCTNNNAGATCGGAAG |
| 47568_2 | TATGAGTGTGGAGTCGTTGCGGACTCAAGTGATTCTTCTGCCTNNNAGATCGGAAG |
| 35751_1 | TATGAGTGTGGAGTCGTTGCCGTGTCTCAGCTCACTGCAAGNNNAGATCGGAAGAG |
| 35751_2 | TATGAGTGTGGAGTCGTTGCCGTGTCTCAGCTCACTGCAAGNNNAGATCGGAAGAG |
| 38976_1 | TATGAGTGTGGAGTCGTTGCGTGAAACCCCGTCTCTACTGAAANNNAGATCGGAAG |
| 38976_2 | TATGAGTGTGGAGTCGTTGCGTGAAACCCCGTCTCTACTGAAANNNAGATCGGAAG |
| 46874_1 | TATGAGTGTGGAGTCGTTGCGACACCACGTCTGGCTAATTTTTNNNAGATCGGAAG |
| 46874_2 | TATGAGTGTGGAGTCGTTGCGACACCACGTCTGGCTAATTTTTNNNAGATCGGAAG |
| 45172_1 | TATGAGTGTGGAGTCGTTGCACCCCTGCTCATCTAAATCTTGGNNNAGATCGGAAG |
| 45172_2 | TATGAGTGTGGAGTCGTTGCACCCCTGCTCATCTAAATCTTGGNNNAGATCGGAAG |
| 28933_1 | TATGAGTGTGGAGTCGTTGCTCCACTGATGTCCGTTGTACATTNNNAGATCGGAAG |
| 28933_2 | TATGAGTGTGGAGTCGTTGCTCCACTGATGTCCGTTGTACATTNNNAGATCGGAAG |
| 41504_1 | TATGAGTGTGGAGTCGTTGCTCCAAAGAGGTGAACATATCCCCNNNAGATCGGAAG |
| 41504_2 | TATGAGTGTGGAGTCGTTGCTCCAAAGAGGTGAACATATCCCCNNNAGATCGGAAG |

| | |
|---|---|
| 36097_1 | TATGAGTGTGGAGTCGTTGCGGTATACACACGCCATAGTGGTTNNNAGATCGGAAG |
| 36097_2 | TATGAGTGTGGAGTCGTTGCGGTATACACACGCCATAGTGGTTNNNAGATCGGAAG |
| 35737_1 | TATGAGTGTGGAGTCGTTGCACAGAAACGAGTTCTCACCATGTNNNAGATCGGAAG |
| 35737_2 | TATGAGTGTGGAGTCGTTGCACAGAAACGAGTTCTCACCATGTNNNAGATCGGAAG |
| 48564_1 | TATGAGTGTGGAGTCGTTGCCAAATTGCTGGGATTACAGGCATNNNAGATCGGAAG |
| 48564_2 | TATGAGTGTGGAGTCGTTGCCAAATTGCTGGGATTACAGGCATNNNAGATCGGAAG |
| 35321_1 | TATGAGTGTGGAGTCGTTGCTTCATGCCATTCTCCTTCCTCAGNNNAGATCGGAAGA |
| 35321_2 | TATGAGTGTGGAGTCGTTGCTTCATGCCATTCTCCTTCCTCAGNNNAGATCGGAAGA |
| 42568_1 | TATGAGTGTGGAGTCGTTGCGCTCTCAGCTTGGATGTTGGTATNNNAGATCGGAAG |
| 42568_2 | TATGAGTGTGGAGTCGTTGCGCTCTCAGCTTGGATGTTGGTATNNNAGATCGGAAG |
| 12964_1 | TATGAGTGTGGAGTCGTTGCCTATGTACATGTTCCTCCAGNNNAGATCGGAAGAGC |
| 12964_2 | TATGAGTGTGGAGTCGTTGCCTATGTACATGTTCCTCCAGNNNAGATCGGAAGAGC |
| 43775_1 | TATGAGTGTGGAGTCGTTGCGGAAGTCCAAGATCAAAGTGTGCNNNAGATCGGAA |
| 43775_2 | TATGAGTGTGGAGTCGTTGCGGAAGTCCAAGATCAAAGTGTGCNNNAGATCGGAA |
| 48599_1 | TATGAGTGTGGAGTCGTTGCGTTCAAACAATTCTCGTGCCTCANNNAGATCGGAAG |
| 48599_2 | TATGAGTGTGGAGTCGTTGCGTTCAAACAATTCTCGTGCCTCANNNAGATCGGAAG |
| 40832_1 | TATGAGTGTGGAGTCGTTGCGACATACAAAAGGCCAACTGACANNNAGATCGGAA |
| 40832_2 | TATGAGTGTGGAGTCGTTGCGACATACAAAAGGCCAACTGACANNNAGATCGGAA |
| 37342_1 | TATGAGTGTGGAGTCGTTGCAGATTATTTGTCCCCAGGTTGCANNNAGATCGGAAG |
| 37342_2 | TATGAGTGTGGAGTCGTTGCAGATTATTTGTCCCCAGGTTGCANNNAGATCGGAAG |
| 49545_1 | TATGAGTGTGGAGTCGTTGCTAGCTGGGAATACAGGTATGTGCNNNAGATCGGAA |
| 49545_2 | TATGAGTGTGGAGTCGTTGCTAGCTGGGAATACAGGTATGTGCNNNAGATCGGAA |
| 35949_1 | TATGAGTGTGGAGTCGTTGCCTGTCATCCCAGCACTTTGAGAGNNNAGATCGGAAG |
| 35949_2 | TATGAGTGTGGAGTCGTTGCCTGTCATCCCAGCACTTTGAGAGNNNAGATCGGAAG |
| 7803_1 | TATGAGTGTGGAGTCGTTGCCAGTATAAGCCAGTGTGCCTAGCNNNAGATCGGAAG |
| 7803_2 | TATGAGTGTGGAGTCGTTGCCAGTATAAGCCAGTGTGCCTAGCNNNAGATCGGAAG |
| 30607_1 | TATGAGTGTGGAGTCGTTGCGTCCCAGTGATTAAACTGAGGTCANNNAGATCGGAA |
| 30607_2 | TATGAGTGTGGAGTCGTTGCGTCCCAGTGATTAAACTGAGGTCANNNAGATCGGAA |
| 45602_1 | TATGAGTGTGGAGTCGTTGCCTATGATCAGGCCACTGGACTTCNNNAGATCGGAAG |
| 45602_2 | TATGAGTGTGGAGTCGTTGCCTATGATCAGGCCACTGGACTTCNNNAGATCGGAAG |
| 37012_1 | TATGAGTGTGGAGTCGTTGCTGTTTTGAGACAGAGTCTTGCTCNNNAGATCGGAAG |
| 37012_2 | TATGAGTGTGGAGTCGTTGCTGTTTTGAGACAGAGTCTTGCTCNNNAGATCGGAAG |
| 12936_1 | TATGAGTGTGGAGTCGTTGCCAACATTTTAGGAACTTTCTAGATNNNAGATCGGAA |
| 12936_2 | TATGAGTGTGGAGTCGTTGCCAACATTTTAGGAACTTTCTAGATNNNAGATCGGAA |
| 52547_1 | TATGAGTGTGGAGTCGTTGCGTTGCAAATGTCGAAACGGAAAGNNNAGATCGGAA |
| 52547_2 | TATGAGTGTGGAGTCGTTGCGTTGCAAATGTCGAAACGGAAAGNNNAGATCGGAA |
| 2258_1 | TATGAGTGTGGAGTCGTTGCATACATCTTGGCTAGGCTCGGTANNNAGATCGGAAG |
| 2258_2 | TATGAGTGTGGAGTCGTTGCATACATCTTGGCTAGGCTCGGTANNNAGATCGGAAG |
| 45137_1 | TATGAGTGTGGAGTCGTTGCGCCAGCCCAGTACTCTACAAATANNNAGATCGGAAG |
| 45137_2 | TATGAGTGTGGAGTCGTTGCGCCAGCCCAGTACTCTACAAATANNNAGATCGGAAG |
| 50431_1 | TATGAGTGTGGAGTCGTTGCACTGTTCAAACCCAGGCAGATATNNNAGATCGGAAG |
| 50431_2 | TATGAGTGTGGAGTCGTTGCACTGTTCAAACCCAGGCAGATATNNNAGATCGGAAG |
| 33422_1 | TATGAGTGTGGAGTCGTTGCTTGTCCCACACAGAATCTTCCTTNNNAGATCGGAAGA |

| | |
|---|---|
| 33422_2 | TATGAGTGTGGAGTCGTTGCTTGTCCCACACAGAATCTTCCTTNNNAGATCGGAAGA |
| 44040_1 | TATGAGTGTGGAGTCGTTGCCTGAGGTTCAGCTTTGTAGGGACNNNAGATCGGAA |
| 44040_2 | TATGAGTGTGGAGTCGTTGCCTGAGGTTCAGCTTTGTAGGGACNNNAGATCGGAA |
| 48753_1 | TATGAGTGTGGAGTCGTTGCAGGTCTCAACAGTGCACCTAAAANNNAGATCGGAAG |
| 48753_2 | TATGAGTGTGGAGTCGTTGCAGGTCTCAACAGTGCACCTAAAANNNAGATCGGAAG |
| 53033_1 | TATGAGTGTGGAGTCGTTGCTTTGACCAGAACTTGACAGGGAANNNAGATCGGAA |
| 53033_2 | TATGAGTGTGGAGTCGTTGCTTTGACCAGAACTTGACAGGGAANNNAGATCGGAA |
| 10403_1 | TATGAGTGTGGAGTCGTTGCATTTCTCCCCACTGTCCTTTCCTNNNAGATCGGAAGA |
| 10403_2 | TATGAGTGTGGAGTCGTTGCATTTCTCCCCACTGTCCTTTCCTNNNAGATCGGAAGA |
| 29769_1 | TATGAGTGTGGAGTCGTTGCTCGCACTCATTTTCAATAGCAGTNNNAGATCGGAAG |
| 29769_2 | TATGAGTGTGGAGTCGTTGCTCGCACTCATTTTCAATAGCAGTNNNAGATCGGAAG |
| 51174_1 | TATGAGTGTGGAGTCGTTGCGACCCCATCCCTCCTATACTCTGNNNAGATCGGAAGA |
| 51174_2 | TATGAGTGTGGAGTCGTTGCGACCCCATCCCTCCTATACTCTGNNNAGATCGGAAGA |
| 43131_1 | TATGAGTGTGGAGTCGTTGCAATTTCAAGCACACTCCACTGACNNNAGATCGGAAG |
| 43131_2 | TATGAGTGTGGAGTCGTTGCAATTTCAAGCACACTCCACTGACNNNAGATCGGAAG |
| 39848_1 | TATGAGTGTGGAGTCGTTGCATTGTTTACTGCCGTGTGACTTCNNNAGATCGGAAG |
| 39848_2 | TATGAGTGTGGAGTCGTTGCATTGTTTACTGCCGTGTGACTTCNNNAGATCGGAAG |
| 39118_1 | TATGAGTGTGGAGTCGTTGCGTGGTACGATCCTAGTTCACTGGNNNAGATCGGAA |
| 39118_2 | TATGAGTGTGGAGTCGTTGCGTGGTACGATCCTAGTTCACTGGNNNAGATCGGAA |
| 45535_1 | TATGAGTGTGGAGTCGTTGCCTTTGGCCTAAAGCTGACACTTCNNNAGATCGGAAG |
| 45535_2 | TATGAGTGTGGAGTCGTTGCCTTTGGCCTAAAGCTGACACTTCNNNAGATCGGAAG |
| 13193_1 | TATGAGTGTGGAGTCGTTGCTAATTTTCATGTAATTGTGTGGTNNNAGATCGGAAG |
| 13193_2 | TATGAGTGTGGAGTCGTTGCTAATTTTCATGTAATTGTGTGGTNNNAGATCGGAAG |
| 45819_1 | TATGAGTGTGGAGTCGTTGCGAGTAGCTGGCACTACAGATGTGNNNAGATCGGAA |
| 45819_2 | TATGAGTGTGGAGTCGTTGCGAGTAGCTGGCACTACAGATGTGNNNAGATCGGAA |
| 41161_1 | TATGAGTGTGGAGTCGTTGCACTCTGGATACCATGCTGAATCCNNNAGATCGGAAG |
| 41161_2 | TATGAGTGTGGAGTCGTTGCACTCTGGATACCATGCTGAATCCNNNAGATCGGAAG |
| 41339_1 | TATGAGTGTGGAGTCGTTGCTTTCACAGGTTGGTGTCAGTTTGNNNAGATCGGAAG |
| 41339_2 | TATGAGTGTGGAGTCGTTGCTTTCACAGGTTGGTGTCAGTTTGNNNAGATCGGAAG |
| 51745_1 | TATGAGTGTGGAGTCGTTGCTACCCACAAGGATCCAGGTTTTCNNNAGATCGGAAG |
| 51745_2 | TATGAGTGTGGAGTCGTTGCTACCCACAAGGATCCAGGTTTTCNNNAGATCGGAAG |
| 38190_1 | TATGAGTGTGGAGTCGTTGCACTAGATGAAAAGCAGAGGTGGANNNAGATCGGA |
| 38190_2 | TATGAGTGTGGAGTCGTTGCACTAGATGAAAAGCAGAGGTGGANNNAGATCGGA |
| 48423_1 | TATGAGTGTGGAGTCGTTGCTCTCCCAAAATACAGCTCTGTGGNNNAGATCGGAAG |
| 48423_2 | TATGAGTGTGGAGTCGTTGCTCTCCCAAAATACAGCTCTGTGGNNNAGATCGGAAG |
| 44666_1 | TATGAGTGTGGAGTCGTTGCCAGGGAACCTAACCTCACATTCANNNAGATCGGAAG |
| 44666_2 | TATGAGTGTGGAGTCGTTGCCAGGGAACCTAACCTCACATTCANNNAGATCGGAAG |
| 39893_1 | TATGAGTGTGGAGTCGTTGCCAAATGAATTCCCCAGCCTTCTGNNNAGATCGGAAG |
| 39893_2 | TATGAGTGTGGAGTCGTTGCCAAATGAATTCCCCAGCCTTCTGNNNAGATCGGAAG |
| 40896_1 | TATGAGTGTGGAGTCGTTGCACGAACCATGAAGTAAACATCACCNNNAGATCGGAA |
| 40896_2 | TATGAGTGTGGAGTCGTTGCACGAACCATGAAGTAAACATCACCNNNAGATCGGAA |
| 2530_1 | TATGAGTGTGGAGTCGTTGCAAGCACGCATATAACAGGACAGGNNNAGATCGGAA |
| 2530_2 | TATGAGTGTGGAGTCGTTGCAAGCACGCATATAACAGGACAGGNNNAGATCGGAA |

| | |
|---|---|
| 13331_1 | TATGAGTGTGGAGTCGTTGCTCTTTTTTGGTTCCATATGAATTNNNAGATCGGAAGA |
| 13331_2 | TATGAGTGTGGAGTCGTTGCTCTTTTTTGGTTCCATATGAATTNNNAGATCGGAAGA |
| 10336_1 | TATGAGTGTGGAGTCGTTGCCATAGGCCAAGTCATCAGGAAGGNNNAGATCGGAA |
| 10336_2 | TATGAGTGTGGAGTCGTTGCCATAGGCCAAGTCATCAGGAAGGNNNAGATCGGAA |
| 32177_1 | TATGAGTGTGGAGTCGTTGCGGTTCACTTGAGACCCTGTAACANNNAGATCGGAAG |
| 32177_2 | TATGAGTGTGGAGTCGTTGCGGTTCACTTGAGACCCTGTAACANNNAGATCGGAAG |
| 51714_1 | TATGAGTGTGGAGTCGTTGCGCTGGATCAACAACAATGCAGTANNNAGATCGGAAG |
| 51714_2 | TATGAGTGTGGAGTCGTTGCGCTGGATCAACAACAATGCAGTANNNAGATCGGAAG |
| 38805_1 | TATGAGTGTGGAGTCGTTGCAAAAAGGGAGAGAAGAACCAAGTNNNAGATCGGA |
| 38805_2 | TATGAGTGTGGAGTCGTTGCAAAAAGGGAGAGAAGAACCAAGTNNNAGATCGGA |
| 47269_1 | TATGAGTGTGGAGTCGTTGCCAAGTAGCTGGGACTACAGGCNNNAGATCGGAAGA |
| 47269_2 | TATGAGTGTGGAGTCGTTGCCAAGTAGCTGGGACTACAGGCNNNAGATCGGAAGA |
| 13171_1 | TATGAGTGTGGAGTCGTTGCTATCATACAAATAAGCCAGTCACNNNAGATCGGAAG |
| 13171_2 | TATGAGTGTGGAGTCGTTGCTATCATACAAATAAGCCAGTCACNNNAGATCGGAAG |
| 36809_1 | TATGAGTGTGGAGTCGTTGCCTCATTCCTGTAATCCCAGCACTNNNAGATCGGAAGA |
| 36809_2 | TATGAGTGTGGAGTCGTTGCCTCATTCCTGTAATCCCAGCACTNNNAGATCGGAAGA |
| 5626_1 | TATGAGTGTGGAGTCGTTGCGAGCCTTTGGAGCAGCTTTACTTNNNAGATCGGAAG |
| 5626_2 | TATGAGTGTGGAGTCGTTGCGAGCCTTTGGAGCAGCTTTACTTNNNAGATCGGAAG |
| 31873_1 | TATGAGTGTGGAGTCGTTGCTTCTCTCTGGATTTTCAGGAGGCNNNAGATCGGAAG |
| 31873_2 | TATGAGTGTGGAGTCGTTGCTTCTCTCTGGATTTTCAGGAGGCNNNAGATCGGAAG |
| 51197_1 | TATGAGTGTGGAGTCGTTGCACACACACACACACACACACAAATTNNNAGATCGGAAG |
| 51197_2 | TATGAGTGTGGAGTCGTTGCACACACACACACACACACACAAATTNNNAGATCGGAAG |
| 46748_1 | TATGAGTGTGGAGTCGTTGCCCCAACTAGCTGGAATTACAGGTNNNAGATCGGAAG |
| 46748_2 | TATGAGTGTGGAGTCGTTGCCCCAACTAGCTGGAATTACAGGTNNNAGATCGGAAG |
| 45416_1 | TATGAGTGTGGAGTCGTTGCTTGGCCAAACTTGCTAAGGAATGNNNAGATCGGAAG |
| 45416_2 | TATGAGTGTGGAGTCGTTGCTTGGCCAAACTTGCTAAGGAATGNNNAGATCGGAAG |
| 37774_1 | TATGAGTGTGGAGTCGTTGCCAGACTACAAAGGCCTTCATCCTNNNAGATCGGAAG |
| 37774_2 | TATGAGTGTGGAGTCGTTGCCAGACTACAAAGGCCTTCATCCTNNNAGATCGGAAG |
| 29532_1 | TATGAGTGTGGAGTCGTTGCAAGAAATGGTTGGTTGGTGGTTTNNNAGATCGGAA |
| 29532_2 | TATGAGTGTGGAGTCGTTGCAAGAAATGGTTGGTTGGTGGTTTNNNAGATCGGAA |
| 44821_1 | TATGAGTGTGGAGTCGTTGCTCAGGAACACTAATGTGGCATGANNNAGATCGGAA |
| 44821_2 | TATGAGTGTGGAGTCGTTGCTCAGGAACACTAATGTGGCATGANNNAGATCGGAA |
| 28773_1 | TATGAGTGTGGAGTCGTTGCAAGAATCACTTGAACCTGGGAGGNNNAGATCGGAA |
| 28773_2 | TATGAGTGTGGAGTCGTTGCAAGAATCACTTGAACCTGGGAGGNNNAGATCGGAA |
| 39072_1 | TATGAGTGTGGAGTCGTTGCACACAGGATAAAGTGGTAACTTGGANNNAGATCGG |
| 39072_2 | TATGAGTGTGGAGTCGTTGCACACAGGATAAAGTGGTAACTTGGANNNAGATCGG |
| 41421_1 | TATGAGTGTGGAGTCGTTGCCAGATGCTGGTGAGGTTTCAAAGNNNAGATCGGAA |
| 41421_2 | TATGAGTGTGGAGTCGTTGCCAGATGCTGGTGAGGTTTCAAAGNNNAGATCGGAA |
| 29092_1 | TATGAGTGTGGAGTCGTTGCTGCTCTCTGTGTGTCATTCTCTCNNNAGATCGGAAGA |
| 29092_2 | TATGAGTGTGGAGTCGTTGCTGCTCTCTGTGTGTCATTCTCTCNNNAGATCGGAAGA |
| 49570_1 | TATGAGTGTGGAGTCGTTGCAGAAACTACAACCAAGAGAAGCTNNNAGATCGGAA |
| 49570_2 | TATGAGTGTGGAGTCGTTGCAGAAACTACAACCAAGAGAAGCTNNNAGATCGGAA |
| 9291_1 | TATGAGTGTGGAGTCGTTGCACAGTAGCAGTGGCAAGAGAATGNNNAGATCGGAA |

| | |
|---|---|
| 9291_2 | TATGAGTGTGGAGTCGTTGCACAGTAGCAGTGGCAAGAGAATGNNNAGATCGGA |
| 52284_1 | TATGAGTGTGGAGTCGTTGCTCCTGTAGTCATTCTTCATGGCANNNAGATCGGAAG |
| 52284_2 | TATGAGTGTGGAGTCGTTGCTCCTGTAGTCATTCTTCATGGCANNNAGATCGGAAG |
| 38220_1 | TATGAGTGTGGAGTCGTTGCGATCACCTGAGGTCAGCAGTTTNNNAGATCGGAAG |
| 38220_2 | TATGAGTGTGGAGTCGTTGCGATCACCTGAGGTCAGCAGTTTNNNAGATCGGAAG |
| 52429_1 | TATGAGTGTGGAGTCGTTGCAAGGAGCATTTCAGCAGCATTTTNNNAGATCGGAAG |
| 52429_2 | TATGAGTGTGGAGTCGTTGCAAGGAGCATTTCAGCAGCATTTTNNNAGATCGGAAG |
| 80_1 | TATGAGTGTGGAGTCGTTGCGACTACATCTCCCGTCCTGCNNNAGATCGGAAGAGC |
| 80_2 | TATGAGTGTGGAGTCGTTGCGACTACATCTCCCGTCCTGCNNNAGATCGGAAGAGC |
| 3087_1 | TATGAGTGTGGAGTCGTTGCCTGGGGTCTCATATACAGGGACCNNNAGATCGGAA |
| 3087_2 | TATGAGTGTGGAGTCGTTGCCTGGGGTCTCATATACAGGGACCNNNAGATCGGAA |
| 33714_1 | TATGAGTGTGGAGTCGTTGCTCAGGAACATCACTAGTGGAAGCNNNAGATCGGAA |
| 33714_2 | TATGAGTGTGGAGTCGTTGCTCAGGAACATCACTAGTGGAAGCNNNAGATCGGAA |
| 28763_1 | TATGAGTGTGGAGTCGTTGCCCTCCTGCATTCAAGCAATTCTCNNNAGATCGGAAGA |
| 28763_2 | TATGAGTGTGGAGTCGTTGCCCTCCTGCATTCAAGCAATTCTCNNNAGATCGGAAGA |
| 31521_1 | TATGAGTGTGGAGTCGTTGCTGTTAGGCAGGTTATTGAGCTGTNNNAGATCGGAA |
| 31521_2 | TATGAGTGTGGAGTCGTTGCTGTTAGGCAGGTTATTGAGCTGTNNNAGATCGGAA |
| 39090_1 | TATGAGTGTGGAGTCGTTGCGGTCAAGTGGGATTTATCATAGGGANNNAGATCGG |
| 39090_2 | TATGAGTGTGGAGTCGTTGCGGTCAAGTGGGATTTATCATAGGGANNNAGATCGG |
| 35824_1 | TATGAGTGTGGAGTCGTTGCTTCAAGCAATTCTCCTGCCTCAGNNNAGATCGGAAG |
| 35824_2 | TATGAGTGTGGAGTCGTTGCTTCAAGCAATTCTCCTGCCTCAGNNNAGATCGGAAG |
| 42349_1 | TATGAGTGTGGAGTCGTTGCCAGTTGCTTAGGAGAGTGTGACTNNNAGATCGGAA |
| 42349_2 | TATGAGTGTGGAGTCGTTGCCAGTTGCTTAGGAGAGTGTGACTNNNAGATCGGAA |
| 39455_1 | TATGAGTGTGGAGTCGTTGCAAAGGCGGGACTCTTTCTTTTTCNNNAGATCGGAAG |
| 39455_2 | TATGAGTGTGGAGTCGTTGCAAAGGCGGGACTCTTTCTTTTTCNNNAGATCGGAAG |
| 51272_1 | TATGAGTGTGGAGTCGTTGCGTGGTTCTGAGAACTGAGAGGAGNNNAGATCGGA |
| 51272_2 | TATGAGTGTGGAGTCGTTGCGTGGTTCTGAGAACTGAGAGGAGNNNAGATCGGA |
| 31146_1 | TATGAGTGTGGAGTCGTTGCCACACACACACTCACACACATACNNNAGATCGGAAGA |
| 31146_2 | TATGAGTGTGGAGTCGTTGCCACACACACACTCACACACATACNNNAGATCGGAAGA |
| 41077_1 | TATGAGTGTGGAGTCGTTGCTAAACATGAGGAAAAACGTGCCCNNNAGATCGGAA |
| 41077_2 | TATGAGTGTGGAGTCGTTGCTAAACATGAGGAAAAACGTGCCCNNNAGATCGGAA |
| 49585_1 | TATGAGTGTGGAGTCGTTGCGTCAAACTCCTGACCTCAACTGANNNAGATCGGAAG |
| 49585_2 | TATGAGTGTGGAGTCGTTGCGTCAAACTCCTGACCTCAACTGANNNAGATCGGAAG |
| 13164_1 | TATGAGTGTGGAGTCGTTGCCCAGGGAAGTATTATACTTACGNNNAGATCGGAAG |
| 13164_2 | TATGAGTGTGGAGTCGTTGCCCAGGGAAGTATTATACTTACGNNNAGATCGGAAG |
| 13187_1 | TATGAGTGTGGAGTCGTTGCTGTTCGTAGACATGAAAGAATNNNAGATCGGAAGA |
| 13187_2 | TATGAGTGTGGAGTCGTTGCTGTTCGTAGACATGAAAGAATNNNAGATCGGAAGA |
| 39703_1 | TATGAGTGTGGAGTCGTTGCCAGCGGCTACTACTCTATTCCAGNNNAGATCGGAAG |
| 39703_2 | TATGAGTGTGGAGTCGTTGCCAGCGGCTACTACTCTATTCCAGNNNAGATCGGAAG |
| 41839_1 | TATGAGTGTGGAGTCGTTGCCCTCTTTACTGGGTGAGACTTCCNNNAGATCGGAAG |
| 41839_2 | TATGAGTGTGGAGTCGTTGCCCTCTTTACTGGGTGAGACTTCCNNNAGATCGGAAG |
| 38459_1 | TATGAGTGTGGAGTCGTTGCGGGGGGCCCTATCATTTTTATCANNNAGATCGGAAG |
| 38459_2 | TATGAGTGTGGAGTCGTTGCGGGGGGCCCTATCATTTTTATCANNNAGATCGGAAG |

| | |
|---|---|
| 37890_1 | TATGAGTGTGGAGTCGTTGCCAGCACCTATGGAGCACTAAGAANNNAGATCGGAA |
| 37890_2 | TATGAGTGTGGAGTCGTTGCCAGCACCTATGGAGCACTAAGAANNNAGATCGGAA |
| 43460_1 | TATGAGTGTGGAGTCGTTGCTCACCATGCCCAGCTAATTTTTGNNNAGATCGGAAG |
| 43460_2 | TATGAGTGTGGAGTCGTTGCTCACCATGCCCAGCTAATTTTTGNNNAGATCGGAAG |
| 39054_1 | TATGAGTGTGGAGTCGTTGCTGCAGACAAGAAAAATGAACCCANNNAGATCGGAA |
| 39054_2 | TATGAGTGTGGAGTCGTTGCTGCAGACAAGAAAAATGAACCCANNNAGATCGGAA |
| 40091_1 | TATGAGTGTGGAGTCGTTGCTACAGCATCATACCCAGTTAGGCNNNAGATCGGAAG |
| 40091_2 | TATGAGTGTGGAGTCGTTGCTACAGCATCATACCCAGTTAGGCNNNAGATCGGAAG |
| 29270_1 | TATGAGTGTGGAGTCGTTGCATGACCTGCTAATGTGTGATGGTNNNAGATCGGAA |
| 29270_2 | TATGAGTGTGGAGTCGTTGCATGACCTGCTAATGTGTGATGGTNNNAGATCGGAA |
| 42781_1 | TATGAGTGTGGAGTCGTTGCGGACTGGACTTAGAGTAACACCGNNNAGATCGGAA |
| 42781_2 | TATGAGTGTGGAGTCGTTGCGGACTGGACTTAGAGTAACACCGNNNAGATCGGAA |
| 29001_1 | TATGAGTGTGGAGTCGTTGCAGAGAGAGAGAGAGAGAGAGAGAGTNNNAGATCGG |
| 29001_2 | TATGAGTGTGGAGTCGTTGCAGAGAGAGAGAGAGAGAGAGAGAGTNNNAGATCGG |
| 42822_1 | TATGAGTGTGGAGTCGTTGCTTTCTCTCTCTTTTGAGCCCTCCNNNAGATCGGAAGA |
| 42822_2 | TATGAGTGTGGAGTCGTTGCTTTCTCTCTCTTTTGAGCCCTCCNNNAGATCGGAAGA |
| 30372_1 | TATGAGTGTGGAGTCGTTGCCTGGCTTGGTTCTCAACTGTCTTNNNAGATCGGAAG |
| 30372_2 | TATGAGTGTGGAGTCGTTGCCTGGCTTGGTTCTCAACTGTCTTNNNAGATCGGAAG |
| 37566_1 | TATGAGTGTGGAGTCGTTGCTGAAACCATTTTCCAGGTGTCTGNNNAGATCGGAAG |
| 37566_2 | TATGAGTGTGGAGTCGTTGCTGAAACCATTTTCCAGGTGTCTGNNNAGATCGGAAG |
| 3444_1 | TATGAGTGTGGAGTCGTTGCACTTCCATCCCCATGAGGCAANNNAGATCGGAAGAG |
| 3444_2 | TATGAGTGTGGAGTCGTTGCACTTCCATCCCCATGAGGCAANNNAGATCGGAAGAG |
| 36066_1 | TATGAGTGTGGAGTCGTTGCGACTCCCATACCCTTACTGTCTGNNNAGATCGGAAGA |
| 36066_2 | TATGAGTGTGGAGTCGTTGCGACTCCCATACCCTTACTGTCTGNNNAGATCGGAAGA |
| 41180_1 | TATGAGTGTGGAGTCGTTGCGGTTAGGATGCAGAGAAACTGGANNNAGATCGGA |
| 41180_2 | TATGAGTGTGGAGTCGTTGCGGTTAGGATGCAGAGAAACTGGANNNAGATCGGA |
| 34852_1 | TATGAGTGTGGAGTCGTTGCCCATGACACCCGGCTAATTTTTTNNNAGATCGGAAG |
| 34852_2 | TATGAGTGTGGAGTCGTTGCCCATGACACCCGGCTAATTTTTTNNNAGATCGGAAG |
| 12537_1 | TATGAGTGTGGAGTCGTTGCGAGATTTTAGCAGATCCGATATNNNAGATCGGAAG |
| 12537_2 | TATGAGTGTGGAGTCGTTGCGAGATTTTAGCAGATCCGATATNNNAGATCGGAAG |
| 36145_1 | TATGAGTGTGGAGTCGTTGCACCTATCCATCCAAGGATCAAGGNNNAGATCGGAAG |
| 36145_2 | TATGAGTGTGGAGTCGTTGCACCTATCCATCCAAGGATCAAGGNNNAGATCGGAAG |
| 43246_1 | TATGAGTGTGGAGTCGTTGCCCACTTTCAGCATTGGACAGATCNNNAGATCGGAAG |
| 43246_2 | TATGAGTGTGGAGTCGTTGCCCACTTTCAGCATTGGACAGATCNNNAGATCGGAAG |
| 48252_1 | TATGAGTGTGGAGTCGTTGCTCACTCATTATTAGTCTGTGCAGGTNNNAGATCGGA |
| 48252_2 | TATGAGTGTGGAGTCGTTGCTCACTCATTATTAGTCTGTGCAGGTNNNAGATCGGA |
| 32486_1 | TATGAGTGTGGAGTCGTTGCCCGAGTAGCTGGGACTATAGGTNNNAGATCGGAAG |
| 32486_2 | TATGAGTGTGGAGTCGTTGCCCGAGTAGCTGGGACTATAGGTNNNAGATCGGAAG |
| 51439_1 | TATGAGTGTGGAGTCGTTGCGGCATGGTGGTGATTTCAAAGTGNNNAGATCGGAA |
| 51439_2 | TATGAGTGTGGAGTCGTTGCGGCATGGTGGTGATTTCAAAGTGNNNAGATCGGAA |
| 46374_1 | TATGAGTGTGGAGTCGTTGCAAACGTGAAACCTGAATACGCTCNNNAGATCGGAAG |
| 46374_2 | TATGAGTGTGGAGTCGTTGCAAACGTGAAACCTGAATACGCTCNNNAGATCGGAAG |
| 46302_1 | TATGAGTGTGGAGTCGTTGCTGGGGGTCTCTCTATATTGTCCANNNAGATCGGAAG |

| | |
|---|---|
| 46302_2 | TATGAGTGTGGAGTCGTTGCTGGGGGTCTCTCTATATTGTCCANNNAGATCGGAAG |
| 40145_1 | TATGAGTGTGGAGTCGTTGCATAGCTTGCAGTGAGCCGAGATCNNNAGATCGGAA |
| 40145_2 | TATGAGTGTGGAGTCGTTGCATAGCTTGCAGTGAGCCGAGATCNNNAGATCGGAA |
| 38907_1 | TATGAGTGTGGAGTCGTTGCCAGCAAACTGAACAGATCTGCAANNNAGATCGGAAG |
| 38907_2 | TATGAGTGTGGAGTCGTTGCCAGCAAACTGAACAGATCTGCAANNNAGATCGGAAG |
| 32671_1 | TATGAGTGTGGAGTCGTTGCGAGTTCATGCAATTCTCCTGCCNNNAGATCGGAAGA |
| 32671_2 | TATGAGTGTGGAGTCGTTGCGAGTTCATGCAATTCTCCTGCCNNNAGATCGGAAGA |
| 52768_1 | TATGAGTGTGGAGTCGTTGCTTCAAGTTTGAACACAGAACGGGNNNAGATCGGAA |
| 52768_2 | TATGAGTGTGGAGTCGTTGCTTCAAGTTTGAACACAGAACGGGNNNAGATCGGAA |
| 29391_1 | TATGAGTGTGGAGTCGTTGCTTTTTTGTGTTAGTTTGTTCTTTNNNAGATCGGAAGA |
| 29391_2 | TATGAGTGTGGAGTCGTTGCTTTTTTGTGTTAGTTTGTTCTTTNNNAGATCGGAAGA |
| 47606_1 | TATGAGTGTGGAGTCGTTGCTCTCGATCTCCTGACCTCTTGATNNNAGATCGGAAGA |
| 47606_2 | TATGAGTGTGGAGTCGTTGCTCTCGATCTCCTGACCTCTTGATNNNAGATCGGAAGA |
| 32189_1 | TATGAGTGTGGAGTCGTTGCCCCTAGTTGCTTTCTCACTCACANNNAGATCGGAAGA |
| 32189_2 | TATGAGTGTGGAGTCGTTGCCCCTAGTTGCTTTCTCACTCACANNNAGATCGGAAGA |
| 48326_1 | TATGAGTGTGGAGTCGTTGCCTGTTTCAGGAGAGATCTCGACCNNNAGATCGGAAG |
| 48326_2 | TATGAGTGTGGAGTCGTTGCCTGTTTCAGGAGAGATCTCGACCNNNAGATCGGAAG |
| 33405_1 | TATGAGTGTGGAGTCGTTGCCCTGTAGTCCCAGCTACTCGGNNNAGATCGGAAGAG |
| 33405_2 | TATGAGTGTGGAGTCGTTGCCCTGTAGTCCCAGCTACTCGGNNNAGATCGGAAGAG |
| 46124_1 | TATGAGTGTGGAGTCGTTGCCATCTCAGCTCACTGCAACATCNNNAGATCGGAAGA |
| 46124_2 | TATGAGTGTGGAGTCGTTGCCATCTCAGCTCACTGCAACATCNNNAGATCGGAAGA |
| 53264_1 | TATGAGTGTGGAGTCGTTGCAATTGCATGTGTTCAAAGGAGCANNNAGATCGGAA |
| 53264_2 | TATGAGTGTGGAGTCGTTGCAATTGCATGTGTTCAAAGGAGCANNNAGATCGGAA |
| 36203_1 | TATGAGTGTGGAGTCGTTGCCTAGAATTCTGAGGGCCTGAGGANNNAGATCGGAA |
| 36203_2 | TATGAGTGTGGAGTCGTTGCCTAGAATTCTGAGGGCCTGAGGANNNAGATCGGAA |
| 33101_1 | TATGAGTGTGGAGTCGTTGCCTCCCGAGTAGCTGAGATTACAGNNNAGATCGGAAG |
| 33101_2 | TATGAGTGTGGAGTCGTTGCCTCCCGAGTAGCTGAGATTACAGNNNAGATCGGAAG |
| 47756_1 | TATGAGTGTGGAGTCGTTGCCCTCCTGAGTGGTTGGAATTACANNNAGATCGGAAG |
| 47756_2 | TATGAGTGTGGAGTCGTTGCCCTCCTGAGTGGTTGGAATTACANNNAGATCGGAAG |
| 45995_1 | TATGAGTGTGGAGTCGTTGCCGAATAGCAGGGACTACAGGTGNNNAGATCGGAAG |
| 45995_2 | TATGAGTGTGGAGTCGTTGCCGAATAGCAGGGACTACAGGTGNNNAGATCGGAAG |
| 33950_1 | TATGAGTGTGGAGTCGTTGCGATCTCCTGACCTCATGATCCACNNNAGATCGGAAG |
| 33950_2 | TATGAGTGTGGAGTCGTTGCGATCTCCTGACCTCATGATCCACNNNAGATCGGAAG |
| 42514_1 | TATGAGTGTGGAGTCGTTGCTGTCAGATTGAAGACACCAAAAGCNNNAGATCGGA |
| 42514_2 | TATGAGTGTGGAGTCGTTGCTGTCAGATTGAAGACACCAAAAGCNNNAGATCGGA |
| 38388_1 | TATGAGTGTGGAGTCGTTGCCTCAAGTGATCCTCCTGCCTTAGNNNAGATCGGAAG |
| 38388_2 | TATGAGTGTGGAGTCGTTGCCTCAAGTGATCCTCCTGCCTTAGNNNAGATCGGAAG |
| 39716_1 | TATGAGTGTGGAGTCGTTGCAGTCAGGATTGTTTCCGAAAGGTNNNAGATCGGAA |
| 39716_2 | TATGAGTGTGGAGTCGTTGCAGTCAGGATTGTTTCCGAAAGGTNNNAGATCGGAA |
| 46169_1 | TATGAGTGTGGAGTCGTTGCTTTTGGTAGACACGAGGTTTCACNNNAGATCGGAAG |
| 46169_2 | TATGAGTGTGGAGTCGTTGCTTTTGGTAGACACGAGGTTTCACNNNAGATCGGAAG |
| 46529_1 | TATGAGTGTGGAGTCGTTGCTCCTGAGTAGCTGGGATTACAGANNNAGATCGGAA |
| 46529_2 | TATGAGTGTGGAGTCGTTGCTCCTGAGTAGCTGGGATTACAGANNNAGATCGGAA |

| | |
|---|---|
| 36726_1 | TATGAGTGTGGAGTCGTTGCTTTGGGAGACGGCTTGATTTTTCNNNAGATCGGAAG |
| 36726_2 | TATGAGTGTGGAGTCGTTGCTTTGGGAGACGGCTTGATTTTTCNNNAGATCGGAAG |
| 50363_1 | TATGAGTGTGGAGTCGTTGCCAACACCAAGCAGATCTAATGCANNNAGATCGGAAG |
| 50363_2 | TATGAGTGTGGAGTCGTTGCCAACACCAAGCAGATCTAATGCANNNAGATCGGAAG |
| 5583_1 | TATGAGTGTGGAGTCGTTGCAACTGGAAATGTTGCACTAGGCANNNAGATCGGAA |
| 5583_2 | TATGAGTGTGGAGTCGTTGCAACTGGAAATGTTGCACTAGGCANNNAGATCGGAA |
| 44805_1 | TATGAGTGTGGAGTCGTTGCTGAGTGCTATGCTCTGGATTCTGNNNAGATCGGAAG |
| 44805_2 | TATGAGTGTGGAGTCGTTGCTGAGTGCTATGCTCTGGATTCTGNNNAGATCGGAAG |
| 42174_1 | TATGAGTGTGGAGTCGTTGCCAGCTCTCTTCTCCTCACACTTTNNNAGATCGGAAGA |
| 42174_2 | TATGAGTGTGGAGTCGTTGCCAGCTCTCTTCTCCTCACACTTTNNNAGATCGGAAGA |
| 36991_1 | TATGAGTGTGGAGTCGTTGCTTTGTGGTGATGGTACAGTCACTNNNAGATCGGAA |
| 36991_2 | TATGAGTGTGGAGTCGTTGCTTTGTGGTGATGGTACAGTCACTNNNAGATCGGAA |
| 42147_1 | TATGAGTGTGGAGTCGTTGCTGATCTTGTCTCACTGCAACCTCNNNAGATCGGAAG |
| 42147_2 | TATGAGTGTGGAGTCGTTGCTGATCTTGTCTCACTGCAACCTCNNNAGATCGGAAG |
| 9970_1 | TATGAGTGTGGAGTCGTTGCTTCTGCAGTTTTTGAACTTGGCTGNNNAGATCGGAA |
| 9970_2 | TATGAGTGTGGAGTCGTTGCTTCTGCAGTTTTTGAACTTGGCTGNNNAGATCGGAA |
| 36605_1 | TATGAGTGTGGAGTCGTTGCATGAATTCCAAGGTCTCTCTCGGNNNAGATCGGAAG |
| 36605_2 | TATGAGTGTGGAGTCGTTGCATGAATTCCAAGGTCTCTCTCGGNNNAGATCGGAAG |
| 52977_1 | TATGAGTGTGGAGTCGTTGCCAGTAGCACAATCACAGCTCACTNNNAGATCGGAAG |
| 52977_2 | TATGAGTGTGGAGTCGTTGCCAGTAGCACAATCACAGCTCACTNNNAGATCGGAAG |
| 45207_1 | TATGAGTGTGGAGTCGTTGCTTGCTAGCATCTTTATTGCCCCTNNNAGATCGGAAGA |
| 45207_2 | TATGAGTGTGGAGTCGTTGCTTGCTAGCATCTTTATTGCCCCTNNNAGATCGGAAGA |
| 45083_1 | TATGAGTGTGGAGTCGTTGCTGAGGACAAAGAACAGCTTCAGTNNNAGATCGGAA |
| 45083_2 | TATGAGTGTGGAGTCGTTGCTGAGGACAAAGAACAGCTTCAGTNNNAGATCGGAA |
| 30262_1 | TATGAGTGTGGAGTCGTTGCACTCCTACAATTGAGGGTTGTGTNNNAGATCGGAAG |
| 30262_2 | TATGAGTGTGGAGTCGTTGCACTCCTACAATTGAGGGTTGTGTNNNAGATCGGAAG |
| 29991_1 | TATGAGTGTGGAGTCGTTGCTCAGATTGGCTGCTGATTCTGATNNNAGATCGGAAG |
| 29991_2 | TATGAGTGTGGAGTCGTTGCTCAGATTGGCTGCTGATTCTGATNNNAGATCGGAAG |
| 2454_1 | TATGAGTGTGGAGTCGTTGCCCATCCTCTTGTCTCTGCAGTACNNNAGATCGGAAGA |
| 2454_2 | TATGAGTGTGGAGTCGTTGCCCATCCTCTTGTCTCTGCAGTACNNNAGATCGGAAGA |
| 48547_1 | TATGAGTGTGGAGTCGTTGCCCATCAGAAGCTCCACTATGGTCNNNAGATCGGAAG |
| 48547_2 | TATGAGTGTGGAGTCGTTGCCCATCAGAAGCTCCACTATGGTCNNNAGATCGGAAG |
| 45135_1 | TATGAGTGTGGAGTCGTTGCCTCTGGTCTTCATCATCACCACCNNNAGATCGGAAGA |
| 45135_2 | TATGAGTGTGGAGTCGTTGCCTCTGGTCTTCATCATCACCACCNNNAGATCGGAAGA |
| 43357_1 | TATGAGTGTGGAGTCGTTGCAGAAAACAGGCATTTGAGCTTCCNNNAGATCGGAA |
| 43357_2 | TATGAGTGTGGAGTCGTTGCAGAAAACAGGCATTTGAGCTTCCNNNAGATCGGAA |
| 51984_1 | TATGAGTGTGGAGTCGTTGCGCATGTCGCATTCCATTTGTTTCNNNAGATCGGAAG |
| 51984_2 | TATGAGTGTGGAGTCGTTGCGCATGTCGCATTCCATTTGTTTCNNNAGATCGGAAG |
| 31840_1 | TATGAGTGTGGAGTCGTTGCTGGAGTGCAGTGTTGTCATCATANNNAGATCGGAA |
| 31840_2 | TATGAGTGTGGAGTCGTTGCTGGAGTGCAGTGTTGTCATCATANNNAGATCGGAA |
| 45631_1 | TATGAGTGTGGAGTCGTTGCATGATCTCGGCTCACTGCAATCNNNAGATCGGAAGA |
| 45631_2 | TATGAGTGTGGAGTCGTTGCATGATCTCGGCTCACTGCAATCNNNAGATCGGAAGA |
| 43855_1 | TATGAGTGTGGAGTCGTTGCGCAACATCAATGAGGCTTCTAGGNNNAGATCGGAA |

| | |
|---|---|
| 43855_2 | TATGAGTGTGGAGTCGTTGCGCAACATCAATGAGGCTTCTAGGNNNAGATCGGAA |
| 47856_1 | TATGAGTGTGGAGTCGTTGCGGAATGAAGTGGCGCAATCATAGNNNAGATCGGAA |
| 47856_2 | TATGAGTGTGGAGTCGTTGCGGAATGAAGTGGCGCAATCATAGNNNAGATCGGAA |
| 51017_1 | TATGAGTGTGGAGTCGTTGCATTCCCCTTCTCCACATTCGTTCNNNAGATCGGAAGA |
| 51017_2 | TATGAGTGTGGAGTCGTTGCATTCCCCTTCTCCACATTCGTTCNNNAGATCGGAAGA |
| 38073_1 | TATGAGTGTGGAGTCGTTGCTCAGCACTGGATTGTAGACTTGTNNNAGATCGGAAG |
| 38073_2 | TATGAGTGTGGAGTCGTTGCTCAGCACTGGATTGTAGACTTGTNNNAGATCGGAAG |
| 35687_1 | TATGAGTGTGGAGTCGTTGCTCACTGCAGATTCATGATCCACANNNAGATCGGAAG |
| 35687_2 | TATGAGTGTGGAGTCGTTGCTCACTGCAGATTCATGATCCACANNNAGATCGGAAG |
| 48804_1 | TATGAGTGTGGAGTCGTTGCGAGTAGCTGGGATTATAGGTGCCNNNAGATCGGAA |
| 48804_2 | TATGAGTGTGGAGTCGTTGCGAGTAGCTGGGATTATAGGTGCCNNNAGATCGGAA |
| 53149_1 | TATGAGTGTGGAGTCGTTGCTTTTGTTTCTGAAAATGTAGGGGTNNNAGATCGGAA |
| 53149_2 | TATGAGTGTGGAGTCGTTGCTTTTGTTTCTGAAAATGTAGGGGTNNNAGATCGGAA |
| 41222_1 | TATGAGTGTGGAGTCGTTGCATCTTTGAAAGAGGAGGCTGAGGNNNAGATCGGAA |
| 41222_2 | TATGAGTGTGGAGTCGTTGCATCTTTGAAAGAGGAGGCTGAGGNNNAGATCGGAA |
| 46995_1 | TATGAGTGTGGAGTCGTTGCCCACAGTACCCGAGATTACATGTNNNAGATCGGAAG |
| 46995_2 | TATGAGTGTGGAGTCGTTGCCCACAGTACCCGAGATTACATGTNNNAGATCGGAAG |
| 13048_1 | TATGAGTGTGGAGTCGTTGCCTTTCTCCTGCTGCTTATTTNNNAGATCGGAAGAGCA |
| 13048_2 | TATGAGTGTGGAGTCGTTGCCTTTCTCCTGCTGCTTATTTNNNAGATCGGAAGAGCA |
| 9952_1 | TATGAGTGTGGAGTCGTTGCTCTGATCATTCACCATGGCAGTGNNNAGATCGGAAG |
| 9952_2 | TATGAGTGTGGAGTCGTTGCTCTGATCATTCACCATGGCAGTGNNNAGATCGGAAG |
| 36746_1 | TATGAGTGTGGAGTCGTTGCTGGATTTTCAGGTCCTGGAGATGNNNAGATCGGAA |
| 36746_2 | TATGAGTGTGGAGTCGTTGCTGGATTTTCAGGTCCTGGAGATGNNNAGATCGGAA |
| 13521_1 | TATGAGTGTGGAGTCGTTGCAAAACAAAACAAGGGTCACTTNNNAGATCGGAAGA |
| 13521_2 | TATGAGTGTGGAGTCGTTGCAAAACAAAACAAGGGTCACTTNNNAGATCGGAAGA |
| 29676_1 | TATGAGTGTGGAGTCGTTGCCCCCACTTCCTGATTTTCCTTGANNNAGATCGGAAGA |
| 29676_2 | TATGAGTGTGGAGTCGTTGCCCCCACTTCCTGATTTTCCTTGANNNAGATCGGAAGA |
| 32444_1 | TATGAGTGTGGAGTCGTTGCTTTGAAGTTGGGTAGCATGATGCNNNAGATCGGAA |
| 32444_2 | TATGAGTGTGGAGTCGTTGCTTTGAAGTTGGGTAGCATGATGCNNNAGATCGGAA |
| 34222_1 | TATGAGTGTGGAGTCGTTGCGATGTGGATTCCAAGGTCCTTCTNNNAGATCGGAAG |
| 34222_2 | TATGAGTGTGGAGTCGTTGCGATGTGGATTCCAAGGTCCTTCTNNNAGATCGGAAG |
| 33912_1 | TATGAGTGTGGAGTCGTTGCCCAAGTAGCTGGGACGACAGNNNAGATCGGAAGAG |
| 33912_2 | TATGAGTGTGGAGTCGTTGCCCAAGTAGCTGGGACGACAGNNNAGATCGGAAGAG |
| 42080_1 | TATGAGTGTGGAGTCGTTGCTCCAGAAGAGGCAAATTCATGGANNNAGATCGGAA |
| 42080_2 | TATGAGTGTGGAGTCGTTGCTCCAGAAGAGGCAAATTCATGGANNNAGATCGGAA |
| 32519_1 | TATGAGTGTGGAGTCGTTGCTTAGCGTGTGTATTTCTGTGGGTNNNAGATCGGAA |
| 32519_2 | TATGAGTGTGGAGTCGTTGCTTAGCGTGTGTATTTCTGTGGGTNNNAGATCGGAA |
| 46711_1 | TATGAGTGTGGAGTCGTTGCTTCACACCATTCTCCTGCCTTAGNNNAGATCGGAAGA |
| 46711_2 | TATGAGTGTGGAGTCGTTGCTTCACACCATTCTCCTGCCTTAGNNNAGATCGGAAGA |
| 53126_1 | TATGAGTGTGGAGTCGTTGCAACGTTTTGTGTAGTGCTTTTCTNNNAGATCGGAAG |
| 53126_2 | TATGAGTGTGGAGTCGTTGCAACGTTTTGTGTAGTGCTTTTCTNNNAGATCGGAAG |
| 36519_1 | TATGAGTGTGGAGTCGTTGCTGTGGAAGGGAACATCCATCTTCNNNAGATCGGAA |
| 36519_2 | TATGAGTGTGGAGTCGTTGCTGTGGAAGGGAACATCCATCTTCNNNAGATCGGAA |

| | |
|---|---|
| 36237_1 | TATGAGTGTGGAGTCGTTGCAAAAACCACAAAGTCCCACTGTGNNNAGATCGGAAG |
| 36237_2 | TATGAGTGTGGAGTCGTTGCAAAAACCACAAAGTCCCACTGTGNNNAGATCGGAAG |
| 30222_1 | TATGAGTGTGGAGTCGTTGCCCTAAAGCACTCTTCCCTGTGGNNNAGATCGGAAGA |
| 30222_2 | TATGAGTGTGGAGTCGTTGCCCTAAAGCACTCTTCCCTGTGGNNNAGATCGGAAGA |
| 43225_1 | TATGAGTGTGGAGTCGTTGCCTAGGATTATAGACGTGGGCCACNNNAGATCGGAA |
| 43225_2 | TATGAGTGTGGAGTCGTTGCCTAGGATTATAGACGTGGGCCACNNNAGATCGGAA |
| 44848_1 | TATGAGTGTGGAGTCGTTGCCCACCAAAACTCATGTTGATGCTNNNAGATCGGAAG |
| 44848_2 | TATGAGTGTGGAGTCGTTGCCCACCAAAACTCATGTTGATGCTNNNAGATCGGAAG |
| 12525_1 | TATGAGTGTGGAGTCGTTGCGAGTTATTTGGCAGTAGATTTTTNNNAGATCGGAAG |
| 12525_2 | TATGAGTGTGGAGTCGTTGCGAGTTATTTGGCAGTAGATTTTTNNNAGATCGGAAG |
| 46186_1 | TATGAGTGTGGAGTCGTTGCGAGAGAGGATCTGGCTCTGTTGNNNAGATCGGAAG |
| 46186_2 | TATGAGTGTGGAGTCGTTGCGAGAGAGGATCTGGCTCTGTTGNNNAGATCGGAAG |
| 43660_1 | TATGAGTGTGGAGTCGTTGCCAAGGAGGCATACATCTTGCAAGNNNAGATCGGAA |
| 43660_2 | TATGAGTGTGGAGTCGTTGCCAAGGAGGCATACATCTTGCAAGNNNAGATCGGAA |
| 51708_1 | TATGAGTGTGGAGTCGTTGCAACCCTTTTTGCTGTAAGTGAGCNNNAGATCGGAAG |
| 51708_2 | TATGAGTGTGGAGTCGTTGCAACCCTTTTTGCTGTAAGTGAGCNNNAGATCGGAAG |
| 37821_1 | TATGAGTGTGGAGTCGTTGCGATGGACGAAATCGCTTGAAGAGNNNAGATCGGAA |
| 37821_2 | TATGAGTGTGGAGTCGTTGCGATGGACGAAATCGCTTGAAGAGNNNAGATCGGAA |
| 44264_1 | TATGAGTGTGGAGTCGTTGCAAGCAAGGACATGGCAAGAATTCNNNAGATCGGAA |
| 44264_2 | TATGAGTGTGGAGTCGTTGCAAGCAAGGACATGGCAAGAATTCNNNAGATCGGAA |
| 30483_1 | TATGAGTGTGGAGTCGTTGCACTCCTAGTAAACAAGTATGCACANNNAGATCGGAA |
| 30483_2 | TATGAGTGTGGAGTCGTTGCACTCCTAGTAAACAAGTATGCACANNNAGATCGGAA |
| 13462_1 | TATGAGTGTGGAGTCGTTGCTGCTATGGTAAGATAACCTCTTNNNAGATCGGAAGA |
| 13462_2 | TATGAGTGTGGAGTCGTTGCTGCTATGGTAAGATAACCTCTTNNNAGATCGGAAGA |
| 48343_1 | TATGAGTGTGGAGTCGTTGCATTGAGTTTTGCTCTTGTTGCCCNNNAGATCGGAAG |
| 48343_2 | TATGAGTGTGGAGTCGTTGCATTGAGTTTTGCTCTTGTTGCCCNNNAGATCGGAAG |
| 38317_1 | TATGAGTGTGGAGTCGTTGCAGAAGGGAACTTCCTTGAGCTAANNNAGATCGGAA |
| 38317_2 | TATGAGTGTGGAGTCGTTGCAGAAGGGAACTTCCTTGAGCTAANNNAGATCGGAA |
| 41365_1 | TATGAGTGTGGAGTCGTTGCTAGAGTTAGCCCCATAGACAGCTNNNAGATCGGAAG |
| 41365_2 | TATGAGTGTGGAGTCGTTGCTAGAGTTAGCCCCATAGACAGCTNNNAGATCGGAAG |
| 46281_1 | TATGAGTGTGGAGTCGTTGCGTTCCCAGCCTATTTCCCTTCTTNNNAGATCGGAAGA |
| 46281_2 | TATGAGTGTGGAGTCGTTGCGTTCCCAGCCTATTTCCCTTCTTNNNAGATCGGAAGA |
| 45639_1 | TATGAGTGTGGAGTCGTTGCGGGAGGACTGGAAAAGGAGTAAGNNNAGATCGGA |
| 45639_2 | TATGAGTGTGGAGTCGTTGCGGGAGGACTGGAAAAGGAGTAAGNNNAGATCGGA |
| 48438_1 | TATGAGTGTGGAGTCGTTGCAAGAGCTGGGATTACAGATCACGNNNAGATCGGAA |
| 48438_2 | TATGAGTGTGGAGTCGTTGCAAGAGCTGGGATTACAGATCACGNNNAGATCGGAA |
| 41267_1 | TATGAGTGTGGAGTCGTTGCCTATGCGTCTTCTGCAAAGCAATNNNAGATCGGAAG |
| 41267_2 | TATGAGTGTGGAGTCGTTGCCTATGCGTCTTCTGCAAAGCAATNNNAGATCGGAAG |
| 13515_1 | TATGAGTGTGGAGTCGTTGCGAAAAATAAACATGCACGATTATNNNAGATCGGAA |
| 13515_2 | TATGAGTGTGGAGTCGTTGCGAAAAATAAACATGCACGATTATNNNAGATCGGAA |
| 115_1 | TATGAGTGTGGAGTCGTTGCGCGGTTGAGTTTTGCATATGAACCNNNAGATCGGA |
| 115_2 | TATGAGTGTGGAGTCGTTGCGCGGTTGAGTTTTGCATATGAACCNNNAGATCGGA |
| 7892_1 | TATGAGTGTGGAGTCGTTGCTAACAATTTGTGCTCATAGGCAAGNNNAGATCGGAA |

| | |
|---|---|
| 7892_2 | TATGAGTGTGGAGTCGTTGCTAACAATTTGTGCTCATAGGCAAGNNNAGATCGGAA |
| 44802_1 | TATGAGTGTGGAGTCGTTGCAGAGACTATGAAGTGGAGGTGGANNNAGATCGGA |
| 44802_2 | TATGAGTGTGGAGTCGTTGCAGAGACTATGAAGTGGAGGTGGANNNAGATCGGA |
| 32766_1 | TATGAGTGTGGAGTCGTTGCAACCAAACCAATAATATGCATGCTNNNAGATCGGAA |
| 32766_2 | TATGAGTGTGGAGTCGTTGCAACCAAACCAATAATATGCATGCTNNNAGATCGGAA |
| 52018_1 | TATGAGTGTGGAGTCGTTGCACTAGCACCCCTATTATTCCCCTNNNAGATCGGAAGA |
| 52018_2 | TATGAGTGTGGAGTCGTTGCACTAGCACCCCTATTATTCCCCTNNNAGATCGGAAGA |
| 45012_1 | TATGAGTGTGGAGTCGTTGCCTCTGTTTCTTCCTAAGCCCTGTNNNAGATCGGAAGA |
| 45012_2 | TATGAGTGTGGAGTCGTTGCCTCTGTTTCTTCCTAAGCCCTGTNNNAGATCGGAAGA |
| 50726_1 | TATGAGTGTGGAGTCGTTGCGAACCCTCACACCTTTGCTCAGNNNAGATCGGAAGA |
| 50726_2 | TATGAGTGTGGAGTCGTTGCGAACCCTCACACCTTTGCTCAGNNNAGATCGGAAGA |
| 39806_1 | TATGAGTGTGGAGTCGTTGCGAGCTTTGGAACACACTGGTTTTNNNAGATCGGAAG |
| 39806_2 | TATGAGTGTGGAGTCGTTGCGAGCTTTGGAACACACTGGTTTTNNNAGATCGGAAG |
| 41973_1 | TATGAGTGTGGAGTCGTTGCGAGTAGCTGGGATTACAGGCACNNNAGATCGGAAG |
| 41973_2 | TATGAGTGTGGAGTCGTTGCGAGTAGCTGGGATTACAGGCACNNNAGATCGGAAG |
| 43381_1 | TATGAGTGTGGAGTCGTTGCTTTTTTTTGACGGAGTCTTGCTCNNNAGATCGGAAG |
| 43381_2 | TATGAGTGTGGAGTCGTTGCTTTTTTTTGACGGAGTCTTGCTCNNNAGATCGGAAG |
| 9213_1 | TATGAGTGTGGAGTCGTTGCTTCAGAGAGCCATCCAACCATTGNNNAGATCGGAAG |
| 9213_2 | TATGAGTGTGGAGTCGTTGCTTCAGAGAGCCATCCAACCATTGNNNAGATCGGAAG |
| 38996_1 | TATGAGTGTGGAGTCGTTGCGAACCCAAGGAAAGAAGATAGGCNNNAGATCGGAA |
| 38996_2 | TATGAGTGTGGAGTCGTTGCGAACCCAAGGAAAGAAGATAGGCNNNAGATCGGAA |
| 41768_1 | TATGAGTGTGGAGTCGTTGCGGAGAGGGCATCTTCTTCTTTTTNNNAGATCGGAAG |
| 41768_2 | TATGAGTGTGGAGTCGTTGCGGAGAGGGCATCTTCTTCTTTTTNNNAGATCGGAAG |
| 48412_1 | TATGAGTGTGGAGTCGTTGCGTAGAACAGTACTTGCTTTGGGCNNNAGATCGGAA |
| 48412_2 | TATGAGTGTGGAGTCGTTGCGTAGAACAGTACTTGCTTTGGGCNNNAGATCGGAA |
| 39812_1 | TATGAGTGTGGAGTCGTTGCACAGGCCTATCCATTTTTCTCGTNNNAGATCGGAAGA |
| 39812_2 | TATGAGTGTGGAGTCGTTGCACAGGCCTATCCATTTTTCTCGTNNNAGATCGGAAGA |
| 42800_1 | TATGAGTGTGGAGTCGTTGCCAAGTGAGATTTACCCCTGGGATNNNAGATCGGAA |
| 42800_2 | TATGAGTGTGGAGTCGTTGCCAAGTGAGATTTACCCCTGGGATNNNAGATCGGAA |
| 32157_1 | TATGAGTGTGGAGTCGTTGCAAAAAAGAGGGGCTTTGTACCACNNNAGATCGGAA |
| 32157_2 | TATGAGTGTGGAGTCGTTGCAAAAAAGAGGGGCTTTGTACCACNNNAGATCGGAA |
| 43456_1 | TATGAGTGTGGAGTCGTTGCAATCTCTCAGGCTCAATGGATCCNNNAGATCGGAAG |
| 43456_2 | TATGAGTGTGGAGTCGTTGCAATCTCTCAGGCTCAATGGATCCNNNAGATCGGAAG |
| 39170_1 | TATGAGTGTGGAGTCGTTGCCAGGAGTGATAGCTTATGCCTGTNNNAGATCGGAA |
| 39170_2 | TATGAGTGTGGAGTCGTTGCCAGGAGTGATAGCTTATGCCTGTNNNAGATCGGAA |
| 33712_1 | TATGAGTGTGGAGTCGTTGCCTGGGAAACATAGTGAGAACCCANNNAGATCGGAA |
| 33712_2 | TATGAGTGTGGAGTCGTTGCCTGGGAAACATAGTGAGAACCCANNNAGATCGGAA |
| 37490_1 | TATGAGTGTGGAGTCGTTGCATAAACGTGGCTTAGTGTGGCTANNNAGATCGGAA |
| 37490_2 | TATGAGTGTGGAGTCGTTGCATAAACGTGGCTTAGTGTGGCTANNNAGATCGGAA |
| 50493_1 | TATGAGTGTGGAGTCGTTGCCACCAGGCTAAACAAAAGACTCGNNNAGATCGGAA |
| 50493_2 | TATGAGTGTGGAGTCGTTGCCACCAGGCTAAACAAAAGACTCGNNNAGATCGGAA |
| 49973_1 | TATGAGTGTGGAGTCGTTGCAACCTCCCAATACTCATCGAGTGNNNAGATCGGAAG |
| 49973_2 | TATGAGTGTGGAGTCGTTGCAACCTCCCAATACTCATCGAGTGNNNAGATCGGAAG |

| | |
|---|---|
| 47863_1 | TATGAGTGTGGAGTCGTTGCAGATGGAGTCTCGTTCTGTTGCNNNAGATCGGAAG |
| 47863_2 | TATGAGTGTGGAGTCGTTGCAGATGGAGTCTCGTTCTGTTGCNNNAGATCGGAAG |
| 37526_1 | TATGAGTGTGGAGTCGTTGCTGTTCATGCCACTGTACTCTAGCNNNAGATCGGAAG |
| 37526_2 | TATGAGTGTGGAGTCGTTGCTGTTCATGCCACTGTACTCTAGCNNNAGATCGGAAG |
| 12708_1 | TATGAGTGTGGAGTCGTTGCTTTTTGACCATACCTGAGTATAANNNAGATCGGAAG |
| 12708_2 | TATGAGTGTGGAGTCGTTGCTTTTTGACCATACCTGAGTATAANNNAGATCGGAAG |
| 31574_1 | TATGAGTGTGGAGTCGTTGCGAAGAAAGACAGAGAGCATGCACNNNAGATCGGAA |
| 31574_2 | TATGAGTGTGGAGTCGTTGCGAAGAAAGACAGAGAGCATGCACNNNAGATCGGAA |
| 31250_1 | TATGAGTGTGGAGTCGTTGCTTTTGCAAATGTTTTACCTCGCTNNNAGATCGGAAGA |
| 31250_2 | TATGAGTGTGGAGTCGTTGCTTTTGCAAATGTTTTACCTCGCTNNNAGATCGGAAGA |
| 43554_1 | TATGAGTGTGGAGTCGTTGCCCTCCTGGGTTCAAGGAATTCTCNNNAGATCGGAAG |
| 43554_2 | TATGAGTGTGGAGTCGTTGCCCTCCTGGGTTCAAGGAATTCTCNNNAGATCGGAAG |
| 44294_1 | TATGAGTGTGGAGTCGTTGCCTGGGTATTGGTTGTCTCTTGGANNNAGATCGGAAG |
| 44294_2 | TATGAGTGTGGAGTCGTTGCCTGGGTATTGGTTGTCTCTTGGANNNAGATCGGAAG |
| 52567_1 | TATGAGTGTGGAGTCGTTGCGAGCACTGTGCAGATCCTGTATANNNAGATCGGAAG |
| 52567_2 | TATGAGTGTGGAGTCGTTGCGAGCACTGTGCAGATCCTGTATANNNAGATCGGAAG |
| 45884_1 | TATGAGTGTGGAGTCGTTGCTCGGCCTTCAAAAGTTCTAGGATNNNAGATCGGAAG |
| 45884_2 | TATGAGTGTGGAGTCGTTGCTCGGCCTTCAAAAGTTCTAGGATNNNAGATCGGAAG |
| 46378_1 | TATGAGTGTGGAGTCGTTGCAAAGCAGGCACTACCACTTTCTANNNAGATCGGAAG |
| 46378_2 | TATGAGTGTGGAGTCGTTGCAAAGCAGGCACTACCACTTTCTANNNAGATCGGAAG |
| 47716_1 | TATGAGTGTGGAGTCGTTGCGCTGGCCTAAGGTCAAATGAATCNNNAGATCGGAAG |
| 47716_2 | TATGAGTGTGGAGTCGTTGCGCTGGCCTAAGGTCAAATGAATCNNNAGATCGGAAG |
| 47574_1 | TATGAGTGTGGAGTCGTTGCGGCAGTTGGAGAAGAATTTGGAANNNAGATCGGAA |
| 47574_2 | TATGAGTGTGGAGTCGTTGCGGCAGTTGGAGAAGAATTTGGAANNNAGATCGGAA |
| 49999_1 | TATGAGTGTGGAGTCGTTGCGACAACACCTTCCCTAGTTTCCTNNNAGATCGGAAGA |
| 49999_2 | TATGAGTGTGGAGTCGTTGCGACAACACCTTCCCTAGTTTCCTNNNAGATCGGAAGA |
| 46195_1 | TATGAGTGTGGAGTCGTTGCCTCCTGAGTAGCTGGGACTACAGNNNAGATCGGAAG |
| 46195_2 | TATGAGTGTGGAGTCGTTGCCTCCTGAGTAGCTGGGACTACAGNNNAGATCGGAAG |
| 53183_1 | TATGAGTGTGGAGTCGTTGCTATGCTATGTCCCCTTTTCTGCANNNAGATCGGAAGA |
| 53183_2 | TATGAGTGTGGAGTCGTTGCTATGCTATGTCCCCTTTTCTGCANNNAGATCGGAAGA |
| 13708_1 | TATGAGTGTGGAGTCGTTGCTCATGCTCTGATTAGATAATCTTNNNAGATCGGAAG |
| 13708_2 | TATGAGTGTGGAGTCGTTGCTCATGCTCTGATTAGATAATCTTNNNAGATCGGAAG |
| 10286_1 | TATGAGTGTGGAGTCGTTGCTGGGTCCTTGTTTGTTTACATTCANNNAGATCGGAAG |
| 10286_2 | TATGAGTGTGGAGTCGTTGCTGGGTCCTTGTTTGTTTACATTCANNNAGATCGGAAG |
| 45267_1 | TATGAGTGTGGAGTCGTTGCACTCAGAGATACTACCAGGGGACNNNAGATCGGAAG |
| 45267_2 | TATGAGTGTGGAGTCGTTGCACTCAGAGATACTACCAGGGGACNNNAGATCGGAAG |
| 43622_1 | TATGAGTGTGGAGTCGTTGCGTGTTTTCTCTGAAGCCGTTTGANNNAGATCGGAAG |
| 43622_2 | TATGAGTGTGGAGTCGTTGCGTGTTTTCTCTGAAGCCGTTTGANNNAGATCGGAAG |
| 47252_1 | TATGAGTGTGGAGTCGTTGCCACTTGTTTCCCAGGTCAAAGTGNNNAGATCGGAAG |
| 47252_2 | TATGAGTGTGGAGTCGTTGCCACTTGTTTCCCAGGTCAAAGTGNNNAGATCGGAAG |
| 42410_1 | TATGAGTGTGGAGTCGTTGCCTAAAGCTGGGGTGGAGTTACAGNNNAGATCGGAA |
| 42410_2 | TATGAGTGTGGAGTCGTTGCCTAAAGCTGGGGTGGAGTTACAGNNNAGATCGGAA |
| 11293_1 | TATGAGTGTGGAGTCGTTGCCTCCCAGAGCCAGATTATAGGGGGNNNAGATCGGA |

| | |
|---|---|
| 11293_2 | TATGAGTGTGGAGTCGTTGCCTCCCAGAGCCAGATTATAGGGGGNNNAGATCGGA |
| 32517_1 | TATGAGTGTGGAGTCGTTGCGCCATATATGACAGACCCACAGTNNNAGATCGGAAG |
| 32517_2 | TATGAGTGTGGAGTCGTTGCGCCATATATGACAGACCCACAGTNNNAGATCGGAAG |
| 39629_1 | TATGAGTGTGGAGTCGTTGCGCAAGGTAGAGCCATACTAGTCCNNNAGATCGGAA |
| 39629_2 | TATGAGTGTGGAGTCGTTGCGCAAGGTAGAGCCATACTAGTCCNNNAGATCGGAA |
| 45803_1 | TATGAGTGTGGAGTCGTTGCTCTCCTGGACTCAAGTGATCCTCNNNAGATCGGAAG |
| 45803_2 | TATGAGTGTGGAGTCGTTGCTCTCCTGGACTCAAGTGATCCTCNNNAGATCGGAAG |
| 37963_1 | TATGAGTGTGGAGTCGTTGCAATGCTTGAGGAGATGCATACCTNNNAGATCGGAA |
| 37963_2 | TATGAGTGTGGAGTCGTTGCAATGCTTGAGGAGATGCATACCTNNNAGATCGGAA |
| 52859_1 | TATGAGTGTGGAGTCGTTGCATAGTCTCTGCTGTGCTGTTGAANNNAGATCGGAAG |
| 52859_2 | TATGAGTGTGGAGTCGTTGCATAGTCTCTGCTGTGCTGTTGAANNNAGATCGGAAG |
| 35928_1 | TATGAGTGTGGAGTCGTTGCTACTCAGAAAACGGTCCAACAGTNNNAGATCGGAAG |
| 35928_2 | TATGAGTGTGGAGTCGTTGCTACTCAGAAAACGGTCCAACAGTNNNAGATCGGAAG |
| 36617_1 | TATGAGTGTGGAGTCGTTGCGAAGGATGGGAAAAGAGCCAAACNNNAGATCGGA |
| 36617_2 | TATGAGTGTGGAGTCGTTGCGAAGGATGGGAAAAGAGCCAAACNNNAGATCGGA |
| 31455_1 | TATGAGTGTGGAGTCGTTGCAGTCAGCTAATGATGAGGTCCACNNNAGATCGGAA |
| 31455_2 | TATGAGTGTGGAGTCGTTGCAGTCAGCTAATGATGAGGTCCACNNNAGATCGGAA |
| 37860_1 | TATGAGTGTGGAGTCGTTGCCCTGTAATCCCAACACCTTGAGANNNAGATCGGAAG |
| 37860_2 | TATGAGTGTGGAGTCGTTGCCCTGTAATCCCAACACCTTGAGANNNAGATCGGAAG |
| 48600_1 | TATGAGTGTGGAGTCGTTGCCCAAGCAATCCTCCTACCTCAGNNNAGATCGGAAGA |
| 48600_2 | TATGAGTGTGGAGTCGTTGCCCAAGCAATCCTCCTACCTCAGNNNAGATCGGAAGA |
| 40183_1 | TATGAGTGTGGAGTCGTTGCCTATTATCAGACCAGCATGGGGGNNNAGATCGGAA |
| 40183_2 | TATGAGTGTGGAGTCGTTGCCTATTATCAGACCAGCATGGGGGNNNAGATCGGAA |
| 52300_1 | TATGAGTGTGGAGTCGTTGCCCTCCCACCTCCCCTAATGGNNNAGATCGGAAGAGC |
| 52300_2 | TATGAGTGTGGAGTCGTTGCCCTCCCACCTCCCCTAATGGNNNAGATCGGAAGAGC |
| 49052_1 | TATGAGTGTGGAGTCGTTGCGCAGAGGCTTCAGGGTACTAATTNNNAGATCGGAA |
| 49052_2 | TATGAGTGTGGAGTCGTTGCGCAGAGGCTTCAGGGTACTAATTNNNAGATCGGAA |
| 13633_1 | TATGAGTGTGGAGTCGTTGCCAAAATGGAATTTGTTTTCAGAGNNNAGATCGGAA |
| 13633_2 | TATGAGTGTGGAGTCGTTGCCAAAATGGAATTTGTTTTCAGAGNNNAGATCGGAA |
| 51447_1 | TATGAGTGTGGAGTCGTTGCGGAGGAAAGGAAGGAAGAGTGAGNNNAGATCGG |
| 51447_2 | TATGAGTGTGGAGTCGTTGCGGAGGAAAGGAAGGAAGAGTGAGNNNAGATCGG |
| 53021_1 | TATGAGTGTGGAGTCGTTGCTAAGGACAGCAGCTCTCATTCAGNNNAGATCGGAAG |
| 53021_2 | TATGAGTGTGGAGTCGTTGCTAAGGACAGCAGCTCTCATTCAGNNNAGATCGGAAG |
| 10261_1 | TATGAGTGTGGAGTCGTTGCTTCTGTTCAAGAGCCTGTGGAAGNNNAGATCGGAA |
| 10261_2 | TATGAGTGTGGAGTCGTTGCTTCTGTTCAAGAGCCTGTGGAAGNNNAGATCGGAA |
| 40423_1 | TATGAGTGTGGAGTCGTTGCCCAGTTTTGTGCTTGAAACCCAGNNNAGATCGGAAG |
| 40423_2 | TATGAGTGTGGAGTCGTTGCCCAGTTTTGTGCTTGAAACCCAGNNNAGATCGGAAG |
| 47534_1 | TATGAGTGTGGAGTCGTTGCCTCAATCTTGGCTCACTTCAACCNNNAGATCGGAAGA |
| 47534_2 | TATGAGTGTGGAGTCGTTGCCTCAATCTTGGCTCACTTCAACCNNNAGATCGGAAGA |
| 45978_1 | TATGAGTGTGGAGTCGTTGCCAAGTAGCTGGGACTATGGGCNNNAGATCGGAAGA |
| 45978_2 | TATGAGTGTGGAGTCGTTGCCAAGTAGCTGGGACTATGGGCNNNAGATCGGAAGA |
| 10476_1 | TATGAGTGTGGAGTCGTTGCATGCAAGAGAAAGACAAGCACCANNNAGATCGGAA |
| 10476_2 | TATGAGTGTGGAGTCGTTGCATGCAAGAGAAAGACAAGCACCANNNAGATCGGAA |

| | |
|---|---|
| 49742_1 | TATGAGTGTGGAGTCGTTGCGAGATGGGAGTCTCACTCTGTTGNNNAGATCGGAA |
| 49742_2 | TATGAGTGTGGAGTCGTTGCGAGATGGGAGTCTCACTCTGTTGNNNAGATCGGAA |
| 45143_1 | TATGAGTGTGGAGTCGTTGCGGTAGGTTCTCACTGTCCAATGTNNNAGATCGGAAG |
| 45143_2 | TATGAGTGTGGAGTCGTTGCGGTAGGTTCTCACTGTCCAATGTNNNAGATCGGAAG |
| 29803_1 | TATGAGTGTGGAGTCGTTGCGCCATTTCTGCATTTTTCTCCAANNNAGATCGGAAGA |
| 29803_2 | TATGAGTGTGGAGTCGTTGCGCCATTTCTGCATTTTTCTCCAANNNAGATCGGAAGA |
| 38294_1 | TATGAGTGTGGAGTCGTTGCTTTCCTACTCCCCCCACTAAGTANNNAGATCGGAAGA |
| 38294_2 | TATGAGTGTGGAGTCGTTGCTTTCCTACTCCCCCCACTAAGTANNNAGATCGGAAGA |
| 46832_1 | TATGAGTGTGGAGTCGTTGCGCTGGAGTACAGTGACATGATCTNNNAGATCGGAA |
| 46832_2 | TATGAGTGTGGAGTCGTTGCGCTGGAGTACAGTGACATGATCTNNNAGATCGGAA |
| 37453_1 | TATGAGTGTGGAGTCGTTGCACGTTTTATCCTGCACTTTGGTGNNNAGATCGGAAG |
| 37453_2 | TATGAGTGTGGAGTCGTTGCACGTTTTATCCTGCACTTTGGTGNNNAGATCGGAAG |
| 33944_1 | TATGAGTGTGGAGTCGTTGCCCGCCTGGCTAATTTTTGTACTTNNNAGATCGGAAG |
| 33944_2 | TATGAGTGTGGAGTCGTTGCCCGCCTGGCTAATTTTTGTACTTNNNAGATCGGAAG |
| 49812_1 | TATGAGTGTGGAGTCGTTGCGCGTATAAGCTGACCTAGACCATNNNAGATCGGAAG |
| 49812_2 | TATGAGTGTGGAGTCGTTGCGCGTATAAGCTGACCTAGACCATNNNAGATCGGAAG |
| 38158_1 | TATGAGTGTGGAGTCGTTGCACCGAGTGCAAGAAGTTTTACACNNNAGATCGGAA |
| 38158_2 | TATGAGTGTGGAGTCGTTGCACCGAGTGCAAGAAGTTTTACACNNNAGATCGGAA |
| 39338_1 | TATGAGTGTGGAGTCGTTGCCCCACTCCTTGTTATACACCCAANNNAGATCGGAAGA |
| 39338_2 | TATGAGTGTGGAGTCGTTGCCCCACTCCTTGTTATACACCCAANNNAGATCGGAAGA |
| 33946_1 | TATGAGTGTGGAGTCGTTGCCCTCCTAGGTTCAAGCGATTCTCNNNAGATCGGAAG |
| 33946_2 | TATGAGTGTGGAGTCGTTGCCCTCCTAGGTTCAAGCGATTCTCNNNAGATCGGAAG |
| 40005_1 | TATGAGTGTGGAGTCGTTGCCAGATCATGAGGTCAGGAGATCGNNNAGATCGGAA |
| 40005_2 | TATGAGTGTGGAGTCGTTGCCAGATCATGAGGTCAGGAGATCGNNNAGATCGGAA |
| 29877_1 | TATGAGTGTGGAGTCGTTGCCCACAACACGTTACTGCAATTCTNNNAGATCGGAAG |
| 29877_2 | TATGAGTGTGGAGTCGTTGCCCACAACACGTTACTGCAATTCTNNNAGATCGGAAG |
| 2158_1 | TATGAGTGTGGAGTCGTTGCACTTCCAAAATCCAGAAGGAAGANNNAGATCGGAA |
| 2158_2 | TATGAGTGTGGAGTCGTTGCACTTCCAAAATCCAGAAGGAAGANNNAGATCGGAA |
| 40155_1 | TATGAGTGTGGAGTCGTTGCGAAGTCAGGAGTTCTAGACCAGCNNNAGATCGGAA |
| 40155_2 | TATGAGTGTGGAGTCGTTGCGAAGTCAGGAGTTCTAGACCAGCNNNAGATCGGAA |
| 51365_1 | TATGAGTGTGGAGTCGTTGCTTCCATCTTGCAAGGTGTGAAACNNNAGATCGGAAG |
| 51365_2 | TATGAGTGTGGAGTCGTTGCTTCCATCTTGCAAGGTGTGAAACNNNAGATCGGAAG |
| 43455_1 | TATGAGTGTGGAGTCGTTGCATTCTTCTGCTTCAGCTTCCTGANNNAGATCGGAAGA |
| 43455_2 | TATGAGTGTGGAGTCGTTGCATTCTTCTGCTTCAGCTTCCTGANNNAGATCGGAAGA |
| 48569_1 | TATGAGTGTGGAGTCGTTGCATTTTAGCGATTCTCCTGCCTCANNNAGATCGGAAG |
| 48569_2 | TATGAGTGTGGAGTCGTTGCATTTTAGCGATTCTCCTGCCTCANNNAGATCGGAAG |
| 43323_1 | TATGAGTGTGGAGTCGTTGCCCTACCAAAGTGCTGGGATTACANNNAGATCGGAAG |
| 43323_2 | TATGAGTGTGGAGTCGTTGCCCTACCAAAGTGCTGGGATTACANNNAGATCGGAAG |
| 35000_1 | TATGAGTGTGGAGTCGTTGCTTGAGACGTAGTCTTGCTCTGTCNNNAGATCGGAAG |
| 35000_2 | TATGAGTGTGGAGTCGTTGCTTGAGACGTAGTCTTGCTCTGTCNNNAGATCGGAAG |
| 29545_1 | TATGAGTGTGGAGTCGTTGCGCTATTTGCAGAGGGGACCTAACNNNAGATCGGAA |
| 29545_2 | TATGAGTGTGGAGTCGTTGCGCTATTTGCAGAGGGGACCTAACNNNAGATCGGAA |
| 30144_1 | TATGAGTGTGGAGTCGTTGCCTTGCAGTTGTTGTTGCTCACATNNNAGATCGGAAG |

| | |
|---|---|
| 30144_2 | TATGAGTGTGGAGTCGTTGCCTTGCAGTTGTTGTTGCTCACATNNNAGATCGGAAG |
| 37933_1 | TATGAGTGTGGAGTCGTTGCCCTGGGCAACAAAAGTGAAACTCNNNAGATCGGAA |
| 37933_2 | TATGAGTGTGGAGTCGTTGCCCTGGGCAACAAAAGTGAAACTCNNNAGATCGGAA |
| 43349_1 | TATGAGTGTGGAGTCGTTGCCAAGTAGCTGGGATTACAAGTGCNNNAGATCGGAA |
| 43349_2 | TATGAGTGTGGAGTCGTTGCCAAGTAGCTGGGATTACAAGTGCNNNAGATCGGAA |
| 43849_1 | TATGAGTGTGGAGTCGTTGCCCCAGGTTTTCAAAGCAGGAAATNNNAGATCGGAA |
| 43849_2 | TATGAGTGTGGAGTCGTTGCCCCAGGTTTTCAAAGCAGGAAATNNNAGATCGGAA |
| 40368_1 | TATGAGTGTGGAGTCGTTGCAAGATGAAGGCTGGTCTGGTATGNNNAGATCGGAA |
| 40368_2 | TATGAGTGTGGAGTCGTTGCAAGATGAAGGCTGGTCTGGTATGNNNAGATCGGAA |
| 33625_1 | TATGAGTGTGGAGTCGTTGCCTGGGTTCACGCCATTCTCTTNNNAGATCGGAAGAG |
| 33625_2 | TATGAGTGTGGAGTCGTTGCCTGGGTTCACGCCATTCTCTTNNNAGATCGGAAGAG |
| 39406_1 | TATGAGTGTGGAGTCGTTGCTTTGAATCCAAACCCCATTCAGANNNAGATCGGAAG |
| 39406_2 | TATGAGTGTGGAGTCGTTGCTTTGAATCCAAACCCCATTCAGANNNAGATCGGAAG |
| 45272_1 | TATGAGTGTGGAGTCGTTGCCCCTCTCTCATACCTACTCCCAANNNAGATCGGAAGA |
| 45272_2 | TATGAGTGTGGAGTCGTTGCCCCTCTCTCATACCTACTCCCAANNNAGATCGGAAGA |
| 35586_1 | TATGAGTGTGGAGTCGTTGCGTTCAAGCAATTCTCTGCCTCAGNNNAGATCGGAAG |
| 35586_2 | TATGAGTGTGGAGTCGTTGCGTTCAAGCAATTCTCTGCCTCAGNNNAGATCGGAAG |
| 52754_1 | TATGAGTGTGGAGTCGTTGCCACAAAGTTCACTGGGGAAACTCNNNAGATCGGAA |
| 52754_2 | TATGAGTGTGGAGTCGTTGCCACAAAGTTCACTGGGGAAACTCNNNAGATCGGAA |
| 30277_1 | TATGAGTGTGGAGTCGTTGCGGGAAAAGCTGAGTACCTGAAGANNNAGATCGGA |
| 30277_2 | TATGAGTGTGGAGTCGTTGCGGGAAAAGCTGAGTACCTGAAGANNNAGATCGGA |
| 32030_1 | TATGAGTGTGGAGTCGTTGCGAGACAAGGTCTTGCTCTGTCANNNAGATCGGAAG |
| 32030_2 | TATGAGTGTGGAGTCGTTGCGAGACAAGGTCTTGCTCTGTCANNNAGATCGGAAG |
| 47518_1 | TATGAGTGTGGAGTCGTTGCTGTTTCAAGTGATTCTCCTGCCTNNNAGATCGGAAG |
| 47518_2 | TATGAGTGTGGAGTCGTTGCTGTTTCAAGTGATTCTCCTGCCTNNNAGATCGGAAG |
| 34006_1 | TATGAGTGTGGAGTCGTTGCGAGATCTCGGTTCACTGCAAACTNNNAGATCGGAAG |
| 34006_2 | TATGAGTGTGGAGTCGTTGCGAGATCTCGGTTCACTGCAAACTNNNAGATCGGAAG |
| 37027_1 | TATGAGTGTGGAGTCGTTGCACACAGAGCCACATCACATATGTNNNAGATCGGAAG |
| 37027_2 | TATGAGTGTGGAGTCGTTGCACACAGAGCCACATCACATATGTNNNAGATCGGAAG |
| 37803_1 | TATGAGTGTGGAGTCGTTGCCCTGTCTCTCTGGTTCTGATGACNNNAGATCGGAAG |
| 37803_2 | TATGAGTGTGGAGTCGTTGCCCTGTCTCTCTGGTTCTGATGACNNNAGATCGGAAG |
| 44355_1 | TATGAGTGTGGAGTCGTTGCTAAAGGCGCTTCTGTGCTGTANNNAGATCGGAAGA |
| 44355_2 | TATGAGTGTGGAGTCGTTGCTAAAGGCGCTTCTGTGCTGTANNNAGATCGGAAGA |
| 49995_1 | TATGAGTGTGGAGTCGTTGCTTCTGTTGGGCACAAAGTCTAGGNNNAGATCGGAA |
| 49995_2 | TATGAGTGTGGAGTCGTTGCTTCTGTTGGGCACAAAGTCTAGGNNNAGATCGGAA |
| 42544_1 | TATGAGTGTGGAGTCGTTGCGTCTTGAACTCCTGGTTTCAAGCNNNAGATCGGAAG |
| 42544_2 | TATGAGTGTGGAGTCGTTGCGTCTTGAACTCCTGGTTTCAAGCNNNAGATCGGAAG |
| 32918_1 | TATGAGTGTGGAGTCGTTGCAGGGGGAAAAGCAAGAGAAGAATNNNAGATCGGA |
| 32918_2 | TATGAGTGTGGAGTCGTTGCAGGGGGAAAAGCAAGAGAAGAATNNNAGATCGGA |
| 36328_1 | TATGAGTGTGGAGTCGTTGCTGCTATGTGATCCAGCAATTCCANNNAGATCGGAAG |
| 36328_2 | TATGAGTGTGGAGTCGTTGCTGCTATGTGATCCAGCAATTCCANNNAGATCGGAAG |
| 3708_1 | TATGAGTGTGGAGTCGTTGCCCAGGGATCCTCATCTGTTGGTANNNAGATCGGAAG |
| 3708_2 | TATGAGTGTGGAGTCGTTGCCCAGGGATCCTCATCTGTTGGTANNNAGATCGGAAG |

| | |
|---|---|
| 43263_1 | TATGAGTGTGGAGTCGTTGCCTCCTGGGTTCAAGGGATTCTCNNNAGATCGGAAGA |
| 43263_2 | TATGAGTGTGGAGTCGTTGCCTCCTGGGTTCAAGGGATTCTCNNNAGATCGGAAGA |
| 13253_1 | TATGAGTGTGGAGTCGTTGCAAACAAACAAATAAAAAGATCTCANNNAGATCGGAA |
| 13253_2 | TATGAGTGTGGAGTCGTTGCAAACAAACAAATAAAAAGATCTCANNNAGATCGGAA |
| 50131_1 | TATGAGTGTGGAGTCGTTGCTAGGTGTCCATTTGGAACCTGAGNNNAGATCGGAA |
| 50131_2 | TATGAGTGTGGAGTCGTTGCTAGGTGTCCATTTGGAACCTGAGNNNAGATCGGAA |
| 40519_1 | TATGAGTGTGGAGTCGTTGCGCGGGTTGCCTTTTACTTTCATTNNNAGATCGGAAG |
| 40519_2 | TATGAGTGTGGAGTCGTTGCGCGGGTTGCCTTTTACTTTCATTNNNAGATCGGAAG |
| 10037_1 | TATGAGTGTGGAGTCGTTGCCTCTCTCCAGTTCTTGCAGCAATNNNAGATCGGAAGA |
| 10037_2 | TATGAGTGTGGAGTCGTTGCCTCTCTCCAGTTCTTGCAGCAATNNNAGATCGGAAGA |
| 33286_1 | TATGAGTGTGGAGTCGTTGCCTGAGTAGGTGGGATTACAGGTGNNNAGATCGGAA |
| 33286_2 | TATGAGTGTGGAGTCGTTGCCTGAGTAGGTGGGATTACAGGTGNNNAGATCGGAA |
| 52016_1 | TATGAGTGTGGAGTCGTTGCACGACAGCATAGATTTGGTTCCTNNNAGATCGGAAG |
| 52016_2 | TATGAGTGTGGAGTCGTTGCACGACAGCATAGATTTGGTTCCTNNNAGATCGGAAG |
| 42156_1 | TATGAGTGTGGAGTCGTTGCGCAAAACATCAGGCCTATGTAGGNNNAGATCGGAA |
| 42156_2 | TATGAGTGTGGAGTCGTTGCGCAAAACATCAGGCCTATGTAGGNNNAGATCGGAA |
| 34001_1 | TATGAGTGTGGAGTCGTTGCCTTCTTCACCATCTTGCATTCCCNNNAGATCGGAAGA |
| 34001_2 | TATGAGTGTGGAGTCGTTGCCTTCTTCACCATCTTGCATTCCCNNNAGATCGGAAGA |
| 39621_1 | TATGAGTGTGGAGTCGTTGCGGAGTTGCAATCCTAGTCTCTGANNNAGATCGGAAG |
| 39621_2 | TATGAGTGTGGAGTCGTTGCGGAGTTGCAATCCTAGTCTCTGANNNAGATCGGAAG |
| 12668_1 | TATGAGTGTGGAGTCGTTGCTTGTCTATGCCTCAGATTTTATNNNAGATCGGAAGA |
| 12668_2 | TATGAGTGTGGAGTCGTTGCTTGTCTATGCCTCAGATTTTATNNNAGATCGGAAGA |
| 42931_1 | TATGAGTGTGGAGTCGTTGCACAGAGAGAGAGGAAGAGAGCTTNNNAGATCGGA |
| 42931_2 | TATGAGTGTGGAGTCGTTGCACAGAGAGAGAGGAAGAGAGCTTNNNAGATCGGA |
| 39180_1 | TATGAGTGTGGAGTCGTTGCGAGAGGTAGAGAAAAGTCAGCGANNNAGATCGGA |
| 39180_2 | TATGAGTGTGGAGTCGTTGCGAGAGGTAGAGAAAAGTCAGCGANNNAGATCGGA |
| 46747_1 | TATGAGTGTGGAGTCGTTGCTGTAGTGGCGTGATCACAGTTTANNNAGATCGGAA |
| 46747_2 | TATGAGTGTGGAGTCGTTGCTGTAGTGGCGTGATCACAGTTTANNNAGATCGGAA |
| 36297_1 | TATGAGTGTGGAGTCGTTGCAGTCCATATTGGGCGCTTTTCTANNNAGATCGGAAG |
| 36297_2 | TATGAGTGTGGAGTCGTTGCAGTCCATATTGGGCGCTTTTCTANNNAGATCGGAAG |
| 28805_1 | TATGAGTGTGGAGTCGTTGCTTGTGTACATGCTTTTCCTCTGCNNNAGATCGGAAGA |
| 28805_2 | TATGAGTGTGGAGTCGTTGCTTGTGTACATGCTTTTCCTCTGCNNNAGATCGGAAGA |
| 36342_1 | TATGAGTGTGGAGTCGTTGCACCAATCTTCCTCTGTCTATGCANNNAGATCGGAAGA |
| 36342_2 | TATGAGTGTGGAGTCGTTGCACCAATCTTCCTCTGTCTATGCANNNAGATCGGAAGA |
| 28982_1 | TATGAGTGTGGAGTCGTTGCAATGAATGGGAGTGCAGATGTCTNNNAGATCGGAA |
| 28982_2 | TATGAGTGTGGAGTCGTTGCAATGAATGGGAGTGCAGATGTCTNNNAGATCGGAA |
| 47257_1 | TATGAGTGTGGAGTCGTTGCATCTCAGCCTACCACAACCTCTANNNAGATCGGAAGA |
| 47257_2 | TATGAGTGTGGAGTCGTTGCATCTCAGCCTACCACAACCTCTANNNAGATCGGAAGA |
| 40537_1 | TATGAGTGTGGAGTCGTTGCATGTTAGCGAGGAGAAGGAAGAGNNNAGATCGGA |
| 40537_2 | TATGAGTGTGGAGTCGTTGCATGTTAGCGAGGAGAAGGAAGAGNNNAGATCGGA |
| 46736_1 | TATGAGTGTGGAGTCGTTGCGATCTTGGCTCATTGCAACCTCNNNAGATCGGAAGA |
| 46736_2 | TATGAGTGTGGAGTCGTTGCGATCTTGGCTCATTGCAACCTCNNNAGATCGGAAGA |
| 29655_1 | TATGAGTGTGGAGTCGTTGCGCAACTAACGAAACTGCCCATACNNNAGATCGGAAG |

| | |
|---|---|
| 29655_2 | TATGAGTGTGGAGTCGTTGCGCAACTAACGAAACTGCCCATACNNNAGATCGGAAG |
| 10357_1 | TATGAGTGTGGAGTCGTTGCCCCATGAAGGTAAACGCTGACTANNNAGATCGGAAG |
| 10357_2 | TATGAGTGTGGAGTCGTTGCCCCATGAAGGTAAACGCTGACTANNNAGATCGGAAG |
| 35819_1 | TATGAGTGTGGAGTCGTTGCACTGCAACATCTACCTCCCAAATNNNAGATCGGAAGA |
| 35819_2 | TATGAGTGTGGAGTCGTTGCACTGCAACATCTACCTCCCAAATNNNAGATCGGAAGA |
| 37791_1 | TATGAGTGTGGAGTCGTTGCAAGAGATTGAGATCATCCTGGCCNNNAGATCGGAAG |
| 37791_2 | TATGAGTGTGGAGTCGTTGCAAGAGATTGAGATCATCCTGGCCNNNAGATCGGAAG |
| 38450_1 | TATGAGTGTGGAGTCGTTGCGGTGAAGAAATGGGGAGCTACTGNNNAGATCGGAA |
| 38450_2 | TATGAGTGTGGAGTCGTTGCGGTGAAGAAATGGGGAGCTACTGNNNAGATCGGAA |
| 31126_1 | TATGAGTGTGGAGTCGTTGCAGGTCTTGGGAACTGAAAGCTTANNNAGATCGGAAG |
| 31126_2 | TATGAGTGTGGAGTCGTTGCAGGTCTTGGGAACTGAAAGCTTANNNAGATCGGAAG |
| 12962_1 | TATGAGTGTGGAGTCGTTGCTAATTGCATTTTAGTTCAAGGGNNNAGATCGGAAGA |
| 12962_2 | TATGAGTGTGGAGTCGTTGCTAATTGCATTTTAGTTCAAGGGNNNAGATCGGAAGA |
| 43770_1 | TATGAGTGTGGAGTCGTTGCAAAAAAAACAGAGGGTTGGGTGCNNNAGATCGGAA |
| 43770_2 | TATGAGTGTGGAGTCGTTGCAAAAAAAACAGAGGGTTGGGTGCNNNAGATCGGAA |
| 34839_1 | TATGAGTGTGGAGTCGTTGCGCTACCATGCCCAGCTAATTTTTNNNAGATCGGAAGA |
| 34839_2 | TATGAGTGTGGAGTCGTTGCGCTACCATGCCCAGCTAATTTTTNNNAGATCGGAAGA |
| 52894_1 | TATGAGTGTGGAGTCGTTGCGCCCCACTCCCTCTACTCTATATNNNAGATCGGAAGAG |
| 52894_2 | TATGAGTGTGGAGTCGTTGCGCCCCACTCCCTCTACTCTATATNNNAGATCGGAAGAG |
| 42211_1 | TATGAGTGTGGAGTCGTTGCACTTATTGCCCTGGCTAGAAACTNNNAGATCGGAAGA |
| 42211_2 | TATGAGTGTGGAGTCGTTGCACTTATTGCCCTGGCTAGAAACTNNNAGATCGGAAGA |
| 32744_1 | TATGAGTGTGGAGTCGTTGCGCCAACATAGCAAAACCTGGTTTNNNAGATCGGAAG |
| 32744_2 | TATGAGTGTGGAGTCGTTGCGCCAACATAGCAAAACCTGGTTTNNNAGATCGGAAG |
| 50064_1 | TATGAGTGTGGAGTCGTTGCCCTTCTAGTTGTGGGTTGCTAGTNNNAGATCGGAAG |
| 50064_2 | TATGAGTGTGGAGTCGTTGCCCTTCTAGTTGTGGGTTGCTAGTNNNAGATCGGAAG |
| 38296_1 | TATGAGTGTGGAGTCGTTGCGTGGTCTTAGCTACTTGAGAGGGNNNAGATCGGAAG |
| 38296_2 | TATGAGTGTGGAGTCGTTGCGTGGTCTTAGCTACTTGAGAGGGNNNAGATCGGAAG |
| 51448_1 | TATGAGTGTGGAGTCGTTGCGAGCAAGAGGTTTGTAGGTGGTANNNAGATCGGAA |
| 51448_2 | TATGAGTGTGGAGTCGTTGCGAGCAAGAGGTTTGTAGGTGGTANNNAGATCGGAA |
| 33477_1 | TATGAGTGTGGAGTCGTTGCCAAGGCAGGAGGATTACTTGAGTNNNAGATCGGAA |
| 33477_2 | TATGAGTGTGGAGTCGTTGCCAAGGCAGGAGGATTACTTGAGTNNNAGATCGGAA |
| 31132_1 | TATGAGTGTGGAGTCGTTGCTGTTTGTGTATTGTCTGCCTTTCCNNNAGATCGGAAG |
| 31132_2 | TATGAGTGTGGAGTCGTTGCTGTTTGTGTATTGTCTGCCTTTCCNNNAGATCGGAAG |
| 31416_1 | TATGAGTGTGGAGTCGTTGCAGCAACTGGTGAATCTAGGTGAGNNNAGATCGGAA |
| 31416_2 | TATGAGTGTGGAGTCGTTGCAGCAACTGGTGAATCTAGGTGAGNNNAGATCGGAA |
| 31453_1 | TATGAGTGTGGAGTCGTTGCTTGTGCTGGATGTCAGTGATTCTNNNAGATCGGAAG |
| 31453_2 | TATGAGTGTGGAGTCGTTGCTTGTGCTGGATGTCAGTGATTCTNNNAGATCGGAAG |
| 48193_1 | TATGAGTGTGGAGTCGTTGCTTCAAGCTATTCTCCTGCCTCAGNNNAGATCGGAAGA |
| 48193_2 | TATGAGTGTGGAGTCGTTGCTTCAAGCTATTCTCCTGCCTCAGNNNAGATCGGAAGA |
| 12198_1 | TATGAGTGTGGAGTCGTTGCCCAACACAAGCCTGGAAATGCNNNAGATCGGAAGAG |
| 12198_2 | TATGAGTGTGGAGTCGTTGCCCAACACAAGCCTGGAAATGCNNNAGATCGGAAGAG |
| 51210_1 | TATGAGTGTGGAGTCGTTGCAGACCCTGTCTCAAAATCGATCANNNAGATCGGAAGA |
| 51210_2 | TATGAGTGTGGAGTCGTTGCAGACCCTGTCTCAAAATCGATCANNNAGATCGGAAGA |

| | |
|---|---|
| 13281_1 | TATGAGTGTGGAGTCGTTGCTATGTATTCGCCAATTTGTAATTNNNAGATCGGAAG |
| 13281_2 | TATGAGTGTGGAGTCGTTGCTATGTATTCGCCAATTTGTAATTNNNAGATCGGAAG |
| 39557_1 | TATGAGTGTGGAGTCGTTGCACCGTCCTGTTCATTTTACCTGANNNAGATCGGAAGA |
| 39557_2 | TATGAGTGTGGAGTCGTTGCACCGTCCTGTTCATTTTACCTGANNNAGATCGGAAGA |
| 52213_1 | TATGAGTGTGGAGTCGTTGCTTCTTCACCAGAGAGCAGAAACANNNAGATCGGAAG |
| 52213_2 | TATGAGTGTGGAGTCGTTGCTTCTTCACCAGAGAGCAGAAACANNNAGATCGGAAG |
| 37334_1 | TATGAGTGTGGAGTCGTTGCGGTAGGCAAATGACATGAACAGANNNAGATCGGAA |
| 37334_2 | TATGAGTGTGGAGTCGTTGCGGTAGGCAAATGACATGAACAGANNNAGATCGGAA |
| 10421_1 | TATGAGTGTGGAGTCGTTGCATGAGTATGTGGGGTGTAGGGGNNNAGATCGGAA |
| 10421_2 | TATGAGTGTGGAGTCGTTGCATGAGTATGTGGGGTGTAGGGGNNNAGATCGGAA |
| 49985_1 | TATGAGTGTGGAGTCGTTGCGATGGACTTGAGAAGCACGTAGTNNNAGATCGGAA |
| 49985_2 | TATGAGTGTGGAGTCGTTGCGATGGACTTGAGAAGCACGTAGTNNNAGATCGGAA |
| 45733_1 | TATGAGTGTGGAGTCGTTGCCCTCCTGATAGCTGGGACTACAGNNNAGATCGGAAG |
| 45733_2 | TATGAGTGTGGAGTCGTTGCCCTCCTGATAGCTGGGACTACAGNNNAGATCGGAAG |
| 39844_1 | TATGAGTGTGGAGTCGTTGCAGAGAGGTGGTTCAATTAGAGGCNNNAGATCGGAA |
| 39844_2 | TATGAGTGTGGAGTCGTTGCAGAGAGGTGGTTCAATTAGAGGCNNNAGATCGGAA |
| 41780_1 | TATGAGTGTGGAGTCGTTGCCTCCCGGGTTCAATCAATTCCTTNNNAGATCGGAAGA |
| 41780_2 | TATGAGTGTGGAGTCGTTGCCTCCCGGGTTCAATCAATTCCTTNNNAGATCGGAAGA |
| 38488_1 | TATGAGTGTGGAGTCGTTGCAGAAATGCTATCTCCAACAGGCTNNNAGATCGGAAG |
| 38488_2 | TATGAGTGTGGAGTCGTTGCAGAAATGCTATCTCCAACAGGCTNNNAGATCGGAAG |
| 49016_1 | TATGAGTGTGGAGTCGTTGCTTTTTGAGACAGAGTCTTGCCCTNNNAGATCGGAAG |
| 49016_2 | TATGAGTGTGGAGTCGTTGCTTTTTGAGACAGAGTCTTGCCCTNNNAGATCGGAAG |
| 40360_1 | TATGAGTGTGGAGTCGTTGCCTGAGGTCAGGAGTTCGAGACNNNAGATCGGAAGA |
| 40360_2 | TATGAGTGTGGAGTCGTTGCCTGAGGTCAGGAGTTCGAGACNNNAGATCGGAAGA |
| 43284_1 | TATGAGTGTGGAGTCGTTGCAGAATGAGTGTGTGTTTACAGAAACNNNAGATCGG |
| 43284_2 | TATGAGTGTGGAGTCGTTGCAGAATGAGTGTGTGTTTACAGAAACNNNAGATCGG |
| 49563_1 | TATGAGTGTGGAGTCGTTGCAAGTAGCTGAGATTATAGGCGCCNNNAGATCGGAA |
| 49563_2 | TATGAGTGTGGAGTCGTTGCAAGTAGCTGAGATTATAGGCGCCNNNAGATCGGAA |
| 41220_1 | TATGAGTGTGGAGTCGTTGCGAGTTGGAGTCTCACTGGGTGNNNAGATCGGAAGA |
| 41220_2 | TATGAGTGTGGAGTCGTTGCGAGTTGGAGTCTCACTGGGTGNNNAGATCGGAAGA |
| 39561_1 | TATGAGTGTGGAGTCGTTGCAAAAGAACTACCCTGAGACTGGGNNNAGATCGGAA |
| 39561_2 | TATGAGTGTGGAGTCGTTGCAAAAGAACTACCCTGAGACTGGGNNNAGATCGGAA |
| 44371_1 | TATGAGTGTGGAGTCGTTGCCTGGAGTGGATGAGAGACTTTCTNNNAGATCGGAA |
| 44371_2 | TATGAGTGTGGAGTCGTTGCCTGGAGTGGATGAGAGACTTTCTNNNAGATCGGAA |
| 36530_1 | TATGAGTGTGGAGTCGTTGCATCTGCTGACCCAGAATATGCATNNNAGATCGGAAG |
| 36530_2 | TATGAGTGTGGAGTCGTTGCATCTGCTGACCCAGAATATGCATNNNAGATCGGAAG |
| 36426_1 | TATGAGTGTGGAGTCGTTGCGCTGGTGTGCAGTTATTATCAGCNNNAGATCGGAAG |
| 36426_2 | TATGAGTGTGGAGTCGTTGCGCTGGTGTGCAGTTATTATCAGCNNNAGATCGGAAG |
| 40523_1 | TATGAGTGTGGAGTCGTTGCAAAACGGGCAAAAGACTTTGACANNNAGATCGGAA |
| 40523_2 | TATGAGTGTGGAGTCGTTGCAAAACGGGCAAAAGACTTTGACANNNAGATCGGAA |
| 32201_1 | TATGAGTGTGGAGTCGTTGCCAGTATTTGGGACTACAGGGGACNNNAGATCGGAA |
| 32201_2 | TATGAGTGTGGAGTCGTTGCCAGTATTTGGGACTACAGGGGACNNNAGATCGGAA |
| 40078_1 | TATGAGTGTGGAGTCGTTGCTCAGGAGTTGAGGAGGGATAAGTNNNAGATCGGA |

| | |
|---|---|
| 40078_2 | TATGAGTGTGGAGTCGTTGCTCAGGAGTTGAGGAGGGATAAGTNNNAGATCGGA |
| 44334_1 | TATGAGTGTGGAGTCGTTGCCTAAGATTAGGGCTGCCGTGGNNNAGATCGGAAGA |
| 44334_2 | TATGAGTGTGGAGTCGTTGCCTAAGATTAGGGCTGCCGTGGNNNAGATCGGAAGA |
| 53347_1 | TATGAGTGTGGAGTCGTTGCAAAGGGGATATCACCACTGTTCCNNNAGATCGGAAG |
| 53347_2 | TATGAGTGTGGAGTCGTTGCAAAGGGGATATCACCACTGTTCCNNNAGATCGGAAG |
| 48947_1 | TATGAGTGTGGAGTCGTTGCGCAATCTCGGTTCACTGCAAGNNNAGATCGGAAGA |
| 48947_2 | TATGAGTGTGGAGTCGTTGCGCAATCTCGGTTCACTGCAAGNNNAGATCGGAAGA |
| 48023_1 | TATGAGTGTGGAGTCGTTGCCCTGCCTGAACCCAGAAATCTTANNNAGATCGGAAG |
| 48023_2 | TATGAGTGTGGAGTCGTTGCCCTGCCTGAACCCAGAAATCTTANNNAGATCGGAAG |
| 31631_1 | TATGAGTGTGGAGTCGTTGCGCGTGAGAGAGAGAAAGAGAGAGNNNAGATCGG |
| 31631_2 | TATGAGTGTGGAGTCGTTGCGCGTGAGAGAGAGAAAGAGAGAGNNNAGATCGG |
| 44791_1 | TATGAGTGTGGAGTCGTTGCGCTTCGACTTCTAATCCCAAAGCNNNAGATCGGAAG |
| 44791_2 | TATGAGTGTGGAGTCGTTGCGCTTCGACTTCTAATCCCAAAGCNNNAGATCGGAAG |
| 12595_1 | TATGAGTGTGGAGTCGTTGCTACAACCTCAAAGAACGTTATNNNAGATCGGAAGAG |
| 12595_2 | TATGAGTGTGGAGTCGTTGCTACAACCTCAAAGAACGTTATNNNAGATCGGAAGAG |
| 45007_1 | TATGAGTGTGGAGTCGTTGCTTCACTTGGAGTTTGGGGAAGAANNNAGATCGGAA |
| 45007_2 | TATGAGTGTGGAGTCGTTGCTTCACTTGGAGTTTGGGGAAGAANNNAGATCGGAA |
| 673_1 | TATGAGTGTGGAGTCGTTGCGAGACAGAGACTGAATTGCTTGAGANNNAGATCGG |
| 673_2 | TATGAGTGTGGAGTCGTTGCGAGACAGAGACTGAATTGCTTGAGANNNAGATCGG |
| 31112_1 | TATGAGTGTGGAGTCGTTGCAGAAACACAATGCTAGAGCCAGANNNAGATCGGAA |
| 31112_2 | TATGAGTGTGGAGTCGTTGCAGAAACACAATGCTAGAGCCAGANNNAGATCGGAA |
| 41607_1 | TATGAGTGTGGAGTCGTTGCTCCTGGGTTCAAATGATTCTCGTNNNAGATCGGAAG |
| 41607_2 | TATGAGTGTGGAGTCGTTGCTCCTGGGTTCAAATGATTCTCGTNNNAGATCGGAAG |
| 42808_1 | TATGAGTGTGGAGTCGTTGCAAACTCCTGACCTCAAGTGATCCNNNAGATCGGAAG |
| 42808_2 | TATGAGTGTGGAGTCGTTGCAAACTCCTGACCTCAAGTGATCCNNNAGATCGGAAG |
| 52261_1 | TATGAGTGTGGAGTCGTTGCTTTCCTGGAACCAGAAGTCTCAGNNNAGATCGGAAG |
| 52261_2 | TATGAGTGTGGAGTCGTTGCTTTCCTGGAACCAGAAGTCTCAGNNNAGATCGGAAG |
| 41568_1 | TATGAGTGTGGAGTCGTTGCGTTCCTAGTACCCCTGGAGAAGANNNAGATCGGAA |
| 41568_2 | TATGAGTGTGGAGTCGTTGCGTTCCTAGTACCCCTGGAGAAGANNNAGATCGGAA |
| 40219_1 | TATGAGTGTGGAGTCGTTGCTAGCACTGATTTTCCCAAGGACANNNAGATCGGAAG |
| 40219_2 | TATGAGTGTGGAGTCGTTGCTAGCACTGATTTTCCCAAGGACANNNAGATCGGAAG |
| 32289_1 | TATGAGTGTGGAGTCGTTGCTGATCTCGGCTCACTGAAACATCNNNAGATCGGAAG |
| 32289_2 | TATGAGTGTGGAGTCGTTGCTGATCTCGGCTCACTGAAACATCNNNAGATCGGAAG |
| 53402_1 | TATGAGTGTGGAGTCGTTGCGGAGGGAAGGAAGGAAAGAAAGANNNAGATCGG |
| 53402_2 | TATGAGTGTGGAGTCGTTGCGGAGGGAAGGAAGGAAAGAAAGANNNAGATCGG |
| 30396_1 | TATGAGTGTGGAGTCGTTGCAGAGCAGTGGTAAAGAACTAGAGTTGANNNAGATC |
| 30396_2 | TATGAGTGTGGAGTCGTTGCAGAGCAGTGGTAAAGAACTAGAGTTGANNNAGATC |
| 30275_1 | TATGAGTGTGGAGTCGTTGCGAGAGACACACCTGCTCCAATTTNNNAGATCGGAAG |
| 30275_2 | TATGAGTGTGGAGTCGTTGCGAGAGACACACCTGCTCCAATTTNNNAGATCGGAAG |
| 50575_1 | TATGAGTGTGGAGTCGTTGCGCAGGGAGAATAATCTGTTCCCANNNAGATCGGAA |
| 50575_2 | TATGAGTGTGGAGTCGTTGCGCAGGGAGAATAATCTGTTCCCANNNAGATCGGAA |
| 39314_1 | TATGAGTGTGGAGTCGTTGCTCAGATGACTTCTCGATAGCAGCNNNAGATCGGAAG |
| 39314_2 | TATGAGTGTGGAGTCGTTGCTCAGATGACTTCTCGATAGCAGCNNNAGATCGGAAG |

| | |
|---|---|
| 47507_1 | TATGAGTGTGGAGTCGTTGCCATGCGAAATGGAAAGGGAGAAGNNNAGATCGGA |
| 47507_2 | TATGAGTGTGGAGTCGTTGCCATGCGAAATGGAAAGGGAGAAGNNNAGATCGGA |
| 35183_1 | TATGAGTGTGGAGTCGTTGCTTTGTTCAATGAAGAGGTTGGGCNNNAGATCGGAA |
| 35183_2 | TATGAGTGTGGAGTCGTTGCTTTGTTCAATGAAGAGGTTGGGCNNNAGATCGGAA |
| 30569_1 | TATGAGTGTGGAGTCGTTGCACATCTTTCAATCCTCTCAAAGCCTTNNNAGATCGGA |
| 30569_2 | TATGAGTGTGGAGTCGTTGCACATCTTTCAATCCTCTCAAAGCCTTNNNAGATCGGA |
| 53002_1 | TATGAGTGTGGAGTCGTTGCACAGGGGAAACTACCAAACACTTNNNAGATCGGAA |
| 53002_2 | TATGAGTGTGGAGTCGTTGCACAGGGGAAACTACCAAACACTTNNNAGATCGGAA |
| 45448_1 | TATGAGTGTGGAGTCGTTGCAGACTTAGCAATCTGACAGTCTTNNNAGATCGGAAG |
| 45448_2 | TATGAGTGTGGAGTCGTTGCAGACTTAGCAATCTGACAGTCTTNNNAGATCGGAAG |
| 12814_1 | TATGAGTGTGGAGTCGTTGCAATTTCCTCCAGCTATTAGTAATNNNAGATCGGAAG |
| 12814_2 | TATGAGTGTGGAGTCGTTGCAATTTCCTCCAGCTATTAGTAATNNNAGATCGGAAG |
| 38391_1 | TATGAGTGTGGAGTCGTTGCCACGCCTATAATCCCACCACTTTNNNAGATCGGAAGA |
| 38391_2 | TATGAGTGTGGAGTCGTTGCCACGCCTATAATCCCACCACTTTNNNAGATCGGAAGA |
| 42492_1 | TATGAGTGTGGAGTCGTTGCTCCACACTCTACTGTCTCAAAGCNNNAGATCGGAAG |
| 42492_2 | TATGAGTGTGGAGTCGTTGCTCCACACTCTACTGTCTCAAAGCNNNAGATCGGAAG |
| 50295_1 | TATGAGTGTGGAGTCGTTGCCTGAAAGCGTCTCCAAAAATCCANNNAGATCGGAAG |
| 50295_2 | TATGAGTGTGGAGTCGTTGCCTGAAAGCGTCTCCAAAAATCCANNNAGATCGGAAG |
| 50066_1 | TATGAGTGTGGAGTCGTTGCTTCCTCTTGGTTTGGAGTACCTGNNNAGATCGGAAG |
| 50066_2 | TATGAGTGTGGAGTCGTTGCTTCCTCTTGGTTTGGAGTACCTGNNNAGATCGGAAG |
| 32606_1 | TATGAGTGTGGAGTCGTTGCCATCCTCTCTGACAACAGTGGAANNNAGATCGGAAG |
| 32606_2 | TATGAGTGTGGAGTCGTTGCCATCCTCTCTGACAACAGTGGAANNNAGATCGGAAG |
| 46204_1 | TATGAGTGTGGAGTCGTTGCCTCCTGGGTTGACGTCATTCTCNNNAGATCGGAAGA |
| 46204_2 | TATGAGTGTGGAGTCGTTGCCTCCTGGGTTGACGTCATTCTCNNNAGATCGGAAGA |
| 28908_1 | TATGAGTGTGGAGTCGTTGCCTCTCATCAGGCGTGTCACTTNNNAGATCGGAAGAG |
| 28908_2 | TATGAGTGTGGAGTCGTTGCCTCTCATCAGGCGTGTCACTTNNNAGATCGGAAGAG |
| 33171_1 | TATGAGTGTGGAGTCGTTGCGGATTCCTGGCTAGTGTTCTCTCNNNAGATCGGAAG |
| 33171_2 | TATGAGTGTGGAGTCGTTGCGGATTCCTGGCTAGTGTTCTCTCNNNAGATCGGAAG |
| 35889_1 | TATGAGTGTGGAGTCGTTGCAACAGAAGTCAGATGTCCTCAGGNNNAGATCGGAA |
| 35889_2 | TATGAGTGTGGAGTCGTTGCAACAGAAGTCAGATGTCCTCAGGNNNAGATCGGAA |
| 38045_1 | TATGAGTGTGGAGTCGTTGCCTGTAGAGAAGGCGGTACTTGAANNNAGATCGGAA |
| 38045_2 | TATGAGTGTGGAGTCGTTGCCTGTAGAGAAGGCGGTACTTGAANNNAGATCGGAA |
| 33579_1 | TATGAGTGTGGAGTCGTTGCATGAGAGACAGGACTAGCTGGATNNNAGATCGGAA |
| 33579_2 | TATGAGTGTGGAGTCGTTGCATGAGAGACAGGACTAGCTGGATNNNAGATCGGAA |
| 45471_1 | TATGAGTGTGGAGTCGTTGCATTGTCTAAATGCCGCATCCTTGNNNAGATCGGAAG |
| 45471_2 | TATGAGTGTGGAGTCGTTGCATTGTCTAAATGCCGCATCCTTGNNNAGATCGGAAG |
| 12805_1 | TATGAGTGTGGAGTCGTTGCTATAAGTTATTATCGCATGCCTANNNAGATCGGAAG |
| 12805_2 | TATGAGTGTGGAGTCGTTGCTATAAGTTATTATCGCATGCCTANNNAGATCGGAAG |
| 34118_1 | TATGAGTGTGGAGTCGTTGCCCGAGGACAGAGTGATGTTAGACNNNAGATCGGAA |
| 34118_2 | TATGAGTGTGGAGTCGTTGCCCGAGGACAGAGTGATGTTAGACNNNAGATCGGAA |
| 51983_1 | TATGAGTGTGGAGTCGTTGCCCCCAAACCATAAGAATCCTGGANNNAGATCGGAAG |
| 51983_2 | TATGAGTGTGGAGTCGTTGCCCCCAAACCATAAGAATCCTGGANNNAGATCGGAAG |
| 31278_1 | TATGAGTGTGGAGTCGTTGCGTAATCCAAAAGCTTTAGCCGCANNNAGATCGGAAG |

| | |
|---|---|
| 31278_2 | TATGAGTGTGGAGTCGTTGCGTAATCCAAAAGCTTTAGCCGCANNNAGATCGGAAG |
| 48541_1 | TATGAGTGTGGAGTCGTTGCTGGGATTACAAGTGTGAGCTACCNNNAGATCGGAA |
| 48541_2 | TATGAGTGTGGAGTCGTTGCTGGGATTACAAGTGTGAGCTACCNNNAGATCGGAA |
| 31802_1 | TATGAGTGTGGAGTCGTTGCTCACAAGTGCACATGGAATGTTCNNNAGATCGGAAG |
| 31802_2 | TATGAGTGTGGAGTCGTTGCTCACAAGTGCACATGGAATGTTCNNNAGATCGGAAG |
| 41565_1 | TATGAGTGTGGAGTCGTTGCCAAGTAGCTGGGACTATAGGCGNNNAGATCGGAAG |
| 41565_2 | TATGAGTGTGGAGTCGTTGCCAAGTAGCTGGGACTATAGGCGNNNAGATCGGAAG |
| 38337_1 | TATGAGTGTGGAGTCGTTGCCTCAGCTCACTGGTGGTAGATACNNNAGATCGGAAG |
| 38337_2 | TATGAGTGTGGAGTCGTTGCCTCAGCTCACTGGTGGTAGATACNNNAGATCGGAAG |
| 30861_1 | TATGAGTGTGGAGTCGTTGCTGTTGTTTTCAGATGAGTGGGGANNNAGATCGGAA |
| 30861_2 | TATGAGTGTGGAGTCGTTGCTGTTGTTTTCAGATGAGTGGGGANNNAGATCGGAA |
| 34038_1 | TATGAGTGTGGAGTCGTTGCGAGATGGAGTCTTGCTTTGTTGCNNNAGATCGGAA |
| 34038_2 | TATGAGTGTGGAGTCGTTGCGAGATGGAGTCTTGCTTTGTTGCNNNAGATCGGAA |
| 47430_1 | TATGAGTGTGGAGTCGTTGCTGTTAGCCAGGATGGTCTTGATCNNNAGATCGGAA |
| 47430_2 | TATGAGTGTGGAGTCGTTGCTGTTAGCCAGGATGGTCTTGATCNNNAGATCGGAA |
| 46192_1 | TATGAGTGTGGAGTCGTTGCCCTCCTGGGTTCAAGTGATTCTCNNNAGATCGGAAG |
| 46192_2 | TATGAGTGTGGAGTCGTTGCCCTCCTGGGTTCAAGTGATTCTCNNNAGATCGGAAG |
| 53096_1 | TATGAGTGTGGAGTCGTTGCCAGCTCCAACTCTATGCAGTTTANNNAGATCGGAAG |
| 53096_2 | TATGAGTGTGGAGTCGTTGCCAGCTCCAACTCTATGCAGTTTANNNAGATCGGAAG |
| 30721_1 | TATGAGTGTGGAGTCGTTGCCAAGTGCTCAATCTCTACCAGGGNNNAGATCGGAAG |
| 30721_2 | TATGAGTGTGGAGTCGTTGCCAAGTGCTCAATCTCTACCAGGGNNNAGATCGGAAG |
| 42010_1 | TATGAGTGTGGAGTCGTTGCTAGGCTGAAGCAATTTTTCTGCCNNNAGATCGGAAG |
| 42010_2 | TATGAGTGTGGAGTCGTTGCTAGGCTGAAGCAATTTTTCTGCCNNNAGATCGGAAG |
| 44136_1 | TATGAGTGTGGAGTCGTTGCGTCTCTTCTTTGCACAGTGTCACNNNAGATCGGAAG |
| 44136_2 | TATGAGTGTGGAGTCGTTGCGTCTCTTCTTTGCACAGTGTCACNNNAGATCGGAAG |
| 32697_1 | TATGAGTGTGGAGTCGTTGCTCAATCTCTGGGACTGAAGCAATNNNAGATCGGAAG |
| 32697_2 | TATGAGTGTGGAGTCGTTGCTCAATCTCTGGGACTGAAGCAATNNNAGATCGGAAG |
| 46704_1 | TATGAGTGTGGAGTCGTTGCGTTCAAACGATTCTTGTGCCTCANNNAGATCGGAAG |
| 46704_2 | TATGAGTGTGGAGTCGTTGCGTTCAAACGATTCTTGTGCCTCANNNAGATCGGAAG |
| 42975_1 | TATGAGTGTGGAGTCGTTGCTAAGTTTGTGTGTGTAGGGGGAGNNNAGATCGGA |
| 42975_2 | TATGAGTGTGGAGTCGTTGCTAAGTTTGTGTGTGTAGGGGGAGNNNAGATCGGA |
| 50382_1 | TATGAGTGTGGAGTCGTTGCAGCCATGATCAGCCTCACACNNNAGATCGGAAGAGC |
| 50382_2 | TATGAGTGTGGAGTCGTTGCAGCCATGATCAGCCTCACACNNNAGATCGGAAGAGC |
| 2761_1 | TATGAGTGTGGAGTCGTTGCTCTTCACTTCCATAAGGTTAGCCTNNNAGATCGGAAG |
| 2761_2 | TATGAGTGTGGAGTCGTTGCTCTTCACTTCCATAAGGTTAGCCTNNNAGATCGGAAG |
| 2882_1 | TATGAGTGTGGAGTCGTTGCTGTAAACGAAGCCCTCAAACTGTNNNAGATCGGAAG |
| 2882_2 | TATGAGTGTGGAGTCGTTGCTGTAAACGAAGCCCTCAAACTGTNNNAGATCGGAAG |
| 46925_1 | TATGAGTGTGGAGTCGTTGCTCAACCTCCTGAGTACTGGGATTNNNAGATCGGAAG |
| 46925_2 | TATGAGTGTGGAGTCGTTGCTCAACCTCCTGAGTACTGGGATTNNNAGATCGGAAG |
| 44010_1 | TATGAGTGTGGAGTCGTTGCCAGAGAAATCTTGACCCTCACCANNNAGATCGGAAG |
| 44010_2 | TATGAGTGTGGAGTCGTTGCCAGAGAAATCTTGACCCTCACCANNNAGATCGGAAG |
| 13202_1 | TATGAGTGTGGAGTCGTTGCTTCTAACTGGAGTGAGATAATACNNNAGATCGGAA |
| 13202_2 | TATGAGTGTGGAGTCGTTGCTTCTAACTGGAGTGAGATAATACNNNAGATCGGAA |

| | |
|---|---|
| 32097_1 | TATGAGTGTGGAGTCGTTGCGGAAGCTAGGGTCTTTGCATGTANNNAGATCGGAA |
| 32097_2 | TATGAGTGTGGAGTCGTTGCGGAAGCTAGGGTCTTTGCATGTANNNAGATCGGAA |
| 40511_1 | TATGAGTGTGGAGTCGTTGCGTCCATTGGTGAGTGCAATTTCANNNAGATCGGAAG |
| 40511_2 | TATGAGTGTGGAGTCGTTGCGTCCATTGGTGAGTGCAATTTCANNNAGATCGGAAG |
| 48605_1 | TATGAGTGTGGAGTCGTTGCTTCGAACGATTCTCCTGCCTAAGNNNAGATCGGAAG |
| 48605_2 | TATGAGTGTGGAGTCGTTGCTTCGAACGATTCTCCTGCCTAAGNNNAGATCGGAAG |
| 36001_1 | TATGAGTGTGGAGTCGTTGCAGAAAGGAGGAGAGACAGAGACANNNAGATCGGA |
| 36001_2 | TATGAGTGTGGAGTCGTTGCAGAAAGGAGGAGAGACAGAGACANNNAGATCGGA |
| 51452_1 | TATGAGTGTGGAGTCGTTGCCTGGGAGTGAAATTCCTGGAGAANNNAGATCGGAA |
| 51452_2 | TATGAGTGTGGAGTCGTTGCCTGGGAGTGAAATTCCTGGAGAANNNAGATCGGAA |
| 28804_1 | TATGAGTGTGGAGTCGTTGCAGCTTGCAAATACTTTCTCCCATNNNAGATCGGAAGA |
| 28804_2 | TATGAGTGTGGAGTCGTTGCAGCTTGCAAATACTTTCTCCCATNNNAGATCGGAAGA |
| 50241_1 | TATGAGTGTGGAGTCGTTGCGTAGGAGACATAGCCTGAGGGAGNNNAGATCGGA |
| 50241_2 | TATGAGTGTGGAGTCGTTGCGTAGGAGACATAGCCTGAGGGAGNNNAGATCGGA |
| 46690_1 | TATGAGTGTGGAGTCGTTGCTGGAATTTCAGAACACCCTCCATNNNAGATCGGAAG |
| 46690_2 | TATGAGTGTGGAGTCGTTGCTGGAATTTCAGAACACCCTCCATNNNAGATCGGAAG |
| 32618_1 | TATGAGTGTGGAGTCGTTGCGGAGCCAAAGACCTGAACAGATANNNAGATCGGAA |
| 32618_2 | TATGAGTGTGGAGTCGTTGCGGAGCCAAAGACCTGAACAGATANNNAGATCGGAA |
| 52065_1 | TATGAGTGTGGAGTCGTTGCCTTGCACAGGATTAGGCATTGAGNNNAGATCGGAA |
| 52065_2 | TATGAGTGTGGAGTCGTTGCCTTGCACAGGATTAGGCATTGAGNNNAGATCGGAA |
| 10031_1 | TATGAGTGTGGAGTCGTTGCATGCCGCAAAAACATTTGCATCANNNAGATCGGAAG |
| 10031_2 | TATGAGTGTGGAGTCGTTGCATGCCGCAAAAACATTTGCATCANNNAGATCGGAAG |
| 53059_1 | TATGAGTGTGGAGTCGTTGCCATAAGCTACGATCATGCCACTGNNNAGATCGGAAG |
| 53059_2 | TATGAGTGTGGAGTCGTTGCCATAAGCTACGATCATGCCACTGNNNAGATCGGAAG |
| 44826_1 | TATGAGTGTGGAGTCGTTGCTTCAGGGTATTGTGTAGGGCTTCNNNAGATCGGAA |
| 44826_2 | TATGAGTGTGGAGTCGTTGCTTCAGGGTATTGTGTAGGGCTTCNNNAGATCGGAA |
| 43764_1 | TATGAGTGTGGAGTCGTTGCAGCAACTTCAGGGAAGCATTTTCNNNAGATCGGAAG |
| 43764_2 | TATGAGTGTGGAGTCGTTGCAGCAACTTCAGGGAAGCATTTTCNNNAGATCGGAAG |
| 35616_1 | TATGAGTGTGGAGTCGTTGCCACAGTACCAGGCGTTATCAGTANNNAGATCGGAAG |
| 35616_2 | TATGAGTGTGGAGTCGTTGCCACAGTACCAGGCGTTATCAGTANNNAGATCGGAAG |
| 13494_1 | TATGAGTGTGGAGTCGTTGCAAAAAGTTCAGGATGTGTAGTNNNAGATCGGAAGA |
| 13494_2 | TATGAGTGTGGAGTCGTTGCAAAAAGTTCAGGATGTGTAGTNNNAGATCGGAAGA |
| 42712_1 | TATGAGTGTGGAGTCGTTGCTTTGGTCTTCAGTCTCTGGAAGANNNAGATCGGAAG |
| 42712_2 | TATGAGTGTGGAGTCGTTGCTTTGGTCTTCAGTCTCTGGAAGANNNAGATCGGAAG |
| 32208_1 | TATGAGTGTGGAGTCGTTGCTATCCAGAACTGAACACCTTGCTNNNAGATCGGAAG |
| 32208_2 | TATGAGTGTGGAGTCGTTGCTATCCAGAACTGAACACCTTGCTNNNAGATCGGAAG |
| 10309_1 | TATGAGTGTGGAGTCGTTGCCGCATTGCCTTCCACTTCAAATTNNNAGATCGGAAGA |
| 10309_2 | TATGAGTGTGGAGTCGTTGCCGCATTGCCTTCCACTTCAAATTNNNAGATCGGAAGA |
| 52035_1 | TATGAGTGTGGAGTCGTTGCCCAAGTAGCTCGGATTATAGGCANNNAGATCGGAA |
| 52035_2 | TATGAGTGTGGAGTCGTTGCCCAAGTAGCTCGGATTATAGGCANNNAGATCGGAA |
| 12676_1 | TATGAGTGTGGAGTCGTTGCATGTCAGATTGAAGATATACAGTNNNAGATCGGAA |
| 12676_2 | TATGAGTGTGGAGTCGTTGCATGTCAGATTGAAGATATACAGTNNNAGATCGGAA |
| 28832_1 | TATGAGTGTGGAGTCGTTGCGAAAACCTGGGAGCTAAGTGAGANNNAGATCGGAA |

| | |
|---|---|
| 28832_2 | TATGAGTGTGGAGTCGTTGCGAAAACCTGGGAGCTAAGTGAGANNNAGATCGGAA |
| 12500_1 | TATGAGTGTGGAGTCGTTGCGCATGTACAGAGTGAGTAATGNNNAGATCGGAAGA |
| 12500_2 | TATGAGTGTGGAGTCGTTGCGCATGTACAGAGTGAGTAATGNNNAGATCGGAAGA |
| 49406_1 | TATGAGTGTGGAGTCGTTGCGAGTTTTGCTCTTGTTGCCCAGNNNAGATCGGAAGA |
| 49406_2 | TATGAGTGTGGAGTCGTTGCGAGTTTTGCTCTTGTTGCCCAGNNNAGATCGGAAGA |
| 39901_1 | TATGAGTGTGGAGTCGTTGCAAGGGAAAGGGAAAAGGAAAGGANNNAGATCGG |
| 39901_2 | TATGAGTGTGGAGTCGTTGCAAGGGAAAGGGAAAAGGAAAGGANNNAGATCGG |
| 52244_1 | TATGAGTGTGGAGTCGTTGCGACACACACACACAGGAGAGTATNNNAGATCGGAA |
| 52244_2 | TATGAGTGTGGAGTCGTTGCGACACACACACACAGGAGAGTATNNNAGATCGGAA |
| 49195_1 | TATGAGTGTGGAGTCGTTGCTCACTCTTCTCCTCCTCTGTTGANNNAGATCGGAAGA |
| 49195_2 | TATGAGTGTGGAGTCGTTGCTCACTCTTCTCCTCCTCTGTTGANNNAGATCGGAAGA |
| 29035_1 | TATGAGTGTGGAGTCGTTGCATACGTACTGTGTGTGTGTGTGTNNNAGATCGGAA |
| 29035_2 | TATGAGTGTGGAGTCGTTGCATACGTACTGTGTGTGTGTGTGTNNNAGATCGGAA |
| 41098_1 | TATGAGTGTGGAGTCGTTGCCTGAGATTACAGGAGTGAGCCACNNNAGATCGGAA |
| 41098_2 | TATGAGTGTGGAGTCGTTGCCTGAGATTACAGGAGTGAGCCACNNNAGATCGGAA |
| 40667_1 | TATGAGTGTGGAGTCGTTGCGACAGAGCGAGACACCATCTAAANNNAGATCGGAA |
| 40667_2 | TATGAGTGTGGAGTCGTTGCGACAGAGCGAGACACCATCTAAANNNAGATCGGAA |
| 50018_1 | TATGAGTGTGGAGTCGTTGCTCACTTTGGGAGGTTCTCTAAGCNNNAGATCGGAAG |
| 50018_2 | TATGAGTGTGGAGTCGTTGCTCACTTTGGGAGGTTCTCTAAGCNNNAGATCGGAAG |
| 51516_1 | TATGAGTGTGGAGTCGTTGCCTTGGGTCAAATGCTCAGATGTGNNNAGATCGGAA |
| 51516_2 | TATGAGTGTGGAGTCGTTGCCTTGGGTCAAATGCTCAGATGTGNNNAGATCGGAA |
| 35998_1 | TATGAGTGTGGAGTCGTTGCCCCACAGGACAAGCTGTAAACTTNNNAGATCGGAAG |
| 35998_2 | TATGAGTGTGGAGTCGTTGCCCCACAGGACAAGCTGTAAACTTNNNAGATCGGAAG |
| 2249_1 | TATGAGTGTGGAGTCGTTGCAAGCCATTTGCAACTTGTCCCTANNNAGATCGGAAG |
| 2249_2 | TATGAGTGTGGAGTCGTTGCAAGCCATTTGCAACTTGTCCCTANNNAGATCGGAAG |
| 10326_1 | TATGAGTGTGGAGTCGTTGCAGAGCCTGTGACAATGGTGAAATNNNAGATCGGAA |
| 10326_2 | TATGAGTGTGGAGTCGTTGCAGAGCCTGTGACAATGGTGAAATNNNAGATCGGAA |
| 44257_1 | TATGAGTGTGGAGTCGTTGCCCACATTAGGGAGGGCTATCATCNNNAGATCGGAAG |
| 44257_2 | TATGAGTGTGGAGTCGTTGCCCACATTAGGGAGGGCTATCATCNNNAGATCGGAAG |
| 28973_1 | TATGAGTGTGGAGTCGTTGCTTTCTTCCTTTTTGTTGCCGTGTNNNAGATCGGAAGA |
| 28973_2 | TATGAGTGTGGAGTCGTTGCTTTCTTCCTTTTTGTTGCCGTGTNNNAGATCGGAAGA |
| 35213_1 | TATGAGTGTGGAGTCGTTGCATCATGTTGGCCAGGCTAGTTTANNNAGATCGGAAG |
| 35213_2 | TATGAGTGTGGAGTCGTTGCATCATGTTGGCCAGGCTAGTTTANNNAGATCGGAAG |
| 50272_1 | TATGAGTGTGGAGTCGTTGCTCTGAAACTCCTGGAGGTTTGAGNNNAGATCGGAA |
| 50272_2 | TATGAGTGTGGAGTCGTTGCTCTGAAACTCCTGGAGGTTTGAGNNNAGATCGGAA |
| 12750_1 | TATGAGTGTGGAGTCGTTGCATTTTGCCAATGCTATTCTATTTNNNAGATCGGAAGA |
| 12750_2 | TATGAGTGTGGAGTCGTTGCATTTTGCCAATGCTATTCTATTTNNNAGATCGGAAGA |
| 39393_1 | TATGAGTGTGGAGTCGTTGCATTTGGAGACCTTTTGCATGTCANNNAGATCGGAAG |
| 39393_2 | TATGAGTGTGGAGTCGTTGCATTTGGAGACCTTTTGCATGTCANNNAGATCGGAAG |
| 52210_1 | TATGAGTGTGGAGTCGTTGCGGCTCAGGTCATTTCAGTAGGAANNNAGATCGGAA |
| 52210_2 | TATGAGTGTGGAGTCGTTGCGGCTCAGGTCATTTCAGTAGGAANNNAGATCGGAA |
| 45502_1 | TATGAGTGTGGAGTCGTTGCCCATCTTAGGAGTGCTGTCAGAGNNNAGATCGGAA |
| 45502_2 | TATGAGTGTGGAGTCGTTGCCCATCTTAGGAGTGCTGTCAGAGNNNAGATCGGAA |

| | |
|---|---|
| 31835_1 | TATGAGTGTGGAGTCGTTGCGGCCCCCACTTAGAAATACCATANNNAGATCGGAAG |
| 31835_2 | TATGAGTGTGGAGTCGTTGCGGCCCCCACTTAGAAATACCATANNNAGATCGGAAG |
| 45289_1 | TATGAGTGTGGAGTCGTTGCCTGAGGAAAAACTGCCGAGCNNNAGATCGGAAGAG |
| 45289_2 | TATGAGTGTGGAGTCGTTGCCTGAGGAAAAACTGCCGAGCNNNAGATCGGAAGAG |
| 52231_1 | TATGAGTGTGGAGTCGTTGCTTTGTCCTTGCCATAGTTTGCTGNNNAGATCGGAAG |
| 52231_2 | TATGAGTGTGGAGTCGTTGCTTTGTCCTTGCCATAGTTTGCTGNNNAGATCGGAAG |
| 50624_1 | TATGAGTGTGGAGTCGTTGCGCCTGACTTTCCATCTCTATCCANNNAGATCGGAAGA |
| 50624_2 | TATGAGTGTGGAGTCGTTGCGCCTGACTTTCCATCTCTATCCANNNAGATCGGAAGA |
| 50741_1 | TATGAGTGTGGAGTCGTTGCTTTTTTGAGACAGAGCCTCACTCNNNAGATCGGAAG |
| 50741_2 | TATGAGTGTGGAGTCGTTGCTTTTTTGAGACAGAGCCTCACTCNNNAGATCGGAAG |
| 35792_1 | TATGAGTGTGGAGTCGTTGCTTCAAACGGCTCTCCTTTCCTTANNNAGATCGGAAGA |
| 35792_2 | TATGAGTGTGGAGTCGTTGCTTCAAACGGCTCTCCTTTCCTTANNNAGATCGGAAGA |
| 9122_1 | TATGAGTGTGGAGTCGTTGCCGTGTAGGTATATTTTGGGGGTATNNNAGATCGGA |
| 9122_2 | TATGAGTGTGGAGTCGTTGCCGTGTAGGTATATTTTGGGGGTATNNNAGATCGGA |
| 45307_1 | TATGAGTGTGGAGTCGTTGCCAGCCTGAGTCTCTGGTTTACATNNNAGATCGGAAG |
| 45307_2 | TATGAGTGTGGAGTCGTTGCCAGCCTGAGTCTCTGGTTTACATNNNAGATCGGAAG |
| 49905_1 | TATGAGTGTGGAGTCGTTGCCACAGACTTTACCCCGAAGCTNNNAGATCGGAAGAG |
| 49905_2 | TATGAGTGTGGAGTCGTTGCCACAGACTTTACCCCGAAGCTNNNAGATCGGAAGAG |
| 37570_1 | TATGAGTGTGGAGTCGTTGCATGTCAAAAGCCAAAAGCCACTTNNNAGATCGGAAG |
| 37570_2 | TATGAGTGTGGAGTCGTTGCATGTCAAAAGCCAAAAGCCACTTNNNAGATCGGAAG |
| 46849_1 | TATGAGTGTGGAGTCGTTGCCTCAGATGCCACAGTTGTTTGTTNNNAGATCGGAAG |
| 46849_2 | TATGAGTGTGGAGTCGTTGCCTCAGATGCCACAGTTGTTTGTTNNNAGATCGGAAG |
| 30135_1 | TATGAGTGTGGAGTCGTTGCCAGCGCTTTAGAACCACATTTGTNNNAGATCGGAAG |
| 30135_2 | TATGAGTGTGGAGTCGTTGCCAGCGCTTTAGAACCACATTTGTNNNAGATCGGAAG |
| 2593_1 | TATGAGTGTGGAGTCGTTGCTGGCTGCTTTTCCTGACCATTAGNNNAGATCGGAAG |
| 2593_2 | TATGAGTGTGGAGTCGTTGCTGGCTGCTTTTCCTGACCATTAGNNNAGATCGGAAG |
| 38275_1 | TATGAGTGTGGAGTCGTTGCCACTTGTGATTGGGTTTCCTGTCNNNAGATCGGAAG |
| 38275_2 | TATGAGTGTGGAGTCGTTGCCACTTGTGATTGGGTTTCCTGTCNNNAGATCGGAAG |
| 35374_1 | TATGAGTGTGGAGTCGTTGCGGCTCTTTTTTCAATCCAGGGACNNNAGATCGGAAG |
| 35374_2 | TATGAGTGTGGAGTCGTTGCGGCTCTTTTTTCAATCCAGGGACNNNAGATCGGAAG |
| 30877_1 | TATGAGTGTGGAGTCGTTGCAATCCACCACAATCAAGTAGGCTNNNAGATCGGAAG |
| 30877_2 | TATGAGTGTGGAGTCGTTGCAATCCACCACAATCAAGTAGGCTNNNAGATCGGAAG |
| 29598_1 | TATGAGTGTGGAGTCGTTGCCAAATGAAGCCAAAAAGAACAGCANNNAGATCGGA |
| 29598_2 | TATGAGTGTGGAGTCGTTGCCAAATGAAGCCAAAAAGAACAGCANNNAGATCGGA |
| 52159_1 | TATGAGTGTGGAGTCGTTGCGTGCAAGCTGTCAATGGATCTACNNNAGATCGGAA |
| 52159_2 | TATGAGTGTGGAGTCGTTGCGTGCAAGCTGTCAATGGATCTACNNNAGATCGGAA |
| 38389_1 | TATGAGTGTGGAGTCGTTGCTGACTTTTGCCGAAGTTCTTTCTNNNAGATCGGAAG |
| 38389_2 | TATGAGTGTGGAGTCGTTGCTGACTTTTGCCGAAGTTCTTTCTNNNAGATCGGAAG |
| 50622_1 | TATGAGTGTGGAGTCGTTGCTTCTGGAAGGAAGGGACCCTAGNNNAGATCGGAAG |
| 50622_2 | TATGAGTGTGGAGTCGTTGCTTCTGGAAGGAAGGGACCCTAGNNNAGATCGGAAG |
| 48694_1 | TATGAGTGTGGAGTCGTTGCTCCCCAGTGATTAGAAGTTGAGCNNNAGATCGGAA |
| 48694_2 | TATGAGTGTGGAGTCGTTGCTCCCCAGTGATTAGAAGTTGAGCNNNAGATCGGAA |
| 44906_1 | TATGAGTGTGGAGTCGTTGCCTGGGAACTGCTTTTCAGGTTCNNNAGATCGGAAGA |

| | |
|---|---|
| 44906_2 | TATGAGTGTGGAGTCGTTGCCTGGGAACTGCTTTTCAGGTTCNNNAGATCGGAAGA |
| 30986_1 | TATGAGTGTGGAGTCGTTGCCAGGCAACAGAAACAGACATTCANNNAGATCGGAAG |
| 30986_2 | TATGAGTGTGGAGTCGTTGCCAGGCAACAGAAACAGACATTCANNNAGATCGGAAG |
| 29578_1 | TATGAGTGTGGAGTCGTTGCGAGAAGTTCTCCTACATCCCCGANNNAGATCGGAAG |
| 29578_2 | TATGAGTGTGGAGTCGTTGCGAGAAGTTCTCCTACATCCCCGANNNAGATCGGAAG |
| 51715_1 | TATGAGTGTGGAGTCGTTGCCTGTACCTTGGCTAGGATTACCCNNNAGATCGGAAG |
| 51715_2 | TATGAGTGTGGAGTCGTTGCCTGTACCTTGGCTAGGATTACCCNNNAGATCGGAAG |
| 31641_1 | TATGAGTGTGGAGTCGTTGCGGTTCTGTTGTGCAAACCTACTCNNNAGATCGGAAG |
| 31641_2 | TATGAGTGTGGAGTCGTTGCGGTTCTGTTGTGCAAACCTACTCNNNAGATCGGAAG |
| 42233_1 | TATGAGTGTGGAGTCGTTGCGAGACAGAGTCACACTGTTTTGCNNNAGATCGGAAG |
| 42233_2 | TATGAGTGTGGAGTCGTTGCGAGACAGAGTCACACTGTTTTGCNNNAGATCGGAAG |
| 8988_1 | TATGAGTGTGGAGTCGTTGCCTGCCCACAGTAGAACCTAGTCTNNNAGATCGGAAG |
| 8988_2 | TATGAGTGTGGAGTCGTTGCCTGCCCACAGTAGAACCTAGTCTNNNAGATCGGAAG |
| 39743_1 | TATGAGTGTGGAGTCGTTGCGTTTCAGTTCCTTTCACATGGAGTNNNAGATCGGAA |
| 39743_2 | TATGAGTGTGGAGTCGTTGCGTTTCAGTTCCTTTCACATGGAGTNNNAGATCGGAA |
| 42288_1 | TATGAGTGTGGAGTCGTTGCAAGTGATGGGATTTCAAGGGTGANNNAGATCGGA |
| 42288_2 | TATGAGTGTGGAGTCGTTGCAAGTGATGGGATTTCAAGGGTGANNNAGATCGGA |
| 43360_1 | TATGAGTGTGGAGTCGTTGCCAATCTCAGTTCACTGCAAGCTCNNNAGATCGGAAG |
| 43360_2 | TATGAGTGTGGAGTCGTTGCCAATCTCAGTTCACTGCAAGCTCNNNAGATCGGAAG |
| 31546_1 | TATGAGTGTGGAGTCGTTGCAGCAAGCTTCTACCTATTGTGGTNNNAGATCGGAAG |
| 31546_2 | TATGAGTGTGGAGTCGTTGCAGCAAGCTTCTACCTATTGTGGTNNNAGATCGGAAG |
| 28776_1 | TATGAGTGTGGAGTCGTTGCCCCCTTCCCATTTCAATTTGCTTNNNAGATCGGAAGA |
| 28776_2 | TATGAGTGTGGAGTCGTTGCCCCCTTCCCATTTCAATTTGCTTNNNAGATCGGAAGA |
| 43787_1 | TATGAGTGTGGAGTCGTTGCGGATGAAAATGCCTTCCCATCAGNNNAGATCGGAAG |
| 43787_2 | TATGAGTGTGGAGTCGTTGCGGATGAAAATGCCTTCCCATCAGNNNAGATCGGAAG |
| 51306_1 | TATGAGTGTGGAGTCGTTGCTCACCCTATACCAGTGCTTTTCCNNNAGATCGGAAGA |
| 51306_2 | TATGAGTGTGGAGTCGTTGCTCACCCTATACCAGTGCTTTTCCNNNAGATCGGAAGA |
| 29599_1 | TATGAGTGTGGAGTCGTTGCCTAAGACCATGCAAACACACACCNNNAGATCGGAAG |
| 29599_2 | TATGAGTGTGGAGTCGTTGCCTAAGACCATGCAAACACACACCNNNAGATCGGAAG |
| 46242_1 | TATGAGTGTGGAGTCGTTGCGCTGGAGGCCATTATCCTAAGAGNNNAGATCGGAA |
| 46242_2 | TATGAGTGTGGAGTCGTTGCGCTGGAGGCCATTATCCTAAGAGNNNAGATCGGAA |
| 29070_1 | TATGAGTGTGGAGTCGTTGCACACACCTAGTGGGGAGATCTAANNNAGATCGGAA |
| 29070_2 | TATGAGTGTGGAGTCGTTGCACACACCTAGTGGGGAGATCTAANNNAGATCGGAA |
| 51783_1 | TATGAGTGTGGAGTCGTTGCATGCAGCTGACTGTTGATCACTANNNAGATCGGAAG |
| 51783_2 | TATGAGTGTGGAGTCGTTGCATGCAGCTGACTGTTGATCACTANNNAGATCGGAAG |
| 31_1 | TATGAGTGTGGAGTCGTTGCGGAGATGGAGGGGAAAAAGGAGANNNAGATCGG |
| 31_2 | TATGAGTGTGGAGTCGTTGCGGAGATGGAGGGGAAAAAGGAGANNNAGATCGG |
| 29339_1 | TATGAGTGTGGAGTCGTTGCTCCTGTGCATAGTCTGATGTCTGNNNAGATCGGAAG |
| 29339_2 | TATGAGTGTGGAGTCGTTGCTCCTGTGCATAGTCTGATGTCTGNNNAGATCGGAAG |
| 37813_1 | TATGAGTGTGGAGTCGTTGCTAGTCTCAGCTACTCAGGAAGCTNNNAGATCGGAAG |
| 37813_2 | TATGAGTGTGGAGTCGTTGCTAGTCTCAGCTACTCAGGAAGCTNNNAGATCGGAAG |
| 36648_1 | TATGAGTGTGGAGTCGTTGCGAGGGATGTGATATTGACCCTGGNNNAGATCGGAA |
| 36648_2 | TATGAGTGTGGAGTCGTTGCGAGGGATGTGATATTGACCCTGGNNNAGATCGGAA |

| | |
|---|---|
| 1972_1 | TATGAGTGTGGAGTCGTTGCTTCTGGGAAGTAAACCACAAGTCANNNAGATCGGAA |
| 1972_2 | TATGAGTGTGGAGTCGTTGCTTCTGGGAAGTAAACCACAAGTCANNNAGATCGGAA |
| 9927_1 | TATGAGTGTGGAGTCGTTGCACACTTCCTACCTTCCTTACCCCNNNAGATCGGAAGAG |
| 9927_2 | TATGAGTGTGGAGTCGTTGCACACTTCCTACCTTCCTTACCCCNNNAGATCGGAAGAG |
| 38445_1 | TATGAGTGTGGAGTCGTTGCCTCAAGGATGGTGCACAACAATCNNNAGATCGGAAG |
| 38445_2 | TATGAGTGTGGAGTCGTTGCCTCAAGGATGGTGCACAACAATCNNNAGATCGGAAG |
| 40610_1 | TATGAGTGTGGAGTCGTTGCTCTACGGGGTGGAATATAAAAAAGTNNNAGATCGGA |
| 40610_2 | TATGAGTGTGGAGTCGTTGCTCTACGGGGTGGAATATAAAAAAGTNNNAGATCGGA |
| 51937_1 | TATGAGTGTGGAGTCGTTGCGGGCTCTTCACTTCCCAGTAGNNNAGATCGGAAGAG |
| 51937_2 | TATGAGTGTGGAGTCGTTGCGGGCTCTTCACTTCCCAGTAGNNNAGATCGGAAGAG |
| 29315_1 | TATGAGTGTGGAGTCGTTGCATAATGGCATTTGCAGTGACCTGNNNAGATCGGAAG |
| 29315_2 | TATGAGTGTGGAGTCGTTGCATAATGGCATTTGCAGTGACCTGNNNAGATCGGAAG |
| 44903_1 | TATGAGTGTGGAGTCGTTGCGCATCTCTCTCTCACCCTCTTTCNNNAGATCGGAAGAG |
| 44903_2 | TATGAGTGTGGAGTCGTTGCGCATCTCTCTCTCACCCTCTTTCNNNAGATCGGAAGAG |
| 34368_1 | TATGAGTGTGGAGTCGTTGCTCAAAACACAATCATGCCCTTCCNNNAGATCGGAAGA |
| 34368_2 | TATGAGTGTGGAGTCGTTGCTCAAAACACAATCATGCCCTTCCNNNAGATCGGAAGA |
| 34833_1 | TATGAGTGTGGAGTCGTTGCTTGGGCAACTTCCTGACATAGTANNNAGATCGGAAG |
| 34833_2 | TATGAGTGTGGAGTCGTTGCTTGGGCAACTTCCTGACATAGTANNNAGATCGGAAG |
| 46838_1 | TATGAGTGTGGAGTCGTTGCGCCTCCCAAAGTGCTAGGATTATNNNAGATCGGAAG |
| 46838_2 | TATGAGTGTGGAGTCGTTGCGCCTCCCAAAGTGCTAGGATTATNNNAGATCGGAAG |
| 45917_1 | TATGAGTGTGGAGTCGTTGCTTCAAGTGATTCTCCTGCCTCAGNNNAGATCGGAAGA |
| 45917_2 | TATGAGTGTGGAGTCGTTGCTTCAAGTGATTCTCCTGCCTCAGNNNAGATCGGAAGA |
| 39662_1 | TATGAGTGTGGAGTCGTTGCACAGCTTTACACAAAAATGCACCNNNAGATCGGAAGA |
| 39662_2 | TATGAGTGTGGAGTCGTTGCACAGCTTTACACAAAAATGCACCNNNAGATCGGAAGA |
| 13372_1 | TATGAGTGTGGAGTCGTTGCTTTTTAAAATGAGATGAAGCCCNNNAGATCGGAAGA |
| 13372_2 | TATGAGTGTGGAGTCGTTGCTTTTTAAAATGAGATGAAGCCCNNNAGATCGGAAGA |
| 48778_1 | TATGAGTGTGGAGTCGTTGCCCACGCCCAGCTAATTTTTGTATNNNAGATCGGAAGA |
| 48778_2 | TATGAGTGTGGAGTCGTTGCCCACGCCCAGCTAATTTTTGTATNNNAGATCGGAAGA |
| 42714_1 | TATGAGTGTGGAGTCGTTGCCATGCCTGTAGTCCCAGTTACTTNNNAGATCGGAAGA |
| 42714_2 | TATGAGTGTGGAGTCGTTGCCATGCCTGTAGTCCCAGTTACTTNNNAGATCGGAAGA |
| 40855_1 | TATGAGTGTGGAGTCGTTGCTGTTATGTGAATCAGAACTTCCCANNNAGATCGGAAG |
| 40855_2 | TATGAGTGTGGAGTCGTTGCTGTTATGTGAATCAGAACTTCCCANNNAGATCGGAAG |
| 39171_1 | TATGAGTGTGGAGTCGTTGCACAGGGTGAAAAGACAACCTACANNNAGATCGGAAG |
| 39171_2 | TATGAGTGTGGAGTCGTTGCACAGGGTGAAAAGACAACCTACANNNAGATCGGAAG |
| 42436_1 | TATGAGTGTGGAGTCGTTGCCAAAAACGAGTTTCTGCAGAGGTNNNAGATCGGAAG |
| 42436_2 | TATGAGTGTGGAGTCGTTGCCAAAAACGAGTTTCTGCAGAGGTNNNAGATCGGAAG |
| 30270_1 | TATGAGTGTGGAGTCGTTGCTTGCTCTCAGATCTTTTCGCTCTNNNAGATCGGAAGA |
| 30270_2 | TATGAGTGTGGAGTCGTTGCTTGCTCTCAGATCTTTTCGCTCTNNNAGATCGGAAGA |
| 51360_1 | TATGAGTGTGGAGTCGTTGCCAATAGCAGTAGTGGGACATGGTNNNAGATCGGAA |
| 51360_2 | TATGAGTGTGGAGTCGTTGCCAATAGCAGTAGTGGGACATGGTNNNAGATCGGAA |
| 13640_1 | TATGAGTGTGGAGTCGTTGCTTGGAACTTGAATTCTGTTTTAANNNAGATCGGAAGA |
| 13640_2 | TATGAGTGTGGAGTCGTTGCTTGGAACTTGAATTCTGTTTTAANNNAGATCGGAAGA |
| 35655_1 | TATGAGTGTGGAGTCGTTGCGTTTCACCATGTTAGCCAGGATGNNNAGATCGGAAG |

| | |
|---|---|
| 35655_2 | TATGAGTGTGGAGTCGTTGCGTTTCACCATGTTAGCCAGGATGNNNAGATCGGAAG |
| 48247_1 | TATGAGTGTGGAGTCGTTGCCAAGTGATTCTCTTGCATCAGCCNNNAGATCGGAAG |
| 48247_2 | TATGAGTGTGGAGTCGTTGCCAAGTGATTCTCTTGCATCAGCCNNNAGATCGGAAG |
| 13784_1 | TATGAGTGTGGAGTCGTTGCATAAGTTTTAAACACCATAACGTNNNAGATCGGAAG |
| 13784_2 | TATGAGTGTGGAGTCGTTGCATAAGTTTTAAACACCATAACGTNNNAGATCGGAAG |
| 32152_1 | TATGAGTGTGGAGTCGTTGCACTGCAAGGCCATTGTCCTAATANNNAGATCGGAAG |
| 32152_2 | TATGAGTGTGGAGTCGTTGCACTGCAAGGCCATTGTCCTAATANNNAGATCGGAAG |
| 42320_1 | TATGAGTGTGGAGTCGTTGCATATACATAGACACACCCACACATNNNAGATCGGAA |
| 42320_2 | TATGAGTGTGGAGTCGTTGCATATACATAGACACACCCACACATNNNAGATCGGAA |
| 31530_1 | TATGAGTGTGGAGTCGTTGCGTCACCAAGCTCCATCTAGTCAANNNAGATCGGAAG |
| 31530_2 | TATGAGTGTGGAGTCGTTGCGTCACCAAGCTCCATCTAGTCAANNNAGATCGGAAG |
| 34198_1 | TATGAGTGTGGAGTCGTTGCCAGAGCTTCTATGTCAGTCCCAANNNAGATCGGAAG |
| 34198_2 | TATGAGTGTGGAGTCGTTGCCAGAGCTTCTATGTCAGTCCCAANNNAGATCGGAAG |
| 39335_1 | TATGAGTGTGGAGTCGTTGCAGAGGTTGTAGAGAGCCGAGATCNNNAGATCGGAA |
| 39335_2 | TATGAGTGTGGAGTCGTTGCAGAGGTTGTAGAGAGCCGAGATCNNNAGATCGGAA |
| 49972_1 | TATGAGTGTGGAGTCGTTGCGAACCTCCCAAGACTGTAGTGTTNNNAGATCGGAAG |
| 49972_2 | TATGAGTGTGGAGTCGTTGCGAACCTCCCAAGACTGTAGTGTTNNNAGATCGGAAG |
| 3144_1 | TATGAGTGTGGAGTCGTTGCAAAATGGAGTGGATTGGGGTGTCNNNAGATCGGA |
| 3144_2 | TATGAGTGTGGAGTCGTTGCAAAATGGAGTGGATTGGGGTGTCNNNAGATCGGA |
| 50003_1 | TATGAGTGTGGAGTCGTTGCTCTCTGGGAATCTCTCTCAGGTTNNNAGATCGGAAG |
| 50003_2 | TATGAGTGTGGAGTCGTTGCTCTCTGGGAATCTCTCTCAGGTTNNNAGATCGGAAG |
| 48807_1 | TATGAGTGTGGAGTCGTTGCTAAGAGGGGACATTGTTTGGTCCNNNAGATCGGAA |
| 48807_2 | TATGAGTGTGGAGTCGTTGCTAAGAGGGGACATTGTTTGGTCCNNNAGATCGGAA |
| 31960_1 | TATGAGTGTGGAGTCGTTGCGTATATGGCTCCACACAAATGGCNNNAGATCGGAAG |
| 31960_2 | TATGAGTGTGGAGTCGTTGCGTATATGGCTCCACACAAATGGCNNNAGATCGGAAG |
| 31029_1 | TATGAGTGTGGAGTCGTTGCTCAAATGCAGCATGTCCAAAACTNNNAGATCGGAAG |
| 31029_2 | TATGAGTGTGGAGTCGTTGCTCAAATGCAGCATGTCCAAAACTNNNAGATCGGAAG |
| 46108_1 | TATGAGTGTGGAGTCGTTGCATAGATTCTCACTCTGTCACCCANNNAGATCGGAAG |
| 46108_2 | TATGAGTGTGGAGTCGTTGCATAGATTCTCACTCTGTCACCCANNNAGATCGGAAG |
| 4283_1 | TATGAGTGTGGAGTCGTTGCTCCCCTGATTTCCATCTGGACATNNNAGATCGGAAG |
| 4283_2 | TATGAGTGTGGAGTCGTTGCTCCCCTGATTTCCATCTGGACATNNNAGATCGGAAG |
| 46929_1 | TATGAGTGTGGAGTCGTTGCCTCACAGTAGGCTAAAGGGATCCNNNAGATCGGAA |
| 46929_2 | TATGAGTGTGGAGTCGTTGCCTCACAGTAGGCTAAAGGGATCCNNNAGATCGGAA |
| 52996_1 | TATGAGTGTGGAGTCGTTGCCCGGTTTCAAGCGATTCTCATACNNNAGATCGGAAG |
| 52996_2 | TATGAGTGTGGAGTCGTTGCCCGGTTTCAAGCGATTCTCATACNNNAGATCGGAAG |
| 52122_1 | TATGAGTGTGGAGTCGTTGCCATCAGGTGACTGTCAGGTAGTTNNNAGATCGGAA |
| 52122_2 | TATGAGTGTGGAGTCGTTGCCATCAGGTGACTGTCAGGTAGTTNNNAGATCGGAA |
| 30715_1 | TATGAGTGTGGAGTCGTTGCTTGAGAGGCTAAAGGGCTACACTNNNAGATCGGAA |
| 30715_2 | TATGAGTGTGGAGTCGTTGCTTGAGAGGCTAAAGGGCTACACTNNNAGATCGGAA |
| 30625_1 | ATGAGTGTGGAGTCGTTGCACATACCACATTGTTCTGGTTACAATNNNAGATCGGA |
| 30625_2 | ATGAGTGTGGAGTCGTTGCACATACCACATTGTTCTGGTTACAATNNNAGATCGGA |
| 50217_1 | TATGAGTGTGGAGTCGTTGCTTGGCTAGGACATGCTATGATCCNNNAGATCGGAAG |
| 50217_2 | TATGAGTGTGGAGTCGTTGCTTGGCTAGGACATGCTATGATCCNNNAGATCGGAAG |

| | |
|---|---|
| 51906_1 | TATGAGTGTGGAGTCGTTGCCCTTGCTTTGCATTTCCTACTGGNNNAGATCGGAAGA |
| 51906_2 | TATGAGTGTGGAGTCGTTGCCCTTGCTTTGCATTTCCTACTGGNNNAGATCGGAAGA |
| 29063_1 | TATGAGTGTGGAGTCGTTGCATGCACACACACAATAAAGGTGGNNNAGATCGGAA |
| 29063_2 | TATGAGTGTGGAGTCGTTGCATGCACACACACAATAAAGGTGGNNNAGATCGGAA |
| 38609_1 | TATGAGTGTGGAGTCGTTGCCAGAAGAATTGCTTGAACCCGAGNNNAGATCGGAA |
| 38609_2 | TATGAGTGTGGAGTCGTTGCCAGAAGAATTGCTTGAACCCGAGNNNAGATCGGAA |
| 35566_1 | TATGAGTGTGGAGTCGTTGCGGATTACTGGCATAAACCACTGCNNNAGATCGGAAG |
| 35566_2 | TATGAGTGTGGAGTCGTTGCGGATTACTGGCATAAACCACTGCNNNAGATCGGAAG |
| 11473_1 | TATGAGTGTGGAGTCGTTGCGCCATCTTCATTTCTGGCTGCAGGAATNNNAGATCG |
| 11473_2 | TATGAGTGTGGAGTCGTTGCGCCATCTTCATTTCTGGCTGCAGGAATNNNAGATCG |
| 42136_1 | TATGAGTGTGGAGTCGTTGCTGATCTCGGCTCACTGCAAGNNNAGATCGGAAGAG |
| 42136_2 | TATGAGTGTGGAGTCGTTGCTGATCTCGGCTCACTGCAAGNNNAGATCGGAAGAG |
| 40356_1 | TATGAGTGTGGAGTCGTTGCAGAGGGATACTTCAGTTCAGGGANNNAGATCGGAA |
| 40356_2 | TATGAGTGTGGAGTCGTTGCAGAGGGATACTTCAGTTCAGGGANNNAGATCGGAA |
| 11163_1 | TATGAGTGTGGAGTCGTTGCATGTTCTTACACTGGCTTTCTGGTNNNAGATCGGAA |
| 11163_2 | TATGAGTGTGGAGTCGTTGCATGTTCTTACACTGGCTTTCTGGTNNNAGATCGGAA |
| 2535_1 | TATGAGTGTGGAGTCGTTGCTCTGCCATTTACAAGCTAGCAGGNNNAGATCGGAAG |
| 2535_2 | TATGAGTGTGGAGTCGTTGCTCTGCCATTTACAAGCTAGCAGGNNNAGATCGGAAG |
| 43539_1 | TATGAGTGTGGAGTCGTTGCCTTCATTTGAGGTCCAGGAAGGGNNNAGATCGGAA |
| 43539_2 | TATGAGTGTGGAGTCGTTGCCTTCATTTGAGGTCCAGGAAGGGNNNAGATCGGAA |
| 48224_1 | TATGAGTGTGGAGTCGTTGCGTTCTTCTGCCTCAATCTCCTGANNNAGATCGGAAGA |
| 48224_2 | TATGAGTGTGGAGTCGTTGCGTTCTTCTGCCTCAATCTCCTGANNNAGATCGGAAGA |
| 35400_1 | TATGAGTGTGGAGTCGTTGCTACCGCCACACCTAGCTAATTTTNNNAGATCGGAAGA |
| 35400_2 | TATGAGTGTGGAGTCGTTGCTACCGCCACACCTAGCTAATTTTNNNAGATCGGAAGA |
| 36998_1 | TATGAGTGTGGAGTCGTTGCAGGAGAATCACTGGATCCTGAGANNNAGATCGGAA |
| 36998_2 | TATGAGTGTGGAGTCGTTGCAGGAGAATCACTGGATCCTGAGANNNAGATCGGAA |
| 46530_1 | TATGAGTGTGGAGTCGTTGCCACTGTGCCCAGCTAATTTTTGTNNNAGATCGGAAG |
| 46530_2 | TATGAGTGTGGAGTCGTTGCCACTGTGCCCAGCTAATTTTTGTNNNAGATCGGAAG |
| 32256_1 | TATGAGTGTGGAGTCGTTGCGATGTTCAAAGAACTGGTGCCAANNNAGATCGGAA |
| 32256_2 | TATGAGTGTGGAGTCGTTGCGATGTTCAAAGAACTGGTGCCAANNNAGATCGGAA |
| 29453_1 | TATGAGTGTGGAGTCGTTGCGGGGAAATCTGGACAGACACATNNNAGATCGGAAG |
| 29453_2 | TATGAGTGTGGAGTCGTTGCGGGGAAATCTGGACAGACACATNNNAGATCGGAAG |
| 32185_1 | TATGAGTGTGGAGTCGTTGCAGGGCCTATAGTCCCAACTACTTNNNAGATCGGAAG |
| 32185_2 | TATGAGTGTGGAGTCGTTGCAGGGCCTATAGTCCCAACTACTTNNNAGATCGGAAG |
| 10627_1 | TATGAGTGTGGAGTCGTTGCTGGCAGTGATGTTCCAAGTTCATNNNAGATCGGAAG |
| 10627_2 | TATGAGTGTGGAGTCGTTGCTGGCAGTGATGTTCCAAGTTCATNNNAGATCGGAAG |
| 48056_1 | TATGAGTGTGGAGTCGTTGCAGATGGAGTCTCTCTCTGTCACCNNNAGATCGGAAG |
| 48056_2 | TATGAGTGTGGAGTCGTTGCAGATGGAGTCTCTCTCTGTCACCNNNAGATCGGAAG |
| 35548_1 | TATGAGTGTGGAGTCGTTGCAGTCTCAACTAACTGCAAGCTCTNNNAGATCGGAAG |
| 35548_2 | TATGAGTGTGGAGTCGTTGCAGTCTCAACTAACTGCAAGCTCTNNNAGATCGGAAG |
| 43211_1 | TATGAGTGTGGAGTCGTTGCATTGCTTTGGGGAGTATGGTCATNNNAGATCGGAA |
| 43211_2 | TATGAGTGTGGAGTCGTTGCATTGCTTTGGGGAGTATGGTCATNNNAGATCGGAA |
| 46903_1 | TATGAGTGTGGAGTCGTTGCTGTGTAACTTTTTCTCTGCCACANNNAGATCGGAAG |

| | |
|---|---|
| 46903_2 | TATGAGTGTGGAGTCGTTGCTGTGTAACTTTTTCTCTGCCACANNNAGATCGGAAG |
| 29415_1 | TATGAGTGTGGAGTCGTTGCGCTGAAGGAGTATTTTGCGTGTGNNNAGATCGGAA |
| 29415_2 | TATGAGTGTGGAGTCGTTGCGCTGAAGGAGTATTTTGCGTGTGNNNAGATCGGAA |
| 41237_1 | TATGAGTGTGGAGTCGTTGCAGATCGAGACCATCTTGGCTAACNNNAGATCGGAAG |
| 41237_2 | TATGAGTGTGGAGTCGTTGCAGATCGAGACCATCTTGGCTAACNNNAGATCGGAAG |
| 39469_1 | TATGAGTGTGGAGTCGTTGCGGCCTTCCCAGATCATTTAGTCTNNNAGATCGGAAG |
| 39469_2 | TATGAGTGTGGAGTCGTTGCGGCCTTCCCAGATCATTTAGTCTNNNAGATCGGAAG |
| 51525_1 | TATGAGTGTGGAGTCGTTGCTGGTGGTAGAAAAGAGGCTTCTCNNNAGATCGGAA |
| 51525_2 | TATGAGTGTGGAGTCGTTGCTGGTGGTAGAAAAGAGGCTTCTCNNNAGATCGGAA |
| 45146_1 | TATGAGTGTGGAGTCGTTGCAATGTCACCCACTCCCAAATGTANNNAGATCGGAAG |
| 45146_2 | TATGAGTGTGGAGTCGTTGCAATGTCACCCACTCCCAAATGTANNNAGATCGGAAG |
| 36952_1 | TATGAGTGTGGAGTCGTTGCCATCACACCCAGCCTTCTTTTTCNNNAGATCGGAAGA |
| 36952_2 | TATGAGTGTGGAGTCGTTGCCATCACACCCAGCCTTCTTTTTCNNNAGATCGGAAGA |
| 44857_1 | TATGAGTGTGGAGTCGTTGCAAACCTGCTGTCCCTAAATGGATNNNAGATCGGAAG |
| 44857_2 | TATGAGTGTGGAGTCGTTGCAAACCTGCTGTCCCTAAATGGATNNNAGATCGGAAG |
| 44444_1 | TATGAGTGTGGAGTCGTTGCCTGGCATGTAGTGGCATGATCTTNNNAGATCGGAAG |
| 44444_2 | TATGAGTGTGGAGTCGTTGCCTGGCATGTAGTGGCATGATCTTNNNAGATCGGAAG |
| 47583_1 | TATGAGTGTGGAGTCGTTGCCAGGTTCACGCCATTCTCCTNNNAGATCGGAAGAGC |
| 47583_2 | TATGAGTGTGGAGTCGTTGCCAGGTTCACGCCATTCTCCTNNNAGATCGGAAGAGC |
| 12194_1 | TATGAGTGTGGAGTCGTTGCAACTAAGGTAGCAACAGAGCTTTNNNAGATCGGAAG |
| 12194_2 | TATGAGTGTGGAGTCGTTGCAACTAAGGTAGCAACAGAGCTTTNNNAGATCGGAAG |
| 48408_1 | TATGAGTGTGGAGTCGTTGCGCTATGCTGTGAGGATGAAATCCNNNAGATCGGAA |
| 48408_2 | TATGAGTGTGGAGTCGTTGCGCTATGCTGTGAGGATGAAATCCNNNAGATCGGAA |
| 39283_1 | TATGAGTGTGGAGTCGTTGCGTGCCTGTAAGTCCCACTACTCNNNAGATCGGAAGA |
| 39283_2 | TATGAGTGTGGAGTCGTTGCGTGCCTGTAAGTCCCACTACTCNNNAGATCGGAAGA |
| 51237_1 | TATGAGTGTGGAGTCGTTGCGATGCAGGAGATCAGACCCTAGANNNAGATCGGAA |
| 51237_2 | TATGAGTGTGGAGTCGTTGCGATGCAGGAGATCAGACCCTAGANNNAGATCGGAA |
| 44996_1 | TATGAGTGTGGAGTCGTTGCAGGGACATAATGGCTCCAGAAAGNNNAGATCGGAA |
| 44996_2 | TATGAGTGTGGAGTCGTTGCAGGGACATAATGGCTCCAGAAAGNNNAGATCGGAA |
| 29778_1 | TATGAGTGTGGAGTCGTTGCTAACTTGCGATAACAACCCAGTCNNNAGATCGGAAG |
| 29778_2 | TATGAGTGTGGAGTCGTTGCTAACTTGCGATAACAACCCAGTCNNNAGATCGGAAG |
| 13094_1 | TATGAGTGTGGAGTCGTTGCATCAAAGATATCAACAGAAAACCNNNAGATCGGAAG |
| 13094_2 | TATGAGTGTGGAGTCGTTGCATCAAAGATATCAACAGAAAACCNNNAGATCGGAAG |
| 37783_1 | TATGAGTGTGGAGTCGTTGCAGTTGAGAGGAACTCTAGCAAATNNNAGATCGGAA |
| 37783_2 | TATGAGTGTGGAGTCGTTGCAGTTGAGAGGAACTCTAGCAAATNNNAGATCGGAA |
| 52386_1 | TATGAGTGTGGAGTCGTTGCGCGCCCGCATTTTTTTTTTTTTTNNNAGATCGGAAGA |
| 52386_2 | TATGAGTGTGGAGTCGTTGCGCGCCCGCATTTTTTTTTTTTTTNNNAGATCGGAAGA |
| 40778_1 | TATGAGTGTGGAGTCGTTGCATAAATGGCACATGGGAAAGCTGNNNAGATCGGAA |
| 40778_2 | TATGAGTGTGGAGTCGTTGCATAAATGGCACATGGGAAAGCTGNNNAGATCGGAA |
| 30191_1 | TATGAGTGTGGAGTCGTTGCGCTTTTGGCTTCTATTGGGTCTCNNNAGATCGGAAG |
| 30191_2 | TATGAGTGTGGAGTCGTTGCGCTTTTGGCTTCTATTGGGTCTCNNNAGATCGGAAG |
| 12613_1 | TATGAGTGTGGAGTCGTTGCCTAAATTGGGAAACAGTTGTTAGNNNAGATCGGAA |
| 12613_2 | TATGAGTGTGGAGTCGTTGCCTAAATTGGGAAACAGTTGTTAGNNNAGATCGGAA |

| | |
|---|---|
| 31476_1 | TATGAGTGTGGAGTCGTTGCGGGTGGTTGTTCATCCTGTATCTNNNAGATCGGAAG |
| 31476_2 | TATGAGTGTGGAGTCGTTGCGGGTGGTTGTTCATCCTGTATCTNNNAGATCGGAAG |
| 31141_1 | TATGAGTGTGGAGTCGTTGCGTTCAAGTGATTCTTCTGCCTCGNNNAGATCGGAAG |
| 31141_2 | TATGAGTGTGGAGTCGTTGCGTTCAAGTGATTCTTCTGCCTCGNNNAGATCGGAAG |
| 34775_1 | TATGAGTGTGGAGTCGTTGCTTGGCTAAAGGCGACGTAAAATGNNNAGATCGGAA |
| 34775_2 | TATGAGTGTGGAGTCGTTGCTTGGCTAAAGGCGACGTAAAATGNNNAGATCGGAA |
| 41018_1 | TATGAGTGTGGAGTCGTTGCCAAAAGAAGACGTACAAGTGGCCNNNAGATCGGAA |
| 41018_2 | TATGAGTGTGGAGTCGTTGCCAAAAGAAGACGTACAAGTGGCCNNNAGATCGGAA |
| 45328_1 | TATGAGTGTGGAGTCGTTGCTACATGGTTTCTAGCCAGCTCAANNNAGATCGGAAG |
| 45328_2 | TATGAGTGTGGAGTCGTTGCTACATGGTTTCTAGCCAGCTCAANNNAGATCGGAAG |
| 31118_1 | TATGAGTGTGGAGTCGTTGCTGGGTAAACAAGAGCTCAGCTAGNNNAGATCGGAA |
| 31118_2 | TATGAGTGTGGAGTCGTTGCTGGGTAAACAAGAGCTCAGCTAGNNNAGATCGGAA |
| 49491_1 | TATGAGTGTGGAGTCGTTGCATGCCATGAGTCAAATCCTTCCTNNNAGATCGGAAG |
| 49491_2 | TATGAGTGTGGAGTCGTTGCATGCCATGAGTCAAATCCTTCCTNNNAGATCGGAAG |
| 43688_1 | TATGAGTGTGGAGTCGTTGCGATGAGGGATCCCATGACTCATGNNNAGATCGGAA |
| 43688_2 | TATGAGTGTGGAGTCGTTGCGATGAGGGATCCCATGACTCATGNNNAGATCGGAA |
| 33209_1 | TATGAGTGTGGAGTCGTTGCGCTGCTAACAAACTGAATCCCTGNNNAGATCGGAAG |
| 33209_2 | TATGAGTGTGGAGTCGTTGCGCTGCTAACAAACTGAATCCCTGNNNAGATCGGAAG |
| 36957_1 | TATGAGTGTGGAGTCGTTGCTGAAAAAGCAGAGAGTTCCTGGANNNAGATCGGAA |
| 36957_2 | TATGAGTGTGGAGTCGTTGCTGAAAAAGCAGAGAGTTCCTGGANNNAGATCGGAA |
| 31536_1 | TATGAGTGTGGAGTCGTTGCACTCCTTACACAAAGACAGCACTNNNAGATCGGAAG |
| 31536_2 | TATGAGTGTGGAGTCGTTGCACTCCTTACACAAAGACAGCACTNNNAGATCGGAAG |
| 43087_1 | TATGAGTGTGGAGTCGTTGCTATGGCAGTGGTCCCATAAGATTNNNAGATCGGAA |
| 43087_2 | TATGAGTGTGGAGTCGTTGCTATGGCAGTGGTCCCATAAGATTNNNAGATCGGAA |
| 31519_1 | TATGAGTGTGGAGTCGTTGCTACCCATACATGCACATTGTCCANNNAGATCGGAAG |
| 31519_2 | TATGAGTGTGGAGTCGTTGCTACCCATACATGCACATTGTCCANNNAGATCGGAAG |
| 36018_1 | TATGAGTGTGGAGTCGTTGCATAAAGACATGGCTGAGACTGGGNNNAGATCGGAA |
| 36018_2 | TATGAGTGTGGAGTCGTTGCATAAAGACATGGCTGAGACTGGGNNNAGATCGGAA |
| 36070_1 | TATGAGTGTGGAGTCGTTGCTGCAATCAGCTATAATCCTTCCCTNNNAGATCGGAAG |
| 36070_2 | TATGAGTGTGGAGTCGTTGCTGCAATCAGCTATAATCCTTCCCTNNNAGATCGGAAG |
| 43361_1 | TATGAGTGTGGAGTCGTTGCTGTAGAGAGGGGGTTTCAACATGNNNAGATCGGAA |
| 43361_2 | TATGAGTGTGGAGTCGTTGCTGTAGAGAGGGGGTTTCAACATGNNNAGATCGGAA |
| 41512_1 | TATGAGTGTGGAGTCGTTGCTAGGTAGTTAGGCAGACATGAGCNNNAGATCGGAA |
| 41512_2 | TATGAGTGTGGAGTCGTTGCTAGGTAGTTAGGCAGACATGAGCNNNAGATCGGAA |
| 48042_1 | TATGAGTGTGGAGTCGTTGCTAGCTAGGGATTACAGGCATGTGNNNAGATCGGAA |
| 48042_2 | TATGAGTGTGGAGTCGTTGCTAGCTAGGGATTACAGGCATGTGNNNAGATCGGAA |
| 47781_1 | TATGAGTGTGGAGTCGTTGCCAGACACAGAGAGGCTTCCTGNNNAGATCGGAAGA |
| 47781_2 | TATGAGTGTGGAGTCGTTGCCAGACACAGAGAGGCTTCCTGNNNAGATCGGAAGA |
| 11153_1 | TATGAGTGTGGAGTCGTTGCGCTACCTGGAACTATTGCAGCCNNNAGATCGGAAGA |
| 11153_2 | TATGAGTGTGGAGTCGTTGCGCTACCTGGAACTATTGCAGCCNNNAGATCGGAAGA |
| 46045_1 | TATGAGTGTGGAGTCGTTGCAAAGCAGAAGCCATCCTATTCCANNNAGATCGGAAG |
| 46045_2 | TATGAGTGTGGAGTCGTTGCAAAGCAGAAGCCATCCTATTCCANNNAGATCGGAAG |
| 51620_1 | TATGAGTGTGGAGTCGTTGCCATTAGAAAATTAGCCCGGCACANNNAGATCGGAAG |

| | |
|---|---|
| 51620_2 | TATGAGTGTGGAGTCGTTGCCATTAGAAAATTAGCCCGGCACANNNAGATCGGAAG |
| 39694_1 | TATGAGTGTGGAGTCGTTGCAAATCGTCCAACAACACAAAGCANNNAGATCGGAAG |
| 39694_2 | TATGAGTGTGGAGTCGTTGCAAATCGTCCAACAACACAAAGCANNNAGATCGGAAG |
| 50283_1 | TATGAGTGTGGAGTCGTTGCCGAGACCAGCCTGACTAACATAGNNNAGATCGGAAG |
| 50283_2 | TATGAGTGTGGAGTCGTTGCCGAGACCAGCCTGACTAACATAGNNNAGATCGGAAG |
| 36565_1 | TATGAGTGTGGAGTCGTTGCAGCAATGTGGCTATGAGAAGAGANNNAGATCGGAA |
| 36565_2 | TATGAGTGTGGAGTCGTTGCAGCAATGTGGCTATGAGAAGAGANNNAGATCGGAA |
| 13402_1 | TATGAGTGTGGAGTCGTTGCTTTTGGTCTTGTTTCAATTTCTTNNNAGATCGGAAGA |
| 13402_2 | TATGAGTGTGGAGTCGTTGCTTTTGGTCTTGTTTCAATTTCTTNNNAGATCGGAAGA |
| 31807_1 | TATGAGTGTGGAGTCGTTGCGATCTTGGCTCACTGCAACCTNNNAGATCGGAAGAG |
| 31807_2 | TATGAGTGTGGAGTCGTTGCGATCTTGGCTCACTGCAACCTNNNAGATCGGAAGAG |
| 46457_1 | TATGAGTGTGGAGTCGTTGCTTTGTAAGGTTACGTGTTGTTTGANNNAGATCGGAA |
| 46457_2 | TATGAGTGTGGAGTCGTTGCTTTGTAAGGTTACGTGTTGTTTGANNNAGATCGGAA |
| 42953_1 | TATGAGTGTGGAGTCGTTGCGGGATTGCATTCTTGATTTGGCTNNNAGATCGGAAG |
| 42953_2 | TATGAGTGTGGAGTCGTTGCGGGATTGCATTCTTGATTTGGCTNNNAGATCGGAAG |
| 31107_1 | TATGAGTGTGGAGTCGTTGCTGTCAGGAGTGAGAATTTCCCAGNNNAGATCGGAAG |
| 31107_2 | TATGAGTGTGGAGTCGTTGCTGTCAGGAGTGAGAATTTCCCAGNNNAGATCGGAAG |
| 42129_1 | TATGAGTGTGGAGTCGTTGCGATCTTGGTTCACTGCAAGCTCNNNAGATCGGAAGA |
| 42129_2 | TATGAGTGTGGAGTCGTTGCGATCTTGGTTCACTGCAAGCTCNNNAGATCGGAAGA |
| 49385_1 | TATGAGTGTGGAGTCGTTGCACCACCCCTGGCTAATTTTTGTANNNAGATCGGAAGA |
| 49385_2 | TATGAGTGTGGAGTCGTTGCACCACCCCTGGCTAATTTTTGTANNNAGATCGGAAGA |
| 30501_1 | TATGAGTGTGGAGTCGTTGCGGATCTGGGACAGTCTTTGTGCTNNNAGATCGGAAG |
| 30501_2 | TATGAGTGTGGAGTCGTTGCGGATCTGGGACAGTCTTTGTGCTNNNAGATCGGAAG |
| 37178_1 | TATGAGTGTGGAGTCGTTGCATGGCAAAATGGGAATTCAGACCNNNAGATCGGAAG |
| 37178_2 | TATGAGTGTGGAGTCGTTGCATGGCAAAATGGGAATTCAGACCNNNAGATCGGAAG |
| 43958_1 | TATGAGTGTGGAGTCGTTGCCAGGGAGAAAAGAGAAGCAGAGANNNAGATCGGA |
| 43958_2 | TATGAGTGTGGAGTCGTTGCCAGGGAGAAAAGAGAAGCAGAGANNNAGATCGGA |
| 29136_1 | TATGAGTGTGGAGTCGTTGCGCTAACGCCAGGAAAGAAAGAAANNNAGATCGGAA |
| 29136_2 | TATGAGTGTGGAGTCGTTGCGCTAACGCCAGGAAAGAAAGAAANNNAGATCGGAA |
| 32798_1 | TATGAGTGTGGAGTCGTTGCGAGATGGAGTTTCACCATGTTGGNNNAGATCGGAAG |
| 32798_2 | TATGAGTGTGGAGTCGTTGCGAGATGGAGTTTCACCATGTTGGNNNAGATCGGAAG |
| 48950_1 | TATGAGTGTGGAGTCGTTGCCCTCCCAGATTCAAGCCATTCTTNNNAGATCGGAAGA |
| 48950_2 | TATGAGTGTGGAGTCGTTGCCCTCCCAGATTCAAGCCATTCTTNNNAGATCGGAAGA |
| 33942_1 | TATGAGTGTGGAGTCGTTGCCCTCCCAGATTCAAGCCATTCTCNNNAGATCGGAAGA |
| 33942_2 | TATGAGTGTGGAGTCGTTGCCCTCCCAGATTCAAGCCATTCTCNNNAGATCGGAAGA |
| 36234_1 | TATGAGTGTGGAGTCGTTGCCAGGGACAGTGCTATTGTTACCTNNNAGATCGGAAG |
| 36234_2 | TATGAGTGTGGAGTCGTTGCCAGGGACAGTGCTATTGTTACCTNNNAGATCGGAAG |
| 10159_1 | TATGAGTGTGGAGTCGTTGCTTGTGAATGTGGCTTTTTGTACANNNAGATCGGAAG |
| 10159_2 | TATGAGTGTGGAGTCGTTGCTTGTGAATGTGGCTTTTTGTACANNNAGATCGGAAG |
| 44321_1 | TATGAGTGTGGAGTCGTTGCATGAAAGGAGGGCTGTGAGATTTNNNAGATCGGAA |
| 44321_2 | TATGAGTGTGGAGTCGTTGCATGAAAGGAGGGCTGTGAGATTTNNNAGATCGGAA |
| 3097_1 | TATGAGTGTGGAGTCGTTGCCACACATTGGAGAGGTGTTTTGCNNNAGATCGGAAG |
| 3097_2 | TATGAGTGTGGAGTCGTTGCCACACATTGGAGAGGTGTTTTGCNNNAGATCGGAAG |

| | |
|---|---|
| 39284_1 | TATGAGTGTGGAGTCGTTGCTGCACTGGTCATGAACTACTTGANNNAGATCGGAAG |
| 39284_2 | TATGAGTGTGGAGTCGTTGCTGCACTGGTCATGAACTACTTGANNNAGATCGGAAG |
| 47696_1 | TATGAGTGTGGAGTCGTTGCGATCAAGCAATCAATCCTCCTGCNNNAGATCGGAAG |
| 47696_2 | TATGAGTGTGGAGTCGTTGCGATCAAGCAATCAATCCTCCTGCNNNAGATCGGAAG |
| 40703_1 | TATGAGTGTGGAGTCGTTGCGGAGGGACTGAAGATAGCTGATGNNNAGATCGGA |
| 40703_2 | TATGAGTGTGGAGTCGTTGCGGAGGGACTGAAGATAGCTGATGNNNAGATCGGA |
| 46390_1 | TATGAGTGTGGAGTCGTTGCTAGAGTGCAGTGGTGAGATCTTGNNNAGATCGGAA |
| 46390_2 | TATGAGTGTGGAGTCGTTGCTAGAGTGCAGTGGTGAGATCTTGNNNAGATCGGAA |
| 29413_1 | TATGAGTGTGGAGTCGTTGCGCATATTACTGGTGCAGGACCATNNNAGATCGGAA |
| 29413_2 | TATGAGTGTGGAGTCGTTGCGCATATTACTGGTGCAGGACCATNNNAGATCGGAA |
| 33451_1 | TATGAGTGTGGAGTCGTTGCCAACAAAAAAACCCAGCTTGGTGNNNAGATCGGAA |
| 33451_2 | TATGAGTGTGGAGTCGTTGCCAACAAAAAAACCCAGCTTGGTGNNNAGATCGGAA |
| 43805_1 | TATGAGTGTGGAGTCGTTGCCCCTAGAATCTCAATGCCACCTTNNNAGATCGGAAGA |
| 43805_2 | TATGAGTGTGGAGTCGTTGCCCCTAGAATCTCAATGCCACCTTNNNAGATCGGAAGA |
| 10049_1 | TATGAGTGTGGAGTCGTTGCCTTCGCTCGGGATAGAGGGAGNNNAGATCGGAAGA |
| 10049_2 | TATGAGTGTGGAGTCGTTGCCTTCGCTCGGGATAGAGGGAGNNNAGATCGGAAGA |
| 37090_1 | TATGAGTGTGGAGTCGTTGCTGTGAGCAACATTATTCTGGGGANNNAGATCGGAA |
| 37090_2 | TATGAGTGTGGAGTCGTTGCTGTGAGCAACATTATTCTGGGGANNNAGATCGGAA |
| 29326_1 | TATGAGTGTGGAGTCGTTGCAGACAGACACAGAGACAGAGAGANNNAGATCGGA |
| 29326_2 | TATGAGTGTGGAGTCGTTGCAGACAGACACAGAGACAGAGAGANNNAGATCGGA |
| 40384_1 | TATGAGTGTGGAGTCGTTGCAAGGTCAGAGAAGAAGACGTGACNNNAGATCGGA |
| 40384_2 | TATGAGTGTGGAGTCGTTGCAAGGTCAGAGAAGAAGACGTGACNNNAGATCGGA |
| 52039_1 | TATGAGTGTGGAGTCGTTGCGAAAGAGGCCAAACAAGCTCAATNNNAGATCGGAA |
| 52039_2 | TATGAGTGTGGAGTCGTTGCGAAAGAGGCCAAACAAGCTCAATNNNAGATCGGAA |
| 41119_1 | TATGAGTGTGGAGTCGTTGCGACCTATTTCTGTCTTCCCTGCTNNNAGATCGGAAGA |
| 41119_2 | TATGAGTGTGGAGTCGTTGCGACCTATTTCTGTCTTCCCTGCTNNNAGATCGGAAGA |
| 47160_1 | TATGAGTGTGGAGTCGTTGCGCTCAAGTGATTCTCCTTCCTCANNNAGATCGGAAGA |
| 47160_2 | TATGAGTGTGGAGTCGTTGCGCTCAAGTGATTCTCCTTCCTCANNNAGATCGGAAGA |
| 30687_1 | TATGAGTGTGGAGTCGTTGCATTATGCTTTCATGCCTCCTCCCNNNAGATCGGAAGA |
| 30687_2 | TATGAGTGTGGAGTCGTTGCATTATGCTTTCATGCCTCCTCCCNNNAGATCGGAAGA |
| 34370_1 | TATGAGTGTGGAGTCGTTGCTCCTGGGGAGTTCTGAACAATTTNNNAGATCGGAAG |
| 34370_2 | TATGAGTGTGGAGTCGTTGCTCCTGGGGAGTTCTGAACAATTTNNNAGATCGGAAG |
| 10196_1 | TATGAGTGTGGAGTCGTTGCGGCATCTTCTAAGCCACAAAAGGNNNAGATCGGAA |
| 10196_2 | TATGAGTGTGGAGTCGTTGCGGCATCTTCTAAGCCACAAAAGGNNNAGATCGGAA |
| 47825_1 | TATGAGTGTGGAGTCGTTGCCAATCTCAGCTCACTGCAACTTCNNNAGATCGGAAGA |
| 47825_2 | TATGAGTGTGGAGTCGTTGCCAATCTCAGCTCACTGCAACTTCNNNAGATCGGAAGA |
| 39303_1 | TATGAGTGTGGAGTCGTTGCGAGGCAGGAGAATGGTGTGAANNNAGATCGGAAG |
| 39303_2 | TATGAGTGTGGAGTCGTTGCGAGGCAGGAGAATGGTGTGAANNNAGATCGGAAG |
| 51207_1 | TATGAGTGTGGAGTCGTTGCATCACTTTGATACCAGCCCAAGTNNNAGATCGGAAG |
| 51207_2 | TATGAGTGTGGAGTCGTTGCATCACTTTGATACCAGCCCAAGTNNNAGATCGGAAG |
| 47646_1 | TATGAGTGTGGAGTCGTTGCCACCCCCGGCTAATTTTTTGTATNNNAGATCGGAAGA |
| 47646_2 | TATGAGTGTGGAGTCGTTGCCACCCCCGGCTAATTTTTTGTATNNNAGATCGGAAGA |
| 52407_1 | TATGAGTGTGGAGTCGTTGCGACAGGATAGCATGCACATCTCANNNAGATCGGAA |

| | |
|---|---|
| 52407_2 | TATGAGTGTGGAGTCGTTGCGACAGGATAGCATGCACATCTCANNNAGATCGGAAG |
| 30697_1 | TATGAGTGTGGAGTCGTTGCGATGGCAAAGAGAAGTGAGGTGANNNAGATCGGA |
| 30697_2 | TATGAGTGTGGAGTCGTTGCGATGGCAAAGAGAAGTGAGGTGANNNAGATCGGA |
| 35523_1 | TATGAGTGTGGAGTCGTTGCCCCAGGTTCAAGCAATTCTCTTGNNNAGATCGGAAG |
| 35523_2 | TATGAGTGTGGAGTCGTTGCCCCAGGTTCAAGCAATTCTCTTGNNNAGATCGGAAG |
| 30691_1 | TATGAGTGTGGAGTCGTTGCAGAGAGATGCTGAGGTTTCTGAGTNNNAGATCGGA |
| 30691_2 | TATGAGTGTGGAGTCGTTGCAGAGAGATGCTGAGGTTTCTGAGTNNNAGATCGGA |
| 45303_1 | TATGAGTGTGGAGTCGTTGCAGCAAGAGAAGGAATTTGAGGGANNNAGATCGGA |
| 45303_2 | TATGAGTGTGGAGTCGTTGCAGCAAGAGAAGGAATTTGAGGGANNNAGATCGGA |
| 53426_1 | TATGAGTGTGGAGTCGTTGCAAGAAAAGGAAGGAAGGGAAGGGNNNAGATCGG |
| 53426_2 | TATGAGTGTGGAGTCGTTGCAAGAAAAGGAAGGAAGGGAAGGGNNNAGATCGG |
| 37356_1 | TATGAGTGTGGAGTCGTTGCTGACAAAGGGAAAACTCCATGTGNNNAGATCGGAA |
| 37356_2 | TATGAGTGTGGAGTCGTTGCTGACAAAGGGAAAACTCCATGTGNNNAGATCGGAA |
| 46249_1 | TATGAGTGTGGAGTCGTTGCTGTTCATTGCAACCATTGTCTCCNNNAGATCGGAAGA |
| 46249_2 | TATGAGTGTGGAGTCGTTGCTGTTCATTGCAACCATTGTCTCCNNNAGATCGGAAGA |
| 30674_1 | TATGAGTGTGGAGTCGTTGCTCACCTTCTTTCCCTCCACTATGTTNNNAGATCGGAAG |
| 30674_2 | TATGAGTGTGGAGTCGTTGCTCACCTTCTTTCCCTCCACTATGTTNNNAGATCGGAAG |
| 36302_1 | TATGAGTGTGGAGTCGTTGCTTATTGAAAAAGAGGCCGCAACANNNAGATCGGAA |
| 36302_2 | TATGAGTGTGGAGTCGTTGCTTATTGAAAAAGAGGCCGCAACANNNAGATCGGAA |
| 47099_1 | TATGAGTGTGGAGTCGTTGCGCCCGAGATAACTACTGTCTCTGNNNAGATCGGAAG |
| 47099_2 | TATGAGTGTGGAGTCGTTGCGCCCGAGATAACTACTGTCTCTGNNNAGATCGGAAG |
| 51222_1 | TATGAGTGTGGAGTCGTTGCGGAGTAGTCGGTTCGAAAAGTCTNNNAGATCGGAA |
| 51222_2 | TATGAGTGTGGAGTCGTTGCGGAGTAGTCGGTTCGAAAAGTCTNNNAGATCGGAA |
| 12614_1 | TATGAGTGTGGAGTCGTTGCGCAGTAAGAGTCATACGAATAANNNAGATCGGAAG |
| 12614_2 | TATGAGTGTGGAGTCGTTGCGCAGTAAGAGTCATACGAATAANNNAGATCGGAAG |
| 39093_1 | TATGAGTGTGGAGTCGTTGCCCATCAAAAAGGGGGCAAATGATNNNAGATCGGAA |
| 39093_2 | TATGAGTGTGGAGTCGTTGCCCATCAAAAAGGGGGCAAATGATNNNAGATCGGAA |
| 50890_1 | TATGAGTGTGGAGTCGTTGCCCTGTTTTACACACATGTGCTCANNNAGATCGGAAGA |
| 50890_2 | TATGAGTGTGGAGTCGTTGCCCTGTTTTACACACATGTGCTCANNNAGATCGGAAGA |
| 51855_1 | TATGAGTGTGGAGTCGTTGCGCTCCACGTAAGTGATAAAAGCCNNNAGATCGGAAG |
| 51855_2 | TATGAGTGTGGAGTCGTTGCGCTCCACGTAAGTGATAAAAGCCNNNAGATCGGAAG |
| 43606_1 | TATGAGTGTGGAGTCGTTGCGAGGACAGGAACAGTGGATCTGNNNAGATCGGAAG |
| 43606_2 | TATGAGTGTGGAGTCGTTGCGAGGACAGGAACAGTGGATCTGNNNAGATCGGAAG |
| 53097_1 | TATGAGTGTGGAGTCGTTGCGAGGCAGAAAAACAGGGTCTAGANNNAGATCGGAA |
| 53097_2 | TATGAGTGTGGAGTCGTTGCGAGGCAGAAAAACAGGGTCTAGANNNAGATCGGAA |
| 44671_1 | TATGAGTGTGGAGTCGTTGCAAAAAGAGATTTGTGGCTGCTCCNNNAGATCGGAAG |
| 44671_2 | TATGAGTGTGGAGTCGTTGCAAAAAGAGATTTGTGGCTGCTCCNNNAGATCGGAAG |
| 29799_1 | TATGAGTGTGGAGTCGTTGCAGGGATCATGTGAGAGCATTGTTNNNAGATCGGAA |
| 29799_2 | TATGAGTGTGGAGTCGTTGCAGGGATCATGTGAGAGCATTGTTNNNAGATCGGAA |
| 31759_1 | TATGAGTGTGGAGTCGTTGCGACACAAGCACTACCACTTTGACNNNAGATCGGAAG |
| 31759_2 | TATGAGTGTGGAGTCGTTGCGACACAAGCACTACCACTTTGACNNNAGATCGGAAG |
| 45734_1 | TATGAGTGTGGAGTCGTTGCCTGAGGAGAAGCAGGAAGTGAGNNNAGATCGGAA |
| 45734_2 | TATGAGTGTGGAGTCGTTGCCTGAGGAGAAGCAGGAAGTGAGNNNAGATCGGAA |

| | |
|---|---|
| 28855_1 | TATGAGTGTGGAGTCGTTGCATGGAGGAGGCTGAAGAAAACTTNNNAGATCGGAA |
| 28855_2 | TATGAGTGTGGAGTCGTTGCATGGAGGAGGCTGAAGAAAACTTNNNAGATCGGAA |
| 40718_1 | TATGAGTGTGGAGTCGTTGCTACAAAAAATTAGCCGGGCATGGNNNAGATCGGAA |
| 40718_2 | TATGAGTGTGGAGTCGTTGCTACAAAAAATTAGCCGGGCATGGNNNAGATCGGAA |
| 46201_1 | TATGAGTGTGGAGTCGTTGCGAGTATACCAACATGCCCAGCTANNNAGATCGGAAG |
| 46201_2 | TATGAGTGTGGAGTCGTTGCGAGTATACCAACATGCCCAGCTANNNAGATCGGAAG |
| 45766_1 | TATGAGTGTGGAGTCGTTGCGCTTTCCAAATAGCTGGGGTTACNNNAGATCGGAAG |
| 45766_2 | TATGAGTGTGGAGTCGTTGCGCTTTCCAAATAGCTGGGGTTACNNNAGATCGGAAG |
| 37413_1 | TATGAGTGTGGAGTCGTTGCGAGCAATGGGAAATCACTGGATGNNNAGATCGGAA |
| 37413_2 | TATGAGTGTGGAGTCGTTGCGAGCAATGGGAAATCACTGGATGNNNAGATCGGAA |
| 43479_1 | TATGAGTGTGGAGTCGTTGCGCCACCACATCAGGCTAATTTTTNNNAGATCGGAAG |
| 43479_2 | TATGAGTGTGGAGTCGTTGCGCCACCACATCAGGCTAATTTTTNNNAGATCGGAAG |
| 39220_1 | TATGAGTGTGGAGTCGTTGCTTTGCACCTTTTCAACCTCAACCNNNAGATCGGAAGA |
| 39220_2 | TATGAGTGTGGAGTCGTTGCTTTGCACCTTTTCAACCTCAACCNNNAGATCGGAAGA |
| 47729_1 | TATGAGTGTGGAGTCGTTGCCACTTCAGTCCCCAGAGTAACTGNNNAGATCGGAAG |
| 47729_2 | TATGAGTGTGGAGTCGTTGCCACTTCAGTCCCCAGAGTAACTGNNNAGATCGGAAG |
| 47954_1 | TATGAGTGTGGAGTCGTTGCATGCCCAGCCACTTTTTTTCTTTNNNAGATCGGAAGA |
| 47954_2 | TATGAGTGTGGAGTCGTTGCATGCCCAGCCACTTTTTTTCTTTNNNAGATCGGAAGA |
| 41770_1 | TATGAGTGTGGAGTCGTTGCTAGCTGGGATTACAGGTATGTGCNNNAGATCGGAA |
| 41770_2 | TATGAGTGTGGAGTCGTTGCTAGCTGGGATTACAGGTATGTGCNNNAGATCGGAA |
| 44601_1 | TATGAGTGTGGAGTCGTTGCAACCAGCTCTACCTCATGTTCAGNNNAGATCGGAAG |
| 44601_2 | TATGAGTGTGGAGTCGTTGCAACCAGCTCTACCTCATGTTCAGNNNAGATCGGAAG |
| 49321_1 | TATGAGTGTGGAGTCGTTGCATCACGAATCATCACAGCCTCAANNNAGATCGGAAG |
| 49321_2 | TATGAGTGTGGAGTCGTTGCATCACGAATCATCACAGCCTCAANNNAGATCGGAAG |
| 50756_1 | TATGAGTGTGGAGTCGTTGCTGTGGACATGAAGCTGAGAGAAANNNAGATCGGAA |
| 50756_2 | TATGAGTGTGGAGTCGTTGCTGTGGACATGAAGCTGAGAGAAANNNAGATCGGAA |
| 30851_1 | TATGAGTGTGGAGTCGTTGCAATACGACTTTGCCAACTGAGTCNNNAGATCGGAAG |
| 30851_2 | TATGAGTGTGGAGTCGTTGCAATACGACTTTGCCAACTGAGTCNNNAGATCGGAAG |
| 30223_1 | TATGAGTGTGGAGTCGTTGCGACAAAACTATGCTTTGCCCCATNNNAGATCGGAAG |
| 30223_2 | TATGAGTGTGGAGTCGTTGCGACAAAACTATGCTTTGCCCCATNNNAGATCGGAAG |
| 9271_1 | TATGAGTGTGGAGTCGTTGCCAGCCAAAGAAGAAACCACTGGANNNAGATCGGAA |
| 9271_2 | TATGAGTGTGGAGTCGTTGCCAGCCAAAGAAGAAACCACTGGANNNAGATCGGAA |
| 37141_1 | TATGAGTGTGGAGTCGTTGCTGAACATCAGGAATCAGGTGTGTNNNAGATCGGAA |
| 37141_2 | TATGAGTGTGGAGTCGTTGCTGAACATCAGGAATCAGGTGTGTNNNAGATCGGAA |
| 44662_1 | TATGAGTGTGGAGTCGTTGCTTCGATGTGAAGGCATTGAGGGNNNAGATCGGAAG |
| 44662_2 | TATGAGTGTGGAGTCGTTGCTTCGATGTGAAGGCATTGAGGGNNNAGATCGGAAG |
| 13389_1 | TATGAGTGTGGAGTCGTTGCAGAACTCACAGATTCTTTCTTATNNNAGATCGGAAGA |
| 13389_2 | TATGAGTGTGGAGTCGTTGCAGAACTCACAGATTCTTTCTTATNNNAGATCGGAAGA |
| 41265_1 | TATGAGTGTGGAGTCGTTGCATTGCTTGAATCTGAGAGACGGANNNAGATCGGAA |
| 41265_2 | TATGAGTGTGGAGTCGTTGCATTGCTTGAATCTGAGAGACGGANNNAGATCGGAA |
| 12620_1 | TATGAGTGTGGAGTCGTTGCTACCTACTACCATTTTTCAATGTNNNAGATCGGAAGA |
| 12620_2 | TATGAGTGTGGAGTCGTTGCTACCTACTACCATTTTTCAATGTNNNAGATCGGAAGA |
| 30948_1 | TATGAGTGTGGAGTCGTTGCGCTAGGATGGTGGCTAAGATACCNNNAGATCGGAA |

| | |
|---|---|
| 30948_2 | TATGAGTGTGGAGTCGTTGCGCTAGGATGGTGGCTAAGATACCNNNAGATCGGAA |
| 44632_1 | TATGAGTGTGGAGTCGTTGCTTCTTGGCCAAATCCATGAGAGTNNNAGATCGGAAG |
| 44632_2 | TATGAGTGTGGAGTCGTTGCTTCTTGGCCAAATCCATGAGAGTNNNAGATCGGAAG |
| 34461_1 | TATGAGTGTGGAGTCGTTGCGAGTAGCGAGGGCTGAAAAATTGNNNAGATCGGA |
| 34461_2 | TATGAGTGTGGAGTCGTTGCGAGTAGCGAGGGCTGAAAAATTGNNNAGATCGGA |
| 31950_1 | TATGAGTGTGGAGTCGTTGCCCTTATGTGTGTTTAGGCCTAGANNNAGATCGGAAG |
| 31950_2 | TATGAGTGTGGAGTCGTTGCCCTTATGTGTGTTTAGGCCTAGANNNAGATCGGAAG |
| 33124_1 | TATGAGTGTGGAGTCGTTGCGTGGGAAAAAGGTAGGGAAGCTANNNAGATCGGA |
| 33124_2 | TATGAGTGTGGAGTCGTTGCGTGGGAAAAAGGTAGGGAAGCTANNNAGATCGGA |
| 39375_1 | TATGAGTGTGGAGTCGTTGCTCTTCTAGAGACTGGCAACACACNNNAGATCGGAAG |
| 39375_2 | TATGAGTGTGGAGTCGTTGCTCTTCTAGAGACTGGCAACACACNNNAGATCGGAAG |
| 32203_1 | TATGAGTGTGGAGTCGTTGCCTTCAAGCAATACTCCTGCCTCANNNAGATCGGAAG |
| 32203_2 | TATGAGTGTGGAGTCGTTGCCTTCAAGCAATACTCCTGCCTCANNNAGATCGGAAG |
| 10963_1 | TATGAGTGTGGAGTCGTTGCTCTGTCTTCCAGTAGCTGGGATTNNNAGATCGGAAG |
| 10963_2 | TATGAGTGTGGAGTCGTTGCTCTGTCTTCCAGTAGCTGGGATTNNNAGATCGGAAG |
| 35350_1 | TATGAGTGTGGAGTCGTTGCGATCTCGGCTCACTGTAAGCTCNNNAGATCGGAAGA |
| 35350_2 | TATGAGTGTGGAGTCGTTGCGATCTCGGCTCACTGTAAGCTCNNNAGATCGGAAGA |
| 47918_1 | TATGAGTGTGGAGTCGTTGCCAGAGTAGCTGGGATTACAGGTGNNNAGATCGGAA |
| 47918_2 | TATGAGTGTGGAGTCGTTGCCAGAGTAGCTGGGATTACAGGTGNNNAGATCGGAA |
| 47250_1 | TATGAGTGTGGAGTCGTTGCCTCCTGGGTTCAAGCCATTCTCNNNAGATCGGAAGA |
| 47250_2 | TATGAGTGTGGAGTCGTTGCCTCCTGGGTTCAAGCCATTCTCNNNAGATCGGAAGA |
| 35115_1 | TATGAGTGTGGAGTCGTTGCGATCTCGGATCACTGCAACCTCNNNAGATCGGAAGA |
| 35115_2 | TATGAGTGTGGAGTCGTTGCGATCTCGGATCACTGCAACCTCNNNAGATCGGAAGA |
| 34716_1 | TATGAGTGTGGAGTCGTTGCCGAGACAGAATCTTGCAACCTCTNNNAGATCGGAAG |
| 34716_2 | TATGAGTGTGGAGTCGTTGCCGAGACAGAATCTTGCAACCTCTNNNAGATCGGAAG |
| 12650_1 | TATGAGTGTGGAGTCGTTGCGACTGGAGAAAAGGTTTCAGNNNAGATCGGAAGAG |
| 12650_2 | TATGAGTGTGGAGTCGTTGCGACTGGAGAAAAGGTTTCAGNNNAGATCGGAAGAG |
| 52548_1 | TATGAGTGTGGAGTCGTTGCCCAGACCTGATCCACTGCTTCNNNAGATCGGAAGAG |
| 52548_2 | TATGAGTGTGGAGTCGTTGCCCAGACCTGATCCACTGCTTCNNNAGATCGGAAGAG |
| 30079_1 | TATGAGTGTGGAGTCGTTGCTGTGTAACTCAGTCAGAACGAGGNNNAGATCGGAA |
| 30079_2 | TATGAGTGTGGAGTCGTTGCTGTGTAACTCAGTCAGAACGAGGNNNAGATCGGAA |
| 10954_1 | TATGAGTGTGGAGTCGTTGCTCAAGAGAAAGCAGAGCCCCAANNNAGATCGGAAG |
| 10954_2 | TATGAGTGTGGAGTCGTTGCTCAAGAGAAAGCAGAGCCCCAANNNAGATCGGAAG |
| 13068_1 | TATGAGTGTGGAGTCGTTGCCTGGTTGATTTTCTTGATACTTTNNNAGATCGGAAG |
| 13068_2 | TATGAGTGTGGAGTCGTTGCCTGGTTGATTTTCTTGATACTTTNNNAGATCGGAAG |
| 34538_1 | TATGAGTGTGGAGTCGTTGCGTGCTTGCTTCCCATAGAAATGGNNNAGATCGGAAG |
| 34538_2 | TATGAGTGTGGAGTCGTTGCGTGCTTGCTTCCCATAGAAATGGNNNAGATCGGAAG |
| 47478_1 | TATGAGTGTGGAGTCGTTGCCAGTGGCATAGTCATAGCTCACTNNNAGATCGGAAG |
| 47478_2 | TATGAGTGTGGAGTCGTTGCCAGTGGCATAGTCATAGCTCACTNNNAGATCGGAAG |
| 10657_1 | TATGAGTGTGGAGTCGTTGCCATAATAGGCTCTCAGGCAGCAGNNNAGATCGGAA |
| 10657_2 | TATGAGTGTGGAGTCGTTGCCATAATAGGCTCTCAGGCAGCAGNNNAGATCGGAA |
| 36142_1 | TATGAGTGTGGAGTCGTTGCGGATCACGAGGTCACGAGATTATNNNAGATCGGAA |
| 36142_2 | TATGAGTGTGGAGTCGTTGCGGATCACGAGGTCACGAGATTATNNNAGATCGGAA |

| | |
|---|---|
| 29279_1 | TATGAGTGTGGAGTCGTTGCCCGGACTACAACTCCCAGAATGNNNAGATCGGAAGA |
| 29279_2 | TATGAGTGTGGAGTCGTTGCCCGGACTACAACTCCCAGAATGNNNAGATCGGAAGA |
| 41543_1 | TATGAGTGTGGAGTCGTTGCGTCTCCCGAGTAGCTGGAATTACNNNAGATCGGAAG |
| 41543_2 | TATGAGTGTGGAGTCGTTGCGTCTCCCGAGTAGCTGGAATTACNNNAGATCGGAAG |
| 43368_1 | TATGAGTGTGGAGTCGTTGCTTTAGTAGAGGCAGGGGTTTCACNNNAGATCGGAA |
| 43368_2 | TATGAGTGTGGAGTCGTTGCTTTAGTAGAGGCAGGGGTTTCACNNNAGATCGGAA |
| 33243_1 | TATGAGTGTGGAGTCGTTGCGTAGCTGGGACTACAGTTTCACANNNAGATCGGAAG |
| 33243_2 | TATGAGTGTGGAGTCGTTGCGTAGCTGGGACTACAGTTTCACANNNAGATCGGAAG |
| 33626_1 | TATGAGTGTGGAGTCGTTGCGTTCACGCCATTCTCCTACCTCNNNAGATCGGAAGAG |
| 33626_2 | TATGAGTGTGGAGTCGTTGCGTTCACGCCATTCTCCTACCTCNNNAGATCGGAAGAG |
| 9047_1 | TATGAGTGTGGAGTCGTTGCACACTGGGTTTGTGGTATCTCCTNNNAGATCGGAAG |
| 9047_2 | TATGAGTGTGGAGTCGTTGCACACTGGGTTTGTGGTATCTCCTNNNAGATCGGAAG |
| 53061_1 | TATGAGTGTGGAGTCGTTGCGTCACCAAAATCAGAAAGCTCCGNNNAGATCGGAAG |
| 53061_2 | TATGAGTGTGGAGTCGTTGCGTCACCAAAATCAGAAAGCTCCGNNNAGATCGGAAG |
| 52130_1 | TATGAGTGTGGAGTCGTTGCGAGTGAGAATATGCGGTGTTTGGNNNAGATCGGAA |
| 52130_2 | TATGAGTGTGGAGTCGTTGCGAGTGAGAATATGCGGTGTTTGGNNNAGATCGGAA |
| 39580_1 | TATGAGTGTGGAGTCGTTGCCCACTTCCAAGAACATTATGGCANNNAGATCGGAAG |
| 39580_2 | TATGAGTGTGGAGTCGTTGCCCACTTCCAAGAACATTATGGCANNNAGATCGGAAG |
| 34795_1 | TATGAGTGTGGAGTCGTTGCCAGGTAGCTGGGATTACAGGTACNNNAGATCGGAA |
| 34795_2 | TATGAGTGTGGAGTCGTTGCCAGGTAGCTGGGATTACAGGTACNNNAGATCGGAA |
| 39771_1 | TATGAGTGTGGAGTCGTTGCCTAGGTGACAGAACGAGACTCTGNNNAGATCGGAA |
| 39771_2 | TATGAGTGTGGAGTCGTTGCCTAGGTGACAGAACGAGACTCTGNNNAGATCGGAA |
| 49336_1 | TATGAGTGTGGAGTCGTTGCTGAAGTGCAGTGATGTGATCTCANNNAGATCGGAA |
| 49336_2 | TATGAGTGTGGAGTCGTTGCTGAAGTGCAGTGATGTGATCTCANNNAGATCGGAA |
| 43814_1 | TATGAGTGTGGAGTCGTTGCCCAAGAGCCACATAAGAAATGCANNNAGATCGGAAG |
| 43814_2 | TATGAGTGTGGAGTCGTTGCCCAAGAGCCACATAAGAAATGCANNNAGATCGGAAG |
| 50877_1 | TATGAGTGTGGAGTCGTTGCCCGGCCGCTTTTTTTTTTTTTTTNNNAGATCGGAAGA |
| 50877_2 | TATGAGTGTGGAGTCGTTGCCCGGCCGCTTTTTTTTTTTTTTTNNNAGATCGGAAGA |
| 35243_1 | TATGAGTGTGGAGTCGTTGCTGGAGTACAGTGGTGTGATCTTGNNNAGATCGGAA |
| 35243_2 | TATGAGTGTGGAGTCGTTGCTGGAGTACAGTGGTGTGATCTTGNNNAGATCGGAA |
| 50819_1 | TATGAGTGTGGAGTCGTTGCCTGAAGACCACTTACCCAGTTCANNNAGATCGGAAG |
| 50819_2 | TATGAGTGTGGAGTCGTTGCCTGAAGACCACTTACCCAGTTCANNNAGATCGGAAG |
| 36710_1 | TATGAGTGTGGAGTCGTTGCGTGACATACACCTGTAGTCCCAGNNNAGATCGGAAG |
| 36710_2 | TATGAGTGTGGAGTCGTTGCGTGACATACACCTGTAGTCCCAGNNNAGATCGGAAG |
| 29487_1 | TATGAGTGTGGAGTCGTTGCAAGAAGCCACTGTGTTAGCATCANNNAGATCGGAAG |
| 29487_2 | TATGAGTGTGGAGTCGTTGCAAGAAGCCACTGTGTTAGCATCANNNAGATCGGAAG |
| 48067_1 | TATGAGTGTGGAGTCGTTGCGCAAAAGTTCACAGTGTGAGTGTNNNAGATCGGAA |
| 48067_2 | TATGAGTGTGGAGTCGTTGCGCAAAAGTTCACAGTGTGAGTGTNNNAGATCGGAA |
| 39924_1 | TATGAGTGTGGAGTCGTTGCGAGGAAGGAAGGAAGGAAGGAAGNNNAGATCGG |
| 39924_2 | TATGAGTGTGGAGTCGTTGCGAGGAAGGAAGGAAGGAAGGAAGNNNAGATCGG |
| 2238_1 | TATGAGTGTGGAGTCGTTGCGCCCAATCCCCTGTTTTCACATANNNAGATCGGAAGA |
| 2238_2 | TATGAGTGTGGAGTCGTTGCGCCCAATCCCCTGTTTTCACATANNNAGATCGGAAGA |
| 35963_1 | TATGAGTGTGGAGTCGTTGCGCCTGTAATCCCAGCACTTTGNNNAGATCGGAAGAG |

| | |
|---|---|
| 35963_2 | TATGAGTGTGGAGTCGTTGCGCCTGTAATCCCAGCACTTTGNNNAGATCGGAAGAG |
| 31889_1 | TATGAGTGTGGAGTCGTTGCTGACATTATTCTGACCCATGGGGNNNAGATCGGAAG |
| 31889_2 | TATGAGTGTGGAGTCGTTGCTGACATTATTCTGACCCATGGGGNNNAGATCGGAAG |
| 37942_1 | TATGAGTGTGGAGTCGTTGCTTCTTGTAGAGACAGGGACTTGCNNNAGATCGGAA |
| 37942_2 | TATGAGTGTGGAGTCGTTGCTTCTTGTAGAGACAGGGACTTGCNNNAGATCGGAA |
| 52669_1 | TATGAGTGTGGAGTCGTTGCCCCGATCCAACCCTTTTCTGAGNNNAGATCGGAAGA |
| 52669_2 | TATGAGTGTGGAGTCGTTGCCCCGATCCAACCCTTTTCTGAGNNNAGATCGGAAGA |
| 41669_1 | TATGAGTGTGGAGTCGTTGCCACCACACCTGGCTAATTTTTGTNNNAGATCGGAAGA |
| 41669_2 | TATGAGTGTGGAGTCGTTGCCACCACACCTGGCTAATTTTTGTNNNAGATCGGAAGA |
| 34783_1 | TATGAGTGTGGAGTCGTTGCTATGAAGCAGTCACTCCCCTTTCNNNAGATCGGAAGA |
| 34783_2 | TATGAGTGTGGAGTCGTTGCTATGAAGCAGTCACTCCCCTTTCNNNAGATCGGAAGA |
| 8813_1 | TATGAGTGTGGAGTCGTTGCCCAGGCAGTCTGTGATGTATGATTNNNAGATCGGAA |
| 8813_2 | TATGAGTGTGGAGTCGTTGCCCAGGCAGTCTGTGATGTATGATTNNNAGATCGGAA |
| 36208_1 | TATGAGTGTGGAGTCGTTGCTGGGGAGGGGTTTTACAATGAAANNNAGATCGGAA |
| 36208_2 | TATGAGTGTGGAGTCGTTGCTGGGGAGGGGTTTTACAATGAAANNNAGATCGGAA |
| 41125_1 | TATGAGTGTGGAGTCGTTGCGGTAACTTGAACATGCCACCATGNNNAGATCGGAAG |
| 41125_2 | TATGAGTGTGGAGTCGTTGCGGTAACTTGAACATGCCACCATGNNNAGATCGGAAG |
| 41852_1 | TATGAGTGTGGAGTCGTTGCCTGGGTTCACGCCATTCTTCTNNNAGATCGGAAGAG |
| 41852_2 | TATGAGTGTGGAGTCGTTGCCTGGGTTCACGCCATTCTTCTNNNAGATCGGAAGAG |
| 44637_1 | TATGAGTGTGGAGTCGTTGCCTTATTTTGTTAGGGGGACCGGGNNNAGATCGGAA |
| 44637_2 | TATGAGTGTGGAGTCGTTGCCTTATTTTGTTAGGGGGACCGGGNNNAGATCGGAA |
| 35164_1 | TATGAGTGTGGAGTCGTTGCCAATAGTGCGAAAGCAGCTCACNNNAGATCGGAAGA |
| 35164_2 | TATGAGTGTGGAGTCGTTGCCAATAGTGCGAAAGCAGCTCACNNNAGATCGGAAGA |
| 52684_1 | TATGAGTGTGGAGTCGTTGCTGCCTTGCTTGATGAGTTCTTTGNNNAGATCGGAAG |
| 52684_2 | TATGAGTGTGGAGTCGTTGCTGCCTTGCTTGATGAGTTCTTTGNNNAGATCGGAAG |
| 40672_1 | TATGAGTGTGGAGTCGTTGCGCCTCTTTTACTTCAGCCCAATGNNNAGATCGGAAGA |
| 40672_2 | TATGAGTGTGGAGTCGTTGCGCCTCTTTTACTTCAGCCCAATGNNNAGATCGGAAGA |
| 53417_1 | TATGAGTGTGGAGTCGTTGCAGAGGGAGGAGAAAAGAAGAGGANNNAGATCGGA |
| 53417_2 | TATGAGTGTGGAGTCGTTGCAGAGGGAGGAGAAAAGAAGAGGANNNAGATCGGA |
| 30119_1 | TATGAGTGTGGAGTCGTTGCTGGATACTTTACTTTGCATTCTGGANNNAGATCGGA |
| 30119_2 | TATGAGTGTGGAGTCGTTGCTGGATACTTTACTTTGCATTCTGGANNNAGATCGGA |
| 51996_1 | TATGAGTGTGGAGTCGTTGCTAGGACAATAGTGGAGGGAAGGTNNNAGATCGGA |
| 51996_2 | TATGAGTGTGGAGTCGTTGCTAGGACAATAGTGGAGGGAAGGTNNNAGATCGGA |
| 30647_1 | TATGAGTGTGGAGTCGTTGCGGCATGAGTGTGCAGATCAGAGNNNAGATCGGAAG |
| 30647_2 | TATGAGTGTGGAGTCGTTGCGGCATGAGTGTGCAGATCAGAGNNNAGATCGGAAG |
| 44419_1 | TATGAGTGTGGAGTCGTTGCGTACATGACTGCACAGTACCAGANNNAGATCGGAAG |
| 44419_2 | TATGAGTGTGGAGTCGTTGCGTACATGACTGCACAGTACCAGANNNAGATCGGAAG |
| 31749_1 | TATGAGTGTGGAGTCGTTGCTCCAGATGGAGAGCTATAGGAGTNNNAGATCGGAA |
| 31749_2 | TATGAGTGTGGAGTCGTTGCTCCAGATGGAGAGCTATAGGAGTNNNAGATCGGAA |
| 50989_1 | TATGAGTGTGGAGTCGTTGCGGGTGAGAACAGTTGGGGATATTNNNAGATCGGAA |
| 50989_2 | TATGAGTGTGGAGTCGTTGCGGGTGAGAACAGTTGGGGATATTNNNAGATCGGAA |
| 46558_1 | TATGAGTGTGGAGTCGTTGCATGTGCCCATACATTTGTGTGTGNNNAGATCGGAAG |
| 46558_2 | TATGAGTGTGGAGTCGTTGCATGTGCCCATACATTTGTGTGTGNNNAGATCGGAAG |

| | |
|---|---|
| 43091_1 | TATGAGTGTGGAGTCGTTGCAGGAGATTGCTTTGGGTCATCATNNNAGATCGGAA |
| 43091_2 | TATGAGTGTGGAGTCGTTGCAGGAGATTGCTTTGGGTCATCATNNNAGATCGGAA |
| 45994_1 | TATGAGTGTGGAGTCGTTGCCTGACAGAACTGGCTTCCCAAAANNNAGATCGGAAG |
| 45994_2 | TATGAGTGTGGAGTCGTTGCCTGACAGAACTGGCTTCCCAAAANNNAGATCGGAAG |
| 39699_1 | TATGAGTGTGGAGTCGTTGCGGTTGTGAAGACCATGGGAAAATNNNAGATCGGAA |
| 39699_2 | TATGAGTGTGGAGTCGTTGCGGTTGTGAAGACCATGGGAAAATNNNAGATCGGAA |
| 41735_1 | TATGAGTGTGGAGTCGTTGCCACAATTTCGGCTCACTGCAGNNNAGATCGGAAGAG |
| 41735_2 | TATGAGTGTGGAGTCGTTGCCACAATTTCGGCTCACTGCAGNNNAGATCGGAAGAG |
| 41825_1 | TATGAGTGTGGAGTCGTTGCCTTTTGGACCTTCAGCCTTGAACNNNAGATCGGAAG |
| 41825_2 | TATGAGTGTGGAGTCGTTGCCTTTTGGACCTTCAGCCTTGAACNNNAGATCGGAAG |
| 29666_1 | TATGAGTGTGGAGTCGTTGCCAGCTCAGCCCTAGTACTTCTGNNNAGATCGGAAGA |
| 29666_2 | TATGAGTGTGGAGTCGTTGCCAGCTCAGCCCTAGTACTTCTGNNNAGATCGGAAGA |
| 46836_1 | TATGAGTGTGGAGTCGTTGCCTGCTGAGCATGGACTAAGATGANNNAGATCGGAA |
| 46836_2 | TATGAGTGTGGAGTCGTTGCCTGCTGAGCATGGACTAAGATGANNNAGATCGGAA |
| 51713_1 | TATGAGTGTGGAGTCGTTGCCTCAGGGTTCCAAACGTTCCANNNAGATCGGAAGAG |
| 51713_2 | TATGAGTGTGGAGTCGTTGCCTCAGGGTTCCAAACGTTCCANNNAGATCGGAAGAG |
| 51161_1 | TATGAGTGTGGAGTCGTTGCCCACTCAAACTTGCACTTCTCACNNNAGATCGGAAGA |
| 51161_2 | TATGAGTGTGGAGTCGTTGCCCACTCAAACTTGCACTTCTCACNNNAGATCGGAAGA |
| 51069_1 | TATGAGTGTGGAGTCGTTGCTGGCTAGGAAAGACTGAAGTCACNNNAGATCGGAA |
| 51069_2 | TATGAGTGTGGAGTCGTTGCTGGCTAGGAAAGACTGAAGTCACNNNAGATCGGAA |
| 907_1 | TATGAGTGTGGAGTCGTTGCGGGAAGCTAGGTGAAAGAACCCANNNAGATCGGAA |
| 907_2 | TATGAGTGTGGAGTCGTTGCGGGAAGCTAGGTGAAAGAACCCANNNAGATCGGAA |
| 45625_1 | TATGAGTGTGGAGTCGTTGCTGTTAGCCAGGATGGTCTTGATCNNNAGATCGGAA |
| 45625_2 | TATGAGTGTGGAGTCGTTGCTGTTAGCCAGGATGGTCTTGATCNNNAGATCGGAA |
| 29860_1 | TATGAGTGTGGAGTCGTTGCACAAAACAAAAAGACGCTTGACTCNNNAGATCGGAA |
| 29860_2 | TATGAGTGTGGAGTCGTTGCACAAAACAAAAAGACGCTTGACTCNNNAGATCGGAA |
| 46216_1 | TATGAGTGTGGAGTCGTTGCAGGTTTGAGTGATTCTTCTGCCTNNNAGATCGGAAG |
| 46216_2 | TATGAGTGTGGAGTCGTTGCAGGTTTGAGTGATTCTTCTGCCTNNNAGATCGGAAG |
| 52243_1 | TATGAGTGTGGAGTCGTTGCCACCACCAATGGATTCAAACCAANNNAGATCGGAAG |
| 52243_2 | TATGAGTGTGGAGTCGTTGCCACCACCAATGGATTCAAACCAANNNAGATCGGAAG |
| 44929_1 | TATGAGTGTGGAGTCGTTGCTCCCTGTGGTGAATTGACAGAATNNNAGATCGGAA |
| 44929_2 | TATGAGTGTGGAGTCGTTGCTCCCTGTGGTGAATTGACAGAATNNNAGATCGGAA |
| 46804_1 | TATGAGTGTGGAGTCGTTGCTTCCTCAGGTTAGTCTGTTTGGTNNNAGATCGGAAG |
| 46804_2 | TATGAGTGTGGAGTCGTTGCTTCCTCAGGTTAGTCTGTTTGGTNNNAGATCGGAAG |
| 45580_1 | TATGAGTGTGGAGTCGTTGCCTGAGTAGCTGGGACTACAGATGNNNAGATCGGAA |
| 45580_2 | TATGAGTGTGGAGTCGTTGCCTGAGTAGCTGGGACTACAGATGNNNAGATCGGAA |
| 53107_1 | TATGAGTGTGGAGTCGTTGCGCCTGTCAGTAATTCCAGCTACTNNNAGATCGGAAG |
| 53107_2 | TATGAGTGTGGAGTCGTTGCGCCTGTCAGTAATTCCAGCTACTNNNAGATCGGAAG |
| 50577_1 | TATGAGTGTGGAGTCGTTGCGAATTGTGCAGGATGGCAAGAAGNNNAGATCGGA |
| 50577_2 | TATGAGTGTGGAGTCGTTGCGAATTGTGCAGGATGGCAAGAAGNNNAGATCGGA |
| 12483_1 | TATGAGTGTGGAGTCGTTGCGTAATCCAGATCCGTAAGCTNNNAGATCGGAAGAG |
| 12483_2 | TATGAGTGTGGAGTCGTTGCGTAATCCAGATCCGTAAGCTNNNAGATCGGAAGAG |
| 45777_1 | TATGAGTGTGGAGTCGTTGCATGATTCTCTCACTTCAGCCTCCNNNAGATCGGAAGA |

| | |
|---|---|
| 45777_2 | TATGAGTGTGGAGTCGTTGCATGATTCTCTCACTTCAGCCTCCNNNAGATCGGAAGA |
| 12953_1 | TATGAGTGTGGAGTCGTTGCAACCAAATGTTCTTTATCCATCTNNNAGATCGGAAGA |
| 12953_2 | TATGAGTGTGGAGTCGTTGCAACCAAATGTTCTTTATCCATCTNNNAGATCGGAAGA |
| 12585_1 | TATGAGTGTGGAGTCGTTGCGCAAGTTCATTCTCATAACAAAANNNAGATCGGAAG |
| 12585_2 | TATGAGTGTGGAGTCGTTGCGCAAGTTCATTCTCATAACAAAANNNAGATCGGAAG |
| 51635_1 | TATGAGTGTGGAGTCGTTGCATATCACCGAAGGCAGAACACTTNNNAGATCGGAAG |
| 51635_2 | TATGAGTGTGGAGTCGTTGCATATCACCGAAGGCAGAACACTTNNNAGATCGGAAG |
| 51923_1 | TATGAGTGTGGAGTCGTTGCCTGTCCCTTAGTGAGCAGTGTAGNNNAGATCGGAAG |
| 51923_2 | TATGAGTGTGGAGTCGTTGCCTGTCCCTTAGTGAGCAGTGTAGNNNAGATCGGAAG |
| 41120_1 | TATGAGTGTGGAGTCGTTGCCTCCTGTAATCCCAGCACTTTGGNNNAGATCGGAAG |
| 41120_2 | TATGAGTGTGGAGTCGTTGCCTCCTGTAATCCCAGCACTTTGGNNNAGATCGGAAG |
| 37318_1 | TATGAGTGTGGAGTCGTTGCAAAAGTTCCTGCAAGCTACTGTGNNNAGATCGGAAG |
| 37318_2 | TATGAGTGTGGAGTCGTTGCAAAAGTTCCTGCAAGCTACTGTGNNNAGATCGGAAG |
| 49123_1 | TATGAGTGTGGAGTCGTTGCGTTCAAGCAATTTTCCCACCTCANNNAGATCGGAAGA |
| 49123_2 | TATGAGTGTGGAGTCGTTGCGTTCAAGCAATTTTCCCACCTCANNNAGATCGGAAGA |
| 3100_1 | TATGAGTGTGGAGTCGTTGCAAATGCTCTTACAGCAGAGAGCCNNNAGATCGGAAG |
| 3100_2 | TATGAGTGTGGAGTCGTTGCAAATGCTCTTACAGCAGAGAGCCNNNAGATCGGAAG |
| 47657_1 | TATGAGTGTGGAGTCGTTGCAAATGTGGAGTTCTTTGTTGCCCNNNAGATCGGAAG |
| 47657_2 | TATGAGTGTGGAGTCGTTGCAAATGTGGAGTTCTTTGTTGCCCNNNAGATCGGAAG |
| 40404_1 | TATGAGTGTGGAGTCGTTGCGGTTGGGGGTTCAATGTGTAATGNNNAGATCGGAA |
| 40404_2 | TATGAGTGTGGAGTCGTTGCGGTTGGGGGTTCAATGTGTAATGNNNAGATCGGAA |
| 37895_1 | TATGAGTGTGGAGTCGTTGCACTGTAGTGAGCTGTGATTGTGTNNNAGATCGGAA |
| 37895_2 | TATGAGTGTGGAGTCGTTGCACTGTAGTGAGCTGTGATTGTGTNNNAGATCGGAA |
| 40526_1 | TATGAGTGTGGAGTCGTTGCCAGCTTCTGAAAACACTGGCATTNNNAGATCGGAAG |
| 40526_2 | TATGAGTGTGGAGTCGTTGCCAGCTTCTGAAAACACTGGCATTNNNAGATCGGAAG |
| 32141_1 | TATGAGTGTGGAGTCGTTGCCAATCTCGGCTCACTGTAACCTCNNNAGATCGGAAG |
| 32141_2 | TATGAGTGTGGAGTCGTTGCCAATCTCGGCTCACTGTAACCTCNNNAGATCGGAAG |
| 39307_1 | TATGAGTGTGGAGTCGTTGCAACCCAAATCCTTACAGCTGACTNNNAGATCGGAAG |
| 39307_2 | TATGAGTGTGGAGTCGTTGCAACCCAAATCCTTACAGCTGACTNNNAGATCGGAAG |
| 39804_1 | TATGAGTGTGGAGTCGTTGCACACCTGACTTACACCACTAAGGNNNAGATCGGAAG |
| 39804_2 | TATGAGTGTGGAGTCGTTGCACACCTGACTTACACCACTAAGGNNNAGATCGGAAG |
| 50545_1 | TATGAGTGTGGAGTCGTTGCCCAGGATCTGAAAGAGAAGCTGANNNAGATCGGAA |
| 50545_2 | TATGAGTGTGGAGTCGTTGCCCAGGATCTGAAAGAGAAGCTGANNNAGATCGGAA |
| 12957_1 | TATGAGTGTGGAGTCGTTGCCCTCTCCTTCTACCATGTTATNNNAGATCGGAAGAGC |
| 12957_2 | TATGAGTGTGGAGTCGTTGCCCTCTCCTTCTACCATGTTATNNNAGATCGGAAGAGC |
| 42690_1 | TATGAGTGTGGAGTCGTTGCAGAAGTCCAGCAACTGTAAGGACNNNAGATCGGAA |
| 42690_2 | TATGAGTGTGGAGTCGTTGCAGAAGTCCAGCAACTGTAAGGACNNNAGATCGGAA |
| 52921_1 | TATGAGTGTGGAGTCGTTGCCTCAGCCTTTCCTCGCTCTAGNNNAGATCGGAAGAG |
| 52921_2 | TATGAGTGTGGAGTCGTTGCCTCAGCCTTTCCTCGCTCTAGNNNAGATCGGAAGAG |
| 44270_1 | TATGAGTGTGGAGTCGTTGCGCTTTGGCATCAACACCCTAAATNNNAGATCGGAAG |
| 44270_2 | TATGAGTGTGGAGTCGTTGCGCTTTGGCATCAACACCCTAAATNNNAGATCGGAAG |
| 48596_1 | TATGAGTGTGGAGTCGTTGCTAGGCCTTCGATGATATTGCCAANNNAGATCGGAAG |
| 48596_2 | TATGAGTGTGGAGTCGTTGCTAGGCCTTCGATGATATTGCCAANNNAGATCGGAAG |

| | |
|---|---|
| 38417_1 | TATGAGTGTGGAGTCGTTGCATACAGAATATGGATGCAGGGGGNNNAGATCGGAA |
| 38417_2 | TATGAGTGTGGAGTCGTTGCATACAGAATATGGATGCAGGGGGNNNAGATCGGAA |
| 39299_1 | TATGAGTGTGGAGTCGTTGCAGGCTGCTACAAATTCCATGAGANNNAGATCGGAAG |
| 39299_2 | TATGAGTGTGGAGTCGTTGCAGGCTGCTACAAATTCCATGAGANNNAGATCGGAAG |
| 51268_1 | TATGAGTGTGGAGTCGTTGCCTTGCTCTCATAGGCTTGGGAGNNNAGATCGGAAGA |
| 51268_2 | TATGAGTGTGGAGTCGTTGCCTTGCTCTCATAGGCTTGGGAGNNNAGATCGGAAGA |
| 48780_1 | TATGAGTGTGGAGTCGTTGCGCTTCCCAAAGTGCTGAGATTACNNNAGATCGGAAG |
| 48780_2 | TATGAGTGTGGAGTCGTTGCGCTTCCCAAAGTGCTGAGATTACNNNAGATCGGAAG |
| 38511_1 | TATGAGTGTGGAGTCGTTGCTGAAGGAATTATTGTTTATGTGTGCTNNNAGATCGG |
| 38511_2 | TATGAGTGTGGAGTCGTTGCTGAAGGAATTATTGTTTATGTGTGCTNNNAGATCGG |
| 49988_1 | TATGAGTGTGGAGTCGTTGCCTTCCCCTTCTTCATGGCAAAAGNNNAGATCGGAAGA |
| 49988_2 | TATGAGTGTGGAGTCGTTGCCTTCCCCTTCTTCATGGCAAAAGNNNAGATCGGAAGA |
| 30690_1 | TATGAGTGTGGAGTCGTTGCTCTTCAGTTCGTGTGTGGAGACANNNAGATCGGAAG |
| 30690_2 | TATGAGTGTGGAGTCGTTGCTCTTCAGTTCGTGTGTGGAGACANNNAGATCGGAAG |
| 29723_1 | TATGAGTGTGGAGTCGTTGCGCTGAGGCACATCTGAATTCAAANNNAGATCGGAAG |
| 29723_2 | TATGAGTGTGGAGTCGTTGCGCTGAGGCACATCTGAATTCAAANNNAGATCGGAAG |
| 50574_1 | TATGAGTGTGGAGTCGTTGCTTTTTACCCTTGCACTCCCTGTANNNAGATCGGAAGA[illegible] |
| 50574_2 | TATGAGTGTGGAGTCGTTGCTTTTTACCCTTGCACTCCCTGTANNNAGATCGGAAGA[illegible] |
| 49654_1 | TATGAGTGTGGAGTCGTTGCCTCAGCGTCCAAAAGTGTTGAAANNNAGATCGGAAG |
| 49654_2 | TATGAGTGTGGAGTCGTTGCCTCAGCGTCCAAAAGTGTTGAAANNNAGATCGGAAG |
| 32086_1 | TATGAGTGTGGAGTCGTTGCGCATCATAGGCAATCAGCTAGGANNNAGATCGGAA[illegible] |
| 32086_2 | TATGAGTGTGGAGTCGTTGCGCATCATAGGCAATCAGCTAGGANNNAGATCGGAA[illegible] |
| 34549_1 | TATGAGTGTGGAGTCGTTGCCCCCCCAAGAAAAAAAACACACANNNAGATCGGAAG |
| 34549_2 | TATGAGTGTGGAGTCGTTGCCCCCCCAAGAAAAAAAACACACANNNAGATCGGAAG |
| 48495_1 | TATGAGTGTGGAGTCGTTGCATAAAGGTGGAGGCCAGATAGGANNNAGATCGGAA |
| 48495_2 | TATGAGTGTGGAGTCGTTGCATAAAGGTGGAGGCCAGATAGGANNNAGATCGGAA |
| 40880_1 | TATGAGTGTGGAGTCGTTGCGAAGTGATGGGGATGCAAGTCTANNNAGATCGGAA |
| 40880_2 | TATGAGTGTGGAGTCGTTGCGAAGTGATGGGGATGCAAGTCTANNNAGATCGGAA |
| 13709_1 | TATGAGTGTGGAGTCGTTGCGTTAACTGAAGATATTCCAACTCNNNAGATCGGAAGA |
| 13709_2 | TATGAGTGTGGAGTCGTTGCGTTAACTGAAGATATTCCAACTCNNNAGATCGGAAGA |
| 46565_1 | TATGAGTGTGGAGTCGTTGCCTCATTGGATTCTGAGGGGTTGANNNAGATCGGAA[illegible] |
| 46565_2 | TATGAGTGTGGAGTCGTTGCCTCATTGGATTCTGAGGGGTTGANNNAGATCGGAA[illegible] |
| 30631_1 | TATGAGTGTGGAGTCGTTGCTCCAGACTCTACAACGAACTCAANNNAGATCGGAAGA |
| 30631_2 | TATGAGTGTGGAGTCGTTGCTCCAGACTCTACAACGAACTCAANNNAGATCGGAAGA |
| 31378_1 | TATGAGTGTGGAGTCGTTGCATTGCTGGAAGTGGAAAAACACANNNAGATCGGAA[illegible] |
| 31378_2 | TATGAGTGTGGAGTCGTTGCATTGCTGGAAGTGGAAAAACACANNNAGATCGGAA[illegible] |
| 39900_1 | TATGAGTGTGGAGTCGTTGCAGCCCCAGTTGAAGTCTCAATAGNNNAGATCGGAAG |
| 39900_2 | TATGAGTGTGGAGTCGTTGCAGCCCCAGTTGAAGTCTCAATAGNNNAGATCGGAAG |
| 50933_1 | TATGAGTGTGGAGTCGTTGCCTGCTATTTGGAGATTGCTCAGCNNNAGATCGGAAG |
| 50933_2 | TATGAGTGTGGAGTCGTTGCCTGCTATTTGGAGATTGCTCAGCNNNAGATCGGAAG |
| 43851_1 | TATGAGTGTGGAGTCGTTGCAGTGGACTAATGGACTAGGGGAANNNAGATCGGAA |
| 43851_2 | TATGAGTGTGGAGTCGTTGCAGTGGACTAATGGACTAGGGGAANNNAGATCGGAA |
| 36732_1 | TATGAGTGTGGAGTCGTTGCAACAGAGCAAGATTCCGTCTCAANNNAGATCGGAAG |

| | |
|---|---|
| 36732_2 | TATGAGTGTGGAGTCGTTGCAACAGAGCAAGATTCCGTCTCAANNNAGATCGGAAG |
| 44687_1 | TATGAGTGTGGAGTCGTTGCTCTCGTTTGTCACTGCATCCTTANNNAGATCGGAAGA |
| 44687_2 | TATGAGTGTGGAGTCGTTGCTCTCGTTTGTCACTGCATCCTTANNNAGATCGGAAGA |
| 10420_1 | TATGAGTGTGGAGTCGTTGCGGAAGAATCCATCTGGAGCATGGNNNAGATCGGAA |
| 10420_2 | TATGAGTGTGGAGTCGTTGCGGAAGAATCCATCTGGAGCATGGNNNAGATCGGAA |
| 32892_1 | TATGAGTGTGGAGTCGTTGCGCTGGGGATACGGGATTCTATTTNNNAGATCGGAA |
| 32892_2 | TATGAGTGTGGAGTCGTTGCGCTGGGGATACGGGATTCTATTTNNNAGATCGGAA |
| 51775_1 | TATGAGTGTGGAGTCGTTGCAACTTCCAGAGGAACGATCAGACNNNAGATCGGAA |
| 51775_2 | TATGAGTGTGGAGTCGTTGCAACTTCCAGAGGAACGATCAGACNNNAGATCGGAA |
| 34276_1 | TATGAGTGTGGAGTCGTTGCCATCCTGGCTAACAAGGTGAAACNNNAGATCGGAA |
| 34276_2 | TATGAGTGTGGAGTCGTTGCCATCCTGGCTAACAAGGTGAAACNNNAGATCGGAA |
| 41145_1 | TATGAGTGTGGAGTCGTTGCAATAATGGGAGAGGATTGCGAGGNNNAGATCGGA |
| 41145_2 | TATGAGTGTGGAGTCGTTGCAATAATGGGAGAGGATTGCGAGGNNNAGATCGGA |
| 12910_1 | TATGAGTGTGGAGTCGTTGCTATCATGAGTGATTCCAACTATGNNNAGATCGGAAG |
| 12910_2 | TATGAGTGTGGAGTCGTTGCTATCATGAGTGATTCCAACTATGNNNAGATCGGAAG |
| 2640_1 | TATGAGTGTGGAGTCGTTGCTGTCCCCGTGGATCTATTTTGTGNNNAGATCGGAAG |
| 2640_2 | TATGAGTGTGGAGTCGTTGCTGTCCCCGTGGATCTATTTTGTGNNNAGATCGGAAG |
| 36742_1 | TATGAGTGTGGAGTCGTTGCTGTTGTTAGATATTGCCAGCCCTNNNAGATCGGAAG |
| 36742_2 | TATGAGTGTGGAGTCGTTGCTGTTGTTAGATATTGCCAGCCCTNNNAGATCGGAAG |
| 50136_1 | TATGAGTGTGGAGTCGTTGCTTCTTACTCTCCCAACCTTGTGGNNNAGATCGGAAGA |
| 50136_2 | TATGAGTGTGGAGTCGTTGCTTCTTACTCTCCCAACCTTGTGGNNNAGATCGGAAGA |
| 38395_1 | TATGAGTGTGGAGTCGTTGCATACAAGCCAAGAGAACCCCAATNNNAGATCGGAAG |
| 38395_2 | TATGAGTGTGGAGTCGTTGCATACAAGCCAAGAGAACCCCAATNNNAGATCGGAAG |
| 43158_1 | TATGAGTGTGGAGTCGTTGCCACCACGCCAGGCTAATTTTTTNNNAGATCGGAAGA |
| 43158_2 | TATGAGTGTGGAGTCGTTGCCACCACGCCAGGCTAATTTTTTNNNAGATCGGAAGA |
| 34278_1 | TATGAGTGTGGAGTCGTTGCCACTTGAGGAGGGGAGAAAGAAANNNAGATCGGA |
| 34278_2 | TATGAGTGTGGAGTCGTTGCCACTTGAGGAGGGGAGAAAGAAANNNAGATCGGA |
| 33118_1 | TATGAGTGTGGAGTCGTTGCGCTCCAGAATTTCTGCCTCATTCNNNAGATCGGAAGA |
| 33118_2 | TATGAGTGTGGAGTCGTTGCGCTCCAGAATTTCTGCCTCATTCNNNAGATCGGAAGA |
| 51081_1 | TATGAGTGTGGAGTCGTTGCAGGAAGGAAAGCTCTTGGTTCAANNNAGATCGGAA |
| 51081_2 | TATGAGTGTGGAGTCGTTGCAGGAAGGAAAGCTCTTGGTTCAANNNAGATCGGAA |
| 33191_1 | TATGAGTGTGGAGTCGTTGCCCTGGCCAACATGGTAAAACCNNNAGATCGGAAGAG |
| 33191_2 | TATGAGTGTGGAGTCGTTGCCCTGGCCAACATGGTAAAACCNNNAGATCGGAAGAG |
| 10036_1 | TATGAGTGTGGAGTCGTTGCAGAGGGAATTCGCAAACTTACCCNNNAGATCGGAAG |
| 10036_2 | TATGAGTGTGGAGTCGTTGCAGAGGGAATTCGCAAACTTACCCNNNAGATCGGAAG |
| 12720_1 | TATGAGTGTGGAGTCGTTGCCCTGGTTATATGGTGGAAAGNNNAGATCGGAAGAG |
| 12720_2 | TATGAGTGTGGAGTCGTTGCCCTGGTTATATGGTGGAAAGNNNAGATCGGAAGAG |
| 49733_1 | TATGAGTGTGGAGTCGTTGCCAATTTTTGGCTTTTGCCACCCNNNAGATCGGAAGAG |
| 49733_2 | TATGAGTGTGGAGTCGTTGCCAATTTTTGGCTTTTGCCACCCNNNAGATCGGAAGAG |
| 52322_1 | TATGAGTGTGGAGTCGTTGCTGGCTGTTAGCTGGCTTGTATTANNNAGATCGGAAG |
| 52322_2 | TATGAGTGTGGAGTCGTTGCTGGCTGTTAGCTGGCTTGTATTANNNAGATCGGAAG |
| 50588_1 | TATGAGTGTGGAGTCGTTGCGCCTCTGCCTACACACACACAGNNNAGATCGGAAGAGC |
| 50588_2 | TATGAGTGTGGAGTCGTTGCGCCTCTGCCTACACACACACAGNNNAGATCGGAAGAGC |

| | |
|---|---|
| 32621_1 | TATGAGTGTGGAGTCGTTGCAACGTACCAAGATTGAACCATGANNNAGATCGGAA |
| 32621_2 | TATGAGTGTGGAGTCGTTGCAACGTACCAAGATTGAACCATGANNNAGATCGGAA |
| 45283_1 | TATGAGTGTGGAGTCGTTGCTGAATTGGGTGAGAGAGAAAGGGNNNAGATCGGA |
| 45283_2 | TATGAGTGTGGAGTCGTTGCTGAATTGGGTGAGAGAGAAAGGGNNNAGATCGGA |
| 38015_1 | TATGAGTGTGGAGTCGTTGCTACACTTGTACTGGGCACTTACCNNNAGATCGGAAG |
| 38015_2 | TATGAGTGTGGAGTCGTTGCTACACTTGTACTGGGCACTTACCNNNAGATCGGAAG |
| 45812_1 | TATGAGTGTGGAGTCGTTGCATTCTCCCTCCTCAGCGTTCTAANNNAGATCGGAAGA |
| 45812_2 | TATGAGTGTGGAGTCGTTGCATTCTCCCTCCTCAGCGTTCTAANNNAGATCGGAAGA |
| 53391_1 | TATGAGTGTGGAGTCGTTGCTTTTTTTTTTCCCCAGGCTGGAGNNNAGATCGGAAG |
| 53391_2 | TATGAGTGTGGAGTCGTTGCTTTTTTTTTTCCCCAGGCTGGAGNNNAGATCGGAAG |
| 12611_1 | TATGAGTGTGGAGTCGTTGCCAAAATACATCTATTCCCATCAGNNNAGATCGGAAG |
| 12611_2 | TATGAGTGTGGAGTCGTTGCCAAAATACATCTATTCCCATCAGNNNAGATCGGAAG |
| 47385_1 | TATGAGTGTGGAGTCGTTGCAGATGAAGTCTTGCTCTGTCTCCNNNAGATCGGAAG |
| 47385_2 | TATGAGTGTGGAGTCGTTGCAGATGAAGTCTTGCTCTGTCTCCNNNAGATCGGAAG |
| 39133_1 | TATGAGTGTGGAGTCGTTGCCTCCTGCAACTAGTTCAGTGTCTNNNAGATCGGAAG |
| 39133_2 | TATGAGTGTGGAGTCGTTGCCTCCTGCAACTAGTTCAGTGTCTNNNAGATCGGAAG |
| 45264_1 | TATGAGTGTGGAGTCGTTGCTGTGCCACCTATCATCACTTCTTNNNAGATCGGAAGA |
| 45264_2 | TATGAGTGTGGAGTCGTTGCTGTGCCACCTATCATCACTTCTTNNNAGATCGGAAGA |
| 35716_1 | TATGAGTGTGGAGTCGTTGCACGCCCGGCTAATTTTTTGTATTNNNAGATCGGAAG |
| 35716_2 | TATGAGTGTGGAGTCGTTGCACGCCCGGCTAATTTTTTGTATTNNNAGATCGGAAG |
| 44809_1 | TATGAGTGTGGAGTCGTTGCAAACTCCCCTGTTCTCAGAATCCNNNAGATCGGAAGA |
| 44809_2 | TATGAGTGTGGAGTCGTTGCAAACTCCCCTGTTCTCAGAATCCNNNAGATCGGAAGA |
| 2665_1 | TATGAGTGTGGAGTCGTTGCTTGGGGAGGGGGATACAATGAATNNNAGATCGGA |
| 2665_2 | TATGAGTGTGGAGTCGTTGCTTGGGGAGGGGGATACAATGAATNNNAGATCGGA |
| 40813_1 | TATGAGTGTGGAGTCGTTGCTAGCCAGCACTACAAGAACTGTTNNNAGATCGGAAG |
| 40813_2 | TATGAGTGTGGAGTCGTTGCTAGCCAGCACTACAAGAACTGTTNNNAGATCGGAAG |
| 43745_1 | TATGAGTGTGGAGTCGTTGCTGGAAGTTGAGAAGTGCAGTCTTNNNAGATCGGAA |
| 43745_2 | TATGAGTGTGGAGTCGTTGCTGGAAGTTGAGAAGTGCAGTCTTNNNAGATCGGAA |
| 31576_1 | TATGAGTGTGGAGTCGTTGCTGGACACTTAGATTGATTCCACANNNAGATCGGAAG |
| 31576_2 | TATGAGTGTGGAGTCGTTGCTGGACACTTAGATTGATTCCACANNNAGATCGGAAG |
| 37146_1 | TATGAGTGTGGAGTCGTTGCTGGCATTTCTTCCCTCCACTAAANNNAGATCGGAAGA |
| 37146_2 | TATGAGTGTGGAGTCGTTGCTGGCATTTCTTCCCTCCACTAAANNNAGATCGGAAGA |
| 45608_1 | TATGAGTGTGGAGTCGTTGCCATTCTCCTGCCTCAGCCTCNNNAGATCGGAAGAGCA |
| 45608_2 | TATGAGTGTGGAGTCGTTGCCATTCTCCTGCCTCAGCCTCNNNAGATCGGAAGAGCA |
| 12934_1 | TATGAGTGTGGAGTCGTTGCATGATGAAGAAGGAGACCATNNNAGATCGGAAGAG |
| 12934_2 | TATGAGTGTGGAGTCGTTGCATGATGAAGAAGGAGACCATNNNAGATCGGAAGAG |
| 29044_1 | TATGAGTGTGGAGTCGTTGCGCAACAGAAAATCTGCATGTAGTNNNAGATCGGAA |
| 29044_2 | TATGAGTGTGGAGTCGTTGCGCAACAGAAAATCTGCATGTAGTNNNAGATCGGAA |
| 37747_1 | TATGAGTGTGGAGTCGTTGCCAGTGTGTGTGTTGCTCTTACAANNNAGATCGGAAG |
| 37747_2 | TATGAGTGTGGAGTCGTTGCCAGTGTGTGTGTTGCTCTTACAANNNAGATCGGAAG |
| 52135_1 | TATGAGTGTGGAGTCGTTGCATTCCTTTCTCCAGCTACCATCCNNNAGATCGGAAGA |
| 52135_2 | TATGAGTGTGGAGTCGTTGCATTCCTTTCTCCAGCTACCATCCNNNAGATCGGAAGA |
| 47020_1 | TATGAGTGTGGAGTCGTTGCTTCAAGTGATTCTCCTGCCTCAANNNAGATCGGAAG |

| | |
|---|---|
| 47020_2 | TATGAGTGTGGAGTCGTTGCTTCAAGTGATTCTCCTGCCTCAANNNAGATCGGAAG |
| 52590_1 | TATGAGTGTGGAGTCGTTGCTGGGGCCTAGGAAGAAAACAAATNNNAGATCGGAA |
| 52590_2 | TATGAGTGTGGAGTCGTTGCTGGGGCCTAGGAAGAAAACAAATNNNAGATCGGAA |
| 40964_1 | TATGAGTGTGGAGTCGTTGCCCTAAGCGACAAGTACAGTGACTNNNAGATCGGAA |
| 40964_2 | TATGAGTGTGGAGTCGTTGCCCTAAGCGACAAGTACAGTGACTNNNAGATCGGAA |
| 13802_1 | TATGAGTGTGGAGTCGTTGCCAGAAAGCAATAAGCACCAGNNNAGATCGGAAGAG |
| 13802_2 | TATGAGTGTGGAGTCGTTGCCAGAAAGCAATAAGCACCAGNNNAGATCGGAAGAG |
| 12969_1 | TATGAGTGTGGAGTCGTTGCTAAGATGGCACCTATATGAGANNNAGATCGGAAGA |
| 12969_2 | TATGAGTGTGGAGTCGTTGCTAAGATGGCACCTATATGAGANNNAGATCGGAAGA |
| 32046_1 | TATGAGTGTGGAGTCGTTGCCTATCTTGGCTCACTGCAACCTCNNNAGATCGGAAG |
| 32046_2 | TATGAGTGTGGAGTCGTTGCCTATCTTGGCTCACTGCAACCTCNNNAGATCGGAAG |
| 43935_1 | TATGAGTGTGGAGTCGTTGCGGTCTCCAAGTCATGATTCCTGTNNNAGATCGGAAG |
| 43935_2 | TATGAGTGTGGAGTCGTTGCGGTCTCCAAGTCATGATTCCTGTNNNAGATCGGAAG |
| 43376_1 | TATGAGTGTGGAGTCGTTGCTGGTTTCAAGCGATTCTCCTGTANNNAGATCGGAAG |
| 43376_2 | TATGAGTGTGGAGTCGTTGCTGGTTTCAAGCGATTCTCCTGTANNNAGATCGGAAG |
| 2488_1 | TATGAGTGTGGAGTCGTTGCCCAACTTCTGCTTTTGACTGCAGNNNAGATCGGAAG |
| 2488_2 | TATGAGTGTGGAGTCGTTGCCCAACTTCTGCTTTTGACTGCAGNNNAGATCGGAAG |
| 35286_1 | TATGAGTGTGGAGTCGTTGCATCTAGTTGTCCTTGTGCCAGTTNNNAGATCGGAAG |
| 35286_2 | TATGAGTGTGGAGTCGTTGCATCTAGTTGTCCTTGTGCCAGTTNNNAGATCGGAAG |
| 45854_1 | TATGAGTGTGGAGTCGTTGCCTTCCAGGGTTCAACCGATTCTCNNNAGATCGGAAG |
| 45854_2 | TATGAGTGTGGAGTCGTTGCCTTCCAGGGTTCAACCGATTCTCNNNAGATCGGAAG |
| 35513_1 | TATGAGTGTGGAGTCGTTGCGATCCACCCACATCCAAAGTACTNNNAGATCGGAAG |
| 35513_2 | TATGAGTGTGGAGTCGTTGCGATCCACCCACATCCAAAGTACTNNNAGATCGGAAG |
| 29815_1 | TATGAGTGTGGAGTCGTTGCATGATGCTAGTGCCCCATAAGAGNNNAGATCGGAA |
| 29815_2 | TATGAGTGTGGAGTCGTTGCATGATGCTAGTGCCCCATAAGAGNNNAGATCGGAA |
| 37677_1 | TATGAGTGTGGAGTCGTTGCAGCCCCGGGTAGAATAACTTTACNNNAGATCGGAAG |
| 37677_2 | TATGAGTGTGGAGTCGTTGCAGCCCCGGGTAGAATAACTTTACNNNAGATCGGAAG |
| 28783_1 | TATGAGTGTGGAGTCGTTGCGGTGGTTTGCCTAGGTTTTCTTCNNNAGATCGGAAG |
| 28783_2 | TATGAGTGTGGAGTCGTTGCGGTGGTTTGCCTAGGTTTTCTTCNNNAGATCGGAAG |
| 53330_1 | TATGAGTGTGGAGTCGTTGCTCCTGTTGGTTCAGTTTCTCCTTNNNAGATCGGAAGA |
| 53330_2 | TATGAGTGTGGAGTCGTTGCTCCTGTTGGTTCAGTTTCTCCTTNNNAGATCGGAAGA |
| 49435_1 | TATGAGTGTGGAGTCGTTGCGAAGCTTCACTTTCTGTCCACACNNNAGATCGGAAG |
| 49435_2 | TATGAGTGTGGAGTCGTTGCGAAGCTTCACTTTCTGTCCACACNNNAGATCGGAAG |
| 41784_1 | TATGAGTGTGGAGTCGTTGCCCTCCTGGTTTCAGTGATTCTCANNNAGATCGGAAG |
| 41784_2 | TATGAGTGTGGAGTCGTTGCCCTCCTGGTTTCAGTGATTCTCANNNAGATCGGAAG |
| 33417_1 | TATGAGTGTGGAGTCGTTGCTGCTTATGCCTATAATCCCAGCANNNAGATCGGAAG |
| 33417_2 | TATGAGTGTGGAGTCGTTGCTGCTTATGCCTATAATCCCAGCANNNAGATCGGAAG |
| 36840_1 | TATGAGTGTGGAGTCGTTGCTGATCTTTTGCTTTGCTTTGTCCTNNNAGATCGGAAG |
| 36840_2 | TATGAGTGTGGAGTCGTTGCTGATCTTTTGCTTTGCTTTGTCCTNNNAGATCGGAAG |
| 13339_1 | TATGAGTGTGGAGTCGTTGCTTTGTAACGAAGTGAAGGAGNNNAGATCGGAAGA |
| 13339_2 | TATGAGTGTGGAGTCGTTGCTTTGTAACGAAGTGAAGGAGNNNAGATCGGAAGA |
| 9111_1 | TATGAGTGTGGAGTCGTTGCAGCCAGGTAGGTATATAGACAACTNNNAGATCGGA |
| 9111_2 | TATGAGTGTGGAGTCGTTGCAGCCAGGTAGGTATATAGACAACTNNNAGATCGGA |

| | |
|---|---|
| 36498_1 | TATGAGTGTGGAGTCGTTGCGAGGCAGGAGAATTGTTTGAACCNNNAGATCGGAA |
| 36498_2 | TATGAGTGTGGAGTCGTTGCGAGGCAGGAGAATTGTTTGAACCNNNAGATCGGAA |
| 47648_1 | TATGAGTGTGGAGTCGTTGCCTGTTCTTAGCACTCCAGAGAGGNNNAGATCGGAAG |
| 47648_2 | TATGAGTGTGGAGTCGTTGCCTGTTCTTAGCACTCCAGAGAGGNNNAGATCGGAAG |
| 34944_1 | TATGAGTGTGGAGTCGTTGCCTTTCACGGAGTCTCACTCTGTTNNNAGATCGGAAGA |
| 34944_2 | TATGAGTGTGGAGTCGTTGCCTTTCACGGAGTCTCACTCTGTTNNNAGATCGGAAGA |
| 8312_1 | TATGAGTGTGGAGTCGTTGCCACCCCTCAGCCCAACTACTTGNNNAGATCGGAAGAG |
| 8312_2 | TATGAGTGTGGAGTCGTTGCCACCCCTCAGCCCAACTACTTGNNNAGATCGGAAGAG |
| 10569_1 | TATGAGTGTGGAGTCGTTGCTGAACTGATGGTATTCGCAGCAANNNAGATCGGAAG |
| 10569_2 | TATGAGTGTGGAGTCGTTGCTGAACTGATGGTATTCGCAGCAANNNAGATCGGAAG |
| 42584_1 | TATGAGTGTGGAGTCGTTGCCAGACCTTACACCCTTCACAGAANNNAGATCGGAAGA |
| 42584_2 | TATGAGTGTGGAGTCGTTGCCAGACCTTACACCCTTCACAGAANNNAGATCGGAAGA |
| 50596_1 | TATGAGTGTGGAGTCGTTGCAACAAGGTTTATCACGTAGGCCTNNNAGATCGGAAG |
| 50596_2 | TATGAGTGTGGAGTCGTTGCAACAAGGTTTATCACGTAGGCCTNNNAGATCGGAAG |
| 51193_1 | TATGAGTGTGGAGTCGTTGCCCACAGTTCTGAATAGCCTCCTTNNNAGATCGGAAGA |
| 51193_2 | TATGAGTGTGGAGTCGTTGCCCACAGTTCTGAATAGCCTCCTTNNNAGATCGGAAGA |
| 37878_1 | TATGAGTGTGGAGTCGTTGCTCTCATCATTCAGCCTCAATGCTNNNAGATCGGAAGA |
| 37878_2 | TATGAGTGTGGAGTCGTTGCTCTCATCATTCAGCCTCAATGCTNNNAGATCGGAAGA |
| 34875_1 | TATGAGTGTGGAGTCGTTGCCTTGGCTCACTGCAACCTCTANNNAGATCGGAAGAG |
| 34875_2 | TATGAGTGTGGAGTCGTTGCCTTGGCTCACTGCAACCTCTANNNAGATCGGAAGAG |
| 53300_1 | TATGAGTGTGGAGTCGTTGCAAACTTCTGACCTCAGGTGATCCNNNAGATCGGAAG |
| 53300_2 | TATGAGTGTGGAGTCGTTGCAAACTTCTGACCTCAGGTGATCCNNNAGATCGGAAG |
| 30999_1 | TATGAGTGTGGAGTCGTTGCGTCCTGCTGGGTAGTCTTTGTTANNNAGATCGGAAG |
| 30999_2 | TATGAGTGTGGAGTCGTTGCGTCCTGCTGGGTAGTCTTTGTTANNNAGATCGGAAG |
| 51754_1 | TATGAGTGTGGAGTCGTTGCGCCATGTGGAAATGTGAATCCATNNNAGATCGGAAG |
| 51754_2 | TATGAGTGTGGAGTCGTTGCGCCATGTGGAAATGTGAATCCATNNNAGATCGGAAG |
| 43379_1 | TATGAGTGTGGAGTCGTTGCGAGGTCTTGTTGCTCTGCATTTTNNNAGATCGGAAG |
| 43379_2 | TATGAGTGTGGAGTCGTTGCGAGGTCTTGTTGCTCTGCATTTTNNNAGATCGGAAG |
| 46272_1 | TATGAGTGTGGAGTCGTTGCCATGGTTTTTTGTGGGGCATTTGNNNAGATCGGAAG |
| 46272_2 | TATGAGTGTGGAGTCGTTGCCATGGTTTTTTGTGGGGCATTTGNNNAGATCGGAAG |
| 39310_1 | TATGAGTGTGGAGTCGTTGCAAAAAGAGAAATCAGGCCACGTCNNNAGATCGGAA |
| 39310_2 | TATGAGTGTGGAGTCGTTGCAAAAAGAGAAATCAGGCCACGTCNNNAGATCGGAA |
| 8925_1 | TATGAGTGTGGAGTCGTTGCGCCCAAAATGACAGCTTATGCAANNNAGATCGGAAG |
| 8925_2 | TATGAGTGTGGAGTCGTTGCGCCCAAAATGACAGCTTATGCAANNNAGATCGGAAG |
| 31326_1 | TATGAGTGTGGAGTCGTTGCACAGTGCAAGAAAGTCCAGTTTTNNNAGATCGGAAG |
| 31326_2 | TATGAGTGTGGAGTCGTTGCACAGTGCAAGAAAGTCCAGTTTTNNNAGATCGGAAG |
| 5861_1 | TATGAGTGTGGAGTCGTTGCTCGTGTCTCAGCTACTCAGTGAGNNNAGATCGGAAG |
| 5861_2 | TATGAGTGTGGAGTCGTTGCTCGTGTCTCAGCTACTCAGTGAGNNNAGATCGGAAG |
| 52476_1 | TATGAGTGTGGAGTCGTTGCAAGAGAGAATCAGACCTGGGGTANNNAGATCGGAA |
| 52476_2 | TATGAGTGTGGAGTCGTTGCAAGAGAGAATCAGACCTGGGGTANNNAGATCGGAA |
| 37653_1 | TATGAGTGTGGAGTCGTTGCCCAGTCAGAACTAAAGGGCTTCTNNNAGATCGGAAG |
| 37653_2 | TATGAGTGTGGAGTCGTTGCCCAGTCAGAACTAAAGGGCTTCTNNNAGATCGGAAG |
| 38588_1 | TATGAGTGTGGAGTCGTTGCCTCTTGCCTATAATCCCAGCACTNNNAGATCGGAAGA |

| | |
|---|---|
| 38588_2 | TATGAGTGTGGAGTCGTTGCCTCTTGCCTATAATCCCAGCACTNNNAGATCGGAAGA |
| 31129_1 | TATGAGTGTGGAGTCGTTGCGCATGTTTGCATTTGGTTTCCAGNNNAGATCGGAAG |
| 31129_2 | TATGAGTGTGGAGTCGTTGCGCATGTTTGCATTTGGTTTCCAGNNNAGATCGGAAG |
| 52528_1 | TATGAGTGTGGAGTCGTTGCCAGATGGAGAAGAGCATCCTACCNNNAGATCGGAA |
| 52528_2 | TATGAGTGTGGAGTCGTTGCCAGATGGAGAAGAGCATCCTACCNNNAGATCGGAA |
| 37448_1 | TATGAGTGTGGAGTCGTTGCAGTACAACCTCCACGGAAAACATNNNAGATCGGAAG |
| 37448_2 | TATGAGTGTGGAGTCGTTGCAGTACAACCTCCACGGAAAACATNNNAGATCGGAAG |
| 37032_1 | TATGAGTGTGGAGTCGTTGCGGAGGCTAAGACAGGAGAATCACNNNAGATCGGAA |
| 37032_2 | TATGAGTGTGGAGTCGTTGCGGAGGCTAAGACAGGAGAATCACNNNAGATCGGAA |
| 50467_1 | TATGAGTGTGGAGTCGTTGCTAGGGAAATACCTCATTTGGCCCNNNAGATCGGAAG |
| 50467_2 | TATGAGTGTGGAGTCGTTGCTAGGGAAATACCTCATTTGGCCCNNNAGATCGGAAG |
| 40459_1 | TATGAGTGTGGAGTCGTTGCCCCTGTCTCTATACACACACACANNNAGATCGGAAGA |
| 40459_2 | TATGAGTGTGGAGTCGTTGCCCCTGTCTCTATACACACACACANNNAGATCGGAAGA |
| 31017_1 | TATGAGTGTGGAGTCGTTGCGTATGTATTTAGAGGTGGGGGCCNNNAGATCGGAA |
| 31017_2 | TATGAGTGTGGAGTCGTTGCGTATGTATTTAGAGGTGGGGGCCNNNAGATCGGAA |
| 38994_1 | TATGAGTGTGGAGTCGTTGCTGCATTCTTGATATGGTTCTCACCNNNAGATCGGAA |
| 38994_2 | TATGAGTGTGGAGTCGTTGCTGCATTCTTGATATGGTTCTCACCNNNAGATCGGAA |
| 30287_1 | TATGAGTGTGGAGTCGTTGCAAGTAAGTAAAAAACAACGCATATCGNNNAGATCGG |
| 30287_2 | TATGAGTGTGGAGTCGTTGCAAGTAAGTAAAAAACAACGCATATCGNNNAGATCGG |
| 32829_1 | TATGAGTGTGGAGTCGTTGCCTGAGTAGCTGGGATTACAGGTGNNNAGATCGGAA |
| 32829_2 | TATGAGTGTGGAGTCGTTGCCTGAGTAGCTGGGATTACAGGTGNNNAGATCGGAA |
| 46676_1 | TATGAGTGTGGAGTCGTTGCCTCTTGTTGACCAGGCTAGAGTGNNNAGATCGGAA |
| 46676_2 | TATGAGTGTGGAGTCGTTGCCTCTTGTTGACCAGGCTAGAGTGNNNAGATCGGAA |
| 51790_1 | TATGAGTGTGGAGTCGTTGCGGAGCTATCAAACACGCAAATGANNNAGATCGGAA |
| 51790_2 | TATGAGTGTGGAGTCGTTGCGGAGCTATCAAACACGCAAATGANNNAGATCGGAA |
| 34397_1 | TATGAGTGTGGAGTCGTTGCCAAAACACCAAAAACAGGGGTCTNNNAGATCGGAA |
| 34397_2 | TATGAGTGTGGAGTCGTTGCCAAAACACCAAAAACAGGGGTCTNNNAGATCGGAA |
| 33014_1 | TATGAGTGTGGAGTCGTTGCCCATTTCCCAGGTTCAAGCAATTNNNAGATCGGAAG |
| 33014_2 | TATGAGTGTGGAGTCGTTGCCCATTTCCCAGGTTCAAGCAATTNNNAGATCGGAAG |
| 35796_1 | TATGAGTGTGGAGTCGTTGCATTCTGTAGGTTGCCTGTTCACTNNNAGATCGGAAG |
| 35796_2 | TATGAGTGTGGAGTCGTTGCATTCTGTAGGTTGCCTGTTCACTNNNAGATCGGAAG |
| 37393_1 | TATGAGTGTGGAGTCGTTGCGAAAAATGAGGCAGGGCATAGTGNNNAGATCGGA |
| 37393_2 | TATGAGTGTGGAGTCGTTGCGAAAAATGAGGCAGGGCATAGTGNNNAGATCGGA |
| 48316_1 | TATGAGTGTGGAGTCGTTGCCTTGGAACAGTCTGAAGGCATTGNNNAGATCGGAA |
| 48316_2 | TATGAGTGTGGAGTCGTTGCCTTGGAACAGTCTGAAGGCATTGNNNAGATCGGAA |
| 10222_1 | TATGAGTGTGGAGTCGTTGCCTGAAGATTCCTGGGGTAGGGAANNNAGATCGGAA |
| 10222_2 | TATGAGTGTGGAGTCGTTGCCTGAAGATTCCTGGGGTAGGGAANNNAGATCGGAA |
| 44051_1 | TATGAGTGTGGAGTCGTTGCTCCTGGCATTGAAAGAACAAACCNNNAGATCGGAAG |
| 44051_2 | TATGAGTGTGGAGTCGTTGCTCCTGGCATTGAAAGAACAAACCNNNAGATCGGAAG |
| 37077_1 | TATGAGTGTGGAGTCGTTGCTGGCTCCTGCTTATTATCCTAGCNNNAGATCGGAAG |
| 37077_2 | TATGAGTGTGGAGTCGTTGCTGGCTCCTGCTTATTATCCTAGCNNNAGATCGGAAG |
| 53346_1 | TATGAGTGTGGAGTCGTTGCACTACCATACGATCCAGCAATCCNNNAGATCGGAAG |
| 53346_2 | TATGAGTGTGGAGTCGTTGCACTACCATACGATCCAGCAATCCNNNAGATCGGAAG |

| | |
|---|---|
| 31855_1 | TATGAGTGTGGAGTCGTTGCTGGCTTCACTGTTGAATTCTACCNNNAGATCGGAAG |
| 31855_2 | TATGAGTGTGGAGTCGTTGCTGGCTTCACTGTTGAATTCTACCNNNAGATCGGAAG |
| 36056_1 | TATGAGTGTGGAGTCGTTGCAGTGTATCAGCTCCACCATTCAGNNNAGATCGGAAG |
| 36056_2 | TATGAGTGTGGAGTCGTTGCAGTGTATCAGCTCCACCATTCAGNNNAGATCGGAAG |
| 45891_1 | TATGAGTGTGGAGTCGTTGCGAGTAGCTGGGACTATAGGCACNNNAGATCGGAAG |
| 45891_2 | TATGAGTGTGGAGTCGTTGCGAGTAGCTGGGACTATAGGCACNNNAGATCGGAAG |
| 50108_1 | TATGAGTGTGGAGTCGTTGCGTTTAGCAAGCACCTTCCAGAAANNNAGATCGGAAG |
| 50108_2 | TATGAGTGTGGAGTCGTTGCGTTTAGCAAGCACCTTCCAGAAANNNAGATCGGAAG |
| 49871_1 | TATGAGTGTGGAGTCGTTGCAGTTGTTCTGATCGCTTCCTTCTNNNAGATCGGAAG |
| 49871_2 | TATGAGTGTGGAGTCGTTGCAGTTGTTCTGATCGCTTCCTTCTNNNAGATCGGAAG |
| 2513_1 | TATGAGTGTGGAGTCGTTGCTGCCTCTCTCTGATCACAGGTAGNNNAGATCGGAAG |
| 2513_2 | TATGAGTGTGGAGTCGTTGCTGCCTCTCTCTGATCACAGGTAGNNNAGATCGGAAG |
| 49199_1 | TATGAGTGTGGAGTCGTTGCCATGATCTTGGCTCACTGCAACNNNAGATCGGAAGA |
| 49199_2 | TATGAGTGTGGAGTCGTTGCCATGATCTTGGCTCACTGCAACNNNAGATCGGAAGA |
| 43867_1 | TATGAGTGTGGAGTCGTTGCGGGTCTTCATGATCCCTGTTCTTNNNAGATCGGAAG |
| 43867_2 | TATGAGTGTGGAGTCGTTGCGGGTCTTCATGATCCCTGTTCTTNNNAGATCGGAAG |
| 29853_1 | TATGAGTGTGGAGTCGTTGCGCCTCTGACTGGGAATAGTTACTNNNAGATCGGAAG |
| 29853_2 | TATGAGTGTGGAGTCGTTGCGCCTCTGACTGGGAATAGTTACTNNNAGATCGGAAG |
| 44070_1 | TATGAGTGTGGAGTCGTTGCGGTTGTTTGTTTGGTTGGTTGGTNNNAGATCGGAA |
| 44070_2 | TATGAGTGTGGAGTCGTTGCGGTTGTTTGTTTGGTTGGTTGGTNNNAGATCGGAA |
| 13441_1 | TATGAGTGTGGAGTCGTTGCTAGACTGAGCTTAGACAGAGNNNAGATCGGAAGAG |
| 13441_2 | TATGAGTGTGGAGTCGTTGCTAGACTGAGCTTAGACAGAGNNNAGATCGGAAGAG |
| 38796_1 | TATGAGTGTGGAGTCGTTGCTAGTGGCTACTATGAGCAACTGCNNNAGATCGGAA |
| 38796_2 | TATGAGTGTGGAGTCGTTGCTAGTGGCTACTATGAGCAACTGCNNNAGATCGGAA |
| 47731_1 | TATGAGTGTGGAGTCGTTGCTCTCCTCTCTTCCCACTTCCTTCNNNAGATCGGAAGAG |
| 47731_2 | TATGAGTGTGGAGTCGTTGCTCTCCTCTCTTCCCACTTCCTTCNNNAGATCGGAAGAG |
| 50322_1 | TATGAGTGTGGAGTCGTTGCGGTATACACACGCCATAGTGGTTNNNAGATCGGAA |
| 50322_2 | TATGAGTGTGGAGTCGTTGCGGTATACACACGCCATAGTGGTTNNNAGATCGGAA |
| 34950_1 | TATGAGTGTGGAGTCGTTGCCTCCTGAGTTGCTGGGATTACAGNNNAGATCGGAA |
| 34950_2 | TATGAGTGTGGAGTCGTTGCCTCCTGAGTTGCTGGGATTACAGNNNAGATCGGAA |
| 52553_1 | TATGAGTGTGGAGTCGTTGCAAGAGGGCTGACTTTTCCGTTTNNNAGATCGGAAG |
| 52553_2 | TATGAGTGTGGAGTCGTTGCAAGAGGGCTGACTTTTCCGTTTNNNAGATCGGAAG |
| 47093_1 | TATGAGTGTGGAGTCGTTGCCTCCCAAAGTGCTGTGATTACAGNNNAGATCGGAAG |
| 47093_2 | TATGAGTGTGGAGTCGTTGCCTCCCAAAGTGCTGTGATTACAGNNNAGATCGGAAG |
| 47314_1 | TATGAGTGTGGAGTCGTTGCTGGAGGACTAGAGCCAATACTGANNNAGATCGGAA |
| 47314_2 | TATGAGTGTGGAGTCGTTGCTGGAGGACTAGAGCCAATACTGANNNAGATCGGAA |
| 38026_1 | TATGAGTGTGGAGTCGTTGCGAGGCAAGAGAATGGCATGAACNNNAGATCGGAA |
| 38026_2 | TATGAGTGTGGAGTCGTTGCGAGGCAAGAGAATGGCATGAACNNNAGATCGGAA |
| 37456_1 | TATGAGTGTGGAGTCGTTGCCGCCTATAATCCCAGCACTTTTGNNNAGATCGGAAG |
| 37456_2 | TATGAGTGTGGAGTCGTTGCCGCCTATAATCCCAGCACTTTTGNNNAGATCGGAAG |
| 45823_1 | TATGAGTGTGGAGTCGTTGCCTTGGGAGCTCTGTCTATGTTCANNNAGATCGGAAG |
| 45823_2 | TATGAGTGTGGAGTCGTTGCCTTGGGAGCTCTGTCTATGTTCANNNAGATCGGAAG |
| 40081_1 | TATGAGTGTGGAGTCGTTGCACTAGACCCCTTTCTTACACTGTNNNAGATCGGAAGA |

| | |
|---|---|
| 40081_2 | TATGAGTGTGGAGTCGTTGCACTAGACCCCTTTCTTACACTGTNNNAGATCGGAAGA |
| 28852_1 | TATGAGTGTGGAGTCGTTGCACTTTGGGGCTCTTCATCTTGAANNNAGATCGGAAG |
| 28852_2 | TATGAGTGTGGAGTCGTTGCACTTTGGGGCTCTTCATCTTGAANNNAGATCGGAAG |
| 34499_1 | TATGAGTGTGGAGTCGTTGCTTCTCTTCTCCTCTGCTCTCTGANNNAGATCGGAAGA |
| 34499_2 | TATGAGTGTGGAGTCGTTGCTTCTCTTCTCCTCTGCTCTCTGANNNAGATCGGAAGA |
| 33894_1 | TATGAGTGTGGAGTCGTTGCAATACGGTGACAAGATCTTGGCTNNNAGATCGGAA |
| 33894_2 | TATGAGTGTGGAGTCGTTGCAATACGGTGACAAGATCTTGGCTNNNAGATCGGAA |
| 8153_1 | TATGAGTGTGGAGTCGTTGCAGTATTGCTCTGTTGCCCTACGNNNAGATCGGAAGA |
| 8153_2 | TATGAGTGTGGAGTCGTTGCAGTATTGCTCTGTTGCCCTACGNNNAGATCGGAAGA |
| 49885_1 | TATGAGTGTGGAGTCGTTGCTGCTGTATTCTCAGAGATGTGCANNNAGATCGGAAG |
| 49885_2 | TATGAGTGTGGAGTCGTTGCTGCTGTATTCTCAGAGATGTGCANNNAGATCGGAAG |
| 34633_1 | TATGAGTGTGGAGTCGTTGCGAGCTTTGCCTGCAGACTCCNNNAGATCGGAAGAG |
| 34633_2 | TATGAGTGTGGAGTCGTTGCGAGCTTTGCCTGCAGACTCCNNNAGATCGGAAGAG |
| 43420_1 | TATGAGTGTGGAGTCGTTGCCTGAGATGGACTGTCACTCACTCNNNAGATCGGAAG |
| 43420_2 | TATGAGTGTGGAGTCGTTGCCTGAGATGGACTGTCACTCACTCNNNAGATCGGAAG |
| 47350_1 | TATGAGTGTGGAGTCGTTGCATGAGACTGGAAACACATCAGCTNNNAGATCGGAA |
| 47350_2 | TATGAGTGTGGAGTCGTTGCATGAGACTGGAAACACATCAGCTNNNAGATCGGAA |
| 43436_1 | TATGAGTGTGGAGTCGTTGCGCCTTCTGAGTAGCTGGAACTACNNNAGATCGGAAG |
| 43436_2 | TATGAGTGTGGAGTCGTTGCGCCTTCTGAGTAGCTGGAACTACNNNAGATCGGAAG |
| 52438_1 | TATGAGTGTGGAGTCGTTGCATCCCTATCAATATGCCACCACCNNNAGATCGGAAGA |
| 52438_2 | TATGAGTGTGGAGTCGTTGCATCCCTATCAATATGCCACCACCNNNAGATCGGAAGA |
| 31325_1 | TATGAGTGTGGAGTCGTTGCCTCCTTCCTTTTGAGACCTCGAANNNAGATCGGAAGA |
| 31325_2 | TATGAGTGTGGAGTCGTTGCCTCCTTCCTTTTGAGACCTCGAANNNAGATCGGAAGA |
| 34231_1 | TATGAGTGTGGAGTCGTTGCTCCATCAGGAAAATACCACATTCCTNNNAGATCGGAA |
| 34231_2 | TATGAGTGTGGAGTCGTTGCTCCATCAGGAAAATACCACATTCCTNNNAGATCGGAA |
| 42229_1 | TATGAGTGTGGAGTCGTTGCAGTTGGAGTCTCACTGTGTCATCNNNAGATCGGAAG |
| 42229_2 | TATGAGTGTGGAGTCGTTGCAGTTGGAGTCTCACTGTGTCATCNNNAGATCGGAAG |
| 52695_1 | TATGAGTGTGGAGTCGTTGCAATACTGTGTCTAAAGAGCCCGGNNNAGATCGGAA |
| 52695_2 | TATGAGTGTGGAGTCGTTGCAATACTGTGTCTAAAGAGCCCGGNNNAGATCGGAA |
| 35021_1 | TATGAGTGTGGAGTCGTTGCCCTCCCAAGTTGCTGTTACTACANNNAGATCGGAAGA |
| 35021_2 | TATGAGTGTGGAGTCGTTGCCCTCCCAAGTTGCTGTTACTACANNNAGATCGGAAGA |
| 33538_1 | TATGAGTGTGGAGTCGTTGCAGTAGAAACAGGCTTTCACCGTGNNNAGATCGGAA |
| 33538_2 | TATGAGTGTGGAGTCGTTGCAGTAGAAACAGGCTTTCACCGTGNNNAGATCGGAA |
| 13230_1 | TATGAGTGTGGAGTCGTTGCTTGTTTCTTTGCCTATAATTGTTNNNAGATCGGAAGA |
| 13230_2 | TATGAGTGTGGAGTCGTTGCTTGTTTCTTTGCCTATAATTGTTNNNAGATCGGAAGA |
| 12573_1 | TATGAGTGTGGAGTCGTTGCCAACAAGATGATTTCTGGAATCNNNAGATCGGAAGA |
| 12573_2 | TATGAGTGTGGAGTCGTTGCCAACAAGATGATTTCTGGAATCNNNAGATCGGAAGA |
| 38645_1 | TATGAGTGTGGAGTCGTTGCCATCACAATCAGCTGGGTTTCATNNNAGATCGGAAG |
| 38645_2 | TATGAGTGTGGAGTCGTTGCCATCACAATCAGCTGGGTTTCATNNNAGATCGGAAG |
| 30420_1 | TATGAGTGTGGAGTCGTTGCATAATGGCAATCTAGTGGCCTGGNNNAGATCGGAA |
| 30420_2 | TATGAGTGTGGAGTCGTTGCATAATGGCAATCTAGTGGCCTGGNNNAGATCGGAA |
| 36244_1 | TATGAGTGTGGAGTCGTTGCGGCCTTAGGTGTATCTATGCTGGNNNAGATCGGAA |
| 36244_2 | TATGAGTGTGGAGTCGTTGCGGCCTTAGGTGTATCTATGCTGGNNNAGATCGGAA |

| | |
|---|---|
| 53224_1 | TATGAGTGTGGAGTCGTTGCTTTGGCCTCAAACTTCAGGTGANNNAGATCGGAAGA |
| 53224_2 | TATGAGTGTGGAGTCGTTGCTTTGGCCTCAAACTTCAGGTGANNNAGATCGGAAGA |
| 48204_1 | TATGAGTGTGGAGTCGTTGCCTTCCTGGGTTGAAGCAATTCTCNNNAGATCGGAAG |
| 48204_2 | TATGAGTGTGGAGTCGTTGCCTTCCTGGGTTGAAGCAATTCTCNNNAGATCGGAAG |
| 32967_1 | TATGAGTGTGGAGTCGTTGCCTACCATAGGGGGCAGTTCAATTNNNAGATCGGAA |
| 32967_2 | TATGAGTGTGGAGTCGTTGCCTACCATAGGGGGCAGTTCAATTNNNAGATCGGAA |
| 46851_1 | TATGAGTGTGGAGTCGTTGCCCGTGTTCAAGCAATTCTCCTGNNNAGATCGGAAGA |
| 46851_2 | TATGAGTGTGGAGTCGTTGCCCGTGTTCAAGCAATTCTCCTGNNNAGATCGGAAGA |
| 44983_1 | TATGAGTGTGGAGTCGTTGCGTTCAAGTACTTCTCCTGCCTCANNNAGATCGGAAG |
| 44983_2 | TATGAGTGTGGAGTCGTTGCGTTCAAGTACTTCTCCTGCCTCANNNAGATCGGAAG |
| 40729_1 | TATGAGTGTGGAGTCGTTGCGATTTGGACAGAGGTCAGTGTGANNNAGATCGGAA |
| 40729_2 | TATGAGTGTGGAGTCGTTGCGATTTGGACAGAGGTCAGTGTGANNNAGATCGGAA |
| 51660_1 | TATGAGTGTGGAGTCGTTGCAGGTAGCCTAGGATCACCTTCATNNNAGATCGGAAG |
| 51660_2 | TATGAGTGTGGAGTCGTTGCAGGTAGCCTAGGATCACCTTCATNNNAGATCGGAAG |
| 51861_1 | TATGAGTGTGGAGTCGTTGCTGCGGAGTAAATGGAACTACAGGNNNAGATCGGAA |
| 51861_2 | TATGAGTGTGGAGTCGTTGCTGCGGAGTAAATGGAACTACAGGNNNAGATCGGAA |
| 33535_1 | TATGAGTGTGGAGTCGTTGCCTTGAACTCCTGACCTCGTGATCNNNAGATCGGAAG |
| 33535_2 | TATGAGTGTGGAGTCGTTGCCTTGAACTCCTGACCTCGTGATCNNNAGATCGGAAG |
| 41695_1 | TATGAGTGTGGAGTCGTTGCCACCACGCCTGGCTAATTTTTANNNAGATCGGAAG |
| 41695_2 | TATGAGTGTGGAGTCGTTGCCACCACGCCTGGCTAATTTTTANNNAGATCGGAAG |
| 33355_1 | TATGAGTGTGGAGTCGTTGCATGGTGACTGAGAGCTCCAAAAANNNAGATCGGAA |
| 33355_2 | TATGAGTGTGGAGTCGTTGCATGGTGACTGAGAGCTCCAAAAANNNAGATCGGAA |
| 31337_1 | TATGAGTGTGGAGTCGTTGCCAACGTGGGCAATTGGATCATAANNNAGATCGGAA |
| 31337_2 | TATGAGTGTGGAGTCGTTGCCAACGTGGGCAATTGGATCATAANNNAGATCGGAA |
| 42354_1 | TATGAGTGTGGAGTCGTTGCTGTTGACTGTGGACATACAACCANNNAGATCGGAAG |
| 42354_2 | TATGAGTGTGGAGTCGTTGCTGTTGACTGTGGACATACAACCANNNAGATCGGAAG |
| 48243_1 | TATGAGTGTGGAGTCGTTGCCTTCCTTCTCCTCTGGATCCTTTNNNAGATCGGAAGA |
| 48243_2 | TATGAGTGTGGAGTCGTTGCCTTCCTTCTCCTCTGGATCCTTTNNNAGATCGGAAGA |
| 39912_1 | TATGAGTGTGGAGTCGTTGCTCAAACACATTGTGGTCAGAGGANNNAGATCGGAA |
| 39912_2 | TATGAGTGTGGAGTCGTTGCTCAAACACATTGTGGTCAGAGGANNNAGATCGGAA |
| 52051_1 | TATGAGTGTGGAGTCGTTGCACTTTTACACGTTTCCAGTGCACNNNAGATCGGAAG |
| 52051_2 | TATGAGTGTGGAGTCGTTGCACTTTTACACGTTTCCAGTGCACNNNAGATCGGAAG |
| 32768_1 | TATGAGTGTGGAGTCGTTGCTGCCTGTAATCTCAGCTACTTGGNNNAGATCGGAAG |
| 32768_2 | TATGAGTGTGGAGTCGTTGCTGCCTGTAATCTCAGCTACTTGGNNNAGATCGGAAG |
| 50470_1 | TATGAGTGTGGAGTCGTTGCCTTTCAGCTTGTGGTTCCTCTTGNNNAGATCGGAAG |
| 50470_2 | TATGAGTGTGGAGTCGTTGCCTTTCAGCTTGTGGTTCCTCTTGNNNAGATCGGAAG |
| 38918_1 | TATGAGTGTGGAGTCGTTGCGCCCTGTGGTTACCAAGACAATANNNAGATCGGAAG |
| 38918_2 | TATGAGTGTGGAGTCGTTGCGCCCTGTGGTTACCAAGACAATANNNAGATCGGAAG |
| 52918_1 | TATGAGTGTGGAGTCGTTGCGGGCAAATACGTCATCTGTGTGNNNAGATCGGAAG |
| 52918_2 | TATGAGTGTGGAGTCGTTGCGGGCAAATACGTCATCTGTGTGNNNAGATCGGAAG |
| 36377_1 | TATGAGTGTGGAGTCGTTGCGCAATGCACTAGTCAGATGCAATNNNAGATCGGAA |
| 36377_2 | TATGAGTGTGGAGTCGTTGCGCAATGCACTAGTCAGATGCAATNNNAGATCGGAA |
| 30080_1 | TATGAGTGTGGAGTCGTTGCCATCCTTTTGAGTGTGTGTGTGTNNNAGATCGGAAG |

| | |
|---|---|
| 30080_2 | TATGAGTGTGGAGTCGTTGCCATCCTTTTGAGTGTGTGTGTGTNNNAGATCGGAAG |
| 40585_1 | TATGAGTGTGGAGTCGTTGCACTGCAATTTTCATTTGAGGGGGNNNAGATCGGAA |
| 40585_2 | TATGAGTGTGGAGTCGTTGCACTGCAATTTTCATTTGAGGGGGNNNAGATCGGAA |
| 32546_1 | TATGAGTGTGGAGTCGTTGCCTGAGGTGGGAGAATTGCTTGAGNNNAGATCGGA |
| 32546_2 | TATGAGTGTGGAGTCGTTGCCTGAGGTGGGAGAATTGCTTGAGNNNAGATCGGA |
| 33736_1 | TATGAGTGTGGAGTCGTTGCTGTGAAACAGTTGCTCCATGAACNNNAGATCGGAAG |
| 33736_2 | TATGAGTGTGGAGTCGTTGCTGTGAAACAGTTGCTCCATGAACNNNAGATCGGAAG |
| 41790_1 | TATGAGTGTGGAGTCGTTGCTATCTCCCAATGCTATCCCTCCCNNNAGATCGGAAGA |
| 41790_2 | TATGAGTGTGGAGTCGTTGCTATCTCCCAATGCTATCCCTCCCNNNAGATCGGAAGA |
| 12507_1 | TATGAGTGTGGAGTCGTTGCAGTAGCAAGATTATATATGTGCANNNAGATCGGAA |
| 12507_2 | TATGAGTGTGGAGTCGTTGCAGTAGCAAGATTATATATGTGCANNNAGATCGGAA |
| 34588_1 | TATGAGTGTGGAGTCGTTGCCCTCCCCTTTTTAGACCGTAGAGNNNAGATCGGAAG |
| 34588_2 | TATGAGTGTGGAGTCGTTGCCCTCCCCTTTTTAGACCGTAGAGNNNAGATCGGAAG |
| 13129_1 | TATGAGTGTGGAGTCGTTGCAAGAGGAACAAAACACCAGTNNNAGATCGGAAGAG |
| 13129_2 | TATGAGTGTGGAGTCGTTGCAAGAGGAACAAAACACCAGTNNNAGATCGGAAGAG |
| 46335_1 | TATGAGTGTGGAGTCGTTGCTAATTCACTTTGTGACCCTGGGTNNNAGATCGGAAG |
| 46335_2 | TATGAGTGTGGAGTCGTTGCTAATTCACTTTGTGACCCTGGGTNNNAGATCGGAAG |
| 28079_1 | TATGAGTGTGGAGTCGTTGCAACAACCTATGAATTCTGAAAAANNNAGATCGGAAG |
| 28079_2 | TATGAGTGTGGAGTCGTTGCAACAACCTATGAATTCTGAAAAANNNAGATCGGAAG |
| 13329_1 | TATGAGTGTGGAGTCGTTGCCAAAGGAATGACTCATACAATGNNNAGATCGGAAG |
| 13329_2 | TATGAGTGTGGAGTCGTTGCCAAAGGAATGACTCATACAATGNNNAGATCGGAAG |
| 13071_1 | TATGAGTGTGGAGTCGTTGCCTGGATCCTGATACAGATAAATGNNNAGATCGGAA |
| 13071_2 | TATGAGTGTGGAGTCGTTGCCTGGATCCTGATACAGATAAATGNNNAGATCGGAA |
| 32921_1 | TATGAGTGTGGAGTCGTTGCGCTTCATTAGTTGCCACACATGTNNNAGATCGGAAG |
| 32921_2 | TATGAGTGTGGAGTCGTTGCGCTTCATTAGTTGCCACACATGTNNNAGATCGGAAG |
| 47246_1 | TATGAGTGTGGAGTCGTTGCCAATGGCATGATCTCAGCTCACTNNNAGATCGGAAG |
| 47246_2 | TATGAGTGTGGAGTCGTTGCCAATGGCATGATCTCAGCTCACTNNNAGATCGGAAG |
| 35355_1 | TATGAGTGTGGAGTCGTTGCCAATCTCAACTCACTGCAAGCTCNNNAGATCGGAAG |
| 35355_2 | TATGAGTGTGGAGTCGTTGCCAATCTCAACTCACTGCAAGCTCNNNAGATCGGAAG |
| 39958_1 | TATGAGTGTGGAGTCGTTGCCATAACACATTGGCACCTGGAACNNNAGATCGGAAG |
| 39958_2 | TATGAGTGTGGAGTCGTTGCCATAACACATTGGCACCTGGAACNNNAGATCGGAAG |
| 46307_1 | TATGAGTGTGGAGTCGTTGCTGCACCCAGACAGGTAACAAAATNNNAGATCGGAA |
| 46307_2 | TATGAGTGTGGAGTCGTTGCTGCACCCAGACAGGTAACAAAATNNNAGATCGGAA |
| 51631_1 | TATGAGTGTGGAGTCGTTGCAGGCATGCAAAGGATTGAATGTCNNNAGATCGGAA |
| 51631_2 | TATGAGTGTGGAGTCGTTGCAGGCATGCAAAGGATTGAATGTCNNNAGATCGGAA |
| 48357_1 | TATGAGTGTGGAGTCGTTGCTTCAAGCAATTCTCCTGCCTCAGNNNAGATCGGAAG |
| 48357_2 | TATGAGTGTGGAGTCGTTGCTTCAAGCAATTCTCCTGCCTCAGNNNAGATCGGAAG |
| 48093_1 | TATGAGTGTGGAGTCGTTGCCCACAACACCCAGCTAAGTTTTTNNNAGATCGGAAG |
| 48093_2 | TATGAGTGTGGAGTCGTTGCCCACAACACCCAGCTAAGTTTTTNNNAGATCGGAAG |
| 50539_1 | TATGAGTGTGGAGTCGTTGCGGTGAAGGTACAGACTCATCCAGNNNAGATCGGAA |
| 50539_2 | TATGAGTGTGGAGTCGTTGCGGTGAAGGTACAGACTCATCCAGNNNAGATCGGAA |
| 42446_1 | TATGAGTGTGGAGTCGTTGCTTTTCCTCATTCCTTCTCTGGCANNNAGATCGGAAGA |
| 42446_2 | TATGAGTGTGGAGTCGTTGCTTTTCCTCATTCCTTCTCTGGCANNNAGATCGGAAGA |

| | |
|---|---|
| 13555_1 | TATGAGTGTGGAGTCGTTGCACTTACTGTCCATAAAATTTTCTNNNAGATCGGAAGA |
| 13555_2 | TATGAGTGTGGAGTCGTTGCACTTACTGTCCATAAAATTTTCTNNNAGATCGGAAGA |
| 34634_1 | TATGAGTGTGGAGTCGTTGCCTCCCCCTCTGTCTCTCCAAGNNNAGATCGGAAGAGC |
| 34634_2 | TATGAGTGTGGAGTCGTTGCCTCCCCCTCTGTCTCTCCAAGNNNAGATCGGAAGAGC |
| 32411_1 | TATGAGTGTGGAGTCGTTGCTCTGGTTACTAGTAAAGCCACGTNNNAGATCGGAAG |
| 32411_2 | TATGAGTGTGGAGTCGTTGCTCTGGTTACTAGTAAAGCCACGTNNNAGATCGGAAG |
| 40608_1 | TATGAGTGTGGAGTCGTTGCACAGTATCAGTATGGACCCATGANNNAGATCGGAA |
| 40608_2 | TATGAGTGTGGAGTCGTTGCACAGTATCAGTATGGACCCATGANNNAGATCGGAA |
| 49099_1 | TATGAGTGTGGAGTCGTTGCGCCTCCTGAGTAACTGGGATTACNNNAGATCGGAAG |
| 49099_2 | TATGAGTGTGGAGTCGTTGCGCCTCCTGAGTAACTGGGATTACNNNAGATCGGAAG |
| 35485_1 | TATGAGTGTGGAGTCGTTGCTCAGTAGCATGATCTCAGCTCACNNNAGATCGGAAG |
| 35485_2 | TATGAGTGTGGAGTCGTTGCTCAGTAGCATGATCTCAGCTCACNNNAGATCGGAAG |
| 38052_1 | TATGAGTGTGGAGTCGTTGCAAACAAAACCTGATAGCCTGGGTNNNAGATCGGAA |
| 38052_2 | TATGAGTGTGGAGTCGTTGCAAACAAAACCTGATAGCCTGGGTNNNAGATCGGAA |
| 46705_1 | TATGAGTGTGGAGTCGTTGCCCAGGTTCAAGCAATTCTCGTGNNNAGATCGGAAGA |
| 46705_2 | TATGAGTGTGGAGTCGTTGCCCAGGTTCAAGCAATTCTCGTGNNNAGATCGGAAGA |
| 46576_1 | TATGAGTGTGGAGTCGTTGCATAACAACAGCAAAGGTGATGCCNNNAGATCGGAA |
| 46576_2 | TATGAGTGTGGAGTCGTTGCATAACAACAGCAAAGGTGATGCCNNNAGATCGGAA |
| 48923_1 | TATGAGTGTGGAGTCGTTGCTCCCAATTAGCAGTTCTTAGTGGTNNNAGATCGGAA |
| 48923_2 | TATGAGTGTGGAGTCGTTGCTCCCAATTAGCAGTTCTTAGTGGTNNNAGATCGGAA |
| 35090_1 | TATGAGTGTGGAGTCGTTGCATGCAATCTCAGCTTACTGCAACNNNAGATCGGAAG |
| 35090_2 | TATGAGTGTGGAGTCGTTGCATGCAATCTCAGCTTACTGCAACNNNAGATCGGAAG |
| 45556_1 | TATGAGTGTGGAGTCGTTGCTTGGACTTCTGCTCTACTTGGACNNNAGATCGGAAG |
| 45556_2 | TATGAGTGTGGAGTCGTTGCTTGGACTTCTGCTCTACTTGGACNNNAGATCGGAAG |
| 49500_1 | TATGAGTGTGGAGTCGTTGCCGAGTAGCTGGGACTACAGGTNNNAGATCGGAAGA |
| 49500_2 | TATGAGTGTGGAGTCGTTGCCGAGTAGCTGGGACTACAGGTNNNAGATCGGAAGA |
| 37574_1 | TATGAGTGTGGAGTCGTTGCTGTACTCCAGTCTGAGTGACAGANNNAGATCGGAA |
| 37574_2 | TATGAGTGTGGAGTCGTTGCTGTACTCCAGTCTGAGTGACAGANNNAGATCGGAA |
| 40018_1 | TATGAGTGTGGAGTCGTTGCAACAGGACTTTGGGAGAGAATGTNNNAGATCGGAA |
| 40018_2 | TATGAGTGTGGAGTCGTTGCAACAGGACTTTGGGAGAGAATGTNNNAGATCGGAA |
| 404_1 | TATGAGTGTGGAGTCGTTGCAAACAAAAGATTGGCTAGGTGTGGNNNAGATCGGA |
| 404_2 | TATGAGTGTGGAGTCGTTGCAAACAAAAGATTGGCTAGGTGTGGNNNAGATCGGA |
| 50438_1 | TATGAGTGTGGAGTCGTTGCTTTTATCCCAAGCATGCTTCCTGNNNAGATCGGAAG |
| 50438_2 | TATGAGTGTGGAGTCGTTGCTTTTATCCCAAGCATGCTTCCTGNNNAGATCGGAAG |
| 50732_1 | TATGAGTGTGGAGTCGTTGCCCACACTGAGACAAACCCAATTCNNNAGATCGGAAG |
| 50732_2 | TATGAGTGTGGAGTCGTTGCCCACACTGAGACAAACCCAATTCNNNAGATCGGAAG |
| 45798_1 | TATGAGTGTGGAGTCGTTGCCCATTTTGACTCACTGCAAGCTCNNNAGATCGGAAG |
| 45798_2 | TATGAGTGTGGAGTCGTTGCCCATTTTGACTCACTGCAAGCTCNNNAGATCGGAAG |
| 33294_1 | TATGAGTGTGGAGTCGTTGCTTTTAGTAGAGACGGGGTTTCGCNNNAGATCGGAA |
| 33294_2 | TATGAGTGTGGAGTCGTTGCTTTTAGTAGAGACGGGGTTTCGCNNNAGATCGGAA |
| 31186_1 | TATGAGTGTGGAGTCGTTGCGAATCCTAGCTCATTGTTTGGCCNNNAGATCGGAAG |
| 31186_2 | TATGAGTGTGGAGTCGTTGCGAATCCTAGCTCATTGTTTGGCCNNNAGATCGGAAG |
| 35089_1 | TATGAGTGTGGAGTCGTTGCAAATGCTAGGATTACAGGCGTGANNNAGATCGGAA |

| | |
|---|---|
| 35089_2 | TATGAGTGTGGAGTCGTTGCAAATGCTAGGATTACAGGCGTGANNNAGATCGGAA |
| 36000_1 | TATGAGTGTGGAGTCGTTGCCTGGGCAATATAGCAAGACTCCANNNAGATCGGAAG |
| 36000_2 | TATGAGTGTGGAGTCGTTGCCTGGGCAATATAGCAAGACTCCANNNAGATCGGAAG |
| 45314_1 | TATGAGTGTGGAGTCGTTGCGTGAGAGGAGAAAGAAAGAGCCANNNAGATCGGA |
| 45314_2 | TATGAGTGTGGAGTCGTTGCGTGAGAGGAGAAAGAAAGAGCCANNNAGATCGGA |
| 29766_1 | ATGAGTGTGGAGTCGTTGCAGTGAAAAAGCTAAACGGTTAATTTTNNNAGATCGG |
| 29766_2 | ATGAGTGTGGAGTCGTTGCAGTGAAAAAGCTAAACGGTTAATTTTNNNAGATCGG |
| 47005_1 | TATGAGTGTGGAGTCGTTGCAGACAGTTTCCTTTTTGTTGCCCNNNAGATCGGAAG |
| 47005_2 | TATGAGTGTGGAGTCGTTGCAGACAGTTTCCTTTTTGTTGCCCNNNAGATCGGAAG |
| 33777_1 | TATGAGTGTGGAGTCGTTGCTCAGCTCTCCTCCTTTTCTCATGNNNAGATCGGAAGA |
| 33777_2 | TATGAGTGTGGAGTCGTTGCTCAGCTCTCCTCCTTTTCTCATGNNNAGATCGGAAGA |
| 45959_1 | TATGAGTGTGGAGTCGTTGCCTGGGTTCAACTGATTCTTCTGCNNNAGATCGGAAG |
| 45959_2 | TATGAGTGTGGAGTCGTTGCCTGGGTTCAACTGATTCTTCTGCNNNAGATCGGAAG |
| 49547_1 | TATGAGTGTGGAGTCGTTGCCTCAAAAGAAAGCAGATGGCACANNNAGATCGGAA |
| 49547_2 | TATGAGTGTGGAGTCGTTGCCTCAAAAGAAAGCAGATGGCACANNNAGATCGGAA |
| 41857_1 | TATGAGTGTGGAGTCGTTGCCTCAAGTGATTCTCCTGCCTCAGNNNAGATCGGAAG |
| 41857_2 | TATGAGTGTGGAGTCGTTGCCTCAAGTGATTCTCCTGCCTCAGNNNAGATCGGAAG |
| 33205_1 | TATGAGTGTGGAGTCGTTGCAGAAATCACAAAACTCAGAGCCANNNAGATCGGAAG |
| 33205_2 | TATGAGTGTGGAGTCGTTGCAGAAATCACAAAACTCAGAGCCANNNAGATCGGAAG |
| 49264_1 | TATGAGTGTGGAGTCGTTGCAAAATCTCGGCTCACTGCGAGNNNAGATCGGAAGAG |
| 49264_2 | TATGAGTGTGGAGTCGTTGCAAAATCTCGGCTCACTGCGAGNNNAGATCGGAAGAG |
| 33079_1 | TATGAGTGTGGAGTCGTTGCTTCAAACGATTCTCCTGCCTCAGNNNAGATCGGAAGA |
| 33079_2 | TATGAGTGTGGAGTCGTTGCTTCAAACGATTCTCCTGCCTCAGNNNAGATCGGAAGA |
| 33572_1 | TATGAGTGTGGAGTCGTTGCAAGTTCAAGTGATTCTCCTGCCTNNNAGATCGGAAG |
| 33572_2 | TATGAGTGTGGAGTCGTTGCAAGTTCAAGTGATTCTCCTGCCTNNNAGATCGGAAG |
| 35592_1 | TATGAGTGTGGAGTCGTTGCTCCTGAGTAGCTGGGATTACAGANNNAGATCGGAA |
| 35592_2 | TATGAGTGTGGAGTCGTTGCTCCTGAGTAGCTGGGATTACAGANNNAGATCGGAA |
| 11277_1 | TATGAGTGTGGAGTCGTTGCTCCTGTTCTGACTTTGGGGATAGANNNAGATCGGAA |
| 11277_2 | TATGAGTGTGGAGTCGTTGCTCCTGTTCTGACTTTGGGGATAGANNNAGATCGGAA |
| 53313_1 | TATGAGTGTGGAGTCGTTGCCTAGCAGGGTTTTTCTGAGGGACNNNAGATCGGAA |
| 53313_2 | TATGAGTGTGGAGTCGTTGCCTAGCAGGGTTTTTCTGAGGGACNNNAGATCGGAA |
| 32591_1 | TATGAGTGTGGAGTCGTTGCGGCTCAAACCTCTCCTTTTTCACNNNAGATCGGAAGA |
| 32591_2 | TATGAGTGTGGAGTCGTTGCGGCTCAAACCTCTCCTTTTTCACNNNAGATCGGAAGA |
| 43439_1 | TATGAGTGTGGAGTCGTTGCCACCACGTCTGGCTAATTTTTGTNNNAGATCGGAAG |
| 43439_2 | TATGAGTGTGGAGTCGTTGCCACCACGTCTGGCTAATTTTTGTNNNAGATCGGAAG |
| 47917_1 | TATGAGTGTGGAGTCGTTGCGTGCTCAGATCTGCTCTCCATCNNNAGATCGGAAGA |
| 47917_2 | TATGAGTGTGGAGTCGTTGCGTGCTCAGATCTGCTCTCCATCNNNAGATCGGAAGA |
| 38979_1 | TATGAGTGTGGAGTCGTTGCTTTCACTTGTCTCTTGAGGCTCCNNNAGATCGGAAGA |
| 38979_2 | TATGAGTGTGGAGTCGTTGCTTTCACTTGTCTCTTGAGGCTCCNNNAGATCGGAAGA |
| 29756_1 | TATGAGTGTGGAGTCGTTGCTTTCCCACGTCTCTTTACCAGTTNNNAGATCGGAAGA |
| 29756_2 | TATGAGTGTGGAGTCGTTGCTTTCCCACGTCTCTTTACCAGTTNNNAGATCGGAAGA |
| 51669_1 | TATGAGTGTGGAGTCGTTGCTTTCCTTCAAATCTTTCCGCCTGNNNAGATCGGAAGA |
| 51669_2 | TATGAGTGTGGAGTCGTTGCTTTCCTTCAAATCTTTCCGCCTGNNNAGATCGGAAGA |

| | |
|---|---|
| 35016_1 | TATGAGTGTGGAGTCGTTGCAGAATCTCACTATGTTGCCCAGGNNNAGATCGGAAG |
| 35016_2 | TATGAGTGTGGAGTCGTTGCAGAATCTCACTATGTTGCCCAGGNNNAGATCGGAAG |
| 34983_1 | TATGAGTGTGGAGTCGTTGCGGTTCAAGTGATTCTTCTGCCTCNNNAGATCGGAAG |
| 34983_2 | TATGAGTGTGGAGTCGTTGCGGTTCAAGTGATTCTTCTGCCTCNNNAGATCGGAAG |
| 30596_1 | TATGAGTGTGGAGTCGTTGCACGCAGGCAAGAAGGAAATACAATANNNAGATCGG |
| 30596_2 | TATGAGTGTGGAGTCGTTGCACGCAGGCAAGAAGGAAATACAATANNNAGATCGG |
| 44423_1 | TATGAGTGTGGAGTCGTTGCCTCCGTGTGATGTCCTCCATAATNNNAGATCGGAAG |
| 44423_2 | TATGAGTGTGGAGTCGTTGCCTCCGTGTGATGTCCTCCATAATNNNAGATCGGAAG |
| 44453_1 | TATGAGTGTGGAGTCGTTGCTTCTCTGGAGAACCCTGACTAATNNNAGATCGGAAG |
| 44453_2 | TATGAGTGTGGAGTCGTTGCTTCTCTGGAGAACCCTGACTAATNNNAGATCGGAAG |
| 30864_1 | TATGAGTGTGGAGTCGTTGCATCAGTGCTGTTGCATAGTGTTCNNNAGATCGGAAG |
| 30864_2 | TATGAGTGTGGAGTCGTTGCATCAGTGCTGTTGCATAGTGTTCNNNAGATCGGAAG |
| 35393_1 | TATGAGTGTGGAGTCGTTGCCATTCTCCTGCCTCAGCCTCNNNAGATCGGAAGAGCA |
| 35393_2 | TATGAGTGTGGAGTCGTTGCCATTCTCCTGCCTCAGCCTCNNNAGATCGGAAGAGCA |
| 10789_1 | TATGAGTGTGGAGTCGTTGCAACAGCTGGCCATTCTCTGAAAGNNNAGATCGGAAG |
| 10789_2 | TATGAGTGTGGAGTCGTTGCAACAGCTGGCCATTCTCTGAAAGNNNAGATCGGAAG |
| 52108_1 | TATGAGTGTGGAGTCGTTGCTTTCTGTGCCAACTCCAGATGATNNNAGATCGGAAG |
| 52108_2 | TATGAGTGTGGAGTCGTTGCTTTCTGTGCCAACTCCAGATGATNNNAGATCGGAAG |
| 34471_1 | TATGAGTGTGGAGTCGTTGCCTCTCTCTCTCTGTCTCCCTGTCNNNAGATCGGAAGA |
| 34471_2 | TATGAGTGTGGAGTCGTTGCCTCTCTCTCTCTGTCTCCCTGTCNNNAGATCGGAAGA |
| 33835_1 | TATGAGTGTGGAGTCGTTGCAATCAGAACACATGGACACAGGANNNAGATCGGAA |
| 33835_2 | TATGAGTGTGGAGTCGTTGCAATCAGAACACATGGACACAGGANNNAGATCGGAA |
| 30282_1 | TATGAGTGTGGAGTCGTTGCAATGTTATCACCATCATCATTCTTAGTNNNAGATCGG |
| 30282_2 | TATGAGTGTGGAGTCGTTGCAATGTTATCACCATCATCATTCTTAGTNNNAGATCGG |
| 7989_1 | TATGAGTGTGGAGTCGTTGCGTCTACCCTATCCACTTGGTACTNNNAGATCGGAAGA |
| 7989_2 | TATGAGTGTGGAGTCGTTGCGTCTACCCTATCCACTTGGTACTNNNAGATCGGAAGA |
| 42829_1 | TATGAGTGTGGAGTCGTTGCGATCATGGCTCACTGTAAGCTCTNNNAGATCGGAAG |
| 42829_2 | TATGAGTGTGGAGTCGTTGCGATCATGGCTCACTGTAAGCTCTNNNAGATCGGAAG |
| 39363_1 | TATGAGTGTGGAGTCGTTGCTTGCTAACCCCCTCATTATGCTTNNNAGATCGGAAGA |
| 39363_2 | TATGAGTGTGGAGTCGTTGCTTGCTAACCCCCTCATTATGCTTNNNAGATCGGAAGA |
| 44403_1 | TATGAGTGTGGAGTCGTTGCGGGCTCTAGTGGGTCCAGAGNNNAGATCGGAAGA |
| 44403_2 | TATGAGTGTGGAGTCGTTGCGGGCTCTAGTGGGTCCAGAGNNNAGATCGGAAGA |
| 30993_1 | TATGAGTGTGGAGTCGTTGCTATTTAGCTTACAGGTCCGTGGGNNNAGATCGGAA |
| 30993_2 | TATGAGTGTGGAGTCGTTGCTATTTAGCTTACAGGTCCGTGGGNNNAGATCGGAA |
| 51472_1 | TATGAGTGTGGAGTCGTTGCGTTGTCTACAGCTCATTCTTCGCNNNAGATCGGAAG |
| 51472_2 | TATGAGTGTGGAGTCGTTGCGTTGTCTACAGCTCATTCTTCGCNNNAGATCGGAAG |
| 50344_1 | TATGAGTGTGGAGTCGTTGCGGATACAACTCAGAGACAGCCAANNNAGATCGGAA |
| 50344_2 | TATGAGTGTGGAGTCGTTGCGGATACAACTCAGAGACAGCCAANNNAGATCGGAA |
| 37640_1 | TATGAGTGTGGAGTCGTTGCGTGGCTCACTCCTGTAATCCTAGNNNAGATCGGAAG |
| 37640_2 | TATGAGTGTGGAGTCGTTGCGTGGCTCACTCCTGTAATCCTAGNNNAGATCGGAAG |
| 37346_1 | TATGAGTGTGGAGTCGTTGCCTGATTGTGAAGACTGTGGGAGANNNAGATCGGAA |
| 37346_2 | TATGAGTGTGGAGTCGTTGCCTGATTGTGAAGACTGTGGGAGANNNAGATCGGAA |
| 48607_1 | TATGAGTGTGGAGTCGTTGCCTGGGATTATAGGAGTGAGCCACNNNAGATCGGAA |

| | |
|---|---|
| 48607_2 | TATGAGTGTGGAGTCGTTGCCTGGGATTATAGGAGTGAGCCACNNNAGATCGGAA |
| 32052_1 | TATGAGTGTGGAGTCGTTGCCCTAGTCTCGGATAAAACAGACGTNNNAGATCGGAA |
| 32052_2 | TATGAGTGTGGAGTCGTTGCCCTAGTCTCGGATAAAACAGACGTNNNAGATCGGAA |
| 43052_1 | TATGAGTGTGGAGTCGTTGCAGGAGAATCACTAGAACCCCAGANNNAGATCGGAA |
| 43052_2 | TATGAGTGTGGAGTCGTTGCAGGAGAATCACTAGAACCCCAGANNNAGATCGGAA |
| 32713_1 | TATGAGTGTGGAGTCGTTGCCTCACATCACGTCTGTCATTCCANNNAGATCGGAAGA |
| 32713_2 | TATGAGTGTGGAGTCGTTGCCTCACATCACGTCTGTCATTCCANNNAGATCGGAAGA |
| 12939_1 | TATGAGTGTGGAGTCGTTGCATTCTTTTCAAAATAGCCTTTGANNNAGATCGGAAG |
| 12939_2 | TATGAGTGTGGAGTCGTTGCATTCTTTTCAAAATAGCCTTTGANNNAGATCGGAAG |
| 35825_1 | TATGAGTGTGGAGTCGTTGCCAATTCTCCTGCCTCAGCCTCNNNAGATCGGAAGAG |
| 35825_2 | TATGAGTGTGGAGTCGTTGCCAATTCTCCTGCCTCAGCCTCNNNAGATCGGAAGAG |
| 44725_1 | TATGAGTGTGGAGTCGTTGCGAAAACCATGGACAGAAGAAGGCNNNAGATCGGA |
| 44725_2 | TATGAGTGTGGAGTCGTTGCGAAAACCATGGACAGAAGAAGGCNNNAGATCGGA |
| 43535_1 | TATGAGTGTGGAGTCGTTGCATTGCTGGGATTACAGATGGGAGNNNAGATCGGAA |
| 43535_2 | TATGAGTGTGGAGTCGTTGCATTGCTGGGATTACAGATGGGAGNNNAGATCGGAA |
| 34376_1 | TATGAGTGTGGAGTCGTTGCTTGAGGCTATGGAAGTTCAAGGGNNNAGATCGGAA |
| 34376_2 | TATGAGTGTGGAGTCGTTGCTTGAGGCTATGGAAGTTCAAGGGNNNAGATCGGAA |
| 38583_1 | TATGAGTGTGGAGTCGTTGCCCCAAAGTGCTAGGATTACAGGTNNNAGATCGGAA |
| 38583_2 | TATGAGTGTGGAGTCGTTGCCCCAAAGTGCTAGGATTACAGGTNNNAGATCGGAA |
| 10605_1 | TATGAGTGTGGAGTCGTTGCTAGGATCATGGGGTGGTCACTTTNNNAGATCGGAA |
| 10605_2 | TATGAGTGTGGAGTCGTTGCTAGGATCATGGGGTGGTCACTTTNNNAGATCGGAA |
| 4509_1 | TATGAGTGTGGAGTCGTTGCCCCACAAGAAATACTCAGCAGCANNNAGATCGGAAG |
| 4509_2 | TATGAGTGTGGAGTCGTTGCCCCACAAGAAATACTCAGCAGCANNNAGATCGGAAG |
| 52123_1 | TATGAGTGTGGAGTCGTTGCAGGAAACTGACTTACTGGCAACANNNAGATCGGAA |
| 52123_2 | TATGAGTGTGGAGTCGTTGCAGGAAACTGACTTACTGGCAACANNNAGATCGGAA |
| 49103_1 | TATGAGTGTGGAGTCGTTGCCGAGTAACTGGGATTGTAGGTGTNNNAGATCGGAA |
| 49103_2 | TATGAGTGTGGAGTCGTTGCCGAGTAACTGGGATTGTAGGTGTNNNAGATCGGAA |
| 13148_1 | TATGAGTGTGGAGTCGTTGCTCTTTAATGATTTGGTCATACTGNNNAGATCGGAAG |
| 13148_2 | TATGAGTGTGGAGTCGTTGCTCTTTAATGATTTGGTCATACTGNNNAGATCGGAAG |
| 50395_1 | TATGAGTGTGGAGTCGTTGCCAGCCTGTGAAGGGAAGGAGNNNAGATCGGAAGA |
| 50395_2 | TATGAGTGTGGAGTCGTTGCCAGCCTGTGAAGGGAAGGAGNNNAGATCGGAAGA |
| 50540_1 | TATGAGTGTGGAGTCGTTGCAATGCTCTTTGGATATGAGGCCANNNAGATCGGAA |
| 50540_2 | TATGAGTGTGGAGTCGTTGCAATGCTCTTTGGATATGAGGCCANNNAGATCGGAA |
| 47407_1 | TATGAGTGTGGAGTCGTTGCCCCAAGTTCAAGCAATTCTCCTGNNNAGATCGGAAG |
| 47407_2 | TATGAGTGTGGAGTCGTTGCCCCAAGTTCAAGCAATTCTCCTGNNNAGATCGGAAG |
| 42103_1 | TATGAGTGTGGAGTCGTTGCCTCCCAAGTAGCCGAGATTACAGNNNAGATCGGAAG |
| 42103_2 | TATGAGTGTGGAGTCGTTGCCTCCCAAGTAGCCGAGATTACAGNNNAGATCGGAAG |
| 36205_1 | TATGAGTGTGGAGTCGTTGCGCTTCAAGAAATTTGGCTGCATGNNNAGATCGGAA |
| 36205_2 | TATGAGTGTGGAGTCGTTGCGCTTCAAGAAATTTGGCTGCATGNNNAGATCGGAA |
| 3143_1 | TATGAGTGTGGAGTCGTTGCGACGTGAGAATTGGAACTCCCTTNNNAGATCGGAA |
| 3143_2 | TATGAGTGTGGAGTCGTTGCGACGTGAGAATTGGAACTCCCTTNNNAGATCGGAA |
| 30888_1 | TATGAGTGTGGAGTCGTTGCCAAGTAGATTTCCTGCAGGGAGTNNNAGATCGGAA |
| 30888_2 | TATGAGTGTGGAGTCGTTGCCAAGTAGATTTCCTGCAGGGAGTNNNAGATCGGAA |

| | |
|---|---|
| 42928_1 | TATGAGTGTGGAGTCGTTGCTCTTTGCGATTGTGAATTGTGCTNNNAGATCGGAAG |
| 42928_2 | TATGAGTGTGGAGTCGTTGCTCTTTGCGATTGTGAATTGTGCTNNNAGATCGGAAG |
| 33223_1 | TATGAGTGTGGAGTCGTTGCGATCTCTGATCACTGCAACCTCCNNNAGATCGGAAG |
| 33223_2 | TATGAGTGTGGAGTCGTTGCGATCTCTGATCACTGCAACCTCCNNNAGATCGGAAG |
| 10363_1 | TATGAGTGTGGAGTCGTTGCCTGTTGATAGCAATGTCAATGTCANNNAGATCGGAA |
| 10363_2 | TATGAGTGTGGAGTCGTTGCCTGTTGATAGCAATGTCAATGTCANNNAGATCGGAA |
| 46837_1 | TATGAGTGTGGAGTCGTTGCGCCTCCTAGGTTCATGTCATTCTNNNAGATCGGAAG |
| 46837_2 | TATGAGTGTGGAGTCGTTGCGCCTCCTAGGTTCATGTCATTCTNNNAGATCGGAAG |
| 48699_1 | TATGAGTGTGGAGTCGTTGCCCTAATAGCTGAGACCACAGGTGNNNAGATCGGAA |
| 48699_2 | TATGAGTGTGGAGTCGTTGCCCTAATAGCTGAGACCACAGGTGNNNAGATCGGAA |
| 53158_1 | TATGAGTGTGGAGTCGTTGCCAATCAGTTCACTTGGCTGACTGNNNAGATCGGAAG |
| 53158_2 | TATGAGTGTGGAGTCGTTGCCAATCAGTTCACTTGGCTGACTGNNNAGATCGGAAG |
| 29847_1 | TATGAGTGTGGAGTCGTTGCAACACCACATACGGATGTAACCTNNNAGATCGGAAG |
| 29847_2 | TATGAGTGTGGAGTCGTTGCAACACCACATACGGATGTAACCTNNNAGATCGGAAG |
| 7918_1 | TATGAGTGTGGAGTCGTTGCGCAGAGCAAGAGTATGAGCATGTTTNNNAGATCGG |
| 7918_2 | TATGAGTGTGGAGTCGTTGCGCAGAGCAAGAGTATGAGCATGTTTNNNAGATCGG |
| 38705_1 | TATGAGTGTGGAGTCGTTGCCTGAGGCAAGAAAATGGCATGAANNNAGATCGGAA |
| 38705_2 | TATGAGTGTGGAGTCGTTGCCTGAGGCAAGAAAATGGCATGAANNNAGATCGGAA |
| 10060_1 | TATGAGTGTGGAGTCGTTGCTGATGGGAGCTGTATTAGGGGAGNNNAGATCGGA |
| 10060_2 | TATGAGTGTGGAGTCGTTGCTGATGGGAGCTGTATTAGGGGAGNNNAGATCGGA |
| 49244_1 | TATGAGTGTGGAGTCGTTGCTTCAAACAATTCTCATGCCTCGGNNNAGATCGGAAG |
| 49244_2 | TATGAGTGTGGAGTCGTTGCTTCAAACAATTCTCATGCCTCGGNNNAGATCGGAAG |
| 52914_1 | TATGAGTGTGGAGTCGTTGCGATGCTGTTCACCCTCAGTTAGANNNAGATCGGAAG |
| 52914_2 | TATGAGTGTGGAGTCGTTGCGATGCTGTTCACCCTCAGTTAGANNNAGATCGGAAG |
| 53077_1 | TATGAGTGTGGAGTCGTTGCGTCTCATTCCTGAACTCAGCCTANNNAGATCGGAAG |
| 53077_2 | TATGAGTGTGGAGTCGTTGCGTCTCATTCCTGAACTCAGCCTANNNAGATCGGAAG |
| 49877_1 | TATGAGTGTGGAGTCGTTGCTCAGTCAGCTATAGTTTAGTTTGGTNNNAGATCGGA |
| 49877_2 | TATGAGTGTGGAGTCGTTGCTCAGTCAGCTATAGTTTAGTTTGGTNNNAGATCGGA |
| 29117_1 | TATGAGTGTGGAGTCGTTGCTGTGAACATTGTCCTTCTTGTGCNNNAGATCGGAAG |
| 29117_2 | TATGAGTGTGGAGTCGTTGCTGTGAACATTGTCCTTCTTGTGCNNNAGATCGGAAG |
| 40794_1 | TATGAGTGTGGAGTCGTTGCTTCTACCATTCTCTCCACCCTCTNNNAGATCGGAAGA |
| 40794_2 | TATGAGTGTGGAGTCGTTGCTTCTACCATTCTCTCCACCCTCTNNNAGATCGGAAGA |
| 52773_1 | TATGAGTGTGGAGTCGTTGCCTGCCTACATGTCCCAAGAGGNNNAGATCGGAAGA |
| 52773_2 | TATGAGTGTGGAGTCGTTGCCTGCCTACATGTCCCAAGAGGNNNAGATCGGAAGA |
| 44674_1 | TATGAGTGTGGAGTCGTTGCCCCCATTTGGAGACAGAGACTACNNNAGATCGGAAG |
| 44674_2 | TATGAGTGTGGAGTCGTTGCCCCCATTTGGAGACAGAGACTACNNNAGATCGGAAG |
| 38274_1 | TATGAGTGTGGAGTCGTTGCACAAAAATTAGCCTGGTGTGGTGNNNAGATCGGAA |
| 38274_2 | TATGAGTGTGGAGTCGTTGCACAAAAATTAGCCTGGTGTGGTGNNNAGATCGGAA |
| 49843_1 | TATGAGTGTGGAGTCGTTGCCCTGAACTCACTGCCAAGAAAAGNNNAGATCGGAAG |
| 49843_2 | TATGAGTGTGGAGTCGTTGCCCTGAACTCACTGCCAAGAAAAGNNNAGATCGGAAG |
| 50843_1 | TATGAGTGTGGAGTCGTTGCACCTAGGGTCACATTAGCTGAAGNNNAGATCGGAA |
| 50843_2 | TATGAGTGTGGAGTCGTTGCACCTAGGGTCACATTAGCTGAAGNNNAGATCGGAA |
| 51536_1 | TATGAGTGTGGAGTCGTTGCCCTAGGAAAGTTGGGGGAATCATNNNAGATCGGAA |

| | |
|---|---|
| 51536_2 | TATGAGTGTGGAGTCGTTGCCCTAGGAAAGTTGGGGGAATCATNNNAGATCGGAA |
| 35615_1 | TATGAGTGTGGAGTCGTTGCTCCTGGGTTCAATGATTCTCCTGNNNAGATCGGAAG |
| 35615_2 | TATGAGTGTGGAGTCGTTGCTCCTGGGTTCAATGATTCTCCTGNNNAGATCGGAAG |
| 50980_1 | TATGAGTGTGGAGTCGTTGCCAAGTCCCCAAACCAAATTCCTCNNNAGATCGGAAGA |
| 50980_2 | TATGAGTGTGGAGTCGTTGCCAAGTCCCCAAACCAAATTCCTCNNNAGATCGGAAGA |
| 52564_1 | TATGAGTGTGGAGTCGTTGCGGGGCTTCCAGATCCAGAAATAANNNAGATCGGAA |
| 52564_2 | TATGAGTGTGGAGTCGTTGCGGGGCTTCCAGATCCAGAAATAANNNAGATCGGAA |
| 39818_1 | TATGAGTGTGGAGTCGTTGCGGTTCTCTCATCTCTTCACACTGTNNNAGATCGGAAG |
| 39818_2 | TATGAGTGTGGAGTCGTTGCGGTTCTCTCATCTCTTCACACTGTNNNAGATCGGAAG |
| 30692_1 | TATGAGTGTGGAGTCGTTGCCCCTCACAAGGTTCTTCAAGTTATNNNAGATCGGAAG |
| 30692_2 | TATGAGTGTGGAGTCGTTGCCCCTCACAAGGTTCTTCAAGTTATNNNAGATCGGAAG |
| 37408_1 | TATGAGTGTGGAGTCGTTGCCCAGCTTGCTTGCTCAGTTTTTANNNAGATCGGAAGA |
| 37408_2 | TATGAGTGTGGAGTCGTTGCCCAGCTTGCTTGCTCAGTTTTTANNNAGATCGGAAGA |
| 44308_1 | TATGAGTGTGGAGTCGTTGCTCTTCCCCAGCCATTAGATAGGANNNAGATCGGAAG |
| 44308_2 | TATGAGTGTGGAGTCGTTGCTCTTCCCCAGCCATTAGATAGGANNNAGATCGGAAG |
| 33916_1 | TATGAGTGTGGAGTCGTTGCCTCCCAGCTTCAAGCAATTCTTCNNNAGATCGGAAGA |
| 33916_2 | TATGAGTGTGGAGTCGTTGCCTCCCAGCTTCAAGCAATTCTTCNNNAGATCGGAAGA |
| 43276_1 | TATGAGTGTGGAGTCGTTGCCCACTTGTTGATTGATGGGCATTNNNAGATCGGAAG |
| 43276_2 | TATGAGTGTGGAGTCGTTGCCCACTTGTTGATTGATGGGCATTNNNAGATCGGAAG |
| 29597_1 | TATGAGTGTGGAGTCGTTGCAGAGCTTTTCGTAGACTGTCCAGNNNAGATCGGAAG |
| 29597_2 | TATGAGTGTGGAGTCGTTGCAGAGCTTTTCGTAGACTGTCCAGNNNAGATCGGAAG |
| 45453_1 | TATGAGTGTGGAGTCGTTGCGGGAAAGGATTTGAAGCAGGAGANNNAGATCGGA |
| 45453_2 | TATGAGTGTGGAGTCGTTGCGGGAAAGGATTTGAAGCAGGAGANNNAGATCGGA |
| 45870_1 | TATGAGTGTGGAGTCGTTGCCAGGTTCACACCATTCTCCTGTNNNAGATCGGAAGA |
| 45870_2 | TATGAGTGTGGAGTCGTTGCCAGGTTCACACCATTCTCCTGTNNNAGATCGGAAGA |
| 29418_1 | TATGAGTGTGGAGTCGTTGCCTCAATCCAGGTTGCCGTCAAAANNNAGATCGGAAG |
| 29418_2 | TATGAGTGTGGAGTCGTTGCCTCAATCCAGGTTGCCGTCAAAANNNAGATCGGAAG |
| 35048_1 | TATGAGTGTGGAGTCGTTGCCCATGCCCGGCTAATTTTTTGTANNNAGATCGGAAG |
| 35048_2 | TATGAGTGTGGAGTCGTTGCCCATGCCCGGCTAATTTTTTGTANNNAGATCGGAAG |
| 47416_1 | TATGAGTGTGGAGTCGTTGCCCTAGTGACAAAAGGCCTTCCAGNNNAGATCGGAAG |
| 47416_2 | TATGAGTGTGGAGTCGTTGCCCTAGTGACAAAAGGCCTTCCAGNNNAGATCGGAAG |
| 42149_1 | TATGAGTGTGGAGTCGTTGCGCAATGGCACTATCTCAACTCACNNNAGATCGGAAG |
| 42149_2 | TATGAGTGTGGAGTCGTTGCGCAATGGCACTATCTCAACTCACNNNAGATCGGAAG |
| 38502_1 | TATGAGTGTGGAGTCGTTGCGAGTCTTGCTCTATCGCCCAGNNNAGATCGGAAGAG |
| 38502_2 | TATGAGTGTGGAGTCGTTGCGAGTCTTGCTCTATCGCCCAGNNNAGATCGGAAGAG |
| 38424_1 | TATGAGTGTGGAGTCGTTGCTGGACAGGAAATGGGAGAGAATGNNNAGATCGGA |
| 38424_2 | TATGAGTGTGGAGTCGTTGCTGGACAGGAAATGGGAGAGAATGNNNAGATCGGA |
| 31133_1 | TATGAGTGTGGAGTCGTTGCGTGTGTCTGTATGTGTGTTCGTGNNNAGATCGGAA |
| 31133_2 | TATGAGTGTGGAGTCGTTGCGTGTGTCTGTATGTGTGTTCGTGNNNAGATCGGAA |
| 49265_1 | TATGAGTGTGGAGTCGTTGCAGTTATGAAGAAGCCTGGAGACGNNNAGATCGGAA |
| 49265_2 | TATGAGTGTGGAGTCGTTGCAGTTATGAAGAAGCCTGGAGACGNNNAGATCGGAA |
| 35678_1 | TATGAGTGTGGAGTCGTTGCCAATGTTTGCTTAGGCTGGTCTCNNNAGATCGGAAG |
| 35678_2 | TATGAGTGTGGAGTCGTTGCCAATGTTTGCTTAGGCTGGTCTCNNNAGATCGGAAG |

| | |
|---|---|
| 37449_1 | TATGAGTGTGGAGTCGTTGCGAGAAGCTTCAAAAAAGGGTCCCNNNAGATCGGAA |
| 37449_2 | TATGAGTGTGGAGTCGTTGCGAGAAGCTTCAAAAAAGGGTCCCNNNAGATCGGAA |
| 28842_1 | TATGAGTGTGGAGTCGTTGCCCCACCACCCCCTTATAGTTTTTNNNAGATCGGAAGA |
| 28842_2 | TATGAGTGTGGAGTCGTTGCCCCACCACCCCCTTATAGTTTTTNNNAGATCGGAAGA |
| 35469_1 | TATGAGTGTGGAGTCGTTGCATGGCAAAATCTCAACTCACAGCNNNAGATCGGAAG |
| 35469_2 | TATGAGTGTGGAGTCGTTGCATGGCAAAATCTCAACTCACAGCNNNAGATCGGAAG |
| 43234_1 | TATGAGTGTGGAGTCGTTGCCACCTGGGGATTACAACTTGAGANNNAGATCGGAA |
| 43234_2 | TATGAGTGTGGAGTCGTTGCCACCTGGGGATTACAACTTGAGANNNAGATCGGAA |
| 51171_1 | TATGAGTGTGGAGTCGTTGCGTGAGAAGTCCTACAATCTGCCANNNAGATCGGAAG |
| 51171_2 | TATGAGTGTGGAGTCGTTGCGTGAGAAGTCCTACAATCTGCCANNNAGATCGGAAG |
| 43403_1 | TATGAGTGTGGAGTCGTTGCAGCCTCCTGAGTAGCTGAGATTANNNAGATCGGAA |
| 43403_2 | TATGAGTGTGGAGTCGTTGCAGCCTCCTGAGTAGCTGAGATTANNNAGATCGGAA |
| 37887_1 | TATGAGTGTGGAGTCGTTGCGGGAAGAAAGCAGCCTTCTTACTNNNAGATCGGAA |
| 37887_2 | TATGAGTGTGGAGTCGTTGCGGGAAGAAAGCAGCCTTCTTACTNNNAGATCGGAA |
| 49404_1 | TATGAGTGTGGAGTCGTTGCTTCTGGAACTTCTACATTCACAGANNNAGATCGGAA |
| 49404_2 | TATGAGTGTGGAGTCGTTGCTTCTGGAACTTCTACATTCACAGANNNAGATCGGAA |
| 36984_1 | TATGAGTGTGGAGTCGTTGCAAACCACCACGAACATAACTGTTNNNAGATCGGAAG |
| 36984_2 | TATGAGTGTGGAGTCGTTGCAAACCACCACGAACATAACTGTTNNNAGATCGGAAG |
| 35410_1 | TATGAGTGTGGAGTCGTTGCTCATGATATTCTCCTGCCTCAGCNNNAGATCGGAAGA |
| 35410_2 | TATGAGTGTGGAGTCGTTGCTCATGATATTCTCCTGCCTCAGCNNNAGATCGGAAGA |
| 31144_1 | TATGAGTGTGGAGTCGTTGCTCTCCTTATCTGACCTCGTCCTTNNNAGATCGGAAGA |
| 31144_2 | TATGAGTGTGGAGTCGTTGCTCTCCTTATCTGACCTCGTCCTTNNNAGATCGGAAGA |
| 45570_1 | TATGAGTGTGGAGTCGTTGCCCTTCCAAGTTCAAGCGATTCTCNNNAGATCGGAAG |
| 45570_2 | TATGAGTGTGGAGTCGTTGCCCTTCCAAGTTCAAGCGATTCTCNNNAGATCGGAAG |
| 29266_1 | TATGAGTGTGGAGTCGTTGCTGTACAGATTTCCCAGCTGACAANNNAGATCGGAAG |
| 29266_2 | TATGAGTGTGGAGTCGTTGCTGTACAGATTTCCCAGCTGACAANNNAGATCGGAAG |
| 50513_1 | TATGAGTGTGGAGTCGTTGCAGGACTGTGTAGTGCAAATCAGGNNNAGATCGGAA |
| 50513_2 | TATGAGTGTGGAGTCGTTGCAGGACTGTGTAGTGCAAATCAGGNNNAGATCGGAA |
| 50903_1 | TATGAGTGTGGAGTCGTTGCAGTATAAAGCCTGGCAATGCAATNNNAGATCGGAA |
| 50903_2 | TATGAGTGTGGAGTCGTTGCAGTATAAAGCCTGGCAATGCAATNNNAGATCGGAA |
| 51926_1 | TATGAGTGTGGAGTCGTTGCAACCCACCTAACCATTGGAGTACNNNAGATCGGAAG |
| 51926_2 | TATGAGTGTGGAGTCGTTGCAACCCACCTAACCATTGGAGTACNNNAGATCGGAAG |
| 43346_1 | TATGAGTGTGGAGTCGTTGCCCATCATGCCTGGCTAATTTTTGNNNAGATCGGAAG |
| 43346_2 | TATGAGTGTGGAGTCGTTGCCCATCATGCCTGGCTAATTTTTGNNNAGATCGGAAG |
| 37404_1 | TATGAGTGTGGAGTCGTTGCTGCAAATTGAACTGGATAGCCTGNNNAGATCGGAA |
| 37404_2 | TATGAGTGTGGAGTCGTTGCTGCAAATTGAACTGGATAGCCTGNNNAGATCGGAA |
| 46191_1 | TATGAGTGTGGAGTCGTTGCGAATTTTGGGAACCTCAGACTGCNNNAGATCGGAA |
| 46191_2 | TATGAGTGTGGAGTCGTTGCGAATTTTGGGAACCTCAGACTGCNNNAGATCGGAA |
| 33374_1 | TATGAGTGTGGAGTCGTTGCGGCCCTTCTCGTCATATCCTAACNNNAGATCGGAAGA |
| 33374_2 | TATGAGTGTGGAGTCGTTGCGGCCCTTCTCGTCATATCCTAACNNNAGATCGGAAGA |
| 42600_1 | TATGAGTGTGGAGTCGTTGCCAGAGTTTCATCATGTTGGCCAGNNNAGATCGGAAG |
| 42600_2 | TATGAGTGTGGAGTCGTTGCCAGAGTTTCATCATGTTGGCCAGNNNAGATCGGAAG |
| 29835_1 | TATGAGTGTGGAGTCGTTGCCCCTGTAGCAGAATGGGAGAAATNNNAGATCGGAA |

| | |
|---|---|
| 29835_2 | TATGAGTGTGGAGTCGTTGCCCCTGTAGCAGAATGGGAGAAATNNNAGATCGGAA |
| 34965_1 | TATGAGTGTGGAGTCGTTGCCATCTCGCCCGGCTAATTTTTTANNNAGATCGGAAGA |
| 34965_2 | TATGAGTGTGGAGTCGTTGCCATCTCGCCCGGCTAATTTTTTANNNAGATCGGAAGA |
| 30188_1 | TATGAGTGTGGAGTCGTTGCACAACCTTTTAGTTCAGTTGCTGTNNNAGATCGGAA |
| 30188_2 | TATGAGTGTGGAGTCGTTGCACAACCTTTTAGTTCAGTTGCTGTNNNAGATCGGAA |
| 10622_1 | TATGAGTGTGGAGTCGTTGCTCATAGCGCTCTTTTCCTTTCCTNNNAGATCGGAAGA |
| 10622_2 | TATGAGTGTGGAGTCGTTGCTCATAGCGCTCTTTTCCTTTCCTNNNAGATCGGAAGA |
| 49474_1 | TATGAGTGTGGAGTCGTTGCACAGTCTCAGAGTCCTTTGCAAANNNAGATCGGAAG |
| 49474_2 | TATGAGTGTGGAGTCGTTGCACAGTCTCAGAGTCCTTTGCAAANNNAGATCGGAAG |
| 31353_1 | TATGAGTGTGGAGTCGTTGCAAACCTCTGACCTATTTGGGAGTNNNAGATCGGAAG |
| 31353_2 | TATGAGTGTGGAGTCGTTGCAAACCTCTGACCTATTTGGGAGTNNNAGATCGGAAG |
| 29776_1 | TATGAGTGTGGAGTCGTTGCAAGCAGCAGTATGCTGAAGAANNNAGATCGGAAGA |
| 29776_2 | TATGAGTGTGGAGTCGTTGCAAGCAGCAGTATGCTGAAGAANNNAGATCGGAAGA |
| 48114_1 | TATGAGTGTGGAGTCGTTGCTTCAAGCAATTCTCCTGCCTCAGNNNAGATCGGAAG |
| 48114_2 | TATGAGTGTGGAGTCGTTGCTTCAAGCAATTCTCCTGCCTCAGNNNAGATCGGAAG |
| 34096_1 | TATGAGTGTGGAGTCGTTGCTAACATAGCAAGACCACCCATCCNNNAGATCGGAAG |
| 34096_2 | TATGAGTGTGGAGTCGTTGCTAACATAGCAAGACCACCCATCCNNNAGATCGGAAG |
| 50457_1 | TATGAGTGTGGAGTCGTTGCCAAGTTCTCTCCCAAGGTGAGTCNNNAGATCGGAAG |
| 50457_2 | TATGAGTGTGGAGTCGTTGCCAAGTTCTCTCCCAAGGTGAGTCNNNAGATCGGAAG |
| 38497_1 | TATGAGTGTGGAGTCGTTGCGATAACAAAGTTGCTTTGGGGGANNNAGATCGGAA |
| 38497_2 | TATGAGTGTGGAGTCGTTGCGATAACAAAGTTGCTTTGGGGGANNNAGATCGGAA |
| 41774_1 | TATGAGTGTGGAGTCGTTGCTCAAGTGATTCTCCTGCCTCATCNNNAGATCGGAAGA |
| 41774_2 | TATGAGTGTGGAGTCGTTGCTCAAGTGATTCTCCTGCCTCATCNNNAGATCGGAAGA |
| 37008_1 | TATGAGTGTGGAGTCGTTGCGCCATTGAGACAATGATCTTGCANNNAGATCGGAAG |
| 37008_2 | TATGAGTGTGGAGTCGTTGCGCCATTGAGACAATGATCTTGCANNNAGATCGGAAG |
| 48396_1 | TATGAGTGTGGAGTCGTTGCATCCTCCTGAGTAGCTGGGATTANNNAGATCGGAAG |
| 48396_2 | TATGAGTGTGGAGTCGTTGCATCCTCCTGAGTAGCTGGGATTANNNAGATCGGAAG |
| 46855_1 | TATGAGTGTGGAGTCGTTGCGGCAAAAGCGTTATGCATTATGCNNNAGATCGGAA |
| 46855_2 | TATGAGTGTGGAGTCGTTGCGGCAAAAGCGTTATGCATTATGCNNNAGATCGGAA |
| 43577_1 | TATGAGTGTGGAGTCGTTGCACATTTGCAGGGAATATCTGGGTNNNAGATCGGAA |
| 43577_2 | TATGAGTGTGGAGTCGTTGCACATTTGCAGGGAATATCTGGGTNNNAGATCGGAA |
| 38213_1 | TATGAGTGTGGAGTCGTTGCTCTTTTGTAGGAAGACTGAGCGCNNNAGATCGGAA |
| 38213_2 | TATGAGTGTGGAGTCGTTGCTCTTTTGTAGGAAGACTGAGCGCNNNAGATCGGAA |
| 52925_1 | TATGAGTGTGGAGTCGTTGCTCTGGGAATGAAGTCACTGACTGNNNAGATCGGAA |
| 52925_2 | TATGAGTGTGGAGTCGTTGCTCTGGGAATGAAGTCACTGACTGNNNAGATCGGAA |
| 46966_1 | TATGAGTGTGGAGTCGTTGCCTCCCAAGTAGCTGAGACTACAGNNNAGATCGGAAG |
| 46966_2 | TATGAGTGTGGAGTCGTTGCCTCCCAAGTAGCTGAGACTACAGNNNAGATCGGAAG |
| 40057_1 | TATGAGTGTGGAGTCGTTGCATTCAACAGGTTACAGGAGGAGCNNNAGATCGGAA |
| 40057_2 | TATGAGTGTGGAGTCGTTGCATTCAACAGGTTACAGGAGGAGCNNNAGATCGGAA |
| 13556_1 | TATGAGTGTGGAGTCGTTGCGAATTATATTCTGCCACTTAAGGNNNAGATCGGAAG |
| 13556_2 | TATGAGTGTGGAGTCGTTGCGAATTATATTCTGCCACTTAAGGNNNAGATCGGAAG |
| 48068_1 | TATGAGTGTGGAGTCGTTGCTCCTGTAGTAGTCTGGAAGTCCANNNAGATCGGAAG |
| 48068_2 | TATGAGTGTGGAGTCGTTGCTCCTGTAGTAGTCTGGAAGTCCANNNAGATCGGAAG |

| | |
|---|---|
| 42032_1 | TATGAGTGTGGAGTCGTTGCTGAGGGAGTAGATTCGCTTGAACNNNAGATCGGAA |
| 42032_2 | TATGAGTGTGGAGTCGTTGCTGAGGGAGTAGATTCGCTTGAACNNNAGATCGGAA |
| 12866_1 | TATGAGTGTGGAGTCGTTGCTATTCCACTGTTCTTTGTGTAANNNAGATCGGAAGA |
| 12866_2 | TATGAGTGTGGAGTCGTTGCTATTCCACTGTTCTTTGTGTAANNNAGATCGGAAGA |
| 44187_1 | TATGAGTGTGGAGTCGTTGCTTAGGGGACTGAAAGGCAAATCANNNAGATCGGAA |
| 44187_2 | TATGAGTGTGGAGTCGTTGCTTAGGGGACTGAAAGGCAAATCANNNAGATCGGAA |
| 38139_1 | TATGAGTGTGGAGTCGTTGCAGCAGCTTTGGAACTGGGTAATANNNAGATCGGAA |
| 38139_2 | TATGAGTGTGGAGTCGTTGCAGCAGCTTTGGAACTGGGTAATANNNAGATCGGAA |
| 37381_1 | TATGAGTGTGGAGTCGTTGCAGGCATTCCAAGCAGAGTGAATANNNAGATCGGAA |
| 37381_2 | TATGAGTGTGGAGTCGTTGCAGGCATTCCAAGCAGAGTGAATANNNAGATCGGAA |
| 29019_1 | TATGAGTGTGGAGTCGTTGCTTGTGCATAGGAACAAGTCCAGANNNAGATCGGAA |
| 29019_2 | TATGAGTGTGGAGTCGTTGCTTGTGCATAGGAACAAGTCCAGANNNAGATCGGAA |
| 43944_1 | TATGAGTGTGGAGTCGTTGCTCCTCTCTCTCTCTCTCTGTGTGNNNAGATCGGAAGA |
| 43944_2 | TATGAGTGTGGAGTCGTTGCTCCTCTCTCTCTCTCTCTGTGTGNNNAGATCGGAAGA |
| 39416_1 | TATGAGTGTGGAGTCGTTGCACACCATGGAAGTCAACAAACACNNNAGATCGGAAG |
| 39416_2 | TATGAGTGTGGAGTCGTTGCACACCATGGAAGTCAACAAACACNNNAGATCGGAAG |
| 41450_1 | TATGAGTGTGGAGTCGTTGCAATTGCATTTCCGTGTCACACAANNNAGATCGGAAG |
| 41450_2 | TATGAGTGTGGAGTCGTTGCAATTGCATTTCCGTGTCACACAANNNAGATCGGAAG |
| 49058_1 | TATGAGTGTGGAGTCGTTGCACCAAATTTATCCCCAGCCTGATNNNAGATCGGAAGA |
| 49058_2 | TATGAGTGTGGAGTCGTTGCACCAAATTTATCCCCAGCCTGATNNNAGATCGGAAGA |
| 7838_1 | TATGAGTGTGGAGTCGTTGCAAGCAATGCAGAAAAGTTTCACTNNNAGATCGGAAG |
| 7838_2 | TATGAGTGTGGAGTCGTTGCAAGCAATGCAGAAAAGTTTCACTNNNAGATCGGAAG |
| 37452_1 | TATGAGTGTGGAGTCGTTGCGACAGAGCAAGACTCCATCTCAANNNAGATCGGAAG |
| 37452_2 | TATGAGTGTGGAGTCGTTGCGACAGAGCAAGACTCCATCTCAANNNAGATCGGAAG |
| 43680_1 | TATGAGTGTGGAGTCGTTGCAGATTTGGAAAAGACCCTCCTCCNNNAGATCGGAAG |
| 43680_2 | TATGAGTGTGGAGTCGTTGCAGATTTGGAAAAGACCCTCCTCCNNNAGATCGGAAG |
| 30876_1 | TATGAGTGTGGAGTCGTTGCTCCCTAGAACCTCCAAAAGAAGCNNNAGATCGGAAG |
| 30876_2 | TATGAGTGTGGAGTCGTTGCTCCCTAGAACCTCCAAAAGAAGCNNNAGATCGGAAG |
| 45908_1 | TATGAGTGTGGAGTCGTTGCTAAGTAGCTGGGATTACAGGCACNNNAGATCGGAA |
| 45908_2 | TATGAGTGTGGAGTCGTTGCTAAGTAGCTGGGATTACAGGCACNNNAGATCGGAA |
| 37250_1 | TATGAGTGTGGAGTCGTTGCCACACACACACACACACACATATNNNAGATCGGAAGA |
| 37250_2 | TATGAGTGTGGAGTCGTTGCCACACACACACACACACACATATNNNAGATCGGAAGA |
| 40107_1 | TATGAGTGTGGAGTCGTTGCTTTTTGCTCAGGATAGCTTTGGCNNNAGATCGGAAG |
| 40107_2 | TATGAGTGTGGAGTCGTTGCTTTTTGCTCAGGATAGCTTTGGCNNNAGATCGGAAG |
| 49550_1 | TATGAGTGTGGAGTCGTTGCGATACCCAGGGTTCCAGTAACAANNNAGATCGGAA |
| 49550_2 | TATGAGTGTGGAGTCGTTGCGATACCCAGGGTTCCAGTAACAANNNAGATCGGAA |
| 44417_1 | TATGAGTGTGGAGTCGTTGCCTGCAACTTGAGATCACCTGTTGNNNAGATCGGAAG |
| 44417_2 | TATGAGTGTGGAGTCGTTGCCTGCAACTTGAGATCACCTGTTGNNNAGATCGGAAG |
| 52932_1 | TATGAGTGTGGAGTCGTTGCGGGCAATGGAGTTATGTTCCAAGNNNAGATCGGAA |
| 52932_2 | TATGAGTGTGGAGTCGTTGCGGGCAATGGAGTTATGTTCCAAGNNNAGATCGGAA |
| 48520_1 | TATGAGTGTGGAGTCGTTGCTTCAAGTGATTCTCCTGCCTCAGNNNAGATCGGAAG |
| 48520_2 | TATGAGTGTGGAGTCGTTGCTTCAAGTGATTCTCCTGCCTCAGNNNAGATCGGAAG |
| 29282_1 | TATGAGTGTGGAGTCGTTGCTATCTTCGTGGGACATGACTTCCNNNAGATCGGAAG |

| | |
|---|---|
| 29282_2 | TATGAGTGTGGAGTCGTTGCTATCTTCGTGGGACATGACTTCCNNNAGATCGGAAG |
| 41725_1 | TATGAGTGTGGAGTCGTTGCTCCTCCCGAGTTCAAGTGATTACNNNAGATCGGAAG |
| 41725_2 | TATGAGTGTGGAGTCGTTGCTCCTCCCGAGTTCAAGTGATTACNNNAGATCGGAAG |
| 48481_1 | TATGAGTGTGGAGTCGTTGCCTCACTTGCCATTCAATTCCACANNNAGATCGGAAGA |
| 48481_2 | TATGAGTGTGGAGTCGTTGCCTCACTTGCCATTCAATTCCACANNNAGATCGGAAGA |
| 33959_1 | TATGAGTGTGGAGTCGTTGCCTCCCAGGTTCATGCCATTCTCNNNAGATCGGAAGAG |
| 33959_2 | TATGAGTGTGGAGTCGTTGCCTCCCAGGTTCATGCCATTCTCNNNAGATCGGAAGAG |
| 13213_1 | TATGAGTGTGGAGTCGTTGCGGGTAAAAAAATAATTTCCCTTCNNNAGATCGGAAG |
| 13213_2 | TATGAGTGTGGAGTCGTTGCGGGTAAAAAAATAATTTCCCTTCNNNAGATCGGAAG |
| 32894_1 | TATGAGTGTGGAGTCGTTGCGTTCCTCAAGTTCCCATCCTCTTNNNAGATCGGAAGA |
| 32894_2 | TATGAGTGTGGAGTCGTTGCGTTCCTCAAGTTCCCATCCTCTTNNNAGATCGGAAGA |
| 35624_1 | TATGAGTGTGGAGTCGTTGCTGTTAGCCAGGATGGTCTTGATCNNNAGATCGGAAG |
| 35624_2 | TATGAGTGTGGAGTCGTTGCTGTTAGCCAGGATGGTCTTGATCNNNAGATCGGAAG |
| 34866_1 | TATGAGTGTGGAGTCGTTGCTTCAAGTGATTCTCCTGCCTCAGNNNAGATCGGAAG |
| 34866_2 | TATGAGTGTGGAGTCGTTGCTTCAAGTGATTCTCCTGCCTCAGNNNAGATCGGAAG |
| 51208_1 | TATGAGTGTGGAGTCGTTGCGTTTTGGCTGCATTCAGAGAGTTNNNAGATCGGAAG |
| 51208_2 | TATGAGTGTGGAGTCGTTGCGTTTTGGCTGCATTCAGAGAGTTNNNAGATCGGAAG |
| 31804_1 | TATGAGTGTGGAGTCGTTGCTCACAAGTGCACATGGAATGTTCNNNAGATCGGAAG |
| 31804_2 | TATGAGTGTGGAGTCGTTGCTCACAAGTGCACATGGAATGTTCNNNAGATCGGAAG |
| 37796_1 | TATGAGTGTGGAGTCGTTGCTGCCTGTAATTGCAGCTATTTGGNNNAGATCGGAAG |
| 37796_2 | TATGAGTGTGGAGTCGTTGCTGCCTGTAATTGCAGCTATTTGGNNNAGATCGGAAG |
| 10483_1 | TATGAGTGTGGAGTCGTTGCGGTAGGAACCAGGGACTGAAGAANNNAGATCGGA |
| 10483_2 | TATGAGTGTGGAGTCGTTGCGGTAGGAACCAGGGACTGAAGAANNNAGATCGGA |
| 33010_1 | TATGAGTGTGGAGTCGTTGCGGAATTTCACTCTTGTTGCCCAGNNNAGATCGGAAG |
| 33010_2 | TATGAGTGTGGAGTCGTTGCGGAATTTCACTCTTGTTGCCCAGNNNAGATCGGAAG |
| 9638_1 | TATGAGTGTGGAGTCGTTGCTCTCTTCCCCTCCTCAATTCTCCNNNAGATCGGAAGAG |
| 9638_2 | TATGAGTGTGGAGTCGTTGCTCTCTTCCCCTCCTCAATTCTCCNNNAGATCGGAAGAG |
| 46675_1 | TATGAGTGTGGAGTCGTTGCGTAACTGGGATTACAGGAGCACANNNAGATCGGAA |
| 46675_2 | TATGAGTGTGGAGTCGTTGCGTAACTGGGATTACAGGAGCACANNNAGATCGGAA |
| 49314_1 | TATGAGTGTGGAGTCGTTGCCCAAAATGCTGGAATTACAGGCANNNAGATCGGAAG |
| 49314_2 | TATGAGTGTGGAGTCGTTGCCCAAAATGCTGGAATTACAGGCANNNAGATCGGAAG |
| 37225_1 | TATGAGTGTGGAGTCGTTGCACAGTCTGGTAGCTTGTGAACATNNNAGATCGGAAG |
| 37225_2 | TATGAGTGTGGAGTCGTTGCACAGTCTGGTAGCTTGTGAACATNNNAGATCGGAAG |
| 37744_1 | TATGAGTGTGGAGTCGTTGCTCCCAGAGATCAGTTTGGTTCTGNNNAGATCGGAAG |
| 37744_2 | TATGAGTGTGGAGTCGTTGCTCCCAGAGATCAGTTTGGTTCTGNNNAGATCGGAAG |
| 33573_1 | TATGAGTGTGGAGTCGTTGCTTCTTGTTCAGTCATGGGAAGGTNNNAGATCGGAAG |
| 33573_2 | TATGAGTGTGGAGTCGTTGCTTCTTGTTCAGTCATGGGAAGGTNNNAGATCGGAAG |
| 51119_1 | TATGAGTGTGGAGTCGTTGCAGCCTGCCATCACTTAGAGAAAANNNAGATCGGAAG |
| 51119_2 | TATGAGTGTGGAGTCGTTGCAGCCTGCCATCACTTAGAGAAAANNNAGATCGGAAG |
| 36443_1 | TATGAGTGTGGAGTCGTTGCCAGGAAGTCAAGGCTGTAGTGAGNNNAGATCGGAA |
| 36443_2 | TATGAGTGTGGAGTCGTTGCCAGGAAGTCAAGGCTGTAGTGAGNNNAGATCGGAA |
| 31962_1 | TATGAGTGTGGAGTCGTTGCGCCTAGGTGCTCATATGGACTTANNNAGATCGGAAG |
| 31962_2 | TATGAGTGTGGAGTCGTTGCGCCTAGGTGCTCATATGGACTTANNNAGATCGGAAG |

| | |
|---|---|
| 35367_1 | TATGAGTGTGGAGTCGTTGCCAGTGGTGCAATTTTGGCTCANNNAGATCGGAAGA |
| 35367_2 | TATGAGTGTGGAGTCGTTGCCAGTGGTGCAATTTTGGCTCANNNAGATCGGAAGA |
| 13079_1 | TATGAGTGTGGAGTCGTTGCAAGTAAAATATTTAAAAGCAACCANNNAGATCGGAA |
| 13079_2 | TATGAGTGTGGAGTCGTTGCAAGTAAAATATTTAAAAGCAACCANNNAGATCGGAA |
| 36620_1 | TATGAGTGTGGAGTCGTTGCGGGGGGGAATAAAAGACTTGTCANNNAGATCGGA |
| 36620_2 | TATGAGTGTGGAGTCGTTGCGGGGGGGAATAAAAGACTTGTCANNNAGATCGGA |
| 32167_1 | TATGAGTGTGGAGTCGTTGCTTTGCTTTTCTGTTGGGGGAAAGNNNAGATCGGAA |
| 32167_2 | TATGAGTGTGGAGTCGTTGCTTTGCTTTTCTGTTGGGGGAAAGNNNAGATCGGAA |
| 30159_1 | TATGAGTGTGGAGTCGTTGCGTTGCCAATGACACCAATAGCTGNNNAGATCGGAAG |
| 30159_2 | TATGAGTGTGGAGTCGTTGCGTTGCCAATGACACCAATAGCTGNNNAGATCGGAAG |
| 40926_1 | TATGAGTGTGGAGTCGTTGCCCAGGCTCCATACACAGTTCTAANNNAGATCGGAAG |
| 40926_2 | TATGAGTGTGGAGTCGTTGCCCAGGCTCCATACACAGTTCTAANNNAGATCGGAAG |
| 39384_1 | TATGAGTGTGGAGTCGTTGCGGTGCTGGGGTCTCAATTATGTANNNAGATCGGAA |
| 39384_2 | TATGAGTGTGGAGTCGTTGCGGTGCTGGGGTCTCAATTATGTANNNAGATCGGAA |
| 2308_1 | TATGAGTGTGGAGTCGTTGCAGTTGTAGAGATGGTTGCACGACNNNAGATCGGAA |
| 2308_2 | TATGAGTGTGGAGTCGTTGCAGTTGTAGAGATGGTTGCACGACNNNAGATCGGAA |
| 40986_1 | TATGAGTGTGGAGTCGTTGCTTCTGTAAGTGGATGTGGAGTGGNNNAGATCGGAA |
| 40986_2 | TATGAGTGTGGAGTCGTTGCTTCTGTAAGTGGATGTGGAGTGGNNNAGATCGGAA |
| 33768_1 | TATGAGTGTGGAGTCGTTGCAATGCAGAAATAACCTGGGGTCANNNAGATCGGAA |
| 33768_2 | TATGAGTGTGGAGTCGTTGCAATGCAGAAATAACCTGGGGTCANNNAGATCGGAA |
| 13100_1 | TATGAGTGTGGAGTCGTTGCGACAAGAACACAGAGACAAGNNNAGATCGGAAGAG |
| 13100_2 | TATGAGTGTGGAGTCGTTGCGACAAGAACACAGAGACAAGNNNAGATCGGAAGAG |
| 33187_1 | TATGAGTGTGGAGTCGTTGCGGAGGGGCCAATATGACAAACTANNNAGATCGGAA |
| 33187_2 | TATGAGTGTGGAGTCGTTGCGGAGGGGCCAATATGACAAACTANNNAGATCGGAA |
| 20463_1 | TATGAGTGTGGAGTCGTTGCTTTTAAAAGGAAAGAATGTCTCTNNNAGATCGGAAG |
| 20463_2 | TATGAGTGTGGAGTCGTTGCTTTTAAAAGGAAAGAATGTCTCTNNNAGATCGGAAG |
| 44387_1 | TATGAGTGTGGAGTCGTTGCCTGGAACGTAACACACTTGCATTNNNAGATCGGAAG |
| 44387_2 | TATGAGTGTGGAGTCGTTGCCTGGAACGTAACACACTTGCATTNNNAGATCGGAAG |
| 39435_1 | TATGAGTGTGGAGTCGTTGCATGTGGCACATATACACCATGGANNNAGATCGGAAG |
| 39435_2 | TATGAGTGTGGAGTCGTTGCATGTGGCACATATACACCATGGANNNAGATCGGAAG |
| 31911_1 | TATGAGTGTGGAGTCGTTGCACTGTACATATTTTTGTGGTACATGTGNNNAGATCG |
| 31911_2 | TATGAGTGTGGAGTCGTTGCACTGTACATATTTTTGTGGTACATGTGNNNAGATCG |
| 30931_1 | TATGAGTGTGGAGTCGTTGCATGAGAGCAAGAACCTTGTCTGTNNNAGATCGGAA |
| 30931_2 | TATGAGTGTGGAGTCGTTGCATGAGAGCAAGAACCTTGTCTGTNNNAGATCGGAA |
| 46776_1 | TATGAGTGTGGAGTCGTTGCCTGGAGTGCAATGGTGTGATTATNNNAGATCGGAA |
| 46776_2 | TATGAGTGTGGAGTCGTTGCCTGGAGTGCAATGGTGTGATTATNNNAGATCGGAA |
| 13099_1 | TATGAGTGTGGAGTCGTTGCTCTCCTTTTATTAATAGGAAGGCNNNAGATCGGAAG |
| 13099_2 | TATGAGTGTGGAGTCGTTGCTCTCCTTTTATTAATAGGAAGGCNNNAGATCGGAAG |
| 37496_1 | TATGAGTGTGGAGTCGTTGCATGCCTGTAATCCCACCACTTTNNNAGATCGGAAGAG |
| 37496_2 | TATGAGTGTGGAGTCGTTGCATGCCTGTAATCCCACCACTTTNNNAGATCGGAAGAG |
| 51710_1 | TATGAGTGTGGAGTCGTTGCATCCATTCTACGTGCTGAGAGTCNNNAGATCGGAAG |
| 51710_2 | TATGAGTGTGGAGTCGTTGCATCCATTCTACGTGCTGAGAGTCNNNAGATCGGAAG |
| 49432_1 | TATGAGTGTGGAGTCGTTGCAACAACTTCAAAAGGCAGCTTGTNNNAGATCGGAAG |

| | |
|---|---|
| 49432_2 | TATGAGTGTGGAGTCGTTGCAACAACTTCAAAAGGCAGCTTGTNNNAGATCGGAAG |
| 37406_1 | TATGAGTGTGGAGTCGTTGCAGGAAGAAATAACCACAGGCACTNNNAGATCGGAA |
| 37406_2 | TATGAGTGTGGAGTCGTTGCAGGAAGAAATAACCACAGGCACTNNNAGATCGGAA |
| 46617_1 | TATGAGTGTGGAGTCGTTGCGAGGGAGCACATTTATCAATGGCNNNAGATCGGAA |
| 46617_2 | TATGAGTGTGGAGTCGTTGCGAGGGAGCACATTTATCAATGGCNNNAGATCGGAA |
| 37948_1 | TATGAGTGTGGAGTCGTTGCCAAGTTTCTCCATCAAATGTGGGANNNAGATCGGAA |
| 37948_2 | TATGAGTGTGGAGTCGTTGCCAAGTTTCTCCATCAAATGTGGGANNNAGATCGGAA |
| 2116_1 | TATGAGTGTGGAGTCGTTGCACAGTAATAAACACTCATGGGCANNNAGATCGGAA |
| 2116_2 | TATGAGTGTGGAGTCGTTGCACAGTAATAAACACTCATGGGCANNNAGATCGGAA |
| 48144_1 | TATGAGTGTGGAGTCGTTGCCCTCCTGGGTTCAAGTGATTCTTNNNAGATCGGAAG |
| 48144_2 | TATGAGTGTGGAGTCGTTGCCCTCCTGGGTTCAAGTGATTCTTNNNAGATCGGAAG |
| 10500_1 | TATGAGTGTGGAGTCGTTGCTTGCTGTGTCTAAGCATCTTTGTNNNAGATCGGAAG |
| 10500_2 | TATGAGTGTGGAGTCGTTGCTTGCTGTGTCTAAGCATCTTTGTNNNAGATCGGAAG |
| 45815_1 | TATGAGTGTGGAGTCGTTGCACTATCTGTGCTCATGGAACGTGNNNAGATCGGAAG |
| 45815_2 | TATGAGTGTGGAGTCGTTGCACTATCTGTGCTCATGGAACGTGNNNAGATCGGAAG |
| 32906_1 | TATGAGTGTGGAGTCGTTGCCTATAGCACAGCAAACCTCAGACNNNAGATCGGAAG |
| 32906_2 | TATGAGTGTGGAGTCGTTGCCTATAGCACAGCAAACCTCAGACNNNAGATCGGAAG |
| 43662_1 | TATGAGTGTGGAGTCGTTGCAGCTGGTTATTCTACTGTTGGGGNNNAGATCGGAA |
| 43662_2 | TATGAGTGTGGAGTCGTTGCAGCTGGTTATTCTACTGTTGGGGNNNAGATCGGAA |
| 36818_1 | TATGAGTGTGGAGTCGTTGCACATATGTGGACACAGAGACACTNNNAGATCGGAA |
| 36818_2 | TATGAGTGTGGAGTCGTTGCACATATGTGGACACAGAGACACTNNNAGATCGGAA |
| 39491_1 | TATGAGTGTGGAGTCGTTGCACCAGAGGAGGATAAGGAGACATNNNAGATCGGA |
| 39491_2 | TATGAGTGTGGAGTCGTTGCACCAGAGGAGGATAAGGAGACATNNNAGATCGGA |
| 13095_1 | TATGAGTGTGGAGTCGTTGCCTTTAGGAATGCTGTGTGTCNNNAGATCGGAAGAG |
| 13095_2 | TATGAGTGTGGAGTCGTTGCCTTTAGGAATGCTGTGTGTCNNNAGATCGGAAGAG |
| 52029_1 | TATGAGTGTGGAGTCGTTGCTTTCAAGCCAGTGGATCTTAGCTNNNAGATCGGAAG |
| 52029_2 | TATGAGTGTGGAGTCGTTGCTTTCAAGCCAGTGGATCTTAGCTNNNAGATCGGAAG |
| 51124_1 | TATGAGTGTGGAGTCGTTGCTAGGACTTCTGGGGGTTATCCTTNNNAGATCGGAAG |
| 51124_2 | TATGAGTGTGGAGTCGTTGCTAGGACTTCTGGGGGTTATCCTTNNNAGATCGGAAG |
| 45915_1 | TATGAGTGTGGAGTCGTTGCCATGATCTCGGCTCACTGCAANNNAGATCGGAAGAG |
| 45915_2 | TATGAGTGTGGAGTCGTTGCCATGATCTCGGCTCACTGCAANNNAGATCGGAAGAG |
| 43609_1 | TATGAGTGTGGAGTCGTTGCCGTCTGGTGTGGTCTGGAAAATNNNAGATCGGAAG |
| 43609_2 | TATGAGTGTGGAGTCGTTGCCGTCTGGTGTGGTCTGGAAAATNNNAGATCGGAAG |
| 42974_1 | TATGAGTGTGGAGTCGTTGCTCACATGACCTGTTCAGAAAGGTNNNAGATCGGAAG |
| 42974_2 | TATGAGTGTGGAGTCGTTGCTCACATGACCTGTTCAGAAAGGTNNNAGATCGGAAG |
| 52986_1 | TATGAGTGTGGAGTCGTTGCAACAAAATGGCAGGAGTAAGTCCNNNAGATCGGAA |
| 52986_2 | TATGAGTGTGGAGTCGTTGCAACAAAATGGCAGGAGTAAGTCCNNNAGATCGGAA |
| 41244_1 | TATGAGTGTGGAGTCGTTGCTAACATAGCAAGACCACCCATCCNNNAGATCGGAAG |
| 41244_2 | TATGAGTGTGGAGTCGTTGCTAACATAGCAAGACCACCCATCCNNNAGATCGGAAG |
| 41330_1 | TATGAGTGTGGAGTCGTTGCAGTCTTGGGTGAGATTTTGGACANNNAGATCGGAA |
| 41330_2 | TATGAGTGTGGAGTCGTTGCAGTCTTGGGTGAGATTTTGGACANNNAGATCGGAA |
| 13486_1 | TATGAGTGTGGAGTCGTTGCCTGTTTATGAGGCTTATGTTGNNNAGATCGGAAGA |
| 13486_2 | TATGAGTGTGGAGTCGTTGCCTGTTTATGAGGCTTATGTTGNNNAGATCGGAAGA |

| | |
|---|---|
| 30799_1 | TATGAGTGTGGAGTCGTTGCAAGACCCAGTGACAAAACTTTGCNNNAGATCGGAAG |
| 30799_2 | TATGAGTGTGGAGTCGTTGCAAGACCCAGTGACAAAACTTTGCNNNAGATCGGAAG |
| 41980_1 | TATGAGTGTGGAGTCGTTGCCTCCTGAGTAGCTGGGATTACAGNNNAGATCGGAA |
| 41980_2 | TATGAGTGTGGAGTCGTTGCCTCCTGAGTAGCTGGGATTACAGNNNAGATCGGAA |
| 922_1 | TATGAGTGTGGAGTCGTTGCTTAGCATCACGCAATATGCCCATANNNAGATCGGAA |
| 922_2 | TATGAGTGTGGAGTCGTTGCTTAGCATCACGCAATATGCCCATANNNAGATCGGAA |
| 12760_1 | TATGAGTGTGGAGTCGTTGCGTGTTCAACAAGAGTTCTTTGNNNAGATCGGAAGA |
| 12760_2 | TATGAGTGTGGAGTCGTTGCGTGTTCAACAAGAGTTCTTTGNNNAGATCGGAAGA |
| 30895_1 | TATGAGTGTGGAGTCGTTGCTCAGTGATGTGGTTATGCGTGTANNNAGATCGGAA |
| 30895_2 | TATGAGTGTGGAGTCGTTGCTCAGTGATGTGGTTATGCGTGTANNNAGATCGGAA |
| 29322_1 | TATGAGTGTGGAGTCGTTGCGTACAGTCCAGGGAAACAGACATNNNAGATCGGAA |
| 29322_2 | TATGAGTGTGGAGTCGTTGCGTACAGTCCAGGGAAACAGACATNNNAGATCGGAA |
| 47222_1 | TATGAGTGTGGAGTCGTTGCCTCATTGAAACCTCCACCTCCTGNNNAGATCGGAAGA |
| 47222_2 | TATGAGTGTGGAGTCGTTGCCTCATTGAAACCTCCACCTCCTGNNNAGATCGGAAGA |
| 2146_1 | TATGAGTGTGGAGTCGTTGCAGCCATAGCCTAATCCAATTGTCANNNAGATCGGAA |
| 2146_2 | TATGAGTGTGGAGTCGTTGCAGCCATAGCCTAATCCAATTGTCANNNAGATCGGAA |
| 32402_1 | TATGAGTGTGGAGTCGTTGCTGATGAACTAGGTCTACATTGCANNNAGATCGGAAG |
| 32402_2 | TATGAGTGTGGAGTCGTTGCTGATGAACTAGGTCTACATTGCANNNAGATCGGAAG |
| 33578_1 | TATGAGTGTGGAGTCGTTGCGGAGTCTCAATCTATCGCCCAGNNNAGATCGGAAGA |
| 33578_2 | TATGAGTGTGGAGTCGTTGCGGAGTCTCAATCTATCGCCCAGNNNAGATCGGAAGA |
| 50751_1 | TATGAGTGTGGAGTCGTTGCCAAGGTCATCAAAGCTGTTCCTGNNNAGATCGGAAG |
| 50751_2 | TATGAGTGTGGAGTCGTTGCCAAGGTCATCAAAGCTGTTCCTGNNNAGATCGGAAG |
| 29403_1 | TATGAGTGTGGAGTCGTTGCTTTTTTCTTTTTCTTTGAGACTGNNNAGATCGGAAGA |
| 29403_2 | TATGAGTGTGGAGTCGTTGCTTTTTTCTTTTTCTTTGAGACTGNNNAGATCGGAAGA |
| 49175_1 | TATGAGTGTGGAGTCGTTGCTTATTTCGGGCACTCCACAGATTNNNAGATCGGAAG |
| 49175_2 | TATGAGTGTGGAGTCGTTGCTTATTTCGGGCACTCCACAGATTNNNAGATCGGAAG |
| 42375_1 | TATGAGTGTGGAGTCGTTGCATTCGATAATGCAGGAGCTCCATNNNAGATCGGAAG |
| 42375_2 | TATGAGTGTGGAGTCGTTGCATTCGATAATGCAGGAGCTCCATNNNAGATCGGAAG |
| 44988_1 | TATGAGTGTGGAGTCGTTGCTTCGAGTCTGAGAGCGAATTGATNNNAGATCGGAA |
| 44988_2 | TATGAGTGTGGAGTCGTTGCTTCGAGTCTGAGAGCGAATTGATNNNAGATCGGAA |
| 31591_1 | TATGAGTGTGGAGTCGTTGCTACAAAGATCATCACGACAGCCANNNAGATCGGAAG |
| 31591_2 | TATGAGTGTGGAGTCGTTGCTACAAAGATCATCACGACAGCCANNNAGATCGGAAG |
| 53013_1 | TATGAGTGTGGAGTCGTTGCCTCAAGTGATTCTCCTGCCTCAGNNNAGATCGGAAG |
| 53013_2 | TATGAGTGTGGAGTCGTTGCCTCAAGTGATTCTCCTGCCTCAGNNNAGATCGGAAG |
| 38376_1 | TATGAGTGTGGAGTCGTTGCAAAAAAAATTAGCCAGGCGTGGTNNNAGATCGGAA |
| 38376_2 | TATGAGTGTGGAGTCGTTGCAAAAAAAATTAGCCAGGCGTGGTNNNAGATCGGAA |
| 48539_1 | TATGAGTGTGGAGTCGTTGCCTGAGTACATCCAGAAAACCCCANNNAGATCGGAAG |
| 48539_2 | TATGAGTGTGGAGTCGTTGCCTGAGTACATCCAGAAAACCCCANNNAGATCGGAAG |
| 34514_1 | TATGAGTGTGGAGTCGTTGCCCCTCCTCAGACCATGTTGTTATNNNAGATCGGAAGA |
| 34514_2 | TATGAGTGTGGAGTCGTTGCCCCTCCTCAGACCATGTTGTTATNNNAGATCGGAAGA |
| 45253_1 | TATGAGTGTGGAGTCGTTGCGTAAAAACACTGGTTGCCTCTGGNNNAGATCGGAA |
| 45253_2 | TATGAGTGTGGAGTCGTTGCGTAAAAACACTGGTTGCCTCTGGNNNAGATCGGAA |
| 13598_1 | TATGAGTGTGGAGTCGTTGCCTTCCATCCCTCTCTTCATTNNNAGATCGGAAGAGCA |

| | |
|---|---|
| 13598_2 | TATGAGTGTGGAGTCGTTGCCTTCCATCCCTCTCTTCATTNNNAGATCGGAAGAGCA |
| 52682_1 | TATGAGTGTGGAGTCGTTGCTAAGCAGCGAACTCTCTGGATACNNNAGATCGGAAG |
| 52682_2 | TATGAGTGTGGAGTCGTTGCTAAGCAGCGAACTCTCTGGATACNNNAGATCGGAAG |
| 6295_1 | TATGAGTGTGGAGTCGTTGCCTCAGATAATGGTGCCACTGTCCNNNAGATCGGAAG |
| 6295_2 | TATGAGTGTGGAGTCGTTGCCTCAGATAATGGTGCCACTGTCCNNNAGATCGGAAG |
| 3700_1 | TATGAGTGTGGAGTCGTTGCGGGTATAGAGGAGTGCACCTGTANNNAGATCGGAA |
| 3700_2 | TATGAGTGTGGAGTCGTTGCGGGTATAGAGGAGTGCACCTGTANNNAGATCGGAA |
| 44189_1 | TATGAGTGTGGAGTCGTTGCCTCCCCATCCCAAGCCTTTGNNNAGATCGGAAGAGC |
| 44189_2 | TATGAGTGTGGAGTCGTTGCCTCCCCATCCCAAGCCTTTGNNNAGATCGGAAGAGC |
| 51858_1 | TATGAGTGTGGAGTCGTTGCCTCAGCCAGACATTACCAGTCATNNNAGATCGGAAG |
| 51858_2 | TATGAGTGTGGAGTCGTTGCCTCAGCCAGACATTACCAGTCATNNNAGATCGGAAG |
| 36007_1 | TATGAGTGTGGAGTCGTTGCACTGAATAAGTGTGGCCTCTTTCNNNAGATCGGAAG |
| 36007_2 | TATGAGTGTGGAGTCGTTGCACTGAATAAGTGTGGCCTCTTTCNNNAGATCGGAAG |
| 33931_1 | TATGAGTGTGGAGTCGTTGCCCCTAGTAACTGGGATTACAGGCNNNAGATCGGAA |
| 33931_2 | TATGAGTGTGGAGTCGTTGCCCCTAGTAACTGGGATTACAGGCNNNAGATCGGAA |
| 48976_1 | TATGAGTGTGGAGTCGTTGCTTTTGCCAGGAAAAAAGATGCCANNNAGATCGGAA |
| 48976_2 | TATGAGTGTGGAGTCGTTGCTTTTGCCAGGAAAAAAGATGCCANNNAGATCGGAA |
| 38809_1 | TATGAGTGTGGAGTCGTTGCTGTGTAGGCTACCCTACCTAACANNNAGATCGGAAG |
| 38809_2 | TATGAGTGTGGAGTCGTTGCTGTGTAGGCTACCCTACCTAACANNNAGATCGGAAG |
| 13503_1 | TATGAGTGTGGAGTCGTTGCTTACCTTTTTCCATGTTTTGTTTNNNAGATCGGAAGA |
| 13503_2 | TATGAGTGTGGAGTCGTTGCTTACCTTTTTCCATGTTTTGTTTNNNAGATCGGAAGA |
| 43732_1 | TATGAGTGTGGAGTCGTTGCGGCGACCAGAAAAGGAAATGAAANNNAGATCGGA |
| 43732_2 | TATGAGTGTGGAGTCGTTGCGGCGACCAGAAAAGGAAATGAAANNNAGATCGGA |
| 9645_1 | TATGAGTGTGGAGTCGTTGCATCCAAAGGCAGTTTCTCCATCCNNNAGATCGGAAG |
| 9645_2 | TATGAGTGTGGAGTCGTTGCATCCAAAGGCAGTTTCTCCATCCNNNAGATCGGAAG |
| 35795_1 | TATGAGTGTGGAGTCGTTGCTTAGGGTACATGTGCACATTGTGNNNAGATCGGAA |
| 35795_2 | TATGAGTGTGGAGTCGTTGCTTAGGGTACATGTGCACATTGTGNNNAGATCGGAA |
| 45767_1 | TATGAGTGTGGAGTCGTTGCTGAGAACAGATATGGGGCTCAACNNNAGATCGGAA |
| 45767_2 | TATGAGTGTGGAGTCGTTGCTGAGAACAGATATGGGGCTCAACNNNAGATCGGAA |
| 51203_1 | TATGAGTGTGGAGTCGTTGCTCTTACAACAGAGTCCTTGGGTTNNNAGATCGGAAG |
| 51203_2 | TATGAGTGTGGAGTCGTTGCTCTTACAACAGAGTCCTTGGGTTNNNAGATCGGAAG |
| 47737_1 | TATGAGTGTGGAGTCGTTGCCAGAAAGGAGGCTTTGCTTTGTTNNNAGATCGGAA |
| 47737_2 | TATGAGTGTGGAGTCGTTGCCAGAAAGGAGGCTTTGCTTTGTTNNNAGATCGGAA |
| 52520_1 | TATGAGTGTGGAGTCGTTGCACATGGTAATAGCTCAGCGAACTNNNAGATCGGAA |
| 52520_2 | TATGAGTGTGGAGTCGTTGCACATGGTAATAGCTCAGCGAACTNNNAGATCGGAA |
| 48896_1 | TATGAGTGTGGAGTCGTTGCAACACACACACACTAGTTTGGATNNNAGATCGGAAG |
| 48896_2 | TATGAGTGTGGAGTCGTTGCAACACACACACACTAGTTTGGATNNNAGATCGGAAG |
| 51065_1 | TATGAGTGTGGAGTCGTTGCCATTGCCAAAAATGTTGCAGGACNNNAGATCGGAA |
| 51065_2 | TATGAGTGTGGAGTCGTTGCCATTGCCAAAAATGTTGCAGGACNNNAGATCGGAA |
| 40382_1 | TATGAGTGTGGAGTCGTTGCTGAAAAAGGCCATCAAGGAGAGANNNAGATCGGA |
| 40382_2 | TATGAGTGTGGAGTCGTTGCTGAAAAAGGCCATCAAGGAGAGANNNAGATCGGA |
| 46221_1 | TATGAGTGTGGAGTCGTTGCGCAGTGGCACAATCTTAGCTTACNNNAGATCGGAAG |
| 46221_2 | TATGAGTGTGGAGTCGTTGCGCAGTGGCACAATCTTAGCTTACNNNAGATCGGAAG |

| | |
|---|---|
| 30946_1 | TATGAGTGTGGAGTCGTTGCTCTGGATATACCCCACCATTCCTNNNAGATCGGAAGA |
| 30946_2 | TATGAGTGTGGAGTCGTTGCTCTGGATATACCCCACCATTCCTNNNAGATCGGAAGA |
| 33160_1 | TATGAGTGTGGAGTCGTTGCTATGGACAGAGCATGGTTTCCTCNNNAGATCGGAAG |
| 33160_2 | TATGAGTGTGGAGTCGTTGCTATGGACAGAGCATGGTTTCCTCNNNAGATCGGAAG |
| 33993_1 | TATGAGTGTGGAGTCGTTGCTAGGCTCAAGTGATTCTAGTGCCNNNAGATCGGAAG |
| 33993_2 | TATGAGTGTGGAGTCGTTGCTAGGCTCAAGTGATTCTAGTGCCNNNAGATCGGAAG |
| 45793_1 | TATGAGTGTGGAGTCGTTGCTTCAAGTGATTCTCCTGCCTCAGNNNAGATCGGAAG |
| 45793_2 | TATGAGTGTGGAGTCGTTGCTTCAAGTGATTCTCCTGCCTCAGNNNAGATCGGAAG |
| 31436_1 | TATGAGTGTGGAGTCGTTGCGCTCTTTGCCACAACAGAGAAAGNNNAGATCGGAA |
| 31436_2 | TATGAGTGTGGAGTCGTTGCGCTCTTTGCCACAACAGAGAAAGNNNAGATCGGAA |
| 51426_1 | TATGAGTGTGGAGTCGTTGCACACAGCCACACCATATCATAGTNNNAGATCGGAAG |
| 51426_2 | TATGAGTGTGGAGTCGTTGCACACAGCCACACCATATCATAGTNNNAGATCGGAAG |
| 33639_1 | TATGAGTGTGGAGTCGTTGCTTCAAGTGATTCTCCTGCCTCAGNNNAGATCGGAAG |
| 33639_2 | TATGAGTGTGGAGTCGTTGCTTCAAGTGATTCTCCTGCCTCAGNNNAGATCGGAAG |
| 30020_1 | TATGAGTGTGGAGTCGTTGCTGGAGCATTGTCATTGATGTCTGNNNAGATCGGAA |
| 30020_2 | TATGAGTGTGGAGTCGTTGCTGGAGCATTGTCATTGATGTCTGNNNAGATCGGAA |
| 37973_1 | TATGAGTGTGGAGTCGTTGCTCCATGTGAGAGAAGTCCAAGTCNNNAGATCGGAA |
| 37973_2 | TATGAGTGTGGAGTCGTTGCTCCATGTGAGAGAAGTCCAAGTCNNNAGATCGGAA |
| 47235_1 | TATGAGTGTGGAGTCGTTGCACGCCCGGGTAATTTTTTTGTANNNAGATCGGAAG |
| 47235_2 | TATGAGTGTGGAGTCGTTGCACGCCCGGGTAATTTTTTTGTANNNAGATCGGAAG |
| 39346_1 | TATGAGTGTGGAGTCGTTGCCTGTGCTTCTGTTACAACCTGACNNNAGATCGGAAG |
| 39346_2 | TATGAGTGTGGAGTCGTTGCCTGTGCTTCTGTTACAACCTGACNNNAGATCGGAAG |
| 36124_1 | TATGAGTGTGGAGTCGTTGCAGATTGGATAGTGGAACAGAATCANNNAGATCGGA |
| 36124_2 | TATGAGTGTGGAGTCGTTGCAGATTGGATAGTGGAACAGAATCANNNAGATCGGA |
| 32290_1 | TATGAGTGTGGAGTCGTTGCAAAGTGGGTTGAAGCAAGATTGGNNNAGATCGGA |
| 32290_2 | TATGAGTGTGGAGTCGTTGCAAAGTGGGTTGAAGCAAGATTGGNNNAGATCGGA |
| 3940_1 | TATGAGTGTGGAGTCGTTGCCAACAGCAACTCCGCTAAGGATNNNAGATCGGAAG |
| 3940_2 | TATGAGTGTGGAGTCGTTGCCAACAGCAACTCCGCTAAGGATNNNAGATCGGAAG |
| 33821_1 | TATGAGTGTGGAGTCGTTGCCCTGTAATCCCAGCACTTTAGGANNNAGATCGGAAG |
| 33821_2 | TATGAGTGTGGAGTCGTTGCCCTGTAATCCCAGCACTTTAGGANNNAGATCGGAAG |
| 30576_1 | TATGAGTGTGGAGTCGTTGCAGCCACCAAGATGCCCCAATAATNNNAGATCGGAAG |
| 30576_2 | TATGAGTGTGGAGTCGTTGCAGCCACCAAGATGCCCCAATAATNNNAGATCGGAAG |
| 49239_1 | TATGAGTGTGGAGTCGTTGCCTCCTCCCAGGTTCAAGCAATTNNNAGATCGGAAGA |
| 49239_2 | TATGAGTGTGGAGTCGTTGCCTCCTCCCAGGTTCAAGCAATTNNNAGATCGGAAGA |
| 34865_1 | TATGAGTGTGGAGTCGTTGCCCTGGGTTCAAGCAATTCTCCTANNNAGATCGGAAG |
| 34865_2 | TATGAGTGTGGAGTCGTTGCCCTGGGTTCAAGCAATTCTCCTANNNAGATCGGAAG |
| 47050_1 | TATGAGTGTGGAGTCGTTGCCCTCTCAAAGTGCTGGGATTACANNNAGATCGGAAG |
| 47050_2 | TATGAGTGTGGAGTCGTTGCCCTCTCAAAGTGCTGGGATTACANNNAGATCGGAAG |
| 34938_1 | TATGAGTGTGGAGTCGTTGCACAATCTCGGCTCACTGCAAGNNNAGATCGGAAGAG |
| 34938_2 | TATGAGTGTGGAGTCGTTGCACAATCTCGGCTCACTGCAAGNNNAGATCGGAAGAG |
| 37482_1 | TATGAGTGTGGAGTCGTTGCAAAAACACCAAAATTAGCCGGCCNNNAGATCGGAAG |
| 37482_2 | TATGAGTGTGGAGTCGTTGCAAAAACACCAAAATTAGCCGGCCNNNAGATCGGAAG |
| 47936_1 | TATGAGTGTGGAGTCGTTGCCCTCTACCCCCAGGAAGGAGNNNAGATCGGAAGAG |

| | |
|---|---|
| 47936_2 | TATGAGTGTGGAGTCGTTGCCCTCTACCCCCAGGAAGGAGNNNAGATCGGAAGAG |
| 44801_1 | TATGAGTGTGGAGTCGTTGCGTCTCCCTTTCTGTTACCTCCAGNNNAGATCGGAAGA |
| 44801_2 | TATGAGTGTGGAGTCGTTGCGTCTCCCTTTCTGTTACCTCCAGNNNAGATCGGAAGA |
| 46060_1 | TATGAGTGTGGAGTCGTTGCCCATCATGCCCGGCTAATTTTTNNNAGATCGGAAGA |
| 46060_2 | TATGAGTGTGGAGTCGTTGCCCATCATGCCCGGCTAATTTTTNNNAGATCGGAAGA |
| 31041_1 | TATGAGTGTGGAGTCGTTGCCAGGGCTAGACAGGAATCAGATCNNNAGATCGGAA |
| 31041_2 | TATGAGTGTGGAGTCGTTGCCAGGGCTAGACAGGAATCAGATCNNNAGATCGGAA |
| 49420_1 | TATGAGTGTGGAGTCGTTGCTGTTAGTAATGACTGCACAAGCANNNAGATCGGAA |
| 49420_2 | TATGAGTGTGGAGTCGTTGCTGTTAGTAATGACTGCACAAGCANNNAGATCGGAA |
| 35927_1 | TATGAGTGTGGAGTCGTTGCGACCACAAGATCAGGAGATCGAGNNNAGATCGGAA |
| 35927_2 | TATGAGTGTGGAGTCGTTGCGACCACAAGATCAGGAGATCGAGNNNAGATCGGAA |
| 52960_1 | TATGAGTGTGGAGTCGTTGCGTGAGCAAGTGGAGACAAAAGTCNNNAGATCGGA |
| 52960_2 | TATGAGTGTGGAGTCGTTGCGTGAGCAAGTGGAGACAAAAGTCNNNAGATCGGA |
| 39757_1 | TATGAGTGTGGAGTCGTTGCGATTGCTTGAGTCCAGGAGGTTNNNAGATCGGAAG |
| 39757_2 | TATGAGTGTGGAGTCGTTGCGATTGCTTGAGTCCAGGAGGTTNNNAGATCGGAAG |
| 35462_1 | TATGAGTGTGGAGTCGTTGCCTGGATCTCGGCTTACTGCAAGNNNAGATCGGAAG |
| 35462_2 | TATGAGTGTGGAGTCGTTGCCTGGATCTCGGCTTACTGCAAGNNNAGATCGGAAG |
| 42038_1 | TATGAGTGTGGAGTCGTTGCAGAGAGAGAGAGAGGCAAAGAGANNNAGATCGG |
| 42038_2 | TATGAGTGTGGAGTCGTTGCAGAGAGAGAGAGAGGCAAAGAGANNNAGATCGG |
| 42056_1 | TATGAGTGTGGAGTCGTTGCCGGCCCATGTCCAACTAATTTTTNNNAGATCGGAAG |
| 42056_2 | TATGAGTGTGGAGTCGTTGCCGGCCCATGTCCAACTAATTTTTNNNAGATCGGAAG |
| 52250_1 | TATGAGTGTGGAGTCGTTGCTGGGCAGCTGTAGATGATTATGGNNNAGATCGGAA |
| 52250_2 | TATGAGTGTGGAGTCGTTGCTGGGCAGCTGTAGATGATTATGGNNNAGATCGGAA |
| 46063_1 | TATGAGTGTGGAGTCGTTGCCCACCACACTCGGCTAATTTTTTNNNAGATCGGAAGA |
| 46063_2 | TATGAGTGTGGAGTCGTTGCCCACCACACTCGGCTAATTTTTTNNNAGATCGGAAGA |
| 49276_1 | TATGAGTGTGGAGTCGTTGCTGGAGTGCAGTGATGTGATTACANNNAGATCGGAA |
| 49276_2 | TATGAGTGTGGAGTCGTTGCTGGAGTGCAGTGATGTGATTACANNNAGATCGGAA |
| 42123_1 | TATGAGTGTGGAGTCGTTGCTCCAACTCCCTGGTTCAAATGATNNNAGATCGGAAG |
| 42123_2 | TATGAGTGTGGAGTCGTTGCTCCAACTCCCTGGTTCAAATGATNNNAGATCGGAAG |
| 37800_1 | TATGAGTGTGGAGTCGTTGCGCCAACAGGTTGCTTATGAAAGANNNAGATCGGAA |
| 37800_2 | TATGAGTGTGGAGTCGTTGCGCCAACAGGTTGCTTATGAAAGANNNAGATCGGAA |
| 47638_1 | TATGAGTGTGGAGTCGTTGCAGACGGAGTCTCTCTCTGTCCNNNAGATCGGAAGAG |
| 47638_2 | TATGAGTGTGGAGTCGTTGCAGACGGAGTCTCTCTCTGTCCNNNAGATCGGAAGAG |
| 35407_1 | TATGAGTGTGGAGTCGTTGCATGATCTCGGCTCACTGCAAGNNNAGATCGGAAGA |
| 35407_2 | TATGAGTGTGGAGTCGTTGCATGATCTCGGCTCACTGCAAGNNNAGATCGGAAGA |
| 30580_1 | TATGAGTGTGGAGTCGTTGCCACCAAGACAAAGAGCTTAGGGCANNNAGATCGGA |
| 30580_2 | TATGAGTGTGGAGTCGTTGCCACCAAGACAAAGAGCTTAGGGCANNNAGATCGGA |
| 33927_1 | TATGAGTGTGGAGTCGTTGCTCTCCGAGTAGCTGGGACTACNNNAGATCGGAAGA |
| 33927_2 | TATGAGTGTGGAGTCGTTGCTCTCCGAGTAGCTGGGACTACNNNAGATCGGAAGA |
| 37851_1 | TATGAGTGTGGAGTCGTTGCACAGAGATGTAACTGCAGATCCTNNNAGATCGGAA |
| 37851_2 | TATGAGTGTGGAGTCGTTGCACAGAGATGTAACTGCAGATCCTNNNAGATCGGAA |
| 52025_1 | TATGAGTGTGGAGTCGTTGCCTTCCTGGCAAAAGTAACTGCAANNNAGATCGGAAG |
| 52025_2 | TATGAGTGTGGAGTCGTTGCCTTCCTGGCAAAAGTAACTGCAANNNAGATCGGAAG |

| | |
|---|---|
| 30402_1 | TATGAGTGTGGAGTCGTTGCTAACTTCAAATGGGCCAGGGGTANNNAGATCGGAA |
| 30402_2 | TATGAGTGTGGAGTCGTTGCTAACTTCAAATGGGCCAGGGGTANNNAGATCGGAA |
| 9929_1 | TATGAGTGTGGAGTCGTTGCCCTTCATCTCTGTCCTTTTACCCCNNNAGATCGGAAG |
| 9929_2 | TATGAGTGTGGAGTCGTTGCCCTTCATCTCTGTCCTTTTACCCCNNNAGATCGGAAG |
| 11454_1 | TATGAGTGTGGAGTCGTTGCACTGCTGCTAAAATTCCTACAGCNNNAGATCGGAAG |
| 11454_2 | TATGAGTGTGGAGTCGTTGCACTGCTGCTAAAATTCCTACAGCNNNAGATCGGAAG |
| 45168_1 | TATGAGTGTGGAGTCGTTGCACCCATCTTCTAAAACGGAAGCTNNNAGATCGGAAG |
| 45168_2 | TATGAGTGTGGAGTCGTTGCACCCATCTTCTAAAACGGAAGCTNNNAGATCGGAAG |
| 44669_1 | TATGAGTGTGGAGTCGTTGCGAGGACCTCTGTAAAAGCCACAGNNNAGATCGGAA |
| 44669_2 | TATGAGTGTGGAGTCGTTGCGAGGACCTCTGTAAAAGCCACAGNNNAGATCGGAA |
| 50142_1 | TATGAGTGTGGAGTCGTTGCCTCTTACCTCACAGCTGAAACCTNNNAGATCGGAAG |
| 50142_2 | TATGAGTGTGGAGTCGTTGCCTCTTACCTCACAGCTGAAACCTNNNAGATCGGAAG |
| 49049_1 | TATGAGTGTGGAGTCGTTGCCCTCCCAAGTAGCTGGAACTATGNNNAGATCGGAAG |
| 49049_2 | TATGAGTGTGGAGTCGTTGCCCTCCCAAGTAGCTGGAACTATGNNNAGATCGGAAG |
| 44844_1 | TATGAGTGTGGAGTCGTTGCAGATAAGAAGTGTGTGCTGGGAGNNNAGATCGGA |
| 44844_2 | TATGAGTGTGGAGTCGTTGCAGATAAGAAGTGTGTGCTGGGAGNNNAGATCGGA |
| 40783_1 | TATGAGTGTGGAGTCGTTGCACCAAAAAGTCTCTCTGAACCAANNNAGATCGGAAG |
| 40783_2 | TATGAGTGTGGAGTCGTTGCACCAAAAAGTCTCTCTGAACCAANNNAGATCGGAAG |
| 31018_1 | TATGAGTGTGGAGTCGTTGCTGCAACACCCAGTGACAAAAATTNNNAGATCGGAAG |
| 31018_2 | TATGAGTGTGGAGTCGTTGCTGCAACACCCAGTGACAAAAATTNNNAGATCGGAAG |
| 34296_1 | TATGAGTGTGGAGTCGTTGCGTGTGTGTTAGTGTGCACAGATGNNNAGATCGGAA |
| 34296_2 | TATGAGTGTGGAGTCGTTGCGTGTGTGTTAGTGTGCACAGATGNNNAGATCGGAA |
| 5525_1 | TATGAGTGTGGAGTCGTTGCACACTCCTGACATGTTTTTCCCCNNNAGATCGGAAGA |
| 5525_2 | TATGAGTGTGGAGTCGTTGCACACTCCTGACATGTTTTTCCCCNNNAGATCGGAAGA |
| 37988_1 | TATGAGTGTGGAGTCGTTGCTGGTCTGTACTACTAGTGTTACCGNNNAGATCGGAA |
| 37988_2 | TATGAGTGTGGAGTCGTTGCTGGTCTGTACTACTAGTGTTACCGNNNAGATCGGAA |
| 42963_1 | TATGAGTGTGGAGTCGTTGCGTTTCGGGAAATGCATCCTTAGGNNNAGATCGGAA |
| 42963_2 | TATGAGTGTGGAGTCGTTGCGTTTCGGGAAATGCATCCTTAGGNNNAGATCGGAA |
| 34114_1 | TATGAGTGTGGAGTCGTTGCAGGAATCGGTAACATTTCACCCANNNAGATCGGAAG |
| 34114_2 | TATGAGTGTGGAGTCGTTGCAGGAATCGGTAACATTTCACCCANNNAGATCGGAAG |
| 46498_1 | TATGAGTGTGGAGTCGTTGCGAATCTGGGCTCACTACCTCTGNNNAGATCGGAAGA |
| 46498_2 | TATGAGTGTGGAGTCGTTGCGAATCTGGGCTCACTACCTCTGNNNAGATCGGAAGA |
| 29575_1 | TATGAGTGTGGAGTCGTTGCAGCATGGATCATCGATAGTAACANNNAGATCGGAA |
| 29575_2 | TATGAGTGTGGAGTCGTTGCAGCATGGATCATCGATAGTAACANNNAGATCGGAA |
| 46103_1 | TATGAGTGTGGAGTCGTTGCTAGTAGCTGGGATTACAAGTGCCNNNAGATCGGAA |
| 46103_2 | TATGAGTGTGGAGTCGTTGCTAGTAGCTGGGATTACAAGTGCCNNNAGATCGGAA |
| 35952_1 | TATGAGTGTGGAGTCGTTGCCCAAGATGAGCAGATCACCTGAGNNNAGATCGGAA |
| 35952_2 | TATGAGTGTGGAGTCGTTGCCCAAGATGAGCAGATCACCTGAGNNNAGATCGGAA |
| 41233_1 | TATGAGTGTGGAGTCGTTGCCAACAGAAAAATCCGTGACCCAANNNAGATCGGAAG |
| 41233_2 | TATGAGTGTGGAGTCGTTGCCAACAGAAAAATCCGTGACCCAANNNAGATCGGAAG |
| 49377_1 | TATGAGTGTGGAGTCGTTGCACGCCCGGCTAATTTTTTGTATTNNNAGATCGGAAG |
| 49377_2 | TATGAGTGTGGAGTCGTTGCACGCCCGGCTAATTTTTTGTATTNNNAGATCGGAAG |
| 48566_1 | TATGAGTGTGGAGTCGTTGCGAGTAGCTGGGATTACAGGCATGNNNAGATCGGAA |

| | |
|---|---|
| 48566_2 | TATGAGTGTGGAGTCGTTGCGAGTAGCTGGGATTACAGGCATGNNNAGATCGGAA |
| 41705_1 | TATGAGTGTGGAGTCGTTGCCAGAGTAGCTGGGACTACAGGTNNNAGATCGGAAG |
| 41705_2 | TATGAGTGTGGAGTCGTTGCCAGAGTAGCTGGGACTACAGGTNNNAGATCGGAAG |
| 43551_1 | TATGAGTGTGGAGTCGTTGCCCACTATGCCTGGCTAATTTTTTNNNAGATCGGAAGA |
| 43551_2 | TATGAGTGTGGAGTCGTTGCCCACTATGCCTGGCTAATTTTTTNNNAGATCGGAAGA |
| 31624_1 | TATGAGTGTGGAGTCGTTGCTCTCACTCAGGGGAACCATCTATNNNAGATCGGAAG |
| 31624_2 | TATGAGTGTGGAGTCGTTGCTCTCACTCAGGGGAACCATCTATNNNAGATCGGAAG |
| 46112_1 | TATGAGTGTGGAGTCGTTGCCTGTGATCTAGGGCAGAGGTTTTNNNAGATCGGAA |
| 46112_2 | TATGAGTGTGGAGTCGTTGCCTGTGATCTAGGGCAGAGGTTTTNNNAGATCGGAA |
| 48701_1 | TATGAGTGTGGAGTCGTTGCTTCAAGTGATTCTCCTGCCTCAGNNNAGATCGGAAGA |
| 48701_2 | TATGAGTGTGGAGTCGTTGCTTCAAGTGATTCTCCTGCCTCAGNNNAGATCGGAAGA |
| 49864_1 | TATGAGTGTGGAGTCGTTGCAGTCTTTATGCCTGCTTCTGACANNNAGATCGGAAGA |
| 49864_2 | TATGAGTGTGGAGTCGTTGCAGTCTTTATGCCTGCTTCTGACANNNAGATCGGAAGA |
| 13746_1 | TATGAGTGTGGAGTCGTTGCCTTCCAAATTCAGATTCCAGTANNNAGATCGGAAGA |
| 13746_2 | TATGAGTGTGGAGTCGTTGCCTTCCAAATTCAGATTCCAGTANNNAGATCGGAAGA |
| 41527_1 | TATGAGTGTGGAGTCGTTGCGATTGCTGACTCTTCATTGAGGGNNNAGATCGGAAG |
| 41527_2 | TATGAGTGTGGAGTCGTTGCGATTGCTGACTCTTCATTGAGGGNNNAGATCGGAAG |
| 31715_1 | TATGAGTGTGGAGTCGTTGCTGTCTTGGTTCAGGCCTAAATCANNNAGATCGGAAG |
| 31715_2 | TATGAGTGTGGAGTCGTTGCTGTCTTGGTTCAGGCCTAAATCANNNAGATCGGAAG |
| 10559_1 | TATGAGTGTGGAGTCGTTGCTCAGTTATGAGGTAGGGTGGCAANNNAGATCGGAA |
| 10559_2 | TATGAGTGTGGAGTCGTTGCTCAGTTATGAGGTAGGGTGGCAANNNAGATCGGAA |
| 35978_1 | TATGAGTGTGGAGTCGTTGCACAAAAGAATGAGGCTGCATTGTNNNAGATCGGAA |
| 35978_2 | TATGAGTGTGGAGTCGTTGCACAAAAGAATGAGGCTGCATTGTNNNAGATCGGAA |
| 52952_1 | TATGAGTGTGGAGTCGTTGCCAAGAATTCGAGCACTGCATGGNNNAGATCGGAAGA |
| 52952_2 | TATGAGTGTGGAGTCGTTGCCAAGAATTCGAGCACTGCATGGNNNAGATCGGAAGA |
| 51271_1 | TATGAGTGTGGAGTCGTTGCACGTGACCACTAGGTGTCATAACNNNAGATCGGAAG |
| 51271_2 | TATGAGTGTGGAGTCGTTGCACGTGACCACTAGGTGTCATAACNNNAGATCGGAAG |
| 32854_1 | TATGAGTGTGGAGTCGTTGCTGCTCCCAGCCATTTTTTGTTTTNNNAGATCGGAAGA |
| 32854_2 | TATGAGTGTGGAGTCGTTGCTGCTCCCAGCCATTTTTTGTTTTNNNAGATCGGAAGA |
| 10780_1 | TATGAGTGTGGAGTCGTTGCAATGCAAAGCCAAAGACACAAGGNNNAGATCGGAA |
| 10780_2 | TATGAGTGTGGAGTCGTTGCAATGCAAAGCCAAAGACACAAGGNNNAGATCGGAA |
| 33801_1 | TATGAGTGTGGAGTCGTTGCAACGTAGTGATCTTGCCAGCTTANNNAGATCGGAAG |
| 33801_2 | TATGAGTGTGGAGTCGTTGCAACGTAGTGATCTTGCCAGCTTANNNAGATCGGAAG |
| 39006_1 | TATGAGTGTGGAGTCGTTGCCATCCTGGCTAACATGGTGAAACNNNAGATCGGAAG |
| 39006_2 | TATGAGTGTGGAGTCGTTGCCATCCTGGCTAACATGGTGAAACNNNAGATCGGAAG |
| 32147_1 | TATGAGTGTGGAGTCGTTGCGCTCAAGGGATCCTTTTACCTCANNNAGATCGGAAGA |
| 32147_2 | TATGAGTGTGGAGTCGTTGCGCTCAAGGGATCCTTTTACCTCANNNAGATCGGAAGA |
| 36725_1 | TATGAGTGTGGAGTCGTTGCGGGGATTTCTAGCCTTGCTCTTANNNAGATCGGAAG |
| 36725_2 | TATGAGTGTGGAGTCGTTGCGGGGATTTCTAGCCTTGCTCTTANNNAGATCGGAAG |
| 37756_1 | TATGAGTGTGGAGTCGTTGCACCTTTGAATCTCCTACACCTGGNNNAGATCGGAAGA |
| 37756_2 | TATGAGTGTGGAGTCGTTGCACCTTTGAATCTCCTACACCTGGNNNAGATCGGAAGA |
| 13600_1 | TATGAGTGTGGAGTCGTTGCACTAAACTCAAGAAGTCAGTATTNNNAGATCGGAAG |
| 13600_2 | TATGAGTGTGGAGTCGTTGCACTAAACTCAAGAAGTCAGTATTNNNAGATCGGAAG |

| | |
|---|---|
| 48344_1 | TATGAGTGTGGAGTCGTTGCTTCAAGTGATTCTCCTGCCTCAGNNNAGATCGGAAG |
| 48344_2 | TATGAGTGTGGAGTCGTTGCTTCAAGTGATTCTCCTGCCTCAGNNNAGATCGGAAG |
| 47752_1 | TATGAGTGTGGAGTCGTTGCCCTCCCAGGTTCAAGTGATTCTCNNNAGATCGGAAG |
| 47752_2 | TATGAGTGTGGAGTCGTTGCCCTCCCAGGTTCAAGTGATTCTCNNNAGATCGGAAG |
| 42005_1 | TATGAGTGTGGAGTCGTTGCTTCAAGTGATTCTCCTGCCTCAGNNNAGATCGGAAG |
| 42005_2 | TATGAGTGTGGAGTCGTTGCTTCAAGTGATTCTCCTGCCTCAGNNNAGATCGGAAG |
| 38963_1 | TATGAGTGTGGAGTCGTTGCGCACTTGGCATTGAAAGTGGTAGNNNAGATCGGAA |
| 38963_2 | TATGAGTGTGGAGTCGTTGCGCACTTGGCATTGAAAGTGGTAGNNNAGATCGGAA |
| 40216_1 | TATGAGTGTGGAGTCGTTGCAACACCTATAATCCCAGTGCGTTNNNAGATCGGAAG |
| 40216_2 | TATGAGTGTGGAGTCGTTGCAACACCTATAATCCCAGTGCGTTNNNAGATCGGAAG |
| 36835_1 | TATGAGTGTGGAGTCGTTGCCTTGCAGTGAGCTGAGATTGTGNNNAGATCGGAAG |
| 36835_2 | TATGAGTGTGGAGTCGTTGCCTTGCAGTGAGCTGAGATTGTGNNNAGATCGGAAG |
| 51934_1 | TATGAGTGTGGAGTCGTTGCACCAACAAACATCTGCAAGTACGNNNAGATCGGAAG |
| 51934_2 | TATGAGTGTGGAGTCGTTGCACCAACAAACATCTGCAAGTACGNNNAGATCGGAAG |
| 44379_1 | TATGAGTGTGGAGTCGTTGCGACAGCACTTCCAGGTATCCATGNNNAGATCGGAAG |
| 44379_2 | TATGAGTGTGGAGTCGTTGCGACAGCACTTCCAGGTATCCATGNNNAGATCGGAAG |
| 30001_1 | TATGAGTGTGGAGTCGTTGCCACGTGTCTGCAAGATACCAATTNNNAGATCGGAAG |
| 30001_2 | TATGAGTGTGGAGTCGTTGCCACGTGTCTGCAAGATACCAATTNNNAGATCGGAAG |
| 43793_1 | TATGAGTGTGGAGTCGTTGCAGTCCCCAAATTAGTGCCATACANNNAGATCGGAAG |
| 43793_2 | TATGAGTGTGGAGTCGTTGCAGTCCCCAAATTAGTGCCATACANNNAGATCGGAAG |
| 49740_1 | TATGAGTGTGGAGTCGTTGCTCATTATGAGGGCTGGCTTTCTTNNNAGATCGGAAG |
| 49740_2 | TATGAGTGTGGAGTCGTTGCTCATTATGAGGGCTGGCTTTCTTNNNAGATCGGAAG |
| 49844_1 | TATGAGTGTGGAGTCGTTGCACTGTATATCAGAGTGTACCTGTGANNNAGATCGGA |
| 49844_2 | TATGAGTGTGGAGTCGTTGCACTGTATATCAGAGTGTACCTGTGANNNAGATCGGA |
| 49993_1 | TATGAGTGTGGAGTCGTTGCGGGGCCCAGGTTATTTATTACCANNNAGATCGGAAG |
| 49993_2 | TATGAGTGTGGAGTCGTTGCGGGGCCCAGGTTATTTATTACCANNNAGATCGGAAG |
| 38541_1 | TATGAGTGTGGAGTCGTTGCTAGCTAGGATTACAGGCATGTGCNNNAGATCGGAA |
| 38541_2 | TATGAGTGTGGAGTCGTTGCTAGCTAGGATTACAGGCATGTGCNNNAGATCGGAA |
| 30581_1 | TATGAGTGTGGAGTCGTTGCGCACCCTGCTAGAAGTGACAAATNNNAGATCGGAAG |
| 30581_2 | TATGAGTGTGGAGTCGTTGCGCACCCTGCTAGAAGTGACAAATNNNAGATCGGAAG |
| 32155_1 | TATGAGTGTGGAGTCGTTGCAGATCTGCCTGCAAAATTCCATTNNNAGATCGGAAG |
| 32155_2 | TATGAGTGTGGAGTCGTTGCAGATCTGCCTGCAAAATTCCATTNNNAGATCGGAAG |
| 53135_1 | TATGAGTGTGGAGTCGTTGCGCCATTTGTGTGGAGCCATATACNNNAGATCGGAAG |
| 53135_2 | TATGAGTGTGGAGTCGTTGCGCCATTTGTGTGGAGCCATATACNNNAGATCGGAAG |
| 52679_1 | TATGAGTGTGGAGTCGTTGCTAAAAGAGCCTCACTTGGGAAGGNNNAGATCGGAA |
| 52679_2 | TATGAGTGTGGAGTCGTTGCTAAAAGAGCCTCACTTGGGAAGGNNNAGATCGGAA |
| 40564_1 | TATGAGTGTGGAGTCGTTGCAACAGCACAAGCAGCCTAAAATTNNNAGATCGGAAG |
| 40564_2 | TATGAGTGTGGAGTCGTTGCAACAGCACAAGCAGCCTAAAATTNNNAGATCGGAAG |
| 43092_1 | TATGAGTGTGGAGTCGTTGCACATAGTCATAAAGGGTTTTCCCTNNNAGATCGGAA |
| 43092_2 | TATGAGTGTGGAGTCGTTGCACATAGTCATAAAGGGTTTTCCCTNNNAGATCGGAA |
| 40236_1 | TATGAGTGTGGAGTCGTTGCAACAACCTTTCTCTCAGGGCTACNNNAGATCGGAAG |
| 40236_2 | TATGAGTGTGGAGTCGTTGCAACAACCTTTCTCTCAGGGCTACNNNAGATCGGAAG |
| 46453_1 | TATGAGTGTGGAGTCGTTGCTGTTAGCCAGGATGGTCTCAATCNNNAGATCGGAAG |

| | |
|---|---|
| 46453_2 | TATGAGTGTGGAGTCGTTGCTGTTAGCCAGGATGGTCTCAATCNNNAGATCGGAAG |
| 43864_1 | TATGAGTGTGGAGTCGTTGCTTAGGCTAGAGCAGTGACTCTCTNNNAGATCGGAAG |
| 43864_2 | TATGAGTGTGGAGTCGTTGCTTAGGCTAGAGCAGTGACTCTCTNNNAGATCGGAAG |
| 31727_1 | TATGAGTGTGGAGTCGTTGCGACCACCAGGAGAGACATTTAGGNNNAGATCGGAA |
| 31727_2 | TATGAGTGTGGAGTCGTTGCGACCACCAGGAGAGACATTTAGGNNNAGATCGGAA |
| 44600_1 | TATGAGTGTGGAGTCGTTGCATCTCACTGAGGAAAAAGGTGCTNNNAGATCGGAA |
| 44600_2 | TATGAGTGTGGAGTCGTTGCATCTCACTGAGGAAAAAGGTGCTNNNAGATCGGAA |
| 31409_1 | TATGAGTGTGGAGTCGTTGCTGTGTGTGTGTGTGTGATGAATGNNNAGATCGGAA |
| 31409_2 | TATGAGTGTGGAGTCGTTGCTGTGTGTGTGTGTGTGATGAATGNNNAGATCGGAA |
| 31313_1 | TATGAGTGTGGAGTCGTTGCCTCTGTGATGGAAGAGTGGTCAGNNNAGATCGGAA |
| 31313_2 | TATGAGTGTGGAGTCGTTGCCTCTGTGATGGAAGAGTGGTCAGNNNAGATCGGAA |
| 44584_1 | TATGAGTGTGGAGTCGTTGCACCCAAACTGTTATCTGAAGCCTNNNAGATCGGAAG |
| 44584_2 | TATGAGTGTGGAGTCGTTGCACCCAAACTGTTATCTGAAGCCTNNNAGATCGGAAG |
| 30249_1 | TATGAGTGTGGAGTCGTTGCAACAAGTCAAACTGCCCATGATGNNNAGATCGGAAG |
| 30249_2 | TATGAGTGTGGAGTCGTTGCAACAAGTCAAACTGCCCATGATGNNNAGATCGGAAG |
| 13053_1 | TATGAGTGTGGAGTCGTTGCTATTGTGGAATGAAATTGTGATGNNNAGATCGGAA |
| 13053_2 | TATGAGTGTGGAGTCGTTGCTATTGTGGAATGAAATTGTGATGNNNAGATCGGAA |
| 40041_1 | TATGAGTGTGGAGTCGTTGCATGCAGTTTCTTCCTAGCATCGANNNAGATCGGAAG |
| 40041_2 | TATGAGTGTGGAGTCGTTGCATGCAGTTTCTTCCTAGCATCGANNNAGATCGGAAG |
| 34106_1 | TATGAGTGTGGAGTCGTTGCTCTCACAGAGAAAGTGAAATCCCTNNNAGATCGGAA |
| 34106_2 | TATGAGTGTGGAGTCGTTGCTCTCACAGAGAAAGTGAAATCCCTNNNAGATCGGAA |
| 29347_1 | TATGAGTGTGGAGTCGTTGCGGAGCTTCTGCAGGCTATTCTGNNNAGATCGGAAG |
| 29347_2 | TATGAGTGTGGAGTCGTTGCGGAGCTTCTGCAGGCTATTCTGNNNAGATCGGAAG |
| 33090_1 | TATGAGTGTGGAGTCGTTGCTCACTATCTCACTCTACTGCCCANNNAGATCGGAAGA |
| 33090_2 | TATGAGTGTGGAGTCGTTGCTCACTATCTCACTCTACTGCCCANNNAGATCGGAAGA |
| 39541_1 | TATGAGTGTGGAGTCGTTGCAGACCCAGCAGCAGAAATAATCANNNAGATCGGAA |
| 39541_2 | TATGAGTGTGGAGTCGTTGCAGACCCAGCAGCAGAAATAATCANNNAGATCGGAA |
| 32789_1 | TATGAGTGTGGAGTCGTTGCCTTGGCTTCATCTTCACCAGGAANNNAGATCGGAAG |
| 32789_2 | TATGAGTGTGGAGTCGTTGCCTTGGCTTCATCTTCACCAGGAANNNAGATCGGAAG |
| 35005_1 | TATGAGTGTGGAGTCGTTGCGAGTTTTGCTCTTGTTGCCCAGNNNAGATCGGAAGA |
| 35005_2 | TATGAGTGTGGAGTCGTTGCGAGTTTTGCTCTTGTTGCCCAGNNNAGATCGGAAGA |
| 35781_1 | TATGAGTGTGGAGTCGTTGCCGAGTAGCTGGTATTACAGGCATNNNAGATCGGAA |
| 35781_2 | TATGAGTGTGGAGTCGTTGCCGAGTAGCTGGTATTACAGGCATNNNAGATCGGAA |
| 42315_1 | TATGAGTGTGGAGTCGTTGCAGCTACTTCTGCTGGTAGGAATGNNNAGATCGGAA |
| 42315_2 | TATGAGTGTGGAGTCGTTGCAGCTACTTCTGCTGGTAGGAATGNNNAGATCGGAA |
| 35327_1 | TATGAGTGTGGAGTCGTTGCAAGTAACACCACACCTGGCTAATNNNAGATCGGAAG |
| 35327_2 | TATGAGTGTGGAGTCGTTGCAAGTAACACCACACCTGGCTAATNNNAGATCGGAAG |
| 32931_1 | TATGAGTGTGGAGTCGTTGCGAGGCAAGATAATCGCTTGAACCNNNAGATCGGAA |
| 32931_2 | TATGAGTGTGGAGTCGTTGCGAGGCAAGATAATCGCTTGAACCNNNAGATCGGAA |
| 44290_1 | TATGAGTGTGGAGTCGTTGCGAACAACACCTGGAGCCTTTTTTNNNAGATCGGAAG |
| 44290_2 | TATGAGTGTGGAGTCGTTGCGAACAACACCTGGAGCCTTTTTTNNNAGATCGGAAG |
| 30026_1 | TATGAGTGTGGAGTCGTTGCGCAAAGCTACACCTTTCCTCATCNNNAGATCGGAAGA |
| 30026_2 | TATGAGTGTGGAGTCGTTGCGCAAAGCTACACCTTTCCTCATCNNNAGATCGGAAGA |

| | |
|---|---|
| 43294_1 | TATGAGTGTGGAGTCGTTGCAGACAATACTGCTGGAAGTGGAGNNNAGATCGGAA |
| 43294_2 | TATGAGTGTGGAGTCGTTGCAGACAATACTGCTGGAAGTGGAGNNNAGATCGGAA |
| 41467_1 | TATGAGTGTGGAGTCGTTGCAGAAACACCAGGAAGAGATGCTCNNNAGATCGGAA |
| 41467_2 | TATGAGTGTGGAGTCGTTGCAGAAACACCAGGAAGAGATGCTCNNNAGATCGGAA |
| 2723_1 | TATGAGTGTGGAGTCGTTGCTGAGCTACTCTGTGTCAGGAACTNNNAGATCGGAAG |
| 2723_2 | TATGAGTGTGGAGTCGTTGCTGAGCTACTCTGTGTCAGGAACTNNNAGATCGGAAG |
| 49722_1 | TATGAGTGTGGAGTCGTTGCGCAGGAGCATATACCTTTGAGGANNNAGATCGGAA |
| 49722_2 | TATGAGTGTGGAGTCGTTGCGCAGGAGCATATACCTTTGAGGANNNAGATCGGAA |
| 38904_1 | TATGAGTGTGGAGTCGTTGCTGCAAAAACCATGGGAAAAGTGTNNNAGATCGGAA |
| 38904_2 | TATGAGTGTGGAGTCGTTGCTGCAAAAACCATGGGAAAAGTGTNNNAGATCGGAA |
| 49364_1 | TATGAGTGTGGAGTCGTTGCCACAATCTTGGCTCACTGCAACNNNAGATCGGAAGA |
| 49364_2 | TATGAGTGTGGAGTCGTTGCCACAATCTTGGCTCACTGCAACNNNAGATCGGAAGA |
| 33397_1 | TATGAGTGTGGAGTCGTTGCACAACTCAAATGTTCATCAGCTGANNNAGATCGGAA |
| 33397_2 | TATGAGTGTGGAGTCGTTGCACAACTCAAATGTTCATCAGCTGANNNAGATCGGAA |
| 31923_1 | TATGAGTGTGGAGTCGTTGCAACTCTGCCTTTATTTGATGCCGNNNAGATCGGAAG |
| 31923_2 | TATGAGTGTGGAGTCGTTGCAACTCTGCCTTTATTTGATGCCGNNNAGATCGGAAG |
| 5552_1 | TATGAGTGTGGAGTCGTTGCGACTTACGAGAGACGACCGAAAANNNAGATCGGAA |
| 5552_2 | TATGAGTGTGGAGTCGTTGCGACTTACGAGAGACGACCGAAAANNNAGATCGGAA |
| 44633_1 | TATGAGTGTGGAGTCGTTGCCTCACTCTGCTGGGATTAGAAGGNNNAGATCGGAA |
| 44633_2 | TATGAGTGTGGAGTCGTTGCCTCACTCTGCTGGGATTAGAAGGNNNAGATCGGAA |
| 49204_1 | TATGAGTGTGGAGTCGTTGCACAGGTTCAAGCAATTCTTCTGCNNNAGATCGGAAG |
| 49204_2 | TATGAGTGTGGAGTCGTTGCACAGGTTCAAGCAATTCTTCTGCNNNAGATCGGAAG |
| 13650_1 | TATGAGTGTGGAGTCGTTGCGTTTCCAATGATAGTAGGAAAACNNNAGATCGGAA |
| 13650_2 | TATGAGTGTGGAGTCGTTGCGTTTCCAATGATAGTAGGAAAACNNNAGATCGGAA |
| 13091_1 | TATGAGTGTGGAGTCGTTGCATCTGACGTACACTTTCCAGNNNAGATCGGAAGAGC |
| 13091_2 | TATGAGTGTGGAGTCGTTGCATCTGACGTACACTTTCCAGNNNAGATCGGAAGAGC |
| 44625_1 | TATGAGTGTGGAGTCGTTGCGGTATCCGTCATGAGGGTCTTGNNNAGATCGGAAG |
| 44625_2 | TATGAGTGTGGAGTCGTTGCGGTATCCGTCATGAGGGTCTTGNNNAGATCGGAAG |
| 31559_1 | TATGAGTGTGGAGTCGTTGCCTCGCTACCTACCTCCTTTCCTANNNAGATCGGAAGA |
| 31559_2 | TATGAGTGTGGAGTCGTTGCCTCGCTACCTACCTCCTTTCCTANNNAGATCGGAAGA |
| 34158_1 | TATGAGTGTGGAGTCGTTGCAGTTCTCCAGCAAGATACCACTGNNNAGATCGGAAG |
| 34158_2 | TATGAGTGTGGAGTCGTTGCAGTTCTCCAGCAAGATACCACTGNNNAGATCGGAAG |
| 39512_1 | TATGAGTGTGGAGTCGTTGCAGTTGCAGTGAGCTGAGATTGTANNNAGATCGGAA |
| 39512_2 | TATGAGTGTGGAGTCGTTGCAGTTGCAGTGAGCTGAGATTGTANNNAGATCGGAA |
| 51569_1 | TATGAGTGTGGAGTCGTTGCACATCATTACACAGTCTGAGGCANNNAGATCGGAAG |
| 51569_2 | TATGAGTGTGGAGTCGTTGCACATCATTACACAGTCTGAGGCANNNAGATCGGAAG |
| 31365_1 | TATGAGTGTGGAGTCGTTGCGCTGTAAGAGGCCTTCGTATTCTNNNAGATCGGAAG |
| 31365_2 | TATGAGTGTGGAGTCGTTGCGCTGTAAGAGGCCTTCGTATTCTNNNAGATCGGAAG |
| 12871_1 | TATGAGTGTGGAGTCGTTGCCAGGTCCTTTTCCAAACTTANNNAGATCGGAAGAGC |
| 12871_2 | TATGAGTGTGGAGTCGTTGCCAGGTCCTTTTCCAAACTTANNNAGATCGGAAGAGC |
| 35298_1 | TATGAGTGTGGAGTCGTTGCCTCCTGGGTTCACGTCATTCTCNNNAGATCGGAAGA |
| 35298_2 | TATGAGTGTGGAGTCGTTGCCTCCTGGGTTCACGTCATTCTCNNNAGATCGGAAGA |
| 46227_1 | TATGAGTGTGGAGTCGTTGCAGACGGAGTCTCGCTCTGTCNNNAGATCGGAAGAG |

| | |
|---|---|
| 46227_2 | TATGAGTGTGGAGTCGTTGCAGACGGAGTCTCGCTCTGTCNNNAGATCGGAAGAG |
| 29435_1 | TATGAGTGTGGAGTCGTTGCGTGGTTAGGTGTCTTTTCCTCCCNNNAGATCGGAAG |
| 29435_2 | TATGAGTGTGGAGTCGTTGCGTGGTTAGGTGTCTTTTCCTCCCNNNAGATCGGAAG |
| 31240_1 | TATGAGTGTGGAGTCGTTGCGGATATACAGAAGCTGACCCCTGNNNAGATCGGAA |
| 31240_2 | TATGAGTGTGGAGTCGTTGCGGATATACAGAAGCTGACCCCTGNNNAGATCGGAA |
| 34601_1 | TATGAGTGTGGAGTCGTTGCTGCGTCACAGAAAACTTCTTTCCNNNAGATCGGAAG |
| 34601_2 | TATGAGTGTGGAGTCGTTGCTGCGTCACAGAAAACTTCTTTCCNNNAGATCGGAAG |
| 49356_1 | TATGAGTGTGGAGTCGTTGCCCTCCCCACAAGTCTATCAACTCNNNAGATCGGAAGA |
| 49356_2 | TATGAGTGTGGAGTCGTTGCCCTCCCCACAAGTCTATCAACTCNNNAGATCGGAAGA |
| 47015_1 | TATGAGTGTGGAGTCGTTGCATAGTGGCATGATCTCAGCTCACNNNAGATCGGAAG |
| 47015_2 | TATGAGTGTGGAGTCGTTGCATAGTGGCATGATCTCAGCTCACNNNAGATCGGAAG |
| 44661_1 | TATGAGTGTGGAGTCGTTGCTCTTTGGGATACTCATTGTGGGGNNNAGATCGGAA |
| 44661_2 | TATGAGTGTGGAGTCGTTGCTCTTTGGGATACTCATTGTGGGGNNNAGATCGGAA |
| 13214_1 | TATGAGTGTGGAGTCGTTGCGCAAGGAAAGAGTATTTAAAAACNNNAGATCGGAA |
| 13214_2 | TATGAGTGTGGAGTCGTTGCGCAAGGAAAGAGTATTTAAAAACNNNAGATCGGAA |
| 46508_1 | TATGAGTGTGGAGTCGTTGCCAGTGACACGATCTTAGCTCACTNNNAGATCGGAAG |
| 46508_2 | TATGAGTGTGGAGTCGTTGCCAGTGACACGATCTTAGCTCACTNNNAGATCGGAAG |
| 30777_1 | TATGAGTGTGGAGTCGTTGCCACACACACATAGACACCTGAGANNNAGATCGGAAG |
| 30777_2 | TATGAGTGTGGAGTCGTTGCCACACACACATAGACACCTGAGANNNAGATCGGAAG |
| 32419_1 | TATGAGTGTGGAGTCGTTGCTCTGCCTTCTAGGTTCAAGCAATNNNAGATCGGAAG |
| 32419_2 | TATGAGTGTGGAGTCGTTGCTCTGCCTTCTAGGTTCAAGCAATNNNAGATCGGAAG |
| 52676_1 | TATGAGTGTGGAGTCGTTGCCGATCTGCACTGATGTACTCTGANNNAGATCGGAAG |
| 52676_2 | TATGAGTGTGGAGTCGTTGCCGATCTGCACTGATGTACTCTGANNNAGATCGGAAG |
| 47831_1 | TATGAGTGTGGAGTCGTTGCGGACGGGAGTTCTAGACCAAAGNNNAGATCGGAAG |
| 47831_2 | TATGAGTGTGGAGTCGTTGCGGACGGGAGTTCTAGACCAAAGNNNAGATCGGAAG |
| 42960_1 | TATGAGTGTGGAGTCGTTGCGAGCAGTCCGAGTGTCATCATTANNNAGATCGGAA |
| 42960_2 | TATGAGTGTGGAGTCGTTGCGAGCAGTCCGAGTGTCATCATTANNNAGATCGGAA |
| 8870_1 | TATGAGTGTGGAGTCGTTGCGGTTTCTGTGGCTAACTGGATGTNNNAGATCGGAA |
| 8870_2 | TATGAGTGTGGAGTCGTTGCGGTTTCTGTGGCTAACTGGATGTNNNAGATCGGAA |
| 49179_1 | TATGAGTGTGGAGTCGTTGCAATTGCAATAGTCTCCCCACCTTNNNAGATCGGAAGA |
| 49179_2 | TATGAGTGTGGAGTCGTTGCAATTGCAATAGTCTCCCCACCTTNNNAGATCGGAAGA |
| 41036_1 | TATGAGTGTGGAGTCGTTGCTGTACACCTATGCTCACAGAAGCNNNAGATCGGAAG |
| 41036_2 | TATGAGTGTGGAGTCGTTGCTGTACACCTATGCTCACAGAAGCNNNAGATCGGAAG |
| 43607_1 | TATGAGTGTGGAGTCGTTGCCGTCTGGTGTGGTCTGGAAAATNNNAGATCGGAAG |
| 43607_2 | TATGAGTGTGGAGTCGTTGCCGTCTGGTGTGGTCTGGAAAATNNNAGATCGGAAG |
| 45174_1 | TATGAGTGTGGAGTCGTTGCGCCCACCATTTTTTTTCCCCTAANNNAGATCGGAAGA |
| 45174_2 | TATGAGTGTGGAGTCGTTGCGCCCACCATTTTTTTTCCCCTAANNNAGATCGGAAGA |
| 45217_1 | TATGAGTGTGGAGTCGTTGCCAGAGGAAAGGGAAACTGCTCTANNNAGATCGGAA |
| 45217_2 | TATGAGTGTGGAGTCGTTGCCAGAGGAAAGGGAAACTGCTCTANNNAGATCGGAA |
| 52088_1 | TATGAGTGTGGAGTCGTTGCTTCATGCACTCTGACATTAGGCTNNNAGATCGGAAG |
| 52088_2 | TATGAGTGTGGAGTCGTTGCTTCATGCACTCTGACATTAGGCTNNNAGATCGGAAG |
| 29585_1 | TATGAGTGTGGAGTCGTTGCCAAAGACACAAGTCCAGCTGAAGNNNAGATCGGAA |
| 29585_2 | TATGAGTGTGGAGTCGTTGCCAAAGACACAAGTCCAGCTGAAGNNNAGATCGGAA |

| | |
|---|---|
| 8284_1 | TATGAGTGTGGAGTCGTTGCGGAGAGAGAATGAGTGGTTTCTANNNAGATCGGAA |
| 8284_2 | TATGAGTGTGGAGTCGTTGCGGAGAGAGAATGAGTGGTTTCTANNNAGATCGGAA |
| 37280_1 | TATGAGTGTGGAGTCGTTGCTCGTGTCTACAAGTATGGCAGTCNNNAGATCGGAAG |
| 37280_2 | TATGAGTGTGGAGTCGTTGCTCGTGTCTACAAGTATGGCAGTCNNNAGATCGGAAG |
| 33421_1 | TATGAGTGTGGAGTCGTTGCAAACTGTGAGGGAAGAGAAGGACNNNAGATCGGAA |
| 33421_2 | TATGAGTGTGGAGTCGTTGCAAACTGTGAGGGAAGAGAAGGACNNNAGATCGGAA |
| 46663_1 | TATGAGTGTGGAGTCGTTGCGTGTGATCTCAGCTTACTGCAACNNNAGATCGGAAG |
| 46663_2 | TATGAGTGTGGAGTCGTTGCGTGTGATCTCAGCTTACTGCAACNNNAGATCGGAAG |
| 41225_1 | TATGAGTGTGGAGTCGTTGCCCACTCCTGCGTATGACATTTTTNNNAGATCGGAAGA |
| 41225_2 | TATGAGTGTGGAGTCGTTGCCCACTCCTGCGTATGACATTTTTNNNAGATCGGAAGA |
| 10594_1 | TATGAGTGTGGAGTCGTTGCAATAAAATCCACCCTGCCAGCATNNNAGATCGGAAGA |
| 10594_2 | TATGAGTGTGGAGTCGTTGCAATAAAATCCACCCTGCCAGCATNNNAGATCGGAAGA |
| 46784_1 | TATGAGTGTGGAGTCGTTGCCTCCCAAAGTGCAGGAATTACAGNNNAGATCGGAAG |
| 46784_2 | TATGAGTGTGGAGTCGTTGCCTCCCAAAGTGCAGGAATTACAGNNNAGATCGGAAG |
| 50035_1 | TATGAGTGTGGAGTCGTTGCAACGTGTTCATCAGGAAGGGATANNNAGATCGGAA |
| 50035_2 | TATGAGTGTGGAGTCGTTGCAACGTGTTCATCAGGAAGGGATANNNAGATCGGAA |
| 49259_1 | TATGAGTGTGGAGTCGTTGCGCCAAATGCAATGTTCTCCTCATNNNAGATCGGAAGA |
| 49259_2 | TATGAGTGTGGAGTCGTTGCGCCAAATGCAATGTTCTCCTCATNNNAGATCGGAAGA |
| 52874_1 | TATGAGTGTGGAGTCGTTGCGATGTTTGAGGGGACAAGCAGTNNNAGATCGGAAG |
| 52874_2 | TATGAGTGTGGAGTCGTTGCGATGTTTGAGGGGACAAGCAGTNNNAGATCGGAAG |
| 353_1 | TATGAGTGTGGAGTCGTTGCATGCTCACATCTTGGGAGTTACTNNNAGATCGGAAG |
| 353_2 | TATGAGTGTGGAGTCGTTGCATGCTCACATCTTGGGAGTTACTNNNAGATCGGAAG |
| 40139_1 | TATGAGTGTGGAGTCGTTGCGGTGAAACCCCGTCTCTACTAAANNNAGATCGGAAG |
| 40139_2 | TATGAGTGTGGAGTCGTTGCGGTGAAACCCCGTCTCTACTAAANNNAGATCGGAAG |
| 48230_1 | TATGAGTGTGGAGTCGTTGCTTCCTCATGTCCTCGATCCTTTGNNNAGATCGGAAGA |
| 48230_2 | TATGAGTGTGGAGTCGTTGCTTCCTCATGTCCTCGATCCTTTGNNNAGATCGGAAGA |
| 42902_1 | TATGAGTGTGGAGTCGTTGCGCACAATCTTGGCTCACTACAGNNNAGATCGGAAGA |
| 42902_2 | TATGAGTGTGGAGTCGTTGCGCACAATCTTGGCTCACTACAGNNNAGATCGGAAGA |
| 44577_1 | TATGAGTGTGGAGTCGTTGCCACACACGCACACACACATATATNNNAGATCGGAAGA |
| 44577_2 | TATGAGTGTGGAGTCGTTGCCACACACGCACACACACATATATNNNAGATCGGAAGA |
| 42638_1 | TATGAGTGTGGAGTCGTTGCTTTGGGGATACATGTGAAGGTGTNNNAGATCGGAA |
| 42638_2 | TATGAGTGTGGAGTCGTTGCTTTGGGGATACATGTGAAGGTGTNNNAGATCGGAA |
| 41689_1 | TATGAGTGTGGAGTCGTTGCGATCTCGGCTCATTGCAACCTCNNNAGATCGGAAGA |
| 41689_2 | TATGAGTGTGGAGTCGTTGCGATCTCGGCTCATTGCAACCTCNNNAGATCGGAAGA |
| 44910_1 | TATGAGTGTGGAGTCGTTGCCAGACAGCCAGATACTCCTCATGNNNAGATCGGAAG |
| 44910_2 | TATGAGTGTGGAGTCGTTGCCAGACAGCCAGATACTCCTCATGNNNAGATCGGAAG |
| 41136_1 | TATGAGTGTGGAGTCGTTGCATCCTAGCTACTCAGAAGGCTGANNNAGATCGGAAG |
| 41136_2 | TATGAGTGTGGAGTCGTTGCATCCTAGCTACTCAGAAGGCTGANNNAGATCGGAAG |
| 37762_1 | TATGAGTGTGGAGTCGTTGCTCAAATTCCTGGCCTCAAGCTATNNNAGATCGGAAGA |
| 37762_2 | TATGAGTGTGGAGTCGTTGCTCAAATTCCTGGCCTCAAGCTATNNNAGATCGGAAGA |
| 36978_1 | TATGAGTGTGGAGTCGTTGCCCAAACATCTCATGTACCCCGTANNNAGATCGGAAGA |
| 36978_2 | TATGAGTGTGGAGTCGTTGCCCAAACATCTCATGTACCCCGTANNNAGATCGGAAGA |
| 39325_1 | TATGAGTGTGGAGTCGTTGCACACACACACACACACACAAAATNNNAGATCGGAAGA |

| | |
|---|---|
| 39325_2 | TATGAGTGTGGAGTCGTTGCACACACACACACACACACACAAAATNNNAGATCGGAAG |
| 46824_1 | TATGAGTGTGGAGTCGTTGCCTTCCAGTTTTTGCCCATTCAGTNNNAGATCGGAAGA |
| 46824_2 | TATGAGTGTGGAGTCGTTGCCTTCCAGTTTTTGCCCATTCAGTNNNAGATCGGAAGA |
| 39741_1 | TATGAGTGTGGAGTCGTTGCGAATCGCTTGAACCTGGAAGTTGNNNAGATCGGAA |
| 39741_2 | TATGAGTGTGGAGTCGTTGCGAATCGCTTGAACCTGGAAGTTGNNNAGATCGGAA |
| 37085_1 | TATGAGTGTGGAGTCGTTGCTTCTACTGGCTTCCTTGTTCCTCNNNAGATCGGAAGA |
| 37085_2 | TATGAGTGTGGAGTCGTTGCTTCTACTGGCTTCCTTGTTCCTCNNNAGATCGGAAGA |
| 50119_1 | TATGAGTGTGGAGTCGTTGCTCCTGGATCACAGTCACTAACACNNNAGATCGGAAG |
| 50119_2 | TATGAGTGTGGAGTCGTTGCTCCTGGATCACAGTCACTAACACNNNAGATCGGAAG |
| 32131_1 | TATGAGTGTGGAGTCGTTGCGGTAGGTATGGGACGGTTACTTCNNNAGATCGGAA |
| 32131_2 | TATGAGTGTGGAGTCGTTGCGGTAGGTATGGGACGGTTACTTCNNNAGATCGGAA |
| 32884_1 | TATGAGTGTGGAGTCGTTGCAGATCGAGACCATCTTGGCTAACNNNAGATCGGAAG |
| 32884_2 | TATGAGTGTGGAGTCGTTGCAGATCGAGACCATCTTGGCTAACNNNAGATCGGAAG |
| 50690_1 | TATGAGTGTGGAGTCGTTGCCACTTCTGCAGTCTCAGTGATCTNNNAGATCGGAAG |
| 50690_2 | TATGAGTGTGGAGTCGTTGCCACTTCTGCAGTCTCAGTGATCTNNNAGATCGGAAG |
| 52372_1 | TATGAGTGTGGAGTCGTTGCGCTCAAAGCAAGCTATCGAACATNNNAGATCGGAAG |
| 52372_2 | TATGAGTGTGGAGTCGTTGCGCTCAAAGCAAGCTATCGAACATNNNAGATCGGAAG |
| 33321_1 | TATGAGTGTGGAGTCGTTGCGAATGGCAATATCTGGCTTCGACNNNAGATCGGAAG |
| 33321_2 | TATGAGTGTGGAGTCGTTGCGAATGGCAATATCTGGCTTCGACNNNAGATCGGAAG |
| 52997_1 | TATGAGTGTGGAGTCGTTGCCTGTCTCTCTGCTGGTCTTTCTTNNNAGATCGGAAGA |
| 52997_2 | TATGAGTGTGGAGTCGTTGCCTGTCTCTCTGCTGGTCTTTCTTNNNAGATCGGAAGA |
| 51004_1 | TATGAGTGTGGAGTCGTTGCGATCTCAGCTCACTGCAACCTCNNNAGATCGGAAGA |
| 51004_2 | TATGAGTGTGGAGTCGTTGCGATCTCAGCTCACTGCAACCTCNNNAGATCGGAAGA |
| 51551_1 | TATGAGTGTGGAGTCGTTGCTGAGCCATCTCTGTGTACATCTCNNNAGATCGGAAG |
| 51551_2 | TATGAGTGTGGAGTCGTTGCTGAGCCATCTCTGTGTACATCTCNNNAGATCGGAAG |
| 5519_1 | TATGAGTGTGGAGTCGTTGCGTCTACATTCGCTGAGAGGCAGNNNAGATCGGAAG |
| 5519_2 | TATGAGTGTGGAGTCGTTGCGTCTACATTCGCTGAGAGGCAGNNNAGATCGGAAG |
| 32031_1 | TATGAGTGTGGAGTCGTTGCCCTCCCACCATGCTAAGATTACANNNAGATCGGAAG |
| 32031_2 | TATGAGTGTGGAGTCGTTGCCCTCCCACCATGCTAAGATTACANNNAGATCGGAAG |
| 52452_1 | TATGAGTGTGGAGTCGTTGCCCCTTTCTTTTCCTTGAGAGGGANNNAGATCGGAAG |
| 52452_2 | TATGAGTGTGGAGTCGTTGCCCCTTTCTTTTCCTTGAGAGGGANNNAGATCGGAAG |
| 39502_1 | TATGAGTGTGGAGTCGTTGCCTTAGTTACTGATGGGGAAGGGGNNNAGATCGGAA |
| 39502_2 | TATGAGTGTGGAGTCGTTGCCTTAGTTACTGATGGGGAAGGGGNNNAGATCGGAA |
| 28986_1 | TATGAGTGTGGAGTCGTTGCCATGTGCATATGTGTGTGTGTGTNNNAGATCGGAA |
| 28986_2 | TATGAGTGTGGAGTCGTTGCCATGTGCATATGTGTGTGTGTGTNNNAGATCGGAA |
| 41803_1 | TATGAGTGTGGAGTCGTTGCCCTCCTGGGTTCAAGTGATTCTCNNNAGATCGGAAG |
| 41803_2 | TATGAGTGTGGAGTCGTTGCCCTCCTGGGTTCAAGTGATTCTCNNNAGATCGGAAG |
| 47536_1 | TATGAGTGTGGAGTCGTTGCTTGCCAATAAAGCCTGAAGTTGGNNNAGATCGGAA |
| 47536_2 | TATGAGTGTGGAGTCGTTGCTTGCCAATAAAGCCTGAAGTTGGNNNAGATCGGAA |
| 49702_1 | TATGAGTGTGGAGTCGTTGCGCTTTTAGCATTCGAGAAAGGCANNNAGATCGGAA |
| 49702_2 | TATGAGTGTGGAGTCGTTGCGCTTTTAGCATTCGAGAAAGGCANNNAGATCGGAA |
| 47386_1 | TATGAGTGTGGAGTCGTTGCCGATCTCCTGACCTCATGATCTGNNNAGATCGGAAG |
| 47386_2 | TATGAGTGTGGAGTCGTTGCCGATCTCCTGACCTCATGATCTGNNNAGATCGGAAG |

| | |
|---|---|
| 39177_1 | TATGAGTGTGGAGTCGTTGCAGGGCAACCAATATAGCCAGAAANNNAGATCGGAA |
| 39177_2 | TATGAGTGTGGAGTCGTTGCAGGGCAACCAATATAGCCAGAAANNNAGATCGGAA |
| 44196_1 | TATGAGTGTGGAGTCGTTGCCATCAGTCTGTGTGAGAAGGGATNNNAGATCGGAA |
| 44196_2 | TATGAGTGTGGAGTCGTTGCCATCAGTCTGTGTGAGAAGGGATNNNAGATCGGAA |
| 34548_1 | TATGAGTGTGGAGTCGTTGCGATACCAGGACCTAGCCTACTGTNNNAGATCGGAAG |
| 34548_2 | TATGAGTGTGGAGTCGTTGCGATACCAGGACCTAGCCTACTGTNNNAGATCGGAAG |
| 13188_1 | TATGAGTGTGGAGTCGTTGCAAAGTTAAAAACGGAGAGAAAATNNNAGATCGGAA |
| 13188_2 | TATGAGTGTGGAGTCGTTGCAAAGTTAAAAACGGAGAGAAAATNNNAGATCGGAA |
| 4445_1 | TATGAGTGTGGAGTCGTTGCTTTGAAGGCATGGTTAGCTGAGGNNNAGATCGGAA |
| 4445_2 | TATGAGTGTGGAGTCGTTGCTTTGAAGGCATGGTTAGCTGAGGNNNAGATCGGAA |
| 44169_1 | TATGAGTGTGGAGTCGTTGCCCAACATGGATGGAGCGTAATTCNNNAGATCGGAAG |
| 44169_2 | TATGAGTGTGGAGTCGTTGCCCAACATGGATGGAGCGTAATTCNNNAGATCGGAAG |
| 13074_1 | TATGAGTGTGGAGTCGTTGCCAATCTACACAAAATAGAAGCAANNNAGATCGGAAG |
| 13074_2 | TATGAGTGTGGAGTCGTTGCCAATCTACACAAAATAGAAGCAANNNAGATCGGAAG |
| 35880_1 | TATGAGTGTGGAGTCGTTGCTGAGAGCAAGACTCCATCTCAAANNNAGATCGGAAG |
| 35880_2 | TATGAGTGTGGAGTCGTTGCTGAGAGCAAGACTCCATCTCAAANNNAGATCGGAAG |
| 41984_1 | TATGAGTGTGGAGTCGTTGCGAATCTCACTTTGTTGCCCAGACNNNAGATCGGAAG |
| 41984_2 | TATGAGTGTGGAGTCGTTGCGAATCTCACTTTGTTGCCCAGACNNNAGATCGGAAG |
| 51471_1 | TATGAGTGTGGAGTCGTTGCCAACTTTGGTCCCTTCAGTGTGNNNAGATCGGAAGA |
| 51471_2 | TATGAGTGTGGAGTCGTTGCCAACTTTGGTCCCTTCAGTGTGNNNAGATCGGAAGA |
| 9301_1 | TATGAGTGTGGAGTCGTTGCGGTAAATTTGGTCCCCAACTCANNNAGATCGGAAG |
| 9301_2 | TATGAGTGTGGAGTCGTTGCGGTAAATTTGGTCCCCAACTCANNNAGATCGGAAG |
| 42751_1 | TATGAGTGTGGAGTCGTTGCAGAAAATGGCAGGAGTAAGTTCCTNNNAGATCGGA |
| 42751_2 | TATGAGTGTGGAGTCGTTGCAGAAAATGGCAGGAGTAAGTTCCTNNNAGATCGGA |
| 46897_1 | TATGAGTGTGGAGTCGTTGCAGAAAACCATCAAACTTGGCTGGNNNAGATCGGAA |
| 46897_2 | TATGAGTGTGGAGTCGTTGCAGAAAACCATCAAACTTGGCTGGNNNAGATCGGAA |
| 33891_1 | TATGAGTGTGGAGTCGTTGCTACAATCTCGGCTCACTGCAAGNNNAGATCGGAAGA |
| 33891_2 | TATGAGTGTGGAGTCGTTGCTACAATCTCGGCTCACTGCAAGNNNAGATCGGAAGA |
| 41976_1 | TATGAGTGTGGAGTCGTTGCCTCCCAGGTTCACACCATTCTCNNNAGATCGGAAGAG |
| 41976_2 | TATGAGTGTGGAGTCGTTGCCTCCCAGGTTCACACCATTCTCNNNAGATCGGAAGAG |
| 40288_1 | TATGAGTGTGGAGTCGTTGCAACAAGAGTGAAAGGCCATCTCANNNAGATCGGAA |
| 40288_2 | TATGAGTGTGGAGTCGTTGCAACAAGAGTGAAAGGCCATCTCANNNAGATCGGAA |
| 39770_1 | TATGAGTGTGGAGTCGTTGCGGCCTCTTCAAGGAGAACTACAANNNAGATCGGAAG |
| 39770_2 | TATGAGTGTGGAGTCGTTGCGGCCTCTTCAAGGAGAACTACAANNNAGATCGGAAG |
| 34109_1 | TATGAGTGTGGAGTCGTTGCAAAGAATGGCTACTACACAGGCANNNAGATCGGAA |
| 34109_2 | TATGAGTGTGGAGTCGTTGCAAAGAATGGCTACTACACAGGCANNNAGATCGGAA |
| 33351_1 | TATGAGTGTGGAGTCGTTGCGGGTTCAAGTGATTCTCCTGTCTNNNAGATCGGAAG |
| 33351_2 | TATGAGTGTGGAGTCGTTGCGGGTTCAAGTGATTCTCCTGTCTNNNAGATCGGAAG |
| 42181_1 | TATGAGTGTGGAGTCGTTGCGACTATGGGAGTAAGCCACCATGNNNAGATCGGAA |
| 42181_2 | TATGAGTGTGGAGTCGTTGCGACTATGGGAGTAAGCCACCATGNNNAGATCGGAA |
| 52664_1 | TATGAGTGTGGAGTCGTTGCGCCTTCGAGCATGTTTTGTTTTCNNNAGATCGGAAG |
| 52664_2 | TATGAGTGTGGAGTCGTTGCGCCTTCGAGCATGTTTTGTTTTCNNNAGATCGGAAG |
| 42369_1 | TATGAGTGTGGAGTCGTTGCCTCTAAGCTAAGGACAGCAGACCNNNAGATCGGAAG |

| | |
|---|---|
| 42369_2 | TATGAGTGTGGAGTCGTTGCCTCTAAGCTAAGGACAGCAGACCNNNAGATCGGAA |
| 37959_1 | TATGAGTGTGGAGTCGTTGCCCAAACACTGTGATCCTGAGTTGNNNAGATCGGAAG |
| 37959_2 | TATGAGTGTGGAGTCGTTGCCCAAACACTGTGATCCTGAGTTGNNNAGATCGGAAG |
| 13778_1 | TATGAGTGTGGAGTCGTTGCTTACAGGAATTCACCAACATGNNNAGATCGGAAGAG |
| 13778_2 | TATGAGTGTGGAGTCGTTGCTTACAGGAATTCACCAACATGNNNAGATCGGAAGAG |
| 48724_1 | TATGAGTGTGGAGTCGTTGCGTAGCTGGGATTACAGGCATACANNNAGATCGGAA |
| 48724_2 | TATGAGTGTGGAGTCGTTGCGTAGCTGGGATTACAGGCATACANNNAGATCGGAA |
| 31370_1 | TATGAGTGTGGAGTCGTTGCCAGTTCGTTGCTCAAGCCTAAAANNNAGATCGGAAG |
| 31370_2 | TATGAGTGTGGAGTCGTTGCCAGTTCGTTGCTCAAGCCTAAAANNNAGATCGGAAG |
| 51066_1 | TATGAGTGTGGAGTCGTTGCCAGGCTAGGAGTTGATGGTTCTTNNNAGATCGGAA |
| 51066_2 | TATGAGTGTGGAGTCGTTGCCAGGCTAGGAGTTGATGGTTCTTNNNAGATCGGAA |
| 32231_1 | TATGAGTGTGGAGTCGTTGCGAGAAAGGGTGAAGGCAAACATCNNNAGATCGGA |
| 32231_2 | TATGAGTGTGGAGTCGTTGCGAGAAAGGGTGAAGGCAAACATCNNNAGATCGGA |
| 46471_1 | TATGAGTGTGGAGTCGTTGCCTTCTGGAACTCCTAGCCTCAAGNNNAGATCGGAAG |
| 46471_2 | TATGAGTGTGGAGTCGTTGCCTTCTGGAACTCCTAGCCTCAAGNNNAGATCGGAAG |
| 53383_1 | TATGAGTGTGGAGTCGTTGCCTTTCCACCTCTTCACTGTCTCANNNAGATCGGAAGA |
| 53383_2 | TATGAGTGTGGAGTCGTTGCCTTTCCACCTCTTCACTGTCTCANNNAGATCGGAAGA |
| 13238_1 | TATGAGTGTGGAGTCGTTGCAAGGCAGAATCAAAAAATAGAAANNNAGATCGGAA |
| 13238_2 | TATGAGTGTGGAGTCGTTGCAAGGCAGAATCAAAAAATAGAAANNNAGATCGGAA |
| 43562_1 | TATGAGTGTGGAGTCGTTGCCTTCATCTCCCAGGTTCAAGTGANNNAGATCGGAAG |
| 43562_2 | TATGAGTGTGGAGTCGTTGCCTTCATCTCCCAGGTTCAAGTGANNNAGATCGGAAG |
| 50804_1 | TATGAGTGTGGAGTCGTTGCCTTGCCATCCCAGGTGACTTNNNAGATCGGAAGAGC |
| 50804_2 | TATGAGTGTGGAGTCGTTGCCTTGCCATCCCAGGTGACTTNNNAGATCGGAAGAGC |
| 39747_1 | TATGAGTGTGGAGTCGTTGCTGAAAGAAGTGAGAGTCAAAAGGCNNNAGATCGG |
| 39747_2 | TATGAGTGTGGAGTCGTTGCTGAAAGAAGTGAGAGTCAAAAGGCNNNAGATCGG |
| 11687_1 | TATGAGTGTGGAGTCGTTGCCGGCAGCCAGCAAAGGAAATNNNAGATCGGAAGAG |
| 11687_2 | TATGAGTGTGGAGTCGTTGCCGGCAGCCAGCAAAGGAAATNNNAGATCGGAAGAG |
| 40320_1 | TATGAGTGTGGAGTCGTTGCAGATAGAGTTTGGAGACAGCTGCNNNAGATCGGAA |
| 40320_2 | TATGAGTGTGGAGTCGTTGCAGATAGAGTTTGGAGACAGCTGCNNNAGATCGGAA |
| 9713_1 | TATGAGTGTGGAGTCGTTGCCGGATTCTGGAAAGGGGCCCNNNAGATCGGAAGAG |
| 9713_2 | TATGAGTGTGGAGTCGTTGCCGGATTCTGGAAAGGGGCCCNNNAGATCGGAAGAG |
| 37155_1 | TATGAGTGTGGAGTCGTTGCTTTCGTGTGTAGTCTCCTTACCGNNNAGATCGGAAG |
| 37155_2 | TATGAGTGTGGAGTCGTTGCTTTCGTGTGTAGTCTCCTTACCGNNNAGATCGGAAG |
| 34252_1 | TATGAGTGTGGAGTCGTTGCTCCAGAACTAGTGGCTCATTCTGNNNAGATCGGAAG |
| 34252_2 | TATGAGTGTGGAGTCGTTGCTCCAGAACTAGTGGCTCATTCTGNNNAGATCGGAAG |
| 38545_1 | TATGAGTGTGGAGTCGTTGCTTCCATTCATGTGTGTGTGTGTGNNNAGATCGGAAG |
| 38545_2 | TATGAGTGTGGAGTCGTTGCTTCCATTCATGTGTGTGTGTGTGNNNAGATCGGAAG |
| 28906_1 | TATGAGTGTGGAGTCGTTGCTTTGTTCTTCCCAGCTACCACTTNNNAGATCGGAAGA |
| 28906_2 | TATGAGTGTGGAGTCGTTGCTTTGTTCTTCCCAGCTACCACTTNNNAGATCGGAAGA |
| 32847_1 | TATGAGTGTGGAGTCGTTGCTTTCAGTAGAGACGGGGTTTCACNNNAGATCGGAA |
| 32847_2 | TATGAGTGTGGAGTCGTTGCTTTCAGTAGAGACGGGGTTTCACNNNAGATCGGAA |
| 37133_1 | TATGAGTGTGGAGTCGTTGCAAAGAAACGGAGGGTAAAATGGCNNNAGATCGGA |
| 37133_2 | TATGAGTGTGGAGTCGTTGCAAAGAAACGGAGGGTAAAATGGCNNNAGATCGGA |

| | |
|---|---|
| 52346_1 | TATGAGTGTGGAGTCGTTGCCTTCATTTCAAAGTGGGCCACATNNNAGATCGGAAG |
| 52346_2 | TATGAGTGTGGAGTCGTTGCCTTCATTTCAAAGTGGGCCACATNNNAGATCGGAAG |
| 52374_1 | TATGAGTGTGGAGTCGTTGCGAGTGTCACTCTCTGGATACTGCNNNAGATCGGAAG |
| 52374_2 | TATGAGTGTGGAGTCGTTGCGAGTGTCACTCTCTGGATACTGCNNNAGATCGGAAG |
| 44443_1 | TATGAGTGTGGAGTCGTTGCGAGGGGGAGAAGGGATGAAAGNNNAGATCGGAA |
| 44443_2 | TATGAGTGTGGAGTCGTTGCGAGGGGGAGAAGGGATGAAAGNNNAGATCGGAA |
| 53055_1 | TATGAGTGTGGAGTCGTTGCTTGGAGACAGAGAGACACAGAGANNNAGATCGGA |
| 53055_2 | TATGAGTGTGGAGTCGTTGCTTGGAGACAGAGAGACACAGAGANNNAGATCGGA |
| 42344_1 | TATGAGTGTGGAGTCGTTGCCTCATGAGCAGCTGGAACTACTGNNNAGATCGGAAG |
| 42344_2 | TATGAGTGTGGAGTCGTTGCCTCATGAGCAGCTGGAACTACTGNNNAGATCGGAAG |
| 37305_1 | TATGAGTGTGGAGTCGTTGCCAGGACACTCTCTGAGCACATAANNNAGATCGGAAG |
| 37305_2 | TATGAGTGTGGAGTCGTTGCCAGGACACTCTCTGAGCACATAANNNAGATCGGAAG |
| 12251_1 | TATGAGTGTGGAGTCGTTGCCTCGGGTGGAGCATCTCCATTNNNAGATCGGAAGAG |
| 12251_2 | TATGAGTGTGGAGTCGTTGCCTCGGGTGGAGCATCTCCATTNNNAGATCGGAAGAG |
| 10969_1 | TATGAGTGTGGAGTCGTTGCTAGCAGAATGGAAGGAAAAGAGGNNNAGATCGGA |
| 10969_2 | TATGAGTGTGGAGTCGTTGCTAGCAGAATGGAAGGAAAAGAGGNNNAGATCGGA |
| 52954_1 | TATGAGTGTGGAGTCGTTGCGGAGTCAAATATTGGGAGAGGGGNNNAGATCGGA |
| 52954_2 | TATGAGTGTGGAGTCGTTGCGGAGTCAAATATTGGGAGAGGGGNNNAGATCGGA |
| 44176_1 | TATGAGTGTGGAGTCGTTGCCGGGGAGGAAGTAACAAAGAACANNNAGATCGGA |
| 44176_2 | TATGAGTGTGGAGTCGTTGCCGGGGAGGAAGTAACAAAGAACANNNAGATCGGA |
| 35638_1 | TATGAGTGTGGAGTCGTTGCCAGTAGCTGGTACTACAGGCGNNNAGATCGGAAGA |
| 35638_2 | TATGAGTGTGGAGTCGTTGCCAGTAGCTGGTACTACAGGCGNNNAGATCGGAAGA |
| 37940_1 | TATGAGTGTGGAGTCGTTGCAAGTGTGCAACAATCGCTATAGCNNNAGATCGGAAG |
| 37940_2 | TATGAGTGTGGAGTCGTTGCAAGTGTGCAACAATCGCTATAGCNNNAGATCGGAAG |
| 29423_1 | TATGAGTGTGGAGTCGTTGCACTTACAGATGCACAGCAGGAAGNNNAGATCGGAA |
| 29423_2 | TATGAGTGTGGAGTCGTTGCACTTACAGATGCACAGCAGGAAGNNNAGATCGGAA |
| 45846_1 | TATGAGTGTGGAGTCGTTGCCCCAAACCAGAATGACGCATAAANNNAGATCGGAAG |
| 45846_2 | TATGAGTGTGGAGTCGTTGCCCCAAACCAGAATGACGCATAAANNNAGATCGGAAG |
| 37156_1 | TATGAGTGTGGAGTCGTTGCGAACAACAGACACCACGAAACATNNNAGATCGGAAG |
| 37156_2 | TATGAGTGTGGAGTCGTTGCGAACAACAGACACCACGAAACATNNNAGATCGGAAG |
| 33788_1 | TATGAGTGTGGAGTCGTTGCAGCAGCAACAACAACAAAACTCTNNNAGATCGGAAG |
| 33788_2 | TATGAGTGTGGAGTCGTTGCAGCAGCAACAACAACAAAACTCTNNNAGATCGGAAG |
| 47265_1 | TATGAGTGTGGAGTCGTTGCTATTTCTTGGTTCGGTCTTGGGANNNAGATCGGAAG |
| 47265_2 | TATGAGTGTGGAGTCGTTGCTATTTCTTGGTTCGGTCTTGGGANNNAGATCGGAAG |
| 30065_1 | TATGAGTGTGGAGTCGTTGCCTCTCTCAGTCTGGATTTTGCTGANNNAGATCGGAA |
| 30065_2 | TATGAGTGTGGAGTCGTTGCCTCTCTCAGTCTGGATTTTGCTGANNNAGATCGGAA |
| 35246_1 | TATGAGTGTGGAGTCGTTGCAGGTTCAAGCAATTCTCCTGTCTNNNAGATCGGAAG |
| 35246_2 | TATGAGTGTGGAGTCGTTGCAGGTTCAAGCAATTCTCCTGTCTNNNAGATCGGAAG |
| 36663_1 | TATGAGTGTGGAGTCGTTGCTGGGATTAGGGAAGATCAGGGAANNNAGATCGGA |
| 36663_2 | TATGAGTGTGGAGTCGTTGCTGGGATTAGGGAAGATCAGGGAANNNAGATCGGA |
| 51188_1 | TATGAGTGTGGAGTCGTTGCCCAAAGCAACAAGAAGTACCCAGNNNAGATCGGAA |
| 51188_2 | TATGAGTGTGGAGTCGTTGCCCAAAGCAACAAGAAGTACCCAGNNNAGATCGGAA |
| 13475_1 | TATGAGTGTGGAGTCGTTGCGGCTATTTTTCCACTGATTAAATNNNAGATCGGAAG |

| | |
|---|---|
| 13475_2 | TATGAGTGTGGAGTCGTTGCGGCTATTTTTCCACTGATTAAATNNNAGATCGGAAG |
| 45051_1 | TATGAGTGTGGAGTCGTTGCGATGATCCAGATGTAAGGGCAGTNNNAGATCGGAA |
| 45051_2 | TATGAGTGTGGAGTCGTTGCGATGATCCAGATGTAAGGGCAGTNNNAGATCGGAA |
| 50996_1 | TATGAGTGTGGAGTCGTTGCAGCCTCCTGAGTAGCTAGGATTANNNAGATCGGAAG |
| 50996_2 | TATGAGTGTGGAGTCGTTGCAGCCTCCTGAGTAGCTAGGATTANNNAGATCGGAAG |
| 41434_1 | TATGAGTGTGGAGTCGTTGCAAAGGGAGAAACAGGAAGTGTGTNNNAGATCGGA |
| 41434_2 | TATGAGTGTGGAGTCGTTGCAAAGGGAGAAACAGGAAGTGTGTNNNAGATCGGA |
| 53207_1 | TATGAGTGTGGAGTCGTTGCGCCTCTGGACAGGATATCACAAANNNAGATCGGAAG |
| 53207_2 | TATGAGTGTGGAGTCGTTGCGCCTCTGGACAGGATATCACAAANNNAGATCGGAAG |
| 13391_1 | TATGAGTGTGGAGTCGTTGCTACTCAATTTAATTGCAACCTTTNNNAGATCGGAAGA |
| 13391_2 | TATGAGTGTGGAGTCGTTGCTACTCAATTTAATTGCAACCTTTNNNAGATCGGAAGA |
| 2317_1 | TATGAGTGTGGAGTCGTTGCTGGCCTCCATGAAACAGTTCTACNNNAGATCGGAAG |
| 2317_2 | TATGAGTGTGGAGTCGTTGCTGGCCTCCATGAAACAGTTCTACNNNAGATCGGAAG |
| 48759_1 | TATGAGTGTGGAGTCGTTGCACACTATTCTCCTACCTCAGCCTNNNAGATCGGAAGA |
| 48759_2 | TATGAGTGTGGAGTCGTTGCACACTATTCTCCTACCTCAGCCTNNNAGATCGGAAGA |
| 32527_1 | TATGAGTGTGGAGTCGTTGCGGAGCATGAGTAGCCAATACAATNNNAGATCGGAA |
| 32527_2 | TATGAGTGTGGAGTCGTTGCGGAGCATGAGTAGCCAATACAATNNNAGATCGGAA |
| 53202_1 | TATGAGTGTGGAGTCGTTGCTAGAGTCTCTGCCTGGTAATCCANNNAGATCGGAAG |
| 53202_2 | TATGAGTGTGGAGTCGTTGCTAGAGTCTCTGCCTGGTAATCCANNNAGATCGGAAG |
| 40547_1 | TATGAGTGTGGAGTCGTTGCAGAAAACTGGAGCTCTACATGCANNNAGATCGGAAG |
| 40547_2 | TATGAGTGTGGAGTCGTTGCAGAAAACTGGAGCTCTACATGCANNNAGATCGGAAG |
| 48464_1 | TATGAGTGTGGAGTCGTTGCCGAGTAGCTGAGATTACAGGCATNNNAGATCGGAA |
| 48464_2 | TATGAGTGTGGAGTCGTTGCCGAGTAGCTGAGATTACAGGCATNNNAGATCGGAA |
| 51786_1 | TATGAGTGTGGAGTCGTTGCGGGCTCCTCACTTCCCAGTANNNAGATCGGAAGAGC |
| 51786_2 | TATGAGTGTGGAGTCGTTGCGGGCTCCTCACTTCCCAGTANNNAGATCGGAAGAGC |
| 38009_1 | TATGAGTGTGGAGTCGTTGCACCTCACTAGTGACCCTGAAAACNNNAGATCGGAAG |
| 38009_2 | TATGAGTGTGGAGTCGTTGCACCTCACTAGTGACCCTGAAAACNNNAGATCGGAAG |
| 30393_1 | TATGAGTGTGGAGTCGTTGCTGGAAACAAGTCACATTTATCTGGGANNNAGATCGG |
| 30393_2 | TATGAGTGTGGAGTCGTTGCTGGAAACAAGTCACATTTATCTGGGANNNAGATCGG |
| 36173_1 | TATGAGTGTGGAGTCGTTGCTACACACACACACACAAATGCAANNNAGATCGGAAG |
| 36173_2 | TATGAGTGTGGAGTCGTTGCTACACACACACACACAAATGCAANNNAGATCGGAAG |
| 30052_1 | TATGAGTGTGGAGTCGTTGCGGGCCTTGGTGCTTATTTTCTCNNNAGATCGGAAGA |
| 30052_2 | TATGAGTGTGGAGTCGTTGCGGGCCTTGGTGCTTATTTTCTCNNNAGATCGGAAGA |
| 35767_1 | TATGAGTGTGGAGTCGTTGCTCAGACGCATTATGATCAAGCTGNNNAGATCGGAAG |
| 35767_2 | TATGAGTGTGGAGTCGTTGCTCAGACGCATTATGATCAAGCTGNNNAGATCGGAAG |
| 29014_1 | TATGAGTGTGGAGTCGTTGCGAGGAGGACTAGAGAGAAAGGGANNNAGATCGGA |
| 29014_2 | TATGAGTGTGGAGTCGTTGCGAGGAGGACTAGAGAGAAAGGGANNNAGATCGGA |
| 45046_1 | TATGAGTGTGGAGTCGTTGCCCACTGCTGCTAGGGAAAAAAAANNNAGATCGGAA |
| 45046_2 | TATGAGTGTGGAGTCGTTGCCCACTGCTGCTAGGGAAAAAAAANNNAGATCGGAA |
| 30059_1 | TGAGTGTGGAGTCGTTGCTGAAGTTGCTAAGGTAGTAGTTTCCANNNAGATCGGA |
| 30059_2 | TGAGTGTGGAGTCGTTGCTGAAGTTGCTAAGGTAGTAGTTTCCANNNAGATCGGA |
| 32704_1 | TATGAGTGTGGAGTCGTTGCTTCAAGCAATTCCCCTGTCTCAGNNNAGATCGGAAG |
| 32704_2 | TATGAGTGTGGAGTCGTTGCTTCAAGCAATTCCCCTGTCTCAGNNNAGATCGGAAG |

| | |
|---|---|
| 36947_1 | TATGAGTGTGGAGTCGTTGCTTAGCTGGGCATGGTAGTATGTGNNNAGATCGGAA |
| 36947_2 | TATGAGTGTGGAGTCGTTGCTTAGCTGGGCATGGTAGTATGTGNNNAGATCGGAA |
| 39319_1 | TATGAGTGTGGAGTCGTTGCATTGTGTCCTACCTCTGTTGGTGNNNAGATCGGAAG |
| 39319_2 | TATGAGTGTGGAGTCGTTGCATTGTGTCCTACCTCTGTTGGTGNNNAGATCGGAAG |
| 13046_1 | TATGAGTGTGGAGTCGTTGCAAAACAAAACAAGGGTCACTTNNNAGATCGGAAGA |
| 13046_2 | TATGAGTGTGGAGTCGTTGCAAAACAAAACAAGGGTCACTTNNNAGATCGGAAGA |
| 33896_1 | TATGAGTGTGGAGTCGTTGCTTCACACCATTCTCCTGCCTCNNNAGATCGGAAGAGC |
| 33896_2 | TATGAGTGTGGAGTCGTTGCTTCACACCATTCTCCTGCCTCNNNAGATCGGAAGAGC |
| 44081_1 | TATGAGTGTGGAGTCGTTGCGAGTGGCAGCAAAAATGACAGATNNNAGATCGGAA |
| 44081_2 | TATGAGTGTGGAGTCGTTGCGAGTGGCAGCAAAAATGACAGATNNNAGATCGGAA |
| 51854_1 | TATGAGTGTGGAGTCGTTGCCATTCCAGGTGATCCTTCTCCTCNNNAGATCGGAAGA |
| 51854_2 | TATGAGTGTGGAGTCGTTGCCATTCCAGGTGATCCTTCTCCTCNNNAGATCGGAAGA |
| 41022_1 | TATGAGTGTGGAGTCGTTGCAGAGAGAACTGAAGCAAATGTGCNNNAGATCGGAA |
| 41022_2 | TATGAGTGTGGAGTCGTTGCAGAGAGAACTGAAGCAAATGTGCNNNAGATCGGAA |
| 47066_1 | TATGAGTGTGGAGTCGTTGCGGATGTCAGGAAGCCTCAAACTANNNAGATCGGAA |
| 47066_2 | TATGAGTGTGGAGTCGTTGCGGATGTCAGGAAGCCTCAAACTANNNAGATCGGAA |
| 35096_1 | TATGAGTGTGGAGTCGTTGCCCTCCAGTGTTCACTCCATTCTCNNNAGATCGGAAGA |
| 35096_2 | TATGAGTGTGGAGTCGTTGCCCTCCAGTGTTCACTCCATTCTCNNNAGATCGGAAGA |
| 12967_1 | TATGAGTGTGGAGTCGTTGCGATAAAACTATGGCGAAGAAAANNNAGATCGGAAG |
| 12967_2 | TATGAGTGTGGAGTCGTTGCGATAAAACTATGGCGAAGAAAANNNAGATCGGAAG |
| 33202_1 | TATGAGTGTGGAGTCGTTGCCAGAAGAAAGGCATTTGTTTTGCNNNAGATCGGAA |
| 33202_2 | TATGAGTGTGGAGTCGTTGCCAGAAGAAAGGCATTTGTTTTGCNNNAGATCGGAA |
| 50379_1 | TATGAGTGTGGAGTCGTTGCCAGCGGACACTCAATGAGATCTANNNAGATCGGAA |
| 50379_2 | TATGAGTGTGGAGTCGTTGCCAGCGGACACTCAATGAGATCTANNNAGATCGGAA |
| 50699_1 | TATGAGTGTGGAGTCGTTGCGCATTGTCTCTGCATTGTCATCCNNNAGATCGGAAG |
| 50699_2 | TATGAGTGTGGAGTCGTTGCGCATTGTCTCTGCATTGTCATCCNNNAGATCGGAAG |
| 37645_1 | TATGAGTGTGGAGTCGTTGCTTCAGACAGTAAAGCTCAGAGCANNNAGATCGGAA |
| 37645_2 | TATGAGTGTGGAGTCGTTGCTTCAGACAGTAAAGCTCAGAGCANNNAGATCGGAA |
| 13419_1 | TATGAGTGTGGAGTCGTTGCGAGAAAGAAGCAATTGTGATGNNNAGATCGGAAG |
| 13419_2 | TATGAGTGTGGAGTCGTTGCGAGAAAGAAGCAATTGTGATGNNNAGATCGGAAG |
| 49392_1 | TATGAGTGTGGAGTCGTTGCCTTGCAGTTGAGTCTCCAGATCTNNNAGATCGGAAG |
| 49392_2 | TATGAGTGTGGAGTCGTTGCCTTGCAGTTGAGTCTCCAGATCTNNNAGATCGGAAG |
| 53064_1 | TATGAGTGTGGAGTCGTTGCCTCCTGCAATATACTGCCAGACANNNAGATCGGAAG |
| 53064_2 | TATGAGTGTGGAGTCGTTGCCTCCTGCAATATACTGCCAGACANNNAGATCGGAAG |
| 49151_1 | TATGAGTGTGGAGTCGTTGCGTAGCTGGGATTACAGGCACATNNNAGATCGGAAG |
| 49151_2 | TATGAGTGTGGAGTCGTTGCGTAGCTGGGATTACAGGCACATNNNAGATCGGAAG |
| 12859_1 | TATGAGTGTGGAGTCGTTGCGAAAATAAGCTGGTTTCTATGTANNNAGATCGGAA |
| 12859_2 | TATGAGTGTGGAGTCGTTGCGAAAATAAGCTGGTTTCTATGTANNNAGATCGGAA |
| 37748_1 | TATGAGTGTGGAGTCGTTGCCTGAGGCAGGAGAATAGCATGAANNNAGATCGGAA |
| 37748_2 | TATGAGTGTGGAGTCGTTGCCTGAGGCAGGAGAATAGCATGAANNNAGATCGGAA |
| 31904_1 | TATGAGTGTGGAGTCGTTGCGAGGCTGAGGCTGGAGAATTGNNNAGATCGGAAG |
| 31904_2 | TATGAGTGTGGAGTCGTTGCGAGGCTGAGGCTGGAGAATTGNNNAGATCGGAAG |
| 35874_1 | TATGAGTGTGGAGTCGTTGCAAATAGACTGTGGTGATGGTGGCNNNAGATCGGAA |

| | |
|---|---|
| 35874_2 | TATGAGTGTGGAGTCGTTGCAAATAGACTGTGGTGATGGTGGCNNNAGATCGGAA |
| 32271_1 | TATGAGTGTGGAGTCGTTGCAGGTGAGAGCAAGAGTAAAATCANNNAGATCGGAA |
| 32271_2 | TATGAGTGTGGAGTCGTTGCAGGTGAGAGCAAGAGTAAAATCANNNAGATCGGAA |
| 44614_1 | TATGAGTGTGGAGTCGTTGCATTCCATTCCTTACGCGATCTGANNNAGATCGGAAG |
| 44614_2 | TATGAGTGTGGAGTCGTTGCATTCCATTCCTTACGCGATCTGANNNAGATCGGAAG |
| 52146_1 | TATGAGTGTGGAGTCGTTGCATGAGAAAAGTCACCTCTGTCCCNNNAGATCGGAAG |
| 52146_2 | TATGAGTGTGGAGTCGTTGCATGAGAAAAGTCACCTCTGTCCCNNNAGATCGGAAG |
| 10157_1 | TATGAGTGTGGAGTCGTTGCAGGCTTGGGGAGCTATGATAAAGANNNAGATCGGA |
| 10157_2 | TATGAGTGTGGAGTCGTTGCAGGCTTGGGGAGCTATGATAAAGANNNAGATCGGA |
| 32959_1 | TATGAGTGTGGAGTCGTTGCCACAGTGGCTCAAGCTTGTAATCNNNAGATCGGAAG |
| 32959_2 | TATGAGTGTGGAGTCGTTGCCACAGTGGCTCAAGCTTGTAATCNNNAGATCGGAAG |
| 33650_1 | TATGAGTGTGGAGTCGTTGCCTTCCTGGGTTCAAGCAATTCTCNNNAGATCGGAAG |
| 33650_2 | TATGAGTGTGGAGTCGTTGCCTTCCTGGGTTCAAGCAATTCTCNNNAGATCGGAAG |
| 52971_1 | TATGAGTGTGGAGTCGTTGCCACACATGCACACACAAAAACACNNNAGATCGGAAG |
| 52971_2 | TATGAGTGTGGAGTCGTTGCCACACATGCACACACAAAAACACNNNAGATCGGAAG |
| 29376_1 | TATGAGTGTGGAGTCGTTGCCTTCCGGTTTCAAGCAATTGGATNNNAGATCGGAAG |
| 29376_2 | TATGAGTGTGGAGTCGTTGCCTTCCGGTTTCAAGCAATTGGATNNNAGATCGGAAG |
| 41188_1 | TATGAGTGTGGAGTCGTTGCCCATCCCCACTCCTTTTCAGTTANNNAGATCGGAAGA |
| 41188_2 | TATGAGTGTGGAGTCGTTGCCCATCCCCACTCCTTTTCAGTTANNNAGATCGGAAGA |
| 46037_1 | TATGAGTGTGGAGTCGTTGCTATACAAGCCTGCAAGAACCACANNNAGATCGGAAG |
| 46037_2 | TATGAGTGTGGAGTCGTTGCTATACAAGCCTGCAAGAACCACANNNAGATCGGAAG |
| 52575_1 | TATGAGTGTGGAGTCGTTGCGGCCTTCACAGTGTTTTTCAAGTNNNAGATCGGAAG |
| 52575_2 | TATGAGTGTGGAGTCGTTGCGGCCTTCACAGTGTTTTTCAAGTNNNAGATCGGAAG |
| 32145_1 | TATGAGTGTGGAGTCGTTGCCTCCCAGGTTCAAGCGATTCTTNNNAGATCGGAAGA |
| 32145_2 | TATGAGTGTGGAGTCGTTGCCTCCCAGGTTCAAGCGATTCTTNNNAGATCGGAAGA |
| 50904_1 | TATGAGTGTGGAGTCGTTGCCAAGTTCTGGGTCTCCTTATCCCNNNAGATCGGAAG |
| 50904_2 | TATGAGTGTGGAGTCGTTGCCAAGTTCTGGGTCTCCTTATCCCNNNAGATCGGAAG |
| 46758_1 | TATGAGTGTGGAGTCGTTGCCTACCAGGTGATTCTACTCTGCCNNNAGATCGGAAG |
| 46758_2 | TATGAGTGTGGAGTCGTTGCCTACCAGGTGATTCTACTCTGCCNNNAGATCGGAAG |
| 34360_1 | TATGAGTGTGGAGTCGTTGCTCTCAAACTCCTGGGTTCAAGTTNNNAGATCGGAAG |
| 34360_2 | TATGAGTGTGGAGTCGTTGCTCTCAAACTCCTGGGTTCAAGTTNNNAGATCGGAAG |
| 53359_1 | TATGAGTGTGGAGTCGTTGCACTTTTTCCAAACCATCAAGGACANNNAGATCGGAA |
| 53359_2 | TATGAGTGTGGAGTCGTTGCACTTTTTCCAAACCATCAAGGACANNNAGATCGGAA |
| 33143_1 | TATGAGTGTGGAGTCGTTGCCATATGTCAAACAGGCACACGAANNNAGATCGGAA |
| 33143_2 | TATGAGTGTGGAGTCGTTGCCATATGTCAAACAGGCACACGAANNNAGATCGGAA |
| 12031_1 | TATGAGTGTGGAGTCGTTGCTGATGAGCAGCGTCAAGGACAACNNNAGATCGGAA |
| 12031_2 | TATGAGTGTGGAGTCGTTGCTGATGAGCAGCGTCAAGGACAACNNNAGATCGGAA |
| 52156_1 | TATGAGTGTGGAGTCGTTGCGAGACGCTCTTCACCTCTCAGNNNAGATCGGAAGAG |
| 52156_2 | TATGAGTGTGGAGTCGTTGCGAGACGCTCTTCACCTCTCAGNNNAGATCGGAAGAG |
| 38768_1 | TATGAGTGTGGAGTCGTTGCCATGCCTGTAATCCCAGCATTTTNNNAGATCGGAAG |
| 38768_2 | TATGAGTGTGGAGTCGTTGCCATGCCTGTAATCCCAGCATTTTNNNAGATCGGAAG |
| 34597_1 | TATGAGTGTGGAGTCGTTGCCCCTACCATGTTCCAGGCAATATNNNAGATCGGAAG |
| 34597_2 | TATGAGTGTGGAGTCGTTGCCCCTACCATGTTCCAGGCAATATNNNAGATCGGAAG |

| | |
|---|---|
| 34184_1 | TATGAGTGTGGAGTCGTTGCTATGTTGACAGACCTCACACCACNNNAGATCGGAAG |
| 34184_2 | TATGAGTGTGGAGTCGTTGCTATGTTGACAGACCTCACACCACNNNAGATCGGAAG |
| 38734_1 | TATGAGTGTGGAGTCGTTGCAGCTCCAGAGTTTCTGATTGATTNNNAGATCGGAAG |
| 38734_2 | TATGAGTGTGGAGTCGTTGCAGCTCCAGAGTTTCTGATTGATTNNNAGATCGGAAG |
| 33713_1 | TATGAGTGTGGAGTCGTTGCATGCCTGTAATCCCAGCACTTTNNNAGATCGGAAGA |
| 33713_2 | TATGAGTGTGGAGTCGTTGCATGCCTGTAATCCCAGCACTTTNNNAGATCGGAAGA |
| 39990_1 | TATGAGTGTGGAGTCGTTGCTCACTTGAGTTCAGGAGTTCGAGNNNAGATCGGAA |
| 39990_2 | TATGAGTGTGGAGTCGTTGCTCACTTGAGTTCAGGAGTTCGAGNNNAGATCGGAA |
| 33382_1 | TATGAGTGTGGAGTCGTTGCTGGATGATAAAACAAAGGCCAACNNNAGATCGGAA |
| 33382_2 | TATGAGTGTGGAGTCGTTGCTGGATGATAAAACAAAGGCCAACNNNAGATCGGAA |
| 33502_1 | TATGAGTGTGGAGTCGTTGCTTGCAACATATAAGGCAAAGGGCNNNAGATCGGAA |
| 33502_2 | TATGAGTGTGGAGTCGTTGCTTGCAACATATAAGGCAAAGGGCNNNAGATCGGAA |
| 51037_1 | TATGAGTGTGGAGTCGTTGCCAATGATAGTGCCCTGCTAGCTANNNAGATCGGAAG |
| 51037_2 | TATGAGTGTGGAGTCGTTGCCAATGATAGTGCCCTGCTAGCTANNNAGATCGGAAG |
| 31756_1 | TATGAGTGTGGAGTCGTTGCACTTCTCTGGAATTTCATGCCCTNNNAGATCGGAAGA |
| 31756_2 | TATGAGTGTGGAGTCGTTGCACTTCTCTGGAATTTCATGCCCTNNNAGATCGGAAGA |
| 33754_1 | TATGAGTGTGGAGTCGTTGCCACGCCTATAATCCCAGCACTTTNNNAGATCGGAAGA |
| 33754_2 | TATGAGTGTGGAGTCGTTGCCACGCCTATAATCCCAGCACTTTNNNAGATCGGAAGA |
| 47988_1 | TATGAGTGTGGAGTCGTTGCCTCCCGAGTAGCTGGAACTACAGNNNAGATCGGAA |
| 47988_2 | TATGAGTGTGGAGTCGTTGCCTCCCGAGTAGCTGGAACTACAGNNNAGATCGGAA |
| 29086_1 | TATGAGTGTGGAGTCGTTGCTTGCCTTGTTCCAGTGAGTCTAANNNAGATCGGAAG |
| 29086_2 | TATGAGTGTGGAGTCGTTGCTTGCCTTGTTCCAGTGAGTCTAANNNAGATCGGAAG |
| 49928_1 | TATGAGTGTGGAGTCGTTGCGCTAGTGGAACTCTGGCTGAAATNNNAGATCGGAA |
| 49928_2 | TATGAGTGTGGAGTCGTTGCGCTAGTGGAACTCTGGCTGAAATNNNAGATCGGAA |
| 12881_1 | TATGAGTGTGGAGTCGTTGCACATAAAATGTGGTAACATACACNNNAGATCGGAAG |
| 12881_2 | TATGAGTGTGGAGTCGTTGCACATAAAATGTGGTAACATACACNNNAGATCGGAAG |
| 13244_1 | TATGAGTGTGGAGTCGTTGCTTTCACATCCTTACATTTCAAAANNNAGATCGGAAGA |
| 13244_2 | TATGAGTGTGGAGTCGTTGCTTTCACATCCTTACATTTCAAAANNNAGATCGGAAGA |
| 36172_1 | TATGAGTGTGGAGTCGTTGCGTCAGGAGATCCAAGTTAGTCCCNNNAGATCGGAA |
| 36172_2 | TATGAGTGTGGAGTCGTTGCGTCAGGAGATCCAAGTTAGTCCCNNNAGATCGGAA |
| 12506_1 | TATGAGTGTGGAGTCGTTGCGAGAAAAGCAAGAGAAGTTAAATNNNAGATCGGA |
| 12506_2 | TATGAGTGTGGAGTCGTTGCGAGAAAAGCAAGAGAAGTTAAATNNNAGATCGGA |
| 34442_1 | TATGAGTGTGGAGTCGTTGCGAGGCACTTCCGCTTTCTGTNNNAGATCGGAAGAGC |
| 34442_2 | TATGAGTGTGGAGTCGTTGCGAGGCACTTCCGCTTTCTGTNNNAGATCGGAAGAGC |
| 29763_1 | TATGAGTGTGGAGTCGTTGCCAACAATGTGTCCACTGCTATGGNNNAGATCGGAAG |
| 29763_2 | TATGAGTGTGGAGTCGTTGCCAACAATGTGTCCACTGCTATGGNNNAGATCGGAAG |
| 46757_1 | TATGAGTGTGGAGTCGTTGCCCACCACACCCAGCTAATTTTTTNNNAGATCGGAAGA |
| 46757_2 | TATGAGTGTGGAGTCGTTGCCCACCACACCCAGCTAATTTTTTNNNAGATCGGAAGA |
| 12891_1 | TATGAGTGTGGAGTCGTTGCGCAGATTAAAAGACAAAACAAAANNNAGATCGGAA |
| 12891_2 | TATGAGTGTGGAGTCGTTGCGCAGATTAAAAGACAAAACAAAANNNAGATCGGAA |
| 13750_1 | TATGAGTGTGGAGTCGTTGCATTGAGACATGAAGAACAGACNNNAGATCGGAAGA |
| 13750_2 | TATGAGTGTGGAGTCGTTGCATTGAGACATGAAGAACAGACNNNAGATCGGAAGA |
| 34089_1 | TATGAGTGTGGAGTCGTTGCCTCCTGAATAGCTGGGACTACAGNNNAGATCGGAA |

| | |
|---|---|
| 34089_2 | TATGAGTGTGGAGTCGTTGCCTCCTGAATAGCTGGGACTACAGNNNAGATCGGAAG |
| 33597_1 | TATGAGTGTGGAGTCGTTGCCCTCCTCAGTTCAAGCGATTTTGNNNAGATCGGAAGA |
| 33597_2 | TATGAGTGTGGAGTCGTTGCCCTCCTCAGTTCAAGCGATTTTGNNNAGATCGGAAGA |
| 44370_1 | TATGAGTGTGGAGTCGTTGCTTGCTGGGTTTGAATTCTGGTTCNNNAGATCGGAAG |
| 44370_2 | TATGAGTGTGGAGTCGTTGCTTGCTGGGTTTGAATTCTGGTTCNNNAGATCGGAAG |
| 38126_1 | TATGAGTGTGGAGTCGTTGCACAAAAATTAGCTGGGCTTGGTGNNNAGATCGGAA |
| 38126_2 | TATGAGTGTGGAGTCGTTGCACAAAAATTAGCTGGGCTTGGTGNNNAGATCGGAA |
| 34934_1 | TATGAGTGTGGAGTCGTTGCCTGGGATTACAGGTGTGAGTCACNNNAGATCGGAA |
| 34934_2 | TATGAGTGTGGAGTCGTTGCCTGGGATTACAGGTGTGAGTCACNNNAGATCGGAA |
| 36510_1 | TATGAGTGTGGAGTCGTTGCAGGGAAAATGTTGCCCAAGTAGANNNAGATCGGAA |
| 36510_2 | TATGAGTGTGGAGTCGTTGCAGGGAAAATGTTGCCCAAGTAGANNNAGATCGGAA |
| 29857_1 | TATGAGTGTGGAGTCGTTGCAAAGACCCCATTCAGTTATGTGTNNNAGATCGGAAG |
| 29857_2 | TATGAGTGTGGAGTCGTTGCAAAGACCCCATTCAGTTATGTGTNNNAGATCGGAAG |
| 32912_1 | TATGAGTGTGGAGTCGTTGCGCAGGAGAATTGCTTGAACCTGNNNAGATCGGAAG |
| 32912_2 | TATGAGTGTGGAGTCGTTGCGCAGGAGAATTGCTTGAACCTGNNNAGATCGGAAG |
| 31762_1 | TATGAGTGTGGAGTCGTTGCGTGTGTGTGTGAGAGAGAGAGAGNNNAGATCGGA |
| 31762_2 | TATGAGTGTGGAGTCGTTGCGTGTGTGTGTGAGAGAGAGAGAGNNNAGATCGGA |
| 29793_1 | TATGAGTGTGGAGTCGTTGCACATACCATTAACATTTGTGTGTAGGANNNAGATCG |
| 29793_2 | TATGAGTGTGGAGTCGTTGCACATACCATTAACATTTGTGTGTAGGANNNAGATCG |
| 46479_1 | TATGAGTGTGGAGTCGTTGCGTGCCCAGCCAGCATAAATATTTNNNAGATCGGAAG |
| 46479_2 | TATGAGTGTGGAGTCGTTGCGTGCCCAGCCAGCATAAATATTTNNNAGATCGGAAG |
| 30232_1 | TATGAGTGTGGAGTCGTTGCTGGTTTTGTGTTTTTGCTTTTCCANNNAGATCGGAAG |
| 30232_2 | TATGAGTGTGGAGTCGTTGCTGGTTTTGTGTTTTTGCTTTTCCANNNAGATCGGAAG |
| 12882_1 | TATGAGTGTGGAGTCGTTGCAATGAATGGCAAATTCTATTTCTNNNAGATCGGAAG |
| 12882_2 | TATGAGTGTGGAGTCGTTGCAATGAATGGCAAATTCTATTTCTNNNAGATCGGAAG |
| 29180_1 | TATGAGTGTGGAGTCGTTGCCTCTCAGACCCTCCCTCTAGAAANNNAGATCGGAAGA |
| 29180_2 | TATGAGTGTGGAGTCGTTGCCTCTCAGACCCTCCCTCTAGAAANNNAGATCGGAAGA |
| 29691_1 | TATGAGTGTGGAGTCGTTGCTCTACGTGAAGGTGAATGGTGGNNNAGATCGGAAG |
| 29691_2 | TATGAGTGTGGAGTCGTTGCTCTACGTGAAGGTGAATGGTGGNNNAGATCGGAAG |
| 36846_1 | TATGAGTGTGGAGTCGTTGCGGGAGACTAAAGTGGAGGATCACNNNAGATCGGAA |
| 36846_2 | TATGAGTGTGGAGTCGTTGCGGGAGACTAAAGTGGAGGATCACNNNAGATCGGAA |
| 31172_1 | TATGAGTGTGGAGTCGTTGCTAATCTCAGCACTTTGGGAGACCNNNAGATCGGAAG |
| 31172_2 | TATGAGTGTGGAGTCGTTGCTAATCTCAGCACTTTGGGAGACCNNNAGATCGGAAG |
| 37936_1 | TATGAGTGTGGAGTCGTTGCGTGGATCACAAGGTCAGGAGATTNNNAGATCGGAA |
| 37936_2 | TATGAGTGTGGAGTCGTTGCGTGGATCACAAGGTCAGGAGATTNNNAGATCGGAA |
| 46215_1 | TATGAGTGTGGAGTCGTTGCGTTCTGTGTTCTCAATGGCCATTNNNAGATCGGAAG |
| 46215_2 | TATGAGTGTGGAGTCGTTGCGTTCTGTGTTCTCAATGGCCATTNNNAGATCGGAAG |
| 44958_1 | TATGAGTGTGGAGTCGTTGCCGGGTGGGAGATAACGTGTAATANNNAGATCGGAA |
| 44958_2 | TATGAGTGTGGAGTCGTTGCCGGGTGGGAGATAACGTGTAATANNNAGATCGGAA |
| 42255_1 | TATGAGTGTGGAGTCGTTGCCTGGTGTCTAATTCCAGACCTCGNNNAGATCGGAAG |
| 42255_2 | TATGAGTGTGGAGTCGTTGCCTGGTGTCTAATTCCAGACCTCGNNNAGATCGGAAG |
| 40158_1 | TATGAGTGTGGAGTCGTTGCACCAGCAAAGGATGAGAGTACATNNNAGATCGGAA |
| 40158_2 | TATGAGTGTGGAGTCGTTGCACCAGCAAAGGATGAGAGTACATNNNAGATCGGAA |

| | |
|---|---|
| 41126_1 | TATGAGTGTGGAGTCGTTGCTCAATTGAGGTCAGGAGTTGGAGNNNAGATCGGAA |
| 41126_2 | TATGAGTGTGGAGTCGTTGCTCAATTGAGGTCAGGAGTTGGAGNNNAGATCGGAA |
| 49829_1 | TATGAGTGTGGAGTCGTTGCCTCTATTACCCAGGCTGGAGTGNNNAGATCGGAAGA |
| 49829_2 | TATGAGTGTGGAGTCGTTGCCTCTATTACCCAGGCTGGAGTGNNNAGATCGGAAGA |
| 29660_1 | TATGAGTGTGGAGTCGTTGCCTCTCTTAACTGCAAGGCTTGTCNNNAGATCGGAAG |
| 29660_2 | TATGAGTGTGGAGTCGTTGCCTCTCTTAACTGCAAGGCTTGTCNNNAGATCGGAAG |
| 47114_1 | TATGAGTGTGGAGTCGTTGCTTCTGGGTTCAAGCGATTCTCATNNNAGATCGGAAG |
| 47114_2 | TATGAGTGTGGAGTCGTTGCTTCTGGGTTCAAGCGATTCTCATNNNAGATCGGAAG |
| 39870_1 | TATGAGTGTGGAGTCGTTGCCTTTCCATCTTGCTTGTCCACTGNNNAGATCGGAAGA |
| 39870_2 | TATGAGTGTGGAGTCGTTGCCTTTCCATCTTGCTTGTCCACTGNNNAGATCGGAAGA |
| 49028_1 | TATGAGTGTGGAGTCGTTGCAGGAAGAGTGGAAACGTTTCTCTNNNAGATCGGAA |
| 49028_2 | TATGAGTGTGGAGTCGTTGCAGGAAGAGTGGAAACGTTTCTCTNNNAGATCGGAA |
| 40256_1 | TATGAGTGTGGAGTCGTTGCCTCTATCCTGAGCTGTTGGATCTNNNAGATCGGAAG |
| 40256_2 | TATGAGTGTGGAGTCGTTGCCTCTATCCTGAGCTGTTGGATCTNNNAGATCGGAAG |
| 49586_1 | TATGAGTGTGGAGTCGTTGCGTAGTTTGATCACAGCTCACTGCNNNAGATCGGAAG |
| 49586_2 | TATGAGTGTGGAGTCGTTGCGTAGTTTGATCACAGCTCACTGCNNNAGATCGGAAG |
| 49222_1 | TATGAGTGTGGAGTCGTTGCTGGCCTGAATCCTCTCTCTACTTNNNAGATCGGAAGA |
| 49222_2 | TATGAGTGTGGAGTCGTTGCTGGCCTGAATCCTCTCTCTACTTNNNAGATCGGAAGA |
| 30730_1 | TATGAGTGTGGAGTCGTTGCTTTTCCAGACAGGCTGATCTTGNNNAGATCGGAAGA |
| 30730_2 | TATGAGTGTGGAGTCGTTGCTTTTCCAGACAGGCTGATCTTGNNNAGATCGGAAGA |
| 37521_1 | TATGAGTGTGGAGTCGTTGCCAGAACCCAACGTTTGATGTCTGNNNAGATCGGAAG |
| 37521_2 | TATGAGTGTGGAGTCGTTGCCAGAACCCAACGTTTGATGTCTGNNNAGATCGGAAG |
| 36192_1 | TATGAGTGTGGAGTCGTTGCAAAGCTTTCCAACTGTGCTCATCNNNAGATCGGAAG |
| 36192_2 | TATGAGTGTGGAGTCGTTGCAAAGCTTTCCAACTGTGCTCATCNNNAGATCGGAAG |
| 39710_1 | TATGAGTGTGGAGTCGTTGCCTCTGAGTTCCATGGCATACACTNNNAGATCGGAAG |
| 39710_2 | TATGAGTGTGGAGTCGTTGCCTCTGAGTTCCATGGCATACACTNNNAGATCGGAAG |
| 13349_1 | TATGAGTGTGGAGTCGTTGCTATTTATTTTTGAGACAGGGTCNNNAGATCGGAAG |
| 13349_2 | TATGAGTGTGGAGTCGTTGCTATTTATTTTTGAGACAGGGTCNNNAGATCGGAAG |
| 29690_1 | TATGAGTGTGGAGTCGTTGCATACACCTTTGCTCTCCCTACAGNNNAGATCGGAAGA |
| 29690_2 | TATGAGTGTGGAGTCGTTGCATACACCTTTGCTCTCCCTACAGNNNAGATCGGAAGA |
| 31166_1 | TATGAGTGTGGAGTCGTTGCTATTTATCTGTCTGAGCCCTGCCNNNAGATCGGAAG |
| 31166_2 | TATGAGTGTGGAGTCGTTGCTATTTATCTGTCTGAGCCCTGCCNNNAGATCGGAAG |
| 40025_1 | TATGAGTGTGGAGTCGTTGCGTTCCAGTGAGCAGAGATCTCAANNNAGATCGGAA |
| 40025_2 | TATGAGTGTGGAGTCGTTGCGTTCCAGTGAGCAGAGATCTCAANNNAGATCGGAA |
| 51339_1 | TATGAGTGTGGAGTCGTTGCGCGACCAGAAGTGGAAAATCCANNNAGATCGGAAG |
| 51339_2 | TATGAGTGTGGAGTCGTTGCGCGACCAGAAGTGGAAAATCCANNNAGATCGGAAG |
| 29710_1 | TATGAGTGTGGAGTCGTTGCCAATCAGTAGCCAGCATCTGCCNNNAGATCGGAAGA |
| 29710_2 | TATGAGTGTGGAGTCGTTGCCAATCAGTAGCCAGCATCTGCCNNNAGATCGGAAGA |
| 42860_1 | TATGAGTGTGGAGTCGTTGCGGCTGATCTTCAACTCCTGAGATNNNAGATCGGAAG |
| 42860_2 | TATGAGTGTGGAGTCGTTGCGGCTGATCTTCAACTCCTGAGATNNNAGATCGGAAG |
| 34212_1 | TATGAGTGTGGAGTCGTTGCGAACAGAGCCTCAAAGACCTGTANNNAGATCGGAA |
| 34212_2 | TATGAGTGTGGAGTCGTTGCGAACAGAGCCTCAAAGACCTGTANNNAGATCGGAA |
| 32911_1 | TATGAGTGTGGAGTCGTTGCGCCACGGAAACCTTTTGTTGATANNNAGATCGGAAG |

| | |
|---|---|
| 32911_2 | TATGAGTGTGGAGTCGTTGCGCCACGGAAACCTTTTGTTGATANNNAGATCGGAAG |
| 31103_1 | TATGAGTGTGGAGTCGTTGCATCGCGATCCACTGGAATACTTNNNAGATCGGAAG |
| 31103_2 | TATGAGTGTGGAGTCGTTGCATCGCGATCCACTGGAATACTTNNNAGATCGGAAG |
| 46348_1 | TATGAGTGTGGAGTCGTTGCCAAGTAGCAGGGATTATAGGCGTNNNAGATCGGAA |
| 46348_2 | TATGAGTGTGGAGTCGTTGCCAAGTAGCAGGGATTATAGGCGTNNNAGATCGGAA |
| 46394_1 | TATGAGTGTGGAGTCGTTGCCTCCTGAGTAGCTGGGACTACAGNNNAGATCGGAA |
| 46394_2 | TATGAGTGTGGAGTCGTTGCCTCCTGAGTAGCTGGGACTACAGNNNAGATCGGAA |
| 41107_1 | TATGAGTGTGGAGTCGTTGCGACTGGGTGAAAGCTGGTTACTANNNAGATCGGAA |
| 41107_2 | TATGAGTGTGGAGTCGTTGCGACTGGGTGAAAGCTGGTTACTANNNAGATCGGAA |
| 13424_1 | TATGAGTGTGGAGTCGTTGCTGTATGTCATTCCTGTTCACNNNAGATCGGAAGAGC |
| 13424_2 | TATGAGTGTGGAGTCGTTGCTGTATGTCATTCCTGTTCACNNNAGATCGGAAGAGC |
| 48319_1 | TATGAGTGTGGAGTCGTTGCCTGAGTTCAAGCAATTCTCCTGCNNNAGATCGGAAG |
| 48319_2 | TATGAGTGTGGAGTCGTTGCCTGAGTTCAAGCAATTCTCCTGCNNNAGATCGGAAG |
| 39065_1 | TATGAGTGTGGAGTCGTTGCTGAGGTAGGAGAATGGTGTGAACNNNAGATCGGA |
| 39065_2 | TATGAGTGTGGAGTCGTTGCTGAGGTAGGAGAATGGTGTGAACNNNAGATCGGA |
| 46350_1 | TATGAGTGTGGAGTCGTTGCCCAGTGAGACAGAACTATGCACTNNNAGATCGGAAG |
| 46350_2 | TATGAGTGTGGAGTCGTTGCCCAGTGAGACAGAACTATGCACTNNNAGATCGGAAG |
| 13449_1 | TATGAGTGTGGAGTCGTTGCCTGACCATTAGCATCACTTAATNNNAGATCGGAAGA |
| 13449_2 | TATGAGTGTGGAGTCGTTGCCTGACCATTAGCATCACTTAATNNNAGATCGGAAGA |
| 39257_1 | TATGAGTGTGGAGTCGTTGCTCAGGAGGTAGAAGGAGCAGTTANNNAGATCGGA |
| 39257_2 | TATGAGTGTGGAGTCGTTGCTCAGGAGGTAGAAGGAGCAGTTANNNAGATCGGA |
| 41797_1 | TATGAGTGTGGAGTCGTTGCTCCAGGTTCAAGTGATTCTCCTGNNNAGATCGGAAG |
| 41797_2 | TATGAGTGTGGAGTCGTTGCTCCAGGTTCAAGTGATTCTCCTGNNNAGATCGGAAG |
| 40014_1 | TATGAGTGTGGAGTCGTTGCGGAGACTGAGGCAGGAGAATTGNNNAGATCGGAA |
| 40014_2 | TATGAGTGTGGAGTCGTTGCGGAGACTGAGGCAGGAGAATTGNNNAGATCGGAA |
| 47724_1 | TATGAGTGTGGAGTCGTTGCGTTTCACTGTGTTAGCCAGGATGNNNAGATCGGAAG |
| 47724_2 | TATGAGTGTGGAGTCGTTGCGTTTCACTGTGTTAGCCAGGATGNNNAGATCGGAAG |
| 37175_1 | TATGAGTGTGGAGTCGTTGCTGATTCCAAGCCCAAACAAATCCNNNAGATCGGAAG |
| 37175_2 | TATGAGTGTGGAGTCGTTGCTGATTCCAAGCCCAAACAAATCCNNNAGATCGGAAG |
| 43072_1 | TATGAGTGTGGAGTCGTTGCTTTTTCAGAGACAGGGTCTTGCTNNNAGATCGGAAG |
| 43072_2 | TATGAGTGTGGAGTCGTTGCTTTTTCAGAGACAGGGTCTTGCTNNNAGATCGGAAG |
| 52421_1 | TATGAGTGTGGAGTCGTTGCTGCCTCTGAAGAAGTCTGAATCCNNNAGATCGGAAG |
| 52421_2 | TATGAGTGTGGAGTCGTTGCTGCCTCTGAAGAAGTCTGAATCCNNNAGATCGGAAG |
| 42278_1 | TATGAGTGTGGAGTCGTTGCTCCTGTCTATGTAGGAGGAAGCANNNAGATCGGAA |
| 42278_2 | TATGAGTGTGGAGTCGTTGCTCCTGTCTATGTAGGAGGAAGCANNNAGATCGGAA |
| 45946_1 | TATGAGTGTGGAGTCGTTGCTTTTTAGTAGAGACGGGGTTGCANNNAGATCGGAA |
| 45946_2 | TATGAGTGTGGAGTCGTTGCTTTTTAGTAGAGACGGGGTTGCANNNAGATCGGAA |
| 37842_1 | TATGAGTGTGGAGTCGTTGCAGAGGAAGAAGAGGAAGGAAGGANNNAGATCGG |
| 37842_2 | TATGAGTGTGGAGTCGTTGCAGAGGAAGAAGAGGAAGGAAGGANNNAGATCGG |
| 41150_1 | TATGAGTGTGGAGTCGTTGCAAGTTAGTCGCCATGCTCTCTACNNNAGATCGGAAG |
| 41150_2 | TATGAGTGTGGAGTCGTTGCAAGTTAGTCGCCATGCTCTCTACNNNAGATCGGAAG |
| 34917_1 | TATGAGTGTGGAGTCGTTGCTTTCACGCCATTCTCCTGTCTCNNNAGATCGGAAGAG |
| 34917_2 | TATGAGTGTGGAGTCGTTGCTTTCACGCCATTCTCCTGTCTCNNNAGATCGGAAGAG |

| | |
|---|---|
| 32449_1 | TATGAGTGTGGAGTCGTTGCCAAGTAGCTGGGACTACAGGCNNNAGATCGGAAGA |
| 32449_2 | TATGAGTGTGGAGTCGTTGCCAAGTAGCTGGGACTACAGGCNNNAGATCGGAAGA |
| 52291_1 | TATGAGTGTGGAGTCGTTGCTAGTCGATATCTTGTGCCAGCTCNNNAGATCGGAAG |
| 52291_2 | TATGAGTGTGGAGTCGTTGCTAGTCGATATCTTGTGCCAGCTCNNNAGATCGGAAG |
| 51322_1 | TATGAGTGTGGAGTCGTTGCCAACAGAGCAAGACTCCATCTCANNNAGATCGGAAG |
| 51322_2 | TATGAGTGTGGAGTCGTTGCCAACAGAGCAAGACTCCATCTCANNNAGATCGGAAG |
| 31942_1 | TATGAGTGTGGAGTCGTTGCAGTTGTCACCACTCTTTCTGTGTNNNAGATCGGAAG |
| 31942_2 | TATGAGTGTGGAGTCGTTGCAGTTGTCACCACTCTTTCTGTGTNNNAGATCGGAAG |
| 44006_1 | TATGAGTGTGGAGTCGTTGCATGCAGGGTCACTATGAAAGAGGNNNAGATCGGAA |
| 44006_2 | TATGAGTGTGGAGTCGTTGCATGCAGGGTCACTATGAAAGAGGNNNAGATCGGAA |
| 35791_1 | TATGAGTGTGGAGTCGTTGCTGGTAGTTTCTTTTGCTGTGCAGNNNAGATCGGAAG |
| 35791_2 | TATGAGTGTGGAGTCGTTGCTGGTAGTTTCTTTTGCTGTGCAGNNNAGATCGGAAG |
| 29369_1 | TATGAGTGTGGAGTCGTTGCGATGTCAGGATGAAGTGGGAAGTNNNAGATCGGA |
| 29369_2 | TATGAGTGTGGAGTCGTTGCGATGTCAGGATGAAGTGGGAAGTNNNAGATCGGA |
| 44815_1 | TATGAGTGTGGAGTCGTTGCATTCCTGAAAGGTGGAAAGCAGANNNAGATCGGAA |
| 44815_2 | TATGAGTGTGGAGTCGTTGCATTCCTGAAAGGTGGAAAGCAGANNNAGATCGGAA |
| 38237_1 | TATGAGTGTGGAGTCGTTGCCAACTGCCCTTTCTTACATGCAGNNNAGATCGGAAG |
| 38237_2 | TATGAGTGTGGAGTCGTTGCCAACTGCCCTTTCTTACATGCAGNNNAGATCGGAAG |
| 29170_1 | TATGAGTGTGGAGTCGTTGCTGCTCAATCTCTCTCTCTCTCTCNNNAGATCGGAAGA |
| 29170_2 | TATGAGTGTGGAGTCGTTGCTGCTCAATCTCTCTCTCTCTCTCNNNAGATCGGAAGA |
| 31640_1 | TATGAGTGTGGAGTCGTTGCACATCCAGTCTTCTCAGTTGCAANNNAGATCGGAAG |
| 31640_2 | TATGAGTGTGGAGTCGTTGCACATCCAGTCTTCTCAGTTGCAANNNAGATCGGAAG |
| 44412_1 | TATGAGTGTGGAGTCGTTGCCCCTCATTTTCGCTTTTGCCTTANNNAGATCGGAAGA |
| 44412_2 | TATGAGTGTGGAGTCGTTGCCCCTCATTTTCGCTTTTGCCTTANNNAGATCGGAAGA |
| 10674_1 | TATGAGTGTGGAGTCGTTGCGCCAGGAAGGAAAAAGTTCTTGCNNNAGATCGGAA |
| 10674_2 | TATGAGTGTGGAGTCGTTGCGCCAGGAAGGAAAAAGTTCTTGCNNNAGATCGGAA |
| 28873_1 | TATGAGTGTGGAGTCGTTGCTAGGCATGAAGAGTAGCACTGTGNNNAGATCGGAA |
| 28873_2 | TATGAGTGTGGAGTCGTTGCTAGGCATGAAGAGTAGCACTGTGNNNAGATCGGAA |
| 43616_1 | TATGAGTGTGGAGTCGTTGCTCCTCAAGAAAACCCTCAAGGAGNNNAGATCGGAAG |
| 43616_2 | TATGAGTGTGGAGTCGTTGCTCCTCAAGAAAACCCTCAAGGAGNNNAGATCGGAAG |
| 53319_1 | TATGAGTGTGGAGTCGTTGCTTCAAGTGATTCTCCTGCCTCAGNNNAGATCGGAAG |
| 53319_2 | TATGAGTGTGGAGTCGTTGCTTCAAGTGATTCTCCTGCCTCAGNNNAGATCGGAAG |
| 8513_1 | TATGAGTGTGGAGTCGTTGCCAGAGACTTGACATGGAAACCTNNNAGATCGGAAG |
| 8513_2 | TATGAGTGTGGAGTCGTTGCCAGAGACTTGACATGGAAACCTNNNAGATCGGAAG |
| 31672_1 | TATGAGTGTGGAGTCGTTGCTCCAGTTTTAGCCCCTTACTTCCNNNAGATCGGAAGA |
| 31672_2 | TATGAGTGTGGAGTCGTTGCTCCAGTTTTAGCCCCTTACTTCCNNNAGATCGGAAGA |
| 48208_1 | TATGAGTGTGGAGTCGTTGCCCTACAGGGTTCAAGCGATTCTCNNNAGATCGGAAG |
| 48208_2 | TATGAGTGTGGAGTCGTTGCCCTACAGGGTTCAAGCGATTCTCNNNAGATCGGAAG |
| 37476_1 | TATGAGTGTGGAGTCGTTGCGACAACCCAAATGATGTCCATGGNNNAGATCGGAAG |
| 37476_2 | TATGAGTGTGGAGTCGTTGCGACAACCCAAATGATGTCCATGGNNNAGATCGGAAG |
| 81_1 | TATGAGTGTGGAGTCGTTGCTGCCACACACACTTCCACATAANNNAGATCGGAAGA |
| 81_2 | TATGAGTGTGGAGTCGTTGCTGCCACACACACTTCCACATAANNNAGATCGGAAGA |
| 52608_1 | TATGAGTGTGGAGTCGTTGCAAAGATTCTGAAAAGCAACGGCCNNNAGATCGGAA |

| | |
|---|---|
| 52608_2 | TATGAGTGTGGAGTCGTTGCAAAGATTCTGAAAAGCAACGGCCNNNAGATCGGAA |
| 8525_1 | TATGAGTGTGGAGTCGTTGCCACCTCAAGGAAGCAATCATCTAANNNAGATCGGAA |
| 8525_2 | TATGAGTGTGGAGTCGTTGCCACCTCAAGGAAGCAATCATCTAANNNAGATCGGAA |
| 51146_1 | TATGAGTGTGGAGTCGTTGCGAGGAGAGAGACAGGAGGTTGTANNNAGATCGGA |
| 51146_2 | TATGAGTGTGGAGTCGTTGCGAGGAGAGAGACAGGAGGTTGTANNNAGATCGGA |
| 12043_1 | TATGAGTGTGGAGTCGTTGCGGTGGTCCTGCTTGTGTCATNNNAGATCGGAAGAG |
| 12043_2 | TATGAGTGTGGAGTCGTTGCGGTGGTCCTGCTTGTGTCATNNNAGATCGGAAGAG |
| 52223_1 | TATGAGTGTGGAGTCGTTGCGAGATGGGGGTTCACTATCTTGGNNNAGATCGGAA |
| 52223_2 | TATGAGTGTGGAGTCGTTGCGAGATGGGGGTTCACTATCTTGGNNNAGATCGGAA |
| 3155_1 | TATGAGTGTGGAGTCGTTGCACAATATTCATGGTGGCTGGCAGNNNAGATCGGAA |
| 3155_2 | TATGAGTGTGGAGTCGTTGCACAATATTCATGGTGGCTGGCAGNNNAGATCGGAA |
| 32247_1 | TATGAGTGTGGAGTCGTTGCCTGAGGTAGTGTTGGAGACAGTCNNNAGATCGGAA |
| 32247_2 | TATGAGTGTGGAGTCGTTGCCTGAGGTAGTGTTGGAGACAGTCNNNAGATCGGAA |
| 53203_1 | TATGAGTGTGGAGTCGTTGCGTGAAGGCCCAGAGAAGATGATANNNAGATCGGAA |
| 53203_2 | TATGAGTGTGGAGTCGTTGCGTGAAGGCCCAGAGAAGATGATANNNAGATCGGAA |
| 36871_1 | TATGAGTGTGGAGTCGTTGCGGCAAGAAAATGAAACAAAGGGCNNNAGATCGGA |
| 36871_2 | TATGAGTGTGGAGTCGTTGCGGCAAGAAAATGAAACAAAGGGCNNNAGATCGGA |
| 36319_1 | TATGAGTGTGGAGTCGTTGCCATGGTTTCATGTGCCTGTAGTTNNNAGATCGGAAG |
| 36319_2 | TATGAGTGTGGAGTCGTTGCCATGGTTTCATGTGCCTGTAGTTNNNAGATCGGAAG |
| 12782_1 | TATGAGTGTGGAGTCGTTGCAAAATCACAGTGAAATTGAAATGNNNAGATCGGAA |
| 12782_2 | TATGAGTGTGGAGTCGTTGCAAAATCACAGTGAAATTGAAATGNNNAGATCGGAA |
| 31685_1 | TATGAGTGTGGAGTCGTTGCATTCTCCTCTGCTTTTAGGGTGGNNNAGATCGGAAG |
| 31685_2 | TATGAGTGTGGAGTCGTTGCATTCTCCTCTGCTTTTAGGGTGGNNNAGATCGGAAG |
| 3050_1 | TATGAGTGTGGAGTCGTTGCTGCTTCTTGGCAGGACTTTGAAANNNAGATCGGAAG |
| 3050_2 | TATGAGTGTGGAGTCGTTGCTGCTTCTTGGCAGGACTTTGAAANNNAGATCGGAAG |
| 37912_1 | TATGAGTGTGGAGTCGTTGCCCTGTAATCCCAGCTACTAGGGANNNAGATCGGAAG |
| 37912_2 | TATGAGTGTGGAGTCGTTGCCCTGTAATCCCAGCTACTAGGGANNNAGATCGGAAG |
| 40736_1 | TATGAGTGTGGAGTCGTTGCATACCCTTTACCAAGTTTCCCCCNNNAGATCGGAAGA |
| 40736_2 | TATGAGTGTGGAGTCGTTGCATACCCTTTACCAAGTTTCCCCCNNNAGATCGGAAGA |
| 10047_1 | TATGAGTGTGGAGTCGTTGCCGTGACACCCATTGCTCTAGATTNNNAGATCGGAAG |
| 10047_2 | TATGAGTGTGGAGTCGTTGCCGTGACACCCATTGCTCTAGATTNNNAGATCGGAAG |
| 47321_1 | TATGAGTGTGGAGTCGTTGCCATGAGCAAATGTGGTGTCCAACNNNAGATCGGAA |
| 47321_2 | TATGAGTGTGGAGTCGTTGCCATGAGCAAATGTGGTGTCCAACNNNAGATCGGAA |
| 38061_1 | TATGAGTGTGGAGTCGTTGCTGGGGGATTCTGTATACAGCATGNNNAGATCGGAA |
| 38061_2 | TATGAGTGTGGAGTCGTTGCTGGGGGATTCTGTATACAGCATGNNNAGATCGGAA |
| 13470_1 | TATGAGTGTGGAGTCGTTGCGCTATTAAAGACACCATTTTCTTNNNAGATCGGAAG |
| 13470_2 | TATGAGTGTGGAGTCGTTGCGCTATTAAAGACACCATTTTCTTNNNAGATCGGAAG |
| 46734_1 | TATGAGTGTGGAGTCGTTGCTTCACACCATTCTCCTGCCTCNNNAGATCGGAAGAGC |
| 46734_2 | TATGAGTGTGGAGTCGTTGCTTCACACCATTCTCCTGCCTCNNNAGATCGGAAGAGC |
| 36101_1 | TATGAGTGTGGAGTCGTTGCCCCGAAACATTTCACATCAGCAANNNAGATCGGAAG |
| 36101_2 | TATGAGTGTGGAGTCGTTGCCCCGAAACATTTCACATCAGCAANNNAGATCGGAAG |
| 36643_1 | TATGAGTGTGGAGTCGTTGCGACTGAACAAACTCCTTCTTGGCNNNAGATCGGAAG |
| 36643_2 | TATGAGTGTGGAGTCGTTGCGACTGAACAAACTCCTTCTTGGCNNNAGATCGGAAG |

33424_1 TATGAGTGTGGAGTCGTTGCAGAGTGATAGAGCAAATGTGGCANNNAGATCGGAA
33424_2 TATGAGTGTGGAGTCGTTGCAGAGTGATAGAGCAAATGTGGCANNNAGATCGGAA
48914_1 TATGAGTGTGGAGTCGTTGCGATCTCAGCTCACTGCAACCTCNNNAGATCGGAAGAC
48914_2 TATGAGTGTGGAGTCGTTGCGATCTCAGCTCACTGCAACCTCNNNAGATCGGAAGAC
34532_1 TATGAGTGTGGAGTCGTTGCCCTGAGGAAAGCCTTGCTCTTTANNNAGATCGGAAG
34532_2 TATGAGTGTGGAGTCGTTGCCCTGAGGAAAGCCTTGCTCTTTANNNAGATCGGAAG
48425_1 TATGAGTGTGGAGTCGTTGCGTTCATACCATTCTCCTGCCTCANNNAGATCGGAAGA
48425_2 TATGAGTGTGGAGTCGTTGCGTTCATACCATTCTCCTGCCTCANNNAGATCGGAAGA
35602_1 TATGAGTGTGGAGTCGTTGCGGCCCTTATCCTGTAACCACTACNNNAGATCGGAAGA
35602_2 TATGAGTGTGGAGTCGTTGCGGCCCTTATCCTGTAACCACTACNNNAGATCGGAAGA
13796_1 TATGAGTGTGGAGTCGTTGCATTCTTTTGAAGTTAATTCTCCCNNNAGATCGGAAGA
13796_2 TATGAGTGTGGAGTCGTTGCATTCTTTTGAAGTTAATTCTCCCNNNAGATCGGAAGA
10272_1 TATGAGTGTGGAGTCGTTGCTGCTTGGCAATGGATGGAATAACNNNAGATCGGAAG
10272_2 TATGAGTGTGGAGTCGTTGCTGCTTGGCAATGGATGGAATAACNNNAGATCGGAAG
40280_1 TATGAGTGTGGAGTCGTTGCACAAAATATGATGCTCCAAAGGAAANNNAGATCGGA
40280_2 TATGAGTGTGGAGTCGTTGCACAAAATATGATGCTCCAAAGGAAANNNAGATCGGA
52062_1 TATGAGTGTGGAGTCGTTGCCTTGAGCTTTCTCCGGCTTTCTCNNNAGATCGGAAGA
52062_2 TATGAGTGTGGAGTCGTTGCCTTGAGCTTTCTCCGGCTTTCTCNNNAGATCGGAAGA
32267_1 TATGAGTGTGGAGTCGTTGCGTACAACAGTCAACACTCCTTGCNNNAGATCGGAAGA
32267_2 TATGAGTGTGGAGTCGTTGCGTACAACAGTCAACACTCCTTGCNNNAGATCGGAAGA
44513_1 TATGAGTGTGGAGTCGTTGCCTTCTTCTCGTCGCTGTCCAANNNAGATCGGAAGAGC
44513_2 TATGAGTGTGGAGTCGTTGCCTTCTTCTCGTCGCTGTCCAANNNAGATCGGAAGAGC
28847_1 TATGAGTGTGGAGTCGTTGCTTTTTTCCAGGGAGGGATTCCATNNNAGATCGGAAG
28847_2 TATGAGTGTGGAGTCGTTGCTTTTTTCCAGGGAGGGATTCCATNNNAGATCGGAAG
41567_1 TATGAGTGTGGAGTCGTTGCATTCTCCTGCCTCAGTCTCCATANNNAGATCGGAAGA
41567_2 TATGAGTGTGGAGTCGTTGCATTCTCCTGCCTCAGTCTCCATANNNAGATCGGAAGA
51779_1 TATGAGTGTGGAGTCGTTGCAGATGGCACATGGAGTGAAAGATNNNAGATCGGAA
51779_2 TATGAGTGTGGAGTCGTTGCAGATGGCACATGGAGTGAAAGATNNNAGATCGGAA
10512_1 TATGAGTGTGGAGTCGTTGCAAAAACAACCTTCTTCCCCCAGTNNNAGATCGGAAGA
10512_2 TATGAGTGTGGAGTCGTTGCAAAAACAACCTTCTTCCCCCAGTNNNAGATCGGAAGA
50123_1 TATGAGTGTGGAGTCGTTGCGGGGCTTTAGATGGGAGAGTTTANNNAGATCGGAA
50123_2 TATGAGTGTGGAGTCGTTGCGGGGCTTTAGATGGGAGAGTTTANNNAGATCGGAA
12188_1 TATGAGTGTGGAGTCGTTGCCCCTCATCCCCCTTCCATGATAANNNAGATCGGAAGAC
12188_2 TATGAGTGTGGAGTCGTTGCCCCTCATCCCCCTTCCATGATAANNNAGATCGGAAGAC
34873_1 TATGAGTGTGGAGTCGTTGCCAGTGGTGCAATCTCAATCTCAGNNNAGATCGGAAG
34873_2 TATGAGTGTGGAGTCGTTGCCAGTGGTGCAATCTCAATCTCAGNNNAGATCGGAAG
13498_1 TATGAGTGTGGAGTCGTTGCAAAAACATTTGCATGGTTATGTANNNAGATCGGAAG
13498_2 TATGAGTGTGGAGTCGTTGCAAAAACATTTGCATGGTTATGTANNNAGATCGGAAG
52573_1 TATGAGTGTGGAGTCGTTGCAAGCCTGTACACAGATGGAGAAGNNNAGATCGGAA
52573_2 TATGAGTGTGGAGTCGTTGCAAGCCTGTACACAGATGGAGAAGNNNAGATCGGAA
10553_1 TATGAGTGTGGAGTCGTTGCTGGCTACAGTGGAAGTTTCTTTTNNNAGATCGGAAG
10553_2 TATGAGTGTGGAGTCGTTGCTGGCTACAGTGGAAGTTTCTTTTNNNAGATCGGAAG
32207_1 TATGAGTGTGGAGTCGTTGCGTAAATTCTTCACACCCCTGTGCNNNAGATCGGAAGA

| | |
|---|---|
| 32207_2 | TATGAGTGTGGAGTCGTTGCGTAAATTCTTCACACCCCTGTGCNNNAGATCGGAAGA |
| 41640_1 | TATGAGTGTGGAGTCGTTGCCTCCTGGGCTGAAGTGATTCTCNNNAGATCGGAAGA |
| 41640_2 | TATGAGTGTGGAGTCGTTGCCTCCTGGGCTGAAGTGATTCTCNNNAGATCGGAAGA |
| 32791_1 | TATGAGTGTGGAGTCGTTGCCTGAGTAGCTCGGATTACTGGTGNNNAGATCGGAA |
| 32791_2 | TATGAGTGTGGAGTCGTTGCCTGAGTAGCTCGGATTACTGGTGNNNAGATCGGAA |
| 5512_1 | TATGAGTGTGGAGTCGTTGCATACTGCTTGATGCTCTCTGCCNNNAGATCGGAAGA |
| 5512_2 | TATGAGTGTGGAGTCGTTGCATACTGCTTGATGCTCTCTGCCNNNAGATCGGAAGA |
| 39824_1 | TATGAGTGTGGAGTCGTTGCCTCATCACACCCGAGTCAAGAAANNNAGATCGGAAG |
| 39824_2 | TATGAGTGTGGAGTCGTTGCCTCATCACACCCGAGTCAAGAAANNNAGATCGGAAG |
| 7818_1 | TATGAGTGTGGAGTCGTTGCCTATCTGGGCTCATTGCTGGTTNNNAGATCGGAAGA |
| 7818_2 | TATGAGTGTGGAGTCGTTGCCTATCTGGGCTCATTGCTGGTTNNNAGATCGGAAGA |
| 50827_1 | TATGAGTGTGGAGTCGTTGCGTGGCATGTTGAAGTCTGGTTACNNNAGATCGGAA |
| 50827_2 | TATGAGTGTGGAGTCGTTGCGTGGCATGTTGAAGTCTGGTTACNNNAGATCGGAA |
| 33662_1 | TATGAGTGTGGAGTCGTTGCCTGAGTAGCTGGGACTACAGATGNNNAGATCGGAA |
| 33662_2 | TATGAGTGTGGAGTCGTTGCCTGAGTAGCTGGGACTACAGATGNNNAGATCGGAA |
| 39752_1 | TATGAGTGTGGAGTCGTTGCAACCACTAAGATGGGGAGAAACCNNNAGATCGGAA |
| 39752_2 | TATGAGTGTGGAGTCGTTGCAACCACTAAGATGGGGAGAAACCNNNAGATCGGAA |
| 51370_1 | TATGAGTGTGGAGTCGTTGCCCTGGACAGAAATAGTGGCTGATNNNAGATCGGAA |
| 51370_2 | TATGAGTGTGGAGTCGTTGCCCTGGACAGAAATAGTGGCTGATNNNAGATCGGAA |
| 43618_1 | TATGAGTGTGGAGTCGTTGCGTCTCTGTCACGTGAATCCCTAGNNNAGATCGGAAG |
| 43618_2 | TATGAGTGTGGAGTCGTTGCGTCTCTGTCACGTGAATCCCTAGNNNAGATCGGAAG |
| 42817_1 | TATGAGTGTGGAGTCGTTGCCCTCACAGGAGCCACAATATACANNNAGATCGGAAG |
| 42817_2 | TATGAGTGTGGAGTCGTTGCCCTCACAGGAGCCACAATATACANNNAGATCGGAAG |
| 47950_1 | TATGAGTGTGGAGTCGTTGCCAATCTCGGCTCACTGTAACCTCNNNAGATCGGAAGA |
| 47950_2 | TATGAGTGTGGAGTCGTTGCCAATCTCGGCTCACTGTAACCTCNNNAGATCGGAAGA |
| 13726_1 | TATGAGTGTGGAGTCGTTGCATGATAAAAGTGGCCTTGTTANNNAGATCGGAAGA |
| 13726_2 | TATGAGTGTGGAGTCGTTGCATGATAAAAGTGGCCTTGTTANNNAGATCGGAAGA |
| 329_1 | TATGAGTGTGGAGTCGTTGCCGGTTTGTAGGTTTCTGCAGAGAGNNNAGATCGGA |
| 329_2 | TATGAGTGTGGAGTCGTTGCCGGTTTGTAGGTTTCTGCAGAGAGNNNAGATCGGA |
| 40725_1 | TATGAGTGTGGAGTCGTTGCTGGTGAAGATGTGTAGAAAGGGANNNAGATCGGA |
| 40725_2 | TATGAGTGTGGAGTCGTTGCTGGTGAAGATGTGTAGAAAGGGANNNAGATCGGA |
| 47417_1 | TATGAGTGTGGAGTCGTTGCGTGGGGGAGAAGTGGCATTATAANNNAGATCGGA |
| 47417_2 | TATGAGTGTGGAGTCGTTGCGTGGGGGAGAAGTGGCATTATAANNNAGATCGGA |
| 44102_1 | TATGAGTGTGGAGTCGTTGCAGTCTGGAGAGATAGACCATCCGNNNAGATCGGAA |
| 44102_2 | TATGAGTGTGGAGTCGTTGCAGTCTGGAGAGATAGACCATCCGNNNAGATCGGAA |
| 50489_1 | TATGAGTGTGGAGTCGTTGCATAAACGGTCTACTGCTGTGGACNNNAGATCGGAAG |
| 50489_2 | TATGAGTGTGGAGTCGTTGCATAAACGGTCTACTGCTGTGGACNNNAGATCGGAAG |
| 51610_1 | TATGAGTGTGGAGTCGTTGCCAGATACTCCCTGCTTCCTCTCTNNNAGATCGGAAGA |
| 51610_2 | TATGAGTGTGGAGTCGTTGCCAGATACTCCCTGCTTCCTCTCTNNNAGATCGGAAGA |
| 47548_1 | TATGAGTGTGGAGTCGTTGCGAAGAACGCTGGAGGTTGATTGNNNAGATCGGAAG |
| 47548_2 | TATGAGTGTGGAGTCGTTGCGAAGAACGCTGGAGGTTGATTGNNNAGATCGGAAG |
| 40656_1 | TATGAGTGTGGAGTCGTTGCACAGAGGCAGAGTTTGGAATGATNNNAGATCGGAA |
| 40656_2 | TATGAGTGTGGAGTCGTTGCACAGAGGCAGAGTTTGGAATGATNNNAGATCGGAA |

| | |
|---|---|
| 46268_1 | TATGAGTGTGGAGTCGTTGCCAAGGCTAGTTGAAGTGGGGTATNNNAGATCGGAA |
| 46268_2 | TATGAGTGTGGAGTCGTTGCCAAGGCTAGTTGAAGTGGGGTATNNNAGATCGGAA |
| 35422_1 | TATGAGTGTGGAGTCGTTGCAGTCTCAAACTCCTGAGCTCAAGNNNAGATCGGAAG |
| 35422_2 | TATGAGTGTGGAGTCGTTGCAGTCTCAAACTCCTGAGCTCAAGNNNAGATCGGAAG |
| 43573_1 | TATGAGTGTGGAGTCGTTGCCTAGTAGCTGGGACTACTGGTGTNNNAGATCGGAA |
| 43573_2 | TATGAGTGTGGAGTCGTTGCCTAGTAGCTGGGACTACTGGTGTNNNAGATCGGAA |
| 44973_1 | TATGAGTGTGGAGTCGTTGCTGCTCATGAGAAATCCAAGTGGANNNAGATCGGAA |
| 44973_2 | TATGAGTGTGGAGTCGTTGCTGCTCATGAGAAATCCAAGTGGANNNAGATCGGAA |
| 46279_1 | TATGAGTGTGGAGTCGTTGCCGAGTAGCTGGGACTACAGGTNNNAGATCGGAAGA |
| 46279_2 | TATGAGTGTGGAGTCGTTGCCGAGTAGCTGGGACTACAGGTNNNAGATCGGAAGA |
| 30272_1 | TATGAGTGTGGAGTCGTTGCGCACAGAAGTAAGTAGACAAAGCANNNAGATCGGA |
| 30272_2 | TATGAGTGTGGAGTCGTTGCGCACAGAAGTAAGTAGACAAAGCANNNAGATCGGA |
| 40549_1 | TATGAGTGTGGAGTCGTTGCTTGAGAAGTGGGATCCAGGATTGNNNAGATCGGAA |
| 40549_2 | TATGAGTGTGGAGTCGTTGCTTGAGAAGTGGGATCCAGGATTGNNNAGATCGGAA |
| 9352_1 | TATGAGTGTGGAGTCGTTGCTACTCTTTGGCTCTGAGGAAGGGNNNAGATCGGAA |
| 9352_2 | TATGAGTGTGGAGTCGTTGCTACTCTTTGGCTCTGAGGAAGGGNNNAGATCGGAA |
| 36985_1 | TATGAGTGTGGAGTCGTTGCGCTTATGACTCCCATGACTATTGCNNNAGATCGGAA |
| 36985_2 | TATGAGTGTGGAGTCGTTGCGCTTATGACTCCCATGACTATTGCNNNAGATCGGAA |
| 38362_1 | TATGAGTGTGGAGTCGTTGCGAGGTTAGGGCAAGAGGTGTTATNNNAGATCGGA |
| 38362_2 | TATGAGTGTGGAGTCGTTGCGAGGTTAGGGCAAGAGGTGTTATNNNAGATCGGA |
| 45691_1 | TATGAGTGTGGAGTCGTTGCCAAGTGATTCTTCTGCTTCAGCCNNNAGATCGGAAG |
| 45691_2 | TATGAGTGTGGAGTCGTTGCCAAGTGATTCTTCTGCTTCAGCCNNNAGATCGGAAG |
| 48489_1 | TATGAGTGTGGAGTCGTTGCTTCCCTGAATTCCTCCCCAACTCNNNAGATCGGAAGA |
| 48489_2 | TATGAGTGTGGAGTCGTTGCTTCCCTGAATTCCTCCCCAACTCNNNAGATCGGAAGA |
| 51948_1 | TATGAGTGTGGAGTCGTTGCTTTGCCCAAGTACCCTCAACATANNNAGATCGGAAG |
| 51948_2 | TATGAGTGTGGAGTCGTTGCTTTGCCCAAGTACCCTCAACATANNNAGATCGGAAG |
| 31557_1 | TATGAGTGTGGAGTCGTTGCAAAGCTCTGGAGGCTACAAGTACNNNAGATCGGAA |
| 31557_2 | TATGAGTGTGGAGTCGTTGCAAAGCTCTGGAGGCTACAAGTACNNNAGATCGGAA |
| 51211_1 | TATGAGTGTGGAGTCGTTGCTGTAGATGATGGGTTGATGGGTGNNNAGATCGGA |
| 51211_2 | TATGAGTGTGGAGTCGTTGCTGTAGATGATGGGTTGATGGGTGNNNAGATCGGA |
| 40707_1 | TATGAGTGTGGAGTCGTTGCACCTAACAAGGGTCTAGTATCCAGANNNAGATCGGA |
| 40707_2 | TATGAGTGTGGAGTCGTTGCACCTAACAAGGGTCTAGTATCCAGANNNAGATCGGA |
| 42789_1 | TATGAGTGTGGAGTCGTTGCTTTGTGTGTGTGTGTGAGAGAGANNNAGATCGGAA |
| 42789_2 | TATGAGTGTGGAGTCGTTGCTTTGTGTGTGTGTGTGAGAGAGANNNAGATCGGAA |
| 47744_1 | TATGAGTGTGGAGTCGTTGCCCATGCCAGGCACATGACTGNNNAGATCGGAAGAG |
| 47744_2 | TATGAGTGTGGAGTCGTTGCCCATGCCAGGCACATGACTGNNNAGATCGGAAGAG |
| 43494_1 | TATGAGTGTGGAGTCGTTGCCTCTGGAGTAGCTGGGATTACAGNNNAGATCGGAA |
| 43494_2 | TATGAGTGTGGAGTCGTTGCCTCTGGAGTAGCTGGGATTACAGNNNAGATCGGAA |
| 41586_1 | TATGAGTGTGGAGTCGTTGCCTGGTCTTGAACTCTTGGACTCANNNAGATCGGAAG |
| 41586_2 | TATGAGTGTGGAGTCGTTGCCTGGTCTTGAACTCTTGGACTCANNNAGATCGGAAG |
| 43815_1 | TATGAGTGTGGAGTCGTTGCGGAACCGATAGATTCCGGATGAANNNAGATCGGAA |
| 43815_2 | TATGAGTGTGGAGTCGTTGCGGAACCGATAGATTCCGGATGAANNNAGATCGGAA |
| 37814_1 | TATGAGTGTGGAGTCGTTGCCTCACTGGATGGGCTCAATAGTANNNAGATCGGAA |

| | |
|---|---|
| 37814_2 | TATGAGTGTGGAGTCGTTGCCTCACTGGATGGGCTCAATAGTANNNAGATCGGAAG |
| 31722_1 | TATGAGTGTGGAGTCGTTGCATTGCAGTTGGAGGTTAGCCAAGNNNAGATCGGAA |
| 31722_2 | TATGAGTGTGGAGTCGTTGCATTGCAGTTGGAGGTTAGCCAAGNNNAGATCGGAA |
| 39396_1 | TATGAGTGTGGAGTCGTTGCAACTGGACTCCTAACTTTTCCCCNNNAGATCGGAAGA |
| 39396_2 | TATGAGTGTGGAGTCGTTGCAACTGGACTCCTAACTTTTCCCCNNNAGATCGGAAGA |
| 41753_1 | TATGAGTGTGGAGTCGTTGCAAGTGATTCTCCTGTCTCAACCCNNNAGATCGGAAGA |
| 41753_2 | TATGAGTGTGGAGTCGTTGCAAGTGATTCTCCTGTCTCAACCCNNNAGATCGGAAGA |
| 52804_1 | TATGAGTGTGGAGTCGTTGCAGCCCTCAGTTTATGGTCACTTTNNNAGATCGGAAGA |
| 52804_2 | TATGAGTGTGGAGTCGTTGCAGCCCTCAGTTTATGGTCACTTTNNNAGATCGGAAGA |
| 2719_1 | TATGAGTGTGGAGTCGTTGCAGGTGCTCTGTTCTATCCCAACTNNNAGATCGGAAGA |
| 2719_2 | TATGAGTGTGGAGTCGTTGCAGGTGCTCTGTTCTATCCCAACTNNNAGATCGGAAGA |
| 53166_1 | TATGAGTGTGGAGTCGTTGCTCTTTAGACCTCCCTCCAAAAGCNNNAGATCGGAAGA |
| 53166_2 | TATGAGTGTGGAGTCGTTGCTCTTTAGACCTCCCTCCAAAAGCNNNAGATCGGAAGA |
| 44050_1 | TATGAGTGTGGAGTCGTTGCTTCAGTGATGAAGGTGTTTCTGTNNNAGATCGGAAG |
| 44050_2 | TATGAGTGTGGAGTCGTTGCTTCAGTGATGAAGGTGTTTCTGTNNNAGATCGGAAG |
| 49786_1 | TATGAGTGTGGAGTCGTTGCTCCTCCACAACACTGTTCTTTCANNNAGATCGGAAGA |
| 49786_2 | TATGAGTGTGGAGTCGTTGCTCCTCCACAACACTGTTCTTTCANNNAGATCGGAAGA |
| 52328_1 | TATGAGTGTGGAGTCGTTGCAAGGATACATGCTCACCCTTTGTNNNAGATCGGAAG |
| 52328_2 | TATGAGTGTGGAGTCGTTGCAAGGATACATGCTCACCCTTTGTNNNAGATCGGAAG |
| 13654_1 | TATGAGTGTGGAGTCGTTGCTAAGTAAAGAATCTGGTCAATTGNNNAGATCGGAAG |
| 13654_2 | TATGAGTGTGGAGTCGTTGCTAAGTAAAGAATCTGGTCAATTGNNNAGATCGGAAG |
| 38530_1 | TATGAGTGTGGAGTCGTTGCGAGGCAAGAGAATGGCATGAACNNNAGATCGGAAG |
| 38530_2 | TATGAGTGTGGAGTCGTTGCGAGGCAAGAGAATGGCATGAACNNNAGATCGGAAG |
| 46142_1 | TATGAGTGTGGAGTCGTTGCTGTCACCCAGGCTAGAGTACAGNNNAGATCGGAAGA |
| 46142_2 | TATGAGTGTGGAGTCGTTGCTGTCACCCAGGCTAGAGTACAGNNNAGATCGGAAGA |
| 31963_1 | TATGAGTGTGGAGTCGTTGCTGAGAGAATGGGTATCAGTGGTTNNNAGATCGGAA |
| 31963_2 | TATGAGTGTGGAGTCGTTGCTGAGAGAATGGGTATCAGTGGTTNNNAGATCGGAA |
| 36112_1 | TATGAGTGTGGAGTCGTTGCCCGGACTGAACCAATGTACTTCTNNNAGATCGGAAG |
| 36112_2 | TATGAGTGTGGAGTCGTTGCCCGGACTGAACCAATGTACTTCTNNNAGATCGGAAG |
| 52506_1 | TATGAGTGTGGAGTCGTTGCTGGCTTCTGCTGTGGCTATATTGNNNAGATCGGAAG |
| 52506_2 | TATGAGTGTGGAGTCGTTGCTGGCTTCTGCTGTGGCTATATTGNNNAGATCGGAAG |
| 29343_1 | TATGAGTGTGGAGTCGTTGCTGTCAGCAATTTCCAGACCCTATNNNAGATCGGAAGA |
| 29343_2 | TATGAGTGTGGAGTCGTTGCTGTCAGCAATTTCCAGACCCTATNNNAGATCGGAAGA |
| 10129_1 | TATGAGTGTGGAGTCGTTGCAGGTATGGCAGCTTACCACTGTANNNAGATCGGAAG |
| 10129_2 | TATGAGTGTGGAGTCGTTGCAGGTATGGCAGCTTACCACTGTANNNAGATCGGAAG |
| 52072_1 | TATGAGTGTGGAGTCGTTGCGGCATTTGTTGGATTTCTCACCANNNAGATCGGAAG |
| 52072_2 | TATGAGTGTGGAGTCGTTGCGGCATTTGTTGGATTTCTCACCANNNAGATCGGAAG |
| 34793_1 | TATGAGTGTGGAGTCGTTGCTCAGTCTCTTGAGTAGCTGGGATNNNAGATCGGAAG |
| 34793_2 | TATGAGTGTGGAGTCGTTGCTCAGTCTCTTGAGTAGCTGGGATNNNAGATCGGAAG |
| 40087_1 | TATGAGTGTGGAGTCGTTGCTACAAAATTAGCTGGGCTGGTCGNNNAGATCGGAA |
| 40087_2 | TATGAGTGTGGAGTCGTTGCTACAAAATTAGCTGGGCTGGTCGNNNAGATCGGAA |
| 41836_1 | TATGAGTGTGGAGTCGTTGCTCAAGCGATTCTTCTGTTTCAGCNNNAGATCGGAAGA |
| 41836_2 | TATGAGTGTGGAGTCGTTGCTCAAGCGATTCTTCTGTTTCAGCNNNAGATCGGAAGA |

| | |
|---|---|
| 38913_1 | TATGAGTGTGGAGTCGTTGCCTCCTCAACAACCCATTCCATGANNNAGATCGGAAGA |
| 38913_2 | TATGAGTGTGGAGTCGTTGCCTCCTCAACAACCCATTCCATGANNNAGATCGGAAGA |
| 48354_1 | TATGAGTGTGGAGTCGTTGCCGTCTCCCAAGTTGAAGCAATTCNNNAGATCGGAAG |
| 48354_2 | TATGAGTGTGGAGTCGTTGCCGTCTCCCAAGTTGAAGCAATTCNNNAGATCGGAAG |
| 2922_1 | TATGAGTGTGGAGTCGTTGCTATTCCCTGTTGCTCAAGCGATTNNNAGATCGGAAG |
| 2922_2 | TATGAGTGTGGAGTCGTTGCTATTCCCTGTTGCTCAAGCGATTNNNAGATCGGAAG |
| 48869_1 | TATGAGTGTGGAGTCGTTGCCCAATAGAGCGAGATCATCACCANNNAGATCGGAAG |
| 48869_2 | TATGAGTGTGGAGTCGTTGCCCAATAGAGCGAGATCATCACCANNNAGATCGGAAG |
| 33751_1 | TATGAGTGTGGAGTCGTTGCCCTTTAGGCAGGGGAAACTCAAANNNAGATCGGAA |
| 33751_2 | TATGAGTGTGGAGTCGTTGCCCTTTAGGCAGGGGAAACTCAAANNNAGATCGGAA |
| 31255_1 | TATGAGTGTGGAGTCGTTGCATCACTTAGGTCAGCAGACACAGNNNAGATCGGAAG |
| 31255_2 | TATGAGTGTGGAGTCGTTGCATCACTTAGGTCAGCAGACACAGNNNAGATCGGAAG |
| 50767_1 | TATGAGTGTGGAGTCGTTGCTTGCTCAAGGGTTGTTATCGGAANNNAGATCGGAA |
| 50767_2 | TATGAGTGTGGAGTCGTTGCTTGCTCAAGGGTTGTTATCGGAANNNAGATCGGAA |
| 43316_1 | TATGAGTGTGGAGTCGTTGCTGAAATCCAGTGGCGTGATCTCNNNAGATCGGAAG |
| 43316_2 | TATGAGTGTGGAGTCGTTGCTGAAATCCAGTGGCGTGATCTCNNNAGATCGGAAG |
| 47930_1 | TATGAGTGTGGAGTCGTTGCTTCAAGTGATTCTCCTGCCTCAGNNNAGATCGGAAG |
| 47930_2 | TATGAGTGTGGAGTCGTTGCTTCAAGTGATTCTCCTGCCTCAGNNNAGATCGGAAG |
| 47329_1 | TATGAGTGTGGAGTCGTTGCGCTTCTCACCCCTTAGGAGTTTTNNNAGATCGGAAG |
| 47329_2 | TATGAGTGTGGAGTCGTTGCGCTTCTCACCCCTTAGGAGTTTTNNNAGATCGGAAG |
| 52286_1 | TATGAGTGTGGAGTCGTTGCCCTGTGGCTTTTCTAGATGCATGNNNAGATCGGAAG |
| 52286_2 | TATGAGTGTGGAGTCGTTGCCCTGTGGCTTTTCTAGATGCATGNNNAGATCGGAAG |
| 40311_1 | TATGAGTGTGGAGTCGTTGCTCAAACACTTCAACCAAAAGACAGANNNAGATCGGA |
| 40311_2 | TATGAGTGTGGAGTCGTTGCTCAAACACTTCAACCAAAAGACAGANNNAGATCGGA |
| 9373_1 | TATGAGTGTGGAGTCGTTGCAGAGTAGAGACAAAGCAGGGAGGNNNAGATCGGA |
| 9373_2 | TATGAGTGTGGAGTCGTTGCAGAGTAGAGACAAAGCAGGGAGGNNNAGATCGGA |
| 50019_1 | TATGAGTGTGGAGTCGTTGCTAGAAATGCGCTTTGAGCAGATGNNNAGATCGGAA |
| 50019_2 | TATGAGTGTGGAGTCGTTGCTAGAAATGCGCTTTGAGCAGATGNNNAGATCGGAA |
| 43188_1 | TATGAGTGTGGAGTCGTTGCAATTTGCCTCACTCTCATGACCTNNNAGATCGGAAGA |
| 43188_2 | TATGAGTGTGGAGTCGTTGCAATTTGCCTCACTCTCATGACCTNNNAGATCGGAAGA |
| 2070_1 | TATGAGTGTGGAGTCGTTGCTGACCACTTTCAAAGCAAAGACANNNAGATCGGAAG |
| 2070_2 | TATGAGTGTGGAGTCGTTGCTGACCACTTTCAAAGCAAAGACANNNAGATCGGAAG |
| 48913_1 | TATGAGTGTGGAGTCGTTGCAGATGGAGTCTCCTTCTTTTGCCNNNAGATCGGAAG |
| 48913_2 | TATGAGTGTGGAGTCGTTGCAGATGGAGTCTCCTTCTTTTGCCNNNAGATCGGAAG |
| 39570_1 | TATGAGTGTGGAGTCGTTGCATGGAATAGTGATTACCAGGGGCNNNAGATCGGAA |
| 39570_2 | TATGAGTGTGGAGTCGTTGCATGGAATAGTGATTACCAGGGGCNNNAGATCGGAA |
| 31984_1 | TATGAGTGTGGAGTCGTTGCACAAGTTGTATTGGTGGAGGACANNNAGATCGGAA |
| 31984_2 | TATGAGTGTGGAGTCGTTGCACAAGTTGTATTGGTGGAGGACANNNAGATCGGAA |
| 5549_1 | TATGAGTGTGGAGTCGTTGCCAGGTGTGGGGTAATTCTGTTCCNNNAGATCGGAA |
| 5549_2 | TATGAGTGTGGAGTCGTTGCCAGGTGTGGGGTAATTCTGTTCCNNNAGATCGGAA |
| 50357_1 | TATGAGTGTGGAGTCGTTGCTGTCCCTAAAGTCTTTGGTAATGGTNNNAGATCGGA |
| 50357_2 | TATGAGTGTGGAGTCGTTGCTGTCCCTAAAGTCTTTGGTAATGGTNNNAGATCGGA |
| 35542_1 | TATGAGTGTGGAGTCGTTGCGTTACACCGTGTTAGCCAGGATANNNAGATCGGAA |

| | |
|---|---|
| 35542_2 | TATGAGTGTGGAGTCGTTGCGTTACACCGTGTTAGCCAGGATANNNAGATCGGAA |
| 44864_1 | TATGAGTGTGGAGTCGTTGCCTTTGAGATGGGCCAGTAAGGTCNNNAGATCGGAA |
| 44864_2 | TATGAGTGTGGAGTCGTTGCCTTTGAGATGGGCCAGTAAGGTCNNNAGATCGGAA |
| 49942_1 | TATGAGTGTGGAGTCGTTGCCAAATGCAGAAGCGGGACTTAGNNNAGATCGGAAG |
| 49942_2 | TATGAGTGTGGAGTCGTTGCCAAATGCAGAAGCGGGACTTAGNNNAGATCGGAAG |
| 49549_1 | TATGAGTGTGGAGTCGTTGCCAGTAACACAATCTCGGCTCACTNNNAGATCGGAAG |
| 49549_2 | TATGAGTGTGGAGTCGTTGCCAGTAACACAATCTCGGCTCACTNNNAGATCGGAAG |
| 11018_1 | TATGAGTGTGGAGTCGTTGCCCCACAAGCAGGAAACCTGAAANNNAGATCGGAAG |
| 11018_2 | TATGAGTGTGGAGTCGTTGCCCCACAAGCAGGAAACCTGAAANNNAGATCGGAAG |
| 33225_1 | TATGAGTGTGGAGTCGTTGCAGTAGATGGGATTACAGGCACACNNNAGATCGGAA |
| 33225_2 | TATGAGTGTGGAGTCGTTGCAGTAGATGGGATTACAGGCACACNNNAGATCGGAA |
| 12811_1 | TATGAGTGTGGAGTCGTTGCAACAGATTTTACAGCAAAGATTTNNNAGATCGGAAG |
| 12811_2 | TATGAGTGTGGAGTCGTTGCAACAGATTTTACAGCAAAGATTTNNNAGATCGGAAG |
| 44259_1 | TATGAGTGTGGAGTCGTTGCGTCTGGTTTTGAAAAGAGGGCAANNNAGATCGGAA |
| 44259_2 | TATGAGTGTGGAGTCGTTGCGTCTGGTTTTGAAAAGAGGGCAANNNAGATCGGAA |
| 10449_1 | TATGAGTGTGGAGTCGTTGCCATACCTCACCTTGCCTCCTCTANNNAGATCGGAAGA |
| 10449_2 | TATGAGTGTGGAGTCGTTGCCATACCTCACCTTGCCTCCTCTANNNAGATCGGAAGA |
| 29986_1 | TATGAGTGTGGAGTCGTTGCTGAGTGTCCAATACAGTGATGGTNNNAGATCGGAA |
| 29986_2 | TATGAGTGTGGAGTCGTTGCTGAGTGTCCAATACAGTGATGGTNNNAGATCGGAA |
| 29523_1 | TATGAGTGTGGAGTCGTTGCTTTCCAGGACCCTACCAAACAGANNNAGATCGGAAG |
| 29523_2 | TATGAGTGTGGAGTCGTTGCTTTCCAGGACCCTACCAAACAGANNNAGATCGGAAG |
| 12547_1 | TATGAGTGTGGAGTCGTTGCGTTATGAAGGTAAATGGATTCACNNNAGATCGGAA |
| 12547_2 | TATGAGTGTGGAGTCGTTGCGTTATGAAGGTAAATGGATTCACNNNAGATCGGAA |
| 29976_1 | TATGAGTGTGGAGTCGTTGCTAGCTGCCTAAAGTCGTGTGTAANNNAGATCGGAA |
| 29976_2 | TATGAGTGTGGAGTCGTTGCTAGCTGCCTAAAGTCGTGTGTAANNNAGATCGGAA |
| 45632_1 | TATGAGTGTGGAGTCGTTGCCCTTAGCATCCTAAGTAGCTGGGNNNAGATCGGAAG |
| 45632_2 | TATGAGTGTGGAGTCGTTGCCCTTAGCATCCTAAGTAGCTGGGNNNAGATCGGAAG |
| 43750_1 | TATGAGTGTGGAGTCGTTGCGAAAACAGCAGAGGGACTAAGGANNNAGATCGGA |
| 43750_2 | TATGAGTGTGGAGTCGTTGCGAAAACAGCAGAGGGACTAAGGANNNAGATCGGA |
| 46887_1 | TATGAGTGTGGAGTCGTTGCCATCACCATGCCCTGCTAATTTTNNNAGATCGGAAGA |
| 46887_2 | TATGAGTGTGGAGTCGTTGCCATCACCATGCCCTGCTAATTTTNNNAGATCGGAAGA |
| 46524_1 | TATGAGTGTGGAGTCGTTGCTTCACACCATTCTCCTGCCTCNNNAGATCGGAAGAGC |
| 46524_2 | TATGAGTGTGGAGTCGTTGCTTCACACCATTCTCCTGCCTCNNNAGATCGGAAGAGC |
| 46780_1 | TATGAGTGTGGAGTCGTTGCCGCCCGGCTAATTTTAGTAGAGANNNAGATCGGAA |
| 46780_2 | TATGAGTGTGGAGTCGTTGCCGCCCGGCTAATTTTAGTAGAGANNNAGATCGGAA |
| 45457_1 | TATGAGTGTGGAGTCGTTGCAGAAGCAGATATCTGGAAACGCTNNNAGATCGGAA |
| 45457_2 | TATGAGTGTGGAGTCGTTGCAGAAGCAGATATCTGGAAACGCTNNNAGATCGGAA |
| 28951_1 | TATGAGTGTGGAGTCGTTGCTGAAAACACAATTCCCTGCTCACNNNAGATCGGAAG |
| 28951_2 | TATGAGTGTGGAGTCGTTGCTGAAAACACAATTCCCTGCTCACNNNAGATCGGAAG |
| 30433_1 | TATGAGTGTGGAGTCGTTGCGGACTGTAGCGGATGGTGATAAANNNAGATCGGAA |
| 30433_2 | TATGAGTGTGGAGTCGTTGCGGACTGTAGCGGATGGTGATAAANNNAGATCGGAA |
| 10584_1 | TATGAGTGTGGAGTCGTTGCTCAGGGAGATTCATTCGGCTTTGNNNAGATCGGAA |
| 10584_2 | TATGAGTGTGGAGTCGTTGCTCAGGGAGATTCATTCGGCTTTGNNNAGATCGGAA |

| | |
|---|---|
| 48383_1 | TATGAGTGTGGAGTCGTTGCCTTCCATCTTAGCCTCCCAAAGTNNNAGATCGGAAGA |
| 48383_2 | TATGAGTGTGGAGTCGTTGCCTTCCATCTTAGCCTCCCAAAGTNNNAGATCGGAAGA |
| 44202_1 | TATGAGTGTGGAGTCGTTGCAATGTATACCTGGGCAGATGAGGNNNAGATCGGAA |
| 44202_2 | TATGAGTGTGGAGTCGTTGCAATGTATACCTGGGCAGATGAGGNNNAGATCGGAA |
| 45184_1 | TATGAGTGTGGAGTCGTTGCTTAGGGTGTAAAGCTGTTACCCCNNNAGATCGGAAG |
| 45184_2 | TATGAGTGTGGAGTCGTTGCTTAGGGTGTAAAGCTGTTACCCCNNNAGATCGGAAG |
| 32357_1 | TATGAGTGTGGAGTCGTTGCCACTGTTTTCGGACCTTCCATGNNNAGATCGGAAGA |
| 32357_2 | TATGAGTGTGGAGTCGTTGCCACTGTTTTCGGACCTTCCATGNNNAGATCGGAAGA |
| 49061_1 | TATGAGTGTGGAGTCGTTGCTGTTTCCTGATTTTAGGGGAGCTNNNAGATCGGAAG |
| 49061_2 | TATGAGTGTGGAGTCGTTGCTGTTTCCTGATTTTAGGGGAGCTNNNAGATCGGAAG |
| 53009_1 | TATGAGTGTGGAGTCGTTGCCCTGTGAGCAAGAGGGTGTAGNNNAGATCGGAAGA |
| 53009_2 | TATGAGTGTGGAGTCGTTGCCCTGTGAGCAAGAGGGTGTAGNNNAGATCGGAAGA |
| 44322_1 | TATGAGTGTGGAGTCGTTGCGGCACTAGATTGGAGATCTAGGCNNNAGATCGGAA |
| 44322_2 | TATGAGTGTGGAGTCGTTGCGGCACTAGATTGGAGATCTAGGCNNNAGATCGGAA |
| 33023_1 | TATGAGTGTGGAGTCGTTGCCATTCTCCTGCCTCAGCCTCNNNAGATCGGAAGAGCA |
| 33023_2 | TATGAGTGTGGAGTCGTTGCCATTCTCCTGCCTCAGCCTCNNNAGATCGGAAGAGCA |
| 41594_1 | TATGAGTGTGGAGTCGTTGCGATTTGGGATAGCTTTGAGGCACNNNAGATCGGAA |
| 41594_2 | TATGAGTGTGGAGTCGTTGCGATTTGGGATAGCTTTGAGGCACNNNAGATCGGAA |
| 47784_1 | TATGAGTGTGGAGTCGTTGCGATTCAAGCAATTCTCCTGCCTCNNNAGATCGGAAGA |
| 47784_2 | TATGAGTGTGGAGTCGTTGCGATTCAAGCAATTCTCCTGCCTCNNNAGATCGGAAGA |
| 31502_1 | TATGAGTGTGGAGTCGTTGCCTCATGTTGACATTTCGAGGCAGNNNAGATCGGAAG |
| 31502_2 | TATGAGTGTGGAGTCGTTGCCTCATGTTGACATTTCGAGGCAGNNNAGATCGGAAG |
| 13516_1 | TATGAGTGTGGAGTCGTTGCGAAAAATAAACATGCACGATTATNNNAGATCGGAAG |
| 13516_2 | TATGAGTGTGGAGTCGTTGCGAAAAATAAACATGCACGATTATNNNAGATCGGAAG |
| 43929_1 | TATGAGTGTGGAGTCGTTGCACACACACACACACACACAAATTNNNAGATCGGAAG |
| 43929_2 | TATGAGTGTGGAGTCGTTGCACACACACACACACACACAAATTNNNAGATCGGAAG |
| 41865_1 | TATGAGTGTGGAGTCGTTGCCCTGTGGCTTCTCTACTCAGAAANNNAGATCGGAAG |
| 41865_2 | TATGAGTGTGGAGTCGTTGCCCTGTGGCTTCTCTACTCAGAAANNNAGATCGGAAG |
| 10866_1 | TATGAGTGTGGAGTCGTTGCTGTTCCAGGATCCTTCTGGTCTTNNNAGATCGGAAG |
| 10866_2 | TATGAGTGTGGAGTCGTTGCTGTTCCAGGATCCTTCTGGTCTTNNNAGATCGGAAG |
| 29297_1 | TATGAGTGTGGAGTCGTTGCTCTGTGCCCACTAAGCTACTTTCNNNAGATCGGAAGA |
| 29297_2 | TATGAGTGTGGAGTCGTTGCTCTGTGCCCACTAAGCTACTTTCNNNAGATCGGAAGA |
| 835_1 | TATGAGTGTGGAGTCGTTGCAAAAGGGAGCCCAAACATCCCNNNAGATCGGAAGA |
| 835_2 | TATGAGTGTGGAGTCGTTGCAAAAGGGAGCCCAAACATCCCNNNAGATCGGAAGA |
| 3041_1 | TATGAGTGTGGAGTCGTTGCCGAAGCTTTTCCCTCATTTCTGCNNNAGATCGGAAGA |
| 3041_2 | TATGAGTGTGGAGTCGTTGCCGAAGCTTTTCCCTCATTTCTGCNNNAGATCGGAAGA |
| 12504_1 | TATGAGTGTGGAGTCGTTGCAACAGTTATTTCTAGTGTTGTTGNNNAGATCGGAAG |
| 12504_2 | TATGAGTGTGGAGTCGTTGCAACAGTTATTTCTAGTGTTGTTGNNNAGATCGGAAG |
| 39441_1 | TATGAGTGTGGAGTCGTTGCCCCCCCACCATTATACTTCTCAGNNNAGATCGGAAGA |
| 39441_2 | TATGAGTGTGGAGTCGTTGCCCCCCCACCATTATACTTCTCAGNNNAGATCGGAAGA |
| 44550_1 | TATGAGTGTGGAGTCGTTGCTACACACACACACACACAAATGTNNNAGATCGGAAG |
| 44550_2 | TATGAGTGTGGAGTCGTTGCTACACACACACACACACAAATGTNNNAGATCGGAAG |
| 29742_1 | TATGAGTGTGGAGTCGTTGCCTCACACAGTGGGTACTAAGCTANNNAGATCGGAAG |

| | |
|---|---|
| 29742_2 | TATGAGTGTGGAGTCGTTGCCTCACACAGTGGGTACTAAGCTANNNAGATCGGAAG |
| 13454_1 | TATGAGTGTGGAGTCGTTGCGGATTCTTTAAAGGGGTTGTNNNAGATCGGAAGAG |
| 13454_2 | TATGAGTGTGGAGTCGTTGCGGATTCTTTAAAGGGGTTGTNNNAGATCGGAAGAG |
| 34992_1 | TATGAGTGTGGAGTCGTTGCCTCAAGTGATTCTCCTGCCTCAGNNNAGATCGGAAG |
| 34992_2 | TATGAGTGTGGAGTCGTTGCCTCAAGTGATTCTCCTGCCTCAGNNNAGATCGGAAG |
| 47793_1 | TATGAGTGTGGAGTCGTTGCCTCTACCTCCTGACCTTGTGATCNNNAGATCGGAAGA |
| 47793_2 | TATGAGTGTGGAGTCGTTGCCTCTACCTCCTGACCTTGTGATCNNNAGATCGGAAGA |
| 41912_1 | TATGAGTGTGGAGTCGTTGCGAATTGCTGGGTCATCAGTTAGCNNNAGATCGGAA |
| 41912_2 | TATGAGTGTGGAGTCGTTGCGAATTGCTGGGTCATCAGTTAGCNNNAGATCGGAA |
| 41885_1 | TATGAGTGTGGAGTCGTTGCTCCCTCTTTTTGCTTTGGAAACCNNNAGATCGGAAGA |
| 41885_2 | TATGAGTGTGGAGTCGTTGCTCCCTCTTTTTGCTTTGGAAACCNNNAGATCGGAAGA |
| 41667_1 | TATGAGTGTGGAGTCGTTGCTCGAACTTTTGACCTCAGGTGATNNNAGATCGGAAG |
| 41667_2 | TATGAGTGTGGAGTCGTTGCTCGAACTTTTGACCTCAGGTGATNNNAGATCGGAAG |
| 30417_1 | TATGAGTGTGGAGTCGTTGCTTCAGTAGTTCACATTGCTGTGTTTNNNAGATCGGA |
| 30417_2 | TATGAGTGTGGAGTCGTTGCTTCAGTAGTTCACATTGCTGTGTTTNNNAGATCGGA |
| 43059_1 | TATGAGTGTGGAGTCGTTGCTTGGGTTCAACTAACTTGCTGGANNNAGATCGGAAG |
| 43059_2 | TATGAGTGTGGAGTCGTTGCTTGGGTTCAACTAACTTGCTGGANNNAGATCGGAAG |
| 10513_1 | TATGAGTGTGGAGTCGTTGCATGCAGCACTTGGTAAAACCAGTNNNAGATCGGAA |
| 10513_2 | TATGAGTGTGGAGTCGTTGCATGCAGCACTTGGTAAAACCAGTNNNAGATCGGAA |
| 35125_1 | TATGAGTGTGGAGTCGTTGCCATGTAGCTGGGATTACAGGCATNNNAGATCGGAA |
| 35125_2 | TATGAGTGTGGAGTCGTTGCCATGTAGCTGGGATTACAGGCATNNNAGATCGGAA |
| 52470_1 | TATGAGTGTGGAGTCGTTGCGTACACTAGAGGGAGAAATGGGCNNNAGATCGGA |
| 52470_2 | TATGAGTGTGGAGTCGTTGCGTACACTAGAGGGAGAAATGGGCNNNAGATCGGA |
| 36897_1 | TATGAGTGTGGAGTCGTTGCACCCCAAAGAAGTAGAAAGATGGGNNNAGATCGGA |
| 36897_2 | TATGAGTGTGGAGTCGTTGCACCCCAAAGAAGTAGAAAGATGGGNNNAGATCGGA |
| 29344_1 | TATGAGTGTGGAGTCGTTGCACCAATGATGACAGCTCTGGAATNNNAGATCGGAA |
| 29344_2 | TATGAGTGTGGAGTCGTTGCACCAATGATGACAGCTCTGGAATNNNAGATCGGAA |
| 39443_1 | TATGAGTGTGGAGTCGTTGCACAAAGTCTTGCTCAGTCACTCANNNAGATCGGAAG |
| 39443_2 | TATGAGTGTGGAGTCGTTGCACAAAGTCTTGCTCAGTCACTCANNNAGATCGGAAG |
| 44108_1 | TATGAGTGTGGAGTCGTTGCTCCGAAATACCCTGTGTCAGATGNNNAGATCGGAAG |
| 44108_2 | TATGAGTGTGGAGTCGTTGCTCCGAAATACCCTGTGTCAGATGNNNAGATCGGAAG |
| 52311_1 | TATGAGTGTGGAGTCGTTGCAAACTTCGCACCCCGTCGAANNNAGATCGGAAGAGC |
| 52311_2 | TATGAGTGTGGAGTCGTTGCAAACTTCGCACCCCGTCGAANNNAGATCGGAAGAGC |
| 45687_1 | TATGAGTGTGGAGTCGTTGCAAGTAGCTGGGATTACAGGCACNNNAGATCGGAAG |
| 45687_2 | TATGAGTGTGGAGTCGTTGCAAGTAGCTGGGATTACAGGCACNNNAGATCGGAAG |
| 50354_1 | TATGAGTGTGGAGTCGTTGCATAGTCCTTCCTTCATGCACTGGNNNAGATCGGAAG |
| 50354_2 | TATGAGTGTGGAGTCGTTGCATAGTCCTTCCTTCATGCACTGGNNNAGATCGGAAG |
| 8878_1 | TATGAGTGTGGAGTCGTTGCAACTCGCCTGTGTAAAGCATACANNNAGATCGGAAG |
| 8878_2 | TATGAGTGTGGAGTCGTTGCAACTCGCCTGTGTAAAGCATACANNNAGATCGGAAG |
| 31798_1 | TATGAGTGTGGAGTCGTTGCATATGTGTGTGTGTGTGTGTGNNNAGATCGGAA |
| 31798_2 | TATGAGTGTGGAGTCGTTGCATATGTGTGTGTGTGTGTGTGNNNAGATCGGAA |
| 52576_1 | TATGAGTGTGGAGTCGTTGCCAACTAAGCCATGCTCCACAATGNNNAGATCGGAAG |
| 52576_2 | TATGAGTGTGGAGTCGTTGCCAACTAAGCCATGCTCCACAATGNNNAGATCGGAAG |

| | |
|---|---|
| 37520_1 | TATGAGTGTGGAGTCGTTGCAAAACATGATTTTGTGGGTGGGGNNNAGATCGGAA |
| 37520_2 | TATGAGTGTGGAGTCGTTGCAAAACATGATTTTGTGGGTGGGGNNNAGATCGGAA |
| 37519_1 | TATGAGTGTGGAGTCGTTGCGGGCATTGTTCAAGTTCTTTTCCNNNAGATCGGAAG |
| 37519_2 | TATGAGTGTGGAGTCGTTGCGGGCATTGTTCAAGTTCTTTTCCNNNAGATCGGAAG |
| 52178_1 | TATGAGTGTGGAGTCGTTGCAGGAGACATCCCATGGACAAAAANNNAGATCGGAA |
| 52178_2 | TATGAGTGTGGAGTCGTTGCAGGAGACATCCCATGGACAAAAANNNAGATCGGAA |
| 6621_1 | TATGAGTGTGGAGTCGTTGCGGGACAACAGTTGGACACCATGNNNAGATCGGAAG |
| 6621_2 | TATGAGTGTGGAGTCGTTGCGGGACAACAGTTGGACACCATGNNNAGATCGGAAG |
| 34969_1 | TATGAGTGTGGAGTCGTTGCTAGCTGGGATTATAGGTGTGTGCNNNAGATCGGAA |
| 34969_2 | TATGAGTGTGGAGTCGTTGCTAGCTGGGATTATAGGTGTGTGCNNNAGATCGGAA |
| 12793_1 | TATGAGTGTGGAGTCGTTGCATTTTACCATGATATACCCTGAANNNAGATCGGAAG |
| 12793_2 | TATGAGTGTGGAGTCGTTGCATTTTACCATGATATACCCTGAANNNAGATCGGAAG |
| 34760_1 | TATGAGTGTGGAGTCGTTGCCGGCCTCTCTCCTTCTTTCTATGNNNAGATCGGAAGA |
| 34760_2 | TATGAGTGTGGAGTCGTTGCCGGCCTCTCTCCTTCTTTCTATGNNNAGATCGGAAGA |
| 43817_1 | TATGAGTGTGGAGTCGTTGCCACTTTTGGATGGGGTTGTTTGTNNNAGATCGGAA |
| 43817_2 | TATGAGTGTGGAGTCGTTGCCACTTTTGGATGGGGTTGTTTGTNNNAGATCGGAA |
| 29610_1 | TATGAGTGTGGAGTCGTTGCTCAGGTTAGTGCAAACAATCCCANNNAGATCGGAAG |
| 29610_2 | TATGAGTGTGGAGTCGTTGCTCAGGTTAGTGCAAACAATCCCANNNAGATCGGAAG |
| 12168_1 | TATGAGTGTGGAGTCGTTGCGACTGACATTGGACACGGAACAAGNNNAGATCGGA |
| 12168_2 | TATGAGTGTGGAGTCGTTGCGACTGACATTGGACACGGAACAAGNNNAGATCGGA |
| 42659_1 | TATGAGTGTGGAGTCGTTGCAGGATGATTTCTCAAGTGGGTCTNNNAGATCGGAA |
| 42659_2 | TATGAGTGTGGAGTCGTTGCAGGATGATTTCTCAAGTGGGTCTNNNAGATCGGAA |
| 34796_1 | TATGAGTGTGGAGTCGTTGCCTAACGCCTATAATCCCAGCACTNNNAGATCGGAAG |
| 34796_2 | TATGAGTGTGGAGTCGTTGCCTAACGCCTATAATCCCAGCACTNNNAGATCGGAAG |
| 12923_1 | TATGAGTGTGGAGTCGTTGCCACTGATACTAAGTTCAGAAACTNNNAGATCGGAAG |
| 12923_2 | TATGAGTGTGGAGTCGTTGCCACTGATACTAAGTTCAGAAACTNNNAGATCGGAAG |
| 40056_1 | TATGAGTGTGGAGTCGTTGCGACCTCACCACTCTGATGTTCTTNNNAGATCGGAAGA |
| 40056_2 | TATGAGTGTGGAGTCGTTGCGACCTCACCACTCTGATGTTCTTNNNAGATCGGAAGA |
| 30412_1 | TATGAGTGTGGAGTCGTTGCGAGAGAGAGAAAGTGTCAATCNNNAGATCGGAAG |
| 30412_2 | TATGAGTGTGGAGTCGTTGCGAGAGAGAGAAAGTGTCAATCNNNAGATCGGAAG |
| 43948_1 | TATGAGTGTGGAGTCGTTGCTTTTCTCTCTCTCTCTCCCCCTTNNNAGATCGGAAGAG |
| 43948_2 | TATGAGTGTGGAGTCGTTGCTTTTCTCTCTCTCTCTCCCCCTTNNNAGATCGGAAGAG |
| 38142_1 | TATGAGTGTGGAGTCGTTGCCCTCTAGTCCCACCTACTTGTGNNNAGATCGGAAGA |
| 38142_2 | TATGAGTGTGGAGTCGTTGCCCTCTAGTCCCACCTACTTGTGNNNAGATCGGAAGA |
| 12711_1 | TATGAGTGTGGAGTCGTTGCGATAGTTTGTCTTTCGAATAGTCNNNAGATCGGAAG |
| 12711_2 | TATGAGTGTGGAGTCGTTGCGATAGTTTGTCTTTCGAATAGTCNNNAGATCGGAAG |
| 44078_1 | TATGAGTGTGGAGTCGTTGCGAGGCTCTCATGTTTTTGGCTTTNNNAGATCGGAAG |
| 44078_2 | TATGAGTGTGGAGTCGTTGCGAGGCTCTCATGTTTTTGGCTTTNNNAGATCGGAAG |
| 38439_1 | TATGAGTGTGGAGTCGTTGCAAAGGACCAAAAAAGGAGCAACCNNNAGATCGGAA |
| 38439_2 | TATGAGTGTGGAGTCGTTGCAAAGGACCAAAAAAGGAGCAACCNNNAGATCGGAA |
| 10282_1 | TATGAGTGTGGAGTCGTTGCGTCTGAGTTCTAGGGAGCAAGGANNNAGATCGGAA |
| 10282_2 | TATGAGTGTGGAGTCGTTGCGTCTGAGTTCTAGGGAGCAAGGANNNAGATCGGAA |
| 8302_1 | TATGAGTGTGGAGTCGTTGCTGCTGTGGCTAGAACTTCCAAAAANNNAGATCGGAA |

| | |
|---|---|
| 8302_2 | TATGAGTGTGGAGTCGTTGCTGCTGTGGCTAGAACTTCCAAAAANNNAGATCGGAA |
| 32242_1 | TATGAGTGTGGAGTCGTTGCAGCTGAAGGATTTACGTTACCCANNNAGATCGGAAG |
| 32242_2 | TATGAGTGTGGAGTCGTTGCAGCTGAAGGATTTACGTTACCCANNNAGATCGGAAG |
| 52532_1 | TATGAGTGTGGAGTCGTTGCAGGGAGAAGGGAAAATGAGTGACNNNAGATCGGA |
| 52532_2 | TATGAGTGTGGAGTCGTTGCAGGGAGAAGGGAAAATGAGTGACNNNAGATCGGA |
| 10654_1 | TATGAGTGTGGAGTCGTTGCAACCCTCGGTAACTGTGAATGAGNNNAGATCGGAA |
| 10654_2 | TATGAGTGTGGAGTCGTTGCAACCCTCGGTAACTGTGAATGAGNNNAGATCGGAA |
| 46904_1 | TATGAGTGTGGAGTCGTTGCCTCCTGAGTAGCTGGGATTTCAGNNNAGATCGGAA |
| 46904_2 | TATGAGTGTGGAGTCGTTGCCTCCTGAGTAGCTGGGATTTCAGNNNAGATCGGAA |
| 29589_1 | TATGAGTGTGGAGTCGTTGCTTTTATCTCCTCCCACACCTTCGNNNAGATCGGAAGA |
| 29589_2 | TATGAGTGTGGAGTCGTTGCTTTTATCTCCTCCCACACCTTCGNNNAGATCGGAAGA |
| 13620_1 | TATGAGTGTGGAGTCGTTGCCTAATAGATTACCTTTATCGCCTNNNAGATCGGAAGA |
| 13620_2 | TATGAGTGTGGAGTCGTTGCCTAATAGATTACCTTTATCGCCTNNNAGATCGGAAGA |
| 52169_1 | TATGAGTGTGGAGTCGTTGCCCCTAGCAGAAGTTCTCCATGAGNNNAGATCGGAAG |
| 52169_2 | TATGAGTGTGGAGTCGTTGCCCCTAGCAGAAGTTCTCCATGAGNNNAGATCGGAAG |
| 37511_1 | TATGAGTGTGGAGTCGTTGCCATCTACTGGGAGTCTTAGCCAGNNNAGATCGGAAG |
| 37511_2 | TATGAGTGTGGAGTCGTTGCCATCTACTGGGAGTCTTAGCCAGNNNAGATCGGAAG |
| 51129_1 | TATGAGTGTGGAGTCGTTGCACTCCTTCAACCTCCCTTTTCTCNNNAGATCGGAAGAG |
| 51129_2 | TATGAGTGTGGAGTCGTTGCACTCCTTCAACCTCCCTTTTCTCNNNAGATCGGAAGAG |
| 32558_1 | TATGAGTGTGGAGTCGTTGCGAGACCAAGGCTCACTCTGTTGNNNAGATCGGAAGA |
| 32558_2 | TATGAGTGTGGAGTCGTTGCGAGACCAAGGCTCACTCTGTTGNNNAGATCGGAAGA |
| 35167_1 | TATGAGTGTGGAGTCGTTGCGGAATATTCCCTATTTCCACGGANNNAGATCGGAAG |
| 35167_2 | TATGAGTGTGGAGTCGTTGCGGAATATTCCCTATTTCCACGGANNNAGATCGGAAG |
| 10858_1 | TATGAGTGTGGAGTCGTTGCTGCTCAACACACAGAAGGATAAANNNAGATCGGAA |
| 10858_2 | TATGAGTGTGGAGTCGTTGCTGCTCAACACACAGAAGGATAAANNNAGATCGGAA |
| 827_1 | TATGAGTGTGGAGTCGTTGCTTTTTCTTCCCCAGTCCCTCANNNAGATCGGAAGAG |
| 827_2 | TATGAGTGTGGAGTCGTTGCTTTTTCTTCCCCAGTCCCTCANNNAGATCGGAAGAG |
| 47586_1 | TATGAGTGTGGAGTCGTTGCGCTGTTAGTCTGATGGGGACTTTNNNAGATCGGAA |
| 47586_2 | TATGAGTGTGGAGTCGTTGCGCTGTTAGTCTGATGGGGACTTTNNNAGATCGGAA |
| 38092_1 | TATGAGTGTGGAGTCGTTGCCAACTCGAGAACATGGTCAGAGANNNAGATCGGAA |
| 38092_2 | TATGAGTGTGGAGTCGTTGCCAACTCGAGAACATGGTCAGAGANNNAGATCGGAA |
| 41065_1 | TATGAGTGTGGAGTCGTTGCCTGCAACCAAGAACTTCCAACAGNNNAGATCGGAAG |
| 41065_2 | TATGAGTGTGGAGTCGTTGCCTGCAACCAAGAACTTCCAACAGNNNAGATCGGAAG |
| 9664_1 | TATGAGTGTGGAGTCGTTGCTGTTATGGATCAGCAAGAAGGCCNNNAGATCGGAA |
| 9664_2 | TATGAGTGTGGAGTCGTTGCTGTTATGGATCAGCAAGAAGGCCNNNAGATCGGAA |
| 36467_1 | TATGAGTGTGGAGTCGTTGCGATGAAAGTGCCTCCCCATCAGNNNAGATCGGAAGA |
| 36467_2 | TATGAGTGTGGAGTCGTTGCGATGAAAGTGCCTCCCCATCAGNNNAGATCGGAAGA |
| 12276_1 | TATGAGTGTGGAGTCGTTGCTGCTAGGCTCAGCTCTTGTTTCNNNAGATCGGAAGA |
| 12276_2 | TATGAGTGTGGAGTCGTTGCTGCTAGGCTCAGCTCTTGTTTCNNNAGATCGGAAGA |
| 45055_1 | TATGAGTGTGGAGTCGTTGCGAATCCAAGGAAACACTGGCATGNNNAGATCGGAA |
| 45055_2 | TATGAGTGTGGAGTCGTTGCGAATCCAAGGAAACACTGGCATGNNNAGATCGGAA |
| 43465_1 | TATGAGTGTGGAGTCGTTGCGAGTAGCTGGGATTACAGGCATGNNNAGATCGGAA |
| 43465_2 | TATGAGTGTGGAGTCGTTGCGAGTAGCTGGGATTACAGGCATGNNNAGATCGGAA |

| | |
|---|---|
| 39845_1 | TATGAGTGTGGAGTCGTTGCTGGAAACACATCTCATGCTCATGNNNAGATCGGAAG |
| 39845_2 | TATGAGTGTGGAGTCGTTGCTGGAAACACATCTCATGCTCATGNNNAGATCGGAAG |
| 42692_1 | TATGAGTGTGGAGTCGTTGCGGAGGTGAAAGATCTCTGCAAGANNNAGATCGGAA |
| 42692_2 | TATGAGTGTGGAGTCGTTGCGGAGGTGAAAGATCTCTGCAAGANNNAGATCGGAA |
| 36298_1 | TATGAGTGTGGAGTCGTTGCACAGATCATCCAGAGGCAAAACTNNNAGATCGGAA |
| 36298_2 | TATGAGTGTGGAGTCGTTGCACAGATCATCCAGAGGCAAAACTNNNAGATCGGAA |
| 39991_1 | TATGAGTGTGGAGTCGTTGCAAATTAGCTGCCTGTAGTCCCAANNNAGATCGGAAG |
| 39991_2 | TATGAGTGTGGAGTCGTTGCAAATTAGCTGCCTGTAGTCCCAANNNAGATCGGAAG |
| 48557_1 | TATGAGTGTGGAGTCGTTGCCTGTAGAGTGTTGGTGTATCCCCNNNAGATCGGAAG |
| 48557_2 | TATGAGTGTGGAGTCGTTGCCTGTAGAGTGTTGGTGTATCCCCNNNAGATCGGAAG |
| 32354_1 | TATGAGTGTGGAGTCGTTGCTTACTGCAACTTCCACCTCCTGNNNAGATCGGAAGAG |
| 32354_2 | TATGAGTGTGGAGTCGTTGCTTACTGCAACTTCCACCTCCTGNNNAGATCGGAAGAG |
| 5412_1 | TATGAGTGTGGAGTCGTTGCACTTGATCCCTCTCCTCCATCTTNNNAGATCGGAAGA |
| 5412_2 | TATGAGTGTGGAGTCGTTGCACTTGATCCCTCTCCTCCATCTTNNNAGATCGGAAGA |
| 4778_1 | TATGAGTGTGGAGTCGTTGCAACTGGCTCAGGCATGAAAACAANNNAGATCGGAA |
| 4778_2 | TATGAGTGTGGAGTCGTTGCAACTGGCTCAGGCATGAAAACAANNNAGATCGGAA |
| 34039_1 | TATGAGTGTGGAGTCGTTGCAGTGGTACAATCTCAGCTCACTGNNNAGATCGGAAG |
| 34039_2 | TATGAGTGTGGAGTCGTTGCAGTGGTACAATCTCAGCTCACTGNNNAGATCGGAAG |
| 30896_1 | TATGAGTGTGGAGTCGTTGCCTGAACAATGGAGGGAGACAGAANNNAGATCGGA |
| 30896_2 | TATGAGTGTGGAGTCGTTGCCTGAACAATGGAGGGAGACAGAANNNAGATCGGA |
| 43983_1 | TATGAGTGTGGAGTCGTTGCAGTTTGCCTCAGTACCCATAAGGNNNAGATCGGAAG |
| 43983_2 | TATGAGTGTGGAGTCGTTGCAGTTTGCCTCAGTACCCATAAGGNNNAGATCGGAAG |
| 41459_1 | TATGAGTGTGGAGTCGTTGCTACGTACTCGAACAAGTGACAGGNNNAGATCGGAA |
| 41459_2 | TATGAGTGTGGAGTCGTTGCTACGTACTCGAACAAGTGACAGGNNNAGATCGGAA |
| 30069_1 | TATGAGTGTGGAGTCGTTGCACCAACATAACTTTAGTCCAAGATGTNNNAGATCGG |
| 30069_2 | TATGAGTGTGGAGTCGTTGCACCAACATAACTTTAGTCCAAGATGTNNNAGATCGG |
| 52670_1 | TATGAGTGTGGAGTCGTTGCCTGGGAATGTTTACTAGCATGCGNNNAGATCGGAA |
| 52670_2 | TATGAGTGTGGAGTCGTTGCCTGGGAATGTTTACTAGCATGCGNNNAGATCGGAA |
| 34656_1 | TATGAGTGTGGAGTCGTTGCATACAAAAATTAGCCGGGCATGGNNNAGATCGGAA |
| 34656_2 | TATGAGTGTGGAGTCGTTGCATACAAAAATTAGCCGGGCATGGNNNAGATCGGAA |
| 43707_1 | TATGAGTGTGGAGTCGTTGCACAGGACTATGAGAGGGTGAAGANNNAGATCGGA |
| 43707_2 | TATGAGTGTGGAGTCGTTGCACAGGACTATGAGAGGGTGAAGANNNAGATCGGA |
| 41448_1 | TATGAGTGTGGAGTCGTTGCAAAGAAAGAGAGTGGTTCACCCTNNNAGATCGGAA |
| 41448_2 | TATGAGTGTGGAGTCGTTGCAAAGAAAGAGAGTGGTTCACCCTNNNAGATCGGAA |
| 8978_1 | TATGAGTGTGGAGTCGTTGCTACCCACACACCTCATGGATTCTNNNAGATCGGAAGA |
| 8978_2 | TATGAGTGTGGAGTCGTTGCTACCCACACACCTCATGGATTCTNNNAGATCGGAAGA |
| 43956_1 | TATGAGTGTGGAGTCGTTGCAGGGAGGAAGGGAGACAGAAATTNNNAGATCGGA |
| 43956_2 | TATGAGTGTGGAGTCGTTGCAGGGAGGAAGGGAGACAGAAATTNNNAGATCGGA |
| 40035_1 | TATGAGTGTGGAGTCGTTGCTCCAGGTTCAAGTGATTCTCCTGNNNAGATCGGAAG |
| 40035_2 | TATGAGTGTGGAGTCGTTGCTCCAGGTTCAAGTGATTCTCCTGNNNAGATCGGAAG |
| 39615_1 | TATGAGTGTGGAGTCGTTGCCAATGATGCTATCTTGGCTCACTGNNNAGATCGGAA |
| 39615_2 | TATGAGTGTGGAGTCGTTGCCAATGATGCTATCTTGGCTCACTGNNNAGATCGGAA |
| 33318_1 | TATGAGTGTGGAGTCGTTGCGCAATCATAGCTCACTGTACCCTNNNAGATCGGAAG |

| | |
|---|---|
| 33318_2 | TATGAGTGTGGAGTCGTTGCGCAATCATAGCTCACTGTACCCTNNNAGATCGGAAG |
| 46835_1 | TATGAGTGTGGAGTCGTTGCTCTCACTGGTTCATTCTTGGAGGNNNAGATCGGAAG |
| 46835_2 | TATGAGTGTGGAGTCGTTGCTCTCACTGGTTCATTCTTGGAGGNNNAGATCGGAAG |
| 29119_1 | TATGAGTGTGGAGTCGTTGCCAAGGACTGAGGAATACTGCAGTNNNAGATCGGAA |
| 29119_2 | TATGAGTGTGGAGTCGTTGCCAAGGACTGAGGAATACTGCAGTNNNAGATCGGAA |
| 42371_1 | TATGAGTGTGGAGTCGTTGCAAACACAGCACATCTGGAAAGTGNNNAGATCGGAA |
| 42371_2 | TATGAGTGTGGAGTCGTTGCAAACACAGCACATCTGGAAAGTGNNNAGATCGGAA |
| 6455_1 | TATGAGTGTGGAGTCGTTGCGCCCAGAGGTGTATAGCCAAAATNNNAGATCGGAA |
| 6455_2 | TATGAGTGTGGAGTCGTTGCGCCCAGAGGTGTATAGCCAAAATNNNAGATCGGAA |
| 45682_1 | TATGAGTGTGGAGTCGTTGCTTCAAGTGATTCTCCTGCCTCAGNNNAGATCGGAAG |
| 45682_2 | TATGAGTGTGGAGTCGTTGCTTCAAGTGATTCTCCTGCCTCAGNNNAGATCGGAAG |
| 42214_1 | TATGAGTGTGGAGTCGTTGCGTGCAGTTTTCTGTGTTTACCGANNNAGATCGGAAG |
| 42214_2 | TATGAGTGTGGAGTCGTTGCGTGCAGTTTTCTGTGTTTACCGANNNAGATCGGAAG |
| 44383_1 | TATGAGTGTGGAGTCGTTGCAAACCCAGCTACCTTTCATTTGCNNNAGATCGGAAGA |
| 44383_2 | TATGAGTGTGGAGTCGTTGCAAACCCAGCTACCTTTCATTTGCNNNAGATCGGAAGA |
| 33613_1 | TATGAGTGTGGAGTCGTTGCATGGATTCTCATTCTGTCACCCANNNAGATCGGAAG |
| 33613_2 | TATGAGTGTGGAGTCGTTGCATGGATTCTCATTCTGTCACCCANNNAGATCGGAAG |
| 41352_1 | TATGAGTGTGGAGTCGTTGCTACTCAATTCCTTGGTCTGCACANNNAGATCGGAAG |
| 41352_2 | TATGAGTGTGGAGTCGTTGCTACTCAATTCCTTGGTCTGCACANNNAGATCGGAAG |
| 38514_1 | TATGAGTGTGGAGTCGTTGCTCCAGGTTCAAGTGATTCTCCTGNNNAGATCGGAAG |
| 38514_2 | TATGAGTGTGGAGTCGTTGCTCCAGGTTCAAGTGATTCTCCTGNNNAGATCGGAAG |
| 41840_1 | TATGAGTGTGGAGTCGTTGCTTCAAGTGATTCTCCTGCCTCAGNNNAGATCGGAAG |
| 41840_2 | TATGAGTGTGGAGTCGTTGCTTCAAGTGATTCTCCTGCCTCAGNNNAGATCGGAAG |
| 53179_1 | TATGAGTGTGGAGTCGTTGCGCTACCATTTACAATGGAGCTGCNNNAGATCGGAAG |
| 53179_2 | TATGAGTGTGGAGTCGTTGCGCTACCATTTACAATGGAGCTGCNNNAGATCGGAAG |
| 13174_1 | TATGAGTGTGGAGTCGTTGCCTTTCTGGGAGTTTTCTAAATGNNNAGATCGGAAGA |
| 13174_2 | TATGAGTGTGGAGTCGTTGCCTTTCTGGGAGTTTTCTAAATGNNNAGATCGGAAGA |
| 31316_1 | TATGAGTGTGGAGTCGTTGCAGGCACAGCTAGTATTCCATTGTNNNAGATCGGAAG |
| 31316_2 | TATGAGTGTGGAGTCGTTGCAGGCACAGCTAGTATTCCATTGTNNNAGATCGGAAG |
| 42556_1 | TATGAGTGTGGAGTCGTTGCCTCCAGGGTTCATGCCATTCTCNNNAGATCGGAAGA |
| 42556_2 | TATGAGTGTGGAGTCGTTGCCTCCAGGGTTCATGCCATTCTCNNNAGATCGGAAGA |
| 36551_1 | TATGAGTGTGGAGTCGTTGCAAAAGAAATGATGCCACGTCTGGNNNAGATCGGAA |
| 36551_2 | TATGAGTGTGGAGTCGTTGCAAAAGAAATGATGCCACGTCTGGNNNAGATCGGAA |
| 44890_1 | TATGAGTGTGGAGTCGTTGCGACAGGTACCAAGGCACTAGAAANNNAGATCGGAA |
| 44890_2 | TATGAGTGTGGAGTCGTTGCGACAGGTACCAAGGCACTAGAAANNNAGATCGGAA |
| 41461_1 | TATGAGTGTGGAGTCGTTGCTTTCACAACCTACTCATCTGCCANNNAGATCGGAAGA |
| 41461_2 | TATGAGTGTGGAGTCGTTGCTTTCACAACCTACTCATCTGCCANNNAGATCGGAAGA |
| 30848_1 | TATGAGTGTGGAGTCGTTGCTTTTTGCGTTCCTTTAGAGCTGGNNNAGATCGGAAG |
| 30848_2 | TATGAGTGTGGAGTCGTTGCTTTTTGCGTTCCTTTAGAGCTGGNNNAGATCGGAAG |
| 2494_1 | TATGAGTGTGGAGTCGTTGCGGTCTGTTTCAGGTGGTTGAGAGNNNAGATCGGAA |
| 2494_2 | TATGAGTGTGGAGTCGTTGCGGTCTGTTTCAGGTGGTTGAGAGNNNAGATCGGAA |
| 33727_1 | TATGAGTGTGGAGTCGTTGCGAAACAGGAGTTCGGTTTGGAACNNNAGATCGGAA |
| 33727_2 | TATGAGTGTGGAGTCGTTGCGAAACAGGAGTTCGGTTTGGAACNNNAGATCGGAA |

| | |
|---|---|
| 2646_1 | TATGAGTGTGGAGTCGTTGCATCTTACGCAGTTATGGGGCTTGNNNAGATCGGAAG |
| 2646_2 | TATGAGTGTGGAGTCGTTGCATCTTACGCAGTTATGGGGCTTGNNNAGATCGGAAG |
| 31005_1 | TATGAGTGTGGAGTCGTTGCTTTCTCTCTGTCTTTCCGGAGTCNNNAGATCGGAAGA |
| 31005_2 | TATGAGTGTGGAGTCGTTGCTTTCTCTCTGTCTTTCCGGAGTCNNNAGATCGGAAGA |
| 33690_1 | TATGAGTGTGGAGTCGTTGCCTGTAATCCCAGCTACTCAGGAGNNNAGATCGGAAG |
| 33690_2 | TATGAGTGTGGAGTCGTTGCCTGTAATCCCAGCTACTCAGGAGNNNAGATCGGAAG |
| 37987_1 | TATGAGTGTGGAGTCGTTGCCTAGTTGAAGGGACGAATTTGCCNNNAGATCGGAA |
| 37987_2 | TATGAGTGTGGAGTCGTTGCCTAGTTGAAGGGACGAATTTGCCNNNAGATCGGAA |
| 45693_1 | TATGAGTGTGGAGTCGTTGCGGCTCAGTCATGGTCTACTGTAGNNNAGATCGGAAG |
| 45693_2 | TATGAGTGTGGAGTCGTTGCGGCTCAGTCATGGTCTACTGTAGNNNAGATCGGAAG |
| 34970_1 | TATGAGTGTGGAGTCGTTGCGAAGAGGAGTCTTGCTGTGTCCNNNAGATCGGAAG |
| 34970_2 | TATGAGTGTGGAGTCGTTGCGAAGAGGAGTCTTGCTGTGTCCNNNAGATCGGAAG |
| 2899_1 | TATGAGTGTGGAGTCGTTGCACACTGCTTCAGAGCTAGAAGGANNNAGATCGGAA |
| 2899_2 | TATGAGTGTGGAGTCGTTGCACACTGCTTCAGAGCTAGAAGGANNNAGATCGGAA |
| 31615_1 | TATGAGTGTGGAGTCGTTGCACCCACTCAGGTATGAACACATCNNNAGATCGGAAG |
| 31615_2 | TATGAGTGTGGAGTCGTTGCACCCACTCAGGTATGAACACATCNNNAGATCGGAAG |
| 43046_1 | TATGAGTGTGGAGTCGTTGCAAATAATGGCATTCGCAGCTACCNNNAGATCGGAAG |
| 43046_2 | TATGAGTGTGGAGTCGTTGCAAATAATGGCATTCGCAGCTACCNNNAGATCGGAAG |
| 47566_1 | TATGAGTGTGGAGTCGTTGCCATGATCTTGGCTCACTGCAACNNNAGATCGGAAGA |
| 47566_2 | TATGAGTGTGGAGTCGTTGCCATGATCTTGGCTCACTGCAACNNNAGATCGGAAGA |
| 37106_1 | TATGAGTGTGGAGTCGTTGCAAAAAAGCAAGGGAGCTGTGTTTNNNAGATCGGAA |
| 37106_2 | TATGAGTGTGGAGTCGTTGCAAAAAAGCAAGGGAGCTGTGTTTNNNAGATCGGAA |
| 53083_1 | TATGAGTGTGGAGTCGTTGCAAACTGTCTCTCAGACCACAGTGNNNAGATCGGAAG |
| 53083_2 | TATGAGTGTGGAGTCGTTGCAAACTGTCTCTCAGACCACAGTGNNNAGATCGGAAG |
| 36601_1 | TATGAGTGTGGAGTCGTTGCCTGATCCCGGTGTCACTATTCATNNNAGATCGGAAG |
| 36601_2 | TATGAGTGTGGAGTCGTTGCCTGATCCCGGTGTCACTATTCATNNNAGATCGGAAG |
| 47369_1 | TATGAGTGTGGAGTCGTTGCCTCCTCAGTTCACGCCATTCTCNNNAGATCGGAAGAG |
| 47369_2 | TATGAGTGTGGAGTCGTTGCCTCCTCAGTTCACGCCATTCTCNNNAGATCGGAAGAG |
| 39960_1 | TATGAGTGTGGAGTCGTTGCCCCTGTCTGCTGTTTTTCTTTCCNNNAGATCGGAAGA |
| 39960_2 | TATGAGTGTGGAGTCGTTGCCCCTGTCTGCTGTTTTTCTTTCCNNNAGATCGGAAGA |
| 39401_1 | TATGAGTGTGGAGTCGTTGCAGAAAAACGCATGCAATGGATCANNNAGATCGGAA |
| 39401_2 | TATGAGTGTGGAGTCGTTGCAGAAAAACGCATGCAATGGATCANNNAGATCGGAA |
| 45829_1 | TATGAGTGTGGAGTCGTTGCGTTCAAGAGATTCTCCTGCCTCANNNAGATCGGAAG |
| 45829_2 | TATGAGTGTGGAGTCGTTGCGTTCAAGAGATTCTCCTGCCTCANNNAGATCGGAAG |
| 36762_1 | TATGAGTGTGGAGTCGTTGCTCAGGAGACTGAGACAGGAGAATNNNAGATCGGAA |
| 36762_2 | TATGAGTGTGGAGTCGTTGCTCAGGAGACTGAGACAGGAGAATNNNAGATCGGAA |
| 37888_1 | TATGAGTGTGGAGTCGTTGCACAGCAGTAGACAAATAGAGCCCNNNAGATCGGAA |
| 37888_2 | TATGAGTGTGGAGTCGTTGCACAGCAGTAGACAAATAGAGCCCNNNAGATCGGAA |
| 42540_1 | TATGAGTGTGGAGTCGTTGCGAGAGATTCCTGACTCTGTGCTTNNNAGATCGGAAG |
| 42540_2 | TATGAGTGTGGAGTCGTTGCGAGAGATTCCTGACTCTGTGCTTNNNAGATCGGAAG |
| 8160_1 | TATGAGTGTGGAGTCGTTGCTCTTGACTGTGCTTCTTTACTCCNNNAGATCGGAAGA |
| 8160_2 | TATGAGTGTGGAGTCGTTGCTCTTGACTGTGCTTCTTTACTCCNNNAGATCGGAAGA |
| 39733_1 | TATGAGTGTGGAGTCGTTGCGCACTTTGGGAGATCTAGGTGAANNNAGATCGGAA |

| | |
|---|---|
| 39733_2 | TATGAGTGTGGAGTCGTTGCGCACTTTGGGAGATCTAGGTGAANNNAGATCGGAA |
| 44684_1 | TATGAGTGTGGAGTCGTTGCCCACATACAGAGCACGTAAGAGANNNAGATCGGAA |
| 44684_2 | TATGAGTGTGGAGTCGTTGCCCACATACAGAGCACGTAAGAGANNNAGATCGGAA |
| 50207_1 | TATGAGTGTGGAGTCGTTGCCCCTGGGATCTCATTTACCTCAGNNNAGATCGGAAG |
| 50207_2 | TATGAGTGTGGAGTCGTTGCCCCTGGGATCTCATTTACCTCAGNNNAGATCGGAAG |
| 28817_1 | TATGAGTGTGGAGTCGTTGCGGTGCAATCATGGCTCACTAGANNNAGATCGGAAG |
| 28817_2 | TATGAGTGTGGAGTCGTTGCGGTGCAATCATGGCTCACTAGANNNAGATCGGAAG |
| 31537_1 | TATGAGTGTGGAGTCGTTGCCCTTATAGGGAGGTCTCTTGGGANNNAGATCGGAA |
| 31537_2 | TATGAGTGTGGAGTCGTTGCCCTTATAGGGAGGTCTCTTGGGANNNAGATCGGAA |
| 53221_1 | TATGAGTGTGGAGTCGTTGCTTTGGCCTCAAACTTCAGGTGANNNAGATCGGAAGA |
| 53221_2 | TATGAGTGTGGAGTCGTTGCTTTGGCCTCAAACTTCAGGTGANNNAGATCGGAAGA |
| 36430_1 | TATGAGTGTGGAGTCGTTGCATATGCACGTTGTTCAACCCAACNNNAGATCGGAAG |
| 36430_2 | TATGAGTGTGGAGTCGTTGCATATGCACGTTGTTCAACCCAACNNNAGATCGGAAG |
| 37918_1 | TATGAGTGTGGAGTCGTTGCTGAGGTCAGGAATTTGAGACCAGNNNAGATCGGAA |
| 37918_2 | TATGAGTGTGGAGTCGTTGCTGAGGTCAGGAATTTGAGACCAGNNNAGATCGGAA |
| 52791_1 | TATGAGTGTGGAGTCGTTGCGGAGGGGAGGGGAAGGTTTCNNNAGATCGGAAGA |
| 52791_2 | TATGAGTGTGGAGTCGTTGCGGAGGGGAGGGGAAGGTTTCNNNAGATCGGAAGA |
| 39362_1 | TATGAGTGTGGAGTCGTTGCACAGACCAAAGAAAGCCAAACACNNNAGATCGGAA |
| 39362_2 | TATGAGTGTGGAGTCGTTGCACAGACCAAAGAAAGCCAAACACNNNAGATCGGAA |
| 44985_1 | TATGAGTGTGGAGTCGTTGCGCAATTTTTTTCAGCACCCCCTANNNAGATCGGAAGA |
| 44985_2 | TATGAGTGTGGAGTCGTTGCGCAATTTTTTTCAGCACCCCCTANNNAGATCGGAAGA |
| 30812_1 | TATGAGTGTGGAGTCGTTGCGTTGTTCTAGTGGGTGGCATCTANNNAGATCGGAA |
| 30812_2 | TATGAGTGTGGAGTCGTTGCGTTGTTCTAGTGGGTGGCATCTANNNAGATCGGAA |
| 43392_1 | TATGAGTGTGGAGTCGTTGCTCAGGTTCAAGCAATTCTTCTGCNNNAGATCGGAAG |
| 43392_2 | TATGAGTGTGGAGTCGTTGCTCAGGTTCAAGCAATTCTTCTGCNNNAGATCGGAAG |
| 7499_1 | TATGAGTGTGGAGTCGTTGCCAGATACGTTACACGTTGGCGTNNNAGATCGGAAGA |
| 7499_2 | TATGAGTGTGGAGTCGTTGCCAGATACGTTACACGTTGGCGTNNNAGATCGGAAGA |
| 9765_1 | TATGAGTGTGGAGTCGTTGCCGTGTGTGTGTGTATGCCTTTTANNNAGATCGGAAG |
| 9765_2 | TATGAGTGTGGAGTCGTTGCCGTGTGTGTGTGTATGCCTTTTANNNAGATCGGAAG |
| 36505_1 | TATGAGTGTGGAGTCGTTGCTGGGCCATACCACTCAATTTACANNNAGATCGGAAG |
| 36505_2 | TATGAGTGTGGAGTCGTTGCTGGGCCATACCACTCAATTTACANNNAGATCGGAAG |
| 51412_1 | TATGAGTGTGGAGTCGTTGCATATTCTGCAGGCAATGGGAAAANNNAGATCGGAA |
| 51412_2 | TATGAGTGTGGAGTCGTTGCATATTCTGCAGGCAATGGGAAAANNNAGATCGGAA |
| 10585_1 | TATGAGTGTGGAGTCGTTGCTTTCCTTTCTTCTCCACCTCCCTNNNAGATCGGAAGAG |
| 10585_2 | TATGAGTGTGGAGTCGTTGCTTTCCTTTCTTCTCCACCTCCCTNNNAGATCGGAAGAG |
| 41305_1 | TATGAGTGTGGAGTCGTTGCTCTTGTCCTATCTCCACAGTTGCNNNAGATCGGAAGA |
| 41305_2 | TATGAGTGTGGAGTCGTTGCTCTTGTCCTATCTCCACAGTTGCNNNAGATCGGAAGA |
| 44145_1 | TATGAGTGTGGAGTCGTTGCGTAGGGCATACCTTGGAGGAAATNNNAGATCGGAA |
| 44145_2 | TATGAGTGTGGAGTCGTTGCGTAGGGCATACCTTGGAGGAAATNNNAGATCGGAA |
| 13399_1 | TATGAGTGTGGAGTCGTTGCCTTCATACAGGGTATTCCATAATNNNAGATCGGAAG |
| 13399_2 | TATGAGTGTGGAGTCGTTGCCTTCATACAGGGTATTCCATAATNNNAGATCGGAAG |
| 34845_1 | TATGAGTGTGGAGTCGTTGCGTAGAGACAGGGTTTCACTGTGTNNNAGATCGGAA |
| 34845_2 | TATGAGTGTGGAGTCGTTGCGTAGAGACAGGGTTTCACTGTGTNNNAGATCGGAA |

| | |
|---|---|
| 43640_1 | TATGAGTGTGGAGTCGTTGCCATTTTCATCCACACCTTCGTCCNNNAGATCGGAAGA |
| 43640_2 | TATGAGTGTGGAGTCGTTGCCATTTTCATCCACACCTTCGTCCNNNAGATCGGAAGA |
| 31681_1 | TATGAGTGTGGAGTCGTTGCCATCAAGTGTCCACCAAGGAAAGNNNAGATCGGAAG |
| 31681_2 | TATGAGTGTGGAGTCGTTGCCATCAAGTGTCCACCAAGGAAAGNNNAGATCGGAAG |
| 53324_1 | TATGAGTGTGGAGTCGTTGCGTGTGTGTGTGTGTGTGTATTNNNAGATCGGAA |
| 53324_2 | TATGAGTGTGGAGTCGTTGCGTGTGTGTGTGTGTGTGTATTNNNAGATCGGAA |
| 36888_1 | TATGAGTGTGGAGTCGTTGCGTAGTCCCAGCTACTCCGGAGNNNAGATCGGAAGAG |
| 36888_2 | TATGAGTGTGGAGTCGTTGCGTAGTCCCAGCTACTCCGGAGNNNAGATCGGAAGAG |
| 41660_1 | TATGAGTGTGGAGTCGTTGCAGTTCTGGGAATACAGGCTTGAGNNNAGATCGGAA |
| 41660_2 | TATGAGTGTGGAGTCGTTGCAGTTCTGGGAATACAGGCTTGAGNNNAGATCGGAA |
| 28820_1 | TATGAGTGTGGAGTCGTTGCGTAATGGCATTCTGAGCAACCTGNNNAGATCGGAAG |
| 28820_2 | TATGAGTGTGGAGTCGTTGCGTAATGGCATTCTGAGCAACCTGNNNAGATCGGAAG |
| 39269_1 | TATGAGTGTGGAGTCGTTGCCCCTGAGAAGGTGCATGTTTTTTNNNAGATCGGAAG |
| 39269_2 | TATGAGTGTGGAGTCGTTGCCCCTGAGAAGGTGCATGTTTTTTNNNAGATCGGAAG |
| 53271_1 | TATGAGTGTGGAGTCGTTGCTTCCTTACTATTGTGTGCCTGCTNNNAGATCGGAAGA |
| 53271_2 | TATGAGTGTGGAGTCGTTGCTTCCTTACTATTGTGTGCCTGCTNNNAGATCGGAAGA |
| 43243_1 | TATGAGTGTGGAGTCGTTGCCAGGTTTGATTGGTGGTCAGAACNNNAGATCGGAA |
| 43243_2 | TATGAGTGTGGAGTCGTTGCCAGGTTTGATTGGTGGTCAGAACNNNAGATCGGAA |
| 34240_1 | TATGAGTGTGGAGTCGTTGCACCCTAATACCATACCCTTCCCANNNAGATCGGAAGA |
| 34240_2 | TATGAGTGTGGAGTCGTTGCACCCTAATACCATACCCTTCCCANNNAGATCGGAAGA |
| 50712_1 | TATGAGTGTGGAGTCGTTGCTGTATCTCTAGAGGACAGGCACTNNNAGATCGGAAG |
| 50712_2 | TATGAGTGTGGAGTCGTTGCTGTATCTCTAGAGGACAGGCACTNNNAGATCGGAAG |
| 30062_1 | TATGAGTGTGGAGTCGTTGCAGTGGTGGTCCAAATGGTATTGANNNAGATCGGAA |
| 30062_2 | TATGAGTGTGGAGTCGTTGCAGTGGTGGTCCAAATGGTATTGANNNAGATCGGAA |
| 10783_1 | TATGAGTGTGGAGTCGTTGCCACCCCTTTTCACACCATCCTAGNNNAGATCGGAAGA |
| 10783_2 | TATGAGTGTGGAGTCGTTGCCACCCCTTTTCACACCATCCTAGNNNAGATCGGAAGA |
| 13109_1 | TATGAGTGTGGAGTCGTTGCAATCTCTATGATCGCATCAGTANNNAGATCGGAAGA |
| 13109_2 | TATGAGTGTGGAGTCGTTGCAATCTCTATGATCGCATCAGTANNNAGATCGGAAGA |
| 49236_1 | TATGAGTGTGGAGTCGTTGCCCACGCCCAGCTAATTTTTGTATNNNAGATCGGAAGA |
| 49236_2 | TATGAGTGTGGAGTCGTTGCCCACGCCCAGCTAATTTTTGTATNNNAGATCGGAAGA |
| 31298_1 | TATGAGTGTGGAGTCGTTGCACACATACATGCACCCAAATTGGNNNAGATCGGAAG |
| 31298_2 | TATGAGTGTGGAGTCGTTGCACACATACATGCACCCAAATTGGNNNAGATCGGAAG |
| 51468_1 | TATGAGTGTGGAGTCGTTGCAAAGTGAAGAGGGAAGCATAGGGNNNAGATCGGA |
| 51468_2 | TATGAGTGTGGAGTCGTTGCAAAGTGAAGAGGGAAGCATAGGGNNNAGATCGGA |
| 53043_1 | TATGAGTGTGGAGTCGTTGCGGTGAATCACAAGGTCAGGAGATNNNAGATCGGAA |
| 53043_2 | TATGAGTGTGGAGTCGTTGCGGTGAATCACAAGGTCAGGAGATNNNAGATCGGAA |
| 32839_1 | TATGAGTGTGGAGTCGTTGCCCTCCTGAGTATCTGGGACTACANNNAGATCGGAAG |
| 32839_2 | TATGAGTGTGGAGTCGTTGCCCTCCTGAGTATCTGGGACTACANNNAGATCGGAAG |
| 43127_1 | TATGAGTGTGGAGTCGTTGCCAGAAGATTGAAACTGGACCCCTNNNAGATCGGAAG |
| 43127_2 | TATGAGTGTGGAGTCGTTGCCAGAAGATTGAAACTGGACCCCTNNNAGATCGGAAG |
| 39874_1 | TATGAGTGTGGAGTCGTTGCCACATATGTGGTCCCAGCTACTTNNNAGATCGGAAG |
| 39874_2 | TATGAGTGTGGAGTCGTTGCCACATATGTGGTCCCAGCTACTTNNNAGATCGGAAG |
| 50494_1 | TATGAGTGTGGAGTCGTTGCCGTCTTTATGGCTTCTCACCGTANNNAGATCGGAAG |

| | |
|---|---|
| 50494_2 | TATGAGTGTGGAGTCGTTGCCGTCTTTATGGCTTCTCACCGTANNNAGATCGGAAGA |
| 51753_1 | TATGAGTGTGGAGTCGTTGCTATGGTTTTGTCTTCTGCTCCCANNNAGATCGGAAGA |
| 51753_2 | TATGAGTGTGGAGTCGTTGCTATGGTTTTGTCTTCTGCTCCCANNNAGATCGGAAGA |
| 33982_1 | TATGAGTGTGGAGTCGTTGCGGAGTTCAGTGTTGCAATCTCAGNNNAGATCGGAAG |
| 33982_2 | TATGAGTGTGGAGTCGTTGCGGAGTTCAGTGTTGCAATCTCAGNNNAGATCGGAAG |
| 47785_1 | TATGAGTGTGGAGTCGTTGCCCTCCTGAGTAGCTGGGACTATANNNAGATCGGAAG |
| 47785_2 | TATGAGTGTGGAGTCGTTGCCCTCCTGAGTAGCTGGGACTATANNNAGATCGGAAG |
| 29065_1 | TATGAGTGTGGAGTCGTTGCGTAGCAGAAACCTGTCCTTCTGANNNAGATCGGAAG |
| 29065_2 | TATGAGTGTGGAGTCGTTGCGTAGCAGAAACCTGTCCTTCTGANNNAGATCGGAAG |
| 49096_1 | TATGAGTGTGGAGTCGTTGCCTGCTAGGCTCAAGTGACACTCNNNAGATCGGAAGA |
| 49096_2 | TATGAGTGTGGAGTCGTTGCCTGCTAGGCTCAAGTGACACTCNNNAGATCGGAAGA |
| 32233_1 | TATGAGTGTGGAGTCGTTGCAGTGCTGGGATTACAGGTGTAAGNNNAGATCGGAA |
| 32233_2 | TATGAGTGTGGAGTCGTTGCAGTGCTGGGATTACAGGTGTAAGNNNAGATCGGAA |
| 41668_1 | TATGAGTGTGGAGTCGTTGCGTGATCCTCCCACTTTAGCCTACNNNAGATCGGAAGA |
| 41668_2 | TATGAGTGTGGAGTCGTTGCGTGATCCTCCCACTTTAGCCTACNNNAGATCGGAAGA |
| 42393_1 | TATGAGTGTGGAGTCGTTGCCCTCTGGTATCATCTGAAGGCTCNNNAGATCGGAAGA |
| 42393_2 | TATGAGTGTGGAGTCGTTGCCCTCTGGTATCATCTGAAGGCTCNNNAGATCGGAAGA |
| 45898_1 | TATGAGTGTGGAGTCGTTGCAGATTACAGGCATAAACCACCGTNNNAGATCGGAAG |
| 45898_2 | TATGAGTGTGGAGTCGTTGCAGATTACAGGCATAAACCACCGTNNNAGATCGGAAG |
| 8036_1 | TATGAGTGTGGAGTCGTTGCATAATCTCCAGGCCAGTGAGTGGNNNAGATCGGAAG |
| 8036_2 | TATGAGTGTGGAGTCGTTGCATAATCTCCAGGCCAGTGAGTGGNNNAGATCGGAAG |
| 36970_1 | TATGAGTGTGGAGTCGTTGCCTCTCTTTTGGTGTGAGCCTCTNNNAGATCGGAAGAG |
| 36970_2 | TATGAGTGTGGAGTCGTTGCCTCTCTTTTGGTGTGAGCCTCTNNNAGATCGGAAGAG |
| 41043_1 | TATGAGTGTGGAGTCGTTGCGTGCTAGAACAATGGTTGCTTCCNNNAGATCGGAAG |
| 41043_2 | TATGAGTGTGGAGTCGTTGCGTGCTAGAACAATGGTTGCTTCCNNNAGATCGGAAG |
| 45176_1 | TATGAGTGTGGAGTCGTTGCAGTTAGAGGTTGGACAGAACACCNNNAGATCGGAAG |
| 45176_2 | TATGAGTGTGGAGTCGTTGCAGTTAGAGGTTGGACAGAACACCNNNAGATCGGAAG |
| 35664_1 | TATGAGTGTGGAGTCGTTGCCAAAGGGAGGTGATTTGTATGCCNNNAGATCGGAAG |
| 35664_2 | TATGAGTGTGGAGTCGTTGCCAAAGGGAGGTGATTTGTATGCCNNNAGATCGGAAG |
| 35841_1 | TATGAGTGTGGAGTCGTTGCAGATCCTCCTACCTTGGACTCTCNNNAGATCGGAAGA |
| 35841_2 | TATGAGTGTGGAGTCGTTGCAGATCCTCCTACCTTGGACTCTCNNNAGATCGGAAGA |
| 39786_1 | TATGAGTGTGGAGTCGTTGCTCCTTGTGGACAAAGGATGGATANNNAGATCGGAAG |
| 39786_2 | TATGAGTGTGGAGTCGTTGCTCCTTGTGGACAAAGGATGGATANNNAGATCGGAAG |
| 52168_1 | TATGAGTGTGGAGTCGTTGCATAATGCTTTTGAGGACCAAGCCNNNAGATCGGAAG |
| 52168_2 | TATGAGTGTGGAGTCGTTGCATAATGCTTTTGAGGACCAAGCCNNNAGATCGGAAG |
| 46550_1 | TATGAGTGTGGAGTCGTTGCCTCACATTGCATCTTTGAGCCAANNNAGATCGGAAGA |
| 46550_2 | TATGAGTGTGGAGTCGTTGCCTCACATTGCATCTTTGAGCCAANNNAGATCGGAAGA |
| 47130_1 | TATGAGTGTGGAGTCGTTGCCGGGTTCAAGCCATTCTCCTNNNAGATCGGAAGAGCA |
| 47130_2 | TATGAGTGTGGAGTCGTTGCCGGGTTCAAGCCATTCTCCTNNNAGATCGGAAGAGCA |
| 37591_1 | TATGAGTGTGGAGTCGTTGCGTTCACAACCTTACTTATGCAGTNNNAGATCGGAAGA |
| 37591_2 | TATGAGTGTGGAGTCGTTGCGTTCACAACCTTACTTATGCAGTNNNAGATCGGAAGA |
| 38062_1 | TATGAGTGTGGAGTCGTTGCGAATTCCCTGAGTGGTCCATCTTNNNAGATCGGAAGA |
| 38062_2 | TATGAGTGTGGAGTCGTTGCGAATTCCCTGAGTGGTCCATCTTNNNAGATCGGAAGA |

| | |
|---|---|
| 32330_1 | TATGAGTGTGGAGTCGTTGCCAAAATGTTGGGAGTACAGGCGNNNAGATCGGAAG |
| 32330_2 | TATGAGTGTGGAGTCGTTGCCAAAATGTTGGGAGTACAGGCGNNNAGATCGGAAG |
| 48754_1 | TATGAGTGTGGAGTCGTTGCTTTTGTGTCTAGGTTCCTCAGGTNNNAGATCGGAAG |
| 48754_2 | TATGAGTGTGGAGTCGTTGCTTTTGTGTCTAGGTTCCTCAGGTNNNAGATCGGAAG |
| 50480_1 | TATGAGTGTGGAGTCGTTGCTACACAGGTACAGGGACAAAGACNNNAGATCGGAA |
| 50480_2 | TATGAGTGTGGAGTCGTTGCTACACAGGTACAGGGACAAAGACNNNAGATCGGAA |
| 44133_1 | TATGAGTGTGGAGTCGTTGCTTTCCCTTGAATGGAAGGACAGTNNNAGATCGGAAG |
| 44133_2 | TATGAGTGTGGAGTCGTTGCTTTCCCTTGAATGGAAGGACAGTNNNAGATCGGAAG |
| 48551_1 | TATGAGTGTGGAGTCGTTGCATAATTTACCCCCCAGAAGCTCCNNNAGATCGGAAGA |
| 48551_2 | TATGAGTGTGGAGTCGTTGCATAATTTACCCCCCAGAAGCTCCNNNAGATCGGAAGA |
| 40136_1 | TATGAGTGTGGAGTCGTTGCTTCCTCTTTTGAACCGCTTTTGGNNNAGATCGGAAGA |
| 40136_2 | TATGAGTGTGGAGTCGTTGCTTCCTCTTTTGAACCGCTTTTGGNNNAGATCGGAAGA |
| 44219_1 | TATGAGTGTGGAGTCGTTGCATTCCTCTCTCAGGTGAAATGGGNNNAGATCGGAAG |
| 44219_2 | TATGAGTGTGGAGTCGTTGCATTCCTCTCTCAGGTGAAATGGGNNNAGATCGGAAG |
| 38329_1 | TATGAGTGTGGAGTCGTTGCATGGTGGTCCTAGCTACTCAGAANNNAGATCGGAAG |
| 38329_2 | TATGAGTGTGGAGTCGTTGCATGGTGGTCCTAGCTACTCAGAANNNAGATCGGAAG |
| 9993_1 | TATGAGTGTGGAGTCGTTGCGCTAGAAGCCAAGGGAAGAGAACNNNAGATCGGAA |
| 9993_2 | TATGAGTGTGGAGTCGTTGCGCTAGAAGCCAAGGGAAGAGAACNNNAGATCGGAA |
| 49370_1 | TATGAGTGTGGAGTCGTTGCGTCTCCCGAGTTCAAGTGATTCTNNNAGATCGGAAG |
| 49370_2 | TATGAGTGTGGAGTCGTTGCGTCTCCCGAGTTCAAGTGATTCTNNNAGATCGGAAG |
| 8400_1 | TATGAGTGTGGAGTCGTTGCATAGGCCACTGGGGTCAAAATTGNNNAGATCGGAA |
| 8400_2 | TATGAGTGTGGAGTCGTTGCATAGGCCACTGGGGTCAAAATTGNNNAGATCGGAA |
| 31127_1 | TATGAGTGTGGAGTCGTTGCACTGATGAGCAATTCCTTCTCCCNNNAGATCGGAAGA |
| 31127_2 | TATGAGTGTGGAGTCGTTGCACTGATGAGCAATTCCTTCTCCCNNNAGATCGGAAGA |
| 32522_1 | TATGAGTGTGGAGTCGTTGCAAAGAAATAAAACTCCCCTGGCCNNNAGATCGGAAG |
| 32522_2 | TATGAGTGTGGAGTCGTTGCAAAGAAATAAAACTCCCCTGGCCNNNAGATCGGAAG |
| 7912_1 | TATGAGTGTGGAGTCGTTGCAAACTCAGAGGGCCAGGATGGAANNNAGATCGGAA |
| 7912_2 | TATGAGTGTGGAGTCGTTGCAAACTCAGAGGGCCAGGATGGAANNNAGATCGGAA |
| 48456_1 | TATGAGTGTGGAGTCGTTGCACAAAGGGAAAAAGAAAAGCCCCNNNAGATCGGAA |
| 48456_2 | TATGAGTGTGGAGTCGTTGCACAAAGGGAAAAAGAAAAGCCCCNNNAGATCGGAA |
| 52738_1 | TATGAGTGTGGAGTCGTTGCCTCAGCCTGGAGACTCTGGAGNNNAGATCGGAAGA |
| 52738_2 | TATGAGTGTGGAGTCGTTGCCTCAGCCTGGAGACTCTGGAGNNNAGATCGGAAGA |
| 48751_1 | TATGAGTGTGGAGTCGTTGCCAACATGGCTTTGTTTTCTGTGCNNNAGATCGGAAG |
| 48751_2 | TATGAGTGTGGAGTCGTTGCCAACATGGCTTTGTTTTCTGTGCNNNAGATCGGAAG |
| 48401_1 | TATGAGTGTGGAGTCGTTGCGCCACCATACCCAGCTAATTTTTNNNAGATCGGAAGA |
| 48401_2 | TATGAGTGTGGAGTCGTTGCGCCACCATACCCAGCTAATTTTTNNNAGATCGGAAGA |
| 38759_1 | TATGAGTGTGGAGTCGTTGCGGAGGCTGAGGCAGAATAATCATNNNAGATCGGAA |
| 38759_2 | TATGAGTGTGGAGTCGTTGCGGAGGCTGAGGCAGAATAATCATNNNAGATCGGAA |
| 51338_1 | TATGAGTGTGGAGTCGTTGCTCTTGAACCTCCAAGCTAACTCCNNNAGATCGGAAGA |
| 51338_2 | TATGAGTGTGGAGTCGTTGCTCTTGAACCTCCAAGCTAACTCCNNNAGATCGGAAGA |
| 31837_1 | TATGAGTGTGGAGTCGTTGCCCAAGATGAGCAGATCACCTGAGNNNAGATCGGAA |
| 31837_2 | TATGAGTGTGGAGTCGTTGCCCAAGATGAGCAGATCACCTGAGNNNAGATCGGAA |
| 29694_1 | TATGAGTGTGGAGTCGTTGCAAATTGAGCCTTTCCCCTTCCTGNNNAGATCGGAAGA |

| | |
|---|---|
| 29694_2 | TATGAGTGTGGAGTCGTTGCAAATTGAGCCTTTCCCCTTCCTGNNNAGATCGGAAGA |
| 10320_1 | TATGAGTGTGGAGTCGTTGCTATGCCCAGTGGTGAAAAATGCTNNNAGATCGGAAG |
| 10320_2 | TATGAGTGTGGAGTCGTTGCTATGCCCAGTGGTGAAAAATGCTNNNAGATCGGAAG |
| 47132_1 | TATGAGTGTGGAGTCGTTGCTCATCAGACCCACCACTTTACTGNNNAGATCGGAAGA |
| 47132_2 | TATGAGTGTGGAGTCGTTGCTCATCAGACCCACCACTTTACTGNNNAGATCGGAAGA |
| 51184_1 | TATGAGTGTGGAGTCGTTGCCTGCTCTGGAAACCCTACATAGGNNNAGATCGGAAG |
| 51184_2 | TATGAGTGTGGAGTCGTTGCCTGCTCTGGAAACCCTACATAGGNNNAGATCGGAAG |
| 41227_1 | TATGAGTGTGGAGTCGTTGCATCACAAGGTCAGGAATTCGAGTNNNAGATCGGAAG |
| 41227_2 | TATGAGTGTGGAGTCGTTGCATCACAAGGTCAGGAATTCGAGTNNNAGATCGGAAG |
| 52190_1 | TATGAGTGTGGAGTCGTTGCTTGGTGAAGGTGAGACTCATGAGNNNAGATCGGAA |
| 52190_2 | TATGAGTGTGGAGTCGTTGCTTGGTGAAGGTGAGACTCATGAGNNNAGATCGGAA |
| 43513_1 | TATGAGTGTGGAGTCGTTGCAAAGAGGGCATGGGAGGTAAAATNNNAGATCGGAA |
| 43513_2 | TATGAGTGTGGAGTCGTTGCAAAGAGGGCATGGGAGGTAAAATNNNAGATCGGAA |
| 31768_1 | TATGAGTGTGGAGTCGTTGCTCCCTCTCTCTCTCTCTCTCTCTNNNAGATCGGAAGAG |
| 31768_2 | TATGAGTGTGGAGTCGTTGCTCCCTCTCTCTCTCTCTCTCTCTNNNAGATCGGAAGAG |
| 45472_1 | TATGAGTGTGGAGTCGTTGCTTGGAAGAGGGCCTTAGTAGAGANNNAGATCGGAA |
| 45472_2 | TATGAGTGTGGAGTCGTTGCTTGGAAGAGGGCCTTAGTAGAGANNNAGATCGGAA |
| 38760_1 | TATGAGTGTGGAGTCGTTGCTATATAAAGGCCAGGTGTGGTGGNNNAGATCGGAA |
| 38760_2 | TATGAGTGTGGAGTCGTTGCTATATAAAGGCCAGGTGTGGTGGNNNAGATCGGAA |
| 30694_1 | TATGAGTGTGGAGTCGTTGCTCCTTTCACTACGTGTTTTCTGAACNNNAGATCGGAAG |
| 30694_2 | TATGAGTGTGGAGTCGTTGCTCCTTTCACTACGTGTTTTCTGAACNNNAGATCGGAAG |
| 43584_1 | TATGAGTGTGGAGTCGTTGCAGCTAAGGCCTGTCGAGAAATCNNNAGATCGGAAGA |
| 43584_2 | TATGAGTGTGGAGTCGTTGCAGCTAAGGCCTGTCGAGAAATCNNNAGATCGGAAGA |
| 38005_1 | TATGAGTGTGGAGTCGTTGCTGTTTGGATGCGTAGTTGAATGANNNAGATCGGAAG |
| 38005_2 | TATGAGTGTGGAGTCGTTGCTGTTTGGATGCGTAGTTGAATGANNNAGATCGGAAG |
| 31564_1 | TATGAGTGTGGAGTCGTTGCTTGTAGACAAAGCTGCCCTCCNNNAGATCGGAAGAG |
| 31564_2 | TATGAGTGTGGAGTCGTTGCTTGTAGACAAAGCTGCCCTCCNNNAGATCGGAAGAG |
| 4595_1 | TATGAGTGTGGAGTCGTTGCTTGAGTTCAGTGAGGGCACAAATNNNAGATCGGAAG |
| 4595_2 | TATGAGTGTGGAGTCGTTGCTTGAGTTCAGTGAGGGCACAAATNNNAGATCGGAAG |
| 50014_1 | TATGAGTGTGGAGTCGTTGCCGGCTCCAGTTCTTGAAAAATGTNNNAGATCGGAAG |
| 50014_2 | TATGAGTGTGGAGTCGTTGCCGGCTCCAGTTCTTGAAAAATGTNNNAGATCGGAAG |
| 46208_1 | TATGAGTGTGGAGTCGTTGCTTTTAGTAGAGACGGGGTTTCACNNNAGATCGGAAG |
| 46208_2 | TATGAGTGTGGAGTCGTTGCTTTTAGTAGAGACGGGGTTTCACNNNAGATCGGAAG |
| 42234_1 | TATGAGTGTGGAGTCGTTGCATGAGTTATGGAGGAGTCCCTGTNNNAGATCGGAAG |
| 42234_2 | TATGAGTGTGGAGTCGTTGCATGAGTTATGGAGGAGTCCCTGTNNNAGATCGGAAG |
| 36086_1 | TATGAGTGTGGAGTCGTTGCGAAAGGAATCGTGGTGTATTGGCNNNAGATCGGAA |
| 36086_2 | TATGAGTGTGGAGTCGTTGCGAAAGGAATCGTGGTGTATTGGCNNNAGATCGGAA |
| 41580_1 | TATGAGTGTGGAGTCGTTGCTCATTGTGGTTTTGACATGCACTNNNAGATCGGAAGA |
| 41580_2 | TATGAGTGTGGAGTCGTTGCTCATTGTGGTTTTGACATGCACTNNNAGATCGGAAGA |
| 43432_1 | TATGAGTGTGGAGTCGTTGCTCCCCCAAAACTGAGATTACTGGNNNAGATCGGAAG |
| 43432_2 | TATGAGTGTGGAGTCGTTGCTCCCCCAAAACTGAGATTACTGGNNNAGATCGGAAG |
| 43921_1 | TATGAGTGTGGAGTCGTTGCTCTTTCTTGGGATGGAGTTTTGCNNNAGATCGGAAG |
| 43921_2 | TATGAGTGTGGAGTCGTTGCTCTTTCTTGGGATGGAGTTTTGCNNNAGATCGGAAG |

| | |
|---|---|
| 39317_1 | TATGAGTGTGGAGTCGTTGCACTTACAAGGGATGTGAAGGACCNNNAGATCGGAA |
| 39317_2 | TATGAGTGTGGAGTCGTTGCACTTACAAGGGATGTGAAGGACCNNNAGATCGGAA |
| 43345_1 | TATGAGTGTGGAGTCGTTGCTGGAGTGCAGTGGTACAATCTCNNNAGATCGGAAG |
| 43345_2 | TATGAGTGTGGAGTCGTTGCTGGAGTGCAGTGGTACAATCTCNNNAGATCGGAAG |
| 12528_1 | TATGAGTGTGGAGTCGTTGCGAGAAGATTTATGAGCCAAGTANNNAGATCGGAAG |
| 12528_2 | TATGAGTGTGGAGTCGTTGCGAGAAGATTTATGAGCCAAGTANNNAGATCGGAAG |
| 47248_1 | TATGAGTGTGGAGTCGTTGCATCAAGTGATTCTCCTGCCTCAGNNNAGATCGGAAG |
| 47248_2 | TATGAGTGTGGAGTCGTTGCATCAAGTGATTCTCCTGCCTCAGNNNAGATCGGAAG |
| 34600_1 | TATGAGTGTGGAGTCGTTGCAGGATTGCTGGTTTGTTATTGCCNNNAGATCGGAAG |
| 34600_2 | TATGAGTGTGGAGTCGTTGCAGGATTGCTGGTTTGTTATTGCCNNNAGATCGGAAG |
| 36642_1 | TATGAGTGTGGAGTCGTTGCACAAAAATTAGCCAGGTGTGGTGNNNAGATCGGAA |
| 36642_2 | TATGAGTGTGGAGTCGTTGCACAAAAATTAGCCAGGTGTGGTGNNNAGATCGGAA |
| 42488_1 | TATGAGTGTGGAGTCGTTGCACAAGACTCTCTGCCTGGTAATCNNNAGATCGGAAG |
| 42488_2 | TATGAGTGTGGAGTCGTTGCACAAGACTCTCTGCCTGGTAATCNNNAGATCGGAAG |
| 46667_1 | TATGAGTGTGGAGTCGTTGCTTGCTAGTTATCCCCAAACCTCCNNNAGATCGGAAGA |
| 46667_2 | TATGAGTGTGGAGTCGTTGCTTGCTAGTTATCCCCAAACCTCCNNNAGATCGGAAGA |
| 13042_1 | TATGAGTGTGGAGTCGTTGCTATCAGGAATGGACTTAGTCTNNNAGATCGGAAGA |
| 13042_2 | TATGAGTGTGGAGTCGTTGCTATCAGGAATGGACTTAGTCTNNNAGATCGGAAGA |
| 44105_1 | TATGAGTGTGGAGTCGTTGCTGATCTGAAGGCCTACAGCATTTNNNAGATCGGAAG |
| 44105_2 | TATGAGTGTGGAGTCGTTGCTGATCTGAAGGCCTACAGCATTTNNNAGATCGGAAG |
| 29126_1 | TATGAGTGTGGAGTCGTTGCCATCCTCCTAGTTTCTCTGGCTGNNNAGATCGGAAGA |
| 29126_2 | TATGAGTGTGGAGTCGTTGCCATCCTCCTAGTTTCTCTGGCTGNNNAGATCGGAAGA |
| 9566_1 | TATGAGTGTGGAGTCGTTGCAGGATGAAACCACCTAATGGTGCNNNAGATCGGAA |
| 9566_2 | TATGAGTGTGGAGTCGTTGCAGGATGAAACCACCTAATGGTGCNNNAGATCGGAA |
| 42098_1 | TATGAGTGTGGAGTCGTTGCGGACAATGTCATCACTGGCTAGTNNNAGATCGGAAG |
| 42098_2 | TATGAGTGTGGAGTCGTTGCGGACAATGTCATCACTGGCTAGTNNNAGATCGGAAG |
| 42236_1 | TATGAGTGTGGAGTCGTTGCTCCTGAGACTTTGCTGAAATTGCNNNAGATCGGAAG |
| 42236_2 | TATGAGTGTGGAGTCGTTGCTCCTGAGACTTTGCTGAAATTGCNNNAGATCGGAAG |
| 46332_1 | TATGAGTGTGGAGTCGTTGCTCCTGGGTAACTGGGACTATAGGNNNAGATCGGAA |
| 46332_2 | TATGAGTGTGGAGTCGTTGCTCCTGGGTAACTGGGACTATAGGNNNAGATCGGAA |
| 49254_1 | TATGAGTGTGGAGTCGTTGCCCACGAAAAAGACACTGTCCTTCNNNAGATCGGAAG |
| 49254_2 | TATGAGTGTGGAGTCGTTGCCCACGAAAAAGACACTGTCCTTCNNNAGATCGGAAG |
| 35756_1 | TATGAGTGTGGAGTCGTTGCCAAAACTGCTGGGATTACAGGTGNNNAGATCGGAA |
| 35756_2 | TATGAGTGTGGAGTCGTTGCCAAAACTGCTGGGATTACAGGTGNNNAGATCGGAA |
| 40244_1 | TATGAGTGTGGAGTCGTTGCCCTAACTCCAGTGAGCTGAACTTNNNAGATCGGAAG |
| 40244_2 | TATGAGTGTGGAGTCGTTGCCCTAACTCCAGTGAGCTGAACTTNNNAGATCGGAAG |
| 38143_1 | TATGAGTGTGGAGTCGTTGCAAAAAGAAAAACAGGCCAGGTGCNNNAGATCGGAA |
| 38143_2 | TATGAGTGTGGAGTCGTTGCAAAAAGAAAAACAGGCCAGGTGCNNNAGATCGGAA |
| 2369_1 | TATGAGTGTGGAGTCGTTGCCCCAGAAGGAAACATAGTCCAGCNNNAGATCGGAA |
| 2369_2 | TATGAGTGTGGAGTCGTTGCCCCAGAAGGAAACATAGTCCAGCNNNAGATCGGAA |
| 47038_1 | TATGAGTGTGGAGTCGTTGCCTGTTCCTTGAGGAGAGTCATCCNNNAGATCGGAAG |
| 47038_2 | TATGAGTGTGGAGTCGTTGCCTGTTCCTTGAGGAGAGTCATCCNNNAGATCGGAAG |
| 9431_1 | TATGAGTGTGGAGTCGTTGCTCCCACAGGTCGGCCTTATATAGNNNAGATCGGAAG |

| | |
|---|---|
| 9431_2 | TATGAGTGTGGAGTCGTTGCTCCCACAGGTCGGCCTTATATAGNNNAGATCGGAAG |
| 45438_1 | TATGAGTGTGGAGTCGTTGCATCAAGGTCAGTGGGCAAAAATGNNNAGATCGGAA |
| 45438_2 | TATGAGTGTGGAGTCGTTGCATCAAGGTCAGTGGGCAAAAATGNNNAGATCGGAA |
| 47181_1 | TATGAGTGTGGAGTCGTTGCGCCTCCTGAGTTCAAGTGATTCTNNNAGATCGGAAG |
| 47181_2 | TATGAGTGTGGAGTCGTTGCGCCTCCTGAGTTCAAGTGATTCTNNNAGATCGGAAG |
| 13257_1 | TATGAGTGTGGAGTCGTTGCAGCCTGTATAAAAATAAAAACAGANNNAGATCGGA |
| 13257_2 | TATGAGTGTGGAGTCGTTGCAGCCTGTATAAAAATAAAAACAGANNNAGATCGGA |
| 2240_1 | TATGAGTGTGGAGTCGTTGCTCCACAAAGAGTGATGGGGGATANNNAGATCGGAA |
| 2240_2 | TATGAGTGTGGAGTCGTTGCTCCACAAAGAGTGATGGGGGATANNNAGATCGGAA |
| 44745_1 | TATGAGTGTGGAGTCGTTGCAGATCCTGAGAGAGTCCTTGTGANNNAGATCGGAA |
| 44745_2 | TATGAGTGTGGAGTCGTTGCAGATCCTGAGAGAGTCCTTGTGANNNAGATCGGAA |
| 37530_1 | TATGAGTGTGGAGTCGTTGCGAGTAGCGTGTGTGATATAGCCANNNAGATCGGAA |
| 37530_2 | TATGAGTGTGGAGTCGTTGCGAGTAGCGTGTGTGATATAGCCANNNAGATCGGAA |
| 53039_1 | TATGAGTGTGGAGTCGTTGCACACAAACTCACCATTCAATCCANNNAGATCGGAAGA |
| 53039_2 | TATGAGTGTGGAGTCGTTGCACACAAACTCACCATTCAATCCANNNAGATCGGAAGA |
| 30868_1 | TATGAGTGTGGAGTCGTTGCAGTAGCTGGGACTATAGGTGTGTNNNAGATCGGAA |
| 30868_2 | TATGAGTGTGGAGTCGTTGCAGTAGCTGGGACTATAGGTGTGTNNNAGATCGGAA |
| 49350_1 | TATGAGTGTGGAGTCGTTGCCAGCTATTGTAAAAGTGGTTGTGTNNNAGATCGGA |
| 49350_2 | TATGAGTGTGGAGTCGTTGCCAGCTATTGTAAAAGTGGTTGTGTNNNAGATCGGA |
| 9446_1 | TATGAGTGTGGAGTCGTTGCTTCTGCATTGTAACAGGGTGGAGNNNAGATCGGAA |
| 9446_2 | TATGAGTGTGGAGTCGTTGCTTCTGCATTGTAACAGGGTGGAGNNNAGATCGGAA |
| 12728_1 | TATGAGTGTGGAGTCGTTGCAGTATCAGGGGAAATCTAAATTTNNNAGATCGGAAG |
| 12728_2 | TATGAGTGTGGAGTCGTTGCAGTATCAGGGGAAATCTAAATTTNNNAGATCGGAAG |
| 40137_1 | TATGAGTGTGGAGTCGTTGCGGTTGCAGTGAGCCAAAATCATNNNAGATCGGAAG |
| 40137_2 | TATGAGTGTGGAGTCGTTGCGGTTGCAGTGAGCCAAAATCATNNNAGATCGGAAG |
| 47989_1 | TATGAGTGTGGAGTCGTTGCCTAATTTCAAAAAGGGCCTGGGGNNNAGATCGGAA |
| 47989_2 | TATGAGTGTGGAGTCGTTGCCTAATTTCAAAAAGGGCCTGGGGNNNAGATCGGAA |
| 32377_1 | TATGAGTGTGGAGTCGTTGCCACCACCACCATTTTTCACAGAANNNAGATCGGAAGA |
| 32377_2 | TATGAGTGTGGAGTCGTTGCCACCACCACCATTTTTCACAGAANNNAGATCGGAAGA |
| 30350_1 | TATGAGTGTGGAGTCGTTGCGTCTAACTGCTGGAAACCGTCAANNNAGATCGGAAG |
| 30350_2 | TATGAGTGTGGAGTCGTTGCGTCTAACTGCTGGAAACCGTCAANNNAGATCGGAAG |
| 50842_1 | TATGAGTGTGGAGTCGTTGCTGCAAGCGTTTTATAAGCCTCTGNNNAGATCGGAAG |
| 50842_2 | TATGAGTGTGGAGTCGTTGCTGCAAGCGTTTTATAAGCCTCTGNNNAGATCGGAAG |
| 36845_1 | TATGAGTGTGGAGTCGTTGCGCAGGCCTTGATTGAGTAGACTTNNNAGATCGGAA |
| 36845_2 | TATGAGTGTGGAGTCGTTGCGCAGGCCTTGATTGAGTAGACTTNNNAGATCGGAA |
| 30865_1 | TATGAGTGTGGAGTCGTTGCAACTGGAATACCTGCCTTCACTTNNNAGATCGGAAG |
| 30865_2 | TATGAGTGTGGAGTCGTTGCAACTGGAATACCTGCCTTCACTTNNNAGATCGGAAG |
| 48295_1 | TATGAGTGTGGAGTCGTTGCCCAACCAATCTACCTCTACCCACNNNAGATCGGAAGA |
| 48295_2 | TATGAGTGTGGAGTCGTTGCCCAACCAATCTACCTCTACCCACNNNAGATCGGAAGA |
| 13022_1 | TATGAGTGTGGAGTCGTTGCAATCTCTATGATCGCATCAGTANNNAGATCGGAAGA |
| 13022_2 | TATGAGTGTGGAGTCGTTGCAATCTCTATGATCGCATCAGTANNNAGATCGGAAGA |
| 13149_1 | TATGAGTGTGGAGTCGTTGCTTTATCACTAAAGAAATGGTTGGNNNAGATCGGAAG |
| 13149_2 | TATGAGTGTGGAGTCGTTGCTTTATCACTAAAGAAATGGTTGGNNNAGATCGGAAG |

| | |
|---|---|
| 34027_1 | TATGAGTGTGGAGTCGTTGCCTCAAGCAATCTTCCCACTTCAGNNNAGATCGGAAGA |
| 34027_2 | TATGAGTGTGGAGTCGTTGCCTCAAGCAATCTTCCCACTTCAGNNNAGATCGGAAGA |
| 44817_1 | TATGAGTGTGGAGTCGTTGCGATAAAAGACAAGCTCCTCGTGCNNNAGATCGGAAG |
| 44817_2 | TATGAGTGTGGAGTCGTTGCGATAAAAGACAAGCTCCTCGTGCNNNAGATCGGAAG |
| 30614_1 | TATGAGTGTGGAGTCGTTGCCATCAGTGCTACCCTCCCATCTANNNAGATCGGAAGA |
| 30614_2 | TATGAGTGTGGAGTCGTTGCCATCAGTGCTACCCTCCCATCTANNNAGATCGGAAGA |
| 37451_1 | TATGAGTGTGGAGTCGTTGCCCGTAGTCCTAGCTACTTGGGAGNNNAGATCGGAAG |
| 37451_2 | TATGAGTGTGGAGTCGTTGCCCGTAGTCCTAGCTACTTGGGAGNNNAGATCGGAAG |
| 35847_1 | TATGAGTGTGGAGTCGTTGCGCATGATCTTGGCTCACTACAACNNNAGATCGGAAG |
| 35847_2 | TATGAGTGTGGAGTCGTTGCGCATGATCTTGGCTCACTACAACNNNAGATCGGAAG |
| 52502_1 | TATGAGTGTGGAGTCGTTGCTGGTACCTTTCTCCCACATTTCANNNAGATCGGAAGA |
| 52502_2 | TATGAGTGTGGAGTCGTTGCTGGTACCTTTCTCCCACATTTCANNNAGATCGGAAGA |
| 32537_1 | TATGAGTGTGGAGTCGTTGCTGTGTCACCTGTATCATGCACATNNNAGATCGGAAG |
| 32537_2 | TATGAGTGTGGAGTCGTTGCTGTGTCACCTGTATCATGCACATNNNAGATCGGAAG |
| 31356_1 | TATGAGTGTGGAGTCGTTGCACATATTCGGAGAAGGGGTGATGNNNAGATCGGAA |
| 31356_2 | TATGAGTGTGGAGTCGTTGCACATATTCGGAGAAGGGGTGATGNNNAGATCGGAA |
| 39687_1 | TATGAGTGTGGAGTCGTTGCAAACTTCTAGGTAGAATGACATGGANNNAGATCGG |
| 39687_2 | TATGAGTGTGGAGTCGTTGCAAACTTCTAGGTAGAATGACATGGANNNAGATCGG |
| 45061_1 | TATGAGTGTGGAGTCGTTGCTGGACTGCTGATTCTGATGAGTCNNNAGATCGGAAG |
| 45061_2 | TATGAGTGTGGAGTCGTTGCTGGACTGCTGATTCTGATGAGTCNNNAGATCGGAAG |
| 46934_1 | TATGAGTGTGGAGTCGTTGCTCGGTGACACTATCTGCTTACTGNNNAGATCGGAAG |
| 46934_2 | TATGAGTGTGGAGTCGTTGCTCGGTGACACTATCTGCTTACTGNNNAGATCGGAAG |
| 48986_1 | TATGAGTGTGGAGTCGTTGCTCTTTGAGACAGAGTTTCGCTGTNNNAGATCGGAAG |
| 48986_2 | TATGAGTGTGGAGTCGTTGCTCTTTGAGACAGAGTTTCGCTGTNNNAGATCGGAAG |
| 48583_1 | TATGAGTGTGGAGTCGTTGCTGCAGTGGAATGATCTCAGTTCANNNAGATCGGAAG |
| 48583_2 | TATGAGTGTGGAGTCGTTGCTGCAGTGGAATGATCTCAGTTCANNNAGATCGGAAG |
| 44596_1 | TATGAGTGTGGAGTCGTTGCGTGGGTGGAGTGTACTTGTACTCNNNAGATCGGAA |
| 44596_2 | TATGAGTGTGGAGTCGTTGCGTGGGTGGAGTGTACTTGTACTCNNNAGATCGGAA |
| 48908_1 | TATGAGTGTGGAGTCGTTGCGGTAGAGACAGGGTTTCCTCATGNNNAGATCGGAA |
| 48908_2 | TATGAGTGTGGAGTCGTTGCGGTAGAGACAGGGTTTCCTCATGNNNAGATCGGAA |
| 35526_1 | TATGAGTGTGGAGTCGTTGCCTGAACCGAAGCAGATCAGAGATNNNAGATCGGAA |
| 35526_2 | TATGAGTGTGGAGTCGTTGCCTGAACCGAAGCAGATCAGAGATNNNAGATCGGAA |
| 29133_1 | TATGAGTGTGGAGTCGTTGCACTGTTCTAGTTTAGCCCTGTGGNNNAGATCGGAAG |
| 29133_2 | TATGAGTGTGGAGTCGTTGCACTGTTCTAGTTTAGCCCTGTGGNNNAGATCGGAAG |
| 5410_1 | TATGAGTGTGGAGTCGTTGCGGGTGTTTGTTACCCCAAGACATNNNAGATCGGAAG |
| 5410_2 | TATGAGTGTGGAGTCGTTGCGGGTGTTTGTTACCCCAAGACATNNNAGATCGGAAG |
| 35725_1 | TATGAGTGTGGAGTCGTTGCATCAAGCAATTCTCCTGCCTCANNNAGATCGGAAGA |
| 35725_2 | TATGAGTGTGGAGTCGTTGCATCAAGCAATTCTCCTGCCTCANNNAGATCGGAAGA |
| 9797_1 | TATGAGTGTGGAGTCGTTGCCAAGCAAGCAAATAACTGGGCTANNNAGATCGGAA |
| 9797_2 | TATGAGTGTGGAGTCGTTGCCAAGCAAGCAAATAACTGGGCTANNNAGATCGGAA |
| 13151_1 | TATGAGTGTGGAGTCGTTGCAAAATATCATCAACTGCATTGAGNNNAGATCGGAAG |
| 13151_2 | TATGAGTGTGGAGTCGTTGCAAAATATCATCAACTGCATTGAGNNNAGATCGGAAG |
| 37792_1 | TATGAGTGTGGAGTCGTTGCTCACTTTATCACCCAGGATAGCGNNNAGATCGGAAG |

| | |
|---|---|
| 37792_2 | TATGAGTGTGGAGTCGTTGCTCACTTTATCACCCAGGATAGCGNNNAGATCGGAAG |
| 35580_1 | TATGAGTGTGGAGTCGTTGCCCTCCCAAGCTGTAGGAGATTACNNNAGATCGGAAG |
| 35580_2 | TATGAGTGTGGAGTCGTTGCCCTCCCAAGCTGTAGGAGATTACNNNAGATCGGAAG |
| 33253_1 | TATGAGTGTGGAGTCGTTGCCGTCTTGTACTACAGTGAGACCCNNNAGATCGGAAG |
| 33253_2 | TATGAGTGTGGAGTCGTTGCCGTCTTGTACTACAGTGAGACCCNNNAGATCGGAAG |
| 52662_1 | TATGAGTGTGGAGTCGTTGCATGAACAGCACTCTGAAGCACTANNNAGATCGGAAG |
| 52662_2 | TATGAGTGTGGAGTCGTTGCATGAACAGCACTCTGAAGCACTANNNAGATCGGAAG |
| 51200_1 | TATGAGTGTGGAGTCGTTGCAACCAAATACCACCTGTACCACANNNAGATCGGAAG |
| 51200_2 | TATGAGTGTGGAGTCGTTGCAACCAAATACCACCTGTACCACANNNAGATCGGAAG |
| 50317_1 | TATGAGTGTGGAGTCGTTGCATCGGGCAAGAACTAGTCAACATNNNAGATCGGAA |
| 50317_2 | TATGAGTGTGGAGTCGTTGCATCGGGCAAGAACTAGTCAACATNNNAGATCGGAA |
| 13235_1 | TATGAGTGTGGAGTCGTTGCATTCTGAAGCTTTTGAAAATACTNNNAGATCGGAAG |
| 13235_2 | TATGAGTGTGGAGTCGTTGCATTCTGAAGCTTTTGAAAATACTNNNAGATCGGAAG |
| 29214_1 | TATGAGTGTGGAGTCGTTGCCGTGTGTGTTTGTGTGTGTATCTNNNAGATCGGAAG |
| 29214_2 | TATGAGTGTGGAGTCGTTGCCGTGTGTGTTTGTGTGTGTATCTNNNAGATCGGAAG |
| 51842_1 | TATGAGTGTGGAGTCGTTGCCATCATTTCCCCTTACGCCAATCNNNAGATCGGAAGA |
| 51842_2 | TATGAGTGTGGAGTCGTTGCCATCATTTCCCCTTACGCCAATCNNNAGATCGGAAGA |
| 51588_1 | TATGAGTGTGGAGTCGTTGCCTAGGACATGGAGAAAGAGCGAGNNNAGATCGGA |
| 51588_2 | TATGAGTGTGGAGTCGTTGCCTAGGACATGGAGAAAGAGCGAGNNNAGATCGGA |
| 36925_1 | TATGAGTGTGGAGTCGTTGCCCAGAAAGCCTCCAACAACATTTNNNAGATCGGAAG |
| 36925_2 | TATGAGTGTGGAGTCGTTGCCCAGAAAGCCTCCAACAACATTTNNNAGATCGGAAG |
| 38765_1 | TATGAGTGTGGAGTCGTTGCTGCCTGAACAATACTGTCAAATGTNNNAGATCGGAA |
| 38765_2 | TATGAGTGTGGAGTCGTTGCTGCCTGAACAATACTGTCAAATGTNNNAGATCGGAA |
| 53304_1 | TATGAGTGTGGAGTCGTTGCTAGGCTCTGTGCTAAAAGCTTGANNNAGATCGGAA |
| 53304_2 | TATGAGTGTGGAGTCGTTGCTAGGCTCTGTGCTAAAAGCTTGANNNAGATCGGAA |
| 13298_1 | TATGAGTGTGGAGTCGTTGCCAATTTCATTATCTTCTTCACTCTNNNAGATCGGAAG |
| 13298_2 | TATGAGTGTGGAGTCGTTGCCAATTTCATTATCTTCTTCACTCTNNNAGATCGGAAG |
| 41414_1 | TATGAGTGTGGAGTCGTTGCAAACTCGTGTCATGGAAGTTTGCNNNAGATCGGAA |
| 41414_2 | TATGAGTGTGGAGTCGTTGCAAACTCGTGTCATGGAAGTTTGCNNNAGATCGGAA |
| 52964_1 | TATGAGTGTGGAGTCGTTGCCAGGCACACACACAGGTATCTATNNNAGATCGGAAG |
| 52964_2 | TATGAGTGTGGAGTCGTTGCCAGGCACACACACAGGTATCTATNNNAGATCGGAAG |
| 43586_1 | TATGAGTGTGGAGTCGTTGCCAAAGGGAACTGGAGAGAAGAGGNNNAGATCGGA |
| 43586_2 | TATGAGTGTGGAGTCGTTGCCAAAGGGAACTGGAGAGAAGAGGNNNAGATCGGA |
| 13674_1 | TATGAGTGTGGAGTCGTTGCACACACTAGGAAGATACCTCNNNAGATCGGAAGAG |
| 13674_2 | TATGAGTGTGGAGTCGTTGCACACACTAGGAAGATACCTCNNNAGATCGGAAGAG |
| 9252_1 | TATGAGTGTGGAGTCGTTGCCCCACAAAAGGCAAAGGAATCTGNNNAGATCGGAA |
| 9252_2 | TATGAGTGTGGAGTCGTTGCCCCACAAAAGGCAAAGGAATCTGNNNAGATCGGAA |
| 37311_1 | TATGAGTGTGGAGTCGTTGCACTTGAGCCATACGTTCATGAGANNNAGATCGGAAG |
| 37311_2 | TATGAGTGTGGAGTCGTTGCACTTGAGCCATACGTTCATGAGANNNAGATCGGAAG |
| 33490_1 | TATGAGTGTGGAGTCGTTGCTCTGGACATAAACCATTCCTGCCNNNAGATCGGAAG |
| 33490_2 | TATGAGTGTGGAGTCGTTGCTCTGGACATAAACCATTCCTGCCNNNAGATCGGAAG |
| 35960_1 | TATGAGTGTGGAGTCGTTGCTTGGCAACATAGCAAGATCTCCTNNNAGATCGGAAG |
| 35960_2 | TATGAGTGTGGAGTCGTTGCTTGGCAACATAGCAAGATCTCCTNNNAGATCGGAAG |

| | |
|---|---|
| 43986_1 | TATGAGTGTGGAGTCGTTGCTGCTGCTAATAAAGACATACCCCANNNAGATCGGAA |
| 43986_2 | TATGAGTGTGGAGTCGTTGCTGCTGCTAATAAAGACATACCCCANNNAGATCGGAA |
| 50723_1 | TATGAGTGTGGAGTCGTTGCGCTCTACTCCTAGGGCAGATAGANNNAGATCGGAAG |
| 50723_2 | TATGAGTGTGGAGTCGTTGCGCTCTACTCCTAGGGCAGATAGANNNAGATCGGAAG |
| 29524_1 | TATGAGTGTGGAGTCGTTGCTTTTTTTTTTTTTTGAGACGGAGNNNAGATCGGAAG |
| 29524_2 | TATGAGTGTGGAGTCGTTGCTTTTTTTTTTTTTTGAGACGGAGNNNAGATCGGAAG |
| 31938_1 | TATGAGTGTGGAGTCGTTGCAGAAACCCTCTTACCCAGAAACCNNNAGATCGGAAG |
| 31938_2 | TATGAGTGTGGAGTCGTTGCAGAAACCCTCTTACCCAGAAACCNNNAGATCGGAAG |
| 50975_1 | TATGAGTGTGGAGTCGTTGCATCAGCCACAACTAAATCGCTCTNNNAGATCGGAAG |
| 50975_2 | TATGAGTGTGGAGTCGTTGCATCAGCCACAACTAAATCGCTCTNNNAGATCGGAAG |
| 36420_1 | TATGAGTGTGGAGTCGTTGCGCGGAGAAAATGTTGATAGGCATNNNAGATCGGAA |
| 36420_2 | TATGAGTGTGGAGTCGTTGCGCGGAGAAAATGTTGATAGGCATNNNAGATCGGAA |
| 37522_1 | TATGAGTGTGGAGTCGTTGCTCTGTAATTGCTCTGGCTTGGATNNNAGATCGGAAG |
| 37522_2 | TATGAGTGTGGAGTCGTTGCTCTGTAATTGCTCTGGCTTGGATNNNAGATCGGAAG |
| 36405_1 | TATGAGTGTGGAGTCGTTGCCTAGACCAAACCTTAGGCTGTGTNNNAGATCGGAAG |
| 36405_2 | TATGAGTGTGGAGTCGTTGCCTAGACCAAACCTTAGGCTGTGTNNNAGATCGGAAG |
| 38282_1 | TATGAGTGTGGAGTCGTTGCCGAAAAAAATCCATGCCAGTCCTNNNAGATCGGAAG |
| 38282_2 | TATGAGTGTGGAGTCGTTGCCGAAAAAAATCCATGCCAGTCCTNNNAGATCGGAAG |
| 48479_1 | TATGAGTGTGGAGTCGTTGCCTCTTTCTCTTCCGCTTCCTTCTNNNAGATCGGAAGAG |
| 48479_2 | TATGAGTGTGGAGTCGTTGCCTCTTTCTCTTCCGCTTCCTTCTNNNAGATCGGAAGAG |
| 51751_1 | TATGAGTGTGGAGTCGTTGCTTGAACTCCTGACCTCATGATCCNNNAGATCGGAAG |
| 51751_2 | TATGAGTGTGGAGTCGTTGCTTGAACTCCTGACCTCATGATCCNNNAGATCGGAAG |
| 49306_1 | TATGAGTGTGGAGTCGTTGCAAGCGATTCTATTCTCCTGCCTCNNNAGATCGGAAGA |
| 49306_2 | TATGAGTGTGGAGTCGTTGCAAGCGATTCTATTCTCCTGCCTCNNNAGATCGGAAGA |
| 32220_1 | TATGAGTGTGGAGTCGTTGCGGAATTAGGGTCACGAGTTTCCTNNNAGATCGGAA |
| 32220_2 | TATGAGTGTGGAGTCGTTGCGGAATTAGGGTCACGAGTTTCCTNNNAGATCGGAA |
| 4295_1 | TATGAGTGTGGAGTCGTTGCTTGTAGGAGGGAGTGTCTCACTCNNNAGATCGGAA |
| 4295_2 | TATGAGTGTGGAGTCGTTGCTTGTAGGAGGGAGTGTCTCACTCNNNAGATCGGAA |
| 31511_1 | TATGAGTGTGGAGTCGTTGCTTTATTAGGCAGAGCAAGGTGGANNNAGATCGGAA |
| 31511_2 | TATGAGTGTGGAGTCGTTGCTTTATTAGGCAGAGCAAGGTGGANNNAGATCGGAA |
| 37372_1 | TATGAGTGTGGAGTCGTTGCTGAGGCTGGCTTCTTTTCATTTGNNNAGATCGGAAG |
| 37372_2 | TATGAGTGTGGAGTCGTTGCTGAGGCTGGCTTCTTTTCATTTGNNNAGATCGGAAG |
| 52410_1 | TATGAGTGTGGAGTCGTTGCCCTAGTCCTCCCCTTCCATAAGANNNAGATCGGAAGA |
| 52410_2 | TATGAGTGTGGAGTCGTTGCCCTAGTCCTCCCCTTCCATAAGANNNAGATCGGAAGA |
| 13287_1 | TATGAGTGTGGAGTCGTTGCAGAACTAATGAGATCATAATCAGANNNAGATCGGA |
| 13287_2 | TATGAGTGTGGAGTCGTTGCAGAACTAATGAGATCATAATCAGANNNAGATCGGA |
| 43430_1 | TATGAGTGTGGAGTCGTTGCGCCCTGTTATCCCCATTCTTTTTNNNAGATCGGAAGA |
| 43430_2 | TATGAGTGTGGAGTCGTTGCGCCCTGTTATCCCCATTCTTTTTNNNAGATCGGAAGA |
| 49088_1 | TATGAGTGTGGAGTCGTTGCCCTCTGAAGATTGCCCCTTACTANNNAGATCGGAAG |
| 49088_2 | TATGAGTGTGGAGTCGTTGCCCTCTGAAGATTGCCCCTTACTANNNAGATCGGAAG |
| 39074_1 | TATGAGTGTGGAGTCGTTGCAAGAGCCAGACCCCTTTTCATAGNNNAGATCGGAAG |
| 39074_2 | TATGAGTGTGGAGTCGTTGCAAGAGCCAGACCCCTTTTCATAGNNNAGATCGGAAG |
| 40214_1 | TATGAGTGTGGAGTCGTTGCATCAATGAGCCAACTATTGCTGCNNNAGATCGGAAG |

| | |
|---|---|
| 40214_2 | TATGAGTGTGGAGTCGTTGCATCAATGAGCCAACTATTGCTGCNNNAGATCGGAAG |
| 45818_1 | TATGAGTGTGGAGTCGTTGCCCAAGTAGCTGGGATTACAGGTGNNNAGATCGGAA |
| 45818_2 | TATGAGTGTGGAGTCGTTGCCCAAGTAGCTGGGATTACAGGTGNNNAGATCGGAA |
| 48081_1 | TATGAGTGTGGAGTCGTTGCGAAGTGTGAGGCATAAATCCTGCNNNAGATCGGAA |
| 48081_2 | TATGAGTGTGGAGTCGTTGCGAAGTGTGAGGCATAAATCCTGCNNNAGATCGGAA |
| 51285_1 | TATGAGTGTGGAGTCGTTGCTGGCTAAGATTGCAGGAATCCATNNNAGATCGGAA |
| 51285_2 | TATGAGTGTGGAGTCGTTGCTGGCTAAGATTGCAGGAATCCATNNNAGATCGGAA |
| 2397_1 | TATGAGTGTGGAGTCGTTGCGAGGCATGAATGCTGAGAAATGCNNNAGATCGGAA |
| 2397_2 | TATGAGTGTGGAGTCGTTGCGAGGCATGAATGCTGAGAAATGCNNNAGATCGGAA |
| 39005_1 | TATGAGTGTGGAGTCGTTGCCTTGCAGTGAGCTGAGATCGTNNNAGATCGGAAGA |
| 39005_2 | TATGAGTGTGGAGTCGTTGCCTTGCAGTGAGCTGAGATCGTNNNAGATCGGAAGA |
| 48890_1 | TATGAGTGTGGAGTCGTTGCCATTCTCTTGCCTCAGCCTCANNNAGATCGGAAGAG |
| 48890_2 | TATGAGTGTGGAGTCGTTGCCATTCTCTTGCCTCAGCCTCANNNAGATCGGAAGAG |
| 36706_1 | TATGAGTGTGGAGTCGTTGCCGCCTGTAATCCTAGCATTTTGGNNNAGATCGGAAG |
| 36706_2 | TATGAGTGTGGAGTCGTTGCCGCCTGTAATCCTAGCATTTTGGNNNAGATCGGAAG |
| 43790_1 | TATGAGTGTGGAGTCGTTGCGAGTTTCACCAGACATCCTCCAGNNNAGATCGGAAG |
| 43790_2 | TATGAGTGTGGAGTCGTTGCGAGTTTCACCAGACATCCTCCAGNNNAGATCGGAAG |
| 39712_1 | TATGAGTGTGGAGTCGTTGCAACAAGATGAATAGCAGAGGCCANNNAGATCGGAA |
| 39712_2 | TATGAGTGTGGAGTCGTTGCAACAAGATGAATAGCAGAGGCCANNNAGATCGGAA |
| 52552_1 | TATGAGTGTGGAGTCGTTGCTGTCCAAATGAGTTGCTTCACTGNNNAGATCGGAAG |
| 52552_2 | TATGAGTGTGGAGTCGTTGCTGTCCAAATGAGTTGCTTCACTGNNNAGATCGGAAG |
| 28868_1 | TATGAGTGTGGAGTCGTTGCCTCTCAACAGGTAGCCAATCAGTNNNAGATCGGAAG |
| 28868_2 | TATGAGTGTGGAGTCGTTGCCTCTCAACAGGTAGCCAATCAGTNNNAGATCGGAAG |
| 45872_1 | TATGAGTGTGGAGTCGTTGCTCTTCCCTCATCCCAGTAGAGAGNNNAGATCGGAAG |
| 45872_2 | TATGAGTGTGGAGTCGTTGCTCTTCCCTCATCCCAGTAGAGAGNNNAGATCGGAAG |
| 32456_1 | TATGAGTGTGGAGTCGTTGCAGTGCAAAAAGGCTGAGAATTCCNNNAGATCGGAA |
| 32456_2 | TATGAGTGTGGAGTCGTTGCAGTGCAAAAAGGCTGAGAATTCCNNNAGATCGGAA |
| 51172_1 | TATGAGTGTGGAGTCGTTGCGTTGGAGTAAGTTGGGCAGTTTCNNNAGATCGGAA |
| 51172_2 | TATGAGTGTGGAGTCGTTGCGTTGGAGTAAGTTGGGCAGTTTCNNNAGATCGGAA |
| 35278_1 | TATGAGTGTGGAGTCGTTGCGAAACTTCTTTTTGACGTGTGCGNNNAGATCGGAAG |
| 35278_2 | TATGAGTGTGGAGTCGTTGCGAAACTTCTTTTTGACGTGTGCGNNNAGATCGGAAG |
| 49577_1 | TATGAGTGTGGAGTCGTTGCTGTCACTGGTACTTTGATAGGGANNNAGATCGGAA |
| 49577_2 | TATGAGTGTGGAGTCGTTGCTGTCACTGGTACTTTGATAGGGANNNAGATCGGAA |
| 34119_1 | TATGAGTGTGGAGTCGTTGCTAGTTGCGGTACGATTTGAGTGANNNAGATCGGAA |
| 34119_2 | TATGAGTGTGGAGTCGTTGCTAGTTGCGGTACGATTTGAGTGANNNAGATCGGAA |
| 32043_1 | TATGAGTGTGGAGTCGTTGCCAAGTAGCTGGGACTACAGGCNNNAGATCGGAAGA |
| 32043_2 | TATGAGTGTGGAGTCGTTGCCAAGTAGCTGGGACTACAGGCNNNAGATCGGAAGA |
| 31957_1 | TATGAGTGTGGAGTCGTTGCAAATGCATAGTGAAGAGAGGGCANNNAGATCGGA |
| 31957_2 | TATGAGTGTGGAGTCGTTGCAAATGCATAGTGAAGAGAGGGCANNNAGATCGGA |
| 42955_1 | TATGAGTGTGGAGTCGTTGCCTCCTGAGTAGCTGGGATTACAGNNNAGATCGGAA |
| 42955_2 | TATGAGTGTGGAGTCGTTGCCTCCTGAGTAGCTGGGATTACAGNNNAGATCGGAA |
| 37050_1 | TATGAGTGTGGAGTCGTTGCCCAGCACCCTTTCATGATAAAGCNNNAGATCGGAAG |
| 37050_2 | TATGAGTGTGGAGTCGTTGCCCAGCACCCTTTCATGATAAAGCNNNAGATCGGAAG |

| | |
|---|---|
| 53238_1 | TATGAGTGTGGAGTCGTTGCGCTGGTAAAACTGCTTGCTGTAANNNAGATCGGAAG |
| 53238_2 | TATGAGTGTGGAGTCGTTGCGCTGGTAAAACTGCTTGCTGTAANNNAGATCGGAAG |
| 38964_1 | TATGAGTGTGGAGTCGTTGCGACGGATGCATTCTTCAGTATGCNNNAGATCGGAAG |
| 38964_2 | TATGAGTGTGGAGTCGTTGCGACGGATGCATTCTTCAGTATGCNNNAGATCGGAAG |
| 13645_1 | TATGAGTGTGGAGTCGTTGCAGAACTCTGATGTTTGTAAAATCNNNAGATCGGAAG |
| 13645_2 | TATGAGTGTGGAGTCGTTGCAGAACTCTGATGTTTGTAAAATCNNNAGATCGGAAG |
| 52384_1 | TATGAGTGTGGAGTCGTTGCGCTCTGTCATTGCCCTCATACTANNNAGATCGGAAGA |
| 52384_2 | TATGAGTGTGGAGTCGTTGCGCTCTGTCATTGCCCTCATACTANNNAGATCGGAAGA |
| 34596_1 | TATGAGTGTGGAGTCGTTGCCCAGAGAAACGACCACTGAAATGNNNAGATCGGAA |
| 34596_2 | TATGAGTGTGGAGTCGTTGCCCAGAGAAACGACCACTGAAATGNNNAGATCGGAA |
| 13411_1 | TATGAGTGTGGAGTCGTTGCCATTAGTAGCCTGACTGTTAATNNNAGATCGGAAGA |
| 13411_2 | TATGAGTGTGGAGTCGTTGCCATTAGTAGCCTGACTGTTAATNNNAGATCGGAAGA |
| 47212_1 | TATGAGTGTGGAGTCGTTGCCTCTTGCATGACTGACTCCTTCTNNNAGATCGGAAGA |
| 47212_2 | TATGAGTGTGGAGTCGTTGCCTCTTGCATGACTGACTCCTTCTNNNAGATCGGAAGA |
| 3180_1 | TATGAGTGTGGAGTCGTTGCACTGCTACCAAAAATTGGTGGGTNNNAGATCGGAA |
| 3180_2 | TATGAGTGTGGAGTCGTTGCACTGCTACCAAAAATTGGTGGGTNNNAGATCGGAA |
| 35360_1 | TATGAGTGTGGAGTCGTTGCCTCCCAGGTTCCAGTGATTCTCNNNAGATCGGAAGA |
| 35360_2 | TATGAGTGTGGAGTCGTTGCCTCCCAGGTTCCAGTGATTCTCNNNAGATCGGAAGA |
| 2093_1 | TATGAGTGTGGAGTCGTTGCTGTAACGACCAGTGTCTTGAACCNNNAGATCGGAAG |
| 2093_2 | TATGAGTGTGGAGTCGTTGCTGTAACGACCAGTGTCTTGAACCNNNAGATCGGAAG |
| 40980_1 | TATGAGTGTGGAGTCGTTGCAAATCACCGAGTATGGTCTCTGGNNNAGATCGGAAG |
| 40980_2 | TATGAGTGTGGAGTCGTTGCAAATCACCGAGTATGGTCTCTGGNNNAGATCGGAAG |
| 32048_1 | TATGAGTGTGGAGTCGTTGCCGCTCAAACAATCTTCCCACTTCNNNAGATCGGAAGA |
| 32048_2 | TATGAGTGTGGAGTCGTTGCCGCTCAAACAATCTTCCCACTTCNNNAGATCGGAAGA |
| 33781_1 | TATGAGTGTGGAGTCGTTGCTTGCATGCCTGTAGTAGTCTCAGNNNAGATCGGAAG |
| 33781_2 | TATGAGTGTGGAGTCGTTGCTTGCATGCCTGTAGTAGTCTCAGNNNAGATCGGAAG |
| 44183_1 | TATGAGTGTGGAGTCGTTGCCTGAGCGCAATGTTTTGTGTTTTNNNAGATCGGAAG |
| 44183_2 | TATGAGTGTGGAGTCGTTGCCTGAGCGCAATGTTTTGTGTTTTNNNAGATCGGAAG |
| 51910_1 | TATGAGTGTGGAGTCGTTGCGAGCTTTCTACCTAACACTGGCTNNNAGATCGGAAGA |
| 51910_2 | TATGAGTGTGGAGTCGTTGCGAGCTTTCTACCTAACACTGGCTNNNAGATCGGAAGA |
| 35821_1 | TATGAGTGTGGAGTCGTTGCCTGGACATTACAGGGCATCTTCTNNNAGATCGGAAG |
| 35821_2 | TATGAGTGTGGAGTCGTTGCCTGGACATTACAGGGCATCTTCTNNNAGATCGGAAG |
| 32084_1 | TATGAGTGTGGAGTCGTTGCGGCCCTGTCTTTGACAATTTCTCNNNAGATCGGAAGA |
| 32084_2 | TATGAGTGTGGAGTCGTTGCGGCCCTGTCTTTGACAATTTCTCNNNAGATCGGAAGA |
| 6206_1 | TATGAGTGTGGAGTCGTTGCTTGGCCTTCACATTAGACTGTAGAGNNNAGATCGGA |
| 6206_2 | TATGAGTGTGGAGTCGTTGCTTGGCCTTCACATTAGACTGTAGAGNNNAGATCGGA |
| 10075_1 | TATGAGTGTGGAGTCGTTGCAGTAGGTGCAGACTAGAAAGCCANNNAGATCGGAA |
| 10075_2 | TATGAGTGTGGAGTCGTTGCAGTAGGTGCAGACTAGAAAGCCANNNAGATCGGAA |
| 35210_1 | TATGAGTGTGGAGTCGTTGCCAGGCTCCCAAATAGCTGGAATANNNAGATCGGAAG |
| 35210_2 | TATGAGTGTGGAGTCGTTGCCAGGCTCCCAAATAGCTGGAATANNNAGATCGGAAG |
| 40632_1 | TATGAGTGTGGAGTCGTTGCCCTGGGAGACAGAGTGAGACTCNNNAGATCGGAAG |
| 40632_2 | TATGAGTGTGGAGTCGTTGCCCTGGGAGACAGAGTGAGACTCNNNAGATCGGAAG |
| 43536_1 | TATGAGTGTGGAGTCGTTGCACGCCCGGCTAATTTTTTGTATTNNNAGATCGGAAGA |

| | |
|---|---|
| 43536_2 | TATGAGTGTGGAGTCGTTGCACGCCCGGCTAATTTTTTGTATTNNNAGATCGGAAG |
| 44975_1 | TATGAGTGTGGAGTCGTTGCGAATGGGAGTTGCAGTCGAGTAGNNNAGATCGGA |
| 44975_2 | TATGAGTGTGGAGTCGTTGCGAATGGGAGTTGCAGTCGAGTAGNNNAGATCGGA |
| 47117_1 | TATGAGTGTGGAGTCGTTGCCTGGAGTGCAATCATGGCTAAAGNNNAGATCGGAA |
| 47117_2 | TATGAGTGTGGAGTCGTTGCCTGGAGTGCAATCATGGCTAAAGNNNAGATCGGAA |
| 30043_1 | TATGAGTGTGGAGTCGTTGCCTACAGGTGAAGCCGGAGTGNNNAGATCGGAAGAG |
| 30043_2 | TATGAGTGTGGAGTCGTTGCCTACAGGTGAAGCCGGAGTGNNNAGATCGGAAGAG |
| 39154_1 | TATGAGTGTGGAGTCGTTGCCACATCAACCAGTGGAACACAATNNNAGATCGGAAG |
| 39154_2 | TATGAGTGTGGAGTCGTTGCCACATCAACCAGTGGAACACAATNNNAGATCGGAAG |
| 34563_1 | TATGAGTGTGGAGTCGTTGCAGAGAAGGGGTCTTACCATGTTGNNNAGATCGGAA |
| 34563_2 | TATGAGTGTGGAGTCGTTGCAGAGAAGGGGTCTTACCATGTTGNNNAGATCGGAA |
| 10693_1 | TATGAGTGTGGAGTCGTTGCCCCGTGATCATCTCTGACCTGATNNNAGATCGGAAG |
| 10693_2 | TATGAGTGTGGAGTCGTTGCCCCGTGATCATCTCTGACCTGATNNNAGATCGGAAG |
| 29863_1 | TATGAGTGTGGAGTCGTTGCGTGAGATGACACAGTTACAGGGANNNAGATCGGAA |
| 29863_2 | TATGAGTGTGGAGTCGTTGCGTGAGATGACACAGTTACAGGGANNNAGATCGGAA |
| 32291_1 | TATGAGTGTGGAGTCGTTGCTGTCTGTTGGTGTATCTGTGTGTNNNAGATCGGAAG |
| 32291_2 | TATGAGTGTGGAGTCGTTGCTGTCTGTTGGTGTATCTGTGTGTNNNAGATCGGAAG |
| 40680_1 | TATGAGTGTGGAGTCGTTGCCCCTCATAGAGCAGATGTGAGACNNNAGATCGGAA |
| 40680_2 | TATGAGTGTGGAGTCGTTGCCCCTCATAGAGCAGATGTGAGACNNNAGATCGGAA |
| 47860_1 | TATGAGTGTGGAGTCGTTGCTCCCAAAGTGCTGAGATTACAGGNNNAGATCGGAA |
| 47860_2 | TATGAGTGTGGAGTCGTTGCTCCCAAAGTGCTGAGATTACAGGNNNAGATCGGAA |
| 13011_1 | TATGAGTGTGGAGTCGTTGCCATACATCTTGATTCTTTTCCATNNNAGATCGGAAGA |
| 13011_2 | TATGAGTGTGGAGTCGTTGCCATACATCTTGATTCTTTTCCATNNNAGATCGGAAGA |
| 47124_1 | TATGAGTGTGGAGTCGTTGCCACCATGCTCGGCTAATTTGTTTNNNAGATCGGAAG |
| 47124_2 | TATGAGTGTGGAGTCGTTGCCACCATGCTCGGCTAATTTGTTTNNNAGATCGGAAG |
| 1998_1 | TATGAGTGTGGAGTCGTTGCCCTGGATGAACACAGTTGCAAAANNNAGATCGGAA |
| 1998_2 | TATGAGTGTGGAGTCGTTGCCCTGGATGAACACAGTTGCAAAANNNAGATCGGAA |
| 43245_1 | TATGAGTGTGGAGTCGTTGCAGAGAGAGGCAGAGAGAGAGAAGNNNAGATCGG |
| 43245_2 | TATGAGTGTGGAGTCGTTGCAGAGAGAGGCAGAGAGAGAGAAGNNNAGATCGG |
| 10906_1 | TATGAGTGTGGAGTCGTTGCGACTAAACTCGACGAAGGCGAANNNAGATCGGAAG |
| 10906_2 | TATGAGTGTGGAGTCGTTGCGACTAAACTCGACGAAGGCGAANNNAGATCGGAAG |
| 45612_1 | TATGAGTGTGGAGTCGTTGCCCAAGTAGCTGGGATTACACGTNNNAGATCGGAAG |
| 45612_2 | TATGAGTGTGGAGTCGTTGCCCAAGTAGCTGGGATTACACGTNNNAGATCGGAAG |
| 44325_1 | TATGAGTGTGGAGTCGTTGCCATGAAGTCGCTGAAGGAAAGTGNNNAGATCGGAA |
| 44325_2 | TATGAGTGTGGAGTCGTTGCCATGAAGTCGCTGAAGGAAAGTGNNNAGATCGGAA |
| 38377_1 | TATGAGTGTGGAGTCGTTGCATGGAACCACTGTTGCATATGTGNNNAGATCGGAA |
| 38377_2 | TATGAGTGTGGAGTCGTTGCATGGAACCACTGTTGCATATGTGNNNAGATCGGAA |
| 34128_1 | TATGAGTGTGGAGTCGTTGCACCTAGGGAAGACTAAGCATCTCTNNNAGATCGGAA |
| 34128_2 | TATGAGTGTGGAGTCGTTGCACCTAGGGAAGACTAAGCATCTCTNNNAGATCGGAA |
| 32597_1 | TATGAGTGTGGAGTCGTTGCATGTCAGATTCAATGCAGTCCCTNNNAGATCGGAAG |
| 32597_2 | TATGAGTGTGGAGTCGTTGCATGTCAGATTCAATGCAGTCCCTNNNAGATCGGAAG |
| 34920_1 | TATGAGTGTGGAGTCGTTGCTAAAAATGCATATATCGGCCGGGNNNAGATCGGAA |
| 34920_2 | TATGAGTGTGGAGTCGTTGCTAAAAATGCATATATCGGCCGGGNNNAGATCGGAA |

| | |
|---|---|
| 32629_1 | TATGAGTGTGGAGTCGTTGCGAGTTTTGCTCTTGTTGCCCAGNNNAGATCGGAAGA |
| 32629_2 | TATGAGTGTGGAGTCGTTGCGAGTTTTGCTCTTGTTGCCCAGNNNAGATCGGAAGA |
| 46881_1 | TATGAGTGTGGAGTCGTTGCAATTGCCCACACAGTTTTTGTCANNNAGATCGGAAG |
| 46881_2 | TATGAGTGTGGAGTCGTTGCAATTGCCCACACAGTTTTTGTCANNNAGATCGGAAG |
| 53375_1 | TATGAGTGTGGAGTCGTTGCCCTCCCAGTCATGTTTCCTGTTANNNAGATCGGAAGA |
| 53375_2 | TATGAGTGTGGAGTCGTTGCCCTCCCAGTCATGTTTCCTGTTANNNAGATCGGAAGA |
| 28843_1 | TATGAGTGTGGAGTCGTTGCCTCCAGACTTTTTCATCTTCCCANNNAGATCGGAAGA |
| 28843_2 | TATGAGTGTGGAGTCGTTGCCTCCAGACTTTTTCATCTTCCCANNNAGATCGGAAGA |
| 39651_1 | TATGAGTGTGGAGTCGTTGCCCAAACAAACAAAAAACTGGCCCNNNAGATCGGAAG |
| 39651_2 | TATGAGTGTGGAGTCGTTGCCCAAACAAACAAAAAACTGGCCCNNNAGATCGGAAG |
| 34746_1 | TATGAGTGTGGAGTCGTTGCTTAGAGTGCAATGGTGGGATCTCNNNAGATCGGAA |
| 34746_2 | TATGAGTGTGGAGTCGTTGCTTAGAGTGCAATGGTGGGATCTCNNNAGATCGGAA |
| 41596_1 | TATGAGTGTGGAGTCGTTGCTCAAGTAGCTGGAATTACAGGCANNNAGATCGGAA |
| 41596_2 | TATGAGTGTGGAGTCGTTGCTCAAGTAGCTGGAATTACAGGCANNNAGATCGGAA |
| 13554_1 | TATGAGTGTGGAGTCGTTGCACAGTTGGTATTTTTAATTCTCCNNNAGATCGGAAG |
| 13554_2 | TATGAGTGTGGAGTCGTTGCACAGTTGGTATTTTTAATTCTCCNNNAGATCGGAAG |
| 12938_1 | TATGAGTGTGGAGTCGTTGCCTTCTAAAAGTTCTTCATCTGAANNNAGATCGGAAG |
| 12938_2 | TATGAGTGTGGAGTCGTTGCCTTCTAAAAGTTCTTCATCTGAANNNAGATCGGAAG |
| 45640_1 | TATGAGTGTGGAGTCGTTGCTAGTAGAGACGGGGTTTCTCCATNNNAGATCGGAA |
| 45640_2 | TATGAGTGTGGAGTCGTTGCTAGTAGAGACGGGGTTTCTCCATNNNAGATCGGAA |
| 36039_1 | TATGAGTGTGGAGTCGTTGCGGGCACTGAGAAACAGAAACAAANNNAGATCGGAA |
| 36039_2 | TATGAGTGTGGAGTCGTTGCGGGCACTGAGAAACAGAAACAAANNNAGATCGGAA |
| 45992_1 | TATGAGTGTGGAGTCGTTGCCCTCCCAGGTTGAATTCGTCTTTNNNAGATCGGAAG |
| 45992_2 | TATGAGTGTGGAGTCGTTGCCCTCCCAGGTTGAATTCGTCTTTNNNAGATCGGAAG |
| 5692_1 | TATGAGTGTGGAGTCGTTGCACATACTGAGTTGGAGTTGCCTGNNNAGATCGGAA |
| 5692_2 | TATGAGTGTGGAGTCGTTGCACATACTGAGTTGGAGTTGCCTGNNNAGATCGGAA |
| 36051_1 | TATGAGTGTGGAGTCGTTGCTGCTTTGTGTATTTGGGAACTCTGNNNAGATCGGAA |
| 36051_2 | TATGAGTGTGGAGTCGTTGCTGCTTTGTGTATTTGGGAACTCTGNNNAGATCGGAA |
| 53248_1 | TATGAGTGTGGAGTCGTTGCGACCAGAAGAGAGCAGAGCTGNNNAGATCGGAAG |
| 53248_2 | TATGAGTGTGGAGTCGTTGCGACCAGAAGAGAGCAGAGCTGNNNAGATCGGAAG |
| 34407_1 | TATGAGTGTGGAGTCGTTGCTTTTTCCCTGACAGTGCTAAGGANNNAGATCGGAAG |
| 34407_2 | TATGAGTGTGGAGTCGTTGCTTTTTCCCTGACAGTGCTAAGGANNNAGATCGGAAG |
| 47829_1 | TATGAGTGTGGAGTCGTTGCCTCCTGAGTAGCTGGGACTACANNNAGATCGGAAG |
| 47829_2 | TATGAGTGTGGAGTCGTTGCCTCCTGAGTAGCTGGGACTACANNNAGATCGGAAG |
| 30659_1 | TATGAGTGTGGAGTCGTTGCCCTTAGCCAGCTCTTTGTCGGNNNAGATCGGAAGAG |
| 30659_2 | TATGAGTGTGGAGTCGTTGCCCTTAGCCAGCTCTTTGTCGGNNNAGATCGGAAGAG |
| 13097_1 | TATGAGTGTGGAGTCGTTGCGGCTATTTTTCCACTGATTAAATNNNAGATCGGAAG |
| 13097_2 | TATGAGTGTGGAGTCGTTGCGGCTATTTTTCCACTGATTAAATNNNAGATCGGAAG |
| 50479_1 | TATGAGTGTGGAGTCGTTGCGCTCTGTTACCTAGTCTGGTGTGNNNAGATCGGAAG |
| 50479_2 | TATGAGTGTGGAGTCGTTGCGCTCTGTTACCTAGTCTGGTGTGNNNAGATCGGAAG |
| 32634_1 | TATGAGTGTGGAGTCGTTGCGTTCAAGCAATTCTCTGCCTCAGNNNAGATCGGAAG |
| 32634_2 | TATGAGTGTGGAGTCGTTGCGTTCAAGCAATTCTCTGCCTCAGNNNAGATCGGAAG |
| 32953_1 | TATGAGTGTGGAGTCGTTGCGCTTGTAATCCCAGCACTTTGAGNNNAGATCGGAAG |

| | |
|---|---|
| 32953_2 | TATGAGTGTGGAGTCGTTGCGCTTGTAATCCCAGCACTTTGAGNNNAGATCGGAAG |
| 50402_1 | TATGAGTGTGGAGTCGTTGCCTGACTGTTCTGTCTCTGCCTACNNNAGATCGGAAGA |
| 50402_2 | TATGAGTGTGGAGTCGTTGCCTGACTGTTCTGTCTCTGCCTACNNNAGATCGGAAGA |
| 13367_1 | TATGAGTGTGGAGTCGTTGCTTGAAGTTTTCCATCAATTAACANNNAGATCGGAAGA |
| 13367_2 | TATGAGTGTGGAGTCGTTGCTTGAAGTTTTCCATCAATTAACANNNAGATCGGAAGA |
| 37110_1 | TATGAGTGTGGAGTCGTTGCTGGTGTGAACCCAGGTCATATTTNNNAGATCGGAAG |
| 37110_2 | TATGAGTGTGGAGTCGTTGCTGGTGTGAACCCAGGTCATATTTNNNAGATCGGAAG |
| 31383_1 | TATGAGTGTGGAGTCGTTGCTTCAAGTCTGAAATCTGCAGGGANNNAGATCGGAA |
| 31383_2 | TATGAGTGTGGAGTCGTTGCTTCAAGTCTGAAATCTGCAGGGANNNAGATCGGAA |
| 34459_1 | TATGAGTGTGGAGTCGTTGCGTCACAGACACGGGTGTTCTTANNNAGATCGGAAGA |
| 34459_2 | TATGAGTGTGGAGTCGTTGCGTCACAGACACGGGTGTTCTTANNNAGATCGGAAGA |
| 51777_1 | TATGAGTGTGGAGTCGTTGCTCTGAATTCTGCCCATGGTACAANNNAGATCGGAAGA |
| 51777_2 | TATGAGTGTGGAGTCGTTGCTCTGAATTCTGCCCATGGTACAANNNAGATCGGAAGA |
| 4147_1 | TATGAGTGTGGAGTCGTTGCCACCACTACTACCGCCAATGAAGNNNAGATCGGAAGA |
| 4147_2 | TATGAGTGTGGAGTCGTTGCCACCACTACTACCGCCAATGAAGNNNAGATCGGAAGA |
| 37856_1 | TATGAGTGTGGAGTCGTTGCTTCTTTTCCCACATGAACCTCCANNNAGATCGGAAGA |
| 37856_2 | TATGAGTGTGGAGTCGTTGCTTCTTTTCCCACATGAACCTCCANNNAGATCGGAAGA |
| 38130_1 | TATGAGTGTGGAGTCGTTGCGAGGTCAGGAGATTGAGACCATCNNNAGATCGGAA |
| 38130_2 | TATGAGTGTGGAGTCGTTGCGAGGTCAGGAGATTGAGACCATCNNNAGATCGGAA |
| 2775_1 | TATGAGTGTGGAGTCGTTGCTTTCCATTTCACATTGGTTTTCCTNNNAGATCGGAAGA |
| 2775_2 | TATGAGTGTGGAGTCGTTGCTTTCCATTTCACATTGGTTTTCCTNNNAGATCGGAAGA |
| 36246_1 | TATGAGTGTGGAGTCGTTGCTGCTAAAGATAATGGCCTCCTCCNNNAGATCGGAAGA |
| 36246_2 | TATGAGTGTGGAGTCGTTGCTGCTAAAGATAATGGCCTCCTCCNNNAGATCGGAAGA |
| 40602_1 | TATGAGTGTGGAGTCGTTGCCTGAGGCAGGAGAATAGCTTGAANNNAGATCGGAA |
| 40602_2 | TATGAGTGTGGAGTCGTTGCCTGAGGCAGGAGAATAGCTTGAANNNAGATCGGAA |
| 29774_1 | TATGAGTGTGGAGTCGTTGCGCCCTATTCTTCAGTACACGAGANNNAGATCGGAAGA |
| 29774_2 | TATGAGTGTGGAGTCGTTGCGCCCTATTCTTCAGTACACGAGANNNAGATCGGAAGA |
| 44827_1 | TATGAGTGTGGAGTCGTTGCACAATAGCCTTTATTGCTGTGGCNNNAGATCGGAAG |
| 44827_2 | TATGAGTGTGGAGTCGTTGCACAATAGCCTTTATTGCTGTGGCNNNAGATCGGAAG |
| 30959_1 | TATGAGTGTGGAGTCGTTGCCACGTATGCTATGCCCACTTTTCNNNAGATCGGAAGA |
| 30959_2 | TATGAGTGTGGAGTCGTTGCCACGTATGCTATGCCCACTTTTCNNNAGATCGGAAGA |
| 30817_1 | TATGAGTGTGGAGTCGTTGCGATTCACAGGGAAATTGGCTCAGNNNAGATCGGAA |
| 30817_2 | TATGAGTGTGGAGTCGTTGCGATTCACAGGGAAATTGGCTCAGNNNAGATCGGAA |
| 29223_1 | TATGAGTGTGGAGTCGTTGCCTCCCTTCCTTTCCCTCTCCTNNNAGATCGGAAGAGCA |
| 29223_2 | TATGAGTGTGGAGTCGTTGCCTCCCTTCCTTTCCCTCTCCTNNNAGATCGGAAGAGCA |
| 47661_1 | TATGAGTGTGGAGTCGTTGCTTCAAGCCATTCTCCTGTCTCAGNNNAGATCGGAAGA |
| 47661_2 | TATGAGTGTGGAGTCGTTGCTTCAAGCCATTCTCCTGTCTCAGNNNAGATCGGAAGA |
| 46914_1 | TATGAGTGTGGAGTCGTTGCTAAAGACAAGAAGAGGTGGGAGGNNNAGATCGGA |
| 46914_2 | TATGAGTGTGGAGTCGTTGCTAAAGACAAGAAGAGGTGGGAGGNNNAGATCGGA |
| 35621_1 | TATGAGTGTGGAGTCGTTGCAATCCAGCTGTGAACACATTTGGNNNAGATCGGAAG |
| 35621_2 | TATGAGTGTGGAGTCGTTGCAATCCAGCTGTGAACACATTTGGNNNAGATCGGAAG |
| 40187_1 | TATGAGTGTGGAGTCGTTGCACACATGTATTTGCTTGCATATGCNNNAGATCGGAA |
| 40187_2 | TATGAGTGTGGAGTCGTTGCACACATGTATTTGCTTGCATATGCNNNAGATCGGAA |

| | |
|---|---|
| 41872_1 | TATGAGTGTGGAGTCGTTGCCAAAATGTTGGGATTACAGGCGTNNNAGATCGGAA |
| 41872_2 | TATGAGTGTGGAGTCGTTGCCAAAATGTTGGGATTACAGGCGTNNNAGATCGGAA |
| 2630_1 | TATGAGTGTGGAGTCGTTGCTTTCATCAGAAGTGAGCCAGGTGNNNAGATCGGAA |
| 2630_2 | TATGAGTGTGGAGTCGTTGCTTTCATCAGAAGTGAGCCAGGTGNNNAGATCGGAA |
| 6884_1 | TATGAGTGTGGAGTCGTTGCGCATAGACAAAGGTGAAGAGGGCNNNAGATCGGA |
| 6884_2 | TATGAGTGTGGAGTCGTTGCGCATAGACAAAGGTGAAGAGGGCNNNAGATCGGA |
| 40402_1 | TATGAGTGTGGAGTCGTTGCGGCCTGCCAAAAACTATTAGGACNNNAGATCGGAA |
| 40402_2 | TATGAGTGTGGAGTCGTTGCGGCCTGCCAAAAACTATTAGGACNNNAGATCGGAA |
| 50165_1 | TATGAGTGTGGAGTCGTTGCCTCTCTAGGGAAAATCCGAGCTCNNNAGATCGGAAG |
| 50165_2 | TATGAGTGTGGAGTCGTTGCCTCTCTAGGGAAAATCCGAGCTCNNNAGATCGGAAG |
| 48470_1 | TATGAGTGTGGAGTCGTTGCCTCAAGCCATACTCCTACCTCAGNNNAGATCGGAAG |
| 48470_2 | TATGAGTGTGGAGTCGTTGCCTCAAGCCATACTCCTACCTCAGNNNAGATCGGAAG |
| 43567_1 | TATGAGTGTGGAGTCGTTGCGCCCAGCCTGTATGTCTTTTTTTNNNAGATCGGAAG |
| 43567_2 | TATGAGTGTGGAGTCGTTGCGCCCAGCCTGTATGTCTTTTTTTNNNAGATCGGAAG |
| 37871_1 | TATGAGTGTGGAGTCGTTGCCTGTAGAGAAGGCGGTACTTGAANNNAGATCGGAA |
| 37871_2 | TATGAGTGTGGAGTCGTTGCCTGTAGAGAAGGCGGTACTTGAANNNAGATCGGAA |
| 29397_1 | TATGAGTGTGGAGTCGTTGCTTGCTTGTCAGATGCCTATGACANNNAGATCGGAAG |
| 29397_2 | TATGAGTGTGGAGTCGTTGCTTGCTTGTCAGATGCCTATGACANNNAGATCGGAAG |
| 36453_1 | TATGAGTGTGGAGTCGTTGCACCAAAAAATGAGGGAGACTGGTNNNAGATCGGAA |
| 36453_2 | TATGAGTGTGGAGTCGTTGCACCAAAAAATGAGGGAGACTGGTNNNAGATCGGAA |
| 42293_1 | TATGAGTGTGGAGTCGTTGCCCCAGGAGATTGATGGTTACACANNNAGATCGGAA |
| 42293_2 | TATGAGTGTGGAGTCGTTGCCCCAGGAGATTGATGGTTACACANNNAGATCGGAA |
| 30593_1 | TATGAGTGTGGAGTCGTTGCAAGCTCCTGCATTTGTGTGAGTTNNNAGATCGGAAG |
| 30593_2 | TATGAGTGTGGAGTCGTTGCAAGCTCCTGCATTTGTGTGAGTTNNNAGATCGGAAG |
| 30416_1 | TATGAGTGTGGAGTCGTTGCGCGGCTTTTTGCCCCTCACGNNNAGATCGGAAGAGC |
| 30416_2 | TATGAGTGTGGAGTCGTTGCGCGGCTTTTTGCCCCTCACGNNNAGATCGGAAGAGC |
| 36941_1 | TATGAGTGTGGAGTCGTTGCTAGCTACTCTTTGAGCCAGCAATNNNAGATCGGAAG |
| 36941_2 | TATGAGTGTGGAGTCGTTGCTAGCTACTCTTTGAGCCAGCAATNNNAGATCGGAAG |
| 45938_1 | TATGAGTGTGGAGTCGTTGCGGAAGAAGAAGGCCAAGAACAAGNNNAGATCGGA |
| 45938_2 | TATGAGTGTGGAGTCGTTGCGGAAGAAGAAGGCCAAGAACAAGNNNAGATCGGA |
| 46853_1 | TATGAGTGTGGAGTCGTTGCCACAATCTCGGCTCATTGCAAGNNNAGATCGGAAGA |
| 46853_2 | TATGAGTGTGGAGTCGTTGCCACAATCTCGGCTCATTGCAAGNNNAGATCGGAAGA |
| 29765_1 | TATGAGTGTGGAGTCGTTGCAGATCATGACCCAGGTATCTTGCNNNAGATCGGAAG |
| 29765_2 | TATGAGTGTGGAGTCGTTGCAGATCATGACCCAGGTATCTTGCNNNAGATCGGAAG |
| 49078_1 | TATGAGTGTGGAGTCGTTGCTTTTCTCTTCTACTGGCTTAGACANNNAGATCGGAAG |
| 49078_2 | TATGAGTGTGGAGTCGTTGCTTTTCTCTTCTACTGGCTTAGACANNNAGATCGGAAG |
| 33212_1 | TATGAGTGTGGAGTCGTTGCCAGACAGAAGTGAAGGCAAACAGNNNAGATCGGAA |
| 33212_2 | TATGAGTGTGGAGTCGTTGCCAGACAGAAGTGAAGGCAAACAGNNNAGATCGGAA |
| 51808_1 | TATGAGTGTGGAGTCGTTGCATGTGTGCTGATTGGTTTGTGAGNNNAGATCGGAA |
| 51808_2 | TATGAGTGTGGAGTCGTTGCATGTGTGCTGATTGGTTTGTGAGNNNAGATCGGAA |
| 32333_1 | TATGAGTGTGGAGTCGTTGCTTGGGTTCTGGTTCTCTTTGGTTNNNAGATCGGAAG |
| 32333_2 | TATGAGTGTGGAGTCGTTGCTTGGGTTCTGGTTCTCTTTGGTTNNNAGATCGGAAG |
| 34058_1 | TATGAGTGTGGAGTCGTTGCCCATTCTCCTGCCTTAGTCTCTCNNNAGATCGGAAGA |

| | |
|---|---|
| 34058_2 | TATGAGTGTGGAGTCGTTGCCCATTCTCCTGCCTTAGTCTCTCNNNAGATCGGAAGA |
| 36442_1 | TATGAGTGTGGAGTCGTTGCATTGTAGTGACTGCTTTGCCTTGNNNAGATCGGAAG |
| 36442_2 | TATGAGTGTGGAGTCGTTGCATTGTAGTGACTGCTTTGCCTTGNNNAGATCGGAAG |
| 42588_1 | TATGAGTGTGGAGTCGTTGCTGTGTATGTGTCTGTGACCTTCANNNAGATCGGAAG |
| 42588_2 | TATGAGTGTGGAGTCGTTGCTGTGTATGTGTCTGTGACCTTCANNNAGATCGGAAG |
| 36418_1 | TATGAGTGTGGAGTCGTTGCAGGTTTATGAAAAAGGAGGGCCANNNAGATCGGAA |
| 36418_2 | TATGAGTGTGGAGTCGTTGCAGGTTTATGAAAAAGGAGGGCCANNNAGATCGGAA |
| 30789_1 | TATGAGTGTGGAGTCGTTGCGCAGGGACCTTCTCTTTCTTGTTNNNAGATCGGAAG |
| 30789_2 | TATGAGTGTGGAGTCGTTGCGCAGGGACCTTCTCTTTCTTGTTNNNAGATCGGAAG |
| 13404_1 | TATGAGTGTGGAGTCGTTGCGAGCAATTAAAACAGATTCCTAANNNAGATCGGAAG |
| 13404_2 | TATGAGTGTGGAGTCGTTGCGAGCAATTAAAACAGATTCCTAANNNAGATCGGAAG |
| 43647_1 | TATGAGTGTGGAGTCGTTGCAGAGACAGGGAGAGAGAGAGATGNNNAGATCGGA |
| 43647_2 | TATGAGTGTGGAGTCGTTGCAGAGACAGGGAGAGAGAGAGATGNNNAGATCGGA |
| 49285_1 | TATGAGTGTGGAGTCGTTGCCAGCCTCCTGAATAGTTGGGATTNNNAGATCGGAAG |
| 49285_2 | TATGAGTGTGGAGTCGTTGCCAGCCTCCTGAATAGTTGGGATTNNNAGATCGGAAG |
| 31051_1 | TATGAGTGTGGAGTCGTTGCCCCCCCACCATTTCTGAGAAATANNNAGATCGGAAG |
| 31051_2 | TATGAGTGTGGAGTCGTTGCCCCCCCACCATTTCTGAGAAATANNNAGATCGGAAG |
| 34890_1 | TATGAGTGTGGAGTCGTTGCCTCTCGAGTGGCTGGGACTATAGNNNAGATCGGAA |
| 34890_2 | TATGAGTGTGGAGTCGTTGCCTCTCGAGTGGCTGGGACTATAGNNNAGATCGGAA |
| 10687_1 | TATGAGTGTGGAGTCGTTGCTACTTTTCCTGCAGTGTACAGCCNNNAGATCGGAAG |
| 10687_2 | TATGAGTGTGGAGTCGTTGCTACTTTTCCTGCAGTGTACAGCCNNNAGATCGGAAG |
| 10562_1 | TATGAGTGTGGAGTCGTTGCTAGCTCCAACCCTGGTTTTGTTTNNNAGATCGGAAG |
| 10562_2 | TATGAGTGTGGAGTCGTTGCTAGCTCCAACCCTGGTTTTGTTTNNNAGATCGGAAG |
| 32454_1 | TATGAGTGTGGAGTCGTTGCGTGTGTGGTGAATTGGAAGGAAGNNNAGATCGGA |
| 32454_2 | TATGAGTGTGGAGTCGTTGCGTGTGTGGTGAATTGGAAGGAAGNNNAGATCGGA |
| 52038_1 | TATGAGTGTGGAGTCGTTGCTTAGTTTGCTGAGGATGATGGCTNNNAGATCGGAA |
| 52038_2 | TATGAGTGTGGAGTCGTTGCTTAGTTTGCTGAGGATGATGGCTNNNAGATCGGAA |
| 43767_1 | TATGAGTGTGGAGTCGTTGCGAACAGAGAGGTGAAGGACCATTNNNAGATCGGAA |
| 43767_2 | TATGAGTGTGGAGTCGTTGCGAACAGAGAGGTGAAGGACCATTNNNAGATCGGAA |
| 50940_1 | TATGAGTGTGGAGTCGTTGCTGGTGTCAGGAAATCTCTTCTGGNNNAGATCGGAA |
| 50940_2 | TATGAGTGTGGAGTCGTTGCTGGTGTCAGGAAATCTCTTCTGGNNNAGATCGGAA |
| 47041_1 | TATGAGTGTGGAGTCGTTGCACCACGCCCAGCTAATTTTGTATNNNAGATCGGAAG |
| 47041_2 | TATGAGTGTGGAGTCGTTGCACCACGCCCAGCTAATTTTGTATNNNAGATCGGAAG |
| 39473_1 | TATGAGTGTGGAGTCGTTGCGCATTCATTGACCTAGGCATGTCNNNAGATCGGAAG |
| 39473_2 | TATGAGTGTGGAGTCGTTGCGCATTCATTGACCTAGGCATGTCNNNAGATCGGAAG |
| 42999_1 | TATGAGTGTGGAGTCGTTGCAGCAGACGTTCTTCTTCCCTTTCNNNAGATCGGAAGA |
| 42999_2 | TATGAGTGTGGAGTCGTTGCAGCAGACGTTCTTCTTCCCTTTCNNNAGATCGGAAGA |
| 47403_1 | TATGAGTGTGGAGTCGTTGCGTTCAAGCGATTCTCATGCCTCNNNAGATCGGAAGA |
| 47403_2 | TATGAGTGTGGAGTCGTTGCGTTCAAGCGATTCTCATGCCTCNNNAGATCGGAAGA |
| 28940_1 | TATGAGTGTGGAGTCGTTGCACACACACACACACAGGTGATTANNNAGATCGGAAG |
| 28940_2 | TATGAGTGTGGAGTCGTTGCACACACACACACACAGGTGATTANNNAGATCGGAAG |
| 29768_1 | TATGAGTGTGGAGTCGTTGCCCAAAAAAGCACTGTGGGAGTTANNNAGATCGGAA |
| 29768_2 | TATGAGTGTGGAGTCGTTGCCCAAAAAAGCACTGTGGGAGTTANNNAGATCGGAA |

| | |
|---|---|
| 31709_1 | TATGAGTGTGGAGTCGTTGCGAAAAGTACCAGAACACACAGGCNNNAGATCGGAA |
| 31709_2 | TATGAGTGTGGAGTCGTTGCGAAAAGTACCAGAACACACAGGCNNNAGATCGGAA |
| 48638_1 | TATGAGTGTGGAGTCGTTGCTAGCTGAGATTATAGGTGCACGCNNNAGATCGGAA |
| 48638_2 | TATGAGTGTGGAGTCGTTGCTAGCTGAGATTATAGGTGCACGCNNNAGATCGGAA |
| 43579_1 | TATGAGTGTGGAGTCGTTGCACATTTGCAGGGAATATCTGGGTNNNAGATCGGAA |
| 43579_2 | TATGAGTGTGGAGTCGTTGCACATTTGCAGGGAATATCTGGGTNNNAGATCGGAA |
| 31114_1 | TATGAGTGTGGAGTCGTTGCTAGCACTATTTCCTGGATGGCAGNNNAGATCGGAAG |
| 31114_2 | TATGAGTGTGGAGTCGTTGCTAGCACTATTTCCTGGATGGCAGNNNAGATCGGAAG |
| 34888_1 | TATGAGTGTGGAGTCGTTGCCCACCACAGCCAGCTAATTTTTTNNNAGATCGGAAG |
| 34888_2 | TATGAGTGTGGAGTCGTTGCCCACCACAGCCAGCTAATTTTTTNNNAGATCGGAAG |
| 47818_1 | TATGAGTGTGGAGTCGTTGCGACAGGGAGGAACAGAGACTCAGNNNAGATCGGA |
| 47818_2 | TATGAGTGTGGAGTCGTTGCGACAGGGAGGAACAGAGACTCAGNNNAGATCGGA |
| 44603_1 | TATGAGTGTGGAGTCGTTGCCCAAGTAGCTGAGACTATAGGCGNNNAGATCGGAA |
| 44603_2 | TATGAGTGTGGAGTCGTTGCCCAAGTAGCTGAGACTATAGGCGNNNAGATCGGAA |
| 29789_1 | TATGAGTGTGGAGTCGTTGCAACTCAATTTCTCCACGTGTAGTNNNAGATCGGAAG |
| 29789_2 | TATGAGTGTGGAGTCGTTGCAACTCAATTTCTCCACGTGTAGTNNNAGATCGGAAG |
| 29336_1 | TATGAGTGTGGAGTCGTTGCGACAGAGATTGAGTGCGCATGNNNAGATCGGAAG |
| 29336_2 | TATGAGTGTGGAGTCGTTGCGACAGAGATTGAGTGCGCATGNNNAGATCGGAAG |
| 47242_1 | TATGAGTGTGGAGTCGTTGCCTTCCGGGTTCAAGTGATTCTCANNNAGATCGGAAG |
| 47242_2 | TATGAGTGTGGAGTCGTTGCCTTCCGGGTTCAAGTGATTCTCANNNAGATCGGAAG |
| 51853_1 | TATGAGTGTGGAGTCGTTGCTTCTGAAGTTAGGCCTGGGATTCNNNAGATCGGAA |
| 51853_2 | TATGAGTGTGGAGTCGTTGCTTCTGAAGTTAGGCCTGGGATTCNNNAGATCGGAA |
| 34127_1 | TATGAGTGTGGAGTCGTTGCTAACCTCAGGCAGGAAAGACATCNNNAGATCGGAA |
| 34127_2 | TATGAGTGTGGAGTCGTTGCTAACCTCAGGCAGGAAAGACATCNNNAGATCGGAA |
| 53182_1 | TATGAGTGTGGAGTCGTTGCGGCCAACCTGGAGATTACTCTTTNNNAGATCGGAAG |
| 53182_2 | TATGAGTGTGGAGTCGTTGCGGCCAACCTGGAGATTACTCTTTNNNAGATCGGAAG |
| 33201_1 | TATGAGTGTGGAGTCGTTGCGGGGTCCTTGGAAAGTTTTTGTCNNNAGATCGGAA |
| 33201_2 | TATGAGTGTGGAGTCGTTGCGGGGTCCTTGGAAAGTTTTTGTCNNNAGATCGGAA |
| 33003_1 | TATGAGTGTGGAGTCGTTGCCCTCCTGAGTAGCTGGGATTAGANNNAGATCGGAA |
| 33003_2 | TATGAGTGTGGAGTCGTTGCCCTCCTGAGTAGCTGGGATTAGANNNAGATCGGAA |
| 49004_1 | TATGAGTGTGGAGTCGTTGCCCATTCTCCTGCCTCAAGCTCNNNAGATCGGAAGAG |
| 49004_2 | TATGAGTGTGGAGTCGTTGCCCATTCTCCTGCCTCAAGCTCNNNAGATCGGAAGAG |
| 46315_1 | TATGAGTGTGGAGTCGTTGCCCTCCTGAGTAGCTAGGACTACANNNAGATCGGAAG |
| 46315_2 | TATGAGTGTGGAGTCGTTGCCCTCCTGAGTAGCTAGGACTACANNNAGATCGGAAG |
| 45028_1 | TATGAGTGTGGAGTCGTTGCAGTACCCCGGTTTGAATTTCAGTNNNAGATCGGAAG |
| 45028_2 | TATGAGTGTGGAGTCGTTGCAGTACCCCGGTTTGAATTTCAGTNNNAGATCGGAAG |
| 47937_1 | TATGAGTGTGGAGTCGTTGCAAGTGCTAGGTTTACAGCTGTGANNNAGATCGGAA |
| 47937_2 | TATGAGTGTGGAGTCGTTGCAAGTGCTAGGTTTACAGCTGTGANNNAGATCGGAA |
| 51402_1 | TATGAGTGTGGAGTCGTTGCTTTACCTGTCTGCCTCCTCATTCNNNAGATCGGAAGA |
| 51402_2 | TATGAGTGTGGAGTCGTTGCTTTACCTGTCTGCCTCCTCATTCNNNAGATCGGAAGA |
| 50619_1 | TATGAGTGTGGAGTCGTTGCTTGCACATTTTGGACTTCGTTGTNNNAGATCGGAAG |
| 50619_2 | TATGAGTGTGGAGTCGTTGCTTGCACATTTTGGACTTCGTTGTNNNAGATCGGAAG |
| 36769_1 | TATGAGTGTGGAGTCGTTGCAGGCAGTGACATAATCGAACTGTNNNAGATCGGAA |

| | |
|---|---|
| 36769_2 | TATGAGTGTGGAGTCGTTGCAGGCAGTGACATAATCGAACTGTNNNAGATCGGAA |
| 39987_1 | TATGAGTGTGGAGTCGTTGCCTGCTCAGATACTCAGCCAGTAGNNNAGATCGGAAG |
| 39987_2 | TATGAGTGTGGAGTCGTTGCCTGCTCAGATACTCAGCCAGTAGNNNAGATCGGAAG |
| 48632_1 | TATGAGTGTGGAGTCGTTGCCTGTCCCATCTCAGATCTGCTCTNNNAGATCGGAAGA |
| 48632_2 | TATGAGTGTGGAGTCGTTGCCTGTCCCATCTCAGATCTGCTCTNNNAGATCGGAAGA |
| 42112_1 | TATGAGTGTGGAGTCGTTGCCTCGAACTCCTGACACACAACTANNNAGATCGGAAG |
| 42112_2 | TATGAGTGTGGAGTCGTTGCCTCGAACTCCTGACACACAACTANNNAGATCGGAAG |
| 47409_1 | TATGAGTGTGGAGTCGTTGCCACCATGCCCGGCTAATTTTTANNNAGATCGGAAG |
| 47409_2 | TATGAGTGTGGAGTCGTTGCCACCATGCCCGGCTAATTTTTANNNAGATCGGAAG |
| 51694_1 | TATGAGTGTGGAGTCGTTGCCATGCCTGTAGTCCCCATTACTTNNNAGATCGGAAG |
| 51694_2 | TATGAGTGTGGAGTCGTTGCCATGCCTGTAGTCCCCATTACTTNNNAGATCGGAAG |
| 52411_1 | TATGAGTGTGGAGTCGTTGCAAACACCTGTCCTTCTCTAGCTGNNNAGATCGGAAG |
| 52411_2 | TATGAGTGTGGAGTCGTTGCAAACACCTGTCCTTCTCTAGCTGNNNAGATCGGAAG |
| 38484_1 | TATGAGTGTGGAGTCGTTGCAAAGTGGTAAGTTCAGGACAAGTNNNAGATCGGAA |
| 38484_2 | TATGAGTGTGGAGTCGTTGCAAAGTGGTAAGTTCAGGACAAGTNNNAGATCGGAA |
| 43367_1 | TATGAGTGTGGAGTCGTTGCTCAGCAAGACAGAGCCATTTACTNNNAGATCGGAAG |
| 43367_2 | TATGAGTGTGGAGTCGTTGCTCAGCAAGACAGAGCCATTTACTNNNAGATCGGAAG |
| 42018_1 | TATGAGTGTGGAGTCGTTGCAAAGAAAGGTGTTGTCTCTGCTGNNNAGATCGGAA |
| 42018_2 | TATGAGTGTGGAGTCGTTGCAAAGAAAGGTGTTGTCTCTGCTGNNNAGATCGGAA |
| 37304_1 | TATGAGTGTGGAGTCGTTGCATCCTGGGTGACAGAGTGAGANNNAGATCGGAAG |
| 37304_2 | TATGAGTGTGGAGTCGTTGCATCCTGGGTGACAGAGTGAGANNNAGATCGGAAG |
| 10741_1 | TATGAGTGTGGAGTCGTTGCTGACCACAAATTGCCCAAGACTTNNNAGATCGGAAG |
| 10741_2 | TATGAGTGTGGAGTCGTTGCTGACCACAAATTGCCCAAGACTTNNNAGATCGGAAG |
| 37197_1 | TATGAGTGTGGAGTCGTTGCAGCCTGAACACATCTGATATCCTNNNAGATCGGAAG |
| 37197_2 | TATGAGTGTGGAGTCGTTGCAGCCTGAACACATCTGATATCCTNNNAGATCGGAAG |
| 40495_1 | TATGAGTGTGGAGTCGTTGCGCTTAGGGCTCCCAACAAATTAGNNNAGATCGGAAG |
| 40495_2 | TATGAGTGTGGAGTCGTTGCGCTTAGGGCTCCCAACAAATTAGNNNAGATCGGAAG |
| 29141_1 | TATGAGTGTGGAGTCGTTGCGGCTCTAGTAGGCAAAGCATCAANNNAGATCGGAA |
| 29141_2 | TATGAGTGTGGAGTCGTTGCGGCTCTAGTAGGCAAAGCATCAANNNAGATCGGAA |
| 32984_1 | TATGAGTGTGGAGTCGTTGCCAAGTTTGTTCTCTCCTGCTCCTNNNAGATCGGAAGA |
| 32984_2 | TATGAGTGTGGAGTCGTTGCCAAGTTTGTTCTCTCCTGCTCCTNNNAGATCGGAAGA |
| 48795_1 | TATGAGTGTGGAGTCGTTGCAGCCTGAGCACAAATCTTTTGTCNNNAGATCGGAAG |
| 48795_2 | TATGAGTGTGGAGTCGTTGCAGCCTGAGCACAAATCTTTTGTCNNNAGATCGGAAG |
| 49536_1 | TATGAGTGTGGAGTCGTTGCGTGCCATTTCTGAAACTGTGACANNNAGATCGGAAG |
| 49536_2 | TATGAGTGTGGAGTCGTTGCGTGCCATTTCTGAAACTGTGACANNNAGATCGGAAG |
| 49930_1 | TATGAGTGTGGAGTCGTTGCTCTCAACGTATTTTTCCTCTGGGTNNNAGATCGGAAG |
| 49930_2 | TATGAGTGTGGAGTCGTTGCTCTCAACGTATTTTTCCTCTGGGTNNNAGATCGGAAG |
| 34352_1 | TATGAGTGTGGAGTCGTTGCGGTGGGAAGGAAGTGAGTATCTCNNNAGATCGGA |
| 34352_2 | TATGAGTGTGGAGTCGTTGCGGTGGGAAGGAAGTGAGTATCTCNNNAGATCGGA |
| 52987_1 | TATGAGTGTGGAGTCGTTGCGATGGTAAGACAAAGGCAAACCANNNAGATCGGAA |
| 52987_2 | TATGAGTGTGGAGTCGTTGCGATGGTAAGACAAAGGCAAACCANNNAGATCGGAA |
| 10616_1 | TATGAGTGTGGAGTCGTTGCTGCTTCACTAACACCCCAAACAANNNAGATCGGAAG |
| 10616_2 | TATGAGTGTGGAGTCGTTGCTGCTTCACTAACACCCCAAACAANNNAGATCGGAAG |

| | |
|---|---|
| 29692_1 | TATGAGTGTGGAGTCGTTGCTCCTGATAGCTAGACCTGAGAATNNNAGATCGGAAG |
| 29692_2 | TATGAGTGTGGAGTCGTTGCTCCTGATAGCTAGACCTGAGAATNNNAGATCGGAAG |
| 47251_1 | TATGAGTGTGGAGTCGTTGCCTCCTGAGTAGCTGGGACTACAGNNNAGATCGGAA |
| 47251_2 | TATGAGTGTGGAGTCGTTGCCTCCTGAGTAGCTGGGACTACAGNNNAGATCGGAA |
| 46241_1 | TATGAGTGTGGAGTCGTTGCTCTTGACTTACCACAACCTCCACNNNAGATCGGAAGA |
| 46241_2 | TATGAGTGTGGAGTCGTTGCTCTTGACTTACCACAACCTCCACNNNAGATCGGAAGA |
| 40670_1 | TATGAGTGTGGAGTCGTTGCCCCTGGAGTGTATTAGGAACTGGNNNAGATCGGAA |
| 40670_2 | TATGAGTGTGGAGTCGTTGCCCCTGGAGTGTATTAGGAACTGGNNNAGATCGGAA |
| 32721_1 | TATGAGTGTGGAGTCGTTGCAAGCCTGTAATCCCAGCACTTTNNNAGATCGGAAGA |
| 32721_2 | TATGAGTGTGGAGTCGTTGCAAGCCTGTAATCCCAGCACTTTNNNAGATCGGAAGA |
| 35111_1 | TATGAGTGTGGAGTCGTTGCTATGCCTGGTTGTGTGATCATCANNNAGATCGGAAG |
| 35111_2 | TATGAGTGTGGAGTCGTTGCTATGCCTGGTTGTGTGATCATCANNNAGATCGGAAG |
| 40458_1 | TATGAGTGTGGAGTCGTTGCCAGGAAGCTGATTCAGGAGGATTNNNAGATCGGAA |
| 40458_2 | TATGAGTGTGGAGTCGTTGCCAGGAAGCTGATTCAGGAGGATTNNNAGATCGGAA |
| 31268_1 | TATGAGTGTGGAGTCGTTGCCACACAATTGGCCAACCTAACATNNNAGATCGGAAG |
| 31268_2 | TATGAGTGTGGAGTCGTTGCCACACAATTGGCCAACCTAACATNNNAGATCGGAAG |
| 30924_1 | TATGAGTGTGGAGTCGTTGCTTGGACTTCTTTAGATGGCAGCTNNNAGATCGGAAG |
| 30924_2 | TATGAGTGTGGAGTCGTTGCTTGGACTTCTTTAGATGGCAGCTNNNAGATCGGAAG |
| 2189_1 | TATGAGTGTGGAGTCGTTGCATTCTAGCCTGGGTATTTGGGGANNNAGATCGGAA |
| 2189_2 | TATGAGTGTGGAGTCGTTGCATTCTAGCCTGGGTATTTGGGGANNNAGATCGGAA |
| 34664_1 | TATGAGTGTGGAGTCGTTGCCTCCTGAGTCACTGGGATTACAGNNNAGATCGGAAG |
| 34664_2 | TATGAGTGTGGAGTCGTTGCCTCCTGAGTCACTGGGATTACAGNNNAGATCGGAAG |
| 37460_1 | TATGAGTGTGGAGTCGTTGCGGTAGAGGCATAAGGGAACCTTCNNNAGATCGGAA |
| 37460_2 | TATGAGTGTGGAGTCGTTGCGGTAGAGGCATAAGGGAACCTTCNNNAGATCGGAA |
| 47551_1 | TATGAGTGTGGAGTCGTTGCCACACCATGCCCAGCTAATTTTTNNNAGATCGGAAGA |
| 47551_2 | TATGAGTGTGGAGTCGTTGCCACACCATGCCCAGCTAATTTTTNNNAGATCGGAAGA |
| 50759_1 | TATGAGTGTGGAGTCGTTGCAAAAGCATTTCACTATGGCTGGGNNNAGATCGGAA |
| 50759_2 | TATGAGTGTGGAGTCGTTGCAAAAGCATTTCACTATGGCTGGGNNNAGATCGGAA |
| 52785_1 | TATGAGTGTGGAGTCGTTGCCTCACCCAAGTCTCGATACCAGNNNAGATCGGAAGA |
| 52785_2 | TATGAGTGTGGAGTCGTTGCCTCACCCAAGTCTCGATACCAGNNNAGATCGGAAGA |
| 47003_1 | TATGAGTGTGGAGTCGTTGCCTTTCCATGGCCTCAACTCTACTNNNAGATCGGAAGA |
| 47003_2 | TATGAGTGTGGAGTCGTTGCCTTTCCATGGCCTCAACTCTACTNNNAGATCGGAAGA |
| 43738_1 | TATGAGTGTGGAGTCGTTGCTGCATTCAACAGCTCTTCATGTGNNNAGATCGGAAG |
| 43738_2 | TATGAGTGTGGAGTCGTTGCTGCATTCAACAGCTCTTCATGTGNNNAGATCGGAAG |
| 46933_1 | TATGAGTGTGGAGTCGTTGCTGAAATTTGGCACCAAGTCACAANNNAGATCGGAAG |
| 46933_2 | TATGAGTGTGGAGTCGTTGCTGAAATTTGGCACCAAGTCACAANNNAGATCGGAAG |
| 38461_1 | TATGAGTGTGGAGTCGTTGCCAAATCAAACTCATGACGCAGCTNNNAGATCGGAAG |
| 38461_2 | TATGAGTGTGGAGTCGTTGCCAAATCAAACTCATGACGCAGCTNNNAGATCGGAAG |
| 53360_1 | TATGAGTGTGGAGTCGTTGCTTTTGAGACAGGGTTTTGCCATGNNNAGATCGGAA |
| 53360_2 | TATGAGTGTGGAGTCGTTGCTTTTGAGACAGGGTTTTGCCATGNNNAGATCGGAA |
| 33854_1 | TATGAGTGTGGAGTCGTTGCGAGGCAGGAGAGTCACTTGAACNNNAGATCGGAAG |
| 33854_2 | TATGAGTGTGGAGTCGTTGCGAGGCAGGAGAGTCACTTGAACNNNAGATCGGAAG |
| 41859_1 | TATGAGTGTGGAGTCGTTGCCATTCTCCTGCCTCAGCCTCNNNAGATCGGAAGAGCA |

| | |
|---|---|
| 41859_2 | TATGAGTGTGGAGTCGTTGCCATTCTCCTGCCTCAGCCTCNNNAGATCGGAAGAGCA |
| 49974_1 | TATGAGTGTGGAGTCGTTGCCTCGCGTATCTCATCTCTGCCNNNAGATCGGAAGAG |
| 49974_2 | TATGAGTGTGGAGTCGTTGCCTCGCGTATCTCATCTCTGCCNNNAGATCGGAAGAG |
| 29659_1 | TATGAGTGTGGAGTCGTTGCTGCACTCTCGAACATCTCTTTGTNNNAGATCGGAAGA |
| 29659_2 | TATGAGTGTGGAGTCGTTGCTGCACTCTCGAACATCTCTTTGTNNNAGATCGGAAGA |
| 31078_1 | TATGAGTGTGGAGTCGTTGCCAAGCATGCATGGTGAGTGTATTNNNAGATCGGAA |
| 31078_2 | TATGAGTGTGGAGTCGTTGCCAAGCATGCATGGTGAGTGTATTNNNAGATCGGAA |
| 50129_1 | TATGAGTGTGGAGTCGTTGCTCCTAAAAGCAAAGAATTGGCCGNNNAGATCGGAA |
| 50129_2 | TATGAGTGTGGAGTCGTTGCTCCTAAAAGCAAAGAATTGGCCGNNNAGATCGGAA |
| 46343_1 | TATGAGTGTGGAGTCGTTGCCGATCTCGGCTCGCTACAAGNNNAGATCGGAAGAG |
| 46343_2 | TATGAGTGTGGAGTCGTTGCCGATCTCGGCTCGCTACAAGNNNAGATCGGAAGAG |
| 40928_1 | TATGAGTGTGGAGTCGTTGCAACACTGACCAGAATGTCTTACANNNAGATCGGAAG |
| 40928_2 | TATGAGTGTGGAGTCGTTGCAACACTGACCAGAATGTCTTACANNNAGATCGGAAG |
| 33486_1 | TATGAGTGTGGAGTCGTTGCCCAAGACCATTCCACTGCATTACNNNAGATCGGAAGA |
| 33486_2 | TATGAGTGTGGAGTCGTTGCCCAAGACCATTCCACTGCATTACNNNAGATCGGAAGA |
| 39300_1 | TATGAGTGTGGAGTCGTTGCCAAGTAGCTGGGATTACAGGCATNNNAGATCGGAA |
| 39300_2 | TATGAGTGTGGAGTCGTTGCCAAGTAGCTGGGATTACAGGCATNNNAGATCGGAA |
| 46710_1 | TATGAGTGTGGAGTCGTTGCTAGCTGGGATTATAGGCATGTGCNNNAGATCGGAA |
| 46710_2 | TATGAGTGTGGAGTCGTTGCTAGCTGGGATTATAGGCATGTGCNNNAGATCGGAA |
| 41805_1 | TATGAGTGTGGAGTCGTTGCGCCTCCTGAATAGCTGAGACTACNNNAGATCGGAAG |
| 41805_2 | TATGAGTGTGGAGTCGTTGCGCCTCCTGAATAGCTGAGACTACNNNAGATCGGAAG |
| 35482_1 | TATGAGTGTGGAGTCGTTGCCTCCTGAGTAGCTGGGACTACAGNNNAGATCGGAA |
| 35482_2 | TATGAGTGTGGAGTCGTTGCCTCCTGAGTAGCTGGGACTACAGNNNAGATCGGAA |
| 50867_1 | TATGAGTGTGGAGTCGTTGCCTGTCTTTTGTGACGTCACGATCNNNAGATCGGAAG |
| 50867_2 | TATGAGTGTGGAGTCGTTGCCTGTCTTTTGTGACGTCACGATCNNNAGATCGGAAG |
| 34692_1 | TATGAGTGTGGAGTCGTTGCCACGATCTCAGCTCAATGCAACNNNAGATCGGAAGA |
| 34692_2 | TATGAGTGTGGAGTCGTTGCCACGATCTCAGCTCAATGCAACNNNAGATCGGAAGA |
| 34755_1 | TATGAGTGTGGAGTCGTTGCGTTCAAGAGATTCTCCTGCCTCANNNAGATCGGAAG |
| 34755_2 | TATGAGTGTGGAGTCGTTGCGTTCAAGAGATTCTCCTGCCTCANNNAGATCGGAAG |
| 50667_1 | TATGAGTGTGGAGTCGTTGCGTAACAGCAAAGAGGGAGGAGAANNNAGATCGGA |
| 50667_2 | TATGAGTGTGGAGTCGTTGCGTAACAGCAAAGAGGGAGGAGAANNNAGATCGGA |
| 32274_1 | TATGAGTGTGGAGTCGTTGCGTCCCTCCCATAACACATGAGAANNNAGATCGGAAG |
| 32274_2 | TATGAGTGTGGAGTCGTTGCGTCCCTCCCATAACACATGAGAANNNAGATCGGAAG |
| 44064_1 | TATGAGTGTGGAGTCGTTGCTAAAGAACAGGAACAGCTAGGGCNNNAGATCGGAA |
| 44064_2 | TATGAGTGTGGAGTCGTTGCTAAAGAACAGGAACAGCTAGGGCNNNAGATCGGAA |
| 47603_1 | TATGAGTGTGGAGTCGTTGCCTCCAGGGTTCACACCATTCTCNNNAGATCGGAAGA |
| 47603_2 | TATGAGTGTGGAGTCGTTGCCTCCAGGGTTCACACCATTCTCNNNAGATCGGAAGA |
| 37488_1 | TATGAGTGTGGAGTCGTTGCGCTGACAAATTGGACTACATCAANNNAGATCGGAAG |
| 37488_2 | TATGAGTGTGGAGTCGTTGCGCTGACAAATTGGACTACATCAANNNAGATCGGAAG |
| 44299_1 | TATGAGTGTGGAGTCGTTGCGACATCTCTTTGCCCAGACCTATNNNAGATCGGAAGA |
| 44299_2 | TATGAGTGTGGAGTCGTTGCGACATCTCTTTGCCCAGACCTATNNNAGATCGGAAGA |
| 51362_1 | TATGAGTGTGGAGTCGTTGCCACCATAGTGGCACCTGAGTAATNNNAGATCGGAAG |
| 51362_2 | TATGAGTGTGGAGTCGTTGCCACCATAGTGGCACCTGAGTAATNNNAGATCGGAAG |

| | |
|---|---|
| 41005_1 | TATGAGTGTGGAGTCGTTGCTATGTACATGAGGCCAGACATGGNNNAGATCGGAA |
| 41005_2 | TATGAGTGTGGAGTCGTTGCTATGTACATGAGGCCAGACATGGNNNAGATCGGAA |
| 52586_1 | TATGAGTGTGGAGTCGTTGCACCCAGAATGGCATATAAGGTCCNNNAGATCGGAAG |
| 52586_2 | TATGAGTGTGGAGTCGTTGCACCCAGAATGGCATATAAGGTCCNNNAGATCGGAAG |
| 40381_1 | TATGAGTGTGGAGTCGTTGCATCACAACCTTATAGTCCTGCCANNNAGATCGGAAG |
| 40381_2 | TATGAGTGTGGAGTCGTTGCATCACAACCTTATAGTCCTGCCANNNAGATCGGAAG |
| 30778_1 | TATGAGTGTGGAGTCGTTGCAAAAGGTCAGGTTCAGAGGCTAGNNNAGATCGGAA |
| 30778_2 | TATGAGTGTGGAGTCGTTGCAAAAGGTCAGGTTCAGAGGCTAGNNNAGATCGGAA |
| 43731_1 | TATGAGTGTGGAGTCGTTGCCTTTGAGACACACTGCGGATACCNNNAGATCGGAAG |
| 43731_2 | TATGAGTGTGGAGTCGTTGCCTTTGAGACACACTGCGGATACCNNNAGATCGGAAG |
| 29090_1 | TATGAGTGTGGAGTCGTTGCTTGCCTCTTTTCCTCTCTGTCTCNNNAGATCGGAAGA |
| 29090_2 | TATGAGTGTGGAGTCGTTGCTTGCCTCTTTTCCTCTCTGTCTCNNNAGATCGGAAGA |
| 42224_1 | TATGAGTGTGGAGTCGTTGCAGTTGGAGTCTCACTGTGTCATCNNNAGATCGGAAG |
| 42224_2 | TATGAGTGTGGAGTCGTTGCAGTTGGAGTCTCACTGTGTCATCNNNAGATCGGAAG |
| 41375_1 | TATGAGTGTGGAGTCGTTGCTCTTCTTTTCTCCTCCCTCTCCTNNNAGATCGGAAGAG |
| 41375_2 | TATGAGTGTGGAGTCGTTGCTCTTCTTTTCTCCTCCCTCTCCTNNNAGATCGGAAGAG |
| 30642_1 | TATGAGTGTGGAGTCGTTGCAGAGTCAACCTCCTCTTTTTGTTNNNAGATCGGAAGA |
| 30642_2 | TATGAGTGTGGAGTCGTTGCAGAGTCAACCTCCTCTTTTTGTTNNNAGATCGGAAGA |
| 30515_1 | TATGAGTGTGGAGTCGTTGCGGTGTTTTAGGGATCCGAAGCTTNNNAGATCGGAA |
| 30515_2 | TATGAGTGTGGAGTCGTTGCGGTGTTTTAGGGATCCGAAGCTTNNNAGATCGGAA |
| 51893_1 | TATGAGTGTGGAGTCGTTGCCCCATGTGCTCTCTCTCTCTTTTNNNAGATCGGAAGA |
| 51893_2 | TATGAGTGTGGAGTCGTTGCCCCATGTGCTCTCTCTCTCTTTTNNNAGATCGGAAGA |
| 33587_1 | TATGAGTGTGGAGTCGTTGCCTCCCAGATCCAAGCGATTCTCNNNAGATCGGAAGA |
| 33587_2 | TATGAGTGTGGAGTCGTTGCCTCCCAGATCCAAGCGATTCTCNNNAGATCGGAAGA |
| 37617_1 | TATGAGTGTGGAGTCGTTGCTTATGAGCCTCAAGATGACCCTGNNNAGATCGGAAG |
| 37617_2 | TATGAGTGTGGAGTCGTTGCTTATGAGCCTCAAGATGACCCTGNNNAGATCGGAAG |
| 37797_1 | TATGAGTGTGGAGTCGTTGCAACTGGATCACCAAGGTTTAGCTNNNAGATCGGAAG |
| 37797_2 | TATGAGTGTGGAGTCGTTGCAACTGGATCACCAAGGTTTAGCTNNNAGATCGGAAG |
| 29810_1 | TATGAGTGTGGAGTCGTTGCCTGTTTTGCTAGAAGGGACATGCNNNAGATCGGAA |
| 29810_2 | TATGAGTGTGGAGTCGTTGCCTGTTTTGCTAGAAGGGACATGCNNNAGATCGGAA |
| 33262_1 | TATGAGTGTGGAGTCGTTGCGGACTAAACTGAAGGCAAAACCCNNNAGATCGGAA |
| 33262_2 | TATGAGTGTGGAGTCGTTGCGGACTAAACTGAAGGCAAAACCCNNNAGATCGGAA |
| 50617_1 | TATGAGTGTGGAGTCGTTGCGCCAGAACCAAAGCCAGATATTGNNNAGATCGGAA |
| 50617_2 | TATGAGTGTGGAGTCGTTGCGCCAGAACCAAAGCCAGATATTGNNNAGATCGGAA |
| 51851_1 | TATGAGTGTGGAGTCGTTGCCTATTCCACAAAACCGCCATTGTNNNAGATCGGAAG |
| 51851_2 | TATGAGTGTGGAGTCGTTGCCTATTCCACAAAACCGCCATTGTNNNAGATCGGAAG |
| 46075_1 | TATGAGTGTGGAGTCGTTGCCACAATCTCAGCTCACTGCAACNNNAGATCGGAAGA |
| 46075_2 | TATGAGTGTGGAGTCGTTGCCACAATCTCAGCTCACTGCAACNNNAGATCGGAAGA |
| 35011_1 | TATGAGTGTGGAGTCGTTGCCCACCATGCCCAGCTAATTTTTTNNNAGATCGGAAGA |
| 35011_2 | TATGAGTGTGGAGTCGTTGCCCACCATGCCCAGCTAATTTTTTNNNAGATCGGAAGA |
| 48712_1 | TATGAGTGTGGAGTCGTTGCGAAGAGTTTCCATTGCTTTGCCTNNNAGATCGGAAG |
| 48712_2 | TATGAGTGTGGAGTCGTTGCGAAGAGTTTCCATTGCTTTGCCTNNNAGATCGGAAG |
| 31115_1 | TATGAGTGTGGAGTCGTTGCAAACAGCAAAAATGTAGCCACGANNNAGATCGGAA |

| | |
|---|---|
| 31115_2 | TATGAGTGTGGAGTCGTTGCAAACAGCAAAAATGTAGCCACGANNNAGATCGGAA |
| 51078_1 | TATGAGTGTGGAGTCGTTGCCTTATGGTGGTAGCAGAGGGACNNNAGATCGGAAG |
| 51078_2 | TATGAGTGTGGAGTCGTTGCCTTATGGTGGTAGCAGAGGGACNNNAGATCGGAAG |
| 13601_1 | TATGAGTGTGGAGTCGTTGCATGGCAGACAGTGAAAAAACNNNAGATCGGAAGAG |
| 13601_2 | TATGAGTGTGGAGTCGTTGCATGGCAGACAGTGAAAAAACNNNAGATCGGAAGAG |
| 31253_1 | TATGAGTGTGGAGTCGTTGCAAGCTGTGATTGATTCTGAGTGCNNNAGATCGGAA |
| 31253_2 | TATGAGTGTGGAGTCGTTGCAAGCTGTGATTGATTCTGAGTGCNNNAGATCGGAA |
| 28769_1 | TATGAGTGTGGAGTCGTTGCGGATGGAGCTGGAGTCCATTATTNNNAGATCGGAA |
| 28769_2 | TATGAGTGTGGAGTCGTTGCGGATGGAGCTGGAGTCCATTATTNNNAGATCGGAA |
| 39064_1 | TATGAGTGTGGAGTCGTTGCAGATACAAACAAGCCCAGACAGTNNNAGATCGGAA |
| 39064_2 | TATGAGTGTGGAGTCGTTGCAGATACAAACAAGCCCAGACAGTNNNAGATCGGAA |
| 35369_1 | TATGAGTGTGGAGTCGTTGCTTCTGGACACCTGGTCATTTGATNNNAGATCGGAAG |
| 35369_2 | TATGAGTGTGGAGTCGTTGCTTCTGGACACCTGGTCATTTGATNNNAGATCGGAAG |
| 47320_1 | TATGAGTGTGGAGTCGTTGCCAATCTTGGCTCACTGCAACTTCNNNAGATCGGAAG |
| 47320_2 | TATGAGTGTGGAGTCGTTGCCAATCTTGGCTCACTGCAACTTCNNNAGATCGGAAG |
| 46662_1 | TATGAGTGTGGAGTCGTTGCGCCTCCCAAGTACCTGAGATTACNNNAGATCGGAAG |
| 46662_2 | TATGAGTGTGGAGTCGTTGCGCCTCCCAAGTACCTGAGATTACNNNAGATCGGAAG |
| 46767_1 | TATGAGTGTGGAGTCGTTGCTGCCTCCAGAGTAGATGGGATTANNNAGATCGGAA |
| 46767_2 | TATGAGTGTGGAGTCGTTGCTGCCTCCAGAGTAGATGGGATTANNNAGATCGGAA |
| 42863_1 | TATGAGTGTGGAGTCGTTGCGCTCCTCTTTGTACCTCCGATAGNNNAGATCGGAAG |
| 42863_2 | TATGAGTGTGGAGTCGTTGCGCTCCTCTTTGTACCTCCGATAGNNNAGATCGGAAG |
| 38787_1 | TATGAGTGTGGAGTCGTTGCGATATCCGCACTTCTGTTTGTGGNNNAGATCGGAAG |
| 38787_2 | TATGAGTGTGGAGTCGTTGCGATATCCGCACTTCTGTTTGTGGNNNAGATCGGAAG |
| 37173_1 | TATGAGTGTGGAGTCGTTGCGTCTGATAGTTCCCAGAAACCCANNNAGATCGGAAG |
| 37173_2 | TATGAGTGTGGAGTCGTTGCGTCTGATAGTTCCCAGAAACCCANNNAGATCGGAAG |
| 35578_1 | TATGAGTGTGGAGTCGTTGCTACACATATGCTGAGTAGGGCACNNNAGATCGGAA |
| 35578_2 | TATGAGTGTGGAGTCGTTGCTACACATATGCTGAGTAGGGCACNNNAGATCGGAA |
| 45395_1 | TATGAGTGTGGAGTCGTTGCTATTGTCAAGACACAACCGCAACNNNAGATCGGAAG |
| 45395_2 | TATGAGTGTGGAGTCGTTGCTATTGTCAAGACACAACCGCAACNNNAGATCGGAAG |
| 43772_1 | TATGAGTGTGGAGTCGTTGCCTTGCAGTAAGCTGAGATCATGCNNNAGATCGGAA |
| 43772_2 | TATGAGTGTGGAGTCGTTGCCTTGCAGTAAGCTGAGATCATGCNNNAGATCGGAA |
| 13665_1 | TATGAGTGTGGAGTCGTTGCTTTTCTTTTTCTTTGAGACTGGNNNAGATCGGAAGA |
| 13665_2 | TATGAGTGTGGAGTCGTTGCTTTTCTTTTTCTTTGAGACTGGNNNAGATCGGAAGA |
| 41747_1 | TATGAGTGTGGAGTCGTTGCTCTCTAGGCCTCAGTTTTTCCACNNNAGATCGGAAGA |
| 41747_2 | TATGAGTGTGGAGTCGTTGCTCTCTAGGCCTCAGTTTTTCCACNNNAGATCGGAAGA |
| 29543_1 | TATGAGTGTGGAGTCGTTGCATGTGTCTAGAATGCAGTCGTGTNNNAGATCGGAA |
| 29543_2 | TATGAGTGTGGAGTCGTTGCATGTGTCTAGAATGCAGTCGTGTNNNAGATCGGAA |
| 30804_1 | TATGAGTGTGGAGTCGTTGCGTTACAATGCGGTGTATGAGCTCNNNAGATCGGAA |
| 30804_2 | TATGAGTGTGGAGTCGTTGCGTTACAATGCGGTGTATGAGCTCNNNAGATCGGAA |
| 41224_1 | TATGAGTGTGGAGTCGTTGCTCTGAACAAAAACACCCACATCCNNNAGATCGGAAG |
| 41224_2 | TATGAGTGTGGAGTCGTTGCTCTGAACAAAAACACCCACATCCNNNAGATCGGAAG |
| 2160_1 | TATGAGTGTGGAGTCGTTGCTGTGAGAGAGAGCCACTTGAAGANNNAGATCGGAA |
| 2160_2 | TATGAGTGTGGAGTCGTTGCTGTGAGAGAGAGCCACTTGAAGANNNAGATCGGAA |

| | |
|---|---|
| 29544_1 | TATGAGTGTGGAGTCGTTGCCTTCAAGAGCCACGTAAGGTTCANNNAGATCGGAAG |
| 29544_2 | TATGAGTGTGGAGTCGTTGCCTTCAAGAGCCACGTAAGGTTCANNNAGATCGGAAG |
| 29697_1 | TATGAGTGTGGAGTCGTTGCACTCTCACCACATATTTGTTTTCCTNNNAGATCGGAAG |
| 29697_2 | TATGAGTGTGGAGTCGTTGCACTCTCACCACATATTTGTTTTCCTNNNAGATCGGAAG |
| 51899_1 | TATGAGTGTGGAGTCGTTGCCCTCCACAGCACTAGTCATTAGGNNNAGATCGGAAGA |
| 51899_2 | TATGAGTGTGGAGTCGTTGCCCTCCACAGCACTAGTCATTAGGNNNAGATCGGAAGA |
| 51968_1 | TATGAGTGTGGAGTCGTTGCCCCGGCCCAAACTTTATTTTTGANNNAGATCGGAAGA |
| 51968_2 | TATGAGTGTGGAGTCGTTGCCCCGGCCCAAACTTTATTTTTGANNNAGATCGGAAGA |
| 51417_1 | TATGAGTGTGGAGTCGTTGCGAGCCTTCTAGGAAGCATCAAGTNNNAGATCGGAAG |
| 51417_2 | TATGAGTGTGGAGTCGTTGCGAGCCTTCTAGGAAGCATCAAGTNNNAGATCGGAAG |
| 38431_1 | TATGAGTGTGGAGTCGTTGCGCAGAGGGGTTTAGTAGAGAAGGNNNAGATCGGAA |
| 38431_2 | TATGAGTGTGGAGTCGTTGCGCAGAGGGGTTTAGTAGAGAAGGNNNAGATCGGAA |
| 36580_1 | TATGAGTGTGGAGTCGTTGCATGTGGATGAGTCATGGACACAANNNAGATCGGAAG |
| 36580_2 | TATGAGTGTGGAGTCGTTGCATGTGGATGAGTCATGGACACAANNNAGATCGGAAG |
| 32561_1 | TATGAGTGTGGAGTCGTTGCCTCTCGGGTTCAAGCAATTCTCNNNAGATCGGAAGAG |
| 32561_2 | TATGAGTGTGGAGTCGTTGCCTCTCGGGTTCAAGCAATTCTCNNNAGATCGGAAGAG |
| 5546_1 | TATGAGTGTGGAGTCGTTGCTGGGAAGGGGAAAACAAAGACTANNNAGATCGGAA |
| 5546_2 | TATGAGTGTGGAGTCGTTGCTGGGAAGGGGAAAACAAAGACTANNNAGATCGGAA |
| 39251_1 | TATGAGTGTGGAGTCGTTGCCTGGGCTGGATAATCTAGAGAGCNNNAGATCGGAAG |
| 39251_2 | TATGAGTGTGGAGTCGTTGCCTGGGCTGGATAATCTAGAGAGCNNNAGATCGGAAG |
| 34694_1 | TATGAGTGTGGAGTCGTTGCAATGGGGTCTTACCATATTGCCCNNNAGATCGGAAG |
| 34694_2 | TATGAGTGTGGAGTCGTTGCAATGGGGTCTTACCATATTGCCCNNNAGATCGGAAG |
| 37827_1 | TATGAGTGTGGAGTCGTTGCTTGTGGGCTTGGAAATAACACTGNNNAGATCGGAAG |
| 37827_2 | TATGAGTGTGGAGTCGTTGCTTGTGGGCTTGGAAATAACACTGNNNAGATCGGAAG |
| 52317_1 | TATGAGTGTGGAGTCGTTGCAAAAAAAAAAAAGTGGCCAGGCGNNNAGATCGGAA |
| 52317_2 | TATGAGTGTGGAGTCGTTGCAAAAAAAAAAAAGTGGCCAGGCGNNNAGATCGGAA |
| 35417_1 | TATGAGTGTGGAGTCGTTGCGATTACAAGTGCACACCACCATGNNNAGATCGGAAG |
| 35417_2 | TATGAGTGTGGAGTCGTTGCGATTACAAGTGCACACCACCATGNNNAGATCGGAAG |
| 29621_1 | TATGAGTGTGGAGTCGTTGCGGTATGGGTAAGGTCCACTACAANNNAGATCGGAAG |
| 29621_2 | TATGAGTGTGGAGTCGTTGCGGTATGGGTAAGGTCCACTACAANNNAGATCGGAAG |
| 48368_1 | TATGAGTGTGGAGTCGTTGCACAATCTCGGCTCACTGCAAGNNNAGATCGGAAGAG |
| 48368_2 | TATGAGTGTGGAGTCGTTGCACAATCTCGGCTCACTGCAAGNNNAGATCGGAAGAG |
| 41701_1 | TATGAGTGTGGAGTCGTTGCAAAGTGCTGGGATTATAGGCGAGNNNAGATCGGAA |
| 41701_2 | TATGAGTGTGGAGTCGTTGCAAAGTGCTGGGATTATAGGCGAGNNNAGATCGGAA |
| 13574_1 | TATGAGTGTGGAGTCGTTGCGTATTCTTGGTTTCTGATGAAATNNNAGATCGGAAG |
| 13574_2 | TATGAGTGTGGAGTCGTTGCGTATTCTTGGTTTCTGATGAAATNNNAGATCGGAAG |
| 37737_1 | TATGAGTGTGGAGTCGTTGCTGACCCCTGAAAGCATAGTCTTTNNNAGATCGGAAGA |
| 37737_2 | TATGAGTGTGGAGTCGTTGCTGACCCCTGAAAGCATAGTCTTTNNNAGATCGGAAGA |
| 51907_1 | TATGAGTGTGGAGTCGTTGCACTCCCTCTTTCCTACCTCAAGANNNAGATCGGAAGAG |
| 51907_2 | TATGAGTGTGGAGTCGTTGCACTCCCTCTTTCCTACCTCAAGANNNAGATCGGAAGAG |
| 8487_1 | TATGAGTGTGGAGTCGTTGCTCAAGCTCCTTGGTAAACAGGCTNNNAGATCGGAAG |
| 8487_2 | TATGAGTGTGGAGTCGTTGCTCAAGCTCCTTGGTAAACAGGCTNNNAGATCGGAAG |
| 43646_1 | TATGAGTGTGGAGTCGTTGCTAGTAGGAAACTCCGCTCTACCANNNAGATCGGAAG |

| | |
|---|---|
| 43646_2 | TATGAGTGTGGAGTCGTTGCTAGTAGGAAACTCCGCTCTACCANNNAGATCGGAAG |
| 43905_1 | TATGAGTGTGGAGTCGTTGCCTTCTGGCAGAGGACGTGATGNNNAGATCGGAAGA |
| 43905_2 | TATGAGTGTGGAGTCGTTGCCTTCTGGCAGAGGACGTGATGNNNAGATCGGAAGA |
| 37604_1 | TATGAGTGTGGAGTCGTTGCCTACTCTCCGGACAGTGGAAAAANNNAGATCGGAAG |
| 37604_2 | TATGAGTGTGGAGTCGTTGCCTACTCTCCGGACAGTGGAAAAANNNAGATCGGAAG |
| 28819_1 | TATGAGTGTGGAGTCGTTGCCCACAGGTATCAAGGGTAATGCTNNNAGATCGGAAG |
| 28819_2 | TATGAGTGTGGAGTCGTTGCCCACAGGTATCAAGGGTAATGCTNNNAGATCGGAAG |
| 42292_1 | TATGAGTGTGGAGTCGTTGCCGCCCTCATTCTGTCTCTATGAANNNAGATCGGAAGA |
| 42292_2 | TATGAGTGTGGAGTCGTTGCCGCCCTCATTCTGTCTCTATGAANNNAGATCGGAAGA |
| 38512_1 | TATGAGTGTGGAGTCGTTGCTGAGGTCATGAGTTTGAGACCAGNNNAGATCGGAA |
| 38512_2 | TATGAGTGTGGAGTCGTTGCTGAGGTCATGAGTTTGAGACCAGNNNAGATCGGAA |
| 32193_1 | TATGAGTGTGGAGTCGTTGCTCAGGCCGTAGCTTAGATGTTACNNNAGATCGGAAG |
| 32193_2 | TATGAGTGTGGAGTCGTTGCTCAGGCCGTAGCTTAGATGTTACNNNAGATCGGAAG |
| 31218_1 | TATGAGTGTGGAGTCGTTGCTACAGCTGCCACCTTATTCACTTNNNAGATCGGAAGA |
| 31218_2 | TATGAGTGTGGAGTCGTTGCTACAGCTGCCACCTTATTCACTTNNNAGATCGGAAGA |
| 39150_1 | TATGAGTGTGGAGTCGTTGCCTAGGAAAACCCAAAGGCAAAGGNNNAGATCGGAA |
| 39150_2 | TATGAGTGTGGAGTCGTTGCCTAGGAAAACCCAAAGGCAAAGGNNNAGATCGGAA |
| 32313_1 | TATGAGTGTGGAGTCGTTGCCACTCTGTTTCCTGTTCAGCATGNNNAGATCGGAAGA |
| 32313_2 | TATGAGTGTGGAGTCGTTGCCACTCTGTTTCCTGTTCAGCATGNNNAGATCGGAAGA |
| 47686_1 | TATGAGTGTGGAGTCGTTGCACGCCCGGCTAATTTTTTGTATTNNNAGATCGGAAGA |
| 47686_2 | TATGAGTGTGGAGTCGTTGCACGCCCGGCTAATTTTTTGTATTNNNAGATCGGAAGA |
| 37412_1 | TATGAGTGTGGAGTCGTTGCGGGGTCCATTACAGAAGCTAGAANNNAGATCGGAA |
| 37412_2 | TATGAGTGTGGAGTCGTTGCGGGGTCCATTACAGAAGCTAGAANNNAGATCGGAA |
| 40477_1 | TATGAGTGTGGAGTCGTTGCCTGAGGCAGAAGGATCACTTGAGNNNAGATCGGAA |
| 40477_2 | TATGAGTGTGGAGTCGTTGCCTGAGGCAGAAGGATCACTTGAGNNNAGATCGGAA |
| 40284_1 | TATGAGTGTGGAGTCGTTGCACTTTCAAGGTCATTACTTTGGGCNNNAGATCGGAA |
| 40284_2 | TATGAGTGTGGAGTCGTTGCACTTTCAAGGTCATTACTTTGGGCNNNAGATCGGAA |
| 31006_1 | TATGAGTGTGGAGTCGTTGCGAGAGAGAGAGAGAGAGAGAGGTNNNAGATCGGA |
| 31006_2 | TATGAGTGTGGAGTCGTTGCGAGAGAGAGAGAGAGAGAGAGGTNNNAGATCGGA |
| 40199_1 | TATGAGTGTGGAGTCGTTGCCTTGTACATTGCTGGTGGGAATGNNNAGATCGGAA |
| 40199_2 | TATGAGTGTGGAGTCGTTGCCTTGTACATTGCTGGTGGGAATGNNNAGATCGGAA |
| 13792_1 | TATGAGTGTGGAGTCGTTGCGAAAAACTGTAAACTGAGTTCTTNNNAGATCGGAAG |
| 13792_2 | TATGAGTGTGGAGTCGTTGCGAAAAACTGTAAACTGAGTTCTTNNNAGATCGGAAG |
| 30083_1 | TATGAGTGTGGAGTCGTTGCTACAATCGTTCTATGAGCCTGGGNNNAGATCGGAAG |
| 30083_2 | TATGAGTGTGGAGTCGTTGCTACAATCGTTCTATGAGCCTGGGNNNAGATCGGAAG |
| 50835_1 | TATGAGTGTGGAGTCGTTGCAGTCAGGACTCGTTTCTGATCTGNNNAGATCGGAAG |
| 50835_2 | TATGAGTGTGGAGTCGTTGCAGTCAGGACTCGTTTCTGATCTGNNNAGATCGGAAG |
| 38906_1 | TATGAGTGTGGAGTCGTTGCTTGCCACAACTTACCTACCTCTTNNNAGATCGGAAGA |
| 38906_2 | TATGAGTGTGGAGTCGTTGCTTGCCACAACTTACCTACCTCTTNNNAGATCGGAAGA |
| 36583_1 | TATGAGTGTGGAGTCGTTGCATTCACAGACCCCTGAACTAACCNNNAGATCGGAAGA |
| 36583_2 | TATGAGTGTGGAGTCGTTGCATTCACAGACCCCTGAACTAACCNNNAGATCGGAAGA |
| 37325_1 | TATGAGTGTGGAGTCGTTGCTCTAGGGACAGAGTGAGATTCCANNNAGATCGGAA |
| 37325_2 | TATGAGTGTGGAGTCGTTGCTCTAGGGACAGAGTGAGATTCCANNNAGATCGGAA |

| | |
|---|---|
| 33917_1 | TATGAGTGTGGAGTCGTTGCTGTAGTGCAGTGGTGTGATCTTGNNNAGATCGGAA |
| 33917_2 | TATGAGTGTGGAGTCGTTGCTGTAGTGCAGTGGTGTGATCTTGNNNAGATCGGAA |
| 44477_1 | TATGAGTGTGGAGTCGTTGCCACAGTTTGAGAGAAGCACCAAGNNNAGATCGGAA |
| 44477_2 | TATGAGTGTGGAGTCGTTGCCACAGTTTGAGAGAAGCACCAAGNNNAGATCGGAA |
| 8082_1 | TATGAGTGTGGAGTCGTTGCTCTAATGAGGGTAGGCTGTTGAANNNAGATCGGAA |
| 8082_2 | TATGAGTGTGGAGTCGTTGCTCTAATGAGGGTAGGCTGTTGAANNNAGATCGGAA |
| 39731_1 | TATGAGTGTGGAGTCGTTGCAACATAAAGCCAGGAAGCTAGGANNNAGATCGGAA |
| 39731_2 | TATGAGTGTGGAGTCGTTGCAACATAAAGCCAGGAAGCTAGGANNNAGATCGGAA |
| 36669_1 | TATGAGTGTGGAGTCGTTGCCCTGATAAGGCCACCCAACAGNNNAGATCGGAAGAG |
| 36669_2 | TATGAGTGTGGAGTCGTTGCCCTGATAAGGCCACCCAACAGNNNAGATCGGAAGAG |
| 35609_1 | TATGAGTGTGGAGTCGTTGCGGAGTACAGTGGCATGATCTCCNNNAGATCGGAAG |
| 35609_2 | TATGAGTGTGGAGTCGTTGCGGAGTACAGTGGCATGATCTCCNNNAGATCGGAAG |
| 29983_1 | TATGAGTGTGGAGTCGTTGCCAATGCTGTAACGCAGATTAGCANNNAGATCGGAAG |
| 29983_2 | TATGAGTGTGGAGTCGTTGCCAATGCTGTAACGCAGATTAGCANNNAGATCGGAAG |
| 33508_1 | TATGAGTGTGGAGTCGTTGCCCTTCAAGGTATGGCTTCTCTCANNNAGATCGGAAG |
| 33508_2 | TATGAGTGTGGAGTCGTTGCCCTTCAAGGTATGGCTTCTCTCANNNAGATCGGAAG |
| 47880_1 | TATGAGTGTGGAGTCGTTGCCTCCTGAGTAGCTGGGATTACACNNNAGATCGGAAG |
| 47880_2 | TATGAGTGTGGAGTCGTTGCCTCCTGAGTAGCTGGGATTACACNNNAGATCGGAAG |
| 52397_1 | TATGAGTGTGGAGTCGTTGCGCCCCAGGTTATTTCTTGCATTTNNNAGATCGGAAG |
| 52397_2 | TATGAGTGTGGAGTCGTTGCGCCCCAGGTTATTTCTTGCATTTNNNAGATCGGAAG |
| 32603_1 | TATGAGTGTGGAGTCGTTGCTCTTCTGAGCTTTGGACCTGTACNNNAGATCGGAAG |
| 32603_2 | TATGAGTGTGGAGTCGTTGCTCTTCTGAGCTTTGGACCTGTACNNNAGATCGGAAG |
| 41845_1 | TATGAGTGTGGAGTCGTTGCTCAAGTGATTCTCCTGTCTGAGCNNNAGATCGGAAG |
| 41845_2 | TATGAGTGTGGAGTCGTTGCTCAAGTGATTCTCCTGTCTGAGCNNNAGATCGGAAG |
| 43556_1 | TATGAGTGTGGAGTCGTTGCTGCCAAGTAAGTTGTTTGCAGAANNNAGATCGGAAG |
| 43556_2 | TATGAGTGTGGAGTCGTTGCTGCCAAGTAAGTTGTTTGCAGAANNNAGATCGGAAG |
| 38819_1 | TATGAGTGTGGAGTCGTTGCATAAATAAAGCCACCACCCTCGANNNAGATCGGAAG |
| 38819_2 | TATGAGTGTGGAGTCGTTGCATAAATAAAGCCACCACCCTCGANNNAGATCGGAAG |
| 38199_1 | TATGAGTGTGGAGTCGTTGCAGCTCAACTCAGGATCAGAAAGGNNNAGATCGGAA |
| 38199_2 | TATGAGTGTGGAGTCGTTGCAGCTCAACTCAGGATCAGAAAGGNNNAGATCGGAA |
| 36856_1 | TATGAGTGTGGAGTCGTTGCTCAGGAGACTCAGTCAGGAGAATNNNAGATCGGAA |
| 36856_2 | TATGAGTGTGGAGTCGTTGCTCAGGAGACTCAGTCAGGAGAATNNNAGATCGGAA |
| 36495_1 | TATGAGTGTGGAGTCGTTGCTAATGTGAACCCAGATTTGCCTGNNNAGATCGGAAG |
| 36495_2 | TATGAGTGTGGAGTCGTTGCTAATGTGAACCCAGATTTGCCTGNNNAGATCGGAAG |
| 36994_1 | TATGAGTGTGGAGTCGTTGCCATGGAATCAACCAAAGTGTCCANNNAGATCGGAAG |
| 36994_2 | TATGAGTGTGGAGTCGTTGCCATGGAATCAACCAAAGTGTCCANNNAGATCGGAAG |
| 45126_1 | TATGAGTGTGGAGTCGTTGCCAATTCTTGAGAAGCCCAGCTTGNNNAGATCGGAAG |
| 45126_2 | TATGAGTGTGGAGTCGTTGCCAATTCTTGAGAAGCCCAGCTTGNNNAGATCGGAAG |
| 45134_1 | TATGAGTGTGGAGTCGTTGCTGCTGGTGCCATCATTTCAATTTNNNAGATCGGAAG |
| 45134_2 | TATGAGTGTGGAGTCGTTGCTGCTGGTGCCATCATTTCAATTTNNNAGATCGGAAG |
| 52937_1 | TATGAGTGTGGAGTCGTTGCTCTCTATCAGATGCCCCCAAAACNNNAGATCGGAAG |
| 52937_2 | TATGAGTGTGGAGTCGTTGCTCTCTATCAGATGCCCCCAAAACNNNAGATCGGAAG |
| 41886_1 | TATGAGTGTGGAGTCGTTGCGAGACATGGACTTGCTCTGTCAANNNAGATCGGAAG |

| | |
|---|---|
| 41886_2 | TATGAGTGTGGAGTCGTTGCGAGACATGGACTTGCTCTGTCAANNNAGATCGGAAG |
| 49030_1 | TATGAGTGTGGAGTCGTTGCGCAACTACGCCCAGCTAATTTTTNNNAGATCGGAAG |
| 49030_2 | TATGAGTGTGGAGTCGTTGCGCAACTACGCCCAGCTAATTTTTNNNAGATCGGAAG |
| 43009_1 | TATGAGTGTGGAGTCGTTGCAATCCATCTGGTCCTGAGCATTTNNNAGATCGGAAG |
| 43009_2 | TATGAGTGTGGAGTCGTTGCAATCCATCTGGTCCTGAGCATTTNNNAGATCGGAAG |
| 51089_1 | TATGAGTGTGGAGTCGTTGCAAAAAAAAAATTAGCCGGGCACGNNNAGATCGGAA |
| 51089_2 | TATGAGTGTGGAGTCGTTGCAAAAAAAAAATTAGCCGGGCACGNNNAGATCGGAA |
| 48217_1 | TATGAGTGTGGAGTCGTTGCCTCCAGGGTTCAAGCGATTCTCNNNAGATCGGAAGA |
| 48217_2 | TATGAGTGTGGAGTCGTTGCCTCCAGGGTTCAAGCGATTCTCNNNAGATCGGAAGA |
| 31271_1 | TATGAGTGTGGAGTCGTTGCCGTGGGTGAGAGATTGAGCAANNNAGATCGGAAG |
| 31271_2 | TATGAGTGTGGAGTCGTTGCCGTGGGTGAGAGATTGAGCAANNNAGATCGGAAG |
| 47703_1 | TATGAGTGTGGAGTCGTTGCCCTGTGCAGCGATTGATTTCTGNNNAGATCGGAAGA |
| 47703_2 | TATGAGTGTGGAGTCGTTGCCCTGTGCAGCGATTGATTTCTGNNNAGATCGGAAGA |
| 10316_1 | TATGAGTGTGGAGTCGTTGCAACTCTGGTTACCTTGGGGAAGTNNNAGATCGGAA |
| 10316_2 | TATGAGTGTGGAGTCGTTGCAACTCTGGTTACCTTGGGGAAGTNNNAGATCGGAA |
| 29967_1 | TATGAGTGTGGAGTCGTTGCACAGAGGTTCAGGATTAATTTCTTCTNNNAGATCGG |
| 29967_2 | TATGAGTGTGGAGTCGTTGCACAGAGGTTCAGGATTAATTTCTTCTNNNAGATCGG |
| 29934_1 | TATGAGTGTGGAGTCGTTGCGGACACAGAGTAATGCTGACAGANNNAGATCGGAA |
| 29934_2 | TATGAGTGTGGAGTCGTTGCGGACACAGAGTAATGCTGACAGANNNAGATCGGAA |
| 51328_1 | TATGAGTGTGGAGTCGTTGCAGGGAAGGGATAGCATTAGGAGANNNAGATCGGA |
| 51328_2 | TATGAGTGTGGAGTCGTTGCAGGGAAGGGATAGCATTAGGAGANNNAGATCGGA |
| 31522_1 | TATGAGTGTGGAGTCGTTGCTGATGGCTACAAGGCATTCCATANNNAGATCGGAAG |
| 31522_2 | TATGAGTGTGGAGTCGTTGCTGATGGCTACAAGGCATTCCATANNNAGATCGGAAG |
| 51438_1 | TATGAGTGTGGAGTCGTTGCCACACCAACCTGAGCTACAAAAGNNNAGATCGGAAG |
| 51438_2 | TATGAGTGTGGAGTCGTTGCCACACCAACCTGAGCTACAAAAGNNNAGATCGGAAG |
| 44875_1 | TATGAGTGTGGAGTCGTTGCCTCTCTGACCTTAGAGTCTGTGCNNNAGATCGGAAG |
| 44875_2 | TATGAGTGTGGAGTCGTTGCCTCTCTGACCTTAGAGTCTGTGCNNNAGATCGGAAG |
| 12880_1 | TATGAGTGTGGAGTCGTTGCAATTGCCCAGAGAGTTTCTTNNNAGATCGGAAGAG |
| 12880_2 | TATGAGTGTGGAGTCGTTGCAATTGCCCAGAGAGTTTCTTNNNAGATCGGAAGAG |
| 45923_1 | TATGAGTGTGGAGTCGTTGCTTCAAGTGATTCTCCTGCCTCAGNNNAGATCGGAAG |
| 45923_2 | TATGAGTGTGGAGTCGTTGCTTCAAGTGATTCTCCTGCCTCAGNNNAGATCGGAAG |
| 13218_1 | TATGAGTGTGGAGTCGTTGCACTCCACTGCTTATTTCATCNNNAGATCGGAAGAGCA |
| 13218_2 | TATGAGTGTGGAGTCGTTGCACTCCACTGCTTATTTCATCNNNAGATCGGAAGAGCA |
| 41301_1 | TATGAGTGTGGAGTCGTTGCCCACTCATCGGAGCTGTATCAATNNNAGATCGGAAG |
| 41301_2 | TATGAGTGTGGAGTCGTTGCCCACTCATCGGAGCTGTATCAATNNNAGATCGGAAG |
| 47498_1 | TATGAGTGTGGAGTCGTTGCGACGAAGTCTTACTCTGTCTCCCNNNAGATCGGAAG |
| 47498_2 | TATGAGTGTGGAGTCGTTGCGACGAAGTCTTACTCTGTCTCCCNNNAGATCGGAAG |
| 35066_1 | TATGAGTGTGGAGTCGTTGCCTCAAGCAATTCTCATGCCTCAGNNNAGATCGGAAG |
| 35066_2 | TATGAGTGTGGAGTCGTTGCCTCAAGCAATTCTCATGCCTCAGNNNAGATCGGAAG |
| 46327_1 | TATGAGTGTGGAGTCGTTGCAGTAGCTCATGCCCAGCTAATTTNNNAGATCGGAAG |
| 46327_2 | TATGAGTGTGGAGTCGTTGCAGTAGCTCATGCCCAGCTAATTTNNNAGATCGGAAG |
| 38633_1 | TATGAGTGTGGAGTCGTTGCCCCAAAAACTCCTAGGCCTGATANNNAGATCGGAAG |
| 38633_2 | TATGAGTGTGGAGTCGTTGCCCCAAAAACTCCTAGGCCTGATANNNAGATCGGAAG |

| | |
|---|---|
| 39974_1 | TATGAGTGTGGAGTCGTTGCAGGAGATGATGAGTTTTGGGACANNNAGATCGGAA |
| 39974_2 | TATGAGTGTGGAGTCGTTGCAGGAGATGATGAGTTTTGGGACANNNAGATCGGAA |
| 2491_1 | TATGAGTGTGGAGTCGTTGCTCGCTCTTCGGTATCACAAAGAGNNNAGATCGGAAG |
| 2491_2 | TATGAGTGTGGAGTCGTTGCTCGCTCTTCGGTATCACAAAGAGNNNAGATCGGAAG |
| 51918_1 | TATGAGTGTGGAGTCGTTGCTGTGACTTCCATCTTGTTTGCACNNNAGATCGGAAGA |
| 51918_2 | TATGAGTGTGGAGTCGTTGCTGTGACTTCCATCTTGTTTGCACNNNAGATCGGAAGA |
| 40795_1 | TATGAGTGTGGAGTCGTTGCCTCACAGATGGAGCAATGGTGAGNNNAGATCGGAA |
| 40795_2 | TATGAGTGTGGAGTCGTTGCCTCACAGATGGAGCAATGGTGAGNNNAGATCGGAA |
| 37909_1 | TATGAGTGTGGAGTCGTTGCATGTCCCAATTTGTGAGATCCCANNNAGATCGGAAG |
| 37909_2 | TATGAGTGTGGAGTCGTTGCATGTCCCAATTTGTGAGATCCCANNNAGATCGGAAG |
| 47061_1 | TATGAGTGTGGAGTCGTTGCCTGACTTCAAGCAATTCTCCTGCNNNAGATCGGAAGA |
| 47061_2 | TATGAGTGTGGAGTCGTTGCCTGACTTCAAGCAATTCTCCTGCNNNAGATCGGAAGA |
| 4404_1 | TATGAGTGTGGAGTCGTTGCTAAGGCAGCACACTTCTACCTCTNNNAGATCGGAAGA |
| 4404_2 | TATGAGTGTGGAGTCGTTGCTAAGGCAGCACACTTCTACCTCTNNNAGATCGGAAGA |
| 46313_1 | TATGAGTGTGGAGTCGTTGCAACCTTAGCTCACTGCAACCTCNNNAGATCGGAAGAG |
| 46313_2 | TATGAGTGTGGAGTCGTTGCAACCTTAGCTCACTGCAACCTCNNNAGATCGGAAGAG |
| 29178_1 | TATGAGTGTGGAGTCGTTGCAAAGATAGATCCTGTTGCCTGCANNNAGATCGGAAG |
| 29178_2 | TATGAGTGTGGAGTCGTTGCAAAGATAGATCCTGTTGCCTGCANNNAGATCGGAAG |
| 49020_1 | TATGAGTGTGGAGTCGTTGCCGCCTGATCCAAACGATAATTGTNNNAGATCGGAAG |
| 49020_2 | TATGAGTGTGGAGTCGTTGCCGCCTGATCCAAACGATAATTGTNNNAGATCGGAAG |
| 42618_1 | TATGAGTGTGGAGTCGTTGCTGGAGTGCAGTGGTACAATCTCNNNAGATCGGAAGA |
| 42618_2 | TATGAGTGTGGAGTCGTTGCTGGAGTGCAGTGGTACAATCTCNNNAGATCGGAAGA |
| 38358_1 | TATGAGTGTGGAGTCGTTGCCCAGAGGTACAAAGAGGAGTTGGNNNAGATCGGAA |
| 38358_2 | TATGAGTGTGGAGTCGTTGCCCAGAGGTACAAAGAGGAGTTGGNNNAGATCGGAA |
| 47427_1 | TATGAGTGTGGAGTCGTTGCTTCAAGCTATTCTCCTGCCTCAGNNNAGATCGGAAGA |
| 47427_2 | TATGAGTGTGGAGTCGTTGCTTCAAGCTATTCTCCTGCCTCAGNNNAGATCGGAAGA |
| 47322_1 | TATGAGTGTGGAGTCGTTGCCTTCCTGGGTAGCTGGGATTACNNNAGATCGGAAGA |
| 47322_2 | TATGAGTGTGGAGTCGTTGCCTTCCTGGGTAGCTGGGATTACNNNAGATCGGAAGA |
| 48214_1 | TATGAGTGTGGAGTCGTTGCAGCCTCCTGAGTAGCTGAGATTANNNAGATCGGAAG |
| 48214_2 | TATGAGTGTGGAGTCGTTGCAGCCTCCTGAGTAGCTGAGATTANNNAGATCGGAAG |
| 34023_1 | TATGAGTGTGGAGTCGTTGCTCAATCTCCTGACCTCGTGATTCNNNAGATCGGAAGA |
| 34023_2 | TATGAGTGTGGAGTCGTTGCTCAATCTCCTGACCTCGTGATTCNNNAGATCGGAAGA |
| 30575_1 | TATGAGTGTGGAGTCGTTGCAAGAAAGGTCGATTCAGCTGGTTNNNAGATCGGAAG |
| 30575_2 | TATGAGTGTGGAGTCGTTGCAAGAAAGGTCGATTCAGCTGGTTNNNAGATCGGAAG |
| 40345_1 | TATGAGTGTGGAGTCGTTGCGAATTTCACCACTAAAGCTGGGCNNNAGATCGGAAG |
| 40345_2 | TATGAGTGTGGAGTCGTTGCGAATTTCACCACTAAAGCTGGGCNNNAGATCGGAAG |
| 43470_1 | TATGAGTGTGGAGTCGTTGCACGCCCGGCTAATTTTTGTATTTNNNAGATCGGAAGA |
| 43470_2 | TATGAGTGTGGAGTCGTTGCACGCCCGGCTAATTTTTGTATTTNNNAGATCGGAAGA |
| 13226_1 | TATGAGTGTGGAGTCGTTGCGTAACAGATGACAAGATCAAAGNNNAGATCGGAAG |
| 13226_2 | TATGAGTGTGGAGTCGTTGCGTAACAGATGACAAGATCAAAGNNNAGATCGGAAG |
| 34287_1 | TATGAGTGTGGAGTCGTTGCTCATCAAAAACCCTGTTTCCTGCNNNAGATCGGAAGA |
| 34287_2 | TATGAGTGTGGAGTCGTTGCTCATCAAAAACCCTGTTTCCTGCNNNAGATCGGAAGA |
| 11601_1 | TATGAGTGTGGAGTCGTTGCACCACACACAGGGAGATATCCAANNNAGATCGGAAG |

| | |
|---|---|
| 11601_2 | TATGAGTGTGGAGTCGTTGCACCACACACAGGGAGATATCCAANNNAGATCGGAA |
| 9401_1 | TATGAGTGTGGAGTCGTTGCTCTAGTGGTGAGGAAAGACCTCGNNNAGATCGGAA |
| 9401_2 | TATGAGTGTGGAGTCGTTGCTCTAGTGGTGAGGAAAGACCTCGNNNAGATCGGAA |
| 40010_1 | TATGAGTGTGGAGTCGTTGCCATGCTTGTAATCCCAGCACTTTNNNAGATCGGAAG |
| 40010_2 | TATGAGTGTGGAGTCGTTGCCATGCTTGTAATCCCAGCACTTTNNNAGATCGGAAG |
| 47174_1 | TATGAGTGTGGAGTCGTTGCCCACGCCCTGCTAATTTTTGTATNNNAGATCGGAAG |
| 47174_2 | TATGAGTGTGGAGTCGTTGCCCACGCCCTGCTAATTTTTGTATNNNAGATCGGAAG |
| 52094_1 | TATGAGTGTGGAGTCGTTGCCATGACCCTGTGTACATCCTTCANNNAGATCGGAAG |
| 52094_2 | TATGAGTGTGGAGTCGTTGCCATGACCCTGTGTACATCCTTCANNNAGATCGGAAG |
| 48272_1 | TATGAGTGTGGAGTCGTTGCGCTACCACACCCAGCTAATTTTTNNNAGATCGGAAG |
| 48272_2 | TATGAGTGTGGAGTCGTTGCGCTACCACACCCAGCTAATTTTTNNNAGATCGGAAG |
| 46421_1 | TATGAGTGTGGAGTCGTTGCCTCCCGGGTTCAAATGATTTTCCNNNAGATCGGAAG |
| 46421_2 | TATGAGTGTGGAGTCGTTGCCTCCCGGGTTCAAATGATTTTCCNNNAGATCGGAAG |
| 39285_1 | TATGAGTGTGGAGTCGTTGCTGACACAGGAAGACAAAACTTGCNNNAGATCGGAA |
| 39285_2 | TATGAGTGTGGAGTCGTTGCTGACACAGGAAGACAAAACTTGCNNNAGATCGGAA |
| 52634_1 | TATGAGTGTGGAGTCGTTGCTAGGAAAGGGGGAATAAGCCATCNNNAGATCGGA |
| 52634_2 | TATGAGTGTGGAGTCGTTGCTAGGAAAGGGGGAATAAGCCATCNNNAGATCGGA |
| 32460_1 | TATGAGTGTGGAGTCGTTGCGTCCAGACACATAGACCAAAGGANNNAGATCGGAA |
| 32460_2 | TATGAGTGTGGAGTCGTTGCGTCCAGACACATAGACCAAAGGANNNAGATCGGAA |
| 43249_1 | TATGAGTGTGGAGTCGTTGCTAGCTAGGATTACAGGTGTGTGCNNNAGATCGGAA |
| 43249_2 | TATGAGTGTGGAGTCGTTGCTAGCTAGGATTACAGGTGTGTGCNNNAGATCGGAA |
| 36849_1 | TATGAGTGTGGAGTCGTTGCGAAACTTCTTGATCGAGCTCTGCNNNAGATCGGAAG |
| 36849_2 | TATGAGTGTGGAGTCGTTGCGAAACTTCTTGATCGAGCTCTGCNNNAGATCGGAAG |
| 38065_1 | TATGAGTGTGGAGTCGTTGCTTTGTCTTCCCAGCAGATCTACCNNNAGATCGGAAG |
| 38065_2 | TATGAGTGTGGAGTCGTTGCTTTGTCTTCCCAGCAGATCTACCNNNAGATCGGAAG |
| 51859_1 | TATGAGTGTGGAGTCGTTGCCTTTACAACCTCCTGGATCCACANNNAGATCGGAAG |
| 51859_2 | TATGAGTGTGGAGTCGTTGCCTTTACAACCTCCTGGATCCACANNNAGATCGGAAG |
| 2437_1 | TATGAGTGTGGAGTCGTTGCGGGCAGTCTGGCAGAATTATCTCNNNAGATCGGAA |
| 2437_2 | TATGAGTGTGGAGTCGTTGCGGGCAGTCTGGCAGAATTATCTCNNNAGATCGGAA |
| 33170_1 | TATGAGTGTGGAGTCGTTGCCAACACTAAACCCTAAGGCTTGCNNNAGATCGGAAG |
| 33170_2 | TATGAGTGTGGAGTCGTTGCCAACACTAAACCCTAAGGCTTGCNNNAGATCGGAAG |
| 51905_1 | TATGAGTGTGGAGTCGTTGCAGTCCTGCCACCTAGAGTGATATNNNAGATCGGAAG |
| 51905_2 | TATGAGTGTGGAGTCGTTGCAGTCCTGCCACCTAGAGTGATATNNNAGATCGGAAG |
| 39931_1 | TATGAGTGTGGAGTCGTTGCAGGCATGGCTACAAAAGATTCCTNNNAGATCGGAA |
| 39931_2 | TATGAGTGTGGAGTCGTTGCAGGCATGGCTACAAAAGATTCCTNNNAGATCGGAA |
| 30905_1 | TATGAGTGTGGAGTCGTTGCACCCAGCTAATACTTGTTGCAGANNNAGATCGGAAG |
| 30905_2 | TATGAGTGTGGAGTCGTTGCACCCAGCTAATACTTGTTGCAGANNNAGATCGGAAG |
| 45467_1 | TATGAGTGTGGAGTCGTTGCTGAGCTGGAAGTGACAGAAAACTNNNAGATCGGAA |
| 45467_2 | TATGAGTGTGGAGTCGTTGCTGAGCTGGAAGTGACAGAAAACTNNNAGATCGGAA |
| 40240_1 | TATGAGTGTGGAGTCGTTGCCCCTACACATTCCCTTGACAGTTNNNAGATCGGAAGA |
| 40240_2 | TATGAGTGTGGAGTCGTTGCCCCTACACATTCCCTTGACAGTTNNNAGATCGGAAGA |
| 46308_1 | TATGAGTGTGGAGTCGTTGCGTGACTTTAGAAGGGTGGCTACANNNAGATCGGAA |
| 46308_2 | TATGAGTGTGGAGTCGTTGCGTGACTTTAGAAGGGTGGCTACANNNAGATCGGAA |

| | |
|---|---|
| 33046_1 | TATGAGTGTGGAGTCGTTGCCTCCTGTGTAGCTGGGACTACAGNNNAGATCGGAAG |
| 33046_2 | TATGAGTGTGGAGTCGTTGCCTCCTGTGTAGCTGGGACTACAGNNNAGATCGGAAG |
| 53425_1 | TATGAGTGTGGAGTCGTTGCTACAAATGTGCCCCTGAACCTAANNNAGATCGGAAG |
| 53425_2 | TATGAGTGTGGAGTCGTTGCTACAAATGTGCCCCTGAACCTAANNNAGATCGGAAG |
| 52647_1 | TATGAGTGTGGAGTCGTTGCGGCCACCTTGAACTTGATAGAGANNNAGATCGGAAG |
| 52647_2 | TATGAGTGTGGAGTCGTTGCGGCCACCTTGAACTTGATAGAGANNNAGATCGGAAG |
| 49835_1 | TATGAGTGTGGAGTCGTTGCTGCTGTCATGATTCCTAACACCANNNAGATCGGAAG |
| 49835_2 | TATGAGTGTGGAGTCGTTGCTGCTGTCATGATTCCTAACACCANNNAGATCGGAAG |
| 51880_1 | TATGAGTGTGGAGTCGTTGCAACCTGCTCATGTGTATGTCTGTNNNAGATCGGAAG |
| 51880_2 | TATGAGTGTGGAGTCGTTGCAACCTGCTCATGTGTATGTCTGTNNNAGATCGGAAG |
| 12856_1 | TATGAGTGTGGAGTCGTTGCAAAAATGTTTCTGTGTGTACTTANNNAGATCGGAAG |
| 12856_2 | TATGAGTGTGGAGTCGTTGCAAAAATGTTTCTGTGTGTACTTANNNAGATCGGAAG |
| 41928_1 | TATGAGTGTGGAGTCGTTGCGATCTTGGCTCACTGCAAGCTCNNNAGATCGGAAGA |
| 41928_2 | TATGAGTGTGGAGTCGTTGCGATCTTGGCTCACTGCAAGCTCNNNAGATCGGAAGA |
| 34124_1 | TATGAGTGTGGAGTCGTTGCACCAAGGAGGCTTTCTAAAGGAGNNNAGATCGGAA |
| 34124_2 | TATGAGTGTGGAGTCGTTGCACCAAGGAGGCTTTCTAAAGGAGNNNAGATCGGAA |
| 10911_1 | TATGAGTGTGGAGTCGTTGCTGAACACCACCACAAAAATCAGANNNAGATCGGAAG |
| 10911_2 | TATGAGTGTGGAGTCGTTGCTGAACACCACCACAAAAATCAGANNNAGATCGGAAG |
| 38186_1 | TATGAGTGTGGAGTCGTTGCACACGGTAAAACCCCATCTCTACNNNAGATCGGAAG |
| 38186_2 | TATGAGTGTGGAGTCGTTGCACACGGTAAAACCCCATCTCTACNNNAGATCGGAAG |
| 38153_1 | TATGAGTGTGGAGTCGTTGCCTGTGCTTCACCCTATCTAGCAANNNAGATCGGAAG |
| 38153_2 | TATGAGTGTGGAGTCGTTGCCTGTGCTTCACCCTATCTAGCAANNNAGATCGGAAG |
| 52607_1 | TATGAGTGTGGAGTCGTTGCACCCAGGACTATGACAATATGCCNNNAGATCGGAAG |
| 52607_2 | TATGAGTGTGGAGTCGTTGCACCCAGGACTATGACAATATGCCNNNAGATCGGAAG |
| 31359_1 | TATGAGTGTGGAGTCGTTGCGCATAACCTTCTGGGCTATCTGANNNAGATCGGAAG |
| 31359_2 | TATGAGTGTGGAGTCGTTGCGCATAACCTTCTGGGCTATCTGANNNAGATCGGAAG |
| 36511_1 | TATGAGTGTGGAGTCGTTGCTTGCAAGTAATGGCACCTTCAAGNNNAGATCGGAAG |
| 36511_2 | TATGAGTGTGGAGTCGTTGCTTGCAAGTAATGGCACCTTCAAGNNNAGATCGGAAG |
| 36626_1 | TATGAGTGTGGAGTCGTTGCGGCCTATCAATCAACTGCAACATNNNAGATCGGAAG |
| 36626_2 | TATGAGTGTGGAGTCGTTGCGGCCTATCAATCAACTGCAACATNNNAGATCGGAAG |
| 47031_1 | TATGAGTGTGGAGTCGTTGCCATGATCTTGGCTCACTGCAACNNNAGATCGGAAGA |
| 47031_2 | TATGAGTGTGGAGTCGTTGCCATGATCTTGGCTCACTGCAACNNNAGATCGGAAGA |
| 12771_1 | TATGAGTGTGGAGTCGTTGCTCATCTAAACATTTAAAAGTGCANNNAGATCGGAAG |
| 12771_2 | TATGAGTGTGGAGTCGTTGCTCATCTAAACATTTAAAAGTGCANNNAGATCGGAAG |
| 51566_1 | TATGAGTGTGGAGTCGTTGCACCCCGAAGAACCCTCCATATATNNNAGATCGGAAG |
| 51566_2 | TATGAGTGTGGAGTCGTTGCACCCCGAAGAACCCTCCATATATNNNAGATCGGAAG |
| 34844_1 | TATGAGTGTGGAGTCGTTGCTAGCTGGATTACTGGCACATACCNNNAGATCGGAAG |
| 34844_2 | TATGAGTGTGGAGTCGTTGCTAGCTGGATTACTGGCACATACCNNNAGATCGGAAG |
| 5622_1 | TATGAGTGTGGAGTCGTTGCCCTCCAGAGCACTGAAAAGTCTTNNNAGATCGGAAG |
| 5622_2 | TATGAGTGTGGAGTCGTTGCCCTCCAGAGCACTGAAAAGTCTTNNNAGATCGGAAG |
| 31021_1 | TATGAGTGTGGAGTCGTTGCTGTAACTACCTCCCACTGTTGTCNNNAGATCGGAAGA |
| 31021_2 | TATGAGTGTGGAGTCGTTGCTGTAACTACCTCCCACTGTTGTCNNNAGATCGGAAGA |
| 44215_1 | TATGAGTGTGGAGTCGTTGCGCTGAAACCCCGTCTCTACTAAANNNAGATCGGAAG |

| | |
|---|---|
| 44215_2 | TATGAGTGTGGAGTCGTTGCGCTGAAACCCCGTCTCTACTAAANNNAGATCGGAAG |
| 42675_1 | TATGAGTGTGGAGTCGTTGCACACTAAGAAATTCTGAAAGGGTGANNNAGATCGG |
| 42675_2 | TATGAGTGTGGAGTCGTTGCACACTAAGAAATTCTGAAAGGGTGANNNAGATCGG |
| 35835_1 | TATGAGTGTGGAGTCGTTGCCTGGGGTTCACGCCATTCTCNNNAGATCGGAAGAGC |
| 35835_2 | TATGAGTGTGGAGTCGTTGCCTGGGGTTCACGCCATTCTCNNNAGATCGGAAGAGC |
| 34046_1 | TATGAGTGTGGAGTCGTTGCCTTGAAATCCCAAAGTTAGCCCCNNNAGATCGGAAG |
| 34046_2 | TATGAGTGTGGAGTCGTTGCCTTGAAATCCCAAAGTTAGCCCCNNNAGATCGGAAG |
| 48967_1 | TATGAGTGTGGAGTCGTTGCGGCGTAATCTCGGCTCACTATAANNNAGATCGGAAG |
| 48967_2 | TATGAGTGTGGAGTCGTTGCGGCGTAATCTCGGCTCACTATAANNNAGATCGGAAG |
| 45049_1 | TATGAGTGTGGAGTCGTTGCAATTCGTATGGATCGACAGCAGTNNNAGATCGGAA |
| 45049_2 | TATGAGTGTGGAGTCGTTGCAATTCGTATGGATCGACAGCAGTNNNAGATCGGAA |
| 39012_1 | TATGAGTGTGGAGTCGTTGCTATCACTCAGTCATCTTCCCTGCNNNAGATCGGAAGA |
| 39012_2 | TATGAGTGTGGAGTCGTTGCTATCACTCAGTCATCTTCCCTGCNNNAGATCGGAAGA |
| 34031_1 | TATGAGTGTGGAGTCGTTGCTGTATGGGTGCAAGAAAAGGAGANNNAGATCGGA |
| 34031_2 | TATGAGTGTGGAGTCGTTGCTGTATGGGTGCAAGAAAAGGAGANNNAGATCGGA |
| 46648_1 | TATGAGTGTGGAGTCGTTGCCAGAGTAGCTGGGATTACAGGTGNNNAGATCGGAA |
| 46648_2 | TATGAGTGTGGAGTCGTTGCCAGAGTAGCTGGGATTACAGGTGNNNAGATCGGAA |
| 35409_1 | TATGAGTGTGGAGTCGTTGCGTCACCATCTCTGCTCACCATAANNNAGATCGGAAGA |
| 35409_2 | TATGAGTGTGGAGTCGTTGCGTCACCATCTCTGCTCACCATAANNNAGATCGGAAGA |
| 51495_1 | TATGAGTGTGGAGTCGTTGCCTTGTATTCTCCAAAGCCAGCAGNNNAGATCGGAAG |
| 51495_2 | TATGAGTGTGGAGTCGTTGCCTTGTATTCTCCAAAGCCAGCAGNNNAGATCGGAAG |
| 41958_1 | TATGAGTGTGGAGTCGTTGCCAGAGTAGCTGGGATTACAGGCNNNAGATCGGAAG |
| 41958_2 | TATGAGTGTGGAGTCGTTGCCAGAGTAGCTGGGATTACAGGCNNNAGATCGGAAG |
| 13695_1 | TATGAGTGTGGAGTCGTTGCTTCACATAAAGGAACATGGTTNNNAGATCGGAAGA |
| 13695_2 | TATGAGTGTGGAGTCGTTGCTTCACATAAAGGAACATGGTTNNNAGATCGGAAGA |
| 45684_1 | TATGAGTGTGGAGTCGTTGCTGGTGTGATCTCAGCTCATTGAANNNAGATCGGAAG |
| 45684_2 | TATGAGTGTGGAGTCGTTGCTGGTGTGATCTCAGCTCATTGAANNNAGATCGGAAG |
| 52393_1 | TATGAGTGTGGAGTCGTTGCCATCTCTGAGCTACTCCCCTGANNNAGATCGGAAGA |
| 52393_2 | TATGAGTGTGGAGTCGTTGCCATCTCTGAGCTACTCCCCTGANNNAGATCGGAAGA |
| 30583_1 | TATGAGTGTGGAGTCGTTGCCTGGGAGATAAAGGCTGCAGTTANNNAGATCGGAA |
| 30583_2 | TATGAGTGTGGAGTCGTTGCCTGGGAGATAAAGGCTGCAGTTANNNAGATCGGAA |
| 38967_1 | TATGAGTGTGGAGTCGTTGCGAAAAACTGTGAGACCCATTGCANNNAGATCGGAA |
| 38967_2 | TATGAGTGTGGAGTCGTTGCGAAAAACTGTGAGACCCATTGCANNNAGATCGGAA |
| 46323_1 | TATGAGTGTGGAGTCGTTGCGCCTCCTGGGTATAAGTCCTCTTNNNAGATCGGAAG |
| 46323_2 | TATGAGTGTGGAGTCGTTGCGCCTCCTGGGTATAAGTCCTCTTNNNAGATCGGAAG |
| 10865_1 | TATGAGTGTGGAGTCGTTGCACGGGCATGTGTATGTGTTTCTTNNNAGATCGGAAG |
| 10865_2 | TATGAGTGTGGAGTCGTTGCACGGGCATGTGTATGTGTTTCTTNNNAGATCGGAAG |
| 44337_1 | TATGAGTGTGGAGTCGTTGCGGACTGAGACTTTTCCCTCAACTNNNAGATCGGAAG |
| 44337_2 | TATGAGTGTGGAGTCGTTGCGGACTGAGACTTTTCCCTCAACTNNNAGATCGGAAG |
| 36364_1 | TATGAGTGTGGAGTCGTTGCGAGGTTGCAGTAAGCTGAGATTGNNNAGATCGGAA |
| 36364_2 | TATGAGTGTGGAGTCGTTGCGAGGTTGCAGTAAGCTGAGATTGNNNAGATCGGAA |
| 48793_1 | TATGAGTGTGGAGTCGTTGCAAATAGTGACTCTGCCAACCCATNNNAGATCGGAAG |
| 48793_2 | TATGAGTGTGGAGTCGTTGCAAATAGTGACTCTGCCAACCCATNNNAGATCGGAAG |

| | |
|---|---|
| 38990_1 | TATGAGTGTGGAGTCGTTGCAGAGGTCTTAGCCAATGCAATCANNNAGATCGGAA |
| 38990_2 | TATGAGTGTGGAGTCGTTGCAGAGGTCTTAGCCAATGCAATCANNNAGATCGGAA |
| 51836_1 | TATGAGTGTGGAGTCGTTGCCGACCCCCTCTTTGAGTTTCATTNNNAGATCGGAAGA |
| 51836_2 | TATGAGTGTGGAGTCGTTGCCGACCCCCTCTTTGAGTTTCATTNNNAGATCGGAAGA |
| 3234_1 | TATGAGTGTGGAGTCGTTGCCAAGATGTATGTTACAGGATTAGTGGANNNAGATC |
| 3234_2 | TATGAGTGTGGAGTCGTTGCCAAGATGTATGTTACAGGATTAGTGGANNNAGATC |
| 32897_1 | TATGAGTGTGGAGTCGTTGCCAAGTCTCTGACTCCCACATGAANNNAGATCGGAAG |
| 32897_2 | TATGAGTGTGGAGTCGTTGCCAAGTCTCTGACTCCCACATGAANNNAGATCGGAAG |
| 42565_1 | TATGAGTGTGGAGTCGTTGCATGCCTGGCCTGATTAGTGATTANNNAGATCGGAAG |
| 42565_2 | TATGAGTGTGGAGTCGTTGCATGCCTGGCCTGATTAGTGATTANNNAGATCGGAAG |
| 49746_1 | TATGAGTGTGGAGTCGTTGCTTCAAGTGATTCTCCTGCCTCAGNNNAGATCGGAAG |
| 49746_2 | TATGAGTGTGGAGTCGTTGCTTCAAGTGATTCTCCTGCCTCAGNNNAGATCGGAAG |
| 37583_1 | TATGAGTGTGGAGTCGTTGCTGTACAACTGCCAGCACAGATATNNNAGATCGGAAG |
| 37583_2 | TATGAGTGTGGAGTCGTTGCTGTACAACTGCCAGCACAGATATNNNAGATCGGAAG |
| 41712_1 | TATGAGTGTGGAGTCGTTGCGACAGGGTCTCGCTCTGTTGNNNAGATCGGAAGAG |
| 41712_2 | TATGAGTGTGGAGTCGTTGCGACAGGGTCTCGCTCTGTTGNNNAGATCGGAAGAG |
| 30727_1 | TATGAGTGTGGAGTCGTTGCGGCTCTATGAAGCAAGTGGAACANNNAGATCGGAA |
| 30727_2 | TATGAGTGTGGAGTCGTTGCGGCTCTATGAAGCAAGTGGAACANNNAGATCGGAA |
| 42859_1 | TATGAGTGTGGAGTCGTTGCTTTTGACATGATCGCACTCTGTGNNNAGATCGGAAG |
| 42859_2 | TATGAGTGTGGAGTCGTTGCTTTTGACATGATCGCACTCTGTGNNNAGATCGGAAG |
| 31171_1 | TATGAGTGTGGAGTCGTTGCATGTACAAGATGGCCCAACAGATNNNAGATCGGAA |
| 31171_2 | TATGAGTGTGGAGTCGTTGCATGTACAAGATGGCCCAACAGATNNNAGATCGGAA |
| 33689_1 | TATGAGTGTGGAGTCGTTGCCATTTCTGAAACTGAGAGGGGGANNNAGATCGGAA |
| 33689_2 | TATGAGTGTGGAGTCGTTGCCATTTCTGAAACTGAGAGGGGGANNNAGATCGGAA |
| 30744_1 | TATGAGTGTGGAGTCGTTGCAGAGACAGAGAGAGACAACGAGANNNAGATCGGA |
| 30744_2 | TATGAGTGTGGAGTCGTTGCAGAGACAGAGAGAGACAACGAGANNNAGATCGGA |
| 3131_1 | TATGAGTGTGGAGTCGTTGCATGTGGCAGAGTGTTCAGTTAGCNNNAGATCGGAA |
| 3131_2 | TATGAGTGTGGAGTCGTTGCATGTGGCAGAGTGTTCAGTTAGCNNNAGATCGGAA |
| 38096_1 | TATGAGTGTGGAGTCGTTGCGTTTTGGGCCAAAATGTGCAAATNNNAGATCGGAA |
| 38096_2 | TATGAGTGTGGAGTCGTTGCGTTTTGGGCCAAAATGTGCAAATNNNAGATCGGAA |
| 52776_1 | TATGAGTGTGGAGTCGTTGCCGGATGACCACGTTTGATACAAANNNAGATCGGAA |
| 52776_2 | TATGAGTGTGGAGTCGTTGCCGGATGACCACGTTTGATACAAANNNAGATCGGAA |
| 30757_1 | TATGAGTGTGGAGTCGTTGCTGAATGGTGCTGGAAATACTGGANNNAGATCGGAA |
| 30757_2 | TATGAGTGTGGAGTCGTTGCTGAATGGTGCTGGAAATACTGGANNNAGATCGGAA |
| 41532_1 | TATGAGTGTGGAGTCGTTGCCACGATCTCAGCTCAATGCAACNNNAGATCGGAAGA |
| 41532_2 | TATGAGTGTGGAGTCGTTGCCACGATCTCAGCTCAATGCAACNNNAGATCGGAAGA |
| 45341_1 | TATGAGTGTGGAGTCGTTGCATCAATGACCAGCACTTCATCCTNNNAGATCGGAAG |
| 45341_2 | TATGAGTGTGGAGTCGTTGCATCAATGACCAGCACTTCATCCTNNNAGATCGGAAG |
| 42768_1 | TATGAGTGTGGAGTCGTTGCCCATGAGGAGAGGAAGTGTTGAANNNAGATCGGA |
| 42768_2 | TATGAGTGTGGAGTCGTTGCCCATGAGGAGAGGAAGTGTTGAANNNAGATCGGA |
| 48266_1 | TATGAGTGTGGAGTCGTTGCCAGGTTCAAGCAATTCTCCTGCNNNAGATCGGAAGA |
| 48266_2 | TATGAGTGTGGAGTCGTTGCCAGGTTCAAGCAATTCTCCTGCNNNAGATCGGAAGA |
| 31136_1 | TATGAGTGTGGAGTCGTTGCGTGGATGTCAAAGTTCTGGCTTCNNNAGATCGGAA |

| | |
|---|---|
| 31136_2 | TATGAGTGTGGAGTCGTTGCGTGGATGTCAAAGTTCTGGCTTCNNNAGATCGGAA |
| 10722_1 | TATGAGTGTGGAGTCGTTGCATCAGGCCTTCCCTCTTCTGATTNNNAGATCGGAAGA |
| 10722_2 | TATGAGTGTGGAGTCGTTGCATCAGGCCTTCCCTCTTCTGATTNNNAGATCGGAAGA |
| 38268_1 | TATGAGTGTGGAGTCGTTGCTCTCATTGCTCTTGGAACTTTCCNNNAGATCGGAAGA |
| 38268_2 | TATGAGTGTGGAGTCGTTGCTCTCATTGCTCTTGGAACTTTCCNNNAGATCGGAAGA |
| 49566_1 | TATGAGTGTGGAGTCGTTGCTCTCCTACCTTTTCTTTTGAGACTNNNAGATCGGAAG |
| 49566_2 | TATGAGTGTGGAGTCGTTGCTCTCCTACCTTTTCTTTTGAGACTNNNAGATCGGAAG |
| 32770_1 | TATGAGTGTGGAGTCGTTGCGTACAGCAACAGTCCCACATCTANNNAGATCGGAAG |
| 32770_2 | TATGAGTGTGGAGTCGTTGCGTACAGCAACAGTCCCACATCTANNNAGATCGGAAG |
| 50170_1 | TATGAGTGTGGAGTCGTTGCCAGTACAGAGTGTTCAGTCAGCTNNNAGATCGGAA |
| 50170_2 | TATGAGTGTGGAGTCGTTGCCAGTACAGAGTGTTCAGTCAGCTNNNAGATCGGAA |
| 30513_1 | TATGAGTGTGGAGTCGTTGCTGTGGGCGTTAGTACTTGGGNNNAGATCGGAAGAG |
| 30513_2 | TATGAGTGTGGAGTCGTTGCTGTGGGCGTTAGTACTTGGGNNNAGATCGGAAGAG |
| 53191_1 | TATGAGTGTGGAGTCGTTGCGATCTCAAACTCCTGGCTTCGANNNAGATCGGAAGA |
| 53191_2 | TATGAGTGTGGAGTCGTTGCGATCTCAAACTCCTGGCTTCGANNNAGATCGGAAGA |
| 37830_1 | TATGAGTGTGGAGTCGTTGCTGACCAAAGGATTATTCTCGTTGCNNNAGATCGGAA |
| 37830_2 | TATGAGTGTGGAGTCGTTGCTGACCAAAGGATTATTCTCGTTGCNNNAGATCGGAA |
| 34374_1 | TATGAGTGTGGAGTCGTTGCATCTTGGGAGGTGGAATGTTTCANNNAGATCGGAA |
| 34374_2 | TATGAGTGTGGAGTCGTTGCATCTTGGGAGGTGGAATGTTTCANNNAGATCGGAA |
| 41463_1 | TATGAGTGTGGAGTCGTTGCGCTGGAGAGGATGTGGAGAAATANNNAGATCGGA |
| 41463_2 | TATGAGTGTGGAGTCGTTGCGCTGGAGAGGATGTGGAGAAATANNNAGATCGGA |
| 52546_1 | TATGAGTGTGGAGTCGTTGCTGTTCCTTTTCCTCCAAACTCCTNNNAGATCGGAAGA |
| 52546_2 | TATGAGTGTGGAGTCGTTGCTGTTCCTTTTCCTCCAAACTCCTNNNAGATCGGAAGA |
| 45597_1 | TATGAGTGTGGAGTCGTTGCTACAATCATGTCTTGACTGCAGCNNNAGATCGGAAG |
| 45597_2 | TATGAGTGTGGAGTCGTTGCTACAATCATGTCTTGACTGCAGCNNNAGATCGGAAG |
| 39680_1 | TATGAGTGTGGAGTCGTTGCTGCTGTTCTTTTCCTTCACAACANNNAGATCGGAAGA |
| 39680_2 | TATGAGTGTGGAGTCGTTGCTGCTGTTCTTTTCCTTCACAACANNNAGATCGGAAGA |
| 35848_1 | TATGAGTGTGGAGTCGTTGCCAATTCTCCTATCTCAGCCTCCCNNNAGATCGGAAGA |
| 35848_2 | TATGAGTGTGGAGTCGTTGCCAATTCTCCTATCTCAGCCTCCCNNNAGATCGGAAGA |
| 39166_1 | TATGAGTGTGGAGTCGTTGCAATCTCTGGAACACAGCTAAGGCNNNAGATCGGAA |
| 39166_2 | TATGAGTGTGGAGTCGTTGCAATCTCTGGAACACAGCTAAGGCNNNAGATCGGAA |
| 32372_1 | TATGAGTGTGGAGTCGTTGCGGAGTACAGTGGTGTGATCTCAGNNNAGATCGGAA |
| 32372_2 | TATGAGTGTGGAGTCGTTGCGGAGTACAGTGGTGTGATCTCAGNNNAGATCGGAA |
| 50437_1 | TATGAGTGTGGAGTCGTTGCAAGAAGAAAGAAGGCTCCAGGTTNNNAGATCGGAA |
| 50437_2 | TATGAGTGTGGAGTCGTTGCAAGAAGAAAGAAGGCTCCAGGTTNNNAGATCGGAA |
| 36709_1 | TATGAGTGTGGAGTCGTTGCAGCAGACTAGGGAATCATTGGATNNNAGATCGGAA |
| 36709_2 | TATGAGTGTGGAGTCGTTGCAGCAGACTAGGGAATCATTGGATNNNAGATCGGAA |
| 13260_1 | TATGAGTGTGGAGTCGTTGCCCAAATAAGGGTAGTAGATTTACNNNAGATCGGAA |
| 13260_2 | TATGAGTGTGGAGTCGTTGCCCAAATAAGGGTAGTAGATTTACNNNAGATCGGAA |
| 33403_1 | TATGAGTGTGGAGTCGTTGCAAGAGGAAGGAACATGACAAGGANNNAGATCGGA |
| 33403_2 | TATGAGTGTGGAGTCGTTGCAAGAGGAAGGAACATGACAAGGANNNAGATCGGA |
| 42596_1 | TATGAGTGTGGAGTCGTTGCCATATGTCTGCGTGTGTGTGTGNNNAGATCGGAAG |
| 42596_2 | TATGAGTGTGGAGTCGTTGCCATATGTCTGCGTGTGTGTGTGNNNAGATCGGAAG |

| | |
|---|---|
| 49635_1 | TATGAGTGTGGAGTCGTTGCCACCACACCCGGATACTTTTTTTNNNAGATCGGAAGA |
| 49635_2 | TATGAGTGTGGAGTCGTTGCCACCACACCCGGATACTTTTTTTNNNAGATCGGAAGA |
| 34677_1 | TATGAGTGTGGAGTCGTTGCCCAAAGTGTTGAGATTACAGGCGNNNAGATCGGAA |
| 34677_2 | TATGAGTGTGGAGTCGTTGCCCAAAGTGTTGAGATTACAGGCGNNNAGATCGGAA |
| 47052_1 | TATGAGTGTGGAGTCGTTGCCAGTGACACGATCATAGCTCACTNNNAGATCGGAAG |
| 47052_2 | TATGAGTGTGGAGTCGTTGCCAGTGACACGATCATAGCTCACTNNNAGATCGGAAG |
| 33599_1 | TATGAGTGTGGAGTCGTTGCTGTGTGTGTGCTTTCTCCTCATANNNAGATCGGAAG |
| 33599_2 | TATGAGTGTGGAGTCGTTGCTGTGTGTGTGCTTTCTCCTCATANNNAGATCGGAAG |
| 52040_1 | TATGAGTGTGGAGTCGTTGCCAGTAGCCCCCAGTGTCTATTTTNNNAGATCGGAAG |
| 52040_2 | TATGAGTGTGGAGTCGTTGCCAGTAGCCCCCAGTGTCTATTTTNNNAGATCGGAAG |
| 44054_1 | TATGAGTGTGGAGTCGTTGCGGGGAAAAGCATTTTTGGTGGATNNNAGATCGGAA |
| 44054_2 | TATGAGTGTGGAGTCGTTGCGGGGAAAAGCATTTTTGGTGGATNNNAGATCGGAA |
| 28853_1 | TATGAGTGTGGAGTCGTTGCGAGGCAGTCACTCCAGGTCTAGNNNAGATCGGAAG |
| 28853_2 | TATGAGTGTGGAGTCGTTGCGAGGCAGTCACTCCAGGTCTAGNNNAGATCGGAAG |
| 46385_1 | TATGAGTGTGGAGTCGTTGCCCATCGTTGCATCCCTAGGATAANNNAGATCGGAAG |
| 46385_2 | TATGAGTGTGGAGTCGTTGCCCATCGTTGCATCCCTAGGATAANNNAGATCGGAAG |
| 31053_1 | TATGAGTGTGGAGTCGTTGCGACGATGTGTACTCTTTGTGCAGNNNAGATCGGAAG |
| 31053_2 | TATGAGTGTGGAGTCGTTGCGACGATGTGTACTCTTTGTGCAGNNNAGATCGGAAG |
| 39962_1 | TATGAGTGTGGAGTCGTTGCGAAATGAGCACTGACTCTGGAGANNNAGATCGGAA |
| 39962_2 | TATGAGTGTGGAGTCGTTGCGAAATGAGCACTGACTCTGGAGANNNAGATCGGAA |
| 39326_1 | TATGAGTGTGGAGTCGTTGCAATGGATGACAGGTAGGGTGTTCNNNAGATCGGAA |
| 39326_2 | TATGAGTGTGGAGTCGTTGCAATGGATGACAGGTAGGGTGTTCNNNAGATCGGAA |
| 40186_1 | TATGAGTGTGGAGTCGTTGCACTGAAAAAATTCTCTGGTTAGTGGANNNAGATCGG |
| 40186_2 | TATGAGTGTGGAGTCGTTGCACTGAAAAAATTCTCTGGTTAGTGGANNNAGATCGG |
| 36744_1 | TATGAGTGTGGAGTCGTTGCCTGTAATCCCAGCTACTCAGGAGNNNAGATCGGAAG |
| 36744_2 | TATGAGTGTGGAGTCGTTGCCTGTAATCCCAGCTACTCAGGAGNNNAGATCGGAAG |
| 51683_1 | TATGAGTGTGGAGTCGTTGCCCCTCTATCAATGTGCTCAGCTANNNAGATCGGAAG |
| 51683_2 | TATGAGTGTGGAGTCGTTGCCCCTCTATCAATGTGCTCAGCTANNNAGATCGGAAG |
| 50982_1 | TATGAGTGTGGAGTCGTTGCTGTGTCTCACTGAGATTTTGTGCNNNAGATCGGAAG |
| 50982_2 | TATGAGTGTGGAGTCGTTGCTGTGTCTCACTGAGATTTTGTGCNNNAGATCGGAAG |
| 32099_1 | TATGAGTGTGGAGTCGTTGCAGTGTTCCTGAGGAAGAAGAGAANNNAGATCGGA |
| 32099_2 | TATGAGTGTGGAGTCGTTGCAGTGTTCCTGAGGAAGAAGAGAANNNAGATCGGA |
| 37619_1 | TATGAGTGTGGAGTCGTTGCTGAGGTCGATCTTTATGGTTCCANNNAGATCGGAAG |
| 37619_2 | TATGAGTGTGGAGTCGTTGCTGAGGTCGATCTTTATGGTTCCANNNAGATCGGAAG |
| 36258_1 | TATGAGTGTGGAGTCGTTGCAGAGAGACTGGGAAGTAGAAGCTNNNAGATCGGA |
| 36258_2 | TATGAGTGTGGAGTCGTTGCAGAGAGACTGGGAAGTAGAAGCTNNNAGATCGGA |
| 35138_1 | TATGAGTGTGGAGTCGTTGCGGAGTCTCACTCTTTCACCCAAGNNNAGATCGGAAG |
| 35138_2 | TATGAGTGTGGAGTCGTTGCGGAGTCTCACTCTTTCACCCAAGNNNAGATCGGAAG |
| 52194_1 | TATGAGTGTGGAGTCGTTGCGGCATGATCATGGCTCACTACAGNNNAGATCGGAAG |
| 52194_2 | TATGAGTGTGGAGTCGTTGCGGCATGATCATGGCTCACTACAGNNNAGATCGGAAG |
| 47216_1 | TATGAGTGTGGAGTCGTTGCGTTCAAGAGATTCTCCTGCCTCANNNAGATCGGAAG |
| 47216_2 | TATGAGTGTGGAGTCGTTGCGTTCAAGAGATTCTCCTGCCTCANNNAGATCGGAAG |
| 37843_1 | TATGAGTGTGGAGTCGTTGCATACATTATGGTTGGGTTGGGCANNNAGATCGGAA |

| | |
|---|---|
| 37843_2 | TATGAGTGTGGAGTCGTTGCATACATTATGGTTGGGTTGGGCANNNAGATCGGAA |
| 30030_1 | TATGAGTGTGGAGTCGTTGCCCATGACTTCTCCTTCACTGACANNNAGATCGGAAGA |
| 30030_2 | TATGAGTGTGGAGTCGTTGCCCATGACTTCTCCTTCACTGACANNNAGATCGGAAGA |
| 37092_1 | TATGAGTGTGGAGTCGTTGCACAAAGGCTACCTCACCTCAAAANNNAGATCGGAAG |
| 37092_2 | TATGAGTGTGGAGTCGTTGCACAAAGGCTACCTCACCTCAAAANNNAGATCGGAAG |
| 4305_1 | TATGAGTGTGGAGTCGTTGCTGAAGCATGATTTTTAGTGGCTGCNNNAGATCGGA |
| 4305_2 | TATGAGTGTGGAGTCGTTGCTGAAGCATGATTTTTAGTGGCTGCNNNAGATCGGA |
| 10972_1 | TATGAGTGTGGAGTCGTTGCACGGCCAAAAGTGACCTAAAGAGNNNAGATCGGAA |
| 10972_2 | TATGAGTGTGGAGTCGTTGCACGGCCAAAAGTGACCTAAAGAGNNNAGATCGGAA |
| 47484_1 | TATGAGTGTGGAGTCGTTGCGAGGTACAGGTTCTCACTCTGTCNNNAGATCGGAAG |
| 47484_2 | TATGAGTGTGGAGTCGTTGCGAGGTACAGGTTCTCACTCTGTCNNNAGATCGGAAG |
| 32443_1 | TATGAGTGTGGAGTCGTTGCCTCACACCATTCTCCTGCCTCNNNAGATCGGAAGAGC |
| 32443_2 | TATGAGTGTGGAGTCGTTGCCTCACACCATTCTCCTGCCTCNNNAGATCGGAAGAGC |
| 11134_1 | TATGAGTGTGGAGTCGTTGCCAGCATCATGCCCGGATAATTTTCNNNAGATCGGAA |
| 11134_2 | TATGAGTGTGGAGTCGTTGCCAGCATCATGCCCGGATAATTTTCNNNAGATCGGAA |
| 41464_1 | TATGAGTGTGGAGTCGTTGCGCTACTCAGGAGACATGTGAGAGNNNAGATCGGAA |
| 41464_2 | TATGAGTGTGGAGTCGTTGCGCTACTCAGGAGACATGTGAGAGNNNAGATCGGAA |
| 40032_1 | TATGAGTGTGGAGTCGTTGCAGACCCAAACCAGTAAGCTGAATNNNAGATCGGAA |
| 40032_2 | TATGAGTGTGGAGTCGTTGCAGACCCAAACCAGTAAGCTGAATNNNAGATCGGAA |
| 48706_1 | TATGAGTGTGGAGTCGTTGCCAAGTAGCTGGGACTACATGTGTNNNAGATCGGAA |
| 48706_2 | TATGAGTGTGGAGTCGTTGCCAAGTAGCTGGGACTACATGTGTNNNAGATCGGAA |
| 43518_1 | TATGAGTGTGGAGTCGTTGCCACCACGCCTGGCTAATTTTTNNNAGATCGGAAGAG |
| 43518_2 | TATGAGTGTGGAGTCGTTGCCACCACGCCTGGCTAATTTTTNNNAGATCGGAAGAG |
| 42539_1 | TATGAGTGTGGAGTCGTTGCAGCTAGCTGTTCTCCCCAAATACNNNAGATCGGAAG |
| 42539_2 | TATGAGTGTGGAGTCGTTGCAGCTAGCTGTTCTCCCCAAATACNNNAGATCGGAAG |
| 45039_1 | TATGAGTGTGGAGTCGTTGCAAGATCACTGCCTTCAGAGGAAGNNNAGATCGGAA |
| 45039_2 | TATGAGTGTGGAGTCGTTGCAAGATCACTGCCTTCAGAGGAAGNNNAGATCGGAA |
| 37802_1 | TATGAGTGTGGAGTCGTTGCGTCAACCCCAACAAAATCCAGAGNNNAGATCGGAAG |
| 37802_2 | TATGAGTGTGGAGTCGTTGCGTCAACCCCAACAAAATCCAGAGNNNAGATCGGAAG |
| 29083_1 | TATGAGTGTGGAGTCGTTGCGGGAAAGGAAGAGAAGAAGAGCANNNAGATCGG |
| 29083_2 | TATGAGTGTGGAGTCGTTGCGGGAAAGGAAGAGAAGAAGAGCANNNAGATCGG |
| 46901_1 | TATGAGTGTGGAGTCGTTGCTCAAGTGATTCTCCTTCCTCAGCNNNAGATCGGAAG |
| 46901_2 | TATGAGTGTGGAGTCGTTGCTCAAGTGATTCTCCTTCCTCAGCNNNAGATCGGAAG |
| 49186_1 | TATGAGTGTGGAGTCGTTGCGCACCAACATTCCCAGCTAATTANNNAGATCGGAAG |
| 49186_2 | TATGAGTGTGGAGTCGTTGCGCACCAACATTCCCAGCTAATTANNNAGATCGGAAG |
| 42703_1 | TATGAGTGTGGAGTCGTTGCTGGTTGATGGTTACTTAGACTGGTNNNAGATCGGA |
| 42703_2 | TATGAGTGTGGAGTCGTTGCTGGTTGATGGTTACTTAGACTGGTNNNAGATCGGA |
| 48222_1 | TATGAGTGTGGAGTCGTTGCCAAGTAGCTGGGATTACAGGTGTNNNAGATCGGAA |
| 48222_2 | TATGAGTGTGGAGTCGTTGCCAAGTAGCTGGGATTACAGGTGTNNNAGATCGGAA |
| 45907_1 | TATGAGTGTGGAGTCGTTGCCTTCCTGGGTTCAAGCAATTCTCNNNAGATCGGAAG |
| 45907_2 | TATGAGTGTGGAGTCGTTGCCTTCCTGGGTTCAAGCAATTCTCNNNAGATCGGAAG |
| 32880_1 | TATGAGTGTGGAGTCGTTGCAGACGGAGTCTCACTCTGTCGNNNAGATCGGAAGA |
| 32880_2 | TATGAGTGTGGAGTCGTTGCAGACGGAGTCTCACTCTGTCGNNNAGATCGGAAGA |

| | |
|---|---|
| 49098_1 | TATGAGTGTGGAGTCGTTGCTTTCACATGATGCTACCACCTCANNNAGATCGGAAG |
| 49098_2 | TATGAGTGTGGAGTCGTTGCTTTCACATGATGCTACCACCTCANNNAGATCGGAAG |
| 5535_1 | TATGAGTGTGGAGTCGTTGCGGCAAAGGCCCCAAATTATGAACNNNAGATCGGAA |
| 5535_2 | TATGAGTGTGGAGTCGTTGCGGCAAAGGCCCCAAATTATGAACNNNAGATCGGAA |
| 12877_1 | TATGAGTGTGGAGTCGTTGCGATTTTATGCTTTCTTTACATGGNNNAGATCGGAAG |
| 12877_2 | TATGAGTGTGGAGTCGTTGCGATTTTATGCTTTCTTTACATGGNNNAGATCGGAAG |
| 49836_1 | TATGAGTGTGGAGTCGTTGCCTCGGTAAATAGTGCCACCTTCTNNNAGATCGGAAG |
| 49836_2 | TATGAGTGTGGAGTCGTTGCCTCGGTAAATAGTGCCACCTTCTNNNAGATCGGAAG |
| 31092_1 | TATGAGTGTGGAGTCGTTGCAAAGTGTCGACAGGATAGTGACCNNNAGATCGGAA |
| 31092_2 | TATGAGTGTGGAGTCGTTGCAAAGTGTCGACAGGATAGTGACCNNNAGATCGGAA |
| 39929_1 | TATGAGTGTGGAGTCGTTGCCATCAAATGCTTTTTCTGTGCCANNNAGATCGGAAG |
| 39929_2 | TATGAGTGTGGAGTCGTTGCCATCAAATGCTTTTTCTGTGCCANNNAGATCGGAAG |
| 2979_1 | TATGAGTGTGGAGTCGTTGCGCAATGACAAAGGTAACAATTCCAGNNNAGATCGG |
| 2979_2 | TATGAGTGTGGAGTCGTTGCGCAATGACAAAGGTAACAATTCCAGNNNAGATCGG |
| 33793_1 | TATGAGTGTGGAGTCGTTGCACTCCTGATCTCATGTGATCTGCNNNAGATCGGAAG |
| 33793_2 | TATGAGTGTGGAGTCGTTGCACTCCTGATCTCATGTGATCTGCNNNAGATCGGAAG |
| 49345_1 | TATGAGTGTGGAGTCGTTGCAACGAAGGGTGCAATAGAAAACCNNNAGATCGGAA |
| 49345_2 | TATGAGTGTGGAGTCGTTGCAACGAAGGGTGCAATAGAAAACCNNNAGATCGGAA |
| 34104_1 | TATGAGTGTGGAGTCGTTGCTAATCCTAACACTTTGGGAGGCCNNNAGATCGGAAG |
| 34104_2 | TATGAGTGTGGAGTCGTTGCTAATCCTAACACTTTGGGAGGCCNNNAGATCGGAAG |
| 38156_1 | TATGAGTGTGGAGTCGTTGCAGATTACCTTGAAGGCAGCAAGANNNAGATCGGAA |
| 38156_2 | TATGAGTGTGGAGTCGTTGCAGATTACCTTGAAGGCAGCAAGANNNAGATCGGAA |
| 53431_1 | TATGAGTGTGGAGTCGTTGCAGGGAGAGAGACAGAGGAAGAAANNNAGATCGG |
| 53431_2 | TATGAGTGTGGAGTCGTTGCAGGGAGAGAGACAGAGGAAGAAANNNAGATCGG |
| 2687_1 | TATGAGTGTGGAGTCGTTGCGCTCAGGTGGTCGAGTATCTCATNNNAGATCGGAAG |
| 2687_2 | TATGAGTGTGGAGTCGTTGCGCTCAGGTGGTCGAGTATCTCATNNNAGATCGGAAG |
| 51545_1 | TATGAGTGTGGAGTCGTTGCAACAAGAAGACAGCCCACTCTTTNNNAGATCGGAAG |
| 51545_2 | TATGAGTGTGGAGTCGTTGCAACAAGAAGACAGCCCACTCTTTNNNAGATCGGAAG |
| 47693_1 | TATGAGTGTGGAGTCGTTGCCCAAGTAGCTGGTACTACAGGTGNNNAGATCGGAA |
| 47693_2 | TATGAGTGTGGAGTCGTTGCCCAAGTAGCTGGTACTACAGGTGNNNAGATCGGAA |
| 39507_1 | TATGAGTGTGGAGTCGTTGCCGTTTTCTCTTCCTCTTCCCTGANNNAGATCGGAAGA |
| 39507_2 | TATGAGTGTGGAGTCGTTGCCGTTTTCTCTTCCTCTTCCCTGANNNAGATCGGAAGA |
| 12685_1 | TATGAGTGTGGAGTCGTTGCAATTTTATGGAATCCACACCTNNNAGATCGGAAGAG |
| 12685_2 | TATGAGTGTGGAGTCGTTGCAATTTTATGGAATCCACACCTNNNAGATCGGAAGAG |
| 49931_1 | TATGAGTGTGGAGTCGTTGCATCCTTGGAGATTCATCAGCTGGNNNAGATCGGAAG |
| 49931_2 | TATGAGTGTGGAGTCGTTGCATCCTTGGAGATTCATCAGCTGGNNNAGATCGGAAG |
| 34972_1 | TATGAGTGTGGAGTCGTTGCATCTCCTGGGTTCAAGCAATTCTNNNAGATCGGAAG |
| 34972_2 | TATGAGTGTGGAGTCGTTGCATCTCCTGGGTTCAAGCAATTCTNNNAGATCGGAAG |
| 46305_1 | TATGAGTGTGGAGTCGTTGCATAATCTCAGCTCACTGCAACCTNNNAGATCGGAAG |
| 46305_2 | TATGAGTGTGGAGTCGTTGCATAATCTCAGCTCACTGCAACCTNNNAGATCGGAAG |
| 10291_1 | TATGAGTGTGGAGTCGTTGCTTGCCATGTATCTCCAGTGTACANNNAGATCGGAAG |
| 10291_2 | TATGAGTGTGGAGTCGTTGCTTGCCATGTATCTCCAGTGTACANNNAGATCGGAAG |
| 34535_1 | TATGAGTGTGGAGTCGTTGCTTCTCCTGCCTTTCACAAACTCTNNNAGATCGGAAGA |

| | |
|---|---|
| 34535_2 | TATGAGTGTGGAGTCGTTGCTTCTCCTGCCTTTCACAAACTCTNNNAGATCGGAAGA |
| 32665_1 | TATGAGTGTGGAGTCGTTGCGAGATCTTGGCTCACTGTAACCCNNNAGATCGGAAG |
| 32665_2 | TATGAGTGTGGAGTCGTTGCGAGATCTTGGCTCACTGTAACCCNNNAGATCGGAAG |
| 46504_1 | TATGAGTGTGGAGTCGTTGCCACAACCACACCCAGCTAATTTTNNNAGATCGGAAGA |
| 46504_2 | TATGAGTGTGGAGTCGTTGCCACAACCACACCCAGCTAATTTTNNNAGATCGGAAGA |
| 29299_1 | TATGAGTGTGGAGTCGTTGCCCATTCACAGTAACTGATCTGCTNNNAGATCGGAAG |
| 29299_2 | TATGAGTGTGGAGTCGTTGCCCATTCACAGTAACTGATCTGCTNNNAGATCGGAAG |
| 35502_1 | TATGAGTGTGGAGTCGTTGCCCAAGTAGCTGCGATTACAGAGANNNAGATCGGAA |
| 35502_2 | TATGAGTGTGGAGTCGTTGCCCAAGTAGCTGCGATTACAGAGANNNAGATCGGAA |
| 29079_1 | TATGAGTGTGGAGTCGTTGCAATGGCTGGGTTAGGTACTTTGTNNNAGATCGGAA |
| 29079_2 | TATGAGTGTGGAGTCGTTGCAATGGCTGGGTTAGGTACTTTGTNNNAGATCGGAA |
| 36161_1 | TATGAGTGTGGAGTCGTTGCAACTCACTATGCAGCTGTCTGAANNNAGATCGGAAG |
| 36161_2 | TATGAGTGTGGAGTCGTTGCAACTCACTATGCAGCTGTCTGAANNNAGATCGGAAG |
| 43048_1 | TATGAGTGTGGAGTCGTTGCACACACAAGAGAGAATGAACACTNNNAGATCGGAA |
| 43048_2 | TATGAGTGTGGAGTCGTTGCACACACAAGAGAGAATGAACACTNNNAGATCGGAA |
| 45169_1 | TATGAGTGTGGAGTCGTTGCGACTACAAGACGACCACCATCCNNNAGATCGGAAGA |
| 45169_2 | TATGAGTGTGGAGTCGTTGCGACTACAAGACGACCACCATCCNNNAGATCGGAAGA |
| 33252_1 | TATGAGTGTGGAGTCGTTGCCGAGTAGCTGGAATTACAGGTGTNNNAGATCGGAA |
| 33252_2 | TATGAGTGTGGAGTCGTTGCCGAGTAGCTGGAATTACAGGTGTNNNAGATCGGAA |
| 13710_1 | TATGAGTGTGGAGTCGTTGCGTTAACTGAAGATATTCCAACTCNNNAGATCGGAAG |
| 13710_2 | TATGAGTGTGGAGTCGTTGCGTTAACTGAAGATATTCCAACTCNNNAGATCGGAAG |
| 38613_1 | TATGAGTGTGGAGTCGTTGCCTTTGATCCCACCAACAATCAGGNNNAGATCGGAAG |
| 38613_2 | TATGAGTGTGGAGTCGTTGCCTTTGATCCCACCAACAATCAGGNNNAGATCGGAAG |
| 49133_1 | TATGAGTGTGGAGTCGTTGCCTTTGTAGAGGCAGGGTTTCGCNNNAGATCGGAAG |
| 49133_2 | TATGAGTGTGGAGTCGTTGCCTTTGTAGAGGCAGGGTTTCGCNNNAGATCGGAAG |
| 37140_1 | TATGAGTGTGGAGTCGTTGCTCTGTCACTCCTCTCTGTACTCANNNAGATCGGAAGA |
| 37140_2 | TATGAGTGTGGAGTCGTTGCTCTGTCACTCCTCTCTGTACTCANNNAGATCGGAAGA |
| 51243_1 | TATGAGTGTGGAGTCGTTGCACCTGCAACTGAAAGACATCAGANNNAGATCGGAA |
| 51243_2 | TATGAGTGTGGAGTCGTTGCACCTGCAACTGAAAGACATCAGANNNAGATCGGAA |
| 39841_1 | TATGAGTGTGGAGTCGTTGCTTCTTGCTTTTCCACCTTGATGTNNNAGATCGGAAGA |
| 39841_2 | TATGAGTGTGGAGTCGTTGCTTCTTGCTTTTCCACCTTGATGTNNNAGATCGGAAGA |
| 4703_1 | TATGAGTGTGGAGTCGTTGCGGCAGGTTTCCTCTCTAAAGCAANNNAGATCGGAAG |
| 4703_2 | TATGAGTGTGGAGTCGTTGCGGCAGGTTTCCTCTCTAAAGCAANNNAGATCGGAAG |
| 39212_1 | TATGAGTGTGGAGTCGTTGCGTTTGGCCACCTTTTGTCTCTTTNNNAGATCGGAAGA |
| 39212_2 | TATGAGTGTGGAGTCGTTGCGTTTGGCCACCTTTTGTCTCTTTNNNAGATCGGAAGA |
| 47576_1 | TATGAGTGTGGAGTCGTTGCCAGCCTCTTGAGAAACTGGGATTNNNAGATCGGAA |
| 47576_2 | TATGAGTGTGGAGTCGTTGCCAGCCTCTTGAGAAACTGGGATTNNNAGATCGGAA |
| 29399_1 | TATGAGTGTGGAGTCGTTGCACTTTTCTTTGCTTGTCAGATGCCNNNAGATCGGAAG |
| 29399_2 | TATGAGTGTGGAGTCGTTGCACTTTTCTTTGCTTGTCAGATGCCNNNAGATCGGAAG |
| 45371_1 | TATGAGTGTGGAGTCGTTGCCTGGGGACTTAGGGATGTAAAGGNNNAGATCGGA |
| 45371_2 | TATGAGTGTGGAGTCGTTGCCTGGGGACTTAGGGATGTAAAGGNNNAGATCGGA |
| 50274_1 | TATGAGTGTGGAGTCGTTGCCTCCCAACTCCCTACTTTCTGTCNNNAGATCGGAAGA |
| 50274_2 | TATGAGTGTGGAGTCGTTGCCTCCCAACTCCCTACTTTCTGTCNNNAGATCGGAAGA |

| | |
|---|---|
| 34164_1 | TATGAGTGTGGAGTCGTTGCAACTCTGATGGAGGAAATGGGACNNNAGATCGGAA |
| 34164_2 | TATGAGTGTGGAGTCGTTGCAACTCTGATGGAGGAAATGGGACNNNAGATCGGAA |
| 47090_1 | TATGAGTGTGGAGTCGTTGCGCTGGAGTGCAGAGGTATAATCANNNAGATCGGAA |
| 47090_2 | TATGAGTGTGGAGTCGTTGCGCTGGAGTGCAGAGGTATAATCANNNAGATCGGAA |
| 52839_1 | TATGAGTGTGGAGTCGTTGCCTCTCCCACACTTGCTCATACCNNNAGATCGGAAGAG |
| 52839_2 | TATGAGTGTGGAGTCGTTGCCTCTCCCACACTTGCTCATACCNNNAGATCGGAAGAG |
| 38361_1 | TATGAGTGTGGAGTCGTTGCGAAGTCAAGAAAGTGGTCCTCCTNNNAGATCGGAA |
| 38361_2 | TATGAGTGTGGAGTCGTTGCGAAGTCAAGAAAGTGGTCCTCCTNNNAGATCGGAA |
| 48029_1 | TATGAGTGTGGAGTCGTTGCCTTGAAAGCCCAGGGAGACTATTNNNAGATCGGAA |
| 48029_2 | TATGAGTGTGGAGTCGTTGCCTTGAAAGCCCAGGGAGACTATTNNNAGATCGGAA |
| 40821_1 | TATGAGTGTGGAGTCGTTGCTTCAAGCAACTCTCTAGCCTCAGNNNAGATCGGAAG |
| 40821_2 | TATGAGTGTGGAGTCGTTGCTTCAAGCAACTCTCTAGCCTCAGNNNAGATCGGAAG |
| 69_1 | TATGAGTGTGGAGTCGTTGCCCTCCCTTATACTGGGCCCCTNNNAGATCGGAAGAG |
| 69_2 | TATGAGTGTGGAGTCGTTGCCCTCCCTTATACTGGGCCCCTNNNAGATCGGAAGAG |
| 35100_1 | TATGAGTGTGGAGTCGTTGCATTCTCAAGCCATCCTCCCATTTNNNAGATCGGAAGA |
| 35100_2 | TATGAGTGTGGAGTCGTTGCATTCTCAAGCCATCCTCCCATTTNNNAGATCGGAAGA |
| 36120_1 | TATGAGTGTGGAGTCGTTGCGGGAGGATCATTTGAGTCAGGAGNNNAGATCGGA |
| 36120_2 | TATGAGTGTGGAGTCGTTGCGGGAGGATCATTTGAGTCAGGAGNNNAGATCGGA |
| 44200_1 | TATGAGTGTGGAGTCGTTGCTGCTAGGATCTGATTACCCGTTGNNNAGATCGGAAG |
| 44200_2 | TATGAGTGTGGAGTCGTTGCTGCTAGGATCTGATTACCCGTTGNNNAGATCGGAAG |
| 46451_1 | TATGAGTGTGGAGTCGTTGCCTTTCCCAGGTTGGAGTGCTATNNNAGATCGGAAGA |
| 46451_2 | TATGAGTGTGGAGTCGTTGCCTTTCCCAGGTTGGAGTGCTATNNNAGATCGGAAGA |
| 52189_1 | TATGAGTGTGGAGTCGTTGCCAGCATGAGAAGACATCCCCTAGNNNAGATCGGAAG |
| 52189_2 | TATGAGTGTGGAGTCGTTGCCAGCATGAGAAGACATCCCCTAGNNNAGATCGGAAG |
| 40551_1 | TATGAGTGTGGAGTCGTTGCTTGGAGGACAATGGTGTGATCTTNNNAGATCGGAA |
| 40551_2 | TATGAGTGTGGAGTCGTTGCTTGGAGGACAATGGTGTGATCTTNNNAGATCGGAA |
| 48167_1 | TATGAGTGTGGAGTCGTTGCTTCAAGTGATTCTCCTGCCTCAGNNNAGATCGGAAG |
| 48167_2 | TATGAGTGTGGAGTCGTTGCTTCAAGTGATTCTCCTGCCTCAGNNNAGATCGGAAG |
| 44392_1 | TATGAGTGTGGAGTCGTTGCCAAGGAAAGGAAATCAACCCCACNNNAGATCGGAAG |
| 44392_2 | TATGAGTGTGGAGTCGTTGCCAAGGAAAGGAAATCAACCCCACNNNAGATCGGAAG |
| 29283_1 | TATGAGTGTGGAGTCGTTGCGCTAAAAAGCTTCTGCACAGTGANNNAGATCGGAAG |
| 29283_2 | TATGAGTGTGGAGTCGTTGCGCTAAAAAGCTTCTGCACAGTGANNNAGATCGGAAG |
| 45722_1 | TATGAGTGTGGAGTCGTTGCTAGTTTGACAGGTTCAGGTTGCTNNNAGATCGGAAG |
| 45722_2 | TATGAGTGTGGAGTCGTTGCTAGTTTGACAGGTTCAGGTTGCTNNNAGATCGGAAG |
| 31801_1 | TATGAGTGTGGAGTCGTTGCTGAGAATCACTTGAACCCAGGAGNNNAGATCGGAA |
| 31801_2 | TATGAGTGTGGAGTCGTTGCTGAGAATCACTTGAACCCAGGAGNNNAGATCGGAA |
| 30029_1 | TATGAGTGTGGAGTCGTTGCGATCTGTCCTGTCTGCGGATTAANNNAGATCGGAAG |
| 30029_2 | TATGAGTGTGGAGTCGTTGCGATCTGTCCTGTCTGCGGATTAANNNAGATCGGAAG |
| 13220_1 | TATGAGTGTGGAGTCGTTGCGTATTCTTGGTTTCTGATGAAATNNNAGATCGGAAG |
| 13220_2 | TATGAGTGTGGAGTCGTTGCGTATTCTTGGTTTCTGATGAAATNNNAGATCGGAAG |
| 49101_1 | TATGAGTGTGGAGTCGTTGCCCACTGGCCACAGGGATATAAAANNNAGATCGGAAG |
| 49101_2 | TATGAGTGTGGAGTCGTTGCCCACTGGCCACAGGGATATAAAANNNAGATCGGAAG |
| 32502_1 | TATGAGTGTGGAGTCGTTGCGAGTCTCACTCTGTCACCCAAGNNNAGATCGGAAGA |

| | |
|---|---|
| 32502_2 | TATGAGTGTGGAGTCGTTGCGAGTCTCACTCTGTCACCCAAGNNNAGATCGGAAGA |
| 31198_1 | TATGAGTGTGGAGTCGTTGCGTGTGTATGTGTAACTTCCCCCTNNNAGATCGGAAG |
| 31198_2 | TATGAGTGTGGAGTCGTTGCGTGTGTATGTGTAACTTCCCCCTNNNAGATCGGAAG |
| 41256_1 | TATGAGTGTGGAGTCGTTGCGTGCCTGTAATCCCAGCTACTCNNNAGATCGGAAGA |
| 41256_2 | TATGAGTGTGGAGTCGTTGCGTGCCTGTAATCCCAGCTACTCNNNAGATCGGAAGA |
| 10344_1 | TATGAGTGTGGAGTCGTTGCTGCTGTGAGGAAACTCAACAGTCNNNAGATCGGAA |
| 10344_2 | TATGAGTGTGGAGTCGTTGCTGCTGTGAGGAAACTCAACAGTCNNNAGATCGGAA |
| 42470_1 | TATGAGTGTGGAGTCGTTGCGAGGGAAGGAGAGAGAGAGAGAGNNNAGATCGG |
| 42470_2 | TATGAGTGTGGAGTCGTTGCGAGGGAAGGAGAGAGAGAGAGAGNNNAGATCGG |
| 39322_1 | TATGAGTGTGGAGTCGTTGCCTTTGAAAGGTGGGCCAATTCATNNNAGATCGGAA |
| 39322_2 | TATGAGTGTGGAGTCGTTGCCTTTGAAAGGTGGGCCAATTCATNNNAGATCGGAA |
| 44707_1 | TATGAGTGTGGAGTCGTTGCGCAGATTGCCCTGACTACTAACANNNAGATCGGAAG |
| 44707_2 | TATGAGTGTGGAGTCGTTGCGCAGATTGCCCTGACTACTAACANNNAGATCGGAAG |
| 33747_1 | TATGAGTGTGGAGTCGTTGCCCAGACCTACAGAGTCTCTGAGTNNNAGATCGGAAG |
| 33747_2 | TATGAGTGTGGAGTCGTTGCCCAGACCTACAGAGTCTCTGAGTNNNAGATCGGAAG |
| 40527_1 | TATGAGTGTGGAGTCGTTGCCCCTGCCCCTTTTCTGATTTAGTNNNAGATCGGAAGA |
| 40527_2 | TATGAGTGTGGAGTCGTTGCCCCTGCCCCTTTTCTGATTTAGTNNNAGATCGGAAGA |
| 12941_1 | TATGAGTGTGGAGTCGTTGCGTCTATGAAGTGAAAAACAACANNNAGATCGGAAG |
| 12941_2 | TATGAGTGTGGAGTCGTTGCGTCTATGAAGTGAAAAACAACANNNAGATCGGAAG |
| 41017_1 | TATGAGTGTGGAGTCGTTGCCTCACGAGTAGCTGGGACTACAGNNNAGATCGGAA |
| 41017_2 | TATGAGTGTGGAGTCGTTGCCTCACGAGTAGCTGGGACTACAGNNNAGATCGGAA |
| 38203_1 | TATGAGTGTGGAGTCGTTGCCTTACAGTGAGCTGAGATCGCGNNNAGATCGGAAG |
| 38203_2 | TATGAGTGTGGAGTCGTTGCCTTACAGTGAGCTGAGATCGCGNNNAGATCGGAAG |
| 48271_1 | TATGAGTGTGGAGTCGTTGCAGACAATCTCTCACTCTGTTGCCNNNAGATCGGAAG |
| 48271_2 | TATGAGTGTGGAGTCGTTGCAGACAATCTCTCACTCTGTTGCCNNNAGATCGGAAG |
| 13161_1 | TATGAGTGTGGAGTCGTTGCAAATTTCCTGTATTCTGAACAAGNNNAGATCGGAAG |
| 13161_2 | TATGAGTGTGGAGTCGTTGCAAATTTCCTGTATTCTGAACAAGNNNAGATCGGAAG |
| 5274_1 | TATGAGTGTGGAGTCGTTGCTCTCTGGCCTTTCCATCTCTTCANNNAGATCGGAAGA |
| 5274_2 | TATGAGTGTGGAGTCGTTGCTCTCTGGCCTTTCCATCTCTTCANNNAGATCGGAAGA |
| 43475_1 | TATGAGTGTGGAGTCGTTGCGTCTTGAACTCCTCAGCTCAGACNNNAGATCGGAAG |
| 43475_2 | TATGAGTGTGGAGTCGTTGCGTCTTGAACTCCTCAGCTCAGACNNNAGATCGGAAG |
| 52128_1 | TATGAGTGTGGAGTCGTTGCCTGTCATGCTCACTCTCACTCTTNNNAGATCGGAAGA |
| 52128_2 | TATGAGTGTGGAGTCGTTGCCTGTCATGCTCACTCTCACTCTTNNNAGATCGGAAGA |
| 49700_1 | TATGAGTGTGGAGTCGTTGCTAGAGATGTGCTGGTTTCACAGTNNNAGATCGGAA |
| 49700_2 | TATGAGTGTGGAGTCGTTGCTAGAGATGTGCTGGTTTCACAGTNNNAGATCGGAA |
| 46177_1 | TATGAGTGTGGAGTCGTTGCGCCTGGCCAAATCTAGACATTTCNNNAGATCGGAAG |
| 46177_2 | TATGAGTGTGGAGTCGTTGCGCCTGGCCAAATCTAGACATTTCNNNAGATCGGAAG |
| 47711_1 | TATGAGTGTGGAGTCGTTGCGGTGTCGAAGGCTATTTGTGATGNNNAGATCGGAA |
| 47711_2 | TATGAGTGTGGAGTCGTTGCGGTGTCGAAGGCTATTTGTGATGNNNAGATCGGAA |
| 2400_1 | TATGAGTGTGGAGTCGTTGCTTTAGAGCCGAATACCCACATGCNNNAGATCGGAAG |
| 2400_2 | TATGAGTGTGGAGTCGTTGCTTTAGAGCCGAATACCCACATGCNNNAGATCGGAAG |
| 36232_1 | TATGAGTGTGGAGTCGTTGCTGTAGTCCCAGCTATTCAGGAGANNNAGATCGGAA |
| 36232_2 | TATGAGTGTGGAGTCGTTGCTGTAGTCCCAGCTATTCAGGAGANNNAGATCGGAA |

| | |
|---|---|
| 49366_1 | TATGAGTGTGGAGTCGTTGCGGCTGGAATGCAGTTGTATGATCNNNAGATCGGAA |
| 49366_2 | TATGAGTGTGGAGTCGTTGCGGCTGGAATGCAGTTGTATGATCNNNAGATCGGAA |
| 42710_1 | TATGAGTGTGGAGTCGTTGCTCCTCTTACATAGTCCCCTGAGNNNAGATCGGAAG |
| 42710_2 | TATGAGTGTGGAGTCGTTGCTCCTCTTACATAGTCCCCTGAGNNNAGATCGGAAG |
| 39342_1 | TATGAGTGTGGAGTCGTTGCGGAGATTGTGAAGCTGGGTCTCNNNAGATCGGAAG |
| 39342_2 | TATGAGTGTGGAGTCGTTGCGGAGATTGTGAAGCTGGGTCTCNNNAGATCGGAAG |
| 3174_1 | TATGAGTGTGGAGTCGTTGCTGATGTAAACTGGCTTCCTCGACNNNAGATCGGAAG |
| 3174_2 | TATGAGTGTGGAGTCGTTGCTGATGTAAACTGGCTTCCTCGACNNNAGATCGGAAG |
| 49153_1 | TATGAGTGTGGAGTCGTTGCTGGAGCTCACTGTCTTTTGTCATNNNAGATCGGAAG |
| 49153_2 | TATGAGTGTGGAGTCGTTGCTGGAGCTCACTGTCTTTTGTCATNNNAGATCGGAAG |
| 45674_1 | TATGAGTGTGGAGTCGTTGCGTTCAAGCAATTCTCTTGCCTCANNNAGATCGGAAG |
| 45674_2 | TATGAGTGTGGAGTCGTTGCGTTCAAGCAATTCTCTTGCCTCANNNAGATCGGAAG |
| 45623_1 | TATGAGTGTGGAGTCGTTGCCCAAAGTGCTGTGATTATAGGCGNNNAGATCGGAA |
| 45623_2 | TATGAGTGTGGAGTCGTTGCCCAAAGTGCTGTGATTATAGGCGNNNAGATCGGAA |
| 41798_1 | TATGAGTGTGGAGTCGTTGCCTCCTGTGTAGCTGGGACTACAGNNNAGATCGGAA |
| 41798_2 | TATGAGTGTGGAGTCGTTGCCTCCTGTGTAGCTGGGACTACAGNNNAGATCGGAA |
| 46942_1 | TATGAGTGTGGAGTCGTTGCGAGATGGGGTTTCATCATGTTGGNNNAGATCGGAA |
| 46942_2 | TATGAGTGTGGAGTCGTTGCGAGATGGGGTTTCATCATGTTGGNNNAGATCGGAA |
| 34412_1 | TATGAGTGTGGAGTCGTTGCACCTCTGAAACGTGGGTGAATAANNNAGATCGGAA |
| 34412_2 | TATGAGTGTGGAGTCGTTGCACCTCTGAAACGTGGGTGAATAANNNAGATCGGAA |
| 32885_1 | TATGAGTGTGGAGTCGTTGCCATATGGGGAAGATCCAACAGCTNNNAGATCGGAA |
| 32885_2 | TATGAGTGTGGAGTCGTTGCCATATGGGGAAGATCCAACAGCTNNNAGATCGGAA |
| 38925_1 | TATGAGTGTGGAGTCGTTGCTCACTCTTGGCTATGGGGAAAAGNNNAGATCGGAA |
| 38925_2 | TATGAGTGTGGAGTCGTTGCTCACTCTTGGCTATGGGGAAAAGNNNAGATCGGAA |
| 34579_1 | TATGAGTGTGGAGTCGTTGCCAAGTGTGTCCGAAAGTAACCTGNNNAGATCGGAA |
| 34579_2 | TATGAGTGTGGAGTCGTTGCCAAGTGTGTCCGAAAGTAACCTGNNNAGATCGGAA |
| 47273_1 | TATGAGTGTGGAGTCGTTGCCACCCAGCCGAGTATTCATTTTGNNNAGATCGGAAG |
| 47273_2 | TATGAGTGTGGAGTCGTTGCCACCCAGCCGAGTATTCATTTTGNNNAGATCGGAAG |
| 52831_1 | TATGAGTGTGGAGTCGTTGCAATGAATGTGATTGTTCCCTGCCNNNAGATCGGAAG |
| 52831_2 | TATGAGTGTGGAGTCGTTGCAATGAATGTGATTGTTCCCTGCCNNNAGATCGGAAG |
| 38051_1 | TATGAGTGTGGAGTCGTTGCCAGGATCCAGGTATAAGCTCCACNNNAGATCGGAAG |
| 38051_2 | TATGAGTGTGGAGTCGTTGCCAGGATCCAGGTATAAGCTCCACNNNAGATCGGAAG |
| 2209_1 | TATGAGTGTGGAGTCGTTGCAAGTTCCCTTGTTAGCCGAATGANNNAGATCGGAAG |
| 2209_2 | TATGAGTGTGGAGTCGTTGCAAGTTCCCTTGTTAGCCGAATGANNNAGATCGGAAG |
| 36457_1 | TATGAGTGTGGAGTCGTTGCAGAATTGCTTGAACCTGGGAGAANNNAGATCGGAA |
| 36457_2 | TATGAGTGTGGAGTCGTTGCAGAATTGCTTGAACCTGGGAGAANNNAGATCGGAA |
| 36893_1 | TATGAGTGTGGAGTCGTTGCCACACCTGTAATCCCAGCACTTTNNNAGATCGGAAG |
| 36893_2 | TATGAGTGTGGAGTCGTTGCCACACCTGTAATCCCAGCACTTTNNNAGATCGGAAG |
| 13657_1 | TATGAGTGTGGAGTCGTTGCTAAAAATTATGTTAGACCTGGTCNNNAGATCGGAAG |
| 13657_2 | TATGAGTGTGGAGTCGTTGCTAAAAATTATGTTAGACCTGGTCNNNAGATCGGAAG |
| 40721_1 | TATGAGTGTGGAGTCGTTGCCAGGAGAATGGCATGAACCTTGNNNAGATCGGAAG |
| 40721_2 | TATGAGTGTGGAGTCGTTGCCAGGAGAATGGCATGAACCTTGNNNAGATCGGAAG |
| 36265_1 | TATGAGTGTGGAGTCGTTGCTCTCTTTCCTCATCCTCTCCTTTNNNAGATCGGAAGAG |

| | |
|---|---|
| 36265_2 | TATGAGTGTGGAGTCGTTGCTCTCTTTCCTCATCCTCTCCTTTNNNAGATCGGAAGAG |
| 36916_1 | TATGAGTGTGGAGTCGTTGCTGCAGGAATCATTTTCAACTGCTNNNAGATCGGAAG |
| 36916_2 | TATGAGTGTGGAGTCGTTGCTGCAGGAATCATTTTCAACTGCTNNNAGATCGGAAG |
| 35789_1 | TATGAGTGTGGAGTCGTTGCATTGTCTGCTTCCTTTCTCAGCTNNNAGATCGGAAGA |
| 35789_2 | TATGAGTGTGGAGTCGTTGCATTGTCTGCTTCCTTTCTCAGCTNNNAGATCGGAAGA |
| 32728_1 | TATGAGTGTGGAGTCGTTGCCTCAAGTGATTCTCCCACCATGANNNAGATCGGAAG |
| 32728_2 | TATGAGTGTGGAGTCGTTGCCTCAAGTGATTCTCCCACCATGANNNAGATCGGAAG |
| 38394_1 | TATGAGTGTGGAGTCGTTGCGCAAACCAAAAGAAAATGGAGGGNNNAGATCGGAA |
| 38394_2 | TATGAGTGTGGAGTCGTTGCGCAAACCAAAAGAAAATGGAGGGNNNAGATCGGAA |
| 32401_1 | TATGAGTGTGGAGTCGTTGCAGGGGATGACTGCTGGGTATATANNNAGATCGGAA |
| 32401_2 | TATGAGTGTGGAGTCGTTGCAGGGGATGACTGCTGGGTATATANNNAGATCGGAA |
| 35276_1 | TATGAGTGTGGAGTCGTTGCAGATTTGTAGAGTTCACCCGTCCNNNAGATCGGAAG |
| 35276_2 | TATGAGTGTGGAGTCGTTGCAGATTTGTAGAGTTCACCCGTCCNNNAGATCGGAAG |
| 34995_1 | TATGAGTGTGGAGTCGTTGCCCACGCCCAGCCAATTTTTATATNNNAGATCGGAAGA |
| 34995_2 | TATGAGTGTGGAGTCGTTGCCCACGCCCAGCCAATTTTTATATNNNAGATCGGAAGA |
| 38943_1 | TATGAGTGTGGAGTCGTTGCAAACTTCAGCCTCCCAGTTGTATNNNAGATCGGAAG |
| 38943_2 | TATGAGTGTGGAGTCGTTGCAAACTTCAGCCTCCCAGTTGTATNNNAGATCGGAAG |
| 35384_1 | TATGAGTGTGGAGTCGTTGCAGACAGAGTCTCGCTTTGTCATCNNNAGATCGGAAG |
| 35384_2 | TATGAGTGTGGAGTCGTTGCAGACAGAGTCTCGCTTTGTCATCNNNAGATCGGAAG |
| 12998_1 | TATGAGTGTGGAGTCGTTGCAATCTTTAACTGCCATGTAAACNNNAGATCGGAAGA |
| 12998_2 | TATGAGTGTGGAGTCGTTGCAATCTTTAACTGCCATGTAAACNNNAGATCGGAAGA |
| 41820_1 | TATGAGTGTGGAGTCGTTGCTTCAAACCATTCTCCTGCCTCAGNNNAGATCGGAAGA |
| 41820_2 | TATGAGTGTGGAGTCGTTGCTTCAAACCATTCTCCTGCCTCAGNNNAGATCGGAAGA |
| 52247_1 | TATGAGTGTGGAGTCGTTGCATAAACAGGCCCAGAGAACTCAGNNNAGATCGGAA |
| 52247_2 | TATGAGTGTGGAGTCGTTGCATAAACAGGCCCAGAGAACTCAGNNNAGATCGGAA |
| 42702_1 | TATGAGTGTGGAGTCGTTGCAGGAGGTGATTGGATCATGAAGGNNNAGATCGGA |
| 42702_2 | TATGAGTGTGGAGTCGTTGCAGGAGGTGATTGGATCATGAAGGNNNAGATCGGA |
| 33089_1 | TATGAGTGTGGAGTCGTTGCGGGTTCAAGTAATTCTCCTGCCTNNNAGATCGGAAG |
| 33089_2 | TATGAGTGTGGAGTCGTTGCGGGTTCAAGTAATTCTCCTGCCTNNNAGATCGGAAG |
| 33161_1 | TATGAGTGTGGAGTCGTTGCCACAGTGGCTCACTCCTGTAATCNNNAGATCGGAAG |
| 33161_2 | TATGAGTGTGGAGTCGTTGCCACAGTGGCTCACTCCTGTAATCNNNAGATCGGAAG |
| 2583_1 | TATGAGTGTGGAGTCGTTGCTAAAAGGCTTACTCCAGTGCTGGNNNAGATCGGAAG |
| 2583_2 | TATGAGTGTGGAGTCGTTGCTAAAAGGCTTACTCCAGTGCTGGNNNAGATCGGAAG |
| 49398_1 | TATGAGTGTGGAGTCGTTGCCCACCAAGACTGGCTAATTTTTGTNNNAGATCGGAA |
| 49398_2 | TATGAGTGTGGAGTCGTTGCCCACCAAGACTGGCTAATTTTTGTNNNAGATCGGAA |
| 38448_1 | TATGAGTGTGGAGTCGTTGCAGTCTTTTAGTGAGTTGCTGACANNNAGATCGGAAG |
| 38448_2 | TATGAGTGTGGAGTCGTTGCAGTCTTTTAGTGAGTTGCTGACANNNAGATCGGAAG |
| 53072_1 | TATGAGTGTGGAGTCGTTGCGATCATTTGAGGCCAGGAGTTTGNNNAGATCGGAA |
| 53072_2 | TATGAGTGTGGAGTCGTTGCGATCATTTGAGGCCAGGAGTTTGNNNAGATCGGAA |
| 43946_1 | TATGAGTGTGGAGTCGTTGCTTCCTTCCTTCCTTCCTTCCTTCNNNAGATCGGAAGAG |
| 43946_2 | TATGAGTGTGGAGTCGTTGCTTCCTTCCTTCCTTCCTTCCTTCNNNAGATCGGAAGAG |
| 12527_1 | TATGAGTGTGGAGTCGTTGCAAACATATCTACTTCCACGTAAGNNNAGATCGGAAG |
| 12527_2 | TATGAGTGTGGAGTCGTTGCAAACATATCTACTTCCACGTAAGNNNAGATCGGAAG |

| | |
|---|---|
| 2411_1 | TATGAGTGTGGAGTCGTTGCTTCCTAGGCCAATTGACAAGTGCNNNAGATCGGAAG |
| 2411_2 | TATGAGTGTGGAGTCGTTGCTTCCTAGGCCAATTGACAAGTGCNNNAGATCGGAAG |
| 40322_1 | TATGAGTGTGGAGTCGTTGCATTCACTCAGAAAAACGCTGACCNNNAGATCGGAAG |
| 40322_2 | TATGAGTGTGGAGTCGTTGCATTCACTCAGAAAAACGCTGACCNNNAGATCGGAAG |
| 42873_1 | TATGAGTGTGGAGTCGTTGCTGTCACATTTTCTTTATCCACTCGTNNNAGATCGGAA |
| 42873_2 | TATGAGTGTGGAGTCGTTGCTGTCACATTTTCTTTATCCACTCGTNNNAGATCGGAA |
| 30592_1 | TATGAGTGTGGAGTCGTTGCCCGAGAAGCAGAGGCACTTTNNNAGATCGGAAGAG |
| 30592_2 | TATGAGTGTGGAGTCGTTGCCCGAGAAGCAGAGGCACTTTNNNAGATCGGAAGAG |
| 41004_1 | TATGAGTGTGGAGTCGTTGCAGACAGTCTCTCACTTTGTCACCNNNAGATCGGAAG |
| 41004_2 | TATGAGTGTGGAGTCGTTGCAGACAGTCTCTCACTTTGTCACCNNNAGATCGGAAG |
| 48331_1 | TATGAGTGTGGAGTCGTTGCGTGCAACAAAAGCTCCCCATAAANNNAGATCGGAAG |
| 48331_2 | TATGAGTGTGGAGTCGTTGCGTGCAACAAAAGCTCCCCATAAANNNAGATCGGAAG |
| 2615_1 | TATGAGTGTGGAGTCGTTGCCCATCTTTCCTCCAATCAGGCTGNNNAGATCGGAAG |
| 2615_2 | TATGAGTGTGGAGTCGTTGCCCATCTTTCCTCCAATCAGGCTGNNNAGATCGGAAG |
| 38340_1 | TATGAGTGTGGAGTCGTTGCACTTCCTTAGGCTCTAGGGCTTANNNAGATCGGAAG |
| 38340_2 | TATGAGTGTGGAGTCGTTGCACTTCCTTAGGCTCTAGGGCTTANNNAGATCGGAAG |
| 51347_1 | TATGAGTGTGGAGTCGTTGCATTGGCTCACACAATCACAAAGGNNNAGATCGGAAG |
| 51347_2 | TATGAGTGTGGAGTCGTTGCATTGGCTCACACAATCACAAAGGNNNAGATCGGAAG |
| 42073_1 | TATGAGTGTGGAGTCGTTGCACGTCTGTAATCCCAGCACTTTNNNAGATCGGAAGA |
| 42073_2 | TATGAGTGTGGAGTCGTTGCACGTCTGTAATCCCAGCACTTTNNNAGATCGGAAGA |
| 30544_1 | TATGAGTGTGGAGTCGTTGCCAGCCATGTGACTCTGCCTTTTTNNNAGATCGGAAG |
| 30544_2 | TATGAGTGTGGAGTCGTTGCCAGCCATGTGACTCTGCCTTTTTNNNAGATCGGAAG |
| 48043_1 | TATGAGTGTGGAGTCGTTGCTATCTCAGCCTTCCAAGTACCTGNNNAGATCGGAAG |
| 48043_2 | TATGAGTGTGGAGTCGTTGCTATCTCAGCCTTCCAAGTACCTGNNNAGATCGGAAG |
| 37992_1 | TATGAGTGTGGAGTCGTTGCACAGAGCCAGTAATAACCCCTTGNNNAGATCGGAAG |
| 37992_2 | TATGAGTGTGGAGTCGTTGCACAGAGCCAGTAATAACCCCTTGNNNAGATCGGAAG |
| 52206_1 | TATGAGTGTGGAGTCGTTGCGATGCTCCTCACTTCCCAGACNNNAGATCGGAAGAG |
| 52206_2 | TATGAGTGTGGAGTCGTTGCGATGCTCCTCACTTCCCAGACNNNAGATCGGAAGAG |
| 35587_1 | TATGAGTGTGGAGTCGTTGCTACAGAACACAAGAGTCATGGGGNNNAGATCGGAA |
| 35587_2 | TATGAGTGTGGAGTCGTTGCTACAGAACACAAGAGTCATGGGGNNNAGATCGGAA |
| 45895_1 | TATGAGTGTGGAGTCGTTGCCCACACCCGGCTAATTTTTTGTANNNAGATCGGAAG |
| 45895_2 | TATGAGTGTGGAGTCGTTGCCCACACCCGGCTAATTTTTTGTANNNAGATCGGAAG |
| 32279_1 | TATGAGTGTGGAGTCGTTGCACCCCTATGTCCCTAGGTGAATTNNNAGATCGGAAG |
| 32279_2 | TATGAGTGTGGAGTCGTTGCACCCCTATGTCCCTAGGTGAATTNNNAGATCGGAAG |
| 35035_1 | TATGAGTGTGGAGTCGTTGCTGTTAGCCAGGATGGTCTCAATCNNNAGATCGGAAG |
| 35035_2 | TATGAGTGTGGAGTCGTTGCTGTTAGCCAGGATGGTCTCAATCNNNAGATCGGAAG |
| 48655_1 | TATGAGTGTGGAGTCGTTGCTTCACATCATTCTCCTGCCTCTGNNNAGATCGGAAGA |
| 48655_2 | TATGAGTGTGGAGTCGTTGCTTCACATCATTCTCCTGCCTCTGNNNAGATCGGAAGA |
| 43043_1 | TATGAGTGTGGAGTCGTTGCTGCAATAATGGCATTCACAGCAANNNAGATCGGAAG |
| 43043_2 | TATGAGTGTGGAGTCGTTGCTGCAATAATGGCATTCACAGCAANNNAGATCGGAAG |
| 32015_1 | TATGAGTGTGGAGTCGTTGCCTCTCGGGTTCAAGCAATTCTGNNNAGATCGGAAGA |
| 32015_2 | TATGAGTGTGGAGTCGTTGCCTCTCGGGTTCAAGCAATTCTGNNNAGATCGGAAGA |
| 31965_1 | TATGAGTGTGGAGTCGTTGCGTATATGGCTCCACACAAATGGCNNNAGATCGGAAG |

| | |
|---|---|
| 31965_2 | TATGAGTGTGGAGTCGTTGCGTATATGGCTCCACACAAATGGCNNNAGATCGGAAG |
| 40419_1 | TATGAGTGTGGAGTCGTTGCTTGGTATATTCAGTGGTGTGCCANNNAGATCGGAA |
| 40419_2 | TATGAGTGTGGAGTCGTTGCTTGGTATATTCAGTGGTGTGCCANNNAGATCGGAA |
| 51828_1 | TATGAGTGTGGAGTCGTTGCTTGGAGATGAGATTTGGAGCCAANNNAGATCGGAA |
| 51828_2 | TATGAGTGTGGAGTCGTTGCTTGGAGATGAGATTTGGAGCCAANNNAGATCGGAA |
| 50606_1 | TATGAGTGTGGAGTCGTTGCCACACATACACACACACACAGACNNNAGATCGGAAGA |
| 50606_2 | TATGAGTGTGGAGTCGTTGCCACACATACACACACACACAGACNNNAGATCGGAAGA |
| 29970_1 | TATGAGTGTGGAGTCGTTGCACACAGAGAACCAGTTTCCAGTTNNNAGATCGGAAG |
| 29970_2 | TATGAGTGTGGAGTCGTTGCACACAGAGAACCAGTTTCCAGTTNNNAGATCGGAAG |
| 50505_1 | TATGAGTGTGGAGTCGTTGCAAGCAGGAGGAGGAGGAGATAAANNNAGATCGGA |
| 50505_2 | TATGAGTGTGGAGTCGTTGCAAGCAGGAGGAGGAGGAGATAAANNNAGATCGGA |
| 48407_1 | TATGAGTGTGGAGTCGTTGCCCGTGCCCAGCTAATTTTTGTATNNNAGATCGGAAGA |
| 48407_2 | TATGAGTGTGGAGTCGTTGCCCGTGCCCAGCTAATTTTTGTATNNNAGATCGGAAGA |
| 12523_1 | TATGAGTGTGGAGTCGTTGCACTAATCTGTCTTACTGTTCAATNNNAGATCGGAAGA |
| 12523_2 | TATGAGTGTGGAGTCGTTGCACTAATCTGTCTTACTGTTCAATNNNAGATCGGAAGA |
| 36144_1 | TATGAGTGTGGAGTCGTTGCCAAGGTGAATGGTTTTTGAGGGTNNNAGATCGGAA |
| 36144_2 | TATGAGTGTGGAGTCGTTGCCAAGGTGAATGGTTTTTGAGGGTNNNAGATCGGAA |
| 30621_1 | ATGAGTGTGGAGTCGTTGCTGGGTATACAAGTATCTCTTCAAGGCTNNNAGATCGG |
| 30621_2 | ATGAGTGTGGAGTCGTTGCTGGGTATACAAGTATCTCTTCAAGGCTNNNAGATCGG |
| 31970_1 | TATGAGTGTGGAGTCGTTGCGCTGTCTGTATCAGATTGGCCTTNNNAGATCGGAAG |
| 31970_2 | TATGAGTGTGGAGTCGTTGCGCTGTCTGTATCAGATTGGCCTTNNNAGATCGGAAG |
| 51509_1 | TATGAGTGTGGAGTCGTTGCGATATGGTGTTTTGCCATGTTGCNNNAGATCGGAAG |
| 51509_2 | TATGAGTGTGGAGTCGTTGCGATATGGTGTTTTGCCATGTTGCNNNAGATCGGAAG |
| 35444_1 | TATGAGTGTGGAGTCGTTGCAGAGTGCAATGTCAGGATCTCAGNNNAGATCGGAA |
| 35444_2 | TATGAGTGTGGAGTCGTTGCAGAGTGCAATGTCAGGATCTCAGNNNAGATCGGAA |
| 43785_1 | TATGAGTGTGGAGTCGTTGCACCCAGCCATCAAAACATGAAAGNNNAGATCGGAAG |
| 43785_2 | TATGAGTGTGGAGTCGTTGCACCCAGCCATCAAAACATGAAAGNNNAGATCGGAAG |
| 2083_1 | TATGAGTGTGGAGTCGTTGCTTCCAAGACCCTCAGTTGGTACANNNAGATCGGAAG |
| 2083_2 | TATGAGTGTGGAGTCGTTGCTTCCAAGACCCTCAGTTGGTACANNNAGATCGGAAG |
| 42298_1 | TATGAGTGTGGAGTCGTTGCCTCAAAATAGCATTTCCTGGGCANNNAGATCGGAAG |
| 42298_2 | TATGAGTGTGGAGTCGTTGCCTCAAAATAGCATTTCCTGGGCANNNAGATCGGAAG |
| 36341_1 | TATGAGTGTGGAGTCGTTGCTGGAAGAGTCATTGTGAGACAGGNNNAGATCGGAA |
| 36341_2 | TATGAGTGTGGAGTCGTTGCTGGAAGAGTCATTGTGAGACAGGNNNAGATCGGAA |
| 44212_1 | TATGAGTGTGGAGTCGTTGCAGAGGAAAACAGGAGGACTCAACNNNAGATCGGAA |
| 44212_2 | TATGAGTGTGGAGTCGTTGCAGAGGAAAACAGGAGGACTCAACNNNAGATCGGAA |
| 5938_1 | TATGAGTGTGGAGTCGTTGCATTCCTCCACATCGTTCTGACAGNNNAGATCGGAAGA |
| 5938_2 | TATGAGTGTGGAGTCGTTGCATTCCTCCACATCGTTCTGACAGNNNAGATCGGAAGA |
| 10122_1 | TATGAGTGTGGAGTCGTTGCACAGAAACATGCCAGCTACAGAGNNNAGATCGGAA |
| 10122_2 | TATGAGTGTGGAGTCGTTGCACAGAAACATGCCAGCTACAGAGNNNAGATCGGAA |
| 44142_1 | TATGAGTGTGGAGTCGTTGCGAATTGCTCCCACTCACACATTCNNNAGATCGGAAGA |
| 44142_2 | TATGAGTGTGGAGTCGTTGCGAATTGCTCCCACTCACACATTCNNNAGATCGGAAGA |
| 47258_1 | TATGAGTGTGGAGTCGTTGCAGATGGAGTCTCTCTCTGTCACCNNNAGATCGGAAG |
| 47258_2 | TATGAGTGTGGAGTCGTTGCAGATGGAGTCTCTCTCTGTCACCNNNAGATCGGAAG |

| | |
|---|---|
| 47743_1 | TATGAGTGTGGAGTCGTTGCCTCAAGCAATCCTCTCACCTCAGNNNAGATCGGAAG |
| 47743_2 | TATGAGTGTGGAGTCGTTGCCTCAAGCAATCCTCTCACCTCAGNNNAGATCGGAAG |
| 38926_1 | TATGAGTGTGGAGTCGTTGCAGTATGTTCAGATCGTGGTACCANNNAGATCGGAA |
| 38926_2 | TATGAGTGTGGAGTCGTTGCAGTATGTTCAGATCGTGGTACCANNNAGATCGGAA |
| 36431_1 | TATGAGTGTGGAGTCGTTGCGGTTCACAGGCCAGTTTTTGTANNNAGATCGGAAG |
| 36431_2 | TATGAGTGTGGAGTCGTTGCGGTTCACAGGCCAGTTTTTGTANNNAGATCGGAAG |
| 10002_1 | TATGAGTGTGGAGTCGTTGCTTTGATGTTGAGGTGACCAAGCANNNAGATCGGAA |
| 10002_2 | TATGAGTGTGGAGTCGTTGCTTTGATGTTGAGGTGACCAAGCANNNAGATCGGAA |
| 29901_1 | TATGAGTGTGGAGTCGTTGCGTTGATTTAAACAGGCAGCCACTNNNAGATCGGAAG |
| 29901_2 | TATGAGTGTGGAGTCGTTGCGTTGATTTAAACAGGCAGCCACTNNNAGATCGGAAG |
| 13699_1 | TATGAGTGTGGAGTCGTTGCCAGTTCTTTCTTTTTCTCACTATNNNAGATCGGAAGA |
| 13699_2 | TATGAGTGTGGAGTCGTTGCCAGTTCTTTCTTTTTCTCACTATNNNAGATCGGAAGA |
| 49531_1 | TATGAGTGTGGAGTCGTTGCTTCTGCAGAAGTCATTTGGTGTGNNNAGATCGGAA |
| 49531_2 | TATGAGTGTGGAGTCGTTGCTTCTGCAGAAGTCATTTGGTGTGNNNAGATCGGAA |
| 44841_1 | TATGAGTGTGGAGTCGTTGCCGATGGAAGGGAACAGATGATGANNNAGATCGGA |
| 44841_2 | TATGAGTGTGGAGTCGTTGCCGATGGAAGGGAACAGATGATGANNNAGATCGGA |
| 53413_1 | TATGAGTGTGGAGTCGTTGCCAGCACTAGGGGAATGATGCTAANNNAGATCGGAA |
| 53413_2 | TATGAGTGTGGAGTCGTTGCCAGCACTAGGGGAATGATGCTAANNNAGATCGGAA |
| 8026_1 | TATGAGTGTGGAGTCGTTGCTATCTTCAAAGCCAGCAATCTGANNNAGATCGGAAG |
| 8026_2 | TATGAGTGTGGAGTCGTTGCTATCTTCAAAGCCAGCAATCTGANNNAGATCGGAAG |
| 47012_1 | TATGAGTGTGGAGTCGTTGCCCATCTCCTGACCTCATGATCTGNNNAGATCGGAAG |
| 47012_2 | TATGAGTGTGGAGTCGTTGCCCATCTCCTGACCTCATGATCTGNNNAGATCGGAAG |
| 36127_1 | TATGAGTGTGGAGTCGTTGCAGAGAATCGCTTGAACTCAGGAGNNNAGATCGGAA |
| 36127_2 | TATGAGTGTGGAGTCGTTGCAGAGAATCGCTTGAACTCAGGAGNNNAGATCGGAA |
| 30796_1 | TATGAGTGTGGAGTCGTTGCGGAACTAGAGAGAAGGGCAAAGTNNNAGATCGGA |
| 30796_2 | TATGAGTGTGGAGTCGTTGCGGAACTAGAGAGAAGGGCAAAGTNNNAGATCGGA |
| 35261_1 | TATGAGTGTGGAGTCGTTGCAGTGAGATCTTGGCTTACTGCAANNNAGATCGGAA |
| 35261_2 | TATGAGTGTGGAGTCGTTGCAGTGAGATCTTGGCTTACTGCAANNNAGATCGGAA |
| 7143_1 | TATGAGTGTGGAGTCGTTGCGAGAGAGAGACAGAAACAGTCTGTNNNAGATCGG |
| 7143_2 | TATGAGTGTGGAGTCGTTGCGAGAGAGAGACAGAAACAGTCTGTNNNAGATCGG |
| 33991_1 | TATGAGTGTGGAGTCGTTGCAGAAAAGCCATTGCCCTTCTTTCNNNAGATCGGAAG |
| 33991_2 | TATGAGTGTGGAGTCGTTGCAGAAAAGCCATTGCCCTTCTTTCNNNAGATCGGAAG |
| 34526_1 | TATGAGTGTGGAGTCGTTGCTGGTGGAGAGCTTTCTTCTTTCANNNAGATCGGAAG |
| 34526_2 | TATGAGTGTGGAGTCGTTGCTGGTGGAGAGCTTTCTTCTTTCANNNAGATCGGAAG |
| 42043_1 | TATGAGTGTGGAGTCGTTGCTGGAAGGTGGAAGCTAAAATCCANNNAGATCGGAA |
| 42043_2 | TATGAGTGTGGAGTCGTTGCTGGAAGGTGGAAGCTAAAATCCANNNAGATCGGAA |
| 31908_1 | TATGAGTGTGGAGTCGTTGCGAAAGAGACAGGGAAAGGGAGAANNNAGATCGG |
| 31908_2 | TATGAGTGTGGAGTCGTTGCGAAAGAGACAGGGAAAGGGAGAANNNAGATCGG |
| 47555_1 | TATGAGTGTGGAGTCGTTGCAGACACACTTTCACTCTTGTTGCNNNAGATCGGAAG |
| 47555_2 | TATGAGTGTGGAGTCGTTGCAGACACACTTTCACTCTTGTTGCNNNAGATCGGAAG |
| 51077_1 | TATGAGTGTGGAGTCGTTGCACACATAGCCTGCTGGAAATCATNNNAGATCGGAAG |
| 51077_2 | TATGAGTGTGGAGTCGTTGCACACATAGCCTGCTGGAAATCATNNNAGATCGGAAG |
| 47123_1 | TATGAGTGTGGAGTCGTTGCTGAGTAGCTGAGATTACAGGTGCNNNAGATCGGAA |

| | |
|---|---|
| 47123_2 | TATGAGTGTGGAGTCGTTGCTGAGTAGCTGAGATTACAGGTGCNNNAGATCGGAA |
| 50711_1 | TATGAGTGTGGAGTCGTTGCCCTCTAGCCTGACTCAACAGTTTNNNAGATCGGAAGA |
| 50711_2 | TATGAGTGTGGAGTCGTTGCCCTCTAGCCTGACTCAACAGTTTNNNAGATCGGAAGA |
| 45205_1 | TATGAGTGTGGAGTCGTTGCCCTCCCTTGAATCCAGACATTCANNNAGATCGGAAGA |
| 45205_2 | TATGAGTGTGGAGTCGTTGCCCTCCCTTGAATCCAGACATTCANNNAGATCGGAAGA |
| 49447_1 | TATGAGTGTGGAGTCGTTGCCAACAAGCCTATCAAGAGCACACNNNAGATCGGAAG |
| 49447_2 | TATGAGTGTGGAGTCGTTGCCAACAAGCCTATCAAGAGCACACNNNAGATCGGAAG |
| 48323_1 | TATGAGTGTGGAGTCGTTGCCCTTCCTGAGTAGCTGGGATTACNNNAGATCGGAAG |
| 48323_2 | TATGAGTGTGGAGTCGTTGCCCTTCCTGAGTAGCTGGGATTACNNNAGATCGGAAG |
| 32156_1 | TATGAGTGTGGAGTCGTTGCAAAAAACAACAGTTCTCTGCCCCNNNAGATCGGAAG |
| 32156_2 | TATGAGTGTGGAGTCGTTGCAAAAAACAACAGTTCTCTGCCCCNNNAGATCGGAAG |
| 34418_1 | TATGAGTGTGGAGTCGTTGCCTCTGGGATCCGTCATCGAAGNNNAGATCGGAAGA |
| 34418_2 | TATGAGTGTGGAGTCGTTGCCTCTGGGATCCGTCATCGAAGNNNAGATCGGAAGA |
| 28872_1 | TATGAGTGTGGAGTCGTTGCAGGGCTCTTCACTAATGTTGAGGNNNAGATCGGAA |
| 28872_2 | TATGAGTGTGGAGTCGTTGCAGGGCTCTTCACTAATGTTGAGGNNNAGATCGGAA |
| 48683_1 | TATGAGTGTGGAGTCGTTGCACATTTGCAGGGAATATCTGGGTNNNAGATCGGAA |
| 48683_2 | TATGAGTGTGGAGTCGTTGCACATTTGCAGGGAATATCTGGGTNNNAGATCGGAA |
| 9380_1 | TATGAGTGTGGAGTCGTTGCGAGACCCCAGATAAATGTGCAGGNNNAGATCGGAA |
| 9380_2 | TATGAGTGTGGAGTCGTTGCGAGACCCCAGATAAATGTGCAGGNNNAGATCGGAA |
| 29419_1 | TATGAGTGTGGAGTCGTTGCACTGACTGGGTGATCTGACCTAGNNNAGATCGGAA |
| 29419_2 | TATGAGTGTGGAGTCGTTGCACTGACTGGGTGATCTGACCTAGNNNAGATCGGAA |
| 29171_1 | TATGAGTGTGGAGTCGTTGCAAAATCATGCTGCTGTTACTGCCNNNAGATCGGAAG |
| 29171_2 | TATGAGTGTGGAGTCGTTGCAAAATCATGCTGCTGTTACTGCCNNNAGATCGGAAG |
| 37955_1 | TATGAGTGTGGAGTCGTTGCCCTGTAATCTCAGCTACTCAGGGNNNAGATCGGAAG |
| 37955_2 | TATGAGTGTGGAGTCGTTGCCCTGTAATCTCAGCTACTCAGGGNNNAGATCGGAAG |
| 51796_1 | TATGAGTGTGGAGTCGTTGCCCGGGCCTATAATCTCATTAGCANNNAGATCGGAAG |
| 51796_2 | TATGAGTGTGGAGTCGTTGCCCGGGCCTATAATCTCATTAGCANNNAGATCGGAAG |
| 43588_1 | TATGAGTGTGGAGTCGTTGCACAAGCTGCTCCTAGGAGAAAAANNNAGATCGGAA |
| 43588_2 | TATGAGTGTGGAGTCGTTGCACAAGCTGCTCCTAGGAGAAAAANNNAGATCGGAA |
| 49209_1 | TATGAGTGTGGAGTCGTTGCAAGTGTTGGGATTACAGGTGTGANNNAGATCGGAA |
| 49209_2 | TATGAGTGTGGAGTCGTTGCAAGTGTTGGGATTACAGGTGTGANNNAGATCGGAA |
| 44409_1 | TATGAGTGTGGAGTCGTTGCAAGCAAAGGGTATTGAGGGATGTNNNAGATCGGA |
| 44409_2 | TATGAGTGTGGAGTCGTTGCAAGCAAAGGGTATTGAGGGATGTNNNAGATCGGA |
| 42261_1 | TATGAGTGTGGAGTCGTTGCGCATCCTCATCACTATTTTGGGTGNNNAGATCGGAA |
| 42261_2 | TATGAGTGTGGAGTCGTTGCGCATCCTCATCACTATTTTGGGTGNNNAGATCGGAA |
| 43445_1 | TATGAGTGTGGAGTCGTTGCGTAGTTGGGATTACAGGTGTCCANNNAGATCGGAA |
| 43445_2 | TATGAGTGTGGAGTCGTTGCGTAGTTGGGATTACAGGTGTCCANNNAGATCGGAA |
| 2072_1 | TATGAGTGTGGAGTCGTTGCCGAATGCCAGAAGGTAAAGGACANNNAGATCGGAA |
| 2072_2 | TATGAGTGTGGAGTCGTTGCCGAATGCCAGAAGGTAAAGGACANNNAGATCGGAA |
| 45478_1 | TATGAGTGTGGAGTCGTTGCATTGTTAGCTCTGTGCCTAACCANNNAGATCGGAAG |
| 45478_2 | TATGAGTGTGGAGTCGTTGCATTGTTAGCTCTGTGCCTAACCANNNAGATCGGAAG |
| 13141_1 | TATGAGTGTGGAGTCGTTGCAATGACAATATTAGGACTTCTCANNNAGATCGGAAG |
| 13141_2 | TATGAGTGTGGAGTCGTTGCAATGACAATATTAGGACTTCTCANNNAGATCGGAAG |

| | |
|---|---|
| 44939_1 | TATGAGTGTGGAGTCGTTGCTATGAGCACAATTGGCCTCCTTANNNAGATCGGAAG |
| 44939_2 | TATGAGTGTGGAGTCGTTGCTATGAGCACAATTGGCCTCCTTANNNAGATCGGAAG |
| 33491_1 | TATGAGTGTGGAGTCGTTGCGGCGAAAATACACTGTGGCATAANNNAGATCGGAA |
| 33491_2 | TATGAGTGTGGAGTCGTTGCGGCGAAAATACACTGTGGCATAANNNAGATCGGAA |
| 38877_1 | TATGAGTGTGGAGTCGTTGCTCCAACAATATGACACTCTGGAGANNNAGATCGGAA |
| 38877_2 | TATGAGTGTGGAGTCGTTGCTCCAACAATATGACACTCTGGAGANNNAGATCGGAA |
| 38667_1 | TATGAGTGTGGAGTCGTTGCCAGTGAACCATGATCACACCTCTNNNAGATCGGAAG |
| 38667_2 | TATGAGTGTGGAGTCGTTGCCAGTGAACCATGATCACACCTCTNNNAGATCGGAAG |
| 13007_1 | TATGAGTGTGGAGTCGTTGCATTCCTGTATTTCTTTAGTTGTCNNNAGATCGGAAGA |
| 13007_2 | TATGAGTGTGGAGTCGTTGCATTCCTGTATTTCTTTAGTTGTCNNNAGATCGGAAGA |
| 34188_1 | TATGAGTGTGGAGTCGTTGCTAATAGGGCAGGTGGTAGAGACTNNNAGATCGGA |
| 34188_2 | TATGAGTGTGGAGTCGTTGCTAATAGGGCAGGTGGTAGAGACTNNNAGATCGGA |
| 44681_1 | TATGAGTGTGGAGTCGTTGCAAGACACTAGGAGATGAGAGGGTNNNAGATCGGA |
| 44681_2 | TATGAGTGTGGAGTCGTTGCAAGACACTAGGAGATGAGAGGGTNNNAGATCGGA |
| 5800_1 | TATGAGTGTGGAGTCGTTGCTACAGGTGTGAGACACTGCGAGCNNNAGATCGGAA |
| 5800_2 | TATGAGTGTGGAGTCGTTGCTACAGGTGTGAGACACTGCGAGCNNNAGATCGGAA |
| 40336_1 | TATGAGTGTGGAGTCGTTGCCTCAGGGGAGCAATAAAAAAGCANNNAGATCGGAA |
| 40336_2 | TATGAGTGTGGAGTCGTTGCCTCAGGGGAGCAATAAAAAAGCANNNAGATCGGAA |
| 13016_1 | TATGAGTGTGGAGTCGTTGCAACTCCTCATATTCAAATTTGAGNNNAGATCGGAAG |
| 13016_2 | TATGAGTGTGGAGTCGTTGCAACTCCTCATATTCAAATTTGAGNNNAGATCGGAAG |
| 2881_1 | TATGAGTGTGGAGTCGTTGCATGAAACACACACAGCCAGATGTNNNAGATCGGAA |
| 2881_2 | TATGAGTGTGGAGTCGTTGCATGAAACACACACAGCCAGATGTNNNAGATCGGAA |
| 32809_1 | TATGAGTGTGGAGTCGTTGCCAAGTAGCTGGGACTACAGGCNNNAGATCGGAAGA |
| 32809_2 | TATGAGTGTGGAGTCGTTGCCAAGTAGCTGGGACTACAGGCNNNAGATCGGAAGA |
| 33398_1 | TATGAGTGTGGAGTCGTTGCCTCTGAGAGGCACCATCTGAAGNNNAGATCGGAAG |
| 33398_2 | TATGAGTGTGGAGTCGTTGCCTCTGAGAGGCACCATCTGAAGNNNAGATCGGAAG |
| 51734_1 | TATGAGTGTGGAGTCGTTGCGCTGTTTTAGACATCTTTCCACGTNNNAGATCGGAA |
| 51734_2 | TATGAGTGTGGAGTCGTTGCGCTGTTTTAGACATCTTTCCACGTNNNAGATCGGAA |
| 45054_1 | TATGAGTGTGGAGTCGTTGCAAGGAGATGCTTTGTCCGTCAGNNNAGATCGGAAG |
| 45054_2 | TATGAGTGTGGAGTCGTTGCAAGGAGATGCTTTGTCCGTCAGNNNAGATCGGAAG |
| 42526_1 | TATGAGTGTGGAGTCGTTGCTGCTTTCCACAGTGACTGAACTANNNAGATCGGAAG |
| 42526_2 | TATGAGTGTGGAGTCGTTGCTGCTTTCCACAGTGACTGAACTANNNAGATCGGAAG |
| 31315_1 | TATGAGTGTGGAGTCGTTGCAAAATGGGAGAAGGGGAGCTAAGNNNAGATCGGA |
| 31315_2 | TATGAGTGTGGAGTCGTTGCAAAATGGGAGAAGGGGAGCTAAGNNNAGATCGGA |
| 36408_1 | TATGAGTGTGGAGTCGTTGCGTGACAGAGAGAGACCCCATCTNNNAGATCGGAAG |
| 36408_2 | TATGAGTGTGGAGTCGTTGCGTGACAGAGAGAGACCCCATCTNNNAGATCGGAAG |
| 32849_1 | TATGAGTGTGGAGTCGTTGCTCTCAAACTCCTGACCTCAAGTGNNNAGATCGGAAG |
| 32849_2 | TATGAGTGTGGAGTCGTTGCTCTCAAACTCCTGACCTCAAGTGNNNAGATCGGAAG |
| 51515_1 | TATGAGTGTGGAGTCGTTGCCTCACTGAAACCTCTGCCTTTCANNNAGATCGGAAGA |
| 51515_2 | TATGAGTGTGGAGTCGTTGCCTCACTGAAACCTCTGCCTTTCANNNAGATCGGAAGA |
| 31323_1 | TATGAGTGTGGAGTCGTTGCTCTGTTTACCTGTCTGTCTTCCCNNNAGATCGGAAGA |
| 31323_2 | TATGAGTGTGGAGTCGTTGCTCTGTTTACCTGTCTGTCTTCCCNNNAGATCGGAAGA |
| 50674_1 | TATGAGTGTGGAGTCGTTGCCAGACAGACTGAAGATGCCAATGNNNAGATCGGAA |

| | |
|---|---|
| 50674_2 | TATGAGTGTGGAGTCGTTGCCAGACAGACTGAAGATGCCAATGNNNAGATCGGAA |
| 31010_1 | TATGAGTGTGGAGTCGTTGCTCTGCTCTGTTTTTTTCCCCATCNNNAGATCGGAAGA |
| 31010_2 | TATGAGTGTGGAGTCGTTGCTCTGCTCTGTTTTTTTCCCCATCNNNAGATCGGAAGA |
| 31368_1 | TATGAGTGTGGAGTCGTTGCACTGTGTGTATGGAAGCGGAATANNNAGATCGGAA |
| 31368_2 | TATGAGTGTGGAGTCGTTGCACTGTGTGTATGGAAGCGGAATANNNAGATCGGAA |
| 10529_1 | TATGAGTGTGGAGTCGTTGCAGCCAAACGATTCAGTGCATTTTNNNAGATCGGAAG |
| 10529_2 | TATGAGTGTGGAGTCGTTGCAGCCAAACGATTCAGTGCATTTTNNNAGATCGGAAG |
| 36671_1 | TATGAGTGTGGAGTCGTTGCTGTTTAGGGCTAGACCTGTTGTCNNNAGATCGGAAG |
| 36671_2 | TATGAGTGTGGAGTCGTTGCTGTTTAGGGCTAGACCTGTTGTCNNNAGATCGGAAG |
| 44611_1 | TATGAGTGTGGAGTCGTTGCACACACTGAGTTGTAGACCTTCCNNNAGATCGGAAG |
| 44611_2 | TATGAGTGTGGAGTCGTTGCACACACTGAGTTGTAGACCTTCCNNNAGATCGGAAG |
| 10599_1 | TATGAGTGTGGAGTCGTTGCCCAACCCCCGTTATCTGTGATTTNNNAGATCGGAAGA |
| 10599_2 | TATGAGTGTGGAGTCGTTGCCCAACCCCCGTTATCTGTGATTTNNNAGATCGGAAGA |
| 35969_1 | TATGAGTGTGGAGTCGTTGCGATGTGTTTTCGCTGGGAAAGTTNNNAGATCGGAA |
| 35969_2 | TATGAGTGTGGAGTCGTTGCGATGTGTTTTCGCTGGGAAAGTTNNNAGATCGGAA |
| 23849_1 | TATGAGTGTGGAGTCGTTGCAGACAGAGAGGATCCATAATATANNNAGATCGGAA |
| 23849_2 | TATGAGTGTGGAGTCGTTGCAGACAGAGAGGATCCATAATATANNNAGATCGGAA |
| 41363_1 | TATGAGTGTGGAGTCGTTGCTGGCATGGCTTAGCTGCTTATAGNNNAGATCGGAAG |
| 41363_2 | TATGAGTGTGGAGTCGTTGCTGGCATGGCTTAGCTGCTTATAGNNNAGATCGGAAG |
| 35973_1 | TATGAGTGTGGAGTCGTTGCTACAAAAAGTAGCTGGGCATGGTNNNAGATCGGAA |
| 35973_2 | TATGAGTGTGGAGTCGTTGCTACAAAAAGTAGCTGGGCATGGTNNNAGATCGGAA |
| 30662_1 | TATGAGTGTGGAGTCGTTGCTTTTCTACTTGGATGGCCTCCANNNAGATCGGAAGA |
| 30662_2 | TATGAGTGTGGAGTCGTTGCTTTTCTACTTGGATGGCCTCCANNNAGATCGGAAGA |
| 47183_1 | TATGAGTGTGGAGTCGTTGCCCCAGTCCTAGAATCAGCCATTTNNNAGATCGGAAG |
| 47183_2 | TATGAGTGTGGAGTCGTTGCCCCAGTCCTAGAATCAGCCATTTNNNAGATCGGAAG |
| 42604_1 | TATGAGTGTGGAGTCGTTGCCCTCAGCAGTGTACATTTGCTTCNNNAGATCGGAAG |
| 42604_2 | TATGAGTGTGGAGTCGTTGCCCTCAGCAGTGTACATTTGCTTCNNNAGATCGGAAG |
| 46510_1 | TATGAGTGTGGAGTCGTTGCCTCCCAAGTAGCTGGAACTACAGNNNAGATCGGAAG |
| 46510_2 | TATGAGTGTGGAGTCGTTGCCTCCCAAGTAGCTGGAACTACAGNNNAGATCGGAAG |
| 41336_1 | TATGAGTGTGGAGTCGTTGCCACAGTTTCCTTCCTTCCTTCCTNNNAGATCGGAAGA |
| 41336_2 | TATGAGTGTGGAGTCGTTGCCACAGTTTCCTTCCTTCCTTCCTNNNAGATCGGAAGA |
| 38437_1 | TATGAGTGTGGAGTCGTTGCTTCATCACCAGTACTTGCCATCANNNAGATCGGAAGA |
| 38437_2 | TATGAGTGTGGAGTCGTTGCTTCATCACCAGTACTTGCCATCANNNAGATCGGAAGA |
| 47773_1 | TATGAGTGTGGAGTCGTTGCCTGAGATTACAAGTGCACACCACNNNAGATCGGAAG |
| 47773_2 | TATGAGTGTGGAGTCGTTGCCTGAGATTACAAGTGCACACCACNNNAGATCGGAAG |
| 49899_1 | TATGAGTGTGGAGTCGTTGCGCTGTCTTCTTTCCCAGGACTTANNNAGATCGGAAGA |
| 49899_2 | TATGAGTGTGGAGTCGTTGCGCTGTCTTCTTTCCCAGGACTTANNNAGATCGGAAGA |
| 31482_1 | TATGAGTGTGGAGTCGTTGCACCATATTCAGTCAGTGCAGGAANNNAGATCGGAAG |
| 31482_2 | TATGAGTGTGGAGTCGTTGCACCATATTCAGTCAGTGCAGGAANNNAGATCGGAAG |
| 29816_1 | TATGAGTGTGGAGTCGTTGCTGGATCTGTGAGGACAAGTATGACNNNAGATCGGA |
| 29816_2 | TATGAGTGTGGAGTCGTTGCTGGATCTGTGAGGACAAGTATGACNNNAGATCGGA |
| 37461_1 | TATGAGTGTGGAGTCGTTGCTTCAAAAATGAATTCAGGGCCGGNNNAGATCGGAA |
| 37461_2 | TATGAGTGTGGAGTCGTTGCTTCAAAAATGAATTCAGGGCCGGNNNAGATCGGAA |

| | |
|---|---|
| 38648_1 | TATGAGTGTGGAGTCGTTGCGGGACAAGAACGAGACTTCATCTNNNAGATCGGAA |
| 38648_2 | TATGAGTGTGGAGTCGTTGCGGGACAAGAACGAGACTTCATCTNNNAGATCGGAA |
| 51969_1 | TATGAGTGTGGAGTCGTTGCGAATGGGATCTACCTGGGTCATGNNNAGATCGGAA |
| 51969_2 | TATGAGTGTGGAGTCGTTGCGAATGGGATCTACCTGGGTCATGNNNAGATCGGAA |
| 36718_1 | TATGAGTGTGGAGTCGTTGCATTTCCCAGCATCCTTTGAGTCTNNNAGATCGGAAGA |
| 36718_2 | TATGAGTGTGGAGTCGTTGCATTTCCCAGCATCCTTTGAGTCTNNNAGATCGGAAGA |
| 50275_1 | TATGAGTGTGGAGTCGTTGCTCCCACCTCAATTTTCCCTTTGANNNAGATCGGAAGA |
| 50275_2 | TATGAGTGTGGAGTCGTTGCTCCCACCTCAATTTTCCCTTTGANNNAGATCGGAAGA |
| 46951_1 | TATGAGTGTGGAGTCGTTGCCTTCCGAGTAGCTGGGATTACAGNNNAGATCGGAA |
| 46951_2 | TATGAGTGTGGAGTCGTTGCCTTCCGAGTAGCTGGGATTACAGNNNAGATCGGAA |
| 36169_1 | TATGAGTGTGGAGTCGTTGCCCCTGGAGAAACTCACACATGTANNNAGATCGGAAG |
| 36169_2 | TATGAGTGTGGAGTCGTTGCCCCTGGAGAAACTCACACATGTANNNAGATCGGAAG |
| 35707_1 | TATGAGTGTGGAGTCGTTGCACAGAAAGAGGGGAAAACCTGTTNNNAGATCGGAA |
| 35707_2 | TATGAGTGTGGAGTCGTTGCACAGAAAGAGGGGAAAACCTGTTNNNAGATCGGAA |
| 43601_1 | TATGAGTGTGGAGTCGTTGCACCCCTACTAGCCTATTGCAAATNNNAGATCGGAAGA |
| 43601_2 | TATGAGTGTGGAGTCGTTGCACCCCTACTAGCCTATTGCAAATNNNAGATCGGAAGA |
| 36832_1 | TATGAGTGTGGAGTCGTTGCTTTGCTTAGTCTGTGAGCTGTCTNNNAGATCGGAAG |
| 36832_2 | TATGAGTGTGGAGTCGTTGCTTTGCTTAGTCTGTGAGCTGTCTNNNAGATCGGAAG |
| 35632_1 | TATGAGTGTGGAGTCGTTGCGTTCAAGCGATTCTTCTGCCTCNNNAGATCGGAAGA |
| 35632_2 | TATGAGTGTGGAGTCGTTGCGTTCAAGCGATTCTTCTGCCTCNNNAGATCGGAAGA |
| 41479_1 | TATGAGTGTGGAGTCGTTGCTTCGAATTTGTTTGCTCTTGCCTNNNAGATCGGAAGA |
| 41479_2 | TATGAGTGTGGAGTCGTTGCTTCGAATTTGTTTGCTCTTGCCTNNNAGATCGGAAGA |
| 40960_1 | TATGAGTGTGGAGTCGTTGCGTGAAGAGAACCTACAGAATGGGANNNAGATCGGA |
| 40960_2 | TATGAGTGTGGAGTCGTTGCGTGAAGAGAACCTACAGAATGGGANNNAGATCGGA |
| 45679_1 | TATGAGTGTGGAGTCGTTGCGCAATCTTCCTGCCTACTGAGTANNNAGATCGGAAG |
| 45679_2 | TATGAGTGTGGAGTCGTTGCGCAATCTTCCTGCCTACTGAGTANNNAGATCGGAAG |
| 12569_1 | TATGAGTGTGGAGTCGTTGCTTTCTCTTTATAGCAGTCGTTAANNNAGATCGGAAGA |
| 12569_2 | TATGAGTGTGGAGTCGTTGCTTTCTCTTTATAGCAGTCGTTAANNNAGATCGGAAGA |
| 33874_1 | TATGAGTGTGGAGTCGTTGCAAACATGGGAAAGTTGCATACCCNNNAGATCGGAA |
| 33874_2 | TATGAGTGTGGAGTCGTTGCAAACATGGGAAAGTTGCATACCCNNNAGATCGGAA |
| 50509_1 | TATGAGTGTGGAGTCGTTGCGAGGAGCAGCAGGTAGGAACNNNAGATCGGAAGA |
| 50509_2 | TATGAGTGTGGAGTCGTTGCGAGGAGCAGCAGGTAGGAACNNNAGATCGGAAGA |
| 51911_1 | TATGAGTGTGGAGTCGTTGCTCTGCTGGAAAATGATGTCTCCANNNAGATCGGAAG |
| 51911_2 | TATGAGTGTGGAGTCGTTGCTCTGCTGGAAAATGATGTCTCCANNNAGATCGGAAG |
| 44227_1 | TATGAGTGTGGAGTCGTTGCTCCTCCTCCTGATCTGAGTCATTNNNAGATCGGAAGA |
| 44227_2 | TATGAGTGTGGAGTCGTTGCTCCTCCTCCTGATCTGAGTCATTNNNAGATCGGAAGA |
| 46633_1 | TATGAGTGTGGAGTCGTTGCTTCTTCAGATGTCCAGTGATGGGNNNAGATCGGAA |
| 46633_2 | TATGAGTGTGGAGTCGTTGCTTCTTCAGATGTCCAGTGATGGGNNNAGATCGGAA |
| 3177_1 | TATGAGTGTGGAGTCGTTGCTCAAGAGATGCCAAGTACTTCAGANNNAGATCGGAA |
| 3177_2 | TATGAGTGTGGAGTCGTTGCTCAAGAGATGCCAAGTACTTCAGANNNAGATCGGAA |
| 40943_1 | TATGAGTGTGGAGTCGTTGCTGAGACATTGCCATCATAACTTCCNNNAGATCGGAA |
| 40943_2 | TATGAGTGTGGAGTCGTTGCTGAGACATTGCCATCATAACTTCCNNNAGATCGGAA |
| 50914_1 | TATGAGTGTGGAGTCGTTGCCACAGACATCCCCGTTCTCTGNNNAGATCGGAAGAG |

| | |
|---|---|
| 50914_2 | TATGAGTGTGGAGTCGTTGCCACAGACATCCCCGTTCTCTGNNNAGATCGGAAGAG |
| 9865_1 | TATGAGTGTGGAGTCGTTGCTGTATGCTACTGCTCTAGCTGGANNNAGATCGGAAG |
| 9865_2 | TATGAGTGTGGAGTCGTTGCTGTATGCTACTGCTCTAGCTGGANNNAGATCGGAAG |
| 34773_1 | TATGAGTGTGGAGTCGTTGCCCTGGAGTAGCTGGGATTACAGNNNAGATCGGAAG |
| 34773_2 | TATGAGTGTGGAGTCGTTGCCCTGGAGTAGCTGGGATTACAGNNNAGATCGGAAG |
| 43708_1 | TATGAGTGTGGAGTCGTTGCGTGTCCAGGAAGTGAGGATGAATNNNAGATCGGA |
| 43708_2 | TATGAGTGTGGAGTCGTTGCGTGTCCAGGAAGTGAGGATGAATNNNAGATCGGA |
| 51095_1 | TATGAGTGTGGAGTCGTTGCGAAAACCCCTTTGAGAGCTTCACNNNAGATCGGAAG |
| 51095_2 | TATGAGTGTGGAGTCGTTGCGAAAACCCCTTTGAGAGCTTCACNNNAGATCGGAAG |
| 34540_1 | TATGAGTGTGGAGTCGTTGCCACACACACACACACACACAATTNNNAGATCGGAAG |
| 34540_2 | TATGAGTGTGGAGTCGTTGCCACACACACACACACACACAATTNNNAGATCGGAAG |
| 43675_1 | TATGAGTGTGGAGTCGTTGCGATGCCACGTCCATTTGCTTCNNNAGATCGGAAGAG |
| 43675_2 | TATGAGTGTGGAGTCGTTGCGATGCCACGTCCATTTGCTTCNNNAGATCGGAAGAG |
| 51895_1 | TATGAGTGTGGAGTCGTTGCCATTCCCTGCTTCTGACCCCNNNAGATCGGAAGAGCA |
| 51895_2 | TATGAGTGTGGAGTCGTTGCCATTCCCTGCTTCTGACCCCNNNAGATCGGAAGAGCA |
| 10475_1 | TATGAGTGTGGAGTCGTTGCTGCTGTTTCTTCAGCTATCACCANNNAGATCGGAAG |
| 10475_2 | TATGAGTGTGGAGTCGTTGCTGCTGTTTCTTCAGCTATCACCANNNAGATCGGAAG |
| 46755_1 | TATGAGTGTGGAGTCGTTGCCAGTGGCATGATCATGGCATACTNNNAGATCGGAA |
| 46755_2 | TATGAGTGTGGAGTCGTTGCCAGTGGCATGATCATGGCATACTNNNAGATCGGAA |
| 33760_1 | TATGAGTGTGGAGTCGTTGCCTGTAATGCCAGCACTTTGGGAGNNNAGATCGGAA |
| 33760_2 | TATGAGTGTGGAGTCGTTGCCTGTAATGCCAGCACTTTGGGAGNNNAGATCGGAA |
| 45329_1 | TATGAGTGTGGAGTCGTTGCTTCCATTTCCTAGCTAGGGCAAGNNNAGATCGGAAG |
| 45329_2 | TATGAGTGTGGAGTCGTTGCTTCCATTTCCTAGCTAGGGCAAGNNNAGATCGGAAG |
| 42534_1 | TATGAGTGTGGAGTCGTTGCTAATGCTGCAATGAACATGTGCANNNAGATCGGAA |
| 42534_2 | TATGAGTGTGGAGTCGTTGCTAATGCTGCAATGAACATGTGCANNNAGATCGGAA |
| 33372_1 | TATGAGTGTGGAGTCGTTGCAGCGAGAGAGAAAGAGAAAGACANNNAGATCGGA |
| 33372_2 | TATGAGTGTGGAGTCGTTGCAGCGAGAGAGAAAGAGAAAGACANNNAGATCGGA |
| 39808_1 | TATGAGTGTGGAGTCGTTGCTTTGGGACTAGAGATGCTCAACCNNNAGATCGGAA |
| 39808_2 | TATGAGTGTGGAGTCGTTGCTTTGGGACTAGAGATGCTCAACCNNNAGATCGGAA |
| 34791_1 | TATGAGTGTGGAGTCGTTGCCTGAGTAGCTGGGATTACAGGTGNNNAGATCGGAA |
| 34791_2 | TATGAGTGTGGAGTCGTTGCCTGAGTAGCTGGGATTACAGGTGNNNAGATCGGAA |
| 38249_1 | TATGAGTGTGGAGTCGTTGCCCTCTCTGCACACAACCTAGAAANNNAGATCGGAAG |
| 38249_2 | TATGAGTGTGGAGTCGTTGCCCTCTCTGCACACAACCTAGAAANNNAGATCGGAAG |
| 50242_1 | TATGAGTGTGGAGTCGTTGCAAAACCAATGCCAGTCTCCATTGNNNAGATCGGAAG |
| 50242_2 | TATGAGTGTGGAGTCGTTGCAAAACCAATGCCAGTCTCCATTGNNNAGATCGGAAG |
| 35413_1 | TATGAGTGTGGAGTCGTTGCGGCATGATCTCGGATCACTGTAANNNAGATCGGAA |
| 35413_2 | TATGAGTGTGGAGTCGTTGCGGCATGATCTCGGATCACTGTAANNNAGATCGGAA |
| 50378_1 | TATGAGTGTGGAGTCGTTGCCAGCTTAGACTGCAGAGGAACTANNNAGATCGGAA |
| 50378_2 | TATGAGTGTGGAGTCGTTGCCAGCTTAGACTGCAGAGGAACTANNNAGATCGGAA |
| 52811_1 | TATGAGTGTGGAGTCGTTGCCCTGCCCAGACATTTGAAACAAANNNAGATCGGAAG |
| 52811_2 | TATGAGTGTGGAGTCGTTGCCCTGCCCAGACATTTGAAACAAANNNAGATCGGAAG |
| 12911_1 | TATGAGTGTGGAGTCGTTGCTCTTTCCTATGTTTTTCTCCATANNNAGATCGGAAGA |
| 12911_2 | TATGAGTGTGGAGTCGTTGCTCTTTCCTATGTTTTTCTCCATANNNAGATCGGAAGA |

| | |
|---|---|
| 41323_1 | TATGAGTGTGGAGTCGTTGCAGCAAACTGAATCCATTAGCACANNNAGATCGGAAG |
| 41323_2 | TATGAGTGTGGAGTCGTTGCAGCAAACTGAATCCATTAGCACANNNAGATCGGAAG |
| 9604_1 | TATGAGTGTGGAGTCGTTGCAACTGGAGAAAGAACTGCAGTGGNNNAGATCGGA |
| 9604_2 | TATGAGTGTGGAGTCGTTGCAACTGGAGAAAGAACTGCAGTGGNNNAGATCGGA |
| 29574_1 | TATGAGTGTGGAGTCGTTGCTTCATCCCCTAGGCTCTTACTCANNNAGATCGGAAGA |
| 29574_2 | TATGAGTGTGGAGTCGTTGCTTCATCCCCTAGGCTCTTACTCANNNAGATCGGAAGA |
| 38647_1 | TATGAGTGTGGAGTCGTTGCTCCCTATCAAAATAACAGCAGGCTNNNAGATCGGAA |
| 38647_2 | TATGAGTGTGGAGTCGTTGCTCCCTATCAAAATAACAGCAGGCTNNNAGATCGGAA |
| 51501_1 | TATGAGTGTGGAGTCGTTGCCTCCACCCCAAATCAACAGGATANNNAGATCGGAAG |
| 51501_2 | TATGAGTGTGGAGTCGTTGCCTCCACCCCAAATCAACAGGATANNNAGATCGGAAG |
| 52432_1 | TATGAGTGTGGAGTCGTTGCGAGGTTTGGGGTTTGATTGAACCNNNAGATCGGAA |
| 52432_2 | TATGAGTGTGGAGTCGTTGCGAGGTTTGGGGTTTGATTGAACCNNNAGATCGGAA |
| 42813_1 | TATGAGTGTGGAGTCGTTGCATGGGGATCAGTTCTACTTGCTGNNNAGATCGGAAG |
| 42813_2 | TATGAGTGTGGAGTCGTTGCATGGGGATCAGTTCTACTTGCTGNNNAGATCGGAAG |
| 32083_1 | TATGAGTGTGGAGTCGTTGCGGTGGGTGTACTTGAGAGAAGAGNNNAGATCGGA |
| 32083_2 | TATGAGTGTGGAGTCGTTGCGGTGGGTGTACTTGAGAGAAGAGNNNAGATCGGA |
| 49388_1 | TATGAGTGTGGAGTCGTTGCCTCCAGGGTACAAGCGATTCTCNNNAGATCGGAAGA |
| 49388_2 | TATGAGTGTGGAGTCGTTGCCTCCAGGGTACAAGCGATTCTCNNNAGATCGGAAGA |
| 30153_1 | TATGAGTGTGGAGTCGTTGCACTTCATGAAGGAGATTCCTAGGCNNNAGATCGGA |
| 30153_2 | TATGAGTGTGGAGTCGTTGCACTTCATGAAGGAGATTCCTAGGCNNNAGATCGGA |
| 31633_1 | TATGAGTGTGGAGTCGTTGCCAAGTAAAGTGACCATGCCAGTGNNNAGATCGGAA |
| 31633_2 | TATGAGTGTGGAGTCGTTGCCAAGTAAAGTGACCATGCCAGTGNNNAGATCGGAA |
| 33869_1 | TATGAGTGTGGAGTCGTTGCTGAGGCAGGAGAATCTCTTGAACNNNAGATCGGAA |
| 33869_2 | TATGAGTGTGGAGTCGTTGCTGAGGCAGGAGAATCTCTTGAACNNNAGATCGGAA |
| 35200_1 | TATGAGTGTGGAGTCGTTGCCAGGTAGCTGGGACTATAGGCANNNAGATCGGAAG |
| 35200_2 | TATGAGTGTGGAGTCGTTGCCAGGTAGCTGGGACTATAGGCANNNAGATCGGAAG |
| 12629_1 | TATGAGTGTGGAGTCGTTGCATGAGCCAAATAAGAAGTTCTTNNNAGATCGGAAG |
| 12629_2 | TATGAGTGTGGAGTCGTTGCATGAGCCAAATAAGAAGTTCTTNNNAGATCGGAAG |
| 32542_1 | TATGAGTGTGGAGTCGTTGCTCCTGTTTCCTTGAATCTCAGGGNNNAGATCGGAAG |
| 32542_2 | TATGAGTGTGGAGTCGTTGCTCCTGTTTCCTTGAATCTCAGGGNNNAGATCGGAAG |
| 51632_1 | TATGAGTGTGGAGTCGTTGCTTCTCAGTGACCCTCCTTTTCTCNNNAGATCGGAAGA |
| 51632_2 | TATGAGTGTGGAGTCGTTGCTTCTCAGTGACCCTCCTTTTCTCNNNAGATCGGAAGA |
| 31216_1 | TATGAGTGTGGAGTCGTTGCCGAGTAGCTGGGATTATAAGCGTNNNAGATCGGAA |
| 31216_2 | TATGAGTGTGGAGTCGTTGCCGAGTAGCTGGGATTATAAGCGTNNNAGATCGGAA |
| 36474_1 | TATGAGTGTGGAGTCGTTGCCACCTCCAACACACATCACATTTNNNAGATCGGAAGA |
| 36474_2 | TATGAGTGTGGAGTCGTTGCCACCTCCAACACACATCACATTTNNNAGATCGGAAGA |
| 3011_1 | TATGAGTGTGGAGTCGTTGCAGATGTAAGTAGGTGCGGGTTTGNNNAGATCGGA |
| 3011_2 | TATGAGTGTGGAGTCGTTGCAGATGTAAGTAGGTGCGGGTTTGNNNAGATCGGA |
| 51290_1 | TATGAGTGTGGAGTCGTTGCTGAGTGGAGGGAACTTAGAGGATNNNAGATCGGA |
| 51290_2 | TATGAGTGTGGAGTCGTTGCTGAGTGGAGGGAACTTAGAGGATNNNAGATCGGA |
| 48486_1 | TATGAGTGTGGAGTCGTTGCTTCAAGTGATTCTCCTGCCTCAGNNNAGATCGGAAG |
| 48486_2 | TATGAGTGTGGAGTCGTTGCTTCAAGTGATTCTCCTGCCTCAGNNNAGATCGGAAG |
| 45128_1 | TATGAGTGTGGAGTCGTTGCCACAGACAGTGGAGTATGTGACANNNAGATCGGAA |

| | |
|---|---|
| 45128_2 | TATGAGTGTGGAGTCGTTGCCACAGACAGTGGAGTATGTGACANNNAGATCGGAA |
| 48419_1 | TATGAGTGTGGAGTCGTTGCTTCAAGCTATTCTCCTGCCTCAGNNNAGATCGGAAGA |
| 48419_2 | TATGAGTGTGGAGTCGTTGCTTCAAGCTATTCTCCTGCCTCAGNNNAGATCGGAAGA |
| 41021_1 | TATGAGTGTGGAGTCGTTGCCACACATGAGTTTGAGAACACCCNNNAGATCGGAAG |
| 41021_2 | TATGAGTGTGGAGTCGTTGCCACACATGAGTTTGAGAACACCCNNNAGATCGGAAG |
| 31839_1 | TATGAGTGTGGAGTCGTTGCAGAGCGACACTCCATCTCAAAAANNNAGATCGGAAG |
| 31839_2 | TATGAGTGTGGAGTCGTTGCAGAGCGACACTCCATCTCAAAAANNNAGATCGGAAG |
| 39350_1 | TATGAGTGTGGAGTCGTTGCCTGTAGATTCCATAGGCCCTTCCNNNAGATCGGAAG |
| 39350_2 | TATGAGTGTGGAGTCGTTGCCTGTAGATTCCATAGGCCCTTCCNNNAGATCGGAAG |
| 53329_1 | TATGAGTGTGGAGTCGTTGCACGTGTATATATGCGTGTACACACNNNAGATCGGAA |
| 53329_2 | TATGAGTGTGGAGTCGTTGCACGTGTATATATGCGTGTACACACNNNAGATCGGAA |
| 48779_1 | TATGAGTGTGGAGTCGTTGCCGTCTCCCTAAAAAGCACTGGGNNNAGATCGGAAGA |
| 48779_2 | TATGAGTGTGGAGTCGTTGCCGTCTCCCTAAAAAGCACTGGGNNNAGATCGGAAGA |
| 50264_1 | TATGAGTGTGGAGTCGTTGCCGCCAGGTGAGTACATCCTCNNNAGATCGGAAGAG |
| 50264_2 | TATGAGTGTGGAGTCGTTGCCGCCAGGTGAGTACATCCTCNNNAGATCGGAAGAG |
| 44488_1 | TATGAGTGTGGAGTCGTTGCCTCAAAGGATCCTTCCACCTCAGNNNAGATCGGAAG |
| 44488_2 | TATGAGTGTGGAGTCGTTGCCTCAAAGGATCCTTCCACCTCAGNNNAGATCGGAAG |
| 35698_1 | TATGAGTGTGGAGTCGTTGCCGTGTCTCAGCTCACTGCAAGNNNAGATCGGAAGAG |
| 35698_2 | TATGAGTGTGGAGTCGTTGCCGTGTCTCAGCTCACTGCAAGNNNAGATCGGAAGAG |
| 44781_1 | TATGAGTGTGGAGTCGTTGCATTTGTGCCTAGCTATCGTCCAGNNNAGATCGGAAG |
| 44781_2 | TATGAGTGTGGAGTCGTTGCATTTGTGCCTAGCTATCGTCCAGNNNAGATCGGAAG |
| 43723_1 | TATGAGTGTGGAGTCGTTGCGATGGAGGTGGAGGTCCTAGAAGNNNAGATCGGA |
| 43723_2 | TATGAGTGTGGAGTCGTTGCGATGGAGGTGGAGGTCCTAGAAGNNNAGATCGGA |
| 39047_1 | TATGAGTGTGGAGTCGTTGCTCAAGGTATATAAACACATGCAGGANNNAGATCGG |
| 39047_2 | TATGAGTGTGGAGTCGTTGCTCAAGGTATATAAACACATGCAGGANNNAGATCGG |
| 45336_1 | TATGAGTGTGGAGTCGTTGCATCTGGGCTGCTTCTTTTTTTCCNNNAGATCGGAAGA |
| 45336_2 | TATGAGTGTGGAGTCGTTGCATCTGGGCTGCTTCTTTTTTTCCNNNAGATCGGAAGA |
| 52175_1 | TATGAGTGTGGAGTCGTTGCGCTAACAGTGAAAGCCAAGAAGGNNNAGATCGGAA |
| 52175_2 | TATGAGTGTGGAGTCGTTGCGCTAACAGTGAAAGCCAAGAAGGNNNAGATCGGAA |
| 5658_1 | TATGAGTGTGGAGTCGTTGCAGGGAGAAAGAGAGAGATCACGCNNNAGATCGGA |
| 5658_2 | TATGAGTGTGGAGTCGTTGCAGGGAGAAAGAGAGAGATCACGCNNNAGATCGGA |
| 36134_1 | TATGAGTGTGGAGTCGTTGCGGTAAAGAGAGGTACAGGAATCGGNNNAGATCGG |
| 36134_2 | TATGAGTGTGGAGTCGTTGCGGTAAAGAGAGGTACAGGAATCGGNNNAGATCGG |
| 10262_1 | TATGAGTGTGGAGTCGTTGCGGGAGCAGGAGAGGAGAATTTTCNNNAGATCGGA |
| 10262_2 | TATGAGTGTGGAGTCGTTGCGGGAGCAGGAGAGGAGAATTTTCNNNAGATCGGA |
| 44749_1 | TATGAGTGTGGAGTCGTTGCTGGGGAAACTCAGGATGATTGTTNNNAGATCGGAA |
| 44749_2 | TATGAGTGTGGAGTCGTTGCTGGGGAAACTCAGGATGATTGTTNNNAGATCGGAA |
| 36125_1 | TATGAGTGTGGAGTCGTTGCATGAAAATGGTACACCTGGTCCANNNAGATCGGAAG |
| 36125_2 | TATGAGTGTGGAGTCGTTGCATGAAAATGGTACACCTGGTCCANNNAGATCGGAAG |
| 49880_1 | TATGAGTGTGGAGTCGTTGCTAGTATGACCCTGGGGAGGTATCNNNAGATCGGAA |
| 49880_2 | TATGAGTGTGGAGTCGTTGCTAGTATGACCCTGGGGAGGTATCNNNAGATCGGAA |
| 45337_1 | TATGAGTGTGGAGTCGTTGCGGGGAAGATAGTAGCTTGCAAGANNNAGATCGGA |
| 45337_2 | TATGAGTGTGGAGTCGTTGCGGGGAAGATAGTAGCTTGCAAGANNNAGATCGGA |

| | |
|---|---|
| 33750_1 | TATGAGTGTGGAGTCGTTGCAGAGAAAGAAAACCCTCCAAGCTNNNAGATCGGAA |
| 33750_2 | TATGAGTGTGGAGTCGTTGCAGAGAAAGAAAACCCTCCAAGCTNNNAGATCGGAA |
| 48493_1 | TATGAGTGTGGAGTCGTTGCCTCCTGCCTTCCTTCCTTTCTTANNNAGATCGGAAGA |
| 48493_2 | TATGAGTGTGGAGTCGTTGCCTCCTGCCTTCCTTCCTTTCTTANNNAGATCGGAAGA |
| 13435_1 | TATGAGTGTGGAGTCGTTGCATAGATATCTGAAGTGTGTGAAGNNNAGATCGGAA |
| 13435_2 | TATGAGTGTGGAGTCGTTGCATAGATATCTGAAGTGTGTGAAGNNNAGATCGGAA |
| 41756_1 | TATGAGTGTGGAGTCGTTGCCCTCTGTGTTCAAGCGATTCTCCNNNAGATCGGAAG |
| 41756_2 | TATGAGTGTGGAGTCGTTGCCCTCTGTGTTCAAGCGATTCTCCNNNAGATCGGAAG |
| 51296_1 | TATGAGTGTGGAGTCGTTGCGAACACAGAGAAGAGTGAGGTGTNNNAGATCGGA |
| 51296_2 | TATGAGTGTGGAGTCGTTGCGAACACAGAGAAGAGTGAGGTGTNNNAGATCGGA |
| 49041_1 | TATGAGTGTGGAGTCGTTGCTGGAATGGGTTCTGCGATGATTANNNAGATCGGAA |
| 49041_2 | TATGAGTGTGGAGTCGTTGCTGGAATGGGTTCTGCGATGATTANNNAGATCGGAA |
| 45059_1 | TATGAGTGTGGAGTCGTTGCTTTAGGTTGCTGGAGATGGGAAGNNNAGATCGGA |
| 45059_2 | TATGAGTGTGGAGTCGTTGCTTTAGGTTGCTGGAGATGGGAAGNNNAGATCGGA |
| 35338_1 | TATGAGTGTGGAGTCGTTGCGCAGGGAGGTAACTTGCATTTTTNNNAGATCGGAA |
| 35338_2 | TATGAGTGTGGAGTCGTTGCGCAGGGAGGTAACTTGCATTTTTNNNAGATCGGAA |
| 37309_1 | TATGAGTGTGGAGTCGTTGCTAACCTTCCATACCTTCCATGCCNNNAGATCGGAAGA |
| 37309_2 | TATGAGTGTGGAGTCGTTGCTAACCTTCCATACCTTCCATGCCNNNAGATCGGAAGA |
| 29100_1 | TATGAGTGTGGAGTCGTTGCTATACACACACACACACACACACNNNAGATCGGAAG |
| 29100_2 | TATGAGTGTGGAGTCGTTGCTATACACACACACACACACACACNNNAGATCGGAAG |
| 13294_1 | TATGAGTGTGGAGTCGTTGCAAGGGGATGATTCTAAATCATTNNNAGATCGGAAG |
| 13294_2 | TATGAGTGTGGAGTCGTTGCAAGGGGATGATTCTAAATCATTNNNAGATCGGAAG |
| 43560_1 | TATGAGTGTGGAGTCGTTGCTGCACCCAGCCTAGAACATTATTNNNAGATCGGAAG |
| 43560_2 | TATGAGTGTGGAGTCGTTGCTGCACCCAGCCTAGAACATTATTNNNAGATCGGAAG |
| 30762_1 | TATGAGTGTGGAGTCGTTGCTGGGTCAGAACTGAAGGTGATACNNNAGATCGGAA |
| 30762_2 | TATGAGTGTGGAGTCGTTGCTGGGTCAGAACTGAAGGTGATACNNNAGATCGGAA |
| 36084_1 | TATGAGTGTGGAGTCGTTGCTACTGTGGCTCACACCTGTAATCNNNAGATCGGAAG |
| 36084_2 | TATGAGTGTGGAGTCGTTGCTACTGTGGCTCACACCTGTAATCNNNAGATCGGAAG |
| 37227_1 | TATGAGTGTGGAGTCGTTGCTAATTCAGTTGGGGATGACAGGGNNNAGATCGGAA |
| 37227_2 | TATGAGTGTGGAGTCGTTGCTAATTCAGTTGGGGATGACAGGGNNNAGATCGGAA |
| 42679_1 | TATGAGTGTGGAGTCGTTGCCCATAGGGTTCTTGTGGGAGAGNNNAGATCGGAAG |
| 42679_2 | TATGAGTGTGGAGTCGTTGCCCATAGGGTTCTTGTGGGAGAGNNNAGATCGGAAG |
| 32549_1 | TATGAGTGTGGAGTCGTTGCTGTGTCTCAAAAACAACAAACAGCNNNAGATCGGAA |
| 32549_2 | TATGAGTGTGGAGTCGTTGCTGTGTCTCAAAAACAACAAACAGCNNNAGATCGGAA |
| 5596_1 | TATGAGTGTGGAGTCGTTGCGCTCTGAAAACTCCCTTGCTACANNNAGATCGGAAG |
| 5596_2 | TATGAGTGTGGAGTCGTTGCGCTCTGAAAACTCCCTTGCTACANNNAGATCGGAAG |
| 2047_1 | TATGAGTGTGGAGTCGTTGCCTCCAGCTCTCATCTGGGTCTAANNNAGATCGGAAG |
| 2047_2 | TATGAGTGTGGAGTCGTTGCCTCCAGCTCTCATCTGGGTCTAANNNAGATCGGAAG |
| 36340_1 | TATGAGTGTGGAGTCGTTGCACATAGTCTCTGACCACAATGGANNNAGATCGGAAG |
| 36340_2 | TATGAGTGTGGAGTCGTTGCACATAGTCTCTGACCACAATGGANNNAGATCGGAAG |
| 49013_1 | TATGAGTGTGGAGTCGTTGCTTCAAGCAATTCTCCTGCCTCAGNNNAGATCGGAAG |
| 49013_2 | TATGAGTGTGGAGTCGTTGCTTCAAGCAATTCTCCTGCCTCAGNNNAGATCGGAAG |
| 49796_1 | TATGAGTGTGGAGTCGTTGCAAACTCCCAGACATACCGAAGTGNNNAGATCGGAAG |

| | |
|---|---|
| 49796_2 | TATGAGTGTGGAGTCGTTGCAAACTCCCAGACATACCGAAGTGNNNAGATCGGAAG |
| 48527_1 | TATGAGTGTGGAGTCGTTGCCCACCAGGCCCAGATAATTTTTGNNNAGATCGGAAG |
| 48527_2 | TATGAGTGTGGAGTCGTTGCCCACCAGGCCCAGATAATTTTTGNNNAGATCGGAAG |
| 30933_1 | TATGAGTGTGGAGTCGTTGCCGAGCCCTATAACCCAAACAGTANNNAGATCGGAAG |
| 30933_2 | TATGAGTGTGGAGTCGTTGCCGAGCCCTATAACCCAAACAGTANNNAGATCGGAAG |
| 40444_1 | TATGAGTGTGGAGTCGTTGCCCTACTAGATCCCTCACATGTGCNNNAGATCGGAAG |
| 40444_2 | TATGAGTGTGGAGTCGTTGCCCTACTAGATCCCTCACATGTGCNNNAGATCGGAAG |
| 53397_1 | TATGAGTGTGGAGTCGTTGCGTGGAGACCCTTCATCTCTTCAGNNNAGATCGGAAG |
| 53397_2 | TATGAGTGTGGAGTCGTTGCGTGGAGACCCTTCATCTCTTCAGNNNAGATCGGAAG |
| 51389_1 | TATGAGTGTGGAGTCGTTGCGCCTGAAGACGTATTTCCCATTGNNNAGATCGGAAG |
| 51389_2 | TATGAGTGTGGAGTCGTTGCGCCTGAAGACGTATTTCCCATTGNNNAGATCGGAAG |
| 47429_1 | TATGAGTGTGGAGTCGTTGCGAGACGAAGTCATGCTCTGTCANNNAGATCGGAAG |
| 47429_2 | TATGAGTGTGGAGTCGTTGCGAGACGAAGTCATGCTCTGTCANNNAGATCGGAAG |
| 48775_1 | TATGAGTGTGGAGTCGTTGCGACCAGGAAGGAGATTTGAGAGGNNNAGATCGGA |
| 48775_2 | TATGAGTGTGGAGTCGTTGCGACCAGGAAGGAGATTTGAGAGGNNNAGATCGGA |
| 405_1 | TATGAGTGTGGAGTCGTTGCCGCTTGAGCAGGAGATCGAGNNNAGATCGGAAGA |
| 405_2 | TATGAGTGTGGAGTCGTTGCCGCTTGAGCAGGAGATCGAGNNNAGATCGGAAGA |
| 41929_1 | TATGAGTGTGGAGTCGTTGCCACGATCTTAGCTCACTGCAAGNNNAGATCGGAAGA |
| 41929_2 | TATGAGTGTGGAGTCGTTGCCACGATCTTAGCTCACTGCAAGNNNAGATCGGAAGA |
| 48972_1 | TATGAGTGTGGAGTCGTTGCACATGGGGTTTTATATTTCCACCTNNNAGATCGGAA |
| 48972_2 | TATGAGTGTGGAGTCGTTGCACATGGGGTTTTATATTTCCACCTNNNAGATCGGAA |
| 52022_1 | TATGAGTGTGGAGTCGTTGCTGTGGTGTTTGGCTTTTTGTTCTNNNAGATCGGAAG |
| 52022_2 | TATGAGTGTGGAGTCGTTGCTGTGGTGTTTGGCTTTTTGTTCTNNNAGATCGGAAG |
| 33515_1 | TATGAGTGTGGAGTCGTTGCAGATACTAGAGGTTGCACAGTGCNNNAGATCGGAA |
| 33515_2 | TATGAGTGTGGAGTCGTTGCAGATACTAGAGGTTGCACAGTGCNNNAGATCGGAA |
| 10231_1 | TATGAGTGTGGAGTCGTTGCAAGGGTGTAAGAGTGTTCTTTGTNNNAGATCGGAA |
| 10231_2 | TATGAGTGTGGAGTCGTTGCAAGGGTGTAAGAGTGTTCTTTGTNNNAGATCGGAA |
| 30815_1 | TATGAGTGTGGAGTCGTTGCCTGGTTCTGAGGAATGGAACTGANNNAGATCGGAA |
| 30815_2 | TATGAGTGTGGAGTCGTTGCCTGGTTCTGAGGAATGGAACTGANNNAGATCGGAA |
| 47397_1 | TATGAGTGTGGAGTCGTTGCCATTCTCCTGCCTCAGCCTCNNNAGATCGGAAGAGCA |
| 47397_2 | TATGAGTGTGGAGTCGTTGCCATTCTCCTGCCTCAGCCTCNNNAGATCGGAAGAGCA |
| 49642_1 | TATGAGTGTGGAGTCGTTGCTTTCCTCCTCCTCCTCCTTTTTGNNNAGATCGGAAGAG |
| 49642_2 | TATGAGTGTGGAGTCGTTGCTTTCCTCCTCCTCCTCCTTTTTGNNNAGATCGGAAGAG |
| 44920_1 | TATGAGTGTGGAGTCGTTGCTCACTCACAGAAAGAAGGCTCTCNNNAGATCGGAAG |
| 44920_2 | TATGAGTGTGGAGTCGTTGCTCACTCACAGAAAGAAGGCTCTCNNNAGATCGGAAG |
| 29472_1 | TATGAGTGTGGAGTCGTTGCATGATCCCACCATAGAGGTGAGGNNNAGATCGGAA |
| 29472_2 | TATGAGTGTGGAGTCGTTGCATGATCCCACCATAGAGGTGAGGNNNAGATCGGAA |
| 34973_1 | TATGAGTGTGGAGTCGTTGCTTCAAGCGATTCTCCTGAGTGGNNNAGATCGGAAGA |
| 34973_2 | TATGAGTGTGGAGTCGTTGCTTCAAGCGATTCTCCTGAGTGGNNNAGATCGGAAGA |
| 47775_1 | TATGAGTGTGGAGTCGTTGCGAGTCAGAGTCTTGCTTTGTCACNNNAGATCGGAAG |
| 47775_2 | TATGAGTGTGGAGTCGTTGCGAGTCAGAGTCTTGCTTTGTCACNNNAGATCGGAAG |
| 31646_1 | TATGAGTGTGGAGTCGTTGCTGTGCGTTCGACAAATGGAAAAGNNNAGATCGGAA |
| 31646_2 | TATGAGTGTGGAGTCGTTGCTGTGCGTTCGACAAATGGAAAAGNNNAGATCGGAA |

| | |
|---|---|
| 50445_1 | TATGAGTGTGGAGTCGTTGCAAAGGTCCCTGGGGTTCTGTTTNNNAGATCGGAAG |
| 50445_2 | TATGAGTGTGGAGTCGTTGCAAAGGTCCCTGGGGTTCTGTTTNNNAGATCGGAAG |
| 29950_1 | TATGAGTGTGGAGTCGTTGCCTCACTCTTTCTTCTGCACTTGCNNNAGATCGGAAGA |
| 29950_2 | TATGAGTGTGGAGTCGTTGCCTCACTCTTTCTTCTGCACTTGCNNNAGATCGGAAGA |
| 34541_1 | TATGAGTGTGGAGTCGTTGCCCCGGCCGAAAAGTTCTTTATTTNNNAGATCGGAAG |
| 34541_2 | TATGAGTGTGGAGTCGTTGCCCCGGCCGAAAAGTTCTTTATTTNNNAGATCGGAAG |
| 46709_1 | TATGAGTGTGGAGTCGTTGCTTCAAGTGATTCTCCCACCTCAGNNNAGATCGGAAG |
| 46709_2 | TATGAGTGTGGAGTCGTTGCTTCAAGTGATTCTCCCACCTCAGNNNAGATCGGAAG |
| 38114_1 | TATGAGTGTGGAGTCGTTGCAACAGGAGAGTTAGCCTATCCCTNNNAGATCGGAAG |
| 38114_2 | TATGAGTGTGGAGTCGTTGCAACAGGAGAGTTAGCCTATCCCTNNNAGATCGGAAG |
| 30863_1 | TATGAGTGTGGAGTCGTTGCCTGGTGCTCTGTCCATATCCTTTNNNAGATCGGAAG |
| 30863_2 | TATGAGTGTGGAGTCGTTGCCTGGTGCTCTGTCCATATCCTTTNNNAGATCGGAAG |
| 40045_1 | TATGAGTGTGGAGTCGTTGCGCCTAGAAACTCTCAGGTGGAAANNNAGATCGGAA |
| 40045_2 | TATGAGTGTGGAGTCGTTGCGCCTAGAAACTCTCAGGTGGAAANNNAGATCGGAA |
| 33377_1 | TATGAGTGTGGAGTCGTTGCACCCTGGAGATATGACTTTTCCTNNNAGATCGGAAG |
| 33377_2 | TATGAGTGTGGAGTCGTTGCACCCTGGAGATATGACTTTTCCTNNNAGATCGGAAG |
| 44508_1 | TATGAGTGTGGAGTCGTTGCCACACACACACACACATATGGACNNNAGATCGGAAG |
| 44508_2 | TATGAGTGTGGAGTCGTTGCCACACACACACACACATATGGACNNNAGATCGGAAG |
| 50527_1 | TATGAGTGTGGAGTCGTTGCCAACTCCTCTTATGCCGCCTAGNNNAGATCGGAAGA |
| 50527_2 | TATGAGTGTGGAGTCGTTGCCAACTCCTCTTATGCCGCCTAGNNNAGATCGGAAGA |
| 42086_1 | TATGAGTGTGGAGTCGTTGCGCTGAGGTAGGAGAATCACTGAGNNNAGATCGGAA |
| 42086_2 | TATGAGTGTGGAGTCGTTGCGCTGAGGTAGGAGAATCACTGAGNNNAGATCGGAA |
| 39241_1 | TATGAGTGTGGAGTCGTTGCTCTGGGCTGAGATTGTATTAGCANNNAGATCGGAA |
| 39241_2 | TATGAGTGTGGAGTCGTTGCTCTGGGCTGAGATTGTATTAGCANNNAGATCGGAA |
| 44457_1 | TATGAGTGTGGAGTCGTTGCGGTACTTTGACCTGACTGACGATNNNAGATCGGAAG |
| 44457_2 | TATGAGTGTGGAGTCGTTGCGGTACTTTGACCTGACTGACGATNNNAGATCGGAAG |
| 13103_1 | TATGAGTGTGGAGTCGTTGCAACTCCTCATATTCAAATTTGAGNNNAGATCGGAAG |
| 13103_2 | TATGAGTGTGGAGTCGTTGCAACTCCTCATATTCAAATTTGAGNNNAGATCGGAAG |
| 37472_1 | TATGAGTGTGGAGTCGTTGCCTCACGCCTGTAATCCTAACACTNNNAGATCGGAAG |
| 37472_2 | TATGAGTGTGGAGTCGTTGCCTCACGCCTGTAATCCTAACACTNNNAGATCGGAAG |
| 2268_1 | TATGAGTGTGGAGTCGTTGCCCTGGCACATAGCAGCTGATTTANNNAGATCGGAAG |
| 2268_2 | TATGAGTGTGGAGTCGTTGCCCTGGCACATAGCAGCTGATTTANNNAGATCGGAAG |
| 31882_1 | TATGAGTGTGGAGTCGTTGCGCAGAATCATTTGAACCTGGGAGNNNAGATCGGAA |
| 31882_2 | TATGAGTGTGGAGTCGTTGCGCAGAATCATTTGAACCTGGGAGNNNAGATCGGAA |
| 43590_1 | TATGAGTGTGGAGTCGTTGCTGTGGGATCAATGGTGATATCCCNNNAGATCGGAA |
| 43590_2 | TATGAGTGTGGAGTCGTTGCTGTGGGATCAATGGTGATATCCCNNNAGATCGGAA |
| 12559_1 | TATGAGTGTGGAGTCGTTGCATATAGAATGTATATCTCAGGGCNNNAGATCGGAA |
| 12559_2 | TATGAGTGTGGAGTCGTTGCATATAGAATGTATATCTCAGGGCNNNAGATCGGAA |
| 13622_1 | TATGAGTGTGGAGTCGTTGCTTTCATAATTTATAAGTGTGGGGNNNAGATCGGAA |
| 13622_2 | TATGAGTGTGGAGTCGTTGCTTTCATAATTTATAAGTGTGGGGNNNAGATCGGAA |
| 42678_1 | TATGAGTGTGGAGTCGTTGCCATGCCTGTAGTCCCATGTACTTNNNAGATCGGAAG |
| 42678_2 | TATGAGTGTGGAGTCGTTGCCATGCCTGTAGTCCCATGTACTTNNNAGATCGGAAG |
| 48335_1 | TATGAGTGTGGAGTCGTTGCTTCAAGTGATTCTCCTGCCTCAGNNNAGATCGGAAG |

| | |
|---|---|
| 48335_2 | TATGAGTGTGGAGTCGTTGCTTCAAGTGATTCTCCTGCCTCAGNNNAGATCGGAAG |
| 30913_1 | TATGAGTGTGGAGTCGTTGCTACTGAGAGACCAAAATCCTGGCNNNAGATCGGAA |
| 30913_2 | TATGAGTGTGGAGTCGTTGCTACTGAGAGACCAAAATCCTGGCNNNAGATCGGAA |
| 29015_1 | TATGAGTGTGGAGTCGTTGCTGCATATGTCAGTCAAGTCACCANNNAGATCGGAAG |
| 29015_2 | TATGAGTGTGGAGTCGTTGCTGCATATGTCAGTCAAGTCACCANNNAGATCGGAAG |
| 13780_1 | TATGAGTGTGGAGTCGTTGCCAATTCTTTTGGATATGTACTCANNNAGATCGGAAG |
| 13780_2 | TATGAGTGTGGAGTCGTTGCCAATTCTTTTGGATATGTACTCANNNAGATCGGAAG |
| 35801_1 | TATGAGTGTGGAGTCGTTGCCTGGAGTGGTATGATCTCAGCTCNNNAGATCGGAA |
| 35801_2 | TATGAGTGTGGAGTCGTTGCCTGGAGTGGTATGATCTCAGCTCNNNAGATCGGAA |
| 31657_1 | TATGAGTGTGGAGTCGTTGCCAAAGGCTGTGAAAGTACTGTGGNNNAGATCGGAA |
| 31657_2 | TATGAGTGTGGAGTCGTTGCCAAAGGCTGTGAAAGTACTGTGGNNNAGATCGGAA |
| 36497_1 | TATGAGTGTGGAGTCGTTGCCAAATGGAAACACATCCCATGCTNNNAGATCGGAAG |
| 36497_2 | TATGAGTGTGGAGTCGTTGCCAAATGGAAACACATCCCATGCTNNNAGATCGGAAG |
| 29852_1 | TATGAGTGTGGAGTCGTTGCTGGGTCGTAGCCATTTAAGAACANNNAGATCGGAA |
| 29852_2 | TATGAGTGTGGAGTCGTTGCTGGGTCGTAGCCATTTAAGAACANNNAGATCGGAA |
| 40652_1 | TATGAGTGTGGAGTCGTTGCCATCTCCTGGGGATGAAGCTATTNNNAGATCGGAAG |
| 40652_2 | TATGAGTGTGGAGTCGTTGCCATCTCCTGGGGATGAAGCTATTNNNAGATCGGAAG |
| 45833_1 | TATGAGTGTGGAGTCGTTGCATACAAGACCAAATCCTCCCCACNNNAGATCGGAAG |
| 45833_2 | TATGAGTGTGGAGTCGTTGCATACAAGACCAAATCCTCCCCACNNNAGATCGGAAG |
| 48588_1 | TATGAGTGTGGAGTCGTTGCAAGTAGCTGAGATTACAGGCACCNNNAGATCGGAA |
| 48588_2 | TATGAGTGTGGAGTCGTTGCAAGTAGCTGAGATTACAGGCACCNNNAGATCGGAA |
| 40412_1 | TATGAGTGTGGAGTCGTTGCAATTGTTACAGGGTGATGACGGANNNAGATCGGAA |
| 40412_2 | TATGAGTGTGGAGTCGTTGCAATTGTTACAGGGTGATGACGGANNNAGATCGGAA |
| 50809_1 | TATGAGTGTGGAGTCGTTGCGTTCAGTATGGCCTCCTTGTGTANNNAGATCGGAAG |
| 50809_2 | TATGAGTGTGGAGTCGTTGCGTTCAGTATGGCCTCCTTGTGTANNNAGATCGGAAG |
| 32893_1 | TATGAGTGTGGAGTCGTTGCGGGATTTTCGCTCAATGGATACANNNAGATCGGAA |
| 32893_2 | TATGAGTGTGGAGTCGTTGCGGGATTTTCGCTCAATGGATACANNNAGATCGGAA |
| 40861_1 | TATGAGTGTGGAGTCGTTGCTGGCTTGGACATATTTTTCTGTAGCNNNAGATCGGA |
| 40861_2 | TATGAGTGTGGAGTCGTTGCTGGCTTGGACATATTTTTCTGTAGCNNNAGATCGGA |
| 29018_1 | TATGAGTGTGGAGTCGTTGCAAATGGTATATAGCCCTCGCAGGNNNAGATCGGAA |
| 29018_2 | TATGAGTGTGGAGTCGTTGCAAATGGTATATAGCCCTCGCAGGNNNAGATCGGAA |
| 44340_1 | TATGAGTGTGGAGTCGTTGCGATGGACAGGAGATGTGGATGTGNNNAGATCGGA |
| 44340_2 | TATGAGTGTGGAGTCGTTGCGATGGACAGGAGATGTGGATGTGNNNAGATCGGA |
| 48225_1 | TATGAGTGTGGAGTCGTTGCCTGTCTTTGGGGAAAAGGACCANNNAGATCGGAAG |
| 48225_2 | TATGAGTGTGGAGTCGTTGCCTGTCTTTGGGGAAAAGGACCANNNAGATCGGAAG |
| 42980_1 | TATGAGTGTGGAGTCGTTGCTGTACACACACAACAACAACAAGANNNAGATCGGAA |
| 42980_2 | TATGAGTGTGGAGTCGTTGCTGTACACACACAACAACAACAAGANNNAGATCGGAA |
| 49690_1 | TATGAGTGTGGAGTCGTTGCATGGAGTTTCACTTTTGTTGCCCNNNAGATCGGAAG |
| 49690_2 | TATGAGTGTGGAGTCGTTGCATGGAGTTTCACTTTTGTTGCCCNNNAGATCGGAAG |
| 43624_1 | TATGAGTGTGGAGTCGTTGCATGCACCTGTGAGGACTGTTTATNNNAGATCGGAAG |
| 43624_2 | TATGAGTGTGGAGTCGTTGCATGCACCTGTGAGGACTGTTTATNNNAGATCGGAAG |
| 28845_1 | TATGAGTGTGGAGTCGTTGCTGCCTAATTCCTCCTACCTCAGANNNAGATCGGAAGA |
| 28845_2 | TATGAGTGTGGAGTCGTTGCTGCCTAATTCCTCCTACCTCAGANNNAGATCGGAAGA |

| | |
|---|---|
| 48831_1 | TATGAGTGTGGAGTCGTTGCTCTTGTCTTCTGCTAGCTTTGGGNNNAGATCGGAAG |
| 48831_2 | TATGAGTGTGGAGTCGTTGCTCTTGTCTTCTGCTAGCTTTGGGNNNAGATCGGAAG |
| 42561_1 | TATGAGTGTGGAGTCGTTGCAGGTAACTCCAATTCCACTCAGCNNNAGATCGGAAG |
| 42561_2 | TATGAGTGTGGAGTCGTTGCAGGTAACTCCAATTCCACTCAGCNNNAGATCGGAAG |
| 50424_1 | TATGAGTGTGGAGTCGTTGCACTGAGATCCTTGAGAGAGGACANNNAGATCGGAA |
| 50424_2 | TATGAGTGTGGAGTCGTTGCACTGAGATCCTTGAGAGAGGACANNNAGATCGGAA |
| 33346_1 | TATGAGTGTGGAGTCGTTGCGCCTCCCAAAGTTGTGATTACAGNNNAGATCGGAAG |
| 33346_2 | TATGAGTGTGGAGTCGTTGCGCCTCCCAAAGTTGTGATTACAGNNNAGATCGGAAG |
| 51168_1 | TATGAGTGTGGAGTCGTTGCTAGAACGAATGCGCTGGACATTNNNAGATCGGAAG |
| 51168_2 | TATGAGTGTGGAGTCGTTGCTAGAACGAATGCGCTGGACATTNNNAGATCGGAAG |
| 47331_1 | TATGAGTGTGGAGTCGTTGCACTCCATTGGACTCTATCAGTGCNNNAGATCGGAAG |
| 47331_2 | TATGAGTGTGGAGTCGTTGCACTCCATTGGACTCTATCAGTGCNNNAGATCGGAAG |
| 33004_1 | TATGAGTGTGGAGTCGTTGCGATCTCGGCTCACTGTAAGCTCNNNAGATCGGAAGA |
| 33004_2 | TATGAGTGTGGAGTCGTTGCGATCTCGGCTCACTGTAAGCTCNNNAGATCGGAAGA |
| 41592_1 | TATGAGTGTGGAGTCGTTGCCCTGTAATCCCAGCACTTTAGGANNNAGATCGGAAG |
| 41592_2 | TATGAGTGTGGAGTCGTTGCCCTGTAATCCCAGCACTTTAGGANNNAGATCGGAAG |
| 47058_1 | TATGAGTGTGGAGTCGTTGCCCATGCCCGGCTAATTTTGTATTNNNAGATCGGAAG |
| 47058_2 | TATGAGTGTGGAGTCGTTGCCCATGCCCGGCTAATTTTGTATTNNNAGATCGGAAG |
| 43982_1 | TATGAGTGTGGAGTCGTTGCAGAGAGAGGAAGGAAGGAAGGAANNNAGATCGG |
| 43982_2 | TATGAGTGTGGAGTCGTTGCAGAGAGAGGAAGGAAGGAAGGAANNNAGATCGG |
| 38985_1 | TATGAGTGTGGAGTCGTTGCAGGAAATCATAGAGCTTTGGGAANNNAGATCGGAA |
| 38985_2 | TATGAGTGTGGAGTCGTTGCAGGAAATCATAGAGCTTTGGGAANNNAGATCGGAA |
| 34261_1 | TATGAGTGTGGAGTCGTTGCTGTACACTGTACCCAACGTGTAGNNNAGATCGGAAG |
| 34261_2 | TATGAGTGTGGAGTCGTTGCTGTACACTGTACCCAACGTGTAGNNNAGATCGGAAG |
| 33051_1 | TATGAGTGTGGAGTCGTTGCCATTCTCCTGCCTCAGCCTCNNNAGATCGGAAGAGCA |
| 33051_2 | TATGAGTGTGGAGTCGTTGCCATTCTCCTGCCTCAGCCTCNNNAGATCGGAAGAGCA |
| 31345_1 | TATGAGTGTGGAGTCGTTGCTCTGCAAGCATTCCAAAGGAAAGNNNAGATCGGAA |
| 31345_2 | TATGAGTGTGGAGTCGTTGCTCTGCAAGCATTCCAAAGGAAAGNNNAGATCGGAA |
| 34797_1 | TATGAGTGTGGAGTCGTTGCCTGAGTAGCTGGGATTACAGGTGNNNAGATCGGAA |
| 34797_2 | TATGAGTGTGGAGTCGTTGCCTGAGTAGCTGGGATTACAGGTGNNNAGATCGGAA |
| 35230_1 | TATGAGTGTGGAGTCGTTGCGTATACATGTGTCATGCTGGTGCNNNAGATCGGAA |
| 35230_2 | TATGAGTGTGGAGTCGTTGCGTATACATGTGTCATGCTGGTGCNNNAGATCGGAA |
| 34072_1 | TATGAGTGTGGAGTCGTTGCTTCAAGCAATTCTCCTGCCTCAGNNNAGATCGGAAG |
| 34072_2 | TATGAGTGTGGAGTCGTTGCTTCAAGCAATTCTCCTGCCTCAGNNNAGATCGGAAG |
| 36416_1 | TATGAGTGTGGAGTCGTTGCGGAAAGAAAGAATGGGCATGGTGNNNAGATCGGA |
| 36416_2 | TATGAGTGTGGAGTCGTTGCGGAAAGAAAGAATGGGCATGGTGNNNAGATCGGA |
| 31344_1 | TATGAGTGTGGAGTCGTTGCAGCAGTGTGTCTTTGATTATGGCNNNAGATCGGAA |
| 31344_2 | TATGAGTGTGGAGTCGTTGCAGCAGTGTGTCTTTGATTATGGCNNNAGATCGGAA |
| 43155_1 | TATGAGTGTGGAGTCGTTGCTTTTTCTGCTGGACACTCTACGTNNNAGATCGGAAG |
| 43155_2 | TATGAGTGTGGAGTCGTTGCTTTTTCTGCTGGACACTCTACGTNNNAGATCGGAAG |
| 13348_1 | TATGAGTGTGGAGTCGTTGCTACATTTTATAGGTTTGTGCAAANNNAGATCGGAAG |
| 13348_2 | TATGAGTGTGGAGTCGTTGCTACATTTTATAGGTTTGTGCAAANNNAGATCGGAAG |
| 37621_1 | TATGAGTGTGGAGTCGTTGCTGTTCATCAGTGTATCCCAAGGGNNNAGATCGGAAG |

| | |
|---|---|
| 37621_2 | TATGAGTGTGGAGTCGTTGCTGTTCATCAGTGTATCCCAAGGGNNNAGATCGGAAG |
| 48182_1 | TATGAGTGTGGAGTCGTTGCACAGGGTTTTGCTCTGTCACCNNNAGATCGGAAGAG |
| 48182_2 | TATGAGTGTGGAGTCGTTGCACAGGGTTTTGCTCTGTCACCNNNAGATCGGAAGAG |
| 30054_1 | TGAGTGTGGAGTCGTTGCTTTTAGGATATGCAATCTAACAAGAGTNNNAGATCGGA |
| 30054_2 | TGAGTGTGGAGTCGTTGCTTTTAGGATATGCAATCTAACAAGAGTNNNAGATCGGA |
| 34787_1 | TATGAGTGTGGAGTCGTTGCCGCCCGGCTAATCTTTGTATTTTNNNAGATCGGAAG |
| 34787_2 | TATGAGTGTGGAGTCGTTGCCGCCCGGCTAATCTTTGTATTTTNNNAGATCGGAAG |
| 47569_1 | TATGAGTGTGGAGTCGTTGCGGTTATGATCCAGGAGCAGCTCNNNAGATCGGAAG |
| 47569_2 | TATGAGTGTGGAGTCGTTGCGGTTATGATCCAGGAGCAGCTCNNNAGATCGGAAG |
| 53272_1 | TATGAGTGTGGAGTCGTTGCCCCCCTTTCTAAGCTCTACTGTGNNNAGATCGGAAGA |
| 53272_2 | TATGAGTGTGGAGTCGTTGCCCCCCTTTCTAAGCTCTACTGTGNNNAGATCGGAAGA |
| 47096_1 | TATGAGTGTGGAGTCGTTGCACTCCTGACTTCAAGTGATCCACNNNAGATCGGAAG |
| 47096_2 | TATGAGTGTGGAGTCGTTGCACTCCTGACTTCAAGTGATCCACNNNAGATCGGAAG |
| 37386_1 | TATGAGTGTGGAGTCGTTGCTAAAGCAGGAAGTGAAAGTCCGANNNAGATCGGAA |
| 37386_2 | TATGAGTGTGGAGTCGTTGCTAAAGCAGGAAGTGAAAGTCCGANNNAGATCGGAA |
| 34428_1 | TATGAGTGTGGAGTCGTTGCATGAGGCCAGATGGTGTCAATTANNNAGATCGGAA |
| 34428_2 | TATGAGTGTGGAGTCGTTGCATGAGGCCAGATGGTGTCAATTANNNAGATCGGAA |
| 48324_1 | TATGAGTGTGGAGTCGTTGCCCAAAATGCTGGGATTACAGGTGNNNAGATCGGAA |
| 48324_2 | TATGAGTGTGGAGTCGTTGCCCAAAATGCTGGGATTACAGGTGNNNAGATCGGAA |
| 32930_1 | TATGAGTGTGGAGTCGTTGCCTGAGGTGGGAGGATTACTTGAGNNNAGATCGGAA |
| 32930_2 | TATGAGTGTGGAGTCGTTGCCTGAGGTGGGAGGATTACTTGAGNNNAGATCGGAA |
| 50298_1 | TATGAGTGTGGAGTCGTTGCCTCATGTCTGTAATCCCAGCACTNNNAGATCGGAAG |
| 50298_2 | TATGAGTGTGGAGTCGTTGCCTCATGTCTGTAATCCCAGCACTNNNAGATCGGAAG |
| 44962_1 | TATGAGTGTGGAGTCGTTGCGGAAGAAGGAGAGAGGGAATCAGNNNAGATCGGA |
| 44962_2 | TATGAGTGTGGAGTCGTTGCGGAAGAAGGAGAGAGGGAATCAGNNNAGATCGGA |
| 49869_1 | TATGAGTGTGGAGTCGTTGCTCTGGTTAGTGTGAAGAAGTGGGNNNAGATCGGAA |
| 49869_2 | TATGAGTGTGGAGTCGTTGCTCTGGTTAGTGTGAAGAAGTGGGNNNAGATCGGAA |
| 39205_1 | TATGAGTGTGGAGTCGTTGCCACCACACCCAGCTAATTTTTGTNNNAGATCGGAAGA |
| 39205_2 | TATGAGTGTGGAGTCGTTGCCACCACACCCAGCTAATTTTTGTNNNAGATCGGAAGA |
| 29254_1 | TATGAGTGTGGAGTCGTTGCTGGCATACAGCAGAACGATCATANNNAGATCGGAA |
| 29254_2 | TATGAGTGTGGAGTCGTTGCTGGCATACAGCAGAACGATCATANNNAGATCGGAA |
| 32968_1 | TATGAGTGTGGAGTCGTTGCGGCAAGACTGGAATCTAACAACANNNAGATCGGAA |
| 32968_2 | TATGAGTGTGGAGTCGTTGCGGCAAGACTGGAATCTAACAACANNNAGATCGGAA |
| 45769_1 | TATGAGTGTGGAGTCGTTGCTGGCCTAATTGCTGGAGAATTCTNNNAGATCGGAAG |
| 45769_2 | TATGAGTGTGGAGTCGTTGCTGGCCTAATTGCTGGAGAATTCTNNNAGATCGGAAG |
| 40270_1 | TATGAGTGTGGAGTCGTTGCCGGGAAGAAGAAGTTGCAATGAGNNNAGATCGGA |
| 40270_2 | TATGAGTGTGGAGTCGTTGCCGGGAAGAAGAAGTTGCAATGAGNNNAGATCGGA |
| 39027_1 | TATGAGTGTGGAGTCGTTGCCCCATGAAAGTGAGGAGGATACANNNAGATCGGAA |
| 39027_2 | TATGAGTGTGGAGTCGTTGCCCCATGAAAGTGAGGAGGATACANNNAGATCGGAA |
| 36578_1 | TATGAGTGTGGAGTCGTTGCCGTGTACCTGTAGTCCTAGCTACNNNAGATCGGAAG |
| 36578_2 | TATGAGTGTGGAGTCGTTGCCGTGTACCTGTAGTCCTAGCTACNNNAGATCGGAAG |
| 33485_1 | TATGAGTGTGGAGTCGTTGCTTTTTCCTTTGCAGACCTCATGGNNNAGATCGGAAG |
| 33485_2 | TATGAGTGTGGAGTCGTTGCTTTTTCCTTTGCAGACCTCATGGNNNAGATCGGAAG |

| | |
|---|---|
| 34078_1 | TATGAGTGTGGAGTCGTTGCGATTCAAGTGGTTCTCCTGCCTCNNNAGATCGGAAG |
| 34078_2 | TATGAGTGTGGAGTCGTTGCGATTCAAGTGGTTCTCCTGCCTCNNNAGATCGGAAG |
| 35916_1 | TATGAGTGTGGAGTCGTTGCGGCTTTTACGCGAAAGGTAAAGTNNNAGATCGGAA |
| 35916_2 | TATGAGTGTGGAGTCGTTGCGGCTTTTACGCGAAAGGTAAAGTNNNAGATCGGAA |
| 35202_1 | TATGAGTGTGGAGTCGTTGCGACGGAGTCTCGTTCTGTCGNNNAGATCGGAAGAG |
| 35202_2 | TATGAGTGTGGAGTCGTTGCGACGGAGTCTCGTTCTGTCGNNNAGATCGGAAGAG |
| 42126_1 | TATGAGTGTGGAGTCGTTGCCAATCTCAGTTCACTGCAACCTCNNNAGATCGGAAGA |
| 42126_2 | TATGAGTGTGGAGTCGTTGCCAATCTCAGTTCACTGCAACCTCNNNAGATCGGAAGA |
| 33757_1 | TATGAGTGTGGAGTCGTTGCGAAAGAGACAAGCTGAGAGGAGGNNNAGATCGGA |
| 33757_2 | TATGAGTGTGGAGTCGTTGCGAAAGAGACAAGCTGAGAGGAGGNNNAGATCGGA |
| 39726_1 | TATGAGTGTGGAGTCGTTGCCTCACTGCAACCTTCACCTCCNNNAGATCGGAAGAGC |
| 39726_2 | TATGAGTGTGGAGTCGTTGCCTCACTGCAACCTTCACCTCCNNNAGATCGGAAGAGC |
| 30733_1 | TATGAGTGTGGAGTCGTTGCAACTTGTCAACCCCTTCGATGTNNNAGATCGGAAGA |
| 30733_2 | TATGAGTGTGGAGTCGTTGCAACTTGTCAACCCCTTCGATGTNNNAGATCGGAAGA |
| 43571_1 | TATGAGTGTGGAGTCGTTGCCTCCTGAGTAGCTGGTATCACAGNNNAGATCGGAAG |
| 43571_2 | TATGAGTGTGGAGTCGTTGCCTCCTGAGTAGCTGGTATCACAGNNNAGATCGGAAG |
| 39117_1 | TATGAGTGTGGAGTCGTTGCTCCACAGTCCAAAGTCTGATCTGNNNAGATCGGAAG |
| 39117_2 | TATGAGTGTGGAGTCGTTGCTCCACAGTCCAAAGTCTGATCTGNNNAGATCGGAAG |
| 39826_1 | TATGAGTGTGGAGTCGTTGCGATTTCCCCACATTTGCTTGGTTNNNAGATCGGAAG |
| 39826_2 | TATGAGTGTGGAGTCGTTGCGATTTCCCCACATTTGCTTGGTTNNNAGATCGGAAG |
| 47520_1 | TATGAGTGTGGAGTCGTTGCTGTAAGACAGAAAAACAGGCAGCNNNAGATCGGAA |
| 47520_2 | TATGAGTGTGGAGTCGTTGCTGTAAGACAGAAAAACAGGCAGCNNNAGATCGGAA |
| 13408_1 | TATGAGTGTGGAGTCGTTGCAGAGAAGTATTAAGAATGACTCCNNNAGATCGGAA |
| 13408_2 | TATGAGTGTGGAGTCGTTGCAGAGAAGTATTAAGAATGACTCCNNNAGATCGGAA |
| 46896_1 | TATGAGTGTGGAGTCGTTGCGAGTAGCTGGGATTACAGGCATGNNNAGATCGGAA |
| 46896_2 | TATGAGTGTGGAGTCGTTGCGAGTAGCTGGGATTACAGGCATGNNNAGATCGGAA |
| 28782_1 | TATGAGTGTGGAGTCGTTGCCCAGGTTCAAGCAATTTTCCTGTNNNAGATCGGAAG |
| 28782_2 | TATGAGTGTGGAGTCGTTGCCCAGGTTCAAGCAATTTTCCTGTNNNAGATCGGAAG |
| 38352_1 | TATGAGTGTGGAGTCGTTGCACCGAAGCTAAATCAACAGAGGTNNNAGATCGGAA |
| 38352_2 | TATGAGTGTGGAGTCGTTGCACCGAAGCTAAATCAACAGAGGTNNNAGATCGGAA |
| 44166_1 | TATGAGTGTGGAGTCGTTGCTGGGTGCCATTTTTTTGAGGATCNNNAGATCGGAAG |
| 44166_2 | TATGAGTGTGGAGTCGTTGCTGGGTGCCATTTTTTTGAGGATCNNNAGATCGGAAG |
| 33526_1 | TATGAGTGTGGAGTCGTTGCCAAGCCCAGAGTGTGAAGACTATNNNAGATCGGAA |
| 33526_2 | TATGAGTGTGGAGTCGTTGCCAAGCCCAGAGTGTGAAGACTATNNNAGATCGGAA |
| 34294_1 | TATGAGTGTGGAGTCGTTGCCTCCTGAGTAGCTGGGATTACAGNNNAGATCGGAAG |
| 34294_2 | TATGAGTGTGGAGTCGTTGCCTCCTGAGTAGCTGGGATTACAGNNNAGATCGGAAG |
| 30426_1 | TATGAGTGTGGAGTCGTTGCTTCAAGGATTCTCCCGCCTTAACCNNNAGATCGGAAG |
| 30426_2 | TATGAGTGTGGAGTCGTTGCTTCAAGGATTCTCCCGCCTTAACCNNNAGATCGGAAG |
| 35131_1 | TATGAGTGTGGAGTCGTTGCCCAAAGTACTGGGATTACAGGCANNNAGATCGGAA |
| 35131_2 | TATGAGTGTGGAGTCGTTGCCCAAAGTACTGGGATTACAGGCANNNAGATCGGAA |
| 49279_1 | TATGAGTGTGGAGTCGTTGCCCCAAGTTCAAGCGATTCTCTTGNNNAGATCGGAAG |
| 49279_2 | TATGAGTGTGGAGTCGTTGCCCCAAGTTCAAGCGATTCTCTTGNNNAGATCGGAAG |
| 13031_1 | TATGAGTGTGGAGTCGTTGCTAGCGACACAAAGTTAAACAANNNAGATCGGAAGA |

| | |
|---|---|
| 13031_2 | TATGAGTGTGGAGTCGTTGCTAGCGACACAAAGTTAAACAANNNAGATCGGAAGA |
| 32871_1 | TATGAGTGTGGAGTCGTTGCATCCAATTTGCCAGTCTGAGTCTNNNAGATCGGAAG |
| 32871_2 | TATGAGTGTGGAGTCGTTGCATCCAATTTGCCAGTCTGAGTCTNNNAGATCGGAAG |
| 47324_1 | TATGAGTGTGGAGTCGTTGCGCTTGGTTGCTGTTGGTGTATAGNNNAGATCGGAA |
| 47324_2 | TATGAGTGTGGAGTCGTTGCGCTTGGTTGCTGTTGGTGTATAGNNNAGATCGGAA |
| 42101_1 | TATGAGTGTGGAGTCGTTGCATACTACAGGCATGCATCACCATNNNAGATCGGAAG |
| 42101_2 | TATGAGTGTGGAGTCGTTGCATACTACAGGCATGCATCACCATNNNAGATCGGAAG |
| 40706_1 | TATGAGTGTGGAGTCGTTGCCTCCAGTATACAGTGGCTAGCAGNNNAGATCGGAA |
| 40706_2 | TATGAGTGTGGAGTCGTTGCCTCCAGTATACAGTGGCTAGCAGNNNAGATCGGAA |
| 9280_1 | TATGAGTGTGGAGTCGTTGCAAGACTGTGTGTATGTGCATATATGNNNAGATCGG |
| 9280_2 | TATGAGTGTGGAGTCGTTGCAAGACTGTGTGTATGTGCATATATGNNNAGATCGG |
| 32181_1 | TATGAGTGTGGAGTCGTTGCACCATGCGATATTTTAGATCCCCANNNAGATCGGAA |
| 32181_2 | TATGAGTGTGGAGTCGTTGCACCATGCGATATTTTAGATCCCCANNNAGATCGGAA |
| 31208_1 | TATGAGTGTGGAGTCGTTGCAGTGCAGCTCTTTTTACCTACCTNNNAGATCGGAAG |
| 31208_2 | TATGAGTGTGGAGTCGTTGCAGTGCAGCTCTTTTTACCTACCTNNNAGATCGGAAG |
| 43943_1 | TATGAGTGTGGAGTCGTTGCCACACACACACACAATTGGTTCTNNNAGATCGGAAG |
| 43943_2 | TATGAGTGTGGAGTCGTTGCCACACACACACACAATTGGTTCTNNNAGATCGGAAG |
| 34931_1 | TATGAGTGTGGAGTCGTTGCAATCTCCACTCACTGCAAGCTCNNNAGATCGGAAGA |
| 34931_2 | TATGAGTGTGGAGTCGTTGCAATCTCCACTCACTGCAAGCTCNNNAGATCGGAAGA |
| 34291_1 | TATGAGTGTGGAGTCGTTGCGTGTAAAGGGAGATGTCTGCTCANNNAGATCGGAA |
| 34291_2 | TATGAGTGTGGAGTCGTTGCGTGTAAAGGGAGATGTCTGCTCANNNAGATCGGAA |
| 46533_1 | TATGAGTGTGGAGTCGTTGCCGATCTTTGCTCACTACAACCTCNNNAGATCGGAAGA |
| 46533_2 | TATGAGTGTGGAGTCGTTGCCGATCTTTGCTCACTACAACCTCNNNAGATCGGAAGA |
| 52216_1 | TATGAGTGTGGAGTCGTTGCCCCGACCCCATCATTGTTTTTTTNNNAGATCGGAAGA |
| 52216_2 | TATGAGTGTGGAGTCGTTGCCCCGACCCCATCATTGTTTTTTTNNNAGATCGGAAGA |
| 35173_1 | TATGAGTGTGGAGTCGTTGCGTTGGTTAGGCTTGTCTTGAACCNNNAGATCGGAAG |
| 35173_2 | TATGAGTGTGGAGTCGTTGCGTTGGTTAGGCTTGTCTTGAACCNNNAGATCGGAAG |
| 47390_1 | TATGAGTGTGGAGTCGTTGCGAGACGGAGTCTTGCTCTGTCNNNAGATCGGAAGA |
| 47390_2 | TATGAGTGTGGAGTCGTTGCGAGACGGAGTCTTGCTCTGTCNNNAGATCGGAAGA |
| 48284_1 | TATGAGTGTGGAGTCGTTGCCAATCTCAGCTCACTGCAACTTCNNNAGATCGGAAGA |
| 48284_2 | TATGAGTGTGGAGTCGTTGCCAATCTCAGCTCACTGCAACTTCNNNAGATCGGAAGA |
| 43821_1 | TATGAGTGTGGAGTCGTTGCTCTCATCCTTCAGTCAGTCTCCTNNNAGATCGGAAGA |
| 43821_2 | TATGAGTGTGGAGTCGTTGCTCTCATCCTTCAGTCAGTCTCCTNNNAGATCGGAAGA |
| 31841_1 | TATGAGTGTGGAGTCGTTGCGTTGCTTTCTGGTGAGTCCTTTCNNNAGATCGGAAG |
| 31841_2 | TATGAGTGTGGAGTCGTTGCGTTGCTTTCTGGTGAGTCCTTTCNNNAGATCGGAAG |
| 43469_1 | TATGAGTGTGGAGTCGTTGCCATATTGGCTCACTGCAACCTGNNNAGATCGGAAGA |
| 43469_2 | TATGAGTGTGGAGTCGTTGCCATATTGGCTCACTGCAACCTGNNNAGATCGGAAGA |
| 41817_1 | TATGAGTGTGGAGTCGTTGCGTTCAAGCAATTCTCTTGCCTCANNNAGATCGGAAGA |
| 41817_2 | TATGAGTGTGGAGTCGTTGCGTTCAAGCAATTCTCTTGCCTCANNNAGATCGGAAGA |
| 52666_1 | TATGAGTGTGGAGTCGTTGCTTGAGTTTCAGTCCCTCGGATTCNNNAGATCGGAAG |
| 52666_2 | TATGAGTGTGGAGTCGTTGCTTGAGTTTCAGTCCCTCGGATTCNNNAGATCGGAAG |
| 49031_1 | TATGAGTGTGGAGTCGTTGCCACAATCTCAGCTCACTGCAACNNNAGATCGGAAGA |
| 49031_2 | TATGAGTGTGGAGTCGTTGCCACAATCTCAGCTCACTGCAACNNNAGATCGGAAGA |

| | |
|---|---|
| 10778_1 | TATGAGTGTGGAGTCGTTGCGGCTCAGTAACTTTTCCTGGAGGNNNAGATCGGAAG |
| 10778_2 | TATGAGTGTGGAGTCGTTGCGGCTCAGTAACTTTTCCTGGAGGNNNAGATCGGAAG |
| 30586_1 | TATGAGTGTGGAGTCGTTGCCAATTCAAGGCCAAAGACCTGTTNNNAGATCGGAAG |
| 30586_2 | TATGAGTGTGGAGTCGTTGCCAATTCAAGGCCAAAGACCTGTTNNNAGATCGGAAG |
| 29340_1 | TATGAGTGTGGAGTCGTTGCACCTTAGGGACTGGTCTGTCATANNNAGATCGGAAG |
| 29340_2 | TATGAGTGTGGAGTCGTTGCACCTTAGGGACTGGTCTGTCATANNNAGATCGGAAG |
| 40359_1 | TATGAGTGTGGAGTCGTTGCTTGGAGACCAACATGGCCAATATNNNAGATCGGAAG |
| 40359_2 | TATGAGTGTGGAGTCGTTGCTTGGAGACCAACATGGCCAATATNNNAGATCGGAAG |
| 36745_1 | TATGAGTGTGGAGTCGTTGCAACTCCTAGTTCTGAAGGTGCTTNNNAGATCGGAAG |
| 36745_2 | TATGAGTGTGGAGTCGTTGCAACTCCTAGTTCTGAAGGTGCTTNNNAGATCGGAAG |
| 37857_1 | TATGAGTGTGGAGTCGTTGCTTGCATACAGTTGGTTGATGGTGNNNAGATCGGAA |
| 37857_2 | TATGAGTGTGGAGTCGTTGCTTGCATACAGTTGGTTGATGGTGNNNAGATCGGAA |
| 49990_1 | TATGAGTGTGGAGTCGTTGCTATGTCTCTCCCTACTCTTCCCCNNNAGATCGGAAGA |
| 49990_2 | TATGAGTGTGGAGTCGTTGCTATGTCTCTCCCTACTCTTCCCCNNNAGATCGGAAGA |
| 42145_1 | TATGAGTGTGGAGTCGTTGCGAGGAGCTGGGACTACAGGANNNAGATCGGAAGA |
| 42145_2 | TATGAGTGTGGAGTCGTTGCGAGGAGCTGGGACTACAGGANNNAGATCGGAAGA |
| 36839_1 | TATGAGTGTGGAGTCGTTGCGGATCACTTGAGCACAGGAGTTCNNNAGATCGGAA |
| 36839_2 | TATGAGTGTGGAGTCGTTGCGGATCACTTGAGCACAGGAGTTCNNNAGATCGGAA |
| 49620_1 | TATGAGTGTGGAGTCGTTGCTGCTATGTTGTTCAGGCTGATCTNNNAGATCGGAAG |
| 49620_2 | TATGAGTGTGGAGTCGTTGCTGCTATGTTGTTCAGGCTGATCTNNNAGATCGGAAG |
| 32555_1 | TATGAGTGTGGAGTCGTTGCCTCACTGTAGCCTCAACTTCTCANNNAGATCGGAAGA |
| 32555_2 | TATGAGTGTGGAGTCGTTGCCTCACTGTAGCCTCAACTTCTCANNNAGATCGGAAGA |
| 29013_1 | TATGAGTGTGGAGTCGTTGCAAGCAAAGATGAAGAAACACGGGNNNAGATCGGAA |
| 29013_2 | TATGAGTGTGGAGTCGTTGCAAGCAAAGATGAAGAAACACGGGNNNAGATCGGAA |
| 45296_1 | TATGAGTGTGGAGTCGTTGCGTTGACAGCTAAGTACTTGGGGANNNAGATCGGAA |
| 45296_2 | TATGAGTGTGGAGTCGTTGCGTTGACAGCTAAGTACTTGGGGANNNAGATCGGAA |
| 38873_1 | TATGAGTGTGGAGTCGTTGCGAGAAGAGCTTTAGAGAAGGCTATNNNAGATCGGA |
| 38873_2 | TATGAGTGTGGAGTCGTTGCGAGAAGAGCTTTAGAGAAGGCTATNNNAGATCGGA |
| 48798_1 | TATGAGTGTGGAGTCGTTGCTTCAAGTGATTCTCCTGCCTCAGNNNAGATCGGAAG |
| 48798_2 | TATGAGTGTGGAGTCGTTGCTTCAAGTGATTCTCCTGCCTCAGNNNAGATCGGAAG |
| 52375_1 | TATGAGTGTGGAGTCGTTGCTCGGGAAGAGAAAGAGCAGATTCNNNAGATCGGAA |
| 52375_2 | TATGAGTGTGGAGTCGTTGCTCGGGAAGAGAAAGAGCAGATTCNNNAGATCGGAA |
| 33232_1 | TATGAGTGTGGAGTCGTTGCACTTCTCGGTTCAAGCTATCCTCNNNAGATCGGAAGA |
| 33232_2 | TATGAGTGTGGAGTCGTTGCACTTCTCGGTTCAAGCTATCCTCNNNAGATCGGAAGA |
| 8865_1 | TATGAGTGTGGAGTCGTTGCATTGTGGGTGGCAAAGAAACTGTNNNAGATCGGAA |
| 8865_2 | TATGAGTGTGGAGTCGTTGCATTGTGGGTGGCAAAGAAACTGTNNNAGATCGGAA |
| 52055_1 | TATGAGTGTGGAGTCGTTGCCAAATGCCTGGAAATCTGCCTTANNNAGATCGGAAG |
| 52055_2 | TATGAGTGTGGAGTCGTTGCCAAATGCCTGGAAATCTGCCTTANNNAGATCGGAAG |
| 38415_1 | TATGAGTGTGGAGTCGTTGCGCCTGTCCTGAAATGATCTCTGTNNNAGATCGGAAG |
| 38415_2 | TATGAGTGTGGAGTCGTTGCGCCTGTCCTGAAATGATCTCTGTNNNAGATCGGAAG |
| 38295_1 | TATGAGTGTGGAGTCGTTGCAAAGTAGGCGAAGGATATGCACANNNAGATCGGAA |
| 38295_2 | TATGAGTGTGGAGTCGTTGCAAAGTAGGCGAAGGATATGCACANNNAGATCGGAA |
| 49641_1 | TATGAGTGTGGAGTCGTTGCCCAAGTAACTGGGACTACATGCANNNAGATCGGAAG |

| | |
|---|---|
| 49641_2 | TATGAGTGTGGAGTCGTTGCCCAAGTAACTGGGACTACATGCANNNAGATCGGAAG |
| 35741_1 | TATGAGTGTGGAGTCGTTGCATCAGTTTCGTGCAGTTCAACAGNNNAGATCGGAAG |
| 35741_2 | TATGAGTGTGGAGTCGTTGCATCAGTTTCGTGCAGTTCAACAGNNNAGATCGGAAG |
| 36503_1 | TATGAGTGTGGAGTCGTTGCAGTATAATGCAGTGCTCCAGACANNNAGATCGGAAG |
| 36503_2 | TATGAGTGTGGAGTCGTTGCAGTATAATGCAGTGCTCCAGACANNNAGATCGGAAG |
| 41879_1 | TATGAGTGTGGAGTCGTTGCCAATTCTTCTGTCTCAGCCTCCTNNNAGATCGGAAGA |
| 41879_2 | TATGAGTGTGGAGTCGTTGCCAATTCTTCTGTCTCAGCCTCCTNNNAGATCGGAAGA |
| 13024_1 | TATGAGTGTGGAGTCGTTGCCCGGTATGTAATTTAACATTCAANNNAGATCGGAAG |
| 13024_2 | TATGAGTGTGGAGTCGTTGCCCGGTATGTAATTTAACATTCAANNNAGATCGGAAG |
| 37274_1 | TATGAGTGTGGAGTCGTTGCTGTGAAAGGGGAGACCATTGTAGNNNAGATCGGAA |
| 37274_2 | TATGAGTGTGGAGTCGTTGCTGTGAAAGGGGAGACCATTGTAGNNNAGATCGGAA |
| 51529_1 | TATGAGTGTGGAGTCGTTGCGGGATCTGGGAAAGGAACAAGAANNNAGATCGGA |
| 51529_2 | TATGAGTGTGGAGTCGTTGCGGGATCTGGGAAAGGAACAAGAANNNAGATCGGA |
| 44974_1 | TATGAGTGTGGAGTCGTTGCGTATTGGGGGGAGTGGTCATTAGNNNAGATCGGA |
| 44974_2 | TATGAGTGTGGAGTCGTTGCGTATTGGGGGGAGTGGTCATTAGNNNAGATCGGA |
| 38977_1 | TATGAGTGTGGAGTCGTTGCAGGAAGGAAGAAGGAAAGAAGGANNNAGATCGG |
| 38977_2 | TATGAGTGTGGAGTCGTTGCAGGAAGGAAGAAGGAAAGAAGGANNNAGATCGG |
| 47826_1 | TATGAGTGTGGAGTCGTTGCGGAGTCTCACTCTGTAACTCAGGNNNAGATCGGAAG |
| 47826_2 | TATGAGTGTGGAGTCGTTGCGGAGTCTCACTCTGTAACTCAGGNNNAGATCGGAAG |
| 46763_1 | TATGAGTGTGGAGTCGTTGCCCTTCGAAGTAGCTGGGATTACANNNAGATCGGAAG |
| 46763_2 | TATGAGTGTGGAGTCGTTGCCCTTCGAAGTAGCTGGGATTACANNNAGATCGGAAG |
| 51651_1 | TATGAGTGTGGAGTCGTTGCACAGCTTCAACCCCCTAAGATTTNNNAGATCGGAAG |
| 51651_2 | TATGAGTGTGGAGTCGTTGCACAGCTTCAACCCCCTAAGATTTNNNAGATCGGAAG |
| 33567_1 | TATGAGTGTGGAGTCGTTGCCTCCCGAGTTCAAGTGATTCTCANNNAGATCGGAAG |
| 33567_2 | TATGAGTGTGGAGTCGTTGCCTCCCGAGTTCAAGTGATTCTCANNNAGATCGGAAG |
| 29351_1 | TATGAGTGTGGAGTCGTTGCAGAGAAGCGTGGTCAATAGTGTTNNNAGATCGGAA |
| 29351_2 | TATGAGTGTGGAGTCGTTGCAGAGAAGCGTGGTCAATAGTGTTNNNAGATCGGAA |
| 44810_1 | TATGAGTGTGGAGTCGTTGCTGAAAGGAGAGGTCAACTGGTTCNNNAGATCGGAA |
| 44810_2 | TATGAGTGTGGAGTCGTTGCTGAAAGGAGAGGTCAACTGGTTCNNNAGATCGGAA |
| 13044_1 | TATGAGTGTGGAGTCGTTGCCTAATGGTACAACTGCTGAAANNNAGATCGGAAGAG |
| 13044_2 | TATGAGTGTGGAGTCGTTGCCTAATGGTACAACTGCTGAAANNNAGATCGGAAGAG |
| 51602_1 | TATGAGTGTGGAGTCGTTGCGCTGGGAAAAGAGAAGAGTCAGANNNAGATCGGA |
| 51602_2 | TATGAGTGTGGAGTCGTTGCGCTGGGAAAAGAGAAGAGTCAGANNNAGATCGGA |
| 35810_1 | TATGAGTGTGGAGTCGTTGCATTATCAGGTCACCACAACCACCNNNAGATCGGAAG |
| 35810_2 | TATGAGTGTGGAGTCGTTGCATTATCAGGTCACCACAACCACCNNNAGATCGGAAG |
| 34963_1 | TATGAGTGTGGAGTCGTTGCCTGCCGGTGAGTTTTCATGAATTNNNAGATCGGAAG |
| 34963_2 | TATGAGTGTGGAGTCGTTGCCTGCCGGTGAGTTTTCATGAATTNNNAGATCGGAAG |
| 46359_1 | TATGAGTGTGGAGTCGTTGCGCCTCCCAAGTAGCTAGGATTACNNNAGATCGGAAG |
| 46359_2 | TATGAGTGTGGAGTCGTTGCGCCTCCCAAGTAGCTAGGATTACNNNAGATCGGAAG |
| 33642_1 | TATGAGTGTGGAGTCGTTGCGGTATAGTGGCATGTTCTCAGCTNNNAGATCGGAAG |
| 33642_2 | TATGAGTGTGGAGTCGTTGCGGTATAGTGGCATGTTCTCAGCTNNNAGATCGGAAG |
| 2365_1 | TATGAGTGTGGAGTCGTTGCAAACTGACGAGGTCGCTATAAGCNNNAGATCGGAAG |
| 2365_2 | TATGAGTGTGGAGTCGTTGCAAACTGACGAGGTCGCTATAAGCNNNAGATCGGAAG |

| | |
|---|---|
| 44835_1 | TATGAGTGTGGAGTCGTTGCATTCAGCGACGGATGAGTAAAGTNNNAGATCGGAA |
| 44835_2 | TATGAGTGTGGAGTCGTTGCATTCAGCGACGGATGAGTAAAGTNNNAGATCGGAA |
| 35310_1 | TATGAGTGTGGAGTCGTTGCGGAGTACAGTGGTGTGATCTCAGNNNAGATCGGAA |
| 35310_2 | TATGAGTGTGGAGTCGTTGCGGAGTACAGTGGTGTGATCTCAGNNNAGATCGGAA |
| 35221_1 | TATGAGTGTGGAGTCGTTGCTGTGTGATAGATCCAGCTCTGTGNNNAGATCGGAA |
| 35221_2 | TATGAGTGTGGAGTCGTTGCTGTGTGATAGATCCAGCTCTGTGNNNAGATCGGAA |
| 41855_1 | TATGAGTGTGGAGTCGTTGCGATATCAGCTCACTGCAACCTCTNNNAGATCGGAAG |
| 41855_2 | TATGAGTGTGGAGTCGTTGCGATATCAGCTCACTGCAACCTCTNNNAGATCGGAAG |
| 12809_1 | TATGAGTGTGGAGTCGTTGCTTGAAAATTGTTGCATTTGTAAANNNAGATCGGAAG |
| 12809_2 | TATGAGTGTGGAGTCGTTGCTTGAAAATTGTTGCATTTGTAAANNNAGATCGGAAG |
| 31880_1 | TATGAGTGTGGAGTCGTTGCATACATCTATGACCTCACCCCCANNNAGATCGGAAGA |
| 31880_2 | TATGAGTGTGGAGTCGTTGCATACATCTATGACCTCACCCCCANNNAGATCGGAAGA |
| 32487_1 | TATGAGTGTGGAGTCGTTGCGGAGTACAGTGGCGCAATCTCNNNAGATCGGAAGA |
| 32487_2 | TATGAGTGTGGAGTCGTTGCGGAGTACAGTGGCGCAATCTCNNNAGATCGGAAGA |
| 38248_1 | TATGAGTGTGGAGTCGTTGCTCACTGTCCATCAGAGAAATGCANNNAGATCGGAAG |
| 38248_2 | TATGAGTGTGGAGTCGTTGCTCACTGTCCATCAGAGAAATGCANNNAGATCGGAAG |
| 29493_1 | TATGAGTGTGGAGTCGTTGCTTCCCACTATCCTCCTTTCTGCANNNAGATCGGAAGA |
| 29493_2 | TATGAGTGTGGAGTCGTTGCTTCCCACTATCCTCCTTTCTGCANNNAGATCGGAAGA |
| 16576_1 | TATGAGTGTGGAGTCGTTGCTCCTATAACACTACCCTAGTTATNNNAGATCGGAAGA |
| 16576_2 | TATGAGTGTGGAGTCGTTGCTCCTATAACACTACCCTAGTTATNNNAGATCGGAAGA |
| 51156_1 | TATGAGTGTGGAGTCGTTGCACCATGTTTGTGAAGTAGGGGTTNNNAGATCGGAA |
| 51156_2 | TATGAGTGTGGAGTCGTTGCACCATGTTTGTGAAGTAGGGGTTNNNAGATCGGAA |
| 33436_1 | TATGAGTGTGGAGTCGTTGCGCCTTCTGGTAGAAAATGCACTCNNNAGATCGGAAG |
| 33436_2 | TATGAGTGTGGAGTCGTTGCGCCTTCTGGTAGAAAATGCACTCNNNAGATCGGAAG |
| 10376_1 | TATGAGTGTGGAGTCGTTGCTAGTTGGCAGTAGTTCCCTCACANNNAGATCGGAAG |
| 10376_2 | TATGAGTGTGGAGTCGTTGCTAGTTGGCAGTAGTTCCCTCACANNNAGATCGGAAG |
| 36988_1 | TATGAGTGTGGAGTCGTTGCAAGTGGATTTTCCCGTAGTCTCCNNNAGATCGGAAG |
| 36988_2 | TATGAGTGTGGAGTCGTTGCAAGTGGATTTTCCCGTAGTCTCCNNNAGATCGGAAG |
| 42772_1 | TATGAGTGTGGAGTCGTTGCAGGGATGCATATTTGAGTGACAANNNAGATCGGAA |
| 42772_2 | TATGAGTGTGGAGTCGTTGCAGGGATGCATATTTGAGTGACAANNNAGATCGGAA |
| 35311_1 | TATGAGTGTGGAGTCGTTGCTTCAAGTGATTCTCCTGCCTCAGNNNAGATCGGAAG |
| 35311_2 | TATGAGTGTGGAGTCGTTGCTTCAAGTGATTCTCCTGCCTCAGNNNAGATCGGAAG |
| 47165_1 | TATGAGTGTGGAGTCGTTGCGAGTAGCTGGGACTACAGGCANNNAGATCGGAAGA |
| 47165_2 | TATGAGTGTGGAGTCGTTGCGAGTAGCTGGGACTACAGGCANNNAGATCGGAAGA |
| 13787_1 | TATGAGTGTGGAGTCGTTGCACGATTAAGACAAGGAGGATNNNAGATCGGAAGA |
| 13787_2 | TATGAGTGTGGAGTCGTTGCACGATTAAGACAAGGAGGATNNNAGATCGGAAGA |
| 36352_1 | TATGAGTGTGGAGTCGTTGCGTGTGTGTGTGTGTGTGTATCNNNAGATCGGAA |
| 36352_2 | TATGAGTGTGGAGTCGTTGCGTGTGTGTGTGTGTGTGTATCNNNAGATCGGAA |
| 49723_1 | TATGAGTGTGGAGTCGTTGCGATTATCCTGTCTCAGCCTCCTGNNNAGATCGGAAG |
| 49723_2 | TATGAGTGTGGAGTCGTTGCGATTATCCTGTCTCAGCCTCCTGNNNAGATCGGAAG |
| 41418_1 | TATGAGTGTGGAGTCGTTGCGACTCTGAGGGAAGAACATGGAGNNNAGATCGGA |
| 41418_2 | TATGAGTGTGGAGTCGTTGCGACTCTGAGGGAAGAACATGGAGNNNAGATCGGA |
| 44876_1 | TATGAGTGTGGAGTCGTTGCAAAAAGCCGGAGACAAAATGAGGNNNAGATCGGA |

| | |
|---|---|
| 44876_2 | TATGAGTGTGGAGTCGTTGCAAAAAGCCGGAGACAAAATGAGGNNNAGATCGGA |
| 34627_1 | TATGAGTGTGGAGTCGTTGCAGCATTTTCAGTCTTGTTTGGGGNNNAGATCGGAAG |
| 34627_2 | TATGAGTGTGGAGTCGTTGCAGCATTTTCAGTCTTGTTTGGGGNNNAGATCGGAAG |
| 39014_1 | TATGAGTGTGGAGTCGTTGCGCCTGTGCTCTATACATGGAACANNNAGATCGGAAG |
| 39014_2 | TATGAGTGTGGAGTCGTTGCGCCTGTGCTCTATACATGGAACANNNAGATCGGAAG |
| 13669_1 | TATGAGTGTGGAGTCGTTGCATAGAGTGTAGATGCATCAAAATNNNAGATCGGAA |
| 13669_2 | TATGAGTGTGGAGTCGTTGCATAGAGTGTAGATGCATCAAAATNNNAGATCGGAA |
| 49749_1 | TATGAGTGTGGAGTCGTTGCTAGTCTTGACCTCCTGACCTCAANNNAGATCGGAAG |
| 49749_2 | TATGAGTGTGGAGTCGTTGCTAGTCTTGACCTCCTGACCTCAANNNAGATCGGAAG |
| 33117_1 | TATGAGTGTGGAGTCGTTGCGAATGTACCAGACAGGACACTCANNNAGATCGGAA |
| 33117_2 | TATGAGTGTGGAGTCGTTGCGAATGTACCAGACAGGACACTCANNNAGATCGGAA |
| 29606_1 | TATGAGTGTGGAGTCGTTGCTAGCATTGGATACAGAGCAGGACNNNAGATCGGAA |
| 29606_2 | TATGAGTGTGGAGTCGTTGCTAGCATTGGATACAGAGCAGGACNNNAGATCGGAA |
| 35356_1 | TATGAGTGTGGAGTCGTTGCCTCCATTACGGTTGAGGTGATCCNNNAGATCGGAAG |
| 35356_2 | TATGAGTGTGGAGTCGTTGCCTCCATTACGGTTGAGGTGATCCNNNAGATCGGAAG |
| 38183_1 | TATGAGTGTGGAGTCGTTGCACAAAGTGGGGTTTATCACAGGANNNAGATCGGAA |
| 38183_2 | TATGAGTGTGGAGTCGTTGCACAAAGTGGGGTTTATCACAGGANNNAGATCGGAA |
| 34439_1 | TATGAGTGTGGAGTCGTTGCCCAATTCAATGCCCTTCATCCAANNNAGATCGGAAGA |
| 34439_2 | TATGAGTGTGGAGTCGTTGCCCAATTCAATGCCCTTCATCCAANNNAGATCGGAAGA |
| 46259_1 | TATGAGTGTGGAGTCGTTGCTCCCATGGGAACCCTTGATTTTANNNAGATCGGAAG |
| 46259_2 | TATGAGTGTGGAGTCGTTGCTCCCATGGGAACCCTTGATTTTANNNAGATCGGAAG |
| 9098_1 | TATGAGTGTGGAGTCGTTGCGTGTGTGTGTCTGTACTAAGCACTNNNAGATCGGAA |
| 9098_2 | TATGAGTGTGGAGTCGTTGCGTGTGTGTGTCTGTACTAAGCACTNNNAGATCGGAA |
| 39701_1 | TATGAGTGTGGAGTCGTTGCGAAAGTAACCACAGACATGCACANNNAGATCGGAA |
| 39701_2 | TATGAGTGTGGAGTCGTTGCGAAAGTAACCACAGACATGCACANNNAGATCGGAA |
| 52735_1 | TATGAGTGTGGAGTCGTTGCAACCTATGTTTCCAGTGGAGCATNNNAGATCGGAAG |
| 52735_2 | TATGAGTGTGGAGTCGTTGCAACCTATGTTTCCAGTGGAGCATNNNAGATCGGAAG |
| 40824_1 | TATGAGTGTGGAGTCGTTGCCATGCACATGTACACACACTTCTNNNAGATCGGAAG |
| 40824_2 | TATGAGTGTGGAGTCGTTGCCATGCACATGTACACACACTTCTNNNAGATCGGAAG |
| 8083_1 | TATGAGTGTGGAGTCGTTGCAGTTCTTCAATGGGTCTGGCTTTNNNAGATCGGAAG |
| 8083_2 | TATGAGTGTGGAGTCGTTGCAGTTCTTCAATGGGTCTGGCTTTNNNAGATCGGAAG |
| 13316_1 | TATGAGTGTGGAGTCGTTGCCAGAAGAATCTTCCTCTTTTTCNNNAGATCGGAAGA |
| 13316_2 | TATGAGTGTGGAGTCGTTGCCAGAAGAATCTTCCTCTTTTTCNNNAGATCGGAAGA |
| 48403_1 | TATGAGTGTGGAGTCGTTGCGGTTCACCCCCATTCTCCTGNNNAGATCGGAAGAGC |
| 48403_2 | TATGAGTGTGGAGTCGTTGCGGTTCACCCCCATTCTCCTGNNNAGATCGGAAGAGC |
| 31228_1 | TATGAGTGTGGAGTCGTTGCAAAGGAGAGTGAGAGAAAGAGCGNNNAGATCGGA |
| 31228_2 | TATGAGTGTGGAGTCGTTGCAAAGGAGAGTGAGAGAAAGAGCGNNNAGATCGGA |
| 50198_1 | TATGAGTGTGGAGTCGTTGCGTGTGGAGATGTGATAGGAAGGGNNNAGATCGGA |
| 50198_2 | TATGAGTGTGGAGTCGTTGCGTGTGGAGATGTGATAGGAAGGGNNNAGATCGGA |
| 13457_1 | TATGAGTGTGGAGTCGTTGCGATTTCTGAGGCACATAGTCNNNAGATCGGAAGAG |
| 13457_2 | TATGAGTGTGGAGTCGTTGCGATTTCTGAGGCACATAGTCNNNAGATCGGAAGAG |
| 38814_1 | TATGAGTGTGGAGTCGTTGCTTCCTTCTGCTTCTTTCTCTCCTNNNAGATCGGAAGAG |
| 38814_2 | TATGAGTGTGGAGTCGTTGCTTCCTTCTGCTTCTTTCTCTCCTNNNAGATCGGAAGAG |

| | |
|---|---|
| 45747_1 | TATGAGTGTGGAGTCGTTGCTGGGATTGCTGGATCAAATGGTANNNAGATCGGAA |
| 45747_2 | TATGAGTGTGGAGTCGTTGCTGGGATTGCTGGATCAAATGGTANNNAGATCGGAA |
| 44629_1 | TATGAGTGTGGAGTCGTTGCTCTCAGTCTCTTTCAAGCACCTCNNNAGATCGGAAGA |
| 44629_2 | TATGAGTGTGGAGTCGTTGCTCTCAGTCTCTTTCAAGCACCTCNNNAGATCGGAAGA |
| 31387_1 | TATGAGTGTGGAGTCGTTGCAAGGGCTGCTGATGATTTTTCACNNNAGATCGGAAG |
| 31387_2 | TATGAGTGTGGAGTCGTTGCAAGGGCTGCTGATGATTTTTCACNNNAGATCGGAAG |
| 37502_1 | TATGAGTGTGGAGTCGTTGCTCTCACTCTATCTCTCAGGCTGGNNNAGATCGGAAG |
| 37502_2 | TATGAGTGTGGAGTCGTTGCTCTCACTCTATCTCTCAGGCTGGNNNAGATCGGAAG |
| 30418_1 | TATGAGTGTGGAGTCGTTGCTTTTTTCACTGAGGTATGAATTGNNNAGATCGGAAG |
| 30418_2 | TATGAGTGTGGAGTCGTTGCTTTTTTCACTGAGGTATGAATTGNNNAGATCGGAAG |
| 49917_1 | TATGAGTGTGGAGTCGTTGCCAGACGCGCCTCTAGCATAGNNNAGATCGGAAGAG |
| 49917_2 | TATGAGTGTGGAGTCGTTGCCAGACGCGCCTCTAGCATAGNNNAGATCGGAAGAG |
| 28883_1 | TATGAGTGTGGAGTCGTTGCGCAAGTTAGCCACAGAATTAGGGNNNAGATCGGAA |
| 28883_2 | TATGAGTGTGGAGTCGTTGCGCAAGTTAGCCACAGAATTAGGGNNNAGATCGGAA |
| 33242_1 | TATGAGTGTGGAGTCGTTGCTCTATCACCTAGACGAGAGTGCANNNAGATCGGAAG |
| 33242_2 | TATGAGTGTGGAGTCGTTGCTCTATCACCTAGACGAGAGTGCANNNAGATCGGAAG |
| 50834_1 | TATGAGTGTGGAGTCGTTGCATTTCCAGCTCAAGTCCTGTCTTNNNAGATCGGAAG |
| 50834_2 | TATGAGTGTGGAGTCGTTGCATTTCCAGCTCAAGTCCTGTCTTNNNAGATCGGAAG |
| 43183_1 | TATGAGTGTGGAGTCGTTGCGCAGAGCAGCTTCATTTTTCAGANNNAGATCGGAAG |
| 43183_2 | TATGAGTGTGGAGTCGTTGCGCAGAGCAGCTTCATTTTTCAGANNNAGATCGGAAG |
| 29311_1 | TATGAGTGTGGAGTCGTTGCTTGATTGAGATCTCCAGCCAACANNNAGATCGGAAG |
| 29311_2 | TATGAGTGTGGAGTCGTTGCTTGATTGAGATCTCCAGCCAACANNNAGATCGGAAG |
| 36400_1 | TATGAGTGTGGAGTCGTTGCGTGCTCTCTTGAGGTCACTGTTANNNAGATCGGAAG |
| 36400_2 | TATGAGTGTGGAGTCGTTGCGTGCTCTCTTGAGGTCACTGTTANNNAGATCGGAAG |
| 45180_1 | TATGAGTGTGGAGTCGTTGCCACAGCCGACTAGCTCTAAGAGNNNAGATCGGAAG |
| 45180_2 | TATGAGTGTGGAGTCGTTGCCACAGCCGACTAGCTCTAAGAGNNNAGATCGGAAG |
| 30434_1 | TATGAGTGTGGAGTCGTTGCGGACTGTAGCGGATGGTGATAAANNNAGATCGGA |
| 30434_2 | TATGAGTGTGGAGTCGTTGCGGACTGTAGCGGATGGTGATAAANNNAGATCGGA |
| 51849_1 | TATGAGTGTGGAGTCGTTGCCTTCTCTCGGCTTGTCTCTTTCTNNNAGATCGGAAGA |
| 51849_2 | TATGAGTGTGGAGTCGTTGCCTTCTCTCGGCTTGTCTCTTTCTNNNAGATCGGAAGA |
| 30646_1 | TATGAGTGTGGAGTCGTTGCGGCATGAGTGTGCAGATCAGAGNNNAGATCGGAA |
| 30646_2 | TATGAGTGTGGAGTCGTTGCGGCATGAGTGTGCAGATCAGAGNNNAGATCGGAA |
| 47452_1 | TATGAGTGTGGAGTCGTTGCCTGGTCTTGAACTCCTGACTTCANNNAGATCGGAAG |
| 47452_2 | TATGAGTGTGGAGTCGTTGCCTGGTCTTGAACTCCTGACTTCANNNAGATCGGAAG |
| 42116_1 | TATGAGTGTGGAGTCGTTGCCCTCCTGAGTAGCTGGGACTATANNNAGATCGGAAG |
| 42116_2 | TATGAGTGTGGAGTCGTTGCCCTCCTGAGTAGCTGGGACTATANNNAGATCGGAAG |
| 13240_1 | TATGAGTGTGGAGTCGTTGCCACCTACTAGTTGTTTATTACCTNNNAGATCGGAAGA |
| 13240_2 | TATGAGTGTGGAGTCGTTGCCACCTACTAGTTGTTTATTACCTNNNAGATCGGAAGA |
| 36063_1 | TATGAGTGTGGAGTCGTTGCTTGCTTACAGGAACAAGGTGGATNNNAGATCGGAA |
| 36063_2 | TATGAGTGTGGAGTCGTTGCTTGCTTACAGGAACAAGGTGGATNNNAGATCGGAA |
| 50968_1 | TATGAGTGTGGAGTCGTTGCCATCTGTCCTCTTTCCCCTTTGANNNAGATCGGAAGA |
| 50968_2 | TATGAGTGTGGAGTCGTTGCCATCTGTCCTCTTTCCCCTTTGANNNAGATCGGAAGA |
| 49626_1 | TATGAGTGTGGAGTCGTTGCCATCCATCCATCCATCCATCCATNNNAGATCGGAAGA |

| | |
|---|---|
| 49626_2 | TATGAGTGTGGAGTCGTTGCCATCCATCCATCCATCCATCCATNNNAGATCGGAAGA |
| 46032_1 | TATGAGTGTGGAGTCGTTGCATACAAGGAGGGTTCACTGCAATNNNAGATCGGAA |
| 46032_2 | TATGAGTGTGGAGTCGTTGCATACAAGGAGGGTTCACTGCAATNNNAGATCGGAA |
| 30472_1 | TATGAGTGTGGAGTCGTTGCTGTAACCCTGATTTGACTTTGGTNNNAGATCGGAAG |
| 30472_2 | TATGAGTGTGGAGTCGTTGCTGTAACCCTGATTTGACTTTGGTNNNAGATCGGAAG |
| 45968_1 | TATGAGTGTGGAGTCGTTGCGGGTTCAAGTGATTCTCCTGTCTNNNAGATCGGAAG |
| 45968_2 | TATGAGTGTGGAGTCGTTGCGGGTTCAAGTGATTCTCCTGTCTNNNAGATCGGAAG |
| 29280_1 | TATGAGTGTGGAGTCGTTGCGTACTCACTCTTGCTGTAGTCCCNNNAGATCGGAAG |
| 29280_2 | TATGAGTGTGGAGTCGTTGCGTACTCACTCTTGCTGTAGTCCCNNNAGATCGGAAG |
| 47794_1 | TATGAGTGTGGAGTCGTTGCCAAGTAGCTGGAACTACAGGTGTNNNAGATCGGAA |
| 47794_2 | TATGAGTGTGGAGTCGTTGCCAAGTAGCTGGAACTACAGGTGTNNNAGATCGGAA |
| 34654_1 | TATGAGTGTGGAGTCGTTGCGCCTTCCAGACATGTCTATTGAGNNNAGATCGGAAG |
| 34654_2 | TATGAGTGTGGAGTCGTTGCGCCTTCCAGACATGTCTATTGAGNNNAGATCGGAAG |
| 44819_1 | TATGAGTGTGGAGTCGTTGCCATGAGGTTCCAGCACTCTCTCNNNAGATCGGAAGA |
| 44819_2 | TATGAGTGTGGAGTCGTTGCCATGAGGTTCCAGCACTCTCTCNNNAGATCGGAAGA |
| 43169_1 | TATGAGTGTGGAGTCGTTGCTCCAGGTTCAAGTGATTCTCCAGNNNAGATCGGAAG |
| 43169_2 | TATGAGTGTGGAGTCGTTGCTCCAGGTTCAAGTGATTCTCCAGNNNAGATCGGAAG |
| 12535_1 | TATGAGTGTGGAGTCGTTGCTGGCTTATTTAGTATCATGTATGNNNAGATCGGAAG |
| 12535_2 | TATGAGTGTGGAGTCGTTGCTGGCTTATTTAGTATCATGTATGNNNAGATCGGAAG |
| 34557_1 | TATGAGTGTGGAGTCGTTGCTCTGGGGATGGTCCTAAGAGGNNNAGATCGGAAGA |
| 34557_2 | TATGAGTGTGGAGTCGTTGCTCTGGGGATGGTCCTAAGAGGNNNAGATCGGAAGA |
| 33665_1 | TATGAGTGTGGAGTCGTTGCATGCTGGGATTACAGGTATGAGGNNNAGATCGGAA |
| 33665_2 | TATGAGTGTGGAGTCGTTGCATGCTGGGATTACAGGTATGAGGNNNAGATCGGAA |
| 50067_1 | TATGAGTGTGGAGTCGTTGCGTTTTTCTTCGAGACGTGGGAAGNNNAGATCGGAA |
| 50067_2 | TATGAGTGTGGAGTCGTTGCGTTTTTCTTCGAGACGTGGGAAGNNNAGATCGGAA |
| 1501_1 | TATGAGTGTGGAGTCGTTGCCTGGTACGTGTGAACCTGTTCATNNNAGATCGGAAG |
| 1501_2 | TATGAGTGTGGAGTCGTTGCCTGGTACGTGTGAACCTGTTCATNNNAGATCGGAAG |
| 28860_1 | TATGAGTGTGGAGTCGTTGCTAGACTCAGTGTGAAAGGGGCTANNNAGATCGGAA |
| 28860_2 | TATGAGTGTGGAGTCGTTGCTAGACTCAGTGTGAAAGGGGCTANNNAGATCGGAA |
| 32326_1 | TATGAGTGTGGAGTCGTTGCGGGTTCAAGTGATTCTCCTGTCTNNNAGATCGGAAG |
| 32326_2 | TATGAGTGTGGAGTCGTTGCGGGTTCAAGTGATTCTCCTGTCTNNNAGATCGGAAG |
| 37998_1 | TATGAGTGTGGAGTCGTTGCATTCCAAAGGCCCTAGAGAAGTGNNNAGATCGGAA |
| 37998_2 | TATGAGTGTGGAGTCGTTGCATTCCAAAGGCCCTAGAGAAGTGNNNAGATCGGAA |
| 33583_1 | TATGAGTGTGGAGTCGTTGCACTACACCATGCCCAGCTAATTTNNNAGATCGGAAG |
| 33583_2 | TATGAGTGTGGAGTCGTTGCACTACACCATGCCCAGCTAATTTNNNAGATCGGAAG |
| 39595_1 | TATGAGTGTGGAGTCGTTGCTGGACAGTAACAAGTGTTGGTGANNNAGATCGGAA |
| 39595_2 | TATGAGTGTGGAGTCGTTGCTGGACAGTAACAAGTGTTGGTGANNNAGATCGGAA |
| 50870_1 | TATGAGTGTGGAGTCGTTGCACCTGGATTCTCAAGAATGGTCTNNNAGATCGGAAG |
| 50870_2 | TATGAGTGTGGAGTCGTTGCACCTGGATTCTCAAGAATGGTCTNNNAGATCGGAAG |
| 29354_1 | TATGAGTGTGGAGTCGTTGCATAACTGAGCTGGGTCAGACTTGNNNAGATCGGAA |
| 29354_2 | TATGAGTGTGGAGTCGTTGCATAACTGAGCTGGGTCAGACTTGNNNAGATCGGAA |
| 45066_1 | TATGAGTGTGGAGTCGTTGCCAAGAAGTTACCCTGGAGTCTGGNNNAGATCGGAA |
| 45066_2 | TATGAGTGTGGAGTCGTTGCCAAGAAGTTACCCTGGAGTCTGGNNNAGATCGGAA |

| | |
|---|---|
| 51278_1 | TATGAGTGTGGAGTCGTTGCCAGTTCTCCGCATTCTGTTTTGTNNNAGATCGGAAG |
| 51278_2 | TATGAGTGTGGAGTCGTTGCCAGTTCTCCGCATTCTGTTTTGTNNNAGATCGGAAG |
| 41194_1 | TATGAGTGTGGAGTCGTTGCGCACCCCTAAAGATAGTTCCCAANNNAGATCGGAAG |
| 41194_2 | TATGAGTGTGGAGTCGTTGCGCACCCCTAAAGATAGTTCCCAANNNAGATCGGAAG |
| 40021_1 | TATGAGTGTGGAGTCGTTGCCACAATCTTGGCTCACTGCAACNNNAGATCGGAAGA |
| 40021_2 | TATGAGTGTGGAGTCGTTGCCACAATCTTGGCTCACTGCAACNNNAGATCGGAAGA |
| 51826_1 | TATGAGTGTGGAGTCGTTGCTCTACATCCCTGTAGCCCTGTTANNNAGATCGGAAG |
| 51826_2 | TATGAGTGTGGAGTCGTTGCTCTACATCCCTGTAGCCCTGTTANNNAGATCGGAAG |
| 38798_1 | TATGAGTGTGGAGTCGTTGCAAGGAAGAATGTTTGGGGGATGANNNAGATCGGA |
| 38798_2 | TATGAGTGTGGAGTCGTTGCAAGGAAGAATGTTTGGGGGATGANNNAGATCGGA |
| 39103_1 | TATGAGTGTGGAGTCGTTGCATCCCAGTGATACCTGATGTCATNNNAGATCGGAAG |
| 39103_2 | TATGAGTGTGGAGTCGTTGCATCCCAGTGATACCTGATGTCATNNNAGATCGGAAG |
| 51679_1 | TATGAGTGTGGAGTCGTTGCCTTCCTGTCTCTTGGGAGTTTCCNNNAGATCGGAAG |
| 51679_2 | TATGAGTGTGGAGTCGTTGCCTTCCTGTCTCTTGGGAGTTTCCNNNAGATCGGAAG |
| 29975_1 | TATGAGTGTGGAGTCGTTGCTGTTTGTGTTGATAATGAAGGGACANNNAGATCGG |
| 29975_2 | TATGAGTGTGGAGTCGTTGCTGTTTGTGTTGATAATGAAGGGACANNNAGATCGG |
| 2756_1 | TATGAGTGTGGAGTCGTTGCAGAACCCTGCACTGAATATCCCTNNNAGATCGGAAG |
| 2756_2 | TATGAGTGTGGAGTCGTTGCAGAACCCTGCACTGAATATCCCTNNNAGATCGGAAG |
| 45216_1 | TATGAGTGTGGAGTCGTTGCTGCAGGGACACTGTAGTAACAAGNNNAGATCGGAA |
| 45216_2 | TATGAGTGTGGAGTCGTTGCTGCAGGGACACTGTAGTAACAAGNNNAGATCGGAA |
| 31918_1 | TATGAGTGTGGAGTCGTTGCTCACATTTTCACTACCTGAAGAACANNNAGATCGGA |
| 31918_2 | TATGAGTGTGGAGTCGTTGCTCACATTTTCACTACCTGAAGAACANNNAGATCGGA |
| 50336_1 | TATGAGTGTGGAGTCGTTGCGTAAGCCTTTCAAATGTGTGGCTNNNAGATCGGAAG |
| 50336_2 | TATGAGTGTGGAGTCGTTGCGTAAGCCTTTCAAATGTGTGGCTNNNAGATCGGAAG |
| 37001_1 | TATGAGTGTGGAGTCGTTGCAGAAAGCTGGAGGCATCACATTANNNAGATCGGAA |
| 37001_2 | TATGAGTGTGGAGTCGTTGCAGAAAGCTGGAGGCATCACATTANNNAGATCGGAA |
| 30639_1 | TATGAGTGTGGAGTCGTTGCCTAACCTGCCAGTTCTTTCCAGNNNAGATCGGAAGA |
| 30639_2 | TATGAGTGTGGAGTCGTTGCCTAACCTGCCAGTTCTTTCCAGNNNAGATCGGAAGA |
| 47692_1 | TATGAGTGTGGAGTCGTTGCCCTTCACACTCCAGCTCTTTCTCNNNAGATCGGAAGA |
| 47692_2 | TATGAGTGTGGAGTCGTTGCCCTTCACACTCCAGCTCTTTCTCNNNAGATCGGAAGA |
| 49372_1 | TATGAGTGTGGAGTCGTTGCCCTCCACCCCCTTACCTGTTATANNNAGATCGGAAGA |
| 49372_2 | TATGAGTGTGGAGTCGTTGCCCTCCACCCCCTTACCTGTTATANNNAGATCGGAAGA |
| 41868_1 | TATGAGTGTGGAGTCGTTGCTTCAAGTGATTCTCCTGCCTCAGNNNAGATCGGAAG |
| 41868_2 | TATGAGTGTGGAGTCGTTGCTTCAAGTGATTCTCCTGCCTCAGNNNAGATCGGAAG |
| 29561_1 | TATGAGTGTGGAGTCGTTGCAGCCCTAGGTATCTTCCAGAAGANNNAGATCGGAAG |
| 29561_2 | TATGAGTGTGGAGTCGTTGCAGCCCTAGGTATCTTCCAGAAGANNNAGATCGGAAG |
| 42645_1 | TATGAGTGTGGAGTCGTTGCTTTCCCAGGCTAGTCTCGAATTCNNNAGATCGGAAG |
| 42645_2 | TATGAGTGTGGAGTCGTTGCTTTCCCAGGCTAGTCTCGAATTCNNNAGATCGGAAG |
| 33069_1 | TATGAGTGTGGAGTCGTTGCAATCTCTCCTCACTGCAAGCTCNNNAGATCGGAAGA |
| 33069_2 | TATGAGTGTGGAGTCGTTGCAATCTCTCCTCACTGCAAGCTCNNNAGATCGGAAGA |
| 34183_1 | TATGAGTGTGGAGTCGTTGCTCTCATCATTCAGCCTCAATGCTNNNAGATCGGAAGA |
| 34183_2 | TATGAGTGTGGAGTCGTTGCTCTCATCATTCAGCCTCAATGCTNNNAGATCGGAAGA |
| 50729_1 | TATGAGTGTGGAGTCGTTGCCCCCATCCTCACTATCTTCACTCNNNAGATCGGAAGA |

| | |
|---|---|
| 50729_2 | TATGAGTGTGGAGTCGTTGCCCCCATCCTCACTATCTTCACTCNNNAGATCGGAAGA |
| 41495_1 | TATGAGTGTGGAGTCGTTGCTACTCAATTCCTTGGTCTGCACANNNAGATCGGAAG |
| 41495_2 | TATGAGTGTGGAGTCGTTGCTACTCAATTCCTTGGTCTGCACANNNAGATCGGAAG |
| 40340_1 | TATGAGTGTGGAGTCGTTGCGTAGCATATTGGAAGCAGACGTCNNNAGATCGGAA |
| 40340_2 | TATGAGTGTGGAGTCGTTGCGTAGCATATTGGAAGCAGACGTCNNNAGATCGGAA |
| 31786_1 | TATGAGTGTGGAGTCGTTGCTCTACTTTGTTCCAGGCTTGACANNNAGATCGGAAG |
| 31786_2 | TATGAGTGTGGAGTCGTTGCTCTACTTTGTTCCAGGCTTGACANNNAGATCGGAAG |
| 44868_1 | TATGAGTGTGGAGTCGTTGCTAGAGGGGGTGAAACAAGAGTTGNNNAGATCGGA |
| 44868_2 | TATGAGTGTGGAGTCGTTGCTAGAGGGGGTGAAACAAGAGTTGNNNAGATCGGA |
| 32029_1 | TATGAGTGTGGAGTCGTTGCCCTCTCAAAGTGCTGGGATTACANNNAGATCGGAAG |
| 32029_2 | TATGAGTGTGGAGTCGTTGCCCTCTCAAAGTGCTGGGATTACANNNAGATCGGAAG |
| 33037_1 | TATGAGTGTGGAGTCGTTGCGAGTAGCTGGGACTACAGGCANNNAGATCGGAAGA |
| 33037_2 | TATGAGTGTGGAGTCGTTGCGAGTAGCTGGGACTACAGGCANNNAGATCGGAAGA |
| 46535_1 | TATGAGTGTGGAGTCGTTGCAATCTCCTGACCTTGTGATCCACNNNAGATCGGAAG |
| 46535_2 | TATGAGTGTGGAGTCGTTGCAATCTCCTGACCTTGTGATCCACNNNAGATCGGAAG |
| 42430_1 | TATGAGTGTGGAGTCGTTGCACTGGATCATGATGGCAGTTTCTNNNAGATCGGAAG |
| 42430_2 | TATGAGTGTGGAGTCGTTGCACTGGATCATGATGGCAGTTTCTNNNAGATCGGAAG |
| 34476_1 | TATGAGTGTGGAGTCGTTGCTCGCTGCTACATTCCTGTAAGTTNNNAGATCGGAAG |
| 34476_2 | TATGAGTGTGGAGTCGTTGCTCGCTGCTACATTCCTGTAAGTTNNNAGATCGGAAG |
| 38684_1 | TATGAGTGTGGAGTCGTTGCAGAGACAGAGAGAAGAAAGAAGCANNNAGATCGG |
| 38684_2 | TATGAGTGTGGAGTCGTTGCAGAGACAGAGAGAAGAAAGAAGCANNNAGATCGG |
| 29130_1 | TATGAGTGTGGAGTCGTTGCACATCTTTGAGGGTACACGCATANNNAGATCGGAAG |
| 29130_2 | TATGAGTGTGGAGTCGTTGCACATCTTTGAGGGTACACGCATANNNAGATCGGAAG |
| 52871_1 | TATGAGTGTGGAGTCGTTGCCAAGAAGTGTGCCTTTGTGCTGNNNAGATCGGAAG |
| 52871_2 | TATGAGTGTGGAGTCGTTGCCAAGAAGTGTGCCTTTGTGCTGNNNAGATCGGAAG |
| 39875_1 | TATGAGTGTGGAGTCGTTGCAGAATGCTTACCTACAGAACACACTNNNAGATCGGA |
| 39875_2 | TATGAGTGTGGAGTCGTTGCAGAATGCTTACCTACAGAACACACTNNNAGATCGGA |
| 33780_1 | TATGAGTGTGGAGTCGTTGCAAGGTCAGGAAATCGAGACCATCNNNAGATCGGAA |
| 33780_2 | TATGAGTGTGGAGTCGTTGCAAGGTCAGGAAATCGAGACCATCNNNAGATCGGAA |
| 44207_1 | TATGAGTGTGGAGTCGTTGCAGATGCTAGTGCTGGTAACAACANNNAGATCGGAA |
| 44207_2 | TATGAGTGTGGAGTCGTTGCAGATGCTAGTGCTGGTAACAACANNNAGATCGGAA |
| 36311_1 | TATGAGTGTGGAGTCGTTGCATTGCTTGAGCCCTAGAGTTCAANNNAGATCGGAAG |
| 36311_2 | TATGAGTGTGGAGTCGTTGCATTGCTTGAGCCCTAGAGTTCAANNNAGATCGGAAG |
| 37294_1 | TATGAGTGTGGAGTCGTTGCCCATCTGCCTGCTGAATTACTTTNNNAGATCGGAAGA |
| 37294_2 | TATGAGTGTGGAGTCGTTGCCCATCTGCCTGCTGAATTACTTTNNNAGATCGGAAGA |
| 48644_1 | TATGAGTGTGGAGTCGTTGCCACACCTGTAATCCCAGCACTATNNNAGATCGGAAG |
| 48644_2 | TATGAGTGTGGAGTCGTTGCCACACCTGTAATCCCAGCACTATNNNAGATCGGAAG |
| 43051_1 | TATGAGTGTGGAGTCGTTGCTGATATGAGCAGCCCATTCAGAANNNAGATCGGAA |
| 43051_2 | TATGAGTGTGGAGTCGTTGCTGATATGAGCAGCCCATTCAGAANNNAGATCGGAA |
| 37043_1 | TATGAGTGTGGAGTCGTTGCCCTCTTAGATTCTGTAGCTGGGCNNNAGATCGGAAG |
| 37043_2 | TATGAGTGTGGAGTCGTTGCCCTCTTAGATTCTGTAGCTGGGCNNNAGATCGGAAG |
| 39516_1 | TATGAGTGTGGAGTCGTTGCGCCATAGACTTCTTCACAGACCANNNAGATCGGAAG |
| 39516_2 | TATGAGTGTGGAGTCGTTGCGCCATAGACTTCTTCACAGACCANNNAGATCGGAAG |

| | |
|---|---|
| 30385_1 | TATGAGTGTGGAGTCGTTGCCCGGTATGTAATTTAACATTCAANNNAGATCGGAAG |
| 30385_2 | TATGAGTGTGGAGTCGTTGCCCGGTATGTAATTTAACATTCAANNNAGATCGGAAG |
| 39552_1 | TATGAGTGTGGAGTCGTTGCCCCAGGATCTTACAAGACTGAGGNNNAGATCGGAA |
| 39552_2 | TATGAGTGTGGAGTCGTTGCCCCAGGATCTTACAAGACTGAGGNNNAGATCGGAA |
| 40815_1 | TATGAGTGTGGAGTCGTTGCTAGAAGCGTTTCCCCAGATATGGNNNAGATCGGAA |
| 40815_2 | TATGAGTGTGGAGTCGTTGCTAGAAGCGTTTCCCCAGATATGGNNNAGATCGGAA |
| 35629_1 | TATGAGTGTGGAGTCGTTGCCAACAAGGCTGGGAGTATAACCTNNNAGATCGGAA |
| 35629_2 | TATGAGTGTGGAGTCGTTGCCAACAAGGCTGGGAGTATAACCTNNNAGATCGGAA |
| 49755_1 | TATGAGTGTGGAGTCGTTGCGAATTCAAGGAACAGATGCCTGGNNNAGATCGGAA |
| 49755_2 | TATGAGTGTGGAGTCGTTGCGAATTCAAGGAACAGATGCCTGGNNNAGATCGGAA |
| 46331_1 | TATGAGTGTGGAGTCGTTGCATTCACGTGAATCTCCTGCCTCNNNAGATCGGAAGA |
| 46331_2 | TATGAGTGTGGAGTCGTTGCATTCACGTGAATCTCCTGCCTCNNNAGATCGGAAGA |
| 34159_1 | TATGAGTGTGGAGTCGTTGCACAATTCTGGAGGCTGAGAAGTTNNNAGATCGGAA |
| 34159_2 | TATGAGTGTGGAGTCGTTGCACAATTCTGGAGGCTGAGAAGTTNNNAGATCGGAA |
| 35553_1 | TATGAGTGTGGAGTCGTTGCGAGATCTCGGCTCACTGCAAGNNNAGATCGGAAGA |
| 35553_2 | TATGAGTGTGGAGTCGTTGCGAGATCTCGGCTCACTGCAAGNNNAGATCGGAAGA |
| 47306_1 | TATGAGTGTGGAGTCGTTGCGATCTCGGCTCTTGGCTTACTGNNNAGATCGGAAGA |
| 47306_2 | TATGAGTGTGGAGTCGTTGCGATCTCGGCTCTTGGCTTACTGNNNAGATCGGAAGA |
| 34322_1 | TATGAGTGTGGAGTCGTTGCTGGCAACATAGATGAACCTGGAANNNAGATCGGAA |
| 34322_2 | TATGAGTGTGGAGTCGTTGCTGGCAACATAGATGAACCTGGAANNNAGATCGGAA |
| 12870_1 | TATGAGTGTGGAGTCGTTGCCACTGAAAATTGATGCTTGTANNNAGATCGGAAGA |
| 12870_2 | TATGAGTGTGGAGTCGTTGCCACTGAAAATTGATGCTTGTANNNAGATCGGAAGA |
| 35267_1 | TATGAGTGTGGAGTCGTTGCATCCCAGACTGAAATCCCAAGACNNNAGATCGGAAG |
| 35267_2 | TATGAGTGTGGAGTCGTTGCATCCCAGACTGAAATCCCAAGACNNNAGATCGGAAG |
| 12149_1 | TATGAGTGTGGAGTCGTTGCGAATGGGCCCTCACTTGACATGANNNAGATCGGAA |
| 12149_2 | TATGAGTGTGGAGTCGTTGCGAATGGGCCCTCACTTGACATGANNNAGATCGGAA |
| 40580_1 | TATGAGTGTGGAGTCGTTGCGGCAAGCAGAGAGACAAATCATGNNNAGATCGGAA |
| 40580_2 | TATGAGTGTGGAGTCGTTGCGGCAAGCAGAGAGACAAATCATGNNNAGATCGGAA |
| 35459_1 | TATGAGTGTGGAGTCGTTGCTAGAGACAGGGTTTGTGTTAGCCNNNAGATCGGAA |
| 35459_2 | TATGAGTGTGGAGTCGTTGCTAGAGACAGGGTTTGTGTTAGCCNNNAGATCGGAA |
| 38640_1 | TATGAGTGTGGAGTCGTTGCAGCCTGTGGTGTTACTAACTCATNNNAGATCGGAAG |
| 38640_2 | TATGAGTGTGGAGTCGTTGCAGCCTGTGGTGTTACTAACTCATNNNAGATCGGAAG |
| 37849_1 | TATGAGTGTGGAGTCGTTGCGAAGAAGAAGAGGTCCAAGGTCANNNAGATCGGA |
| 37849_2 | TATGAGTGTGGAGTCGTTGCGAAGAAGAAGAGGTCCAAGGTCANNNAGATCGGA |
| 32514_1 | TATGAGTGTGGAGTCGTTGCAGTGTGATTTCTGCAATGTAAACANNNAGATCGGAA |
| 32514_2 | TATGAGTGTGGAGTCGTTGCAGTGTGATTTCTGCAATGTAAACANNNAGATCGGAA |
| 44823_1 | TATGAGTGTGGAGTCGTTGCATAGGAAAAGGTTAGCTGGCACANNNAGATCGGAA |
| 44823_2 | TATGAGTGTGGAGTCGTTGCATAGGAAAAGGTTAGCTGGCACANNNAGATCGGAA |
| 38527_1 | TATGAGTGTGGAGTCGTTGCTGGAAGGAGACTTCAGGTCCTATNNNAGATCGGAA |
| 38527_2 | TATGAGTGTGGAGTCGTTGCTGGAAGGAGACTTCAGGTCCTATNNNAGATCGGAA |
| 13649_1 | TATGAGTGTGGAGTCGTTGCGCTAGGATCTTTAGAGAAGTTTANNNAGATCGGAA |
| 13649_2 | TATGAGTGTGGAGTCGTTGCGCTAGGATCTTTAGAGAAGTTTANNNAGATCGGAA |
| 42396_1 | TATGAGTGTGGAGTCGTTGCGAGTGTGAAGAACGGACCAGATNNNAGATCGGAA |

| | |
|---|---|
| 42396_2 | TATGAGTGTGGAGTCGTTGCGAGTGTGAAGAACGGACCAGATNNNAGATCGGAA |
| 39329_1 | TATGAGTGTGGAGTCGTTGCAGGAGAATGGCGTGAACTCGNNNAGATCGGAAGA |
| 39329_2 | TATGAGTGTGGAGTCGTTGCAGGAGAATGGCGTGAACTCGNNNAGATCGGAAGA |
| 12493_1 | TATGAGTGTGGAGTCGTTGCCCGAGATACTTGTTAAAAATTTGNNNAGATCGGAAG |
| 12493_2 | TATGAGTGTGGAGTCGTTGCCCGAGATACTTGTTAAAAATTTGNNNAGATCGGAAG |
| 48772_1 | TATGAGTGTGGAGTCGTTGCGCCTGTTTTCAAGCTCACTCATTNNNAGATCGGAAG |
| 48772_2 | TATGAGTGTGGAGTCGTTGCGCCTGTTTTCAAGCTCACTCATTNNNAGATCGGAAG |
| 37569_1 | TATGAGTGTGGAGTCGTTGCACAAGTTGTCCCTGTTCAGCTAANNNAGATCGGAAG |
| 37569_2 | TATGAGTGTGGAGTCGTTGCACAAGTTGTCCCTGTTCAGCTAANNNAGATCGGAAG |
| 35538_1 | TATGAGTGTGGAGTCGTTGCCTGCAGGTCATTTTCAGCTGAAANNNAGATCGGAAG |
| 35538_2 | TATGAGTGTGGAGTCGTTGCCTGCAGGTCATTTTCAGCTGAAANNNAGATCGGAAG |
| 10571_1 | TATGAGTGTGGAGTCGTTGCGGCTTCTTGGTAAAAGAAATGGTTNNNAGATCGGA |
| 10571_2 | TATGAGTGTGGAGTCGTTGCGGCTTCTTGGTAAAAGAAATGGTTNNNAGATCGGA |
| 36943_1 | TATGAGTGTGGAGTCGTTGCAACTGTTTGACTATCTTCCCCCCNNNAGATCGGAAGA |
| 36943_2 | TATGAGTGTGGAGTCGTTGCAACTGTTTGACTATCTTCCCCCCNNNAGATCGGAAGA |
| 39358_1 | TATGAGTGTGGAGTCGTTGCCCTGTAATCTCAGCACTTTGGGANNNAGATCGGAAG |
| 39358_2 | TATGAGTGTGGAGTCGTTGCCCTGTAATCTCAGCACTTTGGGANNNAGATCGGAAG |
| 46435_1 | TATGAGTGTGGAGTCGTTGCGTGATCTCAGCTCACTGTAACCTNNNAGATCGGAAG |
| 46435_2 | TATGAGTGTGGAGTCGTTGCGTGATCTCAGCTCACTGTAACCTNNNAGATCGGAAG |
| 50096_1 | TATGAGTGTGGAGTCGTTGCCTGTTCTACTTTGGTCACCTCCANNNAGATCGGAAGA |
| 50096_2 | TATGAGTGTGGAGTCGTTGCCTGTTCTACTTTGGTCACCTCCANNNAGATCGGAAGA |
| 51957_1 | TATGAGTGTGGAGTCGTTGCCAGATGACTACTTTCCAGCTGGANNNAGATCGGAAG |
| 51957_2 | TATGAGTGTGGAGTCGTTGCCAGATGACTACTTTCCAGCTGGANNNAGATCGGAAG |
| 36721_1 | TATGAGTGTGGAGTCGTTGCTGAATAGGGAGTAAACGGATGCCNNNAGATCGGAA |
| 36721_2 | TATGAGTGTGGAGTCGTTGCTGAATAGGGAGTAAACGGATGCCNNNAGATCGGAA |
| 32192_1 | TATGAGTGTGGAGTCGTTGCCACAGGTTCAAGCAATTCTCCTGNNNAGATCGGAAG |
| 32192_2 | TATGAGTGTGGAGTCGTTGCCACAGGTTCAAGCAATTCTCCTGNNNAGATCGGAAG |
| 51346_1 | TATGAGTGTGGAGTCGTTGCTCAGATTGAAGAACTTGCCCTCANNNAGATCGGAAG |
| 51346_2 | TATGAGTGTGGAGTCGTTGCTCAGATTGAAGAACTTGCCCTCANNNAGATCGGAAG |
| 47649_1 | TATGAGTGTGGAGTCGTTGCCACGCCCAGCTAATTTTTTTTCGNNNAGATCGGAAG |
| 47649_2 | TATGAGTGTGGAGTCGTTGCCACGCCCAGCTAATTTTTTTTCGNNNAGATCGGAAG |
| 29920_1 | TATGAGTGTGGAGTCGTTGCTTGGCCTCCAAGATTTTGATCCANNNAGATCGGAAG |
| 29920_2 | TATGAGTGTGGAGTCGTTGCTTGGCCTCCAAGATTTTGATCCANNNAGATCGGAAG |
| 31203_1 | TATGAGTGTGGAGTCGTTGCAGAGTGAAACCCCATCTCAAACANNNAGATCGGAAG |
| 31203_2 | TATGAGTGTGGAGTCGTTGCAGAGTGAAACCCCATCTCAAACANNNAGATCGGAAG |
| 52688_1 | TATGAGTGTGGAGTCGTTGCAATGGTGAGTGCAAAACCAAACANNNAGATCGGAA |
| 52688_2 | TATGAGTGTGGAGTCGTTGCAATGGTGAGTGCAAAACCAAACANNNAGATCGGAA |
| 31039_1 | TATGAGTGTGGAGTCGTTGCAAGAGAACAAGAGTCTCTGCCTGNNNAGATCGGAA |
| 31039_2 | TATGAGTGTGGAGTCGTTGCAAGAGAACAAGAGTCTCTGCCTGNNNAGATCGGAA |
| 48196_1 | TATGAGTGTGGAGTCGTTGCTATAAATAGCAGGGAGGGTGGGANNNAGATCGGA |
| 48196_2 | TATGAGTGTGGAGTCGTTGCTATAAATAGCAGGGAGGGTGGGANNNAGATCGGA |
| 2931_1 | TATGAGTGTGGAGTCGTTGCGCATTCCGGACTAAATCCCACTTNNNAGATCGGAAG |
| 2931_2 | TATGAGTGTGGAGTCGTTGCGCATTCCGGACTAAATCCCACTTNNNAGATCGGAAG |

| | |
|---|---|
| 51723_1 | TATGAGTGTGGAGTCGTTGCTGTCATCTAGAGCCTCACCAGTANNNAGATCGGAAG |
| 51723_2 | TATGAGTGTGGAGTCGTTGCTGTCATCTAGAGCCTCACCAGTANNNAGATCGGAAG |
| 45844_1 | TATGAGTGTGGAGTCGTTGCCTCCTGAGTAGCTGGGATTACAGNNNAGATCGGAA |
| 45844_2 | TATGAGTGTGGAGTCGTTGCCTCCTGAGTAGCTGGGATTACAGNNNAGATCGGAA |
| 51181_1 | TATGAGTGTGGAGTCGTTGCTCCACACATTGTAGTGATCGTGTNNNAGATCGGAAG |
| 51181_2 | TATGAGTGTGGAGTCGTTGCTCCACACATTGTAGTGATCGTGTNNNAGATCGGAAG |
| 45204_1 | TATGAGTGTGGAGTCGTTGCGGATGAAGAGTTGAAGGGAGGAGNNNAGATCGG |
| 45204_2 | TATGAGTGTGGAGTCGTTGCGGATGAAGAGTTGAAGGGAGGAGNNNAGATCGG |
| 10959_1 | TATGAGTGTGGAGTCGTTGCTTGTGGCCCCTTTTCAGAGAACNNNAGATCGGAAGA |
| 10959_2 | TATGAGTGTGGAGTCGTTGCTTGTGGCCCCTTTTCAGAGAACNNNAGATCGGAAGA |
| 11549_1 | TATGAGTGTGGAGTCGTTGCCCGTATGCTTTTGTGTCCTCACNNNAGATCGGAAGA |
| 11549_2 | TATGAGTGTGGAGTCGTTGCCCGTATGCTTTTGTGTCCTCACNNNAGATCGGAAGA |
| 44708_1 | TATGAGTGTGGAGTCGTTGCGTTCTGAGTGGTCACTGGAACTGNNNAGATCGGAA |
| 44708_2 | TATGAGTGTGGAGTCGTTGCGTTCTGAGTGGTCACTGGAACTGNNNAGATCGGAA |
| 52675_1 | TATGAGTGTGGAGTCGTTGCTGGAAAGGTGCAGACAACATCTANNNAGATCGGAA |
| 52675_2 | TATGAGTGTGGAGTCGTTGCTGGAAAGGTGCAGACAACATCTANNNAGATCGGAA |
| 48274_1 | TATGAGTGTGGAGTCGTTGCCTCCTGAGTAGCTGGGACTACAGNNNAGATCGGAA |
| 48274_2 | TATGAGTGTGGAGTCGTTGCCTCCTGAGTAGCTGGGACTACAGNNNAGATCGGAA |
| 30769_1 | TATGAGTGTGGAGTCGTTGCTGATTCAAATGCCAGAGAGACGANNNAGATCGGAA |
| 30769_2 | TATGAGTGTGGAGTCGTTGCTGATTCAAATGCCAGAGAGACGANNNAGATCGGAA |
| 2623_1 | TATGAGTGTGGAGTCGTTGCTGACCCCTACAGACTGACTTCTGNNNAGATCGGAAG |
| 2623_2 | TATGAGTGTGGAGTCGTTGCTGACCCCTACAGACTGACTTCTGNNNAGATCGGAAG |
| 51636_1 | TATGAGTGTGGAGTCGTTGCGGGGCTACGTAAAGAACACCANNNAGATCGGAAGA |
| 51636_2 | TATGAGTGTGGAGTCGTTGCGGGGCTACGTAAAGAACACCANNNAGATCGGAAGA |
| 46746_1 | TATGAGTGTGGAGTCGTTGCACTTTTGTGGATATGGGGTCTCCNNNAGATCGGAAG |
| 46746_2 | TATGAGTGTGGAGTCGTTGCACTTTTGTGGATATGGGGTCTCCNNNAGATCGGAAG |
| 37181_1 | TATGAGTGTGGAGTCGTTGCGTATGCCTGCTGCGTAGTAAAAGNNNAGATCGGAA |
| 37181_2 | TATGAGTGTGGAGTCGTTGCGTATGCCTGCTGCGTAGTAAAAGNNNAGATCGGAA |
| 28911_1 | TATGAGTGTGGAGTCGTTGCGACAACACTTTTGCCACTTCTGANNNAGATCGGAAG |
| 28911_2 | TATGAGTGTGGAGTCGTTGCGACAACACTTTTGCCACTTCTGANNNAGATCGGAAG |
| 44095_1 | TATGAGTGTGGAGTCGTTGCGAAGTTAGCTCAGGATAGCACCANNNAGATCGGAA |
| 44095_2 | TATGAGTGTGGAGTCGTTGCGAAGTTAGCTCAGGATAGCACCANNNAGATCGGAA |
| 48818_1 | TATGAGTGTGGAGTCGTTGCCTACATAGCAGCCCCTGGTAATTNNNAGATCGGAAG |
| 48818_2 | TATGAGTGTGGAGTCGTTGCCTACATAGCAGCCCCTGGTAATTNNNAGATCGGAAG |
| 39461_1 | TATGAGTGTGGAGTCGTTGCGTGACGAGAAAAACCCCACAAAANNNAGATCGGAA |
| 39461_2 | TATGAGTGTGGAGTCGTTGCGTGACGAGAAAAACCCCACAAAANNNAGATCGGAA |
| 48695_1 | TATGAGTGTGGAGTCGTTGCCCTGGGCAACATACTTGTGTTTTNNNAGATCGGAAG |
| 48695_2 | TATGAGTGTGGAGTCGTTGCCCTGGGCAACATACTTGTGTTTTNNNAGATCGGAAG |
| 49492_1 | TATGAGTGTGGAGTCGTTGCACCATGCTCGGCTAATTTTTTGTNNNAGATCGGAAG |
| 49492_2 | TATGAGTGTGGAGTCGTTGCACCATGCTCGGCTAATTTTTTGTNNNAGATCGGAAG |
| 38954_1 | TATGAGTGTGGAGTCGTTGCTGAACACATACTCATACTGCACCANNNAGATCGGAA |
| 38954_2 | TATGAGTGTGGAGTCGTTGCTGAACACATACTCATACTGCACCANNNAGATCGGAA |
| 46278_1 | TATGAGTGTGGAGTCGTTGCGATCAATCTCAGCTCACTGCAACNNNAGATCGGAAG |

| | |
|---|---|
| 46278_2 | TATGAGTGTGGAGTCGTTGCGATCAATCTCAGCTCACTGCAACNNNAGATCGGAAG |
| 38398_1 | TATGAGTGTGGAGTCGTTGCTTCTAGGATGGCTCAGTCACATGNNNAGATCGGAAG |
| 38398_2 | TATGAGTGTGGAGTCGTTGCTTCTAGGATGGCTCAGTCACATGNNNAGATCGGAAG |
| 46251_1 | TATGAGTGTGGAGTCGTTGCATACAGGGTCTCACTATGTTGCCNNNAGATCGGAAG |
| 46251_2 | TATGAGTGTGGAGTCGTTGCATACAGGGTCTCACTATGTTGCCNNNAGATCGGAAG |
| 52938_1 | TATGAGTGTGGAGTCGTTGCATTTGCACCATGAGCTTTCTGTCNNNAGATCGGAAG |
| 52938_2 | TATGAGTGTGGAGTCGTTGCATTTGCACCATGAGCTTTCTGTCNNNAGATCGGAAG |
| 41342_1 | TATGAGTGTGGAGTCGTTGCCCAGTGACTGTGGTATCTCATCCNNNAGATCGGAAG |
| 41342_2 | TATGAGTGTGGAGTCGTTGCCCAGTGACTGTGGTATCTCATCCNNNAGATCGGAAG |
| 30761_1 | TATGAGTGTGGAGTCGTTGCAGATCACTGGGAACCCTTGATTTNNNAGATCGGAAG |
| 30761_2 | TATGAGTGTGGAGTCGTTGCAGATCACTGGGAACCCTTGATTTNNNAGATCGGAAG |
| 12599_1 | TATGAGTGTGGAGTCGTTGCGTTTTGACAAACTCAAGCTTANNNAGATCGGAAGAG |
| 12599_2 | TATGAGTGTGGAGTCGTTGCGTTTTGACAAACTCAAGCTTANNNAGATCGGAAGAG |
| 39795_1 | TATGAGTGTGGAGTCGTTGCAAAAGGAGTCTCTGGTCTCATGGNNNAGATCGGAA |
| 39795_2 | TATGAGTGTGGAGTCGTTGCAAAAGGAGTCTCTGGTCTCATGGNNNAGATCGGAA |
| 50620_1 | TATGAGTGTGGAGTCGTTGCCCGTATCAGTGCATAAGACTGGANNNAGATCGGAA |
| 50620_2 | TATGAGTGTGGAGTCGTTGCCCGTATCAGTGCATAAGACTGGANNNAGATCGGAA |
| 34867_1 | TATGAGTGTGGAGTCGTTGCCTGAGTAGCTGGGATTACAGGTGNNNAGATCGGAA |
| 34867_2 | TATGAGTGTGGAGTCGTTGCCTGAGTAGCTGGGATTACAGGTGNNNAGATCGGAA |
| 48473_1 | TATGAGTGTGGAGTCGTTGCCCGTGCCCAGCTAATTTTTGTATNNNAGATCGGAAG |
| 48473_2 | TATGAGTGTGGAGTCGTTGCCCGTGCCCAGCTAATTTTTGTATNNNAGATCGGAAG |
| 46509_1 | TATGAGTGTGGAGTCGTTGCCCAACTCTCAGGTGGACTTGTAANNNAGATCGGAAG |
| 46509_2 | TATGAGTGTGGAGTCGTTGCCCAACTCTCAGGTGGACTTGTAANNNAGATCGGAAG |
| 2580_1 | TATGAGTGTGGAGTCGTTGCAGTGGAGGAGTAGAAAGCACTGGNNNAGATCGGA |
| 2580_2 | TATGAGTGTGGAGTCGTTGCAGTGGAGGAGTAGAAAGCACTGGNNNAGATCGGA |
| 49849_1 | TATGAGTGTGGAGTCGTTGCATACAAAAGCCTGTCTCCCTCAGNNNAGATCGGAAG |
| 49849_2 | TATGAGTGTGGAGTCGTTGCATACAAAAGCCTGTCTCCCTCAGNNNAGATCGGAAG |
| 13689_1 | TATGAGTGTGGAGTCGTTGCATGACATGTTTGGGTCTTTTATANNNAGATCGGAAG |
| 13689_2 | TATGAGTGTGGAGTCGTTGCATGACATGTTTGGGTCTTTTATANNNAGATCGGAAG |
| 28978_1 | TATGAGTGTGGAGTCGTTGCAACTGTCAGGTGAGAAATGGGAANNNAGATCGGAA |
| 28978_2 | TATGAGTGTGGAGTCGTTGCAACTGTCAGGTGAGAAATGGGAANNNAGATCGGAA |
| 34582_1 | TATGAGTGTGGAGTCGTTGCGGAGGGAGAGCATTAGGACAAATNNNAGATCGGA |
| 34582_2 | TATGAGTGTGGAGTCGTTGCGGAGGGAGAGCATTAGGACAAATNNNAGATCGGA |
| 48725_1 | TATGAGTGTGGAGTCGTTGCTCTGTGCCTAGCTTATTTCACTCANNNAGATCGGAAG |
| 48725_2 | TATGAGTGTGGAGTCGTTGCTCTGTGCCTAGCTTATTTCACTCANNNAGATCGGAAG |
| 30185_1 | TATGAGTGTGGAGTCGTTGCACTAATAGCAGGAGCCATGGAACNNNAGATCGGAA |
| 30185_2 | TATGAGTGTGGAGTCGTTGCACTAATAGCAGGAGCCATGGAACNNNAGATCGGAA |
| 34640_1 | TATGAGTGTGGAGTCGTTGCGTTGGGCCTTCCAGTAGACACNNNAGATCGGAAGA |
| 34640_2 | TATGAGTGTGGAGTCGTTGCGTTGGGCCTTCCAGTAGACACNNNAGATCGGAAGA |
| 30693_1 | TATGAGTGTGGAGTCGTTGCCCTCCTATTTGCCGTGGACAACCTGNNNAGATCGGA |
| 30693_2 | TATGAGTGTGGAGTCGTTGCCCTCCTATTTGCCGTGGACAACCTGNNNAGATCGGA |
| 45014_1 | TATGAGTGTGGAGTCGTTGCCAGAACACACCCTTCTTTTGCATNNNAGATCGGAAGA |
| 45014_2 | TATGAGTGTGGAGTCGTTGCCAGAACACACCCTTCTTTTGCATNNNAGATCGGAAGA |

| | |
|---|---|
| 45443_1 | TATGAGTGTGGAGTCGTTGCTCCCTTATCCAAAATGCTTGGGANNNAGATCGGAAG |
| 45443_2 | TATGAGTGTGGAGTCGTTGCTCCCTTATCCAAAATGCTTGGGANNNAGATCGGAAG |
| 5555_1 | TATGAGTGTGGAGTCGTTGCTCCTCCAAAGACATGAGGACTGANNNAGATCGGAAG |
| 5555_2 | TATGAGTGTGGAGTCGTTGCTCCTCCAAAGACATGAGGACTGANNNAGATCGGAAG |
| 45534_1 | TATGAGTGTGGAGTCGTTGCACTTTGGTCAGGGAGAATGGAAANNNAGATCGGAA |
| 45534_2 | TATGAGTGTGGAGTCGTTGCACTTTGGTCAGGGAGAATGGAAANNNAGATCGGAA |
| 39774_1 | TATGAGTGTGGAGTCGTTGCCTGAAATCATACCAACCACGCTCNNNAGATCGGAAG |
| 39774_2 | TATGAGTGTGGAGTCGTTGCCTGAAATCATACCAACCACGCTCNNNAGATCGGAAG |
| 39083_1 | TATGAGTGTGGAGTCGTTGCCGGTTCTCTCACCCCTTTGTATANNNAGATCGGAAGA |
| 39083_2 | TATGAGTGTGGAGTCGTTGCCGGTTCTCTCACCCCTTTGTATANNNAGATCGGAAGA |
| 34419_1 | TATGAGTGTGGAGTCGTTGCCTCAGGAAGTGGGAAAGGATCCNNNAGATCGGAAG |
| 34419_2 | TATGAGTGTGGAGTCGTTGCCTCAGGAAGTGGGAAAGGATCCNNNAGATCGGAAG |
| 38755_1 | TATGAGTGTGGAGTCGTTGCTCCTGTAGAGTTGTTTGAGCTCCNNNAGATCGGAAG |
| 38755_2 | TATGAGTGTGGAGTCGTTGCTCCTGTAGAGTTGTTTGAGCTCCNNNAGATCGGAAG |
| 44065_1 | TATGAGTGTGGAGTCGTTGCTTCCTTGAGTCTGGACACAAACANNNAGATCGGAAG |
| 44065_2 | TATGAGTGTGGAGTCGTTGCTTCCTTGAGTCTGGACACAAACANNNAGATCGGAAG |
| 38733_1 | TATGAGTGTGGAGTCGTTGCAGGCAGATCGGAATCAAACTACANNNAGATCGGAA |
| 38733_2 | TATGAGTGTGGAGTCGTTGCAGGCAGATCGGAATCAAACTACANNNAGATCGGAA |
| 29389_1 | TATGAGTGTGGAGTCGTTGCTTTTTTTTTCTTTGAGATGAAGTNNNAGATCGGAAG |
| 29389_2 | TATGAGTGTGGAGTCGTTGCTTTTTTTTTCTTTGAGATGAAGTNNNAGATCGGAAG |
| 41894_1 | TATGAGTGTGGAGTCGTTGCTTCAAGTGATTCTCCTGCCTCAGNNNAGATCGGAAG |
| 41894_2 | TATGAGTGTGGAGTCGTTGCTTCAAGTGATTCTCCTGCCTCAGNNNAGATCGGAAG |
| 48063_1 | TATGAGTGTGGAGTCGTTGCATGAAAAGGACAAGGGTACTCCGNNNAGATCGGAA |
| 48063_2 | TATGAGTGTGGAGTCGTTGCATGAAAAGGACAAGGGTACTCCGNNNAGATCGGAA |
| 41944_1 | TATGAGTGTGGAGTCGTTGCTCCTGATCTCCTGACCTCATGATNNNAGATCGGAAGA |
| 41944_2 | TATGAGTGTGGAGTCGTTGCTCCTGATCTCCTGACCTCATGATNNNAGATCGGAAGA |
| 53295_1 | TATGAGTGTGGAGTCGTTGCCTGATCCAGGAGGCCGCATCNNNAGATCGGAAGAG |
| 53295_2 | TATGAGTGTGGAGTCGTTGCCTGATCCAGGAGGCCGCATCNNNAGATCGGAAGAG |
| 50846_1 | TATGAGTGTGGAGTCGTTGCTCAGCCCTATCGAAACTTCAAGGNNNAGATCGGAAG |
| 50846_2 | TATGAGTGTGGAGTCGTTGCTCAGCCCTATCGAAACTTCAAGGNNNAGATCGGAAG |
| 44746_1 | TATGAGTGTGGAGTCGTTGCCAGCTACTTCAGGCTCCACTTTANNNAGATCGGAAG |
| 44746_2 | TATGAGTGTGGAGTCGTTGCCAGCTACTTCAGGCTCCACTTTANNNAGATCGGAAG |
| 45576_1 | TATGAGTGTGGAGTCGTTGCTAAAAATGCATATATCGGCCGGGNNNAGATCGGAA |
| 45576_2 | TATGAGTGTGGAGTCGTTGCTAAAAATGCATATATCGGCCGGGNNNAGATCGGAA |
| 48098_1 | TATGAGTGTGGAGTCGTTGCGTGACAACAGACTCTCCAGCAGNNNAGATCGGAAG |
| 48098_2 | TATGAGTGTGGAGTCGTTGCGTGACAACAGACTCTCCAGCAGNNNAGATCGGAAG |
| 50686_1 | TATGAGTGTGGAGTCGTTGCCATTGGTTGTCACAAAGAAGCCTNNNAGATCGGAAG |
| 50686_2 | TATGAGTGTGGAGTCGTTGCCATTGGTTGTCACAAAGAAGCCTNNNAGATCGGAAG |
| 40723_1 | TATGAGTGTGGAGTCGTTGCGTGAGGTCTTGGCTGTACAAAACNNNAGATCGGAA |
| 40723_2 | TATGAGTGTGGAGTCGTTGCGTGAGGTCTTGGCTGTACAAAACNNNAGATCGGAA |
| 31765_1 | TATGAGTGTGGAGTCGTTGCTCTTTTTCAGGGTCTCACTCTGTNNNAGATCGGAAGA |
| 31765_2 | TATGAGTGTGGAGTCGTTGCTCTTTTTCAGGGTCTCACTCTGTNNNAGATCGGAAGA |
| 32170_1 | TATGAGTGTGGAGTCGTTGCTTGACAAGGTCTCACTCTGTTGCNNNAGATCGGAAG |

| | |
|---|---|
| 32170_2 | TATGAGTGTGGAGTCGTTGCTTGACAAGGTCTCACTCTGTTGCNNNAGATCGGAAG |
| 49439_1 | TATGAGTGTGGAGTCGTTGCCTCCCAAAGTGCTGTGATTACAGNNNAGATCGGAAG |
| 49439_2 | TATGAGTGTGGAGTCGTTGCCTCCCAAAGTGCTGTGATTACAGNNNAGATCGGAAG |
| 48824_1 | TATGAGTGTGGAGTCGTTGCGGGTACAAGTGATTCACCTGCTANNNAGATCGGAA |
| 48824_2 | TATGAGTGTGGAGTCGTTGCGGGTACAAGTGATTCACCTGCTANNNAGATCGGAA |
| 46078_1 | TATGAGTGTGGAGTCGTTGCGAGTGCAGTGGCTCAAACAAGNNNAGATCGGAAGA |
| 46078_2 | TATGAGTGTGGAGTCGTTGCGAGTGCAGTGGCTCAAACAAGNNNAGATCGGAAGA |
| 51625_1 | TATGAGTGTGGAGTCGTTGCTGATCAGAGATTGAGCTCATGGGNNNAGATCGGAA |
| 51625_2 | TATGAGTGTGGAGTCGTTGCTGATCAGAGATTGAGCTCATGGGNNNAGATCGGAA |
| 51109_1 | TATGAGTGTGGAGTCGTTGCCCTATGGGCAATTTCCTCAGGACNNNAGATCGGAAG |
| 51109_2 | TATGAGTGTGGAGTCGTTGCCCTATGGGCAATTTCCTCAGGACNNNAGATCGGAAG |
| 28876_1 | TATGAGTGTGGAGTCGTTGCGGAATGACTGTTGTTGTTGAGCANNNAGATCGGAA |
| 28876_2 | TATGAGTGTGGAGTCGTTGCGGAATGACTGTTGTTGTTGAGCANNNAGATCGGAA |
| 48852_1 | TATGAGTGTGGAGTCGTTGCCCAAAGTTCTGGGATTACAGGCANNNAGATCGGAA |
| 48852_2 | TATGAGTGTGGAGTCGTTGCCCAAAGTTCTGGGATTACAGGCANNNAGATCGGAA |
| 45500_1 | TATGAGTGTGGAGTCGTTGCGGAAAAGAAGGAACAGGGTTGGANNNAGATCGGA |
| 45500_2 | TATGAGTGTGGAGTCGTTGCGGAAAAGAAGGAACAGGGTTGGANNNAGATCGGA |
| 34178_1 | TATGAGTGTGGAGTCGTTGCTACTCCTACCACCAGTTTCCTCANNNAGATCGGAAGA |
| 34178_2 | TATGAGTGTGGAGTCGTTGCTACTCCTACCACCAGTTTCCTCANNNAGATCGGAAGA |
| 34700_1 | TATGAGTGTGGAGTCGTTGCCGCCCGGCTTTTCTTTTTTTTTTNNNAGATCGGAAGA |
| 34700_2 | TATGAGTGTGGAGTCGTTGCCGCCCGGCTTTTCTTTTTTTTTTNNNAGATCGGAAGA |
| 33466_1 | TATGAGTGTGGAGTCGTTGCGATTGGAGGACAATGGCAAATGTNNNAGATCGGAA |
| 33466_2 | TATGAGTGTGGAGTCGTTGCGATTGGAGGACAATGGCAAATGTNNNAGATCGGAA |
| 50663_1 | TATGAGTGTGGAGTCGTTGCATGGAAAGAATGAAGTCCCTGCTNNNAGATCGGAA |
| 50663_2 | TATGAGTGTGGAGTCGTTGCATGGAAAGAATGAAGTCCCTGCTNNNAGATCGGAA |
| 35071_1 | TATGAGTGTGGAGTCGTTGCAGCCAATGGTCTAAATGGGTTCTNNNAGATCGGAA |
| 35071_2 | TATGAGTGTGGAGTCGTTGCAGCCAATGGTCTAAATGGGTTCTNNNAGATCGGAA |
| 13437_1 | TATGAGTGTGGAGTCGTTGCTTCATGGGATTAGAAGAATTCAANNNAGATCGGAA |
| 13437_2 | TATGAGTGTGGAGTCGTTGCTTCATGGGATTAGAAGAATTCAANNNAGATCGGAA |
| 38032_1 | TATGAGTGTGGAGTCGTTGCACCCAAAACAACAAGAAGGAAAANNNAGATCGGAA |
| 38032_2 | TATGAGTGTGGAGTCGTTGCACCCAAAACAACAAGAAGGAAAANNNAGATCGGAA |
| 47658_1 | TATGAGTGTGGAGTCGTTGCGATGGAGTCCTACTCTGTTGTCCNNNAGATCGGAAG |
| 47658_2 | TATGAGTGTGGAGTCGTTGCGATGGAGTCCTACTCTGTTGTCCNNNAGATCGGAAG |
| 46049_1 | TATGAGTGTGGAGTCGTTGCGATGGGACTCTTTACCAGTGAGGNNNAGATCGGAA |
| 46049_2 | TATGAGTGTGGAGTCGTTGCGATGGGACTCTTTACCAGTGAGGNNNAGATCGGAA |
| 12560_1 | TATGAGTGTGGAGTCGTTGCAAAATCGAGAAGAGAGGAGANNNAGATCGGAAGA |
| 12560_2 | TATGAGTGTGGAGTCGTTGCAAAATCGAGAAGAGAGGAGANNNAGATCGGAAGA |
| 44811_1 | TATGAGTGTGGAGTCGTTGCCTGATGGCTTCACTGTCGAATTCNNNAGATCGGAAG |
| 44811_2 | TATGAGTGTGGAGTCGTTGCCTGATGGCTTCACTGTCGAATTCNNNAGATCGGAAG |
| 33000_1 | TATGAGTGTGGAGTCGTTGCAAGCGATTCTTATGTCTCAGCCTNNNAGATCGGAAG |
| 33000_2 | TATGAGTGTGGAGTCGTTGCAAGCGATTCTTATGTCTCAGCCTNNNAGATCGGAAG |
| 33136_1 | TATGAGTGTGGAGTCGTTGCGCCCGCCACACAAACATATATATNNNAGATCGGAAG |
| 33136_2 | TATGAGTGTGGAGTCGTTGCGCCCGCCACACAAACATATATATNNNAGATCGGAAG |

| | |
|---|---|
| 51196_1 | TATGAGTGTGGAGTCGTTGCTGTAGATGACAGGTTGATGGGTGNNNAGATCGGA |
| 51196_2 | TATGAGTGTGGAGTCGTTGCTGTAGATGACAGGTTGATGGGTGNNNAGATCGGA |
| 31398_1 | TATGAGTGTGGAGTCGTTGCAGAGGGGAAAGGAGATGAGGTTANNNAGATCGGA |
| 31398_2 | TATGAGTGTGGAGTCGTTGCAGAGGGGAAAGGAGATGAGGTTANNNAGATCGGA |
| 47614_1 | TATGAGTGTGGAGTCGTTGCCTACAGGTACGTGACACTACACCNNNAGATCGGAAG |
| 47614_2 | TATGAGTGTGGAGTCGTTGCCTACAGGTACGTGACACTACACCNNNAGATCGGAAG |
| 50157_1 | TATGAGTGTGGAGTCGTTGCTCAGCACAAATAAATCCCGAGACNNNAGATCGGAAG |
| 50157_2 | TATGAGTGTGGAGTCGTTGCTCAGCACAAATAAATCCCGAGACNNNAGATCGGAAG |
| 43975_1 | TATGAGTGTGGAGTCGTTGCTTCCTCCGTGAAGTACATGATGGNNNAGATCGGAAG |
| 43975_2 | TATGAGTGTGGAGTCGTTGCTTCCTCCGTGAAGTACATGATGGNNNAGATCGGAAG |
| 39559_1 | TATGAGTGTGGAGTCGTTGCACAACCATCTTCAGCAACTACCANNNAGATCGGAAG |
| 39559_2 | TATGAGTGTGGAGTCGTTGCACAACCATCTTCAGCAACTACCANNNAGATCGGAAG |
| 51384_1 | TATGAGTGTGGAGTCGTTGCAGCGATTCAGGACTGTTTTCTCTNNNAGATCGGAAG |
| 51384_2 | TATGAGTGTGGAGTCGTTGCAGCGATTCAGGACTGTTTTCTCTNNNAGATCGGAAG |
| 37357_1 | TATGAGTGTGGAGTCGTTGCCCTACAGGACAATGAACACGAGANNNAGATCGGAA |
| 37357_2 | TATGAGTGTGGAGTCGTTGCCCTACAGGACAATGAACACGAGANNNAGATCGGAA |
| 47615_1 | TATGAGTGTGGAGTCGTTGCCTGGTCCAACCTGGAGAAAGGNNNAGATCGGAAGA |
| 47615_2 | TATGAGTGTGGAGTCGTTGCCTGGTCCAACCTGGAGAAAGGNNNAGATCGGAAGA |
| 41919_1 | TATGAGTGTGGAGTCGTTGCATTTTGGCTCACTGGAACCTACANNNAGATCGGAAG |
| 41919_2 | TATGAGTGTGGAGTCGTTGCATTTTGGCTCACTGGAACCTACANNNAGATCGGAAG |
| 34161_1 | TATGAGTGTGGAGTCGTTGCGTTTCTGGCTCACACCTGTAATCNNNAGATCGGAAG |
| 34161_2 | TATGAGTGTGGAGTCGTTGCGTTTCTGGCTCACACCTGTAATCNNNAGATCGGAAG |
| 10148_1 | TATGAGTGTGGAGTCGTTGCAGAAGTTTGCTTTCCCTAGCTGCNNNAGATCGGAAG |
| 10148_2 | TATGAGTGTGGAGTCGTTGCAGAAGTTTGCTTTCCCTAGCTGCNNNAGATCGGAAG |
| 47381_1 | TATGAGTGTGGAGTCGTTGCCAGGATAGTCTCGATCACCTGACNNNAGATCGGAAG |
| 47381_2 | TATGAGTGTGGAGTCGTTGCCAGGATAGTCTCGATCACCTGACNNNAGATCGGAAG |
| 9094_1 | TATGAGTGTGGAGTCGTTGCTTGTCTCATGTGTCCCTCTCACTNNNAGATCGGAAGA |
| 9094_2 | TATGAGTGTGGAGTCGTTGCTTGTCTCATGTGTCCCTCTCACTNNNAGATCGGAAGA |
| 51942_1 | TATGAGTGTGGAGTCGTTGCCAAAAGGGCAGTCTTATCTCCCTNNNAGATCGGAAG |
| 51942_2 | TATGAGTGTGGAGTCGTTGCCAAAAGGGCAGTCTTATCTCCCTNNNAGATCGGAAG |
| 29637_1 | TATGAGTGTGGAGTCGTTGCACACACAACATTTTTACTTTGTGGCNNNAGATCGGA |
| 29637_2 | TATGAGTGTGGAGTCGTTGCACACACAACATTTTTACTTTGTGGCNNNAGATCGGA |
| 50072_1 | TATGAGTGTGGAGTCGTTGCGTGTGTGTGTGTGTGTGTGTATANNNAGATCGGAA |
| 50072_2 | TATGAGTGTGGAGTCGTTGCGTGTGTGTGTGTGTGTGTGTATANNNAGATCGGAA |
| 43711_1 | TATGAGTGTGGAGTCGTTGCCTTCCGGAATGGTCCACATGTTNNNAGATCGGAAGA |
| 43711_2 | TATGAGTGTGGAGTCGTTGCCTTCCGGAATGGTCCACATGTTNNNAGATCGGAAGA |
| 50225_1 | TATGAGTGTGGAGTCGTTGCCTCTGCAGGTAAGGACTGGAATANNNAGATCGGAA |
| 50225_2 | TATGAGTGTGGAGTCGTTGCCTCTGCAGGTAAGGACTGGAATANNNAGATCGGAA |
| 42513_1 | TATGAGTGTGGAGTCGTTGCCAATTGGCTCACACATTCACACANNNAGATCGGAAG |
| 42513_2 | TATGAGTGTGGAGTCGTTGCCAATTGGCTCACACATTCACACANNNAGATCGGAAG |
| 43164_1 | TATGAGTGTGGAGTCGTTGCAGATGGAGTCTCACTCTGTCACCNNNAGATCGGAAG |
| 43164_2 | TATGAGTGTGGAGTCGTTGCAGATGGAGTCTCACTCTGTCACCNNNAGATCGGAAG |
| 35649_1 | TATGAGTGTGGAGTCGTTGCAGTCTTGACAGGCTGTATGTGTCNNNAGATCGGAA |

| | |
|---|---|
| 35649_2 | TATGAGTGTGGAGTCGTTGCAGTCTTGACAGGCTGTATGTGTCNNNAGATCGGAA |
| 41101_1 | TATGAGTGTGGAGTCGTTGCCTCTACGATGCTTCATGTGCCTANNNAGATCGGAAG |
| 41101_2 | TATGAGTGTGGAGTCGTTGCCTCTACGATGCTTCATGTGCCTANNNAGATCGGAAG |
| 42188_1 | TATGAGTGTGGAGTCGTTGCGTTCAAGAGATTCTCCTGCCTCANNNAGATCGGAAG |
| 42188_2 | TATGAGTGTGGAGTCGTTGCGTTCAAGAGATTCTCCTGCCTCANNNAGATCGGAAG |
| 12044_1 | TATGAGTGTGGAGTCGTTGCAACTAGGCTGGGCCTGGGCTANNNAGATCGGAAGA |
| 12044_2 | TATGAGTGTGGAGTCGTTGCAACTAGGCTGGGCCTGGGCTANNNAGATCGGAAGA |
| 36344_1 | TATGAGTGTGGAGTCGTTGCAGTTCTTCTCAGGCTGTTGGAATNNNAGATCGGAAG |
| 36344_2 | TATGAGTGTGGAGTCGTTGCAGTTCTTCTCAGGCTGTTGGAATNNNAGATCGGAAG |
| 33813_1 | TATGAGTGTGGAGTCGTTGCAAGGGAAAAAGGACAAAGGGGAANNNAGATCGGA |
| 33813_2 | TATGAGTGTGGAGTCGTTGCAAGGGAAAAAGGACAAAGGGGAANNNAGATCGGA |
| 52459_1 | TATGAGTGTGGAGTCGTTGCCCTCCCGAGTAGCTAGGATTACANNNAGATCGGAAG |
| 52459_2 | TATGAGTGTGGAGTCGTTGCCCTCCCGAGTAGCTAGGATTACANNNAGATCGGAAG |
| 48925_1 | TATGAGTGTGGAGTCGTTGCCCACACCCGGCTAATTTTTTGTANNNAGATCGGAAG |
| 48925_2 | TATGAGTGTGGAGTCGTTGCCCACACCCGGCTAATTTTTTGTANNNAGATCGGAAG |
| 13376_1 | TATGAGTGTGGAGTCGTTGCACATATATTTATCATGTGCAACANNNAGATCGGAAG |
| 13376_2 | TATGAGTGTGGAGTCGTTGCACATATATTTATCATGTGCAACANNNAGATCGGAAG |
| 29111_1 | TATGAGTGTGGAGTCGTTGCTCTGAGGAGCATTGAAACCTACCNNNAGATCGGAAG |
| 29111_2 | TATGAGTGTGGAGTCGTTGCTCTGAGGAGCATTGAAACCTACCNNNAGATCGGAAG |
| 49806_1 | TATGAGTGTGGAGTCGTTGCTGGAGTGTCTAAAAGGTCTCTGCNNNAGATCGGAA |
| 49806_2 | TATGAGTGTGGAGTCGTTGCTGGAGTGTCTAAAAGGTCTCTGCNNNAGATCGGAA |
| 36374_1 | TATGAGTGTGGAGTCGTTGCTTCCAATGCAGCAGTCAAATGTANNNAGATCGGAAG |
| 36374_2 | TATGAGTGTGGAGTCGTTGCTTCCAATGCAGCAGTCAAATGTANNNAGATCGGAAG |
| 10843_1 | TATGAGTGTGGAGTCGTTGCCCTGTCAAATCCTGATTCTGCCANNNAGATCGGAAG |
| 10843_2 | TATGAGTGTGGAGTCGTTGCCCTGTCAAATCCTGATTCTGCCANNNAGATCGGAAG |
| 36489_1 | TATGAGTGTGGAGTCGTTGCCTCCTTCCTTGCTCCTATGCTAGNNNAGATCGGAAGA |
| 36489_2 | TATGAGTGTGGAGTCGTTGCCTCCTTCCTTGCTCCTATGCTAGNNNAGATCGGAAGA |
| 41877_1 | TATGAGTGTGGAGTCGTTGCGAAACCTCCACTTCCCATCTCATNNNAGATCGGAAGA |
| 41877_2 | TATGAGTGTGGAGTCGTTGCGAAACCTCCACTTCCCATCTCATNNNAGATCGGAAGA |
| 2981_1 | TATGAGTGTGGAGTCGTTGCATTACCTCAAAGGAAAACTGGAAGAANNNAGATCG |
| 2981_2 | TATGAGTGTGGAGTCGTTGCATTACCTCAAAGGAAAACTGGAAGAANNNAGATCG |
| 48161_1 | TATGAGTGTGGAGTCGTTGCCATACAATCATGGCTCACTGCAGNNNAGATCGGAAG |
| 48161_2 | TATGAGTGTGGAGTCGTTGCCATACAATCATGGCTCACTGCAGNNNAGATCGGAAG |
| 33314_1 | TATGAGTGTGGAGTCGTTGCATTTTCTTCTCAGAGCCTCCCTGNNNAGATCGGAAGA |
| 33314_2 | TATGAGTGTGGAGTCGTTGCATTTTCTTCTCAGAGCCTCCCTGNNNAGATCGGAAGA |
| 33082_1 | TATGAGTGTGGAGTCGTTGCGAGACGGAGTCTTGCTCTGTCNNNAGATCGGAAGA |
| 33082_2 | TATGAGTGTGGAGTCGTTGCGAGACGGAGTCTTGCTCTGTCNNNAGATCGGAAGA |
| 38853_1 | TATGAGTGTGGAGTCGTTGCAGGATAGAGAGAAAACCCTCCCTNNNAGATCGGAA |
| 38853_2 | TATGAGTGTGGAGTCGTTGCAGGATAGAGAGAAAACCCTCCCTNNNAGATCGGAA |
| 43532_1 | TATGAGTGTGGAGTCGTTGCATGATCTCGGCTCACTGCAAGNNNAGATCGGAAGAG |
| 43532_2 | TATGAGTGTGGAGTCGTTGCATGATCTCGGCTCACTGCAAGNNNAGATCGGAAGAG |
| 34889_1 | TATGAGTGTGGAGTCGTTGCGTTCAAGAGATTCTCCTGCCTCANNNAGATCGGAAG |
| 34889_2 | TATGAGTGTGGAGTCGTTGCGTTCAAGAGATTCTCCTGCCTCANNNAGATCGGAAG |

42096_1 TATGAGTGTGGAGTCGTTGCAAAAGTACTCCTGTTTTGGGGGTNNNAGATCGGAAG
42096_2 TATGAGTGTGGAGTCGTTGCAAAAGTACTCCTGTTTTGGGGGTNNNAGATCGGAAG
48307_1 TATGAGTGTGGAGTCGTTGCCACTGCATGTACTCCAGGTCTTTNNNAGATCGGAAGA
48307_2 TATGAGTGTGGAGTCGTTGCCACTGCATGTACTCCAGGTCTTTNNNAGATCGGAAGA
41788_1 TATGAGTGTGGAGTCGTTGCTTGATTCTCTGCTTGGTCACTGTNNNAGATCGGAAGA
41788_2 TATGAGTGTGGAGTCGTTGCTTGATTCTCTGCTTGGTCACTGTNNNAGATCGGAAGA
42493_1 TATGAGTGTGGAGTCGTTGCATAACCCAGAGACTTCTCCTGGANNNAGATCGGAAG
42493_2 TATGAGTGTGGAGTCGTTGCATAACCCAGAGACTTCTCCTGGANNNAGATCGGAAG
10297_1 TATGAGTGTGGAGTCGTTGCCCGACCATATCTGCACAGAACAANNNAGATCGGAAG
10297_2 TATGAGTGTGGAGTCGTTGCCCGACCATATCTGCACAGAACAANNNAGATCGGAAG
38774_1 TATGAGTGTGGAGTCGTTGCCGGGAGGCTAAGGTAAGAGAATCNNNAGATCGGAA
38774_2 TATGAGTGTGGAGTCGTTGCCGGGAGGCTAAGGTAAGAGAATCNNNAGATCGGAA
46682_1 TATGAGTGTGGAGTCGTTGCCCCAGTCCTGTGTTTTAGTCACTNNNAGATCGGAAGA
46682_2 TATGAGTGTGGAGTCGTTGCCCCAGTCCTGTGTTTTAGTCACTNNNAGATCGGAAGA
10767_1 TATGAGTGTGGAGTCGTTGCAGAGACCTGCTATGAGATGGCATNNNAGATCGGAAG
10767_2 TATGAGTGTGGAGTCGTTGCAGAGACCTGCTATGAGATGGCATNNNAGATCGGAAG
51488_1 TATGAGTGTGGAGTCGTTGCCATCCAGGAGGGAGGTGGAGNNNAGATCGGAAGAG
51488_2 TATGAGTGTGGAGTCGTTGCCATCCAGGAGGGAGGTGGAGNNNAGATCGGAAGAG
53410_1 TATGAGTGTGGAGTCGTTGCTGCTACTGCTATCTAGAAGGCAGNNNAGATCGGAAG
53410_2 TATGAGTGTGGAGTCGTTGCTGCTACTGCTATCTAGAAGGCAGNNNAGATCGGAAG
9533_1 TATGAGTGTGGAGTCGTTGCTGGGATCCTCTTCAATGGAAAGCNNNAGATCGGAAG
9533_2 TATGAGTGTGGAGTCGTTGCTGGGATCCTCTTCAATGGAAAGCNNNAGATCGGAAG
39983_1 TATGAGTGTGGAGTCGTTGCCAAAAAGTAGCTGAGTGTGGTGGNNNAGATCGGAA
39983_2 TATGAGTGTGGAGTCGTTGCCAAAAAGTAGCTGAGTGTGGTGGNNNAGATCGGAA
36934_1 TATGAGTGTGGAGTCGTTGCTACCCCAATAAACTCTCTCCTGCNNNAGATCGGAAGA
36934_2 TATGAGTGTGGAGTCGTTGCTACCCCAATAAACTCTCTCCTGCNNNAGATCGGAAGA
604_1 TATGAGTGTGGAGTCGTTGCATTCCTAGACAGTTGTAGCAGGGTNNNAGATCGGAA
604_2 TATGAGTGTGGAGTCGTTGCATTCCTAGACAGTTGTAGCAGGGTNNNAGATCGGAA
45673_1 TATGAGTGTGGAGTCGTTGCTACAGGCATATGCTACCATGACCNNNAGATCGGAAG
45673_2 TATGAGTGTGGAGTCGTTGCTACAGGCATATGCTACCATGACCNNNAGATCGGAAG
42532_1 TATGAGTGTGGAGTCGTTGCTGTGTCTGTGTGTGTTTAGTGGANNNAGATCGGAAG
42532_2 TATGAGTGTGGAGTCGTTGCTGTGTCTGTGTGTGTTTAGTGGANNNAGATCGGAAG
53285_1 TATGAGTGTGGAGTCGTTGCCTGATCCAGGAGGCCGCATCNNNAGATCGGAAGAGC
53285_2 TATGAGTGTGGAGTCGTTGCCTGATCCAGGAGGCCGCATCNNNAGATCGGAAGAGC
37168_1 TATGAGTGTGGAGTCGTTGCTGATACTGTCACCCAGGTACTGANNNAGATCGGAAG
37168_2 TATGAGTGTGGAGTCGTTGCTGATACTGTCACCCAGGTACTGANNNAGATCGGAAG
30155_1 TATGAGTGTGGAGTCGTTGCGCTTTGCCTTGGAGATTTTGAGGNNNAGATCGGAAG
30155_2 TATGAGTGTGGAGTCGTTGCGCTTTGCCTTGGAGATTTTGAGGNNNAGATCGGAAG
33134_1 TATGAGTGTGGAGTCGTTGCATTTCTACACTGCTGGTGGGAATNNNAGATCGGAAG
33134_2 TATGAGTGTGGAGTCGTTGCATTTCTACACTGCTGGTGGGAATNNNAGATCGGAAG
52465_1 TATGAGTGTGGAGTCGTTGCAAGGGAGATCTCATTTTCCCCACNNNAGATCGGAAGA
52465_2 TATGAGTGTGGAGTCGTTGCAAGGGAGATCTCATTTTCCCCACNNNAGATCGGAAGA
35634_1 TATGAGTGTGGAGTCGTTGCCTCAAACCCCTGACCTAGTGATCNNNAGATCGGAAGA

| | |
|---|---|
| 35634_2 | TATGAGTGTGGAGTCGTTGCCTCAAACCCCTGACCTAGTGATCNNNAGATCGGAAG |
| 40575_1 | TATGAGTGTGGAGTCGTTGCCTCTTCCCTGTTCCCTTGGTATGNNNAGATCGGAAGA |
| 40575_2 | TATGAGTGTGGAGTCGTTGCCTCTTCCCTGTTCCCTTGGTATGNNNAGATCGGAAGA |
| 43119_1 | TATGAGTGTGGAGTCGTTGCCCCATGAAAGTGAGGACGATACANNNAGATCGGAA |
| 43119_2 | TATGAGTGTGGAGTCGTTGCCCCATGAAAGTGAGGACGATACANNNAGATCGGAA |
| 50374_1 | TATGAGTGTGGAGTCGTTGCAACGAGATCACCTTGGAGTCACNNNAGATCGGAAG |
| 50374_2 | TATGAGTGTGGAGTCGTTGCAACGAGATCACCTTGGAGTCACNNNAGATCGGAAG |
| 48560_1 | TATGAGTGTGGAGTCGTTGCAGTGTAGAGTGTAGGAGGATCCCNNNAGATCGGAA |
| 48560_2 | TATGAGTGTGGAGTCGTTGCAGTGTAGAGTGTAGGAGGATCCCNNNAGATCGGAA |
| 8308_1 | TATGAGTGTGGAGTCGTTGCTCTGCTCTCTCCTGGTCTGTATGNNNAGATCGGAAG |
| 8308_2 | TATGAGTGTGGAGTCGTTGCTCTGCTCTCTCCTGGTCTGTATGNNNAGATCGGAAG |
| 48663_1 | TATGAGTGTGGAGTCGTTGCCTTTCTGAGGATTGCTTTGGGTANNNAGATCGGAAG |
| 48663_2 | TATGAGTGTGGAGTCGTTGCCTTTCTGAGGATTGCTTTGGGTANNNAGATCGGAAG |
| 29226_1 | TATGAGTGTGGAGTCGTTGCATCAGCTTCTCCACCATCTTGTTNNNAGATCGGAAGA |
| 29226_2 | TATGAGTGTGGAGTCGTTGCATCAGCTTCTCCACCATCTTGTTNNNAGATCGGAAGA |
| 47912_1 | TATGAGTGTGGAGTCGTTGCCGCCCAGCCAGAATAAACATTTTNNNAGATCGGAAG |
| 47912_2 | TATGAGTGTGGAGTCGTTGCCGCCCAGCCAGAATAAACATTTTNNNAGATCGGAAG |
| 2629_1 | TATGAGTGTGGAGTCGTTGCTGAGGTTGAGTTGGAATACCGTGNNNAGATCGGAA |
| 2629_2 | TATGAGTGTGGAGTCGTTGCTGAGGTTGAGTTGGAATACCGTGNNNAGATCGGAA |
| 44807_1 | TATGAGTGTGGAGTCGTTGCGGAGAGGCCTATATGTGCATCATNNNAGATCGGAA |
| 44807_2 | TATGAGTGTGGAGTCGTTGCGGAGAGGCCTATATGTGCATCATNNNAGATCGGAA |
| 44233_1 | TATGAGTGTGGAGTCGTTGCCAGGGGATCAAGGAAACTAGGTGNNNAGATCGGA |
| 44233_2 | TATGAGTGTGGAGTCGTTGCCAGGGGATCAAGGAAACTAGGTGNNNAGATCGGA |
| 29885_1 | TATGAGTGTGGAGTCGTTGCTCACAACTTCCTCTTCTCAGAGCNNNAGATCGGAAGA |
| 29885_2 | TATGAGTGTGGAGTCGTTGCTCACAACTTCCTCTTCTCAGAGCNNNAGATCGGAAGA |
| 37078_1 | TATGAGTGTGGAGTCGTTGCGCCACCATTATCAAGTGGGTTTCNNNAGATCGGAAG |
| 37078_2 | TATGAGTGTGGAGTCGTTGCGCCACCATTATCAAGTGGGTTTCNNNAGATCGGAAG |
| 48847_1 | TATGAGTGTGGAGTCGTTGCCCAAGACTTCTCTGTCCCGTCNNNAGATCGGAAGAG |
| 48847_2 | TATGAGTGTGGAGTCGTTGCCCAAGACTTCTCTGTCCCGTCNNNAGATCGGAAGAG |
| 46209_1 | TATGAGTGTGGAGTCGTTGCATTCTTCTACCTCAGCCTCCTGANNNAGATCGGAAGA |
| 46209_2 | TATGAGTGTGGAGTCGTTGCATTCTTCTACCTCAGCCTCCTGANNNAGATCGGAAGA |
| 50451_1 | TATGAGTGTGGAGTCGTTGCCCCATCCCTCCAGTTGAAGATTTNNNAGATCGGAAG |
| 50451_2 | TATGAGTGTGGAGTCGTTGCCCCATCCCTCCAGTTGAAGATTTNNNAGATCGGAAG |
| 51782_1 | TATGAGTGTGGAGTCGTTGCACAAGGGTTTTCCAGAGATGTCTNNNAGATCGGAAG |
| 51782_2 | TATGAGTGTGGAGTCGTTGCACAAGGGTTTTCCAGAGATGTCTNNNAGATCGGAAG |
| 13455_1 | TATGAGTGTGGAGTCGTTGCCTGACTTGTTTTCCACAAACNNNAGATCGGAAGAGC |
| 13455_2 | TATGAGTGTGGAGTCGTTGCCTGACTTGTTTTCCACAAACNNNAGATCGGAAGAGC |
| 45064_1 | TATGAGTGTGGAGTCGTTGCCACCACACCCAGCTAATTTTTGTNNNAGATCGGAAGA |
| 45064_2 | TATGAGTGTGGAGTCGTTGCCACCACACCCAGCTAATTTTTGTNNNAGATCGGAAGA |
| 37586_1 | TATGAGTGTGGAGTCGTTGCTTTGGGAAATGGCATTCTCCAACNNNAGATCGGAAG |
| 37586_2 | TATGAGTGTGGAGTCGTTGCTTTGGGAAATGGCATTCTCCAACNNNAGATCGGAAG |
| 33497_1 | TATGAGTGTGGAGTCGTTGCCAGCTCTGGTTTCACAAATGAGGNNNAGATCGGAAG |
| 33497_2 | TATGAGTGTGGAGTCGTTGCCAGCTCTGGTTTCACAAATGAGGNNNAGATCGGAAG |

| | |
|---|---|
| 52154_1 | TATGAGTGTGGAGTCGTTGCGTAAGGTGGATGAAGGGTACCTGNNNAGATCGGA |
| 52154_2 | TATGAGTGTGGAGTCGTTGCGTAAGGTGGATGAAGGGTACCTGNNNAGATCGGA |
| 48946_1 | TATGAGTGTGGAGTCGTTGCATTTTTGCAAAGACAGGGTCTCGNNNAGATCGGAA |
| 48946_2 | TATGAGTGTGGAGTCGTTGCATTTTTGCAAAGACAGGGTCTCGNNNAGATCGGAA |
| 33190_1 | TATGAGTGTGGAGTCGTTGCTGGTTTCACTGATTTTCTTCCCANNNAGATCGGAAGA |
| 33190_2 | TATGAGTGTGGAGTCGTTGCTGGTTTCACTGATTTTCTTCCCANNNAGATCGGAAGA |
| 44704_1 | TATGAGTGTGGAGTCGTTGCCCTTTCCCTTGCTGATTTTGGTTNNNAGATCGGAAGA |
| 44704_2 | TATGAGTGTGGAGTCGTTGCCCTTTCCCTTGCTGATTTTGGTTNNNAGATCGGAAGA |
| 47370_1 | TATGAGTGTGGAGTCGTTGCTTGGGTTGATTCCATGTCTTTGCNNNAGATCGGAAG |
| 47370_2 | TATGAGTGTGGAGTCGTTGCTTGGGTTGATTCCATGTCTTTGCNNNAGATCGGAAG |
| 34174_1 | TATGAGTGTGGAGTCGTTGCTGGATCCTTACCAAGTTTCACCANNNAGATCGGAAG |
| 34174_2 | TATGAGTGTGGAGTCGTTGCTGGATCCTTACCAAGTTTCACCANNNAGATCGGAAG |
| 6001_1 | TATGAGTGTGGAGTCGTTGCAGGAGTAGGAGGTAGGGAGGAANNNAGATCGGA |
| 6001_2 | TATGAGTGTGGAGTCGTTGCAGGAGTAGGAGGTAGGGAGGAANNNAGATCGGA |
| 50181_1 | TATGAGTGTGGAGTCGTTGCTCTTTCCCTTTCTCTCTGCTGTCNNNAGATCGGAAGA |
| 50181_2 | TATGAGTGTGGAGTCGTTGCTCTTTCCCTTTCTCTCTGCTGTCNNNAGATCGGAAGA |
| 31210_1 | TATGAGTGTGGAGTCGTTGCTGGTAGAGTGGAAGCATACCCTANNNAGATCGGAA |
| 31210_2 | TATGAGTGTGGAGTCGTTGCTGGTAGAGTGGAAGCATACCCTANNNAGATCGGAA |
| 29745_1 | TATGAGTGTGGAGTCGTTGCTGCCATCTTGCATTTCTTACAGCNNNAGATCGGAAGA |
| 29745_2 | TATGAGTGTGGAGTCGTTGCTGCCATCTTGCATTTCTTACAGCNNNAGATCGGAAGA |
| 38938_1 | TATGAGTGTGGAGTCGTTGCTCCCAAAAGTTTGAAGGTTGCTGNNNAGATCGGAAG |
| 38938_2 | TATGAGTGTGGAGTCGTTGCTCCCAAAAGTTTGAAGGTTGCTGNNNAGATCGGAAG |
| 37102_1 | TATGAGTGTGGAGTCGTTGCTGCAGTGTGTAATTGTTGTTCCCNNNAGATCGGAAG |
| 37102_2 | TATGAGTGTGGAGTCGTTGCTGCAGTGTGTAATTGTTGTTCCCNNNAGATCGGAAG |
| 29946_1 | TATGAGTGTGGAGTCGTTGCCGTAATCAGCTTGTTCTCCCAATNNNAGATCGGAAG |
| 29946_2 | TATGAGTGTGGAGTCGTTGCCGTAATCAGCTTGTTCTCCCAATNNNAGATCGGAAG |
| 37815_1 | TATGAGTGTGGAGTCGTTGCAAGCTGAGGTAGGAGGATCACTTNNNAGATCGGAA |
| 37815_2 | TATGAGTGTGGAGTCGTTGCAAGCTGAGGTAGGAGGATCACTTNNNAGATCGGAA |
| 42704_1 | TATGAGTGTGGAGTCGTTGCCTCCTGGGTTCACACCATTCTCNNNAGATCGGAAGA |
| 42704_2 | TATGAGTGTGGAGTCGTTGCCTCCTGGGTTCACACCATTCTCNNNAGATCGGAAGA |
| 31944_1 | TATGAGTGTGGAGTCGTTGCATATGGTTTCTTGTCTGGCGAGTNNNAGATCGGAAG |
| 31944_2 | TATGAGTGTGGAGTCGTTGCATATGGTTTCTTGTCTGGCGAGTNNNAGATCGGAAG |
| 138_1 | TATGAGTGTGGAGTCGTTGCCCGACTTTTCCACCATTCCTTACANNNAGATCGGAAG |
| 138_2 | TATGAGTGTGGAGTCGTTGCCCGACTTTTCCACCATTCCTTACANNNAGATCGGAAG |
| 30810_1 | TATGAGTGTGGAGTCGTTGCTGTCTAAAGTCTCCATTTCATGCANNNAGATCGGAA |
| 30810_2 | TATGAGTGTGGAGTCGTTGCTGTCTAAAGTCTCCATTTCATGCANNNAGATCGGAA |
| 38122_1 | TATGAGTGTGGAGTCGTTGCTAGAAGTAGGGTGACTGGGGATTNNNAGATCGGA |
| 38122_2 | TATGAGTGTGGAGTCGTTGCTAGAAGTAGGGTGACTGGGGATTNNNAGATCGGA |
| 35078_1 | TATGAGTGTGGAGTCGTTGCGTACACAACGTGCAGGTTTGTTANNNAGATCGGAA |
| 35078_2 | TATGAGTGTGGAGTCGTTGCGTACACAACGTGCAGGTTTGTTANNNAGATCGGAA |
| 13659_1 | TATGAGTGTGGAGTCGTTGCTGGAAGAAGAAATGTTTTTAGTTNNNAGATCGGAA |
| 13659_2 | TATGAGTGTGGAGTCGTTGCTGGAAGAAGAAATGTTTTTAGTTNNNAGATCGGAA |
| 50591_1 | TATGAGTGTGGAGTCGTTGCAGTTCAGGGAAGGGATAGAGGAANNNAGATCGGA |

| | |
|---|---|
| 50591_2 | TATGAGTGTGGAGTCGTTGCAGTTCAGGGAAGGGATAGAGGAANNNAGATCGGA |
| 13772_1 | TATGAGTGTGGAGTCGTTGCTCCTTATCCGTATACTTTGTATCNNNAGATCGGAAGA |
| 13772_2 | TATGAGTGTGGAGTCGTTGCTCCTTATCCGTATACTTTGTATCNNNAGATCGGAAGA |
| 38968_1 | TATGAGTGTGGAGTCGTTGCAGGTTTTCTTAGCTGCTTGGAGANNNAGATCGGAA |
| 38968_2 | TATGAGTGTGGAGTCGTTGCAGGTTTTCTTAGCTGCTTGGAGANNNAGATCGGAA |
| 38349_1 | TATGAGTGTGGAGTCGTTGCTTCCTGACTCTGAGGGGAATTTGNNNAGATCGGAA |
| 38349_2 | TATGAGTGTGGAGTCGTTGCTTCCTGACTCTGAGGGGAATTTGNNNAGATCGGAA |
| 35010_1 | TATGAGTGTGGAGTCGTTGCCACAATCTTGGCTCACTGCAACNNNAGATCGGAAGA |
| 35010_2 | TATGAGTGTGGAGTCGTTGCCACAATCTTGGCTCACTGCAACNNNAGATCGGAAGA |
| 51302_1 | TATGAGTGTGGAGTCGTTGCCACATTTTACTTGGATGGCAGCANNNAGATCGGAAG |
| 51302_2 | TATGAGTGTGGAGTCGTTGCCACATTTTACTTGGATGGCAGCANNNAGATCGGAAG |
| 38385_1 | TATGAGTGTGGAGTCGTTGCGAGAATCACTTGAACCCAGGAGANNNAGATCGGAA |
| 38385_2 | TATGAGTGTGGAGTCGTTGCGAGAATCACTTGAACCCAGGAGANNNAGATCGGAA |
| 34691_1 | TATGAGTGTGGAGTCGTTGCTTCAAGCAATTCTCCTGCCTCAGNNNAGATCGGAAG |
| 34691_2 | TATGAGTGTGGAGTCGTTGCTTCAAGCAATTCTCCTGCCTCAGNNNAGATCGGAAG |
| 32942_1 | TATGAGTGTGGAGTCGTTGCACCTATAATCCCAGCACTTCAGGNNNAGATCGGAAG |
| 32942_2 | TATGAGTGTGGAGTCGTTGCACCTATAATCCCAGCACTTCAGGNNNAGATCGGAAG |
| 43740_1 | TATGAGTGTGGAGTCGTTGCCCATAGACTTTTTCTGCACAGCCNNNAGATCGGAAG |
| 43740_2 | TATGAGTGTGGAGTCGTTGCCCATAGACTTTTTCTGCACAGCCNNNAGATCGGAAG |
| 31494_1 | TATGAGTGTGGAGTCGTTGCAGAGAAGAGAGAGAGATCTGGCANNNAGATCGGA |
| 31494_2 | TATGAGTGTGGAGTCGTTGCAGAGAAGAGAGAGAGATCTGGCANNNAGATCGGA |
| 46604_1 | TATGAGTGTGGAGTCGTTGCCCTCCCAAAGTGCTGAGATTACANNNAGATCGGAAG |
| 46604_2 | TATGAGTGTGGAGTCGTTGCCCTCCCAAAGTGCTGAGATTACANNNAGATCGGAAG |
| 43324_1 | TATGAGTGTGGAGTCGTTGCAGGCACGCTGTAATTTTTGTACTNNNAGATCGGAAG |
| 43324_2 | TATGAGTGTGGAGTCGTTGCAGGCACGCTGTAATTTTTGTACTNNNAGATCGGAAG |
| 39596_1 | TATGAGTGTGGAGTCGTTGCAGATACCATGTCTTGCCAGTCAGNNNAGATCGGAAG |
| 39596_2 | TATGAGTGTGGAGTCGTTGCAGATACCATGTCTTGCCAGTCAGNNNAGATCGGAAG |
| 41277_1 | TATGAGTGTGGAGTCGTTGCCTCACATCACGTCTGTCATTCCANNNAGATCGGAAGA |
| 41277_2 | TATGAGTGTGGAGTCGTTGCCTCACATCACGTCTGTCATTCCANNNAGATCGGAAGA |
| 40235_1 | TATGAGTGTGGAGTCGTTGCAGTTAGTCTTCATGGCCTCTTGGNNNAGATCGGAAG |
| 40235_2 | TATGAGTGTGGAGTCGTTGCAGTTAGTCTTCATGGCCTCTTGGNNNAGATCGGAAG |
| 46922_1 | TATGAGTGTGGAGTCGTTGCGTTGAATCTGCCTGGGAAAACTCNNNAGATCGGAA |
| 46922_2 | TATGAGTGTGGAGTCGTTGCGTTGAATCTGCCTGGGAAAACTCNNNAGATCGGAA |
| 41492_1 | TATGAGTGTGGAGTCGTTGCTAGGTAGTTAGGCAGACATGAGCNNNAGATCGGAA |
| 41492_2 | TATGAGTGTGGAGTCGTTGCTAGGTAGTTAGGCAGACATGAGCNNNAGATCGGAA |
| 53424_1 | TATGAGTGTGGAGTCGTTGCCCCCAGATCAAATCATCTCCCATNNNAGATCGGAAGA |
| 53424_2 | TATGAGTGTGGAGTCGTTGCCCCCAGATCAAATCATCTCCCATNNNAGATCGGAAGA |
| 49697_1 | TATGAGTGTGGAGTCGTTGCACTGCGACTCCCAAAGTTATGTTNNNAGATCGGAAG |
| 49697_2 | TATGAGTGTGGAGTCGTTGCACTGCGACTCCCAAAGTTATGTTNNNAGATCGGAAG |
| 46975_1 | TATGAGTGTGGAGTCGTTGCTTCAAGTGATTCTCCTGCCTCAGNNNAGATCGGAAG |
| 46975_2 | TATGAGTGTGGAGTCGTTGCTTCAAGTGATTCTCCTGCCTCAGNNNAGATCGGAAG |
| 42299_1 | TATGAGTGTGGAGTCGTTGCTTCCTAACTTGTGTCCAGTCCTGNNNAGATCGGAAG |
| 42299_2 | TATGAGTGTGGAGTCGTTGCTTCCTAACTTGTGTCCAGTCCTGNNNAGATCGGAAG |

| | |
|---|---|
| 44165_1 | TATGAGTGTGGAGTCGTTGCAGAAAAGAATGTCCATTGCCACCNNNAGATCGGAAG |
| 44165_2 | TATGAGTGTGGAGTCGTTGCAGAAAAGAATGTCCATTGCCACCNNNAGATCGGAAG |
| 51980_1 | TATGAGTGTGGAGTCGTTGCCTCCTGAGTAGCTGGGACTACAGNNNAGATCGGAA |
| 51980_2 | TATGAGTGTGGAGTCGTTGCCTCCTGAGTAGCTGGGACTACAGNNNAGATCGGAA |
| 10685_1 | TATGAGTGTGGAGTCGTTGCAACAATGGCCAGTCCGACTTCNNNAGATCGGAAGAG |
| 10685_2 | TATGAGTGTGGAGTCGTTGCAACAATGGCCAGTCCGACTTCNNNAGATCGGAAGAG |
| 50717_1 | TATGAGTGTGGAGTCGTTGCCTCCACACCATATGGCACTTCTTNNNAGATCGGAAGA |
| 50717_2 | TATGAGTGTGGAGTCGTTGCCTCCACACCATATGGCACTTCTTNNNAGATCGGAAGA |
| 48673_1 | TATGAGTGTGGAGTCGTTGCATGATCTGGGCTCACTGCAAGNNNAGATCGGAAGA |
| 48673_2 | TATGAGTGTGGAGTCGTTGCATGATCTGGGCTCACTGCAAGNNNAGATCGGAAGA |
| 37056_1 | TATGAGTGTGGAGTCGTTGCAGGGGAGTCAAAGGACACAAAATNNNAGATCGGA |
| 37056_2 | TATGAGTGTGGAGTCGTTGCAGGGGAGTCAAAGGACACAAAATNNNAGATCGGA |
| 41935_1 | TATGAGTGTGGAGTCGTTGCGTTCACGCCATTCTCCTACCTCNNNAGATCGGAAGAG |
| 41935_2 | TATGAGTGTGGAGTCGTTGCGTTCACGCCATTCTCCTACCTCNNNAGATCGGAAGAG |
| 29833_1 | TATGAGTGTGGAGTCGTTGCGCAGATACAGCTTCCACTCTCTCNNNAGATCGGAAG |
| 29833_2 | TATGAGTGTGGAGTCGTTGCGCAGATACAGCTTCCACTCTCTCNNNAGATCGGAAG |
| 30756_1 | TATGAGTGTGGAGTCGTTGCGAATTTTCTCCCAGAGAAGCAGCNNNAGATCGGAAG |
| 30756_2 | TATGAGTGTGGAGTCGTTGCGAATTTTCTCCCAGAGAAGCAGCNNNAGATCGGAAG |
| 47227_1 | TATGAGTGTGGAGTCGTTGCTAGTGAGGTCTCATGACACTACTNNNAGATCGGAAG |
| 47227_2 | TATGAGTGTGGAGTCGTTGCTAGTGAGGTCTCATGACACTACTNNNAGATCGGAAG |
| 12867_1 | TATGAGTGTGGAGTCGTTGCCTCATAGTTCAGAAGAGTTCAGNNNAGATCGGAAG |
| 12867_2 | TATGAGTGTGGAGTCGTTGCCTCATAGTTCAGAAGAGTTCAGNNNAGATCGGAAG |
| 36067_1 | TATGAGTGTGGAGTCGTTGCGCTTTGGGCATACTTCAAAATGCNNNAGATCGGAAG |
| 36067_2 | TATGAGTGTGGAGTCGTTGCGCTTTGGGCATACTTCAAAATGCNNNAGATCGGAAG |
| 13610_1 | TATGAGTGTGGAGTCGTTGCCCATTCTTGAGAGTTCCTTTTTNNNAGATCGGAAGA |
| 13610_2 | TATGAGTGTGGAGTCGTTGCCCATTCTTGAGAGTTCCTTTTTNNNAGATCGGAAGA |
| 52926_1 | TATGAGTGTGGAGTCGTTGCCGGTCCCTCTGCAAATGTTTGNNNAGATCGGAAGAG |
| 52926_2 | TATGAGTGTGGAGTCGTTGCCGGTCCCTCTGCAAATGTTTGNNNAGATCGGAAGAG |
| 32935_1 | TATGAGTGTGGAGTCGTTGCTGACATTCCTCCTTGTGAGAGTGNNNAGATCGGAAG |
| 32935_2 | TATGAGTGTGGAGTCGTTGCTGACATTCCTCCTTGTGAGAGTGNNNAGATCGGAAG |
| 44063_1 | TATGAGTGTGGAGTCGTTGCCAAACTCCTCACACCCATGGTTANNNAGATCGGAAG |
| 44063_2 | TATGAGTGTGGAGTCGTTGCCAAACTCCTCACACCCATGGTTANNNAGATCGGAAG |
| 39229_1 | TATGAGTGTGGAGTCGTTGCTGCTTTATTCAGAGCTGTGGACTNNNAGATCGGAAG |
| 39229_2 | TATGAGTGTGGAGTCGTTGCTGCTTTATTCAGAGCTGTGGACTNNNAGATCGGAAG |
| 32004_1 | TATGAGTGTGGAGTCGTTGCATGTTTAGGAGAGCATGGAAGGGNNNAGATCGGA |
| 32004_2 | TATGAGTGTGGAGTCGTTGCATGTTTAGGAGAGCATGGAAGGGNNNAGATCGGA |
| 11345_1 | TATGAGTGTGGAGTCGTTGCAGATTCCAATAGACCTTGAGGCCCNNNAGATCGGAA |
| 11345_2 | TATGAGTGTGGAGTCGTTGCAGATTCCAATAGACCTTGAGGCCCNNNAGATCGGAA |
| 46424_1 | TATGAGTGTGGAGTCGTTGCACCACGCCTGGCAAATTTTTTTANNNAGATCGGAAG |
| 46424_2 | TATGAGTGTGGAGTCGTTGCACCACGCCTGGCAAATTTTTTTANNNAGATCGGAAG |
| 39288_1 | TATGAGTGTGGAGTCGTTGCTGTCACAACTACTCAACACTGCTNNNAGATCGGAAG |
| 39288_2 | TATGAGTGTGGAGTCGTTGCTGTCACAACTACTCAACACTGCTNNNAGATCGGAAG |
| 33684_1 | TATGAGTGTGGAGTCGTTGCTTTTAGTAGAGACGGGGTTTCACNNNAGATCGGAA |

| | |
|---|---|
| 33684_2 | TATGAGTGTGGAGTCGTTGCTTTTAGTAGAGACGGGGTTTCACNNNAGATCGGAA |
| 32222_1 | TATGAGTGTGGAGTCGTTGCGAGGGAAGGAAGGAAAGAAGGAANNNAGATCGG |
| 32222_2 | TATGAGTGTGGAGTCGTTGCGAGGGAAGGAAGGAAAGAAGGAANNNAGATCGG |
| 38961_1 | TATGAGTGTGGAGTCGTTGCTTCTATGGAGAGGAATTTGGGGCNNNAGATCGGAA |
| 38961_2 | TATGAGTGTGGAGTCGTTGCTTCTATGGAGAGGAATTTGGGGCNNNAGATCGGAA |
| 40231_1 | TATGAGTGTGGAGTCGTTGCCAATGGCTTGATCTCAGCTCACTNNNAGATCGGAAG |
| 40231_2 | TATGAGTGTGGAGTCGTTGCCAATGGCTTGATCTCAGCTCACTNNNAGATCGGAAG |
| 46155_1 | TATGAGTGTGGAGTCGTTGCCTCCTGAGTAGCTGGGATTACAGNNNAGATCGGAA |
| 46155_2 | TATGAGTGTGGAGTCGTTGCCTCCTGAGTAGCTGGGATTACAGNNNAGATCGGAA |
| 33386_1 | TATGAGTGTGGAGTCGTTGCAACCCAAACGTCCATCAGTGATANNNAGATCGGAAG |
| 33386_2 | TATGAGTGTGGAGTCGTTGCAACCCAAACGTCCATCAGTGATANNNAGATCGGAAG |
| 30489_1 | TATGAGTGTGGAGTCGTTGCGATGTGCTTTTAGAAGGCCCAGANNNAGATCGGAA |
| 30489_2 | TATGAGTGTGGAGTCGTTGCGATGTGCTTTTAGAAGGCCCAGANNNAGATCGGAA |
| 53306_1 | TATGAGTGTGGAGTCGTTGCAGCAGACGTTCTTCTTCCCTTTCNNNAGATCGGAAGA |
| 53306_2 | TATGAGTGTGGAGTCGTTGCAGCAGACGTTCTTCTTCCCTTTCNNNAGATCGGAAGA |
| 48036_1 | TATGAGTGTGGAGTCGTTGCCTTCCCCCATGATCAAGCAATTCNNNAGATCGGAAGA |
| 48036_2 | TATGAGTGTGGAGTCGTTGCCTTCCCCCATGATCAAGCAATTCNNNAGATCGGAAGA |
| 41829_1 | TATGAGTGTGGAGTCGTTGCTGATCTCGGCTCACAGCAAGNNNAGATCGGAAGAG |
| 41829_2 | TATGAGTGTGGAGTCGTTGCTGATCTCGGCTCACAGCAAGNNNAGATCGGAAGAG |
| 48180_1 | TATGAGTGTGGAGTCGTTGCTAAGAGCTCTGTGGCTTTGGAAANNNAGATCGGAA |
| 48180_2 | TATGAGTGTGGAGTCGTTGCTAAGAGCTCTGTGGCTTTGGAAANNNAGATCGGAA |
| 29051_1 | TATGAGTGTGGAGTCGTTGCACTGCTATAGCCTTCATGCTCTGNNNAGATCGGAAG |
| 29051_2 | TATGAGTGTGGAGTCGTTGCACTGCTATAGCCTTCATGCTCTGNNNAGATCGGAAG |
| 43884_1 | TATGAGTGTGGAGTCGTTGCTGGGGGTGAATAAATTACCAGCTNNNAGATCGGAA |
| 43884_2 | TATGAGTGTGGAGTCGTTGCTGGGGGTGAATAAATTACCAGCTNNNAGATCGGAA |
| 42210_1 | TATGAGTGTGGAGTCGTTGCAGCCTGCTAAAATCTGTTTGAAANNNAGATCGGAAG |
| 42210_2 | TATGAGTGTGGAGTCGTTGCAGCCTGCTAAAATCTGTTTGAAANNNAGATCGGAAG |
| 28961_1 | TATGAGTGTGGAGTCGTTGCTGTGTGTATATATCGATGCGTGTNNNAGATCGGAA |
| 28961_2 | TATGAGTGTGGAGTCGTTGCTGTGTGTATATATCGATGCGTGTNNNAGATCGGAA |
| 9317_1 | TATGAGTGTGGAGTCGTTGCCCTAATACAACTGTGGGAGCTGGNNNAGATCGGAA |
| 9317_2 | TATGAGTGTGGAGTCGTTGCCCTAATACAACTGTGGGAGCTGGNNNAGATCGGAA |
| 34762_1 | TATGAGTGTGGAGTCGTTGCGATCTCTGATCACTGCAACCTCCNNNAGATCGGAAG |
| 34762_2 | TATGAGTGTGGAGTCGTTGCGATCTCTGATCACTGCAACCTCCNNNAGATCGGAAG |
| 46121_1 | TATGAGTGTGGAGTCGTTGCTGAGTAACTGGGATTACAGGTGCNNNAGATCGGAA |
| 46121_2 | TATGAGTGTGGAGTCGTTGCTGAGTAACTGGGATTACAGGTGCNNNAGATCGGAA |
| 29696_1 | TATGAGTGTGGAGTCGTTGCGAGAAAAGTGTCACCGATCACAAATTTNNNAGATCG |
| 29696_2 | TATGAGTGTGGAGTCGTTGCGAGAAAAGTGTCACCGATCACAAATTTNNNAGATCG |
| 30278_1 | TATGAGTGTGGAGTCGTTGCCTATGCGAGACAAACCCAAACAANNNAGATCGGAAG |
| 30278_2 | TATGAGTGTGGAGTCGTTGCCTATGCGAGACAAACCCAAACAANNNAGATCGGAAG |
| 49109_1 | TATGAGTGTGGAGTCGTTGCAGTGCTGGGATTATAGGCATGAGNNNAGATCGGAA |
| 49109_2 | TATGAGTGTGGAGTCGTTGCAGTGCTGGGATTATAGGCATGAGNNNAGATCGGAA |
| 48087_1 | TATGAGTGTGGAGTCGTTGCTCCCTGGTCTCTCACTTTATCCTNNNAGATCGGAAGA |
| 48087_2 | TATGAGTGTGGAGTCGTTGCTCCCTGGTCTCTCACTTTATCCTNNNAGATCGGAAGA |

| | |
|---|---|
| 48241_1 | TATGAGTGTGGAGTCGTTGCGGTAGCTGGAACTGCATGTCTNNNAGATCGGAAGA |
| 48241_2 | TATGAGTGTGGAGTCGTTGCGGTAGCTGGAACTGCATGTCTNNNAGATCGGAAGA |
| 36196_1 | TATGAGTGTGGAGTCGTTGCGTCCATGATTGAACCACTGCACNNNAGATCGGAAGA |
| 36196_2 | TATGAGTGTGGAGTCGTTGCGTCCATGATTGAACCACTGCACNNNAGATCGGAAGA |
| 38743_1 | TATGAGTGTGGAGTCGTTGCACAAGAGCGAAACTCCATCTAGGNNNAGATCGGAA |
| 38743_2 | TATGAGTGTGGAGTCGTTGCACAAGAGCGAAACTCCATCTAGGNNNAGATCGGAA |
| 52833_1 | TATGAGTGTGGAGTCGTTGCGAGCTGGATACAAGCCTGGGNNNAGATCGGAAGA |
| 52833_2 | TATGAGTGTGGAGTCGTTGCGAGCTGGATACAAGCCTGGGNNNAGATCGGAAGA |
| 38063_1 | TATGAGTGTGGAGTCGTTGCTCTCTTCACTTCTCCTCTCCCCNNNAGATCGGAAGAG |
| 38063_2 | TATGAGTGTGGAGTCGTTGCTCTCTTCACTTCTCCTCTCCCCNNNAGATCGGAAGAG |
| 34836_1 | TATGAGTGTGGAGTCGTTGCCACGCCCAGCTAATTTTTTTTCGNNNAGATCGGAAG |
| 34836_2 | TATGAGTGTGGAGTCGTTGCCACGCCCAGCTAATTTTTTTTCGNNNAGATCGGAAG |
| 49640_1 | TATGAGTGTGGAGTCGTTGCCTCCTGAGTAGCTGGGACTACAGNNNAGATCGGAA |
| 49640_2 | TATGAGTGTGGAGTCGTTGCCTCCTGAGTAGCTGGGACTACAGNNNAGATCGGAA |
| 52909_1 | TATGAGTGTGGAGTCGTTGCTCAGGGTATGTTTGACTCAAGGGNNNAGATCGGAA |
| 52909_2 | TATGAGTGTGGAGTCGTTGCTCAGGGTATGTTTGACTCAAGGGNNNAGATCGGAA |
| 40515_1 | TATGAGTGTGGAGTCGTTGCTCTACTGCCAAGGCCTTTGATACNNNAGATCGGAAG |
| 40515_2 | TATGAGTGTGGAGTCGTTGCTCTACTGCCAAGGCCTTTGATACNNNAGATCGGAAG |
| 30966_1 | TATGAGTGTGGAGTCGTTGCCTCTCAACTTCCCCCTCTTCTTCNNNAGATCGGAAGAG |
| 30966_2 | TATGAGTGTGGAGTCGTTGCCTCTCAACTTCCCCCTCTTCTTCNNNAGATCGGAAGAG |
| 47933_1 | TATGAGTGTGGAGTCGTTGCTCCCCCAGATTCAAGCAATTCTTNNNAGATCGGAAG |
| 47933_2 | TATGAGTGTGGAGTCGTTGCTCCCCCAGATTCAAGCAATTCTTNNNAGATCGGAAG |
| 49381_1 | TATGAGTGTGGAGTCGTTGCAATATTGACTCACTGCAACCCCTNNNAGATCGGAAG |
| 49381_2 | TATGAGTGTGGAGTCGTTGCAATATTGACTCACTGCAACCCCTNNNAGATCGGAAG |
| 29813_1 | TATGAGTGTGGAGTCGTTGCGTGAGAGAGGCTGATGGTAAGTTNNNAGATCGGA |
| 29813_2 | TATGAGTGTGGAGTCGTTGCGTGAGAGAGGCTGATGGTAAGTTNNNAGATCGGA |
| 46104_1 | TATGAGTGTGGAGTCGTTGCCTCCTGGGTAGCTGAGATTACAGNNNAGATCGGAA |
| 46104_2 | TATGAGTGTGGAGTCGTTGCCTCCTGGGTAGCTGAGATTACAGNNNAGATCGGAA |
| 44301_1 | TATGAGTGTGGAGTCGTTGCTTACCATAAATCTGCATCCCCCCNNNAGATCGGAAGA |
| 44301_2 | TATGAGTGTGGAGTCGTTGCTTACCATAAATCTGCATCCCCCCNNNAGATCGGAAGA |
| 13172_1 | TATGAGTGTGGAGTCGTTGCCCATCTGGTTGATCTAAAATTTTNNNAGATCGGAAG |
| 13172_2 | TATGAGTGTGGAGTCGTTGCCCATCTGGTTGATCTAAAATTTTNNNAGATCGGAAG |
| 43720_1 | TATGAGTGTGGAGTCGTTGCGCGGGGAGGGATTATGGTAATAGNNNAGATCGGA |
| 43720_2 | TATGAGTGTGGAGTCGTTGCGCGGGGAGGGATTATGGTAATAGNNNAGATCGGA |
| 37019_1 | TATGAGTGTGGAGTCGTTGCTTCCTGCTCAATTGAATGAAGGTNNNAGATCGGAAG |
| 37019_2 | TATGAGTGTGGAGTCGTTGCTTCCTGCTCAATTGAATGAAGGTNNNAGATCGGAAG |
| 10103_1 | TATGAGTGTGGAGTCGTTGCAGCACAGAAGCATCTCCCAAAAANNNAGATCGGAAG |
| 10103_2 | TATGAGTGTGGAGTCGTTGCAGCACAGAAGCATCTCCCAAAAANNNAGATCGGAAG |
| 49521_1 | TATGAGTGTGGAGTCGTTGCGTCTCCTGGGTTCAAGTGATTCTNNNAGATCGGAAG |
| 49521_2 | TATGAGTGTGGAGTCGTTGCGTCTCCTGGGTTCAAGTGATTCTNNNAGATCGGAAG |
| 42524_1 | TATGAGTGTGGAGTCGTTGCGATCTCAGTTCACTGCAACCTCTNNNAGATCGGAAG |
| 42524_2 | TATGAGTGTGGAGTCGTTGCGATCTCAGTTCACTGCAACCTCTNNNAGATCGGAAG |
| 44159_1 | TATGAGTGTGGAGTCGTTGCCAGGTGTGTTTGGGATGGAGGNNNAGATCGGAAG |

| | |
|---|---|
| 44159_2 | TATGAGTGTGGAGTCGTTGCCAGGTGTGTTTGGGATGGAGGNNNAGATCGGAAG |
| 34545_1 | TATGAGTGTGGAGTCGTTGCCCTCCCACTGGTTTCTTTCAGNNNAGATCGGAAGAG |
| 34545_2 | TATGAGTGTGGAGTCGTTGCCCTCCCACTGGTTTCTTTCAGNNNAGATCGGAAGAG |
| 40717_1 | TATGAGTGTGGAGTCGTTGCACATGCAGTATTTGTCTCCCTGTNNNAGATCGGAAG |
| 40717_2 | TATGAGTGTGGAGTCGTTGCACATGCAGTATTTGTCTCCCTGTNNNAGATCGGAAG |
| 50695_1 | TATGAGTGTGGAGTCGTTGCGACATCATTCTGTGGAGCCTCTGNNNAGATCGGAAG |
| 50695_2 | TATGAGTGTGGAGTCGTTGCGACATCATTCTGTGGAGCCTCTGNNNAGATCGGAAG |
| 42720_1 | TATGAGTGTGGAGTCGTTGCTTTTTTTTTTTTGCCCAGGCTGGNNNAGATCGGAAG |
| 42720_2 | TATGAGTGTGGAGTCGTTGCTTTTTTTTTTTTGCCCAGGCTGGNNNAGATCGGAAG |
| 30998_1 | TATGAGTGTGGAGTCGTTGCCTTAGAGGGAATCTGTATGGGGCNNNAGATCGGAA |
| 30998_2 | TATGAGTGTGGAGTCGTTGCCTTAGAGGGAATCTGTATGGGGCNNNAGATCGGAA |
| 37199_1 | TATGAGTGTGGAGTCGTTGCTTGTACGTTTTCGTTTCTGTGCANNNAGATCGGAAG |
| 37199_2 | TATGAGTGTGGAGTCGTTGCTTGTACGTTTTCGTTTCTGTGCANNNAGATCGGAAG |
| 7874_1 | TATGAGTGTGGAGTCGTTGCTGGGAAGGCAGTACAAAAACTGTNNNAGATCGGAA |
| 7874_2 | TATGAGTGTGGAGTCGTTGCTGGGAAGGCAGTACAAAAACTGTNNNAGATCGGAA |
| 37150_1 | TATGAGTGTGGAGTCGTTGCGCATGCCAGGGAGTGAAAAATAGNNNAGATCGGA |
| 37150_2 | TATGAGTGTGGAGTCGTTGCGCATGCCAGGGAGTGAAAAATAGNNNAGATCGGA |
| 42109_1 | TATGAGTGTGGAGTCGTTGCACCTAGAAGTGGAATTGCTAGACTNNNAGATCGGA |
| 42109_2 | TATGAGTGTGGAGTCGTTGCACCTAGAAGTGGAATTGCTAGACTNNNAGATCGGA |
| 41156_1 | TATGAGTGTGGAGTCGTTGCACCACCACCAAAAAACCAAAAGANNNAGATCGGAAG |
| 41156_2 | TATGAGTGTGGAGTCGTTGCACCACCACCAAAAAACCAAAAGANNNAGATCGGAAG |
| 31618_1 | TATGAGTGTGGAGTCGTTGCGTAGAGAGATTCCCAACAGGTGGNNNAGATCGGAA |
| 31618_2 | TATGAGTGTGGAGTCGTTGCGTAGAGAGATTCCCAACAGGTGGNNNAGATCGGAA |
| 50512_1 | TATGAGTGTGGAGTCGTTGCGCCTCACAGTATTTTGTTCCCAGNNNAGATCGGAAG |
| 50512_2 | TATGAGTGTGGAGTCGTTGCGCCTCACAGTATTTTGTTCCCAGNNNAGATCGGAAG |
| 49992_1 | TATGAGTGTGGAGTCGTTGCAAAGGGCCAGTAGGAAAATCTGTNNNAGATCGGAA |
| 49992_2 | TATGAGTGTGGAGTCGTTGCAAAGGGCCAGTAGGAAAATCTGTNNNAGATCGGAA |
| 33902_1 | TATGAGTGTGGAGTCGTTGCACAACACCACACCCAGCTAATTTNNNAGATCGGAAG |
| 33902_2 | TATGAGTGTGGAGTCGTTGCACAACACCACACCCAGCTAATTTNNNAGATCGGAAG |
| 46653_1 | TATGAGTGTGGAGTCGTTGCACCACGCCCAGCTAATTTTTTTTNNNAGATCGGAAGA |
| 46653_2 | TATGAGTGTGGAGTCGTTGCACCACGCCCAGCTAATTTTTTTTNNNAGATCGGAAGA |
| 47339_1 | TATGAGTGTGGAGTCGTTGCGCCACCACACCAGACTAATTTTTNNNAGATCGGAAG |
| 47339_2 | TATGAGTGTGGAGTCGTTGCGCCACCACACCAGACTAATTTTTNNNAGATCGGAAG |
| 49330_1 | TATGAGTGTGGAGTCGTTGCCAGCAGCATTTCTTACAGTCACCNNNAGATCGGAAG |
| 49330_2 | TATGAGTGTGGAGTCGTTGCCAGCAGCATTTCTTACAGTCACCNNNAGATCGGAAG |
| 33607_1 | TATGAGTGTGGAGTCGTTGCGTAGCTGGGATTACAGGCACATNNNAGATCGGAAG |
| 33607_2 | TATGAGTGTGGAGTCGTTGCGTAGCTGGGATTACAGGCACATNNNAGATCGGAAG |
| 41726_1 | TATGAGTGTGGAGTCGTTGCCTCCTGAGTAGCTGGGATTACAGNNNAGATCGGAA |
| 41726_2 | TATGAGTGTGGAGTCGTTGCCTCCTGAGTAGCTGGGATTACAGNNNAGATCGGAA |
| 50270_1 | TATGAGTGTGGAGTCGTTGCGGTGGAGCTAGGAAAGTCAGAAANNNAGATCGGA |
| 50270_2 | TATGAGTGTGGAGTCGTTGCGGTGGAGCTAGGAAAGTCAGAAANNNAGATCGGA |
| 50024_1 | TATGAGTGTGGAGTCGTTGCGATCTCCACCTCTCCTTCAGTTGNNNAGATCGGAAGA |
| 50024_2 | TATGAGTGTGGAGTCGTTGCGATCTCCACCTCTCCTTCAGTTGNNNAGATCGGAAGA |

| | |
|---|---|
| 35531_1 | TATGAGTGTGGAGTCGTTGCAGGTTCAAGTGATTCTTCTGCCTNNNAGATCGGAAG |
| 35531_2 | TATGAGTGTGGAGTCGTTGCAGGTTCAAGTGATTCTTCTGCCTNNNAGATCGGAAG |
| 48928_1 | TATGAGTGTGGAGTCGTTGCCAAGTAGCTGGGACTACAGGCNNNAGATCGGAAGA |
| 48928_2 | TATGAGTGTGGAGTCGTTGCCAAGTAGCTGGGACTACAGGCNNNAGATCGGAAGA |
| 37539_1 | TATGAGTGTGGAGTCGTTGCAATGATCCTCCCATCTCAGCTTCNNNAGATCGGAAGA |
| 37539_2 | TATGAGTGTGGAGTCGTTGCAATGATCCTCCCATCTCAGCTTCNNNAGATCGGAAGA |
| 47127_1 | TATGAGTGTGGAGTCGTTGCCAATCTTGGCTCACTGCAATCTCNNNAGATCGGAAG |
| 47127_2 | TATGAGTGTGGAGTCGTTGCCAATCTTGGCTCACTGCAATCTCNNNAGATCGGAAG |
| 30074_1 | TATGAGTGTGGAGTCGTTGCGATGCCCAAGTATGCCTAAAACCNNNAGATCGGAAG |
| 30074_2 | TATGAGTGTGGAGTCGTTGCGATGCCCAAGTATGCCTAAAACCNNNAGATCGGAAG |
| 44129_1 | TATGAGTGTGGAGTCGTTGCCCAAATGGGTCTACCTCCTTTCANNNAGATCGGAAG |
| 44129_2 | TATGAGTGTGGAGTCGTTGCCCAAATGGGTCTACCTCCTTTCANNNAGATCGGAAG |
| 43086_1 | TATGAGTGTGGAGTCGTTGCGATATGCAAATGGACACAGGAGCNNNAGATCGGAA |
| 43086_2 | TATGAGTGTGGAGTCGTTGCGATATGCAAATGGACACAGGAGCNNNAGATCGGAA |
| 52544_1 | TATGAGTGTGGAGTCGTTGCCAGACACCCAAAGCCTTTTCTTTNNNAGATCGGAAG |
| 52544_2 | TATGAGTGTGGAGTCGTTGCCAGACACCCAAAGCCTTTTCTTTNNNAGATCGGAAG |
| 51567_1 | TATGAGTGTGGAGTCGTTGCTTCCCATCCTGCTGATTAGTTCCNNNAGATCGGAAGA |
| 51567_2 | TATGAGTGTGGAGTCGTTGCTTCCCATCCTGCTGATTAGTTCCNNNAGATCGGAAGA |
| 39772_1 | TATGAGTGTGGAGTCGTTGCGAGGAGATATCTGCTCTGCCTTCNNNAGATCGGAAG |
| 39772_2 | TATGAGTGTGGAGTCGTTGCGAGGAGATATCTGCTCTGCCTTCNNNAGATCGGAAG |
| 10652_1 | TATGAGTGTGGAGTCGTTGCAATGAACACTTGGTTCTCCAGCGNNNAGATCGGAAG |
| 10652_2 | TATGAGTGTGGAGTCGTTGCAATGAACACTTGGTTCTCCAGCGNNNAGATCGGAAG |
| 13492_1 | TATGAGTGTGGAGTCGTTGCTGATTTCATGGTCTGACTATAAGNNNAGATCGGAAG |
| 13492_2 | TATGAGTGTGGAGTCGTTGCTGATTTCATGGTCTGACTATAAGNNNAGATCGGAAG |
| 46065_1 | TATGAGTGTGGAGTCGTTGCATAACGGGGCTTTGTTATGTTGCNNNAGATCGGAA |
| 46065_2 | TATGAGTGTGGAGTCGTTGCATAACGGGGCTTTGTTATGTTGCNNNAGATCGGAA |
| 44022_1 | TATGAGTGTGGAGTCGTTGCCATTGCTCCTCTGTTCTCCATCCNNNAGATCGGAAGA |
| 44022_2 | TATGAGTGTGGAGTCGTTGCCATTGCTCCTCTGTTCTCCATCCNNNAGATCGGAAGA |
| 47415_1 | TATGAGTGTGGAGTCGTTGCCTCCATCTCCTGACCTTGTGATCNNNAGATCGGAAGA |
| 47415_2 | TATGAGTGTGGAGTCGTTGCCTCCATCTCCTGACCTTGTGATCNNNAGATCGGAAGA |
| 34560_1 | TATGAGTGTGGAGTCGTTGCATGGTCCTAAGAGGAGGAGCAGNNNAGATCGGAA |
| 34560_2 | TATGAGTGTGGAGTCGTTGCATGGTCCTAAGAGGAGGAGCAGNNNAGATCGGAA |
| 8514_1 | TATGAGTGTGGAGTCGTTGCTCCAGTGTAACTTATTTGGTTGAACCNNNAGATCGG |
| 8514_2 | TATGAGTGTGGAGTCGTTGCTCCAGTGTAACTTATTTGGTTGAACCNNNAGATCGG |
| 41012_1 | TATGAGTGTGGAGTCGTTGCAGGTATCGAGTCTCACTATAGCCANNNAGATCGGAA |
| 41012_2 | TATGAGTGTGGAGTCGTTGCAGGTATCGAGTCTCACTATAGCCANNNAGATCGGAA |
| 49596_1 | TATGAGTGTGGAGTCGTTGCACGCCCGGCTAATTTTTGTATTTNNNAGATCGGAAG |
| 49596_2 | TATGAGTGTGGAGTCGTTGCACGCCCGGCTAATTTTTGTATTTNNNAGATCGGAAG |
| 30183_1 | TATGAGTGTGGAGTCGTTGCAATAAGTGGACAGGGCAGAATGTNNNAGATCGGAA |
| 30183_2 | TATGAGTGTGGAGTCGTTGCAATAAGTGGACAGGGCAGAATGTNNNAGATCGGAA |
| 12988_1 | TATGAGTGTGGAGTCGTTGCCTTTTCTCATACCTTTGACAGANNNAGATCGGAAGA |
| 12988_2 | TATGAGTGTGGAGTCGTTGCCTTTTCTCATACCTTTGACAGANNNAGATCGGAAGA |
| 40577_1 | TATGAGTGTGGAGTCGTTGCAGAACACTCAGGTAACTCACTGCNNNAGATCGGAAG |

| | |
|---|---|
| 40577_2 | TATGAGTGTGGAGTCGTTGCAGAACACTCAGGTAACTCACTGCNNNAGATCGGAAG |
| 44331_1 | TATGAGTGTGGAGTCGTTGCGATGACCGTACTGCATCTCAGAGNNNAGATCGGAAG |
| 44331_2 | TATGAGTGTGGAGTCGTTGCGATGACCGTACTGCATCTCAGAGNNNAGATCGGAAG |
| 31079_1 | TATGAGTGTGGAGTCGTTGCCTCTAAAGCAATGAGCAGAGTGGNNNAGATCGGAA |
| 31079_2 | TATGAGTGTGGAGTCGTTGCCTCTAAAGCAATGAGCAGAGTGGNNNAGATCGGAA |
| 28985_1 | TATGAGTGTGGAGTCGTTGCAGACGGAGTCTCTCTCTCTGTCNNNAGATCGGAAGA |
| 28985_2 | TATGAGTGTGGAGTCGTTGCAGACGGAGTCTCTCTCTCTGTCNNNAGATCGGAAGA |
| 46772_1 | TATGAGTGTGGAGTCGTTGCTAGGGACCACCATCTCCTAGAAGNNNAGATCGGAAG |
| 46772_2 | TATGAGTGTGGAGTCGTTGCTAGGGACCACCATCTCCTAGAAGNNNAGATCGGAAG |
| 46376_1 | TATGAGTGTGGAGTCGTTGCAGCCAGAGCAGTTAGACAAGAAANNNAGATCGGAA |
| 46376_2 | TATGAGTGTGGAGTCGTTGCAGCCAGAGCAGTTAGACAAGAAANNNAGATCGGAA |
| 48249_1 | TATGAGTGTGGAGTCGTTGCATAAGGAATGCCACCTGGGATTTNNNAGATCGGAAG |
| 48249_2 | TATGAGTGTGGAGTCGTTGCATAAGGAATGCCACCTGGGATTTNNNAGATCGGAAG |
| 37187_1 | TATGAGTGTGGAGTCGTTGCAGATCTAGTGTTTGGTAGCACCCNNNAGATCGGAAG |
| 37187_2 | TATGAGTGTGGAGTCGTTGCAGATCTAGTGTTTGGTAGCACCCNNNAGATCGGAAG |
| 52710_1 | TATGAGTGTGGAGTCGTTGCGAGTGGAGGAAGGAGGGGTGNNNAGATCGGAAG |
| 52710_2 | TATGAGTGTGGAGTCGTTGCGAGTGGAGGAAGGAGGGGTGNNNAGATCGGAAG |
| 49105_1 | TATGAGTGTGGAGTCGTTGCCCTAGTAGCCAGGATTACAGGTGNNNAGATCGGAA |
| 49105_2 | TATGAGTGTGGAGTCGTTGCCCTAGTAGCCAGGATTACAGGTGNNNAGATCGGAA |
| 52091_1 | TATGAGTGTGGAGTCGTTGCAAATGACATGGCATTCTCTCCCTNNNAGATCGGAAG |
| 52091_2 | TATGAGTGTGGAGTCGTTGCAAATGACATGGCATTCTCTCCCTNNNAGATCGGAAG |
| 52152_1 | TATGAGTGTGGAGTCGTTGCGGGAGGAACCATGTGGAGATAATNNNAGATCGGA |
| 52152_2 | TATGAGTGTGGAGTCGTTGCGGGAGGAACCATGTGGAGATAATNNNAGATCGGA |
| 10270_1 | TATGAGTGTGGAGTCGTTGCGGTATCAGGGTAATGCTCCTGACNNNAGATCGGAAG |
| 10270_2 | TATGAGTGTGGAGTCGTTGCGGTATCAGGGTAATGCTCCTGACNNNAGATCGGAAG |
| 44278_1 | TATGAGTGTGGAGTCGTTGCGTCAGAAACTGATTTTGCCAGAGTNNNAGATCGGA |
| 44278_2 | TATGAGTGTGGAGTCGTTGCGTCAGAAACTGATTTTGCCAGAGTNNNAGATCGGA |
| 12765_1 | TATGAGTGTGGAGTCGTTGCCCAATCGTCTCTTAAGTTTTTATNNNAGATCGGAAGA |
| 12765_2 | TATGAGTGTGGAGTCGTTGCCCAATCGTCTCTTAAGTTTTTATNNNAGATCGGAAGA |
| 44869_1 | TATGAGTGTGGAGTCGTTGCAGGGGAGGAAAGTGAGAAAAAGGNNNAGATCGG |
| 44869_2 | TATGAGTGTGGAGTCGTTGCAGGGGAGGAAAGTGAGAAAAAGGNNNAGATCGG |
| 53029_1 | TATGAGTGTGGAGTCGTTGCTCCCCAGAAATTCCCACCATTAGNNNAGATCGGAAG |
| 53029_2 | TATGAGTGTGGAGTCGTTGCTCCCCAGAAATTCCCACCATTAGNNNAGATCGGAAG |
| 34431_1 | TATGAGTGTGGAGTCGTTGCGCTCAGGTGACGAGTACACTAAANNNAGATCGGAA |
| 34431_2 | TATGAGTGTGGAGTCGTTGCGCTCAGGTGACGAGTACACTAAANNNAGATCGGAA |
| 53320_1 | TATGAGTGTGGAGTCGTTGCCTCTGTGGTGGCTTTTCTCTTTGNNNAGATCGGAAG |
| 53320_2 | TATGAGTGTGGAGTCGTTGCCTCTGTGGTGGCTTTTCTCTTTGNNNAGATCGGAAG |
| 45429_1 | TATGAGTGTGGAGTCGTTGCATTTTCTCTTTTTCCCCTCCCCANNNAGATCGGAAGAG |
| 45429_2 | TATGAGTGTGGAGTCGTTGCATTTTCTCTTTTTCCCCTCCCCANNNAGATCGGAAGAG |
| 30122_1 | TATGAGTGTGGAGTCGTTGCTTAGCTGCCTTGGTAGAGATTGCNNNAGATCGGAAG |
| 30122_2 | TATGAGTGTGGAGTCGTTGCTTAGCTGCCTTGGTAGAGATTGCNNNAGATCGGAAG |
| 41074_1 | TATGAGTGTGGAGTCGTTGCTTCACTATATTGCCCAGGCTTGTNNNAGATCGGAAG |
| 41074_2 | TATGAGTGTGGAGTCGTTGCTTCACTATATTGCCCAGGCTTGTNNNAGATCGGAAG |

| | |
|---|---|
| 12862_1 | TATGAGTGTGGAGTCGTTGCCTGGACTCTATATTGTCAAACTANNNAGATCGGAAG |
| 12862_2 | TATGAGTGTGGAGTCGTTGCCTGGACTCTATATTGTCAAACTANNNAGATCGGAAG |
| 39573_1 | TATGAGTGTGGAGTCGTTGCTCGATGACCTAACCACAATCCTGNNNAGATCGGAAG |
| 39573_2 | TATGAGTGTGGAGTCGTTGCTCGATGACCTAACCACAATCCTGNNNAGATCGGAAG |
| 35449_1 | TATGAGTGTGGAGTCGTTGCAGACGGAGTCTCGCTCTGTCNNNAGATCGGAAGAG |
| 35449_2 | TATGAGTGTGGAGTCGTTGCAGACGGAGTCTCGCTCTGTCNNNAGATCGGAAGAG |
| 46139_1 | TATGAGTGTGGAGTCGTTGCATACTGACACACTTCAGGCTGAGNNNAGATCGGAAG |
| 46139_2 | TATGAGTGTGGAGTCGTTGCATACTGACACACTTCAGGCTGAGNNNAGATCGGAAG |
| 51291_1 | TATGAGTGTGGAGTCGTTGCTCAAGAGGGCAAGCATTCTTACTNNNAGATCGGAAG |
| 51291_2 | TATGAGTGTGGAGTCGTTGCTCAAGAGGGCAAGCATTCTTACTNNNAGATCGGAAG |
| 41749_1 | TATGAGTGTGGAGTCGTTGCGCTAGTTCCTACTGGCATCATGANNNAGATCGGAAG |
| 41749_2 | TATGAGTGTGGAGTCGTTGCGCTAGTTCCTACTGGCATCATGANNNAGATCGGAAG |
| 30430_1 | TATGAGTGTGGAGTCGTTGCAATACTCACAGACCCCCATGTGANNNAGATCGGAAG |
| 30430_2 | TATGAGTGTGGAGTCGTTGCAATACTCACAGACCCCCATGTGANNNAGATCGGAAG |
| 10386_1 | TATGAGTGTGGAGTCGTTGCACCAGTTTGGGCTTGAATACGTTNNNAGATCGGAAG |
| 10386_2 | TATGAGTGTGGAGTCGTTGCACCAGTTTGGGCTTGAATACGTTNNNAGATCGGAAG |
| 52441_1 | TATGAGTGTGGAGTCGTTGCCGATCCTGTCCTTTCCCTGTTTANNNAGATCGGAAGA |
| 52441_2 | TATGAGTGTGGAGTCGTTGCCGATCCTGTCCTTTCCCTGTTTANNNAGATCGGAAGA |
| 45214_1 | TATGAGTGTGGAGTCGTTGCGGTTTCATGTACCGCATCAGATGNNNAGATCGGAAG |
| 45214_2 | TATGAGTGTGGAGTCGTTGCGGTTTCATGTACCGCATCAGATGNNNAGATCGGAAG |
| 50249_1 | TATGAGTGTGGAGTCGTTGCCTGTGGAGATGTTCATGACATGCNNNAGATCGGAA |
| 50249_2 | TATGAGTGTGGAGTCGTTGCCTGTGGAGATGTTCATGACATGCNNNAGATCGGAA |
| 42862_1 | TATGAGTGTGGAGTCGTTGCTGCTCCTATAAACATGTGTGTGCNNNAGATCGGAAG |
| 42862_2 | TATGAGTGTGGAGTCGTTGCTGCTCCTATAAACATGTGTGTGCNNNAGATCGGAAG |
| 36356_1 | TATGAGTGTGGAGTCGTTGCACTCCAATTCCATTTCGCCAAAGNNNAGATCGGAAG |
| 36356_2 | TATGAGTGTGGAGTCGTTGCACTCCAATTCCATTTCGCCAAAGNNNAGATCGGAAG |
| 29197_1 | TATGAGTGTGGAGTCGTTGCGATCTCTCGGCTCATTGTAACCTNNNAGATCGGAAG |
| 29197_2 | TATGAGTGTGGAGTCGTTGCGATCTCTCGGCTCATTGTAACCTNNNAGATCGGAAG |
| 43520_1 | TATGAGTGTGGAGTCGTTGCCCACGCCCAGCTAATTTTGTATTNNNAGATCGGAAG |
| 43520_2 | TATGAGTGTGGAGTCGTTGCCCACGCCCAGCTAATTTTGTATTNNNAGATCGGAAG |
| 32477_1 | TATGAGTGTGGAGTCGTTGCCTTGATCTCCTGACCTCGTGATCNNNAGATCGGAAG |
| 32477_2 | TATGAGTGTGGAGTCGTTGCCTTGATCTCCTGACCTCGTGATCNNNAGATCGGAAG |
| 49998_1 | TATGAGTGTGGAGTCGTTGCCTTGGCCTCCTCTTTCTCACAAANNNAGATCGGAAGA |
| 49998_2 | TATGAGTGTGGAGTCGTTGCCTTGGCCTCCTCTTTCTCACAAANNNAGATCGGAAGA |
| 47148_1 | TATGAGTGTGGAGTCGTTGCGATCTCGGCTCATTACAACCTCTNNNAGATCGGAAG |
| 47148_2 | TATGAGTGTGGAGTCGTTGCGATCTCGGCTCATTACAACCTCTNNNAGATCGGAAG |
| 48459_1 | TATGAGTGTGGAGTCGTTGCCTAGGTCTTGAACTCCTTGGCTTNNNAGATCGGAAG |
| 48459_2 | TATGAGTGTGGAGTCGTTGCCTAGGTCTTGAACTCCTTGGCTTNNNAGATCGGAAG |
| 48139_1 | TATGAGTGTGGAGTCGTTGCAAACTCCTGGCCTCAAGTGATCNNNAGATCGGAAGA |
| 48139_2 | TATGAGTGTGGAGTCGTTGCAAACTCCTGGCCTCAAGTGATCNNNAGATCGGAAGA |
| 13019_1 | TATGAGTGTGGAGTCGTTGCACAAATTAATATCCAGTAGAGAGANNNAGATCGGA |
| 13019_2 | TATGAGTGTGGAGTCGTTGCACAAATTAATATCCAGTAGAGAGANNNAGATCGGA |
| 40647_1 | TATGAGTGTGGAGTCGTTGCTCACATTTCTGGAGGCTAGAAGCNNNAGATCGGAA |

| | |
|---|---|
| 40647_2 | TATGAGTGTGGAGTCGTTGCTCACATTTCTGGAGGCTAGAAGCNNNAGATCGGAAG |
| 29206_1 | TATGAGTGTGGAGTCGTTGCTAGATCTGGCATGTGAAACACCANNNAGATCGGAAG |
| 29206_2 | TATGAGTGTGGAGTCGTTGCTAGATCTGGCATGTGAAACACCANNNAGATCGGAAG |
| 46428_1 | TATGAGTGTGGAGTCGTTGCTCCAGGTTCAAGTGATTCTCCTGNNNAGATCGGAAG |
| 46428_2 | TATGAGTGTGGAGTCGTTGCTCCAGGTTCAAGTGATTCTCCTGNNNAGATCGGAAG |
| 9680_1 | TATGAGTGTGGAGTCGTTGCCGTTCTGTTTACCTGGAGATCCCNNNAGATCGGAAGA |
| 9680_2 | TATGAGTGTGGAGTCGTTGCCGTTCTGTTTACCTGGAGATCCCNNNAGATCGGAAGA |
| 34936_1 | TATGAGTGTGGAGTCGTTGCCACCACAGTCGCCTAATTTTTGTNNNAGATCGGAAGA |
| 34936_2 | TATGAGTGTGGAGTCGTTGCCACCACAGTCGCCTAATTTTTGTNNNAGATCGGAAGA |
| 10160_1 | TATGAGTGTGGAGTCGTTGCGGCTGGGATGGCATAATTTGGAANNNAGATCGGAA |
| 10160_2 | TATGAGTGTGGAGTCGTTGCGGCTGGGATGGCATAATTTGGAANNNAGATCGGAA |
| 12825_1 | TATGAGTGTGGAGTCGTTGCCTAACCAATAAAACAAGAGATCCNNNAGATCGGAAG |
| 12825_2 | TATGAGTGTGGAGTCGTTGCCTAACCAATAAAACAAGAGATCCNNNAGATCGGAAG |
| 36738_1 | TATGAGTGTGGAGTCGTTGCCATCATGGGAGAAAACACTGCAGNNNAGATCGGAA |
| 36738_2 | TATGAGTGTGGAGTCGTTGCCATCATGGGAGAAAACACTGCAGNNNAGATCGGAA |
| 32346_1 | TATGAGTGTGGAGTCGTTGCTCACCAATATCCACTGTTCTGCANNNAGATCGGAAGA |
| 32346_2 | TATGAGTGTGGAGTCGTTGCTCACCAATATCCACTGTTCTGCANNNAGATCGGAAGA |
| 49309_1 | TATGAGTGTGGAGTCGTTGCTTCAAGTGATTCTCCTGCCTCAGNNNAGATCGGAAGA |
| 49309_2 | TATGAGTGTGGAGTCGTTGCTTCAAGTGATTCTCCTGCCTCAGNNNAGATCGGAAGA |
| 35196_1 | TATGAGTGTGGAGTCGTTGCTCCAAGTGATTTGTCTGCCTCAGNNNAGATCGGAAG |
| 35196_2 | TATGAGTGTGGAGTCGTTGCTCCAAGTGATTTGTCTGCCTCAGNNNAGATCGGAAG |
| 39330_1 | TATGAGTGTGGAGTCGTTGCTTTGAAGAAGGCAAGATCAGGGANNNAGATCGGAA |
| 39330_2 | TATGAGTGTGGAGTCGTTGCTTTGAAGAAGGCAAGATCAGGGANNNAGATCGGAA |
| 2254_1 | TATGAGTGTGGAGTCGTTGCGTGAGGCAATGGATGTGTTAGCTANNNAGATCGGA |
| 2254_2 | TATGAGTGTGGAGTCGTTGCGTGAGGCAATGGATGTGTTAGCTANNNAGATCGGA |
| 29600_1 | TATGAGTGTGGAGTCGTTGCAGCAGCATGACAGGAAAACAAGANNNAGATCGGAA |
| 29600_2 | TATGAGTGTGGAGTCGTTGCAGCAGCATGACAGGAAAACAAGANNNAGATCGGAA |
| 42508_1 | TATGAGTGTGGAGTCGTTGCGGTGAGTCTGTGGTCATTTTGTGNNNAGATCGGAA |
| 42508_2 | TATGAGTGTGGAGTCGTTGCGGTGAGTCTGTGGTCATTTTGTGNNNAGATCGGAA |
| 41273_1 | TATGAGTGTGGAGTCGTTGCGGCAGAGACTGGAGTGATGTAGNNNAGATCGGAA |
| 41273_2 | TATGAGTGTGGAGTCGTTGCGGCAGAGACTGGAGTGATGTAGNNNAGATCGGAA |
| 46882_1 | TATGAGTGTGGAGTCGTTGCTAGGTATGGGTCTGTGGATGTGTNNNAGATCGGAA |
| 46882_2 | TATGAGTGTGGAGTCGTTGCTAGGTATGGGTCTGTGGATGTGTNNNAGATCGGAA |
| 39519_1 | TATGAGTGTGGAGTCGTTGCGTAGGAGGATCACTTGAGCTCTGNNNAGATCGGAA |
| 39519_2 | TATGAGTGTGGAGTCGTTGCGTAGGAGGATCACTTGAGCTCTGNNNAGATCGGAA |
| 34658_1 | TATGAGTGTGGAGTCGTTGCTGTACTACAGCCTAGGCAACAAGNNNAGATCGGAAG |
| 34658_2 | TATGAGTGTGGAGTCGTTGCTGTACTACAGCCTAGGCAACAAGNNNAGATCGGAAG |
| 51369_1 | TATGAGTGTGGAGTCGTTGCCATTCTCAGATCTCCAGCTGTGTNNNAGATCGGAAGA |
| 51369_2 | TATGAGTGTGGAGTCGTTGCCATTCTCAGATCTCCAGCTGTGTNNNAGATCGGAAGA |
| 37174_1 | TATGAGTGTGGAGTCGTTGCGCCTTGACTCTATTGTACATGGCNNNAGATCGGAAG |
| 37174_2 | TATGAGTGTGGAGTCGTTGCGCCTTGACTCTATTGTACATGGCNNNAGATCGGAAG |
| 51289_1 | TATGAGTGTGGAGTCGTTGCCAAAAAACTGGTGCGGATCCTAGNNNAGATCGGAA |
| 51289_2 | TATGAGTGTGGAGTCGTTGCCAAAAAACTGGTGCGGATCCTAGNNNAGATCGGAA |

| | |
|---|---|
| 3201_1 | TATGAGTGTGGAGTCGTTGCCAAGCACCTTGAACTCAAAAGNNNAGATCGGAAGAG |
| 3201_2 | TATGAGTGTGGAGTCGTTGCCAAGCACCTTGAACTCAAAAGNNNAGATCGGAAGAG |
| 31157_1 | TATGAGTGTGGAGTCGTTGCCTAAAGAGCCAAATCTGTTGCCCNNNAGATCGGAAG |
| 31157_2 | TATGAGTGTGGAGTCGTTGCCTAAAGAGCCAAATCTGTTGCCCNNNAGATCGGAAG |
| 30356_1 | TATGAGTGTGGAGTCGTTGCTCTTTTGAAGGTATCATGGTCCAGAGANNNAGATCG |
| 30356_2 | TATGAGTGTGGAGTCGTTGCTCTTTTGAAGGTATCATGGTCCAGAGANNNAGATCG |
| 42634_1 | TATGAGTGTGGAGTCGTTGCACAGAGAGAGAGAGATGGGGAAANNNAGATCGGA |
| 42634_2 | TATGAGTGTGGAGTCGTTGCACAGAGAGAGAGAGATGGGGAAANNNAGATCGGA |
| 30024_1 | TATGAGTGTGGAGTCGTTGCACCCATCCTCCATCTATAGCACTNNNAGATCGGAAGA |
| 30024_2 | TATGAGTGTGGAGTCGTTGCACCCATCCTCCATCTATAGCACTNNNAGATCGGAAGA |
| 50993_1 | TATGAGTGTGGAGTCGTTGCCAGATAGAGACAGAGGAAGGGGANNNAGATCGGA |
| 50993_2 | TATGAGTGTGGAGTCGTTGCCAGATAGAGACAGAGGAAGGGGANNNAGATCGGA |
| 47418_1 | TATGAGTGTGGAGTCGTTGCCCTCCCAGGTACAAGCTATTCTTNNNAGATCGGAAGA |
| 47418_2 | TATGAGTGTGGAGTCGTTGCCCTCCCAGGTACAAGCTATTCTTNNNAGATCGGAAGA |
| 34061_1 | TATGAGTGTGGAGTCGTTGCCAAGTAGCTGGGACTATAGGCGNNNAGATCGGAAG |
| 34061_2 | TATGAGTGTGGAGTCGTTGCCAAGTAGCTGGGACTATAGGCGNNNAGATCGGAAG |
| 46391_1 | TATGAGTGTGGAGTCGTTGCCTCACAGGCATCTACCAAACTCTNNNAGATCGGAAGA |
| 46391_2 | TATGAGTGTGGAGTCGTTGCCTCACAGGCATCTACCAAACTCTNNNAGATCGGAAGA |
| 42319_1 | TATGAGTGTGGAGTCGTTGCTGTAATGGAGCCAATATCCCCAANNNAGATCGGAAG |
| 42319_2 | TATGAGTGTGGAGTCGTTGCTGTAATGGAGCCAATATCCCCAANNNAGATCGGAAG |
| 29634_1 | TATGAGTGTGGAGTCGTTGCATACAACCATCGTTTGCTCCTTCNNNAGATCGGAAGA |
| 29634_2 | TATGAGTGTGGAGTCGTTGCATACAACCATCGTTTGCTCCTTCNNNAGATCGGAAGA |
| 45514_1 | TATGAGTGTGGAGTCGTTGCTGTCCAGCTCGAAGACAATTGAANNNAGATCGGAAG |
| 45514_2 | TATGAGTGTGGAGTCGTTGCTGTCCAGCTCGAAGACAATTGAANNNAGATCGGAAG |
| 49965_1 | TATGAGTGTGGAGTCGTTGCTTCATTGCTCTTACCTGTTCCCANNNAGATCGGAAGA |
| 49965_2 | TATGAGTGTGGAGTCGTTGCTTCATTGCTCTTACCTGTTCCCANNNAGATCGGAAGA |
| 39366_1 | TATGAGTGTGGAGTCGTTGCACAGAACTTGGTGTCTCATTGGANNNAGATCGGAAG |
| 39366_2 | TATGAGTGTGGAGTCGTTGCACAGAACTTGGTGTCTCATTGGANNNAGATCGGAAG |
| 40533_1 | TATGAGTGTGGAGTCGTTGCTTACCTCAAGAGGTTGCTGTGAANNNAGATCGGAAG |
| 40533_2 | TATGAGTGTGGAGTCGTTGCTTACCTCAAGAGGTTGCTGTGAANNNAGATCGGAAG |
| 48521_1 | TATGAGTGTGGAGTCGTTGCCAAGTAGCTGGGACTACAGGTGNNNAGATCGGAAG |
| 48521_2 | TATGAGTGTGGAGTCGTTGCCAAGTAGCTGGGACTACAGGTGNNNAGATCGGAAG |
| 51553_1 | TATGAGTGTGGAGTCGTTGCATGAGGTGATATTGGCCAAAAGANNNAGATCGGAA |
| 51553_2 | TATGAGTGTGGAGTCGTTGCATGAGGTGATATTGGCCAAAAGANNNAGATCGGAA |
| 51572_1 | TATGAGTGTGGAGTCGTTGCGGTGTCAAGGGAGAGGCAAAGNNNAGATCGGAAG |
| 51572_2 | TATGAGTGTGGAGTCGTTGCGGTGTCAAGGGAGAGGCAAAGNNNAGATCGGAAG |
| 37463_1 | TATGAGTGTGGAGTCGTTGCTGCAGACCTCAGACCTCAATCTANNNAGATCGGAAGA |
| 37463_2 | TATGAGTGTGGAGTCGTTGCTGCAGACCTCAGACCTCAATCTANNNAGATCGGAAGA |
| 48358_1 | TATGAGTGTGGAGTCGTTGCCCCCTACAGTCACAGAAACTACANNNAGATCGGAAGA |
| 48358_2 | TATGAGTGTGGAGTCGTTGCCCCCTACAGTCACAGAAACTACANNNAGATCGGAAGA |
| 37686_1 | TATGAGTGTGGAGTCGTTGCTGTGAATGAAGCTAAAGGAGTGCNNNAGATCGGAA |
| 37686_2 | TATGAGTGTGGAGTCGTTGCTGTGAATGAAGCTAAAGGAGTGCNNNAGATCGGAA |
| 13406_1 | TATGAGTGTGGAGTCGTTGCCATGGTAATTAGCACTTTTCCNNNAGATCGGAAGAG |

| | |
|---|---|
| 13406_2 | TATGAGTGTGGAGTCGTTGCCATGGTAATTAGCACTTTTCCNNNAGATCGGAAGAG |
| 38247_1 | TATGAGTGTGGAGTCGTTGCTGGAGTTTCATGTTCAGCCAAACNNNAGATCGGAAG |
| 38247_2 | TATGAGTGTGGAGTCGTTGCTGGAGTTTCATGTTCAGCCAAACNNNAGATCGGAAG |
| 51891_1 | TATGAGTGTGGAGTCGTTGCTTGTGGGTTCACTGCCTTTTTTTNNNAGATCGGAAGA |
| 51891_2 | TATGAGTGTGGAGTCGTTGCTTGTGGGTTCACTGCCTTTTTTTNNNAGATCGGAAGA |
| 46176_1 | TATGAGTGTGGAGTCGTTGCTGAAGTTTCCCAGGCTATAGTGGNNNAGATCGGAAG |
| 46176_2 | TATGAGTGTGGAGTCGTTGCTGAAGTTTCCCAGGCTATAGTGGNNNAGATCGGAAG |
| 30834_1 | TATGAGTGTGGAGTCGTTGCTTGGATAGAAGGCTGGAATGCTTNNNAGATCGGAA |
| 30834_2 | TATGAGTGTGGAGTCGTTGCTTGGATAGAAGGCTGGAATGCTTNNNAGATCGGAA |
| 41740_1 | TATGAGTGTGGAGTCGTTGCGAGTGCAGTGGTGCGATCTTNNNAGATCGGAAGAG |
| 41740_2 | TATGAGTGTGGAGTCGTTGCGAGTGCAGTGGTGCGATCTTNNNAGATCGGAAGAG |
| 38539_1 | TATGAGTGTGGAGTCGTTGCTTTAGTACATGTGCAGGATGTGCNNNAGATCGGAAG |
| 38539_2 | TATGAGTGTGGAGTCGTTGCTTTAGTACATGTGCAGGATGTGCNNNAGATCGGAAG |
| 30668_1 | ATGAGTGTGGAGTCGTTGCAATCATACTGACCACTCTCTCAGACTNNNAGATCGGAA |
| 30668_2 | ATGAGTGTGGAGTCGTTGCAATCATACTGACCACTCTCTCAGACTNNNAGATCGGAA |
| 50345_1 | TATGAGTGTGGAGTCGTTGCTGAGGCAGGAGAATCACTTCAATNNNAGATCGGAA |
| 50345_2 | TATGAGTGTGGAGTCGTTGCTGAGGCAGGAGAATCACTTCAATNNNAGATCGGAA |
| 12937_1 | TATGAGTGTGGAGTCGTTGCAATTCTCCATCCTTTCCAAAGNNNAGATCGGAAGAG |
| 12937_2 | TATGAGTGTGGAGTCGTTGCAATTCTCCATCCTTTCCAAAGNNNAGATCGGAAGAG |
| 2594_1 | TATGAGTGTGGAGTCGTTGCGCTGAGGTATTTGTTTGCTCTTGGNNNAGATCGGAA |
| 2594_2 | TATGAGTGTGGAGTCGTTGCGCTGAGGTATTTGTTTGCTCTTGGNNNAGATCGGAA |
| 29713_1 | TATGAGTGTGGAGTCGTTGCAGAACAATTTGGTGAATGGCTGGNNNAGATCGGAA |
| 29713_2 | TATGAGTGTGGAGTCGTTGCAGAACAATTTGGTGAATGGCTGGNNNAGATCGGAA |
| 46156_1 | TATGAGTGTGGAGTCGTTGCTTCACGCCATTCTCCTGTCTCNNNAGATCGGAAGAGC |
| 46156_2 | TATGAGTGTGGAGTCGTTGCTTCACGCCATTCTCCTGTCTCNNNAGATCGGAAGAGC |
| 50892_1 | TATGAGTGTGGAGTCGTTGCAGTCAGATGAAATTCCCTGGGATNNNAGATCGGAAG |
| 50892_2 | TATGAGTGTGGAGTCGTTGCAGTCAGATGAAATTCCCTGGGATNNNAGATCGGAAG |
| 51464_1 | TATGAGTGTGGAGTCGTTGCCTTCCCTCACCTCCTTTTCCAGNNNAGATCGGAAGAG |
| 51464_2 | TATGAGTGTGGAGTCGTTGCCTTCCCTCACCTCCTTTTCCAGNNNAGATCGGAAGAG |
| 49622_1 | TATGAGTGTGGAGTCGTTGCCGATTATGGCTCACTGCAAACTCNNNAGATCGGAAG |
| 49622_2 | TATGAGTGTGGAGTCGTTGCCGATTATGGCTCACTGCAAACTCNNNAGATCGGAAG |
| 46943_1 | TATGAGTGTGGAGTCGTTGCAAATAGCTGGGATTACAGGTGCANNNAGATCGGAA |
| 46943_2 | TATGAGTGTGGAGTCGTTGCAAATAGCTGGGATTACAGGTGCANNNAGATCGGAA |
| 29658_1 | TATGAGTGTGGAGTCGTTGCAGTCACCCTGATTGAAACAGAGCNNNAGATCGGAAG |
| 29658_2 | TATGAGTGTGGAGTCGTTGCAGTCACCCTGATTGAAACAGAGCNNNAGATCGGAAG |
| 49767_1 | TATGAGTGTGGAGTCGTTGCACCCCATTCTCACCAACACATAANNNAGATCGGAAGA |
| 49767_2 | TATGAGTGTGGAGTCGTTGCACCCCATTCTCACCAACACATAANNNAGATCGGAAGA |
| 33593_1 | TATGAGTGTGGAGTCGTTGCTATGTTGCCCAGGTTGATCTTGANNNAGATCGGAAG |
| 33593_2 | TATGAGTGTGGAGTCGTTGCTATGTTGCCCAGGTTGATCTTGANNNAGATCGGAAG |
| 41626_1 | TATGAGTGTGGAGTCGTTGCCAAGTAGCTGGGACTACAGGCNNNAGATCGGAAGA |
| 41626_2 | TATGAGTGTGGAGTCGTTGCCAAGTAGCTGGGACTACAGGCNNNAGATCGGAAGA |
| 39669_1 | TATGAGTGTGGAGTCGTTGCAAGAACTGACACCCTAGCTGCATNNNAGATCGGAAG |
| 39669_2 | TATGAGTGTGGAGTCGTTGCAAGAACTGACACCCTAGCTGCATNNNAGATCGGAAG |

| | |
|---|---|
| 35039_1 | TATGAGTGTGGAGTCGTTGCAATCTCTGCCTCCTGAGTTCAAGNNNAGATCGGAAG |
| 35039_2 | TATGAGTGTGGAGTCGTTGCAATCTCTGCCTCCTGAGTTCAAGNNNAGATCGGAAG |
| 30461_1 | TATGAGTGTGGAGTCGTTGCTCTGCCTATCTCTAGCCCACCTNNNAGATCGGAAGAG |
| 30461_2 | TATGAGTGTGGAGTCGTTGCTCTGCCTATCTCTAGCCCACCTNNNAGATCGGAAGAG |
| 45881_1 | TATGAGTGTGGAGTCGTTGCTTTCAAGCGATTCTCCTGTCTCANNNAGATCGGAAGA |
| 45881_2 | TATGAGTGTGGAGTCGTTGCTTTCAAGCGATTCTCCTGTCTCANNNAGATCGGAAGA |
| 12895_1 | TATGAGTGTGGAGTCGTTGCTTTCCTTGAGCTGTTTTCTGNNNAGATCGGAAGAGC |
| 12895_2 | TATGAGTGTGGAGTCGTTGCTTTCCTTGAGCTGTTTTCTGNNNAGATCGGAAGAGC |
| 35670_1 | TATGAGTGTGGAGTCGTTGCTTCAGAGTTGCCATCACCACATANNNAGATCGGAAG |
| 35670_2 | TATGAGTGTGGAGTCGTTGCTTCAGAGTTGCCATCACCACATANNNAGATCGGAAG |
| 53286_1 | TATGAGTGTGGAGTCGTTGCCTGATCCAGGAGGCCGCATCNNNAGATCGGAAGAG |
| 53286_2 | TATGAGTGTGGAGTCGTTGCCTGATCCAGGAGGCCGCATCNNNAGATCGGAAGAG |
| 29602_1 | TATGAGTGTGGAGTCGTTGCTGACAGGAACTTCTATCTGCCTGNNNAGATCGGAAG |
| 29602_2 | TATGAGTGTGGAGTCGTTGCTGACAGGAACTTCTATCTGCCTGNNNAGATCGGAAG |
| 35415_1 | TATGAGTGTGGAGTCGTTGCGCTGGCTGCCTAAAATGAGTTAGNNNAGATCGGAA |
| 35415_2 | TATGAGTGTGGAGTCGTTGCGCTGGCTGCCTAAAATGAGTTAGNNNAGATCGGAA |
| 30800_1 | TATGAGTGTGGAGTCGTTGCAGAGAGAGAGAGAGAGATCAGCCNNNAGATCGGA |
| 30800_2 | TATGAGTGTGGAGTCGTTGCAGAGAGAGAGAGAGAGATCAGCCNNNAGATCGGA |
| 43013_1 | TATGAGTGTGGAGTCGTTGCAGTAGAGCCTCTTTCTCAACACANNNAGATCGGAAG |
| 43013_2 | TATGAGTGTGGAGTCGTTGCAGTAGAGCCTCTTTCTCAACACANNNAGATCGGAAG |
| 11203_1 | TATGAGTGTGGAGTCGTTGCTTTTCCCCAACCCCACTAATTCCNNNAGATCGGAAGA |
| 11203_2 | TATGAGTGTGGAGTCGTTGCTTTTCCCCAACCCCACTAATTCCNNNAGATCGGAAGA |
| 50922_1 | TATGAGTGTGGAGTCGTTGCCCTGACCAACTCTGGGAAACCNNNAGATCGGAAGAG |
| 50922_2 | TATGAGTGTGGAGTCGTTGCCCTGACCAACTCTGGGAAACCNNNAGATCGGAAGAG |
| 29417_1 | TATGAGTGTGGAGTCGTTGCAGCAGAGTAACAGGAATTTTCTCTNNNAGATCGGAA |
| 29417_2 | TATGAGTGTGGAGTCGTTGCAGCAGAGTAACAGGAATTTTCTCTNNNAGATCGGAA |
| 46974_1 | TATGAGTGTGGAGTCGTTGCGAGGTGAAGTCTCAGTCTGTCACNNNAGATCGGAA |
| 46974_2 | TATGAGTGTGGAGTCGTTGCGAGGTGAAGTCTCAGTCTGTCACNNNAGATCGGAA |
| 46906_1 | TATGAGTGTGGAGTCGTTGCCAAGTAGCTGGGACTACAGGCNNNAGATCGGAAGA |
| 46906_2 | TATGAGTGTGGAGTCGTTGCCAAGTAGCTGGGACTACAGGCNNNAGATCGGAAGA |
| 43024_1 | TATGAGTGTGGAGTCGTTGCGAGAATCGCTTGAACCCAGATTGNNNAGATCGGAA |
| 43024_2 | TATGAGTGTGGAGTCGTTGCGAGAATCGCTTGAACCCAGATTGNNNAGATCGGAA |
| 48273_1 | TATGAGTGTGGAGTCGTTGCAGTTGGAAATCAGGAGAGGCAGNNNAGATCGGAA |
| 48273_2 | TATGAGTGTGGAGTCGTTGCAGTTGGAAATCAGGAGAGGCAGNNNAGATCGGAA |
| 50966_1 | TATGAGTGTGGAGTCGTTGCGGAGCTCTGACCCTGAAATAGAGNNNAGATCGGAA |
| 50966_2 | TATGAGTGTGGAGTCGTTGCGGAGCTCTGACCCTGAAATAGAGNNNAGATCGGAA |
| 42383_1 | TATGAGTGTGGAGTCGTTGCGGAATACCAAGGCAGAGAAAGGANNNAGATCGGA |
| 42383_2 | TATGAGTGTGGAGTCGTTGCGGAATACCAAGGCAGAGAAAGGANNNAGATCGGA |
| 46875_1 | TATGAGTGTGGAGTCGTTGCCCCTAGTAGCTGGGACTACAGGNNNAGATCGGAAG |
| 46875_2 | TATGAGTGTGGAGTCGTTGCCCCTAGTAGCTGGGACTACAGGNNNAGATCGGAAG |
| 37964_1 | TATGAGTGTGGAGTCGTTGCTGAGGAGAGGGAGTGACTCTAAGNNNAGATCGGA |
| 37964_2 | TATGAGTGTGGAGTCGTTGCTGAGGAGAGGGAGTGACTCTAAGNNNAGATCGGA |
| 30950_1 | TATGAGTGTGGAGTCGTTGCCACACACACACTCAAGTCACAAGNNNAGATCGGAAG |

| | |
|---|---|
| 30950_2 | TATGAGTGTGGAGTCGTTGCCACACACACACTCAAGTCACAAGNNNAGATCGGAAG |
| 45941_1 | TATGAGTGTGGAGTCGTTGCCACTACGCCCGGCTAATTTTTTNNNAGATCGGAAGA |
| 45941_2 | TATGAGTGTGGAGTCGTTGCCACTACGCCCGGCTAATTTTTTNNNAGATCGGAAGA |
| 42449_1 | TATGAGTGTGGAGTCGTTGCCACACAGTATCACTCTCTCCTGGNNNAGATCGGAAG |
| 42449_2 | TATGAGTGTGGAGTCGTTGCCACACAGTATCACTCTCTCCTGGNNNAGATCGGAAG |
| 47080_1 | TATGAGTGTGGAGTCGTTGCCATGATCTCGGCTCATTGCAACNNNAGATCGGAAGA |
| 47080_2 | TATGAGTGTGGAGTCGTTGCCATGATCTCGGCTCATTGCAACNNNAGATCGGAAGA |
| 44572_1 | TATGAGTGTGGAGTCGTTGCTTCCATGACTTTGCAATTGTGAANNNAGATCGGAAG |
| 44572_2 | TATGAGTGTGGAGTCGTTGCTTCCATGACTTTGCAATTGTGAANNNAGATCGGAAG |
| 38178_1 | TATGAGTGTGGAGTCGTTGCATGTTTGTTGTCCCAGCTACTCANNNAGATCGGAAG |
| 38178_2 | TATGAGTGTGGAGTCGTTGCATGTTTGTTGTCCCAGCTACTCANNNAGATCGGAAG |
| 52719_1 | TATGAGTGTGGAGTCGTTGCGTGTGTGTCCCTATGTACAGGAGNNNAGATCGGAA |
| 52719_2 | TATGAGTGTGGAGTCGTTGCGTGTGTGTCCCTATGTACAGGAGNNNAGATCGGAA |
| 47791_1 | TATGAGTGTGGAGTCGTTGCGATGTTCTCGATCTCCTGACCTCNNNAGATCGGAAG |
| 47791_2 | TATGAGTGTGGAGTCGTTGCGATGTTCTCGATCTCCTGACCTCNNNAGATCGGAAG |
| 48864_1 | TATGAGTGTGGAGTCGTTGCCCGATTCGCAAATCATTCATTGCNNNAGATCGGAAG |
| 48864_2 | TATGAGTGTGGAGTCGTTGCCCGATTCGCAAATCATTCATTGCNNNAGATCGGAAG |
| 45645_1 | TATGAGTGTGGAGTCGTTGCTTCAAGTGATTCTCCTGCCTCAGNNNAGATCGGAAG |
| 45645_2 | TATGAGTGTGGAGTCGTTGCTTCAAGTGATTCTCCTGCCTCAGNNNAGATCGGAAG |
| 37760_1 | TATGAGTGTGGAGTCGTTGCGTCTCGAATTCCTGACCTCAAGTNNNAGATCGGAAG |
| 37760_2 | TATGAGTGTGGAGTCGTTGCGTCTCGAATTCCTGACCTCAAGTNNNAGATCGGAAG |
| 29667_1 | TATGAGTGTGGAGTCGTTGCCCGCAATGATGAAGGCTTCTCNNNAGATCGGAAGAG |
| 29667_2 | TATGAGTGTGGAGTCGTTGCCCGCAATGATGAAGGCTTCTCNNNAGATCGGAAGAG |
| 4361_1 | TATGAGTGTGGAGTCGTTGCTGGGCAAAGGTTGGGAGAATTACNNNAGATCGGAA |
| 4361_2 | TATGAGTGTGGAGTCGTTGCTGGGCAAAGGTTGGGAGAATTACNNNAGATCGGAA |
| 42830_1 | TATGAGTGTGGAGTCGTTGCAATAGCTGGGATTACAGGTGCATNNNAGATCGGAA |
| 42830_2 | TATGAGTGTGGAGTCGTTGCAATAGCTGGGATTACAGGTGCATNNNAGATCGGAA |
| 29334_1 | TATGAGTGTGGAGTCGTTGCAGTATCTGGGTCACCAAAAGCTTNNNAGATCGGAAG |
| 29334_2 | TATGAGTGTGGAGTCGTTGCAGTATCTGGGTCACCAAAAGCTTNNNAGATCGGAAG |
| 33084_1 | TATGAGTGTGGAGTCGTTGCACAGGATGGTCTCGATCTCCTTANNNAGATCGGAAG |
| 33084_2 | TATGAGTGTGGAGTCGTTGCACAGGATGGTCTCGATCTCCTTANNNAGATCGGAAG |
| 2625_1 | TATGAGTGTGGAGTCGTTGCGTGTGCTCCTTGTCCTCTAAGTGNNNAGATCGGAAG |
| 2625_2 | TATGAGTGTGGAGTCGTTGCGTGTGCTCCTTGTCCTCTAAGTGNNNAGATCGGAAG |
| 32241_1 | TATGAGTGTGGAGTCGTTGCGTGGTGGTGACTGGAGATTAGTTNNNAGATCGGAA |
| 32241_2 | TATGAGTGTGGAGTCGTTGCGTGGTGGTGACTGGAGATTAGTTNNNAGATCGGAA |
| 51060_1 | TATGAGTGTGGAGTCGTTGCAAGGTTTCCAAGTCTCCTCAAGTNNNAGATCGGAAG |
| 51060_2 | TATGAGTGTGGAGTCGTTGCAAGGTTTCCAAGTCTCCTCAAGTNNNAGATCGGAAG |
| 44790_1 | TATGAGTGTGGAGTCGTTGCTAGGAAGTAGTGTGTGCTTGGACNNNAGATCGGAA |
| 44790_2 | TATGAGTGTGGAGTCGTTGCTAGGAAGTAGTGTGTGCTTGGACNNNAGATCGGAA |
| 34135_1 | TATGAGTGTGGAGTCGTTGCCCCACCAAAATTTAGCTGAGCATNNNAGATCGGAAG |
| 34135_2 | TATGAGTGTGGAGTCGTTGCCCCACCAAAATTTAGCTGAGCATNNNAGATCGGAAG |
| 12578_1 | TATGAGTGTGGAGTCGTTGCTCACCGATAAGATCATCTATTTTNNNAGATCGGAAG |
| 12578_2 | TATGAGTGTGGAGTCGTTGCTCACCGATAAGATCATCTATTTTNNNAGATCGGAAG |

| | |
|---|---|
| 36080_1 | TATGAGTGTGGAGTCGTTGCACAAAAAATTTACCGGGCATGGTNNNAGATCGGAAG |
| 36080_2 | TATGAGTGTGGAGTCGTTGCACAAAAAATTTACCGGGCATGGTNNNAGATCGGAAG |
| 13199_1 | TATGAGTGTGGAGTCGTTGCCAACTGAAGCCTTCTTTTAAGNNNAGATCGGAAGAG |
| 13199_2 | TATGAGTGTGGAGTCGTTGCCAACTGAAGCCTTCTTTTAAGNNNAGATCGGAAGAG |
| 32408_1 | TATGAGTGTGGAGTCGTTGCCAGGAAGTACATGTGCAGGTTTGNNNAGATCGGAA |
| 32408_2 | TATGAGTGTGGAGTCGTTGCCAGGAAGTACATGTGCAGGTTTGNNNAGATCGGAA |
| 36698_1 | TATGAGTGTGGAGTCGTTGCAAAGGATGGAGAAGAAAAGGCCANNNAGATCGGA |
| 36698_2 | TATGAGTGTGGAGTCGTTGCAAAGGATGGAGAAGAAAAGGCCANNNAGATCGGA |
| 42192_1 | TATGAGTGTGGAGTCGTTGCCTCCTGAGTAGCTGGGTTTACAGNNNAGATCGGAAG |
| 42192_2 | TATGAGTGTGGAGTCGTTGCCTCCTGAGTAGCTGGGTTTACAGNNNAGATCGGAAG |
| 39642_1 | TATGAGTGTGGAGTCGTTGCTTCAAGTGATTCTCCTGCCTCAGNNNAGATCGGAAG |
| 39642_2 | TATGAGTGTGGAGTCGTTGCTTCAAGTGATTCTCCTGCCTCAGNNNAGATCGGAAG |
| 29502_1 | TATGAGTGTGGAGTCGTTGCACACATTCACTCACACAAAAAGNNNAGATCGGAAGA |
| 29502_2 | TATGAGTGTGGAGTCGTTGCACACATTCACTCACACAAAAAGNNNAGATCGGAAGA |
| 53045_1 | TATGAGTGTGGAGTCGTTGCGAAAGTGTGCACATTCTTTGGCNNNAGATCGGAAGA |
| 53045_2 | TATGAGTGTGGAGTCGTTGCGAAAGTGTGCACATTCTTTGGCNNNAGATCGGAAGA |
| 12985_1 | TATGAGTGTGGAGTCGTTGCGAATCACCTATTACATTTGACTTNNNAGATCGGAAG |
| 12985_2 | TATGAGTGTGGAGTCGTTGCGAATCACCTATTACATTTGACTTNNNAGATCGGAAG |
| 37112_1 | TATGAGTGTGGAGTCGTTGCGCTGATACGCTGTCTTTCTTGTCNNNAGATCGGAAG |
| 37112_2 | TATGAGTGTGGAGTCGTTGCGCTGATACGCTGTCTTTCTTGTCNNNAGATCGGAAG |
| 51627_1 | TATGAGTGTGGAGTCGTTGCGCTCAGTATCAGACTCTCAGACGNNNAGATCGGAAG |
| 51627_2 | TATGAGTGTGGAGTCGTTGCGCTCAGTATCAGACTCTCAGACGNNNAGATCGGAAG |
| 9965_1 | TATGAGTGTGGAGTCGTTGCACCAACTTTTTCAGAGCTTCGGTNNNAGATCGGAAG |
| 9965_2 | TATGAGTGTGGAGTCGTTGCACCAACTTTTTCAGAGCTTCGGTNNNAGATCGGAAG |
| 39689_1 | TATGAGTGTGGAGTCGTTGCGGGCTGAAAAAACTGTGAACAGANNNAGATCGGAA |
| 39689_2 | TATGAGTGTGGAGTCGTTGCGGGCTGAAAAAACTGTGAACAGANNNAGATCGGAA |
| 49910_1 | TATGAGTGTGGAGTCGTTGCTCCCATCGATGTCATGTACTGTGNNNAGATCGGAAG |
| 49910_2 | TATGAGTGTGGAGTCGTTGCTCCCATCGATGTCATGTACTGTGNNNAGATCGGAAG |
| 34832_1 | TATGAGTGTGGAGTCGTTGCGCTTTGCTTGCTCTGTCATCCNNNAGATCGGAAGAG |
| 34832_2 | TATGAGTGTGGAGTCGTTGCGCTTTGCTTGCTCTGTCATCCNNNAGATCGGAAGAG |
| 13676_1 | TATGAGTGTGGAGTCGTTGCATAGTCTGGGCCATACTTTTNNNAGATCGGAAGAGC |
| 13676_2 | TATGAGTGTGGAGTCGTTGCATAGTCTGGGCCATACTTTTNNNAGATCGGAAGAGC |
| 37588_1 | TATGAGTGTGGAGTCGTTGCGCTGCCATGAACATTCGTATACANNNAGATCGGAAG |
| 37588_2 | TATGAGTGTGGAGTCGTTGCGCTGCCATGAACATTCGTATACANNNAGATCGGAAG |
| 9100_1 | TATGAGTGTGGAGTCGTTGCTGCTCTTCACAATGCAAAATGACANNNAGATCGGAA |
| 9100_2 | TATGAGTGTGGAGTCGTTGCTGCTCTTCACAATGCAAAATGACANNNAGATCGGAA |
| 45905_1 | TATGAGTGTGGAGTCGTTGCCCACACCCGGCTTTTTTTTTTTTNNNAGATCGGAAGA |
| 45905_2 | TATGAGTGTGGAGTCGTTGCCCACACCCGGCTTTTTTTTTTTTNNNAGATCGGAAGA |
| 48011_1 | TATGAGTGTGGAGTCGTTGCCATGTAGCTGGGATTACAGGTGTNNNAGATCGGAA |
| 48011_2 | TATGAGTGTGGAGTCGTTGCCATGTAGCTGGGATTACAGGTGTNNNAGATCGGAA |
| 10567_1 | TATGAGTGTGGAGTCGTTGCTAGCCAGCTTCCACTGTTCAAACNNNAGATCGGAAG |
| 10567_2 | TATGAGTGTGGAGTCGTTGCTAGCCAGCTTCCACTGTTCAAACNNNAGATCGGAAG |
| 32107_1 | TATGAGTGTGGAGTCGTTGCGGAGTACAATGGCACAATCTTGGNNNAGATCGGAA |

| | |
|---|---|
| 32107_2 | TATGAGTGTGGAGTCGTTGCGGAGTACAATGGCACAATCTTGGNNNAGATCGGAA |
| 46250_1 | TATGAGTGTGGAGTCGTTGCCCGGATTCACGCCATTCTACTNNNAGATCGGAAGAG |
| 46250_2 | TATGAGTGTGGAGTCGTTGCCCGGATTCACGCCATTCTACTNNNAGATCGGAAGAG |
| 48698_1 | TATGAGTGTGGAGTCGTTGCATCTTAGCTCACTGCAACTTCCANNNAGATCGGAAG |
| 48698_2 | TATGAGTGTGGAGTCGTTGCATCTTAGCTCACTGCAACTTCCANNNAGATCGGAAG |
| 36004_1 | TATGAGTGTGGAGTCGTTGCACGTTCTGGAGAGAATCACCTTCNNNAGATCGGAAG |
| 36004_2 | TATGAGTGTGGAGTCGTTGCACGTTCTGGAGAGAATCACCTTCNNNAGATCGGAAG |
| 48522_1 | TATGAGTGTGGAGTCGTTGCTTCAAGCAATTCTCCTGCCTCAGNNNAGATCGGAAG |
| 48522_2 | TATGAGTGTGGAGTCGTTGCTTCAAGCAATTCTCCTGCCTCAGNNNAGATCGGAAG |
| 13050_1 | TATGAGTGTGGAGTCGTTGCCCATCAATAATAGCAAGTCATTTNNNAGATCGGAAG |
| 13050_2 | TATGAGTGTGGAGTCGTTGCCCATCAATAATAGCAAGTCATTTNNNAGATCGGAAG |
| 13119_1 | TATGAGTGTGGAGTCGTTGCCCATCAATAATAGCAAGTCATTTNNNAGATCGGAAG |
| 13119_2 | TATGAGTGTGGAGTCGTTGCCCATCAATAATAGCAAGTCATTTNNNAGATCGGAAG |
| 3193_1 | TATGAGTGTGGAGTCGTTGCTATGATGTCTGTGTGGAAAGCGGNNNAGATCGGAA |
| 3193_2 | TATGAGTGTGGAGTCGTTGCTATGATGTCTGTGTGGAAAGCGGNNNAGATCGGAA |
| 49246_1 | TATGAGTGTGGAGTCGTTGCCATAATCTTGGCTCACTGCAACCNNNAGATCGGAAG |
| 49246_2 | TATGAGTGTGGAGTCGTTGCCATAATCTTGGCTCACTGCAACCNNNAGATCGGAAG |
| 46769_1 | TATGAGTGTGGAGTCGTTGCCTGACCTCAGGTGATCCATCTGNNNAGATCGGAAGA |
| 46769_2 | TATGAGTGTGGAGTCGTTGCCTGACCTCAGGTGATCCATCTGNNNAGATCGGAAGA |
| 45869_1 | TATGAGTGTGGAGTCGTTGCAGAAGGAGTTTTGCTGTTGTTGCNNNAGATCGGAA |
| 45869_2 | TATGAGTGTGGAGTCGTTGCAGAAGGAGTTTTGCTGTTGTTGCNNNAGATCGGAA |
| 35954_1 | TATGAGTGTGGAGTCGTTGCGAGGCTGAGGCAGGAGAATGNNNAGATCGGAAGA |
| 35954_2 | TATGAGTGTGGAGTCGTTGCGAGGCTGAGGCAGGAGAATGNNNAGATCGGAAGA |
| 30806_1 | TATGAGTGTGGAGTCGTTGCGAGCTCTGACCAGAAACCAACTANNNAGATCGGAAG |
| 30806_2 | TATGAGTGTGGAGTCGTTGCGAGCTCTGACCAGAAACCAACTANNNAGATCGGAAG |
| 46507_1 | TATGAGTGTGGAGTCGTTGCCGCCATGATAAGAGAGCAGAGAANNNAGATCGGAA |
| 46507_2 | TATGAGTGTGGAGTCGTTGCCGCCATGATAAGAGAGCAGAGAANNNAGATCGGAA |
| 1018_1 | TATGAGTGTGGAGTCGTTGCGTTACCATGCCAGGTCTCCTATNNNAGATCGGAAGA |
| 1018_2 | TATGAGTGTGGAGTCGTTGCGTTACCATGCCAGGTCTCCTATNNNAGATCGGAAGA |
| 44213_1 | TATGAGTGTGGAGTCGTTGCTTCAGATCTCAGGTTTACCGTGGNNNAGATCGGAAG |
| 44213_2 | TATGAGTGTGGAGTCGTTGCTTCAGATCTCAGGTTTACCGTGGNNNAGATCGGAAG |
| 1634_1 | TATGAGTGTGGAGTCGTTGCGGAGATGCACGTATGCTGAGAAANNNAGATCGGAA |
| 1634_2 | TATGAGTGTGGAGTCGTTGCGGAGATGCACGTATGCTGAGAAANNNAGATCGGAA |
| 33910_1 | TATGAGTGTGGAGTCGTTGCCACAATCTCAGCTCACTGCAAGNNNAGATCGGAAGA |
| 33910_2 | TATGAGTGTGGAGTCGTTGCCACAATCTCAGCTCACTGCAAGNNNAGATCGGAAGA |
| 35859_1 | TATGAGTGTGGAGTCGTTGCCTGTAGTCCCAGCTACTCAGGAGNNNAGATCGGAAG |
| 35859_2 | TATGAGTGTGGAGTCGTTGCCTGTAGTCCCAGCTACTCAGGAGNNNAGATCGGAAG |
| 41084_1 | TATGAGTGTGGAGTCGTTGCGCATACAGGATTGTGTGATCTGCNNNAGATCGGAA |
| 41084_2 | TATGAGTGTGGAGTCGTTGCGCATACAGGATTGTGTGATCTGCNNNAGATCGGAA |
| 43510_1 | TATGAGTGTGGAGTCGTTGCGAGTAGCTGGCATTACAGGCATANNNAGATCGGAA |
| 43510_2 | TATGAGTGTGGAGTCGTTGCGAGTAGCTGGCATTACAGGCATANNNAGATCGGAA |
| 34574_1 | TATGAGTGTGGAGTCGTTGCAGGACCATGCACATGTCTGAATANNNAGATCGGAA |
| 34574_2 | TATGAGTGTGGAGTCGTTGCAGGACCATGCACATGTCTGAATANNNAGATCGGAA |

| | |
|---|---|
| 42700_1 | TATGAGTGTGGAGTCGTTGCTGCATATCACATTACTTTATCCACTCANNNAGATCGGA |
| 42700_2 | TATGAGTGTGGAGTCGTTGCTGCATATCACATTACTTTATCCACTCANNNAGATCGGA |
| 33440_1 | TATGAGTGTGGAGTCGTTGCGAAGGAGTAGAAGGTGCTGAGAGNNNAGATCGGA |
| 33440_2 | TATGAGTGTGGAGTCGTTGCGAAGGAGTAGAAGGTGCTGAGAGNNNAGATCGGA |
| 39511_1 | TATGAGTGTGGAGTCGTTGCAATCGAAGAACCTAAGGCCATGTNNNAGATCGGAAG |
| 39511_2 | TATGAGTGTGGAGTCGTTGCAATCGAAGAACCTAAGGCCATGTNNNAGATCGGAAG |
| 44087_1 | TATGAGTGTGGAGTCGTTGCCCCAATCTTCACAAAGAACCCACNNNAGATCGGAAGA |
| 44087_2 | TATGAGTGTGGAGTCGTTGCCCCAATCTTCACAAAGAACCCACNNNAGATCGGAAGA |
| 10023_1 | TATGAGTGTGGAGTCGTTGCATCAGTGGAGTGGCATCCTATCANNNAGATCGGAAG |
| 10023_2 | TATGAGTGTGGAGTCGTTGCATCAGTGGAGTGGCATCCTATCANNNAGATCGGAAG |
| 35912_1 | TATGAGTGTGGAGTCGTTGCATTCCTCCAACACCAGTTAGACCNNNAGATCGGAAGA |
| 35912_2 | TATGAGTGTGGAGTCGTTGCATTCCTCCAACACCAGTTAGACCNNNAGATCGGAAGA |
| 33790_1 | TATGAGTGTGGAGTCGTTGCGAGGTGAGGGGGACAAATTGTNNNAGATCGGAAGA |
| 33790_2 | TATGAGTGTGGAGTCGTTGCGAGGTGAGGGGGACAAATTGTNNNAGATCGGAAGA |
| 50677_1 | TATGAGTGTGGAGTCGTTGCTGACAGTAGCTTGGGGAATAAGGNNNAGATCGGAA |
| 50677_2 | TATGAGTGTGGAGTCGTTGCTGACAGTAGCTTGGGGAATAAGGNNNAGATCGGAA |
| 52377_1 | TATGAGTGTGGAGTCGTTGCTCCTTACACTTTACATCCGAGCCNNNAGATCGGAAGA |
| 52377_2 | TATGAGTGTGGAGTCGTTGCTCCTTACACTTTACATCCGAGCCNNNAGATCGGAAGA |
| 52974_1 | TATGAGTGTGGAGTCGTTGCCTTGAGTGAAGACAAAGTGCAGGNNNAGATCGGAA |
| 52974_2 | TATGAGTGTGGAGTCGTTGCCTTGAGTGAAGACAAAGTGCAGGNNNAGATCGGAA |
| 34067_1 | TATGAGTGTGGAGTCGTTGCTACAGTGACATGAACATGGCTCANNNAGATCGGAAG |
| 34067_2 | TATGAGTGTGGAGTCGTTGCTACAGTGACATGAACATGGCTCANNNAGATCGGAAG |
| 46076_1 | TATGAGTGTGGAGTCGTTGCCCCAGGTTCAAGCAATTCTCATGNNNAGATCGGAAGA |
| 46076_2 | TATGAGTGTGGAGTCGTTGCCCCAGGTTCAAGCAATTCTCATGNNNAGATCGGAAGA |
| 47969_1 | TATGAGTGTGGAGTCGTTGCTGCCCGGCTGATTTTTGTATTTTNNNAGATCGGAAGA |
| 47969_2 | TATGAGTGTGGAGTCGTTGCTGCCCGGCTGATTTTTGTATTTTNNNAGATCGGAAGA |
| 36166_1 | TATGAGTGTGGAGTCGTTGCACAATCAGCAGGCATTTACATAGGNNNAGATCGGAA |
| 36166_2 | TATGAGTGTGGAGTCGTTGCACAATCAGCAGGCATTTACATAGGNNNAGATCGGAA |
| 31213_1 | TATGAGTGTGGAGTCGTTGCTGATCCTCTTGGAACCCAAAGAGNNNAGATCGGAAG |
| 31213_2 | TATGAGTGTGGAGTCGTTGCTGATCCTCTTGGAACCCAAAGAGNNNAGATCGGAAG |
| 41443_1 | TATGAGTGTGGAGTCGTTGCAGAGACATTTTGGAGCCCATTGANNNAGATCGGAAG |
| 41443_2 | TATGAGTGTGGAGTCGTTGCAGAGACATTTTGGAGCCCATTGANNNAGATCGGAAG |
| 46346_1 | TATGAGTGTGGAGTCGTTGCCTGGGTTCAAGTAATTCTGCTGCNNNAGATCGGAAG |
| 46346_2 | TATGAGTGTGGAGTCGTTGCCTGGGTTCAAGTAATTCTGCTGCNNNAGATCGGAAG |
| 13182_1 | TATGAGTGTGGAGTCGTTGCTTTGTGATGTGAATACTTTCAAANNNAGATCGGAAG |
| 13182_2 | TATGAGTGTGGAGTCGTTGCTTTGTGATGTGAATACTTTCAAANNNAGATCGGAAG |
| 13366_1 | TATGAGTGTGGAGTCGTTGCTTGAAGTTTTCCATCAATTAACANNNAGATCGGAAGA |
| 13366_2 | TATGAGTGTGGAGTCGTTGCTTGAAGTTTTCCATCAATTAACANNNAGATCGGAAGA |
| 52717_1 | TATGAGTGTGGAGTCGTTGCAATCTCTGCCTGTTTCTGCCATANNNAGATCGGAAGA |
| 52717_2 | TATGAGTGTGGAGTCGTTGCAATCTCTGCCTGTTTCTGCCATANNNAGATCGGAAGA |
| 50491_1 | TATGAGTGTGGAGTCGTTGCCCATCACTATCGCTGAGAGAACANNNAGATCGGAAG |
| 50491_2 | TATGAGTGTGGAGTCGTTGCCCATCACTATCGCTGAGAGAACANNNAGATCGGAAG |
| 2506_1 | TATGAGTGTGGAGTCGTTGCACTTCTGGGCTTCTACTCCTGAGNNNAGATCGGAAGA |

| | |
|---|---|
| 2506_2 | TATGAGTGTGGAGTCGTTGCACTTCTGGGCTTCTACTCCTGAGNNNAGATCGGAAG |
| 36507_1 | TATGAGTGTGGAGTCGTTGCGAAAGAAAAATTGAGTGGGGGGANNNAGATCGGA |
| 36507_2 | TATGAGTGTGGAGTCGTTGCGAAAGAAAAATTGAGTGGGGGGANNNAGATCGGA |
| 39302_1 | TATGAGTGTGGAGTCGTTGCCTAGCAAAAACGTGGCCTTGTAANNNAGATCGGAA |
| 39302_2 | TATGAGTGTGGAGTCGTTGCCTAGCAAAAACGTGGCCTTGTAANNNAGATCGGAA |
| 13227_1 | TATGAGTGTGGAGTCGTTGCTTGCCTTTCATGAATACCAGNNNAGATCGGAAGAGC |
| 13227_2 | TATGAGTGTGGAGTCGTTGCTTGCCTTTCATGAATACCAGNNNAGATCGGAAGAGC |
| 36734_1 | TATGAGTGTGGAGTCGTTGCGTAGTCTCAGCTACTTGAGAGGCNNNAGATCGGAA |
| 36734_2 | TATGAGTGTGGAGTCGTTGCGTAGTCTCAGCTACTTGAGAGGCNNNAGATCGGAA |
| 13623_1 | TATGAGTGTGGAGTCGTTGCTTTCATAATTTATAAGTGTGGGGNNNAGATCGGAA |
| 13623_2 | TATGAGTGTGGAGTCGTTGCTTTCATAATTTATAAGTGTGGGGNNNAGATCGGAA |
| 29131_1 | TATGAGTGTGGAGTCGTTGCGAAACTTCGCAGTAACACTAGGCNNNAGATCGGAA |
| 29131_2 | TATGAGTGTGGAGTCGTTGCGAAACTTCGCAGTAACACTAGGCNNNAGATCGGAA |
| 44188_1 | TATGAGTGTGGAGTCGTTGCTCCCCGGTGATCATCTCCATATANNNAGATCGGAAG |
| 44188_2 | TATGAGTGTGGAGTCGTTGCTCCCCGGTGATCATCTCCATATANNNAGATCGGAAG |
| 40890_1 | TATGAGTGTGGAGTCGTTGCCAGCCTGACAATGCAGAAGAAAANNNAGATCGGAA |
| 40890_2 | TATGAGTGTGGAGTCGTTGCCAGCCTGACAATGCAGAAGAAAANNNAGATCGGAA |
| 36263_1 | TATGAGTGTGGAGTCGTTGCATTGGCCCCTGAATAAAGTCAGTNNNAGATCGGAAG |
| 36263_2 | TATGAGTGTGGAGTCGTTGCATTGGCCCCTGAATAAAGTCAGTNNNAGATCGGAAG |
| 50284_1 | TATGAGTGTGGAGTCGTTGCTGAGAGCAAGACTCCATCTCAAANNNAGATCGGAAG |
| 50284_2 | TATGAGTGTGGAGTCGTTGCTGAGAGCAAGACTCCATCTCAAANNNAGATCGGAAG |
| 2843_1 | TATGAGTGTGGAGTCGTTGCTGGTTTGGCCTTCTATCTACCGANNNAGATCGGAAG |
| 2843_2 | TATGAGTGTGGAGTCGTTGCTGGTTTGGCCTTCTATCTACCGANNNAGATCGGAAG |
| 41138_1 | TATGAGTGTGGAGTCGTTGCGTTGAGACTGTAGTGAGCCATGANNNAGATCGGAA |
| 41138_2 | TATGAGTGTGGAGTCGTTGCGTTGAGACTGTAGTGAGCCATGANNNAGATCGGAA |
| 44471_1 | TATGAGTGTGGAGTCGTTGCAAAAACCAAGACTCTCCCAGGAGNNNAGATCGGAA |
| 44471_2 | TATGAGTGTGGAGTCGTTGCAAAAACCAAGACTCTCCCAGGAGNNNAGATCGGAA |
| 39950_1 | TATGAGTGTGGAGTCGTTGCGTCCAGGTATTCATGCATTGGTGNNNAGATCGGAA |
| 39950_2 | TATGAGTGTGGAGTCGTTGCGTCCAGGTATTCATGCATTGGTGNNNAGATCGGAA |
| 39158_1 | TATGAGTGTGGAGTCGTTGCTCCAAATTGGAAAGAAAGCAATCCANNNAGATCGG |
| 39158_2 | TATGAGTGTGGAGTCGTTGCTCCAAATTGGAAAGAAAGCAATCCANNNAGATCGG |
| 48833_1 | TATGAGTGTGGAGTCGTTGCAATTCAGAGTCCTTGTTCCCCATNNNAGATCGGAAG |
| 48833_2 | TATGAGTGTGGAGTCGTTGCAATTCAGAGTCCTTGTTCCCCATNNNAGATCGGAAG |
| 10431_1 | TATGAGTGTGGAGTCGTTGCAAGAGACACAGCCAACACTCATCNNNAGATCGGAAG |
| 10431_2 | TATGAGTGTGGAGTCGTTGCAAGAGACACAGCCAACACTCATCNNNAGATCGGAAG |
| 47668_1 | TATGAGTGTGGAGTCGTTGCTTCAAGCAATTCTCCTGCCTCAGNNNAGATCGGAAG |
| 47668_2 | TATGAGTGTGGAGTCGTTGCTTCAAGCAATTCTCCTGCCTCAGNNNAGATCGGAAG |
| 46760_1 | TATGAGTGTGGAGTCGTTGCGGTTCAAGTGATTCTTCTGCCTCNNNAGATCGGAAG |
| 46760_2 | TATGAGTGTGGAGTCGTTGCGGTTCAAGTGATTCTTCTGCCTCNNNAGATCGGAAG |
| 31820_1 | TATGAGTGTGGAGTCGTTGCACACACACACAGACACACACATANNNAGATCGGAAG |
| 31820_2 | TATGAGTGTGGAGTCGTTGCACACACACACAGACACACACATANNNAGATCGGAAG |
| 48637_1 | TATGAGTGTGGAGTCGTTGCTAGCTGGGAATACAGAAACCACCNNNAGATCGGAA |
| 48637_2 | TATGAGTGTGGAGTCGTTGCTAGCTGGGAATACAGAAACCACCNNNAGATCGGAA |

| | |
|---|---|
| 35660_1 | TATGAGTGTGGAGTCGTTGCGATGGAGTTTTACTCTTGGTGCCNNNAGATCGGAAG |
| 35660_2 | TATGAGTGTGGAGTCGTTGCGATGGAGTTTTACTCTTGGTGCCNNNAGATCGGAAG |
| 31283_1 | TATGAGTGTGGAGTCGTTGCTCTCACGCCAACATTACCACCNNNAGATCGGAAGAG |
| 31283_2 | TATGAGTGTGGAGTCGTTGCTCTCACGCCAACATTACCACCNNNAGATCGGAAGAG |
| 6105_1 | TATGAGTGTGGAGTCGTTGCTCTGGACTACACTGGCGTGATNNNAGATCGGAAGA |
| 6105_2 | TATGAGTGTGGAGTCGTTGCTCTGGACTACACTGGCGTGATNNNAGATCGGAAGA |
| 29195_1 | TATGAGTGTGGAGTCGTTGCAGGTTTTTTCCATGTAGAGTCGTNNNAGATCGGAAG |
| 29195_2 | TATGAGTGTGGAGTCGTTGCAGGTTTTTTCCATGTAGAGTCGTNNNAGATCGGAAG |
| 12532_1 | TATGAGTGTGGAGTCGTTGCGAAAAGCTACTATAGTCACAGANNNAGATCGGAAG |
| 12532_2 | TATGAGTGTGGAGTCGTTGCGAAAAGCTACTATAGTCACAGANNNAGATCGGAAG |
| 42318_1 | TATGAGTGTGGAGTCGTTGCCTGGAGAATTGCTTGAACTTGGGNNNAGATCGGAA |
| 42318_2 | TATGAGTGTGGAGTCGTTGCCTGGAGAATTGCTTGAACTTGGGNNNAGATCGGAA |
| 35659_1 | TATGAGTGTGGAGTCGTTGCCACCTCTGCCATCTCCTAGATTCNNNAGATCGGAAGA |
| 35659_2 | TATGAGTGTGGAGTCGTTGCCACCTCTGCCATCTCCTAGATTCNNNAGATCGGAAGA |
| 29706_1 | TATGAGTGTGGAGTCGTTGCTGACTGAAAATCCCAAGTAGGAANNNAGATCGGAA |
| 29706_2 | TATGAGTGTGGAGTCGTTGCTGACTGAAAATCCCAAGTAGGAANNNAGATCGGAA |
| 51975_1 | TATGAGTGTGGAGTCGTTGCACTTAGAGACAGTCTTGCTCTGTNNNAGATCGGAAG |
| 51975_2 | TATGAGTGTGGAGTCGTTGCACTTAGAGACAGTCTTGCTCTGTNNNAGATCGGAAG |
| 36221_1 | TATGAGTGTGGAGTCGTTGCCAACCACAAAGTGCATCAGTACCNNNAGATCGGAAG |
| 36221_2 | TATGAGTGTGGAGTCGTTGCCAACCACAAAGTGCATCAGTACCNNNAGATCGGAAG |
| 34690_1 | TATGAGTGTGGAGTCGTTGCGTTCACACCATTCTCCTACCTCANNNAGATCGGAAGA |
| 34690_2 | TATGAGTGTGGAGTCGTTGCGTTCACACCATTCTCCTACCTCANNNAGATCGGAAGA |
| 37419_1 | TATGAGTGTGGAGTCGTTGCATGATCACTTGAAGCCAGGAGTTNNNAGATCGGAA |
| 37419_2 | TATGAGTGTGGAGTCGTTGCATGATCACTTGAAGCCAGGAGTTNNNAGATCGGAA |
| 13401_1 | TATGAGTGTGGAGTCGTTGCAGAGAAGGTTCACTAACTGTNNNAGATCGGAAGAG |
| 13401_2 | TATGAGTGTGGAGTCGTTGCAGAGAAGGTTCACTAACTGTNNNAGATCGGAAGAG |
| 42479_1 | TATGAGTGTGGAGTCGTTGCGCTGAGATGGGTAGAAACAATGCNNNAGATCGGAA |
| 42479_2 | TATGAGTGTGGAGTCGTTGCGCTGAGATGGGTAGAAACAATGCNNNAGATCGGAA |
| 38044_1 | TATGAGTGTGGAGTCGTTGCTCACTCAAATCTTTCCTCCTGGGNNNAGATCGGAAGA |
| 38044_2 | TATGAGTGTGGAGTCGTTGCTCACTCAAATCTTTCCTCCTGGGNNNAGATCGGAAGA |
| 2827_1 | TATGAGTGTGGAGTCGTTGCAACAGTGCATCCCAAGTGAAAGTNNNAGATCGGAA |
| 2827_2 | TATGAGTGTGGAGTCGTTGCAACAGTGCATCCCAAGTGAAAGTNNNAGATCGGAA |
| 40938_1 | TATGAGTGTGGAGTCGTTGCAAATTCTAAGGGCCTCCAACCATNNNAGATCGGAAG |
| 40938_2 | TATGAGTGTGGAGTCGTTGCAAATTCTAAGGGCCTCCAACCATNNNAGATCGGAAG |
| 47694_1 | TATGAGTGTGGAGTCGTTGCCTCACTTCCCATGGACAGCTTNNNAGATCGGAAGAG |
| 47694_2 | TATGAGTGTGGAGTCGTTGCCTCACTTCCCATGGACAGCTTNNNAGATCGGAAGAG |
| 42241_1 | TATGAGTGTGGAGTCGTTGCACATACACACACACCTGTTAGCTNNNAGATCGGAAG |
| 42241_2 | TATGAGTGTGGAGTCGTTGCACATACACACACACCTGTTAGCTNNNAGATCGGAAG |
| 53105_1 | TATGAGTGTGGAGTCGTTGCCCAGGGTGGTCAATACTACAGTGNNNAGATCGGAA |
| 53105_2 | TATGAGTGTGGAGTCGTTGCCCAGGGTGGTCAATACTACAGTGNNNAGATCGGAA |
| 9720_1 | TATGAGTGTGGAGTCGTTGCGCACAGGTCATTTTGGGAATGACNNNAGATCGGAA |
| 9720_2 | TATGAGTGTGGAGTCGTTGCGCACAGGTCATTTTGGGAATGACNNNAGATCGGAA |
| 36102_1 | TATGAGTGTGGAGTCGTTGCCAGCAAGCACTGAGAACAAATGANNNAGATCGGAA |

| | |
|---|---|
| 36102_2 | TATGAGTGTGGAGTCGTTGCCAGCAAGCACTGAGAACAAATGANNNAGATCGGAA |
| 52323_1 | TATGAGTGTGGAGTCGTTGCCCCCCGCTCTTTTTTTTTGAGANNNAGATCGGAAGA |
| 52323_2 | TATGAGTGTGGAGTCGTTGCCCCCCGCTCTTTTTTTTTGAGANNNAGATCGGAAGA |
| 35522_1 | TATGAGTGTGGAGTCGTTGCCTGGTCTTGAACTCCTGAACTCANNNAGATCGGAAGA |
| 35522_2 | TATGAGTGTGGAGTCGTTGCCTGGTCTTGAACTCCTGAACTCANNNAGATCGGAAGA |
| 45324_1 | TATGAGTGTGGAGTCGTTGCCTCAAGGCTTGAAAGGGTAGACTNNNAGATCGGAA |
| 45324_2 | TATGAGTGTGGAGTCGTTGCCTCAAGGCTTGAAAGGGTAGACTNNNAGATCGGAA |
| 43628_1 | TATGAGTGTGGAGTCGTTGCCTCTGTGCTGTTGAACAACCATGNNNAGATCGGAAG |
| 43628_2 | TATGAGTGTGGAGTCGTTGCCTCTGTGCTGTTGAACAACCATGNNNAGATCGGAAG |
| 28789_1 | TATGAGTGTGGAGTCGTTGCGTGTGTGTGTGTATGTGTGTGTGNNNAGATCGGAA |
| 28789_2 | TATGAGTGTGGAGTCGTTGCGTGTGTGTGTGTATGTGTGTGTGNNNAGATCGGAA |
| 12883_1 | TATGAGTGTGGAGTCGTTGCCAGCATATGGACAAAAAACATNNNAGATCGGAAGA |
| 12883_2 | TATGAGTGTGGAGTCGTTGCCAGCATATGGACAAAAAACATNNNAGATCGGAAGA |
| 3167_1 | TATGAGTGTGGAGTCGTTGCTTACTTCCAGGCCTGTTCTTGTGNNNAGATCGGAAGA |
| 3167_2 | TATGAGTGTGGAGTCGTTGCTTACTTCCAGGCCTGTTCTTGTGNNNAGATCGGAAGA |
| 31267_1 | TATGAGTGTGGAGTCGTTGCAGGGCACAGAGAGAAAGTACTTGNNNAGATCGGAA |
| 31267_2 | TATGAGTGTGGAGTCGTTGCAGGGCACAGAGAGAAAGTACTTGNNNAGATCGGAA |
| 42082_1 | TATGAGTGTGGAGTCGTTGCCGTCATGGAGCACTTTATCAACCNNNAGATCGGAAGA |
| 42082_2 | TATGAGTGTGGAGTCGTTGCCGTCATGGAGCACTTTATCAACCNNNAGATCGGAAGA |
| 46096_1 | TATGAGTGTGGAGTCGTTGCGGTTTGAGTGATTCTTCTGCCTCNNNAGATCGGAAGA |
| 46096_2 | TATGAGTGTGGAGTCGTTGCGGTTTGAGTGATTCTTCTGCCTCNNNAGATCGGAAGA |
| 51316_1 | TATGAGTGTGGAGTCGTTGCGAGATTGGACGATAGCATCTCCANNNAGATCGGAAG |
| 51316_2 | TATGAGTGTGGAGTCGTTGCGAGATTGGACGATAGCATCTCCANNNAGATCGGAAG |
| 30504_1 | TATGAGTGTGGAGTCGTTGCTGATTTTCTCCTGTGTGGCCANNNAGATCGGAAGAG |
| 30504_2 | TATGAGTGTGGAGTCGTTGCTGATTTTCTCCTGTGTGGCCANNNAGATCGGAAGAG |
| 45956_1 | TATGAGTGTGGAGTCGTTGCGACTGTGGTGTATGTTGTGTGTGNNNAGATCGGAA |
| 45956_2 | TATGAGTGTGGAGTCGTTGCGACTGTGGTGTATGTTGTGTGTGNNNAGATCGGAA |
| 35346_1 | TATGAGTGTGGAGTCGTTGCATCACAGTTCACTACAGCCTTGANNNAGATCGGAAGA |
| 35346_2 | TATGAGTGTGGAGTCGTTGCATCACAGTTCACTACAGCCTTGANNNAGATCGGAAGA |
| 53041_1 | TATGAGTGTGGAGTCGTTGCATTCCTCCAACACCAGTTAGACCNNNAGATCGGAAGA |
| 53041_2 | TATGAGTGTGGAGTCGTTGCATTCCTCCAACACCAGTTAGACCNNNAGATCGGAAGA |
| 32182_1 | TATGAGTGTGGAGTCGTTGCAAATAATATGGCCAGGCACAGTGNNNAGATCGGAA |
| 32182_2 | TATGAGTGTGGAGTCGTTGCAAATAATATGGCCAGGCACAGTGNNNAGATCGGAA |
| 34430_1 | TATGAGTGTGGAGTCGTTGCTGGGTAGCTTCACACTTTAGTCCNNNAGATCGGAAGA |
| 34430_2 | TATGAGTGTGGAGTCGTTGCTGGGTAGCTTCACACTTTAGTCCNNNAGATCGGAAGA |
| 31364_1 | TATGAGTGTGGAGTCGTTGCGTGGTCAGATGAGCAAGTTGTTCNNNAGATCGGAA |
| 31364_2 | TATGAGTGTGGAGTCGTTGCGTGGTCAGATGAGCAAGTTGTTCNNNAGATCGGAA |
| 43347_1 | TATGAGTGTGGAGTCGTTGCTGTCAACCCCAGTCAGTAAAGAGNNNAGATCGGAAG |
| 43347_2 | TATGAGTGTGGAGTCGTTGCTGTCAACCCCAGTCAGTAAAGAGNNNAGATCGGAAG |
| 46091_1 | TATGAGTGTGGAGTCGTTGCCTCAGCCTCCTAAGTAGCTAGGANNNAGATCGGAAG |
| 46091_2 | TATGAGTGTGGAGTCGTTGCCTCAGCCTCCTAAGTAGCTAGGANNNAGATCGGAAG |
| 34032_1 | TATGAGTGTGGAGTCGTTGCCATTCTCCTGCCTCAGCCTCNNNAGATCGGAAGAGCA |
| 34032_2 | TATGAGTGTGGAGTCGTTGCCATTCTCCTGCCTCAGCCTCNNNAGATCGGAAGAGCA |

| | |
|---|---|
| 49252_1 | TATGAGTGTGGAGTCGTTGCAATTCTCCTACCTCAGTCTCCCANNNAGATCGGAAGA |
| 49252_2 | TATGAGTGTGGAGTCGTTGCAATTCTCCTACCTCAGTCTCCCANNNAGATCGGAAGA |
| 50789_1 | TATGAGTGTGGAGTCGTTGCCCTGTTGATGGCTTCCATTTCTCNNNAGATCGGAAGA |
| 50789_2 | TATGAGTGTGGAGTCGTTGCCCTGTTGATGGCTTCCATTTCTCNNNAGATCGGAAGA |
| 52743_1 | TATGAGTGTGGAGTCGTTGCAATCATGGCTTTGCTATCTCCCANNNAGATCGGAAG |
| 52743_2 | TATGAGTGTGGAGTCGTTGCAATCATGGCTTTGCTATCTCCCANNNAGATCGGAAG |
| 47983_1 | TATGAGTGTGGAGTCGTTGCTGTACCCAGCCAGGATTTTCTTTNNNAGATCGGAAG |
| 47983_2 | TATGAGTGTGGAGTCGTTGCTGTACCCAGCCAGGATTTTCTTTNNNAGATCGGAAG |
| 36976_1 | TATGAGTGTGGAGTCGTTGCGATGTAGGGAGAAGCTGGAGAAGNNNAGATCGGA |
| 36976_2 | TATGAGTGTGGAGTCGTTGCGATGTAGGGAGAAGCTGGAGAAGNNNAGATCGGA |
| 46364_1 | TATGAGTGTGGAGTCGTTGCACCTCCCACCATAGAGAGAAGAANNNAGATCGGAAG |
| 46364_2 | TATGAGTGTGGAGTCGTTGCACCTCCCACCATAGAGAGAAGAANNNAGATCGGAAG |
| 32480_1 | TATGAGTGTGGAGTCGTTGCCTCCGGAGTAGCTGGGACTACNNNAGATCGGAAGA |
| 32480_2 | TATGAGTGTGGAGTCGTTGCCTCCGGAGTAGCTGGGACTACNNNAGATCGGAAGA |
| 32922_1 | TATGAGTGTGGAGTCGTTGCTGAAGCTAGAAGCCATCATCCTCNNNAGATCGGAAG |
| 32922_2 | TATGAGTGTGGAGTCGTTGCTGAAGCTAGAAGCCATCATCCTCNNNAGATCGGAAG |
| 36290_1 | TATGAGTGTGGAGTCGTTGCGCATACCTCTTGTCCTTGCTACTNNNAGATCGGAAGA |
| 36290_2 | TATGAGTGTGGAGTCGTTGCGCATACCTCTTGTCCTTGCTACTNNNAGATCGGAAGA |
| 34343_1 | TATGAGTGTGGAGTCGTTGCGTGTGGTTTTCAGGGGTGGAGNNNAGATCGGAAG |
| 34343_2 | TATGAGTGTGGAGTCGTTGCGTGTGGTTTTCAGGGGTGGAGNNNAGATCGGAAG |
| 39291_1 | TATGAGTGTGGAGTCGTTGCGGGGGTCAATTCAGCAAAAGAATNNNAGATCGGAA |
| 39291_2 | TATGAGTGTGGAGTCGTTGCGGGGGTCAATTCAGCAAAAGAATNNNAGATCGGAA |
| 51321_1 | TATGAGTGTGGAGTCGTTGCACTGAAGAAAGGTAGTGTGTCCCNNNAGATCGGAA |
| 51321_2 | TATGAGTGTGGAGTCGTTGCACTGAAGAAAGGTAGTGTGTCCCNNNAGATCGGAA |
| 46551_1 | TATGAGTGTGGAGTCGTTGCCCTCCTGGGTTCAAGTGATTCTCNNNAGATCGGAAG |
| 46551_2 | TATGAGTGTGGAGTCGTTGCCCTCCTGGGTTCAAGTGATTCTCNNNAGATCGGAAG |
| 51793_1 | TATGAGTGTGGAGTCGTTGCTTAGTATGGGGCACAATCAAGGGNNNAGATCGGAA |
| 51793_2 | TATGAGTGTGGAGTCGTTGCTTAGTATGGGGCACAATCAAGGGNNNAGATCGGAA |
| 39683_1 | TATGAGTGTGGAGTCGTTGCGCCAGCATCACCTTGATAACAAANNNAGATCGGAAG |
| 39683_2 | TATGAGTGTGGAGTCGTTGCGCCAGCATCACCTTGATAACAAANNNAGATCGGAAG |
| 48679_1 | TATGAGTGTGGAGTCGTTGCTGGAATACAGGCACACACCATTANNNAGATCGGAAG |
| 48679_2 | TATGAGTGTGGAGTCGTTGCTGGAATACAGGCACACACCATTANNNAGATCGGAAG |
| 29936_1 | TATGAGTGTGGAGTCGTTGCACAGGGTTATTACGGATATTGGCANNNAGATCGGA |
| 29936_2 | TATGAGTGTGGAGTCGTTGCACAGGGTTATTACGGATATTGGCANNNAGATCGGA |
| 29167_1 | TATGAGTGTGGAGTCGTTGCGAAGGGGAAGGGAAGAAATCCTTNNNAGATCGGA |
| 29167_2 | TATGAGTGTGGAGTCGTTGCGAAGGGGAAGGGAAGAAATCCTTNNNAGATCGGA |
| 32806_1 | TATGAGTGTGGAGTCGTTGCCTCCCAGGTTCAAGCAATTCTCTNNNAGATCGGAAG |
| 32806_2 | TATGAGTGTGGAGTCGTTGCCTCCCAGGTTCAAGCAATTCTCTNNNAGATCGGAAG |
| 12604_1 | TATGAGTGTGGAGTCGTTGCGGAATTCCTTATGTGTGAAATTTNNNAGATCGGAAG |
| 12604_2 | TATGAGTGTGGAGTCGTTGCGGAATTCCTTATGTGTGAAATTTNNNAGATCGGAAG |
| 50461_1 | TATGAGTGTGGAGTCGTTGCACCTTTCTTTCTGATTCCACACANNNAGATCGGAAGA |
| 50461_2 | TATGAGTGTGGAGTCGTTGCACCTTTCTTTCTGATTCCACACANNNAGATCGGAAGA |
| 29244_1 | TATGAGTGTGGAGTCGTTGCGGGGGAGAGAGATAGGGTGATAANNNAGATCGGA |

| | |
|---|---|
| 29244_2 | TATGAGTGTGGAGTCGTTGCGGGGGAGAGAGATAGGGTGATAANNNAGATCGG |
| 36496_1 | TATGAGTGTGGAGTCGTTGCTATTGTTATGGAGTGAGGGTGGGNNNAGATCGGA |
| 36496_2 | TATGAGTGTGGAGTCGTTGCTATTGTTATGGAGTGAGGGTGGGNNNAGATCGGA |
| 10161_1 | TATGAGTGTGGAGTCGTTGCAAGCATGTGTGTATGCCGAGAAANNNAGATCGGAA |
| 10161_2 | TATGAGTGTGGAGTCGTTGCAAGCATGTGTGTATGCCGAGAAANNNAGATCGGAA |
| 31735_1 | TATGAGTGTGGAGTCGTTGCTCATTGTCAGTGGAGTCCTTCAGNNNAGATCGGAAG |
| 31735_2 | TATGAGTGTGGAGTCGTTGCTCATTGTCAGTGGAGTCCTTCAGNNNAGATCGGAAG |
| 38471_1 | TATGAGTGTGGAGTCGTTGCTAGAGGGATCCAGTTTTTCCACCNNNAGATCGGAAG |
| 38471_2 | TATGAGTGTGGAGTCGTTGCTAGAGGGATCCAGTTTTTCCACCNNNAGATCGGAAG |
| 40259_1 | TATGAGTGTGGAGTCGTTGCATTTGAGGTCAGGAGTTCGAGACNNNAGATCGGAA |
| 40259_2 | TATGAGTGTGGAGTCGTTGCATTTGAGGTCAGGAGTTCGAGACNNNAGATCGGAA |
| 36492_1 | TATGAGTGTGGAGTCGTTGCCAGAAGAATGGCGTGAACCTGNNNAGATCGGAAGA |
| 36492_2 | TATGAGTGTGGAGTCGTTGCCAGAAGAATGGCGTGAACCTGNNNAGATCGGAAGA |
| 51824_1 | TATGAGTGTGGAGTCGTTGCAGTGGAGCTCAAAGGGATTTAGGNNNAGATCGGA |
| 51824_2 | TATGAGTGTGGAGTCGTTGCAGTGGAGCTCAAAGGGATTTAGGNNNAGATCGGA |
| 36724_1 | TATGAGTGTGGAGTCGTTGCGGGAGGGAGGGAAAGAAAAGAAANNNAGATCGG |
| 36724_2 | TATGAGTGTGGAGTCGTTGCGGGAGGGAGGGAAAGAAAAGAAANNNAGATCGG |
| 50959_1 | TATGAGTGTGGAGTCGTTGCAAAAAACAGCCTCCCTACTCCCNNNAGATCGGAAGA |
| 50959_2 | TATGAGTGTGGAGTCGTTGCAAAAAACAGCCTCCCTACTCCCNNNAGATCGGAAGA |
| 30597_1 | TATGAGTGTGGAGTCGTTGCAGAGGAGAGAGGCCAAGCTCAATNNNAGATCGGA |
| 30597_2 | TATGAGTGTGGAGTCGTTGCAGAGGAGAGAGGCCAAGCTCAATNNNAGATCGGA |
| 44310_1 | TATGAGTGTGGAGTCGTTGCGCATACCAGACATATGACATGCGNNNAGATCGGAAG |
| 44310_2 | TATGAGTGTGGAGTCGTTGCGCATACCAGACATATGACATGCGNNNAGATCGGAAG |
| 44959_1 | TATGAGTGTGGAGTCGTTGCCGTTAGAAACTCTCGACCATTGCNNNAGATCGGAAG |
| 44959_2 | TATGAGTGTGGAGTCGTTGCCGTTAGAAACTCTCGACCATTGCNNNAGATCGGAAG |
| 45024_1 | TATGAGTGTGGAGTCGTTGCCTCTGGAGTAAGAGGCAACTGAGNNNAGATCGGAA |
| 45024_2 | TATGAGTGTGGAGTCGTTGCCTCTGGAGTAAGAGGCAACTGAGNNNAGATCGGAA |
| 29426_1 | TATGAGTGTGGAGTCGTTGCATTCATTGACTTCAGGTAGATTANNNAGATCGGAAG |
| 29426_2 | TATGAGTGTGGAGTCGTTGCATTCATTGACTTCAGGTAGATTANNNAGATCGGAAG |
| 50637_1 | TATGAGTGTGGAGTCGTTGCTGCCATCATCTTTGCCTATACCANNNAGATCGGAAGA |
| 50637_2 | TATGAGTGTGGAGTCGTTGCTGCCATCATCTTTGCCTATACCANNNAGATCGGAAGA |
| 35065_1 | TATGAGTGTGGAGTCGTTGCTTCAAGCGATTCTCCTGACTCAGNNNAGATCGGAAG |
| 35065_2 | TATGAGTGTGGAGTCGTTGCTTCAAGCGATTCTCCTGACTCAGNNNAGATCGGAAG |
| 41337_1 | TATGAGTGTGGAGTCGTTGCGCAGAGAAAGAAAATGAGGCCTGNNNAGATCGGA |
| 41337_2 | TATGAGTGTGGAGTCGTTGCGCAGAGAAAGAAAATGAGGCCTGNNNAGATCGGA |
| 30546_1 | TATGAGTGTGGAGTCGTTGCGTTCCAGAAAACAGCAGAACTCANNNAGATCGGAAG |
| 30546_2 | TATGAGTGTGGAGTCGTTGCGTTCCAGAAAACAGCAGAACTCANNNAGATCGGAAG |
| 32003_1 | TATGAGTGTGGAGTCGTTGCTCAGACCAGCATCCTTTCACTTANNNAGATCGGAAGA |
| 32003_2 | TATGAGTGTGGAGTCGTTGCTCAGACCAGCATCCTTTCACTTANNNAGATCGGAAGA |
| 10139_1 | TATGAGTGTGGAGTCGTTGCGCTGCAATGAATACAAGAGAGCANNNAGATCGGAA |
| 10139_2 | TATGAGTGTGGAGTCGTTGCGCTGCAATGAATACAAGAGAGCANNNAGATCGGAA |
| 33598_1 | TATGAGTGTGGAGTCGTTGCCGAATAGCTGGGACTACAGGTACNNNAGATCGGAA |
| 33598_2 | TATGAGTGTGGAGTCGTTGCCGAATAGCTGGGACTACAGGTACNNNAGATCGGAA |

| | |
|---|---|
| 45909_1 | TATGAGTGTGGAGTCGTTGCAGATTACAGGCATGAACCACCATNNNAGATCGGAAG |
| 45909_2 | TATGAGTGTGGAGTCGTTGCAGATTACAGGCATGAACCACCATNNNAGATCGGAAG |
| 42155_1 | TATGAGTGTGGAGTCGTTGCCTCCTGGGTTCATGCCATTCTCNNNAGATCGGAAGA |
| 42155_2 | TATGAGTGTGGAGTCGTTGCCTCCTGGGTTCATGCCATTCTCNNNAGATCGGAAGA |
| 50866_1 | TATGAGTGTGGAGTCGTTGCTTTTCACCCATGCCCTAGAGTTTNNNAGATCGGAAG |
| 50866_2 | TATGAGTGTGGAGTCGTTGCTTTTCACCCATGCCCTAGAGTTTNNNAGATCGGAAG |
| 42335_1 | TATGAGTGTGGAGTCGTTGCGTTTCTCCTTCTCTGAGGAACCCNNNAGATCGGAAG |
| 42335_2 | TATGAGTGTGGAGTCGTTGCGTTTCTCCTTCTCTGAGGAACCCNNNAGATCGGAAG |
| 39402_1 | TATGAGTGTGGAGTCGTTGCAATCCCTGCTCAGTAATACAGCCNNNAGATCGGAAG |
| 39402_2 | TATGAGTGTGGAGTCGTTGCAATCCCTGCTCAGTAATACAGCCNNNAGATCGGAAG |
| 46354_1 | TATGAGTGTGGAGTCGTTGCCACTTTGTTCTCAGCAAGAAGGGNNNAGATCGGAA |
| 46354_2 | TATGAGTGTGGAGTCGTTGCCACTTTGTTCTCAGCAAGAAGGGNNNAGATCGGAA |
| 43422_1 | TATGAGTGTGGAGTCGTTGCGCCTCCTGAGTAGCTGGAATTACNNNAGATCGGAAG |
| 43422_2 | TATGAGTGTGGAGTCGTTGCGCCTCCTGAGTAGCTGGAATTACNNNAGATCGGAAG |
| 49166_1 | TATGAGTGTGGAGTCGTTGCGGTTCAAACAATTCTCCTGCCTCNNNAGATCGGAAG |
| 49166_2 | TATGAGTGTGGAGTCGTTGCGGTTCAAACAATTCTCCTGCCTCNNNAGATCGGAAG |
| 31421_1 | TATGAGTGTGGAGTCGTTGCGAGAGACGTGGTGGGAAGTAATTNNNAGATCGGA |
| 31421_2 | TATGAGTGTGGAGTCGTTGCGAGAGACGTGGTGGGAAGTAATTNNNAGATCGGA |
| 30755_1 | TATGAGTGTGGAGTCGTTGCATGTATTTGGAAAAGGGAGGAGTNNNAGATCGGA |
| 30755_2 | TATGAGTGTGGAGTCGTTGCATGTATTTGGAAAAGGGAGGAGTNNNAGATCGGA |
| 44558_1 | TATGAGTGTGGAGTCGTTGCCTGCTGGTAGCGAAATTCAGATGNNNAGATCGGAA |
| 44558_2 | TATGAGTGTGGAGTCGTTGCCTGCTGGTAGCGAAATTCAGATGNNNAGATCGGAA |
| 45072_1 | TATGAGTGTGGAGTCGTTGCCAGAGCTGTATTACCTGATCGCTNNNAGATCGGAAG |
| 45072_2 | TATGAGTGTGGAGTCGTTGCCAGAGCTGTATTACCTGATCGCTNNNAGATCGGAAG |
| 44991_1 | TATGAGTGTGGAGTCGTTGCTTTTTGCCACCGGAGACATCTAGNNNAGATCGGAAG |
| 44991_2 | TATGAGTGTGGAGTCGTTGCTTTTTGCCACCGGAGACATCTAGNNNAGATCGGAAG |
| 53163_1 | TATGAGTGTGGAGTCGTTGCTACCTGACCAGTATAGTGGAGCANNNAGATCGGAA |
| 53163_2 | TATGAGTGTGGAGTCGTTGCTACCTGACCAGTATAGTGGAGCANNNAGATCGGAA |
| 48364_1 | TATGAGTGTGGAGTCGTTGCCTCCTGAGTAGCTGGGATTACAGNNNAGATCGGAA |
| 48364_2 | TATGAGTGTGGAGTCGTTGCCTCCTGAGTAGCTGGGATTACAGNNNAGATCGGAA |
| 51396_1 | TATGAGTGTGGAGTCGTTGCGCAAAAGTCTGACTGGAATGGACNNNAGATCGGAA |
| 51396_2 | TATGAGTGTGGAGTCGTTGCGCAAAAGTCTGACTGGAATGGACNNNAGATCGGAA |
| 50625_1 | TATGAGTGTGGAGTCGTTGCTTGAAATTGCCCTTTCCACTGACNNNAGATCGGAAG |
| 50625_2 | TATGAGTGTGGAGTCGTTGCTTGAAATTGCCCTTTCCACTGACNNNAGATCGGAAG |
| 13544_1 | TATGAGTGTGGAGTCGTTGCGGAAATAGCCTTAAAAAAAAAACTNNNAGATCGGA |
| 13544_2 | TATGAGTGTGGAGTCGTTGCGGAAATAGCCTTAAAAAAAAAACTNNNAGATCGGA |
| 32962_1 | TATGAGTGTGGAGTCGTTGCTCCTAAAGGTGGAAGCTGAAACANNNAGATCGGAA |
| 32962_2 | TATGAGTGTGGAGTCGTTGCTCCTAAAGGTGGAAGCTGAAACANNNAGATCGGAA |
| 31592_1 | TATGAGTGTGGAGTCGTTGCTGAACAGCTAATTCCAACCAACANNNAGATCGGAAG |
| 31592_2 | TATGAGTGTGGAGTCGTTGCTGAACAGCTAATTCCAACCAACANNNAGATCGGAAG |
| 43352_1 | TATGAGTGTGGAGTCGTTGCGTTCAAGAGATTCTCCTGCCTCANNNAGATCGGAAG |
| 43352_2 | TATGAGTGTGGAGTCGTTGCGTTCAAGAGATTCTCCTGCCTCANNNAGATCGGAAG |
| 47097_1 | TATGAGTGTGGAGTCGTTGCTTCAAGTGATTCTCCTGCCTCAGNNNAGATCGGAAG |

| | |
|---|---|
| 47097_2 | TATGAGTGTGGAGTCGTTGCTTCAAGTGATTCTCCTGCCTCAGNNNAGATCGGAAG |
| 12861_1 | TATGAGTGTGGAGTCGTTGCTATATCATTTGAAGGTAGACACCNNNAGATCGGAAG |
| 12861_2 | TATGAGTGTGGAGTCGTTGCTATATCATTTGAAGGTAGACACCNNNAGATCGGAAG |
| 32362_1 | TATGAGTGTGGAGTCGTTGCCCTACTGGGAAGTGAGGAGCCNNNAGATCGGAAGA |
| 32362_2 | TATGAGTGTGGAGTCGTTGCCCTACTGGGAAGTGAGGAGCCNNNAGATCGGAAGA |
| 31471_1 | TATGAGTGTGGAGTCGTTGCCCCAGCCAGATTCTATCATCACANNNAGATCGGAAG |
| 31471_2 | TATGAGTGTGGAGTCGTTGCCCCAGCCAGATTCTATCATCACANNNAGATCGGAAG |
| 48765_1 | TATGAGTGTGGAGTCGTTGCCTGCCTCAACCTCCTGAATAGTTNNNAGATCGGAAG |
| 48765_2 | TATGAGTGTGGAGTCGTTGCCTGCCTCAACCTCCTGAATAGTTNNNAGATCGGAAG |
| 30731_1 | TATGAGTGTGGAGTCGTTGCATGATTTTGGCCTTTTCTGTCATGNNNAGATCGGAA |
| 30731_2 | TATGAGTGTGGAGTCGTTGCATGATTTTGGCCTTTTCTGTCATGNNNAGATCGGAA |
| 31949_1 | TATGAGTGTGGAGTCGTTGCGAGGAGAAAGTACTCAACCAGGGNNNAGATCGGA |
| 31949_2 | TATGAGTGTGGAGTCGTTGCGAGGAGAAAGTACTCAACCAGGGNNNAGATCGGA |
| 43079_1 | TATGAGTGTGGAGTCGTTGCGCACATACACACACAGAAATCCANNNAGATCGGAAG |
| 43079_2 | TATGAGTGTGGAGTCGTTGCGCACATACACACACAGAAATCCANNNAGATCGGAAG |
| 8790_1 | TATGAGTGTGGAGTCGTTGCTCTGAGGAGAATGTCCAGAGCTTNNNAGATCGGAA |
| 8790_2 | TATGAGTGTGGAGTCGTTGCTCTGAGGAGAATGTCCAGAGCTTNNNAGATCGGAA |
| 52847_1 | TATGAGTGTGGAGTCGTTGCCTGGGGGTCTTGCTCACACCNNNAGATCGGAAGAG |
| 52847_2 | TATGAGTGTGGAGTCGTTGCCTGGGGGTCTTGCTCACACCNNNAGATCGGAAGAG |
| 8963_1 | TATGAGTGTGGAGTCGTTGCACTCCTGTACCTCTTGGCAGAATNNNAGATCGGAAG |
| 8963_2 | TATGAGTGTGGAGTCGTTGCACTCCTGTACCTCTTGGCAGAATNNNAGATCGGAAG |
| 40953_1 | TATGAGTGTGGAGTCGTTGCTTTTCCTTAGGTCTCACAGAGGCNNNAGATCGGAAG |
| 40953_2 | TATGAGTGTGGAGTCGTTGCTTTTCCTTAGGTCTCACAGAGGCNNNAGATCGGAAG |
| 34997_1 | TATGAGTGTGGAGTCGTTGCATGGAGTTTTGTCCTTTTGCCTGNNNAGATCGGAAG |
| 34997_2 | TATGAGTGTGGAGTCGTTGCATGGAGTTTTGTCCTTTTGCCTGNNNAGATCGGAAG |
| 41846_1 | TATGAGTGTGGAGTCGTTGCCAATTCTCCTACCTCAGCCTACCNNNAGATCGGAAGA |
| 41846_2 | TATGAGTGTGGAGTCGTTGCCAATTCTCCTACCTCAGCCTACCNNNAGATCGGAAGA |
| 39691_1 | TATGAGTGTGGAGTCGTTGCGTTTATGGAAACTTGGGGCTGTTNNNAGATCGGAA |
| 39691_2 | TATGAGTGTGGAGTCGTTGCGTTTATGGAAACTTGGGGCTGTTNNNAGATCGGAA |
| 38108_1 | TATGAGTGTGGAGTCGTTGCACTACAGAAGAGCGTGCAAGTAANNNAGATCGGAA |
| 38108_2 | TATGAGTGTGGAGTCGTTGCACTACAGAAGAGCGTGCAAGTAANNNAGATCGGAA |
| 38571_1 | TATGAGTGTGGAGTCGTTGCCAAAAAAGAAGTGGGGACTTGCTNNNAGATCGGAA |
| 38571_2 | TATGAGTGTGGAGTCGTTGCCAAAAAAGAAGTGGGGACTTGCTNNNAGATCGGAA |
| 39905_1 | TATGAGTGTGGAGTCGTTGCACAGATTACACTGAAAGGGGACANNNAGATCGGAA |
| 39905_2 | TATGAGTGTGGAGTCGTTGCACAGATTACACTGAAAGGGGACANNNAGATCGGAA |
| 46987_1 | TATGAGTGTGGAGTCGTTGCCCTGGAGTAGCTGGGATTACAGNNNAGATCGGAAG |
| 46987_2 | TATGAGTGTGGAGTCGTTGCCCTGGAGTAGCTGGGATTACAGNNNAGATCGGAAG |
| 33810_1 | TATGAGTGTGGAGTCGTTGCAAATGCTGGGATTACAGACGTGANNNAGATCGGAA |
| 33810_2 | TATGAGTGTGGAGTCGTTGCAAATGCTGGGATTACAGACGTGANNNAGATCGGAA |
| 50190_1 | TATGAGTGTGGAGTCGTTGCCTTTCATCTGCTGACCTGGTACTNNNAGATCGGAAG |
| 50190_2 | TATGAGTGTGGAGTCGTTGCCTTTCATCTGCTGACCTGGTACTNNNAGATCGGAAG |
| 12733_1 | TATGAGTGTGGAGTCGTTGCGTTTTGTGCAAAAGTGTTTGNNNAGATCGGAAGAG |
| 12733_2 | TATGAGTGTGGAGTCGTTGCGTTTTGTGCAAAAGTGTTTGNNNAGATCGGAAGAG |

| | |
|---|---|
| 31175_1 | TATGAGTGTGGAGTCGTTGCTTCTCTCTCTCTCTTTCCCCNNNAGATCGGAAGAG |
| 31175_2 | TATGAGTGTGGAGTCGTTGCTTCTCTCTCTCTCTTTCCCCNNNAGATCGGAAGAG |
| 53015_1 | TATGAGTGTGGAGTCGTTGCTTCAATGTGAGAGATGAGTGGGGNNNAGATCGGA |
| 53015_2 | TATGAGTGTGGAGTCGTTGCTTCAATGTGAGAGATGAGTGGGGNNNAGATCGGA |
| 45417_1 | TATGAGTGTGGAGTCGTTGCTCCAAATGAAAGGAGGGGTACAGNNNAGATCGGA |
| 45417_2 | TATGAGTGTGGAGTCGTTGCTCCAAATGAAAGGAGGGGTACAGNNNAGATCGGA |
| 51884_1 | TATGAGTGTGGAGTCGTTGCAGACTCTCCCCAAATAATGCCAGNNNAGATCGGAAG |
| 51884_2 | TATGAGTGTGGAGTCGTTGCAGACTCTCCCCAAATAATGCCAGNNNAGATCGGAAG |
| 44291_1 | TATGAGTGTGGAGTCGTTGCAGGCCAGTTTACATGATGGTCTTNNNAGATCGGAAG |
| 44291_2 | TATGAGTGTGGAGTCGTTGCAGGCCAGTTTACATGATGGTCTTNNNAGATCGGAAG |
| 49968_1 | TATGAGTGTGGAGTCGTTGCCTTCTGGTCCCTTCACAGAATCANNNAGATCGGAAG |
| 49968_2 | TATGAGTGTGGAGTCGTTGCCTTCTGGTCCCTTCACAGAATCANNNAGATCGGAAG |
| 44726_1 | TATGAGTGTGGAGTCGTTGCAGGTGAGCATGGAGTGAGTTTTTNNNAGATCGGAA |
| 44726_2 | TATGAGTGTGGAGTCGTTGCAGGTGAGCATGGAGTGAGTTTTTNNNAGATCGGAA |
| 43493_1 | TATGAGTGTGGAGTCGTTGCACGATCTTGATCTCCTGACATCGNNNAGATCGGAAG |
| 43493_2 | TATGAGTGTGGAGTCGTTGCACGATCTTGATCTCCTGACATCGNNNAGATCGGAAG |
| 10841_1 | TATGAGTGTGGAGTCGTTGCAGGTCTAACCAGCTCTCTAGGTGNNNAGATCGGAAG |
| 10841_2 | TATGAGTGTGGAGTCGTTGCAGGTCTAACCAGCTCTCTAGGTGNNNAGATCGGAAG |
| 37096_1 | TATGAGTGTGGAGTCGTTGCGCTTCCCATCTTGCTGGTTTATCNNNAGATCGGAAGA |
| 37096_2 | TATGAGTGTGGAGTCGTTGCGCTTCCCATCTTGCTGGTTTATCNNNAGATCGGAAGA |
| 9942_1 | TATGAGTGTGGAGTCGTTGCCCTCTCAGTTCTCTCATGCTGCNNNAGATCGGAAGAG |
| 9942_2 | TATGAGTGTGGAGTCGTTGCCCTCTCAGTTCTCTCATGCTGCNNNAGATCGGAAGAG |
| 10785_1 | TATGAGTGTGGAGTCGTTGCGAAACAGTTCTTGGCCAGAGGAGNNNAGATCGGAA |
| 10785_2 | TATGAGTGTGGAGTCGTTGCGAAACAGTTCTTGGCCAGAGGAGNNNAGATCGGAA |
| 49411_1 | TATGAGTGTGGAGTCGTTGCGTCTTTGGCATTCACACTGGAAANNNAGATCGGAAG |
| 49411_2 | TATGAGTGTGGAGTCGTTGCGTCTTTGGCATTCACACTGGAAANNNAGATCGGAAG |
| 34608_1 | TATGAGTGTGGAGTCGTTGCATCCAGGAGGGAGGTGGGAGNNNAGATCGGAAGA |
| 34608_2 | TATGAGTGTGGAGTCGTTGCATCCAGGAGGGAGGTGGGAGNNNAGATCGGAAGA |
| 48691_1 | TATGAGTGTGGAGTCGTTGCTAGCTGGGATTATAGGCATGTGCNNNAGATCGGAA |
| 48691_2 | TATGAGTGTGGAGTCGTTGCTAGCTGGGATTATAGGCATGTGCNNNAGATCGGAA |
| 41566_1 | TATGAGTGTGGAGTCGTTGCCTGAAGTGTAGTGGTGCGATCATNNNAGATCGGAA |
| 41566_2 | TATGAGTGTGGAGTCGTTGCCTGAAGTGTAGTGGTGCGATCATNNNAGATCGGAA |
| 13035_1 | TATGAGTGTGGAGTCGTTGCAAATTTCCAACTTGTATGTGTTTNNNAGATCGGAAG |
| 13035_2 | TATGAGTGTGGAGTCGTTGCAAATTTCCAACTTGTATGTGTTTNNNAGATCGGAAG |
| 30107_1 | TATGAGTGTGGAGTCGTTGCGAAGTCTAAGTCCTTATCCCCTTTNNNAGATCGGAAG |
| 30107_2 | TATGAGTGTGGAGTCGTTGCGAAGTCTAAGTCCTTATCCCCTTTNNNAGATCGGAAG |
| 49396_1 | TATGAGTGTGGAGTCGTTGCCTCCTGAGTAGCTGGGATTACAGNNNAGATCGGAA |
| 49396_2 | TATGAGTGTGGAGTCGTTGCCTCCTGAGTAGCTGGGATTACAGNNNAGATCGGAA |
| 50182_1 | TATGAGTGTGGAGTCGTTGCCGACCACTTTGTCAAGCTCATTTNNNAGATCGGAAGA |
| 50182_2 | TATGAGTGTGGAGTCGTTGCCGACCACTTTGTCAAGCTCATTTNNNAGATCGGAAGA |
| 33759_1 | TATGAGTGTGGAGTCGTTGCCTGGAAACAACCCCAAATGTTCTNNNAGATCGGAAG |
| 33759_2 | TATGAGTGTGGAGTCGTTGCCTGGAAACAACCCCAAATGTTCTNNNAGATCGGAAG |
| 52184_1 | TATGAGTGTGGAGTCGTTGCAACGCCACCTACATACAGCTAAANNNAGATCGGAAG |

| | |
|---|---|
| 52184_2 | TATGAGTGTGGAGTCGTTGCAACGCCACCTACATACAGCTAAANNNAGATCGGAAG |
| 47142_1 | TATGAGTGTGGAGTCGTTGCAGGTTCAAATGATTCTCCTGCCTNNNAGATCGGAAG |
| 47142_2 | TATGAGTGTGGAGTCGTTGCAGGTTCAAATGATTCTCCTGCCTNNNAGATCGGAAG |
| 2832_1 | TATGAGTGTGGAGTCGTTGCGATGAACCTGCATGACCTGAAGTNNNAGATCGGAA |
| 2832_2 | TATGAGTGTGGAGTCGTTGCGATGAACCTGCATGACCTGAAGTNNNAGATCGGAA |
| 29613_1 | TATGAGTGTGGAGTCGTTGCACAAAAATCACTCCTTAAAAGAGACTCNNNAGATCG |
| 29613_2 | TATGAGTGTGGAGTCGTTGCACAAAAATCACTCCTTAAAAGAGACTCNNNAGATCG |
| 41371_1 | TATGAGTGTGGAGTCGTTGCGCCCCAGCTTCTTGAATTGTTTTNNNAGATCGGAAG |
| 41371_2 | TATGAGTGTGGAGTCGTTGCGCCCCAGCTTCTTGAATTGTTTTNNNAGATCGGAAG |
| 52118_1 | TATGAGTGTGGAGTCGTTGCAAGCTCAGAGAAGCCGTTTTCTANNNAGATCGGAAG |
| 52118_2 | TATGAGTGTGGAGTCGTTGCAAGCTCAGAGAAGCCGTTTTCTANNNAGATCGGAAG |
| 39724_1 | TATGAGTGTGGAGTCGTTGCGAGGTCAGGAGTTCAAGACCAAGNNNAGATCGGAA |
| 39724_2 | TATGAGTGTGGAGTCGTTGCGAGGTCAGGAGTTCAAGACCAAGNNNAGATCGGAA |
| 46005_1 | TATGAGTGTGGAGTCGTTGCACAGACCTGCAGGAATGAAGTAGNNNAGATCGGAA |
| 46005_2 | TATGAGTGTGGAGTCGTTGCACAGACCTGCAGGAATGAAGTAGNNNAGATCGGAA |
| 38852_1 | TATGAGTGTGGAGTCGTTGCAAGTAATGACCCTGTCTTTGCCTNNNAGATCGGAAG |
| 38852_2 | TATGAGTGTGGAGTCGTTGCAAGTAATGACCCTGTCTTTGCCTNNNAGATCGGAAG |
| 46252_1 | TATGAGTGTGGAGTCGTTGCTGCAGTGACAAGATCATGACTCANNNAGATCGGAAG |
| 46252_2 | TATGAGTGTGGAGTCGTTGCTGCAGTGACAAGATCATGACTCANNNAGATCGGAAG |
| 46181_1 | TATGAGTGTGGAGTCGTTGCTTAGTAAAGACAGGGTTTCGCCANNNAGATCGGAA |
| 46181_2 | TATGAGTGTGGAGTCGTTGCTTAGTAAAGACAGGGTTTCGCCANNNAGATCGGAA |
| 8000_1 | TATGAGTGTGGAGTCGTTGCCCTGGGTTCAAAGCCTAATAGCANNNAGATCGGAAG |
| 8000_2 | TATGAGTGTGGAGTCGTTGCCCTGGGTTCAAAGCCTAATAGCANNNAGATCGGAAG |
| 32216_1 | TATGAGTGTGGAGTCGTTGCCATCATGCCCCTCCATAGTGTACNNNAGATCGGAAG |
| 32216_2 | TATGAGTGTGGAGTCGTTGCCATCATGCCCCTCCATAGTGTACNNNAGATCGGAAG |
| 33476_1 | TATGAGTGTGGAGTCGTTGCTGTACCTCAGGGAAATACAAGGCNNNAGATCGGAA |
| 33476_2 | TATGAGTGTGGAGTCGTTGCTGTACCTCAGGGAAATACAAGGCNNNAGATCGGAA |
| 32567_1 | TATGAGTGTGGAGTCGTTGCGATGGAGTCTCATTCTCTCGCCNNNAGATCGGAAGA |
| 32567_2 | TATGAGTGTGGAGTCGTTGCGATGGAGTCTCATTCTCTCGCCNNNAGATCGGAAGA |
| 50459_1 | TATGAGTGTGGAGTCGTTGCTGGCATGAACCAGAAGAAGAAGANNNAGATCGGAA |
| 50459_2 | TATGAGTGTGGAGTCGTTGCTGGCATGAACCAGAAGAAGAAGANNNAGATCGGAA |
| 13655_1 | TATGAGTGTGGAGTCGTTGCAAGAAACATTTGGAAATAGAACANNNAGATCGGAA |
| 13655_2 | TATGAGTGTGGAGTCGTTGCAAGAAACATTTGGAAATAGAACANNNAGATCGGAA |
| 217_1 | TATGAGTGTGGAGTCGTTGCACGATCAGAAAGCAGCATTTCCGNNNAGATCGGAAG |
| 217_2 | TATGAGTGTGGAGTCGTTGCACGATCAGAAAGCAGCATTTCCGNNNAGATCGGAAG |
| 49462_1 | TATGAGTGTGGAGTCGTTGCCCACACCCGGCTAATTTTTTGTANNNAGATCGGAAG |
| 49462_2 | TATGAGTGTGGAGTCGTTGCCCACACCCGGCTAATTTTTTGTANNNAGATCGGAAG |
| 10404_1 | TATGAGTGTGGAGTCGTTGCATTTGAAGAGGACATGTGCCTNNNAGATCGGAAGA |
| 10404_2 | TATGAGTGTGGAGTCGTTGCATTTGAAGAGGACATGTGCCTNNNAGATCGGAAGA |
| 39139_1 | TATGAGTGTGGAGTCGTTGCCCACTCTCCATTTAGCCAGAAGTNNNAGATCGGAAG |
| 39139_2 | TATGAGTGTGGAGTCGTTGCCCACTCTCCATTTAGCCAGAAGTNNNAGATCGGAAG |
| 13476_1 | TATGAGTGTGGAGTCGTTGCGTTTCAGATAGTGCTGTGTTNNNAGATCGGAAGAG |
| 13476_2 | TATGAGTGTGGAGTCGTTGCGTTTCAGATAGTGCTGTGTTNNNAGATCGGAAGAG |

| | |
|---|---|
| 45117_1 | TATGAGTGTGGAGTCGTTGCTTACCTTCGCCTCAAGTACTGTCNNNAGATCGGAAGA |
| 45117_2 | TATGAGTGTGGAGTCGTTGCTTACCTTCGCCTCAAGTACTGTCNNNAGATCGGAAGA |
| 52614_1 | TATGAGTGTGGAGTCGTTGCGGATTTGTTTTGGTCGGGCTAGNNNAGATCGGAA |
| 52614_2 | TATGAGTGTGGAGTCGTTGCGGATTTGTTTTGGTCGGGCTAGNNNAGATCGGAA |
| 10724_1 | TATGAGTGTGGAGTCGTTGCCTTAGACCTGGACGCCATGTTGNNNAGATCGGAAGA |
| 10724_2 | TATGAGTGTGGAGTCGTTGCCTTAGACCTGGACGCCATGTTGNNNAGATCGGAAGA |
| 35703_1 | TATGAGTGTGGAGTCGTTGCTCCATTCTGCTAGCCTAAGTCTTNNNAGATCGGAAGA |
| 35703_2 | TATGAGTGTGGAGTCGTTGCTCCATTCTGCTAGCCTAAGTCTTNNNAGATCGGAAGA |
| 42981_1 | TATGAGTGTGGAGTCGTTGCTACTTTCCTTCTACGCCTTGCATNNNAGATCGGAAGA |
| 42981_2 | TATGAGTGTGGAGTCGTTGCTACTTTCCTTCTACGCCTTGCATNNNAGATCGGAAGA |
| 2154_1 | TATGAGTGTGGAGTCGTTGCAGGAGGGCAGCAATTTGAAAAGANNNAGATCGGAA |
| 2154_2 | TATGAGTGTGGAGTCGTTGCAGGAGGGCAGCAATTTGAAAAGANNNAGATCGGAA |
| 34543_1 | TATGAGTGTGGAGTCGTTGCTCATCTTAGTCCATGCTCAGCAGNNNAGATCGGAAG |
| 34543_2 | TATGAGTGTGGAGTCGTTGCTCATCTTAGTCCATGCTCAGCAGNNNAGATCGGAAG |
| 29643_1 | TATGAGTGTGGAGTCGTTGCGAGTATCCCAAGGACTGTTCTCCNNNAGATCGGAAG |
| 29643_2 | TATGAGTGTGGAGTCGTTGCGAGTATCCCAAGGACTGTTCTCCNNNAGATCGGAAG |
| 31663_1 | TATGAGTGTGGAGTCGTTGCGACTCTGTTTGGGAGAAAGCAACNNNAGATCGGAA |
| 31663_2 | TATGAGTGTGGAGTCGTTGCGACTCTGTTTGGGAGAAAGCAACNNNAGATCGGAA |
| 13276_1 | TATGAGTGTGGAGTCGTTGCCATGCTAGCAATTAACAATCAANNNAGATCGGAAGA |
| 13276_2 | TATGAGTGTGGAGTCGTTGCCATGCTAGCAATTAACAATCAANNNAGATCGGAAGA |
| 49095_1 | TATGAGTGTGGAGTCGTTGCAACCACCTCTTTTCCAGAATGGANNNAGATCGGAAG |
| 49095_2 | TATGAGTGTGGAGTCGTTGCAACCACCTCTTTTCCAGAATGGANNNAGATCGGAAG |
| 52513_1 | TATGAGTGTGGAGTCGTTGCATTAGAAGAAGCCCAGCATGGAANNNAGATCGGAA |
| 52513_2 | TATGAGTGTGGAGTCGTTGCATTAGAAGAAGCCCAGCATGGAANNNAGATCGGAA |
| 32649_1 | TATGAGTGTGGAGTCGTTGCTTCCAGGTTCAAGCAATTCTCCTNNNAGATCGGAAGA |
| 32649_2 | TATGAGTGTGGAGTCGTTGCTTCCAGGTTCAAGCAATTCTCCTNNNAGATCGGAAGA |
| 5337_1 | TATGAGTGTGGAGTCGTTGCCTCCAACCACATTTTTCCCCCTTNNNAGATCGGAAGA |
| 5337_2 | TATGAGTGTGGAGTCGTTGCCTCCAACCACATTTTTCCCCCTTNNNAGATCGGAAGA |
| 39417_1 | TATGAGTGTGGAGTCGTTGCTACAAAAACAAATAGCTGGGCGTNNNAGATCGGAA |
| 39417_2 | TATGAGTGTGGAGTCGTTGCTACAAAAACAAATAGCTGGGCGTNNNAGATCGGAA |
| 1997_1 | TATGAGTGTGGAGTCGTTGCCAGGTGGAACAGTTTGCACAAAANNNAGATCGGAA |
| 1997_2 | TATGAGTGTGGAGTCGTTGCCAGGTGGAACAGTTTGCACAAAANNNAGATCGGAA |
| 33898_1 | TATGAGTGTGGAGTCGTTGCCTCCTGGGTTCATGCCATTCTCNNNAGATCGGAAGA |
| 33898_2 | TATGAGTGTGGAGTCGTTGCCTCCTGGGTTCATGCCATTCTCNNNAGATCGGAAGA |
| 43964_1 | TATGAGTGTGGAGTCGTTGCACCTGCAATATGGGCTAACAAGANNNAGATCGGAA |
| 43964_2 | TATGAGTGTGGAGTCGTTGCACCTGCAATATGGGCTAACAAGANNNAGATCGGAA |
| 34897_1 | TATGAGTGTGGAGTCGTTGCCCTCCCACGTTCAAATGATTCTCNNNAGATCGGAAGA |
| 34897_2 | TATGAGTGTGGAGTCGTTGCCCTCCCACGTTCAAATGATTCTCNNNAGATCGGAAGA |
| 46889_1 | TATGAGTGTGGAGTCGTTGCCAGCAAGACACAACCATTCACTTNNNAGATCGGAAG |
| 46889_2 | TATGAGTGTGGAGTCGTTGCCAGCAAGACACAACCATTCACTTNNNAGATCGGAAG |
| 33701_1 | TATGAGTGTGGAGTCGTTGCAATGGGGACATAACCACCTTCTCNNNAGATCGGAAG |
| 33701_2 | TATGAGTGTGGAGTCGTTGCAATGGGGACATAACCACCTTCTCNNNAGATCGGAAG |
| 33260_1 | TATGAGTGTGGAGTCGTTGCGATCTCGACTCACTGCAAGCTCNNNAGATCGGAAGA |

| | |
|---|---|
| 33260_2 | TATGAGTGTGGAGTCGTTGCGATCTCGACTCACTGCAAGCTCNNNAGATCGGAAGA |
| 48386_1 | TATGAGTGTGGAGTCGTTGCTCTGGATTCAAAGCGATTCTCCTNNNAGATCGGAAG |
| 48386_2 | TATGAGTGTGGAGTCGTTGCTCTGGATTCAAAGCGATTCTCCTNNNAGATCGGAAG |
| 12534_1 | TATGAGTGTGGAGTCGTTGCCTTAATAGGCACATGAGATCTCNNNAGATCGGAAGA |
| 12534_2 | TATGAGTGTGGAGTCGTTGCCTTAATAGGCACATGAGATCTCNNNAGATCGGAAGA |
| 46569_1 | TATGAGTGTGGAGTCGTTGCTTCAAGTGATTCTCCCACCTCAGNNNAGATCGGAAG |
| 46569_2 | TATGAGTGTGGAGTCGTTGCTTCAAGTGATTCTCCCACCTCAGNNNAGATCGGAAG |
| 32287_1 | TATGAGTGTGGAGTCGTTGCTACGTTGCCCAGACTAGTTTCAANNNAGATCGGAAG |
| 32287_2 | TATGAGTGTGGAGTCGTTGCTACGTTGCCCAGACTAGTTTCAANNNAGATCGGAAG |
| 47162_1 | TATGAGTGTGGAGTCGTTGCACCTCCTAGGCTCAGGTAATTCTNNNAGATCGGAAG |
| 47162_2 | TATGAGTGTGGAGTCGTTGCACCTCCTAGGCTCAGGTAATTCTNNNAGATCGGAAG |
| 45381_1 | TATGAGTGTGGAGTCGTTGCGCACATACATGCACATACACACANNNAGATCGGAAG |
| 45381_2 | TATGAGTGTGGAGTCGTTGCGCACATACATGCACATACACACANNNAGATCGGAAG |
| 13733_1 | TATGAGTGTGGAGTCGTTGCACAGTAAATTAATCCTGGTTTTCNNNAGATCGGAAG |
| 13733_2 | TATGAGTGTGGAGTCGTTGCACAGTAAATTAATCCTGGTTTTCNNNAGATCGGAAG |
| 37438_1 | TATGAGTGTGGAGTCGTTGCCTCTTGGTGTCCTATCCACACACNNNAGATCGGAAGA |
| 37438_2 | TATGAGTGTGGAGTCGTTGCCTCTTGGTGTCCTATCCACACACNNNAGATCGGAAGA |
| 33442_1 | TATGAGTGTGGAGTCGTTGCGTGACATGGATGAAGCTGGAAAGNNNAGATCGGA |
| 33442_2 | TATGAGTGTGGAGTCGTTGCGTGACATGGATGAAGCTGGAAAGNNNAGATCGGA |
| 13346_1 | TATGAGTGTGGAGTCGTTGCTTTCATTTGTCTAAGGTTTTTCTNNNAGATCGGAAGA |
| 13346_2 | TATGAGTGTGGAGTCGTTGCTTTCATTTGTCTAAGGTTTTTCTNNNAGATCGGAAGA |
| 38911_1 | TATGAGTGTGGAGTCGTTGCAAATGGTGCTGGGATAACTGGATNNNAGATCGGAA |
| 38911_2 | TATGAGTGTGGAGTCGTTGCAAATGGTGCTGGGATAACTGGATNNNAGATCGGAA |
| 37866_1 | TATGAGTGTGGAGTCGTTGCGTGGCCATAAACTACCCAGAGAANNNAGATCGGAA |
| 37866_2 | TATGAGTGTGGAGTCGTTGCGTGGCCATAAACTACCCAGAGAANNNAGATCGGAA |
| 43582_1 | TATGAGTGTGGAGTCGTTGCAGTGATGATGATGACAATGCTCCNNNAGATCGGAA |
| 43582_2 | TATGAGTGTGGAGTCGTTGCAGTGATGATGATGACAATGCTCCNNNAGATCGGAA |
| 42000_1 | TATGAGTGTGGAGTCGTTGCCATGATCTTGGCTCACTGCAACNNNAGATCGGAAGA |
| 42000_2 | TATGAGTGTGGAGTCGTTGCCATGATCTTGGCTCACTGCAACNNNAGATCGGAAGA |
| 42122_1 | TATGAGTGTGGAGTCGTTGCACCACACCCAGCTAACTTTTGTANNNAGATCGGAAG |
| 42122_2 | TATGAGTGTGGAGTCGTTGCACCACACCCAGCTAACTTTTGTANNNAGATCGGAAG |
| 47045_1 | TATGAGTGTGGAGTCGTTGCAACTAACAACTGTGGGATCCAGGNNNAGATCGGAA |
| 47045_2 | TATGAGTGTGGAGTCGTTGCAACTAACAACTGTGGGATCCAGGNNNAGATCGGAA |
| 45224_1 | TATGAGTGTGGAGTCGTTGCACAAAAAAACCTCTCAGCTCACGNNNAGATCGGAAG |
| 45224_2 | TATGAGTGTGGAGTCGTTGCACAAAAAAACCTCTCAGCTCACGNNNAGATCGGAAG |
| 32503_1 | TATGAGTGTGGAGTCGTTGCTTCAAGCAACTCTCCTGTCTCAGNNNAGATCGGAAG |
| 32503_2 | TATGAGTGTGGAGTCGTTGCTTCAAGCAACTCTCCTGTCTCAGNNNAGATCGGAAG |
| 29632_1 | TATGAGTGTGGAGTCGTTGCCATCTGCTGAGGTCTTCTTCCCNNNAGATCGGAAGA |
| 29632_2 | TATGAGTGTGGAGTCGTTGCCATCTGCTGAGGTCTTCTTCCCNNNAGATCGGAAGA |
| 52471_1 | TATGAGTGTGGAGTCGTTGCAGGTGAGAGGACAAGTCATGAACNNNAGATCGGA |
| 52471_2 | TATGAGTGTGGAGTCGTTGCAGGTGAGAGGACAAGTCATGAACNNNAGATCGGA |
| 12362_1 | TATGAGTGTGGAGTCGTTGCAACCAAATGACCTGGCTTGGAATNNNAGATCGGAA |
| 12362_2 | TATGAGTGTGGAGTCGTTGCAACCAAATGACCTGGCTTGGAATNNNAGATCGGAA |

| | |
|---|---|
| 44468_1 | TATGAGTGTGGAGTCGTTGCGCCTTCCAGGTTGAAGCAATTCTNNNAGATCGGAAG |
| 44468_2 | TATGAGTGTGGAGTCGTTGCGCCTTCCAGGTTGAAGCAATTCTNNNAGATCGGAAG |
| 31161_1 | TATGAGTGTGGAGTCGTTGCAGCCTCACAAAAGATAGTGCCATNNNAGATCGGAAG |
| 31161_2 | TATGAGTGTGGAGTCGTTGCAGCCTCACAAAAGATAGTGCCATNNNAGATCGGAAG |
| 48722_1 | TATGAGTGTGGAGTCGTTGCTGTCCACTCTTCAATCCACATGANNNAGATCGGAAGA |
| 48722_2 | TATGAGTGTGGAGTCGTTGCTGTCCACTCTTCAATCCACATGANNNAGATCGGAAGA |
| 37862_1 | TATGAGTGTGGAGTCGTTGCCAGAACTTGATGGAGTTTCAGCCNNNAGATCGGAAG |
| 37862_2 | TATGAGTGTGGAGTCGTTGCCAGAACTTGATGGAGTTTCAGCCNNNAGATCGGAAG |
| 44654_1 | TATGAGTGTGGAGTCGTTGCCAGAGGGATCTCAGTTGGGAAAANNNAGATCGGAA |
| 44654_2 | TATGAGTGTGGAGTCGTTGCCAGAGGGATCTCAGTTGGGAAAANNNAGATCGGAA |
| 37818_1 | TATGAGTGTGGAGTCGTTGCAGAGATACATAGGGCAAGGCATGNNNAGATCGGAA |
| 37818_2 | TATGAGTGTGGAGTCGTTGCAGAGATACATAGGGCAAGGCATGNNNAGATCGGAA |
| 33290_1 | TATGAGTGTGGAGTCGTTGCAGAAGGAGTCTCACTCTGTCACCNNNAGATCGGAAG |
| 33290_2 | TATGAGTGTGGAGTCGTTGCAGAAGGAGTCTCACTCTGTCACCNNNAGATCGGAAG |
| 33586_1 | TATGAGTGTGGAGTCGTTGCCTCGATCTCCTGACCTTGTGATCNNNAGATCGGAAG |
| 33586_2 | TATGAGTGTGGAGTCGTTGCCTCGATCTCCTGACCTTGTGATCNNNAGATCGGAAG |
| 30688_1 | TATGAGTGTGGAGTCGTTGCGATGTACCTTTGAGAATCCGTCATNNNAGATCGGAA |
| 30688_2 | TATGAGTGTGGAGTCGTTGCGATGTACCTTTGAGAATCCGTCATNNNAGATCGGAA |
| 42759_1 | TATGAGTGTGGAGTCGTTGCACTACTCAGCCATAAAAAGGAACANNNAGATCGGAA |
| 42759_2 | TATGAGTGTGGAGTCGTTGCACTACTCAGCCATAAAAAGGAACANNNAGATCGGAA |
| 41249_1 | TATGAGTGTGGAGTCGTTGCCAAATCTCAAAGACTCAGCCCCANNNAGATCGGAAG |
| 41249_2 | TATGAGTGTGGAGTCGTTGCCAAATCTCAAAGACTCAGCCCCANNNAGATCGGAAG |
| 38702_1 | TATGAGTGTGGAGTCGTTGCGAGACAGAGGCTCAACTTCAGTANNNAGATCGGAA |
| 38702_2 | TATGAGTGTGGAGTCGTTGCGAGACAGAGGCTCAACTTCAGTANNNAGATCGGAA |
| 33063_1 | TATGAGTGTGGAGTCGTTGCGGCTTGAGCAATCAGCAAAACTANNNAGATCGGAAG |
| 33063_2 | TATGAGTGTGGAGTCGTTGCGGCTTGAGCAATCAGCAAAACTANNNAGATCGGAAG |
| 44565_1 | TATGAGTGTGGAGTCGTTGCCTATCACACAGTCTTTGGGTCCANNNAGATCGGAAG |
| 44565_2 | TATGAGTGTGGAGTCGTTGCCTATCACACAGTCTTTGGGTCCANNNAGATCGGAAG |
| 50838_1 | TATGAGTGTGGAGTCGTTGCCGTCATAAGTATCCCAGACCTCGNNNAGATCGGAAG |
| 50838_2 | TATGAGTGTGGAGTCGTTGCCGTCATAAGTATCCCAGACCTCGNNNAGATCGGAAG |
| 37084_1 | TATGAGTGTGGAGTCGTTGCAGATTTCAACATTGCACTCCAGCNNNAGATCGGAAG |
| 37084_2 | TATGAGTGTGGAGTCGTTGCAGATTTCAACATTGCACTCCAGCNNNAGATCGGAAG |
| 43404_1 | TATGAGTGTGGAGTCGTTGCCCACCACACCCAGCTAATTTTTTNNNAGATCGGAAGA |
| 43404_2 | TATGAGTGTGGAGTCGTTGCCCACCACACCCAGCTAATTTTTTNNNAGATCGGAAGA |
| 8032_1 | TATGAGTGTGGAGTCGTTGCGGAGCCTTGCTCTTATCGCTCNNNAGATCGGAAGAG |
| 8032_2 | TATGAGTGTGGAGTCGTTGCGGAGCCTTGCTCTTATCGCTCNNNAGATCGGAAGAG |
| 48301_1 | TATGAGTGTGGAGTCGTTGCGCTGTGCCAATTTCAGATCCTTTNNNAGATCGGAAG |
| 48301_2 | TATGAGTGTGGAGTCGTTGCGCTGTGCCAATTTCAGATCCTTTNNNAGATCGGAAG |
| 48830_1 | TATGAGTGTGGAGTCGTTGCCCCTCCCTCGTCACTCCAGTNNNAGATCGGAAGAGCA |
| 48830_2 | TATGAGTGTGGAGTCGTTGCCCCTCCCTCGTCACTCCAGTNNNAGATCGGAAGAGCA |
| 4144_1 | TATGAGTGTGGAGTCGTTGCCTCTGTTACCCAGATTGAAGGGCNNNAGATCGGAAG |
| 4144_2 | TATGAGTGTGGAGTCGTTGCCTCTGTTACCCAGATTGAAGGGCNNNAGATCGGAAG |
| 44828_1 | TATGAGTGTGGAGTCGTTGCTGACTACTCTGTGGACCTAGAGGNNNAGATCGGAA |

| | |
|---|---|
| 44828_2 | TATGAGTGTGGAGTCGTTGCTGACTACTCTGTGGACCTAGAGGNNNAGATCGGAA[illegible] |
| 50938_1 | TATGAGTGTGGAGTCGTTGCAACCTGGGTTTCCTACGTCAAGNNNAGATCGGAAGA |
| 50938_2 | TATGAGTGTGGAGTCGTTGCAACCTGGGTTTCCTACGTCAAGNNNAGATCGGAAGA |
| 30842_1 | TATGAGTGTGGAGTCGTTGCTGACACCATTTCAGGTTCATTGGNNNAGATCGGAAG |
| 30842_2 | TATGAGTGTGGAGTCGTTGCTGACACCATTTCAGGTTCATTGGNNNAGATCGGAAG |
| 40774_1 | TATGAGTGTGGAGTCGTTGCGACAGGGACAGAGGATCATTCAANNNAGATCGGAA |
| 40774_2 | TATGAGTGTGGAGTCGTTGCGACAGGGACAGAGGATCATTCAANNNAGATCGGAA |
| 41183_1 | TATGAGTGTGGAGTCGTTGCTTCCATTGTGTGTATATGCCACANNNAGATCGGAAG[illegible] |
| 41183_2 | TATGAGTGTGGAGTCGTTGCTTCCATTGTGTGTATATGCCACANNNAGATCGGAAG[illegible] |
| 46854_1 | TATGAGTGTGGAGTCGTTGCCAGGTTCAAGCAATTCTCCTGCNNNAGATCGGAAGA[illegible] |
| 46854_2 | TATGAGTGTGGAGTCGTTGCCAGGTTCAAGCAATTCTCCTGCNNNAGATCGGAAGA[illegible] |
| 30922_1 | TATGAGTGTGGAGTCGTTGCGTAACCAGCACCCAAATCAAGACNNNAGATCGGAAG |
| 30922_2 | TATGAGTGTGGAGTCGTTGCGTAACCAGCACCCAAATCAAGACNNNAGATCGGAAG |
| 49379_1 | TATGAGTGTGGAGTCGTTGCGAGACAAGCTGCCAGTAGTTTTGNNNAGATCGGAA[illegible] |
| 49379_2 | TATGAGTGTGGAGTCGTTGCGAGACAAGCTGCCAGTAGTTTTGNNNAGATCGGAA[illegible] |
| 41523_1 | TATGAGTGTGGAGTCGTTGCAGGGAAAAGGGAAAGGAGAAAGGNNNAGATCGG[illegible] |
| 41523_2 | TATGAGTGTGGAGTCGTTGCAGGGAAAAGGGAAAGGAGAAAGGNNNAGATCGG[illegible] |
| 47627_1 | TATGAGTGTGGAGTCGTTGCCCAAGTAGCTGGGATTACAGGTANNNAGATCGGAA[illegible] |
| 47627_2 | TATGAGTGTGGAGTCGTTGCCCAAGTAGCTGGGATTACAGGTANNNAGATCGGAA[illegible] |
| 2257_1 | TATGAGTGTGGAGTCGTTGCACCAAAACATTGGAGCCACAGTANNNAGATCGGAAG |
| 2257_2 | TATGAGTGTGGAGTCGTTGCACCAAAACATTGGAGCCACAGTANNNAGATCGGAAG |
| 40849_1 | TATGAGTGTGGAGTCGTTGCTCTTGTGGACCAGAAGAAGGAAGNNNAGATCGGAA |
| 40849_2 | TATGAGTGTGGAGTCGTTGCTCTTGTGGACCAGAAGAAGGAAGNNNAGATCGGAA |
| 41951_1 | TATGAGTGTGGAGTCGTTGCGATCTTGGCTTACTGCACCCTNNNAGATCGGAAGAG[illegible] |
| 41951_2 | TATGAGTGTGGAGTCGTTGCGATCTTGGCTTACTGCACCCTNNNAGATCGGAAGAG[illegible] |
| 36493_1 | TATGAGTGTGGAGTCGTTGCATAGCAAGAAGGCATTCACCAGANNNAGATCGGAA[illegible] |
| 36493_2 | TATGAGTGTGGAGTCGTTGCATAGCAAGAAGGCATTCACCAGANNNAGATCGGAA[illegible] |
| 52659_1 | TATGAGTGTGGAGTCGTTGCGGCTCGCTGTGAAGAAAGAAATTNNNAGATCGGAA[illegible] |
| 52659_2 | TATGAGTGTGGAGTCGTTGCGGCTCGCTGTGAAGAAAGAAATTNNNAGATCGGAA[illegible] |
| 46511_1 | TATGAGTGTGGAGTCGTTGCCTCCTGAGTAGCTGGGATTACAGNNNAGATCGGAA[illegible] |
| 46511_2 | TATGAGTGTGGAGTCGTTGCCTCCTGAGTAGCTGGGATTACAGNNNAGATCGGAA[illegible] |
| 5048_1 | TATGAGTGTGGAGTCGTTGCCCCCAGCACACATTTTGTAACACNNNAGATCGGAAG[illegible] |
| 5048_2 | TATGAGTGTGGAGTCGTTGCCCCCAGCACACATTTTGTAACACNNNAGATCGGAAG[illegible] |
| 41943_1 | TATGAGTGTGGAGTCGTTGCAGTAGCTGAGATTACAGGTGCACNNNAGATCGGAA[illegible] |
| 41943_2 | TATGAGTGTGGAGTCGTTGCAGTAGCTGAGATTACAGGTGCACNNNAGATCGGAA[illegible] |
| 32331_1 | TATGAGTGTGGAGTCGTTGCCATCTCCAAACCCCTCCCACNNNAGATCGGAAGAGCA |
| 32331_2 | TATGAGTGTGGAGTCGTTGCCATCTCCAAACCCCTCCCACNNNAGATCGGAAGAGCA |
| 12714_1 | TATGAGTGTGGAGTCGTTGCCTTTTCATCGAAGTATGTGTAAANNNAGATCGGAAG |
| 12714_2 | TATGAGTGTGGAGTCGTTGCCTTTTCATCGAAGTATGTGTAAANNNAGATCGGAAG |
| 33304_1 | TATGAGTGTGGAGTCGTTGCGATTCAAGCAATTCTCCTGCCTCNNNAGATCGGAAG[illegible] |
| 33304_2 | TATGAGTGTGGAGTCGTTGCGATTCAAGCAATTCTCCTGCCTCNNNAGATCGGAAG[illegible] |
| 44378_1 | TATGAGTGTGGAGTCGTTGCTACTCTGCACTGTTACAGGAACANNNAGATCGGAAG |
| 44378_2 | TATGAGTGTGGAGTCGTTGCTACTCTGCACTGTTACAGGAACANNNAGATCGGAAG |

| | |
|---|---|
| 29943_1 | TATGAGTGTGGAGTCGTTGCATTGTCATTCTCAAGTGCAAGGCNNNAGATCGGAAG |
| 29943_2 | TATGAGTGTGGAGTCGTTGCATTGTCATTCTCAAGTGCAAGGCNNNAGATCGGAAG |
| 36309_1 | TATGAGTGTGGAGTCGTTGCCACCATTTGTTTCCACTTCTGCANNNAGATCGGAAGA |
| 36309_2 | TATGAGTGTGGAGTCGTTGCCACCATTTGTTTCCACTTCTGCANNNAGATCGGAAGA |
| 46547_1 | TATGAGTGTGGAGTCGTTGCTGTGGTTTGTTTTGCATGTCAGANNNAGATCGGAAG |
| 46547_2 | TATGAGTGTGGAGTCGTTGCTGTGGTTTGTTTTGCATGTCAGANNNAGATCGGAAG |
| 10313_1 | TATGAGTGTGGAGTCGTTGCAGACAGTGTACCATACTGAGGGTNNNAGATCGGAA |
| 10313_2 | TATGAGTGTGGAGTCGTTGCAGACAGTGTACCATACTGAGGGTNNNAGATCGGAA |
| 9278_1 | TATGAGTGTGGAGTCGTTGCGGAGACCACAGCACAGTCTTATGNNNAGATCGGAA |
| 9278_2 | TATGAGTGTGGAGTCGTTGCGGAGACCACAGCACAGTCTTATGNNNAGATCGGAA |
| 51593_1 | TATGAGTGTGGAGTCGTTGCCTTGGCTCCCCTTCTACACTATCNNNAGATCGGAAGA |
| 51593_2 | TATGAGTGTGGAGTCGTTGCCTTGGCTCCCCTTCTACACTATCNNNAGATCGGAAGA |
| 31265_1 | TATGAGTGTGGAGTCGTTGCGAGGAATAGCATGGTAGGAAGCANNNAGATCGGA |
| 31265_2 | TATGAGTGTGGAGTCGTTGCGAGGAATAGCATGGTAGGAAGCANNNAGATCGGA |
| 13033_1 | TATGAGTGTGGAGTCGTTGCTTTTCCTGTTTAAGCATTCATAANNNAGATCGGAAGA |
| 13033_2 | TATGAGTGTGGAGTCGTTGCTTTTCCTGTTTAAGCATTCATAANNNAGATCGGAAGA |
| 37828_1 | TATGAGTGTGGAGTCGTTGCACAAATCTCAAAATGGCAGATGTNNNAGATCGGAAG |
| 37828_2 | TATGAGTGTGGAGTCGTTGCACAAATCTCAAAATGGCAGATGTNNNAGATCGGAAG |
| 38353_1 | TATGAGTGTGGAGTCGTTGCAGAAAGAATGAGACAATGGGAACTNNNAGATCGGA |
| 38353_2 | TATGAGTGTGGAGTCGTTGCAGAAAGAATGAGACAATGGGAACTNNNAGATCGGA |
| 46725_1 | TATGAGTGTGGAGTCGTTGCAATTAGCTGTGAATTCCTCGGGTNNNAGATCGGAAG |
| 46725_2 | TATGAGTGTGGAGTCGTTGCAATTAGCTGTGAATTCCTCGGGTNNNAGATCGGAAG |
| 52104_1 | TATGAGTGTGGAGTCGTTGCACTAGGACTTGCTTAGGAGTTGCNNNAGATCGGAA |
| 52104_2 | TATGAGTGTGGAGTCGTTGCACTAGGACTTGCTTAGGAGTTGCNNNAGATCGGAA |
| 8823_1 | TATGAGTGTGGAGTCGTTGCGCTTTTGTGTCTGGCTTACTCTTNNNAGATCGGAAG |
| 8823_2 | TATGAGTGTGGAGTCGTTGCGCTTTTGTGTCTGGCTTACTCTTNNNAGATCGGAAG |
| 37682_1 | TATGAGTGTGGAGTCGTTGCCCTCCCACAACAAATGAATTCACTNNNAGATCGGAAG |
| 37682_2 | TATGAGTGTGGAGTCGTTGCCCTCCCACAACAAATGAATTCACTNNNAGATCGGAAG |
| 32680_1 | TATGAGTGTGGAGTCGTTGCCTCACTGAAATCTCCACCTCACANNNAGATCGGAAGA |
| 32680_2 | TATGAGTGTGGAGTCGTTGCCTCACTGAAATCTCCACCTCACANNNAGATCGGAAGA |
| 13429_1 | TATGAGTGTGGAGTCGTTGCCTAGTTGATCCAGATCAGTGANNNAGATCGGAAGA |
| 13429_2 | TATGAGTGTGGAGTCGTTGCCTAGTTGATCCAGATCAGTGANNNAGATCGGAAGA |
| 44993_1 | TATGAGTGTGGAGTCGTTGCCGGTTCCATATTCAGACCACCTTNNNAGATCGGAAG |
| 44993_2 | TATGAGTGTGGAGTCGTTGCCGGTTCCATATTCAGACCACCTTNNNAGATCGGAAG |
| 52368_1 | TATGAGTGTGGAGTCGTTGCGGTACTCCATGCTCAAAAAAGGGNNNAGATCGGAA |
| 52368_2 | TATGAGTGTGGAGTCGTTGCGGTACTCCATGCTCAAAAAAGGGNNNAGATCGGAA |
| 45659_1 | TATGAGTGTGGAGTCGTTGCGATACTCCTACCTCAGCCTCTCANNNAGATCGGAAGA |
| 45659_2 | TATGAGTGTGGAGTCGTTGCGATACTCCTACCTCAGCCTCTCANNNAGATCGGAAGA |
| 39025_1 | TATGAGTGTGGAGTCGTTGCTCAGTCTCTACCATGTGAACAGANNNAGATCGGAAG |
| 39025_2 | TATGAGTGTGGAGTCGTTGCTCAGTCTCTACCATGTGAACAGANNNAGATCGGAAG |
| 37434_1 | TATGAGTGTGGAGTCGTTGCGAAAACTCTGAAGCACTCCACACNNNAGATCGGAAG |
| 37434_2 | TATGAGTGTGGAGTCGTTGCGAAAACTCTGAAGCACTCCACACNNNAGATCGGAAG |
| 53036_1 | TATGAGTGTGGAGTCGTTGCCATGCCTGTAATCCCAGCATTTTNNNAGATCGGAAG |

| | |
|---|---|
| 53036_2 | TATGAGTGTGGAGTCGTTGCCATGCCTGTAATCCCAGCATTTTNNNAGATCGGAAG |
| 45901_1 | TATGAGTGTGGAGTCGTTGCGGTTATAAAGATAGGGGCGGCTANNNAGATCGGA |
| 45901_2 | TATGAGTGTGGAGTCGTTGCGGTTATAAAGATAGGGGCGGCTANNNAGATCGGA |
| 48877_1 | TATGAGTGTGGAGTCGTTGCCTGCCAATTGGAAAACACTCACTNNNAGATCGGAAG |
| 48877_2 | TATGAGTGTGGAGTCGTTGCCTGCCAATTGGAAAACACTCACTNNNAGATCGGAAG |
| 13428_1 | TATGAGTGTGGAGTCGTTGCGTCTATATGCAAAATGAGTTTGANNNAGATCGGAA |
| 13428_2 | TATGAGTGTGGAGTCGTTGCGTCTATATGCAAAATGAGTTTGANNNAGATCGGAA |
| 41154_1 | TATGAGTGTGGAGTCGTTGCTGTAGTCACAGCAACTCAGGAGNNNAGATCGGAAG |
| 41154_2 | TATGAGTGTGGAGTCGTTGCTGTAGTCACAGCAACTCAGGAGNNNAGATCGGAAG |
| 30231_1 | TATGAGTGTGGAGTCGTTGCTTTCCCTTTACACTACGCCAATTNNNAGATCGGAAGA |
| 30231_2 | TATGAGTGTGGAGTCGTTGCTTTCCCTTTACACTACGCCAATTNNNAGATCGGAAGA |
| 30276_1 | TATGAGTGTGGAGTCGTTGCTACCTCTCTGAGTTCTCCCTGAANNNAGATCGGAAGA |
| 30276_2 | TATGAGTGTGGAGTCGTTGCTACCTCTCTGAGTTCTCCCTGAANNNAGATCGGAAGA |
| 51215_1 | TATGAGTGTGGAGTCGTTGCTATTAGTTCTCTCTTCCCCGGGANNNAGATCGGAAGA |
| 51215_2 | TATGAGTGTGGAGTCGTTGCTATTAGTTCTCTCTTCCCCGGGANNNAGATCGGAAGA |
| 8057_1 | TATGAGTGTGGAGTCGTTGCTCCAACTCTCATGATCCTCCCANNNAGATCGGAAGAG |
| 8057_2 | TATGAGTGTGGAGTCGTTGCTCCAACTCTCATGATCCTCCCANNNAGATCGGAAGAG |
| 10179_1 | TATGAGTGTGGAGTCGTTGCAAAACGTAAGGAGCGCACATACTNNNAGATCGGAA |
| 10179_2 | TATGAGTGTGGAGTCGTTGCAAAACGTAAGGAGCGCACATACTNNNAGATCGGAA |
| 53040_1 | TATGAGTGTGGAGTCGTTGCCTGAGGCAGGAGAATCACTTGAANNNAGATCGGAA |
| 53040_2 | TATGAGTGTGGAGTCGTTGCCTGAGGCAGGAGAATCACTTGAANNNAGATCGGAA |
| 52257_1 | TATGAGTGTGGAGTCGTTGCAACTTCGGCTTTGATGACCTAGTNNNAGATCGGAAG |
| 52257_2 | TATGAGTGTGGAGTCGTTGCAACTTCGGCTTTGATGACCTAGTNNNAGATCGGAAG |
| 41832_1 | TATGAGTGTGGAGTCGTTGCCCATATTGGTCAGGCTAGTCTCGNNNAGATCGGAAG |
| 41832_2 | TATGAGTGTGGAGTCGTTGCCCATATTGGTCAGGCTAGTCTCGNNNAGATCGGAAG |
| 41723_1 | TATGAGTGTGGAGTCGTTGCCTCCTGAGTTGCTGAGACTACAGNNNAGATCGGAAG |
| 41723_2 | TATGAGTGTGGAGTCGTTGCCTCCTGAGTTGCTGAGACTACAGNNNAGATCGGAAG |
| 43920_1 | TATGAGTGTGGAGTCGTTGCACACACACTCACACACACATATTNNNAGATCGGAAGA |
| 43920_2 | TATGAGTGTGGAGTCGTTGCACACACACTCACACACACATATTNNNAGATCGGAAGA |
| 31833_1 | TATGAGTGTGGAGTCGTTGCTATTTTCAGTGGTGCTTCCTCCANNNAGATCGGAAG |
| 31833_2 | TATGAGTGTGGAGTCGTTGCTATTTTCAGTGGTGCTTCCTCCANNNAGATCGGAAG |
| 1532_1 | TATGAGTGTGGAGTCGTTGCAGAAGGGGCAAAAGGTTTTTGTTNNNAGATCGGAA |
| 1532_2 | TATGAGTGTGGAGTCGTTGCAGAAGGGGCAAAAGGTTTTTGTTNNNAGATCGGAA |
| 49716_1 | TATGAGTGTGGAGTCGTTGCGTTCTGTGAAGAATGCCAATGGTNNNAGATCGGAA |
| 49716_2 | TATGAGTGTGGAGTCGTTGCGTTCTGTGAAGAATGCCAATGGTNNNAGATCGGAA |
| 43166_1 | TATGAGTGTGGAGTCGTTGCAACGTGCAGAGATTGGAAAAGTGNNNAGATCGGA |
| 43166_2 | TATGAGTGTGGAGTCGTTGCAACGTGCAGAGATTGGAAAAGTGNNNAGATCGGA |
| 45650_1 | TATGAGTGTGGAGTCGTTGCTTCAAGCGATTCTCCTGCCTTAGNNNAGATCGGAAG |
| 45650_2 | TATGAGTGTGGAGTCGTTGCTTCAAGCGATTCTCCTGCCTTAGNNNAGATCGGAAG |
| 34444_1 | TATGAGTGTGGAGTCGTTGCTGTGTTGGATATGGAAGTGAGGGNNNAGATCGGA |
| 34444_2 | TATGAGTGTGGAGTCGTTGCTGTGTTGGATATGGAAGTGAGGGNNNAGATCGGA |
| 38222_1 | TATGAGTGTGGAGTCGTTGCTAGACGTGAAAGCCATACACACANNNAGATCGGAA |
| 38222_2 | TATGAGTGTGGAGTCGTTGCTAGACGTGAAAGCCATACACACANNNAGATCGGAA |

| | |
|---|---|
| 2908_1 | TATGAGTGTGGAGTCGTTGCACACTGATAGACACTTGGGTTGTNNNAGATCGGAAG |
| 2908_2 | TATGAGTGTGGAGTCGTTGCACACTGATAGACACTTGGGTTGTNNNAGATCGGAAG |
| 40903_1 | TATGAGTGTGGAGTCGTTGCAGGAATGATGAGACCCATGATGGNNNAGATCGGAA |
| 40903_2 | TATGAGTGTGGAGTCGTTGCAGGAATGATGAGACCCATGATGGNNNAGATCGGAA |
| 44622_1 | TATGAGTGTGGAGTCGTTGCCACAAGAAGAAATGAAGCCCCTGNNNAGATCGGAA |
| 44622_2 | TATGAGTGTGGAGTCGTTGCCACAAGAAGAAATGAAGCCCCTGNNNAGATCGGAA |
| 28880_1 | TATGAGTGTGGAGTCGTTGCCCATTGTAGGCCAGAGTTAGTGTNNNAGATCGGAAG |
| 28880_2 | TATGAGTGTGGAGTCGTTGCCCATTGTAGGCCAGAGTTAGTGTNNNAGATCGGAAG |
| 50875_1 | TATGAGTGTGGAGTCGTTGCTCCCAAGTCAAATTTCACCTTTCANNNAGATCGGAAG |
| 50875_2 | TATGAGTGTGGAGTCGTTGCTCCCAAGTCAAATTTCACCTTTCANNNAGATCGGAAG |
| 49782_1 | TATGAGTGTGGAGTCGTTGCAGTGAGCATGCAAATCAGGAAACNNNAGATCGGAA |
| 49782_2 | TATGAGTGTGGAGTCGTTGCAGTGAGCATGCAAATCAGGAAACNNNAGATCGGAA |
| 34000_1 | TATGAGTGTGGAGTCGTTGCCTGAGTAGCTGGGATTACAGGTGNNNAGATCGGAA |
| 34000_2 | TATGAGTGTGGAGTCGTTGCCTGAGTAGCTGGGATTACAGGTGNNNAGATCGGAA |
| 38519_1 | TATGAGTGTGGAGTCGTTGCACACTCTAAGAGAATCAAGAGATCCTNNNAGATCGG |
| 38519_2 | TATGAGTGTGGAGTCGTTGCACACTCTAAGAGAATCAAGAGATCCTNNNAGATCGG |
| 53019_1 | TATGAGTGTGGAGTCGTTGCCCACTCCAGCAAATGCATATGAGNNNAGATCGGAAG |
| 53019_2 | TATGAGTGTGGAGTCGTTGCCCACTCCAGCAAATGCATATGAGNNNAGATCGGAAG |
| 52988_1 | TATGAGTGTGGAGTCGTTGCCTGTAGTCCCAGCTACTCAGGAGNNNAGATCGGAAG |
| 52988_2 | TATGAGTGTGGAGTCGTTGCCTGTAGTCCCAGCTACTCAGGAGNNNAGATCGGAAG |
| 32448_1 | TATGAGTGTGGAGTCGTTGCCATTCTCCTGCCTCAGCCTCNNNAGATCGGAAGAGCA |
| 32448_2 | TATGAGTGTGGAGTCGTTGCCATTCTCCTGCCTCAGCCTCNNNAGATCGGAAGAGCA |
| 53281_1 | TATGAGTGTGGAGTCGTTGCTGGTATATGGCAGTTTCCGTTCANNNAGATCGGAAG |
| 53281_2 | TATGAGTGTGGAGTCGTTGCTGGTATATGGCAGTTTCCGTTCANNNAGATCGGAAG |
| 39482_1 | TATGAGTGTGGAGTCGTTGCGCAAAAAGAACAAAGCTGAAGGCNNNAGATCGGAA |
| 39482_2 | TATGAGTGTGGAGTCGTTGCGCAAAAAGAACAAAGCTGAAGGCNNNAGATCGGAA |
| 44724_1 | TATGAGTGTGGAGTCGTTGCAGAAAGAGCAAGCAATAGGGAGANNNAGATCGGA |
| 44724_2 | TATGAGTGTGGAGTCGTTGCAGAAAGAGCAAGCAATAGGGAGANNNAGATCGGA |
| 35382_1 | TATGAGTGTGGAGTCGTTGCATGATCTCAGTTCACTGCAACCTNNNAGATCGGAAG |
| 35382_2 | TATGAGTGTGGAGTCGTTGCATGATCTCAGTTCACTGCAACCTNNNAGATCGGAAG |
| 43896_1 | TATGAGTGTGGAGTCGTTGCGGGGGTGTTGGTGGTATTACATANNNAGATCGGAA |
| 43896_2 | TATGAGTGTGGAGTCGTTGCGGGGGTGTTGGTGGTATTACATANNNAGATCGGAA |
| 43978_1 | TATGAGTGTGGAGTCGTTGCGAGAATTCTCCTGGATCTTGCCANNNAGATCGGAAG |
| 43978_2 | TATGAGTGTGGAGTCGTTGCGAGAATTCTCCTGGATCTTGCCANNNAGATCGGAAG |
| 46406_1 | TATGAGTGTGGAGTCGTTGCTCTTAGAGGCCACTGCATTTTCTNNNAGATCGGAAG |
| 46406_2 | TATGAGTGTGGAGTCGTTGCTCTTAGAGGCCACTGCATTTTCTNNNAGATCGGAAG |
| 48512_1 | TATGAGTGTGGAGTCGTTGCCCCACCTCTTGATATCCACATGTNNNAGATCGGAAGA |
| 48512_2 | TATGAGTGTGGAGTCGTTGCCCCACCTCTTGATATCCACATGTNNNAGATCGGAAGA |
| 36532_1 | TATGAGTGTGGAGTCGTTGCGTGTGAGCAACAAGGCTGTTTATNNNAGATCGGAA |
| 36532_2 | TATGAGTGTGGAGTCGTTGCGTGTGAGCAACAAGGCTGTTTATNNNAGATCGGAA |
| 43304_1 | TATGAGTGTGGAGTCGTTGCCACCTTAGCCTTCTGAGTAGCTGNNNAGATCGGAAG |
| 43304_2 | TATGAGTGTGGAGTCGTTGCCACCTTAGCCTTCTGAGTAGCTGNNNAGATCGGAAG |
| 47374_1 | TATGAGTGTGGAGTCGTTGCAAGCAATCCTTCCATCTCAACCTNNNAGATCGGAAGA |

| | |
|---|---|
| 47374_2 | TATGAGTGTGGAGTCGTTGCAAGCAATCCTTCCATCTCAACCTNNNAGATCGGAAGA |
| 13209_1 | TATGAGTGTGGAGTCGTTGCCCCTATCTTTTCAACCTAATTTTNNNAGATCGGAAGA |
| 13209_2 | TATGAGTGTGGAGTCGTTGCCCCTATCTTTTCAACCTAATTTTNNNAGATCGGAAGA |
| 34978_1 | TATGAGTGTGGAGTCGTTGCGGTAACACACGTGAGGTTTTTGTNNNAGATCGGAA |
| 34978_2 | TATGAGTGTGGAGTCGTTGCGGTAACACACGTGAGGTTTTTGTNNNAGATCGGAA |
| 33391_1 | TATGAGTGTGGAGTCGTTGCGAGTGTCCTTCATGCTTTCTTTTNNNAGATCGGAAGA |
| 33391_2 | TATGAGTGTGGAGTCGTTGCGAGTGTCCTTCATGCTTTCTTTTNNNAGATCGGAAGA |
| 49671_1 | TATGAGTGTGGAGTCGTTGCCACTGTGCCCAGCTAATTTTGTANNNAGATCGGAAG |
| 49671_2 | TATGAGTGTGGAGTCGTTGCCACTGTGCCCAGCTAATTTTGTANNNAGATCGGAAG |
| 46718_1 | TATGAGTGTGGAGTCGTTGCGAGTGGAGAGGCGTATGAGTATGNNNAGATCGGA |
| 46718_2 | TATGAGTGTGGAGTCGTTGCGAGTGGAGAGGCGTATGAGTATGNNNAGATCGGA |
| 52985_1 | TATGAGTGTGGAGTCGTTGCACACACACACACACACACAGTATNNNAGATCGGAAG |
| 52985_2 | TATGAGTGTGGAGTCGTTGCACACACACACACACACACAGTATNNNAGATCGGAAG |
| 50310_1 | TATGAGTGTGGAGTCGTTGCGAGACACAGACACAGAGGAGAAGNNNAGATCGGA |
| 50310_2 | TATGAGTGTGGAGTCGTTGCGAGACACAGACACAGAGGAGAAGNNNAGATCGGA |
| 37822_1 | TATGAGTGTGGAGTCGTTGCCAGGTTCAGTATCCACAGTCCAANNNAGATCGGAAG |
| 37822_2 | TATGAGTGTGGAGTCGTTGCCAGGTTCAGTATCCACAGTCCAANNNAGATCGGAAG |
| 38006_1 | TATGAGTGTGGAGTCGTTGCTGCTCTTGCCACAAAAACAGAAANNNAGATCGGAAG |
| 38006_2 | TATGAGTGTGGAGTCGTTGCTGCTCTTGCCACAAAAACAGAAANNNAGATCGGAAG |
| 10593_1 | TATGAGTGTGGAGTCGTTGCTCAAAAAGGCTCCTGTCTCTCCTNNNAGATCGGAAG |
| 10593_2 | TATGAGTGTGGAGTCGTTGCTCAAAAAGGCTCCTGTCTCTCCTNNNAGATCGGAAG |
| 38644_1 | TATGAGTGTGGAGTCGTTGCAAAATTAGCTGGGTGTGGTGATGNNNAGATCGGAA |
| 38644_2 | TATGAGTGTGGAGTCGTTGCAAAATTAGCTGGGTGTGGTGATGNNNAGATCGGAA |
| 37946_1 | TATGAGTGTGGAGTCGTTGCTAAAGCACATACAGTAGGCCCTGNNNAGATCGGAA |
| 37946_2 | TATGAGTGTGGAGTCGTTGCTAAAGCACATACAGTAGGCCCTGNNNAGATCGGAA |
| 49756_1 | TATGAGTGTGGAGTCGTTGCACTACCCATTTCTCAATGTGCCTNNNAGATCGGAAGA |
| 49756_2 | TATGAGTGTGGAGTCGTTGCACTACCCATTTCTCAATGTGCCTNNNAGATCGGAAGA |
| 35106_1 | TATGAGTGTGGAGTCGTTGCTTTGGAGAGGACCTCTTTTTTTTTNNNAGATCGGAA |
| 35106_2 | TATGAGTGTGGAGTCGTTGCTTTGGAGAGGACCTCTTTTTTTTTNNNAGATCGGAA |
| 48460_1 | TATGAGTGTGGAGTCGTTGCCTGGGTTCAAATGAGTCTCCTGANNNAGATCGGAAG |
| 48460_2 | TATGAGTGTGGAGTCGTTGCCTGGGTTCAAATGAGTCTCCTGANNNAGATCGGAAG |
| 10933_1 | TATGAGTGTGGAGTCGTTGCAGAAGGAAGTGCTCAATTTCAGTTNNNAGATCGGA |
| 10933_2 | TATGAGTGTGGAGTCGTTGCAGAAGGAAGTGCTCAATTTCAGTTNNNAGATCGGA |
| 166_1 | TATGAGTGTGGAGTCGTTGCTTAGTCCAGCCAGTCTCTTCCTCNNNAGATCGGAAGA |
| 166_2 | TATGAGTGTGGAGTCGTTGCTTAGTCCAGCCAGTCTCTTCCTCNNNAGATCGGAAGA |
| 29904_1 | TATGAGTGTGGAGTCGTTGCCCCAAGTCATATGCAAACAAGCANNNAGATCGGAAG |
| 29904_2 | TATGAGTGTGGAGTCGTTGCCCCAAGTCATATGCAAACAAGCANNNAGATCGGAAG |
| 50094_1 | TATGAGTGTGGAGTCGTTGCAAGTACAGGACAAGGACCCATTCNNNAGATCGGAA |
| 50094_2 | TATGAGTGTGGAGTCGTTGCAAGTACAGGACAAGGACCCATTCNNNAGATCGGAA |
| 38107_1 | TATGAGTGTGGAGTCGTTGCGAGGTTGTAGTGAGCCGAGATTGNNNAGATCGGAA |
| 38107_2 | TATGAGTGTGGAGTCGTTGCGAGGTTGTAGTGAGCCGAGATTGNNNAGATCGGAA |
| 29661_1 | TATGAGTGTGGAGTCGTTGCCTCAAGATCTGTGGTTTACCCCANNNAGATCGGAAG |
| 29661_2 | TATGAGTGTGGAGTCGTTGCCTCAAGATCTGTGGTTTACCCCANNNAGATCGGAAG |

| | |
|---|---|
| 50312_1 | TATGAGTGTGGAGTCGTTGCCTCTCATCTAATCTAGCCCTCGCNNNAGATCGGAAGA |
| 50312_2 | TATGAGTGTGGAGTCGTTGCCTCTCATCTAATCTAGCCCTCGCNNNAGATCGGAAGA |
| 31064_1 | TATGAGTGTGGAGTCGTTGCAAGTGCATATGACACATCCATGANNNAGATCGGAAG |
| 31064_2 | TATGAGTGTGGAGTCGTTGCAAGTGCATATGACACATCCATGANNNAGATCGGAAG |
| 53067_1 | TATGAGTGTGGAGTCGTTGCGAAAGCAGAATTTGGAGGCTGAGNNNAGATCGGA |
| 53067_2 | TATGAGTGTGGAGTCGTTGCGAAAGCAGAATTTGGAGGCTGAGNNNAGATCGGA |
| 30429_1 | TATGAGTGTGGAGTCGTTGCAATACTCACAGACCCCCATGTGANNNAGATCGGAAG |
| 30429_2 | TATGAGTGTGGAGTCGTTGCAATACTCACAGACCCCCATGTGANNNAGATCGGAAG |
| 30161_1 | TATGAGTGTGGAGTCGTTGCTGTAAGTTAATCAGGAATTTCTTGGTNNNAGATCGG |
| 30161_2 | TATGAGTGTGGAGTCGTTGCTGTAAGTTAATCAGGAATTTCTTGGTNNNAGATCGG |
| 47841_1 | TATGAGTGTGGAGTCGTTGCCCTCCCAAGTTCAAGCGATTTTCNNNAGATCGGAAG |
| 47841_2 | TATGAGTGTGGAGTCGTTGCCCTCCCAAGTTCAAGCGATTTTCNNNAGATCGGAAG |
| 43626_1 | TATGAGTGTGGAGTCGTTGCCAGGATTTTGAGGCTGTTTTCCANNNAGATCGGAAG |
| 43626_2 | TATGAGTGTGGAGTCGTTGCCAGGATTTTGAGGCTGTTTTCCANNNAGATCGGAAG |
| 53176_1 | TATGAGTGTGGAGTCGTTGCTCCATCAGTTCACCTTGTGTCATNNNAGATCGGAAGA |
| 53176_2 | TATGAGTGTGGAGTCGTTGCTCCATCAGTTCACCTTGTGTCATNNNAGATCGGAAGA |
| 43393_1 | TATGAGTGTGGAGTCGTTGCGAGACAAGTCTCACTCTGTCACCNNNAGATCGGAAG |
| 43393_2 | TATGAGTGTGGAGTCGTTGCGAGACAAGTCTCACTCTGTCACCNNNAGATCGGAAG |
| 47233_1 | TATGAGTGTGGAGTCGTTGCACCATGCCCGGCTAATTTTTGTANNNAGATCGGAAG |
| 47233_2 | TATGAGTGTGGAGTCGTTGCACCATGCCCGGCTAATTTTTGTANNNAGATCGGAAG |
| 33977_1 | TATGAGTGTGGAGTCGTTGCCCATCGCGTCCAGCTAATTTTTNNNAGATCGGAAGA |
| 33977_2 | TATGAGTGTGGAGTCGTTGCCCATCGCGTCCAGCTAATTTTTNNNAGATCGGAAGA |
| 53393_1 | TATGAGTGTGGAGTCGTTGCTGTAAGATGTGACTTGCTCCTCCNNNAGATCGGAAG |
| 53393_2 | TATGAGTGTGGAGTCGTTGCTGTAAGATGTGACTTGCTCCTCCNNNAGATCGGAAG |
| 29937_1 | TATGAGTGTGGAGTCGTTGCTCCGCTTGACTTATTTTTAACACCCNNNAGATCGGAA |
| 29937_2 | TATGAGTGTGGAGTCGTTGCTCCGCTTGACTTATTTTTAACACCCNNNAGATCGGAA |
| 37575_1 | TATGAGTGTGGAGTCGTTGCAAGAGAAAGAAAAAGACAGGGGTNNNAGATCGGA |
| 37575_2 | TATGAGTGTGGAGTCGTTGCAAGAGAAAGAAAAAGACAGGGGTNNNAGATCGGA |
| 32562_1 | TATGAGTGTGGAGTCGTTGCCTCAAGTGATTCTCCTGCCTCAGNNNAGATCGGAAG |
| 32562_2 | TATGAGTGTGGAGTCGTTGCCTCAAGTGATTCTCCTGCCTCAGNNNAGATCGGAAG |
| 39590_1 | TATGAGTGTGGAGTCGTTGCAGCCAGAAAAGTGACTCAGACTTNNNAGATCGGAA |
| 39590_2 | TATGAGTGTGGAGTCGTTGCAGCCAGAAAAGTGACTCAGACTTNNNAGATCGGAA |
| 52699_1 | TATGAGTGTGGAGTCGTTGCGGCATTTGTCGTGACTATGAGTGNNNAGATCGGAA |
| 52699_2 | TATGAGTGTGGAGTCGTTGCGGCATTTGTCGTGACTATGAGTGNNNAGATCGGAA |
| 42567_1 | TATGAGTGTGGAGTCGTTGCAACAGTGCTAATGGATTAGGGCTNNNAGATCGGAA |
| 42567_2 | TATGAGTGTGGAGTCGTTGCAACAGTGCTAATGGATTAGGGCTNNNAGATCGGAA |
| 46024_1 | TATGAGTGTGGAGTCGTTGCGTCTCCTGGGTTCCAATGATTCTNNNAGATCGGAAG |
| 46024_2 | TATGAGTGTGGAGTCGTTGCGTCTCCTGGGTTCCAATGATTCTNNNAGATCGGAAG |
| 9523_1 | TATGAGTGTGGAGTCGTTGCGCAGTAGGTGCTCCACAAAAATGNNNAGATCGGAA |
| 9523_2 | TATGAGTGTGGAGTCGTTGCGCAGTAGGTGCTCCACAAAAATGNNNAGATCGGAA |
| 45494_1 | TATGAGTGTGGAGTCGTTGCTCCTGGGGATGGACAGTCAGNNNAGATCGGAAGAG |
| 45494_2 | TATGAGTGTGGAGTCGTTGCTCCTGGGGATGGACAGTCAGNNNAGATCGGAAGAG |
| 38112_1 | TATGAGTGTGGAGTCGTTGCTGCTGAATGAAAAAAGCCAGACANNNAGATCGGAA |

| | |
|---|---|
| 38112_2 | TATGAGTGTGGAGTCGTTGCTGCTGAATGAAAAAAGCCAGACANNNAGATCGGAA |
| 43108_1 | TATGAGTGTGGAGTCGTTGCGGTGGATCATGAGGTCAAGAGATNNNAGATCGGA |
| 43108_2 | TATGAGTGTGGAGTCGTTGCGGTGGATCATGAGGTCAAGAGATNNNAGATCGGA |
| 52262_1 | TATGAGTGTGGAGTCGTTGCTTGTGGGATGGAATGAACATTGCNNNAGATCGGAA |
| 52262_2 | TATGAGTGTGGAGTCGTTGCTTGTGGGATGGAATGAACATTGCNNNAGATCGGAA |
| 2481_1 | TATGAGTGTGGAGTCGTTGCTTATTCAAACGATCCCTGCCCAGNNNAGATCGGAAG |
| 2481_2 | TATGAGTGTGGAGTCGTTGCTTATTCAAACGATCCCTGCCCAGNNNAGATCGGAAG |
| 37044_1 | TATGAGTGTGGAGTCGTTGCTTGCATAAGTAACCAGGAGCTGANNNAGATCGGAA |
| 37044_2 | TATGAGTGTGGAGTCGTTGCTTGCATAAGTAACCAGGAGCTGANNNAGATCGGAA |
| 32352_1 | TATGAGTGTGGAGTCGTTGCCCTCACACACACATACTCAGTCANNNAGATCGGAAG |
| 32352_2 | TATGAGTGTGGAGTCGTTGCCCTCACACACACATACTCAGTCANNNAGATCGGAAG |
| 8212_1 | TATGAGTGTGGAGTCGTTGCTAGGGTCTACTCTGTCACCCAGGTNNNAGATCGGAA |
| 8212_2 | TATGAGTGTGGAGTCGTTGCTAGGGTCTACTCTGTCACCCAGGTNNNAGATCGGAA |
| 29850_1 | TATGAGTGTGGAGTCGTTGCCTGTGGTAAAAAGATAACCTCAGAANNNAGATCGG |
| 29850_2 | TATGAGTGTGGAGTCGTTGCCTGTGGTAAAAAGATAACCTCAGAANNNAGATCGG |
| 29830_1 | TATGAGTGTGGAGTCGTTGCTCAGAGAATGTTCCTTCAGCTCCNNNAGATCGGAAG |
| 29830_2 | TATGAGTGTGGAGTCGTTGCTCAGAGAATGTTCCTTCAGCTCCNNNAGATCGGAAG |
| 13017_1 | TATGAGTGTGGAGTCGTTGCAGCATCAAAGTGGTAAATAAAATNNNAGATCGGAA |
| 13017_2 | TATGAGTGTGGAGTCGTTGCAGCATCAAAGTGGTAAATAAAATNNNAGATCGGAA |
| 50839_1 | TATGAGTGTGGAGTCGTTGCGTCTCATCTCTGTTCCCCACATTNNNAGATCGGAAGA |
| 50839_2 | TATGAGTGTGGAGTCGTTGCGTCTCATCTCTGTTCCCCACATTNNNAGATCGGAAGA |
| 32696_1 | TATGAGTGTGGAGTCGTTGCTTCACACCATTCTCCTGCCTCNNNAGATCGGAAGAGC |
| 32696_2 | TATGAGTGTGGAGTCGTTGCTTCACACCATTCTCCTGCCTCNNNAGATCGGAAGAGC |
| 50475_1 | TATGAGTGTGGAGTCGTTGCTTTGCTGAGGAGTTGAGTTCAGTNNNAGATCGGAA |
| 50475_2 | TATGAGTGTGGAGTCGTTGCTTTGCTGAGGAGTTGAGTTCAGTNNNAGATCGGAA |
| 52539_1 | TATGAGTGTGGAGTCGTTGCCTACTTAGTCAGGAAACCCAGGCNNNAGATCGGAAG |
| 52539_2 | TATGAGTGTGGAGTCGTTGCCTACTTAGTCAGGAAACCCAGGCNNNAGATCGGAAG |
| 44448_1 | TATGAGTGTGGAGTCGTTGCCTCCTGAGTAGCTGGGACTACAGNNNAGATCGGAA |
| 44448_2 | TATGAGTGTGGAGTCGTTGCCTCCTGAGTAGCTGGGACTACAGNNNAGATCGGAA |
| 46314_1 | TATGAGTGTGGAGTCGTTGCAGTGGCACAATTACAGCTCACTANNNAGATCGGAAG |
| 46314_2 | TATGAGTGTGGAGTCGTTGCAGTGGCACAATTACAGCTCACTANNNAGATCGGAAG |
| 47659_1 | TATGAGTGTGGAGTCGTTGCAAGACAGTCTATGAAGCTGGAGCNNNAGATCGGAA |
| 47659_2 | TATGAGTGTGGAGTCGTTGCAAGACAGTCTATGAAGCTGGAGCNNNAGATCGGAA |
| 31793_1 | TATGAGTGTGGAGTCGTTGCTGCTCCAAATTTTCATCACAGGANNNAGATCGGAAG |
| 31793_2 | TATGAGTGTGGAGTCGTTGCTGCTCCAAATTTTCATCACAGGANNNAGATCGGAAG |
| 53411_1 | TATGAGTGTGGAGTCGTTGCGGAGGAAGAGAGAGAGAAGGAGANNNAGATCGG |
| 53411_2 | TATGAGTGTGGAGTCGTTGCGGAGGAAGAGAGAGAGAAGGAGANNNAGATCGG |
| 42823_1 | TATGAGTGTGGAGTCGTTGCGCTCTTCCCTTTAGCTCCATGAANNNAGATCGGAAG |
| 42823_2 | TATGAGTGTGGAGTCGTTGCGCTCTTCCCTTTAGCTCCATGAANNNAGATCGGAAG |
| 40548_1 | TATGAGTGTGGAGTCGTTGCCAATGGAACAGAATAGAGCGCACNNNAGATCGGAA |
| 40548_2 | TATGAGTGTGGAGTCGTTGCCAATGGAACAGAATAGAGCGCACNNNAGATCGGAA |
| 30493_1 | TATGAGTGTGGAGTCGTTGCCTCTGGTGAGGATGGTTTCAGGTNNNAGATCGGAA |
| 30493_2 | TATGAGTGTGGAGTCGTTGCCTCTGGTGAGGATGGTTTCAGGTNNNAGATCGGAA |

| | |
|---|---|
| 43264_1 | TATGAGTGTGGAGTCGTTGCAAACAGTCTTCTCACGTTAGCCTNNNAGATCGGAAG |
| 43264_2 | TATGAGTGTGGAGTCGTTGCAAACAGTCTTCTCACGTTAGCCTNNNAGATCGGAAG |
| 47105_1 | TATGAGTGTGGAGTCGTTGCCGCCTGGCTGGTTTTTGTATTTTNNNAGATCGGAAG |
| 47105_2 | TATGAGTGTGGAGTCGTTGCCGCCTGGCTGGTTTTTGTATTTTNNNAGATCGGAAG |
| 39835_1 | TATGAGTGTGGAGTCGTTGCGAATGGAGACACCTGTAGCGTATNNNAGATCGGAA |
| 39835_2 | TATGAGTGTGGAGTCGTTGCGAATGGAGACACCTGTAGCGTATNNNAGATCGGAA |
| 38196_1 | TATGAGTGTGGAGTCGTTGCCTTACACCTGTAATCCCAGCACTNNNAGATCGGAAGA |
| 38196_2 | TATGAGTGTGGAGTCGTTGCCTTACACCTGTAATCCCAGCACTNNNAGATCGGAAGA |
| 18202_1 | TATGAGTGTGGAGTCGTTGCTTCTCTCTCACATCTCACACNNNAGATCGGAAGAGCA |
| 18202_2 | TATGAGTGTGGAGTCGTTGCTTCTCTCTCACATCTCACACNNNAGATCGGAAGAGCA |
| 39023_1 | TATGAGTGTGGAGTCGTTGCCCCAGGGGTCCAAACATACATATNNNAGATCGGAAG |
| 39023_2 | TATGAGTGTGGAGTCGTTGCCCCAGGGGTCCAAACATACATATNNNAGATCGGAAG |
| 30203_1 | TATGAGTGTGGAGTCGTTGCCCCACAGGCTTCTTTTTTTCACANNNAGATCGGAAGA |
| 30203_2 | TATGAGTGTGGAGTCGTTGCCCCACAGGCTTCTTTTTTTCACANNNAGATCGGAAGA |
| 46440_1 | TATGAGTGTGGAGTCGTTGCGTGACACGATTGCAGTTCACTACNNNAGATCGGAAG |
| 46440_2 | TATGAGTGTGGAGTCGTTGCGTGACACGATTGCAGTTCACTACNNNAGATCGGAAG |
| 47439_1 | TATGAGTGTGGAGTCGTTGCCTCCCGGATTCGAGCAATTTTCNNNAGATCGGAAGA |
| 47439_2 | TATGAGTGTGGAGTCGTTGCCTCCCGGATTCGAGCAATTTTCNNNAGATCGGAAGA |
| 32858_1 | TATGAGTGTGGAGTCGTTGCCTCCACCACGCCAAGCTAATTTTNNNAGATCGGAAGA |
| 32858_2 | TATGAGTGTGGAGTCGTTGCCTCCACCACGCCAAGCTAATTTTNNNAGATCGGAAGA |
| 50854_1 | TATGAGTGTGGAGTCGTTGCTATTGTTGGTGACTGACTCAGGGNNNAGATCGGAA |
| 50854_2 | TATGAGTGTGGAGTCGTTGCTATTGTTGGTGACTGACTCAGGGNNNAGATCGGAA |
| 49952_1 | TATGAGTGTGGAGTCGTTGCATCACTAAGTGACAGGCTAAGCCNNNAGATCGGAA |
| 49952_2 | TATGAGTGTGGAGTCGTTGCATCACTAAGTGACAGGCTAAGCCNNNAGATCGGAA |
| 31653_1 | TATGAGTGTGGAGTCGTTGCGGCTTTGTTTCCTATCACGAAGGNNNAGATCGGAAG |
| 31653_2 | TATGAGTGTGGAGTCGTTGCGGCTTTGTTTCCTATCACGAAGGNNNAGATCGGAAG |
| 52144_1 | TATGAGTGTGGAGTCGTTGCCATAAGGGCCTAACCAAGACAGTNNNAGATCGGAA |
| 52144_2 | TATGAGTGTGGAGTCGTTGCCATAAGGGCCTAACCAAGACAGTNNNAGATCGGAA |
| 50015_1 | TATGAGTGTGGAGTCGTTGCCTTGGTGAAAACATGAAGGGGACNNNAGATCGGAA |
| 50015_2 | TATGAGTGTGGAGTCGTTGCCTTGGTGAAAACATGAAGGGGACNNNAGATCGGAA |
| 37671_1 | TATGAGTGTGGAGTCGTTGCGATTCTCCTCTGCAGGTTTCAGANNNAGATCGGAAG |
| 37671_2 | TATGAGTGTGGAGTCGTTGCGATTCTCCTCTGCAGGTTTCAGANNNAGATCGGAAG |
| 12831_1 | TATGAGTGTGGAGTCGTTGCATGTTTGGAATGTTTTTAAAACCNNNAGATCGGAAG |
| 12831_2 | TATGAGTGTGGAGTCGTTGCATGTTTGGAATGTTTTTAAAACCNNNAGATCGGAAG |
| 43237_1 | TATGAGTGTGGAGTCGTTGCTGAGTTCAGAAGTATGAGCCCACNNNAGATCGGAA |
| 43237_2 | TATGAGTGTGGAGTCGTTGCTGAGTTCAGAAGTATGAGCCCACNNNAGATCGGAA |
| 36636_1 | TATGAGTGTGGAGTCGTTGCAAAGAAATTAGCCAGGTGCAGTGNNNAGATCGGAA |
| 36636_2 | TATGAGTGTGGAGTCGTTGCAAAGAAATTAGCCAGGTGCAGTGNNNAGATCGGAA |
| 53421_1 | TATGAGTGTGGAGTCGTTGCGGGAGAGAGAGAGAGAGAGGTTTNNNAGATCGGA |
| 53421_2 | TATGAGTGTGGAGTCGTTGCGGGAGAGAGAGAGAGAGAGGTTTNNNAGATCGGA |
| 48390_1 | TATGAGTGTGGAGTCGTTGCACTGTTGTACATATGCCCTGACANNNAGATCGGAAG |
| 48390_2 | TATGAGTGTGGAGTCGTTGCACTGTTGTACATATGCCCTGACANNNAGATCGGAAG |
| 40866_1 | TATGAGTGTGGAGTCGTTGCGGATTCCCTTGGGTCTTTCACTANNNAGATCGGAAG |

| | |
|---|---|
| 40866_2 | TATGAGTGTGGAGTCGTTGCGGATTCCCTTGGGTCTTTCACTANNNAGATCGGAAG |
| 31744_1 | TATGAGTGTGGAGTCGTTGCCCACAGCCTAAATTACTAGCCCANNNAGATCGGAAG |
| 31744_2 | TATGAGTGTGGAGTCGTTGCCCACAGCCTAAATTACTAGCCCANNNAGATCGGAAG |
| 53326_1 | TATGAGTGTGGAGTCGTTGCGTTTGTCTGGATCCACAGAAAGCNNNAGATCGGAA |
| 53326_2 | TATGAGTGTGGAGTCGTTGCGTTTGTCTGGATCCACAGAAAGCNNNAGATCGGAA |
| 48517_1 | TATGAGTGTGGAGTCGTTGCCCTAATGAACGCCTACCTGACATNNNAGATCGGAAG |
| 48517_2 | TATGAGTGTGGAGTCGTTGCCCTAATGAACGCCTACCTGACATNNNAGATCGGAAG |
| 34635_1 | TATGAGTGTGGAGTCGTTGCGTTTCACACGTGTAACCCAGAACNNNAGATCGGAAG |
| 34635_2 | TATGAGTGTGGAGTCGTTGCGTTTCACACGTGTAACCCAGAACNNNAGATCGGAAG |
| 37347_1 | TATGAGTGTGGAGTCGTTGCTCCACCTTCTGTTTCCCTTCATCNNNAGATCGGAAGA |
| 37347_2 | TATGAGTGTGGAGTCGTTGCTCCACCTTCTGTTTCCCTTCATCNNNAGATCGGAAGA |
| 52863_1 | TATGAGTGTGGAGTCGTTGCATGAATCTCCCGGAAGATGAGCNNNAGATCGGAAG |
| 52863_2 | TATGAGTGTGGAGTCGTTGCATGAATCTCCCGGAAGATGAGCNNNAGATCGGAAG |
| 32716_1 | TATGAGTGTGGAGTCGTTGCCCCAGTTTTGCACTTGAAACTCANNNAGATCGGAAG |
| 32716_2 | TATGAGTGTGGAGTCGTTGCCCCAGTTTTGCACTTGAAACTCANNNAGATCGGAAG |
| 38332_1 | TATGAGTGTGGAGTCGTTGCTGCCTAGCTTCATCACTGCTATTNNNAGATCGGAAGA |
| 38332_2 | TATGAGTGTGGAGTCGTTGCTGCCTAGCTTCATCACTGCTATTNNNAGATCGGAAGA |
| 33271_1 | TATGAGTGTGGAGTCGTTGCGCATAATCATAGCTCACTGCAGCNNNAGATCGGAAG |
| 33271_2 | TATGAGTGTGGAGTCGTTGCGCATAATCATAGCTCACTGCAGCNNNAGATCGGAAG |
| 43187_1 | TATGAGTGTGGAGTCGTTGCGGTTCAAGTGATTTTCCTGCCTCNNNAGATCGGAAG |
| 43187_2 | TATGAGTGTGGAGTCGTTGCGGTTCAAGTGATTTTCCTGCCTCNNNAGATCGGAAG |
| 37450_1 | TATGAGTGTGGAGTCGTTGCTCGCCAAGAAAAGTCTCAACTCTNNNAGATCGGAAG |
| 37450_2 | TATGAGTGTGGAGTCGTTGCTCGCCAAGAAAAGTCTCAACTCTNNNAGATCGGAAG |
| 40127_1 | TATGAGTGTGGAGTCGTTGCATGGAACAGATCAGAGAACCTGGNNNAGATCGGAA |
| 40127_2 | TATGAGTGTGGAGTCGTTGCATGGAACAGATCAGAGAACCTGGNNNAGATCGGAA |
| 48915_1 | TATGAGTGTGGAGTCGTTGCCCACCACACCCAGCTAATTTTTTNNNAGATCGGAAGA |
| 48915_2 | TATGAGTGTGGAGTCGTTGCCCACCACACCCAGCTAATTTTTTNNNAGATCGGAAGA |
| 49201_1 | TATGAGTGTGGAGTCGTTGCCCTGTCTCCAAGTTCAGAGAGTGNNNAGATCGGAAG |
| 49201_2 | TATGAGTGTGGAGTCGTTGCCCTGTCTCCAAGTTCAGAGAGTGNNNAGATCGGAAG |
| 32214_1 | TATGAGTGTGGAGTCGTTGCGAGCTTCTGACTGACAGCTCACNNNAGATCGGAAGA |
| 32214_2 | TATGAGTGTGGAGTCGTTGCGAGCTTCTGACTGACAGCTCACNNNAGATCGGAAGA |
| 50958_1 | TATGAGTGTGGAGTCGTTGCATACATTTGGCCCTGTCAGTCTTNNNAGATCGGAAG |
| 50958_2 | TATGAGTGTGGAGTCGTTGCATACATTTGGCCCTGTCAGTCTTNNNAGATCGGAAG |
| 32462_1 | TATGAGTGTGGAGTCGTTGCGCAGGCATCTGTAATCCCATCTANNNAGATCGGAAG |
| 32462_2 | TATGAGTGTGGAGTCGTTGCGCAGGCATCTGTAATCCCATCTANNNAGATCGGAAG |
| 12658_1 | TATGAGTGTGGAGTCGTTGCCCAGTGTATATGCTAAGCAGNNNAGATCGGAAGAG |
| 12658_2 | TATGAGTGTGGAGTCGTTGCCCAGTGTATATGCTAAGCAGNNNAGATCGGAAGAG |
| 30233_1 | TATGAGTGTGGAGTCGTTGCGTGAGGGTGAGTCATTTAGAGCANNNAGATCGGAA |
| 30233_2 | TATGAGTGTGGAGTCGTTGCGTGAGGGTGAGTCATTTAGAGCANNNAGATCGGAA |
| 51963_1 | TATGAGTGTGGAGTCGTTGCACATTAGTCCAATGCCATCACCTNNNAGATCGGAAG |
| 51963_2 | TATGAGTGTGGAGTCGTTGCACATTAGTCCAATGCCATCACCTNNNAGATCGGAAG |
| 39593_1 | TATGAGTGTGGAGTCGTTGCGTGAATCACTTGAGGTCAGGAGTNNNAGATCGGAA |
| 39593_2 | TATGAGTGTGGAGTCGTTGCGTGAATCACTTGAGGTCAGGAGTNNNAGATCGGAA |

| | |
|---|---|
| 46518_1 | TATGAGTGTGGAGTCGTTGCGAGGCGGAGTCTTACTCTGTCNNNAGATCGGAAGA |
| 46518_2 | TATGAGTGTGGAGTCGTTGCGAGGCGGAGTCTTACTCTGTCNNNAGATCGGAAGA |
| 46810_1 | TATGAGTGTGGAGTCGTTGCGTAGGACCTGTGAGAGAAGATGCNNNAGATCGGA |
| 46810_2 | TATGAGTGTGGAGTCGTTGCGTAGGACCTGTGAGAGAAGATGCNNNAGATCGGA |
| 34469_1 | TATGAGTGTGGAGTCGTTGCGTTCCTGTGTTAGAGTGCTGAGANNNAGATCGGAA |
| 34469_2 | TATGAGTGTGGAGTCGTTGCGTTCCTGTGTTAGAGTGCTGAGANNNAGATCGGAA |
| 13559_1 | TATGAGTGTGGAGTCGTTGCCATACTGTAGGGTAATAAATTTGTNNNAGATCGGAA |
| 13559_2 | TATGAGTGTGGAGTCGTTGCCATACTGTAGGGTAATAAATTTGTNNNAGATCGGAA |
| 10624_1 | TATGAGTGTGGAGTCGTTGCTGTCCTTCAAAGTCTAGTCTCCCTNNNAGATCGGAAG |
| 10624_2 | TATGAGTGTGGAGTCGTTGCTGTCCTTCAAAGTCTAGTCTCCCTNNNAGATCGGAAG |
| 36539_1 | TATGAGTGTGGAGTCGTTGCCTTTGTAGCCTGAAGGGGAGAATNNNAGATCGGAA |
| 36539_2 | TATGAGTGTGGAGTCGTTGCCTTTGTAGCCTGAAGGGGAGAATNNNAGATCGGAA |
| 30259_1 | TATGAGTGTGGAGTCGTTGCCCAAAGGGTAAATCACACTGACTNNNAGATCGGAAG |
| 30259_2 | TATGAGTGTGGAGTCGTTGCCCAAAGGGTAAATCACACTGACTNNNAGATCGGAAG |
| 41925_1 | TATGAGTGTGGAGTCGTTGCCCTCCTGGGTTCAAGCTATTCTCNNNAGATCGGAAG |
| 41925_2 | TATGAGTGTGGAGTCGTTGCCCTCCTGGGTTCAAGCTATTCTCNNNAGATCGGAAG |
| 30470_1 | TATGAGTGTGGAGTCGTTGCGTTCATGCCATTCTCCTACCTCANNNAGATCGGAAGA |
| 30470_2 | TATGAGTGTGGAGTCGTTGCGTTCATGCCATTCTCCTACCTCANNNAGATCGGAAGA |
| 13758_1 | TATGAGTGTGGAGTCGTTGCTTTGAGAGATTTCCTTTTTCAAANNNAGATCGGAAG |
| 13758_2 | TATGAGTGTGGAGTCGTTGCTTTGAGAGATTTCCTTTTTCAAANNNAGATCGGAAG |
| 33151_1 | TATGAGTGTGGAGTCGTTGCTGCCTACATCAACAAAAGAGGAANNNAGATCGGAA |
| 33151_2 | TATGAGTGTGGAGTCGTTGCTGCCTACATCAACAAAAGAGGAANNNAGATCGGAA |
| 10069_1 | TATGAGTGTGGAGTCGTTGCCTGCGGTGAGCACTTACAGTATTNNNAGATCGGAAG |
| 10069_2 | TATGAGTGTGGAGTCGTTGCCTGCGGTGAGCACTTACAGTATTNNNAGATCGGAAG |
| 41643_1 | TATGAGTGTGGAGTCGTTGCCTCCTGAGTAGCTGGGACTACAGNNNAGATCGGAA |
| 41643_2 | TATGAGTGTGGAGTCGTTGCCTCCTGAGTAGCTGGGACTACAGNNNAGATCGGAA |
| 42079_1 | TATGAGTGTGGAGTCGTTGCGAGGGGGTAACCTTGGATGAATANNNAGATCGGA |
| 42079_2 | TATGAGTGTGGAGTCGTTGCGAGGGGGTAACCTTGGATGAATANNNAGATCGGA |
| 5564_1 | TATGAGTGTGGAGTCGTTGCTGATATTGGTTTCTGGGCACTGGNNNAGATCGGAA |
| 5564_2 | TATGAGTGTGGAGTCGTTGCTGATATTGGTTTCTGGGCACTGGNNNAGATCGGAA |
| 34074_1 | TATGAGTGTGGAGTCGTTGCTCTTCTCGAACTGACAGTGGATGNNNAGATCGGAAG |
| 34074_2 | TATGAGTGTGGAGTCGTTGCTCTTCTCGAACTGACAGTGGATGNNNAGATCGGAAG |
| 52745_1 | TATGAGTGTGGAGTCGTTGCGCATGTGACAGAGACAGACAAACNNNAGATCGGAA |
| 52745_2 | TATGAGTGTGGAGTCGTTGCGCATGTGACAGAGACAGACAAACNNNAGATCGGAA |
| 12779_1 | TATGAGTGTGGAGTCGTTGCGGAGGTAAGAATAACAAATCATCNNNAGATCGGAA |
| 12779_2 | TATGAGTGTGGAGTCGTTGCGGAGGTAAGAATAACAAATCATCNNNAGATCGGAA |
| 47441_1 | TATGAGTGTGGAGTCGTTGCTGTCTTTTCCACTTGATTCCATGCNNNAGATCGGAAG |
| 47441_2 | TATGAGTGTGGAGTCGTTGCTGTCTTTTCCACTTGATTCCATGCNNNAGATCGGAAG |
| 30114_1 | TATGAGTGTGGAGTCGTTGCTGAATCGAAAATGCTTAATGTTACAAANNNAGATCG |
| 30114_2 | TATGAGTGTGGAGTCGTTGCTGAATCGAAAATGCTTAATGTTACAAANNNAGATCG |
| 41379_1 | TATGAGTGTGGAGTCGTTGCTGAGAGGCCTGAGAGAAAGAGAANNNAGATCGGA |
| 41379_2 | TATGAGTGTGGAGTCGTTGCTGAGAGGCCTGAGAGAAAGAGAANNNAGATCGGA |
| 10107_1 | TATGAGTGTGGAGTCGTTGCGCCCAGCCAGGAAATAAACCTAGNNNAGATCGGAA |

| | |
|---|---|
| 10107_2 | TATGAGTGTGGAGTCGTTGCGCCCAGCCAGGAAATAAACCTAGNNNAGATCGGAA |
| 30808_1 | TATGAGTGTGGAGTCGTTGCCACCACTGGGACTACATTAGGTCNNNAGATCGGAAG |
| 30808_2 | TATGAGTGTGGAGTCGTTGCCACCACTGGGACTACATTAGGTCNNNAGATCGGAAG |
| 48797_1 | TATGAGTGTGGAGTCGTTGCTTAGTAGAGACGGGATTTCACCGNNNAGATCGGAA |
| 48797_2 | TATGAGTGTGGAGTCGTTGCTTAGTAGAGACGGGATTTCACCGNNNAGATCGGAA |
| 49821_1 | TATGAGTGTGGAGTCGTTGCGGAAGCCTTTACTCCAAAGCAAGNNNAGATCGGAAG |
| 49821_2 | TATGAGTGTGGAGTCGTTGCGGAAGCCTTTACTCCAAAGCAAGNNNAGATCGGAAG |
| 32869_1 | TATGAGTGTGGAGTCGTTGCCTCCCGAGTAGCTAGGACTACAGNNNAGATCGGAAG |
| 32869_2 | TATGAGTGTGGAGTCGTTGCCTCCCGAGTAGCTAGGACTACAGNNNAGATCGGAAG |
| 39864_1 | TATGAGTGTGGAGTCGTTGCAGTAAGCTGAGATCGCGCTACNNNAGATCGGAAGA |
| 39864_2 | TATGAGTGTGGAGTCGTTGCAGTAAGCTGAGATCGCGCTACNNNAGATCGGAAGA |
| 330_1 | TATGAGTGTGGAGTCGTTGCACTACGTCCAGTTAGGTTTCCAAGNNNAGATCGGAA |
| 330_2 | TATGAGTGTGGAGTCGTTGCACTACGTCCAGTTAGGTTTCCAAGNNNAGATCGGAA |
| 29228_1 | TATGAGTGTGGAGTCGTTGCACACACACAGAGAGAGAGAGAGANNNAGATCGGA |
| 29228_2 | TATGAGTGTGGAGTCGTTGCACACACACAGAGAGAGAGAGAGANNNAGATCGGA |
| 13718_1 | TATGAGTGTGGAGTCGTTGCGAAACCCTAATTCAATGTTTGAANNNAGATCGGAAG |
| 13718_2 | TATGAGTGTGGAGTCGTTGCGAAACCCTAATTCAATGTTTGAANNNAGATCGGAAG |
| 32777_1 | TATGAGTGTGGAGTCGTTGCCACTTGTAATCCCAGCACTTTGGNNNAGATCGGAAG |
| 32777_2 | TATGAGTGTGGAGTCGTTGCCACTTGTAATCCCAGCACTTTGGNNNAGATCGGAAG |
| 50762_1 | TATGAGTGTGGAGTCGTTGCGAAGACAGCGCCATTGAGGATCNNNAGATCGGAAG |
| 50762_2 | TATGAGTGTGGAGTCGTTGCGAAGACAGCGCCATTGAGGATCNNNAGATCGGAAG |
| 39061_1 | TATGAGTGTGGAGTCGTTGCTGTGATTGGATTGTTTGCAAGTCANNNAGATCGGA |
| 39061_2 | TATGAGTGTGGAGTCGTTGCTGTGATTGGATTGTTTGCAAGTCANNNAGATCGGA |
| 33877_1 | TATGAGTGTGGAGTCGTTGCGACATGGAGAAGTGAGGAGAAGGNNNAGATCGGA |
| 33877_2 | TATGAGTGTGGAGTCGTTGCGACATGGAGAAGTGAGGAGAAGGNNNAGATCGGA |
| 13317_1 | TATGAGTGTGGAGTCGTTGCCAGAAGAATCTTCCTCTTTTTCNNNAGATCGGAAGA |
| 13317_2 | TATGAGTGTGGAGTCGTTGCCAGAAGAATCTTCCTCTTTTTCNNNAGATCGGAAGA |
| 41319_1 | TATGAGTGTGGAGTCGTTGCGTAGTGACCTCAGTGTCTCCAAGNNNAGATCGGAAG |
| 41319_2 | TATGAGTGTGGAGTCGTTGCGTAGTGACCTCAGTGTCTCCAAGNNNAGATCGGAAG |
| 44268_1 | TATGAGTGTGGAGTCGTTGCGAGATGGGACACTCCAGTATTGGNNNAGATCGGAA |
| 44268_2 | TATGAGTGTGGAGTCGTTGCGAGATGGGACACTCCAGTATTGGNNNAGATCGGAA |
| 32450_1 | TATGAGTGTGGAGTCGTTGCGATCTCAGCTCACCACAACCTCNNNAGATCGGAAGA |
| 32450_2 | TATGAGTGTGGAGTCGTTGCGATCTCAGCTCACCACAACCTCNNNAGATCGGAAGA |
| 36664_1 | TATGAGTGTGGAGTCGTTGCCTCACACCTGTAATCTCAGCACTNNNAGATCGGAAGA |
| 36664_2 | TATGAGTGTGGAGTCGTTGCCTCACACCTGTAATCTCAGCACTNNNAGATCGGAAGA |
| 299_1 | TATGAGTGTGGAGTCGTTGCCATCCGTGCAACAAACAACAACTNNNAGATCGGAAG |
| 299_2 | TATGAGTGTGGAGTCGTTGCCATCCGTGCAACAAACAACAACTNNNAGATCGGAAG |
| 2373_1 | TATGAGTGTGGAGTCGTTGCACTTTTCCCACACAACATTCAGCNNNAGATCGGAAGA |
| 2373_2 | TATGAGTGTGGAGTCGTTGCACTTTTCCCACACAACATTCAGCNNNAGATCGGAAGA |
| 10551_1 | TATGAGTGTGGAGTCGTTGCTGCGTGTTTGTTTTGCCTTCTTTNNNAGATCGGAAGA |
| 10551_2 | TATGAGTGTGGAGTCGTTGCTGCGTGTTTGTTTTGCCTTCTTTNNNAGATCGGAAGA |
| 32564_1 | TATGAGTGTGGAGTCGTTGCCATTCTCCTGCCTCAGCCTCNNNAGATCGGAAGAGCA |
| 32564_2 | TATGAGTGTGGAGTCGTTGCCATTCTCCTGCCTCAGCCTCNNNAGATCGGAAGAGCA |

| | |
|---|---|
| 43326_1 | TATGAGTGTGGAGTCGTTGCCCAAAGTGCTGGGGAGTATTTTNNNAGATCGGAA |
| 43326_2 | TATGAGTGTGGAGTCGTTGCCCAAAGTGCTGGGGAGTATTTTNNNAGATCGGAA |
| 391_1 | TATGAGTGTGGAGTCGTTGCGACCACACAACTTTGCAACTTGNNNAGATCGGAAGA |
| 391_2 | TATGAGTGTGGAGTCGTTGCGACCACACAACTTTGCAACTTGNNNAGATCGGAAGA |
| 48741_1 | TATGAGTGTGGAGTCGTTGCTTATAACACCCCTGAGCTTTGCTNNNAGATCGGAAG |
| 48741_2 | TATGAGTGTGGAGTCGTTGCTTATAACACCCCTGAGCTTTGCTNNNAGATCGGAAG |
| 47063_1 | TATGAGTGTGGAGTCGTTGCATCTCCACTAACTGCAACCTCTGNNNAGATCGGAAGA |
| 47063_2 | TATGAGTGTGGAGTCGTTGCATCTCCACTAACTGCAACCTCTGNNNAGATCGGAAGA |
| 34903_1 | TATGAGTGTGGAGTCGTTGCGGGTTCAAGTAATTCTCCTGCCTNNNAGATCGGAAG |
| 34903_2 | TATGAGTGTGGAGTCGTTGCGGGTTCAAGTAATTCTCCTGCCTNNNAGATCGGAAG |
| 37148_1 | TATGAGTGTGGAGTCGTTGCACAGTTTTTTTTTCTCCAAGGGTNNNAGATCGGAAG |
| 37148_2 | TATGAGTGTGGAGTCGTTGCACAGTTTTTTTTTCTCCAAGGGTNNNAGATCGGAAG |
| 51096_1 | TATGAGTGTGGAGTCGTTGCTTCTTCACACATACCCTGTCAGGNNNAGATCGGAAG |
| 51096_2 | TATGAGTGTGGAGTCGTTGCTTCTTCACACATACCCTGTCAGGNNNAGATCGGAAG |
| 51730_1 | TATGAGTGTGGAGTCGTTGCCTTTCTAGAATGTTCGCCTTGCCNNNAGATCGGAAG |
| 51730_2 | TATGAGTGTGGAGTCGTTGCCTTTCTAGAATGTTCGCCTTGCCNNNAGATCGGAAG |
| 49391_1 | TATGAGTGTGGAGTCGTTGCAATACTGCAGCATGTATCGCCTANNNAGATCGGAAG |
| 49391_2 | TATGAGTGTGGAGTCGTTGCAATACTGCAGCATGTATCGCCTANNNAGATCGGAAG |
| 44151_1 | TATGAGTGTGGAGTCGTTGCATTATGACTGCAGGACAAAGGCTNNNAGATCGGAA |
| 44151_2 | TATGAGTGTGGAGTCGTTGCATTATGACTGCAGGACAAAGGCTNNNAGATCGGAA |
| 38861_1 | TATGAGTGTGGAGTCGTTGCTGGGTTGCAAATAAACTCCAAAGTNNNAGATCGGA |
| 38861_2 | TATGAGTGTGGAGTCGTTGCTGGGTTGCAAATAAACTCCAAAGTNNNAGATCGGA |
| 36917_1 | TATGAGTGTGGAGTCGTTGCATCATCCTTCTCACTGGCACATTNNNAGATCGGAAGA |
| 36917_2 | TATGAGTGTGGAGTCGTTGCATCATCCTTCTCACTGGCACATTNNNAGATCGGAAGA |
| 35929_1 | TATGAGTGTGGAGTCGTTGCGGTGAATCACAAGGTCAGGAGATNNNAGATCGGAA |
| 35929_2 | TATGAGTGTGGAGTCGTTGCGGTGAATCACAAGGTCAGGAGATNNNAGATCGGAA |
| 41205_1 | TATGAGTGTGGAGTCGTTGCTTTGTCATGAACCATCTGTTGCCNNNAGATCGGAAG |
| 41205_2 | TATGAGTGTGGAGTCGTTGCTTTGTCATGAACCATCTGTTGCCNNNAGATCGGAAG |
| 48584_1 | TATGAGTGTGGAGTCGTTGCCAATCACGGCTTTCGGAATCAGNNNAGATCGGAAGA |
| 48584_2 | TATGAGTGTGGAGTCGTTGCCAATCACGGCTTTCGGAATCAGNNNAGATCGGAAGA |
| 43633_1 | TATGAGTGTGGAGTCGTTGCCGGGATGAACCATGACCTCAGNNNAGATCGGAAGA |
| 43633_2 | TATGAGTGTGGAGTCGTTGCCGGGATGAACCATGACCTCAGNNNAGATCGGAAGA |
| 41505_1 | TATGAGTGTGGAGTCGTTGCATCACTTGAGACCAGGAGTTCACNNNAGATCGGAAG |
| 41505_2 | TATGAGTGTGGAGTCGTTGCATCACTTGAGACCAGGAGTTCACNNNAGATCGGAAG |
| 31680_1 | TATGAGTGTGGAGTCGTTGCAGAAAGACTATGAAAGCCTGCCANNNAGATCGGAA |
| 31680_2 | TATGAGTGTGGAGTCGTTGCAGAAAGACTATGAAAGCCTGCCANNNAGATCGGAA |
| 42034_1 | TATGAGTGTGGAGTCGTTGCAGAAAGGAGGAGAGACAGAGACANNNAGATCGGA |
| 42034_2 | TATGAGTGTGGAGTCGTTGCAGAAAGGAGGAGAGACAGAGACANNNAGATCGGA |
| 49168_1 | TATGAGTGTGGAGTCGTTGCAACACTGTAATGCAAGTCCATGCNNNAGATCGGAAG |
| 49168_2 | TATGAGTGTGGAGTCGTTGCAACACTGTAATGCAAGTCCATGCNNNAGATCGGAAG |
| 12570_1 | TATGAGTGTGGAGTCGTTGCTAAAGGTGTTAGTCAAACAAAACNNNAGATCGGAA |
| 12570_2 | TATGAGTGTGGAGTCGTTGCTAAAGGTGTTAGTCAAACAAAACNNNAGATCGGAA |
| 40536_1 | TATGAGTGTGGAGTCGTTGCAAAGAAAGACCATTTCCACTGCCNNNAGATCGGAAG |

| | |
|---|---|
| 40536_2 | TATGAGTGTGGAGTCGTTGCAAAGAAAGACCATTTCCACTGCCNNNAGATCGGAAG |
| 52767_1 | TATGAGTGTGGAGTCGTTGCCTCCCCTTCAGAAGCCACTTAGNNNAGATCGGAAGA |
| 52767_2 | TATGAGTGTGGAGTCGTTGCCTCCCCTTCAGAAGCCACTTAGNNNAGATCGGAAGA |
| 36042_1 | TATGAGTGTGGAGTCGTTGCGACACAGAGACAGAGAGACACAGNNNAGATCGGA |
| 36042_2 | TATGAGTGTGGAGTCGTTGCGACACAGAGACAGAGAGACACAGNNNAGATCGGA |
| 40757_1 | TATGAGTGTGGAGTCGTTGCGGTTCAAGTGATTGTCCTGCTTCNNNAGATCGGAAG |
| 40757_2 | TATGAGTGTGGAGTCGTTGCGGTTCAAGTGATTGTCCTGCTTCNNNAGATCGGAAG |
| 12876_1 | TATGAGTGTGGAGTCGTTGCGACAAGACACAAATCTAAGAACNNNAGATCGGAAG |
| 12876_2 | TATGAGTGTGGAGTCGTTGCGACAAGACACAAATCTAAGAACNNNAGATCGGAAG |
| 41473_1 | TATGAGTGTGGAGTCGTTGCGTGAACATGTTCTCAGTGTCTGCNNNAGATCGGAAG |
| 41473_2 | TATGAGTGTGGAGTCGTTGCGTGAACATGTTCTCAGTGTCTGCNNNAGATCGGAAG |
| 50342_1 | TATGAGTGTGGAGTCGTTGCAGCTAACAAAGTGCCGACAGATANNNAGATCGGAA |
| 50342_2 | TATGAGTGTGGAGTCGTTGCAGCTAACAAAGTGCCGACAGATANNNAGATCGGAA |
| 34446_1 | TATGAGTGTGGAGTCGTTGCTAGCTATCCCCACTCTCTCTAGCNNNAGATCGGAAGA |
| 34446_2 | TATGAGTGTGGAGTCGTTGCTAGCTATCCCCACTCTCTCTAGCNNNAGATCGGAAGA |
| 50634_1 | TATGAGTGTGGAGTCGTTGCTCACAAGGAAACCCAGGATGAAANNNAGATCGGAA |
| 50634_2 | TATGAGTGTGGAGTCGTTGCTCACAAGGAAACCCAGGATGAAANNNAGATCGGAA |
| 36069_1 | TATGAGTGTGGAGTCGTTGCTGGACCTCCTTTCTTTGCAAAACNNNAGATCGGAAG |
| 36069_2 | TATGAGTGTGGAGTCGTTGCTGGACCTCCTTTCTTTGCAAAACNNNAGATCGGAAG |
| 41557_1 | TATGAGTGTGGAGTCGTTGCTTCTGTTCACTACAACCTCCACCNNNAGATCGGAAGA |
| 41557_2 | TATGAGTGTGGAGTCGTTGCTTCTGTTCACTACAACCTCCACCNNNAGATCGGAAGA |
| 41003_1 | TATGAGTGTGGAGTCGTTGCGATCACCTGAGGCTAGGAGTTTGNNNAGATCGGAA |
| 41003_2 | TATGAGTGTGGAGTCGTTGCGATCACCTGAGGCTAGGAGTTTGNNNAGATCGGAA |
| 51152_1 | TATGAGTGTGGAGTCGTTGCTGGAAACGTGACGGGTATAGTTTNNNAGATCGGAA |
| 51152_2 | TATGAGTGTGGAGTCGTTGCTGGAAACGTGACGGGTATAGTTTNNNAGATCGGAA |
| 32209_1 | TATGAGTGTGGAGTCGTTGCGGTAGGACTGCCACCAGAAATTANNNAGATCGGAA |
| 32209_2 | TATGAGTGTGGAGTCGTTGCGGTAGGACTGCCACCAGAAATTANNNAGATCGGAA |
| 40362_1 | TATGAGTGTGGAGTCGTTGCACCCCATTTACCCTGATGTGATTNNNAGATCGGAAG |
| 40362_2 | TATGAGTGTGGAGTCGTTGCACCCCATTTACCCTGATGTGATTNNNAGATCGGAAG |
| 40616_1 | TATGAGTGTGGAGTCGTTGCTTGGCGAAGGACTTTTCATGTTCNNNAGATCGGAAG |
| 40616_2 | TATGAGTGTGGAGTCGTTGCTTGGCGAAGGACTTTTCATGTTCNNNAGATCGGAAG |
| 31412_1 | TATGAGTGTGGAGTCGTTGCCACACACACACACACTCATTTNNNAGATCGGAAGA |
| 31412_2 | TATGAGTGTGGAGTCGTTGCCACACACACACACACTCATTTNNNAGATCGGAAGA |
| 9788_1 | TATGAGTGTGGAGTCGTTGCACACCACTCCTGTTGGTTCTGTANNNAGATCGGAAG |
| 9788_2 | TATGAGTGTGGAGTCGTTGCACACCACTCCTGTTGGTTCTGTANNNAGATCGGAAG |
| 33798_1 | TATGAGTGTGGAGTCGTTGCGGCTGAATGGATGGAAGAACAAGNNNAGATCGGA |
| 33798_2 | TATGAGTGTGGAGTCGTTGCGGCTGAATGGATGGAAGAACAAGNNNAGATCGGA |
| 43806_1 | TATGAGTGTGGAGTCGTTGCCACTTTCGGTTCATCTTCTGCTCNNNAGATCGGAAGA |
| 43806_2 | TATGAGTGTGGAGTCGTTGCCACTTTCGGTTCATCTTCTGCTCNNNAGATCGGAAGA |
| 41057_1 | TATGAGTGTGGAGTCGTTGCCGTAGGCCCTCATGATTGGATTANNNAGATCGGAAG |
| 41057_2 | TATGAGTGTGGAGTCGTTGCCGTAGGCCCTCATGATTGGATTANNNAGATCGGAAG |
| 10712_1 | TATGAGTGTGGAGTCGTTGCTTTTCCCTTTGTTTCCTGCCTCANNNAGATCGGAAGA |
| 10712_2 | TATGAGTGTGGAGTCGTTGCTTTTCCCTTTGTTTCCTGCCTCANNNAGATCGGAAGA |

| | |
|---|---|
| 31439_1 | TATGAGTGTGGAGTCGTTGCCTGTATCTACTTCGTGCTCCTCCNNNAGATCGGAAGA |
| 31439_2 | TATGAGTGTGGAGTCGTTGCCTGTATCTACTTCGTGCTCCTCCNNNAGATCGGAAGA |
| 49093_1 | TATGAGTGTGGAGTCGTTGCCTTGAGGCTTTTGTTTCCCCATNNNAGATCGGAAGA |
| 49093_2 | TATGAGTGTGGAGTCGTTGCCTTGAGGCTTTTGTTTCCCCATNNNAGATCGGAAGA |
| 40806_1 | TATGAGTGTGGAGTCGTTGCGTTTGACCTTCACACTAGCCCTANNNAGATCGGAAGA |
| 40806_2 | TATGAGTGTGGAGTCGTTGCGTTTGACCTTCACACTAGCCCTANNNAGATCGGAAGA |
| 30118_1 | TATGAGTGTGGAGTCGTTGCACAAAAGGTGTTTTGATATCTCATCGTNNNAGATCG |
| 30118_2 | TATGAGTGTGGAGTCGTTGCACAAAAGGTGTTTTGATATCTCATCGTNNNAGATCG |
| 37321_1 | TATGAGTGTGGAGTCGTTGCGGGCAACAGAGCAATATTCCATCNNNAGATCGGAAG |
| 37321_2 | TATGAGTGTGGAGTCGTTGCGGGCAACAGAGCAATATTCCATCNNNAGATCGGAAG |
| 50874_1 | TATGAGTGTGGAGTCGTTGCAGTGTGTGGGTCAAAGTATTCCANNNAGATCGGAA |
| 50874_2 | TATGAGTGTGGAGTCGTTGCAGTGTGTGGGTCAAAGTATTCCANNNAGATCGGAA |
| 33973_1 | TATGAGTGTGGAGTCGTTGCTCGTGAACTCCTGAACTTGTGATNNNAGATCGGAAG |
| 33973_2 | TATGAGTGTGGAGTCGTTGCTCGTGAACTCCTGAACTTGTGATNNNAGATCGGAAG |
| 49249_1 | TATGAGTGTGGAGTCGTTGCTTCAAGCGATTTTCATGCCTCAGNNNAGATCGGAAGA |
| 49249_2 | TATGAGTGTGGAGTCGTTGCTTCAAGCGATTTTCATGCCTCAGNNNAGATCGGAAGA |
| 41170_1 | TATGAGTGTGGAGTCGTTGCGTTCCCACTGACCAGCCTATTTANNNAGATCGGAAGA |
| 41170_2 | TATGAGTGTGGAGTCGTTGCGTTCCCACTGACCAGCCTATTTANNNAGATCGGAAGA |
| 32820_1 | TATGAGTGTGGAGTCGTTGCGCACCTTGGTAAAGACAGAAACCNNNAGATCGGAA |
| 32820_2 | TATGAGTGTGGAGTCGTTGCGCACCTTGGTAAAGACAGAAACCNNNAGATCGGAA |
| 3161_1 | TATGAGTGTGGAGTCGTTGCTGGAATCCCTAGAAACTCGGTCTNNNAGATCGGAAG |
| 3161_2 | TATGAGTGTGGAGTCGTTGCTGGAATCCCTAGAAACTCGGTCTNNNAGATCGGAAG |
| 47517_1 | TATGAGTGTGGAGTCGTTGCTTCCATTTTGTTGGTCTCCAAGCNNNAGATCGGAAGA |
| 47517_2 | TATGAGTGTGGAGTCGTTGCTTCCATTTTGTTGGTCTCCAAGCNNNAGATCGGAAGA |
| 45677_1 | TATGAGTGTGGAGTCGTTGCCTGCGGAGTAACTAGGACTACAGNNNAGATCGGAA |
| 45677_2 | TATGAGTGTGGAGTCGTTGCCTGCGGAGTAACTAGGACTACAGNNNAGATCGGAA |
| 47422_1 | TATGAGTGTGGAGTCGTTGCCACCACCAAACCTGGCTAATTTTNNNAGATCGGAAGA |
| 47422_2 | TATGAGTGTGGAGTCGTTGCCACCACCAAACCTGGCTAATTTTNNNAGATCGGAAGA |
| 47401_1 | TATGAGTGTGGAGTCGTTGCATAATCTCGGCTTACTGCAACCTNNNAGATCGGAAGA |
| 47401_2 | TATGAGTGTGGAGTCGTTGCATAATCTCGGCTTACTGCAACCTNNNAGATCGGAAGA |
| 47782_1 | TATGAGTGTGGAGTCGTTGCACAGTGGCTCACACCTATAATCCNNNAGATCGGAAGA |
| 47782_2 | TATGAGTGTGGAGTCGTTGCACAGTGGCTCACACCTATAATCCNNNAGATCGGAAGA |
| 10143_1 | TATGAGTGTGGAGTCGTTGCTTTCCTGATTACCAGGCTGCAAGNNNAGATCGGAAG |
| 10143_2 | TATGAGTGTGGAGTCGTTGCTTTCCTGATTACCAGGCTGCAAGNNNAGATCGGAAG |
| 38973_1 | TATGAGTGTGGAGTCGTTGCTGGAGATTGCTTTGGGGAGTATTNNNAGATCGGAA |
| 38973_2 | TATGAGTGTGGAGTCGTTGCTGGAGATTGCTTTGGGGAGTATTNNNAGATCGGAA |
| 39108_1 | TATGAGTGTGGAGTCGTTGCGTAGTGACCTCAGTGTCTCCAAGNNNAGATCGGAAG |
| 39108_2 | TATGAGTGTGGAGTCGTTGCGTAGTGACCTCAGTGTCTCCAAGNNNAGATCGGAAG |
| 48935_1 | TATGAGTGTGGAGTCGTTGCGTGATCTCAACTCACTGCAACTTNNNAGATCGGAAGA |
| 48935_2 | TATGAGTGTGGAGTCGTTGCGTGATCTCAACTCACTGCAACTTNNNAGATCGGAAGA |
| 44475_1 | TATGAGTGTGGAGTCGTTGCTCCTTTGGGGGAAATGCTAATGTNNNAGATCGGAA |
| 44475_2 | TATGAGTGTGGAGTCGTTGCTCCTTTGGGGGAAATGCTAATGTNNNAGATCGGAA |
| 9631_1 | TATGAGTGTGGAGTCGTTGCCCTTTTGTCAGAAATCATGCAAGCNNNAGATCGGAA |

| | |
|---|---|
| 9631_2 | TATGAGTGTGGAGTCGTTGCCCTTTTGTCAGAAATCATGCAAGCNNNAGATCGGAA |
| 35730_1 | TATGAGTGTGGAGTCGTTGCTCTGACCTTTTCCACTAAGTGCANNNAGATCGGAAGA |
| 35730_2 | TATGAGTGTGGAGTCGTTGCTCTGACCTTTTCCACTAAGTGCANNNAGATCGGAAGA |
| 45075_1 | TATGAGTGTGGAGTCGTTGCTGTTAGAAGGTGGGACTTTGGAGNNNAGATCGGAA |
| 45075_2 | TATGAGTGTGGAGTCGTTGCTGTTAGAAGGTGGGACTTTGGAGNNNAGATCGGAA |
| 47749_1 | TATGAGTGTGGAGTCGTTGCGGCAGGTATGTTCAGAGGTCCNNNAGATCGGAAGA |
| 47749_2 | TATGAGTGTGGAGTCGTTGCGGCAGGTATGTTCAGAGGTCCNNNAGATCGGAAGA |
| 34202_1 | TATGAGTGTGGAGTCGTTGCCACTAGAAAAGGAGCCTGGGAATNNNAGATCGGAA |
| 34202_2 | TATGAGTGTGGAGTCGTTGCCACTAGAAAAGGAGCCTGGGAATNNNAGATCGGAA |
| 47864_1 | TATGAGTGTGGAGTCGTTGCCAATCTCAGCTTATTGCAGCCTCNNNAGATCGGAAGA |
| 47864_2 | TATGAGTGTGGAGTCGTTGCCAATCTCAGCTTATTGCAGCCTCNNNAGATCGGAAGA |
| 44715_1 | TATGAGTGTGGAGTCGTTGCCCTGCTCGGAAGAAGAAGATTTGNNNAGATCGGAA |
| 44715_2 | TATGAGTGTGGAGTCGTTGCCCTGCTCGGAAGAAGAAGATTTGNNNAGATCGGAA |
| 2335_1 | TATGAGTGTGGAGTCGTTGCAATGCAGGTTGTTGAGGATTTGANNNAGATCGGAA |
| 2335_2 | TATGAGTGTGGAGTCGTTGCAATGCAGGTTGTTGAGGATTTGANNNAGATCGGAA |
| 41232_1 | TATGAGTGTGGAGTCGTTGCATGGATGGAAGAGGAGGTCACTANNNAGATCGGA |
| 41232_2 | TATGAGTGTGGAGTCGTTGCATGGATGGAAGAGGAGGTCACTANNNAGATCGGA |
| 45755_1 | TATGAGTGTGGAGTCGTTGCCTCCCGGGTTCAAGTGATTCTCNNNAGATCGGAAGA |
| 45755_2 | TATGAGTGTGGAGTCGTTGCCTCCCGGGTTCAAGTGATTCTCNNNAGATCGGAAGA |
| 33551_1 | TATGAGTGTGGAGTCGTTGCGGTTCAAGCAATTCTCCTGTCTCNNNAGATCGGAAG |
| 33551_2 | TATGAGTGTGGAGTCGTTGCGGTTCAAGCAATTCTCCTGTCTCNNNAGATCGGAAG |
| 37994_1 | TATGAGTGTGGAGTCGTTGCCCCAGTGTCCATCAGCATATGAANNNAGATCGGAAG |
| 37994_2 | TATGAGTGTGGAGTCGTTGCCCCAGTGTCCATCAGCATATGAANNNAGATCGGAAG |
| 48264_1 | TATGAGTGTGGAGTCGTTGCTTCAAGCTATTCTCCTGCCTCAGNNNAGATCGGAAGA |
| 48264_2 | TATGAGTGTGGAGTCGTTGCTTCAAGCTATTCTCCTGCCTCAGNNNAGATCGGAAGA |
| 48138_1 | TATGAGTGTGGAGTCGTTGCCGAGTAACTGGGATTACAGGCATNNNAGATCGGAA |
| 48138_2 | TATGAGTGTGGAGTCGTTGCCGAGTAACTGGGATTACAGGCATNNNAGATCGGAA |
| 10111_1 | TATGAGTGTGGAGTCGTTGCAGTTGCCAGATAAATACAGGGAGCNNNAGATCGGA |
| 10111_2 | TATGAGTGTGGAGTCGTTGCAGTTGCCAGATAAATACAGGGAGCNNNAGATCGGA |
| 49883_1 | TATGAGTGTGGAGTCGTTGCTGAGCTGTGTTTTAGGAATTGCCNNNAGATCGGAA |
| 49883_2 | TATGAGTGTGGAGTCGTTGCTGAGCTGTGTTTTAGGAATTGCCNNNAGATCGGAA |
| 31464_1 | TATGAGTGTGGAGTCGTTGCAAAGAGAGAGACAGAGAGGGGAANNNAGATCGG |
| 31464_2 | TATGAGTGTGGAGTCGTTGCAAAGAGAGAGACAGAGAGGGGAANNNAGATCGG |
| 34097_1 | TATGAGTGTGGAGTCGTTGCCAAATCTCAAAGACTCAGCCCCANNNAGATCGGAAG |
| 34097_2 | TATGAGTGTGGAGTCGTTGCCAAATCTCAAAGACTCAGCCCCANNNAGATCGGAAG |
| 51985_1 | TATGAGTGTGGAGTCGTTGCCCTCCCCTCCTGTAGATGAGTATNNNAGATCGGAAG |
| 51985_2 | TATGAGTGTGGAGTCGTTGCCCTCCCCTCCTGTAGATGAGTATNNNAGATCGGAAG |
| 52125_1 | TATGAGTGTGGAGTCGTTGCTTATAGACATCTACCACCACGCCNNNAGATCGGAAG |
| 52125_2 | TATGAGTGTGGAGTCGTTGCTTATAGACATCTACCACCACGCCNNNAGATCGGAAG |
| 50576_1 | TATGAGTGTGGAGTCGTTGCTGATGGCAGAAATAGGGATGCATNNNAGATCGGAA |
| 50576_2 | TATGAGTGTGGAGTCGTTGCTGATGGCAGAAATAGGGATGCATNNNAGATCGGAA |
| 49416_1 | TATGAGTGTGGAGTCGTTGCACGTTAGTGTTCAACAAGCAAGCNNNAGATCGGAA |
| 49416_2 | TATGAGTGTGGAGTCGTTGCACGTTAGTGTTCAACAAGCAAGCNNNAGATCGGAA |

| | |
|---|---|
| 40413_1 | TATGAGTGTGGAGTCGTTGCAGTGCCTGCTTATAACTGGATGTNNNAGATCGGAAG |
| 40413_2 | TATGAGTGTGGAGTCGTTGCAGTGCCTGCTTATAACTGGATGTNNNAGATCGGAAG |
| 44642_1 | TATGAGTGTGGAGTCGTTGCGACACAGAGGGAGCAAAGATCAGNNNAGATCGGAA |
| 44642_2 | TATGAGTGTGGAGTCGTTGCGACACAGAGGGAGCAAAGATCAGNNNAGATCGGAA |
| 43625_1 | TATGAGTGTGGAGTCGTTGCCCAAGTCTAGGACACTCGGAAAANNNAGATCGGAAG |
| 43625_2 | TATGAGTGTGGAGTCGTTGCCCAAGTCTAGGACACTCGGAAAANNNAGATCGGAAG |
| 52652_1 | TATGAGTGTGGAGTCGTTGCTTGGCAGATGACTCCAGAACTACNNNAGATCGGAAG |
| 52652_2 | TATGAGTGTGGAGTCGTTGCTTGGCAGATGACTCCAGAACTACNNNAGATCGGAAG |
| 37778_1 | TATGAGTGTGGAGTCGTTGCTGAGGAGAAAAATGAAGCAGGGANNNAGATCGGAA |
| 37778_2 | TATGAGTGTGGAGTCGTTGCTGAGGAGAAAAATGAAGCAGGGANNNAGATCGGAA |
| 46866_1 | TATGAGTGTGGAGTCGTTGCCTCCTGAGTAGCTGGGACTACAGNNNAGATCGGAAG |
| 46866_2 | TATGAGTGTGGAGTCGTTGCCTCCTGAGTAGCTGGGACTACAGNNNAGATCGGAAG |
| 49841_1 | TATGAGTGTGGAGTCGTTGCGTCTTTTTCATGTCAACTGGGCANNNAGATCGGAAGA |
| 49841_2 | TATGAGTGTGGAGTCGTTGCGTCTTTTTCATGTCAACTGGGCANNNAGATCGGAAGA |
| 40802_1 | TATGAGTGTGGAGTCGTTGCCTCAAGCAGAAGGAGTTTTGTCCNNNAGATCGGAAG |
| 40802_2 | TATGAGTGTGGAGTCGTTGCCTCAAGCAGAAGGAGTTTTGTCCNNNAGATCGGAAG |
| 29053_1 | TATGAGTGTGGAGTCGTTGCGGAATAGGGAGATTTGGCAGAGTNNNAGATCGGAA |
| 29053_2 | TATGAGTGTGGAGTCGTTGCGGAATAGGGAGATTTGGCAGAGTNNNAGATCGGAA |
| 30392_1 | TATGAGTGTGGAGTCGTTGCCTGTGGGTCATTCGTGGAAGATANNNAGATCGGAAG |
| 30392_2 | TATGAGTGTGGAGTCGTTGCCTGTGGGTCATTCGTGGAAGATANNNAGATCGGAAG |
| 46431_1 | TATGAGTGTGGAGTCGTTGCTGGAGACAGGATTTCACCATGTTNNNAGATCGGAAG |
| 46431_2 | TATGAGTGTGGAGTCGTTGCTGGAGACAGGATTTCACCATGTTNNNAGATCGGAAG |
| 33122_1 | TATGAGTGTGGAGTCGTTGCCAGGAGAATGGCGTGAACCTNNNAGATCGGAAGAG |
| 33122_2 | TATGAGTGTGGAGTCGTTGCCAGGAGAATGGCGTGAACCTNNNAGATCGGAAGAG |
| 9972_1 | TATGAGTGTGGAGTCGTTGCAGGGGGTAAAAGGGAGGAGTTTTNNNAGATCGGAA |
| 9972_2 | TATGAGTGTGGAGTCGTTGCAGGGGGTAAAAGGGAGGAGTTTTNNNAGATCGGAA |
| 2844_1 | TATGAGTGTGGAGTCGTTGCAGGCTGGGAAAGATATACCGAGTNNNAGATCGGAA |
| 2844_2 | TATGAGTGTGGAGTCGTTGCAGGCTGGGAAAGATATACCGAGTNNNAGATCGGAA |
| 48132_1 | TATGAGTGTGGAGTCGTTGCCAAGTAGCTGGGATTACAGGCATNNNAGATCGGAAG |
| 48132_2 | TATGAGTGTGGAGTCGTTGCCAAGTAGCTGGGATTACAGGCATNNNAGATCGGAAG |
| 37379_1 | TATGAGTGTGGAGTCGTTGCACATGGACATCATCATTTTACCCANNNAGATCGGAAG |
| 37379_2 | TATGAGTGTGGAGTCGTTGCACATGGACATCATCATTTTACCCANNNAGATCGGAAG |
| 39430_1 | TATGAGTGTGGAGTCGTTGCCCTATAGTCCCAGCTACTCAGGANNNAGATCGGAAGA |
| 39430_2 | TATGAGTGTGGAGTCGTTGCCCTATAGTCCCAGCTACTCAGGANNNAGATCGGAAGA |
| 37620_1 | TATGAGTGTGGAGTCGTTGCGAGGGAAAAAAACCTCTTACCCTNNNAGATCGGAAG |
| 37620_2 | TATGAGTGTGGAGTCGTTGCGAGGGAAAAAAACCTCTTACCCTNNNAGATCGGAAG |
| 44248_1 | TATGAGTGTGGAGTCGTTGCTCTCCTCTATCAGTAGTCCCCACNNNAGATCGGAAGA |
| 44248_2 | TATGAGTGTGGAGTCGTTGCTCTCCTCTATCAGTAGTCCCCACNNNAGATCGGAAGA |
| 32801_1 | TATGAGTGTGGAGTCGTTGCCCCAGGTTCAAGCAATTCTCCTANNNAGATCGGAAGA |
| 32801_2 | TATGAGTGTGGAGTCGTTGCCCCAGGTTCAAGCAATTCTCCTANNNAGATCGGAAGA |
| 48126_1 | TATGAGTGTGGAGTCGTTGCAGTGGTTCTCAAAGTGTGGTCTTNNNAGATCGGAAG |
| 48126_2 | TATGAGTGTGGAGTCGTTGCAGTGGTTCTCAAAGTGTGGTCTTNNNAGATCGGAAG |
| 41932_1 | TATGAGTGTGGAGTCGTTGCAATCTTGGCTCACTGAAACCTCTNNNAGATCGGAAGA |

| | |
|---|---|
| 41932_2 | TATGAGTGTGGAGTCGTTGCAATCTTGGCTCACTGAAACCTCTNNNAGATCGGAAG |
| 53075_1 | TATGAGTGTGGAGTCGTTGCGTGGTGGCTCACACATGTAATTCNNNAGATCGGAAG |
| 53075_2 | TATGAGTGTGGAGTCGTTGCGTGGTGGCTCACACATGTAATTCNNNAGATCGGAAG |
| 31217_1 | TATGAGTGTGGAGTCGTTGCTGAACTCACTGAATAGCTCCTGGNNNAGATCGGAAG |
| 31217_2 | TATGAGTGTGGAGTCGTTGCTGAACTCACTGAATAGCTCCTGGNNNAGATCGGAAG |
| 48059_1 | TATGAGTGTGGAGTCGTTGCTCAAGTAGCTGGGATTACAAGCANNNAGATCGGAA |
| 48059_2 | TATGAGTGTGGAGTCGTTGCTCAAGTAGCTGGGATTACAAGCANNNAGATCGGAA |
| 46580_1 | TATGAGTGTGGAGTCGTTGCATCTAAGCTCACCACAACCTCTGNNNAGATCGGAAG |
| 46580_2 | TATGAGTGTGGAGTCGTTGCATCTAAGCTCACCACAACCTCTGNNNAGATCGGAAG |
| 3081_1 | TATGAGTGTGGAGTCGTTGCATAAATAAGCTGCAGGCAGGGTCNNNAGATCGGAA |
| 3081_2 | TATGAGTGTGGAGTCGTTGCATAAATAAGCTGCAGGCAGGGTCNNNAGATCGGAA |
| 33769_1 | TATGAGTGTGGAGTCGTTGCAAAAAAAACCACCACGAAACCACNNNAGATCGGAAG |
| 33769_2 | TATGAGTGTGGAGTCGTTGCAAAAAAAACCACCACGAAACCACNNNAGATCGGAAG |
| 6359_1 | TATGAGTGTGGAGTCGTTGCCAGACCAAGATGTCACACCAGTCNNNAGATCGGAAG |
| 6359_2 | TATGAGTGTGGAGTCGTTGCCAGACCAAGATGTCACACCAGTCNNNAGATCGGAAG |
| 52668_1 | TATGAGTGTGGAGTCGTTGCTCCTTCTAAGCGCGTGTACCNNNAGATCGGAAGAGC |
| 52668_2 | TATGAGTGTGGAGTCGTTGCTCCTTCTAAGCGCGTGTACCNNNAGATCGGAAGAGC |
| 43853_1 | TATGAGTGTGGAGTCGTTGCGGGGGAGAAGTGGGAGTGAGNNNAGATCGGAAG |
| 43853_2 | TATGAGTGTGGAGTCGTTGCGGGGGAGAAGTGGGAGTGAGNNNAGATCGGAAG |
| 52556_1 | TATGAGTGTGGAGTCGTTGCTGCCTGGAATTCCCTTTTTACCTNNNAGATCGGAAGA |
| 52556_2 | TATGAGTGTGGAGTCGTTGCTGCCTGGAATTCCCTTTTTACCTNNNAGATCGGAAGA |
| 37442_1 | TATGAGTGTGGAGTCGTTGCTGCTGGGAATTTGATAGTATTTGCANNNAGATCGGA |
| 37442_2 | TATGAGTGTGGAGTCGTTGCTGCTGGGAATTTGATAGTATTTGCANNNAGATCGGA |
| 38276_1 | TATGAGTGTGGAGTCGTTGCTCAGCATGGAAATGTTTCAACCCNNNAGATCGGAAG |
| 38276_2 | TATGAGTGTGGAGTCGTTGCTCAGCATGGAAATGTTTCAACCCNNNAGATCGGAAG |
| 29656_1 | TATGAGTGTGGAGTCGTTGCCAGATCACGAGGTCAGGAGTTCNNNAGATCGGAAG |
| 29656_2 | TATGAGTGTGGAGTCGTTGCCAGATCACGAGGTCAGGAGTTCNNNAGATCGGAAG |
| 617_1 | TATGAGTGTGGAGTCGTTGCCTCTTCCAGCCATCATTGCATCANNNAGATCGGAAGA |
| 617_2 | TATGAGTGTGGAGTCGTTGCCTCTTCCAGCCATCATTGCATCANNNAGATCGGAAGA |
| 39709_1 | TATGAGTGTGGAGTCGTTGCCAAAGTGACAGTAACCAAGTGGCNNNAGATCGGAA |
| 39709_2 | TATGAGTGTGGAGTCGTTGCCAAAGTGACAGTAACCAAGTGGCNNNAGATCGGAA |
| 49960_1 | TATGAGTGTGGAGTCGTTGCTGACATTTTTCACCCGGAATTGGNNNAGATCGGAAG |
| 49960_2 | TATGAGTGTGGAGTCGTTGCTGACATTTTTCACCCGGAATTGGNNNAGATCGGAAG |
| 43706_1 | TATGAGTGTGGAGTCGTTGCACCCTGAGTACAAGACCCAATTCNNNAGATCGGAAG |
| 43706_2 | TATGAGTGTGGAGTCGTTGCACCCTGAGTACAAGACCCAATTCNNNAGATCGGAAG |
| 50945_1 | TATGAGTGTGGAGTCGTTGCCTTGCTAGAAGACCCTCAAGCTCNNNAGATCGGAAG |
| 50945_2 | TATGAGTGTGGAGTCGTTGCCTTGCTAGAAGACCCTCAAGCTCNNNAGATCGGAAG |
| 36061_1 | TATGAGTGTGGAGTCGTTGCAGAATCTTACTGTGTTCTGGGGGNNNAGATCGGAA |
| 36061_2 | TATGAGTGTGGAGTCGTTGCAGAATCTTACTGTGTTCTGGGGGNNNAGATCGGAA |
| 38935_1 | TATGAGTGTGGAGTCGTTGCTACCTGACTGTTTTGCTTCACCTNNNAGATCGGAAGA |
| 38935_2 | TATGAGTGTGGAGTCGTTGCTACCTGACTGTTTTGCTTCACCTNNNAGATCGGAAGA |
| 40948_1 | TATGAGTGTGGAGTCGTTGCCTTTTCCCCACCCAGCTTTATTGNNNAGATCGGAAGA |
| 40948_2 | TATGAGTGTGGAGTCGTTGCCTTTTCCCCACCCAGCTTTATTGNNNAGATCGGAAGA |

| | |
|---|---|
| 34239_1 | TATGAGTGTGGAGTCGTTGCGCAGACAGACTCAAATCATGAAGANNNAGATCGGA |
| 34239_2 | TATGAGTGTGGAGTCGTTGCGCAGACAGACTCAAATCATGAAGANNNAGATCGGA |
| 33581_1 | TATGAGTGTGGAGTCGTTGCTGGTATGATTTCGGCTCACTGAANNNAGATCGGAAG |
| 33581_2 | TATGAGTGTGGAGTCGTTGCTGGTATGATTTCGGCTCACTGAANNNAGATCGGAAG |
| 34094_1 | TATGAGTGTGGAGTCGTTGCTCTGAACAAAAACACCCACATCCNNNAGATCGGAAG |
| 34094_2 | TATGAGTGTGGAGTCGTTGCTCTGAACAAAAACACCCACATCCNNNAGATCGGAAG |
| 41678_1 | TATGAGTGTGGAGTCGTTGCGATCTCAGCTCACTGCAACCTCNNNAGATCGGAAGA |
| 41678_2 | TATGAGTGTGGAGTCGTTGCGATCTCAGCTCACTGCAACCTCNNNAGATCGGAAGA |
| 41252_1 | TATGAGTGTGGAGTCGTTGCAGCGAGAGAGAAAGAGAAAGACANNNAGATCGGA |
| 41252_2 | TATGAGTGTGGAGTCGTTGCAGCGAGAGAGAAAGAGAAAGACANNNAGATCGGA |
| 10038_1 | TATGAGTGTGGAGTCGTTGCAGAGGCCTAGTGTAACTGAAGGGNNNAGATCGGAA |
| 10038_2 | TATGAGTGTGGAGTCGTTGCAGAGGCCTAGTGTAACTGAAGGGNNNAGATCGGAA |
| 32779_1 | TATGAGTGTGGAGTCGTTGCAAAGCTGCACACTTACAACCATTNNNAGATCGGAAG |
| 32779_2 | TATGAGTGTGGAGTCGTTGCAAAGCTGCACACTTACAACCATTNNNAGATCGGAAG |
| 47454_1 | TATGAGTGTGGAGTCGTTGCGCTCCTACCATGCATTGTGTTTTNNNAGATCGGAAG |
| 47454_2 | TATGAGTGTGGAGTCGTTGCGCTCCTACCATGCATTGTGTTTTNNNAGATCGGAAG |
| 37018_1 | TATGAGTGTGGAGTCGTTGCAGATCAGCAAGCGGTAACAAAACNNNAGATCGGAA |
| 37018_2 | TATGAGTGTGGAGTCGTTGCAGATCAGCAAGCGGTAACAAAACNNNAGATCGGAA |
| 48328_1 | TATGAGTGTGGAGTCGTTGCGGGATTATAGGCACACATCACCANNNAGATCGGAAG |
| 48328_2 | TATGAGTGTGGAGTCGTTGCGGGATTATAGGCACACATCACCANNNAGATCGGAAG |
| 13443_1 | TATGAGTGTGGAGTCGTTGCATTTCCCTACATGATGTACATTTNNNAGATCGGAAGA |
| 13443_2 | TATGAGTGTGGAGTCGTTGCATTTCCCTACATGATGTACATTTNNNAGATCGGAAGA |
| 33383_1 | TATGAGTGTGGAGTCGTTGCGTCAATCACGAGGTCAGGAGTTCNNNAGATCGGAA |
| 33383_2 | TATGAGTGTGGAGTCGTTGCGTCAATCACGAGGTCAGGAGTTCNNNAGATCGGAA |
| 29510_1 | TATGAGTGTGGAGTCGTTGCTTTTTCTCTTTCAGCTGTTGTCNNNAGATCGGAAGAG |
| 29510_2 | TATGAGTGTGGAGTCGTTGCTTTTTCTCTTTCAGCTGTTGTCNNNAGATCGGAAGAG |
| 36805_1 | TATGAGTGTGGAGTCGTTGCCCCTTTCACCACTCATTGTCAACNNNAGATCGGAAGA |
| 36805_2 | TATGAGTGTGGAGTCGTTGCCCCTTTCACCACTCATTGTCAACNNNAGATCGGAAGA |
| 40388_1 | TATGAGTGTGGAGTCGTTGCTGAGACACTGTAGAACAACTAGCANNNAGATCGGAA |
| 40388_2 | TATGAGTGTGGAGTCGTTGCTGAGACACTGTAGAACAACTAGCANNNAGATCGGAA |
| 33194_1 | TATGAGTGTGGAGTCGTTGCTTGCCCTATATCTGAGGTTGCTCNNNAGATCGGAAG |
| 33194_2 | TATGAGTGTGGAGTCGTTGCTTGCCCTATATCTGAGGTTGCTCNNNAGATCGGAAG |
| 13216_1 | TATGAGTGTGGAGTCGTTGCTTAGTGTCTGGCTTAAATTCCNNNAGATCGGAAGAG |
| 13216_2 | TATGAGTGTGGAGTCGTTGCTTAGTGTCTGGCTTAAATTCCNNNAGATCGGAAGAG |
| 34185_1 | TATGAGTGTGGAGTCGTTGCGGTATGGTAGCTCATGCCTGTAANNNAGATCGGAA |
| 34185_2 | TATGAGTGTGGAGTCGTTGCGGTATGGTAGCTCATGCCTGTAANNNAGATCGGAA |
| 47014_1 | TATGAGTGTGGAGTCGTTGCCAAAGTGCTGGGATTACAGGTGTNNNAGATCGGAA |
| 47014_2 | TATGAGTGTGGAGTCGTTGCCAAAGTGCTGGGATTACAGGTGTNNNAGATCGGAA |
| 32250_1 | TATGAGTGTGGAGTCGTTGCGCTTTGTGACGTATAAGGCCTTCNNNAGATCGGAAG |
| 32250_2 | TATGAGTGTGGAGTCGTTGCGCTTTGTGACGTATAAGGCCTTCNNNAGATCGGAAG |
| 52950_1 | TATGAGTGTGGAGTCGTTGCACAGTGTAACTGAAAGGGGGATGNNNAGATCGGAA |
| 52950_2 | TATGAGTGTGGAGTCGTTGCACAGTGTAACTGAAAGGGGGATGNNNAGATCGGAA |
| 10530_1 | TATGAGTGTGGAGTCGTTGCTGTCAGTGTATTGCTTCTGCTTTNNNAGATCGGAAG |

| | |
|---|---|
| 10530_2 | TATGAGTGTGGAGTCGTTGCTGTCAGTGTATTGCTTCTGCTTTNNNAGATCGGAAG |
| 37642_1 | TATGAGTGTGGAGTCGTTGCTACCTACCACAGTGCTGGATAGANNNAGATCGGAAG |
| 37642_2 | TATGAGTGTGGAGTCGTTGCTACCTACCACAGTGCTGGATAGANNNAGATCGGAAG |
| 30829_1 | TATGAGTGTGGAGTCGTTGCCTCTCTCTCTCTCACACACACACNNNAGATCGGAAGA |
| 30829_2 | TATGAGTGTGGAGTCGTTGCCTCTCTCTCTCTCACACACACACNNNAGATCGGAAGA |
| 40788_1 | TATGAGTGTGGAGTCGTTGCTTAGAGAAGAAGCTTCCAGGCAGNNNAGATCGGAA |
| 40788_2 | TATGAGTGTGGAGTCGTTGCTTAGAGAAGAAGCTTCCAGGCAGNNNAGATCGGAA |
| 51090_1 | TATGAGTGTGGAGTCGTTGCCAAAAAATTAGCCAGGCATTGGGNNNAGATCGGAA |
| 51090_2 | TATGAGTGTGGAGTCGTTGCCAAAAAATTAGCCAGGCATTGGGNNNAGATCGGAA |
| 29591_1 | TATGAGTGTGGAGTCGTTGCTCCCCCCGCGCTAAATATTGNNNAGATCGGAAGAGC |
| 29591_2 | TATGAGTGTGGAGTCGTTGCTCCCCCCGCGCTAAATATTGNNNAGATCGGAAGAGC |
| 34616_1 | TATGAGTGTGGAGTCGTTGCCCCTGACATTCCCCTAGAGGTNNNAGATCGGAAGAG |
| 34616_2 | TATGAGTGTGGAGTCGTTGCCCCTGACATTCCCCTAGAGGTNNNAGATCGGAAGAG |
| 40126_1 | TATGAGTGTGGAGTCGTTGCATTTCAGCTCACTGCAACTTCTGNNNAGATCGGAAG |
| 40126_2 | TATGAGTGTGGAGTCGTTGCATTTCAGCTCACTGCAACTTCTGNNNAGATCGGAAG |
| 49156_1 | TATGAGTGTGGAGTCGTTGCGATCTAGGCTCACTGCAAGCTCNNNAGATCGGAAGA |
| 49156_2 | TATGAGTGTGGAGTCGTTGCGATCTAGGCTCACTGCAAGCTCNNNAGATCGGAAGA |
| 42226_1 | TATGAGTGTGGAGTCGTTGCAGTTGGAGTCTCACTGTGTCATCNNNAGATCGGAAG |
| 42226_2 | TATGAGTGTGGAGTCGTTGCAGTTGGAGTCTCACTGTGTCATCNNNAGATCGGAAG |
| 42948_1 | TATGAGTGTGGAGTCGTTGCTGTCTTTGTCTGGCTAGGATATCNNNAGATCGGAAG |
| 42948_2 | TATGAGTGTGGAGTCGTTGCTGTCTTTGTCTGGCTAGGATATCNNNAGATCGGAAG |
| 50204_1 | TATGAGTGTGGAGTCGTTGCCATATGCACATTATGGCTCCTCGNNNAGATCGGAAG |
| 50204_2 | TATGAGTGTGGAGTCGTTGCCATATGCACATTATGGCTCCTCGNNNAGATCGGAAG |
| 13084_1 | TATGAGTGTGGAGTCGTTGCTTGAAAAGAAATCGAAACTGTTANNNAGATCGGAA |
| 13084_2 | TATGAGTGTGGAGTCGTTGCTTGAAAAGAAATCGAAACTGTTANNNAGATCGGAA |
| 29364_1 | TATGAGTGTGGAGTCGTTGCGGGCATTATCAAGGCCAGTTTTTNNNAGATCGGAAG |
| 29364_2 | TATGAGTGTGGAGTCGTTGCGGGCATTATCAAGGCCAGTTTTTNNNAGATCGGAAG |
| 31997_1 | TATGAGTGTGGAGTCGTTGCGTAATCCCAGCTACTTGAGAGGCNNNAGATCGGAA |
| 31997_2 | TATGAGTGTGGAGTCGTTGCGTAATCCCAGCTACTTGAGAGGCNNNAGATCGGAA |
| 8792_1 | TATGAGTGTGGAGTCGTTGCTGTCTGTGTAATTTGGCAAGCTTNNNAGATCGGAAG |
| 8792_2 | TATGAGTGTGGAGTCGTTGCTGTCTGTGTAATTTGGCAAGCTTNNNAGATCGGAAG |
| 29874_1 | TATGAGTGTGGAGTCGTTGCTTCGAGCCATTTTCTTCACTGTGNNNAGATCGGAAG |
| 29874_2 | TATGAGTGTGGAGTCGTTGCTTCGAGCCATTTTCTTCACTGTGNNNAGATCGGAAG |
| 41890_1 | TATGAGTGTGGAGTCGTTGCTAGAGGGTAGTGGCATGATCTCANNNAGATCGGAA |
| 41890_2 | TATGAGTGTGGAGTCGTTGCTAGAGGGTAGTGGCATGATCTCANNNAGATCGGAA |
| 28800_1 | TATGAGTGTGGAGTCGTTGCTTTTGCAGCAACTTAGATGGAGCNNNAGATCGGAA |
| 28800_2 | TATGAGTGTGGAGTCGTTGCTTTTGCAGCAACTTAGATGGAGCNNNAGATCGGAA |
| 47728_1 | TATGAGTGTGGAGTCGTTGCAATAAACAGGATGCTCAGTTCCTNNNAGATCGGAAG |
| 47728_2 | TATGAGTGTGGAGTCGTTGCAATAAACAGGATGCTCAGTTCCTNNNAGATCGGAAG |
| 32810_1 | TATGAGTGTGGAGTCGTTGCCTCCTGGGTTCACACCATTCTCNNNAGATCGGAAGA |
| 32810_2 | TATGAGTGTGGAGTCGTTGCCTCCTGGGTTCACACCATTCTCNNNAGATCGGAAGA |
| 9973_1 | TATGAGTGTGGAGTCGTTGCCTGGAAAAGTGGTGGCTAGGTTTNNNAGATCGGAA |
| 9973_2 | TATGAGTGTGGAGTCGTTGCCTGGAAAAGTGGTGGCTAGGTTTNNNAGATCGGAA |

| | |
|---|---|
| 40557_1 | TATGAGTGTGGAGTCGTTGCAAACAAAACTCAAACCTGGTGGCNNNAGATCGGAAG |
| 40557_2 | TATGAGTGTGGAGTCGTTGCAAACAAAACTCAAACCTGGTGGCNNNAGATCGGAAG |
| 42525_1 | TATGAGTGTGGAGTCGTTGCTTCTTCATCCACTCGTTGACTGANNNAGATCGGAAGA |
| 42525_2 | TATGAGTGTGGAGTCGTTGCTTCTTCATCCACTCGTTGACTGANNNAGATCGGAAGA |
| 45874_1 | TATGAGTGTGGAGTCGTTGCCAGTAGCATGATCATGGCTCTCTNNNAGATCGGAAG |
| 45874_2 | TATGAGTGTGGAGTCGTTGCCAGTAGCATGATCATGGCTCTCTNNNAGATCGGAAG |
| 50303_1 | TATGAGTGTGGAGTCGTTGCAAAATCTGAAAGTCACGGAAGGGNNNAGATCGGAA |
| 50303_2 | TATGAGTGTGGAGTCGTTGCAAAATCTGAAAGTCACGGAAGGGNNNAGATCGGAA |
| 34035_1 | TATGAGTGTGGAGTCGTTGCGAACTCCTGAGCTCAAGCAATCTNNNAGATCGGAAG |
| 34035_2 | TATGAGTGTGGAGTCGTTGCGAACTCCTGAGCTCAAGCAATCTNNNAGATCGGAAG |
| 39769_1 | TATGAGTGTGGAGTCGTTGCAGAGATGACATGAAGTTAGGACTGANNNAGATCGG |
| 39769_2 | TATGAGTGTGGAGTCGTTGCAGAGATGACATGAAGTTAGGACTGANNNAGATCGG |
| 2685_1 | TATGAGTGTGGAGTCGTTGCGCCCATGTATGTGTGATTGGCATNNNAGATCGGAAG |
| 2685_2 | TATGAGTGTGGAGTCGTTGCGCCCATGTATGTGTGATTGGCATNNNAGATCGGAAG |
| 7910_1 | TATGAGTGTGGAGTCGTTGCTTGCAGAGGGGAGATGTCACCTTNNNAGATCGGAA[illegible] |
| 7910_2 | TATGAGTGTGGAGTCGTTGCTTGCAGAGGGGAGATGTCACCTTNNNAGATCGGAA[illegible] |
| 9911_1 | TATGAGTGTGGAGTCGTTGCACTTACCTAGCAACCCACAGAGANNNAGATCGGAAG |
| 9911_2 | TATGAGTGTGGAGTCGTTGCACTTACCTAGCAACCCACAGAGANNNAGATCGGAAG |
| 49813_1 | TATGAGTGTGGAGTCGTTGCCTTTCTGCAGTCCAAATCCACTGNNNAGATCGGAAGA |
| 49813_2 | TATGAGTGTGGAGTCGTTGCCTTTCTGCAGTCCAAATCCACTGNNNAGATCGGAAGA |
| 36329_1 | TATGAGTGTGGAGTCGTTGCGAGATTACACCACTGCACTCCAGNNNAGATCGGAAG |
| 36329_2 | TATGAGTGTGGAGTCGTTGCGAGATTACACCACTGCACTCCAGNNNAGATCGGAAG |
| 48462_1 | TATGAGTGTGGAGTCGTTGCGAGACAGTCTTGCTCTGTTGCCNNNAGATCGGAAGA |
| 48462_2 | TATGAGTGTGGAGTCGTTGCGAGACAGTCTTGCTCTGTTGCCNNNAGATCGGAAGA |
| 42778_1 | TATGAGTGTGGAGTCGTTGCATACAGCTAATCTAACACCCCCCNNNAGATCGGAAGA |
| 42778_2 | TATGAGTGTGGAGTCGTTGCATACAGCTAATCTAACACCCCCCNNNAGATCGGAAGA |
| 39884_1 | TATGAGTGTGGAGTCGTTGCCTATACAAGTGGCCCTGTGCTATNNNAGATCGGAAG |
| 39884_2 | TATGAGTGTGGAGTCGTTGCCTATACAAGTGGCCCTGTGCTATNNNAGATCGGAAG |
| 34705_1 | TATGAGTGTGGAGTCGTTGCGACAGATTCTCACTCTCACCCAGNNNAGATCGGAAGA |
| 34705_2 | TATGAGTGTGGAGTCGTTGCGACAGATTCTCACTCTCACCCAGNNNAGATCGGAAGA |
| 9038_1 | TATGAGTGTGGAGTCGTTGCCAGTGACTATCCAAAGGCAAGCTNNNAGATCGGAAG |
| 9038_2 | TATGAGTGTGGAGTCGTTGCCAGTGACTATCCAAAGGCAAGCTNNNAGATCGGAAG |
| 39937_1 | TATGAGTGTGGAGTCGTTGCCAGAGCCAGACTCCATCTCAAAANNNAGATCGGAAG |
| 39937_2 | TATGAGTGTGGAGTCGTTGCCAGAGCCAGACTCCATCTCAAAANNNAGATCGGAAG |
| 39979_1 | TATGAGTGTGGAGTCGTTGCCTTTTTGGGAGGCAGAATGGAAGNNNAGATCGGAA |
| 39979_2 | TATGAGTGTGGAGTCGTTGCCTTTTTGGGAGGCAGAATGGAAGNNNAGATCGGAA |
| 44354_1 | TATGAGTGTGGAGTCGTTGCCTGTACAAGCAAGACAGCAGTTGNNNAGATCGGAA[illegible] |
| 44354_2 | TATGAGTGTGGAGTCGTTGCCTGTACAAGCAAGACAGCAGTTGNNNAGATCGGAA[illegible] |
| 44008_1 | TATGAGTGTGGAGTCGTTGCATGCTGTAGTCTGCCTTTTACCANNNAGATCGGAAGA |
| 44008_2 | TATGAGTGTGGAGTCGTTGCATGCTGTAGTCTGCCTTTTACCANNNAGATCGGAAGA |
| 44156_1 | TATGAGTGTGGAGTCGTTGCATGCTAAAAGGCCACTTCCCATANNNAGATCGGAAG[illegible] |
| 44156_2 | TATGAGTGTGGAGTCGTTGCATGCTAAAAGGCCACTTCCCATANNNAGATCGGAAG[illegible] |
| 33154_1 | TATGAGTGTGGAGTCGTTGCTGACATTCCTCCTTGTGAGAGTGNNNAGATCGGAAG |

| | |
|---|---|
| 33154_2 | TATGAGTGTGGAGTCGTTGCTGACATTCCTCCTTGTGAGAGTGNNNAGATCGGAAG |
| 35174_1 | TATGAGTGTGGAGTCGTTGCAAGTGATTGTCCTTCCTCAGCTTNNNAGATCGGAAG |
| 35174_2 | TATGAGTGTGGAGTCGTTGCAAGTGATTGTCCTTCCTCAGCTTNNNAGATCGGAAG |
| 8881_1 | TATGAGTGTGGAGTCGTTGCCCAAACAGTGGGTGAGTGAATGTNNNAGATCGGAA |
| 8881_2 | TATGAGTGTGGAGTCGTTGCCCAAACAGTGGGTGAGTGAATGTNNNAGATCGGAA |
| 30338_1 | TATGAGTGTGGAGTCGTTGCCTCTTCCTGCCCGTCCTCTTTTNNNAGATCGGAAGAG |
| 30338_2 | TATGAGTGTGGAGTCGTTGCCTCTTCCTGCCCGTCCTCTTTTNNNAGATCGGAAGAG |
| 50653_1 | TATGAGTGTGGAGTCGTTGCATAAAGATGTGCAGCTGGACAGANNNAGATCGGAA |
| 50653_2 | TATGAGTGTGGAGTCGTTGCATAAAGATGTGCAGCTGGACAGANNNAGATCGGAA |
| 45078_1 | TATGAGTGTGGAGTCGTTGCGGAGGGGGTTCAGGAAATAAGTTNNNAGATCGGA |
| 45078_2 | TATGAGTGTGGAGTCGTTGCGGAGGGGGTTCAGGAAATAAGTTNNNAGATCGGA |
| 40427_1 | TATGAGTGTGGAGTCGTTGCGACCTGGACATTGGTAGTGTCTTNNNAGATCGGAA |
| 40427_2 | TATGAGTGTGGAGTCGTTGCGACCTGGACATTGGTAGTGTCTTNNNAGATCGGAA |
| 45349_1 | TATGAGTGTGGAGTCGTTGCTGCTTTTCTCTGTGTGCATATGCNNNAGATCGGAAG |
| 45349_2 | TATGAGTGTGGAGTCGTTGCTGCTTTTCTCTGTGTGCATATGCNNNAGATCGGAAG |
| 10812_1 | TATGAGTGTGGAGTCGTTGCTTCCCACACTCCTCTGTCTGTGNNNAGATCGGAAGAG |
| 10812_2 | TATGAGTGTGGAGTCGTTGCTTCCCACACTCCTCTGTCTGTGNNNAGATCGGAAGAG |
| 10138_1 | TATGAGTGTGGAGTCGTTGCCTTCTAAAACCTGCTCCAGTGCTNNNAGATCGGAAG |
| 10138_2 | TATGAGTGTGGAGTCGTTGCCTTCTAAAACCTGCTCCAGTGCTNNNAGATCGGAAG |
| 29648_1 | TATGAGTGTGGAGTCGTTGCACTGATGGATGTAAGCCGAGCNNNAGATCGGAAGA |
| 29648_2 | TATGAGTGTGGAGTCGTTGCACTGATGGATGTAAGCCGAGCNNNAGATCGGAAGA |
| 47360_1 | TATGAGTGTGGAGTCGTTGCCCATAGACGTGTGGTGGTCGNNNAGATCGGAAGAG |
| 47360_2 | TATGAGTGTGGAGTCGTTGCCCATAGACGTGTGGTGGTCGNNNAGATCGGAAGAG |
| 29469_1 | TATGAGTGTGGAGTCGTTGCAAACTAACCTACAGGCTTTAAGTAGANNNAGATCGG |
| 29469_2 | TATGAGTGTGGAGTCGTTGCAAACTAACCTACAGGCTTTAAGTAGANNNAGATCGG |
| 12499_1 | TATGAGTGTGGAGTCGTTGCTGAACCAGCAATAATGTTTTCNNNAGATCGGAAGAG |
| 12499_2 | TATGAGTGTGGAGTCGTTGCTGAACCAGCAATAATGTTTTCNNNAGATCGGAAGAG |
| 10628_1 | TATGAGTGTGGAGTCGTTGCCAACTGAACCAGAGCACTGTCATNNNAGATCGGAAG |
| 10628_2 | TATGAGTGTGGAGTCGTTGCCAACTGAACCAGAGCACTGTCATNNNAGATCGGAAG |
| 39169_1 | TATGAGTGTGGAGTCGTTGCGGTGTTTCCAGAGGAGATAAGCANNNAGATCGGAA |
| 39169_2 | TATGAGTGTGGAGTCGTTGCGGTGTTTCCAGAGGAGATAAGCANNNAGATCGGAA |
| 7970_1 | TATGAGTGTGGAGTCGTTGCTCTCCACTCACAGCCACTTCTANNNAGATCGGAAGAG |
| 7970_2 | TATGAGTGTGGAGTCGTTGCTCTCCACTCACAGCCACTTCTANNNAGATCGGAAGAG |
| 30568_1 | TATGAGTGTGGAGTCGTTGCAACCTTGGAAAAATGTCTAGGTCTGNNNAGATCGG |
| 30568_2 | TATGAGTGTGGAGTCGTTGCAACCTTGGAAAAATGTCTAGGTCTGNNNAGATCGG |
| 49612_1 | TATGAGTGTGGAGTCGTTGCTCTAATGGTCAGGTCCCTCTTCTNNNAGATCGGAAG |
| 49612_2 | TATGAGTGTGGAGTCGTTGCTCTAATGGTCAGGTCCCTCTTCTNNNAGATCGGAAG |
| 3069_1 | TATGAGTGTGGAGTCGTTGCCTCGTAGGTTCAGTACCATTGGTGNNNAGATCGGAA |
| 3069_2 | TATGAGTGTGGAGTCGTTGCCTCGTAGGTTCAGTACCATTGGTGNNNAGATCGGAA |
| 48071_1 | TATGAGTGTGGAGTCGTTGCAGAGAAGGGGAGACTGAGAGAATNNNAGATCGGA |
| 48071_2 | TATGAGTGTGGAGTCGTTGCAGAGAAGGGGAGACTGAGAGAATNNNAGATCGGA |
| 49468_1 | TATGAGTGTGGAGTCGTTGCATACAAACAGGTGCATTTCGGAGNNNAGATCGGAA |
| 49468_2 | TATGAGTGTGGAGTCGTTGCATACAAACAGGTGCATTTCGGAGNNNAGATCGGAA |

| | |
|---|---|
| 51137_1 | TATGAGTGTGGAGTCGTTGCGTGGAACTGATTGAACCACACAGNNNAGATCGGAA |
| 51137_2 | TATGAGTGTGGAGTCGTTGCGTGGAACTGATTGAACCACACAGNNNAGATCGGAA |
| 50122_1 | TATGAGTGTGGAGTCGTTGCAACAACTTGAGAGTGACAGAGGGNNNAGATCGGA |
| 50122_2 | TATGAGTGTGGAGTCGTTGCAACAACTTGAGAGTGACAGAGGGNNNAGATCGGA |
| 40033_1 | TATGAGTGTGGAGTCGTTGCATTGCCCATTTGGTAACTTGGACNNNAGATCGGAAG |
| 40033_2 | TATGAGTGTGGAGTCGTTGCATTGCCCATTTGGTAACTTGGACNNNAGATCGGAAG |
| 31754_1 | TATGAGTGTGGAGTCGTTGCCCCTCTCTCCTCTCTCTCTTCTCNNNAGATCGGAAGAG |
| 31754_2 | TATGAGTGTGGAGTCGTTGCCCCTCTCTCCTCTCTCTCTTCTCNNNAGATCGGAAGAG |
| 53091_1 | TATGAGTGTGGAGTCGTTGCACAATCTTCATCAGCAAGACCCTNNNAGATCGGAAG |
| 53091_2 | TATGAGTGTGGAGTCGTTGCACAATCTTCATCAGCAAGACCCTNNNAGATCGGAAG |
| 29221_1 | TATGAGTGTGGAGTCGTTGCCAAGTTTGCAGTATACAAGGTCANNNAGATCGGAA |
| 29221_2 | TATGAGTGTGGAGTCGTTGCCAAGTTTGCAGTATACAAGGTCANNNAGATCGGAA |
| 39988_1 | TATGAGTGTGGAGTCGTTGCGTGTGTGTGGAGTGTATGAGAGANNNAGATCGGA |
| 39988_2 | TATGAGTGTGGAGTCGTTGCGTGTGTGTGGAGTGTATGAGAGANNNAGATCGGA |
| 39828_1 | TATGAGTGTGGAGTCGTTGCCCACCACGTCTGGCTAATTTTTTNNNAGATCGGAAGA |
| 39828_2 | TATGAGTGTGGAGTCGTTGCCCACCACGTCTGGCTAATTTTTTNNNAGATCGGAAGA |
| 34733_1 | TATGAGTGTGGAGTCGTTGCCTCAGGGGAGAGGTTGAGAATTTNNNAGATCGGAA |
| 34733_2 | TATGAGTGTGGAGTCGTTGCCTCAGGGGAGAGGTTGAGAATTTNNNAGATCGGAA |
| 37232_1 | TATGAGTGTGGAGTCGTTGCTGAATTTACACTTTCCAGTTTGCTNNNAGATCGGAAG |
| 37232_2 | TATGAGTGTGGAGTCGTTGCTGAATTTACACTTTCCAGTTTGCTNNNAGATCGGAAG |
| 35939_1 | TATGAGTGTGGAGTCGTTGCGGAGGTTGTAGTGAGCCAAGATGNNNAGATCGGA |
| 35939_2 | TATGAGTGTGGAGTCGTTGCGGAGGTTGTAGTGAGCCAAGATGNNNAGATCGGA |
| 31548_1 | TATGAGTGTGGAGTCGTTGCCCCCTCTGTTGCCTTTAGAAAGANNNAGATCGGAAG |
| 31548_2 | TATGAGTGTGGAGTCGTTGCCCCCTCTGTTGCCTTTAGAAAGANNNAGATCGGAAG |
| 42460_1 | TATGAGTGTGGAGTCGTTGCAGTTACACAAGCAACCATGAGGANNNAGATCGGAA |
| 42460_2 | TATGAGTGTGGAGTCGTTGCAGTTACACAAGCAACCATGAGGANNNAGATCGGAA |
| 31135_1 | TATGAGTGTGGAGTCGTTGCCTGTCATTCACCCAATTGCTCAGNNNAGATCGGAAG |
| 31135_2 | TATGAGTGTGGAGTCGTTGCCTGTCATTCACCCAATTGCTCAGNNNAGATCGGAAG |
| 34990_1 | TATGAGTGTGGAGTCGTTGCCCACGCACGGCTAATTTTTGTATNNNAGATCGGAAG |
| 34990_2 | TATGAGTGTGGAGTCGTTGCCCACGCACGGCTAATTTTTGTATNNNAGATCGGAAG |
| 44554_1 | TATGAGTGTGGAGTCGTTGCCACACACACACCCCAAAGTAAAANNNAGATCGGAAG |
| 44554_2 | TATGAGTGTGGAGTCGTTGCCACACACACACCCCAAAGTAAAANNNAGATCGGAAG |
| 33086_1 | TATGAGTGTGGAGTCGTTGCTGATCTCAGCTCACTGCAACTCNNNAGATCGGAAGA |
| 33086_2 | TATGAGTGTGGAGTCGTTGCTGATCTCAGCTCACTGCAACTCNNNAGATCGGAAGA |
| 50038_1 | TATGAGTGTGGAGTCGTTGCGAGCGAGGTACACACTTTCAAACNNNAGATCGGAA |
| 50038_2 | TATGAGTGTGGAGTCGTTGCGAGCGAGGTACACACTTTCAAACNNNAGATCGGAA |
| 29669_1 | TATGAGTGTGGAGTCGTTGCAGGTGACCTCAACACTGCTGNNNAGATCGGAAGAG |
| 29669_2 | TATGAGTGTGGAGTCGTTGCAGGTGACCTCAACACTGCTGNNNAGATCGGAAGAG |
| 13591_1 | TATGAGTGTGGAGTCGTTGCTAATCCTGGTTTTCAAGAGTATTNNNAGATCGGAAG |
| 13591_2 | TATGAGTGTGGAGTCGTTGCTAATCCTGGTTTTCAAGAGTATTNNNAGATCGGAAG |
| 30365_1 | TATGAGTGTGGAGTCGTTGCTGTAAACCTCATGGCACCTATCANNNAGATCGGAAG |
| 30365_2 | TATGAGTGTGGAGTCGTTGCTGTAAACCTCATGGCACCTATCANNNAGATCGGAAG |
| 52756_1 | TATGAGTGTGGAGTCGTTGCCTGTGACACGTCAACTGTCTTTCNNNAGATCGGAAG |

| | |
|---|---|
| 52756_2 | TATGAGTGTGGAGTCGTTGCCTGTGACACGTCAACTGTCTTTCNNNAGATCGGAAG |
| 41802_1 | TATGAGTGTGGAGTCGTTGCCTGCCTCCTAGGTTCAAGTGATTNNNAGATCGGAAG |
| 41802_2 | TATGAGTGTGGAGTCGTTGCCTGCCTCCTAGGTTCAAGTGATTNNNAGATCGGAAG |
| 30039_1 | TATGAGTGTGGAGTCGTTGCTCATTTGGCATTTCTTGGAGTGTNNNAGATCGGAAG |
| 30039_2 | TATGAGTGTGGAGTCGTTGCTCATTTGGCATTTCTTGGAGTGTNNNAGATCGGAAG |
| 9005_1 | TATGAGTGTGGAGTCGTTGCTCTCTCCTCTCCTCTTTTGATCTNNNAGATCGGAAGA |
| 9005_2 | TATGAGTGTGGAGTCGTTGCTCTCTCCTCTCCTCTTTTGATCTNNNAGATCGGAAGA |
| 36759_1 | TATGAGTGTGGAGTCGTTGCTGGGAGGAAAGGAGAAAATGAGGNNNAGATCGG |
| 36759_2 | TATGAGTGTGGAGTCGTTGCTGGGAGGAAAGGAGAAAATGAGGNNNAGATCGG |
| 1194_1 | TATGAGTGTGGAGTCGTTGCAACTATCCTCCCAACCTCAGCTTNNNAGATCGGAAGA |
| 1194_2 | TATGAGTGTGGAGTCGTTGCAACTATCCTCCCAACCTCAGCTTNNNAGATCGGAAGA |
| 45077_1 | TATGAGTGTGGAGTCGTTGCAGGGGGGGTACCTATAACATGATNNNAGATCGGAA |
| 45077_2 | TATGAGTGTGGAGTCGTTGCAGGGGGGGTACCTATAACATGATNNNAGATCGGAA |
| 43006_1 | TATGAGTGTGGAGTCGTTGCCTTCTGAGGCCTAAAGGAACTGTNNNAGATCGGAA |
| 43006_2 | TATGAGTGTGGAGTCGTTGCCTTCTGAGGCCTAAAGGAACTGTNNNAGATCGGAA |
| 40432_1 | TATGAGTGTGGAGTCGTTGCTTGGTGAAAACTTGCTGTGTTGANNNAGATCGGAA |
| 40432_2 | TATGAGTGTGGAGTCGTTGCTTGGTGAAAACTTGCTGTGTTGANNNAGATCGGAA |
| 46027_1 | TATGAGTGTGGAGTCGTTGCCATTCTCCTGCCTCAGCCTCNNNAGATCGGAAGAGC |
| 46027_2 | TATGAGTGTGGAGTCGTTGCCATTCTCCTGCCTCAGCCTCNNNAGATCGGAAGAGC |
| 43608_1 | TATGAGTGTGGAGTCGTTGCAAGGTCTTGGTTCTGCTTTACCANNNAGATCGGAAG |
| 43608_2 | TATGAGTGTGGAGTCGTTGCAAGGTCTTGGTTCTGCTTTACCANNNAGATCGGAAG |
| 39940_1 | TATGAGTGTGGAGTCGTTGCAATCCCAGCTACTTGGAAGTCTGNNNAGATCGGAAG |
| 39940_2 | TATGAGTGTGGAGTCGTTGCAATCCCAGCTACTTGGAAGTCTGNNNAGATCGGAAG |
| 47460_1 | TATGAGTGTGGAGTCGTTGCATGTGCCACCATTCCTAGCTAAANNNAGATCGGAAG |
| 47460_2 | TATGAGTGTGGAGTCGTTGCATGTGCCACCATTCCTAGCTAAANNNAGATCGGAAG |
| 46239_1 | TATGAGTGTGGAGTCGTTGCCTCAAGCTATTCTCCTGCCTCAANNNAGATCGGAAGA |
| 46239_2 | TATGAGTGTGGAGTCGTTGCCTCAAGCTATTCTCCTGCCTCAANNNAGATCGGAAGA |
| 45018_1 | TATGAGTGTGGAGTCGTTGCTGCATACTCCTCAGGAACAGATGNNNAGATCGGAAG |
| 45018_2 | TATGAGTGTGGAGTCGTTGCTGCATACTCCTCAGGAACAGATGNNNAGATCGGAAG |
| 31334_1 | TATGAGTGTGGAGTCGTTGCGCTCTTCCACATCTATTGGGTCANNNAGATCGGAAG |
| 31334_2 | TATGAGTGTGGAGTCGTTGCGCTCTTCCACATCTATTGGGTCANNNAGATCGGAAG |
| 52495_1 | TATGAGTGTGGAGTCGTTGCGAAAGAAAGGAAAACGCTGTCCANNNAGATCGGAA |
| 52495_2 | TATGAGTGTGGAGTCGTTGCGAAAGAAAGGAAAACGCTGTCCANNNAGATCGGAA |
| 46035_1 | TATGAGTGTGGAGTCGTTGCCACCACGCCCAGCTAATTTTTNNNAGATCGGAAGAG |
| 46035_2 | TATGAGTGTGGAGTCGTTGCCACCACGCCCAGCTAATTTTTNNNAGATCGGAAGAG |
| 51165_1 | TATGAGTGTGGAGTCGTTGCCTTAGTTTCGGGAGTGTGAGGAANNNAGATCGGAA |
| 51165_2 | TATGAGTGTGGAGTCGTTGCCTTAGTTTCGGGAGTGTGAGGAANNNAGATCGGAA |
| 35446_1 | TATGAGTGTGGAGTCGTTGCTCCAAGATGGACTTCAACTCCTGNNNAGATCGGAAG |
| 35446_2 | TATGAGTGTGGAGTCGTTGCTCCAAGATGGACTTCAACTCCTGNNNAGATCGGAAG |
| 51401_1 | TATGAGTGTGGAGTCGTTGCCTGTCAGAAAAATGGCCAAACCTNNNAGATCGGAAG |
| 51401_2 | TATGAGTGTGGAGTCGTTGCCTGTCAGAAAAATGGCCAAACCTNNNAGATCGGAAG |
| 45491_1 | TATGAGTGTGGAGTCGTTGCTTTCTGGAGACGGTTCATAACGTNNNAGATCGGAAG |
| 45491_2 | TATGAGTGTGGAGTCGTTGCTTTCTGGAGACGGTTCATAACGTNNNAGATCGGAAG |

| | |
|---|---|
| 13120_1 | TATGAGTGTGGAGTCGTTGCCTAGGTTAGTTTGTGTTAACATGNNNAGATCGGAAG |
| 13120_2 | TATGAGTGTGGAGTCGTTGCCTAGGTTAGTTTGTGTTAACATGNNNAGATCGGAAG |
| 10589_1 | TATGAGTGTGGAGTCGTTGCGTCGAAACGTGATGCGATGAATTNNNAGATCGGAA |
| 10589_2 | TATGAGTGTGGAGTCGTTGCGTCGAAACGTGATGCGATGAATTNNNAGATCGGAA |
| 29037_1 | TATGAGTGTGGAGTCGTTGCTCTATCATGAAATCTGTCAAGCATCNNNAGATCGGA |
| 29037_2 | TATGAGTGTGGAGTCGTTGCTCTATCATGAAATCTGTCAAGCATCNNNAGATCGGA |
| 35121_1 | TATGAGTGTGGAGTCGTTGCATTCGTGTCATGGAAGTTTGCTGNNNAGATCGGAAG |
| 35121_2 | TATGAGTGTGGAGTCGTTGCATTCGTGTCATGGAAGTTTGCTGNNNAGATCGGAAG |
| 12586_1 | TATGAGTGTGGAGTCGTTGCTTTCTCTGGATGCAAAATAAAATNNNAGATCGGAAG |
| 12586_2 | TATGAGTGTGGAGTCGTTGCTTTCTCTGGATGCAAAATAAAATNNNAGATCGGAAG |
| 35739_1 | TATGAGTGTGGAGTCGTTGCATTGTCTGCTTCCTTTCTCAGCTNNNAGATCGGAAGA |
| 35739_2 | TATGAGTGTGGAGTCGTTGCATTGTCTGCTTCCTTTCTCAGCTNNNAGATCGGAAGA |
| 43666_1 | TATGAGTGTGGAGTCGTTGCAATGTGGATGCAAGCAGGAAATCNNNAGATCGGAA |
| 43666_2 | TATGAGTGTGGAGTCGTTGCAATGTGGATGCAAGCAGGAAATCNNNAGATCGGAA |
| 38154_1 | TATGAGTGTGGAGTCGTTGCGGATCACGAAGTCAGGAGATTGANNNAGATCGGAA |
| 38154_2 | TATGAGTGTGGAGTCGTTGCGGATCACGAAGTCAGGAGATTGANNNAGATCGGAA |
| 37765_1 | TATGAGTGTGGAGTCGTTGCCTGTAGTCTCAGCTACTTGGGAGNNNAGATCGGAAG |
| 37765_2 | TATGAGTGTGGAGTCGTTGCCTGTAGTCTCAGCTACTTGGGAGNNNAGATCGGAAG |
| 49482_1 | TATGAGTGTGGAGTCGTTGCGAGAGAGAGAGAGCAAGCTGATGNNNAGATCGGA |
| 49482_2 | TATGAGTGTGGAGTCGTTGCGAGAGAGAGAGAGCAAGCTGATGNNNAGATCGGA |
| 36271_1 | TATGAGTGTGGAGTCGTTGCATGAGGACAACTGGCAAACTTTGNNNAGATCGGAA |
| 36271_2 | TATGAGTGTGGAGTCGTTGCATGAGGACAACTGGCAAACTTTGNNNAGATCGGAA |
| 50742_1 | TATGAGTGTGGAGTCGTTGCGATTTTCGAACGCACCTCCTTCNNNAGATCGGAAGA |
| 50742_2 | TATGAGTGTGGAGTCGTTGCGATTTTCGAACGCACCTCCTTCNNNAGATCGGAAGA |
| 31945_1 | TATGAGTGTGGAGTCGTTGCTTTGGACTTGTGTGGGTTCTGTANNNAGATCGGAAG |
| 31945_2 | TATGAGTGTGGAGTCGTTGCTTTGGACTTGTGTGGGTTCTGTANNNAGATCGGAAG |
| 36214_1 | TATGAGTGTGGAGTCGTTGCGGAGTGCTTTTTCTCTGAGTTGCNNNAGATCGGAAG |
| 36214_2 | TATGAGTGTGGAGTCGTTGCGGAGTGCTTTTTCTCTGAGTTGCNNNAGATCGGAAG |
| 42257_1 | TATGAGTGTGGAGTCGTTGCCTGAGTTCAGGCTATCTGCTCATNNNAGATCGGAAG |
| 42257_2 | TATGAGTGTGGAGTCGTTGCCTGAGTTCAGGCTATCTGCTCATNNNAGATCGGAAG |
| 37366_1 | TATGAGTGTGGAGTCGTTGCACTAAAAAAAAACAGCTGGGCACANNNAGATCGGAA |
| 37366_2 | TATGAGTGTGGAGTCGTTGCACTAAAAAAAAACAGCTGGGCACANNNAGATCGGAA |
| 32378_1 | TATGAGTGTGGAGTCGTTGCCAGAATAGCTTGAACCCAGGAGGNNNAGATCGGAA |
| 32378_2 | TATGAGTGTGGAGTCGTTGCCAGAATAGCTTGAACCCAGGAGGNNNAGATCGGAA |
| 32755_1 | TATGAGTGTGGAGTCGTTGCAGGCTCTTCTGTTCCTTTTACCTNNNAGATCGGAAGA |
| 32755_2 | TATGAGTGTGGAGTCGTTGCAGGCTCTTCTGTTCCTTTTACCTNNNAGATCGGAAGA |
| 33129_1 | TATGAGTGTGGAGTCGTTGCGTCAGCAATCACAGCAACACATANNNAGATCGGAAG |
| 33129_2 | TATGAGTGTGGAGTCGTTGCGTCAGCAATCACAGCAACACATANNNAGATCGGAAG |
| 33356_1 | TATGAGTGTGGAGTCGTTGCACAGTCTTGCTCTATTGCCTAGGNNNAGATCGGAAG |
| 33356_2 | TATGAGTGTGGAGTCGTTGCACAGTCTTGCTCTATTGCCTAGGNNNAGATCGGAAG |
| 29994_1 | TATGAGTGTGGAGTCGTTGCCAGTAACAGAGGACACGAGTTCANNNAGATCGGAA |
| 29994_2 | TATGAGTGTGGAGTCGTTGCCAGTAACAGAGGACACGAGTTCANNNAGATCGGAA |
| 49544_1 | TATGAGTGTGGAGTCGTTGCGAGACAGGGTTTTGCCATATTGGNNNAGATCGGAA |

| | |
|---|---|
| 49544_2 | TATGAGTGTGGAGTCGTTGCGAGACAGGGTTTTGCCATATTGGNNNAGATCGGAA |
| 34906_1 | TATGAGTGTGGAGTCGTTGCCACCATGCCCAGCTAATTTTTGTNNNAGATCGGAAG |
| 34906_2 | TATGAGTGTGGAGTCGTTGCCACCATGCCCAGCTAATTTTTGTNNNAGATCGGAAG |
| 37254_1 | TATGAGTGTGGAGTCGTTGCGAACTGGAAGGGGAAAGTTGTCTNNNAGATCGGAA |
| 37254_2 | TATGAGTGTGGAGTCGTTGCGAACTGGAAGGGGAAAGTTGTCTNNNAGATCGGAA |
| 13456_1 | TATGAGTGTGGAGTCGTTGCCTTTAAGATAAAACACAGGCTTANNNAGATCGGAAG |
| 13456_2 | TATGAGTGTGGAGTCGTTGCCTTTAAGATAAAACACAGGCTTANNNAGATCGGAAG |
| 38412_1 | TATGAGTGTGGAGTCGTTGCAGCAAGGGCATCAAATAGGGAATNNNAGATCGGAA |
| 38412_2 | TATGAGTGTGGAGTCGTTGCAGCAAGGGCATCAAATAGGGAATNNNAGATCGGAA |
| 49657_1 | TATGAGTGTGGAGTCGTTGCGTATACCTCCTCCAGCCATGAAGNNNAGATCGGAAG |
| 49657_2 | TATGAGTGTGGAGTCGTTGCGTATACCTCCTCCAGCCATGAAGNNNAGATCGGAAG |
| 33275_1 | TATGAGTGTGGAGTCGTTGCCGGGATCTCAGCTCACTGTAAGNNNAGATCGGAAG |
| 33275_2 | TATGAGTGTGGAGTCGTTGCCGGGATCTCAGCTCACTGTAAGNNNAGATCGGAAG |
| 53292_1 | TATGAGTGTGGAGTCGTTGCCTGATCCAGGAGGCCGCATCNNNAGATCGGAAGAG |
| 53292_2 | TATGAGTGTGGAGTCGTTGCCTGATCCAGGAGGCCGCATCNNNAGATCGGAAGAG |
| 36103_1 | TATGAGTGTGGAGTCGTTGCACATGAACATGGGCATTACACTGNNNAGATCGGAA |
| 36103_2 | TATGAGTGTGGAGTCGTTGCACATGAACATGGGCATTACACTGNNNAGATCGGAA |
| 34529_1 | TATGAGTGTGGAGTCGTTGCGCCAAAGGGTTCTCTCTCTTTCTNNNAGATCGGAAG |
| 34529_2 | TATGAGTGTGGAGTCGTTGCGCCAAAGGGTTCTCTCTCTTTCTNNNAGATCGGAAG |
| 7399_1 | TATGAGTGTGGAGTCGTTGCGAAACACAGCCAAATACCAGTCGNNNAGATCGGAA |
| 7399_2 | TATGAGTGTGGAGTCGTTGCGAAACACAGCCAAATACCAGTCGNNNAGATCGGAA |
| 42117_1 | TATGAGTGTGGAGTCGTTGCGGTCTCGAACTCCTAACCTCAAGNNNAGATCGGAAG |
| 42117_2 | TATGAGTGTGGAGTCGTTGCGGTCTCGAACTCCTAACCTCAAGNNNAGATCGGAAG |
| 42141_1 | TATGAGTGTGGAGTCGTTGCCCAAGTAGCTGGGATTACAGGTANNNAGATCGGAA |
| 42141_2 | TATGAGTGTGGAGTCGTTGCCCAAGTAGCTGGGATTACAGGTANNNAGATCGGAA |
| 32178_1 | TATGAGTGTGGAGTCGTTGCTTATCATGCTTCACCTTCCCACANNNAGATCGGAAGA |
| 32178_2 | TATGAGTGTGGAGTCGTTGCTTATCATGCTTCACCTTCCCACANNNAGATCGGAAGA |
| 9253_1 | TATGAGTGTGGAGTCGTTGCACCTAAAGCAAAGGCTTTTTGCTGNNNAGATCGGAA |
| 9253_2 | TATGAGTGTGGAGTCGTTGCACCTAAAGCAAAGGCTTTTTGCTGNNNAGATCGGAA |
| 45156_1 | TATGAGTGTGGAGTCGTTGCACCTTTTTGGAGCAAGAGAAGGANNNAGATCGGAA |
| 45156_2 | TATGAGTGTGGAGTCGTTGCACCTTTTTGGAGCAAGAGAAGGANNNAGATCGGAA |
| 43071_1 | TATGAGTGTGGAGTCGTTGCTGAGTTGGTTCCACATTTTTGCANNNAGATCGGAAG |
| 43071_2 | TATGAGTGTGGAGTCGTTGCTGAGTTGGTTCCACATTTTTGCANNNAGATCGGAAG |
| 42370_1 | TATGAGTGTGGAGTCGTTGCTTTTTGAGACAGGGTCTTGCTCTNNNAGATCGGAAG |
| 42370_2 | TATGAGTGTGGAGTCGTTGCTTTTTGAGACAGGGTCTTGCTCTNNNAGATCGGAAG |
| 32308_1 | TATGAGTGTGGAGTCGTTGCCACTTGTGTGGGTATGTGTATCTNNNAGATCGGAAG |
| 32308_2 | TATGAGTGTGGAGTCGTTGCCACTTGTGTGGGTATGTGTATCTNNNAGATCGGAAG |
| 10539_1 | TATGAGTGTGGAGTCGTTGCTGGCAGAAGTATCAGCAGCATTTNNNAGATCGGAA |
| 10539_2 | TATGAGTGTGGAGTCGTTGCTGGCAGAAGTATCAGCAGCATTTNNNAGATCGGAA |
| 34847_1 | TATGAGTGTGGAGTCGTTGCCCAAGTTCAAGTGATCCTTCTGCNNNAGATCGGAAG |
| 34847_2 | TATGAGTGTGGAGTCGTTGCCCAAGTTCAAGTGATCCTTCTGCNNNAGATCGGAAG |
| 12663_1 | TATGAGTGTGGAGTCGTTGCATCAAGAGGATAAACTGCATTNNNAGATCGGAAGA |
| 12663_2 | TATGAGTGTGGAGTCGTTGCATCAAGAGGATAAACTGCATTNNNAGATCGGAAGA |

| | |
|---|---|
| 30075_1 | TATGAGTGTGGAGTCGTTGCACCACCAATATCTCAGGTATGCANNNAGATCGGAAG |
| 30075_2 | TATGAGTGTGGAGTCGTTGCACCACCAATATCTCAGGTATGCANNNAGATCGGAAG |
| 50353_1 | TATGAGTGTGGAGTCGTTGCGGCTTACGAGTCATGTAACCAGANNNAGATCGGAA |
| 50353_2 | TATGAGTGTGGAGTCGTTGCGGCTTACGAGTCATGTAACCAGANNNAGATCGGAA |
| 37134_1 | TATGAGTGTGGAGTCGTTGCACAACAAAGAAAAATCAGCAGTGANNNAGATCGGA |
| 37134_2 | TATGAGTGTGGAGTCGTTGCACAACAAAGAAAAATCAGCAGTGANNNAGATCGGA |
| 3188_1 | TATGAGTGTGGAGTCGTTGCCCCACCTCTTACCGATTTCTTCCNNNAGATCGGAAGA |
| 3188_2 | TATGAGTGTGGAGTCGTTGCCCCACCTCTTACCGATTTCTTCCNNNAGATCGGAAGA |
| 51191_1 | TATGAGTGTGGAGTCGTTGCTGGAGCACTTCAAGCTTCCTTATNNNAGATCGGAAG |
| 51191_2 | TATGAGTGTGGAGTCGTTGCTGGAGCACTTCAAGCTTCCTTATNNNAGATCGGAAG |
| 40516_1 | TATGAGTGTGGAGTCGTTGCATGTGTAATGAAGGAAGACCCCCNNNAGATCGGAA |
| 40516_2 | TATGAGTGTGGAGTCGTTGCATGTGTAATGAAGGAAGACCCCCNNNAGATCGGAA |
| 44566_1 | TATGAGTGTGGAGTCGTTGCCACACACACCCCACATTTTCTTTNNNAGATCGGAAGA |
| 44566_2 | TATGAGTGTGGAGTCGTTGCCACACACACCCCACATTTTCTTTNNNAGATCGGAAGA |
| 38728_1 | TATGAGTGTGGAGTCGTTGCACCTGCAATTCTAGCTACTCGAGNNNAGATCGGAAG |
| 38728_2 | TATGAGTGTGGAGTCGTTGCACCTGCAATTCTAGCTACTCGAGNNNAGATCGGAAG |
| 41051_1 | TATGAGTGTGGAGTCGTTGCTGCACATTCCAGGTACTAGACACNNNAGATCGGAAG |
| 41051_2 | TATGAGTGTGGAGTCGTTGCTGCACATTCCAGGTACTAGACACNNNAGATCGGAAG |
| 50282_1 | TATGAGTGTGGAGTCGTTGCAATACAAAATTAGCCGGGCATGGNNNAGATCGGAA |
| 50282_2 | TATGAGTGTGGAGTCGTTGCAATACAAAATTAGCCGGGCATGGNNNAGATCGGAA |
| 13725_1 | TATGAGTGTGGAGTCGTTGCTCGCATAAATATTTTAGGAAGTGNNNAGATCGGAAG |
| 13725_2 | TATGAGTGTGGAGTCGTTGCTCGCATAAATATTTTAGGAAGTGNNNAGATCGGAAG |
| 35898_1 | TATGAGTGTGGAGTCGTTGCAACATTTCTGCAGAGGTTTGACCNNNAGATCGGAAG |
| 35898_2 | TATGAGTGTGGAGTCGTTGCAACATTTCTGCAGAGGTTTGACCNNNAGATCGGAAG |
| 35359_1 | TATGAGTGTGGAGTCGTTGCTTCAAGCAATTCTCCTGCCTCAGNNNAGATCGGAAG |
| 35359_2 | TATGAGTGTGGAGTCGTTGCTTCAAGCAATTCTCCTGCCTCAGNNNAGATCGGAAG |
| 33641_1 | TATGAGTGTGGAGTCGTTGCGACAGAGTATCGATCGGTCTTGTNNNAGATCGGAA |
| 33641_2 | TATGAGTGTGGAGTCGTTGCGACAGAGTATCGATCGGTCTTGTNNNAGATCGGAA |
| 48176_1 | TATGAGTGTGGAGTCGTTGCACTCAGGAGTTCAAGTGATCCACNNNAGATCGGAAG |
| 48176_2 | TATGAGTGTGGAGTCGTTGCACTCAGGAGTTCAAGTGATCCACNNNAGATCGGAAG |
| 11459_1 | TATGAGTGTGGAGTCGTTGCACCCAAACTGATACACATGTTTTCTNNNAGATCGGAA |
| 11459_2 | TATGAGTGTGGAGTCGTTGCACCCAAACTGATACACATGTTTTCTNNNAGATCGGAA |
| 47155_1 | TATGAGTGTGGAGTCGTTGCCCTTCCAAGTAGCTGGGATTACANNNAGATCGGAAG |
| 47155_2 | TATGAGTGTGGAGTCGTTGCCCTTCCAAGTAGCTGGGATTACANNNAGATCGGAAG |
| 44366_1 | TATGAGTGTGGAGTCGTTGCGAAGAATGCCCATGTCAGGGAAGNNNAGATCGGAA |
| 44366_2 | TATGAGTGTGGAGTCGTTGCGAAGAATGCCCATGTCAGGGAAGNNNAGATCGGAA |
| 43972_1 | TATGAGTGTGGAGTCGTTGCTATTTAGGAGTGTGGGATTCCCGNNNAGATCGGAA |
| 43972_2 | TATGAGTGTGGAGTCGTTGCTATTTAGGAGTGTGGGATTCCCGNNNAGATCGGAA |
| 41624_1 | TATGAGTGTGGAGTCGTTGCCTCCTGAGTAGCTGGGACTACAGNNNAGATCGGAA |
| 41624_2 | TATGAGTGTGGAGTCGTTGCCTCCTGAGTAGCTGGGACTACAGNNNAGATCGGAA |
| 29890_1 | TATGAGTGTGGAGTCGTTGCTGTGAACTGGGTTACTGACTGAGNNNAGATCGGAA |
| 29890_2 | TATGAGTGTGGAGTCGTTGCTGTGAACTGGGTTACTGACTGAGNNNAGATCGGAA |
| 39971_1 | TATGAGTGTGGAGTCGTTGCATCTCAGCTCACTGTAACCTCTGNNNAGATCGGAAG |

| | |
|---|---|
| 39971_2 | TATGAGTGTGGAGTCGTTGCATCTCAGCTCACTGTAACCTCTGNNNAGATCGGAAG |
| 30456_1 | TATGAGTGTGGAGTCGTTGCGAGAAGGGATCTAGGACTCCTGGNNNAGATCGGAA |
| 30456_2 | TATGAGTGTGGAGTCGTTGCGAGAAGGGATCTAGGACTCCTGGNNNAGATCGGAA |
| 34025_1 | TATGAGTGTGGAGTCGTTGCTAATGTCGCGATCTCGGCTCNNNAGATCGGAAGAG |
| 34025_2 | TATGAGTGTGGAGTCGTTGCTAATGTCGCGATCTCGGCTCNNNAGATCGGAAGAG |
| 36491_1 | TATGAGTGTGGAGTCGTTGCGCAGTGAGCTATGATCACAACACNNNAGATCGGAA |
| 36491_2 | TATGAGTGTGGAGTCGTTGCGCAGTGAGCTATGATCACAACACNNNAGATCGGAA |
| 43878_1 | TATGAGTGTGGAGTCGTTGCACAAGCCAGAATGCAAAACTTCANNNAGATCGGAAG |
| 43878_2 | TATGAGTGTGGAGTCGTTGCACAAGCCAGAATGCAAAACTTCANNNAGATCGGAAG |
| 38829_1 | TATGAGTGTGGAGTCGTTGCGTCATTGCTATGCTGCTTACCTGNNNAGATCGGAAG |
| 38829_2 | TATGAGTGTGGAGTCGTTGCGTCATTGCTATGCTGCTTACCTGNNNAGATCGGAAG |
| 46190_1 | TATGAGTGTGGAGTCGTTGCCTGAGTAGCTGGGACTACAGTCANNNAGATCGGAA |
| 46190_2 | TATGAGTGTGGAGTCGTTGCCTGAGTAGCTGGGACTACAGTCANNNAGATCGGAA |
| 35830_1 | TATGAGTGTGGAGTCGTTGCCAATCTTGTCTCACTGCAACCTCNNNAGATCGGAAGA |
| 35830_2 | TATGAGTGTGGAGTCGTTGCCAATCTTGTCTCACTGCAACCTCNNNAGATCGGAAGA |
| 46047_1 | TATGAGTGTGGAGTCGTTGCGCCTCCTGAGTAGCTGGTATTACNNNAGATCGGAAG |
| 46047_2 | TATGAGTGTGGAGTCGTTGCGCCTCCTGAGTAGCTGGTATTACNNNAGATCGGAAG |
| 50403_1 | TATGAGTGTGGAGTCGTTGCATAGAAGGTCCGGAAACAATGGGNNNAGATCGGA |
| 50403_2 | TATGAGTGTGGAGTCGTTGCATAGAAGGTCCGGAAACAATGGGNNNAGATCGGA |
| 34326_1 | TATGAGTGTGGAGTCGTTGCGGTTACTATTAGCTCGGTCAGGGNNNAGATCGGAA |
| 34326_2 | TATGAGTGTGGAGTCGTTGCGGTTACTATTAGCTCGGTCAGGGNNNAGATCGGAA |
| 38380_1 | TATGAGTGTGGAGTCGTTGCTGTGCTTATAGTCCCAGCAACTTNNNAGATCGGAAG |
| 38380_2 | TATGAGTGTGGAGTCGTTGCTGTGCTTATAGTCCCAGCAACTTNNNAGATCGGAAG |
| 36085_1 | TATGAGTGTGGAGTCGTTGCTCCACAAAAGCAATTGAAAACTCTNNNAGATCGGAA |
| 36085_2 | TATGAGTGTGGAGTCGTTGCTCCACAAAAGCAATTGAAAACTCTNNNAGATCGGAA |
| 49452_1 | TATGAGTGTGGAGTCGTTGCACATGGATGGACCCAAAGTCATTNNNAGATCGGAA |
| 49452_2 | TATGAGTGTGGAGTCGTTGCACATGGATGGACCCAAAGTCATTNNNAGATCGGAA |
| 1949_1 | TATGAGTGTGGAGTCGTTGCCAAGCAATGTGGATGATGACAGCNNNAGATCGGAA |
| 1949_2 | TATGAGTGTGGAGTCGTTGCCAAGCAATGTGGATGATGACAGCNNNAGATCGGAA |
| 51989_1 | TATGAGTGTGGAGTCGTTGCCTCTTCTGGGTCTTGCTTTCTCTNNNAGATCGGAAGA |
| 51989_2 | TATGAGTGTGGAGTCGTTGCCTCTTCTGGGTCTTGCTTTCTCTNNNAGATCGGAAGA |
| 40927_1 | TATGAGTGTGGAGTCGTTGCGCATTTCTGATTGCTTCCTTTGGNNNAGATCGGAAG |
| 40927_2 | TATGAGTGTGGAGTCGTTGCGCATTTCTGATTGCTTCCTTTGGNNNAGATCGGAAG |
| 34662_1 | TATGAGTGTGGAGTCGTTGCGGGAGCCCAGAAATAGAGTACTGNNNAGATCGGAA |
| 34662_2 | TATGAGTGTGGAGTCGTTGCGGGAGCCCAGAAATAGAGTACTGNNNAGATCGGAA |
| 32337_1 | TATGAGTGTGGAGTCGTTGCTGACAAGAAGGACAGAAGAGCTGNNNAGATCGGA |
| 32337_2 | TATGAGTGTGGAGTCGTTGCTGACAAGAAGGACAGAAGAGCTGNNNAGATCGGA |
| 40539_1 | TATGAGTGTGGAGTCGTTGCTCTTCCACTGATGCACAATGTCTNNNAGATCGGAAG |
| 40539_2 | TATGAGTGTGGAGTCGTTGCTCTTCCACTGATGCACAATGTCTNNNAGATCGGAAG |
| 30042_1 | TATGAGTGTGGAGTCGTTGCGAGAAGAAGAAGGAGTAAGGCTTNNNAGATCGGA |
| 30042_2 | TATGAGTGTGGAGTCGTTGCGAGAAGAAGAAGGAGTAAGGCTTNNNAGATCGGA |
| 41208_1 | TATGAGTGTGGAGTCGTTGCACTGAGAATGATGGTTTCCAGCTNNNAGATCGGAA |
| 41208_2 | TATGAGTGTGGAGTCGTTGCACTGAGAATGATGGTTTCCAGCTNNNAGATCGGAA |

| | |
|---|---|
| 32373_1 | TATGAGTGTGGAGTCGTTGCGGATAGCTTGAGATCAGGAGGTGNNNAGATCGGA |
| 32373_2 | TATGAGTGTGGAGTCGTTGCGGATAGCTTGAGATCAGGAGGTGNNNAGATCGGA |
| 2472_1 | TATGAGTGTGGAGTCGTTGCTGGAGAGAGGAATGTGGAGGAAGNNNAGATCGG |
| 2472_2 | TATGAGTGTGGAGTCGTTGCTGGAGAGAGGAATGTGGAGGAAGNNNAGATCGG |
| 46441_1 | TATGAGTGTGGAGTCGTTGCCTCCTGGGTAGCTGGAATTACAGNNNAGATCGGAA |
| 46441_2 | TATGAGTGTGGAGTCGTTGCCTCCTGGGTAGCTGGAATTACAGNNNAGATCGGAA |
| 33364_1 | TATGAGTGTGGAGTCGTTGCAAGTAGCTGGGACTACAAGCACNNNAGATCGGAAG |
| 33364_2 | TATGAGTGTGGAGTCGTTGCAAGTAGCTGGGACTACAAGCACNNNAGATCGGAAG |
| 51480_1 | TATGAGTGTGGAGTCGTTGCGGCACCCAGTAAATACCTGTAGANNNAGATCGGAA |
| 51480_2 | TATGAGTGTGGAGTCGTTGCGGCACCCAGTAAATACCTGTAGANNNAGATCGGAA |
| 41031_1 | TATGAGTGTGGAGTCGTTGCAAAAAGAAAGCCACTTCCCACTGNNNAGATCGGAAG |
| 41031_2 | TATGAGTGTGGAGTCGTTGCAAAAAGAAAGCCACTTCCCACTGNNNAGATCGGAAG |
| 36821_1 | TATGAGTGTGGAGTCGTTGCGGTGAAACCCCGTCTCTACTAAANNNAGATCGGAAG |
| 36821_2 | TATGAGTGTGGAGTCGTTGCGGTGAAACCCCGTCTCTACTAAANNNAGATCGGAAG |
| 10368_1 | TATGAGTGTGGAGTCGTTGCAGAGCACATGGCATATTATTGGCANNNAGATCGGA |
| 10368_2 | TATGAGTGTGGAGTCGTTGCAGAGCACATGGCATATTATTGGCANNNAGATCGGA |
| 13146_1 | TATGAGTGTGGAGTCGTTGCTTACAATCTGCAAAGTACAAATTNNNAGATCGGAAG |
| 13146_2 | TATGAGTGTGGAGTCGTTGCTTACAATCTGCAAAGTACAAATTNNNAGATCGGAAG |
| 31473_1 | TATGAGTGTGGAGTCGTTGCCACACACACACACCATGAAATATNNNAGATCGGAAG |
| 31473_2 | TATGAGTGTGGAGTCGTTGCCACACACACACACCATGAAATATNNNAGATCGGAAG |
| 40509_1 | TATGAGTGTGGAGTCGTTGCACTTGTAATCCCAGCAATCCCAGNNNAGATCGGAAG |
| 40509_2 | TATGAGTGTGGAGTCGTTGCACTTGTAATCCCAGCAATCCCAGNNNAGATCGGAAG |
| 41562_1 | TATGAGTGTGGAGTCGTTGCGCACCACACCCAGCTAAATTTTTNNNAGATCGGAAG |
| 41562_2 | TATGAGTGTGGAGTCGTTGCGCACCACACCCAGCTAAATTTTTNNNAGATCGGAAG |
| 34321_1 | TATGAGTGTGGAGTCGTTGCGTCAGAAGAGACAGAGGGAACAGNNNAGATCGG |
| 34321_2 | TATGAGTGTGGAGTCGTTGCGTCAGAAGAGACAGAGGGAACAGNNNAGATCGG |
| 34621_1 | TATGAGTGTGGAGTCGTTGCGTCAGAAATGCGAACACCTGAAANNNAGATCGGAA |
| 34621_2 | TATGAGTGTGGAGTCGTTGCGTCAGAAATGCGAACACCTGAAANNNAGATCGGAA |
| 36368_1 | TATGAGTGTGGAGTCGTTGCTGAGATAGCACCTCTGTACTCCANNNAGATCGGAAG |
| 36368_2 | TATGAGTGTGGAGTCGTTGCTGAGATAGCACCTCTGTACTCCANNNAGATCGGAAG |
| 12772_1 | TATGAGTGTGGAGTCGTTGCCACCTGTTGATCTCTCCTTTNNNAGATCGGAAGAGCA |
| 12772_2 | TATGAGTGTGGAGTCGTTGCCACCTGTTGATCTCTCCTTTNNNAGATCGGAAGAGCA |
| 13341_1 | TATGAGTGTGGAGTCGTTGCACTGTTAACTACTGTACAACCNNNAGATCGGAAGAG |
| 13341_2 | TATGAGTGTGGAGTCGTTGCACTGTTAACTACTGTACAACCNNNAGATCGGAAGAG |
| 32172_1 | TATGAGTGTGGAGTCGTTGCGTTCAAGTGATTCTCCTGCCTCANNNAGATCGGAAG |
| 32172_2 | TATGAGTGTGGAGTCGTTGCGTTCAAGTGATTCTCCTGCCTCANNNAGATCGGAAG |
| 48747_1 | TATGAGTGTGGAGTCGTTGCCATAGAGTCAGCAAACAGCAACCNNNAGATCGGAA |
| 48747_2 | TATGAGTGTGGAGTCGTTGCCATAGAGTCAGCAAACAGCAACCNNNAGATCGGAA |
| 3077_1 | TATGAGTGTGGAGTCGTTGCGGCAGAAGAAGAGGATGTTCAGGNNNAGATCGG |
| 3077_2 | TATGAGTGTGGAGTCGTTGCGGCAGAAGAAGAGGATGTTCAGGNNNAGATCGG |
| 12519_1 | TATGAGTGTGGAGTCGTTGCCTTTTATAGCCTAATGAACACACNNNAGATCGGAAG |
| 12519_2 | TATGAGTGTGGAGTCGTTGCCTTTTATAGCCTAATGAACACACNNNAGATCGGAAG |
| 40435_1 | TATGAGTGTGGAGTCGTTGCTCACCGCAAAAATTAGTGTCACCNNNAGATCGGAAG |

| | |
|---|---|
| 40435_2 | TATGAGTGTGGAGTCGTTGCTCACCGCAAAAATTAGTGTCACCNNNAGATCGGAAGA |
| 37237_1 | TATGAGTGTGGAGTCGTTGCAGGAGCTGGTACCATTTCTTCTGNNNAGATCGGAAG |
| 37237_2 | TATGAGTGTGGAGTCGTTGCAGGAGCTGGTACCATTTCTTCTGNNNAGATCGGAAG |
| 13279_1 | TATGAGTGTGGAGTCGTTGCCACTAGTACCTCCATGTTATTTANNNAGATCGGAAGA |
| 13279_2 | TATGAGTGTGGAGTCGTTGCCACTAGTACCTCCATGTTATTTANNNAGATCGGAAGA |
| 44862_1 | TATGAGTGTGGAGTCGTTGCGCACCCCAACCAGAATTCAATAGNNNAGATCGGAAG |
| 44862_2 | TATGAGTGTGGAGTCGTTGCGCACCCCAACCAGAATTCAATAGNNNAGATCGGAAG |
| 34851_1 | TATGAGTGTGGAGTCGTTGCCACAATCTCAGCTCACTGCAACNNNAGATCGGAAGAC |
| 34851_2 | TATGAGTGTGGAGTCGTTGCCACAATCTCAGCTCACTGCAACNNNAGATCGGAAGAC |
| 51936_1 | TATGAGTGTGGAGTCGTTGCAGGTAGAGGATGGAACTGGGTTANNNAGATCGGAA |
| 51936_2 | TATGAGTGTGGAGTCGTTGCAGGTAGAGGATGGAACTGGGTTANNNAGATCGGAA |
| 35626_1 | TATGAGTGTGGAGTCGTTGCGAAGGTGAGTCTCGTTCTGTCACNNNAGATCGGAAG |
| 35626_2 | TATGAGTGTGGAGTCGTTGCGAAGGTGAGTCTCGTTCTGTCACNNNAGATCGGAAG |
| 44969_1 | TATGAGTGTGGAGTCGTTGCAGATAAAACACAGGCAGGTGGATNNNAGATCGGAA |
| 44969_2 | TATGAGTGTGGAGTCGTTGCAGATAAAACACAGGCAGGTGGATNNNAGATCGGAA |
| 43996_1 | TATGAGTGTGGAGTCGTTGCATAAAGCAGCTGTGGCAAAATGTNNNAGATCGGAAG |
| 43996_2 | TATGAGTGTGGAGTCGTTGCATAAAGCAGCTGTGGCAAAATGTNNNAGATCGGAAG |
| 2133_1 | TATGAGTGTGGAGTCGTTGCAGCACATAGCTTGTAGGGTTATCANNNAGATCGGAA |
| 2133_2 | TATGAGTGTGGAGTCGTTGCAGCACATAGCTTGTAGGGTTATCANNNAGATCGGAA |
| 40474_1 | TATGAGTGTGGAGTCGTTGCTAAAGGCTACTCAACATGTGCCANNNAGATCGGAAG |
| 40474_2 | TATGAGTGTGGAGTCGTTGCTAAAGGCTACTCAACATGTGCCANNNAGATCGGAAG |
| 38419_1 | TATGAGTGTGGAGTCGTTGCTATGGAACCAACCCAAATGCCTANNNAGATCGGAAG |
| 38419_2 | TATGAGTGTGGAGTCGTTGCTATGGAACCAACCCAAATGCCTANNNAGATCGGAAG |
| 49351_1 | TATGAGTGTGGAGTCGTTGCCGCTTGGATGTTCATTGGGAATTNNNAGATCGGAAG |
| 49351_2 | TATGAGTGTGGAGTCGTTGCCGCTTGGATGTTCATTGGGAATTNNNAGATCGGAAG |
| 38278_1 | TATGAGTGTGGAGTCGTTGCATATTGCTCAGCTTTCTGTCACCNNNAGATCGGAAGA |
| 38278_2 | TATGAGTGTGGAGTCGTTGCATATTGCTCAGCTTTCTGTCACCNNNAGATCGGAAGA |
| 52218_1 | TATGAGTGTGGAGTCGTTGCTAGAAGGCAAGAAGGCAATCAGTNNNAGATCGGAA |
| 52218_2 | TATGAGTGTGGAGTCGTTGCTAGAAGGCAAGAAGGCAATCAGTNNNAGATCGGAA |
| 47186_1 | TATGAGTGTGGAGTCGTTGCCTGAATAGCTGGGATTACAGGCANNNAGATCGGAAG |
| 47186_2 | TATGAGTGTGGAGTCGTTGCCTGAATAGCTGGGATTACAGGCANNNAGATCGGAAG |
| 41931_1 | TATGAGTGTGGAGTCGTTGCAGAAAAGGTCTCACTCTGTCACCNNNAGATCGGAAG |
| 41931_2 | TATGAGTGTGGAGTCGTTGCAGAAAAGGTCTCACTCTGTCACCNNNAGATCGGAAG |
| 30424_1 | TATGAGTGTGGAGTCGTTGCTCAATCTTGATTACCAGGTGGCANNNAGATCGGAAG |
| 30424_2 | TATGAGTGTGGAGTCGTTGCTCAATCTTGATTACCAGGTGGCANNNAGATCGGAAG |
| 44583_1 | TATGAGTGTGGAGTCGTTGCGCCATAGTTCCAGAGGTCATTGANNNAGATCGGAAG |
| 44583_2 | TATGAGTGTGGAGTCGTTGCGCCATAGTTCCAGAGGTCATTGANNNAGATCGGAAG |
| 52466_1 | TATGAGTGTGGAGTCGTTGCACTTTTTGGATGTCCAGGTTTGGNNNAGATCGGAAG |
| 52466_2 | TATGAGTGTGGAGTCGTTGCACTTTTTGGATGTCCAGGTTTGGNNNAGATCGGAAG |
| 53118_1 | TATGAGTGTGGAGTCGTTGCCCAATCCTCCTGCAAACATTTCANNNAGATCGGAAGA |
| 53118_2 | TATGAGTGTGGAGTCGTTGCCCAATCCTCCTGCAAACATTTCANNNAGATCGGAAGA |
| 34763_1 | TATGAGTGTGGAGTCGTTGCCTCCTGAGTAGCTGGGACTACAGNNNAGATCGGAAG |
| 34763_2 | TATGAGTGTGGAGTCGTTGCCTCCTGAGTAGCTGGGACTACAGNNNAGATCGGAAG |

| | |
|---|---|
| 33885_1 | TATGAGTGTGGAGTCGTTGCTTGAGACGTAGTCTTGCTCTGTCNNNAGATCGGAAG |
| 33885_2 | TATGAGTGTGGAGTCGTTGCTTGAGACGTAGTCTTGCTCTGTCNNNAGATCGGAAG |
| 1481_1 | TATGAGTGTGGAGTCGTTGCCGAAGTTCACTCTGTCGCCCNNNAGATCGGAAGAGCA |
| 1481_2 | TATGAGTGTGGAGTCGTTGCCGAAGTTCACTCTGTCGCCCNNNAGATCGGAAGAGCA |
| 49407_1 | TATGAGTGTGGAGTCGTTGCTGTTAGCCAGGATGGTCTCAATCNNNAGATCGGAAG |
| 49407_2 | TATGAGTGTGGAGTCGTTGCTGTTAGCCAGGATGGTCTCAATCNNNAGATCGGAAG |
| 50246_1 | TATGAGTGTGGAGTCGTTGCGGTGGTCAACAGCATCTTCAAAGNNNAGATCGGAAG |
| 50246_2 | TATGAGTGTGGAGTCGTTGCGGTGGTCAACAGCATCTTCAAAGNNNAGATCGGAAG |
| 42805_1 | TATGAGTGTGGAGTCGTTGCACACACACACAAATTTAGCTGGGNNNAGATCGGAAG |
| 42805_2 | TATGAGTGTGGAGTCGTTGCACACACACACAAATTTAGCTGGGNNNAGATCGGAAG |
| 52044_1 | TATGAGTGTGGAGTCGTTGCCCAAACACTCTTCCCAGTCTTTGNNNAGATCGGAAGA |
| 52044_2 | TATGAGTGTGGAGTCGTTGCCCAAACACTCTTCCCAGTCTTTGNNNAGATCGGAAGA |
| 32072_1 | TATGAGTGTGGAGTCGTTGCCGCCCGGCTAACTTTTGTATTTTNNNAGATCGGAAGA |
| 32072_2 | TATGAGTGTGGAGTCGTTGCCGCCCGGCTAACTTTTGTATTTTNNNAGATCGGAAGA |
| 34050_1 | TATGAGTGTGGAGTCGTTGCTAGCAGAGCTACTGATTTGTGCANNNAGATCGGAAG |
| 34050_2 | TATGAGTGTGGAGTCGTTGCTAGCAGAGCTACTGATTTGTGCANNNAGATCGGAAG |
| 32500_1 | TATGAGTGTGGAGTCGTTGCGATCTTGGCTCGCTGCAAATTCNNNAGATCGGAAGA |
| 32500_2 | TATGAGTGTGGAGTCGTTGCGATCTTGGCTCGCTGCAAATTCNNNAGATCGGAAGA |
| 36776_1 | TATGAGTGTGGAGTCGTTGCGGAATAAAGTAAGCCACACAGCCNNNAGATCGGAAG |
| 36776_2 | TATGAGTGTGGAGTCGTTGCGGAATAAAGTAAGCCACACAGCCNNNAGATCGGAAG |
| 35636_1 | TATGAGTGTGGAGTCGTTGCTTTTTGGAGACTGAGTCCCACTCNNNAGATCGGAAGA |
| 35636_2 | TATGAGTGTGGAGTCGTTGCTTTTTGGAGACTGAGTCCCACTCNNNAGATCGGAAGA |
| 52772_1 | TATGAGTGTGGAGTCGTTGCTACACAAAGTTGTATGCTGTGCGNNNAGATCGGAAG |
| 52772_2 | TATGAGTGTGGAGTCGTTGCTACACAAAGTTGTATGCTGTGCGNNNAGATCGGAAG |
| 40913_1 | TATGAGTGTGGAGTCGTTGCAAACAAAACAAAAGACTAGGAAGGTNNNAGATCGG |
| 40913_2 | TATGAGTGTGGAGTCGTTGCAAACAAAACAAAAGACTAGGAAGGTNNNAGATCGG |
| 10299_1 | TATGAGTGTGGAGTCGTTGCTCAGCAATCCCTGTTGACCTTTTNNNAGATCGGAAGA |
| 10299_2 | TATGAGTGTGGAGTCGTTGCTCAGCAATCCCTGTTGACCTTTTNNNAGATCGGAAGA |
| 41506_1 | TATGAGTGTGGAGTCGTTGCAACCACTGCTTTTGACTTTCTGGNNNAGATCGGAAGA |
| 41506_2 | TATGAGTGTGGAGTCGTTGCAACCACTGCTTTTGACTTTCTGGNNNAGATCGGAAGA |
| 36052_1 | TATGAGTGTGGAGTCGTTGCAGAGCTCAATCCCAAGATAGCTCNNNAGATCGGAAG |
| 36052_2 | TATGAGTGTGGAGTCGTTGCAGAGCTCAATCCCAAGATAGCTCNNNAGATCGGAAG |
| 29327_1 | TATGAGTGTGGAGTCGTTGCTAAAATGGATCCAAGGGCCAAGANNNAGATCGGAAG |
| 29327_2 | TATGAGTGTGGAGTCGTTGCTAAAATGGATCCAAGGGCCAAGANNNAGATCGGAAG |
| 41496_1 | TATGAGTGTGGAGTCGTTGCTGTGATTGGGACCACAGTCTAAANNNAGATCGGAAG |
| 41496_2 | TATGAGTGTGGAGTCGTTGCTGTGATTGGGACCACAGTCTAAANNNAGATCGGAAG |
| 50308_1 | TATGAGTGTGGAGTCGTTGCGCCCATGTTCTCAGAACCTTCTANNNAGATCGGAAGA |
| 50308_2 | TATGAGTGTGGAGTCGTTGCGCCCATGTTCTCAGAACCTTCTANNNAGATCGGAAGA |
| 42463_1 | TATGAGTGTGGAGTCGTTGCGTAAGATATGCCTTGCTTCCCCTNNNAGATCGGAAGA |
| 42463_2 | TATGAGTGTGGAGTCGTTGCGTAAGATATGCCTTGCTTCCCCTNNNAGATCGGAAGA |
| 49856_1 | TATGAGTGTGGAGTCGTTGCAACTCATTCCAGATTGCCTCGATNNNAGATCGGAAGA |
| 49856_2 | TATGAGTGTGGAGTCGTTGCAACTCATTCCAGATTGCCTCGATNNNAGATCGGAAGA |
| 50359_1 | TATGAGTGTGGAGTCGTTGCCAGCATTACTTCACTGATGCCTGNNNAGATCGGAAGA |

| | |
|---|---|
| 50359_2 | TATGAGTGTGGAGTCGTTGCCAGCATTACTTCACTGATGCCTGNNNAGATCGGAAG |
| 36385_1 | TATGAGTGTGGAGTCGTTGCAGAGAGCTCTGATTGTGCTACTGNNNAGATCGGAAG |
| 36385_2 | TATGAGTGTGGAGTCGTTGCAGAGAGCTCTGATTGTGCTACTGNNNAGATCGGAAG |
| 36334_1 | TATGAGTGTGGAGTCGTTGCCACCCTGGACCTACAAAATCAGANNNAGATCGGAAG |
| 36334_2 | TATGAGTGTGGAGTCGTTGCCACCCTGGACCTACAAAATCAGANNNAGATCGGAAG |
| 42131_1 | TATGAGTGTGGAGTCGTTGCACGCCCGGCTAATTTTTGTATTTNNNAGATCGGAAG |
| 42131_2 | TATGAGTGTGGAGTCGTTGCACGCCCGGCTAATTTTTGTATTTNNNAGATCGGAAG |
| 39630_1 | TATGAGTGTGGAGTCGTTGCGGAACAAACTAAGGATGCTTGGTNNNAGATCGGAA |
| 39630_2 | TATGAGTGTGGAGTCGTTGCGGAACAAACTAAGGATGCTTGGTNNNAGATCGGAA |
| 28765_1 | TATGAGTGTGGAGTCGTTGCATATGTGTGTGTGTGTGTGTGTGNNNAGATCGGAA |
| 28765_2 | TATGAGTGTGGAGTCGTTGCATATGTGTGTGTGTGTGTGTGTGNNNAGATCGGAA |
| 43503_1 | TATGAGTGTGGAGTCGTTGCGAGTAGCTGCGATTACAGGAGTGNNNAGATCGGAA |
| 43503_2 | TATGAGTGTGGAGTCGTTGCGAGTAGCTGCGATTACAGGAGTGNNNAGATCGGAA |
| 46297_1 | TATGAGTGTGGAGTCGTTGCCATGCCTCAGTAGCTGGGATTACNNNAGATCGGAAG |
| 46297_2 | TATGAGTGTGGAGTCGTTGCCATGCCTCAGTAGCTGGGATTACNNNAGATCGGAAG |
| 47487_1 | TATGAGTGTGGAGTCGTTGCCCTCCCAAAGTGCTAGGATTACANNNAGATCGGAAG |
| 47487_2 | TATGAGTGTGGAGTCGTTGCCCTCCCAAAGTGCTAGGATTACANNNAGATCGGAAG |
| 32541_1 | TATGAGTGTGGAGTCGTTGCAAAAAAGACAGACACAGGAGCAGNNNAGATCGGAA |
| 32541_2 | TATGAGTGTGGAGTCGTTGCAAAAAAGACAGACACAGGAGCAGNNNAGATCGGAA |
| 13651_1 | TATGAGTGTGGAGTCGTTGCCAAACAGATTGCATATTATCTGANNNAGATCGGAAG |
| 13651_2 | TATGAGTGTGGAGTCGTTGCCAAACAGATTGCATATTATCTGANNNAGATCGGAAG |
| 53270_1 | TATGAGTGTGGAGTCGTTGCATTCCACTTCCCCTTCTTCCAAANNNAGATCGGAAGA |
| 53270_2 | TATGAGTGTGGAGTCGTTGCATTCCACTTCCCCTTCTTCCAAANNNAGATCGGAAGA |
| 46859_1 | TATGAGTGTGGAGTCGTTGCTGAGTAGATGAGATTACAGGCGCNNNAGATCGGAA |
| 46859_2 | TATGAGTGTGGAGTCGTTGCTGAGTAGATGAGATTACAGGCGCNNNAGATCGGAA |
| 42089_1 | TATGAGTGTGGAGTCGTTGCCTTTGTTCAGGTGTGCTCTCAAGNNNAGATCGGAAG |
| 42089_2 | TATGAGTGTGGAGTCGTTGCCTTTGTTCAGGTGTGCTCTCAAGNNNAGATCGGAAG |
| 35995_1 | TATGAGTGTGGAGTCGTTGCGTTGCTTTCTGGTGAGTCCTTTCNNNAGATCGGAAG |
| 35995_2 | TATGAGTGTGGAGTCGTTGCGTTGCTTTCTGGTGAGTCCTTTCNNNAGATCGGAAG |
| 47564_1 | TATGAGTGTGGAGTCGTTGCCTTGAGCTTGAGGGTCCCTGNNNAGATCGGAAGAG |
| 47564_2 | TATGAGTGTGGAGTCGTTGCCTTGAGCTTGAGGGTCCCTGNNNAGATCGGAAGAG |
| 39016_1 | TATGAGTGTGGAGTCGTTGCAAACCGGATCTGTAAAACTGGGTNNNAGATCGGAA |
| 39016_2 | TATGAGTGTGGAGTCGTTGCAAACCGGATCTGTAAAACTGGGTNNNAGATCGGAA |
| 36355_1 | TATGAGTGTGGAGTCGTTGCGCCAAGTCTCCACATGTTTGTTCNNNAGATCGGAAG |
| 36355_2 | TATGAGTGTGGAGTCGTTGCGCCAAGTCTCCACATGTTTGTTCNNNAGATCGGAAG |
| 47630_1 | TATGAGTGTGGAGTCGTTGCATGAAGTTCTGCCCGTAGATCTGNNNAGATCGGAAG |
| 47630_2 | TATGAGTGTGGAGTCGTTGCATGAAGTTCTGCCCGTAGATCTGNNNAGATCGGAAG |
| 38375_1 | TATGAGTGTGGAGTCGTTGCAGAAAACTTCCCTGCCAACATTGNNNAGATCGGAAG |
| 38375_2 | TATGAGTGTGGAGTCGTTGCAGAAAACTTCCCTGCCAACATTGNNNAGATCGGAAG |
| 32964_1 | TATGAGTGTGGAGTCGTTGCAGAAGTTGCAGTGAGCCAAGATCNNNAGATCGGAA |
| 32964_2 | TATGAGTGTGGAGTCGTTGCAGAAGTTGCAGTGAGCCAAGATCNNNAGATCGGAA |
| 9817_1 | TATGAGTGTGGAGTCGTTGCTCCAAGACCATCACTCAGCCATANNNAGATCGGAAG |
| 9817_2 | TATGAGTGTGGAGTCGTTGCTCCAAGACCATCACTCAGCCATANNNAGATCGGAAG |

| | |
|---|---|
| 8943_1 | TATGAGTGTGGAGTCGTTGCTGTGGGAGTACAAGTACATAGAGTNNNAGATCGGA |
| 8943_2 | TATGAGTGTGGAGTCGTTGCTGTGGGAGTACAAGTACATAGAGTNNNAGATCGGA |
| 38368_1 | TATGAGTGTGGAGTCGTTGCGTTGGAAGGGTTATGAGGGTTGANNNAGATCGGA |
| 38368_2 | TATGAGTGTGGAGTCGTTGCGTTGGAAGGGTTATGAGGGTTGANNNAGATCGGA |
| 2805_1 | TATGAGTGTGGAGTCGTTGCGCTGACATCACAGTCTTCTCTCTNNNAGATCGGAAGA |
| 2805_2 | TATGAGTGTGGAGTCGTTGCGCTGACATCACAGTCTTCTCTCTNNNAGATCGGAAGA |
| 37930_1 | TATGAGTGTGGAGTCGTTGCTGTAATCCTAGCACTTTGGGAGGNNNAGATCGGAA |
| 37930_2 | TATGAGTGTGGAGTCGTTGCTGTAATCCTAGCACTTTGGGAGGNNNAGATCGGAA |
| 228_1 | TATGAGTGTGGAGTCGTTGCCAAGATGGCAGTGCGATGACCGNNNAGATCGGAAG |
| 228_2 | TATGAGTGTGGAGTCGTTGCCAAGATGGCAGTGCGATGACCGNNNAGATCGGAAG |
| 38475_1 | TATGAGTGTGGAGTCGTTGCAAAACAGATGCATTGACCAGTGGNNNAGATCGGAA |
| 38475_2 | TATGAGTGTGGAGTCGTTGCAAAACAGATGCATTGACCAGTGGNNNAGATCGGAA |
| 10874_1 | TATGAGTGTGGAGTCGTTGCAGGCAAATGGGATTGCATCAATCANNNAGATCGGA |
| 10874_2 | TATGAGTGTGGAGTCGTTGCAGGCAAATGGGATTGCATCAATCANNNAGATCGGA |
| 47596_1 | TATGAGTGTGGAGTCGTTGCCAAGGTTTGGCTGGTTTTCCATGNNNAGATCGGAAG |
| 47596_2 | TATGAGTGTGGAGTCGTTGCCAAGGTTTGGCTGGTTTTCCATGNNNAGATCGGAAG |
| 33654_1 | TATGAGTGTGGAGTCGTTGCCCTCCCAGGTTCAAGTGATTTTCNNNAGATCGGAAG |
| 33654_2 | TATGAGTGTGGAGTCGTTGCCCTCCCAGGTTCAAGTGATTTTCNNNAGATCGGAAG |
| 33441_1 | TATGAGTGTGGAGTCGTTGCGCATGGGCTTTGCAATTGAGTANNNAGATCGGAAG |
| 33441_2 | TATGAGTGTGGAGTCGTTGCGCATGGGCTTTGCAATTGAGTANNNAGATCGGAAG |
| 53423_1 | TATGAGTGTGGAGTCGTTGCAGACACAGAGGGAAAGAGAACACNNNAGATCGGA |
| 53423_2 | TATGAGTGTGGAGTCGTTGCAGACACAGAGGGAAAGAGAACACNNNAGATCGGA |
| 33937_1 | TATGAGTGTGGAGTCGTTGCGCTGGAACTACAGGTACATGCTANNNAGATCGGAA |
| 33937_2 | TATGAGTGTGGAGTCGTTGCGCTGGAACTACAGGTACATGCTANNNAGATCGGAA |
| 6623_1 | TATGAGTGTGGAGTCGTTGCACTACAGCACTACTTCCCCACTTNNNAGATCGGAAGA |
| 6623_2 | TATGAGTGTGGAGTCGTTGCACTACAGCACTACTTCCCCACTTNNNAGATCGGAAGA |
| 32963_1 | TATGAGTGTGGAGTCGTTGCTCCTCAGCAAACTAACACAGGAANNNAGATCGGAAG |
| 32963_2 | TATGAGTGTGGAGTCGTTGCTCCTCAGCAAACTAACACAGGAANNNAGATCGGAAG |
| 33524_1 | TATGAGTGTGGAGTCGTTGCGGGTCACCTTTATTCCAGATCCCNNNAGATCGGAAG |
| 33524_2 | TATGAGTGTGGAGTCGTTGCGGGTCACCTTTATTCCAGATCCCNNNAGATCGGAAG |
| 32678_1 | TATGAGTGTGGAGTCGTTGCTTCAAGTGATTCTCCTGCCTCAGNNNAGATCGGAAG |
| 32678_2 | TATGAGTGTGGAGTCGTTGCTTCAAGTGATTCTCCTGCCTCAGNNNAGATCGGAAG |
| 50817_1 | TATGAGTGTGGAGTCGTTGCAGCTAGAGGACAACAGACTTTGGNNNAGATCGGAA |
| 50817_2 | TATGAGTGTGGAGTCGTTGCAGCTAGAGGACAACAGACTTTGGNNNAGATCGGAA |
| 38266_1 | TATGAGTGTGGAGTCGTTGCCGAGGCTATGGAGAAAAGGGAATNNNAGATCGGA |
| 38266_2 | TATGAGTGTGGAGTCGTTGCCGAGGCTATGGAGAAAAGGGAATNNNAGATCGGA |
| 13383_1 | TATGAGTGTGGAGTCGTTGCTTCCAGAAGTAGAAGACGAGNNNAGATCGGAAGAG |
| 13383_2 | TATGAGTGTGGAGTCGTTGCTTCCAGAAGTAGAAGACGAGNNNAGATCGGAAGAG |
| 38288_1 | TATGAGTGTGGAGTCGTTGCACTCTGCAGAAGCTCTTTAGTTTNNNAGATCGGAAG |
| 38288_2 | TATGAGTGTGGAGTCGTTGCACTCTGCAGAAGCTCTTTAGTTTNNNAGATCGGAAG |
| 41280_1 | TATGAGTGTGGAGTCGTTGCGGCCCTTCTCGTCATATCCTAACNNNAGATCGGAAGA |
| 41280_2 | TATGAGTGTGGAGTCGTTGCGGCCCTTCTCGTCATATCCTAACNNNAGATCGGAAGA |
| 32992_1 | TATGAGTGTGGAGTCGTTGCGTAGTGCAGTGGAGTGATCTCAANNNAGATCGGAA |

| | |
|---|---|
| 32992_2 | TATGAGTGTGGAGTCGTTGCGTAGTGCAGTGGAGTGATCTCAANNNAGATCGGAA |
| 47680_1 | TATGAGTGTGGAGTCGTTGCAGATCTAAGCTCACTGCAACCTCNNNAGATCGGAAG |
| 47680_2 | TATGAGTGTGGAGTCGTTGCAGATCTAAGCTCACTGCAACCTCNNNAGATCGGAAG |
| 41899_1 | TATGAGTGTGGAGTCGTTGCCAATCATGGCTCCTGGAGACTTANNNAGATCGGAAG |
| 41899_2 | TATGAGTGTGGAGTCGTTGCCAATCATGGCTCCTGGAGACTTANNNAGATCGGAAG |
| 45400_1 | TATGAGTGTGGAGTCGTTGCCACACATTTGCAAGGCCAAATTTNNNAGATCGGAAG |
| 45400_2 | TATGAGTGTGGAGTCGTTGCCACACATTTGCAAGGCCAAATTTNNNAGATCGGAAG |
| 36359_1 | TATGAGTGTGGAGTCGTTGCGGTGTGGACATAAGTTTTGGTGTNNNAGATCGGAA |
| 36359_2 | TATGAGTGTGGAGTCGTTGCGGTGTGGACATAAGTTTTGGTGTNNNAGATCGGAA |
| 33624_1 | TATGAGTGTGGAGTCGTTGCAACTCCTGAACTTGTGATCCACANNNAGATCGGAAG |
| 33624_2 | TATGAGTGTGGAGTCGTTGCAACTCCTGAACTTGTGATCCACANNNAGATCGGAAG |
| 30334_1 | TATGAGTGTGGAGTCGTTGCCCACACCTGCTTAAGGGGGTTCNNNAGATCGGAAGA |
| 30334_2 | TATGAGTGTGGAGTCGTTGCCCACACCTGCTTAAGGGGGTTCNNNAGATCGGAAGA |
| 51992_1 | TATGAGTGTGGAGTCGTTGCGGAGGAAAATACCAGGCACCTATNNNAGATCGGAA |
| 51992_2 | TATGAGTGTGGAGTCGTTGCGGAGGAAAATACCAGGCACCTATNNNAGATCGGAA |
| 52153_1 | TATGAGTGTGGAGTCGTTGCATAGCTTAGGCCATTGCTTCAGANNNAGATCGGAAG |
| 52153_2 | TATGAGTGTGGAGTCGTTGCATAGCTTAGGCCATTGCTTCAGANNNAGATCGGAAG |
| 39668_1 | TATGAGTGTGGAGTCGTTGCTAGGTTGCAATGAGAGGAGATCGNNNAGATCGGAA |
| 39668_2 | TATGAGTGTGGAGTCGTTGCTAGGTTGCAATGAGAGGAGATCGNNNAGATCGGAA |
| 32364_1 | TATGAGTGTGGAGTCGTTGCACACTATGCAAAGCACAAATCCCNNNAGATCGGAAG |
| 32364_2 | TATGAGTGTGGAGTCGTTGCACACTATGCAAAGCACAAATCCCNNNAGATCGGAAG |
| 44074_1 | TATGAGTGTGGAGTCGTTGCTACCTTGAACCATCTCCTCTCCTNNNAGATCGGAAGA |
| 44074_2 | TATGAGTGTGGAGTCGTTGCTACCTTGAACCATCTCCTCTCCTNNNAGATCGGAAGA |
| 51817_1 | TATGAGTGTGGAGTCGTTGCGTATGTGATGTTCCCCTTCCTGTNNNAGATCGGAAG |
| 51817_2 | TATGAGTGTGGAGTCGTTGCGTATGTGATGTTCCCCTTCCTGTNNNAGATCGGAAG |
| 30068_1 | TATGAGTGTGGAGTCGTTGCTCACTTTGAATTCTACACGATACCTNNNAGATCGGAA |
| 30068_2 | TATGAGTGTGGAGTCGTTGCTCACTTTGAATTCTACACGATACCTNNNAGATCGGAA |
| 38279_1 | TATGAGTGTGGAGTCGTTGCTACCTGGAAGTCTCATTGCTGACNNNAGATCGGAAG |
| 38279_2 | TATGAGTGTGGAGTCGTTGCTACCTGGAAGTCTCATTGCTGACNNNAGATCGGAAG |
| 46890_1 | TATGAGTGTGGAGTCGTTGCAAGTAGCTGGGACTAAAGGCACNNNAGATCGGAAG |
| 46890_2 | TATGAGTGTGGAGTCGTTGCAAGTAGCTGGGACTAAAGGCACNNNAGATCGGAAG |
| 44512_1 | TATGAGTGTGGAGTCGTTGCATATCTCATCTTGACCCAGGGCTNNNAGATCGGAAG |
| 44512_2 | TATGAGTGTGGAGTCGTTGCATATCTCATCTTGACCCAGGGCTNNNAGATCGGAAG |
| 39602_1 | TATGAGTGTGGAGTCGTTGCAACATTCTTAGACCACAGTGGCTNNNAGATCGGAAG |
| 39602_2 | TATGAGTGTGGAGTCGTTGCAACATTCTTAGACCACAGTGGCTNNNAGATCGGAAG |
| 37396_1 | TATGAGTGTGGAGTCGTTGCCAATCTGGAGAAACCCCAACAACNNNAGATCGGAAG |
| 37396_2 | TATGAGTGTGGAGTCGTTGCCAATCTGGAGAAACCCCAACAACNNNAGATCGGAAG |
| 36778_1 | TATGAGTGTGGAGTCGTTGCCATGCCTGTAGTCTCAGCTACTTNNNAGATCGGAAG |
| 36778_2 | TATGAGTGTGGAGTCGTTGCCATGCCTGTAGTCTCAGCTACTTNNNAGATCGGAAG |
| 45042_1 | TATGAGTGTGGAGTCGTTGCCAGAGAGACAGAGAGGGAATGCNNNAGATCGGAA |
| 45042_2 | TATGAGTGTGGAGTCGTTGCCAGAGAGACAGAGAGGGAATGCNNNAGATCGGAA |
| 33002_1 | TATGAGTGTGGAGTCGTTGCGATCTTGGCTCACTACAACCTCCNNNAGATCGGAAG |
| 33002_2 | TATGAGTGTGGAGTCGTTGCGATCTTGGCTCACTACAACCTCCNNNAGATCGGAAG |

| | |
|---|---|
| 40077_1 | TATGAGTGTGGAGTCGTTGCTGATGTACACCTGTAGTTCCAGCNNNAGATCGGAAG |
| 40077_2 | TATGAGTGTGGAGTCGTTGCTGATGTACACCTGTAGTTCCAGCNNNAGATCGGAAG |
| 29871_1 | TATGAGTGTGGAGTCGTTGCATGGGCAGGTACGGCAAAAGNNNAGATCGGAAGA |
| 29871_2 | TATGAGTGTGGAGTCGTTGCATGGGCAGGTACGGCAAAAGNNNAGATCGGAAGA |
| 42709_1 | TATGAGTGTGGAGTCGTTGCCACTGGCTTGTGAAAAACAAACANNNAGATCGGAA |
| 42709_2 | TATGAGTGTGGAGTCGTTGCCACTGGCTTGTGAAAAACAAACANNNAGATCGGAA |
| 52390_1 | TATGAGTGTGGAGTCGTTGCACATCTCATCTGAGGACAGCTTGNNNAGATCGGAAG |
| 52390_2 | TATGAGTGTGGAGTCGTTGCACATCTCATCTGAGGACAGCTTGNNNAGATCGGAAG |
| 37732_1 | TATGAGTGTGGAGTCGTTGCGTCACATCTGTAATCCCAGCACTNNNAGATCGGAAG |
| 37732_2 | TATGAGTGTGGAGTCGTTGCGTCACATCTGTAATCCCAGCACTNNNAGATCGGAAG |
| 47152_1 | TATGAGTGTGGAGTCGTTGCTGCTGAATCCTGGAGGTAAGTTCNNNAGATCGGAA |
| 47152_2 | TATGAGTGTGGAGTCGTTGCTGCTGAATCCTGGAGGTAAGTTCNNNAGATCGGAA |
| 44887_1 | TATGAGTGTGGAGTCGTTGCACAGGAGTTCTACTGGCTTTCTGNNNAGATCGGAAG |
| 44887_2 | TATGAGTGTGGAGTCGTTGCACAGGAGTTCTACTGGCTTTCTGNNNAGATCGGAAG |
| 42551_1 | TATGAGTGTGGAGTCGTTGCTCTTGGAATGTCTTTGTCTGGCTNNNAGATCGGAAG |
| 42551_2 | TATGAGTGTGGAGTCGTTGCTCTTGGAATGTCTTTGTCTGGCTNNNAGATCGGAAG |
| 41236_1 | TATGAGTGTGGAGTCGTTGCGCAAATGTATTCTTCACGGGAGGNNNAGATCGGAA |
| 41236_2 | TATGAGTGTGGAGTCGTTGCGCAAATGTATTCTTCACGGGAGGNNNAGATCGGAA |
| 44080_1 | TATGAGTGTGGAGTCGTTGCGCATACCTGATTCCAATGTCTGCNNNAGATCGGAAG |
| 44080_2 | TATGAGTGTGGAGTCGTTGCGCATACCTGATTCCAATGTCTGCNNNAGATCGGAAG |
| 47923_1 | TATGAGTGTGGAGTCGTTGCGGCTAGAGTACAGTGGTGTGATCNNNAGATCGGAA |
| 47923_2 | TATGAGTGTGGAGTCGTTGCGGCTAGAGTACAGTGGTGTGATCNNNAGATCGGAA |
| 50757_1 | TATGAGTGTGGAGTCGTTGCCGAAAAGGAAACCCGCTAAAGAANNNAGATCGGAA |
| 50757_2 | TATGAGTGTGGAGTCGTTGCCGAAAAGGAAACCCGCTAAAGAANNNAGATCGGAA |
| 35130_1 | TATGAGTGTGGAGTCGTTGCCACCGTGCCTAGTCTCAAACTAANNNAGATCGGAAG |
| 35130_2 | TATGAGTGTGGAGTCGTTGCCACCGTGCCTAGTCTCAAACTAANNNAGATCGGAAG |
| 51057_1 | TATGAGTGTGGAGTCGTTGCTTCTCAATTCCATCCAGGAGGTGNNNAGATCGGAAG |
| 51057_2 | TATGAGTGTGGAGTCGTTGCTTCTCAATTCCATCCAGGAGGTGNNNAGATCGGAAG |
| 36425_1 | TATGAGTGTGGAGTCGTTGCTTTGTGGTCAGCATTCTCTCTCCNNNAGATCGGAAGA |
| 36425_2 | TATGAGTGTGGAGTCGTTGCTTTGTGGTCAGCATTCTCTCTCCNNNAGATCGGAAGA |
| 39253_1 | TATGAGTGTGGAGTCGTTGCTCTTGACTACTCCTCTCACGCTANNNAGATCGGAAGA |
| 39253_2 | TATGAGTGTGGAGTCGTTGCTCTTGACTACTCCTCTCACGCTANNNAGATCGGAAGA |
| 44657_1 | TATGAGTGTGGAGTCGTTGCCTGCAGTCTCCGTGCATAGAGNNNAGATCGGAAGA |
| 44657_2 | TATGAGTGTGGAGTCGTTGCCTGCAGTCTCCGTGCATAGAGNNNAGATCGGAAGA |
| 44534_1 | TATGAGTGTGGAGTCGTTGCGTCTTTTGGAGCAACATGAATGGANNNAGATCGGA |
| 44534_2 | TATGAGTGTGGAGTCGTTGCGTCTTTTGGAGCAACATGAATGGANNNAGATCGGA |
| 49281_1 | TATGAGTGTGGAGTCGTTGCAATAGCTGGGATTACAGGTACGCNNNAGATCGGAA |
| 49281_2 | TATGAGTGTGGAGTCGTTGCAATAGCTGGGATTACAGGTACGCNNNAGATCGGAA |
| 10214_1 | TATGAGTGTGGAGTCGTTGCGGTCATGGAGCTCCTATTCAGGANNNAGATCGGAA |
| 10214_2 | TATGAGTGTGGAGTCGTTGCGGTCATGGAGCTCCTATTCAGGANNNAGATCGGAA |
| 9500_1 | TATGAGTGTGGAGTCGTTGCGTCTCTTTGTCTCTCTGGCCATGNNNAGATCGGAAG |
| 9500_2 | TATGAGTGTGGAGTCGTTGCGTCTCTTTGTCTCTCTGGCCATGNNNAGATCGGAAG |
| 10307_1 | TATGAGTGTGGAGTCGTTGCAGAAGAAACAGGGAGTGGGGATTNNNAGATCGGA |

| | |
|---|---|
| 10307_2 | TATGAGTGTGGAGTCGTTGCAGAAGAAACAGGGAGTGGGGATTNNNAGATCGGA |
| 49445_1 | TATGAGTGTGGAGTCGTTGCTTCACACCATTCTCCTGCCTCNNNAGATCGGAAGAGC |
| 49445_2 | TATGAGTGTGGAGTCGTTGCTTCACACCATTCTCCTGCCTCNNNAGATCGGAAGAGC |
| 48418_1 | TATGAGTGTGGAGTCGTTGCGATTCTCCTGTCAGCCTCCTAAGNNNAGATCGGAAG |
| 48418_2 | TATGAGTGTGGAGTCGTTGCGATTCTCCTGTCAGCCTCCTAAGNNNAGATCGGAAG |
| 13154_1 | TATGAGTGTGGAGTCGTTGCTAAAACCAGCTCTGAAAAGAAANNNAGATCGGAAG |
| 13154_2 | TATGAGTGTGGAGTCGTTGCTAAAACCAGCTCTGAAAAGAAANNNAGATCGGAAG |
| 46692_1 | TATGAGTGTGGAGTCGTTGCTCTGTTGGTCATGATGGTTAGCANNNAGATCGGAA |
| 46692_2 | TATGAGTGTGGAGTCGTTGCTCTGTTGGTCATGATGGTTAGCANNNAGATCGGAA |
| 46152_1 | TATGAGTGTGGAGTCGTTGCCTGAATGAGATGGAATGACCCCANNNAGATCGGAA |
| 46152_2 | TATGAGTGTGGAGTCGTTGCCTGAATGAGATGGAATGACCCCANNNAGATCGGAA |
| 44234_1 | TATGAGTGTGGAGTCGTTGCTCTCCAGATTTTGATGGTGCTGANNNAGATCGGAAG |
| 44234_2 | TATGAGTGTGGAGTCGTTGCTCTCCAGATTTTGATGGTGCTGANNNAGATCGGAAG |
| 32531_1 | TATGAGTGTGGAGTCGTTGCGCAGGGTTTGCAATCCTAGTTTCNNNAGATCGGAAG |
| 32531_2 | TATGAGTGTGGAGTCGTTGCGCAGGGTTTGCAATCCTAGTTTCNNNAGATCGGAAG |
| 52116_1 | TATGAGTGTGGAGTCGTTGCCTAGTAAGGCAGTCTGGTCACAGNNNAGATCGGAA |
| 52116_2 | TATGAGTGTGGAGTCGTTGCCTAGTAAGGCAGTCTGGTCACAGNNNAGATCGGAA |
| 44523_1 | TATGAGTGTGGAGTCGTTGCTTTGAGAGATCATAGGCATGGCCNNNAGATCGGAA |
| 44523_2 | TATGAGTGTGGAGTCGTTGCTTTGAGAGATCATAGGCATGGCCNNNAGATCGGAA |
| 49386_1 | TATGAGTGTGGAGTCGTTGCGCTCACTGAAACCTCTCTCTCTCNNNAGATCGGAAGA |
| 49386_2 | TATGAGTGTGGAGTCGTTGCGCTCACTGAAACCTCTCTCTCTCNNNAGATCGGAAGA |
| 9923_1 | TATGAGTGTGGAGTCGTTGCTCCCCATCAATTTGCATGACAGANNNAGATCGGAAG |
| 9923_2 | TATGAGTGTGGAGTCGTTGCTCCCCATCAATTTGCATGACAGANNNAGATCGGAAG |
| 35871_1 | TATGAGTGTGGAGTCGTTGCGAGAGAGAGAGAGTCTTCTTGCGNNNAGATCGGAA |
| 35871_2 | TATGAGTGTGGAGTCGTTGCGAGAGAGAGAGAGTCTTCTTGCGNNNAGATCGGAA |
| 4696_1 | TATGAGTGTGGAGTCGTTGCTGGCTATGGGGAAATAACTGAGCNNNAGATCGGAA |
| 4696_2 | TATGAGTGTGGAGTCGTTGCTGGCTATGGGGAAATAACTGAGCNNNAGATCGGAA |
| 39385_1 | TATGAGTGTGGAGTCGTTGCGATCACAAGGTCAGGGGATCAAGNNNAGATCGGAA |
| 39385_2 | TATGAGTGTGGAGTCGTTGCGATCACAAGGTCAGGGGATCAAGNNNAGATCGGAA |
| 41471_1 | TATGAGTGTGGAGTCGTTGCAAAAGTTGTTGCCTGTTCATGGANNNAGATCGGAA |
| 41471_2 | TATGAGTGTGGAGTCGTTGCAAAAGTTGTTGCCTGTTCATGGANNNAGATCGGAA |
| 11010_1 | TATGAGTGTGGAGTCGTTGCTGGTATTCTCCCCTTGTGTCTGTNNNAGATCGGAAG |
| 11010_2 | TATGAGTGTGGAGTCGTTGCTGGTATTCTCCCCTTGTGTCTGTNNNAGATCGGAAG |
| 33787_1 | TATGAGTGTGGAGTCGTTGCGCCAGAGTACCAGTGTTCAAAACNNNAGATCGGAA |
| 33787_2 | TATGAGTGTGGAGTCGTTGCGCCAGAGTACCAGTGTTCAAAACNNNAGATCGGAA |
| 37198_1 | TATGAGTGTGGAGTCGTTGCGCTGGCACAGATTTGAGATTTGANNNAGATCGGAA |
| 37198_2 | TATGAGTGTGGAGTCGTTGCGCTGGCACAGATTTGAGATTTGANNNAGATCGGAA |
| 36797_1 | TATGAGTGTGGAGTCGTTGCACTCTTGTATGAGCCAGACTTGGNNNAGATCGGAAG |
| 36797_2 | TATGAGTGTGGAGTCGTTGCACTCTTGTATGAGCCAGACTTGGNNNAGATCGGAAG |
| 36911_1 | TATGAGTGTGGAGTCGTTGCAAAATGATGGGGCTGTGTCAAAGNNNAGATCGGAA |
| 36911_2 | TATGAGTGTGGAGTCGTTGCAAAATGATGGGGCTGTGTCAAAGNNNAGATCGGAA |
| 39474_1 | TATGAGTGTGGAGTCGTTGCATCCTGTATCTAGCACTTTGCCANNNAGATCGGAAG |
| 39474_2 | TATGAGTGTGGAGTCGTTGCATCCTGTATCTAGCACTTTGCCANNNAGATCGGAAG |

| | |
|---|---|
| 52177_1 | TATGAGTGTGGAGTCGTTGCGCAGATCACTTGAGGTCAGGAGNNNAGATCGGAAG |
| 52177_2 | TATGAGTGTGGAGTCGTTGCGCAGATCACTTGAGGTCAGGAGNNNAGATCGGAAG |
| 29650_1 | TATGAGTGTGGAGTCGTTGCCTCGGTGACCATCTTAGTCAGTTNNNAGATCGGAAG |
| 29650_2 | TATGAGTGTGGAGTCGTTGCCTCGGTGACCATCTTAGTCAGTTNNNAGATCGGAAG |
| 37028_1 | TATGAGTGTGGAGTCGTTGCAGAGGACTTGAACAGATGCTTCTNNNAGATCGGAA |
| 37028_2 | TATGAGTGTGGAGTCGTTGCAGAGGACTTGAACAGATGCTTCTNNNAGATCGGAA |
| 36791_1 | TATGAGTGTGGAGTCGTTGCATAAATTAGCTGGGCATGGTGGTNNNAGATCGGAA |
| 36791_2 | TATGAGTGTGGAGTCGTTGCATAAATTAGCTGGGCATGGTGGTNNNAGATCGGAA |
| 42055_1 | TATGAGTGTGGAGTCGTTGCCCACCAAAAGGCAACTAGAACTGNNNAGATCGGAA |
| 42055_2 | TATGAGTGTGGAGTCGTTGCCCACCAAAAGGCAACTAGAACTGNNNAGATCGGAA |
| 53042_1 | TATGAGTGTGGAGTCGTTGCTAGGTAAAAGGAGCTGTCATCCCNNNAGATCGGAA |
| 53042_2 | TATGAGTGTGGAGTCGTTGCTAGGTAAAAGGAGCTGTCATCCCNNNAGATCGGAA |
| 31639_1 | TATGAGTGTGGAGTCGTTGCAGTGTATGCTACTGAAGGAGCTGNNNAGATCGGAA |
| 31639_2 | TATGAGTGTGGAGTCGTTGCAGTGTATGCTACTGAAGGAGCTGNNNAGATCGGAA |
| 29729_1 | TATGAGTGTGGAGTCGTTGCCTGTATATGGCCTAACATCCCCNNNAGATCGGAAG |
| 29729_2 | TATGAGTGTGGAGTCGTTGCCTGTATATGGCCTAACATCCCCNNNAGATCGGAAG |
| 33553_1 | TATGAGTGTGGAGTCGTTGCGTTCAAGTGATTCTCTTGCCTCGNNNAGATCGGAAG |
| 33553_2 | TATGAGTGTGGAGTCGTTGCGTTCAAGTGATTCTCTTGCCTCGNNNAGATCGGAAG |
| 44255_1 | TATGAGTGTGGAGTCGTTGCCATCCCCCTGAGAGGAAAGAATCNNNAGATCGGAA |
| 44255_2 | TATGAGTGTGGAGTCGTTGCCATCCCCCTGAGAGGAAAGAATCNNNAGATCGGAA |
| 43667_1 | TATGAGTGTGGAGTCGTTGCGATGCCACGTCCATTTGCTTCNNNAGATCGGAAGAG |
| 43667_2 | TATGAGTGTGGAGTCGTTGCGATGCCACGTCCATTTGCTTCNNNAGATCGGAAGAG |
| 352_1 | TATGAGTGTGGAGTCGTTGCTCCCAGTTCCACCTTACTAGTGTNNNAGATCGGAAGA |
| 352_2 | TATGAGTGTGGAGTCGTTGCTCCCAGTTCCACCTTACTAGTGTNNNAGATCGGAAGA |
| 46650_1 | TATGAGTGTGGAGTCGTTGCCTCCTGAGTAGCTTGGACTACAGNNNAGATCGGAAG |
| 46650_2 | TATGAGTGTGGAGTCGTTGCCTCCTGAGTAGCTTGGACTACAGNNNAGATCGGAAG |
| 13753_1 | TATGAGTGTGGAGTCGTTGCATGATATGTGCTTTCCTGTTTNNNAGATCGGAAGAG |
| 13753_2 | TATGAGTGTGGAGTCGTTGCATGATATGTGCTTTCCTGTTTNNNAGATCGGAAGAG |
| 35818_1 | TATGAGTGTGGAGTCGTTGCCATTCTGACTGGTGTGAGATGGTNNNAGATCGGAA |
| 35818_2 | TATGAGTGTGGAGTCGTTGCCATTCTGACTGGTGTGAGATGGTNNNAGATCGGAA |
| 47078_1 | TATGAGTGTGGAGTCGTTGCTGGCTGCAGTATATGACACAGAANNNAGATCGGAA |
| 47078_2 | TATGAGTGTGGAGTCGTTGCTGGCTGCAGTATATGACACAGAANNNAGATCGGAA |
| 52256_1 | TATGAGTGTGGAGTCGTTGCGGTGCATGCCTATAGTCTCAGTTNNNAGATCGGAAG |
| 52256_2 | TATGAGTGTGGAGTCGTTGCGGTGCATGCCTATAGTCTCAGTTNNNAGATCGGAAG |
| 50609_1 | TATGAGTGTGGAGTCGTTGCTTGTTTTTTTGGAACACAGGCCTNNNAGATCGGAAG |
| 50609_2 | TATGAGTGTGGAGTCGTTGCTTGTTTTTTTGGAACACAGGCCTNNNAGATCGGAAG |
| 43498_1 | TATGAGTGTGGAGTCGTTGCGGCGGGATTGAGTGAGTTTCTTANNNAGATCGGAA |
| 43498_2 | TATGAGTGTGGAGTCGTTGCGGCGGGATTGAGTGAGTTTCTTANNNAGATCGGAA |
| 13354_1 | TATGAGTGTGGAGTCGTTGCTAGAGAGAACACAAAGAAATCATNNNAGATCGGAA |
| 13354_2 | TATGAGTGTGGAGTCGTTGCTAGAGAGAACACAAAGAAATCATNNNAGATCGGAA |
| 38586_1 | TATGAGTGTGGAGTCGTTGCCCACCATGCCCAGCTAATTTTTTNNNAGATCGGAAGA |
| 38586_2 | TATGAGTGTGGAGTCGTTGCCCACCATGCCCAGCTAATTTTTTNNNAGATCGGAAGA |
| 42095_1 | TATGAGTGTGGAGTCGTTGCCCAGCCACAGCCATAGATAGAAANNNAGATCGGAA |

| | |
|---|---|
| 42095_2 | TATGAGTGTGGAGTCGTTGCCCAGCCACAGCCATAGATAGAAANNNAGATCGGAAG |
| 37135_1 | TATGAGTGTGGAGTCGTTGCCTAGGTGACAGAGTGAGACCTTGNNNAGATCGGAAG |
| 37135_2 | TATGAGTGTGGAGTCGTTGCCTAGGTGACAGAGTGAGACCTTGNNNAGATCGGAAG |
| 42385_1 | TATGAGTGTGGAGTCGTTGCGCCACTAGTCCATGTACAGATGANNNAGATCGGAAG |
| 42385_2 | TATGAGTGTGGAGTCGTTGCGCCACTAGTCCATGTACAGATGANNNAGATCGGAAG |
| 36332_1 | TATGAGTGTGGAGTCGTTGCTGTTGTTTTTCTCTTTCCCGACANNNAGATCGGAAGA |
| 36332_2 | TATGAGTGTGGAGTCGTTGCTGTTGTTTTTCTCTTTCCCGACANNNAGATCGGAAGA |
| 45124_1 | TATGAGTGTGGAGTCGTTGCAGAATATACAACAAGAGTGGGCCTNNNAGATCGGAA |
| 45124_2 | TATGAGTGTGGAGTCGTTGCAGAATATACAACAAGAGTGGGCCTNNNAGATCGGAA |
| 2307_1 | TATGAGTGTGGAGTCGTTGCTGCTGTTGGAACGTGTGATAGACNNNAGATCGGAAG |
| 2307_2 | TATGAGTGTGGAGTCGTTGCTGCTGTTGGAACGTGTGATAGACNNNAGATCGGAAG |
| 51987_1 | TATGAGTGTGGAGTCGTTGCACCCAAAAAAAACCCACCCAATANNNAGATCGGAAGA |
| 51987_2 | TATGAGTGTGGAGTCGTTGCACCCAAAAAAAACCCACCCAATANNNAGATCGGAAGA |
| 35047_1 | TATGAGTGTGGAGTCGTTGCCCTCCTGAGTAGCTGCAATTACANNNAGATCGGAAGA |
| 35047_2 | TATGAGTGTGGAGTCGTTGCCCTCCTGAGTAGCTGCAATTACANNNAGATCGGAAGA |
| 41675_1 | TATGAGTGTGGAGTCGTTGCCAGTAGCACAATCATGGCTCACTNNNAGATCGGAAG |
| 41675_2 | TATGAGTGTGGAGTCGTTGCCAGTAGCACAATCATGGCTCACTNNNAGATCGGAAG |
| 51628_1 | TATGAGTGTGGAGTCGTTGCGATGGGGGAGAGGAGAGTAACCNNNAGATCGGAA |
| 51628_2 | TATGAGTGTGGAGTCGTTGCGATGGGGGAGAGGAGAGTAACCNNNAGATCGGAA |
| 50979_1 | TATGAGTGTGGAGTCGTTGCTTTGCACCAAAAGCATAGACTGGNNNAGATCGGAAG |
| 50979_2 | TATGAGTGTGGAGTCGTTGCTTTGCACCAAAAGCATAGACTGGNNNAGATCGGAAG |
| 49197_1 | TATGAGTGTGGAGTCGTTGCCTACATTTTATGCGCAGAGGAGCNNNAGATCGGAAG |
| 49197_2 | TATGAGTGTGGAGTCGTTGCCTACATTTTATGCGCAGAGGAGCNNNAGATCGGAAG |
| 39821_1 | TATGAGTGTGGAGTCGTTGCGGATTGGAGAGAGGAAAGCTCAANNNAGATCGGAA |
| 39821_2 | TATGAGTGTGGAGTCGTTGCGGATTGGAGAGAGGAAAGCTCAANNNAGATCGGAA |
| 51123_1 | TATGAGTGTGGAGTCGTTGCGCGTCACATTTTACAGTATGGGTNNNAGATCGGAAG |
| 51123_2 | TATGAGTGTGGAGTCGTTGCGCGTCACATTTTACAGTATGGGTNNNAGATCGGAAG |
| 36960_1 | TATGAGTGTGGAGTCGTTGCGTGCTCTGTGGAGGAAGTAAGAANNNAGATCGGAA |
| 36960_2 | TATGAGTGTGGAGTCGTTGCGTGCTCTGTGGAGGAAGTAAGAANNNAGATCGGAA |
| 43521_1 | TATGAGTGTGGAGTCGTTGCCAAACTCCTGACCTCAAATGAGCNNNAGATCGGAAGA |
| 43521_2 | TATGAGTGTGGAGTCGTTGCCAAACTCCTGACCTCAAATGAGCNNNAGATCGGAAGA |
| 32890_1 | TATGAGTGTGGAGTCGTTGCGCATCTCAACAGCTGAAACACTTNNNAGATCGGAAGA |
| 32890_2 | TATGAGTGTGGAGTCGTTGCGCATCTCAACAGCTGAAACACTTNNNAGATCGGAAGA |
| 13271_1 | TATGAGTGTGGAGTCGTTGCAGTCCAAGGTATTATTCTTATGCNNNAGATCGGAAGA |
| 13271_2 | TATGAGTGTGGAGTCGTTGCAGTCCAAGGTATTATTCTTATGCNNNAGATCGGAAGA |
| 30887_1 | TATGAGTGTGGAGTCGTTGCGCTGGGACAGGTATTTGATGGTANNNAGATCGGAA |
| 30887_2 | TATGAGTGTGGAGTCGTTGCGCTGGGACAGGTATTTGATGGTANNNAGATCGGAA |
| 44434_1 | TATGAGTGTGGAGTCGTTGCACATGTATGGGTGTTATATGTACACANNNAGATCGG |
| 44434_2 | TATGAGTGTGGAGTCGTTGCACATGTATGGGTGTTATATGTACACANNNAGATCGG |
| 46577_1 | TATGAGTGTGGAGTCGTTGCCCAGGTTCAAGCGATTCTCCTNNNAGATCGGAAGAG |
| 46577_2 | TATGAGTGTGGAGTCGTTGCCCAGGTTCAAGCGATTCTCCTNNNAGATCGGAAGAG |
| 48580_1 | TATGAGTGTGGAGTCGTTGCGAGATCTCGCTATGTTGCCCAGNNNAGATCGGAAGA |
| 48580_2 | TATGAGTGTGGAGTCGTTGCGAGATCTCGCTATGTTGCCCAGNNNAGATCGGAAGA |

| | |
|---|---|
| 44396_1 | TATGAGTGTGGAGTCGTTGCGAGAGTGGGGGAGTAGACGTNNNAGATCGGAAGA |
| 44396_2 | TATGAGTGTGGAGTCGTTGCGAGAGTGGGGGAGTAGACGTNNNAGATCGGAAGA |
| 38013_1 | TATGAGTGTGGAGTCGTTGCCACAGCAGCAGACTATCCACATANNNAGATCGGAAG |
| 38013_2 | TATGAGTGTGGAGTCGTTGCCACAGCAGCAGACTATCCACATANNNAGATCGGAAG |
| 34279_1 | TATGAGTGTGGAGTCGTTGCAGGTCTGTGAGAGATGCAGGATANNNAGATCGGA |
| 34279_2 | TATGAGTGTGGAGTCGTTGCAGGTCTGTGAGAGATGCAGGATANNNAGATCGGA |
| 51264_1 | TATGAGTGTGGAGTCGTTGCTGTGGGTGAAGAGAAAGGGTATGNNNAGATCGGA |
| 51264_2 | TATGAGTGTGGAGTCGTTGCTGTGGGTGAAGAGAAAGGGTATGNNNAGATCGGA |
| 41933_1 | TATGAGTGTGGAGTCGTTGCTGAAGTGCAGTGGTGTGATCTTNNNAGATCGGAAG |
| 41933_2 | TATGAGTGTGGAGTCGTTGCTGAAGTGCAGTGGTGTGATCTTNNNAGATCGGAAG |
| 41177_1 | TATGAGTGTGGAGTCGTTGCAGACAAGGTCCTTGCATTGTTTGNNNAGATCGGAAG |
| 41177_2 | TATGAGTGTGGAGTCGTTGCAGACAAGGTCCTTGCATTGTTTGNNNAGATCGGAAG |
| 42231_1 | TATGAGTGTGGAGTCGTTGCGAATGTGTTAGGAAGGGAGTCCCNNNAGATCGGAA |
| 42231_2 | TATGAGTGTGGAGTCGTTGCGAATGTGTTAGGAAGGGAGTCCCNNNAGATCGGAA |
| 10323_1 | TATGAGTGTGGAGTCGTTGCGGGCTAGGGGGTTGAGTATGTATNNNAGATCGGA |
| 10323_2 | TATGAGTGTGGAGTCGTTGCGGGCTAGGGGGTTGAGTATGTATNNNAGATCGGA |
| 38491_1 | TATGAGTGTGGAGTCGTTGCACCCTTCCTACCTCATACCTCATNNNAGATCGGAAGA |
| 38491_2 | TATGAGTGTGGAGTCGTTGCACCCTTCCTACCTCATACCTCATNNNAGATCGGAAGA |
| 52143_1 | TATGAGTGTGGAGTCGTTGCTCAGTGGCTCCCCAAATTTCTTANNNAGATCGGAAG |
| 52143_2 | TATGAGTGTGGAGTCGTTGCTCAGTGGCTCCCCAAATTTCTTANNNAGATCGGAAG |
| 50656_1 | TATGAGTGTGGAGTCGTTGCTTCTGCCTCTGGTTGATAGCAATNNNAGATCGGAAG |
| 50656_2 | TATGAGTGTGGAGTCGTTGCTTCTGCCTCTGGTTGATAGCAATNNNAGATCGGAAG |
| 28867_1 | TATGAGTGTGGAGTCGTTGCAATGATTGGCAGCATGACAAGAGNNNAGATCGGAA |
| 28867_2 | TATGAGTGTGGAGTCGTTGCAATGATTGGCAGCATGACAAGAGNNNAGATCGGAA |
| 31814_1 | TATGAGTGTGGAGTCGTTGCGCTAGGCTATCTTGAAGGGTACCNNNAGATCGGAA |
| 31814_2 | TATGAGTGTGGAGTCGTTGCGCTAGGCTATCTTGAAGGGTACCNNNAGATCGGAA |
| 40768_1 | TATGAGTGTGGAGTCGTTGCAGCTTGGAAGATTGCTTTTAGGCNNNAGATCGGAA |
| 40768_2 | TATGAGTGTGGAGTCGTTGCAGCTTGGAAGATTGCTTTTAGGCNNNAGATCGGAA |
| 47854_1 | TATGAGTGTGGAGTCGTTGCGATCCTCCCTAGTAGCTGTGAGANNNAGATCGGAAG |
| 47854_2 | TATGAGTGTGGAGTCGTTGCGATCCTCCCTAGTAGCTGTGAGANNNAGATCGGAAG |
| 13426_1 | TATGAGTGTGGAGTCGTTGCATCATGTTTAGATCACAATTAGGNNNAGATCGGAAG |
| 13426_2 | TATGAGTGTGGAGTCGTTGCATCATGTTTAGATCACAATTAGGNNNAGATCGGAAG |
| 48428_1 | TATGAGTGTGGAGTCGTTGCCGCGTTTCATGTCATTTTCCTTCNNNAGATCGGAAGA |
| 48428_2 | TATGAGTGTGGAGTCGTTGCCGCGTTTCATGTCATTTTCCTTCNNNAGATCGGAAGA |
| 36631_1 | TATGAGTGTGGAGTCGTTGCGAGGCCCTGTCTCCAAAAAAAAANNNAGATCGGAA |
| 36631_2 | TATGAGTGTGGAGTCGTTGCGAGGCCCTGTCTCCAAAAAAAAANNNAGATCGGAA |
| 45696_1 | TATGAGTGTGGAGTCGTTGCGATTTTGGCTCACTGCAACCTTTNNNAGATCGGAAG |
| 45696_2 | TATGAGTGTGGAGTCGTTGCGATTTTGGCTCACTGCAACCTTTNNNAGATCGGAAG |
| 30103_1 | TATGAGTGTGGAGTCGTTGCAGAGACAGTGGCATAATCAGTGANNNAGATCGGAA |
| 30103_2 | TATGAGTGTGGAGTCGTTGCAGAGACAGTGGCATAATCAGTGANNNAGATCGGAA |
| 42368_1 | TATGAGTGTGGAGTCGTTGCAGAGAGAGGGAATTGGAGAATGCNNNAGATCGGA |
| 42368_2 | TATGAGTGTGGAGTCGTTGCAGAGAGAGGGAATTGGAGAATGCNNNAGATCGGA |
| 45218_1 | TATGAGTGTGGAGTCGTTGCTCAGGAGGGTACAAAGCTTTTGTNNNAGATCGGAA |

| | |
|---|---|
| 45218_2 | TATGAGTGTGGAGTCGTTGCTCAGGAGGGTACAAAGCTTTTGTNNNAGATCGGAA |
| 31797_1 | TATGAGTGTGGAGTCGTTGCGAGTGCAGTAGTGTGATCTCAGTNNNAGATCGGAA |
| 31797_2 | TATGAGTGTGGAGTCGTTGCGAGTGCAGTAGTGTGATCTCAGTNNNAGATCGGAA |
| 46693_1 | TATGAGTGTGGAGTCGTTGCAGGTTCAAGTGATTCTTCTGCCTNNNAGATCGGAAG |
| 46693_2 | TATGAGTGTGGAGTCGTTGCAGGTTCAAGTGATTCTTCTGCCTNNNAGATCGGAAG |
| 50484_1 | TATGAGTGTGGAGTCGTTGCCTGAACTCTGGGTCTAGAAGCTCNNNAGATCGGAAG |
| 50484_2 | TATGAGTGTGGAGTCGTTGCCTGAACTCTGGGTCTAGAAGCTCNNNAGATCGGAAG |
| 35351_1 | TATGAGTGTGGAGTCGTTGCCTCCTGAGTAGCTGGGACTACAGNNNAGATCGGAAG |
| 35351_2 | TATGAGTGTGGAGTCGTTGCCTCCTGAGTAGCTGGGACTACAGNNNAGATCGGAAG |
| 46542_1 | TATGAGTGTGGAGTCGTTGCCGTCTCCCAAGTTGAAGCAATTCNNNAGATCGGAAG |
| 46542_2 | TATGAGTGTGGAGTCGTTGCCGTCTCCCAAGTTGAAGCAATTCNNNAGATCGGAAG |
| 13553_1 | TATGAGTGTGGAGTCGTTGCACAGTTGGTATTTTTAATTCTCCNNNAGATCGGAAG |
| 13553_2 | TATGAGTGTGGAGTCGTTGCACAGTTGGTATTTTTAATTCTCCNNNAGATCGGAAG |
| 10717_1 | TATGAGTGTGGAGTCGTTGCTGGCCCTCTTTTGCCTACTATTGNNNAGATCGGAAGA |
| 10717_2 | TATGAGTGTGGAGTCGTTGCTGGCCCTCTTTTGCCTACTATTGNNNAGATCGGAAGA |
| 47358_1 | TATGAGTGTGGAGTCGTTGCTCTGGAACTCCTGACCTCAAATGNNNAGATCGGAAG |
| 47358_2 | TATGAGTGTGGAGTCGTTGCTCTGGAACTCCTGACCTCAAATGNNNAGATCGGAAG |
| 51729_1 | TATGAGTGTGGAGTCGTTGCGCGAAAGAAAAGGGCTTGGTTATNNNAGATCGGAA |
| 51729_2 | TATGAGTGTGGAGTCGTTGCGCGAAAGAAAAGGGCTTGGTTATNNNAGATCGGAA |
| 41185_1 | TATGAGTGTGGAGTCGTTGCACATGTGCAGGTTTGTTACATGGNNNAGATCGGAA |
| 41185_2 | TATGAGTGTGGAGTCGTTGCACATGTGCAGGTTTGTTACATGGNNNAGATCGGAA |
| 37661_1 | TATGAGTGTGGAGTCGTTGCGGGCAAGGGACTTCCTGTAATAANNNAGATCGGAA |
| 37661_2 | TATGAGTGTGGAGTCGTTGCGGGCAAGGGACTTCCTGTAATAANNNAGATCGGAA |
| 49745_1 | TATGAGTGTGGAGTCGTTGCACCCAGCTTTTTTAGGTGTTGTGNNNAGATCGGAAG |
| 49745_2 | TATGAGTGTGGAGTCGTTGCACCCAGCTTTTTTAGGTGTTGTGNNNAGATCGGAAG |
| 13524_1 | TATGAGTGTGGAGTCGTTGCGGTATGCCTGTATCATATTTTTTNNNAGATCGGAAG |
| 13524_2 | TATGAGTGTGGAGTCGTTGCGGTATGCCTGTATCATATTTTTTNNNAGATCGGAAG |
| 12609_1 | TATGAGTGTGGAGTCGTTGCAAAGTTCTTCAGAAGCATCAANNNAGATCGGAAGA |
| 12609_2 | TATGAGTGTGGAGTCGTTGCAAAGTTCTTCAGAAGCATCAANNNAGATCGGAAGA |
| 42716_1 | TATGAGTGTGGAGTCGTTGCACTCTAGCCTGGGAAGCAGAGNNNAGATCGGAAGA |
| 42716_2 | TATGAGTGTGGAGTCGTTGCACTCTAGCCTGGGAAGCAGAGNNNAGATCGGAAGA |
| 10469_1 | TATGAGTGTGGAGTCGTTGCAATCACAAAACCAGCTATGGGCANNNAGATCGGAA |
| 10469_2 | TATGAGTGTGGAGTCGTTGCAATCACAAAACCAGCTATGGGCANNNAGATCGGAA |
| 31937_1 | TATGAGTGTGGAGTCGTTGCATTTCATGAGGCCAGCATCATTCNNNAGATCGGAAG |
| 31937_2 | TATGAGTGTGGAGTCGTTGCATTTCATGAGGCCAGCATCATTCNNNAGATCGGAAG |
| 43401_1 | TATGAGTGTGGAGTCGTTGCCCTCCCAAGTCCAAGTGATTCTCNNNAGATCGGAAG |
| 43401_2 | TATGAGTGTGGAGTCGTTGCCCTCCCAAGTCCAAGTGATTCTCNNNAGATCGGAAG |
| 38836_1 | TATGAGTGTGGAGTCGTTGCCAGACACACACACACACATATGCNNNAGATCGGAAG |
| 38836_2 | TATGAGTGTGGAGTCGTTGCCAGACACACACACACACATATGCNNNAGATCGGAAG |
| 50048_1 | TATGAGTGTGGAGTCGTTGCGACAAGATGTCCTCCCTTTCAGTNNNAGATCGGAAG |
| 50048_2 | TATGAGTGTGGAGTCGTTGCGACAAGATGTCCTCCCTTTCAGTNNNAGATCGGAAG |
| 41116_1 | TATGAGTGTGGAGTCGTTGCCCGACATTTGAAAAGAGTGCACTNNNAGATCGGAA |
| 41116_2 | TATGAGTGTGGAGTCGTTGCCCGACATTTGAAAAGAGTGCACTNNNAGATCGGAA |

| | |
|---|---|
| 32927_1 | TATGAGTGTGGAGTCGTTGCAAAAAATTAGCTGGGCATGGTGGNNNAGATCGGAA |
| 32927_2 | TATGAGTGTGGAGTCGTTGCAAAAAATTAGCTGGGCATGGTGGNNNAGATCGGAA |
| 44038_1 | TATGAGTGTGGAGTCGTTGCACTTCCTGGATCAGAACTTGCATNNNAGATCGGAAG |
| 44038_2 | TATGAGTGTGGAGTCGTTGCACTTCCTGGATCAGAACTTGCATNNNAGATCGGAAG |
| 47645_1 | TATGAGTGTGGAGTCGTTGCGGTTGAACTCCTTCCCTTCTAGGNNNAGATCGGAAG |
| 47645_2 | TATGAGTGTGGAGTCGTTGCGGTTGAACTCCTTCCCTTCTAGGNNNAGATCGGAAG |
| 29672_1 | TATGAGTGTGGAGTCGTTGCCCCAAGACCCGAAGTTCTCCNNNAGATCGGAAGAGC |
| 29672_2 | TATGAGTGTGGAGTCGTTGCCCCAAGACCCGAAGTTCTCCNNNAGATCGGAAGAGC |
| 32002_1 | TATGAGTGTGGAGTCGTTGCAGCATGAACAGAGGCTCAGAATTNNNAGATCGGAA |
| 32002_2 | TATGAGTGTGGAGTCGTTGCAGCATGAACAGAGGCTCAGAATTNNNAGATCGGAA |
| 13765_1 | TATGAGTGTGGAGTCGTTGCAAAAACTTTCCCAGTGAAGTATNNNAGATCGGAAGA |
| 13765_2 | TATGAGTGTGGAGTCGTTGCAAAAACTTTCCCAGTGAAGTATNNNAGATCGGAAGA |
| 28794_1 | TATGAGTGTGGAGTCGTTGCAGGCATATATAGGTGCTCAACGTNNNAGATCGGAA |
| 28794_2 | TATGAGTGTGGAGTCGTTGCAGGCATATATAGGTGCTCAACGTNNNAGATCGGAA |
| 35187_1 | TATGAGTGTGGAGTCGTTGCCAATTTTCTCCCTTACCCTCCCANNNAGATCGGAAGA |
| 35187_2 | TATGAGTGTGGAGTCGTTGCCAATTTTCTCCCTTACCCTCCCANNNAGATCGGAAGA |
| 48025_1 | TATGAGTGTGGAGTCGTTGCAGCCTCCTGAGTAGCTAGGATCNNNAGATCGGAAG |
| 48025_2 | TATGAGTGTGGAGTCGTTGCAGCCTCCTGAGTAGCTAGGATCNNNAGATCGGAAG |
| 41915_1 | TATGAGTGTGGAGTCGTTGCTGTAATCTCAGCTCACTGCAACTNNNAGATCGGAAG |
| 41915_2 | TATGAGTGTGGAGTCGTTGCTGTAATCTCAGCTCACTGCAACTNNNAGATCGGAAG |
| 46940_1 | TATGAGTGTGGAGTCGTTGCCACTAAAGAAATTCTGGCTGGGCNNNAGATCGGAA |
| 46940_2 | TATGAGTGTGGAGTCGTTGCCACTAAAGAAATTCTGGCTGGGCNNNAGATCGGAA |
| 47219_1 | TATGAGTGTGGAGTCGTTGCTAGAGACGAGGTTTTGCCATGTTNNNAGATCGGAA |
| 47219_2 | TATGAGTGTGGAGTCGTTGCTAGAGACGAGGTTTTGCCATGTTNNNAGATCGGAA |
| 31299_1 | TATGAGTGTGGAGTCGTTGCTGTATGAAAGACAGACTGGCCAANNNAGATCGGAA |
| 31299_2 | TATGAGTGTGGAGTCGTTGCTGTATGAAAGACAGACTGGCCAANNNAGATCGGAA |
| 51542_1 | TATGAGTGTGGAGTCGTTGCTGCAAGGCTATAAGGCTTAGCATNNNAGATCGGAA |
| 51542_2 | TATGAGTGTGGAGTCGTTGCTGCAAGGCTATAAGGCTTAGCATNNNAGATCGGAA |
| 41683_1 | TATGAGTGTGGAGTCGTTGCGATGAAAGCAATTCTCCTGCCTCNNNAGATCGGAAG |
| 41683_2 | TATGAGTGTGGAGTCGTTGCGATGAAAGCAATTCTCCTGCCTCNNNAGATCGGAAG |
| 45771_1 | TATGAGTGTGGAGTCGTTGCAAAGAGCTTTCCTCCAGCCTTCNNNAGATCGGAAGA |
| 45771_2 | TATGAGTGTGGAGTCGTTGCAAAGAGCTTTCCTCCAGCCTTCNNNAGATCGGAAGA |
| 36523_1 | TATGAGTGTGGAGTCGTTGCTCCTCTTGTAGTTCTAGTCCCAGTNNNAGATCGGAAG |
| 36523_2 | TATGAGTGTGGAGTCGTTGCTCCTCTTGTAGTTCTAGTCCCAGTNNNAGATCGGAAG |
| 31308_1 | TATGAGTGTGGAGTCGTTGCACCATTTGTTTTATGTGGCCTGGNNNAGATCGGAAG |
| 31308_2 | TATGAGTGTGGAGTCGTTGCACCATTTGTTTTATGTGGCCTGGNNNAGATCGGAAG |
| 34399_1 | TATGAGTGTGGAGTCGTTGCCCAGACTCAAGTAATCCTCCCACNNNAGATCGGAAG |
| 34399_2 | TATGAGTGTGGAGTCGTTGCCCAGACTCAAGTAATCCTCCCACNNNAGATCGGAAG |
| 49018_1 | TATGAGTGTGGAGTCGTTGCAAGAGTTTTTGCTCTGTCCTCCANNNAGATCGGAAG |
| 49018_2 | TATGAGTGTGGAGTCGTTGCAAGAGTTTTTGCTCTGTCCTCCANNNAGATCGGAAG |
| 38318_1 | TATGAGTGTGGAGTCGTTGCTGAAAAGATAGCTGCACTCCTGTNNNAGATCGGAAG |
| 38318_2 | TATGAGTGTGGAGTCGTTGCTGAAAAGATAGCTGCACTCCTGTNNNAGATCGGAAG |
| 50501_1 | TATGAGTGTGGAGTCGTTGCACATACTCTGAGATCCTTCCCCANNNAGATCGGAAGA |

| | |
|---|---|
| 50501_2 | TATGAGTGTGGAGTCGTTGCACATACTCTGAGATCCTTCCCCANNNAGATCGGAAGA |
| 47809_1 | TATGAGTGTGGAGTCGTTGCCCAAGTAGCTGGGATTACAGGTGNNNAGATCGGAA |
| 47809_2 | TATGAGTGTGGAGTCGTTGCCCAAGTAGCTGGGATTACAGGTGNNNAGATCGGAA |
| 42747_1 | TATGAGTGTGGAGTCGTTGCATCTGACATTTCCACAACCTCCANNNAGATCGGAAGA |
| 42747_2 | TATGAGTGTGGAGTCGTTGCATCTGACATTTCCACAACCTCCANNNAGATCGGAAGA |
| 45587_1 | TATGAGTGTGGAGTCGTTGCCATGGGTTCAAGCAATTCTCCTGNNNAGATCGGAAG |
| 45587_2 | TATGAGTGTGGAGTCGTTGCCATGGGTTCAAGCAATTCTCCTGNNNAGATCGGAAG |
| 32611_1 | TATGAGTGTGGAGTCGTTGCAAAGAAAGCAAAGAAACCCTGGCNNNAGATCGGAA |
| 32611_2 | TATGAGTGTGGAGTCGTTGCAAAGAAAGCAAAGAAACCCTGGCNNNAGATCGGAA |
| 42425_1 | TATGAGTGTGGAGTCGTTGCTCAACAGAAGAAAGGAGGAGAGCNNNAGATCGGA |
| 42425_2 | TATGAGTGTGGAGTCGTTGCTCAACAGAAGAAAGGAGGAGAGCNNNAGATCGGA |
| 41474_1 | TATGAGTGTGGAGTCGTTGCAGGAGGTTACAGTGAGCCAAAATNNNAGATCGGAA |
| 41474_2 | TATGAGTGTGGAGTCGTTGCAGGAGGTTACAGTGAGCCAAAATNNNAGATCGGAA |
| 30723_1 | TATGAGTGTGGAGTCGTTGCTATGGGCTTTTAATCTGCTAGCTNNNAGATCGGAA |
| 30723_2 | TATGAGTGTGGAGTCGTTGCTATGGGCTTTTAATCTGCTAGCTNNNAGATCGGAA |
| 45820_1 | TATGAGTGTGGAGTCGTTGCACTATCTGTGCTCATGGAACGTGNNNAGATCGGAAG |
| 45820_2 | TATGAGTGTGGAGTCGTTGCACTATCTGTGCTCATGGAACGTGNNNAGATCGGAAG |
| 51818_1 | TATGAGTGTGGAGTCGTTGCATCTACCCTACCAGTCTTCCCTCNNNAGATCGGAAGA |
| 51818_2 | TATGAGTGTGGAGTCGTTGCATCTACCCTACCAGTCTTCCCTCNNNAGATCGGAAGA |
| 29135_1 | TATGAGTGTGGAGTCGTTGCAATAGTGACTGCATGCTGTAGCTNNNAGATCGGAAG |
| 29135_2 | TATGAGTGTGGAGTCGTTGCAATAGTGACTGCATGCTGTAGCTNNNAGATCGGAAG |
| 36433_1 | TATGAGTGTGGAGTCGTTGCGTCTCACTCTGTTGCTGGGATTANNNAGATCGGAAG |
| 36433_2 | TATGAGTGTGGAGTCGTTGCGTCTCACTCTGTTGCTGGGATTANNNAGATCGGAAG |
| 1613_1 | TATGAGTGTGGAGTCGTTGCTTTCCCAGAACATGGCCTACCTANNNAGATCGGAAG |
| 1613_2 | TATGAGTGTGGAGTCGTTGCTTTCCCAGAACATGGCCTACCTANNNAGATCGGAAG |
| 39720_1 | TATGAGTGTGGAGTCGTTGCTTCATGACATTCTCCTGCCTCAGNNNAGATCGGAAGA |
| 39720_2 | TATGAGTGTGGAGTCGTTGCTTCATGACATTCTCCTGCCTCAGNNNAGATCGGAAGA |
| 35688_1 | TATGAGTGTGGAGTCGTTGCGAAAGACCCCTAGCAAGGCTAATNNNAGATCGGAA |
| 35688_2 | TATGAGTGTGGAGTCGTTGCGAAAGACCCCTAGCAAGGCTAATNNNAGATCGGAA |
| 12982_1 | TATGAGTGTGGAGTCGTTGCGTTAAATTCAAGGTCAATCCAGNNNAGATCGGAAGA |
| 12982_2 | TATGAGTGTGGAGTCGTTGCGTTAAATTCAAGGTCAATCCAGNNNAGATCGGAAGA |
| 39232_1 | TATGAGTGTGGAGTCGTTGCTTCCTAGAGCATGTCTTGGCTTTNNNAGATCGGAAG |
| 39232_2 | TATGAGTGTGGAGTCGTTGCTTCCTAGAGCATGTCTTGGCTTTNNNAGATCGGAAG |
| 34808_1 | TATGAGTGTGGAGTCGTTGCTGGAATAGCTGGAATTACAGGCGNNNAGATCGGAA |
| 34808_2 | TATGAGTGTGGAGTCGTTGCTGGAATAGCTGGAATTACAGGCGNNNAGATCGGAA |
| 31400_1 | TATGAGTGTGGAGTCGTTGCAGTACAGTGTTCACTACTGCCATNNNAGATCGGAAG |
| 31400_2 | TATGAGTGTGGAGTCGTTGCAGTACAGTGTTCACTACTGCCATNNNAGATCGGAAG |
| 29285_1 | TATGAGTGTGGAGTCGTTGCAATCCATTCATGAAGGTGGAGCTNNNAGATCGGAA |
| 29285_2 | TATGAGTGTGGAGTCGTTGCAATCCATTCATGAAGGTGGAGCTNNNAGATCGGAA |
| 36854_1 | TATGAGTGTGGAGTCGTTGCTTATGAAAAAAAGCCAGGCGTGGNNNAGATCGGAA |
| 36854_2 | TATGAGTGTGGAGTCGTTGCTTATGAAAAAAAGCCAGGCGTGGNNNAGATCGGAA |
| 2505_1 | TATGAGTGTGGAGTCGTTGCCGTGTGTGTAATAGGGAGTGGAGNNNAGATCGGA |
| 2505_2 | TATGAGTGTGGAGTCGTTGCCGTGTGTGTAATAGGGAGTGGAGNNNAGATCGGA |

| | |
|---|---|
| 45053_1 | TATGAGTGTGGAGTCGTTGCAGGGGTCAAGTTACTTCTTAGTACANNNAGATCGGA |
| 45053_2 | TATGAGTGTGGAGTCGTTGCAGGGGTCAAGTTACTTCTTAGTACANNNAGATCGGA |
| 3139_1 | TATGAGTGTGGAGTCGTTGCGCAACTCTGGCAAAACGAAATGANNNAGATCGGAA |
| 3139_2 | TATGAGTGTGGAGTCGTTGCGCAACTCTGGCAAAACGAAATGANNNAGATCGGAA |
| 13417_1 | TATGAGTGTGGAGTCGTTGCCACCATTCATTTCATTTTGTTTTNNNAGATCGGAAGA |
| 13417_2 | TATGAGTGTGGAGTCGTTGCCACCATTCATTTCATTTTGTTTTNNNAGATCGGAAGA |
| 9836_1 | TATGAGTGTGGAGTCGTTGCGAAACCGGGAGAAATGATGGTGANNNAGATCGGA |
| 9836_2 | TATGAGTGTGGAGTCGTTGCGAAACCGGGAGAAATGATGGTGANNNAGATCGGA |
| 45192_1 | TATGAGTGTGGAGTCGTTGCGACTGAGGATACAGATGGAGCAGNNNAGATCGGA |
| 45192_2 | TATGAGTGTGGAGTCGTTGCGACTGAGGATACAGATGGAGCAGNNNAGATCGGA |
| 35491_1 | TATGAGTGTGGAGTCGTTGCTGGAATGAAATGGCACGATCTTGNNNAGATCGGAA |
| 35491_2 | TATGAGTGTGGAGTCGTTGCTGGAATGAAATGGCACGATCTTGNNNAGATCGGAA |
| 48870_1 | TATGAGTGTGGAGTCGTTGCCCACCAAGCCTGGCTAATTTTTTNNNAGATCGGAAG |
| 48870_2 | TATGAGTGTGGAGTCGTTGCCCACCAAGCCTGGCTAATTTTTTNNNAGATCGGAAG |
| 46834_1 | TATGAGTGTGGAGTCGTTGCAGCGATCTTCCCACTTCCATATCNNNAGATCGGAAGA |
| 46834_2 | TATGAGTGTGGAGTCGTTGCAGCGATCTTCCCACTTCCATATCNNNAGATCGGAAGA |
| 41535_1 | TATGAGTGTGGAGTCGTTGCCTTCCCAGGTTCAAGCAATTCTCNNNAGATCGGAAG |
| 41535_2 | TATGAGTGTGGAGTCGTTGCCTTCCCAGGTTCAAGCAATTCTCNNNAGATCGGAAG |
| 46132_1 | TATGAGTGTGGAGTCGTTGCTCAGCGGCCTAAATACACAATCTNNNAGATCGGAAG |
| 46132_2 | TATGAGTGTGGAGTCGTTGCTCAGCGGCCTAAATACACAATCTNNNAGATCGGAAG |
| 40445_1 | TATGAGTGTGGAGTCGTTGCGCCACCATGCCAGTTTACTTTTTNNNAGATCGGAAGA |
| 40445_2 | TATGAGTGTGGAGTCGTTGCGCCACCATGCCAGTTTACTTTTTNNNAGATCGGAAGA |
| 37241_1 | TATGAGTGTGGAGTCGTTGCATGTGGAGAAAAGGGGAACACTTNNNAGATCGGAA |
| 37241_2 | TATGAGTGTGGAGTCGTTGCATGTGGAGAAAAGGGGAACACTTNNNAGATCGGAA |
| 51762_1 | TATGAGTGTGGAGTCGTTGCGGCTTGGGAACATCCATCTAGATNNNAGATCGGAAG |
| 51762_2 | TATGAGTGTGGAGTCGTTGCGGCTTGGGAACATCCATCTAGATNNNAGATCGGAAG |
| 36345_1 | TATGAGTGTGGAGTCGTTGCGTCACACGTGAGTGGTTTTGAAANNNAGATCGGAA |
| 36345_2 | TATGAGTGTGGAGTCGTTGCGTCACACGTGAGTGGTTTTGAAANNNAGATCGGAA |
| 31379_1 | TATGAGTGTGGAGTCGTTGCAGATGAATGAAGGAGAGGAGGGANNNAGATCGGA |
| 31379_2 | TATGAGTGTGGAGTCGTTGCAGATGAATGAAGGAGAGGAGGGANNNAGATCGGA |
| 35375_1 | TATGAGTGTGGAGTCGTTGCTGCCTTACACAACTAAGTCACCTNNNAGATCGGAAG |
| 35375_2 | TATGAGTGTGGAGTCGTTGCTGCCTTACACAACTAAGTCACCTNNNAGATCGGAAG |
| 49060_1 | TATGAGTGTGGAGTCGTTGCCGAGTAGCTGGGATTACAGGCNNNAGATCGGAAGA |
| 49060_2 | TATGAGTGTGGAGTCGTTGCCGAGTAGCTGGGATTACAGGCNNNAGATCGGAAGA |
| 40118_1 | TATGAGTGTGGAGTCGTTGCTGATTGATGTATGCATTTTGGTGTNNNAGATCGGAA |
| 40118_2 | TATGAGTGTGGAGTCGTTGCTGATTGATGTATGCATTTTGGTGTNNNAGATCGGAA |
| 32124_1 | TATGAGTGTGGAGTCGTTGCGAGTTTTGCTCTTGTTGCCCAGNNNAGATCGGAAGA |
| 32124_2 | TATGAGTGTGGAGTCGTTGCGAGTTTTGCTCTTGTTGCCCAGNNNAGATCGGAAGA |
| 49426_1 | TATGAGTGTGGAGTCGTTGCCCTCACAAGTAGCTGGGATTACANNNAGATCGGAAG |
| 49426_2 | TATGAGTGTGGAGTCGTTGCCCTCACAAGTAGCTGGGATTACANNNAGATCGGAAG |
| 32383_1 | TATGAGTGTGGAGTCGTTGCAAGTGCTGATGTAAAAGCTGCAGNNNAGATCGGAA |
| 32383_2 | TATGAGTGTGGAGTCGTTGCAAGTGCTGATGTAAAAGCTGCAGNNNAGATCGGAA |
| 29958_1 | TATGAGTGTGGAGTCGTTGCCAGCCTTGGTTTTCTGGGAAAAANNNAGATCGGAAG |

| | |
|---|---|
| 29958_2 | TATGAGTGTGGAGTCGTTGCCAGCCTTGGTTTTCTGGGAAAAANNNAGATCGGAAG |
| 48078_1 | TATGAGTGTGGAGTCGTTGCCTCCCGGGTTCATGCCATTATCNNNAGATCGGAAGA |
| 48078_2 | TATGAGTGTGGAGTCGTTGCCTCCCGGGTTCATGCCATTATCNNNAGATCGGAAGA |
| 31766_1 | TATGAGTGTGGAGTCGTTGCCATGATCTCAGCTCACTGCAACNNNAGATCGGAAGA |
| 31766_2 | TATGAGTGTGGAGTCGTTGCCATGATCTCAGCTCACTGCAACNNNAGATCGGAAGA |
| 41653_1 | TATGAGTGTGGAGTCGTTGCCACAATCTCAGCTCACTGCAAGNNNAGATCGGAAGA |
| 41653_2 | TATGAGTGTGGAGTCGTTGCCACAATCTCAGCTCACTGCAAGNNNAGATCGGAAGA |
| 5714_1 | TATGAGTGTGGAGTCGTTGCTCTTGTGTGTCTGTTCTTGCTCANNNAGATCGGAAGA |
| 5714_2 | TATGAGTGTGGAGTCGTTGCTCTTGTGTGTCTGTTCTTGCTCANNNAGATCGGAAGA |
| 30923_1 | TATGAGTGTGGAGTCGTTGCTTACATCCTCTGTTTGATGGGCANNNAGATCGGAAG |
| 30923_2 | TATGAGTGTGGAGTCGTTGCTTACATCCTCTGTTTGATGGGCANNNAGATCGGAAG |
| 12908_1 | TATGAGTGTGGAGTCGTTGCACTTGGATATCTGCATGAATCNNNAGATCGGAAGAG |
| 12908_2 | TATGAGTGTGGAGTCGTTGCACTTGGATATCTGCATGAATCNNNAGATCGGAAGAG |
| 40751_1 | TATGAGTGTGGAGTCGTTGCGCCTCAGCAGGAAGTTTAGAAGANNNAGATCGGAA |
| 40751_2 | TATGAGTGTGGAGTCGTTGCGCCTCAGCAGGAAGTTTAGAAGANNNAGATCGGAA |
| 35590_1 | TATGAGTGTGGAGTCGTTGCCATGATCTTGGCTCACTGCAATGNNNAGATCGGAAG |
| 35590_2 | TATGAGTGTGGAGTCGTTGCCATGATCTTGGCTCACTGCAATGNNNAGATCGGAAG |
| 47158_1 | TATGAGTGTGGAGTCGTTGCCCTGGCTCTTTGTTGAAGTGATGNNNAGATCGGAAG |
| 47158_2 | TATGAGTGTGGAGTCGTTGCCCTGGCTCTTTGTTGAAGTGATGNNNAGATCGGAAG |
| 43312_1 | TATGAGTGTGGAGTCGTTGCGACAGGGTCTCACCTTATTACCCNNNAGATCGGAAG |
| 43312_2 | TATGAGTGTGGAGTCGTTGCGACAGGGTCTCACCTTATTACCCNNNAGATCGGAAG |
| 39427_1 | TATGAGTGTGGAGTCGTTGCGCTTACAGTGAGCTGAGATCGTNNNAGATCGGAAG |
| 39427_2 | TATGAGTGTGGAGTCGTTGCGCTTACAGTGAGCTGAGATCGTNNNAGATCGGAAG |
| 37769_1 | TATGAGTGTGGAGTCGTTGCCCTGTAGTCCCAGCTACTTAGGANNNAGATCGGAAG |
| 37769_2 | TATGAGTGTGGAGTCGTTGCCCTGTAGTCCCAGCTACTTAGGANNNAGATCGGAAG |
| 40898_1 | TATGAGTGTGGAGTCGTTGCTGTAGACCCAGCTACTTGAGAGANNNAGATCGGAA |
| 40898_2 | TATGAGTGTGGAGTCGTTGCTGTAGACCCAGCTACTTGAGAGANNNAGATCGGAA |
| 30738_1 | TATGAGTGTGGAGTCGTTGCCTCTCTCTCTCTCTCACACCANNNAGATCGGAAGAG |
| 30738_2 | TATGAGTGTGGAGTCGTTGCCTCTCTCTCTCTCTCACACCANNNAGATCGGAAGAG |
| 41777_1 | TATGAGTGTGGAGTCGTTGCCCATGTTTCCCACACTGATCTTGNNNAGATCGGAAGA |
| 41777_2 | TATGAGTGTGGAGTCGTTGCCCATGTTTCCCACACTGATCTTGNNNAGATCGGAAGA |
| 41137_1 | TATGAGTGTGGAGTCGTTGCAAAACCAGCAAGACTCCATCTCTNNNAGATCGGAAG |
| 41137_2 | TATGAGTGTGGAGTCGTTGCAAAACCAGCAAGACTCCATCTCTNNNAGATCGGAAG |
| 2127_1 | TATGAGTGTGGAGTCGTTGCGGGACACAGTGAGAAAGCAATCANNNAGATCGGAA |
| 2127_2 | TATGAGTGTGGAGTCGTTGCGGGACACAGTGAGAAAGCAATCANNNAGATCGGAA |
| 13431_1 | TATGAGTGTGGAGTCGTTGCTCGATTGTCTTCTCTAGTGANNNAGATCGGAAGAGC |
| 13431_2 | TATGAGTGTGGAGTCGTTGCTCGATTGTCTTCTCTAGTGANNNAGATCGGAAGAGC |
| 34288_1 | TATGAGTGTGGAGTCGTTGCAAAGAGGGAAGGAAAGGAAAGGGNNNAGATCGG |
| 34288_2 | TATGAGTGTGGAGTCGTTGCAAAGAGGGAAGGAAAGGAAAGGGNNNAGATCGG |
| 12832_1 | TATGAGTGTGGAGTCGTTGCAAGGGTTTTTTACATCATAAGACNNNAGATCGGAAG |
| 12832_2 | TATGAGTGTGGAGTCGTTGCAAGGGTTTTTTACATCATAAGACNNNAGATCGGAAG |
| 43865_1 | TATGAGTGTGGAGTCGTTGCTGCAATGAGATTTGTCAACAGCTNNNAGATCGGAAG |
| 43865_2 | TATGAGTGTGGAGTCGTTGCTGCAATGAGATTTGTCAACAGCTNNNAGATCGGAAG |

| | |
|---|---|
| 51380_1 | TATGAGTGTGGAGTCGTTGCCCACATGACATACTGAGACACCANNNAGATCGGAAG |
| 51380_2 | TATGAGTGTGGAGTCGTTGCCCACATGACATACTGAGACACCANNNAGATCGGAAG |
| 49289_1 | TATGAGTGTGGAGTCGTTGCTGGAGTGCAGTGGTACAATCTCNNNAGATCGGAAG |
| 49289_2 | TATGAGTGTGGAGTCGTTGCTGGAGTGCAGTGGTACAATCTCNNNAGATCGGAAG |
| 8067_1 | TATGAGTGTGGAGTCGTTGCGTCAGGAGTTTGACACCATCGCNNNAGATCGGAAG |
| 8067_2 | TATGAGTGTGGAGTCGTTGCGTCAGGAGTTTGACACCATCGCNNNAGATCGGAAG |
| 40277_1 | TATGAGTGTGGAGTCGTTGCGAAGCAGGAGAATCACTTGAACCNNNAGATCGGAA |
| 40277_2 | TATGAGTGTGGAGTCGTTGCGAAGCAGGAGAATCACTTGAACCNNNAGATCGGAA |
| 45212_1 | TATGAGTGTGGAGTCGTTGCTTTTACTCCTGGTTCTATGCCCCNNNAGATCGGAAGA |
| 45212_2 | TATGAGTGTGGAGTCGTTGCTTTTACTCCTGGTTCTATGCCCCNNNAGATCGGAAGA |
| 40625_1 | TATGAGTGTGGAGTCGTTGCCCTCCCGAGTAGCTGAAATTACANNNAGATCGGAAG |
| 40625_2 | TATGAGTGTGGAGTCGTTGCCCTCCCGAGTAGCTGAAATTACANNNAGATCGGAAG |
| 40159_1 | TATGAGTGTGGAGTCGTTGCCGGCTGTAATCCTAGCATTTTGGNNNAGATCGGAAG |
| 40159_2 | TATGAGTGTGGAGTCGTTGCCGGCTGTAATCCTAGCATTTTGGNNNAGATCGGAAG |
| 41315_1 | TATGAGTGTGGAGTCGTTGCTCAGTGGATTTCTGGCAATACCANNNAGATCGGAAG |
| 41315_2 | TATGAGTGTGGAGTCGTTGCTCAGTGGATTTCTGGCAATACCANNNAGATCGGAAG |
| 44793_1 | TATGAGTGTGGAGTCGTTGCTGAGTTTCCCTAACTGAGTGAGCNNNAGATCGGAAG |
| 44793_2 | TATGAGTGTGGAGTCGTTGCTGAGTTTCCCTAACTGAGTGAGCNNNAGATCGGAAG |
| 29868_1 | TATGAGTGTGGAGTCGTTGCCAGGCGATCAAGGAGATATTCCANNNAGATCGGAA |
| 29868_2 | TATGAGTGTGGAGTCGTTGCCAGGCGATCAAGGAGATATTCCANNNAGATCGGAA |
| 33107_1 | TATGAGTGTGGAGTCGTTGCATCACAAGGTCAGGAATTCGAGTNNNAGATCGGAA |
| 33107_2 | TATGAGTGTGGAGTCGTTGCATCACAAGGTCAGGAATTCGAGTNNNAGATCGGAA |
| 51718_1 | TATGAGTGTGGAGTCGTTGCCGCCAAAAAAAAGGAGTCTCAGTNNNAGATCGGAA |
| 51718_2 | TATGAGTGTGGAGTCGTTGCCGCCAAAAAAAAGGAGTCTCAGTNNNAGATCGGAA |
| 48032_1 | TATGAGTGTGGAGTCGTTGCTGTGCAAGAAAGAGACCTGGAATNNNAGATCGGAA |
| 48032_2 | TATGAGTGTGGAGTCGTTGCTGTGCAAGAAAGAGACCTGGAATNNNAGATCGGAA |
| 41585_1 | TATGAGTGTGGAGTCGTTGCTTCAAGTGATTCTCCTGCCTCAGNNNAGATCGGAAG |
| 41585_2 | TATGAGTGTGGAGTCGTTGCTTCAAGTGATTCTCCTGCCTCAGNNNAGATCGGAAG |
| 51040_1 | TATGAGTGTGGAGTCGTTGCTCCTTTCCATTTCTCTCTTGAACCTNNNAGATCGGAAG |
| 51040_2 | TATGAGTGTGGAGTCGTTGCTCCTTTCCATTTCTCTCTTGAACCTNNNAGATCGGAAG |
| 13551_1 | TATGAGTGTGGAGTCGTTGCTAACTTTGCAAAATACTTGAGTCNNNAGATCGGAAG |
| 13551_2 | TATGAGTGTGGAGTCGTTGCTAACTTTGCAAAATACTTGAGTCNNNAGATCGGAAG |
| 33846_1 | TATGAGTGTGGAGTCGTTGCGCAGAGATTGTAGTGAGCTGAGANNNAGATCGGAA |
| 33846_2 | TATGAGTGTGGAGTCGTTGCGCAGAGATTGTAGTGAGCTGAGANNNAGATCGGAA |
| 34033_1 | TATGAGTGTGGAGTCGTTGCAAGTAGCTGAGATTACAGGTGCCNNNAGATCGGAA |
| 34033_2 | TATGAGTGTGGAGTCGTTGCAAGTAGCTGAGATTACAGGTGCCNNNAGATCGGAA |
| 50061_1 | TATGAGTGTGGAGTCGTTGCAGCCTAAGTGTCCATCAATTGATGNNNAGATCGGAA |
| 50061_2 | TATGAGTGTGGAGTCGTTGCAGCCTAAGTGTCCATCAATTGATGNNNAGATCGGAA |
| 43886_1 | TATGAGTGTGGAGTCGTTGCTTGGAAAAAACAGTTGAGGAGCGNNNAGATCGGAA |
| 43886_2 | TATGAGTGTGGAGTCGTTGCTTGGAAAAAACAGTTGAGGAGCGNNNAGATCGGAA |
| 35427_1 | TATGAGTGTGGAGTCGTTGCCTTACTGTAAGCTCCACCTCCCNNNAGATCGGAAGAG |
| 35427_2 | TATGAGTGTGGAGTCGTTGCCTTACTGTAAGCTCCACCTCCCNNNAGATCGGAAGAG |
| 52638_1 | TATGAGTGTGGAGTCGTTGCACTGTCAAAATCAGCTAGTGGGANNNAGATCGGAA |

| | |
|---|---|
| 52638_2 | TATGAGTGTGGAGTCGTTGCACTGTCAAAATCAGCTAGTGGGANNNAGATCGGAA |
| 30200_1 | TATGAGTGTGGAGTCGTTGCGCATGAAAGACCACCCCTATTTGNNNAGATCGGAAG |
| 30200_2 | TATGAGTGTGGAGTCGTTGCGCATGAAAGACCACCCCTATTTGNNNAGATCGGAAG |
| 38905_1 | TATGAGTGTGGAGTCGTTGCTTTAGGCACTCAAGGGAATCTCCNNNAGATCGGAAG |
| 38905_2 | TATGAGTGTGGAGTCGTTGCTTTAGGCACTCAAGGGAATCTCCNNNAGATCGGAAG |
| 44053_1 | TATGAGTGTGGAGTCGTTGCCATTACCCTGGTGTCCTTGATGANNNAGATCGGAAG |
| 44053_2 | TATGAGTGTGGAGTCGTTGCCATTACCCTGGTGTCCTTGATGANNNAGATCGGAAG |
| 12051_1 | TATGAGTGTGGAGTCGTTGCTGAAACAACAACCTGATGAGGTAGNNNAGATCGGA |
| 12051_2 | TATGAGTGTGGAGTCGTTGCTGAAACAACAACCTGATGAGGTAGNNNAGATCGGA |
| 37290_1 | TATGAGTGTGGAGTCGTTGCTTGTGTGCAGAGAGGTGATTGTANNNAGATCGGAA |
| 37290_2 | TATGAGTGTGGAGTCGTTGCTTGTGTGCAGAGAGGTGATTGTANNNAGATCGGAA |
| 33116_1 | TATGAGTGTGGAGTCGTTGCTGTTGTCCCAGGCATTTCTGATANNNAGATCGGAAG |
| 33116_2 | TATGAGTGTGGAGTCGTTGCTGTTGTCCCAGGCATTTCTGATANNNAGATCGGAAG |
| 46687_1 | TATGAGTGTGGAGTCGTTGCCACCACACCCAGCTAATTTTTGTNNNAGATCGGAAGA |
| 46687_2 | TATGAGTGTGGAGTCGTTGCCACCACACCCAGCTAATTTTTGTNNNAGATCGGAAGA |
| 46349_1 | TATGAGTGTGGAGTCGTTGCACCAAGTAATGGGATTGCTGGATNNNAGATCGGAA |
| 46349_2 | TATGAGTGTGGAGTCGTTGCACCAAGTAATGGGATTGCTGGATNNNAGATCGGAA |
| 40687_1 | TATGAGTGTGGAGTCGTTGCCAGTGAAATTACCTGGGTCTGGANNNAGATCGGAA |
| 40687_2 | TATGAGTGTGGAGTCGTTGCCAGTGAAATTACCTGGGTCTGGANNNAGATCGGAA |
| 35582_1 | TATGAGTGTGGAGTCGTTGCCCCAAACCCATGAGGACATTTTCNNNAGATCGGAAG |
| 35582_2 | TATGAGTGTGGAGTCGTTGCCCCAAACCCATGAGGACATTTTCNNNAGATCGGAAG |
| 48251_1 | TATGAGTGTGGAGTCGTTGCTTCAAGCGATTCTCCTGTCTCAGNNNAGATCGGAAG |
| 48251_2 | TATGAGTGTGGAGTCGTTGCTTCAAGCGATTCTCCTGTCTCAGNNNAGATCGGAAG |
| 37074_1 | TATGAGTGTGGAGTCGTTGCTTTAGAACAGATCCTGGCACACGNNNAGATCGGAAG |
| 37074_2 | TATGAGTGTGGAGTCGTTGCTTTAGAACAGATCCTGGCACACGNNNAGATCGGAAG |
| 44759_1 | TATGAGTGTGGAGTCGTTGCTTGGGTCCCTATATCTGTGAGGANNNAGATCGGAA |
| 44759_2 | TATGAGTGTGGAGTCGTTGCTTGGGTCCCTATATCTGTGAGGANNNAGATCGGAA |
| 43900_1 | TATGAGTGTGGAGTCGTTGCGGACTGTACTGGCCATACATGAANNNAGATCGGAA |
| 43900_2 | TATGAGTGTGGAGTCGTTGCGGACTGTACTGGCCATACATGAANNNAGATCGGAA |
| 44957_1 | TATGAGTGTGGAGTCGTTGCCAAGAACTAGGTTCAGGTGGGAANNNAGATCGGAA |
| 44957_2 | TATGAGTGTGGAGTCGTTGCCAAGAACTAGGTTCAGGTGGGAANNNAGATCGGAA |
| 42017_1 | TATGAGTGTGGAGTCGTTGCGAGTAGCTGGGATTACAGGCATGNNNAGATCGGAA |
| 42017_2 | TATGAGTGTGGAGTCGTTGCGAGTAGCTGGGATTACAGGCATGNNNAGATCGGAA |
| 40691_1 | TATGAGTGTGGAGTCGTTGCCTGAGTAGCTGGGATTACAGGTGNNNAGATCGGAA |
| 40691_2 | TATGAGTGTGGAGTCGTTGCCTGAGTAGCTGGGATTACAGGTGNNNAGATCGGAA |
| 37501_1 | TATGAGTGTGGAGTCGTTGCGCTTTTCTACACTTTCACCCCACNNNAGATCGGAAGA |
| 37501_2 | TATGAGTGTGGAGTCGTTGCGCTTTTCTACACTTTCACCCCACNNNAGATCGGAAGA |
| 52239_1 | TATGAGTGTGGAGTCGTTGCCCCCCAACAATTGCTTTGAGAATNNNAGATCGGAAG |
| 52239_2 | TATGAGTGTGGAGTCGTTGCCCCCCAACAATTGCTTTGAGAATNNNAGATCGGAAG |
| 51684_1 | TATGAGTGTGGAGTCGTTGCTGTGGTCTTTTGTGACTGACTTCNNNAGATCGGAAG |
| 51684_2 | TATGAGTGTGGAGTCGTTGCTGTGGTCTTTTGTGACTGACTTCNNNAGATCGGAAG |
| 46097_1 | TATGAGTGTGGAGTCGTTGCGTTCAAGAGATTCTCCTGCCTCANNNAGATCGGAAG |
| 46097_2 | TATGAGTGTGGAGTCGTTGCGTTCAAGAGATTCTCCTGCCTCANNNAGATCGGAAG |

| | |
|---|---|
| 35166_1 | TATGAGTGTGGAGTCGTTGCATCTCGGCTCACTGAAAGCTTTANNNAGATCGGAAG |
| 35166_2 | TATGAGTGTGGAGTCGTTGCATCTCGGCTCACTGAAAGCTTTANNNAGATCGGAAG |
| 9467_1 | TATGAGTGTGGAGTCGTTGCAACACCAAAGACAGAAGGCTCAGNNNAGATCGGAA |
| 9467_2 | TATGAGTGTGGAGTCGTTGCAACACCAAAGACAGAAGGCTCAGNNNAGATCGGAA |
| 31702_1 | TATGAGTGTGGAGTCGTTGCCAAGACCCCTTTGCAAAAGAGAGNNNAGATCGGAA |
| 31702_2 | TATGAGTGTGGAGTCGTTGCCAAGACCCCTTTGCAAAAGAGAGNNNAGATCGGAA |
| 37348_1 | TATGAGTGTGGAGTCGTTGCCCATTTCATGAGGCTATAGCTGCNNNAGATCGGAAG |
| 37348_2 | TATGAGTGTGGAGTCGTTGCCCATTTCATGAGGCTATAGCTGCNNNAGATCGGAAG |
| 49989_1 | TATGAGTGTGGAGTCGTTGCCGTGCAGACATTCAGCAGTAATCNNNAGATCGGAAG |
| 49989_2 | TATGAGTGTGGAGTCGTTGCCGTGCAGACATTCAGCAGTAATCNNNAGATCGGAAG |
| 35146_1 | TATGAGTGTGGAGTCGTTGCATACAATCTCTGCCACCCATGTTNNNAGATCGGAAG |
| 35146_2 | TATGAGTGTGGAGTCGTTGCATACAATCTCTGCCACCCATGTTNNNAGATCGGAAG |
| 42870_1 | TATGAGTGTGGAGTCGTTGCGGGGGGTGTATTTAGGAGCTAAGNNNAGATCGGA |
| 42870_2 | TATGAGTGTGGAGTCGTTGCGGGGGGTGTATTTAGGAGCTAAGNNNAGATCGGA |
| 32139_1 | TATGAGTGTGGAGTCGTTGCCATTCTCCTGCCTCAGCCTCNNNAGATCGGAAGAGCA |
| 32139_2 | TATGAGTGTGGAGTCGTTGCCATTCTCCTGCCTCAGCCTCNNNAGATCGGAAGAGCA |
| 36362_1 | TATGAGTGTGGAGTCGTTGCTGGACTCTATTTTGTTTCCTTGGTNNNAGATCGGAA |
| 36362_2 | TATGAGTGTGGAGTCGTTGCTGGACTCTATTTTGTTTCCTTGGTNNNAGATCGGAA |
| 38227_1 | TATGAGTGTGGAGTCGTTGCCACACGCAGCTTTGTCAAATACTNNNAGATCGGAAG |
| 38227_2 | TATGAGTGTGGAGTCGTTGCCACACGCAGCTTTGTCAAATACTNNNAGATCGGAAG |
| 38727_1 | TATGAGTGTGGAGTCGTTGCGGGTTGCAATCCTAGTCTCTGATNNNAGATCGGAAG |
| 38727_2 | TATGAGTGTGGAGTCGTTGCGGGTTGCAATCCTAGTCTCTGATNNNAGATCGGAAG |
| 31993_1 | TATGAGTGTGGAGTCGTTGCGTACTGTGCTAGGCAAGGGTATTNNNAGATCGGAA |
| 31993_2 | TATGAGTGTGGAGTCGTTGCGTACTGTGCTAGGCAAGGGTATTNNNAGATCGGAA |
| 45463_1 | TATGAGTGTGGAGTCGTTGCGGGATGAATCTGTGACACTCCTTNNNAGATCGGAAG |
| 45463_2 | TATGAGTGTGGAGTCGTTGCGGGATGAATCTGTGACACTCCTTNNNAGATCGGAAG |
| 13611_1 | TATGAGTGTGGAGTCGTTGCCCATTCTTGAGAGTTCCTTTTTNNNAGATCGGAAGA |
| 13611_2 | TATGAGTGTGGAGTCGTTGCCCATTCTTGAGAGTTCCTTTTTNNNAGATCGGAAGA |
| 34871_1 | TATGAGTGTGGAGTCGTTGCAGTAGCTGGGACTACAGGTATGTNNNAGATCGGAA |
| 34871_2 | TATGAGTGTGGAGTCGTTGCAGTAGCTGGGACTACAGGTATGTNNNAGATCGGAA |
| 43186_1 | TATGAGTGTGGAGTCGTTGCTTCAAGTGATTCTCCTGCCTCAGNNNAGATCGGAAG |
| 43186_2 | TATGAGTGTGGAGTCGTTGCTTCAAGTGATTCTCCTGCCTCAGNNNAGATCGGAAG |
| 31913_1 | TATGAGTGTGGAGTCGTTGCAAGAATTCAGTTCACTGCCAAGCNNNAGATCGGAAG |
| 31913_2 | TATGAGTGTGGAGTCGTTGCAAGAATTCAGTTCACTGCCAAGCNNNAGATCGGAAG |
| 40688_1 | TATGAGTGTGGAGTCGTTGCGGAATCAATCCAAATGCCCATCANNNAGATCGGAAG |
| 40688_2 | TATGAGTGTGGAGTCGTTGCGGAATCAATCCAAATGCCCATCANNNAGATCGGAAG |
| 10045_1 | TATGAGTGTGGAGTCGTTGCCAGTTCCTACTTCTGCGATCCAGNNNAGATCGGAAG |
| 10045_2 | TATGAGTGTGGAGTCGTTGCCAGTTCCTACTTCTGCGATCCAGNNNAGATCGGAAG |
| 43125_1 | TATGAGTGTGGAGTCGTTGCGAGTCAAGACCTGTTAGTGTGCTNNNAGATCGGAA |
| 43125_2 | TATGAGTGTGGAGTCGTTGCGAGTCAAGACCTGTTAGTGTGCTNNNAGATCGGAA |
| 49460_1 | TATGAGTGTGGAGTCGTTGCAGCTTTATGTGGTACTGATGGCANNNAGATCGGAA |
| 49460_2 | TATGAGTGTGGAGTCGTTGCAGCTTTATGTGGTACTGATGGCANNNAGATCGGAA |
| 40538_1 | TATGAGTGTGGAGTCGTTGCTTTACAGTGTTCTCCAACCACCANNNAGATCGGAAG |

| | |
|---|---|
| 40538_2 | TATGAGTGTGGAGTCGTTGCTTTACAGTGTTCTCCAACCACCANNNAGATCGGAAGA |
| 13156_1 | TATGAGTGTGGAGTCGTTGCGATGATCTTCAGTATCAGGACNNNAGATCGGAAGAG |
| 13156_2 | TATGAGTGTGGAGTCGTTGCGATGATCTTCAGTATCAGGACNNNAGATCGGAAGAG |
| 30358_1 | TATGAGTGTGGAGTCGTTGCCCCAAAAGTCTCCCATAGACTTCCNNNAGATCGGAAG |
| 30358_2 | TATGAGTGTGGAGTCGTTGCCCCAAAAGTCTCCCATAGACTTCCNNNAGATCGGAAG |
| 12916_1 | TATGAGTGTGGAGTCGTTGCTCTGCATCTCTTGTTCTTTGNNNAGATCGGAAGAGCA |
| 12916_2 | TATGAGTGTGGAGTCGTTGCTCTGCATCTCTTGTTCTTTGNNNAGATCGGAAGAGCA |
| 32405_1 | TATGAGTGTGGAGTCGTTGCGAGAATGGCATGAACCTGGGAGNNNAGATCGGAAG |
| 32405_2 | TATGAGTGTGGAGTCGTTGCGAGAATGGCATGAACCTGGGAGNNNAGATCGGAAG |
| 45982_1 | TATGAGTGTGGAGTCGTTGCCGATCTCCTGACCTCATGATCTGNNNAGATCGGAAGA |
| 45982_2 | TATGAGTGTGGAGTCGTTGCCGATCTCCTGACCTCATGATCTGNNNAGATCGGAAGA |
| 31898_1 | TATGAGTGTGGAGTCGTTGCGCATTGACTTCTGGCTCTGATTCNNNAGATCGGAAGA |
| 31898_2 | TATGAGTGTGGAGTCGTTGCGCATTGACTTCTGGCTCTGATTCNNNAGATCGGAAGA |
| 39308_1 | TATGAGTGTGGAGTCGTTGCTGGCAATTTCAGAGGCGAACTATNNNAGATCGGAAG |
| 39308_2 | TATGAGTGTGGAGTCGTTGCTGGCAATTTCAGAGGCGAACTATNNNAGATCGGAAG |
| 37385_1 | TATGAGTGTGGAGTCGTTGCGCAACATAGTGAAACCCCATCTCNNNAGATCGGAAGA |
| 37385_2 | TATGAGTGTGGAGTCGTTGCGCAACATAGTGAAACCCCATCTCNNNAGATCGGAAGA |
| 49286_1 | TATGAGTGTGGAGTCGTTGCCCACCATGCCCAGCTAATTTTTTNNNAGATCGGAAGA |
| 49286_2 | TATGAGTGTGGAGTCGTTGCCCACCATGCCCAGCTAATTTTTTNNNAGATCGGAAGA |
| 12475_1 | TATGAGTGTGGAGTCGTTGCTACACTCTGCTAAACTTATTTCANNNAGATCGGAAGA |
| 12475_2 | TATGAGTGTGGAGTCGTTGCTACACTCTGCTAAACTTATTTCANNNAGATCGGAAGA |
| 47836_1 | TATGAGTGTGGAGTCGTTGCCCTGAGTCATTTCAGAGGTGTCANNNAGATCGGAAG |
| 47836_2 | TATGAGTGTGGAGTCGTTGCCCTGAGTCATTTCAGAGGTGTCANNNAGATCGGAAG |
| 35855_1 | TATGAGTGTGGAGTCGTTGCTAGTCTTGACCTCCTGACCTCAANNNAGATCGGAAGA |
| 35855_2 | TATGAGTGTGGAGTCGTTGCTAGTCTTGACCTCCTGACCTCAANNNAGATCGGAAGA |
| 8510_1 | TATGAGTGTGGAGTCGTTGCTTCCAAGCACAGCTATCTCCTTTNNNAGATCGGAAGA |
| 8510_2 | TATGAGTGTGGAGTCGTTGCTTCCAAGCACAGCTATCTCCTTTNNNAGATCGGAAGA |
| 13715_1 | TATGAGTGTGGAGTCGTTGCAAAACACATTCTTTGCAGCTNNNAGATCGGAAGAGCA |
| 13715_2 | TATGAGTGTGGAGTCGTTGCAAAACACATTCTTTGCAGCTNNNAGATCGGAAGAGCA |
| 38669_1 | TATGAGTGTGGAGTCGTTGCCGTTACACATGCCTTTTTCCTCANNNAGATCGGAAGA |
| 38669_2 | TATGAGTGTGGAGTCGTTGCCGTTACACATGCCTTTTTCCTCANNNAGATCGGAAGA |
| 10410_1 | TATGAGTGTGGAGTCGTTGCGCCACCTTTCATTGTGGACCTATNNNAGATCGGAAGA |
| 10410_2 | TATGAGTGTGGAGTCGTTGCGCCACCTTTCATTGTGGACCTATNNNAGATCGGAAGA |
| 34804_1 | TATGAGTGTGGAGTCGTTGCTAGAGTGCAGTGACATGATCTCGNNNAGATCGGAAG |
| 34804_2 | TATGAGTGTGGAGTCGTTGCTAGAGTGCAGTGACATGATCTCGNNNAGATCGGAAG |
| 43171_1 | TATGAGTGTGGAGTCGTTGCATCTGTGAAATCTGTGCCTGTGANNNAGATCGGAAG |
| 43171_2 | TATGAGTGTGGAGTCGTTGCATCTGTGAAATCTGTGCCTGTGANNNAGATCGGAAG |
| 31579_1 | TATGAGTGTGGAGTCGTTGCTCAACACCTCACTTTCAGCATTGNNNAGATCGGAAGA |
| 31579_2 | TATGAGTGTGGAGTCGTTGCTCAACACCTCACTTTCAGCATTGNNNAGATCGGAAGA |
| 30066_1 | TATGAGTGTGGAGTCGTTGCCTCTCTCAGTCTGGATTTTGCTGANNNAGATCGGAAG |
| 30066_2 | TATGAGTGTGGAGTCGTTGCCTCTCTCAGTCTGGATTTTGCTGANNNAGATCGGAAG |
| 46486_1 | TATGAGTGTGGAGTCGTTGCGGGTTCAAGTGATTCTCCTGTCTNNNAGATCGGAAG |
| 46486_2 | TATGAGTGTGGAGTCGTTGCGGGTTCAAGTGATTCTCCTGTCTNNNAGATCGGAAG |

| | |
|---|---|
| 30551_1 | TATGAGTGTGGAGTCGTTGCGTGAGTGAAAATATCAGCTAGGTGCANNNAGATCG |
| 30551_2 | TATGAGTGTGGAGTCGTTGCGTGAGTGAAAATATCAGCTAGGTGCANNNAGATCG |
| 32428_1 | TATGAGTGTGGAGTCGTTGCGAGAAGGAATCTTGCTCTGTTGCNNNAGATCGGAA |
| 32428_2 | TATGAGTGTGGAGTCGTTGCGAGAAGGAATCTTGCTCTGTTGCNNNAGATCGGAA |
| 38966_1 | TATGAGTGTGGAGTCGTTGCTGATTTGGCTCTTGGTAACTGGTNNNAGATCGGAAG |
| 38966_2 | TATGAGTGTGGAGTCGTTGCTGATTTGGCTCTTGGTAACTGGTNNNAGATCGGAAG |
| 45727_1 | TATGAGTGTGGAGTCGTTGCCCTCCCAGGTTCAAGTGATTCTCNNNAGATCGGAAG |
| 45727_2 | TATGAGTGTGGAGTCGTTGCCCTCCCAGGTTCAAGTGATTCTCNNNAGATCGGAAG |
| 43542_1 | TATGAGTGTGGAGTCGTTGCTTCAGCTTCTGCCTGTATCTCAGNNNAGATCGGAAG |
| 43542_2 | TATGAGTGTGGAGTCGTTGCTTCAGCTTCTGCCTGTATCTCAGNNNAGATCGGAAG |
| 39978_1 | TATGAGTGTGGAGTCGTTGCAGACAAAAATCGTGAAGGGCTTGNNNAGATCGGAA |
| 39978_2 | TATGAGTGTGGAGTCGTTGCAGACAAAAATCGTGAAGGGCTTGNNNAGATCGGAA |
| 48342_1 | TATGAGTGTGGAGTCGTTGCTGTTAGCCAGGATGGTCTCAATCNNNAGATCGGAAG |
| 48342_2 | TATGAGTGTGGAGTCGTTGCTGTTAGCCAGGATGGTCTCAATCNNNAGATCGGAAG |
| 13219_1 | TATGAGTGTGGAGTCGTTGCGATTTAAATCCCTCGCAAGANNNAGATCGGAAGAG |
| 13219_2 | TATGAGTGTGGAGTCGTTGCGATTTAAATCCCTCGCAAGANNNAGATCGGAAGAG |
| 38889_1 | TATGAGTGTGGAGTCGTTGCTTAGACCGATTCCATGCTACTGGNNNAGATCGGAAG |
| 38889_2 | TATGAGTGTGGAGTCGTTGCTTAGACCGATTCCATGCTACTGGNNNAGATCGGAAG |
| 29372_1 | TATGAGTGTGGAGTCGTTGCCCCTCTGTCTCTCTCTCTCTCNNNAGATCGGAAGAG |
| 29372_2 | TATGAGTGTGGAGTCGTTGCCCCTCTGTCTCTCTCTCTCTCNNNAGATCGGAAGAG |
| 30082_1 | TATGAGTGTGGAGTCGTTGCTGGTGTATGTTTTGATGTCATCAGTNNNAGATCGGA |
| 30082_2 | TATGAGTGTGGAGTCGTTGCTGGTGTATGTTTTGATGTCATCAGTNNNAGATCGGA |
| 10992_1 | TATGAGTGTGGAGTCGTTGCTTTATTTTTGCACCCCACTCGCAGNNNAGATCGGAAG |
| 10992_2 | TATGAGTGTGGAGTCGTTGCTTTATTTTTGCACCCCACTCGCAGNNNAGATCGGAAG |
| 39467_1 | TATGAGTGTGGAGTCGTTGCACTTCCAGATCTGCCACAAATCTNNNAGATCGGAAG |
| 39467_2 | TATGAGTGTGGAGTCGTTGCACTTCCAGATCTGCCACAAATCTNNNAGATCGGAAG |
| 39902_1 | TATGAGTGTGGAGTCGTTGCCATGCTTGTAATCCCAGCACTTTNNNAGATCGGAAG |
| 39902_2 | TATGAGTGTGGAGTCGTTGCCATGCTTGTAATCCCAGCACTTTNNNAGATCGGAAG |
| 33758_1 | TATGAGTGTGGAGTCGTTGCGGAAAGACTCAGGCTGTTTCTCTNNNAGATCGGAAG |
| 33758_2 | TATGAGTGTGGAGTCGTTGCGGAAAGACTCAGGCTGTTTCTCTNNNAGATCGGAAG |
| 31059_1 | TATGAGTGTGGAGTCGTTGCACGTGCACCTCAACTGTTACATANNNAGATCGGAAG |
| 31059_2 | TATGAGTGTGGAGTCGTTGCACGTGCACCTCAACTGTTACATANNNAGATCGGAAG |
| 12657_1 | TATGAGTGTGGAGTCGTTGCCTTCTCAAAACATCTTCTTTCCNNNAGATCGGAAGAG |
| 12657_2 | TATGAGTGTGGAGTCGTTGCCTTCTCAAAACATCTTCTTTCCNNNAGATCGGAAGAG |
| 43927_1 | TATGAGTGTGGAGTCGTTGCACACACACAGAGAGAGAGAGAGANNNAGATCGGA |
| 43927_2 | TATGAGTGTGGAGTCGTTGCACACACACAGAGAGAGAGAGAGANNNAGATCGGA |
| 42907_1 | TATGAGTGTGGAGTCGTTGCGCTGGGTCCATAACTTTGCAATTNNNAGATCGGAAG |
| 42907_2 | TATGAGTGTGGAGTCGTTGCGCTGGGTCCATAACTTTGCAATTNNNAGATCGGAAG |
| 32140_1 | TATGAGTGTGGAGTCGTTGCCAGTGGTGTGATCTTGGCTCANNNAGATCGGAAGA |
| 32140_2 | TATGAGTGTGGAGTCGTTGCCAGTGGTGTGATCTTGGCTCANNNAGATCGGAAGA |
| 29580_1 | TATGAGTGTGGAGTCGTTGCTTGTCAGTTCCACTCTGCAACAANNNAGATCGGAAG |
| 29580_2 | TATGAGTGTGGAGTCGTTGCTTGTCAGTTCCACTCTGCAACAANNNAGATCGGAAG |
| 11300_1 | TATGAGTGTGGAGTCGTTGCTCCATTACACAGCCAATAGCCAGNNNAGATCGGAAG |

| | |
|---|---|
| 11300_2 | TATGAGTGTGGAGTCGTTGCTCCATTACACAGCCAATAGCCAGNNNAGATCGGAAG |
| 35899_1 | TATGAGTGTGGAGTCGTTGCTACACAGGAGGCTTCAAAAGGAANNNAGATCGGAA |
| 35899_2 | TATGAGTGTGGAGTCGTTGCTACACAGGAGGCTTCAAAAGGAANNNAGATCGGAA |
| 43410_1 | TATGAGTGTGGAGTCGTTGCCACCACACCTGGCTAATTTTTGTNNNAGATCGGAAG |
| 43410_2 | TATGAGTGTGGAGTCGTTGCCACCACACCTGGCTAATTTTTGTNNNAGATCGGAAG |
| 4271_1 | TATGAGTGTGGAGTCGTTGCTCAGGATTTCTGGTTTCTGGACAGNNNAGATCGGAA |
| 4271_2 | TATGAGTGTGGAGTCGTTGCTCAGGATTTCTGGTTTCTGGACAGNNNAGATCGGAA |
| 37360_1 | TATGAGTGTGGAGTCGTTGCCAAAAACAAACCAAACAACCCCCNNNAGATCGGAAG |
| 37360_2 | TATGAGTGTGGAGTCGTTGCCAAAAACAAACCAAACAACCCCCNNNAGATCGGAAG |
| 35742_1 | TATGAGTGTGGAGTCGTTGCCCTCCCGAGTAGCTAGAACTACANNNAGATCGGAAG |
| 35742_2 | TATGAGTGTGGAGTCGTTGCCCTCCCGAGTAGCTAGAACTACANNNAGATCGGAAG |
| 37435_1 | TATGAGTGTGGAGTCGTTGCATTTCTCACAGTTTTGGAGGCTGNNNAGATCGGAAG |
| 37435_2 | TATGAGTGTGGAGTCGTTGCATTTCTCACAGTTTTGGAGGCTGNNNAGATCGGAAG |
| 52915_1 | TATGAGTGTGGAGTCGTTGCACTTTCGCAGCACAGTGGAGNNNAGATCGGAAGAG |
| 52915_2 | TATGAGTGTGGAGTCGTTGCACTTTCGCAGCACAGTGGAGNNNAGATCGGAAGAG |
| 4474_1 | TATGAGTGTGGAGTCGTTGCTGATGACCACTTCTACTTCTGCCNNNAGATCGGAAGA |
| 4474_2 | TATGAGTGTGGAGTCGTTGCTGATGACCACTTCTACTTCTGCCNNNAGATCGGAAGA |
| 37295_1 | TATGAGTGTGGAGTCGTTGCGCAGTGGCTCACACCTATGATAANNNAGATCGGAAG |
| 37295_2 | TATGAGTGTGGAGTCGTTGCGCAGTGGCTCACACCTATGATAANNNAGATCGGAAG |
| 36513_1 | TATGAGTGTGGAGTCGTTGCTGATCGATTCTGGAAGTTGGGTTNNNAGATCGGAA |
| 36513_2 | TATGAGTGTGGAGTCGTTGCTGATCGATTCTGGAAGTTGGGTTNNNAGATCGGAA |
| 39304_1 | TATGAGTGTGGAGTCGTTGCAGCTAAAGGACCAGAAAAAAGAAGCNNNAGATCGG |
| 39304_2 | TATGAGTGTGGAGTCGTTGCAGCTAAAGGACCAGAAAAAAGAAGCNNNAGATCGG |
| 45763_1 | TATGAGTGTGGAGTCGTTGCCACTGTGCCTAGCTCCTGTATAGNNNAGATCGGAAG |
| 45763_2 | TATGAGTGTGGAGTCGTTGCCACTGTGCCTAGCTCCTGTATAGNNNAGATCGGAAG |
| 29303_1 | TATGAGTGTGGAGTCGTTGCCATAACCTCAGCTCACTGCAACNNNAGATCGGAAGA |
| 29303_2 | TATGAGTGTGGAGTCGTTGCCATAACCTCAGCTCACTGCAACNNNAGATCGGAAGA |
| 44712_1 | TATGAGTGTGGAGTCGTTGCACGGATGTTGTCACTTTGGTTTCNNNAGATCGGAAG |
| 44712_2 | TATGAGTGTGGAGTCGTTGCACGGATGTTGTCACTTTGGTTTCNNNAGATCGGAAG |
| 46081_1 | TATGAGTGTGGAGTCGTTGCGATCTTGACTCACTGCAACCTCTNNNAGATCGGAAGA |
| 46081_2 | TATGAGTGTGGAGTCGTTGCGATCTTGACTCACTGCAACCTCTNNNAGATCGGAAGA |
| 50426_1 | TATGAGTGTGGAGTCGTTGCAAAAAGGTTTCCTGGACCCAAACNNNAGATCGGAAG |
| 50426_2 | TATGAGTGTGGAGTCGTTGCAAAAAGGTTTCCTGGACCCAAACNNNAGATCGGAAG |
| 36581_1 | TATGAGTGTGGAGTCGTTGCGCCTGTTGGATAGTGCTGTCATANNNAGATCGGAAG |
| 36581_2 | TATGAGTGTGGAGTCGTTGCGCCTGTTGGATAGTGCTGTCATANNNAGATCGGAAG |
| 34908_1 | TATGAGTGTGGAGTCGTTGCCCTCCTAGGTTCAAGCGATTCTCNNNAGATCGGAAGA |
| 34908_2 | TATGAGTGTGGAGTCGTTGCCCTCCTAGGTTCAAGCGATTCTCNNNAGATCGGAAGA |
| 46741_1 | TATGAGTGTGGAGTCGTTGCGCCTCCCAAAGTAGCTGAGATTANNNAGATCGGAAG |
| 46741_2 | TATGAGTGTGGAGTCGTTGCGCCTCCCAAAGTAGCTGAGATTANNNAGATCGGAAG |
| 51476_1 | TATGAGTGTGGAGTCGTTGCAGTCCCTGTAAGGCTGTTGTTTCNNNAGATCGGAAG |
| 51476_2 | TATGAGTGTGGAGTCGTTGCAGTCCCTGTAAGGCTGTTGTTTCNNNAGATCGGAAG |
| 47025_1 | TATGAGTGTGGAGTCGTTGCAAGTGCTGGTATTACAGGCATGANNNAGATCGGAA |
| 47025_2 | TATGAGTGTGGAGTCGTTGCAAGTGCTGGTATTACAGGCATGANNNAGATCGGAA |

| | |
|---|---|
| 48123_1 | TATGAGTGTGGAGTCGTTGCGGCTGTCTCATGGAGAAGAGAAANNNAGATCGGAA |
| 48123_2 | TATGAGTGTGGAGTCGTTGCGGCTGTCTCATGGAGAAGAGAAANNNAGATCGGAA |
| 53427_1 | TATGAGTGTGGAGTCGTTGCCTTACTACCATGACGGCACCATANNNAGATCGGAAG |
| 53427_2 | TATGAGTGTGGAGTCGTTGCCTTACTACCATGACGGCACCATANNNAGATCGGAAG |
| 35201_1 | TATGAGTGTGGAGTCGTTGCAACCCCTGCTCAAGAGATTTTCTNNNAGATCGGAAG |
| 35201_2 | TATGAGTGTGGAGTCGTTGCAACCCCTGCTCAAGAGATTTTCTNNNAGATCGGAAG |
| 34726_1 | TATGAGTGTGGAGTCGTTGCCCTTCCAAGTTCAAGCGATTCTCNNNAGATCGGAAG |
| 34726_2 | TATGAGTGTGGAGTCGTTGCCCTTCCAAGTTCAAGCGATTCTCNNNAGATCGGAAG |
| 31543_1 | TATGAGTGTGGAGTCGTTGCTTCTTGCCTGATATTCCCTGACCNNNAGATCGGAAGA |
| 31543_2 | TATGAGTGTGGAGTCGTTGCTTCTTGCCTGATATTCCCTGACCNNNAGATCGGAAGA |
| 30969_1 | TATGAGTGTGGAGTCGTTGCCCCCATAATAAGTGCTTGCTTGGNNNAGATCGGAAG |
| 30969_2 | TATGAGTGTGGAGTCGTTGCCCCCATAATAAGTGCTTGCTTGGNNNAGATCGGAAG |
| 46102_1 | TATGAGTGTGGAGTCGTTGCAGGCTGGAGTATAGTGACTGGATNNNAGATCGGAA |
| 46102_2 | TATGAGTGTGGAGTCGTTGCAGGCTGGAGTATAGTGACTGGATNNNAGATCGGAA |
| 13452_1 | TATGAGTGTGGAGTCGTTGCCAATGTCAGGATCCAGAAAGNNNAGATCGGAAGAG |
| 13452_2 | TATGAGTGTGGAGTCGTTGCCAATGTCAGGATCCAGAAAGNNNAGATCGGAAGAG |
| 32995_1 | TATGAGTGTGGAGTCGTTGCCTGGGTTCAAGCAATTTCTCCTGNNNAGATCGGAAG |
| 32995_2 | TATGAGTGTGGAGTCGTTGCCTGGGTTCAAGCAATTTCTCCTGNNNAGATCGGAAG |
| 53312_1 | TATGAGTGTGGAGTCGTTGCCTAGCAGGGTTTTTCTGAGGGACNNNAGATCGGAA |
| 53312_2 | TATGAGTGTGGAGTCGTTGCCTAGCAGGGTTTTTCTGAGGGACNNNAGATCGGAA |
| 42643_1 | TATGAGTGTGGAGTCGTTGCCTCGGTTGTGCTGTTTTACTGAGNNNAGATCGGAAG |
| 42643_2 | TATGAGTGTGGAGTCGTTGCCTCGGTTGTGCTGTTTTACTGAGNNNAGATCGGAAG |
| 8753_1 | TATGAGTGTGGAGTCGTTGCATTTCCTCTTGCTTGACCCCTTTNNNAGATCGGAAGA |
| 8753_2 | TATGAGTGTGGAGTCGTTGCATTTCCTCTTGCTTGACCCCTTTNNNAGATCGGAAGA |
| 46150_1 | TATGAGTGTGGAGTCGTTGCCTGAGTAGCTGGGATTACAGGTGNNNAGATCGGAA |
| 46150_2 | TATGAGTGTGGAGTCGTTGCCTGAGTAGCTGGGATTACAGGTGNNNAGATCGGAA |
| 31061_1 | TATGAGTGTGGAGTCGTTGCGATTCTAGCAGTGGGAAAAGTGCNNNAGATCGGAA |
| 31061_2 | TATGAGTGTGGAGTCGTTGCGATTCTAGCAGTGGGAAAAGTGCNNNAGATCGGAA |
| 51847_1 | TATGAGTGTGGAGTCGTTGCGAGGGACTGAAGTTTCCACAGTTNNNAGATCGGAA |
| 51847_2 | TATGAGTGTGGAGTCGTTGCGAGGGACTGAAGTTTCCACAGTTNNNAGATCGGAA |
| 37319_1 | TATGAGTGTGGAGTCGTTGCGGGGGGTTATGCTGTTTATACCANNNAGATCGGAA |
| 37319_2 | TATGAGTGTGGAGTCGTTGCGGGGGGTTATGCTGTTTATACCANNNAGATCGGAA |
| 42365_1 | TATGAGTGTGGAGTCGTTGCTTGTAGACCAGCAAGAGAAAGCTNNNAGATCGGAA |
| 42365_2 | TATGAGTGTGGAGTCGTTGCTTGTAGACCAGCAAGAGAAAGCTNNNAGATCGGAA |
| 47671_1 | TATGAGTGTGGAGTCGTTGCGGAGTGTATTGACCACAACCTCTNNNAGATCGGAAG |
| 47671_2 | TATGAGTGTGGAGTCGTTGCGGAGTGTATTGACCACAACCTCTNNNAGATCGGAAG |
| 50228_1 | TATGAGTGTGGAGTCGTTGCTCTCTTCCATCTCAACTCCACAGNNNAGATCGGAAGA |
| 50228_2 | TATGAGTGTGGAGTCGTTGCTCTCTTCCATCTCAACTCCACAGNNNAGATCGGAAGA |
| 38582_1 | TATGAGTGTGGAGTCGTTGCGCAGAAGAGGAGGTCATAGTGACNNNAGATCGGA |
| 38582_2 | TATGAGTGTGGAGTCGTTGCGCAGAAGAGGAGGTCATAGTGACNNNAGATCGGA |
| 12520_1 | TATGAGTGTGGAGTCGTTGCTTGTTTATTCGAGAAAATGATCANNNAGATCGGAAG |
| 12520_2 | TATGAGTGTGGAGTCGTTGCTTGTTTATTCGAGAAAATGATCANNNAGATCGGAAG |
| 388_1 | TATGAGTGTGGAGTCGTTGCCAGAAGATAGTGTAGTGAGGCTACNNNAGATCGGA |

| | |
|---|---|
| 388_2 | TATGAGTGTGGAGTCGTTGCCAGAAGATAGTGTAGTGAGGCTACNNNAGATCGGA |
| 49181_1 | TATGAGTGTGGAGTCGTTGCCGAGTAGCTGGGACTACAGGTNNNAGATCGGAAGA |
| 49181_2 | TATGAGTGTGGAGTCGTTGCCGAGTAGCTGGGACTACAGGTNNNAGATCGGAAGA |
| 1105_1 | TATGAGTGTGGAGTCGTTGCACAGAGACACACTTTTCAAAGCANNNAGATCGGAAG |
| 1105_2 | TATGAGTGTGGAGTCGTTGCACAGAGACACACTTTTCAAAGCANNNAGATCGGAAG |
| 48_1 | TATGAGTGTGGAGTCGTTGCGCCCCGTTACACCGAGTTCTTNNNAGATCGGAAGAG |
| 48_2 | TATGAGTGTGGAGTCGTTGCGCCCCGTTACACCGAGTTCTTNNNAGATCGGAAGAG |
| 41099_1 | TATGAGTGTGGAGTCGTTGCCATGAAAGCTTTGAAGGACCTGGNNNAGATCGGAA |
| 41099_2 | TATGAGTGTGGAGTCGTTGCCATGAAAGCTTTGAAGGACCTGGNNNAGATCGGAA |
| 39963_1 | TATGAGTGTGGAGTCGTTGCCATATGCCCTTTGACCAAAGCAANNNAGATCGGAAG |
| 39963_2 | TATGAGTGTGGAGTCGTTGCCATATGCCCTTTGACCAAAGCAANNNAGATCGGAAG |
| 51562_1 | TATGAGTGTGGAGTCGTTGCTTTCTGTCACCTGTGAGCATTTGNNNAGATCGGAAG |
| 51562_2 | TATGAGTGTGGAGTCGTTGCTTTCTGTCACCTGTGAGCATTTGNNNAGATCGGAAG |
| 38140_1 | TATGAGTGTGGAGTCGTTGCGAGGTGTTTTGGACCTATTACGTNNNAGATCGGAAG |
| 38140_2 | TATGAGTGTGGAGTCGTTGCGAGGTGTTTTGGACCTATTACGTNNNAGATCGGAAG |
| 33287_1 | TATGAGTGTGGAGTCGTTGCGAGTGTAGTGGCACGATCTCAGNNNAGATCGGAAG |
| 33287_2 | TATGAGTGTGGAGTCGTTGCGAGTGTAGTGGCACGATCTCAGNNNAGATCGGAAG |
| 51284_1 | TATGAGTGTGGAGTCGTTGCATATATGTGTGTGTGTGTGCGTGNNNAGATCGGAA |
| 51284_2 | TATGAGTGTGGAGTCGTTGCATATATGTGTGTGTGTGTGCGTGNNNAGATCGGAA |
| 38799_1 | TATGAGTGTGGAGTCGTTGCAGCCTGGAGTACAGTAGTGAGATNNNAGATCGGAA |
| 38799_2 | TATGAGTGTGGAGTCGTTGCAGCCTGGAGTACAGTAGTGAGATNNNAGATCGGAA |
| 2275_1 | TATGAGTGTGGAGTCGTTGCCGAGCAACAGACCAAAGGAAAACNNNAGATCGGAA |
| 2275_2 | TATGAGTGTGGAGTCGTTGCCGAGCAACAGACCAAAGGAAAACNNNAGATCGGAA |
| 30199_1 | TATGAGTGTGGAGTCGTTGCCAAATTAAAATGCTTCTGAGATTCACANNNAGATCG |
| 30199_2 | TATGAGTGTGGAGTCGTTGCCAAATTAAAATGCTTCTGAGATTCACANNNAGATCG |
| 51409_1 | TATGAGTGTGGAGTCGTTGCACAAAGTTCTGACCAAAACACCGNNNAGATCGGAAG |
| 51409_2 | TATGAGTGTGGAGTCGTTGCACAAAGTTCTGACCAAAACACCGNNNAGATCGGAAG |
| 40586_1 | TATGAGTGTGGAGTCGTTGCTGCAGTCTTGAATTTCTCTGAGTGNNNAGATCGGAA |
| 40586_2 | TATGAGTGTGGAGTCGTTGCTGCAGTCTTGAATTTCTCTGAGTGNNNAGATCGGAA |
| 51366_1 | TATGAGTGTGGAGTCGTTGCATTTCCGCATACCCGACTCATAANNNAGATCGGAAG |
| 51366_2 | TATGAGTGTGGAGTCGTTGCATTTCCGCATACCCGACTCATAANNNAGATCGGAAG |
| 32288_1 | TATGAGTGTGGAGTCGTTGCCATCGTGTAATTGCTCACCTCCTNNNAGATCGGAAGA |
| 32288_2 | TATGAGTGTGGAGTCGTTGCCATCGTGTAATTGCTCACCTCCTNNNAGATCGGAAGA |
| 49423_1 | TATGAGTGTGGAGTCGTTGCTTCTTCACAGAGGAAAGGCTTCTNNNAGATCGGAAG |
| 49423_2 | TATGAGTGTGGAGTCGTTGCTTCTTCACAGAGGAAAGGCTTCTNNNAGATCGGAAG |
| 31244_1 | TATGAGTGTGGAGTCGTTGCTGAACAAAAGCCATTGTGTCAGGNNNAGATCGGAA |
| 31244_2 | TATGAGTGTGGAGTCGTTGCTGAACAAAAGCCATTGTGTCAGGNNNAGATCGGAA |
| 49106_1 | TATGAGTGTGGAGTCGTTGCTTCCAGGTTCAAGTGATTCTCGTNNNAGATCGGAAG |
| 49106_2 | TATGAGTGTGGAGTCGTTGCTTCCAGGTTCAAGTGATTCTCGTNNNAGATCGGAAG |
| 33379_1 | TATGAGTGTGGAGTCGTTGCGCTTCAAATGGTGCAAGCTATGANNNAGATCGGAA |
| 33379_2 | TATGAGTGTGGAGTCGTTGCGCTTCAAATGGTGCAAGCTATGANNNAGATCGGAA |
| 30038_1 | TATGAGTGTGGAGTCGTTGCTTTGGCTGAACATCAAGAGTCCTNNNAGATCGGAAG |
| 30038_2 | TATGAGTGTGGAGTCGTTGCTTTGGCTGAACATCAAGAGTCCTNNNAGATCGGAAG |

| | |
|---|---|
| 44131_1 | TATGAGTGTGGAGTCGTTGCCCACTCAGACCCAACATGTAAGTNNNAGATCGGAAG |
| 44131_2 | TATGAGTGTGGAGTCGTTGCCCACTCAGACCCAACATGTAAGTNNNAGATCGGAAG |
| 49895_1 | TATGAGTGTGGAGTCGTTGCTGCTGATAGTGACAACCTGTACCNNNAGATCGGAAG |
| 49895_2 | TATGAGTGTGGAGTCGTTGCTGCTGATAGTGACAACCTGTACCNNNAGATCGGAAG |
| 34406_1 | TATGAGTGTGGAGTCGTTGCCGAAGCCAAAAAAAGAAGCAGGANNNAGATCGGAA |
| 34406_2 | TATGAGTGTGGAGTCGTTGCCGAAGCCAAAAAAAGAAGCAGGANNNAGATCGGAA |
| 33075_1 | TATGAGTGTGGAGTCGTTGCGCAATCTCACCTCACCATAACCTNNNAGATCGGAAGA |
| 33075_2 | TATGAGTGTGGAGTCGTTGCGCAATCTCACCTCACCATAACCTNNNAGATCGGAAGA |
| 13177_1 | TATGAGTGTGGAGTCGTTGCGACAATGTTTTTTAAAGCTTGATNNNAGATCGGAAG |
| 13177_2 | TATGAGTGTGGAGTCGTTGCGACAATGTTTTTTAAAGCTTGATNNNAGATCGGAAG |
| 30330_1 | TATGAGTGTGGAGTCGTTGCCCGCAGAAATTACTTGAGCAGTGNNNAGATCGGAA |
| 30330_2 | TATGAGTGTGGAGTCGTTGCCCGCAGAAATTACTTGAGCAGTGNNNAGATCGGAA |
| 36700_1 | TATGAGTGTGGAGTCGTTGCAGCGAGACACCGTTTCAAAAAAANNNAGATCGGAA |
| 36700_2 | TATGAGTGTGGAGTCGTTGCAGCGAGACACCGTTTCAAAAAAANNNAGATCGGAA |
| 29071_1 | TATGAGTGTGGAGTCGTTGCTCCCTCTCACTTGTTTCCCAAATNNNAGATCGGAAGA |
| 29071_2 | TATGAGTGTGGAGTCGTTGCTCCCTCTCACTTGTTTCCCAAATNNNAGATCGGAAGA |
| 33720_1 | TATGAGTGTGGAGTCGTTGCCTAACATGGTGAAACCGCATCTCNNNAGATCGGAAG |
| 33720_2 | TATGAGTGTGGAGTCGTTGCCTAACATGGTGAAACCGCATCTCNNNAGATCGGAAG |
| 53341_1 | TATGAGTGTGGAGTCGTTGCGGCTTGCACAATTATAGAGGCTGNNNAGATCGGAA |
| 53341_2 | TATGAGTGTGGAGTCGTTGCGGCTTGCACAATTATAGAGGCTGNNNAGATCGGAA |
| 39917_1 | TATGAGTGTGGAGTCGTTGCTTTGCACCACTATATTCCAGCCTNNNAGATCGGAAGA |
| 39917_2 | TATGAGTGTGGAGTCGTTGCTTTGCACCACTATATTCCAGCCTNNNAGATCGGAAGA |
| 42541_1 | TATGAGTGTGGAGTCGTTGCTCTTGAACTCCTGGACTCAAGTGNNNAGATCGGAAG |
| 42541_2 | TATGAGTGTGGAGTCGTTGCTCTTGAACTCCTGGACTCAAGTGNNNAGATCGGAAG |
| 39768_1 | TATGAGTGTGGAGTCGTTGCATGTCCACGGAAAGCTTTGTACANNNAGATCGGAAG |
| 39768_2 | TATGAGTGTGGAGTCGTTGCATGTCCACGGAAAGCTTTGTACANNNAGATCGGAAG |
| 33761_1 | TATGAGTGTGGAGTCGTTGCTGCATCCATCGAGATTATCATGCNNNAGATCGGAAG |
| 33761_2 | TATGAGTGTGGAGTCGTTGCTGCATCCATCGAGATTATCATGCNNNAGATCGGAAG |
| 10759_1 | TATGAGTGTGGAGTCGTTGCAAACATCAGGAGTTGATGTCCCTNNNAGATCGGAAG |
| 10759_2 | TATGAGTGTGGAGTCGTTGCAAACATCAGGAGTTGATGTCCCTNNNAGATCGGAAG |
| 30263_1 | TATGAGTGTGGAGTCGTTGCTACAAGGCTGTCTGTCATGAAGCNNNAGATCGGAA |
| 30263_2 | TATGAGTGTGGAGTCGTTGCTACAAGGCTGTCTGTCATGAAGCNNNAGATCGGAA |
| 8953_1 | TATGAGTGTGGAGTCGTTGCTGACTTCCATTCAGTGCAACAGTNNNAGATCGGAAG |
| 8953_2 | TATGAGTGTGGAGTCGTTGCTGACTTCCATTCAGTGCAACAGTNNNAGATCGGAAG |
| 6721_1 | TATGAGTGTGGAGTCGTTGCATTCACCGAGTGTTTTGTCAGCTCNNNAGATCGGAA |
| 6721_2 | TATGAGTGTGGAGTCGTTGCATTCACCGAGTGTTTTGTCAGCTCNNNAGATCGGAA |
| 52443_1 | TATGAGTGTGGAGTCGTTGCTCTTGTCTCAACAACCAACCAGANNNAGATCGGAAG |
| 52443_2 | TATGAGTGTGGAGTCGTTGCTCTTGTCTCAACAACCAACCAGANNNAGATCGGAAG |
| 47264_1 | TATGAGTGTGGAGTCGTTGCACAAATGGTGGCATACTATACATGCNNNAGATCGGA |
| 47264_2 | TATGAGTGTGGAGTCGTTGCACAAATGGTGGCATACTATACATGCNNNAGATCGGA |
| 49653_1 | TATGAGTGTGGAGTCGTTGCTTGAATAGCTGGGACTACAGGTGNNNAGATCGGAA |
| 49653_2 | TATGAGTGTGGAGTCGTTGCTTGAATAGCTGGGACTACAGGTGNNNAGATCGGAA |
| 36786_1 | TATGAGTGTGGAGTCGTTGCAGCTCTGTCTAGAAAACTTACTGANNNAGATCGGAA |

| | |
|---|---|
| 36786_2 | TATGAGTGTGGAGTCGTTGCAGCTCTGTCTAGAAAACTTACTGANNNAGATCGGAA |
| 43716_1 | TATGAGTGTGGAGTCGTTGCAAATTGAAAGGCCAATGTCCCTGNNNAGATCGGAAG |
| 43716_2 | TATGAGTGTGGAGTCGTTGCAAATTGAAAGGCCAATGTCCCTGNNNAGATCGGAAG |
| 52046_1 | TATGAGTGTGGAGTCGTTGCTGAAACAAAAAGCTGGTTCTTTGANNNAGATCGGAA |
| 52046_2 | TATGAGTGTGGAGTCGTTGCTGAAACAAAAAGCTGGTTCTTTGANNNAGATCGGAA |
| 2033_1 | TATGAGTGTGGAGTCGTTGCCCGAAAAGTGAACTCACCCAGAANNNAGATCGGAAG |
| 2033_2 | TATGAGTGTGGAGTCGTTGCCCGAAAAGTGAACTCACCCAGAANNNAGATCGGAAG |
| 6291_1 | TATGAGTGTGGAGTCGTTGCCAGAGGGATAGTGGAGGCCATANNNAGATCGGAA |
| 6291_2 | TATGAGTGTGGAGTCGTTGCCAGAGGGATAGTGGAGGCCATANNNAGATCGGAA |
| 51214_1 | TATGAGTGTGGAGTCGTTGCGAGAACTCTGACTCCAGCCCTAANNNAGATCGGAAG |
| 51214_2 | TATGAGTGTGGAGTCGTTGCGAGAACTCTGACTCCAGCCCTAANNNAGATCGGAAG |
| 13221_1 | TATGAGTGTGGAGTCGTTGCATACAATAGAAAGTAAACAGCACNNNAGATCGGAA |
| 13221_2 | TATGAGTGTGGAGTCGTTGCATACAATAGAAAGTAAACAGCACNNNAGATCGGAA |
| 29457_1 | TATGAGTGTGGAGTCGTTGCTCTAGTGGTGGAAAAGGAAGTCGNNNAGATCGGAA |
| 29457_2 | TATGAGTGTGGAGTCGTTGCTCTAGTGGTGGAAAAGGAAGTCGNNNAGATCGGAA |
| 9015_1 | TATGAGTGTGGAGTCGTTGCTTCTGCTTTGGTGGTGGAAATCTNNNAGATCGGAAG |
| 9015_2 | TATGAGTGTGGAGTCGTTGCTTCTGCTTTGGTGGTGGAAATCTNNNAGATCGGAAG |
| 9118_1 | TATGAGTGTGGAGTCGTTGCTTGGCAGAGATAAGAGCTAGACANNNAGATCGGAA |
| 9118_2 | TATGAGTGTGGAGTCGTTGCTTGGCAGAGATAAGAGCTAGACANNNAGATCGGAA |
| 36694_1 | TATGAGTGTGGAGTCGTTGCACTTTGCCTCATGGAACTTACCTNNNAGATCGGAAG |
| 36694_2 | TATGAGTGTGGAGTCGTTGCACTTTGCCTCATGGAACTTACCTNNNAGATCGGAAG |
| 30124_1 | TATGAGTGTGGAGTCGTTGCTCCCCACCTGCTTTTTAAACAAANNNAGATCGGAAGA |
| 30124_2 | TATGAGTGTGGAGTCGTTGCTCCCCACCTGCTTTTTAAACAAANNNAGATCGGAAGA |
| 45642_1 | TATGAGTGTGGAGTCGTTGCCAGTGGTATGATCACAGCTCACTNNNAGATCGGAAG |
| 45642_2 | TATGAGTGTGGAGTCGTTGCCAGTGGTATGATCACAGCTCACTNNNAGATCGGAAG |
| 46182_1 | TATGAGTGTGGAGTCGTTGCAGTGCCATGTTTGTCTAACAGGANNNAGATCGGAA |
| 46182_2 | TATGAGTGTGGAGTCGTTGCAGTGCCATGTTTGTCTAACAGGANNNAGATCGGAA |
| 43478_1 | TATGAGTGTGGAGTCGTTGCTTAGTAGAGATGGGGGAAGAGGGNNNAGATCGGA |
| 43478_2 | TATGAGTGTGGAGTCGTTGCTTAGTAGAGATGGGGGAAGAGGGNNNAGATCGGA |
| 32804_1 | TATGAGTGTGGAGTCGTTGCCTCCTGGGTTCAAGCGATTCTCNNNAGATCGGAAGA |
| 32804_2 | TATGAGTGTGGAGTCGTTGCCTCCTGGGTTCAAGCGATTCTCNNNAGATCGGAAGA |
| 29072_1 | TATGAGTGTGGAGTCGTTGCTCATTGGGTTTCTTCCTCCTCAGNNNAGATCGGAAG |
| 29072_2 | TATGAGTGTGGAGTCGTTGCTCATTGGGTTTCTTCCTCCTCAGNNNAGATCGGAAG |
| 48044_1 | TATGAGTGTGGAGTCGTTGCCTCCTGAATAGCTGGGACTACAGNNNAGATCGGAA |
| 48044_2 | TATGAGTGTGGAGTCGTTGCCTCCTGAATAGCTGGGACTACAGNNNAGATCGGAA |
| 37605_1 | TATGAGTGTGGAGTCGTTGCTACAAAAAATTAGCTGGGCGTGGNNNAGATCGGAA |
| 37605_2 | TATGAGTGTGGAGTCGTTGCTACAAAAAATTAGCTGGGCGTGGNNNAGATCGGAA |
| 44486_1 | TATGAGTGTGGAGTCGTTGCGACGGGTTGGTAGCATTCATTTTNNNAGATCGGAA |
| 44486_2 | TATGAGTGTGGAGTCGTTGCGACGGGTTGGTAGCATTCATTTTNNNAGATCGGAA |
| 2619_1 | TATGAGTGTGGAGTCGTTGCTGTGCTGCTTTTTCCGATTATCTGNNNAGATCGGAA |
| 2619_2 | TATGAGTGTGGAGTCGTTGCTGTGCTGCTTTTTCCGATTATCTGNNNAGATCGGAA |
| 40343_1 | TATGAGTGTGGAGTCGTTGCCCAGATCCATGGGAGAAAAATGCNNNAGATCGGAA |
| 40343_2 | TATGAGTGTGGAGTCGTTGCCCAGATCCATGGGAGAAAAATGCNNNAGATCGGAA |

| | |
|---|---|
| 30585_1 | TATGAGTGTGGAGTCGTTGCTTCCCAGAACATTTTTGCCAGGATNNNAGATCGGAA |
| 30585_2 | TATGAGTGTGGAGTCGTTGCTTCCCAGAACATTTTTGCCAGGATNNNAGATCGGAA |
| 10189_1 | TATGAGTGTGGAGTCGTTGCGACTTCTGGAAGGGTGGCAATGNNNAGATCGGAAG |
| 10189_2 | TATGAGTGTGGAGTCGTTGCGACTTCTGGAAGGGTGGCAATGNNNAGATCGGAAG |
| 33717_1 | TATGAGTGTGGAGTCGTTGCAACGAAATGTGCTTTTGTTTGCTNNNAGATCGGAAG |
| 33717_2 | TATGAGTGTGGAGTCGTTGCAACGAAATGTGCTTTTGTTTGCTNNNAGATCGGAAG |
| 48523_1 | TATGAGTGTGGAGTCGTTGCCAGAGCAAGCAATCCAAGAACAANNNAGATCGGAA |
| 48523_2 | TATGAGTGTGGAGTCGTTGCCAGAGCAAGCAATCCAAGAACAANNNAGATCGGAA |
| 47691_1 | TATGAGTGTGGAGTCGTTGCCCAAAGTGCTGGGATTACAAGTGNNNAGATCGGAA |
| 47691_2 | TATGAGTGTGGAGTCGTTGCCCAAAGTGCTGGGATTACAAGTGNNNAGATCGGAA |
| 35357_1 | TATGAGTGTGGAGTCGTTGCATTGCTTTGTACTGGAGGAGACCNNNAGATCGGAA |
| 35357_2 | TATGAGTGTGGAGTCGTTGCATTGCTTTGTACTGGAGGAGACCNNNAGATCGGAA |
| 10372_1 | TATGAGTGTGGAGTCGTTGCCACTGAAGTCCCACTATTACACCANNNAGATCGGAA |
| 10372_2 | TATGAGTGTGGAGTCGTTGCCACTGAAGTCCCACTATTACACCANNNAGATCGGAA |
| 42619_1 | TATGAGTGTGGAGTCGTTGCTTTGTGTGCGTGTGTGTGTAAATNNNAGATCGGAA |
| 42619_2 | TATGAGTGTGGAGTCGTTGCTTTGTGTGCGTGTGTGTGTAAATNNNAGATCGGAA |
| 39271_1 | TATGAGTGTGGAGTCGTTGCCTTGATCTCCTGGCCTCATGATCNNNAGATCGGAAG |
| 39271_2 | TATGAGTGTGGAGTCGTTGCCTTGATCTCCTGGCCTCATGATCNNNAGATCGGAAG |
| 2600_1 | TATGAGTGTGGAGTCGTTGCGCACCTGTGTATGAATGCACATGNNNAGATCGGAAG |
| 2600_2 | TATGAGTGTGGAGTCGTTGCGCACCTGTGTATGAATGCACATGNNNAGATCGGAAG |
| 45480_1 | TATGAGTGTGGAGTCGTTGCAAGCAAGCAGCAGAGATATCAGTNNNAGATCGGAA |
| 45480_2 | TATGAGTGTGGAGTCGTTGCAAGCAAGCAGCAGAGATATCAGTNNNAGATCGGAA |
| 52493_1 | TATGAGTGTGGAGTCGTTGCGTGTTACTGAATGTGGCATCAGGNNNAGATCGGAA |
| 52493_2 | TATGAGTGTGGAGTCGTTGCGTGTTACTGAATGTGGCATCAGGNNNAGATCGGAA |
| 35719_1 | TATGAGTGTGGAGTCGTTGCTGTTCTCAAGACAGATAACACAGTNNNAGATCGGAA |
| 35719_2 | TATGAGTGTGGAGTCGTTGCTGTTCTCAAGACAGATAACACAGTNNNAGATCGGAA |
| 30595_1 | TATGAGTGTGGAGTCGTTGCTTTTGCATGTTGCTTTTGTTTTATAAANNNAGATCGG |
| 30595_2 | TATGAGTGTGGAGTCGTTGCTTTTGCATGTTGCTTTTGTTTTATAAANNNAGATCGG |
| 32057_1 | TATGAGTGTGGAGTCGTTGCTGTCATTGTGGTTTTGATTTGCANNNAGATCGGAAG |
| 32057_2 | TATGAGTGTGGAGTCGTTGCTGTCATTGTGGTTTTGATTTGCANNNAGATCGGAAG |
| 32434_1 | TATGAGTGTGGAGTCGTTGCCCAGGTCAAAGTGATTCTCTTGCNNNAGATCGGAAG |
| 32434_2 | TATGAGTGTGGAGTCGTTGCCCAGGTCAAAGTGATTCTCTTGCNNNAGATCGGAAG |
| 13461_1 | TATGAGTGTGGAGTCGTTGCTTTTCAAGTGAGGTAACCAACNNNAGATCGGAAGA |
| 13461_2 | TATGAGTGTGGAGTCGTTGCTTTTCAAGTGAGGTAACCAACNNNAGATCGGAAGA |
| 33325_1 | TATGAGTGTGGAGTCGTTGCTCAAGTAGCTGGGATTACAGGTGNNNAGATCGGAA |
| 33325_2 | TATGAGTGTGGAGTCGTTGCTCAAGTAGCTGGGATTACAGGTGNNNAGATCGGAA |
| 52227_1 | TATGAGTGTGGAGTCGTTGCTCTGACCCCAAAGCTTTCAAGTANNNAGATCGGAAG |
| 52227_2 | TATGAGTGTGGAGTCGTTGCTCTGACCCCAAAGCTTTCAAGTANNNAGATCGGAAG |
| 43315_1 | TATGAGTGTGGAGTCGTTGCACTTTCACAGTTCTCTGCTCCATNNNAGATCGGAAGA |
| 43315_2 | TATGAGTGTGGAGTCGTTGCACTTTCACAGTTCTCTGCTCCATNNNAGATCGGAAGA |
| 34927_1 | TATGAGTGTGGAGTCGTTGCGGGTTCAAGCGATTCTCTTACCTNNNAGATCGGAAG |
| 34927_2 | TATGAGTGTGGAGTCGTTGCGGGTTCAAGCGATTCTCTTACCTNNNAGATCGGAAG |
| 48997_1 | TATGAGTGTGGAGTCGTTGCCATTCTCCTGCCTCAGCCTCNNNAGATCGGAAGAGCA |

| | |
|---|---|
| 48997_2 | TATGAGTGTGGAGTCGTTGCCATTCTCCTGCCTCAGCCTCNNNAGATCGGAAGAGCA |
| 39700_1 | TATGAGTGTGGAGTCGTTGCCTATTCCTGGTAGGGGCTCTTTCNNNAGATCGGAAG |
| 39700_2 | TATGAGTGTGGAGTCGTTGCCTATTCCTGGTAGGGGCTCTTTCNNNAGATCGGAAG |
| 48900_1 | TATGAGTGTGGAGTCGTTGCCCACCACACCCAGGTAATTTTTGNNNAGATCGGAAG |
| 48900_2 | TATGAGTGTGGAGTCGTTGCCCACCACACCCAGGTAATTTTTGNNNAGATCGGAAG |
| 41718_1 | TATGAGTGTGGAGTCGTTGCCACTACGCCCGGCTAATTTTTTTNNNAGATCGGAAGA |
| 41718_2 | TATGAGTGTGGAGTCGTTGCCACTACGCCCGGCTAATTTTTTTNNNAGATCGGAAGA |
| 49169_1 | TATGAGTGTGGAGTCGTTGCAATTTTAGTACCTCAGCCTCCCGNNNAGATCGGAAG |
| 49169_2 | TATGAGTGTGGAGTCGTTGCAATTTTAGTACCTCAGCCTCCCGNNNAGATCGGAAG |
| 45164_1 | TATGAGTGTGGAGTCGTTGCTAAAACTCAGCATGACCTCAGCTNNNAGATCGGAAG |
| 45164_2 | TATGAGTGTGGAGTCGTTGCTAAAACTCAGCATGACCTCAGCTNNNAGATCGGAAG |
| 43176_1 | TATGAGTGTGGAGTCGTTGCACATCCTGGGTTCAAGTGATTCTNNNAGATCGGAAG |
| 43176_2 | TATGAGTGTGGAGTCGTTGCACATCCTGGGTTCAAGTGATTCTNNNAGATCGGAAG |
| 47326_1 | TATGAGTGTGGAGTCGTTGCATTAGCGGATGTGAACCCCATTTNNNAGATCGGAAG |
| 47326_2 | TATGAGTGTGGAGTCGTTGCATTAGCGGATGTGAACCCCATTTNNNAGATCGGAAG |
| 43858_1 | TATGAGTGTGGAGTCGTTGCGATAGTTCTTTCGGGGGTTCGATNNNAGATCGGAA |
| 43858_2 | TATGAGTGTGGAGTCGTTGCGATAGTTCTTTCGGGGGTTCGATNNNAGATCGGAA |
| 41617_1 | TATGAGTGTGGAGTCGTTGCGGTTCAGTCTTGGTGGGATGTATNNNAGATCGGAA |
| 41617_2 | TATGAGTGTGGAGTCGTTGCGGTTCAGTCTTGGTGGGATGTATNNNAGATCGGAA |
| 36898_1 | TATGAGTGTGGAGTCGTTGCAGGGTGACACATCTTATCTTTGGANNNAGATCGGAA |
| 36898_2 | TATGAGTGTGGAGTCGTTGCAGGGTGACACATCTTATCTTTGGANNNAGATCGGAA |
| 31563_1 | TATGAGTGTGGAGTCGTTGCACTCCTGACAAAGCTCATACAGANNNAGATCGGAAG |
| 31563_2 | TATGAGTGTGGAGTCGTTGCACTCCTGACAAAGCTCATACAGANNNAGATCGGAAG |
| 47475_1 | TATGAGTGTGGAGTCGTTGCCCAAGTAGCTGGGATTACAGGTGNNNAGATCGGAA |
| 47475_2 | TATGAGTGTGGAGTCGTTGCCCAAGTAGCTGGGATTACAGGTGNNNAGATCGGAA |
| 43202_1 | TATGAGTGTGGAGTCGTTGCTATCTAAATCCTTGCTCCAGCCCNNNAGATCGGAAGA |
| 43202_2 | TATGAGTGTGGAGTCGTTGCTATCTAAATCCTTGCTCCAGCCCNNNAGATCGGAAGA |
| 45858_1 | TATGAGTGTGGAGTCGTTGCTTGAAATCTGGGCAACTGTCTCTNNNAGATCGGAAG |
| 45858_2 | TATGAGTGTGGAGTCGTTGCTTGAAATCTGGGCAACTGTCTCTNNNAGATCGGAAG |
| 44895_1 | TATGAGTGTGGAGTCGTTGCAGGAGCAGTGTCCTAGTAGGAATNNNAGATCGGAA |
| 44895_2 | TATGAGTGTGGAGTCGTTGCAGGAGCAGTGTCCTAGTAGGAATNNNAGATCGGAA |
| 50536_1 | TATGAGTGTGGAGTCGTTGCCCTATTACACCCCCTCCTCAAATNNNAGATCGGAAGA |
| 50536_2 | TATGAGTGTGGAGTCGTTGCCCTATTACACCCCCTCCTCAAATNNNAGATCGGAAGA |
| 43203_1 | TATGAGTGTGGAGTCGTTGCCACCACGCCTGGCTAATTTTTTANNNAGATCGGAAG |
| 43203_2 | TATGAGTGTGGAGTCGTTGCCACCACGCCTGGCTAATTTTTTANNNAGATCGGAAG |
| 52059_1 | TATGAGTGTGGAGTCGTTGCGGGAGTGATTTCGGTTCCTAAGTNNNAGATCGGAA |
| 52059_2 | TATGAGTGTGGAGTCGTTGCGGGAGTGATTTCGGTTCCTAAGTNNNAGATCGGAA |
| 50714_1 | TATGAGTGTGGAGTCGTTGCGGCCCATTTTATTGTCTGCCTTTNNNAGATCGGAAGA |
| 50714_2 | TATGAGTGTGGAGTCGTTGCGGCCCATTTTATTGTCTGCCTTTNNNAGATCGGAAGA |
| 30273_1 | TATGAGTGTGGAGTCGTTGCAAATGTTGGAATAGATTAAGACTGCANNNAGATCG |
| 30273_2 | TATGAGTGTGGAGTCGTTGCAAATGTTGGAATAGATTAAGACTGCANNNAGATCG |
| 35383_1 | TATGAGTGTGGAGTCGTTGCCAAGTAGCTGGGAATACAGGTGTNNNAGATCGGAA |
| 35383_2 | TATGAGTGTGGAGTCGTTGCCAAGTAGCTGGGAATACAGGTGTNNNAGATCGGAA |

| | |
|---|---|
| 38002_1 | TATGAGTGTGGAGTCGTTGCGGGACCAGCTGAAATTATTGACCNNNAGATCGGAAG |
| 38002_2 | TATGAGTGTGGAGTCGTTGCGGGACCAGCTGAAATTATTGACCNNNAGATCGGAAG |
| 35486_1 | TATGAGTGTGGAGTCGTTGCTGCCAAAAGAGCAGCATGAAAATNNNAGATCGGAA |
| 35486_2 | TATGAGTGTGGAGTCGTTGCTGCCAAAAGAGCAGCATGAAAATNNNAGATCGGAA |
| 43635_1 | TATGAGTGTGGAGTCGTTGCATTGATTGTGACTCTTGGGGGTTNNNAGATCGGAAG |
| 43635_2 | TATGAGTGTGGAGTCGTTGCATTGATTGTGACTCTTGGGGGTTNNNAGATCGGAAG |
| 31583_1 | TATGAGTGTGGAGTCGTTGCCATGGTAGTACTCTGGGAACACANNNAGATCGGAAG |
| 31583_2 | TATGAGTGTGGAGTCGTTGCCATGGTAGTACTCTGGGAACACANNNAGATCGGAAG |
| 50562_1 | TATGAGTGTGGAGTCGTTGCCCCTTCTCCCCCAAAATACTCAANNNAGATCGGAAGA |
| 50562_2 | TATGAGTGTGGAGTCGTTGCCCCTTCTCCCCCAAAATACTCAANNNAGATCGGAAGA |
| 42799_1 | TATGAGTGTGGAGTCGTTGCATATTGACCAGGTCCCATGGTTCNNNAGATCGGAAG |
| 42799_2 | TATGAGTGTGGAGTCGTTGCATATTGACCAGGTCCCATGGTTCNNNAGATCGGAAG |
| 13679_1 | TATGAGTGTGGAGTCGTTGCTACAAACGTATGGAATGCTTNNNAGATCGGAAGAG |
| 13679_2 | TATGAGTGTGGAGTCGTTGCTACAAACGTATGGAATGCTTNNNAGATCGGAAGAG |
| 10531_1 | TATGAGTGTGGAGTCGTTGCGTATGGATTCTGCGGTGAGCTTTNNNAGATCGGAAG |
| 10531_2 | TATGAGTGTGGAGTCGTTGCGTATGGATTCTGCGGTGAGCTTTNNNAGATCGGAAG |
| 44377_1 | TATGAGTGTGGAGTCGTTGCATCCATTTGTCAGTTGAGGGACANNNAGATCGGAAG |
| 44377_2 | TATGAGTGTGGAGTCGTTGCATCCATTTGTCAGTTGAGGGACANNNAGATCGGAAG |
| 38931_1 | TATGAGTGTGGAGTCGTTGCTGCCAAAAACCAGTTGCTTTGTANNNAGATCGGAAG |
| 38931_2 | TATGAGTGTGGAGTCGTTGCTGCCAAAAACCAGTTGCTTTGTANNNAGATCGGAAG |
| 13782_1 | TATGAGTGTGGAGTCGTTGCGGAAGAGACAGGATTTTCTGNNNAGATCGGAAGAG |
| 13782_2 | TATGAGTGTGGAGTCGTTGCGGAAGAGACAGGATTTTCTGNNNAGATCGGAAGAG |
| 46488_1 | TATGAGTGTGGAGTCGTTGCTAGTAGCTGGGATTACAGGCATGNNNAGATCGGAA |
| 46488_2 | TATGAGTGTGGAGTCGTTGCTAGTAGCTGGGATTACAGGCATGNNNAGATCGGAA |
| 44469_1 | TATGAGTGTGGAGTCGTTGCATTTTGCAGCAAGTTGGATGGAANNNAGATCGGAA |
| 44469_2 | TATGAGTGTGGAGTCGTTGCATTTTGCAGCAAGTTGGATGGAANNNAGATCGGAA |
| 53085_1 | TATGAGTGTGGAGTCGTTGCAATGGATTTGCTGAGTTGTGCACNNNAGATCGGAAG |
| 53085_2 | TATGAGTGTGGAGTCGTTGCAATGGATTTGCTGAGTTGTGCACNNNAGATCGGAAG |
| 41538_1 | TATGAGTGTGGAGTCGTTGCCTCCTGAGTAGCTGGGACTACAGNNNAGATCGGAAG |
| 41538_2 | TATGAGTGTGGAGTCGTTGCCTCCTGAGTAGCTGGGACTACAGNNNAGATCGGAAG |
| 9717_1 | TATGAGTGTGGAGTCGTTGCACCCAATGACCTTTGGCAAGTACNNNAGATCGGAAG |
| 9717_2 | TATGAGTGTGGAGTCGTTGCACCCAATGACCTTTGGCAAGTACNNNAGATCGGAAG |
| 30500_1 | TATGAGTGTGGAGTCGTTGCCCACTTCTAATGCATGCCAGAGCNNNAGATCGGAAG |
| 30500_2 | TATGAGTGTGGAGTCGTTGCCCACTTCTAATGCATGCCAGAGCNNNAGATCGGAAG |
| 31979_1 | TATGAGTGTGGAGTCGTTGCTGTAATTCTCGGAGAGGACACACNNNAGATCGGAAG |
| 31979_2 | TATGAGTGTGGAGTCGTTGCTGTAATTCTCGGAGAGGACACACNNNAGATCGGAAG |
| 29114_1 | TATGAGTGTGGAGTCGTTGCACACACACACGCACATCCTATTANNNAGATCGGAAG |
| 29114_2 | TATGAGTGTGGAGTCGTTGCACACACACACGCACATCCTATTANNNAGATCGGAAG |
| 37213_1 | TATGAGTGTGGAGTCGTTGCACTGGCAAACAGAATTCAACAATNNNAGATCGGAAG |
| 37213_2 | TATGAGTGTGGAGTCGTTGCACTGGCAAACAGAATTCAACAATNNNAGATCGGAAG |
| 46031_1 | TATGAGTGTGGAGTCGTTGCACAAATGATTGCTCAACACAGGANNNAGATCGGAAG |
| 46031_2 | TATGAGTGTGGAGTCGTTGCACAAATGATTGCTCAACACAGGANNNAGATCGGAAG |
| 34028_1 | TATGAGTGTGGAGTCGTTGCGTTCAAGAGATTCTCCTGCCTCANNNAGATCGGAAG |

| | |
|---|---|
| 34028_2 | TATGAGTGTGGAGTCGTTGCGTTCAAGAGATTCTCCTGCCTCANNNAGATCGGAAG |
| 32727_1 | TATGAGTGTGGAGTCGTTGCGCCTGTAATCCCAGCACTTTGNNNAGATCGGAAGAG |
| 32727_2 | TATGAGTGTGGAGTCGTTGCGCCTGTAATCCCAGCACTTTGNNNAGATCGGAAGAG |
| 37729_1 | TATGAGTGTGGAGTCGTTGCGAGGCACAAAAGGAGACATTACANNNAGATCGGAA |
| 37729_2 | TATGAGTGTGGAGTCGTTGCGAGGCACAAAAGGAGACATTACANNNAGATCGGAA |
| 35392_1 | TATGAGTGTGGAGTCGTTGCAATGTCTGGGATTACAGGCATGTNNNAGATCGGAA |
| 35392_2 | TATGAGTGTGGAGTCGTTGCAATGTCTGGGATTACAGGCATGTNNNAGATCGGAA |
| 35654_1 | TATGAGTGTGGAGTCGTTGCAGATGGAGTCTCGCACTGTTTCNNNAGATCGGAAG |
| 35654_2 | TATGAGTGTGGAGTCGTTGCAGATGGAGTCTCGCACTGTTTCNNNAGATCGGAAG |
| 49708_1 | TATGAGTGTGGAGTCGTTGCGATTACAGGCATGCACCATCATGNNNAGATCGGAAG |
| 49708_2 | TATGAGTGTGGAGTCGTTGCGATTACAGGCATGCACCATCATGNNNAGATCGGAAG |
| 32784_1 | TATGAGTGTGGAGTCGTTGCCATGCCTGTAATTCCAGCACTTTNNNAGATCGGAAG |
| 32784_2 | TATGAGTGTGGAGTCGTTGCCATGCCTGTAATTCCAGCACTTTNNNAGATCGGAAG |
| 31483_1 | TATGAGTGTGGAGTCGTTGCTCTAAGGATTCAGGTGTGCAGTCNNNAGATCGGAA |
| 31483_2 | TATGAGTGTGGAGTCGTTGCTCTAAGGATTCAGGTGTGCAGTCNNNAGATCGGAA |
| 48905_1 | TATGAGTGTGGAGTCGTTGCTGGAGCTGGAGGCCATTATTTTANNNAGATCGGAA |
| 48905_2 | TATGAGTGTGGAGTCGTTGCTGGAGCTGGAGGCCATTATTTTANNNAGATCGGAA |
| 30520_1 | TATGAGTGTGGAGTCGTTGCGTGTCTAAGTGAGATCGTGTGGGNNNAGATCGGAA |
| 30520_2 | TATGAGTGTGGAGTCGTTGCGTGTCTAAGTGAGATCGTGTGGGNNNAGATCGGAA |
| 3031_1 | TATGAGTGTGGAGTCGTTGCTCTTCTCACTGAACCAGACCTGTNNNAGATCGGAAG |
| 3031_2 | TATGAGTGTGGAGTCGTTGCTCTTCTCACTGAACCAGACCTGTNNNAGATCGGAAG |
| 50520_1 | TATGAGTGTGGAGTCGTTGCTGTATGTTTGGGGAAAGGGAGAGNNNAGATCGGA |
| 50520_2 | TATGAGTGTGGAGTCGTTGCTGTATGTTTGGGGAAAGGGAGAGNNNAGATCGGA |
| 45669_1 | TATGAGTGTGGAGTCGTTGCTCTGCCCAGGTGTAATAGCTTACNNNAGATCGGAAG |
| 45669_2 | TATGAGTGTGGAGTCGTTGCTCTGCCCAGGTGTAATAGCTTACNNNAGATCGGAAG |
| 29887_1 | TATGAGTGTGGAGTCGTTGCTGCTTATGATCTGTCAAAGAAGAGANNNAGATCGG |
| 29887_2 | TATGAGTGTGGAGTCGTTGCTGCTTATGATCTGTCAAAGAAGAGANNNAGATCGG |
| 32091_1 | TATGAGTGTGGAGTCGTTGCCTTCCAGGTTCAAGCGATTCTCCNNNAGATCGGAAG |
| 32091_2 | TATGAGTGTGGAGTCGTTGCCTTCCAGGTTCAAGCGATTCTCCNNNAGATCGGAAG |
| 42888_1 | TATGAGTGTGGAGTCGTTGCTTGAATTCCTGGGCTCAAGAGATNNNAGATCGGAA |
| 42888_2 | TATGAGTGTGGAGTCGTTGCTTGAATTCCTGGGCTCAAGAGATNNNAGATCGGAA |
| 51927_1 | TATGAGTGTGGAGTCGTTGCGTGCTGTCCTGAGTATGTGTACANNNAGATCGGAA |
| 51927_2 | TATGAGTGTGGAGTCGTTGCGTGCTGTCCTGAGTATGTGTACANNNAGATCGGAA |
| 50026_1 | TATGAGTGTGGAGTCGTTGCGCGAATGACTACTCAGAGTCTGTNNNAGATCGGAA |
| 50026_2 | TATGAGTGTGGAGTCGTTGCGCGAATGACTACTCAGAGTCTGTNNNAGATCGGAA |
| 45161_1 | TATGAGTGTGGAGTCGTTGCCACCTTAGTTCCGACCCAGAGNNNAGATCGGAAGAG |
| 45161_2 | TATGAGTGTGGAGTCGTTGCCACCTTAGTTCCGACCCAGAGNNNAGATCGGAAGAG |
| 38441_1 | TATGAGTGTGGAGTCGTTGCTAGCGTTCTAGCCTCTTTTCTCCNNNAGATCGGAAGA |
| 38441_2 | TATGAGTGTGGAGTCGTTGCTAGCGTTCTAGCCTCTTTTCTCCNNNAGATCGGAAGA |
| 12355_1 | TATGAGTGTGGAGTCGTTGCTTCACTTTCTCCTCCCACCCTGNNNAGATCGGAAGAG |
| 12355_2 | TATGAGTGTGGAGTCGTTGCTTCACTTTCTCCTCCCACCCTGNNNAGATCGGAAGAG |
| 51768_1 | TATGAGTGTGGAGTCGTTGCCTGTTCATGCTGGGGAGATTTTGNNNAGATCGGAA |
| 51768_2 | TATGAGTGTGGAGTCGTTGCCTGTTCATGCTGGGGAGATTTTGNNNAGATCGGAA |

| | |
|---|---|
| 36116_1 | TATGAGTGTGGAGTCGTTGCCCTGCAAAGGAAAACAATCCTGTNNNAGATCGGAAG |
| 36116_2 | TATGAGTGTGGAGTCGTTGCCCTGCAAAGGAAAACAATCCTGTNNNAGATCGGAAG |
| 38546_1 | TATGAGTGTGGAGTCGTTGCGTACGAGGTAGGGATCCCATTTCNNNAGATCGGAAG |
| 38546_2 | TATGAGTGTGGAGTCGTTGCGTACGAGGTAGGGATCCCATTTCNNNAGATCGGAAG |
| 7901_1 | TATGAGTGTGGAGTCGTTGCTAGGCTGAAATGCAGTGTGGCAANNNAGATCGGAA |
| 7901_2 | TATGAGTGTGGAGTCGTTGCTAGGCTGAAATGCAGTGTGGCAANNNAGATCGGAA |
| 47243_1 | TATGAGTGTGGAGTCGTTGCACGCCCGGCTAATTTTTTGTATTNNNAGATCGGAAGA |
| 47243_2 | TATGAGTGTGGAGTCGTTGCACGCCCGGCTAATTTTTTGTATTNNNAGATCGGAAGA |
| 41493_1 | TATGAGTGTGGAGTCGTTGCCCATGGATAACGGGACTATTGTGNNNAGATCGGAAG |
| 41493_2 | TATGAGTGTGGAGTCGTTGCCCATGGATAACGGGACTATTGTGNNNAGATCGGAAG |
| 52435_1 | TATGAGTGTGGAGTCGTTGCGAGCACACAGGGATCATATTTGCNNNAGATCGGAAG |
| 52435_2 | TATGAGTGTGGAGTCGTTGCGAGCACACAGGGATCATATTTGCNNNAGATCGGAAG |
| 36313_1 | TATGAGTGTGGAGTCGTTGCTGATTTGGGATAAGAGCCAGAGCNNNAGATCGGAA |
| 36313_2 | TATGAGTGTGGAGTCGTTGCTGATTTGGGATAAGAGCCAGAGCNNNAGATCGGAA |
| 13794_1 | TATGAGTGTGGAGTCGTTGCGTCAAATTTCTCAACTCTGTGNNNAGATCGGAAGAG |
| 13794_2 | TATGAGTGTGGAGTCGTTGCGTCAAATTTCTCAACTCTGTGNNNAGATCGGAAGAG |
| 13351_1 | TATGAGTGTGGAGTCGTTGCAATTAACAAATGTTAGGGGAAATNNNAGATCGGAAG |
| 13351_2 | TATGAGTGTGGAGTCGTTGCAATTAACAAATGTTAGGGGAAATNNNAGATCGGAAG |
| 44292_1 | TATGAGTGTGGAGTCGTTGCTTGTGGTGGTGTTATCCCTGTAGNNNAGATCGGAAG |
| 44292_2 | TATGAGTGTGGAGTCGTTGCTTGTGGTGGTGTTATCCCTGTAGNNNAGATCGGAAG |
| 45410_1 | TATGAGTGTGGAGTCGTTGCACAAAGTTCTGACCAAAACACCGNNNAGATCGGAAG |
| 45410_2 | TATGAGTGTGGAGTCGTTGCACAAAGTTCTGACCAAAACACCGNNNAGATCGGAAG |
| 37270_1 | TATGAGTGTGGAGTCGTTGCTCTGGAAGTCTATCACAACTGGCNNNAGATCGGAAG |
| 37270_2 | TATGAGTGTGGAGTCGTTGCTCTGGAAGTCTATCACAACTGGCNNNAGATCGGAAG |
| 12497_1 | TATGAGTGTGGAGTCGTTGCGATATTCCACCAGAATCGTAGNNNAGATCGGAAGAG |
| 12497_2 | TATGAGTGTGGAGTCGTTGCGATATTCCACCAGAATCGTAGNNNAGATCGGAAGAG |
| 30320_1 | TATGAGTGTGGAGTCGTTGCCAGCTCATGAAGGGTACCTCATGNNNAGATCGGAAG |
| 30320_2 | TATGAGTGTGGAGTCGTTGCCAGCTCATGAAGGGTACCTCATGNNNAGATCGGAAG |
| 46476_1 | TATGAGTGTGGAGTCGTTGCCTGCAATTACAGGTGTAAGCCACNNNAGATCGGAAG |
| 46476_2 | TATGAGTGTGGAGTCGTTGCCTGCAATTACAGGTGTAAGCCACNNNAGATCGGAAG |
| 10710_1 | TATGAGTGTGGAGTCGTTGCTCAACCCTCCAATATGGCTCTCANNNAGATCGGAAGA |
| 10710_2 | TATGAGTGTGGAGTCGTTGCTCAACCCTCCAATATGGCTCTCANNNAGATCGGAAGA |
| 9780_1 | TATGAGTGTGGAGTCGTTGCCACACACACGACTTTTGCTCTTANNNAGATCGGAAGA |
| 9780_2 | TATGAGTGTGGAGTCGTTGCCACACACACGACTTTTGCTCTTANNNAGATCGGAAGA |
| 47573_1 | TATGAGTGTGGAGTCGTTGCCCCAGGTTCAAGCAATTATCTGCNNNAGATCGGAAGA |
| 47573_2 | TATGAGTGTGGAGTCGTTGCCCCAGGTTCAAGCAATTATCTGCNNNAGATCGGAAGA |
| 40763_1 | TATGAGTGTGGAGTCGTTGCGGTGTAAAAGAGCATGTGGACACNNNAGATCGGAA |
| 40763_2 | TATGAGTGTGGAGTCGTTGCGGTGTAAAAGAGCATGTGGACACNNNAGATCGGAA |
| 46478_1 | TATGAGTGTGGAGTCGTTGCTTCAAGTGATTCTCCTGCCTCAGNNNAGATCGGAAGA |
| 46478_2 | TATGAGTGTGGAGTCGTTGCTTCAAGTGATTCTCCTGCCTCAGNNNAGATCGGAAGA |
| 35873_1 | TATGAGTGTGGAGTCGTTGCAAATCGTACCAATTCCTCCCTCCNNNAGATCGGAAGA |
| 35873_2 | TATGAGTGTGGAGTCGTTGCAAATCGTACCAATTCCTCCCTCCNNNAGATCGGAAGA |
| 29082_1 | TATGAGTGTGGAGTCGTTGCAGATCAGTGAAAACAGGAGCTGTNNNAGATCGGAA |

| | |
|---|---|
| 29082_2 | TATGAGTGTGGAGTCGTTGCAGATCAGTGAAAACAGGAGCTGTNNNAGATCGGAA |
| 12639_1 | TATGAGTGTGGAGTCGTTGCTGCTTATATAGGTTTGATGAGTANNNAGATCGGAAG |
| 12639_2 | TATGAGTGTGGAGTCGTTGCTGCTTATATAGGTTTGATGAGTANNNAGATCGGAAG |
| 35053_1 | TATGAGTGTGGAGTCGTTGCCCACGCCCAGCTAATTTTTTGTANNNAGATCGGAAGA |
| 35053_2 | TATGAGTGTGGAGTCGTTGCCCACGCCCAGCTAATTTTTTGTANNNAGATCGGAAGA |
| 36576_1 | TATGAGTGTGGAGTCGTTGCCACTCAGCTGTTGTAGAGGATGTNNNAGATCGGAAG |
| 36576_2 | TATGAGTGTGGAGTCGTTGCCACTCAGCTGTTGTAGAGGATGTNNNAGATCGGAAG |
| 32835_1 | TATGAGTGTGGAGTCGTTGCCCTCTTGGGTTCATGCCATTCTCNNNAGATCGGAAGA |
| 32835_2 | TATGAGTGTGGAGTCGTTGCCCTCTTGGGTTCATGCCATTCTCNNNAGATCGGAAGA |
| 13384_1 | TATGAGTGTGGAGTCGTTGCATTCTTGTTTTTAAACTTAGCCANNNAGATCGGAAGA |
| 13384_2 | TATGAGTGTGGAGTCGTTGCATTCTTGTTTTTAAACTTAGCCANNNAGATCGGAAGA |
| 39999_1 | TATGAGTGTGGAGTCGTTGCAGATGAGGTCTCACTATGTTGCCNNNAGATCGGAAG |
| 39999_2 | TATGAGTGTGGAGTCGTTGCAGATGAGGTCTCACTATGTTGCCNNNAGATCGGAAG |
| 32807_1 | TATGAGTGTGGAGTCGTTGCAGGTTCAAGGGATTCTTCTGCTTNNNAGATCGGAAG |
| 32807_2 | TATGAGTGTGGAGTCGTTGCAGGTTCAAGGGATTCTTCTGCTTNNNAGATCGGAAG |
| 50921_1 | TATGAGTGTGGAGTCGTTGCCCACCACGATCCCCTTACTTTTANNNAGATCGGAAGA |
| 50921_2 | TATGAGTGTGGAGTCGTTGCCCACCACGATCCCCTTACTTTTANNNAGATCGGAAGA |
| 40279_1 | TATGAGTGTGGAGTCGTTGCGTGCCCCATTGCAAAAGAAAATGNNNAGATCGGAAG |
| 40279_2 | TATGAGTGTGGAGTCGTTGCGTGCCCCATTGCAAAAGAAAATGNNNAGATCGGAAG |
| 42069_1 | TATGAGTGTGGAGTCGTTGCACCCACAGGCAACATAGGATAAGNNNAGATCGGAAG |
| 42069_2 | TATGAGTGTGGAGTCGTTGCACCCACAGGCAACATAGGATAAGNNNAGATCGGAAG |
| 40640_1 | TATGAGTGTGGAGTCGTTGCAACTGAGAAACAGAAGGGACAGANNNAGATCGGAA |
| 40640_2 | TATGAGTGTGGAGTCGTTGCAACTGAGAAACAGAAGGGACAGANNNAGATCGGAA |
| 45929_1 | TATGAGTGTGGAGTCGTTGCCTCACTGCAACCTGGAGTGTAATNNNAGATCGGAAG |
| 45929_2 | TATGAGTGTGGAGTCGTTGCCTCACTGCAACCTGGAGTGTAATNNNAGATCGGAAG |
| 29075_1 | TATGAGTGTGGAGTCGTTGCGGATGACTGGCAATTCCCTTTCTNNNAGATCGGAAGA |
| 29075_2 | TATGAGTGTGGAGTCGTTGCGGATGACTGGCAATTCCCTTTCTNNNAGATCGGAAGA |
| 53106_1 | TATGAGTGTGGAGTCGTTGCGTGACAGAGTACGGTTAGAACGTNNNAGATCGGAA |
| 53106_2 | TATGAGTGTGGAGTCGTTGCGTGACAGAGTACGGTTAGAACGTNNNAGATCGGAA |
| 10048_1 | TATGAGTGTGGAGTCGTTGCCTGAGTGATATGGCAGCCAACTTNNNAGATCGGAAG |
| 10048_2 | TATGAGTGTGGAGTCGTTGCCTGAGTGATATGGCAGCCAACTTNNNAGATCGGAAG |
| 45047_1 | TATGAGTGTGGAGTCGTTGCCTTTCGGACTGGTGCAGGGTAGNNNAGATCGGAAG |
| 45047_2 | TATGAGTGTGGAGTCGTTGCCTTTCGGACTGGTGCAGGGTAGNNNAGATCGGAAG |
| 35799_1 | TATGAGTGTGGAGTCGTTGCCCATTTCAGCCTCCCAAGTATCTNNNAGATCGGAAGA |
| 35799_2 | TATGAGTGTGGAGTCGTTGCCCATTTCAGCCTCCCAAGTATCTNNNAGATCGGAAGA |
| 46347_1 | TATGAGTGTGGAGTCGTTGCCTCCTAAAGTGCTGGGACTACAGNNNAGATCGGAAG |
| 46347_2 | TATGAGTGTGGAGTCGTTGCCTCCTAAAGTGCTGGGACTACAGNNNAGATCGGAAG |
| 32609_1 | TATGAGTGTGGAGTCGTTGCGGAGGGGAGAGAGAAACAGAAAANNNAGATCGGA |
| 32609_2 | TATGAGTGTGGAGTCGTTGCGGAGGGGAGAGAGAAACAGAAAANNNAGATCGGA |
| 53366_1 | TATGAGTGTGGAGTCGTTGCATTATGTTTCCTGAAGCCTCCCCNNNAGATCGGAAGA |
| 53366_2 | TATGAGTGTGGAGTCGTTGCATTATGTTTCCTGAAGCCTCCCCNNNAGATCGGAAGA |
| 43458_1 | TATGAGTGTGGAGTCGTTGCGGTCTCACTTCTGGCAAGTAGAANNNAGATCGGAAG |
| 43458_2 | TATGAGTGTGGAGTCGTTGCGGTCTCACTTCTGGCAAGTAGAANNNAGATCGGAAG |

| | |
|---|---|
| 39850_1 | TATGAGTGTGGAGTCGTTGCCCTTTTCCAGACCACTAGTCACANNNAGATCGGAAGA |
| 39850_2 | TATGAGTGTGGAGTCGTTGCCCTTTTCCAGACCACTAGTCACANNNAGATCGGAAGA |
| 39581_1 | TATGAGTGTGGAGTCGTTGCAGAAAGAAAGAGAGAGAGGGGGANNNAGATCGGA |
| 39581_2 | TATGAGTGTGGAGTCGTTGCAGAAAGAAAGAGAGAGAGGGGGANNNAGATCGGA |
| 9976_1 | TATGAGTGTGGAGTCGTTGCGCCTTTCTCTGCCTCCATTTCTTNNNAGATCGGAAGAC |
| 9976_2 | TATGAGTGTGGAGTCGTTGCGCCTTTCTCTGCCTCCATTTCTTNNNAGATCGGAAGAC |
| 52220_1 | TATGAGTGTGGAGTCGTTGCCACTGCCTCTTAGTCCCCATCNNNAGATCGGAAGAGC |
| 52220_2 | TATGAGTGTGGAGTCGTTGCCACTGCCTCTTAGTCCCCATCNNNAGATCGGAAGAGC |
| 52642_1 | TATGAGTGTGGAGTCGTTGCTTTTATGGGCTGACACTCATGGANNNAGATCGGAAG |
| 52642_2 | TATGAGTGTGGAGTCGTTGCTTTTATGGGCTGACACTCATGGANNNAGATCGGAAG |
| 37499_1 | TATGAGTGTGGAGTCGTTGCACAGTCATGAGTCATGGAAGCATNNNAGATCGGAAC |
| 37499_2 | TATGAGTGTGGAGTCGTTGCACAGTCATGAGTCATGGAAGCATNNNAGATCGGAAC |
| 44778_1 | TATGAGTGTGGAGTCGTTGCTCTAGCCTTCTGTCAGTTCTGTGNNNAGATCGGAAGA |
| 44778_2 | TATGAGTGTGGAGTCGTTGCTCTAGCCTTCTGTCAGTTCTGTGNNNAGATCGGAAGA |
| 32812_1 | TATGAGTGTGGAGTCGTTGCCTCAAGCAATCCTCCTACCTCAGNNNAGATCGGAAGA |
| 32812_2 | TATGAGTGTGGAGTCGTTGCCTCAAGCAATCCTCCTACCTCAGNNNAGATCGGAAGA |
| 39883_1 | TATGAGTGTGGAGTCGTTGCCAACGTTGCAGTGAGCTATGATTNNNAGATCGGAAG |
| 39883_2 | TATGAGTGTGGAGTCGTTGCCAACGTTGCAGTGAGCTATGATTNNNAGATCGGAAG |
| 43627_1 | TATGAGTGTGGAGTCGTTGCGTAGTGTATAGACAGCCCTGTGGNNNAGATCGGAAC |
| 43627_2 | TATGAGTGTGGAGTCGTTGCGTAGTGTATAGACAGCCCTGTGGNNNAGATCGGAAC |
| 48904_1 | TATGAGTGTGGAGTCGTTGCTTCCACGCCCCTTTCAATCATATNNNAGATCGGAAGA |
| 48904_2 | TATGAGTGTGGAGTCGTTGCTTCCACGCCCCTTTCAATCATATNNNAGATCGGAAGA |
| 51965_1 | TATGAGTGTGGAGTCGTTGCCATTACACTCCCATCAGAAGCCTNNNAGATCGGAAGA |
| 51965_2 | TATGAGTGTGGAGTCGTTGCCATTACACTCCCATCAGAAGCCTNNNAGATCGGAAGA |
| 45456_1 | TATGAGTGTGGAGTCGTTGCAGATTAGTTTCACACAGCCAAGCNNNAGATCGGAAG |
| 45456_2 | TATGAGTGTGGAGTCGTTGCAGATTAGTTTCACACAGCCAAGCNNNAGATCGGAAG |
| 46196_1 | TATGAGTGTGGAGTCGTTGCAGATGGAGTCTCACTCTGTCACCNNNAGATCGGAAG |
| 46196_2 | TATGAGTGTGGAGTCGTTGCAGATGGAGTCTCACTCTGTCACCNNNAGATCGGAAG |
| 47109_1 | TATGAGTGTGGAGTCGTTGCCCCTTGATACGCCTCAGGATAAANNNAGATCGGAAG |
| 47109_2 | TATGAGTGTGGAGTCGTTGCCCCTTGATACGCCTCAGGATAAANNNAGATCGGAAG |
| 46229_1 | TATGAGTGTGGAGTCGTTGCTCCCAAGTAGCTGAGACTACAGANNNAGATCGGAAG |
| 46229_2 | TATGAGTGTGGAGTCGTTGCTCCCAAGTAGCTGAGACTACAGANNNAGATCGGAAG |
| 10126_1 | TATGAGTGTGGAGTCGTTGCGCCCAAGATGTTCTGAAAATGGCNNNAGATCGGAAG |
| 10126_2 | TATGAGTGTGGAGTCGTTGCGCCCAAGATGTTCTGAAAATGGCNNNAGATCGGAAG |
| 51502_1 | TATGAGTGTGGAGTCGTTGCAAGGGGCAACAGAAAAGAGAGAANNNAGATCGGA |
| 51502_2 | TATGAGTGTGGAGTCGTTGCAAGGGGCAACAGAAAAGAGAGAANNNAGATCGGA |
| 40323_1 | TATGAGTGTGGAGTCGTTGCAAGTGCCTCTTAGGAAAACTGGTNNNAGATCGGAAG |
| 40323_2 | TATGAGTGTGGAGTCGTTGCAAGTGCCTCTTAGGAAAACTGGTNNNAGATCGGAAG |
| 51720_1 | TATGAGTGTGGAGTCGTTGCTTTTCTGTCTCTCTGCCCACTTCNNNAGATCGGAAGAC |
| 51720_2 | TATGAGTGTGGAGTCGTTGCTTTTCTGTCTCTCTGCCCACTTCNNNAGATCGGAAGAC |
| 41837_1 | TATGAGTGTGGAGTCGTTGCAGATGGAGTCTCACTCTGTTGCNNNAGATCGGAAGA |
| 41837_2 | TATGAGTGTGGAGTCGTTGCAGATGGAGTCTCACTCTGTTGCNNNAGATCGGAAGA |
| 33324_1 | TATGAGTGTGGAGTCGTTGCGTCAACCTCCCAGGCTCAAGNNNAGATCGGAAGAGC |

| | |
|---|---|
| 33324_2 | TATGAGTGTGGAGTCGTTGCGTCAACCTCCCAGGCTCAAGNNNAGATCGGAAGAGC |
| 11939_1 | TATGAGTGTGGAGTCGTTGCCTTGGTCTCTTCGCACTGGACANNNAGATCGGAAGA |
| 11939_2 | TATGAGTGTGGAGTCGTTGCCTTGGTCTCTTCGCACTGGACANNNAGATCGGAAGA |
| 29796_1 | TATGAGTGTGGAGTCGTTGCTAGACTGAAAGTTACAGGCCACCNNNAGATCGGAAG |
| 29796_2 | TATGAGTGTGGAGTCGTTGCTAGACTGAAAGTTACAGGCCACCNNNAGATCGGAAG |
| 50764_1 | TATGAGTGTGGAGTCGTTGCCCACGTATTCCTACATAGCCACTNNNAGATCGGAAGA |
| 50764_2 | TATGAGTGTGGAGTCGTTGCCCACGTATTCCTACATAGCCACTNNNAGATCGGAAGA |
| 50059_1 | TATGAGTGTGGAGTCGTTGCAGCTGCTGTGAACATATTTGTACANNNAGATCGGAA |
| 50059_2 | TATGAGTGTGGAGTCGTTGCAGCTGCTGTGAACATATTTGTACANNNAGATCGGAA |
| 34293_1 | TATGAGTGTGGAGTCGTTGCCAGATTCAGTGTCGGTTTTGGACNNNAGATCGGAAG |
| 34293_2 | TATGAGTGTGGAGTCGTTGCCAGATTCAGTGTCGGTTTTGGACNNNAGATCGGAAG |
| 47103_1 | TATGAGTGTGGAGTCGTTGCTGCAGTACAGTTTGAAGTCAGGTNNNAGATCGGAA |
| 47103_2 | TATGAGTGTGGAGTCGTTGCTGCAGTACAGTTTGAAGTCAGGTNNNAGATCGGAA |
| 29163_1 | TATGAGTGTGGAGTCGTTGCACCCTTTTCGGACAAATTATCCTNNNAGATCGGAAGA |
| 29163_2 | TATGAGTGTGGAGTCGTTGCACCCTTTTCGGACAAATTATCCTNNNAGATCGGAAGA |
| 52865_1 | TATGAGTGTGGAGTCGTTGCAGAACATGATGGACCCCTACAACNNNAGATCGGAAG |
| 52865_2 | TATGAGTGTGGAGTCGTTGCAGAACATGATGGACCCCTACAACNNNAGATCGGAAG |
| 38341_1 | TATGAGTGTGGAGTCGTTGCAGAGAGAGAAGGATCATGGTGACNNNAGATCGGAA |
| 38341_2 | TATGAGTGTGGAGTCGTTGCAGAGAGAGAAGGATCATGGTGACNNNAGATCGGAA |
| 52155_1 | TATGAGTGTGGAGTCGTTGCTCTATCGGAATACCACAAGCTGGNNNAGATCGGAAG |
| 52155_2 | TATGAGTGTGGAGTCGTTGCTCTATCGGAATACCACAAGCTGGNNNAGATCGGAAG |
| 45904_1 | TATGAGTGTGGAGTCGTTGCAACCTCCTGAGTAGCTGGGATTANNNAGATCGGAAG |
| 45904_2 | TATGAGTGTGGAGTCGTTGCAACCTCCTGAGTAGCTGGGATTANNNAGATCGGAAG |
| 40068_1 | TATGAGTGTGGAGTCGTTGCCCTTTTGGTATGGGCTGACTTTGNNNAGATCGGAAG |
| 40068_2 | TATGAGTGTGGAGTCGTTGCCCTTTTGGTATGGGCTGACTTTGNNNAGATCGGAAG |
| 8117_1 | TATGAGTGTGGAGTCGTTGCCTGTGATGCCCCTGTGCATAGANNNAGATCGGAAGA |
| 8117_2 | TATGAGTGTGGAGTCGTTGCCTGTGATGCCCCTGTGCATAGANNNAGATCGGAAGA |
| 40221_1 | TATGAGTGTGGAGTCGTTGCCTGGAAAAGCATTCTCTTTGCCANNNAGATCGGAAG |
| 40221_2 | TATGAGTGTGGAGTCGTTGCCTGGAAAAGCATTCTCTTTGCCANNNAGATCGGAAG |
| 42820_1 | TATGAGTGTGGAGTCGTTGCATTCATACTCAAATGCCCAAGCCNNNAGATCGGAAGA |
| 42820_2 | TATGAGTGTGGAGTCGTTGCATTCATACTCAAATGCCCAAGCCNNNAGATCGGAAGA |
| 44404_1 | TATGAGTGTGGAGTCGTTGCGAACGAGCATACCATCACAGTGANNNAGATCGGAAG |
| 44404_2 | TATGAGTGTGGAGTCGTTGCGAACGAGCATACCATCACAGTGANNNAGATCGGAAG |
| 50514_1 | TATGAGTGTGGAGTCGTTGCTGAGAATCAGAAGACTCGCTTGGNNNAGATCGGAA |
| 50514_2 | TATGAGTGTGGAGTCGTTGCTGAGAATCAGAAGACTCGCTTGGNNNAGATCGGAA |
| 31377_1 | TATGAGTGTGGAGTCGTTGCTGGCAGCAAGTAATATCCACATGNNNAGATCGGAAG |
| 31377_2 | TATGAGTGTGGAGTCGTTGCTGGCAGCAAGTAATATCCACATGNNNAGATCGGAAG |
| 39031_1 | TATGAGTGTGGAGTCGTTGCCAAGGCTGGTTCAACATATGCAANNNAGATCGGAAG |
| 39031_2 | TATGAGTGTGGAGTCGTTGCCAAGGCTGGTTCAACATATGCAANNNAGATCGGAAG |
| 40182_1 | TATGAGTGTGGAGTCGTTGCCTCACACCTGTAATCTCCACACTNNNAGATCGGAAGA |
| 40182_2 | TATGAGTGTGGAGTCGTTGCCTCACACCTGTAATCTCCACACTNNNAGATCGGAAGA |
| 32826_1 | TATGAGTGTGGAGTCGTTGCGAGTAGCTAGGATTACAGGTGCCNNNAGATCGGAA |
| 32826_2 | TATGAGTGTGGAGTCGTTGCGAGTAGCTAGGATTACAGGTGCCNNNAGATCGGAA |

| | |
|---|---|
| 39153_1 | TATGAGTGTGGAGTCGTTGCACATGATAGTTGACCCTATGGAGTNNNAGATCGGAA |
| 39153_2 | TATGAGTGTGGAGTCGTTGCACATGATAGTTGACCCTATGGAGTNNNAGATCGGAA |
| 35134_1 | TATGAGTGTGGAGTCGTTGCCTTTGCTGTGCAGAGGCTTTTANNNAGATCGGAAG |
| 35134_2 | TATGAGTGTGGAGTCGTTGCCTTTGCTGTGCAGAGGCTTTTANNNAGATCGGAAG |
| 33131_1 | TATGAGTGTGGAGTCGTTGCGAGAACAAAGATGTGCTGCTACGNNNAGATCGGAA |
| 33131_2 | TATGAGTGTGGAGTCGTTGCGAGAACAAAGATGTGCTGCTACGNNNAGATCGGAA |
| 30196_1 | TATGAGTGTGGAGTCGTTGCAAGCTGAACTTTCCTAGTGCTGANNNAGATCGGAAG |
| 30196_2 | TATGAGTGTGGAGTCGTTGCAAGCTGAACTTTCCTAGTGCTGANNNAGATCGGAAG |
| 43883_1 | TATGAGTGTGGAGTCGTTGCCAAAGGAGAGATCCCCTCAACTGNNNAGATCGGAAG |
| 43883_2 | TATGAGTGTGGAGTCGTTGCCAAAGGAGAGATCCCCTCAACTGNNNAGATCGGAAG |
| 48299_1 | TATGAGTGTGGAGTCGTTGCCCATCTCAAAGTTTGGCCCAATTNNNAGATCGGAAGA |
| 48299_2 | TATGAGTGTGGAGTCGTTGCCCATCTCAAAGTTTGGCCCAATTNNNAGATCGGAAGA |
| 44599_1 | TATGAGTGTGGAGTCGTTGCTGGAAGTGAGAAGGGAAAGATGGNNNAGATCGGA |
| 44599_2 | TATGAGTGTGGAGTCGTTGCTGGAAGTGAGAAGGGAAAGATGGNNNAGATCGGA |
| 12921_1 | TATGAGTGTGGAGTCGTTGCGGTGAAAGGTTAAAACATATCTTNNNAGATCGGAAG |
| 12921_2 | TATGAGTGTGGAGTCGTTGCGGTGAAAGGTTAAAACATATCTTNNNAGATCGGAAG |
| 50608_1 | TATGAGTGTGGAGTCGTTGCTTGAGTGACTTTCTCATCCTCCGNNNAGATCGGAAGA |
| 50608_2 | TATGAGTGTGGAGTCGTTGCTTGAGTGACTTTCTCATCCTCCGNNNAGATCGGAAGA |
| 36864_1 | TATGAGTGTGGAGTCGTTGCGTAATCCCAGCTACTTGAGAGGCNNNAGATCGGAAG |
| 36864_2 | TATGAGTGTGGAGTCGTTGCGTAATCCCAGCTACTTGAGAGGCNNNAGATCGGAAG |
| 12955_1 | TATGAGTGTGGAGTCGTTGCAATGTCACCTGTATCTGTGANNNAGATCGGAAGAGC |
| 12955_2 | TATGAGTGTGGAGTCGTTGCAATGTCACCTGTATCTGTGANNNAGATCGGAAGAGC |
| 28932_1 | TATGAGTGTGGAGTCGTTGCCAACAAAACACACAAACTGGTGCNNNAGATCGGAAG |
| 28932_2 | TATGAGTGTGGAGTCGTTGCCAACAAAACACACAAACTGGTGCNNNAGATCGGAAG |
| 34131_1 | TATGAGTGTGGAGTCGTTGCTATGTACGTGTGTGTGTGTGTGTNNNAGATCGGAA |
| 34131_2 | TATGAGTGTGGAGTCGTTGCTATGTACGTGTGTGTGTGTGTGTNNNAGATCGGAA |
| 47650_1 | TATGAGTGTGGAGTCGTTGCCTCCTGAGTAGCTGGGATTACAGNNNAGATCGGAAG |
| 47650_2 | TATGAGTGTGGAGTCGTTGCCTCCTGAGTAGCTGGGATTACAGNNNAGATCGGAAG |
| 44104_1 | TATGAGTGTGGAGTCGTTGCTGTGATGCCTTAGAGTGGATGAANNNAGATCGGAA |
| 44104_2 | TATGAGTGTGGAGTCGTTGCTGTGATGCCTTAGAGTGGATGAANNNAGATCGGAA |
| 52595_1 | TATGAGTGTGGAGTCGTTGCAAAGGCGAAGATGTTTCACGAAGNNNAGATCGGAA |
| 52595_2 | TATGAGTGTGGAGTCGTTGCAAAGGCGAAGATGTTTCACGAAGNNNAGATCGGAA |
| 40641_1 | TATGAGTGTGGAGTCGTTGCACAGAGAAGCAACACCATTTCTTNNNAGATCGGAAG |
| 40641_2 | TATGAGTGTGGAGTCGTTGCACAGAGAAGCAACACCATTTCTTNNNAGATCGGAAG |
| 42060_1 | TATGAGTGTGGAGTCGTTGCTCTCTGTACCTTGAAACAGTGGCNNNAGATCGGAAG |
| 42060_2 | TATGAGTGTGGAGTCGTTGCTCTCTGTACCTTGAAACAGTGGCNNNAGATCGGAAG |
| 53046_1 | TATGAGTGTGGAGTCGTTGCAAGAAATCTCAGAGACTCAGCCCNNNAGATCGGAAG |
| 53046_2 | TATGAGTGTGGAGTCGTTGCAAGAAATCTCAGAGACTCAGCCCNNNAGATCGGAAG |
| 42064_1 | TATGAGTGTGGAGTCGTTGCTGAGGAGGCAGAGAAAACGAATTNNNAGATCGGAA |
| 42064_2 | TATGAGTGTGGAGTCGTTGCTGAGGAGGCAGAGAAAACGAATTNNNAGATCGGAA |
| 46084_1 | TATGAGTGTGGAGTCGTTGCCAGAGTCTTGTTCTGTCACCCANNNAGATCGGAAGA |
| 46084_2 | TATGAGTGTGGAGTCGTTGCCAGAGTCTTGTTCTGTCACCCANNNAGATCGGAAGA |
| 29652_1 | TATGAGTGTGGAGTCGTTGCCCCTGACCAAGGAGAAACATGATNNNAGATCGGAA |

| | |
|---|---|
| 29652_2 | TATGAGTGTGGAGTCGTTGCCCCTGACCAAGGAGAAACATGATNNNAGATCGGAA |
| 41260_1 | TATGAGTGTGGAGTCGTTGCGCCTCCCTCTAACCATGAGAAAANNNAGATCGGAAG |
| 41260_2 | TATGAGTGTGGAGTCGTTGCGCCTCCCTCTAACCATGAGAAAANNNAGATCGGAAG |
| 33085_1 | TATGAGTGTGGAGTCGTTGCAGGCTGCAGTTGAGTATCTTGATNNNAGATCGGAA |
| 33085_2 | TATGAGTGTGGAGTCGTTGCAGGCTGCAGTTGAGTATCTTGATNNNAGATCGGAA |
| 37104_1 | TATGAGTGTGGAGTCGTTGCCAAAACAAAACCTGCACCACCTANNNAGATCGGAAG |
| 37104_2 | TATGAGTGTGGAGTCGTTGCCAAAACAAAACCTGCACCACCTANNNAGATCGGAAG |
| 42617_1 | TATGAGTGTGGAGTCGTTGCGTTCAAGAGAGTCTCCTGCCTCNNNAGATCGGAAGA |
| 42617_2 | TATGAGTGTGGAGTCGTTGCGTTCAAGAGAGTCTCCTGCCTCNNNAGATCGGAAGA |
| 32213_1 | TATGAGTGTGGAGTCGTTGCAAAGGAGGAGGGTTCATGGTTACNNNAGATCGGAA |
| 32213_2 | TATGAGTGTGGAGTCGTTGCAAAGGAGGAGGGTTCATGGTTACNNNAGATCGGAA |
| 53132_1 | TATGAGTGTGGAGTCGTTGCCCATATCCCCATTCACTGTCCTTNNNAGATCGGAAGA |
| 53132_2 | TATGAGTGTGGAGTCGTTGCCCATATCCCCATTCACTGTCCTTNNNAGATCGGAAGA |
| 29827_1 | TATGAGTGTGGAGTCGTTGCATTGAGGGCTTCACAGGTAATGTNNNAGATCGGAA |
| 29827_2 | TATGAGTGTGGAGTCGTTGCATTGAGGGCTTCACAGGTAATGTNNNAGATCGGAA |
| 33452_1 | TATGAGTGTGGAGTCGTTGCCAGAGGCTGTGGTGAGAAGATAGNNNAGATCGGA |
| 33452_2 | TATGAGTGTGGAGTCGTTGCCAGAGGCTGTGGTGAGAAGATAGNNNAGATCGGA |
| 13471_1 | TATGAGTGTGGAGTCGTTGCTCCACATTTTAAAAACTAAGCTTNNNAGATCGGAAG |
| 13471_2 | TATGAGTGTGGAGTCGTTGCTCCACATTTTAAAAACTAAGCTTNNNAGATCGGAAG |
| 13290_1 | TATGAGTGTGGAGTCGTTGCTTTATCTTTTTTCTGCCATCTTTNNNAGATCGGAAGA |
| 13290_2 | TATGAGTGTGGAGTCGTTGCTTTATCTTTTTTCTGCCATCTTTNNNAGATCGGAAGA |
| 44422_1 | TATGAGTGTGGAGTCGTTGCCTCAGTAGGGTTCTGTTTCCACANNNAGATCGGAAG |
| 44422_2 | TATGAGTGTGGAGTCGTTGCCTCAGTAGGGTTCTGTTTCCACANNNAGATCGGAAG |
| 43728_1 | TATGAGTGTGGAGTCGTTGCAGATGTGACAAGGATGAGGCTTTNNNAGATCGGAA |
| 43728_2 | TATGAGTGTGGAGTCGTTGCAGATGTGACAAGGATGAGGCTTTNNNAGATCGGAA |
| 48178_1 | TATGAGTGTGGAGTCGTTGCCTGGTCACAAATGATTCTCCTGCNNNAGATCGGAAG |
| 48178_2 | TATGAGTGTGGAGTCGTTGCCTGGTCACAAATGATTCTCCTGCNNNAGATCGGAAG |
| 40299_1 | TATGAGTGTGGAGTCGTTGCATGAGAGGGAAGAGTCAAAGCTGNNNAGATCGGA |
| 40299_2 | TATGAGTGTGGAGTCGTTGCATGAGAGGGAAGAGTCAAAGCTGNNNAGATCGGA |
| 32785_1 | TATGAGTGTGGAGTCGTTGCCAATGGCCTTGAGAAGCAAAAGANNNAGATCGGAA |
| 32785_2 | TATGAGTGTGGAGTCGTTGCCAATGGCCTTGAGAAGCAAAAGANNNAGATCGGAA |
| 34721_1 | TATGAGTGTGGAGTCGTTGCGTGATCTCAGCTCACTACAACCTNNNAGATCGGAAG |
| 34721_2 | TATGAGTGTGGAGTCGTTGCGTGATCTCAGCTCACTACAACCTNNNAGATCGGAAG |
| 31810_1 | TATGAGTGTGGAGTCGTTGCGTTCAAACGATTTTCCTGCCTCANNNAGATCGGAAG |
| 31810_2 | TATGAGTGTGGAGTCGTTGCGTTCAAACGATTTTCCTGCCTCANNNAGATCGGAAG |
| 45541_1 | TATGAGTGTGGAGTCGTTGCCGCCATGTACACCATCATCAACNNNAGATCGGAAGA |
| 45541_2 | TATGAGTGTGGAGTCGTTGCCGCCATGTACACCATCATCAACNNNAGATCGGAAGA |
| 37410_1 | TATGAGTGTGGAGTCGTTGCAGAAATGCCACATGACACACATGNNNAGATCGGAA |
| 37410_2 | TATGAGTGTGGAGTCGTTGCAGAAATGCCACATGACACACATGNNNAGATCGGAA |
| 13072_1 | TATGAGTGTGGAGTCGTTGCGTCATCCTAGAAGGAATGGTNNNAGATCGGAAGAG |
| 13072_2 | TATGAGTGTGGAGTCGTTGCGTCATCCTAGAAGGAATGGTNNNAGATCGGAAGAG |
| 40361_1 | TATGAGTGTGGAGTCGTTGCCACTTGAAATGTGGCTAGTGCAANNNAGATCGGAA |
| 40361_2 | TATGAGTGTGGAGTCGTTGCCACTTGAAATGTGGCTAGTGCAANNNAGATCGGAA |

| | |
|---|---|
| 13627_1 | TATGAGTGTGGAGTCGTTGCGAATAAGCATAACTATTGTTCCANNNAGATCGGAAG |
| 13627_2 | TATGAGTGTGGAGTCGTTGCGAATAAGCATAACTATTGTTCCANNNAGATCGGAAG |
| 46621_1 | TATGAGTGTGGAGTCGTTGCCTGGAGCATGTTGGTATGATCCTNNNAGATCGGAAG |
| 46621_2 | TATGAGTGTGGAGTCGTTGCCTGGAGCATGTTGGTATGATCCTNNNAGATCGGAAG |
| 39927_1 | TATGAGTGTGGAGTCGTTGCTCTGTTGTTTAGTGCAGCTACTCTNNNAGATCGGAAG |
| 39927_2 | TATGAGTGTGGAGTCGTTGCTCTGTTGTTTAGTGCAGCTACTCTNNNAGATCGGAAG |
| 34955_1 | TATGAGTGTGGAGTCGTTGCTGGAATGGGTTCTGCGATGATTANNNAGATCGGAA |
| 34955_2 | TATGAGTGTGGAGTCGTTGCTGGAATGGGTTCTGCGATGATTANNNAGATCGGAA |
| 29229_1 | TATGAGTGTGGAGTCGTTGCGCAGCTTCTACTAACCAAGAGGTNNNAGATCGGAAG |
| 29229_2 | TATGAGTGTGGAGTCGTTGCGCAGCTTCTACTAACCAAGAGGTNNNAGATCGGAAG |
| 39643_1 | TATGAGTGTGGAGTCGTTGCACCAAAAAAGGAGAGGGGGAAATNNNAGATCGGA |
| 39643_2 | TATGAGTGTGGAGTCGTTGCACCAAAAAAGGAGAGGGGGAAATNNNAGATCGGA |
| 44899_1 | TATGAGTGTGGAGTCGTTGCAGATGATATGTGGGAGAAGGTGCNNNAGATCGGA |
| 44899_2 | TATGAGTGTGGAGTCGTTGCAGATGATATGTGGGAGAAGGTGCNNNAGATCGGA |
| 38499_1 | TATGAGTGTGGAGTCGTTGCGCCATCCTCCCACATACATGTTANNNAGATCGGAAGA |
| 38499_2 | TATGAGTGTGGAGTCGTTGCGCCATCCTCCCACATACATGTTANNNAGATCGGAAGA |
| 47032_1 | TATGAGTGTGGAGTCGTTGCACACCATCACACCCAGCTAATTTNNNAGATCGGAAGA |
| 47032_2 | TATGAGTGTGGAGTCGTTGCACACCATCACACCCAGCTAATTTNNNAGATCGGAAGA |
| 44696_1 | TATGAGTGTGGAGTCGTTGCGACCTCTTCACATTTTTGAGGCCNNNAGATCGGAAG |
| 44696_2 | TATGAGTGTGGAGTCGTTGCGACCTCTTCACATTTTTGAGGCCNNNAGATCGGAAG |
| 43553_1 | TATGAGTGTGGAGTCGTTGCCAGTGACATCATCTCAGCTCACTNNNAGATCGGAAG |
| 43553_2 | TATGAGTGTGGAGTCGTTGCCAGTGACATCATCTCAGCTCACTNNNAGATCGGAAG |
| 10351_1 | TATGAGTGTGGAGTCGTTGCTCCAAGCTGTTCTGGCTTATTCCNNNAGATCGGAAG |
| 10351_2 | TATGAGTGTGGAGTCGTTGCTCCAAGCTGTTCTGGCTTATTCCNNNAGATCGGAAG |
| 49789_1 | TATGAGTGTGGAGTCGTTGCTGTGCATTTTGAACACTAAGAACCANNNAGATCGGA |
| 49789_2 | TATGAGTGTGGAGTCGTTGCTGTGCATTTTGAACACTAAGAACCANNNAGATCGGA |
| 37306_1 | TATGAGTGTGGAGTCGTTGCTCCGGCAGAACTTTGAGTTTCTANNNAGATCGGAAG |
| 37306_2 | TATGAGTGTGGAGTCGTTGCTCCGGCAGAACTTTGAGTTTCTANNNAGATCGGAAG |
| 52910_1 | TATGAGTGTGGAGTCGTTGCAAGGGCTGCTGATGATTTTTCACNNNAGATCGGAAG |
| 52910_2 | TATGAGTGTGGAGTCGTTGCAAGGGCTGCTGATGATTTTTCACNNNAGATCGGAAG |
| 13086_1 | TATGAGTGTGGAGTCGTTGCTCCAAATGCTGCAAAATAATAATNNNAGATCGGAAG |
| 13086_2 | TATGAGTGTGGAGTCGTTGCTCCAAATGCTGCAAAATAATAATNNNAGATCGGAAG |
| 50330_1 | TATGAGTGTGGAGTCGTTGCGCCTGTTCACCCCAATCCTATTANNNAGATCGGAAGA |
| 50330_2 | TATGAGTGTGGAGTCGTTGCGCCTGTTCACCCCAATCCTATTANNNAGATCGGAAGA |
| 33174_1 | TATGAGTGTGGAGTCGTTGCCTGTAAGCAGAGGTGACCTTACCNNNAGATCGGAAG |
| 33174_2 | TATGAGTGTGGAGTCGTTGCCTGTAAGCAGAGGTGACCTTACCNNNAGATCGGAAG |
| 40142_1 | TATGAGTGTGGAGTCGTTGCCGGTTCAGACTGCATTTTTGACTNNNAGATCGGAAG |
| 40142_2 | TATGAGTGTGGAGTCGTTGCCGGTTCAGACTGCATTTTTGACTNNNAGATCGGAAG |
| 41092_1 | TATGAGTGTGGAGTCGTTGCAATGGGGCACTAATGAGCTGATTNNNAGATCGGAA |
| 41092_2 | TATGAGTGTGGAGTCGTTGCAATGGGGCACTAATGAGCTGATTNNNAGATCGGAA |
| 31497_1 | TATGAGTGTGGAGTCGTTGCTGGTCTCATAGGTTATTTGGGCANNNAGATCGGAAG |
| 31497_2 | TATGAGTGTGGAGTCGTTGCTGGTCTCATAGGTTATTTGGGCANNNAGATCGGAAG |
| 51376_1 | TATGAGTGTGGAGTCGTTGCCATGACTTACTGAGACACCAGCTNNNAGATCGGAAG |

| | |
|---|---|
| 51376_2 | TATGAGTGTGGAGTCGTTGCCATGACTTACTGAGACACCAGCTNNNAGATCGGAAG |
| 34335_1 | TATGAGTGTGGAGTCGTTGCCCAGAGTTCTTGGCAGGATCTTANNNAGATCGGAAG |
| 34335_2 | TATGAGTGTGGAGTCGTTGCCCAGAGTTCTTGGCAGGATCTTANNNAGATCGGAAG |
| 42473_1 | TATGAGTGTGGAGTCGTTGCGGTGTCTCAGCTACTTGAAAAGCNNNAGATCGGAA |
| 42473_2 | TATGAGTGTGGAGTCGTTGCGGTGTCTCAGCTACTTGAAAAGCNNNAGATCGGAA |
| 29908_1 | TATGAGTGTGGAGTCGTTGCGACAGAGACTTTTTTTCCCCCCTNNNAGATCGGAAG |
| 29908_2 | TATGAGTGTGGAGTCGTTGCGACAGAGACTTTTTTTCCCCCCTNNNAGATCGGAAG |
| 37538_1 | TATGAGTGTGGAGTCGTTGCCACAGCATTGGGGGAACTATTTGNNNAGATCGGAA |
| 37538_2 | TATGAGTGTGGAGTCGTTGCCACAGCATTGGGGGAACTATTTGNNNAGATCGGAA |
| 31886_1 | TATGAGTGTGGAGTCGTTGCCCTTTCCAGGACCAGCAGATAATNNNAGATCGGAAG |
| 31886_2 | TATGAGTGTGGAGTCGTTGCCCTTTCCAGGACCAGCAGATAATNNNAGATCGGAAG |
| 46964_1 | TATGAGTGTGGAGTCGTTGCCTCCTGAGTAGCTGGAACTACAGNNNAGATCGGAA |
| 46964_2 | TATGAGTGTGGAGTCGTTGCCTCCTGAGTAGCTGGAACTACAGNNNAGATCGGAA |
| 36383_1 | TATGAGTGTGGAGTCGTTGCGCTTGGAAGAAAATCTTGGAGGCNNNAGATCGGAA |
| 36383_2 | TATGAGTGTGGAGTCGTTGCGCTTGGAAGAAAATCTTGGAGGCNNNAGATCGGAA |
| 45195_1 | TATGAGTGTGGAGTCGTTGCACAGATCCTCAAACTCCTTCCACNNNAGATCGGAAGA |
| 45195_2 | TATGAGTGTGGAGTCGTTGCACAGATCCTCAAACTCCTTCCACNNNAGATCGGAAGA |
| 38510_1 | TATGAGTGTGGAGTCGTTGCGAGTTTGAGACTAGCCTGACCAANNNAGATCGGAA |
| 38510_2 | TATGAGTGTGGAGTCGTTGCGAGTTTGAGACTAGCCTGACCAANNNAGATCGGAA |
| 10463_1 | TATGAGTGTGGAGTCGTTGCCACACAGGCACTGGGATAATTCANNNAGATCGGAAG |
| 10463_2 | TATGAGTGTGGAGTCGTTGCCACACAGGCACTGGGATAATTCANNNAGATCGGAAG |
| 37208_1 | TATGAGTGTGGAGTCGTTGCCAGAGATACAAAGGGCTGACTGTNNNAGATCGGAA |
| 37208_2 | TATGAGTGTGGAGTCGTTGCCAGAGATACAAAGGGCTGACTGTNNNAGATCGGAA |
| 40094_1 | TATGAGTGTGGAGTCGTTGCGGCAAATCCATGTTCTGTGTTCANNNAGATCGGAAG |
| 40094_2 | TATGAGTGTGGAGTCGTTGCGGCAAATCCATGTTCTGTGTTCANNNAGATCGGAAG |
| 45166_1 | TATGAGTGTGGAGTCGTTGCTACCAGTTGGGAACTCCTAGGAANNNAGATCGGAA |
| 45166_2 | TATGAGTGTGGAGTCGTTGCTACCAGTTGGGAACTCCTAGGAANNNAGATCGGAA |
| 35497_1 | TATGAGTGTGGAGTCGTTGCATTAGAGGTATGAGCCACTGTGCNNNAGATCGGAA |
| 35497_2 | TATGAGTGTGGAGTCGTTGCATTAGAGGTATGAGCCACTGTGCNNNAGATCGGAA |
| 49682_1 | TATGAGTGTGGAGTCGTTGCAGACCATTGTTGAAGCTTGCAAANNNAGATCGGAA |
| 49682_2 | TATGAGTGTGGAGTCGTTGCAGACCATTGTTGAAGCTTGCAAANNNAGATCGGAA |
| 34583_1 | TATGAGTGTGGAGTCGTTGCGAGGAGAGGGGAGGGGAAGTNNNAGATCGGAAG |
| 34583_2 | TATGAGTGTGGAGTCGTTGCGAGGAGAGGGGAGGGGAAGTNNNAGATCGGAAG |
| 37045_1 | TATGAGTGTGGAGTCGTTGCGGATAGGGGAGTGAGTTTTGACANNNAGATCGGA |
| 37045_2 | TATGAGTGTGGAGTCGTTGCGGATAGGGGAGTGAGTTTTGACANNNAGATCGGA |
| 28919_1 | TATGAGTGTGGAGTCGTTGCTCAATCAAATGCAGGCCAAACATNNNAGATCGGAAG |
| 28919_2 | TATGAGTGTGGAGTCGTTGCTCAATCAAATGCAGGCCAAACATNNNAGATCGGAAG |
| 29842_1 | TATGAGTGTGGAGTCGTTGCAGAAATCACAGCAAGACTGAGGCNNNAGATCGGAA |
| 29842_2 | TATGAGTGTGGAGTCGTTGCAGAAATCACAGCAAGACTGAGGCNNNAGATCGGAA |
| 36021_1 | TATGAGTGTGGAGTCGTTGCGGAAAATCTGGCGAAATGCAAACNNNAGATCGGAA |
| 36021_2 | TATGAGTGTGGAGTCGTTGCGGAAAATCTGGCGAAATGCAAACNNNAGATCGGAA |
| 39101_1 | TATGAGTGTGGAGTCGTTGCAGATGGAGTCTCACTCTGTCACCNNNAGATCGGAAG |
| 39101_2 | TATGAGTGTGGAGTCGTTGCAGATGGAGTCTCACTCTGTCACCNNNAGATCGGAAG |

| | |
|---|---|
| 41248_1 | TATGAGTGTGGAGTCGTTGCGATGTGTTTTCGCTGGGAAAGTTNNNAGATCGGAA |
| 41248_2 | TATGAGTGTGGAGTCGTTGCGATGTGTTTTCGCTGGGAAAGTTNNNAGATCGGAA |
| 45309_1 | TATGAGTGTGGAGTCGTTGCAGCTTTATGAAGTCACTGGAGCCNNNAGATCGGAA |
| 45309_2 | TATGAGTGTGGAGTCGTTGCAGCTTTATGAAGTCACTGGAGCCNNNAGATCGGAA |
| 45384_1 | TATGAGTGTGGAGTCGTTGCCAGTTCGTCGAAGAGTCCGAAGNNNAGATCGGAAG |
| 45384_2 | TATGAGTGTGGAGTCGTTGCCAGTTCGTCGAAGAGTCCGAAGNNNAGATCGGAAG |
| 46910_1 | TATGAGTGTGGAGTCGTTGCATTCTCCTTCCTCAGCCTCCTNNNAGATCGGAAGAGC |
| 46910_2 | TATGAGTGTGGAGTCGTTGCATTCTCCTTCCTCAGCCTCCTNNNAGATCGGAAGAGC |
| 41763_1 | TATGAGTGTGGAGTCGTTGCTTGCATTTTTTTTGTCTGTGCCCNNNAGATCGGAAGA |
| 41763_2 | TATGAGTGTGGAGTCGTTGCTTGCATTTTTTTTGTCTGTGCCCNNNAGATCGGAAGA |
| 49430_1 | TATGAGTGTGGAGTCGTTGCATCTAGCCAGAGTCCTCTTAGCANNNAGATCGGAAG |
| 49430_2 | TATGAGTGTGGAGTCGTTGCATCTAGCCAGAGTCCTCTTAGCANNNAGATCGGAAG |
| 10840_1 | TATGAGTGTGGAGTCGTTGCGTGGGGTTTTCTAGAGTTTGGCANNNAGATCGGAA |
| 10840_2 | TATGAGTGTGGAGTCGTTGCGTGGGGTTTTCTAGAGTTTGGCANNNAGATCGGAA |
| 29626_1 | TATGAGTGTGGAGTCGTTGCTGGAGTCATGTATTCCAAACAAAGGNNNAGATCGG |
| 29626_2 | TATGAGTGTGGAGTCGTTGCTGGAGTCATGTATTCCAAACAAAGGNNNAGATCGG |
| 32020_1 | TATGAGTGTGGAGTCGTTGCTTCAAGTGATTCTCCTGCCTCAANNNAGATCGGAAG |
| 32020_2 | TATGAGTGTGGAGTCGTTGCTTCAAGTGATTCTCCTGCCTCAANNNAGATCGGAAG |
| 46735_1 | TATGAGTGTGGAGTCGTTGCGTTCCTAAAGGCAACTCAGGGTANNNAGATCGGAA |
| 46735_2 | TATGAGTGTGGAGTCGTTGCGTTCCTAAAGGCAACTCAGGGTANNNAGATCGGAA |
| 10237_1 | TATGAGTGTGGAGTCGTTGCTTCGCATTTGTTTAGTCCTGCCTNNNAGATCGGAAG |
| 10237_2 | TATGAGTGTGGAGTCGTTGCTTCGCATTTGTTTAGTCCTGCCTNNNAGATCGGAAG |
| 39911_1 | TATGAGTGTGGAGTCGTTGCGAAACTGGTGATGAGCAACTTCCNNNAGATCGGAA |
| 39911_2 | TATGAGTGTGGAGTCGTTGCGAAACTGGTGATGAGCAACTTCCNNNAGATCGGAA |
| 13520_1 | TATGAGTGTGGAGTCGTTGCAATAATTTCCCTATTAGCCAGAANNNAGATCGGAAG |
| 13520_2 | TATGAGTGTGGAGTCGTTGCAATAATTTCCCTATTAGCCAGAANNNAGATCGGAAG |
| 47973_1 | TATGAGTGTGGAGTCGTTGCAAGCTAAGATCCACTGGCTTGAANNNAGATCGGAA |
| 47973_2 | TATGAGTGTGGAGTCGTTGCAAGCTAAGATCCACTGGCTTGAANNNAGATCGGAA |
| 34386_1 | TATGAGTGTGGAGTCGTTGCTTTGAGCCAGGATCTCTCTTAGCNNNAGATCGGAAG |
| 34386_2 | TATGAGTGTGGAGTCGTTGCTTTGAGCCAGGATCTCTCTTAGCNNNAGATCGGAAG |
| 53403_1 | TATGAGTGTGGAGTCGTTGCAAGAATCTAGACACACAGGCCTGNNNAGATCGGAA |
| 53403_2 | TATGAGTGTGGAGTCGTTGCAAGAATCTAGACACACAGGCCTGNNNAGATCGGAA |
| 35672_1 | TATGAGTGTGGAGTCGTTGCGATTGTCTTGGTGGCTCTTCTTGNNNAGATCGGAAG |
| 35672_2 | TATGAGTGTGGAGTCGTTGCGATTGTCTTGGTGGCTCTTCTTGNNNAGATCGGAAG |
| 33687_1 | TATGAGTGTGGAGTCGTTGCATCTTTGAAAGAGGAGGCTGAGGNNNAGATCGGA |
| 33687_2 | TATGAGTGTGGAGTCGTTGCATCTTTGAAAGAGGAGGCTGAGGNNNAGATCGGA |
| 13056_1 | TATGAGTGTGGAGTCGTTGCCCTATTCCCTATAAGAAATACGANNNAGATCGGAAG |
| 13056_2 | TATGAGTGTGGAGTCGTTGCCCTATTCCCTATAAGAAATACGANNNAGATCGGAAG |
| 40800_1 | TATGAGTGTGGAGTCGTTGCTAGCCAGGGATAAAGCATGAACTNNNAGATCGGAA |
| 40800_2 | TATGAGTGTGGAGTCGTTGCTAGCCAGGGATAAAGCATGAACTNNNAGATCGGAA |
| 46490_1 | TATGAGTGTGGAGTCGTTGCGGGTTCAAGTGATTCTCCTGTCTNNNAGATCGGAAG |
| 46490_2 | TATGAGTGTGGAGTCGTTGCGGGTTCAAGTGATTCTCCTGTCTNNNAGATCGGAAG |
| 10525_1 | TATGAGTGTGGAGTCGTTGCTCAGTTGCCATGTGCTATGGTTTNNNAGATCGGAAG |

| | |
|---|---|
| 10525_2 | TATGAGTGTGGAGTCGTTGCTCAGTTGCCATGTGCTATGGTTTNNNAGATCGGAAG |
| 35404_1 | TATGAGTGTGGAGTCGTTGCAATACCTCATGAGCTCAAGCGATNNNAGATCGGAAG |
| 35404_2 | TATGAGTGTGGAGTCGTTGCAATACCTCATGAGCTCAAGCGATNNNAGATCGGAAG |
| 31582_1 | TATGAGTGTGGAGTCGTTGCACACAGACACATACACACACAGANNNAGATCGGAAG |
| 31582_2 | TATGAGTGTGGAGTCGTTGCACACAGACACATACACACACAGANNNAGATCGGAAG |
| 42416_1 | TATGAGTGTGGAGTCGTTGCACACACACAGACACAGAAGTGATNNNAGATCGGAA |
| 42416_2 | TATGAGTGTGGAGTCGTTGCACACACACAGACACAGAAGTGATNNNAGATCGGAA |
| 37576_1 | TATGAGTGTGGAGTCGTTGCCCAGTGCAACATGAATTGTGCTANNNAGATCGGAAG |
| 37576_2 | TATGAGTGTGGAGTCGTTGCCCAGTGCAACATGAATTGTGCTANNNAGATCGGAAG |
| 30410_1 | TATGAGTGTGGAGTCGTTGCTGAAAGGGCATGCAATAGACTCCNNNAGATCGGAA |
| 30410_2 | TATGAGTGTGGAGTCGTTGCTGAAAGGGCATGCAATAGACTCCNNNAGATCGGAA |
| 34932_1 | TATGAGTGTGGAGTCGTTGCTTCAAGCAATTTTCCTGCCTCAGNNNAGATCGGAAG |
| 34932_2 | TATGAGTGTGGAGTCGTTGCTTCAAGCAATTTTCCTGCCTCAGNNNAGATCGGAAG |
| 33015_1 | TATGAGTGTGGAGTCGTTGCCCACCACTCCCTGCTAATTTTTGNNNAGATCGGAAGA |
| 33015_2 | TATGAGTGTGGAGTCGTTGCCCACCACTCCCTGCTAATTTTTGNNNAGATCGGAAGA |
| 37904_1 | TATGAGTGTGGAGTCGTTGCTGCCTGTAATTCCAGCTACTCAGNNNAGATCGGAAG |
| 37904_2 | TATGAGTGTGGAGTCGTTGCTGCCTGTAATTCCAGCTACTCAGNNNAGATCGGAAG |
| 13325_1 | TATGAGTGTGGAGTCGTTGCATGTTAGTAATGAGGACAACTAGNNNAGATCGGAA |
| 13325_2 | TATGAGTGTGGAGTCGTTGCATGTTAGTAATGAGGACAACTAGNNNAGATCGGAA |
| 40609_1 | TATGAGTGTGGAGTCGTTGCTTTCCAAGCTGTTTTTTTGGGGTNNNAGATCGGAAG |
| 40609_2 | TATGAGTGTGGAGTCGTTGCTTTCCAAGCTGTTTTTTTGGGGTNNNAGATCGGAAG |
| 49043_1 | TATGAGTGTGGAGTCGTTGCAGTCTCCTGAGTAGCTGGGATTNNNAGATCGGAAG |
| 49043_2 | TATGAGTGTGGAGTCGTTGCAGTCTCCTGAGTAGCTGGGATTNNNAGATCGGAAG |
| 31486_1 | TATGAGTGTGGAGTCGTTGCACATGCTTACGTGAACATTCTGCNNNAGATCGGAAG |
| 31486_2 | TATGAGTGTGGAGTCGTTGCACATGCTTACGTGAACATTCTGCNNNAGATCGGAAG |
| 34487_1 | TATGAGTGTGGAGTCGTTGCCAAAGTACTGGGATGACAGGTGTNNNAGATCGGAA |
| 34487_2 | TATGAGTGTGGAGTCGTTGCCAAAGTACTGGGATGACAGGTGTNNNAGATCGGAA |
| 34565_1 | TATGAGTGTGGAGTCGTTGCCTTCTAGGCCCCGTCTCTCAGNNNAGATCGGAAGAG |
| 34565_2 | TATGAGTGTGGAGTCGTTGCCTTCTAGGCCCCGTCTCTCAGNNNAGATCGGAAGAG |
| 38921_1 | TATGAGTGTGGAGTCGTTGCACATTCTTCTCAGTGCCACATGGNNNAGATCGGAAG |
| 38921_2 | TATGAGTGTGGAGTCGTTGCACATTCTTCTCAGTGCCACATGGNNNAGATCGGAAG |
| 38011_1 | TATGAGTGTGGAGTCGTTGCCGTTTATACACTACCAAACTGGCANNNAGATCGGAA |
| 38011_2 | TATGAGTGTGGAGTCGTTGCCGTTTATACACTACCAAACTGGCANNNAGATCGGAA |
| 35896_1 | TATGAGTGTGGAGTCGTTGCACACAAACTCACCATTCAATCCANNNAGATCGGAAGA |
| 35896_2 | TATGAGTGTGGAGTCGTTGCACACAAACTCACCATTCAATCCANNNAGATCGGAAGA |
| 37023_1 | TATGAGTGTGGAGTCGTTGCAAGGGTGAGGAGAAAACAGAACCNNNAGATCGGA |
| 37023_2 | TATGAGTGTGGAGTCGTTGCAAGGGTGAGGAGAAAACAGAACCNNNAGATCGGA |
| 10561_1 | TATGAGTGTGGAGTCGTTGCAGCGCATTGTTTTGCATCCTTTTNNNAGATCGGAAG |
| 10561_2 | TATGAGTGTGGAGTCGTTGCAGCGCATTGTTTTGCATCCTTTTNNNAGATCGGAAG |
| 32137_1 | TATGAGTGTGGAGTCGTTGCAAATTGGGAACTGACCTTTTCCANNNAGATCGGAAG |
| 32137_2 | TATGAGTGTGGAGTCGTTGCAAATTGGGAACTGACCTTTTCCANNNAGATCGGAAG |
| 41981_1 | TATGAGTGTGGAGTCGTTGCCAGTAGCACTATCTTGGCTCACTNNNAGATCGGAAG |
| 41981_2 | TATGAGTGTGGAGTCGTTGCCAGTAGCACTATCTTGGCTCACTNNNAGATCGGAAG |

| | |
|---|---|
| 40669_1 | TATGAGTGTGGAGTCGTTGCGCAGGACACAAGCTTGAACTTAGNNNAGATCGGAA |
| 40669_2 | TATGAGTGTGGAGTCGTTGCGCAGGACACAAGCTTGAACTTAGNNNAGATCGGAA |
| 51995_1 | TATGAGTGTGGAGTCGTTGCTGGGTACAAATAAGGCCAGAGAGNNNAGATCGGAA |
| 51995_2 | TATGAGTGTGGAGTCGTTGCTGGGTACAAATAAGGCCAGAGAGNNNAGATCGGAA |
| 34486_1 | TATGAGTGTGGAGTCGTTGCTATTTTCCAGGCCTTTTCCCTGTNNNAGATCGGAAGA |
| 34486_2 | TATGAGTGTGGAGTCGTTGCTATTTTCCAGGCCTTTTCCCTGTNNNAGATCGGAAGA |
| 52034_1 | TATGAGTGTGGAGTCGTTGCATTCCAGATGTGATTTGGGTGGANNNAGATCGGAA |
| 52034_2 | TATGAGTGTGGAGTCGTTGCATTCCAGATGTGATTTGGGTGGANNNAGATCGGAA |
| 32248_1 | TATGAGTGTGGAGTCGTTGCCTGTTGCTCAGGCTATAGTCTCANNNAGATCGGAAG |
| 32248_2 | TATGAGTGTGGAGTCGTTGCCTGTTGCTCAGGCTATAGTCTCANNNAGATCGGAAG |
| 42029_1 | TATGAGTGTGGAGTCGTTGCAGATCAAAGCATGCCACATTCAGNNNAGATCGGAAG |
| 42029_2 | TATGAGTGTGGAGTCGTTGCAGATCAAAGCATGCCACATTCAGNNNAGATCGGAAG |
| 38479_1 | TATGAGTGTGGAGTCGTTGCCCACCACACCCAGCCATAAATTANNNAGATCGGAAG |
| 38479_2 | TATGAGTGTGGAGTCGTTGCCCACCACACCCAGCCATAAATTANNNAGATCGGAAG |
| 37893_1 | TATGAGTGTGGAGTCGTTGCCTGTAGTCCCAGCTACTCAGGAGNNNAGATCGGAAG |
| 37893_2 | TATGAGTGTGGAGTCGTTGCCTGTAGTCCCAGCTACTCAGGAGNNNAGATCGGAAG |
| 52730_1 | TATGAGTGTGGAGTCGTTGCTTCTGCCAGGTTTCAAGGTGAGNNNAGATCGGAAG |
| 52730_2 | TATGAGTGTGGAGTCGTTGCTTCTGCCAGGTTTCAAGGTGAGNNNAGATCGGAAG |
| 48621_1 | TATGAGTGTGGAGTCGTTGCGCCTCCTGAATAGCTGGGATTATNNNAGATCGGAAG |
| 48621_2 | TATGAGTGTGGAGTCGTTGCGCCTCCTGAATAGCTGGGATTATNNNAGATCGGAAG |
| 53409_1 | TATGAGTGTGGAGTCGTTGCCATGGTGGTTCATGCCTGTATTCNNNAGATCGGAAG |
| 53409_2 | TATGAGTGTGGAGTCGTTGCCATGGTGGTTCATGCCTGTATTCNNNAGATCGGAAG |
| 46909_1 | TATGAGTGTGGAGTCGTTGCGGAATTCCAGAGCGATTGAGTTGNNNAGATCGGAA |
| 46909_2 | TATGAGTGTGGAGTCGTTGCGGAATTCCAGAGCGATTGAGTTGNNNAGATCGGAA |
| 32520_1 | TATGAGTGTGGAGTCGTTGCACAGCACATGACCTGTTTAGAGANNNAGATCGGAAG |
| 32520_2 | TATGAGTGTGGAGTCGTTGCACAGCACATGACCTGTTTAGAGANNNAGATCGGAAG |
| 37752_1 | TATGAGTGTGGAGTCGTTGCCATTCCAGGCAATGTGGTAGAACNNNAGATCGGAAG |
| 37752_2 | TATGAGTGTGGAGTCGTTGCCATTCCAGGCAATGTGGTAGAACNNNAGATCGGAAG |
| 31724_1 | TATGAGTGTGGAGTCGTTGCCGTCTTGCTCCTCGATGTGAGNNNAGATCGGAAGAG |
| 31724_2 | TATGAGTGTGGAGTCGTTGCCGTCTTGCTCCTCGATGTGAGNNNAGATCGGAAGAG |
| 53332_1 | TATGAGTGTGGAGTCGTTGCTTTGGATGTGGTTTATGCCCATGNNNAGATCGGAAG |
| 53332_2 | TATGAGTGTGGAGTCGTTGCTTTGGATGTGGTTTATGCCCATGNNNAGATCGGAAG |
| 52442_1 | TATGAGTGTGGAGTCGTTGCAAGAATGAAGCCAGCTTTTGCTTNNNAGATCGGAAG |
| 52442_2 | TATGAGTGTGGAGTCGTTGCAAGAATGAAGCCAGCTTTTGCTTNNNAGATCGGAAG |
| 29258_1 | TATGAGTGTGGAGTCGTTGCAGACTACTGTGCTCCAAGACATGNNNAGATCGGAAG |
| 29258_2 | TATGAGTGTGGAGTCGTTGCAGACTACTGTGCTCCAAGACATGNNNAGATCGGAAG |
| 31125_1 | TATGAGTGTGGAGTCGTTGCACACACACACACACCTCTTCTAANNNAGATCGGAAGA |
| 31125_2 | TATGAGTGTGGAGTCGTTGCACACACACACACACCTCTTCTAANNNAGATCGGAAGA |
| 35043_1 | TATGAGTGTGGAGTCGTTGCCCACGCCCAGCTAATTTTTGTATNNNAGATCGGAAG |
| 35043_2 | TATGAGTGTGGAGTCGTTGCCCACGCCCAGCTAATTTTTGTATNNNAGATCGGAAG |
| 50511_1 | TATGAGTGTGGAGTCGTTGCGAGAAGAGAGCGTATACTGACCGNNNAGATCGGAA |
| 50511_2 | TATGAGTGTGGAGTCGTTGCGAGAAGAGAGCGTATACTGACCGNNNAGATCGGAA |
| 46927_1 | TATGAGTGTGGAGTCGTTGCGTTTCAAGCAATTCTCCCACCTCNNNAGATCGGAAGA |

| | |
|---|---|
| 46927_2 | TATGAGTGTGGAGTCGTTGCGTTTCAAGCAATTCTCCCACCTCNNNAGATCGGAAGA |
| 4824_1 | TATGAGTGTGGAGTCGTTGCGAGCAACAGAAAGAACACAATTGCNNNAGATCGGAA |
| 4824_2 | TATGAGTGTGGAGTCGTTGCGAGCAACAGAAAGAACACAATTGCNNNAGATCGGAA |
| 44624_1 | TATGAGTGTGGAGTCGTTGCGTCTCCTGGGTTCAAGTGATTCTNNNAGATCGGAAG |
| 44624_2 | TATGAGTGTGGAGTCGTTGCGTCTCCTGGGTTCAAGTGATTCTNNNAGATCGGAAG |
| 13628_1 | TATGAGTGTGGAGTCGTTGCCATGAGTTCATTCAAGTAAAGTANNNAGATCGGAAG |
| 13628_2 | TATGAGTGTGGAGTCGTTGCCATGAGTTCATTCAAGTAAAGTANNNAGATCGGAAG |
| 44989_1 | TATGAGTGTGGAGTCGTTGCTCTGTCCGCCAAGTATTCATTGANNNAGATCGGAAGA |
| 44989_2 | TATGAGTGTGGAGTCGTTGCTCTGTCCGCCAAGTATTCATTGANNNAGATCGGAAGA |
| 5751_1 | TATGAGTGTGGAGTCGTTGCAGAGGAGAGAGAGACTTTGGTGCNNNAGATCGGAA |
| 5751_2 | TATGAGTGTGGAGTCGTTGCAGAGGAGAGAGAGACTTTGGTGCNNNAGATCGGAA |
| 12850_1 | TATGAGTGTGGAGTCGTTGCGAGCTTAATAAACATGTAGCAAANNNAGATCGGAAG |
| 12850_2 | TATGAGTGTGGAGTCGTTGCGAGCTTAATAAACATGTAGCAAANNNAGATCGGAAG |
| 50054_1 | TATGAGTGTGGAGTCGTTGCCACCCAGAGGAACCCTAACAAATNNNAGATCGGAAG |
| 50054_2 | TATGAGTGTGGAGTCGTTGCCACCCAGAGGAACCCTAACAAATNNNAGATCGGAAG |
| 29700_1 | TATGAGTGTGGAGTCGTTGCGCCCAGACAGAAACGTTTACCTANNNAGATCGGAAG |
| 29700_2 | TATGAGTGTGGAGTCGTTGCGCCCAGACAGAAACGTTTACCTANNNAGATCGGAAG |
| 34739_1 | TATGAGTGTGGAGTCGTTGCAGACGGAGTCTCTCTCTGTCCNNNAGATCGGAAGAG |
| 34739_2 | TATGAGTGTGGAGTCGTTGCAGACGGAGTCTCTCTCTGTCCNNNAGATCGGAAGAG |
| 51209_1 | TATGAGTGTGGAGTCGTTGCCTACCGCAAAGGGATTTAGGACTNNNAGATCGGAAG |
| 51209_2 | TATGAGTGTGGAGTCGTTGCCTACCGCAAAGGGATTTAGGACTNNNAGATCGGAAG |
| 34949_1 | TATGAGTGTGGAGTCGTTGCGCAAGTAGCTTTGGTTTTTGACTNNNAGATCGGAAG |
| 34949_2 | TATGAGTGTGGAGTCGTTGCGCAAGTAGCTTTGGTTTTTGACTNNNAGATCGGAAG |
| 49463_1 | TATGAGTGTGGAGTCGTTGCTTAGCCATTTCTCTACTGTTGCANNNAGATCGGAAGA |
| 49463_2 | TATGAGTGTGGAGTCGTTGCTTAGCCATTTCTCTACTGTTGCANNNAGATCGGAAGA |
| 2474_1 | TATGAGTGTGGAGTCGTTGCCACTCCGTCACTCTGGTTGAAGNNNAGATCGGAAGA |
| 2474_2 | TATGAGTGTGGAGTCGTTGCCACTCCGTCACTCTGGTTGAAGNNNAGATCGGAAGA |
| 52061_1 | TATGAGTGTGGAGTCGTTGCGTACTCTCTCCGCAGTGAACTGNNNAGATCGGAAGA |
| 52061_2 | TATGAGTGTGGAGTCGTTGCGTACTCTCTCCGCAGTGAACTGNNNAGATCGGAAGA |
| 49629_1 | TATGAGTGTGGAGTCGTTGCTCCCTTTCCTGATCCTTTGACTGNNNAGATCGGAAGA |
| 49629_2 | TATGAGTGTGGAGTCGTTGCTCCCTTTCCTGATCCTTTGACTGNNNAGATCGGAAGA |
| 50571_1 | TATGAGTGTGGAGTCGTTGCGGCCTCCTCCCTCAACACAGNNNAGATCGGAAGAGCA |
| 50571_2 | TATGAGTGTGGAGTCGTTGCGGCCTCCTCCCTCAACACAGNNNAGATCGGAAGAGCA |
| 36842_1 | TATGAGTGTGGAGTCGTTGCTCAACATAAAAAGCCGTTTTGCCNNNAGATCGGAAGA |
| 36842_2 | TATGAGTGTGGAGTCGTTGCTCAACATAAAAAGCCGTTTTGCCNNNAGATCGGAAGA |
| 29541_1 | TATGAGTGTGGAGTCGTTGCCTTCTGCTCACCTCTCTCCTGNNNAGATCGGAAGAGCA |
| 29541_2 | TATGAGTGTGGAGTCGTTGCCTTCTGCTCACCTCTCTCCTGNNNAGATCGGAAGAGCA |
| 30776_1 | TATGAGTGTGGAGTCGTTGCTCCAGTTTCATGTCTTCTCTGGGNNNAGATCGGAAGA |
| 30776_2 | TATGAGTGTGGAGTCGTTGCTCCAGTTTCATGTCTTCTCTGGGNNNAGATCGGAAGA |
| 33111_1 | TATGAGTGTGGAGTCGTTGCCATCCTGGCTAACAAGGTGAAACNNNAGATCGGAAG |
| 33111_2 | TATGAGTGTGGAGTCGTTGCCATCCTGGCTAACAAGGTGAAACNNNAGATCGGAAG |
| 49050_1 | TATGAGTGTGGAGTCGTTGCGGTTCAAACAATTCTCCTGCCTCNNNAGATCGGAAGA |
| 49050_2 | TATGAGTGTGGAGTCGTTGCGGTTCAAACAATTCTCCTGCCTCNNNAGATCGGAAGA |

| | |
|---|---|
| 10697_1 | TATGAGTGTGGAGTCGTTGCCGATTGGTCAGTGAGTTCCTGTTNNNAGATCGGAAG |
| 10697_2 | TATGAGTGTGGAGTCGTTGCCGATTGGTCAGTGAGTTCCTGTTNNNAGATCGGAAG |
| 41959_1 | TATGAGTGTGGAGTCGTTGCGTTGGTCAGGATGGTCTTGATCTNNNAGATCGGAA |
| 41959_2 | TATGAGTGTGGAGTCGTTGCGTTGGTCAGGATGGTCTTGATCTNNNAGATCGGAA |
| 38462_1 | TATGAGTGTGGAGTCGTTGCTGGAAAGTAGCAGCAGAACTAGGNNNAGATCGGAA |
| 38462_2 | TATGAGTGTGGAGTCGTTGCTGGAAAGTAGCAGCAGAACTAGGNNNAGATCGGAA |
| 51624_1 | TATGAGTGTGGAGTCGTTGCGCCCCCATCCTACAATATCACTTNNNAGATCGGAAGA |
| 51624_2 | TATGAGTGTGGAGTCGTTGCGCCCCCATCCTACAATATCACTTNNNAGATCGGAAGA |
| 36750_1 | TATGAGTGTGGAGTCGTTGCTCAGTGTCTGTGTAATGAGCAGTNNNAGATCGGAA |
| 36750_2 | TATGAGTGTGGAGTCGTTGCTCAGTGTCTGTGTAATGAGCAGTNNNAGATCGGAA |
| 37240_1 | TATGAGTGTGGAGTCGTTGCAGATATGTGCACTCCCATGTTCANNNAGATCGGAAG |
| 37240_2 | TATGAGTGTGGAGTCGTTGCAGATATGTGCACTCCCATGTTCANNNAGATCGGAAG |
| 10719_1 | TATGAGTGTGGAGTCGTTGCCAAGCCCTTTCTGGGAGTTTTCTNNNAGATCGGAAG |
| 10719_2 | TATGAGTGTGGAGTCGTTGCCAAGCCCTTTCTGGGAGTTTTCTNNNAGATCGGAAG |
| 48194_1 | TATGAGTGTGGAGTCGTTGCCTCAAGTGATCCTCCTACCTCAGNNNAGATCGGAAG |
| 48194_2 | TATGAGTGTGGAGTCGTTGCCTCAAGTGATCCTCCTACCTCAGNNNAGATCGGAAG |
| 12854_1 | TATGAGTGTGGAGTCGTTGCATAGAGTGTAGATGCATCAAAATNNNAGATCGGAA |
| 12854_2 | TATGAGTGTGGAGTCGTTGCATAGAGTGTAGATGCATCAAAATNNNAGATCGGAA |
| 32718_1 | TATGAGTGTGGAGTCGTTGCACCAAATTGCAGCTGTACCTCTANNNAGATCGGAAG |
| 32718_2 | TATGAGTGTGGAGTCGTTGCACCAAATTGCAGCTGTACCTCTANNNAGATCGGAAG |
| 32677_1 | TATGAGTGTGGAGTCGTTGCGATGGAGTCTCACTCTGTCGCNNNAGATCGGAAGA |
| 32677_2 | TATGAGTGTGGAGTCGTTGCGATGGAGTCTCACTCTGTCGCNNNAGATCGGAAGA |
| 42923_1 | TATGAGTGTGGAGTCGTTGCAGTTCTATGCCTGTAGCCTATGCNNNAGATCGGAAG |
| 42923_2 | TATGAGTGTGGAGTCGTTGCAGTTCTATGCCTGTAGCCTATGCNNNAGATCGGAAG |
| 2263_1 | TATGAGTGTGGAGTCGTTGCGATGGAGCCTGAAACTGCTTGTANNNAGATCGGAA |
| 2263_2 | TATGAGTGTGGAGTCGTTGCGATGGAGCCTGAAACTGCTTGTANNNAGATCGGAA |
| 51216_1 | TATGAGTGTGGAGTCGTTGCGGTAGGGAGACCCAAGCAGGNNNAGATCGGAAGA |
| 51216_2 | TATGAGTGTGGAGTCGTTGCGGTAGGGAGACCCAAGCAGGNNNAGATCGGAAGA |
| 33267_1 | TATGAGTGTGGAGTCGTTGCGCTAGTAGCTGGGATTACAGGTGNNNAGATCGGAA |
| 33267_2 | TATGAGTGTGGAGTCGTTGCGCTAGTAGCTGGGATTACAGGTGNNNAGATCGGAA |
| 39892_1 | TATGAGTGTGGAGTCGTTGCTGGCTCATGACTATAATCCCAGCNNNAGATCGGAAG |
| 39892_2 | TATGAGTGTGGAGTCGTTGCTGGCTCATGACTATAATCCCAGCNNNAGATCGGAAG |
| 52825_1 | TATGAGTGTGGAGTCGTTGCCTTCCAGCCCCTCTGAGAGCNNNAGATCGGAAGAGC |
| 52825_2 | TATGAGTGTGGAGTCGTTGCCTTCCAGCCCCTCTGAGAGCNNNAGATCGGAAGAGC |
| 48750_1 | TATGAGTGTGGAGTCGTTGCTTCTTTTGCCTCAGGACAGTGATNNNAGATCGGAAG |
| 48750_2 | TATGAGTGTGGAGTCGTTGCTTCTTTTGCCTCAGGACAGTGATNNNAGATCGGAAG |
| 32559_1 | TATGAGTGTGGAGTCGTTGCCTTCCAGGTTCACGCCATTCTCNNNAGATCGGAAGA |
| 32559_2 | TATGAGTGTGGAGTCGTTGCCTTCCAGGTTCACGCCATTCTCNNNAGATCGGAAGA |
| 44713_1 | TATGAGTGTGGAGTCGTTGCGTAGACTGACACAAGAGTTCCCCNNNAGATCGGAAG |
| 44713_2 | TATGAGTGTGGAGTCGTTGCGTAGACTGACACAAGAGTTCCCCNNNAGATCGGAAG |
| 36502_1 | TATGAGTGTGGAGTCGTTGCAACATGGCGAAATCCCATCTCTANNNAGATCGGAAG |
| 36502_2 | TATGAGTGTGGAGTCGTTGCAACATGGCGAAATCCCATCTCTANNNAGATCGGAAG |
| 38933_1 | TATGAGTGTGGAGTCGTTGCGGGTGAGGGAGATAAGGAGGTATNNNAGATCGGA |

| | |
|---|---|
| 38933_2 | TATGAGTGTGGAGTCGTTGCGGGTGAGGGAGATAAGGAGGTATNNNAGATCGGA |
| 49399_1 | TATGAGTGTGGAGTCGTTGCATGCTGGGATTACAGGTGTAAGCNNNAGATCGGAA |
| 49399_2 | TATGAGTGTGGAGTCGTTGCATGCTGGGATTACAGGTGTAAGCNNNAGATCGGAA |
| 35151_1 | TATGAGTGTGGAGTCGTTGCCATCTTTGTGTGGCATTGTGTGTNNNAGATCGGAAG |
| 35151_2 | TATGAGTGTGGAGTCGTTGCCATCTTTGTGTGGCATTGTGTGTNNNAGATCGGAAG |
| 45157_1 | TATGAGTGTGGAGTCGTTGCCCCTCACAGAACCACCATAGTTCNNNAGATCGGAAG |
| 45157_2 | TATGAGTGTGGAGTCGTTGCCCCTCACAGAACCACCATAGTTCNNNAGATCGGAAG |
| 35313_1 | TATGAGTGTGGAGTCGTTGCTGCCCAGTGAAATGAATATCAGCNNNAGATCGGAAG |
| 35313_2 | TATGAGTGTGGAGTCGTTGCTGCCCAGTGAAATGAATATCAGCNNNAGATCGGAAG |
| 48128_1 | TATGAGTGTGGAGTCGTTGCTCTTCTTCTCGTAAAGCCTACGANNNAGATCGGAAG |
| 48128_2 | TATGAGTGTGGAGTCGTTGCTCTTCTTCTCGTAAAGCCTACGANNNAGATCGGAAG |
| 12887_1 | TATGAGTGTGGAGTCGTTGCAGTGTTCCTTCATTAATTTAAGCNNNAGATCGGAAG |
| 12887_2 | TATGAGTGTGGAGTCGTTGCAGTGTTCCTTCATTAATTTAAGCNNNAGATCGGAAG |
| 48768_1 | TATGAGTGTGGAGTCGTTGCTGAGATGTTGGTCAAAGGGTACANNNAGATCGGAA |
| 48768_2 | TATGAGTGTGGAGTCGTTGCTGAGATGTTGGTCAAAGGGTACANNNAGATCGGAA |
| 10603_1 | TATGAGTGTGGAGTCGTTGCAGGTCCAGGAAATACTCCAGCTTNNNAGATCGGAAG |
| 10603_2 | TATGAGTGTGGAGTCGTTGCAGGTCCAGGAAATACTCCAGCTTNNNAGATCGGAAG |
| 35877_1 | TATGAGTGTGGAGTCGTTGCCTCATCACAAGCAGCAGTTAGATNNNAGATCGGAAG |
| 35877_2 | TATGAGTGTGGAGTCGTTGCCTCATCACAAGCAGCAGTTAGATNNNAGATCGGAAG |
| 44312_1 | TATGAGTGTGGAGTCGTTGCAATAAAATACCTCAGGCGGGTGGNNNAGATCGGAA |
| 44312_2 | TATGAGTGTGGAGTCGTTGCAATAAAATACCTCAGGCGGGTGGNNNAGATCGGAA |
| 35925_1 | TATGAGTGTGGAGTCGTTGCTCCAGTTCTGTCTACTCCAATTCANNNAGATCGGAAG |
| 35925_2 | TATGAGTGTGGAGTCGTTGCTCCAGTTCTGTCTACTCCAATTCANNNAGATCGGAAG |
| 47895_1 | TATGAGTGTGGAGTCGTTGCCCGCTGTTCTTTCTAGAAGCTTTNNNAGATCGGAAG |
| 47895_2 | TATGAGTGTGGAGTCGTTGCCCGCTGTTCTTTCTAGAAGCTTTNNNAGATCGGAAG |
| 2178_1 | TATGAGTGTGGAGTCGTTGCGACACATTTCTGCAGTGGGTAGANNNAGATCGGAA |
| 2178_2 | TATGAGTGTGGAGTCGTTGCGACACATTTCTGCAGTGGGTAGANNNAGATCGGAA |
| 46358_1 | TATGAGTGTGGAGTCGTTGCCTGCTCTCAGGGATGTGGAATTTNNNAGATCGGAAG |
| 46358_2 | TATGAGTGTGGAGTCGTTGCCTGCTCTCAGGGATGTGGAATTTNNNAGATCGGAAG |
| 51400_1 | TATGAGTGTGGAGTCGTTGCTGTATTGTGCAGGTACTCAGGAGNNNAGATCGGAA |
| 51400_2 | TATGAGTGTGGAGTCGTTGCTGTATTGTGCAGGTACTCAGGAGNNNAGATCGGAA |
| 30326_1 | TATGAGTGTGGAGTCGTTGCTCTTTGGTGAGAAGAGCAAAGCCNNNAGATCGGAA |
| 30326_2 | TATGAGTGTGGAGTCGTTGCTCTTTGGTGAGAAGAGCAAAGCCNNNAGATCGGAA |
| 46515_1 | TATGAGTGTGGAGTCGTTGCCAGGTTCAAGCAATTCTCCTGCNNNAGATCGGAAGA |
| 46515_2 | TATGAGTGTGGAGTCGTTGCCAGGTTCAAGCAATTCTCCTGCNNNAGATCGGAAGA |
| 13722_1 | TATGAGTGTGGAGTCGTTGCAAGATAGTCTCCACTTTTTAATGNNNAGATCGGAAG |
| 13722_2 | TATGAGTGTGGAGTCGTTGCAAGATAGTCTCCACTTTTTAATGNNNAGATCGGAAG |
| 43371_1 | TATGAGTGTGGAGTCGTTGCTTCGAACGATTCTCCTGACTCAGNNNAGATCGGAAG |
| 43371_2 | TATGAGTGTGGAGTCGTTGCTTCGAACGATTCTCCTGACTCAGNNNAGATCGGAAG |
| 38392_1 | TATGAGTGTGGAGTCGTTGCTGAGGCAGAAGAATTGCTTGAACNNNAGATCGGAA |
| 38392_2 | TATGAGTGTGGAGTCGTTGCTGAGGCAGAAGAATTGCTTGAACNNNAGATCGGAA |
| 34312_1 | TATGAGTGTGGAGTCGTTGCGTGTCCCTTGCTGACTCATGATANNNAGATCGGAAG |
| 34312_2 | TATGAGTGTGGAGTCGTTGCGTGTCCCTTGCTGACTCATGATANNNAGATCGGAAG |

| | |
|---|---|
| 53356_1 | TATGAGTGTGGAGTCGTTGCCTCTCTCCCTGGGACATAAATGGNNNAGATCGGAAG |
| 53356_2 | TATGAGTGTGGAGTCGTTGCCTCTCTCCCTGGGACATAAATGGNNNAGATCGGAAG |
| 40407_1 | TATGAGTGTGGAGTCGTTGCTCTAGCCAATCCAGCATTCTGTTNNNAGATCGGAAGA |
| 40407_2 | TATGAGTGTGGAGTCGTTGCTCTAGCCAATCCAGCATTCTGTTNNNAGATCGGAAGA |
| 47153_1 | TATGAGTGTGGAGTCGTTGCCACCATGCCCAGCTAATTTTTGTNNNAGATCGGAAGA |
| 47153_2 | TATGAGTGTGGAGTCGTTGCCACCATGCCCAGCTAATTTTTGTNNNAGATCGGAAGA |
| 44839_1 | TATGAGTGTGGAGTCGTTGCTGCTCTCACTCTCACTTCCTTTCNNNAGATCGGAAGAG |
| 44839_2 | TATGAGTGTGGAGTCGTTGCTGCTCTCACTCTCACTTCCTTTCNNNAGATCGGAAGAG |
| 49441_1 | TATGAGTGTGGAGTCGTTGCACTCACTAGAATCTTTGGGTCCCNNNAGATCGGAAGA |
| 49441_2 | TATGAGTGTGGAGTCGTTGCACTCACTAGAATCTTTGGGTCCCNNNAGATCGGAAGA |
| 49116_1 | TATGAGTGTGGAGTCGTTGCCCCACTGCTTCATTTTGACTAGCNNNAGATCGGAAGA |
| 49116_2 | TATGAGTGTGGAGTCGTTGCCCCACTGCTTCATTTTGACTAGCNNNAGATCGGAAGA |
| 37392_1 | TATGAGTGTGGAGTCGTTGCGGGAGAACAGAGAAACAAAGAGGNNNAGATCGGA |
| 37392_2 | TATGAGTGTGGAGTCGTTGCGGGAGAACAGAGAAACAAAGAGGNNNAGATCGGA |
| 50285_1 | TATGAGTGTGGAGTCGTTGCGATTCCAGAACCTTCCTCCCAAANNNAGATCGGAAGA |
| 50285_2 | TATGAGTGTGGAGTCGTTGCGATTCCAGAACCTTCCTCCCAAANNNAGATCGGAAGA |
| 45866_1 | TATGAGTGTGGAGTCGTTGCCAACTCACGGGCTCTGCTTCNNNAGATCGGAAGAGCA |
| 45866_2 | TATGAGTGTGGAGTCGTTGCCAACTCACGGGCTCTGCTTCNNNAGATCGGAAGAGCA |
| 49937_1 | TATGAGTGTGGAGTCGTTGCGAGAAAGGAGAAAGGAGGGAAGGNNNAGATCGGA |
| 49937_2 | TATGAGTGTGGAGTCGTTGCGAGAAAGGAGAAAGGAGGGAAGGNNNAGATCGGA |
| 40681_1 | TATGAGTGTGGAGTCGTTGCTCTTCAGTTAGTCAAACAGGCCANNNAGATCGGAAG |
| 40681_2 | TATGAGTGTGGAGTCGTTGCTCTTCAGTTAGTCAAACAGGCCANNNAGATCGGAAG |
| 48704_1 | TATGAGTGTGGAGTCGTTGCCAGCCACATTGATCACTCTGTTGNNNAGATCGGAAGA |
| 48704_2 | TATGAGTGTGGAGTCGTTGCCAGCCACATTGATCACTCTGTTGNNNAGATCGGAAGA |
| 49519_1 | TATGAGTGTGGAGTCGTTGCCTGTTCTGAAAGTTGCATGAGGGNNNAGATCGGAAG |
| 49519_2 | TATGAGTGTGGAGTCGTTGCCTGTTCTGAAAGTTGCATGAGGGNNNAGATCGGAAG |
| 29199_1 | TATGAGTGTGGAGTCGTTGCTATTTGCACCCCTTCTTGTCTGTNNNAGATCGGAAGA |
| 29199_2 | TATGAGTGTGGAGTCGTTGCTATTTGCACCCCTTCTTGTCTGTNNNAGATCGGAAGA |
| 51689_1 | TATGAGTGTGGAGTCGTTGCCCCGGCCTGGTAGAAAAAATTTTNNNAGATCGGAAG |
| 51689_2 | TATGAGTGTGGAGTCGTTGCCCCGGCCTGGTAGAAAAAATTTTNNNAGATCGGAAG |
| 50828_1 | TATGAGTGTGGAGTCGTTGCAAAATGGCAAATTACCGAGGAGCNNNAGATCGGAAG |
| 50828_2 | TATGAGTGTGGAGTCGTTGCAAAATGGCAAATTACCGAGGAGCNNNAGATCGGAAG |
| 35027_1 | TATGAGTGTGGAGTCGTTGCCTCAAACTCCTGACCTCGTGATCNNNAGATCGGAAGA |
| 35027_2 | TATGAGTGTGGAGTCGTTGCCTCAAACTCCTGACCTCGTGATCNNNAGATCGGAAGA |
| 10794_1 | TATGAGTGTGGAGTCGTTGCCTCTCTCCTCCTACTGTGTGAGCNNNAGATCGGAAGA |
| 10794_2 | TATGAGTGTGGAGTCGTTGCCTCTCTCCTCCTACTGTGTGAGCNNNAGATCGGAAGA |
| 40017_1 | TATGAGTGTGGAGTCGTTGCACAGTGGTAGAACAGACATTGGGNNNAGATCGGAA |
| 40017_2 | TATGAGTGTGGAGTCGTTGCACAGTGGTAGAACAGACATTGGGNNNAGATCGGAA |
| 39434_1 | TATGAGTGTGGAGTCGTTGCGATGTTCTGATCCCACATCAGGANNNAGATCGGAAG |
| 39434_2 | TATGAGTGTGGAGTCGTTGCGATGTTCTGATCCCACATCAGGANNNAGATCGGAAG |
| 34226_1 | TATGAGTGTGGAGTCGTTGCAAAAGAGCCTGAGTAGCAAAAGCNNNAGATCGGAA |
| 34226_2 | TATGAGTGTGGAGTCGTTGCAAAAGAGCCTGAGTAGCAAAAGCNNNAGATCGGAA |
| 49714_1 | TATGAGTGTGGAGTCGTTGCCCCAAATGCTTTCAGGCTTTACTNNNAGATCGGAAGA |

| | |
|---|---|
| 49714_2 | TATGAGTGTGGAGTCGTTGCCCCAAATGCTTTCAGGCTTTACTNNNAGATCGGAAG |
| 39086_1 | TATGAGTGTGGAGTCGTTGCTGGTGTGCAATATGGTTTTTCTGTNNNAGATCGGAA |
| 39086_2 | TATGAGTGTGGAGTCGTTGCTGGTGTGCAATATGGTTTTTCTGTNNNAGATCGGAA |
| 33550_1 | TATGAGTGTGGAGTCGTTGCCTGGGTTCAATTGATTCTCCTGCNNNAGATCGGAAG |
| 33550_2 | TATGAGTGTGGAGTCGTTGCCTGGGTTCAATTGATTCTCCTGCNNNAGATCGGAAG |
| 12681_1 | TATGAGTGTGGAGTCGTTGCTATGATTTGTTTGTTTGTTTGTGNNNAGATCGGAAG |
| 12681_2 | TATGAGTGTGGAGTCGTTGCTATGATTTGTTTGTTTGTTTGTGNNNAGATCGGAAG |
| 46184_1 | TATGAGTGTGGAGTCGTTGCTACGAACTTTCACTGAACAGCACNNNAGATCGGAAG |
| 46184_2 | TATGAGTGTGGAGTCGTTGCTACGAACTTTCACTGAACAGCACNNNAGATCGGAAG |
| 4247_1 | TATGAGTGTGGAGTCGTTGCGGTGAACATCCACAGGGAAACAGNNNAGATCGGAA |
| 4247_2 | TATGAGTGTGGAGTCGTTGCGGTGAACATCCACAGGGAAACAGNNNAGATCGGAA |
| 39172_1 | TATGAGTGTGGAGTCGTTGCGCCCAGGACTAGAAGGATTCAAANNNAGATCGGAA |
| 39172_2 | TATGAGTGTGGAGTCGTTGCGCCCAGGACTAGAAGGATTCAAANNNAGATCGGAA |
| 52939_1 | TATGAGTGTGGAGTCGTTGCCTCCAGGTGCGTATTCTCTTTCTNNNAGATCGGAAG |
| 52939_2 | TATGAGTGTGGAGTCGTTGCCTCCAGGTGCGTATTCTCTTTCTNNNAGATCGGAAG |
| 28998_1 | TATGAGTGTGGAGTCGTTGCTGTTTTTTGGTTTGCGGTTACCANNNAGATCGGAAG |
| 28998_2 | TATGAGTGTGGAGTCGTTGCTGTTTTTTGGTTTGCGGTTACCANNNAGATCGGAAG |
| 42576_1 | TATGAGTGTGGAGTCGTTGCGGAAGCTTAGGAACTCTCATCCCNNNAGATCGGAAG |
| 42576_2 | TATGAGTGTGGAGTCGTTGCGGAAGCTTAGGAACTCTCATCCCNNNAGATCGGAAG |
| 51512_1 | TATGAGTGTGGAGTCGTTGCTTTCCAGAAAACCAGCTCCTGATNNNAGATCGGAAG |
| 51512_2 | TATGAGTGTGGAGTCGTTGCTTTCCAGAAAACCAGCTCCTGATNNNAGATCGGAAG |
| 52815_1 | TATGAGTGTGGAGTCGTTGCAAGTTGAAGTTGGGGGAGATGTTNNNAGATCGGA |
| 52815_2 | TATGAGTGTGGAGTCGTTGCAAGTTGAAGTTGGGGGAGATGTTNNNAGATCGGA |
| 29618_1 | TATGAGTGTGGAGTCGTTGCGGCTGGACCAGTGTGTATCTGNNNAGATCGGAAGA |
| 29618_2 | TATGAGTGTGGAGTCGTTGCGGCTGGACCAGTGTGTATCTGNNNAGATCGGAAGA |
| 51756_1 | TATGAGTGTGGAGTCGTTGCAAGTCCTGAGCAAACTGGGATAGNNNAGATCGGAA |
| 51756_2 | TATGAGTGTGGAGTCGTTGCAAGTCCTGAGCAAACTGGGATAGNNNAGATCGGAA |
| 39296_1 | TATGAGTGTGGAGTCGTTGCTCTGGGATCTCTATGTTGTTCCANNNAGATCGGAAG |
| 39296_2 | TATGAGTGTGGAGTCGTTGCTCTGGGATCTCTATGTTGTTCCANNNAGATCGGAAG |
| 39429_1 | TATGAGTGTGGAGTCGTTGCCCACCATCTCAGTAATATGGGCANNNAGATCGGAAG |
| 39429_2 | TATGAGTGTGGAGTCGTTGCCCACCATCTCAGTAATATGGGCANNNAGATCGGAAG |
| 42583_1 | TATGAGTGTGGAGTCGTTGCATAGAGAGAGATAGGGTCTCGCTNNNAGATCGGAA |
| 42583_2 | TATGAGTGTGGAGTCGTTGCATAGAGAGAGATAGGGTCTCGCTNNNAGATCGGAA |
| 53222_1 | TATGAGTGTGGAGTCGTTGCTTTGGCCTCAAACTTCAGGTGANNNAGATCGGAAGA |
| 53222_2 | TATGAGTGTGGAGTCGTTGCTTTGGCCTCAAACTTCAGGTGANNNAGATCGGAAGA |
| 48707_1 | TATGAGTGTGGAGTCGTTGCTTCAAGTGATTCTCCTGCCTCAGNNNAGATCGGAAG |
| 48707_2 | TATGAGTGTGGAGTCGTTGCTTCAAGTGATTCTCCTGCCTCAGNNNAGATCGGAAG |
| 30375_1 | TATGAGTGTGGAGTCGTTGCAATGAACTAGCTAGCAGACCCCTNNNAGATCGGAAG |
| 30375_2 | TATGAGTGTGGAGTCGTTGCAATGAACTAGCTAGCAGACCCCTNNNAGATCGGAAG |
| 44720_1 | TATGAGTGTGGAGTCGTTGCCTAGGTCTCCTGTTTGCCATGTNNNAGATCGGAAGA |
| 44720_2 | TATGAGTGTGGAGTCGTTGCCTAGGTCTCCTGTTTGCCATGTNNNAGATCGGAAGA |
| 30795_1 | TATGAGTGTGGAGTCGTTGCCAAAGGCAGGTGAATGTTACAGGNNNAGATCGGAA |
| 30795_2 | TATGAGTGTGGAGTCGTTGCCAAAGGCAGGTGAATGTTACAGGNNNAGATCGGAA |

| | |
|---|---|
| 43718_1 | TATGAGTGTGGAGTCGTTGCACAGAAGGTGCATCATACCCATTNNNAGATCGGAAG |
| 43718_2 | TATGAGTGTGGAGTCGTTGCACAGAAGGTGCATCATACCCATTNNNAGATCGGAAG |
| 37919_1 | TATGAGTGTGGAGTCGTTGCGACAGATGGGGACATGAGACAGNNNAGATCGGAA |
| 37919_2 | TATGAGTGTGGAGTCGTTGCGACAGATGGGGACATGAGACAGNNNAGATCGGAA |
| 45614_1 | TATGAGTGTGGAGTCGTTGCCTCCCAGGTTCAAGTGATACTCCNNNAGATCGGAAG |
| 45614_2 | TATGAGTGTGGAGTCGTTGCCTCCCAGGTTCAAGTGATACTCCNNNAGATCGGAAG |
| 51340_1 | TATGAGTGTGGAGTCGTTGCCTTCTGGAATCAAGAGAGGCCTTNNNAGATCGGAA |
| 51340_2 | TATGAGTGTGGAGTCGTTGCCTTCTGGAATCAAGAGAGGCCTTNNNAGATCGGAA |
| 39904_1 | TATGAGTGTGGAGTCGTTGCGGTGGGAACAGATGTGTTTCATGNNNAGATCGGAA |
| 39904_2 | TATGAGTGTGGAGTCGTTGCGGTGGGAACAGATGTGTTTCATGNNNAGATCGGAA |
| 46779_1 | TATGAGTGTGGAGTCGTTGCTCTCCTGAGTAGCTGGGATTACANNNAGATCGGAAG |
| 46779_2 | TATGAGTGTGGAGTCGTTGCTCTCCTGAGTAGCTGGGATTACANNNAGATCGGAAG |
| 36771_1 | TATGAGTGTGGAGTCGTTGCCGACAGAACAAGACCCCATCTAANNNAGATCGGAAG |
| 36771_2 | TATGAGTGTGGAGTCGTTGCCGACAGAACAAGACCCCATCTAANNNAGATCGGAAG |
| 33414_1 | TATGAGTGTGGAGTCGTTGCGTGCTGACACACACTACAACATGNNNAGATCGGAAG |
| 33414_2 | TATGAGTGTGGAGTCGTTGCGTGCTGACACACACTACAACATGNNNAGATCGGAAG |
| 45011_1 | TATGAGTGTGGAGTCGTTGCCCCTGGGAATCCTAGACTCTTTGNNNAGATCGGAAG |
| 45011_2 | TATGAGTGTGGAGTCGTTGCCCCTGGGAATCCTAGACTCTTTGNNNAGATCGGAAG |
| 39644_1 | TATGAGTGTGGAGTCGTTGCGAGGGTATTGGGGGAGTCAAAATNNNAGATCGGA |
| 39644_2 | TATGAGTGTGGAGTCGTTGCGAGGGTATTGGGGGAGTCAAAATNNNAGATCGGA |
| 35932_1 | TATGAGTGTGGAGTCGTTGCTGTAATCTCAGCACTTTGGGAGGNNNAGATCGGAA |
| 35932_2 | TATGAGTGTGGAGTCGTTGCTGTAATCTCAGCACTTTGGGAGGNNNAGATCGGAA |
| 38946_1 | TATGAGTGTGGAGTCGTTGCAAATGACCATACAGCCAAAAGCANNNAGATCGGAA |
| 38946_2 | TATGAGTGTGGAGTCGTTGCAAATGACCATACAGCCAAAAGCANNNAGATCGGAA |
| 12583_1 | TATGAGTGTGGAGTCGTTGCAGAGAGGTTTTGATTACTCAAATNNNAGATCGGAA |
| 12583_2 | TATGAGTGTGGAGTCGTTGCAGAGAGGTTTTGATTACTCAAATNNNAGATCGGAA |
| 10273_1 | TATGAGTGTGGAGTCGTTGCACACTTTCCTCCCCAAAGGAAACNNNAGATCGGAAG |
| 10273_2 | TATGAGTGTGGAGTCGTTGCACACTTTCCTCCCCAAAGGAAACNNNAGATCGGAAG |
| 33799_1 | TATGAGTGTGGAGTCGTTGCCCTAAGGGACTTATGGGCTACCANNNAGATCGGAA |
| 33799_2 | TATGAGTGTGGAGTCGTTGCCCTAAGGGACTTATGGGCTACCANNNAGATCGGAA |
| 37397_1 | TATGAGTGTGGAGTCGTTGCCTGTAATCCCAGCTCCAGAACTTNNNAGATCGGAAG |
| 37397_2 | TATGAGTGTGGAGTCGTTGCCTGTAATCCCAGCTCCAGAACTTNNNAGATCGGAAG |
| 43040_1 | TATGAGTGTGGAGTCGTTGCACACACACACACACACAAAACAANNNAGATCGGAAG |
| 43040_2 | TATGAGTGTGGAGTCGTTGCACACACACACACACACAAAACAANNNAGATCGGAAG |
| 42744_1 | TATGAGTGTGGAGTCGTTGCAATGTTGAGCGAGGAAAGCTAGANNNAGATCGGAA |
| 42744_2 | TATGAGTGTGGAGTCGTTGCAATGTTGAGCGAGGAAAGCTAGANNNAGATCGGAA |
| 38058_1 | TATGAGTGTGGAGTCGTTGCGCCCCCCGTCTCTACTAAAATACNNNAGATCGGAAG |
| 38058_2 | TATGAGTGTGGAGTCGTTGCGCCCCCCGTCTCTACTAAAATACNNNAGATCGGAAG |
| 44069_1 | TATGAGTGTGGAGTCGTTGCGAAGGGAACAGGATGGAGTCATTNNNAGATCGGA |
| 44069_2 | TATGAGTGTGGAGTCGTTGCGAAGGGAACAGGATGGAGTCATTNNNAGATCGGA |
| 35884_1 | TATGAGTGTGGAGTCGTTGCGCTGAGGAGGGAAGATTTCTTGANNNAGATCGGAA |
| 35884_2 | TATGAGTGTGGAGTCGTTGCGCTGAGGAGGGAAGATTTCTTGANNNAGATCGGAA |
| 48639_1 | TATGAGTGTGGAGTCGTTGCTTCAAGCTATTCTCCTGCCTCAGNNNAGATCGGAAG |

| | |
|---|---|
| 48639_2 | TATGAGTGTGGAGTCGTTGCTTCAAGCTATTCTCCTGCCTCAGNNNAGATCGGAAGA |
| 31581_1 | TATGAGTGTGGAGTCGTTGCTGTGTGTGTGTGTTTGTATCTGTNNNAGATCGGAAG |
| 31581_2 | TATGAGTGTGGAGTCGTTGCTGTGTGTGTGTGTTTGTATCTGTNNNAGATCGGAAG |
| 49591_1 | TATGAGTGTGGAGTCGTTGCCTCCTGGGTTCACTCCATTCTCNNNAGATCGGAAGAG |
| 49591_2 | TATGAGTGTGGAGTCGTTGCCTCCTGGGTTCACTCCATTCTCNNNAGATCGGAAGAG |
| 41801_1 | TATGAGTGTGGAGTCGTTGCCTGAGTAGCTGGGATTACAGGTGNNNAGATCGGAA |
| 41801_2 | TATGAGTGTGGAGTCGTTGCCTGAGTAGCTGGGATTACAGGTGNNNAGATCGGAA |
| 38259_1 | TATGAGTGTGGAGTCGTTGCATTCTGTTTTCCACCCTAGGCATNNNAGATCGGAAGA |
| 38259_2 | TATGAGTGTGGAGTCGTTGCATTCTGTTTTCCACCCTAGGCATNNNAGATCGGAAGA |
| 43088_1 | TATGAGTGTGGAGTCGTTGCACCACACACATACACACAGATGTNNNAGATCGGAAG |
| 43088_2 | TATGAGTGTGGAGTCGTTGCACCACACACATACACACAGATGTNNNAGATCGGAAG |
| 41358_1 | TATGAGTGTGGAGTCGTTGCCTATGGCTAACACAGGGAGAGTCNNNAGATCGGAA |
| 41358_2 | TATGAGTGTGGAGTCGTTGCCTATGGCTAACACAGGGAGAGTCNNNAGATCGGAA |
| 31732_1 | TATGAGTGTGGAGTCGTTGCAAAAACACTGTGACTTAGGGGGTNNNAGATCGGAA |
| 31732_2 | TATGAGTGTGGAGTCGTTGCAAAAACACTGTGACTTAGGGGGTNNNAGATCGGAA |
| 49975_1 | TATGAGTGTGGAGTCGTTGCCAGGGGTCACCTTCCTAATTTGANNNAGATCGGAAG |
| 49975_2 | TATGAGTGTGGAGTCGTTGCCAGGGGTCACCTTCCTAATTTGANNNAGATCGGAAG |
| 45675_1 | TATGAGTGTGGAGTCGTTGCGTCTGTCGGTGCAGTTCATCTATNNNAGATCGGAAG |
| 45675_2 | TATGAGTGTGGAGTCGTTGCGTCTGTCGGTGCAGTTCATCTATNNNAGATCGGAAG |
| 29411_1 | TATGAGTGTGGAGTCGTTGCGGGAATGTTGTGGGAAAAGGGATNNNAGATCGGA |
| 29411_2 | TATGAGTGTGGAGTCGTTGCGGGAATGTTGTGGGAAAAGGGATNNNAGATCGGA |
| 51780_1 | TATGAGTGTGGAGTCGTTGCAGAGCTGGAAGTAGAACCTAGGTNNNAGATCGGAA |
| 51780_2 | TATGAGTGTGGAGTCGTTGCAGAGCTGGAAGTAGAACCTAGGTNNNAGATCGGAA |
| 48415_1 | TATGAGTGTGGAGTCGTTGCGAGTCAGGGTCTCACTCTGTTACNNNAGATCGGAAG |
| 48415_2 | TATGAGTGTGGAGTCGTTGCGAGTCAGGGTCTCACTCTGTTACNNNAGATCGGAAG |
| 13490_1 | TATGAGTGTGGAGTCGTTGCGCTTGATTTTGAATACTGATTTTNNNAGATCGGAAG |
| 13490_2 | TATGAGTGTGGAGTCGTTGCGCTTGATTTTGAATACTGATTTTNNNAGATCGGAAG |
| 31034_1 | TATGAGTGTGGAGTCGTTGCCACCCTGCCCTTACTTATGATCTNNNAGATCGGAAGA |
| 31034_2 | TATGAGTGTGGAGTCGTTGCCACCCTGCCCTTACTTATGATCTNNNAGATCGGAAGA |
| 40869_1 | TATGAGTGTGGAGTCGTTGCCCCAGCATGTAGGCACGTATTATNNNAGATCGGAAG |
| 40869_2 | TATGAGTGTGGAGTCGTTGCCCCAGCATGTAGGCACGTATTATNNNAGATCGGAAG |
| 13801_1 | TATGAGTGTGGAGTCGTTGCCAAACTATGGCTTACAACATCNNNAGATCGGAAGAG |
| 13801_2 | TATGAGTGTGGAGTCGTTGCCAAACTATGGCTTACAACATCNNNAGATCGGAAGAG |
| 47074_1 | TATGAGTGTGGAGTCGTTGCGGTGTGCAATAGCGTGATCTTGNNNAGATCGGAAG |
| 47074_2 | TATGAGTGTGGAGTCGTTGCGGTGTGCAATAGCGTGATCTTGNNNAGATCGGAAG |
| 12348_1 | TATGAGTGTGGAGTCGTTGCCACCAGGAAAGTGGAAAATGGGTANNNAGATCGGA |
| 12348_2 | TATGAGTGTGGAGTCGTTGCCACCAGGAAAGTGGAAAATGGGTANNNAGATCGGA |
| 50469_1 | TATGAGTGTGGAGTCGTTGCTCCCTTTGTTCAGACTCCAACAANNNAGATCGGAAGA |
| 50469_2 | TATGAGTGTGGAGTCGTTGCTCCCTTTGTTCAGACTCCAACAANNNAGATCGGAAGA |
| 48873_1 | TATGAGTGTGGAGTCGTTGCCTCCTGAGTAGCTGGGATTACAGNNNAGATCGGAA |
| 48873_2 | TATGAGTGTGGAGTCGTTGCCTCCTGAGTAGCTGGGATTACAGNNNAGATCGGAA |
| 40653_1 | TATGAGTGTGGAGTCGTTGCTTCAGTCATTCCACAATGCAAGCNNNAGATCGGAAG |
| 40653_2 | TATGAGTGTGGAGTCGTTGCTTCAGTCATTCCACAATGCAAGCNNNAGATCGGAAG |

| | |
|---|---|
| 29749_1 | TATGAGTGTGGAGTCGTTGCATTCTGTGCCTTGACAAGTGAGANNNAGATCGGAAG |
| 29749_2 | TATGAGTGTGGAGTCGTTGCATTCTGTGCCTTGACAAGTGAGANNNAGATCGGAAG |
| 43794_1 | TATGAGTGTGGAGTCGTTGCCTGGGATGGTAACTCTCCTTCACNNNAGATCGGAAG |
| 43794_2 | TATGAGTGTGGAGTCGTTGCCTGGGATGGTAACTCTCCTTCACNNNAGATCGGAAG |
| 46475_1 | TATGAGTGTGGAGTCGTTGCTTACCTCTTTCAGTCTTGGGAGCNNNAGATCGGAAG |
| 46475_2 | TATGAGTGTGGAGTCGTTGCTTACCTCTTTCAGTCTTGGGAGCNNNAGATCGGAAG |
| 36247_1 | TATGAGTGTGGAGTCGTTGCTCTTGTGTTGCTTTGGGAAATCCNNNAGATCGGAAG |
| 36247_2 | TATGAGTGTGGAGTCGTTGCTCTTGTGTTGCTTTGGGAAATCCNNNAGATCGGAAG |
| 453_1 | TATGAGTGTGGAGTCGTTGCAATCCCTCATCTTGCTCCACCTANNNAGATCGGAAGA |
| 453_2 | TATGAGTGTGGAGTCGTTGCAATCCCTCATCTTGCTCCACCTANNNAGATCGGAAGA |
| 48995_1 | TATGAGTGTGGAGTCGTTGCAATGATTCTTTTGCCTCAGCCTCNNNAGATCGGAAGA |
| 48995_2 | TATGAGTGTGGAGTCGTTGCAATGATTCTTTTGCCTCAGCCTCNNNAGATCGGAAGA |
| 48262_1 | TATGAGTGTGGAGTCGTTGCCGAGTAGCTGGGATTACAGGTACNNNAGATCGGAA |
| 48262_2 | TATGAGTGTGGAGTCGTTGCCGAGTAGCTGGGATTACAGGTACNNNAGATCGGAA |
| 50871_1 | TATGAGTGTGGAGTCGTTGCTGACTCTGTCAGAAAGCACTCTCNNNAGATCGGAAG |
| 50871_2 | TATGAGTGTGGAGTCGTTGCTGACTCTGTCAGAAAGCACTCTCNNNAGATCGGAAG |
| 38333_1 | TATGAGTGTGGAGTCGTTGCAGATGGGAGGTAGGTGTGACTATNNNAGATCGGA |
| 38333_2 | TATGAGTGTGGAGTCGTTGCAGATGGGAGGTAGGTGTGACTATNNNAGATCGGA |
| 41091_1 | TATGAGTGTGGAGTCGTTGCCACACCTGTAATCCCAGCACTTTNNNAGATCGGAAGA |
| 41091_2 | TATGAGTGTGGAGTCGTTGCCACACCTGTAATCCCAGCACTTTNNNAGATCGGAAGA |
| 47924_1 | TATGAGTGTGGAGTCGTTGCCCACGCCTGGAGAATTTTTTGTANNNAGATCGGAAG |
| 47924_2 | TATGAGTGTGGAGTCGTTGCCCACGCCTGGAGAATTTTTTGTANNNAGATCGGAAG |
| 30194_1 | TATGAGTGTGGAGTCGTTGCCACTAATTTGATCGGGGGCTTGNNNAGATCGGAAG |
| 30194_2 | TATGAGTGTGGAGTCGTTGCCACTAATTTGATCGGGGGCTTGNNNAGATCGGAAG |
| 34709_1 | TATGAGTGTGGAGTCGTTGCACCATGCCCGGCTATTTTTTTTTNNNAGATCGGAAGA |
| 34709_2 | TATGAGTGTGGAGTCGTTGCACCATGCCCGGCTATTTTTTTTTNNNAGATCGGAAGA |
| 33506_1 | TATGAGTGTGGAGTCGTTGCGCCTGTAAACAGAATCTGCCCTANNNAGATCGGAAG |
| 33506_2 | TATGAGTGTGGAGTCGTTGCGCCTGTAAACAGAATCTGCCCTANNNAGATCGGAAG |
| 41141_1 | TATGAGTGTGGAGTCGTTGCGAGGCAGGAGAATCACTTGAACTNNNAGATCGGAA |
| 41141_2 | TATGAGTGTGGAGTCGTTGCGAGGCAGGAGAATCACTTGAACTNNNAGATCGGAA |
| 42289_1 | TATGAGTGTGGAGTCGTTGCTTCAAGTGATTCTCCTGCCTCAGNNNAGATCGGAAG |
| 42289_2 | TATGAGTGTGGAGTCGTTGCTTCAAGTGATTCTCCTGCCTCAGNNNAGATCGGAAG |
| 9161_1 | TATGAGTGTGGAGTCGTTGCGCCCTATGTAATGAAGCCTGCATNNNAGATCGGAAG |
| 9161_2 | TATGAGTGTGGAGTCGTTGCGCCCTATGTAATGAAGCCTGCATNNNAGATCGGAAG |
| 29604_1 | TATGAGTGTGGAGTCGTTGCGAAACAGGACATGGCAACCAGAANNNAGATCGGAA |
| 29604_2 | TATGAGTGTGGAGTCGTTGCGAAACAGGACATGGCAACCAGAANNNAGATCGGAA |
| 30873_1 | TATGAGTGTGGAGTCGTTGCTCTCTCTCTCTCTCTGTGTGTGTNNNAGATCGGAAGA |
| 30873_2 | TATGAGTGTGGAGTCGTTGCTCTCTCTCTCTCTCTGTGTGTGTNNNAGATCGGAAGA |
| 52208_1 | TATGAGTGTGGAGTCGTTGCTTGAGTTCAAGGTAAAGTCCCCCNNNAGATCGGAAG |
| 52208_2 | TATGAGTGTGGAGTCGTTGCTTGAGTTCAAGGTAAAGTCCCCCNNNAGATCGGAAG |
| 41591_1 | TATGAGTGTGGAGTCGTTGCTTCAAGCAATTCTCCTGCCTCAGNNNAGATCGGAAG |
| 41591_2 | TATGAGTGTGGAGTCGTTGCTTCAAGCAATTCTCCTGCCTCAGNNNAGATCGGAAG |
| 7980_1 | TATGAGTGTGGAGTCGTTGCCAGCCTACCTCCCAATTCACAGTNNNAGATCGGAAG |

| | |
|---|---|
| 7980_2 | TATGAGTGTGGAGTCGTTGCCAGCCTACCTCCCAATTCACAGTNNNAGATCGGAAGA |
| 35967_1 | TATGAGTGTGGAGTCGTTGCCCTGGGTGACAGAGTAACACTCNNNAGATCGGAAG |
| 35967_2 | TATGAGTGTGGAGTCGTTGCCCTGGGTGACAGAGTAACACTCNNNAGATCGGAAG |
| 42601_1 | TATGAGTGTGGAGTCGTTGCGCTGAGAATTCCCAAAATCTGCANNNAGATCGGAAG |
| 42601_2 | TATGAGTGTGGAGTCGTTGCGCTGAGAATTCCCAAAATCTGCANNNAGATCGGAAG |
| 52741_1 | TATGAGTGTGGAGTCGTTGCTAAACTTGGGAGACAGTCAGAGCNNNAGATCGGAA |
| 52741_2 | TATGAGTGTGGAGTCGTTGCTAAACTTGGGAGACAGTCAGAGCNNNAGATCGGAA |
| 32914_1 | TATGAGTGTGGAGTCGTTGCCACACACACATACACACACACTCNNNAGATCGGAAGA |
| 32914_2 | TATGAGTGTGGAGTCGTTGCCACACACACATACACACACACTCNNNAGATCGGAAGA |
| 44367_1 | TATGAGTGTGGAGTCGTTGCTGAGCAAATAGTGTTGGCCTACANNNAGATCGGAAG |
| 44367_2 | TATGAGTGTGGAGTCGTTGCTGAGCAAATAGTGTTGGCCTACANNNAGATCGGAAG |
| 52353_1 | TATGAGTGTGGAGTCGTTGCTCTGAGATGTACAGAGCTTGCTTNNNAGATCGGAAG |
| 52353_2 | TATGAGTGTGGAGTCGTTGCTCTGAGATGTACAGAGCTTGCTTNNNAGATCGGAAG |
| 2721_1 | TATGAGTGTGGAGTCGTTGCAGAAACAGAACTACACAGCAACTNNNAGATCGGAAG |
| 2721_2 | TATGAGTGTGGAGTCGTTGCAGAAACAGAACTACACAGCAACTNNNAGATCGGAAG |
| 50221_1 | TATGAGTGTGGAGTCGTTGCCCCCAAAAAATGCCTTCTGAGTTNNNAGATCGGAAG |
| 50221_2 | TATGAGTGTGGAGTCGTTGCCCCCAAAAAATGCCTTCTGAGTTNNNAGATCGGAAG |
| 44559_1 | TATGAGTGTGGAGTCGTTGCCACACACACACACACACATATNNNAGATCGGAAGA |
| 44559_2 | TATGAGTGTGGAGTCGTTGCCACACACACACACACACATATNNNAGATCGGAAGA |
| 37854_1 | TATGAGTGTGGAGTCGTTGCGAACAACAAAGCCTGGATGACAGNNNAGATCGGAA |
| 37854_2 | TATGAGTGTGGAGTCGTTGCGAACAACAAAGCCTGGATGACAGNNNAGATCGGAA |
| 51759_1 | TATGAGTGTGGAGTCGTTGCATTCACCCTGTAAGGAAGGCTTCNNNAGATCGGAAG |
| 51759_2 | TATGAGTGTGGAGTCGTTGCATTCACCCTGTAAGGAAGGCTTCNNNAGATCGGAAG |
| 38411_1 | TATGAGTGTGGAGTCGTTGCACTAACCGAGAGGCTCATAGGTNNNAGATCGGAAG |
| 38411_2 | TATGAGTGTGGAGTCGTTGCACTAACCGAGAGGCTCATAGGTNNNAGATCGGAAG |
| 40218_1 | TATGAGTGTGGAGTCGTTGCTGCCAAGGAAACACAGGTAGTTANNNAGATCGGAA |
| 40218_2 | TATGAGTGTGGAGTCGTTGCTGCCAAGGAAACACAGGTAGTTANNNAGATCGGAA |
| 2483_1 | TATGAGTGTGGAGTCGTTGCTAATTTCCCCAGTCTTGCTCCAGNNNAGATCGGAAGA |
| 2483_2 | TATGAGTGTGGAGTCGTTGCTAATTTCCCCAGTCTTGCTCCAGNNNAGATCGGAAGA |
| 11213_1 | TATGAGTGTGGAGTCGTTGCCGACCCCCTGGCCAAGGTCTANNNAGATCGGAAGAG |
| 11213_2 | TATGAGTGTGGAGTCGTTGCCGACCCCCTGGCCAAGGTCTANNNAGATCGGAAGAG |
| 42165_1 | TATGAGTGTGGAGTCGTTGCGGCATGATCATGGCTCACTGTAGNNNAGATCGGAAG |
| 42165_2 | TATGAGTGTGGAGTCGTTGCGGCATGATCATGGCTCACTGTAGNNNAGATCGGAAG |
| 34734_1 | TATGAGTGTGGAGTCGTTGCCTCCTGAGTAGCTGGGACTACANNNAGATCGGAAGA |
| 34734_2 | TATGAGTGTGGAGTCGTTGCCTCCTGAGTAGCTGGGACTACANNNAGATCGGAAGA |
| 29242_1 | TATGAGTGTGGAGTCGTTGCTTTTGAGTTGGAGTCTCGCTTTGNNNAGATCGGAAG |
| 29242_2 | TATGAGTGTGGAGTCGTTGCTTTTGAGTTGGAGTCTCGCTTTGNNNAGATCGGAAG |
| 45350_1 | TATGAGTGTGGAGTCGTTGCATGGAAAGCAGGCTGAAGACATANNNAGATCGGAA |
| 45350_2 | TATGAGTGTGGAGTCGTTGCATGGAAAGCAGGCTGAAGACATANNNAGATCGGAA |
| 36454_1 | TATGAGTGTGGAGTCGTTGCAAAAAAGAGGGAGAGAGCTTCCCNNNAGATCGGAA |
| 36454_2 | TATGAGTGTGGAGTCGTTGCAAAAAAGAGGGAGAGAGCTTCCCNNNAGATCGGAA |
| 48322_1 | TATGAGTGTGGAGTCGTTGCGGGCAAGGTGATGTGATGAAATCNNNAGATCGGAA |
| 48322_2 | TATGAGTGTGGAGTCGTTGCGGGCAAGGTGATGTGATGAAATCNNNAGATCGGAA |

| | |
|---|---|
| 46623_1 | TATGAGTGTGGAGTCGTTGCGCGTCCGGCTAAGTTTTTGTATTNNNAGATCGGAAG |
| 46623_2 | TATGAGTGTGGAGTCGTTGCGCGTCCGGCTAAGTTTTTGTATTNNNAGATCGGAAG |
| 47125_1 | TATGAGTGTGGAGTCGTTGCAGTGGTACAATCTCAGCTCACTGNNNAGATCGGAAG |
| 47125_2 | TATGAGTGTGGAGTCGTTGCAGTGGTACAATCTCAGCTCACTGNNNAGATCGGAAG |
| 11892_1 | TATGAGTGTGGAGTCGTTGCGACAGAGCCGTTCAGGAGAAANNNAGATCGGAAGA |
| 11892_2 | TATGAGTGTGGAGTCGTTGCGACAGAGCCGTTCAGGAGAAANNNAGATCGGAAGA |
| 37727_1 | TATGAGTGTGGAGTCGTTGCTTTATTTTGTGATGTGCATGGCANNNAGATCGGAAG |
| 37727_2 | TATGAGTGTGGAGTCGTTGCTTTATTTTGTGATGTGCATGGCANNNAGATCGGAAG |
| 2699_1 | TATGAGTGTGGAGTCGTTGCCTTGTGGCCATCACCATTGAGATNNNAGATCGGAAG |
| 2699_2 | TATGAGTGTGGAGTCGTTGCCTTGTGGCCATCACCATTGAGATNNNAGATCGGAAG |
| 13502_1 | TATGAGTGTGGAGTCGTTGCTTACCTTTTTCCATGTTTTGTTTNNNAGATCGGAAGA |
| 13502_2 | TATGAGTGTGGAGTCGTTGCTTACCTTTTTCCATGTTTTGTTTNNNAGATCGGAAGA |
| 35192_1 | TATGAGTGTGGAGTCGTTGCTTCAAGTGATTCTCCTGCCTCAGNNNAGATCGGAAG |
| 35192_2 | TATGAGTGTGGAGTCGTTGCTTCAAGTGATTCTCCTGCCTCAGNNNAGATCGGAAG |
| 42837_1 | TATGAGTGTGGAGTCGTTGCCCATAATTTCGGCTCACTGCAGNNNAGATCGGAAGA |
| 42837_2 | TATGAGTGTGGAGTCGTTGCCCATAATTTCGGCTCACTGCAGNNNAGATCGGAAGA |
| 35918_1 | TATGAGTGTGGAGTCGTTGCATAAGAGGAGAGGTTAGTGCCCTNNNAGATCGGAA |
| 35918_2 | TATGAGTGTGGAGTCGTTGCATAAGAGGAGAGGTTAGTGCCCTNNNAGATCGGAA |
| 10391_1 | TATGAGTGTGGAGTCGTTGCGCTGGAGTGCAGTAGAATTCTGTNNNAGATCGGAA |
| 10391_2 | TATGAGTGTGGAGTCGTTGCGCTGGAGTGCAGTAGAATTCTGTNNNAGATCGGAA |
| 45873_1 | TATGAGTGTGGAGTCGTTGCGATCTTGGCTCACTGAAACCTCTNNNAGATCGGAAG |
| 45873_2 | TATGAGTGTGGAGTCGTTGCGATCTTGGCTCACTGAAACCTCTNNNAGATCGGAAG |
| 43760_1 | TATGAGTGTGGAGTCGTTGCTGTGATTTAGAATCATGGGGCGANNNAGATCGGAA |
| 43760_2 | TATGAGTGTGGAGTCGTTGCTGTGATTTAGAATCATGGGGCGANNNAGATCGGAA |
| 36153_1 | TATGAGTGTGGAGTCGTTGCCCTTCCTCATATAGTGTCCCTGCNNNAGATCGGAAGA |
| 36153_2 | TATGAGTGTGGAGTCGTTGCCCTTCCTCATATAGTGTCCCTGCNNNAGATCGGAAGA |
| 39403_1 | TATGAGTGTGGAGTCGTTGCAAGTACCCACAGAAGCTTTACCCNNNAGATCGGAAG |
| 39403_2 | TATGAGTGTGGAGTCGTTGCAAGTACCCACAGAAGCTTTACCCNNNAGATCGGAAG |
| 33995_1 | TATGAGTGTGGAGTCGTTGCACCTCTCAAGTTCAAGCGATTCTNNNAGATCGGAAG |
| 33995_2 | TATGAGTGTGGAGTCGTTGCACCTCTCAAGTTCAAGCGATTCTNNNAGATCGGAAG |
| 32985_1 | TATGAGTGTGGAGTCGTTGCTTTTGTACTCTATTGCTGCAGTCTNNNAGATCGGAAG |
| 32985_2 | TATGAGTGTGGAGTCGTTGCTTTTGTACTCTATTGCTGCAGTCTNNNAGATCGGAAG |
| 39059_1 | TATGAGTGTGGAGTCGTTGCGGCTAAGGAAGGACAATTGCTTGNNNAGATCGGAA |
| 39059_2 | TATGAGTGTGGAGTCGTTGCGGCTAAGGAAGGACAATTGCTTGNNNAGATCGGAA |
| 34854_1 | TATGAGTGTGGAGTCGTTGCCACCACACCCGGCTAATTTTTNNNAGATCGGAAGAG |
| 34854_2 | TATGAGTGTGGAGTCGTTGCCACCACACCCGGCTAATTTTTNNNAGATCGGAAGAG |
| 44656_1 | TATGAGTGTGGAGTCGTTGCAGACAGGAGGTTCTTTCACTTCCNNNAGATCGGAAG |
| 44656_2 | TATGAGTGTGGAGTCGTTGCAGACAGGAGGTTCTTTCACTTCCNNNAGATCGGAAG |
| 39323_1 | TATGAGTGTGGAGTCGTTGCTAGAGATGAACCCTGGGTGTTTGNNNAGATCGGAA |
| 39323_2 | TATGAGTGTGGAGTCGTTGCTAGAGATGAACCCTGGGTGTTTGNNNAGATCGGAA |
| 13518_1 | TATGAGTGTGGAGTCGTTGCCTATGGTCGTTATTTTCTCATTTNNNAGATCGGAAGA |
| 13518_2 | TATGAGTGTGGAGTCGTTGCCTATGGTCGTTATTTTCTCATTTNNNAGATCGGAAGA |
| 45185_1 | TATGAGTGTGGAGTCGTTGCTGCTCCAAGGAAAAATAGTGGGTNNNAGATCGGAA |

| | |
|---|---|
| 45185_2 | TATGAGTGTGGAGTCGTTGCTGCTCCAAGGAAAAATAGTGGGTNNNAGATCGGAA |
| 42125_1 | TATGAGTGTGGAGTCGTTGCGGGTTTGGAATTCCCTGTCCTATNNNAGATCGGAAG |
| 42125_2 | TATGAGTGTGGAGTCGTTGCGGGTTTGGAATTCCCTGTCCTATNNNAGATCGGAAG |
| 48444_1 | TATGAGTGTGGAGTCGTTGCGTGGCTTCTAAACACGATGCACNNNAGATCGGAAG |
| 48444_2 | TATGAGTGTGGAGTCGTTGCGTGGCTTCTAAACACGATGCACNNNAGATCGGAAG |
| 40198_1 | TATGAGTGTGGAGTCGTTGCAGAGGAGAAGAGAAAACACTCTCANNNAGATCGGA |
| 40198_2 | TATGAGTGTGGAGTCGTTGCAGAGGAGAAGAGAAAACACTCTCANNNAGATCGGA |
| 32053_1 | TATGAGTGTGGAGTCGTTGCCGCCCCAACTGATTTTCATCATANNNAGATCGGAAG |
| 32053_2 | TATGAGTGTGGAGTCGTTGCCGCCCCAACTGATTTTCATCATANNNAGATCGGAAG |
| 32596_1 | TATGAGTGTGGAGTCGTTGCAGATGGAGTCTTTCTCTGTTGCCNNNAGATCGGAAG |
| 32596_2 | TATGAGTGTGGAGTCGTTGCAGATGGAGTCTTTCTCTGTTGCCNNNAGATCGGAAG |
| 48559_1 | TATGAGTGTGGAGTCGTTGCTGGGTGTGTCTATTGTAGAGTGCNNNAGATCGGAA |
| 48559_2 | TATGAGTGTGGAGTCGTTGCTGGGTGTGTCTATTGTAGAGTGCNNNAGATCGGAA |
| 44949_1 | TATGAGTGTGGAGTCGTTGCCAGTGCATTTCTGATTTCACCCANNNAGATCGGAAG |
| 44949_2 | TATGAGTGTGGAGTCGTTGCCAGTGCATTTCTGATTTCACCCANNNAGATCGGAAG |
| 50658_1 | TATGAGTGTGGAGTCGTTGCTGAGGGAGAAGCCAAAAGATCAANNNAGATCGGAA |
| 50658_2 | TATGAGTGTGGAGTCGTTGCTGAGGGAGAAGCCAAAAGATCAANNNAGATCGGAA |
| 30901_1 | TATGAGTGTGGAGTCGTTGCCCTGTGTGAGTGAAAGACAGAGANNNAGATCGGAA |
| 30901_2 | TATGAGTGTGGAGTCGTTGCCCTGTGTGAGTGAAAGACAGAGANNNAGATCGGAA |
| 35794_1 | TATGAGTGTGGAGTCGTTGCAGCTCATGTATTCTGCCCGTTTANNNAGATCGGAAG |
| 35794_2 | TATGAGTGTGGAGTCGTTGCAGCTCATGTATTCTGCCCGTTTANNNAGATCGGAAG |
| 49744_1 | TATGAGTGTGGAGTCGTTGCTTTCCTCAGCCATGTCTCATCAANNNAGATCGGAAGA |
| 49744_2 | TATGAGTGTGGAGTCGTTGCTTTCCTCAGCCATGTCTCATCAANNNAGATCGGAAGA |
| 52628_1 | TATGAGTGTGGAGTCGTTGCACCTGAGAACTAAGCTCTGATGTNNNAGATCGGAAG |
| 52628_2 | TATGAGTGTGGAGTCGTTGCACCTGAGAACTAAGCTCTGATGTNNNAGATCGGAAG |
| 44350_1 | TATGAGTGTGGAGTCGTTGCGCAGGAGTTTCAAAACATGGAGTNNNAGATCGGAA |
| 44350_2 | TATGAGTGTGGAGTCGTTGCGCAGGAGTTTCAAAACATGGAGTNNNAGATCGGAA |
| 48603_1 | TATGAGTGTGGAGTCGTTGCCCCAAAGTGCTAGGATTACAGGTNNNAGATCGGAAG |
| 48603_2 | TATGAGTGTGGAGTCGTTGCCCCAAAGTGCTAGGATTACAGGTNNNAGATCGGAAG |
| 31185_1 | TATGAGTGTGGAGTCGTTGCAAGTGCTGAGATTACAGGCATGANNNAGATCGGAA |
| 31185_2 | TATGAGTGTGGAGTCGTTGCAAGTGCTGAGATTACAGGCATGANNNAGATCGGAA |
| 37498_1 | TATGAGTGTGGAGTCGTTGCAAGTTTTGCCCCGGTTTTACAAANNNAGATCGGAAG |
| 37498_2 | TATGAGTGTGGAGTCGTTGCAAGTTTTGCCCCGGTTTTACAAANNNAGATCGGAAG |
| 46369_1 | TATGAGTGTGGAGTCGTTGCGAGAGAAGTCTCGTTCTTGTCCCNNNAGATCGGAAG |
| 46369_2 | TATGAGTGTGGAGTCGTTGCGAGAGAAGTCTCGTTCTTGTCCCNNNAGATCGGAAG |
| 29538_1 | TATGAGTGTGGAGTCGTTGCAGAGCGACTACTCCAGGTGAGNNNAGATCGGAAGA |
| 29538_2 | TATGAGTGTGGAGTCGTTGCAGAGCGACTACTCCAGGTGAGNNNAGATCGGAAGA |
| 48211_1 | TATGAGTGTGGAGTCGTTGCCAGCTGATGCCATGTTTCCCTNNNAGATCGGAAGAG |
| 48211_2 | TATGAGTGTGGAGTCGTTGCCAGCTGATGCCATGTTTCCCTNNNAGATCGGAAGAG |
| 45368_1 | TATGAGTGTGGAGTCGTTGCCTCATTGGTTGGTGGGCATTTAGNNNAGATCGGAA |
| 45368_2 | TATGAGTGTGGAGTCGTTGCCTCATTGGTTGGTGGGCATTTAGNNNAGATCGGAA |
| 36922_1 | TATGAGTGTGGAGTCGTTGCTAAGGATGAGGTAGGAGGATCCCNNNAGATCGGAA |
| 36922_2 | TATGAGTGTGGAGTCGTTGCTAAGGATGAGGTAGGAGGATCCCNNNAGATCGGAA |

| | |
|---|---|
| 13189_1 | TATGAGTGTGGAGTCGTTGCAGGTAGGGTGATTTTAATGATATNNNAGATCGGAA |
| 13189_2 | TATGAGTGTGGAGTCGTTGCAGGTAGGGTGATTTTAATGATATNNNAGATCGGAA |
| 29267_1 | TATGAGTGTGGAGTCGTTGCTTCTCCCAACAGAAGCAGAAAGGNNNAGATCGGAA |
| 29267_2 | TATGAGTGTGGAGTCGTTGCTTCTCCCAACAGAAGCAGAAAGGNNNAGATCGGAA |
| 49903_1 | TATGAGTGTGGAGTCGTTGCGCCAGGGCTATCTAAAAGAGTCANNNAGATCGGAA |
| 49903_2 | TATGAGTGTGGAGTCGTTGCGCCAGGGCTATCTAAAAGAGTCANNNAGATCGGAA |
| 48359_1 | TATGAGTGTGGAGTCGTTGCTCTGCTTCCTGAGTTCAAGTGATNNNAGATCGGAAG |
| 48359_2 | TATGAGTGTGGAGTCGTTGCTCTGCTTCCTGAGTTCAAGTGATNNNAGATCGGAAG |
| 8094_1 | TATGAGTGTGGAGTCGTTGCTAGTGGCTAATTGACATTCCCAACNNNAGATCGGAA |
| 8094_2 | TATGAGTGTGGAGTCGTTGCTAGTGGCTAATTGACATTCCCAACNNNAGATCGGAA |
| 48118_1 | TATGAGTGTGGAGTCGTTGCTTCCAGGTTCATCCATGTTGTCANNNAGATCGGAAG |
| 48118_2 | TATGAGTGTGGAGTCGTTGCTTCCAGGTTCATCCATGTTGTCANNNAGATCGGAAG |
| 35617_1 | TATGAGTGTGGAGTCGTTGCTCAAGAAATTCTCCCTCCTCAGCNNNAGATCGGAAGA |
| 35617_2 | TATGAGTGTGGAGTCGTTGCTCAAGAAATTCTCCCTCCTCAGCNNNAGATCGGAAGA |
| 45931_1 | TATGAGTGTGGAGTCGTTGCGATTCTCTTACCTCAGCCTCCAGNNNAGATCGGAAGA |
| 45931_2 | TATGAGTGTGGAGTCGTTGCGATTCTCTTACCTCAGCCTCCAGNNNAGATCGGAAGA |
| 38730_1 | TATGAGTGTGGAGTCGTTGCAGCAAAATATTGAAGGAGCTGCANNNAGATCGGAA |
| 38730_2 | TATGAGTGTGGAGTCGTTGCAGCAAAATATTGAAGGAGCTGCANNNAGATCGGAA |
| 30673_1 | TATGAGTGTGGAGTCGTTGCGATATTAGGCAGGCTTGCAGTTACNNNAGATCGGA |
| 30673_2 | TATGAGTGTGGAGTCGTTGCGATATTAGGCAGGCTTGCAGTTACNNNAGATCGGA |
| 32939_1 | TATGAGTGTGGAGTCGTTGCGCTCAAGCCTGTCTCAAAAAAAANNNAGATCGGAAG |
| 32939_2 | TATGAGTGTGGAGTCGTTGCGCTCAAGCCTGTCTCAAAAAAAANNNAGATCGGAAG |
| 47851_1 | TATGAGTGTGGAGTCGTTGCCTCCTTGGCACTCCTTTCCTATGNNNAGATCGGAAGA |
| 47851_2 | TATGAGTGTGGAGTCGTTGCCTCCTTGGCACTCCTTTCCTATGNNNAGATCGGAAGA |
| 32863_1 | TATGAGTGTGGAGTCGTTGCAGAGTCTCACTCTATCCCTCAGGNNNAGATCGGAAG |
| 32863_2 | TATGAGTGTGGAGTCGTTGCAGAGTCTCACTCTATCCCTCAGGNNNAGATCGGAAG |
| 40628_1 | TATGAGTGTGGAGTCGTTGCGAGTGGGGACAAAAAAAGAAGCANNNAGATCGGA |
| 40628_2 | TATGAGTGTGGAGTCGTTGCGAGTGGGGACAAAAAAAGAAGCANNNAGATCGGA |
| 40128_1 | TATGAGTGTGGAGTCGTTGCGAAGGAGGATCGCTTGAGACTAGNNNAGATCGGA |
| 40128_2 | TATGAGTGTGGAGTCGTTGCGAAGGAGGATCGCTTGAGACTAGNNNAGATCGGA |
| 50974_1 | TATGAGTGTGGAGTCGTTGCTTTCTGGGACCAAGAGAGAGATGNNNAGATCGGAA |
| 50974_2 | TATGAGTGTGGAGTCGTTGCTTTCTGGGACCAAGAGAGAGATGNNNAGATCGGAA |
| 48729_1 | TATGAGTGTGGAGTCGTTGCTCTCTGGTTTTTCTCCAACTGTTCNNNAGATCGGAAG |
| 48729_2 | TATGAGTGTGGAGTCGTTGCTCTCTGGTTTTTCTCCAACTGTTCNNNAGATCGGAAG |
| 45864_1 | TATGAGTGTGGAGTCGTTGCAGATGGAGTCTCTCTCTGTCACCNNNAGATCGGAAG |
| 45864_2 | TATGAGTGTGGAGTCGTTGCAGATGGAGTCTCTCTCTGTCACCNNNAGATCGGAAG |
| 39996_1 | TATGAGTGTGGAGTCGTTGCCCTTTAGGAATTGATTGGGCCTGNNNAGATCGGAA |
| 39996_2 | TATGAGTGTGGAGTCGTTGCCCTTTAGGAATTGATTGGGCCTGNNNAGATCGGAA |
| 37984_1 | TATGAGTGTGGAGTCGTTGCAAAACGCACACAAAGTATCCCAGNNNAGATCGGAAG |
| 37984_2 | TATGAGTGTGGAGTCGTTGCAAAACGCACACAAAGTATCCCAGNNNAGATCGGAAG |
| 41553_1 | TATGAGTGTGGAGTCGTTGCCAATCTTGTCTCACTGCAGCATCNNNAGATCGGAAG |
| 41553_2 | TATGAGTGTGGAGTCGTTGCCAATCTTGTCTCACTGCAGCATCNNNAGATCGGAAG |
| 29564_1 | TATGAGTGTGGAGTCGTTGCTTAGAAAAGCGCTTGGGTTCTTGNNNAGATCGGAA |

| | |
|---|---|
| 29564_2 | TATGAGTGTGGAGTCGTTGCTTAGAAAAGCGCTTGGGTTCTTGNNNAGATCGGAA |
| 33244_1 | TATGAGTGTGGAGTCGTTGCGATGGAGGCTCGCTCTATCATCNNNAGATCGGAAGA |
| 33244_2 | TATGAGTGTGGAGTCGTTGCGATGGAGGCTCGCTCTATCATCNNNAGATCGGAAGA |
| 30089_1 | TATGAGTGTGGAGTCGTTGCTTTTTGAAGACCATTGAGGATGTTNNNAGATCGGAA |
| 30089_2 | TATGAGTGTGGAGTCGTTGCTTTTTGAAGACCATTGAGGATGTTNNNAGATCGGAA |
| 48586_1 | TATGAGTGTGGAGTCGTTGCGATCTCGGCTCACTGCAACCNNNAGATCGGAAGAGC |
| 48586_2 | TATGAGTGTGGAGTCGTTGCGATCTCGGCTCACTGCAACCNNNAGATCGGAAGAGC |
| 47838_1 | TATGAGTGTGGAGTCGTTGCGAGATGGGGGTCTCACTATGTTGNNNAGATCGGAA |
| 47838_2 | TATGAGTGTGGAGTCGTTGCGAGATGGGGGTCTCACTATGTTGNNNAGATCGGAA |
| 32210_1 | TATGAGTGTGGAGTCGTTGCACGGGACAGAAGAAAATTTTGACTNNNAGATCGGA |
| 32210_2 | TATGAGTGTGGAGTCGTTGCACGGGACAGAAGAAAATTTTGACTNNNAGATCGGA |
| 29328_1 | TATGAGTGTGGAGTCGTTGCTCGAGGGAGAAAAAAGGAGGAAGNNNAGATCGGA |
| 29328_2 | TATGAGTGTGGAGTCGTTGCTCGAGGGAGAAAAAAGGAGGAAGNNNAGATCGGA |
| 28809_1 | TATGAGTGTGGAGTCGTTGCCTCCCACTGATTACATGAGGCTTNNNAGATCGGAAG |
| 28809_2 | TATGAGTGTGGAGTCGTTGCCTCCCACTGATTACATGAGGCTTNNNAGATCGGAAG |
| 29324_1 | TATGAGTGTGGAGTCGTTGCGTGGTTACATGTGGTCACAAGTGNNNAGATCGGAA |
| 29324_2 | TATGAGTGTGGAGTCGTTGCGTGGTTACATGTGGTCACAAGTGNNNAGATCGGAA |
| 42398_1 | TATGAGTGTGGAGTCGTTGCTTGATGAACATTTGGGCTGGTTCNNNAGATCGGAAG |
| 42398_2 | TATGAGTGTGGAGTCGTTGCTTGATGAACATTTGGGCTGGTTCNNNAGATCGGAAG |
| 50559_1 | TATGAGTGTGGAGTCGTTGCCTCATGGAGAAGCAGCAACTCTANNNAGATCGGAAG |
| 50559_2 | TATGAGTGTGGAGTCGTTGCCTCATGGAGAAGCAGCAACTCTANNNAGATCGGAAG |
| 37618_1 | TATGAGTGTGGAGTCGTTGCAGTGAGCTGTGATTGTGTCCTACNNNAGATCGGAAG |
| 37618_2 | TATGAGTGTGGAGTCGTTGCAGTGAGCTGTGATTGTGTCCTACNNNAGATCGGAAG |
| 33933_1 | TATGAGTGTGGAGTCGTTGCAGGTTCAAATGATTCTCCTGCCTNNNAGATCGGAAG |
| 33933_2 | TATGAGTGTGGAGTCGTTGCAGGTTCAAATGATTCTCCTGCCTNNNAGATCGGAAG |
| 30145_1 | TATGAGTGTGGAGTCGTTGCCTTGCAGTTGTTGTTGCTCACATNNNAGATCGGAAG |
| 30145_2 | TATGAGTGTGGAGTCGTTGCCTTGCAGTTGTTGTTGCTCACATNNNAGATCGGAAG |
| 30011_1 | TATGAGTGTGGAGTCGTTGCAAAGTGTGCAGTTATACTCACGTNNNAGATCGGAAG |
| 30011_2 | TATGAGTGTGGAGTCGTTGCAAAGTGTGCAGTTATACTCACGTNNNAGATCGGAAG |
| 53060_1 | TATGAGTGTGGAGTCGTTGCAAGGACACTAGTGTAGCCATAGCNNNAGATCGGAA |
| 53060_2 | TATGAGTGTGGAGTCGTTGCAAGGACACTAGTGTAGCCATAGCNNNAGATCGGAA |
| 13409_1 | TATGAGTGTGGAGTCGTTGCACACAATCTGGATAATATTGAGANNNAGATCGGAAG |
| 13409_2 | TATGAGTGTGGAGTCGTTGCACACAATCTGGATAATATTGAGANNNAGATCGGAAG |
| 52339_1 | TATGAGTGTGGAGTCGTTGCATGATGTTCCTCTTTTGCTTGCCNNNAGATCGGAAGA |
| 52339_2 | TATGAGTGTGGAGTCGTTGCATGATGTTCCTCTTTTGCTTGCCNNNAGATCGGAAGA |
| 31234_1 | TATGAGTGTGGAGTCGTTGCGGCTCTGCATACCAGGTTTTTTTNNNAGATCGGAAG |
| 31234_2 | TATGAGTGTGGAGTCGTTGCGGCTCTGCATACCAGGTTTTTTTNNNAGATCGGAAG |
| 44579_1 | TATGAGTGTGGAGTCGTTGCGATCTGGGGAACCTCTGTTTCTCNNNAGATCGGAAG |
| 44579_2 | TATGAGTGTGGAGTCGTTGCGATCTGGGGAACCTCTGTTTCTCNNNAGATCGGAAG |
| 53401_1 | TATGAGTGTGGAGTCGTTGCGTACCAAAAAGCAGAGGGAGAGANNNAGATCGGA |
| 53401_2 | TATGAGTGTGGAGTCGTTGCGTACCAAAAAGCAGAGGGAGAGANNNAGATCGGA |
| 32619_1 | TATGAGTGTGGAGTCGTTGCCCAGTGACAGTCATTTTAGCAGCNNNAGATCGGAAG |
| 32619_2 | TATGAGTGTGGAGTCGTTGCCCAGTGACAGTCATTTTAGCAGCNNNAGATCGGAAG |

| | |
|---|---|
| 29693_1 | TATGAGTGTGGAGTCGTTGCCCATCATGTTTGGGGCTGTAATGNNNAGATCGGAAG |
| 29693_2 | TATGAGTGTGGAGTCGTTGCCCATCATGTTTGGGGCTGTAATGNNNAGATCGGAAG |
| 46173_1 | TATGAGTGTGGAGTCGTTGCGGTTCAAGTGATTCTTCTGCCTCNNNAGATCGGAAGA |
| 46173_2 | TATGAGTGTGGAGTCGTTGCGGTTCAAGTGATTCTTCTGCCTCNNNAGATCGGAAGA |
| 30651_1 | TATGAGTGTGGAGTCGTTGCCTCACCTTGGGATCGTGCAAGAANNNAGATCGGAAG |
| 30651_2 | TATGAGTGTGGAGTCGTTGCCTCACCTTGGGATCGTGCAAGAANNNAGATCGGAAG |
| 33623_1 | TATGAGTGTGGAGTCGTTGCCAAGTAGCTGGGACTACAGGCNNNAGATCGGAAGA |
| 33623_2 | TATGAGTGTGGAGTCGTTGCCAAGTAGCTGGGACTACAGGCNNNAGATCGGAAGA |
| 46640_1 | TATGAGTGTGGAGTCGTTGCCTCAGTTCAAGCAATTCTCCTGCNNNAGATCGGAAGA |
| 46640_2 | TATGAGTGTGGAGTCGTTGCCTCAGTTCAAGCAATTCTCCTGCNNNAGATCGGAAGA |
| 41554_1 | TATGAGTGTGGAGTCGTTGCAGCTCACTGCAAACTTGAACTTCNNNAGATCGGAAGA |
| 41554_2 | TATGAGTGTGGAGTCGTTGCAGCTCACTGCAAACTTGAACTTCNNNAGATCGGAAGA |
| 52110_1 | TATGAGTGTGGAGTCGTTGCGAGGATACCATTTCCAGCTCCATNNNAGATCGGAAGA |
| 52110_2 | TATGAGTGTGGAGTCGTTGCGAGGATACCATTTCCAGCTCCATNNNAGATCGGAAGA |
| 41729_1 | TATGAGTGTGGAGTCGTTGCGGAGTGCAATGGCTTGATCTTGNNNAGATCGGAAGA |
| 41729_2 | TATGAGTGTGGAGTCGTTGCGGAGTGCAATGGCTTGATCTTGNNNAGATCGGAAGA |
| 34789_1 | TATGAGTGTGGAGTCGTTGCTTTTTGAGACAGGGTCTTGCTCTNNNAGATCGGAAG |
| 34789_2 | TATGAGTGTGGAGTCGTTGCTTTTTGAGACAGGGTCTTGCTCTNNNAGATCGGAAG |
| 46829_1 | TATGAGTGTGGAGTCGTTGCCTCCTGAGTAGCTGGGACTACAGNNNAGATCGGAAG |
| 46829_2 | TATGAGTGTGGAGTCGTTGCCTCCTGAGTAGCTGGGACTACAGNNNAGATCGGAAG |
| 30406_1 | TATGAGTGTGGAGTCGTTGCGGAATGGCACTCTAGATAGCAGCNNNAGATCGGAAG |
| 30406_2 | TATGAGTGTGGAGTCGTTGCGGAATGGCACTCTAGATAGCAGCNNNAGATCGGAAG |
| 29828_1 | TATGAGTGTGGAGTCGTTGCAGTCCTAAGGGGCATCCTAAGTCNNNAGATCGGAAG |
| 29828_2 | TATGAGTGTGGAGTCGTTGCAGTCCTAAGGGGCATCCTAAGTCNNNAGATCGGAAG |
| 13124_1 | TATGAGTGTGGAGTCGTTGCGAAGTGTTCGTTCAGATATTTCNNNAGATCGGAAGA |
| 13124_2 | TATGAGTGTGGAGTCGTTGCGAAGTGTTCGTTCAGATATTTCNNNAGATCGGAAGA |
| 46245_1 | TATGAGTGTGGAGTCGTTGCCCACCACACCCAGCTAATTTTTTNNNAGATCGGAAGA |
| 46245_2 | TATGAGTGTGGAGTCGTTGCCCACCACACCCAGCTAATTTTTTNNNAGATCGGAAGA |
| 39754_1 | TATGAGTGTGGAGTCGTTGCAGGAGCTCACAATAAAGTGCAGANNNAGATCGGAAG |
| 39754_2 | TATGAGTGTGGAGTCGTTGCAGGAGCTCACAATAAAGTGCAGANNNAGATCGGAAG |
| 40685_1 | TATGAGTGTGGAGTCGTTGCATATGCTGGCATCTTGATGTTGCNNNAGATCGGAAG |
| 40685_2 | TATGAGTGTGGAGTCGTTGCATATGCTGGCATCTTGATGTTGCNNNAGATCGGAAG |
| 42608_1 | TATGAGTGTGGAGTCGTTGCGGTGAGTCTGTGGTCATTTTGTGNNNAGATCGGAAG |
| 42608_2 | TATGAGTGTGGAGTCGTTGCGGTGAGTCTGTGGTCATTTTGTGNNNAGATCGGAAG |
| 31545_1 | TATGAGTGTGGAGTCGTTGCAAGCAGAAGTTCAAAGCATGACCNNNAGATCGGAAG |
| 31545_2 | TATGAGTGTGGAGTCGTTGCAAGCAGAAGTTCAAAGCATGACCNNNAGATCGGAAG |
| 49117_1 | TATGAGTGTGGAGTCGTTGCTGTGTCTTTTTTTGTGGCTTGGTNNNAGATCGGAAGA |
| 49117_2 | TATGAGTGTGGAGTCGTTGCTGTGTCTTTTTTTGTGGCTTGGTNNNAGATCGGAAGA |
| 35352_1 | TATGAGTGTGGAGTCGTTGCCAATAGCAGCACAATCTCAGCTCNNNAGATCGGAAGA |
| 35352_2 | TATGAGTGTGGAGTCGTTGCCAATAGCAGCACAATCTCAGCTCNNNAGATCGGAAGA |
| 31520_1 | TATGAGTGTGGAGTCGTTGCCTTTGTCTGGAAGAACAGAGGGANNNAGATCGGAA |
| 31520_2 | TATGAGTGTGGAGTCGTTGCCTTTGTCTGGAAGAACAGAGGGANNNAGATCGGAA |
| 44282_1 | TATGAGTGTGGAGTCGTTGCTGAACAACCTTCTGAAGCTCTGTNNNAGATCGGAAGA |

| | |
|---|---|
| 44282_2 | TATGAGTGTGGAGTCGTTGCTGAACAACCTTCTGAAGCTCTGTNNNAGATCGGAAG |
| 10826_1 | TATGAGTGTGGAGTCGTTGCTCAGGTCCATCTTTCAAGCCCTANNNAGATCGGAAG |
| 10826_2 | TATGAGTGTGGAGTCGTTGCTCAGGTCCATCTTTCAAGCCCTANNNAGATCGGAAG |
| 1795_1 | TATGAGTGTGGAGTCGTTGCAGAGCCCCATTCAAAGGCTCCGNNNAGATCGGAAGA |
| 1795_2 | TATGAGTGTGGAGTCGTTGCAGAGCCCCATTCAAAGGCTCCGNNNAGATCGGAAGA |
| 40428_1 | TATGAGTGTGGAGTCGTTGCAAAACCAAGTAACCAGGCCTAGTNNNAGATCGGAA |
| 40428_2 | TATGAGTGTGGAGTCGTTGCAAAACCAAGTAACCAGGCCTAGTNNNAGATCGGAA |
| 46267_1 | TATGAGTGTGGAGTCGTTGCACTATCTCGGCTCACTGCAACNNNAGATCGGAAGAG |
| 46267_2 | TATGAGTGTGGAGTCGTTGCACTATCTCGGCTCACTGCAACNNNAGATCGGAAGAG |
| 34663_1 | TATGAGTGTGGAGTCGTTGCAGAGCTGAGATAAACGCCCTTTCNNNAGATCGGAAG |
| 34663_2 | TATGAGTGTGGAGTCGTTGCAGAGCTGAGATAAACGCCCTTTCNNNAGATCGGAAG |
| 46172_1 | TATGAGTGTGGAGTCGTTGCGAAGAGGAGTCTTGCTGTGTCCNNNAGATCGGAAG |
| 46172_2 | TATGAGTGTGGAGTCGTTGCGAAGAGGAGTCTTGCTGTGTCCNNNAGATCGGAAG |
| 39145_1 | TATGAGTGTGGAGTCGTTGCTCTCAGAATTAGCAGACAAGAGCANNNAGATCGGA |
| 39145_2 | TATGAGTGTGGAGTCGTTGCTCTCAGAATTAGCAGACAAGAGCANNNAGATCGGA |
| 8174_1 | TATGAGTGTGGAGTCGTTGCAAGTGGCATGATCTTTGCTCACGANNNAGATCGGAA |
| 8174_2 | TATGAGTGTGGAGTCGTTGCAAGTGGCATGATCTTTGCTCACGANNNAGATCGGAA |
| 35907_1 | TATGAGTGTGGAGTCGTTGCGGCCCCCACTTAGAAATACCATANNNAGATCGGAAG |
| 35907_2 | TATGAGTGTGGAGTCGTTGCGGCCCCCACTTAGAAATACCATANNNAGATCGGAAG |
| 32800_1 | TATGAGTGTGGAGTCGTTGCCTCACTGCAAGCTCTGTCTCTCNNNAGATCGGAAGA |
| 32800_2 | TATGAGTGTGGAGTCGTTGCCTCACTGCAAGCTCTGTCTCTCNNNAGATCGGAAGA |
| 36350_1 | TATGAGTGTGGAGTCGTTGCAAACCTTGTTTGCAAAGGACCAGNNNAGATCGGAA |
| 36350_2 | TATGAGTGTGGAGTCGTTGCAAACCTTGTTTGCAAAGGACCAGNNNAGATCGGAA |
| 51446_1 | TATGAGTGTGGAGTCGTTGCTTCTCAGTGACCAAAGCTGGAATNNNAGATCGGAAG |
| 51446_2 | TATGAGTGTGGAGTCGTTGCTTCTCAGTGACCAAAGCTGGAATNNNAGATCGGAAG |
| 47541_1 | TATGAGTGTGGAGTCGTTGCGATCTCCACTCACTGCAAGCTCNNNAGATCGGAAGA |
| 47541_2 | TATGAGTGTGGAGTCGTTGCGATCTCCACTCACTGCAAGCTCNNNAGATCGGAAGA |
| 48687_1 | TATGAGTGTGGAGTCGTTGCTCATTCATTCATTCCCCCCACATNNNAGATCGGAAGA |
| 48687_2 | TATGAGTGTGGAGTCGTTGCTCATTCATTCATTCCCCCCACATNNNAGATCGGAAGA |
| 43774_1 | TATGAGTGTGGAGTCGTTGCTAGCTGAGCATTATGTTGTGTGCNNNAGATCGGAA |
| 43774_2 | TATGAGTGTGGAGTCGTTGCTAGCTGAGCATTATGTTGTGTGCNNNAGATCGGAA |
| 42110_1 | TATGAGTGTGGAGTCGTTGCCTGATGTCAGGAGTTTGAGACCANNNAGATCGGAA |
| 42110_2 | TATGAGTGTGGAGTCGTTGCCTGATGTCAGGAGTTTGAGACCANNNAGATCGGAA |
| 29506_1 | TATGAGTGTGGAGTCGTTGCATGATTAATGAAAACTGACTGGGNNNAGATCGGAA |
| 29506_2 | TATGAGTGTGGAGTCGTTGCATGATTAATGAAAACTGACTGGGNNNAGATCGGAA |
| 40467_1 | TATGAGTGTGGAGTCGTTGCAAAGGAGGATGGTAAAGGCAGAGNNNAGATCGGA |
| 40467_2 | TATGAGTGTGGAGTCGTTGCAAAGGAGGATGGTAAAGGCAGAGNNNAGATCGGA |
| 34144_1 | TATGAGTGTGGAGTCGTTGCATTTTAGCCCACTGAGACCCATTNNNAGATCGGAAG |
| 34144_2 | TATGAGTGTGGAGTCGTTGCATTTTAGCCCACTGAGACCCATTNNNAGATCGGAAG |
| 51164_1 | TATGAGTGTGGAGTCGTTGCAGGTGGATATAAGCGGAACTGACNNNAGATCGGAA |
| 51164_2 | TATGAGTGTGGAGTCGTTGCAGGTGGATATAAGCGGAACTGACNNNAGATCGGAA |
| 10275_1 | TATGAGTGTGGAGTCGTTGCACTGGTCACAGGTCTGGATTGTANNNAGATCGGAA |
| 10275_2 | TATGAGTGTGGAGTCGTTGCACTGGTCACAGGTCTGGATTGTANNNAGATCGGAA |

| | |
|---|---|
| 31154_1 | TATGAGTGTGGAGTCGTTGCCATCTGAGAGCCACAAAACCATCNNNAGATCGGAAG |
| 31154_2 | TATGAGTGTGGAGTCGTTGCCATCTGAGAGCCACAAAACCATCNNNAGATCGGAAG |
| 45170_1 | TATGAGTGTGGAGTCGTTGCAAGAGATGGAACCTCATTGCCATNNNAGATCGGAAG |
| 45170_2 | TATGAGTGTGGAGTCGTTGCAAGAGATGGAACCTCATTGCCATNNNAGATCGGAAG |
| 45566_1 | TATGAGTGTGGAGTCGTTGCTTCTTGCCTAAAACCTCCACAGTNNNAGATCGGAAGA |
| 45566_2 | TATGAGTGTGGAGTCGTTGCTTCTTGCCTAAAACCTCCACAGTNNNAGATCGGAAGA |
| 39144_1 | TATGAGTGTGGAGTCGTTGCTAGGGAGTTGTAGTTTTCCGTGGNNNAGATCGGAA |
| 39144_2 | TATGAGTGTGGAGTCGTTGCTAGGGAGTTGTAGTTTTCCGTGGNNNAGATCGGAA |
| 33901_1 | TATGAGTGTGGAGTCGTTGCGATTCAAGCGATTCTCCTGCCTCNNNAGATCGGAAGA |
| 33901_2 | TATGAGTGTGGAGTCGTTGCGATTCAAGCGATTCTCCTGCCTCNNNAGATCGGAAGA |
| 45969_1 | TATGAGTGTGGAGTCGTTGCGGAATAGAGGTGTGAGCCATCATNNNAGATCGGAA |
| 45969_2 | TATGAGTGTGGAGTCGTTGCGGAATAGAGGTGTGAGCCATCATNNNAGATCGGAA |
| 13583_1 | TATGAGTGTGGAGTCGTTGCCTAAACGAATAAAGCAGTTACATNNNAGATCGGAAG |
| 13583_2 | TATGAGTGTGGAGTCGTTGCCTAAACGAATAAAGCAGTTACATNNNAGATCGGAAG |
| 50160_1 | TATGAGTGTGGAGTCGTTGCCTCTTCTGGGACCCTGGATTTTCNNNAGATCGGAAGA |
| 50160_2 | TATGAGTGTGGAGTCGTTGCCTCTTCTGGGACCCTGGATTTTCNNNAGATCGGAAGA |
| 41582_1 | TATGAGTGTGGAGTCGTTGCGTCTCACTCTGTCGCCCATGNNNAGATCGGAAGAGCA |
| 41582_2 | TATGAGTGTGGAGTCGTTGCGTCTCACTCTGTCGCCCATGNNNAGATCGGAAGAGCA |
| 39717_1 | TATGAGTGTGGAGTCGTTGCGAATCTGTGTCCTTGATGGCTTTNNNAGATCGGAAG |
| 39717_2 | TATGAGTGTGGAGTCGTTGCGAATCTGTGTCCTTGATGGCTTTNNNAGATCGGAAG |
| 30242_1 | TATGAGTGTGGAGTCGTTGCCAAAGACAGATGCAAGCCATGATNNNAGATCGGAA |
| 30242_2 | TATGAGTGTGGAGTCGTTGCCAAAGACAGATGCAAGCCATGATNNNAGATCGGAA |
| 13423_1 | TATGAGTGTGGAGTCGTTGCTGTATGTCATTCCTGTTCACNNNAGATCGGAAGAGCA |
| 13423_2 | TATGAGTGTGGAGTCGTTGCTGTATGTCATTCCTGTTCACNNNAGATCGGAAGAGCA |
| 33299_1 | TATGAGTGTGGAGTCGTTGCCAAGTAGCTGGGACTACAGGCNNNAGATCGGAAGA |
| 33299_2 | TATGAGTGTGGAGTCGTTGCCAAGTAGCTGGGACTACAGGCNNNAGATCGGAAGA |
| 3090_1 | TATGAGTGTGGAGTCGTTGCTGTGTAGCTTGAACCCAGGAGGTNNNAGATCGGAA |
| 3090_2 | TATGAGTGTGGAGTCGTTGCTGTGTAGCTTGAACCCAGGAGGTNNNAGATCGGAA |
| 51835_1 | TATGAGTGTGGAGTCGTTGCAGCTGTTCCAATAGGTATGTGGGNNNAGATCGGAA |
| 51835_2 | TATGAGTGTGGAGTCGTTGCAGCTGTTCCAATAGGTATGTGGGNNNAGATCGGAA |
| 47254_1 | TATGAGTGTGGAGTCGTTGCACCACACCCGGCTAATTTTTTTTNNNAGATCGGAAGA |
| 47254_2 | TATGAGTGTGGAGTCGTTGCACCACACCCGGCTAATTTTTTTTNNNAGATCGGAAGA |
| 10287_1 | TATGAGTGTGGAGTCGTTGCGAGTGGACCTGGTAAGAAGGTCANNNAGATCGGAA |
| 10287_2 | TATGAGTGTGGAGTCGTTGCGAGTGGACCTGGTAAGAAGGTCANNNAGATCGGAA |
| 37335_1 | TATGAGTGTGGAGTCGTTGCCCAGGGGCTTTACACAAGGTATANNNAGATCGGAAG |
| 37335_2 | TATGAGTGTGGAGTCGTTGCCCAGGGGCTTTACACAAGGTATANNNAGATCGGAAG |
| 32276_1 | TATGAGTGTGGAGTCGTTGCAAACTAGGAATGTTGCCCCTCATNNNAGATCGGAAG |
| 32276_2 | TATGAGTGTGGAGTCGTTGCAAACTAGGAATGTTGCCCCTCATNNNAGATCGGAAG |
| 42346_1 | TATGAGTGTGGAGTCGTTGCGGAAAGTTTGCTGCATCTTCCTTNNNAGATCGGAAGA |
| 42346_2 | TATGAGTGTGGAGTCGTTGCGGAAAGTTTGCTGCATCTTCCTTNNNAGATCGGAAGA |
| 53307_1 | TATGAGTGTGGAGTCGTTGCAGCAGACGTTCTTCTTCCCTTTCNNNAGATCGGAAGA |
| 53307_2 | TATGAGTGTGGAGTCGTTGCAGCAGACGTTCTTCTTCCCTTTCNNNAGATCGGAAGA |
| 42171_1 | TATGAGTGTGGAGTCGTTGCGTTCAAGAGATTCTCCTGCCTCANNNAGATCGGAAGA |

| | |
|---|---|
| 42171_2 | TATGAGTGTGGAGTCGTTGCGTTCAAGAGATTCTCCTGCCTCANNNAGATCGGAAG |
| 46342_1 | TATGAGTGTGGAGTCGTTGCTCAAGAAACCTAGGGCTCAGAACNNNAGATCGGAAG |
| 46342_2 | TATGAGTGTGGAGTCGTTGCTCAAGAAACCTAGGGCTCAGAACNNNAGATCGGAAG |
| 36424_1 | TATGAGTGTGGAGTCGTTGCAGGCAAACTTGTGAAATCTCAGCNNNAGATCGGAAG |
| 36424_2 | TATGAGTGTGGAGTCGTTGCAGGCAAACTTGTGAAATCTCAGCNNNAGATCGGAAG |
| 51717_1 | TATGAGTGTGGAGTCGTTGCCCCTCTCTCCCCCCTCTCTCNNNAGATCGGAAGAGCAC |
| 51717_2 | TATGAGTGTGGAGTCGTTGCCCCTCTCTCCCCCCTCTCTCNNNAGATCGGAAGAGCAC |
| 48101_1 | TATGAGTGTGGAGTCGTTGCGTTCAAGCAATTCTCATGCCTCANNNAGATCGGAAG |
| 48101_2 | TATGAGTGTGGAGTCGTTGCGTTCAAGCAATTCTCATGCCTCANNNAGATCGGAAG |
| 51666_1 | TATGAGTGTGGAGTCGTTGCAAGGCACATACACTCACAGAGTTNNNAGATCGGAAG |
| 51666_2 | TATGAGTGTGGAGTCGTTGCAAGGCACATACACTCACAGAGTTNNNAGATCGGAAG |
| 37696_1 | TATGAGTGTGGAGTCGTTGCAGAAAGAGGAAGTGCTCCAAACANNNAGATCGGAA |
| 37696_2 | TATGAGTGTGGAGTCGTTGCAGAAAGAGGAAGTGCTCCAAACANNNAGATCGGAA |
| 31434_1 | TATGAGTGTGGAGTCGTTGCACATCTAGGGAATGTCTATGTGTTNNNAGATCGGAA |
| 31434_2 | TATGAGTGTGGAGTCGTTGCACATCTAGGGAATGTCTATGTGTTNNNAGATCGGAA |
| 46930_1 | TATGAGTGTGGAGTCGTTGCTTCCAGGTTCAAGCAATTCTCCTNNNAGATCGGAAG |
| 46930_2 | TATGAGTGTGGAGTCGTTGCTTCCAGGTTCAAGCAATTCTCCTNNNAGATCGGAAG |
| 48202_1 | TATGAGTGTGGAGTCGTTGCGGCAATTCTGGTCTTTTGTGGTTNNNAGATCGGAAG |
| 48202_2 | TATGAGTGTGGAGTCGTTGCGGCAATTCTGGTCTTTTGTGGTTNNNAGATCGGAAG |
| 45237_1 | TATGAGTGTGGAGTCGTTGCTCCATGCTACTCTTTCTTCCGAGNNNAGATCGGAAGA |
| 45237_2 | TATGAGTGTGGAGTCGTTGCTCCATGCTACTCTTTCTTCCGAGNNNAGATCGGAAGA |
| 29802_1 | TATGAGTGTGGAGTCGTTGCCCACCTCGGTTCCATATTCAGACNNNAGATCGGAAG |
| 29802_2 | TATGAGTGTGGAGTCGTTGCCCACCTCGGTTCCATATTCAGACNNNAGATCGGAAG |
| 32447_1 | TATGAGTGTGGAGTCGTTGCCTCATGTGATCCCCCACCTTAGNNNAGATCGGAAGA |
| 32447_2 | TATGAGTGTGGAGTCGTTGCCTCATGTGATCCCCCACCTTAGNNNAGATCGGAAGA |
| 33825_1 | TATGAGTGTGGAGTCGTTGCGTAGTATGTGCCTGTAGTCCCAGNNNAGATCGGAAG |
| 33825_2 | TATGAGTGTGGAGTCGTTGCGTAGTATGTGCCTGTAGTCCCAGNNNAGATCGGAAG |
| 13228_1 | TATGAGTGTGGAGTCGTTGCTTGTTTCTTTGCCTATAATTGTTNNNAGATCGGAAGA |
| 13228_2 | TATGAGTGTGGAGTCGTTGCTTGTTTCTTTGCCTATAATTGTTNNNAGATCGGAAGA |
| 33062_1 | TATGAGTGTGGAGTCGTTGCCAGGAGAGTAGCTGGGATTACAGNNNAGATCGGAA |
| 33062_2 | TATGAGTGTGGAGTCGTTGCCAGGAGAGTAGCTGGGATTACAGNNNAGATCGGAA |
| 33999_1 | TATGAGTGTGGAGTCGTTGCTGATCTTGGCTCACTACAACCTCNNNAGATCGGAAG |
| 33999_2 | TATGAGTGTGGAGTCGTTGCTGATCTTGGCTCACTACAACCTCNNNAGATCGGAAG |
| 46019_1 | TATGAGTGTGGAGTCGTTGCCACGCCCGGCTAATTTTTTGTANNNAGATCGGAAGA |
| 46019_2 | TATGAGTGTGGAGTCGTTGCCACGCCCGGCTAATTTTTTGTANNNAGATCGGAAGA |
| 38575_1 | TATGAGTGTGGAGTCGTTGCGGAAAGAGATGAAAGTGGGGACANNNAGATCGGA |
| 38575_2 | TATGAGTGTGGAGTCGTTGCGGAAAGAGATGAAAGTGGGGACANNNAGATCGGA |
| 31075_1 | TATGAGTGTGGAGTCGTTGCTGCTACAACATGGATGAACCTCANNNAGATCGGAAG |
| 31075_2 | TATGAGTGTGGAGTCGTTGCTGCTACAACATGGATGAACCTCANNNAGATCGGAAG |
| 48834_1 | TATGAGTGTGGAGTCGTTGCGAGTTGAGTCTTCTCTTCTGGCANNNAGATCGGAAG |
| 48834_2 | TATGAGTGTGGAGTCGTTGCGAGTTGAGTCTTCTCTTCTGGCANNNAGATCGGAAG |
| 52884_1 | TATGAGTGTGGAGTCGTTGCCCTACCACGTTGACTTTGAAAGGNNNAGATCGGAAG |
| 52884_2 | TATGAGTGTGGAGTCGTTGCCCTACCACGTTGACTTTGAAAGGNNNAGATCGGAAG |

| | |
|---|---|
| 12965_1 | TATGAGTGTGGAGTCGTTGCCAGTACTATTATGAACCACCATANNNAGATCGGAAG |
| 12965_2 | TATGAGTGTGGAGTCGTTGCCAGTACTATTATGAACCACCATANNNAGATCGGAAG |
| 42358_1 | TATGAGTGTGGAGTCGTTGCCCCCACACATACACACATCTGTTNNNAGATCGGAAGA |
| 42358_2 | TATGAGTGTGGAGTCGTTGCCCCCACACATACACACATCTGTTNNNAGATCGGAAGA |
| 33329_1 | TATGAGTGTGGAGTCGTTGCTCTCCCAGGTTCATGTGATTCTCNNNAGATCGGAAG |
| 33329_2 | TATGAGTGTGGAGTCGTTGCTCTCCCAGGTTCATGTGATTCTCNNNAGATCGGAAG |
| 43426_1 | TATGAGTGTGGAGTCGTTGCACTTTTGTCTTCCGGGAGATACCNNNAGATCGGAAG |
| 43426_2 | TATGAGTGTGGAGTCGTTGCACTTTTGTCTTCCGGGAGATACCNNNAGATCGGAAG |
| 38250_1 | TATGAGTGTGGAGTCGTTGCCATTCTGTGAAGCCAGTATCACCNNNAGATCGGAAG |
| 38250_2 | TATGAGTGTGGAGTCGTTGCCATTCTGTGAAGCCAGTATCACCNNNAGATCGGAAG |
| 31263_1 | TATGAGTGTGGAGTCGTTGCACACACACACACACACACATTTTNNNAGATCGGAAGA |
| 31263_2 | TATGAGTGTGGAGTCGTTGCACACACACACACACACACATTTTNNNAGATCGGAAGA |
| 45947_1 | TATGAGTGTGGAGTCGTTGCCAGACAGGTGCTGAAATCAAAGGNNNAGATCGGAA |
| 45947_2 | TATGAGTGTGGAGTCGTTGCCAGACAGGTGCTGAAATCAAAGGNNNAGATCGGAA |
| 41641_1 | TATGAGTGTGGAGTCGTTGCTTCAAGTGATTCTCCTGCCTCAGNNNAGATCGGAAG |
| 41641_2 | TATGAGTGTGGAGTCGTTGCTTCAAGTGATTCTCCTGCCTCAGNNNAGATCGGAAG |
| 11450_1 | TATGAGTGTGGAGTCGTTGCGGGTTTATGTTGATGGGGAAAAGNNNAGATCGGA |
| 11450_2 | TATGAGTGTGGAGTCGTTGCGGGTTTATGTTGATGGGGAAAAGNNNAGATCGGA |
| 48822_1 | TATGAGTGTGGAGTCGTTGCGAGACGGAGTCTTGCTGTCTCNNNAGATCGGAAGA |
| 48822_2 | TATGAGTGTGGAGTCGTTGCGAGACGGAGTCTTGCTGTCTCNNNAGATCGGAAGA |
| 51229_1 | TATGAGTGTGGAGTCGTTGCTAGCAACAATCACCCAACAGTCTNNNAGATCGGAAG |
| 51229_2 | TATGAGTGTGGAGTCGTTGCTAGCAACAATCACCCAACAGTCTNNNAGATCGGAAG |
| 42994_1 | TATGAGTGTGGAGTCGTTGCCACACATACACACACACACACATNNNAGATCGGAAG |
| 42994_2 | TATGAGTGTGGAGTCGTTGCCACACATACACACACACACACATNNNAGATCGGAAG |
| 2649_1 | TATGAGTGTGGAGTCGTTGCTCCCAAGATCTGTACTGCAGTTGNNNAGATCGGAAG |
| 2649_2 | TATGAGTGTGGAGTCGTTGCTCCCAAGATCTGTACTGCAGTTGNNNAGATCGGAAG |
| 43309_1 | TATGAGTGTGGAGTCGTTGCCATTCTCCTGCCTCAGCCTCNNNAGATCGGAAGAGCA |
| 43309_2 | TATGAGTGTGGAGTCGTTGCCATTCTCCTGCCTCAGCCTCNNNAGATCGGAAGAGCA |
| 50104_1 | TATGAGTGTGGAGTCGTTGCAAGCTTTGTATCTGGGGTCTTCCNNNAGATCGGAAG |
| 50104_2 | TATGAGTGTGGAGTCGTTGCAAGCTTTGTATCTGGGGTCTTCCNNNAGATCGGAAG |
| 43969_1 | TATGAGTGTGGAGTCGTTGCCAGAGGCATCATCATGGCTCANNNAGATCGGAAGA |
| 43969_2 | TATGAGTGTGGAGTCGTTGCCAGAGGCATCATCATGGCTCANNNAGATCGGAAGA |
| 38070_1 | TATGAGTGTGGAGTCGTTGCCAATCAGGGAAAGGGAAACACACNNNAGATCGGAA |
| 38070_2 | TATGAGTGTGGAGTCGTTGCCAATCAGGGAAAGGGAAACACACNNNAGATCGGAA |
| 12524_1 | TATGAGTGTGGAGTCGTTGCCTTTCAAAAGATCCAGTTCTCTNNNAGATCGGAAGA |
| 12524_2 | TATGAGTGTGGAGTCGTTGCCTTTCAAAAGATCCAGTTCTCTNNNAGATCGGAAGA |
| 52565_1 | TATGAGTGTGGAGTCGTTGCATTAGAATCTGTGGCAGGCAAGANNNAGATCGGAA |
| 52565_2 | TATGAGTGTGGAGTCGTTGCATTAGAATCTGTGGCAGGCAAGANNNAGATCGGAA |
| 12533_1 | TATGAGTGTGGAGTCGTTGCTATACTCACAGTACTCACACTNNNAGATCGGAAGAG |
| 12533_2 | TATGAGTGTGGAGTCGTTGCTATACTCACAGTACTCACACTNNNAGATCGGAAGAG |
| 52314_1 | TATGAGTGTGGAGTCGTTGCCTTGTCTGAGTGGAAGGCTGTATNNNAGATCGGAA |
| 52314_2 | TATGAGTGTGGAGTCGTTGCCTTGTCTGAGTGGAAGGCTGTATNNNAGATCGGAA |
| 12617_1 | TATGAGTGTGGAGTCGTTGCAATTCCCAGTCTTCATTTCCNNNAGATCGGAAGAGCA |

| | |
|---|---|
| 12617_2 | TATGAGTGTGGAGTCGTTGCAATTCCCAGTCTTCATTTCCNNNAGATCGGAAGAGCA |
| 13102_1 | TATGAGTGTGGAGTCGTTGCGAACAACCCCTTATACTTAATTGNNNAGATCGGAAG |
| 13102_2 | TATGAGTGTGGAGTCGTTGCGAACAACCCCTTATACTTAATTGNNNAGATCGGAAG |
| 30207_1 | TATGAGTGTGGAGTCGTTGCGACACCACCAGTTTCCTAATTGGNNNAGATCGGAAG |
| 30207_2 | TATGAGTGTGGAGTCGTTGCGACACCACCAGTTTCCTAATTGGNNNAGATCGGAAG |
| 33418_1 | TATGAGTGTGGAGTCGTTGCGCCTGGGTTTGAATTCTGAACTCNNNAGATCGGAAG |
| 33418_2 | TATGAGTGTGGAGTCGTTGCGCCTGGGTTTGAATTCTGAACTCNNNAGATCGGAAG |
| 29464_1 | TATGAGTGTGGAGTCGTTGCACACCCATTAAGCAAAGATAAGAGCANNNAGATCGG |
| 29464_2 | TATGAGTGTGGAGTCGTTGCACACCCATTAAGCAAAGATAAGAGCANNNAGATCGG |
| 50745_1 | TATGAGTGTGGAGTCGTTGCCAAAGGCCAGAGTTTCAGGAAAGNNNAGATCGGAA |
| 50745_2 | TATGAGTGTGGAGTCGTTGCCAAAGGCCAGAGTTTCAGGAAAGNNNAGATCGGAA |
| 42471_1 | TATGAGTGTGGAGTCGTTGCTGCGAGAGAGAGAGAGAGAAAGANNNAGATCGGA |
| 42471_2 | TATGAGTGTGGAGTCGTTGCTGCGAGAGAGAGAGAGAGAAAGANNNAGATCGGA |
| 47211_1 | TATGAGTGTGGAGTCGTTGCCTGGAGTACAATGACGGGATCTCNNNAGATCGGAA |
| 47211_2 | TATGAGTGTGGAGTCGTTGCCTGGAGTACAATGACGGGATCTCNNNAGATCGGAA |
| 29404_1 | TATGAGTGTGGAGTCGTTGCCCTACTCCCCTGCCTTCCTAGNNNAGATCGGAAGAGC |
| 29404_2 | TATGAGTGTGGAGTCGTTGCCCTACTCCCCTGCCTTCCTAGNNNAGATCGGAAGAGC |
| 36229_1 | TATGAGTGTGGAGTCGTTGCAAACGCATCATGGCAAAATCTCTNNNAGATCGGAAG |
| 36229_2 | TATGAGTGTGGAGTCGTTGCAAACGCATCATGGCAAAATCTCTNNNAGATCGGAAG |
| 33047_1 | TATGAGTGTGGAGTCGTTGCCTCACTGCAGTCTTGAATTCCTGNNNAGATCGGAAG |
| 33047_2 | TATGAGTGTGGAGTCGTTGCCTCACTGCAGTCTTGAATTCCTGNNNAGATCGGAAG |
| 36099_1 | TATGAGTGTGGAGTCGTTGCGAGGGACTGCTAGATCAATTCGTNNNAGATCGGAA |
| 36099_2 | TATGAGTGTGGAGTCGTTGCGAGGGACTGCTAGATCAATTCGTNNNAGATCGGAA |
| 32110_1 | TATGAGTGTGGAGTCGTTGCCAAGTAGCTGGGACTACAGGTGNNNAGATCGGAAG |
| 32110_2 | TATGAGTGTGGAGTCGTTGCCAAGTAGCTGGGACTACAGGTGNNNAGATCGGAAG |
| 40042_1 | TATGAGTGTGGAGTCGTTGCACCACTCTAACCTCTGGTTTCTTNNNAGATCGGAAGA |
| 40042_2 | TATGAGTGTGGAGTCGTTGCACCACTCTAACCTCTGGTTTCTTNNNAGATCGGAAGA |
| 46713_1 | TATGAGTGTGGAGTCGTTGCCAGTTGACATTTCAAGGCCCTTTNNNAGATCGGAAG |
| 46713_2 | TATGAGTGTGGAGTCGTTGCCAGTTGACATTTCAAGGCCCTTTNNNAGATCGGAAG |
| 48553_1 | TATGAGTGTGGAGTCGTTGCATGATCTTGGCTCACTGAAACCTNNNAGATCGGAAG |
| 48553_2 | TATGAGTGTGGAGTCGTTGCATGATCTTGGCTCACTGAAACCTNNNAGATCGGAAG |
| 51706_1 | TATGAGTGTGGAGTCGTTGCTTTGTCTCTACATCAACTGCCCANNNAGATCGGAAGA |
| 51706_2 | TATGAGTGTGGAGTCGTTGCTTTGTCTCTACATCAACTGCCCANNNAGATCGGAAGA |
| 34591_1 | TATGAGTGTGGAGTCGTTGCCATGCTAGGTGGATATCTGCTGTNNNAGATCGGAA |
| 34591_2 | TATGAGTGTGGAGTCGTTGCCATGCTAGGTGGATATCTGCTGTNNNAGATCGGAA |
| 32205_1 | TATGAGTGTGGAGTCGTTGCGCTGTGTGATAGTTCCAGGAATTNNNAGATCGGAA |
| 32205_2 | TATGAGTGTGGAGTCGTTGCGCTGTGTGATAGTTCCAGGAATTNNNAGATCGGAA |
| 46880_1 | TATGAGTGTGGAGTCGTTGCCAACCTCTATGTCCTGGGTTCAANNNAGATCGGAAG |
| 46880_2 | TATGAGTGTGGAGTCGTTGCCAACCTCTATGTCCTGGGTTCAANNNAGATCGGAAG |
| 30819_1 | TATGAGTGTGGAGTCGTTGCACAAGATCCACCTACACGTACACNNNAGATCGGAAG |
| 30819_2 | TATGAGTGTGGAGTCGTTGCACAAGATCCACCTACACGTACACNNNAGATCGGAAG |
| 40758_1 | TATGAGTGTGGAGTCGTTGCAAAGAAAAGGGAGTACAGGGAGCNNNAGATCGGA |
| 40758_2 | TATGAGTGTGGAGTCGTTGCAAAGAAAAGGGAGTACAGGGAGCNNNAGATCGGA |

| | |
|---|---|
| 50857_1 | TATGAGTGTGGAGTCGTTGCTGGTCTCTATTACTGGCTCCTCANNNAGATCGGAAGA |
| 50857_2 | TATGAGTGTGGAGTCGTTGCTGGTCTCTATTACTGGCTCCTCANNNAGATCGGAAGA |
| 41497_1 | TATGAGTGTGGAGTCGTTGCAAACTCTGAACAATGCCTGAAGCNNNAGATCGGAAG |
| 41497_2 | TATGAGTGTGGAGTCGTTGCAAACTCTGAACAATGCCTGAAGCNNNAGATCGGAAG |
| 37712_1 | TATGAGTGTGGAGTCGTTGCCTTGAACTCAGAAGGCAGAGGTTNNNAGATCGGAA |
| 37712_2 | TATGAGTGTGGAGTCGTTGCCTTGAACTCAGAAGGCAGAGGTTNNNAGATCGGAA |
| 31111_1 | TATGAGTGTGGAGTCGTTGCACTGGCTCCACATTTTTGCAATTNNNAGATCGGAAGA |
| 31111_2 | TATGAGTGTGGAGTCGTTGCACTGGCTCCACATTTTTGCAATTNNNAGATCGGAAGA |
| 37616_1 | TATGAGTGTGGAGTCGTTGCGAGATGGATTTTCACCCTTGTCGNNNAGATCGGAAG |
| 37616_2 | TATGAGTGTGGAGTCGTTGCGAGATGGATTTTCACCCTTGTCGNNNAGATCGGAAG |
| 37823_1 | TATGAGTGTGGAGTCGTTGCGCTCATGCCTGTAATACCAACACNNNAGATCGGAAGA |
| 37823_2 | TATGAGTGTGGAGTCGTTGCGCTCATGCCTGTAATACCAACACNNNAGATCGGAAGA |
| 42030_1 | TATGAGTGTGGAGTCGTTGCTACAAAAAGTAGCTGGGCATGGTNNNAGATCGGAA |
| 42030_2 | TATGAGTGTGGAGTCGTTGCTACAAAAAGTAGCTGGGCATGGTNNNAGATCGGAA |
| 10123_1 | TATGAGTGTGGAGTCGTTGCATACAGTGAAGGATTTCACAGCTNNNAGATCGGAAG |
| 10123_2 | TATGAGTGTGGAGTCGTTGCATACAGTGAAGGATTTCACAGCTNNNAGATCGGAAG |
| 44610_1 | TATGAGTGTGGAGTCGTTGCCATTGAGGATACCGGAGGAAGAGNNNAGATCGGAA |
| 44610_2 | TATGAGTGTGGAGTCGTTGCCATTGAGGATACCGGAGGAAGAGNNNAGATCGGAA |
| 43437_1 | TATGAGTGTGGAGTCGTTGCCTCGAACTCCTGACCTCATGATCNNNAGATCGGAAGA |
| 43437_2 | TATGAGTGTGGAGTCGTTGCCTCGAACTCCTGACCTCATGATCNNNAGATCGGAAGA |
| 52075_1 | TATGAGTGTGGAGTCGTTGCCTGAACAGAACTTGGCTCCTTTGNNNAGATCGGAAG |
| 52075_2 | TATGAGTGTGGAGTCGTTGCCTGAACAGAACTTGGCTCCTTTGNNNAGATCGGAAG |
| 29839_1 | TATGAGTGTGGAGTCGTTGCTGTGGACAAACATGCTACTTCCTNNNAGATCGGAAG |
| 29839_2 | TATGAGTGTGGAGTCGTTGCTGTGGACAAACATGCTACTTCCTNNNAGATCGGAAG |
| 30976_1 | TATGAGTGTGGAGTCGTTGCAAGCCTGATAAAACCTTCCAACTNNNAGATCGGAAGA |
| 30976_2 | TATGAGTGTGGAGTCGTTGCAAGCCTGATAAAACCTTCCAACTNNNAGATCGGAAGA |
| 28887_1 | TATGAGTGTGGAGTCGTTGCTGCCTGGCTAATTACTACTCACCNNNAGATCGGAAGA |
| 28887_2 | TATGAGTGTGGAGTCGTTGCTGCCTGGCTAATTACTACTCACCNNNAGATCGGAAGA |
| 43113_1 | TATGAGTGTGGAGTCGTTGCACACAAAACTCTCCCTCCCAAATNNNAGATCGGAAGA |
| 43113_2 | TATGAGTGTGGAGTCGTTGCACACAAAACTCTCCCTCCCAAATNNNAGATCGGAAGA |
| 3186_1 | TATGAGTGTGGAGTCGTTGCCTTGCAGCAGATAAGGGCACTTANNNAGATCGGAA |
| 3186_2 | TATGAGTGTGGAGTCGTTGCCTTGCAGCAGATAAGGGCACTTANNNAGATCGGAA |
| 50666_1 | TATGAGTGTGGAGTCGTTGCCCTTTGTCTCTGACCACAGCTATNNNAGATCGGAAGA |
| 50666_2 | TATGAGTGTGGAGTCGTTGCCCTTTGTCTCTGACCACAGCTATNNNAGATCGGAAGA |
| 53350_1 | TATGAGTGTGGAGTCGTTGCTGTCTCTCTCTGTGTCTCTACGTNNNAGATCGGAAGA |
| 53350_2 | TATGAGTGTGGAGTCGTTGCTGTCTCTCTCTGTGTCTCTACGTNNNAGATCGGAAGA |
| 51544_1 | TATGAGTGTGGAGTCGTTGCCAGGACTATGCACCTCCAATCTTNNNAGATCGGAAGA |
| 51544_2 | TATGAGTGTGGAGTCGTTGCCAGGACTATGCACCTCCAATCTTNNNAGATCGGAAGA |
| 35895_1 | TATGAGTGTGGAGTCGTTGCAGATGCCAAGACCTCTGCTAAAANNNAGATCGGAAG |
| 35895_2 | TATGAGTGTGGAGTCGTTGCAGATGCCAAGACCTCTGCTAAAANNNAGATCGGAAG |
| 51257_1 | TATGAGTGTGGAGTCGTTGCGCGGACTCTGACACTTGTTAGTNNNAGATCGGAAGA |
| 51257_2 | TATGAGTGTGGAGTCGTTGCGCGGACTCTGACACTTGTTAGTNNNAGATCGGAAGA |
| 43734_1 | TATGAGTGTGGAGTCGTTGCACCAGAGATGAAAAAGGATCCCCNNNAGATCGGAA |

| | |
|---|---|
| 43734_2 | TATGAGTGTGGAGTCGTTGCACCAGAGATGAAAAAGGATCCCCNNNAGATCGGAA |
| 43869_1 | TATGAGTGTGGAGTCGTTGCAAGTGCTTTATGTTGCAATGGCTNNNAGATCGGAAG |
| 43869_2 | TATGAGTGTGGAGTCGTTGCAAGTGCTTTATGTTGCAATGGCTNNNAGATCGGAAG |
| 29771_1 | TATGAGTGTGGAGTCGTTGCGCTGCAGCCACAGTTCCTATAAANNNAGATCGGAAG |
| 29771_2 | TATGAGTGTGGAGTCGTTGCGCTGCAGCCACAGTTCCTATAAANNNAGATCGGAAG |
| 37971_1 | TATGAGTGTGGAGTCGTTGCAGACTGAGACTCTTGTTGAAGCCNNNAGATCGGAAG |
| 37971_2 | TATGAGTGTGGAGTCGTTGCAGACTGAGACTCTTGTTGAAGCCNNNAGATCGGAAG |
| 30619_1 | TATGAGTGTGGAGTCGTTGCACACTTTGCTGACATTCCAAAGCNNNAGATCGGAAG |
| 30619_2 | TATGAGTGTGGAGTCGTTGCACACTTTGCTGACATTCCAAAGCNNNAGATCGGAAG |
| 41026_1 | TATGAGTGTGGAGTCGTTGCGGGGCTTTGTACCACACTGAATANNNAGATCGGAAG |
| 41026_2 | TATGAGTGTGGAGTCGTTGCGGGGCTTTGTACCACACTGAATANNNAGATCGGAAG |
| 50642_1 | TATGAGTGTGGAGTCGTTGCTGCTGTTAGGAGAGTGTTGTACCNNNAGATCGGAA |
| 50642_2 | TATGAGTGTGGAGTCGTTGCTGCTGTTAGGAGAGTGTTGTACCNNNAGATCGGAA |
| 43321_1 | TATGAGTGTGGAGTCGTTGCCAGAGACAGGTTTTTGCCATGTTNNNAGATCGGAAG |
| 43321_2 | TATGAGTGTGGAGTCGTTGCCAGAGACAGGTTTTTGCCATGTTNNNAGATCGGAAG |
| 10068_1 | TATGAGTGTGGAGTCGTTGCCCGCCAGGGTTTGTTTGTTTTAGNNNAGATCGGAAG |
| 10068_2 | TATGAGTGTGGAGTCGTTGCCCGCCAGGGTTTGTTTGTTTTAGNNNAGATCGGAAG |
| 13790_1 | TATGAGTGTGGAGTCGTTGCCTACCATCCTTAGTTTGATAGAGNNNAGATCGGAAG |
| 13790_2 | TATGAGTGTGGAGTCGTTGCCTACCATCCTTAGTTTGATAGAGNNNAGATCGGAAG |
| 32882_1 | TATGAGTGTGGAGTCGTTGCCCATGATGAGACTATCAGCACCANNNAGATCGGAAG |
| 32882_2 | TATGAGTGTGGAGTCGTTGCCCATGATGAGACTATCAGCACCANNNAGATCGGAAG |
| 39458_1 | TATGAGTGTGGAGTCGTTGCAGGACACAATCCCATTCCCAATTNNNAGATCGGAAG |
| 39458_2 | TATGAGTGTGGAGTCGTTGCAGGACACAATCCCATTCCCAATTNNNAGATCGGAAG |
| 43875_1 | TATGAGTGTGGAGTCGTTGCCTTAGTATTATGGCCGGCAAAGCNNNAGATCGGAAG |
| 43875_2 | TATGAGTGTGGAGTCGTTGCCTTAGTATTATGGCCGGCAAAGCNNNAGATCGGAAG |
| 4784_1 | TATGAGTGTGGAGTCGTTGCGCAGAGCTTTTTCGACACTAACCNNNAGATCGGAAG |
| 4784_2 | TATGAGTGTGGAGTCGTTGCGCAGAGCTTTTTCGACACTAACCNNNAGATCGGAAG |
| 36868_1 | TATGAGTGTGGAGTCGTTGCAAGAACCCCCAAATTCCTTAGCTNNNAGATCGGAAG |
| 36868_2 | TATGAGTGTGGAGTCGTTGCAAGAACCCCCAAATTCCTTAGCTNNNAGATCGGAAG |
| 13728_1 | TATGAGTGTGGAGTCGTTGCATTGTTGTATCACCAGTTAGTTNNNAGATCGGAAGA |
| 13728_2 | TATGAGTGTGGAGTCGTTGCATTGTTGTATCACCAGTTAGTTNNNAGATCGGAAGA |
| 36973_1 | TATGAGTGTGGAGTCGTTGCTCTGCAAAAACAGAAACACCTGANNNAGATCGGAAG |
| 36973_2 | TATGAGTGTGGAGTCGTTGCTCTGCAAAAACAGAAACACCTGANNNAGATCGGAAG |
| 52947_1 | TATGAGTGTGGAGTCGTTGCACCTGCAGTTTCTCCACATGTATNNNAGATCGGAAG |
| 52947_2 | TATGAGTGTGGAGTCGTTGCACCTGCAGTTTCTCCACATGTATNNNAGATCGGAAG |
| 41776_1 | TATGAGTGTGGAGTCGTTGCCTCCTGGGTTCAAGCGATTCTCNNNAGATCGGAAGA |
| 41776_2 | TATGAGTGTGGAGTCGTTGCCTCCTGGGTTCAAGCGATTCTCNNNAGATCGGAAGA |
| 37945_1 | TATGAGTGTGGAGTCGTTGCAGAACTCTAGCAACTAGCAGCAGNNNAGATCGGAA |
| 37945_2 | TATGAGTGTGGAGTCGTTGCAGAACTCTAGCAACTAGCAGCAGNNNAGATCGGAA |
| 9843_1 | TATGAGTGTGGAGTCGTTGCGGAGAATGACAAAAGACAGCTGANNNAGATCGGAA |
| 9843_2 | TATGAGTGTGGAGTCGTTGCGGAGAATGACAAAAGACAGCTGANNNAGATCGGAA |
| 34829_1 | TATGAGTGTGGAGTCGTTGCCCAAGTAGCTGGGATGACAGGNNNAGATCGGAAGA |
| 34829_2 | TATGAGTGTGGAGTCGTTGCCCAAGTAGCTGGGATGACAGGNNNAGATCGGAAGA |

42402_1 TATGAGTGTGGAGTCGTTGCATGGCTAGAAATGTCACTCCTTTNNNAGATCGGAAG
42402_2 TATGAGTGTGGAGTCGTTGCATGGCTAGAAATGTCACTCCTTTNNNAGATCGGAAG
31740_1 TATGAGTGTGGAGTCGTTGCGTGGCCACTGAGTTTTCTGTTTTNNNAGATCGGAAG
31740_2 TATGAGTGTGGAGTCGTTGCGTGGCCACTGAGTTTTCTGTTTTNNNAGATCGGAAG
40875_1 TATGAGTGTGGAGTCGTTGCGTGGTAAGCTGAAGACGTAGGAANNNAGATCGGA
40875_2 TATGAGTGTGGAGTCGTTGCGTGGTAAGCTGAAGACGTAGGAANNNAGATCGGA
10354_1 TATGAGTGTGGAGTCGTTGCGGAGGAATGAAAGCCATGCAAACNNNAGATCGGAA
10354_2 TATGAGTGTGGAGTCGTTGCGGAGGAATGAAAGCCATGCAAACNNNAGATCGGAA
52715_1 TATGAGTGTGGAGTCGTTGCCTTCAGGACAAGCTTTGTGTGGNNNAGATCGGAAG
52715_2 TATGAGTGTGGAGTCGTTGCCTTCAGGACAAGCTTTGTGTGGNNNAGATCGGAAG
32569_1 TATGAGTGTGGAGTCGTTGCCTGAGTAGCTGGGATTACAGGTGNNNAGATCGGAA
32569_2 TATGAGTGTGGAGTCGTTGCCTGAGTAGCTGGGATTACAGGTGNNNAGATCGGAA
13403_1 TATGAGTGTGGAGTCGTTGCTTTTGGTCTTGTTTCAATTTCTTNNNAGATCGGAAGA
13403_2 TATGAGTGTGGAGTCGTTGCTTTTGGTCTTGTTTCAATTTCTTNNNAGATCGGAAGA
35438_1 TATGAGTGTGGAGTCGTTGCCTCGCTCTCCTGAGTAATTGGAANNNAGATCGGAAG
35438_2 TATGAGTGTGGAGTCGTTGCCTCGCTCTCCTGAGTAATTGGAANNNAGATCGGAAG
36283_1 TATGAGTGTGGAGTCGTTGCAGTAGTTACACCTGGAGAGGGAANNNAGATCGGAA
36283_2 TATGAGTGTGGAGTCGTTGCAGTAGTTACACCTGGAGAGGGAANNNAGATCGGAA
29620_1 TATGAGTGTGGAGTCGTTGCCAGCAAGGAGGCATTTGAATGTGNNNAGATCGGAA
29620_2 TATGAGTGTGGAGTCGTTGCCAGCAAGGAGGCATTTGAATGTGNNNAGATCGGAA
33408_1 TATGAGTGTGGAGTCGTTGCCTGTAATCCCAGCACTTTCAGGANNNAGATCGGAAG
33408_2 TATGAGTGTGGAGTCGTTGCCTGTAATCCCAGCACTTTCAGGANNNAGATCGGAAG
49961_1 TATGAGTGTGGAGTCGTTGCAGGGAAAATCCATCCTCAGACACNNNAGATCGGAAG
49961_2 TATGAGTGTGGAGTCGTTGCAGGGAAAATCCATCCTCAGACACNNNAGATCGGAAG
49618_1 TATGAGTGTGGAGTCGTTGCGCCACTCTGTATCTTTTGATTGGTNNNAGATCGGAA
49618_2 TATGAGTGTGGAGTCGTTGCGCCACTCTGTATCTTTTGATTGGTNNNAGATCGGAA
35966_1 TATGAGTGTGGAGTCGTTGCTCTCTCTCTGGGGAAGTTTTGTGNNNAGATCGGAAG
35966_2 TATGAGTGTGGAGTCGTTGCTCTCTCTCTGGGGAAGTTTTGTGNNNAGATCGGAAG
40453_1 TATGAGTGTGGAGTCGTTGCATAATTTCAAGGCAGAGCAGCTGNNNAGATCGGAA
40453_2 TATGAGTGTGGAGTCGTTGCATAATTTCAAGGCAGAGCAGCTGNNNAGATCGGAA
52431_1 TATGAGTGTGGAGTCGTTGCTGGACTCTAATTCTTACCGCAGGNNNAGATCGGAAG
52431_2 TATGAGTGTGGAGTCGTTGCTGGACTCTAATTCTTACCGCAGGNNNAGATCGGAAG
13067_1 TATGAGTGTGGAGTCGTTGCGAAGAAGAGAGTCAGAGTCANNNAGATCGGAAGA
13067_2 TATGAGTGTGGAGTCGTTGCGAAGAAGAGAGTCAGAGTCANNNAGATCGGAAGA
12170_1 TATGAGTGTGGAGTCGTTGCTTCTCTCCGGCGGATACGGCANNNAGATCGGAAGAG
12170_2 TATGAGTGTGGAGTCGTTGCTTCTCTCCGGCGGATACGGCANNNAGATCGGAAGAG
13061_1 TATGAGTGTGGAGTCGTTGCCTAATTCAGAATCATTTTGTGGANNNAGATCGGAAG
13061_2 TATGAGTGTGGAGTCGTTGCCTAATTCAGAATCATTTTGTGGANNNAGATCGGAAG
29490_1 TATGAGTGTGGAGTCGTTGCTGACTGCTGATGTTCTTTCTGCNNNAGATCGGAAGA
29490_2 TATGAGTGTGGAGTCGTTGCTGACTGCTGATGTTCTTTCTGCNNNAGATCGGAAGA
50887_1 TATGAGTGTGGAGTCGTTGCGGTTACTTCATGTATGCACAGCCNNNAGATCGGAAG
50887_2 TATGAGTGTGGAGTCGTTGCGGTTACTTCATGTATGCACAGCCNNNAGATCGGAAG
37164_1 TATGAGTGTGGAGTCGTTGCGGACCTGAGGTGAGGGAATAAAGNNNAGATCGGA

| | |
|---|---|
| 37164_2 | TATGAGTGTGGAGTCGTTGCGGACCTGAGGTGAGGGAATAAAGNNNAGATCGGA |
| 49092_1 | TATGAGTGTGGAGTCGTTGCAACTTCTGCCTCCTGGATTCAAGNNNAGATCGGAAG |
| 49092_2 | TATGAGTGTGGAGTCGTTGCAACTTCTGCCTCCTGGATTCAAGNNNAGATCGGAAG |
| 32005_1 | TATGAGTGTGGAGTCGTTGCGAGACAGAGATACCGTTCCCATCNNNAGATCGGAAG |
| 32005_2 | TATGAGTGTGGAGTCGTTGCGAGACAGAGATACCGTTCCCATCNNNAGATCGGAAG |
| 43629_1 | TATGAGTGTGGAGTCGTTGCAAGCAATTCCTCCTGTAGACCTCNNNAGATCGGAAG |
| 43629_2 | TATGAGTGTGGAGTCGTTGCAAGCAATTCCTCCTGTAGACCTCNNNAGATCGGAAG |
| 40558_1 | TATGAGTGTGGAGTCGTTGCGGGTTGTGAAGACTATGGGACAANNNAGATCGGAA |
| 40558_2 | TATGAGTGTGGAGTCGTTGCGGGTTGTGAAGACTATGGGACAANNNAGATCGGAA |
| 48999_1 | TATGAGTGTGGAGTCGTTGCCATTCTCCTGCCTCAGCCTCNNNAGATCGGAAGAGCA |
| 48999_2 | TATGAGTGTGGAGTCGTTGCCATTCTCCTGCCTCAGCCTCNNNAGATCGGAAGAGCA |
| 39033_1 | TATGAGTGTGGAGTCGTTGCTCAAACTCCTAGGCTCAAGCAATNNNAGATCGGAAG |
| 39033_2 | TATGAGTGTGGAGTCGTTGCTCAAACTCCTAGGCTCAAGCAATNNNAGATCGGAAG |
| 45694_1 | TATGAGTGTGGAGTCGTTGCTTCCAGTGATTCTTGTGCCTCAGNNNAGATCGGAAG |
| 45694_2 | TATGAGTGTGGAGTCGTTGCTTCCAGTGATTCTTGTGCCTCAGNNNAGATCGGAAG |
| 40148_1 | TATGAGTGTGGAGTCGTTGCCTCCCGGGTTCAAGTGATTCTCNNNAGATCGGAAGA |
| 40148_2 | TATGAGTGTGGAGTCGTTGCCTCCCGGGTTCAAGTGATTCTCNNNAGATCGGAAGA |
| 52503_1 | TATGAGTGTGGAGTCGTTGCGATCCTTTCCTGTCCCGTGTGNNNAGATCGGAAGAG |
| 52503_2 | TATGAGTGTGGAGTCGTTGCGATCCTTTCCTGTCCCGTGTGNNNAGATCGGAAGAG |
| 41102_1 | TATGAGTGTGGAGTCGTTGCGAGAGAGAGAGAGACAGGGTCTTNNNAGATCGGA |
| 41102_2 | TATGAGTGTGGAGTCGTTGCGAGAGAGAGAGAGACAGGGTCTTNNNAGATCGGA |
| 29468_1 | TATGAGTGTGGAGTCGTTGCGTCATTTTAAAGTCCTTGCCCATNNNAGATCGGAAGA |
| 29468_2 | TATGAGTGTGGAGTCGTTGCGTCATTTTAAAGTCCTTGCCCATNNNAGATCGGAAGA |
| 31320_1 | TATGAGTGTGGAGTCGTTGCGATGCCTTTGCTTTTTGTCTTGCNNNAGATCGGAAGA |
| 31320_2 | TATGAGTGTGGAGTCGTTGCGATGCCTTTGCTTTTTGTCTTGCNNNAGATCGGAAGA |
| 50599_1 | TATGAGTGTGGAGTCGTTGCCTGTTCAGTGGTGATAGCAATGCNNNAGATCGGAA |
| 50599_2 | TATGAGTGTGGAGTCGTTGCCTGTTCAGTGGTGATAGCAATGCNNNAGATCGGAA |
| 44932_1 | TATGAGTGTGGAGTCGTTGCCAACTCCTGTTCCCCATTCTGTANNNAGATCGGAAGA |
| 44932_2 | TATGAGTGTGGAGTCGTTGCCAACTCCTGTTCCCCATTCTGTANNNAGATCGGAAGA |
| 10941_1 | TATGAGTGTGGAGTCGTTGCCCGTGACAAACTAAACGGGAATNNNAGATCGGAAG |
| 10941_2 | TATGAGTGTGGAGTCGTTGCCCGTGACAAACTAAACGGGAATNNNAGATCGGAAG |
| 30725_1 | TATGAGTGTGGAGTCGTTGCCAGCCTTTACCTCACGGTCTCGNNNAGATCGGAAGA |
| 30725_2 | TATGAGTGTGGAGTCGTTGCCAGCCTTTACCTCACGGTCTCGNNNAGATCGGAAGA |
| 34666_1 | TATGAGTGTGGAGTCGTTGCGGTGACAGAGTGAAACTCCATCTNNNAGATCGGAA |
| 34666_2 | TATGAGTGTGGAGTCGTTGCGGTGACAGAGTGAAACTCCATCTNNNAGATCGGAA |
| 30468_1 | TATGAGTGTGGAGTCGTTGCTGCTAAACAAAGAAGGGAGGGANNNAGATCGGAA |
| 30468_2 | TATGAGTGTGGAGTCGTTGCTGCTAAACAAAGAAGGGAGGGANNNAGATCGGAA |
| 29466_1 | TATGAGTGTGGAGTCGTTGCGGCTGGTAAAGTGTATGGATGGGNNNAGATCGGA |
| 29466_2 | TATGAGTGTGGAGTCGTTGCGGCTGGTAAAGTGTATGGATGGGNNNAGATCGGA |
| 31661_1 | TATGAGTGTGGAGTCGTTGCTCCTTTAGTCCCAGGACCAATTGNNNAGATCGGAAG |
| 31661_2 | TATGAGTGTGGAGTCGTTGCTCCTTTAGTCCCAGGACCAATTGNNNAGATCGGAAG |
| 41058_1 | TATGAGTGTGGAGTCGTTGCTAACAGAATAGTCTAGGGCGCTGNNNAGATCGGAA |
| 41058_2 | TATGAGTGTGGAGTCGTTGCTAACAGAATAGTCTAGGGCGCTGNNNAGATCGGAA |

| | |
|---|---|
| 51085_1 | TATGAGTGTGGAGTCGTTGCCGTTCCTGGGTTATCATGTGAGANNNAGATCGGAAG |
| 51085_2 | TATGAGTGTGGAGTCGTTGCCGTTCCTGGGTTATCATGTGAGANNNAGATCGGAAG |
| 53276_1 | TATGAGTGTGGAGTCGTTGCGGAAACTTTGCCTTCCTGAGAGANNNAGATCGGAAG |
| 53276_2 | TATGAGTGTGGAGTCGTTGCGGAAACTTTGCCTTCCTGAGAGANNNAGATCGGAAG |
| 36139_1 | TATGAGTGTGGAGTCGTTGCCAGATATGGCATTCAAGGAAGCGNNNAGATCGGAA |
| 36139_2 | TATGAGTGTGGAGTCGTTGCCAGATATGGCATTCAAGGAAGCGNNNAGATCGGAA |
| 32726_1 | TATGAGTGTGGAGTCGTTGCGGATCACTTGAGGTCAGGAGTTTNNNAGATCGGAA |
| 32726_2 | TATGAGTGTGGAGTCGTTGCGGATCACTTGAGGTCAGGAGTTTNNNAGATCGGAA |
| 39592_1 | TATGAGTGTGGAGTCGTTGCGGAAAAAAGAAAGCTCATCCGGGNNNAGATCGGAA |
| 39592_2 | TATGAGTGTGGAGTCGTTGCGGAAAAAAGAAAGCTCATCCGGGNNNAGATCGGAA |
| 40371_1 | TATGAGTGTGGAGTCGTTGCGGACAGACTATCTAGGCTTGTCCNNNAGATCGGAAG |
| 40371_2 | TATGAGTGTGGAGTCGTTGCGGACAGACTATCTAGGCTTGTCCNNNAGATCGGAAG |
| 47133_1 | TATGAGTGTGGAGTCGTTGCTTTGTGACATTGAGAACATGCCANNNAGATCGGAAG |
| 47133_2 | TATGAGTGTGGAGTCGTTGCTTTGTGACATTGAGAACATGCCANNNAGATCGGAAG |
| 32175_1 | TATGAGTGTGGAGTCGTTGCAGAAGAGTTCTCACTGTGTTGCCNNNAGATCGGAAG |
| 32175_2 | TATGAGTGTGGAGTCGTTGCAGAAGAGTTCTCACTGTGTTGCCNNNAGATCGGAAG |
| 37966_1 | TATGAGTGTGGAGTCGTTGCCTGGAGAACTGGCGAAATTTGAGNNNAGATCGGAA |
| 37966_2 | TATGAGTGTGGAGTCGTTGCCTGGAGAACTGGCGAAATTTGAGNNNAGATCGGAA |
| 45743_1 | TATGAGTGTGGAGTCGTTGCGGTTGCAAAAACCTTAGAGGGACNNNAGATCGGAA |
| 45743_2 | TATGAGTGTGGAGTCGTTGCGGTTGCAAAAACCTTAGAGGGACNNNAGATCGGAA |
| 37968_1 | TATGAGTGTGGAGTCGTTGCATTCCCTGGCAAAGAAAATTCCCNNNAGATCGGAAG |
| 37968_2 | TATGAGTGTGGAGTCGTTGCATTCCCTGGCAAAGAAAATTCCCNNNAGATCGGAAG |
| 41975_1 | TATGAGTGTGGAGTCGTTGCCCTCCTGAGTAGCTGGGACTANNNAGATCGGAAGA |
| 41975_2 | TATGAGTGTGGAGTCGTTGCCCTCCTGAGTAGCTGGGACTANNNAGATCGGAAGA |
| 28761_1 | TATGAGTGTGGAGTCGTTGCTTAGTACAGACGAGGTTTTGCCANNNAGATCGGAA |
| 28761_2 | TATGAGTGTGGAGTCGTTGCTTAGTACAGACGAGGTTTTGCCANNNAGATCGGAA |
| 30527_1 | TATGAGTGTGGAGTCGTTGCGTTATCTGTCTCTAGCTCCAGAGGTTCNNNAGATCG |
| 30527_2 | TATGAGTGTGGAGTCGTTGCGTTATCTGTCTCTAGCTCCAGAGGTTCNNNAGATCG |
| 12829_1 | TATGAGTGTGGAGTCGTTGCACTCCACTGCTTATTTCATCNNNAGATCGGAAGAGCA |
| 12829_2 | TATGAGTGTGGAGTCGTTGCACTCCACTGCTTATTTCATCNNNAGATCGGAAGAGCA |
| 47878_1 | TATGAGTGTGGAGTCGTTGCGAGTAGCTGGGATTACAGGAGTGNNNAGATCGGA |
| 47878_2 | TATGAGTGTGGAGTCGTTGCGAGTAGCTGGGATTACAGGAGTGNNNAGATCGGA |
| 50579_1 | TATGAGTGTGGAGTCGTTGCAAAGCTAGGATTTTGCCTTTCCCNNNAGATCGGAAG |
| 50579_2 | TATGAGTGTGGAGTCGTTGCAAAGCTAGGATTTTGCCTTTCCCNNNAGATCGGAAG |
| 52101_1 | TATGAGTGTGGAGTCGTTGCTGCTAATCCCTTGTCTCAATTGANNNAGATCGGAAG |
| 52101_2 | TATGAGTGTGGAGTCGTTGCTGCTAATCCCTTGTCTCAATTGANNNAGATCGGAAG |
| 32855_1 | TATGAGTGTGGAGTCGTTGCCTTGATCTCGGCTCACTGCAANNNAGATCGGAAGAG |
| 32855_2 | TATGAGTGTGGAGTCGTTGCCTTGATCTCGGCTCACTGCAANNNAGATCGGAAGAG |
| 42985_1 | TATGAGTGTGGAGTCGTTGCACCTTGGACCATAACTTGATGCTNNNAGATCGGAAG |
| 42985_2 | TATGAGTGTGGAGTCGTTGCACCTTGGACCATAACTTGATGCTNNNAGATCGGAAG |
| 35405_1 | TATGAGTGTGGAGTCGTTGCCATTCTCCTGCCTCAGCCTCNNNAGATCGGAAGAGCA |
| 35405_2 | TATGAGTGTGGAGTCGTTGCCATTCTCCTGCCTCAGCCTCNNNAGATCGGAAGAGCA |
| 45799_1 | TATGAGTGTGGAGTCGTTGCTCTTGCAGCTCAACCTCTTTACTNNNAGATCGGAAGA |

| | |
|---|---|
| 45799_2 | TATGAGTGTGGAGTCGTTGCTCTTGCAGCTCAACCTCTTTACTNNNAGATCGGAAGA |
| 50984_1 | TATGAGTGTGGAGTCGTTGCTGCAGGATGGGAGTCTAGTAAGANNNAGATCGGAA |
| 50984_2 | TATGAGTGTGGAGTCGTTGCTGCAGGATGGGAGTCTAGTAAGANNNAGATCGGAA |
| 42760_1 | TATGAGTGTGGAGTCGTTGCAAAGCATCACAGTCTTCCAGTCANNNAGATCGGAAG |
| 42760_2 | TATGAGTGTGGAGTCGTTGCAAAGCATCACAGTCTTCCAGTCANNNAGATCGGAAG |
| 31812_1 | TATGAGTGTGGAGTCGTTGCCATTCCTGCCAGCTGATGAAATTNNNAGATCGGAAG |
| 31812_2 | TATGAGTGTGGAGTCGTTGCCATTCCTGCCAGCTGATGAAATTNNNAGATCGGAAG |
| 47547_1 | TATGAGTGTGGAGTCGTTGCTGGCAACCATAAAACCTGGAAACNNNAGATCGGAA |
| 47547_2 | TATGAGTGTGGAGTCGTTGCTGGCAACCATAAAACCTGGAAACNNNAGATCGGAA |
| 51088_1 | TATGAGTGTGGAGTCGTTGCGAGACTCCATCCTCCTACTCGGNNNAGATCGGAAGA |
| 51088_2 | TATGAGTGTGGAGTCGTTGCGAGACTCCATCCTCCTACTCGGNNNAGATCGGAAGA |
| 30113_1 | TATGAGTGTGGAGTCGTTGCTGGAAACCAGCCTTACTGCCNNNAGATCGGAAGAG |
| 30113_2 | TATGAGTGTGGAGTCGTTGCTGGAAACCAGCCTTACTGCCNNNAGATCGGAAGAG |
| 10300_1 | TATGAGTGTGGAGTCGTTGCAGGTGGAGGATAACCAAGTGTTTNNNAGATCGGAA |
| 10300_2 | TATGAGTGTGGAGTCGTTGCAGGTGGAGGATAACCAAGTGTTTNNNAGATCGGAA |
| 48966_1 | TATGAGTGTGGAGTCGTTGCATCAGTGCTGTTGCATAGTGTTCNNNAGATCGGAAG |
| 48966_2 | TATGAGTGTGGAGTCGTTGCATCAGTGCTGTTGCATAGTGTTCNNNAGATCGGAAG |
| 36011_1 | TATGAGTGTGGAGTCGTTGCGGCAAAATGAAACAGCAGACTCTNNNAGATCGGAA |
| 36011_2 | TATGAGTGTGGAGTCGTTGCGGCAAAATGAAACAGCAGACTCTNNNAGATCGGAA |
| 12489_1 | TATGAGTGTGGAGTCGTTGCCCCATGTTGCCTATTTATTTTTTNNNAGATCGGAAGA |
| 12489_2 | TATGAGTGTGGAGTCGTTGCCCCATGTTGCCTATTTATTTTTTNNNAGATCGGAAGA |
| 40499_1 | TATGAGTGTGGAGTCGTTGCACCTGTAATCCTAGCACTTTGGGNNNAGATCGGAAG |
| 40499_2 | TATGAGTGTGGAGTCGTTGCACCTGTAATCCTAGCACTTTGGGNNNAGATCGGAAG |
| 12705_1 | TATGAGTGTGGAGTCGTTGCATTCATTCAGTGTAGGTGTGNNNAGATCGGAAGAG |
| 12705_2 | TATGAGTGTGGAGTCGTTGCATTCATTCAGTGTAGGTGTGNNNAGATCGGAAGAG |
| 49506_1 | TATGAGTGTGGAGTCGTTGCCCACGCCCAGCTAATTTTTGTATNNNAGATCGGAAG |
| 49506_2 | TATGAGTGTGGAGTCGTTGCCCACGCCCAGCTAATTTTTGTATNNNAGATCGGAAG |
| 41738_1 | TATGAGTGTGGAGTCGTTGCCCCTGGTTCAAGCAATTCTTCTGNNNAGATCGGAAG |
| 41738_2 | TATGAGTGTGGAGTCGTTGCCCCTGGTTCAAGCAATTCTTCTGNNNAGATCGGAAG |
| 10566_1 | TATGAGTGTGGAGTCGTTGCTGCTTTACTGATGACCTTCTCAGCNNNAGATCGGAA |
| 10566_2 | TATGAGTGTGGAGTCGTTGCTGCTTTACTGATGACCTTCTCAGCNNNAGATCGGAA |
| 39941_1 | TATGAGTGTGGAGTCGTTGCGAGGGACACATTGGAAAACTTGGNNNAGATCGGAA |
| 39941_2 | TATGAGTGTGGAGTCGTTGCGAGGGACACATTGGAAAACTTGGNNNAGATCGGAA |
| 44431_1 | TATGAGTGTGGAGTCGTTGCACGGATTTCCTTGCAACTCAAAGNNNAGATCGGAAG |
| 44431_2 | TATGAGTGTGGAGTCGTTGCACGGATTTCCTTGCAACTCAAAGNNNAGATCGGAAG |
| 46293_1 | TATGAGTGTGGAGTCGTTGCGAGATGGAGTATCGCCCTGTCTCNNNAGATCGGAA |
| 46293_2 | TATGAGTGTGGAGTCGTTGCGAGATGGAGTATCGCCCTGTCTCNNNAGATCGGAA |
| 42001_1 | TATGAGTGTGGAGTCGTTGCAGTGCAGTGGTGTGATCTCGNNNAGATCGGAAGAG |
| 42001_2 | TATGAGTGTGGAGTCGTTGCAGTGCAGTGGTGTGATCTCGNNNAGATCGGAAGAG |
| 5518_1 | TATGAGTGTGGAGTCGTTGCCCCCCTGCTCCATTCATACATTCNNNAGATCGGAAGA |
| 5518_2 | TATGAGTGTGGAGTCGTTGCCCCCCTGCTCCATTCATACATTCNNNAGATCGGAAGA |
| 48860_1 | TATGAGTGTGGAGTCGTTGCCTGGAGTGAAATGGCATGATCTCNNNAGATCGGAA |
| 48860_2 | TATGAGTGTGGAGTCGTTGCCTGGAGTGAAATGGCATGATCTCNNNAGATCGGAA |

| | |
|---|---|
| 46069_1 | TATGAGTGTGGAGTCGTTGCAGCAATGAGGGCTAATTCTGTCANNNAGATCGGAAG |
| 46069_2 | TATGAGTGTGGAGTCGTTGCAGCAATGAGGGCTAATTCTGTCANNNAGATCGGAAG |
| 49567_1 | TATGAGTGTGGAGTCGTTGCCCTCGGTTCTCTTGAGAGCAGNNNAGATCGGAAGAG |
| 49567_2 | TATGAGTGTGGAGTCGTTGCCCTCGGTTCTCTTGAGAGCAGNNNAGATCGGAAGAG |
| 42158_1 | TATGAGTGTGGAGTCGTTGCCAAGTCTTGCTCTGTTACCCAAGNNNAGATCGGAAG |
| 42158_2 | TATGAGTGTGGAGTCGTTGCCAAGTCTTGCTCTGTTACCCAAGNNNAGATCGGAAG |
| 46960_1 | TATGAGTGTGGAGTCGTTGCACTACTGATCAGCACACGAGTTTNNNAGATCGGAAG |
| 46960_2 | TATGAGTGTGGAGTCGTTGCACTACTGATCAGCACACGAGTTTNNNAGATCGGAAG |
| 13089_1 | TATGAGTGTGGAGTCGTTGCATTTATTTTACGACAACAACAACNNNAGATCGGAAG |
| 13089_2 | TATGAGTGTGGAGTCGTTGCATTTATTTTACGACAACAACAACNNNAGATCGGAAG |
| 35842_1 | TATGAGTGTGGAGTCGTTGCGGTTCAAGCAATTATCCTGCCTCNNNAGATCGGAAG |
| 35842_2 | TATGAGTGTGGAGTCGTTGCGGTTCAAGCAATTATCCTGCCTCNNNAGATCGGAAG |
| 51503_1 | TATGAGTGTGGAGTCGTTGCTTGCTGGGTTGAATAGTGGTTCTNNNAGATCGGAAG |
| 51503_2 | TATGAGTGTGGAGTCGTTGCTTGCTGGGTTGAATAGTGGTTCTNNNAGATCGGAAG |
| 10310_1 | TATGAGTGTGGAGTCGTTGCGCCCTAGAGAAGTCTGCATTTGTNNNAGATCGGAAG |
| 10310_2 | TATGAGTGTGGAGTCGTTGCGCCCTAGAGAAGTCTGCATTTGTNNNAGATCGGAAG |
| 12852_1 | TATGAGTGTGGAGTCGTTGCATCTGACAGGTGATATTCAAACNNNAGATCGGAAGA |
| 12852_2 | TATGAGTGTGGAGTCGTTGCATCTGACAGGTGATATTCAAACNNNAGATCGGAAGA |
| 53433_1 | TATGAGTGTGGAGTCGTTGCAACAGAGGAAGAACACATTCACANNNAGATCGGAA |
| 53433_2 | TATGAGTGTGGAGTCGTTGCAACAGAGGAAGAACACATTCACANNNAGATCGGAA |
| 42788_1 | TATGAGTGTGGAGTCGTTGCTAATGGCATTTTCAGCAACCTGGNNNAGATCGGAAG |
| 42788_2 | TATGAGTGTGGAGTCGTTGCTAATGGCATTTTCAGCAACCTGGNNNAGATCGGAAG |
| 46423_1 | TATGAGTGTGGAGTCGTTGCGACAGAGTCTTGTTCTTGTTGCCNNNAGATCGGAAG |
| 46423_2 | TATGAGTGTGGAGTCGTTGCGACAGAGTCTTGTTCTTGTTGCCNNNAGATCGGAAG |
| 13385_1 | TATGAGTGTGGAGTCGTTGCATAAAACCAAAAAACAGGTAAGTNNNAGATCGGAA |
| 13385_2 | TATGAGTGTGGAGTCGTTGCATAAAACCAAAAAACAGGTAAGTNNNAGATCGGAA |
| 44972_1 | TATGAGTGTGGAGTCGTTGCCACTGTCTGCTGTGAAGGTCTAGNNNAGATCGGAAG |
| 44972_2 | TATGAGTGTGGAGTCGTTGCCACTGTCTGCTGTGAAGGTCTAGNNNAGATCGGAAG |
| 43580_1 | TATGAGTGTGGAGTCGTTGCATGTCATTCATTCATTCCCCCCANNNAGATCGGAAGA |
| 43580_2 | TATGAGTGTGGAGTCGTTGCATGTCATTCATTCATTCCCCCCANNNAGATCGGAAGA |
| 13580_1 | TATGAGTGTGGAGTCGTTGCTTTATTTTTTTGAGACAGAGTCTNNNAGATCGGAAG |
| 13580_2 | TATGAGTGTGGAGTCGTTGCTTTATTTTTTTGAGACAGAGTCTNNNAGATCGGAAG |
| 39348_1 | TATGAGTGTGGAGTCGTTGCTTTTCCTTTTTTGCAGGCAGTCANNNAGATCGGAAG |
| 39348_2 | TATGAGTGTGGAGTCGTTGCTTTTCCTTTTTTGCAGGCAGTCANNNAGATCGGAAG |
| 40745_1 | TATGAGTGTGGAGTCGTTGCGCTTATGCTTCCCTATGGTGAGANNNAGATCGGAAG |
| 40745_2 | TATGAGTGTGGAGTCGTTGCGCTTATGCTTCCCTATGGTGAGANNNAGATCGGAAG |
| 50294_1 | TATGAGTGTGGAGTCGTTGCGGATCACGAGGTCAAGAGATCAANNNAGATCGGAA |
| 50294_2 | TATGAGTGTGGAGTCGTTGCGGATCACGAGGTCAAGAGATCAANNNAGATCGGAA |
| 47017_1 | TATGAGTGTGGAGTCGTTGCAGAGCTACTGTCATGAAATGCCANNNAGATCGGAAG |
| 47017_2 | TATGAGTGTGGAGTCGTTGCAGAGCTACTGTCATGAAATGCCANNNAGATCGGAAG |
| 39745_1 | TATGAGTGTGGAGTCGTTGCAGGATTCTCAAGAGGGGTAAACCNNNAGATCGGAA |
| 39745_2 | TATGAGTGTGGAGTCGTTGCAGGATTCTCAAGAGGGGTAAACCNNNAGATCGGAA |
| 52540_1 | TATGAGTGTGGAGTCGTTGCTCTCCTTCAACAAGTCCTCCAAGNNNAGATCGGAAGA |

| | |
|---|---|
| 52540_2 | TATGAGTGTGGAGTCGTTGCTCTCCTTCAACAAGTCCTCCAAGNNNAGATCGGAAGA |
| 35336_1 | TATGAGTGTGGAGTCGTTGCCTCCTCAGCTGGGATTACAGATGNNNAGATCGGAAG |
| 35336_2 | TATGAGTGTGGAGTCGTTGCCTCCTCAGCTGGGATTACAGATGNNNAGATCGGAAG |
| 11718_1 | TATGAGTGTGGAGTCGTTGCTCAGAGTGAAGGACCTTTCAGCTNNNAGATCGGAAG |
| 11718_2 | TATGAGTGTGGAGTCGTTGCTCAGAGTGAAGGACCTTTCAGCTNNNAGATCGGAAG |
| 40576_1 | TATGAGTGTGGAGTCGTTGCACCTGACTAAGGGGCAAGAAAATNNNAGATCGGAA |
| 40576_2 | TATGAGTGTGGAGTCGTTGCACCTGACTAAGGGGCAAGAAAATNNNAGATCGGAA |
| 51890_1 | TATGAGTGTGGAGTCGTTGCTTAGTTTGCTGAGGATGATGGCTNNNAGATCGGAA |
| 51890_2 | TATGAGTGTGGAGTCGTTGCTTAGTTTGCTGAGGATGATGGCTNNNAGATCGGAA |
| 47766_1 | TATGAGTGTGGAGTCGTTGCCAGATGGAATCTTACCCTGTCGTNNNAGATCGGAAG |
| 47766_2 | TATGAGTGTGGAGTCGTTGCCAGATGGAATCTTACCCTGTCGTNNNAGATCGGAAG |
| 10681_1 | TATGAGTGTGGAGTCGTTGCCTCCCTCATGCCTCTTGATTTCCNNNAGATCGGAAGA |
| 10681_2 | TATGAGTGTGGAGTCGTTGCCTCCCTCATGCCTCTTGATTTCCNNNAGATCGGAAGA |
| 42929_1 | TATGAGTGTGGAGTCGTTGCCGTGCGTACAACACATTCTCTTTNNNAGATCGGAAG |
| 42929_2 | TATGAGTGTGGAGTCGTTGCCGTGCGTACAACACATTCTCTTTNNNAGATCGGAAG |
| 48083_1 | TATGAGTGTGGAGTCGTTGCGAGAGTCATGCCCTACAAACCATNNNAGATCGGAAG |
| 48083_2 | TATGAGTGTGGAGTCGTTGCGAGAGTCATGCCCTACAAACCATNNNAGATCGGAAG |
| 48253_1 | TATGAGTGTGGAGTCGTTGCCTCCTGGTGTTCTGGGTTAGGNNNAGATCGGAAGA |
| 48253_2 | TATGAGTGTGGAGTCGTTGCCTCCTGGTGTTCTGGGTTAGGNNNAGATCGGAAGA |
| 45736_1 | TATGAGTGTGGAGTCGTTGCGAGGCTGAGGAGAAGCAGGAGNNNAGATCGGAAG |
| 45736_2 | TATGAGTGTGGAGTCGTTGCGAGGCTGAGGAGAAGCAGGAGNNNAGATCGGAAG |
| 9488_1 | TATGAGTGTGGAGTCGTTGCCCTACTTTGCTGTACCACCACAGNNNAGATCGGAAG |
| 9488_2 | TATGAGTGTGGAGTCGTTGCCCTACTTTGCTGTACCACCACAGNNNAGATCGGAAG |
| 8846_1 | TATGAGTGTGGAGTCGTTGCTCTCCTCTCCTTGAGCTTTCTGTNNNAGATCGGAAGA |
| 8846_2 | TATGAGTGTGGAGTCGTTGCTCTCCTCTCCTTGAGCTTTCTGTNNNAGATCGGAAGA |
| 51820_1 | TATGAGTGTGGAGTCGTTGCCCTCCTGGGTTCAAGTGATTCTCNNNAGATCGGAAG |
| 51820_2 | TATGAGTGTGGAGTCGTTGCCCTCCTGGGTTCAAGTGATTCTCNNNAGATCGGAAG |
| 47748_1 | TATGAGTGTGGAGTCGTTGCGAGTAGCAGGGATTACAGGCATGNNNAGATCGGAA |
| 47748_2 | TATGAGTGTGGAGTCGTTGCGAGTAGCAGGGATTACAGGCATGNNNAGATCGGAA |
| 34432_1 | TATGAGTGTGGAGTCGTTGCCATGTTGGATCACGCCTGTAATCNNNAGATCGGAAG |
| 34432_2 | TATGAGTGTGGAGTCGTTGCCATGTTGGATCACGCCTGTAATCNNNAGATCGGAAG |
| 33292_1 | TATGAGTGTGGAGTCGTTGCTGTTAGAACTCCTGGGCTTATGGNNNAGATCGGAAG |
| 33292_2 | TATGAGTGTGGAGTCGTTGCTGTTAGAACTCCTGGGCTTATGGNNNAGATCGGAAG |
| 13440_1 | TATGAGTGTGGAGTCGTTGCGAGAATAGGAACCCAGATATGNNNAGATCGGAAGA |
| 13440_2 | TATGAGTGTGGAGTCGTTGCGAGAATAGGAACCCAGATATGNNNAGATCGGAAGA |
| 41466_1 | TATGAGTGTGGAGTCGTTGCTTCATCCAGTGTGTGTGTTTGTGNNNAGATCGGAAG |
| 41466_2 | TATGAGTGTGGAGTCGTTGCTTCATCCAGTGTGTGTGTTTGTGNNNAGATCGGAAG |
| 2962_1 | TATGAGTGTGGAGTCGTTGCACCATGTCTTCTGCCTCACATTTNNNAGATCGGAAGA |
| 2962_2 | TATGAGTGTGGAGTCGTTGCACCATGTCTTCTGCCTCACATTTNNNAGATCGGAAGA |
| 46593_1 | TATGAGTGTGGAGTCGTTGCTGATCTCGGCTCACTGCAATATCNNNAGATCGGAAG |
| 46593_2 | TATGAGTGTGGAGTCGTTGCTGATCTCGGCTCACTGCAATATCNNNAGATCGGAAG |
| 31067_1 | TATGAGTGTGGAGTCGTTGCGAGCTGCAATAAACTCACTAGCGNNNAGATCGGAAG |
| 31067_2 | TATGAGTGTGGAGTCGTTGCGAGCTGCAATAAACTCACTAGCGNNNAGATCGGAAG |

| | |
|---|---|
| 10064_1 | TATGAGTGTGGAGTCGTTGCAGGAGGATAAGATGACATCCGGGNNNAGATCGGAA |
| 10064_2 | TATGAGTGTGGAGTCGTTGCAGGAGGATAAGATGACATCCGGGNNNAGATCGGAA |
| 47557_1 | TATGAGTGTGGAGTCGTTGCCTCCTGAGTAGCTGGGACTACAGNNNAGATCGGAAG |
| 47557_2 | TATGAGTGTGGAGTCGTTGCCTCCTGAGTAGCTGGGACTACAGNNNAGATCGGAAG |
| 47474_1 | TATGAGTGTGGAGTCGTTGCGGATTACCCCCAAATACCCACTTNNNAGATCGGAAGA |
| 47474_2 | TATGAGTGTGGAGTCGTTGCGGATTACCCCCAAATACCCACTTNNNAGATCGGAAGA |
| 43160_1 | TATGAGTGTGGAGTCGTTGCCTGGGTTCAAGTGCTTTTCCTGNNNAGATCGGAAGA |
| 43160_2 | TATGAGTGTGGAGTCGTTGCCTGGGTTCAAGTGCTTTTCCTGNNNAGATCGGAAGA |
| 52351_1 | TATGAGTGTGGAGTCGTTGCCTTGAAGGCAGTCGCAGTTCTAGNNNAGATCGGAAG |
| 52351_2 | TATGAGTGTGGAGTCGTTGCCTTGAAGGCAGTCGCAGTTCTAGNNNAGATCGGAAG |
| 9703_1 | TATGAGTGTGGAGTCGTTGCATTCCTTGGGTTCAACCTTTCCCNNNAGATCGGAAGA |
| 9703_2 | TATGAGTGTGGAGTCGTTGCATTCCTTGGGTTCAACCTTTCCCNNNAGATCGGAAGA |
| 35337_1 | TATGAGTGTGGAGTCGTTGCTGTTGGGATTACAGGAGTAAGCCNNNAGATCGGAA |
| 35337_2 | TATGAGTGTGGAGTCGTTGCTGTTGGGATTACAGGAGTAAGCCNNNAGATCGGAA |
| 47196_1 | TATGAGTGTGGAGTCGTTGCACTACGCCCGGCTAATTTTTTTTNNNAGATCGGAAGA |
| 47196_2 | TATGAGTGTGGAGTCGTTGCACTACGCCCGGCTAATTTTTTTTNNNAGATCGGAAGA |
| 13700_1 | TATGAGTGTGGAGTCGTTGCCAGTTCTTTCTTTTTCTCACTATNNNAGATCGGAAGA |
| 13700_2 | TATGAGTGTGGAGTCGTTGCCAGTTCTTTCTTTTTCTCACTATNNNAGATCGGAAGA |
| 44497_1 | TATGAGTGTGGAGTCGTTGCCTTCCTGGGTTCAAGCAATTCTCNNNAGATCGGAAGA |
| 44497_2 | TATGAGTGTGGAGTCGTTGCCTTCCTGGGTTCAAGCAATTCTCNNNAGATCGGAAGA |
| 45726_1 | TATGAGTGTGGAGTCGTTGCCAGTCATGATTGCAGGATAGGGTNNNAGATCGGAA |
| 45726_2 | TATGAGTGTGGAGTCGTTGCCAGTCATGATTGCAGGATAGGGTNNNAGATCGGAA |
| 45897_1 | TATGAGTGTGGAGTCGTTGCTCTTACCTGGCTCTAGGAAGACANNNAGATCGGAAG |
| 45897_2 | TATGAGTGTGGAGTCGTTGCTCTTACCTGGCTCTAGGAAGACANNNAGATCGGAAG |
| 47925_1 | TATGAGTGTGGAGTCGTTGCGCCTCATCCCCCACAAGTATATCNNNAGATCGGAAGA |
| 47925_2 | TATGAGTGTGGAGTCGTTGCGCCTCATCCCCCACAAGTATATCNNNAGATCGGAAGA |
| 38748_1 | TATGAGTGTGGAGTCGTTGCTCATTCAAAAAAGGCTCCCCTCTNNNAGATCGGAAGA |
| 38748_2 | TATGAGTGTGGAGTCGTTGCTCATTCAAAAAAGGCTCCCCTCTNNNAGATCGGAAGA |
| 41721_1 | TATGAGTGTGGAGTCGTTGCGGATCAAGCGATTCTCCTACCTCNNNAGATCGGAAGA |
| 41721_2 | TATGAGTGTGGAGTCGTTGCGGATCAAGCGATTCTCCTACCTCNNNAGATCGGAAGA |
| 40567_1 | TATGAGTGTGGAGTCGTTGCGCAAGCCCACCTTAGGATAATCTNNNAGATCGGAAG |
| 40567_2 | TATGAGTGTGGAGTCGTTGCGCAAGCCCACCTTAGGATAATCTNNNAGATCGGAAG |
| 30930_1 | TATGAGTGTGGAGTCGTTGCATAGCTTGTGAGGCTCTTGATCCNNNAGATCGGAAG |
| 30930_2 | TATGAGTGTGGAGTCGTTGCATAGCTTGTGAGGCTCTTGATCCNNNAGATCGGAAG |
| 36748_1 | TATGAGTGTGGAGTCGTTGCGAAAGAAACCAGTTGCAAAAGGCNNNAGATCGGAA |
| 36748_2 | TATGAGTGTGGAGTCGTTGCGAAAGAAACCAGTTGCAAAAGGCNNNAGATCGGAA |
| 44195_1 | TATGAGTGTGGAGTCGTTGCTGTATGAGCTTGTTTGGGCTTTGNNNAGATCGGAAG |
| 44195_2 | TATGAGTGTGGAGTCGTTGCTGTATGAGCTTGTTTGGGCTTTGNNNAGATCGGAAG |
| 39094_1 | TATGAGTGTGGAGTCGTTGCTTCTTTGAAAGGGAGTCTTGCTCNNNAGATCGGAAG |
| 39094_2 | TATGAGTGTGGAGTCGTTGCTTCTTTGAAAGGGAGTCTTGCTCNNNAGATCGGAAG |
| 30793_1 | TATGAGTGTGGAGTCGTTGCCACACACACACACACACACATATNNNAGATCGGAAGA |
| 30793_2 | TATGAGTGTGGAGTCGTTGCCACACACACACACACACACATATNNNAGATCGGAAGA |
| 50140_1 | TATGAGTGTGGAGTCGTTGCTCTAAACCCCTGTCTCCCTGTATNNNAGATCGGAAGA |

| | |
|---|---|
| 50140_2 | TATGAGTGTGGAGTCGTTGCTCTAAACCCCTGTCTCCCTGTATNNNAGATCGGAAGA |
| 31890_1 | TATGAGTGTGGAGTCGTTGCGACACAGAGACAGAGAGACACAGNNNAGATCGGAA |
| 31890_2 | TATGAGTGTGGAGTCGTTGCGACACAGAGACAGAGAGACACAGNNNAGATCGGAA |
| 51511_1 | TATGAGTGTGGAGTCGTTGCTATGAAGCTCATGCGTACTCCTCNNNAGATCGGAAG |
| 51511_2 | TATGAGTGTGGAGTCGTTGCTATGAAGCTCATGCGTACTCCTCNNNAGATCGGAAG |
| 31920_1 | TATGAGTGTGGAGTCGTTGCAACAAAATACCACACACTGGGTGNNNAGATCGGAA |
| 31920_2 | TATGAGTGTGGAGTCGTTGCAACAAAATACCACACACTGGGTGNNNAGATCGGAA |
| 40332_1 | TATGAGTGTGGAGTCGTTGCGCCTGATGGTTGTCCTTTCTTTCNNNAGATCGGAAG |
| 40332_2 | TATGAGTGTGGAGTCGTTGCGCCTGATGGTTGTCCTTTCTTTCNNNAGATCGGAAG |
| 41223_1 | TATGAGTGTGGAGTCGTTGCCCAAAGCAAGCAGATCTCTTCAGNNNAGATCGGAAG |
| 41223_2 | TATGAGTGTGGAGTCGTTGCCCAAAGCAAGCAGATCTCTTCAGNNNAGATCGGAAG |
| 40452_1 | TATGAGTGTGGAGTCGTTGCTGGCTTGACCTAAGAAGCACTAANNNAGATCGGAA |
| 40452_2 | TATGAGTGTGGAGTCGTTGCTGGCTTGACCTAAGAAGCACTAANNNAGATCGGAA |
| 31447_1 | TATGAGTGTGGAGTCGTTGCAAGCTTCATCTCTAACATTGGCTNNNAGATCGGAAG |
| 31447_2 | TATGAGTGTGGAGTCGTTGCAAGCTTCATCTCTAACATTGGCTNNNAGATCGGAAG |
| 48157_1 | TATGAGTGTGGAGTCGTTGCAAGCTCAGGCAGAAGGAAGACNNNAGATCGGAAG |
| 48157_2 | TATGAGTGTGGAGTCGTTGCAAGCTCAGGCAGAAGGAAGACNNNAGATCGGAAG |
| 12526_1 | TATGAGTGTGGAGTCGTTGCAAACACTGATCTATTCACCTATTNNNAGATCGGAAGA |
| 12526_2 | TATGAGTGTGGAGTCGTTGCAAACACTGATCTATTCACCTATTNNNAGATCGGAAGA |
| 31565_1 | TATGAGTGTGGAGTCGTTGCACTCAAGTGTACACACAGAAGCANNNAGATCGGAA |
| 31565_2 | TATGAGTGTGGAGTCGTTGCACTCAAGTGTACACACAGAAGCANNNAGATCGGAA |
| 38029_1 | TATGAGTGTGGAGTCGTTGCACTGAATTCCGAAGGAGGAAGACNNNAGATCGGAA |
| 38029_2 | TATGAGTGTGGAGTCGTTGCACTGAATTCCGAAGGAGGAAGACNNNAGATCGGAA |
| 13397_1 | TATGAGTGTGGAGTCGTTGCTATCAGGAATGGACTTAGTCTNNNAGATCGGAAGA |
| 13397_2 | TATGAGTGTGGAGTCGTTGCTATCAGGAATGGACTTAGTCTNNNAGATCGGAAGA |
| 48507_1 | TATGAGTGTGGAGTCGTTGCCCTCTCAGGATCACACCATTCTCNNNAGATCGGAAGA |
| 48507_2 | TATGAGTGTGGAGTCGTTGCCCTCTCAGGATCACACCATTCTCNNNAGATCGGAAGA |
| 34193_1 | TATGAGTGTGGAGTCGTTGCGAGGTTGCATTGAGCTGTGATTGNNNAGATCGGAA |
| 34193_2 | TATGAGTGTGGAGTCGTTGCGAGGTTGCATTGAGCTGTGATTGNNNAGATCGGAA |
| 31953_1 | TATGAGTGTGGAGTCGTTGCCTCCCACAGGAAGGAAAGAGAAANNNAGATCGGAA |
| 31953_2 | TATGAGTGTGGAGTCGTTGCCTCCCACAGGAAGGAAAGAGAAANNNAGATCGGAA |
| 41923_1 | TATGAGTGTGGAGTCGTTGCGGTTCAAGCGATTCTTTTCCCTCNNNAGATCGGAAG |
| 41923_2 | TATGAGTGTGGAGTCGTTGCGGTTCAAGCGATTCTTTTCCCTCNNNAGATCGGAAG |
| 10092_1 | TATGAGTGTGGAGTCGTTGCTCCCAGGTCAACTTAGTCATGGTNNNAGATCGGAAG |
| 10092_2 | TATGAGTGTGGAGTCGTTGCTCCCAGGTCAACTTAGTCATGGTNNNAGATCGGAAG |
| 44653_1 | TATGAGTGTGGAGTCGTTGCCCAAACCAAGTGGAAGCTCTTGNNNAGATCGGAAG |
| 44653_2 | TATGAGTGTGGAGTCGTTGCCCAAACCAAGTGGAAGCTCTTGNNNAGATCGGAAG |
| 30157_1 | TATGAGTGTGGAGTCGTTGCTCAATCTCAGAATATTGGTGTGCTNNNAGATCGGAA |
| 30157_2 | TATGAGTGTGGAGTCGTTGCTCAATCTCAGAATATTGGTGTGCTNNNAGATCGGAA |
| 42065_1 | TATGAGTGTGGAGTCGTTGCGTGTAAAACTTTGGGGTGCTTCANNNAGATCGGAA |
| 42065_2 | TATGAGTGTGGAGTCGTTGCGTGTAAAACTTTGGGGTGCTTCANNNAGATCGGAA |
| 34295_1 | TATGAGTGTGGAGTCGTTGCCACAATTCAGCCATGACCCTGNNNAGATCGGAAGAG |
| 34295_2 | TATGAGTGTGGAGTCGTTGCCACAATTCAGCCATGACCCTGNNNAGATCGGAAGAG |

| | |
|---|---|
| 52723_1 | TATGAGTGTGGAGTCGTTGCCACACTCAAAGGTGACTTGCAATNNNAGATCGGAAG |
| 52723_2 | TATGAGTGTGGAGTCGTTGCCACACTCAAAGGTGACTTGCAATNNNAGATCGGAAG |
| 49701_1 | TATGAGTGTGGAGTCGTTGCGCATGGGGATGTCTAGCTTTTCTNNNAGATCGGAAG |
| 49701_2 | TATGAGTGTGGAGTCGTTGCGCATGGGGATGTCTAGCTTTTCTNNNAGATCGGAAG |
| 50939_1 | TATGAGTGTGGAGTCGTTGCATTTATGGCCATCGAGGACTCTCNNNAGATCGGAAG |
| 50939_2 | TATGAGTGTGGAGTCGTTGCATTTATGGCCATCGAGGACTCTCNNNAGATCGGAAG |
| 46586_1 | TATGAGTGTGGAGTCGTTGCAGACAGTGCTTCTCACCTGAAATNNNAGATCGGAAG |
| 46586_2 | TATGAGTGTGGAGTCGTTGCAGACAGTGCTTCTCACCTGAAATNNNAGATCGGAAG |
| 43414_1 | TATGAGTGTGGAGTCGTTGCCAAGTAGCTGGGACTATAGGCACNNNAGATCGGAA |
| 43414_2 | TATGAGTGTGGAGTCGTTGCCAAGTAGCTGGGACTATAGGCACNNNAGATCGGAA |
| 52330_1 | TATGAGTGTGGAGTCGTTGCCTCCCAAGGAATTGCATCCCGNNNAGATCGGAAGAG |
| 52330_2 | TATGAGTGTGGAGTCGTTGCCTCCCAAGGAATTGCATCCCGNNNAGATCGGAAGAG |
| 38628_1 | TATGAGTGTGGAGTCGTTGCAGATAACTGTATCTACTAGCCAGGANNNAGATCGGA |
| 38628_2 | TATGAGTGTGGAGTCGTTGCAGATAACTGTATCTACTAGCCAGGANNNAGATCGGA |
| 47542_1 | TATGAGTGTGGAGTCGTTGCAACCATGCCCAGCTAATTTTTGTNNNAGATCGGAAG |
| 47542_2 | TATGAGTGTGGAGTCGTTGCAACCATGCCCAGCTAATTTTTGTNNNAGATCGGAAG |
| 43576_1 | TATGAGTGTGGAGTCGTTGCATCATTCATTCATTCCCCCCACANNNAGATCGGAAGA |
| 43576_2 | TATGAGTGTGGAGTCGTTGCATCATTCATTCATTCCCCCCACANNNAGATCGGAAGA |
| 35774_1 | TATGAGTGTGGAGTCGTTGCCACCCAGATGATGTTCCTCTTCANNNAGATCGGAAG |
| 35774_2 | TATGAGTGTGGAGTCGTTGCCACCCAGATGATGTTCCTCTTCANNNAGATCGGAAG |
| 29113_1 | TATGAGTGTGGAGTCGTTGCGTGTCTGCTCCCAGTTCAATTTCNNNAGATCGGAAG |
| 29113_2 | TATGAGTGTGGAGTCGTTGCGTGTCTGCTCCCAGTTCAATTTCNNNAGATCGGAAG |
| 35419_1 | TATGAGTGTGGAGTCGTTGCCAGGAGTGCAGTGGTATGATCATNNNAGATCGGAA |
| 35419_2 | TATGAGTGTGGAGTCGTTGCCAGGAGTGCAGTGGTATGATCATNNNAGATCGGAA |
| 33337_1 | TATGAGTGTGGAGTCGTTGCGTATTTTTGGTAGAGGGGGGGAGNNNAGATCGGA |
| 33337_2 | TATGAGTGTGGAGTCGTTGCGTATTTTTGGTAGAGGGGGGGAGNNNAGATCGGA |
| 32071_1 | TATGAGTGTGGAGTCGTTGCGAGCTTACAGTGAGCCGAGATAGNNNAGATCGGAA |
| 32071_2 | TATGAGTGTGGAGTCGTTGCGAGCTTACAGTGAGCCGAGATAGNNNAGATCGGAA |
| 52578_1 | TATGAGTGTGGAGTCGTTGCCTGCCCACCACAAGATTGTAAACNNNAGATCGGAAG |
| 52578_2 | TATGAGTGTGGAGTCGTTGCCTGCCCACCACAAGATTGTAAACNNNAGATCGGAAG |
| 37180_1 | TATGAGTGTGGAGTCGTTGCGCAGGAGAATTGAGCCAAGATTGNNNAGATCGGAA |
| 37180_2 | TATGAGTGTGGAGTCGTTGCGCAGGAGAATTGAGCCAAGATTGNNNAGATCGGAA |
| 52531_1 | TATGAGTGTGGAGTCGTTGCGAATGAAGCAATGCCTGGAAGAANNNAGATCGGAA |
| 52531_2 | TATGAGTGTGGAGTCGTTGCGAATGAAGCAATGCCTGGAAGAANNNAGATCGGAA |
| 44587_1 | TATGAGTGTGGAGTCGTTGCTTTGATTGATCAGGCTAAGTCCANNNAGATCGGAAG |
| 44587_2 | TATGAGTGTGGAGTCGTTGCTTTGATTGATCAGGCTAAGTCCANNNAGATCGGAAG |
| 29906_1 | TATGAGTGTGGAGTCGTTGCGCTCAGAAGACACCCATGATCTNNNAGATCGGAAGA |
| 29906_2 | TATGAGTGTGGAGTCGTTGCGCTCAGAAGACACCCATGATCTNNNAGATCGGAAGA |
| 38102_1 | TATGAGTGTGGAGTCGTTGCGGTGGATTGCTAGAGCTTAGGAGNNNAGATCGGAA |
| 38102_2 | TATGAGTGTGGAGTCGTTGCGGTGGATTGCTAGAGCTTAGGAGNNNAGATCGGAA |
| 34364_1 | TATGAGTGTGGAGTCGTTGCGTTTTGACTTCCCTCCCAAACTGNNNAGATCGGAAGA |
| 34364_2 | TATGAGTGTGGAGTCGTTGCGTTTTGACTTCCCTCCCAAACTGNNNAGATCGGAAGA |
| 30041_1 | TATGAGTGTGGAGTCGTTGCGCCGTCTGCAACATATACTTCAGNNNAGATCGGAAG |

| | |
|---|---|
| 30041_2 | TATGAGTGTGGAGTCGTTGCGCCGTCTGCAACATATACTTCAGNNNAGATCGGAAG |
| 34581_1 | TATGAGTGTGGAGTCGTTGCCAGGGAACGTTCATATGCAAGTTNNNAGATCGGAA |
| 34581_2 | TATGAGTGTGGAGTCGTTGCCAGGGAACGTTCATATGCAAGTTNNNAGATCGGAA |
| 29631_1 | TATGAGTGTGGAGTCGTTGCAGATGGTTGATGAGGTTCACACANNNAGATCGGAA |
| 29631_2 | TATGAGTGTGGAGTCGTTGCAGATGGTTGATGAGGTTCACACANNNAGATCGGAA |
| 47040_1 | TATGAGTGTGGAGTCGTTGCAAGAGATTCTCCTACCTCAGCCTNNNAGATCGGAAG |
| 47040_2 | TATGAGTGTGGAGTCGTTGCAAGAGATTCTCCTACCTCAGCCTNNNAGATCGGAAG |
| 29080_1 | TATGAGTGTGGAGTCGTTGCTGACATCCTGGTTTTTTTCCTTTNNNAGATCGGAAGA |
| 29080_2 | TATGAGTGTGGAGTCGTTGCTGACATCCTGGTTTTTTTCCTTTNNNAGATCGGAAGA |
| 40545_1 | TATGAGTGTGGAGTCGTTGCTGAGCTCATTTGGCAAACGAATTNNNAGATCGGAAG |
| 40545_2 | TATGAGTGTGGAGTCGTTGCTGAGCTCATTTGGCAAACGAATTNNNAGATCGGAAG |
| 41411_1 | TATGAGTGTGGAGTCGTTGCTTACAAAACCAGATGTTGGGCTGNNNAGATCGGAA |
| 41411_2 | TATGAGTGTGGAGTCGTTGCTTACAAAACCAGATGTTGGGCTGNNNAGATCGGAA |
| 39357_1 | TATGAGTGTGGAGTCGTTGCCCATCTTTCTGCCACCATCAATGNNNAGATCGGAAGA |
| 39357_2 | TATGAGTGTGGAGTCGTTGCCCATCTTTCTGCCACCATCAATGNNNAGATCGGAAGA |
| 3098_1 | TATGAGTGTGGAGTCGTTGCTTGACTGCACTGGAAATCCTCTGNNNAGATCGGAAG |
| 3098_2 | TATGAGTGTGGAGTCGTTGCTTGACTGCACTGGAAATCCTCTGNNNAGATCGGAAG |
| 43269_1 | TATGAGTGTGGAGTCGTTGCCCAGGTTCAAGCAATTCTACTGCNNNAGATCGGAAG |
| 43269_2 | TATGAGTGTGGAGTCGTTGCCCAGGTTCAAGCAATTCTACTGCNNNAGATCGGAAG |
| 32040_1 | TATGAGTGTGGAGTCGTTGCCTGAGGCAGGAGAATCACTTGAANNNAGATCGGAA |
| 32040_2 | TATGAGTGTGGAGTCGTTGCCTGAGGCAGGAGAATCACTTGAANNNAGATCGGAA |
| 42119_1 | TATGAGTGTGGAGTCGTTGCCACCTGGCTCAAAGTGAAACTTTNNNAGATCGGAAG |
| 42119_2 | TATGAGTGTGGAGTCGTTGCCACCTGGCTCAAAGTGAAACTTTNNNAGATCGGAAG |
| 51177_1 | TATGAGTGTGGAGTCGTTGCGGCTCCAATATCTGTGGCTCTAANNNAGATCGGAAG |
| 51177_2 | TATGAGTGTGGAGTCGTTGCGGCTCCAATATCTGTGGCTCTAANNNAGATCGGAAG |
| 40939_1 | TATGAGTGTGGAGTCGTTGCGTCACTACAATCTCCACCTCCTGNNNAGATCGGAAGA |
| 40939_2 | TATGAGTGTGGAGTCGTTGCGTCACTACAATCTCCACCTCCTGNNNAGATCGGAAGA |
| 50541_1 | TATGAGTGTGGAGTCGTTGCCATCATGTGTGTGAACAAGCCTTNNNAGATCGGAAG |
| 50541_2 | TATGAGTGTGGAGTCGTTGCCATCATGTGTGTGAACAAGCCTTNNNAGATCGGAAG |
| 50636_1 | TATGAGTGTGGAGTCGTTGCCACTATCTTGGAGCTGGAGATCCNNNAGATCGGAAG |
| 50636_2 | TATGAGTGTGGAGTCGTTGCCACTATCTTGGAGCTGGAGATCCNNNAGATCGGAAG |
| 13370_1 | TATGAGTGTGGAGTCGTTGCCTCAAAACCATACTATTACATGGNNNAGATCGGAAG |
| 13370_2 | TATGAGTGTGGAGTCGTTGCCTCAAAACCATACTATTACATGGNNNAGATCGGAAG |
| 8221_1 | TATGAGTGTGGAGTCGTTGCGGGGTTTTCACTCTTGTCACCANNNAGATCGGAAGA |
| 8221_2 | TATGAGTGTGGAGTCGTTGCGGGGTTTTCACTCTTGTCACCANNNAGATCGGAAGA |
| 44247_1 | TATGAGTGTGGAGTCGTTGCGTGCTGCCTCTTTTTTTAGGGAANNNAGATCGGAAG |
| 44247_2 | TATGAGTGTGGAGTCGTTGCGTGCTGCCTCTTTTTTTAGGGAANNNAGATCGGAAG |
| 36304_1 | TATGAGTGTGGAGTCGTTGCCTCATGCCTGTAATCCCAGAACTNNNAGATCGGAAG |
| 36304_2 | TATGAGTGTGGAGTCGTTGCCTCATGCCTGTAATCCCAGAACTNNNAGATCGGAAG |
| 44927_1 | TATGAGTGTGGAGTCGTTGCGAACAGTGCATTAGGAAGGAAGCNNNAGATCGGAA |
| 44927_2 | TATGAGTGTGGAGTCGTTGCGAACAGTGCATTAGGAAGGAAGCNNNAGATCGGAA |
| 35685_1 | TATGAGTGTGGAGTCGTTGCTCCCAAAATGATGGAGGAGTGTTNNNAGATCGGAA |
| 35685_2 | TATGAGTGTGGAGTCGTTGCTCCCAAAATGATGGAGGAGTGTTNNNAGATCGGAA |

| | |
|---|---|
| 50851_1 | TATGAGTGTGGAGTCGTTGCCAGAGGCATTTCCATTCTGCTTTNNNAGATCGGAAGA |
| 50851_2 | TATGAGTGTGGAGTCGTTGCCAGAGGCATTTCCATTCTGCTTTNNNAGATCGGAAGA |
| 45575_1 | TATGAGTGTGGAGTCGTTGCCCTCCTAGGTTCAAGCGATTCTCNNNAGATCGGAAGA |
| 45575_2 | TATGAGTGTGGAGTCGTTGCCCTCCTAGGTTCAAGCGATTCTCNNNAGATCGGAAGA |
| 11673_1 | TATGAGTGTGGAGTCGTTGCTCCGCGTCAAGACGACAGCGNNNAGATCGGAAGAG |
| 11673_2 | TATGAGTGTGGAGTCGTTGCTCCGCGTCAAGACGACAGCGNNNAGATCGGAAGAG |
| 48608_1 | TATGAGTGTGGAGTCGTTGCCATAGCTCACTACAGCCTCAGTCNNNAGATCGGAAGA |
| 48608_2 | TATGAGTGTGGAGTCGTTGCCATAGCTCACTACAGCCTCAGTCNNNAGATCGGAAGA |
| 10311_1 | TATGAGTGTGGAGTCGTTGCAAAAAGTGCTTGGAGGAAAATGTNNNAGATCGGAA |
| 10311_2 | TATGAGTGTGGAGTCGTTGCAAAAAGTGCTTGGAGGAAAATGTNNNAGATCGGAA |
| 45005_1 | TATGAGTGTGGAGTCGTTGCTTCCCAAACACAGGAAGATGACTNNNAGATCGGAAG |
| 45005_2 | TATGAGTGTGGAGTCGTTGCTTCCCAAACACAGGAAGATGACTNNNAGATCGGAAG |
| 31314_1 | TATGAGTGTGGAGTCGTTGCACTAAAGCTGCAAAACTAAGGCCNNNAGATCGGAAG |
| 31314_2 | TATGAGTGTGGAGTCGTTGCACTAAAGCTGCAAAACTAAGGCCNNNAGATCGGAAG |
| 35613_1 | TATGAGTGTGGAGTCGTTGCAAGCTTCTTTTCTGTGATGGCTGNNNAGATCGGAAG |
| 35613_2 | TATGAGTGTGGAGTCGTTGCAAGCTTCTTTTCTGTGATGGCTGNNNAGATCGGAAG |
| 49763_1 | TATGAGTGTGGAGTCGTTGCCATACCATCCAATGTGAAGGGGANNNAGATCGGAA |
| 49763_2 | TATGAGTGTGGAGTCGTTGCCATACCATCCAATGTGAAGGGGANNNAGATCGGAA |
| 44197_1 | TATGAGTGTGGAGTCGTTGCAGAGGGCTTCTGAGAATGACTTGNNNAGATCGGAA |
| 44197_2 | TATGAGTGTGGAGTCGTTGCAGAGGGCTTCTGAGAATGACTTGNNNAGATCGGAA |
| 33133_1 | TATGAGTGTGGAGTCGTTGCTGGTCAGGAGAAGTAATGAGACCNNNAGATCGGAA |
| 33133_2 | TATGAGTGTGGAGTCGTTGCTGGTCAGGAGAAGTAATGAGACCNNNAGATCGGAA |
| 2573_1 | TATGAGTGTGGAGTCGTTGCGAAAAGAGAGAAAGGGGGTTGGGNNNAGATCGGA |
| 2573_2 | TATGAGTGTGGAGTCGTTGCGAAAAGAGAGAAAGGGGGTTGGGNNNAGATCGGA |
| 10384_1 | TATGAGTGTGGAGTCGTTGCCCTGCTGTTCTGGTATCTGCTTTNNNAGATCGGAAGA |
| 10384_2 | TATGAGTGTGGAGTCGTTGCCCTGCTGTTCTGGTATCTGCTTTNNNAGATCGGAAGA |
| 13615_1 | TATGAGTGTGGAGTCGTTGCTGTATAATAAAACTTCGAAGCTCNNNAGATCGGAAG |
| 13615_2 | TATGAGTGTGGAGTCGTTGCTGTATAATAAAACTTCGAAGCTCNNNAGATCGGAAG |
| 35171_1 | TATGAGTGTGGAGTCGTTGCTCAGCTCTTGAACCTCTTCTTCCNNNAGATCGGAAGA |
| 35171_2 | TATGAGTGTGGAGTCGTTGCTCAGCTCTTGAACCTCTTCTTCCNNNAGATCGGAAGA |
| 10506_1 | TATGAGTGTGGAGTCGTTGCGAAGAAGTCCCACTATCCCAGGTNNNAGATCGGAAG |
| 10506_2 | TATGAGTGTGGAGTCGTTGCGAAGAAGTCCCACTATCCCAGGTNNNAGATCGGAAG |
| 33142_1 | TATGAGTGTGGAGTCGTTGCCTCTTGGACAGGAAGGTTCATGANNNAGATCGGAA |
| 33142_2 | TATGAGTGTGGAGTCGTTGCCTCTTGGACAGGAAGGTTCATGANNNAGATCGGAA |
| 49059_1 | TATGAGTGTGGAGTCGTTGCATCTCGGCTCATTGTAACCTCTGNNNAGATCGGAAGA |
| 49059_2 | TATGAGTGTGGAGTCGTTGCATCTCGGCTCATTGTAACCTCTGNNNAGATCGGAAGA |
| 45655_1 | TATGAGTGTGGAGTCGTTGCAGACGGAGTCTCGCTCTGTCNNNAGATCGGAAGAG |
| 45655_2 | TATGAGTGTGGAGTCGTTGCAGACGGAGTCTCGCTCTGTCNNNAGATCGGAAGAG |
| 37002_1 | TATGAGTGTGGAGTCGTTGCACTTTATCAACCAGCCATTGCACNNNAGATCGGAAGA |
| 37002_2 | TATGAGTGTGGAGTCGTTGCACTTTATCAACCAGCCATTGCACNNNAGATCGGAAGA |
| 13055_1 | TATGAGTGTGGAGTCGTTGCTCCATTTCTTCTAAAATAGTCCANNNAGATCGGAAGA |
| 13055_2 | TATGAGTGTGGAGTCGTTGCTCCATTTCTTCTAAAATAGTCCANNNAGATCGGAAGA |
| 52294_1 | TATGAGTGTGGAGTCGTTGCTACAGGCTCTAGAGAGGGAGAAGNNNAGATCGGAA |

| | |
|---|---|
| 52294_2 | TATGAGTGTGGAGTCGTTGCTACAGGCTCTAGAGAGGGAGAAGNNNAGATCGGA |
| 48239_1 | TATGAGTGTGGAGTCGTTGCCCTGGGTTCAAGCAATTCTCTTGNNNAGATCGGAAG |
| 48239_2 | TATGAGTGTGGAGTCGTTGCCCTGGGTTCAAGCAATTCTCTTGNNNAGATCGGAAG |
| 39313_1 | TATGAGTGTGGAGTCGTTGCACATTAGCCAACTTGGTGCTCTANNNAGATCGGAAG |
| 39313_2 | TATGAGTGTGGAGTCGTTGCACATTAGCCAACTTGGTGCTCTANNNAGATCGGAAG |
| 13510_1 | TATGAGTGTGGAGTCGTTGCAAGATGGGAAATCATATTGAGTNNNAGATCGGAAG |
| 13510_2 | TATGAGTGTGGAGTCGTTGCAAGATGGGAAATCATATTGAGTNNNAGATCGGAAG |
| 45308_1 | TATGAGTGTGGAGTCGTTGCCACTCCAGTAGTCATCTGTAGGCNNNAGATCGGAAG |
| 45308_2 | TATGAGTGTGGAGTCGTTGCCACTCCAGTAGTCATCTGTAGGCNNNAGATCGGAAG |
| 51094_1 | TATGAGTGTGGAGTCGTTGCGTCTTCCTACACCCAGTCTCATCNNNAGATCGGAAGA |
| 51094_2 | TATGAGTGTGGAGTCGTTGCGTCTTCCTACACCCAGTCTCATCNNNAGATCGGAAGA |
| 35489_1 | TATGAGTGTGGAGTCGTTGCCAAGTAGCTGGGACTACAGGCNNNAGATCGGAAGA |
| 35489_2 | TATGAGTGTGGAGTCGTTGCCAAGTAGCTGGGACTACAGGCNNNAGATCGGAAGA |
| 40737_1 | TATGAGTGTGGAGTCGTTGCCTCACATGAGATGGGCATGAAATNNNAGATCGGAA |
| 40737_2 | TATGAGTGTGGAGTCGTTGCCTCACATGAGATGGGCATGAAATNNNAGATCGGAA |
| 13386_1 | TATGAGTGTGGAGTCGTTGCGAAACCATAAATGTAACACTCATNNNAGATCGGAAG |
| 13386_2 | TATGAGTGTGGAGTCGTTGCGAAACCATAAATGTAACACTCATNNNAGATCGGAAG |
| 45339_1 | TATGAGTGTGGAGTCGTTGCCAAGCCAAGAATGGAGACATGTGNNNAGATCGGAA |
| 45339_2 | TATGAGTGTGGAGTCGTTGCCAAGCCAAGAATGGAGACATGTGNNNAGATCGGAA |
| 45985_1 | TATGAGTGTGGAGTCGTTGCATCTCCCAGGTTCAAGCGATTCNNNAGATCGGAAGA |
| 45985_2 | TATGAGTGTGGAGTCGTTGCATCTCCCAGGTTCAAGCGATTCNNNAGATCGGAAGA |
| 33345_1 | TATGAGTGTGGAGTCGTTGCGCCTACTGAGTAGCTGGGATTACNNNAGATCGGAA |
| 33345_2 | TATGAGTGTGGAGTCGTTGCGCCTACTGAGTAGCTGGGATTACNNNAGATCGGAA |
| 46629_1 | TATGAGTGTGGAGTCGTTGCACCACGCCTGGCTAATTTTTTTTNNNAGATCGGAAG |
| 46629_2 | TATGAGTGTGGAGTCGTTGCACCACGCCTGGCTAATTTTTTTTNNNAGATCGGAAG |
| 34314_1 | TATGAGTGTGGAGTCGTTGCCACCCCCTCCCTACTTCCAGNNNAGATCGGAAGAGCA |
| 34314_2 | TATGAGTGTGGAGTCGTTGCCACCCCCTCCCTACTTCCAGNNNAGATCGGAAGAGCA |
| 40253_1 | TATGAGTGTGGAGTCGTTGCCACCACTGTGGAAAGCAATTTGANNNAGATCGGAA |
| 40253_2 | TATGAGTGTGGAGTCGTTGCCACCACTGTGGAAAGCAATTTGANNNAGATCGGAA |
| 42057_1 | TATGAGTGTGGAGTCGTTGCAGAGAGAGAGAGAGAGAGCATCCNNNAGATCGGA |
| 42057_2 | TATGAGTGTGGAGTCGTTGCAGAGAGAGAGAGAGAGAGCATCCNNNAGATCGGA |
| 44789_1 | TATGAGTGTGGAGTCGTTGCCTCTTCTGTTCTCCCCTCCCTTCNNNAGATCGGAAGAG |
| 44789_2 | TATGAGTGTGGAGTCGTTGCCTCTTCTGTTCTCCCCTCCCTTCNNNAGATCGGAAGAG |
| 12587_1 | TATGAGTGTGGAGTCGTTGCTTAGTCTTCATTCCCATCCTGNNNAGATCGGAAGAG |
| 12587_2 | TATGAGTGTGGAGTCGTTGCTTAGTCTTCATTCCCATCCTGNNNAGATCGGAAGAG |
| 49904_1 | TATGAGTGTGGAGTCGTTGCTTCTCCGCTGCCGAAAGGTAGNNNAGATCGGAAGA |
| 49904_2 | TATGAGTGTGGAGTCGTTGCTTCTCCGCTGCCGAAAGGTAGNNNAGATCGGAAGA |
| 48012_1 | TATGAGTGTGGAGTCGTTGCTGGTTTGGAGAAGTTCTGTGGAGNNNAGATCGGAA |
| 48012_2 | TATGAGTGTGGAGTCGTTGCTGGTTTGGAGAAGTTCTGTGGAGNNNAGATCGGAA |
| 43882_1 | TATGAGTGTGGAGTCGTTGCCTTCTCCTGATCTGCTGTGGATTNNNAGATCGGAAG |
| 43882_2 | TATGAGTGTGGAGTCGTTGCCTTCTCCTGATCTGCTGTGGATTNNNAGATCGGAAG |
| 38658_1 | TATGAGTGTGGAGTCGTTGCTTGTCCACTTGTTTCACCTGGTANNNAGATCGGAAG |
| 38658_2 | TATGAGTGTGGAGTCGTTGCTTGTCCACTTGTTTCACCTGGTANNNAGATCGGAAG |

| | |
|---|---|
| 12298_1 | TATGAGTGTGGAGTCGTTGCACTGCCAGATGTCACCTCTCTGNNNAGATCGGAAGA |
| 12298_2 | TATGAGTGTGGAGTCGTTGCACTGCCAGATGTCACCTCTCTGNNNAGATCGGAAGA |
| 39773_1 | TATGAGTGTGGAGTCGTTGCTGAAGGCCACAAATAGTATTGCANNNAGATCGGAA |
| 39773_2 | TATGAGTGTGGAGTCGTTGCTGAAGGCCACAAATAGTATTGCANNNAGATCGGAA |
| 33514_1 | TATGAGTGTGGAGTCGTTGCAGGTAGGAGAATTGCTTGAACCCNNNAGATCGGAA |
| 33514_2 | TATGAGTGTGGAGTCGTTGCAGGTAGGAGAATTGCTTGAACCCNNNAGATCGGAA |
| 28945_1 | TATGAGTGTGGAGTCGTTGCTGTTTTCTTCTGTCATCCCACCANNNAGATCGGAAGA |
| 28945_2 | TATGAGTGTGGAGTCGTTGCTGTTTTCTTCTGTCATCCCACCANNNAGATCGGAAGA |
| 39961_1 | TATGAGTGTGGAGTCGTTGCGAAATTTTTGACTGGAGGGGCGNNNAGATCGGAAG |
| 39961_2 | TATGAGTGTGGAGTCGTTGCGAAATTTTTGACTGGAGGGGCGNNNAGATCGGAAG |
| 50547_1 | TATGAGTGTGGAGTCGTTGCTCTTCTCCACTCCCAAATGATCGNNNAGATCGGAAGA |
| 50547_2 | TATGAGTGTGGAGTCGTTGCTCTTCTCCACTCCCAAATGATCGNNNAGATCGGAAGA |
| 42286_1 | TATGAGTGTGGAGTCGTTGCTGTTCATGTGCTTTTTGGGCATTNNNAGATCGGAAG |
| 42286_2 | TATGAGTGTGGAGTCGTTGCTGTTCATGTGCTTTTTGGGCATTNNNAGATCGGAAG |
| 31423_1 | TATGAGTGTGGAGTCGTTGCTGGTATCAAAGTTGGGACCAGAGNNNAGATCGGAA |
| 31423_2 | TATGAGTGTGGAGTCGTTGCTGGTATCAAAGTTGGGACCAGAGNNNAGATCGGAA |
| 45539_1 | TATGAGTGTGGAGTCGTTGCCGAGAATGATGATGACACCCTCTNNNAGATCGGAAG |
| 45539_2 | TATGAGTGTGGAGTCGTTGCCGAGAATGATGATGACACCCTCTNNNAGATCGGAAG |
| 48827_1 | TATGAGTGTGGAGTCGTTGCGTAGAGTAGCTGGGATTACAGGCNNNAGATCGGAA |
| 48827_2 | TATGAGTGTGGAGTCGTTGCGTAGAGTAGCTGGGATTACAGGCNNNAGATCGGAA |
| 50352_1 | TATGAGTGTGGAGTCGTTGCAGGTTCTTCCTATTGGCTCATGCNNNAGATCGGAAG |
| 50352_2 | TATGAGTGTGGAGTCGTTGCAGGTTCTTCCTATTGGCTCATGCNNNAGATCGGAAG |
| 42239_1 | TATGAGTGTGGAGTCGTTGCAATCGTATATGCACAGCACTCCANNNAGATCGGAAG |
| 42239_2 | TATGAGTGTGGAGTCGTTGCAATCGTATATGCACAGCACTCCANNNAGATCGGAAG |
| 13450_1 | TATGAGTGTGGAGTCGTTGCCTGACCATTAGCATCACTTAATNNNAGATCGGAAGA |
| 13450_2 | TATGAGTGTGGAGTCGTTGCCTGACCATTAGCATCACTTAATNNNAGATCGGAAGA |
| 47367_1 | TATGAGTGTGGAGTCGTTGCTTAGTCTGTTCTCACACTGCTGTNNNAGATCGGAAGA |
| 47367_2 | TATGAGTGTGGAGTCGTTGCTTAGTCTGTTCTCACACTGCTGTNNNAGATCGGAAGA |
| 32245_1 | TATGAGTGTGGAGTCGTTGCTTCCCCCCAGAAAAATGCAGATANNNAGATCGGAAG |
| 32245_2 | TATGAGTGTGGAGTCGTTGCTTCCCCCCAGAAAAATGCAGATANNNAGATCGGAAG |
| 45567_1 | TATGAGTGTGGAGTCGTTGCGAGAGGGAGGGAAGGAGAAAAAGNNNAGATCGG |
| 45567_2 | TATGAGTGTGGAGTCGTTGCGAGAGGGAGGGAAGGAGAAAAAGNNNAGATCGG |
| 41706_1 | TATGAGTGTGGAGTCGTTGCTTCAAGCGATTGTCCTTCCTCAGNNNAGATCGGAAG |
| 41706_2 | TATGAGTGTGGAGTCGTTGCTTCAAGCGATTGTCCTTCCTCAGNNNAGATCGGAAG |
| 43267_1 | TATGAGTGTGGAGTCGTTGCATTACAGGAGTTACTGCACCAGGNNNAGATCGGAA |
| 43267_2 | TATGAGTGTGGAGTCGTTGCATTACAGGAGTTACTGCACCAGGNNNAGATCGGAA |
| 33500_1 | TATGAGTGTGGAGTCGTTGCCACACCTGTAATCCCAGCACTTTNNNAGATCGGAAGA |
| 33500_2 | TATGAGTGTGGAGTCGTTGCCACACCTGTAATCCCAGCACTTTNNNAGATCGGAAGA |
| 12565_1 | TATGAGTGTGGAGTCGTTGCATTCTCCTTTAGTCATTCAACAANNNAGATCGGAAGA |
| 12565_2 | TATGAGTGTGGAGTCGTTGCATTCTCCTTTAGTCATTCAACAANNNAGATCGGAAGA |
| 53129_1 | TATGAGTGTGGAGTCGTTGCAGACTTTGTGCTCAGTGGTATGTNNNAGATCGGAA |
| 53129_2 | TATGAGTGTGGAGTCGTTGCAGACTTTGTGCTCAGTGGTATGTNNNAGATCGGAA |
| 44260_1 | TATGAGTGTGGAGTCGTTGCGAGGTTTTGGGCATTTTCAAGGANNNAGATCGGAA |

| | |
|---|---|
| 44260_2 | TATGAGTGTGGAGTCGTTGCGAGGTTTTGGGCATTTTCAAGGANNNAGATCGGAA |
| 40116_1 | TATGAGTGTGGAGTCGTTGCAGGAATGTTGCAGAGAGGATTCANNNAGATCGGAA |
| 40116_2 | TATGAGTGTGGAGTCGTTGCAGGAATGTTGCAGAGAGGATTCANNNAGATCGGAA |
| 29068_1 | TATGAGTGTGGAGTCGTTGCACACATACACACACATATATATACACGNNNAGATCGG |
| 29068_2 | TATGAGTGTGGAGTCGTTGCACACATACACACACATATATATACACGNNNAGATCGG |
| 34228_1 | TATGAGTGTGGAGTCGTTGCGTGGAGATTATTTCCTGGCAAGGNNNAGATCGGAA |
| 34228_2 | TATGAGTGTGGAGTCGTTGCGTGGAGATTATTTCCTGGCAAGGNNNAGATCGGAA |
| 29880_1 | TATGAGTGTGGAGTCGTTGCTTTGGTAGAGAAGGTGTCTTGCTNNNAGATCGGAA |
| 29880_2 | TATGAGTGTGGAGTCGTTGCTTTGGTAGAGAAGGTGTCTTGCTNNNAGATCGGAA |
| 44439_1 | TATGAGTGTGGAGTCGTTGCAGCAGCTGAGATTACAGGCATAANNNAGATCGGAA |
| 44439_2 | TATGAGTGTGGAGTCGTTGCAGCAGCTGAGATTACAGGCATAANNNAGATCGGAA |
| 11689_1 | TATGAGTGTGGAGTCGTTGCATTATGGGCATCTACAACCATGCNNNAGATCGGAAG |
| 11689_2 | TATGAGTGTGGAGTCGTTGCATTATGGGCATCTACAACCATGCNNNAGATCGGAAG |
| 52418_1 | TATGAGTGTGGAGTCGTTGCGAGGAGAGAGGGAGGAGAGTGNNNAGATCGGAA |
| 52418_2 | TATGAGTGTGGAGTCGTTGCGAGGAGAGAGGGAGGAGAGTGNNNAGATCGGAA |
| 45559_1 | TATGAGTGTGGAGTCGTTGCCATTCCAGAGTAGAGGCCGAATTNNNAGATCGGAA |
| 45559_2 | TATGAGTGTGGAGTCGTTGCCATTCCAGAGTAGAGGCCGAATTNNNAGATCGGAA |
| 40245_1 | TATGAGTGTGGAGTCGTTGCTCTAAAACTAACCCCCTCCTTGCNNNAGATCGGAAGA |
| 40245_2 | TATGAGTGTGGAGTCGTTGCTCTAAAACTAACCCCCTCCTTGCNNNAGATCGGAAGA |
| 39152_1 | TATGAGTGTGGAGTCGTTGCAAACAAGGAAGCAATAGCCTCTGNNNAGATCGGAA |
| 39152_2 | TATGAGTGTGGAGTCGTTGCAAACAAGGAAGCAATAGCCTCTGNNNAGATCGGAA |
| 10292_1 | TATGAGTGTGGAGTCGTTGCGGTAACAAGGTCAAATCTGCACANNNAGATCGGAA |
| 10292_2 | TATGAGTGTGGAGTCGTTGCGGTAACAAGGTCAAATCTGCACANNNAGATCGGAA |
| 39582_1 | TATGAGTGTGGAGTCGTTGCCTGAGGCAGGAGAATCACTTGAANNNAGATCGGAA |
| 39582_2 | TATGAGTGTGGAGTCGTTGCCTGAGGCAGGAGAATCACTTGAANNNAGATCGGAA |
| 12396_1 | TATGAGTGTGGAGTCGTTGCAGATACAGTCTCACTTTGTGGCCANNNAGATCGGAA |
| 12396_2 | TATGAGTGTGGAGTCGTTGCAGATACAGTCTCACTTTGTGGCCANNNAGATCGGAA |
| 13724_1 | TATGAGTGTGGAGTCGTTGCTCGCATAAATATTTTAGGAAGTGNNNAGATCGGAA |
| 13724_2 | TATGAGTGTGGAGTCGTTGCTCGCATAAATATTTTAGGAAGTGNNNAGATCGGAA |
| 3156_1 | TATGAGTGTGGAGTCGTTGCTTGGAGTTGCAGTTGATGTGTGANNNAGATCGGAA |
| 3156_2 | TATGAGTGTGGAGTCGTTGCTTGGAGTTGCAGTTGATGTGTGANNNAGATCGGAA |
| 12691_1 | TATGAGTGTGGAGTCGTTGCAGTTGTTACTCATAGAGGTATTGNNNAGATCGGAA |
| 12691_2 | TATGAGTGTGGAGTCGTTGCAGTTGTTACTCATAGAGGTATTGNNNAGATCGGAA |
| 33961_1 | TATGAGTGTGGAGTCGTTGCCAATCTTGGCTCACTGCAACCNNNAGATCGGAAGAG |
| 33961_2 | TATGAGTGTGGAGTCGTTGCCAATCTTGGCTCACTGCAACCNNNAGATCGGAAGAG |
| 531_1 | TATGAGTGTGGAGTCGTTGCTCCCAGCCCTGAGCAAGACATNNNAGATCGGAAGAG |
| 531_2 | TATGAGTGTGGAGTCGTTGCTCCCAGCCCTGAGCAAGACATNNNAGATCGGAAGAG |
| 29402_1 | TATGAGTGTGGAGTCGTTGCGGTCAACATCTTTTACATTCTGNNNAGATCGGAAGA |
| 29402_2 | TATGAGTGTGGAGTCGTTGCGGTCAACATCTTTTACATTCTGNNNAGATCGGAAGA |
| 34648_1 | TATGAGTGTGGAGTCGTTGCTGGTGCCAGTTGTAAAGAAGTCTNNNAGATCGGAA |
| 34648_2 | TATGAGTGTGGAGTCGTTGCTGGTGCCAGTTGTAAAGAAGTCTNNNAGATCGGAA |
| 35980_1 | TATGAGTGTGGAGTCGTTGCCTCCCTTCATCCTGCAGCTTTTCNNNAGATCGGAAGA |
| 35980_2 | TATGAGTGTGGAGTCGTTGCCTCCCTTCATCCTGCAGCTTTTCNNNAGATCGGAAGA |

| | |
|---|---|
| 30331_1 | TATGAGTGTGGAGTCGTTGCTTCCTCAGATGTGCTTTTCCTGGNNNAGATCGGAAG |
| 30331_2 | TATGAGTGTGGAGTCGTTGCTTCCTCAGATGTGCTTTTCCTGGNNNAGATCGGAAG |
| 49172_1 | TATGAGTGTGGAGTCGTTGCGAGTTTTGCTCTTATTGCCCAGGNNNAGATCGGAAG |
| 49172_2 | TATGAGTGTGGAGTCGTTGCGAGTTTTGCTCTTATTGCCCAGGNNNAGATCGGAAG |
| 45189_1 | TATGAGTGTGGAGTCGTTGCATATGGATGGAGCAGGGGGTATANNNAGATCGGA |
| 45189_2 | TATGAGTGTGGAGTCGTTGCATATGGATGGAGCAGGGGGTATANNNAGATCGGA |
| 12975_1 | TATGAGTGTGGAGTCGTTGCTTTGGTCGTGTGTACATATTTATNNNAGATCGGAAG |
| 12975_2 | TATGAGTGTGGAGTCGTTGCTTTGGTCGTGTGTACATATTTATNNNAGATCGGAAG |
| 50856_1 | TATGAGTGTGGAGTCGTTGCCCATCCACACTCCCATTACAAGANNNAGATCGGAAG |
| 50856_2 | TATGAGTGTGGAGTCGTTGCCCATCCACACTCCCATTACAAGANNNAGATCGGAAG |
| 48936_1 | TATGAGTGTGGAGTCGTTGCCAACCCCTTTTCATGCATCCATGNNNAGATCGGAAGA |
| 48936_2 | TATGAGTGTGGAGTCGTTGCCAACCCCTTTTCATGCATCCATGNNNAGATCGGAAGA |
| 45301_1 | TATGAGTGTGGAGTCGTTGCGAGGAAGAAAGGAGGGTGTCTCNNNAGATCGGAA |
| 45301_2 | TATGAGTGTGGAGTCGTTGCGAGGAAGAAAGGAGGGTGTCTCNNNAGATCGGAA |
| 30689_1 | TATGAGTGTGGAGTCGTTGCAGAGAGATGCTGAGGTTTCTGAGTNNNAGATCGGA |
| 30689_2 | TATGAGTGTGGAGTCGTTGCAGAGAGATGCTGAGGTTTCTGAGTNNNAGATCGGA |
| 44710_1 | TATGAGTGTGGAGTCGTTGCCCTTAGGTACCTTAGGGGGCATANNNAGATCGGAA |
| 44710_2 | TATGAGTGTGGAGTCGTTGCCCTTAGGTACCTTAGGGGGCATANNNAGATCGGAA |
| 49787_1 | TATGAGTGTGGAGTCGTTGCACCTTTGTTGCAGTTCTCTCACTNNNAGATCGGAAGA |
| 49787_2 | TATGAGTGTGGAGTCGTTGCACCTTTGTTGCAGTTCTCTCACTNNNAGATCGGAAGA |
| 34603_1 | TATGAGTGTGGAGTCGTTGCGAAGGAGCGGAAAAGTTGGAAGNNNAGATCGGAA |
| 34603_2 | TATGAGTGTGGAGTCGTTGCGAAGGAGCGGAAAAGTTGGAAGNNNAGATCGGAA |
| 35159_1 | TATGAGTGTGGAGTCGTTGCCTGGGACTATAGGCATGTACCACNNNAGATCGGAA |
| 35159_2 | TATGAGTGTGGAGTCGTTGCCTGGGACTATAGGCATGTACCACNNNAGATCGGAA |
| 45523_1 | TATGAGTGTGGAGTCGTTGCACATCTCAGCAGGGAAAATAGCANNNAGATCGGAA |
| 45523_2 | TATGAGTGTGGAGTCGTTGCACATCTCAGCAGGGAAAATAGCANNNAGATCGGAA |
| 36231_1 | TATGAGTGTGGAGTCGTTGCAGCCAACTTCATCACTCAGTCATNNNAGATCGGAAG |
| 36231_2 | TATGAGTGTGGAGTCGTTGCAGCCAACTTCATCACTCAGTCATNNNAGATCGGAAG |
| 35162_1 | TATGAGTGTGGAGTCGTTGCTTCTGCCGATTCAAGCAATTCTGNNNAGATCGGAAG |
| 35162_2 | TATGAGTGTGGAGTCGTTGCTTCTGCCGATTCAAGCAATTCTGNNNAGATCGGAAG |
| 23238_1 | TATGAGTGTGGAGTCGTTGCTTTGAATGATAAACTTTCCTCTCNNNAGATCGGAAG |
| 23238_2 | TATGAGTGTGGAGTCGTTGCTTTGAATGATAAACTTTCCTCTCNNNAGATCGGAAG |
| 53343_1 | TATGAGTGTGGAGTCGTTGCACACACACACACACAAGCTATCTNNNAGATCGGAAG |
| 53343_2 | TATGAGTGTGGAGTCGTTGCACACACACACACACAAGCTATCTNNNAGATCGGAAG |
| 50222_1 | TATGAGTGTGGAGTCGTTGCTGTGTGGATGTGGGAGACTAATCNNNAGATCGGAA |
| 50222_2 | TATGAGTGTGGAGTCGTTGCTGTGTGGATGTGGGAGACTAATCNNNAGATCGGAA |
| 28802_1 | TATGAGTGTGGAGTCGTTGCAGCCACTGGAGAAACATAAGTGANNNAGATCGGAA |
| 28802_2 | TATGAGTGTGGAGTCGTTGCAGCCACTGGAGAAACATAAGTGANNNAGATCGGAA |
| 29784_1 | TATGAGTGTGGAGTCGTTGCAGGCTTCAAAACCTTCTTCTGTCNNNAGATCGGAAG |
| 29784_2 | TATGAGTGTGGAGTCGTTGCAGGCTTCAAAACCTTCTTCTGTCNNNAGATCGGAAG |
| 50992_1 | TATGAGTGTGGAGTCGTTGCCAGAACTTCTGGCCTCTGAGAATNNNAGATCGGAAG |
| 50992_2 | TATGAGTGTGGAGTCGTTGCCAGAACTTCTGGCCTCTGAGAATNNNAGATCGGAAG |
| 48075_1 | TATGAGTGTGGAGTCGTTGCGATGGAGTCTCACTCTAGTTGCCNNNAGATCGGAAG |

| | |
|---|---|
| 48075_2 | TATGAGTGTGGAGTCGTTGCGATGGAGTCTCACTCTAGTTGCCNNNAGATCGGAAG |
| 34718_1 | TATGAGTGTGGAGTCGTTGCTAGAACCATCACGCTATTAGGGCNNNAGATCGGAAG |
| 34718_2 | TATGAGTGTGGAGTCGTTGCTAGAACCATCACGCTATTAGGGCNNNAGATCGGAAG |
| 46523_1 | TATGAGTGTGGAGTCGTTGCTGGTTTTGATTTGCATTTCCCAGANNNAGATCGGAA |
| 46523_2 | TATGAGTGTGGAGTCGTTGCTGGTTTTGATTTGCATTTCCCAGANNNAGATCGGAA |
| 52355_1 | TATGAGTGTGGAGTCGTTGCCCCTTCCTCTTCTTTACACCCTCNNNAGATCGGAAGAG |
| 52355_2 | TATGAGTGTGGAGTCGTTGCCCCTTCCTCTTCTTTACACCCTCNNNAGATCGGAAGAG |
| 51724_1 | TATGAGTGTGGAGTCGTTGCTGCAAAAGAGACCTTCCGTTTTCNNNAGATCGGAAG |
| 51724_2 | TATGAGTGTGGAGTCGTTGCTGCAAAAGAGACCTTCCGTTTTCNNNAGATCGGAAG |
| 13735_1 | TATGAGTGTGGAGTCGTTGCATTTTCTTAGCACCATTTCTTAGNNNAGATCGGAAGA |
| 13735_2 | TATGAGTGTGGAGTCGTTGCATTTTCTTAGCACCATTTCTTAGNNNAGATCGGAAGA |
| 49548_1 | TATGAGTGTGGAGTCGTTGCACACACATTATCAGAGCCACCTTNNNAGATCGGAAG |
| 49548_2 | TATGAGTGTGGAGTCGTTGCACACACATTATCAGAGCCACCTTNNNAGATCGGAAG |
| 41783_1 | TATGAGTGTGGAGTCGTTGCAGACGGAGTCTCACTCTGTCGNNNAGATCGGAAGA |
| 41783_2 | TATGAGTGTGGAGTCGTTGCAGACGGAGTCTCACTCTGTCGNNNAGATCGGAAGA |
| 35765_1 | TATGAGTGTGGAGTCGTTGCTGCGATCTCAGCTCACTACAAGNNNAGATCGGAAGA |
| 35765_2 | TATGAGTGTGGAGTCGTTGCTGCGATCTCAGCTCACTACAAGNNNAGATCGGAAGA |
| 40705_1 | TATGAGTGTGGAGTCGTTGCCATGTGCAGGTGGATTGGATTTTNNNAGATCGGAA |
| 40705_2 | TATGAGTGTGGAGTCGTTGCCATGTGCAGGTGGATTGGATTTTNNNAGATCGGAA |
| 35072_1 | TATGAGTGTGGAGTCGTTGCCTCAGCCACCTGAATAGTTGAGANNNAGATCGGAAG |
| 35072_2 | TATGAGTGTGGAGTCGTTGCCTCAGCCACCTGAATAGTTGAGANNNAGATCGGAAG |
| 40696_1 | TATGAGTGTGGAGTCGTTGCATGTACATGCCTCTTTGTCGGAANNNAGATCGGAAG |
| 40696_2 | TATGAGTGTGGAGTCGTTGCATGTACATGCCTCTTTGTCGGAANNNAGATCGGAAG |
| 32316_1 | TATGAGTGTGGAGTCGTTGCTGTTCAGATTCTTTGGTCCCACANNNAGATCGGAAG |
| 32316_2 | TATGAGTGTGGAGTCGTTGCTGTTCAGATTCTTTGGTCCCACANNNAGATCGGAAG |
| 29961_1 | TATGAGTGTGGAGTCGTTGCCTGAATGTACCTGAGCTGCAGAGNNNAGATCGGAA |
| 29961_2 | TATGAGTGTGGAGTCGTTGCCTGAATGTACCTGAGCTGCAGAGNNNAGATCGGAA |
| 31413_1 | TATGAGTGTGGAGTCGTTGCACATTTCATAGCTCAGTGCAGTGNNNAGATCGGAAG |
| 31413_2 | TATGAGTGTGGAGTCGTTGCACATTTCATAGCTCAGTGCAGTGNNNAGATCGGAAG |
| 33475_1 | TATGAGTGTGGAGTCGTTGCCAGACGCACAAAAGGTCACATAGNNNAGATCGGAA |
| 33475_2 | TATGAGTGTGGAGTCGTTGCCAGACGCACAAAAGGTCACATAGNNNAGATCGGAA |
| 13312_1 | TATGAGTGTGGAGTCGTTGCAATCTGAATAACGGCCAAATNNNAGATCGGAAGAG |
| 13312_2 | TATGAGTGTGGAGTCGTTGCAATCTGAATAACGGCCAAATNNNAGATCGGAAGAG |
| 48455_1 | TATGAGTGTGGAGTCGTTGCGGACGATCGCTGAGTAACATTTGNNNAGATCGGAA |
| 48455_2 | TATGAGTGTGGAGTCGTTGCGGACGATCGCTGAGTAACATTTGNNNAGATCGGAA |
| 5429_1 | TATGAGTGTGGAGTCGTTGCGGGGGTTGTCTGTGAATGAGAAANNNAGATCGGA |
| 5429_2 | TATGAGTGTGGAGTCGTTGCGGGGGTTGTCTGTGAATGAGAAANNNAGATCGGA |
| 12554_1 | TATGAGTGTGGAGTCGTTGCTGACCTGGTTAATAAATATGCTANNNAGATCGGAAG |
| 12554_2 | TATGAGTGTGGAGTCGTTGCTGACCTGGTTAATAAATATGCTANNNAGATCGGAAG |
| 10015_1 | TATGAGTGTGGAGTCGTTGCATAAAAGAGAAACCTTGGGCCGGNNNAGATCGGAA |
| 10015_2 | TATGAGTGTGGAGTCGTTGCATAAAAGAGAAACCTTGGGCCGGNNNAGATCGGAA |
| 32074_1 | TATGAGTGTGGAGTCGTTGCCAGGTTCAAGCGATTGTCCTGNNNAGATCGGAAGA |
| 32074_2 | TATGAGTGTGGAGTCGTTGCCAGGTTCAAGCGATTGTCCTGNNNAGATCGGAAGA |

| | |
|---|---|
| 6137_1 | TATGAGTGTGGAGTCGTTGCAACTGGAGGAACAATGTGTGTGTNNNAGATCGGAA |
| 6137_2 | TATGAGTGTGGAGTCGTTGCAACTGGAGGAACAATGTGTGTGTNNNAGATCGGAA |
| 48416_1 | TATGAGTGTGGAGTCGTTGCCTCCCAGGTTCATGCCATTCTCNNNAGATCGGAAGA |
| 48416_2 | TATGAGTGTGGAGTCGTTGCCTCCCAGGTTCATGCCATTCTCNNNAGATCGGAAGA |
| 32051_1 | TATGAGTGTGGAGTCGTTGCTAAGAAAAGAGCGGGGAAACAGANNNAGATCGGA |
| 32051_2 | TATGAGTGTGGAGTCGTTGCTAAGAAAAGAGCGGGGAAACAGANNNAGATCGGA |
| 36378_1 | TATGAGTGTGGAGTCGTTGCCCCAGGGTAAGAGGTTGAAACAANNNAGATCGGAA |
| 36378_2 | TATGAGTGTGGAGTCGTTGCCCCAGGGTAAGAGGTTGAAACAANNNAGATCGGAA |
| 42352_1 | TATGAGTGTGGAGTCGTTGCTAATATTTCACAACGCACCCGTGNNNAGATCGGAAG |
| 42352_2 | TATGAGTGTGGAGTCGTTGCTAATATTTCACAACGCACCCGTGNNNAGATCGGAAG |
| 45883_1 | TATGAGTGTGGAGTCGTTGCCCAGACTAGTGTCCTCAAGCATTNNNAGATCGGAAG |
| 45883_2 | TATGAGTGTGGAGTCGTTGCCCAGACTAGTGTCCTCAAGCATTNNNAGATCGGAAG |
| 43133_1 | TATGAGTGTGGAGTCGTTGCTGTTCCAGAATGTTCTCTCCCAGNNNAGATCGGAAG |
| 43133_2 | TATGAGTGTGGAGTCGTTGCTGTTCCAGAATGTTCTCTCCCAGNNNAGATCGGAAG |
| 31080_1 | TATGAGTGTGGAGTCGTTGCCACACACATGCACAGACACGNNNAGATCGGAAGAG |
| 31080_2 | TATGAGTGTGGAGTCGTTGCCACACACATGCACAGACACGNNNAGATCGGAAGAG |
| 52167_1 | TATGAGTGTGGAGTCGTTGCTCTACCAACTGCTTGCACTATGTNNNAGATCGGAAG |
| 52167_2 | TATGAGTGTGGAGTCGTTGCTCTACCAACTGCTTGCACTATGTNNNAGATCGGAAG |
| 43997_1 | TATGAGTGTGGAGTCGTTGCTTTAGTCACCTTGCACTGTTGGANNNAGATCGGAAG |
| 43997_2 | TATGAGTGTGGAGTCGTTGCTTTAGTCACCTTGCACTGTTGGANNNAGATCGGAAG |
| 40440_1 | TATGAGTGTGGAGTCGTTGCCAGGAGTTGCTCACTGTTACAGTNNNAGATCGGAAG |
| 40440_2 | TATGAGTGTGGAGTCGTTGCCAGGAGTTGCTCACTGTTACAGTNNNAGATCGGAAG |
| 13786_1 | TATGAGTGTGGAGTCGTTGCGAGACCTACATTACACCACTNNNAGATCGGAAGAGC |
| 13786_2 | TATGAGTGTGGAGTCGTTGCGAGACCTACATTACACCACTNNNAGATCGGAAGAGC |
| 3130_1 | TATGAGTGTGGAGTCGTTGCCTCTCTCCAGTGTGAATCCTCCTNNNAGATCGGAAGA |
| 3130_2 | TATGAGTGTGGAGTCGTTGCCTCTCTCCAGTGTGAATCCTCCTNNNAGATCGGAAGA |
| 40974_1 | TATGAGTGTGGAGTCGTTGCCTGAGTGCTAATTGTGCTGCTAGNNNAGATCGGAA |
| 40974_2 | TATGAGTGTGGAGTCGTTGCCTGAGTGCTAATTGTGCTGCTAGNNNAGATCGGAA |
| 41386_1 | TATGAGTGTGGAGTCGTTGCACACAAGCAAAAATCCTTCTGGGNNNAGATCGGAAG |
| 41386_2 | TATGAGTGTGGAGTCGTTGCACACAAGCAAAAATCCTTCTGGGNNNAGATCGGAAG |
| 48267_1 | TATGAGTGTGGAGTCGTTGCGTCTTGATCTCCTGACCTTGTGANNNAGATCGGAAG |
| 48267_2 | TATGAGTGTGGAGTCGTTGCGTCTTGATCTCCTGACCTTGTGANNNAGATCGGAAG |
| 47372_1 | TATGAGTGTGGAGTCGTTGCGAGTCTAGCTCTGTCGCCCANNNAGATCGGAAGAG |
| 47372_2 | TATGAGTGTGGAGTCGTTGCGAGTCTAGCTCTGTCGCCCANNNAGATCGGAAGAG |
| 40364_1 | TATGAGTGTGGAGTCGTTGCGCAGGAAAACATGATGGGCTATCNNNAGATCGGAA |
| 40364_2 | TATGAGTGTGGAGTCGTTGCGCAGGAAAACATGATGGGCTATCNNNAGATCGGAA |
| 12623_1 | TATGAGTGTGGAGTCGTTGCCAGGCTCTCAAGATAAAAGATANNNAGATCGGAAG |
| 12623_2 | TATGAGTGTGGAGTCGTTGCCAGGCTCTCAAGATAAAAGATANNNAGATCGGAAG |
| 12477_1 | TATGAGTGTGGAGTCGTTGCTGCTAAATAATTTAAATTGCAGCNNNAGATCGGAAG |
| 12477_2 | TATGAGTGTGGAGTCGTTGCTGCTAAATAATTTAAATTGCAGCNNNAGATCGGAAG |
| 34504_1 | TATGAGTGTGGAGTCGTTGCGTCTCTTCTGAGTCTAGTGACGCNNNAGATCGGAAG |
| 34504_2 | TATGAGTGTGGAGTCGTTGCGTCTCTTCTGAGTCTAGTGACGCNNNAGATCGGAAG |
| 44243_1 | TATGAGTGTGGAGTCGTTGCATAAGAACATTCCTGTCACCCCCNNNAGATCGGAAG |

| | |
|---|---|
| 44243_2 | TATGAGTGTGGAGTCGTTGCATAAGAACATTCCTGTCACCCCCNNNAGATCGGAAGA |
| 50216_1 | TATGAGTGTGGAGTCGTTGCCAGGCTACCCGTCAGACATCNNNAGATCGGAAGAGC |
| 50216_2 | TATGAGTGTGGAGTCGTTGCCAGGCTACCCGTCAGACATCNNNAGATCGGAAGAGC |
| 43634_1 | TATGAGTGTGGAGTCGTTGCGATAAACCATGACCTCAGCCGANNNAGATCGGAAGA |
| 43634_2 | TATGAGTGTGGAGTCGTTGCGATAAACCATGACCTCAGCCGANNNAGATCGGAAGA |
| 29439_1 | TATGAGTGTGGAGTCGTTGCCAATTTCATTATCTTCTTCACTCTNNNAGATCGGAAGA |
| 29439_2 | TATGAGTGTGGAGTCGTTGCCAATTTCATTATCTTCTTCACTCTNNNAGATCGGAAGA |
| 45403_1 | TATGAGTGTGGAGTCGTTGCGTACATTTTTGTGCCCGTGTGTANNNAGATCGGAAG |
| 45403_2 | TATGAGTGTGGAGTCGTTGCGTACATTTTTGTGCCCGTGTGTANNNAGATCGGAAG |
| 37563_1 | TATGAGTGTGGAGTCGTTGCTGCCATAAGGGACAAAGGTTCTTNNNAGATCGGAAG |
| 37563_2 | TATGAGTGTGGAGTCGTTGCTGCCATAAGGGACAAAGGTTCTTNNNAGATCGGAAG |
| 44194_1 | TATGAGTGTGGAGTCGTTGCTCAGTTTAGGATCTGCTATGGGCNNNAGATCGGAAG |
| 44194_2 | TATGAGTGTGGAGTCGTTGCTCAGTTTAGGATCTGCTATGGGCNNNAGATCGGAAG |
| 43167_1 | TATGAGTGTGGAGTCGTTGCGGCATAGACAATCTGCAAAGACCNNNAGATCGGAAG |
| 43167_2 | TATGAGTGTGGAGTCGTTGCGGCATAGACAATCTGCAAAGACCNNNAGATCGGAAG |
| 29582_1 | TATGAGTGTGGAGTCGTTGCTTTCCATCGGGACTCATTTCTGTNNNAGATCGGAAGA |
| 29582_2 | TATGAGTGTGGAGTCGTTGCTTTCCATCGGGACTCATTTCTGTNNNAGATCGGAAGA |
| 37549_1 | TATGAGTGTGGAGTCGTTGCTCCTGTGGGAAATATGCCATACANNNAGATCGGAAG |
| 37549_2 | TATGAGTGTGGAGTCGTTGCTCCTGTGGGAAATATGCCATACANNNAGATCGGAAG |
| 52233_1 | TATGAGTGTGGAGTCGTTGCCAGGAGGTGAGATCAGGAGCTNNNAGATCGGAAGA |
| 52233_2 | TATGAGTGTGGAGTCGTTGCCAGGAGGTGAGATCAGGAGCTNNNAGATCGGAAGA |
| 52749_1 | TATGAGTGTGGAGTCGTTGCAGCGGTAGAGTGTAATGAAAGCANNNAGATCGGAA |
| 52749_2 | TATGAGTGTGGAGTCGTTGCAGCGGTAGAGTGTAATGAAAGCANNNAGATCGGAA |
| 36632_1 | TATGAGTGTGGAGTCGTTGCCCTCTCATTGCAATAGCTTTCCGNNNAGATCGGAAGA |
| 36632_2 | TATGAGTGTGGAGTCGTTGCCCTCTCATTGCAATAGCTTTCCGNNNAGATCGGAAGA |
| 34136_1 | TATGAGTGTGGAGTCGTTGCCACATCACGAGGTCAGGAGATCNNNAGATCGGAAGA |
| 34136_2 | TATGAGTGTGGAGTCGTTGCCACATCACGAGGTCAGGAGATCNNNAGATCGGAAGA |
| 45638_1 | TATGAGTGTGGAGTCGTTGCACTCCTGGGTTCAAGTGATTCTCNNNAGATCGGAAGA |
| 45638_2 | TATGAGTGTGGAGTCGTTGCACTCCTGGGTTCAAGTGATTCTCNNNAGATCGGAAGA |
| 10957_1 | TATGAGTGTGGAGTCGTTGCGGGTGGGAGAGGCTCAGGAATGNNNAGATCGGAA |
| 10957_2 | TATGAGTGTGGAGTCGTTGCGGGTGGGAGAGGCTCAGGAATGNNNAGATCGGAA |
| 13250_1 | TATGAGTGTGGAGTCGTTGCATAAAATAGAAAGAATGGCAAGCNNNAGATCGGAA |
| 13250_2 | TATGAGTGTGGAGTCGTTGCATAAAATAGAAAGAATGGCAAGCNNNAGATCGGAA |
| 13125_1 | TATGAGTGTGGAGTCGTTGCTATGTTACAGGATTAGTGGATTTNNNAGATCGGAAG |
| 13125_2 | TATGAGTGTGGAGTCGTTGCTATGTTACAGGATTAGTGGATTTNNNAGATCGGAAG |
| 13274_1 | TATGAGTGTGGAGTCGTTGCTTACATGAACTCTATTAGGTGTCNNNAGATCGGAAGA |
| 13274_2 | TATGAGTGTGGAGTCGTTGCTTACATGAACTCTATTAGGTGTCNNNAGATCGGAAGA |
| 47591_1 | TATGAGTGTGGAGTCGTTGCCATGGGAAGACATTTCCGACAAGNNNAGATCGGAAG |
| 47591_2 | TATGAGTGTGGAGTCGTTGCCATGGGAAGACATTTCCGACAAGNNNAGATCGGAAG |
| 31123_1 | TATGAGTGTGGAGTCGTTGCCTGTTATCATCTCCCCCTCACTGNNNAGATCGGAAGAG |
| 31123_2 | TATGAGTGTGGAGTCGTTGCCTGTTATCATCTCCCCCTCACTGNNNAGATCGGAAGAG |
| 47943_1 | TATGAGTGTGGAGTCGTTGCTAGTTTGCTGACACATCACTCCANNNAGATCGGAAGA |
| 47943_2 | TATGAGTGTGGAGTCGTTGCTAGTTTGCTGACACATCACTCCANNNAGATCGGAAGA |

| | |
|---|---|
| 31697_1 | TATGAGTGTGGAGTCGTTGCATGGTCATGATGGGAAAAGTGGANNNAGATCGGA |
| 31697_2 | TATGAGTGTGGAGTCGTTGCATGGTCATGATGGGAAAAGTGGANNNAGATCGGA |
| 39872_1 | TATGAGTGTGGAGTCGTTGCAAGTGAAGATCATCTGCCACTGTNNNAGATCGGAAG |
| 39872_2 | TATGAGTGTGGAGTCGTTGCAAGTGAAGATCATCTGCCACTGTNNNAGATCGGAAG |
| 39301_1 | TATGAGTGTGGAGTCGTTGCAAGGAGGATCAAGATGAGGAACANNNAGATCGGA |
| 39301_2 | TATGAGTGTGGAGTCGTTGCAAGGAGGATCAAGATGAGGAACANNNAGATCGGA |
| 46064_1 | TATGAGTGTGGAGTCGTTGCGTATTGGGATTACAGGCATGAGCNNNAGATCGGAA |
| 46064_2 | TATGAGTGTGGAGTCGTTGCGTATTGGGATTACAGGCATGAGCNNNAGATCGGAA |
| 31084_1 | TATGAGTGTGGAGTCGTTGCTTTTTTGTGCATGTTGTCCTGGANNNAGATCGGAAG |
| 31084_2 | TATGAGTGTGGAGTCGTTGCTTTTTTGTGCATGTTGTCCTGGANNNAGATCGGAAG |
| 31009_1 | TATGAGTGTGGAGTCGTTGCGATGGAGGGGTCACAACAAGTATNNNAGATCGGAA |
| 31009_2 | TATGAGTGTGGAGTCGTTGCGATGGAGGGGTCACAACAAGTATNNNAGATCGGAA |
| 50782_1 | TATGAGTGTGGAGTCGTTGCCAGACACCTTCGTTCAAACTTGTNNNAGATCGGAAG |
| 50782_2 | TATGAGTGTGGAGTCGTTGCCAGACACCTTCGTTCAAACTTGTNNNAGATCGGAAG |
| 30058_1 | TATGAGTGTGGAGTCGTTGCTGGTCCAGACAGAGGTATGTAGANNNAGATCGGAA |
| 30058_2 | TATGAGTGTGGAGTCGTTGCTGGTCCAGACAGAGGTATGTAGANNNAGATCGGAA |
| 34801_1 | TATGAGTGTGGAGTCGTTGCTGTAGTGCAGTGGTGTGATCTCNNNAGATCGGAAG |
| 34801_2 | TATGAGTGTGGAGTCGTTGCTGTAGTGCAGTGGTGTGATCTCNNNAGATCGGAAG |
| 35152_1 | TATGAGTGTGGAGTCGTTGCTGATCTCGGCTCACTGCAAGNNNAGATCGGAAGAG |
| 35152_2 | TATGAGTGTGGAGTCGTTGCTGATCTCGGCTCACTGCAAGNNNAGATCGGAAGAG |
| 35314_1 | TATGAGTGTGGAGTCGTTGCCAGCCTCCTAAAGTACTGGGAAANNNAGATCGGAAG |
| 35314_2 | TATGAGTGTGGAGTCGTTGCCAGCCTCCTAAAGTACTGGGAAANNNAGATCGGAAG |
| 40123_1 | TATGAGTGTGGAGTCGTTGCACTGATTCTGTCATTCAAATTTGGTNNNAGATCGGA |
| 40123_2 | TATGAGTGTGGAGTCGTTGCACTGATTCTGTCATTCAAATTTGGTNNNAGATCGGA |
| 47726_1 | TATGAGTGTGGAGTCGTTGCAGAAGAGTTCAACGGGCATTACTNNNAGATCGGAA |
| 47726_2 | TATGAGTGTGGAGTCGTTGCAGAAGAGTTCAACGGGCATTACTNNNAGATCGGAA |
| 46092_1 | TATGAGTGTGGAGTCGTTGCCACAGGTTCAAGCAATTCTCCTGNNNAGATCGGAAG |
| 46092_2 | TATGAGTGTGGAGTCGTTGCCACAGGTTCAAGCAATTCTCCTGNNNAGATCGGAAG |
| 30228_1 | TATGAGTGTGGAGTCGTTGCGGTACCTCCAGTCCTCATGAAAGNNNAGATCGGAAG |
| 30228_2 | TATGAGTGTGGAGTCGTTGCGGTACCTCCAGTCCTCATGAAAGNNNAGATCGGAAG |
| 3047_1 | TATGAGTGTGGAGTCGTTGCCCACATCAGCCTAGTTTCGCTCNNNAGATCGGAAGA |
| 3047_2 | TATGAGTGTGGAGTCGTTGCCCACATCAGCCTAGTTTCGCTCNNNAGATCGGAAGA |
| 34902_1 | TATGAGTGTGGAGTCGTTGCCAGATGGAGTCTCACTCTGTCACNNNAGATCGGAAG |
| 34902_2 | TATGAGTGTGGAGTCGTTGCCAGATGGAGTCTCACTCTGTCACNNNAGATCGGAAG |
| 52021_1 | TATGAGTGTGGAGTCGTTGCGAGGAAAGGCAGGATAGTGTCTTNNNAGATCGGA |
| 52021_2 | TATGAGTGTGGAGTCGTTGCGAGGAAAGGCAGGATAGTGTCTTNNNAGATCGGA |
| 11254_1 | TATGAGTGTGGAGTCGTTGCGTGGAGTATTCATTCCAGCACAGNNNAGATCGGAA |
| 11254_2 | TATGAGTGTGGAGTCGTTGCGTGGAGTATTCATTCCAGCACAGNNNAGATCGGAA |
| 47232_1 | TATGAGTGTGGAGTCGTTGCAAAACAGATTGAGTCTTGTGCCANNNAGATCGGAA |
| 47232_2 | TATGAGTGTGGAGTCGTTGCAAAACAGATTGAGTCTTGTGCCANNNAGATCGGAA |
| 48394_1 | TATGAGTGTGGAGTCGTTGCCTAGGAAAGACCACGCTGATCAANNNAGATCGGAA |
| 48394_2 | TATGAGTGTGGAGTCGTTGCCTAGGAAAGACCACGCTGATCAANNNAGATCGGAA |
| 30837_1 | TATGAGTGTGGAGTCGTTGCGACTCAACTTATCCCCAGGTGTCNNNAGATCGGAAG |

| | |
|---|---|
| 30837_2 | TATGAGTGTGGAGTCGTTGCGACTCAACTTATCCCCAGGTGTCNNNAGATCGGAAG |
| 28846_1 | TATGAGTGTGGAGTCGTTGCCTTTCTCTCACCCACTTGCTTTGNNNAGATCGGAAGA |
| 28846_2 | TATGAGTGTGGAGTCGTTGCCTTTCTCTCACCCACTTGCTTTGNNNAGATCGGAAGA |
| 48290_1 | TATGAGTGTGGAGTCGTTGCTTCCTTTGTTTCCCTTCCCTACCNNNAGATCGGAAGAG |
| 48290_2 | TATGAGTGTGGAGTCGTTGCTTCCTTTGTTTCCCTTCCCTACCNNNAGATCGGAAGAG |
| 33401_1 | TATGAGTGTGGAGTCGTTGCCGAATTCATTTGGGCCTCATTCANNNAGATCGGAAG |
| 33401_2 | TATGAGTGTGGAGTCGTTGCCGAATTCATTTGGGCCTCATTCANNNAGATCGGAAG |
| 34910_1 | TATGAGTGTGGAGTCGTTGCCTCCACATCTCAGGTTCAAGTGANNNAGATCGGAAG |
| 34910_2 | TATGAGTGTGGAGTCGTTGCCTCCACATCTCAGGTTCAAGTGANNNAGATCGGAAG |
| 46571_1 | TATGAGTGTGGAGTCGTTGCAGCCTCCTGAGTAGCTGAGATTANNNAGATCGGAAG |
| 46571_2 | TATGAGTGTGGAGTCGTTGCAGCCTCCTGAGTAGCTGAGATTANNNAGATCGGAAG |
| 37716_1 | TATGAGTGTGGAGTCGTTGCGTGCTTGCCTCCTTCTTTTCATGNNNAGATCGGAAGA |
| 37716_2 | TATGAGTGTGGAGTCGTTGCGTGCTTGCCTCCTTCTTTTCATGNNNAGATCGGAAGA |
| 12865_1 | TATGAGTGTGGAGTCGTTGCGAGCATGTTATACTTCAGAAACNNNAGATCGGAAGA |
| 12865_2 | TATGAGTGTGGAGTCGTTGCGAGCATGTTATACTTCAGAAACNNNAGATCGGAAGA |
| 46122_1 | TATGAGTGTGGAGTCGTTGCCCACCATGCCCAGCTAATTTTTTNNNAGATCGGAAGA |
| 46122_2 | TATGAGTGTGGAGTCGTTGCCCACCATGCCCAGCTAATTTTTTNNNAGATCGGAAGA |
| 47904_1 | TATGAGTGTGGAGTCGTTGCTGTCTGAGAAAGGAAGCCAGATCNNNAGATCGGAA |
| 47904_2 | TATGAGTGTGGAGTCGTTGCTGTCTGAGAAAGGAAGCCAGATCNNNAGATCGGAA |
| 32045_1 | TATGAGTGTGGAGTCGTTGCGCTTAGGGTAGGCAACTAAAGGTNNNAGATCGGAA |
| 32045_2 | TATGAGTGTGGAGTCGTTGCGCTTAGGGTAGGCAACTAAAGGTNNNAGATCGGAA |
| 13169_1 | TATGAGTGTGGAGTCGTTGCGAAGGGGTTAACATGACAGTNNNAGATCGGAAGAG |
| 13169_2 | TATGAGTGTGGAGTCGTTGCGAAGGGGTTAACATGACAGTNNNAGATCGGAAGAG |
| 46785_1 | TATGAGTGTGGAGTCGTTGCCTGGGTTCAAATGATTCTCCTGCNNNAGATCGGAAG |
| 46785_2 | TATGAGTGTGGAGTCGTTGCCTGGGTTCAAATGATTCTCCTGCNNNAGATCGGAAG |
| 29281_1 | TATGAGTGTGGAGTCGTTGCAAACTCCCCCTGTCTAATCCCTANNNAGATCGGAAGA |
| 29281_2 | TATGAGTGTGGAGTCGTTGCAAACTCCCCCTGTCTAATCCCTANNNAGATCGGAAGA |
| 53384_1 | TATGAGTGTGGAGTCGTTGCCTTTCCCTCTCACTCCTTCTGTCNNNAGATCGGAAGAG |
| 53384_2 | TATGAGTGTGGAGTCGTTGCCTTTCCCTCTCACTCCTTCTGTCNNNAGATCGGAAGAG |
| 29988_1 | TATGAGTGTGGAGTCGTTGCAGATGACACTAGAGGAGAGCTGTNNNAGATCGGAA |
| 29988_2 | TATGAGTGTGGAGTCGTTGCAGATGACACTAGAGGAGAGCTGTNNNAGATCGGAA |
| 48548_1 | TATGAGTGTGGAGTCGTTGCGAAGCTCCAGCATAATTTACCCCNNNAGATCGGAAG |
| 48548_2 | TATGAGTGTGGAGTCGTTGCGAAGCTCCAGCATAATTTACCCCNNNAGATCGGAAG |
| 2484_1 | TATGAGTGTGGAGTCGTTGCAAATGCCAATAGATGTGCCCAGANNNAGATCGGAAG |
| 2484_2 | TATGAGTGTGGAGTCGTTGCAAATGCCAATAGATGTGCCCAGANNNAGATCGGAAG |
| 30110_1 | TATGAGTGTGGAGTCGTTGCGACACCAATGGACATTGCAGTTCNNNAGATCGGAAG |
| 30110_2 | TATGAGTGTGGAGTCGTTGCGACACCAATGGACATTGCAGTTCNNNAGATCGGAAG |
| 50925_1 | TATGAGTGTGGAGTCGTTGCACCCCATTCAGGACTCATTTTCANNNAGATCGGAAGA |
| 50925_2 | TATGAGTGTGGAGTCGTTGCACCCCATTCAGGACTCATTTTCANNNAGATCGGAAGA |
| 50101_1 | TATGAGTGTGGAGTCGTTGCGACAAAGTCTCATTCTGTTGCCCNNNAGATCGGAAG |
| 50101_2 | TATGAGTGTGGAGTCGTTGCGACAAAGTCTCATTCTGTTGCCCNNNAGATCGGAAG |
| 31943_1 | TATGAGTGTGGAGTCGTTGCGAAGGCAACAAAACTGGATGGAGNNNAGATCGGAA |
| 31943_2 | TATGAGTGTGGAGTCGTTGCGAAGGCAACAAAACTGGATGGAGNNNAGATCGGAA |

| | |
|---|---|
| 32466_1 | TATGAGTGTGGAGTCGTTGCTCAGTCCAGGATACAATGAAGGGNNNAGATCGGAA |
| 32466_2 | TATGAGTGTGGAGTCGTTGCTCAGTCCAGGATACAATGAAGGGNNNAGATCGGAA |
| 38050_1 | TATGAGTGTGGAGTCGTTGCTAGATCTGGGAACTGGGGATGATNNNAGATCGGAA |
| 38050_2 | TATGAGTGTGGAGTCGTTGCTAGATCTGGGAACTGGGGATGATNNNAGATCGGAA |
| 51850_1 | TATGAGTGTGGAGTCGTTGCCGCTCCTCACTTCCCAGATGNNNAGATCGGAAGAGC |
| 51850_2 | TATGAGTGTGGAGTCGTTGCCGCTCCTCACTTCCCAGATGNNNAGATCGGAAGAGC |
| 40968_1 | TATGAGTGTGGAGTCGTTGCACGCAAGGTTATTCATAGGCTCTNNNAGATCGGAAG |
| 40968_2 | TATGAGTGTGGAGTCGTTGCACGCAAGGTTATTCATAGGCTCTNNNAGATCGGAAG |
| 50347_1 | TATGAGTGTGGAGTCGTTGCTCTTCCATGACAAAATGAAGCAGANNNAGATCGGAA |
| 50347_2 | TATGAGTGTGGAGTCGTTGCTCTTCCATGACAAAATGAAGCAGANNNAGATCGGAA |
| 37064_1 | TATGAGTGTGGAGTCGTTGCCTACGGTATCAGAAACTCCAGGGNNNAGATCGGAA |
| 37064_2 | TATGAGTGTGGAGTCGTTGCCTACGGTATCAGAAACTCCAGGGNNNAGATCGGAA |
| 50681_1 | TATGAGTGTGGAGTCGTTGCGTGTATGGGGAACACAAACCTTGNNNAGATCGGAA |
| 50681_2 | TATGAGTGTGGAGTCGTTGCGTGTATGGGGAACACAAACCTTGNNNAGATCGGAA |
| 1641_1 | TATGAGTGTGGAGTCGTTGCGGGAGATTCTTCAAGTAAAGGGCANNNAGATCGGA |
| 1641_2 | TATGAGTGTGGAGTCGTTGCGGGAGATTCTTCAAGTAAAGGGCANNNAGATCGGA |
| 32891_1 | TATGAGTGTGGAGTCGTTGCATCAGTCTGCAGACAACTCATGANNNAGATCGGAAG |
| 32891_2 | TATGAGTGTGGAGTCGTTGCATCAGTCTGCAGACAACTCATGANNNAGATCGGAAG |
| 48744_1 | TATGAGTGTGGAGTCGTTGCTGACCCAGGTGTGTTTGAAAATGNNNAGATCGGAA |
| 48744_2 | TATGAGTGTGGAGTCGTTGCTGACCCAGGTGTGTTTGAAAATGNNNAGATCGGAA |
| 34440_1 | TATGAGTGTGGAGTCGTTGCTAGATGACAGTTTGATGGGTGCANNNAGATCGGAA |
| 34440_2 | TATGAGTGTGGAGTCGTTGCTAGATGACAGTTTGATGGGTGCANNNAGATCGGAA |
| 49389_1 | TATGAGTGTGGAGTCGTTGCGTAGTTGGGATTACAAGCGTGTGNNNAGATCGGAA |
| 49389_2 | TATGAGTGTGGAGTCGTTGCGTAGTTGGGATTACAAGCGTGTGNNNAGATCGGAA |
| 13720_1 | TATGAGTGTGGAGTCGTTGCGACTGTTAGTAACTTAGTCTCCNNNAGATCGGAAGA |
| 13720_2 | TATGAGTGTGGAGTCGTTGCGACTGTTAGTAACTTAGTCTCCNNNAGATCGGAAGA |
| 13333_1 | TATGAGTGTGGAGTCGTTGCCCTTTTGTGTGTTCTTTTATGANNNAGATCGGAAGA |
| 13333_2 | TATGAGTGTGGAGTCGTTGCCCTTTTGTGTGTTCTTTTATGANNNAGATCGGAAGA |
| 42777_1 | TATGAGTGTGGAGTCGTTGCGCACCTATTTCTTTGCCCTTGTTNNNAGATCGGAAGA |
| 42777_2 | TATGAGTGTGGAGTCGTTGCGCACCTATTTCTTTGCCCTTGTTNNNAGATCGGAAGA |
| 41055_1 | TATGAGTGTGGAGTCGTTGCGTCAACTAGGTTGCTGTAAAGGCNNNAGATCGGAA |
| 41055_2 | TATGAGTGTGGAGTCGTTGCGTCAACTAGGTTGCTGTAAAGGCNNNAGATCGGAA |
| 39260_1 | TATGAGTGTGGAGTCGTTGCGGGTTCAAGCAATTCTCCTACCTNNNAGATCGGAAG |
| 39260_2 | TATGAGTGTGGAGTCGTTGCGGGTTCAAGCAATTCTCCTACCTNNNAGATCGGAAG |
| 44933_1 | TATGAGTGTGGAGTCGTTGCCTTGGCATATCCCACTTGACCTANNNAGATCGGAAG |
| 44933_2 | TATGAGTGTGGAGTCGTTGCCTTGGCATATCCCACTTGACCTANNNAGATCGGAAG |
| 32501_1 | TATGAGTGTGGAGTCGTTGCGGCTCCTGAGTAGCTGAGATTACNNNAGATCGGAA |
| 32501_2 | TATGAGTGTGGAGTCGTTGCGGCTCCTGAGTAGCTGAGATTACNNNAGATCGGAA |
| 42184_1 | TATGAGTGTGGAGTCGTTGCCTAGTAGCTGGGATTACAGGCACNNNAGATCGGAA |
| 42184_2 | TATGAGTGTGGAGTCGTTGCCTAGTAGCTGGGATTACAGGCACNNNAGATCGGAA |
| 43350_1 | TATGAGTGTGGAGTCGTTGCCTAGTAGGTGGGATTACAGGTGCNNNAGATCGGAA |
| 43350_2 | TATGAGTGTGGAGTCGTTGCCTAGTAGGTGGGATTACAGGTGCNNNAGATCGGAA |
| 37551_1 | TATGAGTGTGGAGTCGTTGCTCACCTCACTCAGAGTAAAAGCCNNNAGATCGGAAG |

| | |
|---|---|
| 37551_2 | TATGAGTGTGGAGTCGTTGCTCACCTCACTCAGAGTAAAAGCCNNNAGATCGGAAG |
| 43913_1 | TATGAGTGTGGAGTCGTTGCTGGTAAGGGCGAAGACTGAAAATNNNAGATCGGA |
| 43913_2 | TATGAGTGTGGAGTCGTTGCTGGTAAGGGCGAAGACTGAAAATNNNAGATCGGA |
| 39038_1 | TATGAGTGTGGAGTCGTTGCAAATGGATAGTTGGACCCAGAGGNNNAGATCGGAA |
| 39038_2 | TATGAGTGTGGAGTCGTTGCAAATGGATAGTTGGACCCAGAGGNNNAGATCGGAA |
| 50696_1 | TATGAGTGTGGAGTCGTTGCCTCCTGAGTAGCTGGGATTACAGNNNAGATCGGAA |
| 50696_2 | TATGAGTGTGGAGTCGTTGCCTCCTGAGTAGCTGGGATTACAGNNNAGATCGGAA |
| 39684_1 | TATGAGTGTGGAGTCGTTGCGAGTCTCACTCTATCACCCATGCNNNAGATCGGAAG |
| 39684_2 | TATGAGTGTGGAGTCGTTGCGAGTCTCACTCTATCACCCATGCNNNAGATCGGAAG |
| 51220_1 | TATGAGTGTGGAGTCGTTGCTTCAAATATGGCGGTCTCTTGGANNNAGATCGGAAG |
| 51220_2 | TATGAGTGTGGAGTCGTTGCTTCAAATATGGCGGTCTCTTGGANNNAGATCGGAAG |
| 40780_1 | TATGAGTGTGGAGTCGTTGCGCATTTATTGGCCAGTGCAATTGNNNAGATCGGAAG |
| 40780_2 | TATGAGTGTGGAGTCGTTGCGCATTTATTGGCCAGTGCAATTGNNNAGATCGGAAG |
| 30891_1 | TATGAGTGTGGAGTCGTTGCAAGGGGAAATGTGTGGAACAGATNNNAGATCGGA |
| 30891_2 | TATGAGTGTGGAGTCGTTGCAAGGGGAAATGTGTGGAACAGATNNNAGATCGGA |
| 38773_1 | TATGAGTGTGGAGTCGTTGCCCATGTGTGACAGAGTAGGATCCNNNAGATCGGAA |
| 38773_2 | TATGAGTGTGGAGTCGTTGCCCATGTGTGACAGAGTAGGATCCNNNAGATCGGAA |
| 32759_1 | TATGAGTGTGGAGTCGTTGCTCTAGCCCAGAGTCATCACCTTNNNAGATCGGAAGA |
| 32759_2 | TATGAGTGTGGAGTCGTTGCTCTAGCCCAGAGTCATCACCTTNNNAGATCGGAAGA |
| 28774_1 | TATGAGTGTGGAGTCGTTGCGTAATCCCAGCTACTCAGTAGGCNNNAGATCGGAAG |
| 28774_2 | TATGAGTGTGGAGTCGTTGCGTAATCCCAGCTACTCAGTAGGCNNNAGATCGGAAG |
| 47277_1 | TATGAGTGTGGAGTCGTTGCCTAGAGTAGCTGGGATTACAGGCNNNAGATCGGAA |
| 47277_2 | TATGAGTGTGGAGTCGTTGCCTAGAGTAGCTGGGATTACAGGCNNNAGATCGGAA |
| 33378_1 | TATGAGTGTGGAGTCGTTGCAACATCATTCTCCTCCTCCTCCTNNNAGATCGGAAGA |
| 33378_2 | TATGAGTGTGGAGTCGTTGCAACATCATTCTCCTCCTCCTCCTNNNAGATCGGAAGA |
| 44445_1 | TATGAGTGTGGAGTCGTTGCGAGCCATTTCGAACAGATGGAAGNNNAGATCGGAA |
| 44445_2 | TATGAGTGTGGAGTCGTTGCGAGCCATTTCGAACAGATGGAAGNNNAGATCGGAA |
| 9376_1 | TATGAGTGTGGAGTCGTTGCGGATCATTCAAAGGCTACCGAGGNNNAGATCGGAA |
| 9376_2 | TATGAGTGTGGAGTCGTTGCGGATCATTCAAAGGCTACCGAGGNNNAGATCGGAA |
| 37132_1 | TATGAGTGTGGAGTCGTTGCATCCCCTTCCTCTGCCCTATTATNNNAGATCGGAAGA |
| 37132_2 | TATGAGTGTGGAGTCGTTGCATCCCCTTCCTCTGCCCTATTATNNNAGATCGGAAGA |
| 38997_1 | TATGAGTGTGGAGTCGTTGCCAAGATGGTCTCGATCTCCTGACNNNAGATCGGAAG |
| 38997_2 | TATGAGTGTGGAGTCGTTGCCAAGATGGTCTCGATCTCCTGACNNNAGATCGGAAG |
| 12730_1 | TATGAGTGTGGAGTCGTTGCAGAACACTTGGATACCTTAAATANNNAGATCGGAAG |
| 12730_2 | TATGAGTGTGGAGTCGTTGCAGAACACTTGGATACCTTAAATANNNAGATCGGAAG |
| 12900_1 | TATGAGTGTGGAGTCGTTGCTAAATTCAATACACAGTTCAGTGNNNAGATCGGAAG |
| 12900_2 | TATGAGTGTGGAGTCGTTGCTAAATTCAATACACAGTTCAGTGNNNAGATCGGAAG |
| 43963_1 | TATGAGTGTGGAGTCGTTGCGAGAGAGAGAGAGAGCAAATGGGNNNAGATCGG |
| 43963_2 | TATGAGTGTGGAGTCGTTGCGAGAGAGAGAGAGAGCAAATGGGNNNAGATCGG |
| 50341_1 | TATGAGTGTGGAGTCGTTGCACAGGCCAGGTTGTTCTTATAGGNNNAGATCGGAA |
| 50341_2 | TATGAGTGTGGAGTCGTTGCACAGGCCAGGTTGTTCTTATAGGNNNAGATCGGAA |
| 40981_1 | TATGAGTGTGGAGTCGTTGCCGACAACGACAACAACAACAAAANNNAGATCGGAA |
| 40981_2 | TATGAGTGTGGAGTCGTTGCCGACAACGACAACAACAACAAAANNNAGATCGGAA |

| | |
|---|---|
| 42914_1 | TATGAGTGTGGAGTCGTTGCATAGAGAGAGAGACGGAGTTTCGNNNAGATCGGA |
| 42914_2 | TATGAGTGTGGAGTCGTTGCATAGAGAGAGAGACGGAGTTTCGNNNAGATCGGA |
| 13178_1 | TATGAGTGTGGAGTCGTTGCCTCACCTGGTACAAACAATATNNNAGATCGGAAGAG |
| 13178_2 | TATGAGTGTGGAGTCGTTGCCTCACCTGGTACAAACAATATNNNAGATCGGAAGAG |
| 12707_1 | TATGAGTGTGGAGTCGTTGCGAGAAGAACCATATGTAAGTGTNNNAGATCGGAAG |
| 12707_2 | TATGAGTGTGGAGTCGTTGCGAGAAGAACCATATGTAAGTGTNNNAGATCGGAAG |
| 30970_1 | TATGAGTGTGGAGTCGTTGCAAAATAAAACCCACCTCGAGCCTNNNAGATCGGAAG |
| 30970_2 | TATGAGTGTGGAGTCGTTGCAAAATAAAACCCACCTCGAGCCTNNNAGATCGGAAG |
| 36213_1 | TATGAGTGTGGAGTCGTTGCACTTTTGATGGCTACAGCTCTGTNNNAGATCGGAAG |
| 36213_2 | TATGAGTGTGGAGTCGTTGCACTTTTGATGGCTACAGCTCTGTNNNAGATCGGAAG |
| 46520_1 | TATGAGTGTGGAGTCGTTGCCTCCCGGATTCATGCCATTCTCNNNAGATCGGAAGA |
| 46520_2 | TATGAGTGTGGAGTCGTTGCCTCCCGGATTCATGCCATTCTCNNNAGATCGGAAGA |
| 39000_1 | TATGAGTGTGGAGTCGTTGCTGGAATGTAACAAGGTGACTTTGCNNNAGATCGGA |
| 39000_2 | TATGAGTGTGGAGTCGTTGCTGGAATGTAACAAGGTGACTTTGCNNNAGATCGGA |
| 13400_1 | TATGAGTGTGGAGTCGTTGCCTTCATACAGGGTATTCCATAATNNNAGATCGGAAG |
| 13400_2 | TATGAGTGTGGAGTCGTTGCCTTCATACAGGGTATTCCATAATNNNAGATCGGAAG |
| 13144_1 | TATGAGTGTGGAGTCGTTGCAGAGAAGAAAACAAAGGTAGGNNNAGATCGGAAG |
| 13144_2 | TATGAGTGTGGAGTCGTTGCAGAGAAGAAAACAAAGGTAGGNNNAGATCGGAAG |
| 50578_1 | TATGAGTGTGGAGTCGTTGCTGATTGTAGTCCTCCTGCTTGTCNNNAGATCGGAAG |
| 50578_2 | TATGAGTGTGGAGTCGTTGCTGATTGTAGTCCTCCTGCTTGTCNNNAGATCGGAAG |
| 53087_1 | TATGAGTGTGGAGTCGTTGCAGAGTGTGACCAAAATGTAGGGCNNNAGATCGGAA |
| 53087_2 | TATGAGTGTGGAGTCGTTGCAGAGTGTGACCAAAATGTAGGGCNNNAGATCGGAA |
| 32594_1 | TATGAGTGTGGAGTCGTTGCCAGAGGGCTCCTGTATTTATGCANNNAGATCGGAAG |
| 32594_2 | TATGAGTGTGGAGTCGTTGCCAGAGGGCTCCTGTATTTATGCANNNAGATCGGAAG |
| 45469_1 | TATGAGTGTGGAGTCGTTGCCAAGAGCCTAGACCAATGGTAGGNNNAGATCGGAA |
| 45469_2 | TATGAGTGTGGAGTCGTTGCCAAGAGCCTAGACCAATGGTAGGNNNAGATCGGAA |
| 33831_1 | TATGAGTGTGGAGTCGTTGCCATGCAAGACAAAACCACAATGCNNNAGATCGGAAG |
| 33831_2 | TATGAGTGTGGAGTCGTTGCCATGCAAGACAAAACCACAATGCNNNAGATCGGAAG |
| 50956_1 | TATGAGTGTGGAGTCGTTGCCCCCCTCCGCTTCCTAATTTCNNNAGATCGGAAGAGC |
| 50956_2 | TATGAGTGTGGAGTCGTTGCCCCCCTCCGCTTCCTAATTTCNNNAGATCGGAAGAGC |
| 10318_1 | TATGAGTGTGGAGTCGTTGCGCAGCACGTTAGGTTTCATTTTCTNNNAGATCGGAA |
| 10318_2 | TATGAGTGTGGAGTCGTTGCGCAGCACGTTAGGTTTCATTTTCTNNNAGATCGGAA |
| 33879_1 | TATGAGTGTGGAGTCGTTGCGACCTATTTCTGTCTTCCCTGCTNNNAGATCGGAAGA |
| 33879_2 | TATGAGTGTGGAGTCGTTGCGACCTATTTCTGTCTTCCCTGCTNNNAGATCGGAAGA |
| 33948_1 | TATGAGTGTGGAGTCGTTGCGTCTTCCGAGTAGCTGGGACTANNNAGATCGGAAG |
| 33948_2 | TATGAGTGTGGAGTCGTTGCGTCTTCCGAGTAGCTGGGACTANNNAGATCGGAAG |
| 38817_1 | TATGAGTGTGGAGTCGTTGCCGTTGTGAAGACCATGGGAAAAGNNNAGATCGGAA |
| 38817_2 | TATGAGTGTGGAGTCGTTGCCGTTGTGAAGACCATGGGAAAAGNNNAGATCGGAA |
| 31534_1 | TATGAGTGTGGAGTCGTTGCGGGTAATAAGAAGAGAGAGCCGGNNNAGATCGGA |
| 31534_2 | TATGAGTGTGGAGTCGTTGCGGGTAATAAGAAGAGAGAGCCGGNNNAGATCGGA |
| 37585_1 | TATGAGTGTGGAGTCGTTGCACTGCCTCGATTACCCATGAAATNNNAGATCGGAAG |
| 37585_2 | TATGAGTGTGGAGTCGTTGCACTGCCTCGATTACCCATGAAATNNNAGATCGGAAG |
| 40375_1 | TATGAGTGTGGAGTCGTTGCGACTACCAGGACTGCCTACCTANNNAGATCGGAAGA |

| | |
|---|---|
| 40375_2 | TATGAGTGTGGAGTCGTTGCGACTACCAGGACTGCCTACCTANNNAGATCGGAAGA |
| 8568_1 | TATGAGTGTGGAGTCGTTGCTGGTGTCTCTGATTTGCTCAGTGNNNAGATCGGAAG |
| 8568_2 | TATGAGTGTGGAGTCGTTGCTGGTGTCTCTGATTTGCTCAGTGNNNAGATCGGAAG |
| 32647_1 | TATGAGTGTGGAGTCGTTGCGATCTTGGCTCACTGCAACCTCNNNAGATCGGAAGA |
| 32647_2 | TATGAGTGTGGAGTCGTTGCGATCTTGGCTCACTGCAACCTCNNNAGATCGGAAGA |
| 44307_1 | TATGAGTGTGGAGTCGTTGCTCTGTTATCTTCGCTGACACACANNNAGATCGGAAG |
| 44307_2 | TATGAGTGTGGAGTCGTTGCTCTGTTATCTTCGCTGACACACANNNAGATCGGAAG |
| 40265_1 | TATGAGTGTGGAGTCGTTGCTTCCAGATCCTTTGCCTATGGACNNNAGATCGGAAG |
| 40265_2 | TATGAGTGTGGAGTCGTTGCTTCCAGATCCTTTGCCTATGGACNNNAGATCGGAAG |
| 40921_1 | TATGAGTGTGGAGTCGTTGCGATTTCCACCAGCAGCATATAGCNNNAGATCGGAAG |
| 40921_2 | TATGAGTGTGGAGTCGTTGCGATTTCCACCAGCAGCATATAGCNNNAGATCGGAAG |
| 42730_1 | TATGAGTGTGGAGTCGTTGCACACTGCTCAAAGAAATCAGAGANNNAGATCGGAA |
| 42730_2 | TATGAGTGTGGAGTCGTTGCACACTGCTCAAAGAAATCAGAGANNNAGATCGGAA |
| 977_1 | TATGAGTGTGGAGTCGTTGCATAGCAGAATGTCACTAGACCCATNNNAGATCGGAA |
| 977_2 | TATGAGTGTGGAGTCGTTGCATAGCAGAATGTCACTAGACCCATNNNAGATCGGAA |
| 45300_1 | TATGAGTGTGGAGTCGTTGCTTTAGGGCCATTTGCAGTAGAGTNNNAGATCGGAA |
| 45300_2 | TATGAGTGTGGAGTCGTTGCTTTAGGGCCATTTGCAGTAGAGTNNNAGATCGGAA |
| 38141_1 | TATGAGTGTGGAGTCGTTGCTGTTGTCTTCCATATCAGGTCCTNNNAGATCGGAAG |
| 38141_2 | TATGAGTGTGGAGTCGTTGCTGTTGTCTTCCATATCAGGTCCTNNNAGATCGGAAG |
| 44226_1 | TATGAGTGTGGAGTCGTTGCAAAAGGAGGGTGGACTCTGTTAGNNNAGATCGGAA |
| 44226_2 | TATGAGTGTGGAGTCGTTGCAAAAGGAGGGTGGACTCTGTTAGNNNAGATCGGAA |
| 45065_1 | TATGAGTGTGGAGTCGTTGCAGAGTGTTACTGAGTCAGCACTGNNNAGATCGGAA |
| 45065_2 | TATGAGTGTGGAGTCGTTGCAGAGTGTTACTGAGTCAGCACTGNNNAGATCGGAA |
| 45837_1 | TATGAGTGTGGAGTCGTTGCCAAGCCTAGGTTAGGGTTCCATTNNNAGATCGGAAG |
| 45837_2 | TATGAGTGTGGAGTCGTTGCCAAGCCTAGGTTAGGGTTCCATTNNNAGATCGGAAG |
| 39333_1 | TATGAGTGTGGAGTCGTTGCAGACTGAAGGCAAAGAGATGAGGNNNAGATCGGA |
| 39333_2 | TATGAGTGTGGAGTCGTTGCAGACTGAAGGCAAAGAGATGAGGNNNAGATCGGA |
| 45006_1 | TATGAGTGTGGAGTCGTTGCTGTAGACAGAGGAGTATGGCAGANNNAGATCGGA |
| 45006_2 | TATGAGTGTGGAGTCGTTGCTGTAGACAGAGGAGTATGGCAGANNNAGATCGGA |
| 32639_1 | TATGAGTGTGGAGTCGTTGCCCTCCTAGGTTCAAGCGATTCTTNNNAGATCGGAAG |
| 32639_2 | TATGAGTGTGGAGTCGTTGCCCTCCTAGGTTCAAGCGATTCTTNNNAGATCGGAAG |
| 46292_1 | TATGAGTGTGGAGTCGTTGCCTCCCGGATTCAAGCAATTCTTCNNNAGATCGGAAG |
| 46292_2 | TATGAGTGTGGAGTCGTTGCCTCCCGGATTCAAGCAATTCTTCNNNAGATCGGAAG |
| 41082_1 | TATGAGTGTGGAGTCGTTGCGTGGATAACAGCAAACAACCTCCNNNAGATCGGAAG |
| 41082_2 | TATGAGTGTGGAGTCGTTGCGTGGATAACAGCAAACAACCTCCNNNAGATCGGAAG |
| 49346_1 | TATGAGTGTGGAGTCGTTGCTCCTTGCCTCATTTCCCAAGTTANNNAGATCGGAAGA |
| 49346_2 | TATGAGTGTGGAGTCGTTGCTCCTTGCCTCATTTCCCAAGTTANNNAGATCGGAAGA |
| 53434_1 | TATGAGTGTGGAGTCGTTGCCAAACCATCCCCATGATCCAAACNNNAGATCGGAAG |
| 53434_2 | TATGAGTGTGGAGTCGTTGCCAAACCATCCCCATGATCCAAACNNNAGATCGGAAG |
| 29741_1 | TATGAGTGTGGAGTCGTTGCCAGGACCAAGGAGACTATGTCTGNNNAGATCGGAA |
| 29741_2 | TATGAGTGTGGAGTCGTTGCCAGGACCAAGGAGACTATGTCTGNNNAGATCGGAA |
| 42138_1 | TATGAGTGTGGAGTCGTTGCCAGTGAGCCAAGATAGAACCACTNNNAGATCGGAA |
| 42138_2 | TATGAGTGTGGAGTCGTTGCCAGTGAGCCAAGATAGAACCACTNNNAGATCGGAA |

| | |
|---|---|
| 51770_1 | TATGAGTGTGGAGTCGTTGCGGGGGAAATTCCAAACTTCAACANNNAGATCGGAA |
| 51770_2 | TATGAGTGTGGAGTCGTTGCGGGGGAAATTCCAAACTTCAACANNNAGATCGGAA |
| 49450_1 | TATGAGTGTGGAGTCGTTGCGCCTTCCAAAACATCACACTCATNNNAGATCGGAAG |
| 49450_2 | TATGAGTGTGGAGTCGTTGCGCCTTCCAAAACATCACACTCATNNNAGATCGGAAG |
| 35645_1 | TATGAGTGTGGAGTCGTTGCCCCGAGTAGCTGGGACTACAGNNNAGATCGGAAGA |
| 35645_2 | TATGAGTGTGGAGTCGTTGCCCCGAGTAGCTGGGACTACAGNNNAGATCGGAAGA |
| 39887_1 | TATGAGTGTGGAGTCGTTGCTCTTTGTACAGAAGGCAAGACCANNNAGATCGGAA |
| 39887_2 | TATGAGTGTGGAGTCGTTGCTCTTTGTACAGAAGGCAAGACCANNNAGATCGGAA |
| 35857_1 | TATGAGTGTGGAGTCGTTGCTTTTTAGTAGAGACGGGGGCTTCNNNAGATCGGAA |
| 35857_2 | TATGAGTGTGGAGTCGTTGCTTTTTAGTAGAGACGGGGGCTTCNNNAGATCGGAA |
| 35697_1 | TATGAGTGTGGAGTCGTTGCGGCATGATTTTGGCTCACTACAANNNAGATCGGAAG |
| 35697_2 | TATGAGTGTGGAGTCGTTGCGGCATGATTTTGGCTCACTACAANNNAGATCGGAAG |
| 49091_1 | TATGAGTGTGGAGTCGTTGCGCTGTTTGAATTCTGTCCTTCCCNNNAGATCGGAAG |
| 49091_2 | TATGAGTGTGGAGTCGTTGCGCTGTTTGAATTCTGTCCTTCCCNNNAGATCGGAAG |
| 29002_1 | TATGAGTGTGGAGTCGTTGCTGCTATTGTGTAAGGAAATCTGGTNNNAGATCGGA |
| 29002_2 | TATGAGTGTGGAGTCGTTGCTGCTATTGTGTAAGGAAATCTGGTNNNAGATCGGA |
| 30318_1 | TATGAGTGTGGAGTCGTTGCGACAGAAGAGGGAGGTCAGTCTTNNNAGATCGGA |
| 30318_2 | TATGAGTGTGGAGTCGTTGCGACAGAAGAGGGAGGTCAGTCTTNNNAGATCGGA |
| 43648_1 | TATGAGTGTGGAGTCGTTGCGTTTTCCTGCAGAGAGAGACAGANNNAGATCGGAA |
| 43648_2 | TATGAGTGTGGAGTCGTTGCGTTTTCCTGCAGAGAGAGACAGANNNAGATCGGAA |
| 10040_1 | TATGAGTGTGGAGTCGTTGCGTGACCCTGGATACTTGAGACCTNNNAGATCGGAAG |
| 10040_2 | TATGAGTGTGGAGTCGTTGCGTGACCCTGGATACTTGAGACCTNNNAGATCGGAAG |
| 35530_1 | TATGAGTGTGGAGTCGTTGCCATTCTCCTGCCTCAGCCTCNNNAGATCGGAAGAGCA |
| 35530_2 | TATGAGTGTGGAGTCGTTGCCATTCTCCTGCCTCAGCCTCNNNAGATCGGAAGAGCA |
| 43862_1 | TATGAGTGTGGAGTCGTTGCAGGCAAATCTCCAAGGAGTCTTTNNNAGATCGGAAG |
| 43862_2 | TATGAGTGTGGAGTCGTTGCAGGCAAATCTCCAAGGAGTCTTTNNNAGATCGGAAG |
| 41861_1 | TATGAGTGTGGAGTCGTTGCCCTGTGTTCAAGCAATTCTCCTGNNNAGATCGGAAG |
| 41861_2 | TATGAGTGTGGAGTCGTTGCCCTGTGTTCAAGCAATTCTCCTGNNNAGATCGGAAG |
| 30655_1 | TATGAGTGTGGAGTCGTTGCACTATTCTATTGATGGGCATGCANNNAGATCGGAAG |
| 30655_2 | TATGAGTGTGGAGTCGTTGCACTATTCTATTGATGGGCATGCANNNAGATCGGAAG |
| 31787_1 | TATGAGTGTGGAGTCGTTGCAGAGAAGAGAAGAGAAGGCTCCTNNNAGATCGGA |
| 31787_2 | TATGAGTGTGGAGTCGTTGCAGAGAAGAGAAGAGAAGGCTCCTNNNAGATCGGA |
| 50134_1 | TATGAGTGTGGAGTCGTTGCTGGAAGATACTATCCCCACCCTTNNNAGATCGGAAG |
| 50134_2 | TATGAGTGTGGAGTCGTTGCTGGAAGATACTATCCCCACCCTTNNNAGATCGGAAG |
| 37610_1 | TATGAGTGTGGAGTCGTTGCAGGTCAGGAGATAGAGACCATCCNNNAGATCGGAA |
| 37610_2 | TATGAGTGTGGAGTCGTTGCAGGTCAGGAGATAGAGACCATCCNNNAGATCGGAA |
| 48320_1 | TATGAGTGTGGAGTCGTTGCCAAGGGACTGCAATGTGATCTTGNNNAGATCGGAA |
| 48320_2 | TATGAGTGTGGAGTCGTTGCCAAGGGACTGCAATGTGATCTTGNNNAGATCGGAA |
| 43474_1 | TATGAGTGTGGAGTCGTTGCCCACGCCCAGCTAATTTTTTGTANNNAGATCGGAAG |
| 43474_2 | TATGAGTGTGGAGTCGTTGCCCACGCCCAGCTAATTTTTTGTANNNAGATCGGAAG |
| 39446_1 | TATGAGTGTGGAGTCGTTGCGCAGGAAAACAAAGGGCAGATATNNNAGATCGGA |
| 39446_2 | TATGAGTGTGGAGTCGTTGCGCAGGAAAACAAAGGGCAGATATNNNAGATCGGA |
| 51134_1 | TATGAGTGTGGAGTCGTTGCGATTCCAGCCAGGTAAGGTCTTTNNNAGATCGGAAG |

| | |
|---|---|
| 51134_2 | TATGAGTGTGGAGTCGTTGCGATTCCAGCCAGGTAAGGTCTTTNNNAGATCGGAAG |
| 41782_1 | TATGAGTGTGGAGTCGTTGCCGCCATTCTTACGTGAGTCTTTTNNNAGATCGGAAG |
| 41782_2 | TATGAGTGTGGAGTCGTTGCCGCCATTCTTACGTGAGTCTTTTNNNAGATCGGAAG |
| 2306_1 | TATGAGTGTGGAGTCGTTGCAGGGACACACCCTTACCATAGACNNNAGATCGGAAG |
| 2306_2 | TATGAGTGTGGAGTCGTTGCAGGGACACACCCTTACCATAGACNNNAGATCGGAAG |
| 37754_1 | TATGAGTGTGGAGTCGTTGCGGTCCAAAAAGAATTCCAGGCTGNNNAGATCGGAA |
| 37754_2 | TATGAGTGTGGAGTCGTTGCGGTCCAAAAAGAATTCCAGGCTGNNNAGATCGGAA |
| 42549_1 | TATGAGTGTGGAGTCGTTGCGTCTAGATCACTAGCAATGCTGCNNNAGATCGGAAG |
| 42549_2 | TATGAGTGTGGAGTCGTTGCGTCTAGATCACTAGCAATGCTGCNNNAGATCGGAAG |
| 45555_1 | TATGAGTGTGGAGTCGTTGCTGGTAAGTCTCAAGTTTCTCCCANNNAGATCGGAAG |
| 45555_2 | TATGAGTGTGGAGTCGTTGCTGGTAAGTCTCAAGTTTCTCCCANNNAGATCGGAAG |
| 49431_1 | TATGAGTGTGGAGTCGTTGCGCAATCTCAGTTCACTGCAACTTNNNAGATCGGAAG |
| 49431_2 | TATGAGTGTGGAGTCGTTGCGCAATCTCAGTTCACTGCAACTTNNNAGATCGGAAG |
| 45013_1 | TATGAGTGTGGAGTCGTTGCGAAGCTGCTCTGTATGCTAAAGCNNNAGATCGGAAG |
| 45013_2 | TATGAGTGTGGAGTCGTTGCGAAGCTGCTCTGTATGCTAAAGCNNNAGATCGGAAG |
| 34505_1 | TATGAGTGTGGAGTCGTTGCTCTCATTTGTGCCTGAAATTCCGNNNAGATCGGAAG |
| 34505_2 | TATGAGTGTGGAGTCGTTGCTCTCATTTGTGCCTGAAATTCCGNNNAGATCGGAAG |
| 52598_1 | TATGAGTGTGGAGTCGTTGCATCTGCCACAGGTCAAAATTGTGNNNAGATCGGAAG |
| 52598_2 | TATGAGTGTGGAGTCGTTGCATCTGCCACAGGTCAAAATTGTGNNNAGATCGGAAG |
| 33109_1 | TATGAGTGTGGAGTCGTTGCAAAACCGCTTGAATGAGAACAGGNNNAGATCGGAA |
| 33109_2 | TATGAGTGTGGAGTCGTTGCAAAACCGCTTGAATGAGAACAGGNNNAGATCGGAA |
| 36547_1 | TATGAGTGTGGAGTCGTTGCACTGCATGTTGAATGAACTCCACNNNAGATCGGAAG |
| 36547_2 | TATGAGTGTGGAGTCGTTGCACTGCATGTTGAATGAACTCCACNNNAGATCGGAAG |
| 42306_1 | TATGAGTGTGGAGTCGTTGCAGTGGGTGGTTGAATACTGATGTNNNAGATCGGAA |
| 42306_2 | TATGAGTGTGGAGTCGTTGCAGTGGGTGGTTGAATACTGATGTNNNAGATCGGAA |
| 33911_1 | TATGAGTGTGGAGTCGTTGCGTAGTTCAATGGCACAGTCTTGGNNNAGATCGGAA |
| 33911_2 | TATGAGTGTGGAGTCGTTGCGTAGTTCAATGGCACAGTCTTGGNNNAGATCGGAA |
| 50069_1 | TATGAGTGTGGAGTCGTTGCTTTCCCTTCCTCATTCTGTCTCGNNNAGATCGGAAGA |
| 50069_2 | TATGAGTGTGGAGTCGTTGCTTTCCCTTCCTCATTCTGTCTCGNNNAGATCGGAAGA |
| 41490_1 | TATGAGTGTGGAGTCGTTGCAGCCTCCCATCAAATAACTGAGGNNNAGATCGGAAG |
| 41490_2 | TATGAGTGTGGAGTCGTTGCAGCCTCCCATCAAATAACTGAGGNNNAGATCGGAAG |
| 43654_1 | TATGAGTGTGGAGTCGTTGCATGTCCTTTTCTTCCTGGGTACCNNNAGATCGGAAGA |
| 43654_2 | TATGAGTGTGGAGTCGTTGCATGTCCTTTTCTTCCTGGGTACCNNNAGATCGGAAGA |
| 53215_1 | TATGAGTGTGGAGTCGTTGCTTTGGCCTCAAACTTCAGGTGANNNAGATCGGAAGA |
| 53215_2 | TATGAGTGTGGAGTCGTTGCTTTGGCCTCAAACTTCAGGTGANNNAGATCGGAAGA |
| 43053_1 | TATGAGTGTGGAGTCGTTGCCGCCCTCATTCTGTCTCTATGAANNNAGATCGGAAGA |
| 43053_2 | TATGAGTGTGGAGTCGTTGCCGCCCTCATTCTGTCTCTATGAANNNAGATCGGAAGA |
| 47852_1 | TATGAGTGTGGAGTCGTTGCCAAAGTGCTGGGATTACATGCATNNNAGATCGGAA |
| 47852_2 | TATGAGTGTGGAGTCGTTGCCAAAGTGCTGGGATTACATGCATNNNAGATCGGAA |
| 52658_1 | TATGAGTGTGGAGTCGTTGCATAGAGACTTCAGCCTAACACGCNNNAGATCGGAAG |
| 52658_2 | TATGAGTGTGGAGTCGTTGCATAGAGACTTCAGCCTAACACGCNNNAGATCGGAAG |
| 9996_1 | TATGAGTGTGGAGTCGTTGCCCAAGGCATGTGTCTTACTGACANNNAGATCGGAAG |
| 9996_2 | TATGAGTGTGGAGTCGTTGCCCAAGGCATGTGTCTTACTGACANNNAGATCGGAAG |

| | |
|---|---|
| 29102_1 | TATGAGTGTGGAGTCGTTGCCACACACACACACACATATATNNNAGATCGGAAG |
| 29102_2 | TATGAGTGTGGAGTCGTTGCCACACACACACACACATATATNNNAGATCGGAAG |
| 46636_1 | TATGAGTGTGGAGTCGTTGCTAGAGTACAGTGGCGTGATCTTGNNNAGATCGGAA |
| 46636_2 | TATGAGTGTGGAGTCGTTGCTAGAGTACAGTGGCGTGATCTTGNNNAGATCGGAA |
| 10034_1 | TATGAGTGTGGAGTCGTTGCTCATCACCTCCGGAAAGTGTAGTNNNAGATCGGAAG |
| 10034_2 | TATGAGTGTGGAGTCGTTGCTCATCACCTCCGGAAAGTGTAGTNNNAGATCGGAAG |
| 47892_1 | TATGAGTGTGGAGTCGTTGCCTGCCTCCTAGGTTCAAGTGATTNNNAGATCGGAAG |
| 47892_2 | TATGAGTGTGGAGTCGTTGCCTGCCTCCTAGGTTCAAGTGATTNNNAGATCGGAAG |
| 34951_1 | TATGAGTGTGGAGTCGTTGCGTTCAAGCGATTCTCTTGCCTCNNNAGATCGGAAGA |
| 34951_2 | TATGAGTGTGGAGTCGTTGCGTTCAAGCGATTCTCTTGCCTCNNNAGATCGGAAGA |
| 44045_1 | TATGAGTGTGGAGTCGTTGCTCGTTCAGTTCTACTTGAAGGCANNNAGATCGGAAG |
| 44045_2 | TATGAGTGTGGAGTCGTTGCTCGTTCAGTTCTACTTGAAGGCANNNAGATCGGAAG |
| 52639_1 | TATGAGTGTGGAGTCGTTGCGTCGAAGAAAACTGGGGAAAGTGNNNAGATCGGA |
| 52639_2 | TATGAGTGTGGAGTCGTTGCGTCGAAGAAAACTGGGGAAAGTGNNNAGATCGGA |
| 50704_1 | TATGAGTGTGGAGTCGTTGCATGAGGTCATTGCAGGCTTGATTNNNAGATCGGAA |
| 50704_2 | TATGAGTGTGGAGTCGTTGCATGAGGTCATTGCAGGCTTGATTNNNAGATCGGAA |
| 29062_1 | TATGAGTGTGGAGTCGTTGCTGAATCCCCAGCTCATAAAACCANNNAGATCGGAAG |
| 29062_2 | TATGAGTGTGGAGTCGTTGCTGAATCCCCAGCTCATAAAACCANNNAGATCGGAAG |
| 41444_1 | TATGAGTGTGGAGTCGTTGCGCAGGGCTTGAAATCGTAGTCTANNNAGATCGGAA |
| 41444_2 | TATGAGTGTGGAGTCGTTGCGCAGGGCTTGAAATCGTAGTCTANNNAGATCGGAA |
| 13013_1 | TATGAGTGTGGAGTCGTTGCGACAAGAACACAGAGACAAGNNNAGATCGGAAGA |
| 13013_2 | TATGAGTGTGGAGTCGTTGCGACAAGAACACAGAGACAAGNNNAGATCGGAAGA |
| 51914_1 | TATGAGTGTGGAGTCGTTGCAAAAAATTAGACGGGCATGGTGGNNNAGATCGGA |
| 51914_2 | TATGAGTGTGGAGTCGTTGCAAAAAATTAGACGGGCATGGTGGNNNAGATCGGA |
| 43523_1 | TATGAGTGTGGAGTCGTTGCCACAATCTTGGCTCACTGCAAGNNNAGATCGGAAGA |
| 43523_2 | TATGAGTGTGGAGTCGTTGCCACAATCTTGGCTCACTGCAAGNNNAGATCGGAAGA |
| 49461_1 | TATGAGTGTGGAGTCGTTGCATATTGCTCAGGCTGGTCTTGAANNNAGATCGGAAG |
| 49461_2 | TATGAGTGTGGAGTCGTTGCATATTGCTCAGGCTGGTCTTGAANNNAGATCGGAAG |
| 48499_1 | TATGAGTGTGGAGTCGTTGCATGTTCAAGTGATTCTCCTGCCTNNNAGATCGGAAG |
| 48499_2 | TATGAGTGTGGAGTCGTTGCATGTTCAAGTGATTCTCCTGCCTNNNAGATCGGAAG |
| 5465_1 | TATGAGTGTGGAGTCGTTGCTAGAGTGTAGGGTTGGGCACAAANNNAGATCGGA |
| 5465_2 | TATGAGTGTGGAGTCGTTGCTAGAGTGTAGGGTTGGGCACAAANNNAGATCGGA |
| 13393_1 | TATGAGTGTGGAGTCGTTGCATATCTTAAGGAATAATTGTGGATNNNAGATCGGA |
| 13393_2 | TATGAGTGTGGAGTCGTTGCATATCTTAAGGAATAATTGTGGATNNNAGATCGGA |
| 52654_1 | TATGAGTGTGGAGTCGTTGCCCATACATACTCAGGACCCCAACNNNAGATCGGAAG |
| 52654_2 | TATGAGTGTGGAGTCGTTGCCCATACATACTCAGGACCCCAACNNNAGATCGGAAG |
| 52490_1 | TATGAGTGTGGAGTCGTTGCCTGACATCACCCTACTTGTCCAGNNNAGATCGGAAG |
| 52490_2 | TATGAGTGTGGAGTCGTTGCCTGACATCACCCTACTTGTCCAGNNNAGATCGGAAG |
| 36212_1 | TATGAGTGTGGAGTCGTTGCCTGTAAACCCAACACTTGACGAGNNNAGATCGGAAG |
| 36212_2 | TATGAGTGTGGAGTCGTTGCCTGTAAACCCAACACTTGACGAGNNNAGATCGGAAG |
| 2539_1 | TATGAGTGTGGAGTCGTTGCTGGTTGTCAATTATGATGGCAGANNNAGATCGGAA |
| 2539_2 | TATGAGTGTGGAGTCGTTGCTGGTTGTCAATTATGATGGCAGANNNAGATCGGAA |
| 12920_1 | TATGAGTGTGGAGTCGTTGCTGCTTCGTAAGGGAAACTATNNNAGATCGGAAGAG |

| | |
|---|---|
| 12920_2 | TATGAGTGTGGAGTCGTTGCTGCTTCGTAAGGGAAACTATNNNAGATCGGAAGAG |
| 2795_1 | TATGAGTGTGGAGTCGTTGCTTACTGCCACAAAATGGAGTGCTNNNAGATCGGAAG |
| 2795_2 | TATGAGTGTGGAGTCGTTGCTTACTGCCACAAAATGGAGTGCTNNNAGATCGGAAG |
| 50499_1 | TATGAGTGTGGAGTCGTTGCTAGTGGTGAAGGAGGAGTTAGGTNNNAGATCGGA |
| 50499_2 | TATGAGTGTGGAGTCGTTGCTAGTGGTGAAGGAGGAGTTAGGTNNNAGATCGGA |
| 9149_1 | TATGAGTGTGGAGTCGTTGCGCATACACACACCTAGAAGGCATNNNAGATCGGAAG |
| 9149_2 | TATGAGTGTGGAGTCGTTGCGCATACACACACCTAGAAGGCATNNNAGATCGGAAG |
| 51521_1 | TATGAGTGTGGAGTCGTTGCAGCAATGGCAGTGATTTCCTCTANNNAGATCGGAAG |
| 51521_2 | TATGAGTGTGGAGTCGTTGCAGCAATGGCAGTGATTTCCTCTANNNAGATCGGAAG |
| 37103_1 | TATGAGTGTGGAGTCGTTGCAATATTGGGGAATCTCTGCAGGANNNAGATCGGAA |
| 37103_2 | TATGAGTGTGGAGTCGTTGCAATATTGGGGAATCTCTGCAGGANNNAGATCGGAA |
| 38643_1 | TATGAGTGTGGAGTCGTTGCGGGGGTAGCAGTATCCAGAAATCNNNAGATCGGAA |
| 38643_2 | TATGAGTGTGGAGTCGTTGCGGGGGTAGCAGTATCCAGAAATCNNNAGATCGGAA |
| 45616_1 | TATGAGTGTGGAGTCGTTGCCTCCTGGGTTCAAGGGATTCACNNNAGATCGGAAGA |
| 45616_2 | TATGAGTGTGGAGTCGTTGCCTCCTGGGTTCAAGGGATTCACNNNAGATCGGAAGA |
| 49341_1 | TATGAGTGTGGAGTCGTTGCATGGCCCAACTGTTCTTTTCATGNNNAGATCGGAAG |
| 49341_2 | TATGAGTGTGGAGTCGTTGCATGGCCCAACTGTTCTTTTCATGNNNAGATCGGAAG |
| 49737_1 | TATGAGTGTGGAGTCGTTGCAGTATATGTCCTGGTTTGGCCANNNAGATCGGAAGA |
| 49737_2 | TATGAGTGTGGAGTCGTTGCAGTATATGTCCTGGTTTGGCCANNNAGATCGGAAGA |
| 35064_1 | TATGAGTGTGGAGTCGTTGCTCCTTTTTTGGTGTCCTCTAAGCTNNNAGATCGGAAG |
| 35064_2 | TATGAGTGTGGAGTCGTTGCTCCTTTTTTGGTGTCCTCTAAGCTNNNAGATCGGAAG |
| 36108_1 | TATGAGTGTGGAGTCGTTGCTCAAAGGGACAAGCACAGACATANNNAGATCGGAA |
| 36108_2 | TATGAGTGTGGAGTCGTTGCTCAAAGGGACAAGCACAGACATANNNAGATCGGAA |
| 45397_1 | TATGAGTGTGGAGTCGTTGCTTCTCCAGAGAACCCTGACTAATNNNAGATCGGAAG |
| 45397_2 | TATGAGTGTGGAGTCGTTGCTTCTCCAGAGAACCCTGACTAATNNNAGATCGGAAG |
| 41906_1 | TATGAGTGTGGAGTCGTTGCCACCATGCCCAGCTAATTTTTGTNNNAGATCGGAAG |
| 41906_2 | TATGAGTGTGGAGTCGTTGCCACCATGCCCAGCTAATTTTTGTNNNAGATCGGAAG |
| 32566_1 | TATGAGTGTGGAGTCGTTGCTCAAGTAATTCTCCTGCCTCCACNNNAGATCGGAAGA |
| 32566_2 | TATGAGTGTGGAGTCGTTGCTCAAGTAATTCTCCTGCCTCCACNNNAGATCGGAAGA |
| 53023_1 | TATGAGTGTGGAGTCGTTGCCAGCTAGGCACAACTACCTACTCNNNAGATCGGAAG |
| 53023_2 | TATGAGTGTGGAGTCGTTGCCAGCTAGGCACAACTACCTACTCNNNAGATCGGAAG |
| 30040_1 | TATGAGTGTGGAGTCGTTGCACTTTGCCATTCTTAATCTGTAAAACANNNAGATCGG |
| 30040_2 | TATGAGTGTGGAGTCGTTGCACTTTGCCATTCTTAATCTGTAAAACANNNAGATCGG |
| 36639_1 | TATGAGTGTGGAGTCGTTGCAGATTCTCCAACCCTAGAGAGGANNNAGATCGGAA |
| 36639_2 | TATGAGTGTGGAGTCGTTGCAGATTCTCCAACCCTAGAGAGGANNNAGATCGGAA |
| 48399_1 | TATGAGTGTGGAGTCGTTGCGCCACCACACACAGCTAATTTTTNNNAGATCGGAAG |
| 48399_2 | TATGAGTGTGGAGTCGTTGCGCCACCACACACAGCTAATTTTTNNNAGATCGGAAG |
| 30260_1 | TATGAGTGTGGAGTCGTTGCTGGCATGGGTAAACCAGCTATTANNNAGATCGGAA |
| 30260_2 | TATGAGTGTGGAGTCGTTGCTGGCATGGGTAAACCAGCTATTANNNAGATCGGAA |
| 2987_1 | TATGAGTGTGGAGTCGTTGCGCTGACCAAAAATGTCTCAAAGTCAANNNAGATCGG |
| 2987_2 | TATGAGTGTGGAGTCGTTGCGCTGACCAAAAATGTCTCAAAGTCAANNNAGATCGG |
| 41539_1 | TATGAGTGTGGAGTCGTTGCTCAGTCTCTTGAGTAGCTGGGATNNNAGATCGGAA |
| 41539_2 | TATGAGTGTGGAGTCGTTGCTCAGTCTCTTGAGTAGCTGGGATNNNAGATCGGAA |

| | |
|---|---|
| 53180_1 | TATGAGTGTGGAGTCGTTGCCCTTTTCTTCCATGGTAGCCTCTNNNAGATCGGAAGA |
| 53180_2 | TATGAGTGTGGAGTCGTTGCCCTTTTCTTCCATGGTAGCCTCTNNNAGATCGGAAGA |
| 36693_1 | TATGAGTGTGGAGTCGTTGCATACCTACTATGGCATGGTTGGTNNNAGATCGGAAG |
| 36693_2 | TATGAGTGTGGAGTCGTTGCATACCTACTATGGCATGGTTGGTNNNAGATCGGAAG |
| 50584_1 | TATGAGTGTGGAGTCGTTGCTTCTTTTGAACCCTTCTTCCCCANNNAGATCGGAAGA |
| 50584_2 | TATGAGTGTGGAGTCGTTGCTTCTTTTGAACCCTTCTTCCCCANNNAGATCGGAAGA |
| 53016_1 | TATGAGTGTGGAGTCGTTGCGACACAGACAAGAGAAGGCCATANNNAGATCGGAA |
| 53016_2 | TATGAGTGTGGAGTCGTTGCGACACAGACAAGAGAAGGCCATANNNAGATCGGAA |
| 43029_1 | TATGAGTGTGGAGTCGTTGCGCTGAAGTGTAGTGGTACGATCTNNNAGATCGGAA |
| 43029_2 | TATGAGTGTGGAGTCGTTGCGCTGAAGTGTAGTGGTACGATCTNNNAGATCGGAA |
| 33678_1 | TATGAGTGTGGAGTCGTTGCCCTTGGGCATTTTGGTCATTTTANNNAGATCGGAAG |
| 33678_2 | TATGAGTGTGGAGTCGTTGCCCTTGGGCATTTTGGTCATTTTANNNAGATCGGAAG |
| 30036_1 | TATGAGTGTGGAGTCGTTGCGACACCTGTTGGAGGGAAAATTGNNNAGATCGGAA |
| 30036_2 | TATGAGTGTGGAGTCGTTGCGACACCTGTTGGAGGGAAAATTGNNNAGATCGGAA |
| 36685_1 | TATGAGTGTGGAGTCGTTGCTCTGACACATGGTGAGAACTCAGNNNAGATCGGAA |
| 36685_2 | TATGAGTGTGGAGTCGTTGCTCTGACACATGGTGAGAACTCAGNNNAGATCGGAA |
| 39930_1 | TATGAGTGTGGAGTCGTTGCAACAAACATGCTTGCCTTGTTCTNNNAGATCGGAAG |
| 39930_2 | TATGAGTGTGGAGTCGTTGCAACAAACATGCTTGCCTTGTTCTNNNAGATCGGAAG |
| 31322_1 | TATGAGTGTGGAGTCGTTGCGTGATTCCACATGCTTTGCAGTANNNAGATCGGAAG |
| 31322_2 | TATGAGTGTGGAGTCGTTGCGTGATTCCACATGCTTTGCAGTANNNAGATCGGAAG |
| 30411_1 | TATGAGTGTGGAGTCGTTGCAGCCTCCTAAGTAGCTGGGATTANNNAGATCGGAAG |
| 30411_2 | TATGAGTGTGGAGTCGTTGCAGCCTCCTAAGTAGCTGGGATTANNNAGATCGGAAG |
| 49090_1 | TATGAGTGTGGAGTCGTTGCTTCAAGCGATTCTCCTGACTCAGNNNAGATCGGAAG |
| 49090_2 | TATGAGTGTGGAGTCGTTGCTTCAAGCGATTCTCCTGACTCAGNNNAGATCGGAAG |
| 35103_1 | TATGAGTGTGGAGTCGTTGCAGAAAAACATGAAAAGCGAAGCTNNNAGATCGGAA |
| 35103_2 | TATGAGTGTGGAGTCGTTGCAGAAAAACATGAAAAGCGAAGCTNNNAGATCGGAA |
| 40304_1 | TATGAGTGTGGAGTCGTTGCGCATTGGGAGATATGCCTAATGCNNNAGATCGGAA |
| 40304_2 | TATGAGTGTGGAGTCGTTGCGCATTGGGAGATATGCCTAATGCNNNAGATCGGAA |
| 44774_1 | TATGAGTGTGGAGTCGTTGCAAACATACCGAGAAGCATGCAAGNNNAGATCGGAA |
| 44774_2 | TATGAGTGTGGAGTCGTTGCAAACATACCGAGAAGCATGCAAGNNNAGATCGGAA |
| 50630_1 | TATGAGTGTGGAGTCGTTGCGACCTTACTGGCCCATCTCAAAGNNNAGATCGGAAG |
| 50630_2 | TATGAGTGTGGAGTCGTTGCGACCTTACTGGCCCATCTCAAAGNNNAGATCGGAAG |
| 48574_1 | TATGAGTGTGGAGTCGTTGCTTGCTGGTGACCTTTACTCTACCNNNAGATCGGAAG |
| 48574_2 | TATGAGTGTGGAGTCGTTGCTTGCTGGTGACCTTTACTCTACCNNNAGATCGGAAG |
| 43583_1 | TATGAGTGTGGAGTCGTTGCGATCTCAAGCTGTATCCTGGGAGNNNAGATCGGAA |
| 43583_2 | TATGAGTGTGGAGTCGTTGCGATCTCAAGCTGTATCCTGGGAGNNNAGATCGGAA |
| 8252_1 | TATGAGTGTGGAGTCGTTGCATGGGCTCTCACTTTGATATGCANNNAGATCGGAAG |
| 8252_2 | TATGAGTGTGGAGTCGTTGCATGGGCTCTCACTTTGATATGCANNNAGATCGGAAG |
| 32996_1 | TATGAGTGTGGAGTCGTTGCTTCAAGTGATTCTCCTGCCTCAGNNNAGATCGGAAG |
| 32996_2 | TATGAGTGTGGAGTCGTTGCTTCAAGTGATTCTCCTGCCTCAGNNNAGATCGGAAG |
| 29287_1 | TATGAGTGTGGAGTCGTTGCGGAGGCTGAACACTTGGAAATTGNNNAGATCGGAA |
| 29287_2 | TATGAGTGTGGAGTCGTTGCGGAGGCTGAACACTTGGAAATTGNNNAGATCGGAA |
| 5673_1 | TATGAGTGTGGAGTCGTTGCTCGTGAGCCAATATCTCATCACTNNNAGATCGGAAG |

| | |
|---|---|
| 5673_2 | TATGAGTGTGGAGTCGTTGCTCGTGAGCCAATATCTCATCACTNNNAGATCGGAAG |
| 39316_1 | TATGAGTGTGGAGTCGTTGCTACACATGGGGTGAGTCAGTTTTNNNAGATCGGAAG |
| 39316_2 | TATGAGTGTGGAGTCGTTGCTACACATGGGGTGAGTCAGTTTTNNNAGATCGGAAG |
| 41000_1 | TATGAGTGTGGAGTCGTTGCGAGACAGAGTTTCGCTGTTGTTGNNNAGATCGGAA |
| 41000_2 | TATGAGTGTGGAGTCGTTGCGAGACAGAGTTTCGCTGTTGTTGNNNAGATCGGAA |
| 33473_1 | TATGAGTGTGGAGTCGTTGCCACTTGACCAGCGTTATGGAATCNNNAGATCGGAAG |
| 33473_2 | TATGAGTGTGGAGTCGTTGCCACTTGACCAGCGTTATGGAATCNNNAGATCGGAAG |
| 39830_1 | TATGAGTGTGGAGTCGTTGCAGCCAATGTAATCCACTCCATCCNNNAGATCGGAAG |
| 39830_2 | TATGAGTGTGGAGTCGTTGCAGCCAATGTAATCCACTCCATCCNNNAGATCGGAAG |
| 37094_1 | TATGAGTGTGGAGTCGTTGCAAAACATTTCACGGTTGAGGCAANNNAGATCGGAAG |
| 37094_2 | TATGAGTGTGGAGTCGTTGCAAAACATTTCACGGTTGAGGCAANNNAGATCGGAAG |
| 42336_1 | TATGAGTGTGGAGTCGTTGCTTTGGTTCAGCAATCCCAGTAGTNNNAGATCGGAAG |
| 42336_2 | TATGAGTGTGGAGTCGTTGCTTTGGTTCAGCAATCCCAGTAGTNNNAGATCGGAAG |
| 43443_1 | TATGAGTGTGGAGTCGTTGCTTGTTGTGTTTCTGCCAGGTTTTNNNAGATCGGAAG |
| 43443_2 | TATGAGTGTGGAGTCGTTGCTTGTTGTGTTTCTGCCAGGTTTTNNNAGATCGGAAG |
| 38783_1 | TATGAGTGTGGAGTCGTTGCGAGATGGCCAAATAGGAACAGCTNNNAGATCGGAA |
| 38783_2 | TATGAGTGTGGAGTCGTTGCGAGATGGCCAAATAGGAACAGCTNNNAGATCGGAA |
| 31089_1 | TATGAGTGTGGAGTCGTTGCCACAGGATTATAGGGGCCAAGAANNNAGATCGGAA |
| 31089_2 | TATGAGTGTGGAGTCGTTGCCACAGGATTATAGGGGCCAAGAANNNAGATCGGAA |
| 40007_1 | TATGAGTGTGGAGTCGTTGCAGTCAGTCCTTATTTCCCTTCTTTCANNNAGATCGGA |
| 40007_2 | TATGAGTGTGGAGTCGTTGCAGTCAGTCCTTATTTCCCTTCTTTCANNNAGATCGGA |
| 40931_1 | TATGAGTGTGGAGTCGTTGCGAGAATGGCATGAACCCAGGAGNNNAGATCGGAAG |
| 40931_2 | TATGAGTGTGGAGTCGTTGCGAGAATGGCATGAACCCAGGAGNNNAGATCGGAAG |
| 42593_1 | TATGAGTGTGGAGTCGTTGCTTCTCTGGACACGTTTTGAGAGTNNNAGATCGGAAG |
| 42593_2 | TATGAGTGTGGAGTCGTTGCTTCTCTGGACACGTTTTGAGAGTNNNAGATCGGAAG |
| 2380_1 | TATGAGTGTGGAGTCGTTGCTGATGATGAGGGTGACAAGGACTNNNAGATCGGA |
| 2380_2 | TATGAGTGTGGAGTCGTTGCTGATGATGAGGGTGACAAGGACTNNNAGATCGGA |
| 43653_1 | TATGAGTGTGGAGTCGTTGCAAAAATACAAACACTAGCCGGGCNNNAGATCGGAAG |
| 43653_2 | TATGAGTGTGGAGTCGTTGCAAAAATACAAACACTAGCCGGGCNNNAGATCGGAAG |
| 31764_1 | TATGAGTGTGGAGTCGTTGCGGGATGGGGTGATGTAAGCATGNNNAGATCGGAA |
| 31764_2 | TATGAGTGTGGAGTCGTTGCGGGATGGGGTGATGTAAGCATGNNNAGATCGGAA |
| 52524_1 | TATGAGTGTGGAGTCGTTGCGTTGCTTAGGACAGTTGTGTCACNNNAGATCGGAAG |
| 52524_2 | TATGAGTGTGGAGTCGTTGCGTTGCTTAGGACAGTTGTGTCACNNNAGATCGGAAG |
| 47525_1 | TATGAGTGTGGAGTCGTTGCGGAGTGAATCCCTGATACTCTGANNNAGATCGGAAG |
| 47525_2 | TATGAGTGTGGAGTCGTTGCGGAGTGAATCCCTGATACTCTGANNNAGATCGGAAG |
| 45419_1 | TATGAGTGTGGAGTCGTTGCCGCCTTACCTCGCTTACTAACTANNNAGATCGGAAGA |
| 45419_2 | TATGAGTGTGGAGTCGTTGCCGCCTTACCTCGCTTACTAACTANNNAGATCGGAAGA |
| 35614_1 | TATGAGTGTGGAGTCGTTGCCTCAAAGTGCTGGGATTACAAGCNNNAGATCGGAAG |
| 35614_2 | TATGAGTGTGGAGTCGTTGCCTCAAAGTGCTGGGATTACAAGCNNNAGATCGGAAG |
| 47893_1 | TATGAGTGTGGAGTCGTTGCTTGAGATTCATCCATGTTGTTTGTNNNAGATCGGAA |
| 47893_2 | TATGAGTGTGGAGTCGTTGCTTGAGATTCATCCATGTTGTTTGTNNNAGATCGGAA |
| 40812_1 | TATGAGTGTGGAGTCGTTGCCCCAATGATCTGTTTGTTGCCTANNNAGATCGGAAG |
| 40812_2 | TATGAGTGTGGAGTCGTTGCCCCAATGATCTGTTTGTTGCCTANNNAGATCGGAAG |

| | |
|---|---|
| 43257_1 | TATGAGTGTGGAGTCGTTGCACTTATGAGTTTTAGTAGCTTTTTGGTNNNAGATCG |
| 43257_2 | TATGAGTGTGGAGTCGTTGCACTTATGAGTTTTAGTAGCTTTTTGGTNNNAGATCG |
| 32123_1 | TATGAGTGTGGAGTCGTTGCTACAACTTGGAGAAGTCAGGGTGNNNAGATCGGAA |
| 32123_2 | TATGAGTGTGGAGTCGTTGCTACAACTTGGAGAAGTCAGGGTGNNNAGATCGGAA |
| 44113_1 | TATGAGTGTGGAGTCGTTGCGGTTCTCCCTAGCCTACCTCCNNNAGATCGGAAGAG |
| 44113_2 | TATGAGTGTGGAGTCGTTGCGGTTCTCCCTAGCCTACCTCCNNNAGATCGGAAGAG |
| 18711_1 | TATGAGTGTGGAGTCGTTGCATTTTGTTGGCTGTGAAATCNNNAGATCGGAAGAG |
| 18711_2 | TATGAGTGTGGAGTCGTTGCATTTTGTTGGCTGTGAAATCNNNAGATCGGAAGAG |
| 36432_1 | TATGAGTGTGGAGTCGTTGCGACCCTTGGAGGAAAAATGTTGGNNNAGATCGGAA |
| 36432_2 | TATGAGTGTGGAGTCGTTGCGACCCTTGGAGGAAAAATGTTGGNNNAGATCGGAA |
| 2536_1 | TATGAGTGTGGAGTCGTTGCGTTTCTGATCTCATCCTGGCAGGNNNAGATCGGAAG |
| 2536_2 | TATGAGTGTGGAGTCGTTGCGTTTCTGATCTCATCCTGGCAGGNNNAGATCGGAAG |
| 52085_1 | TATGAGTGTGGAGTCGTTGCCATTGTCTGTTCTTGCCATCCAGNNNAGATCGGAAG |
| 52085_2 | TATGAGTGTGGAGTCGTTGCCATTGTCTGTTCTTGCCATCCAGNNNAGATCGGAAG |
| 39652_1 | TATGAGTGTGGAGTCGTTGCTCAGAGCAGCAATAAATGAAGTTGANNNAGATCGG |
| 39652_2 | TATGAGTGTGGAGTCGTTGCTCAGAGCAGCAATAAATGAAGTTGANNNAGATCGG |
| 43104_1 | TATGAGTGTGGAGTCGTTGCACGCCCGGCTAAATTTTTGTATTNNNAGATCGGAAG |
| 43104_2 | TATGAGTGTGGAGTCGTTGCACGCCCGGCTAAATTTTTGTATTNNNAGATCGGAAG |
| 40357_1 | TATGAGTGTGGAGTCGTTGCTAACAAATTAGCTGGGCATGGTGNNNAGATCGGAA |
| 40357_2 | TATGAGTGTGGAGTCGTTGCTAACAAATTAGCTGGGCATGGTGNNNAGATCGGAA |
| 49212_1 | TATGAGTGTGGAGTCGTTGCAAACTCCTGACCTCAAGTGATCCNNNAGATCGGAAG |
| 49212_2 | TATGAGTGTGGAGTCGTTGCAAACTCCTGACCTCAAGTGATCCNNNAGATCGGAAG |
| 52759_1 | TATGAGTGTGGAGTCGTTGCCCTCTCAAGGGTGTCAGCTGNNNAGATCGGAAGAG |
| 52759_2 | TATGAGTGTGGAGTCGTTGCCCTCTCAAGGGTGTCAGCTGNNNAGATCGGAAGAG |
| 29712_1 | TATGAGTGTGGAGTCGTTGCACCATTTGTCCTGTAGCTACCTGNNNAGATCGGAAG |
| 29712_2 | TATGAGTGTGGAGTCGTTGCACCATTTGTCCTGTAGCTACCTGNNNAGATCGGAAG |
| 50971_1 | TATGAGTGTGGAGTCGTTGCGAGCTCTGACACCTATTCTTGGTNNNAGATCGGAAG |
| 50971_2 | TATGAGTGTGGAGTCGTTGCGAGCTCTGACACCTATTCTTGGTNNNAGATCGGAAG |
| 30149_1 | TATGAGTGTGGAGTCGTTGCCTGATAACCCAAATGGCTGCAAANNNAGATCGGAAG |
| 30149_2 | TATGAGTGTGGAGTCGTTGCCTGATAACCCAAATGGCTGCAAANNNAGATCGGAAG |
| 30266_1 | TATGAGTGTGGAGTCGTTGCTTTTGTTTTTGTAATGTTGTGCCANNNAGATCGGAA |
| 30266_2 | TATGAGTGTGGAGTCGTTGCTTTTGTTTTTGTAATGTTGTGCCANNNAGATCGGAA |
| 51054_1 | TATGAGTGTGGAGTCGTTGCCTACACAGGTTGTTCGGAAACACNNNAGATCGGAAG |
| 51054_2 | TATGAGTGTGGAGTCGTTGCCTACACAGGTTGTTCGGAAACACNNNAGATCGGAAG |
| 29335_1 | TATGAGTGTGGAGTCGTTGCAGGCATTAGACATTCTCACGTGANNNAGATCGGAAG |
| 29335_2 | TATGAGTGTGGAGTCGTTGCAGGCATTAGACATTCTCACGTGANNNAGATCGGAAG |
| 43208_1 | TATGAGTGTGGAGTCGTTGCTCATCTGTCTACCTATCATCTAGCTNNNAGATCGGAA |
| 43208_2 | TATGAGTGTGGAGTCGTTGCTCATCTGTCTACCTATCATCTAGCTNNNAGATCGGAA |
| 45788_1 | TATGAGTGTGGAGTCGTTGCCGGGTTTACGCCATTCTCCTNNNAGATCGGAAGAGC |
| 45788_2 | TATGAGTGTGGAGTCGTTGCCGGGTTTACGCCATTCTCCTNNNAGATCGGAAGAGC |
| 45058_1 | TATGAGTGTGGAGTCGTTGCTGGAAAGAAGGATGCTAGTTGGGNNNAGATCGGA |
| 45058_2 | TATGAGTGTGGAGTCGTTGCTGGAAAGAAGGATGCTAGTTGGGNNNAGATCGGA |
| 44813_1 | TATGAGTGTGGAGTCGTTGCAAGCAGAAAGAAGGAGAAGGAGGNNNAGATCGG |

| | |
|---|---|
| 44813_2 | TATGAGTGTGGAGTCGTTGCAAGCAGAAAGAAGGAGAAGGAGGNNNAGATCGGA |
| 29261_1 | TATGAGTGTGGAGTCGTTGCCCTGTGGGGATGATATCTCCATGNNNAGATCGGAAG |
| 29261_2 | TATGAGTGTGGAGTCGTTGCCCTGTGGGGATGATATCTCCATGNNNAGATCGGAAG |
| 9062_1 | TATGAGTGTGGAGTCGTTGCTCCTATATTTGTCCCAAGGGCCTNNNAGATCGGAAG |
| 9062_2 | TATGAGTGTGGAGTCGTTGCTCCTATATTTGTCCCAAGGGCCTNNNAGATCGGAAG |
| 42198_1 | TATGAGTGTGGAGTCGTTGCGGGGTGCTGGAACATGGTAAATANNNAGATCGGAA |
| 42198_2 | TATGAGTGTGGAGTCGTTGCGGGGTGCTGGAACATGGTAAATANNNAGATCGGAA |
| 52750_1 | TATGAGTGTGGAGTCGTTGCCATGTTCAGCTCAAAGTCCACAGNNNAGATCGGAAG |
| 52750_2 | TATGAGTGTGGAGTCGTTGCCATGTTCAGCTCAAAGTCCACAGNNNAGATCGGAAG |
| 35987_1 | TATGAGTGTGGAGTCGTTGCGGGAAACTCTGACTCAGCCAATANNNAGATCGGAAG |
| 35987_2 | TATGAGTGTGGAGTCGTTGCGGGAAACTCTGACTCAGCCAATANNNAGATCGGAAG |
| 36548_1 | TATGAGTGTGGAGTCGTTGCCTGTAGTCCCAGCTACTCAGGAGNNNAGATCGGAAG |
| 36548_2 | TATGAGTGTGGAGTCGTTGCCTGTAGTCCCAGCTACTCAGGAGNNNAGATCGGAAG |
| 35677_1 | TATGAGTGTGGAGTCGTTGCAGTAGCTGGGACTACAGGCANNNAGATCGGAAGAG |
| 35677_2 | TATGAGTGTGGAGTCGTTGCAGTAGCTGGGACTACAGGCANNNAGATCGGAAGAG |
| 38309_1 | TATGAGTGTGGAGTCGTTGCGTCACAGCTCAGTGTTTGCATTANNNAGATCGGAAG |
| 38309_2 | TATGAGTGTGGAGTCGTTGCGTCACAGCTCAGTGTTTGCATTANNNAGATCGGAAG |
| 47185_1 | TATGAGTGTGGAGTCGTTGCGTAGAGATGGGGTTTTACCACGTNNNAGATCGGAA |
| 47185_2 | TATGAGTGTGGAGTCGTTGCGTAGAGATGGGGTTTTACCACGTNNNAGATCGGAA |
| 37283_1 | TATGAGTGTGGAGTCGTTGCGCTCTGCACTCCTTTTGACTTTTNNNAGATCGGAAGA |
| 37283_2 | TATGAGTGTGGAGTCGTTGCGCTCTGCACTCCTTTTGACTTTTNNNAGATCGGAAGA |
| 51297_1 | TATGAGTGTGGAGTCGTTGCATGTCCGTATACCACTTGCCAATNNNAGATCGGAAG |
| 51297_2 | TATGAGTGTGGAGTCGTTGCATGTCCGTATACCACTTGCCAATNNNAGATCGGAAG |
| 41642_1 | TATGAGTGTGGAGTCGTTGCCCATTCTCCTACCTCAGTCTCCCNNNAGATCGGAAGA |
| 41642_2 | TATGAGTGTGGAGTCGTTGCCCATTCTCCTACCTCAGTCTCCCNNNAGATCGGAAGA |
| 46727_1 | TATGAGTGTGGAGTCGTTGCCTAGAGTGCAGTGCCACAATCANNNAGATCGGAAGA |
| 46727_2 | TATGAGTGTGGAGTCGTTGCCTAGAGTGCAGTGCCACAATCANNNAGATCGGAAGA |
| 33919_1 | TATGAGTGTGGAGTCGTTGCGAGTCAGTGTCTCACTCTGTCACNNNAGATCGGAAG |
| 33919_2 | TATGAGTGTGGAGTCGTTGCGAGTCAGTGTCTCACTCTGTCACNNNAGATCGGAAG |
| 9375_1 | TATGAGTGTGGAGTCGTTGCACAGTAGTCCCCTTTATCCGAGGNNNAGATCGGAAG |
| 9375_2 | TATGAGTGTGGAGTCGTTGCACAGTAGTCCCCTTTATCCGAGGNNNAGATCGGAAG |
| 35024_1 | TATGAGTGTGGAGTCGTTGCGAGTGCAGTGGTGCTATCTCTGNNNAGATCGGAAG |
| 35024_2 | TATGAGTGTGGAGTCGTTGCGAGTGCAGTGGTGCTATCTCTGNNNAGATCGGAAG |
| 35511_1 | TATGAGTGTGGAGTCGTTGCGGAATAGGTGGGGTTTGGTTACTNNNAGATCGGAA |
| 35511_2 | TATGAGTGTGGAGTCGTTGCGGAATAGGTGGGGTTTGGTTACTNNNAGATCGGAA |
| 50977_1 | TATGAGTGTGGAGTCGTTGCCTAGTAATGTGCCCCTGGTTCTTNNNAGATCGGAAG |
| 50977_2 | TATGAGTGTGGAGTCGTTGCCTAGTAATGTGCCCCTGGTTCTTNNNAGATCGGAAG |
| 41372_1 | TATGAGTGTGGAGTCGTTGCCCTGCAGTGGGGAAATCTATTTANNNAGATCGGAAG |
| 41372_2 | TATGAGTGTGGAGTCGTTGCCCTGCAGTGGGGAAATCTATTTANNNAGATCGGAAG |
| 31245_1 | TATGAGTGTGGAGTCGTTGCCGCAGACCTCTCATTGACATTTCNNNAGATCGGAAG |
| 31245_2 | TATGAGTGTGGAGTCGTTGCCGCAGACCTCTCATTGACATTTCNNNAGATCGGAAG |
| 29978_1 | TATGAGTGTGGAGTCGTTGCCCCCGGAAGCAATGACAGTANNNAGATCGGAAGAG |
| 29978_2 | TATGAGTGTGGAGTCGTTGCCCCCGGAAGCAATGACAGTANNNAGATCGGAAGAG |

| | |
|---|---|
| 48757_1 | TATGAGTGTGGAGTCGTTGCCCACCATGCTCCGTCTGATTTATNNNAGATCGGAAGA |
| 48757_2 | TATGAGTGTGGAGTCGTTGCCCACCATGCTCCGTCTGATTTATNNNAGATCGGAAGA |
| 33059_1 | TATGAGTGTGGAGTCGTTGCCTCCCAGGTTCCAGTGATTCTTCNNNAGATCGGAAGA |
| 33059_2 | TATGAGTGTGGAGTCGTTGCCTCCCAGGTTCCAGTGATTCTTCNNNAGATCGGAAGA |
| 35139_1 | TATGAGTGTGGAGTCGTTGCCTCACTGTAACCTCGAACTCCAANNNAGATCGGAAGA |
| 35139_2 | TATGAGTGTGGAGTCGTTGCCTCACTGTAACCTCGAACTCCAANNNAGATCGGAAGA |
| 51281_1 | TATGAGTGTGGAGTCGTTGCAGATAACGGGCTCACCAAAATCTNNNAGATCGGAAG |
| 51281_2 | TATGAGTGTGGAGTCGTTGCAGATAACGGGCTCACCAAAATCTNNNAGATCGGAAG |
| 37481_1 | TATGAGTGTGGAGTCGTTGCACAGTCCAATTAGAAAAAGGGCANNNAGATCGGAA |
| 37481_2 | TATGAGTGTGGAGTCGTTGCACAGTCCAATTAGAAAAAGGGCANNNAGATCGGAA |
| 45511_1 | TATGAGTGTGGAGTCGTTGCCTGTAGTGACGAAAGAGCAGGTANNNAGATCGGAA |
| 45511_2 | TATGAGTGTGGAGTCGTTGCCTGTAGTGACGAAAGAGCAGGTANNNAGATCGGAA |
| 49159_1 | TATGAGTGTGGAGTCGTTGCCTCCCCAGAAGCTTGGATTACAGNNNAGATCGGAAG |
| 49159_2 | TATGAGTGTGGAGTCGTTGCCTCCCCAGAAGCTTGGATTACAGNNNAGATCGGAAG |
| 48933_1 | TATGAGTGTGGAGTCGTTGCCCTTCCAAGTAGCTGGGATTACANNNAGATCGGAAG |
| 48933_2 | TATGAGTGTGGAGTCGTTGCCCTTCCAAGTAGCTGGGATTACANNNAGATCGGAAG |
| 44863_1 | TATGAGTGTGGAGTCGTTGCTGTTGATTGAGGGGAGATGACAGNNNAGATCGGAA |
| 44863_2 | TATGAGTGTGGAGTCGTTGCTGTTGATTGAGGGGAGATGACAGNNNAGATCGGAA |
| 33361_1 | TATGAGTGTGGAGTCGTTGCCTCAATCTTGGCTCACTGCAAGNNNAGATCGGAAGA |
| 33361_2 | TATGAGTGTGGAGTCGTTGCCTCAATCTTGGCTCACTGCAAGNNNAGATCGGAAGA |
| 40503_1 | TATGAGTGTGGAGTCGTTGCGAGGTAGGAGAATGGCGTGAACNNNAGATCGGAA |
| 40503_2 | TATGAGTGTGGAGTCGTTGCGAGGTAGGAGAATGGCGTGAACNNNAGATCGGAA |
| 38170_1 | TATGAGTGTGGAGTCGTTGCGTGGGGTCCTAAAAGCCATAAGANNNAGATCGGAA |
| 38170_2 | TATGAGTGTGGAGTCGTTGCGTGGGGTCCTAAAAGCCATAAGANNNAGATCGGAA |
| 30400_1 | TATGAGTGTGGAGTCGTTGCAAGTTGTGTTTTGGCTGAGGATCNNNAGATCGGAA |
| 30400_2 | TATGAGTGTGGAGTCGTTGCAAGTTGTGTTTTGGCTGAGGATCNNNAGATCGGAA |
| 39223_1 | TATGAGTGTGGAGTCGTTGCCTGTAGTCCCAGCTACTCAGGAGNNNAGATCGGAAG |
| 39223_2 | TATGAGTGTGGAGTCGTTGCCTGTAGTCCCAGCTACTCAGGAGNNNAGATCGGAAG |
| 43482_1 | TATGAGTGTGGAGTCGTTGCCTCACTGCAACTTCAAACTCCTGNNNAGATCGGAAGA |
| 43482_2 | TATGAGTGTGGAGTCGTTGCCTCACTGCAACTTCAAACTCCTGNNNAGATCGGAAGA |
| 48420_1 | TATGAGTGTGGAGTCGTTGCCCAGAATGGTCTCATGAGGTCAANNNAGATCGGAA |
| 48420_2 | TATGAGTGTGGAGTCGTTGCCCAGAATGGTCTCATGAGGTCAANNNAGATCGGAA |
| 13332_1 | TATGAGTGTGGAGTCGTTGCAAAGGCTGCAGTTTTATACTNNNAGATCGGAAGAG |
| 13332_2 | TATGAGTGTGGAGTCGTTGCAAAGGCTGCAGTTTTATACTNNNAGATCGGAAGAG |
| 38189_1 | TATGAGTGTGGAGTCGTTGCGGGTTTGAGGTTACAGTGAGCTANNNAGATCGGAA |
| 38189_2 | TATGAGTGTGGAGTCGTTGCGGGTTTGAGGTTACAGTGAGCTANNNAGATCGGAA |
| 50944_1 | TATGAGTGTGGAGTCGTTGCCTCCCGTTTGGCCATTTCTCCNNNAGATCGGAAGAG |
| 50944_2 | TATGAGTGTGGAGTCGTTGCCTCCCGTTTGGCCATTTCTCCNNNAGATCGGAAGAG |
| 47138_1 | TATGAGTGTGGAGTCGTTGCGATCTCCACTCACTGCAACCTNNNAGATCGGAAGAG |
| 47138_2 | TATGAGTGTGGAGTCGTTGCGATCTCCACTCACTGCAACCTNNNAGATCGGAAGAG |
| 12851_1 | TATGAGTGTGGAGTCGTTGCTGATTAATGCAGATGGAAAGATANNNAGATCGGAA |
| 12851_2 | TATGAGTGTGGAGTCGTTGCTGATTAATGCAGATGGAAAGATANNNAGATCGGAA |
| 32949_1 | TATGAGTGTGGAGTCGTTGCGGGGCCTTATGACCTTATTTTGCNNNAGATCGGAAG |

| | |
|---|---|
| 32949_2 | TATGAGTGTGGAGTCGTTGCGGGGCCTTATGACCTTATTTTGCNNNAGATCGGAAG |
| 42049_1 | TATGAGTGTGGAGTCGTTGCCAACATAGCAAGACCCTGTCTCTNNNAGATCGGAAG |
| 42049_2 | TATGAGTGTGGAGTCGTTGCCAACATAGCAAGACCCTGTCTCTNNNAGATCGGAAG |
| 43089_1 | TATGAGTGTGGAGTCGTTGCAAGATCATTTGAGCCCAAGAGGTNNNAGATCGGAA |
| 43089_2 | TATGAGTGTGGAGTCGTTGCAAGATCATTTGAGCCCAAGAGGTNNNAGATCGGAA |
| 13267_1 | TATGAGTGTGGAGTCGTTGCGTTAATAACTGCATTGGATTTTCNNNAGATCGGAAG |
| 13267_2 | TATGAGTGTGGAGTCGTTGCGTTAATAACTGCATTGGATTTTCNNNAGATCGGAAG |
| 9593_1 | TATGAGTGTGGAGTCGTTGCTTCGTACTTTACCTTTGGAGGGCNNNAGATCGGAAG |
| 9593_2 | TATGAGTGTGGAGTCGTTGCTTCGTACTTTACCTTTGGAGGGCNNNAGATCGGAAG |
| 41751_1 | TATGAGTGTGGAGTCGTTGCCATTCTCCTGCCTCAGCCTCNNNAGATCGGAAGAGCA |
| 41751_2 | TATGAGTGTGGAGTCGTTGCCATTCTCCTGCCTCAGCCTCNNNAGATCGGAAGAGCA |
| 50518_1 | TATGAGTGTGGAGTCGTTGCGGAAATATCCCTACCACCCATCCNNNAGATCGGAAG |
| 50518_2 | TATGAGTGTGGAGTCGTTGCGGAAATATCCCTACCACCCATCCNNNAGATCGGAAG |
| 29209_1 | TATGAGTGTGGAGTCGTTGCTCACACATGTAGCCTTGCATAACNNNAGATCGGAAG |
| 29209_2 | TATGAGTGTGGAGTCGTTGCTCACACATGTAGCCTTGCATAACNNNAGATCGGAAG |
| 51864_1 | TATGAGTGTGGAGTCGTTGCTCACCACCTCTGGAACTCAATTCNNNAGATCGGAAG |
| 51864_2 | TATGAGTGTGGAGTCGTTGCTCACCACCTCTGGAACTCAATTCNNNAGATCGGAAG |
| 33525_1 | TATGAGTGTGGAGTCGTTGCTTTGGGGTGTCACATGATCAGATNNNAGATCGGAA |
| 33525_2 | TATGAGTGTGGAGTCGTTGCTTTGGGGTGTCACATGATCAGATNNNAGATCGGAA |
| 13137_1 | TATGAGTGTGGAGTCGTTGCATTGATTCAGATGAAGAAGATCANNNAGATCGGAA |
| 13137_2 | TATGAGTGTGGAGTCGTTGCATTGATTCAGATGAAGAAGATCANNNAGATCGGAA |
| 52037_1 | TATGAGTGTGGAGTCGTTGCTATTACAAGCATGAGCCATTGCGNNNAGATCGGAAG |
| 52037_2 | TATGAGTGTGGAGTCGTTGCTATTACAAGCATGAGCCATTGCGNNNAGATCGGAAG |
| 10076_1 | TATGAGTGTGGAGTCGTTGCATACCAAACTCCTGTCCTCAAGCNNNAGATCGGAAG |
| 10076_2 | TATGAGTGTGGAGTCGTTGCATACCAAACTCCTGTCCTCAAGCNNNAGATCGGAAG |
| 34220_1 | TATGAGTGTGGAGTCGTTGCAAATGCTTAGGGGAAATCTGCCTNNNAGATCGGAA |
| 34220_2 | TATGAGTGTGGAGTCGTTGCAAATGCTTAGGGGAAATCTGCCTNNNAGATCGGAA |
| 10399_1 | TATGAGTGTGGAGTCGTTGCTCAGTAGGAACGTTATCGGCTCTNNNAGATCGGAAG |
| 10399_2 | TATGAGTGTGGAGTCGTTGCTCAGTAGGAACGTTATCGGCTCTNNNAGATCGGAAG |
| 34807_1 | TATGAGTGTGGAGTCGTTGCTGACAGTATCTCACTCTTTCGCCNNNAGATCGGAAG |
| 34807_2 | TATGAGTGTGGAGTCGTTGCTGACAGTATCTCACTCTTTCGCCNNNAGATCGGAAG |
| 30742_1 | TATGAGTGTGGAGTCGTTGCAAAAGATGTTGCATATCCAGGCCNNNAGATCGGAA |
| 30742_2 | TATGAGTGTGGAGTCGTTGCAAAAGATGTTGCATATCCAGGCCNNNAGATCGGAA |
| 36049_1 | TATGAGTGTGGAGTCGTTGCGAGACAATGAGGACAAGGGTAGGNNNAGATCGGA |
| 36049_2 | TATGAGTGTGGAGTCGTTGCGAGACAATGAGGACAAGGGTAGGNNNAGATCGGA |
| 50602_1 | TATGAGTGTGGAGTCGTTGCGACTGACATGGGTAGAGGTGATCNNNAGATCGGAA |
| 50602_2 | TATGAGTGTGGAGTCGTTGCGACTGACATGGGTAGAGGTGATCNNNAGATCGGAA |
| 45610_1 | TATGAGTGTGGAGTCGTTGCTTCAAGTGATTCTCCTGCCTCAGNNNAGATCGGAAG |
| 45610_2 | TATGAGTGTGGAGTCGTTGCTTCAAGTGATTCTCCTGCCTCAGNNNAGATCGGAAG |
| 29353_1 | TATGAGTGTGGAGTCGTTGCCTGGAGACAGGAGAGAAAAGCAANNNAGATCGGA |
| 29353_2 | TATGAGTGTGGAGTCGTTGCCTGGAGACAGGAGAGAAAAGCAANNNAGATCGGA |
| 50787_1 | TATGAGTGTGGAGTCGTTGCGCCCAGTGCTATGAACATTCATGNNNAGATCGGAAG |
| 50787_2 | TATGAGTGTGGAGTCGTTGCGCCCAGTGCTATGAACATTCATGNNNAGATCGGAAG |

| | |
|---|---|
| 10333_1 | TATGAGTGTGGAGTCGTTGCAAACTCCTCCAAGAAACTGGACANNNAGATCGGAAG |
| 10333_2 | TATGAGTGTGGAGTCGTTGCAAACTCCTCCAAGAAACTGGACANNNAGATCGGAAG |
| 33339_1 | TATGAGTGTGGAGTCGTTGCGTTCAAGCCATTCTCCTACCTCANNNAGATCGGAAGA |
| 33339_2 | TATGAGTGTGGAGTCGTTGCGTTCAAGCCATTCTCCTACCTCANNNAGATCGGAAGA |
| 9063_1 | TATGAGTGTGGAGTCGTTGCAGTCCAGTGTAAGTACTTCGCCTNNNAGATCGGAAG |
| 9063_2 | TATGAGTGTGGAGTCGTTGCAGTCCAGTGTAAGTACTTCGCCTNNNAGATCGGAAG |
| 5588_1 | TATGAGTGTGGAGTCGTTGCTGAGTGCAGACGTCATTTTAGCANNNAGATCGGAAG |
| 5588_2 | TATGAGTGTGGAGTCGTTGCTGAGTGCAGACGTCATTTTAGCANNNAGATCGGAAG |
| 49005_1 | TATGAGTGTGGAGTCGTTGCCTCCTGAATAGCTGGGACTACAGNNNAGATCGGAAG |
| 49005_2 | TATGAGTGTGGAGTCGTTGCCTCCTGAATAGCTGGGACTACAGNNNAGATCGGAAG |
| 49187_1 | TATGAGTGTGGAGTCGTTGCGAAGAAGAGCCTGAAAATGCCAGNNNAGATCGGAA |
| 49187_2 | TATGAGTGTGGAGTCGTTGCGAAGAAGAGCCTGAAAATGCCAGNNNAGATCGGAA |
| 48748_1 | TATGAGTGTGGAGTCGTTGCGGGCCTTTTTTGGTTGGTATGTTNNNAGATCGGAAG |
| 48748_2 | TATGAGTGTGGAGTCGTTGCGGGCCTTTTTTGGTTGGTATGTTNNNAGATCGGAAG |
| 29185_1 | TATGAGTGTGGAGTCGTTGCAACATGTGGTAGGAGCAAGAAGTNNNAGATCGGAA |
| 29185_2 | TATGAGTGTGGAGTCGTTGCAACATGTGGTAGGAGCAAGAAGTNNNAGATCGGAA |
| 45113_1 | TATGAGTGTGGAGTCGTTGCTGTGACTGTGATTTGTTTTATGGTNNNAGATCGGAA |
| 45113_2 | TATGAGTGTGGAGTCGTTGCTGTGACTGTGATTTGTTTTATGGTNNNAGATCGGAA |
| 49954_1 | TATGAGTGTGGAGTCGTTGCTTTGAGCAATAGTGTCAGTGGGANNNAGATCGGAA |
| 49954_2 | TATGAGTGTGGAGTCGTTGCTTTGAGCAATAGTGTCAGTGGGANNNAGATCGGAA |
| 50196_1 | TATGAGTGTGGAGTCGTTGCGCACTAGAACAAAAGCTCACCTGNNNAGATCGGAAG |
| 50196_2 | TATGAGTGTGGAGTCGTTGCGCACTAGAACAAAAGCTCACCTGNNNAGATCGGAAG |
| 31892_1 | TATGAGTGTGGAGTCGTTGCAGAATCTTACTGTGTTCTGGGGGNNNAGATCGGAA |
| 31892_2 | TATGAGTGTGGAGTCGTTGCAGAATCTTACTGTGTTCTGGGGGNNNAGATCGGAA |
| 32310_1 | TATGAGTGTGGAGTCGTTGCTTGATGACAGGGAGCACGAATTANNNAGATCGGAA |
| 32310_2 | TATGAGTGTGGAGTCGTTGCTTGATGACAGGGAGCACGAATTANNNAGATCGGAA |
| 38762_1 | TATGAGTGTGGAGTCGTTGCATGTTGCCCATTCACTAGCAATGNNNAGATCGGAAG |
| 38762_2 | TATGAGTGTGGAGTCGTTGCATGTTGCCCATTCACTAGCAATGNNNAGATCGGAAG |
| 41654_1 | TATGAGTGTGGAGTCGTTGCGAGACAGAGCTTGGCTTTTGTTGNNNAGATCGGAA |
| 41654_2 | TATGAGTGTGGAGTCGTTGCGAGACAGAGCTTGGCTTTTGTTGNNNAGATCGGAA |
| 43486_1 | TATGAGTGTGGAGTCGTTGCACAGAACAACAACAACAACAGCANNNAGATCGGAAG |
| 43486_2 | TATGAGTGTGGAGTCGTTGCACAGAACAACAACAACAACAGCANNNAGATCGGAAG |
| 51195_1 | TATGAGTGTGGAGTCGTTGCCTTTTAGCTGAGTTTGACCTGGGNNNAGATCGGAAG |
| 51195_2 | TATGAGTGTGGAGTCGTTGCCTTTTAGCTGAGTTTGACCTGGGNNNAGATCGGAAG |
| 49773_1 | TATGAGTGTGGAGTCGTTGCGTCTCTCCAGATCACGAACCTAGNNNAGATCGGAAG |
| 49773_2 | TATGAGTGTGGAGTCGTTGCGTCTCTCCAGATCACGAACCTAGNNNAGATCGGAAG |
| 29046_1 | TATGAGTGTGGAGTCGTTGCGCATGCACACGTGTATACTCAGANNNAGATCGGAAG |
| 29046_2 | TATGAGTGTGGAGTCGTTGCGCATGCACACGTGTATACTCAGANNNAGATCGGAAG |
| 748_1 | TATGAGTGTGGAGTCGTTGCTTCACCTCTCCATGGGTCTCATTNNNAGATCGGAAGA |
| 748_2 | TATGAGTGTGGAGTCGTTGCTTCACCTCTCCATGGGTCTCATTNNNAGATCGGAAGA |
| 41522_1 | TATGAGTGTGGAGTCGTTGCATCCTGCAGGTACAGTGAAGAAGNNNAGATCGGAA |
| 41522_2 | TATGAGTGTGGAGTCGTTGCATCCTGCAGGTACAGTGAAGAAGNNNAGATCGGAA |
| 39175_1 | TATGAGTGTGGAGTCGTTGCAGTAAAAAGAGCCCACATAGCCANNNAGATCGGAAG |

| | |
|---|---|
| 39175_2 | TATGAGTGTGGAGTCGTTGCAGTAAAAAGAGCCCACATAGCCANNNAGATCGGAA |
| 40099_1 | TATGAGTGTGGAGTCGTTGCACCACTGACACCACAGAAATAAANNNAGATCGGAAG |
| 40099_2 | TATGAGTGTGGAGTCGTTGCACCACTGACACCACAGAAATAAANNNAGATCGGAAG |
| 52858_1 | TATGAGTGTGGAGTCGTTGCAGTGTGCCATTCATCCTGTCTCNNNAGATCGGAAGA |
| 52858_2 | TATGAGTGTGGAGTCGTTGCAGTGTGCCATTCATCCTGTCTCNNNAGATCGGAAGA |
| 35081_1 | TATGAGTGTGGAGTCGTTGCTATCTCAGTGAGCAGAGGGATGANNNAGATCGGAA |
| 35081_2 | TATGAGTGTGGAGTCGTTGCTATCTCAGTGAGCAGAGGGATGANNNAGATCGGAA |
| 32151_1 | TATGAGTGTGGAGTCGTTGCCTGGCAGCTACAAATCCCAATATNNNAGATCGGAAG |
| 32151_2 | TATGAGTGTGGAGTCGTTGCCTGGCAGCTACAAATCCCAATATNNNAGATCGGAAG |
| 45724_1 | TATGAGTGTGGAGTCGTTGCATAAGAGGGTCACGGATGTCCATNNNAGATCGGAA |
| 45724_2 | TATGAGTGTGGAGTCGTTGCATAAGAGGGTCACGGATGTCCATNNNAGATCGGAA |
| 43011_1 | TATGAGTGTGGAGTCGTTGCTCCAGCCAGAATGAAGAAACTGANNNAGATCGGAA |
| 43011_2 | TATGAGTGTGGAGTCGTTGCTCCAGCCAGAATGAAGAAACTGANNNAGATCGGAA |
| 43291_1 | TATGAGTGTGGAGTCGTTGCCCTCCAGAGTAGCTAGGACTACANNNAGATCGGAAG |
| 43291_2 | TATGAGTGTGGAGTCGTTGCCCTCCAGAGTAGCTAGGACTACANNNAGATCGGAAG |
| 47098_1 | TATGAGTGTGGAGTCGTTGCATCTCGGCTAACTGCAAGCTCNNNAGATCGGAAGAG |
| 47098_2 | TATGAGTGTGGAGTCGTTGCATCTCGGCTAACTGCAAGCTCNNNAGATCGGAAGAG |
| 36465_1 | TATGAGTGTGGAGTCGTTGCGGGAAAGGTCCAGATATTTGGGTNNNAGATCGGAA |
| 36465_2 | TATGAGTGTGGAGTCGTTGCGGGAAAGGTCCAGATATTTGGGTNNNAGATCGGAA |
| 32945_1 | TATGAGTGTGGAGTCGTTGCACCCAATAATCAAAAGCCTCCCANNNAGATCGGAAG |
| 32945_2 | TATGAGTGTGGAGTCGTTGCACCCAATAATCAAAAGCCTCCCANNNAGATCGGAAG |
| 32342_1 | TATGAGTGTGGAGTCGTTGCCAGACACCAGGGAAAAGAGGTAANNNAGATCGGA |
| 32342_2 | TATGAGTGTGGAGTCGTTGCCAGACACCAGGGAAAAGAGGTAANNNAGATCGGA |
| 40132_1 | TATGAGTGTGGAGTCGTTGCTTGATTCCACCCAGGTTTCATCTNNNAGATCGGAAG |
| 40132_2 | TATGAGTGTGGAGTCGTTGCTTGATTCCACCCAGGTTTCATCTNNNAGATCGGAAG |
| 42487_1 | TATGAGTGTGGAGTCGTTGCAAATAGTCACTGCCTGGTAACCCNNNAGATCGGAAG |
| 42487_2 | TATGAGTGTGGAGTCGTTGCAAATAGTCACTGCCTGGTAACCCNNNAGATCGGAAG |
| 39555_1 | TATGAGTGTGGAGTCGTTGCACACGTGGGAATTGAAGATGAGANNNAGATCGGA |
| 39555_2 | TATGAGTGTGGAGTCGTTGCACACGTGGGAATTGAAGATGAGANNNAGATCGGA |
| 51470_1 | TATGAGTGTGGAGTCGTTGCAGTCTTTTTGCAAAATGGGACCCNNNAGATCGGAAG |
| 51470_2 | TATGAGTGTGGAGTCGTTGCAGTCTTTTTGCAAAATGGGACCCNNNAGATCGGAAG |
| 32708_1 | TATGAGTGTGGAGTCGTTGCGTTCAAGCATCCTCCTACCTCAGNNNAGATCGGAAG |
| 32708_2 | TATGAGTGTGGAGTCGTTGCGTTCAAGCATCCTCCTACCTCAGNNNAGATCGGAAG |
| 41173_1 | TATGAGTGTGGAGTCGTTGCATGGTAAGCTCTAAAGAACCACTNNNAGATCGGAAG |
| 41173_2 | TATGAGTGTGGAGTCGTTGCATGGTAAGCTCTAAAGAACCACTNNNAGATCGGAAG |
| 49425_1 | TATGAGTGTGGAGTCGTTGCTTCAAGTGATTCTCCTGCCTGAGNNNAGATCGGAAG |
| 49425_2 | TATGAGTGTGGAGTCGTTGCTTCAAGTGATTCTCCTGCCTGAGNNNAGATCGGAAG |
| 45148_1 | TATGAGTGTGGAGTCGTTGCGAGGAGAGACAAGGGAGAAACAGNNNAGATCGGA |
| 45148_2 | TATGAGTGTGGAGTCGTTGCGAGGAGAGACAAGGGAGAAACAGNNNAGATCGGA |
| 38360_1 | TATGAGTGTGGAGTCGTTGCACACTGGAAAGTCTCAGCAATAGANNNAGATCGGA |
| 38360_2 | TATGAGTGTGGAGTCGTTGCACACTGGAAAGTCTCAGCAATAGANNNAGATCGGA |
| 40664_1 | TATGAGTGTGGAGTCGTTGCCACCATCGCCATCTCTTTGTACTNNNAGATCGGAAGA |
| 40664_2 | TATGAGTGTGGAGTCGTTGCCACCATCGCCATCTCTTTGTACTNNNAGATCGGAAGA |

| | |
|---|---|
| 51413_1 | TATGAGTGTGGAGTCGTTGCTCCCCCTTTTGAATTCTTTCCCCNNNAGATCGGAAGA |
| 51413_2 | TATGAGTGTGGAGTCGTTGCTCCCCCTTTTGAATTCTTTCCCCNNNAGATCGGAAGA |
| 31435_1 | TATGAGTGTGGAGTCGTTGCACAGGGACAATGCATAGAACAGANNNAGATCGGAA |
| 31435_2 | TATGAGTGTGGAGTCGTTGCACAGGGACAATGCATAGAACAGANNNAGATCGGAA |
| 31636_1 | TATGAGTGTGGAGTCGTTGCTATCAAATGGAGGAGAGGGGACTNNNAGATCGGAA |
| 31636_2 | TATGAGTGTGGAGTCGTTGCTATCAAATGGAGGAGAGGGGACTNNNAGATCGGAA |
| 33235_1 | TATGAGTGTGGAGTCGTTGCCCTATAAGCATGATCCTCCCACCNNNAGATCGGAAGA |
| 33235_2 | TATGAGTGTGGAGTCGTTGCCCTATAAGCATGATCCTCCCACCNNNAGATCGGAAGA |
| 31257_1 | TATGAGTGTGGAGTCGTTGCACGTTTTTCAGGTTGAGGTTTTTNNNAGATCGGAAG |
| 31257_2 | TATGAGTGTGGAGTCGTTGCACGTTTTTCAGGTTGAGGTTTTTNNNAGATCGGAAG |
| 53283_1 | TATGAGTGTGGAGTCGTTGCTAACTACTTCCTTGTGCGTGTCANNNAGATCGGAAGA |
| 53283_2 | TATGAGTGTGGAGTCGTTGCTAACTACTTCCTTGTGCGTGTCANNNAGATCGGAAGA |
| 50408_1 | TATGAGTGTGGAGTCGTTGCCTTTCAGCCTTTTCCAGAACCAGNNNAGATCGGAAGA |
| 50408_2 | TATGAGTGTGGAGTCGTTGCCTTTCAGCCTTTTCCAGAACCAGNNNAGATCGGAAGA |
| 53101_1 | TATGAGTGTGGAGTCGTTGCACACCCAAGGGAATATTCTGAGGNNNAGATCGGAA |
| 53101_2 | TATGAGTGTGGAGTCGTTGCACACCCAAGGGAATATTCTGAGGNNNAGATCGGAA |
| 37687_1 | TATGAGTGTGGAGTCGTTGCAGTCTCATAGAACCACTCCAGGTNNNAGATCGGAAG |
| 37687_2 | TATGAGTGTGGAGTCGTTGCAGTCTCATAGAACCACTCCAGGTNNNAGATCGGAAG |
| 33786_1 | TATGAGTGTGGAGTCGTTGCTGACCTATCTCCTACACTCCCTGNNNAGATCGGAAGA |
| 33786_2 | TATGAGTGTGGAGTCGTTGCTGACCTATCTCCTACACTCCCTGNNNAGATCGGAAGA |
| 43791_1 | TATGAGTGTGGAGTCGTTGCAGAGAACCACTTGAACCCAGAAGNNNAGATCGGAA |
| 43791_2 | TATGAGTGTGGAGTCGTTGCAGAGAACCACTTGAACCCAGAAGNNNAGATCGGAA |
| 34079_1 | TATGAGTGTGGAGTCGTTGCGGAGTGCAGTGGTGTGATCTCNNNAGATCGGAAGA |
| 34079_2 | TATGAGTGTGGAGTCGTTGCGGAGTGCAGTGGTGTGATCTCNNNAGATCGGAAGA |
| 36469_1 | TATGAGTGTGGAGTCGTTGCTGGGACAGGGTAGGCATTCTATANNNAGATCGGAA |
| 36469_2 | TATGAGTGTGGAGTCGTTGCTGGGACAGGGTAGGCATTCTATANNNAGATCGGAA |
| 44461_1 | TATGAGTGTGGAGTCGTTGCGGGAGAGTGGGGGTATATACAGANNNAGATCGGA |
| 44461_2 | TATGAGTGTGGAGTCGTTGCGGGAGAGTGGGGGTATATACAGANNNAGATCGGA |
| 38217_1 | TATGAGTGTGGAGTCGTTGCGAGGTCAGGAGATTGAGACCATCNNNAGATCGGAA |
| 38217_2 | TATGAGTGTGGAGTCGTTGCGAGGTCAGGAGATTGAGACCATCNNNAGATCGGAA |
| 28862_1 | TATGAGTGTGGAGTCGTTGCGCTGTATAGAACTTGCAACAGGGNNNAGATCGGAA |
| 28862_2 | TATGAGTGTGGAGTCGTTGCGCTGTATAGAACTTGCAACAGGGNNNAGATCGGAA |
| 39112_1 | TATGAGTGTGGAGTCGTTGCCTGTAGTCCCAGCTACTCGAGAGNNNAGATCGGAAG |
| 39112_2 | TATGAGTGTGGAGTCGTTGCCTGTAGTCCCAGCTACTCGAGAGNNNAGATCGGAAG |
| 38118_1 | TATGAGTGTGGAGTCGTTGCAAAGAAAAGAAAACTGCAGGCCANNNAGATCGGAA |
| 38118_2 | TATGAGTGTGGAGTCGTTGCAAAGAAAAGAAAACTGCAGGCCANNNAGATCGGAA |
| 47354_1 | TATGAGTGTGGAGTCGTTGCCTCCCTGGTTCAAGCGATTCTCNNNAGATCGGAAGA |
| 47354_2 | TATGAGTGTGGAGTCGTTGCCTCCCTGGTTCAAGCGATTCTCNNNAGATCGGAAGA |
| 44287_1 | TATGAGTGTGGAGTCGTTGCAAGACCTGGACCAATGTGTATCCNNNAGATCGGAAG |
| 44287_2 | TATGAGTGTGGAGTCGTTGCAAGACCTGGACCAATGTGTATCCNNNAGATCGGAAG |
| 44485_1 | TATGAGTGTGGAGTCGTTGCACCTGAACGGGTACTTTTCTCAANNNAGATCGGAAG |
| 44485_2 | TATGAGTGTGGAGTCGTTGCACCTGAACGGGTACTTTTCTCAANNNAGATCGGAAG |
| 44328_1 | TATGAGTGTGGAGTCGTTGCCTCTCGGAGAGCTTAGTGATGTCNNNAGATCGGAAG |

| | |
|---|---|
| 44328_2 | TATGAGTGTGGAGTCGTTGCCTCTCGGAGAGCTTAGTGATGTCNNNAGATCGGAAG |
| 34919_1 | TATGAGTGTGGAGTCGTTGCCTCACATTGCATCTTTGAGCCAANNNAGATCGGAAG |
| 34919_2 | TATGAGTGTGGAGTCGTTGCCTCACATTGCATCTTTGAGCCAANNNAGATCGGAAG |
| 41759_1 | TATGAGTGTGGAGTCGTTGCGTCAGGAGCATTTTGAAGTGGACNNNAGATCGGAA |
| 41759_2 | TATGAGTGTGGAGTCGTTGCGTCAGGAGCATTTTGAAGTGGACNNNAGATCGGAA |
| 33096_1 | TATGAGTGTGGAGTCGTTGCCAGGTTCAAGCGGTTCTCCTNNNAGATCGGAAGAGC |
| 33096_2 | TATGAGTGTGGAGTCGTTGCCAGGTTCAAGCGGTTCTCCTNNNAGATCGGAAGAGC |
| 36863_1 | TATGAGTGTGGAGTCGTTGCCTACCAGAATGGAAGCCCTTTCANNNAGATCGGAAG |
| 36863_2 | TATGAGTGTGGAGTCGTTGCCTACCAGAATGGAAGCCCTTTCANNNAGATCGGAAG |
| 48325_1 | TATGAGTGTGGAGTCGTTGCGGCTGAACTGGAAAAAAGAATGGANNNAGATCGG |
| 48325_2 | TATGAGTGTGGAGTCGTTGCGGCTGAACTGGAAAAAAGAATGGANNNAGATCGG |
| 49219_1 | TATGAGTGTGGAGTCGTTGCTGTAGATGAAACAAGCCCCATGANNNAGATCGGAA |
| 49219_2 | TATGAGTGTGGAGTCGTTGCTGTAGATGAAACAAGCCCCATGANNNAGATCGGAA |
| 12993_1 | TATGAGTGTGGAGTCGTTGCGGAACTCCTGAATAATTTTTTCANNNAGATCGGAAG |
| 12993_2 | TATGAGTGTGGAGTCGTTGCGGAACTCCTGAATAATTTTTTCANNNAGATCGGAAG |
| 43926_1 | TATGAGTGTGGAGTCGTTGCGCTGCTTCCTCAGATAAGAAGGANNNAGATCGGAA |
| 43926_2 | TATGAGTGTGGAGTCGTTGCGCTGCTTCCTCAGATAAGAAGGANNNAGATCGGAA |
| 33826_1 | TATGAGTGTGGAGTCGTTGCCCCCAGCTCATCTGATCTAACTTNNNAGATCGGAAGA |
| 33826_2 | TATGAGTGTGGAGTCGTTGCCCCCAGCTCATCTGATCTAACTTNNNAGATCGGAAGA |
| 31465_1 | TATGAGTGTGGAGTCGTTGCTAGCTGGGACAACAGACACATGNNNAGATCGGAAG |
| 31465_2 | TATGAGTGTGGAGTCGTTGCTAGCTGGGACAACAGACACATGNNNAGATCGGAAG |
| 43825_1 | TATGAGTGTGGAGTCGTTGCCTCTACAAGGCTGCCTTAGTTGANNNAGATCGGAAG |
| 43825_2 | TATGAGTGTGGAGTCGTTGCCTCTACAAGGCTGCCTTAGTTGANNNAGATCGGAAG |
| 31808_1 | TATGAGTGTGGAGTCGTTGCTTGGATGAAACTGGAGGCCATTANNNAGATCGGAA |
| 31808_2 | TATGAGTGTGGAGTCGTTGCTTGGATGAAACTGGAGGCCATTANNNAGATCGGAA |
| 10788_1 | TATGAGTGTGGAGTCGTTGCTACATTGTTGCTGCTGTGGGAAGNNNAGATCGGAA |
| 10788_2 | TATGAGTGTGGAGTCGTTGCTACATTGTTGCTGCTGTGGGAAGNNNAGATCGGAA |
| 42026_1 | TATGAGTGTGGAGTCGTTGCTCCAGTTCTGTCTACTCCAATTCANNNAGATCGGAAG |
| 42026_2 | TATGAGTGTGGAGTCGTTGCTCCAGTTCTGTCTACTCCAATTCANNNAGATCGGAAG |
| 10058_1 | TATGAGTGTGGAGTCGTTGCAGTTCAAAAGCAGTGACATTGCANNNAGATCGGAA |
| 10058_2 | TATGAGTGTGGAGTCGTTGCAGTTCAAAAGCAGTGACATTGCANNNAGATCGGAA |
| 37819_1 | TATGAGTGTGGAGTCGTTGCCACCTGTAATCCCAGCATTTTGGNNNAGATCGGAAG |
| 37819_2 | TATGAGTGTGGAGTCGTTGCCACCTGTAATCCCAGCATTTTGGNNNAGATCGGAAG |
| 3066_1 | TATGAGTGTGGAGTCGTTGCATGTTGATGGGATTCCAGTAGCCNNNAGATCGGAA |
| 3066_2 | TATGAGTGTGGAGTCGTTGCATGTTGATGGGATTCCAGTAGCCNNNAGATCGGAA |
| 230_1 | TATGAGTGTGGAGTCGTTGCGGTAAAATAGGGGTGGAGGTCAAANNNAGATCGG |
| 230_2 | TATGAGTGTGGAGTCGTTGCGGTAAAATAGGGGTGGAGGTCAAANNNAGATCGG |
| 44214_1 | TATGAGTGTGGAGTCGTTGCTGGGACATCTAGGTACCTCAACTNNNAGATCGGAAG |
| 44214_2 | TATGAGTGTGGAGTCGTTGCTGGGACATCTAGGTACCTCAACTNNNAGATCGGAAG |
| 44595_1 | TATGAGTGTGGAGTCGTTGCGTCTGCCAACTGGATGCATATTCNNNAGATCGGAAG |
| 44595_2 | TATGAGTGTGGAGTCGTTGCGTCTGCCAACTGGATGCATATTCNNNAGATCGGAAG |
| 35290_1 | TATGAGTGTGGAGTCGTTGCCTGGCCAAGTTTCAGTCTTCTTGNNNAGATCGGAAG |
| 35290_2 | TATGAGTGTGGAGTCGTTGCCTGGCCAAGTTTCAGTCTTCTTGNNNAGATCGGAAG |

| | |
|---|---|
| 34506_1 | TATGAGTGTGGAGTCGTTGCTTAGCAAGGAACAACGCATTCTCNNNAGATCGGAAG |
| 34506_2 | TATGAGTGTGGAGTCGTTGCTTAGCAAGGAACAACGCATTCTCNNNAGATCGGAAG |
| 49387_1 | TATGAGTGTGGAGTCGTTGCCCAAGGTACTAGGATCACAGGTGNNNAGATCGGAAG |
| 49387_2 | TATGAGTGTGGAGTCGTTGCCCAAGGTACTAGGATCACAGGTGNNNAGATCGGAAG |
| 41152_1 | TATGAGTGTGGAGTCGTTGCACATTGTAGCCTCAAACTCCTGGNNNAGATCGGAAGA |
| 41152_2 | TATGAGTGTGGAGTCGTTGCACATTGTAGCCTCAAACTCCTGGNNNAGATCGGAAGA |
| 40848_1 | TATGAGTGTGGAGTCGTTGCTTTGGGGTACAAGTGGTTTCTCANNNAGATCGGAAG |
| 40848_2 | TATGAGTGTGGAGTCGTTGCTTTGGGGTACAAGTGGTTTCTCANNNAGATCGGAAG |
| 50405_1 | TATGAGTGTGGAGTCGTTGCACTGAAGGTTCTCCTGGGAATTCNNNAGATCGGAAG |
| 50405_2 | TATGAGTGTGGAGTCGTTGCACTGAAGGTTCTCCTGGGAATTCNNNAGATCGGAAG |
| 35546_1 | TATGAGTGTGGAGTCGTTGCGCTGGGATTACAGATGTACACCANNNAGATCGGAAG |
| 35546_2 | TATGAGTGTGGAGTCGTTGCGCTGGGATTACAGATGTACACCANNNAGATCGGAAG |
| 31507_1 | TATGAGTGTGGAGTCGTTGCGGGAGCCCTCATCTCTACAAAAANNNAGATCGGAAG |
| 31507_2 | TATGAGTGTGGAGTCGTTGCGGGAGCCCTCATCTCTACAAAAANNNAGATCGGAAG |
| 52312_1 | TATGAGTGTGGAGTCGTTGCTTTGTACTTGGTTGAATCGTGGCNNNAGATCGGAAG |
| 52312_2 | TATGAGTGTGGAGTCGTTGCTTTGTACTTGGTTGAATCGTGGCNNNAGATCGGAAG |
| 34043_1 | TATGAGTGTGGAGTCGTTGCATCTTCTGACCTCATGATCCACCNNNAGATCGGAAGA |
| 34043_2 | TATGAGTGTGGAGTCGTTGCATCTTCTGACCTCATGATCCACCNNNAGATCGGAAGA |
| 31896_1 | TATGAGTGTGGAGTCGTTGCATCTGAACTTTTGGGGGTCAACTNNNAGATCGGAAG |
| 31896_2 | TATGAGTGTGGAGTCGTTGCATCTGAACTTTTGGGGGTCAACTNNNAGATCGGAAG |
| 33126_1 | TATGAGTGTGGAGTCGTTGCTAGATATACCCACTCACCCCACCNNNAGATCGGAAGA |
| 33126_2 | TATGAGTGTGGAGTCGTTGCTAGATATACCCACTCACCCCACCNNNAGATCGGAAGA |
| 46454_1 | TATGAGTGTGGAGTCGTTGCCTCCCTGGTTCAAGCGATTCTCNNNAGATCGGAAGAG |
| 46454_2 | TATGAGTGTGGAGTCGTTGCCTCCCTGGTTCAAGCGATTCTCNNNAGATCGGAAGAG |
| 40534_1 | TATGAGTGTGGAGTCGTTGCGTAGTCCCAGCTACTTGAGAAGCNNNAGATCGGAAG |
| 40534_2 | TATGAGTGTGGAGTCGTTGCGTAGTCCCAGCTACTTGAGAAGCNNNAGATCGGAAG |
| 48472_1 | TATGAGTGTGGAGTCGTTGCCTGTAATCCCAGCTACTCAGGTGNNNAGATCGGAAG |
| 48472_2 | TATGAGTGTGGAGTCGTTGCCTGTAATCCCAGCTACTCAGGTGNNNAGATCGGAAG |
| 38206_1 | TATGAGTGTGGAGTCGTTGCCAAGATTGTGTCATTGCACTCCANNNAGATCGGAAGA |
| 38206_2 | TATGAGTGTGGAGTCGTTGCCAAGATTGTGTCATTGCACTCCANNNAGATCGGAAGA |
| 45705_1 | TATGAGTGTGGAGTCGTTGCGTGACGGAGTCTCGTTCTGTCNNNAGATCGGAAGAG |
| 45705_2 | TATGAGTGTGGAGTCGTTGCGTGACGGAGTCTCGTTCTGTCNNNAGATCGGAAGAG |
| 33297_1 | TATGAGTGTGGAGTCGTTGCCAAGTAGCTGGGACTACAGGCNNNAGATCGGAAGA |
| 33297_2 | TATGAGTGTGGAGTCGTTGCCAAGTAGCTGGGACTACAGGCNNNAGATCGGAAGA |
| 30839_1 | TATGAGTGTGGAGTCGTTGCGCACAAATACCTGGGGTAGCTATNNNAGATCGGAAG |
| 30839_2 | TATGAGTGTGGAGTCGTTGCGCACAAATACCTGGGGTAGCTATNNNAGATCGGAAG |
| 51313_1 | TATGAGTGTGGAGTCGTTGCTTCAAACTCTTGGCTACAGCTCTNNNAGATCGGAAGA |
| 51313_2 | TATGAGTGTGGAGTCGTTGCTTCAAACTCTTGGCTACAGCTCTNNNAGATCGGAAGA |
| 12787_1 | TATGAGTGTGGAGTCGTTGCTTCGATATTCAAAGACTGTTTTGNNNAGATCGGAAGA |
| 12787_2 | TATGAGTGTGGAGTCGTTGCTTCGATATTCAAAGACTGTTTTGNNNAGATCGGAAGA |
| 48505_1 | TATGAGTGTGGAGTCGTTGCTTGCTCTTCCTTTCCCATCTCATNNNAGATCGGAAGAG |
| 48505_2 | TATGAGTGTGGAGTCGTTGCTTGCTCTTCCTTTCCCATCTCATNNNAGATCGGAAGAG |
| 29709_1 | TATGAGTGTGGAGTCGTTGCGTCCTCCTCCAACAGCAATACTANNNAGATCGGAAGA |

| | |
|---|---|
| 29709_2 | TATGAGTGTGGAGTCGTTGCGTCCTCCTCCAACAGCAATACTANNNAGATCGGAAG |
| 36481_1 | TATGAGTGTGGAGTCGTTGCAGATAGAGCTACAACCCCACAGANNNAGATCGGAAG |
| 36481_2 | TATGAGTGTGGAGTCGTTGCAGATAGAGCTACAACCCCACAGANNNAGATCGGAAG |
| 37711_1 | TATGAGTGTGGAGTCGTTGCTAAATGAGCCCTGTCTAACCACCNNNAGATCGGAAG |
| 37711_2 | TATGAGTGTGGAGTCGTTGCTAAATGAGCCCTGTCTAACCACCNNNAGATCGGAAG |
| 43112_1 | TATGAGTGTGGAGTCGTTGCGTAATCCTTACCAGCGAAAGCACNNNAGATCGGAAG |
| 43112_2 | TATGAGTGTGGAGTCGTTGCGTAATCCTTACCAGCGAAAGCACNNNAGATCGGAAG |
| 10088_1 | TATGAGTGTGGAGTCGTTGCTCTTCTGTTCTTCCTTGTGTGGGNNNAGATCGGAAG |
| 10088_2 | TATGAGTGTGGAGTCGTTGCTCTTCTGTTCTTCCTTGTGTGGGNNNAGATCGGAAG |
| 31238_1 | TATGAGTGTGGAGTCGTTGCTTACACAGGCTATGCTGGGAAATNNNAGATCGGAA |
| 31238_2 | TATGAGTGTGGAGTCGTTGCTTACACAGGCTATGCTGGGAAATNNNAGATCGGAA |
| 34537_1 | TATGAGTGTGGAGTCGTTGCTTGAGATTTGGGTGGGGACATAGNNNAGATCGGA |
| 34537_2 | TATGAGTGTGGAGTCGTTGCTTGAGATTTGGGTGGGGACATAGNNNAGATCGGA |
| 10501_1 | TATGAGTGTGGAGTCGTTGCTGCTTGGATTTGCCGATAGTTGTNNNAGATCGGAAG |
| 10501_2 | TATGAGTGTGGAGTCGTTGCTGCTTGGATTTGCCGATAGTTGTNNNAGATCGGAAG |
| 44229_1 | TATGAGTGTGGAGTCGTTGCCCTGTCTTTGTTCTATCGGAGCTNNNAGATCGGAAG |
| 44229_2 | TATGAGTGTGGAGTCGTTGCCCTGTCTTTGTTCTATCGGAGCTNNNAGATCGGAAG |
| 52973_1 | TATGAGTGTGGAGTCGTTGCACAGAGACACATCGAGATTGATGNNNAGATCGGAA |
| 52973_2 | TATGAGTGTGGAGTCGTTGCACAGAGACACATCGAGATTGATGNNNAGATCGGAA |
| 41455_1 | TATGAGTGTGGAGTCGTTGCTCCAACCTCAGAGTTTGAATCTTNNNAGATCGGAAG |
| 41455_2 | TATGAGTGTGGAGTCGTTGCTCCAACCTCAGAGTTTGAATCTTNNNAGATCGGAAG |
| 51563_1 | TATGAGTGTGGAGTCGTTGCGCATGAAGATGGTGTTGGTGTAGNNNAGATCGGA |
| 51563_2 | TATGAGTGTGGAGTCGTTGCGCATGAAGATGGTGTTGGTGTAGNNNAGATCGGA |
| 50369_1 | TATGAGTGTGGAGTCGTTGCTGCAATGGTTTGTGCCTGTAATCNNNAGATCGGAAG |
| 50369_2 | TATGAGTGTGGAGTCGTTGCTGCAATGGTTTGTGCCTGTAATCNNNAGATCGGAAG |
| 51534_1 | TATGAGTGTGGAGTCGTTGCTAAGAGGCAACTCCCCAACATACNNNAGATCGGAAG |
| 51534_2 | TATGAGTGTGGAGTCGTTGCTAAGAGGCAACTCCCCAACATACNNNAGATCGGAAG |
| 33978_1 | TATGAGTGTGGAGTCGTTGCATCTCCCAGGTTCAAGCAATTCTNNNAGATCGGAAG |
| 33978_2 | TATGAGTGTGGAGTCGTTGCATCTCCCAGGTTCAAGCAATTCTNNNAGATCGGAAG |
| 49077_1 | TATGAGTGTGGAGTCGTTGCCTCTCAAGTGGCTAGGACTACCANNNAGATCGGAAG |
| 49077_2 | TATGAGTGTGGAGTCGTTGCCTCTCAAGTGGCTAGGACTACCANNNAGATCGGAAG |
| 49042_1 | TATGAGTGTGGAGTCGTTGCGTTCCAGTGATTCTCCTGCCTTANNNAGATCGGAAG |
| 49042_2 | TATGAGTGTGGAGTCGTTGCGTTCCAGTGATTCTCCTGCCTTANNNAGATCGGAAG |
| 39920_1 | TATGAGTGTGGAGTCGTTGCACCATATGTTAGGCCAGAAAGCANNNAGATCGGAA |
| 39920_2 | TATGAGTGTGGAGTCGTTGCACCATATGTTAGGCCAGAAAGCANNNAGATCGGAA |
| 51993_1 | TATGAGTGTGGAGTCGTTGCTGAGGAACTCCAGGACAGATACTNNNAGATCGGAA |
| 51993_2 | TATGAGTGTGGAGTCGTTGCTGAGGAACTCCAGGACAGATACTNNNAGATCGGAA |
| 40909_1 | TATGAGTGTGGAGTCGTTGCGTGCCTTGGAGAAATCAAAGAGCNNNAGATCGGAA |
| 40909_2 | TATGAGTGTGGAGTCGTTGCGTGCCTTGGAGAAATCAAAGAGCNNNAGATCGGAA |
| 51422_1 | TATGAGTGTGGAGTCGTTGCAAGGAAGCAAAATGGGTTGTCTGNNNAGATCGGAA |
| 51422_2 | TATGAGTGTGGAGTCGTTGCAAGGAAGCAAAATGGGTTGTCTGNNNAGATCGGAA |
| 43631_1 | TATGAGTGTGGAGTCGTTGCTTTGCAGACGTTTCTCGGTAATGNNNAGATCGGAAG |
| 43631_2 | TATGAGTGTGGAGTCGTTGCTTTGCAGACGTTTCTCGGTAATGNNNAGATCGGAAG |

| | |
|---|---|
| 52777_1 | TATGAGTGTGGAGTCGTTGCCAGAAAAGTGCTGTAGATGGGGTNNNAGATCGGAA |
| 52777_2 | TATGAGTGTGGAGTCGTTGCCAGAAAAGTGCTGTAGATGGGGTNNNAGATCGGAA |
| 4_1 | TATGAGTGTGGAGTCGTTGCAGGTAAACGTGCTATTCTGCCTGNNNAGATCGGAAG |
| 4_2 | TATGAGTGTGGAGTCGTTGCAGGTAAACGTGCTATTCTGCCTGNNNAGATCGGAAG |
| 2983_1 | TATGAGTGTGGAGTCGTTGCGCGGGGTCCTTCCCATGATACATNNNAGATCGGAAG |
| 2983_2 | TATGAGTGTGGAGTCGTTGCGCGGGGTCCTTCCCATGATACATNNNAGATCGGAAG |
| 33512_1 | TATGAGTGTGGAGTCGTTGCTCTTGCTAGTGGATAAGCTGACCNNNAGATCGGAAG |
| 33512_2 | TATGAGTGTGGAGTCGTTGCTCTTGCTAGTGGATAAGCTGACCNNNAGATCGGAAG |
| 41761_1 | TATGAGTGTGGAGTCGTTGCTGTAGGAAAGCATCACATGGTGANNNAGATCGGAA |
| 41761_2 | TATGAGTGTGGAGTCGTTGCTGTAGGAAAGCATCACATGGTGANNNAGATCGGAA |
| 33565_1 | TATGAGTGTGGAGTCGTTGCTAGAGTGCAATGGTGTGATCTCGNNNAGATCGGAA |
| 33565_2 | TATGAGTGTGGAGTCGTTGCTAGAGTGCAATGGTGTGATCTCGNNNAGATCGGAA |
| 45412_1 | TATGAGTGTGGAGTCGTTGCACAAAGTTCTGACCAAAACACCGNNNAGATCGGAAG |
| 45412_2 | TATGAGTGTGGAGTCGTTGCACAAAGTTCTGACCAAAACACCGNNNAGATCGGAAG |
| 38802_1 | TATGAGTGTGGAGTCGTTGCTTCAGCAGAGTTTCCGGATACAANNNAGATCGGAAG |
| 38802_2 | TATGAGTGTGGAGTCGTTGCTTCAGCAGAGTTTCCGGATACAANNNAGATCGGAAG |
| 47047_1 | TATGAGTGTGGAGTCGTTGCCATCTCGGCTCACTGCAACTTNNNAGATCGGAAGAG |
| 47047_2 | TATGAGTGTGGAGTCGTTGCCATCTCGGCTCACTGCAACTTNNNAGATCGGAAGAG |
| 51230_1 | TATGAGTGTGGAGTCGTTGCCCTACTGGGAAGTGAGGAGCCNNNAGATCGGAAGA |
| 51230_2 | TATGAGTGTGGAGTCGTTGCCCTACTGGGAAGTGAGGAGCCNNNAGATCGGAAGA |
| 37163_1 | TATGAGTGTGGAGTCGTTGCCATGTCATATGGGCATGGAAAGGNNNAGATCGGAA |
| 37163_2 | TATGAGTGTGGAGTCGTTGCCATGTCATATGGGCATGGAAAGGNNNAGATCGGAA |
| 51218_1 | TATGAGTGTGGAGTCGTTGCGAGATACAGGGGAGTTGACCAACNNNAGATCGGAA |
| 51218_2 | TATGAGTGTGGAGTCGTTGCGAGATACAGGGGAGTTGACCAACNNNAGATCGGAA |
| 37838_1 | TATGAGTGTGGAGTCGTTGCCGGCATATAGCAGGCACTCAATANNNAGATCGGAA |
| 37838_2 | TATGAGTGTGGAGTCGTTGCCGGCATATAGCAGGCACTCAATANNNAGATCGGAA |
| 12827_1 | TATGAGTGTGGAGTCGTTGCCGTTCTGCTTTATTCTTGATAGNNNAGATCGGAAGA |
| 12827_2 | TATGAGTGTGGAGTCGTTGCCGTTCTGCTTTATTCTTGATAGNNNAGATCGGAAGA |
| 40383_1 | TATGAGTGTGGAGTCGTTGCAATGCAGATTCTCAGACTCACCANNNAGATCGGAAG |
| 40383_2 | TATGAGTGTGGAGTCGTTGCAATGCAGATTCTCAGACTCACCANNNAGATCGGAAG |
| 32638_1 | TATGAGTGTGGAGTCGTTGCCTCTTGAGTAGCTGGGACTACAGNNNAGATCGGAA |
| 32638_2 | TATGAGTGTGGAGTCGTTGCCTCTTGAGTAGCTGGGACTACAGNNNAGATCGGAA |
| 52511_1 | TATGAGTGTGGAGTCGTTGCCTGTTTCTGTTTTGGTTGGGGAGNNNAGATCGGAA |
| 52511_2 | TATGAGTGTGGAGTCGTTGCCTGTTTCTGTTTTGGTTGGGGAGNNNAGATCGGAA |
| 12073_1 | TATGAGTGTGGAGTCGTTGCCTCCATGTCTCCCTCCTCTTGTANNNAGATCGGAAGA |
| 12073_2 | TATGAGTGTGGAGTCGTTGCCTCCATGTCTCCCTCCTCTTGTANNNAGATCGGAAGA |
| 40799_1 | TATGAGTGTGGAGTCGTTGCATAGAACACTTCACATGCCACCTNNNAGATCGGAAG |
| 40799_2 | TATGAGTGTGGAGTCGTTGCATAGAACACTTCACATGCCACCTNNNAGATCGGAAG |
| 51227_1 | TATGAGTGTGGAGTCGTTGCGAGGTATACCCAACAGCTCACTGNNNAGATCGGAAG |
| 51227_2 | TATGAGTGTGGAGTCGTTGCGAGGTATACCCAACAGCTCACTGNNNAGATCGGAAG |
| 12873_1 | TATGAGTGTGGAGTCGTTGCGAAAGAGAATGATTTTGCTCTTANNNAGATCGGAA |
| 12873_2 | TATGAGTGTGGAGTCGTTGCGAAAGAGAATGATTTTGCTCTTANNNAGATCGGAA |
| 41235_1 | TATGAGTGTGGAGTCGTTGCTACTCAGAAAACGGTCCAACAGTNNNAGATCGGAAG |

| | |
|---|---|
| 41235_2 | TATGAGTGTGGAGTCGTTGCTACTCAGAAAACGGTCCAACAGTNNNAGATCGGAAG |
| 52006_1 | TATGAGTGTGGAGTCGTTGCGTTACTTTGGTCAGCACCTTTCCNNNAGATCGGAAG |
| 52006_2 | TATGAGTGTGGAGTCGTTGCGTTACTTTGGTCAGCACCTTTCCNNNAGATCGGAAG |
| 33696_1 | TATGAGTGTGGAGTCGTTGCTTATAGGATAAGGACCTGGCACGNNNAGATCGGAA |
| 33696_2 | TATGAGTGTGGAGTCGTTGCTTATAGGATAAGGACCTGGCACGNNNAGATCGGAA |
| 31662_1 | TATGAGTGTGGAGTCGTTGCCCTTCCGACCTTCTTTCTACACANNNAGATCGGAAGA |
| 31662_2 | TATGAGTGTGGAGTCGTTGCCCTTCCGACCTTCTTTCTACACANNNAGATCGGAAGA |
| 30990_1 | TATGAGTGTGGAGTCGTTGCGGCTTCCTGAGTCCAATTTTCACNNNAGATCGGAAG |
| 30990_2 | TATGAGTGTGGAGTCGTTGCGGCTTCCTGAGTCCAATTTTCACNNNAGATCGGAAG |
| 46141_1 | TATGAGTGTGGAGTCGTTGCTCCCCATTCAAACTGTCTGCATANNNAGATCGGAAGA |
| 46141_2 | TATGAGTGTGGAGTCGTTGCTCCCCATTCAAACTGTCTGCATANNNAGATCGGAAGA |
| 51141_1 | TATGAGTGTGGAGTCGTTGCAAGACACCTTAGAAACCTGCCTTNNNAGATCGGAAG |
| 51141_2 | TATGAGTGTGGAGTCGTTGCAAGACACCTTAGAAACCTGCCTTNNNAGATCGGAAG |
| 44139_1 | TATGAGTGTGGAGTCGTTGCGAGCAAAAAGAATGTTGGGGACANNNAGATCGGAA |
| 44139_2 | TATGAGTGTGGAGTCGTTGCGAGCAAAAAGAATGTTGGGGACANNNAGATCGGAA |
| 32269_1 | TATGAGTGTGGAGTCGTTGCGATGCTAATCCTGAGCGAAAACCNNNAGATCGGAAG |
| 32269_2 | TATGAGTGTGGAGTCGTTGCGATGCTAATCCTGAGCGAAAACCNNNAGATCGGAAG |
| 33264_1 | TATGAGTGTGGAGTCGTTGCCATTCTCCTGCCTCAGCCTCNNNAGATCGGAAGAGCA |
| 33264_2 | TATGAGTGTGGAGTCGTTGCCATTCTCCTGCCTCAGCCTCNNNAGATCGGAAGAGCA |
| 38642_1 | TATGAGTGTGGAGTCGTTGCGATCACGAGGTCAGGAGTTCAAGNNNAGATCGGAA |
| 38642_2 | TATGAGTGTGGAGTCGTTGCGATCACGAGGTCAGGAGTTCAAGNNNAGATCGGAA |
| 2540_1 | TATGAGTGTGGAGTCGTTGCTACAGCAAGGGCAGAGATAGAGGNNNAGATCGGA |
| 2540_2 | TATGAGTGTGGAGTCGTTGCTACAGCAAGGGCAGAGATAGAGGNNNAGATCGGA |
| 46671_1 | TATGAGTGTGGAGTCGTTGCTCCAGGTTCAAGTGATTCTCCTGNNNAGATCGGAAG |
| 46671_2 | TATGAGTGTGGAGTCGTTGCTCCAGGTTCAAGTGATTCTCCTGNNNAGATCGGAAG |
| 32049_1 | TATGAGTGTGGAGTCGTTGCTGCCTGTAATACCAGCTACTTGGNNNAGATCGGAAG |
| 32049_2 | TATGAGTGTGGAGTCGTTGCTGCCTGTAATACCAGCTACTTGGNNNAGATCGGAAG |
| 43364_1 | TATGAGTGTGGAGTCGTTGCGGTTGGTGGTAATGTCCCTTTTGNNNAGATCGGAAG |
| 43364_2 | TATGAGTGTGGAGTCGTTGCGGTTGGTGGTAATGTCCCTTTTGNNNAGATCGGAAG |
| 12666_1 | TATGAGTGTGGAGTCGTTGCTTCTAAAAGTCATGTTAGACAGTNNNAGATCGGAAG |
| 12666_2 | TATGAGTGTGGAGTCGTTGCTTCTAAAAGTCATGTTAGACAGTNNNAGATCGGAAG |
| 39264_1 | TATGAGTGTGGAGTCGTTGCGACGCAGCATACACTCAAAAAGANNNAGATCGGAA |
| 39264_2 | TATGAGTGTGGAGTCGTTGCGACGCAGCATACACTCAAAAAGANNNAGATCGGAA |
| 33830_1 | TATGAGTGTGGAGTCGTTGCAAAGCATAAGGGGTTAGACTGGGNNNAGATCGGA |
| 33830_2 | TATGAGTGTGGAGTCGTTGCAAAGCATAAGGGGTTAGACTGGGNNNAGATCGGA |
| 13497_1 | TATGAGTGTGGAGTCGTTGCAAAAACATTTGCATGGTTATGTANNNAGATCGGAAG |
| 13497_2 | TATGAGTGTGGAGTCGTTGCAAAAACATTTGCATGGTTATGTANNNAGATCGGAAG |
| 32277_1 | TATGAGTGTGGAGTCGTTGCTTTTTTTCGAGACGGAGTTTCGCNNNAGATCGGAAG |
| 32277_2 | TATGAGTGTGGAGTCGTTGCTTTTTTTCGAGACGGAGTTTCGCNNNAGATCGGAAG |
| 991_1 | TATGAGTGTGGAGTCGTTGCGGTACTACGGTCCTTGGTAGAAGNNNAGATCGGAA |
| 991_2 | TATGAGTGTGGAGTCGTTGCGGTACTACGGTCCTTGGTAGAAGNNNAGATCGGAA |
| 29494_1 | TATGAGTGTGGAGTCGTTGCGCAATACCGCGAGGAAGAGGAGCNNNAGATCGGA |
| 29494_2 | TATGAGTGTGGAGTCGTTGCGCAATACCGCGAGGAAGAGGAGCNNNAGATCGGA |

| | |
|---|---|
| 29957_1 | TATGAGTGTGGAGTCGTTGCTTATCCACTGTGTTGCAGGCATTNNNAGATCGGAAG |
| 29957_2 | TATGAGTGTGGAGTCGTTGCTTATCCACTGTGTTGCAGGCATTNNNAGATCGGAAG |
| 13668_1 | TATGAGTGTGGAGTCGTTGCCTTATTTGACCTCCCTATAACCNNNAGATCGGAAGAG |
| 13668_2 | TATGAGTGTGGAGTCGTTGCCTTATTTGACCTCCCTATAACCNNNAGATCGGAAGAG |
| 35576_1 | TATGAGTGTGGAGTCGTTGCGGATCTCGGCTCACTGCAAGNNNAGATCGGAAGAG |
| 35576_2 | TATGAGTGTGGAGTCGTTGCGGATCTCGGCTCACTGCAAGNNNAGATCGGAAGAG |
| 34438_1 | TATGAGTGTGGAGTCGTTGCTCTCATTTCCAACTGAAGCCTGANNNAGATCGGAAG |
| 34438_2 | TATGAGTGTGGAGTCGTTGCTCTCATTTCCAACTGAAGCCTGANNNAGATCGGAAG |
| 34665_1 | TATGAGTGTGGAGTCGTTGCGAGTCTTCAAGGTTACAGTGAGCNNNAGATCGGAA |
| 34665_2 | TATGAGTGTGGAGTCGTTGCGAGTCTTCAAGGTTACAGTGAGCNNNAGATCGGAA |
| 46652_1 | TATGAGTGTGGAGTCGTTGCCACCACGCCCAGCTAATTTTTTNNNAGATCGGAAGA |
| 46652_2 | TATGAGTGTGGAGTCGTTGCCACCACGCCCAGCTAATTTTTTNNNAGATCGGAAGA |
| 52559_1 | TATGAGTGTGGAGTCGTTGCAATGGACTTCACCTCTCTTCCAGNNNAGATCGGAAG |
| 52559_2 | TATGAGTGTGGAGTCGTTGCAATGGACTTCACCTCTCTTCCAGNNNAGATCGGAAG |
| 31461_1 | TATGAGTGTGGAGTCGTTGCAGGCACAAAAAACAGACAACACANNNAGATCGGAA |
| 31461_2 | TATGAGTGTGGAGTCGTTGCAGGCACAAAAAACAGACAACACANNNAGATCGGAA |
| 33128_1 | TATGAGTGTGGAGTCGTTGCTGCTGTGGAAATAAGAGCACTCTNNNAGATCGGAA |
| 33128_2 | TATGAGTGTGGAGTCGTTGCTGCTGTGGAAATAAGAGCACTCTNNNAGATCGGAA |
| 13288_1 | TATGAGTGTGGAGTCGTTGCCTTTAGTAGCCTTCTCTTCATATNNNAGATCGGAAGA |
| 13288_2 | TATGAGTGTGGAGTCGTTGCCTTTAGTAGCCTTCTCTTCATATNNNAGATCGGAAGA |
| 36228_1 | TATGAGTGTGGAGTCGTTGCGTTGGTGAGAATGCCAGGTATTGNNNAGATCGGAA |
| 36228_2 | TATGAGTGTGGAGTCGTTGCGTTGGTGAGAATGCCAGGTATTGNNNAGATCGGAA |
| 2224_1 | TATGAGTGTGGAGTCGTTGCTGCTGTTTGCCTAAGAACTGTGANNNAGATCGGAAG |
| 2224_2 | TATGAGTGTGGAGTCGTTGCTGCTGTTTGCCTAAGAACTGTGANNNAGATCGGAAG |
| 50144_1 | TATGAGTGTGGAGTCGTTGCAGCTGTTTCAGAAGAGAGTGAGGNNNAGATCGGAA |
| 50144_2 | TATGAGTGTGGAGTCGTTGCAGCTGTTTCAGAAGAGAGTGAGGNNNAGATCGGAA |
| 44609_1 | TATGAGTGTGGAGTCGTTGCGCGTGAAGTCTCTGTTCCTTTTCNNNAGATCGGAAG |
| 44609_2 | TATGAGTGTGGAGTCGTTGCGCGTGAAGTCTCTGTTCCTTTTCNNNAGATCGGAAG |
| 40700_1 | TATGAGTGTGGAGTCGTTGCGATCTGTCACTGTCTCCCATCACNNNAGATCGGAAG |
| 40700_2 | TATGAGTGTGGAGTCGTTGCGATCTGTCACTGTCTCCCATCACNNNAGATCGGAAG |
| 32251_1 | TATGAGTGTGGAGTCGTTGCTTGTATTCCTCACACTTCCCTGGNNNAGATCGGAAGA |
| 32251_2 | TATGAGTGTGGAGTCGTTGCTTGTATTCCTCACACTTCCCTGGNNNAGATCGGAAGA |
| 39452_1 | TATGAGTGTGGAGTCGTTGCATTCATCTGTGGTTTTTGGCTCANNNAGATCGGAAG |
| 39452_2 | TATGAGTGTGGAGTCGTTGCATTCATCTGTGGTTTTTGGCTCANNNAGATCGGAAG |
| 40542_1 | TATGAGTGTGGAGTCGTTGCCAAAGATGTCCCTCATGCTACCTNNNAGATCGGAAG |
| 40542_2 | TATGAGTGTGGAGTCGTTGCCAAAGATGTCCCTCATGCTACCTNNNAGATCGGAAG |
| 42989_1 | TATGAGTGTGGAGTCGTTGCAGCAAAAGTTAGTGGACGGAAGANNNAGATCGGA |
| 42989_2 | TATGAGTGTGGAGTCGTTGCAGCAAAAGTTAGTGGACGGAAGANNNAGATCGGA |
| 45768_1 | TATGAGTGTGGAGTCGTTGCCTTGAGGAGAGAGAATTGGGGTCNNNAGATCGGA |
| 45768_2 | TATGAGTGTGGAGTCGTTGCCTTGAGGAGAGAGAATTGGGGTCNNNAGATCGGA |
| 33024_1 | TATGAGTGTGGAGTCGTTGCGAGTAGCTGGGACTACAAGCATGNNNAGATCGGAA |
| 33024_2 | TATGAGTGTGGAGTCGTTGCGAGTAGCTGGGACTACAAGCATGNNNAGATCGGAA |
| 43120_1 | TATGAGTGTGGAGTCGTTGCTTGAGGATGTGTGTCCATGACATNNNAGATCGGAA |

| | |
|---|---|
| 43120_2 | TATGAGTGTGGAGTCGTTGCTTGAGGATGTGTGTCCATGACATNNNAGATCGGAA |
| 39373_1 | TATGAGTGTGGAGTCGTTGCACATTGGCAGGTCTTAGGTCTTTNNNAGATCGGAAG |
| 39373_2 | TATGAGTGTGGAGTCGTTGCACATTGGCAGGTCTTAGGTCTTTNNNAGATCGGAAG |
| 50439_1 | TATGAGTGTGGAGTCGTTGCGTCCCATCCTTTTACCTTGAGGGNNNAGATCGGAAG |
| 50439_2 | TATGAGTGTGGAGTCGTTGCGTCCCATCCTTTTACCTTGAGGGNNNAGATCGGAAG |
| 39992_1 | TATGAGTGTGGAGTCGTTGCTTCAAGACCAGCCTGAACAACATNNNAGATCGGAAG |
| 39992_2 | TATGAGTGTGGAGTCGTTGCTTCAAGACCAGCCTGAACAACATNNNAGATCGGAAG |
| 8779_1 | TATGAGTGTGGAGTCGTTGCTCCTGCAATCCTGCAAGTTTCTTNNNAGATCGGAAGA |
| 8779_2 | TATGAGTGTGGAGTCGTTGCTCCTGCAATCCTGCAAGTTTCTTNNNAGATCGGAAGA |
| 8527_1 | TATGAGTGTGGAGTCGTTGCGTGCTCTGATTTAGGTGATCCAAGGNNNAGATCGG |
| 8527_2 | TATGAGTGTGGAGTCGTTGCGTGCTCTGATTTAGGTGATCCAAGGNNNAGATCGG |
| 51726_1 | TATGAGTGTGGAGTCGTTGCCAAGCACCCACGTTTATTTCCTTNNNAGATCGGAAGA |
| 51726_2 | TATGAGTGTGGAGTCGTTGCCAAGCACCCACGTTTATTTCCTTNNNAGATCGGAAGA |
| 30486_1 | TATGAGTGTGGAGTCGTTGCCTGTTAATGTGCGATCCGACAGGNNNAGATCGGAA |
| 30486_2 | TATGAGTGTGGAGTCGTTGCCTGTTAATGTGCGATCCGACAGGNNNAGATCGGAA |
| 47530_1 | TATGAGTGTGGAGTCGTTGCACGCCCGGCTAATTTTTTGTATTNNNAGATCGGAAG |
| 47530_2 | TATGAGTGTGGAGTCGTTGCACGCCCGGCTAATTTTTTGTATTNNNAGATCGGAAG |
| 45504_1 | TATGAGTGTGGAGTCGTTGCTGTACATGGTTCTGTGGGTTTCANNNAGATCGGAAG |
| 45504_2 | TATGAGTGTGGAGTCGTTGCTGTACATGGTTCTGTGGGTTTCANNNAGATCGGAAG |
| 5447_1 | TATGAGTGTGGAGTCGTTGCCTCAGCCGGGACAAGGAGGGNNNAGATCGGAAGA |
| 5447_2 | TATGAGTGTGGAGTCGTTGCCTCAGCCGGGACAAGGAGGGNNNAGATCGGAAGA |
| 47187_1 | TATGAGTGTGGAGTCGTTGCGGATCACCTGAGGTCAGAAGTTTNNNAGATCGGAA |
| 47187_2 | TATGAGTGTGGAGTCGTTGCGGATCACCTGAGGTCAGAAGTTTNNNAGATCGGAA |
| 40881_1 | TATGAGTGTGGAGTCGTTGCAGACGTCTATTATTGGGTGCACANNNAGATCGGAAG |
| 40881_2 | TATGAGTGTGGAGTCGTTGCAGACGTCTATTATTGGGTGCACANNNAGATCGGAAG |
| 34302_1 | TATGAGTGTGGAGTCGTTGCTTGCATGCATGGATTCTTGCTTTNNNAGATCGGAAG |
| 34302_2 | TATGAGTGTGGAGTCGTTGCTTGCATGCATGGATTCTTGCTTTNNNAGATCGGAAG |
| 41263_1 | TATGAGTGTGGAGTCGTTGCAAGCAACCTGAGAAGCTAGTTCANNNAGATCGGAA |
| 41263_2 | TATGAGTGTGGAGTCGTTGCAAGCAACCTGAGAAGCTAGTTCANNNAGATCGGAA |
| 35094_1 | TATGAGTGTGGAGTCGTTGCTGATGCGAACTCAGCTTACTGTANNNAGATCGGAAG |
| 35094_2 | TATGAGTGTGGAGTCGTTGCTGATGCGAACTCAGCTTACTGTANNNAGATCGGAAG |
| 42441_1 | TATGAGTGTGGAGTCGTTGCCCGATTCATGAAACTAACCCTTGGNNNAGATCGGAA |
| 42441_2 | TATGAGTGTGGAGTCGTTGCCCGATTCATGAAACTAACCCTTGGNNNAGATCGGAA |
| 40817_1 | TATGAGTGTGGAGTCGTTGCTGTGGTCCTTAGTTATTTTCTCTGGANNNAGATCGG |
| 40817_2 | TATGAGTGTGGAGTCGTTGCTGTGGTCCTTAGTTATTTTCTCTGGANNNAGATCGG |
| 46868_1 | TATGAGTGTGGAGTCGTTGCTAGTAGCTGGGATTACAGGCATGNNNAGATCGGAA |
| 46868_2 | TATGAGTGTGGAGTCGTTGCTAGTAGCTGGGATTACAGGCATGNNNAGATCGGAA |
| 53094_1 | TATGAGTGTGGAGTCGTTGCAAAGTTTGAAACATGCTACAGGCNNNAGATCGGAA |
| 53094_2 | TATGAGTGTGGAGTCGTTGCAAAGTTTGAAACATGCTACAGGCNNNAGATCGGAA |
| 40369_1 | TATGAGTGTGGAGTCGTTGCGCAGTGTCATGGAGAATATGTGCNNNAGATCGGAA |
| 40369_2 | TATGAGTGTGGAGTCGTTGCGCAGTGTCATGGAGAATATGTGCNNNAGATCGGAA |
| 38848_1 | TATGAGTGTGGAGTCGTTGCAGTTACAAAAGCCGTGGGAAAAGNNNAGATCGGAA |
| 38848_2 | TATGAGTGTGGAGTCGTTGCAGTTACAAAAGCCGTGGGAAAAGNNNAGATCGGAA |

| | |
|---|---|
| 37159_1 | TATGAGTGTGGAGTCGTTGCAAGAACTACCATTTGACCCAGCANNNAGATCGGAAG |
| 37159_2 | TATGAGTGTGGAGTCGTTGCAAGAACTACCATTTGACCCAGCANNNAGATCGGAAG |
| 33990_1 | TATGAGTGTGGAGTCGTTGCCAATGGCGTGATCTCAGGTTACTNNNAGATCGGAAG |
| 33990_2 | TATGAGTGTGGAGTCGTTGCCAATGGCGTGATCTCAGGTTACTNNNAGATCGGAAG |
| 35936_1 | TATGAGTGTGGAGTCGTTGCAGTTTGAAAGCTAAAGTGTGGGCNNNAGATCGGAA |
| 35936_2 | TATGAGTGTGGAGTCGTTGCAGTTTGAAAGCTAAAGTGTGGGCNNNAGATCGGAA |
| 45861_1 | TATGAGTGTGGAGTCGTTGCGAAACCGACGACAGGAAAATGAGNNNAGATCGGAA |
| 45861_2 | TATGAGTGTGGAGTCGTTGCGAAACCGACGACAGGAAAATGAGNNNAGATCGGAA |
| 43704_1 | TATGAGTGTGGAGTCGTTGCGCAATCCACACAACAGGTATGTCNNNAGATCGGAAG |
| 43704_2 | TATGAGTGTGGAGTCGTTGCGCAATCCACACAACAGGTATGTCNNNAGATCGGAAG |
| 42738_1 | TATGAGTGTGGAGTCGTTGCTGACAAAATGGGAGGAAAAAGACCNNNAGATCGGA |
| 42738_2 | TATGAGTGTGGAGTCGTTGCTGACAAAATGGGAGGAAAAAGACCNNNAGATCGGA |
| 42872_1 | TATGAGTGTGGAGTCGTTGCGATGGAGTCTCACTCTGTCGCNNNAGATCGGAAGAG |
| 42872_2 | TATGAGTGTGGAGTCGTTGCGATGGAGTCTCACTCTGTCGCNNNAGATCGGAAGAG |
| 48485_1 | TATGAGTGTGGAGTCGTTGCGATCCCCCAGAGGTTTCTCATCNNNAGATCGGAAGA |
| 48485_2 | TATGAGTGTGGAGTCGTTGCGATCCCCCAGAGGTTTCTCATCNNNAGATCGGAAGA |
| 2521_1 | TATGAGTGTGGAGTCGTTGCAGAACATGAAGGGACACACATCGNNNAGATCGGAA |
| 2521_2 | TATGAGTGTGGAGTCGTTGCAGAACATGAAGGGACACACATCGNNNAGATCGGAA |
| 52707_1 | TATGAGTGTGGAGTCGTTGCAAACTAATGTGGGTGGGAAGAGGNNNAGATCGGA |
| 52707_2 | TATGAGTGTGGAGTCGTTGCAAACTAATGTGGGTGGGAAGAGGNNNAGATCGGA |
| 49693_1 | TATGAGTGTGGAGTCGTTGCACCTTTGAAGCTGTCAGAGGGNNNAGATCGGAAGA |
| 49693_2 | TATGAGTGTGGAGTCGTTGCACCTTTGAAGCTGTCAGAGGGNNNAGATCGGAAGA |
| 45465_1 | TATGAGTGTGGAGTCGTTGCGATCCACAAGGGGTCATAGGAAANNNAGATCGGAA |
| 45465_2 | TATGAGTGTGGAGTCGTTGCGATCCACAAGGGGTCATAGGAAANNNAGATCGGAA |
| 32206_1 | TATGAGTGTGGAGTCGTTGCTCTTGTTCTTTTTTATGGCTGCANNNAGATCGGAAGA |
| 32206_2 | TATGAGTGTGGAGTCGTTGCTCTTGTTCTTTTTTATGGCTGCANNNAGATCGGAAGA |
| 37200_1 | TATGAGTGTGGAGTCGTTGCAAATCCTTGCAATTTATCTGCCANNNAGATCGGAAGA |
| 37200_2 | TATGAGTGTGGAGTCGTTGCAAATCCTTGCAATTTATCTGCCANNNAGATCGGAAGA |
| 33011_1 | TATGAGTGTGGAGTCGTTGCTCTTGACTTACCACAACCTCCACNNNAGATCGGAAGA |
| 33011_2 | TATGAGTGTGGAGTCGTTGCTCTTGACTTACCACAACCTCCACNNNAGATCGGAAGA |
| 49097_1 | TATGAGTGTGGAGTCGTTGCCCTGGGTTCAAGCAATTCTCTTGNNNAGATCGGAAG |
| 49097_2 | TATGAGTGTGGAGTCGTTGCCCTGGGTTCAAGCAATTCTCTTGNNNAGATCGGAAG |
| 49956_1 | TATGAGTGTGGAGTCGTTGCAAATTCTTGCAGTCAAAAGGGCANNNAGATCGGAA |
| 49956_2 | TATGAGTGTGGAGTCGTTGCAAATTCTTGCAGTCAAAAGGGCANNNAGATCGGAA |
| 44686_1 | TATGAGTGTGGAGTCGTTGCGGATGATGGGAAGACTCACTCTCNNNAGATCGGAA |
| 44686_2 | TATGAGTGTGGAGTCGTTGCGGATGATGGGAAGACTCACTCTCNNNAGATCGGAA |
| 33447_1 | TATGAGTGTGGAGTCGTTGCAGAAAAGGGCCTAGGGATTTGAGNNNAGATCGGAA |
| 33447_2 | TATGAGTGTGGAGTCGTTGCAGAAAAGGGCCTAGGGATTTGAGNNNAGATCGGAA |
| 31878_1 | TATGAGTGTGGAGTCGTTGCGAAAGTGTGCACATTCTTTGGCNNNAGATCGGAAGA |
| 31878_2 | TATGAGTGTGGAGTCGTTGCGAAAGTGTGCACATTCTTTGGCNNNAGATCGGAAGA |
| 10413_1 | TATGAGTGTGGAGTCGTTGCGTGTGTGTCAGGGGAGATTGTTGNNNAGATCGGAA |
| 10413_2 | TATGAGTGTGGAGTCGTTGCGTGTGTGTCAGGGGAGATTGTTGNNNAGATCGGAA |
| 42586_1 | TATGAGTGTGGAGTCGTTGCGTTCAAGTGATCCTCCTGTCTCANNNAGATCGGAAGA |

| | |
|---|---|
| 42586_2 | TATGAGTGTGGAGTCGTTGCGTTCAAGTGATCCTCCTGTCTCANNNAGATCGGAAG |
| 45111_1 | TATGAGTGTGGAGTCGTTGCTGAGCACAGTTCAGATTTTTGGTNNNAGATCGGAAG |
| 45111_2 | TATGAGTGTGGAGTCGTTGCTGAGCACAGTTCAGATTTTTGGTNNNAGATCGGAAG |
| 47201_1 | TATGAGTGTGGAGTCGTTGCGAAGAGGGGAGTCTGATGACAAANNNAGATCGGA |
| 47201_2 | TATGAGTGTGGAGTCGTTGCGAAGAGGGGAGTCTGATGACAAANNNAGATCGGA |
| 11834_1 | TATGAGTGTGGAGTCGTTGCCACAGAGAAGGGACAGAAGCAGNNNAGATCGGAA |
| 11834_2 | TATGAGTGTGGAGTCGTTGCCACAGAGAAGGGACAGAAGCAGNNNAGATCGGAA |
| 43887_1 | TATGAGTGTGGAGTCGTTGCCCTATCTGTTGAGCGCTGTCTATNNNAGATCGGAAG |
| 43887_2 | TATGAGTGTGGAGTCGTTGCCCTATCTGTTGAGCGCTGTCTATNNNAGATCGGAAG |
| 35986_1 | TATGAGTGTGGAGTCGTTGCATACAAAAATTAGCTGGGTGCGGNNNAGATCGGAA |
| 35986_2 | TATGAGTGTGGAGTCGTTGCATACAAAAATTAGCTGGGTGCGGNNNAGATCGGAA |
| 46602_1 | TATGAGTGTGGAGTCGTTGCTCTCACTTTGTCACCTACACTGGNNNAGATCGGAAGA |
| 46602_2 | TATGAGTGTGGAGTCGTTGCTCTCACTTTGTCACCTACACTGGNNNAGATCGGAAGA |
| 40313_1 | TATGAGTGTGGAGTCGTTGCACTGCGACGTCAATCAAAACAATNNNAGATCGGAAG |
| 40313_2 | TATGAGTGTGGAGTCGTTGCACTGCGACGTCAATCAAAACAATNNNAGATCGGAAG |
| 51541_1 | TATGAGTGTGGAGTCGTTGCCTGGTGATGAGGAAGGAAGGAAGNNNAGATCGGA |
| 51541_2 | TATGAGTGTGGAGTCGTTGCCTGGTGATGAGGAAGGAAGGAAGNNNAGATCGGA |
| 45893_1 | TATGAGTGTGGAGTCGTTGCCAAGTAGCTGGGACTACAGGCNNNAGATCGGAAGA |
| 45893_2 | TATGAGTGTGGAGTCGTTGCCAAGTAGCTGGGACTACAGGCNNNAGATCGGAAGA |
| 44556_1 | TATGAGTGTGGAGTCGTTGCTACCTAACCAAGGAGGCAAAACANNNAGATCGGAA |
| 44556_2 | TATGAGTGTGGAGTCGTTGCTACCTAACCAAGGAGGCAAAACANNNAGATCGGAA |
| 33856_1 | TATGAGTGTGGAGTCGTTGCGCTCATGTCTGTAATTCCAGCACNNNAGATCGGAAG |
| 33856_2 | TATGAGTGTGGAGTCGTTGCGCTCATGTCTGTAATTCCAGCACNNNAGATCGGAAG |
| 52766_1 | TATGAGTGTGGAGTCGTTGCACGGACCTGAGAACATTATGCTTNNNAGATCGGAAG |
| 52766_2 | TATGAGTGTGGAGTCGTTGCACGGACCTGAGAACATTATGCTTNNNAGATCGGAAG |
| 31528_1 | TATGAGTGTGGAGTCGTTGCATCCTCACTCCAAAAAACCCACTNNNAGATCGGAAGA |
| 31528_2 | TATGAGTGTGGAGTCGTTGCATCCTCACTCCAAAAAACCCACTNNNAGATCGGAAGA |
| 32358_1 | TATGAGTGTGGAGTCGTTGCCACGCCCAGCTAATTTTGTATGTNNNAGATCGGAAG |
| 32358_2 | TATGAGTGTGGAGTCGTTGCCACGCCCAGCTAATTTTGTATGTNNNAGATCGGAAG |
| 40911_1 | TATGAGTGTGGAGTCGTTGCAATCTCGGCTCACTGAAACCTCNNNAGATCGGAAGA |
| 40911_2 | TATGAGTGTGGAGTCGTTGCAATCTCGGCTCACTGAAACCTCNNNAGATCGGAAGA |
| 31954_1 | TATGAGTGTGGAGTCGTTGCTGCCAAACACACAGAACTAGTTTNNNAGATCGGAAG |
| 31954_2 | TATGAGTGTGGAGTCGTTGCTGCCAAACACACAGAACTAGTTTNNNAGATCGGAAG |
| 2689_1 | TATGAGTGTGGAGTCGTTGCCTCCCCTTGCCATTCTTCTTCATNNNAGATCGGAAGA |
| 2689_2 | TATGAGTGTGGAGTCGTTGCCTCCCCTTGCCATTCTTCTTCATNNNAGATCGGAAGA |
| 42168_1 | TATGAGTGTGGAGTCGTTGCCACGCCCGGCTAATTTTTTTGTANNNAGATCGGAAG |
| 42168_2 | TATGAGTGTGGAGTCGTTGCCACGCCCGGCTAATTTTTTTGTANNNAGATCGGAAG |
| 48549_1 | TATGAGTGTGGAGTCGTTGCCACCATGGTCTACTCACCAGAAGNNNAGATCGGAAG |
| 48549_2 | TATGAGTGTGGAGTCGTTGCCACCATGGTCTACTCACCAGAAGNNNAGATCGGAAG |
| 2919_1 | TATGAGTGTGGAGTCGTTGCACTGGGGTGTATGTGAGGTCATTNNNAGATCGGAA |
| 2919_2 | TATGAGTGTGGAGTCGTTGCACTGGGGTGTATGTGAGGTCATTNNNAGATCGGAA |
| 9934_1 | TATGAGTGTGGAGTCGTTGCTTCACCTGGAACAGCATGTCAGGNNNAGATCGGAAG |
| 9934_2 | TATGAGTGTGGAGTCGTTGCTTCACCTGGAACAGCATGTCAGGNNNAGATCGGAAG |

| | |
|---|---|
| 42882_1 | TATGAGTGTGGAGTCGTTGCACAGGCTGGCTATTTACCACAATNNNAGATCGGAAG |
| 42882_2 | TATGAGTGTGGAGTCGTTGCACAGGCTGGCTATTTACCACAATNNNAGATCGGAAG |
| 11330_1 | TATGAGTGTGGAGTCGTTGCGAGGGAGAGTTTGGAATTTCCCAGNNNAGATCGGA |
| 11330_2 | TATGAGTGTGGAGTCGTTGCGAGGGAGAGTTTGGAATTTCCCAGNNNAGATCGGA |
| 11176_1 | TATGAGTGTGGAGTCGTTGCTGGACTGGCTTGTAACTTGCTTTNNNAGATCGGAAG |
| 11176_2 | TATGAGTGTGGAGTCGTTGCTGGACTGGCTTGTAACTTGCTTTNNNAGATCGGAAG |
| 44261_1 | TATGAGTGTGGAGTCGTTGCGAGAGCCTCTCTGTGCATACAATNNNAGATCGGAAG |
| 44261_2 | TATGAGTGTGGAGTCGTTGCGAGAGCCTCTCTGTGCATACAATNNNAGATCGGAAG |
| 35750_1 | TATGAGTGTGGAGTCGTTGCTTCAAGTGATTCTCCCACCTCAGNNNAGATCGGAAG |
| 35750_2 | TATGAGTGTGGAGTCGTTGCTTCAAGTGATTCTCCCACCTCAGNNNAGATCGGAAG |
| 35844_1 | TATGAGTGTGGAGTCGTTGCCTCCTGAGTAGCTGGGATTACAGNNNAGATCGGAA |
| 35844_2 | TATGAGTGTGGAGTCGTTGCCTCCTGAGTAGCTGGGATTACAGNNNAGATCGGAA |
| 50676_1 | TATGAGTGTGGAGTCGTTGCTGATGGGAAGTGGTAACAACTGTNNNAGATCGGAA |
| 50676_2 | TATGAGTGTGGAGTCGTTGCTGATGGGAAGTGGTAACAACTGTNNNAGATCGGAA |
| 37129_1 | TATGAGTGTGGAGTCGTTGCCCTGGGCAACAAAGTGAGACTCNNNAGATCGGAAG |
| 37129_2 | TATGAGTGTGGAGTCGTTGCCCTGGGCAACAAAGTGAGACTCNNNAGATCGGAAG |
| 39082_1 | TATGAGTGTGGAGTCGTTGCTCTTGTCCTATCTCCACAGTTGCNNNAGATCGGAAGA |
| 39082_2 | TATGAGTGTGGAGTCGTTGCTCTTGTCCTATCTCCACAGTTGCNNNAGATCGGAAGA |
| 13677_1 | TATGAGTGTGGAGTCGTTGCTGTTTCTAGATAGCACAGGANNNAGATCGGAAGAG |
| 13677_2 | TATGAGTGTGGAGTCGTTGCTGTTTCTAGATAGCACAGGANNNAGATCGGAAGAG |
| 45688_1 | TATGAGTGTGGAGTCGTTGCTCAAGTGATTCTCCTTCCTCAGCNNNAGATCGGAAG |
| 45688_2 | TATGAGTGTGGAGTCGTTGCTCAAGTGATTCTCCTTCCTCAGCNNNAGATCGGAAG |
| 43325_1 | TATGAGTGTGGAGTCGTTGCGAAGGTTTACTGCCTGCTCTTCTNNNAGATCGGAAG |
| 43325_2 | TATGAGTGTGGAGTCGTTGCGAAGGTTTACTGCCTGCTCTTCTNNNAGATCGGAAG |
| 34946_1 | TATGAGTGTGGAGTCGTTGCAAGACAGTCTATGAAGCTGGAGCNNNAGATCGGAA |
| 34946_2 | TATGAGTGTGGAGTCGTTGCAAGACAGTCTATGAAGCTGGAGCNNNAGATCGGAA |
| 41755_1 | TATGAGTGTGGAGTCGTTGCAATCTTGTCTCACTGCAACCTCTNNNAGATCGGAAGA |
| 41755_2 | TATGAGTGTGGAGTCGTTGCAATCTTGTCTCACTGCAACCTCTNNNAGATCGGAAGA |
| 37131_1 | TATGAGTGTGGAGTCGTTGCCAATGTGGAAAACTGAGGGGTTCNNNAGATCGGAA |
| 37131_2 | TATGAGTGTGGAGTCGTTGCCAATGTGGAAAACTGAGGGGTTCNNNAGATCGGAA |
| 47554_1 | TATGAGTGTGGAGTCGTTGCCACCACGCCTGACTAATTTTTGTNNNAGATCGGAAG |
| 47554_2 | TATGAGTGTGGAGTCGTTGCCACCACGCCTGACTAATTTTTGTNNNAGATCGGAAG |
| 30212_1 | TATGAGTGTGGAGTCGTTGCTGGAGCTTCACAATCCTAAAGCANNNAGATCGGAAG |
| 30212_2 | TATGAGTGTGGAGTCGTTGCTGGAGCTTCACAATCCTAAAGCANNNAGATCGGAAG |
| 44265_1 | TATGAGTGTGGAGTCGTTGCGGTTCGTCAGTTGTCTTCTCCTNNNAGATCGGAAGA |
| 44265_2 | TATGAGTGTGGAGTCGTTGCGGTTCGTCAGTTGTCTTCTCCTNNNAGATCGGAAGA |
| 41250_1 | TATGAGTGTGGAGTCGTTGCATTCGTGAAGTTGAAGTGGCTCANNNAGATCGGAA |
| 41250_2 | TATGAGTGTGGAGTCGTTGCATTCGTGAAGTTGAAGTGGCTCANNNAGATCGGAA |
| 30786_1 | TATGAGTGTGGAGTCGTTGCAAGACACTAGTCATTGGGGTAGGNNNAGATCGGAA |
| 30786_2 | TATGAGTGTGGAGTCGTTGCAAGACACTAGTCATTGGGGTAGGNNNAGATCGGAA |
| 37431_1 | TATGAGTGTGGAGTCGTTGCGCAGGAGAATCAATTGAACCCAGNNNAGATCGGAA |
| 37431_2 | TATGAGTGTGGAGTCGTTGCGCAGGAGAATCAATTGAACCCAGNNNAGATCGGAA |
| 50848_1 | TATGAGTGTGGAGTCGTTGCTGGATGTGGTTATGTTCCAATCACNNNAGATCGGAA |

| | |
|---|---|
| 50848_2 | TATGAGTGTGGAGTCGTTGCTGGATGTGGTTATGTTCCAATCACNNNAGATCGGAA |
| 43118_1 | TATGAGTGTGGAGTCGTTGCCCAGTAACTGCCAGACTACTGTTNNNAGATCGGAAG |
| 43118_2 | TATGAGTGTGGAGTCGTTGCCCAGTAACTGCCAGACTACTGTTNNNAGATCGGAAG |
| 34500_1 | TATGAGTGTGGAGTCGTTGCGGGCGCCTCACTTCCTAGTANNNAGATCGGAAGAGC |
| 34500_2 | TATGAGTGTGGAGTCGTTGCGGGCGCCTCACTTCCTAGTANNNAGATCGGAAGAGC |
| 36796_1 | TATGAGTGTGGAGTCGTTGCTTGTGTCAGGGAAACTGGTGAATNNNAGATCGGAA |
| 36796_2 | TATGAGTGTGGAGTCGTTGCTTGTGTCAGGGAAACTGGTGAATNNNAGATCGGAA |
| 12818_1 | TATGAGTGTGGAGTCGTTGCCTGACTTACCCTAGTACTTAGTANNNAGATCGGAAG |
| 12818_2 | TATGAGTGTGGAGTCGTTGCCTGACTTACCCTAGTACTTAGTANNNAGATCGGAAG |
| 30973_1 | TATGAGTGTGGAGTCGTTGCCCATTCACTTCCTCAACCCACTANNNAGATCGGAAGA |
| 30973_2 | TATGAGTGTGGAGTCGTTGCCCATTCACTTCCTCAACCCACTANNNAGATCGGAAGA |
| 51908_1 | TATGAGTGTGGAGTCGTTGCCCCCCTGACACTTTATTATCCCTNNNAGATCGGAAGA |
| 51908_2 | TATGAGTGTGGAGTCGTTGCCCCCCTGACACTTTATTATCCCTNNNAGATCGGAAGA |
| 47457_1 | TATGAGTGTGGAGTCGTTGCAAGTGCTGGCATTATAGGTGTGANNNAGATCGGAA |
| 47457_2 | TATGAGTGTGGAGTCGTTGCAAGTGCTGGCATTATAGGTGTGANNNAGATCGGAA |
| 39022_1 | TATGAGTGTGGAGTCGTTGCAAGAAGCAGGATGTCTTGGTCATNNNAGATCGGAA |
| 39022_2 | TATGAGTGTGGAGTCGTTGCAAGAAGCAGGATGTCTTGGTCATNNNAGATCGGAA |
| 31399_1 | TATGAGTGTGGAGTCGTTGCTAGAAAAAGGAAGAGAGTGGGGCNNNAGATCGGA |
| 31399_2 | TATGAGTGTGGAGTCGTTGCTAGAAAAAGGAAGAGAGTGGGGCNNNAGATCGGA |
| 51072_1 | TATGAGTGTGGAGTCGTTGCAGATGATTCCTGCTAGTGACAGCNNNAGATCGGAAG |
| 51072_2 | TATGAGTGTGGAGTCGTTGCAGATGATTCCTGCTAGTGACAGCNNNAGATCGGAAG |
| 44872_1 | TATGAGTGTGGAGTCGTTGCTCCCCTTTTTTCTCCCCAAAGAANNNAGATCGGAAGA |
| 44872_2 | TATGAGTGTGGAGTCGTTGCTCCCCTTTTTTCTCCCCAAAGAANNNAGATCGGAAGA |
| 47472_1 | TATGAGTGTGGAGTCGTTGCCCTCCTAGGTTCAAGCGATTCTCNNNAGATCGGAAG |
| 47472_2 | TATGAGTGTGGAGTCGTTGCCCTCCTAGGTTCAAGCGATTCTCNNNAGATCGGAAG |
| 30989_1 | TATGAGTGTGGAGTCGTTGCCATACACATTCTCTCTCTCCCCCNNNAGATCGGAAGAG |
| 30989_2 | TATGAGTGTGGAGTCGTTGCCATACACATTCTCTCTCTCCCCCNNNAGATCGGAAGAG |
| 52012_1 | TATGAGTGTGGAGTCGTTGCGCCCATGTGTTCTCATTGTTCAANNNAGATCGGAAG |
| 52012_2 | TATGAGTGTGGAGTCGTTGCGCCCATGTGTTCTCATTGTTCAANNNAGATCGGAAG |
| 43918_1 | TATGAGTGTGGAGTCGTTGCTCTGTCTGTCCATCCATCCTACTNNNAGATCGGAAGA |
| 43918_2 | TATGAGTGTGGAGTCGTTGCTCTGTCTGTCCATCCATCCTACTNNNAGATCGGAAGA |
| 50873_1 | TATGAGTGTGGAGTCGTTGCCCATGACACGGCAGAAGTAAAAGNNNAGATCGGAA |
| 50873_2 | TATGAGTGTGGAGTCGTTGCCCATGACACGGCAGAAGTAAAAGNNNAGATCGGAA |
| 37809_1 | TATGAGTGTGGAGTCGTTGCTGCCCTCAACTTGATAAAGAGCANNNAGATCGGAAG |
| 37809_2 | TATGAGTGTGGAGTCGTTGCTGCCCTCAACTTGATAAAGAGCANNNAGATCGGAAG |
| 48414_1 | TATGAGTGTGGAGTCGTTGCTGCATGTTGAGGATAACCTTCCANNNAGATCGGAAG |
| 48414_2 | TATGAGTGTGGAGTCGTTGCTGCATGTTGAGGATAACCTTCCANNNAGATCGGAAG |
| 44351_1 | TATGAGTGTGGAGTCGTTGCTTCCCTAACTGGACTGTGCATTCNNNAGATCGGAAG |
| 44351_2 | TATGAGTGTGGAGTCGTTGCTTCCCTAACTGGACTGTGCATTCNNNAGATCGGAAG |
| 36964_1 | TATGAGTGTGGAGTCGTTGCCCCAACAAAGAAAAGCCCATGATNNNAGATCGGAAG |
| 36964_2 | TATGAGTGTGGAGTCGTTGCCCCAACAAAGAAAAGCCCATGATNNNAGATCGGAAG |
| 43643_1 | TATGAGTGTGGAGTCGTTGCTCATTCAGGAGGAGAGCTAGACANNNAGATCGGAA |
| 43643_2 | TATGAGTGTGGAGTCGTTGCTCATTCAGGAGGAGAGCTAGACANNNAGATCGGAA |

| | |
|---|---|
| 30295_1 | TATGAGTGTGGAGTCGTTGCCACTCTTCAGCACAATCAACCAGNNNAGATCGGAAG |
| 30295_2 | TATGAGTGTGGAGTCGTTGCCACTCTTCAGCACAATCAACCAGNNNAGATCGGAAG |
| 52092_1 | TATGAGTGTGGAGTCGTTGCCCTGGACCACTGTCTACATTCAANNNAGATCGGAAG |
| 52092_2 | TATGAGTGTGGAGTCGTTGCCCTGGACCACTGTCTACATTCAANNNAGATCGGAAG |
| 51269_1 | TATGAGTGTGGAGTCGTTGCCCCAGCGGCTCATTGAGAACNNNAGATCGGAAGAG |
| 51269_2 | TATGAGTGTGGAGTCGTTGCCCCAGCGGCTCATTGAGAACNNNAGATCGGAAGAG |
| 30251_1 | TATGAGTGTGGAGTCGTTGCATTTCTGTCCCTTACTCTCAGCCNNNAGATCGGAAGA |
| 30251_2 | TATGAGTGTGGAGTCGTTGCATTTCTGTCCCTTACTCTCAGCCNNNAGATCGGAAGA |
| 43993_1 | TATGAGTGTGGAGTCGTTGCGCTGGGAATATGGTGCAAGAAAANNNAGATCGGAA |
| 43993_2 | TATGAGTGTGGAGTCGTTGCGCTGGGAATATGGTGCAAGAAAANNNAGATCGGAA |
| 30379_1 | TATGAGTGTGGAGTCGTTGCCCGGTATGTAATTTAACATTCAANNNAGATCGGAAG |
| 30379_2 | TATGAGTGTGGAGTCGTTGCCCGGTATGTAATTTAACATTCAANNNAGATCGGAAG |
| 12083_1 | TATGAGTGTGGAGTCGTTGCTTCTTCCACCTCACCAAGGTAGCNNNAGATCGGAAG |
| 12083_2 | TATGAGTGTGGAGTCGTTGCTTCTTCCACCTCACCAAGGTAGCNNNAGATCGGAAG |
| 46226_1 | TATGAGTGTGGAGTCGTTGCGGTTCAAGTGATTTTCCTGCCTCNNNAGATCGGAAG |
| 46226_2 | TATGAGTGTGGAGTCGTTGCGGTTCAAGTGATTTTCCTGCCTCNNNAGATCGGAAG |
| 33725_1 | TATGAGTGTGGAGTCGTTGCGCCTATAATTCCAGCTACTCGGGNNNAGATCGGAAG |
| 33725_2 | TATGAGTGTGGAGTCGTTGCGCCTATAATTCCAGCTACTCGGGNNNAGATCGGAAG |
| 39565_1 | TATGAGTGTGGAGTCGTTGCTTTGCAATTGCTGTCTTCCCTTTNNNAGATCGGAAGA |
| 39565_2 | TATGAGTGTGGAGTCGTTGCTTTGCAATTGCTGTCTTCCCTTTNNNAGATCGGAAGA |
| 34483_1 | TATGAGTGTGGAGTCGTTGCTTCCTAAGGCTAACATGAATGCCNNNAGATCGGAAG |
| 34483_2 | TATGAGTGTGGAGTCGTTGCTTCCTAAGGCTAACATGAATGCCNNNAGATCGGAAG |
| 10770_1 | TATGAGTGTGGAGTCGTTGCTTGTCTTTAGGCTGGGTGTAGTGNNNAGATCGGAA |
| 10770_2 | TATGAGTGTGGAGTCGTTGCTTGTCTTTAGGCTGGGTGTAGTGNNNAGATCGGAA |
| 46839_1 | TATGAGTGTGGAGTCGTTGCAAGTTCAAGTGATTCTCCTGCCTNNNAGATCGGAAG |
| 46839_2 | TATGAGTGTGGAGTCGTTGCAAGTTCAAGTGATTCTCCTGCCTNNNAGATCGGAAG |
| 50333_1 | TATGAGTGTGGAGTCGTTGCCTGCAAAACAAACTGAAGGGGAANNNAGATCGGAA |
| 50333_2 | TATGAGTGTGGAGTCGTTGCCTGCAAAACAAACTGAAGGGGAANNNAGATCGGAA |
| 51061_1 | TATGAGTGTGGAGTCGTTGCTTTCTTTTTCAGAGGGGCTTCCTNNNAGATCGGAAG |
| 51061_2 | TATGAGTGTGGAGTCGTTGCTTTCTTTTTCAGAGGGGCTTCCTNNNAGATCGGAAG |
| 51359_1 | TATGAGTGTGGAGTCGTTGCTGTTCTGCACATGTATCCCAGAANNNAGATCGGAAG |
| 51359_2 | TATGAGTGTGGAGTCGTTGCTGTTCTGCACATGTATCCCAGAANNNAGATCGGAAG |
| 31077_1 | TATGAGTGTGGAGTCGTTGCTTTCACCTTGGCTGGAGTACAATNNNAGATCGGAAG |
| 31077_2 | TATGAGTGTGGAGTCGTTGCTTTCACCTTGGCTGGAGTACAATNNNAGATCGGAAG |
| 37730_1 | TATGAGTGTGGAGTCGTTGCCGTACTCAAAAAGGCCCAATGTTNNNAGATCGGAAG |
| 37730_2 | TATGAGTGTGGAGTCGTTGCCGTACTCAAAAAGGCCCAATGTTNNNAGATCGGAAG |
| 47402_1 | TATGAGTGTGGAGTCGTTGCTTGTGACAGAAGACATGCGTTTCNNNAGATCGGAA |
| 47402_2 | TATGAGTGTGGAGTCGTTGCTTGTGACAGAAGACATGCGTTTCNNNAGATCGGAA |
| 32187_1 | TATGAGTGTGGAGTCGTTGCGGTGAAACCCCGTCTCTACTAAANNNAGATCGGAAG |
| 32187_2 | TATGAGTGTGGAGTCGTTGCGGTGAAACCCCGTCTCTACTAAANNNAGATCGGAAG |
| 35435_1 | TATGAGTGTGGAGTCGTTGCCACCTCACCCGGCTAATTTTTTTNNNAGATCGGAAGA |
| 35435_2 | TATGAGTGTGGAGTCGTTGCCACCTCACCCGGCTAATTTTTTTNNNAGATCGGAAGA |
| 2227_1 | TATGAGTGTGGAGTCGTTGCTAACATGAGCTCAACGCTCTTGANNNAGATCGGAAG |

| | |
|---|---|
| 2227_2 | TATGAGTGTGGAGTCGTTGCTAACATGAGCTCAACGCTCTTGANNNAGATCGGAAG |
| 35864_1 | TATGAGTGTGGAGTCGTTGCTGAAGATCACCTTGAGAGAACCCNNNAGATCGGAAG |
| 35864_2 | TATGAGTGTGGAGTCGTTGCTGAAGATCACCTTGAGAGAACCCNNNAGATCGGAAG |
| 47837_1 | TATGAGTGTGGAGTCGTTGCGTGTCCACCAAGAAGTCACAGAGNNNAGATCGGAA |
| 47837_2 | TATGAGTGTGGAGTCGTTGCGTGTCCACCAAGAAGTCACAGAGNNNAGATCGGAA |
| 38556_1 | TATGAGTGTGGAGTCGTTGCACATGCTTGTAGTCCCAGCTACNNNAGATCGGAAGA |
| 38556_2 | TATGAGTGTGGAGTCGTTGCACATGCTTGTAGTCCCAGCTACNNNAGATCGGAAGA |
| 10524_1 | TATGAGTGTGGAGTCGTTGCCAGCCCCATCTCAATTGGTCTTTNNNAGATCGGAAGA |
| 10524_2 | TATGAGTGTGGAGTCGTTGCCAGCCCCATCTCAATTGGTCTTTNNNAGATCGGAAGA |
| 51589_1 | TATGAGTGTGGAGTCGTTGCGAGTCAATCAGGGGTGTGTCTGNNNAGATCGGAAG |
| 51589_2 | TATGAGTGTGGAGTCGTTGCGAGTCAATCAGGGGTGTGTCTGNNNAGATCGGAAG |
| 41986_1 | TATGAGTGTGGAGTCGTTGCTTATCCCACAGAAAGACCTGGTGNNNAGATCGGAAG |
| 41986_2 | TATGAGTGTGGAGTCGTTGCTTATCCCACAGAAAGACCTGGTGNNNAGATCGGAAG |
| 36542_1 | TATGAGTGTGGAGTCGTTGCGGTCAGTCTAAGCCTCCTGTTTTNNNAGATCGGAAG |
| 36542_2 | TATGAGTGTGGAGTCGTTGCGGTCAGTCTAAGCCTCCTGTTTTNNNAGATCGGAAG |
| 46814_1 | TATGAGTGTGGAGTCGTTGCAGCAGAAACAAAGAAACATAGCCCNNNAGATCGGA |
| 46814_2 | TATGAGTGTGGAGTCGTTGCAGCAGAAACAAAGAAACATAGCCCNNNAGATCGGA |
| 8784_1 | TATGAGTGTGGAGTCGTTGCCCATAAGCCAAGACTGCCATCTTNNNAGATCGGAAG |
| 8784_2 | TATGAGTGTGGAGTCGTTGCCCATAAGCCAAGACTGCCATCTTNNNAGATCGGAAG |
| 52362_1 | TATGAGTGTGGAGTCGTTGCCCATCCAGACAGACACTCTTCCNNNAGATCGGAAGA |
| 52362_2 | TATGAGTGTGGAGTCGTTGCCCATCCAGACAGACACTCTTCCNNNAGATCGGAAGA |
| 51976_1 | TATGAGTGTGGAGTCGTTGCGGAGACAGTGTTTCACCATGTTGNNNAGATCGGAA |
| 51976_2 | TATGAGTGTGGAGTCGTTGCGGAGACAGTGTTTCACCATGTTGNNNAGATCGGAA |
| 37529_1 | TATGAGTGTGGAGTCGTTGCACACACCTGTAGTCCTAGCTACTNNNAGATCGGAAG |
| 37529_2 | TATGAGTGTGGAGTCGTTGCACACACCTGTAGTCCTAGCTACTNNNAGATCGGAAG |
| 45943_1 | TATGAGTGTGGAGTCGTTGCAGATGCTACACATGAGGTGGTTTNNNAGATCGGAA |
| 45943_2 | TATGAGTGTGGAGTCGTTGCAGATGCTACACATGAGGTGGTTTNNNAGATCGGAA |
| 49174_1 | TATGAGTGTGGAGTCGTTGCGGTTCAAGTGATTCTCCTGCCTNNNAGATCGGAAGA |
| 49174_2 | TATGAGTGTGGAGTCGTTGCGGTTCAAGTGATTCTCCTGCCTNNNAGATCGGAAGA |
| 41785_1 | TATGAGTGTGGAGTCGTTGCTGATCTCGGCTCACTGCAAGNNNAGATCGGAAGAG |
| 41785_2 | TATGAGTGTGGAGTCGTTGCTGATCTCGGCTCACTGCAAGNNNAGATCGGAAGAG |
| 44267_1 | TATGAGTGTGGAGTCGTTGCGGAGACTGTACATCCGAGTCTTGNNNAGATCGGAA |
| 44267_2 | TATGAGTGTGGAGTCGTTGCGGAGACTGTACATCCGAGTCTTGNNNAGATCGGAA |
| 40470_1 | TATGAGTGTGGAGTCGTTGCTTGAGCATGTGGAGAAATTGGAGNNNAGATCGGA |
| 40470_2 | TATGAGTGTGGAGTCGTTGCTTGAGCATGTGGAGAAATTGGAGNNNAGATCGGA |
| 51868_1 | TATGAGTGTGGAGTCGTTGCCACTGTTTTCGGACCTTCCATGNNNAGATCGGAAGA |
| 51868_2 | TATGAGTGTGGAGTCGTTGCCACTGTTTTCGGACCTTCCATGNNNAGATCGGAAGA |
| 18929_1 | TATGAGTGTGGAGTCGTTGCATGAATGGATTTTAAATCCTAGAAANNNAGATCGGA |
| 18929_2 | TATGAGTGTGGAGTCGTTGCATGAATGGATTTTAAATCCTAGAAANNNAGATCGGA |
| 49170_1 | TATGAGTGTGGAGTCGTTGCGTTTCACCATGTTAGCCAGGATGNNNAGATCGGAAG |
| 49170_2 | TATGAGTGTGGAGTCGTTGCGTTTCACCATGTTAGCCAGGATGNNNAGATCGGAAG |
| 41550_1 | TATGAGTGTGGAGTCGTTGCGTTCAAGCGATTTCCGGCTAATTNNNAGATCGGAAG |
| 41550_2 | TATGAGTGTGGAGTCGTTGCGTTCAAGCGATTTCCGGCTAATTNNNAGATCGGAAG |

| | |
|---|---|
| 38074_1 | TATGAGTGTGGAGTCGTTGCGCCGACTGAGAAGAACCTTTTTCNNNAGATCGGAAG |
| 38074_2 | TATGAGTGTGGAGTCGTTGCGCCGACTGAGAAGAACCTTTTTCNNNAGATCGGAAG |
| 36913_1 | TATGAGTGTGGAGTCGTTGCCCTGTAATTGCAGCTACTTGGGANNNAGATCGGAAG |
| 36913_2 | TATGAGTGTGGAGTCGTTGCCCTGTAATTGCAGCTACTTGGGANNNAGATCGGAAG |
| 33985_1 | TATGAGTGTGGAGTCGTTGCTTCAAGCAATTCCCCTACCTCAGNNNAGATCGGAAG |
| 33985_2 | TATGAGTGTGGAGTCGTTGCTTCAAGCAATTCCCCTACCTCAGNNNAGATCGGAAG |
| 33029_1 | TATGAGTGTGGAGTCGTTGCAATCTCAGCTCACTGCAACCTCNNNAGATCGGAAGA |
| 33029_2 | TATGAGTGTGGAGTCGTTGCAATCTCAGCTCACTGCAACCTCNNNAGATCGGAAGA |
| 45950_1 | TATGAGTGTGGAGTCGTTGCTTCAAGCGATTCTCCTGCCTCNNNAGATCGGAAGAG |
| 45950_2 | TATGAGTGTGGAGTCGTTGCTTCAAGCGATTCTCCTGCCTCNNNAGATCGGAAGAG |
| 51815_1 | TATGAGTGTGGAGTCGTTGCACACCCCCAAAAAAAGAATTCCTNNNAGATCGGAAG |
| 51815_2 | TATGAGTGTGGAGTCGTTGCACACCCCCAAAAAAAGAATTCCTNNNAGATCGGAAG |
| 48506_1 | TATGAGTGTGGAGTCGTTGCTTCAAGCAATTCTCCTGCCTCAGNNNAGATCGGAAG |
| 48506_2 | TATGAGTGTGGAGTCGTTGCTTCAAGCAATTCTCCTGCCTCAGNNNAGATCGGAAG |
| 49014_1 | TATGAGTGTGGAGTCGTTGCTGTTTTGGGCTGACCTCCTATACNNNAGATCGGAAG |
| 49014_2 | TATGAGTGTGGAGTCGTTGCTGTTTTGGGCTGACCTCCTATACNNNAGATCGGAAG |
| 13396_1 | TATGAGTGTGGAGTCGTTGCTATCAGGAATGGACTTAGTCTNNNAGATCGGAAGA |
| 13396_2 | TATGAGTGTGGAGTCGTTGCTATCAGGAATGGACTTAGTCTNNNAGATCGGAAGA |
| 32169_1 | TATGAGTGTGGAGTCGTTGCAGGCCTAAGACAAAGTCCAATGTNNNAGATCGGAA |
| 32169_2 | TATGAGTGTGGAGTCGTTGCAGGCCTAAGACAAAGTCCAATGTNNNAGATCGGAA |
| 32188_1 | TATGAGTGTGGAGTCGTTGCCAATCTCGGCTCACTGCAAGNNNAGATCGGAAGAG |
| 32188_2 | TATGAGTGTGGAGTCGTTGCCAATCTCGGCTCACTGCAAGNNNAGATCGGAAGAG |
| 10406_1 | TATGAGTGTGGAGTCGTTGCCTTACAACCCAGCAAGTGTCACTNNNAGATCGGAAG |
| 10406_2 | TATGAGTGTGGAGTCGTTGCCTTACAACCCAGCAAGTGTCACTNNNAGATCGGAAG |
| 43405_1 | TATGAGTGTGGAGTCGTTGCGTTGAGCCCATCCAATGAGTTTTNNNAGATCGGAAG |
| 43405_2 | TATGAGTGTGGAGTCGTTGCGTTGAGCCCATCCAATGAGTTTTNNNAGATCGGAAG |
| 31809_1 | TATGAGTGTGGAGTCGTTGCTGGGTCAAATAGGAAGTCTCAAGGNNNAGATCGGA |
| 31809_2 | TATGAGTGTGGAGTCGTTGCTGGGTCAAATAGGAAGTCTCAAGGNNNAGATCGGA |
| 30189_1 | TATGAGTGTGGAGTCGTTGCTGATGGCTCTGATGAAGGCTATCNNNAGATCGGAA |
| 30189_2 | TATGAGTGTGGAGTCGTTGCTGATGGCTCTGATGAAGGCTATCNNNAGATCGGAA |
| 35536_1 | TATGAGTGTGGAGTCGTTGCCATTCTCCTGCCTCAGCCTCNNNAGATCGGAAGAGCA |
| 35536_2 | TATGAGTGTGGAGTCGTTGCCATTCTCCTGCCTCAGCCTCNNNAGATCGGAAGAGCA |
| 51776_1 | TATGAGTGTGGAGTCGTTGCTTCCTATGTCTGTCCTTTGGGTTNNNAGATCGGAAG |
| 51776_2 | TATGAGTGTGGAGTCGTTGCTTCCTATGTCTGTCCTTTGGGTTNNNAGATCGGAAG |
| 34280_1 | TATGAGTGTGGAGTCGTTGCTGGTGGTTATCAAGGCGAAAGTANNNAGATCGGAA |
| 34280_2 | TATGAGTGTGGAGTCGTTGCTGGTGGTTATCAAGGCGAAAGTANNNAGATCGGAA |
| 35547_1 | TATGAGTGTGGAGTCGTTGCTTACCAGGTGGGAAAGACTCTTCNNNAGATCGGAA |
| 35547_2 | TATGAGTGTGGAGTCGTTGCTTACCAGGTGGGAAAGACTCTTCNNNAGATCGGAA |
| 50390_1 | TATGAGTGTGGAGTCGTTGCCAGCCTGTGAAGGGAAGGAGNNNAGATCGGAAGA |
| 50390_2 | TATGAGTGTGGAGTCGTTGCCAGCCTGTGAAGGGAAGGAGNNNAGATCGGAAGA |
| 39376_1 | TATGAGTGTGGAGTCGTTGCAACAAAAATCTTGGCCATGCACANNNAGATCGGAAG |
| 39376_2 | TATGAGTGTGGAGTCGTTGCAACAAAAATCTTGGCCATGCACANNNAGATCGGAAG |
| 45362_1 | TATGAGTGTGGAGTCGTTGCCTCCAGGATGTTACTCAGAGCTCNNNAGATCGGAAG |

| | |
|---|---|
| 45362_2 | TATGAGTGTGGAGTCGTTGCCTCCAGGATGTTACTCAGAGCTCNNNAGATCGGAAG |
| 29085_1 | TATGAGTGTGGAGTCGTTGCTTCTAGGAATGATGCTGTCCGTCNNNAGATCGGAAG |
| 29085_2 | TATGAGTGTGGAGTCGTTGCTTCTAGGAATGATGCTGTCCGTCNNNAGATCGGAAG |
| 10377_1 | TATGAGTGTGGAGTCGTTGCCACCTACAGGCCAAATTGGGTAANNNAGATCGGAA |
| 10377_2 | TATGAGTGTGGAGTCGTTGCCACCTACAGGCCAAATTGGGTAANNNAGATCGGAA |
| 49757_1 | TATGAGTGTGGAGTCGTTGCATGAGCTCCTCAGGCATTTTTCTNNNAGATCGGAAG |
| 49757_2 | TATGAGTGTGGAGTCGTTGCATGAGCTCCTCAGGCATTTTTCTNNNAGATCGGAAG |
| 48650_1 | TATGAGTGTGGAGTCGTTGCGTCTCAAACTCCTGACCTCAAGTNNNAGATCGGAAG |
| 48650_2 | TATGAGTGTGGAGTCGTTGCGTCTCAAACTCCTGACCTCAAGTNNNAGATCGGAAG |
| 29897_1 | TATGAGTGTGGAGTCGTTGCCCACTCACAGAGGAAACAAACACNNNAGATCGGAAG |
| 29897_2 | TATGAGTGTGGAGTCGTTGCCCACTCACAGAGGAAACAAACACNNNAGATCGGAAG |
| 49130_1 | TATGAGTGTGGAGTCGTTGCCCTCTCATGCTCAAGTGATCCTTNNNAGATCGGAAGA |
| 49130_2 | TATGAGTGTGGAGTCGTTGCCCTCTCATGCTCAAGTGATCCTTNNNAGATCGGAAGA |
| 30512_1 | TATGAGTGTGGAGTCGTTGCCCTCTTAAAGCCCCCAGATTCATGNNNAGATCGGAA |
| 30512_2 | TATGAGTGTGGAGTCGTTGCCCTCTTAAAGCCCCCAGATTCATGNNNAGATCGGAA |
| 45627_1 | TATGAGTGTGGAGTCGTTGCGAGTGCAGTGGAGCAATCTTGNNNAGATCGGAAGA |
| 45627_2 | TATGAGTGTGGAGTCGTTGCGAGTGCAGTGGAGCAATCTTGNNNAGATCGGAAGA |
| 30324_1 | TATGAGTGTGGAGTCGTTGCATTAAAGGTGCGATTAGGGAGCGNNNAGATCGGA |
| 30324_2 | TATGAGTGTGGAGTCGTTGCATTAAAGGTGCGATTAGGGAGCGNNNAGATCGGA |
| 29759_1 | TATGAGTGTGGAGTCGTTGCCTCACGCTGTGAAACTTTCTTCCNNNAGATCGGAAGA |
| 29759_2 | TATGAGTGTGGAGTCGTTGCCTCACGCTGTGAAACTTTCTTCCNNNAGATCGGAAGA |
| 10164_1 | TATGAGTGTGGAGTCGTTGCGCTCTGAGTGTCTTGTAGCCTTTNNNAGATCGGAAG |
| 10164_2 | TATGAGTGTGGAGTCGTTGCGCTCTGAGTGTCTTGTAGCCTTTNNNAGATCGGAAG |
| 50318_1 | TATGAGTGTGGAGTCGTTGCGATCTGGTCGTTGGGATCCTCNNNAGATCGGAAGA |
| 50318_2 | TATGAGTGTGGAGTCGTTGCGATCTGGTCGTTGGGATCCTCNNNAGATCGGAAGA |
| 39368_1 | TATGAGTGTGGAGTCGTTGCAAATGAAGTGGAGATGGGAAGCTNNNAGATCGGA |
| 39368_2 | TATGAGTGTGGAGTCGTTGCAAATGAAGTGGAGATGGGAAGCTNNNAGATCGGA |
| 28937_1 | TATGAGTGTGGAGTCGTTGCTGTATGTGTGTGTGTGTTCCTCANNNAGATCGGAAG |
| 28937_2 | TATGAGTGTGGAGTCGTTGCTGTATGTGTGTGTGTGTTCCTCANNNAGATCGGAAG |
| 42413_1 | TATGAGTGTGGAGTCGTTGCTCTCTCTCTCCTAGAAACTGGGGNNNAGATCGGAAG |
| 42413_2 | TATGAGTGTGGAGTCGTTGCTCTCTCTCTCCTAGAAACTGGGGNNNAGATCGGAAG |
| 12563_1 | TATGAGTGTGGAGTCGTTGCCAAATAGTTCTCGCTCAGTGNNNAGATCGGAAGAG |
| 12563_2 | TATGAGTGTGGAGTCGTTGCCAAATAGTTCTCGCTCAGTGNNNAGATCGGAAGAG |
| 11157_1 | TATGAGTGTGGAGTCGTTGCGGGGAAGCTCAACCTCACTAAAANNNAGATCGGAA |
| 11157_2 | TATGAGTGTGGAGTCGTTGCGGGGAAGCTCAACCTCACTAAAANNNAGATCGGAA |
| 52437_1 | TATGAGTGTGGAGTCGTTGCCGAAAAGGCAGTCCAGTATCCANNNAGATCGGAAG |
| 52437_2 | TATGAGTGTGGAGTCGTTGCCGAAAAGGCAGTCCAGTATCCANNNAGATCGGAAG |
| 36827_1 | TATGAGTGTGGAGTCGTTGCTTCAAGTGACTGGCAGTGGATATNNNAGATCGGAA |
| 36827_2 | TATGAGTGTGGAGTCGTTGCTTCAAGTGACTGGCAGTGGATATNNNAGATCGGAA |
| 10106_1 | TATGAGTGTGGAGTCGTTGCGCAGTTGGGAGAAAAGGAGTAGGNNNAGATCGGA |
| 10106_2 | TATGAGTGTGGAGTCGTTGCGCAGTTGGGAGAAAAGGAGTAGGNNNAGATCGGA |
| 47164_1 | TATGAGTGTGGAGTCGTTGCGATCTCGGCTCACTAGAGTTTCCNNNAGATCGGAAG |
| 47164_2 | TATGAGTGTGGAGTCGTTGCGATCTCGGCTCACTAGAGTTTCCNNNAGATCGGAAG |

| | |
|---|---|
| 316_1 | TATGAGTGTGGAGTCGTTGCCTGTTGGTCTCAGGTGCCTATNNNAGATCGGAAGAG |
| 316_2 | TATGAGTGTGGAGTCGTTGCCTGTTGGTCTCAGGTGCCTATNNNAGATCGGAAGAG |
| 37083_1 | TATGAGTGTGGAGTCGTTGCCACTCCTGTAATCCCAGCACTTTNNNAGATCGGAAGA |
| 37083_2 | TATGAGTGTGGAGTCGTTGCCACTCCTGTAATCCCAGCACTTTNNNAGATCGGAAGA |
| 46322_1 | TATGAGTGTGGAGTCGTTGCTTGCAGCCCAGTATGTATTCAGTNNNAGATCGGAAG |
| 46322_2 | TATGAGTGTGGAGTCGTTGCTTGCAGCCCAGTATGTATTCAGTNNNAGATCGGAAG |
| 42465_1 | TATGAGTGTGGAGTCGTTGCTACAGTGGCACAATCATAGCTCANNNAGATCGGAAG |
| 42465_2 | TATGAGTGTGGAGTCGTTGCTACAGTGGCACAATCATAGCTCANNNAGATCGGAAG |
| 50763_1 | TATGAGTGTGGAGTCGTTGCTATGTTTCCTACACCACCATGGGNNNAGATCGGAAG |
| 50763_2 | TATGAGTGTGGAGTCGTTGCTATGTTTCCTACACCACCATGGGNNNAGATCGGAAG |
| 29670_1 | TATGAGTGTGGAGTCGTTGCATGGAGTAAGTATGGCCACATGGNNNAGATCGGAA |
| 29670_2 | TATGAGTGTGGAGTCGTTGCATGGAGTAAGTATGGCCACATGGNNNAGATCGGAA |
| 48437_1 | TATGAGTGTGGAGTCGTTGCAATCTTGTCTCACTGCAACCTCTNNNAGATCGGAAGA |
| 48437_2 | TATGAGTGTGGAGTCGTTGCAATCTTGTCTCACTGCAACCTCTNNNAGATCGGAAGA |
| 10532_1 | TATGAGTGTGGAGTCGTTGCAGAGAGAGAGAGTTACCTCCTTTNNNAGATCGGAA |
| 10532_2 | TATGAGTGTGGAGTCGTTGCAGAGAGAGAGAGTTACCTCCTTTNNNAGATCGGAA |
| 37493_1 | TATGAGTGTGGAGTCGTTGCGCCTATAGTCCCAGCTATGAGGANNNAGATCGGAA |
| 37493_2 | TATGAGTGTGGAGTCGTTGCGCCTATAGTCCCAGCTATGAGGANNNAGATCGGAA |
| 50324_1 | TATGAGTGTGGAGTCGTTGCAGTACAAAGAAGAGCTGGTACCANNNAGATCGGAA |
| 50324_2 | TATGAGTGTGGAGTCGTTGCAGTACAAAGAAGAGCTGGTACCANNNAGATCGGAA |
| 49669_1 | TATGAGTGTGGAGTCGTTGCCATTTCTCCAAGGAGCATGGTTCNNNAGATCGGAAG |
| 49669_2 | TATGAGTGTGGAGTCGTTGCCATTTCTCCAAGGAGCATGGTTCNNNAGATCGGAAG |
| 32661_1 | TATGAGTGTGGAGTCGTTGCCCTCTCTGGTTCGAGTGATTCTTNNNAGATCGGAAG |
| 32661_2 | TATGAGTGTGGAGTCGTTGCCCTCTCTGGTTCGAGTGATTCTTNNNAGATCGGAAG |
| 34704_1 | TATGAGTGTGGAGTCGTTGCCATTCTCCTGCCTCAGCCTCNNNAGATCGGAAGAGCA |
| 34704_2 | TATGAGTGTGGAGTCGTTGCCATTCTCCTGCCTCAGCCTCNNNAGATCGGAAGAGCA |
| 12748_1 | TATGAGTGTGGAGTCGTTGCATACTGAGAGTTAGAAGTTAAGGNNNAGATCGGAA |
| 12748_2 | TATGAGTGTGGAGTCGTTGCATACTGAGAGTTAGAAGTTAAGGNNNAGATCGGAA |
| 13639_1 | TATGAGTGTGGAGTCGTTGCTGTATGTTATTTAACAACAAGCANNNAGATCGGAAG |
| 13639_2 | TATGAGTGTGGAGTCGTTGCTGTATGTTATTTAACAACAAGCANNNAGATCGGAAG |
| 29104_1 | TATGAGTGTGGAGTCGTTGCATGAGAGCCTCCAAGTTCTCAAGNNNAGATCGGAAG |
| 29104_2 | TATGAGTGTGGAGTCGTTGCATGAGAGCCTCCAAGTTCTCAAGNNNAGATCGGAAG |
| 34318_1 | TATGAGTGTGGAGTCGTTGCTTTTGTCCTCAGAAGACACGGTCNNNAGATCGGAAG |
| 34318_2 | TATGAGTGTGGAGTCGTTGCTTTTGTCCTCAGAAGACACGGTCNNNAGATCGGAAG |
| 31328_1 | TATGAGTGTGGAGTCGTTGCGGTATACAAAACAGGCTCTGGGANNNAGATCGGAA |
| 31328_2 | TATGAGTGTGGAGTCGTTGCGGTATACAAAACAGGCTCTGGGANNNAGATCGGAA |
| 35627_1 | TATGAGTGTGGAGTCGTTGCTTCACACCATTCTCCTGCCTCNNNAGATCGGAAGAGC |
| 35627_2 | TATGAGTGTGGAGTCGTTGCTTCACACCATTCTCCTGCCTCNNNAGATCGGAAGAGC |
| 53062_1 | TATGAGTGTGGAGTCGTTGCAGAGTTGCAGTCACTCGAATCTTNNNAGATCGGAAG |
| 53062_2 | TATGAGTGTGGAGTCGTTGCAGAGTTGCAGTCACTCGAATCTTNNNAGATCGGAAG |
| 45863_1 | TATGAGTGTGGAGTCGTTGCACGCCCGGCTAATTTTTTGTATTNNNAGATCGGAAG |
| 45863_2 | TATGAGTGTGGAGTCGTTGCACGCCCGGCTAATTTTTTGTATTNNNAGATCGGAAG |
| 31654_1 | TATGAGTGTGGAGTCGTTGCCTTTCCCCCTCACTCTGAATGAANNNAGATCGGAAGA |

| | |
|---|---|
| 31654_2 | TATGAGTGTGGAGTCGTTGCCTTTCCCCCTCACTCTGAATGAANNNAGATCGGAAGA |
| 45101_1 | TATGAGTGTGGAGTCGTTGCTCCTGATACCTACCACCTCCAAANNNAGATCGGAAGA |
| 45101_2 | TATGAGTGTGGAGTCGTTGCTCCTGATACCTACCACCTCCAAANNNAGATCGGAAGA |
| 5875_1 | TATGAGTGTGGAGTCGTTGCACACAATCAAAGGACTGCCTGTTNNNAGATCGGAAG |
| 5875_2 | TATGAGTGTGGAGTCGTTGCACACAATCAAAGGACTGCCTGTTNNNAGATCGGAAG |
| 32526_1 | TATGAGTGTGGAGTCGTTGCGTGGGATCACCAAATGCTAAGAANNNAGATCGGAA |
| 32526_2 | TATGAGTGTGGAGTCGTTGCGTGGGATCACCAAATGCTAAGAANNNAGATCGGAA |
| 52944_1 | TATGAGTGTGGAGTCGTTGCCTCGCTGCTCTCATACTCATCATNNNAGATCGGAAGA |
| 52944_2 | TATGAGTGTGGAGTCGTTGCCTCGCTGCTCTCATACTCATCATNNNAGATCGGAAGA |
| 30445_1 | TATGAGTGTGGAGTCGTTGCCTCAAGTAATCCTCCCACCTCAGNNNAGATCGGAAGA |
| 30445_2 | TATGAGTGTGGAGTCGTTGCCTCAAGTAATCCTCCCACCTCAGNNNAGATCGGAAGA |
| 40820_1 | TATGAGTGTGGAGTCGTTGCACCACCTGAATTCAGAAGGTCAANNNAGATCGGAAG |
| 40820_2 | TATGAGTGTGGAGTCGTTGCACCACCTGAATTCAGAAGGTCAANNNAGATCGGAAG |
| 35254_1 | TATGAGTGTGGAGTCGTTGCGTAGTGCAGTGGTTTGATCTTGGNNNAGATCGGAA |
| 35254_2 | TATGAGTGTGGAGTCGTTGCGTAGTGCAGTGGTTTGATCTTGGNNNAGATCGGAA |
| 10260_1 | TATGAGTGTGGAGTCGTTGCCCCCATTCTGTGGAAGTAACCAGNNNAGATCGGAAG |
| 10260_2 | TATGAGTGTGGAGTCGTTGCCCCCATTCTGTGGAAGTAACCAGNNNAGATCGGAAG |
| 50385_1 | TATGAGTGTGGAGTCGTTGCGAGGAGGGGGGAATCCTGATNNNAGATCGGAAGA |
| 50385_2 | TATGAGTGTGGAGTCGTTGCGAGGAGGGGGGAATCCTGATNNNAGATCGGAAGA |
| 44541_1 | TATGAGTGTGGAGTCGTTGCCTGAGGAGCTCATATCCATGTCCNNNAGATCGGAAG |
| 44541_2 | TATGAGTGTGGAGTCGTTGCCTGAGGAGCTCATATCCATGTCCNNNAGATCGGAAG |
| 38030_1 | TATGAGTGTGGAGTCGTTGCACTGCCTGAAAGATCCATCAACTNNNAGATCGGAAG |
| 38030_2 | TATGAGTGTGGAGTCGTTGCACTGCCTGAAAGATCCATCAACTNNNAGATCGGAAG |
| 49405_1 | TATGAGTGTGGAGTCGTTGCGTAGAGATGAGGCCTTGCTATGTNNNAGATCGGAA |
| 49405_2 | TATGAGTGTGGAGTCGTTGCGTAGAGATGAGGCCTTGCTATGTNNNAGATCGGAA |
| 51574_1 | TATGAGTGTGGAGTCGTTGCCCTTGTTCCTCAGACACGGATGNNNAGATCGGAAGA |
| 51574_2 | TATGAGTGTGGAGTCGTTGCCCTTGTTCCTCAGACACGGATGNNNAGATCGGAAGA |
| 41197_1 | TATGAGTGTGGAGTCGTTGCCCTTTTTGAGCTGTAACACTCCCNNNAGATCGGAAGA |
| 41197_2 | TATGAGTGTGGAGTCGTTGCCCTTTTTGAGCTGTAACACTCCCNNNAGATCGGAAGA |
| 13308_1 | TATGAGTGTGGAGTCGTTGCTTATAACTTGATTCTTCCCATTGNNNAGATCGGAAGA |
| 13308_2 | TATGAGTGTGGAGTCGTTGCTTATAACTTGATTCTTCCCATTGNNNAGATCGGAAGA |
| 30366_1 | TATGAGTGTGGAGTCGTTGCGTCCAGTCCTGTGTGTGGATTANNNAGATCGGAAGA |
| 30366_2 | TATGAGTGTGGAGTCGTTGCGTCCAGTCCTGTGTGTGGATTANNNAGATCGGAAGA |
| 40043_1 | TATGAGTGTGGAGTCGTTGCGCCCATTGGTGTGTAGTTGAAAANNNAGATCGGAA |
| 40043_2 | TATGAGTGTGGAGTCGTTGCGCCCATTGGTGTGTAGTTGAAAANNNAGATCGGAA |
| 43761_1 | TATGAGTGTGGAGTCGTTGCTCATATGACATGCTCCAGGGTTCNNNAGATCGGAAG |
| 43761_2 | TATGAGTGTGGAGTCGTTGCTCATATGACATGCTCCAGGGTTCNNNAGATCGGAAG |
| 35535_1 | TATGAGTGTGGAGTCGTTGCTGCCTAGGTTTTCTTCTAGGGTTNNNAGATCGGAAG |
| 35535_2 | TATGAGTGTGGAGTCGTTGCTGCCTAGGTTTTCTTCTAGGGTTNNNAGATCGGAAG |
| 30308_1 | TATGAGTGTGGAGTCGTTGCTGAATAAGGAAACCTCATGGGGGNNNAGATCGGAA |
| 30308_2 | TATGAGTGTGGAGTCGTTGCTGAATAAGGAAACCTCATGGGGGNNNAGATCGGAA |
| 30475_1 | TATGAGTGTGGAGTCGTTGCAGACAGAAGCATAAGAGGCAACATNNNAGATCGGA |
| 30475_2 | TATGAGTGTGGAGTCGTTGCAGACAGAAGCATAAGAGGCAACATNNNAGATCGGA |

| | |
|---|---|
| 8016_1 | TATGAGTGTGGAGTCGTTGCAAACTAACTCTTCGATGCCCAGANNNAGATCGGAAG |
| 8016_2 | TATGAGTGTGGAGTCGTTGCAAACTAACTCTTCGATGCCCAGANNNAGATCGGAAG |
| 52961_1 | TATGAGTGTGGAGTCGTTGCGGACACTTGTGATTGGATTTGGGNNNAGATCGGAA |
| 52961_2 | TATGAGTGTGGAGTCGTTGCGGACACTTGTGATTGGATTTGGGNNNAGATCGGAA |
| 37394_1 | TATGAGTGTGGAGTCGTTGCTCATGCAGCTATGGAGAAGGATGNNNAGATCGGAA |
| 37394_2 | TATGAGTGTGGAGTCGTTGCTCATGCAGCTATGGAGAAGGATGNNNAGATCGGAA |
| 39247_1 | TATGAGTGTGGAGTCGTTGCGCATAACCCACATGAGGAAAGGANNNAGATCGGAA |
| 39247_2 | TATGAGTGTGGAGTCGTTGCGCATAACCCACATGAGGAAAGGANNNAGATCGGAA |
| 42326_1 | TATGAGTGTGGAGTCGTTGCGAGACAGAGACTCAGTTTGTTGCNNNAGATCGGAA |
| 42326_2 | TATGAGTGTGGAGTCGTTGCGAGACAGAGACTCAGTTTGTTGCNNNAGATCGGAA |
| 12958_1 | TATGAGTGTGGAGTCGTTGCACCTCCTATGAAAATTGCAGNNNAGATCGGAAGAG |
| 12958_2 | TATGAGTGTGGAGTCGTTGCACCTCCTATGAAAATTGCAGNNNAGATCGGAAGAG |
| 43714_1 | TATGAGTGTGGAGTCGTTGCCAGAGGCTCAGAAAAAGAACGTGNNNAGATCGGAA |
| 43714_2 | TATGAGTGTGGAGTCGTTGCCAGAGGCTCAGAAAAAGAACGTGNNNAGATCGGAA |
| 38089_1 | TATGAGTGTGGAGTCGTTGCCGCTCTACCTATTGACCTTCCTCNNNAGATCGGAAGA |
| 38089_2 | TATGAGTGTGGAGTCGTTGCCGCTCTACCTATTGACCTTCCTCNNNAGATCGGAAGA |
| 12619_1 | TATGAGTGTGGAGTCGTTGCATAGGAACATGAAATAGCAATGANNNAGATCGGAA |
| 12619_2 | TATGAGTGTGGAGTCGTTGCATAGGAACATGAAATAGCAATGANNNAGATCGGAA |
| 45132_1 | TATGAGTGTGGAGTCGTTGCCCCAACATACACATGCACACATTNNNAGATCGGAAGA |
| 45132_2 | TATGAGTGTGGAGTCGTTGCCCCAACATACACATGCACACATTNNNAGATCGGAAGA |
| 32990_1 | TATGAGTGTGGAGTCGTTGCTTCAAGTGATTCTCCTGCCTCAGNNNAGATCGGAAGA |
| 32990_2 | TATGAGTGTGGAGTCGTTGCTTCAAGTGATTCTCCTGCCTCAGNNNAGATCGGAAGA |
| 48531_1 | TATGAGTGTGGAGTCGTTGCAGAGCAGGTTTCACAGCATCTTANNNAGATCGGAAG |
| 48531_2 | TATGAGTGTGGAGTCGTTGCAGAGCAGGTTTCACAGCATCTTANNNAGATCGGAAG |
| 48957_1 | TATGAGTGTGGAGTCGTTGCCTTCTCTCTATCCCCACAGCATCNNNAGATCGGAAGA |
| 48957_2 | TATGAGTGTGGAGTCGTTGCCTTCTCTCTATCCCCACAGCATCNNNAGATCGGAAGA |
| 35901_1 | TATGAGTGTGGAGTCGTTGCATCCTAATATTGCATGCAACTGCNNNAGATCGGAAG |
| 35901_2 | TATGAGTGTGGAGTCGTTGCATCCTAATATTGCATGCAACTGCNNNAGATCGGAAG |
| 43813_1 | TATGAGTGTGGAGTCGTTGCATGCCACAAATGTGCATGGTAAANNNAGATCGGAA |
| 43813_2 | TATGAGTGTGGAGTCGTTGCATGCCACAAATGTGCATGGTAAANNNAGATCGGAA |
| 35601_1 | TATGAGTGTGGAGTCGTTGCCTCTGGGGATCCTTTGCTTTCTANNNAGATCGGAAG |
| 35601_2 | TATGAGTGTGGAGTCGTTGCCTCTGGGGATCCTTTGCTTTCTANNNAGATCGGAAG |
| 49270_1 | TATGAGTGTGGAGTCGTTGCGGTTCAAGTGATTCTCTTGCCTCNNNAGATCGGAAG |
| 49270_2 | TATGAGTGTGGAGTCGTTGCGGTTCAAGTGATTCTCTTGCCTCNNNAGATCGGAAG |
| 42883_1 | TATGAGTGTGGAGTCGTTGCTAAGTAGCTGGGACTACAGGTGTNNNAGATCGGAA |
| 42883_2 | TATGAGTGTGGAGTCGTTGCTAAGTAGCTGGGACTACAGGTGTNNNAGATCGGAA |
| 52200_1 | TATGAGTGTGGAGTCGTTGCGCAAATTCAGCTGAGCTCCATAGNNNAGATCGGAAG |
| 52200_2 | TATGAGTGTGGAGTCGTTGCGCAAATTCAGCTGAGCTCCATAGNNNAGATCGGAAG |
| 34636_1 | TATGAGTGTGGAGTCGTTGCTGGATAGTTCTCAGTGACCCAACNNNAGATCGGAAG |
| 34636_2 | TATGAGTGTGGAGTCGTTGCTGGATAGTTCTCAGTGACCCAACNNNAGATCGGAAG |
| 52413_1 | TATGAGTGTGGAGTCGTTGCGGGTAGTCTCCACACTCTGTTTTNNNAGATCGGAAG |
| 52413_2 | TATGAGTGTGGAGTCGTTGCGGGTAGTCTCCACACTCTGTTTTNNNAGATCGGAAG |
| 32065_1 | TATGAGTGTGGAGTCGTTGCATGGTAACTTCACTGGGCTAGACNNNAGATCGGAA |

| | |
|---|---|
| 32065_2 | TATGAGTGTGGAGTCGTTGCATGGTAACTTCACTGGGCTAGACNNNAGATCGGAAG |
| 44775_1 | TATGAGTGTGGAGTCGTTGCAGCCCTGAAAGGATGATGTACAGNNNAGATCGGAA |
| 44775_2 | TATGAGTGTGGAGTCGTTGCAGCCCTGAAAGGATGATGTACAGNNNAGATCGGAA |
| 43126_1 | TATGAGTGTGGAGTCGTTGCACCTGAAGGCAATGGATTACAGTNNNAGATCGGAA |
| 43126_2 | TATGAGTGTGGAGTCGTTGCACCTGAAGGCAATGGATTACAGTNNNAGATCGGAA |
| 50852_1 | TATGAGTGTGGAGTCGTTGCAGCAGTCAAGAAAGAAAGCCAACNNNAGATCGGAA |
| 50852_2 | TATGAGTGTGGAGTCGTTGCAGCAGTCAAGAAAGAAAGCCAACNNNAGATCGGAA |
| 34255_1 | TATGAGTGTGGAGTCGTTGCCACACCTGTAATCCCACCACTATNNNAGATCGGAAGA |
| 34255_2 | TATGAGTGTGGAGTCGTTGCCACACCTGTAATCCCACCACTATNNNAGATCGGAAGA |
| 36737_1 | TATGAGTGTGGAGTCGTTGCTGTGTAGTTGTGGACTCATCTCTNNNAGATCGGAAG |
| 36737_2 | TATGAGTGTGGAGTCGTTGCTGTGTAGTTGTGGACTCATCTCTNNNAGATCGGAAG |
| 30380_1 | TATGAGTGTGGAGTCGTTGCCCGGTATGTAATTTAACATTCAANNNAGATCGGAAG |
| 30380_2 | TATGAGTGTGGAGTCGTTGCCCGGTATGTAATTTAACATTCAANNNAGATCGGAAG |
| 34843_1 | TATGAGTGTGGAGTCGTTGCGGGAGGCCTAGATGAAACAAGAANNNAGATCGGA |
| 34843_2 | TATGAGTGTGGAGTCGTTGCGGGAGGCCTAGATGAAACAAGAANNNAGATCGGA |
| 50337_1 | TATGAGTGTGGAGTCGTTGCCCACAGCCACATATCACAATCACNNNAGATCGGAAG |
| 50337_2 | TATGAGTGTGGAGTCGTTGCCCACAGCCACATATCACAATCACNNNAGATCGGAAG |
| 42769_1 | TATGAGTGTGGAGTCGTTGCAGGCCATCATGCTAAGCAGATTANNNAGATCGGAAG |
| 42769_2 | TATGAGTGTGGAGTCGTTGCAGGCCATCATGCTAAGCAGATTANNNAGATCGGAAG |
| 42553_1 | TATGAGTGTGGAGTCGTTGCTTCCCTCCTCCTCTACCTTTTGANNNAGATCGGAAGA |
| 42553_2 | TATGAGTGTGGAGTCGTTGCTTCCCTCCTCCTCTACCTTTTGANNNAGATCGGAAGA |
| 43928_1 | TATGAGTGTGGAGTCGTTGCCCTCTTTCTTCTCCCTCTCCTTCNNNAGATCGGAAGAG |
| 43928_2 | TATGAGTGTGGAGTCGTTGCCCTCTTTCTTCTCCCTCTCCTTCNNNAGATCGGAAGAG |
| 32461_1 | TATGAGTGTGGAGTCGTTGCGTTCCCTTCAAGTCTGGTAGGTTNNNAGATCGGAAG |
| 32461_2 | TATGAGTGTGGAGTCGTTGCGTTCCCTTCAAGTCTGGTAGGTTNNNAGATCGGAAG |
| 35260_1 | TATGAGTGTGGAGTCGTTGCATACAATCTCTGCCACCCATGTTNNNAGATCGGAAGA |
| 35260_2 | TATGAGTGTGGAGTCGTTGCATACAATCTCTGCCACCCATGTTNNNAGATCGGAAGA |
| 44706_1 | TATGAGTGTGGAGTCGTTGCGATCGGAGCGGTAAAAGTGAATGNNNAGATCGGA |
| 44706_2 | TATGAGTGTGGAGTCGTTGCGATCGGAGCGGTAAAAGTGAATGNNNAGATCGGA |
| 36076_1 | TATGAGTGTGGAGTCGTTGCGGATGGATACTTGCTGCTTTTCCNNNAGATCGGAAG |
| 36076_2 | TATGAGTGTGGAGTCGTTGCGGATGGATACTTGCTGCTTTTCCNNNAGATCGGAAG |
| 42364_1 | TATGAGTGTGGAGTCGTTGCATCCTAGCTACTCAGAAGGCTGANNNAGATCGGAAG |
| 42364_2 | TATGAGTGTGGAGTCGTTGCATCCTAGCTACTCAGAAGGCTGANNNAGATCGGAAG |
| 6247_1 | TATGAGTGTGGAGTCGTTGCCCTTGTATGCTGTGTACTGCCTGANNNAGATCGGAA |
| 6247_2 | TATGAGTGTGGAGTCGTTGCCCTTGTATGCTGTGTACTGCCTGANNNAGATCGGAA |
| 34989_1 | TATGAGTGTGGAGTCGTTGCCGCATCCCATCAACTCATGTCTTNNNAGATCGGAAGA |
| 34989_2 | TATGAGTGTGGAGTCGTTGCCGCATCCCATCAACTCATGTCTTNNNAGATCGGAAGA |
| 41483_1 | TATGAGTGTGGAGTCGTTGCTGTTTAGTTCTTATTCTACCTGTGCTNNNAGATCGGA |
| 41483_2 | TATGAGTGTGGAGTCGTTGCTGTTTAGTTCTTATTCTACCTGTGCTNNNAGATCGGA |
| 35303_1 | TATGAGTGTGGAGTCGTTGCGATCTTAGCTCACTGCAACCTCTNNNAGATCGGAAG |
| 35303_2 | TATGAGTGTGGAGTCGTTGCGATCTTAGCTCACTGCAACCTCTNNNAGATCGGAAG |
| 50009_1 | TATGAGTGTGGAGTCGTTGCACTTTTCCAGGCCTCATCTAGTGNNNAGATCGGAAG |
| 50009_2 | TATGAGTGTGGAGTCGTTGCACTTTTCCAGGCCTCATCTAGTGNNNAGATCGGAAG |

| | |
|---|---|
| 29879_1 | TATGAGTGTGGAGTCGTTGCTGGTCTTTGCCATAGCGACTGNNNAGATCGGAAGAG |
| 29879_2 | TATGAGTGTGGAGTCGTTGCTGGTCTTTGCCATAGCGACTGNNNAGATCGGAAGAG |
| 45129_1 | TATGAGTGTGGAGTCGTTGCGTATGAGAGAGCATGTGCCAGTANNNAGATCGGAA |
| 45129_2 | TATGAGTGTGGAGTCGTTGCGTATGAGAGAGCATGTGCCAGTANNNAGATCGGAA |
| 10192_1 | TATGAGTGTGGAGTCGTTGCAGAGAGGTTAGGAATTGGGGAGGNNNAGATCGGA |
| 10192_2 | TATGAGTGTGGAGTCGTTGCAGAGAGGTTAGGAATTGGGGAGGNNNAGATCGGA |
| 3178_1 | TATGAGTGTGGAGTCGTTGCAGCAGAATTCCCCAGTATTTTGGGNNNAGATCGGAA |
| 3178_2 | TATGAGTGTGGAGTCGTTGCAGCAGAATTCCCCAGTATTTTGGGNNNAGATCGGAA |
| 35802_1 | TATGAGTGTGGAGTCGTTGCTCATGTGCCTGCCTTGACTTATANNNAGATCGGAAG |
| 35802_2 | TATGAGTGTGGAGTCGTTGCTCATGTGCCTGCCTTGACTTATANNNAGATCGGAAG |
| 41433_1 | TATGAGTGTGGAGTCGTTGCGGGAGAAACAGGAAGTGCAAAAANNNAGATCGGA |
| 41433_2 | TATGAGTGTGGAGTCGTTGCGGGAGAAACAGGAAGTGCAAAAANNNAGATCGGA |
| 43031_1 | TATGAGTGTGGAGTCGTTGCTACTGGTGGGAATGTGAAATGGTNNNAGATCGGAA |
| 43031_2 | TATGAGTGTGGAGTCGTTGCTACTGGTGGGAATGTGAAATGGTNNNAGATCGGAA |
| 39933_1 | TATGAGTGTGGAGTCGTTGCACTGGTTTTGGAGCTGAGACAATNNNAGATCGGAA |
| 39933_2 | TATGAGTGTGGAGTCGTTGCACTGGTTTTGGAGCTGAGACAATNNNAGATCGGAA |
| 43273_1 | TATGAGTGTGGAGTCGTTGCGGAGTGCAATGGGATGATTTCAGNNNAGATCGGAA |
| 43273_2 | TATGAGTGTGGAGTCGTTGCGGAGTGCAATGGGATGATTTCAGNNNAGATCGGAA |
| 35552_1 | TATGAGTGTGGAGTCGTTGCTCATGACTCTGTTGGGAATGACANNNAGATCGGAAG |
| 35552_2 | TATGAGTGTGGAGTCGTTGCTCATGACTCTGTTGGGAATGACANNNAGATCGGAAG |
| 3020_1 | TATGAGTGTGGAGTCGTTGCACTGCTGCAACTACATCACAGAGNNNAGATCGGAAG |
| 3020_2 | TATGAGTGTGGAGTCGTTGCACTGCTGCAACTACATCACAGAGNNNAGATCGGAAG |
| 41167_1 | TATGAGTGTGGAGTCGTTGCTGTACATGTTCACCCTGTATGCANNNAGATCGGAAG |
| 41167_2 | TATGAGTGTGGAGTCGTTGCTGTACATGTTCACCCTGTATGCANNNAGATCGGAAG |
| 48746_1 | TATGAGTGTGGAGTCGTTGCGCGCCCGGCTAATTTTTTGTANNNAGATCGGAAGAG |
| 48746_2 | TATGAGTGTGGAGTCGTTGCGCGCCCGGCTAATTTTTTGTANNNAGATCGGAAGAG |
| 29739_1 | TATGAGTGTGGAGTCGTTGCATACACCATGGAATGCTATGCAGNNNAGATCGGAAG |
| 29739_2 | TATGAGTGTGGAGTCGTTGCATACACCATGGAATGCTATGCAGNNNAGATCGGAAG |
| 32622_1 | TATGAGTGTGGAGTCGTTGCAAAGTTCTAACTGAGCTGGAGCANNNAGATCGGAA |
| 32622_2 | TATGAGTGTGGAGTCGTTGCAAAGTTCTAACTGAGCTGGAGCANNNAGATCGGAA |
| 30405_1 | TATGAGTGTGGAGTCGTTGCAGAGTTTTGTTTGTTTCTAATCCNNNAGATCGGAAG |
| 30405_2 | TATGAGTGTGGAGTCGTTGCAGAGTTTTGTTTGTTTCTAATCCNNNAGATCGGAAG |
| 35777_1 | TATGAGTGTGGAGTCGTTGCACTATCTCAGCTCACTGCAAGCNNNAGATCGGAAGA |
| 35777_2 | TATGAGTGTGGAGTCGTTGCACTATCTCAGCTCACTGCAAGCNNNAGATCGGAAGA |
| 47957_1 | TATGAGTGTGGAGTCGTTGCTTCAGTCAACGAAGAAGCCTCTGNNNAGATCGGAAG |
| 47957_2 | TATGAGTGTGGAGTCGTTGCTTCAGTCAACGAAGAAGCCTCTGNNNAGATCGGAAG |
| 48799_1 | TATGAGTGTGGAGTCGTTGCCAGGTGCAGTGCCAAAAACATATNNNAGATCGGAA |
| 48799_2 | TATGAGTGTGGAGTCGTTGCCAGGTGCAGTGCCAAAAACATATNNNAGATCGGAA |
| 48398_1 | TATGAGTGTGGAGTCGTTGCCGGGGACCAGGTACTAAAGGATANNNAGATCGGAA |
| 48398_2 | TATGAGTGTGGAGTCGTTGCCGGGGACCAGGTACTAAAGGATANNNAGATCGGAA |
| 7886_1 | TATGAGTGTGGAGTCGTTGCAGATAGAGTGGTGGGGCAAGAAGNNNAGATCGGA |
| 7886_2 | TATGAGTGTGGAGTCGTTGCAGATAGAGTGGTGGGGCAAGAAGNNNAGATCGGA |
| 49608_1 | TATGAGTGTGGAGTCGTTGCCCACCCAAGTAGCTGAGATTACANNNAGATCGGAAG |

| | |
|---|---|
| 49608_2 | TATGAGTGTGGAGTCGTTGCCCACCCAAGTAGCTGAGATTACANNNAGATCGGAAG |
| 52060_1 | TATGAGTGTGGAGTCGTTGCTGCAGGAATATGAGAGCTTCTCCNNNAGATCGGAAG |
| 52060_2 | TATGAGTGTGGAGTCGTTGCTGCAGGAATATGAGAGCTTCTCCNNNAGATCGGAAG |
| 34528_1 | TATGAGTGTGGAGTCGTTGCTCCTCTCTTCTCAATAGTCCCCNNNAGATCGGAAGA |
| 34528_2 | TATGAGTGTGGAGTCGTTGCTCCTCTCTTCTCAATAGTCCCCNNNAGATCGGAAGA |
| 39259_1 | TATGAGTGTGGAGTCGTTGCTTCCAGACCAGCCTGACCAATATNNNAGATCGGAAG |
| 39259_2 | TATGAGTGTGGAGTCGTTGCTTCCAGACCAGCCTGACCAATATNNNAGATCGGAAG |
| 36398_1 | TATGAGTGTGGAGTCGTTGCCATTCAATCTGCATGAAGGGCATNNNAGATCGGAAG |
| 36398_2 | TATGAGTGTGGAGTCGTTGCCATTCAATCTGCATGAAGGGCATNNNAGATCGGAAG |
| 5592_1 | TATGAGTGTGGAGTCGTTGCATCCCCAAATACAGATGGGTCCANNNAGATCGGAAG |
| 5592_2 | TATGAGTGTGGAGTCGTTGCATCCCCAAATACAGATGGGTCCANNNAGATCGGAAG |
| 30022_1 | TATGAGTGTGGAGTCGTTGCAACAGTTGCTTGGGAAGAAAGTGNNNAGATCGGAA |
| 30022_2 | TATGAGTGTGGAGTCGTTGCAACAGTTGCTTGGGAAGAAAGTGNNNAGATCGGAA |
| 31357_1 | TATGAGTGTGGAGTCGTTGCTTCCTGGCTACTGCTTATGTTCANNNAGATCGGAAG |
| 31357_2 | TATGAGTGTGGAGTCGTTGCTTCCTGGCTACTGCTTATGTTCANNNAGATCGGAAG |
| 44950_1 | TATGAGTGTGGAGTCGTTGCAATCTTTAGATGTCGAAGCCCCGNNNAGATCGGAAG |
| 44950_2 | TATGAGTGTGGAGTCGTTGCAATCTTTAGATGTCGAAGCCCCGNNNAGATCGGAAG |
| 12740_1 | TATGAGTGTGGAGTCGTTGCCTTTCTTGACATCCCTTTAAAAANNNAGATCGGAAGA |
| 12740_2 | TATGAGTGTGGAGTCGTTGCCTTTCTTGACATCCCTTTAAAAANNNAGATCGGAAGA |
| 35434_1 | TATGAGTGTGGAGTCGTTGCCAATGGTGCAATCATAGCTCACANNNAGATCGGAAG |
| 35434_2 | TATGAGTGTGGAGTCGTTGCCAATGGTGCAATCATAGCTCACANNNAGATCGGAAG |
| 50972_1 | TATGAGTGTGGAGTCGTTGCTTCTCCCACACTGATCTCCTTTGNNNAGATCGGAAGA |
| 50972_2 | TATGAGTGTGGAGTCGTTGCTTCTCCCACACTGATCTCCTTTGNNNAGATCGGAAGA |
| 40424_1 | TATGAGTGTGGAGTCGTTGCCAAGTGGACAGAAAACCAGCAAANNNAGATCGGAA |
| 40424_2 | TATGAGTGTGGAGTCGTTGCCAAGTGGACAGAAAACCAGCAAANNNAGATCGGAA |
| 39862_1 | TATGAGTGTGGAGTCGTTGCTTTGCATATGAATGTCTTGGGCANNNAGATCGGAAG |
| 39862_2 | TATGAGTGTGGAGTCGTTGCTTTGCATATGAATGTCTTGGGCANNNAGATCGGAAG |
| 39432_1 | TATGAGTGTGGAGTCGTTGCATTACTTCCTACCAAGGGGCAAGNNNAGATCGGAAG |
| 39432_2 | TATGAGTGTGGAGTCGTTGCATTACTTCCTACCAAGGGGCAAGNNNAGATCGGAAG |
| 47081_1 | TATGAGTGTGGAGTCGTTGCTTATTTTCTGGGTGGAGCCATCANNNAGATCGGAAG |
| 47081_2 | TATGAGTGTGGAGTCGTTGCTTATTTTCTGGGTGGAGCCATCANNNAGATCGGAAG |
| 40084_1 | TATGAGTGTGGAGTCGTTGCACAAGAACATTTGAGATGGTGGTNNNAGATCGGAA |
| 40084_2 | TATGAGTGTGGAGTCGTTGCACAAGAACATTTGAGATGGTGGTNNNAGATCGGAA |
| 44323_1 | TATGAGTGTGGAGTCGTTGCCTCTTCCTGTGTGGTATTTGCACNNNAGATCGGAAG |
| 44323_2 | TATGAGTGTGGAGTCGTTGCCTCTTCCTGTGTGGTATTTGCACNNNAGATCGGAAG |
| 45160_1 | TATGAGTGTGGAGTCGTTGCGAGAGGTGAAGATTAGGAGGTGCNNNAGATCGGA |
| 45160_2 | TATGAGTGTGGAGTCGTTGCGAGAGGTGAAGATTAGGAGGTGCNNNAGATCGGA |
| 47146_1 | TATGAGTGTGGAGTCGTTGCAGGAGATTTTTCAGCTCACGAGTNNNAGATCGGAA |
| 47146_2 | TATGAGTGTGGAGTCGTTGCAGGAGATTTTTCAGCTCACGAGTNNNAGATCGGAA |
| 44537_1 | TATGAGTGTGGAGTCGTTGCAGTATCTTTCCTTCAGGGCTGTGNNNAGATCGGAAG |
| 44537_2 | TATGAGTGTGGAGTCGTTGCAGTATCTTTCCTTCAGGGCTGTGNNNAGATCGGAAG |
| 49941_1 | TATGAGTGTGGAGTCGTTGCTCCTTCAGACAGAGGACCTTACTNNNAGATCGGAAG |
| 49941_2 | TATGAGTGTGGAGTCGTTGCTCCTTCAGACAGAGGACCTTACTNNNAGATCGGAAG |

| | |
|---|---|
| 48441_1 | TATGAGTGTGGAGTCGTTGCGCTGGTACTGGGAAGACAAAGATNNNAGATCGGAA |
| 48441_2 | TATGAGTGTGGAGTCGTTGCGCTGGTACTGGGAAGACAAAGATNNNAGATCGGAA |
| 41996_1 | TATGAGTGTGGAGTCGTTGCGATTCAAGCAATTCTCCTGCCTCNNNAGATCGGAAGA |
| 41996_2 | TATGAGTGTGGAGTCGTTGCGATTCAAGCAATTCTCCTGCCTCNNNAGATCGGAAGA |
| 28838_1 | TATGAGTGTGGAGTCGTTGCAATCACTGGAGAAGGAAGAGCAGNNNAGATCGGA |
| 28838_2 | TATGAGTGTGGAGTCGTTGCAATCACTGGAGAAGGAAGAGCAGNNNAGATCGGA |
| 36282_1 | TATGAGTGTGGAGTCGTTGCAGAAGGCAAATGGTATAGGCACANNNAGATCGGAA |
| 36282_2 | TATGAGTGTGGAGTCGTTGCAGAAGGCAAATGGTATAGGCACANNNAGATCGGAA |
| 50515_1 | TATGAGTGTGGAGTCGTTGCTTCCTCGGGTTTTAGTGTAGTGGNNNAGATCGGAA |
| 50515_2 | TATGAGTGTGGAGTCGTTGCTTCCTCGGGTTTTAGTGTAGTGGNNNAGATCGGAA |
| 32426_1 | TATGAGTGTGGAGTCGTTGCCAATCTCGGCTCACTGCAAGNNNAGATCGGAAGAG |
| 32426_2 | TATGAGTGTGGAGTCGTTGCCAATCTCGGCTCACTGCAAGNNNAGATCGGAAGAG |
| 35749_1 | TATGAGTGTGGAGTCGTTGCGCCTCCTGAGTAGCTGAGATTACNNNAGATCGGAAG |
| 35749_2 | TATGAGTGTGGAGTCGTTGCGCCTCCTGAGTAGCTGAGATTACNNNAGATCGGAAG |
| 29635_1 | TATGAGTGTGGAGTCGTTGCTGCAAGTTCATTAACTACAGTGTCTNNNAGATCGGA |
| 29635_2 | TATGAGTGTGGAGTCGTTGCTGCAAGTTCATTAACTACAGTGTCTNNNAGATCGGA |
| 40915_1 | TATGAGTGTGGAGTCGTTGCACTCCCTTGATTGAGTCACCTTCNNNAGATCGGAAGA |
| 40915_2 | TATGAGTGTGGAGTCGTTGCACTCCCTTGATTGAGTCACCTTCNNNAGATCGGAAGA |
| 53298_1 | TATGAGTGTGGAGTCGTTGCGTTTATCCAGCCAGCCTCTCATCNNNAGATCGGAAGA |
| 53298_2 | TATGAGTGTGGAGTCGTTGCGTTTATCCAGCCAGCCTCTCATCNNNAGATCGGAAGA |
| 39063_1 | TATGAGTGTGGAGTCGTTGCTCCACATGCTTTTCTTTCTCCCTNNNAGATCGGAAGA |
| 39063_2 | TATGAGTGTGGAGTCGTTGCTCCACATGCTTTTCTTTCTCCCTNNNAGATCGGAAGA |
| 41589_1 | TATGAGTGTGGAGTCGTTGCTTCAAGCGATTTTTCTGCCTCAGNNNAGATCGGAAG |
| 41589_2 | TATGAGTGTGGAGTCGTTGCTTCAAGCGATTTTTCTGCCTCAGNNNAGATCGGAAG |
| 48216_1 | TATGAGTGTGGAGTCGTTGCAGACTCAAGCAATCCTTCCACTTNNNAGATCGGAAG |
| 48216_2 | TATGAGTGTGGAGTCGTTGCAGACTCAAGCAATCCTTCCACTTNNNAGATCGGAAG |
| 12845_1 | TATGAGTGTGGAGTCGTTGCCTCCTTAAGCCATTCTAAAGTTNNNAGATCGGAAGA |
| 12845_2 | TATGAGTGTGGAGTCGTTGCCTCCTTAAGCCATTCTAAAGTTNNNAGATCGGAAGA |
| 33144_1 | TATGAGTGTGGAGTCGTTGCGGATCACGAGGTCAAGAGATTGANNNAGATCGGAA |
| 33144_2 | TATGAGTGTGGAGTCGTTGCGGATCACGAGGTCAAGAGATTGANNNAGATCGGAA |
| 52383_1 | TATGAGTGTGGAGTCGTTGCGAGAGTGTGGAGGAGTTGACACNNNAGATCGGAA |
| 52383_2 | TATGAGTGTGGAGTCGTTGCGAGAGTGTGGAGGAGTTGACACNNNAGATCGGAA |
| 44618_1 | TATGAGTGTGGAGTCGTTGCTGGACATAGGCCTGACAGTAGTANNNAGATCGGAA |
| 44618_2 | TATGAGTGTGGAGTCGTTGCTGGACATAGGCCTGACAGTAGTANNNAGATCGGAA |
| 41814_1 | TATGAGTGTGGAGTCGTTGCCTCGGACCTTGTTTTCTGCTTTTNNNAGATCGGAAGA |
| 41814_2 | TATGAGTGTGGAGTCGTTGCCTCGGACCTTGTTTTCTGCTTTTNNNAGATCGGAAGA |
| 39783_1 | TATGAGTGTGGAGTCGTTGCCCTTCAAGTCCTAGTAAGGCTGGNNNAGATCGGAAG |
| 39783_2 | TATGAGTGTGGAGTCGTTGCCCTTCAAGTCCTAGTAAGGCTGGNNNAGATCGGAAG |
| 42058_1 | TATGAGTGTGGAGTCGTTGCCCATACTACCTCTATCATTTGCCCANNNAGATCGGAA |
| 42058_2 | TATGAGTGTGGAGTCGTTGCCCATACTACCTCTATCATTTGCCCANNNAGATCGGAA |
| 28812_1 | TATGAGTGTGGAGTCGTTGCACACAGCCTCAGAAACTTGTAAGNNNAGATCGGAAG |
| 28812_2 | TATGAGTGTGGAGTCGTTGCACACAGCCTCAGAAACTTGTAAGNNNAGATCGGAAG |
| 52722_1 | TATGAGTGTGGAGTCGTTGCCCAGGAGACAAGGTAAGAACGAANNNAGATCGGAA |

| | |
|---|---|
| 52722_2 | TATGAGTGTGGAGTCGTTGCCCAGGAGACAAGGTAAGAACGAANNNAGATCGGA |
| 29592_1 | TATGAGTGTGGAGTCGTTGCCCCCGCTTTCGTAGTTTTCATTTNNNAGATCGGAAGA |
| 29592_2 | TATGAGTGTGGAGTCGTTGCCCCCGCTTTCGTAGTTTTCATTTNNNAGATCGGAAGA |
| 33651_1 | TATGAGTGTGGAGTCGTTGCCAAGTAGCTGGGACTACAGGCNNNAGATCGGAAGA |
| 33651_2 | TATGAGTGTGGAGTCGTTGCCAAGTAGCTGGGACTACAGGCNNNAGATCGGAAGA |
| 47799_1 | TATGAGTGTGGAGTCGTTGCACAATCTCGGCTCACTGTAAACTNNNAGATCGGAAG |
| 47799_2 | TATGAGTGTGGAGTCGTTGCACAATCTCGGCTCACTGTAAACTNNNAGATCGGAAG |
| 31082_1 | TATGAGTGTGGAGTCGTTGCGTGTGTGTGTGTGTGTGTGTATCNNNAGATCGGAA |
| 31082_2 | TATGAGTGTGGAGTCGTTGCGTGTGTGTGTGTGTGTGTGTATCNNNAGATCGGAA |
| 46801_1 | TATGAGTGTGGAGTCGTTGCCTTTTTGAGACGGAGTCTTGCTCNNNAGATCGGAAG |
| 46801_2 | TATGAGTGTGGAGTCGTTGCCTTTTTGAGACGGAGTCTTGCTCNNNAGATCGGAAG |
| 53217_1 | TATGAGTGTGGAGTCGTTGCTTTGGCCTCAAACTTCAGGTGANNNAGATCGGAAGA |
| 53217_2 | TATGAGTGTGGAGTCGTTGCTTTGGCCTCAAACTTCAGGTGANNNAGATCGGAAGA |
| 29550_1 | TATGAGTGTGGAGTCGTTGCTTTGGGTTTTGGGTCTTTGAACCNNNAGATCGGAAG |
| 29550_2 | TATGAGTGTGGAGTCGTTGCTTTGGGTTTTGGGTCTTTGAACCNNNAGATCGGAAG |
| 5516_1 | TATGAGTGTGGAGTCGTTGCGTTGTTGTACGGAATGTGCAAGGNNNAGATCGGAA |
| 5516_2 | TATGAGTGTGGAGTCGTTGCGTTGTTGTACGGAATGTGCAAGGNNNAGATCGGAA |
| 32554_1 | TATGAGTGTGGAGTCGTTGCGAGTAGCTGGTACTACAGGTGTGNNNAGATCGGAA |
| 32554_2 | TATGAGTGTGGAGTCGTTGCGAGTAGCTGGTACTACAGGTGTGNNNAGATCGGAA |
| 51630_1 | TATGAGTGTGGAGTCGTTGCCCCCAAATCTCTATCCCTCACAGNNNAGATCGGAAGA |
| 51630_2 | TATGAGTGTGGAGTCGTTGCCCCCAAATCTCTATCCCTCACAGNNNAGATCGGAAGA |
| 32017_1 | TATGAGTGTGGAGTCGTTGCATTCAGAAGAGAGGTCAGGGGTANNNAGATCGGA |
| 32017_2 | TATGAGTGTGGAGTCGTTGCATTCAGAAGAGAGGTCAGGGGTANNNAGATCGGA |
| 41634_1 | TATGAGTGTGGAGTCGTTGCTCCCAAGTTCAACCAATTCTCCTNNNAGATCGGAAGA |
| 41634_2 | TATGAGTGTGGAGTCGTTGCTCCCAAGTTCAACCAATTCTCCTNNNAGATCGGAAGA |
| 45840_1 | TATGAGTGTGGAGTCGTTGCATGAGTAGGGGAGAGACTTTGGANNNAGATCGGA |
| 45840_2 | TATGAGTGTGGAGTCGTTGCATGAGTAGGGGAGAGACTTTGGANNNAGATCGGA |
| 43543_1 | TATGAGTGTGGAGTCGTTGCGTTTCACCATGTTAGCCAGGATGNNNAGATCGGAAG |
| 43543_2 | TATGAGTGTGGAGTCGTTGCGTTTCACCATGTTAGCCAGGATGNNNAGATCGGAAG |
| 38401_1 | TATGAGTGTGGAGTCGTTGCAAGAGTCAGTGTCATTGCTAGGANNNAGATCGGAA |
| 38401_2 | TATGAGTGTGGAGTCGTTGCAAGAGTCAGTGTCATTGCTAGGANNNAGATCGGAA |
| 45322_1 | TATGAGTGTGGAGTCGTTGCGGATGAGTTTGACTGTGAAGTGGNNNAGATCGGA |
| 45322_2 | TATGAGTGTGGAGTCGTTGCGGATGAGTTTGACTGTGAAGTGGNNNAGATCGGA |
| 46799_1 | TATGAGTGTGGAGTCGTTGCCTATCTTGGCTCACTGCAAGCTCNNNAGATCGGAAG |
| 46799_2 | TATGAGTGTGGAGTCGTTGCCTATCTTGGCTCACTGCAAGCTCNNNAGATCGGAAG |
| 52899_1 | TATGAGTGTGGAGTCGTTGCATGAGCTGTAGGTCCTTTGGCNNNAGATCGGAAGA |
| 52899_2 | TATGAGTGTGGAGTCGTTGCATGAGCTGTAGGTCCTTTGGCNNNAGATCGGAAGA |
| 13337_1 | TATGAGTGTGGAGTCGTTGCAGAAAAAAGCAAATTAAAACACCNNNAGATCGGAA |
| 13337_2 | TATGAGTGTGGAGTCGTTGCAGAAAAAAGCAAATTAAAACACCNNNAGATCGGAA |
| 40595_1 | TATGAGTGTGGAGTCGTTGCACTGAGTTGTAACCTACAGTGCCNNNAGATCGGAAG |
| 40595_2 | TATGAGTGTGGAGTCGTTGCACTGAGTTGTAACCTACAGTGCCNNNAGATCGGAAG |
| 49225_1 | TATGAGTGTGGAGTCGTTGCCTTCACGGGTTCAAGCAATTCTCNNNAGATCGGAAG |
| 49225_2 | TATGAGTGTGGAGTCGTTGCCTTCACGGGTTCAAGCAATTCTCNNNAGATCGGAAG |

| | |
|---|---|
| 36050_1 | TATGAGTGTGGAGTCGTTGCTGGTCCAGTAGGATACTCACCTTNNNAGATCGGAAG |
| 36050_2 | TATGAGTGTGGAGTCGTTGCTGGTCCAGTAGGATACTCACCTTNNNAGATCGGAAG |
| 50746_1 | TATGAGTGTGGAGTCGTTGCACACACCTACCACCTTTCGAAGNNNAGATCGGAAGA |
| 50746_2 | TATGAGTGTGGAGTCGTTGCACACACCTACCACCTTTCGAAGNNNAGATCGGAAGA |
| 51755_1 | TATGAGTGTGGAGTCGTTGCGTAATCTCTGGCAGACCTGGAAANNNAGATCGGAA |
| 51755_2 | TATGAGTGTGGAGTCGTTGCGTAATCTCTGGCAGACCTGGAAANNNAGATCGGAA |
| 31607_1 | TATGAGTGTGGAGTCGTTGCATGAGCTGTCATTCACACTCCAGNNNAGATCGGAAG |
| 31607_2 | TATGAGTGTGGAGTCGTTGCATGAGCTGTCATTCACACTCCAGNNNAGATCGGAAG |
| 51839_1 | TATGAGTGTGGAGTCGTTGCACCCATAACACAGAAGAGGAGTGNNNAGATCGGAA |
| 51839_2 | TATGAGTGTGGAGTCGTTGCACCCATAACACAGAAGAGGAGTGNNNAGATCGGAA |
| 30359_1 | TATGAGTGTGGAGTCGTTGCCCCAAAAGTCTCCCATAGACTTCCNNNAGATCGGAAG |
| 30359_2 | TATGAGTGTGGAGTCGTTGCCCCAAAAGTCTCCCATAGACTTCCNNNAGATCGGAAG |
| 2350_1 | TATGAGTGTGGAGTCGTTGCAAACGATCAACCTCTTTGCCTCCNNNAGATCGGAAGA |
| 2350_2 | TATGAGTGTGGAGTCGTTGCAAACGATCAACCTCTTTGCCTCCNNNAGATCGGAAGA |
| 31088_1 | TATGAGTGTGGAGTCGTTGCCCACATCGGACTAAAAACCACACNNNAGATCGGAAG |
| 31088_2 | TATGAGTGTGGAGTCGTTGCCCACATCGGACTAAAAACCACACNNNAGATCGGAAG |
| 9995_1 | TATGAGTGTGGAGTCGTTGCTGGGTTAGCTATCTGTGGAACCANNNAGATCGGAA |
| 9995_2 | TATGAGTGTGGAGTCGTTGCTGGGTTAGCTATCTGTGGAACCANNNAGATCGGAA |
| 31606_1 | TATGAGTGTGGAGTCGTTGCCCCCGACACTTTTACTCCCTAANNNAGATCGGAAGA |
| 31606_2 | TATGAGTGTGGAGTCGTTGCCCCCGACACTTTTACTCCCTAANNNAGATCGGAAGA |
| 49347_1 | TATGAGTGTGGAGTCGTTGCTGCCCAGGCTAGAAGAATTTTCTNNNAGATCGGAAG |
| 49347_2 | TATGAGTGTGGAGTCGTTGCTGCCCAGGCTAGAAGAATTTTCTNNNAGATCGGAAG |
| 31327_1 | TATGAGTGTGGAGTCGTTGCCTCACAGGAGCACTAATCCTACCNNNAGATCGGAAG |
| 31327_2 | TATGAGTGTGGAGTCGTTGCCTCACAGGAGCACTAATCCTACCNNNAGATCGGAAG |
| 35204_1 | TATGAGTGTGGAGTCGTTGCGTAGTTTTAGTGGAGACGGGGTTNNNAGATCGGAA |
| 35204_2 | TATGAGTGTGGAGTCGTTGCGTAGTTTTAGTGGAGACGGGGTTNNNAGATCGGAA |
| 46624_1 | TATGAGTGTGGAGTCGTTGCCAAGTAGCTGGGATTACACATGCNNNAGATCGGAA |
| 46624_2 | TATGAGTGTGGAGTCGTTGCCAAGTAGCTGGGATTACACATGCNNNAGATCGGAA |
| 29236_1 | TATGAGTGTGGAGTCGTTGCCACCATGGAATACTACTCAGCCANNNAGATCGGAAG |
| 29236_2 | TATGAGTGTGGAGTCGTTGCCACCATGGAATACTACTCAGCCANNNAGATCGGAAG |
| 51373_1 | TATGAGTGTGGAGTCGTTGCCGTCCCCTTATACTGTCCTCATGNNNAGATCGGAAGA |
| 51373_2 | TATGAGTGTGGAGTCGTTGCCGTCCCCTTATACTGTCCTCATGNNNAGATCGGAAGA |
| 35233_1 | TATGAGTGTGGAGTCGTTGCCAGCCAGTGTCCAAATTCACTTCNNNAGATCGGAAG |
| 35233_2 | TATGAGTGTGGAGTCGTTGCCAGCCAGTGTCCAAATTCACTTCNNNAGATCGGAAG |
| 32263_1 | TATGAGTGTGGAGTCGTTGCATGACCCATGATGAAACCCATGANNNAGATCGGAAG |
| 32263_2 | TATGAGTGTGGAGTCGTTGCATGACCCATGATGAAACCCATGANNNAGATCGGAAG |
| 46029_1 | TATGAGTGTGGAGTCGTTGCTTTACCACAGATGACAAGGGTGGNNNAGATCGGAA |
| 46029_2 | TATGAGTGTGGAGTCGTTGCTTTACCACAGATGACAAGGGTGGNNNAGATCGGAA |
| 35018_1 | TATGAGTGTGGAGTCGTTGCCCTCCTGGGTTCAAGTGATTCTCNNNAGATCGGAAG |
| 35018_2 | TATGAGTGTGGAGTCGTTGCCCTCCTGGGTTCAAGTGATTCTCNNNAGATCGGAAG |
| 47335_1 | TATGAGTGTGGAGTCGTTGCATCTCAGCTTACTGCAACCTCCNNNAGATCGGAAGA |
| 47335_2 | TATGAGTGTGGAGTCGTTGCATCTCAGCTTACTGCAACCTCCNNNAGATCGGAAGA |
| 13262_1 | TATGAGTGTGGAGTCGTTGCTCCGGCTAATTTTTGTATTTTTANNNAGATCGGAAGA |

| | |
|---|---|
| 13262_2 | TATGAGTGTGGAGTCGTTGCTCCGGCTAATTTTTGTATTTTTANNNAGATCGGAAG |
| 11291_1 | TATGAGTGTGGAGTCGTTGCGGCTCTCTTTAGGCTGGGTCTCNNNAGATCGGAAGA |
| 11291_2 | TATGAGTGTGGAGTCGTTGCGGCTCTCTTTAGGCTGGGTCTCNNNAGATCGGAAGA |
| 48843_1 | TATGAGTGTGGAGTCGTTGCCTGCGGTTTCTTCTCTGTCTCCNNNAGATCGGAAGAG |
| 48843_2 | TATGAGTGTGGAGTCGTTGCCTGCGGTTTCTTCTCTGTCTCCNNNAGATCGGAAGAG |
| 36961_1 | TATGAGTGTGGAGTCGTTGCATGAAGGTTTTGATGTGCTGACTNNNAGATCGGAA |
| 36961_2 | TATGAGTGTGGAGTCGTTGCATGAAGGTTTTGATGTGCTGACTNNNAGATCGGAA |
| 33666_1 | TATGAGTGTGGAGTCGTTGCGTTTCACCATGTTAGCCAGGATGNNNAGATCGGAAG |
| 33666_2 | TATGAGTGTGGAGTCGTTGCGTTTCACCATGTTAGCCAGGATGNNNAGATCGGAAG |
| 42162_1 | TATGAGTGTGGAGTCGTTGCAGGAAGTAAGTTTGCATTGTAGCANNNAGATCGGA |
| 42162_2 | TATGAGTGTGGAGTCGTTGCAGGAAGTAAGTTTGCATTGTAGCANNNAGATCGGA |
| 10016_1 | TATGAGTGTGGAGTCGTTGCAGTGAGACCCAGTCACCACATTANNNAGATCGGAAG |
| 10016_2 | TATGAGTGTGGAGTCGTTGCAGTGAGACCCAGTCACCACATTANNNAGATCGGAAG |
| 52816_1 | TATGAGTGTGGAGTCGTTGCCTCACAAAGAGCCCCAGTGATCNNNAGATCGGAAGA |
| 52816_2 | TATGAGTGTGGAGTCGTTGCCTCACAAAGAGCCCCAGTGATCNNNAGATCGGAAGA |
| 12715_1 | TATGAGTGTGGAGTCGTTGCAAGGATCTTCTTCAGACAGCNNNAGATCGGAAGAG |
| 12715_2 | TATGAGTGTGGAGTCGTTGCAAGGATCTTCTTCAGACAGCNNNAGATCGGAAGAG |
| 12670_1 | TATGAGTGTGGAGTCGTTGCTAATTTAGACCCTGAAAAAGAGANNNAGATCGGAA |
| 12670_2 | TATGAGTGTGGAGTCGTTGCTAATTTAGACCCTGAAAAAGAGANNNAGATCGGAA |
| 38173_1 | TATGAGTGTGGAGTCGTTGCATGAGAGAGAGCAGTTTCAAGCANNNAGATCGGAA |
| 38173_2 | TATGAGTGTGGAGTCGTTGCATGAGAGAGAGCAGTTTCAAGCANNNAGATCGGAA |
| 48952_1 | TATGAGTGTGGAGTCGTTGCCAAACTTCCTGTGGGTCACCTCNNNAGATCGGAAGA |
| 48952_2 | TATGAGTGTGGAGTCGTTGCCAAACTTCCTGTGGGTCACCTCNNNAGATCGGAAGA |
| 39185_1 | TATGAGTGTGGAGTCGTTGCCTGATGGGGAAGGATCACTTGAANNNAGATCGGAA |
| 39185_2 | TATGAGTGTGGAGTCGTTGCCTGATGGGGAAGGATCACTTGAANNNAGATCGGAA |
| 47092_1 | TATGAGTGTGGAGTCGTTGCTTCAAGTGATTCTCCTGCCTCAGNNNAGATCGGAAG |
| 47092_2 | TATGAGTGTGGAGTCGTTGCTTCAAGTGATTCTCCTGCCTCAGNNNAGATCGGAAG |
| 32577_1 | TATGAGTGTGGAGTCGTTGCCAATCTCAACTCACTGCAACCTCNNNAGATCGGAAGA |
| 32577_2 | TATGAGTGTGGAGTCGTTGCCAATCTCAACTCACTGCAACCTCNNNAGATCGGAAGA |
| 12644_1 | TATGAGTGTGGAGTCGTTGCCTTCATGTTCAGAAAAACTCCNNNAGATCGGAAGAG |
| 12644_2 | TATGAGTGTGGAGTCGTTGCCTTCATGTTCAGAAAAACTCCNNNAGATCGGAAGAG |
| 29330_1 | TATGAGTGTGGAGTCGTTGCAAGGGAAGGAGGACCATAGAGAANNNAGATCGGA |
| 29330_2 | TATGAGTGTGGAGTCGTTGCAAGGGAAGGAGGACCATAGAGAANNNAGATCGGA |
| 31281_1 | TATGAGTGTGGAGTCGTTGCATAGCACTTAGTGGTCATGGTGGNNNAGATCGGAA |
| 31281_2 | TATGAGTGTGGAGTCGTTGCATAGCACTTAGTGGTCATGGTGGNNNAGATCGGAA |
| 39504_1 | TATGAGTGTGGAGTCGTTGCTTGAGGTCAGGAGTTCAAAACCANNNAGATCGGAA |
| 39504_2 | TATGAGTGTGGAGTCGTTGCTTGAGGTCAGGAGTTCAAAACCANNNAGATCGGAA |
| 32738_1 | TATGAGTGTGGAGTCGTTGCGAGGCAGACCTTGTCTCTAAACANNNAGATCGGAAG |
| 32738_2 | TATGAGTGTGGAGTCGTTGCGAGGCAGACCTTGTCTCTAAACANNNAGATCGGAAG |
| 33974_1 | TATGAGTGTGGAGTCGTTGCGATCTCGGCTCACTGCAAACTCNNNAGATCGGAAGA |
| 33974_2 | TATGAGTGTGGAGTCGTTGCGATCTCGGCTCACTGCAAACTCNNNAGATCGGAAGA |
| 42697_1 | TATGAGTGTGGAGTCGTTGCATACTAGCAAACCGAATCCAGCANNNAGATCGGAAG |
| 42697_2 | TATGAGTGTGGAGTCGTTGCATACTAGCAAACCGAATCCAGCANNNAGATCGGAAG |

| | |
|---|---|
| 12574_1 | TATGAGTGTGGAGTCGTTGCCTGTTCACTTTCTCTTACAAGNNNAGATCGGAAGAG |
| 12574_2 | TATGAGTGTGGAGTCGTTGCCTGTTCACTTTCTCTTACAAGNNNAGATCGGAAGAG |
| 38129_1 | TATGAGTGTGGAGTCGTTGCTGTGGAATTCCAGTTGCTCTACANNNAGATCGGAAG |
| 38129_2 | TATGAGTGTGGAGTCGTTGCTGTGGAATTCCAGTTGCTCTACANNNAGATCGGAAG |
| 49258_1 | TATGAGTGTGGAGTCGTTGCTAGATTACAATGGCGCGATCTCANNNAGATCGGAAG |
| 49258_2 | TATGAGTGTGGAGTCGTTGCTAGATTACAATGGCGCGATCTCANNNAGATCGGAAG |
| 49008_1 | TATGAGTGTGGAGTCGTTGCCCACGCCCAGCTAATTTTTGTATNNNAGATCGGAAG |
| 49008_2 | TATGAGTGTGGAGTCGTTGCCCACGCCCAGCTAATTTTTGTATNNNAGATCGGAAG |
| 45452_1 | TATGAGTGTGGAGTCGTTGCCTCAAGTTATCCTCCCACCTCAGNNNAGATCGGAAGA |
| 45452_2 | TATGAGTGTGGAGTCGTTGCCTCAAGTTATCCTCCCACCTCAGNNNAGATCGGAAGA |
| 12647_1 | TATGAGTGTGGAGTCGTTGCTAATTGCATTTTAGTTCAAGGGNNNAGATCGGAAGA |
| 12647_2 | TATGAGTGTGGAGTCGTTGCTAATTGCATTTTAGTTCAAGGGNNNAGATCGGAAGA |
| 44843_1 | TATGAGTGTGGAGTCGTTGCAAGAGTTTGCCTAAACCTCCCATNNNAGATCGGAAG |
| 44843_2 | TATGAGTGTGGAGTCGTTGCAAGAGTTTGCCTAAACCTCCCATNNNAGATCGGAAG |
| 29031_1 | TATGAGTGTGGAGTCGTTGCATAAGGTGGATTCAGCCAACTGTNNNAGATCGGAA |
| 29031_2 | TATGAGTGTGGAGTCGTTGCATAAGGTGGATTCAGCCAACTGTNNNAGATCGGAA |
| 35308_1 | TATGAGTGTGGAGTCGTTGCCTGACCTCAGGTGATCCATCCANNNAGATCGGAAGA |
| 35308_2 | TATGAGTGTGGAGTCGTTGCCTGACCTCAGGTGATCCATCCANNNAGATCGGAAGA |
| 36891_1 | TATGAGTGTGGAGTCGTTGCAAACGTCGAACAATGCACATCAANNNAGATCGGAAG |
| 36891_2 | TATGAGTGTGGAGTCGTTGCAAACGTCGAACAATGCACATCAANNNAGATCGGAAG |
| 36927_1 | TATGAGTGTGGAGTCGTTGCGGACACAGGCACAGAAGGTATAANNNAGATCGGAA |
| 36927_2 | TATGAGTGTGGAGTCGTTGCGGACACAGGCACAGAAGGTATAANNNAGATCGGAA |
| 43555_1 | TATGAGTGTGGAGTCGTTGCCTCCTGAGTAGCTGGGATTACAGNNNAGATCGGAA |
| 43555_2 | TATGAGTGTGGAGTCGTTGCCTCCTGAGTAGCTGGGATTACAGNNNAGATCGGAA |
| 50598_1 | TATGAGTGTGGAGTCGTTGCCTGTCCCCTCACTTTCCATATCCNNNAGATCGGAAGA |
| 50598_2 | TATGAGTGTGGAGTCGTTGCCTGTCCCCTCACTTTCCATATCCNNNAGATCGGAAGA |
| 45565_1 | TATGAGTGTGGAGTCGTTGCGATGATAGGAACCCTGGAACGAANNNAGATCGGAA |
| 45565_2 | TATGAGTGTGGAGTCGTTGCGATGATAGGAACCCTGGAACGAANNNAGATCGGAA |
| 50580_1 | TATGAGTGTGGAGTCGTTGCCCCCCAACTAGATTTTCTGCAACNNNAGATCGGAAGA |
| 50580_2 | TATGAGTGTGGAGTCGTTGCCCCCCAACTAGATTTTCTGCAACNNNAGATCGGAAGA |
| 51922_1 | TATGAGTGTGGAGTCGTTGCTCAGTCTGAAGCTCTTCCAACTCNNNAGATCGGAAGA |
| 51922_2 | TATGAGTGTGGAGTCGTTGCTCAGTCTGAAGCTCTTCCAACTCNNNAGATCGGAAGA |
| 42050_1 | TATGAGTGTGGAGTCGTTGCACCAGGTTAGGTAAAACAGCTGTNNNAGATCGGAA |
| 42050_2 | TATGAGTGTGGAGTCGTTGCACCAGGTTAGGTAAAACAGCTGTNNNAGATCGGAA |
| 12944_1 | TATGAGTGTGGAGTCGTTGCTAGTGACACTGACAACCATCNNNAGATCGGAAGAG |
| 12944_2 | TATGAGTGTGGAGTCGTTGCTAGTGACACTGACAACCATCNNNAGATCGGAAGAG |
| 35611_1 | TATGAGTGTGGAGTCGTTGCTCATCCTCATGGTCTGTTCATGGNNNAGATCGGAAG |
| 35611_2 | TATGAGTGTGGAGTCGTTGCTCATCCTCATGGTCTGTTCATGGNNNAGATCGGAAG |
| 43902_1 | TATGAGTGTGGAGTCGTTGCGTGTGTGTGTGTGTGTGTGTGTATTNNNAGATCGGAA |
| 43902_2 | TATGAGTGTGGAGTCGTTGCGTGTGTGTGTGTGTGTGTGTGTATTNNNAGATCGGAA |
| 41903_1 | TATGAGTGTGGAGTCGTTGCGGTTCAAGCGAGTCTCCTATCTCNNNAGATCGGAAG |
| 41903_2 | TATGAGTGTGGAGTCGTTGCGGTTCAAGCGAGTCTCCTATCTCNNNAGATCGGAAG |
| 48234_1 | TATGAGTGTGGAGTCGTTGCTGATGATGATGTCTATCACATGGANNNAGATCGGAA |

| | |
|---|---|
| 48234_2 | TATGAGTGTGGAGTCGTTGCTGATGATGATGTCTATCACATGGANNNAGATCGGAA |
| 46422_1 | TATGAGTGTGGAGTCGTTGCTTTCAGTGATTCTCCTGCCTCAGNNNAGATCGGAAGA |
| 46422_2 | TATGAGTGTGGAGTCGTTGCTTTCAGTGATTCTCCTGCCTCAGNNNAGATCGGAAGA |
| 43093_1 | TATGAGTGTGGAGTCGTTGCTTCTCTCTCACAAAACCTAGGGCNNNAGATCGGAAGA |
| 43093_2 | TATGAGTGTGGAGTCGTTGCTTCTCTCTCACAAAACCTAGGGCNNNAGATCGGAAGA |
| 42324_1 | TATGAGTGTGGAGTCGTTGCCACAGAGTCACACATCCTTGAGANNNAGATCGGAAG |
| 42324_2 | TATGAGTGTGGAGTCGTTGCCACAGAGTCACACATCCTTGAGANNNAGATCGGAAG |
| 51397_1 | TATGAGTGTGGAGTCGTTGCTTGATACATCTGAAAGCCACCCANNNAGATCGGAAGA |
| 51397_2 | TATGAGTGTGGAGTCGTTGCTTGATACATCTGAAAGCCACCCANNNAGATCGGAAGA |
| 53117_1 | TATGAGTGTGGAGTCGTTGCCACCTCCTGGATTCGTTGATTTTNNNAGATCGGAAGA |
| 53117_2 | TATGAGTGTGGAGTCGTTGCCACCTCCTGGATTCGTTGATTTTNNNAGATCGGAAGA |
| 34883_1 | TATGAGTGTGGAGTCGTTGCGGCAAAGTCTGTTTCTGACTCTGNNNAGATCGGAAG |
| 34883_2 | TATGAGTGTGGAGTCGTTGCGGCAAAGTCTGTTTCTGACTCTGNNNAGATCGGAAG |
| 37702_1 | TATGAGTGTGGAGTCGTTGCTGAGCTCCAAAATTGAATCAGCANNNAGATCGGAAG |
| 37702_2 | TATGAGTGTGGAGTCGTTGCTGAGCTCCAAAATTGAATCAGCANNNAGATCGGAAG |
| 40117_1 | TATGAGTGTGGAGTCGTTGCAACTGGACGGAGAATGAGTTTGANNNAGATCGGAA |
| 40117_2 | TATGAGTGTGGAGTCGTTGCAACTGGACGGAGAATGAGTTTGANNNAGATCGGAA |
| 45513_1 | TATGAGTGTGGAGTCGTTGCGGAAGGTGGGAACTTATGGGATTNNNAGATCGGAA |
| 45513_2 | TATGAGTGTGGAGTCGTTGCGGAAGGTGGGAACTTATGGGATTNNNAGATCGGAA |
| 49753_1 | TATGAGTGTGGAGTCGTTGCGCAGCCAATCTCTCTCTCTGAATNNNAGATCGGAAGA |
| 49753_2 | TATGAGTGTGGAGTCGTTGCGCAGCCAATCTCTCTCTCTGAATNNNAGATCGGAAGA |
| 48743_1 | TATGAGTGTGGAGTCGTTGCTGCCCCCAACTTGTTTCTATCTTNNNAGATCGGAAGA |
| 48743_2 | TATGAGTGTGGAGTCGTTGCTGCCCCCAACTTGTTTCTATCTTNNNAGATCGGAAGA |
| 29949_1 | TATGAGTGTGGAGTCGTTGCAGGAAGTGAAGTATTTGCATATGACACNNNAGATCG |
| 29949_2 | TATGAGTGTGGAGTCGTTGCAGGAAGTGAAGTATTTGCATATGACACNNNAGATCG |
| 34346_1 | TATGAGTGTGGAGTCGTTGCGCTCTATTTGCCTTCAGTGCTCTNNNAGATCGGAAGA |
| 34346_2 | TATGAGTGTGGAGTCGTTGCGCTCTATTTGCCTTCAGTGCTCTNNNAGATCGGAAGA |
| 50661_1 | TATGAGTGTGGAGTCGTTGCCTCCTTTGATCACACACACACACNNNAGATCGGAAGA |
| 50661_2 | TATGAGTGTGGAGTCGTTGCCTCCTTTGATCACACACACACACNNNAGATCGGAAGA |
| 50560_1 | TATGAGTGTGGAGTCGTTGCCCCTTCTCCCCCAAAATACTCAANNNAGATCGGAAGA |
| 50560_2 | TATGAGTGTGGAGTCGTTGCCCCTTCTCCCCCAAAATACTCAANNNAGATCGGAAGA |
| 32296_1 | TATGAGTGTGGAGTCGTTGCAGGGCACAGACAAACAAAAAGACNNNAGATCGGAA |
| 32296_2 | TATGAGTGTGGAGTCGTTGCAGGGCACAGACAAACAAAAAGACNNNAGATCGGAA |
| 13159_1 | TATGAGTGTGGAGTCGTTGCCAGAGAAACCAAACTAACTTTAGNNNAGATCGGAAG |
| 13159_2 | TATGAGTGTGGAGTCGTTGCCAGAGAAACCAAACTAACTTTAGNNNAGATCGGAAG |
| 44612_1 | TATGAGTGTGGAGTCGTTGCCTTGCGGTTTCGAGTCTGAATACNNNAGATCGGAAG |
| 44612_2 | TATGAGTGTGGAGTCGTTGCCTTGCGGTTTCGAGTCTGAATACNNNAGATCGGAAG |
| 44702_1 | TATGAGTGTGGAGTCGTTGCTACCCAACTCCACTTTTTGTCCANNNAGATCGGAAGA |
| 44702_2 | TATGAGTGTGGAGTCGTTGCTACCCAACTCCACTTTTTGTCCANNNAGATCGGAAGA |
| 49191_1 | TATGAGTGTGGAGTCGTTGCCTTCCGAGTAGCTGGGATTACAGNNNAGATCGGAAG |
| 49191_2 | TATGAGTGTGGAGTCGTTGCCTTCCGAGTAGCTGGGATTACAGNNNAGATCGGAAG |
| 49300_1 | TATGAGTGTGGAGTCGTTGCAAACTGCTGGGATTACAGGCTTNNNAGATCGGAAGA |
| 49300_2 | TATGAGTGTGGAGTCGTTGCAAACTGCTGGGATTACAGGCTTNNNAGATCGGAAGA |

| | |
|---|---|
| 30600_1 | TATGAGTGTGGAGTCGTTGCAGCCCTATGGGAACTAACTTTGAGNNNAGATCGGAA |
| 30600_2 | TATGAGTGTGGAGTCGTTGCAGCCCTATGGGAACTAACTTTGAGNNNAGATCGGAA |
| 29775_1 | TATGAGTGTGGAGTCGTTGCTGTGCAGTGGGGAGATTTTAGATNNNAGATCGGAA |
| 29775_2 | TATGAGTGTGGAGTCGTTGCTGTGCAGTGGGGAGATTTTAGATNNNAGATCGGAA |
| 33559_1 | TATGAGTGTGGAGTCGTTGCAGTAGCTGGGACTATAGGAGCCNNNAGATCGGAAG |
| 33559_2 | TATGAGTGTGGAGTCGTTGCAGTAGCTGGGACTATAGGAGCCNNNAGATCGGAAG |
| 47345_1 | TATGAGTGTGGAGTCGTTGCGGGCCAGGAGAATCCAAAATTTCNNNAGATCGGAA |
| 47345_2 | TATGAGTGTGGAGTCGTTGCGGGCCAGGAGAATCCAAAATTTCNNNAGATCGGAA |
| 44049_1 | TATGAGTGTGGAGTCGTTGCTTCACTAGGAGGATGGGACCTAGNNNAGATCGGAA |
| 44049_2 | TATGAGTGTGGAGTCGTTGCTTCACTAGGAGGATGGGACCTAGNNNAGATCGGAA |
| 42782_1 | TATGAGTGTGGAGTCGTTGCATTCACATCTGCCTCCATCTCAGNNNAGATCGGAAGA |
| 42782_2 | TATGAGTGTGGAGTCGTTGCATTCACATCTGCCTCCATCTCAGNNNAGATCGGAAGA |
| 48671_1 | TATGAGTGTGGAGTCGTTGCCTGTTGCTGACCAAGGTGATTTTNNNAGATCGGAAG |
| 48671_2 | TATGAGTGTGGAGTCGTTGCCTGTTGCTGACCAAGGTGATTTTNNNAGATCGGAAG |
| 225_1 | TATGAGTGTGGAGTCGTTGCCAGTTGCTCGAAGGGGCTTCCCNNNAGATCGGAAGA |
| 225_2 | TATGAGTGTGGAGTCGTTGCCAGTTGCTCGAAGGGGCTTCCCNNNAGATCGGAAGA |
| 48652_1 | TATGAGTGTGGAGTCGTTGCTGAGACAGAGTCTTGCTGTGTCNNNAGATCGGAAGA |
| 48652_2 | TATGAGTGTGGAGTCGTTGCTGAGACAGAGTCTTGCTGTGTCNNNAGATCGGAAGA |
| 41283_1 | TATGAGTGTGGAGTCGTTGCGAGGGACTGCTAGATCAATTCGTNNNAGATCGGAA |
| 41283_2 | TATGAGTGTGGAGTCGTTGCGAGGGACTGCTAGATCAATTCGTNNNAGATCGGAA |
| 42517_1 | TATGAGTGTGGAGTCGTTGCGCTCCCAAATTGCAGAGTACATGNNNAGATCGGAAG |
| 42517_2 | TATGAGTGTGGAGTCGTTGCGCTCCCAAATTGCAGAGTACATGNNNAGATCGGAAG |
| 50205_1 | TATGAGTGTGGAGTCGTTGCGGCTAGGTGGGGTAAAATCTTCANNNAGATCGGAA |
| 50205_2 | TATGAGTGTGGAGTCGTTGCGGCTAGGTGGGGTAAAATCTTCANNNAGATCGGAA |
| 49573_1 | TATGAGTGTGGAGTCGTTGCCCAGACTCGAGGGATTCTTGTGNNNAGATCGGAAGA |
| 49573_2 | TATGAGTGTGGAGTCGTTGCCCAGACTCGAGGGATTCTTGTGNNNAGATCGGAAGA |
| 39588_1 | TATGAGTGTGGAGTCGTTGCACTATCATTCGATCCAGCAACCCNNNAGATCGGAAGA |
| 39588_2 | TATGAGTGTGGAGTCGTTGCACTATCATTCGATCCAGCAACCCNNNAGATCGGAAGA |
| 40596_1 | TATGAGTGTGGAGTCGTTGCGAGGCTGTGTTTAGAGGATCACTNNNAGATCGGAA |
| 40596_2 | TATGAGTGTGGAGTCGTTGCGAGGCTGTGTTTAGAGGATCACTNNNAGATCGGAA |
| 49792_1 | TATGAGTGTGGAGTCGTTGCAGGGGAAAACTGGGCTATATTGGNNNAGATCGGAA |
| 49792_2 | TATGAGTGTGGAGTCGTTGCAGGGGAAAACTGGGCTATATTGGNNNAGATCGGAA |
| 42748_1 | TATGAGTGTGGAGTCGTTGCACACACACACACACACACAAATTNNNAGATCGGAAGA |
| 42748_2 | TATGAGTGTGGAGTCGTTGCACACACACACACACACACAAATTNNNAGATCGGAAGA |
| 30177_1 | TATGAGTGTGGAGTCGTTGCAAAAGCTACAGCCCTAACCATCANNNAGATCGGAAGA |
| 30177_2 | TATGAGTGTGGAGTCGTTGCAAAAGCTACAGCCCTAACCATCANNNAGATCGGAAGA |
| 42910_1 | TATGAGTGTGGAGTCGTTGCCCTGGTTTCAAGCAATTCTCCTGNNNAGATCGGAAGA |
| 42910_2 | TATGAGTGTGGAGTCGTTGCCCTGGTTTCAAGCAATTCTCCTGNNNAGATCGGAAGA |
| 30894_1 | TATGAGTGTGGAGTCGTTGCCGGAGAGAGAGAGAGAGAGAGAGNNNAGATCGGA |
| 30894_2 | TATGAGTGTGGAGTCGTTGCCGGAGAGAGAGAGAGAGAGAGAGNNNAGATCGGA |
| 30720_1 | TATGAGTGTGGAGTCGTTGCGGTAACAACAGCAAGACCCTGTGNNNAGATCGGAA |
| 30720_2 | TATGAGTGTGGAGTCGTTGCGGTAACAACAGCAAGACCCTGTGNNNAGATCGGAA |
| 47968_1 | TATGAGTGTGGAGTCGTTGCCTCTGGAGTAGCTGGGATTACAGNNNAGATCGGAA |

| | |
|---|---|
| 47968_2 | TATGAGTGTGGAGTCGTTGCCTCTGGAGTAGCTGGGATTACAGNNNAGATCGGAA |
| 36705_1 | TATGAGTGTGGAGTCGTTGCAAATAGTCACGGGGGACAAAGAANNNAGATCGGA |
| 36705_2 | TATGAGTGTGGAGTCGTTGCAAATAGTCACGGGGGACAAAGAANNNAGATCGGA |
| 40923_1 | TATGAGTGTGGAGTCGTTGCCAATGCCATACTGTGCAACTCTGNNNAGATCGGAAG |
| 40923_2 | TATGAGTGTGGAGTCGTTGCCAATGCCATACTGTGCAACTCTGNNNAGATCGGAAG |
| 30803_1 | TATGAGTGTGGAGTCGTTGCGCAAGTCACTGAAAATGCCTCTTNNNAGATCGGAAG |
| 30803_2 | TATGAGTGTGGAGTCGTTGCGCAAGTCACTGAAAATGCCTCTTNNNAGATCGGAAG |
| 49639_1 | TATGAGTGTGGAGTCGTTGCTCAAGCGATTCTCCTATCTCAGCNNNAGATCGGAAG |
| 49639_2 | TATGAGTGTGGAGTCGTTGCTCAAGCGATTCTCCTATCTCAGCNNNAGATCGGAAG |
| 33009_1 | TATGAGTGTGGAGTCGTTGCGATCTCGGCTCACTACAAGCTCNNNAGATCGGAAGA |
| 33009_2 | TATGAGTGTGGAGTCGTTGCGATCTCGGCTCACTACAAGCTCNNNAGATCGGAAGA |
| 39485_1 | TATGAGTGTGGAGTCGTTGCTTTCTTTTTAGTTGAGCGTGTGCNNNAGATCGGAAG |
| 39485_2 | TATGAGTGTGGAGTCGTTGCTTTCTTTTTAGTTGAGCGTGTGCNNNAGATCGGAAG |
| 31259_1 | TATGAGTGTGGAGTCGTTGCTACACACACACACACAAACACACNNNAGATCGGAAG |
| 31259_2 | TATGAGTGTGGAGTCGTTGCTACACACACACACACAAACACACNNNAGATCGGAAG |
| 42958_1 | TATGAGTGTGGAGTCGTTGCCCTCCTGAGTTCACACCATTCTCNNNAGATCGGAAGA |
| 42958_2 | TATGAGTGTGGAGTCGTTGCCCTCCTGAGTTCACACCATTCTCNNNAGATCGGAAGA |
| 46891_1 | TATGAGTGTGGAGTCGTTGCAGAGATTCTCTTGTGTGCCATCANNNAGATCGGAAG |
| 46891_2 | TATGAGTGTGGAGTCGTTGCAGAGATTCTCTTGTGTGCCATCANNNAGATCGGAAG |
| 8151_1 | TATGAGTGTGGAGTCGTTGCGTAGCACATACACACCTGTGGTNNNAGATCGGAAGA |
| 8151_2 | TATGAGTGTGGAGTCGTTGCGTAGCACATACACACCTGTGGTNNNAGATCGGAAGA |
| 44772_1 | TATGAGTGTGGAGTCGTTGCACATAAAGCAGTTCTGAGGCTGANNNAGATCGGAA |
| 44772_2 | TATGAGTGTGGAGTCGTTGCACATAAAGCAGTTCTGAGGCTGANNNAGATCGGAA |
| 50080_1 | TATGAGTGTGGAGTCGTTGCTATAAAACAGACCCACAGCCCTCNNNAGATCGGAAG |
| 50080_2 | TATGAGTGTGGAGTCGTTGCTATAAAACAGACCCACAGCCCTCNNNAGATCGGAAG |
| 29730_1 | TATGAGTGTGGAGTCGTTGCCTTTCCTACTGCCTGAACTGTCTNNNAGATCGGAAGA |
| 29730_2 | TATGAGTGTGGAGTCGTTGCCTTTCCTACTGCCTGAACTGTCTNNNAGATCGGAAGA |
| 42609_1 | TATGAGTGTGGAGTCGTTGCATTTTGTGTGTGTGTGTGTGTNNNAGATCGGAA |
| 42609_2 | TATGAGTGTGGAGTCGTTGCATTTTGTGTGTGTGTGTGTGTNNNAGATCGGAA |
| 9455_1 | TATGAGTGTGGAGTCGTTGCGGTAGGCGCCATTTCCTAGAAGNNNAGATCGGAAG |
| 9455_2 | TATGAGTGTGGAGTCGTTGCGGTAGGCGCCATTTCCTAGAAGNNNAGATCGGAAG |
| 52802_1 | TATGAGTGTGGAGTCGTTGCCAGCGGTGAACAAAATCCCTTTANNNAGATCGGAAG |
| 52802_2 | TATGAGTGTGGAGTCGTTGCCAGCGGTGAACAAAATCCCTTTANNNAGATCGGAAG |
| 3165_1 | TATGAGTGTGGAGTCGTTGCCTGTTCTCCTTCCCTGTCCCTAANNNAGATCGGAAGA |
| 3165_2 | TATGAGTGTGGAGTCGTTGCCTGTTCTCCTTCCCTGTCCCTAANNNAGATCGGAAGA |
| 42632_1 | TATGAGTGTGGAGTCGTTGCCTAGTTGTGTGTGTGTGTGTGTGNNNAGATCGGAA |
| 42632_2 | TATGAGTGTGGAGTCGTTGCCTAGTTGTGTGTGTGTGTGTGTGNNNAGATCGGAA |
| 44782_1 | TATGAGTGTGGAGTCGTTGCGTGAAAGAAATTCTAAGGGCCGGNNNAGATCGGAA |
| 44782_2 | TATGAGTGTGGAGTCGTTGCGTGAAAGAAATTCTAAGGGCCGGNNNAGATCGGAA |
| 51317_1 | TATGAGTGTGGAGTCGTTGCTGAACCACCATGGCATGTGTATANNNAGATCGGAAG |
| 51317_2 | TATGAGTGTGGAGTCGTTGCTGAACCACCATGGCATGTGTATANNNAGATCGGAAG |
| 35577_1 | TATGAGTGTGGAGTCGTTGCTGACACACAATCTCAGCATCTCANNNAGATCGGAAG |
| 35577_2 | TATGAGTGTGGAGTCGTTGCTGACACACAATCTCAGCATCTCANNNAGATCGGAAG |

| | |
|---|---|
| 52716_1 | TATGAGTGTGGAGTCGTTGCAGGACTTGATTTGTTGAGACCCANNNAGATCGGAAG |
| 52716_2 | TATGAGTGTGGAGTCGTTGCAGGACTTGATTTGTTGAGACCCANNNAGATCGGAAG |
| 45839_1 | TATGAGTGTGGAGTCGTTGCTATGCAAGATGGGGACAATTTGCNNNAGATCGGAA |
| 45839_2 | TATGAGTGTGGAGTCGTTGCTATGCAAGATGGGGACAATTTGCNNNAGATCGGAA |
| 32642_1 | TATGAGTGTGGAGTCGTTGCGAGATTACAGGCGCCTAGCTAATNNNAGATCGGAA |
| 32642_2 | TATGAGTGTGGAGTCGTTGCGAGATTACAGGCGCCTAGCTAATNNNAGATCGGAA |
| 12479_1 | TATGAGTGTGGAGTCGTTGCAATAGTCCAATGAGTGTGTGNNNAGATCGGAAGAG |
| 12479_2 | TATGAGTGTGGAGTCGTTGCAATAGTCCAATGAGTGTGTGNNNAGATCGGAAGAG |
| 48669_1 | TATGAGTGTGGAGTCGTTGCGTTCTCAGATCTGACCCCAAACTNNNAGATCGGAAG |
| 48669_2 | TATGAGTGTGGAGTCGTTGCGTTCTCAGATCTGACCCCAAACTNNNAGATCGGAAG |
| 40627_1 | TATGAGTGTGGAGTCGTTGCAATCACCAATATCAGACCAGGCANNNAGATCGGAAG |
| 40627_2 | TATGAGTGTGGAGTCGTTGCAATCACCAATATCAGACCAGGCANNNAGATCGGAAG |
| 47413_1 | TATGAGTGTGGAGTCGTTGCTAGGAGGCTGGACAAACTTGTTTNNNAGATCGGAA |
| 47413_2 | TATGAGTGTGGAGTCGTTGCTAGGAGGCTGGACAAACTTGTTTNNNAGATCGGAA |
| 36285_1 | TATGAGTGTGGAGTCGTTGCGCCCAAAGTCATATAGCTACCGANNNAGATCGGAAG |
| 36285_2 | TATGAGTGTGGAGTCGTTGCGCCCAAAGTCATATAGCTACCGANNNAGATCGGAAG |
| 9008_1 | TATGAGTGTGGAGTCGTTGCTTCGGCTATTTCCTACCCGATCTNNNAGATCGGAAGA |
| 9008_2 | TATGAGTGTGGAGTCGTTGCTTCGGCTATTTCCTACCCGATCTNNNAGATCGGAAGA |
| 51044_1 | TATGAGTGTGGAGTCGTTGCTCCACCTTGCCAATGTATAAGCTNNNAGATCGGAAG |
| 51044_2 | TATGAGTGTGGAGTCGTTGCTCCACCTTGCCAATGTATAAGCTNNNAGATCGGAAG |
| 43644_1 | TATGAGTGTGGAGTCGTTGCGGATCTGGGGGAGGGATTTGNNNAGATCGGAAGA |
| 43644_2 | TATGAGTGTGGAGTCGTTGCGGATCTGGGGGAGGGATTTGNNNAGATCGGAAGA |
| 45903_1 | TATGAGTGTGGAGTCGTTGCCCTCAAGTAGCTGGGACTACAGNNNAGATCGGAAG |
| 45903_2 | TATGAGTGTGGAGTCGTTGCCCTCAAGTAGCTGGGACTACAGNNNAGATCGGAAG |
| 29219_1 | TATGAGTGTGGAGTCGTTGCGGGGTGACAATACAGAGCTCTAGNNNAGATCGGAA |
| 29219_2 | TATGAGTGTGGAGTCGTTGCGGGGTGACAATACAGAGCTCTAGNNNAGATCGGAA |
| 31275_1 | TATGAGTGTGGAGTCGTTGCTCTTGCCACTAACTAGCTTCCTGNNNAGATCGGAAGA |
| 31275_2 | TATGAGTGTGGAGTCGTTGCTCTTGCCACTAACTAGCTTCCTGNNNAGATCGGAAGA |
| 51046_1 | TATGAGTGTGGAGTCGTTGCAAGCCATTGAGTTTGACCTCCTANNNAGATCGGAAG |
| 51046_2 | TATGAGTGTGGAGTCGTTGCAAGCCATTGAGTTTGACCTCCTANNNAGATCGGAAG |
| 47072_1 | TATGAGTGTGGAGTCGTTGCGGTTCAAGTGATTTTCCTGCCTCNNNAGATCGGAAG |
| 47072_2 | TATGAGTGTGGAGTCGTTGCGGTTCAAGTGATTTTCCTGCCTCNNNAGATCGGAAG |
| 12919_1 | TATGAGTGTGGAGTCGTTGCCTAGTAAGTACGGATAATGGATCNNNAGATCGGAA |
| 12919_2 | TATGAGTGTGGAGTCGTTGCCTAGTAAGTACGGATAATGGATCNNNAGATCGGAA |
| 9969_1 | TATGAGTGTGGAGTCGTTGCACCCAGAATCAACAGAAACAAACANNNAGATCGGAA |
| 9969_2 | TATGAGTGTGGAGTCGTTGCACCCAGAATCAACAGAAACAAACANNNAGATCGGAA |
| 36859_1 | TATGAGTGTGGAGTCGTTGCGGGAGTTGAGGAATGACTGCTAANNNAGATCGGAA |
| 36859_2 | TATGAGTGTGGAGTCGTTGCGGGAGTTGAGGAATGACTGCTAANNNAGATCGGAA |
| 34862_1 | TATGAGTGTGGAGTCGTTGCGTTCAAGCAATTCTCCTGCCTCNNNAGATCGGAAGA |
| 34862_2 | TATGAGTGTGGAGTCGTTGCGTTCAAGCAATTCTCCTGCCTCNNNAGATCGGAAGA |
| 31785_1 | TATGAGTGTGGAGTCGTTGCTCTCACCTCTAAGGATCTCACGTNNNAGATCGGAAG |
| 31785_2 | TATGAGTGTGGAGTCGTTGCTCTCACCTCTAAGGATCTCACGTNNNAGATCGGAAG |
| 33860_1 | TATGAGTGTGGAGTCGTTGCAGGAAAGCTGAACTCCAGAAGAGNNNAGATCGGAA |

| | |
|---|---|
| 33860_2 | TATGAGTGTGGAGTCGTTGCAGGAAAGCTGAACTCCAGAAGAGNNNAGATCGGAA |
| 41666_1 | TATGAGTGTGGAGTCGTTGCGGGTTCAAGTGATTCTCCTGTCTNNNAGATCGGAAG |
| 41666_2 | TATGAGTGTGGAGTCGTTGCGGGTTCAAGTGATTCTCCTGTCTNNNAGATCGGAAG |
| 46590_1 | TATGAGTGTGGAGTCGTTGCAAATGTCTCAGCAAGCAGAAACCNNNAGATCGGAAG |
| 46590_2 | TATGAGTGTGGAGTCGTTGCAAATGTCTCAGCAAGCAGAAACCNNNAGATCGGAAG |
| 8975_1 | TATGAGTGTGGAGTCGTTGCGCCATGCTACAAGCCCTTATTCTNNNAGATCGGAAGA |
| 8975_2 | TATGAGTGTGGAGTCGTTGCGCCATGCTACAAGCCCTTATTCTNNNAGATCGGAAGA |
| 37206_1 | TATGAGTGTGGAGTCGTTGCAGTTTCAGTAGGAATGGGACCAGNNNAGATCGGAA |
| 37206_2 | TATGAGTGTGGAGTCGTTGCAGTTTCAGTAGGAATGGGACCAGNNNAGATCGGAA |
| 35012_1 | TATGAGTGTGGAGTCGTTGCCCGAGTAGCTGGGATTACAAGTGNNNAGATCGGAAG |
| 35012_2 | TATGAGTGTGGAGTCGTTGCCCGAGTAGCTGGGATTACAAGTGNNNAGATCGGAAG |
| 41989_1 | TATGAGTGTGGAGTCGTTGCCACTTCCAGGGTTCAAGCAATTCNNNAGATCGGAAGA |
| 41989_2 | TATGAGTGTGGAGTCGTTGCCACTTCCAGGGTTCAAGCAATTCNNNAGATCGGAAGA |
| 29911_1 | TATGAGTGTGGAGTCGTTGCGACTGTGCCAAGCCCTTGAATATNNNAGATCGGAAG |
| 29911_2 | TATGAGTGTGGAGTCGTTGCGACTGTGCCAAGCCCTTGAATATNNNAGATCGGAAG |
| 10633_1 | TATGAGTGTGGAGTCGTTGCCAGGGGCACACTCACACATAAATNNNAGATCGGAAG |
| 10633_2 | TATGAGTGTGGAGTCGTTGCCAGGGGCACACTCACACATAAATNNNAGATCGGAAG |
| 40298_1 | TATGAGTGTGGAGTCGTTGCTACACCAACAAGAGACAAGCAGANNNAGATCGGAAG |
| 40298_2 | TATGAGTGTGGAGTCGTTGCTACACCAACAAGAGACAAGCAGANNNAGATCGGAAG |
| 46105_1 | TATGAGTGTGGAGTCGTTGCCTCCTGAGTAGCTGGGATTACAGNNNAGATCGGAAG |
| 46105_2 | TATGAGTGTGGAGTCGTTGCCTCCTGAGTAGCTGGGATTACAGNNNAGATCGGAAG |
| 51560_1 | TATGAGTGTGGAGTCGTTGCTTGGCAAAAACATTCCACTGTGTNNNAGATCGGAAG |
| 51560_2 | TATGAGTGTGGAGTCGTTGCTTGGCAAAAACATTCCACTGTGTNNNAGATCGGAAG |
| 29806_1 | TATGAGTGTGGAGTCGTTGCCAGAGGCATAAACTGAGGTAGAAGNNNAGATCGGA |
| 29806_2 | TATGAGTGTGGAGTCGTTGCCAGAGGCATAAACTGAGGTAGAAGNNNAGATCGGA |
| 51736_1 | TATGAGTGTGGAGTCGTTGCGACCAGGGGGAAGGTAAACACNNNAGATCGGAAGA |
| 51736_2 | TATGAGTGTGGAGTCGTTGCGACCAGGGGGAAGGTAAACACNNNAGATCGGAAGA |
| 31396_1 | TATGAGTGTGGAGTCGTTGCGGCTGTGTGAAAATATGTGCCAANNNAGATCGGAAG |
| 31396_2 | TATGAGTGTGGAGTCGTTGCGGCTGTGTGAAAATATGTGCCAANNNAGATCGGAAG |
| 51042_1 | TATGAGTGTGGAGTCGTTGCGCTTCAGTCCCTAATCACCTAGGNNNAGATCGGAAGA |
| 51042_2 | TATGAGTGTGGAGTCGTTGCGCTTCAGTCCCTAATCACCTAGGNNNAGATCGGAAGA |
| 46932_1 | TATGAGTGTGGAGTCGTTGCCTTCATTCCTGAGAGGTCTGGACNNNAGATCGGAAG |
| 46932_2 | TATGAGTGTGGAGTCGTTGCCTTCATTCCTGAGAGGTCTGGACNNNAGATCGGAAG |
| 34788_1 | TATGAGTGTGGAGTCGTTGCCCCCACCTGGCTAATTTTTGTTTNNNAGATCGGAAGA |
| 34788_2 | TATGAGTGTGGAGTCGTTGCCCCCACCTGGCTAATTTTTGTTTNNNAGATCGGAAGA |
| 38233_1 | TATGAGTGTGGAGTCGTTGCAGAAAACCTCAGGCCAATATCCTNNNAGATCGGAAG |
| 38233_2 | TATGAGTGTGGAGTCGTTGCAGAAAACCTCAGGCCAATATCCTNNNAGATCGGAAG |
| 49302_1 | TATGAGTGTGGAGTCGTTGCGCTCACTATAACCTCCGACTCTCNNNAGATCGGAAGA |
| 49302_2 | TATGAGTGTGGAGTCGTTGCGCTCACTATAACCTCCGACTCTCNNNAGATCGGAAGA |
| 9554_1 | TATGAGTGTGGAGTCGTTGCGGAGGTGATGGGTCTTTGTCATCNNNAGATCGGAAG |
| 9554_2 | TATGAGTGTGGAGTCGTTGCGGAGGTGATGGGTCTTTGTCATCNNNAGATCGGAAG |
| 48717_1 | TATGAGTGTGGAGTCGTTGCTTCCAGGCTCAAGTGTAAAGACANNNAGATCGGAAG |
| 48717_2 | TATGAGTGTGGAGTCGTTGCTTCCAGGCTCAAGTGTAAAGACANNNAGATCGGAAG |

| | |
|---|---|
| 38455_1 | TATGAGTGTGGAGTCGTTGCCTCCCGTAATCCCAGCACTTTNNNAGATCGGAAGAG |
| 38455_2 | TATGAGTGTGGAGTCGTTGCCTCCCGTAATCCCAGCACTTTNNNAGATCGGAAGAG |
| 34409_1 | TATGAGTGTGGAGTCGTTGCCAGAAGAAAGGTCTAGGGCTGAANNNAGATCGGAA |
| 34409_2 | TATGAGTGTGGAGTCGTTGCCAGAAGAAAGGTCTAGGGCTGAANNNAGATCGGAA |
| 41014_1 | TATGAGTGTGGAGTCGTTGCTCAGATGAAAGAAAAAGCTGAGACANNNAGATCGG |
| 41014_2 | TATGAGTGTGGAGTCGTTGCTCAGATGAAAGAAAAAGCTGAGACANNNAGATCGG |
| 12977_1 | TATGAGTGTGGAGTCGTTGCAAAGACCAACAGTTCTTTTTGNNNAGATCGGAAGAG |
| 12977_2 | TATGAGTGTGGAGTCGTTGCAAAGACCAACAGTTCTTTTTGNNNAGATCGGAAGAG |
| 40019_1 | TATGAGTGTGGAGTCGTTGCAAAAAAACCCTCCAGTAACCCCANNNAGATCGGAAG |
| 40019_2 | TATGAGTGTGGAGTCGTTGCAAAAAAACCCTCCAGTAACCCCANNNAGATCGGAAG |
| 31586_1 | TATGAGTGTGGAGTCGTTGCACATGGTTGATCAAGATCGGAGTNNNAGATCGGAA |
| 31586_2 | TATGAGTGTGGAGTCGTTGCACATGGTTGATCAAGATCGGAGTNNNAGATCGGAA |
| 29954_1 | TATGAGTGTGGAGTCGTTGCAAGCCAAGAAAGTTCAATGGCAANNNAGATCGGAA |
| 29954_2 | TATGAGTGTGGAGTCGTTGCAAGCCAAGAAAGTTCAATGGCAANNNAGATCGGAA |
| 31783_1 | TATGAGTGTGGAGTCGTTGCAGAAAGAAAGAAGGAAGGGAGGGNNNAGATCGG |
| 31783_2 | TATGAGTGTGGAGTCGTTGCAGAAAGAAAGAAGGAAGGGAGGGNNNAGATCGG |
| 29722_1 | TATGAGTGTGGAGTCGTTGCTGGAATGTTTTTGCTTAAATGAGAACCNNNAGATCG |
| 29722_2 | TATGAGTGTGGAGTCGTTGCTGGAATGTTTTTGCTTAAATGAGAACCNNNAGATCG |
| 33620_1 | TATGAGTGTGGAGTCGTTGCATCTCGGCTCACTGCAAACTCNNNAGATCGGAAGAG |
| 33620_2 | TATGAGTGTGGAGTCGTTGCATCTCGGCTCACTGCAAACTCNNNAGATCGGAAGAG |
| 44630_1 | TATGAGTGTGGAGTCGTTGCGTCCCCCTCTACATTCCCTTTACNNNAGATCGGAAGA |
| 44630_2 | TATGAGTGTGGAGTCGTTGCGTCCCCCTCTACATTCCCTTTACNNNAGATCGGAAGA |
| 36914_1 | TATGAGTGTGGAGTCGTTGCCTGAGATGGGAGGATCACTTGAGNNNAGATCGGAA |
| 36914_2 | TATGAGTGTGGAGTCGTTGCCTGAGATGGGAGGATCACTTGAGNNNAGATCGGAA |
| 35408_1 | TATGAGTGTGGAGTCGTTGCCACTATAACATCCACCTCCTCGGNNNAGATCGGAAGA |
| 35408_2 | TATGAGTGTGGAGTCGTTGCCACTATAACATCCACCTCCTCGGNNNAGATCGGAAGA |
| 51913_1 | TATGAGTGTGGAGTCGTTGCAAAACAATTAGCCAGGTGTGGTGNNNAGATCGGAA |
| 51913_2 | TATGAGTGTGGAGTCGTTGCAAAACAATTAGCCAGGTGTGGTGNNNAGATCGGAA |
| 41939_1 | TATGAGTGTGGAGTCGTTGCTCCTGGTTCAAGTGATTCTCCTGNNNAGATCGGAAG |
| 41939_2 | TATGAGTGTGGAGTCGTTGCTCCTGGTTCAAGTGATTCTCCTGNNNAGATCGGAAG |
| 32343_1 | TATGAGTGTGGAGTCGTTGCCACAATCATGACTCATTGCAGCTNNNAGATCGGAAG |
| 32343_2 | TATGAGTGTGGAGTCGTTGCCACAATCATGACTCATTGCAGCTNNNAGATCGGAAG |
| 33934_1 | TATGAGTGTGGAGTCGTTGCGATCTCAGCTCACTGCAACCTCNNNAGATCGGAAGA |
| 33934_2 | TATGAGTGTGGAGTCGTTGCGATCTCAGCTCACTGCAACCTCNNNAGATCGGAAGA |
| 35699_1 | TATGAGTGTGGAGTCGTTGCGCTCGTGATCACAGAGGACATATNNNAGATCGGAA |
| 35699_2 | TATGAGTGTGGAGTCGTTGCGCTCGTGATCACAGAGGACATATNNNAGATCGGAA |
| 52729_1 | TATGAGTGTGGAGTCGTTGCCCATTTGAGACACACAGACACACNNNAGATCGGAAG |
| 52729_2 | TATGAGTGTGGAGTCGTTGCCCATTTGAGACACACAGACACACNNNAGATCGGAAG |
| 44836_1 | TATGAGTGTGGAGTCGTTGCTGGGATTGTGGGAACACTTAGGNNNAGATCGGAAG |
| 44836_2 | TATGAGTGTGGAGTCGTTGCTGGGATTGTGGGAACACTTAGGNNNAGATCGGAAG |
| 30179_1 | TATGAGTGTGGAGTCGTTGCCTTCATCACGGCATTTACAGTCCNNNAGATCGGAAGA |
| 30179_2 | TATGAGTGTGGAGTCGTTGCCTTCATCACGGCATTTACAGTCCNNNAGATCGGAAGA |
| 47849_1 | TATGAGTGTGGAGTCGTTGCGCAGTAGCACAATCACAACTCTGNNNAGATCGGAAG |

| | |
|---|---|
| 47849_2 | TATGAGTGTGGAGTCGTTGCGCAGTAGCACAATCACAACTCTGNNNAGATCGGAAG |
| 41376_1 | TATGAGTGTGGAGTCGTTGCCCAGCTAGTCAGGAGTTTGAAGTNNNAGATCGGAAC |
| 41376_2 | TATGAGTGTGGAGTCGTTGCCCAGCTAGTCAGGAGTTTGAAGTNNNAGATCGGAAC |
| 49872_1 | TATGAGTGTGGAGTCGTTGCACAGAAATACCAGGCTCTAGGAGNNNAGATCGGAAC |
| 49872_2 | TATGAGTGTGGAGTCGTTGCACAGAAATACCAGGCTCTAGGAGNNNAGATCGGAAC |
| 13420_1 | TATGAGTGTGGAGTCGTTGCCTTAAAAGTGTTGTGTTACTTTGNNNAGATCGGAAG |
| 13420_2 | TATGAGTGTGGAGTCGTTGCCTTAAAAGTGTTGTGTTACTTTGNNNAGATCGGAAG |
| 33570_1 | TATGAGTGTGGAGTCGTTGCCAGTGGTATGATCTTGGCTCAGTNNNAGATCGGAAG |
| 33570_2 | TATGAGTGTGGAGTCGTTGCCAGTGGTATGATCTTGGCTCAGTNNNAGATCGGAAG |
| 10758_1 | TATGAGTGTGGAGTCGTTGCTCTGATGAAGGGTGAAAGAAGCTNNNAGATCGGAA |
| 10758_2 | TATGAGTGTGGAGTCGTTGCTCTGATGAAGGGTGAAAGAAGCTNNNAGATCGGAA |
| 44474_1 | TATGAGTGTGGAGTCGTTGCTGAATTTTTGGCATTGAGCATGANNNAGATCGGAAG |
| 44474_2 | TATGAGTGTGGAGTCGTTGCTGAATTTTTGGCATTGAGCATGANNNAGATCGGAAG |
| 13562_1 | TATGAGTGTGGAGTCGTTGCGAAATAAAGTGCACAATAAATGCNNNAGATCGGAAC |
| 13562_2 | TATGAGTGTGGAGTCGTTGCGAAATAAAGTGCACAATAAATGCNNNAGATCGGAAC |
| 31214_1 | TATGAGTGTGGAGTCGTTGCGGCTTGAAAATTACAGTGGTGGTNNNAGATCGGAAC |
| 31214_2 | TATGAGTGTGGAGTCGTTGCGGCTTGAAAATTACAGTGGTGGTNNNAGATCGGAAC |
| 34473_1 | TATGAGTGTGGAGTCGTTGCCCCACAGGAGACAGCACTATTTANNNAGATCGGAAG |
| 34473_2 | TATGAGTGTGGAGTCGTTGCCCCACAGGAGACAGCACTATTTANNNAGATCGGAAG |
| 29829_1 | TATGAGTGTGGAGTCGTTGCAAGAACCAATTCACATGTGGCTCNNNAGATCGGAAG |
| 29829_2 | TATGAGTGTGGAGTCGTTGCAAGAACCAATTCACATGTGGCTCNNNAGATCGGAAG |
| 48992_1 | TATGAGTGTGGAGTCGTTGCCTGCTTCTAGGGTGTTAGAGAGCNNNAGATCGGAAC |
| 48992_2 | TATGAGTGTGGAGTCGTTGCCTGCTTCTAGGGTGTTAGAGAGCNNNAGATCGGAAC |
| 40015_1 | TATGAGTGTGGAGTCGTTGCTGACTACTCTAAGACACCCCTGANNNAGATCGGAAGA |
| 40015_2 | TATGAGTGTGGAGTCGTTGCTGACTACTCTAAGACACCCCTGANNNAGATCGGAAGA |
| 30543_1 | TATGAGTGTGGAGTCGTTGCTGGGGGACTGAGATGTTTTGCNNNAGATCGGAAGA |
| 30543_2 | TATGAGTGTGGAGTCGTTGCTGGGGGACTGAGATGTTTTGCNNNAGATCGGAAGA |
| 50962_1 | TATGAGTGTGGAGTCGTTGCCCCAAAGACCACCATATTTTGCANNNAGATCGGAAGA |
| 50962_2 | TATGAGTGTGGAGTCGTTGCCCCAAAGACCACCATATTTTGCANNNAGATCGGAAGA |
| 42858_1 | TATGAGTGTGGAGTCGTTGCAGCATATCCACCACCTCAAACATNNNAGATCGGAAGA |
| 42858_2 | TATGAGTGTGGAGTCGTTGCAGCATATCCACCACCTCAAACATNNNAGATCGGAAGA |
| 13357_1 | TATGAGTGTGGAGTCGTTGCGGGGGAAAAAAACATCTAATAAANNNAGATCGGAA |
| 13357_2 | TATGAGTGTGGAGTCGTTGCGGGGGAAAAAAACATCTAATAAANNNAGATCGGAA |
| 28850_1 | TATGAGTGTGGAGTCGTTGCAGAGAGAGGGAGAGAGAGAGAGANNNAGATCGGA |
| 28850_2 | TATGAGTGTGGAGTCGTTGCAGAGAGAGGGAGAGAGAGAGAGANNNAGATCGGA |
| 44613_1 | TATGAGTGTGGAGTCGTTGCGAAAGGCATGAGGTGGCTAACAGNNNAGATCGGAA |
| 44613_2 | TATGAGTGTGGAGTCGTTGCGAAAGGCATGAGGTGGCTAACAGNNNAGATCGGAA |
| 30941_1 | TATGAGTGTGGAGTCGTTGCAAGTTGTTGTTGTTTGCTTTGGCNNNAGATCGGAAG |
| 30941_2 | TATGAGTGTGGAGTCGTTGCAAGTTGTTGTTGTTTGCTTTGGCNNNAGATCGGAAG |
| 38671_1 | TATGAGTGTGGAGTCGTTGCTGAGAACACTGAAGCTCAGAGAGNNNAGATCGGAA |
| 38671_2 | TATGAGTGTGGAGTCGTTGCTGAGAACACTGAAGCTCAGAGAGNNNAGATCGGAA |
| 40734_1 | TATGAGTGTGGAGTCGTTGCTGGATAACTTTGGGAAGTAGGAGANNNAGATCGGA |
| 40734_2 | TATGAGTGTGGAGTCGTTGCTGGATAACTTTGGGAAGTAGGAGANNNAGATCGGA |

| | |
|---|---|
| 47684_1 | TATGAGTGTGGAGTCGTTGCGGGTCTGCTTCCCAGTCCTCNNNAGATCGGAAGAGC |
| 47684_2 | TATGAGTGTGGAGTCGTTGCGGGTCTGCTTCCCAGTCCTCNNNAGATCGGAAGAGC |
| 32581_1 | TATGAGTGTGGAGTCGTTGCTGATTCCAGCTCACTACAACCTCNNNAGATCGGAAG |
| 32581_2 | TATGAGTGTGGAGTCGTTGCTGATTCCAGCTCACTACAACCTCNNNAGATCGGAAG |
| 40271_1 | TATGAGTGTGGAGTCGTTGCAGAAAAAGTAATGTGCATCCCACTNNNAGATCGGAA |
| 40271_2 | TATGAGTGTGGAGTCGTTGCAGAAAAAGTAATGTGCATCCCACTNNNAGATCGGAA |
| 9497_1 | TATGAGTGTGGAGTCGTTGCGGAGTAGAGCAACCAAGGTCAAGNNNAGATCGGA |
| 9497_2 | TATGAGTGTGGAGTCGTTGCGGAGTAGAGCAACCAAGGTCAAGNNNAGATCGGA |
| 42740_1 | TATGAGTGTGGAGTCGTTGCCTTACGCCTGTAATCCCAGAACTNNNAGATCGGAAG |
| 42740_2 | TATGAGTGTGGAGTCGTTGCCTTACGCCTGTAATCCCAGAACTNNNAGATCGGAAG |
| 41343_1 | TATGAGTGTGGAGTCGTTGCCTGTTTGGATGGCAAGTACAACANNNAGATCGGAA |
| 41343_2 | TATGAGTGTGGAGTCGTTGCCTGTTTGGATGGCAAGTACAACANNNAGATCGGAA |
| 42356_1 | TATGAGTGTGGAGTCGTTGCTTTGGTACAGACAGGGATTCACCNNNAGATCGGAA |
| 42356_2 | TATGAGTGTGGAGTCGTTGCTTTGGTACAGACAGGGATTCACCNNNAGATCGGAA |
| 39560_1 | TATGAGTGTGGAGTCGTTGCTGTAGGGACATGGATGAAACTGGNNNAGATCGGA |
| 39560_2 | TATGAGTGTGGAGTCGTTGCTGTAGGGACATGGATGAAACTGGNNNAGATCGGA |
| 48250_1 | TATGAGTGTGGAGTCGTTGCGGATGGATAGATCATGTTGGCCTNNNAGATCGGAA |
| 48250_2 | TATGAGTGTGGAGTCGTTGCGGATGGATAGATCATGTTGGCCTNNNAGATCGGAA |
| 37784_1 | TATGAGTGTGGAGTCGTTGCATTTTGCTAGGTGAGATGGCTCANNNAGATCGGAA |
| 37784_2 | TATGAGTGTGGAGTCGTTGCATTTTGCTAGGTGAGATGGCTCANNNAGATCGGAA |
| 1651_1 | TATGAGTGTGGAGTCGTTGCCACCTGACCTGTCCTGTCCTTTTNNNAGATCGGAAGA |
| 1651_2 | TATGAGTGTGGAGTCGTTGCCACCTGACCTGTCCTGTCCTTTTNNNAGATCGGAAGA |
| 39609_1 | TATGAGTGTGGAGTCGTTGCCATGCCTGTAATCCCAGCACTATNNNAGATCGGAAG |
| 39609_2 | TATGAGTGTGGAGTCGTTGCCATGCCTGTAATCCCAGCACTATNNNAGATCGGAAG |
| 33939_1 | TATGAGTGTGGAGTCGTTGCCTTCCCAGGTTCAAGCAATTCTCNNNAGATCGGAAG |
| 33939_2 | TATGAGTGTGGAGTCGTTGCCTTCCCAGGTTCAAGCAATTCTCNNNAGATCGGAAG |
| 41423_1 | TATGAGTGTGGAGTCGTTGCCCATAATCCCAGCTAGTCAGGAGNNNAGATCGGAAG |
| 41423_2 | TATGAGTGTGGAGTCGTTGCCCATAATCCCAGCTAGTCAGGAGNNNAGATCGGAAG |
| 46503_1 | TATGAGTGTGGAGTCGTTGCCTTTCGGCGTTTCAAGACAAAGANNNAGATCGGAAG |
| 46503_2 | TATGAGTGTGGAGTCGTTGCCTTTCGGCGTTTCAAGACAAAGANNNAGATCGGAAG |
| 38780_1 | TATGAGTGTGGAGTCGTTGCACTGCTTTTCTCCCATGGATTCTNNNAGATCGGAAGA |
| 38780_2 | TATGAGTGTGGAGTCGTTGCACTGCTTTTCTCCCATGGATTCTNNNAGATCGGAAGA |
| 12631_1 | TATGAGTGTGGAGTCGTTGCAGATGAGTAAAAACATCAGCTNNNAGATCGGAAGA |
| 12631_2 | TATGAGTGTGGAGTCGTTGCAGATGAGTAAAAACATCAGCTNNNAGATCGGAAGA |
| 33080_1 | TATGAGTGTGGAGTCGTTGCCTCCCAGGTTCACACCATTCTCNNNAGATCGGAAGAG |
| 33080_2 | TATGAGTGTGGAGTCGTTGCCTCCCAGGTTCACACCATTCTCNNNAGATCGGAAGAG |
| 41509_1 | TATGAGTGTGGAGTCGTTGCATTTTTTCCTGAAGTGGCCACTGNNNAGATCGGAAG |
| 41509_2 | TATGAGTGTGGAGTCGTTGCATTTTTTCCTGAAGTGGCCACTGNNNAGATCGGAAG |
| 38751_1 | TATGAGTGTGGAGTCGTTGCAAAACCCTCAGCAAAATTGGCATNNNAGATCGGAAG |
| 38751_2 | TATGAGTGTGGAGTCGTTGCAAAACCCTCAGCAAAATTGGCATNNNAGATCGGAAG |
| 28886_1 | TATGAGTGTGGAGTCGTTGCTGGCTTGGTTCTCAACTGTCTTANNNAGATCGGAAG |
| 28886_2 | TATGAGTGTGGAGTCGTTGCTGGCTTGGTTCTCAACTGTCTTANNNAGATCGGAAG |
| 37210_1 | TATGAGTGTGGAGTCGTTGCATCATGAGGTCAGGAGATCGAGANNNAGATCGGAA |

| | |
|---|---|
| 37210_2 | TATGAGTGTGGAGTCGTTGCATCATGAGGTCAGGAGATCGAGANNNAGATCGGAA |
| 46413_1 | TATGAGTGTGGAGTCGTTGCGGTTCAAGCAATTCTCCCACTTCNNNAGATCGGAAGA |
| 46413_2 | TATGAGTGTGGAGTCGTTGCGGTTCAAGCAATTCTCCCACTTCNNNAGATCGGAAGA |
| 49935_1 | TATGAGTGTGGAGTCGTTGCAGATTTGCCGTGTCTTGTGATTGNNNAGATCGGAAG |
| 49935_2 | TATGAGTGTGGAGTCGTTGCAGATTTGCCGTGTCTTGTGATTGNNNAGATCGGAAG |
| 38449_1 | TATGAGTGTGGAGTCGTTGCAGCTGCAGAATGATAGGTACAGTNNNAGATCGGAA |
| 38449_2 | TATGAGTGTGGAGTCGTTGCAGCTGCAGAATGATAGGTACAGTNNNAGATCGGAA |
| 30579_1 | TATGAGTGTGGAGTCGTTGCGGATTAAGGCAATCATGGATTGGANNNAGATCGGA |
| 30579_2 | TATGAGTGTGGAGTCGTTGCGGATTAAGGCAATCATGGATTGGANNNAGATCGGA |
| 13658_1 | TATGAGTGTGGAGTCGTTGCTGGAAGAAGAAATGTTTTTAGTTNNNAGATCGGAA |
| 13658_2 | TATGAGTGTGGAGTCGTTGCTGGAAGAAGAAATGTTTTTAGTTNNNAGATCGGAA |
| 48851_1 | TATGAGTGTGGAGTCGTTGCCACCACGCCTGGCTAATTTTTTNNNAGATCGGAAGA |
| 48851_2 | TATGAGTGTGGAGTCGTTGCCACCACGCCTGGCTAATTTTTTNNNAGATCGGAAGA |
| 13229_1 | TATGAGTGTGGAGTCGTTGCAATCTGAATAACGGCCAAATNNNAGATCGGAAGAG |
| 13229_2 | TATGAGTGTGGAGTCGTTGCAATCTGAATAACGGCCAAATNNNAGATCGGAAGAG |
| 31599_1 | TATGAGTGTGGAGTCGTTGCTTTGTGTATGGTGAGAGACAGGGNNNAGATCGGAA |
| 31599_2 | TATGAGTGTGGAGTCGTTGCTTTGTGTATGGTGAGAGACAGGGNNNAGATCGGAA |
| 33969_1 | TATGAGTGTGGAGTCGTTGCCTGAGTAGCTGGAACTACAGGTGNNNAGATCGGAA |
| 33969_2 | TATGAGTGTGGAGTCGTTGCCTGAGTAGCTGGAACTACAGGTGNNNAGATCGGAA |
| 9772_1 | TATGAGTGTGGAGTCGTTGCAACATGCACATACACATACCCCTNNNAGATCGGAAGA |
| 9772_2 | TATGAGTGTGGAGTCGTTGCAACATGCACATACACATACCCCTNNNAGATCGGAAGA |
| 42867_1 | TATGAGTGTGGAGTCGTTGCTGTCCTTTTCCCTGTTGTCAGAANNNAGATCGGAAGA |
| 42867_2 | TATGAGTGTGGAGTCGTTGCTGTCCTTTTCCCTGTTGTCAGAANNNAGATCGGAAGA |
| 38965_1 | TATGAGTGTGGAGTCGTTGCGATATTCCCTTTCACCACCCCTTNNNAGATCGGAAGA |
| 38965_2 | TATGAGTGTGGAGTCGTTGCGATATTCCCTTTCACCACCCCTTNNNAGATCGGAAGA |
| 32266_1 | TATGAGTGTGGAGTCGTTGCTATTACCTGGGCTGGTCTCAAACNNNAGATCGGAAG |
| 32266_2 | TATGAGTGTGGAGTCGTTGCTATTACCTGGGCTGGTCTCAAACNNNAGATCGGAAG |
| 44854_1 | TATGAGTGTGGAGTCGTTGCACTGTCCAGCGGTATGTTACTTTNNNAGATCGGAAG |
| 44854_2 | TATGAGTGTGGAGTCGTTGCACTGTCCAGCGGTATGTTACTTTNNNAGATCGGAAG |
| 44277_1 | TATGAGTGTGGAGTCGTTGCCTGGACAGTCTCTGAGATGATGGNNNAGATCGGAA |
| 44277_2 | TATGAGTGTGGAGTCGTTGCCTGGACAGTCTCTGAGATGATGGNNNAGATCGGAA |
| 36810_1 | TATGAGTGTGGAGTCGTTGCGAGTGAAAAGGCAACCCATGAAANNNAGATCGGAA |
| 36810_2 | TATGAGTGTGGAGTCGTTGCGAGTGAAAAGGCAACCCATGAAANNNAGATCGGAA |
| 42922_1 | TATGAGTGTGGAGTCGTTGCCCTCCTGCGTTCAACCTATTCTCNNNAGATCGGAAGA |
| 42922_2 | TATGAGTGTGGAGTCGTTGCCCTCCTGCGTTCAACCTATTCTCNNNAGATCGGAAGA |
| 36106_1 | TATGAGTGTGGAGTCGTTGCCAATTCCTGGGCTCAAGCAATCNNNAGATCGGAAGA |
| 36106_2 | TATGAGTGTGGAGTCGTTGCCAATTCCTGGGCTCAAGCAATCNNNAGATCGGAAGA |
| 29184_1 | TATGAGTGTGGAGTCGTTGCTACATGGGCTGTTATCTGTCTGGNNNAGATCGGAAG |
| 29184_2 | TATGAGTGTGGAGTCGTTGCTACATGGGCTGTTATCTGTCTGGNNNAGATCGGAAG |
| 46739_1 | TATGAGTGTGGAGTCGTTGCACCACCACGCCTGACTAATTATTNNNAGATCGGAAGA |
| 46739_2 | TATGAGTGTGGAGTCGTTGCACCACCACGCCTGACTAATTATTNNNAGATCGGAAGA |
| 47272_1 | TATGAGTGTGGAGTCGTTGCTTCAAGTGATTCTCCTGCCTCAGNNNAGATCGGAAGA |
| 47272_2 | TATGAGTGTGGAGTCGTTGCTTCAAGTGATTCTCCTGCCTCAGNNNAGATCGGAAGA |

36630_1 TATGAGTGTGGAGTCGTTGCCTCCCTGGGTGCTAGAATTACAGNNNAGATCGGAAG
36630_2 TATGAGTGTGGAGTCGTTGCCTCCCTGGGTGCTAGAATTACAGNNNAGATCGGAAG
36013_1 TATGAGTGTGGAGTCGTTGCGAGACACAGACACAGAGGAGAAGNNNAGATCGGA
36013_2 TATGAGTGTGGAGTCGTTGCGAGACACAGACACAGAGGAGAAGNNNAGATCGGA
53188_1 TATGAGTGTGGAGTCGTTGCCACTTGAGCGAGCAGAGAGTCNNNAGATCGGAAGA
53188_2 TATGAGTGTGGAGTCGTTGCCACTTGAGCGAGCAGAGAGTCNNNAGATCGGAAGA
50615_1 TATGAGTGTGGAGTCGTTGCCCACCAGAGAAACAGCATTTCAGNNNAGATCGGAAG
50615_2 TATGAGTGTGGAGTCGTTGCCCACCAGAGAAACAGCATTTCAGNNNAGATCGGAAG
30206_1 TATGAGTGTGGAGTCGTTGCGATGGCACAGAGAGAGAAACCATNNNAGATCGGAA
30206_2 TATGAGTGTGGAGTCGTTGCGATGGCACAGAGAGAGAAACCATNNNAGATCGGAA
38521_1 TATGAGTGTGGAGTCGTTGCTTTGGGAACAGGGGAACAGAGNNNAGATCGGAAG
38521_2 TATGAGTGTGGAGTCGTTGCTTTGGGAACAGGGGAACAGAGNNNAGATCGGAAG
51274_1 TATGAGTGTGGAGTCGTTGCTGAGATTTGGATTGGGACACAGANNNAGATCGGAA
51274_2 TATGAGTGTGGAGTCGTTGCTGAGATTTGGATTGGGACACAGANNNAGATCGGAA
44540_1 TATGAGTGTGGAGTCGTTGCTGGGAACATGGATGGAGTTGTAGNNNAGATCGGA
44540_2 TATGAGTGTGGAGTCGTTGCTGGGAACATGGATGGAGTTGTAGNNNAGATCGGA
47934_1 TATGAGTGTGGAGTCGTTGCCAGCACACCCAGCTAATTTTTGTNNNAGATCGGAAG
47934_2 TATGAGTGTGGAGTCGTTGCCAGCACACCCAGCTAATTTTTGTNNNAGATCGGAAG
49449_1 TATGAGTGTGGAGTCGTTGCAATTTCTCAGCTCACCGTAACCTNNNAGATCGGAAGA
49449_2 TATGAGTGTGGAGTCGTTGCAATTTCTCAGCTCACCGTAACCTNNNAGATCGGAAGA
36594_1 TATGAGTGTGGAGTCGTTGCTGCCTTTAGAGAGATAAGTGGCANNNAGATCGGAA
36594_2 TATGAGTGTGGAGTCGTTGCTGCCTTTAGAGAGATAAGTGGCANNNAGATCGGAA
46971_1 TATGAGTGTGGAGTCGTTGCGCCTCCTGAGTAGCTGAGATAACNNNAGATCGGAAG
46971_2 TATGAGTGTGGAGTCGTTGCGCCTCCTGAGTAGCTGAGATAACNNNAGATCGGAAG
33936_1 TATGAGTGTGGAGTCGTTGCCTCTTGTTACCCAGACTGGAGTGNNNAGATCGGAAG
33936_2 TATGAGTGTGGAGTCGTTGCCTCTTGTTACCCAGACTGGAGTGNNNAGATCGGAAG
35214_1 TATGAGTGTGGAGTCGTTGCTTCAGTGGCATGATCTTGTCTCANNNAGATCGGAAG
35214_2 TATGAGTGTGGAGTCGTTGCTTCAGTGGCATGATCTTGTCTCANNNAGATCGGAAG
41114_1 TATGAGTGTGGAGTCGTTGCTGTTTTCCGACATGGCTATACCANNNAGATCGGAAG
41114_2 TATGAGTGTGGAGTCGTTGCTGTTTTCCGACATGGCTATACCANNNAGATCGGAAG
9694_1 TATGAGTGTGGAGTCGTTGCAAAACTCTAGCAGGACCAAACCCNNNAGATCGGAAG
9694_2 TATGAGTGTGGAGTCGTTGCAAAACTCTAGCAGGACCAAACCCNNNAGATCGGAAG
38603_1 TATGAGTGTGGAGTCGTTGCACCAAGGGCTAAGAGAAAGGAAGNNNAGATCGGA
38603_2 TATGAGTGTGGAGTCGTTGCACCAAGGGCTAAGAGAAAGGAAGNNNAGATCGGA
30121_1 TATGAGTGTGGAGTCGTTGCCTATCAACGTAGGGCAGCTGAATNNNAGATCGGAA
30121_2 TATGAGTGTGGAGTCGTTGCCTATCAACGTAGGGCAGCTGAATNNNAGATCGGAA
48654_1 TATGAGTGTGGAGTCGTTGCAATACAGCACACTGATGGGTCTTNNNAGATCGGAAG
48654_2 TATGAGTGTGGAGTCGTTGCAATACAGCACACTGATGGGTCTTNNNAGATCGGAAG
2994_1 TATGAGTGTGGAGTCGTTGCTTTGCCTAGAGAAGTCACAGTGNNNAGATCGGAAG
2994_2 TATGAGTGTGGAGTCGTTGCTTTGCCTAGAGAAGTCACAGTGNNNAGATCGGAAG
45320_1 TATGAGTGTGGAGTCGTTGCGCTTCCCTCCATTGGTGTGTATANNNAGATCGGAAG
45320_2 TATGAGTGTGGAGTCGTTGCGCTTCCCTCCATTGGTGTGTATANNNAGATCGGAAG
29318_1 TATGAGTGTGGAGTCGTTGCACTCTCCCTTGCAGATTACATGANNNAGATCGGAAG

29318_2 TATGAGTGTGGAGTCGTTGCACTCTCCCTTGCAGATTACATGANNNAGATCGGAAG
12074_1 TATGAGTGTGGAGTCGTTGCCCATTCAACTCACTACAGCCTTNNNAGATCGGAAGAG
12074_2 TATGAGTGTGGAGTCGTTGCCCATTCAACTCACTACAGCCTTNNNAGATCGGAAGAG
53168_1 TATGAGTGTGGAGTCGTTGCTTCAATTGCTTGCAAAACCCAGTNNNAGATCGGAAG
53168_2 TATGAGTGTGGAGTCGTTGCTTCAATTGCTTGCAAAACCCAGTNNNAGATCGGAAG
46656_1 TATGAGTGTGGAGTCGTTGCTGTTTCTGGGCTATTCTGTTCCANNNAGATCGGAAG
46656_2 TATGAGTGTGGAGTCGTTGCTGTTTCTGGGCTATTCTGTTCCANNNAGATCGGAAG
45414_1 TATGAGTGTGGAGTCGTTGCACCTTGGCTATTGTGAACAATGCNNNAGATCGGAAG
45414_2 TATGAGTGTGGAGTCGTTGCACCTTGGCTATTGTGAACAATGCNNNAGATCGGAAG
38071_1 TATGAGTGTGGAGTCGTTGCCATCCCAAAAACGACATCTGAGGNNNAGATCGGAAG
38071_2 TATGAGTGTGGAGTCGTTGCCATCCCAAAAACGACATCTGAGGNNNAGATCGGAAG
36121_1 TATGAGTGTGGAGTCGTTGCGTGTAATCCCAGCTACTCAGGAGNNNAGATCGGAA
36121_2 TATGAGTGTGGAGTCGTTGCGTGTAATCCCAGCTACTCAGGAGNNNAGATCGGAA
38373_1 TATGAGTGTGGAGTCGTTGCTCATGTGTTTGTGAATTTTCCTGTNNNAGATCGGAA
38373_2 TATGAGTGTGGAGTCGTTGCTCATGTGTTTGTGAATTTTCCTGTNNNAGATCGGAA
42722_1 TATGAGTGTGGAGTCGTTGCGGCAAAAAACAAAACCCTGTTGGNNNAGATCGGAA
42722_2 TATGAGTGTGGAGTCGTTGCGGCAAAAAACAAAACCCTGTTGGNNNAGATCGGAA
29477_1 TATGAGTGTGGAGTCGTTGCCCCTTGCGGATTAGGCTCTTTAANNNAGATCGGAAG
29477_2 TATGAGTGTGGAGTCGTTGCCCCTTGCGGATTAGGCTCTTTAANNNAGATCGGAAG
51234_1 TATGAGTGTGGAGTCGTTGCCAGTTGTAATGCTGCCAGCTCNNNAGATCGGAAGAG
51234_2 TATGAGTGTGGAGTCGTTGCCAGTTGTAATGCTGCCAGCTCNNNAGATCGGAAGAG
39377_1 TATGAGTGTGGAGTCGTTGCGAGGTCAGGAGTTCGAGACAAGNNNAGATCGGAA
39377_2 TATGAGTGTGGAGTCGTTGCGAGGTCAGGAGTTCGAGACAAGNNNAGATCGGAA
40177_1 TATGAGTGTGGAGTCGTTGCTCCCTCATTCTTTATCCCACCAGNNNAGATCGGAAGA
40177_2 TATGAGTGTGGAGTCGTTGCTCCCTCATTCTTTATCCCACCAGNNNAGATCGGAAGA
39347_1 TATGAGTGTGGAGTCGTTGCTGCTGCAACTTTCTTATGAGTACCNNNAGATCGGAA
39347_2 TATGAGTGTGGAGTCGTTGCTGCTGCAACTTTCTTATGAGTACCNNNAGATCGGAA
31726_1 TATGAGTGTGGAGTCGTTGCCGTCTCTCTCTCTCTCTCTCTNNNAGATCGGAAGAG
31726_2 TATGAGTGTGGAGTCGTTGCCGTCTCTCTCTCTCTCTCTCTNNNAGATCGGAAGAG
12498_1 TATGAGTGTGGAGTCGTTGCTCCATGAGAATCTTCCTCAGNNNAGATCGGAAGAGC
12498_2 TATGAGTGTGGAGTCGTTGCTCCATGAGAATCTTCCTCAGNNNAGATCGGAAGAGC
46765_1 TATGAGTGTGGAGTCGTTGCCTTCTGAGTAGCTGGGACTACAGNNNAGATCGGAA
46765_2 TATGAGTGTGGAGTCGTTGCCTTCTGAGTAGCTGGGACTACAGNNNAGATCGGAA
53334_1 TATGAGTGTGGAGTCGTTGCGTGGAAGCCATTATCCTCAATGCNNNAGATCGGAAG
53334_2 TATGAGTGTGGAGTCGTTGCGTGGAAGCCATTATCCTCAATGCNNNAGATCGGAAG
38573_1 TATGAGTGTGGAGTCGTTGCTAGCCTTTCAAAATGAACAGCCCNNNAGATCGGAAG
38573_2 TATGAGTGTGGAGTCGTTGCTAGCCTTTCAAAATGAACAGCCCNNNAGATCGGAAG
31755_1 TATGAGTGTGGAGTCGTTGCCATCACCTCGATCTGGCTCATTANNNAGATCGGAAG
31755_2 TATGAGTGTGGAGTCGTTGCCATCACCTCGATCTGGCTCATTANNNAGATCGGAAG
36089_1 TATGAGTGTGGAGTCGTTGCAGATGAAGTTGACTCAAATTTGCTNNNAGATCGGAA
36089_2 TATGAGTGTGGAGTCGTTGCAGATGAAGTTGACTCAAATTTGCTNNNAGATCGGAA
34361_1 TATGAGTGTGGAGTCGTTGCCATAGTCGTATCACTGCACTCCANNNAGATCGGAAG
34361_2 TATGAGTGTGGAGTCGTTGCCATAGTCGTATCACTGCACTCCANNNAGATCGGAAG

| | |
|---|---|
| 41117_1 | TATGAGTGTGGAGTCGTTGCTGTTTTCCGACATGGCTATACCANNNAGATCGGAAG |
| 41117_2 | TATGAGTGTGGAGTCGTTGCTGTTTTCCGACATGGCTATACCANNNAGATCGGAAG |
| 33792_1 | TATGAGTGTGGAGTCGTTGCCCAAAAGAGCTAGCAGAATCCACNNNAGATCGGAAG |
| 33792_2 | TATGAGTGTGGAGTCGTTGCCCAAAAGAGCTAGCAGAATCCACNNNAGATCGGAAG |
| 31179_1 | TATGAGTGTGGAGTCGTTGCATAGAGAGCTCAGAAATAGGCCCNNNAGATCGGAA |
| 31179_2 | TATGAGTGTGGAGTCGTTGCATAGAGAGCTCAGAAATAGGCCCNNNAGATCGGAA |
| 52618_1 | TATGAGTGTGGAGTCGTTGCAAAATCATTTTCCTCCCCAACCGNNNAGATCGGAAGA |
| 52618_2 | TATGAGTGTGGAGTCGTTGCAAAATCATTTTCCTCCCCAACCGNNNAGATCGGAAGA |
| 53280_1 | TATGAGTGTGGAGTCGTTGCTTCAGTGTCTGGAGCATTTCAGANNNAGATCGGAAG |
| 53280_2 | TATGAGTGTGGAGTCGTTGCTTCAGTGTCTGGAGCATTTCAGANNNAGATCGGAAG |
| 44162_1 | TATGAGTGTGGAGTCGTTGCGGTGAGTCCTGGAAATGCATTTTNNNAGATCGGAAG |
| 44162_2 | TATGAGTGTGGAGTCGTTGCGGTGAGTCCTGGAAATGCATTTTNNNAGATCGGAAG |
| 44024_1 | TATGAGTGTGGAGTCGTTGCGACCTGGAAAAAGCACGAATTGANNNAGATCGGAA |
| 44024_2 | TATGAGTGTGGAGTCGTTGCGACCTGGAAAAAGCACGAATTGANNNAGATCGGAA |
| 30255_1 | TATGAGTGTGGAGTCGTTGCAGTCTGCTTAAATGATTGTGTATGATNNNAGATCGG |
| 30255_2 | TATGAGTGTGGAGTCGTTGCAGTCTGCTTAAATGATTGTGTATGATNNNAGATCGG |
| 31958_1 | TATGAGTGTGGAGTCGTTGCTTCTGTGTGAACTGTGTCAGGAANNNAGATCGGAAG |
| 31958_2 | TATGAGTGTGGAGTCGTTGCTTCTGTGTGAACTGTGTCAGGAANNNAGATCGGAAG |
| 48612_1 | TATGAGTGTGGAGTCGTTGCGTCCTAGGCAATTCAGGGATCTTNNNAGATCGGAAG |
| 48612_2 | TATGAGTGTGGAGTCGTTGCGTCCTAGGCAATTCAGGGATCTTNNNAGATCGGAAG |
| 29551_1 | TATGAGTGTGGAGTCGTTGCCTCACTGCAGACTCAATCTTCTTGGNNNAGATCGGAA |
| 29551_2 | TATGAGTGTGGAGTCGTTGCCTCACTGCAGACTCAATCTTCTTGGNNNAGATCGGAA |
| 47828_1 | TATGAGTGTGGAGTCGTTGCCCAGGCTAGATTCAAACACTCCTNNNAGATCGGAAG |
| 47828_2 | TATGAGTGTGGAGTCGTTGCCCAGGCTAGATTCAAACACTCCTNNNAGATCGGAAG |
| 36624_1 | TATGAGTGTGGAGTCGTTGCTACAATGCAGGATGTTTGTGCTGNNNAGATCGGAAG |
| 36624_2 | TATGAGTGTGGAGTCGTTGCTACAATGCAGGATGTTTGTGCTGNNNAGATCGGAAG |
| 43940_1 | TATGAGTGTGGAGTCGTTGCACTCACTCATCTGTGCTTAGAGGNNNAGATCGGAAG |
| 43940_2 | TATGAGTGTGGAGTCGTTGCACTCACTCATCTGTGCTTAGAGGNNNAGATCGGAAG |
| 52686_1 | TATGAGTGTGGAGTCGTTGCCCATGGCAAATGTAACAGGCTAGNNNAGATCGGAA |
| 52686_2 | TATGAGTGTGGAGTCGTTGCCCATGGCAAATGTAACAGGCTAGNNNAGATCGGAA |
| 45426_1 | TATGAGTGTGGAGTCGTTGCGAGCTTTCCCCCTATGTTCTCTTNNNAGATCGGAAGA |
| 45426_2 | TATGAGTGTGGAGTCGTTGCGAGCTTTCCCCCTATGTTCTCTTNNNAGATCGGAAGA |
| 36606_1 | TATGAGTGTGGAGTCGTTGCGTTGCAGTCCTGGTCTCTGATAANNNAGATCGGAAG |
| 36606_2 | TATGAGTGTGGAGTCGTTGCGTTGCAGTCCTGGTCTCTGATAANNNAGATCGGAAG |
| 39339_1 | TATGAGTGTGGAGTCGTTGCCCTGTAATTCCAGCTACTTGGGANNNAGATCGGAAG |
| 39339_2 | TATGAGTGTGGAGTCGTTGCCCTGTAATTCCAGCTACTTGGGANNNAGATCGGAAG |
| 42161_1 | TATGAGTGTGGAGTCGTTGCAGCCTTGGATGGAAACTGATTCTNNNAGATCGGAAG |
| 42161_2 | TATGAGTGTGGAGTCGTTGCAGCCTTGGATGGAAACTGATTCTNNNAGATCGGAAG |
| 10425_1 | TATGAGTGTGGAGTCGTTGCCCTCATTCTCCCCCAAACATGAGNNNAGATCGGAAGA |
| 10425_2 | TATGAGTGTGGAGTCGTTGCCCTCATTCTCCCCCAAACATGAGNNNAGATCGGAAGA |
| 33871_1 | TATGAGTGTGGAGTCGTTGCGGAAAGAAGTGAGCTATCCATGCNNNAGATCGGAA |
| 33871_2 | TATGAGTGTGGAGTCGTTGCGGAAAGAAGTGAGCTATCCATGCNNNAGATCGGAA |
| 29509_1 | TATGAGTGTGGAGTCGTTGCCCAGAATCCCAGTAAGTAAAGTNNNAGATCGGAAG |

| | |
|---|---|
| 29509_2 | TATGAGTGTGGAGTCGTTGCCCAGAATCCCAGTAAGTAAAGTNNNAGATCGGAAG |
| 39457_1 | TATGAGTGTGGAGTCGTTGCGACAATGGCTGTATGATCCTTGCNNNAGATCGGAAG |
| 39457_2 | TATGAGTGTGGAGTCGTTGCGACAATGGCTGTATGATCCTTGCNNNAGATCGGAAG |
| 35539_1 | TATGAGTGTGGAGTCGTTGCTTTTGCCTCAGCCTTCTCAGTATNNNAGATCGGAAGA |
| 35539_2 | TATGAGTGTGGAGTCGTTGCTTTTGCCTCAGCCTTCTCAGTATNNNAGATCGGAAGA |
| 32823_1 | TATGAGTGTGGAGTCGTTGCCTCCCGAGTAGCTGGAACTACAGNNNAGATCGGAAG |
| 32823_2 | TATGAGTGTGGAGTCGTTGCCTCCCGAGTAGCTGGAACTACAGNNNAGATCGGAAG |
| 30537_1 | TATGAGTGTGGAGTCGTTGCAGCCTGACTCCATCCTTATCAATNNNAGATCGGAAG |
| 30537_2 | TATGAGTGTGGAGTCGTTGCAGCCTGACTCCATCCTTATCAATNNNAGATCGGAAG |
| 38563_1 | TATGAGTGTGGAGTCGTTGCAAAAAATTAGCTGGGCATGGTGGNNNAGATCGGAA |
| 38563_2 | TATGAGTGTGGAGTCGTTGCAAAAAATTAGCTGGGCATGGTGGNNNAGATCGGAA |
| 52746_1 | TATGAGTGTGGAGTCGTTGCCCATTTCCAGGCAGAACTTCAGNNNAGATCGGAAGA |
| 52746_2 | TATGAGTGTGGAGTCGTTGCCCATTTCCAGGCAGAACTTCAGNNNAGATCGGAAGA |
| 42148_1 | TATGAGTGTGGAGTCGTTGCACAGGGATACAGTTACAGCTCACNNNAGATCGGAAG |
| 42148_2 | TATGAGTGTGGAGTCGTTGCACAGGGATACAGTTACAGCTCACNNNAGATCGGAAG |
| 13147_1 | TATGAGTGTGGAGTCGTTGCTATCATAGAATAATGCAAGTGGANNNAGATCGGAA |
| 13147_2 | TATGAGTGTGGAGTCGTTGCTATCATAGAATAATGCAAGTGGANNNAGATCGGAA |
| 44948_1 | TATGAGTGTGGAGTCGTTGCAAACAAACAAACCTCACAACCGTNNNAGATCGGAAG |
| 44948_2 | TATGAGTGTGGAGTCGTTGCAAACAAACAAACCTCACAACCGTNNNAGATCGGAAG |
| 49673_1 | TATGAGTGTGGAGTCGTTGCCTGCCACTAAGCTGAGAACCATANNNAGATCGGAAG |
| 49673_2 | TATGAGTGTGGAGTCGTTGCCTGCCACTAAGCTGAGAACCATANNNAGATCGGAAG |
| 39725_1 | TATGAGTGTGGAGTCGTTGCAGAACATAAACTCCAGCACCAGANNNAGATCGGAAG |
| 39725_2 | TATGAGTGTGGAGTCGTTGCAGAACATAAACTCCAGCACCAGANNNAGATCGGAAG |
| 12622_1 | TATGAGTGTGGAGTCGTTGCATGCTATCCTGGACTTTGAGNNNAGATCGGAAGAG |
| 12622_2 | TATGAGTGTGGAGTCGTTGCATGCTATCCTGGACTTTGAGNNNAGATCGGAAGAG |
| 49809_1 | TATGAGTGTGGAGTCGTTGCATGCAAAAGTCAACTCCTACCCANNNAGATCGGAAG |
| 49809_2 | TATGAGTGTGGAGTCGTTGCATGCAAAAGTCAACTCCTACCCANNNAGATCGGAAG |
| 13140_1 | TATGAGTGTGGAGTCGTTGCCCTTTGGAAAGTAAAACAAAAAANNNAGATCGGAAG |
| 13140_2 | TATGAGTGTGGAGTCGTTGCCCTTTGGAAAGTAAAACAAAAAANNNAGATCGGAAG |
| 40204_1 | TATGAGTGTGGAGTCGTTGCCTCCTGAGTAGCTGGGACTACAGNNNAGATCGGAAG |
| 40204_2 | TATGAGTGTGGAGTCGTTGCCTCCTGAGTAGCTGGGACTACAGNNNAGATCGGAAG |
| 38551_1 | TATGAGTGTGGAGTCGTTGCGTTTTCGTTGGCAGACATGTGTANNNAGATCGGAAG |
| 38551_2 | TATGAGTGTGGAGTCGTTGCGTTTTCGTTGGCAGACATGTGTANNNAGATCGGAAG |
| 9127_1 | TATGAGTGTGGAGTCGTTGCGGTGAATGGTGTGGAAGGTTGATNNNAGATCGGA |
| 9127_2 | TATGAGTGTGGAGTCGTTGCGGTGAATGGTGTGGAAGGTTGATNNNAGATCGGA |
| 46178_1 | TATGAGTGTGGAGTCGTTGCTCCTCTTGCTACTTGGTGGTAACNNNAGATCGGAAG |
| 46178_2 | TATGAGTGTGGAGTCGTTGCTCCTCTTGCTACTTGGTGGTAACNNNAGATCGGAAG |
| 50592_1 | TATGAGTGTGGAGTCGTTGCGATTGAACCCGGACAGAAACATGNNNAGATCGGAA |
| 50592_2 | TATGAGTGTGGAGTCGTTGCGATTGAACCCGGACAGAAACATGNNNAGATCGGAA |
| 37789_1 | TATGAGTGTGGAGTCGTTGCAAGACACATACCGGCTGTAACTTNNNAGATCGGAAG |
| 37789_2 | TATGAGTGTGGAGTCGTTGCAAGACACATACCGGCTGTAACTTNNNAGATCGGAAG |
| 38690_1 | TATGAGTGTGGAGTCGTTGCACATGATGACTCTTGGGATGGACNNNAGATCGGAA |
| 38690_2 | TATGAGTGTGGAGTCGTTGCACATGATGACTCTTGGGATGGACNNNAGATCGGAA |

| | |
|---|---|
| 38496_1 | TATGAGTGTGGAGTCGTTGCAATACAACAATTAGCTGGGCGTGNNNAGATCGGAA |
| 38496_2 | TATGAGTGTGGAGTCGTTGCAATACAACAATTAGCTGGGCGTGNNNAGATCGGAA |
| 29081_1 | TATGAGTGTGGAGTCGTTGCGAGAGAGAGAGAGAGAGACCCAANNNAGATCGGA |
| 29081_2 | TATGAGTGTGGAGTCGTTGCGAGAGAGAGAGAGAGAGACCCAANNNAGATCGGA |
| 40965_1 | TATGAGTGTGGAGTCGTTGCGTATGTGACGTGATGGATATGTAAANNNAGATCGG |
| 40965_2 | TATGAGTGTGGAGTCGTTGCGTATGTGACGTGATGGATATGTAAANNNAGATCGG |
| 35128_1 | TATGAGTGTGGAGTCGTTGCCTCTGCATCTTGGTTTCAAGCAANNNAGATCGGAAG |
| 35128_2 | TATGAGTGTGGAGTCGTTGCCTCTGCATCTTGGTTTCAAGCAANNNAGATCGGAAG |
| 49902_1 | TATGAGTGTGGAGTCGTTGCATGAGTTAGAAGTGGGGTAGCACNNNAGATCGGAA |
| 49902_2 | TATGAGTGTGGAGTCGTTGCATGAGTTAGAAGTGGGGTAGCACNNNAGATCGGAA |
| 46587_1 | TATGAGTGTGGAGTCGTTGCCCAGTATCTCCTGCTAAGGTTTGANNNAGATCGGAA |
| 46587_2 | TATGAGTGTGGAGTCGTTGCCCAGTATCTCCTGCTAAGGTTTGANNNAGATCGGAA |
| 48108_1 | TATGAGTGTGGAGTCGTTGCTATTGGCCGCATGTATGTCTTCTNNNAGATCGGAAG |
| 48108_2 | TATGAGTGTGGAGTCGTTGCTATTGGCCGCATGTATGTCTTCTNNNAGATCGGAAG |
| 12181_1 | TATGAGTGTGGAGTCGTTGCAGTCATTGGGGAGAATCTGAGAGTNNNAGATCGGA |
| 12181_2 | TATGAGTGTGGAGTCGTTGCAGTCATTGGGGAGAATCTGAGAGTNNNAGATCGGA |
| 32011_1 | TATGAGTGTGGAGTCGTTGCGTGACCATGGCAGGCTATTTTTTNNNAGATCGGAAG |
| 32011_2 | TATGAGTGTGGAGTCGTTGCGTGACCATGGCAGGCTATTTTTTNNNAGATCGGAAG |
| 48374_1 | TATGAGTGTGGAGTCGTTGCCCTACTGAGTAGCTGGGACTACANNNAGATCGGAAG |
| 48374_2 | TATGAGTGTGGAGTCGTTGCCCTACTGAGTAGCTGGGACTACANNNAGATCGGAAG |
| 40248_1 | TATGAGTGTGGAGTCGTTGCGCTACATGTGATTGGCCAAAACTNNNAGATCGGAAG |
| 40248_2 | TATGAGTGTGGAGTCGTTGCGCTACATGTGATTGGCCAAAACTNNNAGATCGGAAG |
| 32174_1 | TATGAGTGTGGAGTCGTTGCCATTCTCCTGCCTCAGCCTCNNNAGATCGGAAGAGCA |
| 32174_2 | TATGAGTGTGGAGTCGTTGCCATTCTCCTGCCTCAGCCTCNNNAGATCGGAAGAGCA |
| 47506_1 | TATGAGTGTGGAGTCGTTGCTAGTAAGGTAGGGGTGTGTGTGTNNNAGATCGGAA |
| 47506_2 | TATGAGTGTGGAGTCGTTGCTAGTAAGGTAGGGGTGTGTGTGTNNNAGATCGGAA |
| 30374_1 | TATGAGTGTGGAGTCGTTGCAATGAACTAGCTAGCAGACCCCTNNNAGATCGGAAG |
| 30374_2 | TATGAGTGTGGAGTCGTTGCAATGAACTAGCTAGCAGACCCCTNNNAGATCGGAAG |
| 50643_1 | TATGAGTGTGGAGTCGTTGCTAAGCCAGTCCAGAGCACTAAAGNNNAGATCGGAA |
| 50643_2 | TATGAGTGTGGAGTCGTTGCTAAGCCAGTCCAGAGCACTAAAGNNNAGATCGGAA |
| 49223_1 | TATGAGTGTGGAGTCGTTGCTCTCTTTCTATCACACAGGCTGGNNNAGATCGGAAG |
| 49223_2 | TATGAGTGTGGAGTCGTTGCTCTCTTTCTATCACACAGGCTGGNNNAGATCGGAAG |
| 31279_1 | TATGAGTGTGGAGTCGTTGCCTGAGAACTTCTCCCTCCACATCNNNAGATCGGAAGA |
| 31279_2 | TATGAGTGTGGAGTCGTTGCCTGAGAACTTCTCCCTCCACATCNNNAGATCGGAAGA |
| 38436_1 | TATGAGTGTGGAGTCGTTGCGGGTTGCAATCCCAGTCTCTAATNNNAGATCGGAAG |
| 38436_2 | TATGAGTGTGGAGTCGTTGCGGGTTGCAATCCCAGTCTCTAATNNNAGATCGGAAG |
| 33994_1 | TATGAGTGTGGAGTCGTTGCCCTCCCGATTAGCAGAGATTACANNNAGATCGGAAG |
| 33994_2 | TATGAGTGTGGAGTCGTTGCCCTCCCGATTAGCAGAGATTACANNNAGATCGGAAG |
| 52627_1 | TATGAGTGTGGAGTCGTTGCGAGATCCCTCAGCGCATGACNNNAGATCGGAAGAG |
| 52627_2 | TATGAGTGTGGAGTCGTTGCGAGATCCCTCAGCGCATGACNNNAGATCGGAAGAG |
| 50990_1 | TATGAGTGTGGAGTCGTTGCCCCTTTGTGTACAGCAAACACTTNNNAGATCGGAAG |
| 50990_2 | TATGAGTGTGGAGTCGTTGCCCCTTTGTGTACAGCAAACACTTNNNAGATCGGAAG |
| 44994_1 | TATGAGTGTGGAGTCGTTGCGAAGACATGTGTCAGGGACTCTCNNNAGATCGGAA |

| | |
|---|---|
| 44994_2 | TATGAGTGTGGAGTCGTTGCGAAGACATGTGTCAGGGACTCTCNNNAGATCGGAA |
| 33884_1 | TATGAGTGTGGAGTCGTTGCGATTTTGAGACGGAGTCTTGCTCNNNAGATCGGAA |
| 33884_2 | TATGAGTGTGGAGTCGTTGCGATTTTGAGACGGAGTCTTGCTCNNNAGATCGGAA |
| 41147_1 | TATGAGTGTGGAGTCGTTGCGAGGTCAGGAGATTGAGACCATCNNNAGATCGGAA |
| 41147_2 | TATGAGTGTGGAGTCGTTGCGAGGTCAGGAGATTGAGACCATCNNNAGATCGGAA |
| 35958_1 | TATGAGTGTGGAGTCGTTGCCTGTAATCCCAGCTACTCAGGAGNNNAGATCGGAAG |
| 35958_2 | TATGAGTGTGGAGTCGTTGCCTGTAATCCCAGCTACTCAGGAGNNNAGATCGGAAG |
| 39646_1 | TATGAGTGTGGAGTCGTTGCATATTCCCCTGAATGTCTGCCAGNNNAGATCGGAAG |
| 39646_2 | TATGAGTGTGGAGTCGTTGCATATTCCCCTGAATGTCTGCCAGNNNAGATCGGAAG |
| 12605_1 | TATGAGTGTGGAGTCGTTGCCCTTTCAAATTAGCTTATTTCACNNNAGATCGGAAGA |
| 12605_2 | TATGAGTGTGGAGTCGTTGCCCTTTCAAATTAGCTTATTTCACNNNAGATCGGAAGA |
| 36415_1 | TATGAGTGTGGAGTCGTTGCCCAATGCCTATACCCCCATTGTANNNAGATCGGAAG |
| 36415_2 | TATGAGTGTGGAGTCGTTGCCCAATGCCTATACCCCCATTGTANNNAGATCGGAAG |
| 45515_1 | TATGAGTGTGGAGTCGTTGCTCCAAAGGATCCTAAACGTTCCCNNNAGATCGGAAG |
| 45515_2 | TATGAGTGTGGAGTCGTTGCTCCAAAGGATCCTAAACGTTCCCNNNAGATCGGAAG |
| 33729_1 | TATGAGTGTGGAGTCGTTGCAGTTGGCATGCTTGTTTATTGATNNNAGATCGGAAG |
| 33729_2 | TATGAGTGTGGAGTCGTTGCAGTTGGCATGCTTGTTTATTGATNNNAGATCGGAAG |
| 44304_1 | TATGAGTGTGGAGTCGTTGCGCAGACACAGAGAAAGAGTGAGANNNAGATCGGA |
| 44304_2 | TATGAGTGTGGAGTCGTTGCGCAGACACAGAGAAAGAGTGAGANNNAGATCGGA |
| 50635_1 | TATGAGTGTGGAGTCGTTGCCTGTCGGTGACATAGTGGTACAGNNNAGATCGGAA |
| 50635_2 | TATGAGTGTGGAGTCGTTGCCTGTCGGTGACATAGTGGTACAGNNNAGATCGGAA |
| 2379_1 | TATGAGTGTGGAGTCGTTGCTTTACCTACTCCGATATGCACGCNNNAGATCGGAAG |
| 2379_2 | TATGAGTGTGGAGTCGTTGCTTTACCTACTCCGATATGCACGCNNNAGATCGGAAG |
| 38970_1 | TATGAGTGTGGAGTCGTTGCCTTAGTAGAGACGGCGTTTCACCNNNAGATCGGAAG |
| 38970_2 | TATGAGTGTGGAGTCGTTGCCTTAGTAGAGACGGCGTTTCACCNNNAGATCGGAAG |
| 47928_1 | TATGAGTGTGGAGTCGTTGCGAGAAGAGCCCCCCTCATCTNNNAGATCGGAAGAG |
| 47928_2 | TATGAGTGTGGAGTCGTTGCGAGAAGAGCCCCCCTCATCTNNNAGATCGGAAGAG |
| 30634_1 | TATGAGTGTGGAGTCGTTGCGACTCCTCCACATGTTGGAAATGNNNAGATCGGAAG |
| 30634_2 | TATGAGTGTGGAGTCGTTGCGACTCCTCCACATGTTGGAAATGNNNAGATCGGAAG |
| 45353_1 | TATGAGTGTGGAGTCGTTGCGAGTGGGAATTCACCAAACATGGNNNAGATCGGAA |
| 45353_2 | TATGAGTGTGGAGTCGTTGCGAGTGGGAATTCACCAAACATGGNNNAGATCGGAA |
| 52487_1 | TATGAGTGTGGAGTCGTTGCCAGCTGATCTCTTTGGCAAGTTCNNNAGATCGGAAG |
| 52487_2 | TATGAGTGTGGAGTCGTTGCCAGCTGATCTCTTTGGCAAGTTCNNNAGATCGGAAG |
| 34553_1 | TATGAGTGTGGAGTCGTTGCGGCAAGTAGGAGCAAAAACAAGANNNAGATCGGA |
| 34553_2 | TATGAGTGTGGAGTCGTTGCGGCAAGTAGGAGCAAAAACAAGANNNAGATCGGA |
| 37370_1 | TATGAGTGTGGAGTCGTTGCGCTGAGAAAGAGGAAGACAGGAANNNAGATCGGA |
| 37370_2 | TATGAGTGTGGAGTCGTTGCGCTGAGAAAGAGGAAGACAGGAANNNAGATCGGA |
| 48530_1 | TATGAGTGTGGAGTCGTTGCGAGTAGCTGGGATTACAGATGCANNNAGATCGGAA |
| 48530_2 | TATGAGTGTGGAGTCGTTGCGAGTAGCTGGGATTACAGATGCANNNAGATCGGAA |
| 41387_1 | TATGAGTGTGGAGTCGTTGCTCACAAGAGGCAAAAAAGGATGTNNNAGATCGGAA |
| 41387_2 | TATGAGTGTGGAGTCGTTGCTCACAAGAGGCAAAAAAGGATGTNNNAGATCGGAA |
| 4725_1 | TATGAGTGTGGAGTCGTTGCAAAGGCATGAAATACAGCAACTCNNNAGATCGGAA |
| 4725_2 | TATGAGTGTGGAGTCGTTGCAAAGGCATGAAATACAGCAACTCNNNAGATCGGAA |

| | |
|---|---|
| 46426_1 | TATGAGTGTGGAGTCGTTGCTTCACGCTATATTCTCCTGCCTCNNNAGATCGGAAGA |
| 46426_2 | TATGAGTGTGGAGTCGTTGCTTCACGCTATATTCTCCTGCCTCNNNAGATCGGAAGA |
| 32916_1 | TATGAGTGTGGAGTCGTTGCACATTTGCATGCTTTTGACTTGTGNNNAGATCGGAA |
| 32916_2 | TATGAGTGTGGAGTCGTTGCACATTTGCATGCTTTTGACTTGTGNNNAGATCGGAA |
| 10470_1 | TATGAGTGTGGAGTCGTTGCCTTTCATTCCAAGTTGGTTGCCANNNAGATCGGAAG |
| 10470_2 | TATGAGTGTGGAGTCGTTGCCTTTCATTCCAAGTTGGTTGCCANNNAGATCGGAAG |
| 39925_1 | TATGAGTGTGGAGTCGTTGCACAACATATGTACCTACCCTCAAGGNNNAGATCGGA |
| 39925_2 | TATGAGTGTGGAGTCGTTGCACAACATATGTACCTACCCTCAAGGNNNAGATCGGA |
| 10195_1 | TATGAGTGTGGAGTCGTTGCGAATTGCAGGTAGCAGGAGAAGGNNNAGATCGGA |
| 10195_2 | TATGAGTGTGGAGTCGTTGCGAATTGCAGGTAGCAGGAGAAGGNNNAGATCGGA |
| 53025_1 | TATGAGTGTGGAGTCGTTGCCAGTAATCCCAGCTGCTTGAAAGNNNAGATCGGAAG |
| 53025_2 | TATGAGTGTGGAGTCGTTGCCAGTAATCCCAGCTGCTTGAAAGNNNAGATCGGAAG |
| 45335_1 | TATGAGTGTGGAGTCGTTGCTGTTCTTGGGAGTCTGATGACAGNNNAGATCGGAA |
| 45335_2 | TATGAGTGTGGAGTCGTTGCTGTTCTTGGGAGTCTGATGACAGNNNAGATCGGAA |
| 35900_1 | TATGAGTGTGGAGTCGTTGCGGTGACTCGAAGACCCCTATTGNNNAGATCGGAAG |
| 35900_2 | TATGAGTGTGGAGTCGTTGCGGTGACTCGAAGACCCCTATTGNNNAGATCGGAAG |
| 32122_1 | TATGAGTGTGGAGTCGTTGCCTCCCATGTTCACACCGTTCTCNNNAGATCGGAAGAG |
| 32122_2 | TATGAGTGTGGAGTCGTTGCCTCCCATGTTCACACCGTTCTCNNNAGATCGGAAGAG |
| 28900_1 | TATGAGTGTGGAGTCGTTGCCTGTGGTTCTTGCACACTTGTAGNNNAGATCGGAAG |
| 28900_2 | TATGAGTGTGGAGTCGTTGCCTGTGGTTCTTGCACACTTGTAGNNNAGATCGGAAG |
| 29903_1 | TATGAGTGTGGAGTCGTTGCTGGCAGTATACAGGTTGAGGTGANNNAGATCGGAA |
| 29903_2 | TATGAGTGTGGAGTCGTTGCTGGCAGTATACAGGTTGAGGTGANNNAGATCGGAA |
| 46809_1 | TATGAGTGTGGAGTCGTTGCAAATCCTGGGCTCAAGTGATCTTNNNAGATCGGAAG |
| 46809_2 | TATGAGTGTGGAGTCGTTGCAAATCCTGGGCTCAAGTGATCTTNNNAGATCGGAAG |
| 50897_1 | TATGAGTGTGGAGTCGTTGCCGTGATCTGGAGCAGTATGTCAANNNAGATCGGAA |
| 50897_2 | TATGAGTGTGGAGTCGTTGCCGTGATCTGGAGCAGTATGTCAANNNAGATCGGAA |
| 51259_1 | TATGAGTGTGGAGTCGTTGCAAACCAGAAGTGCACATACAAGCNNNAGATCGGAA |
| 51259_2 | TATGAGTGTGGAGTCGTTGCAAACCAGAAGTGCACATACAAGCNNNAGATCGGAA |
| 50724_1 | TATGAGTGTGGAGTCGTTGCATCTGAGTGCATTGAGCTGTGTANNNAGATCGGAA |
| 50724_2 | TATGAGTGTGGAGTCGTTGCATCTGAGTGCATTGAGCTGTGTANNNAGATCGGAA |
| 52277_1 | TATGAGTGTGGAGTCGTTGCTCCTCCTGACTCTTTGATTGGTGNNNAGATCGGAAG |
| 52277_2 | TATGAGTGTGGAGTCGTTGCTCCTCCTGACTCTTTGATTGGTGNNNAGATCGGAAG |
| 37118_1 | TATGAGTGTGGAGTCGTTGCTAGGCTCGAAATTGTCACACTGANNNAGATCGGAAG |
| 37118_2 | TATGAGTGTGGAGTCGTTGCTAGGCTCGAAATTGTCACACTGANNNAGATCGGAAG |
| 52965_1 | TATGAGTGTGGAGTCGTTGCAACAGATCCTTCCCTAGAGCCTTNNNAGATCGGAAG |
| 52965_2 | TATGAGTGTGGAGTCGTTGCAACAGATCCTTCCCTAGAGCCTTNNNAGATCGGAAG |
| 47494_1 | TATGAGTGTGGAGTCGTTGCCTCTGTCACCAATTCCCCACCNNNAGATCGGAAGAGC |
| 47494_2 | TATGAGTGTGGAGTCGTTGCCTCTGTCACCAATTCCCCACCNNNAGATCGGAAGAGC |
| 35982_1 | TATGAGTGTGGAGTCGTTGCTGTTGTGAGTCGTGCTATCAAGANNNAGATCGGAA |
| 35982_2 | TATGAGTGTGGAGTCGTTGCTGTTGTGAGTCGTGCTATCAAGANNNAGATCGGAA |
| 38464_1 | TATGAGTGTGGAGTCGTTGCGCATGATGATGGTTCACTGTAGCNNNAGATCGGAA |
| 38464_2 | TATGAGTGTGGAGTCGTTGCGCATGATGATGGTTCACTGTAGCNNNAGATCGGAA |
| 39230_1 | TATGAGTGTGGAGTCGTTGCACCAGCTGCAAATATACACTCTCANNNAGATCGGAA |

| | |
|---|---|
| 39230_2 | TATGAGTGTGGAGTCGTTGCACCAGCTGCAAATATACACTCTCANNNAGATCGGAA |
| 42272_1 | TATGAGTGTGGAGTCGTTGCTTATAGCTCACTGCAGCCTCAAANNNAGATCGGAAG |
| 42272_2 | TATGAGTGTGGAGTCGTTGCTTATAGCTCACTGCAGCCTCAAANNNAGATCGGAAG |
| 47406_1 | TATGAGTGTGGAGTCGTTGCCTCCTGAGTTCACGCCATTCTCNNNAGATCGGAAGA |
| 47406_2 | TATGAGTGTGGAGTCGTTGCCTCCTGAGTTCACGCCATTCTCNNNAGATCGGAAGA |
| 12640_1 | TATGAGTGTGGAGTCGTTGCTCAAGGAAGAATTGAGCTTATGNNNAGATCGGAAG |
| 12640_2 | TATGAGTGTGGAGTCGTTGCTCAAGGAAGAATTGAGCTTATGNNNAGATCGGAAG |
| 31831_1 | TATGAGTGTGGAGTCGTTGCGCTTTGTTGAAGATCAGTTGGCTNNNAGATCGGAA |
| 31831_2 | TATGAGTGTGGAGTCGTTGCGCTTTGTTGAAGATCAGTTGGCTNNNAGATCGGAA |
| 13152_1 | TATGAGTGTGGAGTCGTTGCAAAAATGAGTTATGAATACTGCANNNAGATCGGAA |
| 13152_2 | TATGAGTGTGGAGTCGTTGCAAAAATGAGTTATGAATACTGCANNNAGATCGGAA |
| 47682_1 | TATGAGTGTGGAGTCGTTGCAATAACACATTTAGGGCCCCACTNNNAGATCGGAAG |
| 47682_2 | TATGAGTGTGGAGTCGTTGCAATAACACATTTAGGGCCCCACTNNNAGATCGGAAG |
| 31634_1 | TATGAGTGTGGAGTCGTTGCGTTGAAGAGGCCAAAAGCAGAATNNNAGATCGGAA |
| 31634_2 | TATGAGTGTGGAGTCGTTGCGTTGAAGAGGCCAAAAGCAGAATNNNAGATCGGAA |
| 10925_1 | TATGAGTGTGGAGTCGTTGCGGAAACAATGAACCTGACTTGCAGNNNAGATCGGA |
| 10925_2 | TATGAGTGTGGAGTCGTTGCGGAAACAATGAACCTGACTTGCAGNNNAGATCGGA |
| 36600_1 | TATGAGTGTGGAGTCGTTGCCCAGGACACATTGGTAGCAATTTNNNAGATCGGAAG |
| 36600_2 | TATGAGTGTGGAGTCGTTGCCCAGGACACATTGGTAGCAATTTNNNAGATCGGAAG |
| 31844_1 | TATGAGTGTGGAGTCGTTGCCCTGGATTGCAGTGGATCCTAAANNNAGATCGGAA |
| 31844_2 | TATGAGTGTGGAGTCGTTGCCCTGGATTGCAGTGGATCCTAAANNNAGATCGGAA |
| 46247_1 | TATGAGTGTGGAGTCGTTGCTCTCAGCAGCATCACTGACTATGNNNAGATCGGAAG |
| 46247_2 | TATGAGTGTGGAGTCGTTGCTCTCAGCAGCATCACTGACTATGNNNAGATCGGAAG |
| 12706_1 | TATGAGTGTGGAGTCGTTGCTTGTAAGAATTGTTAAGCATTCCNNNAGATCGGAAG |
| 12706_2 | TATGAGTGTGGAGTCGTTGCTTGTAAGAATTGTTAAGCATTCCNNNAGATCGGAAG |
| 30164_1 | TATGAGTGTGGAGTCGTTGCCATGTGCTAGCCTTGGATGAATGNNNAGATCGGAA |
| 30164_2 | TATGAGTGTGGAGTCGTTGCCATGTGCTAGCCTTGGATGAATGNNNAGATCGGAA |
| 33008_1 | TATGAGTGTGGAGTCGTTGCCAAGTAGATGGGACTACAGGCACNNNAGATCGGAA |
| 33008_2 | TATGAGTGTGGAGTCGTTGCCAAGTAGATGGGACTACAGGCACNNNAGATCGGAA |
| 49173_1 | TATGAGTGTGGAGTCGTTGCTCTTGTTATGTTACCCAGGCTGGNNNAGATCGGAAG |
| 49173_2 | TATGAGTGTGGAGTCGTTGCTCTTGTTATGTTACCCAGGCTGGNNNAGATCGGAAG |
| 47007_1 | TATGAGTGTGGAGTCGTTGCAAGGGGTTCGACCTGTTTATGTTNNNAGATCGGAA |
| 47007_2 | TATGAGTGTGGAGTCGTTGCAAGGGGTTCGACCTGTTTATGTTNNNAGATCGGAA |
| 37303_1 | TATGAGTGTGGAGTCGTTGCAAAGAAATGATAAGTGCCCTGCCNNNAGATCGGAA |
| 37303_2 | TATGAGTGTGGAGTCGTTGCAAAGAAATGATAAGTGCCCTGCCNNNAGATCGGAA |
| 34614_1 | TATGAGTGTGGAGTCGTTGCCTGCACTGACATCTCTCTGTTCTNNNAGATCGGAAGA |
| 34614_2 | TATGAGTGTGGAGTCGTTGCCTGCACTGACATCTCTCTGTTCTNNNAGATCGGAAGA |
| 47118_1 | TATGAGTGTGGAGTCGTTGCCTACCTCAGCCTCCCAAAAGTAGNNNAGATCGGAAG |
| 47118_2 | TATGAGTGTGGAGTCGTTGCCTACCTCAGCCTCCCAAAAGTAGNNNAGATCGGAAG |
| 41662_1 | TATGAGTGTGGAGTCGTTGCCTCCTGAGTAGCTGGGATTACAGNNNAGATCGGAA |
| 41662_2 | TATGAGTGTGGAGTCGTTGCCTCCTGAGTAGCTGGGATTACAGNNNAGATCGGAA |
| 28936_1 | TATGAGTGTGGAGTCGTTGCTGACTGAAACAGCCCCTAGTTACNNNAGATCGGAAG |
| 28936_2 | TATGAGTGTGGAGTCGTTGCTGACTGAAACAGCCCCTAGTTACNNNAGATCGGAAG |

| | |
|---|---|
| 28766_1 | TATGAGTGTGGAGTCGTTGCATGGGCTTGAAATTAGTAGGACATNNNAGATCGGA |
| 28766_2 | TATGAGTGTGGAGTCGTTGCATGGGCTTGAAATTAGTAGGACATNNNAGATCGGA |
| 42100_1 | TATGAGTGTGGAGTCGTTGCGGCATTTCTGGATTGTGAGGTTCNNNAGATCGGAA |
| 42100_2 | TATGAGTGTGGAGTCGTTGCGGCATTTCTGGATTGTGAGGTTCNNNAGATCGGAA |
| 30968_1 | TATGAGTGTGGAGTCGTTGCGTGACATCAGACAAAGAAGGTGCNNNAGATCGGAA |
| 30968_2 | TATGAGTGTGGAGTCGTTGCGTGACATCAGACAAAGAAGGTGCNNNAGATCGGAA |
| 2495_1 | TATGAGTGTGGAGTCGTTGCGAAAGGGAAGGTGGGTTTGAGAGNNNAGATCGGA |
| 2495_2 | TATGAGTGTGGAGTCGTTGCGAAAGGGAAGGTGGGTTTGAGAGNNNAGATCGGA |
| 31001_1 | TATGAGTGTGGAGTCGTTGCTTTGAGCGTACAGTCAGTCACTGNNNAGATCGGAAG |
| 31001_2 | TATGAGTGTGGAGTCGTTGCTTTGAGCGTACAGTCAGTCACTGNNNAGATCGGAAG |
| 37255_1 | TATGAGTGTGGAGTCGTTGCTGTTCCTCTCACTTAGGTTCTCANNNAGATCGGAAGA |
| 37255_2 | TATGAGTGTGGAGTCGTTGCTGTTCCTCTCACTTAGGTTCTCANNNAGATCGGAAGA |
| 30322_1 | TATGAGTGTGGAGTCGTTGCGGATGTTGTTCTGCCTGTGTTTGNNNAGATCGGAAG |
| 30322_2 | TATGAGTGTGGAGTCGTTGCGGATGTTGTTCTGCCTGTGTTTGNNNAGATCGGAAG |
| 4614_1 | TATGAGTGTGGAGTCGTTGCCTGATAACCCTTGGTGCAGTCAGNNNAGATCGGAAG |
| 4614_2 | TATGAGTGTGGAGTCGTTGCCTGATAACCCTTGGTGCAGTCAGNNNAGATCGGAAG |
| 50002_1 | TATGAGTGTGGAGTCGTTGCATGAAGGCAAGCTCCTATTTGGANNNAGATCGGAA |
| 50002_2 | TATGAGTGTGGAGTCGTTGCATGAAGGCAAGCTCCTATTTGGANNNAGATCGGAA |
| 43137_1 | TATGAGTGTGGAGTCGTTGCAGCAAGGAAATGTACATTGTGCCNNNAGATCGGAA |
| 43137_2 | TATGAGTGTGGAGTCGTTGCAGCAAGGAAATGTACATTGTGCCNNNAGATCGGAA |
| 2272_1 | TATGAGTGTGGAGTCGTTGCTTCCTTTGCATACGGTCTCTTCGNNNAGATCGGAAGA |
| 2272_2 | TATGAGTGTGGAGTCGTTGCTTCCTTTGCATACGGTCTCTTCGNNNAGATCGGAAGA |
| 35191_1 | TATGAGTGTGGAGTCGTTGCCGACTAACTGGGATTACAGGCATNNNAGATCGGAA |
| 35191_2 | TATGAGTGTGGAGTCGTTGCCGACTAACTGGGATTACAGGCATNNNAGATCGGAA |
| 38273_1 | TATGAGTGTGGAGTCGTTGCCCACCAAGGAAAATTCTTGAACCANNNAGATCGGAA |
| 38273_2 | TATGAGTGTGGAGTCGTTGCCCACCAAGGAAAATTCTTGAACCANNNAGATCGGAA |
| 30716_1 | TATGAGTGTGGAGTCGTTGCTCCGAATGGCTTAACACCATCCCNNNAGATCGGAAG |
| 30716_2 | TATGAGTGTGGAGTCGTTGCTCCGAATGGCTTAACACCATCCCNNNAGATCGGAAG |
| 35704_1 | TATGAGTGTGGAGTCGTTGCCTCACTGCAAACTCCTCCTTCCNNNAGATCGGAAGAG |
| 35704_2 | TATGAGTGTGGAGTCGTTGCCTCACTGCAAACTCCTCCTTCCNNNAGATCGGAAGAG |
| 13342_1 | TATGAGTGTGGAGTCGTTGCTACTGTTAACTACTGTACAACCNNNAGATCGGAAGA |
| 13342_2 | TATGAGTGTGGAGTCGTTGCTACTGTTAACTACTGTACAACCNNNAGATCGGAAGA |
| 43476_1 | TATGAGTGTGGAGTCGTTGCCTTCCGAGTAGCTGGGATTACAGNNNAGATCGGAA |
| 43476_2 | TATGAGTGTGGAGTCGTTGCCTTCCGAGTAGCTGGGATTACAGNNNAGATCGGAA |
| 30138_1 | TATGAGTGTGGAGTCGTTGCCAAGATGACTGTGGTGATGAGGANNNAGATCGGAA |
| 30138_2 | TATGAGTGTGGAGTCGTTGCCAAGATGACTGTGGTGATGAGGANNNAGATCGGAA |
| 33238_1 | TATGAGTGTGGAGTCGTTGCCTCCTGGGTTCACACCATTCTCNNNAGATCGGAAGA |
| 33238_2 | TATGAGTGTGGAGTCGTTGCCTCCTGGGTTCACACCATTCTCNNNAGATCGGAAGA |
| 37564_1 | TATGAGTGTGGAGTCGTTGCTTGTCAGGGGAATTGGAGAAAGTNNNAGATCGGAA |
| 37564_2 | TATGAGTGTGGAGTCGTTGCTTGTCAGGGGAATTGGAGAAAGTNNNAGATCGGAA |
| 34143_1 | TATGAGTGTGGAGTCGTTGCGGAGCTCCTTACTGTTCATCTCCNNNAGATCGGAAG |
| 34143_2 | TATGAGTGTGGAGTCGTTGCGGAGCTCCTTACTGTTCATCTCCNNNAGATCGGAAG |
| 37763_1 | TATGAGTGTGGAGTCGTTGCGAGAATTGCTTGAATCTGGGAGGNNNAGATCGGAA |

| | |
|---|---|
| 37763_2 | TATGAGTGTGGAGTCGTTGCGAGAATTGCTTGAATCTGGGAGGNNNAGATCGGAA |
| 32430_1 | TATGAGTGTGGAGTCGTTGCTTCAAGTGATTCTCCTGCCTCAGNNNAGATCGGAAG |
| 32430_2 | TATGAGTGTGGAGTCGTTGCTTCAAGTGATTCTCCTGCCTCAGNNNAGATCGGAAG |
| 36764_1 | TATGAGTGTGGAGTCGTTGCAGCCTAGAGGACATCTCACTCTTNNNAGATCGGAAG |
| 36764_2 | TATGAGTGTGGAGTCGTTGCAGCCTAGAGGACATCTCACTCTTNNNAGATCGGAAG |
| 46277_1 | TATGAGTGTGGAGTCGTTGCTCTCCCAGGTTCAAGTGTTTCTCNNNAGATCGGAAG |
| 46277_2 | TATGAGTGTGGAGTCGTTGCTCTCCCAGGTTCAAGTGTTTCTCNNNAGATCGGAAG |
| 44042_1 | TATGAGTGTGGAGTCGTTGCCTCTGGAGGCTGAGCTATATGTGNNNAGATCGGAA |
| 44042_2 | TATGAGTGTGGAGTCGTTGCCTCTGGAGGCTGAGCTATATGTGNNNAGATCGGAA |
| 31692_1 | TATGAGTGTGGAGTCGTTGCAAGGACACCAGTGCTATTGGATTNNNAGATCGGAA |
| 31692_2 | TATGAGTGTGGAGTCGTTGCAAGGACACCAGTGCTATTGGATTNNNAGATCGGAA |
| 12581_1 | TATGAGTGTGGAGTCGTTGCAAATCCCATACAAGTAAGAGTTANNNAGATCGGAAG |
| 12581_2 | TATGAGTGTGGAGTCGTTGCAAATCCCATACAAGTAAGAGTTANNNAGATCGGAAG |
| 8205_1 | TATGAGTGTGGAGTCGTTGCCTCTCACTCACTCACTGTTCCTGNNNAGATCGGAAGA |
| 8205_2 | TATGAGTGTGGAGTCGTTGCCTCTCACTCACTCACTGTTCCTGNNNAGATCGGAAGA |
| 41884_1 | TATGAGTGTGGAGTCGTTGCATGAAATCTTGCTCTGTCACCCANNNAGATCGGAAG |
| 41884_2 | TATGAGTGTGGAGTCGTTGCATGAAATCTTGCTCTGTCACCCANNNAGATCGGAAG |
| 35726_1 | TATGAGTGTGGAGTCGTTGCCCAAGTAGCTGGGATTACAGTCANNNAGATCGGAA |
| 35726_2 | TATGAGTGTGGAGTCGTTGCCCAAGTAGCTGGGATTACAGTCANNNAGATCGGAA |
| 36720_1 | TATGAGTGTGGAGTCGTTGCAATCCTAGAATTTTGGGAGGCCANNNAGATCGGAA |
| 36720_2 | TATGAGTGTGGAGTCGTTGCAATCCTAGAATTTTGGGAGGCCANNNAGATCGGAA |
| 36761_1 | TATGAGTGTGGAGTCGTTGCAGTCACAAAAAGAGAAGCCCTGANNNAGATCGGAA |
| 36761_2 | TATGAGTGTGGAGTCGTTGCAGTCACAAAAAGAGAAGCCCTGANNNAGATCGGAA |
| 34156_1 | TATGAGTGTGGAGTCGTTGCGACCTCGCTTTTTAGGTAGAGCTNNNAGATCGGAAG |
| 34156_2 | TATGAGTGTGGAGTCGTTGCGACCTCGCTTTTTAGGTAGAGCTNNNAGATCGGAAG |
| 32987_1 | TATGAGTGTGGAGTCGTTGCTAACACTGACATTGCATCCGGTANNNAGATCGGAAG |
| 32987_2 | TATGAGTGTGGAGTCGTTGCTAACACTGACATTGCATCCGGTANNNAGATCGGAAG |
| 51357_1 | TATGAGTGTGGAGTCGTTGCATTCTCTTTCTCAGGGACGTTCCNNNAGATCGGAAG |
| 51357_2 | TATGAGTGTGGAGTCGTTGCATTCTCTTTCTCAGGGACGTTCCNNNAGATCGGAAG |
| 34607_1 | TATGAGTGTGGAGTCGTTGCTATCATGAGGAGGTGGCTACAGANNNAGATCGGAA |
| 34607_2 | TATGAGTGTGGAGTCGTTGCTATCATGAGGAGGTGGCTACAGANNNAGATCGGAA |
| 33899_1 | TATGAGTGTGGAGTCGTTGCCATTCTCCTGCCTCAGCCTCNNNAGATCGGAAGAGCA |
| 33899_2 | TATGAGTGTGGAGTCGTTGCCATTCTCCTGCCTCAGCCTCNNNAGATCGGAAGAGCA |
| 41967_1 | TATGAGTGTGGAGTCGTTGCCTCCCAAAGTGCTGGAATTACAGNNNAGATCGGAAG |
| 41967_2 | TATGAGTGTGGAGTCGTTGCCTCCCAAAGTGCTGGAATTACAGNNNAGATCGGAAG |
| 31747_1 | TATGAGTGTGGAGTCGTTGCATGAAAGCTGAGGGAAAAGAAGCNNNAGATCGGA |
| 31747_2 | TATGAGTGTGGAGTCGTTGCATGAAAGCTGAGGGAAAAGAAGCNNNAGATCGGA |
| 42142_1 | TATGAGTGTGGAGTCGTTGCGAGTAGCTGGGATACAGGTGTGNNNAGATCGGAA |
| 42142_2 | TATGAGTGTGGAGTCGTTGCGAGTAGCTGGGATACAGGTGTGNNNAGATCGGAA |
| 41625_1 | TATGAGTGTGGAGTCGTTGCAAGTAGCTGGGACTACAGGCATNNNAGATCGGAAG |
| 41625_2 | TATGAGTGTGGAGTCGTTGCAAGTAGCTGGGACTACAGGCATNNNAGATCGGAAG |
| 32472_1 | TATGAGTGTGGAGTCGTTGCGGTTCAAATGATTCTCCTGCCTCNNNAGATCGGAAG |
| 32472_2 | TATGAGTGTGGAGTCGTTGCGGTTCAAATGATTCTCCTGCCTCNNNAGATCGGAAG |

| | |
|---|---|
| 45732_1 | TATGAGTGTGGAGTCGTTGCGGACCCTCCATACTGTTTTCCATNNNAGATCGGAAGA |
| 45732_2 | TATGAGTGTGGAGTCGTTGCGGACCCTCCATACTGTTTTCCATNNNAGATCGGAAGA |
| 50230_1 | TATGAGTGTGGAGTCGTTGCTAGAATCCCAGAAGTCCAACAGCNNNAGATCGGAAG |
| 50230_2 | TATGAGTGTGGAGTCGTTGCTAGAATCCCAGAAGTCCAACAGCNNNAGATCGGAAG |
| 46907_1 | TATGAGTGTGGAGTCGTTGCCAAGTATCTGGGAGTAGAGGCATNNNAGATCGGAA |
| 46907_2 | TATGAGTGTGGAGTCGTTGCCAAGTATCTGGGAGTAGAGGCATNNNAGATCGGAA |
| 37704_1 | TATGAGTGTGGAGTCGTTGCAGAAAGACCTCAAGTGAGTAACCTNNNAGATCGGAA |
| 37704_2 | TATGAGTGTGGAGTCGTTGCAGAAAGACCTCAAGTGAGTAACCTNNNAGATCGGAA |
| 45203_1 | TATGAGTGTGGAGTCGTTGCGGAGCAGCTCGAAAGGGATTNNNAGATCGGAAGAG |
| 45203_2 | TATGAGTGTGGAGTCGTTGCGGAGCAGCTCGAAAGGGATTNNNAGATCGGAAGAG |
| 42090_1 | TATGAGTGTGGAGTCGTTGCAATTGCTTGAATCTGGAAGGCAGNNNAGATCGGAAG |
| 42090_2 | TATGAGTGTGGAGTCGTTGCAATTGCTTGAATCTGGAAGGCAGNNNAGATCGGAAG |
| 35452_1 | TATGAGTGTGGAGTCGTTGCCTCTTGAGTAGCTGGGACTACAGNNNAGATCGGAAG |
| 35452_2 | TATGAGTGTGGAGTCGTTGCCTCTTGAGTAGCTGGGACTACAGNNNAGATCGGAAG |
| 8298_1 | TATGAGTGTGGAGTCGTTGCAGTACCTCTTTGACATTCCACTTTNNNAGATCGGAAG |
| 8298_2 | TATGAGTGTGGAGTCGTTGCAGTACCTCTTTGACATTCCACTTTNNNAGATCGGAAG |
| 36561_1 | TATGAGTGTGGAGTCGTTGCGCCGTCAAAGAGGCAAATAAAATNNNAGATCGGAAG |
| 36561_2 | TATGAGTGTGGAGTCGTTGCGCCGTCAAAGAGGCAAATAAAATNNNAGATCGGAAG |
| 30918_1 | TATGAGTGTGGAGTCGTTGCTTGCTCTTTCATGTCTCAGTGGTNNNAGATCGGAAGA |
| 30918_2 | TATGAGTGTGGAGTCGTTGCTTGCTCTTTCATGTCTCAGTGGTNNNAGATCGGAAGA |
| 41699_1 | TATGAGTGTGGAGTCGTTGCTTTCACCATGTTAGCCAGGCTNNNAGATCGGAAGAG |
| 41699_2 | TATGAGTGTGGAGTCGTTGCTTTCACCATGTTAGCCAGGCTNNNAGATCGGAAGAG |
| 51410_1 | TATGAGTGTGGAGTCGTTGCACAAAGTTCTGACCAAAACACCGNNNAGATCGGAAG |
| 51410_2 | TATGAGTGTGGAGTCGTTGCACAAAGTTCTGACCAAAACACCGNNNAGATCGGAAG |
| 46002_1 | TATGAGTGTGGAGTCGTTGCTATCTCCCAATGCTATCCCTCCCNNNAGATCGGAAGAG |
| 46002_2 | TATGAGTGTGGAGTCGTTGCTATCTCCCAATGCTATCCCTCCCNNNAGATCGGAAGAG |
| 52215_1 | TATGAGTGTGGAGTCGTTGCCCCGACCCCATCATTGTTTTTTTNNNAGATCGGAAGAG |
| 52215_2 | TATGAGTGTGGAGTCGTTGCCCCGACCCCATCATTGTTTTTTTNNNAGATCGGAAGAG |
| 28798_1 | TATGAGTGTGGAGTCGTTGCGCTAACATGGTGAAACCCCTTCTNNNAGATCGGAAGA |
| 28798_2 | TATGAGTGTGGAGTCGTTGCGCTAACATGGTGAAACCCCTTCTNNNAGATCGGAAGA |
| 40468_1 | TATGAGTGTGGAGTCGTTGCATGTCTTCAAGTGGCATCCTACCNNNAGATCGGAAGA |
| 40468_2 | TATGAGTGTGGAGTCGTTGCATGTCTTCAAGTGGCATCCTACCNNNAGATCGGAAGA |
| 39353_1 | TATGAGTGTGGAGTCGTTGCTTCCATGGTGTATATGTGCCACANNNAGATCGGAAG |
| 39353_2 | TATGAGTGTGGAGTCGTTGCTTCCATGGTGTATATGTGCCACANNNAGATCGGAAG |
| 10302_1 | TATGAGTGTGGAGTCGTTGCTGTGTGTGTGTGAAAAGGAGGTTNNNAGATCGGAA |
| 10302_2 | TATGAGTGTGGAGTCGTTGCTGTGTGTGTGTGAAAAGGAGGTTNNNAGATCGGAA |
| 37637_1 | TATGAGTGTGGAGTCGTTGCCTGAGGCAGGAGAATTGCTTGAANNNAGATCGGAA |
| 37637_2 | TATGAGTGTGGAGTCGTTGCCTGAGGCAGGAGAATTGCTTGAANNNAGATCGGAA |
| 35141_1 | TATGAGTGTGGAGTCGTTGCTTCATGCCATTCTCCTGTCTCAGNNNAGATCGGAAGA |
| 35141_2 | TATGAGTGTGGAGTCGTTGCTTCATGCCATTCTCCTGTCTCAGNNNAGATCGGAAGA |
| 33435_1 | TATGAGTGTGGAGTCGTTGCGGGAGATGACAGGCAGATACAAANNNAGATCGGAA |
| 33435_2 | TATGAGTGTGGAGTCGTTGCGGGAGATGACAGGCAGATACAAANNNAGATCGGAA |
| 51763_1 | TATGAGTGTGGAGTCGTTGCATTTCTGAATGCTCCTGTGTCCTNNNAGATCGGAAGA |

| | |
|---|---|
| 51763_2 | TATGAGTGTGGAGTCGTTGCATTTCTGAATGCTCCTGTGTCCTNNNAGATCGGAAG |
| 38652_1 | TATGAGTGTGGAGTCGTTGCCGATGCAGAAGATGGGTGATTTCNNNAGATCGGAA |
| 38652_2 | TATGAGTGTGGAGTCGTTGCCGATGCAGAAGATGGGTGATTTCNNNAGATCGGAA |
| 34087_1 | TATGAGTGTGGAGTCGTTGCAAGTGTTGGGATTACAGGCATGANNNAGATCGGAA |
| 34087_2 | TATGAGTGTGGAGTCGTTGCAAGTGTTGGGATTACAGGCATGANNNAGATCGGAA |
| 28968_1 | TATGAGTGTGGAGTCGTTGCTATCTGCAGATTTGGTCTTGGGANNNAGATCGGAA |
| 28968_2 | TATGAGTGTGGAGTCGTTGCTATCTGCAGATTTGGTCTTGGGANNNAGATCGGAA |
| 10517_1 | TATGAGTGTGGAGTCGTTGCGGGGTAGGGAGGCTGATTATTCTNNNAGATCGGAA |
| 10517_2 | TATGAGTGTGGAGTCGTTGCGGGGTAGGGAGGCTGATTATTCTNNNAGATCGGAA |
| 38339_1 | TATGAGTGTGGAGTCGTTGCGTGCCAGCCTGACATTTAGAAAANNNAGATCGGAA |
| 38339_2 | TATGAGTGTGGAGTCGTTGCGTGCCAGCCTGACATTTAGAAAANNNAGATCGGAA |
| 13200_1 | TATGAGTGTGGAGTCGTTGCACTTAAATTCTGTCTGAACAAAGNNNAGATCGGAAG |
| 13200_2 | TATGAGTGTGGAGTCGTTGCACTTAAATTCTGTCTGAACAAAGNNNAGATCGGAAG |
| 35477_1 | TATGAGTGTGGAGTCGTTGCTTCATCCATGTCCCTACAAAGGANNNAGATCGGAAG |
| 35477_2 | TATGAGTGTGGAGTCGTTGCTTCATCCATGTCCCTACAAAGGANNNAGATCGGAAG |
| 50533_1 | TATGAGTGTGGAGTCGTTGCTCATTCTAGGTTGGGCCAAATGANNNAGATCGGAA |
| 50533_2 | TATGAGTGTGGAGTCGTTGCTCATTCTAGGTTGGGCCAAATGANNNAGATCGGAA |
| 35662_1 | TATGAGTGTGGAGTCGTTGCACCTAGGATGGTCTCAATCTCCTNNNAGATCGGAAG |
| 35662_2 | TATGAGTGTGGAGTCGTTGCACCTAGGATGGTCTCAATCTCCTNNNAGATCGGAAG |
| 47908_1 | TATGAGTGTGGAGTCGTTGCAGGTTGAAATGATTCTCCTGCCTNNNAGATCGGAAG |
| 47908_2 | TATGAGTGTGGAGTCGTTGCAGGTTGAAATGATTCTCCTGCCTNNNAGATCGGAAG |
| 40894_1 | TATGAGTGTGGAGTCGTTGCTCTGGATAAGGAAGTGTGGCAAANNNAGATCGGAA |
| 40894_2 | TATGAGTGTGGAGTCGTTGCTCTGGATAAGGAAGTGTGGCAAANNNAGATCGGAA |
| 44235_1 | TATGAGTGTGGAGTCGTTGCTCAAATGTCAGTGACCACTAGGGNNNAGATCGGAA |
| 44235_2 | TATGAGTGTGGAGTCGTTGCTCAAATGTCAGTGACCACTAGGGNNNAGATCGGAA |
| 42150_1 | TATGAGTGTGGAGTCGTTGCCATTCTCCTGCCTCAGCCTCNNNAGATCGGAAGAGCA |
| 42150_2 | TATGAGTGTGGAGTCGTTGCCATTCTCCTGCCTCAGCCTCNNNAGATCGGAAGAGCA |
| 32769_1 | TATGAGTGTGGAGTCGTTGCGGCTGATGTGGAAGGATCACTTANNNAGATCGGAA |
| 32769_2 | TATGAGTGTGGAGTCGTTGCGGCTGATGTGGAAGGATCACTTANNNAGATCGGAA |
| 30556_1 | TATGAGTGTGGAGTCGTTGCTTTTCTAAGTTCGGAGTCAGTGTTANNNAGATCGGA |
| 30556_2 | TATGAGTGTGGAGTCGTTGCTTTTCTAAGTTCGGAGTCAGTGTTANNNAGATCGGA |
| 29999_1 | TATGAGTGTGGAGTCGTTGCGCGAGGTGAGAGTAAAGTCCANNNAGATCGGAAG |
| 29999_2 | TATGAGTGTGGAGTCGTTGCGCGAGGTGAGAGTAAAGTCCANNNAGATCGGAAG |
| 42811_1 | TATGAGTGTGGAGTCGTTGCAGAGAGAGAGAAAGACATGAGGTNNNAGATCGGA |
| 42811_2 | TATGAGTGTGGAGTCGTTGCAGAGAGAGAGAAAGACATGAGGTNNNAGATCGGA |
| 52111_1 | TATGAGTGTGGAGTCGTTGCCACTGTCTGGGAAGTGAGAAGAANNNAGATCGGAA |
| 52111_2 | TATGAGTGTGGAGTCGTTGCCACTGTCTGGGAAGTGAGAAGAANNNAGATCGGAA |
| 41275_1 | TATGAGTGTGGAGTCGTTGCGAGATCACACCATTGCATTCCAGNNNAGATCGGAAG |
| 41275_2 | TATGAGTGTGGAGTCGTTGCGAGATCACACCATTGCATTCCAGNNNAGATCGGAAG |
| 44923_1 | TATGAGTGTGGAGTCGTTGCGTTCCCTCTGATTTTTTGCTGCTNNNAGATCGGAAGA |
| 44923_2 | TATGAGTGTGGAGTCGTTGCGTTCCCTCTGATTTTTTGCTGCTNNNAGATCGGAAGA |
| 44107_1 | TATGAGTGTGGAGTCGTTGCCTTCAAAGTCAGCAGGCATCTTCNNNAGATCGGAAG |
| 44107_2 | TATGAGTGTGGAGTCGTTGCCTTCAAAGTCAGCAGGCATCTTCNNNAGATCGGAAG |

| | |
|---|---|
| 42726_1 | TATGAGTGTGGAGTCGTTGCCAGAGGTTGCAGTGAGAAGAGATNNNAGATCGGAA |
| 42726_2 | TATGAGTGTGGAGTCGTTGCCAGAGGTTGCAGTGAGAAGAGATNNNAGATCGGAA |
| 50432_1 | TATGAGTGTGGAGTCGTTGCACTGAAAAGGATGTGGAGGACTCNNNAGATCGGAA |
| 50432_2 | TATGAGTGTGGAGTCGTTGCACTGAAAAGGATGTGGAGGACTCNNNAGATCGGAA |
| 36154_1 | TATGAGTGTGGAGTCGTTGCGCATGGAGCACATTCATGTTTCTNNNAGATCGGAAG |
| 36154_2 | TATGAGTGTGGAGTCGTTGCGCATGGAGCACATTCATGTTTCTNNNAGATCGGAAG |
| 41773_1 | TATGAGTGTGGAGTCGTTGCAGATGGCATCTCACTCTGTCACNNNAGATCGGAAGA |
| 41773_2 | TATGAGTGTGGAGTCGTTGCAGATGGCATCTCACTCTGTCACNNNAGATCGGAAGA |
| 37651_1 | TATGAGTGTGGAGTCGTTGCCTTGAACCAGGGAGTTAGAGCTTNNNAGATCGGAA |
| 37651_2 | TATGAGTGTGGAGTCGTTGCCTTGAACCAGGGAGTTAGAGCTTNNNAGATCGGAA |
| 33229_1 | TATGAGTGTGGAGTCGTTGCCTCAAACTCCTGACCTCGTGATCNNNAGATCGGAAGA |
| 33229_2 | TATGAGTGTGGAGTCGTTGCCTCAAACTCCTGACCTCGTGATCNNNAGATCGGAAGA |
| 30451_1 | TATGAGTGTGGAGTCGTTGCGCCCCAACAGTAATCCAATCACANNNAGATCGGAAG |
| 30451_2 | TATGAGTGTGGAGTCGTTGCGCCCCAACAGTAATCCAATCACANNNAGATCGGAAG |
| 38908_1 | TATGAGTGTGGAGTCGTTGCTCACAGCCAAATACTGCTACACANNNAGATCGGAAG |
| 38908_2 | TATGAGTGTGGAGTCGTTGCTCACAGCCAAATACTGCTACACANNNAGATCGGAAG |
| 2848_1 | TATGAGTGTGGAGTCGTTGCTGGTGTATGTCCAGGATGTGAGTNNNAGATCGGAA |
| 2848_2 | TATGAGTGTGGAGTCGTTGCTGGTGTATGTCCAGGATGTGAGTNNNAGATCGGAA |
| 13380_1 | TATGAGTGTGGAGTCGTTGCAATTATCCTACCTCAGCCTCNNNAGATCGGAAGAGCA |
| 13380_2 | TATGAGTGTGGAGTCGTTGCAATTATCCTACCTCAGCCTCNNNAGATCGGAAGAGCA |
| 41621_1 | TATGAGTGTGGAGTCGTTGCTCTTGAACTCCTGACCTCAAGTGNNNAGATCGGAAG |
| 41621_2 | TATGAGTGTGGAGTCGTTGCTCTTGAACTCCTGACCTCAAGTGNNNAGATCGGAAG |
| 44884_1 | TATGAGTGTGGAGTCGTTGCGACCTTCAGTCACCCAGACATCNNNAGATCGGAAGA |
| 44884_2 | TATGAGTGTGGAGTCGTTGCGACCTTCAGTCACCCAGACATCNNNAGATCGGAAGA |
| 28930_1 | TATGAGTGTGGAGTCGTTGCCTGAGAACCCAAACTGGATGTTGNNNAGATCGGAAG |
| 28930_2 | TATGAGTGTGGAGTCGTTGCCTGAGAACCCAAACTGGATGTTGNNNAGATCGGAAG |
| 41525_1 | TATGAGTGTGGAGTCGTTGCTGGGATTACAGGTGTGAACCAATNNNAGATCGGAA |
| 41525_2 | TATGAGTGTGGAGTCGTTGCTGGGATTACAGGTGTGAACCAATNNNAGATCGGAA |
| 37512_1 | TATGAGTGTGGAGTCGTTGCCAACTGTGCTGTGATACTTAGAAANNNAGATCGGAA |
| 37512_2 | TATGAGTGTGGAGTCGTTGCCAACTGTGCTGTGATACTTAGAAANNNAGATCGGAA |
| 31440_1 | TATGAGTGTGGAGTCGTTGCATCTTGAGGTCAGGAGATCGAGANNNAGATCGGAA |
| 31440_2 | TATGAGTGTGGAGTCGTTGCATCTTGAGGTCAGGAGATCGAGANNNAGATCGGAA |
| 42713_1 | TATGAGTGTGGAGTCGTTGCCACACACACACACCATGGAATACNNNAGATCGGAAG |
| 42713_2 | TATGAGTGTGGAGTCGTTGCCACACACACACACCATGGAATACNNNAGATCGGAAG |
| 29939_1 | TATGAGTGTGGAGTCGTTGCACAGTATGTAGCTTTGGCCATCANNNAGATCGGAAG |
| 29939_2 | TATGAGTGTGGAGTCGTTGCACAGTATGTAGCTTTGGCCATCANNNAGATCGGAAG |
| 36275_1 | TATGAGTGTGGAGTCGTTGCTTTCCAAGAGATGCTGATGTTGCNNNAGATCGGAAG |
| 36275_2 | TATGAGTGTGGAGTCGTTGCTTTCCAAGAGATGCTGATGTTGCNNNAGATCGGAAG |
| 45944_1 | TATGAGTGTGGAGTCGTTGCAGGCAAAGTCTCATTCTTGTTGCNNNAGATCGGAAG |
| 45944_2 | TATGAGTGTGGAGTCGTTGCAGGCAAAGTCTCATTCTTGTTGCNNNAGATCGGAAG |
| 13237_1 | TATGAGTGTGGAGTCGTTGCTTTTATTTTTTTGAGACAGAGGCNNNAGATCGGAAG |
| 13237_2 | TATGAGTGTGGAGTCGTTGCTTTTATTTTTTTGAGACAGAGGCNNNAGATCGGAAG |
| 46499_1 | TATGAGTGTGGAGTCGTTGCCCTACTAAATCCTGCCCATCTCCNNNAGATCGGAAGA |

| | |
|---|---|
| 46499_2 | TATGAGTGTGGAGTCGTTGCCCTACTAAATCCTGCCCATCTCCNNNAGATCGGAAGA |
| 29459_1 | TATGAGTGTGGAGTCGTTGCTATGTGTAGCAGGGGCTTCATTGNNNAGATCGGAA |
| 29459_2 | TATGAGTGTGGAGTCGTTGCTATGTGTAGCAGGGGCTTCATTGNNNAGATCGGAA |
| 37733_1 | TATGAGTGTGGAGTCGTTGCACAACAAAAAGGTAAAATGTGGGGANNNAGATCGG |
| 37733_2 | TATGAGTGTGGAGTCGTTGCACAACAAAAAGGTAAAATGTGGGGANNNAGATCGG |
| 44517_1 | TATGAGTGTGGAGTCGTTGCTGAAGTCCATCAGCTGTCAAGAANNNAGATCGGAAG |
| 44517_2 | TATGAGTGTGGAGTCGTTGCTGAAGTCCATCAGCTGTCAAGAANNNAGATCGGAAG |
| 46857_1 | TATGAGTGTGGAGTCGTTGCCTCAAGCCATTCTCCTGCTTCAGNNNAGATCGGAAGA |
| 46857_2 | TATGAGTGTGGAGTCGTTGCCTCAAGCCATTCTCCTGCTTCAGNNNAGATCGGAAGA |
| 33049_1 | TATGAGTGTGGAGTCGTTGCCATGATCTCAGCTCACTGCAACNNNAGATCGGAAGA |
| 33049_2 | TATGAGTGTGGAGTCGTTGCCATGATCTCAGCTCACTGCAACNNNAGATCGGAAGA |
| 46375_1 | TATGAGTGTGGAGTCGTTGCATGATCTTGACTCACTGCAACCTNNNAGATCGGAAGA |
| 46375_2 | TATGAGTGTGGAGTCGTTGCATGATCTTGACTCACTGCAACCTNNNAGATCGGAAGA |
| 50791_1 | TATGAGTGTGGAGTCGTTGCCCGGGTCACTGAATTGAAAAAGTNNNAGATCGGAA |
| 50791_2 | TATGAGTGTGGAGTCGTTGCCCGGGTCACTGAATTGAAAAAGTNNNAGATCGGAA |
| 39779_1 | TATGAGTGTGGAGTCGTTGCCCCAGAAACAAATCCACACTTCANNNAGATCGGAAGA |
| 39779_2 | TATGAGTGTGGAGTCGTTGCCCCAGAAACAAATCCACACTTCANNNAGATCGGAAGA |
| 12979_1 | TATGAGTGTGGAGTCGTTGCATTTTGGCTACATTAAGCTTTTNNNAGATCGGAAGA |
| 12979_2 | TATGAGTGTGGAGTCGTTGCATTTTGGCTACATTAAGCTTTTNNNAGATCGGAAGA |
| 42762_1 | TATGAGTGTGGAGTCGTTGCGCCTTCAAGCCAAGTCAGTGATANNNAGATCGGAAG |
| 42762_2 | TATGAGTGTGGAGTCGTTGCGCCTTCAAGCCAAGTCAGTGATANNNAGATCGGAAG |
| 46303_1 | TATGAGTGTGGAGTCGTTGCCCTGGGTTCAAGCAATTCTTCTGNNNAGATCGGAAG |
| 46303_2 | TATGAGTGTGGAGTCGTTGCCCTGGGTTCAAGCAATTCTTCTGNNNAGATCGGAAG |
| 44997_1 | TATGAGTGTGGAGTCGTTGCATAACCACGATCTCGATTTGGGGNNNAGATCGGAAG |
| 44997_2 | TATGAGTGTGGAGTCGTTGCATAACCACGATCTCGATTTGGGGNNNAGATCGGAAG |
| 43002_1 | TATGAGTGTGGAGTCGTTGCTTTGTGCCCCTAAAGTTTCAGTGNNNAGATCGGAAG |
| 43002_2 | TATGAGTGTGGAGTCGTTGCTTTGTGCCCCTAAAGTTTCAGTGNNNAGATCGGAAG |
| 31649_1 | TATGAGTGTGGAGTCGTTGCCTCACACACACACACACACAAATNNNAGATCGGAAGA |
| 31649_2 | TATGAGTGTGGAGTCGTTGCCTCACACACACACACACACAAATNNNAGATCGGAAGA |
| 48963_1 | TATGAGTGTGGAGTCGTTGCGCCACTATGCCCAGCTAATTTTTNNNAGATCGGAAGA |
| 48963_2 | TATGAGTGTGGAGTCGTTGCGCCACTATGCCCAGCTAATTTTTNNNAGATCGGAAGA |
| 51068_1 | TATGAGTGTGGAGTCGTTGCGAGAACCTAAGGGAGTCAAGACCNNNAGATCGGAA |
| 51068_2 | TATGAGTGTGGAGTCGTTGCGAGAACCTAAGGGAGTCAAGACCNNNAGATCGGAA |
| 36785_1 | TATGAGTGTGGAGTCGTTGCGCCATGTATGATAGACTCACAGCNNNAGATCGGAAG |
| 36785_2 | TATGAGTGTGGAGTCGTTGCGCCATGTATGATAGACTCACAGCNNNAGATCGGAAG |
| 10213_1 | TATGAGTGTGGAGTCGTTGCCTGAAGGTGATTCTGTCTGGTGANNNAGATCGGAA |
| 10213_2 | TATGAGTGTGGAGTCGTTGCCTGAAGGTGATTCTGTCTGGTGANNNAGATCGGAA |
| 34610_1 | TATGAGTGTGGAGTCGTTGCCTCAGTCCAGGAGGGAGGTGNNNAGATCGGAAGA |
| 34610_2 | TATGAGTGTGGAGTCGTTGCCTCAGTCCAGGAGGGAGGTGNNNAGATCGGAAGA |
| 48263_1 | TATGAGTGTGGAGTCGTTGCCACACCTGCCCAGCTAATTTTTANNNAGATCGGAAGA |
| 48263_2 | TATGAGTGTGGAGTCGTTGCCACACCTGCCCAGCTAATTTTTANNNAGATCGGAAGA |
| 6797_1 | TATGAGTGTGGAGTCGTTGCGTGTGTTGCACCTGGCAATTTTNNNAGATCGGAAGA |
| 6797_2 | TATGAGTGTGGAGTCGTTGCGTGTGTTGCACCTGGCAATTTTNNNAGATCGGAAGA |

| | |
|---|---|
| 46728_1 | TATGAGTGTGGAGTCGTTGCACACCAAAGTGCTAATTGGCAAANNNAGATCGGAA |
| 46728_2 | TATGAGTGTGGAGTCGTTGCACACCAAAGTGCTAATTGGCAAANNNAGATCGGAA |
| 38629_1 | TATGAGTGTGGAGTCGTTGCTTTCCAGCAACAGACTCCAAAGANNNAGATCGGAAG |
| 38629_2 | TATGAGTGTGGAGTCGTTGCTTTCCAGCAACAGACTCCAAAGANNNAGATCGGAAG |
| 44297_1 | TATGAGTGTGGAGTCGTTGCTGTTCTCTCTATGTAGCAGCACCNNNAGATCGGAAG |
| 44297_2 | TATGAGTGTGGAGTCGTTGCTGTTCTCTCTATGTAGCAGCACCNNNAGATCGGAAG |
| 39183_1 | TATGAGTGTGGAGTCGTTGCAAGAAAAAATGCAGAGGGAAGCCNNNAGATCGGA |
| 39183_2 | TATGAGTGTGGAGTCGTTGCAAGAAAAAATGCAGAGGGAAGCCNNNAGATCGGA |
| 45506_1 | TATGAGTGTGGAGTCGTTGCGTCCTCAGACATGGAGTTCAACTNNNAGATCGGAAG |
| 45506_2 | TATGAGTGTGGAGTCGTTGCGTCCTCAGACATGGAGTTCAACTNNNAGATCGGAAG |
| 2361_1 | TATGAGTGTGGAGTCGTTGCGGCATGAAAGGAGATAAGCAAAGCNNNAGATCGG |
| 2361_2 | TATGAGTGTGGAGTCGTTGCGGCATGAAAGGAGATAAGCAAAGCNNNAGATCGG |
| 39815_1 | TATGAGTGTGGAGTCGTTGCTCATTTGGGTTGCTCCTTTCTCTNNNAGATCGGAAGA |
| 39815_2 | TATGAGTGTGGAGTCGTTGCTCATTTGGGTTGCTCCTTTCTCTNNNAGATCGGAAGA |
| 41998_1 | TATGAGTGTGGAGTCGTTGCCAGGTTCAAGCAATTCTCCTGCNNNAGATCGGAAGA |
| 41998_2 | TATGAGTGTGGAGTCGTTGCCAGGTTCAAGCAATTCTCCTGCNNNAGATCGGAAGA |
| 43464_1 | TATGAGTGTGGAGTCGTTGCCCAGATACTTGACTTCCACAGCTNNNAGATCGGAAG |
| 43464_2 | TATGAGTGTGGAGTCGTTGCCCAGATACTTGACTTCCACAGCTNNNAGATCGGAAG |
| 35270_1 | TATGAGTGTGGAGTCGTTGCGTCACAACCAGCCTACCTATTCANNNAGATCGGAAG |
| 35270_2 | TATGAGTGTGGAGTCGTTGCGTCACAACCAGCCTACCTATTCANNNAGATCGGAAG |
| 31717_1 | TATGAGTGTGGAGTCGTTGCAGAGAGGGAGAGAGAGAGAGAGANNNAGATCGG |
| 31717_2 | TATGAGTGTGGAGTCGTTGCAGAGAGGGAGAGAGAGAGAGAGANNNAGATCGG |
| 10102_1 | TATGAGTGTGGAGTCGTTGCAGGACAAGACAAATAGGGGACCANNNAGATCGGA |
| 10102_2 | TATGAGTGTGGAGTCGTTGCAGGACAAGACAAATAGGGGACCANNNAGATCGGA |
| 40098_1 | TATGAGTGTGGAGTCGTTGCGCCTAAACAACATGGAGAAACCCNNNAGATCGGAA |
| 40098_2 | TATGAGTGTGGAGTCGTTGCGCCTAAACAACATGGAGAAACCCNNNAGATCGGAA |
| 52017_1 | TATGAGTGTGGAGTCGTTGCGCACCCTGTGGGACAAAATATTCNNNAGATCGGAAG |
| 52017_2 | TATGAGTGTGGAGTCGTTGCGCACCCTGTGGGACAAAATATTCNNNAGATCGGAAG |
| 36822_1 | TATGAGTGTGGAGTCGTTGCGACATGGAAACAGGAAGTCTTGCNNNAGATCGGAA |
| 36822_2 | TATGAGTGTGGAGTCGTTGCGACATGGAAACAGGAAGTCTTGCNNNAGATCGGAA |
| 34774_1 | TATGAGTGTGGAGTCGTTGCAACCTCTGACTCCCAACTTCAAGNNNAGATCGGAAG |
| 34774_2 | TATGAGTGTGGAGTCGTTGCAACCTCTGACTCCCAACTTCAAGNNNAGATCGGAAG |
| 20550_1 | TATGAGTGTGGAGTCGTTGCCTAGTGAGCTGTAAGTTTCTTNNNAGATCGGAAGAG |
| 20550_2 | TATGAGTGTGGAGTCGTTGCCTAGTGAGCTGTAAGTTTCTTNNNAGATCGGAAGAG |
| 10800_1 | TATGAGTGTGGAGTCGTTGCTACCTCCCCTGACCTCTTTCTTCNNNAGATCGGAAGA |
| 10800_2 | TATGAGTGTGGAGTCGTTGCTACCTCCCCTGACCTCTTTCTTCNNNAGATCGGAAGA |
| 37922_1 | TATGAGTGTGGAGTCGTTGCCAACGACTCCATGACACTCTTTGNNNAGATCGGAAG |
| 37922_2 | TATGAGTGTGGAGTCGTTGCCAACGACTCCATGACACTCTTTGNNNAGATCGGAAG |
| 10853_1 | TATGAGTGTGGAGTCGTTGCACACATTGCTACTATCTGGTACAANNNAGATCGGAA |
| 10853_2 | TATGAGTGTGGAGTCGTTGCACACATTGCTACTATCTGGTACAANNNAGATCGGAA |
| 13065_1 | TATGAGTGTGGAGTCGTTGCTCAGTACTGGAGATTCATTTGNNNAGATCGGAAGA |
| 13065_2 | TATGAGTGTGGAGTCGTTGCTCAGTACTGGAGATTCATTTGNNNAGATCGGAAGA |
| 10271_1 | TATGAGTGTGGAGTCGTTGCTCCAGCGTAAGAAATTGAGGGACNNNAGATCGGAA |

| | |
|---|---|
| 10271_2 | TATGAGTGTGGAGTCGTTGCTCCAGCGTAAGAAATTGAGGGACNNNAGATCGGAA |
| 2149_1 | TATGAGTGTGGAGTCGTTGCCCATTCACCACTGACTGTCTTCANNNAGATCGGAAGA |
| 2149_2 | TATGAGTGTGGAGTCGTTGCCCATTCACCACTGACTGTCTTCANNNAGATCGGAAGA |
| 29356_1 | TATGAGTGTGGAGTCGTTGCTGGAATAAAGGGGACAAGAGCTGNNNAGATCGGA |
| 29356_2 | TATGAGTGTGGAGTCGTTGCTGGAATAAAGGGGACAAGAGCTGNNNAGATCGGA |
| 44100_1 | TATGAGTGTGGAGTCGTTGCGCAGACCTTGAAGGCAGATTTACNNNAGATCGGAA |
| 44100_2 | TATGAGTGTGGAGTCGTTGCGCAGACCTTGAAGGCAGATTTACNNNAGATCGGAA |
| 39628_1 | TATGAGTGTGGAGTCGTTGCTAGGTTTCCACTTTGTTGTTGGTNNNAGATCGGAAG |
| 39628_2 | TATGAGTGTGGAGTCGTTGCTAGGTTTCCACTTTGTTGTTGGTNNNAGATCGGAAG |
| 52439_1 | TATGAGTGTGGAGTCGTTGCGGAAAGGGAGCCTAAATACGTCANNNAGATCGGAA |
| 52439_2 | TATGAGTGTGGAGTCGTTGCGGAAAGGGAGCCTAAATACGTCANNNAGATCGGAA |
| 53243_1 | TATGAGTGTGGAGTCGTTGCTCTAGGAATAGCCCCGGAACTATNNNAGATCGGAAG |
| 53243_2 | TATGAGTGTGGAGTCGTTGCTCTAGGAATAGCCCCGGAACTATNNNAGATCGGAAG |
| 9019_1 | TATGAGTGTGGAGTCGTTGCTGCTAGGTCATGTACTAAGTGCTNNNAGATCGGAAG |
| 9019_2 | TATGAGTGTGGAGTCGTTGCTGCTAGGTCATGTACTAAGTGCTNNNAGATCGGAAG |
| 50474_1 | TATGAGTGTGGAGTCGTTGCGCATGCCCCCACTGAATAAAATTNNNAGATCGGAAG |
| 50474_2 | TATGAGTGTGGAGTCGTTGCGCATGCCCCCACTGAATAAAATTNNNAGATCGGAAG |
| 40978_1 | TATGAGTGTGGAGTCGTTGCACAAGCCAAACTGAAACAGCAATNNNAGATCGGAA |
| 40978_2 | TATGAGTGTGGAGTCGTTGCACAAGCCAAACTGAAACAGCAATNNNAGATCGGAA |
| 46111_1 | TATGAGTGTGGAGTCGTTGCTTATGGAAAGATGGAGGTGCACANNNAGATCGGAA |
| 46111_2 | TATGAGTGTGGAGTCGTTGCTTATGGAAAGATGGAGGTGCACANNNAGATCGGAA |
| 52199_1 | TATGAGTGTGGAGTCGTTGCGCCATAACAAAAAGTCCACAGCTNNNAGATCGGAAG |
| 52199_2 | TATGAGTGTGGAGTCGTTGCGCCATAACAAAAAGTCCACAGCTNNNAGATCGGAAG |
| 49687_1 | TATGAGTGTGGAGTCGTTGCGATCTCAGCTCACTGCAACCTNNNAGATCGGAAGAG |
| 49687_2 | TATGAGTGTGGAGTCGTTGCGATCTCAGCTCACTGCAACCTNNNAGATCGGAAGAG |
| 41564_1 | TATGAGTGTGGAGTCGTTGCCCTCCTGGGTTGAAGTGATTCTTNNNAGATCGGAAG |
| 41564_2 | TATGAGTGTGGAGTCGTTGCCCTCCTGGGTTGAAGTGATTCTTNNNAGATCGGAAG |
| 48184_1 | TATGAGTGTGGAGTCGTTGCGCCGTTGGGGTATAGAAGAGTTANNNAGATCGGAA |
| 48184_2 | TATGAGTGTGGAGTCGTTGCGCCGTTGGGGTATAGAAGAGTTANNNAGATCGGAA |
| 42527_1 | TATGAGTGTGGAGTCGTTGCAAAAAAAGGCTAGGCACAGTGACNNNAGATCGGAA |
| 42527_2 | TATGAGTGTGGAGTCGTTGCAAAAAAAGGCTAGGCACAGTGACNNNAGATCGGAA |
| 37025_1 | TATGAGTGTGGAGTCGTTGCGGTTACATTGAGCTGAGTTTGCANNNAGATCGGAA |
| 37025_2 | TATGAGTGTGGAGTCGTTGCGGTTACATTGAGCTGAGTTTGCANNNAGATCGGAA |
| 31577_1 | TATGAGTGTGGAGTCGTTGCTGAGTCACAGTAGCACATTTCTTNNNAGATCGGAAG |
| 31577_2 | TATGAGTGTGGAGTCGTTGCTGAGTCACAGTAGCACATTTCTTNNNAGATCGGAAG |
| 49320_1 | TATGAGTGTGGAGTCGTTGCACCACGCCCAGCTAATTTTTTTTNNNAGATCGGAAGA |
| 49320_2 | TATGAGTGTGGAGTCGTTGCACCACGCCCAGCTAATTTTTTTTNNNAGATCGGAAGA |
| 32395_1 | TATGAGTGTGGAGTCGTTGCCATTCTCCTGCCTCAGCCTCNNNAGATCGGAAGAGCA |
| 32395_2 | TATGAGTGTGGAGTCGTTGCCATTCTCCTGCCTCAGCCTCNNNAGATCGGAAGAGCA |
| 51371_1 | TATGAGTGTGGAGTCGTTGCTAGGATCCCTCCTCAAAGTAGCANNNAGATCGGAAG |
| 51371_2 | TATGAGTGTGGAGTCGTTGCTAGGATCCCTCCTCAAAGTAGCANNNAGATCGGAAG |
| 32856_1 | TATGAGTGTGGAGTCGTTGCTTCAAGTGATTCTCCTGCCTCAGNNNAGATCGGAAG |
| 32856_2 | TATGAGTGTGGAGTCGTTGCTTCAAGTGATTCTCCTGCCTCAGNNNAGATCGGAAG |

| | |
|---|---|
| 10367_1 | TATGAGTGTGGAGTCGTTGCTGACACCACTGAACTTTTATGCANNNAGATCGGAAG |
| 10367_2 | TATGAGTGTGGAGTCGTTGCTGACACCACTGAACTTTTATGCANNNAGATCGGAAG |
| 10810_1 | TATGAGTGTGGAGTCGTTGCTGGGAAAGTGCATACATGTGTCCNNNAGATCGGAA |
| 10810_2 | TATGAGTGTGGAGTCGTTGCTGGGAAAGTGCATACATGTGTCCNNNAGATCGGAA |
| 50618_1 | TATGAGTGTGGAGTCGTTGCGGCTGCTTCTCCTAGTAGATTCCNNNAGATCGGAAG |
| 50618_2 | TATGAGTGTGGAGTCGTTGCGGCTGCTTCTCCTAGTAGATTCCNNNAGATCGGAAG |
| 32412_1 | TATGAGTGTGGAGTCGTTGCTCACACAAAGACATACCATAGTGANNNAGATCGGAA |
| 32412_2 | TATGAGTGTGGAGTCGTTGCTCACACAAAGACATACCATAGTGANNNAGATCGGAA |
| 36591_1 | TATGAGTGTGGAGTCGTTGCAAACACAAACTCTACAGATGGCANNNAGATCGGAAG |
| 36591_2 | TATGAGTGTGGAGTCGTTGCAAACACAAACTCTACAGATGGCANNNAGATCGGAAG |
| 38505_1 | TATGAGTGTGGAGTCGTTGCACACCTGATGATTGTGGATCACANNNAGATCGGAAG |
| 38505_2 | TATGAGTGTGGAGTCGTTGCACACCTGATGATTGTGGATCACANNNAGATCGGAAG |
| 33763_1 | TATGAGTGTGGAGTCGTTGCGGAGAATCTCTTGAGCCCAGTAGNNNAGATCGGAA |
| 33763_2 | TATGAGTGTGGAGTCGTTGCGGAGAATCTCTTGAGCCCAGTAGNNNAGATCGGAA |
| 47611_1 | TATGAGTGTGGAGTCGTTGCCAATCTCAGCTCAGTGCAACATCNNNAGATCGGAAG |
| 47611_2 | TATGAGTGTGGAGTCGTTGCCAATCTCAGCTCAGTGCAACATCNNNAGATCGGAAG |
| 29956_1 | TATGAGTGTGGAGTCGTTGCCTGTTTTGGAACCAGCCAGATTTNNNAGATCGGAAG |
| 29956_2 | TATGAGTGTGGAGTCGTTGCCTGTTTTGGAACCAGCCAGATTTNNNAGATCGGAAG |
| 41189_1 | TATGAGTGTGGAGTCGTTGCTACGTTGCCCAGACTAGTTTCAANNNAGATCGGAAG |
| 41189_2 | TATGAGTGTGGAGTCGTTGCTACGTTGCCCAGACTAGTTTCAANNNAGATCGGAAG |
| 45389_1 | TATGAGTGTGGAGTCGTTGCTTAGGGAATTGTGAAGGGGAGTCNNNAGATCGGAA |
| 45389_2 | TATGAGTGTGGAGTCGTTGCTTAGGGAATTGTGAAGGGGAGTCNNNAGATCGGAA |
| 35437_1 | TATGAGTGTGGAGTCGTTGCAGTGATTATCTTGCCTCAGCCTCNNNAGATCGGAAG |
| 35437_2 | TATGAGTGTGGAGTCGTTGCAGTGATTATCTTGCCTCAGCCTCNNNAGATCGGAAG |
| 49743_1 | TATGAGTGTGGAGTCGTTGCTCATTATGAGGGCTGGCTTTCTTNNNAGATCGGAAG |
| 49743_2 | TATGAGTGTGGAGTCGTTGCTCATTATGAGGGCTGGCTTTCTTNNNAGATCGGAAG |
| 29848_1 | TATGAGTGTGGAGTCGTTGCCGGAATGGGGAGAAGACGAATAANNNAGATCGGA |
| 29848_2 | TATGAGTGTGGAGTCGTTGCCGGAATGGGGAGAAGACGAATAANNNAGATCGGA |
| 44546_1 | TATGAGTGTGGAGTCGTTGCATGCATATTAGGGCTGGTTCCATNNNAGATCGGAAG |
| 44546_2 | TATGAGTGTGGAGTCGTTGCATGCATATTAGGGCTGGTTCCATNNNAGATCGGAAG |
| 47815_1 | TATGAGTGTGGAGTCGTTGCCCTCAAACTCCTGGGCTCAAGNNNAGATCGGAAGAG |
| 47815_2 | TATGAGTGTGGAGTCGTTGCCCTCAAACTCCTGGGCTCAAGNNNAGATCGGAAGAG |
| 42219_1 | TATGAGTGTGGAGTCGTTGCATGTGTGAGACCAAAGTTTCACANNNAGATCGGAAG |
| 42219_2 | TATGAGTGTGGAGTCGTTGCATGTGTGAGACCAAAGTTTCACANNNAGATCGGAAG |
| 52641_1 | TATGAGTGTGGAGTCGTTGCACACATAAAAAACTGTGCCCCAGNNNAGATCGGAAG |
| 52641_2 | TATGAGTGTGGAGTCGTTGCACACATAAAAAACTGTGCCCCAGNNNAGATCGGAAG |
| 42896_1 | TATGAGTGTGGAGTCGTTGCTGAAATATGTATACATTGTGGAATGGCNNNAGATCG |
| 42896_2 | TATGAGTGTGGAGTCGTTGCTGAAATATGTATACATTGTGGAATGGCNNNAGATCG |
| 13259_1 | TATGAGTGTGGAGTCGTTGCTATGTTGATTTACCTCCCATATTNNNAGATCGGAAGA |
| 13259_2 | TATGAGTGTGGAGTCGTTGCTATGTTGATTTACCTCCCATATTNNNAGATCGGAAGA |
| 47086_1 | TATGAGTGTGGAGTCGTTGCGTTCAAGCAATTCTTCTGCCTCANNNAGATCGGAAG |
| 47086_2 | TATGAGTGTGGAGTCGTTGCGTTCAAGCAATTCTTCTGCCTCANNNAGATCGGAAG |
| 33743_1 | TATGAGTGTGGAGTCGTTGCTGTAGTCCTAGCTACTTGGGAGGNNNAGATCGGAA |

| | |
|---|---|
| 33743_2 | TATGAGTGTGGAGTCGTTGCTGTAGTCCTAGCTACTTGGGAGGNNNAGATCGGAA |
| 35760_1 | TATGAGTGTGGAGTCGTTGCAAACATTTCACCCGCTCCTTTTTNNNAGATCGGAAGA |
| 35760_2 | TATGAGTGTGGAGTCGTTGCAAACATTTCACCCGCTCCTTTTTNNNAGATCGGAAGA |
| 7888_1 | TATGAGTGTGGAGTCGTTGCCTGCAAGCTCTGCCTCCTATGANNNAGATCGGAAGA |
| 7888_2 | TATGAGTGTGGAGTCGTTGCCTGCAAGCTCTGCCTCCTATGANNNAGATCGGAAGA |
| 45746_1 | TATGAGTGTGGAGTCGTTGCGTTTCACTCTTGTTGACCAGGCNNNAGATCGGAAGA |
| 45746_2 | TATGAGTGTGGAGTCGTTGCGTTTCACTCTTGTTGACCAGGCNNNAGATCGGAAGA |
| 51510_1 | TATGAGTGTGGAGTCGTTGCTCCTGGTACATATTCCCCACAAANNNAGATCGGAAGA |
| 51510_2 | TATGAGTGTGGAGTCGTTGCTCCTGGTACATATTCCCCACAAANNNAGATCGGAAGA |
| 43173_1 | TATGAGTGTGGAGTCGTTGCGAGTTTTGCTCTTGTTGCCCAGNNNAGATCGGAAGA |
| 43173_2 | TATGAGTGTGGAGTCGTTGCGAGTTTTGCTCTTGTTGCCCAGNNNAGATCGGAAGA |
| 51657_1 | TATGAGTGTGGAGTCGTTGCAGGGTTCACTCCTGATGTTGTACNNNAGATCGGAAG |
| 51657_2 | TATGAGTGTGGAGTCGTTGCAGGGTTCACTCCTGATGTTGTACNNNAGATCGGAAG |
| 52417_1 | TATGAGTGTGGAGTCGTTGCGAGACAGAGTCTTGCTCCATCATNNNAGATCGGAAG |
| 52417_2 | TATGAGTGTGGAGTCGTTGCGAGACAGAGTCTTGCTCCATCATNNNAGATCGGAAG |
| 36597_1 | TATGAGTGTGGAGTCGTTGCTTACAGATTCAATGCCATCCCCANNNAGATCGGAAGA |
| 36597_2 | TATGAGTGTGGAGTCGTTGCTTACAGATTCAATGCCATCCCCANNNAGATCGGAAGA |
| 44892_1 | TATGAGTGTGGAGTCGTTGCGGTGGGGTGATAGAGTTCTGCNNNAGATCGGAAGA |
| 44892_2 | TATGAGTGTGGAGTCGTTGCGGTGGGGTGATAGAGTTCTGCNNNAGATCGGAAGA |
| 38081_1 | TATGAGTGTGGAGTCGTTGCCCTTGAATGCCAGATGAAAGGTGNNNAGATCGGAA |
| 38081_2 | TATGAGTGTGGAGTCGTTGCCCTTGAATGCCAGATGAAAGGTGNNNAGATCGGAA |
| 39759_1 | TATGAGTGTGGAGTCGTTGCTTGCAGAGTCAGTCTGATTTCCANNNAGATCGGAAG |
| 39759_2 | TATGAGTGTGGAGTCGTTGCTTGCAGAGTCAGTCTGATTTCCANNNAGATCGGAAG |
| 29251_1 | TATGAGTGTGGAGTCGTTGCTACTCTTAGTTTGGGATGGGTCCNNNAGATCGGAAG |
| 29251_2 | TATGAGTGTGGAGTCGTTGCTACTCTTAGTTTGGGATGGGTCCNNNAGATCGGAAG |
| 40261_1 | TATGAGTGTGGAGTCGTTGCCAGACACTCAGAATGTTAGAGCCNNNAGATCGGAAG |
| 40261_2 | TATGAGTGTGGAGTCGTTGCCAGACACTCAGAATGTTAGAGCCNNNAGATCGGAAG |
| 13803_1 | TATGAGTGTGGAGTCGTTGCCTGAAGGTTGCCAAAGAAATNNNAGATCGGAAGAG |
| 13803_2 | TATGAGTGTGGAGTCGTTGCCTGAAGGTTGCCAAAGAAATNNNAGATCGGAAGAG |
| 50161_1 | TATGAGTGTGGAGTCGTTGCACGATCCTTCTCGCCAACCCNNNAGATCGGAAGAGCA |
| 50161_2 | TATGAGTGTGGAGTCGTTGCACGATCCTTCTCGCCAACCCNNNAGATCGGAAGAGCA |
| 49636_1 | TATGAGTGTGGAGTCGTTGCCATCCATGTTGTAGCCGAATGTCNNNAGATCGGAAG |
| 49636_2 | TATGAGTGTGGAGTCGTTGCCATCCATGTTGTAGCCGAATGTCNNNAGATCGGAAG |
| 35940_1 | TATGAGTGTGGAGTCGTTGCAGGAGTTTCAGACCAGCAAAGATNNNAGATCGGAA |
| 35940_2 | TATGAGTGTGGAGTCGTTGCAGGAGTTTCAGACCAGCAAAGATNNNAGATCGGAA |
| 40237_1 | TATGAGTGTGGAGTCGTTGCAAGCTAGGATATGAAGCTCAGGCNNNAGATCGGAA |
| 40237_2 | TATGAGTGTGGAGTCGTTGCAAGCTAGGATATGAAGCTCAGGCNNNAGATCGGAA |
| 47948_1 | TATGAGTGTGGAGTCGTTGCAGGGTAGCTAAGACCTCACTCCNNNAGATCGGAAGA |
| 47948_2 | TATGAGTGTGGAGTCGTTGCAGGGTAGCTAAGACCTCACTCCNNNAGATCGGAAGA |
| 10383_1 | TATGAGTGTGGAGTCGTTGCCACACACACACTGAGCTCTCTTTNNNAGATCGGAAGA |
| 10383_2 | TATGAGTGTGGAGTCGTTGCCACACACACACTGAGCTCTCTTTNNNAGATCGGAAGA |
| 42691_1 | TATGAGTGTGGAGTCGTTGCCTTATGGTTTTGGCTCACTGCAGNNNAGATCGGAAG |
| 42691_2 | TATGAGTGTGGAGTCGTTGCCTTATGGTTTTGGCTCACTGCAGNNNAGATCGGAAG |

| | |
|---|---|
| 51293_1 | TATGAGTGTGGAGTCGTTGCGTCCAGGTGTGAAACTTGTGATGNNNAGATCGGAA |
| 51293_2 | TATGAGTGTGGAGTCGTTGCGTCCAGGTGTGAAACTTGTGATGNNNAGATCGGAA |
| 8498_1 | TATGAGTGTGGAGTCGTTGCCTCCCCATAGCCAGGGATTGTATCNNNAGATCGGAA |
| 8498_2 | TATGAGTGTGGAGTCGTTGCCTCCCCATAGCCAGGGATTGTATCNNNAGATCGGAA |
| 40854_1 | TATGAGTGTGGAGTCGTTGCCTGCAAATGTACCAAGAGTGTCANNNAGATCGGAAG |
| 40854_2 | TATGAGTGTGGAGTCGTTGCCTGCAAATGTACCAAGAGTGTCANNNAGATCGGAAG |
| 42970_1 | TATGAGTGTGGAGTCGTTGCCATGCCCAGAAAACTTTAGCCAANNNAGATCGGAAG |
| 42970_2 | TATGAGTGTGGAGTCGTTGCCATGCCCAGAAAACTTTAGCCAANNNAGATCGGAAG |
| 45351_1 | TATGAGTGTGGAGTCGTTGCTTCAAGTGATTCTCCTGCCTCAGNNNAGATCGGAAG |
| 45351_2 | TATGAGTGTGGAGTCGTTGCTTCAAGTGATTCTCCTGCCTCAGNNNAGATCGGAAG |
| 33893_1 | TATGAGTGTGGAGTCGTTGCAGTAACTGGGACTACAGGGATGANNNAGATCGGAA |
| 33893_2 | TATGAGTGTGGAGTCGTTGCAGTAACTGGGACTACAGGGATGANNNAGATCGGAA |
| 49853_1 | TATGAGTGTGGAGTCGTTGCACTTACCCTTCTGCACCAATCAANNNAGATCGGAAGA |
| 49853_2 | TATGAGTGTGGAGTCGTTGCACTTACCCTTCTGCACCAATCAANNNAGATCGGAAGA |
| 48183_1 | TATGAGTGTGGAGTCGTTGCTTCAAGTGATTCTCCTGCCTCAGNNNAGATCGGAAG |
| 48183_2 | TATGAGTGTGGAGTCGTTGCTTCAAGTGATTCTCCTGCCTCAGNNNAGATCGGAAG |
| 50911_1 | TATGAGTGTGGAGTCGTTGCACAGATCCTGTGTCCACTTTCTCNNNAGATCGGAAGA |
| 50911_2 | TATGAGTGTGGAGTCGTTGCACAGATCCTGTGTCCACTTTCTCNNNAGATCGGAAGA |
| 10565_1 | TATGAGTGTGGAGTCGTTGCATTCTGCTTCCAAGCCCTGTTTTNNNAGATCGGAAGA |
| 10565_2 | TATGAGTGTGGAGTCGTTGCATTCTGCTTCCAAGCCCTGTTTTNNNAGATCGGAAGA |
| 29678_1 | TATGAGTGTGGAGTCGTTGCTTTGGTCAAGTTCAGAGTGTGGANNNAGATCGGAA |
| 29678_2 | TATGAGTGTGGAGTCGTTGCTTTGGTCAAGTTCAGAGTGTGGANNNAGATCGGAA |
| 44697_1 | TATGAGTGTGGAGTCGTTGCCTCCTATCCAGCTCTTGACACTGNNNAGATCGGAAGA |
| 44697_2 | TATGAGTGTGGAGTCGTTGCCTCCTATCCAGCTCTTGACACTGNNNAGATCGGAAGA |
| 43256_1 | TATGAGTGTGGAGTCGTTGCCTCAAGAAGCTGGAGTTACAGGTNNNAGATCGGAA |
| 43256_2 | TATGAGTGTGGAGTCGTTGCCTCAAGAAGCTGGAGTTACAGGTNNNAGATCGGAA |
| 37905_1 | TATGAGTGTGGAGTCGTTGCGTAATCTCAGCTACTCAGGAGGCNNNAGATCGGAAG |
| 37905_2 | TATGAGTGTGGAGTCGTTGCGTAATCTCAGCTACTCAGGAGGCNNNAGATCGGAAG |
| 47411_1 | TATGAGTGTGGAGTCGTTGCTATTACAGCAGTGACGTGTGTGANNNAGATCGGAA |
| 47411_2 | TATGAGTGTGGAGTCGTTGCTATTACAGCAGTGACGTGTGTGANNNAGATCGGAA |
| 3000_1 | TATGAGTGTGGAGTCGTTGCGGGACAGGTAAAGAGACGAGAAAGNNNAGATCGG |
| 3000_2 | TATGAGTGTGGAGTCGTTGCGGGACAGGTAAAGAGACGAGAAAGNNNAGATCGG |
| 6120_1 | TATGAGTGTGGAGTCGTTGCCAAGTCCGCTCTGGAGATAACTNNNAGATCGGAAGA |
| 6120_2 | TATGAGTGTGGAGTCGTTGCCAAGTCCGCTCTGGAGATAACTNNNAGATCGGAAGA |
| 33354_1 | TATGAGTGTGGAGTCGTTGCCCTACCAAAGTGCTGGGATTACANNNAGATCGGAAG |
| 33354_2 | TATGAGTGTGGAGTCGTTGCCCTACCAAAGTGCTGGGATTACANNNAGATCGGAAG |
| 29210_1 | TATGAGTGTGGAGTCGTTGCTGAGATTTTTTGTGGAACCTCCANNNAGATCGGAAG |
| 29210_2 | TATGAGTGTGGAGTCGTTGCTGAGATTTTTTGTGGAACCTCCANNNAGATCGGAAG |
| 47986_1 | TATGAGTGTGGAGTCGTTGCAACGGGGAGCTAGTCTCAAGATNNNAGATCGGAAG |
| 47986_2 | TATGAGTGTGGAGTCGTTGCAACGGGGAGCTAGTCTCAAGATNNNAGATCGGAAG |
| 37567_1 | TATGAGTGTGGAGTCGTTGCTGGTTCCAAGTTACTACAGCGTTNNNAGATCGGAAG |
| 37567_2 | TATGAGTGTGGAGTCGTTGCTGGTTCCAAGTTACTACAGCGTTNNNAGATCGGAAG |
| 47010_1 | TATGAGTGTGGAGTCGTTGCTCAGAGTTCACTGTAGCCTCAACNNNAGATCGGAAG |

| | |
|---|---|
| 47010_2 | TATGAGTGTGGAGTCGTTGCTCAGAGTTCACTGTAGCCTCAACNNNAGATCGGAAG |
| 10352_1 | TATGAGTGTGGAGTCGTTGCGCATGAGAATCTTGAATCCCGGANNNAGATCGGAA |
| 10352_2 | TATGAGTGTGGAGTCGTTGCGCATGAGAATCTTGAATCCCGGANNNAGATCGGAA |
| 30740_1 | TATGAGTGTGGAGTCGTTGCCTGCCAAGTCTCACGTTGAAATTNNNAGATCGGAAG |
| 30740_2 | TATGAGTGTGGAGTCGTTGCCTGCCAAGTCTCACGTTGAAATTNNNAGATCGGAAG |
| 51142_1 | TATGAGTGTGGAGTCGTTGCGGGGTAGAGTGCTGGGTATTATGNNNAGATCGGA |
| 51142_2 | TATGAGTGTGGAGTCGTTGCGGGGTAGAGTGCTGGGTATTATGNNNAGATCGGA |
| 30612_1 | TATGAGTGTGGAGTCGTTGCAAGACTGCAGGTATGATGGCCTNNNAGATCGGAAG |
| 30612_2 | TATGAGTGTGGAGTCGTTGCAAGACTGCAGGTATGATGGCCTNNNAGATCGGAAG |
| 33249_1 | TATGAGTGTGGAGTCGTTGCTATCAAGTGATTCTCCTGCCTCGNNNAGATCGGAAG |
| 33249_2 | TATGAGTGTGGAGTCGTTGCTATCAAGTGATTCTCCTGCCTCGNNNAGATCGGAAG |
| 34916_1 | TATGAGTGTGGAGTCGTTGCCGCTTCCTGAGTTCAAGTGATTCNNNAGATCGGAAG |
| 34916_2 | TATGAGTGTGGAGTCGTTGCCGCTTCCTGAGTTCAAGTGATTCNNNAGATCGGAAG |
| 48829_1 | TATGAGTGTGGAGTCGTTGCGACGGAGTCTCTCTCTGTTGCNNNAGATCGGAAGAG |
| 48829_2 | TATGAGTGTGGAGTCGTTGCGACGGAGTCTCTCTCTGTTGCNNNAGATCGGAAGAG |
| 29503_1 | TATGAGTGTGGAGTCGTTGCGTGGGATAGATGCTCATTTTAAANNNAGATCGGAA |
| 29503_2 | TATGAGTGTGGAGTCGTTGCGTGGGATAGATGCTCATTTTAAANNNAGATCGGAA |
| 47293_1 | TATGAGTGTGGAGTCGTTGCTTTGACTTAGTGATCCTCCTGCCNNNAGATCGGAAG |
| 47293_2 | TATGAGTGTGGAGTCGTTGCTTTGACTTAGTGATCCTCCTGCCNNNAGATCGGAAG |
| 31492_1 | TATGAGTGTGGAGTCGTTGCTTGTGAAGTCAGGTTGCTAATGCNNNAGATCGGAA |
| 31492_2 | TATGAGTGTGGAGTCGTTGCTTGTGAAGTCAGGTTGCTAATGCNNNAGATCGGAA |
| 52969_1 | TATGAGTGTGGAGTCGTTGCAGCAGACGTTCTTCTTCCCTTTCNNNAGATCGGAAGA |
| 52969_2 | TATGAGTGTGGAGTCGTTGCAGCAGACGTTCTTCTTCCCTTTCNNNAGATCGGAAGA |
| 29264_1 | TATGAGTGTGGAGTCGTTGCACCAATGGAATTCAGTCAACCATNNNAGATCGGAAG |
| 29264_2 | TATGAGTGTGGAGTCGTTGCACCAATGGAATTCAGTCAACCATNNNAGATCGGAAG |
| 47544_1 | TATGAGTGTGGAGTCGTTGCCTGAGTAGCTGGGATTACAAGCANNNAGATCGGAA |
| 47544_2 | TATGAGTGTGGAGTCGTTGCCTGAGTAGCTGGGATTACAAGCANNNAGATCGGAA |
| 49486_1 | TATGAGTGTGGAGTCGTTGCTTCACACCATTCTCCTGCCTCNNNAGATCGGAAGAGC |
| 49486_2 | TATGAGTGTGGAGTCGTTGCTTCACACCATTCTCCTGCCTCNNNAGATCGGAAGAGC |
| 9300_1 | TATGAGTGTGGAGTCGTTGCATCAATTTGGAGTTGGGGGTTGGNNNAGATCGGAA |
| 9300_2 | TATGAGTGTGGAGTCGTTGCATCAATTTGGAGTTGGGGGTTGGNNNAGATCGGAA |
| 49162_1 | TATGAGTGTGGAGTCGTTGCGCCTCCCAAGTAGCTAGGATTACNNNAGATCGGAAG |
| 49162_2 | TATGAGTGTGGAGTCGTTGCGCCTCCCAAGTAGCTAGGATTACNNNAGATCGGAAG |
| 46913_1 | TATGAGTGTGGAGTCGTTGCAAACAACAACGACAAGCCACCNNNAGATCGGAAGA |
| 46913_2 | TATGAGTGTGGAGTCGTTGCAAACAACAACGACAAGCCACCNNNAGATCGGAAGA |
| 48287_1 | TATGAGTGTGGAGTCGTTGCCTGAACTACTGGCCTCAAGTGATNNNAGATCGGAAG |
| 48287_2 | TATGAGTGTGGAGTCGTTGCCTGAACTACTGGCCTCAAGTGATNNNAGATCGGAAG |
| 43613_1 | TATGAGTGTGGAGTCGTTGCCCTGAGGTTAGGAGTTTGAGACCNNNAGATCGGAA |
| 43613_2 | TATGAGTGTGGAGTCGTTGCCCTGAGGTTAGGAGTTTGAGACCNNNAGATCGGAA |
| 50693_1 | TATGAGTGTGGAGTCGTTGCTTCACCCCACAAAAAGGGAACTCNNNAGATCGGAAG |
| 50693_2 | TATGAGTGTGGAGTCGTTGCTTCACCCCACAAAAAGGGAACTCNNNAGATCGGAAG |
| 32135_1 | TATGAGTGTGGAGTCGTTGCTCTTTTCCCCATTTTGCTTGGTCNNNAGATCGGAAGA |
| 32135_2 | TATGAGTGTGGAGTCGTTGCTCTTTTCCCCATTTTGCTTGGTCNNNAGATCGGAAGA |

| | |
|---|---|
| 29275_1 | TATGAGTGTGGAGTCGTTGCAATGTCATTTTGCCATGATGCCANNNAGATCGGAAG |
| 29275_2 | TATGAGTGTGGAGTCGTTGCAATGTCATTTTGCCATGATGCCANNNAGATCGGAAG |
| 45219_1 | TATGAGTGTGGAGTCGTTGCGACAGTCACTCGGATGCTGGNNNAGATCGGAAGAG |
| 45219_2 | TATGAGTGTGGAGTCGTTGCGACAGTCACTCGGATGCTGGNNNAGATCGGAAGAG |
| 13743_1 | TATGAGTGTGGAGTCGTTGCTAATAAAATGCACATTGAAACCNNNAGATCGGAAG |
| 13743_2 | TATGAGTGTGGAGTCGTTGCTAATAAAATGCACATTGAAACCNNNAGATCGGAAG |
| 2453_1 | TATGAGTGTGGAGTCGTTGCGGGTGGGTTGTATTAGTCCGTTCNNNAGATCGGAA |
| 2453_2 | TATGAGTGTGGAGTCGTTGCGGGTGGGTTGTATTAGTCCGTTCNNNAGATCGGAA |
| 43327_1 | TATGAGTGTGGAGTCGTTGCGGAGTCTCGCTCTTGTTACCTAGNNNAGATCGGAAG |
| 43327_2 | TATGAGTGTGGAGTCGTTGCGGAGTCTCGCTCTTGTTACCTAGNNNAGATCGGAAG |
| 36028_1 | TATGAGTGTGGAGTCGTTGCGCTTTTATGGGGTTTGCAGGAATNNNAGATCGGAA |
| 36028_2 | TATGAGTGTGGAGTCGTTGCGCTTTTATGGGGTTTGCAGGAATNNNAGATCGGAA |
| 10155_1 | TATGAGTGTGGAGTCGTTGCCACCACCAACAAACACCTTAGGANNNAGATCGGAAG |
| 10155_2 | TATGAGTGTGGAGTCGTTGCCACCACCAACAAACACCTTAGGANNNAGATCGGAAG |
| 30247_1 | TATGAGTGTGGAGTCGTTGCCAGCGTTGTAGAGGCATCTTCTANNNAGATCGGAAG |
| 30247_2 | TATGAGTGTGGAGTCGTTGCCAGCGTTGTAGAGGCATCTTCTANNNAGATCGGAAG |
| 39215_1 | TATGAGTGTGGAGTCGTTGCAGTTTAGAGGAGACTCCACGAGANNNAGATCGGAA |
| 39215_2 | TATGAGTGTGGAGTCGTTGCAGTTTAGAGGAGACTCCACGAGANNNAGATCGGAA |
| 30775_1 | TATGAGTGTGGAGTCGTTGCGTGTGTGTGTGTGTGTGTGTTTANNNAGATCGGAA |
| 30775_2 | TATGAGTGTGGAGTCGTTGCGTGTGTGTGTGTGTGTGTGTTTANNNAGATCGGAA |
| 51941_1 | TATGAGTGTGGAGTCGTTGCCAAGTGGTGTTTGGTTTCCTGTTNNNAGATCGGAAG |
| 51941_2 | TATGAGTGTGGAGTCGTTGCCAAGTGGTGTTTGGTTTCCTGTTNNNAGATCGGAAG |
| 10071_1 | TATGAGTGTGGAGTCGTTGCCTTTCCTTTCCTCCCCACTCTCTNNNAGATCGGAAGAG |
| 10071_2 | TATGAGTGTGGAGTCGTTGCCTTTCCTTTCCTCCCCACTCTCTNNNAGATCGGAAGAG |
| 8531_1 | TATGAGTGTGGAGTCGTTGCGTGGCTATCTCCAACTTGCTGGGNNNAGATCGGAAG |
| 8531_2 | TATGAGTGTGGAGTCGTTGCGTGGCTATCTCCAACTTGCTGGGNNNAGATCGGAAG |
| 44232_1 | TATGAGTGTGGAGTCGTTGCATGGGCACTCTAACTAGGAGGTANNNAGATCGGAA |
| 44232_2 | TATGAGTGTGGAGTCGTTGCATGGGCACTCTAACTAGGAGGTANNNAGATCGGAA |
| 45957_1 | TATGAGTGTGGAGTCGTTGCGTACCTCAGCCTCCCCAGTAGNNNAGATCGGAAGAG |
| 45957_2 | TATGAGTGTGGAGTCGTTGCGTACCTCAGCCTCCCCAGTAGNNNAGATCGGAAGAG |
| 37684_1 | TATGAGTGTGGAGTCGTTGCTCTCAGAGTCCCAGAAGATGAGANNNAGATCGGAA |
| 37684_2 | TATGAGTGTGGAGTCGTTGCTCTCAGAGTCCCAGAAGATGAGANNNAGATCGGAA |
| 49698_1 | TATGAGTGTGGAGTCGTTGCCCTTCTTGCTGTTCTGTCCATTGNNNAGATCGGAAGA |
| 49698_2 | TATGAGTGTGGAGTCGTTGCCCTTCTTGCTGTTCTGTCCATTGNNNAGATCGGAAGA |
| 51947_1 | TATGAGTGTGGAGTCGTTGCCTATCTCAGGACCATCCGATCAANNNAGATCGGAAG |
| 51947_2 | TATGAGTGTGGAGTCGTTGCCTATCTCAGGACCATCCGATCAANNNAGATCGGAAG |
| 40144_1 | TATGAGTGTGGAGTCGTTGCGTTCAAGAGATTCTCCAGCCTCANNNAGATCGGAAG |
| 40144_2 | TATGAGTGTGGAGTCGTTGCGTTCAAGAGATTCTCCAGCCTCANNNAGATCGGAAG |
| 51356_1 | TATGAGTGTGGAGTCGTTGCAGGCATAGCATTGGGAGGTATACNNNAGATCGGAA |
| 51356_2 | TATGAGTGTGGAGTCGTTGCAGGCATAGCATTGGGAGGTATACNNNAGATCGGAA |
| 39837_1 | TATGAGTGTGGAGTCGTTGCCAGTGCACCAGGGAGACTATTATNNNAGATCGGAA |
| 39837_2 | TATGAGTGTGGAGTCGTTGCCAGTGCACCAGGGAGACTATTATNNNAGATCGGAA |
| 35353_1 | TATGAGTGTGGAGTCGTTGCCTGGGATTATAGGAGTGAGCCACNNNAGATCGGAA |

| | |
|---|---|
| 35353_2 | TATGAGTGTGGAGTCGTTGCCTGGGATTATAGGAGTGAGCCACNNNAGATCGGAA |
| 43337_1 | TATGAGTGTGGAGTCGTTGCTTCACTTGGGGGAGTTTTGTTTGNNNAGATCGGAAG |
| 43337_2 | TATGAGTGTGGAGTCGTTGCTTCACTTGGGGGAGTTTTGTTTGNNNAGATCGGAAG |
| 44676_1 | TATGAGTGTGGAGTCGTTGCGAGGGAGGTCGGATTCCATTATCNNNAGATCGGAA |
| 44676_2 | TATGAGTGTGGAGTCGTTGCGAGGGAGGTCGGATTCCATTATCNNNAGATCGGAA |
| 46288_1 | TATGAGTGTGGAGTCGTTGCCTCCCAGGTTCACACCATTCTTCNNNAGATCGGAAGA |
| 46288_2 | TATGAGTGTGGAGTCGTTGCCTCCCAGGTTCACACCATTCTTCNNNAGATCGGAAGA |
| 50289_1 | TATGAGTGTGGAGTCGTTGCACATTTGTTCCTCTGGTGCTTTGNNNAGATCGGAAG |
| 50289_2 | TATGAGTGTGGAGTCGTTGCACATTTGTTCCTCTGGTGCTTTGNNNAGATCGGAAG |
| 32904_1 | TATGAGTGTGGAGTCGTTGCCCTTCTAGGTCCACATCAGCATTNNNAGATCGGAAG |
| 32904_2 | TATGAGTGTGGAGTCGTTGCCCTTCTAGGTCCACATCAGCATTNNNAGATCGGAAG |
| 51603_1 | TATGAGTGTGGAGTCGTTGCACAGGCCAAGTCCTAGGAGAGNNNAGATCGGAAGA |
| 51603_2 | TATGAGTGTGGAGTCGTTGCACAGGCCAAGTCCTAGGAGAGNNNAGATCGGAAGA |
| 42582_1 | TATGAGTGTGGAGTCGTTGCCAGGCTGGAGTACAATGGTATGANNNAGATCGGAA |
| 42582_2 | TATGAGTGTGGAGTCGTTGCCAGGCTGGAGTACAATGGTATGANNNAGATCGGAA |
| 37861_1 | TATGAGTGTGGAGTCGTTGCAAAAAAATTAGCTGGGGGTGGTGNNNAGATCGGA |
| 37861_2 | TATGAGTGTGGAGTCGTTGCAAAAAAATTAGCTGGGGGTGGTGNNNAGATCGGA |
| 42477_1 | TATGAGTGTGGAGTCGTTGCGAGTCTCTCTCCATTGCCCAGNNNAGATCGGAAGAG |
| 42477_2 | TATGAGTGTGGAGTCGTTGCGAGTCTCTCTCCATTGCCCAGNNNAGATCGGAAGAG |
| 30352_1 | TATGAGTGTGGAGTCGTTGCCTGAGTAGCTGGGATTACAGGTGNNNAGATCGGAA |
| 30352_2 | TATGAGTGTGGAGTCGTTGCCTGAGTAGCTGGGATTACAGGTGNNNAGATCGGAA |
| 46043_1 | TATGAGTGTGGAGTCGTTGCGCCTCCTGAGTAGCTGAGATTACNNNAGATCGGAAG |
| 46043_2 | TATGAGTGTGGAGTCGTTGCGCCTCCTGAGTAGCTGAGATTACNNNAGATCGGAAG |
| 50502_1 | TATGAGTGTGGAGTCGTTGCGATGAAGGTTTTCCCAACCCTTGNNNAGATCGGAAG |
| 50502_2 | TATGAGTGTGGAGTCGTTGCGATGAAGGTTTTCCCAACCCTTGNNNAGATCGGAAG |
| 47444_1 | TATGAGTGTGGAGTCGTTGCCTCACTACAACCTCCACTTCCTGNNNAGATCGGAAGA |
| 47444_2 | TATGAGTGTGGAGTCGTTGCCTCACTACAACCTCCACTTCCTGNNNAGATCGGAAGA |
| 12890_1 | TATGAGTGTGGAGTCGTTGCAGATCATCGAACTCTTAGATATGNNNAGATCGGAAG |
| 12890_2 | TATGAGTGTGGAGTCGTTGCAGATCATCGAACTCTTAGATATGNNNAGATCGGAAG |
| 12932_1 | TATGAGTGTGGAGTCGTTGCCCCAAATTACTACAAAGCATAGNNNAGATCGGAAGA |
| 12932_2 | TATGAGTGTGGAGTCGTTGCCCCAAATTACTACAAAGCATAGNNNAGATCGGAAGA |
| 42831_1 | TATGAGTGTGGAGTCGTTGCCTTTTTGTGTGTGCAGGCATTTCNNNAGATCGGAAG |
| 42831_2 | TATGAGTGTGGAGTCGTTGCCTTTTTGTGTGTGCAGGCATTTCNNNAGATCGGAAG |
| 45486_1 | TATGAGTGTGGAGTCGTTGCAAGGTGGCTCTAACTCTGAACTGNNNAGATCGGAAG |
| 45486_2 | TATGAGTGTGGAGTCGTTGCAAGGTGGCTCTAACTCTGAACTGNNNAGATCGGAAG |
| 42810_1 | TATGAGTGTGGAGTCGTTGCGGAGTGCAGTGGTGTGATCTCNNNAGATCGGAAGA |
| 42810_2 | TATGAGTGTGGAGTCGTTGCGGAGTGCAGTGGTGTGATCTCNNNAGATCGGAAGA |
| 35132_1 | TATGAGTGTGGAGTCGTTGCCAGTCTATGCCTTTTGATTGGTGANNNAGATCGGAA |
| 35132_2 | TATGAGTGTGGAGTCGTTGCCAGTCTATGCCTTTTGATTGGTGANNNAGATCGGAA |
| 46370_1 | TATGAGTGTGGAGTCGTTGCATTCAAGTGATTCTCCTGCCTCANNNAGATCGGAAG |
| 46370_2 | TATGAGTGTGGAGTCGTTGCATTCAAGTGATTCTCCTGCCTCANNNAGATCGGAAG |
| 34348_1 | TATGAGTGTGGAGTCGTTGCTTTTGATCTTGTTCCCCAGGCTANNNAGATCGGAAG |
| 34348_2 | TATGAGTGTGGAGTCGTTGCTTTTGATCTTGTTCCCCAGGCTANNNAGATCGGAAG |

| | |
|---|---|
| 37740_1 | TATGAGTGTGGAGTCGTTGCCCCATTCTTTAGCAGTCTGTGTGNNNAGATCGGAAG |
| 37740_2 | TATGAGTGTGGAGTCGTTGCCCCATTCTTTAGCAGTCTGTGTGNNNAGATCGGAAG |
| 5977_1 | TATGAGTGTGGAGTCGTTGCACTCTGGCCTGGGAAATACAGCNNNAGATCGGAAG |
| 5977_2 | TATGAGTGTGGAGTCGTTGCACTCTGGCCTGGGAAATACAGCNNNAGATCGGAAG |
| 5375_1 | TATGAGTGTGGAGTCGTTGCTCCGTTTGGCAGGATATTTACTANNNAGATCGGAAG |
| 5375_2 | TATGAGTGTGGAGTCGTTGCTCCGTTTGGCAGGATATTTACTANNNAGATCGGAAG |
| 2016_1 | TATGAGTGTGGAGTCGTTGCCCAAAGTATGCCAGGACAGACAANNNAGATCGGAA |
| 2016_2 | TATGAGTGTGGAGTCGTTGCCCAAAGTATGCCAGGACAGACAANNNAGATCGGAA |
| 30095_1 | TATGAGTGTGGAGTCGTTGCGAACCGCTTTCTTTTGGATCTCCNNNAGATCGGAAG |
| 30095_2 | TATGAGTGTGGAGTCGTTGCGAACCGCTTTCTTTTGGATCTCCNNNAGATCGGAAG |
| 36466_1 | TATGAGTGTGGAGTCGTTGCTGGCAGACATCTGGTTAGAAACANNNAGATCGGAA |
| 36466_2 | TATGAGTGTGGAGTCGTTGCTGGCAGACATCTGGTTAGAAACANNNAGATCGGAA |
| 32442_1 | TATGAGTGTGGAGTCGTTGCCCTCCCAAGTAGCTGGAATTACANNNAGATCGGAAG |
| 32442_2 | TATGAGTGTGGAGTCGTTGCCCTCCCAAGTAGCTGGAATTACANNNAGATCGGAAG |
| 51151_1 | TATGAGTGTGGAGTCGTTGCCATCAATTGAGTGCTGTGGGAAGNNNAGATCGGAA |
| 51151_2 | TATGAGTGTGGAGTCGTTGCCATCAATTGAGTGCTGTGGGAAGNNNAGATCGGAA |
| 28959_1 | TATGAGTGTGGAGTCGTTGCTGTCTTTGTTTCCTTCAAGGTGTNNNAGATCGGAAG |
| 28959_2 | TATGAGTGTGGAGTCGTTGCTGTCTTTGTTTCCTTCAAGGTGTNNNAGATCGGAAG |
| 48293_1 | TATGAGTGTGGAGTCGTTGCCACCACGCCTGGCTAATTTTTTTNNNAGATCGGAAGA |
| 48293_2 | TATGAGTGTGGAGTCGTTGCCACCACGCCTGGCTAATTTTTTTNNNAGATCGGAAGA |
| 39619_1 | TATGAGTGTGGAGTCGTTGCAACTTGTTCAAGGTCACACTTGTNNNAGATCGGAAG |
| 39619_2 | TATGAGTGTGGAGTCGTTGCAACTTGTTCAAGGTCACACTTGTNNNAGATCGGAAG |
| 42563_1 | TATGAGTGTGGAGTCGTTGCTATGATCAAATCACCTGCCACCANNNAGATCGGAAG |
| 42563_2 | TATGAGTGTGGAGTCGTTGCTATGATCAAATCACCTGCCACCANNNAGATCGGAAG |
| 45999_1 | TATGAGTGTGGAGTCGTTGCTCTTCTAACAGTCAGACCCCTCTNNNAGATCGGAAGA |
| 45999_2 | TATGAGTGTGGAGTCGTTGCTCTTCTAACAGTCAGACCCCTCTNNNAGATCGGAAGA |
| 43135_1 | TATGAGTGTGGAGTCGTTGCAAGAAGGAAGGGAGAGAGGAAGGNNNAGATCGG |
| 43135_2 | TATGAGTGTGGAGTCGTTGCAAGAAGGAAGGGAGAGAGGAAGGNNNAGATCGG |
| 40976_1 | TATGAGTGTGGAGTCGTTGCACAATGCGTGGGAATTCAAGATGNNNAGATCGGAA |
| 40976_2 | TATGAGTGTGGAGTCGTTGCACAATGCGTGGGAATTCAAGATGNNNAGATCGGAA |
| 52714_1 | TATGAGTGTGGAGTCGTTGCTTCCCTCATCCATCTGAATGTGGNNNAGATCGGAAG |
| 52714_2 | TATGAGTGTGGAGTCGTTGCTTCCCTCATCCATCTGAATGTGGNNNAGATCGGAAG |
| 38526_1 | TATGAGTGTGGAGTCGTTGCTGCATGCTACAAAATGAATGTGCNNNAGATCGGAA |
| 38526_2 | TATGAGTGTGGAGTCGTTGCTGCATGCTACAAAATGAATGTGCNNNAGATCGGAA |
| 40716_1 | TATGAGTGTGGAGTCGTTGCGGAAACTAGAAGACCAATGTATCACTNNNAGATCGG |
| 40716_2 | TATGAGTGTGGAGTCGTTGCGGAAACTAGAAGACCAATGTATCACTNNNAGATCGG |
| 31891_1 | TATGAGTGTGGAGTCGTTGCAGTGTATCAGCTCCACCATTCAGNNNAGATCGGAAG |
| 31891_2 | TATGAGTGTGGAGTCGTTGCAGTGTATCAGCTCCACCATTCAGNNNAGATCGGAAG |
| 35180_1 | TATGAGTGTGGAGTCGTTGCGAGTGAGTGTAATGGCACGATCTNNNAGATCGGAA |
| 35180_2 | TATGAGTGTGGAGTCGTTGCGAGTGAGTGTAATGGCACGATCTNNNAGATCGGAA |
| 44769_1 | TATGAGTGTGGAGTCGTTGCTCCTCGTATACGCAACATCAGTCNNNAGATCGGAAG |
| 44769_2 | TATGAGTGTGGAGTCGTTGCTCCTCGTATACGCAACATCAGTCNNNAGATCGGAAG |
| 29698_1 | TATGAGTGTGGAGTCGTTGCTGAAACAGAAAAACAGCACAGCANNNAGATCGGAA |

| | |
|---|---|
| 29698_2 | TATGAGTGTGGAGTCGTTGCTGAAACAGAAAAACAGCACAGCANNNAGATCGGAA |
| 10678_1 | TATGAGTGTGGAGTCGTTGCCTTAGCTACTTTGGCATTGCTGCNNNAGATCGGAAG |
| 10678_2 | TATGAGTGTGGAGTCGTTGCCTTAGCTACTTTGGCATTGCTGCNNNAGATCGGAAG |
| 44881_1 | TATGAGTGTGGAGTCGTTGCAGCACCATATTTTCCTCAGGGTTNNNAGATCGGAAG |
| 44881_2 | TATGAGTGTGGAGTCGTTGCAGCACCATATTTTCCTCAGGGTTNNNAGATCGGAAG |
| 44685_1 | TATGAGTGTGGAGTCGTTGCCACCTGCACAGTGAGTACAAATGNNNAGATCGGAA |
| 44685_2 | TATGAGTGTGGAGTCGTTGCCACCTGCACAGTGAGTACAAATGNNNAGATCGGAA |
| 46199_1 | TATGAGTGTGGAGTCGTTGCCTCCCAGGTTCAAGCAATTCTTCNNNAGATCGGAAG |
| 46199_2 | TATGAGTGTGGAGTCGTTGCCTCCCAGGTTCAAGCAATTCTTCNNNAGATCGGAAG |
| 50736_1 | TATGAGTGTGGAGTCGTTGCACTTTCAAGGACTGCTACAACCANNNAGATCGGAAG |
| 50736_2 | TATGAGTGTGGAGTCGTTGCACTTTCAAGGACTGCTACAACCANNNAGATCGGAAG |
| 7608_1 | TATGAGTGTGGAGTCGTTGCTTTCAGAGTGTGTTGTGCCCCTNNNAGATCGGAAGA |
| 7608_2 | TATGAGTGTGGAGTCGTTGCTTTCAGAGTGTGTTGTGCCCCTNNNAGATCGGAAGA |
| 49395_1 | TATGAGTGTGGAGTCGTTGCGTAGAGACGGGGTTTCACTGTGNNNAGATCGGAAG |
| 49395_2 | TATGAGTGTGGAGTCGTTGCGTAGAGACGGGGTTTCACTGTGNNNAGATCGGAAG |
| 12642_1 | TATGAGTGTGGAGTCGTTGCGCTTTAAATTCCTCTCGAAATTNNNAGATCGGAAGA |
| 12642_2 | TATGAGTGTGGAGTCGTTGCGCTTTAAATTCCTCTCGAAATTNNNAGATCGGAAGA |
| 2711_1 | TATGAGTGTGGAGTCGTTGCGGGCAGGAGGATGGTGTAGATATNNNAGATCGGA |
| 2711_2 | TATGAGTGTGGAGTCGTTGCGGGCAGGAGGATGGTGTAGATATNNNAGATCGGA |
| 47676_1 | TATGAGTGTGGAGTCGTTGCCCGCCATGAACTCTTTCTACAACNNNAGATCGGAAGA |
| 47676_2 | TATGAGTGTGGAGTCGTTGCCCGCCATGAACTCTTTCTACAACNNNAGATCGGAAGA |
| 32535_1 | TATGAGTGTGGAGTCGTTGCGAAAGAAGGAAGGAAGGAGGGAGNNNAGATCGG |
| 32535_2 | TATGAGTGTGGAGTCGTTGCGAAAGAAGGAAGGAAGGAGGGAGNNNAGATCGG |
| 49110_1 | TATGAGTGTGGAGTCGTTGCGCTGGGATTATAGGTGTGTGTCANNNAGATCGGAA |
| 49110_2 | TATGAGTGTGGAGTCGTTGCGCTGGGATTATAGGTGTGTGTCANNNAGATCGGAA |
| 32530_1 | TATGAGTGTGGAGTCGTTGCGAAAAGAAAACTTGGGACAGGGGNNNAGATCGGA |
| 32530_2 | TATGAGTGTGGAGTCGTTGCGAAAAGAAAACTTGGGACAGGGGNNNAGATCGGA |
| 2297_1 | TATGAGTGTGGAGTCGTTGCAGAATGAGCAGTGATGGACAACTNNNAGATCGGAA |
| 2297_2 | TATGAGTGTGGAGTCGTTGCAGAATGAGCAGTGATGGACAACTNNNAGATCGGAA |
| 36300_1 | TATGAGTGTGGAGTCGTTGCTTCAGAAAGGTTGTCTTGCAAGCNNNAGATCGGAA |
| 36300_2 | TATGAGTGTGGAGTCGTTGCTTCAGAAAGGTTGTCTTGCAAGCNNNAGATCGGAA |
| 51616_1 | TATGAGTGTGGAGTCGTTGCAGAGCAGGGTTTCTCAATCTCTGNNNAGATCGGAAG |
| 51616_2 | TATGAGTGTGGAGTCGTTGCAGAGCAGGGTTTCTCAATCTCTGNNNAGATCGGAAG |
| 30216_1 | TATGAGTGTGGAGTCGTTGCACATTGTCAGGGTGAGTCCATTTNNNAGATCGGAAG |
| 30216_2 | TATGAGTGTGGAGTCGTTGCACATTGTCAGGGTGAGTCCATTTNNNAGATCGGAAG |
| 43538_1 | TATGAGTGTGGAGTCGTTGCAGAGAGGGAGTCTCACTCTGTCNNNAGATCGGAAG |
| 43538_2 | TATGAGTGTGGAGTCGTTGCAGAGAGGGAGTCTCACTCTGTCNNNAGATCGGAAG |
| 36014_1 | TATGAGTGTGGAGTCGTTGCGTAATCCTTACCAGCGAAAGCACNNNAGATCGGAAG |
| 36014_2 | TATGAGTGTGGAGTCGTTGCGTAATCCTTACCAGCGAAAGCACNNNAGATCGGAAG |
| 44467_1 | TATGAGTGTGGAGTCGTTGCGCAACTACCAAGACAGAGACAGANNNAGATCGGAA |
| 44467_2 | TATGAGTGTGGAGTCGTTGCGCAACTACCAAGACAGAGACAGANNNAGATCGGAA |
| 33429_1 | TATGAGTGTGGAGTCGTTGCAAAAAATTAGCTGGGTGTGGTGGNNNAGATCGGAA |
| 33429_2 | TATGAGTGTGGAGTCGTTGCAAAAAATTAGCTGGGTGTGGTGGNNNAGATCGGAA |

| | |
|---|---|
| 50538_1 | TATGAGTGTGGAGTCGTTGCAAATACCTGTGGATGGGGAGAACNNNAGATCGGAA |
| 50538_2 | TATGAGTGTGGAGTCGTTGCAAATACCTGTGGATGGGGAGAACNNNAGATCGGAA |
| 51176_1 | TATGAGTGTGGAGTCGTTGCGGACAGTGGAGAAGTGAAGGTAGNNNAGATCGGA |
| 51176_2 | TATGAGTGTGGAGTCGTTGCGGACAGTGGAGAAGTGAAGGTAGNNNAGATCGGA |
| 40941_1 | TATGAGTGTGGAGTCGTTGCAGTACTGGAGAGGATGTGGAGAANNNAGATCGGA |
| 40941_2 | TATGAGTGTGGAGTCGTTGCAGTACTGGAGAGGATGTGGAGAANNNAGATCGGA |
| 37070_1 | TATGAGTGTGGAGTCGTTGCCTGCAAAAGTTACAGCCACAGTANNNAGATCGGAAG |
| 37070_2 | TATGAGTGTGGAGTCGTTGCCTGCAAAAGTTACAGCCACAGTANNNAGATCGGAAG |
| 37670_1 | TATGAGTGTGGAGTCGTTGCCATCAGTGTTTGCCAGAGGTTGNNNAGATCGGAAG |
| 37670_2 | TATGAGTGTGGAGTCGTTGCCATCAGTGTTTGCCAGAGGTTGNNNAGATCGGAAG |
| 49602_1 | TATGAGTGTGGAGTCGTTGCGATCTTGGCTCACTGCAAGGTCNNNAGATCGGAAGA |
| 49602_2 | TATGAGTGTGGAGTCGTTGCGATCTTGGCTCACTGCAAGGTCNNNAGATCGGAAGA |
| 49218_1 | TATGAGTGTGGAGTCGTTGCGATGGACTTTCACTCTTGTTGCCNNNAGATCGGAAG |
| 49218_2 | TATGAGTGTGGAGTCGTTGCGATGGACTTTCACTCTTGTTGCCNNNAGATCGGAAG |
| 38754_1 | TATGAGTGTGGAGTCGTTGCCCAAATGGGTGATTTTCTTGTCGANNNAGATCGGAA |
| 38754_2 | TATGAGTGTGGAGTCGTTGCCCAAATGGGTGATTTTCTTGTCGANNNAGATCGGAA |
| 41979_1 | TATGAGTGTGGAGTCGTTGCTTAGAGATGGGGTTTCACTGTGTNNNAGATCGGAA |
| 41979_2 | TATGAGTGTGGAGTCGTTGCTTAGAGATGGGGTTTCACTGTGTNNNAGATCGGAA |
| 53299_1 | TATGAGTGTGGAGTCGTTGCACACACACTCATATCCACACATTNNNAGATCGGAAGA |
| 53299_2 | TATGAGTGTGGAGTCGTTGCACACACACTCATATCCACACATTNNNAGATCGGAAGA |
| 41587_1 | TATGAGTGTGGAGTCGTTGCGTCTTGCTCTGTCGCCTAGATTNNNAGATCGGAAGA |
| 41587_2 | TATGAGTGTGGAGTCGTTGCGTCTTGCTCTGTCGCCTAGATTNNNAGATCGGAAGA |
| 35706_1 | TATGAGTGTGGAGTCGTTGCAAACATTTCACCCGCTCCTTTTTNNNAGATCGGAAGA |
| 35706_2 | TATGAGTGTGGAGTCGTTGCAAACATTTCACCCGCTCCTTTTTNNNAGATCGGAAGA |
| 49003_1 | TATGAGTGTGGAGTCGTTGCCCCAATCATAAGGTCAGCTCAGTNNNAGATCGGAAG |
| 49003_2 | TATGAGTGTGGAGTCGTTGCCCCAATCATAAGGTCAGCTCAGTNNNAGATCGGAAG |
| 32928_1 | TATGAGTGTGGAGTCGTTGCGGGTTGCAATCCTAGTCTCTGATNNNAGATCGGAAG |
| 32928_2 | TATGAGTGTGGAGTCGTTGCGGGTTGCAATCCTAGTCTCTGATNNNAGATCGGAAG |
| 41936_1 | TATGAGTGTGGAGTCGTTGCTCTTCCTTCCTGAATGATGCCAANNNAGATCGGAAG |
| 41936_2 | TATGAGTGTGGAGTCGTTGCTCTTCCTTCCTGAATGATGCCAANNNAGATCGGAAG |
| 38697_1 | TATGAGTGTGGAGTCGTTGCGGAGAATGGCATGAATCGAGGAGNNNAGATCGGA |
| 38697_2 | TATGAGTGTGGAGTCGTTGCGGAGAATGGCATGAATCGAGGAGNNNAGATCGGA |
| 33035_1 | TATGAGTGTGGAGTCGTTGCGAGTGTTGCTCTGTTGCCCANNNAGATCGGAAGAG |
| 33035_2 | TATGAGTGTGGAGTCGTTGCGAGTGTTGCTCTGTTGCCCANNNAGATCGGAAGAG |
| 42653_1 | TATGAGTGTGGAGTCGTTGCTATTCATGTGCTTGGCTTATGGCNNNAGATCGGAAG |
| 42653_2 | TATGAGTGTGGAGTCGTTGCTATTCATGTGCTTGGCTTATGGCNNNAGATCGGAAG |
| 38507_1 | TATGAGTGTGGAGTCGTTGCCAACCTAGGACAGAAGCCTGAAANNNAGATCGGAA |
| 38507_2 | TATGAGTGTGGAGTCGTTGCCAACCTAGGACAGAAGCCTGAAANNNAGATCGGAA |
| 37026_1 | TATGAGTGTGGAGTCGTTGCAGTTGCAACTCCTTCCCAAAGATNNNAGATCGGAAG |
| 37026_2 | TATGAGTGTGGAGTCGTTGCAGTTGCAACTCCTTCCCAAAGATNNNAGATCGGAAG |
| 48611_1 | TATGAGTGTGGAGTCGTTGCTGGCATAATTGTTACACGTGTGGNNNAGATCGGAA |
| 48611_2 | TATGAGTGTGGAGTCGTTGCTGGCATAATTGTTACACGTGTGGNNNAGATCGGAA |
| 32604_1 | TATGAGTGTGGAGTCGTTGCACACCAGCAATAGAGGGGAATTTNNNAGATCGGAA |

| | |
|---|---|
| 32604_2 | TATGAGTGTGGAGTCGTTGCACACCAGCAATAGAGGGGAATTTNNNAGATCGGAA |
| 4572_1 | TATGAGTGTGGAGTCGTTGCACATCTGGGATCCAGCAAGAGACNNNAGATCGGAA |
| 4572_2 | TATGAGTGTGGAGTCGTTGCACATCTGGGATCCAGCAAGAGACNNNAGATCGGAA |
| 44274_1 | TATGAGTGTGGAGTCGTTGCACCTCCTTCTCAATCTCCTCTGANNNAGATCGGAAGA |
| 44274_2 | TATGAGTGTGGAGTCGTTGCACCTCCTTCTCAATCTCCTCTGANNNAGATCGGAAGA |
| 37722_1 | TATGAGTGTGGAGTCGTTGCGAAACAACCCAGCTGTCTTTCAGNNNAGATCGGAAG |
| 37722_2 | TATGAGTGTGGAGTCGTTGCGAAACAACCCAGCTGTCTTTCAGNNNAGATCGGAAG |
| 30481_1 | TATGAGTGTGGAGTCGTTGCCTTTAACTCTTCCAAAAGCCCTCNNNAGATCGGAAGA |
| 30481_2 | TATGAGTGTGGAGTCGTTGCCTTTAACTCTTCCAAAAGCCCTCNNNAGATCGGAAGA |
| 48602_1 | TATGAGTGTGGAGTCGTTGCGTCTCCTGGGTTCAAGTGATTCTNNNAGATCGGAAG |
| 48602_2 | TATGAGTGTGGAGTCGTTGCGTCTCCTGGGTTCAAGTGATTCTNNNAGATCGGAAG |
| 52709_1 | TATGAGTGTGGAGTCGTTGCAACGAAAAGAGAGGGGGGCTAGNNNAGATCGGAA |
| 52709_2 | TATGAGTGTGGAGTCGTTGCAACGAAAAGAGAGGGGGGCTAGNNNAGATCGGAA |
| 37979_1 | TATGAGTGTGGAGTCGTTGCAAGAAGACAAGCCACAGACTAGGNNNAGATCGGAA |
| 37979_2 | TATGAGTGTGGAGTCGTTGCAAGAAGACAAGCCACAGACTAGGNNNAGATCGGAA |
| 36869_1 | TATGAGTGTGGAGTCGTTGCAGAAAAGAAAAGCAAGCCCATGTNNNAGATCGGAA |
| 36869_2 | TATGAGTGTGGAGTCGTTGCAGAAAAGAAAAGCAAGCCCATGTNNNAGATCGGAA |
| 46893_1 | TATGAGTGTGGAGTCGTTGCCCAAATAGCTGGGACTACAGGTGNNNAGATCGGAA |
| 46893_2 | TATGAGTGTGGAGTCGTTGCCCAAATAGCTGGGACTACAGGTGNNNAGATCGGAA |
| 35653_1 | TATGAGTGTGGAGTCGTTGCTATCTCCTCCCAGTTTCAGCCTANNNAGATCGGAAGA |
| 35653_2 | TATGAGTGTGGAGTCGTTGCTATCTCCTCCCAGTTTCAGCCTANNNAGATCGGAAGA |
| 13083_1 | TATGAGTGTGGAGTCGTTGCGAAACTAATCTAACAGACACAAANNNAGATCGGAAG |
| 13083_2 | TATGAGTGTGGAGTCGTTGCGAAACTAATCTAACAGACACAAANNNAGATCGGAAG |
| 31292_1 | TATGAGTGTGGAGTCGTTGCAATTTCTTGAGGACTGGGACAGCNNNAGATCGGAA |
| 31292_2 | TATGAGTGTGGAGTCGTTGCAATTTCTTGAGGACTGGGACAGCNNNAGATCGGAA |
| 13006_1 | TATGAGTGTGGAGTCGTTGCAGGTGTTAATGAAAAACAAAGAANNNAGATCGGAA |
| 13006_2 | TATGAGTGTGGAGTCGTTGCAGGTGTTAATGAAAAACAAAGAANNNAGATCGGAA |
| 46044_1 | TATGAGTGTGGAGTCGTTGCCTAGTAGCTGGGATTATAGGCGCNNNAGATCGGAA |
| 46044_2 | TATGAGTGTGGAGTCGTTGCCTAGTAGCTGGGATTATAGGCGCNNNAGATCGGAA |
| 42013_1 | TATGAGTGTGGAGTCGTTGCCCAAGTAGCTGGGATTACAGGAGNNNAGATCGGAA |
| 42013_2 | TATGAGTGTGGAGTCGTTGCCCAAGTAGCTGGGATTACAGGAGNNNAGATCGGAA |
| 30407_1 | TATGAGTGTGGAGTCGTTGCAGGAGAGTGGACATGGGTTTCATNNNAGATCGGAA |
| 30407_2 | TATGAGTGTGGAGTCGTTGCAGGAGAGTGGACATGGGTTTCATNNNAGATCGGAA |
| 34299_1 | TATGAGTGTGGAGTCGTTGCGCGCTGTTGTGACTTCCTAATTTNNNAGATCGGAAG |
| 34299_2 | TATGAGTGTGGAGTCGTTGCGCGCTGTTGTGACTTCCTAATTTNNNAGATCGGAAG |
| 9346_1 | TATGAGTGTGGAGTCGTTGCTGTGGGTAGGAAAGACCTTAGGGNNNAGATCGGA |
| 9346_2 | TATGAGTGTGGAGTCGTTGCTGTGGGTAGGAAAGACCTTAGGGNNNAGATCGGA |
| 51825_1 | TATGAGTGTGGAGTCGTTGCCTGCCACGTCTGGGAAGTGANNNAGATCGGAAGAG |
| 51825_2 | TATGAGTGTGGAGTCGTTGCCTGCCACGTCTGGGAAGTGANNNAGATCGGAAGAG |
| 43519_1 | TATGAGTGTGGAGTCGTTGCCTCCCAGGTTCAAGCCATTCTCNNNAGATCGGAAGA |
| 43519_2 | TATGAGTGTGGAGTCGTTGCCTCCCAGGTTCAAGCCATTCTCNNNAGATCGGAAGA |
| 52359_1 | TATGAGTGTGGAGTCGTTGCAGAGGTTTTGTGGAGTTCAGTGTNNNAGATCGGAA |
| 52359_2 | TATGAGTGTGGAGTCGTTGCAGAGGTTTTGTGGAGTTCAGTGTNNNAGATCGGAA |

| | |
|---|---|
| 41247_1 | TATGAGTGTGGAGTCGTTGCCCTGGGTGACAGAGTAACACTCNNNAGATCGGAAG |
| 41247_2 | TATGAGTGTGGAGTCGTTGCCCTGGGTGACAGAGTAACACTCNNNAGATCGGAAG |
| 10909_1 | TATGAGTGTGGAGTCGTTGCTACCTGCTATAACTCCTGGCGNNNAGATCGGAAGAG |
| 10909_2 | TATGAGTGTGGAGTCGTTGCTACCTGCTATAACTCCTGGCGNNNAGATCGGAAGAG |
| 51548_1 | TATGAGTGTGGAGTCGTTGCAAGAAGAGCAGAGAATCAGGGTGNNNAGATCGGA |
| 51548_2 | TATGAGTGTGGAGTCGTTGCAAGAAGAGCAGAGAATCAGGGTGNNNAGATCGGA |
| 10006_1 | TATGAGTGTGGAGTCGTTGCTGAAGTTACGACGTAGAGCCTTTNNNAGATCGGAA |
| 10006_2 | TATGAGTGTGGAGTCGTTGCTGAAGTTACGACGTAGAGCCTTTNNNAGATCGGAA |
| 34478_1 | TATGAGTGTGGAGTCGTTGCTTGGACGACTTTGAGTAAGTGGTNNNAGATCGGAA |
| 34478_2 | TATGAGTGTGGAGTCGTTGCTTGGACGACTTTGAGTAAGTGGTNNNAGATCGGAA |
| 10554_1 | TATGAGTGTGGAGTCGTTGCCACTTGGGACTTGGCAGTAGTTTNNNAGATCGGAA |
| 10554_2 | TATGAGTGTGGAGTCGTTGCCACTTGGGACTTGGCAGTAGTTTNNNAGATCGGAA |
| 33849_1 | TATGAGTGTGGAGTCGTTGCCCTTACTTTCCTGTGTCTCGTCANNNAGATCGGAAGA |
| 33849_2 | TATGAGTGTGGAGTCGTTGCCCTTACTTTCCTGTGTCTCGTCANNNAGATCGGAAGA |
| 34142_1 | TATGAGTGTGGAGTCGTTGCGGTTTACAAAGGGTCACTGAGGANNNAGATCGGAA |
| 34142_2 | TATGAGTGTGGAGTCGTTGCGGTTTACAAAGGGTCACTGAGGANNNAGATCGGAA |
| 31451_1 | TATGAGTGTGGAGTCGTTGCCATGCATGTGGTGACAGTGAATTNNNAGATCGGAA |
| 31451_2 | TATGAGTGTGGAGTCGTTGCCATGCATGTGGTGACAGTGAATTNNNAGATCGGAA |
| 33265_1 | TATGAGTGTGGAGTCGTTGCTTCAAGTGATTCTCCTGCCTCAGNNNAGATCGGAAG |
| 33265_2 | TATGAGTGTGGAGTCGTTGCTTCAAGTGATTCTCCTGCCTCAGNNNAGATCGGAAG |
| 50886_1 | TATGAGTGTGGAGTCGTTGCGCAAGTCTTCTCAGGAACTGCTANNNAGATCGGAAG |
| 50886_2 | TATGAGTGTGGAGTCGTTGCGCAAGTCTTCTCAGGAACTGCTANNNAGATCGGAAG |
| 29960_1 | TATGAGTGTGGAGTCGTTGCGTGCCAGGTGAGAAAATGTTCTGNNNAGATCGGAA |
| 29960_2 | TATGAGTGTGGAGTCGTTGCGTGCCAGGTGAGAAAATGTTCTGNNNAGATCGGAA |
| 44511_1 | TATGAGTGTGGAGTCGTTGCCTGTAATCCCAGCTACTGAGGAGNNNAGATCGGAA |
| 44511_2 | TATGAGTGTGGAGTCGTTGCCTGTAATCCCAGCTACTGAGGAGNNNAGATCGGAA |
| 12912_1 | TATGAGTGTGGAGTCGTTGCTCCTAATAACTCATCAAGAAACANNNAGATCGGAAG |
| 12912_2 | TATGAGTGTGGAGTCGTTGCTCCTAATAACTCATCAAGAAACANNNAGATCGGAAG |
| 35379_1 | TATGAGTGTGGAGTCGTTGCCACGCCCGGCTAATTTTTTGTATNNNAGATCGGAAG |
| 35379_2 | TATGAGTGTGGAGTCGTTGCCACGCCCGGCTAATTTTTTGTATNNNAGATCGGAAG |
| 36353_1 | TATGAGTGTGGAGTCGTTGCAGTTGGAGAACTAACATCAGCTGANNNAGATCGGA |
| 36353_2 | TATGAGTGTGGAGTCGTTGCAGTTGGAGAACTAACATCAGCTGANNNAGATCGGA |
| 40393_1 | TATGAGTGTGGAGTCGTTGCGAGGTCAGGAGATCGAGACCATCNNNAGATCGGAA |
| 40393_2 | TATGAGTGTGGAGTCGTTGCGAGGTCAGGAGATCGAGACCATCNNNAGATCGGAA |
| 43746_1 | TATGAGTGTGGAGTCGTTGCTTCCAAACAAGTATGGTCTCGGANNNAGATCGGAAG |
| 43746_2 | TATGAGTGTGGAGTCGTTGCTTCCAAACAAGTATGGTCTCGGANNNAGATCGGAAG |
| 32583_1 | TATGAGTGTGGAGTCGTTGCTTCAAGTGATTCTCCTGCCTCAGNNNAGATCGGAAG |
| 32583_2 | TATGAGTGTGGAGTCGTTGCTTCAAGTGATTCTCCTGCCTCAGNNNAGATCGGAAG |
| 43081_1 | TATGAGTGTGGAGTCGTTGCAATTTCCCTCAAGGCCTGATGAANNNAGATCGGAAG |
| 43081_2 | TATGAGTGTGGAGTCGTTGCAATTTCCCTCAAGGCCTGATGAANNNAGATCGGAAG |
| 43288_1 | TATGAGTGTGGAGTCGTTGCCACACACACACACACACATACAGNNNAGATCGGAAG |
| 43288_2 | TATGAGTGTGGAGTCGTTGCCACACACACACACACACATACAGNNNAGATCGGAAG |
| 31184_1 | TATGAGTGTGGAGTCGTTGCACTCTCAGTTTCATCATCCCTGANNNAGATCGGAAGA |

| | |
|---|---|
| 31184_2 | TATGAGTGTGGAGTCGTTGCACTCTCAGTTTCATCATCCCTGANNNAGATCGGAAGA |
| 50040_1 | TATGAGTGTGGAGTCGTTGCTTTGGACTCAAGTGATCCTCCTGNNNAGATCGGAAG |
| 50040_2 | TATGAGTGTGGAGTCGTTGCTTTGGACTCAAGTGATCCTCCTGNNNAGATCGGAAG |
| 45312_1 | TATGAGTGTGGAGTCGTTGCGGACCGCCAGACTAACAAAGTNNNAGATCGGAAGA |
| 45312_2 | TATGAGTGTGGAGTCGTTGCGGACCGCCAGACTAACAAAGTNNNAGATCGGAAGA |
| 40058_1 | TATGAGTGTGGAGTCGTTGCGAGGCAGGATAATTGCTTGAACCNNNAGATCGGAA |
| 40058_2 | TATGAGTGTGGAGTCGTTGCGAGGCAGGATAATTGCTTGAACCNNNAGATCGGAA |
| 41356_1 | TATGAGTGTGGAGTCGTTGCAGGTTTATCTTGCAGGTAAGGCANNNAGATCGGAA |
| 41356_2 | TATGAGTGTGGAGTCGTTGCAGGTTTATCTTGCAGGTAAGGCANNNAGATCGGAA |
| 49301_1 | TATGAGTGTGGAGTCGTTGCCCTCAGCAACAATGAATCTGCTTNNNAGATCGGAAG |
| 49301_2 | TATGAGTGTGGAGTCGTTGCCCTCAGCAACAATGAATCTGCTTNNNAGATCGGAAG |
| 45503_1 | TATGAGTGTGGAGTCGTTGCCTTTCAGTGTTGCAGGAGCGTTCNNNAGATCGGAAG |
| 45503_2 | TATGAGTGTGGAGTCGTTGCCTTTCAGTGTTGCAGGAGCGTTCNNNAGATCGGAAG |
| 47060_1 | TATGAGTGTGGAGTCGTTGCAGATGGAGTCTGACTCTGTTGCNNNAGATCGGAAG |
| 47060_2 | TATGAGTGTGGAGTCGTTGCAGATGGAGTCTGACTCTGTTGCNNNAGATCGGAAG |
| 32286_1 | TATGAGTGTGGAGTCGTTGCGTTTGCAGGATGACCTCTGACTANNNAGATCGGAAG |
| 32286_2 | TATGAGTGTGGAGTCGTTGCGTTTGCAGGATGACCTCTGACTANNNAGATCGGAAG |
| 37100_1 | TATGAGTGTGGAGTCGTTGCAAACTACCCGTTATCTGAGTGGGNNNAGATCGGAA |
| 37100_2 | TATGAGTGTGGAGTCGTTGCAAACTACCCGTTATCTGAGTGGGNNNAGATCGGAA |
| 13684_1 | TATGAGTGTGGAGTCGTTGCGTTTCTAAGGACTTTTTCTTAGGNNNAGATCGGAAG |
| 13684_2 | TATGAGTGTGGAGTCGTTGCGTTTCTAAGGACTTTTTCTTAGGNNNAGATCGGAAG |
| 7973_1 | TATGAGTGTGGAGTCGTTGCGCTAGGTGGATCACTTGAGTGCNNNAGATCGGAAG |
| 7973_2 | TATGAGTGTGGAGTCGTTGCGCTAGGTGGATCACTTGAGTGCNNNAGATCGGAAG |
| 36105_1 | TATGAGTGTGGAGTCGTTGCAATATTACGCATCCTGTCCTGGCNNNAGATCGGAAG |
| 36105_2 | TATGAGTGTGGAGTCGTTGCAATATTACGCATCCTGTCCTGGCNNNAGATCGGAAG |
| 29432_1 | TATGAGTGTGGAGTCGTTGCACTTCATGTTCTTTTCAAAGACAGTNNNAGATCGGAA |
| 29432_2 | TATGAGTGTGGAGTCGTTGCACTTCATGTTCTTTTCAAAGACAGTNNNAGATCGGAA |
| 39738_1 | TATGAGTGTGGAGTCGTTGCGGAGAATGGTATGAACCCAGGAGNNNAGATCGGA |
| 39738_2 | TATGAGTGTGGAGTCGTTGCGGAGAATGGTATGAACCCAGGAGNNNAGATCGGA |
| 48203_1 | TATGAGTGTGGAGTCGTTGCCGGCCCTCTACTTCTTTTACGATNNNAGATCGGAAGA |
| 48203_2 | TATGAGTGTGGAGTCGTTGCCGGCCCTCTACTTCTTTTACGATNNNAGATCGGAAGA |
| 36529_1 | TATGAGTGTGGAGTCGTTGCCCTAAGGACACACAGCTACACTTNNNAGATCGGAAG |
| 36529_2 | TATGAGTGTGGAGTCGTTGCCCTAAGGACACACAGCTACACTTNNNAGATCGGAAG |
| 40209_1 | TATGAGTGTGGAGTCGTTGCAAAAGAAAAGTGGGAGATCAGGTNNNAGATCGGA |
| 40209_2 | TATGAGTGTGGAGTCGTTGCAAAAGAAAAGTGGGAGATCAGGTNNNAGATCGGA |
| 32972_1 | TATGAGTGTGGAGTCGTTGCACAAAAAAATTAGCCGGGCGTAATNNNAGATCGGAA |
| 32972_2 | TATGAGTGTGGAGTCGTTGCACAAAAAAATTAGCCGGGCGTAATNNNAGATCGGAA |
| 36371_1 | TATGAGTGTGGAGTCGTTGCGCTTAGTGCCTGGCCTATTTTTCNNNAGATCGGAAG |
| 36371_2 | TATGAGTGTGGAGTCGTTGCGCTTAGTGCCTGGCCTATTTTTCNNNAGATCGGAAG |
| 35606_1 | TATGAGTGTGGAGTCGTTGCTCAAGTGATTCTCCTGCCTTAGCNNNAGATCGGAAG |
| 35606_2 | TATGAGTGTGGAGTCGTTGCTCAAGTGATTCTCCTGCCTTAGCNNNAGATCGGAAG |
| 39879_1 | TATGAGTGTGGAGTCGTTGCGAGCAGGGAGAAAGAACATGAACNNNAGATCGGA |
| 39879_2 | TATGAGTGTGGAGTCGTTGCGAGCAGGGAGAAAGAACATGAACNNNAGATCGGA |

| | |
|---|---|
| 39039_1 | TATGAGTGTGGAGTCGTTGCAGCAAAATGGCAGAGTTTAGCTGNNNAGATCGGAA |
| 39039_2 | TATGAGTGTGGAGTCGTTGCAGCAAAATGGCAGAGTTTAGCTGNNNAGATCGGAA |
| 53340_1 | TATGAGTGTGGAGTCGTTGCCTGACTGTGTTAGTGGTTGCATGNNNAGATCGGAA |
| 53340_2 | TATGAGTGTGGAGTCGTTGCCTGACTGTGTTAGTGGTTGCATGNNNAGATCGGAA |
| 39060_1 | TATGAGTGTGGAGTCGTTGCCATCCTGGACATAGTAACAGGCANNNAGATCGGAA |
| 39060_2 | TATGAGTGTGGAGTCGTTGCCATCCTGGACATAGTAACAGGCANNNAGATCGGAA |
| 32479_1 | TATGAGTGTGGAGTCGTTGCGAGACGGAGTCTTGCTCTGTCNNNAGATCGGAAGA |
| 32479_2 | TATGAGTGTGGAGTCGTTGCGAGACGGAGTCTTGCTCTGTCNNNAGATCGGAAGA |
| 40982_1 | TATGAGTGTGGAGTCGTTGCGAACGTTATCAGGCGATATTGGANNNAGATCGGAA |
| 40982_2 | TATGAGTGTGGAGTCGTTGCGAACGTTATCAGGCGATATTGGANNNAGATCGGAA |
| 38763_1 | TATGAGTGTGGAGTCGTTGCCTGAGGCAGAAGAAGTGCTTGAANNNAGATCGGAA |
| 38763_2 | TATGAGTGTGGAGTCGTTGCCTGAGGCAGAAGAAGTGCTTGAANNNAGATCGGAA |
| 29265_1 | TATGAGTGTGGAGTCGTTGCTGCCTTCTTTGCCTTCCTACATTNNNAGATCGGAAGA |
| 29265_2 | TATGAGTGTGGAGTCGTTGCTGCCTTCTTTGCCTTCCTACATTNNNAGATCGGAAGA |
| 42889_1 | TATGAGTGTGGAGTCGTTGCATCCTGTCATTTGCCACAACATGNNNAGATCGGAAG |
| 42889_2 | TATGAGTGTGGAGTCGTTGCATCCTGTCATTTGCCACAACATGNNNAGATCGGAAG |
| 33621_1 | TATGAGTGTGGAGTCGTTGCCTCCTGAGTAGCTGGGACTACAGNNNAGATCGGAA |
| 33621_2 | TATGAGTGTGGAGTCGTTGCCTCCTGAGTAGCTGGGACTACAGNNNAGATCGGAA |
| 28836_1 | TATGAGTGTGGAGTCGTTGCCACTGATCTCTGCCTGTCTTCAANNNAGATCGGAAGA |
| 28836_2 | TATGAGTGTGGAGTCGTTGCCACTGATCTCTGCCTGTCTTCAANNNAGATCGGAAGA |
| 33426_1 | TATGAGTGTGGAGTCGTTGCGAATTTCACCCTGGCAACGTAAANNNAGATCGGAAG |
| 33426_2 | TATGAGTGTGGAGTCGTTGCGAATTTCACCCTGGCAACGTAAANNNAGATCGGAAG |
| 33172_1 | TATGAGTGTGGAGTCGTTGCGGGACTCTGCATTTTCTGGAGATNNNAGATCGGAA |
| 33172_2 | TATGAGTGTGGAGTCGTTGCGGGACTCTGCATTTTCTGGAGATNNNAGATCGGAA |
| 35495_1 | TATGAGTGTGGAGTCGTTGCACTCCCAAAGTGCTGTGATTACANNNAGATCGGAAG |
| 35495_2 | TATGAGTGTGGAGTCGTTGCACTCCCAAAGTGCTGTGATTACANNNAGATCGGAAG |
| 52649_1 | TATGAGTGTGGAGTCGTTGCTCATCTTCATCACTGCTCAGCTTNNNAGATCGGAAGA |
| 52649_2 | TATGAGTGTGGAGTCGTTGCTCATCTTCATCACTGCTCAGCTTNNNAGATCGGAAGA |
| 41274_1 | TATGAGTGTGGAGTCGTTGCTTATAGGATAAGGACCTGGCACGNNNAGATCGGAA |
| 41274_2 | TATGAGTGTGGAGTCGTTGCTTATAGGATAAGGACCTGGCACGNNNAGATCGGAA |
| 48536_1 | TATGAGTGTGGAGTCGTTGCCTGGTTTCAAGTGATTCTCCTGCNNNAGATCGGAAG |
| 48536_2 | TATGAGTGTGGAGTCGTTGCCTGGTTTCAAGTGATTCTCCTGCNNNAGATCGGAAG |
| 51520_1 | TATGAGTGTGGAGTCGTTGCCCTAGTTGCGAAGCTGGTATTTGNNNAGATCGGAA |
| 51520_2 | TATGAGTGTGGAGTCGTTGCCCTAGTTGCGAAGCTGGTATTTGNNNAGATCGGAA |
| 37492_1 | TATGAGTGTGGAGTCGTTGCGAGTGAGTGTGACACAGCCAANNNAGATCGGAAGA |
| 37492_2 | TATGAGTGTGGAGTCGTTGCGAGTGAGTGTGACACAGCCAANNNAGATCGGAAGA |
| 51006_1 | TATGAGTGTGGAGTCGTTGCAATGAAGACCTGCAGCAGAAGTTNNNAGATCGGAA |
| 51006_2 | TATGAGTGTGGAGTCGTTGCAATGAAGACCTGCAGCAGAAGTTNNNAGATCGGAA |
| 48841_1 | TATGAGTGTGGAGTCGTTGCAGTGCTGGGATTATAGGTGTGAGNNNAGATCGGAA |
| 48841_2 | TATGAGTGTGGAGTCGTTGCAGTGCTGGGATTATAGGTGTGAGNNNAGATCGGAA |
| 51865_1 | TATGAGTGTGGAGTCGTTGCCCTGCCTGGCTTAGATCATACTTNNNAGATCGGAAG |
| 51865_2 | TATGAGTGTGGAGTCGTTGCCCTGCCTGGCTTAGATCATACTTNNNAGATCGGAAG |
| 9538_1 | TATGAGTGTGGAGTCGTTGCTCGTCAACTTCAACAGTTCCCTCNNNAGATCGGAAGA |

| | |
|---|---|
| 9538_2 | TATGAGTGTGGAGTCGTTGCTCGTCAACTTCAACAGTTCCCTCNNNAGATCGGAAGA |
| 35127_1 | TATGAGTGTGGAGTCGTTGCTTGTTGCTGTTGTTTGGAACTGTNNNAGATCGGAAG |
| 35127_2 | TATGAGTGTGGAGTCGTTGCTTGTTGCTGTTGTTTGGAACTGTNNNAGATCGGAAG |
| 12634_1 | TATGAGTGTGGAGTCGTTGCGCAACTTTATATCAGTGAGAATTNNNAGATCGGAAG |
| 12634_2 | TATGAGTGTGGAGTCGTTGCGCAACTTTATATCAGTGAGAATTNNNAGATCGGAAG |
| 48893_1 | TATGAGTGTGGAGTCGTTGCGATCTTGGCTCACTGCAACTTCNNNAGATCGGAAGA |
| 48893_2 | TATGAGTGTGGAGTCGTTGCGATCTTGGCTCACTGCAACTTCNNNAGATCGGAAGA |
| 52285_1 | TATGAGTGTGGAGTCGTTGCCACTGTCTTGCACATTTGAGGTCNNNAGATCGGAAG |
| 52285_2 | TATGAGTGTGGAGTCGTTGCCACTGTCTTGCACATTTGAGGTCNNNAGATCGGAAG |
| 34785_1 | TATGAGTGTGGAGTCGTTGCGATCTCAGCTCACTGCAACCTCNNNAGATCGGAAGA |
| 34785_2 | TATGAGTGTGGAGTCGTTGCGATCTCAGCTCACTGCAACCTCNNNAGATCGGAAGA |
| 32889_1 | TATGAGTGTGGAGTCGTTGCACAATCAGCAGGCATTTACATAGGNNNAGATCGGAA |
| 32889_2 | TATGAGTGTGGAGTCGTTGCACAATCAGCAGGCATTTACATAGGNNNAGATCGGAA |
| 51199_1 | TATGAGTGTGGAGTCGTTGCGTCTCAGATCTGCTTATCCCCAGNNNAGATCGGAAG |
| 51199_2 | TATGAGTGTGGAGTCGTTGCGTCTCAGATCTGCTTATCCCCAGNNNAGATCGGAAG |
| 46894_1 | TATGAGTGTGGAGTCGTTGCGTGTTGTGATTACAGGTGTGAGCNNNAGATCGGAA |
| 46894_2 | TATGAGTGTGGAGTCGTTGCGTGTTGTGATTACAGGTGTGAGCNNNAGATCGGAA |
| 34842_1 | TATGAGTGTGGAGTCGTTGCATAGCTAAAATTACAGGCGCCCANNNAGATCGGAAG |
| 34842_2 | TATGAGTGTGGAGTCGTTGCATAGCTAAAATTACAGGCGCCCANNNAGATCGGAAG |
| 43593_1 | TATGAGTGTGGAGTCGTTGCTCAAGTAGCTGAGATTACAGGCGNNNAGATCGGAA |
| 43593_2 | TATGAGTGTGGAGTCGTTGCTCAAGTAGCTGAGATTACAGGCGNNNAGATCGGAA |
| 44160_1 | TATGAGTGTGGAGTCGTTGCCCTCGCTTTGAGAGACAGGTATCNNNAGATCGGAAG |
| 44160_2 | TATGAGTGTGGAGTCGTTGCCCTCGCTTTGAGAGACAGGTATCNNNAGATCGGAAG |
| 52742_1 | TATGAGTGTGGAGTCGTTGCTCTGTTCATCTGAGGAGAATGGCNNNAGATCGGAA |
| 52742_2 | TATGAGTGTGGAGTCGTTGCTCTGTTCATCTGAGGAGAATGGCNNNAGATCGGAA |
| 32153_1 | TATGAGTGTGGAGTCGTTGCTCGTTAGGTGGTAAAGATCTGACANNNAGATCGGA |
| 32153_2 | TATGAGTGTGGAGTCGTTGCTCGTTAGGTGGTAAAGATCTGACANNNAGATCGGA |
| 32956_1 | TATGAGTGTGGAGTCGTTGCTGACTAGAGGCATCTGGTACTCANNNAGATCGGAA |
| 32956_2 | TATGAGTGTGGAGTCGTTGCTGACTAGAGGCATCTGGTACTCANNNAGATCGGAA |
| 47366_1 | TATGAGTGTGGAGTCGTTGCTCTCTGCCTCCTAAGTTCAAGTGNNNAGATCGGAAG |
| 47366_2 | TATGAGTGTGGAGTCGTTGCTCTCTGCCTCCTAAGTTCAAGTGNNNAGATCGGAAG |
| 40169_1 | TATGAGTGTGGAGTCGTTGCTGAGGTCAGGAATTCAAGACCAGNNNAGATCGGAA |
| 40169_2 | TATGAGTGTGGAGTCGTTGCTGAGGTCAGGAATTCAAGACCAGNNNAGATCGGAA |
| 46337_1 | TATGAGTGTGGAGTCGTTGCAATGAGTACAGCTGCTTCTCACANNNAGATCGGAAG |
| 46337_2 | TATGAGTGTGGAGTCGTTGCAATGAGTACAGCTGCTTCTCACANNNAGATCGGAAG |
| 45220_1 | TATGAGTGTGGAGTCGTTGCCTGACAGTCAAAAAAGGGCAAGTNNNAGATCGGAA |
| 45220_2 | TATGAGTGTGGAGTCGTTGCCTGACAGTCAAAAAAGGGCAAGTNNNAGATCGGAA |
| 40193_1 | TATGAGTGTGGAGTCGTTGCAGTGAGCTATGTGGTCTTGCTACNNNAGATCGGAA |
| 40193_2 | TATGAGTGTGGAGTCGTTGCAGTGAGCTATGTGGTCTTGCTACNNNAGATCGGAA |
| 50297_1 | TATGAGTGTGGAGTCGTTGCTTGTGTACATTTTTTGGCTGGGCNNNAGATCGGAAG |
| 50297_2 | TATGAGTGTGGAGTCGTTGCTTGTGTACATTTTTTGGCTGGGCNNNAGATCGGAAG |
| 44925_1 | TATGAGTGTGGAGTCGTTGCATCTGTTTCAACTCCGAGACTGGNNNAGATCGGAAG |
| 44925_2 | TATGAGTGTGGAGTCGTTGCATCTGTTTCAACTCCGAGACTGGNNNAGATCGGAAG |

| | |
|---|---|
| 33737_1 | TATGAGTGTGGAGTCGTTGCCACACCTGTAATCCCAGCACTTTNNNAGATCGGAAG |
| 33737_2 | TATGAGTGTGGAGTCGTTGCCACACCTGTAATCCCAGCACTTTNNNAGATCGGAAG |
| 30033_1 | TATGAGTGTGGAGTCGTTGCTTGATGCTTTCAAAGAAGGTGGCNNNAGATCGGAA |
| 30033_2 | TATGAGTGTGGAGTCGTTGCTTGATGCTTTCAAAGAAGGTGGCNNNAGATCGGAA |
| 30452_1 | TATGAGTGTGGAGTCGTTGCTATCTCTTGACCTGCCTACCTTCNNNAGATCGGAAGA |
| 30452_2 | TATGAGTGTGGAGTCGTTGCTATCTCTTGACCTGCCTACCTTCNNNAGATCGGAAGA |
| 30774_1 | TATGAGTGTGGAGTCGTTGCACAGACTCCACTTCTCCACTAGANNNAGATCGGAAG |
| 30774_2 | TATGAGTGTGGAGTCGTTGCACAGACTCCACTTCTCCACTAGANNNAGATCGGAAG |
| 44127_1 | TATGAGTGTGGAGTCGTTGCCTATCTCCATGACCCCTTGAAGGNNNAGATCGGAAG |
| 44127_2 | TATGAGTGTGGAGTCGTTGCCTATCTCCATGACCCCTTGAAGGNNNAGATCGGAAG |
| 48110_1 | TATGAGTGTGGAGTCGTTGCCTTCCAGTTCTGTCTTGCATGTCNNNAGATCGGAAG |
| 48110_2 | TATGAGTGTGGAGTCGTTGCCTTCCAGTTCTGTCTTGCATGTCNNNAGATCGGAAG |
| 52406_1 | TATGAGTGTGGAGTCGTTGCTTACCTTTTCCCTGGAGCTTACCNNNAGATCGGAAGA |
| 52406_2 | TATGAGTGTGGAGTCGTTGCTTACCTTTTCCCTGGAGCTTACCNNNAGATCGGAAGA |
| 35600_1 | TATGAGTGTGGAGTCGTTGCGATTACAGTGGCACAATCTTGGCNNNAGATCGGAA |
| 35600_2 | TATGAGTGTGGAGTCGTTGCGATTACAGTGGCACAATCTTGGCNNNAGATCGGAA |
| 36195_1 | TATGAGTGTGGAGTCGTTGCATTGGAGGTGATGGATGTGCTAANNNAGATCGGAA |
| 36195_2 | TATGAGTGTGGAGTCGTTGCATTGGAGGTGATGGATGTGCTAANNNAGATCGGAA |
| 44433_1 | TATGAGTGTGGAGTCGTTGCATACACGGAGGTTACAAAGGCAANNNAGATCGGAA |
| 44433_2 | TATGAGTGTGGAGTCGTTGCATACACGGAGGTTACAAAGGCAANNNAGATCGGAA |
| 38348_1 | TATGAGTGTGGAGTCGTTGCAAGGGGATTCCTAAGGTTGCATTNNNAGATCGGAA |
| 38348_2 | TATGAGTGTGGAGTCGTTGCAAGGGGATTCCTAAGGTTGCATTNNNAGATCGGAA |
| 31260_1 | TATGAGTGTGGAGTCGTTGCATGTAAATGCATGTGTGTGTACANNNAGATCGGAA |
| 31260_2 | TATGAGTGTGGAGTCGTTGCATGTAAATGCATGTGTGTGTACANNNAGATCGGAA |
| 49401_1 | TATGAGTGTGGAGTCGTTGCCTTTTTTTGAGACAGGGTCCCACNNNAGATCGGAAG |
| 49401_2 | TATGAGTGTGGAGTCGTTGCCTTTTTTTGAGACAGGGTCCCACNNNAGATCGGAAG |
| 37201_1 | TATGAGTGTGGAGTCGTTGCAACAAAACATGCACAGAATCTGTNNNAGATCGGAAG |
| 37201_2 | TATGAGTGTGGAGTCGTTGCAACAAAACATGCACAGAATCTGTNNNAGATCGGAAG |
| 35015_1 | TATGAGTGTGGAGTCGTTGCGCACGCCCAGCTAATTTTTTTTNNNAGATCGGAAG |
| 35015_2 | TATGAGTGTGGAGTCGTTGCGCACGCCCAGCTAATTTTTTTTNNNAGATCGGAAG |
| 39081_1 | TATGAGTGTGGAGTCGTTGCTCCTAGAAAAACATGAAAAGCAAAGCNNNAGATCG |
| 39081_2 | TATGAGTGTGGAGTCGTTGCTCCTAGAAAAACATGAAAAGCAAAGCNNNAGATCG |
| 30712_1 | TATGAGTGTGGAGTCGTTGCTGAGCCACCACACTATCTCTTTATCNNNAGATCGGAA |
| 30712_2 | TATGAGTGTGGAGTCGTTGCTGAGCCACCACACTATCTCTTTATCNNNAGATCGGAA |
| 37324_1 | TATGAGTGTGGAGTCGTTGCCTGAGGCAGGAGAATCACTTGAANNNAGATCGGAA |
| 37324_2 | TATGAGTGTGGAGTCGTTGCCTGAGGCAGGAGAATCACTTGAANNNAGATCGGAA |
| 39842_1 | TATGAGTGTGGAGTCGTTGCCACCACGCCTGGCTAATTTTTTNNNAGATCGGAAGA |
| 39842_2 | TATGAGTGTGGAGTCGTTGCCACCACGCCTGGCTAATTTTTTNNNAGATCGGAAGA |
| 40076_1 | TATGAGTGTGGAGTCGTTGCAGAAATGAGGGGGTTTGCCTAATNNNAGATCGGAA |
| 40076_2 | TATGAGTGTGGAGTCGTTGCAGAAATGAGGGGGTTTGCCTAATNNNAGATCGGAA |
| 47344_1 | TATGAGTGTGGAGTCGTTGCCTCAGCGTTTGCTTGTCTGTAAANNNAGATCGGAAG |
| 47344_2 | TATGAGTGTGGAGTCGTTGCCTCAGCGTTTGCTTGTCTGTAAANNNAGATCGGAAG |
| 29689_1 | TATGAGTGTGGAGTCGTTGCCCTGCAATCCCTTCCATTCATTCNNNAGATCGGAAGA |

| | |
|---|---|
| 29689_2 | TATGAGTGTGGAGTCGTTGCCCTGCAATCCCTTCCATTCATTCNNNAGATCGGAAGA |
| 49862_1 | TATGAGTGTGGAGTCGTTGCCATTTTACACCCTGAGCAAGCAANNNAGATCGGAAG |
| 49862_2 | TATGAGTGTGGAGTCGTTGCCATTTTACACCCTGAGCAAGCAANNNAGATCGGAAG |
| 12759_1 | TATGAGTGTGGAGTCGTTGCTATGAAAGGGTTAAGAATTTGTGNNNAGATCGGAA |
| 12759_2 | TATGAGTGTGGAGTCGTTGCTATGAAAGGGTTAAGAATTTGTGNNNAGATCGGAA |
| 34493_1 | TATGAGTGTGGAGTCGTTGCCCCCAGTTCTGCAGCAGTATCNNNAGATCGGAAGAG |
| 34493_2 | TATGAGTGTGGAGTCGTTGCCCCCAGTTCTGCAGCAGTATCNNNAGATCGGAAGAG |
| 45115_1 | TATGAGTGTGGAGTCGTTGCTGTAGCATGTGCATTGTAGCATGNNNAGATCGGAA |
| 45115_2 | TATGAGTGTGGAGTCGTTGCTGTAGCATGTGCATTGTAGCATGNNNAGATCGGAA |
| 39856_1 | TATGAGTGTGGAGTCGTTGCTTGTGTAGGGCTACTATGCTGTGNNNAGATCGGAA |
| 39856_2 | TATGAGTGTGGAGTCGTTGCTTGTGTAGGGCTACTATGCTGTGNNNAGATCGGAA |
| 36147_1 | TATGAGTGTGGAGTCGTTGCGGATTCAGAGTCAATGGTGCTTCNNNAGATCGGAA |
| 36147_2 | TATGAGTGTGGAGTCGTTGCGGATTCAGAGTCAATGGTGCTTCNNNAGATCGGAA |
| 30458_1 | TATGAGTGTGGAGTCGTTGCGAGTCACTGGAATCTATAAGGCTCTNNNAGATCGGA |
| 30458_2 | TATGAGTGTGGAGTCGTTGCGAGTCACTGGAATCTATAAGGCTCTNNNAGATCGGA |
| 40853_1 | TATGAGTGTGGAGTCGTTGCCCTTACAGTGCCTCACAAGAGATNNNAGATCGGAAG |
| 40853_2 | TATGAGTGTGGAGTCGTTGCCCTTACAGTGCCTCACAAGAGATNNNAGATCGGAAG |
| 34723_1 | TATGAGTGTGGAGTCGTTGCAAGTGATCCTCCTGCCTTGATTTNNNAGATCGGAAG |
| 34723_2 | TATGAGTGTGGAGTCGTTGCAAGTGATCCTCCTGCCTTGATTTNNNAGATCGGAAG |
| 36376_1 | TATGAGTGTGGAGTCGTTGCTGATGTCAGGTTTTAGTCTGCCANNNAGATCGGAAG |
| 36376_2 | TATGAGTGTGGAGTCGTTGCTGATGTCAGGTTTTAGTCTGCCANNNAGATCGGAAG |
| 49143_1 | TATGAGTGTGGAGTCGTTGCGTCCCTAATCTGGAAATCCCGAANNNAGATCGGAAG |
| 49143_2 | TATGAGTGTGGAGTCGTTGCGTCCCTAATCTGGAAATCCCGAANNNAGATCGGAAG |
| 30208_1 | TATGAGTGTGGAGTCGTTGCTAAAACATTTGCCAGCAGCTACCNNNAGATCGGAAG |
| 30208_2 | TATGAGTGTGGAGTCGTTGCTAAAACATTTGCCAGCAGCTACCNNNAGATCGGAAG |
| 46946_1 | TATGAGTGTGGAGTCGTTGCGATCTGCCTTCTCAAGCTCTCATNNNAGATCGGAAGA |
| 46946_2 | TATGAGTGTGGAGTCGTTGCGATCTGCCTTCTCAAGCTCTCATNNNAGATCGGAAGA |
| 36253_1 | TATGAGTGTGGAGTCGTTGCGGGTGAGGAGATTGGAATTTCCTNNNAGATCGGAA |
| 36253_2 | TATGAGTGTGGAGTCGTTGCGGGTGAGGAGATTGGAATTTCCTNNNAGATCGGAA |
| 47296_1 | TATGAGTGTGGAGTCGTTGCCCTCCTGAGTAGCTGGGACTATANNNAGATCGGAAG |
| 47296_2 | TATGAGTGTGGAGTCGTTGCCCTCCTGAGTAGCTGGGACTATANNNAGATCGGAAG |
| 34016_1 | TATGAGTGTGGAGTCGTTGCCAAGTAGCTGGGACTACAGGTGNNNAGATCGGAAG |
| 34016_2 | TATGAGTGTGGAGTCGTTGCCAAGTAGCTGGGACTACAGGTGNNNAGATCGGAAG |
| 39968_1 | TATGAGTGTGGAGTCGTTGCACAGGTGGATTGTTTTTTTCTGGANNNAGATCGGAA |
| 39968_2 | TATGAGTGTGGAGTCGTTGCACAGGTGGATTGTTTTTTTCTGGANNNAGATCGGAA |
| 46055_1 | TATGAGTGTGGAGTCGTTGCCAGCCTCCTGAGTAACTGGAATTNNNAGATCGGAAG |
| 46055_2 | TATGAGTGTGGAGTCGTTGCCAGCCTCCTGAGTAACTGGAATTNNNAGATCGGAAG |
| 37332_1 | TATGAGTGTGGAGTCGTTGCTGAGGAATGAGAGAGGGTGAGATNNNAGATCGGA |
| 37332_2 | TATGAGTGTGGAGTCGTTGCTGAGGAATGAGAGAGGGTGAGATNNNAGATCGGA |
| 13101_1 | TATGAGTGTGGAGTCGTTGCATTTAACTCCTTGGGTTTTTTAGNNNAGATCGGAAG |
| 13101_2 | TATGAGTGTGGAGTCGTTGCATTTAACTCCTTGGGTTTTTTAGNNNAGATCGGAAG |
| 36122_1 | TATGAGTGTGGAGTCGTTGCGAAGCAGGGATAATTGCCTGAACNNNAGATCGGAA |
| 36122_2 | TATGAGTGTGGAGTCGTTGCGAAGCAGGGATAATTGCCTGAACNNNAGATCGGAA |

| | |
|---|---|
| 45636_1 | TATGAGTGTGGAGTCGTTGCCCAAAGTGCTGTTCTGGGATTACNNNAGATCGGAAG |
| 45636_2 | TATGAGTGTGGAGTCGTTGCCCAAAGTGCTGTTCTGGGATTACNNNAGATCGGAAG |
| 50816_1 | TATGAGTGTGGAGTCGTTGCAGAAGCTAGCCAACAACAACAACNNNAGATCGGAAG |
| 50816_2 | TATGAGTGTGGAGTCGTTGCAGAAGCTAGCCAACAACAACAACNNNAGATCGGAAG |
| 13087_1 | TATGAGTGTGGAGTCGTTGCCATAATTAGTGCCAATATACCACNNNAGATCGGAAG |
| 13087_2 | TATGAGTGTGGAGTCGTTGCCATAATTAGTGCCAATATACCACNNNAGATCGGAAG |
| 32033_1 | TATGAGTGTGGAGTCGTTGCGGAGAACCCTTACTCTGAGATGCNNNAGATCGGAAG |
| 32033_2 | TATGAGTGTGGAGTCGTTGCGGAGAACCCTTACTCTGAGATGCNNNAGATCGGAAG |
| 52585_1 | TATGAGTGTGGAGTCGTTGCTTTAGGGGCTGAGCGTATCAATTNNNAGATCGGAA |
| 52585_2 | TATGAGTGTGGAGTCGTTGCTTTAGGGGCTGAGCGTATCAATTNNNAGATCGGAA |
| 31201_1 | TATGAGTGTGGAGTCGTTGCTTCAGACCCAGAAGAGTTGATGGNNNAGATCGGAA |
| 31201_2 | TATGAGTGTGGAGTCGTTGCTTCAGACCCAGAAGAGTTGATGGNNNAGATCGGAA |
| 43102_1 | TATGAGTGTGGAGTCGTTGCAAGGAGTGTTGATGTTGATGTGTNNNAGATCGGAA |
| 43102_2 | TATGAGTGTGGAGTCGTTGCAAGGAGTGTTGATGTTGATGTGTNNNAGATCGGAA |
| 32293_1 | TATGAGTGTGGAGTCGTTGCGAATCAAGAATCAGGCTACCACGNNNAGATCGGAA |
| 32293_2 | TATGAGTGTGGAGTCGTTGCGAATCAAGAATCAGGCTACCACGNNNAGATCGGAA |
| 31156_1 | TATGAGTGTGGAGTCGTTGCCCAGGGACATGAAGAGTGAAGTTNNNAGATCGGAA |
| 31156_2 | TATGAGTGTGGAGTCGTTGCCCAGGGACATGAAGAGTGAAGTTNNNAGATCGGAA |
| 35862_1 | TATGAGTGTGGAGTCGTTGCCTACCGGGAAGGCTGAGATAATCNNNAGATCGGAA |
| 35862_2 | TATGAGTGTGGAGTCGTTGCCTACCGGGAAGGCTGAGATAATCNNNAGATCGGAA |
| 42140_1 | TATGAGTGTGGAGTCGTTGCAGTAGCTGGGACTACAGGCANNNAGATCGGAAGAG |
| 42140_2 | TATGAGTGTGGAGTCGTTGCAGTAGCTGGGACTACAGGCANNNAGATCGGAAGAG |
| 43197_1 | TATGAGTGTGGAGTCGTTGCGAAGCAGCCAGACAAAAGATGAANNNAGATCGGAA |
| 43197_2 | TATGAGTGTGGAGTCGTTGCGAAGCAGCCAGACAAAAGATGAANNNAGATCGGAA |
| 6090_1 | TATGAGTGTGGAGTCGTTGCAGGCAAACTGCTAGTCTCCAATTCNNNAGATCGGAA |
| 6090_2 | TATGAGTGTGGAGTCGTTGCAGGCAAACTGCTAGTCTCCAATTCNNNAGATCGGAA |
| 35999_1 | TATGAGTGTGGAGTCGTTGCCTATGGGTGGATCATGAGGTCAGNNNAGATCGGAA |
| 35999_2 | TATGAGTGTGGAGTCGTTGCCTATGGGTGGATCATGAGGTCAGNNNAGATCGGAA |
| 31728_1 | TATGAGTGTGGAGTCGTTGCCCAAGAAGAATCTCCCTCTCTGGNNNAGATCGGAAG |
| 31728_2 | TATGAGTGTGGAGTCGTTGCCCAAGAAGAATCTCCCTCTCTGGNNNAGATCGGAAG |
| 48356_1 | TATGAGTGTGGAGTCGTTGCCCACGCCCGGCTAATTTTTATATNNNAGATCGGAAG |
| 48356_2 | TATGAGTGTGGAGTCGTTGCCCACGCCCGGCTAATTTTTATATNNNAGATCGGAAG |
| 49923_1 | TATGAGTGTGGAGTCGTTGCACCAAGCAGCATATTTCCAACATNNNAGATCGGAAG |
| 49923_2 | TATGAGTGTGGAGTCGTTGCACCAAGCAGCATATTTCCAACATNNNAGATCGGAAG |
| 49543_1 | TATGAGTGTGGAGTCGTTGCCTCCTGTGTAGCTGGGATTACAGNNNAGATCGGAAG |
| 49543_2 | TATGAGTGTGGAGTCGTTGCCTCCTGTGTAGCTGGGATTACAGNNNAGATCGGAAG |
| 31501_1 | TATGAGTGTGGAGTCGTTGCGGGTGAAGGAGAAGGGTTGTTATNNNAGATCGGA |
| 31501_2 | TATGAGTGTGGAGTCGTTGCGGGTGAAGGAGAAGGGTTGTTATNNNAGATCGGA |
| 39481_1 | TATGAGTGTGGAGTCGTTGCGTTTTATTCTTATCGCCCAGGCCNNNAGATCGGAAG |
| 39481_2 | TATGAGTGTGGAGTCGTTGCGTTTTATTCTTATCGCCCAGGCCNNNAGATCGGAAG |
| 36828_1 | TATGAGTGTGGAGTCGTTGCGACTAATACAGAGGCCCAGGATGNNNAGATCGGAA |
| 36828_2 | TATGAGTGTGGAGTCGTTGCGACTAATACAGAGGCCCAGGATGNNNAGATCGGAA |
| 30006_1 | TATGAGTGTGGAGTCGTTGCGCTGATTGCAGAACAAGCTATACCNNNAGATCGGAA |

| | |
|---|---|
| 30006_2 | TATGAGTGTGGAGTCGTTGCGCTGATTGCAGAACAAGCTATACCNNNAGATCGGAA |
| 29237_1 | TATGAGTGTGGAGTCGTTGCGCAATTCTGTGTGTTGTGAGTGANNNAGATCGGAA |
| 29237_2 | TATGAGTGTGGAGTCGTTGCGCAATTCTGTGTGTTGTGAGTGANNNAGATCGGAA |
| 36959_1 | TATGAGTGTGGAGTCGTTGCCAACCCAGGAACCAGGAATCATANNNAGATCGGAAG |
| 36959_2 | TATGAGTGTGGAGTCGTTGCCAACCCAGGAACCAGGAATCATANNNAGATCGGAAG |
| 38845_1 | TATGAGTGTGGAGTCGTTGCACTTCCGCTCATATACTTCAGCANNNAGATCGGAAG |
| 38845_2 | TATGAGTGTGGAGTCGTTGCACTTCCGCTCATATACTTCAGCANNNAGATCGGAAG |
| 42677_1 | TATGAGTGTGGAGTCGTTGCTGATTGACTCCCCAACATCCATTNNNAGATCGGAAG |
| 42677_2 | TATGAGTGTGGAGTCGTTGCTGATTGACTCCCCAACATCCATTNNNAGATCGGAAG |
| 45739_1 | TATGAGTGTGGAGTCGTTGCTGTTAGTGATGGTATTGGCTGCTNNNAGATCGGAA |
| 45739_2 | TATGAGTGTGGAGTCGTTGCTGTTAGTGATGGTATTGGCTGCTNNNAGATCGGAA |
| 32379_1 | TATGAGTGTGGAGTCGTTGCGAAAACATGGGAGCACCTTCTTCNNNAGATCGGAAG |
| 32379_2 | TATGAGTGTGGAGTCGTTGCGAAAACATGGGAGCACCTTCTTCNNNAGATCGGAAG |
| 10258_1 | TATGAGTGTGGAGTCGTTGCCTTTGCTTGCACATGCTGTCAGNNNAGATCGGAAGA |
| 10258_2 | TATGAGTGTGGAGTCGTTGCCTTTGCTTGCACATGCTGTCAGNNNAGATCGGAAGA |
| 52004_1 | TATGAGTGTGGAGTCGTTGCTTTTTGCTTAGGAATGCCTTGGGNNNAGATCGGAAG |
| 52004_2 | TATGAGTGTGGAGTCGTTGCTTTTTGCTTAGGAATGCCTTGGGNNNAGATCGGAAG |
| 40999_1 | TATGAGTGTGGAGTCGTTGCATATGTATGTGTTCTCCCCACCCNNNAGATCGGAAG |
| 40999_2 | TATGAGTGTGGAGTCGTTGCATATGTATGTGTTCTCCCCACCCNNNAGATCGGAAG |
| 29680_1 | TATGAGTGTGGAGTCGTTGCTGAATCTATTCACAGGGCCATGGNNNAGATCGGAAG |
| 29680_2 | TATGAGTGTGGAGTCGTTGCTGAATCTATTCACAGGGCCATGGNNNAGATCGGAAG |
| 50805_1 | TATGAGTGTGGAGTCGTTGCTGTGCTGAGAACACAAGGGTATTNNNAGATCGGAA |
| 50805_2 | TATGAGTGTGGAGTCGTTGCTGTGCTGAGAACACAAGGGTATTNNNAGATCGGAA |
| 50011_1 | TATGAGTGTGGAGTCGTTGCTGCACACTCCTTATTGTTCAGGANNNAGATCGGAAG |
| 50011_2 | TATGAGTGTGGAGTCGTTGCTGCACACTCCTTATTGTTCAGGANNNAGATCGGAAG |
| 49467_1 | TATGAGTGTGGAGTCGTTGCGTTCCTGGTTGTCTGTTGTATGCNNNAGATCGGAAG |
| 49467_2 | TATGAGTGTGGAGTCGTTGCGTTCCTGGTTGTCTGTTGTATGCNNNAGATCGGAAG |
| 32386_1 | TATGAGTGTGGAGTCGTTGCCAGCTCACTGCAGTCTCAATTTCNNNAGATCGGAAG |
| 32386_2 | TATGAGTGTGGAGTCGTTGCCAGCTCACTGCAGTCTCAATTTCNNNAGATCGGAAG |
| 31695_1 | TATGAGTGTGGAGTCGTTGCTCTCTCTCCCTGGATTTTTCTGCNNNAGATCGGAAGA |
| 31695_2 | TATGAGTGTGGAGTCGTTGCTCTCTCTCCCTGGATTTTTCTGCNNNAGATCGGAAGA |
| 36053_1 | TATGAGTGTGGAGTCGTTGCGAGGTCTTGCCACTGTACTCCNNNAGATCGGAAGAG |
| 36053_2 | TATGAGTGTGGAGTCGTTGCGAGGTCTTGCCACTGTACTCCNNNAGATCGGAAGAG |
| 43691_1 | TATGAGTGTGGAGTCGTTGCGATCTCATGTCTCAGCTCTGGAGNNNAGATCGGAAG |
| 43691_2 | TATGAGTGTGGAGTCGTTGCGATCTCATGTCTCAGCTCTGGAGNNNAGATCGGAAG |
| 33045_1 | TATGAGTGTGGAGTCGTTGCACCGTGCCAGGCTATTTTTTTTTNNNAGATCGGAAG |
| 33045_2 | TATGAGTGTGGAGTCGTTGCACCGTGCCAGGCTATTTTTTTTTNNNAGATCGGAAG |
| 30617_1 | TATGAGTGTGGAGTCGTTGCGTCAACTGTGATTGTGCCCCCNNNAGATCGGAAGAG |
| 30617_2 | TATGAGTGTGGAGTCGTTGCGTCAACTGTGATTGTGCCCCCNNNAGATCGGAAGAG |
| 49047_1 | TATGAGTGTGGAGTCGTTGCGATCTAGGCTCACTGCAACCTCNNNAGATCGGAAGA |
| 49047_2 | TATGAGTGTGGAGTCGTTGCGATCTAGGCTCACTGCAACCTCNNNAGATCGGAAGA |
| 45802_1 | TATGAGTGTGGAGTCGTTGCCAGTAGCTGGGATTACAGGCACNNNAGATCGGAAG |
| 45802_2 | TATGAGTGTGGAGTCGTTGCCAGTAGCTGGGATTACAGGCACNNNAGATCGGAAG |

| | |
|---|---|
| 10110_1 | TATGAGTGTGGAGTCGTTGCCTGGATGAGGTTTCCCTTTGGTCNNNAGATCGGAAG |
| 10110_2 | TATGAGTGTGGAGTCGTTGCCTGGATGAGGTTTCCCTTTGGTCNNNAGATCGGAAG |
| 47963_1 | TATGAGTGTGGAGTCGTTGCGGAAAGTGTCCTTCAGAGCAGAGNNNAGATCGGAA |
| 47963_2 | TATGAGTGTGGAGTCGTTGCGGAAAGTGTCCTTCAGAGCAGAGNNNAGATCGGAA |
| 51748_1 | TATGAGTGTGGAGTCGTTGCTGGCATTATACTAGCATTCGCCTNNNAGATCGGAAG |
| 51748_2 | TATGAGTGTGGAGTCGTTGCTGGCATTATACTAGCATTCGCCTNNNAGATCGGAAG |
| 45357_1 | TATGAGTGTGGAGTCGTTGCCTAAAATCCCTCCAACCCAAGGANNNAGATCGGAAG |
| 45357_2 | TATGAGTGTGGAGTCGTTGCCTAAAATCCCTCCAACCCAAGGANNNAGATCGGAAG |
| 49940_1 | TATGAGTGTGGAGTCGTTGCTTGAAATCAGCTTTCGGGGTAGANNNAGATCGGAA |
| 49940_2 | TATGAGTGTGGAGTCGTTGCTTGAAATCAGCTTTCGGGGTAGANNNAGATCGGAA |
| 42076_1 | TATGAGTGTGGAGTCGTTGCTGCCGATCTCTTTCCACATAGAGNNNAGATCGGAAG |
| 42076_2 | TATGAGTGTGGAGTCGTTGCTGCCGATCTCTTTCCACATAGAGNNNAGATCGGAAG |
| 39711_1 | TATGAGTGTGGAGTCGTTGCCTGGGTGACAGGATCATCTGTACNNNAGATCGGAA |
| 39711_2 | TATGAGTGTGGAGTCGTTGCCTGGGTGACAGGATCATCTGTACNNNAGATCGGAA |
| 36407_1 | TATGAGTGTGGAGTCGTTGCACAAAGGACAGACAGAGCTCAAANNNAGATCGGAA |
| 36407_2 | TATGAGTGTGGAGTCGTTGCACAAAGGACAGACAGAGCTCAAANNNAGATCGGAA |
| 9739_1 | TATGAGTGTGGAGTCGTTGCGGGGAGGAGAGAGAATTTTCCACNNNAGATCGGAA |
| 9739_2 | TATGAGTGTGGAGTCGTTGCGGGGAGGAGAGAGAATTTTCCACNNNAGATCGGAA |
| 45376_1 | TATGAGTGTGGAGTCGTTGCTGGGTTCAAGCAATTCTCTCTGTNNNAGATCGGAAG |
| 45376_2 | TATGAGTGTGGAGTCGTTGCTGGGTTCAAGCAATTCTCTCTGTNNNAGATCGGAAG |
| 31875_1 | TATGAGTGTGGAGTCGTTGCTGTAATCTCAGCACTTTGGGAGGNNNAGATCGGAA |
| 31875_2 | TATGAGTGTGGAGTCGTTGCTGTAATCTCAGCACTTTGGGAGGNNNAGATCGGAA |
| 36612_1 | TATGAGTGTGGAGTCGTTGCAGATCATGTGTGTATGGATCGGGNNNAGATCGGAA |
| 36612_2 | TATGAGTGTGGAGTCGTTGCAGATCATGTGTGTATGGATCGGGNNNAGATCGGAA |
| 45198_1 | TATGAGTGTGGAGTCGTTGCGTAAGAAGGATGCCAGGGGTTATNNNAGATCGGAA |
| 45198_2 | TATGAGTGTGGAGTCGTTGCGTAAGAAGGATGCCAGGGGTTATNNNAGATCGGAA |
| 29105_1 | TATGAGTGTGGAGTCGTTGCTTGAGACCACAGCCATCTATCACNNNAGATCGGAAG |
| 29105_2 | TATGAGTGTGGAGTCGTTGCTTGAGACCACAGCCATCTATCACNNNAGATCGGAAG |
| 45585_1 | TATGAGTGTGGAGTCGTTGCTCTCGATCTCCTGACCTCATGATNNNAGATCGGAAGA |
| 45585_2 | TATGAGTGTGGAGTCGTTGCTCTCGATCTCCTGACCTCATGATNNNAGATCGGAAGA |
| 43062_1 | TATGAGTGTGGAGTCGTTGCTTCCTGCAAGAAGGCCTTTATCTNNNAGATCGGAAG |
| 43062_2 | TATGAGTGTGGAGTCGTTGCTTCCTGCAAGAAGGCCTTTATCTNNNAGATCGGAAG |
| 49948_1 | TATGAGTGTGGAGTCGTTGCCCCCCCACTTCTAATTTAGCAGTNNNAGATCGGAAGA |
| 49948_2 | TATGAGTGTGGAGTCGTTGCCCCCCCACTTCTAATTTAGCAGTNNNAGATCGGAAGA |
| 49418_1 | TATGAGTGTGGAGTCGTTGCAGTCGCCATAATTCTGTAGCACTNNNAGATCGGAAG |
| 49418_2 | TATGAGTGTGGAGTCGTTGCAGTCGCCATAATTCTGTAGCACTNNNAGATCGGAAG |
| 45920_1 | TATGAGTGTGGAGTCGTTGCAGGTTTTCTCAGCTCCATTTCCTNNNAGATCGGAAGA |
| 45920_2 | TATGAGTGTGGAGTCGTTGCAGGTTTTCTCAGCTCCATTTCCTNNNAGATCGGAAGA |
| 32925_1 | TATGAGTGTGGAGTCGTTGCGTGTGCTGGAAAAACTTGCTCTANNNAGATCGGAA |
| 32925_2 | TATGAGTGTGGAGTCGTTGCGTGTGCTGGAAAAACTTGCTCTANNNAGATCGGAA |
| 35527_1 | TATGAGTGTGGAGTCGTTGCTGTTAGCCAGGATGGTCTTGATCNNNAGATCGGAA |
| 35527_2 | TATGAGTGTGGAGTCGTTGCTGTTAGCCAGGATGGTCTTGATCNNNAGATCGGAA |
| 38027_1 | TATGAGTGTGGAGTCGTTGCAAGAGGAGAACTGGGAAAACCTCNNNAGATCGGAA |

| | |
|---|---|
| 38027_2 | TATGAGTGTGGAGTCGTTGCAAGAGGAGAACTGGGAAAACCTCNNNAGATCGGAA |
| 12754_1 | TATGAGTGTGGAGTCGTTGCATCGGCATTTTATATTAAAGTGGNNNAGATCGGAAG |
| 12754_2 | TATGAGTGTGGAGTCGTTGCATCGGCATTTTATATTAAAGTGGNNNAGATCGGAAG |
| 34981_1 | TATGAGTGTGGAGTCGTTGCTGGAGTACAGTGACACGATCTTGNNNAGATCGGAA |
| 34981_2 | TATGAGTGTGGAGTCGTTGCTGGAGTACAGTGACACGATCTTGNNNAGATCGGAA |
| 37881_1 | TATGAGTGTGGAGTCGTTGCTTGTCTCGATCCATCACAGTCAGNNNAGATCGGAAG |
| 37881_2 | TATGAGTGTGGAGTCGTTGCTTGTCTCGATCCATCACAGTCAGNNNAGATCGGAAG |
| 30023_1 | TATGAGTGTGGAGTCGTTGCTGATTGCAAAAATGCCTTCTTCANNNAGATCGGAAG |
| 30023_2 | TATGAGTGTGGAGTCGTTGCTGATTGCAAAAATGCCTTCTTCANNNAGATCGGAAG |
| 46258_1 | TATGAGTGTGGAGTCGTTGCAAACATACTTCTGCCACCTTTGCNNNAGATCGGAAGA |
| 46258_2 | TATGAGTGTGGAGTCGTTGCAAACATACTTCTGCCACCTTTGCNNNAGATCGGAAGA |
| 52873_1 | TATGAGTGTGGAGTCGTTGCCACTCCTCTCCCCTTTCTCTTTCNNNAGATCGGAAGAG |
| 52873_2 | TATGAGTGTGGAGTCGTTGCCACTCCTCTCCCCTTTCTCTTTCNNNAGATCGGAAGAG |
| 13204_1 | TATGAGTGTGGAGTCGTTGCCTGAAATCTCTTAGGCAATTTCNNNAGATCGGAAGA |
| 13204_2 | TATGAGTGTGGAGTCGTTGCCTGAAATCTCTTAGGCAATTTCNNNAGATCGGAAGA |
| 35839_1 | TATGAGTGTGGAGTCGTTGCTTCAAGCTATTCTCCTGCCTCAGNNNAGATCGGAAGA |
| 35839_2 | TATGAGTGTGGAGTCGTTGCTTCAAGCTATTCTCCTGCCTCAGNNNAGATCGGAAGA |
| 32100_1 | TATGAGTGTGGAGTCGTTGCGAACTTCAAAAGTGGGGAAGGTGNNNAGATCGGAA |
| 32100_2 | TATGAGTGTGGAGTCGTTGCGAACTTCAAAAGTGGGGAAGGTGNNNAGATCGGAA |
| 49144_1 | TATGAGTGTGGAGTCGTTGCCAATTCTCCTGCCTTAGCCTTCTNNNAGATCGGAAGA |
| 49144_2 | TATGAGTGTGGAGTCGTTGCCAATTCTCCTGCCTTAGCCTTCTNNNAGATCGGAAGA |
| 48270_1 | TATGAGTGTGGAGTCGTTGCCACTATGCCTGGCTAATTTTGCANNNAGATCGGAAG |
| 48270_2 | TATGAGTGTGGAGTCGTTGCCACTATGCCTGGCTAATTTTGCANNNAGATCGGAAG |
| 47255_1 | TATGAGTGTGGAGTCGTTGCTCCTGACCTCAAGTGATCTACCTNNNAGATCGGAAGA |
| 47255_2 | TATGAGTGTGGAGTCGTTGCTCCTGACCTCAAGTGATCTACCTNNNAGATCGGAAGA |
| 50900_1 | TATGAGTGTGGAGTCGTTGCGAAACTTTTTTAGGGGGGGTTGGNNNAGATCGGAA |
| 50900_2 | TATGAGTGTGGAGTCGTTGCGAAACTTTTTTAGGGGGGGTTGGNNNAGATCGGAA |
| 48062_1 | TATGAGTGTGGAGTCGTTGCCAGGTGACCATGAGGACAATCATNNNAGATCGGAA |
| 48062_2 | TATGAGTGTGGAGTCGTTGCCAGGTGACCATGAGGACAATCATNNNAGATCGGAA |
| 39698_1 | TATGAGTGTGGAGTCGTTGCCAACAGAGCAAGACTCCATCTCANNNAGATCGGAAG |
| 39698_2 | TATGAGTGTGGAGTCGTTGCCAACAGAGCAAGACTCCATCTCANNNAGATCGGAAG |
| 36684_1 | TATGAGTGTGGAGTCGTTGCAATTTGTATTGGGCCACATGTGGNNNAGATCGGAA |
| 36684_2 | TATGAGTGTGGAGTCGTTGCAATTTGTATTGGGCCACATGTGGNNNAGATCGGAA |
| 47036_1 | TATGAGTGTGGAGTCGTTGCCCTCCCGAATAGCTAGGACTACANNNAGATCGGAAG |
| 47036_2 | TATGAGTGTGGAGTCGTTGCCCTCCCGAATAGCTAGGACTACANNNAGATCGGAAG |
| 6273_1 | TATGAGTGTGGAGTCGTTGCATATAAGCAATGAGCCACCAAATNNNAGATCGGAAG |
| 6273_2 | TATGAGTGTGGAGTCGTTGCATATAAGCAATGAGCCACCAAATNNNAGATCGGAAG |
| 33851_1 | TATGAGTGTGGAGTCGTTGCGAGAAAAACAGCCAGCAAAAGGANNNAGATCGGAA |
| 33851_2 | TATGAGTGTGGAGTCGTTGCGAGAAAAACAGCCAGCAAAAGGANNNAGATCGGAA |
| 29945_1 | TATGAGTGTGGAGTCGTTGCGTTGTGGGTTTCTCAAGCAGAATNNNAGATCGGAA |
| 29945_2 | TATGAGTGTGGAGTCGTTGCGTTGTGGGTTTCTCAAGCAGAATNNNAGATCGGAA |
| 40450_1 | TATGAGTGTGGAGTCGTTGCGGGTTGCAATCCTAGTCTCTGATNNNAGATCGGAAG |
| 40450_2 | TATGAGTGTGGAGTCGTTGCGGGTTGCAATCCTAGTCTCTGATNNNAGATCGGAAG |

| | |
|---|---|
| 10324_1 | TATGAGTGTGGAGTCGTTGCAGGATGTGGGAGATGTTCTGACANNNAGATCGGAA |
| 10324_2 | TATGAGTGTGGAGTCGTTGCAGGATGTGGGAGATGTTCTGACANNNAGATCGGAA |
| 44870_1 | TATGAGTGTGGAGTCGTTGCAAATAGCCCTGGGTGTGATTCATNNNAGATCGGAA |
| 44870_2 | TATGAGTGTGGAGTCGTTGCAAATAGCCCTGGGTGTGATTCATNNNAGATCGGAA |
| 31602_1 | TATGAGTGTGGAGTCGTTGCCATTGTCAGTCCTCCACCTACTCNNNAGATCGGAAGA |
| 31602_2 | TATGAGTGTGGAGTCGTTGCCATTGTCAGTCCTCCACCTACTCNNNAGATCGGAAGA |
| 46399_1 | TATGAGTGTGGAGTCGTTGCTCTGTCTAGTTTTGGTTATCAGGGTNNNAGATCGGA |
| 46399_2 | TATGAGTGTGGAGTCGTTGCTCTGTCTAGTTTTGGTTATCAGGGTNNNAGATCGGA |
| 46915_1 | TATGAGTGTGGAGTCGTTGCCTCCTGAGTAGCTGGGATTACAGNNNAGATCGGAA |
| 46915_2 | TATGAGTGTGGAGTCGTTGCCTCCTGAGTAGCTGGGATTACAGNNNAGATCGGAA |
| 10466_1 | TATGAGTGTGGAGTCGTTGCATGGGCTGTGCTTTCAATTCTCANNNAGATCGGAAG |
| 10466_2 | TATGAGTGTGGAGTCGTTGCATGGGCTGTGCTTTCAATTCTCANNNAGATCGGAAG |
| 39121_1 | TATGAGTGTGGAGTCGTTGCAAATTGTCGAGTCCAGCAACAACNNNAGATCGGAAG |
| 39121_2 | TATGAGTGTGGAGTCGTTGCAAATTGTCGAGTCCAGCAACAACNNNAGATCGGAAG |
| 47113_1 | TATGAGTGTGGAGTCGTTGCCAACTCCTGGCAGTGTGATTTTTNNNAGATCGGAAG |
| 47113_2 | TATGAGTGTGGAGTCGTTGCCAACTCCTGGCAGTGTGATTTTTNNNAGATCGGAAG |
| 38715_1 | TATGAGTGTGGAGTCGTTGCTAAACACTTGGGTTGCTTTCACCNNNAGATCGGAAG |
| 38715_2 | TATGAGTGTGGAGTCGTTGCTAAACACTTGGGTTGCTTTCACCNNNAGATCGGAAG |
| 13578_1 | TATGAGTGTGGAGTCGTTGCATTAGAACTATAGCGGCTGGNNNAGATCGGAAGAG |
| 13578_2 | TATGAGTGTGGAGTCGTTGCATTAGAACTATAGCGGCTGGNNNAGATCGGAAGAG |
| 49727_1 | TATGAGTGTGGAGTCGTTGCGTTTCACCATGTTAGCCAGGATGNNNAGATCGGAAG |
| 49727_2 | TATGAGTGTGGAGTCGTTGCGTTTCACCATGTTAGCCAGGATGNNNAGATCGGAAG |
| 43823_1 | TATGAGTGTGGAGTCGTTGCAATGTCTCTGAAACCCAGTCTCCNNNAGATCGGAAG |
| 43823_2 | TATGAGTGTGGAGTCGTTGCAATGTCTCTGAAACCCAGTCTCCNNNAGATCGGAAG |
| 32215_1 | TATGAGTGTGGAGTCGTTGCCTGGGCAACAAGAGCAAAACTCNNNAGATCGGAAG |
| 32215_2 | TATGAGTGTGGAGTCGTTGCCTGGGCAACAAGAGCAAAACTCNNNAGATCGGAAG |
| 38820_1 | TATGAGTGTGGAGTCGTTGCGGAGGTTGAGGCAGGAGAATGNNNAGATCGGAAG |
| 38820_2 | TATGAGTGTGGAGTCGTTGCGGAGGTTGAGGCAGGAGAATGNNNAGATCGGAAG |
| 30027_1 | TATGAGTGTGGAGTCGTTGCGCTAGCAAACACTTCTCCAACAANNNAGATCGGAAG |
| 30027_2 | TATGAGTGTGGAGTCGTTGCGCTAGCAAACACTTCTCCAACAANNNAGATCGGAAG |
| 32088_1 | TATGAGTGTGGAGTCGTTGCGAGTGCCTGTAATCCCAGCTACNNNAGATCGGAAGA |
| 32088_2 | TATGAGTGTGGAGTCGTTGCGAGTGCCTGTAATCCCAGCTACNNNAGATCGGAAGA |
| 34233_1 | TATGAGTGTGGAGTCGTTGCGATCACAAGGTCAGGAGATCGAGNNNAGATCGGAA |
| 34233_2 | TATGAGTGTGGAGTCGTTGCGATCACAAGGTCAGGAGATCGAGNNNAGATCGGAA |
| 37906_1 | TATGAGTGTGGAGTCGTTGCATACAGGCTGCTGACCATTATGTNNNAGATCGGAAG |
| 37906_2 | TATGAGTGTGGAGTCGTTGCATACAGGCTGCTGACCATTATGTNNNAGATCGGAAG |
| 49721_1 | TATGAGTGTGGAGTCGTTGCCTGGTCTTGAACTTCTGGACTCANNNAGATCGGAAG |
| 49721_2 | TATGAGTGTGGAGTCGTTGCCTGGTCTTGAACTTCTGGACTCANNNAGATCGGAAG |
| 48381_1 | TATGAGTGTGGAGTCGTTGCAGCACACTCCTAAGCATTTGAGANNNAGATCGGAAG |
| 48381_2 | TATGAGTGTGGAGTCGTTGCAGCACACTCCTAAGCATTTGAGANNNAGATCGGAAG |
| 145_1 | TATGAGTGTGGAGTCGTTGCTTTCATTCTTCTCCGGCTCCCGNNNAGATCGGAAGAG |
| 145_2 | TATGAGTGTGGAGTCGTTGCTTTCATTCTTCTCCGGCTCCCGNNNAGATCGGAAGAG |
| 46028_1 | TATGAGTGTGGAGTCGTTGCTTCCCTGGAGAGAAATGAAGAGCNNNAGATCGGAA |

| | |
|---|---|
| 46028_2 | TATGAGTGTGGAGTCGTTGCTTCCCTGGAGAGAAATGAAGAGCNNNAGATCGGAA |
| 31289_1 | TATGAGTGTGGAGTCGTTGCGTGAGCTATGATCACACCAATGCNNNAGATCGGAAG |
| 31289_2 | TATGAGTGTGGAGTCGTTGCGTGAGCTATGATCACACCAATGCNNNAGATCGGAAG |
| 35442_1 | TATGAGTGTGGAGTCGTTGCCCAATAGGCGTTTTCTTTGAGTGTNNNAGATCGGAA |
| 35442_2 | TATGAGTGTGGAGTCGTTGCCCAATAGGCGTTTTCTTTGAGTGTNNNAGATCGGAA |
| 45356_1 | TATGAGTGTGGAGTCGTTGCTTCTGGGATACATGTGCAGAACANNNAGATCGGAA |
| 45356_2 | TATGAGTGTGGAGTCGTTGCTTCTGGGATACATGTGCAGAACANNNAGATCGGAA |
| 52809_1 | TATGAGTGTGGAGTCGTTGCTCCCGTCTAAATTTAGAGAGCGGNNNAGATCGGAA |
| 52809_2 | TATGAGTGTGGAGTCGTTGCTCCCGTCTAAATTTAGAGAGCGGNNNAGATCGGAA |
| 30665_1 | TATGAGTGTGGAGTCGTTGCTGTGTGTCTCCATCCGAGACATTNNNAGATCGGAAG |
| 30665_2 | TATGAGTGTGGAGTCGTTGCTGTGTGTCTCCATCCGAGACATTNNNAGATCGGAAG |
| 36841_1 | TATGAGTGTGGAGTCGTTGCACTGGTAAACTGGCTTCTATCCANNNAGATCGGAAG |
| 36841_2 | TATGAGTGTGGAGTCGTTGCACTGGTAAACTGGCTTCTATCCANNNAGATCGGAAG |
| 50978_1 | TATGAGTGTGGAGTCGTTGCAAGGCTCCAAGATGTCACTGAATNNNAGATCGGAA |
| 50978_2 | TATGAGTGTGGAGTCGTTGCAAGGCTCCAAGATGTCACTGAATNNNAGATCGGAA |
| 5543_1 | TATGAGTGTGGAGTCGTTGCTGCCAAGGACACAGAAACATTCANNNAGATCGGAA |
| 5543_2 | TATGAGTGTGGAGTCGTTGCTGCCAAGGACACAGAAACATTCANNNAGATCGGAA |
| 2560_1 | TATGAGTGTGGAGTCGTTGCCCTCTAAGTTCTCCTCTCAGGGGNNNAGATCGGAAG |
| 2560_2 | TATGAGTGTGGAGTCGTTGCCCTCTAAGTTCTCCTCTCAGGGGNNNAGATCGGAAG |
| 33878_1 | TATGAGTGTGGAGTCGTTGCCATCTGACAAAATTCAGTGCCCTNNNAGATCGGAAG |
| 33878_2 | TATGAGTGTGGAGTCGTTGCCATCTGACAAAATTCAGTGCCCTNNNAGATCGGAAG |
| 44206_1 | TATGAGTGTGGAGTCGTTGCCAAATCTCGGCTTCCACTTGTTTNNNAGATCGGAAGA |
| 44206_2 | TATGAGTGTGGAGTCGTTGCCAAATCTCGGCTTCCACTTGTTTNNNAGATCGGAAGA |
| 50460_1 | TATGAGTGTGGAGTCGTTGCTAGGGAAATACCTCATTTGGCCCNNNAGATCGGAAG |
| 50460_2 | TATGAGTGTGGAGTCGTTGCTAGGGAAATACCTCATTTGGCCCNNNAGATCGGAAG |
| 40810_1 | TATGAGTGTGGAGTCGTTGCAAAGCCCATTTACACAAGCAGTCNNNAGATCGGAAG |
| 40810_2 | TATGAGTGTGGAGTCGTTGCAAAGCCCATTTACACAAGCAGTCNNNAGATCGGAAG |
| 34211_1 | TATGAGTGTGGAGTCGTTGCCCTTGAGAGCAGAGATCACATCANNNAGATCGGAA |
| 34211_2 | TATGAGTGTGGAGTCGTTGCCCTTGAGAGCAGAGATCACATCANNNAGATCGGAA |
| 50155_1 | TATGAGTGTGGAGTCGTTGCATAGTATGCATGGCCAAGTGTCANNNAGATCGGAA |
| 50155_2 | TATGAGTGTGGAGTCGTTGCATAGTATGCATGGCCAAGTGTCANNNAGATCGGAA |
| 3594_1 | TATGAGTGTGGAGTCGTTGCATCACATTTGGGAGACTAGAATTGANNNAGATCGG |
| 3594_2 | TATGAGTGTGGAGTCGTTGCATCACATTTGGGAGACTAGAATTGANNNAGATCGG |
| 51833_1 | TATGAGTGTGGAGTCGTTGCATGCACCACAGAAACAAAGTTCGNNNAGATCGGAA |
| 51833_2 | TATGAGTGTGGAGTCGTTGCATGCACCACAGAAACAAAGTTCGNNNAGATCGGAA |
| 31120_1 | TATGAGTGTGGAGTCGTTGCCAACTTGCTGTCATGCAGAAGATNNNAGATCGGAAG |
| 31120_2 | TATGAGTGTGGAGTCGTTGCCAACTTGCTGTCATGCAGAAGATNNNAGATCGGAAG |
| 29558_1 | TATGAGTGTGGAGTCGTTGCGGAACCCCTTGCATAATGAAGAGNNNAGATCGGAA |
| 29558_2 | TATGAGTGTGGAGTCGTTGCGGAACCCCTTGCATAATGAAGAGNNNAGATCGGAA |
| 40852_1 | TATGAGTGTGGAGTCGTTGCACTGTGAATACCCCGACTTGATCNNNAGATCGGAAG |
| 40852_2 | TATGAGTGTGGAGTCGTTGCACTGTGAATACCCCGACTTGATCNNNAGATCGGAAG |
| 43701_1 | TATGAGTGTGGAGTCGTTGCAGGGGTCTATATGATCTGGGGTTNNNAGATCGGAA |
| 43701_2 | TATGAGTGTGGAGTCGTTGCAGGGGTCTATATGATCTGGGGTTNNNAGATCGGAA |

| | |
|---|---|
| 49348_1 | TATGAGTGTGGAGTCGTTGCGTATTGTTCAGGCTCCTGGGAGNNNAGATCGGAAG |
| 49348_2 | TATGAGTGTGGAGTCGTTGCGTATTGTTCAGGCTCCTGGGAGNNNAGATCGGAAG |
| 51405_1 | TATGAGTGTGGAGTCGTTGCACAAAGTTCTGACCAAAACACCGNNNAGATCGGAAG |
| 51405_2 | TATGAGTGTGGAGTCGTTGCACAAAGTTCTGACCAAAACACCGNNNAGATCGGAAG |
| 29218_1 | TATGAGTGTGGAGTCGTTGCGTGCATAGAGTTATGTGAGGAGCNNNAGATCGGAA |
| 29218_2 | TATGAGTGTGGAGTCGTTGCGTGCATAGAGTTATGTGAGGAGCNNNAGATCGGAA |
| 45023_1 | TATGAGTGTGGAGTCGTTGCCTCCAGAAGCCGAGATGAACATANNNAGATCGGAA |
| 45023_2 | TATGAGTGTGGAGTCGTTGCCTCCAGAAGCCGAGATGAACATANNNAGATCGGAA |
| 38236_1 | TATGAGTGTGGAGTCGTTGCTGTGAGTTTGGGATGAGGTGTAGNNNAGATCGGA |
| 38236_2 | TATGAGTGTGGAGTCGTTGCTGTGAGTTTGGGATGAGGTGTAGNNNAGATCGGA |
| 52342_1 | TATGAGTGTGGAGTCGTTGCTCTCTCTCCCCATCTCTGTCTTTNNNAGATCGGAAGA |
| 52342_2 | TATGAGTGTGGAGTCGTTGCTCTCTCTCCCCATCTCTGTCTTTNNNAGATCGGAAGA |
| 40521_1 | TATGAGTGTGGAGTCGTTGCCTACCAGGTTCACGCCATTCTCNNNAGATCGGAAGA |
| 40521_2 | TATGAGTGTGGAGTCGTTGCCTACCAGGTTCACGCCATTCTCNNNAGATCGGAAGA |
| 41086_1 | TATGAGTGTGGAGTCGTTGCCTGAGGCAGGAGAATTGCTTGAGNNNAGATCGGAA |
| 41086_2 | TATGAGTGTGGAGTCGTTGCCTGAGGCAGGAGAATTGCTTGAGNNNAGATCGGAA |
| 43255_1 | TATGAGTGTGGAGTCGTTGCGTGTGTGTGTGTGTGTGTGTATTNNNAGATCGGAA |
| 43255_2 | TATGAGTGTGGAGTCGTTGCGTGTGTGTGTGTGTGTGTGTATTNNNAGATCGGAA |
| 42107_1 | TATGAGTGTGGAGTCGTTGCTTCAAGTGATTCTCCTGCCTCAGNNNAGATCGGAAG |
| 42107_2 | TATGAGTGTGGAGTCGTTGCTTCAAGTGATTCTCCTGCCTCAGNNNAGATCGGAAG |
| 34687_1 | TATGAGTGTGGAGTCGTTGCCAATTCTCCTATCTCAGCCTCCCNNNAGATCGGAAGA |
| 34687_2 | TATGAGTGTGGAGTCGTTGCCAATTCTCCTATCTCAGCCTCCCNNNAGATCGGAAGA |
| 52031_1 | TATGAGTGTGGAGTCGTTGCTGTGTTCTCATTTTTCACCTGCCNNNAGATCGGAAGA |
| 52031_2 | TATGAGTGTGGAGTCGTTGCTGTGTTCTCATTTTTCACCTGCCNNNAGATCGGAAGA |
| 34769_1 | TATGAGTGTGGAGTCGTTGCTCCTGAGGAGCTGAGACTACATGNNNAGATCGGAA |
| 34769_2 | TATGAGTGTGGAGTCGTTGCTCCTGAGGAGCTGAGACTACATGNNNAGATCGGAA |
| 31394_1 | TATGAGTGTGGAGTCGTTGCCACAAGTCACTGAAGCTCCTTTGNNNAGATCGGAAG |
| 31394_2 | TATGAGTGTGGAGTCGTTGCCACAAGTCACTGAAGCTCCTTTGNNNAGATCGGAAG |
| 41258_1 | TATGAGTGTGGAGTCGTTGCTGAGGTAGGAGAATCGCTTGAACNNNAGATCGGAA |
| 41258_2 | TATGAGTGTGGAGTCGTTGCTGAGGTAGGAGAATCGCTTGAACNNNAGATCGGAA |
| 43698_1 | TATGAGTGTGGAGTCGTTGCCTGGAGCTGATTGAGGGCTTTATNNNAGATCGGAA |
| 43698_2 | TATGAGTGTGGAGTCGTTGCCTGGAGCTGATTGAGGGCTTTATNNNAGATCGGAA |
| 48085_1 | TATGAGTGTGGAGTCGTTGCAGATTCCTGCTGGTCTGTATCTGNNNAGATCGGAAG |
| 48085_2 | TATGAGTGTGGAGTCGTTGCAGATTCCTGCTGGTCTGTATCTGNNNAGATCGGAAG |
| 52929_1 | TATGAGTGTGGAGTCGTTGCGCAGATGGCTTCCTTGAAACATTNNNAGATCGGAAG |
| 52929_2 | TATGAGTGTGGAGTCGTTGCGCAGATGGCTTCCTTGAAACATTNNNAGATCGGAAG |
| 31504_1 | TATGAGTGTGGAGTCGTTGCTCCTTTGACCAGGCAACTTATCTNNNAGATCGGAAG |
| 31504_2 | TATGAGTGTGGAGTCGTTGCTCCTTTGACCAGGCAACTTATCTNNNAGATCGGAAG |
| 42581_1 | TATGAGTGTGGAGTCGTTGCTCAACCTCCTAGGATCAGGTGATNNNAGATCGGAAG |
| 42581_2 | TATGAGTGTGGAGTCGTTGCTCAACCTCCTAGGATCAGGTGATNNNAGATCGGAAG |
| 48131_1 | TATGAGTGTGGAGTCGTTGCAGCTCTGGAGAACATGTAAGTGGNNNAGATCGGAA |
| 48131_2 | TATGAGTGTGGAGTCGTTGCAGCTCTGGAGAACATGTAAGTGGNNNAGATCGGAA |
| 31072_1 | TATGAGTGTGGAGTCGTTGCATATAGGCAGTTGCATGCTAGCANNNAGATCGGAA |

| | |
|---|---|
| 31072_2 | TATGAGTGTGGAGTCGTTGCATATAGGCAGTTGCATGCTAGCANNNAGATCGGAA |
| 30243_1 | TATGAGTGTGGAGTCGTTGCATGTTTTTTCTCAGGGTGTGCAGNNNAGATCGGAAG |
| 30243_2 | TATGAGTGTGGAGTCGTTGCATGTTTTTTCTCAGGGTGTGCAGNNNAGATCGGAAG |
| 33804_1 | TATGAGTGTGGAGTCGTTGCACCAACCCACTCATGTGATAACANNNAGATCGGAAG |
| 33804_2 | TATGAGTGTGGAGTCGTTGCACCAACCCACTCATGTGATAACANNNAGATCGGAAG |
| 39250_1 | TATGAGTGTGGAGTCGTTGCTCTATCCTAGGCTATTCTGGCTCTNNNAGATCGGAAG |
| 39250_2 | TATGAGTGTGGAGTCGTTGCTCTATCCTAGGCTATTCTGGCTCTNNNAGATCGGAAG |
| 9776_1 | TATGAGTGTGGAGTCGTTGCTGCCCAGAAAGCCAAACCATTTANNNAGATCGGAAG |
| 9776_2 | TATGAGTGTGGAGTCGTTGCTGCCCAGAAAGCCAAACCATTTANNNAGATCGGAAG |
| 30902_1 | TATGAGTGTGGAGTCGTTGCTGTTTATCCCTAAATGGCCAGCTNNNAGATCGGAAG |
| 30902_2 | TATGAGTGTGGAGTCGTTGCTGTTTATCCCTAAATGGCCAGCTNNNAGATCGGAAG |
| 30237_1 | TATGAGTGTGGAGTCGTTGCAGCTAATAGCAGTCTCCCATGTGNNNAGATCGGAAG |
| 30237_2 | TATGAGTGTGGAGTCGTTGCAGCTAATAGCAGTCTCCCATGTGNNNAGATCGGAAG |
| 46923_1 | TATGAGTGTGGAGTCGTTGCACAAGGCTTACCAATGAAAGTGCNNNAGATCGGAA |
| 46923_2 | TATGAGTGTGGAGTCGTTGCACAAGGCTTACCAATGAAAGTGCNNNAGATCGGAA |
| 2694_1 | TATGAGTGTGGAGTCGTTGCAACAACCAATTCCTGACCCAGATNNNAGATCGGAAG |
| 2694_2 | TATGAGTGTGGAGTCGTTGCAACAACCAATTCCTGACCCAGATNNNAGATCGGAAG |
| 44649_1 | TATGAGTGTGGAGTCGTTGCTTCCTTGAGGGTTATCGCAGTAGNNNAGATCGGAA |
| 44649_2 | TATGAGTGTGGAGTCGTTGCTTCCTTGAGGGTTATCGCAGTAGNNNAGATCGGAA |
| 2793_1 | TATGAGTGTGGAGTCGTTGCACAATCATCTGATCCAATTCTGCTNNNAGATCGGAAG |
| 2793_2 | TATGAGTGTGGAGTCGTTGCACAATCATCTGATCCAATTCTGCTNNNAGATCGGAAG |
| 52357_1 | TATGAGTGTGGAGTCGTTGCATTGTTGCCCGAAGAAAACTGACNNNAGATCGGAA |
| 52357_2 | TATGAGTGTGGAGTCGTTGCATTGTTGCCCGAAGAAAACTGACNNNAGATCGGAA |
| 43890_1 | TATGAGTGTGGAGTCGTTGCTCCACCTAGATCTGAGGACAAGTNNNAGATCGGAAG |
| 43890_2 | TATGAGTGTGGAGTCGTTGCTCCACCTAGATCTGAGGACAAGTNNNAGATCGGAAG |
| 52778_1 | TATGAGTGTGGAGTCGTTGCATCCTATGCCAACATTCAGGGAGNNNAGATCGGAAG |
| 52778_2 | TATGAGTGTGGAGTCGTTGCATCCTATGCCAACATTCAGGGAGNNNAGATCGGAAG |
| 50473_1 | TATGAGTGTGGAGTCGTTGCCTGGCGATGTTCAGGTATATGGANNNAGATCGGAA |
| 50473_2 | TATGAGTGTGGAGTCGTTGCCTGGCGATGTTCAGGTATATGGANNNAGATCGGAA |
| 38150_1 | TATGAGTGTGGAGTCGTTGCCTGAGGTGGGAAGATCACTTGAGNNNAGATCGGAA |
| 38150_2 | TATGAGTGTGGAGTCGTTGCCTGAGGTGGGAAGATCACTTGAGNNNAGATCGGAA |
| 43038_1 | TATGAGTGTGGAGTCGTTGCGGAGGTTGAAGTGAGTCCAGATTNNNAGATCGGAA |
| 43038_2 | TATGAGTGTGGAGTCGTTGCGGAGGTTGAAGTGAGTCCAGATTNNNAGATCGGAA |
| 41555_1 | TATGAGTGTGGAGTCGTTGCCACCCAGCCCTAATTCACTTTTGNNNAGATCGGAAGA |
| 41555_2 | TATGAGTGTGGAGTCGTTGCCACCCAGCCCTAATTCACTTTTGNNNAGATCGGAAGA |
| 40772_1 | TATGAGTGTGGAGTCGTTGCTTTGGGCAAAAACGAAAGCAAACNNNAGATCGGAA |
| 40772_2 | TATGAGTGTGGAGTCGTTGCTTTGGGCAAAAACGAAAGCAAACNNNAGATCGGAA |
| 30760_1 | TATGAGTGTGGAGTCGTTGCTCCCTTTGATCTCCTTTCACCTGNNNAGATCGGAAGA |
| 30760_2 | TATGAGTGTGGAGTCGTTGCTCCCTTTGATCTCCTTTCACCTGNNNAGATCGGAAGA |
| 43848_1 | TATGAGTGTGGAGTCGTTGCTTCCAAGGATCCAGAGACATTGGNNNAGATCGGAA |
| 43848_2 | TATGAGTGTGGAGTCGTTGCTTCCAAGGATCCAGAGACATTGGNNNAGATCGGAA |
| 42857_1 | TATGAGTGTGGAGTCGTTGCGGGCCTTTATTTCAAGGACCAACNNNAGATCGGAAG |
| 42857_2 | TATGAGTGTGGAGTCGTTGCGGGCCTTTATTTCAAGGACCAACNNNAGATCGGAAG |

| | |
|---|---|
| 44877_1 | TATGAGTGTGGAGTCGTTGCAGGGAGGAAAACAGGGAAGAAAGNNNAGATCGGA |
| 44877_2 | TATGAGTGTGGAGTCGTTGCAGGGAGGAAAACAGGGAAGAAAGNNNAGATCGGA |
| 45310_1 | TATGAGTGTGGAGTCGTTGCCTCTTGAATGCCGGTTTGATGGNNNAGATCGGAAGA |
| 45310_2 | TATGAGTGTGGAGTCGTTGCCTCTTGAATGCCGGTTTGATGGNNNAGATCGGAAGA |
| 35981_1 | TATGAGTGTGGAGTCGTTGCTGGAGTGCAGTGTTGTCATCATANNNAGATCGGAAG |
| 35981_2 | TATGAGTGTGGAGTCGTTGCTGGAGTGCAGTGTTGTCATCATANNNAGATCGGAAG |
| 39324_1 | TATGAGTGTGGAGTCGTTGCCAGTTGTTGACAGAAACTGGACANNNAGATCGGAAG |
| 39324_2 | TATGAGTGTGGAGTCGTTGCCAGTTGTTGACAGAAACTGGACANNNAGATCGGAAG |
| 50694_1 | TATGAGTGTGGAGTCGTTGCGCAAGAGTTCTTTCATGGGAAGCNNNAGATCGGAAG |
| 50694_2 | TATGAGTGTGGAGTCGTTGCGCAAGAGTTCTTTCATGGGAAGCNNNAGATCGGAAG |
| 47503_1 | TATGAGTGTGGAGTCGTTGCCTCCTGAGTAGCTGGGATTACAGNNNAGATCGGAAG |
| 47503_2 | TATGAGTGTGGAGTCGTTGCCTCCTGAGTAGCTGGGATTACAGNNNAGATCGGAAG |
| 31701_1 | TATGAGTGTGGAGTCGTTGCAGCTCATCTGCAACTTGTCTTTTNNNAGATCGGAAGA |
| 31701_2 | TATGAGTGTGGAGTCGTTGCAGCTCATCTGCAACTTGTCTTTTNNNAGATCGGAAGA |
| 38490_1 | TATGAGTGTGGAGTCGTTGCGAAAAAGGCGAAGCTGGAGAAATNNNAGATCGGAA |
| 38490_2 | TATGAGTGTGGAGTCGTTGCGAAAAAGGCGAAGCTGGAGAAATNNNAGATCGGAA |
| 40883_1 | TATGAGTGTGGAGTCGTTGCTGAAAGCAACAACAACACAAACANNNAGATCGGAAG |
| 40883_2 | TATGAGTGTGGAGTCGTTGCTGAAAGCAACAACAACACAAACANNNAGATCGGAAG |
| 33960_1 | TATGAGTGTGGAGTCGTTGCCTCTCAAGTAGCTGGGACAACAGNNNAGATCGGAAG |
| 33960_2 | TATGAGTGTGGAGTCGTTGCCTCTCAAGTAGCTGGGACAACAGNNNAGATCGGAAG |
| 45794_1 | TATGAGTGTGGAGTCGTTGCCCATGCCCAGCTAACTTTTTGAANNNAGATCGGAAGA |
| 45794_2 | TATGAGTGTGGAGTCGTTGCCCATGCCCAGCTAACTTTTTGAANNNAGATCGGAAGA |
| 49408_1 | TATGAGTGTGGAGTCGTTGCCCTCCTGAGTAACTGGGACTACANNNAGATCGGAAG |
| 49408_2 | TATGAGTGTGGAGTCGTTGCCCTCCTGAGTAACTGGGACTACANNNAGATCGGAAG |
| 41351_1 | TATGAGTGTGGAGTCGTTGCTTTACTTCATGAGGTCAGCACCANNNAGATCGGAAGA |
| 41351_2 | TATGAGTGTGGAGTCGTTGCTTTACTTCATGAGGTCAGCACCANNNAGATCGGAAGA |
| 31000_1 | TATGAGTGTGGAGTCGTTGCTGTGTGTGTGTAAAGGGGAGTTTNNNAGATCGGAA |
| 31000_2 | TATGAGTGTGGAGTCGTTGCTGTGTGTGTGTAAAGGGGAGTTTNNNAGATCGGAA |
| 36044_1 | TATGAGTGTGGAGTCGTTGCCTTGCAGTAAGCTGAGATCATGCNNNAGATCGGAAG |
| 36044_2 | TATGAGTGTGGAGTCGTTGCCTTGCAGTAAGCTGAGATCATGCNNNAGATCGGAAG |
| 13186_1 | TATGAGTGTGGAGTCGTTGCAAAAAGTGCGGAATTTCCATATNNNAGATCGGAAGA |
| 13186_2 | TATGAGTGTGGAGTCGTTGCAAAAAGTGCGGAATTTCCATATNNNAGATCGGAAGA |
| 32457_1 | TATGAGTGTGGAGTCGTTGCCACTGGCAGCATTACAACATGTANNNAGATCGGAAG |
| 32457_2 | TATGAGTGTGGAGTCGTTGCCACTGGCAGCATTACAACATGTANNNAGATCGGAAG |
| 28837_1 | TATGAGTGTGGAGTCGTTGCTTGAAGAAGGGAGCAATCAGACANNNAGATCGGAA |
| 28837_2 | TATGAGTGTGGAGTCGTTGCTTGAAGAAGGGAGCAATCAGACANNNAGATCGGAA |
| 30143_1 | TATGAGTGTGGAGTCGTTGCGAGCAGACAGACTCTCACTGATTNNNAGATCGGAAG |
| 30143_2 | TATGAGTGTGGAGTCGTTGCGAGCAGACAGACTCTCACTGATTNNNAGATCGGAAG |
| 49607_1 | TATGAGTGTGGAGTCGTTGCGATCACAGCTCACTACAACCTCANNNAGATCGGAAGA |
| 49607_2 | TATGAGTGTGGAGTCGTTGCGATCACAGCTCACTACAACCTCANNNAGATCGGAAGA |
| 35721_1 | TATGAGTGTGGAGTCGTTGCCTTGCTCTTTACACAGGGAAAGCNNNAGATCGGAAG |
| 35721_2 | TATGAGTGTGGAGTCGTTGCCTTGCTCTTTACACAGGGAAAGCNNNAGATCGGAAG |
| 50423_1 | TATGAGTGTGGAGTCGTTGCCACCAGCATCAAAGGTATTCCTGNNNAGATCGGAAG |

| | |
|---|---|
| 50423_2 | TATGAGTGTGGAGTCGTTGCCACCAGCATCAAAGGTATTCCTGNNNAGATCGGAAG |
| 41895_1 | TATGAGTGTGGAGTCGTTGCCTCACAGGTTCATGCCATTGTCNNNAGATCGGAAGA |
| 41895_2 | TATGAGTGTGGAGTCGTTGCCTCACAGGTTCATGCCATTGTCNNNAGATCGGAAGA |
| 37831_1 | TATGAGTGTGGAGTCGTTGCGAGAGAGATCCTCCTGCTTTAGCNNNAGATCGGAAG |
| 37831_2 | TATGAGTGTGGAGTCGTTGCGAGAGAGATCCTCCTGCTTTAGCNNNAGATCGGAAG |
| 10427_1 | TATGAGTGTGGAGTCGTTGCGGAACTCGAACCAGGAGCTAGAGNNNAGATCGGAA |
| 10427_2 | TATGAGTGTGGAGTCGTTGCGGAACTCGAACCAGGAGCTAGAGNNNAGATCGGAA |
| 28981_1 | TATGAGTGTGGAGTCGTTGCTTGAGGCTGCAGTGAACTATGATNNNAGATCGGAA |
| 28981_2 | TATGAGTGTGGAGTCGTTGCTTGAGGCTGCAGTGAACTATGATNNNAGATCGGAA |
| 37578_1 | TATGAGTGTGGAGTCGTTGCGGCTGCCATATCTTTCTGTTGTCNNNAGATCGGAAG |
| 37578_2 | TATGAGTGTGGAGTCGTTGCGGCTGCCATATCTTTCTGTTGTCNNNAGATCGGAAG |
| 38880_1 | TATGAGTGTGGAGTCGTTGCATTATCTTGGGAGTTCTAGGGCCNNNAGATCGGAAG |
| 38880_2 | TATGAGTGTGGAGTCGTTGCATTATCTTGGGAGTTCTAGGGCCNNNAGATCGGAAG |
| 37359_1 | TATGAGTGTGGAGTCGTTGCCCCAACTCTTCATTGGAGGAAGANNNAGATCGGAAG |
| 37359_2 | TATGAGTGTGGAGTCGTTGCCCCAACTCTTCATTGGAGGAAGANNNAGATCGGAAG |
| 52275_1 | TATGAGTGTGGAGTCGTTGCGAACCCTTCTCTGCACTCAGTGNNNAGATCGGAAGA |
| 52275_2 | TATGAGTGTGGAGTCGTTGCGAACCCTTCTCTGCACTCAGTGNNNAGATCGGAAGA |
| 13302_1 | TATGAGTGTGGAGTCGTTGCTATAATCATTTGCAATGCATACANNNAGATCGGAAG |
| 13302_2 | TATGAGTGTGGAGTCGTTGCTATAATCATTTGCAATGCATACANNNAGATCGGAAG |
| 51103_1 | TATGAGTGTGGAGTCGTTGCCATAAGGAGCAAGGAACCCAGATNNNAGATCGGAA |
| 51103_2 | TATGAGTGTGGAGTCGTTGCCATAAGGAGCAAGGAACCCAGATNNNAGATCGGAA |
| 41332_1 | TATGAGTGTGGAGTCGTTGCCTCTAGTCCCAGCTACTCAGGAGNNNAGATCGGAAG |
| 41332_2 | TATGAGTGTGGAGTCGTTGCCTCTAGTCCCAGCTACTCAGGAGNNNAGATCGGAAG |
| 46198_1 | TATGAGTGTGGAGTCGTTGCCCAGGTTCAAGCGATTCAATTGTNNNAGATCGGAAG |
| 46198_2 | TATGAGTGTGGAGTCGTTGCCCAGGTTCAAGCGATTCAATTGTNNNAGATCGGAAG |
| 36268_1 | TATGAGTGTGGAGTCGTTGCGGAAAGAAAATTAGCCTGGTGTGGNNNAGATCGGA |
| 36268_2 | TATGAGTGTGGAGTCGTTGCGGAAAGAAAATTAGCCTGGTGTGGNNNAGATCGGA |
| 2635_1 | TATGAGTGTGGAGTCGTTGCGCTTGCGCACATGTGTATATGTGNNNAGATCGGAAG |
| 2635_2 | TATGAGTGTGGAGTCGTTGCGCTTGCGCACATGTGTATATGTGNNNAGATCGGAAG |
| 42519_1 | TATGAGTGTGGAGTCGTTGCATAAAGCAGCTGTGGCAAAATGTNNNAGATCGGAA |
| 42519_2 | TATGAGTGTGGAGTCGTTGCATAAAGCAGCTGTGGCAAAATGTNNNAGATCGGAA |
| 48585_1 | TATGAGTGTGGAGTCGTTGCTTCAAGCGATTTTCCGGTCTCAANNNAGATCGGAAG |
| 48585_2 | TATGAGTGTGGAGTCGTTGCTTCAAGCGATTTTCCGGTCTCAANNNAGATCGGAAG |
| 38482_1 | TATGAGTGTGGAGTCGTTGCCCTTTATAGCCTACTCCTCCCCTNNNAGATCGGAAGA |
| 38482_2 | TATGAGTGTGGAGTCGTTGCCCTTTATAGCCTACTCCTCCCCTNNNAGATCGGAAGA |
| 52141_1 | TATGAGTGTGGAGTCGTTGCGAGCCCACTCCCTCAGCTTGNNNAGATCGGAAGAGC |
| 52141_2 | TATGAGTGTGGAGTCGTTGCGAGCCCACTCCCTCAGCTTGNNNAGATCGGAAGAGC |
| 32126_1 | TATGAGTGTGGAGTCGTTGCCATTCTCCTGCCTCAGCCTCNNNAGATCGGAAGAGCA |
| 32126_2 | TATGAGTGTGGAGTCGTTGCCATTCTCCTGCCTCAGCCTCNNNAGATCGGAAGAGCA |
| 11245_1 | TATGAGTGTGGAGTCGTTGCGGTCTCAGAGGTGGCTAGCTTATCNNNAGATCGGA |
| 11245_2 | TATGAGTGTGGAGTCGTTGCGGTCTCAGAGGTGGCTAGCTTATCNNNAGATCGGA |
| 53328_1 | TATGAGTGTGGAGTCGTTGCTGCTATCCCCATTACACACAGAGNNNAGATCGGAAG |
| 53328_2 | TATGAGTGTGGAGTCGTTGCTGCTATCCCCATTACACACAGAGNNNAGATCGGAAG |

| | |
|---|---|
| 42834_1 | TATGAGTGTGGAGTCGTTGCGGTAAATAGTCTCAGCTCCTCGGNNNAGATCGGAAG |
| 42834_2 | TATGAGTGTGGAGTCGTTGCGGTAAATAGTCTCAGCTCCTCGGNNNAGATCGGAAG |
| 28779_1 | TATGAGTGTGGAGTCGTTGCTGGGTAGATATGCAGTAGTGGGANNNAGATCGGAA |
| 28779_2 | TATGAGTGTGGAGTCGTTGCTGGGTAGATATGCAGTAGTGGGANNNAGATCGGAA |
| 34578_1 | TATGAGTGTGGAGTCGTTGCCACAGAGACCGTGACCACCTNNNAGATCGGAAGAG |
| 34578_2 | TATGAGTGTGGAGTCGTTGCCACAGAGACCGTGACCACCTNNNAGATCGGAAGAG |
| 37691_1 | TATGAGTGTGGAGTCGTTGCACAGGGAACTTACATTGCTTGATNNNAGATCGGAAG |
| 37691_2 | TATGAGTGTGGAGTCGTTGCACAGGGAACTTACATTGCTTGATNNNAGATCGGAAG |
| 51073_1 | TATGAGTGTGGAGTCGTTGCGTTGGTGATTTATGGCGCCAATANNNAGATCGGAA |
| 51073_2 | TATGAGTGTGGAGTCGTTGCGTTGGTGATTTATGGCGCCAATANNNAGATCGGAA |
| 10784_1 | TATGAGTGTGGAGTCGTTGCCCTGCTGATAACCCAAAAGGGAGNNNAGATCGGAA |
| 10784_2 | TATGAGTGTGGAGTCGTTGCCCTGCTGATAACCCAAAAGGGAGNNNAGATCGGAA |
| 44955_1 | TATGAGTGTGGAGTCGTTGCTACCAATCAGTTTCCTCCACTGGNNNAGATCGGAAGA |
| 44955_2 | TATGAGTGTGGAGTCGTTGCTACCAATCAGTTTCCTCCACTGGNNNAGATCGGAAGA |
| 46596_1 | TATGAGTGTGGAGTCGTTGCCACGTCCACCATACCTGAGTATTNNNAGATCGGAAGA |
| 46596_2 | TATGAGTGTGGAGTCGTTGCCACGTCCACCATACCTGAGTATTNNNAGATCGGAAGA |
| 42816_1 | TATGAGTGTGGAGTCGTTGCATTCCTAGTCTGTCCTCTGAAAGNNNAGATCGGAAGA |
| 42816_2 | TATGAGTGTGGAGTCGTTGCATTCCTAGTCTGTCCTCTGAAAGNNNAGATCGGAAGA |
| 34777_1 | TATGAGTGTGGAGTCGTTGCCTAGTAGCTGGGATTACAGGCACNNNAGATCGGAA |
| 34777_2 | TATGAGTGTGGAGTCGTTGCCTAGTAGCTGGGATTACAGGCACNNNAGATCGGAA |
| 30525_1 | TATGAGTGTGGAGTCGTTGCCCTTTATTCTGCCCCATGCTGCNNNAGATCGGAAGAG |
| 30525_2 | TATGAGTGTGGAGTCGTTGCCCTTTATTCTGCCCCATGCTGCNNNAGATCGGAAGAG |
| 39935_1 | TATGAGTGTGGAGTCGTTGCTCCCAAATAGGCAGACTGGAAAANNNAGATCGGAA |
| 39935_2 | TATGAGTGTGGAGTCGTTGCTCCCAAATAGGCAGACTGGAAAANNNAGATCGGAA |
| 50420_1 | TATGAGTGTGGAGTCGTTGCGACTCCCATACCCTTACTGTCTGNNNAGATCGGAAGA |
| 50420_2 | TATGAGTGTGGAGTCGTTGCGACTCCCATACCCTTACTGTCTGNNNAGATCGGAAGA |
| 13192_1 | TATGAGTGTGGAGTCGTTGCGAGTACTGAATGACAGCCATNNNAGATCGGAAGAG |
| 13192_2 | TATGAGTGTGGAGTCGTTGCGAGTACTGAATGACAGCCATNNNAGATCGGAAGAG |
| 52119_1 | TATGAGTGTGGAGTCGTTGCCCAGTGATCAGAGCTTGAGTTCTNNNAGATCGGAAG |
| 52119_2 | TATGAGTGTGGAGTCGTTGCCCAGTGATCAGAGCTTGAGTTCTNNNAGATCGGAAG |
| 45316_1 | TATGAGTGTGGAGTCGTTGCCCTAAGGGTATCGTTAGGGATGCNNNAGATCGGAA |
| 45316_2 | TATGAGTGTGGAGTCGTTGCCCTAAGGGTATCGTTAGGGATGCNNNAGATCGGAA |
| 51341_1 | TATGAGTGTGGAGTCGTTGCTGTAGATGATGGGTTGATGGGTGNNNAGATCGGAA |
| 51341_2 | TATGAGTGTGGAGTCGTTGCTGTAGATGATGGGTTGATGGGTGNNNAGATCGGAA |
| 48150_1 | TATGAGTGTGGAGTCGTTGCTTCAAGTGATTCTCCTGCCTCAGNNNAGATCGGAAGA |
| 48150_2 | TATGAGTGTGGAGTCGTTGCTTCAAGTGATTCTCCTGCCTCAGNNNAGATCGGAAGA |
| 49358_1 | TATGAGTGTGGAGTCGTTGCGGGTTCAGCTAAAGACTCTGTGANNNAGATCGGAA |
| 49358_2 | TATGAGTGTGGAGTCGTTGCGGGTTCAGCTAAAGACTCTGTGANNNAGATCGGAA |
| 3057_1 | TATGAGTGTGGAGTCGTTGCATCAATCAAGACCTCGACGCTTTNNNAGATCGGAAGA |
| 3057_2 | TATGAGTGTGGAGTCGTTGCATCAATCAAGACCTCGACGCTTTNNNAGATCGGAAGA |
| 30261_1 | TATGAGTGTGGAGTCGTTGCTTCAGCTCTTAGTTCCTCTGCTGNNNAGATCGGAAGA |
| 30261_2 | TATGAGTGTGGAGTCGTTGCTTCAGCTCTTAGTTCCTCTGCTGNNNAGATCGGAAGA |
| 35555_1 | TATGAGTGTGGAGTCGTTGCTTCAAACGGCTCTCCTTTCCTTANNNAGATCGGAAGA |

| | |
|---|---|
| 35555_2 | TATGAGTGTGGAGTCGTTGCTTCAAACGGCTCTCCTTTCCTTANNNAGATCGGAAGA |
| 52054_1 | TATGAGTGTGGAGTCGTTGCTGGACAGAATTACTGGACCTGACNNNAGATCGGAA |
| 52054_2 | TATGAGTGTGGAGTCGTTGCTGGACAGAATTACTGGACCTGACNNNAGATCGGAA |
| 30209_1 | TATGAGTGTGGAGTCGTTGCATGGTGTTGAAGTGAGTGAGTCANNNAGATCGGAA |
| 30209_2 | TATGAGTGTGGAGTCGTTGCATGGTGTTGAAGTGAGTGAGTCANNNAGATCGGAA |
| 44860_1 | TATGAGTGTGGAGTCGTTGCACCTTACAGATCTAGCACCAAGGNNNAGATCGGAAG |
| 44860_2 | TATGAGTGTGGAGTCGTTGCACCTTACAGATCTAGCACCAAGGNNNAGATCGGAAG |
| 52067_1 | TATGAGTGTGGAGTCGTTGCGAAGAACAAGGTGGGGAAACTTGNNNAGATCGGA |
| 52067_2 | TATGAGTGTGGAGTCGTTGCGAAGAACAAGGTGGGGAAACTTGNNNAGATCGGA |
| 52497_1 | TATGAGTGTGGAGTCGTTGCCTGAAGCAGTTCCCAAAGTCATGNNNAGATCGGAAG |
| 52497_2 | TATGAGTGTGGAGTCGTTGCCTGAAGCAGTTCCCAAAGTCATGNNNAGATCGGAAG |
| 52492_1 | TATGAGTGTGGAGTCGTTGCTGAGAAATGCTACGAGGTGTTCANNNAGATCGGAA |
| 52492_2 | TATGAGTGTGGAGTCGTTGCTGAGAAATGCTACGAGGTGTTCANNNAGATCGGAA |
| 46673_1 | TATGAGTGTGGAGTCGTTGCAGGTTCAAGTGATTCTTCTGCCTNNNAGATCGGAAG |
| 46673_2 | TATGAGTGTGGAGTCGTTGCAGGTTCAAGTGATTCTTCTGCCTNNNAGATCGGAAG |
| 48617_1 | TATGAGTGTGGAGTCGTTGCCCGGGGAGAGAAAAGTGACTTTANNNAGATCGGAA |
| 48617_2 | TATGAGTGTGGAGTCGTTGCCCGGGGAGAGAAAAGTGACTTTANNNAGATCGGAA |
| 37014_1 | TATGAGTGTGGAGTCGTTGCGGTTGCTTTTTTCCCTGGAGATTNNNAGATCGGAAG |
| 37014_2 | TATGAGTGTGGAGTCGTTGCGGTTGCTTTTTTCCCTGGAGATTNNNAGATCGGAAG |
| 41186_1 | TATGAGTGTGGAGTCGTTGCACCAAATGATACCAAAACATGCTGNNNAGATCGGAA |
| 41186_2 | TATGAGTGTGGAGTCGTTGCACCAAATGATACCAAAACATGCTGNNNAGATCGGAA |
| 52948_1 | TATGAGTGTGGAGTCGTTGCTACTGAAAAGGTGCCTTGATCCANNNAGATCGGAAG |
| 52948_2 | TATGAGTGTGGAGTCGTTGCTACTGAAAAGGTGCCTTGATCCANNNAGATCGGAAG |
| 52113_1 | TATGAGTGTGGAGTCGTTGCTTTCTTTGAAAGTTGGAACGCCTNNNAGATCGGAAG |
| 52113_2 | TATGAGTGTGGAGTCGTTGCTTTCTTTGAAAGTTGGAACGCCTNNNAGATCGGAAG |
| 46833_1 | TATGAGTGTGGAGTCGTTGCTTTTCTCAGCTCAGATTGGGACANNNAGATCGGAAG |
| 46833_2 | TATGAGTGTGGAGTCGTTGCTTTTCTCAGCTCAGATTGGGACANNNAGATCGGAAG |
| 36305_1 | TATGAGTGTGGAGTCGTTGCAAGAAACATGGATCCCCACACTTNNNAGATCGGAAG |
| 36305_2 | TATGAGTGTGGAGTCGTTGCAAGAAACATGGATCCCCACACTTNNNAGATCGGAAG |
| 38019_1 | TATGAGTGTGGAGTCGTTGCAAACGATGGGAATTCGAAAAGCCNNNAGATCGGAA |
| 38019_2 | TATGAGTGTGGAGTCGTTGCAAACGATGGGAATTCGAAAAGCCNNNAGATCGGAA |
| 41779_1 | TATGAGTGTGGAGTCGTTGCGGAGTTTCAGTCTTGTTGCTCAGNNNAGATCGGAAG |
| 41779_2 | TATGAGTGTGGAGTCGTTGCGGAGTTTCAGTCTTGTTGCTCAGNNNAGATCGGAAG |
| 45331_1 | TATGAGTGTGGAGTCGTTGCGGGTGAGAGAGAGAGAGAGAGAGNNNAGATCGG |
| 45331_2 | TATGAGTGTGGAGTCGTTGCGGGTGAGAGAGAGAGAGAGAGAGNNNAGATCGG |
| 43307_1 | TATGAGTGTGGAGTCGTTGCAATTCCCAAAATCTGCAGTCAGCNNNAGATCGGAAG |
| 43307_2 | TATGAGTGTGGAGTCGTTGCAATTCCCAAAATCTGCAGTCAGCNNNAGATCGGAAG |
| 12747_1 | TATGAGTGTGGAGTCGTTGCTAAAAATACAAAAACTAGCCAGGNNNAGATCGGAA |
| 12747_2 | TATGAGTGTGGAGTCGTTGCTAAAAATACAAAAACTAGCCAGGNNNAGATCGGAA |
| 29149_1 | TATGAGTGTGGAGTCGTTGCACACACACACACACACAGAGTNNNAGATCGGAAGAG |
| 29149_2 | TATGAGTGTGGAGTCGTTGCACACACACACACACACAGAGTNNNAGATCGGAAGAG |
| 13233_1 | TATGAGTGTGGAGTCGTTGCCATTCTCATAAACACTCCCAANNNAGATCGGAAGAG |
| 13233_2 | TATGAGTGTGGAGTCGTTGCCATTCTCATAAACACTCCCAANNNAGATCGGAAGAG |

| | |
|---|---|
| 30460_1 | TATGAGTGTGGAGTCGTTGCCCACCTGTACCATCACAGTTACANNNAGATCGGAAGA |
| 30460_2 | TATGAGTGTGGAGTCGTTGCCCACCTGTACCATCACAGTTACANNNAGATCGGAAGA |
| 46884_1 | TATGAGTGTGGAGTCGTTGCGTTCAAGCAATTCTTCTGCCTCANNNAGATCGGAAGA |
| 46884_2 | TATGAGTGTGGAGTCGTTGCGTTCAAGCAATTCTTCTGCCTCANNNAGATCGGAAGA |
| 44381_1 | TATGAGTGTGGAGTCGTTGCTTTTCTCTGGTTCGTCTGCTTTGNNNAGATCGGAAGA |
| 44381_2 | TATGAGTGTGGAGTCGTTGCTTTTCTCTGGTTCGTCTGCTTTGNNNAGATCGGAAGA |
| 10950_1 | TATGAGTGTGGAGTCGTTGCATAACAGGCCCCCACCACTACATNNNAGATCGGAAGA |
| 10950_2 | TATGAGTGTGGAGTCGTTGCATAACAGGCCCCCACCACTACATNNNAGATCGGAAGA |
| 34249_1 | TATGAGTGTGGAGTCGTTGCCACCTCTTTCTAACCTCCAGGACNNNAGATCGGAAGA |
| 34249_2 | TATGAGTGTGGAGTCGTTGCCACCTCTTTCTAACCTCCAGGACNNNAGATCGGAAGA |
| 12843_1 | TATGAGTGTGGAGTCGTTGCCCTTGTCTTATGTTCCAGTATTNNNAGATCGGAAGAG |
| 12843_2 | TATGAGTGTGGAGTCGTTGCCCTTGTCTTATGTTCCAGTATTNNNAGATCGGAAGAG |
| 36837_1 | TATGAGTGTGGAGTCGTTGCACACCTCACACTTTCTTCCTTACTNNNAGATCGGAAGA |
| 36837_2 | TATGAGTGTGGAGTCGTTGCACACCTCACACTTTCTTCCTTACTNNNAGATCGGAAGA |
| 45258_1 | TATGAGTGTGGAGTCGTTGCGATAATCGTTGTAACCCTGACGCNNNAGATCGGAAG |
| 45258_2 | TATGAGTGTGGAGTCGTTGCGATAATCGTTGTAACCCTGACGCNNNAGATCGGAAG |
| 48064_1 | TATGAGTGTGGAGTCGTTGCTGGTATCAGAATGTTGTTGGCCTNNNAGATCGGAAG |
| 48064_2 | TATGAGTGTGGAGTCGTTGCTGGTATCAGAATGTTGTTGGCCTNNNAGATCGGAAG |
| 36077_1 | TATGAGTGTGGAGTCGTTGCCATAAGCTACGATCATGCCACTGNNNAGATCGGAAG |
| 36077_2 | TATGAGTGTGGAGTCGTTGCCATAAGCTACGATCATGCCACTGNNNAGATCGGAAG |
| 34629_1 | TATGAGTGTGGAGTCGTTGCTCCTCCTGAAGAATCCTTGCAAANNNAGATCGGAAGA |
| 34629_2 | TATGAGTGTGGAGTCGTTGCTCCTCCTGAAGAATCCTTGCAAANNNAGATCGGAAGA |
| 41480_1 | TATGAGTGTGGAGTCGTTGCTCCCAGAGATTCATTGTCCTGTGNNNAGATCGGAAGA |
| 41480_2 | TATGAGTGTGGAGTCGTTGCTCCCAGAGATTCATTGTCCTGTGNNNAGATCGGAAGA |
| 30724_1 | TATGAGTGTGGAGTCGTTGCGACGGGGAAAGAGAGAACAGAAGNNNAGATCGGA |
| 30724_2 | TATGAGTGTGGAGTCGTTGCGACGGGGAAAGAGAGAACAGAAGNNNAGATCGGA |
| 31916_1 | TATGAGTGTGGAGTCGTTGCCGAACAGTTAGAGGAGTGGCATANNNAGATCGGAA |
| 31916_2 | TATGAGTGTGGAGTCGTTGCCGAACAGTTAGAGGAGTGGCATANNNAGATCGGAA |
| 34425_1 | TATGAGTGTGGAGTCGTTGCACAGGTGTATGCATGATATCCCCNNNAGATCGGAAG |
| 34425_2 | TATGAGTGTGGAGTCGTTGCACAGGTGTATGCATGATATCCCCNNNAGATCGGAAG |
| 49963_1 | TATGAGTGTGGAGTCGTTGCTTTTAGCTCCTACCTGCATCTCCNNNAGATCGGAAGA |
| 49963_2 | TATGAGTGTGGAGTCGTTGCTTTTAGCTCCTACCTGCATCTCCNNNAGATCGGAAGA |
| 33954_1 | TATGAGTGTGGAGTCGTTGCGAGTGCAGTGGCGAGATCTTAGNNNAGATCGGAAG |
| 33954_2 | TATGAGTGTGGAGTCGTTGCGAGTGCAGTGGCGAGATCTTAGNNNAGATCGGAAG |
| 51554_1 | TATGAGTGTGGAGTCGTTGCTCACTCAAGGCTTGTGATAACCANNNAGATCGGAAG |
| 51554_2 | TATGAGTGTGGAGTCGTTGCTCACTCAAGGCTTGTGATAACCANNNAGATCGGAAG |
| 51372_1 | TATGAGTGTGGAGTCGTTGCAACCAAGAAGCTGTGTTGTTTCCNNNAGATCGGAAG |
| 51372_2 | TATGAGTGTGGAGTCGTTGCAACCAAGAAGCTGTGTTGTTTCCNNNAGATCGGAAG |
| 28969_1 | TATGAGTGTGGAGTCGTTGCTCCTTTTTCTCTTGCTTCAGTCTGNNNAGATCGGAAGA |
| 28969_2 | TATGAGTGTGGAGTCGTTGCTCCTTTTTCTCTTGCTTCAGTCTGNNNAGATCGGAAGA |
| 37846_1 | TATGAGTGTGGAGTCGTTGCAAGATGGATGGAGCAACAGATGTNNNAGATCGGAA |
| 37846_2 | TATGAGTGTGGAGTCGTTGCAAGATGGATGGAGCAACAGATGTNNNAGATCGGAA |
| 36589_1 | TATGAGTGTGGAGTCGTTGCAGATGAAAAATGGTTCACCTTTGTNNNAGATCGGAA |

| | |
|---|---|
| 36589_2 | TATGAGTGTGGAGTCGTTGCAGATGAAAAATGGTTCACCTTTGTNNNAGATCGGAA |
| 35476_1 | TATGAGTGTGGAGTCGTTGCCAATCTCGGCTCACTGCAGGNNNAGATCGGAAGAG |
| 35476_2 | TATGAGTGTGGAGTCGTTGCCAATCTCGGCTCACTGCAGGNNNAGATCGGAAGAG |
| 30257_1 | TATGAGTGTGGAGTCGTTGCTGCTTTACATTATTTTACATGGTTTGTNNNAGATCGG |
| 30257_2 | TATGAGTGTGGAGTCGTTGCTGCTTTACATTATTTTACATGGTTTGTNNNAGATCGG |
| 2679_1 | TATGAGTGTGGAGTCGTTGCGTTCCAGCCTCATAATCAGCCATNNNAGATCGGAAG |
| 2679_2 | TATGAGTGTGGAGTCGTTGCGTTCCAGCCTCATAATCAGCCATNNNAGATCGGAAG |
| 40251_1 | TATGAGTGTGGAGTCGTTGCCCTCTTCACAGATGGAAGCTCTTNNNAGATCGGAAG |
| 40251_2 | TATGAGTGTGGAGTCGTTGCCCTCTTCACAGATGGAAGCTCTTNNNAGATCGGAAG |
| 49959_1 | TATGAGTGTGGAGTCGTTGCTCCAGGCTTAGCTTTTTGTGTTGNNNAGATCGGAAG |
| 49959_2 | TATGAGTGTGGAGTCGTTGCTCCAGGCTTAGCTTTTTGTGTTGNNNAGATCGGAAG |
| 33730_1 | TATGAGTGTGGAGTCGTTGCGGAAACTTGCTACCCCTCAGTAANNNAGATCGGAAG |
| 33730_2 | TATGAGTGTGGAGTCGTTGCGGAAACTTGCTACCCCTCAGTAANNNAGATCGGAAG |
| 40940_1 | TATGAGTGTGGAGTCGTTGCCTATTCACAAAGGCCAGGGTTTGNNNAGATCGGAA |
| 40940_2 | TATGAGTGTGGAGTCGTTGCCTATTCACAAAGGCCAGGGTTTGNNNAGATCGGAA |
| 44395_1 | TATGAGTGTGGAGTCGTTGCCTCTAGAATTCGGGGCTTGGATTNNNAGATCGGAA |
| 44395_2 | TATGAGTGTGGAGTCGTTGCCTCTAGAATTCGGGGCTTGGATTNNNAGATCGGAA |
| 41597_1 | TATGAGTGTGGAGTCGTTGCTTCAAGCAATTCTCCTGCCTCAGNNNAGATCGGAAG |
| 41597_2 | TATGAGTGTGGAGTCGTTGCTTCAAGCAATTCTCCTGCCTCAGNNNAGATCGGAAG |
| 30878_1 | TATGAGTGTGGAGTCGTTGCACACACACACACACACACAAATTNNNAGATCGGAAG |
| 30878_2 | TATGAGTGTGGAGTCGTTGCACACACACACACACACACAAATTNNNAGATCGGAAG |
| 41278_1 | TATGAGTGTGGAGTCGTTGCATGTGCCTGTAATCCCAGCTACNNNAGATCGGAAGA |
| 41278_2 | TATGAGTGTGGAGTCGTTGCATGTGCCTGTAATCCCAGCTACNNNAGATCGGAAGA |
| 29458_1 | ATGAGTGTGGAGTCGTTGCACTGAATGAAACAGAGTCACCAGANNNAGATCGGAA |
| 29458_2 | ATGAGTGTGGAGTCGTTGCACTGAATGAAACAGAGTCACCAGANNNAGATCGGAA |
| 13082_1 | TATGAGTGTGGAGTCGTTGCTGCAAAGGAAAACAAAAAATAATNNNAGATCGGAA |
| 13082_2 | TATGAGTGTGGAGTCGTTGCTGCAAAGGAAAACAAAAAATAATNNNAGATCGGAA |
| 35731_1 | TATGAGTGTGGAGTCGTTGCGAGATCTCGGCTCACTGCAAGNNNAGATCGGAAGA |
| 35731_2 | TATGAGTGTGGAGTCGTTGCGAGATCTCGGCTCACTGCAAGNNNAGATCGGAAGA |
| 36043_1 | TATGAGTGTGGAGTCGTTGCGCTCACCTTTTTCTCTCTACCCTNNNAGATCGGAAGA |
| 36043_2 | TATGAGTGTGGAGTCGTTGCGCTCACCTTTTTCTCTCTACCCTNNNAGATCGGAAGA |
| 31817_1 | TATGAGTGTGGAGTCGTTGCTGCAGTTTGGACAAGTAGAGAGANNNAGATCGGAA |
| 31817_2 | TATGAGTGTGGAGTCGTTGCTGCAGTTTGGACAAGTAGAGAGANNNAGATCGGAA |
| 13382_1 | TATGAGTGTGGAGTCGTTGCATATATAGTGGAGAGACTCAGAANNNAGATCGGAA |
| 13382_2 | TATGAGTGTGGAGTCGTTGCATATATAGTGGAGAGACTCAGAANNNAGATCGGAA |
| 44529_1 | TATGAGTGTGGAGTCGTTGCGATGGTCAGGAATGGGACAAGATNNNAGATCGGA |
| 44529_2 | TATGAGTGTGGAGTCGTTGCGATGGTCAGGAATGGGACAAGATNNNAGATCGGA |
| 35804_1 | TATGAGTGTGGAGTCGTTGCCCAAGAACCTGGGATAATAGCCANNNAGATCGGAA |
| 35804_2 | TATGAGTGTGGAGTCGTTGCCCAAGAACCTGGGATAATAGCCANNNAGATCGGAA |
| 46689_1 | TATGAGTGTGGAGTCGTTGCCTGAAGTGCATTGATGTGATCCCNNNAGATCGGAAG |
| 46689_2 | TATGAGTGTGGAGTCGTTGCCTGAAGTGCATTGATGTGATCCCNNNAGATCGGAAG |
| 45232_1 | TATGAGTGTGGAGTCGTTGCCAGGATCTTGCTCTATCGCCTAGNNNAGATCGGAAG |
| 45232_2 | TATGAGTGTGGAGTCGTTGCCAGGATCTTGCTCTATCGCCTAGNNNAGATCGGAAG |

| | |
|---|---|
| 38457_1 | TATGAGTGTGGAGTCGTTGCGGACACAGGGTCTCACTATGTTGNNNAGATCGGAAG |
| 38457_2 | TATGAGTGTGGAGTCGTTGCGGACACAGGGTCTCACTATGTTGNNNAGATCGGAAG |
| 29834_1 | TATGAGTGTGGAGTCGTTGCACAGTCACCAGTATTAGTGGAGACNNNAGATCGGAA |
| 29834_2 | TATGAGTGTGGAGTCGTTGCACAGTCACCAGTATTAGTGGAGACNNNAGATCGGAA |
| 42977_1 | TATGAGTGTGGAGTCGTTGCTTTCTTCCTGGAGTACAGTGGTGNNNAGATCGGAAG |
| 42977_2 | TATGAGTGTGGAGTCGTTGCTTTCTTCCTGGAGTACAGTGGTGNNNAGATCGGAAG |
| 46164_1 | TATGAGTGTGGAGTCGTTGCGATGGAGTCTCGCTATGTCATCCNNNAGATCGGAAG |
| 46164_2 | TATGAGTGTGGAGTCGTTGCGATGGAGTCTCGCTATGTCATCCNNNAGATCGGAAG |
| 52999_1 | TATGAGTGTGGAGTCGTTGCATAGAAGGTATTTGGGCGATGGGNNNAGATCGGAA |
| 52999_2 | TATGAGTGTGGAGTCGTTGCATAGAAGGTATTTGGGCGATGGGNNNAGATCGGAA |
| 8541_1 | TATGAGTGTGGAGTCGTTGCAGGAAATGTGAAGGAGGCAGGAANNNAGATCGGA |
| 8541_2 | TATGAGTGTGGAGTCGTTGCAGGAAATGTGAAGGAGGCAGGAANNNAGATCGGA |
| 36657_1 | TATGAGTGTGGAGTCGTTGCGCTTTTGCTCAGACATCACCTTTNNNAGATCGGAAGA |
| 36657_2 | TATGAGTGTGGAGTCGTTGCGCTTTTGCTCAGACATCACCTTTNNNAGATCGGAAGA |
| 35038_1 | TATGAGTGTGGAGTCGTTGCGTCGTGTCACAAGAACTTGGTTTNNNAGATCGGAAG |
| 35038_2 | TATGAGTGTGGAGTCGTTGCGTCGTGTCACAAGAACTTGGTTTNNNAGATCGGAAG |
| 10001_1 | TATGAGTGTGGAGTCGTTGCGCACACTGGAACAGAGAACAAGANNNAGATCGGAA |
| 10001_2 | TATGAGTGTGGAGTCGTTGCGCACACTGGAACAGAGAACAAGANNNAGATCGGAA |
| 36269_1 | TATGAGTGTGGAGTCGTTGCTTGACAGCACAACAGAGTGACTANNNAGATCGGAAG |
| 36269_2 | TATGAGTGTGGAGTCGTTGCTTGACAGCACAACAGAGTGACTANNNAGATCGGAAG |
| 30061_1 | TATGAGTGTGGAGTCGTTGCAATGTGTGCTTACGAAGTTCCAGNNNAGATCGGAAG |
| 30061_2 | TATGAGTGTGGAGTCGTTGCAATGTGTGCTTACGAAGTTCCAGNNNAGATCGGAAG |
| 35169_1 | TATGAGTGTGGAGTCGTTGCCTTGCCTCGGCCTCATGAATAGNNNAGATCGGAAGA |
| 35169_2 | TATGAGTGTGGAGTCGTTGCCTTGCCTCGGCCTCATGAATAGNNNAGATCGGAAGA |
| 38923_1 | TATGAGTGTGGAGTCGTTGCGCAAGTCAGTCACTGTGAAACTCNNNAGATCGGAAG |
| 38923_2 | TATGAGTGTGGAGTCGTTGCGCAAGTCAGTCACTGTGAAACTCNNNAGATCGGAAG |
| 30408_1 | TATGAGTGTGGAGTCGTTGCAGGAGAGTGGACATGGGTTTCATNNNAGATCGGAA |
| 30408_2 | TATGAGTGTGGAGTCGTTGCAGGAGAGTGGACATGGGTTTCATNNNAGATCGGAA |
| 45894_1 | TATGAGTGTGGAGTCGTTGCCAGAAAGGCATTGCAAAAGGAGANNNAGATCGGAA |
| 45894_2 | TATGAGTGTGGAGTCGTTGCCAGAAAGGCATTGCAAAAGGAGANNNAGATCGGAA |
| 936_1 | TATGAGTGTGGAGTCGTTGCGGAGCACATGTATAGCAGGACAGNNNAGATCGGAA |
| 936_2 | TATGAGTGTGGAGTCGTTGCGGAGCACATGTATAGCAGGACAGNNNAGATCGGAA |
| 34381_1 | TATGAGTGTGGAGTCGTTGCCCTCCTGGGTTCTAGTGATTCTCNNNAGATCGGAAGA |
| 34381_2 | TATGAGTGTGGAGTCGTTGCCCTCCTGGGTTCTAGTGATTCTCNNNAGATCGGAAGA |
| 52626_1 | TATGAGTGTGGAGTCGTTGCCAGCAGCTGAACACGTCCACNNNAGATCGGAAGAGC |
| 52626_2 | TATGAGTGTGGAGTCGTTGCCAGCAGCTGAACACGTCCACNNNAGATCGGAAGAGC |
| 52663_1 | TATGAGTGTGGAGTCGTTGCTCTCATGAAACCCGTCTTTCCATNNNAGATCGGAAGA |
| 52663_2 | TATGAGTGTGGAGTCGTTGCTCTCATGAAACCCGTCTTTCCATNNNAGATCGGAAGA |
| 34037_1 | TATGAGTGTGGAGTCGTTGCCTCAAGTGATTCTCCCACCTCAGNNNAGATCGGAAGA |
| 34037_2 | TATGAGTGTGGAGTCGTTGCCTCAAGTGATTCTCCCACCTCAGNNNAGATCGGAAGA |
| 29528_1 | TATGAGTGTGGAGTCGTTGCTGGATAAAGTTCCAGAGCCTGGANNNAGATCGGAAG |
| 29528_2 | TATGAGTGTGGAGTCGTTGCTGGATAAAGTTCCAGAGCCTGGANNNAGATCGGAAG |
| 13698_1 | TATGAGTGTGGAGTCGTTGCTTTTTTTCTCTCTTCCTCTATCTNNNAGATCGGAAGAG |

| | |
|---|---|
| 13698_2 | TATGAGTGTGGAGTCGTTGCTTTTTTTCTCTCTTCCTCTATCTNNNAGATCGGAAGAG |
| 40225_1 | TATGAGTGTGGAGTCGTTGCGGGAATATCAACTGGTGACATGGNNNAGATCGGAA |
| 40225_2 | TATGAGTGTGGAGTCGTTGCGGGAATATCAACTGGTGACATGGNNNAGATCGGAA |
| 53150_1 | TATGAGTGTGGAGTCGTTGCATGCTGCCTTTTGTTTGAGAGTCNNNAGATCGGAAG |
| 53150_2 | TATGAGTGTGGAGTCGTTGCATGCTGCCTTTTGTTTGAGAGTCNNNAGATCGGAAG |
| 12974_1 | TATGAGTGTGGAGTCGTTGCAATACTTGCAATTTTCTTAACGTNNNAGATCGGAAGA |
| 12974_2 | TATGAGTGTGGAGTCGTTGCAATACTTGCAATTTTCTTAACGTNNNAGATCGGAAGA |
| 42936_1 | TATGAGTGTGGAGTCGTTGCTCCCTTGTGCATGTACTGTCTTTNNNAGATCGGAAGA |
| 42936_2 | TATGAGTGTGGAGTCGTTGCTCCCTTGTGCATGTACTGTCTTTNNNAGATCGGAAGA |
| 43988_1 | TATGAGTGTGGAGTCGTTGCGAGGGAGGGAAAGGAAAGAAAGANNNAGATCGGA |
| 43988_2 | TATGAGTGTGGAGTCGTTGCGAGGGAGGGAAAGGAAAGAAAGANNNAGATCGGA |
| 53227_1 | TATGAGTGTGGAGTCGTTGCTTTGGCCTCAAACTTCAGGTGANNNAGATCGGAAGA |
| 53227_2 | TATGAGTGTGGAGTCGTTGCTTTGGCCTCAAACTTCAGGTGANNNAGATCGGAAGA |
| 35647_1 | TATGAGTGTGGAGTCGTTGCCTGGAATGCAATGGTGTGATCTCNNNAGATCGGAAG |
| 35647_2 | TATGAGTGTGGAGTCGTTGCCTGGAATGCAATGGTGTGATCTCNNNAGATCGGAAG |
| 13690_1 | TATGAGTGTGGAGTCGTTGCATGACATGTTTGGGTCTTTTATANNNAGATCGGAAG |
| 13690_2 | TATGAGTGTGGAGTCGTTGCATGACATGTTTGGGTCTTTTATANNNAGATCGGAAG |
| 4518_1 | TATGAGTGTGGAGTCGTTGCAGTAGCCTCCACACATTCCTTCTNNNAGATCGGAAGA |
| 4518_2 | TATGAGTGTGGAGTCGTTGCAGTAGCCTCCACACATTCCTTCTNNNAGATCGGAAGA |
| 33272_1 | TATGAGTGTGGAGTCGTTGCCTCCTGGGTTCAAAGGATCCTCNNNAGATCGGAAGA |
| 33272_2 | TATGAGTGTGGAGTCGTTGCCTCCTGGGTTCAAAGGATCCTCNNNAGATCGGAAGA |
| 5617_1 | TATGAGTGTGGAGTCGTTGCTCCACCCTAAAGAGATAAGAGCANNNAGATCGGAAG |
| 5617_2 | TATGAGTGTGGAGTCGTTGCTCCACCCTAAAGAGATAAGAGCANNNAGATCGGAAG |
| 46549_1 | TATGAGTGTGGAGTCGTTGCGCCATCACACCCAGCTAATTTTTNNNAGATCGGAAGA |
| 46549_2 | TATGAGTGTGGAGTCGTTGCGCCATCACACCCAGCTAATTTTTNNNAGATCGGAAGA |
| 45228_1 | TATGAGTGTGGAGTCGTTGCATTAGTGAGAGAGGCAAAGTGGGNNNAGATCGGA |
| 45228_2 | TATGAGTGTGGAGTCGTTGCATTAGTGAGAGAGGCAAAGTGGGNNNAGATCGGA |
| 12473_1 | TATGAGTGTGGAGTCGTTGCGGTACTATGAAGTAAGTATGCATNNNAGATCGGAA |
| 12473_2 | TATGAGTGTGGAGTCGTTGCGGTACTATGAAGTAAGTATGCATNNNAGATCGGAA |
| 44885_1 | TATGAGTGTGGAGTCGTTGCCAAGGAGAACTGAAGTCAGGGAGNNNAGATCGGA |
| 44885_2 | TATGAGTGTGGAGTCGTTGCCAAGGAGAACTGAAGTCAGGGAGNNNAGATCGGA |
| 33087_1 | TATGAGTGTGGAGTCGTTGCCTGGCACAGCAGAAAAATACCATNNNAGATCGGAAG |
| 33087_2 | TATGAGTGTGGAGTCGTTGCCTGGCACAGCAGAAAAATACCATNNNAGATCGGAAG |
| 30682_1 | TATGAGTGTGGAGTCGTTGCATTATATGGCCATCTTGGAGCTCCNNNAGATCGGAA |
| 30682_2 | TATGAGTGTGGAGTCGTTGCATTATATGGCCATCTTGGAGCTCCNNNAGATCGGAA |
| 47228_1 | TATGAGTGTGGAGTCGTTGCGTCACCATGCCCAGCTAATTTTTNNNAGATCGGAAGA |
| 47228_2 | TATGAGTGTGGAGTCGTTGCGTCACCATGCCCAGCTAATTTTTNNNAGATCGGAAGA |
| 10711_1 | TATGAGTGTGGAGTCGTTGCGGTAGTGCAAGGGGTCACAGATNNNAGATCGGAAG |
| 10711_2 | TATGAGTGTGGAGTCGTTGCGGTAGTGCAAGGGGTCACAGATNNNAGATCGGAAG |
| 8522_1 | TATGAGTGTGGAGTCGTTGCGGAAAATCACAGGCTGAGTGAAAGNNNAGATCGGA |
| 8522_2 | TATGAGTGTGGAGTCGTTGCGGAAAATCACAGGCTGAGTGAAAGNNNAGATCGGA |
| 52483_1 | TATGAGTGTGGAGTCGTTGCCTCATTCCGTAAGGCTCTCTCTCNNNAGATCGGAAGA |
| 52483_2 | TATGAGTGTGGAGTCGTTGCCTCATTCCGTAAGGCTCTCTCTCNNNAGATCGGAAGA |

| | |
|---|---|
| 50316_1 | TATGAGTGTGGAGTCGTTGCTGTGCATCCATTCTCCAGTAGTTNNNAGATCGGAAG |
| 50316_2 | TATGAGTGTGGAGTCGTTGCTGTGCATCCATTCTCCAGTAGTTNNNAGATCGGAAG |
| 44963_1 | TATGAGTGTGGAGTCGTTGCTTTAGGGAGGAGGTAGAAGGGAGNNNAGATCGGA |
| 44963_2 | TATGAGTGTGGAGTCGTTGCTTTAGGGAGGAGGTAGAAGGGAGNNNAGATCGGA |
| 32631_1 | TATGAGTGTGGAGTCGTTGCGTCTCAATCTCCTGACCTTGTGTNNNAGATCGGAAG |
| 32631_2 | TATGAGTGTGGAGTCGTTGCGTCTCAATCTCCTGACCTTGTGTNNNAGATCGGAAG |
| 42879_1 | TATGAGTGTGGAGTCGTTGCCTGACCAATACAGACCCTTCCTCNNNAGATCGGAAG |
| 42879_2 | TATGAGTGTGGAGTCGTTGCCTGACCAATACAGACCCTTCCTCNNNAGATCGGAAG |
| 49397_1 | TATGAGTGTGGAGTCGTTGCTGTAGTGCAATGACGTGATCTCANNNAGATCGGAA |
| 49397_2 | TATGAGTGTGGAGTCGTTGCTGTAGTGCAATGACGTGATCTCANNNAGATCGGAA |
| 47310_1 | TATGAGTGTGGAGTCGTTGCTAGGACAATAGTGGAGGGAAGGTNNNAGATCGGA |
| 47310_2 | TATGAGTGTGGAGTCGTTGCTAGGACAATAGTGGAGGGAAGGTNNNAGATCGGA |
| 45630_1 | TATGAGTGTGGAGTCGTTGCGTGTAATGGTGCCATCTTGGTTCNNNAGATCGGAAG |
| 45630_2 | TATGAGTGTGGAGTCGTTGCGTGTAATGGTGCCATCTTGGTTCNNNAGATCGGAAG |
| 39817_1 | TATGAGTGTGGAGTCGTTGCCTGAGTAGCTGGGACTACAACTGNNNAGATCGGAA |
| 39817_2 | TATGAGTGTGGAGTCGTTGCCTGAGTAGCTGGGACTACAACTGNNNAGATCGGAA |
| 32039_1 | TATGAGTGTGGAGTCGTTGCTGACCACCACCATTTAGAAGTTGNNNAGATCGGAAG |
| 32039_2 | TATGAGTGTGGAGTCGTTGCTGACCACCACCATTTAGAAGTTGNNNAGATCGGAAG |
| 37810_1 | TATGAGTGTGGAGTCGTTGCGGTGAAACCCCGTCTCTACTAAANNNAGATCGGAAG |
| 37810_2 | TATGAGTGTGGAGTCGTTGCGGTGAAACCCCGTCTCTACTAAANNNAGATCGGAAG |
| 42804_1 | TATGAGTGTGGAGTCGTTGCGTAGAGGTCTCTTGAAGTGGTCGNNNAGATCGGAA |
| 42804_2 | TATGAGTGTGGAGTCGTTGCGTAGAGGTCTCTTGAAGTGGTCGNNNAGATCGGAA |
| 53336_1 | TATGAGTGTGGAGTCGTTGCGGTGAATCCAGGCAGTTAGAAAANNNAGATCGGAA |
| 53336_2 | TATGAGTGTGGAGTCGTTGCGGTGAATCCAGGCAGTTAGAAAANNNAGATCGGAA |
| 44458_1 | TATGAGTGTGGAGTCGTTGCATGATCTCGGCTCACTGCAANNNAGATCGGAAGAG |
| 44458_2 | TATGAGTGTGGAGTCGTTGCATGATCTCGGCTCACTGCAANNNAGATCGGAAGAG |
| 36802_1 | TATGAGTGTGGAGTCGTTGCGCAGTCTCAACAGTCACTGAGTANNNAGATCGGAAG |
| 36802_2 | TATGAGTGTGGAGTCGTTGCGCAGTCTCAACAGTCACTGAGTANNNAGATCGGAAG |
| 38585_1 | TATGAGTGTGGAGTCGTTGCCAGTGAGCTATGATTGCACCACTNNNAGATCGGAAG |
| 38585_2 | TATGAGTGTGGAGTCGTTGCCAGTGAGCTATGATTGCACCACTNNNAGATCGGAAG |
| 13105_1 | TATGAGTGTGGAGTCGTTGCGTATTGACATCTAATCTGCATTTNNNAGATCGGAAG |
| 13105_2 | TATGAGTGTGGAGTCGTTGCGTATTGACATCTAATCTGCATTTNNNAGATCGGAAG |
| 37840_1 | TATGAGTGTGGAGTCGTTGCCTGGGGAGACAGACTATGTGAACNNNAGATCGGAA |
| 37840_2 | TATGAGTGTGGAGTCGTTGCCTGGGGAGACAGACTATGTGAACNNNAGATCGGAA |
| 50365_1 | TATGAGTGTGGAGTCGTTGCCATGCTGCTGTCATTTTGTGGTANNNAGATCGGAAG |
| 50365_2 | TATGAGTGTGGAGTCGTTGCCATGCTGCTGTCATTTTGTGGTANNNAGATCGGAAG |
| 51668_1 | TATGAGTGTGGAGTCGTTGCTTTTCAAGAAGTGGACTTTGGCGNNNAGATCGGAA |
| 51668_2 | TATGAGTGTGGAGTCGTTGCTTTTCAAGAAGTGGACTTTGGCGNNNAGATCGGAA |
| 36216_1 | TATGAGTGTGGAGTCGTTGCCTATGTCTGGATTTGCCTAGTGCNNNAGATCGGAAG |
| 36216_2 | TATGAGTGTGGAGTCGTTGCCTATGTCTGGATTTGCCTAGTGCNNNAGATCGGAAG |
| 46597_1 | TATGAGTGTGGAGTCGTTGCGATCTCCTGACCTCATGATCCACNNNAGATCGGAAG |
| 46597_2 | TATGAGTGTGGAGTCGTTGCGATCTCCTGACCTCATGATCCACNNNAGATCGGAAG |
| 47456_1 | TATGAGTGTGGAGTCGTTGCCTTCAAGTGATTCTCGTGCCTCANNNAGATCGGAAG |

| | |
|---|---|
| 47456_2 | TATGAGTGTGGAGTCGTTGCCTTCAAGTGATTCTCGTGCCTCANNNAGATCGGAAG |
| 31069_1 | TATGAGTGTGGAGTCGTTGCTTACCATCACTTAGAGGCCAGTGNNNAGATCGGAAG |
| 31069_2 | TATGAGTGTGGAGTCGTTGCTTACCATCACTTAGAGGCCAGTGNNNAGATCGGAAG |
| 44530_1 | TATGAGTGTGGAGTCGTTGCTAGGAAGCTCAGAAGGAAACTGGNNNAGATCGGAA |
| 44530_2 | TATGAGTGTGGAGTCGTTGCTAGGAAGCTCAGAAGGAAACTGGNNNAGATCGGAA |
| 41333_1 | TATGAGTGTGGAGTCGTTGCGCTCATGCTTGTAATCCCAGAACNNNAGATCGGAAG |
| 41333_2 | TATGAGTGTGGAGTCGTTGCGCTCATGCTTGTAATCCCAGAACNNNAGATCGGAAG |
| 32695_1 | TATGAGTGTGGAGTCGTTGCGAGCTGAGGTCTCACTCTTGTTGNNNAGATCGGAAG |
| 32695_2 | TATGAGTGTGGAGTCGTTGCGAGCTGAGGTCTCACTCTTGTTGNNNAGATCGGAAG |
| 3006_1 | TATGAGTGTGGAGTCGTTGCACTGTGATGCTGAGGCATGAAAANNNAGATCGGAA |
| 3006_2 | TATGAGTGTGGAGTCGTTGCACTGTGATGCTGAGGCATGAAAANNNAGATCGGAA |
| 37908_1 | TATGAGTGTGGAGTCGTTGCTTGGAGGTTGCAGTGAACTATCANNNAGATCGGAA |
| 37908_2 | TATGAGTGTGGAGTCGTTGCTTGGAGGTTGCAGTGAACTATCANNNAGATCGGAA |
| 12602_1 | TATGAGTGTGGAGTCGTTGCCTGGTGACCTAAAAAAGAAATAANNNAGATCGGAA |
| 12602_2 | TATGAGTGTGGAGTCGTTGCCTGGTGACCTAAAAAAGAAATAANNNAGATCGGAA |
| 29467_1 | TATGAGTGTGGAGTCGTTGCCAATGTCAGGATCCAGAAAGNNNAGATCGGAAGAG |
| 29467_2 | TATGAGTGTGGAGTCGTTGCCAATGTCAGGATCCAGAAAGNNNAGATCGGAAGAG |
| 32425_1 | TATGAGTGTGGAGTCGTTGCCTCCTGAGTAGCTGGGACTACAGNNNAGATCGGAA |
| 32425_2 | TATGAGTGTGGAGTCGTTGCCTCCTGAGTAGCTGGGACTACAGNNNAGATCGGAA |
| 53309_1 | TATGAGTGTGGAGTCGTTGCCACACACACACACTTCAGTGAATNNNAGATCGGAAG |
| 53309_2 | TATGAGTGTGGAGTCGTTGCCACACACACACACTTCAGTGAATNNNAGATCGGAAG |
| 47271_1 | TATGAGTGTGGAGTCGTTGCTTCACACCATTCTCCTGCCTCNNNAGATCGGAAGAGC |
| 47271_2 | TATGAGTGTGGAGTCGTTGCTTCACACCATTCTCCTGCCTCNNNAGATCGGAAGAGC |
| 50194_1 | TATGAGTGTGGAGTCGTTGCCCAGGAATAGAGACCAGAACCACNNNAGATCGGAA |
| 50194_2 | TATGAGTGTGGAGTCGTTGCCCAGGAATAGAGACCAGAACCACNNNAGATCGGAA |
| 46472_1 | TATGAGTGTGGAGTCGTTGCCATCACGCCCGGCTAATTTTTTTNNNAGATCGGAAGA |
| 46472_2 | TATGAGTGTGGAGTCGTTGCCATCACGCCCGGCTAATTTTTTTNNNAGATCGGAAGA |
| 39906_1 | TATGAGTGTGGAGTCGTTGCTTCCCATAGAAGGCAGCCAAAATNNNAGATCGGAAG |
| 39906_2 | TATGAGTGTGGAGTCGTTGCTTCCCATAGAAGGCAGCCAAAATNNNAGATCGGAAG |
| 38168_1 | TATGAGTGTGGAGTCGTTGCGCAATAGCATAGACAAAGGCCTGNNNAGATCGGAA |
| 38168_2 | TATGAGTGTGGAGTCGTTGCGCAATAGCATAGACAAAGGCCTGNNNAGATCGGAA |
| 46564_1 | TATGAGTGTGGAGTCGTTGCTTCAAGCAATTCTCCTGCCTCAGNNNAGATCGGAAGA |
| 46564_2 | TATGAGTGTGGAGTCGTTGCTTCAAGCAATTCTCCTGCCTCAGNNNAGATCGGAAGA |
| 38093_1 | TATGAGTGTGGAGTCGTTGCGGTTTGAAAGCAAGTGGGTAACANNNAGATCGGAA |
| 38093_2 | TATGAGTGTGGAGTCGTTGCGGTTTGAAAGCAAGTGGGTAACANNNAGATCGGAA |
| 52079_1 | TATGAGTGTGGAGTCGTTGCGGAGAAGGAGGAGACTACAAGGANNNAGATCGGA |
| 52079_2 | TATGAGTGTGGAGTCGTTGCGGAGAAGGAGGAGACTACAAGGANNNAGATCGGA |
| 49371_1 | TATGAGTGTGGAGTCGTTGCTCAATGGTTTCTTTCCCCATCCTNNNAGATCGGAAGA |
| 49371_2 | TATGAGTGTGGAGTCGTTGCTCAATGGTTTCTTTCCCCATCCTNNNAGATCGGAAGA |
| 39066_1 | TATGAGTGTGGAGTCGTTGCAGTGGTTCAAGCAAGATAGCAGANNNAGATCGGAA |
| 39066_2 | TATGAGTGTGGAGTCGTTGCAGTGGTTCAAGCAAGATAGCAGANNNAGATCGGAA |
| 31888_1 | TATGAGTGTGGAGTCGTTGCTACCTTTGGCTGTGACATTGTCTNNNAGATCGGAAG |
| 31888_2 | TATGAGTGTGGAGTCGTTGCTACCTTTGGCTGTGACATTGTCTNNNAGATCGGAAG |

| | |
|---|---|
| 45081_1 | TATGAGTGTGGAGTCGTTGCACAGTCAACTTCCCTTCACAGTTNNNAGATCGGAAGA |
| 45081_2 | TATGAGTGTGGAGTCGTTGCACAGTCAACTTCCCTTCACAGTTNNNAGATCGGAAGA |
| 34727_1 | TATGAGTGTGGAGTCGTTGCTTCAAGTGATTCTCCTGCCTCAGNNNAGATCGGAAG |
| 34727_2 | TATGAGTGTGGAGTCGTTGCTTCAAGTGATTCTCCTGCCTCAGNNNAGATCGGAAG |
| 45973_1 | TATGAGTGTGGAGTCGTTGCCTCTTCCAGGGTTCAACCAACTANNNAGATCGGAAG |
| 45973_2 | TATGAGTGTGGAGTCGTTGCCTCTTCCAGGGTTCAACCAACTANNNAGATCGGAAG |
| 48020_1 | TATGAGTGTGGAGTCGTTGCCCATTCTTCTGCCTCAGACTCTTNNNAGATCGGAAGA |
| 48020_2 | TATGAGTGTGGAGTCGTTGCCCATTCTTCTGCCTCAGACTCTTNNNAGATCGGAAGA |
| 44091_1 | TATGAGTGTGGAGTCGTTGCACTGCTCAAAAGGAACTCAGTGANNNAGATCGGAA |
| 44091_2 | TATGAGTGTGGAGTCGTTGCACTGCTCAAAAGGAACTCAGTGANNNAGATCGGAA |
| 49194_1 | TATGAGTGTGGAGTCGTTGCAGTTTGGTGGGAGTAGTGTTGAANNNAGATCGGAA |
| 49194_2 | TATGAGTGTGGAGTCGTTGCAGTTTGGTGGGAGTAGTGTTGAANNNAGATCGGAA |
| 7994_1 | TATGAGTGTGGAGTCGTTGCGACTGGCATAAGATCCAGCAATCNNNAGATCGGAA |
| 7994_2 | TATGAGTGTGGAGTCGTTGCGACTGGCATAAGATCCAGCAATCNNNAGATCGGAA |
| 13469_1 | TATGAGTGTGGAGTCGTTGCATTTTCTTAGCACCATTTCTTAGNNNAGATCGGAAGA |
| 13469_2 | TATGAGTGTGGAGTCGTTGCATTTTCTTAGCACCATTTCTTAGNNNAGATCGGAAGA |
| 50564_1 | TATGAGTGTGGAGTCGTTGCCCCTTCTCCCCCAAAATACTCAANNNAGATCGGAAGA |
| 50564_2 | TATGAGTGTGGAGTCGTTGCCCCTTCTCCCCCAAAATACTCAANNNAGATCGGAAGA |
| 45070_1 | TATGAGTGTGGAGTCGTTGCTTTATCCTGGTGAAGATGGCCAANNNAGATCGGAA |
| 45070_2 | TATGAGTGTGGAGTCGTTGCTTTATCCTGGTGAAGATGGCCAANNNAGATCGGAA |
| 31475_1 | TATGAGTGTGGAGTCGTTGCCCCCTTCTCCTGTCAGAATCATTNNNAGATCGGAAGA |
| 31475_2 | TATGAGTGTGGAGTCGTTGCCCCCTTCTCCTGTCAGAATCATTNNNAGATCGGAAGA |
| 46919_1 | TATGAGTGTGGAGTCGTTGCGTATGAAGAGCCCACCCTCTTGNNNAGATCGGAAGA |
| 46919_2 | TATGAGTGTGGAGTCGTTGCGTATGAAGAGCCCACCCTCTTGNNNAGATCGGAAGA |
| 28829_1 | TATGAGTGTGGAGTCGTTGCTATAAAACTGTCACCCAGGCTGGNNNAGATCGGAAG |
| 28829_2 | TATGAGTGTGGAGTCGTTGCTATAAAACTGTCACCCAGGCTGGNNNAGATCGGAAG |
| 47021_1 | TATGAGTGTGGAGTCGTTGCGGAGTACATTGGCGCAATCATGNNNAGATCGGAAG |
| 47021_2 | TATGAGTGTGGAGTCGTTGCGGAGTACATTGGCGCAATCATGNNNAGATCGGAAG |
| 44783_1 | TATGAGTGTGGAGTCGTTGCGGACTATCCCAGAATTCGACGCNNNAGATCGGAAGA |
| 44783_2 | TATGAGTGTGGAGTCGTTGCGGACTATCCCAGAATTCGACGCNNNAGATCGGAAGA |
| 37545_1 | TATGAGTGTGGAGTCGTTGCTGGAGTGTAAAACTGTGGGTCTTNNNAGATCGGAA |
| 37545_2 | TATGAGTGTGGAGTCGTTGCTGGAGTGTAAAACTGTGGGTCTTNNNAGATCGGAA |
| 32387_1 | TATGAGTGTGGAGTCGTTGCGGATCTCGGCTCACTGCAAGNNNAGATCGGAAGAG |
| 32387_2 | TATGAGTGTGGAGTCGTTGCGGATCTCGGCTCACTGCAAGNNNAGATCGGAAGAG |
| 43248_1 | TATGAGTGTGGAGTCGTTGCGACACGATCTCAGCTCACTGTAANNNAGATCGGAAG |
| 43248_2 | TATGAGTGTGGAGTCGTTGCGACACGATCTCAGCTCACTGTAANNNAGATCGGAAG |
| 45574_1 | TATGAGTGTGGAGTCGTTGCTGGTTCAAGTCATTCTCTTGCCTNNNAGATCGGAAG |
| 45574_2 | TATGAGTGTGGAGTCGTTGCTGGTTCAAGTCATTCTCTTGCCTNNNAGATCGGAAG |
| 49833_1 | TATGAGTGTGGAGTCGTTGCTTATAACATTTCCTTTGGGGCGCNNNAGATCGGAAG |
| 49833_2 | TATGAGTGTGGAGTCGTTGCTTATAACATTTCCTTTGGGGCGCNNNAGATCGGAAG |
| 48350_1 | TATGAGTGTGGAGTCGTTGCAGTAGATCCCAAAGACAGAAGGCNNNAGATCGGAA |
| 48350_2 | TATGAGTGTGGAGTCGTTGCAGTAGATCCCAAAGACAGAAGGCNNNAGATCGGAA |
| 31571_1 | TATGAGTGTGGAGTCGTTGCCTGCTGTATTTCCACCTTTCACANNNAGATCGGAAGA |

| | |
|---|---|
| 31571_2 | TATGAGTGTGGAGTCGTTGCCTGCTGTATTTCCACCTTTCACANNNAGATCGGAAGA |
| 47267_1 | TATGAGTGTGGAGTCGTTGCTTGTGGCCCAATTGATTTCTTCCNNNAGATCGGAAGA |
| 47267_2 | TATGAGTGTGGAGTCGTTGCTTGTGGCCCAATTGATTTCTTCCNNNAGATCGGAAGA |
| 13582_1 | TATGAGTGTGGAGTCGTTGCAAGAAAAGAGATAAGGTAAAGTTNNNAGATCGGAA |
| 13582_2 | TATGAGTGTGGAGTCGTTGCAAGAAAAGAGATAAGGTAAAGTTNNNAGATCGGAA |
| 36434_1 | TATGAGTGTGGAGTCGTTGCTGACAAAGCAAGACTCCATCTCANNNAGATCGGAAG |
| 36434_2 | TATGAGTGTGGAGTCGTTGCTGACAAAGCAAGACTCCATCTCANNNAGATCGGAAG |
| 34707_1 | TATGAGTGTGGAGTCGTTGCCTGAGTAGCTGGGATTACAGGTGNNNAGATCGGAA |
| 34707_2 | TATGAGTGTGGAGTCGTTGCCTGAGTAGCTGGGATTACAGGTGNNNAGATCGGAA |
| 33048_1 | TATGAGTGTGGAGTCGTTGCGATGGAGTCTCACTCACTCTGTCNNNAGATCGGAAG |
| 33048_2 | TATGAGTGTGGAGTCGTTGCGATGGAGTCTCACTCACTCTGTCNNNAGATCGGAAG |
| 9986_1 | TATGAGTGTGGAGTCGTTGCGGTGAGAGAGCCCAAATCTCCTANNNAGATCGGAA |
| 9986_2 | TATGAGTGTGGAGTCGTTGCGGTGAGAGAGCCCAAATCTCCTANNNAGATCGGAA |
| 28811_1 | TATGAGTGTGGAGTCGTTGCCCTGGATGGAATTGGAGACCATTNNNAGATCGGAA |
| 28811_2 | TATGAGTGTGGAGTCGTTGCCCTGGATGGAATTGGAGACCATTNNNAGATCGGAA |
| 33102_1 | TATGAGTGTGGAGTCGTTGCCATTCTCCTGCCTCAGCCTCNNNAGATCGGAAGAGCA |
| 33102_2 | TATGAGTGTGGAGTCGTTGCCATTCTCCTGCCTCAGCCTCNNNAGATCGGAAGAGCA |
| 13572_1 | TATGAGTGTGGAGTCGTTGCATCTTGCAACAGAGAGATTATCNNNAGATCGGAAGA |
| 13572_2 | TATGAGTGTGGAGTCGTTGCATCTTGCAACAGAGAGATTATCNNNAGATCGGAAGA |
| 38167_1 | TATGAGTGTGGAGTCGTTGCTTTCAGATTAGGACCAAGCCTGGNNNAGATCGGAAG |
| 38167_2 | TATGAGTGTGGAGTCGTTGCTTTCAGATTAGGACCAAGCCTGGNNNAGATCGGAAG |
| 31297_1 | TATGAGTGTGGAGTCGTTGCTAATATTTGGCAAGGAGAGGGGGNNNAGATCGGA |
| 31297_2 | TATGAGTGTGGAGTCGTTGCTAATATTTGGCAAGGAGAGGGGGNNNAGATCGGA |
| 29144_1 | TATGAGTGTGGAGTCGTTGCCTTCACTTGTGTCATGGCAGTTTNNNAGATCGGAAG |
| 29144_2 | TATGAGTGTGGAGTCGTTGCCTTCACTTGTGTCATGGCAGTTTNNNAGATCGGAAG |
| 43661_1 | TATGAGTGTGGAGTCGTTGCAAGGAAAGATTTGGGGTTCCAGANNNAGATCGGAA |
| 43661_2 | TATGAGTGTGGAGTCGTTGCAAGGAAAGATTTGGGGTTCCAGANNNAGATCGGAA |
| 39298_1 | TATGAGTGTGGAGTCGTTGCACACACACACACACAAAACCAATNNNAGATCGGAAG |
| 39298_2 | TATGAGTGTGGAGTCGTTGCACACACACACACACAAAACCAATNNNAGATCGGAAG |
| 53165_1 | TATGAGTGTGGAGTCGTTGCGAGGTCTAACCGGAGGAGGTAGNNNAGATCGGAA |
| 53165_2 | TATGAGTGTGGAGTCGTTGCGAGGTCTAACCGGAGGAGGTAGNNNAGATCGGAA |
| 49969_1 | TATGAGTGTGGAGTCGTTGCGCATAGCACCACTCGAAATGAAANNNAGATCGGAA |
| 49969_2 | TATGAGTGTGGAGTCGTTGCGCATAGCACCACTCGAAATGAAANNNAGATCGGAA |
| 28963_1 | TATGAGTGTGGAGTCGTTGCTGAGTCAGAAACTGCTATGCACTNNNAGATCGGAAG |
| 28963_2 | TATGAGTGTGGAGTCGTTGCTGAGTCAGAAACTGCTATGCACTNNNAGATCGGAAG |
| 41599_1 | TATGAGTGTGGAGTCGTTGCCATTCTCCTGCCTCAGCCTCNNNAGATCGGAAGAGCA |
| 41599_2 | TATGAGTGTGGAGTCGTTGCCATTCTCCTGCCTCAGCCTCNNNAGATCGGAAGAGCA |
| 43028_1 | TATGAGTGTGGAGTCGTTGCATCAAAAAAGGCAAAGAAGGGCANNNAGATCGGAA |
| 43028_2 | TATGAGTGTGGAGTCGTTGCATCAAAAAAGGCAAAGAAGGGCANNNAGATCGGAA |
| 47873_1 | TATGAGTGTGGAGTCGTTGCCCACCAAGCCCAGGTAATTTTTTNNNAGATCGGAAG |
| 47873_2 | TATGAGTGTGGAGTCGTTGCCCACCAAGCCCAGGTAATTTTTTNNNAGATCGGAAG |
| 38012_1 | TATGAGTGTGGAGTCGTTGCAGCCTAATGGAAGATGAAAATGCANNNAGATCGGA |
| 38012_2 | TATGAGTGTGGAGTCGTTGCAGCCTAATGGAAGATGAAAATGCANNNAGATCGGA |

| | |
|---|---|
| 30570_1 | TATGAGTGTGGAGTCGTTGCTCTGTGGTGTTCCCTATGATGTATNNNAGATCGGAA |
| 30570_2 | TATGAGTGTGGAGTCGTTGCTCTGTGGTGTTCCCTATGATGTATNNNAGATCGGAA |
| 48280_1 | TATGAGTGTGGAGTCGTTGCCTTCCCAGGTTCAAGCAATTCTCNNNAGATCGGAAG |
| 48280_2 | TATGAGTGTGGAGTCGTTGCCTTCCCAGGTTCAAGCAATTCTCNNNAGATCGGAAG |
| 28758_1 | TATGAGTGTGGAGTCGTTGCACTGCAGCATCTAAGGTAGTGTCNNNAGATCGGAA |
| 28758_2 | TATGAGTGTGGAGTCGTTGCACTGCAGCATCTAAGGTAGTGTCNNNAGATCGGAA |
| 38538_1 | TATGAGTGTGGAGTCGTTGCCTCACTGTAATCCCAGCACTTTGNNNAGATCGGAAG |
| 38538_2 | TATGAGTGTGGAGTCGTTGCCTCACTGTAATCCCAGCACTTTGNNNAGATCGGAAG |
| 52030_1 | TATGAGTGTGGAGTCGTTGCAATGTTATTCACCTCCCACTGCTNNNAGATCGGAAGA |
| 52030_2 | TATGAGTGTGGAGTCGTTGCAATGTTATTCACCTCCCACTGCTNNNAGATCGGAAGA |
| 41714_1 | TATGAGTGTGGAGTCGTTGCGTTCAACCAATTCTCCTGCCACNNNAGATCGGAAGA |
| 41714_2 | TATGAGTGTGGAGTCGTTGCGTTCAACCAATTCTCCTGCCACNNNAGATCGGAAGA |
| 33076_1 | TATGAGTGTGGAGTCGTTGCCTGGGTTCAATCAATTCTCCTGCNNNAGATCGGAAG |
| 33076_2 | TATGAGTGTGGAGTCGTTGCCTGGGTTCAATCAATTCTCCTGCNNNAGATCGGAAG |
| 35922_1 | TATGAGTGTGGAGTCGTTGCCACAAACACAATAAACGCAGGGANNNAGATCGGAA |
| 35922_2 | TATGAGTGTGGAGTCGTTGCCACAAACACAATAAACGCAGGGANNNAGATCGGAA |
| 42372_1 | TATGAGTGTGGAGTCGTTGCGTTGGAAGCAAGTAAAACTCCCCNNNAGATCGGAA |
| 42372_2 | TATGAGTGTGGAGTCGTTGCGTTGGAAGCAAGTAAAACTCCCCNNNAGATCGGAA |
| 50041_1 | TATGAGTGTGGAGTCGTTGCGTAGCTGGGATTACATATGCCCANNNAGATCGGAA |
| 50041_2 | TATGAGTGTGGAGTCGTTGCGTAGCTGGGATTACATATGCCCANNNAGATCGGAA |
| 41296_1 | TATGAGTGTGGAGTCGTTGCAGCTTTTTCCAGTTCTGTGAAGANNNAGATCGGAAG |
| 41296_2 | TATGAGTGTGGAGTCGTTGCAGCTTTTTCCAGTTCTGTGAAGANNNAGATCGGAAG |
| 50917_1 | TATGAGTGTGGAGTCGTTGCCTCCTTCTCTCTCTCACACACACNNNAGATCGGAAGA |
| 50917_2 | TATGAGTGTGGAGTCGTTGCCTCCTTCTCTCTCTCACACACACNNNAGATCGGAAGA |
| 10395_1 | TATGAGTGTGGAGTCGTTGCAGTTTCTCATAAGGTGGTTGTCGTNNNAGATCGGAA |
| 10395_2 | TATGAGTGTGGAGTCGTTGCAGTTTCTCATAAGGTGGTTGTCGTNNNAGATCGGAA |
| 50411_1 | TATGAGTGTGGAGTCGTTGCGGAGGTTGTAGTGAGCCAAGATGNNNAGATCGGA |
| 50411_2 | TATGAGTGTGGAGTCGTTGCGGAGGTTGTAGTGAGCCAAGATGNNNAGATCGGA |
| 29708_1 | TATGAGTGTGGAGTCGTTGCCAACTTCACAAGGAATTGGAGCTNNNAGATCGGAA |
| 29708_2 | TATGAGTGTGGAGTCGTTGCCAACTTCACAAGGAATTGGAGCTNNNAGATCGGAA |
| 50647_1 | TATGAGTGTGGAGTCGTTGCTAGCTTCAGTAGTAGGACCGTCANNNAGATCGGAA |
| 50647_2 | TATGAGTGTGGAGTCGTTGCTAGCTTCAGTAGTAGGACCGTCANNNAGATCGGAA |
| 39029_1 | TATGAGTGTGGAGTCGTTGCAAGATCACACCACTGCACTCTAGNNNAGATCGGAAG |
| 39029_2 | TATGAGTGTGGAGTCGTTGCAAGATCACACCACTGCACTCTAGNNNAGATCGGAAG |
| 31856_1 | TATGAGTGTGGAGTCGTTGCGAGATTGCAGGCAAGAGTCCTATNNNAGATCGGAA |
| 31856_2 | TATGAGTGTGGAGTCGTTGCGAGATTGCAGGCAAGAGTCCTATNNNAGATCGGAA |
| 39292_1 | TATGAGTGTGGAGTCGTTGCAAAACCTACAGGCCAACATCTCTNNNAGATCGGAAG |
| 39292_2 | TATGAGTGTGGAGTCGTTGCAAAACCTACAGGCCAACATCTCTNNNAGATCGGAAG |
| 31381_1 | TATGAGTGTGGAGTCGTTGCTTGATGCTGAAGTGAATTGTGGANNNAGATCGGAA |
| 31381_2 | TATGAGTGTGGAGTCGTTGCTTGATGCTGAAGTGAATTGTGGANNNAGATCGGAA |
| 49860_1 | TATGAGTGTGGAGTCGTTGCAAGTTGCTAGGTACTTGTGAGGCNNNAGATCGGAA |
| 49860_2 | TATGAGTGTGGAGTCGTTGCAAGTTGCTAGGTACTTGTGAGGCNNNAGATCGGAA |
| 12366_1 | TATGAGTGTGGAGTCGTTGCGAGAGAGAAAGAGAGCGCACATCNNNAGATCGGA |

| | |
|---|---|
| 12366_2 | TATGAGTGTGGAGTCGTTGCGAGAGAGAAAGAGAGCGCACATCNNNAGATCGGAA |
| 32196_1 | TATGAGTGTGGAGTCGTTGCGCTGTCAGACTGGGTGTAAAGTANNNAGATCGGAA |
| 32196_2 | TATGAGTGTGGAGTCGTTGCGCTGTCAGACTGGGTGTAAAGTANNNAGATCGGAA |
| 47327_1 | TATGAGTGTGGAGTCGTTGCGCCTCCCAAGTAGCTAGGATTACNNNAGATCGGAAG |
| 47327_2 | TATGAGTGTGGAGTCGTTGCGCCTCCCAAGTAGCTAGGATTACNNNAGATCGGAAG |
| 39436_1 | TATGAGTGTGGAGTCGTTGCATAATGGAATAGAAGGCCCCACCNNNAGATCGGAA |
| 39436_2 | TATGAGTGTGGAGTCGTTGCATAATGGAATAGAAGGCCCCACCNNNAGATCGGAA |
| 12963_1 | TATGAGTGTGGAGTCGTTGCAGGCAAATCATTTTAATCTTGAANNNAGATCGGAAG |
| 12963_2 | TATGAGTGTGGAGTCGTTGCAGGCAAATCATTTTAATCTTGAANNNAGATCGGAAG |
| 34384_1 | TATGAGTGTGGAGTCGTTGCTACCCAATCTCACCCTATCCCAGNNNAGATCGGAAGA |
| 34384_2 | TATGAGTGTGGAGTCGTTGCTACCCAATCTCACCCTATCCCAGNNNAGATCGGAAGA |
| 53010_1 | TATGAGTGTGGAGTCGTTGCTGGCAAAGGGAGTGAGTTGATATNNNAGATCGGAA |
| 53010_2 | TATGAGTGTGGAGTCGTTGCTGGCAAAGGGAGTGAGTTGATATNNNAGATCGGAA |
| 34819_1 | TATGAGTGTGGAGTCGTTGCCCAAGTAGCTGGGATTATAGGCGNNNAGATCGGAA |
| 34819_2 | TATGAGTGTGGAGTCGTTGCCCAAGTAGCTGGGATTATAGGCGNNNAGATCGGAA |
| 8075_1 | TATGAGTGTGGAGTCGTTGCTTCAAGCCAACTACCCACCTCNNNAGATCGGAAGAG |
| 8075_2 | TATGAGTGTGGAGTCGTTGCTTCAAGCCAACTACCCACCTCNNNAGATCGGAAGAG |
| 45141_1 | TATGAGTGTGGAGTCGTTGCCCTTCTTTCTCCACACCCACTCNNNAGATCGGAAGAG |
| 45141_2 | TATGAGTGTGGAGTCGTTGCCCTTCTTTCTCCACACCCACTCNNNAGATCGGAAGAG |
| 32773_1 | TATGAGTGTGGAGTCGTTGCCATGCTTGTAATCCCAGCACTTTNNNAGATCGGAAG |
| 32773_2 | TATGAGTGTGGAGTCGTTGCCATGCTTGTAATCCCAGCACTTTNNNAGATCGGAAG |
| 10568_1 | TATGAGTGTGGAGTCGTTGCTGAGTAGATGTGGATGCAGCAAGNNNAGATCGGAA |
| 10568_2 | TATGAGTGTGGAGTCGTTGCTGAGTAGATGTGGATGCAGCAAGNNNAGATCGGAA |
| 50779_1 | TATGAGTGTGGAGTCGTTGCCTGAAGAGGATACGGAAGGAACCNNNAGATCGGAA |
| 50779_2 | TATGAGTGTGGAGTCGTTGCCTGAAGAGGATACGGAAGGAACCNNNAGATCGGAA |
| 32307_1 | TATGAGTGTGGAGTCGTTGCACTAGGAAGAATTGGCTCACAGGNNNAGATCGGAA |
| 32307_2 | TATGAGTGTGGAGTCGTTGCACTAGGAAGAATTGGCTCACAGGNNNAGATCGGAA |
| 34108_1 | TATGAGTGTGGAGTCGTTGCACTGTGATTTAGAAGGCTTGCCTNNNAGATCGGAAG |
| 34108_2 | TATGAGTGTGGAGTCGTTGCACTGTGATTTAGAAGGCTTGCCTNNNAGATCGGAAG |
| 42381_1 | TATGAGTGTGGAGTCGTTGCTTACTCCAACACAAAAGCCATGCNNNAGATCGGAAG |
| 42381_2 | TATGAGTGTGGAGTCGTTGCTTACTCCAACACAAAAGCCATGCNNNAGATCGGAAG |
| 10408_1 | TATGAGTGTGGAGTCGTTGCGTGGGTACGTGGAGGTTCATTATNNNAGATCGGAA |
| 10408_2 | TATGAGTGTGGAGTCGTTGCGTGGGTACGTGGAGGTTCATTATNNNAGATCGGAA |
| 49837_1 | TATGAGTGTGGAGTCGTTGCGGCTCCCAGAAGTAGCAGATAAGNNNAGATCGGAA |
| 49837_2 | TATGAGTGTGGAGTCGTTGCGGCTCCCAGAAGTAGCAGATAAGNNNAGATCGGAA |
| 36619_1 | TATGAGTGTGGAGTCGTTGCTTGCAAAACTCTCTAGTCCCACANNNAGATCGGAAG |
| 36619_2 | TATGAGTGTGGAGTCGTTGCTTGCAAAACTCTCTAGTCCCACANNNAGATCGGAAG |
| 49063_1 | TATGAGTGTGGAGTCGTTGCTGCACTCAGACATATGGATGCTTNNNAGATCGGAAG |
| 49063_2 | TATGAGTGTGGAGTCGTTGCTGCACTCAGACATATGGATGCTTNNNAGATCGGAAG |
| 32328_1 | TATGAGTGTGGAGTCGTTGCCACTCCCGACCCCCAAATTAGNNNAGATCGGAAGAG |
| 32328_2 | TATGAGTGTGGAGTCGTTGCCACTCCCGACCCCCAAATTAGNNNAGATCGGAAGAG |
| 48148_1 | TATGAGTGTGGAGTCGTTGCAACTCCTGATCTCAAGTGATCCGNNNAGATCGGAAG |
| 48148_2 | TATGAGTGTGGAGTCGTTGCAACTCCTGATCTCAAGTGATCCGNNNAGATCGGAAG |

| | |
|---|---|
| 13759_1 | TATGAGTGTGGAGTCGTTGCGACAGAATTATGTAGGCTTTATCNNNAGATCGGAAG |
| 13759_2 | TATGAGTGTGGAGTCGTTGCGACAGAATTATGTAGGCTTTATCNNNAGATCGGAAG |
| 51522_1 | TATGAGTGTGGAGTCGTTGCTCTTGCAAATGTCTTTGGCAGTTNNNAGATCGGAAG |
| 51522_2 | TATGAGTGTGGAGTCGTTGCTCTTGCAAATGTCTTTGGCAGTTNNNAGATCGGAAG |
| 43116_1 | TATGAGTGTGGAGTCGTTGCTACTGTGGCTCACACCTGTAATCNNNAGATCGGAAG |
| 43116_2 | TATGAGTGTGGAGTCGTTGCTACTGTGGCTCACACCTGTAATCNNNAGATCGGAAG |
| 45112_1 | TATGAGTGTGGAGTCGTTGCTTACGGTTGCCTAATGACTTCCANNNAGATCGGAAG |
| 45112_2 | TATGAGTGTGGAGTCGTTGCTTACGGTTGCCTAATGACTTCCANNNAGATCGGAAG |
| 12513_1 | TATGAGTGTGGAGTCGTTGCCCAAATAGCTAGGACTCCAGNNNAGATCGGAAGAG |
| 12513_2 | TATGAGTGTGGAGTCGTTGCCCAAATAGCTAGGACTCCAGNNNAGATCGGAAGAG |
| 9386_1 | TATGAGTGTGGAGTCGTTGCATACAATACACTCCACCCCCAGGNNNAGATCGGAAG |
| 9386_2 | TATGAGTGTGGAGTCGTTGCATACAATACACTCCACCCCCAGGNNNAGATCGGAAG |
| 41438_1 | TATGAGTGTGGAGTCGTTGCAAGCTGAACTCATGGGATGGAATNNNAGATCGGAA |
| 41438_2 | TATGAGTGTGGAGTCGTTGCAAGCTGAACTCATGGGATGGAATNNNAGATCGGAA |
| 50894_1 | TATGAGTGTGGAGTCGTTGCCATGCATATCAGAAGGAGTGGGANNNAGATCGGAA |
| 50894_2 | TATGAGTGTGGAGTCGTTGCCATGCATATCAGAAGGAGTGGGANNNAGATCGGAA |
| 42213_1 | TATGAGTGTGGAGTCGTTGCAGCATCCTTAGCAGCTGGAATTANNNAGATCGGAAG |
| 42213_2 | TATGAGTGTGGAGTCGTTGCAGCATCCTTAGCAGCTGGAATTANNNAGATCGGAAG |
| 34605_1 | TATGAGTGTGGAGTCGTTGCAGGTGTTCTTGCTAAACTGACCTNNNAGATCGGAAG |
| 34605_2 | TATGAGTGTGGAGTCGTTGCAGGTGTTCTTGCTAAACTGACCTNNNAGATCGGAAG |
| 53137_1 | TATGAGTGTGGAGTCGTTGCATCACAAACACCCAATCTCTCCANNNAGATCGGAAGA |
| 53137_2 | TATGAGTGTGGAGTCGTTGCATCACAAACACCCAATCTCTCCANNNAGATCGGAAGA |
| 43363_1 | TATGAGTGTGGAGTCGTTGCGTCTCACTTTATTGCCCAACCTGNNNAGATCGGAAGA |
| 43363_2 | TATGAGTGTGGAGTCGTTGCGTCTCACTTTATTGCCCAACCTGNNNAGATCGGAAGA |
| 50829_1 | TATGAGTGTGGAGTCGTTGCTCCCCAAAATAACCCAACCTTGANNNAGATCGGAAG |
| 50829_2 | TATGAGTGTGGAGTCGTTGCTCCCCAAAATAACCCAACCTTGANNNAGATCGGAAG |
| 35562_1 | TATGAGTGTGGAGTCGTTGCCCGAGTAGCTGGGACTACAAATGNNNAGATCGGAA |
| 35562_2 | TATGAGTGTGGAGTCGTTGCCCGAGTAGCTGGGACTACAAATGNNNAGATCGGAA |
| 50706_1 | TATGAGTGTGGAGTCGTTGCTGACTGAGAGTTAGAGAAGCTGCNNNAGATCGGAA |
| 50706_2 | TATGAGTGTGGAGTCGTTGCTGACTGAGAGTTAGAGAAGCTGCNNNAGATCGGAA |
| 32525_1 | TATGAGTGTGGAGTCGTTGCTCTGGAGACATCAACTGACAACCNNNAGATCGGAAG |
| 32525_2 | TATGAGTGTGGAGTCGTTGCTCTGGAGACATCAACTGACAACCNNNAGATCGGAAG |
| 10690_1 | TATGAGTGTGGAGTCGTTGCTACTGAATGACAGCCATCTGGTCNNNAGATCGGAAG |
| 10690_2 | TATGAGTGTGGAGTCGTTGCTACTGAATGACAGCCATCTGGTCNNNAGATCGGAAG |
| 35809_1 | TATGAGTGTGGAGTCGTTGCATGCTAAGTAGGGCACTCTGTTCNNNAGATCGGAAG |
| 35809_2 | TATGAGTGTGGAGTCGTTGCATGCTAAGTAGGGCACTCTGTTCNNNAGATCGGAAG |
| 41663_1 | TATGAGTGTGGAGTCGTTGCCTCCCAGGTTCATGCCATTCTCNNNAGATCGGAAGAG |
| 41663_2 | TATGAGTGTGGAGTCGTTGCCTCCCAGGTTCATGCCATTCTCNNNAGATCGGAAGAG |
| 30254_1 | TATGAGTGTGGAGTCGTTGCACCCCTACGTCTACATCTTCAGANNNAGATCGGAAGA |
| 30254_2 | TATGAGTGTGGAGTCGTTGCACCCCTACGTCTACATCTTCAGANNNAGATCGGAAGA |
| 45497_1 | TATGAGTGTGGAGTCGTTGCGGTAATTTAGGCCCCAAACCAACNNNAGATCGGAAG |
| 45497_2 | TATGAGTGTGGAGTCGTTGCGGTAATTTAGGCCCCAAACCAACNNNAGATCGGAAG |
| 52114_1 | TATGAGTGTGGAGTCGTTGCTTTAGTAGAGATGGGGTTTCGCCNNNAGATCGGAA |

| | |
|---|---|
| 52114_2 | TATGAGTGTGGAGTCGTTGCTTTAGTAGAGATGGGGTTTCGCCNNNAGATCGGAA |
| 32144_1 | TATGAGTGTGGAGTCGTTGCGTGAATGGCATAATTTCGGCTCANNNAGATCGGAAG |
| 32144_2 | TATGAGTGTGGAGTCGTTGCGTGAATGGCATAATTTCGGCTCANNNAGATCGGAAG |
| 38483_1 | TATGAGTGTGGAGTCGTTGCCAGTGAGCTATCATCACACCACTNNNAGATCGGAAG |
| 38483_2 | TATGAGTGTGGAGTCGTTGCCAGTGAGCTATCATCACACCACTNNNAGATCGGAAG |
| 28833_1 | TATGAGTGTGGAGTCGTTGCCAATTTCGGCTCACTGTAACCTCNNNAGATCGGAAG |
| 28833_2 | TATGAGTGTGGAGTCGTTGCCAATTTCGGCTCACTGTAACCTCNNNAGATCGGAAG |
| 31286_1 | TATGAGTGTGGAGTCGTTGCGCAGAACAGTCCACACCATTTACNNNAGATCGGAAG |
| 31286_2 | TATGAGTGTGGAGTCGTTGCGCAGAACAGTCCACACCATTTACNNNAGATCGGAAG |
| 41514_1 | TATGAGTGTGGAGTCGTTGCATCCAATAAGAAACAAGCTGCCCNNNAGATCGGAAG |
| 41514_2 | TATGAGTGTGGAGTCGTTGCATCCAATAAGAAACAAGCTGCCCNNNAGATCGGAAG |
| 47847_1 | TATGAGTGTGGAGTCGTTGCAAAGTGCTGGGATTACAGGCATANNNAGATCGGAA |
| 47847_2 | TATGAGTGTGGAGTCGTTGCAAAGTGCTGGGATTACAGGCATANNNAGATCGGAA |
| 34203_1 | TATGAGTGTGGAGTCGTTGCTAGGAAGAAGGAAGTTCTTGCCCNNNAGATCGGAA |
| 34203_2 | TATGAGTGTGGAGTCGTTGCTAGGAAGAAGGAAGTTCTTGCCCNNNAGATCGGAA |
| 13104_1 | TATGAGTGTGGAGTCGTTGCAGCATCAAAGTGGTAAATAAAATNNNAGATCGGAA |
| 13104_2 | TATGAGTGTGGAGTCGTTGCAGCATCAAAGTGGTAAATAAAATNNNAGATCGGAA |
| 29947_1 | TATGAGTGTGGAGTCGTTGCAGATGCTGATGTATGTGCACAGANNNAGATCGGAA |
| 29947_2 | TATGAGTGTGGAGTCGTTGCAGATGCTGATGTATGTGCACAGANNNAGATCGGAA |
| 45701_1 | TATGAGTGTGGAGTCGTTGCAACAAAATCAAAATGCCAGCCCTNNNAGATCGGAAG |
| 45701_2 | TATGAGTGTGGAGTCGTTGCAACAAAATCAAAATGCCAGCCCTNNNAGATCGGAAG |
| 33708_1 | TATGAGTGTGGAGTCGTTGCAAAACATTTTAGCTGGGGGCAATNNNAGATCGGAA |
| 33708_2 | TATGAGTGTGGAGTCGTTGCAAAACATTTTAGCTGGGGGCAATNNNAGATCGGAA |
| 43992_1 | TATGAGTGTGGAGTCGTTGCCTATCTGCTTGGTGAATCCCTGTNNNAGATCGGAAG |
| 43992_2 | TATGAGTGTGGAGTCGTTGCCTATCTGCTTGGTGAATCCCTGTNNNAGATCGGAAG |
| 46180_1 | TATGAGTGTGGAGTCGTTGCGGCTCTCCAGCGTGAGGATGNNNAGATCGGAAGAG |
| 46180_2 | TATGAGTGTGGAGTCGTTGCGGCTCTCCAGCGTGAGGATGNNNAGATCGGAAGAG |
| 44945_1 | TATGAGTGTGGAGTCGTTGCCGAGATAACCCAGGCTGGAATTANNNAGATCGGAA |
| 44945_2 | TATGAGTGTGGAGTCGTTGCCGAGATAACCCAGGCTGGAATTANNNAGATCGGAA |
| 43239_1 | TATGAGTGTGGAGTCGTTGCCGTGGGGATTACAATTCGACATGNNNAGATCGGAA |
| 43239_2 | TATGAGTGTGGAGTCGTTGCCGTGGGGATTACAATTCGACATGNNNAGATCGGAA |
| 35232_1 | TATGAGTGTGGAGTCGTTGCCCTTCATTGCGGATTCATGATCCNNNAGATCGGAAG |
| 35232_2 | TATGAGTGTGGAGTCGTTGCCCTTCATTGCGGATTCATGATCCNNNAGATCGGAAG |
| 49732_1 | TATGAGTGTGGAGTCGTTGCGCAATTATCCTTCCTCACGCTTCNNNAGATCGGAAGA |
| 49732_2 | TATGAGTGTGGAGTCGTTGCGCAATTATCCTTCCTCACGCTTCNNNAGATCGGAAGA |
| 46474_1 | TATGAGTGTGGAGTCGTTGCCAGCGCAACAAGACTTCATATGTNNNAGATCGGAAG |
| 46474_2 | TATGAGTGTGGAGTCGTTGCCAGCGCAACAAGACTTCATATGTNNNAGATCGGAAG |
| 44245_1 | TATGAGTGTGGAGTCGTTGCAGGGTATGCTGTAATGTGGGAAGNNNAGATCGGA |
| 44245_2 | TATGAGTGTGGAGTCGTTGCAGGGTATGCTGTAATGTGGGAAGNNNAGATCGGA |
| 44548_1 | TATGAGTGTGGAGTCGTTGCGTTTAGTGGTGGTGATGTTGGTGNNNAGATCGGAA |
| 44548_2 | TATGAGTGTGGAGTCGTTGCGTTTAGTGGTGGTGATGTTGGTGNNNAGATCGGAA |
| 9486_1 | TATGAGTGTGGAGTCGTTGCTGACTACCACCAAATGAGCACAGNNNAGATCGGAAG |
| 9486_2 | TATGAGTGTGGAGTCGTTGCTGACTACCACCAAATGAGCACAGNNNAGATCGGAAG |

| | |
|---|---|
| 32850_1 | TATGAGTGTGGAGTCGTTGCGATTCAAGCAATTCTCCTGCCTCNNNAGATCGGAAG |
| 32850_2 | TATGAGTGTGGAGTCGTTGCGATTCAAGCAATTCTCCTGCCTCNNNAGATCGGAAG |
| 35747_1 | TATGAGTGTGGAGTCGTTGCGTTTCACTGTGTTAGCCAGGATGNNNAGATCGGAA |
| 35747_2 | TATGAGTGTGGAGTCGTTGCGTTTCACTGTGTTAGCCAGGATGNNNAGATCGGAA |
| 33688_1 | TATGAGTGTGGAGTCGTTGCGGATCTTGCTATGTTTTCCAGGCNNNAGATCGGAAG |
| 33688_2 | TATGAGTGTGGAGTCGTTGCGGATCTTGCTATGTTTTCCAGGCNNNAGATCGGAAG |
| 36289_1 | TATGAGTGTGGAGTCGTTGCACAGTCCTCCAAATTGCCAAGTANNNAGATCGGAAG |
| 36289_2 | TATGAGTGTGGAGTCGTTGCACAGTCCTCCAAATTGCCAAGTANNNAGATCGGAAG |
| 49120_1 | TATGAGTGTGGAGTCGTTGCGGCCGCATGGAAGATAATGAAAANNNAGATCGGAA |
| 49120_2 | TATGAGTGTGGAGTCGTTGCGGCCGCATGGAAGATAATGAAAANNNAGATCGGAA |
| 43411_1 | TATGAGTGTGGAGTCGTTGCCCACCACTCCCAGCTAATTTTTTNNNAGATCGGAAGA |
| 43411_2 | TATGAGTGTGGAGTCGTTGCCCACCACTCCCAGCTAATTTTTTNNNAGATCGGAAGA |
| 36975_1 | TATGAGTGTGGAGTCGTTGCTGGAAATATTGCCACAGGGAACTNNNAGATCGGAA |
| 36975_2 | TATGAGTGTGGAGTCGTTGCTGGAAATATTGCCACAGGGAACTNNNAGATCGGAA |
| 40760_1 | TATGAGTGTGGAGTCGTTGCACTATCAAGTTGGTTCCATCCCCNNNAGATCGGAAG |
| 40760_2 | TATGAGTGTGGAGTCGTTGCACTATCAAGTTGGTTCCATCCCCNNNAGATCGGAAG |
| 46023_1 | TATGAGTGTGGAGTCGTTGCCTCCTGGGTTCATGGCATTCTCNNNAGATCGGAAGA |
| 46023_2 | TATGAGTGTGGAGTCGTTGCCTCCTGGGTTCATGGCATTCTCNNNAGATCGGAAGA |
| 50017_1 | TATGAGTGTGGAGTCGTTGCCGCGGAATCTGTTGACATTTACTNNNAGATCGGAAG |
| 50017_2 | TATGAGTGTGGAGTCGTTGCCGCGGAATCTGTTGACATTTACTNNNAGATCGGAAG |
| 37120_1 | TATGAGTGTGGAGTCGTTGCCATCCAGCTGAGTACTCTCTGTGNNNAGATCGGAAG |
| 37120_2 | TATGAGTGTGGAGTCGTTGCCATCCAGCTGAGTACTCTCTGTGNNNAGATCGGAAG |
| 36322_1 | TATGAGTGTGGAGTCGTTGCAAAAACCAAGTCAGAAGTTGGGGNNNAGATCGGAA |
| 36322_2 | TATGAGTGTGGAGTCGTTGCAAAAACCAAGTCAGAAGTTGGGGNNNAGATCGGAA |
| 33984_1 | TATGAGTGTGGAGTCGTTGCCAAGTAGCTGGGACTACAGGCNNNAGATCGGAAGA |
| 33984_2 | TATGAGTGTGGAGTCGTTGCCAAGTAGCTGGGACTACAGGCNNNAGATCGGAAGA |
| 32417_1 | TATGAGTGTGGAGTCGTTGCCACCTGCAGTTCAAGCTACTTGNNNAGATCGGAAGA |
| 32417_2 | TATGAGTGTGGAGTCGTTGCCACCTGCAGTTCAAGCTACTTGNNNAGATCGGAAGA |
| 45468_1 | TATGAGTGTGGAGTCGTTGCGGGTCTGTACAAGAGAAGGTTGTNNNAGATCGGAA |
| 45468_2 | TATGAGTGTGGAGTCGTTGCGGGTCTGTACAAGAGAAGGTTGTNNNAGATCGGAA |
| 34806_1 | TATGAGTGTGGAGTCGTTGCTTCACACCATTCTCCTGCCTCNNNAGATCGGAAGAGC |
| 34806_2 | TATGAGTGTGGAGTCGTTGCTTCACACCATTCTCCTGCCTCNNNAGATCGGAAGAGC |
| 53233_1 | TATGAGTGTGGAGTCGTTGCTCGGCTTCTTTTTGTCTTCCTCTNNNAGATCGGAAGA |
| 53233_2 | TATGAGTGTGGAGTCGTTGCTCGGCTTCTTTTTGTCTTCCTCTNNNAGATCGGAAGA |
| 44120_1 | TATGAGTGTGGAGTCGTTGCCACTGCGGAAAAAAGAGGGAATCNNNAGATCGGAA |
| 44120_2 | TATGAGTGTGGAGTCGTTGCCACTGCGGAAAAAAGAGGGAATCNNNAGATCGGAA |
| 36060_1 | TATGAGTGTGGAGTCGTTGCAAGAATCACTTCAACCCGGGAGNNNAGATCGGAAG |
| 36060_2 | TATGAGTGTGGAGTCGTTGCAAGAATCACTTCAACCCGGGAGNNNAGATCGGAAG |
| 10284_1 | TATGAGTGTGGAGTCGTTGCATTCCCAGTCCACCAATACCAGANNNAGATCGGAAG |
| 10284_2 | TATGAGTGTGGAGTCGTTGCATTCCCAGTCCACCAATACCAGANNNAGATCGGAAG |
| 49777_1 | TATGAGTGTGGAGTCGTTGCGTGATTTCCTTTTGCTTGGAGCTNNNAGATCGGAAG |
| 49777_2 | TATGAGTGTGGAGTCGTTGCGTGATTTCCTTTTGCTTGGAGCTNNNAGATCGGAAG |
| 28890_1 | TATGAGTGTGGAGTCGTTGCCCACTTGTTTTGGTACTGCTTGTNNNAGATCGGAAG |

| | |
|---|---|
| 28890_2 | TATGAGTGTGGAGTCGTTGCCCACTTGTTTTGGTACTGCTTGTNNNAGATCGGAAG |
| 37158_1 | TATGAGTGTGGAGTCGTTGCAAGGGCCCTGTATCAGAGTATCANNNAGATCGGAAG |
| 37158_2 | TATGAGTGTGGAGTCGTTGCAAGGGCCCTGTATCAGAGTATCANNNAGATCGGAAG |
| 47486_1 | TATGAGTGTGGAGTCGTTGCACCATGCCCAGCTAATTGTTTTCNNNAGATCGGAAG |
| 47486_2 | TATGAGTGTGGAGTCGTTGCACCATGCCCAGCTAATTGTTTTCNNNAGATCGGAAG |
| 10863_1 | TATGAGTGTGGAGTCGTTGCTCAGGGTAGGTCAAGGAACTGTTNNNAGATCGGAA |
| 10863_2 | TATGAGTGTGGAGTCGTTGCTCAGGGTAGGTCAAGGAACTGTTNNNAGATCGGAA |
| 52399_1 | TATGAGTGTGGAGTCGTTGCTACAGGTTTGATCCTTACACGGGNNNAGATCGGAAG |
| 52399_2 | TATGAGTGTGGAGTCGTTGCTACAGGTTTGATCCTTACACGGGNNNAGATCGGAAG |
| 38301_1 | TATGAGTGTGGAGTCGTTGCACCAATAAAAGAGCAGTAGCAGCNNNAGATCGGAA |
| 38301_2 | TATGAGTGTGGAGTCGTTGCACCAATAAAAGAGCAGTAGCAGCNNNAGATCGGAA |
| 35764_1 | TATGAGTGTGGAGTCGTTGCATTACAGGAGTGAGCCAACACATNNNAGATCGGAAG |
| 35764_2 | TATGAGTGTGGAGTCGTTGCATTACAGGAGTGAGCCAACACATNNNAGATCGGAAG |
| 38721_1 | TATGAGTGTGGAGTCGTTGCCCTAATCAAGGAGGTGAAAGACCNNNAGATCGGAA |
| 38721_2 | TATGAGTGTGGAGTCGTTGCCCTAATCAAGGAGGTGAAAGACCNNNAGATCGGAA |
| 40859_1 | TATGAGTGTGGAGTCGTTGCGATGCGAACACATGGACACATAGNNNAGATCGGAA |
| 40859_2 | TATGAGTGTGGAGTCGTTGCGATGCGAACACATGGACACATAGNNNAGATCGGAA |
| 30743_1 | TATGAGTGTGGAGTCGTTGCGACTGTGGGAAATTTGAGACAGCNNNAGATCGGAA |
| 30743_2 | TATGAGTGTGGAGTCGTTGCGACTGTGGGAAATTTGAGACAGCNNNAGATCGGAA |
| 47675_1 | TATGAGTGTGGAGTCGTTGCGTTCAAGAGATTCTCCTGCCTCANNNAGATCGGAAG |
| 47675_2 | TATGAGTGTGGAGTCGTTGCGTTCAAGAGATTCTCCTGCCTCANNNAGATCGGAAG |
| 38955_1 | TATGAGTGTGGAGTCGTTGCAGCCAAAGAACTGAGTAGACACCNNNAGATCGGAA |
| 38955_2 | TATGAGTGTGGAGTCGTTGCAGCCAAAGAACTGAGTAGACACCNNNAGATCGGAA |
| 36428_1 | TATGAGTGTGGAGTCGTTGCTAATCCCATCAACATCCTCTGCCNNNAGATCGGAAGA |
| 36428_2 | TATGAGTGTGGAGTCGTTGCTAATCCCATCAACATCCTCTGCCNNNAGATCGGAAGA |
| 33834_1 | TATGAGTGTGGAGTCGTTGCGCAGGAATATAAGGAAAGCTGTGGNNNAGATCGG |
| 33834_2 | TATGAGTGTGGAGTCGTTGCGCAGGAATATAAGGAAAGCTGTGGNNNAGATCGG |
| 2017_1 | TATGAGTGTGGAGTCGTTGCACCTGACGATCCATGAGAGACAANNNAGATCGGAAG |
| 2017_2 | TATGAGTGTGGAGTCGTTGCACCTGACGATCCATGAGAGACAANNNAGATCGGAAG |
| 38193_1 | TATGAGTGTGGAGTCGTTGCTGGATTACCAGGGAGAGACTCTTNNNAGATCGGAA |
| 38193_2 | TATGAGTGTGGAGTCGTTGCTGGATTACCAGGGAGAGACTCTTNNNAGATCGGAA |
| 29271_1 | TATGAGTGTGGAGTCGTTGCCACATTCAGTCCATAACAGCCACNNNAGATCGGAAG |
| 29271_2 | TATGAGTGTGGAGTCGTTGCCACATTCAGTCCATAACAGCCACNNNAGATCGGAAG |
| 43452_1 | TATGAGTGTGGAGTCGTTGCCTCCTGAGTAGCTGGGACTACANNNAGATCGGAAG |
| 43452_2 | TATGAGTGTGGAGTCGTTGCCTCCTGAGTAGCTGGGACTACANNNAGATCGGAAGA |
| 41665_1 | TATGAGTGTGGAGTCGTTGCGAGACGGAGTCTTGCTCTGTCNNNAGATCGGAAGA |
| 41665_2 | TATGAGTGTGGAGTCGTTGCGAGACGGAGTCTTGCTCTGTCNNNAGATCGGAAGA |
| 37714_1 | TATGAGTGTGGAGTCGTTGCGCAGAGTCATATTTTGGAGGAGCNNNAGATCGGAA |
| 37714_2 | TATGAGTGTGGAGTCGTTGCGCAGAGTCATATTTTGGAGGAGCNNNAGATCGGAA |
| 42046_1 | TATGAGTGTGGAGTCGTTGCCACCCCATTCTAAAAAGCAGGTGNNNAGATCGGAAG |
| 42046_2 | TATGAGTGTGGAGTCGTTGCCACCCCATTCTAAAAAGCAGGTGNNNAGATCGGAAG |
| 37812_1 | TATGAGTGTGGAGTCGTTGCGTGGATCACTTGAGGTCAGAAGTNNNAGATCGGAA |
| 37812_2 | TATGAGTGTGGAGTCGTTGCGTGGATCACTTGAGGTCAGAAGTNNNAGATCGGAA |

| | |
|---|---|
| 43515_1 | TATGAGTGTGGAGTCGTTGCCAAGTAGCTGGGATTACAGGCATNNNAGATCGGAA |
| 43515_2 | TATGAGTGTGGAGTCGTTGCCAAGTAGCTGGGATTACAGGCATNNNAGATCGGAA |
| 49024_1 | TATGAGTGTGGAGTCGTTGCTGTAGTGCAGTGGTGTGATCTCNNNAGATCGGAAG |
| 49024_2 | TATGAGTGTGGAGTCGTTGCTGTAGTGCAGTGGTGTGATCTCNNNAGATCGGAAG |
| 29193_1 | TATGAGTGTGGAGTCGTTGCAAATGTGGATCATGCGAGAGACTNNNAGATCGGAA |
| 29193_2 | TATGAGTGTGGAGTCGTTGCAAATGTGGATCATGCGAGAGACTNNNAGATCGGAA |
| 49977_1 | TATGAGTGTGGAGTCGTTGCTCTACCCACTTCCCTACCATTCTNNNAGATCGGAAGA |
| 49977_2 | TATGAGTGTGGAGTCGTTGCTCTACCCACTTCCCTACCATTCTNNNAGATCGGAAGA |
| 35309_1 | TATGAGTGTGGAGTCGTTGCAGATGGAATGGAGTGCAGTTGTANNNAGATCGGAA |
| 35309_2 | TATGAGTGTGGAGTCGTTGCAGATGGAATGGAGTGCAGTTGTANNNAGATCGGAA |
| 52234_1 | TATGAGTGTGGAGTCGTTGCTCGGATGTATTCCAAGTGCCTAGNNNAGATCGGAAG |
| 52234_2 | TATGAGTGTGGAGTCGTTGCTCGGATGTATTCCAAGTGCCTAGNNNAGATCGGAAG |
| 43189_1 | TATGAGTGTGGAGTCGTTGCTTCAAGTGATTCTCCTGCCTCAGNNNAGATCGGAAG |
| 43189_2 | TATGAGTGTGGAGTCGTTGCTTCAAGTGATTCTCCTGCCTCAGNNNAGATCGGAAG |
| 31046_1 | TATGAGTGTGGAGTCGTTGCCCTTTCCAAAACTCACCTTTGCTNNNAGATCGGAAGA |
| 31046_2 | TATGAGTGTGGAGTCGTTGCCCTTTCCAAAACTCACCTTTGCTNNNAGATCGGAAGA |
| 43753_1 | TATGAGTGTGGAGTCGTTGCTCCCCCTAAAAATGTGTCCGTTCNNNAGATCGGAAG |
| 43753_2 | TATGAGTGTGGAGTCGTTGCTCCCCCTAAAAATGTGTCCGTTCNNNAGATCGGAAG |
| 42926_1 | TATGAGTGTGGAGTCGTTGCGACTGTCCTTTGTGTCACTAGGTNNNAGATCGGAAG |
| 42926_2 | TATGAGTGTGGAGTCGTTGCGACTGTCCTTTGTGTCACTAGGTNNNAGATCGGAAG |
| 34413_1 | TATGAGTGTGGAGTCGTTGCCTGCCTAACATTCCGACCTCATANNNAGATCGGAAGA |
| 34413_2 | TATGAGTGTGGAGTCGTTGCCTGCCTAACATTCCGACCTCATANNNAGATCGGAAGA |
| 36829_1 | TATGAGTGTGGAGTCGTTGCACAAGTCCCTGTAATGCCTTGAANNNAGATCGGAAG |
| 36829_2 | TATGAGTGTGGAGTCGTTGCACAAGTCCCTGTAATGCCTTGAANNNAGATCGGAAG |
| 33483_1 | TATGAGTGTGGAGTCGTTGCTGGACTCAATGTGAATTTTTGAGANNNAGATCGGAA |
| 33483_2 | TATGAGTGTGGAGTCGTTGCTGGACTCAATGTGAATTTTTGAGANNNAGATCGGAA |
| 48568_1 | TATGAGTGTGGAGTCGTTGCTTCAAGCAATTCTCTTGCCTTGGNNNAGATCGGAAG |
| 48568_2 | TATGAGTGTGGAGTCGTTGCTTCAAGCAATTCTCTTGCCTTGGNNNAGATCGGAAG |
| 31071_1 | TATGAGTGTGGAGTCGTTGCCATTTTTGAGACACGGTCATGCTNNNAGATCGGAAG |
| 31071_2 | TATGAGTGTGGAGTCGTTGCCATTTTTGAGACACGGTCATGCTNNNAGATCGGAAG |
| 51387_1 | TATGAGTGTGGAGTCGTTGCGCCATTCCCTCATTATCTCAGGTNNNAGATCGGAAGA |
| 51387_2 | TATGAGTGTGGAGTCGTTGCGCCATTCCCTCATTATCTCAGGTNNNAGATCGGAAGA |
| 9913_1 | TATGAGTGTGGAGTCGTTGCAATGAGTGTAGGAGGTGCTCAGANNNAGATCGGAA |
| 9913_2 | TATGAGTGTGGAGTCGTTGCAATGAGTGTAGGAGGTGCTCAGANNNAGATCGGAA |
| 44911_1 | TATGAGTGTGGAGTCGTTGCAAGAAAGACTTGCTCACACATGCNNNAGATCGGAAG |
| 44911_2 | TATGAGTGTGGAGTCGTTGCAAGAAAGACTTGCTCACACATGCNNNAGATCGGAAG |
| 41627_1 | TATGAGTGTGGAGTCGTTGCCAAGTAGCTGGGATTACAGGCATNNNAGATCGGAA |
| 41627_2 | TATGAGTGTGGAGTCGTTGCCAAGTAGCTGGGATTACAGGCATNNNAGATCGGAA |
| 41604_1 | TATGAGTGTGGAGTCGTTGCCTCCTGAGCTCAAGGGATTCTGNNNAGATCGGAAGA |
| 41604_2 | TATGAGTGTGGAGTCGTTGCCTCCTGAGCTCAAGGGATTCTGNNNAGATCGGAAGA |
| 33967_1 | TATGAGTGTGGAGTCGTTGCGATCATGGGTCACTACAGCCTCNNNAGATCGGAAGA |
| 33967_2 | TATGAGTGTGGAGTCGTTGCGATCATGGGTCACTACAGCCTCNNNAGATCGGAAGA |
| 49478_1 | TATGAGTGTGGAGTCGTTGCTTACTTCACATGTCACGTGACCTNNNAGATCGGAAG |

| | |
|---|---|
| 49478_2 | TATGAGTGTGGAGTCGTTGCTTACTTCACATGTCACGTGACCTNNNAGATCGGAAGA |
| 31600_1 | TATGAGTGTGGAGTCGTTGCGGAGGGAAGTACAGCGTTGATCNNNAGATCGGAAG |
| 31600_2 | TATGAGTGTGGAGTCGTTGCGGAGGGAAGTACAGCGTTGATCNNNAGATCGGAAG |
| 37725_1 | TATGAGTGTGGAGTCGTTGCGTGGATCACAAGGTCAGGAGATTNNNAGATCGGAA |
| 37725_2 | TATGAGTGTGGAGTCGTTGCGTGGATCACAAGGTCAGGAGATTNNNAGATCGGAA |
| 34772_1 | TATGAGTGTGGAGTCGTTGCTTCAAGCTATCCTCATGCCTCAGNNNAGATCGGAAGA |
| 34772_2 | TATGAGTGTGGAGTCGTTGCTTCAAGCTATCCTCATGCCTCAGNNNAGATCGGAAGA |
| 43457_1 | TATGAGTGTGGAGTCGTTGCGAGTAACTGGGATTACAGGTGCTNNNAGATCGGAAG |
| 43457_2 | TATGAGTGTGGAGTCGTTGCGAGTAACTGGGATTACAGGTGCTNNNAGATCGGAAG |
| 43642_1 | TATGAGTGTGGAGTCGTTGCGGAACATTCCGGTCATTCTCTCTNNNAGATCGGAAGA |
| 43642_2 | TATGAGTGTGGAGTCGTTGCGGAACATTCCGGTCATTCTCTCTNNNAGATCGGAAGA |
| 39975_1 | TATGAGTGTGGAGTCGTTGCGAGTTTATGGCCTAGTGAGAGCANNNAGATCGGAAG |
| 39975_2 | TATGAGTGTGGAGTCGTTGCGAGTTTATGGCCTAGTGAGAGCANNNAGATCGGAAG |
| 33682_1 | TATGAGTGTGGAGTCGTTGCAATGATTCTCCTGCCTCAGTCTCNNNAGATCGGAAGA |
| 33682_2 | TATGAGTGTGGAGTCGTTGCAATGATTCTCCTGCCTCAGTCTCNNNAGATCGGAAGA |
| 42987_1 | TATGAGTGTGGAGTCGTTGCCTCCTGAGTAGCTGGGACTACAGNNNAGATCGGAAG |
| 42987_2 | TATGAGTGTGGAGTCGTTGCCTCCTGAGTAGCTGGGACTACAGNNNAGATCGGAAG |
| 49695_1 | TATGAGTGTGGAGTCGTTGCTTGGTCTCCTCATCATTCACCTGNNNAGATCGGAAGA |
| 49695_2 | TATGAGTGTGGAGTCGTTGCTTGGTCTCCTCATCATTCACCTGNNNAGATCGGAAGA |
| 46193_1 | TATGAGTGTGGAGTCGTTGCCTCAGGGAGGTTGCTTTCTTTTGNNNAGATCGGAAG |
| 46193_2 | TATGAGTGTGGAGTCGTTGCCTCAGGGAGGTTGCTTTCTTTTGNNNAGATCGGAAG |
| 43843_1 | TATGAGTGTGGAGTCGTTGCATCATAAAGGGCCATCAGGACAANNNAGATCGGAAG |
| 43843_2 | TATGAGTGTGGAGTCGTTGCATCATAAAGGGCCATCAGGACAANNNAGATCGGAAG |
| 38072_1 | TATGAGTGTGGAGTCGTTGCAAAATACAAAATCAGCCAGGCGTNNNAGATCGGAAG |
| 38072_2 | TATGAGTGTGGAGTCGTTGCAAAATACAAAATCAGCCAGGCGTNNNAGATCGGAAG |
| 47678_1 | TATGAGTGTGGAGTCGTTGCCAATCATAGCTCACTGCAACCTCNNNAGATCGGAAGA |
| 47678_2 | TATGAGTGTGGAGTCGTTGCCAATCATAGCTCACTGCAACCTCNNNAGATCGGAAGA |
| 38364_1 | TATGAGTGTGGAGTCGTTGCGAGAATGGCATGAACCCAGGAGNNNAGATCGGAAG |
| 38364_2 | TATGAGTGTGGAGTCGTTGCGAGAATGGCATGAACCCAGGAGNNNAGATCGGAAG |
| 46290_1 | TATGAGTGTGGAGTCGTTGCTCTTAGCTCACTACAACCTGTGCNNNAGATCGGAAGA |
| 46290_2 | TATGAGTGTGGAGTCGTTGCTCTTAGCTCACTACAACCTGTGCNNNAGATCGGAAGA |
| 38413_1 | TATGAGTGTGGAGTCGTTGCCAAAAATTACCCAGGTGTGGTGGNNNAGATCGGAAG |
| 38413_2 | TATGAGTGTGGAGTCGTTGCCAAAAATTACCCAGGTGTGGTGGNNNAGATCGGAAG |
| 43454_1 | TATGAGTGTGGAGTCGTTGCAGATCTCATCTCACTGCAACCTCNNNAGATCGGAAGA |
| 43454_2 | TATGAGTGTGGAGTCGTTGCAGATCTCATCTCACTGCAACCTCNNNAGATCGGAAGA |
| 28907_1 | TATGAGTGTGGAGTCGTTGCTGTGAATGCACCTCCTAAATTGCNNNAGATCGGAAGA |
| 28907_2 | TATGAGTGTGGAGTCGTTGCTGTGAATGCACCTCCTAAATTGCNNNAGATCGGAAGA |
| 5844_1 | TATGAGTGTGGAGTCGTTGCGGGCGGTAAGGCTCAGGCATGNNNAGATCGGAAGA |
| 5844_2 | TATGAGTGTGGAGTCGTTGCGGGCGGTAAGGCTCAGGCATGNNNAGATCGGAAGA |
| 44904_1 | TATGAGTGTGGAGTCGTTGCAGGTGATGACATCGGAAAAGTCANNNAGATCGGAA |
| 44904_2 | TATGAGTGTGGAGTCGTTGCAGGTGATGACATCGGAAAAGTCANNNAGATCGGAA |
| 40294_1 | TATGAGTGTGGAGTCGTTGCGAGGCTGAGGCAGGAGAATGNNNAGATCGGAAGA |
| 40294_2 | TATGAGTGTGGAGTCGTTGCGAGGCTGAGGCAGGAGAATGNNNAGATCGGAAGA |

| | |
|---|---|
| 32349_1 | TATGAGTGTGGAGTCGTTGCGTGGTGTAGATCTGTGGGAACTCNNNAGATCGGAA |
| 32349_2 | TATGAGTGTGGAGTCGTTGCGTGGTGTAGATCTGTGGGAACTCNNNAGATCGGAA |
| 43444_1 | TATGAGTGTGGAGTCGTTGCCTGAGTAGCTGGAATTACAGGCANNNAGATCGGAA |
| 43444_2 | TATGAGTGTGGAGTCGTTGCCTGAGTAGCTGGAATTACAGGCANNNAGATCGGAA |
| 49282_1 | TATGAGTGTGGAGTCGTTGCCACCGTTCCTGGCTAATTTTTGTNNNAGATCGGAAG |
| 49282_2 | TATGAGTGTGGAGTCGTTGCCACCGTTCCTGGCTAATTTTTGTNNNAGATCGGAAG |
| 40447_1 | TATGAGTGTGGAGTCGTTGCAAACAAGGTGCTATTGGGCAATCNNNAGATCGGAA |
| 40447_2 | TATGAGTGTGGAGTCGTTGCAAACAAGGTGCTATTGGGCAATCNNNAGATCGGAA |
| 38369_1 | TATGAGTGTGGAGTCGTTGCGCTTATCCACCACGATCAATTCGNNNAGATCGGAAG |
| 38369_2 | TATGAGTGTGGAGTCGTTGCGCTTATCCACCACGATCAATTCGNNNAGATCGGAAG |
| 45315_1 | TATGAGTGTGGAGTCGTTGCATGCTTTATTCTTGTCAGGCAGCNNNAGATCGGAAG |
| 45315_2 | TATGAGTGTGGAGTCGTTGCATGCTTTATTCTTGTCAGGCAGCNNNAGATCGGAAG |
| 31246_1 | TATGAGTGTGGAGTCGTTGCTGTGCCCTTTCTGAAGATTAGCTNNNAGATCGGAAG |
| 31246_2 | TATGAGTGTGGAGTCGTTGCTGTGCCCTTTCTGAAGATTAGCTNNNAGATCGGAAG |
| 13571_1 | TATGAGTGTGGAGTCGTTGCAGGTACCACTAGTCAAGGATNNNAGATCGGAAGAG |
| 13571_2 | TATGAGTGTGGAGTCGTTGCAGGTACCACTAGTCAAGGATNNNAGATCGGAAGAG |
| 35573_1 | TATGAGTGTGGAGTCGTTGCGGTTCACAAAATTCTCCTGCCTCNNNAGATCGGAAG |
| 35573_2 | TATGAGTGTGGAGTCGTTGCGGTTCACAAAATTCTCCTGCCTCNNNAGATCGGAAG |
| 29987_1 | TATGAGTGTGGAGTCGTTGCTGCCAATTCTCCACCACATTTACNNNAGATCGGAAGA |
| 29987_2 | TATGAGTGTGGAGTCGTTGCTGCCAATTCTCCACCACATTTACNNNAGATCGGAAGA |
| 29668_1 | TATGAGTGTGGAGTCGTTGCGAAGGGATCTCAGCTCACTGTAGNNNAGATCGGAA |
| 29668_2 | TATGAGTGTGGAGTCGTTGCGAAGGGATCTCAGCTCACTGTAGNNNAGATCGGAA |
| 39244_1 | TATGAGTGTGGAGTCGTTGCTTACGACATGGATGAGCCTTGAGNNNAGATCGGAA |
| 39244_2 | TATGAGTGTGGAGTCGTTGCTTACGACATGGATGAGCCTTGAGNNNAGATCGGAA |
| 41029_1 | TATGAGTGTGGAGTCGTTGCGAGGAAAGGGCAGTCTTTTCAACNNNAGATCGGAA |
| 41029_2 | TATGAGTGTGGAGTCGTTGCGAGGAAAGGGCAGTCTTTTCAACNNNAGATCGGAA |
| 39723_1 | TATGAGTGTGGAGTCGTTGCAACCTTCAAAAACTACCCACACTNNNAGATCGGAAG |
| 39723_2 | TATGAGTGTGGAGTCGTTGCAACCTTCAAAAACTACCCACACTNNNAGATCGGAAG |
| 13340_1 | TATGAGTGTGGAGTCGTTGCGGATATAGCAGAGAGAGGTTNNNAGATCGGAAGA |
| 13340_2 | TATGAGTGTGGAGTCGTTGCGGATATAGCAGAGAGAGGTTNNNAGATCGGAAGA |
| 40944_1 | TATGAGTGTGGAGTCGTTGCGGAATTGCATATCCATGGGGTTGNNNAGATCGGAA |
| 40944_2 | TATGAGTGTGGAGTCGTTGCGGAATTGCATATCCATGGGGTTGNNNAGATCGGAA |
| 49516_1 | TATGAGTGTGGAGTCGTTGCTGGTAGTGATGAATTCCCTCACCNNNAGATCGGAAG |
| 49516_2 | TATGAGTGTGGAGTCGTTGCTGGTAGTGATGAATTCCCTCACCNNNAGATCGGAAG |
| 48226_1 | TATGAGTGTGGAGTCGTTGCACCATGCCCGGCTATTTTTTTTTNNNAGATCGGAAGA |
| 48226_2 | TATGAGTGTGGAGTCGTTGCACCATGCCCGGCTATTTTTTTTTNNNAGATCGGAAGA |
| 2923_1 | TATGAGTGTGGAGTCGTTGCCGTCTATGGTGTGTGTTGGGATTNNNAGATCGGAA |
| 2923_2 | TATGAGTGTGGAGTCGTTGCCGTCTATGGTGTGTGTTGGGATTNNNAGATCGGAA |
| 52751_1 | TATGAGTGTGGAGTCGTTGCTTCCTCTGATGCTCTCCTTCCTTNNNAGATCGGAAGA |
| 52751_2 | TATGAGTGTGGAGTCGTTGCTTCCTCTGATGCTCTCCTTCCTTNNNAGATCGGAAGA |
| 38342_1 | TATGAGTGTGGAGTCGTTGCCCCAACACACTTAGGGATTGTGANNNAGATCGGAAG |
| 38342_2 | TATGAGTGTGGAGTCGTTGCCCCAACACACTTAGGGATTGTGANNNAGATCGGAAG |
| 34068_1 | TATGAGTGTGGAGTCGTTGCTGTCACCTTGAACCCATACACATNNNAGATCGGAAG |

| | |
|---|---|
| 34068_2 | TATGAGTGTGGAGTCGTTGCTGTCACCTTGAACCCATACACATNNNAGATCGGAAGA |
| 38701_1 | TATGAGTGTGGAGTCGTTGCCACTGACAGCACTAGACAGATCANNNAGATCGGAAG |
| 38701_2 | TATGAGTGTGGAGTCGTTGCCACTGACAGCACTAGACAGATCANNNAGATCGGAAG |
| 34105_1 | TATGAGTGTGGAGTCGTTGCGTCTTCTGCTCATTTAGGCCTCANNNAGATCGGAAGA |
| 34105_2 | TATGAGTGTGGAGTCGTTGCGTCTTCTGCTCATTTAGGCCTCANNNAGATCGGAAGA |
| 9962_1 | TATGAGTGTGGAGTCGTTGCTCCACCTGACAAAAACGTCAGTTNNNAGATCGGAAGA |
| 9962_2 | TATGAGTGTGGAGTCGTTGCTCCACCTGACAAAAACGTCAGTTNNNAGATCGGAAGA |
| 44413_1 | TATGAGTGTGGAGTCGTTGCAAACTCTCTCAGTCTCTGTCTGANNNAGATCGGAAGA |
| 44413_2 | TATGAGTGTGGAGTCGTTGCAAACTCTCTCAGTCTCTGTCTGANNNAGATCGGAAGA |
| 35860_1 | TATGAGTGTGGAGTCGTTGCCAGTGCACACTTTGAAATGGGTANNNAGATCGGAAG |
| 35860_2 | TATGAGTGTGGAGTCGTTGCCAGTGCACACTTTGAAATGGGTANNNAGATCGGAAG |
| 45945_1 | TATGAGTGTGGAGTCGTTGCCCAAAATGCTGGGATTATAGGCGNNNAGATCGGAAG |
| 45945_2 | TATGAGTGTGGAGTCGTTGCCCAAAATGCTGGGATTATAGGCGNNNAGATCGGAAG |
| 39018_1 | TATGAGTGTGGAGTCGTTGCGTGTGATGCAGACAAAAGAGGTCNNNAGATCGGAA |
| 39018_2 | TATGAGTGTGGAGTCGTTGCGTGTGATGCAGACAAAAGAGGTCNNNAGATCGGAA |
| 30988_1 | TATGAGTGTGGAGTCGTTGCAGTGCCAAAGCTGTTTCAAACAANNNAGATCGGAAG |
| 30988_2 | TATGAGTGTGGAGTCGTTGCAGTGCCAAAGCTGTTTCAAACAANNNAGATCGGAAG |
| 45446_1 | TATGAGTGTGGAGTCGTTGCACACAGACTTCACACCTTACACANNNAGATCGGAAGA |
| 45446_2 | TATGAGTGTGGAGTCGTTGCACACAGACTTCACACCTTACACANNNAGATCGGAAGA |
| 46678_1 | TATGAGTGTGGAGTCGTTGCCGCGTCCAGCCTAGTTTTTTATTNNNAGATCGGAAGA |
| 46678_2 | TATGAGTGTGGAGTCGTTGCCGCGTCCAGCCTAGTTTTTTATTNNNAGATCGGAAGA |
| 44365_1 | TATGAGTGTGGAGTCGTTGCGAGATTGGAGAGACCAGGGTTTTNNNAGATCGGAA |
| 44365_2 | TATGAGTGTGGAGTCGTTGCGAGATTGGAGAGACCAGGGTTTTNNNAGATCGGAA |
| 35160_1 | TATGAGTGTGGAGTCGTTGCCTGGCCCATGCTGATTTTACATTNNNAGATCGGAAGA |
| 35160_2 | TATGAGTGTGGAGTCGTTGCCTGGCCCATGCTGATTTTACATTNNNAGATCGGAAGA |
| 43094_1 | TATGAGTGTGGAGTCGTTGCTCATAGCACCACAGATGAGTTCCNNNAGATCGGAAG |
| 43094_2 | TATGAGTGTGGAGTCGTTGCTCATAGCACCACAGATGAGTTCCNNNAGATCGGAAG |
| 45226_1 | TATGAGTGTGGAGTCGTTGCATATCCAAGAAACAAGTCCCCCCNNNAGATCGGAAGA |
| 45226_2 | TATGAGTGTGGAGTCGTTGCATATCCAAGAAACAAGTCCCCCCNNNAGATCGGAAGA |
| 40559_1 | TATGAGTGTGGAGTCGTTGCTTCACAATCCAGGCTTGTCAGTANNNAGATCGGAAGA |
| 40559_2 | TATGAGTGTGGAGTCGTTGCTTCACAATCCAGGCTTGTCAGTANNNAGATCGGAAGA |
| 39293_1 | TATGAGTGTGGAGTCGTTGCCACCACCATGCCACACTAAAATTNNNAGATCGGAAGA |
| 39293_2 | TATGAGTGTGGAGTCGTTGCCACCACCATGCCACACTAAAATTNNNAGATCGGAAGA |
| 33348_1 | TATGAGTGTGGAGTCGTTGCTCCCTTCTCTTGCATTTTTTCAGANNNAGATCGGAAGA |
| 33348_2 | TATGAGTGTGGAGTCGTTGCTCCCTTCTCTTGCATTTTTTCAGANNNAGATCGGAAGA |
| 52713_1 | TATGAGTGTGGAGTCGTTGCCAAAGCAAACCTTTCAGCGTTTTNNNAGATCGGAAGA |
| 52713_2 | TATGAGTGTGGAGTCGTTGCCAAAGCAAACCTTTCAGCGTTTTNNNAGATCGGAAGA |
| 38269_1 | TATGAGTGTGGAGTCGTTGCGTGTAGACACCAGAGGGAATCTCNNNAGATCGGAAG |
| 38269_2 | TATGAGTGTGGAGTCGTTGCGTGTAGACACCAGAGGGAATCTCNNNAGATCGGAAG |
| 48801_1 | TATGAGTGTGGAGTCGTTGCCCTAGCGATTCCAAATTTAGGCCNNNAGATCGGAAGA |
| 48801_2 | TATGAGTGTGGAGTCGTTGCCCTAGCGATTCCAAATTTAGGCCNNNAGATCGGAAGA |
| 36702_1 | TATGAGTGTGGAGTCGTTGCGGACAAGCAGGAATGTAATGACGNNNAGATCGGAA |
| 36702_2 | TATGAGTGTGGAGTCGTTGCGGACAAGCAGGAATGTAATGACGNNNAGATCGGAA |

| | |
|---|---|
| 46377_1 | TATGAGTGTGGAGTCGTTGCGAGGGACTGAAGTAAATAGGCCANNNAGATCGGAA |
| 46377_2 | TATGAGTGTGGAGTCGTTGCGAGGGACTGAAGTAAATAGGCCANNNAGATCGGAA |
| 45723_1 | TATGAGTGTGGAGTCGTTGCTCTTCTCCACATCCTCACCAGTANNNAGATCGGAAGA |
| 45723_2 | TATGAGTGTGGAGTCGTTGCTCTTCTCCACATCCTCACCAGTANNNAGATCGGAAGA |
| 45629_1 | TATGAGTGTGGAGTCGTTGCCCTGGGTTCAAGCAATTCTCCTANNNAGATCGGAAG |
| 45629_2 | TATGAGTGTGGAGTCGTTGCCCTGGGTTCAAGCAATTCTCCTANNNAGATCGGAAG |
| 13355_1 | TATGAGTGTGGAGTCGTTGCAAGTTATAGTCATTGGGGTAAAANNNAGATCGGAA |
| 13355_2 | TATGAGTGTGGAGTCGTTGCAAGTTATAGTCATTGGGGTAAAANNNAGATCGGAA |
| 49354_1 | TATGAGTGTGGAGTCGTTGCCCAATTTGCCAGTCTGTGACTTTNNNAGATCGGAAG |
| 49354_2 | TATGAGTGTGGAGTCGTTGCCCAATTTGCCAGTCTGTGACTTTNNNAGATCGGAAG |
| 45498_1 | TATGAGTGTGGAGTCGTTGCGGGGAAGGAGGAGGAGGAAGNNNAGATCGGAAG |
| 45498_2 | TATGAGTGTGGAGTCGTTGCGGGGAAGGAGGAGGAGGAAGNNNAGATCGGAAG |
| 3153_1 | TATGAGTGTGGAGTCGTTGCCTTTGGGTGCTAATGGTGGAGTTNNNAGATCGGAA |
| 3153_2 | TATGAGTGTGGAGTCGTTGCCTTTGGGTGCTAATGGTGGAGTTNNNAGATCGGAA |
| 41909_1 | TATGAGTGTGGAGTCGTTGCTTCAAGCAATTCTCCTGCCTCAGNNNAGATCGGAAG |
| 41909_2 | TATGAGTGTGGAGTCGTTGCTTCAAGCAATTCTCCTGCCTCAGNNNAGATCGGAAG |
| 35994_1 | TATGAGTGTGGAGTCGTTGCCAAGTAGCTGGGACTACAGGTGNNNAGATCGGAAG |
| 35994_2 | TATGAGTGTGGAGTCGTTGCCAAGTAGCTGGGACTACAGGTGNNNAGATCGGAAG |
| 35955_1 | TATGAGTGTGGAGTCGTTGCCTCATGTCTGTAATCCCAGCACTNNNAGATCGGAAG |
| 35955_2 | TATGAGTGTGGAGTCGTTGCCTCATGTCTGTAATCCCAGCACTNNNAGATCGGAAG |
| 49508_1 | TATGAGTGTGGAGTCGTTGCACTTTGTGCTCTCTTGAGGACAANNNAGATCGGAAG |
| 49508_2 | TATGAGTGTGGAGTCGTTGCACTTTGTGCTCTCTTGAGGACAANNNAGATCGGAAG |
| 44246_1 | TATGAGTGTGGAGTCGTTGCTTCCTGAGGAGTGGACTAGAGAGNNNAGATCGGAA |
| 44246_2 | TATGAGTGTGGAGTCGTTGCTTCCTGAGGAGTGGACTAGAGAGNNNAGATCGGAA |
| 13184_1 | TATGAGTGTGGAGTCGTTGCAAACTGTTATCACACTTAGAAAGNNNAGATCGGAAG |
| 13184_2 | TATGAGTGTGGAGTCGTTGCAAACTGTTATCACACTTAGAAAGNNNAGATCGGAAG |
| 53412_1 | TATGAGTGTGGAGTCGTTGCTGTTTGTCTCTGGATTACCTAGCNNNAGATCGGAAG |
| 53412_2 | TATGAGTGTGGAGTCGTTGCTGTTTGTCTCTGGATTACCTAGCNNNAGATCGGAAG |
| 29262_1 | TATGAGTGTGGAGTCGTTGCTGCAGTCTCTGAAAGTGCATTTGNNNAGATCGGAAG |
| 29262_2 | TATGAGTGTGGAGTCGTTGCTGCAGTCTCTGAAAGTGCATTTGNNNAGATCGGAAG |
| 52331_1 | TATGAGTGTGGAGTCGTTGCCATAATAATCTGGCTCTGGCAGCNNNAGATCGGAAG |
| 52331_2 | TATGAGTGTGGAGTCGTTGCCATAATAATCTGGCTCTGGCAGCNNNAGATCGGAAG |
| 31438_1 | TATGAGTGTGGAGTCGTTGCTGGAGGAAAAAACACAAACCACCNNNAGATCGGAA |
| 31438_2 | TATGAGTGTGGAGTCGTTGCTGGAGGAAAAAACACAAACCACCNNNAGATCGGAA |
| 42775_1 | TATGAGTGTGGAGTCGTTGCAGAAGAAAGAGAAAAGGGAGCCANNNAGATCGGA |
| 42775_2 | TATGAGTGTGGAGTCGTTGCAGAAGAAAGAGAAAAGGGAGCCANNNAGATCGGA |
| 12516_1 | TATGAGTGTGGAGTCGTTGCCAACTGACTGAACATGAAAATATNNNAGATCGGAAG |
| 12516_2 | TATGAGTGTGGAGTCGTTGCCAACTGACTGAACATGAAAATATNNNAGATCGGAAG |
| 37599_1 | TATGAGTGTGGAGTCGTTGCCCAGTTTTGTGCTTGAAACCCAGNNNAGATCGGAAG |
| 37599_2 | TATGAGTGTGGAGTCGTTGCCCAGTTTTGTGCTTGAAACCCAGNNNAGATCGGAAG |
| 51960_1 | TATGAGTGTGGAGTCGTTGCGAGTCTCTCTCTGTTGCTAGGCNNNAGATCGGAAGA |
| 51960_2 | TATGAGTGTGGAGTCGTTGCGAGTCTCTCTCTGTTGCTAGGCNNNAGATCGGAAGA |
| 10115_1 | TATGAGTGTGGAGTCGTTGCTTACTTGGTCTGTGAGGTAGGCANNNAGATCGGAA |

| | |
|---|---|
| 10115_2 | TATGAGTGTGGAGTCGTTGCTTACTTGGTCTGTGAGGTAGGCANNNAGATCGGAA |
| 47319_1 | TATGAGTGTGGAGTCGTTGCAATGCAGTGGTATGATCTCAGCTNNNAGATCGGAA |
| 47319_2 | TATGAGTGTGGAGTCGTTGCAATGCAGTGGTATGATCTCAGCTNNNAGATCGGAA |
| 30343_1 | TATGAGTGTGGAGTCGTTGCGTCCATCCTGCACAGATTGTAGGNNNAGATCGGAAG |
| 30343_2 | TATGAGTGTGGAGTCGTTGCGTCCATCCTGCACAGATTGTAGGNNNAGATCGGAAG |
| 37098_1 | TATGAGTGTGGAGTCGTTGCACTGGGGATGTTAGAACATACCANNNAGATCGGAA |
| 37098_2 | TATGAGTGTGGAGTCGTTGCACTGGGGATGTTAGAACATACCANNNAGATCGGAA |
| 50646_1 | TATGAGTGTGGAGTCGTTGCCCAGACAAAGCCCTTTTCTGAAANNNAGATCGGAAG |
| 50646_2 | TATGAGTGTGGAGTCGTTGCCCAGACAAAGCCCTTTTCTGAAANNNAGATCGGAAG |
| 29306_1 | TATGAGTGTGGAGTCGTTGCGTACGTAAAGTAGCCCAGTCACANNNAGATCGGAA |
| 29306_2 | TATGAGTGTGGAGTCGTTGCGTACGTAAAGTAGCCCAGTCACANNNAGATCGGAA |
| 12553_1 | TATGAGTGTGGAGTCGTTGCAATTTTGCTGAAAATCCAGATATNNNAGATCGGAAG |
| 12553_2 | TATGAGTGTGGAGTCGTTGCAATTTTGCTGAAAATCCAGATATNNNAGATCGGAAG |
| 13652_1 | TATGAGTGTGGAGTCGTTGCCAAACAGATTGCATATTATCTGANNNAGATCGGAAG |
| 13652_2 | TATGAGTGTGGAGTCGTTGCCAAACAGATTGCATATTATCTGANNNAGATCGGAAG |
| 49957_1 | TATGAGTGTGGAGTCGTTGCTAGGCTAGACGTCTTTCATGTGCNNNAGATCGGAAG |
| 49957_2 | TATGAGTGTGGAGTCGTTGCTAGGCTAGACGTCTTTCATGTGCNNNAGATCGGAAG |
| 36549_1 | TATGAGTGTGGAGTCGTTGCGTTTGTGTCCCTTGTTGCTATCCNNNAGATCGGAAG |
| 36549_2 | TATGAGTGTGGAGTCGTTGCGTTTGTGTCCCTTGTTGCTATCCNNNAGATCGGAAG |
| 39422_1 | TATGAGTGTGGAGTCGTTGCTAATCCTAGCTACTCAGGAGGCTNNNAGATCGGAAG |
| 39422_2 | TATGAGTGTGGAGTCGTTGCTAATCCTAGCTACTCAGGAGGCTNNNAGATCGGAAG |
| 35490_1 | TATGAGTGTGGAGTCGTTGCTTGTTGAGATGGGGTTTCACTGTNNNAGATCGGAA |
| 35490_2 | TATGAGTGTGGAGTCGTTGCTTGTTGAGATGGGGTTTCACTGTNNNAGATCGGAA |
| 29819_1 | TATGAGTGTGGAGTCGTTGCATGGCACAAAGAAGGAAAACTGGNNNAGATCGGA |
| 29819_2 | TATGAGTGTGGAGTCGTTGCATGGCACAAAGAAGGAAAACTGGNNNAGATCGGA |
| 53003_1 | TATGAGTGTGGAGTCGTTGCTTTGTTGTTGTTGTGTTTCGTGCNNNAGATCGGAAG |
| 53003_2 | TATGAGTGTGGAGTCGTTGCTTTGTTGTTGTTGTGTTTCGTGCNNNAGATCGGAAG |
| 48994_1 | TATGAGTGTGGAGTCGTTGCCTCCTGAGTAGCTGGGATTACAGNNNAGATCGGAA |
| 48994_2 | TATGAGTGTGGAGTCGTTGCCTCCTGAGTAGCTGGGATTACAGNNNAGATCGGAA |
| 46651_1 | TATGAGTGTGGAGTCGTTGCCAGCAAAGGCCAAGGAATGTTTANNNAGATCGGAA |
| 46651_2 | TATGAGTGTGGAGTCGTTGCCAGCAAAGGCCAAGGAATGTTTANNNAGATCGGAA |
| 47771_1 | TATGAGTGTGGAGTCGTTGCCCTCCTGAGTAGCTGGGACTATANNNAGATCGGAAG |
| 47771_2 | TATGAGTGTGGAGTCGTTGCCCTCCTGAGTAGCTGGGACTATANNNAGATCGGAAG |
| 34956_1 | TATGAGTGTGGAGTCGTTGCTTTTTTTTTTTCCGCCTGCTGTGNNNAGATCGGAAGA |
| 34956_2 | TATGAGTGTGGAGTCGTTGCTTTTTTTTTTTCCGCCTGCTGTGNNNAGATCGGAAGA |
| 48475_1 | TATGAGTGTGGAGTCGTTGCGAGTAGCTAGGATTACAGGCACCNNNAGATCGGAA |
| 48475_2 | TATGAGTGTGGAGTCGTTGCGAGTAGCTAGGATTACAGGCACCNNNAGATCGGAA |
| 34571_1 | TATGAGTGTGGAGTCGTTGCCCATGTGGGGGGAAGAAAAATTCNNNAGATCGGAA |
| 34571_2 | TATGAGTGTGGAGTCGTTGCCCATGTGGGGGGAAGAAAAATTCNNNAGATCGGAA |
| 30097_1 | TATGAGTGTGGAGTCGTTGCGCATACCACTGGATTGAGGATGGNNNAGATCGGAA |
| 30097_2 | TATGAGTGTGGAGTCGTTGCGCATACCACTGGATTGAGGATGGNNNAGATCGGAA |
| 41079_1 | TATGAGTGTGGAGTCGTTGCCTCACTGCAATCTCCAACCTCTNNNAGATCGGAAGAG |
| 41079_2 | TATGAGTGTGGAGTCGTTGCCTCACTGCAATCTCCAACCTCTNNNAGATCGGAAGAG |

| | |
|---|---|
| 30048_1 | TATGAGTGTGGAGTCGTTGCAAGAGGATTCTGGAGCTGGAATGNNNAGATCGGAA |
| 30048_2 | TATGAGTGTGGAGTCGTTGCAAGAGGATTCTGGAGCTGGAATGNNNAGATCGGAA |
| 40334_1 | TATGAGTGTGGAGTCGTTGCAGAGAGAGAGAGAAAAACCCACCNNNAGATCGGAA |
| 40334_2 | TATGAGTGTGGAGTCGTTGCAGAGAGAGAGAGAAAAACCCACCNNNAGATCGGAA |
| 47291_1 | TATGAGTGTGGAGTCGTTGCTTGGTGTAAAGGGGAGATGTCTGNNNAGATCGGAA |
| 47291_2 | TATGAGTGTGGAGTCGTTGCTTGGTGTAAAGGGGAGATGTCTGNNNAGATCGGAA |
| 47364_1 | TATGAGTGTGGAGTCGTTGCGGGCATTTTCACCTTCCCTAATGNNNAGATCGGAAGA |
| 47364_2 | TATGAGTGTGGAGTCGTTGCGGGCATTTTCACCTTCCCTAATGNNNAGATCGGAAGA |
| 50351_1 | TATGAGTGTGGAGTCGTTGCGATGATCGCTCTCCACCACAAAANNNAGATCGGAAGA |
| 50351_2 | TATGAGTGTGGAGTCGTTGCGATGATCGCTCTCCACCACAAAANNNAGATCGGAAGA |
| 38194_1 | TATGAGTGTGGAGTCGTTGCAGCATATGCAAAGAAGCTCAACANNNAGATCGGAAG |
| 38194_2 | TATGAGTGTGGAGTCGTTGCAGCATATGCAAAGAAGCTCAACANNNAGATCGGAAG |
| 29894_1 | TATGAGTGTGGAGTCGTTGCGTGAGCCTTGAGGAGACTATGAANNNAGATCGGAA |
| 29894_2 | TATGAGTGTGGAGTCGTTGCGTGAGCCTTGAGGAGACTATGAANNNAGATCGGAA |
| 41985_1 | TATGAGTGTGGAGTCGTTGCATAGTAGCTGGGACTACAGGTGTNNNAGATCGGAAG |
| 41985_2 | TATGAGTGTGGAGTCGTTGCATAGTAGCTGGGACTACAGGTGTNNNAGATCGGAAG |
| 44921_1 | TATGAGTGTGGAGTCGTTGCTTTGGAAGGTGAGGAAAGAGGTTNNNAGATCGGAA |
| 44921_2 | TATGAGTGTGGAGTCGTTGCTTTGGAAGGTGAGGAAAGAGGTTNNNAGATCGGAA |
| 42362_1 | TATGAGTGTGGAGTCGTTGCTCCTTGACACCTTTCCTTTCAAGANNNAGATCGGAAG |
| 42362_2 | TATGAGTGTGGAGTCGTTGCTCCTTGACACCTTTCCTTTCAAGANNNAGATCGGAAG |
| 36156_1 | TATGAGTGTGGAGTCGTTGCAATGTGATCTACCCTCTAAGCAAANNNAGATCGGAAG |
| 36156_2 | TATGAGTGTGGAGTCGTTGCAATGTGATCTACCCTCTAAGCAAANNNAGATCGGAAG |
| 38240_1 | TATGAGTGTGGAGTCGTTGCAAGGGATCAGTGCAGCAATAAGANNNAGATCGGAA |
| 38240_2 | TATGAGTGTGGAGTCGTTGCAAGGGATCAGTGCAGCAATAAGANNNAGATCGGAA |
| 3137_1 | TATGAGTGTGGAGTCGTTGCCTTGGATATTTCTGCCCTGGGACNNNAGATCGGAAG |
| 3137_2 | TATGAGTGTGGAGTCGTTGCCTTGGATATTTCTGCCCTGGGACNNNAGATCGGAAG |
| 43483_1 | TATGAGTGTGGAGTCGTTGCCCATTCACTGCAACATCCACTTCNNNAGATCGGAAGA |
| 43483_2 | TATGAGTGTGGAGTCGTTGCCCATTCACTGCAACATCCACTTCNNNAGATCGGAAGA |
| 42972_1 | TATGAGTGTGGAGTCGTTGCGTGCCTAACTTCCCTTCCTTGTANNNAGATCGGAAGA |
| 42972_2 | TATGAGTGTGGAGTCGTTGCGTGCCTAACTTCCCTTCCTTGTANNNAGATCGGAAGA |
| 30507_1 | TATGAGTGTGGAGTCGTTGCGCAGTGTGTTAACTGGCATGGGNNNAGATCGGAAG |
| 30507_2 | TATGAGTGTGGAGTCGTTGCGCAGTGTGTTAACTGGCATGGGNNNAGATCGGAAG |
| 30050_1 | TATGAGTGTGGAGTCGTTGCAACCTGTCCATCTATTGAGCCAGNNNAGATCGGAAGA |
| 30050_2 | TATGAGTGTGGAGTCGTTGCAACCTGTCCATCTATTGAGCCAGNNNAGATCGGAAGA |
| 46397_1 | TATGAGTGTGGAGTCGTTGCTTCAACAATCTCACTGCTGGGTANNNAGATCGGAAGA |
| 46397_2 | TATGAGTGTGGAGTCGTTGCTTCAACAATCTCACTGCTGGGTANNNAGATCGGAAGA |
| 41468_1 | TATGAGTGTGGAGTCGTTGCTGGAGCACGGGTCATTGTATTTANNNAGATCGGAAG |
| 41468_2 | TATGAGTGTGGAGTCGTTGCTGGAGCACGGGTCATTGTATTTANNNAGATCGGAAG |
| 12732_1 | TATGAGTGTGGAGTCGTTGCCTTTGCAAACTCTAGAGAAAAAANNNAGATCGGAAG |
| 12732_2 | TATGAGTGTGGAGTCGTTGCCTTTGCAAACTCTAGAGAAAAAANNNAGATCGGAAG |
| 44865_1 | TATGAGTGTGGAGTCGTTGCCAGCCAGCCTGAATTGACATTGNNNAGATCGGAAGA |
| 44865_2 | TATGAGTGTGGAGTCGTTGCCAGCCAGCCTGAATTGACATTGNNNAGATCGGAAGA |
| 48300_1 | TATGAGTGTGGAGTCGTTGCCTTGACTCAGCCTTCCTAGTAGCNNNAGATCGGAAGA |

| | |
|---|---|
| 48300_2 | TATGAGTGTGGAGTCGTTGCCTTGACTCAGCCTTCCTAGTAGCNNNAGATCGGAAG |
| 29166_1 | TATGAGTGTGGAGTCGTTGCTCCTGATCTCATGAGGGTAGACTNNNAGATCGGAAG |
| 29166_2 | TATGAGTGTGGAGTCGTTGCTCCTGATCTCATGAGGGTAGACTNNNAGATCGGAAG |
| 42180_1 | TATGAGTGTGGAGTCGTTGCTGGAGTGCAATGGTACAATCTCANNNAGATCGGAA |
| 42180_2 | TATGAGTGTGGAGTCGTTGCTGGAGTGCAATGGTACAATCTCANNNAGATCGGAA |
| 42280_1 | TATGAGTGTGGAGTCGTTGCTTTCTCCAAGAAGCCCTAGTTCCNNNAGATCGGAAG |
| 42280_2 | TATGAGTGTGGAGTCGTTGCTTTCTCCAAGAAGCCCTAGTTCCNNNAGATCGGAAG |
| 40728_1 | TATGAGTGTGGAGTCGTTGCTGTGGTCTTCCTTCTAGGTTTGGNNNAGATCGGAAG |
| 40728_2 | TATGAGTGTGGAGTCGTTGCTGTGGTCTTCCTTCTAGGTTTGGNNNAGATCGGAAG |
| 43726_1 | TATGAGTGTGGAGTCGTTGCCTGGCTCCAAACAACTTCTTCTGNNNAGATCGGAAG |
| 43726_2 | TATGAGTGTGGAGTCGTTGCCTGGCTCCAAACAACTTCTTCTGNNNAGATCGGAAG |
| 48009_1 | TATGAGTGTGGAGTCGTTGCGCGTGTGTGCATTTTAGTCAGATNNNAGATCGGAA |
| 48009_2 | TATGAGTGTGGAGTCGTTGCGCGTGTGTGCATTTTAGTCAGATNNNAGATCGGAA |
| 43026_1 | TATGAGTGTGGAGTCGTTGCTTGGTTCTTCCCTCTGGAGAATGNNNAGATCGGAAG |
| 43026_2 | TATGAGTGTGGAGTCGTTGCTTGGTTCTTCCCTCTGGAGAATGNNNAGATCGGAAG |
| 13197_1 | TATGAGTGTGGAGTCGTTGCGTTACCTTACGAAGATATTTCAGNNNAGATCGGAAG |
| 13197_2 | TATGAGTGTGGAGTCGTTGCGTTACCTTACGAAGATATTTCAGNNNAGATCGGAAG |
| 13588_1 | TATGAGTGTGGAGTCGTTGCAACTTTTTTTTGACATGAAAACANNNAGATCGGAAG |
| 13588_2 | TATGAGTGTGGAGTCGTTGCAACTTTTTTTTGACATGAAAACANNNAGATCGGAAG |
| 52534_1 | TATGAGTGTGGAGTCGTTGCGGCTGTGGCATGTTACATTACAANNNAGATCGGAA |
| 52534_2 | TATGAGTGTGGAGTCGTTGCGGCTGTGGCATGTTACATTACAANNNAGATCGGAA |
| 31249_1 | TATGAGTGTGGAGTCGTTGCCCCTTCTTTGGAGCAAGACAGTANNNAGATCGGAAG |
| 31249_2 | TATGAGTGTGGAGTCGTTGCCCCTTCTTTGGAGCAAGACAGTANNNAGATCGGAAG |
| 30283_1 | TATGAGTGTGGAGTCGTTGCTCCCATCTGCACTATTGACGTTTNNNAGATCGGAAG |
| 30283_2 | TATGAGTGTGGAGTCGTTGCTCCCATCTGCACTATTGACGTTTNNNAGATCGGAAG |
| 35455_1 | TATGAGTGTGGAGTCGTTGCTTCAAACTCCTGGGTTCAAGTGANNNAGATCGGAAG |
| 35455_2 | TATGAGTGTGGAGTCGTTGCTTCAAACTCCTGGGTTCAAGTGANNNAGATCGGAAG |
| 31336_1 | TATGAGTGTGGAGTCGTTGCATTCTAAAGAGGAGCTGCCAGACNNNAGATCGGAA |
| 31336_2 | TATGAGTGTGGAGTCGTTGCATTCTAAAGAGGAGCTGCCAGACNNNAGATCGGAA |
| 53380_1 | TATGAGTGTGGAGTCGTTGCCTCTCTTTCTTTCCCTCCCTTCCNNNAGATCGGAAGAG |
| 53380_2 | TATGAGTGTGGAGTCGTTGCCTCTCTTTCTTTCCCTCCCTTCCNNNAGATCGGAAGAG |
| 48433_1 | TATGAGTGTGGAGTCGTTGCCCACCATGCTCGGCTAATTTTTTNNNAGATCGGAAG |
| 48433_2 | TATGAGTGTGGAGTCGTTGCCCACCATGCTCGGCTAATTTTTTNNNAGATCGGAAG |
| 41341_1 | TATGAGTGTGGAGTCGTTGCCTCATGTCTGTAATCCCAGCACTNNNAGATCGGAAG |
| 41341_2 | TATGAGTGTGGAGTCGTTGCCTCATGTCTGTAATCCCAGCACTNNNAGATCGGAAG |
| 8900_1 | TATGAGTGTGGAGTCGTTGCTAAGGACATTGCAGCCAAATGGTNNNAGATCGGAA |
| 8900_2 | TATGAGTGTGGAGTCGTTGCTAAGGACATTGCAGCCAAATGGTNNNAGATCGGAA |
| 12906_1 | TATGAGTGTGGAGTCGTTGCATTTGTAAGAAAATACAGAGCAGNNNAGATCGGAA |
| 12906_2 | TATGAGTGTGGAGTCGTTGCATTTGTAAGAAAATACAGAGCAGNNNAGATCGGAA |
| 33533_1 | TATGAGTGTGGAGTCGTTGCGTTCAAGTGATCCTCCTGCCTCNNNAGATCGGAAGA |
| 33533_2 | TATGAGTGTGGAGTCGTTGCGTTCAAGTGATCCTCCTGCCTCNNNAGATCGGAAGA |
| 13799_1 | TATGAGTGTGGAGTCGTTGCCTCCGTCTCAACAACAAAAANNNAGATCGGAAGAGC |
| 13799_2 | TATGAGTGTGGAGTCGTTGCCTCCGTCTCAACAACAAAAANNNAGATCGGAAGAGC |

| | |
|---|---|
| 44928_1 | TATGAGTGTGGAGTCGTTGCGCAGCTGCTCTAAGTCTCAAAAGNNNAGATCGGAAG |
| 44928_2 | TATGAGTGTGGAGTCGTTGCGCAGCTGCTCTAAGTCTCAAAAGNNNAGATCGGAAG |
| 50760_1 | TATGAGTGTGGAGTCGTTGCCAAGAGGAGTATGATGACTGCGANNNAGATCGGAA |
| 50760_2 | TATGAGTGTGGAGTCGTTGCCAAGAGGAGTATGATGACTGCGANNNAGATCGGAA |
| 32334_1 | TATGAGTGTGGAGTCGTTGCAACTGTAATGCCTTTTAGTTCACANNNAGATCGGAA |
| 32334_2 | TATGAGTGTGGAGTCGTTGCAACTGTAATGCCTTTTAGTTCACANNNAGATCGGAA |
| 46712_1 | TATGAGTGTGGAGTCGTTGCCAGTCTTGACCTCATGGACTCAANNNAGATCGGAAG |
| 46712_2 | TATGAGTGTGGAGTCGTTGCCAGTCTTGACCTCATGGACTCAANNNAGATCGGAAG |
| 3062_1 | TATGAGTGTGGAGTCGTTGCCCCAGTGTAGGCAAATGGCTATCNNNAGATCGGAA |
| 3062_2 | TATGAGTGTGGAGTCGTTGCCCCAGTGTAGGCAAATGGCTATCNNNAGATCGGAA |
| 36599_1 | TATGAGTGTGGAGTCGTTGCGCAAGGTCCTGACTCTTTCTACANNNAGATCGGAAG |
| 36599_2 | TATGAGTGTGGAGTCGTTGCGCAAGGTCCTGACTCTTTCTACANNNAGATCGGAAG |
| 5994_1 | TATGAGTGTGGAGTCGTTGCGCTTCAGATTCCGGAGTTCCTTGNNNAGATCGGAAG |
| 5994_2 | TATGAGTGTGGAGTCGTTGCGCTTCAGATTCCGGAGTTCCTTGNNNAGATCGGAAG |
| 40844_1 | TATGAGTGTGGAGTCGTTGCACAGAAAAGCAAACACCACATGTNNNAGATCGGAA |
| 40844_2 | TATGAGTGTGGAGTCGTTGCACAGAAAAGCAAACACCACATGTNNNAGATCGGAA |
| 30745_1 | TATGAGTGTGGAGTCGTTGCCATGCATGCCCATAGTTACACACNNNAGATCGGAAG |
| 30745_2 | TATGAGTGTGGAGTCGTTGCCATGCATGCCCATAGTTACACACNNNAGATCGGAAG |
| 46677_1 | TATGAGTGTGGAGTCGTTGCGAGTCTCACTCTTTTTGCCCAGGNNNAGATCGGAAG |
| 46677_2 | TATGAGTGTGGAGTCGTTGCGAGTCTCACTCTTTTTGCCCAGGNNNAGATCGGAAG |
| 49034_1 | TATGAGTGTGGAGTCGTTGCTTCAAGCAATTTTCCTGCCTCAGNNNAGATCGGAAG |
| 49034_2 | TATGAGTGTGGAGTCGTTGCTTCAAGCAATTTTCCTGCCTCAGNNNAGATCGGAAG |
| 11942_1 | TATGAGTGTGGAGTCGTTGCAGGACAAAGCCTGGGTTATGTTTCNNNAGATCGGA |
| 11942_2 | TATGAGTGTGGAGTCGTTGCAGGACAAAGCCTGGGTTATGTTTCNNNAGATCGGA |
| 1867_1 | TATGAGTGTGGAGTCGTTGCCCATGCCTTTGCAGGTAAGTANNNAGATCGGAAGA |
| 1867_2 | TATGAGTGTGGAGTCGTTGCCCATGCCTTTGCAGGTAAGTANNNAGATCGGAAGA |
| 45347_1 | TATGAGTGTGGAGTCGTTGCACGTAGCAATCAGCAAAACACTTNNNAGATCGGAAG |
| 45347_2 | TATGAGTGTGGAGTCGTTGCACGTAGCAATCAGCAAAACACTTNNNAGATCGGAAG |
| 44688_1 | TATGAGTGTGGAGTCGTTGCGCTTTTACAAAAGAAGCCCCAGTNNNAGATCGGAAG |
| 44688_2 | TATGAGTGTGGAGTCGTTGCGCTTTTACAAAAGAAGCCCCAGTNNNAGATCGGAAG |
| 37119_1 | TATGAGTGTGGAGTCGTTGCCTGTTCTCAGACTTTTGTGTGGGNNNAGATCGGAAG |
| 37119_2 | TATGAGTGTGGAGTCGTTGCCTGTTCTCAGACTTTTGTGTGGGNNNAGATCGGAAG |
| 34686_1 | TATGAGTGTGGAGTCGTTGCTTATTTATCTGCCTCAGCCTCCCNNNAGATCGGAAGA |
| 34686_2 | TATGAGTGTGGAGTCGTTGCTTATTTATCTGCCTCAGCCTCCCNNNAGATCGGAAGA |
| 45275_1 | TATGAGTGTGGAGTCGTTGCTGGGACATTCACAGAGCTGATTANNNAGATCGGAA |
| 45275_2 | TATGAGTGTGGAGTCGTTGCTGGGACATTCACAGAGCTGATTANNNAGATCGGAA |
| 13560_1 | TATGAGTGTGGAGTCGTTGCCATACTGTAGGGTAATAAATTTGTNNNAGATCGGAA |
| 13560_2 | TATGAGTGTGGAGTCGTTGCCATACTGTAGGGTAATAAATTTGTNNNAGATCGGAA |
| 34350_1 | TATGAGTGTGGAGTCGTTGCATGACCCATGATGAAACCCATGANNNAGATCGGAAG |
| 34350_2 | TATGAGTGTGGAGTCGTTGCATGACCCATGATGAAACCCATGANNNAGATCGGAAG |
| 30637_1 | TATGAGTGTGGAGTCGTTGCCAATTTTACAGAGTGCTGCCTGCNNNAGATCGGAAG |
| 30637_2 | TATGAGTGTGGAGTCGTTGCCAATTTTACAGAGTGCTGCCTGCNNNAGATCGGAAG |
| 33175_1 | TATGAGTGTGGAGTCGTTGCAGGTAGTGGCCCTTAGAGAGAATNNNAGATCGGAA |

| | |
|---|---|
| 33175_2 | TATGAGTGTGGAGTCGTTGCAGGTAGTGGCCCTTAGAGAGAATNNNAGATCGGAA |
| 37005_1 | TATGAGTGTGGAGTCGTTGCGCAAGACTCCGACTCGAAAAAAANNNAGATCGGAA |
| 37005_2 | TATGAGTGTGGAGTCGTTGCGCAAGACTCCGACTCGAAAAAAANNNAGATCGGAA |
| 10523_1 | TATGAGTGTGGAGTCGTTGCTTTTCTAGCCTTGGGCTTGGTTTNNNAGATCGGAAG |
| 10523_2 | TATGAGTGTGGAGTCGTTGCTTTTCTAGCCTTGGGCTTGGTTTNNNAGATCGGAAG |
| 10536_1 | TATGAGTGTGGAGTCGTTGCTCTTTCCTAGGGATCCTTGCTTTNNNAGATCGGAAGA |
| 10536_2 | TATGAGTGTGGAGTCGTTGCTCTTTCCTAGGGATCCTTGCTTTNNNAGATCGGAAGA |
| 51711_1 | TATGAGTGTGGAGTCGTTGCCACAGTTGTAGAGTCAGCTCACTNNNAGATCGGAAG |
| 51711_2 | TATGAGTGTGGAGTCGTTGCCACAGTTGTAGAGTCAGCTCACTNNNAGATCGGAAG |
| 48791_1 | TATGAGTGTGGAGTCGTTGCTTCAAGTGATTCTCCTGCCTCAGNNNAGATCGGAAG |
| 48791_2 | TATGAGTGTGGAGTCGTTGCTTCAAGTGATTCTCCTGCCTCAGNNNAGATCGGAAG |
| 48930_1 | TATGAGTGTGGAGTCGTTGCGGCTTCACATTTCTTCCTCTTGGNNNAGATCGGAAGA |
| 48930_2 | TATGAGTGTGGAGTCGTTGCGGCTTCACATTTCTTCCTCTTGGNNNAGATCGGAAGA |
| 42067_1 | TATGAGTGTGGAGTCGTTGCGACGAAAGTGTCTTCCCATCAGCNNNAGATCGGAAG |
| 42067_2 | TATGAGTGTGGAGTCGTTGCGACGAAAGTGTCTTCCCATCAGCNNNAGATCGGAAG |
| 37710_1 | TATGAGTGTGGAGTCGTTGCAGAGTTCCCATATACCCTTTGCCNNNAGATCGGAAG |
| 37710_2 | TATGAGTGTGGAGTCGTTGCAGAGTTCCCATATACCCTTTGCCNNNAGATCGGAAG |
| 32211_1 | TATGAGTGTGGAGTCGTTGCGGAATTTGCTAAGGAGATGGCTGNNNAGATCGGAA |
| 32211_2 | TATGAGTGTGGAGTCGTTGCGGAATTTGCTAAGGAGATGGCTGNNNAGATCGGAA |
| 51741_1 | TATGAGTGTGGAGTCGTTGCCGCTTTCCAATAGTACTTCCTGCNNNAGATCGGAAGA |
| 51741_2 | TATGAGTGTGGAGTCGTTGCCGCTTTCCAATAGTACTTCCTGCNNNAGATCGGAAGA |
| 36277_1 | TATGAGTGTGGAGTCGTTGCTGCTCTCAGTGGTCATCTTTTCANNNAGATCGGAAGA |
| 36277_2 | TATGAGTGTGGAGTCGTTGCTGCTCTCAGTGGTCATCTTTTCANNNAGATCGGAAGA |
| 29873_1 | TATGAGTGTGGAGTCGTTGCAAAGGGCCTAAAAGGAGAGATGGNNNAGATCGGA |
| 29873_2 | TATGAGTGTGGAGTCGTTGCAAAGGGCCTAAAAGGAGAGATGGNNNAGATCGGA |
| 37307_1 | TATGAGTGTGGAGTCGTTGCTGGAGAGAGAGAGAATGACAACGNNNAGATCGGA |
| 37307_2 | TATGAGTGTGGAGTCGTTGCTGGAGAGAGAGAGAATGACAACGNNNAGATCGGA |
| 42856_1 | TATGAGTGTGGAGTCGTTGCCAGGATGCTTTTGTTGGGCAGNNNAGATCGGAAGA |
| 42856_2 | TATGAGTGTGGAGTCGTTGCCAGGATGCTTTTGTTGGGCAGNNNAGATCGGAAGA |
| 29724_1 | TATGAGTGTGGAGTCGTTGCCTGAGTCGGGAACTGCTATGATTNNNAGATCGGAA |
| 29724_2 | TATGAGTGTGGAGTCGTTGCCTGAGTCGGGAACTGCTATGATTNNNAGATCGGAA |
| 1003_1 | TATGAGTGTGGAGTCGTTGCCTGTCTTCTGCGCTTGCTCGTAANNNAGATCGGAAG |
| 1003_2 | TATGAGTGTGGAGTCGTTGCCTGTCTTCTGCGCTTGCTCGTAANNNAGATCGGAAG |
| 31401_1 | TATGAGTGTGGAGTCGTTGCTTCTTCTCCCCAGATACCAATGGNNNAGATCGGAAGA |
| 31401_2 | TATGAGTGTGGAGTCGTTGCTTCTTCTCCCCAGATACCAATGGNNNAGATCGGAAGA |
| 10873_1 | TATGAGTGTGGAGTCGTTGCCGCTTCGGGATTGTTTGGGGTNNNAGATCGGAAGA |
| 10873_2 | TATGAGTGTGGAGTCGTTGCCGCTTCGGGATTGTTTGGGGTNNNAGATCGGAAGA |
| 32197_1 | TATGAGTGTGGAGTCGTTGCTAATCTCGACTCACTGCAACCTCNNNAGATCGGAAGA |
| 32197_2 | TATGAGTGTGGAGTCGTTGCTAATCTCGACTCACTGCAACCTCNNNAGATCGGAAGA |
| 32896_1 | TATGAGTGTGGAGTCGTTGCGAGCCATAGGATTGGTAATGGGTNNNAGATCGGAA |
| 32896_2 | TATGAGTGTGGAGTCGTTGCGAGCCATAGGATTGGTAATGGGTNNNAGATCGGAA |
| 34790_1 | TATGAGTGTGGAGTCGTTGCCAGGTTCAAGCAATTCTCCTGCNNNAGATCGGAAGA |
| 34790_2 | TATGAGTGTGGAGTCGTTGCCAGGTTCAAGCAATTCTCCTGCNNNAGATCGGAAGA |

| | |
|---|---|
| 29033_1 | TATGAGTGTGGAGTCGTTGCTGACTGTATTTGGAGATAGGGCCNNNAGATCGGAA |
| 29033_2 | TATGAGTGTGGAGTCGTTGCTGACTGTATTTGGAGATAGGGCCNNNAGATCGGAA |
| 38932_1 | TATGAGTGTGGAGTCGTTGCCAAAAATTAGCCAGGAGTGGTGGNNNAGATCGGAA |
| 38932_2 | TATGAGTGTGGAGTCGTTGCCAAAAATTAGCCAGGAGTGGTGGNNNAGATCGGAA |
| 48242_1 | TATGAGTGTGGAGTCGTTGCGCCTCCCAGGTTCAAGCAATTNNNAGATCGGAAGAG |
| 48242_2 | TATGAGTGTGGAGTCGTTGCGCCTCCCAGGTTCAAGCAATTNNNAGATCGGAAGAG |
| 50638_1 | TATGAGTGTGGAGTCGTTGCATCAATCTCCATCACATCCAGGGNNNAGATCGGAAG |
| 50638_2 | TATGAGTGTGGAGTCGTTGCATCAATCTCCATCACATCCAGGGNNNAGATCGGAAG |
| 33311_1 | TATGAGTGTGGAGTCGTTGCTTCCATGTTTAGTCTTGGGAGGGNNNAGATCGGAA |
| 33311_2 | TATGAGTGTGGAGTCGTTGCTTCCATGTTTAGTCTTGGGAGGGNNNAGATCGGAA |
| 34822_1 | TATGAGTGTGGAGTCGTTGCTTCAAGTGATTCTCCTGCCTCAGNNNAGATCGGAAG |
| 34822_2 | TATGAGTGTGGAGTCGTTGCTTCAAGTGATTCTCCTGCCTCAGNNNAGATCGGAAG |
| 51105_1 | TATGAGTGTGGAGTCGTTGCGCCAAACTGATGTAGCAGGATTCNNNAGATCGGAA |
| 51105_2 | TATGAGTGTGGAGTCGTTGCGCCAAACTGATGTAGCAGGATTCNNNAGATCGGAA |
| 31791_1 | TATGAGTGTGGAGTCGTTGCAGGCAATTTTTTCTTACAGGCCTNNNAGATCGGAAG |
| 31791_2 | TATGAGTGTGGAGTCGTTGCAGGCAATTTTTTCTTACAGGCCTNNNAGATCGGAAG |
| 45485_1 | TATGAGTGTGGAGTCGTTGCGGTTCAGTCCAGCTTCCTTTTTGNNNAGATCGGAAG |
| 45485_2 | TATGAGTGTGGAGTCGTTGCGGTTCAGTCCAGCTTCCTTTTTGNNNAGATCGGAAG |
| 46847_1 | TATGAGTGTGGAGTCGTTGCGCCTTGCTTTTTTTGCTTTCTTTNNNAGATCGGAAGA |
| 46847_2 | TATGAGTGTGGAGTCGTTGCGCCTTGCTTTTTTTGCTTTCTTTNNNAGATCGGAAGA |
| 33067_1 | TATGAGTGTGGAGTCGTTGCGCTAGAGTACAGTGGCATGATCTNNNAGATCGGAA |
| 33067_2 | TATGAGTGTGGAGTCGTTGCGCTAGAGTACAGTGGCATGATCTNNNAGATCGGAA |
| 1659_1 | TATGAGTGTGGAGTCGTTGCGCTGATACTTGCTACTGCAGTTNNNAGATCGGAAGA |
| 1659_2 | TATGAGTGTGGAGTCGTTGCGCTGATACTTGCTACTGCAGTTNNNAGATCGGAAGA |
| 10162_1 | TATGAGTGTGGAGTCGTTGCTCAAGAGGGAGAATGCTGACAAANNNAGATCGGAA |
| 10162_2 | TATGAGTGTGGAGTCGTTGCTCAAGAGGGAGAATGCTGACAAANNNAGATCGGAA |
| 47959_1 | TATGAGTGTGGAGTCGTTGCCCTCCTGGATTCAAGCGATTCTCNNNAGATCGGAAG |
| 47959_2 | TATGAGTGTGGAGTCGTTGCCCTCCTGGATTCAAGCGATTCTCNNNAGATCGGAAG |
| 52087_1 | TATGAGTGTGGAGTCGTTGCCTTAGGGCTAGACTCCAAAAGCANNNAGATCGGAA |
| 52087_2 | TATGAGTGTGGAGTCGTTGCCTTAGGGCTAGACTCCAAAAGCANNNAGATCGGAA |
| 52691_1 | TATGAGTGTGGAGTCGTTGCACATCTTACACTTGCTCGACACTNNNAGATCGGAAG |
| 52691_2 | TATGAGTGTGGAGTCGTTGCACATCTTACACTTGCTCGACACTNNNAGATCGGAAG |
| 34151_1 | TATGAGTGTGGAGTCGTTGCGATTCCCCTCCCCATCTGTATTCNNNAGATCGGAAGA |
| 34151_2 | TATGAGTGTGGAGTCGTTGCGATTCCCCTCCCCATCTGTATTCNNNAGATCGGAAGA |
| 29321_1 | TATGAGTGTGGAGTCGTTGCTGTCAGAAAGTGTCAAATGCTGTNNNAGATCGGAA |
| 29321_2 | TATGAGTGTGGAGTCGTTGCTGTCAGAAAGTGTCAAATGCTGTNNNAGATCGGAA |
| 35472_1 | TATGAGTGTGGAGTCGTTGCCCTCGGTTCAAGCTATTCTCCTGNNNAGATCGGAAG |
| 35472_2 | TATGAGTGTGGAGTCGTTGCCCTCGGTTCAAGCTATTCTCCTGNNNAGATCGGAAG |
| 44099_1 | TATGAGTGTGGAGTCGTTGCCGAGACTTTTCCTCCAGCTGTATNNNAGATCGGAAG |
| 44099_2 | TATGAGTGTGGAGTCGTTGCCGAGACTTTTCCTCCAGCTGTATNNNAGATCGGAAG |
| 36540_1 | TATGAGTGTGGAGTCGTTGCGAACCTGCCCTTGTTTTATCAGTNNNAGATCGGAAG |
| 36540_2 | TATGAGTGTGGAGTCGTTGCGAACCTGCCCTTGTTTTATCAGTNNNAGATCGGAAG |
| 35084_1 | TATGAGTGTGGAGTCGTTGCCCTCCCAAGTAGCTGTGATTACANNNAGATCGGAAG |

| | |
|---|---|
| 35084_2 | TATGAGTGTGGAGTCGTTGCCCTCCCAAGTAGCTGTGATTACANNNAGATCGGAAG |
| 33676_1 | TATGAGTGTGGAGTCGTTGCAGGTTCAAGTGATTCTTCTGCCTNNNAGATCGGAAG |
| 33676_2 | TATGAGTGTGGAGTCGTTGCAGGTTCAAGTGATTCTTCTGCCTNNNAGATCGGAAG |
| 48711_1 | TATGAGTGTGGAGTCGTTGCGAGGAGTAGGAGGAGGAGGATTCNNNAGATCGGA |
| 48711_2 | TATGAGTGTGGAGTCGTTGCGAGGAGTAGGAGGAGGAGGATTCNNNAGATCGGA |
| 29374_1 | TATGAGTGTGGAGTCGTTGCGTGTCTGTGTGTCTGTGTGTTTTNNNAGATCGGAAG |
| 29374_2 | TATGAGTGTGGAGTCGTTGCGTGTCTGTGTGTCTGTGTGTTTTNNNAGATCGGAAG |
| 46034_1 | TATGAGTGTGGAGTCGTTGCCAATTTTGGCGCTCATTATTGGCNNNAGATCGGAAG |
| 46034_2 | TATGAGTGTGGAGTCGTTGCCAATTTTGGCGCTCATTATTGGCNNNAGATCGGAAG |
| 5634_1 | TATGAGTGTGGAGTCGTTGCAAAGACACACACATCCAGCTAGCNNNAGATCGGAAG |
| 5634_2 | TATGAGTGTGGAGTCGTTGCAAAGACACACACATCCAGCTAGCNNNAGATCGGAAG |
| 48497_1 | TATGAGTGTGGAGTCGTTGCACACCCACCTTTATGCATCTGATNNNAGATCGGAAGA |
| 48497_2 | TATGAGTGTGGAGTCGTTGCACACCCACCTTTATGCATCTGATNNNAGATCGGAAGA |
| 45598_1 | TATGAGTGTGGAGTCGTTGCCATGATCTTGGCTCACTGCAACNNNAGATCGGAAGA |
| 45598_2 | TATGAGTGTGGAGTCGTTGCCATGATCTTGGCTCACTGCAACNNNAGATCGGAAGA |
| 30561_1 | ATGAGTGTGGAGTCGTTGCTGTGGGGTACATGACCTACTATGTACCNNNAGATCGG |
| 30561_2 | ATGAGTGTGGAGTCGTTGCTGTGGGGTACATGACCTACTATGTACCNNNAGATCGG |
| 48173_1 | TATGAGTGTGGAGTCGTTGCGTTTCACCATGTTAGCCAGGATGNNNAGATCGGAAG |
| 48173_2 | TATGAGTGTGGAGTCGTTGCGTTTCACCATGTTAGCCAGGATGNNNAGATCGGAAG |
| 9963_1 | TATGAGTGTGGAGTCGTTGCATTGATCGTAGGACAGGACATGCNNNAGATCGGAA |
| 9963_2 | TATGAGTGTGGAGTCGTTGCATTGATCGTAGGACAGGACATGCNNNAGATCGGAA |
| 10090_1 | TATGAGTGTGGAGTCGTTGCCCTGTTCCATATCTCTGGAGAACCNNNAGATCGGAAG |
| 10090_2 | TATGAGTGTGGAGTCGTTGCCCTGTTCCATATCTCTGGAGAACCNNNAGATCGGAAG |
| 31601_1 | TATGAGTGTGGAGTCGTTGCCCCCACAAATCAGTGCATGTAAANNNAGATCGGAAG |
| 31601_2 | TATGAGTGTGGAGTCGTTGCCCCCACAAATCAGTGCATGTAAANNNAGATCGGAAG |
| 51404_1 | TATGAGTGTGGAGTCGTTGCATCCAAACCATGTAAGTCACCCTNNNAGATCGGAAG |
| 51404_2 | TATGAGTGTGGAGTCGTTGCATCCAAACCATGTAAGTCACCCTNNNAGATCGGAAG |
| 48842_1 | TATGAGTGTGGAGTCGTTGCATGGTGCCATACTGTTGTCTTCTNNNAGATCGGAAG |
| 48842_2 | TATGAGTGTGGAGTCGTTGCATGGTGCCATACTGTTGTCTTCTNNNAGATCGGAAG |
| 33390_1 | TATGAGTGTGGAGTCGTTGCCTGAGACACGAGAATCCCTTGAANNNAGATCGGAAG |
| 33390_2 | TATGAGTGTGGAGTCGTTGCCTGAGACACGAGAATCCCTTGAANNNAGATCGGAAG |
| 40119_1 | TATGAGTGTGGAGTCGTTGCTAAAAAATTAGCTGGGCGTGGTGNNNAGATCGGAA |
| 40119_2 | TATGAGTGTGGAGTCGTTGCTAAAAAATTAGCTGGGCGTGGTGNNNAGATCGGAA |
| 50483_1 | TATGAGTGTGGAGTCGTTGCAACGCTGTCTGTAACTCTGGAATNNNAGATCGGAAG |
| 50483_2 | TATGAGTGTGGAGTCGTTGCAACGCTGTCTGTAACTCTGGAATNNNAGATCGGAAG |
| 28915_1 | TATGAGTGTGGAGTCGTTGCTGCTTGAGACCAGAAGTGTTTCANNNAGATCGGAAG |
| 28915_2 | TATGAGTGTGGAGTCGTTGCTGCTTGAGACCAGAAGTGTTTCANNNAGATCGGAAG |
| 10670_1 | TATGAGTGTGGAGTCGTTGCTCAATGCGATGCAGCAATCATTCNNNAGATCGGAAG |
| 10670_2 | TATGAGTGTGGAGTCGTTGCTCAATGCGATGCAGCAATCATTCNNNAGATCGGAAG |
| 43897_1 | TATGAGTGTGGAGTCGTTGCACTTTGACATTGGACCGAGAGTTNNNAGATCGGAAG |
| 43897_2 | TATGAGTGTGGAGTCGTTGCACTTTGACATTGGACCGAGAGTTNNNAGATCGGAAG |
| 49661_1 | TATGAGTGTGGAGTCGTTGCCATCAGGAAACTCCAAACTGCTGNNNAGATCGGAAG |
| 49661_2 | TATGAGTGTGGAGTCGTTGCCATCAGGAAACTCCAAACTGCTGNNNAGATCGGAAG |

| | |
|---|---|
| 37457_1 | TATGAGTGTGGAGTCGTTGCGAAAGGAAGAGGATTAGAGAAGGANNNAGATCGG |
| 37457_2 | TATGAGTGTGGAGTCGTTGCGAAAGGAAGAGGATTAGAGAAGGANNNAGATCGG |
| 46404_1 | TATGAGTGTGGAGTCGTTGCCTTCCTCCAATCTACTTGCACCTNNNAGATCGGAAGA |
| 46404_2 | TATGAGTGTGGAGTCGTTGCCTTCCTCCAATCTACTTGCACCTNNNAGATCGGAAGA |
| 8250_1 | TATGAGTGTGGAGTCGTTGCACTATTCTCTGCAGCCACAAGCCNNNAGATCGGAAG |
| 8250_2 | TATGAGTGTGGAGTCGTTGCACTATTCTCTGCAGCCACAAGCCNNNAGATCGGAAG |
| 13737_1 | TATGAGTGTGGAGTCGTTGCTTAAGAAAATAGAGGCTTATCCANNNAGATCGGAAG |
| 13737_2 | TATGAGTGTGGAGTCGTTGCTTAAGAAAATAGAGGCTTATCCANNNAGATCGGAAG |
| 36782_1 | TATGAGTGTGGAGTCGTTGCCCTAATCTAGGGTTGCTCTGTTANNNAGATCGGAAG |
| 36782_2 | TATGAGTGTGGAGTCGTTGCCCTAATCTAGGGTTGCTCTGTTANNNAGATCGGAAG |
| 34377_1 | TATGAGTGTGGAGTCGTTGCGTCCTCTCAAAGTGCTGGGATTANNNAGATCGGAAG |
| 34377_2 | TATGAGTGTGGAGTCGTTGCGTCCTCTCAAAGTGCTGGGATTANNNAGATCGGAAG |
| 34117_1 | TATGAGTGTGGAGTCGTTGCATGGGCCTACTAAGTAGTCACCTNNNAGATCGGAAG |
| 34117_2 | TATGAGTGTGGAGTCGTTGCATGGGCCTACTAAGTAGTCACCTNNNAGATCGGAAG |
| 32699_1 | TATGAGTGTGGAGTCGTTGCATAAGCCTGATTCTTCCTTGGCANNNAGATCGGAAG |
| 32699_2 | TATGAGTGTGGAGTCGTTGCATAAGCCTGATTCTTCCTTGGCANNNAGATCGGAAG |
| 29714_1 | TATGAGTGTGGAGTCGTTGCATTTGACTGACAGTAGGAGGTGGNNNAGATCGGAA |
| 29714_2 | TATGAGTGTGGAGTCGTTGCATTTGACTGACAGTAGGAGGTGGNNNAGATCGGAA |
| 39345_1 | TATGAGTGTGGAGTCGTTGCCGGGATCTCGTTATGTAGACTGGNNNAGATCGGAA |
| 39345_2 | TATGAGTGTGGAGTCGTTGCCGGGATCTCGTTATGTAGACTGGNNNAGATCGGAA |
| 52366_1 | TATGAGTGTGGAGTCGTTGCTCCAACCTTCTTCACAGGAGAACNNNAGATCGGAAG |
| 52366_2 | TATGAGTGTGGAGTCGTTGCTCCAACCTTCTTCACAGGAGAACNNNAGATCGGAAG |
| 52983_1 | TATGAGTGTGGAGTCGTTGCCCTCCCTCTCTGTCTCTCTTTCTNNNAGATCGGAAGAG |
| 52983_2 | TATGAGTGTGGAGTCGTTGCCCTCCCTCTCTGTCTCTCTTTCTNNNAGATCGGAAGAG |
| 12485_1 | TATGAGTGTGGAGTCGTTGCCATTAGACTTCTTGTTGGAAAGNNNAGATCGGAAGA |
| 12485_2 | TATGAGTGTGGAGTCGTTGCCATTAGACTTCTTGTTGGAAAGNNNAGATCGGAAGA |
| 37422_1 | TATGAGTGTGGAGTCGTTGCGCTGAGGTAGGAGAATCGTTTGANNNAGATCGGAA |
| 37422_2 | TATGAGTGTGGAGTCGTTGCGCTGAGGTAGGAGAATCGTTTGANNNAGATCGGAA |
| 28913_1 | TATGAGTGTGGAGTCGTTGCACAGAGACAGAGACAGAGAGACANNNAGATCGGA |
| 28913_2 | TATGAGTGTGGAGTCGTTGCACAGAGACAGAGACAGAGAGACANNNAGATCGGA |
| 33387_1 | TATGAGTGTGGAGTCGTTGCTGGTGAAACCCCATCTCTACTAANNNAGATCGGAAG |
| 33387_2 | TATGAGTGTGGAGTCGTTGCTGGTGAAACCCCATCTCTACTAANNNAGATCGGAAG |
| 51901_1 | TATGAGTGTGGAGTCGTTGCCCTTGGATCAGAGGTCAGAAGACNNNAGATCGGAA |
| 51901_2 | TATGAGTGTGGAGTCGTTGCCCTTGGATCAGAGGTCAGAAGACNNNAGATCGGAA |
| 48641_1 | TATGAGTGTGGAGTCGTTGCCTCTTGGGCTCAAGTGATTCTCANNNAGATCGGAAG |
| 48641_2 | TATGAGTGTGGAGTCGTTGCCTCTTGGGCTCAAGTGATTCTCANNNAGATCGGAAG |
| 49983_1 | TATGAGTGTGGAGTCGTTGCTCTGCACAGAGCTTTTACATGGTNNNAGATCGGAAG |
| 49983_2 | TATGAGTGTGGAGTCGTTGCTCTGCACAGAGCTTTTACATGGTNNNAGATCGGAAG |
| 37523_1 | TATGAGTGTGGAGTCGTTGCCACATTGACCCAACAACTTCCTCNNNAGATCGGAAGA |
| 37523_2 | TATGAGTGTGGAGTCGTTGCCACATTGACCCAACAACTTCCTCNNNAGATCGGAAGA |
| 53416_1 | TATGAGTGTGGAGTCGTTGCAGAGGGATGGAGACAGAGAAAGANNNAGATCGGA |
| 53416_2 | TATGAGTGTGGAGTCGTTGCAGAGGGATGGAGACAGAGAAAGANNNAGATCGGA |
| 12501_1 | TATGAGTGTGGAGTCGTTGCCTTGTGAAGGAATCTAGCATAANNNAGATCGGAAG |

| | |
|---|---|
| 12501_2 | TATGAGTGTGGAGTCGTTGCCTTGTGAAGGAATCTAGCATAANNNAGATCGGAAG |
| 51496_1 | TATGAGTGTGGAGTCGTTGCATTAGCAGAAGTCTCAACCCCTGNNNAGATCGGAAG |
| 51496_2 | TATGAGTGTGGAGTCGTTGCATTAGCAGAAGTCTCAACCCCTGNNNAGATCGGAAG |
| 30063_1 | TGAGTGTGGAGTCGTTGCACTGCTTTTTCTCTTAAAATCTTTGCCNNNAGATCGGAA |
| 30063_2 | TGAGTGTGGAGTCGTTGCACTGCTTTTTCTCTTAAAATCTTTGCCNNNAGATCGGAA |
| 40655_1 | TATGAGTGTGGAGTCGTTGCTTGTGTGATCTTTGCCATGTTGTNNNAGATCGGAAG |
| 40655_2 | TATGAGTGTGGAGTCGTTGCTTGTGTGATCTTTGCCATGTTGTNNNAGATCGGAAG |
| 38244_1 | TATGAGTGTGGAGTCGTTGCAAGGACACACTTAGGGAAATGCANNNAGATCGGAA |
| 38244_2 | TATGAGTGTGGAGTCGTTGCAAGGACACACTTAGGGAAATGCANNNAGATCGGAA |
| 46751_1 | TATGAGTGTGGAGTCGTTGCTACAAGCAATTCTCCTGCCTCAGNNNAGATCGGAAG |
| 46751_2 | TATGAGTGTGGAGTCGTTGCTACAAGCAATTCTCCTGCCTCAGNNNAGATCGGAAG |
| 32508_1 | TATGAGTGTGGAGTCGTTGCTTCCTTCTGTGCAGATCATCTCCNNNAGATCGGAAGA |
| 32508_2 | TATGAGTGTGGAGTCGTTGCTTCCTTCTGTGCAGATCATCTCCNNNAGATCGGAAGA |
| 32041_1 | TATGAGTGTGGAGTCGTTGCCAGAATTTCACTCGTTGCCCAGNNNAGATCGGAAGA |
| 32041_2 | TATGAGTGTGGAGTCGTTGCCAGAATTTCACTCGTTGCCCAGNNNAGATCGGAAGA |
| 34313_1 | TATGAGTGTGGAGTCGTTGCTGCCCAAGTCCTATGTTTCTTGANNNAGATCGGAAG |
| 34313_2 | TATGAGTGTGGAGTCGTTGCTGCCCAAGTCCTATGTTTCTTGANNNAGATCGGAAG |
| 10802_1 | TATGAGTGTGGAGTCGTTGCCCACCCTGGGAAATTAGATGCTCNNNAGATCGGAAG |
| 10802_2 | TATGAGTGTGGAGTCGTTGCCCACCCTGGGAAATTAGATGCTCNNNAGATCGGAAG |
| 31187_1 | TATGAGTGTGGAGTCGTTGCGGCCCCAAACATCTGTATCAAAANNNAGATCGGAAG |
| 31187_2 | TATGAGTGTGGAGTCGTTGCGGCCCCAAACATCTGTATCAAAANNNAGATCGGAAG |
| 52657_1 | TATGAGTGTGGAGTCGTTGCACAGATCATCTCCCACACATCACNNNAGATCGGAAGA |
| 52657_2 | TATGAGTGTGGAGTCGTTGCACAGATCATCTCCCACACATCACNNNAGATCGGAAGA |
| 47126_1 | TATGAGTGTGGAGTCGTTGCCTCCCAAGTAGGTGGAACTACAGNNNAGATCGGAA |
| 47126_2 | TATGAGTGTGGAGTCGTTGCCTCCCAAGTAGGTGGAACTACAGNNNAGATCGGAA |
| 34423_1 | TATGAGTGTGGAGTCGTTGCCCGCTACTGGGAAGTGAGGANNNAGATCGGAAGAG |
| 34423_2 | TATGAGTGTGGAGTCGTTGCCCGCTACTGGGAAGTGAGGANNNAGATCGGAAGAG |
| 32255_1 | TATGAGTGTGGAGTCGTTGCAGAAACTGGGAGTCATCGATCACNNNAGATCGGAA |
| 32255_2 | TATGAGTGTGGAGTCGTTGCAGAAACTGGGAGTCATCGATCACNNNAGATCGGAA |
| 13130_1 | TATGAGTGTGGAGTCGTTGCCCTTCTAGCCATAGTAACATATTNNNAGATCGGAAG |
| 13130_2 | TATGAGTGTGGAGTCGTTGCCCTTCTAGCCATAGTAACATATTNNNAGATCGGAAG |
| 39534_1 | TATGAGTGTGGAGTCGTTGCTCCATCTGCTTATATCTCCAGTGANNNAGATCGGAAG |
| 39534_2 | TATGAGTGTGGAGTCGTTGCTCCATCTGCTTATATCTCCAGTGANNNAGATCGGAAG |
| 36490_1 | TATGAGTGTGGAGTCGTTGCCGGTATGGGAGCCAATTTTTGAANNNAGATCGGAA |
| 36490_2 | TATGAGTGTGGAGTCGTTGCCGGTATGGGAGCCAATTTTTGAANNNAGATCGGAA |
| 40109_1 | TATGAGTGTGGAGTCGTTGCGCTTTGCCACCGTGTTTTTTTTTNNNAGATCGGAAGA |
| 40109_2 | TATGAGTGTGGAGTCGTTGCGCTTTGCCACCGTGTTTTTTTTTNNNAGATCGGAAGA |
| 50522_1 | TATGAGTGTGGAGTCGTTGCTGATTTCCTCCCTAGTGCTTTCCNNNAGATCGGAAGA |
| 50522_2 | TATGAGTGTGGAGTCGTTGCTGATTTCCTCCCTAGTGCTTTCCNNNAGATCGGAAGA |
| 41245_1 | TATGAGTGTGGAGTCGTTGCGCCTGTAATCCCAGCACTTTGNNNAGATCGGAAGAG |
| 41245_2 | TATGAGTGTGGAGTCGTTGCGCCTGTAATCCCAGCACTTTGNNNAGATCGGAAGAG |
| 47492_1 | TATGAGTGTGGAGTCGTTGCGCCTCCTGAGTAGCTGAGATTACNNNAGATCGGAAG |
| 47492_2 | TATGAGTGTGGAGTCGTTGCGCCTCCTGAGTAGCTGAGATTACNNNAGATCGGAAG |

| | |
|---|---|
| 31493_1 | TATGAGTGTGGAGTCGTTGCGGCAATCTGTATTATCAAGGGCCNNNAGATCGGAAG |
| 31493_2 | TATGAGTGTGGAGTCGTTGCGGCAATCTGTATTATCAAGGGCCNNNAGATCGGAAG |
| 36908_1 | TATGAGTGTGGAGTCGTTGCGCAGAACCACTGGAACTCTCATANNNAGATCGGAAG |
| 36908_2 | TATGAGTGTGGAGTCGTTGCGCAGAACCACTGGAACTCTCATANNNAGATCGGAAG |
| 41015_1 | TATGAGTGTGGAGTCGTTGCGACAAGGCACTAAGTACCCAAGANNNAGATCGGAA |
| 41015_2 | TATGAGTGTGGAGTCGTTGCGACAAGGCACTAAGTACCCAAGANNNAGATCGGAA |
| 42729_1 | TATGAGTGTGGAGTCGTTGCGTTCCACATTTTTGCAATTGCGANNNAGATCGGAAG |
| 42729_2 | TATGAGTGTGGAGTCGTTGCGTTCCACATTTTTGCAATTGCGANNNAGATCGGAAG |
| 48931_1 | TATGAGTGTGGAGTCGTTGCCCTCCAGAGTAGCTAGGACTACANNNAGATCGGAAG |
| 48931_2 | TATGAGTGTGGAGTCGTTGCCCTCCAGAGTAGCTAGGACTACANNNAGATCGGAAG |
| 39495_1 | TATGAGTGTGGAGTCGTTGCGGGGGGATGATAATGGTGGTTATNNNAGATCGGA |
| 39495_2 | TATGAGTGTGGAGTCGTTGCGGGGGGATGATAATGGTGGTTATNNNAGATCGGA |
| 31235_1 | TATGAGTGTGGAGTCGTTGCCTCCCCATCTACACACATACTTNNNAGATCGGAAGA |
| 31235_2 | TATGAGTGTGGAGTCGTTGCCTCCCCATCTACACACATACTTNNNAGATCGGAAGA |
| 51546_1 | TATGAGTGTGGAGTCGTTGCTTGGCCTCTCAAAGTATTGGGATNNNAGATCGGAAG |
| 51546_2 | TATGAGTGTGGAGTCGTTGCTTGGCCTCTCAAAGTATTGGGATNNNAGATCGGAAG |
| 41921_1 | TATGAGTGTGGAGTCGTTGCTGTTAGCCAGAATGGTCTCCATCNNNAGATCGGAAG |
| 41921_2 | TATGAGTGTGGAGTCGTTGCTGTTAGCCAGAATGGTCTCCATCNNNAGATCGGAAG |
| 51747_1 | TATGAGTGTGGAGTCGTTGCCCCTCCACACACACACATCTATTNNNAGATCGGAAGA |
| 51747_2 | TATGAGTGTGGAGTCGTTGCCCCTCCACACACACACATCTATTNNNAGATCGGAAGA |
| 33664_1 | TATGAGTGTGGAGTCGTTGCGTGTTGGAACTACAAGTGTGAGCNNNAGATCGGAA |
| 33664_2 | TATGAGTGTGGAGTCGTTGCGTGTTGGAACTACAAGTGTGAGCNNNAGATCGGAA |
| 37506_1 | TATGAGTGTGGAGTCGTTGCCTGAGGCAGGAAAATTGCTTGAANNNAGATCGGAA |
| 37506_2 | TATGAGTGTGGAGTCGTTGCCTGAGGCAGGAAAATTGCTTGAANNNAGATCGGAA |
| 13360_1 | TATGAGTGTGGAGTCGTTGCCTCTGTTTACCTGGTTATGTAANNNAGATCGGAAGA |
| 13360_2 | TATGAGTGTGGAGTCGTTGCCTCTGTTTACCTGGTTATGTAANNNAGATCGGAAGA |
| 36242_1 | TATGAGTGTGGAGTCGTTGCTAGATTCATACGAGCCAGCATGTNNNAGATCGGAAG |
| 36242_2 | TATGAGTGTGGAGTCGTTGCTAGATTCATACGAGCCAGCATGTNNNAGATCGGAAG |
| 52161_1 | TATGAGTGTGGAGTCGTTGCGAATTGCATGGGACCCTGAAAAANNNAGATCGGAA |
| 52161_2 | TATGAGTGTGGAGTCGTTGCGAATTGCATGGGACCCTGAAAAANNNAGATCGGAA |
| 47410_1 | TATGAGTGTGGAGTCGTTGCTGATATAGGATGAGCTGGGCAAGNNNAGATCGGAA |
| 47410_2 | TATGAGTGTGGAGTCGTTGCTGATATAGGATGAGCTGGGCAAGNNNAGATCGGAA |
| 48934_1 | TATGAGTGTGGAGTCGTTGCGTTCAAGCGATTCTCATGCCTCNNNAGATCGGAAGA |
| 48934_2 | TATGAGTGTGGAGTCGTTGCGTTCAAGCGATTCTCATGCCTCNNNAGATCGGAAGA |
| 6859_1 | TATGAGTGTGGAGTCGTTGCAGCCATCTGAAGCTGGGGTTGNNNAGATCGGAAGA |
| 6859_2 | TATGAGTGTGGAGTCGTTGCAGCCATCTGAAGCTGGGGTTGNNNAGATCGGAAGA |
| 39647_1 | TATGAGTGTGGAGTCGTTGCAATGCTTATACACTGCTGGTGGANNNAGATCGGAAG |
| 39647_2 | TATGAGTGTGGAGTCGTTGCAATGCTTATACACTGCTGGTGGANNNAGATCGGAAG |
| 31731_1 | TATGAGTGTGGAGTCGTTGCCCAAAGGGCTTTCTTGTGACTTTNNNAGATCGGAAG |
| 31731_2 | TATGAGTGTGGAGTCGTTGCCCAAAGGGCTTTCTTGTGACTTTNNNAGATCGGAAG |
| 30391_1 | TATGAGTGTGGAGTCGTTGCATAAAAGTGTAGAAATCTCTTCAACGTNNNAGATCG |
| 30391_2 | TATGAGTGTGGAGTCGTTGCATAAAAGTGTAGAAATCTCTTCAACGTNNNAGATCG |
| 35500_1 | TATGAGTGTGGAGTCGTTGCGGAGACAGGTTCTTGCTATGTCANNNAGATCGGAA |

| | |
|---|---|
| 35500_2 | TATGAGTGTGGAGTCGTTGCGGAGACAGGTTCTTGCTATGTCANNNAGATCGGAA |
| 34036_1 | TATGAGTGTGGAGTCGTTGCCAGTGACATGATCTTGGCTCACTNNNAGATCGGAAG |
| 34036_2 | TATGAGTGTGGAGTCGTTGCCAGTGACATGATCTTGGCTCACTNNNAGATCGGAAG |
| 44664_1 | TATGAGTGTGGAGTCGTTGCGCAGAGACAGGGAATCAAAGGTANNNAGATCGGA |
| 44664_2 | TATGAGTGTGGAGTCGTTGCGCAGAGACAGGGAATCAAAGGTANNNAGATCGGA |
| 46701_1 | TATGAGTGTGGAGTCGTTGCGAGTGCAGTGGACTCAATCTCAGNNNAGATCGGAA |
| 46701_2 | TATGAGTGTGGAGTCGTTGCGAGTGCAGTGGACTCAATCTCAGNNNAGATCGGAA |
| 18970_1 | TATGAGTGTGGAGTCGTTGCATACATGCTACAAAAAGAAAGAGNNNAGATCGGAA |
| 18970_2 | TATGAGTGTGGAGTCGTTGCATACATGCTACAAAAAGAAAGAGNNNAGATCGGAA |
| 2140_1 | TATGAGTGTGGAGTCGTTGCCCTTCTCCACACCTCTCTTCTCANNNAGATCGGAAGA |
| 2140_2 | TATGAGTGTGGAGTCGTTGCCCTTCTCCACACCTCTCTTCTCANNNAGATCGGAAGA |
| 51582_1 | TATGAGTGTGGAGTCGTTGCTGTTGAAGAGACGAAGTGACCAANNNAGATCGGAA |
| 51582_2 | TATGAGTGTGGAGTCGTTGCTGTTGAAGAGACGAAGTGACCAANNNAGATCGGAA |
| 10736_1 | TATGAGTGTGGAGTCGTTGCGAGCACTTCAGAGCCAAGCTATTNNNAGATCGGAAG |
| 10736_2 | TATGAGTGTGGAGTCGTTGCGAGCACTTCAGAGCCAAGCTATTNNNAGATCGGAAG |
| 40335_1 | TATGAGTGTGGAGTCGTTGCCAACTGAAAGGCTAGTGTGGTTGNNNAGATCGGAA |
| 40335_2 | TATGAGTGTGGAGTCGTTGCCAACTGAAAGGCTAGTGTGGTTGNNNAGATCGGAA |
| 33704_1 | TATGAGTGTGGAGTCGTTGCTCCTGTATGTCAAGACAAATTCCANNNAGATCGGAA |
| 33704_2 | TATGAGTGTGGAGTCGTTGCTCCTGTATGTCAAGACAAATTCCANNNAGATCGGAA |
| 35504_1 | TATGAGTGTGGAGTCGTTGCGTAGAGACAGGATTTCACGGTGTNNNAGATCGGAA |
| 35504_2 | TATGAGTGTGGAGTCGTTGCGTAGAGACAGGATTTCACGGTGTNNNAGATCGGAA |
| 31986_1 | TATGAGTGTGGAGTCGTTGCACGTAAGTGGACTCAACTATCCANNNAGATCGGAAG |
| 31986_2 | TATGAGTGTGGAGTCGTTGCACGTAAGTGGACTCAACTATCCANNNAGATCGGAAG |
| 10334_1 | TATGAGTGTGGAGTCGTTGCGTTTATCTGGCCCACCAGTTAGGNNNAGATCGGAAG |
| 10334_2 | TATGAGTGTGGAGTCGTTGCGTTTATCTGGCCCACCAGTTAGGNNNAGATCGGAAG |
| 32545_1 | TATGAGTGTGGAGTCGTTGCGGTGATGGAAACAGAGAGGGAAANNNAGATCGGA |
| 32545_2 | TATGAGTGTGGAGTCGTTGCGGTGATGGAAACAGAGAGGGAAANNNAGATCGGA |
| 38876_1 | TATGAGTGTGGAGTCGTTGCGGGGGGTCTGTTTATATTATTTGCCNNNAGATCGGA |
| 38876_2 | TATGAGTGTGGAGTCGTTGCGGGGGGTCTGTTTATATTATTTGCCNNNAGATCGGA |
| 8086_1 | TATGAGTGTGGAGTCGTTGCAATGTCTGTCAAAACGGGAAAGGNNNAGATCGGAA |
| 8086_2 | TATGAGTGTGGAGTCGTTGCAATGTCTGTCAAAACGGGAAAGGNNNAGATCGGAA |
| 12723_1 | TATGAGTGTGGAGTCGTTGCAGAGAGGGTATTTAGACCAGNNNAGATCGGAAGA |
| 12723_2 | TATGAGTGTGGAGTCGTTGCAGAGAGGGTATTTAGACCAGNNNAGATCGGAAGA |
| 40603_1 | TATGAGTGTGGAGTCGTTGCGGATGAGGCAGGAGAATCACTTTNNNAGATCGGAA |
| 40603_2 | TATGAGTGTGGAGTCGTTGCGGATGAGGCAGGAGAATCACTTTNNNAGATCGGAA |
| 46401_1 | TATGAGTGTGGAGTCGTTGCCCACGCCCAGCTAATTTTTTGTANNNAGATCGGAAG |
| 46401_2 | TATGAGTGTGGAGTCGTTGCCCACGCCCAGCTAATTTTTTGTANNNAGATCGGAAG |
| 47392_1 | TATGAGTGTGGAGTCGTTGCGCCTCCCAAGTAGCTAGGATTACNNNAGATCGGAAG |
| 47392_2 | TATGAGTGTGGAGTCGTTGCGCCTCCCAAGTAGCTAGGATTACNNNAGATCGGAAG |
| 53267_1 | TATGAGTGTGGAGTCGTTGCGAGAAGATGGCTGGAGGACGNNNAGATCGGAAGA |
| 53267_2 | TATGAGTGTGGAGTCGTTGCGAGAAGATGGCTGGAGGACGNNNAGATCGGAAGA |
| 38851_1 | TATGAGTGTGGAGTCGTTGCGTAGAGATGGGGTTTCGCCATATNNNAGATCGGAA |
| 38851_2 | TATGAGTGTGGAGTCGTTGCGTAGAGATGGGGTTTCGCCATATNNNAGATCGGAA |

| | |
|---|---|
| 471_1 | TATGAGTGTGGAGTCGTTGCAATCCACTCATCCTACCCATCCTNNNAGATCGGAAGA |
| 471_2 | TATGAGTGTGGAGTCGTTGCAATCCACTCATCCTACCCATCCTNNNAGATCGGAAGA |
| 30870_1 | TATGAGTGTGGAGTCGTTGCGAGAGGAGGTAGACAGAGAGGAGNNNAGATCGGA |
| 30870_2 | TATGAGTGTGGAGTCGTTGCGAGAGGAGGTAGACAGAGAGGAGNNNAGATCGGA |
| 9922_1 | TATGAGTGTGGAGTCGTTGCGATTTCCAAGCATGGCCTACAGANNNAGATCGGAAG |
| 9922_2 | TATGAGTGTGGAGTCGTTGCGATTTCCAAGCATGGCCTACAGANNNAGATCGGAAG |
| 49676_1 | TATGAGTGTGGAGTCGTTGCTGTAGAGACAGGGTTTCACTGTGNNNAGATCGGAA |
| 49676_2 | TATGAGTGTGGAGTCGTTGCTGTAGAGACAGGGTTTCACTGTGNNNAGATCGGAA |
| 31274_1 | TATGAGTGTGGAGTCGTTGCAGAGAGAGAGAGAGAGAGTGCAGNNNAGATCGGA |
| 31274_2 | TATGAGTGTGGAGTCGTTGCAGAGAGAGAGAGAGAGAGTGCAGNNNAGATCGGA |
| 46917_1 | TATGAGTGTGGAGTCGTTGCCTCCCACGTAGCTGAGATTACAGNNNAGATCGGAAG |
| 46917_2 | TATGAGTGTGGAGTCGTTGCCTCCCACGTAGCTGAGATTACAGNNNAGATCGGAAG |
| 49393_1 | TATGAGTGTGGAGTCGTTGCCACCACACCTGGCTAATTTTTGTNNNAGATCGGAAGA |
| 49393_2 | TATGAGTGTGGAGTCGTTGCCACCACACCTGGCTAATTTTTGTNNNAGATCGGAAGA |
| 52630_1 | TATGAGTGTGGAGTCGTTGCTTGCAAACCCATGATTTTGGTGTNNNAGATCGGAAG |
| 52630_2 | TATGAGTGTGGAGTCGTTGCTTGCAAACCCATGATTTTGGTGTNNNAGATCGGAAG |
| 51180_1 | TATGAGTGTGGAGTCGTTGCGACGCAGAAGATGGATGATCTCTNNNAGATCGGAA |
| 51180_2 | TATGAGTGTGGAGTCGTTGCGACGCAGAAGATGGATGATCTCTNNNAGATCGGAA |
| 31823_1 | TATGAGTGTGGAGTCGTTGCACACACACACACACACACCTATANNNAGATCGGAAGA |
| 31823_2 | TATGAGTGTGGAGTCGTTGCACACACACACACACACACCTATANNNAGATCGGAAGA |
| 52886_1 | TATGAGTGTGGAGTCGTTGCAGGAGAACTCCTACGTGTCCAGNNNAGATCGGAAGA |
| 52886_2 | TATGAGTGTGGAGTCGTTGCAGGAGAACTCCTACGTGTCCAGNNNAGATCGGAAGA |
| 44061_1 | TATGAGTGTGGAGTCGTTGCCCCCACACTCAGGAAGTTAGAAANNNAGATCGGAAG |
| 44061_2 | TATGAGTGTGGAGTCGTTGCCCCCACACTCAGGAAGTTAGAAANNNAGATCGGAAG |
| 32483_1 | TATGAGTGTGGAGTCGTTGCTTCAGGGATTCAACTTCCTCCTGNNNAGATCGGAAGA |
| 32483_2 | TATGAGTGTGGAGTCGTTGCTTCAGGGATTCAACTTCCTCCTGNNNAGATCGGAAGA |
| 50184_1 | TATGAGTGTGGAGTCGTTGCGGACAGGGATGGGAGGCTACNNNAGATCGGAAGA |
| 50184_2 | TATGAGTGTGGAGTCGTTGCGGACAGGGATGGGAGGCTACNNNAGATCGGAAGA |
| 42698_1 | TATGAGTGTGGAGTCGTTGCAAATGGCATTCACAGAAACCTGGNNNAGATCGGAA |
| 42698_2 | TATGAGTGTGGAGTCGTTGCAAATGGCATTCACAGAAACCTGGNNNAGATCGGAA |
| 35481_1 | TATGAGTGTGGAGTCGTTGCTGATCTCGGCTTACTACAAGCTCNNNAGATCGGAAGA |
| 35481_2 | TATGAGTGTGGAGTCGTTGCTGATCTCGGCTTACTACAAGCTCNNNAGATCGGAAGA |
| 36384_1 | TATGAGTGTGGAGTCGTTGCCCAAGTGGGTGACGTCAATTATCNNNAGATCGGAA |
| 36384_2 | TATGAGTGTGGAGTCGTTGCCCAAGTGGGTGACGTCAATTATCNNNAGATCGGAA |
| 33149_1 | TATGAGTGTGGAGTCGTTGCGATGTATAGGACAACTGGGGAAANNNAGATCGGAA |
| 33149_2 | TATGAGTGTGGAGTCGTTGCGATGTATAGGACAACTGGGGAAANNNAGATCGGAA |
| 44663_1 | TATGAGTGTGGAGTCGTTGCTCAAGGACTGAATCACAGCAAGANNNAGATCGGAA |
| 44663_2 | TATGAGTGTGGAGTCGTTGCTCAAGGACTGAATCACAGCAAGANNNAGATCGGAA |
| 32063_1 | TATGAGTGTGGAGTCGTTGCTTGAGGTTTTCTTGGGACTTACANNNAGATCGGAAG |
| 32063_2 | TATGAGTGTGGAGTCGTTGCTTGAGGTTTTCTTGGGACTTACANNNAGATCGGAAG |
| 46161_1 | TATGAGTGTGGAGTCGTTGCGATCTCAGCTCACTGCAACCTCNNNAGATCGGAAGA |
| 46161_2 | TATGAGTGTGGAGTCGTTGCGATCTCAGCTCACTGCAACCTCNNNAGATCGGAAGA |
| 50882_1 | TATGAGTGTGGAGTCGTTGCTGAAGAGGAAAAGTGTTGGTCCANNNAGATCGGAA |

| | |
|---|---|
| 50882_2 | TATGAGTGTGGAGTCGTTGCTGAAGAGGAAAAGTGTTGGTCCANNNAGATCGGAA |
| 44982_1 | TATGAGTGTGGAGTCGTTGCAGCTCTGATCTCTTCTTCCGTTGNNNAGATCGGAAGA |
| 44982_2 | TATGAGTGTGGAGTCGTTGCAGCTCTGATCTCTTCTTCCGTTGNNNAGATCGGAAGA |
| 10820_1 | TATGAGTGTGGAGTCGTTGCATGACAGAGGAGGGAGAGATCACNNNAGATCGGAA |
| 10820_2 | TATGAGTGTGGAGTCGTTGCATGACAGAGGAGGGAGAGATCACNNNAGATCGGAA |
| 39043_1 | TATGAGTGTGGAGTCGTTGCCTTTGCATGGCTGAAAAAAAGGTNNNAGATCGGAAC |
| 39043_2 | TATGAGTGTGGAGTCGTTGCCTTTGCATGGCTGAAAAAAAGGTNNNAGATCGGAAC |
| 34111_1 | TATGAGTGTGGAGTCGTTGCTTGTTGACCTCTGCCTCTGTTTANNNAGATCGGAAGA |
| 34111_2 | TATGAGTGTGGAGTCGTTGCTTGTTGACCTCTGCCTCTGTTTANNNAGATCGGAAGA |
| 40769_1 | TATGAGTGTGGAGTCGTTGCAATCAAATTGTCCCTCTTTGCGGNNNAGATCGGAAGA |
| 40769_2 | TATGAGTGTGGAGTCGTTGCAATCAAATTGTCCCTCTTTGCGGNNNAGATCGGAAGA |
| 9759_1 | TATGAGTGTGGAGTCGTTGCCCATGCCTTTCAGGAAAACCAACNNNAGATCGGAAGA |
| 9759_2 | TATGAGTGTGGAGTCGTTGCCCATGCCTTTCAGGAAAACCAACNNNAGATCGGAAGA |
| 42143_1 | TATGAGTGTGGAGTCGTTGCTGTCCTTCTGCATCAGGCTATTTNNNAGATCGGAAGA |
| 42143_2 | TATGAGTGTGGAGTCGTTGCTGTCCTTCTGCATCAGGCTATTTNNNAGATCGGAAGA |
| 52740_1 | TATGAGTGTGGAGTCGTTGCCCTCCTACAAGAGCCAGAGTTGNNNAGATCGGAAGA |
| 52740_2 | TATGAGTGTGGAGTCGTTGCCCTCCTACAAGAGCCAGAGTTGNNNAGATCGGAAGA |
| 32212_1 | TATGAGTGTGGAGTCGTTGCTTTGTCATGAACCATCTGTTGCCNNNAGATCGGAAGA |
| 32212_2 | TATGAGTGTGGAGTCGTTGCTTTGTCATGAACCATCTGTTGCCNNNAGATCGGAAGA |
| 45230_1 | TATGAGTGTGGAGTCGTTGCTCTCAAACACCTCGATCGAGATCNNNAGATCGGAAGA |
| 45230_2 | TATGAGTGTGGAGTCGTTGCTCTCAAACACCTCGATCGAGATCNNNAGATCGGAAGA |
| 29038_1 | TATGAGTGTGGAGTCGTTGCCACACAGAGAGAGAGAGAGAGAGANNNAGATCGGA |
| 29038_2 | TATGAGTGTGGAGTCGTTGCCACACAGAGAGAGAGAGAGAGAGANNNAGATCGGA |
| 10117_1 | TATGAGTGTGGAGTCGTTGCGTCACATGAAGGCTGCTATAGCTNNNAGATCGGAAG |
| 10117_2 | TATGAGTGTGGAGTCGTTGCGTCACATGAAGGCTGCTATAGCTNNNAGATCGGAAG |
| 29859_1 | TATGAGTGTGGAGTCGTTGCTCTGAAGACTCACTTTAAAGGCTNNNAGATCGGAAG |
| 29859_2 | TATGAGTGTGGAGTCGTTGCTCTGAAGACTCACTTTAAAGGCTNNNAGATCGGAAG |
| 48546_1 | TATGAGTGTGGAGTCGTTGCGAATCTCCCACCATGGTCTACTCNNNAGATCGGAAGA |
| 48546_2 | TATGAGTGTGGAGTCGTTGCGAATCTCCCACCATGGTCTACTCNNNAGATCGGAAGA |
| 39178_1 | TATGAGTGTGGAGTCGTTGCCTTCCTAAGTGAAGGCACAGAGTNNNAGATCGGAAG |
| 39178_2 | TATGAGTGTGGAGTCGTTGCCTTCCTAAGTGAAGGCACAGAGTNNNAGATCGGAAG |
| 44838_1 | TATGAGTGTGGAGTCGTTGCTCTCTTCTCTCTCAGGACGGAATNNNAGATCGGAAGA |
| 44838_2 | TATGAGTGTGGAGTCGTTGCTCTCTTCTCTCTCAGGACGGAATNNNAGATCGGAAGA |
| 42966_1 | TATGAGTGTGGAGTCGTTGCCTCAAAATAGCATTTCCTGGGCANNNAGATCGGAAG |
| 42966_2 | TATGAGTGTGGAGTCGTTGCCTCAAAATAGCATTTCCTGGGCANNNAGATCGGAAG |
| 29867_1 | TATGAGTGTGGAGTCGTTGCCCCTTCCCCAAAGCAGTTTTAAGNNNAGATCGGAAGA |
| 29867_2 | TATGAGTGTGGAGTCGTTGCCCCTTCCCCAAAGCAGTTTTAAGNNNAGATCGGAAGA |
| 39270_1 | TATGAGTGTGGAGTCGTTGCAAAGTTTTAGGGTCCAAAGTCCTNNNAGATCGGAAG |
| 39270_2 | TATGAGTGTGGAGTCGTTGCAAAGTTTTAGGGTCCAAAGTCCTNNNAGATCGGAAG |
| 37629_1 | TATGAGTGTGGAGTCGTTGCAAACGACAATACATCATGGCCACNNNAGATCGGAAG |
| 37629_2 | TATGAGTGTGGAGTCGTTGCAAACGACAATACATCATGGCCACNNNAGATCGGAAG |
| 43286_1 | TATGAGTGTGGAGTCGTTGCGGCTCTTTTCTCTTTTGCATTGCNNNAGATCGGAAGA |
| 43286_2 | TATGAGTGTGGAGTCGTTGCGGCTCTTTTCTCTTTTGCATTGCNNNAGATCGGAAGA |

| | |
|---|---|
| 45118_1 | TATGAGTGTGGAGTCGTTGCTTGTTTTGCTCTGCTAGGAGACTNNNAGATCGGAAG |
| 45118_2 | TATGAGTGTGGAGTCGTTGCTTGTTTTGCTCTGCTAGGAGACTNNNAGATCGGAAG |
| 39510_1 | TATGAGTGTGGAGTCGTTGCATAGTCTTCATCTCTGCCTTGGCNNNAGATCGGAAGA |
| 39510_2 | TATGAGTGTGGAGTCGTTGCATAGTCTTCATCTCTGCCTTGGCNNNAGATCGGAAGA |
| 38520_1 | TATGAGTGTGGAGTCGTTGCCAATAGTGAAATATGGCCAGGCGNNNAGATCGGAA[illegible] |
| 38520_2 | TATGAGTGTGGAGTCGTTGCCAATAGTGAAATATGGCCAGGCGNNNAGATCGGAA[illegible] |
| 41334_1 | TATGAGTGTGGAGTCGTTGCTGACCCATTTGTACTGAGCAAGANNNAGATCGGAAG |
| 41334_2 | TATGAGTGTGGAGTCGTTGCTGACCCATTTGTACTGAGCAAGANNNAGATCGGAAG |
| 51682_1 | TATGAGTGTGGAGTCGTTGCCTCCTAATGACCAGTCTTGGCTTNNNAGATCGGAAGA |
| 51682_2 | TATGAGTGTGGAGTCGTTGCCTCCTAATGACCAGTCTTGGCTTNNNAGATCGGAAGA |
| 44882_1 | TATGAGTGTGGAGTCGTTGCGGGTGAAACCGCTGGATCTATAANNNAGATCGGAA[illegible] |
| 44882_2 | TATGAGTGTGGAGTCGTTGCGGGTGAAACCGCTGGATCTATAANNNAGATCGGAA[illegible] |
| 28895_1 | TATGAGTGTGGAGTCGTTGCCACTGTGGGTACAAGTAGAACCANNNAGATCGGAA[illegible] |
| 28895_2 | TATGAGTGTGGAGTCGTTGCCACTGTGGGTACAAGTAGAACCANNNAGATCGGAA[illegible] |
| 46916_1 | TATGAGTGTGGAGTCGTTGCCAGTTATAAGGGACAAGGACCCGNNNAGATCGGAA[illegible] |
| 46916_2 | TATGAGTGTGGAGTCGTTGCCAGTTATAAGGGACAAGGACCCGNNNAGATCGGAA[illegible] |
| 39535_1 | TATGAGTGTGGAGTCGTTGCTACAAGAATGGACACTGTGCTCANNNAGATCGGAAG |
| 39535_2 | TATGAGTGTGGAGTCGTTGCTACAAGAATGGACACTGTGCTCANNNAGATCGGAAG |
| 35305_1 | TATGAGTGTGGAGTCGTTGCGGGGACTTGTACAAAAAAGTGGGNNNAGATCGGAA |
| 35305_2 | TATGAGTGTGGAGTCGTTGCGGGGACTTGTACAAAAAAGTGGGNNNAGATCGGAA |
| 41519_1 | TATGAGTGTGGAGTCGTTGCATCACTTGAGCTCAGGAGACAAGNNNAGATCGGAA[illegible] |
| 41519_2 | TATGAGTGTGGAGTCGTTGCATCACTTGAGCTCAGGAGACAAGNNNAGATCGGAA[illegible] |
| 30987_1 | TATGAGTGTGGAGTCGTTGCTCCTACTTTGGGGCATCTAGTCTNNNAGATCGGAAGA |
| 30987_2 | TATGAGTGTGGAGTCGTTGCTCCTACTTTGGGGCATCTAGTCTNNNAGATCGGAAGA |
| 50506_1 | TATGAGTGTGGAGTCGTTGCCACCTCTGTGGCTCTTCACCNNNAGATCGGAAGAGCA |
| 50506_2 | TATGAGTGTGGAGTCGTTGCCACCTCTGTGGCTCTTCACCNNNAGATCGGAAGAGCA |
| 45934_1 | TATGAGTGTGGAGTCGTTGCCATAGACGGGGTTTTTCCATGTTNNNAGATCGGAAG |
| 45934_2 | TATGAGTGTGGAGTCGTTGCCATAGACGGGGTTTTTCCATGTTNNNAGATCGGAAG |
| 32658_1 | TATGAGTGTGGAGTCGTTGCTGTAGAGTCTCACTATGTTGCCCNNNAGATCGGAAGA |
| 32658_2 | TATGAGTGTGGAGTCGTTGCTGTAGAGTCTCACTATGTTGCCCNNNAGATCGGAAGA |
| 31211_1 | TATGAGTGTGGAGTCGTTGCTATTCTTGCTCTTGCTGGTGACTNNNAGATCGGAAGA |
| 31211_2 | TATGAGTGTGGAGTCGTTGCTATTCTTGCTCTTGCTGGTGACTNNNAGATCGGAAGA |
| 38725_1 | TATGAGTGTGGAGTCGTTGCAAAGGAACCCTAAAAACCATGCANNNAGATCGGAAG |
| 38725_2 | TATGAGTGTGGAGTCGTTGCAAAGGAACCCTAAAAACCATGCANNNAGATCGGAAG |
| 44729_1 | TATGAGTGTGGAGTCGTTGCAGAAAGAGCAAGCAATAGGGAGANNNAGATCGGAA |
| 44729_2 | TATGAGTGTGGAGTCGTTGCAGAAAGAGCAAGCAATAGGGAGANNNAGATCGGAA |
| 40639_1 | TATGAGTGTGGAGTCGTTGCTCTTGTCATTTGTAACAGCATGGANNNAGATCGGAA[illegible] |
| 40639_2 | TATGAGTGTGGAGTCGTTGCTCTTGTCATTTGTAACAGCATGGANNNAGATCGGAA[illegible] |
| 10516_1 | TATGAGTGTGGAGTCGTTGCGCGTCTTTTGCTCTTGGTTTTCTNNNAGATCGGAAGA |
| 10516_2 | TATGAGTGTGGAGTCGTTGCGCGTCTTTTGCTCTTGGTTTTCTNNNAGATCGGAAGA |
| 48554_1 | TATGAGTGTGGAGTCGTTGCCACCATGCCCAGCTAATTTTTGTNNNAGATCGGAAGA |
| 48554_2 | TATGAGTGTGGAGTCGTTGCCACCATGCCCAGCTAATTTTTGTNNNAGATCGGAAGA |
| 10966_1 | TATGAGTGTGGAGTCGTTGCGACTGGGAGAGGGTCCATGTCNNNAGATCGGAAGA |

| | |
|---|---|
| 10966_2 | TATGAGTGTGGAGTCGTTGCGACTGGGAGAGGGTCCATGTCNNNAGATCGGAAGA |
| 6237_1 | TATGAGTGTGGAGTCGTTGCATGCCAGCTGATGGTAAGTACCANNNAGATCGGAAG |
| 6237_2 | TATGAGTGTGGAGTCGTTGCATGCCAGCTGATGGTAAGTACCANNNAGATCGGAAG |
| 39426_1 | TATGAGTGTGGAGTCGTTGCATTTTACTAGCAGCCCCAACAGANNNAGATCGGAAGA |
| 39426_2 | TATGAGTGTGGAGTCGTTGCATTTTACTAGCAGCCCCAACAGANNNAGATCGGAAGA |
| 40160_1 | TATGAGTGTGGAGTCGTTGCCAATCTTGGCTCATGGAAACCTCNNNAGATCGGAAGA |
| 40160_2 | TATGAGTGTGGAGTCGTTGCCAATCTTGGCTCATGGAAACCTCNNNAGATCGGAAGA |
| 29089_1 | TATGAGTGTGGAGTCGTTGCAGGGTGTCCACAATAAGAAGAGCNNNAGATCGGAA |
| 29089_2 | TATGAGTGTGGAGTCGTTGCAGGGTGTCCACAATAAGAAGAGCNNNAGATCGGAA |
| 50050_1 | TATGAGTGTGGAGTCGTTGCGCATACGCTAAGTGGCATAACAGNNNAGATCGGAAG |
| 50050_2 | TATGAGTGTGGAGTCGTTGCGCATACGCTAAGTGGCATAACAGNNNAGATCGGAAG |
| 36772_1 | TATGAGTGTGGAGTCGTTGCCTGCCTCATAGTCTCTCAACGTTNNNAGATCGGAAGA |
| 36772_2 | TATGAGTGTGGAGTCGTTGCCTGCCTCATAGTCTCTCAACGTTNNNAGATCGGAAGA |
| 40350_1 | TATGAGTGTGGAGTCGTTGCACACATTTCTGTCCTTGTCCCTTNNNAGATCGGAAGAG |
| 40350_2 | TATGAGTGTGGAGTCGTTGCACACATTTCTGTCCTTGTCCCTTNNNAGATCGGAAGAG |
| 37378_1 | TATGAGTGTGGAGTCGTTGCCCAAAACCAAAACAAAACCGGAANNNAGATCGGAAG |
| 37378_2 | TATGAGTGTGGAGTCGTTGCCCAAAACCAAAACAAAACCGGAANNNAGATCGGAAG |
| 36892_1 | TATGAGTGTGGAGTCGTTGCTGTAATGGACACACCAGCTCAAANNNAGATCGGAAG |
| 36892_2 | TATGAGTGTGGAGTCGTTGCTGTAATGGACACACCAGCTCAAANNNAGATCGGAAG |
| 36260_1 | TATGAGTGTGGAGTCGTTGCTGACTTTCATAGGCAGCTAATTGCNNNAGATCGGAAG |
| 36260_2 | TATGAGTGTGGAGTCGTTGCTGACTTTCATAGGCAGCTAATTGCNNNAGATCGGAAG |
| 49453_1 | TATGAGTGTGGAGTCGTTGCCCTCCGATTCTCTGGGTTCTATGNNNAGATCGGAAGA |
| 49453_2 | TATGAGTGTGGAGTCGTTGCCCTCCGATTCTCTGGGTTCTATGNNNAGATCGGAAGA |
| 41348_1 | TATGAGTGTGGAGTCGTTGCATAACAGAATGGTGGTGCAGGATNNNAGATCGGAA |
| 41348_2 | TATGAGTGTGGAGTCGTTGCATAACAGAATGGTGGTGCAGGATNNNAGATCGGAA |
| 42270_1 | TATGAGTGTGGAGTCGTTGCGGTGAGTCTGTGGTCATTTTGTGNNNAGATCGGAAG |
| 42270_2 | TATGAGTGTGGAGTCGTTGCGGTGAGTCTGTGGTCATTTTGTGNNNAGATCGGAAG |
| 10857_1 | TATGAGTGTGGAGTCGTTGCGCCTGTTGTCTGGGAAGGATAAANNNAGATCGGAAG |
| 10857_2 | TATGAGTGTGGAGTCGTTGCGCCTGTTGTCTGGGAAGGATAAANNNAGATCGGAAG |
| 34672_1 | TATGAGTGTGGAGTCGTTGCTCCTTTTGGGGTGACGATAATGTNNNAGATCGGAAG |
| 34672_2 | TATGAGTGTGGAGTCGTTGCTCCTTTTGGGGTGACGATAATGTNNNAGATCGGAAG |
| 44208_1 | TATGAGTGTGGAGTCGTTGCTCCCCTTCACAAAACCAAGAGAANNNAGATCGGAAGA |
| 44208_2 | TATGAGTGTGGAGTCGTTGCTCCCCTTCACAAAACCAAGAGAANNNAGATCGGAAGA |
| 31603_1 | TATGAGTGTGGAGTCGTTGCAACGTAAGGGAGAGATAGGTGGANNNAGATCGGAA |
| 31603_2 | TATGAGTGTGGAGTCGTTGCAACGTAAGGGAGAGATAGGTGGANNNAGATCGGAA |
| 29101_1 | TATGAGTGTGGAGTCGTTGCCAAAAGGAAGGCCTGTATCCTCTNNNAGATCGGAAG |
| 29101_2 | TATGAGTGTGGAGTCGTTGCCAAAAGGAAGGCCTGTATCCTCTNNNAGATCGGAAG |
| 46310_1 | TATGAGTGTGGAGTCGTTGCCAATTCTCCTGCCTCAGTCTCCNNNAGATCGGAAGAG |
| 46310_2 | TATGAGTGTGGAGTCGTTGCCAATTCTCCTGCCTCAGTCTCCNNNAGATCGGAAGAG |
| 45420_1 | TATGAGTGTGGAGTCGTTGCTTTTGTACTCAGGCCTCAGATCCNNNAGATCGGAAGA |
| 45420_2 | TATGAGTGTGGAGTCGTTGCTTTTGTACTCAGGCCTCAGATCCNNNAGATCGGAAGA |
| 48223_1 | TATGAGTGTGGAGTCGTTGCCTCAAGTGATCCTCCCAACTCAANNNAGATCGGAAGA |
| 48223_2 | TATGAGTGTGGAGTCGTTGCCTCAAGTGATCCTCCCAACTCAANNNAGATCGGAAGA |

| | |
|---|---|
| 44388_1 | TATGAGTGTGGAGTCGTTGCACCCGTAATTGACAGTCATCTCCNNNAGATCGGAAGA |
| 44388_2 | TATGAGTGTGGAGTCGTTGCACCCGTAATTGACAGTCATCTCCNNNAGATCGGAAGA |
| 48336_1 | TATGAGTGTGGAGTCGTTGCTGGCCCAATCATAGCTCACTATCNNNAGATCGGAAGA |
| 48336_2 | TATGAGTGTGGAGTCGTTGCTGGCCCAATCATAGCTCACTATCNNNAGATCGGAAGA |
| 318_1 | TATGAGTGTGGAGTCGTTGCGCTTGTGGGGACAGAATCAAGANNNAGATCGGAAG |
| 318_2 | TATGAGTGTGGAGTCGTTGCGCTTGTGGGGACAGAATCAAGANNNAGATCGGAAG |
| 48565_1 | TATGAGTGTGGAGTCGTTGCCTTCTTGGCCTTCTGAGTAGCTANNNAGATCGGAAGA |
| 48565_2 | TATGAGTGTGGAGTCGTTGCCTTCTTGGCCTTCTGAGTAGCTANNNAGATCGGAAGA |
| 40090_1 | TATGAGTGTGGAGTCGTTGCTAACCGTGCTACTTCCAGATGACNNNAGATCGGAAGA |
| 40090_2 | TATGAGTGTGGAGTCGTTGCTAACCGTGCTACTTCCAGATGACNNNAGATCGGAAGA |
| 46054_1 | TATGAGTGTGGAGTCGTTGCAGATGAGTGGGAAGCAAAGAGAGNNNAGATCGGA |
| 46054_2 | TATGAGTGTGGAGTCGTTGCAGATGAGTGGGAAGCAAAGAGAGNNNAGATCGGA |
| 34433_1 | TATGAGTGTGGAGTCGTTGCCTGGAGTGTAGTGGCATGATCTTNNNAGATCGGAAG |
| 34433_2 | TATGAGTGTGGAGTCGTTGCCTGGAGTGTAGTGGCATGATCTTNNNAGATCGGAAG |
| 35581_1 | TATGAGTGTGGAGTCGTTGCAGGGGATTAGAAAGAAGACAGCCNNNAGATCGGAA |
| 35581_2 | TATGAGTGTGGAGTCGTTGCAGGGGATTAGAAAGAAGACAGCCNNNAGATCGGAA |
| 44373_1 | TATGAGTGTGGAGTCGTTGCCTCCAAATCACTCCACCTCAGTANNNAGATCGGAAGA |
| 44373_2 | TATGAGTGTGGAGTCGTTGCCTCCAAATCACTCCACCTCAGTANNNAGATCGGAAGA |
| 36477_1 | TATGAGTGTGGAGTCGTTGCACTTTACACTCCCACCAACAGTTNNNAGATCGGAAGA |
| 36477_2 | TATGAGTGTGGAGTCGTTGCACTTTACACTCCCACCAACAGTTNNNAGATCGGAAGA |
| 38022_1 | TATGAGTGTGGAGTCGTTGCGAGGCCAGCATCATTTTGATACCNNNAGATCGGAAG |
| 38022_2 | TATGAGTGTGGAGTCGTTGCGAGGCCAGCATCATTTTGATACCNNNAGATCGGAAG |
| 6138_1 | TATGAGTGTGGAGTCGTTGCACAGAGTTTCGCTATTGCCCACNNNAGATCGGAAGA |
| 6138_2 | TATGAGTGTGGAGTCGTTGCACAGAGTTTCGCTATTGCCCACNNNAGATCGGAAGA |
| 43178_1 | TATGAGTGTGGAGTCGTTGCCATGATCTTGGCTCACTGCAACNNNAGATCGGAAGA |
| 43178_2 | TATGAGTGTGGAGTCGTTGCCATGATCTTGGCTCACTGCAACNNNAGATCGGAAGA |
| 47019_1 | TATGAGTGTGGAGTCGTTGCTTCAAGTGATTCTCCTGCCTCAANNNAGATCGGAAGA |
| 47019_2 | TATGAGTGTGGAGTCGTTGCTTCAAGTGATTCTCCTGCCTCAANNNAGATCGGAAGA |
| 37145_1 | TATGAGTGTGGAGTCGTTGCGAGGCTGAGGCAGGAGAATGNNNAGATCGGAAGA |
| 37145_2 | TATGAGTGTGGAGTCGTTGCGAGGCTGAGGCAGGAGAATGNNNAGATCGGAAGA |
| 36672_1 | TATGAGTGTGGAGTCGTTGCTTCTAAGTGCCCACCGTAAATCANNNAGATCGGAAGA |
| 36672_2 | TATGAGTGTGGAGTCGTTGCTTCTAAGTGCCCACCGTAAATCANNNAGATCGGAAGA |
| 29401_1 | TATGAGTGTGGAGTCGTTGCATTTTACAATAAAACTTTGCTGCNNNAGATCGGAAGA |
| 29401_2 | TATGAGTGTGGAGTCGTTGCATTTTACAATAAAACTTTGCTGCNNNAGATCGGAAGA |
| 43973_1 | TATGAGTGTGGAGTCGTTGCAGAAATGGTTGAGAAACAGTGCTNNNAGATCGGAA |
| 43973_2 | TATGAGTGTGGAGTCGTTGCAGAAATGGTTGAGAAACAGTGCTNNNAGATCGGAA |
| 34357_1 | TATGAGTGTGGAGTCGTTGCCTGAGGTACCAGGTTCATCTCACNNNAGATCGGAAG |
| 34357_2 | TATGAGTGTGGAGTCGTTGCCTGAGGTACCAGGTTCATCTCACNNNAGATCGGAAG |
| 33505_1 | TATGAGTGTGGAGTCGTTGCGCAATGATCATGTGTACTCAGCCNNNAGATCGGAAG |
| 33505_2 | TATGAGTGTGGAGTCGTTGCGCAATGATCATGTGTACTCAGCCNNNAGATCGGAAG |
| 49206_1 | TATGAGTGTGGAGTCGTTGCTTCCAAAGCAAACACTGTTGACANNNAGATCGGAAG |
| 49206_2 | TATGAGTGTGGAGTCGTTGCTTCCAAAGCAAACACTGTTGACANNNAGATCGGAAG |
| 29685_1 | TATGAGTGTGGAGTCGTTGCTCGACTCTCCCAACTATAAGCTGNNNAGATCGGAAGA |

| | |
|---|---|
| 29685_2 | TATGAGTGTGGAGTCGTTGCTCGACTCTCCCAACTATAAGCTGNNNAGATCGGAAG |
| 39790_1 | TATGAGTGTGGAGTCGTTGCCATATCTGACAGTCACAGAGGGGNNNAGATCGGAA |
| 39790_2 | TATGAGTGTGGAGTCGTTGCCATATCTGACAGTCACAGAGGGGNNNAGATCGGAA |
| 35009_1 | TATGAGTGTGGAGTCGTTGCTTCAAGCTATTCTCCTGCCTCAGNNNAGATCGGAAGA |
| 35009_2 | TATGAGTGTGGAGTCGTTGCTTCAAGCTATTCTCCTGCCTCAGNNNAGATCGGAAGA |
| 40026_1 | TATGAGTGTGGAGTCGTTGCCTCCATGTACTGTACCCATCACCNNNAGATCGGAAGA |
| 40026_2 | TATGAGTGTGGAGTCGTTGCCTCCATGTACTGTACCCATCACCNNNAGATCGGAAGA |
| 43329_1 | TATGAGTGTGGAGTCGTTGCTTTAGCAACCAAATGGCAGAACCNNNAGATCGGAAG |
| 43329_2 | TATGAGTGTGGAGTCGTTGCTTTAGCAACCAAATGGCAGAACCNNNAGATCGGAAG |
| 39938_1 | TATGAGTGTGGAGTCGTTGCTTTAGTCAGGTCTTTCTGCTGCTNNNAGATCGGAAG |
| 39938_2 | TATGAGTGTGGAGTCGTTGCTTTAGTCAGGTCTTTCTGCTGCTNNNAGATCGGAAG |
| 31341_1 | TATGAGTGTGGAGTCGTTGCCTACCTGGCTACCTCACTTGTTTNNNAGATCGGAAGA |
| 31341_2 | TATGAGTGTGGAGTCGTTGCCTACCTGGCTACCTCACTTGTTTNNNAGATCGGAAGA |
| 46818_1 | TATGAGTGTGGAGTCGTTGCGTTCAAGCGATTCTCCTACCTCANNNAGATCGGAAG |
| 46818_2 | TATGAGTGTGGAGTCGTTGCGTTCAAGCGATTCTCCTACCTCANNNAGATCGGAAG |
| 43418_1 | TATGAGTGTGGAGTCGTTGCTGCCAAGGTGGGGTAAACATATANNNAGATCGGAA |
| 43418_2 | TATGAGTGTGGAGTCGTTGCTGCCAAGGTGGGGTAAACATATANNNAGATCGGAA |
| 43970_1 | TATGAGTGTGGAGTCGTTGCTCTCGAACTGTGTCAGTCCAATTNNNAGATCGGAAG |
| 43970_2 | TATGAGTGTGGAGTCGTTGCTCTCGAACTGTGTCAGTCCAATTNNNAGATCGGAAG |
| 8493_1 | TATGAGTGTGGAGTCGTTGCATGTAATAACTCACTGTGGCCTGGANNNAGATCGGA |
| 8493_2 | TATGAGTGTGGAGTCGTTGCATGTAATAACTCACTGTGGCCTGGANNNAGATCGGA |
| 11690_1 | TATGAGTGTGGAGTCGTTGCGCCCTGAACTCCTGTACTCATGCNNNAGATCGGAAG |
| 11690_2 | TATGAGTGTGGAGTCGTTGCGCCCTGAACTCCTGTACTCATGCNNNAGATCGGAAG |
| 49188_1 | TATGAGTGTGGAGTCGTTGCCCAGGTTAGAGTTCATTGACCCANNNAGATCGGAAG |
| 49188_2 | TATGAGTGTGGAGTCGTTGCCCAGGTTAGAGTTCATTGACCCANNNAGATCGGAAG |
| 31851_1 | TATGAGTGTGGAGTCGTTGCCTGAATCCAGCAGCACATCAAAANNNAGATCGGAAG |
| 31851_2 | TATGAGTGTGGAGTCGTTGCCTGAATCCAGCAGCACATCAAAANNNAGATCGGAAG |
| 46286_1 | TATGAGTGTGGAGTCGTTGCGGGTTCAAGTGATTCTCCTTCCTNNNAGATCGGAAG |
| 46286_2 | TATGAGTGTGGAGTCGTTGCGGGTTCAAGTGATTCTCCTTCCTNNNAGATCGGAAG |
| 44110_1 | TATGAGTGTGGAGTCGTTGCGCCTAAGACCTTTCTTCTAGGGCNNNAGATCGGAAG |
| 44110_2 | TATGAGTGTGGAGTCGTTGCGCCTAAGACCTTTCTTCTAGGGCNNNAGATCGGAAG |
| 47432_1 | TATGAGTGTGGAGTCGTTGCTCTAGGTACTGGCTTCATCTAGTNNNAGATCGGAAG |
| 47432_2 | TATGAGTGTGGAGTCGTTGCTCTAGGTACTGGCTTCATCTAGTNNNAGATCGGAAG |
| 45268_1 | TATGAGTGTGGAGTCGTTGCCACACATGGTCCTTTTTTCCTCCNNNAGATCGGAAGA |
| 45268_2 | TATGAGTGTGGAGTCGTTGCCACACATGGTCCTTTTTTCCTCCNNNAGATCGGAAGA |
| 32852_1 | TATGAGTGTGGAGTCGTTGCTACTGAGTTTCACCATCTTGGCCNNNAGATCGGAAG |
| 32852_2 | TATGAGTGTGGAGTCGTTGCTACTGAGTTTCACCATCTTGGCCNNNAGATCGGAAG |
| 40676_1 | TATGAGTGTGGAGTCGTTGCCTGAGTTTTGCTCTTGTTGCCCNNNAGATCGGAAGA |
| 40676_2 | TATGAGTGTGGAGTCGTTGCCTGAGTTTTGCTCTTGTTGCCCNNNAGATCGGAAGA |
| 34064_1 | TATGAGTGTGGAGTCGTTGCGCAATCTTGGCTCACTGCAAGNNNAGATCGGAAGA |
| 34064_2 | TATGAGTGTGGAGTCGTTGCGCAATCTTGGCTCACTGCAAGNNNAGATCGGAAGA |
| 35378_1 | TATGAGTGTGGAGTCGTTGCTTCTAGCGATTCTTCTGCCTCAGNNNAGATCGGAAG |
| 35378_2 | TATGAGTGTGGAGTCGTTGCTTCTAGCGATTCTTCTGCCTCAGNNNAGATCGGAAG |

| | |
|---|---|
| 13043_1 | TATGAGTGTGGAGTCGTTGCTTGCTTTGGGATCACATTATTTNNNAGATCGGAAGA |
| 13043_2 | TATGAGTGTGGAGTCGTTGCTTGCTTTGGGATCACATTATTTNNNAGATCGGAAGA |
| 36396_1 | TATGAGTGTGGAGTCGTTGCTTGTTCCTAGTCCCACAAAGTCTNNNAGATCGGAAGA |
| 36396_2 | TATGAGTGTGGAGTCGTTGCTTGTTCCTAGTCCCACAAAGTCTNNNAGATCGGAAGA |
| 41157_1 | TATGAGTGTGGAGTCGTTGCTGAGGCAGAATTTCGCTATGTTGNNNAGATCGGAA |
| 41157_2 | TATGAGTGTGGAGTCGTTGCTGAGGCAGAATTTCGCTATGTTGNNNAGATCGGAA |
| 45603_1 | TATGAGTGTGGAGTCGTTGCCTCCAAGTAGCTGGGACTACAGNNNAGATCGGAAGA |
| 45603_2 | TATGAGTGTGGAGTCGTTGCCTCCAAGTAGCTGGGACTACAGNNNAGATCGGAAGA |
| 32760_1 | TATGAGTGTGGAGTCGTTGCTTCTGAGGAAAGAAATTGCCAGCNNNAGATCGGAA |
| 32760_2 | TATGAGTGTGGAGTCGTTGCTTCTGAGGAAAGAAATTGCCAGCNNNAGATCGGAA |
| 36361_1 | TATGAGTGTGGAGTCGTTGCTCTCTGTGTCCCTCTCTATCTCTNNNAGATCGGAAGA |
| 36361_2 | TATGAGTGTGGAGTCGTTGCTCTCTGTGTCCCTCTCTATCTCTNNNAGATCGGAAGA |
| 50305_1 | TATGAGTGTGGAGTCGTTGCCCAGTTCACTAAGACCCTGAGTGNNNAGATCGGAAG |
| 50305_2 | TATGAGTGTGGAGTCGTTGCCCAGTTCACTAAGACCCTGAGTGNNNAGATCGGAAG |
| 31468_1 | TATGAGTGTGGAGTCGTTGCCACTCTCACACTCTTCTTGGGAANNNAGATCGGAAGA |
| 31468_2 | TATGAGTGTGGAGTCGTTGCCACTCTCACACTCTTCTTGGGAANNNAGATCGGAAGA |
| 38067_1 | TATGAGTGTGGAGTCGTTGCACATAAAGAAGTTCTGGCCAGGTNNNAGATCGGAA |
| 38067_2 | TATGAGTGTGGAGTCGTTGCACATAAAGAAGTTCTGGCCAGGTNNNAGATCGGAA |
| 44435_1 | TATGAGTGTGGAGTCGTTGCAGAAATGCTGCTAACCAGAGGAANNNAGATCGGAA |
| 44435_2 | TATGAGTGTGGAGTCGTTGCAGAAATGCTGCTAACCAGAGGAANNNAGATCGGAA |
| 9646_1 | TATGAGTGTGGAGTCGTTGCTAAGGGAAATCCTGCCATTTGCCNNNAGATCGGAAG |
| 9646_2 | TATGAGTGTGGAGTCGTTGCTAAGGGAAATCCTGCCATTTGCCNNNAGATCGGAAG |
| 31858_1 | TATGAGTGTGGAGTCGTTGCAAAGTTGAGGAAGGAGAATCGCTNNNAGATCGGAA |
| 31858_2 | TATGAGTGTGGAGTCGTTGCAAAGTTGAGGAAGGAGAATCGCTNNNAGATCGGAA |
| 28849_1 | TATGAGTGTGGAGTCGTTGCTGGTGTTTCTGACTTCCCTTTCTNNNAGATCGGAAGA |
| 28849_2 | TATGAGTGTGGAGTCGTTGCTGGTGTTTCTGACTTCCCTTTCTNNNAGATCGGAAGA |
| 52687_1 | TATGAGTGTGGAGTCGTTGCGGCAGTTCTATTTCAGCAGCAATNNNAGATCGGAAG |
| 52687_2 | TATGAGTGTGGAGTCGTTGCGGCAGTTCTATTTCAGCAGCAATNNNAGATCGGAAG |
| 39164_1 | TATGAGTGTGGAGTCGTTGCTTGATTTCCGGACCCATCAATCTNNNAGATCGGAAGA |
| 39164_2 | TATGAGTGTGGAGTCGTTGCTTGATTTCCGGACCCATCAATCTNNNAGATCGGAAGA |
| 33479_1 | TATGAGTGTGGAGTCGTTGCTACACTTCCCCTCAGAATCTTGCNNNAGATCGGAAGA |
| 33479_2 | TATGAGTGTGGAGTCGTTGCTACACTTCCCCTCAGAATCTTGCNNNAGATCGGAAGA |
| 49464_1 | TATGAGTGTGGAGTCGTTGCGAGAGGCTCTGAGTGATCATCAGNNNAGATCGGAA |
| 49464_2 | TATGAGTGTGGAGTCGTTGCGAGAGGCTCTGAGTGATCATCAGNNNAGATCGGAA |
| 10575_1 | TATGAGTGTGGAGTCGTTGCGACAGACAAAGAGCCCAAGAGGNNNAGATCGGAAG |
| 10575_2 | TATGAGTGTGGAGTCGTTGCGACAGACAAAGAGCCCAAGAGGNNNAGATCGGAAG |
| 2029_1 | TATGAGTGTGGAGTCGTTGCTCAACCAGAAACATGTGGCTCAANNNAGATCGGAAG |
| 2029_2 | TATGAGTGTGGAGTCGTTGCTCAACCAGAAACATGTGGCTCAANNNAGATCGGAAG |
| 39665_1 | TATGAGTGTGGAGTCGTTGCGTAACCACACATCAGAACAGTGCNNNAGATCGGAAG |
| 39665_2 | TATGAGTGTGGAGTCGTTGCGTAACCACACATCAGAACAGTGCNNNAGATCGGAAG |
| 50159_1 | TATGAGTGTGGAGTCGTTGCCACAAAGGCTGCTAGGAAGTCTANNNAGATCGGAA |
| 50159_2 | TATGAGTGTGGAGTCGTTGCCACAAAGGCTGCTAGGAAGTCTANNNAGATCGGAA |
| 51013_1 | TATGAGTGTGGAGTCGTTGCGACCACAACCAGTCTGACTACATNNNAGATCGGAAG |

| | |
|---|---|
| 51013_2 | TATGAGTGTGGAGTCGTTGCGACCACAACCAGTCTGACTACATNNNAGATCGGAAG |
| 44536_1 | TATGAGTGTGGAGTCGTTGCCAAAGAAAAGACAGTAGCTCGCCNNNAGATCGGAA |
| 44536_2 | TATGAGTGTGGAGTCGTTGCCAAAGAAAAGACAGTAGCTCGCCNNNAGATCGGAA |
| 30248_1 | TATGAGTGTGGAGTCGTTGCTATGGGGATGAGACCTCATGGATNNNAGATCGGAA |
| 30248_2 | TATGAGTGTGGAGTCGTTGCTATGGGGATGAGACCTCATGGATNNNAGATCGGAA |
| 44934_1 | TATGAGTGTGGAGTCGTTGCAGCTGAAGGTTTTGGTCTCAGAANNNAGATCGGAA |
| 44934_2 | TATGAGTGTGGAGTCGTTGCAGCTGAAGGTTTTGGTCTCAGAANNNAGATCGGAA |
| 31408_1 | TATGAGTGTGGAGTCGTTGCCTTGGAGTCTGACATCCCACTAGNNNAGATCGGAAG |
| 31408_2 | TATGAGTGTGGAGTCGTTGCCTTGGAGTCTGACATCCCACTAGNNNAGATCGGAAG |
| 33167_1 | TATGAGTGTGGAGTCGTTGCATTATTCAAGGCCTTGCATATGCNNNAGATCGGAAG |
| 33167_2 | TATGAGTGTGGAGTCGTTGCATTATTCAAGGCCTTGCATATGCNNNAGATCGGAAG |
| 44205_1 | TATGAGTGTGGAGTCGTTGCCTTCCTGACTTTGGGGAATCTGANNNAGATCGGAAG |
| 44205_2 | TATGAGTGTGGAGTCGTTGCCTTCCTGACTTTGGGGAATCTGANNNAGATCGGAAG |
| 36996_1 | TATGAGTGTGGAGTCGTTGCTAGAAAACCCCGTAGTCTTGGTCNNNAGATCGGAAG |
| 36996_2 | TATGAGTGTGGAGTCGTTGCTAGAAAACCCCGTAGTCTTGGTCNNNAGATCGGAAG |
| 46066_1 | TATGAGTGTGGAGTCGTTGCGAGGCTTGTTCTCATCCTACTCCNNNAGATCGGAAG |
| 46066_2 | TATGAGTGTGGAGTCGTTGCGAGGCTTGTTCTCATCCTACTCCNNNAGATCGGAAG |
| 10306_1 | TATGAGTGTGGAGTCGTTGCTCATGGACATTCCTCTCTACCTTNNNAGATCGGAAGA |
| 10306_2 | TATGAGTGTGGAGTCGTTGCTCATGGACATTCCTCTCTACCTTNNNAGATCGGAAGA |
| 47515_1 | TATGAGTGTGGAGTCGTTGCCCTGAAGGGGAGTCTCAGTCTNNNAGATCGGAAGA |
| 47515_2 | TATGAGTGTGGAGTCGTTGCCCTGAAGGGGAGTCTCAGTCTNNNAGATCGGAAGA |
| 12822_1 | TATGAGTGTGGAGTCGTTGCGTATCCAATGTTTAGCTTTTCANNNAGATCGGAAGA |
| 12822_2 | TATGAGTGTGGAGTCGTTGCGTATCCAATGTTTAGCTTTTCANNNAGATCGGAAGA |
| 46594_1 | TATGAGTGTGGAGTCGTTGCATCTCCTGGGTTCAAGCAATTCTNNNAGATCGGAAG |
| 46594_2 | TATGAGTGTGGAGTCGTTGCATCTCCTGGGTTCAAGCAATTCTNNNAGATCGGAAG |
| 37884_1 | TATGAGTGTGGAGTCGTTGCGACAGGGACGCTAGTTCATAGAGNNNAGATCGGAA |
| 37884_2 | TATGAGTGTGGAGTCGTTGCGACAGGGACGCTAGTTCATAGAGNNNAGATCGGAA |
| 31527_1 | TATGAGTGTGGAGTCGTTGCTCCAAAACACAGAAATCACTGCCNNNAGATCGGAAG |
| 31527_2 | TATGAGTGTGGAGTCGTTGCTCCAAAACACAGAAATCACTGCCNNNAGATCGGAAG |
| 39721_1 | TATGAGTGTGGAGTCGTTGCTCCATATTTCCTTTCCTCCTTTCCANNNAGATCGGAAG |
| 39721_2 | TATGAGTGTGGAGTCGTTGCTCCATATTTCCTTTCCTCCTTTCCANNNAGATCGGAAG |
| 53311_1 | TATGAGTGTGGAGTCGTTGCAAATGGTGAGGCAAAGTGTATGCNNNAGATCGGAA |
| 53311_2 | TATGAGTGTGGAGTCGTTGCAAATGGTGAGGCAAAGTGTATGCNNNAGATCGGAA |
| 30772_1 | TATGAGTGTGGAGTCGTTGCTTAGGGGGAACAATGGAGACCTANNNAGATCGGAA |
| 30772_2 | TATGAGTGTGGAGTCGTTGCTTAGGGGGAACAATGGAGACCTANNNAGATCGGAA |
| 10626_1 | TATGAGTGTGGAGTCGTTGCGGTCCTCAACCAGACATTCTGATNNNAGATCGGAAG |
| 10626_2 | TATGAGTGTGGAGTCGTTGCGGTCCTCAACCAGACATTCTGATNNNAGATCGGAAG |
| 50981_1 | TATGAGTGTGGAGTCGTTGCCCATTGCCATAGTTGAGACCCTANNNAGATCGGAAG |
| 50981_2 | TATGAGTGTGGAGTCGTTGCCCATTGCCATAGTTGAGACCCTANNNAGATCGGAAG |
| 39799_1 | TATGAGTGTGGAGTCGTTGCCAGGAGTTGCAATCGTAGTCTCTNNNAGATCGGAAG |
| 39799_2 | TATGAGTGTGGAGTCGTTGCCAGGAGTTGCAATCGTAGTCTCTNNNAGATCGGAAG |
| 30980_1 | TATGAGTGTGGAGTCGTTGCACACAGACATGCACATACTCACTNNNAGATCGGAAG |
| 30980_2 | TATGAGTGTGGAGTCGTTGCACACAGACATGCACATACTCACTNNNAGATCGGAAG |

| | |
|---|---|
| 36907_1 | TATGAGTGTGGAGTCGTTGCCTGTAGTCCCAGCTACTCAGGAGNNNAGATCGGAAG |
| 36907_2 | TATGAGTGTGGAGTCGTTGCCTGTAGTCCCAGCTACTCAGGAGNNNAGATCGGAAG |
| 32672_1 | TATGAGTGTGGAGTCGTTGCTGTTCATGGCTTACTCTCTCACCNNNAGATCGGAAGA |
| 32672_2 | TATGAGTGTGGAGTCGTTGCTGTTCATGGCTTACTCTCTCACCNNNAGATCGGAAGA |
| 40605_1 | TATGAGTGTGGAGTCGTTGCCTAAGGCAGGGGAATCACTTGAANNNAGATCGGAA |
| 40605_2 | TATGAGTGTGGAGTCGTTGCCTAAGGCAGGGGAATCACTTGAANNNAGATCGGAA |
| 33478_1 | TATGAGTGTGGAGTCGTTGCCCTGACGCAACATAAAATGGCTANNNAGATCGGAAG |
| 33478_2 | TATGAGTGTGGAGTCGTTGCCCTGACGCAACATAAAATGGCTANNNAGATCGGAAG |
| 12392_1 | TATGAGTGTGGAGTCGTTGCGTCCATTGCCCTGTCTCCTCTTTNNNAGATCGGAAGA |
| 12392_2 | TATGAGTGTGGAGTCGTTGCGTCCATTGCCCTGTCTCCTCTTTNNNAGATCGGAAGA |
| 47088_1 | TATGAGTGTGGAGTCGTTGCCTGAGTAGCTGGGATTACAGGTGNNNAGATCGGAA |
| 47088_2 | TATGAGTGTGGAGTCGTTGCCTGAGTAGCTGGGATTACAGGTGNNNAGATCGGAA |
| 32198_1 | TATGAGTGTGGAGTCGTTGCGCCTCCAGCATAGCCAGAATTATNNNAGATCGGAAG |
| 32198_2 | TATGAGTGTGGAGTCGTTGCGCCTCCAGCATAGCCAGAATTATNNNAGATCGGAAG |
| 12971_1 | TATGAGTGTGGAGTCGTTGCAATCTGAATCTTGTGGTCTGNNNAGATCGGAAGAG |
| 12971_2 | TATGAGTGTGGAGTCGTTGCAATCTGAATCTTGTGGTCTGNNNAGATCGGAAGAG |
| 42629_1 | TATGAGTGTGGAGTCGTTGCGTGTCTGTCTGTCTGTGTGTGTANNNAGATCGGAAG |
| 42629_2 | TATGAGTGTGGAGTCGTTGCGTGTCTGTCTGTCTGTGTGTGTANNNAGATCGGAAG |
| 51007_1 | TATGAGTGTGGAGTCGTTGCATGGCCCAAATAAGATCCATGCANNNAGATCGGAAG |
| 51007_2 | TATGAGTGTGGAGTCGTTGCATGGCCCAAATAAGATCCATGCANNNAGATCGGAAG |
| 51352_1 | TATGAGTGTGGAGTCGTTGCGGGTGGGGATACTGATCAGTTCNNNAGATCGGAAG |
| 51352_2 | TATGAGTGTGGAGTCGTTGCGGGTGGGGATACTGATCAGTTCNNNAGATCGGAAG |
| 33749_1 | TATGAGTGTGGAGTCGTTGCTCCACAACGAGATACAGCAAAGANNNAGATCGGAA |
| 33749_2 | TATGAGTGTGGAGTCGTTGCTCCACAACGAGATACAGCAAAGANNNAGATCGGAA |
| 51444_1 | TATGAGTGTGGAGTCGTTGCCAATTGGCTTTTGCAAACACCAANNNAGATCGGAAG |
| 51444_2 | TATGAGTGTGGAGTCGTTGCCAATTGGCTTTTGCAAACACCAANNNAGATCGGAAG |
| 30339_1 | TATGAGTGTGGAGTCGTTGCGGAGTCTGTGGAGTGTAGATCGTNNNAGATCGGAA |
| 30339_2 | TATGAGTGTGGAGTCGTTGCGGAGTCTGTGGAGTGTAGATCGTNNNAGATCGGAA |
| 47297_1 | TATGAGTGTGGAGTCGTTGCTCCCAGGCTCAAGCAATTCTTATNNNAGATCGGAAG |
| 47297_2 | TATGAGTGTGGAGTCGTTGCTCCCAGGCTCAAGCAATTCTTATNNNAGATCGGAAG |
| 46708_1 | TATGAGTGTGGAGTCGTTGCTTCAAGTGATTCTCCTGCCTCAGNNNAGATCGGAAG |
| 46708_2 | TATGAGTGTGGAGTCGTTGCTTCAAGTGATTCTCCTGCCTCAGNNNAGATCGGAAG |
| 31925_1 | TATGAGTGTGGAGTCGTTGCCCCTGTTGCAATTCTGATCAAGCNNNAGATCGGAAG |
| 31925_2 | TATGAGTGTGGAGTCGTTGCCCCTGTTGCAATTCTGATCAAGCNNNAGATCGGAAG |
| 37157_1 | TATGAGTGTGGAGTCGTTGCTGGTAAGAGAATGATGACTTGTTGCNNNAGATCGG |
| 37157_2 | TATGAGTGTGGAGTCGTTGCTGGTAAGAGAATGATGACTTGTTGCNNNAGATCGG |
| 2764_1 | TATGAGTGTGGAGTCGTTGCAGGAAGACAATGTACAGAGGCCTNNNAGATCGGAA |
| 2764_2 | TATGAGTGTGGAGTCGTTGCAGGAAGACAATGTACAGAGGCCTNNNAGATCGGAA |
| 48892_1 | TATGAGTGTGGAGTCGTTGCGATTGTCTCGATCTCCTGACCTCNNNAGATCGGAAG |
| 48892_2 | TATGAGTGTGGAGTCGTTGCGATTGTCTCGATCTCCTGACCTCNNNAGATCGGAAG |
| 38397_1 | TATGAGTGTGGAGTCGTTGCCCACTTTTACATGTGCCACCTTTNNNAGATCGGAAGA |
| 38397_2 | TATGAGTGTGGAGTCGTTGCCCACTTTTACATGTGCCACCTTTNNNAGATCGGAAGA |
| 47443_1 | TATGAGTGTGGAGTCGTTGCACAATCTCGGCTCACTGCAAGNNNAGATCGGAAGAG |

| | |
|---|---|
| 47443_2 | TATGAGTGTGGAGTCGTTGCACAATCTCGGCTCACTGCAAGNNNAGATCGGAAGAG |
| 51539_1 | TATGAGTGTGGAGTCGTTGCGTGTAGAGTGGAGCTGAGGATGNNNAGATCGGAA |
| 51539_2 | TATGAGTGTGGAGTCGTTGCGTGTAGAGTGGAGCTGAGGATGNNNAGATCGGAA |
| 39708_1 | TATGAGTGTGGAGTCGTTGCAGATTCTCAGGTCCCACTACAGANNNAGATCGGAAG |
| 39708_2 | TATGAGTGTGGAGTCGTTGCAGATTCTCAGGTCCCACTACAGANNNAGATCGGAAG |
| 38440_1 | TATGAGTGTGGAGTCGTTGCCAGTGGTCTTGAAACAGACACACNNNAGATCGGAAG |
| 38440_2 | TATGAGTGTGGAGTCGTTGCCAGTGGTCTTGAAACAGACACACNNNAGATCGGAAG |
| 39046_1 | TATGAGTGTGGAGTCGTTGCATCCTGCAAGTGACTCAAAGGAANNNAGATCGGAA |
| 39046_2 | TATGAGTGTGGAGTCGTTGCATCCTGCAAGTGACTCAAAGGAANNNAGATCGGAA |
| 39041_1 | TATGAGTGTGGAGTCGTTGCTACAGGAGGCAAAAGATCTCCACNNNAGATCGGAA |
| 39041_2 | TATGAGTGTGGAGTCGTTGCTACAGGAGGCAAAAGATCTCCACNNNAGATCGGAA |
| 37326_1 | TATGAGTGTGGAGTCGTTGCGAAAGGTAAATCTGGCGATGCTGNNNAGATCGGAA |
| 37326_2 | TATGAGTGTGGAGTCGTTGCGAAAGGTAAATCTGGCGATGCTGNNNAGATCGGAA |
| 42941_1 | TATGAGTGTGGAGTCGTTGCTGCAAACAGGGACTGTCTGAATANNNAGATCGGAA |
| 42941_2 | TATGAGTGTGGAGTCGTTGCTGCAAACAGGGACTGTCTGAATANNNAGATCGGAA |
| 10602_1 | TATGAGTGTGGAGTCGTTGCCGGTACAATTAGCCCCTTTCCATNNNAGATCGGAAG |
| 10602_2 | TATGAGTGTGGAGTCGTTGCCGGTACAATTAGCCCCTTTCCATNNNAGATCGGAAG |
| 32908_1 | TATGAGTGTGGAGTCGTTGCGGATCACTTGAGTCCAGGAGTTTNNNAGATCGGAA |
| 32908_2 | TATGAGTGTGGAGTCGTTGCGGATCACTTGAGTCCAGGAGTTTNNNAGATCGGAA |
| 44978_1 | TATGAGTGTGGAGTCGTTGCAAGCTCCAGAGGCTATCTTTTCCNNNAGATCGGAAG |
| 44978_2 | TATGAGTGTGGAGTCGTTGCAAGCTCCAGAGGCTATCTTTTCCNNNAGATCGGAAG |
| 42993_1 | TATGAGTGTGGAGTCGTTGCATTTGAGACGGAGTCACTCTGTTNNNAGATCGGAAG |
| 42993_2 | TATGAGTGTGGAGTCGTTGCATTTGAGACGGAGTCACTCTGTTNNNAGATCGGAAG |
| 38534_1 | TATGAGTGTGGAGTCGTTGCGAATAAATGACAAGCCGGGCATGNNNAGATCGGAA |
| 38534_2 | TATGAGTGTGGAGTCGTTGCGAATAAATGACAAGCCGGGCATGNNNAGATCGGAA |
| 30384_1 | TATGAGTGTGGAGTCGTTGCCCGGTATGTAATTTAACATTCAANNNAGATCGGAAG |
| 30384_2 | TATGAGTGTGGAGTCGTTGCCCGGTATGTAATTTAACATTCAANNNAGATCGGAAG |
| 43035_1 | TATGAGTGTGGAGTCGTTGCAAATGTAAGCGTCATCCCCACTANNNAGATCGGAAG |
| 43035_2 | TATGAGTGTGGAGTCGTTGCAAATGTAAGCGTCATCCCCACTANNNAGATCGGAAG |
| 30297_1 | TATGAGTGTGGAGTCGTTGCCAACTGTTGCGCTATGAACAATANNNAGATCGGAAG |
| 30297_2 | TATGAGTGTGGAGTCGTTGCCAACTGTTGCGCTATGAACAATANNNAGATCGGAAG |
| 12952_1 | TATGAGTGTGGAGTCGTTGCCAAAGACTTAGAGCAGAAGCNNNAGATCGGAAGAG |
| 12952_2 | TATGAGTGTGGAGTCGTTGCCAAAGACTTAGAGCAGAAGCNNNAGATCGGAAGAG |
| 10255_1 | TATGAGTGTGGAGTCGTTGCTACCAGAACCCAGAGATTGAGGGNNNAGATCGGAA |
| 10255_2 | TATGAGTGTGGAGTCGTTGCTACCAGAACCCAGAGATTGAGGGNNNAGATCGGAA |
| 50307_1 | TATGAGTGTGGAGTCGTTGCCTATGGGTGGATCATGAGGTCAGNNNAGATCGGAA |
| 50307_2 | TATGAGTGTGGAGTCGTTGCCTATGGGTGGATCATGAGGTCAGNNNAGATCGGAA |
| 38661_1 | TATGAGTGTGGAGTCGTTGCAGAGGTGTTTAGAGGCAGATTGGNNNAGATCGGA |
| 38661_2 | TATGAGTGTGGAGTCGTTGCAGAGGTGTTTAGAGGCAGATTGGNNNAGATCGGA |
| 51505_1 | TATGAGTGTGGAGTCGTTGCAGGTACGCATTCTCTTTCTCAGGNNNAGATCGGAAG |
| 51505_2 | TATGAGTGTGGAGTCGTTGCAGGTACGCATTCTCTTTCTCAGGNNNAGATCGGAAG |
| 42845_1 | TATGAGTGTGGAGTCGTTGCCCTCTCCAGCTACAAATGTCATGNNNAGATCGGAAG |
| 42845_2 | TATGAGTGTGGAGTCGTTGCCCTCTCCAGCTACAAATGTCATGNNNAGATCGGAAG |

| | |
|---|---|
| 38867_1 | TATGAGTGTGGAGTCGTTGCTTAGTAGCTACGTATGGCCTTGGNNNAGATCGGAAG |
| 38867_2 | TATGAGTGTGGAGTCGTTGCTTAGTAGCTACGTATGGCCTTGGNNNAGATCGGAAG |
| 42457_1 | TATGAGTGTGGAGTCGTTGCCTCCCTCCCTCCCTCTCTCTNNNAGATCGGAAGAGCAC |
| 42457_2 | TATGAGTGTGGAGTCGTTGCCTCCCTCCCTCCCTCTCTCTNNNAGATCGGAAGAGCAC |
| 30653_1 | TATGAGTGTGGAGTCGTTGCTGTGTATGCCTTTGCACCTATGGNNNAGATCGGAAG |
| 30653_2 | TATGAGTGTGGAGTCGTTGCTGTGTATGCCTTTGCACCTATGGNNNAGATCGGAAG |
| 2035_1 | TATGAGTGTGGAGTCGTTGCAGCAGCAAATTCACACAGGAGAANNNAGATCGGAAG |
| 2035_2 | TATGAGTGTGGAGTCGTTGCAGCAGCAAATTCACACAGGAGAANNNAGATCGGAAG |
| 32580_1 | TATGAGTGTGGAGTCGTTGCAATGACTTTGCCCTATGTGCCTANNNAGATCGGAAGA |
| 32580_2 | TATGAGTGTGGAGTCGTTGCAATGACTTTGCCCTATGTGCCTANNNAGATCGGAAGA |
| 48510_1 | TATGAGTGTGGAGTCGTTGCGGGCCTCCATATCACTTTTACCANNNAGATCGGAAGA |
| 48510_2 | TATGAGTGTGGAGTCGTTGCGGGCCTCCATATCACTTTTACCANNNAGATCGGAAGA |
| 8519_1 | TATGAGTGTGGAGTCGTTGCGAGAGGGCTACGAAGTGAGAGTAGNNNAGATCGGA |
| 8519_2 | TATGAGTGTGGAGTCGTTGCGAGAGGGCTACGAAGTGAGAGTAGNNNAGATCGGA |
| 38666_1 | TATGAGTGTGGAGTCGTTGCACAAAAATTAGCTGGGCATGGTGNNNAGATCGGAAG |
| 38666_2 | TATGAGTGTGGAGTCGTTGCACAAAAATTAGCTGGGCATGGTGNNNAGATCGGAAG |
| 50049_1 | TATGAGTGTGGAGTCGTTGCTGTAGAGTCACTAGGAGCTTCCANNNAGATCGGAAG |
| 50049_2 | TATGAGTGTGGAGTCGTTGCTGTAGAGTCACTAGGAGCTTCCANNNAGATCGGAAG |
| 33627_1 | TATGAGTGTGGAGTCGTTGCTTCTGCACTATTCCCGACTGATTNNNAGATCGGAAGA |
| 33627_2 | TATGAGTGTGGAGTCGTTGCTTCTGCACTATTCCCGACTGATTNNNAGATCGGAAGA |
| 52426_1 | TATGAGTGTGGAGTCGTTGCCTGCAGCTCTGATTTCTCATCCTNNNAGATCGGAAGA |
| 52426_2 | TATGAGTGTGGAGTCGTTGCCTGCAGCTCTGATTTCTCATCCTNNNAGATCGGAAGA |
| 53338_1 | TATGAGTGTGGAGTCGTTGCATCAAGGTTTCTTCTGGGTGAGGNNNAGATCGGAAG |
| 53338_2 | TATGAGTGTGGAGTCGTTGCATCAAGGTTTCTTCTGGGTGAGGNNNAGATCGGAAG |
| 42909_1 | TATGAGTGTGGAGTCGTTGCCTGTGTGTGTATATGTGTGTGCANNNAGATCGGAAG |
| 42909_2 | TATGAGTGTGGAGTCGTTGCCTGTGTGTGTATATGTGTGTGCANNNAGATCGGAAG |
| 36003_1 | TATGAGTGTGGAGTCGTTGCAAGAAACCTTCCCTTTCACCTCCNNNAGATCGGAAGA |
| 36003_2 | TATGAGTGTGGAGTCGTTGCAAGAAACCTTCCCTTTCACCTCCNNNAGATCGGAAGA |
| 35572_1 | TATGAGTGTGGAGTCGTTGCATCTCTAACAAGTGGTTGCTCCANNNAGATCGGAAGA |
| 35572_2 | TATGAGTGTGGAGTCGTTGCATCTCTAACAAGTGGTTGCTCCANNNAGATCGGAAGA |
| 30768_1 | TATGAGTGTGGAGTCGTTGCTAATTTGGTGAAAGCATGCCAGGNNNAGATCGGAAG |
| 30768_2 | TATGAGTGTGGAGTCGTTGCTAATTTGGTGAAAGCATGCCAGGNNNAGATCGGAAG |
| 40373_1 | TATGAGTGTGGAGTCGTTGCGAAATAGGGTCTCGCTCTGTCATNNNAGATCGGAAG |
| 40373_2 | TATGAGTGTGGAGTCGTTGCGAAATAGGGTCTCGCTCTGTCATNNNAGATCGGAAG |
| 46961_1 | TATGAGTGTGGAGTCGTTGCTGTAGCTAGGATTACAGGCACACNNNAGATCGGAAG |
| 46961_2 | TATGAGTGTGGAGTCGTTGCTGTAGCTAGGATTACAGGCACACNNNAGATCGGAAG |
| 31721_1 | TATGAGTGTGGAGTCGTTGCCGTGTGAGAGAGAGAGAGAGAGANNNAGATCGGA |
| 31721_2 | TATGAGTGTGGAGTCGTTGCCGTGTGAGAGAGAGAGAGAGAGANNNAGATCGGA |
| 36390_1 | TATGAGTGTGGAGTCGTTGCTGGCAAGAAATACAGTTCTCACGNNNAGATCGGAAG |
| 36390_2 | TATGAGTGTGGAGTCGTTGCTGGCAAGAAATACAGTTCTCACGNNNAGATCGGAAG |
| 33640_1 | TATGAGTGTGGAGTCGTTGCGTTCAAGTAGTTCTCCTGCCTCANNNAGATCGGAAGA |
| 33640_2 | TATGAGTGTGGAGTCGTTGCGTTCAAGTAGTTCTCCTGCCTCANNNAGATCGGAAGA |
| 36245_1 | TATGAGTGTGGAGTCGTTGCCCCATATAAGTATGAGGGAGCCCNNNAGATCGGAAG |

| | |
|---|---|
| 36245_2 | TATGAGTGTGGAGTCGTTGCCCCATATAAGTATGAGGGAGCCCNNNAGATCGGAA |
| 37869_1 | TATGAGTGTGGAGTCGTTGCATCAGGACTTGGTGTATCTTCGGNNNAGATCGGAAG |
| 37869_2 | TATGAGTGTGGAGTCGTTGCATCAGGACTTGGTGTATCTTCGGNNNAGATCGGAAG |
| 38229_1 | TATGAGTGTGGAGTCGTTGCACCACAAATACCGAGAATAGCCTNNNAGATCGGAAG |
| 38229_2 | TATGAGTGTGGAGTCGTTGCACCACAAATACCGAGAATAGCCTNNNAGATCGGAAG |
| 28909_1 | TATGAGTGTGGAGTCGTTGCCATTTTCTTCACCCTGCCACTTTNNNAGATCGGAAGA |
| 28909_2 | TATGAGTGTGGAGTCGTTGCCATTTTCTTCACCCTGCCACTTTNNNAGATCGGAAGA |
| 34248_1 | TATGAGTGTGGAGTCGTTGCACATACAGTGGTTTCCTCCCTTCNNNAGATCGGAAGA |
| 34248_2 | TATGAGTGTGGAGTCGTTGCACATACAGTGGTTTCCTCCCTTCNNNAGATCGGAAGA |
| 33809_1 | TATGAGTGTGGAGTCGTTGCTAGAAGCTATGCACATCCCCCTANNNAGATCGGAAG |
| 33809_2 | TATGAGTGTGGAGTCGTTGCTAGAAGCTATGCACATCCCCCTANNNAGATCGGAAG |
| 37484_1 | TATGAGTGTGGAGTCGTTGCCTGCAATCCTACCACTTTGGGAGNNNAGATCGGAAG |
| 37484_2 | TATGAGTGTGGAGTCGTTGCCTGCAATCCTACCACTTTGGGAGNNNAGATCGGAAG |
| 2760_1 | TATGAGTGTGGAGTCGTTGCGGCATGTTTTATGTTCTGAGCCTNNNAGATCGGAAG |
| 2760_2 | TATGAGTGTGGAGTCGTTGCGGCATGTTTTATGTTCTGAGCCTNNNAGATCGGAAG |
| 10738_1 | TATGAGTGTGGAGTCGTTGCGAAGGGAGTGTCCCTAGAACCTTNNNAGATCGGAA |
| 10738_2 | TATGAGTGTGGAGTCGTTGCGAAGGGAGTGTCCCTAGAACCTTNNNAGATCGGAA |
| 39207_1 | TATGAGTGTGGAGTCGTTGCCCTGAGGGGAGGGGTACAATTATNNNAGATCGGAA |
| 39207_2 | TATGAGTGTGGAGTCGTTGCCCTGAGGGGAGGGGTACAATTATNNNAGATCGGAA |
| 12756_1 | TATGAGTGTGGAGTCGTTGCGGTGATGTTTAAAACAATAAAGCNNNAGATCGGAA |
| 12756_2 | TATGAGTGTGGAGTCGTTGCGGTGATGTTTAAAACAATAAAGCNNNAGATCGGAA |
| 33895_1 | TATGAGTGTGGAGTCGTTGCGCCACCATGCCAGACTATTTTTTNNNAGATCGGAAGA |
| 33895_2 | TATGAGTGTGGAGTCGTTGCGCCACCATGCCAGACTATTTTTTNNNAGATCGGAAGA |
| 46665_1 | TATGAGTGTGGAGTCGTTGCCACAATCTCAGCTCACTGCAAGNNNAGATCGGAAGA |
| 46665_2 | TATGAGTGTGGAGTCGTTGCCACAATCTCAGCTCACTGCAAGNNNAGATCGGAAGA |
| 36595_1 | TATGAGTGTGGAGTCGTTGCTCGTGCGTAGCATCACAATTTTTNNNAGATCGGAAG |
| 36595_2 | TATGAGTGTGGAGTCGTTGCTCGTGCGTAGCATCACAATTTTTNNNAGATCGGAAG |
| 29973_1 | TATGAGTGTGGAGTCGTTGCTGCACCACTCTAGTGACTTGTACNNNAGATCGGAAG |
| 29973_2 | TATGAGTGTGGAGTCGTTGCTGCACCACTCTAGTGACTTGTACNNNAGATCGGAAG |
| 12601_1 | TATGAGTGTGGAGTCGTTGCTTATAAAGCAAATATTTCCTCCCNNNAGATCGGAAGA |
| 12601_2 | TATGAGTGTGGAGTCGTTGCTTATAAAGCAAATATTTCCTCCCNNNAGATCGGAAGA |
| 13561_1 | TATGAGTGTGGAGTCGTTGCTTCTGGAAAAGCTGGAAAAGNNNAGATCGGAAGAG |
| 13561_2 | TATGAGTGTGGAGTCGTTGCTTCTGGAAAAGCTGGAAAAGNNNAGATCGGAAGAG |
| 49231_1 | TATGAGTGTGGAGTCGTTGCGATCTCCTGGTCTCAACTGATCCNNNAGATCGGAAG |
| 49231_2 | TATGAGTGTGGAGTCGTTGCGATCTCCTGGTCTCAACTGATCCNNNAGATCGGAAG |
| 362_1 | TATGAGTGTGGAGTCGTTGCTGAGGCCAAAGAAAGAACAACCTNNNAGATCGGAA |
| 362_2 | TATGAGTGTGGAGTCGTTGCTGAGGCCAAAGAAAGAACAACCTNNNAGATCGGAA |
| 46819_1 | TATGAGTGTGGAGTCGTTGCTGCAAGTAATTCTCCTGCCTCAGNNNAGATCGGAAG |
| 46819_2 | TATGAGTGTGGAGTCGTTGCTGCAAGTAATTCTCCTGCCTCAGNNNAGATCGGAAG |
| 4328_1 | TATGAGTGTGGAGTCGTTGCACATGAACCCAAGCACAGAGATTNNNAGATCGGAAG |
| 4328_2 | TATGAGTGTGGAGTCGTTGCACATGAACCCAAGCACAGAGATTNNNAGATCGGAAG |
| 49412_1 | TATGAGTGTGGAGTCGTTGCGAGAGACACATCCCACTCAACTTNNNAGATCGGAAG |
| 49412_2 | TATGAGTGTGGAGTCGTTGCGAGAGACACATCCCACTCAACTTNNNAGATCGGAAG |

| | |
|---|---|
| 36739_1 | TATGAGTGTGGAGTCGTTGCAAAACGAAACAAACCCCCCAAATNNNAGATCGGAAG |
| 36739_2 | TATGAGTGTGGAGTCGTTGCAAAACGAAACAAACCCCCCAAATNNNAGATCGGAAG |
| 47941_1 | TATGAGTGTGGAGTCGTTGCGTACAAGTGATTCTCCTGCCTCANNNAGATCGGAAG |
| 47941_2 | TATGAGTGTGGAGTCGTTGCGTACAAGTGATTCTCCTGCCTCANNNAGATCGGAAG |
| 41709_1 | TATGAGTGTGGAGTCGTTGCCATTCTCCTGCCTCTCGAGTAGNNNAGATCGGAAGA |
| 41709_2 | TATGAGTGTGGAGTCGTTGCCATTCTCCTGCCTCTCGAGTAGNNNAGATCGGAAGA |
| 51807_1 | TATGAGTGTGGAGTCGTTGCGGGCTCCTCACTTCCCAGTANNNAGATCGGAAGAGC |
| 51807_2 | TATGAGTGTGGAGTCGTTGCGGGCTCCTCACTTCCCAGTANNNAGATCGGAAGAGC |
| 35143_1 | TATGAGTGTGGAGTCGTTGCCTCCTGGGTTCACACCATTCTCNNNAGATCGGAAGA |
| 35143_2 | TATGAGTGTGGAGTCGTTGCCTCCTGGGTTCACACCATTCTCNNNAGATCGGAAGA |
| 45181_1 | TATGAGTGTGGAGTCGTTGCCTCGTAGCTTGAGAAAAGGGACTNNNAGATCGGAA |
| 45181_2 | TATGAGTGTGGAGTCGTTGCCTCGTAGCTTGAGAAAAGGGACTNNNAGATCGGAA |
| 2909_1 | TATGAGTGTGGAGTCGTTGCGAAACCATCTTTGCAATGGTTGTNNNAGATCGGAAG |
| 2909_2 | TATGAGTGTGGAGTCGTTGCGAAACCATCTTTGCAATGGTTGTNNNAGATCGGAAG |
| 30313_1 | TATGAGTGTGGAGTCGTTGCGCAGGGGCATCCTTAATACAGTCNNNAGATCGGAA |
| 30313_2 | TATGAGTGTGGAGTCGTTGCGCAGGGGCATCCTTAATACAGTCNNNAGATCGGAA |
| 13517_1 | TATGAGTGTGGAGTCGTTGCCTCTCCAGGAGATAATATACAAANNNAGATCGGAAG |
| 13517_2 | TATGAGTGTGGAGTCGTTGCCTCTCCAGGAGATAATATACAAANNNAGATCGGAAG |
| 30567_1 | TATGAGTGTGGAGTCGTTGCAAAATTCCTCCACGCTGACTACCNNNAGATCGGAAG |
| 30567_2 | TATGAGTGTGGAGTCGTTGCAAAATTCCTCCACGCTGACTACCNNNAGATCGGAAG |
| 33544_1 | TATGAGTGTGGAGTCGTTGCAGAATCTCACTATGTTGCCCAGGNNNAGATCGGAAG |
| 33544_2 | TATGAGTGTGGAGTCGTTGCAGAATCTCACTATGTTGCCCAGGNNNAGATCGGAAG |
| 46731_1 | TATGAGTGTGGAGTCGTTGCTTGACTGTTGTTGGTGTACAGGANNNAGATCGGAA |
| 46731_2 | TATGAGTGTGGAGTCGTTGCTTGACTGTTGTTGGTGTACAGGANNNAGATCGGAA |
| 43960_1 | TATGAGTGTGGAGTCGTTGCAGTCCAAGTCTGACAACCTGAAANNNAGATCGGAAG |
| 43960_2 | TATGAGTGTGGAGTCGTTGCAGTCCAAGTCTGACAACCTGAAANNNAGATCGGAAG |
| 10141_1 | TATGAGTGTGGAGTCGTTGCCAAACCAAAGTAGGTGGGTCCTGNNNAGATCGGAA |
| 10141_2 | TATGAGTGTGGAGTCGTTGCCAAACCAAAGTAGGTGGGTCCTGNNNAGATCGGAA |
| 50415_1 | TATGAGTGTGGAGTCGTTGCACAAAAGATGTTCCTCACAGCTGNNNAGATCGGAAG |
| 50415_2 | TATGAGTGTGGAGTCGTTGCACAAAAGATGTTCCTCACAGCTGNNNAGATCGGAAG |
| 3520_1 | TATGAGTGTGGAGTCGTTGCGTGATAGAACAGCTCTGCATCTTNNNAGATCGGAAG |
| 3520_2 | TATGAGTGTGGAGTCGTTGCGTGATAGAACAGCTCTGCATCTTNNNAGATCGGAAG |
| 50612_1 | TATGAGTGTGGAGTCGTTGCTCCTGGTTCTGTTGCTTATTGTCNNNAGATCGGAAG |
| 50612_2 | TATGAGTGTGGAGTCGTTGCTCCTGGTTCTGTTGCTTATTGTCNNNAGATCGGAAG |
| 44604_1 | TATGAGTGTGGAGTCGTTGCAGCATAGCATGTGTGAGTGATGTNNNAGATCGGAA |
| 44604_2 | TATGAGTGTGGAGTCGTTGCAGCATAGCATGTGTGAGTGATGTNNNAGATCGGAA |
| 31825_1 | TATGAGTGTGGAGTCGTTGCAGTGTGTGTATATGTGTGTGTGTNNNAGATCGGAA |
| 31825_2 | TATGAGTGTGGAGTCGTTGCAGTGTGTGTATATGTGTGTGTGTNNNAGATCGGAA |
| 44762_1 | TATGAGTGTGGAGTCGTTGCGTCCCTGAGAAGTGTCACAGATCNNNAGATCGGAA |
| 44762_2 | TATGAGTGTGGAGTCGTTGCGTCCCTGAGAAGTGTCACAGATCNNNAGATCGGAA |
| 13145_1 | TATGAGTGTGGAGTCGTTGCCAAGATGGTTGATGAAATAATCANNNAGATCGGAA |
| 13145_2 | TATGAGTGTGGAGTCGTTGCCAAGATGGTTGATGAAATAATCANNNAGATCGGAA |
| 32905_1 | TATGAGTGTGGAGTCGTTGCATGCACGAGTATCTGTTACGGTTNNNAGATCGGAAG |

| | |
|---|---|
| 32905_2 | TATGAGTGTGGAGTCGTTGCATGCACGAGTATCTGTTACGGTTNNNAGATCGGAAG |
| 44138_1 | TATGAGTGTGGAGTCGTTGCCAGATGAGCCAACAACAATGTGTNNNAGATCGGAAG |
| 44138_2 | TATGAGTGTGGAGTCGTTGCCAGATGAGCCAACAACAATGTGTNNNAGATCGGAAG |
| 834_1 | TATGAGTGTGGAGTCGTTGCATCTGTTTCTCAATCCCCGAGTTGNNNAGATCGGAAG |
| 834_2 | TATGAGTGTGGAGTCGTTGCATCTGTTTCTCAATCCCCGAGTTGNNNAGATCGGAAG |
| 44286_1 | TATGAGTGTGGAGTCGTTGCGAGGAAAAGAGGGAGGGAAGAAGNNNAGATCGGA |
| 44286_2 | TATGAGTGTGGAGTCGTTGCGAGGAAAAGAGGGAGGGAAGAAGNNNAGATCGGA |
| 47346_1 | TATGAGTGTGGAGTCGTTGCCTAGTAGCTGGGACTACAGGCANNNAGATCGGAAGA |
| 47346_2 | TATGAGTGTGGAGTCGTTGCCTAGTAGCTGGGACTACAGGCANNNAGATCGGAAGA |
| 42087_1 | TATGAGTGTGGAGTCGTTGCGATCAGAGTTAGTTTTCCACGCGNNNAGATCGGAAG |
| 42087_2 | TATGAGTGTGGAGTCGTTGCGATCAGAGTTAGTTTTCCACGCGNNNAGATCGGAAG |
| 38930_1 | TATGAGTGTGGAGTCGTTGCAGCAGAGGAAAGAATCTCAGAGCNNNAGATCGGAA |
| 38930_2 | TATGAGTGTGGAGTCGTTGCAGCAGAGGAAAGAATCTCAGAGCNNNAGATCGGAA |
| 48988_1 | TATGAGTGTGGAGTCGTTGCCTCTTAGCCTCAAGTGATCCTCCNNNAGATCGGAAGA |
| 48988_2 | TATGAGTGTGGAGTCGTTGCCTCTTAGCCTCAAGTGATCCTCCNNNAGATCGGAAGA |
| 37580_1 | TATGAGTGTGGAGTCGTTGCGGAAGATATCTGGGTGATGGGTCNNNAGATCGGAA |
| 37580_2 | TATGAGTGTGGAGTCGTTGCGGAAGATATCTGGGTGATGGGTCNNNAGATCGGAA |
| 9755_1 | TATGAGTGTGGAGTCGTTGCCTGGGACTTCACTGGGTTAGAACNNNAGATCGGAAG |
| 9755_2 | TATGAGTGTGGAGTCGTTGCCTGGGACTTCACTGGGTTAGAACNNNAGATCGGAAG |
| 51318_1 | TATGAGTGTGGAGTCGTTGCGGGAAGTGGTGACATTTGCAATTNNNAGATCGGAA |
| 51318_2 | TATGAGTGTGGAGTCGTTGCGGGAAGTGGTGACATTTGCAATTNNNAGATCGGAA |
| 40530_1 | TATGAGTGTGGAGTCGTTGCGATCCAAAAGGCAGAGTGGTAGANNNAGATCGGAA |
| 40530_2 | TATGAGTGTGGAGTCGTTGCGATCCAAAAGGCAGAGTGGTAGANNNAGATCGGAA |
| 34182_1 | TATGAGTGTGGAGTCGTTGCGAAAGCTGAAGGTTGTTGGTCTTNNNAGATCGGAA |
| 34182_2 | TATGAGTGTGGAGTCGTTGCGAAAGCTGAAGGTTGTTGGTCTTNNNAGATCGGAA |
| 38145_1 | TATGAGTGTGGAGTCGTTGCTTTTGCCACACTTTCTGGGTTTTNNNAGATCGGAAGA |
| 38145_2 | TATGAGTGTGGAGTCGTTGCTTTTGCCACACTTTCTGGGTTTTNNNAGATCGGAAGA |
| 42647_1 | TATGAGTGTGGAGTCGTTGCAGAGGAAGAGAGGAGAGACTCTGNNNAGATCGGA |
| 42647_2 | TATGAGTGTGGAGTCGTTGCAGAGGAAGAGAGGAGAGACTCTGNNNAGATCGGA |
| 34145_1 | TATGAGTGTGGAGTCGTTGCTTTCCGGTGTAAAACGTTGATGGNNNAGATCGGAAG |
| 34145_2 | TATGAGTGTGGAGTCGTTGCTTTCCGGTGTAAAACGTTGATGGNNNAGATCGGAAG |
| 32409_1 | TATGAGTGTGGAGTCGTTGCAGTAACATTCTCAGGAGGGAGGANNNAGATCGGAA |
| 32409_2 | TATGAGTGTGGAGTCGTTGCAGTAACATTCTCAGGAGGGAGGANNNAGATCGGAA |
| 12824_1 | TATGAGTGTGGAGTCGTTGCTAAAAAGGGAAACTTGGAAAAACNNNAGATCGGAA |
| 12824_2 | TATGAGTGTGGAGTCGTTGCTAAAAAGGGAAACTTGGAAAAACNNNAGATCGGAA |
| 9873_1 | TATGAGTGTGGAGTCGTTGCAAGCTAGGATAGACACACAATGGANNNAGATCGGA |
| 9873_2 | TATGAGTGTGGAGTCGTTGCAAGCTAGGATAGACACACAATGGANNNAGATCGGA |
| 35759_1 | TATGAGTGTGGAGTCGTTGCCTCACTGCAAACTCCTCCTTCCNNNAGATCGGAAGAG |
| 35759_2 | TATGAGTGTGGAGTCGTTGCCTCACTGCAAACTCCTCCTTCCNNNAGATCGGAAGAG |
| 35675_1 | TATGAGTGTGGAGTCGTTGCCCACCACTCTCGGCTAATTTTTGNNNAGATCGGAAGA |
| 35675_2 | TATGAGTGTGGAGTCGTTGCCCACCACTCTCGGCTAATTTTTGNNNAGATCGGAAGA |
| 35890_1 | TATGAGTGTGGAGTCGTTGCGCCTCATATTGTCCTGCTTTTCCNNNAGATCGGAAGA |
| 35890_2 | TATGAGTGTGGAGTCGTTGCGCCTCATATTGTCCTGCTTTTCCNNNAGATCGGAAGA |

| | |
|---|---|
| 10210_1 | TATGAGTGTGGAGTCGTTGCGGGGAAAAGTCAGGAGAGAGAACNNNAGATCGGA |
| 10210_2 | TATGAGTGTGGAGTCGTTGCGGGGAAAAGTCAGGAGAGAGAACNNNAGATCGGA |
| 53200_1 | TATGAGTGTGGAGTCGTTGCCTCCAACTTACCACTAGCGTCCNNNAGATCGGAAGA |
| 53200_2 | TATGAGTGTGGAGTCGTTGCCTCCAACTTACCACTAGCGTCCNNNAGATCGGAAGA |
| 29096_1 | TATGAGTGTGGAGTCGTTGCCATGCCACGTTCAATCAACAGTANNNAGATCGGAAG |
| 29096_2 | TATGAGTGTGGAGTCGTTGCCATGCCACGTTCAATCAACAGTANNNAGATCGGAAG |
| 30577_1 | TATGAGTGTGGAGTCGTTGCAGTCTTATACTGTAGCCTGGGGTNNNAGATCGGAA |
| 30577_2 | TATGAGTGTGGAGTCGTTGCAGTCTTATACTGTAGCCTGGGGTNNNAGATCGGAA |
| 46968_1 | TATGAGTGTGGAGTCGTTGCCACCACACCTGGCCTATATTGAANNNAGATCGGAAG |
| 46968_2 | TATGAGTGTGGAGTCGTTGCCACCACACCTGGCCTATATTGAANNNAGATCGGAAG |
| 30170_1 | TATGAGTGTGGAGTCGTTGCTGCCTTAACTGACAACTTTAATTTACCNNNAGATCGG |
| 30170_2 | TATGAGTGTGGAGTCGTTGCTGCCTTAACTGACAACTTTAATTTACCNNNAGATCGG |
| 31542_1 | TATGAGTGTGGAGTCGTTGCGATGATGGGGTTGAGATGACACTNNNAGATCGGAA |
| 31542_2 | TATGAGTGTGGAGTCGTTGCGATGATGGGGTTGAGATGACACTNNNAGATCGGAA |
| 32463_1 | TATGAGTGTGGAGTCGTTGCTGTTCTTAGCAAAGTGCACAAACTNNNAGATCGGAA |
| 32463_2 | TATGAGTGTGGAGTCGTTGCTGTTCTTAGCAAAGTGCACAAACTNNNAGATCGGAA |
| 31894_1 | TATGAGTGTGGAGTCGTTGCACAAAAAATTTACCGGGCATGGTNNNAGATCGGAA |
| 31894_2 | TATGAGTGTGGAGTCGTTGCACAAAAAATTTACCGGGCATGGTNNNAGATCGGAA |
| 45553_1 | TATGAGTGTGGAGTCGTTGCAAATCAGGAGCTAGCATCACACANNNAGATCGGAA |
| 45553_2 | TATGAGTGTGGAGTCGTTGCAAATCAGGAGCTAGCATCACACANNNAGATCGGAA |
| 9418_1 | TATGAGTGTGGAGTCGTTGCTAAACATTGTTGCCAGTAGCACGNNNAGATCGGAAG |
| 9418_2 | TATGAGTGTGGAGTCGTTGCTAAACATTGTTGCCAGTAGCACGNNNAGATCGGAAG |
| 49518_1 | TATGAGTGTGGAGTCGTTGCTCAAAAAAGCATTTCCTCTGGGGNNNAGATCGGAAG |
| 49518_2 | TATGAGTGTGGAGTCGTTGCTCAAAAAAGCATTTCCTCTGGGGNNNAGATCGGAAG |
| 52842_1 | TATGAGTGTGGAGTCGTTGCAATGAAGACGGTCTCAGTGGAAANNNAGATCGGAA |
| 52842_2 | TATGAGTGTGGAGTCGTTGCAATGAAGACGGTCTCAGTGGAAANNNAGATCGGAA |
| 13605_1 | TATGAGTGTGGAGTCGTTGCGTAATAGGAGCTAGTATCAGAAANNNAGATCGGAA |
| 13605_2 | TATGAGTGTGGAGTCGTTGCGTAATAGGAGCTAGTATCAGAAANNNAGATCGGAA |
| 45385_1 | TATGAGTGTGGAGTCGTTGCCATCACTGCACTCACATGAACTGNNNAGATCGGAAG |
| 45385_2 | TATGAGTGTGGAGTCGTTGCCATCACTGCACTCACATGAACTGNNNAGATCGGAAG |
| 10464_1 | TATGAGTGTGGAGTCGTTGCCAAGCCCACACATGTCTTAGTCANNNAGATCGGAAG |
| 10464_2 | TATGAGTGTGGAGTCGTTGCCAAGCCCACACATGTCTTAGTCANNNAGATCGGAAG |
| 50092_1 | TATGAGTGTGGAGTCGTTGCAGTTATCCATCCTCCAGAAGTGCNNNAGATCGGAAG |
| 50092_2 | TATGAGTGTGGAGTCGTTGCAGTTATCCATCCTCCAGAAGTGCNNNAGATCGGAAG |
| 34502_1 | TATGAGTGTGGAGTCGTTGCGGAATTCGGCTGTCTCCTTTCTANNNAGATCGGAAG |
| 34502_2 | TATGAGTGTGGAGTCGTTGCGGAATTCGGCTGTCTCCTTTCTANNNAGATCGGAAG |
| 12536_1 | TATGAGTGTGGAGTCGTTGCCCTAACAACAGCAATCAATTAATNNNAGATCGGAAG |
| 12536_2 | TATGAGTGTGGAGTCGTTGCCCTAACAACAGCAATCAATTAATNNNAGATCGGAAG |
| 46393_1 | TATGAGTGTGGAGTCGTTGCAAGGTGGAATCATAGGGTAGGGANNNAGATCGGA |
| 46393_2 | TATGAGTGTGGAGTCGTTGCAAGGTGGAATCATAGGGTAGGGANNNAGATCGGA |
| 44023_1 | TATGAGTGTGGAGTCGTTGCTTCAAGAAGTGCTACATGGTGGTNNNAGATCGGAA |
| 44023_2 | TATGAGTGTGGAGTCGTTGCTTCAAGAAGTGCTACATGGTGGTNNNAGATCGGAA |
| 36149_1 | TATGAGTGTGGAGTCGTTGCAAATGTCTCATCCAAGAGGAGCANNNAGATCGGAA |

| | |
|---|---|
| 36149_2 | TATGAGTGTGGAGTCGTTGCAAATGTCTCATCCAAGAGGAGCANNNAGATCGGAAG |
| 13785_1 | TATGAGTGTGGAGTCGTTGCCTTTTTGTTTTTCAGATCCTTCTNNNAGATCGGAAGA |
| 13785_2 | TATGAGTGTGGAGTCGTTGCCTTTTTGTTTTTCAGATCCTTCTNNNAGATCGGAAGA |
| 33340_1 | TATGAGTGTGGAGTCGTTGCCTACCTCAACAATTCTCCTGCCTNNNAGATCGGAAGA |
| 33340_2 | TATGAGTGTGGAGTCGTTGCCTACCTCAACAATTCTCCTGCCTNNNAGATCGGAAGA |
| 34265_1 | TATGAGTGTGGAGTCGTTGCCCATTTTTGCCTCCTAGACCTCTNNNAGATCGGAAGA |
| 34265_2 | TATGAGTGTGGAGTCGTTGCCCATTTTTGCCTCCTAGACCTCTNNNAGATCGGAAGA |
| 29865_1 | TATGAGTGTGGAGTCGTTGCTGTTTGTCTACATAACTGAAGAGCANNNAGATCGGA |
| 29865_2 | TATGAGTGTGGAGTCGTTGCTGTTTGTCTACATAACTGAAGAGCANNNAGATCGGA |
| 46409_1 | TATGAGTGTGGAGTCGTTGCTCCTCCCCAGATCCTACTCTTTTNNNAGATCGGAAGA |
| 46409_2 | TATGAGTGTGGAGTCGTTGCTCCTCCCCAGATCCTACTCTTTTNNNAGATCGGAAGA |
| 37041_1 | TATGAGTGTGGAGTCGTTGCGCATTGAACATCCTCCAGGAAAGNNNAGATCGGAAG |
| 37041_2 | TATGAGTGTGGAGTCGTTGCGCATTGAACATCCTCCAGGAAAGNNNAGATCGGAAG |
| 43737_1 | TATGAGTGTGGAGTCGTTGCGAAGTCCTCCTGTGTGGGAATAGNNNAGATCGGAA |
| 43737_2 | TATGAGTGTGGAGTCGTTGCGAAGTCCTCCTGTGTGGGAATAGNNNAGATCGGAA |
| 46068_1 | TATGAGTGTGGAGTCGTTGCTCAAGTAATTCTCCTGCCTCAGCNNNAGATCGGAAG |
| 46068_2 | TATGAGTGTGGAGTCGTTGCTCAAGTAATTCTCCTGCCTCAGCNNNAGATCGGAAG |
| 34720_1 | TATGAGTGTGGAGTCGTTGCTTCATGCTATTCTCCTGCCTCAGNNNAGATCGGAAGA |
| 34720_2 | TATGAGTGTGGAGTCGTTGCTTCATGCTATTCTCCTGCCTCAGNNNAGATCGGAAGA |
| 10371_1 | TATGAGTGTGGAGTCGTTGCCTCCTATCCTCTACTTGGCACCANNNAGATCGGAAGA |
| 10371_2 | TATGAGTGTGGAGTCGTTGCCTCCTATCCTCTACTTGGCACCANNNAGATCGGAAGA |
| 8494_1 | TATGAGTGTGGAGTCGTTGCGGTCTCACTCTGTCCATGCTGTACNNNAGATCGGAAG |
| 8494_2 | TATGAGTGTGGAGTCGTTGCGGTCTCACTCTGTCCATGCTGTACNNNAGATCGGAAG |
| 28870_1 | TATGAGTGTGGAGTCGTTGCGGTCTGTTTTGTCTGCCTAGAGANNNAGATCGGAAG |
| 28870_2 | TATGAGTGTGGAGTCGTTGCGGTCTGTTTTGTCTGCCTAGAGANNNAGATCGGAAG |
| 42361_1 | TATGAGTGTGGAGTCGTTGCTGGGCTGGAAGGAATCACAATTTNNNAGATCGGAA |
| 42361_2 | TATGAGTGTGGAGTCGTTGCTGGGCTGGAAGGAATCACAATTTNNNAGATCGGAA |
| 32368_1 | TATGAGTGTGGAGTCGTTGCCCCCGCGAAGATATGTTGAAATCNNNAGATCGGAAG |
| 32368_2 | TATGAGTGTGGAGTCGTTGCCCCCGCGAAGATATGTTGAAATCNNNAGATCGGAAG |
| 52186_1 | TATGAGTGTGGAGTCGTTGCGTCCTGGGAAGCTGATTCTGAGNNNAGATCGGAAG |
| 52186_2 | TATGAGTGTGGAGTCGTTGCGTCCTGGGAAGCTGATTCTGAGNNNAGATCGGAAG |
| 47419_1 | TATGAGTGTGGAGTCGTTGCAAACTCCTGACCTCAAGTGATCCNNNAGATCGGAAG |
| 47419_2 | TATGAGTGTGGAGTCGTTGCAAACTCCTGACCTCAAGTGATCCNNNAGATCGGAAG |
| 41088_1 | TATGAGTGTGGAGTCGTTGCAGTGATGAAGATGGTCAACTGCTNNNAGATCGGAA |
| 41088_2 | TATGAGTGTGGAGTCGTTGCAGTGATGAAGATGGTCAACTGCTNNNAGATCGGAA |
| 31449_1 | TATGAGTGTGGAGTCGTTGCTTCTCATGCTTCCTCTCCACTTCNNNAGATCGGAAGAG |
| 31449_2 | TATGAGTGTGGAGTCGTTGCTTCTCATGCTTCCTCTCCACTTCNNNAGATCGGAAGAG |
| 29553_1 | TATGAGTGTGGAGTCGTTGCCACCAGCAGCTCCTACACCGNNNAGATCGGAAGAGC |
| 29553_2 | TATGAGTGTGGAGTCGTTGCCACCAGCAGCTCCTACACCGNNNAGATCGGAAGAGC |
| 41670_1 | TATGAGTGTGGAGTCGTTGCCACCACTACCCCCAGCTAATTTTNNNAGATCGGAAGA |
| 41670_2 | TATGAGTGTGGAGTCGTTGCCACCACTACCCCCAGCTAATTTTNNNAGATCGGAAGA |
| 35618_1 | TATGAGTGTGGAGTCGTTGCAACCTCCTGAGTAACTGGGACTANNNAGATCGGAAG |
| 35618_2 | TATGAGTGTGGAGTCGTTGCAACCTCCTGAGTAACTGGGACTANNNAGATCGGAAG |

| | |
|---|---|
| 13731_1 | TATGAGTGTGGAGTCGTTGCTATCAAATATTTAGCATGGATCCNNNAGATCGGAAG |
| 13731_2 | TATGAGTGTGGAGTCGTTGCTATCAAATATTTAGCATGGATCCNNNAGATCGGAAG |
| 42779_1 | TATGAGTGTGGAGTCGTTGCAGTGGAGCCAAGCATACAACTAANNNAGATCGGAA |
| 42779_2 | TATGAGTGTGGAGTCGTTGCAGTGGAGCCAAGCATACAACTAANNNAGATCGGAA |
| 30983_1 | TATGAGTGTGGAGTCGTTGCCCCTCATTCAGCACTTTGTTCTGNNNAGATCGGAAGA |
| 30983_2 | TATGAGTGTGGAGTCGTTGCCCCTCATTCAGCACTTTGTTCTGNNNAGATCGGAAGA |
| 35461_1 | TATGAGTGTGGAGTCGTTGCCTTGGCCAAGATCAGTACTGGATNNNAGATCGGAAG |
| 35461_2 | TATGAGTGTGGAGTCGTTGCCTTGGCCAAGATCAGTACTGGATNNNAGATCGGAAG |
| 48609_1 | TATGAGTGTGGAGTCGTTGCCAAGTGATTCTCCTACCTCAGCCNNNAGATCGGAAGA |
| 48609_2 | TATGAGTGTGGAGTCGTTGCCAAGTGATTCTCCTACCTCAGCCNNNAGATCGGAAGA |
| 2667_1 | TATGAGTGTGGAGTCGTTGCAAGGCATATCAGCCCCTTTGAATNNNAGATCGGAAG |
| 2667_2 | TATGAGTGTGGAGTCGTTGCAAGGCATATCAGCCCCTTTGAATNNNAGATCGGAAG |
| 44025_1 | TATGAGTGTGGAGTCGTTGCACTTGAGACTGCCCAAATAGGAGNNNAGATCGGAA |
| 44025_2 | TATGAGTGTGGAGTCGTTGCACTTGAGACTGCCCAAATAGGAGNNNAGATCGGAA |
| 39674_1 | TATGAGTGTGGAGTCGTTGCTCCTGGACTTGGAGTTGAAGAAGNNNAGATCGGAA |
| 39674_2 | TATGAGTGTGGAGTCGTTGCTCCTGGACTTGGAGTTGAAGAAGNNNAGATCGGAA |
| 13052_1 | TATGAGTGTGGAGTCGTTGCGGCTGCAATAAGAGATATTTTAANNNAGATCGGAAG |
| 13052_2 | TATGAGTGTGGAGTCGTTGCGGCTGCAATAAGAGATATTTTAANNNAGATCGGAAG |
| 34659_1 | TATGAGTGTGGAGTCGTTGCACCCAACATTCCTAGAAGCTGGNNNAGATCGGAAGA |
| 34659_2 | TATGAGTGTGGAGTCGTTGCACCCAACATTCCTAGAAGCTGGNNNAGATCGGAAGA |
| 39654_1 | TATGAGTGTGGAGTCGTTGCGCCTAGCTTCTCAGAGTTCCATTNNNAGATCGGAAGA |
| 39654_2 | TATGAGTGTGGAGTCGTTGCGCCTAGCTTCTCAGAGTTCCATTNNNAGATCGGAAGA |
| 53066_1 | TATGAGTGTGGAGTCGTTGCGGAACTGAAACAACTGTTGGAGGNNNAGATCGGAA |
| 53066_2 | TATGAGTGTGGAGTCGTTGCGGAACTGAAACAACTGTTGGAGGNNNAGATCGGAA |
| 35460_1 | TATGAGTGTGGAGTCGTTGCCTGGAGTGTAGTTCTGCAGTCTTNNNAGATCGGAAG |
| 35460_2 | TATGAGTGTGGAGTCGTTGCCTGGAGTGTAGTTCTGCAGTCTTNNNAGATCGGAAG |
| 10814_1 | TATGAGTGTGGAGTCGTTGCGTAGCATTTCCACTTCTGAGGCCNNNAGATCGGAAG |
| 10814_2 | TATGAGTGTGGAGTCGTTGCGTAGCATTTCCACTTCTGAGGCCNNNAGATCGGAAG |
| 31637_1 | TATGAGTGTGGAGTCGTTGCTATTGAAACATCCCCCCAACACANNNAGATCGGAAGA |
| 31637_2 | TATGAGTGTGGAGTCGTTGCTATTGAAACATCCCCCCAACACANNNAGATCGGAAGA |
| 39352_1 | TATGAGTGTGGAGTCGTTGCCCCAACTGTCCCCAAAATGTTTTNNNAGATCGGAAGA |
| 39352_2 | TATGAGTGTGGAGTCGTTGCCCCAACTGTCCCCAAAATGTTTTNNNAGATCGGAAGA |
| 29238_1 | TATGAGTGTGGAGTCGTTGCAGCCTCTGGTCACAATATGTTCANNNAGATCGGAAG |
| 29238_2 | TATGAGTGTGGAGTCGTTGCAGCCTCTGGTCACAATATGTTCANNNAGATCGGAAG |
| 47002_1 | TATGAGTGTGGAGTCGTTGCCTGGGATTACAGGTGTGAGTCACNNNAGATCGGAA |
| 47002_2 | TATGAGTGTGGAGTCGTTGCCTGGGATTACAGGTGTGAGTCACNNNAGATCGGAA |
| 45241_1 | TATGAGTGTGGAGTCGTTGCTTTCTTCTTCGGAACTGGCTTCTNNNAGATCGGAAGA |
| 45241_2 | TATGAGTGTGGAGTCGTTGCTTTCTTCTTCGGAACTGGCTTCTNNNAGATCGGAAGA |
| 37464_1 | TATGAGTGTGGAGTCGTTGCTACTCACTGTTCCCTTACCTTGCNNNAGATCGGAAGA |
| 37464_2 | TATGAGTGTGGAGTCGTTGCTACTCACTGTTCCCTTACCTTGCNNNAGATCGGAAGA |
| 40759_1 | TATGAGTGTGGAGTCGTTGCATTGTCTCTTCCCCAAAGTCAGGNNNAGATCGGAAG |
| 40759_2 | TATGAGTGTGGAGTCGTTGCATTGTCTCTTCCCCAAAGTCAGGNNNAGATCGGAAG |
| 36508_1 | TATGAGTGTGGAGTCGTTGCTGTTCAGATGTGTGTAACCTCCTNNNAGATCGGAAG |

| | |
|---|---|
| 36508_2 | TATGAGTGTGGAGTCGTTGCTGTTCAGATGTGTGTAACCTCCTNNNAGATCGGAAG |
| 53123_1 | TATGAGTGTGGAGTCGTTGCACAAAAACTTTCTGAGCATGCCANNNAGATCGGAAG |
| 53123_2 | TATGAGTGTGGAGTCGTTGCACAAAAACTTTCTGAGCATGCCANNNAGATCGGAAG |
| 52889_1 | TATGAGTGTGGAGTCGTTGCTCCCATATCAAAAGACACAGGGGNNNAGATCGGAA |
| 52889_2 | TATGAGTGTGGAGTCGTTGCTCCCATATCAAAAGACACAGGGGNNNAGATCGGAA |
| 48436_1 | TATGAGTGTGGAGTCGTTGCCCTGAGTAGCTGGGATTACAAGGNNNAGATCGGAA |
| 48436_2 | TATGAGTGTGGAGTCGTTGCCCTGAGTAGCTGGGATTACAAGGNNNAGATCGGAA |
| 51573_1 | TATGAGTGTGGAGTCGTTGCCTTGGGGGAGACACATTATGCATNNNAGATCGGAA |
| 51573_2 | TATGAGTGTGGAGTCGTTGCCTTGGGGGAGACACATTATGCATNNNAGATCGGAA |
| 40306_1 | TATGAGTGTGGAGTCGTTGCCATCCCTTCAAACTTCAGCACAANNNAGATCGGAAG |
| 40306_2 | TATGAGTGTGGAGTCGTTGCCATCCCTTCAAACTTCAGCACAANNNAGATCGGAAG |
| 44766_1 | TATGAGTGTGGAGTCGTTGCAGTCAGCCTTGTAGAATTCCCAGNNNAGATCGGAAG |
| 44766_2 | TATGAGTGTGGAGTCGTTGCAGTCAGCCTTGTAGAATTCCCAGNNNAGATCGGAAG |
| 49967_1 | TATGAGTGTGGAGTCGTTGCCCGGCTAGATCTTCATCATGGAANNNAGATCGGAAG |
| 49967_2 | TATGAGTGTGGAGTCGTTGCCCGGCTAGATCTTCATCATGGAANNNAGATCGGAAG |
| 45600_1 | TATGAGTGTGGAGTCGTTGCACTGCTAGATGTCTAAGCTGTCTNNNAGATCGGAAG |
| 45600_2 | TATGAGTGTGGAGTCGTTGCACTGCTAGATGTCTAAGCTGTCTNNNAGATCGGAAG |
| 45760_1 | TATGAGTGTGGAGTCGTTGCCACCACCACGCTTGACTAATTTTNNNAGATCGGAAG |
| 45760_2 | TATGAGTGTGGAGTCGTTGCCACCACCACGCTTGACTAATTTTNNNAGATCGGAAG |
| 41548_1 | TATGAGTGTGGAGTCGTTGCGCCTCCCAAGTAGCTAGGATTACNNNAGATCGGAAG |
| 41548_2 | TATGAGTGTGGAGTCGTTGCGCCTCCCAAGTAGCTAGGATTACNNNAGATCGGAAG |
| 45305_1 | TATGAGTGTGGAGTCGTTGCCTCTCACATAGGCCATTTGTTGCNNNAGATCGGAAG |
| 45305_2 | TATGAGTGTGGAGTCGTTGCCTCTCACATAGGCCATTTGTTGCNNNAGATCGGAAG |
| 42237_1 | TATGAGTGTGGAGTCGTTGCTCCTGTGTCTTTTGGTGTGTTTCNNNAGATCGGAAG |
| 42237_2 | TATGAGTGTGGAGTCGTTGCTCCTGTGTCTTTTGGTGTGTTTCNNNAGATCGGAAG |
| 32786_1 | TATGAGTGTGGAGTCGTTGCGAATCAGACTCACAGGACCCAGNNNAGATCGGAAG |
| 32786_2 | TATGAGTGTGGAGTCGTTGCGAATCAGACTCACAGGACCCAGNNNAGATCGGAAG |
| 40650_1 | TATGAGTGTGGAGTCGTTGCGGTCCACTGCGTTGAATAAAACANNNAGATCGGAA |
| 40650_2 | TATGAGTGTGGAGTCGTTGCGGTCCACTGCGTTGAATAAAACANNNAGATCGGAA |
| 10212_1 | TATGAGTGTGGAGTCGTTGCTGGACTGCAGTTGCTCTAGAGTANNNAGATCGGAA |
| 10212_2 | TATGAGTGTGGAGTCGTTGCTGGACTGCAGTTGCTCTAGAGTANNNAGATCGGAA |
| 51266_1 | TATGAGTGTGGAGTCGTTGCAGTAGGAGATTAGGGATTGGGGCNNNAGATCGGA |
| 51266_2 | TATGAGTGTGGAGTCGTTGCAGTAGGAGATTAGGGATTGGGGCNNNAGATCGGA |
| 50976_1 | TATGAGTGTGGAGTCGTTGCTGGTTCTGAATAGAGCCTTTGGGNNNAGATCGGAA |
| 50976_2 | TATGAGTGTGGAGTCGTTGCTGGTTCTGAATAGAGCCTTTGGGNNNAGATCGGAA |
| 39463_1 | TATGAGTGTGGAGTCGTTGCGAAGCAGGAGAATCACATGAACCNNNAGATCGGAA |
| 39463_2 | TATGAGTGTGGAGTCGTTGCGAAGCAGGAGAATCACATGAACCNNNAGATCGGAA |
| 30415_1 | TATGAGTGTGGAGTCGTTGCGGTTCAAGCAATTCTCATGCCTTNNNAGATCGGAAG |
| 30415_2 | TATGAGTGTGGAGTCGTTGCGGTTCAAGCAATTCTCATGCCTTNNNAGATCGGAAG |
| 50607_1 | TATGAGTGTGGAGTCGTTGCTCCTTCCTTCCTTCTTATTCCCTNNNAGATCGGAAGAG |
| 50607_2 | TATGAGTGTGGAGTCGTTGCTCCTTCCTTCCTTCTTATTCCCTNNNAGATCGGAAGAG |
| 30498_1 | ATGAGTGTGGAGTCGTTGCACTTCTTTCACATTATGATTACCTCATNNNAGATCGGA |
| 30498_2 | ATGAGTGTGGAGTCGTTGCACTTCTTTCACATTATGATTACCTCATNNNAGATCGGA |

| | |
|---|---|
| 33644_1 | TATGAGTGTGGAGTCGTTGCACACCGTTTCACTATGTTAGCCANNNAGATCGGAAG |
| 33644_2 | TATGAGTGTGGAGTCGTTGCACACCGTTTCACTATGTTAGCCANNNAGATCGGAAG |
| 44293_1 | TATGAGTGTGGAGTCGTTGCCTATGCAGGAGACAGTAAAGCCANNNAGATCGGAA |
| 44293_2 | TATGAGTGTGGAGTCGTTGCCTATGCAGGAGACAGTAAAGCCANNNAGATCGGAA |
| 30239_1 | TATGAGTGTGGAGTCGTTGCGGTATGGTGGGCTCTGAGATAAGNNNAGATCGGAAA |
| 30239_2 | TATGAGTGTGGAGTCGTTGCGGTATGGTGGGCTCTGAGATAAGNNNAGATCGGAAA |
| 35503_1 | TATGAGTGTGGAGTCGTTGCTCAGCCTCCCAATTATTTGCAACNNNAGATCGGAAGA |
| 35503_2 | TATGAGTGTGGAGTCGTTGCTCAGCCTCCCAATTATTTGCAACNNNAGATCGGAAGA |
| 49037_1 | TATGAGTGTGGAGTCGTTGCCTGTCAAGCCCAGTGTTCAATTTNNNAGATCGGAAG |
| 49037_2 | TATGAGTGTGGAGTCGTTGCCTGTCAAGCCCAGTGTTCAATTTNNNAGATCGGAAG |
| 50060_1 | TATGAGTGTGGAGTCGTTGCTAGTAGCCTGCATGGAGTTTGAGNNNAGATCGGAA |
| 50060_2 | TATGAGTGTGGAGTCGTTGCTAGTAGCCTGCATGGAGTTTGAGNNNAGATCGGAA |
| 12830_1 | TATGAGTGTGGAGTCGTTGCTTAGAATTCATCAGAGAGTTCATNNNAGATCGGAAG |
| 12830_2 | TATGAGTGTGGAGTCGTTGCTTAGAATTCATCAGAGAGTTCATNNNAGATCGGAAG |
| 45392_1 | TATGAGTGTGGAGTCGTTGCAATAATGGCATTTGCAGCAACCTNNNAGATCGGAAG |
| 45392_2 | TATGAGTGTGGAGTCGTTGCAATAATGGCATTTGCAGCAACCTNNNAGATCGGAAG |
| 42054_1 | TATGAGTGTGGAGTCGTTGCTCCACAGTCCAAAGTCTGATCTGNNNAGATCGGAAG |
| 42054_2 | TATGAGTGTGGAGTCGTTGCTCCACAGTCCAAAGTCTGATCTGNNNAGATCGGAAG |
| 51335_1 | TATGAGTGTGGAGTCGTTGCCTTTTCCACTTTCCTGGGGGACNNNAGATCGGAAGA |
| 51335_2 | TATGAGTGTGGAGTCGTTGCCTTTTCCACTTTCCTGGGGGACNNNAGATCGGAAGA |
| 51307_1 | TATGAGTGTGGAGTCGTTGCACCCTAAAGAGGAGAATCAGTGCNNNAGATCGGAA |
| 51307_2 | TATGAGTGTGGAGTCGTTGCACCCTAAAGAGGAGAATCAGTGCNNNAGATCGGAA |
| 35190_1 | TATGAGTGTGGAGTCGTTGCGCATTCAAGGAATTCTCTGCCTCNNNAGATCGGAAG |
| 35190_2 | TATGAGTGTGGAGTCGTTGCGCATTCAAGGAATTCTCTGCCTCNNNAGATCGGAAG |
| 31174_1 | TATGAGTGTGGAGTCGTTGCCCTCCCCCTCTCTCTTTCTTTTTNNNAGATCGGAAGAG |
| 31174_2 | TATGAGTGTGGAGTCGTTGCCCTCCCCCTCTCTCTTTCTTTTTNNNAGATCGGAAGAG |
| 30480_1 | TATGAGTGTGGAGTCGTTGCTAACTCTTCCAAAAGCCCTCACGNNNAGATCGGAAGA |
| 30480_2 | TATGAGTGTGGAGTCGTTGCTAACTCTTCCAAAAGCCCTCACGNNNAGATCGGAAGA |
| 32737_1 | TATGAGTGTGGAGTCGTTGCGGAGAATCACTTGAACCTGGGATNNNAGATCGGAA |
| 32737_2 | TATGAGTGTGGAGTCGTTGCGGAGAATCACTTGAACCTGGGATNNNAGATCGGAA |
| 30217_1 | TATGAGTGTGGAGTCGTTGCCTCATGCCATTCGGATTTACCTTNNNAGATCGGAAGA |
| 30217_2 | TATGAGTGTGGAGTCGTTGCCTCATGCCATTCGGATTTACCTTNNNAGATCGGAAGA |
| 46263_1 | TATGAGTGTGGAGTCGTTGCATCCCAGGTTCAAGTGATTCTCCNNNAGATCGGAAG |
| 46263_2 | TATGAGTGTGGAGTCGTTGCATCCCAGGTTCAAGTGATTCTCCNNNAGATCGGAAG |
| 43415_1 | TATGAGTGTGGAGTCGTTGCCCGAGTAGCTGGGACTATAGGTNNNAGATCGGAAG |
| 43415_2 | TATGAGTGTGGAGTCGTTGCCCGAGTAGCTGGGACTATAGGTNNNAGATCGGAAG |
| 48463_1 | TATGAGTGTGGAGTCGTTGCGGTTCAAATGATTCTCCTGCCTCNNNAGATCGGAAG |
| 48463_2 | TATGAGTGTGGAGTCGTTGCGGTTCAAATGATTCTCCTGCCTCNNNAGATCGGAAG |
| 36569_1 | TATGAGTGTGGAGTCGTTGCAAATGGGCAAACTGTATGGCATGNNNAGATCGGAA |
| 36569_2 | TATGAGTGTGGAGTCGTTGCAAATGGGCAAACTGTATGGCATGNNNAGATCGGAA |
| 30787_1 | TATGAGTGTGGAGTCGTTGCAGACTTTCCAAACCATACATTGCANNNAGATCGGAAG |
| 30787_2 | TATGAGTGTGGAGTCGTTGCAGACTTTCCAAACCATACATTGCANNNAGATCGGAAG |
| 31628_1 | TATGAGTGTGGAGTCGTTGCTGAGGCAATCTATAGCGAAACCTNNNAGATCGGAAG |

| | |
|---|---|
| 31628_2 | TATGAGTGTGGAGTCGTTGCTGAGGCAATCTATAGCGAAACCTNNNAGATCGGAAG |
| 40006_1 | TATGAGTGTGGAGTCGTTGCCATGCTTGTAATCCCAGCACTTTNNNAGATCGGAAGA |
| 40006_2 | TATGAGTGTGGAGTCGTTGCCATGCTTGTAATCCCAGCACTTTNNNAGATCGGAAGA |
| 40796_1 | TATGAGTGTGGAGTCGTTGCCTGGGGGTTAGAATATCCAGGAANNNAGATCGGAA |
| 40796_2 | TATGAGTGTGGAGTCGTTGCCTGGGGGTTAGAATATCCAGGAANNNAGATCGGAA |
| 36143_1 | TATGAGTGTGGAGTCGTTGCTGGGCAACATAGTGAGACTTCATNNNAGATCGGAAG |
| 36143_2 | TATGAGTGTGGAGTCGTTGCTGGGCAACATAGTGAGACTTCATNNNAGATCGGAAG |
| 36023_1 | TATGAGTGTGGAGTCGTTGCAGTCTTCAAGTCTATACCCCCCANNNAGATCGGAAGA |
| 36023_2 | TATGAGTGTGGAGTCGTTGCAGTCTTCAAGTCTATACCCCCCANNNAGATCGGAAGA |
| 32375_1 | TATGAGTGTGGAGTCGTTGCCACACACACACACACACACATGNNNAGATCGGAAGA |
| 32375_2 | TATGAGTGTGGAGTCGTTGCCACACACACACACACACACATGNNNAGATCGGAAGA |
| 36133_1 | TATGAGTGTGGAGTCGTTGCTGTCTTCTTCCGTCTTCTCAAGGNNNAGATCGGAAGA |
| 36133_2 | TATGAGTGTGGAGTCGTTGCTGTCTTCTTCCGTCTTCTCAAGGNNNAGATCGGAAGA |
| 48024_1 | TATGAGTGTGGAGTCGTTGCCTCCCAGGTTCAAGCGATTCTCNNNAGATCGGAAGA |
| 48024_2 | TATGAGTGTGGAGTCGTTGCCTCCCAGGTTCAAGCGATTCTCNNNAGATCGGAAGA |
| 52694_1 | TATGAGTGTGGAGTCGTTGCCCACACCTCCCTATCACCATTGNNNAGATCGGAAGAG |
| 52694_2 | TATGAGTGTGGAGTCGTTGCCCACACCTCCCTATCACCATTGNNNAGATCGGAAGAG |
| 31311_1 | TATGAGTGTGGAGTCGTTGCCGAGACTGGATATGCGGAACTAANNNAGATCGGAA |
| 31311_2 | TATGAGTGTGGAGTCGTTGCCGAGACTGGATATGCGGAACTAANNNAGATCGGAA |
| 36590_1 | TATGAGTGTGGAGTCGTTGCTAGCAGTGGTTTGTAGTTCTCCCNNNAGATCGGAAG |
| 36590_2 | TATGAGTGTGGAGTCGTTGCTAGCAGTGGTTTGTAGTTCTCCCNNNAGATCGGAAG |
| 44239_1 | TATGAGTGTGGAGTCGTTGCAAAACCTGATATGGGTTGGCAGANNNAGATCGGAA |
| 44239_2 | TATGAGTGTGGAGTCGTTGCAAAACCTGATATGGGTTGGCAGANNNAGATCGGAA |
| 48334_1 | TATGAGTGTGGAGTCGTTGCCTGCCTCAGCCTCCTGATTAGNNNAGATCGGAAGAG |
| 48334_2 | TATGAGTGTGGAGTCGTTGCCTGCCTCAGCCTCCTGATTAGNNNAGATCGGAAGAG |
| 10109_1 | TATGAGTGTGGAGTCGTTGCTGGAAAAGCACTCTGTAGCTCAGNNNAGATCGGAAG |
| 10109_2 | TATGAGTGTGGAGTCGTTGCTGGAAAAGCACTCTGTAGCTCAGNNNAGATCGGAAG |
| 50251_1 | TATGAGTGTGGAGTCGTTGCCCATCTCAGAGTTCCCTTCTGTGNNNAGATCGGAAGA |
| 50251_2 | TATGAGTGTGGAGTCGTTGCCCATCTCAGAGTTCCCTTCTGTGNNNAGATCGGAAGA |
| 45579_1 | TATGAGTGTGGAGTCGTTGCGGTAACACACGTGAGGTTTTTGTNNNAGATCGGAAG |
| 45579_2 | TATGAGTGTGGAGTCGTTGCGGTAACACACGTGAGGTTTTTGTNNNAGATCGGAAG |
| 10207_1 | TATGAGTGTGGAGTCGTTGCACTTGCACACAAATGTTCAACCCNNNAGATCGGAAGA |
| 10207_2 | TATGAGTGTGGAGTCGTTGCACTTGCACACAAATGTTCAACCCNNNAGATCGGAAGA |
| 45211_1 | TATGAGTGTGGAGTCGTTGCTCAGGAAACTTCAGGCAGCATATNNNAGATCGGAAG |
| 45211_2 | TATGAGTGTGGAGTCGTTGCTCAGGAAACTTCAGGCAGCATATNNNAGATCGGAAG |
| 13688_1 | TATGAGTGTGGAGTCGTTGCGTAGCGTATACATTTGTTTTGGNNNAGATCGGAAGA |
| 13688_2 | TATGAGTGTGGAGTCGTTGCGTAGCGTATACATTTGTTTTGGNNNAGATCGGAAGA |
| 32732_1 | TATGAGTGTGGAGTCGTTGCCTGACAGTCAAGAGAGCAAGGAGNNNAGATCGGAA |
| 32732_2 | TATGAGTGTGGAGTCGTTGCCTGACAGTCAAGAGAGCAAGGAGNNNAGATCGGAA |
| 32272_1 | TATGAGTGTGGAGTCGTTGCCCCCAATTTAGAAGCTTGTGTGGNNNAGATCGGAAG |
| 32272_2 | TATGAGTGTGGAGTCGTTGCCCCCAATTTAGAAGCTTGTGTGGNNNAGATCGGAAG |
| 52343_1 | TATGAGTGTGGAGTCGTTGCTTCTGTCTTCACACCAATCTGGANNNAGATCGGAAGA |
| 52343_2 | TATGAGTGTGGAGTCGTTGCTTCTGTCTTCACACCAATCTGGANNNAGATCGGAAGA |

| | |
|---|---|
| 33283_1 | TATGAGTGTGGAGTCGTTGCTCTTGTTCAGTCTTCAGAGGGTGNNNAGATCGGAAG |
| 33283_2 | TATGAGTGTGGAGTCGTTGCTCTTGTTCAGTCTTCAGAGGGTGNNNAGATCGGAAG |
| 47428_1 | TATGAGTGTGGAGTCGTTGCCTTCCTGGTAGCTGGGATTAGAGNNNAGATCGGAA |
| 47428_2 | TATGAGTGTGGAGTCGTTGCCTTCCTGGTAGCTGGGATTAGAGNNNAGATCGGAA |
| 10964_1 | TATGAGTGTGGAGTCGTTGCGCCAGCATCATCCTAATACCCCNNNAGATCGGAAG |
| 10964_2 | TATGAGTGTGGAGTCGTTGCGCCAGCATCATCCTAATACCCCNNNAGATCGGAAG |
| 31976_1 | TATGAGTGTGGAGTCGTTGCCATGCTGCTGTCATTTTGTGGTANNNAGATCGGAAG |
| 31976_2 | TATGAGTGTGGAGTCGTTGCCATGCTGCTGTCATTTTGTGGTANNNAGATCGGAAG |
| 40095_1 | TATGAGTGTGGAGTCGTTGCCTGGGGTATGCTTGCTTTGTTTTNNNAGATCGGAAG |
| 40095_2 | TATGAGTGTGGAGTCGTTGCCTGGGGTATGCTTGCTTTGTTTTNNNAGATCGGAAG |
| 29953_1 | TATGAGTGTGGAGTCGTTGCAAGCCAAGAAAGTTCAATGGCAANNNAGATCGGAA |
| 29953_2 | TATGAGTGTGGAGTCGTTGCAAGCCAAGAAAGTTCAATGGCAANNNAGATCGGAA |
| 35879_1 | TATGAGTGTGGAGTCGTTGCCGAGACCAGCCTGACTAACATAGNNNAGATCGGAAG |
| 35879_2 | TATGAGTGTGGAGTCGTTGCCGAGACCAGCCTGACTAACATAGNNNAGATCGGAAG |
| 39129_1 | TATGAGTGTGGAGTCGTTGCAGTCTTGGGTGAGATTTTGGACANNNAGATCGGAA |
| 39129_2 | TATGAGTGTGGAGTCGTTGCAGTCTTGGGTGAGATTTTGGACANNNAGATCGGAA |
| 37073_1 | TATGAGTGTGGAGTCGTTGCATCCTATTGAGAGGCAAACTGGGNNNAGATCGGAA |
| 37073_2 | TATGAGTGTGGAGTCGTTGCATCCTATTGAGAGGCAAACTGGGNNNAGATCGGAA |
| 36770_1 | TATGAGTGTGGAGTCGTTGCACCCACAAGTCTCTGGAAATTCANNNAGATCGGAAG |
| 36770_2 | TATGAGTGTGGAGTCGTTGCACCCACAAGTCTCTGGAAATTCANNNAGATCGGAAG |
| 33384_1 | TATGAGTGTGGAGTCGTTGCTAAAGCCTGCAGAACCATCCATNNNAGATCGGAAGA |
| 33384_2 | TATGAGTGTGGAGTCGTTGCTAAAGCCTGCAGAACCATCCATNNNAGATCGGAAGA |
| 44479_1 | TATGAGTGTGGAGTCGTTGCTTTTTTTCCCCGAGACAGAGTCTNNNAGATCGGAAG |
| 44479_2 | TATGAGTGTGGAGTCGTTGCTTTTTTTCCCCGAGACAGAGTCTNNNAGATCGGAAG |
| 12739_1 | TATGAGTGTGGAGTCGTTGCGTTTCTCAGTCATGATTCCCNNNAGATCGGAAGAGC |
| 12739_2 | TATGAGTGTGGAGTCGTTGCGTTTCTCAGTCATGATTCCCNNNAGATCGGAAGAGC |
| 52887_1 | TATGAGTGTGGAGTCGTTGCGTTCCTCCTGGTCTGTCTGATTTNNNAGATCGGAAG |
| 52887_2 | TATGAGTGTGGAGTCGTTGCGTTCCTCCTGGTCTGTCTGATTTNNNAGATCGGAAG |
| 34837_1 | TATGAGTGTGGAGTCGTTGCCCTTCTGGGTTTACGCCATTCTCNNNAGATCGGAAG |
| 34837_2 | TATGAGTGTGGAGTCGTTGCCCTTCTGGGTTTACGCCATTCTCNNNAGATCGGAAG |
| 42871_1 | TATGAGTGTGGAGTCGTTGCGATCTTGCTAATGTTCCCCAAGCNNNAGATCGGAAG |
| 42871_2 | TATGAGTGTGGAGTCGTTGCGATCTTGCTAATGTTCCCCAAGCNNNAGATCGGAAG |
| 30962_1 | TATGAGTGTGGAGTCGTTGCATGTTGTGATACCAGGCAGCTAANNNAGATCGGAAG |
| 30962_2 | TATGAGTGTGGAGTCGTTGCATGTTGTGATACCAGGCAGCTAANNNAGATCGGAAG |
| 37272_1 | TATGAGTGTGGAGTCGTTGCACAAATACCCCATCCAAAGGTCANNNAGATCGGAAG |
| 37272_2 | TATGAGTGTGGAGTCGTTGCACAAATACCCCATCCAAAGGTCANNNAGATCGGAAG |
| 46137_1 | TATGAGTGTGGAGTCGTTGCCCTGCTATTCTTAGGCCTTTTCCNNNAGATCGGAAGA |
| 46137_2 | TATGAGTGTGGAGTCGTTGCCCTGCTATTCTTAGGCCTTTTCCNNNAGATCGGAAGA |
| 2490_1 | TATGAGTGTGGAGTCGTTGCCGCAGACCCTGCATATAGTCAGNNNAGATCGGAAG |
| 2490_2 | TATGAGTGTGGAGTCGTTGCCGCAGACCCTGCATATAGTCAGNNNAGATCGGAAG |
| 34362_1 | TATGAGTGTGGAGTCGTTGCTGTCTAACCATCCTTCAGCTGACNNNAGATCGGAAG |
| 34362_2 | TATGAGTGTGGAGTCGTTGCTGTCTAACCATCCTTCAGCTGACNNNAGATCGGAAG |
| 46197_1 | TATGAGTGTGGAGTCGTTGCAAACTCCTAGGCTCAAGTGATCCNNNAGATCGGAAG |

| | |
|---|---|
| 46197_2 | TATGAGTGTGGAGTCGTTGCAAACTCCTAGGCTCAAGTGATCCNNNAGATCGGAAG |
| 52521_1 | TATGAGTGTGGAGTCGTTGCTCCAACTTACTAAACAGGGCTCCNNNAGATCGGAAG |
| 52521_2 | TATGAGTGTGGAGTCGTTGCTCCAACTTACTAAACAGGGCTCCNNNAGATCGGAAG |
| 44384_1 | TATGAGTGTGGAGTCGTTGCTCTCAGGAATAGGTCAAAGCAGGNNNAGATCGGAA |
| 44384_2 | TATGAGTGTGGAGTCGTTGCTCTCAGGAATAGGTCAAAGCAGGNNNAGATCGGAA |
| 30256_1 | TATGAGTGTGGAGTCGTTGCGCCCTATTCTTTAGTAGCCACGTNNNAGATCGGAAG |
| 30256_2 | TATGAGTGTGGAGTCGTTGCGCCCTATTCTTTAGTAGCCACGTNNNAGATCGGAAG |
| 34606_1 | TATGAGTGTGGAGTCGTTGCCCTTCCACCACTATGCTAACCATNNNAGATCGGAAGA |
| 34606_2 | TATGAGTGTGGAGTCGTTGCCCTTCCACCACTATGCTAACCATNNNAGATCGGAAGA |
| 10303_1 | TATGAGTGTGGAGTCGTTGCCGGGTTGCATATATGTCCGAGTTNNNAGATCGGAAG |
| 10303_2 | TATGAGTGTGGAGTCGTTGCCGGGTTGCATATATGTCCGAGTTNNNAGATCGGAAG |
| 31940_1 | TATGAGTGTGGAGTCGTTGCAGTTTCTTCTTGGCCTCTCTGAGNNNAGATCGGAAG |
| 31940_2 | TATGAGTGTGGAGTCGTTGCAGTTTCTTCTTGGCCTCTCTGAGNNNAGATCGGAAG |
| 48498_1 | TATGAGTGTGGAGTCGTTGCCTCACTCCAAAGTAAGTGGGGAGNNNAGATCGGAA |
| 48498_2 | TATGAGTGTGGAGTCGTTGCCTCACTCCAAAGTAAGTGGGGAGNNNAGATCGGAA |
| 30181_1 | TATGAGTGTGGAGTCGTTGCCCACCCAGAAAGTATTATCATCTTAATNNNAGATCGG |
| 30181_2 | TATGAGTGTGGAGTCGTTGCCCACCCAGAAAGTATTATCATCTTAATNNNAGATCGG |
| 47167_1 | TATGAGTGTGGAGTCGTTGCCAGTGGTATGATCATGGCTCACTNNNAGATCGGAAG |
| 47167_2 | TATGAGTGTGGAGTCGTTGCCAGTGGTATGATCATGGCTCACTNNNAGATCGGAAG |
| 10625_1 | TATGAGTGTGGAGTCGTTGCCAGTGTCGTGTTCATAGCTGACTNNNAGATCGGAAG |
| 10625_2 | TATGAGTGTGGAGTCGTTGCCAGTGTCGTGTTCATAGCTGACTNNNAGATCGGAAG |
| 53377_1 | TATGAGTGTGGAGTCGTTGCCCCCCTTCTCTCTCTCTCTCATANNNAGATCGGAAGAG |
| 53377_2 | TATGAGTGTGGAGTCGTTGCCCCCCTTCTCTCTCTCTCTCATANNNAGATCGGAAGAG |
| 44141_1 | TATGAGTGTGGAGTCGTTGCCTTTCATACCTCTCACCTGTGCTNNNAGATCGGAAGA |
| 44141_2 | TATGAGTGTGGAGTCGTTGCCTTTCATACCTCTCACCTGTGCTNNNAGATCGGAAGA |
| 31236_1 | TATGAGTGTGGAGTCGTTGCTCTCTCCCTCTCTCTCTCTCTCTNNNAGATCGGAAGAG |
| 31236_2 | TATGAGTGTGGAGTCGTTGCTCTCTCCCTCTCTCTCTCTCTCTNNNAGATCGGAAGAG |
| 37403_1 | TATGAGTGTGGAGTCGTTGCAAGAAGGATGAGTGCTGGCTTATNNNAGATCGGAA |
| 37403_2 | TATGAGTGTGGAGTCGTTGCAAGAAGGATGAGTGCTGGCTTATNNNAGATCGGAA |
| 37267_1 | TATGAGTGTGGAGTCGTTGCCTGAGATGGGAGAATCACCTGAGNNNAGATCGGAA |
| 37267_2 | TATGAGTGTGGAGTCGTTGCCTGAGATGGGAGAATCACCTGAGNNNAGATCGGAA |
| 45922_1 | TATGAGTGTGGAGTCGTTGCCCCAAGTTCAAGCAATTCTCCTGNNNAGATCGGAAG |
| 45922_2 | TATGAGTGTGGAGTCGTTGCCCCAAGTTCAAGCAATTCTCCTGNNNAGATCGGAAG |
| 29365_1 | TATGAGTGTGGAGTCGTTGCGCTCTGCCCATAACCACTTATCANNNAGATCGGAAGA |
| 29365_2 | TATGAGTGTGGAGTCGTTGCGCTCTGCCCATAACCACTTATCANNNAGATCGGAAGA |
| 44072_1 | TATGAGTGTGGAGTCGTTGCTATCACTATGTGCCCCACTCATGNNNAGATCGGAAG |
| 44072_2 | TATGAGTGTGGAGTCGTTGCTATCACTATGTGCCCCACTCATGNNNAGATCGGAAG |
| 52880_1 | TATGAGTGTGGAGTCGTTGCGTACCACCATCGTGAAGACCATNNNAGATCGGAAGA |
| 52880_2 | TATGAGTGTGGAGTCGTTGCGTACCACCATCGTGAAGACCATNNNAGATCGGAAGA |
| 40710_1 | TATGAGTGTGGAGTCGTTGCTGCTGAATTTCCCTCCAGAAAGANNNAGATCGGAAG |
| 40710_2 | TATGAGTGTGGAGTCGTTGCTGCTGAATTTCCCTCCAGAAAGANNNAGATCGGAAG |
| 44313_1 | TATGAGTGTGGAGTCGTTGCAGAGAGCTGTCCATGTAAGTGTGNNNAGATCGGAA |
| 44313_2 | TATGAGTGTGGAGTCGTTGCAGAGAGCTGTCCATGTAAGTGTGNNNAGATCGGAA |

| 42967_1 | TATGAGTGTGGAGTCGTTGCTTCCTAACTTGTGTCCAGTCCTGNNNAGATCGGAAG |
|---|---|
| 42967_2 | TATGAGTGTGGAGTCGTTGCTTCCTAACTTGTGTCCAGTCCTGNNNAGATCGGAAG |
| 46450_1 | TATGAGTGTGGAGTCGTTGCCCTCTTGGGTTCATGCCATTCTCNNNAGATCGGAAG |
| 46450_2 | TATGAGTGTGGAGTCGTTGCCCTCTTGGGTTCATGCCATTCTCNNNAGATCGGAAG |
| 31419_1 | TATGAGTGTGGAGTCGTTGCGTGAGTTGCATCCTAAGGGAGAANNNAGATCGGAA |
| 31419_2 | TATGAGTGTGGAGTCGTTGCGTGAGTTGCATCCTAAGGGAGAANNNAGATCGGAA |
| 36527_1 | TATGAGTGTGGAGTCGTTGCCTATAGCTCTGCTGAGGACAGTGNNNAGATCGGAA |
| 36527_2 | TATGAGTGTGGAGTCGTTGCCTATAGCTCTGCTGAGGACAGTGNNNAGATCGGAA |
| 42864_1 | TATGAGTGTGGAGTCGTTGCAATGTCTTTTGCAGCAACTTGGANNNAGATCGGAAG |
| 42864_2 | TATGAGTGTGGAGTCGTTGCAATGTCTTTTGCAGCAACTTGGANNNAGATCGGAAG |
| 49527_1 | TATGAGTGTGGAGTCGTTGCGGAGCAGCTTGAGTTACAGGAAANNNAGATCGGAA |
| 49527_2 | TATGAGTGTGGAGTCGTTGCGGAGCAGCTTGAGTTACAGGAAANNNAGATCGGAA |
| 37429_1 | TATGAGTGTGGAGTCGTTGCCCAGAAAACCAGAACATGTGGTCNNNAGATCGGAA |
| 37429_2 | TATGAGTGTGGAGTCGTTGCCCAGAAAACCAGAACATGTGGTCNNNAGATCGGAA |
| 48646_1 | TATGAGTGTGGAGTCGTTGCGTATCTGGGAATACAGGTGCACANNNAGATCGGAA |
| 48646_2 | TATGAGTGTGGAGTCGTTGCGTATCTGGGAATACAGGTGCACANNNAGATCGGAA |
| 32353_1 | TATGAGTGTGGAGTCGTTGCCAACACACACACACACACACATCNNNAGATCGGAAG |
| 32353_2 | TATGAGTGTGGAGTCGTTGCCAACACACACACACACACACATCNNNAGATCGGAAG |
| 43182_1 | TATGAGTGTGGAGTCGTTGCGCACGTACTATTCCGTTTGTCAGNNNAGATCGGAAG |
| 43182_2 | TATGAGTGTGGAGTCGTTGCGCACGTACTATTCCGTTTGTCAGNNNAGATCGGAAG |
| 10042_1 | TATGAGTGTGGAGTCGTTGCCTCTCAGTCCATCTTGTCATGGCNNNAGATCGGAAG |
| 10042_2 | TATGAGTGTGGAGTCGTTGCCTCTCAGTCCATCTTGTCATGGCNNNAGATCGGAAG |
| 48720_1 | TATGAGTGTGGAGTCGTTGCTGTTAGCCAGGATGGTCTTGATCNNNAGATCGGAAG |
| 48720_2 | TATGAGTGTGGAGTCGTTGCTGTTAGCCAGGATGGTCTTGATCNNNAGATCGGAAG |
| 49735_1 | TATGAGTGTGGAGTCGTTGCTTTTAGTAGAGACGGGGCTTCACNNNAGATCGGAA |
| 49735_2 | TATGAGTGTGGAGTCGTTGCTTTTAGTAGAGACGGGGCTTCACNNNAGATCGGAA |
| 49650_1 | TATGAGTGTGGAGTCGTTGCTTTTTCTTGAGACAGGCTCTTGCNNNAGATCGGAAG |
| 49650_2 | TATGAGTGTGGAGTCGTTGCTTTTTCTTGAGACAGGCTCTTGCNNNAGATCGGAAG |
| 29707_1 | TATGAGTGTGGAGTCGTTGCAAGACATTTGTGTTGCAGTCAGANNNAGATCGGAA |
| 29707_2 | TATGAGTGTGGAGTCGTTGCAAGACATTTGTGTTGCAGTCAGANNNAGATCGGAA |
| 48840_1 | TATGAGTGTGGAGTCGTTGCTTGAAGTCACATTGCTTCTGCTGNNNAGATCGGAAG |
| 48840_2 | TATGAGTGTGGAGTCGTTGCTTGAAGTCACATTGCTTCTGCTGNNNAGATCGGAAG |
| 48268_1 | TATGAGTGTGGAGTCGTTGCTGAAAGTGTCTGGGAATGGCATANNNAGATCGGAA |
| 48268_2 | TATGAGTGTGGAGTCGTTGCTGAAAGTGTCTGGGAATGGCATANNNAGATCGGAA |
| 10100_1 | TATGAGTGTGGAGTCGTTGCATTACGGTTTCATCTGCTGAAGCNNNAGATCGGAAG |
| 10100_2 | TATGAGTGTGGAGTCGTTGCATTACGGTTTCATCTGCTGAAGCNNNAGATCGGAAG |
| 36885_1 | TATGAGTGTGGAGTCGTTGCCAGGCCTGTAATCCCAACACTTTNNNAGATCGGAAG |
| 36885_2 | TATGAGTGTGGAGTCGTTGCCAGGCCTGTAATCCCAACACTTTNNNAGATCGGAAG |
| 53235_1 | TATGAGTGTGGAGTCGTTGCATGAAGGTAGTTCTGTTCCCACANNNAGATCGGAAG |
| 53235_2 | TATGAGTGTGGAGTCGTTGCATGAAGGTAGTTCTGTTCCCACANNNAGATCGGAAG |
| 13195_1 | TATGAGTGTGGAGTCGTTGCGATACAATGGTGAGTGAAAATATNNNAGATCGGAA |
| 13195_2 | TATGAGTGTGGAGTCGTTGCGATACAATGGTGAGTGAAAATATNNNAGATCGGAA |
| 31363_1 | TATGAGTGTGGAGTCGTTGCCCAGTGGTCTTTCTGAAGGAGAANNNAGATCGGAA |

| | |
|---|---|
| 31363_2 | TATGAGTGTGGAGTCGTTGCCCAGTGGTCTTTCTGAAGGAGAANNNAGATCGGAA |
| 35439_1 | TATGAGTGTGGAGTCGTTGCGCCTCCCAGGTTCATGTTATTTTNNNAGATCGGAAG |
| 35439_2 | TATGAGTGTGGAGTCGTTGCGCCTCCCAGGTTCATGTTATTTTNNNAGATCGGAAG |
| 48909_1 | TATGAGTGTGGAGTCGTTGCGGAAAGCTGTCCTGGTCTAATCTNNNAGATCGGAAG |
| 48909_2 | TATGAGTGTGGAGTCGTTGCGGAAAGCTGTCCTGGTCTAATCTNNNAGATCGGAAG |
| 50820_1 | TATGAGTGTGGAGTCGTTGCGTCCATGACCCTCTAGAAGACAGNNNAGATCGGAAG |
| 50820_2 | TATGAGTGTGGAGTCGTTGCGTCCATGACCCTCTAGAAGACAGNNNAGATCGGAAG |
| 39885_1 | TATGAGTGTGGAGTCGTTGCGTGGCTCACTCCTATAATCCCAGNNNAGATCGGAAG |
| 39885_2 | TATGAGTGTGGAGTCGTTGCGTGGCTCACTCCTATAATCCCAGNNNAGATCGGAAG |
| 37436_1 | TATGAGTGTGGAGTCGTTGCTCCTGTCATTTGCAACAACATGGNNNAGATCGGAAG |
| 37436_2 | TATGAGTGTGGAGTCGTTGCTCCTGTCATTTGCAACAACATGGNNNAGATCGGAAG |
| 35088_1 | TATGAGTGTGGAGTCGTTGCCTCCTAGGCTCAAGTGATTCTCCNNNAGATCGGAAG |
| 35088_2 | TATGAGTGTGGAGTCGTTGCCTCCTAGGCTCAAGTGATTCTCCNNNAGATCGGAAG |
| 51459_1 | TATGAGTGTGGAGTCGTTGCACCCTTTTCCCAGAAGAAATCGANNNAGATCGGAAG |
| 51459_2 | TATGAGTGTGGAGTCGTTGCACCCTTTTCCCAGAAGAAATCGANNNAGATCGGAAG |
| 31917_1 | TATGAGTGTGGAGTCGTTGCACCTCACTGATGTTTACCTTGGGNNNAGATCGGAAG |
| 31917_2 | TATGAGTGTGGAGTCGTTGCACCTCACTGATGTTTACCTTGGGNNNAGATCGGAAG |
| 45987_1 | TATGAGTGTGGAGTCGTTGCCTCTCAAGTAGCTGGGACTACAGNNNAGATCGGAA |
| 45987_2 | TATGAGTGTGGAGTCGTTGCCTCTCAAGTAGCTGGGACTACAGNNNAGATCGGAA |
| 33402_1 | TATGAGTGTGGAGTCGTTGCAACAACAAACTACCTGAAAAGGAANNNAGATCGGA |
| 33402_2 | TATGAGTGTGGAGTCGTTGCAACAACAAACTACCTGAAAAGGAANNNAGATCGGA |
| 48031_1 | TATGAGTGTGGAGTCGTTGCTTCAAGCAATTCTCCTGCCTCAGNNNAGATCGGAAG |
| 48031_2 | TATGAGTGTGGAGTCGTTGCTTCAAGCAATTCTCCTGCCTCAGNNNAGATCGGAAG |
| 46863_1 | TATGAGTGTGGAGTCGTTGCAGTATCTGGGATTACATGTGCCCNNNAGATCGGAAG |
| 46863_2 | TATGAGTGTGGAGTCGTTGCAGTATCTGGGATTACATGTGCCCNNNAGATCGGAAG |
| 39258_1 | TATGAGTGTGGAGTCGTTGCCCATGGAAGAACCAATCTGTGTGNNNAGATCGGAA |
| 39258_2 | TATGAGTGTGGAGTCGTTGCCCATGGAAGAACCAATCTGTGTGNNNAGATCGGAA |
| 51353_1 | TATGAGTGTGGAGTCGTTGCCCACACACTGTTCTAGGTGATGANNNAGATCGGAAG |
| 51353_2 | TATGAGTGTGGAGTCGTTGCCCACACACTGTTCTAGGTGATGANNNAGATCGGAAG |
| 41716_1 | TATGAGTGTGGAGTCGTTGCGGGTTCAAGTGATTCTCCTACCTNNNAGATCGGAAG |
| 41716_2 | TATGAGTGTGGAGTCGTTGCGGGTTCAAGTGATTCTCCTACCTNNNAGATCGGAAG |
| 28952_1 | TATGAGTGTGGAGTCGTTGCGGTGAGTGAATGAGTGCATTGTTNNNAGATCGGAA |
| 28952_2 | TATGAGTGTGGAGTCGTTGCGGTGAGTGAATGAGTGCATTGTTNNNAGATCGGAA |
| 38974_1 | TATGAGTGTGGAGTCGTTGCAGAGAGAGAGAGACAAAGAGGCTNNNAGATCGGA |
| 38974_2 | TATGAGTGTGGAGTCGTTGCAGAGAGAGAGAGACAAAGAGGCTNNNAGATCGGA |
| 50053_1 | TATGAGTGTGGAGTCGTTGCGGGGAGGGAAAGAAAGAGAAAGANNNAGATCGG |
| 50053_2 | TATGAGTGTGGAGTCGTTGCGGGGAGGGAAAGAAAGAGAAAGANNNAGATCGG |
| 50678_1 | TATGAGTGTGGAGTCGTTGCTCTCACTTTTAGAGGCTTCACCCNNNAGATCGGAAGA |
| 50678_2 | TATGAGTGTGGAGTCGTTGCTCTCACTTTTAGAGGCTTCACCCNNNAGATCGGAAGA |
| 50859_1 | TATGAGTGTGGAGTCGTTGCCCCCTGGTTTGAAGAAAGGAGAANNNAGATCGGAA |
| 50859_2 | TATGAGTGTGGAGTCGTTGCCCCCTGGTTTGAAGAAAGGAGAANNNAGATCGGAA |
| 51524_1 | TATGAGTGTGGAGTCGTTGCGAGGGAGGAAGGAAGGAAACTTCNNNAGATCGGA |
| 51524_2 | TATGAGTGTGGAGTCGTTGCGAGGGAGGAAGGAAGGAAACTTCNNNAGATCGGA |

| | |
|---|---|
| 42222_1 | TATGAGTGTGGAGTCGTTGCTTTTCAGTGGAAGGTGTTTTGGCNNNAGATCGGAA |
| 42222_2 | TATGAGTGTGGAGTCGTTGCTTTTCAGTGGAAGGTGTTTTGGCNNNAGATCGGAA |
| 45759_1 | TATGAGTGTGGAGTCGTTGCGGTTCAAGTGATTTTCCTGCCTCNNNAGATCGGAAG |
| 45759_2 | TATGAGTGTGGAGTCGTTGCGGTTCAAGTGATTTTCCTGCCTCNNNAGATCGGAAG |
| 35306_1 | TATGAGTGTGGAGTCGTTGCTTCCTGGGTTCATGCCATTCTCNNNAGATCGGAAGA |
| 35306_2 | TATGAGTGTGGAGTCGTTGCTTCCTGGGTTCATGCCATTCTCNNNAGATCGGAAGA |
| 49704_1 | TATGAGTGTGGAGTCGTTGCGGTCTTCTGCAGTGACATTTGTCNNNAGATCGGAAG |
| 49704_2 | TATGAGTGTGGAGTCGTTGCGGTCTTCTGCAGTGACATTTGTCNNNAGATCGGAAG |
| 43215_1 | TATGAGTGTGGAGTCGTTGCCGAGTAGCTGGGACTACAGGTNNNAGATCGGAAGA |
| 43215_2 | TATGAGTGTGGAGTCGTTGCCGAGTAGCTGGGACTACAGGTNNNAGATCGGAAGA |
| 31205_1 | TATGAGTGTGGAGTCGTTGCTTTTTGTGGTGACCATTGTGCTTNNNAGATCGGAAG |
| 31205_2 | TATGAGTGTGGAGTCGTTGCTTTTTGTGGTGACCATTGTGCTTNNNAGATCGGAAG |
| 7797_1 | TATGAGTGTGGAGTCGTTGCCAAGCGATACTCCTATCCCAGGNNNAGATCGGAAGA |
| 7797_2 | TATGAGTGTGGAGTCGTTGCCAAGCGATACTCCTATCCCAGGNNNAGATCGGAAGA |
| 36728_1 | TATGAGTGTGGAGTCGTTGCCAATGAGTCTGTGGGGTGTCTAANNNAGATCGGAA |
| 36728_2 | TATGAGTGTGGAGTCGTTGCCAATGAGTCTGTGGGGTGTCTAANNNAGATCGGAA |
| 49272_1 | TATGAGTGTGGAGTCGTTGCCTCCTGAGTAGCTGGGACTACAGNNNAGATCGGAA |
| 49272_2 | TATGAGTGTGGAGTCGTTGCCTCCTGAGTAGCTGGGACTACAGNNNAGATCGGAA |
| 35019_1 | TATGAGTGTGGAGTCGTTGCGAGTGTAATGGCATGATCTTGGCNNNAGATCGGAA |
| 35019_2 | TATGAGTGTGGAGTCGTTGCGAGTGTAATGGCATGATCTTGGCNNNAGATCGGAA |
| 36087_1 | TATGAGTGTGGAGTCGTTGCCACCCCATTCTAAAAAGCAGGTGNNNAGATCGGAAG |
| 36087_2 | TATGAGTGTGGAGTCGTTGCCACCCCATTCTAAAAAGCAGGTGNNNAGATCGGAAG |
| 28902_1 | TATGAGTGTGGAGTCGTTGCGTGTGTGTGTGTGTGTGTATATATNNNAGATCGGA |
| 28902_2 | TATGAGTGTGGAGTCGTTGCGTGTGTGTGTGTGTGTGTATATATNNNAGATCGGA |
| 53371_1 | TATGAGTGTGGAGTCGTTGCTGGGTCAGAACAAAGCCAGAATANNNAGATCGGAA |
| 53371_2 | TATGAGTGTGGAGTCGTTGCTGGGTCAGAACAAAGCCAGAATANNNAGATCGGAA |
| 618_1 | TATGAGTGTGGAGTCGTTGCGGAGAGAGAGAAAATGGGAGCAGNNNAGATCGGA |
| 618_2 | TATGAGTGTGGAGTCGTTGCGGAGAGAGAGAAAATGGGAGCAGNNNAGATCGGA |
| 13179_1 | TATGAGTGTGGAGTCGTTGCTTTCTAAAAGGAGACTCCCTCNNNAGATCGGAAGAG |
| 13179_2 | TATGAGTGTGGAGTCGTTGCTTTCTAAAAGGAGACTCCCTCNNNAGATCGGAAGAG |
| 11171_1 | TATGAGTGTGGAGTCGTTGCATAGGCAGCAGGTATTAGTGACACNNNAGATCGGA |
| 11171_2 | TATGAGTGTGGAGTCGTTGCATAGGCAGCAGGTATTAGTGACACNNNAGATCGGA |
| 39525_1 | TATGAGTGTGGAGTCGTTGCAGAATGATCTGGTCCAATTGAGCTNNNAGATCGGAA |
| 39525_2 | TATGAGTGTGGAGTCGTTGCAGAATGATCTGGTCCAATTGAGCTNNNAGATCGGAA |
| 38689_1 | TATGAGTGTGGAGTCGTTGCGGCTACAGGGGAATCAGATTTGANNNAGATCGGAA |
| 38689_2 | TATGAGTGTGGAGTCGTTGCGGCTACAGGGGAATCAGATTTGANNNAGATCGGAA |
| 33913_1 | TATGAGTGTGGAGTCGTTGCTCAGCCTCCTGAGTAGTTAGGACNNNAGATCGGAAG |
| 33913_2 | TATGAGTGTGGAGTCGTTGCTCAGCCTCCTGAGTAGTTAGGACNNNAGATCGGAAG |
| 38542_1 | TATGAGTGTGGAGTCGTTGCCTTGGAGATGTGTGCCTGTAGTCNNNAGATCGGAA |
| 38542_2 | TATGAGTGTGGAGTCGTTGCCTTGGAGATGTGTGCCTGTAGTCNNNAGATCGGAA |
| 28944_1 | TATGAGTGTGGAGTCGTTGCGCACACGTATGCATTTAGGGAAANNNAGATCGGAA |
| 28944_2 | TATGAGTGTGGAGTCGTTGCGCACACGTATGCATTTAGGGAAANNNAGATCGGAA |
| 50404_1 | TATGAGTGTGGAGTCGTTGCATCCTAATATTGCATGCAACTGCNNNAGATCGGAAG |

| | |
|---|---|
| 50404_2 | TATGAGTGTGGAGTCGTTGCATCCTAATATTGCATGCAACTGCNNNAGATCGGAAG |
| 12508_1 | TATGAGTGTGGAGTCGTTGCGAACGTAAGTGAGGAAAATATAAANNNAGATCGGA |
| 12508_2 | TATGAGTGTGGAGTCGTTGCGAACGTAAGTGAGGAAAATATAAANNNAGATCGGA |
| 45569_1 | TATGAGTGTGGAGTCGTTGCCACAAGCTGCCAAAATAGTGTCANNNAGATCGGAAG |
| 45569_2 | TATGAGTGTGGAGTCGTTGCCACAAGCTGCCAAAATAGTGTCANNNAGATCGGAAG |
| 32301_1 | TATGAGTGTGGAGTCGTTGCTGTGTGTGTTTGTGGTATGTGTGNNNAGATCGGAA |
| 32301_2 | TATGAGTGTGGAGTCGTTGCTGTGTGTGTTTGTGGTATGTGTGNNNAGATCGGAA |
| 10202_1 | TATGAGTGTGGAGTCGTTGCGGGTTCAGGTGAGTGTTGAGATCNNNAGATCGGAA |
| 10202_2 | TATGAGTGTGGAGTCGTTGCGGGTTCAGGTGAGTGTTGAGATCNNNAGATCGGAA |
| 42183_1 | TATGAGTGTGGAGTCGTTGCCTAGAGTGCAGGGTCACAATCTTNNNAGATCGGAAG |
| 42183_2 | TATGAGTGTGGAGTCGTTGCCTAGAGTGCAGGGTCACAATCTTNNNAGATCGGAAG |
| 38458_1 | TATGAGTGTGGAGTCGTTGCCCTAAAACGTACTGTGGTGATGGNNNAGATCGGAA |
| 38458_2 | TATGAGTGTGGAGTCGTTGCCCTAAAACGTACTGTGGTGATGGNNNAGATCGGAA |
| 29250_1 | TATGAGTGTGGAGTCGTTGCCTTGTCTTCCTCATCCCTCTGTCNNNAGATCGGAAGA |
| 29250_2 | TATGAGTGTGGAGTCGTTGCCTTGTCTTCCTCATCCCTCTGTCNNNAGATCGGAAGA |
| 46703_1 | TATGAGTGTGGAGTCGTTGCTTTTAGTAGAGATGGGTGCTGGGNNNAGATCGGAA |
| 46703_2 | TATGAGTGTGGAGTCGTTGCTTTTAGTAGAGATGGGTGCTGGGNNNAGATCGGAA |
| 29169_1 | TATGAGTGTGGAGTCGTTGCTCTCTTGCTTTCTCTCTCTCTGTNNNAGATCGGAAGA |
| 29169_2 | TATGAGTGTGGAGTCGTTGCTCTCTTGCTTTCTCTCTCTCTGTNNNAGATCGGAAGA |
| 39550_1 | TATGAGTGTGGAGTCGTTGCGCCTCCTGAGTAACTGTGACTACNNNAGATCGGAAG |
| 39550_2 | TATGAGTGTGGAGTCGTTGCGCCTCCTGAGTAACTGTGACTACNNNAGATCGGAAG |
| 35083_1 | TATGAGTGTGGAGTCGTTGCCACCACGTCCGGCTAATTTTTTTNNNAGATCGGAAGA |
| 35083_2 | TATGAGTGTGGAGTCGTTGCCACCACGTCCGGCTAATTTTTTTNNNAGATCGGAAGA |
| 33543_1 | TATGAGTGTGGAGTCGTTGCCCTGGGTTCAAGCAATTCTCATGNNNAGATCGGAAG |
| 33543_2 | TATGAGTGTGGAGTCGTTGCCCTGGGTTCAAGCAATTCTCATGNNNAGATCGGAAG |
| 40167_1 | TATGAGTGTGGAGTCGTTGCTGGAACCGGAACTTACTGTACTCNNNAGATCGGAAG |
| 40167_2 | TATGAGTGTGGAGTCGTTGCTGGAACCGGAACTTACTGTACTCNNNAGATCGGAAG |
| 44428_1 | TATGAGTGTGGAGTCGTTGCTCCTGTGTCTGGGGTTTTTGTATNNNAGATCGGAAG |
| 44428_2 | TATGAGTGTGGAGTCGTTGCTCCTGTGTCTGGGGTTTTTGTATNNNAGATCGGAAG |
| 52761_1 | TATGAGTGTGGAGTCGTTGCGATGTGTGCTAAGATCCCAAACGNNNAGATCGGAA |
| 52761_2 | TATGAGTGTGGAGTCGTTGCGATGTGTGCTAAGATCCCAAACGNNNAGATCGGAA |
| 51014_1 | TATGAGTGTGGAGTCGTTGCAAAAGTCAGCTTTGTGGCAACTTNNNAGATCGGAAG |
| 51014_2 | TATGAGTGTGGAGTCGTTGCAAAAGTCAGCTTTGTGGCAACTTNNNAGATCGGAAG |
| 44125_1 | TATGAGTGTGGAGTCGTTGCTTCCTCCCTATTGAGTCTTTGCCNNNAGATCGGAAGA |
| 44125_2 | TATGAGTGTGGAGTCGTTGCTTCCTCCCTATTGAGTCTTTGCCNNNAGATCGGAAGA |
| 50206_1 | TATGAGTGTGGAGTCGTTGCCTCTTCTGCTGACGTCACAAAGNNNAGATCGGAAGA |
| 50206_2 | TATGAGTGTGGAGTCGTTGCCTCTTCTGCTGACGTCACAAAGNNNAGATCGGAAGA |
| 10373_1 | TATGAGTGTGGAGTCGTTGCATCTTGGTGCAGCACACTTTACANNNAGATCGGAAG |
| 10373_2 | TATGAGTGTGGAGTCGTTGCATCTTGGTGCAGCACACTTTACANNNAGATCGGAAG |
| 8542_1 | TATGAGTGTGGAGTCGTTGCAGTAGCAGATGGCGCTAATTTCCTANNNAGATCGGA |
| 8542_2 | TATGAGTGTGGAGTCGTTGCAGTAGCAGATGGCGCTAATTTCCTANNNAGATCGGA |
| 43971_1 | TATGAGTGTGGAGTCGTTGCCAAGGACCCTATCTGTCTTGTCCNNNAGATCGGAAG |
| 43971_2 | TATGAGTGTGGAGTCGTTGCCAAGGACCCTATCTGTCTTGTCCNNNAGATCGGAAG |

| | |
|---|---|
| 30076_1 | TATGAGTGTGGAGTCGTTGCCTTTGCTGTTTAAAAGGGGGGAANNNAGATCGGAA |
| 30076_2 | TATGAGTGTGGAGTCGTTGCCTTTGCTGTTTAAAAGGGGGGAANNNAGATCGGAA |
| 10734_1 | TATGAGTGTGGAGTCGTTGCGGTCTTGGGGAACAAGTGATGTTNNNAGATCGGAA |
| 10734_2 | TATGAGTGTGGAGTCGTTGCGGTCTTGGGGAACAAGTGATGTTNNNAGATCGGAA |
| 51467_1 | TATGAGTGTGGAGTCGTTGCCGCATCAGGAAGAACAGCTAATGNNNAGATCGGAA |
| 51467_2 | TATGAGTGTGGAGTCGTTGCCGCATCAGGAAGAACAGCTAATGNNNAGATCGGAA |
| 13008_1 | TATGAGTGTGGAGTCGTTGCATCAAAGATATCAACAGAAAACCNNNAGATCGGAAG |
| 13008_2 | TATGAGTGTGGAGTCGTTGCATCAAAGATATCAACAGAAAACCNNNAGATCGGAAG |
| 43287_1 | TATGAGTGTGGAGTCGTTGCATTTTGGTGGCATGTCGTACTTGNNNAGATCGGAAG |
| 43287_2 | TATGAGTGTGGAGTCGTTGCATTTTGGTGGCATGTCGTACTTGNNNAGATCGGAAG |
| 29174_1 | TATGAGTGTGGAGTCGTTGCAGCACTGAGATTTACTGCAGTGANNNAGATCGGAA |
| 29174_2 | TATGAGTGTGGAGTCGTTGCAGCACTGAGATTTACTGCAGTGANNNAGATCGGAA |
| 50102_1 | TATGAGTGTGGAGTCGTTGCTCCCCAGTTTTTTTTTGCTGTGTNNNAGATCGGAAGA |
| 50102_2 | TATGAGTGTGGAGTCGTTGCTCCCCAGTTTTTTTTTGCTGTGTNNNAGATCGGAAGA |
| 31718_1 | TATGAGTGTGGAGTCGTTGCGTTCTTTTTTTCCAGCTGGGGAGNNNAGATCGGAAG |
| 31718_2 | TATGAGTGTGGAGTCGTTGCGTTCTTTTTTTCCAGCTGGGGAGNNNAGATCGGAAG |
| 48800_1 | TATGAGTGTGGAGTCGTTGCCAGTGGGAGCAGAGGAAACTCNNNAGATCGGAAG |
| 48800_2 | TATGAGTGTGGAGTCGTTGCCAGTGGGAGCAGAGGAAACTCNNNAGATCGGAAG |
| 42437_1 | TATGAGTGTGGAGTCGTTGCTTCCCGAGACCTGAGTGTTTTAGNNNAGATCGGAAG |
| 42437_2 | TATGAGTGTGGAGTCGTTGCTTCCCGAGACCTGAGTGTTTTAGNNNAGATCGGAAG |
| 29979_1 | TATGAGTGTGGAGTCGTTGCGAAAGTGACCACGATAGAGGGGNNNAGATCGGAA |
| 29979_2 | TATGAGTGTGGAGTCGTTGCGAAAGTGACCACGATAGAGGGGNNNAGATCGGAA |
| 36924_1 | TATGAGTGTGGAGTCGTTGCGAACTTGAACTCAGGGCTCAGTANNNAGATCGGAA |
| 36924_2 | TATGAGTGTGGAGTCGTTGCGAACTTGAACTCAGGGCTCAGTANNNAGATCGGAA |
| 12511_1 | TATGAGTGTGGAGTCGTTGCTAATTTCCCCAAAAAAATGAAACNNNAGATCGGAAG |
| 12511_2 | TATGAGTGTGGAGTCGTTGCTAATTTCCCCAAAAAAATGAAACNNNAGATCGGAAG |
| 38115_1 | TATGAGTGTGGAGTCGTTGCTGTGGTACACTCCTACAATCAAANNNAGATCGGAAG |
| 38115_2 | TATGAGTGTGGAGTCGTTGCTGTGGTACACTCCTACAATCAAANNNAGATCGGAAG |
| 42595_1 | TATGAGTGTGGAGTCGTTGCGCAAATGACAGTTCTTTCCTCTCANNNAGATCGGAA |
| 42595_2 | TATGAGTGTGGAGTCGTTGCGCAAATGACAGTTCTTTCCTCTCANNNAGATCGGAA |
| 43233_1 | TATGAGTGTGGAGTCGTTGCCATCTCCCTTCAGGTAACCAACANNNAGATCGGAAG |
| 43233_2 | TATGAGTGTGGAGTCGTTGCCATCTCCCTTCAGGTAACCAACANNNAGATCGGAAG |
| 13587_1 | TATGAGTGTGGAGTCGTTGCGACATTTGTCAAAAGTTAGTAAANNNAGATCGGAA |
| 13587_2 | TATGAGTGTGGAGTCGTTGCGACATTTGTCAAAAGTTAGTAAANNNAGATCGGAA |
| 36875_1 | TATGAGTGTGGAGTCGTTGCGACTGCTGTGCTAGATTTTGGACNNNAGATCGGAA |
| 36875_2 | TATGAGTGTGGAGTCGTTGCGACTGCTGTGCTAGATTTTGGACNNNAGATCGGAA |
| 33998_1 | TATGAGTGTGGAGTCGTTGCAGACTGGAGTGCAATGATGTGATNNNAGATCGGAA |
| 33998_2 | TATGAGTGTGGAGTCGTTGCAGACTGGAGTGCAATGATGTGATNNNAGATCGGAA |
| 33072_1 | TATGAGTGTGGAGTCGTTGCGCCTCCAGCTACATTCATGTTTCNNNAGATCGGAAGA |
| 33072_2 | TATGAGTGTGGAGTCGTTGCGCCTCCAGCTACATTCATGTTTCNNNAGATCGGAAGA |
| 42598_1 | TATGAGTGTGGAGTCGTTGCAAAAGAATGAGATGGCCAGGTGTNNNAGATCGGAA |
| 42598_2 | TATGAGTGTGGAGTCGTTGCAAAAGAATGAGATGGCCAGGTGTNNNAGATCGGAA |
| 38501_1 | TATGAGTGTGGAGTCGTTGCGCCTGTAATCCCAACACTTTGAGNNNAGATCGGAAG |

| | |
|---|---|
| 38501_2 | TATGAGTGTGGAGTCGTTGCGCCTGTAATCCCAACACTTTGAGNNNAGATCGGAAG |
| 41053_1 | TATGAGTGTGGAGTCGTTGCTGTGATCATGCCACTAAACTCCANNNAGATCGGAAG |
| 41053_2 | TATGAGTGTGGAGTCGTTGCTGTGATCATGCCACTAAACTCCANNNAGATCGGAAG |
| 13472_1 | TATGAGTGTGGAGTCGTTGCTCTTCAAATCCGTATACAATTTGNNNAGATCGGAAG |
| 13472_2 | TATGAGTGTGGAGTCGTTGCTCTTCAAATCCGTATACAATTTGNNNAGATCGGAAG |
| 52563_1 | TATGAGTGTGGAGTCGTTGCTACGCTATGCCATTGTATTCCCTNNNAGATCGGAAG |
| 52563_2 | TATGAGTGTGGAGTCGTTGCTACGCTATGCCATTGTATTCCCTNNNAGATCGGAAG |
| 47180_1 | TATGAGTGTGGAGTCGTTGCCCACGCCCAGCTAATTTTTGTATNNNAGATCGGAAG |
| 47180_2 | TATGAGTGTGGAGTCGTTGCCCACGCCCAGCTAATTTTTGTATNNNAGATCGGAAG |
| 45526_1 | TATGAGTGTGGAGTCGTTGCAAATTCAGGTGTGTGCTTATGCCNNNAGATCGGAAG |
| 45526_2 | TATGAGTGTGGAGTCGTTGCAAATTCAGGTGTGTGCTTATGCCNNNAGATCGGAAG |
| 34828_1 | TATGAGTGTGGAGTCGTTGCACGCCCGGCTAATTTTTTGTATTNNNAGATCGGAAG |
| 34828_2 | TATGAGTGTGGAGTCGTTGCACGCCCGGCTAATTTTTTGTATTNNNAGATCGGAAG |
| 43670_1 | TATGAGTGTGGAGTCGTTGCTCCAGGTTTAGGAGGGGTCTCNNNAGATCGGAAGA |
| 43670_2 | TATGAGTGTGGAGTCGTTGCTCCAGGTTTAGGAGGGGTCTCNNNAGATCGGAAGA |
| 47501_1 | TATGAGTGTGGAGTCGTTGCCCACCATGCCAGGGTAATTTTTTNNNAGATCGGAAG |
| 47501_2 | TATGAGTGTGGAGTCGTTGCCCACCATGCCAGGGTAATTTTTTNNNAGATCGGAAG |
| 51158_1 | TATGAGTGTGGAGTCGTTGCCCCTTTCTTCTGTACTCTGACCCNNNAGATCGGAAGA |
| 51158_2 | TATGAGTGTGGAGTCGTTGCCCCTTTCTTCTGTACTCTGACCCNNNAGATCGGAAGA |
| 10667_1 | TATGAGTGTGGAGTCGTTGCTACTATGTGCCTCCTGCAGAAAGNNNAGATCGGAAG |
| 10667_2 | TATGAGTGTGGAGTCGTTGCTACTATGTGCCTCCTGCAGAAAGNNNAGATCGGAAG |
| 44438_1 | TATGAGTGTGGAGTCGTTGCGAAGGGTTTGGAGGAATAGGTGANNNAGATCGGA |
| 44438_2 | TATGAGTGTGGAGTCGTTGCGAAGGGTTTGGAGGAATAGGTGANNNAGATCGGA |
| 43366_1 | TATGAGTGTGGAGTCGTTGCCACTACGCCCGGCTAATTTTTTANNNAGATCGGAAG |
| 43366_2 | TATGAGTGTGGAGTCGTTGCCACTACGCCCGGCTAATTTTTTANNNAGATCGGAAG |
| 36962_1 | TATGAGTGTGGAGTCGTTGCGGGTGGGGGTTGTTCTACTAATTNNNAGATCGGAA |
| 36962_2 | TATGAGTGTGGAGTCGTTGCGGGTGGGGGTTGTTCTACTAATTNNNAGATCGGAA |
| 44266_1 | TATGAGTGTGGAGTCGTTGCCGGAGATGAGATACGTCCTTCTCNNNAGATCGGAAG |
| 44266_2 | TATGAGTGTGGAGTCGTTGCCGGAGATGAGATACGTCCTTCTCNNNAGATCGGAAG |
| 39653_1 | TATGAGTGTGGAGTCGTTGCTCGCAGACTGAAAGAGACAAAGANNNAGATCGGAA |
| 39653_2 | TATGAGTGTGGAGTCGTTGCTCGCAGACTGAAAGAGACAAAGANNNAGATCGGAA |
| 28892_1 | TATGAGTGTGGAGTCGTTGCTTCAGGCTTGTTTTTGCTTCTGTNNNAGATCGGAAG |
| 28892_2 | TATGAGTGTGGAGTCGTTGCTTCAGGCTTGTTTTTGCTTCTGTNNNAGATCGGAAG |
| 12698_1 | TATGAGTGTGGAGTCGTTGCTAGGAATAGATCCAAACTAGCTANNNAGATCGGAA |
| 12698_2 | TATGAGTGTGGAGTCGTTGCTAGGAATAGATCCAAACTAGCTANNNAGATCGGAA |
| 46443_1 | TATGAGTGTGGAGTCGTTGCATGATCTCGGCTCACTGCAAANNNAGATCGGAAGAG |
| 46443_2 | TATGAGTGTGGAGTCGTTGCATGATCTCGGCTCACTGCAAANNNAGATCGGAAGAG |
| 12844_1 | TATGAGTGTGGAGTCGTTGCCTCAGCTACACCAAGAGATTNNNAGATCGGAAGAG |
| 12844_2 | TATGAGTGTGGAGTCGTTGCCTCAGCTACACCAAGAGATTNNNAGATCGGAAGAG |
| 29094_1 | TATGAGTGTGGAGTCGTTGCTACTGCCTCCCAGTGTTGTTTTTNNNAGATCGGAAG |
| 29094_2 | TATGAGTGTGGAGTCGTTGCTACTGCCTCCCAGTGTTGTTTTTNNNAGATCGGAAG |
| 45408_1 | TATGAGTGTGGAGTCGTTGCACAAAGTTCTGACCAAAACACCGNNNAGATCGGAAG |
| 45408_2 | TATGAGTGTGGAGTCGTTGCACAAAGTTCTGACCAAAACACCGNNNAGATCGGAAG |

| | |
|---|---|
| 41297_1 | TATGAGTGTGGAGTCGTTGCAGTACTGAAAACTGGCTCTGTCANNNAGATCGGAAG |
| 41297_2 | TATGAGTGTGGAGTCGTTGCAGTACTGAAAACTGGCTCTGTCANNNAGATCGGAAG |
| 48696_1 | TATGAGTGTGGAGTCGTTGCCAAGTGATTCTCCTGTCTCAGCTNNNAGATCGGAAG |
| 48696_2 | TATGAGTGTGGAGTCGTTGCCAAGTGATTCTCCTGTCTCAGCTNNNAGATCGGAAG |
| 47757_1 | TATGAGTGTGGAGTCGTTGCCAAGTAGCTGGGACTACAGGTGNNNAGATCGGAAG |
| 47757_2 | TATGAGTGTGGAGTCGTTGCCAAGTAGCTGGGACTACAGGTGNNNAGATCGGAAG |
| 34559_1 | TATGAGTGTGGAGTCGTTGCGGATGGCCCTAAGAGGAGGANNNAGATCGGAAGA |
| 34559_2 | TATGAGTGTGGAGTCGTTGCGGATGGCCCTAAGAGGAGGANNNAGATCGGAAGA |
| 31907_1 | TATGAGTGTGGAGTCGTTGCAGCAAAAAGGTGGCCTTGTTATGNNNAGATCGGAA |
| 31907_2 | TATGAGTGTGGAGTCGTTGCAGCAAAAAGGTGGCCTTGTTATGNNNAGATCGGAA |
| 50325_1 | TATGAGTGTGGAGTCGTTGCGTTCTTCCTTCCAACCCTCTCAANNNAGATCGGAAGA |
| 50325_2 | TATGAGTGTGGAGTCGTTGCGTTCTTCCTTCCAACCCTCTCAANNNAGATCGGAAGA |
| 34467_1 | TATGAGTGTGGAGTCGTTGCGAACTGACTCAGTGCAAAAGGACNNNAGATCGGAA |
| 34467_2 | TATGAGTGTGGAGTCGTTGCGAACTGACTCAGTGCAAAAGGACNNNAGATCGGAA |
| 30701_1 | TATGAGTGTGGAGTCGTTGCTAGTGGTTCTAGAGGTTACATATTTGANNNAGATCG |
| 30701_2 | TATGAGTGTGGAGTCGTTGCTAGTGGTTCTAGAGGTTACATATTTGANNNAGATCG |
| 39040_1 | TATGAGTGTGGAGTCGTTGCCTCCCCACTTTGCTCTCTATCTCNNNAGATCGGAAGA |
| 39040_2 | TATGAGTGTGGAGTCGTTGCCTCCCCACTTTGCTCTCTATCTCNNNAGATCGGAAGA |
| 13513_1 | TATGAGTGTGGAGTCGTTGCTTCAGATTTTTATGAGCCTTTACNNNAGATCGGAAG |
| 13513_2 | TATGAGTGTGGAGTCGTTGCTTCAGATTTTTATGAGCCTTTACNNNAGATCGGAAG |
| 29437_1 | TATGAGTGTGGAGTCGTTGCGCAAGGAAAGAGTATTTAAAAACNNNAGATCGGAA |
| 29437_2 | TATGAGTGTGGAGTCGTTGCGCAAGGAAAGAGTATTTAAAAACNNNAGATCGGAA |
| 40657_1 | TATGAGTGTGGAGTCGTTGCAACTCTTCGGCTGTTCTTTTTCGNNNAGATCGGAAGA |
| 40657_2 | TATGAGTGTGGAGTCGTTGCAACTCTTCGGCTGTTCTTTTTCGNNNAGATCGGAAGA |
| 38291_1 | TATGAGTGTGGAGTCGTTGCTTGCATCCATCAATCACTTTGGGNNNAGATCGGAAG |
| 38291_2 | TATGAGTGTGGAGTCGTTGCTTGCATCCATCAATCACTTTGGGNNNAGATCGGAAG |
| 43132_1 | TATGAGTGTGGAGTCGTTGCGCCAGATCTACAAAAAGCTGCAANNNAGATCGGAAG |
| 43132_2 | TATGAGTGTGGAGTCGTTGCGCCAGATCTACAAAAAGCTGCAANNNAGATCGGAAG |
| 33655_1 | TATGAGTGTGGAGTCGTTGCGAGTAGCTGGGATTACAGGAAGGNNNAGATCGGA |
| 33655_2 | TATGAGTGTGGAGTCGTTGCGAGTAGCTGGGATTACAGGAAGGNNNAGATCGGA |
| 48282_1 | TATGAGTGTGGAGTCGTTGCGATACAGGGTCTCACTGTGTCACNNNAGATCGGAAG |
| 48282_2 | TATGAGTGTGGAGTCGTTGCGATACAGGGTCTCACTGTGTCACNNNAGATCGGAAG |
| 32422_1 | TATGAGTGTGGAGTCGTTGCCTGAGTAGCTGGGATTATAGGCGNNNAGATCGGAA |
| 32422_2 | TATGAGTGTGGAGTCGTTGCCTGAGTAGCTGGGATTATAGGCGNNNAGATCGGAA |
| 46805_1 | TATGAGTGTGGAGTCGTTGCACCCCCAGCAAGTTGTGTAATAANNNAGATCGGAAG |
| 46805_2 | TATGAGTGTGGAGTCGTTGCACCCCCAGCAAGTTGTGTAATAANNNAGATCGGAAG |
| 44272_1 | TATGAGTGTGGAGTCGTTGCACCTGACAACCAAACAGCTTCTANNNAGATCGGAAG |
| 44272_2 | TATGAGTGTGGAGTCGTTGCACCTGACAACCAAACAGCTTCTANNNAGATCGGAAG |
| 28759_1 | TATGAGTGTGGAGTCGTTGCTGCTCAATGAACGAAGGAAGAGTNNNAGATCGGAA |
| 28759_2 | TATGAGTGTGGAGTCGTTGCTGCTCAATGAACGAAGGAAGAGTNNNAGATCGGAA |
| 37560_1 | TATGAGTGTGGAGTCGTTGCTTTAGCCTACAATCAGGGAAGGGNNNAGATCGGAA |
| 37560_2 | TATGAGTGTGGAGTCGTTGCTTTAGCCTACAATCAGGGAAGGGNNNAGATCGGAA |
| 47801_1 | TATGAGTGTGGAGTCGTTGCCTAGTGGGGGCTCCTTTTTAGAGNNNAGATCGGAA |

| | |
|---|---|
| 47801_2 | TATGAGTGTGGAGTCGTTGCCTAGTGGGGGCTCCTTTTTAGAGNNNAGATCGGAA |
| 32149_1 | TATGAGTGTGGAGTCGTTGCGCAAGATCTTAGCTCACTGCAAGNNNAGATCGGAAG |
| 32149_2 | TATGAGTGTGGAGTCGTTGCGCAAGATCTTAGCTCACTGCAAGNNNAGATCGGAAG |
| 36423_1 | TATGAGTGTGGAGTCGTTGCTCCCTAACCTACAAGCTTTGGAANNNAGATCGGAAG |
| 36423_2 | TATGAGTGTGGAGTCGTTGCTCCCTAACCTACAAGCTTTGGAANNNAGATCGGAAG |
| 37745_1 | TATGAGTGTGGAGTCGTTGCGAAAGCATTCGGTCTTGCCTTATNNNAGATCGGAAG |
| 37745_2 | TATGAGTGTGGAGTCGTTGCGAAAGCATTCGGTCTTGCCTTATNNNAGATCGGAAG |
| 43204_1 | TATGAGTGTGGAGTCGTTGCTGAAGAACTGGTCTTTGAGCTGTNNNAGATCGGAA |
| 43204_2 | TATGAGTGTGGAGTCGTTGCTGAAGAACTGGTCTTTGAGCTGTNNNAGATCGGAA |
| 50413_1 | TATGAGTGTGGAGTCGTTGCTTCAAGATCAGCCTGGACAACATNNNAGATCGGAAG |
| 50413_2 | TATGAGTGTGGAGTCGTTGCTTCAAGATCAGCCTGGACAACATNNNAGATCGGAAG |
| 34657_1 | TATGAGTGTGGAGTCGTTGCTACGTGGAAGATGGGCATAACATNNNAGATCGGAA |
| 34657_2 | TATGAGTGTGGAGTCGTTGCTACGTGGAAGATGGGCATAACATNNNAGATCGGAA |
| 51619_1 | TATGAGTGTGGAGTCGTTGCTAAGGCCCACTTATCAGGTCATGNNNAGATCGGAAG |
| 51619_2 | TATGAGTGTGGAGTCGTTGCTAAGGCCCACTTATCAGGTCATGNNNAGATCGGAAG |
| 41409_1 | TATGAGTGTGGAGTCGTTGCTAGAGAGTAGAATGGTGGTTGCCNNNAGATCGGAA |
| 41409_2 | TATGAGTGTGGAGTCGTTGCTAGAGAGTAGAATGGTGGTTGCCNNNAGATCGGAA |
| 36679_1 | TATGAGTGTGGAGTCGTTGCCAGCCCAGTCAACCACTAATCTANNNAGATCGGAAG |
| 36679_2 | TATGAGTGTGGAGTCGTTGCCAGCCCAGTCAACCACTAATCTANNNAGATCGGAAG |
| 43808_1 | TATGAGTGTGGAGTCGTTGCGCTAGCTGCTAGTATGTAGAAAAATNNNAGATCGG |
| 43808_2 | TATGAGTGTGGAGTCGTTGCGCTAGCTGCTAGTATGTAGAAAAATNNNAGATCGG |
| 10764_1 | TATGAGTGTGGAGTCGTTGCTCCTTGTGTTTGGATAGCAAACTNNNAGATCGGAAG |
| 10764_2 | TATGAGTGTGGAGTCGTTGCTCCTTGTGTTTGGATAGCAAACTNNNAGATCGGAAG |
| 42453_1 | TATGAGTGTGGAGTCGTTGCCCTCTTTCTCTCTCTCCCTCTNNNAGATCGGAAGAG |
| 42453_2 | TATGAGTGTGGAGTCGTTGCCCTCTTTCTCTCTCTCCCTCTNNNAGATCGGAAGAG |
| 43240_1 | TATGAGTGTGGAGTCGTTGCTTATGAACACCTCTATGCACACANNNAGATCGGAAG |
| 43240_2 | TATGAGTGTGGAGTCGTTGCTTATGAACACCTCTATGCACACANNNAGATCGGAAG |
| 48406_1 | TATGAGTGTGGAGTCGTTGCTCAACTTCCTGTGGTCAAGTGATNNNAGATCGGAAG |
| 48406_2 | TATGAGTGTGGAGTCGTTGCTCAACTTCCTGTGGTCAAGTGATNNNAGATCGGAAG |
| 49136_1 | TATGAGTGTGGAGTCGTTGCGGCCCTCCTCTCCTGATTTATTTNNNAGATCGGAAGA |
| 49136_2 | TATGAGTGTGGAGTCGTTGCGGCCCTCCTCTCCTGATTTATTTNNNAGATCGGAAGA |
| 39934_1 | TATGAGTGTGGAGTCGTTGCAAATTCTTTTTTGGGGGGGGTTCNNNAGATCGGAA |
| 39934_2 | TATGAGTGTGGAGTCGTTGCAAATTCTTTTTTGGGGGGGGTTCNNNAGATCGGAA |
| 47288_1 | TATGAGTGTGGAGTCGTTGCAATGCAAAATTGGTTCCAGGAGGNNNAGATCGGAA |
| 47288_2 | TATGAGTGTGGAGTCGTTGCAATGCAAAATTGGTTCCAGGAGGNNNAGATCGGAA |
| 11408_1 | TATGAGTGTGGAGTCGTTGCGGAAGGTGGTTGAACTATGTGTTNNNAGATCGGAA |
| 11408_2 | TATGAGTGTGGAGTCGTTGCGGAAGGTGGTTGAACTATGTGTTNNNAGATCGGAA |
| 45857_1 | TATGAGTGTGGAGTCGTTGCTCCTCTGCTTCCTGAGTTCAATCNNNAGATCGGAAGA |
| 45857_2 | TATGAGTGTGGAGTCGTTGCTCCTCTGCTTCCTGAGTTCAATCNNNAGATCGGAAGA |
| 50719_1 | TATGAGTGTGGAGTCGTTGCCCACCACACCCAGCTAATTTTTTNNNAGATCGGAAGA |
| 50719_2 | TATGAGTGTGGAGTCGTTGCCCACCACACCCAGCTAATTTTTTNNNAGATCGGAAGA |
| 35780_1 | TATGAGTGTGGAGTCGTTGCGTTCATACCATTCTCCTGCCTCANNNAGATCGGAAGA |
| 35780_2 | TATGAGTGTGGAGTCGTTGCGTTCATACCATTCTCCTGCCTCANNNAGATCGGAAGA |

| | |
|---|---|
| 30788_1 | TATGAGTGTGGAGTCGTTGCGGTTGGTCTTTCTAGTATGGCCANNNAGATCGGAAG |
| 30788_2 | TATGAGTGTGGAGTCGTTGCGGTTGGTCTTTCTAGTATGGCCANNNAGATCGGAAG |
| 31506_1 | TATGAGTGTGGAGTCGTTGCAATCTGGCAAGAATGGACACTCANNNAGATCGGAAG |
| 31506_2 | TATGAGTGTGGAGTCGTTGCAATCTGGCAAGAATGGACACTCANNNAGATCGGAAG |
| 5765_1 | TATGAGTGTGGAGTCGTTGCTGGTTATGCAGGTATGCAAGTCCNNNAGATCGGAAG |
| 5765_2 | TATGAGTGTGGAGTCGTTGCTGGTTATGCAGGTATGCAAGTCCNNNAGATCGGAAG |
| 40196_1 | TATGAGTGTGGAGTCGTTGCGAAAGAAAAATTGCCAAGCGTGGNNNAGATCGGAA |
| 40196_2 | TATGAGTGTGGAGTCGTTGCGAAAGAAAAATTGCCAAGCGTGGNNNAGATCGGAA |
| 38420_1 | TATGAGTGTGGAGTCGTTGCGGACATACAAATGACCAATAGGCANNNAGATCGGAA |
| 38420_2 | TATGAGTGTGGAGTCGTTGCGGACATACAAATGACCAATAGGCANNNAGATCGGAA |
| 32832_1 | TATGAGTGTGGAGTCGTTGCTCCACCTCCCAGTTCAAATGATTNNNAGATCGGAAGA |
| 32832_2 | TATGAGTGTGGAGTCGTTGCTCCACCTCCCAGTTCAAATGATTNNNAGATCGGAAGA |
| 44597_1 | TATGAGTGTGGAGTCGTTGCCACCTCTCAGGACATGCAGATAGNNNAGATCGGAAG |
| 44597_2 | TATGAGTGTGGAGTCGTTGCCACCTCTCAGGACATGCAGATAGNNNAGATCGGAAG |
| 51147_1 | TATGAGTGTGGAGTCGTTGCTGGAAACGTGACGGGTATAGTTTNNNAGATCGGAAG |
| 51147_2 | TATGAGTGTGGAGTCGTTGCTGGAAACGTGACGGGTATAGTTTNNNAGATCGGAAG |
| 43844_1 | TATGAGTGTGGAGTCGTTGCAGAAAGGGCTTCCATGGAGTTAGNNNAGATCGGAA |
| 43844_2 | TATGAGTGTGGAGTCGTTGCAGAAAGGGCTTCCATGGAGTTAGNNNAGATCGGAA |
| 31699_1 | TATGAGTGTGGAGTCGTTGCTAGGGAAGACAGAAAGAAGGGGANNNAGATCGGA |
| 31699_2 | TATGAGTGTGGAGTCGTTGCTAGGGAAGACAGAAAGAAGGGGANNNAGATCGGA |
| 40141_1 | TATGAGTGTGGAGTCGTTGCAGTAGGCTGCTTCTAAAATGGCTNNNAGATCGGAAG |
| 40141_2 | TATGAGTGTGGAGTCGTTGCAGTAGGCTGCTTCTAAAATGGCTNNNAGATCGGAAG |
| 3880_1 | TATGAGTGTGGAGTCGTTGCCAAGCAATCGGTCCATCTCGTCNNNAGATCGGAAGAG |
| 3880_2 | TATGAGTGTGGAGTCGTTGCCAAGCAATCGGTCCATCTCGTCNNNAGATCGGAAGAG |
| 38359_1 | TATGAGTGTGGAGTCGTTGCTGAGGTCAGGAGTTCAAGATCAGNNNAGATCGGAA |
| 38359_2 | TATGAGTGTGGAGTCGTTGCTGAGGTCAGGAGTTCAAGATCAGNNNAGATCGGAA |
| 46768_1 | TATGAGTGTGGAGTCGTTGCCATCTCATTCTCTTCCAGCCTGTNNNAGATCGGAAGAG |
| 46768_2 | TATGAGTGTGGAGTCGTTGCCATCTCATTCTCTTCCAGCCTGTNNNAGATCGGAAGAG |
| 40178_1 | TATGAGTGTGGAGTCGTTGCTTCATCCTAGCACTTTGGGAGTCNNNAGATCGGAAGA |
| 40178_2 | TATGAGTGTGGAGTCGTTGCTTCATCCTAGCACTTTGGGAGTCNNNAGATCGGAAGA |
| 33604_1 | TATGAGTGTGGAGTCGTTGCTGGCAGTGGTATTGTCATCATTCNNNAGATCGGAAG |
| 33604_2 | TATGAGTGTGGAGTCGTTGCTGGCAGTGGTATTGTCATCATTCNNNAGATCGGAAG |
| 46366_1 | TATGAGTGTGGAGTCGTTGCGTAGCTGGGATTATAGGCACACANNNAGATCGGAAG |
| 46366_2 | TATGAGTGTGGAGTCGTTGCGTAGCTGGGATTATAGGCACACANNNAGATCGGAAG |
| 42015_1 | TATGAGTGTGGAGTCGTTGCATGGAGTCTCCCTTTTTTGTTGCNNNAGATCGGAAGA |
| 42015_2 | TATGAGTGTGGAGTCGTTGCATGGAGTCTCCCTTTTTTGTTGCNNNAGATCGGAAGA |
| 40022_1 | TATGAGTGTGGAGTCGTTGCTCTCTTCCACCTCTTCAGAGTCANNNAGATCGGAAGA |
| 40022_2 | TATGAGTGTGGAGTCGTTGCTCTCTTCCACCTCTTCAGAGTCANNNAGATCGGAAGA |
| 31693_1 | TATGAGTGTGGAGTCGTTGCGAGGATGGGGTTGATGGTAGAAGNNNAGATCGGA |
| 31693_2 | TATGAGTGTGGAGTCGTTGCGAGGATGGGGTTGATGGTAGAAGNNNAGATCGGA |
| 33393_1 | TATGAGTGTGGAGTCGTTGCATTCCATTAGAGGCTGCCCTAAGNNNAGATCGGAAG |
| 33393_2 | TATGAGTGTGGAGTCGTTGCATTCCATTAGAGGCTGCCCTAAGNNNAGATCGGAAG |
| 32650_1 | TATGAGTGTGGAGTCGTTGCACCACATCCCGCTAATATTGTGTNNNAGATCGGAAGA |

| | |
|---|---|
| 32650_2 | TATGAGTGTGGAGTCGTTGCACCACATCCCGCTAATATTGTGTNNNAGATCGGAAG |
| 39820_1 | TATGAGTGTGGAGTCGTTGCGACACTAGGGATGACTTGACTGCNNNAGATCGGAA |
| 39820_2 | TATGAGTGTGGAGTCGTTGCGACACTAGGGATGACTTGACTGCNNNAGATCGGAA |
| 30599_1 | TATGAGTGTGGAGTCGTTGCAGAAAGGCAACTTCATTGGAGCTNNNAGATCGGAA |
| 30599_2 | TATGAGTGTGGAGTCGTTGCAGAAAGGCAACTTCATTGGAGCTNNNAGATCGGAA |
| 30916_1 | TATGAGTGTGGAGTCGTTGCTTCCTACGTCATGAAAGCAGCATNNNAGATCGGAAG |
| 30916_2 | TATGAGTGTGGAGTCGTTGCTTCCTACGTCATGAAAGCAGCATNNNAGATCGGAAG |
| 50471_1 | TATGAGTGTGGAGTCGTTGCGCTATTTTCAGGTGTGTGGACTGNNNAGATCGGAA |
| 50471_2 | TATGAGTGTGGAGTCGTTGCGCTATTTTCAGGTGTGTGGACTGNNNAGATCGGAA |
| 35483_1 | TATGAGTGTGGAGTCGTTGCCTGGTCTCAAACTCCTGACTTGANNNAGATCGGAAG |
| 35483_2 | TATGAGTGTGGAGTCGTTGCCTGGTCTCAAACTCCTGACTTGANNNAGATCGGAAG |
| 10321_1 | TATGAGTGTGGAGTCGTTGCCATTAGTCACATGAAGCTGGCCTNNNAGATCGGAAG |
| 10321_2 | TATGAGTGTGGAGTCGTTGCCATTAGTCACATGAAGCTGGCCTNNNAGATCGGAAG |
| 43341_1 | TATGAGTGTGGAGTCGTTGCTTTCACCGTGTTAGCTAACAGGANNNAGATCGGAAG |
| 43341_2 | TATGAGTGTGGAGTCGTTGCTTTCACCGTGTTAGCTAACAGGANNNAGATCGGAAG |
| 51130_1 | TATGAGTGTGGAGTCGTTGCGGAGGGGGGACTCTTATTTTGTTNNNAGATCGGAA |
| 51130_2 | TATGAGTGTGGAGTCGTTGCGGAGGGGGGACTCTTATTTTGTTNNNAGATCGGAA |
| 38941_1 | TATGAGTGTGGAGTCGTTGCGGGAGGAAAGTCTTTGAAGAGGANNNAGATCGGA |
| 38941_2 | TATGAGTGTGGAGTCGTTGCGGGAGGAAAGTCTTTGAAGAGGANNNAGATCGGA |
| 10488_1 | TATGAGTGTGGAGTCGTTGCAAAACCTATGCCTGGCACTGAAANNNAGATCGGAAG |
| 10488_2 | TATGAGTGTGGAGTCGTTGCAAAACCTATGCCTGGCACTGAAANNNAGATCGGAAG |
| 36038_1 | TATGAGTGTGGAGTCGTTGCTACCTTTGGCTGTGACATTGTCTNNNAGATCGGAAG |
| 36038_2 | TATGAGTGTGGAGTCGTTGCTACCTTTGGCTGTGACATTGTCTNNNAGATCGGAAG |
| 47769_1 | TATGAGTGTGGAGTCGTTGCCCTGTGACTTTGATTCCTCCTCANNNAGATCGGAAGA |
| 47769_2 | TATGAGTGTGGAGTCGTTGCCCTGTGACTTTGATTCCTCCTCANNNAGATCGGAAGA |
| 41132_1 | TATGAGTGTGGAGTCGTTGCACTCACAGCCCTCTGATATAGGTNNNAGATCGGAAG |
| 41132_2 | TATGAGTGTGGAGTCGTTGCACTCACAGCCCTCTGATATAGGTNNNAGATCGGAAG |
| 52967_1 | TATGAGTGTGGAGTCGTTGCTGGAGCTAGAGGCCATTATCCTANNNAGATCGGAA |
| 52967_2 | TATGAGTGTGGAGTCGTTGCTGGAGCTAGAGGCCATTATCCTANNNAGATCGGAA |
| 33742_1 | TATGAGTGTGGAGTCGTTGCATGGTCTGAATGTTTGTGTCCCTNNNAGATCGGAAG |
| 33742_2 | TATGAGTGTGGAGTCGTTGCATGGTCTGAATGTTTGTGTCCCTNNNAGATCGGAAG |
| 34624_1 | TATGAGTGTGGAGTCGTTGCCATTTAGTGGGTAAGGCCAGAGANNNAGATCGGAA |
| 34624_2 | TATGAGTGTGGAGTCGTTGCCATTTAGTGGGTAAGGCCAGAGANNNAGATCGGAA |
| 34730_1 | TATGAGTGTGGAGTCGTTGCAAATACCCCACCATGCAAACTTGNNNAGATCGGAAG |
| 34730_2 | TATGAGTGTGGAGTCGTTGCAAATACCCCACCATGCAAACTTGNNNAGATCGGAAG |
| 34250_1 | TATGAGTGTGGAGTCGTTGCCTTGTCTCTTCCCACTCTCCTTCNNNAGATCGGAAGA |
| 34250_2 | TATGAGTGTGGAGTCGTTGCCTTGTCTCTTCCCACTCTCCTTCNNNAGATCGGAAGA |
| 50893_1 | TATGAGTGTGGAGTCGTTGCCTCTGAGCCTGATTATTCCTGCANNNAGATCGGAAG |
| 50893_2 | TATGAGTGTGGAGTCGTTGCCTCTGAGCCTGATTATTCCTGCANNNAGATCGGAAG |
| 42705_1 | TATGAGTGTGGAGTCGTTGCGGCAACTCTAAAAGCCAACATGTNNNAGATCGGAAG |
| 42705_2 | TATGAGTGTGGAGTCGTTGCGGCAACTCTAAAAGCCAACATGTNNNAGATCGGAAG |
| 42785_1 | TATGAGTGTGGAGTCGTTGCGCCTGTAGTCCCAGCTACTTGNNNAGATCGGAAGAG |
| 42785_2 | TATGAGTGTGGAGTCGTTGCGCCTGTAGTCCCAGCTACTTGNNNAGATCGGAAGAG |

| | |
|---|---|
| 43123_1 | TATGAGTGTGGAGTCGTTGCCCTGAAGCACCTAGTCTACATGGNNNAGATCGGAAG |
| 43123_2 | TATGAGTGTGGAGTCGTTGCCCTGAAGCACCTAGTCTACATGGNNNAGATCGGAAG |
| 33444_1 | TATGAGTGTGGAGTCGTTGCCTAAGGAGAAGGACTAGGAGGCTNNNAGATCGGAA |
| 33444_2 | TATGAGTGTGGAGTCGTTGCCTAAGGAGAAGGACTAGGAGGCTNNNAGATCGGAA |
| 52966_1 | TATGAGTGTGGAGTCGTTGCTTGAAGCAACATGGATGCATCTGNNNAGATCGGAAG |
| 52966_2 | TATGAGTGTGGAGTCGTTGCTTGAAGCAACATGGATGCATCTGNNNAGATCGGAAG |
| 50208_1 | TATGAGTGTGGAGTCGTTGCACCGCGATGAGGTCATATTTGTANNNAGATCGGAAG |
| 50208_2 | TATGAGTGTGGAGTCGTTGCACCGCGATGAGGTCATATTTGTANNNAGATCGGAAG |
| 53260_1 | TATGAGTGTGGAGTCGTTGCCCCAGCAGTCTTCTTAGCTCTACNNNAGATCGGAAGA |
| 53260_2 | TATGAGTGTGGAGTCGTTGCCCCAGCAGTCTTCTTAGCTCTACNNNAGATCGGAAGA |
| 42574_1 | TATGAGTGTGGAGTCGTTGCGTATGTGTATGTGTGGGAGAGGGNNNAGATCGGAA |
| 42574_2 | TATGAGTGTGGAGTCGTTGCGTATGTGTATGTGTGGGAGAGGGNNNAGATCGGAA |
| 50567_1 | TATGAGTGTGGAGTCGTTGCGACACCAGGGAAAAGAGGTAGTCNNNAGATCGGAA |
| 50567_2 | TATGAGTGTGGAGTCGTTGCGACACCAGGGAAAAGAGGTAGTCNNNAGATCGGAA |
| 45588_1 | TATGAGTGTGGAGTCGTTGCCTCCTGAGTAGCTGGGACTACAGNNNAGATCGGAAG |
| 45588_2 | TATGAGTGTGGAGTCGTTGCCTCCTGAGTAGCTGGGACTACAGNNNAGATCGGAAG |
| 44364_1 | TATGAGTGTGGAGTCGTTGCATGCAAAGGATTCTGAGGACCATNNNAGATCGGAAG |
| 44364_2 | TATGAGTGTGGAGTCGTTGCATGCAAAGGATTCTGAGGACCATNNNAGATCGGAAG |
| 40339_1 | TATGAGTGTGGAGTCGTTGCAGGACTAAGGAAAGCCAGAGAGANNNAGATCGGAA |
| 40339_2 | TATGAGTGTGGAGTCGTTGCAGGACTAAGGAAAGCCAGAGAGANNNAGATCGGAA |
| 48058_1 | TATGAGTGTGGAGTCGTTGCGATTTCAGCTCACTGCAAACTCCNNNAGATCGGAAGA |
| 48058_2 | TATGAGTGTGGAGTCGTTGCGATTTCAGCTCACTGCAAACTCCNNNAGATCGGAAGA |
| 38383_1 | TATGAGTGTGGAGTCGTTGCACTGTTTTCCATAATGGCTGCACNNNAGATCGGAAGA |
| 38383_2 | TATGAGTGTGGAGTCGTTGCACTGTTTTCCATAATGGCTGCACNNNAGATCGGAAGA |
| 48640_1 | TATGAGTGTGGAGTCGTTGCCTCCCGAGTAGCTGGAATTACAGNNNAGATCGGAAG |
| 48640_2 | TATGAGTGTGGAGTCGTTGCCTCCCGAGTAGCTGGAATTACAGNNNAGATCGGAAG |
| 30934_1 | TATGAGTGTGGAGTCGTTGCGAAAGACATGGGAACTCCAGAGTNNNAGATCGGAA |
| 30934_2 | TATGAGTGTGGAGTCGTTGCGAAAGACATGGGAACTCCAGAGTNNNAGATCGGAA |
| 42672_1 | TATGAGTGTGGAGTCGTTGCGAGTAGCTGGGACTAAAGGTGTGNNNAGATCGGAA |
| 42672_2 | TATGAGTGTGGAGTCGTTGCGAGTAGCTGGGACTAAAGGTGTGNNNAGATCGGAA |
| 47935_1 | TATGAGTGTGGAGTCGTTGCGAAGACACTTAGGGTGGGTGATGNNNAGATCGGAA |
| 47935_2 | TATGAGTGTGGAGTCGTTGCGAAGACACTTAGGGTGGGTGATGNNNAGATCGGAA |
| 43594_1 | TATGAGTGTGGAGTCGTTGCTTCTGAAGCAACCAGATACTGCTNNNAGATCGGAAG |
| 43594_2 | TATGAGTGTGGAGTCGTTGCTTCTGAAGCAACCAGATACTGCTNNNAGATCGGAAG |
| 52762_1 | TATGAGTGTGGAGTCGTTGCGGAAACTCCTTTCTGTACCTCCCNNNAGATCGGAAGA |
| 52762_2 | TATGAGTGTGGAGTCGTTGCGGAAACTCCTTTCTGTACCTCCCNNNAGATCGGAAGA |
| 43768_1 | TATGAGTGTGGAGTCGTTGCAAGAAAAGAAAGAGTTGCTGCCCNNNAGATCGGAAG |
| 43768_2 | TATGAGTGTGGAGTCGTTGCAAGAAAAGAAAGAGTTGCTGCCCNNNAGATCGGAAG |
| 38580_1 | TATGAGTGTGGAGTCGTTGCAAAAGAGTAGTTTAGCCGACCGTNNNAGATCGGAAG |
| 38580_2 | TATGAGTGTGGAGTCGTTGCAAAAGAGTAGTTTAGCCGACCGTNNNAGATCGGAAG |
| 12540_1 | TATGAGTGTGGAGTCGTTGCGATGCTACTGGACAATCTTTTNNNAGATCGGAAGAG |
| 12540_2 | TATGAGTGTGGAGTCGTTGCGATGCTACTGGACAATCTTTTNNNAGATCGGAAGAG |
| 38547_1 | TATGAGTGTGGAGTCGTTGCACATGTTTATAGTCCCAGCTGCTNNNAGATCGGAAGA |

| | |
|---|---|
| 38547_2 | TATGAGTGTGGAGTCGTTGCACATGTTTATAGTCCCAGCTGCTNNNAGATCGGAAG |
| 50772_1 | TATGAGTGTGGAGTCGTTGCCACCAGAACCTGCCTAGATGTCNNNAGATCGGAAGA |
| 50772_2 | TATGAGTGTGGAGTCGTTGCCACCAGAACCTGCCTAGATGTCNNNAGATCGGAAGA |
| 47451_1 | TATGAGTGTGGAGTCGTTGCTGTGTCTCTGTGCTCACATTACANNNAGATCGGAAG |
| 47451_2 | TATGAGTGTGGAGTCGTTGCTGTGTCTCTGTGCTCACATTACANNNAGATCGGAAG |
| 32390_1 | TATGAGTGTGGAGTCGTTGCCCTCCTGAGTAGCTGGGACTACNNNAGATCGGAAGA |
| 32390_2 | TATGAGTGTGGAGTCGTTGCCCTCCTGAGTAGCTGGGACTACNNNAGATCGGAAGA |
| 30704_1 | TATGAGTGTGGAGTCGTTGCATAATGTGTAAAATGTGGAGGTGCANNNAGATCGG |
| 30704_2 | TATGAGTGTGGAGTCGTTGCATAATGTGTAAAATGTGGAGGTGCANNNAGATCGG |
| 32743_1 | TATGAGTGTGGAGTCGTTGCCCCACAGCAAAACAAAACAAACCNNNAGATCGGAAG |
| 32743_2 | TATGAGTGTGGAGTCGTTGCCCCACAGCAAAACAAAACAAACCNNNAGATCGGAAG |
| 29754_1 | TATGAGTGTGGAGTCGTTGCAAGTTCCTGCTTCTGTCTCTCCTNNNAGATCGGAAGA |
| 29754_2 | TATGAGTGTGGAGTCGTTGCAAGTTCCTGCTTCTGTCTCTCCTNNNAGATCGGAAGA |
| 11669_1 | TATGAGTGTGGAGTCGTTGCAAACAAAGCAGCAAAAGGCCCCNNNAGATCGGAAG |
| 11669_2 | TATGAGTGTGGAGTCGTTGCAAACAAAGCAGCAAAAGGCCCCNNNAGATCGGAAG |
| 32000_1 | TATGAGTGTGGAGTCGTTGCGCAGTGACAAGCAAGGATTGTAANNNAGATCGGAA |
| 32000_2 | TATGAGTGTGGAGTCGTTGCGCAGTGACAAGCAAGGATTGTAANNNAGATCGGAA |
| 46655_1 | TATGAGTGTGGAGTCGTTGCAGAGTCTCACTGTTTTGTCCAGGNNNAGATCGGAAG |
| 46655_2 | TATGAGTGTGGAGTCGTTGCAGAGTCTCACTGTTTTGTCCAGGNNNAGATCGGAAG |
| 49600_1 | TATGAGTGTGGAGTCGTTGCCCACCACGTCCAGCTAATTTTTTNNNAGATCGGAAGA |
| 49600_2 | TATGAGTGTGGAGTCGTTGCCCACCACGTCCAGCTAATTTTTTNNNAGATCGGAAGA |
| 13395_1 | TATGAGTGTGGAGTCGTTGCGCCCTCTAGTTGAATTCTAAANNNAGATCGGAAGAG |
| 13395_2 | TATGAGTGTGGAGTCGTTGCGCCCTCTAGTTGAATTCTAAANNNAGATCGGAAGAG |
| 43617_1 | TATGAGTGTGGAGTCGTTGCCTGTGAAGCTCTCCCTGACATGNNNAGATCGGAAGA |
| 43617_2 | TATGAGTGTGGAGTCGTTGCCTGTGAAGCTCTCCCTGACATGNNNAGATCGGAAGA |
| 47706_1 | TATGAGTGTGGAGTCGTTGCCTATGTCTTCCAGGCTGGTCTTGNNNAGATCGGAAG |
| 47706_2 | TATGAGTGTGGAGTCGTTGCCTATGTCTTCCAGGCTGGTCTTGNNNAGATCGGAAG |
| 31321_1 | TATGAGTGTGGAGTCGTTGCGATCTTTGCACACTCCAGTTTCCNNNAGATCGGAAGA |
| 31321_2 | TATGAGTGTGGAGTCGTTGCGATCTTTGCACACTCCAGTTTCCNNNAGATCGGAAGA |
| 29772_1 | TATGAGTGTGGAGTCGTTGCCTGGAAATGAGGGGGGAAAGAAANNNAGATCGGA |
| 29772_2 | TATGAGTGTGGAGTCGTTGCCTGGAAATGAGGGGGGAAAGAAANNNAGATCGGA |
| 31700_1 | TATGAGTGTGGAGTCGTTGCCTGCTCTCGTACTTCCTTGAGACNNNAGATCGGAAG |
| 31700_2 | TATGAGTGTGGAGTCGTTGCCTGCTCTCGTACTTCCTTGAGACNNNAGATCGGAAG |
| 10019_1 | TATGAGTGTGGAGTCGTTGCGGGAGAGGAAAGGAGAGGTATCANNNAGATCGGA |
| 10019_2 | TATGAGTGTGGAGTCGTTGCGGGAGAGGAAAGGAGAGGTATCANNNAGATCGGA |
| 44146_1 | TATGAGTGTGGAGTCGTTGCATGAAGCATCCGTTCATCCCAATNNNAGATCGGAAG |
| 44146_2 | TATGAGTGTGGAGTCGTTGCATGAAGCATCCGTTCATCCCAATNNNAGATCGGAAG |
| 53011_1 | TATGAGTGTGGAGTCGTTGCCTTACTATGTCCCCTGCCATCTGNNNAGATCGGAAGA |
| 53011_2 | TATGAGTGTGGAGTCGTTGCCTTACTATGTCCCCTGCCATCTGNNNAGATCGGAAGA |
| 50186_1 | TATGAGTGTGGAGTCGTTGCGAAAACCCTGAACCACAAGATGGNNNAGATCGGAA |
| 50186_2 | TATGAGTGTGGAGTCGTTGCGAAAACCCTGAACCACAAGATGGNNNAGATCGGAA |
| 29965_1 | TATGAGTGTGGAGTCGTTGCACATTGTAGTACCAGATGAGCCTNNNAGATCGGAAG |
| 29965_2 | TATGAGTGTGGAGTCGTTGCACATTGTAGTACCAGATGAGCCTNNNAGATCGGAAG |

| | |
|---|---|
| 10043_1 | TATGAGTGTGGAGTCGTTGCGGGAGCTTCAGTTCATAGCTTGANNNAGATCGGAA |
| 10043_2 | TATGAGTGTGGAGTCGTTGCGGGAGCTTCAGTTCATAGCTTGANNNAGATCGGAA |
| 33093_1 | TATGAGTGTGGAGTCGTTGCCGAGTAGCTGGGATTACAGGTGNNNAGATCGGAAG |
| 33093_2 | TATGAGTGTGGAGTCGTTGCCGAGTAGCTGGGATTACAGGTGNNNAGATCGGAAG |
| 29856_1 | TATGAGTGTGGAGTCGTTGCTTTTCTTTTGGGCTGGTTTGCTTNNNAGATCGGAAG |
| 29856_2 | TATGAGTGTGGAGTCGTTGCTTTTCTTTTGGGCTGGTTTGCTTNNNAGATCGGAAG |
| 42607_1 | TATGAGTGTGGAGTCGTTGCGGTAAATAGTCTCAGCTCCTCGGNNNAGATCGGAAG |
| 42607_2 | TATGAGTGTGGAGTCGTTGCGGTAAATAGTCTCAGCTCCTCGGNNNAGATCGGAAG |
| 35441_1 | TATGAGTGTGGAGTCGTTGCAGATGGAGTCTCACTCTTTCGCNNNAGATCGGAAGA |
| 35441_2 | TATGAGTGTGGAGTCGTTGCAGATGGAGTCTCACTCTTTCGCNNNAGATCGGAAGA |
| 50128_1 | TATGAGTGTGGAGTCGTTGCAACAACACGACATTGTAAAAGCANNNAGATCGGAA |
| 50128_2 | TATGAGTGTGGAGTCGTTGCAACAACACGACATTGTAAAAGCANNNAGATCGGAA |
| 37297_1 | TATGAGTGTGGAGTCGTTGCCAGGGGTTGCAATCATAGTCTCTNNNAGATCGGAAG |
| 37297_2 | TATGAGTGTGGAGTCGTTGCCAGGGGTTGCAATCATAGTCTCTNNNAGATCGGAAG |
| 47753_1 | TATGAGTGTGGAGTCGTTGCGTTTCACCATGTTGCCTAGGTTGNNNAGATCGGAAG |
| 47753_2 | TATGAGTGTGGAGTCGTTGCGTTTCACCATGTTGCCTAGGTTGNNNAGATCGGAAG |
| 48450_1 | TATGAGTGTGGAGTCGTTGCTAGAATTATGGGACCAAGCCACANNNAGATCGGAA |
| 48450_2 | TATGAGTGTGGAGTCGTTGCTAGAATTATGGGACCAAGCCACANNNAGATCGGAA |
| 50163_1 | TATGAGTGTGGAGTCGTTGCCCATCCCTCAGTACACTCAGAAGNNNAGATCGGAAG |
| 50163_2 | TATGAGTGTGGAGTCGTTGCCCATCCCTCAGTACACTCAGAAGNNNAGATCGGAAG |
| 35515_1 | TATGAGTGTGGAGTCGTTGCTGGTACATGTGATGTTTTGAGCCNNNAGATCGGAAG |
| 35515_2 | TATGAGTGTGGAGTCGTTGCTGGTACATGTGATGTTTTGAGCCNNNAGATCGGAAG |
| 46646_1 | TATGAGTGTGGAGTCGTTGCCAGCAGCTGAGGCAGAAGATATANNNAGATCGGAA |
| 46646_2 | TATGAGTGTGGAGTCGTTGCCAGCAGCTGAGGCAGAAGATATANNNAGATCGGAA |
| 34896_1 | TATGAGTGTGGAGTCGTTGCCCACCACACTGGGCTAATTTTTTNNNAGATCGGAAG |
| 34896_2 | TATGAGTGTGGAGTCGTTGCCCACCACACTGGGCTAATTTTTTNNNAGATCGGAAG |
| 47198_1 | TATGAGTGTGGAGTCGTTGCGTTCACACTATTCTCCTGCCTCANNNAGATCGGAAGA |
| 47198_2 | TATGAGTGTGGAGTCGTTGCGTTCACACTATTCTCCTGCCTCANNNAGATCGGAAGA |
| 33289_1 | TATGAGTGTGGAGTCGTTGCAAACTCCACCTCATGAGCTCAATNNNAGATCGGAAG |
| 33289_2 | TATGAGTGTGGAGTCGTTGCAAACTCCACCTCATGAGCTCAATNNNAGATCGGAAG |
| 50517_1 | TATGAGTGTGGAGTCGTTGCAATGTAACAGCTGACCCTGACCNNNAGATCGGAAGA |
| 50517_2 | TATGAGTGTGGAGTCGTTGCAATGTAACAGCTGACCCTGACCNNNAGATCGGAAGA |
| 12897_1 | TATGAGTGTGGAGTCGTTGCTGTCTCTAACAGGAGAACTAANNNAGATCGGAAGA |
| 12897_2 | TATGAGTGTGGAGTCGTTGCTGTCTCTAACAGGAGAACTAANNNAGATCGGAAGA |
| 4795_1 | TATGAGTGTGGAGTCGTTGCGTTCCCGTGGAGGAAGAGGTGGNNNAGATCGGAA |
| 4795_2 | TATGAGTGTGGAGTCGTTGCGTTCCCGTGGAGGAAGAGGTGGNNNAGATCGGAA |
| 41130_1 | TATGAGTGTGGAGTCGTTGCACTTCTATGGTGAAACGTAACCCANNNAGATCGGAA |
| 41130_2 | TATGAGTGTGGAGTCGTTGCACTTCTATGGTGAAACGTAACCCANNNAGATCGGAA |
| 44672_1 | TATGAGTGTGGAGTCGTTGCTAGAGCAAGTGGAGATCAGTGTGNNNAGATCGGA |
| 44672_2 | TATGAGTGTGGAGTCGTTGCTAGAGCAAGTGGAGATCAGTGTGNNNAGATCGGA |
| 49597_1 | TATGAGTGTGGAGTCGTTGCGGGTTCAAGTGATTCTCCTGTCTNNNAGATCGGAAG |
| 49597_2 | TATGAGTGTGGAGTCGTTGCGGGTTCAAGTGATTCTCCTGTCTNNNAGATCGGAAG |
| 13258_1 | TATGAGTGTGGAGTCGTTGCTCAGAAAGCAAGATTTTAAACTTNNNAGATCGGAAG |

| | |
|---|---|
| 13258_2 | TATGAGTGTGGAGTCGTTGCTCAGAAAGCAAGATTTTAAACTTNNNAGATCGGAAG |
| 3152_1 | TATGAGTGTGGAGTCGTTGCCTACAAGGACAGGCCCATTTGAGNNNAGATCGGAA |
| 3152_2 | TATGAGTGTGGAGTCGTTGCCTACAAGGACAGGCCCATTTGAGNNNAGATCGGAA |
| 50057_1 | TATGAGTGTGGAGTCGTTGCTCCTCTCTCTTTCTCTCTCTCCCNNNAGATCGGAAGAG |
| 50057_2 | TATGAGTGTGGAGTCGTTGCTCCTCTCTCTTTCTCTCTCTCCCNNNAGATCGGAAGAG |
| 39780_1 | TATGAGTGTGGAGTCGTTGCAACAAAACAGCATGGTACTGACANNNAGATCGGAA |
| 39780_2 | TATGAGTGTGGAGTCGTTGCAACAAAACAGCATGGTACTGACANNNAGATCGGAA |
| 46255_1 | TATGAGTGTGGAGTCGTTGCGAGTAGCTGGGACTATAGGCACNNNAGATCGGAAG |
| 46255_2 | TATGAGTGTGGAGTCGTTGCGAGTAGCTGGGACTATAGGCACNNNAGATCGGAAG |
| 411_1 | TATGAGTGTGGAGTCGTTGCTCTACTGCACTTATCACTATCTGACANNNAGATCGGA |
| 411_2 | TATGAGTGTGGAGTCGTTGCTCTACTGCACTTATCACTATCTGACANNNAGATCGGA |
| 42459_1 | TATGAGTGTGGAGTCGTTGCATGATTTGGGCTCACTACAACCTNNNAGATCGGAAG |
| 42459_2 | TATGAGTGTGGAGTCGTTGCATGATTTGGGCTCACTACAACCTNNNAGATCGGAAG |
| 46585_1 | TATGAGTGTGGAGTCGTTGCTGAGTAGCTGAGATTACAGGCACNNNAGATCGGAA |
| 46585_2 | TATGAGTGTGGAGTCGTTGCTGAGTAGCTGAGATTACAGGCACNNNAGATCGGAA |
| 48643_1 | TATGAGTGTGGAGTCGTTGCCCGAGGAACTGGGATTATAGGTGNNNAGATCGGAA |
| 48643_2 | TATGAGTGTGGAGTCGTTGCCCGAGGAACTGGGATTATAGGTGNNNAGATCGGAA |
| 49457_1 | TATGAGTGTGGAGTCGTTGCTGTGTTGTTTCCCCTGTGTTCTANNNAGATCGGAAG |
| 49457_2 | TATGAGTGTGGAGTCGTTGCTGTGTTGTTTCCCCTGTGTTCTANNNAGATCGGAAG |
| 10247_1 | TATGAGTGTGGAGTCGTTGCCCACTTTCCCACCAAGCAGAAATNNNAGATCGGAAG |
| 10247_2 | TATGAGTGTGGAGTCGTTGCCCACTTTCCCACCAAGCAGAAATNNNAGATCGGAAG |
| 36379_1 | TATGAGTGTGGAGTCGTTGCGAGAATTGCCTGAAAGCACAGAGNNNAGATCGGAA |
| 36379_2 | TATGAGTGTGGAGTCGTTGCGAGAATTGCCTGAAAGCACAGAGNNNAGATCGGAA |
| 37125_1 | TATGAGTGTGGAGTCGTTGCAAAACACCACAGAAAAAGATGTACTNNNAGATCGG |
| 37125_2 | TATGAGTGTGGAGTCGTTGCAAAACACCACAGAAAAAGATGTACTNNNAGATCGG |
| 12616_1 | TATGAGTGTGGAGTCGTTGCCTCAATGGTTTGTGATTTTTTTTNNNAGATCGGAAG |
| 12616_2 | TATGAGTGTGGAGTCGTTGCCTCAATGGTTTGTGATTTTTTTTNNNAGATCGGAAG |
| 50446_1 | TATGAGTGTGGAGTCGTTGCGGAAAGGGCTTTTGTTGGGAAAGNNNAGATCGGA |
| 50446_2 | TATGAGTGTGGAGTCGTTGCGGAAAGGGCTTTTGTTGGGAAAGNNNAGATCGGA |
| 52636_1 | TATGAGTGTGGAGTCGTTGCACTATCTGTAAGTGTGGGTGGTGNNNAGATCGGAA |
| 52636_2 | TATGAGTGTGGAGTCGTTGCACTATCTGTAAGTGTGGGTGGTGNNNAGATCGGAA |
| 36187_1 | TATGAGTGTGGAGTCGTTGCCAGTGAAGACAAGGTGGGGAATANNNAGATCGGA |
| 36187_2 | TATGAGTGTGGAGTCGTTGCCAGTGAAGACAAGGTGGGGAATANNNAGATCGGA |
| 35177_1 | TATGAGTGTGGAGTCGTTGCGTGCCCTCAAGTTCAAGTGATTCNNNAGATCGGAAG |
| 35177_2 | TATGAGTGTGGAGTCGTTGCGTGCCCTCAAGTTCAAGTGATTCNNNAGATCGGAAG |
| 46460_1 | TATGAGTGTGGAGTCGTTGCAGTGTTTCATTCTTGTTGCCCAGNNNAGATCGGAAG |
| 46460_2 | TATGAGTGTGGAGTCGTTGCAGTGTTTCATTCTTGTTGCCCAGNNNAGATCGGAAG |
| 39972_1 | TATGAGTGTGGAGTCGTTGCAAGATTGTACCTCTGCACTCCAGNNNAGATCGGAAG |
| 39972_2 | TATGAGTGTGGAGTCGTTGCAAGATTGTACCTCTGCACTCCAGNNNAGATCGGAAG |
| 43798_1 | TATGAGTGTGGAGTCGTTGCCAACATAGTGAGACCCACCTGAANNNAGATCGGAAG |
| 43798_2 | TATGAGTGTGGAGTCGTTGCCAACATAGTGAGACCCACCTGAANNNAGATCGGAAG |
| 34994_1 | TATGAGTGTGGAGTCGTTGCAGCCTCCTGAGTATCTGGGATTANNNAGATCGGAAG |
| 34994_2 | TATGAGTGTGGAGTCGTTGCAGCCTCCTGAGTATCTGGGATTANNNAGATCGGAAG |

| | |
|---|---|
| 35820_1 | TATGAGTGTGGAGTCGTTGCCAATCTCGGCTCACTGCAATTTCNNNAGATCGGAAG |
| 35820_2 | TATGAGTGTGGAGTCGTTGCCAATCTCGGCTCACTGCAATTTCNNNAGATCGGAAG |
| 32168_1 | TATGAGTGTGGAGTCGTTGCTCCAGAGTAGCCGGGACTACNNNAGATCGGAAGAG |
| 32168_2 | TATGAGTGTGGAGTCGTTGCTCCAGAGTAGCCGGGACTACNNNAGATCGGAAGAG |
| 43587_1 | TATGAGTGTGGAGTCGTTGCATGCCTGTAAAATGGATGAGGGTNNNAGATCGGAA |
| 43587_2 | TATGAGTGTGGAGTCGTTGCATGCCTGTAAAATGGATGAGGGTNNNAGATCGGAA |
| 49027_1 | TATGAGTGTGGAGTCGTTGCGCCCGATGACAGGGAAAAAGATANNNAGATCGGAA |
| 49027_2 | TATGAGTGTGGAGTCGTTGCGCCCGATGACAGGGAAAAAGATANNNAGATCGGAA |
| 51806_1 | TATGAGTGTGGAGTCGTTGCGTCTTCGTCCTGTGAGGTAATGANNNAGATCGGAAG |
| 51806_2 | TATGAGTGTGGAGTCGTTGCGTCTTCGTCCTGTGAGGTAATGANNNAGATCGGAAG |
| 39165_1 | TATGAGTGTGGAGTCGTTGCTGAGGCAGGAGAATAGTGTGAACNNNAGATCGGA |
| 39165_2 | TATGAGTGTGGAGTCGTTGCTGAGGCAGGAGAATAGTGTGAACNNNAGATCGGA |
| 40197_1 | TATGAGTGTGGAGTCGTTGCCTGGCAAAGAGCTTAGACGTTTCNNNAGATCGGAAG |
| 40197_2 | TATGAGTGTGGAGTCGTTGCCTGGCAAAGAGCTTAGACGTTTCNNNAGATCGGAAG |
| 47702_1 | TATGAGTGTGGAGTCGTTGCTTGGCTGCCTGACAATAAAACTTNNNAGATCGGAAG |
| 47702_2 | TATGAGTGTGGAGTCGTTGCTTGGCTGCCTGACAATAAAACTTNNNAGATCGGAAG |
| 36301_1 | TATGAGTGTGGAGTCGTTGCAATAAAAAAGGAGGGAATGGCGGNNNAGATCGGA |
| 36301_2 | TATGAGTGTGGAGTCGTTGCAATAAAAAAGGAGGGAATGGCGGNNNAGATCGGA |
| 12606_1 | TATGAGTGTGGAGTCGTTGCTAAAAGAAAAACTTCAGCTGAATNNNAGATCGGAAG |
| 12606_2 | TATGAGTGTGGAGTCGTTGCTAAAAGAAAAACTTCAGCTGAATNNNAGATCGGAAG |
| 38480_1 | TATGAGTGTGGAGTCGTTGCGGATCCATGAGGTTTGGGTGTTANNNAGATCGGAA |
| 38480_2 | TATGAGTGTGGAGTCGTTGCGGATCCATGAGGTTTGGGTGTTANNNAGATCGGAA |
| 36949_1 | TATGAGTGTGGAGTCGTTGCCACTACTTGCTTCTCCTGGAACTNNNAGATCGGAAGA |
| 36949_2 | TATGAGTGTGGAGTCGTTGCCACTACTTGCTTCTCCTGGAACTNNNAGATCGGAAGA |
| 30954_1 | TATGAGTGTGGAGTCGTTGCGCAAGAGAGACCGAGAAAGAGAANNNAGATCGGA |
| 30954_2 | TATGAGTGTGGAGTCGTTGCGCAAGAGAGACCGAGAAAGAGAANNNAGATCGGA |
| 49839_1 | TATGAGTGTGGAGTCGTTGCAGAGTTTGTCATTGGAACTGGGANNNAGATCGGAA |
| 49839_2 | TATGAGTGTGGAGTCGTTGCAGAGTTTGTCATTGGAACTGGGANNNAGATCGGAA |
| 30672_1 | TATGAGTGTGGAGTCGTTGCCGAACTTTTCCAGAGTGTTTGTNNNAGATCGGAAGA |
| 30672_2 | TATGAGTGTGGAGTCGTTGCCGAACTTTTCCAGAGTGTTTGTNNNAGATCGGAAGA |
| 50261_1 | TATGAGTGTGGAGTCGTTGCCCTCAAAGGAAGCACTCAGGTATNNNAGATCGGAAG |
| 50261_2 | TATGAGTGTGGAGTCGTTGCCCTCAAAGGAAGCACTCAGGTATNNNAGATCGGAAG |
| 29767_1 | TATGAGTGTGGAGTCGTTGCAACGATACTCTAAAAGGGGGCAGNNNAGATCGGAA |
| 29767_2 | TATGAGTGTGGAGTCGTTGCAACGATACTCTAAAAGGGGGCAGNNNAGATCGGAA |
| 44483_1 | TATGAGTGTGGAGTCGTTGCTTTGACTTTGTTGGTCCTCCTGTNNNAGATCGGAAG |
| 44483_2 | TATGAGTGTGGAGTCGTTGCTTTGACTTTGTTGGTCCTCCTGTNNNAGATCGGAAG |
| 34757_1 | TATGAGTGTGGAGTCGTTGCTTCAAGTGATTCTCCTGCCTCAGNNNAGATCGGAAG |
| 34757_2 | TATGAGTGTGGAGTCGTTGCTTCAAGTGATTCTCCTGCCTCAGNNNAGATCGGAAG |
| 44147_1 | TATGAGTGTGGAGTCGTTGCCAGCATTTACTCAGCAGGGTTTGNNNAGATCGGAAG |
| 44147_2 | TATGAGTGTGGAGTCGTTGCCAGCATTTACTCAGCAGGGTTTGNNNAGATCGGAAG |
| 40512_1 | TATGAGTGTGGAGTCGTTGCGTTCTGAAGGCTGAGAAGTCTGANNNAGATCGGAA |
| 40512_2 | TATGAGTGTGGAGTCGTTGCGTTCTGAAGGCTGAGAAGTCTGANNNAGATCGGAA |
| 30658_1 | TATGAGTGTGGAGTCGTTGCACACAGCGGGACTCTATCAAAAANNNAGATCGGAAG |

| | |
|---|---|
| 30658_2 | TATGAGTGTGGAGTCGTTGCACACAGCGGGACTCTATCAAAAANNNAGATCGGAA |
| 40912_1 | TATGAGTGTGGAGTCGTTGCCAAACATGACTTTGAGGCTTCGTNNNAGATCGGAAG |
| 40912_2 | TATGAGTGTGGAGTCGTTGCCAAACATGACTTTGAGGCTTCGTNNNAGATCGGAAG |
| 48080_1 | TATGAGTGTGGAGTCGTTGCAGACGAGGGAAGAGGGAATGATANNNAGATCGGA |
| 48080_2 | TATGAGTGTGGAGTCGTTGCAGACGAGGGAAGAGGGAATGATANNNAGATCGGA |
| 45235_1 | TATGAGTGTGGAGTCGTTGCTAGGACAAAAGGTTGAAGGGGAGNNNAGATCGGA |
| 45235_2 | TATGAGTGTGGAGTCGTTGCTAGGACAAAAGGTTGAAGGGGAGNNNAGATCGGA |
| 40250_1 | TATGAGTGTGGAGTCGTTGCCAGAGCAAGACTCCATCTCCAAANNNAGATCGGAAG |
| 40250_2 | TATGAGTGTGGAGTCGTTGCCAGAGCAAGACTCCATCTCCAAANNNAGATCGGAAG |
| 3017_1 | TATGAGTGTGGAGTCGTTGCTTGCTGTGAAAAAGACGTTCACGNNNAGATCGGAA |
| 3017_2 | TATGAGTGTGGAGTCGTTGCTTGCTGTGAAAAAGACGTTCACGNNNAGATCGGAA |
| 33192_1 | TATGAGTGTGGAGTCGTTGCGCATGTGTCTGTAATCCCAGCTANNNAGATCGGAAG |
| 33192_2 | TATGAGTGTGGAGTCGTTGCGCATGTGTCTGTAATCCCAGCTANNNAGATCGGAAG |
| 36079_1 | TATGAGTGTGGAGTCGTTGCCTAAAGGCACTTGTCAGGTACAGNNNAGATCGGAA |
| 36079_2 | TATGAGTGTGGAGTCGTTGCCTAAAGGCACTTGTCAGGTACAGNNNAGATCGGAA |
| 48532_1 | TATGAGTGTGGAGTCGTTGCCATGCCCAGCCCTAGATATTCTTNNNAGATCGGAAG |
| 48532_2 | TATGAGTGTGGAGTCGTTGCCATGCCCAGCCCTAGATATTCTTNNNAGATCGGAAG |
| 39327_1 | TATGAGTGTGGAGTCGTTGCTTGGATCCACATCTCATGCCATANNNAGATCGGAAG |
| 39327_2 | TATGAGTGTGGAGTCGTTGCTTGGATCCACATCTCATGCCATANNNAGATCGGAAG |
| 37985_1 | TATGAGTGTGGAGTCGTTGCCTCTCAGTCTCCAATTCCTTCCCNNNAGATCGGAAGA |
| 37985_2 | TATGAGTGTGGAGTCGTTGCCTCTCAGTCTCCAATTCCTTCCCNNNAGATCGGAAGA |
| 49785_1 | TATGAGTGTGGAGTCGTTGCTTTACTGGTCTAAGCTCGTTGGGNNNAGATCGGAAG |
| 49785_2 | TATGAGTGTGGAGTCGTTGCTTTACTGGTCTAAGCTCGTTGGGNNNAGATCGGAAG |
| 39351_1 | TATGAGTGTGGAGTCGTTGCCCATGTGACTATCCTGTTTCCCANNNAGATCGGAAGA |
| 39351_2 | TATGAGTGTGGAGTCGTTGCCCATGTGACTATCCTGTTTCCCANNNAGATCGGAAGA |
| 32843_1 | TATGAGTGTGGAGTCGTTGCACTATTCGGCTCACTGCAAGTTNNNAGATCGGAAGA |
| 32843_2 | TATGAGTGTGGAGTCGTTGCACTATTCGGCTCACTGCAAGTTNNNAGATCGGAAGA |
| 39546_1 | TATGAGTGTGGAGTCGTTGCGACAGAGTCTCACTTCTGTCACCNNNAGATCGGAAG |
| 39546_2 | TATGAGTGTGGAGTCGTTGCGACAGAGTCTCACTTCTGTCACCNNNAGATCGGAAG |
| 10651_1 | TATGAGTGTGGAGTCGTTGCGGGCACCTTGACTAATACGAAGGNNNAGATCGGAA |
| 10651_2 | TATGAGTGTGGAGTCGTTGCGGGCACCTTGACTAATACGAAGGNNNAGATCGGAA |
| 10745_1 | TATGAGTGTGGAGTCGTTGCAGCACTTTGGAGAAACTACCTGTNNNAGATCGGAAG |
| 10745_2 | TATGAGTGTGGAGTCGTTGCAGCACTTTGGAGAAACTACCTGTNNNAGATCGGAAG |
| 41930_1 | TATGAGTGTGGAGTCGTTGCCAGACTCCTGGCCTCAATCTATCNNNAGATCGGAAG |
| 41930_2 | TATGAGTGTGGAGTCGTTGCCAGACTCCTGGCCTCAATCTATCNNNAGATCGGAAG |
| 30591_1 | TATGAGTGTGGAGTCGTTGCCCTGGGTCCCTACTAAAACACGTNNNAGATCGGAAG |
| 30591_2 | TATGAGTGTGGAGTCGTTGCCCTGGGTCCCTACTAAAACACGTNNNAGATCGGAAG |
| 46461_1 | TATGAGTGTGGAGTCGTTGCTACGACGCCTGGCTAATTTTTTGNNNAGATCGGAAG |
| 46461_2 | TATGAGTGTGGAGTCGTTGCTACGACGCCTGGCTAATTTTTTGNNNAGATCGGAAG |
| 37367_1 | TATGAGTGTGGAGTCGTTGCTGAATTAGACCCCACCTTTTGTGNNNAGATCGGAAG |
| 37367_2 | TATGAGTGTGGAGTCGTTGCTGAATTAGACCCCACCTTTTGTGNNNAGATCGGAAG |
| 39068_1 | TATGAGTGTGGAGTCGTTGCTGTCAAGGTGATTGTGAAAACCANNNAGATCGGAA |
| 39068_2 | TATGAGTGTGGAGTCGTTGCTGTCAAGGTGATTGTGAAAACCANNNAGATCGGAA |

| | |
|---|---|
| 53148_1 | TATGAGTGTGGAGTCGTTGCGTTTTCAGGTTTCAACTCTCCCGNNNAGATCGGAAGA |
| 53148_2 | TATGAGTGTGGAGTCGTTGCGTTTTCAGGTTTCAACTCTCCCGNNNAGATCGGAAGA |
| 30116_1 | TATGAGTGTGGAGTCGTTGCCCCGTTTCTCTAAAAGCTTTCAAGGNNNAGATCGGAA |
| 30116_2 | TATGAGTGTGGAGTCGTTGCCCCGTTTCTCTAAAAGCTTTCAAGGNNNAGATCGGAA |
| 10498_1 | TATGAGTGTGGAGTCGTTGCTGAGTTGTCTGGCTTTTTGGGAANNNAGATCGGAA |
| 10498_2 | TATGAGTGTGGAGTCGTTGCTGAGTTGTCTGGCTTTTTGGGAANNNAGATCGGAA |
| 42957_1 | TATGAGTGTGGAGTCGTTGCGAGTAGCTGGAAGTACAGGTGTCNNNAGATCGGAA |
| 42957_2 | TATGAGTGTGGAGTCGTTGCGAGTAGCTGGAAGTACAGGTGTCNNNAGATCGGAA |
| 9754_1 | TATGAGTGTGGAGTCGTTGCCTGCACTACCCAAAGGTAGGAACNNNAGATCGGAA |
| 9754_2 | TATGAGTGTGGAGTCGTTGCCTGCACTACCCAAAGGTAGGAACNNNAGATCGGAA |
| 48662_1 | TATGAGTGTGGAGTCGTTGCCTAGGAATGGGGAGGTCAAGTGANNNAGATCGGAA |
| 48662_2 | TATGAGTGTGGAGTCGTTGCCTAGGAATGGGGAGGTCAAGTGANNNAGATCGGAA |
| 30117_1 | TATGAGTGTGGAGTCGTTGCCCCGTTTCTCTAAAAGCTTTCAAGGNNNAGATCGGAA |
| 30117_2 | TATGAGTGTGGAGTCGTTGCCCCGTTTCTCTAAAAGCTTTCAAGGNNNAGATCGGAA |
| 44119_1 | TATGAGTGTGGAGTCGTTGCCTCACAGTGTCTCATGGTCCGNNNAGATCGGAAGAG |
| 44119_2 | TATGAGTGTGGAGTCGTTGCCTCACAGTGTCTCATGGTCCGNNNAGATCGGAAGAG |
| 45477_1 | TATGAGTGTGGAGTCGTTGCTTGACTGGAAACTGAGTCTTGCTNNNAGATCGGAAG |
| 45477_2 | TATGAGTGTGGAGTCGTTGCTTGACTGGAAACTGAGTCTTGCTNNNAGATCGGAAG |
| 2643_1 | TATGAGTGTGGAGTCGTTGCTCATGGAGCAGAAGACAACCTTGNNNAGATCGGAA |
| 2643_2 | TATGAGTGTGGAGTCGTTGCTCATGGAGCAGAAGACAACCTTGNNNAGATCGGAA |
| 29814_1 | TATGAGTGTGGAGTCGTTGCACTTGCATACACTTCACTTTTGCANNNAGATCGGAAG |
| 29814_2 | TATGAGTGTGGAGTCGTTGCACTTGCATACACTTCACTTTTGCANNNAGATCGGAAG |
| 41536_1 | TATGAGTGTGGAGTCGTTGCCTCAGGGGAGAGGTTGAGAATTTNNNAGATCGGAA |
| 41536_2 | TATGAGTGTGGAGTCGTTGCCTCAGGGGAGAGGTTGAGAATTTNNNAGATCGGAA |
| 46959_1 | TATGAGTGTGGAGTCGTTGCTAAAAATGAGGCTGGGATGAGCTNNNAGATCGGAA |
| 46959_2 | TATGAGTGTGGAGTCGTTGCTAAAAATGAGGCTGGGATGAGCTNNNAGATCGGAA |
| 33495_1 | TATGAGTGTGGAGTCGTTGCATCCTACTACCCAGCAATGATCANNNAGATCGGAAGA |
| 33495_2 | TATGAGTGTGGAGTCGTTGCATCCTACTACCCAGCAATGATCANNNAGATCGGAAGA |
| 37641_1 | TATGAGTGTGGAGTCGTTGCGGATCATGAGGTCAGGAGTTCAANNNAGATCGGAA |
| 37641_2 | TATGAGTGTGGAGTCGTTGCGGATCATGAGGTCAGGAGTTCAANNNAGATCGGAA |
| 12529_1 | TATGAGTGTGGAGTCGTTGCAACTACCTCTACCTCCTCTTNNNAGATCGGAAGAGCA |
| 12529_2 | TATGAGTGTGGAGTCGTTGCAACTACCTCTACCTCCTCTTNNNAGATCGGAAGAGCA |
| 47938_1 | TATGAGTGTGGAGTCGTTGCCCATTCTTTCACTTACAAGGCCGNNNAGATCGGAAGA |
| 47938_2 | TATGAGTGTGGAGTCGTTGCCCATTCTTTCACTTACAAGGCCGNNNAGATCGGAAGA |
| 30201_1 | TATGAGTGTGGAGTCGTTGCGTTCAATTTGTGCTTCTCGTGCANNNAGATCGGAAGA |
| 30201_2 | TATGAGTGTGGAGTCGTTGCGTTCAATTTGTGCTTCTCGTGCANNNAGATCGGAAGA |
| 33631_1 | TATGAGTGTGGAGTCGTTGCAAAGTGCTGGGATTACAGGCATNNNAGATCGGAAG |
| 33631_2 | TATGAGTGTGGAGTCGTTGCAAAGTGCTGGGATTACAGGCATNNNAGATCGGAAG |
| 31716_1 | TATGAGTGTGGAGTCGTTGCTACAAAGGCAAAGTCCCACTACANNNAGATCGGAAG |
| 31716_2 | TATGAGTGTGGAGTCGTTGCTACAAAGGCAAAGTCCCACTACANNNAGATCGGAAG |
| 43130_1 | TATGAGTGTGGAGTCGTTGCATGCTGTTTCTAACTGTGGGGTTNNNAGATCGGAAG |
| 43130_2 | TATGAGTGTGGAGTCGTTGCATGCTGTTTCTAACTGTGGGGTTNNNAGATCGGAAG |
| 9752_1 | TATGAGTGTGGAGTCGTTGCTTCAGCTCCTGTTTCCTTGGAACNNNAGATCGGAAGA |

| | |
|---|---|
| 9752_2 | TATGAGTGTGGAGTCGTTGCTTCAGCTCCTGTTTCCTTGGAACNNNAGATCGGAAGA |
| 37594_1 | TATGAGTGTGGAGTCGTTGCGCAAGACACAGCATTGCAAAATTNNNAGATCGGAAG |
| 37594_2 | TATGAGTGTGGAGTCGTTGCGCAAGACACAGCATTGCAAAATTNNNAGATCGGAAG |
| 32979_1 | TATGAGTGTGGAGTCGTTGCGGCCAAAACCTTGAGATACATCANNNAGATCGGAAG |
| 32979_2 | TATGAGTGTGGAGTCGTTGCGGCCAAAACCTTGAGATACATCANNNAGATCGGAAG |
| 38264_1 | TATGAGTGTGGAGTCGTTGCGAAACTGCTTTGTGATGTGTGGANNNAGATCGGAAG |
| 38264_2 | TATGAGTGTGGAGTCGTTGCGAAACTGCTTTGTGATGTGTGGANNNAGATCGGAAG |
| 34176_1 | TATGAGTGTGGAGTCGTTGCGTTGAATTAGAACAACCCCTGCCNNNAGATCGGAAG |
| 34176_2 | TATGAGTGTGGAGTCGTTGCGTTGAATTAGAACAACCCCTGCCNNNAGATCGGAAG |
| 30532_1 | TATGAGTGTGGAGTCGTTGCAGCAAAGCTCTAATGAGCAGGTTTNNNAGATCGGAA |
| 30532_2 | TATGAGTGTGGAGTCGTTGCAGCAAAGCTCTAATGAGCAGGTTTNNNAGATCGGAA |
| 51857_1 | TATGAGTGTGGAGTCGTTGCGTGACACACAGCGATTCAGAACNNNAGATCGGAAGA |
| 51857_2 | TATGAGTGTGGAGTCGTTGCGTGACACACAGCGATTCAGAACNNNAGATCGGAAGA |
| 2271_1 | TATGAGTGTGGAGTCGTTGCGCCTACACAACTGGGGCATTTTANNNAGATCGGAAG |
| 2271_2 | TATGAGTGTGGAGTCGTTGCGCCTACACAACTGGGGCATTTTANNNAGATCGGAAG |
| 47745_1 | TATGAGTGTGGAGTCGTTGCTATCTGGGATTACAGACAAGCGCNNNAGATCGGAAG |
| 47745_2 | TATGAGTGTGGAGTCGTTGCTATCTGGGATTACAGACAAGCGCNNNAGATCGGAAG |
| 40529_1 | TATGAGTGTGGAGTCGTTGCGGGGAGATTGGTCTACAGTTTGTNNNAGATCGGAAG |
| 40529_2 | TATGAGTGTGGAGTCGTTGCGGGGAGATTGGTCTACAGTTTGTNNNAGATCGGAAG |
| 46306_1 | TATGAGTGTGGAGTCGTTGCCACGCCCGGGTAATTTTTTTTGTNNNAGATCGGAAGA |
| 46306_2 | TATGAGTGTGGAGTCGTTGCCACGCCCGGGTAATTTTTTTTGTNNNAGATCGGAAGA |
| 31094_1 | TATGAGTGTGGAGTCGTTGCGGTAGGCCAGTATTCACATGCTANNNAGATCGGAAG |
| 31094_2 | TATGAGTGTGGAGTCGTTGCGGTAGGCCAGTATTCACATGCTANNNAGATCGGAAG |
| 13268_1 | TATGAGTGTGGAGTCGTTGCCACCACCTTAAATATTTGTCTTTNNNAGATCGGAAGA |
| 13268_2 | TATGAGTGTGGAGTCGTTGCCACCACCTTAAATATTTGTCTTTNNNAGATCGGAAGA |
| 33456_1 | TATGAGTGTGGAGTCGTTGCAGAGTTTCAATCCTGGTCTCTGANNNAGATCGGAAG |
| 33456_2 | TATGAGTGTGGAGTCGTTGCAGAGTTTCAATCCTGGTCTCTGANNNAGATCGGAAG |
| 30871_1 | TATGAGTGTGGAGTCGTTGCGTGTGCACATTCCATAGTGGTTCNNNAGATCGGAAG |
| 30871_2 | TATGAGTGTGGAGTCGTTGCGTGTGCACATTCCATAGTGGTTCNNNAGATCGGAAG |
| 5062_1 | TATGAGTGTGGAGTCGTTGCAAGCACGATTGCTAATTGTGGGANNNAGATCGGAAG |
| 5062_2 | TATGAGTGTGGAGTCGTTGCAAGCACGATTGCTAATTGTGGGANNNAGATCGGAAG |
| 50814_1 | TATGAGTGTGGAGTCGTTGCAAAAGAGAGAGAGAGAAAGCGCANNNAGATCGGAA |
| 50814_2 | TATGAGTGTGGAGTCGTTGCAAAAGAGAGAGAGAGAAAGCGCANNNAGATCGGAA |
| 44386_1 | TATGAGTGTGGAGTCGTTGCAAGGAAGGAATTCAGTCCCCATCNNNAGATCGGAAG |
| 44386_2 | TATGAGTGTGGAGTCGTTGCAAGGAAGGAATTCAGTCCCCATCNNNAGATCGGAAG |
| 45935_1 | TATGAGTGTGGAGTCGTTGCGACAGAGTTTTCTCCTTGTTGCCNNNAGATCGGAAGA |
| 45935_2 | TATGAGTGTGGAGTCGTTGCGACAGAGTTTTCTCCTTGTTGCCNNNAGATCGGAAGA |
| 47944_1 | TATGAGTGTGGAGTCGTTGCGCAGTGAGTCTTGGTTGAGAGATNNNAGATCGGAA |
| 47944_2 | TATGAGTGTGGAGTCGTTGCGCAGTGAGTCTTGGTTGAGAGATNNNAGATCGGAA |
| 29622_1 | TATGAGTGTGGAGTCGTTGCGATATAGACTGCGGAGACGTTGTNNNAGATCGGAAG |
| 29622_2 | TATGAGTGTGGAGTCGTTGCGATATAGACTGCGGAGACGTTGTNNNAGATCGGAAG |
| 32455_1 | TATGAGTGTGGAGTCGTTGCGTGCATATCTAGTGGTTCATTGGCNNNAGATCGGAA |
| 32455_2 | TATGAGTGTGGAGTCGTTGCGTGCATATCTAGTGGTTCATTGGCNNNAGATCGGAA |

| | |
|---|---|
| 13263_1 | TATGAGTGTGGAGTCGTTGCTTTGTAACGAAGTGAAGGAGNNNAGATCGGAAGAG |
| 13263_2 | TATGAGTGTGGAGTCGTTGCTTTGTAACGAAGTGAAGGAGNNNAGATCGGAAGAG |
| 33872_1 | TATGAGTGTGGAGTCGTTGCGCAACAATGATGCATAGCCCTTTNNNAGATCGGAAG |
| 33872_2 | TATGAGTGTGGAGTCGTTGCGCAACAATGATGCATAGCCCTTTNNNAGATCGGAAG |
| 37776_1 | TATGAGTGTGGAGTCGTTGCCTTTCTCTCCTGCTACCTTGTGANNNAGATCGGAAGA |
| 37776_2 | TATGAGTGTGGAGTCGTTGCCTTTCTCTCCTGCTACCTTGTGANNNAGATCGGAAGA |
| 45542_1 | TATGAGTGTGGAGTCGTTGCACAGCTAACAAGGCTTCCCATAANNNAGATCGGAAG |
| 45542_2 | TATGAGTGTGGAGTCGTTGCACAGCTAACAAGGCTTCCCATAANNNAGATCGGAAG |
| 50340_1 | TATGAGTGTGGAGTCGTTGCACCCCTGATGAACATTGATGTGANNNAGATCGGAAG |
| 50340_2 | TATGAGTGTGGAGTCGTTGCACCCCTGATGAACATTGATGTGANNNAGATCGGAAG |
| 29657_1 | TATGAGTGTGGAGTCGTTGCTGGTCTCAAGCAATTCTCCTACCNNNAGATCGGAAGA |
| 29657_2 | TATGAGTGTGGAGTCGTTGCTGGTCTCAAGCAATTCTCCTACCNNNAGATCGGAAGA |
| 52706_1 | TATGAGTGTGGAGTCGTTGCTCCCATAAGGTGACTGAAAAGTGTNNNAGATCGGAA |
| 52706_2 | TATGAGTGTGGAGTCGTTGCTCCCATAAGGTGACTGAAAAGTGTNNNAGATCGGAA |
| 29440_1 | TATGAGTGTGGAGTCGTTGCACCCAATCAAAAATGGGCTCTTGNNNAGATCGGAAG |
| 29440_2 | TATGAGTGTGGAGTCGTTGCACCCAATCAAAAATGGGCTCTTGNNNAGATCGGAAG |
| 39036_1 | TATGAGTGTGGAGTCGTTGCTACTCACAATCTCCAGTTGCTGCNNNAGATCGGAAGA |
| 39036_2 | TATGAGTGTGGAGTCGTTGCTACTCACAATCTCCAGTTGCTGCNNNAGATCGGAAGA |
| 38407_1 | TATGAGTGTGGAGTCGTTGCGAGCAACCTTGACCAAACCAAATNNNAGATCGGAAG |
| 38407_2 | TATGAGTGTGGAGTCGTTGCGAGCAACCTTGACCAAACCAAATNNNAGATCGGAAG |
| 52485_1 | TATGAGTGTGGAGTCGTTGCAACATGGTGAAACCTGGTCTCTANNNAGATCGGAAG |
| 52485_2 | TATGAGTGTGGAGTCGTTGCAACATGGTGAAACCTGGTCTCTANNNAGATCGGAAG |
| 31424_1 | TATGAGTGTGGAGTCGTTGCTCTGGAATCTTTAGGCTTCCCAGNNNAGATCGGAAGA |
| 31424_2 | TATGAGTGTGGAGTCGTTGCTCTGGAATCTTTAGGCTTCCCAGNNNAGATCGGAAGA |
| 30530_1 | TATGAGTGTGGAGTCGTTGCTGACAGAGGGAGACGTAGGTGNNNAGATCGGAAGA |
| 30530_2 | TATGAGTGTGGAGTCGTTGCTGACAGAGGGAGACGTAGGTGNNNAGATCGGAAGA |
| 10295_1 | TATGAGTGTGGAGTCGTTGCAAAGGCCTGAAAGAATACAAACCANNNAGATCGGAA |
| 10295_2 | TATGAGTGTGGAGTCGTTGCAAAGGCCTGAAAGAATACAAACCANNNAGATCGGAA |
| 10379_1 | TATGAGTGTGGAGTCGTTGCTCTGGCCTAACACCAAAAGAGAANNNAGATCGGAAG |
| 10379_2 | TATGAGTGTGGAGTCGTTGCTCTGGCCTAACACCAAAAGAGAANNNAGATCGGAAG |
| 50671_1 | TATGAGTGTGGAGTCGTTGCGCCATAGACTCAGCTCTGTTCAANNNAGATCGGAAGA |
| 50671_2 | TATGAGTGTGGAGTCGTTGCGCCATAGACTCAGCTCTGTTCAANNNAGATCGGAAGA |
| 35752_1 | TATGAGTGTGGAGTCGTTGCGCTCGTGATCACAGAGGACATATNNNAGATCGGAAG |
| 35752_2 | TATGAGTGTGGAGTCGTTGCGCTCGTGATCACAGAGGACATATNNNAGATCGGAAG |
| 48784_1 | TATGAGTGTGGAGTCGTTGCCCCACATTACATTGAGTCATGCCNNNAGATCGGAAGA |
| 48784_2 | TATGAGTGTGGAGTCGTTGCCCCACATTACATTGAGTCATGCCNNNAGATCGGAAGA |
| 38723_1 | TATGAGTGTGGAGTCGTTGCTGAAGAGGCAACCCATAGATTGGNNNAGATCGGAAG |
| 38723_2 | TATGAGTGTGGAGTCGTTGCTGAAGAGGCAACCCATAGATTGGNNNAGATCGGAAG |
| 46574_1 | TATGAGTGTGGAGTCGTTGCGGTAGATGGCGGAAAAAAGAACANNNAGATCGGAA |
| 46574_2 | TATGAGTGTGGAGTCGTTGCGGTAGATGGCGGAAAAAAGAACANNNAGATCGGAA |
| 30855_1 | TATGAGTGTGGAGTCGTTGCCAAGCTATAGACTCAGGAACGCTNNNAGATCGGAAG |
| 30855_2 | TATGAGTGTGGAGTCGTTGCCAAGCTATAGACTCAGGAACGCTNNNAGATCGGAAG |
| 35768_1 | TATGAGTGTGGAGTCGTTGCACCTGCATAGTGACACATTGCTANNNAGATCGGAAG |

| | |
|---|---|
| 35768_2 | TATGAGTGTGGAGTCGTTGCACCTGCATAGTGACACATTGCTANNNAGATCGGAAG |
| 39115_1 | TATGAGTGTGGAGTCGTTGCAGCAAACTGAATCCATTAGCACANNNAGATCGGAAG |
| 39115_2 | TATGAGTGTGGAGTCGTTGCAGCAAACTGAATCCATTAGCACANNNAGATCGGAAG |
| 46456_1 | TATGAGTGTGGAGTCGTTGCCTGGAGTGAAGTGGTGCAATCTNNNAGATCGGAAG |
| 46456_2 | TATGAGTGTGGAGTCGTTGCCTGGAGTGAAGTGGTGCAATCTNNNAGATCGGAAG |
| 51648_1 | TATGAGTGTGGAGTCGTTGCCTGGTTATTTAGAAGCGTGTGGCNNNAGATCGGAA |
| 51648_2 | TATGAGTGTGGAGTCGTTGCCTGGTTATTTAGAAGCGTGTGGCNNNAGATCGGAA |
| 48902_1 | TATGAGTGTGGAGTCGTTGCATTGCTCTCTATGCTGGAAAACANNNAGATCGGAAG |
| 48902_2 | TATGAGTGTGGAGTCGTTGCATTGCTCTCTATGCTGGAAAACANNNAGATCGGAAG |
| 6259_1 | TATGAGTGTGGAGTCGTTGCGATATGGTGGCACCCAGCTACTCNNNAGATCGGAAG |
| 6259_2 | TATGAGTGTGGAGTCGTTGCGATATGGTGGCACCCAGCTACTCNNNAGATCGGAAG |
| 33430_1 | TATGAGTGTGGAGTCGTTGCCACCTGTAATCCCAGCACTTTTGNNNAGATCGGAAG |
| 33430_2 | TATGAGTGTGGAGTCGTTGCCACCTGTAATCCCAGCACTTTTGNNNAGATCGGAAG |
| 53247_1 | TATGAGTGTGGAGTCGTTGCAAGGGAATAGGAGTTGGACACACNNNAGATCGGA |
| 53247_2 | TATGAGTGTGGAGTCGTTGCAAGGGAATAGGAGTTGGACACACNNNAGATCGGA |
| 39533_1 | TATGAGTGTGGAGTCGTTGCTACAAACACACACACACACACACNNNAGATCGGAAG |
| 39533_2 | TATGAGTGTGGAGTCGTTGCTACAAACACACACACACACACACNNNAGATCGGAAG |
| 13439_1 | TATGAGTGTGGAGTCGTTGCTTATTTTGTTTGTTTCATTTGGGNNNAGATCGGAAG |
| 13439_2 | TATGAGTGTGGAGTCGTTGCTTATTTTGTTTGTTTCATTTGGGNNNAGATCGGAAG |
| 38443_1 | TATGAGTGTGGAGTCGTTGCGGGAGTCCTTGGTAGATAAGAGACNNNAGATCGGA |
| 38443_2 | TATGAGTGTGGAGTCGTTGCGGGAGTCCTTGGTAGATAAGAGACNNNAGATCGGA |
| 33646_1 | TATGAGTGTGGAGTCGTTGCCCATTCTCCTGCCTCAACCTCNNNAGATCGGAAGAGC |
| 33646_2 | TATGAGTGTGGAGTCGTTGCCCATTCTCCTGCCTCAACCTCNNNAGATCGGAAGAGC |
| 52446_1 | TATGAGTGTGGAGTCGTTGCGTTTTCCACTGGAATCTGAGGGANNNAGATCGGAA |
| 52446_2 | TATGAGTGTGGAGTCGTTGCGTTTTCCACTGGAATCTGAGGGANNNAGATCGGAA |
| 34958_1 | TATGAGTGTGGAGTCGTTGCAACCTCTGACTCCCTGGTTCAAGNNNAGATCGGAAG |
| 34958_2 | TATGAGTGTGGAGTCGTTGCAACCTCTGACTCCCTGGTTCAAGNNNAGATCGGAAG |
| 12042_1 | TATGAGTGTGGAGTCGTTGCAGGCTCACATACGTGGAAAGGAAGNNNAGATCGGA |
| 12042_2 | TATGAGTGTGGAGTCGTTGCAGGCTCACATACGTGGAAAGGAAGNNNAGATCGGA |
| 44650_1 | TATGAGTGTGGAGTCGTTGCAGGATACCTAGAGGGGGAATTGGNNNAGATCGGA |
| 44650_2 | TATGAGTGTGGAGTCGTTGCAGGATACCTAGAGGGGGAATTGGNNNAGATCGGA |
| 39576_1 | TATGAGTGTGGAGTCGTTGCGCAAAAGACTTGAAGAGGCACTTNNNAGATCGGAA |
| 39576_2 | TATGAGTGTGGAGTCGTTGCGCAAAAGACTTGAAGAGGCACTTNNNAGATCGGAA |
| 34311_1 | TATGAGTGTGGAGTCGTTGCTTCCTGTGTCCATGTGTTCTCATNNNAGATCGGAAGA |
| 34311_2 | TATGAGTGTGGAGTCGTTGCTTCCTGTGTCCATGTGTTCTCATNNNAGATCGGAAGA |
| 49747_1 | TATGAGTGTGGAGTCGTTGCCTGAAATGCTTGGGACTAGAAGCNNNAGATCGGAA |
| 49747_2 | TATGAGTGTGGAGTCGTTGCCTGAAATGCTTGGGACTAGAAGCNNNAGATCGGAA |
| 43121_1 | TATGAGTGTGGAGTCGTTGCAACTGAGGTGACAGTCAATGGAGNNNAGATCGGAA |
| 43121_2 | TATGAGTGTGGAGTCGTTGCAACTGAGGTGACAGTCAATGGAGNNNAGATCGGAA |
| 39749_1 | TATGAGTGTGGAGTCGTTGCATGTGACCAAGTGGAGTTTGTCTNNNAGATCGGAA |
| 39749_2 | TATGAGTGTGGAGTCGTTGCATGTGACCAAGTGGAGTTTGTCTNNNAGATCGGAA |
| 44902_1 | TATGAGTGTGGAGTCGTTGCGTGTGTATCTGAGGGTATGCAGTNNNAGATCGGAA |
| 44902_2 | TATGAGTGTGGAGTCGTTGCGTGTGTATCTGAGGGTATGCAGTNNNAGATCGGAA |

| | |
|---|---|
| 45415_1 | TATGAGTGTGGAGTCGTTGCTGATGCTTGAAAATCTCCTGGGANNNAGATCGGAAG |
| 45415_2 | TATGAGTGTGGAGTCGTTGCTGATGCTTGAAAATCTCCTGGGANNNAGATCGGAAG |
| 33498_1 | TATGAGTGTGGAGTCGTTGCCTTTGGTTCTAACAAAGGCCTGGNNNAGATCGGAAG |
| 33498_2 | TATGAGTGTGGAGTCGTTGCCTTTGGTTCTAACAAAGGCCTGGNNNAGATCGGAAG |
| 30327_1 | TATGAGTGTGGAGTCGTTGCGATGCAAAAGTGCTCAGTTTTATTTNNNAGATCGGA |
| 30327_2 | TATGAGTGTGGAGTCGTTGCGATGCAAAAGTGCTCAGTTTTATTTNNNAGATCGGA |
| 34784_1 | TATGAGTGTGGAGTCGTTGCCCCAAGTTCAAGCAATTCTCCTGNNNAGATCGGAAG |
| 34784_2 | TATGAGTGTGGAGTCGTTGCCCCAAGTTCAAGCAATTCTCCTGNNNAGATCGGAAG |
| 36792_1 | TATGAGTGTGGAGTCGTTGCTACGTTTTTACACTGGGCAGAGANNNAGATCGGAAG |
| 36792_2 | TATGAGTGTGGAGTCGTTGCTACGTTTTTACACTGGGCAGAGANNNAGATCGGAAG |
| 32687_1 | TATGAGTGTGGAGTCGTTGCCAATCTCAGCTCACTGCAAGTTCNNNAGATCGGAAG |
| 32687_2 | TATGAGTGTGGAGTCGTTGCCAATCTCAGCTCACTGCAAGTTCNNNAGATCGGAAG |
| 34292_1 | TATGAGTGTGGAGTCGTTGCTCCCCCTAGGTCTTTCTCATGATNNNAGATCGGAAGA |
| 34292_2 | TATGAGTGTGGAGTCGTTGCTCCCCCTAGGTCTTTCTCATGATNNNAGATCGGAAGA |
| 33601_1 | TATGAGTGTGGAGTCGTTGCCTCCTGAGTAGCTGGGACTACAGNNNAGATCGGAA |
| 33601_2 | TATGAGTGTGGAGTCGTTGCCTCCTGAGTAGCTGGGACTACAGNNNAGATCGGAA |
| 48362_1 | TATGAGTGTGGAGTCGTTGCCTCTCGAGTGGCTGGGACTATAGNNNAGATCGGAA |
| 48362_2 | TATGAGTGTGGAGTCGTTGCCTCTCGAGTGGCTGGGACTATAGNNNAGATCGGAA |
| 51425_1 | TATGAGTGTGGAGTCGTTGCGAAGACGGGTGATTTCTGCATTTNNNAGATCGGAA |
| 51425_2 | TATGAGTGTGGAGTCGTTGCGAAGACGGGTGATTTCTGCATTTNNNAGATCGGAA |
| 38622_1 | TATGAGTGTGGAGTCGTTGCTCATGAGGTCAGGAGTTTGAGACNNNAGATCGGAA |
| 38622_2 | TATGAGTGTGGAGTCGTTGCTCATGAGGTCAGGAGTTTGAGACNNNAGATCGGAA |
| 37203_1 | TATGAGTGTGGAGTCGTTGCGCAGAAGTGGCCATACTTTATCACNNNAGATCGGAA |
| 37203_2 | TATGAGTGTGGAGTCGTTGCGCAGAAGTGGCCATACTTTATCACNNNAGATCGGAA |
| 38981_1 | TATGAGTGTGGAGTCGTTGCAACGAACAGGCAAAACATCAACANNNAGATCGGAAG |
| 38981_2 | TATGAGTGTGGAGTCGTTGCAACGAACAGGCAAAACATCAACANNNAGATCGGAAG |
| 36413_1 | TATGAGTGTGGAGTCGTTGCGCGAGTCACTTAGTTTCTCCTGTNNNAGATCGGAAG |
| 36413_2 | TATGAGTGTGGAGTCGTTGCGCGAGTCACTTAGTTTCTCCTGTNNNAGATCGGAAG |
| 31824_1 | TATGAGTGTGGAGTCGTTGCATCAACAGAATTTGGACACAGCANNNAGATCGGAAG |
| 31824_2 | TATGAGTGTGGAGTCGTTGCATCAACAGAATTTGGACACAGCANNNAGATCGGAAG |
| 52808_1 | TATGAGTGTGGAGTCGTTGCCTTCTTGGCTCGGTGTCCTGNNNAGATCGGAAGAGC |
| 52808_2 | TATGAGTGTGGAGTCGTTGCCTTCTTGGCTCGGTGTCCTGNNNAGATCGGAAGAGC |
| 48859_1 | TATGAGTGTGGAGTCGTTGCGGTTCAAGCGATTCTATTGCCTCNNNAGATCGGAAG |
| 48859_2 | TATGAGTGTGGAGTCGTTGCGGTTCAAGCGATTCTATTGCCTCNNNAGATCGGAAG |
| 52032_1 | TATGAGTGTGGAGTCGTTGCAACTCTGCAGTTGTCAGCTAAATNNNAGATCGGAAG |
| 52032_2 | TATGAGTGTGGAGTCGTTGCAACTCTGCAGTTGTCAGCTAAATNNNAGATCGGAAG |
| 12949_1 | TATGAGTGTGGAGTCGTTGCATTGGAATGTTTGTTAATCACAANNNAGATCGGAAG |
| 12949_2 | TATGAGTGTGGAGTCGTTGCATTGGAATGTTTGTTAATCACAANNNAGATCGGAAG |
| 43372_1 | TATGAGTGTGGAGTCGTTGCCACCACAGCCAGCTCATTTTTTTNNNAGATCGGAAGA |
| 43372_2 | TATGAGTGTGGAGTCGTTGCCACCACAGCCAGCTCATTTTTTTNNNAGATCGGAAGA |
| 38234_1 | TATGAGTGTGGAGTCGTTGCTCTATGAGGCCAGCATCATTCTGNNNAGATCGGAAG |
| 38234_2 | TATGAGTGTGGAGTCGTTGCTCTATGAGGCCAGCATCATTCTGNNNAGATCGGAAG |
| 40697_1 | TATGAGTGTGGAGTCGTTGCAAACAAAGGCATGAAAAGGGGAGNNNAGATCGGA |

| | |
|---|---|
| 40697_2 | TATGAGTGTGGAGTCGTTGCAAACAAAGGCATGAAAAGGGGAGNNNAGATCGGA |
| 46188_1 | TATGAGTGTGGAGTCGTTGCGGCGATGAAGAGACTTTTCAAACANNNAGATCGGA |
| 46188_2 | TATGAGTGTGGAGTCGTTGCGGCGATGAAGAGACTTTTCAAACANNNAGATCGGA |
| 47736_1 | TATGAGTGTGGAGTCGTTGCGTTTCACCATGTTAGCCAGGATGNNNAGATCGGAAG |
| 47736_2 | TATGAGTGTGGAGTCGTTGCGTTTCACCATGTTAGCCAGGATGNNNAGATCGGAAG |
| 28962_1 | TATGAGTGTGGAGTCGTTGCGTGTCACACATACATACGCACGNNNAGATCGGAAGA |
| 28962_2 | TATGAGTGTGGAGTCGTTGCGTGTCACACATACATACGCACGNNNAGATCGGAAGA |
| 47834_1 | TATGAGTGTGGAGTCGTTGCGATCTCGGCTCACTGAAAACTCCNNNAGATCGGAAG |
| 47834_2 | TATGAGTGTGGAGTCGTTGCGATCTCGGCTCACTGAAAACTCCNNNAGATCGGAAG |
| 8337_1 | TATGAGTGTGGAGTCGTTGCACAACTATGTCCCTTCCCCTAGGNNNAGATCGGAAG |
| 8337_2 | TATGAGTGTGGAGTCGTTGCACAACTATGTCCCTTCCCCTAGGNNNAGATCGGAAG |
| 13009_1 | TATGAGTGTGGAGTCGTTGCCTTTAGGAATGCTGTGTGTCNNNAGATCGGAAGAG |
| 13009_2 | TATGAGTGTGGAGTCGTTGCCTTTAGGAATGCTGTGTGTCNNNAGATCGGAAGAG |
| 29562_1 | TATGAGTGTGGAGTCGTTGCCCGGCAGAAGGATAACCTCAATANNNAGATCGGAA |
| 29562_2 | TATGAGTGTGGAGTCGTTGCCCGGCAGAAGGATAACCTCAATANNNAGATCGGAA |
| 47526_1 | TATGAGTGTGGAGTCGTTGCCAAGTGATTCTCTTGCTTCAGCCNNNAGATCGGAAG |
| 47526_2 | TATGAGTGTGGAGTCGTTGCCAAGTGATTCTCTTGCTTCAGCCNNNAGATCGGAAG |
| 37625_1 | TATGAGTGTGGAGTCGTTGCCAGGTTTTCAGCCTTCCATATGCNNNAGATCGGAAG |
| 37625_2 | TATGAGTGTGGAGTCGTTGCCAGGTTTTCAGCCTTCCATATGCNNNAGATCGGAAG |
| 45902_1 | TATGAGTGTGGAGTCGTTGCTGCCTTTTGGGTTCAAGTGATTCNNNAGATCGGAAG |
| 45902_2 | TATGAGTGTGGAGTCGTTGCTGCCTTTTGGGTTCAAGTGATTCNNNAGATCGGAAG |
| 34827_1 | TATGAGTGTGGAGTCGTTGCCCATGCCAGGCTCATTTTTTGTANNNAGATCGGAAG |
| 34827_2 | TATGAGTGTGGAGTCGTTGCCCATGCCAGGCTCATTTTTTGTANNNAGATCGGAAG |
| 49619_1 | TATGAGTGTGGAGTCGTTGCTTTCCTGGGCTCAGATGATTTGANNNAGATCGGAAG |
| 49619_2 | TATGAGTGTGGAGTCGTTGCTTTCCTGGGCTCAGATGATTTGANNNAGATCGGAAG |
| 43283_1 | TATGAGTGTGGAGTCGTTGCTGGCCGGGATTTCTAGTATGATGNNNAGATCGGAA |
| 43283_2 | TATGAGTGTGGAGTCGTTGCTGGCCGGGATTTCTAGTATGATGNNNAGATCGGAA |
| 33986_1 | TATGAGTGTGGAGTCGTTGCCTTCCAAGTAGCTGGGACTACGNNNAGATCGGAAGA |
| 33986_2 | TATGAGTGTGGAGTCGTTGCCTTCCAAGTAGCTGGGACTACGNNNAGATCGGAAGA |
| 39638_1 | TATGAGTGTGGAGTCGTTGCTGGAATACCGTTTCACCCATCTTNNNAGATCGGAAG |
| 39638_2 | TATGAGTGTGGAGTCGTTGCTGGAATACCGTTTCACCCATCTTNNNAGATCGGAAG |
| 13121_1 | TATGAGTGTGGAGTCGTTGCTCATTAGCATTAAGAGCATAAACNNNAGATCGGAAG |
| 13121_2 | TATGAGTGTGGAGTCGTTGCTCATTAGCATTAAGAGCATAAACNNNAGATCGGAAG |
| 10499_1 | TATGAGTGTGGAGTCGTTGCTTGTCCCCAGAAAACAACCCATTNNNAGATCGGAAG |
| 10499_2 | TATGAGTGTGGAGTCGTTGCTTGTCCCCAGAAAACAACCCATTNNNAGATCGGAAG |
| 7602_1 | TATGAGTGTGGAGTCGTTGCCTTGAGCCTGGGTGGGAGACNNNAGATCGGAAGA |
| 7602_2 | TATGAGTGTGGAGTCGTTGCCTTGAGCCTGGGTGGGAGACNNNAGATCGGAAGA |
| 32633_1 | TATGAGTGTGGAGTCGTTGCGTTCATGCCATTCTCCTACCTCANNNAGATCGGAAGA |
| 32633_2 | TATGAGTGTGGAGTCGTTGCGTTCATGCCATTCTCCTACCTCANNNAGATCGGAAGA |
| 42297_1 | TATGAGTGTGGAGTCGTTGCATAGCTCACTGCAATCTCCACCNNNAGATCGGAAGA |
| 42297_2 | TATGAGTGTGGAGTCGTTGCATAGCTCACTGCAATCTCCACCNNNAGATCGGAAGA |
| 44109_1 | TATGAGTGTGGAGTCGTTGCTTCTGTGTCTGTGGTTACTAGGCNNNAGATCGGAAG |
| 44109_2 | TATGAGTGTGGAGTCGTTGCTTCTGTGTCTGTGGTTACTAGGCNNNAGATCGGAAG |

| | |
|---|---|
| 48158_1 | TATGAGTGTGGAGTCGTTGCTAGTAGAGACAGGGTTTCGTCCTNNNAGATCGGAA |
| 48158_2 | TATGAGTGTGGAGTCGTTGCTAGTAGAGACAGGGTTTCGTCCTNNNAGATCGGAA |
| 49051_1 | TATGAGTGTGGAGTCGTTGCTACAATCTCGGCTCACTGTAACCNNNAGATCGGAAG |
| 49051_2 | TATGAGTGTGGAGTCGTTGCTACAATCTCGGCTCACTGTAACCNNNAGATCGGAAG |
| 42842_1 | TATGAGTGTGGAGTCGTTGCTTACAGGCTCATGAACCACCTAGNNNAGATCGGAAG |
| 42842_2 | TATGAGTGTGGAGTCGTTGCTTACAGGCTCATGAACCACCTAGNNNAGATCGGAAG |
| 34084_1 | TATGAGTGTGGAGTCGTTGCGCTCAAGCCTTCTTCCTAACTCANNNAGATCGGAAGA |
| 34084_2 | TATGAGTGTGGAGTCGTTGCGCTCAAGCCTTCTTCCTAACTCANNNAGATCGGAAGA |
| 38037_1 | TATGAGTGTGGAGTCGTTGCTGGCAGTATCGCTGAGTTTTAGTNNNAGATCGGAA |
| 38037_2 | TATGAGTGTGGAGTCGTTGCTGGCAGTATCGCTGAGTTTTAGTNNNAGATCGGAA |
| 13519_1 | TATGAGTGTGGAGTCGTTGCCTTAATCCTCAAAACGGTAAATGNNNAGATCGGAAG |
| 13519_2 | TATGAGTGTGGAGTCGTTGCCTTAATCCTCAAAACGGTAAATGNNNAGATCGGAAG |
| 52064_1 | TATGAGTGTGGAGTCGTTGCTTTTTTGTGATGGAGTCTCGCTCNNNAGATCGGAAG |
| 52064_2 | TATGAGTGTGGAGTCGTTGCTTTTTTGTGATGGAGTCTCGCTCNNNAGATCGGAAG |
| 31047_1 | TATGAGTGTGGAGTCGTTGCGGGTTGACTAGAGACTCAGCATTNNNAGATCGGAA |
| 31047_2 | TATGAGTGTGGAGTCGTTGCGGGTTGACTAGAGACTCAGCATTNNNAGATCGGAA |
| 49040_1 | TATGAGTGTGGAGTCGTTGCTTGAATACCCATTGCCCACTCTTNNNAGATCGGAAGA |
| 49040_2 | TATGAGTGTGGAGTCGTTGCTTGAATACCCATTGCCCACTCTTNNNAGATCGGAAGA |
| 31665_1 | TATGAGTGTGGAGTCGTTGCTTAGAAGCAATGACATCCCAGCANNNAGATCGGAAG |
| 31665_2 | TATGAGTGTGGAGTCGTTGCTTAGAAGCAATGACATCCCAGCANNNAGATCGGAAG |
| 46154_1 | TATGAGTGTGGAGTCGTTGCGAGTCTCTATCACCCAGGTTGGANNNAGATCGGAAG |
| 46154_2 | TATGAGTGTGGAGTCGTTGCGAGTCTCTATCACCCAGGTTGGANNNAGATCGGAAG |
| 53392_1 | TATGAGTGTGGAGTCGTTGCCTCCCAGGTTCAAGTGATTCTCANNNAGATCGGAAG |
| 53392_2 | TATGAGTGTGGAGTCGTTGCCTCCCAGGTTCAAGTGATTCTCANNNAGATCGGAAG |
| 38982_1 | TATGAGTGTGGAGTCGTTGCTGGGAGAAAAATGTTTGCAAATCANNNAGATCGGA |
| 38982_2 | TATGAGTGTGGAGTCGTTGCTGGGAGAAAAATGTTTGCAAATCANNNAGATCGGA |
| 31251_1 | TATGAGTGTGGAGTCGTTGCGAGTCATTGTGGCAGAAACAGACNNNAGATCGGAA |
| 31251_2 | TATGAGTGTGGAGTCGTTGCGAGTCATTGTGGCAGAAACAGACNNNAGATCGGAA |
| 50595_1 | TATGAGTGTGGAGTCGTTGCACAGTAAAGAGGCCAACCAGAATNNNAGATCGGAA |
| 50595_2 | TATGAGTGTGGAGTCGTTGCACAGTAAAGAGGCCAACCAGAATNNNAGATCGGAA |
| 43216_1 | TATGAGTGTGGAGTCGTTGCAACTGCACTTTGAGAACCATTGGNNNAGATCGGAAG |
| 43216_2 | TATGAGTGTGGAGTCGTTGCAACTGCACTTTGAGAACCATTGGNNNAGATCGGAAG |
| 2995_1 | TATGAGTGTGGAGTCGTTGCGGGATATTACTCGGTCCAGGTNNNAGATCGGAAGA |
| 2995_2 | TATGAGTGTGGAGTCGTTGCGGGATATTACTCGGTCCAGGTNNNAGATCGGAAGA |
| 12858_1 | TATGAGTGTGGAGTCGTTGCAGATTGTATCTCCAAAGGTTAATNNNAGATCGGAAG |
| 12858_2 | TATGAGTGTGGAGTCGTTGCAGATTGTATCTCCAAAGGTTAATNNNAGATCGGAAG |
| 47608_1 | TATGAGTGTGGAGTCGTTGCCCACCATGCCCAGCTAATTTTTTNNNAGATCGGAAGA |
| 47608_2 | TATGAGTGTGGAGTCGTTGCCCACCATGCCCAGCTAATTTTTTNNNAGATCGGAAGA |
| 43549_1 | TATGAGTGTGGAGTCGTTGCCTAGTCAATATGGTCCAGCCCTCNNNAGATCGGAAG |
| 43549_2 | TATGAGTGTGGAGTCGTTGCCTAGTCAATATGGTCCAGCCCTCNNNAGATCGGAAG |
| 51958_1 | TATGAGTGTGGAGTCGTTGCTTGTCTCATTCACCTTGTCCTGTNNNAGATCGGAAGA |
| 51958_2 | TATGAGTGTGGAGTCGTTGCTTGTCTCATTCACCTTGTCCTGTNNNAGATCGGAAGA |
| 40744_1 | TATGAGTGTGGAGTCGTTGCTGCAAGATGGGTATCCTAGACTCNNNAGATCGGAA |

| | |
|---|---|
| 40744_2 | TATGAGTGTGGAGTCGTTGCTGCAAGATGGGTATCCTAGACTCNNNAGATCGGAAG |
| 40191_1 | TATGAGTGTGGAGTCGTTGCGGATCACTTGAGGTCAGGAGTTCNNNAGATCGGAA |
| 40191_2 | TATGAGTGTGGAGTCGTTGCGGATCACTTGAGGTCAGGAGTTCNNNAGATCGGAA |
| 49080_1 | TATGAGTGTGGAGTCGTTGCTGCCTCAGTTTCAGAACTCGTTANNNAGATCGGAAGA |
| 49080_2 | TATGAGTGTGGAGTCGTTGCTGCCTCAGTTTCAGAACTCGTTANNNAGATCGGAAGA |
| 13078_1 | TATGAGTGTGGAGTCGTTGCTCTAAGAAAAAATGTAAAAGCCCNNNAGATCGGAAG |
| 13078_2 | TATGAGTGTGGAGTCGTTGCTCTAAGAAAAAATGTAAAAGCCCNNNAGATCGGAAG |
| 32125_1 | TATGAGTGTGGAGTCGTTGCCCTCCCAGGTTCAAGCTATTCTTNNNAGATCGGAAGA |
| 32125_2 | TATGAGTGTGGAGTCGTTGCCCTCCCAGGTTCAAGCTATTCTTNNNAGATCGGAAGA |
| 37029_1 | TATGAGTGTGGAGTCGTTGCTCCCTTTTCTCCATATCCTTGCCNNNAGATCGGAAGA |
| 37029_2 | TATGAGTGTGGAGTCGTTGCTCCCTTTTCTCCATATCCTTGCCNNNAGATCGGAAGA |
| 46205_1 | TATGAGTGTGGAGTCGTTGCAGTAGTTGGGAATTACAGGTGCCNNNAGATCGGAA |
| 46205_2 | TATGAGTGTGGAGTCGTTGCAGTAGTTGGGAATTACAGGTGCCNNNAGATCGGAA |
| 35480_1 | TATGAGTGTGGAGTCGTTGCAGCCACCACATCTTTCCTACAATNNNAGATCGGAAGA |
| 35480_2 | TATGAGTGTGGAGTCGTTGCAGCCACCACATCTTTCCTACAATNNNAGATCGGAAGA |
| 47087_1 | TATGAGTGTGGAGTCGTTGCCAGCCATTTTGGTCTTCTTCCTGNNNAGATCGGAAGA |
| 47087_2 | TATGAGTGTGGAGTCGTTGCCAGCCATTTTGGTCTTCTTCCTGNNNAGATCGGAAGA |
| 41752_1 | TATGAGTGTGGAGTCGTTGCCAAAGCTCTGGGATTACAGGCTANNNAGATCGGAAG |
| 41752_2 | TATGAGTGTGGAGTCGTTGCCAAAGCTCTGGGATTACAGGCTANNNAGATCGGAAG |
| 127_1 | TATGAGTGTGGAGTCGTTGCCGGTGGCTGTGGCAGGTGAGNNNAGATCGGAAGA |
| 127_2 | TATGAGTGTGGAGTCGTTGCCGGTGGCTGTGGCAGGTGAGNNNAGATCGGAAGA |
| 32620_1 | TATGAGTGTGGAGTCGTTGCCTCTTCAGGACAGTCAGGGAAAGNNNAGATCGGAA |
| 32620_2 | TATGAGTGTGGAGTCGTTGCCTCTTCAGGACAGTCAGGGAAAGNNNAGATCGGAA |
| 49429_1 | TATGAGTGTGGAGTCGTTGCGAGACAGTTTTGCTCTTGTTGCCNNNAGATCGGAAG |
| 49429_2 | TATGAGTGTGGAGTCGTTGCGAGACAGTTTTGCTCTTGTTGCCNNNAGATCGGAAG |
| 40738_1 | TATGAGTGTGGAGTCGTTGCAACAACAAACAAAGGCTGCCTAANNNAGATCGGAAG |
| 40738_2 | TATGAGTGTGGAGTCGTTGCAACAACAAACAAAGGCTGCCTAANNNAGATCGGAAG |
| 51159_1 | TATGAGTGTGGAGTCGTTGCCCATCATGGCTGAACCTCTATCTNNNAGATCGGAAGA |
| 51159_2 | TATGAGTGTGGAGTCGTTGCCCATCATGGCTGAACCTCTATCTNNNAGATCGGAAGA |
| 33158_1 | TATGAGTGTGGAGTCGTTGCGAATGGGAGTGGTAGTGGTAGTGNNNAGATCGGAA |
| 33158_2 | TATGAGTGTGGAGTCGTTGCGAATGGGAGTGGTAGTGGTAGTGNNNAGATCGGAA |
| 32719_1 | TATGAGTGTGGAGTCGTTGCGAGTTGGGAGTCACTGTACACAANNNAGATCGGAA |
| 32719_2 | TATGAGTGTGGAGTCGTTGCGAGTTGGGAGTCACTGTACACAANNNAGATCGGAA |
| 48790_1 | TATGAGTGTGGAGTCGTTGCTGGGTTCAAGTGATTCTCCTACCNNNAGATCGGAAGA |
| 48790_2 | TATGAGTGTGGAGTCGTTGCTGGGTTCAAGTGATTCTCCTACCNNNAGATCGGAAGA |
| 36747_1 | TATGAGTGTGGAGTCGTTGCGGGGCCATAAGGATGTTCTTACANNNAGATCGGAA |
| 36747_2 | TATGAGTGTGGAGTCGTTGCGGGGCCATAAGGATGTTCTTACANNNAGATCGGAA |
| 52671_1 | TATGAGTGTGGAGTCGTTGCTTCAGACACTATGGCCCAGAAAANNNAGATCGGAAG |
| 52671_2 | TATGAGTGTGGAGTCGTTGCTTCAGACACTATGGCCCAGAAAANNNAGATCGGAAG |
| 49294_1 | TATGAGTGTGGAGTCGTTGCTTTGAGACATGGTCTTGCTCTGTNNNAGATCGGAAG |
| 49294_2 | TATGAGTGTGGAGTCGTTGCTTTGAGACATGGTCTTGCTCTGTNNNAGATCGGAAG |
| 51523_1 | TATGAGTGTGGAGTCGTTGCACCTTGGCTACTGGAAGACATTTNNNAGATCGGAAG |
| 51523_2 | TATGAGTGTGGAGTCGTTGCACCTTGGCTACTGGAAGACATTTNNNAGATCGGAAG |

| | |
|---|---|
| 8708_1 | TATGAGTGTGGAGTCGTTGCCAGCATCCCTTTTAGCCAAAACGNNNAGATCGGAAG |
| 8708_2 | TATGAGTGTGGAGTCGTTGCCAGCATCCCTTTTAGCCAAAACGNNNAGATCGGAAG |
| 36250_1 | TATGAGTGTGGAGTCGTTGCAGAGGATTACTTGTGCCTGGAAGNNNAGATCGGAA |
| 36250_2 | TATGAGTGTGGAGTCGTTGCAGAGGATTACTTGTGCCTGGAAGNNNAGATCGGAA |
| 31222_1 | TATGAGTGTGGAGTCGTTGCCTCTCTCTCTCAGATTGCTCACCNNNAGATCGGAAGA |
| 31222_2 | TATGAGTGTGGAGTCGTTGCCTCTCTCTCTCAGATTGCTCACCNNNAGATCGGAAGA |
| 10973_1 | TATGAGTGTGGAGTCGTTGCTTTATTGCTGGTGGGGACTTTGCNNNAGATCGGAAG |
| 10973_2 | TATGAGTGTGGAGTCGTTGCTTTATTGCTGGTGGGGACTTTGCNNNAGATCGGAAG |
| 49861_1 | TATGAGTGTGGAGTCGTTGCGCAGTGTGGAGACTTATGCAAACNNNAGATCGGAA |
| 49861_2 | TATGAGTGTGGAGTCGTTGCGCAGTGTGGAGACTTATGCAAACNNNAGATCGGAA |
| 10634_1 | TATGAGTGTGGAGTCGTTGCGAACTGTTGAAGCACCTGCTTTGNNNAGATCGGAAG |
| 10634_2 | TATGAGTGTGGAGTCGTTGCGAACTGTTGAAGCACCTGCTTTGNNNAGATCGGAAG |
| 35219_1 | TATGAGTGTGGAGTCGTTGCGTCCAGAACCCAGTTACTGTGATNNNAGATCGGAAG |
| 35219_2 | TATGAGTGTGGAGTCGTTGCGTCCAGAACCCAGTTACTGTGATNNNAGATCGGAAG |
| 46846_1 | TATGAGTGTGGAGTCGTTGCCTCTCAGCAGCTTAGTCCTCAAANNNAGATCGGAAG |
| 46846_2 | TATGAGTGTGGAGTCGTTGCCTCTCAGCAGCTTAGTCCTCAAANNNAGATCGGAAG |
| 36736_1 | TATGAGTGTGGAGTCGTTGCCCTTCCTAGATGGTGAGACCCTANNNAGATCGGAAG |
| 36736_2 | TATGAGTGTGGAGTCGTTGCCCTTCCTAGATGGTGAGACCCTANNNAGATCGGAAG |
| 32036_1 | TATGAGTGTGGAGTCGTTGCTCCCGAGAACAATGGGCATTATTNNNAGATCGGAAG |
| 32036_2 | TATGAGTGTGGAGTCGTTGCTCCCGAGAACAATGGGCATTATTNNNAGATCGGAAG |
| 33007_1 | TATGAGTGTGGAGTCGTTGCGAGACGGAGTTTCATCATGTTGCNNNAGATCGGAA |
| 33007_2 | TATGAGTGTGGAGTCGTTGCGAGACGGAGTTTCATCATGTTGCNNNAGATCGGAA |
| 40492_1 | TATGAGTGTGGAGTCGTTGCCACCTAGAAGGGGGCCAATAAATNNNAGATCGGAA |
| 40492_2 | TATGAGTGTGGAGTCGTTGCCACCTAGAAGGGGGCCAATAAATNNNAGATCGGAA |
| 43754_1 | TATGAGTGTGGAGTCGTTGCCCTCCACCACTAGCACACACNNNAGATCGGAAGAGC |
| 43754_2 | TATGAGTGTGGAGTCGTTGCCCTCCACCACTAGCACACACNNNAGATCGGAAGAGC |
| 32166_1 | TATGAGTGTGGAGTCGTTGCCTCCTGGGTTCACACCATTCTCNNNAGATCGGAAGAG |
| 32166_2 | TATGAGTGTGGAGTCGTTGCCTCCTGGGTTCACACCATTCTCNNNAGATCGGAAGAG |
| 5641_1 | TATGAGTGTGGAGTCGTTGCTCACCCCACTCTGTAACCTAGTTNNNAGATCGGAAGA |
| 5641_2 | TATGAGTGTGGAGTCGTTGCTCACCCCACTCTGTAACCTAGTTNNNAGATCGGAAGA |
| 41419_1 | TATGAGTGTGGAGTCGTTGCTCCTCTGACACAATTCTTGGTGTNNNAGATCGGAAG |
| 41419_2 | TATGAGTGTGGAGTCGTTGCTCCTCTGACACAATTCTTGGTGTNNNAGATCGGAAG |
| 47778_1 | TATGAGTGTGGAGTCGTTGCCTCCTGAGTAGCTGGGACTACAGNNNAGATCGGAA |
| 47778_2 | TATGAGTGTGGAGTCGTTGCCTCCTGAGTAGCTGGGACTACAGNNNAGATCGGAA |
| 31335_1 | TATGAGTGTGGAGTCGTTGCAACCTTGCAATGTGTTAGCTTCCNNNAGATCGGAAG |
| 31335_2 | TATGAGTGTGGAGTCGTTGCAACCTTGCAATGTGTTAGCTTCCNNNAGATCGGAAG |
| 41292_1 | TATGAGTGTGGAGTCGTTGCTGGGATTATGTAAAGCAACCAAACCNNNAGATCGG |
| 41292_2 | TATGAGTGTGGAGTCGTTGCTGGGATTATGTAAAGCAACCAAACCNNNAGATCGG |
| 47634_1 | TATGAGTGTGGAGTCGTTGCTCCACGGAGAATCAAAAGAGAGGNNNAGATCGGAA |
| 47634_2 | TATGAGTGTGGAGTCGTTGCTCCACGGAGAATCAAAAGAGAGGNNNAGATCGGAA |
| 46014_1 | TATGAGTGTGGAGTCGTTGCGCCTCCCAAGTATCTGGAACTACNNNAGATCGGAAG |
| 46014_2 | TATGAGTGTGGAGTCGTTGCGCCTCCCAAGTATCTGGAACTACNNNAGATCGGAAG |
| 51810_1 | TATGAGTGTGGAGTCGTTGCTAGACCGGTACAAGGCTTAGAGANNNAGATCGGAA |

| | |
|---|---|
| 51810_2 | TATGAGTGTGGAGTCGTTGCTAGACCGGTACAAGGCTTAGAGANNNAGATCGGAA |
| 33494_1 | TATGAGTGTGGAGTCGTTGCTTGAAACCACACAGCAACTTTGTNNNAGATCGGAAG |
| 33494_2 | TATGAGTGTGGAGTCGTTGCTTGAAACCACACAGCAACTTTGTNNNAGATCGGAAG |
| 36071_1 | TATGAGTGTGGAGTCGTTGCATGATGGCATATGCTTCTGGTCTNNNAGATCGGAAG |
| 36071_2 | TATGAGTGTGGAGTCGTTGCATGATGGCATATGCTTCTGGTCTNNNAGATCGGAAG |
| 47217_1 | TATGAGTGTGGAGTCGTTGCCTCCTAAGTGGCTGGGATTACAGNNNAGATCGGAA |
| 47217_2 | TATGAGTGTGGAGTCGTTGCCTCCTAAGTGGCTGGGATTACAGNNNAGATCGGAA |
| 48134_1 | TATGAGTGTGGAGTCGTTGCCACGCCCGGCTAATTTTTTTTTTNNNAGATCGGAAGA |
| 48134_2 | TATGAGTGTGGAGTCGTTGCCACGCCCGGCTAATTTTTTTTTTNNNAGATCGGAAGA |
| 10256_1 | TATGAGTGTGGAGTCGTTGCGCATACATCAAGAAAGTGCAACGGNNNAGATCGGA |
| 10256_2 | TATGAGTGTGGAGTCGTTGCGCATACATCAAGAAAGTGCAACGGNNNAGATCGGA |
| 45753_1 | TATGAGTGTGGAGTCGTTGCTTCAAGCGATTCTCCTGTCTCAGNNNAGATCGGAAG |
| 45753_2 | TATGAGTGTGGAGTCGTTGCTTCAAGCGATTCTCCTGTCTCAGNNNAGATCGGAAG |
| 29179_1 | TATGAGTGTGGAGTCGTTGCTCTATTTGTTTGGATGCCCTGGANNNAGATCGGAAG |
| 29179_2 | TATGAGTGTGGAGTCGTTGCTCTATTTGTTTGGATGCCCTGGANNNAGATCGGAAG |
| 33784_1 | TATGAGTGTGGAGTCGTTGCCGAGGCAGGATATAGTGACTCTGNNNAGATCGGAA |
| 33784_2 | TATGAGTGTGGAGTCGTTGCCGAGGCAGGATATAGTGACTCTGNNNAGATCGGAA |
| 52260_1 | TATGAGTGTGGAGTCGTTGCTGAAGCCATCCCAAATTTTGCAANNNAGATCGGAAG |
| 52260_2 | TATGAGTGTGGAGTCGTTGCTGAAGCCATCCCAAATTTTGCAANNNAGATCGGAAG |
| 47619_1 | TATGAGTGTGGAGTCGTTGCTCGAACTCGTCACTCAGGTAATGNNNAGATCGGAAG |
| 47619_2 | TATGAGTGTGGAGTCGTTGCTCGAACTCGTCACTCAGGTAATGNNNAGATCGGAAG |
| 32725_1 | TATGAGTGTGGAGTCGTTGCATAGCCCTTACTCCCCAAAACAGNNNAGATCGGAAG |
| 32725_2 | TATGAGTGTGGAGTCGTTGCATAGCCCTTACTCCCCAAAACAGNNNAGATCGGAAG |
| 38823_1 | TATGAGTGTGGAGTCGTTGCATGCAAACTAAGATGGCAACGAGNNNAGATCGGAA |
| 38823_2 | TATGAGTGTGGAGTCGTTGCATGCAAACTAAGATGGCAACGAGNNNAGATCGGAA |
| 41676_1 | TATGAGTGTGGAGTCGTTGCCCACCACACCTGGCTAATTTTTTNNNAGATCGGAAGA |
| 41676_2 | TATGAGTGTGGAGTCGTTGCCCACCACACCTGGCTAATTTTTTNNNAGATCGGAAGA |
| 37914_1 | TATGAGTGTGGAGTCGTTGCCCTAATGTGTGTGTACTGTGTGCNNNAGATCGGAAG |
| 37914_2 | TATGAGTGTGGAGTCGTTGCCCTAATGTGTGTGTACTGTGTGCNNNAGATCGGAAG |
| 37872_1 | TATGAGTGTGGAGTCGTTGCCAGGAGACAGAGAAAGACCCTTCNNNAGATCGGAA |
| 37872_2 | TATGAGTGTGGAGTCGTTGCCAGGAGACAGAGAAAGACCCTTCNNNAGATCGGAA |
| 31664_1 | TATGAGTGTGGAGTCGTTGCATTTGACAAGGGCACCAAGAAAANNNAGATCGGAA |
| 31664_2 | TATGAGTGTGGAGTCGTTGCATTTGACAAGGGCACCAAGAAAANNNAGATCGGAA |
| 39179_1 | TATGAGTGTGGAGTCGTTGCACATGCTGCTTCCAAGTTTTGTTNNNAGATCGGAAG |
| 39179_2 | TATGAGTGTGGAGTCGTTGCACATGCTGCTTCCAAGTTTTGTTNNNAGATCGGAAG |
| 48102_1 | TATGAGTGTGGAGTCGTTGCTTTCAGCAGCTTGACATGGTTTTNNNAGATCGGAAG |
| 48102_2 | TATGAGTGTGGAGTCGTTGCTTTCAGCAGCTTGACATGGTTTTNNNAGATCGGAAG |
| 31926_1 | TATGAGTGTGGAGTCGTTGCACTGAGCAATGCATAGGTTCAGANNNAGATCGGAA |
| 31926_2 | TATGAGTGTGGAGTCGTTGCACTGAGCAATGCATAGGTTCAGANNNAGATCGGAA |
| 34157_1 | TATGAGTGTGGAGTCGTTGCCTTTCAACGAATCCATGCCATGTNNNAGATCGGAAG |
| 34157_2 | TATGAGTGTGGAGTCGTTGCCTTTCAACGAATCCATGCCATGTNNNAGATCGGAAG |
| 38637_1 | TATGAGTGTGGAGTCGTTGCGAAACCCCATACCCATTAGCAGTNNNAGATCGGAAG |
| 38637_2 | TATGAGTGTGGAGTCGTTGCGAAACCCCATACCCATTAGCAGTNNNAGATCGGAAG |

38825_1 TATGAGTGTGGAGTCGTTGCACAAAATAGACCGCTAGCAAGACNNNAGATCGGAA
38825_2 TATGAGTGTGGAGTCGTTGCACAAAATAGACCGCTAGCAAGACNNNAGATCGGAA
40451_1 TATGAGTGTGGAGTCGTTGCACTGGTAGTTCCCCCACAAAAATNNNAGATCGGAAG
40451_2 TATGAGTGTGGAGTCGTTGCACTGGTAGTTCCCCCACAAAAATNNNAGATCGGAAG
37399_1 TATGAGTGTGGAGTCGTTGCAAAATGGGGAGCTGATGAGAGTTNNNAGATCGGA
37399_2 TATGAGTGTGGAGTCGTTGCAAAATGGGGAGCTGATGAGAGTTNNNAGATCGGA
47373_1 TATGAGTGTGGAGTCGTTGCGTACTTCCTCAGCTTCTGTCTGGNNNAGATCGGAAG
47373_2 TATGAGTGTGGAGTCGTTGCGTACTTCCTCAGCTTCTGTCTGGNNNAGATCGGAAG
31622_1 TATGAGTGTGGAGTCGTTGCACCACCAAGGAATTGTATGGGAANNNAGATCGGAA
31622_2 TATGAGTGTGGAGTCGTTGCACCACCAAGGAATTGTATGGGAANNNAGATCGGAA
30265_1 TATGAGTGTGGAGTCGTTGCTTATCCCAAAGACATTTTTGCGANNNAGATCGGAAG
30265_2 TATGAGTGTGGAGTCGTTGCTTATCCCAAAGACATTTTTGCGANNNAGATCGGAAG
34264_1 TATGAGTGTGGAGTCGTTGCGGAGGTGTACTCAACAGCTCATTNNNAGATCGGAAG
34264_2 TATGAGTGTGGAGTCGTTGCGGAGGTGTACTCAACAGCTCATTNNNAGATCGGAAG
12904_1 TATGAGTGTGGAGTCGTTGCCATTCTCCTCATCAAATTTCTCNNNAGATCGGAAGAG
12904_2 TATGAGTGTGGAGTCGTTGCCATTCTCCTCATCAAATTTCTCNNNAGATCGGAAGAG
37469_1 TATGAGTGTGGAGTCGTTGCAACTGAAAAGCCAACAACACCAANNNAGATCGGAAG
37469_2 TATGAGTGTGGAGTCGTTGCAACTGAAAAGCCAACAACACCAANNNAGATCGGAAG
39631_1 TATGAGTGTGGAGTCGTTGCGCAACCTAGATGAAATGGGCAAANNNAGATCGGAA
39631_2 TATGAGTGTGGAGTCGTTGCGCAACCTAGATGAAATGGGCAAANNNAGATCGGAA
34186_1 TATGAGTGTGGAGTCGTTGCTTGAGACCAGCCTGGCTAATATGNNNAGATCGGAAG
34186_2 TATGAGTGTGGAGTCGTTGCTTGAGACCAGCCTGGCTAATATGNNNAGATCGGAAG
33032_1 TATGAGTGTGGAGTCGTTGCCACCATGCCCGACTAATTTTTGTNNNAGATCGGAAG
33032_2 TATGAGTGTGGAGTCGTTGCCACCATGCCCGACTAATTTTTGTNNNAGATCGGAAG
39290_1 TATGAGTGTGGAGTCGTTGCAGAAAGAAAGGAAGAAGCTTGGCNNNAGATCGGA
39290_2 TATGAGTGTGGAGTCGTTGCAGAAAGAAAGGAAGAAGCTTGGCNNNAGATCGGA
31188_1 TATGAGTGTGGAGTCGTTGCCCAAACTCCTGGGCTCAAGTNNNAGATCGGAAGAGC
31188_2 TATGAGTGTGGAGTCGTTGCCCAAACTCCTGGGCTCAAGTNNNAGATCGGAAGAGC
36422_1 TATGAGTGTGGAGTCGTTGCGCCATTAGCTCTTTCTCCTAGCANNNAGATCGGAAGA
36422_2 TATGAGTGTGGAGTCGTTGCGCCATTAGCTCTTTCTCCTAGCANNNAGATCGGAAGA
44901_1 TATGAGTGTGGAGTCGTTGCCTAGCCAGGGAAATCAGAACCAGNNNAGATCGGAA
44901_2 TATGAGTGTGGAGTCGTTGCCTAGCCAGGGAAATCAGAACCAGNNNAGATCGGAA
31989_1 TATGAGTGTGGAGTCGTTGCAACAGTGCTAATGGATTAGGGCTNNNAGATCGGAA
31989_2 TATGAGTGTGGAGTCGTTGCAACAGTGCTAATGGATTAGGGCTNNNAGATCGGAA
35575_1 TATGAGTGTGGAGTCGTTGCCTAGCAAGCTAAAATCCACTGCCNNNAGATCGGAAG
35575_2 TATGAGTGTGGAGTCGTTGCCTAGCAAGCTAAAATCCACTGCCNNNAGATCGGAAG
3053_1 TATGAGTGTGGAGTCGTTGCCACCTTTCTGATGTGGTTCTGCTNNNAGATCGGAAG
3053_2 TATGAGTGTGGAGTCGTTGCCACCTTTCTGATGTGGTTCTGCTNNNAGATCGGAAG
2348_1 TATGAGTGTGGAGTCGTTGCTCATCTAATCCCCACAGACCTCCNNNAGATCGGAAGA
2348_2 TATGAGTGTGGAGTCGTTGCTCATCTAATCCCCACAGACCTCCNNNAGATCGGAAGA
32481_1 TATGAGTGTGGAGTCGTTGCCAAGTAGCTGGGACTACAGGCNNNAGATCGGAAGA
32481_2 TATGAGTGTGGAGTCGTTGCCAAGTAGCTGGGACTACAGGCNNNAGATCGGAAGA
30553_1 TATGAGTGTGGAGTCGTTGCTTGAGTGAGGAGGGGAGATCAAGNNNAGATCGGA

| | |
|---|---|
| 30553_2 | TATGAGTGTGGAGTCGTTGCTTGAGTGAGGAGGGGAGATCAAGNNNAGATCGGA |
| 39532_1 | TATGAGTGTGGAGTCGTTGCCTAAGGGCAGGAGAATCACTTGANNNAGATCGGAA |
| 39532_2 | TATGAGTGTGGAGTCGTTGCCTAAGGGCAGGAGAATCACTTGANNNAGATCGGAA |
| 12014_1 | TATGAGTGTGGAGTCGTTGCTGCTCCAAACCTTTCTGCTCACTNNNAGATCGGAAGA |
| 12014_2 | TATGAGTGTGGAGTCGTTGCTGCTCCAAACCTTTCTGCTCACTNNNAGATCGGAAGA |
| 53079_1 | TATGAGTGTGGAGTCGTTGCGCAGGTGATTCACATTTCAAGGGNNNAGATCGGAA |
| 53079_2 | TATGAGTGTGGAGTCGTTGCGCAGGTGATTCACATTTCAAGGGNNNAGATCGGAA |
| 52711_1 | TATGAGTGTGGAGTCGTTGCCTGCTTTCCTCTGGGGGCTCNNNAGATCGGAAGAGC |
| 52711_2 | TATGAGTGTGGAGTCGTTGCCTGCTTTCCTCTGGGGGCTCNNNAGATCGGAAGAGC |
| 51517_1 | TATGAGTGTGGAGTCGTTGCGGATAATGGATGGGTGCTTCTCANNNAGATCGGAA |
| 51517_2 | TATGAGTGTGGAGTCGTTGCGGATAATGGATGGGTGCTTCTCANNNAGATCGGAA |
| 9432_1 | TATGAGTGTGGAGTCGTTGCTTTGGGAGGAGAAGGGTCCATAGNNNAGATCGGA |
| 9432_2 | TATGAGTGTGGAGTCGTTGCTTTGGGAGGAGAAGGGTCCATAGNNNAGATCGGA |
| 33317_1 | TATGAGTGTGGAGTCGTTGCATTTGGCAGGGTCATAGGACAATNNNAGATCGGAA |
| 33317_2 | TATGAGTGTGGAGTCGTTGCATTTGGCAGGGTCATAGGACAATNNNAGATCGGAA |
| 37700_1 | TATGAGTGTGGAGTCGTTGCTTATGGGATGCTTACACTGTGCTNNNAGATCGGAAG |
| 37700_2 | TATGAGTGTGGAGTCGTTGCTTATGGGATGCTTACACTGTGCTNNNAGATCGGAAG |
| 3085_1 | TATGAGTGTGGAGTCGTTGCCCGCTGGGAAGAAGAAAACATCTNNNAGATCGGAA |
| 3085_2 | TATGAGTGTGGAGTCGTTGCCCGCTGGGAAGAAGAAAACATCTNNNAGATCGGAA |
| 39243_1 | TATGAGTGTGGAGTCGTTGCTGCCTAGTATCAGGGTATGAGGANNNAGATCGGAA |
| 39243_2 | TATGAGTGTGGAGTCGTTGCTGCCTAGTATCAGGGTATGAGGANNNAGATCGGAA |
| 36586_1 | TATGAGTGTGGAGTCGTTGCTTACTTTTCACTGCTAGGGGGACNNNAGATCGGAAG |
| 36586_2 | TATGAGTGTGGAGTCGTTGCTTACTTTTCACTGCTAGGGGGACNNNAGATCGGAAG |
| 44750_1 | TATGAGTGTGGAGTCGTTGCTTGAGTTTGGGAAGGAGACTGAGNNNAGATCGGA |
| 44750_2 | TATGAGTGTGGAGTCGTTGCTTGAGTTTGGGAAGGAGACTGAGNNNAGATCGGA |
| 39825_1 | TATGAGTGTGGAGTCGTTGCGGAAGGTAGGAAGGTACTCAAATNNNAGATCGGA |
| 39825_2 | TATGAGTGTGGAGTCGTTGCGGAAGGTAGGAAGGTACTCAAATNNNAGATCGGA |
| 34643_1 | TATGAGTGTGGAGTCGTTGCTACTGTCTTCTCCACTGGAATGGNNNAGATCGGAAG |
| 34643_2 | TATGAGTGTGGAGTCGTTGCTACTGTCTTCTCCACTGGAATGGNNNAGATCGGAAG |
| 46264_1 | TATGAGTGTGGAGTCGTTGCCTCCCAGGTTCAAGCGATTCTCNNNAGATCGGAAGA |
| 46264_2 | TATGAGTGTGGAGTCGTTGCCTCCCAGGTTCAAGCGATTCTCNNNAGATCGGAAGA |
| 40297_1 | TATGAGTGTGGAGTCGTTGCCAGTGAGATGCAGCAGGTTCTATNNNAGATCGGAA |
| 40297_2 | TATGAGTGTGGAGTCGTTGCCAGTGAGATGCAGCAGGTTCTATNNNAGATCGGAA |
| 29629_1 | TATGAGTGTGGAGTCGTTGCCTGCAATTAAAAGATCCACAGAACTNNNAGATCGGA |
| 29629_2 | TATGAGTGTGGAGTCGTTGCCTGCAATTAAAAGATCCACAGAACTNNNAGATCGGA |
| 10555_1 | TATGAGTGTGGAGTCGTTGCAGTCTGCTTACAGGAATTTGGCTNNNAGATCGGAAG |
| 10555_2 | TATGAGTGTGGAGTCGTTGCAGTCTGCTTACAGGAATTTGGCTNNNAGATCGGAAG |
| 29647_1 | TATGAGTGTGGAGTCGTTGCCAGGGGGGCCATGCTCTAAANNNAGATCGGAAGAG |
| 29647_2 | TATGAGTGTGGAGTCGTTGCCAGGGGGGCCATGCTCTAAANNNAGATCGGAAGAG |
| 53353_1 | TATGAGTGTGGAGTCGTTGCTTGAAGAGTTAGTGACCGTAGCCNNNAGATCGGAA |
| 53353_2 | TATGAGTGTGGAGTCGTTGCTTGAAGAGTTAGTGACCGTAGCCNNNAGATCGGAA |
| 35816_1 | TATGAGTGTGGAGTCGTTGCGGCATGATTTTGGCTCACTACAANNNAGATCGGAAG |
| 35816_2 | TATGAGTGTGGAGTCGTTGCGGCATGATTTTGGCTCACTACAANNNAGATCGGAAG |

| | |
|---|---|
| 51477_1 | TATGAGTGTGGAGTCGTTGCACAAAAATTAGCTGGGTGTGGTGNNNAGATCGGAA |
| 51477_2 | TATGAGTGTGGAGTCGTTGCACAAAAATTAGCTGGGTGTGGTGNNNAGATCGGAA |
| 12833_1 | TATGAGTGTGGAGTCGTTGCCACCTACATTTTGGAAATTGAANNNAGATCGGAAGA |
| 12833_2 | TATGAGTGTGGAGTCGTTGCCACCTACATTTTGGAAATTGAANNNAGATCGGAAGA |
| 45519_1 | TATGAGTGTGGAGTCGTTGCACAACAAGGAATCACTGACAGGANNNAGATCGGAA |
| 45519_2 | TATGAGTGTGGAGTCGTTGCACAACAAGGAATCACTGACAGGANNNAGATCGGAA |
| 30007_1 | TATGAGTGTGGAGTCGTTGCCTTGAGCAGTGCTTGTGAATCTGNNNAGATCGGAAG |
| 30007_2 | TATGAGTGTGGAGTCGTTGCCTTGAGCAGTGCTTGTGAATCTGNNNAGATCGGAAG |
| 508_1 | TATGAGTGTGGAGTCGTTGCCAAGTGGGTCTCTGTGATGGATTCNNNAGATCGGA |
| 508_2 | TATGAGTGTGGAGTCGTTGCCAAGTGGGTCTCTGTGATGGATTCNNNAGATCGGA |
| 48371_1 | TATGAGTGTGGAGTCGTTGCTCTCCTGAGTAGCTGGGAATACANNNAGATCGGAAG |
| 48371_2 | TATGAGTGTGGAGTCGTTGCTCTCCTGAGTAGCTGGGAATACANNNAGATCGGAAG |
| 48447_1 | TATGAGTGTGGAGTCGTTGCTGACGCTCTTCCATGGATCTAAGNNNAGATCGGAAG |
| 48447_2 | TATGAGTGTGGAGTCGTTGCTGACGCTCTTCCATGGATCTAAGNNNAGATCGGAAG |
| 51591_1 | TATGAGTGTGGAGTCGTTGCACGGGATTTGTTGATTCTCTCCANNNAGATCGGAAG |
| 51591_2 | TATGAGTGTGGAGTCGTTGCACGGGATTTGTTGATTCTCTCCANNNAGATCGGAAG |
| 48434_1 | TATGAGTGTGGAGTCGTTGCCCTGATTCCATACCACAGCCTCNNNAGATCGGAAGAG |
| 48434_2 | TATGAGTGTGGAGTCGTTGCCCTGATTCCATACCACAGCCTCNNNAGATCGGAAGAG |
| 35525_1 | TATGAGTGTGGAGTCGTTGCTCTTGAACTCCAGACCTCAAGTGNNNAGATCGGAAG |
| 35525_2 | TATGAGTGTGGAGTCGTTGCTCTTGAACTCCAGACCTCAAGTGNNNAGATCGGAAG |
| 34195_1 | TATGAGTGTGGAGTCGTTGCTCTACTGTTTCAAGTGACTCACCTNNNAGATCGGAAG |
| 34195_2 | TATGAGTGTGGAGTCGTTGCTCTACTGTTTCAAGTGACTCACCTNNNAGATCGGAAG |
| 35220_1 | TATGAGTGTGGAGTCGTTGCAGAACATTATTGGGGAGCCACATNNNAGATCGGAA |
| 35220_2 | TATGAGTGTGGAGTCGTTGCAGAACATTATTGGGGAGCCACATNNNAGATCGGAA |
| 35776_1 | TATGAGTGTGGAGTCGTTGCGGAGTATCACTCTGTTGCTCAGGNNNAGATCGGAAG |
| 35776_2 | TATGAGTGTGGAGTCGTTGCGGAGTATCACTCTGTTGCTCAGGNNNAGATCGGAAG |
| 13318_1 | TATGAGTGTGGAGTCGTTGCCATGAAAACAAGGCAAAATATTTNNNAGATCGGAAG |
| 13318_2 | TATGAGTGTGGAGTCGTTGCCATGAAAACAAGGCAAAATATTTNNNAGATCGGAAG |
| 13721_1 | TATGAGTGTGGAGTCGTTGCGACTGTTAGTAACTTAGTCTCCNNNAGATCGGAAGA |
| 13721_2 | TATGAGTGTGGAGTCGTTGCGACTGTTAGTAACTTAGTCTCCNNNAGATCGGAAGA |
| 45043_1 | TATGAGTGTGGAGTCGTTGCCCAGGAAAAGACAGACGACGATANNNAGATCGGAA |
| 45043_2 | TATGAGTGTGGAGTCGTTGCCCAGGAAAAGACAGACGACGATANNNAGATCGGAA |
| 39002_1 | TATGAGTGTGGAGTCGTTGCTGAGCTTCATGAGTACATTTCCANNNAGATCGGAAG |
| 39002_2 | TATGAGTGTGGAGTCGTTGCTGAGCTTCATGAGTACATTTCCANNNAGATCGGAAG |
| 44631_1 | TATGAGTGTGGAGTCGTTGCTGGGACTTGTCTTGTAGGAATCCNNNAGATCGGAAG |
| 44631_2 | TATGAGTGTGGAGTCGTTGCTGGGACTTGTCTTGTAGGAATCCNNNAGATCGGAAG |
| 36692_1 | TATGAGTGTGGAGTCGTTGCAATTAGAAACTCTGAGGCTGGGGNNNAGATCGGAA |
| 36692_2 | TATGAGTGTGGAGTCGTTGCAATTAGAAACTCTGAGGCTGGGGNNNAGATCGGAA |
| 37877_1 | TATGAGTGTGGAGTCGTTGCTTTTACTCAGAGGGGGGACTACTNNNAGATCGGAA |
| 37877_2 | TATGAGTGTGGAGTCGTTGCTTTTACTCAGAGGGGGGACTACTNNNAGATCGGAA |
| 25276_1 | TATGAGTGTGGAGTCGTTGCTAATGGATGTGACTGCTATAAAGNNNAGATCGGAA |
| 25276_2 | TATGAGTGTGGAGTCGTTGCTAATGGATGTGACTGCTATAAAGNNNAGATCGGAA |
| 30679_1 | TATGAGTGTGGAGTCGTTGCTGGATGTGCCTACGTTTCTGAAGNNNAGATCGGAAG |

| | |
|---|---|
| 30679_2 | TATGAGTGTGGAGTCGTTGCTGGATGTGCCTACGTTTCTGAAGNNNAGATCGGAAG |
| 38709_1 | TATGAGTGTGGAGTCGTTGCTACCTGAGATTGAGGCAGACAACNNNAGATCGGAA |
| 38709_2 | TATGAGTGTGGAGTCGTTGCTACCTGAGATTGAGGCAGACAACNNNAGATCGGAA |
| 45595_1 | TATGAGTGTGGAGTCGTTGCACGCCCGGCTAATTTTTTGTATTNNNAGATCGGAAG |
| 45595_2 | TATGAGTGTGGAGTCGTTGCACGCCCGGCTAATTTTTTGTATTNNNAGATCGGAAG |
| 48665_1 | TATGAGTGTGGAGTCGTTGCGATTTTCCAGTCTCAGCCTCTCANNNAGATCGGAAGA |
| 48665_2 | TATGAGTGTGGAGTCGTTGCGATTTTCCAGTCTCAGCCTCTCANNNAGATCGGAAGA |
| 10688_1 | TATGAGTGTGGAGTCGTTGCTCAGGGATTTCAAAGGACAAGCCNNNAGATCGGAA |
| 10688_2 | TATGAGTGTGGAGTCGTTGCTCAGGGATTTCAAAGGACAAGCCNNNAGATCGGAA |
| 39479_1 | TATGAGTGTGGAGTCGTTGCAGCCATGGAAGTAACCAGAATGTNNNAGATCGGAA |
| 39479_2 | TATGAGTGTGGAGTCGTTGCAGCCATGGAAGTAACCAGAATGTNNNAGATCGGAA |
| 36550_1 | TATGAGTGTGGAGTCGTTGCGCCATGAACTGACACTTGTTGAANNNAGATCGGAAG |
| 36550_2 | TATGAGTGTGGAGTCGTTGCGCCATGAACTGACACTTGTTGAANNNAGATCGGAAG |
| 32879_1 | TATGAGTGTGGAGTCGTTGCGTTCAAGCGATTCTCTTGCCTCNNNAGATCGGAAGA |
| 32879_2 | TATGAGTGTGGAGTCGTTGCGTTCAAGCGATTCTCTTGCCTCNNNAGATCGGAAGA |
| 31484_1 | TATGAGTGTGGAGTCGTTGCTCACAGACACATATGCACAGACANNNAGATCGGAAG |
| 31484_2 | TATGAGTGTGGAGTCGTTGCTCACAGACACATATGCACAGACANNNAGATCGGAAG |
| 31090_1 | TATGAGTGTGGAGTCGTTGCGGGCATATAGGTTGGAGCTATGTNNNAGATCGGAA |
| 31090_2 | TATGAGTGTGGAGTCGTTGCGGGCATATAGGTTGGAGCTATGTNNNAGATCGGAA |
| 45346_1 | TATGAGTGTGGAGTCGTTGCAACGTGACAATCAAATTTGGGCTNNNAGATCGGAAG |
| 45346_2 | TATGAGTGTGGAGTCGTTGCAACGTGACAATCAAATTTGGGCTNNNAGATCGGAAG |
| 36114_1 | TATGAGTGTGGAGTCGTTGCACGGTAAGACCCTGTCTCAAAAANNNAGATCGGAAG |
| 36114_2 | TATGAGTGTGGAGTCGTTGCACGGTAAGACCCTGTCTCAAAAANNNAGATCGGAAG |
| 52753_1 | TATGAGTGTGGAGTCGTTGCCTGCTTGTTCTCATTCAGGCAAANNNAGATCGGAAG |
| 52753_2 | TATGAGTGTGGAGTCGTTGCCTGCTTGTTCTCATTCAGGCAAANNNAGATCGGAAG |
| 48010_1 | TATGAGTGTGGAGTCGTTGCTCCCAAAGTGCTGGAATTACAGTNNNAGATCGGAAG |
| 48010_2 | TATGAGTGTGGAGTCGTTGCTCCCAAAGTGCTGGAATTACAGTNNNAGATCGGAAG |
| 49476_1 | TATGAGTGTGGAGTCGTTGCATAGGCAACACAAAGGCAAGTTTNNNAGATCGGAA |
| 49476_2 | TATGAGTGTGGAGTCGTTGCATAGGCAACACAAAGGCAAGTTTNNNAGATCGGAA |
| 37777_1 | TATGAGTGTGGAGTCGTTGCCTTGTTATGTTCCCCAGGCTAGTNNNAGATCGGAAG |
| 37777_2 | TATGAGTGTGGAGTCGTTGCCTTGTTATGTTCCCCAGGCTAGTNNNAGATCGGAAG |
| 48981_1 | TATGAGTGTGGAGTCGTTGCCAGTGTAAGAGCAGCTACAGGAANNNAGATCGGAA |
| 48981_2 | TATGAGTGTGGAGTCGTTGCCAGTGTAAGAGCAGCTACAGGAANNNAGATCGGAA |
| 35465_1 | TATGAGTGTGGAGTCGTTGCCCTCCCTTATTTCCTCAAGCAGTNNNAGATCGGAAGA |
| 35465_2 | TATGAGTGTGGAGTCGTTGCCCTCCCTTATTTCCTCAAGCAGTNNNAGATCGGAAGA |
| 48221_1 | TATGAGTGTGGAGTCGTTGCATCTCCCAGGTTCAAGCAATTCTNNNAGATCGGAAG |
| 48221_2 | TATGAGTGTGGAGTCGTTGCATCTCCCAGGTTCAAGCAATTCTNNNAGATCGGAAG |
| 34138_1 | TATGAGTGTGGAGTCGTTGCGGAGATTTCACTTCTTGGGAGGTNNNAGATCGGAA |
| 34138_2 | TATGAGTGTGGAGTCGTTGCGGAGATTTCACTTCTTGGGAGGTNNNAGATCGGAA |
| 49140_1 | TATGAGTGTGGAGTCGTTGCTCTCCTGACCTCATTATCCACCTNNNAGATCGGAAGA |
| 49140_2 | TATGAGTGTGGAGTCGTTGCTCTCCTGACCTCATTATCCACCTNNNAGATCGGAAGA |
| 31922_1 | TATGAGTGTGGAGTCGTTGCGCTGAAGGGCATTAGCTACATTGNNNAGATCGGAA |
| 31922_2 | TATGAGTGTGGAGTCGTTGCGCTGAAGGGCATTAGCTACATTGNNNAGATCGGAA |

| | |
|---|---|
| 35042_1 | TATGAGTGTGGAGTCGTTGCGCTCCCTGAGTCTCAAGAAAAGANNNAGATCGGAAG |
| 35042_2 | TATGAGTGTGGAGTCGTTGCGCTCCCTGAGTCTCAAGAAAAGANNNAGATCGGAAG |
| 47754_1 | TATGAGTGTGGAGTCGTTGCTTCAAGCTATTCTCCTGCCTCAGNNNAGATCGGAAGA |
| 47754_2 | TATGAGTGTGGAGTCGTTGCTTCAAGCTATTCTCCTGCCTCAGNNNAGATCGGAAGA |
| 32692_1 | TATGAGTGTGGAGTCGTTGCGGTTCAAGTGATTTTCCTGCCTCNNNAGATCGGAAGA |
| 32692_2 | TATGAGTGTGGAGTCGTTGCGGTTCAAGTGATTTTCCTGCCTCNNNAGATCGGAAGA |
| 50293_1 | TATGAGTGTGGAGTCGTTGCTGGTAGATGACAGATTGACAGGTGNNNAGATCGGA |
| 50293_2 | TATGAGTGTGGAGTCGTTGCTGGTAGATGACAGATTGACAGGTGNNNAGATCGGA |
| 41193_1 | TATGAGTGTGGAGTCGTTGCTCCTAATATCCAGGTGGGGAGAGNNNAGATCGGAAG |
| 41193_2 | TATGAGTGTGGAGTCGTTGCTCCTAATATCCAGGTGGGGAGAGNNNAGATCGGAAG |
| 43832_1 | TATGAGTGTGGAGTCGTTGCCCTAGCAGGTCAGTTAGTTGGTTNNNAGATCGGAAG |
| 43832_2 | TATGAGTGTGGAGTCGTTGCCCTAGCAGGTCAGTTAGTTGGTTNNNAGATCGGAAG |
| 51499_1 | TATGAGTGTGGAGTCGTTGCGCTTTTTACACATCACTGCGGATNNNAGATCGGAAGA |
| 51499_2 | TATGAGTGTGGAGTCGTTGCGCTTTTTACACATCACTGCGGATNNNAGATCGGAAGA |
| 40122_1 | TATGAGTGTGGAGTCGTTGCGGCAGAAAGTAGGTCTATAGGGCNNNAGATCGGAA |
| 40122_2 | TATGAGTGTGGAGTCGTTGCGGCAGAAAGTAGGTCTATAGGGCNNNAGATCGGAA |
| 37943_1 | TATGAGTGTGGAGTCGTTGCGCAAGCATTATAAACGTATGCACCNNNAGATCGGAA |
| 37943_2 | TATGAGTGTGGAGTCGTTGCGCAAGCATTATAAACGTATGCACCNNNAGATCGGAA |
| 50682_1 | TATGAGTGTGGAGTCGTTGCCCTTCTTGGCCTTTCTCATCTCTNNNAGATCGGAAGAG |
| 50682_2 | TATGAGTGTGGAGTCGTTGCCCTTCTTGGCCTTTCTCATCTCTNNNAGATCGGAAGAG |
| 52979_1 | TATGAGTGTGGAGTCGTTGCGCATGTGCACATTTTTCTCCCTANNNAGATCGGAAGA |
| 52979_2 | TATGAGTGTGGAGTCGTTGCGCATGTGCACATTTTTCTCCCTANNNAGATCGGAAGA |
| 29385_1 | TATGAGTGTGGAGTCGTTGCACAAATGGCCTCTGCTACTTACTNNNAGATCGGAAGA |
| 29385_2 | TATGAGTGTGGAGTCGTTGCACAAATGGCCTCTGCTACTTACTNNNAGATCGGAAGA |
| 31525_1 | TATGAGTGTGGAGTCGTTGCTGTACCCAATGTCTCCAATCCTCNNNAGATCGGAAGA |
| 31525_2 | TATGAGTGTGGAGTCGTTGCTGTACCCAATGTCTCCAATCCTCNNNAGATCGGAAGA |
| 47000_1 | TATGAGTGTGGAGTCGTTGCCACAGGAAGGAGACGATCCAGNNNAGATCGGAAGA |
| 47000_2 | TATGAGTGTGGAGTCGTTGCCACAGGAAGGAGACGATCCAGNNNAGATCGGAAGA |
| 46744_1 | TATGAGTGTGGAGTCGTTGCTAACTGAGACTACAGGTGTGTGCNNNAGATCGGAAG |
| 46744_2 | TATGAGTGTGGAGTCGTTGCTAACTGAGACTACAGGTGTGTGCNNNAGATCGGAAG |
| 50872_1 | TATGAGTGTGGAGTCGTTGCGCTGCATGACACAAATGGACATANNNAGATCGGAAG |
| 50872_2 | TATGAGTGTGGAGTCGTTGCGCTGCATGACACAAATGGACATANNNAGATCGGAAG |
| 48996_1 | TATGAGTGTGGAGTCGTTGCCAGCCTCCTGAGTAAATGGGATTNNNAGATCGGAAG |
| 48996_2 | TATGAGTGTGGAGTCGTTGCCAGCCTCCTGAGTAAATGGGATTNNNAGATCGGAAG |
| 40582_1 | TATGAGTGTGGAGTCGTTGCACTTTCTAGAAACCTGCCCTCTGNNNAGATCGGAAGA |
| 40582_2 | TATGAGTGTGGAGTCGTTGCACTTTCTAGAAACCTGCCCTCTGNNNAGATCGGAAGA |
| 38841_1 | TATGAGTGTGGAGTCGTTGCTGAAAAGCACAGGCAACAATAACANNNAGATCGGAA |
| 38841_2 | TATGAGTGTGGAGTCGTTGCTGAAAAGCACAGGCAACAATAACANNNAGATCGGAA |
| 51275_1 | TATGAGTGTGGAGTCGTTGCTGTACCCAACAGCTCATTGAGAANNNAGATCGGAAG |
| 51275_2 | TATGAGTGTGGAGTCGTTGCTGTACCCAACAGCTCATTGAGAANNNAGATCGGAAG |
| 35589_1 | TATGAGTGTGGAGTCGTTGCTCAGCACTATGTTATGGGTTTTCTNNNAGATCGGAAG |
| 35589_2 | TATGAGTGTGGAGTCGTTGCTCAGCACTATGTTATGGGTTTTCTNNNAGATCGGAAG |
| 36041_1 | TATGAGTGTGGAGTCGTTGCTGACATTATTCTGACCCATGGGGNNNAGATCGGAAG |

| | |
|---|---|
| 36041_2 | TATGAGTGTGGAGTCGTTGCTGACATTATTCTGACCCATGGGGNNNAGATCGGAAG |
| 29645_1 | TATGAGTGTGGAGTCGTTGCCTGTGAGTACTAGGGAGGTGTCANNNAGATCGGAA |
| 29645_2 | TATGAGTGTGGAGTCGTTGCCTGTGAGTACTAGGGAGGTGTCANNNAGATCGGAA |
| 42314_1 | TATGAGTGTGGAGTCGTTGCAGCTACTTCTGCTGGTAGGAATGNNNAGATCGGAA |
| 42314_2 | TATGAGTGTGGAGTCGTTGCAGCTACTTCTGCTGGTAGGAATGNNNAGATCGGAA |
| 34849_1 | TATGAGTGTGGAGTCGTTGCCCTCCTAGGTTCAAGCGATTCTGNNNAGATCGGAAG |
| 34849_2 | TATGAGTGTGGAGTCGTTGCCCTCCTAGGTTCAAGCGATTCTGNNNAGATCGGAAG |
| 35334_1 | TATGAGTGTGGAGTCGTTGCTCATTGCAGCCTCAAATTCTTGGNNNAGATCGGAAG |
| 35334_2 | TATGAGTGTGGAGTCGTTGCTCATTGCAGCCTCAAATTCTTGGNNNAGATCGGAAG |
| 47750_1 | TATGAGTGTGGAGTCGTTGCGTCTCATTCTGTCACCTAGGCTGNNNAGATCGGAAG |
| 47750_2 | TATGAGTGTGGAGTCGTTGCGTCTCATTCTGTCACCTAGGCTGNNNAGATCGGAAG |
| 30862_1 | TATGAGTGTGGAGTCGTTGCGGTGTGGTGCCTCATACTTAGAANNNAGATCGGAA |
| 30862_2 | TATGAGTGTGGAGTCGTTGCGGTGTGGTGCCTCATACTTAGAANNNAGATCGGAA |
| 50888_1 | TATGAGTGTGGAGTCGTTGCTGTCCCTGTGTTTTACTTCAGGANNNAGATCGGAAG |
| 50888_2 | TATGAGTGTGGAGTCGTTGCTGTCCCTGTGTTTTACTTCAGGANNNAGATCGGAAG |
| 13191_1 | TATGAGTGTGGAGTCGTTGCGACAGAATTATGTAGGCTTTATCNNNAGATCGGAAG |
| 13191_2 | TATGAGTGTGGAGTCGTTGCGACAGAATTATGTAGGCTTTATCNNNAGATCGGAAG |
| 44731_1 | TATGAGTGTGGAGTCGTTGCAGGTGAGCATGGAGTGAGTTTTTNNNAGATCGGAA |
| 44731_2 | TATGAGTGTGGAGTCGTTGCAGGTGAGCATGGAGTGAGTTTTTNNNAGATCGGAA |
| 47175_1 | TATGAGTGTGGAGTCGTTGCAGACAGAGTTTTGCTGTTGTTGCNNNAGATCGGAAG |
| 47175_2 | TATGAGTGTGGAGTCGTTGCAGACAGAGTTTTGCTGTTGTTGCNNNAGATCGGAAG |
| 47226_1 | TATGAGTGTGGAGTCGTTGCGCATCTGGTCTAGGATGTATGCANNNAGATCGGAA |
| 47226_2 | TATGAGTGTGGAGTCGTTGCGCATCTGGTCTAGGATGTATGCANNNAGATCGGAA |
| 52327_1 | TATGAGTGTGGAGTCGTTGCCAGAGAAGCTATAAGAGGAGGCGNNNAGATCGGA |
| 52327_2 | TATGAGTGTGGAGTCGTTGCCAGAGAAGCTATAAGAGGAGGCGNNNAGATCGGA |
| 37631_1 | TATGAGTGTGGAGTCGTTGCATTGCTTGAGGTCAGGAGTTTGANNNAGATCGGAA |
| 37631_2 | TATGAGTGTGGAGTCGTTGCATTGCTTGAGGTCAGGAGTTTGANNNAGATCGGAA |
| 51955_1 | TATGAGTGTGGAGTCGTTGCAGGTAGGTAGGGTAACCAACTGTNNNAGATCGGAA |
| 51955_2 | TATGAGTGTGGAGTCGTTGCAGGTAGGTAGGGTAACCAACTGTNNNAGATCGGAA |
| 52112_1 | TATGAGTGTGGAGTCGTTGCCTCTTCTTGGTCAGAGCGAAGGNNNAGATCGGAAG |
| 52112_2 | TATGAGTGTGGAGTCGTTGCCTCTTCTTGGTCAGAGCGAAGGNNNAGATCGGAAG |
| 10935_1 | TATGAGTGTGGAGTCGTTGCGTGCATGAGTGTGAAAAGGGAGNNNAGATCGGAA |
| 10935_2 | TATGAGTGTGGAGTCGTTGCGTGCATGAGTGTGAAAAGGGAGNNNAGATCGGAA |
| 51022_1 | TATGAGTGTGGAGTCGTTGCGAATGTGGTGAGCTACACACATGNNNAGATCGGAA |
| 51022_2 | TATGAGTGTGGAGTCGTTGCGAATGTGGTGAGCTACACACATGNNNAGATCGGAA |
| 51475_1 | TATGAGTGTGGAGTCGTTGCTTGCTGGGTTCAAATGATCCTCTNNNAGATCGGAAG |
| 51475_2 | TATGAGTGTGGAGTCGTTGCTTGCTGGGTTCAAATGATCCTCTNNNAGATCGGAAG |
| 29542_1 | TATGAGTGTGGAGTCGTTGCGGCAATCACTGGGCTCATGANNNAGATCGGAAGAG |
| 29542_2 | TATGAGTGTGGAGTCGTTGCGGCAATCACTGGGCTCATGANNNAGATCGGAAGAG |
| 38017_1 | TATGAGTGTGGAGTCGTTGCGCAGGGGAAAGAACCATTTGAAANNNAGATCGGAA |
| 38017_2 | TATGAGTGTGGAGTCGTTGCGCAGGGGAAAGAACCATTTGAAANNNAGATCGGAA |
| 46179_1 | TATGAGTGTGGAGTCGTTGCTGAAGTTTCCCAGGCTATAGTGGNNNAGATCGGAA |
| 46179_2 | TATGAGTGTGGAGTCGTTGCTGAAGTTTCCCAGGCTATAGTGGNNNAGATCGGAA |

| | |
|---|---|
| 28795_1 | TATGAGTGTGGAGTCGTTGCGTGTGTGTGTGTGTGTGTGTATANNNAGATCGGAA |
| 28795_2 | TATGAGTGTGGAGTCGTTGCGTGTGTGTGTGTGTGTGTGTATANNNAGATCGGAA |
| 40070_1 | TATGAGTGTGGAGTCGTTGCGTAATCCCAGCACTTTGGAATGCNNNAGATCGGAAG |
| 40070_2 | TATGAGTGTGGAGTCGTTGCGTAATCCCAGCACTTTGGAATGCNNNAGATCGGAAG |
| 35942_1 | TATGAGTGTGGAGTCGTTGCATGCAGCTTCCAGAAGAATCAGANNNAGATCGGAA |
| 35942_2 | TATGAGTGTGGAGTCGTTGCATGCAGCTTCCAGAAGAATCAGANNNAGATCGGAA |
| 11097_1 | TATGAGTGTGGAGTCGTTGCGGGTTTTACTTTGTTGCCACCAANNNAGATCGGAAG |
| 11097_2 | TATGAGTGTGGAGTCGTTGCGGGTTTTACTTTGTTGCCACCAANNNAGATCGGAAG |
| 51757_1 | TATGAGTGTGGAGTCGTTGCGAGTGAGAATATGCGGTGTTTGGNNNAGATCGGA |
| 51757_2 | TATGAGTGTGGAGTCGTTGCGAGTGAGAATATGCGGTGTTTGGNNNAGATCGGA |
| 13374_1 | TATGAGTGTGGAGTCGTTGCGTTGTACTACAAGAACATCAGTNNNAGATCGGAAG |
| 13374_2 | TATGAGTGTGGAGTCGTTGCGTTGTACTACAAGAACATCAGTNNNAGATCGGAAG |
| 12753_1 | TATGAGTGTGGAGTCGTTGCGAAAAATACCAGCTAGAGAGAANNNAGATCGGAAG |
| 12753_2 | TATGAGTGTGGAGTCGTTGCGAAAAATACCAGCTAGAGAGAANNNAGATCGGAAG |
| 35161_1 | TATGAGTGTGGAGTCGTTGCCACAATCTCAGCTCACTGCAACNNNAGATCGGAAGA |
| 35161_2 | TATGAGTGTGGAGTCGTTGCCACAATCTCAGCTCACTGCAACNNNAGATCGGAAGA |
| 13430_1 | TATGAGTGTGGAGTCGTTGCTCGATTGTCTTCTCTAGTGANNNAGATCGGAAGAGC |
| 13430_2 | TATGAGTGTGGAGTCGTTGCTCGATTGTCTTCTCTAGTGANNNAGATCGGAAGAGC |
| 45313_1 | TATGAGTGTGGAGTCGTTGCTGATTCAAAATTCACCACAGGCCNNNAGATCGGAAG |
| 45313_2 | TATGAGTGTGGAGTCGTTGCTGATTCAAAATTCACCACAGGCCNNNAGATCGGAAG |
| 40798_1 | TATGAGTGTGGAGTCGTTGCACATAGGAAGGAGACCACAATCGNNNAGATCGGAA |
| 40798_2 | TATGAGTGTGGAGTCGTTGCACATAGGAAGGAGACCACAATCGNNNAGATCGGAA |
| 31526_1 | TATGAGTGTGGAGTCGTTGCGACAATGGTAGAACGTTGCCAAANNNAGATCGGAA |
| 31526_2 | TATGAGTGTGGAGTCGTTGCGACAATGGTAGAACGTTGCCAAANNNAGATCGGAA |
| 50781_1 | TATGAGTGTGGAGTCGTTGCGGCTGATTCCTAAAGTGAATGCGNNNAGATCGGAA |
| 50781_2 | TATGAGTGTGGAGTCGTTGCGGCTGATTCCTAAAGTGAATGCGNNNAGATCGGAA |
| 44001_1 | TATGAGTGTGGAGTCGTTGCTCGTGAGTAAGTAGCTGGGACTANNNAGATCGGAA |
| 44001_2 | TATGAGTGTGGAGTCGTTGCTCGTGAGTAAGTAGCTGGGACTANNNAGATCGGAA |
| 36512_1 | TATGAGTGTGGAGTCGTTGCGTGCACACACAAATTCATGCAAANNNAGATCGGAAG |
| 36512_2 | TATGAGTGTGGAGTCGTTGCGTGCACACACAAATTCATGCAAANNNAGATCGGAAG |
| 29134_1 | TATGAGTGTGGAGTCGTTGCTTAGCTCACTTCTTTCCCTCACCNNNAGATCGGAAGA |
| 29134_2 | TATGAGTGTGGAGTCGTTGCTTAGCTCACTTCTTTCCCTCACCNNNAGATCGGAAGA |
| 33611_1 | TATGAGTGTGGAGTCGTTGCAGCCTCCCAAAATGCTGAGATTANNNAGATCGGAAG |
| 33611_2 | TATGAGTGTGGAGTCGTTGCAGCCTCCCAAAATGCTGAGATTANNNAGATCGGAAG |
| 34120_1 | TATGAGTGTGGAGTCGTTGCAAGCTAAGGCAAGAGAATGGCATNNNAGATCGGAA |
| 34120_2 | TATGAGTGTGGAGTCGTTGCAAGCTAAGGCAAGAGAATGGCATNNNAGATCGGAA |
| 31671_1 | TATGAGTGTGGAGTCGTTGCCAGATTGGGCAAAGCAAACAAACNNNAGATCGGAA |
| 31671_2 | TATGAGTGTGGAGTCGTTGCCAGATTGGGCAAAGCAAACAAACNNNAGATCGGAA |
| 46050_1 | TATGAGTGTGGAGTCGTTGCCTACAACCTCTGCCTCATGAGTTNNNAGATCGGAAG |
| 46050_2 | TATGAGTGTGGAGTCGTTGCCTACAACCTCTGCCTCATGAGTTNNNAGATCGGAAG |
| 49647_1 | TATGAGTGTGGAGTCGTTGCCGAGTAGATGGGATTACAGGCATNNNAGATCGGAA |
| 49647_2 | TATGAGTGTGGAGTCGTTGCCGAGTAGATGGGATTACAGGCATNNNAGATCGGAA |
| 9359_1 | TATGAGTGTGGAGTCGTTGCGAGAGAGAGCAAACAACCTACGGNNNAGATCGGAA |

| | |
|---|---|
| 9359_2 | TATGAGTGTGGAGTCGTTGCGAGAGAGAGCAAACAACCTACGGNNNAGATCGGAA |
| 34630_1 | TATGAGTGTGGAGTCGTTGCTAGAGGTTCTCCATAAGGGTCCCNNNAGATCGGAAG |
| 34630_2 | TATGAGTGTGGAGTCGTTGCTAGAGGTTCTCCATAAGGGTCCCNNNAGATCGGAAG |
| 10721_1 | TATGAGTGTGGAGTCGTTGCAAAGGATCGAATGGCCAAGTACCNNNAGATCGGAA |
| 10721_2 | TATGAGTGTGGAGTCGTTGCAAAGGATCGAATGGCCAAGTACCNNNAGATCGGAA |
| 50169_1 | TATGAGTGTGGAGTCGTTGCCTAACCCCCCACAAGAGCTTAGNNNAGATCGGAAGA |
| 50169_2 | TATGAGTGTGGAGTCGTTGCCTAACCCCCCACAAGAGCTTAGNNNAGATCGGAAGA |
| 47491_1 | TATGAGTGTGGAGTCGTTGCATGAGCAAGACATTGGACTGGAANNNAGATCGGAA |
| 47491_2 | TATGAGTGTGGAGTCGTTGCATGAGCAAGACATTGGACTGGAANNNAGATCGGAA |
| 43526_1 | TATGAGTGTGGAGTCGTTGCTTCAAGTGATTCTCCTGCCTCAGNNNAGATCGGAAG |
| 43526_2 | TATGAGTGTGGAGTCGTTGCTTCAAGTGATTCTCCTGCCTCAGNNNAGATCGGAAG |
| 33571_1 | TATGAGTGTGGAGTCGTTGCCCGTGTTCAAGCAATTCTCCTGNNNAGATCGGAAGA |
| 33571_2 | TATGAGTGTGGAGTCGTTGCCCGTGTTCAAGCAATTCTCCTGNNNAGATCGGAAGA |
| 10080_1 | TATGAGTGTGGAGTCGTTGCAGGTGCTCAAACGTGAGTTGAATNNNAGATCGGAA |
| 10080_2 | TATGAGTGTGGAGTCGTTGCAGGTGCTCAAACGTGAGTTGAATNNNAGATCGGAA |
| 49665_1 | TATGAGTGTGGAGTCGTTGCCACTATAGCCTCAACCTCCAAGGNNNAGATCGGAAG |
| 49665_2 | TATGAGTGTGGAGTCGTTGCCACTATAGCCTCAACCTCCAAGGNNNAGATCGGAAG |
| 13537_1 | TATGAGTGTGGAGTCGTTGCTTATCCATTATCATGAAATGCTGNNNAGATCGGAAG |
| 13537_2 | TATGAGTGTGGAGTCGTTGCTTATCCATTATCATGAAATGCTGNNNAGATCGGAAG |
| 29305_1 | TATGAGTGTGGAGTCGTTGCGCAACCAAGACCTCATAGGAAGANNNAGATCGGAA |
| 29305_2 | TATGAGTGTGGAGTCGTTGCGCAACCAAGACCTCATAGGAAGANNNAGATCGGAA |
| 52882_1 | TATGAGTGTGGAGTCGTTGCCATCTTGCGTCTTGAGTCATGTGNNNAGATCGGAAG |
| 52882_2 | TATGAGTGTGGAGTCGTTGCCATCTTGCGTCTTGAGTCATGTGNNNAGATCGGAAG |
| 29407_1 | TATGAGTGTGGAGTCGTTGCGGGTTAAGGGTGGTTGTCAGTGNNNAGATCGGAA |
| 29407_2 | TATGAGTGTGGAGTCGTTGCGGGTTAAGGGTGGTTGTCAGTGNNNAGATCGGAA |
| 52712_1 | TATGAGTGTGGAGTCGTTGCGTCTCCTCCCTCCCTTCCTCNNNAGATCGGAAGAGCA |
| 52712_2 | TATGAGTGTGGAGTCGTTGCGTCTCCTCCCTCCCTTCCTCNNNAGATCGGAAGAGCA |
| 41482_1 | TATGAGTGTGGAGTCGTTGCGCATGCAACCAGATTTCCAGTTANNNAGATCGGAAG |
| 41482_2 | TATGAGTGTGGAGTCGTTGCGCATGCAACCAGATTTCCAGTTANNNAGATCGGAAG |
| 30518_1 | TATGAGTGTGGAGTCGTTGCCTGTGCTTGGCTTACTTCTGTTATNNNAGATCGGAA |
| 30518_2 | TATGAGTGTGGAGTCGTTGCCTGTGCTTGGCTTACTTCTGTTATNNNAGATCGGAA |
| 8989_1 | TATGAGTGTGGAGTCGTTGCTGTCCCTGCTCCACTATTTCTCTNNNAGATCGGAAGA |
| 8989_2 | TATGAGTGTGGAGTCGTTGCTGTCCCTGCTCCACTATTTCTCTNNNAGATCGGAAGA |
| 53032_1 | TATGAGTGTGGAGTCGTTGCTAGTTTTGCCAAACAAGCATTTGNNNAGATCGGAAG |
| 53032_2 | TATGAGTGTGGAGTCGTTGCTAGTTTTGCCAAACAAGCATTTGNNNAGATCGGAAG |
| 46469_1 | TATGAGTGTGGAGTCGTTGCTAATGCGGAATCACACATGAAGCNNNAGATCGGAA |
| 46469_2 | TATGAGTGTGGAGTCGTTGCTAATGCGGAATCACACATGAAGCNNNAGATCGGAA |
| 37016_1 | TATGAGTGTGGAGTCGTTGCTTGGTAAGTGGCAATACTGGTGANNNAGATCGGAA |
| 37016_2 | TATGAGTGTGGAGTCGTTGCTTGGTAAGTGGCAATACTGGTGANNNAGATCGGAA |
| 33389_1 | TATGAGTGTGGAGTCGTTGCTGTAGTGAGCCTAGAGCAGAGATNNNAGATCGGAA |
| 33389_2 | TATGAGTGTGGAGTCGTTGCTGTAGTGAGCCTAGAGCAGAGATNNNAGATCGGAA |
| 32689_1 | TATGAGTGTGGAGTCGTTGCCTCCCGGGTTCAAGTGATTCTCNNNAGATCGGAAGA |
| 32689_2 | TATGAGTGTGGAGTCGTTGCCTCCCGGGTTCAAGTGATTCTCNNNAGATCGGAAGA |

| | |
|---|---|
| 30147_1 | TATGAGTGTGGAGTCGTTGCTCCTCAAGCTGGGTTTGTAAAAACNNNAGATCGGAA |
| 30147_2 | TATGAGTGTGGAGTCGTTGCTCCTCAAGCTGGGTTTGTAAAAACNNNAGATCGGAA |
| 40733_1 | TATGAGTGTGGAGTCGTTGCTCAAGTGTCTGTCAACAAATGCANNNAGATCGGAAG |
| 40733_2 | TATGAGTGTGGAGTCGTTGCTCAAGTGTCTGTCAACAAATGCANNNAGATCGGAAG |
| 42529_1 | TATGAGTGTGGAGTCGTTGCATAGCACTCTTCTCCCTCCCTTANNNAGATCGGAAGA |
| 42529_2 | TATGAGTGTGGAGTCGTTGCATAGCACTCTTCTCCCTCCCTTANNNAGATCGGAAGA |
| 34628_1 | TATGAGTGTGGAGTCGTTGCTTGTTCCACCTCAGATCATCAGGNNNAGATCGGAAG |
| 34628_2 | TATGAGTGTGGAGTCGTTGCTTGTTCCACCTCAGATCATCAGGNNNAGATCGGAAG |
| 29239_1 | TATGAGTGTGGAGTCGTTGCCTTGACCCATCTGCACCATAGATNNNAGATCGGAAG |
| 29239_2 | TATGAGTGTGGAGTCGTTGCCTTGACCCATCTGCACCATAGATNNNAGATCGGAAG |
| 38303_1 | TATGAGTGTGGAGTCGTTGCACAGAAGAAATGCTAAAGGGAGTNNNAGATCGGAA |
| 38303_2 | TATGAGTGTGGAGTCGTTGCACAGAAGAAATGCTAAAGGGAGTNNNAGATCGGAA |
| 13465_1 | TATGAGTGTGGAGTCGTTGCTTACATATCATCTCGTCCAATAANNNAGATCGGAAGA |
| 13465_2 | TATGAGTGTGGAGTCGTTGCTTACATATCATCTCGTCCAATAANNNAGATCGGAAGA |
| 5620_1 | TATGAGTGTGGAGTCGTTGCACTCATTCTTTCATTGCCCCGTTNNNAGATCGGAAGA |
| 5620_2 | TATGAGTGTGGAGTCGTTGCACTCATTCTTTCATTGCCCCGTTNNNAGATCGGAAGA |
| 39219_1 | TATGAGTGTGGAGTCGTTGCCAGCTATTGTGAAAGGGGTTGTGNNNAGATCGGAA |
| 39219_2 | TATGAGTGTGGAGTCGTTGCCAGCTATTGTGAAAGGGGTTGTGNNNAGATCGGAA |
| 52100_1 | TATGAGTGTGGAGTCGTTGCCAGCTTTCCAGACTGAACCAATGNNNAGATCGGAAG |
| 52100_2 | TATGAGTGTGGAGTCGTTGCCAGCTTTCCAGACTGAACCAATGNNNAGATCGGAAG |
| 39055_1 | TATGAGTGTGGAGTCGTTGCTCCCCAAATCACTAAGCTAAGGGNNNAGATCGGAAG |
| 39055_2 | TATGAGTGTGGAGTCGTTGCTCCCCAAATCACTAAGCTAAGGGNNNAGATCGGAAG |
| 3092_1 | TATGAGTGTGGAGTCGTTGCGCCCACAGCATAAACAAAGTTGGNNNAGATCGGAA |
| 3092_2 | TATGAGTGTGGAGTCGTTGCGCCCACAGCATAAACAAAGTTGGNNNAGATCGGAA |
| 40175_1 | TATGAGTGTGGAGTCGTTGCCTTTCCAGACACTTTCCCTACCANNNAGATCGGAAGA |
| 40175_2 | TATGAGTGTGGAGTCGTTGCCTTTCCAGACACTTTCCCTACCANNNAGATCGGAAGA |
| 30314_1 | TGAGTGTGGAGTCGTTGCACTTCTTGTGGAAGAAACATTAAGTGCNNNAGATCGGA |
| 30314_2 | TGAGTGTGGAGTCGTTGCACTTCTTGTGGAAGAAACATTAAGTGCNNNAGATCGGA |
| 31524_1 | TATGAGTGTGGAGTCGTTGCCCACATATTGTCCCAGTTCTCCANNNAGATCGGAAGA |
| 31524_2 | TATGAGTGTGGAGTCGTTGCCCACATATTGTCCCAGTTCTCCANNNAGATCGGAAGA |
| 40949_1 | TATGAGTGTGGAGTCGTTGCAAGTCTCTTCCCTTTACTTCGGCNNNAGATCGGAAGA |
| 40949_2 | TATGAGTGTGGAGTCGTTGCAAGTCTCTTCCCTTTACTTCGGCNNNAGATCGGAAGA |
| 43295_1 | TATGAGTGTGGAGTCGTTGCGGGTTCAAGTGATTCTCCTGTCTNNNAGATCGGAAG |
| 43295_2 | TATGAGTGTGGAGTCGTTGCGGGTTCAAGTGATTCTCCTGTCTNNNAGATCGGAAG |
| 41484_1 | TATGAGTGTGGAGTCGTTGCACAACTGAAGAAGGTGTCACAGANNNAGATCGGAA |
| 41484_2 | TATGAGTGTGGAGTCGTTGCACAACTGAAGAAGGTGTCACAGANNNAGATCGGAA |
| 49317_1 | TATGAGTGTGGAGTCGTTGCCTCCCAGGTTCATGCCATTCTCNNNAGATCGGAAGA |
| 49317_2 | TATGAGTGTGGAGTCGTTGCCTCCCAGGTTCATGCCATTCTCNNNAGATCGGAAGA |
| 44658_1 | TATGAGTGTGGAGTCGTTGCCACCAAGTGTACTTTGATGCACCNNNAGATCGGAAG |
| 44658_2 | TATGAGTGTGGAGTCGTTGCCACCAAGTGTACTTTGATGCACCNNNAGATCGGAAG |
| 46986_1 | TATGAGTGTGGAGTCGTTGCCTGAGTAGCTGGGATTACAGGTGNNNAGATCGGAA |
| 46986_2 | TATGAGTGTGGAGTCGTTGCCTGAGTAGCTGGGATTACAGGTGNNNAGATCGGAA |
| 44627_1 | TATGAGTGTGGAGTCGTTGCACTGGAGGGGATTTTTCAGATCCNNNAGATCGGAA |

| | |
|---|---|
| 44627_2 | TATGAGTGTGGAGTCGTTGCACTGGAGGGGATTTTTCAGATCCNNNAGATCGGAA |
| 42717_1 | TATGAGTGTGGAGTCGTTGCCAGTGGAGCACAACATAGAATGCNNNAGATCGGAA |
| 42717_2 | TATGAGTGTGGAGTCGTTGCCAGTGGAGCACAACATAGAATGCNNNAGATCGGAA |
| 35920_1 | TATGAGTGTGGAGTCGTTGCGTGTTCAGATTGAGCCATTGCATNNNAGATCGGAAG |
| 35920_2 | TATGAGTGTGGAGTCGTTGCGTGTTCAGATTGAGCCATTGCATNNNAGATCGGAAG |
| 30765_1 | TATGAGTGTGGAGTCGTTGCTGAGGTTTCCTTTTGTGTTGTCCNNNAGATCGGAAG |
| 30765_2 | TATGAGTGTGGAGTCGTTGCTGAGGTTTCCTTTTGTGTTGTCCNNNAGATCGGAAG |
| 34508_1 | TATGAGTGTGGAGTCGTTGCTGTTCAGTTCCCACCTATGAGTGNNNAGATCGGAAG |
| 34508_2 | TATGAGTGTGGAGTCGTTGCTGTTCAGTTCCCACCTATGAGTGNNNAGATCGGAAG |
| 47120_1 | TATGAGTGTGGAGTCGTTGCCTAGTAGCTGGGATTACAGGCGNNNAGATCGGAAG |
| 47120_2 | TATGAGTGTGGAGTCGTTGCCTAGTAGCTGGGATTACAGGCGNNNAGATCGGAAG |
| 2584_1 | TATGAGTGTGGAGTCGTTGCAACCTTCACTGCAATTTTGCTGGNNNAGATCGGAAG |
| 2584_2 | TATGAGTGTGGAGTCGTTGCAACCTTCACTGCAATTTTGCTGGNNNAGATCGGAAG |
| 36645_1 | TATGAGTGTGGAGTCGTTGCAGCATTCCAAAATCAAGCCATGTNNNAGATCGGAAG |
| 36645_2 | TATGAGTGTGGAGTCGTTGCAGCATTCCAAAATCAAGCCATGTNNNAGATCGGAAG |
| 41270_1 | TATGAGTGTGGAGTCGTTGCTTTCACAGGCAATGTCCATCCTANNNAGATCGGAAG |
| 41270_2 | TATGAGTGTGGAGTCGTTGCTTTCACAGGCAATGTCCATCCTANNNAGATCGGAAG |
| 30003_1 | TATGAGTGTGGAGTCGTTGCGCGAAAACTATCATCAGCTCCACNNNAGATCGGAAG |
| 30003_2 | TATGAGTGTGGAGTCGTTGCGCGAAAACTATCATCAGCTCCACNNNAGATCGGAAG |
| 51282_1 | TATGAGTGTGGAGTCGTTGCGTTGTGGTGGCAGTGGTGAGNNNAGATCGGAAGA |
| 51282_2 | TATGAGTGTGGAGTCGTTGCGTTGTGGTGGCAGTGGTGAGNNNAGATCGGAAGA |
| 41863_1 | TATGAGTGTGGAGTCGTTGCCACTACGCCCGGCTAATTTTTTNNNAGATCGGAAGAG |
| 41863_2 | TATGAGTGTGGAGTCGTTGCCACTACGCCCGGCTAATTTTTTNNNAGATCGGAAGAG |
| 640_1 | TATGAGTGTGGAGTCGTTGCCAAGAGGAAGAGGGGGCTTTAGAANNNAGATCGG |
| 640_2 | TATGAGTGTGGAGTCGTTGCCAAGAGGAAGAGGGGGCTTTAGAANNNAGATCGG |
| 53246_1 | TATGAGTGTGGAGTCGTTGCCTCACAGTCACCGATTCTCCAAGNNNAGATCGGAAG |
| 53246_2 | TATGAGTGTGGAGTCGTTGCCTCACAGTCACCGATTCTCCAAGNNNAGATCGGAAG |
| 40620_1 | TATGAGTGTGGAGTCGTTGCCCCAGGCAATGAAAATGTATCCCNNNAGATCGGAAG |
| 40620_2 | TATGAGTGTGGAGTCGTTGCCCCAGGCAATGAAAATGTATCCCNNNAGATCGGAAG |
| 50127_1 | TATGAGTGTGGAGTCGTTGCCTCTTCACCCTTTCTACCACCAANNNAGATCGGAAGA |
| 50127_2 | TATGAGTGTGGAGTCGTTGCCTCTTCACCCTTTCTACCACCAANNNAGATCGGAAGA |
| 31551_1 | TATGAGTGTGGAGTCGTTGCTCCTGCTTCATTTTCGGAAACACNNNAGATCGGAAG |
| 31551_2 | TATGAGTGTGGAGTCGTTGCTCCTGCTTCATTTTCGGAAACACNNNAGATCGGAAG |
| 42247_1 | TATGAGTGTGGAGTCGTTGCAATTCATGTGATAGGTGGCACCANNNAGATCGGAA |
| 42247_2 | TATGAGTGTGGAGTCGTTGCAATTCATGTGATAGGTGGCACCANNNAGATCGGAA |
| 47054_1 | TATGAGTGTGGAGTCGTTGCGGTATATTGTGTAATCCCGGGCTNNNAGATCGGAA |
| 47054_2 | TATGAGTGTGGAGTCGTTGCGGTATATTGTGTAATCCCGGGCTNNNAGATCGGAA |
| 10995_1 | TATGAGTGTGGAGTCGTTGCGCCCTGCAAAGGATTCCATGTTNNNAGATCGGAAGA |
| 10995_2 | TATGAGTGTGGAGTCGTTGCGCCCTGCAAAGGATTCCATGTTNNNAGATCGGAAGA |
| 30163_1 | TATGAGTGTGGAGTCGTTGCGACGGGAAAGAGACTGTGTAAGTNNNAGATCGGA |
| 30163_2 | TATGAGTGTGGAGTCGTTGCGACGGGAAAGAGACTGTGTAAGTNNNAGATCGGA |
| 33816_1 | TATGAGTGTGGAGTCGTTGCGAGCCAGAGCAATCTTGAACAAANNNAGATCGGAA |
| 33816_2 | TATGAGTGTGGAGTCGTTGCGAGCCAGAGCAATCTTGAACAAANNNAGATCGGAA |

| | |
|---|---|
| 45375_1 | TATGAGTGTGGAGTCGTTGCTGGAGGGTAATCAGGAGGGTATTNNNAGATCGGA |
| 45375_2 | TATGAGTGTGGAGTCGTTGCTGGAGGGTAATCAGGAGGGTATTNNNAGATCGGA |
| 51709_1 | TATGAGTGTGGAGTCGTTGCGCCTTCCTTTGTAGCTTGTTCAGNNNAGATCGGAAG |
| 51709_2 | TATGAGTGTGGAGTCGTTGCGCCTTCCTTTGTAGCTTGTTCAGNNNAGATCGGAAG |
| 36065_1 | TATGAGTGTGGAGTCGTTGCCATATGATCGCTACCACAACAGCNNNAGATCGGAAG |
| 36065_2 | TATGAGTGTGGAGTCGTTGCCATATGATCGCTACCACAACAGCNNNAGATCGGAAG |
| 49982_1 | TATGAGTGTGGAGTCGTTGCCTTCTGCATTGAGCTTCAAGGACNNNAGATCGGAAG |
| 49982_2 | TATGAGTGTGGAGTCGTTGCCTTCTGCATTGAGCTTCAAGGACNNNAGATCGGAAG |
| 52440_1 | TATGAGTGTGGAGTCGTTGCTGCACGGAGTATTCAAGGGTAAANNNAGATCGGAA |
| 52440_2 | TATGAGTGTGGAGTCGTTGCTGCACGGAGTATTCAAGGGTAAANNNAGATCGGAA |
| 51489_1 | TATGAGTGTGGAGTCGTTGCGATTGATTGTGTGTGTGTGGTGTNNNAGATCGGAA |
| 51489_2 | TATGAGTGTGGAGTCGTTGCGATTGATTGTGTGTGTGTGGTGTNNNAGATCGGAA |
| 47896_1 | TATGAGTGTGGAGTCGTTGCTGCAATATCGGTTCACTGCAATCNNNAGATCGGAAG |
| 47896_2 | TATGAGTGTGGAGTCGTTGCTGCAATATCGGTTCACTGCAATCNNNAGATCGGAAG |
| 40699_1 | TATGAGTGTGGAGTCGTTGCCATGCCTGTAATCCCAGCTACTNNNAGATCGGAAGA |
| 40699_2 | TATGAGTGTGGAGTCGTTGCCATGCCTGTAATCCCAGCTACTNNNAGATCGGAAGA |
| 44496_1 | TATGAGTGTGGAGTCGTTGCCAAATCCTGGACACTGTGGTAGANNNAGATCGGAA |
| 44496_2 | TATGAGTGTGGAGTCGTTGCCAAATCCTGGACACTGTGGTAGANNNAGATCGGAA |
| 28914_1 | TATGAGTGTGGAGTCGTTGCTGAGCCAAATGCACCCCTTTATANNNAGATCGGAAG |
| 28914_2 | TATGAGTGTGGAGTCGTTGCTGAGCCAAATGCACCCCTTTATANNNAGATCGGAAG |
| 39685_1 | TATGAGTGTGGAGTCGTTGCCTGCGGACACAAAGATAATTCCCNNNAGATCGGAAG |
| 39685_2 | TATGAGTGTGGAGTCGTTGCCTGCGGACACAAAGATAATTCCCNNNAGATCGGAAG |
| 3157_1 | TATGAGTGTGGAGTCGTTGCATAGTCCATTGGAAAGGGAGGCANNNAGATCGGAA |
| 3157_2 | TATGAGTGTGGAGTCGTTGCATAGTCCATTGGAAAGGGAGGCANNNAGATCGGAA |
| 33842_1 | TATGAGTGTGGAGTCGTTGCAAACCCAACCAACCAAACAAACANNNAGATCGGAAG |
| 33842_2 | TATGAGTGTGGAGTCGTTGCAAACCCAACCAACCAAACAAACANNNAGATCGGAAG |
| 12933_1 | TATGAGTGTGGAGTCGTTGCTTTTCTGGAATGGAAAATTAAGTNNNAGATCGGAA |
| 12933_2 | TATGAGTGTGGAGTCGTTGCTTTTCTGGAATGGAAAATTAAGTNNNAGATCGGAA |
| 44358_1 | TATGAGTGTGGAGTCGTTGCAGAGTTATACCAGTCAGGGTCCANNNAGATCGGAA |
| 44358_2 | TATGAGTGTGGAGTCGTTGCAGAGTTATACCAGTCAGGGTCCANNNAGATCGGAA |
| 45646_1 | TATGAGTGTGGAGTCGTTGCCTCCCAAGTTCACGCGATTCTNNNAGATCGGAAGAG |
| 45646_2 | TATGAGTGTGGAGTCGTTGCCTCCCAAGTTCACGCGATTCTNNNAGATCGGAAGAG |
| 45699_1 | TATGAGTGTGGAGTCGTTGCCATGATATCGGCTCACTGCAACNNNAGATCGGAAGA |
| 45699_2 | TATGAGTGTGGAGTCGTTGCCATGATATCGGCTCACTGCAACNNNAGATCGGAAGA |
| 51315_1 | TATGAGTGTGGAGTCGTTGCCAGCTGCTGTGCTCGATTTCNNNAGATCGGAAGAGC |
| 51315_2 | TATGAGTGTGGAGTCGTTGCCAGCTGCTGTGCTCGATTTCNNNAGATCGGAAGAGC |
| 48355_1 | TATGAGTGTGGAGTCGTTGCTTCAAGCAATTCTCCTGCCTCAGNNNAGATCGGAAG |
| 48355_2 | TATGAGTGTGGAGTCGTTGCTTCAAGCAATTCTCCTGCCTCAGNNNAGATCGGAAG |
| 36708_1 | TATGAGTGTGGAGTCGTTGCCTTCTGTCCCCTTCGTTTAGTCCNNNAGATCGGAAGA |
| 36708_2 | TATGAGTGTGGAGTCGTTGCCTTCTGTCCCCTTCGTTTAGTCCNNNAGATCGGAAGA |
| 33741_1 | TATGAGTGTGGAGTCGTTGCACAAAAAAACTGTCCAGTGAGCTNNNAGATCGGAA |
| 33741_2 | TATGAGTGTGGAGTCGTTGCACAAAAAAACTGTCCAGTGAGCTNNNAGATCGGAA |
| 33789_1 | TATGAGTGTGGAGTCGTTGCTTGGCACAACTCTTATAGTGGGTNNNAGATCGGAAG |

| | |
|---|---|
| 33789_2 | TATGAGTGTGGAGTCGTTGCTTGGCACAACTCTTATAGTGGGTNNNAGATCGGAAG |
| 47302_1 | TATGAGTGTGGAGTCGTTGCGGTTCAAGTGATTCTCCTGCCTCNNNAGATCGGAAG |
| 47302_2 | TATGAGTGTGGAGTCGTTGCGGTTCAAGTGATTCTCCTGCCTCNNNAGATCGGAAG |
| 47999_1 | TATGAGTGTGGAGTCGTTGCGACCAGGTTGTCACTGAGAAGATNNNAGATCGGAA |
| 47999_2 | TATGAGTGTGGAGTCGTTGCGACCAGGTTGTCACTGAGAAGATNNNAGATCGGAA |
| 31590_1 | TATGAGTGTGGAGTCGTTGCTTTGGGGCAATCCAGAAATCAACNNNAGATCGGAAG |
| 31590_2 | TATGAGTGTGGAGTCGTTGCTTTGGGGCAATCCAGAAATCAACNNNAGATCGGAAG |
| 33740_1 | TATGAGTGTGGAGTCGTTGCAGCTCAGATCACTTTTCTCACCTNNNAGATCGGAAGA |
| 33740_2 | TATGAGTGTGGAGTCGTTGCAGCTCAGATCACTTTTCTCACCTNNNAGATCGGAAGA |
| 51887_1 | TATGAGTGTGGAGTCGTTGCATTCTTTCTTTCCCTTCCCACCANNNAGATCGGAAGAG |
| 51887_2 | TATGAGTGTGGAGTCGTTGCATTCTTTCTTTCCCTTCCCACCANNNAGATCGGAAGAG |
| 41968_1 | TATGAGTGTGGAGTCGTTGCGGGGAGAGGGGCATTGTAAATATNNNAGATCGGA |
| 41968_2 | TATGAGTGTGGAGTCGTTGCGGGGAGAGGGGCATTGTAAATATNNNAGATCGGA |
| 32406_1 | TATGAGTGTGGAGTCGTTGCTGGCAAGAGGAATGTGGATAACANNNAGATCGGAA |
| 32406_2 | TATGAGTGTGGAGTCGTTGCTGGCAAGAGGAATGTGGATAACANNNAGATCGGAA |
| 48237_1 | TATGAGTGTGGAGTCGTTGCGAGGTCTCACTATATTGCCCAGGNNNAGATCGGAAG |
| 48237_2 | TATGAGTGTGGAGTCGTTGCGAGGTCTCACTATATTGCCCAGGNNNAGATCGGAAG |
| 30558_1 | ATGAGTGTGGAGTCGTTGCTCCTTAAATGTAGAGCTGTTATGATTTNNNAGATCGG |
| 30558_2 | ATGAGTGTGGAGTCGTTGCTCCTTAAATGTAGAGCTGTTATGATTTNNNAGATCGG |
| 12836_1 | TATGAGTGTGGAGTCGTTGCCTATGAAGTCATGGAAGCTCNNNAGATCGGAAGAG |
| 12836_2 | TATGAGTGTGGAGTCGTTGCCTATGAAGTCATGGAAGCTCNNNAGATCGGAAGAG |
| 8346_1 | TATGAGTGTGGAGTCGTTGCGATAACCTCTGCTCACCGCAANNNAGATCGGAAGAG |
| 8346_2 | TATGAGTGTGGAGTCGTTGCGATAACCTCTGCTCACCGCAANNNAGATCGGAAGAG |
| 47479_1 | TATGAGTGTGGAGTCGTTGCGCCTCTCAAGTAGCTGGGATTACNNNAGATCGGAAG |
| 47479_2 | TATGAGTGTGGAGTCGTTGCGCCTCTCAAGTAGCTGGGATTACNNNAGATCGGAAG |
| 48014_1 | TATGAGTGTGGAGTCGTTGCGGAGTTGTGTTCTTATTGCCCAGNNNAGATCGGAAG |
| 48014_2 | TATGAGTGTGGAGTCGTTGCGGAGTTGTGTTCTTATTGCCCAGNNNAGATCGGAAG |
| 37527_1 | TATGAGTGTGGAGTCGTTGCGCAGTGGTTCATGCCTGTAATCNNNAGATCGGAAGA |
| 37527_2 | TATGAGTGTGGAGTCGTTGCGCAGTGGTTCATGCCTGTAATCNNNAGATCGGAAGA |
| 33088_1 | TATGAGTGTGGAGTCGTTGCGAGGGGACAACATCTTTTCTGGANNNAGATCGGAA |
| 33088_2 | TATGAGTGTGGAGTCGTTGCGAGGGGACAACATCTTTTCTGGANNNAGATCGGAA |
| 50845_1 | TATGAGTGTGGAGTCGTTGCAGGAAGCCTGAATGGGAATGAAANNNAGATCGGAA |
| 50845_2 | TATGAGTGTGGAGTCGTTGCAGGAAGCCTGAATGGGAATGAAANNNAGATCGGAA |
| 41285_1 | TATGAGTGTGGAGTCGTTGCACATGAACATGGGCATTACACTGNNNAGATCGGAAG |
| 41285_2 | TATGAGTGTGGAGTCGTTGCACATGAACATGGGCATTACACTGNNNAGATCGGAAG |
| 50950_1 | TATGAGTGTGGAGTCGTTGCGGAACTTGAGCAGGAAGTCGTAGNNNAGATCGGAA |
| 50950_2 | TATGAGTGTGGAGTCGTTGCGGAACTTGAGCAGGAAGTCGTAGNNNAGATCGGAA |
| 35390_1 | TATGAGTGTGGAGTCGTTGCGGCTTGGTAATGGCAAATTTCCTNNNAGATCGGAAG |
| 35390_2 | TATGAGTGTGGAGTCGTTGCGGCTTGGTAATGGCAAATTTCCTNNNAGATCGGAAG |
| 47575_1 | TATGAGTGTGGAGTCGTTGCAGCCACTAGTCCACTTTTGTGATNNNAGATCGGAAG |
| 47575_2 | TATGAGTGTGGAGTCGTTGCAGCCACTAGTCCACTTTTGTGATNNNAGATCGGAAG |
| 35237_1 | TATGAGTGTGGAGTCGTTGCGATTTTGACACTGACGGTCTTGGNNNAGATCGGAAG |
| 35237_2 | TATGAGTGTGGAGTCGTTGCGATTTTGACACTGACGGTCTTGGNNNAGATCGGAAG |

| | |
|---|---|
| 41168_1 | TATGAGTGTGGAGTCGTTGCTGTGAACAGTTTGGGGTGACATANNNAGATCGGAA |
| 41168_2 | TATGAGTGTGGAGTCGTTGCTGTGAACAGTTTGGGGTGACATANNNAGATCGGAA |
| 48821_1 | TATGAGTGTGGAGTCGTTGCATCAGTGCAGACAGTGTGTTAGTNNNAGATCGGAA |
| 48821_2 | TATGAGTGTGGAGTCGTTGCATCAGTGCAGACAGTGTGTTAGTNNNAGATCGGAA |
| 43780_1 | TATGAGTGTGGAGTCGTTGCTAGCCATTGTAACAGGTGTGGAANNNAGATCGGAA |
| 43780_2 | TATGAGTGTGGAGTCGTTGCTAGCCATTGTAACAGGTGTGGAANNNAGATCGGAA |
| 29284_1 | TATGAGTGTGGAGTCGTTGCAGTTTTCCTGTGTTTTTGCCAATNNNAGATCGGAAG |
| 29284_2 | TATGAGTGTGGAGTCGTTGCAGTTTTCCTGTGTTTTTGCCAATNNNAGATCGGAAG |
| 32285_1 | TATGAGTGTGGAGTCGTTGCGGCTCGCAACTAATTTCCTGTTTNNNAGATCGGAAG |
| 32285_2 | TATGAGTGTGGAGTCGTTGCGGCTCGCAACTAATTTCCTGTTTNNNAGATCGGAAG |
| 51800_1 | TATGAGTGTGGAGTCGTTGCGTGATTTGGTGTTGATCGCACANNNAGATCGGAAG |
| 51800_2 | TATGAGTGTGGAGTCGTTGCGTGATTTGGTGTTGATCGCACANNNAGATCGGAAG |
| 51696_1 | TATGAGTGTGGAGTCGTTGCGCATGGCCCTAAGAATTTCACAANNNAGATCGGAAG |
| 51696_2 | TATGAGTGTGGAGTCGTTGCGCATGGCCCTAAGAATTTCACAANNNAGATCGGAAG |
| 35093_1 | TATGAGTGTGGAGTCGTTGCCATTCTCCTGCCTCAGCCTCNNNAGATCGGAAGAGCA |
| 35093_2 | TATGAGTGTGGAGTCGTTGCCATTCTCCTGCCTCAGCCTCNNNAGATCGGAAGAGCA |
| 48610_1 | TATGAGTGTGGAGTCGTTGCGAACTATCAGGCCTAGGATCAGCNNNAGATCGGAA |
| 48610_2 | TATGAGTGTGGAGTCGTTGCGAACTATCAGGCCTAGGATCAGCNNNAGATCGGAA |
| 47844_1 | TATGAGTGTGGAGTCGTTGCTCTCTCTCTCTTACTCAGGCTGGNNNAGATCGGAAG |
| 47844_2 | TATGAGTGTGGAGTCGTTGCTCTCTCTCTCTTACTCAGGCTGGNNNAGATCGGAAG |
| 46072_1 | TATGAGTGTGGAGTCGTTGCAGAGATGGGGTTTCACGTGTTAGNNNAGATCGGAA |
| 46072_2 | TATGAGTGTGGAGTCGTTGCAGAGATGGGGTTTCACGTGTTAGNNNAGATCGGAA |
| 42906_1 | TATGAGTGTGGAGTCGTTGCTGCTATTGTGAATAGTGCTGTGANNNAGATCGGAA |
| 42906_2 | TATGAGTGTGGAGTCGTTGCTGCTATTGTGAATAGTGCTGTGANNNAGATCGGAA |
| 38386_1 | TATGAGTGTGGAGTCGTTGCGAGAGATCTCGAACTGTGTCAGTNNNAGATCGGAA |
| 38386_2 | TATGAGTGTGGAGTCGTTGCGAGAGATCTCGAACTGTGTCAGTNNNAGATCGGAA |
| 13433_1 | TATGAGTGTGGAGTCGTTGCCTGAAAGTCGCAAAAGTTCTNNNAGATCGGAAGAG |
| 13433_2 | TATGAGTGTGGAGTCGTTGCCTGAAAGTCGCAAAAGTTCTNNNAGATCGGAAGAG |
| 34022_1 | TATGAGTGTGGAGTCGTTGCGTTCAAGAGATTCTCCTGCCTCANNNAGATCGGAAG |
| 34022_2 | TATGAGTGTGGAGTCGTTGCGTTCAAGAGATTCTCCTGCCTCANNNAGATCGGAAG |
| 36573_1 | TATGAGTGTGGAGTCGTTGCGAGCAGAAAACTGAAAGCACTGTNNNAGATCGGAA |
| 36573_2 | TATGAGTGTGGAGTCGTTGCGAGCAGAAAACTGAAAGCACTGTNNNAGATCGGAA |
| 47384_1 | TATGAGTGTGGAGTCGTTGCGAACTACTGACCTCAGGTGATCCNNNAGATCGGAAG |
| 47384_2 | TATGAGTGTGGAGTCGTTGCGAACTACTGACCTCAGGTGATCCNNNAGATCGGAAG |
| 44930_1 | TATGAGTGTGGAGTCGTTGCTGCTCATGTTGATGGAGATGTCANNNAGATCGGAA |
| 44930_2 | TATGAGTGTGGAGTCGTTGCTGCTCATGTTGATGGAGATGTCANNNAGATCGGAA |
| 51680_1 | TATGAGTGTGGAGTCGTTGCTGGTTTCACTGACCTTTGGCTATNNNAGATCGGAAG |
| 51680_2 | TATGAGTGTGGAGTCGTTGCTGGTTTCACTGACCTTTGGCTATNNNAGATCGGAAG |
| 35975_1 | TATGAGTGTGGAGTCGTTGCAAGAAAAGAAAGAGTTGCTGCCCNNNAGATCGGAA |
| 35975_2 | TATGAGTGTGGAGTCGTTGCAAGAAAAGAAAGAGTTGCTGCCCNNNAGATCGGAA |
| 13183_1 | TATGAGTGTGGAGTCGTTGCCATCCAAGTTCTTCTTACTACTTNNNAGATCGGAAGA |
| 13183_2 | TATGAGTGTGGAGTCGTTGCCATCCAAGTTCTTCTTACTACTTNNNAGATCGGAAGA |
| 42176_1 | TATGAGTGTGGAGTCGTTGCCTCCTGAGTAGCTGGGACTACAGNNNAGATCGGAA |

| | |
|---|---|
| 42176_2 | TATGAGTGTGGAGTCGTTGCCTCCTGAGTAGCTGGGACTACAGNNNAGATCGGAA |
| 40633_1 | TATGAGTGTGGAGTCGTTGCCTCCTGGGTTCATGCCATTCTCNNNAGATCGGAAGA |
| 40633_2 | TATGAGTGTGGAGTCGTTGCCTCCTGGGTTCATGCCATTCTCNNNAGATCGGAAGA |
| 44400_1 | TATGAGTGTGGAGTCGTTGCCCCCTCTCTGGTGATCTTGTAAGNNNAGATCGGAAG |
| 44400_2 | TATGAGTGTGGAGTCGTTGCCCCCTCTCTGGTGATCTTGTAAGNNNAGATCGGAAG |
| 40028_1 | TATGAGTGTGGAGTCGTTGCTGGCAATGAGTAGCAAAAGAACCNNNAGATCGGAA |
| 40028_2 | TATGAGTGTGGAGTCGTTGCTGGCAATGAGTAGCAAAAGAACCNNNAGATCGGAA |
| 34823_1 | TATGAGTGTGGAGTCGTTGCCTCCTGAGTAGCTGGGATTACAGNNNAGATCGGAA |
| 34823_2 | TATGAGTGTGGAGTCGTTGCCTCCTGAGTAGCTGGGATTACAGNNNAGATCGGAA |
| 34330_1 | TATGAGTGTGGAGTCGTTGCTTCCACTACCCTTTCATTCCTGGNNNAGATCGGAAGA |
| 34330_2 | TATGAGTGTGGAGTCGTTGCTTCCACTACCCTTTCATTCCTGGNNNAGATCGGAAGA |
| 30852_1 | TATGAGTGTGGAGTCGTTGCTGGAGATGGCAGACTCTAAGAGANNNAGATCGGAA |
| 30852_2 | TATGAGTGTGGAGTCGTTGCTGGAGATGGCAGACTCTAAGAGANNNAGATCGGAA |
| 35429_1 | TATGAGTGTGGAGTCGTTGCGCACAATCTCTGCTCACTTCAAGNNNAGATCGGAAG |
| 35429_2 | TATGAGTGTGGAGTCGTTGCGCACAATCTCTGCTCACTTCAAGNNNAGATCGGAAG |
| 40329_1 | TATGAGTGTGGAGTCGTTGCTGGCTTTGGACTCTCTCTTTCTCNNNAGATCGGAAGA |
| 40329_2 | TATGAGTGTGGAGTCGTTGCTGGCTTTGGACTCTCTCTTTCTCNNNAGATCGGAAGA |
| 35671_1 | TATGAGTGTGGAGTCGTTGCCCAGGCCTGATTTTTGAGTTCTGNNNAGATCGGAAG |
| 35671_2 | TATGAGTGTGGAGTCGTTGCCCAGGCCTGATTTTTGAGTTCTGNNNAGATCGGAAG |
| 47629_1 | TATGAGTGTGGAGTCGTTGCCTTTCAAGGAACCAGCCTCATTGNNNAGATCGGAAG |
| 47629_2 | TATGAGTGTGGAGTCGTTGCCTTTCAAGGAACCAGCCTCATTGNNNAGATCGGAAG |
| 41994_1 | TATGAGTGTGGAGTCGTTGCAGATGAAGTCTTGCTCTGTCACCNNNAGATCGGAAG |
| 41994_2 | TATGAGTGTGGAGTCGTTGCAGATGAAGTCTTGCTCTGTCACCNNNAGATCGGAAG |
| 31874_1 | TATGAGTGTGGAGTCGTTGCGACCACAAGATCAGGAGATCGAGNNNAGATCGGAA |
| 31874_2 | TATGAGTGTGGAGTCGTTGCGACCACAAGATCAGGAGATCGAGNNNAGATCGGAA |
| 33450_1 | TATGAGTGTGGAGTCGTTGCTTCCCTTTTGCTACTGTCCACATNNNAGATCGGAAGA |
| 33450_2 | TATGAGTGTGGAGTCGTTGCTTCCCTTTTGCTACTGTCCACATNNNAGATCGGAAGA |
| 48927_1 | TATGAGTGTGGAGTCGTTGCTTTTTCTGCCATTGCTCAGGAAGNNNAGATCGGAAG |
| 48927_2 | TATGAGTGTGGAGTCGTTGCTTTTTCTGCCATTGCTCAGGAAGNNNAGATCGGAAG |
| 33778_1 | TATGAGTGTGGAGTCGTTGCCCTAGTGGTACATGCCAGTAGTCNNNAGATCGGAAG |
| 33778_2 | TATGAGTGTGGAGTCGTTGCCCTAGTGGTACATGCCAGTAGTCNNNAGATCGGAAG |
| 34073_1 | TATGAGTGTGGAGTCGTTGCGATCTCAGCTCACTGCAAACTCTNNNAGATCGGAAG |
| 34073_2 | TATGAGTGTGGAGTCGTTGCGATCTCAGCTCACTGCAAACTCTNNNAGATCGGAAG |
| 33511_1 | TATGAGTGTGGAGTCGTTGCGAGATCATGCCACTACACTCCAGNNNAGATCGGAAG |
| 33511_2 | TATGAGTGTGGAGTCGTTGCGAGATCATGCCACTACACTCCAGNNNAGATCGGAAG |
| 43408_1 | TATGAGTGTGGAGTCGTTGCAGACAGAGTCTCAATCTTGTTGCNNNAGATCGGAAG |
| 43408_2 | TATGAGTGTGGAGTCGTTGCAGACAGAGTCTCAATCTTGTTGCNNNAGATCGGAAG |
| 37121_1 | TATGAGTGTGGAGTCGTTGCTCCTTAGCAAAAGGCATCCTCATNNNAGATCGGAAG |
| 37121_2 | TATGAGTGTGGAGTCGTTGCTCCTTAGCAAAAGGCATCCTCATNNNAGATCGGAAG |
| 13247_1 | TATGAGTGTGGAGTCGTTGCGGAGAAGCCTAAGTAAAAATAAANNNAGATCGGAA |
| 13247_2 | TATGAGTGTGGAGTCGTTGCGGAGAAGCCTAAGTAAAAATAAANNNAGATCGGAA |
| 51001_1 | TATGAGTGTGGAGTCGTTGCAGAAATGGTCTCCATGGAGTCACNNNAGATCGGAA |
| 51001_2 | TATGAGTGTGGAGTCGTTGCAGAAATGGTCTCCATGGAGTCACNNNAGATCGGAA |

| | |
|---|---|
| 35235_1 | TATGAGTGTGGAGTCGTTGCACAGAGATAGGCCACAAAAAGGTNNNAGATCGGAA |
| 35235_2 | TATGAGTGTGGAGTCGTTGCACAGAGATAGGCCACAAAAAGGTNNNAGATCGGAA |
| 51428_1 | TATGAGTGTGGAGTCGTTGCAGCTCATGAGAAGTCACTCAGTGNNNAGATCGGAAG |
| 51428_2 | TATGAGTGTGGAGTCGTTGCAGCTCATGAGAAGTCACTCAGTGNNNAGATCGGAAG |
| 35025_1 | TATGAGTGTGGAGTCGTTGCACCTCTGCCTTCTGAGTTCAAATNNNAGATCGGAAGA |
| 35025_2 | TATGAGTGTGGAGTCGTTGCACCTCTGCCTTCTGAGTTCAAATNNNAGATCGGAAGA |
| 42316_1 | TATGAGTGTGGAGTCGTTGCACAATGTGGCAATTTCTGACTGGNNNAGATCGGAAG |
| 42316_2 | TATGAGTGTGGAGTCGTTGCACAATGTGGCAATTTCTGACTGGNNNAGATCGGAAG |
| 46168_1 | TATGAGTGTGGAGTCGTTGCCCACCACAGCCAGCTAATTTTTTNNNAGATCGGAAGA |
| 46168_2 | TATGAGTGTGGAGTCGTTGCCCACCACAGCCAGCTAATTTTTTNNNAGATCGGAAGA |
| 49365_1 | TATGAGTGTGGAGTCGTTGCTTGCTGGGATTACAGGCATAAGTNNNAGATCGGAAG |
| 49365_2 | TATGAGTGTGGAGTCGTTGCTTGCTGGGATTACAGGCATAAGTNNNAGATCGGAAG |
| 29059_1 | TATGAGTGTGGAGTCGTTGCCACACACACACACACATACACATNNNAGATCGGAAGA |
| 29059_2 | TATGAGTGTGGAGTCGTTGCCACACACACACACACATACACATNNNAGATCGGAAGA |
| 39337_1 | TATGAGTGTGGAGTCGTTGCACTGACACCACAGAAACAGAAGANNNAGATCGGAAG |
| 39337_2 | TATGAGTGTGGAGTCGTTGCACTGACACCACAGAAACAGAAGANNNAGATCGGAAG |
| 29465_1 | TATGAGTGTGGAGTCGTTGCTGGCAGAGGAGGAGCAATTCATTNNNAGATCGGAA |
| 29465_2 | TATGAGTGTGGAGTCGTTGCTGGCAGAGGAGGAGCAATTCATTNNNAGATCGGAA |
| 46997_1 | TATGAGTGTGGAGTCGTTGCACCACGCCTGGCTAATTTGTATTNNNAGATCGGAAGA |
| 46997_2 | TATGAGTGTGGAGTCGTTGCACCACGCCTGGCTAATTTGTATTNNNAGATCGGAAGA |
| 42796_1 | TATGAGTGTGGAGTCGTTGCACACACACACACAAATTAGCTGGNNNAGATCGGAAG |
| 42796_2 | TATGAGTGTGGAGTCGTTGCACACACACACACAAATTAGCTGGNNNAGATCGGAAG |
| 38655_1 | TATGAGTGTGGAGTCGTTGCGGAAGGAAGGTAGGTAGGTAGGTNNNAGATCGGA |
| 38655_2 | TATGAGTGTGGAGTCGTTGCGGAAGGAAGGTAGGTAGGTAGGTNNNAGATCGGA |
| 32421_1 | TATGAGTGTGGAGTCGTTGCTTTTTGTAGAGATGGGGTTTCCANNNAGATCGGAAG |
| 32421_2 | TATGAGTGTGGAGTCGTTGCTTTTTGTAGAGATGGGGTTTCCANNNAGATCGGAAG |
| 10806_1 | TATGAGTGTGGAGTCGTTGCGTGTGTGTGTTTTCTTAGCAGGCNNNAGATCGGAAG |
| 10806_2 | TATGAGTGTGGAGTCGTTGCGTGTGTGTGTTTTCTTAGCAGGCNNNAGATCGGAAG |
| 5597_1 | TATGAGTGTGGAGTCGTTGCCACACGATCTACGTGGAAGGACNNNAGATCGGAAGA |
| 5597_2 | TATGAGTGTGGAGTCGTTGCCACACGATCTACGTGGAAGGACNNNAGATCGGAAGA |
| 37518_1 | TATGAGTGTGGAGTCGTTGCGACGAATGGTCAAACAAACTGGTNNNAGATCGGAAG |
| 37518_2 | TATGAGTGTGGAGTCGTTGCGACGAATGGTCAAACAAACTGGTNNNAGATCGGAAG |
| 39418_1 | TATGAGTGTGGAGTCGTTGCTATTTTGAGACAGAGCCCCACTCNNNAGATCGGAAGA |
| 39418_2 | TATGAGTGTGGAGTCGTTGCTATTTTGAGACAGAGCCCCACTCNNNAGATCGGAAGA |
| 38602_1 | TATGAGTGTGGAGTCGTTGCGGTGGTACTTCAGCTTGGTATCANNNAGATCGGAAG |
| 38602_2 | TATGAGTGTGGAGTCGTTGCGGTGGTACTTCAGCTTGGTATCANNNAGATCGGAAG |
| 40686_1 | TATGAGTGTGGAGTCGTTGCAAAGAAAACCCCATCAGCCAATGNNNAGATCGGAAG |
| 40686_2 | TATGAGTGTGGAGTCGTTGCAAAGAAAACCCCATCAGCCAATGNNNAGATCGGAAG |
| 40801_1 | TATGAGTGTGGAGTCGTTGCCACAACAGAAAAACCAAGCTGTGNNNAGATCGGAAG |
| 40801_2 | TATGAGTGTGGAGTCGTTGCCACAACAGAAAAACCAAGCTGTGNNNAGATCGGAAG |
| 49729_1 | TATGAGTGTGGAGTCGTTGCCCTCTCAGGTTCAAGCGATTCTCNNNAGATCGGAAGA |
| 49729_2 | TATGAGTGTGGAGTCGTTGCCCTCTCAGGTTCAAGCGATTCTCNNNAGATCGGAAGA |
| 43572_1 | TATGAGTGTGGAGTCGTTGCTCATTATGAGGGCTGGCTTTCTTNNNAGATCGGAAG |

| | |
|---|---|
| 43572_2 | TATGAGTGTGGAGTCGTTGCTCATTATGAGGGCTGGCTTTCTTNNNAGATCGGAAG |
| 52591_1 | TATGAGTGTGGAGTCGTTGCTCCCGCCTCCTAAGTAACTCTTANNNAGATCGGAAGA |
| 52591_2 | TATGAGTGTGGAGTCGTTGCTCCCGCCTCCTAAGTAACTCTTANNNAGATCGGAAGA |
| 51308_1 | TATGAGTGTGGAGTCGTTGCATCTAAAACATAAGCAACCGGCCNNNAGATCGGAAG |
| 51308_2 | TATGAGTGTGGAGTCGTTGCATCTAAAACATAAGCAACCGGCCNNNAGATCGGAAG |
| 49044_1 | TATGAGTGTGGAGTCGTTGCCATGCGCCATTATTTCCAGCTAANNNAGATCGGAAG |
| 49044_2 | TATGAGTGTGGAGTCGTTGCCATGCGCCATTATTTCCAGCTAANNNAGATCGGAAG |
| 43835_1 | TATGAGTGTGGAGTCGTTGCAGGGGAAAAGATCAGAAGTGACCNNNAGATCGGA |
| 43835_2 | TATGAGTGTGGAGTCGTTGCAGGGGAAAAGATCAGAAGTGACCNNNAGATCGGA |
| 40110_1 | TATGAGTGTGGAGTCGTTGCCAGTTATTCTGGAAGCTGAGGCANNNAGATCGGAA |
| 40110_2 | TATGAGTGTGGAGTCGTTGCCAGTTATTCTGGAAGCTGAGGCANNNAGATCGGAA |
| 30524_1 | TATGAGTGTGGAGTCGTTGCTTCTGTTTGCAGCCCATTTCTCTNNNAGATCGGAAGA |
| 30524_2 | TATGAGTGTGGAGTCGTTGCTTCTGTTTGCAGCCCATTTCTCTNNNAGATCGGAAGA |
| 40572_1 | TATGAGTGTGGAGTCGTTGCGTAATCATATAGGGAGGTGGGGCNNNAGATCGGA |
| 40572_2 | TATGAGTGTGGAGTCGTTGCGTAATCATATAGGGAGGTGGGGCNNNAGATCGGA |
| 41209_1 | TATGAGTGTGGAGTCGTTGCCACACAGGAATAGGCCTCTTGTANNNAGATCGGAAG |
| 41209_2 | TATGAGTGTGGAGTCGTTGCCACACAGGAATAGGCCTCTTGTANNNAGATCGGAAG |
| 35976_1 | TATGAGTGTGGAGTCGTTGCGGTCTCTCTTAGCCATCACAGAGNNNAGATCGGAAG |
| 35976_2 | TATGAGTGTGGAGTCGTTGCGGTCTCTCTTAGCCATCACAGAGNNNAGATCGGAAG |
| 46162_1 | TATGAGTGTGGAGTCGTTGCAATGGGGTCTTACCATATTGCCCNNNAGATCGGAAG |
| 46162_2 | TATGAGTGTGGAGTCGTTGCAATGGGGTCTTACCATATTGCCCNNNAGATCGGAAG |
| 40111_1 | TATGAGTGTGGAGTCGTTGCCACCTTTGCCCAAGTTTTGCTTNNNAGATCGGAAGAG |
| 40111_2 | TATGAGTGTGGAGTCGTTGCCACCTTTGCCCAAGTTTTGCTTNNNAGATCGGAAGAG |
| 46778_1 | TATGAGTGTGGAGTCGTTGCCTTTCTTCTGCTGGGTTTGGTTTNNNAGATCGGAAG |
| 46778_2 | TATGAGTGTGGAGTCGTTGCCTTTCTTCTGCTGGGTTTGGTTTNNNAGATCGGAAG |
| 36923_1 | TATGAGTGTGGAGTCGTTGCAGAAATGCTATGCTGGTACAAACTNNNAGATCGGA |
| 36923_2 | TATGAGTGTGGAGTCGTTGCAGAAATGCTATGCTGGTACAAACTNNNAGATCGGA |
| 45622_1 | TATGAGTGTGGAGTCGTTGCAGATGAAGTCTCACTCTGTCGCNNNAGATCGGAAGA |
| 45622_2 | TATGAGTGTGGAGTCGTTGCAGATGAAGTCTCACTCTGTCGCNNNAGATCGGAAGA |
| 51485_1 | TATGAGTGTGGAGTCGTTGCGAGAAGTAAACTCAGGGGACACCNNNAGATCGGAA |
| 51485_2 | TATGAGTGTGGAGTCGTTGCGAGAAGTAAACTCAGGGGACACCNNNAGATCGGAA |
| 12942_1 | TATGAGTGTGGAGTCGTTGCGAGAAGGAGGAAAAAAAACAGNNNAGATCGGAAG |
| 12942_2 | TATGAGTGTGGAGTCGTTGCGAGAAGGAGGAAAAAAAACAGNNNAGATCGGAAG |
| 34088_1 | TATGAGTGTGGAGTCGTTGCCAGAATGAACTTGCAGGGGTTTTNNNAGATCGGAA |
| 34088_2 | TATGAGTGTGGAGTCGTTGCCAGAATGAACTTGCAGGGGTTTTNNNAGATCGGAA |
| 36987_1 | TATGAGTGTGGAGTCGTTGCCCCATCTCTTCTGGCTTGTAAGANNNAGATCGGAAG |
| 36987_2 | TATGAGTGTGGAGTCGTTGCCCCATCTCTTCTGGCTTGTAAGANNNAGATCGGAAG |
| 46468_1 | TATGAGTGTGGAGTCGTTGCGTAGCTGGGACTACAGGCATAAGNNNAGATCGGAA |
| 46468_2 | TATGAGTGTGGAGTCGTTGCGTAGCTGGGACTACAGGCATAAGNNNAGATCGGAA |
| 40755_1 | TATGAGTGTGGAGTCGTTGCACCCATCAATCCCATTACTGAGTNNNAGATCGGAAG |
| 40755_2 | TATGAGTGTGGAGTCGTTGCACCCATCAATCCCATTACTGAGTNNNAGATCGGAAG |
| 49713_1 | TATGAGTGTGGAGTCGTTGCTTGAGTTCGTCATAGATGCTGGANNNAGATCGGAA |
| 49713_2 | TATGAGTGTGGAGTCGTTGCTTGAGTTCGTCATAGATGCTGGANNNAGATCGGAA |

| | |
|---|---|
| 41420_1 | TATGAGTGTGGAGTCGTTGCGAGTTAGACCCCTTTATCCCACCNNNAGATCGGAAG |
| 41420_2 | TATGAGTGTGGAGTCGTTGCGAGTTAGACCCCTTTATCCCACCNNNAGATCGGAAG |
| 40389_1 | TATGAGTGTGGAGTCGTTGCCAACAAGAGCGAAACTCCATCTCNNNAGATCGGAAG |
| 40389_2 | TATGAGTGTGGAGTCGTTGCCAACAAGAGCGAAACTCCATCTCNNNAGATCGGAAG |
| 49628_1 | TATGAGTGTGGAGTCGTTGCCTAGCTGAAACTCTGCATGCTTCNNNAGATCGGAAG |
| 49628_2 | TATGAGTGTGGAGTCGTTGCCTAGCTGAAACTCTGCATGCTTCNNNAGATCGGAAG |
| 51874_1 | TATGAGTGTGGAGTCGTTGCCACTGTTTTCGGACCTTCCATGNNNAGATCGGAAGA |
| 51874_2 | TATGAGTGTGGAGTCGTTGCCACTGTTTTCGGACCTTCCATGNNNAGATCGGAAGA |
| 35635_1 | TATGAGTGTGGAGTCGTTGCGTCCAAGTGATTATCCTGCCTCANNNAGATCGGAAG |
| 35635_2 | TATGAGTGTGGAGTCGTTGCGTCCAAGTGATTATCCTGCCTCANNNAGATCGGAAG |
| 38409_1 | TATGAGTGTGGAGTCGTTGCAAGCCAACAAATACAGTTCAGCANNNAGATCGGAAG |
| 38409_2 | TATGAGTGTGGAGTCGTTGCAAGCCAACAAATACAGTTCAGCANNNAGATCGGAAG |
| 13487_1 | TATGAGTGTGGAGTCGTTGCTTTATTTGTAATTTTTGCTGCTGNNNAGATCGGAAG |
| 13487_2 | TATGAGTGTGGAGTCGTTGCTTTATTTGTAATTTTTGCTGCTGNNNAGATCGGAAG |
| 47274_1 | TATGAGTGTGGAGTCGTTGCGTTCCAGTTATTCTCCTGCCTCANNNAGATCGGAAGA |
| 47274_2 | TATGAGTGTGGAGTCGTTGCGTTCCAGTTATTCTCCTGCCTCANNNAGATCGGAAGA |
| 52232_1 | TATGAGTGTGGAGTCGTTGCGAGAGGCACAGAGAGGGAATATGNNNAGATCGGA |
| 52232_2 | TATGAGTGTGGAGTCGTTGCGAGAGGCACAGAGAGGGAATATGNNNAGATCGGA |
| 49696_1 | TATGAGTGTGGAGTCGTTGCTTCATGTCATTCTCCTGCCTCAGNNNAGATCGGAAGA |
| 49696_2 | TATGAGTGTGGAGTCGTTGCTTCATGTCATTCTCCTGCCTCAGNNNAGATCGGAAGA |
| 50433_1 | TATGAGTGTGGAGTCGTTGCTTTGGGCTGGTAGTAAGTTTGGTNNNAGATCGGAA |
| 50433_2 | TATGAGTGTGGAGTCGTTGCTTTGGGCTGGTAGTAAGTTTGGTNNNAGATCGGAA |
| 49284_1 | TATGAGTGTGGAGTCGTTGCCAGTGGTGCAATCATGGCTTACNNNAGATCGGAAG |
| 49284_2 | TATGAGTGTGGAGTCGTTGCCAGTGGTGCAATCATGGCTTACNNNAGATCGGAAG |
| 43336_1 | TATGAGTGTGGAGTCGTTGCATAACCAAAGAGTGGGAAGGTCCNNNAGATCGGAA |
| 43336_2 | TATGAGTGTGGAGTCGTTGCATAACCAAAGAGTGGGAAGGTCCNNNAGATCGGAA |
| 4581_1 | TATGAGTGTGGAGTCGTTGCACCCAAACATTACACCGTAGACTNNNAGATCGGAAG |
| 4581_2 | TATGAGTGTGGAGTCGTTGCACCCAAACATTACACCGTAGACTNNNAGATCGGAAG |
| 35087_1 | TATGAGTGTGGAGTCGTTGCCTCCCAGGTTCACACCATTCTCNNNAGATCGGAAGAG |
| 35087_2 | TATGAGTGTGGAGTCGTTGCCTCCCAGGTTCACACCATTCTCNNNAGATCGGAAGAG |
| 37726_1 | TATGAGTGTGGAGTCGTTGCAGCACGAATATAACTTCCTGGCANNNAGATCGGAAG |
| 37726_2 | TATGAGTGTGGAGTCGTTGCAGCACGAATATAACTTCCTGGCANNNAGATCGGAAG |
| 31467_1 | TATGAGTGTGGAGTCGTTGCTGTTCAACCTCATATCCCCAGTGNNNAGATCGGAAG |
| 31467_2 | TATGAGTGTGGAGTCGTTGCTGTTCAACCTCATATCCCCAGTGNNNAGATCGGAAG |
| 50713_1 | TATGAGTGTGGAGTCGTTGCATGGAGCTCAATTGGAGTGAGTTNNNAGATCGGAA |
| 50713_2 | TATGAGTGTGGAGTCGTTGCATGGAGCTCAATTGGAGTGAGTTNNNAGATCGGAA |
| 50121_1 | TATGAGTGTGGAGTCGTTGCCTTGGAGAGGAGGCTGCTGAANNNAGATCGGAAG |
| 50121_2 | TATGAGTGTGGAGTCGTTGCCTTGGAGAGGAGGCTGCTGAANNNAGATCGGAAG |
| 49253_1 | TATGAGTGTGGAGTCGTTGCTGTGAGTGCAGTGGTGAGATAATNNNAGATCGGAA |
| 49253_2 | TATGAGTGTGGAGTCGTTGCTGTGAGTGCAGTGGTGAGATAATNNNAGATCGGAA |
| 44077_1 | TATGAGTGTGGAGTCGTTGCGGAAATGAAGTTCCCTCAGTCCTNNNAGATCGGAAG |
| 44077_2 | TATGAGTGTGGAGTCGTTGCGGAAATGAAGTTCCCTCAGTCCTNNNAGATCGGAAG |
| 36381_1 | TATGAGTGTGGAGTCGTTGCTTTCCTGGTTGAAAGGTGGTGAANNNAGATCGGAA |

| | |
|---|---|
| 36381_2 | TATGAGTGTGGAGTCGTTGCTTTCCTGGTTGAAAGGTGGTGAANNNAGATCGGAA |
| 45577_1 | TATGAGTGTGGAGTCGTTGCTTTTTTTTTTTCCGCCTGCTGTGNNNAGATCGGAAGA |
| 45577_2 | TATGAGTGTGGAGTCGTTGCTTTTTTTTTTTCCGCCTGCTGTGNNNAGATCGGAAGA |
| 42501_1 | TATGAGTGTGGAGTCGTTGCGGAAGGAAGGAAAGAAGGAAGGANNNAGATCGG |
| 42501_2 | TATGAGTGTGGAGTCGTTGCGGAAGGAAGGAAAGAAGGAAGGANNNAGATCGG |
| 36047_1 | TATGAGTGTGGAGTCGTTGCGCAGGACTGTTCACAATAGCAAANNNAGATCGGAA |
| 36047_2 | TATGAGTGTGGAGTCGTTGCGCAGGACTGTTCACAATAGCAAANNNAGATCGGAA |
| 35625_1 | TATGAGTGTGGAGTCGTTGCCTCAAGTGATTCTCCTGCCTCCNNNAGATCGGAAGA |
| 35625_2 | TATGAGTGTGGAGTCGTTGCCTCAAGTGATTCTCCTGCCTCCNNNAGATCGGAAGA |
| 48974_1 | TATGAGTGTGGAGTCGTTGCAAAGAATGAGAGAGAGGGAGGGANNNAGATCGG |
| 48974_2 | TATGAGTGTGGAGTCGTTGCAAAGAATGAGAGAGAGGGAGGGANNNAGATCGG |
| 12749_1 | TATGAGTGTGGAGTCGTTGCGTAGTGCAGAATCTTTAATTTCANNNAGATCGGAAG |
| 12749_2 | TATGAGTGTGGAGTCGTTGCGTAGTGCAGAATCTTTAATTTCANNNAGATCGGAAG |
| 40839_1 | TATGAGTGTGGAGTCGTTGCCAACTGTGGCTAAGGAAAAAGGGNNNAGATCGGA |
| 40839_2 | TATGAGTGTGGAGTCGTTGCCAACTGTGGCTAAGGAAAAAGGGNNNAGATCGGA |
| 36969_1 | TATGAGTGTGGAGTCGTTGCTCAATAAATGGCGCTGGGTAAACNNNAGATCGGAA |
| 36969_2 | TATGAGTGTGGAGTCGTTGCTCAATAAATGGCGCTGGGTAAACNNNAGATCGGAA |
| 48844_1 | TATGAGTGTGGAGTCGTTGCCTTGAATCAATCCTCCCACCTCANNNAGATCGGAAGA |
| 48844_2 | TATGAGTGTGGAGTCGTTGCCTTGAATCAATCCTCCCACCTCANNNAGATCGGAAGA |
| 53209_1 | TATGAGTGTGGAGTCGTTGCTATGTGGTGGATAAGTGTGTGGGNNNAGATCGGA |
| 53209_2 | TATGAGTGTGGAGTCGTTGCTATGTGGTGGATAAGTGTGTGGGNNNAGATCGGA |
| 31233_1 | TATGAGTGTGGAGTCGTTGCGGATGTACACACAGACAGTCACTNNNAGATCGGAA |
| 31233_2 | TATGAGTGTGGAGTCGTTGCGGATGTACACACAGACAGTCACTNNNAGATCGGAA |
| 34799_1 | TATGAGTGTGGAGTCGTTGCGATCAAGTGATTCTCCTGCCTCANNNAGATCGGAAG |
| 34799_2 | TATGAGTGTGGAGTCGTTGCGATCAAGTGATTCTCCTGCCTCANNNAGATCGGAAG |
| 13234_1 | TATGAGTGTGGAGTCGTTGCCAGAAGAATCTTCCTCTTTTTCNNNAGATCGGAAGA |
| 13234_2 | TATGAGTGTGGAGTCGTTGCCAGAAGAATCTTCCTCTTTTTCNNNAGATCGGAAGA |
| 46211_1 | TATGAGTGTGGAGTCGTTGCTAGGGGAAAGTAGAGGGATGAGGNNNAGATCGG |
| 46211_2 | TATGAGTGTGGAGTCGTTGCTAGGGGAAAGTAGAGGGATGAGGNNNAGATCGG |
| 13313_1 | TATGAGTGTGGAGTCGTTGCCATCATTTCTGGAGTCTCAGNNNAGATCGGAAGAGC |
| 13313_2 | TATGAGTGTGGAGTCGTTGCCATCATTTCTGGAGTCTCAGNNNAGATCGGAAGAGC |
| 48664_1 | TATGAGTGTGGAGTCGTTGCAGAGAGTAACCTGTAACCTGGGANNNAGATCGGAA |
| 48664_2 | TATGAGTGTGGAGTCGTTGCAGAGAGTAACCTGTAACCTGGGANNNAGATCGGAA |
| 30078_1 | TATGAGTGTGGAGTCGTTGCACATCAGGATCCTGGACAGAAACNNNAGATCGGAA |
| 30078_2 | TATGAGTGTGGAGTCGTTGCACATCAGGATCCTGGACAGAAACNNNAGATCGGAA |
| 47972_1 | TATGAGTGTGGAGTCGTTGCCCTCTCAAGCCAGGTGATCAACNNNAGATCGGAAGA |
| 47972_2 | TATGAGTGTGGAGTCGTTGCCCTCTCAAGCCAGGTGATCAACNNNAGATCGGAAGA |
| 31985_1 | TATGAGTGTGGAGTCGTTGCTGTGTGTGTTTTGGTGCTTTGAANNNAGATCGGAA |
| 31985_2 | TATGAGTGTGGAGTCGTTGCTGTGTGTGTTTTGGTGCTTTGAANNNAGATCGGAA |
| 41636_1 | TATGAGTGTGGAGTCGTTGCCTCCCAAAGTGCTAGGATGACAGNNNAGATCGGAA |
| 41636_2 | TATGAGTGTGGAGTCGTTGCCTCCCAAAGTGCTAGGATGACAGNNNAGATCGGAA |
| 32093_1 | TATGAGTGTGGAGTCGTTGCGAGACGGAGTCTTGCTCTGTCNNNAGATCGGAAGA |
| 32093_2 | TATGAGTGTGGAGTCGTTGCGAGACGGAGTCTTGCTCTGTCNNNAGATCGGAAGA |

| | |
|---|---|
| 34872_1 | TATGAGTGTGGAGTCGTTGCTTCTTTTTTGAGACGGAGTCCCANNNAGATCGGAAG |
| 34872_2 | TATGAGTGTGGAGTCGTTGCTTCTTTTTTGAGACGGAGTCCCANNNAGATCGGAAG |
| 38662_1 | TATGAGTGTGGAGTCGTTGCGAAGTCTGCAACCGGTTGTAATCNNNAGATCGGAA |
| 38662_2 | TATGAGTGTGGAGTCGTTGCGAAGTCTGCAACCGGTTGTAATCNNNAGATCGGAA |
| 50032_1 | TATGAGTGTGGAGTCGTTGCCACGCCTATAATCCCAGCACTTTNNNAGATCGGAAG |
| 50032_2 | TATGAGTGTGGAGTCGTTGCCACGCCTATAATCCCAGCACTTTNNNAGATCGGAAG |
| 36035_1 | TATGAGTGTGGAGTCGTTGCAGAATTGCTTGAACCTGGGAGAANNNAGATCGGAA |
| 36035_2 | TATGAGTGTGGAGTCGTTGCAGAATTGCTTGAACCTGGGAGAANNNAGATCGGAA |
| 29840_1 | TATGAGTGTGGAGTCGTTGCGGTAAGACCTGTTTATTTTATAGCTGANNNAGATCG |
| 29840_2 | TATGAGTGTGGAGTCGTTGCGGTAAGACCTGTTTATTTTATAGCTGANNNAGATCG |
| 47857_1 | TATGAGTGTGGAGTCGTTGCCAAAGTGCTGGGATTACAGGAGTNNNAGATCGGAA |
| 47857_2 | TATGAGTGTGGAGTCGTTGCCAAAGTGCTGGGATTACAGGAGTNNNAGATCGGAA |
| 12653_1 | TATGAGTGTGGAGTCGTTGCCTGAGGGTCATGATCTATATTCNNNAGATCGGAAGA |
| 12653_2 | TATGAGTGTGGAGTCGTTGCCTGAGGGTCATGATCTATATTCNNNAGATCGGAAGA |
| 31083_1 | TATGAGTGTGGAGTCGTTGCGTGCAGTATTGTGATCATGGCTCNNNAGATCGGAA |
| 31083_2 | TATGAGTGTGGAGTCGTTGCGTGCAGTATTGTGATCATGGCTCNNNAGATCGGAA |
| 49238_1 | TATGAGTGTGGAGTCGTTGCCAACCTCCCAAGTAGCTAGGATCNNNAGATCGGAAG |
| 49238_2 | TATGAGTGTGGAGTCGTTGCCAACCTCCCAAGTAGCTAGGATCNNNAGATCGGAAG |
| 34110_1 | TATGAGTGTGGAGTCGTTGCGCCCCACCAATAGTACCCATTATNNNAGATCGGAAG |
| 34110_2 | TATGAGTGTGGAGTCGTTGCGCCCCACCAATAGTACCCATTATNNNAGATCGGAAG |
| 30945_1 | TATGAGTGTGGAGTCGTTGCTTGTCACTTGGTAACTGGTCTCCNNNAGATCGGAAG |
| 30945_2 | TATGAGTGTGGAGTCGTTGCTTGTCACTTGGTAACTGGTCTCCNNNAGATCGGAAG |
| 43949_1 | TATGAGTGTGGAGTCGTTGCGCCTCCCGAATTCAAGCAATTCNNNAGATCGGAAGA |
| 43949_2 | TATGAGTGTGGAGTCGTTGCGCCTCCCGAATTCAAGCAATTCNNNAGATCGGAAGA |
| 41671_1 | TATGAGTGTGGAGTCGTTGCAAACTCCTGGGCTCAAGTGATCNNNAGATCGGAAGA |
| 41671_2 | TATGAGTGTGGAGTCGTTGCAAACTCCTGGGCTCAAGTGATCNNNAGATCGGAAGA |
| 34800_1 | TATGAGTGTGGAGTCGTTGCGTCTCAATCTTCTGACCTCGTGANNNAGATCGGAAG |
| 34800_2 | TATGAGTGTGGAGTCGTTGCGTCTCAATCTTCTGACCTCGTGANNNAGATCGGAAG |
| 34840_1 | TATGAGTGTGGAGTCGTTGCCCGGATTCAAACGATTCTCCTCTNNNAGATCGGAAG |
| 34840_2 | TATGAGTGTGGAGTCGTTGCCCGGATTCAAACGATTCTCCTCTNNNAGATCGGAAG |
| 35757_1 | TATGAGTGTGGAGTCGTTGCGCTCAATTGATCCTCCCACCTTANNNAGATCGGAAGA |
| 35757_2 | TATGAGTGTGGAGTCGTTGCGCTCAATTGATCCTCCCACCTTANNNAGATCGGAAGA |
| 38572_1 | TATGAGTGTGGAGTCGTTGCGCAGGAGAATTGATTGAACCTGGNNNAGATCGGAA |
| 38572_2 | TATGAGTGTGGAGTCGTTGCGCAGGAGAATTGATTGAACCTGGNNNAGATCGGAA |
| 42825_1 | TATGAGTGTGGAGTCGTTGCGCCTCTACACATTCCTGCAAATGNNNAGATCGGAAG |
| 42825_2 | TATGAGTGTGGAGTCGTTGCGCCTCTACACATTCCTGCAAATGNNNAGATCGGAAG |
| 35063_1 | TATGAGTGTGGAGTCGTTGCTCCCAAAGTGCTGGAATTACAGANNNAGATCGGAA |
| 35063_2 | TATGAGTGTGGAGTCGTTGCTCCCAAAGTGCTGGAATTACAGANNNAGATCGGAA |
| 40164_1 | TATGAGTGTGGAGTCGTTGCCCTGTAATCCCAGCACTTTAGGANNNAGATCGGAAG |
| 40164_2 | TATGAGTGTGGAGTCGTTGCCCTGTAATCCCAGCACTTTAGGANNNAGATCGGAAG |
| 10337_1 | TATGAGTGTGGAGTCGTTGCTACATTCCTACACCCTCCACTCGNNNAGATCGGAAGA |
| 10337_2 | TATGAGTGTGGAGTCGTTGCTACATTCCTACACCCTCCACTCGNNNAGATCGGAAGA |
| 31028_1 | TATGAGTGTGGAGTCGTTGCTGGAGATAGAGCCATCATGCTTCNNNAGATCGGAA |

| | |
|---|---|
| 31028_2 | TATGAGTGTGGAGTCGTTGCTGGAGATAGAGCCATCATGCTTCNNNAGATCGGAAG |
| 34779_1 | TATGAGTGTGGAGTCGTTGCCTTGAACTCCTGACCTCGTGATCNNNAGATCGGAAG |
| 34779_2 | TATGAGTGTGGAGTCGTTGCCTTGAACTCCTGACCTCGTGATCNNNAGATCGGAAG |
| 29711_1 | TATGAGTGTGGAGTCGTTGCTTGGGGTGATGTGGTGATTAACTNNNAGATCGGAA |
| 29711_2 | TATGAGTGTGGAGTCGTTGCTTGGGGTGATGTGGTGATTAACTNNNAGATCGGAA |
| 10412_1 | TATGAGTGTGGAGTCGTTGCAAGCCTGAGTGAGGTATGGGAATNNNAGATCGGAA |
| 10412_2 | TATGAGTGTGGAGTCGTTGCAAGCCTGAGTGAGGTATGGGAATNNNAGATCGGAA |
| 52526_1 | TATGAGTGTGGAGTCGTTGCGCCGGGGAGTTATTTCTGAACTANNNAGATCGGAAG |
| 52526_2 | TATGAGTGTGGAGTCGTTGCGCCGGGGAGTTATTTCTGAACTANNNAGATCGGAAG |
| 49857_1 | TATGAGTGTGGAGTCGTTGCAAGTGATCTGTGTTCACCTGTGTNNNAGATCGGAAG |
| 49857_2 | TATGAGTGTGGAGTCGTTGCAAGTGATCTGTGTTCACCTGTGTNNNAGATCGGAAG |
| 50795_1 | TATGAGTGTGGAGTCGTTGCTCCAACCAGGTATCAATAGAGCGNNNAGATCGGAAG |
| 50795_2 | TATGAGTGTGGAGTCGTTGCTCCAACCAGGTATCAATAGAGCGNNNAGATCGGAAG |
| 36909_1 | TATGAGTGTGGAGTCGTTGCTGCACGGAAAGAAGGAAGATACANNNAGATCGGAA |
| 36909_2 | TATGAGTGTGGAGTCGTTGCTGCACGGAAAGAAGGAAGATACANNNAGATCGGAA |
| 51611_1 | TATGAGTGTGGAGTCGTTGCCCAGGTCCTTTCCCAGAGATGNNNAGATCGGAAGAG |
| 51611_2 | TATGAGTGTGGAGTCGTTGCCCAGGTCCTTTCCCAGAGATGNNNAGATCGGAAGAG |
| 35463_1 | TATGAGTGTGGAGTCGTTGCAGGATGACACTGGCCTCATAAAANNNAGATCGGAA |
| 35463_2 | TATGAGTGTGGAGTCGTTGCAGGATGACACTGGCCTCATAAAANNNAGATCGGAA |
| 38195_1 | TATGAGTGTGGAGTCGTTGCGGATGGACATTTCAGTTGATCCCNNNAGATCGGAAG |
| 38195_2 | TATGAGTGTGGAGTCGTTGCGGATGGACATTTCAGTTGATCCCNNNAGATCGGAAG |
| 44281_1 | TATGAGTGTGGAGTCGTTGCGGCAGCCTGTCTACTTTGTAGAANNNAGATCGGAAG |
| 44281_2 | TATGAGTGTGGAGTCGTTGCGGCAGCCTGTCTACTTTGTAGAANNNAGATCGGAAG |
| 34531_1 | TATGAGTGTGGAGTCGTTGCCTTTGGACTGCATTGGCTACAAANNNAGATCGGAAG |
| 34531_2 | TATGAGTGTGGAGTCGTTGCCTTTGGACTGCATTGGCTACAAANNNAGATCGGAAG |
| 35304_1 | TATGAGTGTGGAGTCGTTGCAGTCAGATCAGTCACCTTTTGCCNNNAGATCGGAAG |
| 35304_2 | TATGAGTGTGGAGTCGTTGCAGTCAGATCAGTCACCTTTTGCCNNNAGATCGGAAG |
| 42968_1 | TATGAGTGTGGAGTCGTTGCACAATCTTGGCTCACTACAACCTNNNAGATCGGAAG |
| 42968_2 | TATGAGTGTGGAGTCGTTGCACAATCTTGGCTCACTACAACCTNNNAGATCGGAAG |
| 29057_1 | TATGAGTGTGGAGTCGTTGCTTGTCTAGGTGGCAAATATCCCGNNNAGATCGGAAG |
| 29057_2 | TATGAGTGTGGAGTCGTTGCTTGTCTAGGTGGCAAATATCCCGNNNAGATCGGAAG |
| 42597_1 | TATGAGTGTGGAGTCGTTGCGTGTGTGTGTGTGTGTGTGTATANNNAGATCGGAA |
| 42597_2 | TATGAGTGTGGAGTCGTTGCGTGTGTGTGTGTGTGTGTGTATANNNAGATCGGAA |
| 13049_1 | TATGAGTGTGGAGTCGTTGCTTGGTAATCTTTACTTCAAAACCNNNAGATCGGAAG |
| 13049_2 | TATGAGTGTGGAGTCGTTGCTTGGTAATCTTTACTTCAAAACCNNNAGATCGGAAG |
| 46429_1 | TATGAGTGTGGAGTCGTTGCTAGCTGAGATTACAGGTGTGTGCNNNAGATCGGAA |
| 46429_2 | TATGAGTGTGGAGTCGTTGCTAGCTGAGATTACAGGTGTGTGCNNNAGATCGGAA |
| 32261_1 | TATGAGTGTGGAGTCGTTGCCATCTGTGGTTCCAGCTACTCAGNNNAGATCGGAAG |
| 32261_2 | TATGAGTGTGGAGTCGTTGCCATCTGTGGTTCCAGCTACTCAGNNNAGATCGGAAG |
| 48494_1 | TATGAGTGTGGAGTCGTTGCTAATGTTCCAGATGAAGCCTGCTNNNAGATCGGAAG |
| 48494_2 | TATGAGTGTGGAGTCGTTGCTAATGTTCCAGATGAAGCCTGCTNNNAGATCGGAAG |
| 30854_1 | TATGAGTGTGGAGTCGTTGCCCTCTGTTTGCAATCAATCCGTTNNNAGATCGGAAG |
| 30854_2 | TATGAGTGTGGAGTCGTTGCCCTCTGTTTGCAATCAATCCGTTNNNAGATCGGAAG |

| | |
|---|---|
| 35129_1 | TATGAGTGTGGAGTCGTTGCCCTTCAGATTTTGCTGAGGGTTTNNNAGATCGGAAG |
| 35129_2 | TATGAGTGTGGAGTCGTTGCCCTTCAGATTTTGCTGAGGGTTTNNNAGATCGGAAG |
| 43966_1 | TATGAGTGTGGAGTCGTTGCGGAGGAGGAGGAAAGAAAGTGAANNNAGATCGGA |
| 43966_2 | TATGAGTGTGGAGTCGTTGCGGAGGAGGAGGAAAGAAAGTGAANNNAGATCGGA |
| 33277_1 | TATGAGTGTGGAGTCGTTGCCACCATGCCCGACTAATTTTTGTNNNAGATCGGAAG |
| 33277_2 | TATGAGTGTGGAGTCGTTGCCACCATGCCCGACTAATTTTTGTNNNAGATCGGAAG |
| 51052_1 | TATGAGTGTGGAGTCGTTGCCTCCTCTTGTCACCTCCCGTCNNNAGATCGGAAGAGC |
| 51052_2 | TATGAGTGTGGAGTCGTTGCCTCCTCTTGTCACCTCCCGTCNNNAGATCGGAAGAGC |
| 46073_1 | TATGAGTGTGGAGTCGTTGCCAAGTAGCTGGGACTACAGGCNNNAGATCGGAAGA |
| 46073_2 | TATGAGTGTGGAGTCGTTGCCAAGTAGCTGGGACTACAGGCNNNAGATCGGAAGA |
| 34957_1 | TATGAGTGTGGAGTCGTTGCCCTACTGAGTAGCTGGGACTACANNNAGATCGGAA |
| 34957_2 | TATGAGTGTGGAGTCGTTGCCCTACTGAGTAGCTGGGACTACANNNAGATCGGAA |
| 41009_1 | TATGAGTGTGGAGTCGTTGCAAAAGAAAGAAAGAAGGCCAGGCNNNAGATCGGA |
| 41009_2 | TATGAGTGTGGAGTCGTTGCAAAAGAAAGAAAGAAGGCCAGGCNNNAGATCGGA |
| 10601_1 | TATGAGTGTGGAGTCGTTGCTTTTGCATTGCTCCCAAGTAGGTNNNAGATCGGAAG |
| 10601_2 | TATGAGTGTGGAGTCGTTGCTTTTGCATTGCTCCCAAGTAGGTNNNAGATCGGAAG |
| 45277_1 | TATGAGTGTGGAGTCGTTGCAAAGAGATGTGGGGAGATCATGGNNNAGATCGGA |
| 45277_2 | TATGAGTGTGGAGTCGTTGCAAAGAGATGTGGGGAGATCATGGNNNAGATCGGA |
| 2391_1 | TATGAGTGTGGAGTCGTTGCCCATCCCATCTGCCTATTATGGCNNNAGATCGGAAGA |
| 2391_2 | TATGAGTGTGGAGTCGTTGCCCATCCCATCTGCCTATTATGGCNNNAGATCGGAAGA |
| 51451_1 | TATGAGTGTGGAGTCGTTGCACCAACATGGCACATGTATACCTNNNAGATCGGAAG |
| 51451_2 | TATGAGTGTGGAGTCGTTGCACCAACATGGCACATGTATACCTNNNAGATCGGAAG |
| 40394_1 | TATGAGTGTGGAGTCGTTGCAGACATAAGGCAGGAAAACAGGTNNNAGATCGGAA |
| 40394_2 | TATGAGTGTGGAGTCGTTGCAGACATAAGGCAGGAAAACAGGTNNNAGATCGGAA |
| 39556_1 | TATGAGTGTGGAGTCGTTGCGCCACCAACCCAGCTAATTTTTTNNNAGATCGGAAGA |
| 39556_2 | TATGAGTGTGGAGTCGTTGCGCCACCAACCCAGCTAATTTTTTNNNAGATCGGAAGA |
| 50586_1 | TATGAGTGTGGAGTCGTTGCCTCAGCTTGTTCATGTCCTGGAGNNNAGATCGGAAG |
| 50586_2 | TATGAGTGTGGAGTCGTTGCCTCAGCTTGTTCATGTCCTGGAGNNNAGATCGGAAG |
| 12549_1 | TATGAGTGTGGAGTCGTTGCATAACTTTCAATTACAGTCCAACNNNAGATCGGAAGA |
| 12549_2 | TATGAGTGTGGAGTCGTTGCATAACTTTCAATTACAGTCCAACNNNAGATCGGAAGA |
| 42699_1 | TATGAGTGTGGAGTCGTTGCGCTGAATCCATCAACGCATCTAANNNAGATCGGAAG |
| 42699_2 | TATGAGTGTGGAGTCGTTGCGCTGAATCCATCAACGCATCTAANNNAGATCGGAAG |
| 10504_1 | TATGAGTGTGGAGTCGTTGCTCCTATGTGTGATGAGCTCTGGTNNNAGATCGGAAG |
| 10504_2 | TATGAGTGTGGAGTCGTTGCTCCTATGTGTGATGAGCTCTGGTNNNAGATCGGAAG |
| 50665_1 | TATGAGTGTGGAGTCGTTGCCTTGCCTACCTTAGTTGAACCCANNNAGATCGGAAGA |
| 50665_2 | TATGAGTGTGGAGTCGTTGCCTTGCCTACCTTAGTTGAACCCANNNAGATCGGAAGA |
| 49778_1 | TATGAGTGTGGAGTCGTTGCCATACCAATGTGTTTTCGCCACANNNAGATCGGAAG |
| 49778_2 | TATGAGTGTGGAGTCGTTGCCATACCAATGTGTTTTCGCCACANNNAGATCGGAAG |
| 43462_1 | TATGAGTGTGGAGTCGTTGCGTTCAAGAGATTCTCCTGCCTCANNNAGATCGGAAG |
| 43462_2 | TATGAGTGTGGAGTCGTTGCGTTCAAGAGATTCTCCTGCCTCANNNAGATCGGAAG |
| 3030_1 | TATGAGTGTGGAGTCGTTGCAGACTTGCCTCCATATCGAGTGTNNNAGATCGGAAG |
| 3030_2 | TATGAGTGTGGAGTCGTTGCAGACTTGCCTCCATATCGAGTGTNNNAGATCGGAAG |
| 35931_1 | TATGAGTGTGGAGTCGTTGCGGACTGTAAAACCACTTGTCCTANNNAGATCGGAAG |

| | |
|---|---|
| 35931_2 | TATGAGTGTGGAGTCGTTGCGGACTGTAAAACCACTTGTCCTANNNAGATCGGAAG |
| 52992_1 | TATGAGTGTGGAGTCGTTGCGACCATGATGATACTGCCTGTCTNNNAGATCGGAAG |
| 52992_2 | TATGAGTGTGGAGTCGTTGCGACCATGATGATACTGCCTGTCTNNNAGATCGGAAG |
| 45121_1 | TATGAGTGTGGAGTCGTTGCAGCCTCCCATACTTTACCCTCTANNNAGATCGGAAGA |
| 45121_2 | TATGAGTGTGGAGTCGTTGCAGCCTCCCATACTTTACCCTCTANNNAGATCGGAAGA |
| 47368_1 | TATGAGTGTGGAGTCGTTGCACGCCCGGCTAATTTTTTGTATTNNNAGATCGGAAG |
| 47368_2 | TATGAGTGTGGAGTCGTTGCACGCCCGGCTAATTTTTTGTATTNNNAGATCGGAAG |
| 45868_1 | TATGAGTGTGGAGTCGTTGCATCTCCCAGGTTTGAGCAATTCTNNNAGATCGGAAG |
| 45868_2 | TATGAGTGTGGAGTCGTTGCATCTCCCAGGTTTGAGCAATTCTNNNAGATCGGAAG |
| 29725_1 | TATGAGTGTGGAGTCGTTGCTTTGGAATTCCTGCGACTGTTTCNNNAGATCGGAAG |
| 29725_2 | TATGAGTGTGGAGTCGTTGCTTTGGAATTCCTGCGACTGTTTCNNNAGATCGGAAG |
| 4700_1 | TATGAGTGTGGAGTCGTTGCAGTAATGGGAAGGATGACCCACANNNAGATCGGAA |
| 4700_2 | TATGAGTGTGGAGTCGTTGCAGTAATGGGAAGGATGACCCACANNNAGATCGGAA |
| 48170_1 | TATGAGTGTGGAGTCGTTGCGGGAGAGAAGAGCAGCCTATTACNNNAGATCGGAA |
| 48170_2 | TATGAGTGTGGAGTCGTTGCGGGAGAGAAGAGCAGCCTATTACNNNAGATCGGAA |
| 47173_1 | TATGAGTGTGGAGTCGTTGCCTCCTGGGTTCAAGGGATTCTCNNNAGATCGGAAGA |
| 47173_2 | TATGAGTGTGGAGTCGTTGCCTCCTGGGTTCAAGGGATTCTCNNNAGATCGGAAGA |
| 48710_1 | TATGAGTGTGGAGTCGTTGCGATCTCCCCGAACTTCTCAAACANNNAGATCGGAAG |
| 48710_2 | TATGAGTGTGGAGTCGTTGCGATCTCCCCGAACTTCTCAAACANNNAGATCGGAAG |
| 34392_1 | TATGAGTGTGGAGTCGTTGCCTCATTGCAGCCTCAAACTCCTNNNAGATCGGAAGAG |
| 34392_2 | TATGAGTGTGGAGTCGTTGCCTCATTGCAGCCTCAAACTCCTNNNAGATCGGAAGAG |
| 47835_1 | TATGAGTGTGGAGTCGTTGCGAGGGAAAGCCTGGCATATCACNNNAGATCGGAAG |
| 47835_2 | TATGAGTGTGGAGTCGTTGCGAGGGAAAGCCTGGCATATCACNNNAGATCGGAAG |
| 11152_1 | TATGAGTGTGGAGTCGTTGCCCAGCCAGAACATGAACCTAGAGNNNAGATCGGAA |
| 11152_2 | TATGAGTGTGGAGTCGTTGCCCAGCCAGAACATGAACCTAGAGNNNAGATCGGAA |
| 38151_1 | TATGAGTGTGGAGTCGTTGCTGTCTTGGGTATTTGGGAGACTGNNNAGATCGGAA |
| 38151_2 | TATGAGTGTGGAGTCGTTGCTGTCTTGGGTATTTGGGAGACTGNNNAGATCGGAA |
| 12968_1 | TATGAGTGTGGAGTCGTTGCGAGAAGTCATCACAACTCATCNNNAGATCGGAAGAG |
| 12968_2 | TATGAGTGTGGAGTCGTTGCGAGAAGTCATCACAACTCATCNNNAGATCGGAAGAG |
| 29173_1 | TATGAGTGTGGAGTCGTTGCTGTGTGTGTGTGTTTTGTCCAATNNNAGATCGGAAG |
| 29173_2 | TATGAGTGTGGAGTCGTTGCTGTGTGTGTGTGTTTTGTCCAATNNNAGATCGGAAG |
| 43272_1 | TATGAGTGTGGAGTCGTTGCTTGTGCTGCTATAAACGTGTGTGNNNAGATCGGAA |
| 43272_2 | TATGAGTGTGGAGTCGTTGCTTGTGCTGCTATAAACGTGTGTGNNNAGATCGGAA |
| 10650_1 | TATGAGTGTGGAGTCGTTGCTCCGCCTCCTGGTGTATATAAGGNNNAGATCGGAAG |
| 10650_2 | TATGAGTGTGGAGTCGTTGCTCCGCCTCCTGGTGTATATAAGGNNNAGATCGGAAG |
| 28966_1 | TATGAGTGTGGAGTCGTTGCGCATCTTAGAGGAGACCAGTGACNNNAGATCGGAA |
| 28966_2 | TATGAGTGTGGAGTCGTTGCGCATCTTAGAGGAGACCAGTGACNNNAGATCGGAA |
| 48006_1 | TATGAGTGTGGAGTCGTTGCCTCCTCAGAAGCTGGGACTACAGNNNAGATCGGAA |
| 48006_2 | TATGAGTGTGGAGTCGTTGCCTCCTCAGAAGCTGGGACTACAGNNNAGATCGGAA |
| 34054_1 | TATGAGTGTGGAGTCGTTGCCATTCTCCTGCCTCAGCCTCNNNAGATCGGAAGAGCA |
| 34054_2 | TATGAGTGTGGAGTCGTTGCCATTCTCCTGCCTCAGCCTCNNNAGATCGGAAGAGCA |
| 48874_1 | TATGAGTGTGGAGTCGTTGCGAGTTTTGCTCTTGTTGCCCAGNNNAGATCGGAAGA |
| 48874_2 | TATGAGTGTGGAGTCGTTGCGAGTTTTGCTCTTGTTGCCCAGNNNAGATCGGAAGA |

| | |
|---|---|
| 38594_1 | TATGAGTGTGGAGTCGTTGCGGAAGTAGGCAAAAGACAAGACANNNAGATCGGA |
| 38594_2 | TATGAGTGTGGAGTCGTTGCGGAAGTAGGCAAAAGACAAGACANNNAGATCGGA |
| 44682_1 | TATGAGTGTGGAGTCGTTGCGTCAATGGACTTTGGCCAATGAANNNAGATCGGAA |
| 44682_2 | TATGAGTGTGGAGTCGTTGCGTCAATGGACTTTGGCCAATGAANNNAGATCGGAA |
| 41238_1 | TATGAGTGTGGAGTCGTTGCCATGCTTGTAATCCCAGCACTTTNNNAGATCGGAAG |
| 41238_2 | TATGAGTGTGGAGTCGTTGCCATGCTTGTAATCCCAGCACTTTNNNAGATCGGAAG |
| 48195_1 | TATGAGTGTGGAGTCGTTGCTGAGAAGGTCCAGGCACAAATGNNNAGATCGGAAG |
| 48195_2 | TATGAGTGTGGAGTCGTTGCTGAGAAGGTCCAGGCACAAATGNNNAGATCGGAAG |
| 30399_1 | TATGAGTGTGGAGTCGTTGCAGCATAACACTGGTTCAAAATTTACGNNNAGATCGG |
| 30399_2 | TATGAGTGTGGAGTCGTTGCAGCATAACACTGGTTCAAAATTTACGNNNAGATCGG |
| 34644_1 | TATGAGTGTGGAGTCGTTGCAGGCCTATCATTTGGGCAGAAATNNNAGATCGGAA |
| 34644_2 | TATGAGTGTGGAGTCGTTGCAGGCCTATCATTTGGGCAGAAATNNNAGATCGGAA |
| 32320_1 | TATGAGTGTGGAGTCGTTGCACAATACAATGAAACGTTGAGACGNNNAGATCGGA |
| 32320_2 | TATGAGTGTGGAGTCGTTGCACAATACAATGAAACGTTGAGACGNNNAGATCGGA |
| 46163_1 | TATGAGTGTGGAGTCGTTGCTAGAGTGCAGTGACATGATCTCGNNNAGATCGGAA |
| 46163_2 | TATGAGTGTGGAGTCGTTGCTAGAGTGCAGTGACATGATCTCGNNNAGATCGGAA |
| 48702_1 | TATGAGTGTGGAGTCGTTGCAAGTGATTCTTGTGCCTCAGTCTNNNAGATCGGAAG |
| 48702_2 | TATGAGTGTGGAGTCGTTGCAAGTGATTCTTGTGCCTCAGTCTNNNAGATCGGAAG |
| 10173_1 | TATGAGTGTGGAGTCGTTGCTCCCCAATTTATGGTAGCTTGCTNNNAGATCGGAAG |
| 10173_2 | TATGAGTGTGGAGTCGTTGCTCCCCAATTTATGGTAGCTTGCTNNNAGATCGGAAG |
| 30728_1 | TATGAGTGTGGAGTCGTTGCGTGTGTGTGTTTGGGAGAAATAGNNNAGATCGGAA |
| 30728_2 | TATGAGTGTGGAGTCGTTGCGTGTGTGTGTTTGGGAGAAATAGNNNAGATCGGAA |
| 37991_1 | TATGAGTGTGGAGTCGTTGCGGGCCAGCTTTCTAGTTCATAGANNNAGATCGGAAG |
| 37991_2 | TATGAGTGTGGAGTCGTTGCGGGCCAGCTTTCTAGTTCATAGANNNAGATCGGAAG |
| 41510_1 | TATGAGTGTGGAGTCGTTGCAGCCTCCCATCAAATAACTGAGGNNNAGATCGGAAG |
| 41510_2 | TATGAGTGTGGAGTCGTTGCAGCCTCCCATCAAATAACTGAGGNNNAGATCGGAAG |
| 42818_1 | TATGAGTGTGGAGTCGTTGCGGAAGCGTATTGGAACAGGAAACNNNAGATCGGAA |
| 42818_2 | TATGAGTGTGGAGTCGTTGCGGAAGCGTATTGGAACAGGAAACNNNAGATCGGAA |
| 38059_1 | TATGAGTGTGGAGTCGTTGCGCCAGGTATGATTCCAGCTACTTNNNAGATCGGAAG |
| 38059_2 | TATGAGTGTGGAGTCGTTGCGCCAGGTATGATTCCAGCTACTTNNNAGATCGGAAG |
| 34481_1 | TATGAGTGTGGAGTCGTTGCGGGCTCCTCACTTCCCAGTANNNAGATCGGAAGAGC |
| 34481_2 | TATGAGTGTGGAGTCGTTGCGGGCTCCTCACTTCCCAGTANNNAGATCGGAAGAGC |
| 29129_1 | TATGAGTGTGGAGTCGTTGCACACACACAGAGAGAGAGAGAGANNNAGATCGGA |
| 29129_2 | TATGAGTGTGGAGTCGTTGCACACACACAGAGAGAGAGAGAGANNNAGATCGGA |
| 41311_1 | TATGAGTGTGGAGTCGTTGCATGAGAAGGTGCTCAACATCACTNNNAGATCGGAA |
| 41311_2 | TATGAGTGTGGAGTCGTTGCATGAGAAGGTGCTCAACATCACTNNNAGATCGGAA |
| 10074_1 | TATGAGTGTGGAGTCGTTGCACACACACACCAGGGAACAAAATNNNAGATCGGAA |
| 10074_2 | TATGAGTGTGGAGTCGTTGCACACACACACCAGGGAACAAAATNNNAGATCGGAA |
| 52320_1 | TATGAGTGTGGAGTCGTTGCGAGGTCTCTGATCCCATGTGCNNNAGATCGGAAGA |
| 52320_2 | TATGAGTGTGGAGTCGTTGCGAGGTCTCTGATCCCATGTGCNNNAGATCGGAAGA |
| 8486_1 | TATGAGTGTGGAGTCGTTGCACTTTTTGATCAGAGCACGGACANNNAGATCGGAAG |
| 8486_2 | TATGAGTGTGGAGTCGTTGCACTTTTTGATCAGAGCACGGACANNNAGATCGGAAG |
| 45666_1 | TATGAGTGTGGAGTCGTTGCAAAGGGATTACAGACGTGAGCTTNNNAGATCGGAA |

| | |
|---|---|
| 45666_2 | TATGAGTGTGGAGTCGTTGCAAAGGGATTACAGACGTGAGCTTNNNAGATCGGAA |
| 51761_1 | TATGAGTGTGGAGTCGTTGCACCTGAAAATGGAAGTGCTGTTTNNNAGATCGGAA |
| 51761_2 | TATGAGTGTGGAGTCGTTGCACCTGAAAATGGAAGTGCTGTTTNNNAGATCGGAA |
| 29653_1 | TATGAGTGTGGAGTCGTTGCGGTCAACCTGGTACAGACAGAGNNNAGATCGGAAG |
| 29653_2 | TATGAGTGTGGAGTCGTTGCGGTCAACCTGGTACAGACAGAGNNNAGATCGGAAG |
| 46723_1 | TATGAGTGTGGAGTCGTTGCTTTGCAGCTGGGAAGTAGTAGAGNNNAGATCGGAA |
| 46723_2 | TATGAGTGTGGAGTCGTTGCTTTGCAGCTGGGAAGTAGTAGAGNNNAGATCGGAA |
| 2556_1 | TATGAGTGTGGAGTCGTTGCACACATCTTTTCACCACACAGGGNNNAGATCGGAAG |
| 2556_2 | TATGAGTGTGGAGTCGTTGCACACATCTTTTCACCACACAGGGNNNAGATCGGAAG |
| 29073_1 | TATGAGTGTGGAGTCGTTGCGCCCTCATGTTGTCTTTCTGAACNNNAGATCGGAAG |
| 29073_2 | TATGAGTGTGGAGTCGTTGCGCCCTCATGTTGTCTTTCTGAACNNNAGATCGGAAG |
| 29212_1 | TATGAGTGTGGAGTCGTTGCTGGTTGGTCTCACACACCTTATTNNNAGATCGGAAG |
| 29212_2 | TATGAGTGTGGAGTCGTTGCTGGTTGGTCTCACACACCTTATTNNNAGATCGGAAG |
| 43846_1 | TATGAGTGTGGAGTCGTTGCGACAGGAGAGGCTTCTTGTGAGNNNAGATCGGAAG |
| 43846_2 | TATGAGTGTGGAGTCGTTGCGACAGGAGAGGCTTCTTGTGAGNNNAGATCGGAAG |
| 2087_1 | TATGAGTGTGGAGTCGTTGCCCTTTTTGGGTAACCACTACCCANNNAGATCGGAAG |
| 2087_2 | TATGAGTGTGGAGTCGTTGCCCTTTTTGGGTAACCACTACCCANNNAGATCGGAAG |
| 38125_1 | TATGAGTGTGGAGTCGTTGCTCCTTTAGGGAATAACAGCAAGGANNNAGATCGGA |
| 38125_2 | TATGAGTGTGGAGTCGTTGCTCCTTTAGGGAATAACAGCAAGGANNNAGATCGGA |
| 49691_1 | TATGAGTGTGGAGTCGTTGCACAAAAATTGACTGTGCTCTGGGNNNAGATCGGAA |
| 49691_2 | TATGAGTGTGGAGTCGTTGCACAAAAATTGACTGTGCTCTGGGNNNAGATCGGAA |
| 43277_1 | TATGAGTGTGGAGTCGTTGCTGGTTCCACATTGTTGCAATTGTNNNAGATCGGAAG |
| 43277_2 | TATGAGTGTGGAGTCGTTGCTGGTTCCACATTGTTGCAATTGTNNNAGATCGGAAG |
| 41700_1 | TATGAGTGTGGAGTCGTTGCGTTCAAGCGATTCTCCTGCTTCNNNAGATCGGAAGA |
| 41700_2 | TATGAGTGTGGAGTCGTTGCGTTCAAGCGATTCTCCTGCTTCNNNAGATCGGAAGA |
| 30176_1 | TATGAGTGTGGAGTCGTTGCTTGTGCCAGGTCTTGAATACGTANNNAGATCGGAAG |
| 30176_2 | TATGAGTGTGGAGTCGTTGCTTGTGCCAGGTCTTGAATACGTANNNAGATCGGAAG |
| 12764_1 | TATGAGTGTGGAGTCGTTGCGGAGGAGACAGCTTCATAAGNNNAGATCGGAAGA |
| 12764_2 | TATGAGTGTGGAGTCGTTGCGGAGGAGACAGCTTCATAAGNNNAGATCGGAAGA |
| 40606_1 | TATGAGTGTGGAGTCGTTGCCAGATCATGAGGTCAGGAGATGGNNNAGATCGGA |
| 40606_2 | TATGAGTGTGGAGTCGTTGCCAGATCATGAGGTCAGGAGATGGNNNAGATCGGA |
| 40828_1 | TATGAGTGTGGAGTCGTTGCGCCTGGAATCTTACAGGAGAACANNNAGATCGGAA |
| 40828_2 | TATGAGTGTGGAGTCGTTGCGCCTGGAATCTTACAGGAGAACANNNAGATCGGAA |
| 40031_1 | TATGAGTGTGGAGTCGTTGCTGTTTCCCACAGAAGTACAGGAGNNNAGATCGGAA |
| 40031_2 | TATGAGTGTGGAGTCGTTGCTGTTTCCCACAGAAGTACAGGAGNNNAGATCGGAA |
| 44647_1 | TATGAGTGTGGAGTCGTTGCGTGGGGAGAAAGAAAGCGAAATCNNNAGATCGGA |
| 44647_2 | TATGAGTGTGGAGTCGTTGCGTGGGGAGAAAGAAAGCGAAATCNNNAGATCGGA |
| 48625_1 | TATGAGTGTGGAGTCGTTGCCTGGGTTCAAGCAATTATCCTGCNNNAGATCGGAAG |
| 48625_2 | TATGAGTGTGGAGTCGTTGCCTGGGTTCAAGCAATTATCCTGCNNNAGATCGGAAG |
| 3055_1 | TATGAGTGTGGAGTCGTTGCGGAAGAAGACAGTGAAGGGCATGNNNAGATCGGA |
| 3055_2 | TATGAGTGTGGAGTCGTTGCGGAAGAAGACAGTGAAGGGCATGNNNAGATCGGA |
| 36992_1 | TATGAGTGTGGAGTCGTTGCTCTTCTAGTCCTGCTTAGGTCCTNNNAGATCGGAAG |
| 36992_2 | TATGAGTGTGGAGTCGTTGCTCTTCTAGTCCTGCTTAGGTCCTNNNAGATCGGAAG |

| | |
|---|---|
| 36333_1 | TATGAGTGTGGAGTCGTTGCTGGTATCCATCGTCACTGGAATCNNNAGATCGGAAG |
| 36333_2 | TATGAGTGTGGAGTCGTTGCTGGTATCCATCGTCACTGGAATCNNNAGATCGGAAG |
| 46527_1 | TATGAGTGTGGAGTCGTTGCCAAGTGAAAGACTGGTTGGCAAANNNAGATCGGAA |
| 46527_2 | TATGAGTGTGGAGTCGTTGCCAAGTGAAAGACTGGTTGGCAAANNNAGATCGGAA |
| 29583_1 | TATGAGTGTGGAGTCGTTGCTCTCTCATGTGCTGAGGTCGTTANNNAGATCGGAAG |
| 29583_2 | TATGAGTGTGGAGTCGTTGCTCTCTCATGTGCTGAGGTCGTTANNNAGATCGGAAG |
| 51160_1 | TATGAGTGTGGAGTCGTTGCCCACGATCCACATTTTACAGCAGNNNAGATCGGAAG |
| 51160_2 | TATGAGTGTGGAGTCGTTGCCCACGATCCACATTTTACAGCAGNNNAGATCGGAAG |
| 41559_1 | TATGAGTGTGGAGTCGTTGCCCTCAACCTCCCAAAGTACTAGGNNNAGATCGGAAG |
| 41559_2 | TATGAGTGTGGAGTCGTTGCCCTCAACCTCCCAAAGTACTAGGNNNAGATCGGAAG |
| 31304_1 | TATGAGTGTGGAGTCGTTGCCCCACACACACTAACCAGAGTATNNNAGATCGGAAG |
| 31304_2 | TATGAGTGTGGAGTCGTTGCCCCACACACACTAACCAGAGTATNNNAGATCGGAAG |
| 49126_1 | TATGAGTGTGGAGTCGTTGCTTTACTCTTGTCACTCAGGCTGGNNNAGATCGGAAG |
| 49126_2 | TATGAGTGTGGAGTCGTTGCTTTACTCTTGTCACTCAGGCTGGNNNAGATCGGAAG |
| 11665_1 | TATGAGTGTGGAGTCGTTGCCACTTTTCCCTCTCTCCTTCACTTNNNAGATCGGAAGA |
| 11665_2 | TATGAGTGTGGAGTCGTTGCCACTTTTCCCTCTCTCCTTCACTTNNNAGATCGGAAGA |
| 46480_1 | TATGAGTGTGGAGTCGTTGCCTCCTGAGTAGCTGGGACTACAGNNNAGATCGGAA |
| 46480_2 | TATGAGTGTGGAGTCGTTGCCTCCTGAGTAGCTGGGACTACAGNNNAGATCGGAA |
| 31694_1 | TATGAGTGTGGAGTCGTTGCCACCTGCACAGAAAACAAAGTCANNNAGATCGGAAG |
| 31694_2 | TATGAGTGTGGAGTCGTTGCCACCTGCACAGAAAACAAAGTCANNNAGATCGGAAG |
| 49852_1 | TATGAGTGTGGAGTCGTTGCCTTGACTCAGACCATGTTCCTGANNNAGATCGGAAG |
| 49852_2 | TATGAGTGTGGAGTCGTTGCCTTGACTCAGACCATGTTCCTGANNNAGATCGGAAG |
| 31376_1 | TATGAGTGTGGAGTCGTTGCTGACTGAGTAGTTTGAAGACATGCNNNAGATCGGA |
| 31376_2 | TATGAGTGTGGAGTCGTTGCTGACTGAGTAGTTTGAAGACATGCNNNAGATCGGA |
| 42815_1 | TATGAGTGTGGAGTCGTTGCCATCCTGGCTAACATGGTGAAACNNNAGATCGGAAG |
| 42815_2 | TATGAGTGTGGAGTCGTTGCCATCCTGGCTAACATGGTGAAACNNNAGATCGGAAG |
| 51048_1 | TATGAGTGTGGAGTCGTTGCACAAAATTAGCTGGCCATGATGGNNNAGATCGGAA |
| 51048_2 | TATGAGTGTGGAGTCGTTGCACAAAATTAGCTGGCCATGATGGNNNAGATCGGAA |
| 38775_1 | TATGAGTGTGGAGTCGTTGCAACTGAGGAAACTGGAACATGGANNNAGATCGGAA |
| 38775_2 | TATGAGTGTGGAGTCGTTGCAACTGAGGAAACTGGAACATGGANNNAGATCGGAA |
| 47699_1 | TATGAGTGTGGAGTCGTTGCACCACGCCCGGATAATTTTTTTTNNNAGATCGGAAG |
| 47699_2 | TATGAGTGTGGAGTCGTTGCACCACGCCCGGATAATTTTTTTTNNNAGATCGGAAG |
| 31795_1 | TATGAGTGTGGAGTCGTTGCCCCACCCTTATGACCTCTTTCAANNNAGATCGGAAGA |
| 31795_2 | TATGAGTGTGGAGTCGTTGCCCCACCCTTATGACCTCTTTCAANNNAGATCGGAAGA |
| 40765_1 | TATGAGTGTGGAGTCGTTGCGAGGGCGTTGAAGAGATGAAAAGNNNAGATCGGA |
| 40765_2 | TATGAGTGTGGAGTCGTTGCGAGGGCGTTGAAGAGATGAAAAGNNNAGATCGGA |
| 52718_1 | TATGAGTGTGGAGTCGTTGCTTATCTGCGAGCTCCCTTTTTCTNNNAGATCGGAAGA |
| 52718_2 | TATGAGTGTGGAGTCGTTGCTTATCTGCGAGCTCCCTTTTTCTNNNAGATCGGAAGA |
| 33667_1 | TATGAGTGTGGAGTCGTTGCTTTAGTAGAGATGGGGTTTCGCCNNNAGATCGGAA |
| 33667_2 | TATGAGTGTGGAGTCGTTGCTTTAGTAGAGATGGGGTTTCGCCNNNAGATCGGAA |
| 41739_1 | TATGAGTGTGGAGTCGTTGCCATGACCTCCACTCACTGCAANNNAGATCGGAAGAG |
| 41739_2 | TATGAGTGTGGAGTCGTTGCCATGACCTCCACTCACTGCAANNNAGATCGGAAGAG |
| 53153_1 | TATGAGTGTGGAGTCGTTGCGGGTTCTGGCTATTGGTTCCTAANNNAGATCGGAAG |

| | |
|---|---|
| 53153_2 | TATGAGTGTGGAGTCGTTGCGGGTTCTGGCTATTGGTTCCTAANNNAGATCGGAAG |
| 46185_1 | TATGAGTGTGGAGTCGTTGCCTCCTAGGCTCAAGCAATCCTCNNNAGATCGGAAGA |
| 46185_2 | TATGAGTGTGGAGTCGTTGCCTCCTAGGCTCAAGCAATCCTCNNNAGATCGGAAGA |
| 46383_1 | TATGAGTGTGGAGTCGTTGCTGAAGCAGAGGTAGATGGTAAGCNNNAGATCGGA |
| 46383_2 | TATGAGTGTGGAGTCGTTGCTGAAGCAGAGGTAGATGGTAAGCNNNAGATCGGA |
| 45529_1 | TATGAGTGTGGAGTCGTTGCGGTCGGGGATTTCTAATAGCCATNNNAGATCGGAA |
| 45529_2 | TATGAGTGTGGAGTCGTTGCGGTCGGGGATTTCTAATAGCCATNNNAGATCGGAA |
| 2081_1 | TATGAGTGTGGAGTCGTTGCTCATACCCATGTACCCACGTACANNNAGATCGGAAG |
| 2081_2 | TATGAGTGTGGAGTCGTTGCTCATACCCATGTACCCACGTACANNNAGATCGGAAG |
| 13277_1 | TATGAGTGTGGAGTCGTTGCTCTGTACTAAGTTTAAAAAGCACNNNAGATCGGAAG |
| 13277_2 | TATGAGTGTGGAGTCGTTGCTCTGTACTAAGTTTAAAAAGCACNNNAGATCGGAAG |
| 35965_1 | TATGAGTGTGGAGTCGTTGCGTCAAGACCAGTCTGGGTAACATNNNAGATCGGAA |
| 35965_2 | TATGAGTGTGGAGTCGTTGCGTCAAGACCAGTCTGGGTAACATNNNAGATCGGAA |
| 33923_1 | TATGAGTGTGGAGTCGTTGCGATTACAGGCATGCACTACCATGNNNAGATCGGAAG |
| 33923_2 | TATGAGTGTGGAGTCGTTGCGATTACAGGCATGCACTACCATGNNNAGATCGGAAG |
| 34786_1 | TATGAGTGTGGAGTCGTTGCCTCTCTCTCTCCCTTCATCCTCTNNNAGATCGGAAGAG |
| 34786_2 | TATGAGTGTGGAGTCGTTGCCTCTCTCTCTCCCTTCATCCTCTNNNAGATCGGAAGAG |
| 37546_1 | TATGAGTGTGGAGTCGTTGCTGTCATGGAAGTAACACAGTCAGTNNNAGATCGGA |
| 37546_2 | TATGAGTGTGGAGTCGTTGCTGTCATGGAAGTAACACAGTCAGTNNNAGATCGGA |
| 40352_1 | TATGAGTGTGGAGTCGTTGCGAAAATGAGAGAGGCACTGCAGNNNAGATCGGAA |
| 40352_2 | TATGAGTGTGGAGTCGTTGCGAAAATGAGAGAGGCACTGCAGNNNAGATCGGAA |
| 44071_1 | TATGAGTGTGGAGTCGTTGCTCCTTCTCTCCCTGATGCTTCTANNNAGATCGGAAGA |
| 44071_2 | TATGAGTGTGGAGTCGTTGCTCCTTCTCTCCCTGATGCTTCTANNNAGATCGGAAGA |
| 28857_1 | TATGAGTGTGGAGTCGTTGCTGTCCCAAGGAGAGCATCATTTTNNNAGATCGGAAG |
| 28857_2 | TATGAGTGTGGAGTCGTTGCTGTCCCAAGGAGAGCATCATTTTNNNAGATCGGAAG |
| 47581_1 | TATGAGTGTGGAGTCGTTGCAGGAAGGAAGAGCAGGCATTTCNNNAGATCGGAA |
| 47581_2 | TATGAGTGTGGAGTCGTTGCAGGAAGGAAGAGCAGGCATTTCNNNAGATCGGAA |
| 37322_1 | TATGAGTGTGGAGTCGTTGCACATACACTAGGAACTGCAATGCNNNAGATCGGAAG |
| 37322_2 | TATGAGTGTGGAGTCGTTGCACATACACTAGGAACTGCAATGCNNNAGATCGGAAG |
| 51003_1 | TATGAGTGTGGAGTCGTTGCCAGAGAGAGACTCCATCACACACNNNAGATCGGAAG |
| 51003_2 | TATGAGTGTGGAGTCGTTGCCAGAGAGAGACTCCATCACACACNNNAGATCGGAAG |
| 32808_1 | TATGAGTGTGGAGTCGTTGCTGGAGTATAATGGTGCGATCTCGNNNAGATCGGAA |
| 32808_2 | TATGAGTGTGGAGTCGTTGCTGGAGTATAATGGTGCGATCTCGNNNAGATCGGAA |
| 46405_1 | TATGAGTGTGGAGTCGTTGCTCCGAATGATTCTCTTGCCTCAGNNNAGATCGGAAG |
| 46405_2 | TATGAGTGTGGAGTCGTTGCTCCGAATGATTCTCTTGCCTCAGNNNAGATCGGAAG |
| 29483_1 | TATGAGTGTGGAGTCGTTGCCTTGAAACTCTTCTTTGGCTTTTTCANNNAGATCGGA |
| 29483_2 | TATGAGTGTGGAGTCGTTGCCTTGAAACTCTTCTTTGGCTTTTTCANNNAGATCGGA |
| 48000_1 | TATGAGTGTGGAGTCGTTGCCTATCTAGGCTCACTGCAACCTCNNNAGATCGGAAG |
| 48000_2 | TATGAGTGTGGAGTCGTTGCCTATCTAGGCTCACTGCAACCTCNNNAGATCGGAAG |
| 32799_1 | TATGAGTGTGGAGTCGTTGCCGTAGTAGCCCTAGTAGCTGGTANNNAGATCGGAA |
| 32799_2 | TATGAGTGTGGAGTCGTTGCCGTAGTAGCCCTAGTAGCTGGTANNNAGATCGGAA |
| 38300_1 | TATGAGTGTGGAGTCGTTGCAAGTGGGTCTCAGACTAGTCCTTNNNAGATCGGAAG |
| 38300_2 | TATGAGTGTGGAGTCGTTGCAAGTGGGTCTCAGACTAGTCCTTNNNAGATCGGAAG |

| | |
|---|---|
| 53220_1 | TATGAGTGTGGAGTCGTTGCTTTGGCCTCAAACTTCAGGTGANNNAGATCGGAAGA |
| 53220_2 | TATGAGTGTGGAGTCGTTGCTTTGGCCTCAAACTTCAGGTGANNNAGATCGGAAGA |
| 45442_1 | TATGAGTGTGGAGTCGTTGCCACTCCTGTAATCCCAGCACTTTNNNAGATCGGAAGA |
| 45442_2 | TATGAGTGTGGAGTCGTTGCCACTCCTGTAATCCCAGCACTTTNNNAGATCGGAAGA |
| 46607_1 | TATGAGTGTGGAGTCGTTGCCCTCTCCTTTTCCTAGGCTCTTGNNNAGATCGGAAGA |
| 46607_2 | TATGAGTGTGGAGTCGTTGCCCTCTCCTTTTCCTAGGCTCTTGNNNAGATCGGAAGA |
| 29855_1 | TATGAGTGTGGAGTCGTTGCTGAAAACCCACAAATATGACTCATCTNNNAGATCGG |
| 29855_2 | TATGAGTGTGGAGTCGTTGCTGAAAACCCACAAATATGACTCATCTNNNAGATCGG |
| 49035_1 | TATGAGTGTGGAGTCGTTGCTGTGTGTCTCTCTAGGACTTTGCNNNAGATCGGAAG |
| 49035_2 | TATGAGTGTGGAGTCGTTGCTGTGTGTCTCTCTAGGACTTTGCNNNAGATCGGAAG |
| 31682_1 | TATGAGTGTGGAGTCGTTGCTGCATACACTGGGTCATTAGAGGNNNAGATCGGAA |
| 31682_2 | TATGAGTGTGGAGTCGTTGCTGCATACACTGGGTCATTAGAGGNNNAGATCGGAA |
| 31675_1 | TATGAGTGTGGAGTCGTTGCGCTCCAAAGCGACCTAAAGATTTNNNAGATCGGAAG |
| 31675_2 | TATGAGTGTGGAGTCGTTGCGCTCCAAAGCGACCTAAAGATTTNNNAGATCGGAAG |
| 36215_1 | TATGAGTGTGGAGTCGTTGCGCCATCTACAAACCAGGAAGAGANNNAGATCGGAA |
| 36215_2 | TATGAGTGTGGAGTCGTTGCGCCATCTACAAACCAGGAAGAGANNNAGATCGGAA |
| 33301_1 | TATGAGTGTGGAGTCGTTGCGGAGTGCAATGACGCAATCTCNNNAGATCGGAAGA |
| 33301_2 | TATGAGTGTGGAGTCGTTGCGGAGTGCAATGACGCAATCTCNNNAGATCGGAAGA |
| 51050_1 | TATGAGTGTGGAGTCGTTGCAACCACCAAGGGACATGTATACCNNNAGATCGGAAG |
| 51050_2 | TATGAGTGTGGAGTCGTTGCAACCACCAAGGGACATGTATACCNNNAGATCGGAAG |
| 32403_1 | TATGAGTGTGGAGTCGTTGCTGATTGGTTCTCTACGTTCACCTNNNAGATCGGAAG |
| 32403_2 | TATGAGTGTGGAGTCGTTGCTGATTGGTTCTCTACGTTCACCTNNNAGATCGGAAG |
| 49656_1 | TATGAGTGTGGAGTCGTTGCCTCCTGAGTAGCTGGGATTACAGNNNAGATCGGAA |
| 49656_2 | TATGAGTGTGGAGTCGTTGCCTCCTGAGTAGCTGGGATTACAGNNNAGATCGGAA |
| 36312_1 | TATGAGTGTGGAGTCGTTGCGTCTGACTTCATCCAAGGGTTCTNNNAGATCGGAAG |
| 36312_2 | TATGAGTGTGGAGTCGTTGCGTCTGACTTCATCCAAGGGTTCTNNNAGATCGGAAG |
| 32811_1 | TATGAGTGTGGAGTCGTTGCCCTCCATAGTAGCTGGGACTACANNNAGATCGGAAG |
| 32811_2 | TATGAGTGTGGAGTCGTTGCCCTCCATAGTAGCTGGGACTACANNNAGATCGGAAG |
| 45152_1 | TATGAGTGTGGAGTCGTTGCTGTCAGCAGAAGTTGAGTGGTTANNNAGATCGGAA |
| 45152_2 | TATGAGTGTGGAGTCGTTGCTGTCAGCAGAAGTTGAGTGGTTANNNAGATCGGAA |
| 42415_1 | TATGAGTGTGGAGTCGTTGCTTGAGGAACCCCATACTGTTCTCNNNAGATCGGAAG |
| 42415_2 | TATGAGTGTGGAGTCGTTGCTTGAGGAACCCCATACTGTTCTCNNNAGATCGGAAG |
| 32903_1 | TATGAGTGTGGAGTCGTTGCATCCTGGGAAGTTGATGCTGTAGNNNAGATCGGAA |
| 32903_2 | TATGAGTGTGGAGTCGTTGCATCCTGGGAAGTTGATGCTGTAGNNNAGATCGGAA |
| 53022_1 | TATGAGTGTGGAGTCGTTGCTGGTTCAAAAGGCAAAATACACGANNNAGATCGGA |
| 53022_2 | TATGAGTGTGGAGTCGTTGCTGGTTCAAAAGGCAAAATACACGANNNAGATCGGA |
| 41485_1 | TATGAGTGTGGAGTCGTTGCGCATCAATCCATCCTACTCCCATNNNAGATCGGAAGA |
| 41485_2 | TATGAGTGTGGAGTCGTTGCGCATCAATCCATCCTACTCCCATNNNAGATCGGAAGA |
| 30850_1 | TATGAGTGTGGAGTCGTTGCAATCACACAGCTAGGAAGAAGCANNNAGATCGGAA |
| 30850_2 | TATGAGTGTGGAGTCGTTGCAATCACACAGCTAGGAAGAAGCANNNAGATCGGAA |
| 32374_1 | TATGAGTGTGGAGTCGTTGCGAGAGAGAGAGAGAGAGAGACTGCTNNNAGATCGGA |
| 32374_2 | TATGAGTGTGGAGTCGTTGCGAGAGAGAGAGAGAGAGAGACTGCTNNNAGATCGGA |
| 2858_1 | TATGAGTGTGGAGTCGTTGCGTATCCACAACCTCAGTCAAGGTNNNAGATCGGAAG |

| | |
|---|---|
| 2858_2 | TATGAGTGTGGAGTCGTTGCGTATCCACAACCTCAGTCAAGGTNNNAGATCGGAAG |
| 32299_1 | TATGAGTGTGGAGTCGTTGCTATAACAGTGTTGAGAGGTGGGCNNNAGATCGGAA |
| 32299_2 | TATGAGTGTGGAGTCGTTGCTATAACAGTGTTGAGAGGTGGGCNNNAGATCGGAA |
| 46788_1 | TATGAGTGTGGAGTCGTTGCGCCTTGGTTTGAAGCAGGATTAGNNNAGATCGGAA |
| 46788_2 | TATGAGTGTGGAGTCGTTGCGCCTTGGTTTGAAGCAGGATTAGNNNAGATCGGAA |
| 42434_1 | TATGAGTGTGGAGTCGTTGCCATGTGTATTGCAGCACAAGTCANNNAGATCGGAAG |
| 42434_2 | TATGAGTGTGGAGTCGTTGCCATGTGTATTGCAGCACAAGTCANNNAGATCGGAAG |
| 42382_1 | TATGAGTGTGGAGTCGTTGCAATCTCTTTTGGCAGTGCAGAACNNNAGATCGGAAG |
| 42382_2 | TATGAGTGTGGAGTCGTTGCAATCTCTTTTGGCAGTGCAGAACNNNAGATCGGAAG |
| 41990_1 | TATGAGTGTGGAGTCGTTGCGATGGAGTCTCACTCTGTCGCNNNAGATCGGAAGAG |
| 41990_2 | TATGAGTGTGGAGTCGTTGCGATGGAGTCTCACTCTGTCGCNNNAGATCGGAAGAG |
| 51334_1 | TATGAGTGTGGAGTCGTTGCGTGTACCCAACAGCTCATTGAGANNNAGATCGGAAG |
| 51334_2 | TATGAGTGTGGAGTCGTTGCGTGTACCCAACAGCTCATTGAGANNNAGATCGGAAG |
| 39612_1 | TATGAGTGTGGAGTCGTTGCTGGGGTTAGGACAGGAAAATGAGNNNAGATCGGA |
| 39612_2 | TATGAGTGTGGAGTCGTTGCTGGGGTTAGGACAGGAAAATGAGNNNAGATCGGA |
| 31508_1 | TATGAGTGTGGAGTCGTTGCAGTTCTTTGCTTTGTCAAACTGCTNNNAGATCGGAAG |
| 31508_2 | TATGAGTGTGGAGTCGTTGCAGTTCTTTGCTTTGTCAAACTGCTNNNAGATCGGAAG |
| 13371_1 | TATGAGTGTGGAGTCGTTGCGCTCTCATCATAAAATTCAGAAANNNAGATCGGAAG |
| 13371_2 | TATGAGTGTGGAGTCGTTGCGCTCTCATCATAAAATTCAGAAANNNAGATCGGAAG |
| 13168_1 | TATGAGTGTGGAGTCGTTGCGAGAAAAAGAGATATGTGATGTTNNNAGATCGGAA |
| 13168_2 | TATGAGTGTGGAGTCGTTGCGAGAAAAAGAGATATGTGATGTTNNNAGATCGGAA |
| 50233_1 | TATGAGTGTGGAGTCGTTGCAAGGTGGTTGGAGATGACGTTTANNNAGATCGGAA |
| 50233_2 | TATGAGTGTGGAGTCGTTGCAAGGTGGTTGGAGATGACGTTTANNNAGATCGGAA |
| 50262_1 | TATGAGTGTGGAGTCGTTGCTGACATACCAGCTGAAGATCCTGNNNAGATCGGAAG |
| 50262_2 | TATGAGTGTGGAGTCGTTGCTGACATACCAGCTGAAGATCCTGNNNAGATCGGAAG |
| 12798_1 | TATGAGTGTGGAGTCGTTGCATTCTTTGACTTTTGAGAAGATTNNNAGATCGGAAG |
| 12798_2 | TATGAGTGTGGAGTCGTTGCATTCTTTGACTTTTGAGAAGATTNNNAGATCGGAAG |
| 22789_1 | TATGAGTGTGGAGTCGTTGCAAGAGCTGATTGTTAAATATTCCNNNAGATCGGAAG |
| 22789_2 | TATGAGTGTGGAGTCGTTGCAAGAGCTGATTGTTAAATATTCCNNNAGATCGGAAG |
| 48117_1 | TATGAGTGTGGAGTCGTTGCGTTGGTTCTGAAGTGTGGGTTGNNNAGATCGGAAG |
| 48117_2 | TATGAGTGTGGAGTCGTTGCGTTGGTTCTGAAGTGTGGGTTGNNNAGATCGGAAG |
| 52430_1 | TATGAGTGTGGAGTCGTTGCTGTCCAACATGTGCTACACCTTANNNAGATCGGAAG |
| 52430_2 | TATGAGTGTGGAGTCGTTGCTGTCCAACATGTGCTACACCTTANNNAGATCGGAAG |
| 33012_1 | TATGAGTGTGGAGTCGTTGCTTCACACCATTCTCCTGCCTCNNNAGATCGGAAGAGC |
| 33012_2 | TATGAGTGTGGAGTCGTTGCTTCACACCATTCTCCTGCCTCNNNAGATCGGAAGAGC |
| 34297_1 | TATGAGTGTGGAGTCGTTGCCTGAGGCAGGAGAATCACTTGAANNNAGATCGGAA |
| 34297_2 | TATGAGTGTGGAGTCGTTGCCTGAGGCAGGAGAATCACTTGAANNNAGATCGGAA |
| 42357_1 | TATGAGTGTGGAGTCGTTGCACATCAATCTGGAGAACAGGCATNNNAGATCGGAA |
| 42357_2 | TATGAGTGTGGAGTCGTTGCACATCAATCTGGAGAACAGGCATNNNAGATCGGAA |
| 11292_1 | TATGAGTGTGGAGTCGTTGCAGACTTACCAGCTTGGGCCAATGNNNAGATCGGAAG |
| 11292_2 | TATGAGTGTGGAGTCGTTGCAGACTTACCAGCTTGGGCCAATGNNNAGATCGGAAG |
| 30409_1 | TATGAGTGTGGAGTCGTTGCTGAAAGGGCATGCAATAGACTCCNNNAGATCGGAA |
| 30409_2 | TATGAGTGTGGAGTCGTTGCTGAAAGGGCATGCAATAGACTCCNNNAGATCGGAA |

| | |
|---|---|
| 38888_1 | TATGAGTGTGGAGTCGTTGCCCCAACCCAATAAGCAATCACTGNNNAGATCGGAAG |
| 38888_2 | TATGAGTGTGGAGTCGTTGCCCCAACCCAATAAGCAATCACTGNNNAGATCGGAAG |
| 13254_1 | TATGAGTGTGGAGTCGTTGCCCTACTATGTACCCACAAATAATNNNAGATCGGAAG |
| 13254_2 | TATGAGTGTGGAGTCGTTGCCCTACTATGTACCCACAAATAATNNNAGATCGGAAG |
| 51659_1 | TATGAGTGTGGAGTCGTTGCCTGCTCCAGTGAGTCTTTTTTGGNNNAGATCGGAAG |
| 51659_2 | TATGAGTGTGGAGTCGTTGCCTGCTCCAGTGAGTCTTTTTTGGNNNAGATCGGAAG |
| 43899_1 | TATGAGTGTGGAGTCGTTGCCTCTCGTGCCAAACAGTGCAGNNNAGATCGGAAGA |
| 43899_2 | TATGAGTGTGGAGTCGTTGCCTCTCGTGCCAAACAGTGCAGNNNAGATCGGAAGA |
| 37258_1 | TATGAGTGTGGAGTCGTTGCCGTTGACCCCTGTTCTAGATCATNNNAGATCGGAAG |
| 37258_2 | TATGAGTGTGGAGTCGTTGCCGTTGACCCCTGTTCTAGATCATNNNAGATCGGAAG |
| 47034_1 | TATGAGTGTGGAGTCGTTGCCCAAATTGCTGGGATTACAGACGNNNAGATCGGAA |
| 47034_2 | TATGAGTGTGGAGTCGTTGCCCAAATTGCTGGGATTACAGACGNNNAGATCGGAA |
| 30280_1 | TATGAGTGTGGAGTCGTTGCATTTCCTCATTTTGCCTTTGCCANNNAGATCGGAAGA |
| 30280_2 | TATGAGTGTGGAGTCGTTGCATTTCCTCATTTTGCCTTTGCCANNNAGATCGGAAGA |
| 34619_1 | TATGAGTGTGGAGTCGTTGCACGCTAAGTGGCACATTACAGTANNNAGATCGGAA |
| 34619_2 | TATGAGTGTGGAGTCGTTGCACGCTAAGTGGCACATTACAGTANNNAGATCGGAA |
| 36910_1 | TATGAGTGTGGAGTCGTTGCGGTAGCACTGAGTTTTTGGCTACNNNAGATCGGAA |
| 36910_2 | TATGAGTGTGGAGTCGTTGCGGTAGCACTGAGTTTTTGGCTACNNNAGATCGGAA |
| 42191_1 | TATGAGTGTGGAGTCGTTGCCTGATTATCCTGCCTTAGCCTCCNNNAGATCGGAAGA |
| 42191_2 | TATGAGTGTGGAGTCGTTGCCTGATTATCCTGCCTTAGCCTCCNNNAGATCGGAAGA |
| 28828_1 | TATGAGTGTGGAGTCGTTGCAAAGAGAGAGAGAGAAATGGCCANNNAGATCGGA |
| 28828_2 | TATGAGTGTGGAGTCGTTGCAAAGAGAGAGAGAGAAATGGCCANNNAGATCGGA |
| 36227_1 | TATGAGTGTGGAGTCGTTGCTGAGGATTGTTATTTCTGTGGGCNNNAGATCGGAA |
| 36227_2 | TATGAGTGTGGAGTCGTTGCTGAGGATTGTTATTTCTGTGGGCNNNAGATCGGAA |
| 46775_1 | TATGAGTGTGGAGTCGTTGCCGTGCCCGGCTAATTTTTTGTATNNNAGATCGGAAG |
| 46775_2 | TATGAGTGTGGAGTCGTTGCCGTGCCCGGCTAATTTTTTGTATNNNAGATCGGAAG |
| 39601_1 | TATGAGTGTGGAGTCGTTGCCGCTTATAGTCCCAGCTACTCGNNNAGATCGGAAGA |
| 39601_2 | TATGAGTGTGGAGTCGTTGCCGCTTATAGTCCCAGCTACTCGNNNAGATCGGAAGA |
| 35454_1 | TATGAGTGTGGAGTCGTTGCTACCATGCCCGGCTAATTTTTTGNNNAGATCGGAAG |
| 35454_2 | TATGAGTGTGGAGTCGTTGCTACCATGCCCGGCTAATTTTTTGNNNAGATCGGAAG |
| 32191_1 | TATGAGTGTGGAGTCGTTGCCTGCTTCAACCTCCTGAGTAACTNNNAGATCGGAAG |
| 32191_2 | TATGAGTGTGGAGTCGTTGCCTGCTTCAACCTCCTGAGTAACTNNNAGATCGGAAG |
| 42216_1 | TATGAGTGTGGAGTCGTTGCAAGCAGGCAACTTTCCTTCATTCNNNAGATCGGAAG |
| 42216_2 | TATGAGTGTGGAGTCGTTGCAAGCAGGCAACTTTCCTTCATTCNNNAGATCGGAAG |
| 37693_1 | TATGAGTGTGGAGTCGTTGCACCAAAAAACAACGATTAGGCCTNNNAGATCGGAAG |
| 37693_2 | TATGAGTGTGGAGTCGTTGCACCAAAAAACAACGATTAGGCCTNNNAGATCGGAAG |
| 12474_1 | TATGAGTGTGGAGTCGTTGCAACACAACCTTAAAAAGAGAAAANNNAGATCGGAA |
| 12474_2 | TATGAGTGTGGAGTCGTTGCAACACAACCTTAAAAAGAGAAAANNNAGATCGGAA |
| 41867_1 | TATGAGTGTGGAGTCGTTGCATGATCTTGGCTCACTGCAACTTNNNAGATCGGAAG |
| 41867_2 | TATGAGTGTGGAGTCGTTGCATGATCTTGGCTCACTGCAACTTNNNAGATCGGAAG |
| 30640_1 | ATGAGTGTGGAGTCGTTGCGGAAACTAGAAGTTGTTGGCAGTAGTANNNAGATCG |
| 30640_2 | ATGAGTGTGGAGTCGTTGCGGAAACTAGAAGTTGTTGGCAGTAGTANNNAGATCG |
| 31676_1 | TATGAGTGTGGAGTCGTTGCATACCCAGCTTCTTTTGGCCATANNNAGATCGGAAG |

| | |
|---|---|
| 31676_2 | TATGAGTGTGGAGTCGTTGCATACCCAGCTTCTTTTGGCCATANNNAGATCGGAAG |
| 44852_1 | TATGAGTGTGGAGTCGTTGCGGCCCTCTGAACACAAAGGATAANNNAGATCGGAA |
| 44852_2 | TATGAGTGTGGAGTCGTTGCGGCCCTCTGAACACAAAGGATAANNNAGATCGGAA |
| 38639_1 | TATGAGTGTGGAGTCGTTGCAGTCCTTCCATCTTCCCTCTTCTNNNAGATCGGAAGA |
| 38639_2 | TATGAGTGTGGAGTCGTTGCAGTCCTTCCATCTTCCCTCTTCTNNNAGATCGGAAGA |
| 50651_1 | TATGAGTGTGGAGTCGTTGCTCTCTTTGTTCCAGCAGACATCANNNAGATCGGAAG |
| 50651_2 | TATGAGTGTGGAGTCGTTGCTCTCTTTGTTCCAGCAGACATCANNNAGATCGGAAG |
| 50548_1 | TATGAGTGTGGAGTCGTTGCTAGCATCTAGTGACAGTGCCATCNNNAGATCGGAAG |
| 50548_2 | TATGAGTGTGGAGTCGTTGCTAGCATCTAGTGACAGTGCCATCNNNAGATCGGAAG |
| 51744_1 | TATGAGTGTGGAGTCGTTGCCATCAATCCAACTGCACTTTCGANNNAGATCGGAAG |
| 51744_2 | TATGAGTGTGGAGTCGTTGCCATCAATCCAACTGCACTTTCGANNNAGATCGGAAG |
| 39822_1 | TATGAGTGTGGAGTCGTTGCACCTTCCTCAGTACCCAGACTATNNNAGATCGGAAG |
| 39822_2 | TATGAGTGTGGAGTCGTTGCACCTTCCTCAGTACCCAGACTATNNNAGATCGGAAG |
| 46892_1 | TATGAGTGTGGAGTCGTTGCGGTTCAAATGTTTCTCCTGCCTCNNNAGATCGGAAG |
| 46892_2 | TATGAGTGTGGAGTCGTTGCGGTTCAAATGTTTCTCCTGCCTCNNNAGATCGGAAG |
| 10798_1 | TATGAGTGTGGAGTCGTTGCTAGGCATCTCCCAACTGCTTTTCNNNAGATCGGAAG |
| 10798_2 | TATGAGTGTGGAGTCGTTGCTAGGCATCTCCCAACTGCTTTTCNNNAGATCGGAAG |
| 42838_1 | TATGAGTGTGGAGTCGTTGCTCCATCCATGCCCCAAAAGATTANNNAGATCGGAAG |
| 42838_2 | TATGAGTGTGGAGTCGTTGCTCCATCCATGCCCCAAAAGATTANNNAGATCGGAAG |
| 35451_1 | TATGAGTGTGGAGTCGTTGCGCACAATCTCAGCTAACTGCAAANNNAGATCGGAAG |
| 35451_2 | TATGAGTGTGGAGTCGTTGCGCACAATCTCAGCTAACTGCAAANNNAGATCGGAAG |
| 33569_1 | TATGAGTGTGGAGTCGTTGCAGTTTCTCTCTCTCTCTCACCCANNNAGATCGGAAGA |
| 33569_2 | TATGAGTGTGGAGTCGTTGCAGTTTCTCTCTCTCTCTCACCCANNNAGATCGGAAGA |
| 44174_1 | TATGAGTGTGGAGTCGTTGCGTTTGCTTTTCTGGGGCGATACNNNAGATCGGAAG |
| 44174_2 | TATGAGTGTGGAGTCGTTGCGTTTGCTTTTCTGGGGCGATACNNNAGATCGGAAG |
| 34363_1 | TATGAGTGTGGAGTCGTTGCGGTTGCCACAAAGTCTTGGAAATNNNAGATCGGAA |
| 34363_2 | TATGAGTGTGGAGTCGTTGCGGTTGCCACAAAGTCTTGGAAATNNNAGATCGGAA |
| 10249_1 | TATGAGTGTGGAGTCGTTGCCCCAACGAGATAGGTAACAGGTGNNNAGATCGGAA |
| 10249_2 | TATGAGTGTGGAGTCGTTGCCCCAACGAGATAGGTAACAGGTGNNNAGATCGGAA |
| 37923_1 | TATGAGTGTGGAGTCGTTGCTGAGCTGTTGGAGTTTCTTCTGTNNNAGATCGGAAG |
| 37923_2 | TATGAGTGTGGAGTCGTTGCTGAGCTGTTGGAGTTTCTTCTGTNNNAGATCGGAAG |
| 12987_1 | TATGAGTGTGGAGTCGTTGCGGTAAGGTTAAAATGGAAAATCTNNNAGATCGGAA |
| 12987_2 | TATGAGTGTGGAGTCGTTGCGGTAAGGTTAAAATGGAAAATCTNNNAGATCGGAA |
| 2533_1 | TATGAGTGTGGAGTCGTTGCCTGCTAACATGATCACCTCCAGGNNNAGATCGGAAG |
| 2533_2 | TATGAGTGTGGAGTCGTTGCCTGCTAACATGATCACCTCCAGGNNNAGATCGGAAG |
| 38238_1 | TATGAGTGTGGAGTCGTTGCTGTGGTTCATAACTTGGGTCACTNNNAGATCGGAAG |
| 38238_2 | TATGAGTGTGGAGTCGTTGCTGTGGTTCATAACTTGGGTCACTNNNAGATCGGAAG |
| 37087_1 | TATGAGTGTGGAGTCGTTGCTTGACCCATTTGAGCACAATTCCNNNAGATCGGAAG |
| 37087_2 | TATGAGTGTGGAGTCGTTGCTTGACCCATTTGAGCACAATTCCNNNAGATCGGAAG |
| 31478_1 | TATGAGTGTGGAGTCGTTGCACTGAGCACACCTTGGTTTCTAANNNAGATCGGAAG |
| 31478_2 | TATGAGTGTGGAGTCGTTGCACTGAGCACACCTTGGTTTCTAANNNAGATCGGAAG |
| 33873_1 | TATGAGTGTGGAGTCGTTGCCACTGTGGAGACTTTGAGTGAGTNNNAGATCGGAA |
| 33873_2 | TATGAGTGTGGAGTCGTTGCCACTGTGGAGACTTTGAGTGAGTNNNAGATCGGAA |

| | |
|---|---|
| 10620_1 | TATGAGTGTGGAGTCGTTGCACCGAGTGACAATACACAGCANNNAGATCGGAAGA |
| 10620_2 | TATGAGTGTGGAGTCGTTGCACCGAGTGACAATACACAGCANNNAGATCGGAAGA |
| 52555_1 | TATGAGTGTGGAGTCGTTGCGTCTGGCTGGAATGGGAGTAATTNNNAGATCGGAA |
| 52555_2 | TATGAGTGTGGAGTCGTTGCGTCTGGCTGGAATGGGAGTAATTNNNAGATCGGAA |
| 43396_1 | TATGAGTGTGGAGTCGTTGCCATGCCCGGCCTTTTTTTTTTTNNNAGATCGGAAGA |
| 43396_2 | TATGAGTGTGGAGTCGTTGCCATGCCCGGCCTTTTTTTTTTTNNNAGATCGGAAGA |
| 52422_1 | TATGAGTGTGGAGTCGTTGCTCCCTTCTCTGTCACAGAAATCGNNNAGATCGGAAG |
| 52422_2 | TATGAGTGTGGAGTCGTTGCTCCCTTCTCTGTCACAGAAATCGNNNAGATCGGAAG |
| 38995_1 | TATGAGTGTGGAGTCGTTGCCAGCTGAATCAACCAAGTGGAAGNNNAGATCGGAA |
| 38995_2 | TATGAGTGTGGAGTCGTTGCCAGCTGAATCAACCAAGTGGAAGNNNAGATCGGAA |
| 33139_1 | TATGAGTGTGGAGTCGTTGCGCCTATAATCCAGCACTTTGGGANNNAGATCGGAAG |
| 33139_2 | TATGAGTGTGGAGTCGTTGCGCCTATAATCCAGCACTTTGGGANNNAGATCGGAAG |
| 10208_1 | TATGAGTGTGGAGTCGTTGCTGCCCCAAGAAAACTCTTCTCACNNNAGATCGGAAG |
| 10208_2 | TATGAGTGTGGAGTCGTTGCTGCCCCAAGAAAACTCTTCTCACNNNAGATCGGAAG |
| 33649_1 | TATGAGTGTGGAGTCGTTGCTTCAACTGATTCTCCTGCCTCAGNNNAGATCGGAAGA |
| 33649_2 | TATGAGTGTGGAGTCGTTGCTTCAACTGATTCTCCTGCCTCAGNNNAGATCGGAAGA |
| 46949_1 | TATGAGTGTGGAGTCGTTGCGAACCATTCACTCCCCTGGAAAGNNNAGATCGGAAG |
| 46949_2 | TATGAGTGTGGAGTCGTTGCGAACCATTCACTCCCCTGGAAAGNNNAGATCGGAAG |
| 36025_1 | TATGAGTGTGGAGTCGTTGCCCTTTCCAGGACCAGCAGATAATNNNAGATCGGAAG |
| 36025_2 | TATGAGTGTGGAGTCGTTGCCCTTTCCAGGACCAGCAGATAATNNNAGATCGGAAG |
| 46664_1 | TATGAGTGTGGAGTCGTTGCCAATCTTGGCTAACTGCAACCTCNNNAGATCGGAAG |
| 46664_2 | TATGAGTGTGGAGTCGTTGCCAATCTTGGCTAACTGCAACCTCNNNAGATCGGAAG |
| 43605_1 | TATGAGTGTGGAGTCGTTGCGCTGTCATTTTTTGGTGACCGTANNNAGATCGGAAG |
| 43605_2 | TATGAGTGTGGAGTCGTTGCGCTGTCATTTTTTGGTGACCGTANNNAGATCGGAAG |
| 46752_1 | TATGAGTGTGGAGTCGTTGCGTGTTGGGATTACAGGCATAAGCNNNAGATCGGAA |
| 46752_2 | TATGAGTGTGGAGTCGTTGCGTGTTGGGATTACAGGCATAAGCNNNAGATCGGAA |
| 33602_1 | TATGAGTGTGGAGTCGTTGCTCAAGTGATTCTCGTGTCTCAGCNNNAGATCGGAAG |
| 33602_2 | TATGAGTGTGGAGTCGTTGCTCAAGTGATTCTCGTGTCTCAGCNNNAGATCGGAAG |
| 42824_1 | TATGAGTGTGGAGTCGTTGCCAGAATAAACCCTGCAGACTCCTNNNAGATCGGAAG |
| 42824_2 | TATGAGTGTGGAGTCGTTGCCAGAATAAACCCTGCAGACTCCTNNNAGATCGGAAG |
| 8492_1 | TATGAGTGTGGAGTCGTTGCATGTGAAAGCCAGAACTCAGCCNNNAGATCGGAAG |
| 8492_2 | TATGAGTGTGGAGTCGTTGCATGTGAAAGCCAGAACTCAGCCNNNAGATCGGAAG |
| 10774_1 | TATGAGTGTGGAGTCGTTGCCTGGAAACAGTCTCACTCTCTGGNNNAGATCGGAAG |
| 10774_2 | TATGAGTGTGGAGTCGTTGCCTGGAAACAGTCTCACTCTCTGGNNNAGATCGGAAG |
| 47420_1 | TATGAGTGTGGAGTCGTTGCGTGTAGCTGGGACTACAGGTGNNNAGATCGGAAGA |
| 47420_2 | TATGAGTGTGGAGTCGTTGCGTGTAGCTGGGACTACAGGTGNNNAGATCGGAAGA |
| 41165_1 | TATGAGTGTGGAGTCGTTGCGAAGCTAAGACAGGAGAATCGCTNNNAGATCGGAA |
| 41165_2 | TATGAGTGTGGAGTCGTTGCGAAGCTAAGACAGGAGAATCGCTNNNAGATCGGAA |
| 31462_1 | TATGAGTGTGGAGTCGTTGCCCCCACACACTTATACCCTCATTNNNAGATCGGAAGA |
| 31462_2 | TATGAGTGTGGAGTCGTTGCCCCCACACACTTATACCCTCATTNNNAGATCGGAAGA |
| 2864_1 | TATGAGTGTGGAGTCGTTGCTACCATTCTGGGCTTGATGAGGTNNNAGATCGGAA |
| 2864_2 | TATGAGTGTGGAGTCGTTGCTACCATTCTGGGCTTGATGAGGTNNNAGATCGGAA |
| 33925_1 | TATGAGTGTGGAGTCGTTGCTGTTCCAGTTCTCAGAGGGAATGNNNAGATCGGAA |

| | |
|---|---|
| 33925_2 | TATGAGTGTGGAGTCGTTGCTGTTCCAGTTCTCAGAGGGAATGNNNAGATCGGAA |
| 51695_1 | TATGAGTGTGGAGTCGTTGCCTTATCCCTGCAGGCTTAGGAATNNNAGATCGGAAG |
| 51695_2 | TATGAGTGTGGAGTCGTTGCCTTATCCCTGCAGGCTTAGGAATNNNAGATCGGAAG |
| 42965_1 | TATGAGTGTGGAGTCGTTGCATAGCTCACTGCAATCTCCACCNNNAGATCGGAAGA |
| 42965_2 | TATGAGTGTGGAGTCGTTGCATAGCTCACTGCAATCTCCACCNNNAGATCGGAAGA |
| 52214_1 | TATGAGTGTGGAGTCGTTGCCCCGACCCCATCATTGTTTTTTTNNNAGATCGGAAGA |
| 52214_2 | TATGAGTGTGGAGTCGTTGCCCCGACCCCATCATTGTTTTTTTNNNAGATCGGAAGA |
| 40037_1 | TATGAGTGTGGAGTCGTTGCTGTTTTGCTGCTCAATCAGAGAGNNNAGATCGGAAG |
| 40037_2 | TATGAGTGTGGAGTCGTTGCTGTTTTGCTGCTCAATCAGAGAGNNNAGATCGGAAG |
| 51873_1 | TATGAGTGTGGAGTCGTTGCCACTGTTTTCGGACCTTCCATGNNNAGATCGGAAGA |
| 51873_2 | TATGAGTGTGGAGTCGTTGCCACTGTTTTCGGACCTTCCATGNNNAGATCGGAAGA |
| 39374_1 | TATGAGTGTGGAGTCGTTGCGCTCCCTAAACAATGTTTGCTGTNNNAGATCGGAAG |
| 39374_2 | TATGAGTGTGGAGTCGTTGCGCTCCCTAAACAATGTTTGCTGTNNNAGATCGGAAG |
| 45892_1 | TATGAGTGTGGAGTCGTTGCTACCTTGGATTACAGGCACGTGNNNAGATCGGAAG |
| 45892_2 | TATGAGTGTGGAGTCGTTGCTACCTTGGATTACAGGCACGTGNNNAGATCGGAAG |
| 38599_1 | TATGAGTGTGGAGTCGTTGCTGAGCACCTACTATTTACCAGGTNNNAGATCGGAAG |
| 38599_2 | TATGAGTGTGGAGTCGTTGCTGAGCACCTACTATTTACCAGGTNNNAGATCGGAAG |
| 410_1 | TATGAGTGTGGAGTCGTTGCGCAGTTAGAAAAGTGGCAGACACNNNAGATCGGAA |
| 410_2 | TATGAGTGTGGAGTCGTTGCGCAGTTAGAAAAGTGGCAGACACNNNAGATCGGAA |
| 34243_1 | TATGAGTGTGGAGTCGTTGCTGGAGATGGGTGAACAACTTTGANNNAGATCGGAA |
| 34243_2 | TATGAGTGTGGAGTCGTTGCTGGAGATGGGTGAACAACTTTGANNNAGATCGGAA |
| 49638_1 | TATGAGTGTGGAGTCGTTGCCATTCCAAAGTGCTGGGGATTACNNNAGATCGGAAG |
| 49638_2 | TATGAGTGTGGAGTCGTTGCCATTCCAAAGTGCTGGGGATTACNNNAGATCGGAAG |
| 36204_1 | TATGAGTGTGGAGTCGTTGCTGTTCAACCATTGTGGAAGATGGNNNAGATCGGAA |
| 36204_2 | TATGAGTGTGGAGTCGTTGCTGTTCAACCATTGTGGAAGATGGNNNAGATCGGAA |
| 38566_1 | TATGAGTGTGGAGTCGTTGCATCTGGAGGCATCACATTACCTGNNNAGATCGGAAG |
| 38566_2 | TATGAGTGTGGAGTCGTTGCATCTGGAGGCATCACATTACCTGNNNAGATCGGAAG |
| 42685_1 | TATGAGTGTGGAGTCGTTGCTTTTCAGCTCCTAGAGGCCATTTNNNAGATCGGAAG |
| 42685_2 | TATGAGTGTGGAGTCGTTGCTTTTCAGCTCCTAGAGGCCATTTNNNAGATCGGAAG |
| 34225_1 | TATGAGTGTGGAGTCGTTGCCTTGCAGTGAGCCGAGATTGTNNNAGATCGGAAGA |
| 34225_2 | TATGAGTGTGGAGTCGTTGCCTTGCAGTGAGCCGAGATTGTNNNAGATCGGAAGA |
| 44014_1 | TATGAGTGTGGAGTCGTTGCAAGTCAAGGGAAAGTTCTCCTGGNNNAGATCGGAA |
| 44014_2 | TATGAGTGTGGAGTCGTTGCAAGTCAAGGGAAAGTTCTCCTGGNNNAGATCGGAA |
| 30221_1 | TATGAGTGTGGAGTCGTTGCTGTCAATTTTCCATGTTTGTCTCANNNAGATCGGAAG |
| 30221_2 | TATGAGTGTGGAGTCGTTGCTGTCAATTTTCCATGTTTGTCTCANNNAGATCGGAAG |
| 47621_1 | TATGAGTGTGGAGTCGTTGCTTCAAGCAGTTCTCATGCCTCAGNNNAGATCGGAAG |
| 47621_2 | TATGAGTGTGGAGTCGTTGCTTCAAGCAGTTCTCATGCCTCAGNNNAGATCGGAAG |
| 32384_1 | TATGAGTGTGGAGTCGTTGCGTTTGCTCCTCACAAGAACCTTTNNNAGATCGGAAG |
| 32384_2 | TATGAGTGTGGAGTCGTTGCGTTTGCTCCTCACAAGAACCTTTNNNAGATCGGAAG |
| 38862_1 | TATGAGTGTGGAGTCGTTGCCTCAAAAGACTTGCAGCAGTGAANNNAGATCGGAA |
| 38862_2 | TATGAGTGTGGAGTCGTTGCCTCAAAAGACTTGCAGCAGTGAANNNAGATCGGAA |
| 44615_1 | TATGAGTGTGGAGTCGTTGCAGGCCACCTCTACTACTTCTTCTNNNAGATCGGAAGA |
| 44615_2 | TATGAGTGTGGAGTCGTTGCAGGCCACCTCTACTACTTCTTCTNNNAGATCGGAAGA |

| | |
|---|---|
| 51949_1 | TATGAGTGTGGAGTCGTTGCCCTTGGTTTCCTTAGTGGTTCCTNNNAGATCGGAAGA |
| 51949_2 | TATGAGTGTGGAGTCGTTGCCCTTGGTTTCCTTAGTGGTTCCTNNNAGATCGGAAGA |
| 40203_1 | TATGAGTGTGGAGTCGTTGCCACACCTGTAATCCCAGCACTTTNNNAGATCGGAAGA |
| 40203_2 | TATGAGTGTGGAGTCGTTGCCACACCTGTAATCCCAGCACTTTNNNAGATCGGAAGA |
| 33030_1 | TATGAGTGTGGAGTCGTTGCGATCTCAGCTCACTGCAACCTCNNNAGATCGGAAGA |
| 33030_2 | TATGAGTGTGGAGTCGTTGCGATCTCAGCTCACTGCAACCTCNNNAGATCGGAAGA |
| 41673_1 | TATGAGTGTGGAGTCGTTGCTGTCTGGGGACTTGTAACCAAAANNNAGATCGGAA |
| 41673_2 | TATGAGTGTGGAGTCGTTGCTGTCTGGGGACTTGTAACCAAAANNNAGATCGGAA |
| 13612_1 | TATGAGTGTGGAGTCGTTGCTTGAGAGATGAGATGTAAAATGCNNNAGATCGGAA |
| 13612_2 | TATGAGTGTGGAGTCGTTGCTTGAGAGATGAGATGTAAAATGCNNNAGATCGGAA |
| 33857_1 | TATGAGTGTGGAGTCGTTGCGCTGATGATTATAGGCTCTGGGTNNNAGATCGGAA |
| 33857_2 | TATGAGTGTGGAGTCGTTGCGCTGATGATTATAGGCTCTGGGTNNNAGATCGGAA |
| 32303_1 | TATGAGTGTGGAGTCGTTGCTGTCACAACAATCCAATGAGCTGNNNAGATCGGAAG |
| 32303_2 | TATGAGTGTGGAGTCGTTGCTGTCACAACAATCCAATGAGCTGNNNAGATCGGAAG |
| 52850_1 | TATGAGTGTGGAGTCGTTGCCCACAGAATCCCCAGCACACNNNAGATCGGAAGAGCA |
| 52850_2 | TATGAGTGTGGAGTCGTTGCCCACAGAATCCCCAGCACACNNNAGATCGGAAGAGCA |
| 33198_1 | TATGAGTGTGGAGTCGTTGCAATTCAAAAACCCGTGGATGTCCNNNAGATCGGAAG |
| 33198_2 | TATGAGTGTGGAGTCGTTGCAATTCAAAAACCCGTGGATGTCCNNNAGATCGGAAG |
| 43799_1 | TATGAGTGTGGAGTCGTTGCCACTTGTAATCCGAGCACTTTGGNNNAGATCGGAAG |
| 43799_2 | TATGAGTGTGGAGTCGTTGCCACTTGTAATCCGAGCACTTTGGNNNAGATCGGAAG |
| 44520_1 | TATGAGTGTGGAGTCGTTGCCTTTGCCTACCAGAAACGTAGTGNNNAGATCGGAAG |
| 44520_2 | TATGAGTGTGGAGTCGTTGCCTTTGCCTACCAGAAACGTAGTGNNNAGATCGGAAG |
| 40995_1 | TATGAGTGTGGAGTCGTTGCATGAAAACCTCTGATGGCCTTGANNNAGATCGGAAG |
| 40995_2 | TATGAGTGTGGAGTCGTTGCATGAAAACCTCTGATGGCCTTGANNNAGATCGGAAG |
| 53084_1 | TATGAGTGTGGAGTCGTTGCCCTAGTCTGGGCACACAAGTATTNNNAGATCGGAAG |
| 53084_2 | TATGAGTGTGGAGTCGTTGCCCTAGTCTGGGCACACAAGTATTNNNAGATCGGAAG |
| 30974_1 | TATGAGTGTGGAGTCGTTGCGCCTTGAAGGAAAAGACCAAGTCNNNAGATCGGAA |
| 30974_2 | TATGAGTGTGGAGTCGTTGCGCCTTGAAGGAAAAGACCAAGTCNNNAGATCGGAA |
| 41417_1 | TATGAGTGTGGAGTCGTTGCGGGAGGAAAAAAAGACCACAGACNNNAGATCGGAA |
| 41417_2 | TATGAGTGTGGAGTCGTTGCGGGAGGAAAAAAAGACCACAGACNNNAGATCGGAA |
| 35049_1 | TATGAGTGTGGAGTCGTTGCGCCTCCTGAATAGCTGGGAATACNNNAGATCGGAA |
| 35049_2 | TATGAGTGTGGAGTCGTTGCGCCTCCTGAATAGCTGGGAATACNNNAGATCGGAA |
| 30463_1 | TATGAGTGTGGAGTCGTTGCATGGTGCGGGACATAAAATCTGGNNNAGATCGGAA |
| 30463_2 | TATGAGTGTGGAGTCGTTGCATGGTGCGGGACATAAAATCTGGNNNAGATCGGAA |
| 10508_1 | TATGAGTGTGGAGTCGTTGCTAGAGATTGTCTGGCATGTCCGTNNNAGATCGGAA |
| 10508_2 | TATGAGTGTGGAGTCGTTGCTAGAGATTGTCTGGCATGTCCGTNNNAGATCGGAA |
| 39947_1 | TATGAGTGTGGAGTCGTTGCCATCCTGTCTAACACGGTGAAACNNNAGATCGGAAG |
| 39947_2 | TATGAGTGTGGAGTCGTTGCCATCCTGTCTAACACGGTGAAACNNNAGATCGGAAG |
| 33055_1 | TATGAGTGTGGAGTCGTTGCATAGAATCTCACTCTGTTGCCCANNNAGATCGGAAGA |
| 33055_2 | TATGAGTGTGGAGTCGTTGCATAGAATCTCACTCTGTTGCCCANNNAGATCGGAAGA |
| 49827_1 | TATGAGTGTGGAGTCGTTGCCTTCCCAAGTTTTCTGAGCGTGNNNAGATCGGAAGA |
| 49827_2 | TATGAGTGTGGAGTCGTTGCCTTCCCAAGTTTTCTGAGCGTGNNNAGATCGGAAGA |
| 44767_1 | TATGAGTGTGGAGTCGTTGCTAGGGAGAATCATTGGACGTTGGNNNAGATCGGAA |

| | |
|---|---|
| 44767_2 | TATGAGTGTGGAGTCGTTGCTAGGGAGAATCATTGGACGTTGGNNNAGATCGGAA |
| 53169_1 | TATGAGTGTGGAGTCGTTGCTTGACTCCCTCTACCCATTTGTGNNNAGATCGGAAGA |
| 53169_2 | TATGAGTGTGGAGTCGTTGCTTGACTCCCTCTACCCATTTGTGNNNAGATCGGAAGA |
| 10087_1 | TATGAGTGTGGAGTCGTTGCGAAAAAGCATGAGTTGGTGGGTGNNNAGATCGGA |
| 10087_2 | TATGAGTGTGGAGTCGTTGCGAAAAAGCATGAGTTGGTGGGTGNNNAGATCGGA |
| 8998_1 | TATGAGTGTGGAGTCGTTGCTTCAACGGTCTGATGTACCCTCTNNNAGATCGGAAG |
| 8998_2 | TATGAGTGTGGAGTCGTTGCTTCAACGGTCTGATGTACCCTCTNNNAGATCGGAAG |
| 10456_1 | TATGAGTGTGGAGTCGTTGCGCCCCATTATACCTGGAAGAGGANNNAGATCGGAA |
| 10456_2 | TATGAGTGTGGAGTCGTTGCGCCCCATTATACCTGGAAGAGGANNNAGATCGGAA |
| 10701_1 | TATGAGTGTGGAGTCGTTGCCAACGCCAACACCACTACAGATANNNAGATCGGAAG |
| 10701_2 | TATGAGTGTGGAGTCGTTGCCAACGCCAACACCACTACAGATANNNAGATCGGAAG |
| 51875_1 | TATGAGTGTGGAGTCGTTGCTTTGGCCTCAAACTTCAGGTGANNNAGATCGGAAGA |
| 51875_2 | TATGAGTGTGGAGTCGTTGCTTTGGCCTCAAACTTCAGGTGANNNAGATCGGAAGA |
| 50928_1 | TATGAGTGTGGAGTCGTTGCCTCTTCCACCCACTTCTTTTCCTNNNAGATCGGAAGAG |
| 50928_2 | TATGAGTGTGGAGTCGTTGCCTCTTCCACCCACTTCTTTTCCTNNNAGATCGGAAGAG |
| 49150_1 | TATGAGTGTGGAGTCGTTGCTTCAACTGATTCTCCTGCCTCAGNNNAGATCGGAAGA |
| 49150_2 | TATGAGTGTGGAGTCGTTGCTTCAACTGATTCTCCTGCCTCAGNNNAGATCGGAAGA |
| 34040_1 | TATGAGTGTGGAGTCGTTGCTGTTACCCAGATTGGAGTGCAGNNNAGATCGGAAG |
| 34040_2 | TATGAGTGTGGAGTCGTTGCTGTTACCCAGATTGGAGTGCAGNNNAGATCGGAAG |
| 44329_1 | TATGAGTGTGGAGTCGTTGCCTTCTTGCCTAAAGAGAAGGGGGNNNAGATCGGAA |
| 44329_2 | TATGAGTGTGGAGTCGTTGCCTTCTTGCCTAAAGAGAAGGGGGNNNAGATCGGAA |
| 2996_1 | TATGAGTGTGGAGTCGTTGCAACATACAATAAGAACAGCATCAAACNNNAGATCGG |
| 2996_2 | TATGAGTGTGGAGTCGTTGCAACATACAATAAGAACAGCATCAAACNNNAGATCGG |
| 50218_1 | TATGAGTGTGGAGTCGTTGCCTCAATCCACTCCTCCTTATCGGNNNAGATCGGAAGA |
| 50218_2 | TATGAGTGTGGAGTCGTTGCCTCAATCCACTCCTCCTTATCGGNNNAGATCGGAAGA |
| 35797_1 | TATGAGTGTGGAGTCGTTGCCTTGGACCCCCTGTATTCTTTCANNNAGATCGGAAGA |
| 35797_2 | TATGAGTGTGGAGTCGTTGCCTTGGACCCCCTGTATTCTTTCANNNAGATCGGAAGA |
| 51765_1 | TATGAGTGTGGAGTCGTTGCCCTCAACTTTGCTCTGTGATCAGNNNAGATCGGAAG |
| 51765_2 | TATGAGTGTGGAGTCGTTGCCCTCAACTTTGCTCTGTGATCAGNNNAGATCGGAAG |
| 45363_1 | TATGAGTGTGGAGTCGTTGCCACCATTATGTTGTTCTCGGACGNNNAGATCGGAAG |
| 45363_2 | TATGAGTGTGGAGTCGTTGCCACCATTATGTTGTTCTCGGACGNNNAGATCGGAAG |
| 47842_1 | TATGAGTGTGGAGTCGTTGCGCTGGAATGCAGTGACATGATCNNNAGATCGGAAG |
| 47842_2 | TATGAGTGTGGAGTCGTTGCGCTGGAATGCAGTGACATGATCNNNAGATCGGAAG |
| 47499_1 | TATGAGTGTGGAGTCGTTGCTGTACTCCTGTGCTCAAGTGATCNNNAGATCGGAAG |
| 47499_2 | TATGAGTGTGGAGTCGTTGCTGTACTCCTGTGCTCAAGTGATCNNNAGATCGGAAG |
| 49316_1 | TATGAGTGTGGAGTCGTTGCTGAATACCTGGGCTCAAGTGATCNNNAGATCGGAA |
| 49316_2 | TATGAGTGTGGAGTCGTTGCTGAATACCTGGGCTCAAGTGATCNNNAGATCGGAA |
| 49633_1 | TATGAGTGTGGAGTCGTTGCCCCAGCCATAGAGAGACTTTGTTNNNAGATCGGAAG |
| 49633_2 | TATGAGTGTGGAGTCGTTGCCCCAGCCATAGAGAGACTTTGTTNNNAGATCGGAAG |
| 47424_1 | TATGAGTGTGGAGTCGTTGCCTGCCACCACCCGACTAATTTNNNAGATCGGAAGAG |
| 47424_2 | TATGAGTGTGGAGTCGTTGCCTGCCACCACCCGACTAATTTNNNAGATCGGAAGAG |
| 41922_1 | TATGAGTGTGGAGTCGTTGCGATCTCGGCTAACTGATACGTCCNNNAGATCGGAAG |
| 41922_2 | TATGAGTGTGGAGTCGTTGCGATCTCGGCTAACTGATACGTCCNNNAGATCGGAAG |

| | |
|---|---|
| 13634_1 | TATGAGTGTGGAGTCGTTGCCAAAATGGAATTTGTTTTCAGAGNNNAGATCGGAAG |
| 13634_2 | TATGAGTGTGGAGTCGTTGCCAAAATGGAATTTGTTTTCAGAGNNNAGATCGGAAG |
| 40009_1 | TATGAGTGTGGAGTCGTTGCCTACTGGGAATGGCGTGAACCNNNAGATCGGAAGA |
| 40009_2 | TATGAGTGTGGAGTCGTTGCCTACTGGGAATGGCGTGAACCNNNAGATCGGAAGA |
| 29534_1 | TATGAGTGTGGAGTCGTTGCATCCTACGGGAGAAGAGAAAGGGNNNAGATCGGAA |
| 29534_2 | TATGAGTGTGGAGTCGTTGCATCCTACGGGAGAAGAGAAAGGGNNNAGATCGGAA |
| 32273_1 | TATGAGTGTGGAGTCGTTGCGTAGTGGGCCTTTGAGTCTGAGNNNAGATCGGAAG |
| 32273_2 | TATGAGTGTGGAGTCGTTGCGTAGTGGGCCTTTGAGTCTGAGNNNAGATCGGAAG |
| 38696_1 | TATGAGTGTGGAGTCGTTGCACAGTCAGGAACTACATAAAGCCTNNNAGATCGGAA |
| 38696_2 | TATGAGTGTGGAGTCGTTGCACAGTCAGGAACTACATAAAGCCTNNNAGATCGGAA |
| 45125_1 | TATGAGTGTGGAGTCGTTGCAAAAATCAAGTGCTGACTCGTGGNNNAGATCGGAA |
| 45125_2 | TATGAGTGTGGAGTCGTTGCAAAAATCAAGTGCTGACTCGTGGNNNAGATCGGAA |
| 42733_1 | TATGAGTGTGGAGTCGTTGCATGTAAACCACACAGATGGGGTTNNNAGATCGGAA |
| 42733_2 | TATGAGTGTGGAGTCGTTGCATGTAAACCACACAGATGGGGTTNNNAGATCGGAA |
| 37003_1 | TATGAGTGTGGAGTCGTTGCACGATTAGAGGAGTCAGGAGAGTNNNAGATCGGAA |
| 37003_2 | TATGAGTGTGGAGTCGTTGCACGATTAGAGGAGTCAGGAGAGTNNNAGATCGGAA |
| 49260_1 | TATGAGTGTGGAGTCGTTGCCATTCTCCTGCCTCAGCCTCNNNAGATCGGAAGAGCA |
| 49260_2 | TATGAGTGTGGAGTCGTTGCCATTCTCCTGCCTCAGCCTCNNNAGATCGGAAGAGCA |
| 45262_1 | TATGAGTGTGGAGTCGTTGCGTCTGCTGAAGAAACCAGTGTTGNNNAGATCGGAA |
| 45262_2 | TATGAGTGTGGAGTCGTTGCGTCTGCTGAAGAAACCAGTGTTGNNNAGATCGGAA |
| 38101_1 | TATGAGTGTGGAGTCGTTGCATCACTGGTAGCAGCTTTCAGAANNNAGATCGGAAG |
| 38101_2 | TATGAGTGTGGAGTCGTTGCATCACTGGTAGCAGCTTTCAGAANNNAGATCGGAAG |
| 43083_1 | TATGAGTGTGGAGTCGTTGCGCATCTTGCTAAACAGTGGGTACNNNAGATCGGAAG |
| 43083_2 | TATGAGTGTGGAGTCGTTGCGCATCTTGCTAAACAGTGGGTACNNNAGATCGGAAG |
| 31753_1 | TATGAGTGTGGAGTCGTTGCGTCTCTCTCTCTCCCCTCTTTCTNNNAGATCGGAAGAG |
| 31753_2 | TATGAGTGTGGAGTCGTTGCGTCTCTCTCTCTCCCCTCTTTCTNNNAGATCGGAAGAG |
| 34831_1 | TATGAGTGTGGAGTCGTTGCAAGTGATTTTCCTGCCTCAGCTTNNNAGATCGGAAGA |
| 34831_2 | TATGAGTGTGGAGTCGTTGCAAGTGATTTTCCTGCCTCAGCTTNNNAGATCGGAAGA |
| 32670_1 | TATGAGTGTGGAGTCGTTGCCATTCTCCTGCCTCAGCCTCNNNAGATCGGAAGAGCA |
| 32670_2 | TATGAGTGTGGAGTCGTTGCCATTCTCCTGCCTCAGCCTCNNNAGATCGGAAGAGCA |
| 13239_1 | TATGAGTGTGGAGTCGTTGCCAGATCTGATTGAGAGTTGGNNNAGATCGGAAGAG |
| 13239_2 | TATGAGTGTGGAGTCGTTGCCAGATCTGATTGAGAGTTGGNNNAGATCGGAAGAG |
| 30031_1 | TATGAGTGTGGAGTCGTTGCCTGCTCTCTGTCAATCTCTCTGGNNNAGATCGGAAGA |
| 30031_2 | TATGAGTGTGGAGTCGTTGCCTGCTCTCTGTCAATCTCTCTGGNNNAGATCGGAAGA |
| 31668_1 | TATGAGTGTGGAGTCGTTGCTACACACTTATGGACAACAGGCANNNAGATCGGAAG |
| 31668_2 | TATGAGTGTGGAGTCGTTGCTACACACTTATGGACAACAGGCANNNAGATCGGAAG |
| 3168_1 | TATGAGTGTGGAGTCGTTGCCCTGAGCACTTACACTCACTGTCNNNAGATCGGAAGA |
| 3168_2 | TATGAGTGTGGAGTCGTTGCCCTGAGCACTTACACTCACTGTCNNNAGATCGGAAGA |
| 28859_1 | TATGAGTGTGGAGTCGTTGCAGAGGATAGCAGTGGTTGTTTCANNNAGATCGGAA |
| 28859_2 | TATGAGTGTGGAGTCGTTGCAGAGGATAGCAGTGGTTGTTTCANNNAGATCGGAA |
| 36768_1 | TATGAGTGTGGAGTCGTTGCTCTGAGGCAAAGTTTAGACCACANNNAGATCGGAAG |
| 36768_2 | TATGAGTGTGGAGTCGTTGCTCTGAGGCAAAGTTTAGACCACANNNAGATCGGAAG |
| 31515_1 | TATGAGTGTGGAGTCGTTGCACCTGTCAATATCTGCCTGTGAGNNNAGATCGGAAG |

| | |
|---|---|
| 31515_2 | TATGAGTGTGGAGTCGTTGCACCTGTCAATATCTGCCTGTGAGNNNAGATCGGAAG |
| 36488_1 | TATGAGTGTGGAGTCGTTGCAACCTCCTTGGGTCTCTTATGTGNNNAGATCGGAAG |
| 36488_2 | TATGAGTGTGGAGTCGTTGCAACCTCCTTGGGTCTCTTATGTGNNNAGATCGGAAG |
| 43623_1 | TATGAGTGTGGAGTCGTTGCGTGTTACACGTACTCTCACCGTNNNAGATCGGAAGA |
| 43623_2 | TATGAGTGTGGAGTCGTTGCGTGTTACACGTACTCTCACCGTNNNAGATCGGAAGA |
| 45689_1 | TATGAGTGTGGAGTCGTTGCCTGTCAGGTCCTCGGTGCTAGNNNAGATCGGAAGA |
| 45689_2 | TATGAGTGTGGAGTCGTTGCCTGTCAGGTCCTCGGTGCTAGNNNAGATCGGAAGA |
| 43373_1 | TATGAGTGTGGAGTCGTTGCGAGTCTCACTCTGTGTCAACCAGNNNAGATCGGAAG |
| 43373_2 | TATGAGTGTGGAGTCGTTGCGAGTCTCACTCTGTGTCAACCAGNNNAGATCGGAAG |
| 36211_1 | TATGAGTGTGGAGTCGTTGCTCACTTGAGGTCAGGAGTTCAAGNNNAGATCGGAA |
| 36211_2 | TATGAGTGTGGAGTCGTTGCTCACTTGAGGTCAGGAGTTCAAGNNNAGATCGGAA |
| 33342_1 | TATGAGTGTGGAGTCGTTGCCTCCTGAATAGCTGGGACCAGAGNNNAGATCGGAA |
| 33342_2 | TATGAGTGTGGAGTCGTTGCCTCCTGAATAGCTGGGACCAGAGNNNAGATCGGAA |
| 34702_1 | TATGAGTGTGGAGTCGTTGCGTCTTGAACTCCTCAGCTCAGACNNNAGATCGGAAG |
| 34702_2 | TATGAGTGTGGAGTCGTTGCGTCTTGAACTCCTCAGCTCAGACNNNAGATCGGAAG |
| 44531_1 | TATGAGTGTGGAGTCGTTGCTTCCAGTTTCAGCTTGGAGTCATNNNAGATCGGAAG |
| 44531_2 | TATGAGTGTGGAGTCGTTGCTTCCAGTTTCAGCTTGGAGTCATNNNAGATCGGAAG |
| 29181_1 | TATGAGTGTGGAGTCGTTGCTAGAGATCTTCAAGGCCAAGAGCNNNAGATCGGAA |
| 29181_2 | TATGAGTGTGGAGTCGTTGCTAGAGATCTTCAAGGCCAAGAGCNNNAGATCGGAA |
| 30714_1 | TATGAGTGTGGAGTCGTTGCTTGAGAGGCTAAAGGGCTACACTNNNAGATCGGAA |
| 30714_2 | TATGAGTGTGGAGTCGTTGCTTGAGAGGCTAAAGGGCTACACTNNNAGATCGGAA |
| 29409_1 | TATGAGTGTGGAGTCGTTGCTTGGCTTCCTGTCAGTGTTTTTTNNNAGATCGGAAGA |
| 29409_2 | TATGAGTGTGGAGTCGTTGCTTGGCTTCCTGTCAGTGTTTTTTNNNAGATCGGAAGA |
| 51407_1 | TATGAGTGTGGAGTCGTTGCACAAAGTTCTGACCAAAACACCGNNNAGATCGGAAG |
| 51407_2 | TATGAGTGTGGAGTCGTTGCACAAAGTTCTGACCAAAACACCGNNNAGATCGGAAG |
| 35386_1 | TATGAGTGTGGAGTCGTTGCCTGAGTAGCTGGGACTACAAGTGNNNAGATCGGAA |
| 35386_2 | TATGAGTGTGGAGTCGTTGCCTGAGTAGCTGGGACTACAAGTGNNNAGATCGGAA |
| 45939_1 | TATGAGTGTGGAGTCGTTGCCTCCCCAGTAGCTAGGACTACAGNNNAGATCGGAAG |
| 45939_2 | TATGAGTGTGGAGTCGTTGCCTCCCCAGTAGCTAGGACTACAGNNNAGATCGGAAG |
| 28807_1 | TATGAGTGTGGAGTCGTTGCTCATTGATTGATGGGCATTTGGGNNNAGATCGGAA |
| 28807_2 | TATGAGTGTGGAGTCGTTGCTCATTGATTGATGGGCATTTGGGNNNAGATCGGAA |
| 51830_1 | TATGAGTGTGGAGTCGTTGCTGACATTGAGCAACAAATGGCTTNNNAGATCGGAAG |
| 51830_2 | TATGAGTGTGGAGTCGTTGCTGACATTGAGCAACAAATGGCTTNNNAGATCGGAAG |
| 13542_1 | TATGAGTGTGGAGTCGTTGCAAAGAACACAAATAGGAAATCAANNNAGATCGGAA |
| 13542_2 | TATGAGTGTGGAGTCGTTGCAAAGAACACAAATAGGAAATCAANNNAGATCGGAA |
| 31317_1 | TATGAGTGTGGAGTCGTTGCGCTGTATTTGTAACCCATGCAGTNNNAGATCGGAAG |
| 31317_2 | TATGAGTGTGGAGTCGTTGCGCTGTATTTGTAACCCATGCAGTNNNAGATCGGAAG |
| 36414_1 | TATGAGTGTGGAGTCGTTGCATGGGGAAATGAGGCTAAAATGCNNNAGATCGGAA |
| 36414_2 | TATGAGTGTGGAGTCGTTGCATGGGGAAATGAGGCTAAAATGCNNNAGATCGGAA |
| 30921_1 | TATGAGTGTGGAGTCGTTGCGGCACCAGAATTGTGAGTGAAAANNNAGATCGGAA |
| 30921_2 | TATGAGTGTGGAGTCGTTGCGGCACCAGAATTGTGAGTGAAAANNNAGATCGGAA |
| 31300_1 | TATGAGTGTGGAGTCGTTGCATGGAGGAATTAGGGACATTGGTNNNAGATCGGAA |
| 31300_2 | TATGAGTGTGGAGTCGTTGCATGGAGGAATTAGGGACATTGGTNNNAGATCGGAA |

| | |
|---|---|
| 51792_1 | TATGAGTGTGGAGTCGTTGCAACACAATTTGAGGTTCTTGCCCNNNAGATCGGAAG |
| 51792_2 | TATGAGTGTGGAGTCGTTGCAACACAATTTGAGGTTCTTGCCCNNNAGATCGGAAG |
| 42374_1 | TATGAGTGTGGAGTCGTTGCGGACTAAAGTCAGAGCTACAGGGNNNAGATCGGAA |
| 42374_2 | TATGAGTGTGGAGTCGTTGCGGACTAAAGTCAGAGCTACAGGGNNNAGATCGGAA |
| 46330_1 | TATGAGTGTGGAGTCGTTGCTTCACACCATTCTCCTGCCTCNNNAGATCGGAAGAGC |
| 46330_2 | TATGAGTGTGGAGTCGTTGCTTCACACCATTCTCCTGCCTCNNNAGATCGGAAGAGC |
| 36980_1 | TATGAGTGTGGAGTCGTTGCTACTGACCTCCATTCCTCCATCTNNNAGATCGGAAGA |
| 36980_2 | TATGAGTGTGGAGTCGTTGCTACTGACCTCCATTCCTCCATCTNNNAGATCGGAAGA |
| 10691_1 | TATGAGTGTGGAGTCGTTGCCAGAGAGGAAAAACACCCAGGACNNNAGATCGGAA |
| 10691_2 | TATGAGTGTGGAGTCGTTGCCAGAGAGGAAAAACACCCAGGACNNNAGATCGGAA |
| 39859_1 | TATGAGTGTGGAGTCGTTGCTACTGATTCAGATGAGTTGGCCCNNNAGATCGGAAG |
| 39859_2 | TATGAGTGTGGAGTCGTTGCTACTGATTCAGATGAGTTGGCCCNNNAGATCGGAAG |
| 50786_1 | TATGAGTGTGGAGTCGTTGCCTCTGCTCTCAGTGCTGTCATTCNNNAGATCGGAAGA |
| 50786_2 | TATGAGTGTGGAGTCGTTGCCTCTGCTCTCAGTGCTGTCATTCNNNAGATCGGAAGA |
| 13599_1 | TATGAGTGTGGAGTCGTTGCCTTCCATCCCTCTCTTCATTNNNAGATCGGAAGAGCA |
| 13599_2 | TATGAGTGTGGAGTCGTTGCCTTCCATCCCTCTCTTCATTNNNAGATCGGAAGAGCA |
| 9087_1 | TATGAGTGTGGAGTCGTTGCTTTCCTACTCCAAGGTCATGACTNNNAGATCGGAAGA |
| 9087_2 | TATGAGTGTGGAGTCGTTGCTTTCCTACTCCAAGGTCATGACTNNNAGATCGGAAGA |
| 33127_1 | TATGAGTGTGGAGTCGTTGCTCCTTCAAATCCCAGCTGCATTANNNAGATCGGAAGA |
| 33127_2 | TATGAGTGTGGAGTCGTTGCTCCTTCAAATCCCAGCTGCATTANNNAGATCGGAAGA |
| 10817_1 | TATGAGTGTGGAGTCGTTGCTGCTAAACAGACACTTGCAGACCNNNAGATCGGAAG |
| 10817_2 | TATGAGTGTGGAGTCGTTGCTGCTAAACAGACACTTGCAGACCNNNAGATCGGAAG |
| 46864_1 | TATGAGTGTGGAGTCGTTGCCTTCCGAGTAGCTGGGACTACAGNNNAGATCGGAA |
| 46864_2 | TATGAGTGTGGAGTCGTTGCCTTCCGAGTAGCTGGGACTACAGNNNAGATCGGAA |
| 29501_1 | TATGAGTGTGGAGTCGTTGCTTTTCATTGCTAATGCTATTCCANNNAGATCGGAAGA |
| 29501_2 | TATGAGTGTGGAGTCGTTGCTTTTCATTGCTAATGCTATTCCANNNAGATCGGAAGA |
| 13304_1 | TATGAGTGTGGAGTCGTTGCTACTATTGTCTATTAAGTGTGCANNNAGATCGGAAG |
| 13304_2 | TATGAGTGTGGAGTCGTTGCTACTATTGTCTATTAAGTGTGCANNNAGATCGGAAG |
| 45090_1 | TATGAGTGTGGAGTCGTTGCACTTGGGAGGCTTTATCTGTTTGNNNAGATCGGAAG |
| 45090_2 | TATGAGTGTGGAGTCGTTGCACTTGGGAGGCTTTATCTGTTTGNNNAGATCGGAAG |
| 46924_1 | TATGAGTGTGGAGTCGTTGCTAGTTCAAGCAATTCTCCTGCCTNNNAGATCGGAAGA |
| 46924_2 | TATGAGTGTGGAGTCGTTGCTAGTTCAAGCAATTCTCCTGCCTNNNAGATCGGAAGA |
| 1044_1 | TATGAGTGTGGAGTCGTTGCCCTGGCCATAGCCTATTTTGAAAGNNNAGATCGGAA |
| 1044_2 | TATGAGTGTGGAGTCGTTGCCCTGGCCATAGCCTATTTTGAAAGNNNAGATCGGAA |
| 29445_1 | TATGAGTGTGGAGTCGTTGCAGCAACATTCACTTCACTCCCTCNNNAGATCGGAAGA |
| 29445_2 | TATGAGTGTGGAGTCGTTGCAGCAACATTCACTTCACTCCCTCNNNAGATCGGAAGA |
| 45302_1 | TATGAGTGTGGAGTCGTTGCGTCATACTGGGAAGAGGTAGTGTNNNAGATCGGAA |
| 45302_2 | TATGAGTGTGGAGTCGTTGCGTCATACTGGGAAGAGGTAGTGTNNNAGATCGGAA |
| 13343_1 | TATGAGTGTGGAGTCGTTGCAAAGGTACTGTTAACTACTGTACNNNAGATCGGAAG |
| 13343_2 | TATGAGTGTGGAGTCGTTGCAAAGGTACTGTTAACTACTGTACNNNAGATCGGAAG |
| 46143_1 | TATGAGTGTGGAGTCGTTGCGCATGATAATGGCTCGCTGTAGNNNAGATCGGAAG |
| 46143_2 | TATGAGTGTGGAGTCGTTGCGCATGATAATGGCTCGCTGTAGNNNAGATCGGAAG |
| 41830_1 | TATGAGTGTGGAGTCGTTGCTGTGACCAGTAGGAATGATGTCANNNAGATCGGAA |

| | |
|---|---|
| 41830_2 | TATGAGTGTGGAGTCGTTGCTGTGACCAGTAGGAATGATGTCANNNAGATCGGAA |
| 31834_1 | TATGAGTGTGGAGTCGTTGCAGATGCCAAGACCTCTGCTAAAANNNAGATCGGAAG |
| 31834_2 | TATGAGTGTGGAGTCGTTGCAGATGCCAAGACCTCTGCTAAAANNNAGATCGGAAG |
| 47560_1 | TATGAGTGTGGAGTCGTTGCCCATACCCATTAGTGAGCAAGGANNNAGATCGGAAG |
| 47560_2 | TATGAGTGTGGAGTCGTTGCCCATACCCATTAGTGAGCAAGGANNNAGATCGGAAG |
| 31479_1 | TATGAGTGTGGAGTCGTTGCTGTTGACTTTTGTTGGTCTTGCANNNAGATCGGAAG |
| 31479_2 | TATGAGTGTGGAGTCGTTGCTGTTGACTTTTGTTGGTCTTGCANNNAGATCGGAAG |
| 44574_1 | TATGAGTGTGGAGTCGTTGCGTGGACACCTGTAATCCAAGCTANNNAGATCGGAAG |
| 44574_2 | TATGAGTGTGGAGTCGTTGCGTGGACACCTGTAATCCAAGCTANNNAGATCGGAAG |
| 2432_1 | TATGAGTGTGGAGTCGTTGCCCTGAATGTGGCTATGTGTCCTCNNNAGATCGGAAG |
| 2432_2 | TATGAGTGTGGAGTCGTTGCCCTGAATGTGGCTATGTGTCCTCNNNAGATCGGAAG |
| 48614_1 | TATGAGTGTGGAGTCGTTGCGAGTAGCTGGGATTACAGGACTGNNNAGATCGGAA |
| 48614_2 | TATGAGTGTGGAGTCGTTGCGAGTAGCTGGGATTACAGGACTGNNNAGATCGGAA |
| 41046_1 | TATGAGTGTGGAGTCGTTGCGCTTGCAGTGAGCCAAGATAATGNNNAGATCGGAA |
| 41046_2 | TATGAGTGTGGAGTCGTTGCGCTTGCAGTGAGCCAAGATAATGNNNAGATCGGAA |
| 44122_1 | TATGAGTGTGGAGTCGTTGCGTACAACAGTCAACACTCCTTGCNNNAGATCGGAAG |
| 44122_2 | TATGAGTGTGGAGTCGTTGCGTACAACAGTCAACACTCCTTGCNNNAGATCGGAAG |
| 34805_1 | TATGAGTGTGGAGTCGTTGCGAGTGCAGTGACACGATCTCAGNNNAGATCGGAAG |
| 34805_2 | TATGAGTGTGGAGTCGTTGCGAGTGCAGTGACACGATCTCAGNNNAGATCGGAAG |
| 41854_1 | TATGAGTGTGGAGTCGTTGCTTAGGGTACATGTGCACATTGTGNNNAGATCGGAA |
| 41854_2 | TATGAGTGTGGAGTCGTTGCTTAGGGTACATGTGCACATTGTGNNNAGATCGGAA |
| 32983_1 | TATGAGTGTGGAGTCGTTGCAAGCCAGTCTGATTTTATCCCCANNNAGATCGGAAG |
| 32983_2 | TATGAGTGTGGAGTCGTTGCAAGCCAGTCTGATTTTATCCCCANNNAGATCGGAAG |
| 42307_1 | TATGAGTGTGGAGTCGTTGCTTTGTATAGGGCAAGCACAGCTANNNAGATCGGAA |
| 42307_2 | TATGAGTGTGGAGTCGTTGCTTTGTATAGGGCAAGCACAGCTANNNAGATCGGAA |
| 52622_1 | TATGAGTGTGGAGTCGTTGCGACTGGAGCCAAAATGCCAAAGNNNAGATCGGAAG |
| 52622_2 | TATGAGTGTGGAGTCGTTGCGACTGGAGCCAAAATGCCAAAGNNNAGATCGGAAG |
| 41609_1 | TATGAGTGTGGAGTCGTTGCTGTTGGGATTATAGGTGTGAGCCNNNAGATCGGAA |
| 41609_2 | TATGAGTGTGGAGTCGTTGCTGTTGGGATTATAGGTGTGAGCCNNNAGATCGGAA |
| 36358_1 | TATGAGTGTGGAGTCGTTGCGCCACACAAAAAGAGCTAGTCAGNNNAGATCGGAA |
| 36358_2 | TATGAGTGTGGAGTCGTTGCGCCACACAAAAAGAGCTAGTCAGNNNAGATCGGAA |
| 43872_1 | TATGAGTGTGGAGTCGTTGCCCAAAGTTCTGCAGTGTAAGTGGNNNAGATCGGAA |
| 43872_2 | TATGAGTGTGGAGTCGTTGCCCAAAGTTCTGCAGTGTAAGTGGNNNAGATCGGAA |
| 32691_1 | TATGAGTGTGGAGTCGTTGCAAACTCCTGACCTCAAGTGATCCNNNAGATCGGAAG |
| 32691_2 | TATGAGTGTGGAGTCGTTGCAAACTCCTGACCTCAAGTGATCCNNNAGATCGGAAG |
| 38657_1 | TATGAGTGTGGAGTCGTTGCTTACTGAATCCCCAAACTCTGCTNNNAGATCGGAAGA |
| 38657_2 | TATGAGTGTGGAGTCGTTGCTTACTGAATCCCCAAACTCTGCTNNNAGATCGGAAGA |
| 45447_1 | TATGAGTGTGGAGTCGTTGCTGAGGCCAGGAGTTCAATATCAGNNNAGATCGGAA |
| 45447_2 | TATGAGTGTGGAGTCGTTGCTGAGGCCAGGAGTTCAATATCAGNNNAGATCGGAA |
| 51403_1 | TATGAGTGTGGAGTCGTTGCGGGGTACATCAGAGAAACCATCANNNAGATCGGAA |
| 51403_2 | TATGAGTGTGGAGTCGTTGCGGGGTACATCAGAGAAACCATCANNNAGATCGGAA |
| 39520_1 | TATGAGTGTGGAGTCGTTGCGCTCTACCACATATGAACTGGGANNNAGATCGGAAG |
| 39520_2 | TATGAGTGTGGAGTCGTTGCGCTCTACCACATATGAACTGGGANNNAGATCGGAAG |

| | |
|---|---|
| 41163_1 | TATGAGTGTGGAGTCGTTGCCTGTAATCCCAGCTACTGAGGAGNNNAGATCGGAAG |
| 41163_2 | TATGAGTGTGGAGTCGTTGCCTGTAATCCCAGCTACTGAGGAGNNNAGATCGGAAG |
| 50456_1 | TATGAGTGTGGAGTCGTTGCTCACTTTGCATGCTCCTTGAATGNNNAGATCGGAAGA |
| 50456_2 | TATGAGTGTGGAGTCGTTGCTCACTTTGCATGCTCCTTGAATGNNNAGATCGGAAGA |
| 37420_1 | TATGAGTGTGGAGTCGTTGCGGAGGGATTCTGTCATCTCTTCCNNNAGATCGGAAG |
| 37420_2 | TATGAGTGTGGAGTCGTTGCGGAGGGATTCTGTCATCTCTTCCNNNAGATCGGAAG |
| 48528_1 | TATGAGTGTGGAGTCGTTGCGCAATTTTGTGTAGGCCCATGAANNNAGATCGGAAG |
| 48528_2 | TATGAGTGTGGAGTCGTTGCGCAATTTTGTGTAGGCCCATGAANNNAGATCGGAAG |
| 37398_1 | TATGAGTGTGGAGTCGTTGCATAACACAGGGCTCCAAAGTCATNNNAGATCGGAAG |
| 37398_2 | TATGAGTGTGGAGTCGTTGCATAACACAGGGCTCCAAAGTCATNNNAGATCGGAAG |
| 39739_1 | TATGAGTGTGGAGTCGTTGCGGTAGACGGGGCTAGTTATGAAGNNNAGATCGGAA |
| 39739_2 | TATGAGTGTGGAGTCGTTGCGGTAGACGGGGCTAGTTATGAAGNNNAGATCGGAA |
| 30333_1 | TATGAGTGTGGAGTCGTTGCGGCTGGGTATGTGTGATGGAAAANNNAGATCGGAA |
| 30333_2 | TATGAGTGTGGAGTCGTTGCGGCTGGGTATGTGTGATGGAAAANNNAGATCGGAA |
| 30503_1 | TATGAGTGTGGAGTCGTTGCTGTTCACCCGATCCTTTCAAACANNNAGATCGGAAGA |
| 30503_2 | TATGAGTGTGGAGTCGTTGCTGTTCACCCGATCCTTTCAAACANNNAGATCGGAAGA |
| 40961_1 | TATGAGTGTGGAGTCGTTGCCCATCAGGAAGCACAAAGGTAACNNNAGATCGGAA |
| 40961_2 | TATGAGTGTGGAGTCGTTGCCCATCAGGAAGCACAAAGGTAACNNNAGATCGGAA |
| 48351_1 | TATGAGTGTGGAGTCGTTGCCAGCTCCAACCTGAAACTAGAGTNNNAGATCGGAAG |
| 48351_2 | TATGAGTGTGGAGTCGTTGCCAGCTCCAACCTGAAACTAGAGTNNNAGATCGGAAG |
| 42706_1 | TATGAGTGTGGAGTCGTTGCTCTATTTCTCTGGAGAGCCCTGANNNAGATCGGAAG |
| 42706_2 | TATGAGTGTGGAGTCGTTGCTCTATTTCTCTGGAGAGCCCTGANNNAGATCGGAAG |
| 33989_1 | TATGAGTGTGGAGTCGTTGCGTTCAAGCGATTCTTCTGCCTCNNNAGATCGGAAGA |
| 33989_2 | TATGAGTGTGGAGTCGTTGCGTTCAAGCGATTCTTCTGCCTCNNNAGATCGGAAGA |
| 36008_1 | TATGAGTGTGGAGTCGTTGCTCGATTAGTTGAGCCTGTAGTCCNNNAGATCGGAAG |
| 36008_2 | TATGAGTGTGGAGTCGTTGCTCGATTAGTTGAGCCTGTAGTCCNNNAGATCGGAAG |
| 34345_1 | TATGAGTGTGGAGTCGTTGCTGTGTGTGTGTACAGGTGTGTATNNNAGATCGGAA |
| 34345_2 | TATGAGTGTGGAGTCGTTGCTGTGTGTGTGTACAGGTGTGTATNNNAGATCGGAA |
| 13496_1 | TATGAGTGTGGAGTCGTTGCCACCTGTGAAGAAGATCTTGNNNAGATCGGAAGAG |
| 13496_2 | TATGAGTGTGGAGTCGTTGCCACCTGTGAAGAAGATCTTGNNNAGATCGGAAGAG |
| 40355_1 | TATGAGTGTGGAGTCGTTGCTCCAATACTGCCATTCATCCACANNNAGATCGGAAGA |
| 40355_2 | TATGAGTGTGGAGTCGTTGCTCCAATACTGCCATTCATCCACANNNAGATCGGAAGA |
| 43690_1 | TATGAGTGTGGAGTCGTTGCACCAGGGACTGTAAGCTCTCCNNNAGATCGGAAGAG |
| 43690_2 | TATGAGTGTGGAGTCGTTGCACCAGGGACTGTAAGCTCTCCNNNAGATCGGAAGAG |
| 49991_1 | TATGAGTGTGGAGTCGTTGCTCAGTTCTCCATCTCTCCAGAGTNNNAGATCGGAAGA |
| 49991_2 | TATGAGTGTGGAGTCGTTGCTCAGTTCTCCATCTCTCCAGAGTNNNAGATCGGAAGA |
| 29800_1 | TATGAGTGTGGAGTCGTTGCGTAGACAGGATGGCTTCATCGACNNNAGATCGGAA |
| 29800_2 | TATGAGTGTGGAGTCGTTGCGTAGACAGGATGGCTTCATCGACNNNAGATCGGAA |
| 37635_1 | TATGAGTGTGGAGTCGTTGCTAAGGCACGAGAATTGCTTGAACNNNAGATCGGAA |
| 37635_2 | TATGAGTGTGGAGTCGTTGCTAAGGCACGAGAATTGCTTGAACNNNAGATCGGAA |
| 38410_1 | TATGAGTGTGGAGTCGTTGCTGACCTTGTCTAAGTCATTGGCANNNAGATCGGAAG |
| 38410_2 | TATGAGTGTGGAGTCGTTGCTGACCTTGTCTAAGTCATTGGCANNNAGATCGGAAG |
| 42670_1 | TATGAGTGTGGAGTCGTTGCACCCTGTCTCCACAAACAATACANNNAGATCGGAAGA |

| | |
|---|---|
| 42670_2 | TATGAGTGTGGAGTCGTTGCACCCTGTCTCCACAAACAATACANNNAGATCGGAAG |
| 45171_1 | TATGAGTGTGGAGTCGTTGCAGGTCTGTGAGAGATGCAGGATANNNAGATCGGA |
| 45171_2 | TATGAGTGTGGAGTCGTTGCAGGTCTGTGAGAGATGCAGGATANNNAGATCGGA |
| 48103_1 | TATGAGTGTGGAGTCGTTGCTTGTTGCTGTTGTTTGGAACTGTNNNAGATCGGAAG |
| 48103_2 | TATGAGTGTGGAGTCGTTGCTTGTTGCTGTTGTTTGGAACTGTNNNAGATCGGAAG |
| 39919_1 | TATGAGTGTGGAGTCGTTGCAGAGTTGGAGAACTGACACTTCCNNNAGATCGGAA |
| 39919_2 | TATGAGTGTGGAGTCGTTGCAGAGTTGGAGAACTGACACTTCCNNNAGATCGGAA |
| 42340_1 | TATGAGTGTGGAGTCGTTGCGTTGAATTTCACCTGGCACAACTNNNAGATCGGAAG |
| 42340_2 | TATGAGTGTGGAGTCGTTGCGTTGAATTTCACCTGGCACAACTNNNAGATCGGAAG |
| 35885_1 | TATGAGTGTGGAGTCGTTGCGATTCCAGAACCTTCCTCCCAAANNNAGATCGGAAG |
| 35885_2 | TATGAGTGTGGAGTCGTTGCGATTCCAGAACCTTCCTCCCAAANNNAGATCGGAAG |
| 42404_1 | TATGAGTGTGGAGTCGTTGCTCTCAATAATGCCACAACTGGGANNNAGATCGGAAG |
| 42404_2 | TATGAGTGTGGAGTCGTTGCTCTCAATAATGCCACAACTGGGANNNAGATCGGAAG |
| 31964_1 | TATGAGTGTGGAGTCGTTGCAGCTGCATACAAATTGACTCTCCNNNAGATCGGAAG |
| 31964_2 | TATGAGTGTGGAGTCGTTGCAGCTGCATACAAATTGACTCTCCNNNAGATCGGAAG |
| 34242_1 | TATGAGTGTGGAGTCGTTGCTGGTTTAGTTTGCTTCAGGTGTGNNNAGATCGGAA |
| 34242_2 | TATGAGTGTGGAGTCGTTGCTGGTTTAGTTTGCTTCAGGTGTGNNNAGATCGGAA |
| 42911_1 | TATGAGTGTGGAGTCGTTGCTCTCCTGAGTAGCTGGGATTACANNNAGATCGGAAG |
| 42911_2 | TATGAGTGTGGAGTCGTTGCTCTCCTGAGTAGCTGGGATTACANNNAGATCGGAAG |
| 31466_1 | TATGAGTGTGGAGTCGTTGCAGATGACAAAGCTCTAGCCAACANNNAGATCGGAA |
| 31466_2 | TATGAGTGTGGAGTCGTTGCAGATGACAAAGCTCTAGCCAACANNNAGATCGGAA |
| 33221_1 | TATGAGTGTGGAGTCGTTGCTTATTTATCTGCCTCAGCCTCCCNNNAGATCGGAAGA |
| 33221_2 | TATGAGTGTGGAGTCGTTGCTTATTTATCTGCCTCAGCCTCCCNNNAGATCGGAAGA |
| 48719_1 | TATGAGTGTGGAGTCGTTGCCCAAGTAGCTGGGATTACAGGTGNNNAGATCGGAA |
| 48719_2 | TATGAGTGTGGAGTCGTTGCCCAAGTAGCTGGGATTACAGGTGNNNAGATCGGAA |
| 45263_1 | TATGAGTGTGGAGTCGTTGCCTAACAGTCACCAAACAAGCCAGNNNAGATCGGAAG |
| 45263_2 | TATGAGTGTGGAGTCGTTGCCTAACAGTCACCAAACAAGCCAGNNNAGATCGGAAG |
| 44347_1 | TATGAGTGTGGAGTCGTTGCAGGATTTCTAAAACTCCCCGAGGNNNAGATCGGAAG |
| 44347_2 | TATGAGTGTGGAGTCGTTGCAGGATTTCTAAAACTCCCCGAGGNNNAGATCGGAAG |
| 10146_1 | TATGAGTGTGGAGTCGTTGCCATTAGGATGTTGGGGGTTGGTCNNNAGATCGGAA |
| 10146_2 | TATGAGTGTGGAGTCGTTGCCATTAGGATGTTGGGGGTTGGTCNNNAGATCGGAA |
| 52496_1 | TATGAGTGTGGAGTCGTTGCGGTATGATCTCAGCTCACTGCAANNNAGATCGGAAG |
| 52496_2 | TATGAGTGTGGAGTCGTTGCGGTATGATCTCAGCTCACTGCAANNNAGATCGGAAG |
| 34534_1 | TATGAGTGTGGAGTCGTTGCACACACACACACACACAATTCTGNNNAGATCGGAAG |
| 34534_2 | TATGAGTGTGGAGTCGTTGCACACACACACACACACAATTCTGNNNAGATCGGAAG |
| 36480_1 | TATGAGTGTGGAGTCGTTGCGCAAAACAGAGTGTTACATGTGGANNNAGATCGGA |
| 36480_2 | TATGAGTGTGGAGTCGTTGCGCAAAACAGAGTGTTACATGTGGANNNAGATCGGA |
| 36336_1 | TATGAGTGTGGAGTCGTTGCGGTAAACTGCACATGGGTCAATTNNNAGATCGGAA |
| 36336_2 | TATGAGTGTGGAGTCGTTGCGGTAAACTGCACATGGGTCAATTNNNAGATCGGAA |
| 50183_1 | TATGAGTGTGGAGTCGTTGCAACCTGAAGATCACATCTGGCTTNNNAGATCGGAAG |
| 50183_2 | TATGAGTGTGGAGTCGTTGCAACCTGAAGATCACATCTGGCTTNNNAGATCGGAAG |
| 35917_1 | TATGAGTGTGGAGTCGTTGCTAGGTAAAAGGAGCTGTCATCCCNNNAGATCGGAA |
| 35917_2 | TATGAGTGTGGAGTCGTTGCTAGGTAAAAGGAGCTGTCATCCCNNNAGATCGGAA |

| | |
|---|---|
| 32436_1 | TATGAGTGTGGAGTCGTTGCAGGGATTGCATTGCATCTGTAGANNNAGATCGGAA |
| 32436_2 | TATGAGTGTGGAGTCGTTGCAGGGATTGCATTGCATCTGTAGANNNAGATCGGAA |
| 48040_1 | TATGAGTGTGGAGTCGTTGCTTCCAGTGATTCTCCTGCTTCAGNNNAGATCGGAAG |
| 48040_2 | TATGAGTGTGGAGTCGTTGCTTCCAGTGATTCTCCTGCTTCAGNNNAGATCGGAAG |
| 43769_1 | TATGAGTGTGGAGTCGTTGCGTGTTCAGATTGAGCCATTGCATNNNAGATCGGAAG |
| 43769_2 | TATGAGTGTGGAGTCGTTGCGTGTTCAGATTGAGCCATTGCATNNNAGATCGGAAG |
| 38623_1 | TATGAGTGTGGAGTCGTTGCAGAGGTTGCAGTGAACTGAGATCNNNAGATCGGAA |
| 38623_2 | TATGAGTGTGGAGTCGTTGCAGAGGTTGCAGTGAACTGAGATCNNNAGATCGGAA |
| 13566_1 | TATGAGTGTGGAGTCGTTGCATTAGATTCTCATAGCAGCACNNNAGATCGGAAGAG |
| 13566_2 | TATGAGTGTGGAGTCGTTGCATTAGATTCTCATAGCAGCACNNNAGATCGGAAGAG |
| 29628_1 | TATGAGTGTGGAGTCGTTGCTCATCCCACATAAGATGTTGTTCTNNNAGATCGGAA |
| 29628_2 | TATGAGTGTGGAGTCGTTGCTCATCCCACATAAGATGTTGTTCTNNNAGATCGGAA |
| 49066_1 | TATGAGTGTGGAGTCGTTGCGGTGATCTGTCATCTTTGTGCAGNNNAGATCGGAAG |
| 49066_2 | TATGAGTGTGGAGTCGTTGCGGTGATCTGTCATCTTTGTGCAGNNNAGATCGGAAG |
| 33106_1 | TATGAGTGTGGAGTCGTTGCTACAAAAAATTAGCCGGGTGCAGNNNAGATCGGAA |
| 33106_2 | TATGAGTGTGGAGTCGTTGCTACAAAAAATTAGCCGGGTGCAGNNNAGATCGGAA |
| 51743_1 | TATGAGTGTGGAGTCGTTGCGTCATGCGCAAAGTTTTCAGGATNNNAGATCGGAA |
| 51743_2 | TATGAGTGTGGAGTCGTTGCGTCATGCGCAAAGTTTTCAGGATNNNAGATCGGAA |
| 47665_1 | TATGAGTGTGGAGTCGTTGCGATTACAGGCATGCACCATCATGNNNAGATCGGAAG |
| 47665_2 | TATGAGTGTGGAGTCGTTGCGATTACAGGCATGCACCATCATGNNNAGATCGGAAG |
| 48483_1 | TATGAGTGTGGAGTCGTTGCCAAGTAGCTGGGATTACAGGCATNNNAGATCGGAA |
| 48483_2 | TATGAGTGTGGAGTCGTTGCCAAGTAGCTGGGATTACAGGCATNNNAGATCGGAA |
| 48070_1 | TATGAGTGTGGAGTCGTTGCTTGGTCTATAGCAGTGCTACAGANNNAGATCGGAA |
| 48070_2 | TATGAGTGTGGAGTCGTTGCTTGGTCTATAGCAGTGCTACAGANNNAGATCGGAA |
| 40845_1 | TATGAGTGTGGAGTCGTTGCGCTATAAGGTGCATATGGCTTGCNNNAGATCGGAA |
| 40845_2 | TATGAGTGTGGAGTCGTTGCGCTATAAGGTGCATATGGCTTGCNNNAGATCGGAA |
| 31165_1 | TATGAGTGTGGAGTCGTTGCCCTGTCCCCCATTCCAATTTCTANNNAGATCGGAAGA |
| 31165_2 | TATGAGTGTGGAGTCGTTGCCCTGTCCCCCATTCCAATTTCTANNNAGATCGGAAGA |
| 28835_1 | TATGAGTGTGGAGTCGTTGCCAAAGAAAAAGCTGTGACAGGGTNNNAGATCGGAA |
| 28835_2 | TATGAGTGTGGAGTCGTTGCCAAAGAAAAAGCTGTGACAGGGTNNNAGATCGGAA |
| 38111_1 | TATGAGTGTGGAGTCGTTGCTCTCTCTCTCCTTGGCTTACAGANNNAGATCGGAAGA |
| 38111_2 | TATGAGTGTGGAGTCGTTGCTCTCTCTCTCCTTGGCTTACAGANNNAGATCGGAAGA |
| 53197_1 | TATGAGTGTGGAGTCGTTGCTTTTTGAGACAGTCTTGCTCTGTNNNAGATCGGAAG |
| 53197_2 | TATGAGTGTGGAGTCGTTGCTTTTTGAGACAGTCTTGCTCTGTNNNAGATCGGAAG |
| 32571_1 | TATGAGTGTGGAGTCGTTGCCTCCCAGGTTCACACCATTCTCNNNAGATCGGAAGAG |
| 32571_2 | TATGAGTGTGGAGTCGTTGCCTCCCAGGTTCACACCATTCTCNNNAGATCGGAAGAG |
| 52150_1 | TATGAGTGTGGAGTCGTTGCCCAAATATCAATGCGCACCTGAANNNAGATCGGAAG |
| 52150_2 | TATGAGTGTGGAGTCGTTGCCCAAATATCAATGCGCACCTGAANNNAGATCGGAAG |
| 40889_1 | TATGAGTGTGGAGTCGTTGCCCTGTGAGATGCAGATCTGTGTNNNAGATCGGAAG |
| 40889_2 | TATGAGTGTGGAGTCGTTGCCCTGTGAGATGCAGATCTGTGTNNNAGATCGGAAG |
| 45373_1 | TATGAGTGTGGAGTCGTTGCCAGCTCCACTTCCCTGGGTANNNAGATCGGAAGAGC |
| 45373_2 | TATGAGTGTGGAGTCGTTGCCAGCTCCACTTCCCTGGGTANNNAGATCGGAAGAGC |
| 44798_1 | TATGAGTGTGGAGTCGTTGCTAGGTCATCTAACAGCCCCTGTANNNAGATCGGAAG |

| | |
|---|---|
| 44798_2 | TATGAGTGTGGAGTCGTTGCTAGGTCATCTAACAGCCCCTGTANNNAGATCGGAAG |
| 37678_1 | TATGAGTGTGGAGTCGTTGCGGTGAAACCCCGTCTCTACTAAANNNAGATCGGAAG |
| 37678_2 | TATGAGTGTGGAGTCGTTGCGGTGAAACCCCGTCTCTACTAAANNNAGATCGGAAG |
| 41611_1 | TATGAGTGTGGAGTCGTTGCAGATGGACTCTCACTCTGTTGCNNNAGATCGGAAGA |
| 41611_2 | TATGAGTGTGGAGTCGTTGCAGATGGACTCTCACTCTGTTGCNNNAGATCGGAAGA |
| 34572_1 | TATGAGTGTGGAGTCGTTGCGATCTTGGGAGAAGATGAGCTCCNNNAGATCGGAA |
| 34572_2 | TATGAGTGTGGAGTCGTTGCGATCTTGGGAGAAGATGAGCTCCNNNAGATCGGAA |
| 30491_1 | TATGAGTGTGGAGTCGTTGCCTGAGTACAGATTCCCCGTGGTTNNNAGATCGGAAG |
| 30491_2 | TATGAGTGTGGAGTCGTTGCCTGAGTACAGATTCCCCGTGGTTNNNAGATCGGAAG |
| 30377_1 | TATGAGTGTGGAGTCGTTGCCCGGTATGTAATTTAACATTCAANNNAGATCGGAAG |
| 30377_2 | TATGAGTGTGGAGTCGTTGCCCGGTATGTAATTTAACATTCAANNNAGATCGGAAG |
| 40524_1 | TATGAGTGTGGAGTCGTTGCAGCTCCTAGCAGACGTTTTTGTANNNAGATCGGAAG |
| 40524_2 | TATGAGTGTGGAGTCGTTGCAGCTCCTAGCAGACGTTTTTGTANNNAGATCGGAAG |
| 45489_1 | TATGAGTGTGGAGTCGTTGCGTTTTTGAGACGGAGTCTTGCTCNNNAGATCGGAAG |
| 45489_2 | TATGAGTGTGGAGTCGTTGCGTTTTTGAGACGGAGTCTTGCTCNNNAGATCGGAAG |
| 12930_1 | TATGAGTGTGGAGTCGTTGCATGTTAGTAATGAGGACAACTAGNNNAGATCGGAA |
| 12930_2 | TATGAGTGTGGAGTCGTTGCATGTTAGTAATGAGGACAACTAGNNNAGATCGGAA |
| 41335_1 | TATGAGTGTGGAGTCGTTGCTGAATCCCAAGTAATCTGACGACTNNNAGATCGGAA |
| 41335_2 | TATGAGTGTGGAGTCGTTGCTGAATCCCAAGTAATCTGACGACTNNNAGATCGGAA |
| 47949_1 | TATGAGTGTGGAGTCGTTGCGACAAAGCTGACAAAAACAAGCGNNNAGATCGGAA |
| 47949_2 | TATGAGTGTGGAGTCGTTGCGACAAAGCTGACAAAAACAAGCGNNNAGATCGGAA |
| 29590_1 | TATGAGTGTGGAGTCGTTGCCACGTGTGCGACCTCATGTCNNNAGATCGGAAGAGC |
| 29590_2 | TATGAGTGTGGAGTCGTTGCCACGTGTGCGACCTCATGTCNNNAGATCGGAAGAGC |
| 34512_1 | TATGAGTGTGGAGTCGTTGCCACCACCTAGGCCAAGATCTTTANNNAGATCGGAAG |
| 34512_2 | TATGAGTGTGGAGTCGTTGCCACCACCTAGGCCAAGATCTTTANNNAGATCGGAAG |
| 53429_1 | TATGAGTGTGGAGTCGTTGCCAGACAGGAGACCATGAGAAACANNNAGATCGGAA |
| 53429_2 | TATGAGTGTGGAGTCGTTGCCAGACAGGAGACCATGAGAAACANNNAGATCGGAA |
| 46009_1 | TATGAGTGTGGAGTCGTTGCCAAGTAGCGGGGATTACAGGAATNNNAGATCGGAA |
| 46009_2 | TATGAGTGTGGAGTCGTTGCCAAGTAGCGGGGATTACAGGAATNNNAGATCGGAA |
| 38494_1 | TATGAGTGTGGAGTCGTTGCGAGCACGCTTTATTCATCCACTCNNNAGATCGGAAGA |
| 38494_2 | TATGAGTGTGGAGTCGTTGCGAGCACGCTTTATTCATCCACTCNNNAGATCGGAAGA |
| 35869_1 | TATGAGTGTGGAGTCGTTGCGGGGGACCCTAAATGCATCTTTANNNAGATCGGAAG |
| 35869_2 | TATGAGTGTGGAGTCGTTGCGGGGGACCCTAAATGCATCTTTANNNAGATCGGAAG |
| 43781_1 | TATGAGTGTGGAGTCGTTGCAGATGAGACATCACAGGTATGGCNNNAGATCGGAA |
| 43781_2 | TATGAGTGTGGAGTCGTTGCAGATGAGACATCACAGGTATGGCNNNAGATCGGAA |
| 31627_1 | TATGAGTGTGGAGTCGTTGCTATTTGTCTCTCTCCCTCCACCTNNNAGATCGGAAGA |
| 31627_2 | TATGAGTGTGGAGTCGTTGCTATTTGTCTCTCTCCCTCCACCTNNNAGATCGGAAGA |
| 45259_1 | TATGAGTGTGGAGTCGTTGCAGGGGGACTGAGATCTTTCCATANNNAGATCGGAA |
| 45259_2 | TATGAGTGTGGAGTCGTTGCAGGGGGACTGAGATCTTTCCATANNNAGATCGGAA |
| 51192_1 | TATGAGTGTGGAGTCGTTGCTTCATTTCAGGAGCCACTTCTGANNNAGATCGGAAG |
| 51192_2 | TATGAGTGTGGAGTCGTTGCTTCATTTCAGGAGCCACTTCTGANNNAGATCGGAAG |
| 13273_1 | TATGAGTGTGGAGTCGTTGCGCCTAAAGCTAGACTATATTTCTNNNAGATCGGAAG |
| 13273_2 | TATGAGTGTGGAGTCGTTGCGCCTAAAGCTAGACTATATTTCTNNNAGATCGGAAG |

| | |
|---|---|
| 48587_1 | TATGAGTGTGGAGTCGTTGCATAATCCTAGCACTTTGGGAGGCNNNAGATCGGAAG |
| 48587_2 | TATGAGTGTGGAGTCGTTGCATAATCCTAGCACTTTGGGAGGCNNNAGATCGGAAG |
| 49421_1 | TATGAGTGTGGAGTCGTTGCGACGGAAGTCTCGCTCTGTTGNNNAGATCGGAAGA |
| 49421_2 | TATGAGTGTGGAGTCGTTGCGACGGAAGTCTCGCTCTGTTGNNNAGATCGGAAGA |
| 53301_1 | TATGAGTGTGGAGTCGTTGCAATCTTGTGGAGATAGGATGGCCNNNAGATCGGAA |
| 53301_2 | TATGAGTGTGGAGTCGTTGCAATCTTGTGGAGATAGGATGGCCNNNAGATCGGAA |
| 32385_1 | TATGAGTGTGGAGTCGTTGCGATTACAGGCATGCACCATCATGNNNAGATCGGAAG |
| 32385_2 | TATGAGTGTGGAGTCGTTGCGATTACAGGCATGCACCATCATGNNNAGATCGGAAG |
| 43857_1 | TATGAGTGTGGAGTCGTTGCATGCTGCAGAGAAACAAGCAAATNNNAGATCGGAA |
| 43857_2 | TATGAGTGTGGAGTCGTTGCATGCTGCAGAGAAACAAGCAAATNNNAGATCGGAA |
| 33539_1 | TATGAGTGTGGAGTCGTTGCCCATGACACCCGGCTAATTTTTTNNNAGATCGGAAG |
| 33539_2 | TATGAGTGTGGAGTCGTTGCCCATGACACCCGGCTAATTTTTTNNNAGATCGGAAG |
| 34825_1 | TATGAGTGTGGAGTCGTTGCGAGTGCAATGGCGTGATCTTGNNNAGATCGGAAGA |
| 34825_2 | TATGAGTGTGGAGTCGTTGCGAGTGCAATGGCGTGATCTTGNNNAGATCGGAAGA |
| 38998_1 | TATGAGTGTGGAGTCGTTGCCAGAACTACCACTGGACACATCANNNAGATCGGAAG |
| 38998_2 | TATGAGTGTGGAGTCGTTGCCAGAACTACCACTGGACACATCANNNAGATCGGAAG |
| 50614_1 | TATGAGTGTGGAGTCGTTGCTGCAATTTTCACCTTCTCGTCACNNNAGATCGGAAGA |
| 50614_2 | TATGAGTGTGGAGTCGTTGCTGCAATTTTCACCTTCTCGTCACNNNAGATCGGAAGA |
| 36882_1 | TATGAGTGTGGAGTCGTTGCAGTTTCTCCACATCCTCATCCACNNNAGATCGGAAGA |
| 36882_2 | TATGAGTGTGGAGTCGTTGCAGTTTCTCCACATCCTCATCCACNNNAGATCGGAAGA |
| 30453_1 | TATGAGTGTGGAGTCGTTGCTCCGTCTGTTGGTAAAGCTTCTGNNNAGATCGGAAG |
| 30453_2 | TATGAGTGTGGAGTCGTTGCTCCGTCTGTTGGTAAAGCTTCTGNNNAGATCGGAAG |
| 5182_1 | TATGAGTGTGGAGTCGTTGCACACCTTTTCACACTGATACACGTNNNAGATCGGAAG |
| 5182_2 | TATGAGTGTGGAGTCGTTGCACACCTTTTCACACTGATACACGTNNNAGATCGGAAG |
| 31212_1 | TATGAGTGTGGAGTCGTTGCAGTATCAGTTGAAGAGGAACGCANNNAGATCGGAA |
| 31212_2 | TATGAGTGTGGAGTCGTTGCAGTATCAGTTGAAGAGGAACGCANNNAGATCGGAA |
| 49428_1 | TATGAGTGTGGAGTCGTTGCCAGAGTAGCTGGGATTACAGGCNNNAGATCGGAAG |
| 49428_2 | TATGAGTGTGGAGTCGTTGCCAGAGTAGCTGGGATTACAGGCNNNAGATCGGAAG |
| 52095_1 | TATGAGTGTGGAGTCGTTGCTTGGTGGCTTTAGAAGTTCCTCANNNAGATCGGAAG |
| 52095_2 | TATGAGTGTGGAGTCGTTGCTTGGTGGCTTTAGAAGTTCCTCANNNAGATCGGAAG |
| 46324_1 | TATGAGTGTGGAGTCGTTGCCATGATCTTGGCTCACTGCAACNNNAGATCGGAAGA |
| 46324_2 | TATGAGTGTGGAGTCGTTGCCATGATCTTGGCTCACTGCAACNNNAGATCGGAAGA |
| 47561_1 | TATGAGTGTGGAGTCGTTGCCCCTGGGTAGAAAGCCTTGGNNNAGATCGGAAGAG |
| 47561_2 | TATGAGTGTGGAGTCGTTGCCCCTGGGTAGAAAGCCTTGGNNNAGATCGGAAGAG |
| 29525_1 | TATGAGTGTGGAGTCGTTGCCGAGTAGCTGGGACTACAGGTNNNAGATCGGAAGA |
| 29525_2 | TATGAGTGTGGAGTCGTTGCCGAGTAGCTGGGACTACAGGTNNNAGATCGGAAGA |
| 53141_1 | TATGAGTGTGGAGTCGTTGCCTCCCATGAGAAACAGTAGGCTTNNNAGATCGGAAG |
| 53141_2 | TATGAGTGTGGAGTCGTTGCCTCCCATGAGAAACAGTAGGCTTNNNAGATCGGAAG |
| 37899_1 | TATGAGTGTGGAGTCGTTGCGTAATCCCAGCTACTCAGAAGGCNNNAGATCGGAAG |
| 37899_2 | TATGAGTGTGGAGTCGTTGCGTAATCCCAGCTACTCAGAAGGCNNNAGATCGGAAG |
| 37286_1 | TATGAGTGTGGAGTCGTTGCGACACAAAAGATCCCCAGAGAGTNNNAGATCGGAA |
| 37286_2 | TATGAGTGTGGAGTCGTTGCGACACAAAAGATCCCCAGAGAGTNNNAGATCGGAA |
| 31644_1 | TATGAGTGTGGAGTCGTTGCAACTGGTTGAATGGATGGCAAAANNNAGATCGGAA |

| | |
|---|---|
| 31644_2 | TATGAGTGTGGAGTCGTTGCAACTGGTTGAATGGATGGCAAAANNNAGATCGGAA |
| 52278_1 | TATGAGTGTGGAGTCGTTGCGTAGGACAAAGGACATGGCTACANNNAGATCGGAA |
| 52278_2 | TATGAGTGTGGAGTCGTTGCGTAGGACAAAGGACATGGCTACANNNAGATCGGAA |
| 5600_1 | TATGAGTGTGGAGTCGTTGCAAATGCTACACATCCTCCTAGCTNNNAGATCGGAAG |
| 5600_2 | TATGAGTGTGGAGTCGTTGCAAATGCTACACATCCTCCTAGCTNNNAGATCGGAAG |
| 47049_1 | TATGAGTGTGGAGTCGTTGCCCAGATGAGTGACTGAGGAACAANNNAGATCGGAA |
| 47049_2 | TATGAGTGTGGAGTCGTTGCCCAGATGAGTGACTGAGGAACAANNNAGATCGGAA |
| 33521_1 | TATGAGTGTGGAGTCGTTGCAGGCACCATGGGTATCATTTTCANNNAGATCGGAAG |
| 33521_2 | TATGAGTGTGGAGTCGTTGCAGGCACCATGGGTATCATTTTCANNNAGATCGGAAG |
| 36658_1 | TATGAGTGTGGAGTCGTTGCTCTGTAGCACTCTTGAGCATCTGNNNAGATCGGAAG |
| 36658_2 | TATGAGTGTGGAGTCGTTGCTCTGTAGCACTCTTGAGCATCTGNNNAGATCGGAAG |
| 30440_1 | TATGAGTGTGGAGTCGTTGCATGATGAGGCTGTAACTGAGTGANNNAGATCGGAA |
| 30440_2 | TATGAGTGTGGAGTCGTTGCATGATGAGGCTGTAACTGAGTGANNNAGATCGGAA |
| 32439_1 | TATGAGTGTGGAGTCGTTGCCATTCTCCTGCCTCAGCCTCNNNAGATCGGAAGAGCA |
| 32439_2 | TATGAGTGTGGAGTCGTTGCCATTCTCCTGCCTCAGCCTCNNNAGATCGGAAGAGCA |
| 8267_1 | TATGAGTGTGGAGTCGTTGCGAATGTGAACAGGTCCAAACGACANNNAGATCGGA |
| 8267_2 | TATGAGTGTGGAGTCGTTGCGAATGTGAACAGGTCCAAACGACANNNAGATCGGA |
| 36482_1 | TATGAGTGTGGAGTCGTTGCAGAAGAAATGCACGTTCTCCTCANNNAGATCGGAAG |
| 36482_2 | TATGAGTGTGGAGTCGTTGCAGAAGAAATGCACGTTCTCCTCANNNAGATCGGAAG |
| 35041_1 | TATGAGTGTGGAGTCGTTGCCCTCCCAAGTTCATGCTACTCTCNNNAGATCGGAAGA |
| 35041_2 | TATGAGTGTGGAGTCGTTGCCCTCCCAAGTTCATGCTACTCTCNNNAGATCGGAAGA |
| 52780_1 | TATGAGTGTGGAGTCGTTGCGTAGGGAGACGTCAAGCTCTTGNNNAGATCGGAAG |
| 52780_2 | TATGAGTGTGGAGTCGTTGCGTAGGGAGACGTCAAGCTCTTGNNNAGATCGGAAG |
| 13730_1 | TATGAGTGTGGAGTCGTTGCTATCAAATATTTAGCATGGATCCNNNAGATCGGAAG |
| 13730_2 | TATGAGTGTGGAGTCGTTGCTATCAAATATTTAGCATGGATCCNNNAGATCGGAAG |
| 46706_1 | TATGAGTGTGGAGTCGTTGCTTTTGCACTTCCAACATCACTGTNNNAGATCGGAAGA |
| 46706_2 | TATGAGTGTGGAGTCGTTGCTTTTGCACTTCCAACATCACTGTNNNAGATCGGAAGA |
| 32667_1 | TATGAGTGTGGAGTCGTTGCACTCCCAGGTTCAAGTGATTCTCNNNAGATCGGAAG |
| 32667_2 | TATGAGTGTGGAGTCGTTGCACTCCCAGGTTCAAGTGATTCTCNNNAGATCGGAAG |
| 41697_1 | TATGAGTGTGGAGTCGTTGCTGTCTGCTCTAACTTGGACTTCCNNNAGATCGGAAG |
| 41697_2 | TATGAGTGTGGAGTCGTTGCTGTCTGCTCTAACTTGGACTTCCNNNAGATCGGAAG |
| 53229_1 | TATGAGTGTGGAGTCGTTGCGTGAACCCACAATTCCGGAATAGNNNAGATCGGAA |
| 53229_2 | TATGAGTGTGGAGTCGTTGCGTGAACCCACAATTCCGGAATAGNNNAGATCGGAA |
| 50098_1 | TATGAGTGTGGAGTCGTTGCGTCCTCATCTTTACTGTCACCGTNNNAGATCGGAAGA |
| 50098_2 | TATGAGTGTGGAGTCGTTGCGTCCTCATCTTTACTGTCACCGTNNNAGATCGGAAGA |
| 37361_1 | TATGAGTGTGGAGTCGTTGCGCACAGGTGAGTCATTCAGTTTCNNNAGATCGGAAG |
| 37361_2 | TATGAGTGTGGAGTCGTTGCGCACAGGTGAGTCATTCAGTTTCNNNAGATCGGAAG |
| 41251_1 | TATGAGTGTGGAGTCGTTGCATACAAAAATTAGCTGGGTGCGGNNNAGATCGGAA |
| 41251_2 | TATGAGTGTGGAGTCGTTGCATACAAAAATTAGCTGGGTGCGGNNNAGATCGGAA |
| 29556_1 | TATGAGTGTGGAGTCGTTGCTTTTTACGGCAGCCTGTTTAGTGNNNAGATCGGAAG |
| 29556_2 | TATGAGTGTGGAGTCGTTGCTTTTTACGGCAGCCTGTTTAGTGNNNAGATCGGAAG |
| 47998_1 | TATGAGTGTGGAGTCGTTGCCAGCTATTACCCATCCTGCTTCTNNNAGATCGGAAGA |
| 47998_2 | TATGAGTGTGGAGTCGTTGCCAGCTATTACCCATCCTGCTTCTNNNAGATCGGAAGA |

| | |
|---|---|
| 50986_1 | TATGAGTGTGGAGTCGTTGCATTCTGACACCTCCAACATCCAGNNNAGATCGGAAGA |
| 50986_2 | TATGAGTGTGGAGTCGTTGCATTCTGACACCTCCAACATCCAGNNNAGATCGGAAGA |
| 39655_1 | TATGAGTGTGGAGTCGTTGCTCCCATACCTTGTTTGTCACGATNNNAGATCGGAAGA |
| 39655_2 | TATGAGTGTGGAGTCGTTGCTCCCATACCTTGTTTGTCACGATNNNAGATCGGAAGA |
| 41540_1 | TATGAGTGTGGAGTCGTTGCTGGAATAGCTGGAATTACAGGCGNNNAGATCGGAA |
| 41540_2 | TATGAGTGTGGAGTCGTTGCTGGAATAGCTGGAATTACAGGCGNNNAGATCGGAA |
| 41871_1 | TATGAGTGTGGAGTCGTTGCGCATCTATTTGATTCCTCTGTCTTTNNNAGATCGGAAG |
| 41871_2 | TATGAGTGTGGAGTCGTTGCGCATCTATTTGATTCCTCTGTCTTTNNNAGATCGGAAG |
| 42186_1 | TATGAGTGTGGAGTCGTTGCAATCTCAGCTCACTGCAACCTCNNNAGATCGGAAGAG |
| 42186_2 | TATGAGTGTGGAGTCGTTGCAATCTCAGCTCACTGCAACCTCNNNAGATCGGAAGAG |
| 31331_1 | TATGAGTGTGGAGTCGTTGCAGAGACTCAAGGTGTAGCTTACCNNNAGATCGGAAG |
| 31331_2 | TATGAGTGTGGAGTCGTTGCAGAGACTCAAGGTGTAGCTTACCNNNAGATCGGAAG |
| 29931_1 | TATGAGTGTGGAGTCGTTGCGTTATGCAATAAGCTTAACGTGTTTNNNAGATCGGAA |
| 29931_2 | TATGAGTGTGGAGTCGTTGCGTTATGCAATAAGCTTAACGTGTTTNNNAGATCGGAA |
| 50240_1 | TATGAGTGTGGAGTCGTTGCCATTACGATGCCCCATCACTTTCNNNAGATCGGAAGA |
| 50240_2 | TATGAGTGTGGAGTCGTTGCCATTACGATGCCCCATCACTTTCNNNAGATCGGAAGA |
| 50414_1 | TATGAGTGTGGAGTCGTTGCAACAAAACAGAGTGAGCTCCAGANNNAGATCGGAAG |
| 50414_2 | TATGAGTGTGGAGTCGTTGCAACAAAACAGAGTGAGCTCCAGANNNAGATCGGAAG |
| 49817_1 | TATGAGTGTGGAGTCGTTGCAACGTTCTCGCTGCATTTTTGCNNNAGATCGGAAGAG |
| 49817_2 | TATGAGTGTGGAGTCGTTGCAACGTTCTCGCTGCATTTTTGCNNNAGATCGGAAGAG |
| 35853_1 | TATGAGTGTGGAGTCGTTGCGTTCAGATGATTCTCCTGCCTCANNNAGATCGGAAGA |
| 35853_2 | TATGAGTGTGGAGTCGTTGCGTTCAGATGATTCTCCTGCCTCANNNAGATCGGAAGA |
| 46166_1 | TATGAGTGTGGAGTCGTTGCAAAGTGCTGGGATTACAGGCATNNNAGATCGGAAG |
| 46166_2 | TATGAGTGTGGAGTCGTTGCAAAGTGCTGGGATTACAGGCATNNNAGATCGGAAG |
| 45512_1 | TATGAGTGTGGAGTCGTTGCAGCAGAAAAGGAAGAAAAGACGCNNNAGATCGGAA |
| 45512_2 | TATGAGTGTGGAGTCGTTGCAGCAGAAAAGGAAGAAAAGACGCNNNAGATCGGAA |
| 38620_1 | TATGAGTGTGGAGTCGTTGCAAGCAAAGCAAAGCAAAGCAAAGNNNAGATCGGAA |
| 38620_2 | TATGAGTGTGGAGTCGTTGCAAGCAAAGCAAAGCAAAGCAAAGNNNAGATCGGAA |
| 38601_1 | TATGAGTGTGGAGTCGTTGCATAAAGTTGGACCCTTACCTCGCNNNAGATCGGAAGA |
| 38601_2 | TATGAGTGTGGAGTCGTTGCATAAAGTTGGACCCTTACCTCGCNNNAGATCGGAAGA |
| 47077_1 | TATGAGTGTGGAGTCGTTGCTCAGCAGTAAAGCTATCCAGTCCNNNAGATCGGAAG |
| 47077_2 | TATGAGTGTGGAGTCGTTGCTCAGCAGTAAAGCTATCCAGTCCNNNAGATCGGAAG |
| 13138_1 | TATGAGTGTGGAGTCGTTGCTCTCTATAAAAATGGTTCTTTGCNNNAGATCGGAAGA |
| 13138_2 | TATGAGTGTGGAGTCGTTGCTCTCTATAAAAATGGTTCTTTGCNNNAGATCGGAAGA |
| 49791_1 | TATGAGTGTGGAGTCGTTGCGGGGGAAAAACACTCAAGGAATGNNNAGATCGGAA |
| 49791_2 | TATGAGTGTGGAGTCGTTGCGGGGGAAAAACACTCAAGGAATGNNNAGATCGGAA |
| 48959_1 | TATGAGTGTGGAGTCGTTGCCCAAGTAGCTGGGATTACAGGCNNNAGATCGGAAGA |
| 48959_2 | TATGAGTGTGGAGTCGTTGCCCAAGTAGCTGGGATTACAGGCNNNAGATCGGAAGA |
| 41364_1 | TATGAGTGTGGAGTCGTTGCAGGTGCAATATCAGCTCACTTCANNNAGATCGGAAG |
| 41364_2 | TATGAGTGTGGAGTCGTTGCAGGTGCAATATCAGCTCACTTCANNNAGATCGGAAG |
| 43974_1 | TATGAGTGTGGAGTCGTTGCCATTCATGAAGGATCCACCTCCANNNAGATCGGAAGA |
| 43974_2 | TATGAGTGTGGAGTCGTTGCCATTCATGAAGGATCCACCTCCANNNAGATCGGAAGA |
| 31821_1 | TATGAGTGTGGAGTCGTTGCTAAGAAAACCATGCCATTTGCGANNNAGATCGGAAG |

| | |
|---|---|
| 31821_2 | TATGAGTGTGGAGTCGTTGCTAAGAAAACCATGCCATTTGCGANNNAGATCGGAAG |
| 10714_1 | TATGAGTGTGGAGTCGTTGCCTTCCCGCCCATGCTGTTATTATNNNAGATCGGAAGA |
| 10714_2 | TATGAGTGTGGAGTCGTTGCCTTCCCGCCCATGCTGTTATTATNNNAGATCGGAAGA |
| 43724_1 | TATGAGTGTGGAGTCGTTGCGTTCCCAACCAAGGTCTGAGTAGNNNAGATCGGAAG |
| 43724_2 | TATGAGTGTGGAGTCGTTGCGTTCCCAACCAAGGTCTGAGTAGNNNAGATCGGAAG |
| 50076_1 | TATGAGTGTGGAGTCGTTGCTTAGAGTCCAATGTTTGAGGGCANNNAGATCGGAAG |
| 50076_2 | TATGAGTGTGGAGTCGTTGCTTAGAGTCCAATGTTTGAGGGCANNNAGATCGGAAG |
| 50899_1 | TATGAGTGTGGAGTCGTTGCTCCTGTGCAGAAACCTTACTTGTNNNAGATCGGAAGA |
| 50899_2 | TATGAGTGTGGAGTCGTTGCTCCTGTGCAGAAACCTTACTTGTNNNAGATCGGAAGA |
| 48622_1 | TATGAGTGTGGAGTCGTTGCCCCGGATTCAAACAATTCTCTGCNNNAGATCGGAAGA |
| 48622_2 | TATGAGTGTGGAGTCGTTGCCCCGGATTCAAACAATTCTCTGCNNNAGATCGGAAGA |
| 49368_1 | TATGAGTGTGGAGTCGTTGCAGGCCTCTCGAAGTTTTGAGATTNNNAGATCGGAAG |
| 49368_2 | TATGAGTGTGGAGTCGTTGCAGGCCTCTCGAAGTTTTGAGATTNNNAGATCGGAAG |
| 38922_1 | TATGAGTGTGGAGTCGTTGCTCCAGGAGCTGTTTTTTTGAAAANNNAGATCGGAAG |
| 38922_2 | TATGAGTGTGGAGTCGTTGCTCCAGGAGCTGTTTTTTTGAAAANNNAGATCGGAAG |
| 29246_1 | TATGAGTGTGGAGTCGTTGCCTATGAGCTACAGTGAGAGCTGGNNNAGATCGGAAG |
| 29246_2 | TATGAGTGTGGAGTCGTTGCCTATGAGCTACAGTGAGAGCTGGNNNAGATCGGAAG |
| 39200_1 | TATGAGTGTGGAGTCGTTGCTCCAGCCATTCCATCTCTATGTGNNNAGATCGGAAGA |
| 39200_2 | TATGAGTGTGGAGTCGTTGCTCCAGCCATTCCATCTCTATGTGNNNAGATCGGAAGA |
| 43665_1 | TATGAGTGTGGAGTCGTTGCCTTCGGGCCACAAAGGACAATATNNNAGATCGGAAG |
| 43665_2 | TATGAGTGTGGAGTCGTTGCCTTCGGGCCACAAAGGACAATATNNNAGATCGGAAG |
| 966_1 | TATGAGTGTGGAGTCGTTGCACAGATATCACCCACACACACCTNNNAGATCGGAAGA |
| 966_2 | TATGAGTGTGGAGTCGTTGCACAGATATCACCCACACACACCTNNNAGATCGGAAGA |
| 42724_1 | TATGAGTGTGGAGTCGTTGCAACACACAGGAAAATACGGCAAGNNNAGATCGGAAG |
| 42724_2 | TATGAGTGTGGAGTCGTTGCAACACACAGGAAAATACGGCAAGNNNAGATCGGAAG |
| 13788_1 | TATGAGTGTGGAGTCGTTGCAGACTTTCTACATATTAAAGAGGTNNNAGATCGGAA |
| 13788_2 | TATGAGTGTGGAGTCGTTGCAGACTTTCTACATATTAAAGAGGTNNNAGATCGGAA |
| 33066_1 | TATGAGTGTGGAGTCGTTGCACTTCTACTCTCCAAGTCCGAGANNNAGATCGGAAGA |
| 33066_2 | TATGAGTGTGGAGTCGTTGCACTTCTACTCTCCAAGTCCGAGANNNAGATCGGAAGA |
| 48979_1 | TATGAGTGTGGAGTCGTTGCGAGGGGTGACTTCAGCTCAATAANNNAGATCGGAAG |
| 48979_2 | TATGAGTGTGGAGTCGTTGCGAGGGGTGACTTCAGCTCAATAANNNAGATCGGAAG |
| 29341_1 | TATGAGTGTGGAGTCGTTGCGCTGACTGTATCTCTCTCTCTGCNNNAGATCGGAAGA |
| 29341_2 | TATGAGTGTGGAGTCGTTGCGCTGACTGTATCTCTCTCTCTGCNNNAGATCGGAAGA |
| 52515_1 | TATGAGTGTGGAGTCGTTGCTAGGCCTGCTTAGAGGAAACAAGNNNAGATCGGAAG |
| 52515_2 | TATGAGTGTGGAGTCGTTGCTAGGCCTGCTTAGAGGAAACAAGNNNAGATCGGAAG |
| 51012_1 | TATGAGTGTGGAGTCGTTGCGTATTGAAGGTAAAGACGGGGGANNNAGATCGGAA |
| 51012_2 | TATGAGTGTGGAGTCGTTGCGTATTGAAGGTAAAGACGGGGGANNNAGATCGGAA |
| 42616_1 | TATGAGTGTGGAGTCGTTGCGCATGATCATGGCTCACTGTAACNNNAGATCGGAAG |
| 42616_2 | TATGAGTGTGGAGTCGTTGCGCATGATCATGGCTCACTGTAACNNNAGATCGGAAG |
| 38840_1 | TATGAGTGTGGAGTCGTTGCCCAGGAGTAGGAGACTACAGTGANNNAGATCGGAA |
| 38840_2 | TATGAGTGTGGAGTCGTTGCCCAGGAGTAGGAGACTACAGTGANNNAGATCGGAA |
| 29916_1 | TATGAGTGTGGAGTCGTTGCTGTCAAGTCAGAAACAGGTGTCTNNNAGATCGGAAG |
| 29916_2 | TATGAGTGTGGAGTCGTTGCTGTCAAGTCAGAAACAGGTGTCTNNNAGATCGGAAG |

| | |
|---|---|
| 51000_1 | TATGAGTGTGGAGTCGTTGCGGAATAGCTGGGATTACAGGCATNNNAGATCGGAA |
| 51000_2 | TATGAGTGTGGAGTCGTTGCGGAATAGCTGGGATTACAGGCATNNNAGATCGGAA |
| 52089_1 | TATGAGTGTGGAGTCGTTGCCTCAGTGACCTTGATGTCAGACANNNAGATCGGAAG |
| 52089_2 | TATGAGTGTGGAGTCGTTGCCTCAGTGACCTTGATGTCAGACANNNAGATCGGAAG |
| 39553_1 | TATGAGTGTGGAGTCGTTGCTCACCACCACAATCAACATGAAGNNNAGATCGGAAG |
| 39553_2 | TATGAGTGTGGAGTCGTTGCTCACCACCACAATCAACATGAAGNNNAGATCGGAAG |
| 53205_1 | TATGAGTGTGGAGTCGTTGCGAACACAACTGGCCATTCAATGANNNAGATCGGAAG |
| 53205_2 | TATGAGTGTGGAGTCGTTGCGAACACAACTGGCCATTCAATGANNNAGATCGGAAG |
| 41373_1 | TATGAGTGTGGAGTCGTTGCCTGCAGTGAACCATGATCATGTCNNNAGATCGGAAG |
| 41373_2 | TATGAGTGTGGAGTCGTTGCCTGCAGTGAACCATGATCATGTCNNNAGATCGGAAG |
| 10821_1 | TATGAGTGTGGAGTCGTTGCCCCAGTCTGCCTTAGGTAAACACNNNAGATCGGAAG |
| 10821_2 | TATGAGTGTGGAGTCGTTGCCCCAGTCTGCCTTAGGTAAACACNNNAGATCGGAAG |
| 47482_1 | TATGAGTGTGGAGTCGTTGCAGAAATTATCCTGCCTCAGCCTCNNNAGATCGGAAG |
| 47482_2 | TATGAGTGTGGAGTCGTTGCAGAAATTATCCTGCCTCAGCCTCNNNAGATCGGAAG |
| 49720_1 | TATGAGTGTGGAGTCGTTGCGTAGGCACCATGATGCTTTTGTTNNNAGATCGGAAG |
| 49720_2 | TATGAGTGTGGAGTCGTTGCGTAGGCACCATGATGCTTTTGTTNNNAGATCGGAAG |
| 33119_1 | TATGAGTGTGGAGTCGTTGCGAAAAATGTTACTGGACCCCACCNNNAGATCGGAAG |
| 33119_2 | TATGAGTGTGGAGTCGTTGCGAAAAATGTTACTGGACCCCACCNNNAGATCGGAAG |
| 48887_1 | TATGAGTGTGGAGTCGTTGCCCCACATTATTACGCGTAAACCCNNNAGATCGGAAG |
| 48887_2 | TATGAGTGTGGAGTCGTTGCCCCACATTATTACGCGTAAACCCNNNAGATCGGAAG |
| 1994_1 | TATGAGTGTGGAGTCGTTGCGCGGAGGGGAGATAGTCAAAAANNNAGATCGGAA |
| 1994_2 | TATGAGTGTGGAGTCGTTGCGCGGAGGGGAGATAGTCAAAAANNNAGATCGGAA |
| 42274_1 | TATGAGTGTGGAGTCGTTGCGCACTTGTAAATGGAGGTTGCTTNNNAGATCGGAA |
| 42274_2 | TATGAGTGTGGAGTCGTTGCGCACTTGTAAATGGAGGTTGCTTNNNAGATCGGAA |
| 40131_1 | TATGAGTGTGGAGTCGTTGCGGCTCTTGGTTACCTGTGTTGTANNNAGATCGGAAG |
| 40131_2 | TATGAGTGTGGAGTCGTTGCGGCTCTTGGTTACCTGTGTTGTANNNAGATCGGAAG |
| 34855_1 | TATGAGTGTGGAGTCGTTGCGCAACTACGCCCAGCTAATTTTTNNNAGATCGGAAG |
| 34855_2 | TATGAGTGTGGAGTCGTTGCGCAACTACGCCCAGCTAATTTTTNNNAGATCGGAAG |
| 49513_1 | TATGAGTGTGGAGTCGTTGCGTTCAAGCGATTCTCCTACCTCANNNAGATCGGAAG |
| 49513_2 | TATGAGTGTGGAGTCGTTGCGTTCAAGCGATTCTCCTACCTCANNNAGATCGGAAG |
| 1829_1 | TATGAGTGTGGAGTCGTTGCCAGAGTTTTCAGCCAGACTTTGTCNNNAGATCGGAA |
| 1829_2 | TATGAGTGTGGAGTCGTTGCCAGAGTTTTCAGCCAGACTTTGTCNNNAGATCGGAA |
| 31469_1 | TATGAGTGTGGAGTCGTTGCACTTTGTCCTGTTCACTGCTGTANNNAGATCGGAAG |
| 31469_2 | TATGAGTGTGGAGTCGTTGCACTTTGTCCTGTTCACTGCTGTANNNAGATCGGAAG |
| 42304_1 | TATGAGTGTGGAGTCGTTGCAATCCATCTGGTCCTGAGCATTTNNNAGATCGGAAG |
| 42304_2 | TATGAGTGTGGAGTCGTTGCAATCCATCTGGTCCTGAGCATTTNNNAGATCGGAAG |
| 34781_1 | TATGAGTGTGGAGTCGTTGCCCCAAGTTCAAGCAATTCTCCTGNNNAGATCGGAAG |
| 34781_2 | TATGAGTGTGGAGTCGTTGCCCCAAGTTCAAGCAATTCTCCTGNNNAGATCGGAAG |
| 38277_1 | TATGAGTGTGGAGTCGTTGCGTTGCAAGAGAATGAGGGAGGANNNAGATCGGAA |
| 38277_2 | TATGAGTGTGGAGTCGTTGCGTTGCAAGAGAATGAGGGAGGANNNAGATCGGAA |
| 13745_1 | TATGAGTGTGGAGTCGTTGCATTCATTTCAACTTGGTACATTTNNNAGATCGGAAGA |
| 13745_2 | TATGAGTGTGGAGTCGTTGCATTCATTTCAACTTGGTACATTTNNNAGATCGGAAGA |
| 35399_1 | TATGAGTGTGGAGTCGTTGCCACTTCCTCAGCTTCTGGAGTAGNNNAGATCGGAAG |

| | |
|---|---|
| 35399_2 | TATGAGTGTGGAGTCGTTGCCACTTCCTCAGCTTCTGGAGTAGNNNAGATCGGAAG |
| 29663_1 | TATGAGTGTGGAGTCGTTGCCCTAAGCTTCAGGTGTTGAGGATNNNAGATCGGAA |
| 29663_2 | TATGAGTGTGGAGTCGTTGCCCTAAGCTTCAGGTGTTGAGGATNNNAGATCGGAA |
| 2043_1 | TATGAGTGTGGAGTCGTTGCGGAAGGGTGAGAAAAGGCTTGAANNNAGATCGGA |
| 2043_2 | TATGAGTGTGGAGTCGTTGCGGAAGGGTGAGAAAAGGCTTGAANNNAGATCGGA |
| 10315_1 | TATGAGTGTGGAGTCGTTGCTGTGTTACCGAGGACATAGGAGTNNNAGATCGGAA |
| 10315_2 | TATGAGTGTGGAGTCGTTGCTGTGTTACCGAGGACATAGGAGTNNNAGATCGGAA |
| 5615_1 | TATGAGTGTGGAGTCGTTGCCGGGTGGGCCGAACATATTATTTNNNAGATCGGAA |
| 5615_2 | TATGAGTGTGGAGTCGTTGCCGGGTGGGCCGAACATATTATTTNNNAGATCGGAA |
| 48961_1 | TATGAGTGTGGAGTCGTTGCAAAAACCATGTGGCAACAGACATNNNAGATCGGAA |
| 48961_2 | TATGAGTGTGGAGTCGTTGCAAAAACCATGTGGCAACAGACATNNNAGATCGGAA |
| 10659_1 | TATGAGTGTGGAGTCGTTGCTACTTGGCATGCTGCTATGACAGNNNAGATCGGAAG |
| 10659_2 | TATGAGTGTGGAGTCGTTGCTACTTGGCATGCTGCTATGACAGNNNAGATCGGAAG |
| 31195_1 | TATGAGTGTGGAGTCGTTGCACAAATACAAATGACCTTCGCCCNNNAGATCGGAAG |
| 31195_2 | TATGAGTGTGGAGTCGTTGCACAAATACAAATGACCTTCGCCCNNNAGATCGGAAG |
| 3507_1 | TATGAGTGTGGAGTCGTTGCGAAGCAGAAGTGTTTGGGATTTGGNNNAGATCGG |
| 3507_2 | TATGAGTGTGGAGTCGTTGCGAAGCAGAAGTGTTTGGGATTTGGNNNAGATCGG |
| 47905_1 | TATGAGTGTGGAGTCGTTGCCAAAGAGCTTGGACTACAGGCANNNAGATCGGAAG |
| 47905_2 | TATGAGTGTGGAGTCGTTGCCAAAGAGCTTGGACTACAGGCANNNAGATCGGAAG |
| 40124_1 | TATGAGTGTGGAGTCGTTGCTGTCTCAGTGTCCTTCAGTTCAGNNNAGATCGGAAG |
| 40124_2 | TATGAGTGTGGAGTCGTTGCTGTCTCAGTGTCCTTCAGTTCAGNNNAGATCGGAAG |
| 51314_1 | TATGAGTGTGGAGTCGTTGCCCAAATTCTTTCCAGCCACTCACNNNAGATCGGAAGA |
| 51314_2 | TATGAGTGTGGAGTCGTTGCCCAAATTCTTTCCAGCCACTCACNNNAGATCGGAAGA |
| 45710_1 | TATGAGTGTGGAGTCGTTGCGATTACAAATGTGCACCACCACANNNAGATCGGAAG |
| 45710_2 | TATGAGTGTGGAGTCGTTGCGATTACAAATGTGCACCACCACANNNAGATCGGAAG |
| 42206_1 | TATGAGTGTGGAGTCGTTGCCGTAACGTGGATGAGAAGTCTGANNNAGATCGGAA |
| 42206_2 | TATGAGTGTGGAGTCGTTGCCGTAACGTGGATGAGAAGTCTGANNNAGATCGGAA |
| 46223_1 | TATGAGTGTGGAGTCGTTGCGTCACTATAAACTCCACCTCCCGNNNAGATCGGAAGA |
| 46223_2 | TATGAGTGTGGAGTCGTTGCGTCACTATAAACTCCACCTCCCGNNNAGATCGGAAGA |
| 46920_1 | TATGAGTGTGGAGTCGTTGCAGGGAGCAGGGAGTCATTTATTGNNNAGATCGGAA |
| 46920_2 | TATGAGTGTGGAGTCGTTGCAGGGAGCAGGGAGTCATTTATTGNNNAGATCGGAA |
| 13300_1 | TATGAGTGTGGAGTCGTTGCTTAGTGTCTGGCTTAAATTCCNNNAGATCGGAAGAG |
| 13300_2 | TATGAGTGTGGAGTCGTTGCTTAGTGTCTGGCTTAAATTCCNNNAGATCGGAAGAG |
| 36883_1 | TATGAGTGTGGAGTCGTTGCCGCCATGCACTTCTGTACAAATTNNNAGATCGGAAG |
| 36883_2 | TATGAGTGTGGAGTCGTTGCCGCCATGCACTTCTGTACAAATTNNNAGATCGGAAG |
| 41876_1 | TATGAGTGTGGAGTCGTTGCCTCCTGGGTTCACACCATTCTCNNNAGATCGGAAGA |
| 41876_2 | TATGAGTGTGGAGTCGTTGCCTCCTGGGTTCACACCATTCTCNNNAGATCGGAAGA |
| 41870_1 | TATGAGTGTGGAGTCGTTGCCCTTTTGGGTTCAAGCGATTCTCNNNAGATCGGAAG |
| 41870_2 | TATGAGTGTGGAGTCGTTGCCCTTTTGGGTTCAAGCGATTCTCNNNAGATCGGAAG |
| 30626_1 | TATGAGTGTGGAGTCGTTGCGGGAGGAAGTCATTACACTGGGGNNNAGATCGGA |
| 30626_2 | TATGAGTGTGGAGTCGTTGCGGGAGGAAGTCATTACACTGGGGNNNAGATCGGA |
| 51021_1 | TATGAGTGTGGAGTCGTTGCTTTTGCTGATGTGGGTGAGGATANNNAGATCGGAA |
| 51021_2 | TATGAGTGTGGAGTCGTTGCTTTTGCTGATGTGGGTGAGGATANNNAGATCGGAA |

| | |
|---|---|
| 47279_1 | TATGAGTGTGGAGTCGTTGCGCCATCATGCCCAGCTAATTTTTNNNAGATCGGAAGA |
| 47279_2 | TATGAGTGTGGAGTCGTTGCGCCATCATGCCCAGCTAATTTTTNNNAGATCGGAAGA |
| 36286_1 | TATGAGTGTGGAGTCGTTGCGAGAATGGCATGAACTCGGGAGNNNAGATCGGAAG |
| 36286_2 | TATGAGTGTGGAGTCGTTGCGAGAATGGCATGAACTCGGGAGNNNAGATCGGAAG |
| 46109_1 | TATGAGTGTGGAGTCGTTGCAAAAAAGGAGGAGCATTCACAGCNNNAGATCGGAA |
| 46109_2 | TATGAGTGTGGAGTCGTTGCAAAAAAGGAGGAGCATTCACAGCNNNAGATCGGAA |
| 51375_1 | TATGAGTGTGGAGTCGTTGCCACTGGTTTGTGTTTGAGTCGAANNNAGATCGGAAG |
| 51375_2 | TATGAGTGTGGAGTCGTTGCCACTGGTTTGTGTTTGAGTCGAANNNAGATCGGAAG |
| 48782_1 | TATGAGTGTGGAGTCGTTGCTCCTTTTTTGGTGTCCTCTAAGCTNNNAGATCGGAAG |
| 48782_2 | TATGAGTGTGGAGTCGTTGCTCCTTTTTTGGTGTCCTCTAAGCTNNNAGATCGGAAG |
| 37277_1 | TATGAGTGTGGAGTCGTTGCGCAACAGAGCGAGATTCCATATCNNNAGATCGGAAG |
| 37277_2 | TATGAGTGTGGAGTCGTTGCGCAACAGAGCGAGATTCCATATCNNNAGATCGGAAG |
| 46637_1 | TATGAGTGTGGAGTCGTTGCGATTACAGGCACATGCTACCATGNNNAGATCGGAAG |
| 46637_2 | TATGAGTGTGGAGTCGTTGCGATTACAGGCACATGCTACCATGNNNAGATCGGAAG |
| 49826_1 | TATGAGTGTGGAGTCGTTGCACAGGTAACAGCACGAATACTCANNNAGATCGGAAG |
| 49826_2 | TATGAGTGTGGAGTCGTTGCACAGGTAACAGCACGAATACTCANNNAGATCGGAAG |
| 47812_1 | TATGAGTGTGGAGTCGTTGCATCTCCCAGTTTCAAGCGATTCTNNNAGATCGGAAGA |
| 47812_2 | TATGAGTGTGGAGTCGTTGCATCTCCCAGTTTCAAGCGATTCTNNNAGATCGGAAGA |
| 40893_1 | TATGAGTGTGGAGTCGTTGCTTCAACCCTCATTTGCAATGTCCNNNAGATCGGAAGA |
| 40893_2 | TATGAGTGTGGAGTCGTTGCTTCAACCCTCATTTGCAATGTCCNNNAGATCGGAAGA |
| 3065_1 | TATGAGTGTGGAGTCGTTGCTTCATCACCACCAAGTCCAATGGNNNAGATCGGAAGA |
| 3065_2 | TATGAGTGTGGAGTCGTTGCTTCATCACCACCAAGTCCAATGGNNNAGATCGGAAGA |
| 28764_1 | TATGAGTGTGGAGTCGTTGCCTCCCAGGTTCATGCCATTCTCNNNAGATCGGAAGAG |
| 28764_2 | TATGAGTGTGGAGTCGTTGCCTCCCAGGTTCATGCCATTCTCNNNAGATCGGAAGAG |
| 34516_1 | TATGAGTGTGGAGTCGTTGCACCTCAACCTACGCTAATGAAGGNNNAGATCGGAAG |
| 34516_2 | TATGAGTGTGGAGTCGTTGCACCTCAACCTACGCTAATGAAGGNNNAGATCGGAAG |
| 40885_1 | TATGAGTGTGGAGTCGTTGCAACATGGAGGAATCTGGGAAGACNNNAGATCGGAA |
| 40885_2 | TATGAGTGTGGAGTCGTTGCAACATGGAGGAATCTGGGAAGACNNNAGATCGGAA |
| 10672_1 | TATGAGTGTGGAGTCGTTGCAAAAACCATGCCCAGGAGTACTCNNNAGATCGGAAG |
| 10672_2 | TATGAGTGTGGAGTCGTTGCAAAAACCATGCCCAGGAGTACTCNNNAGATCGGAAG |
| 2162_1 | TATGAGTGTGGAGTCGTTGCAGATGAAATGGGTTATGCCACAGANNNAGATCGGA |
| 2162_2 | TATGAGTGTGGAGTCGTTGCAGATGAAATGGGTTATGCCACAGANNNAGATCGGA |
| 44055_1 | TATGAGTGTGGAGTCGTTGCAGACCACTTACAAGCCTATTGCANNNAGATCGGAAGA |
| 44055_2 | TATGAGTGTGGAGTCGTTGCAGACCACTTACAAGCCTATTGCANNNAGATCGGAAGA |
| 48235_1 | TATGAGTGTGGAGTCGTTGCGATCTCGACTCACTGCAAGCTCNNNAGATCGGAAGA |
| 48235_2 | TATGAGTGTGGAGTCGTTGCGATCTCGACTCACTGCAAGCTCNNNAGATCGGAAGA |
| 49944_1 | TATGAGTGTGGAGTCGTTGCACTGAGTAGGTAGATGTGTCCCANNNAGATCGGAA |
| 49944_2 | TATGAGTGTGGAGTCGTTGCACTGAGTAGGTAGATGTGTCCCANNNAGATCGGAA |
| 39057_1 | TATGAGTGTGGAGTCGTTGCTGAGGCAGGAGAATCACTTGAAANNNAGATCGGAA |
| 39057_2 | TATGAGTGTGGAGTCGTTGCTGAGGCAGGAGAATCACTTGAAANNNAGATCGGAA |
| 41507_1 | TATGAGTGTGGAGTCGTTGCTCTCAAAAAGGGAGGGGTGTATGNNNAGATCGGAA |
| 41507_2 | TATGAGTGTGGAGTCGTTGCTCTCAAAAAGGGAGGGGTGTATGNNNAGATCGGAA |
| 42520_1 | TATGAGTGTGGAGTCGTTGCTTCAAGCGATTCTCCTGACTCAGNNNAGATCGGAAGA |

| | |
|---|---|
| 42520_2 | TATGAGTGTGGAGTCGTTGCTTCAAGCGATTCTCCTGACTCAGNNNAGATCGGAAG |
| 34924_1 | TATGAGTGTGGAGTCGTTGCGCAAAACATCAGGCCTATGTAGGNNNAGATCGGAA |
| 34924_2 | TATGAGTGTGGAGTCGTTGCGCAAAACATCAGGCCTATGTAGGNNNAGATCGGAA |
| 53303_1 | TATGAGTGTGGAGTCGTTGCCACACACACACACACACACATATNNNAGATCGGAAGA |
| 53303_2 | TATGAGTGTGGAGTCGTTGCCACACACACACACACACACATATNNNAGATCGGAAGA |
| 32980_1 | TATGAGTGTGGAGTCGTTGCTTCTGCCCAATGTATGACTCCTCNNNAGATCGGAAGA |
| 32980_2 | TATGAGTGTGGAGTCGTTGCTTCTGCCCAATGTATGACTCCTCNNNAGATCGGAAGA |
| 45000_1 | TATGAGTGTGGAGTCGTTGCAGAAACCTCAGGAAGGAACTCACNNNAGATCGGAA |
| 45000_2 | TATGAGTGTGGAGTCGTTGCAGAAACCTCAGGAAGGAACTCACNNNAGATCGGAA |
| 3141_1 | TATGAGTGTGGAGTCGTTGCGCCTCTGCTGAATGACTAGCTGNNNAGATCGGAAGA |
| 3141_2 | TATGAGTGTGGAGTCGTTGCGCCTCTGCTGAATGACTAGCTGNNNAGATCGGAAGA |
| 45991_1 | TATGAGTGTGGAGTCGTTGCCAGGCTTGTCTCGAACTTCTAGTNNNAGATCGGAAG |
| 45991_2 | TATGAGTGTGGAGTCGTTGCCAGGCTTGTCTCGAACTTCTAGTNNNAGATCGGAAG |
| 39666_1 | TATGAGTGTGGAGTCGTTGCGAATTCCCTGAGCACCTCCTAGNNNAGATCGGAAGA |
| 39666_2 | TATGAGTGTGGAGTCGTTGCGAATTCCCTGAGCACCTCCTAGNNNAGATCGGAAGA |
| 13291_1 | TATGAGTGTGGAGTCGTTGCATGGCCCTTTAACTTATTAAAAANNNAGATCGGAAG |
| 13291_2 | TATGAGTGTGGAGTCGTTGCATGGCCCTTTAACTTATTAAAAANNNAGATCGGAAG |
| 39492_1 | TATGAGTGTGGAGTCGTTGCAAGTTGTGTGAAGGCAGAAACTGNNNAGATCGGAA |
| 39492_2 | TATGAGTGTGGAGTCGTTGCAAGTTGTGTGAAGGCAGAAACTGNNNAGATCGGAA |
| 38714_1 | TATGAGTGTGGAGTCGTTGCGTGCCATGACTTTTTGTTGCATGNNNAGATCGGAAG |
| 38714_2 | TATGAGTGTGGAGTCGTTGCGTGCCATGACTTTTTGTTGCATGNNNAGATCGGAAG |
| 37537_1 | TATGAGTGTGGAGTCGTTGCTCAAAAGATACCAAGGTCAGAGGTNNNAGATCGGA |
| 37537_2 | TATGAGTGTGGAGTCGTTGCTCAAAAGATACCAAGGTCAGAGGTNNNAGATCGGA |
| 34219_1 | TATGAGTGTGGAGTCGTTGCTGCCAACAAAAAAAGTCCAGGACNNNAGATCGGAA |
| 34219_2 | TATGAGTGTGGAGTCGTTGCTGCCAACAAAAAAAGTCCAGGACNNNAGATCGGAA |
| 49686_1 | TATGAGTGTGGAGTCGTTGCTTGGAATTGGCACCTTTGATTCCNNNAGATCGGAAG |
| 49686_2 | TATGAGTGTGGAGTCGTTGCTTGGAATTGGCACCTTTGATTCCNNNAGATCGGAAG |
| 51834_1 | TATGAGTGTGGAGTCGTTGCGGGCTCCTCACTTCCCAGTANNNAGATCGGAAGAGC |
| 51834_2 | TATGAGTGTGGAGTCGTTGCGGGCTCCTCACTTCCCAGTANNNAGATCGGAAGAGC |
| 30346_1 | TATGAGTGTGGAGTCGTTGCGAAACCTTCCATGGAGTCCAGACNNNAGATCGGAAG |
| 30346_2 | TATGAGTGTGGAGTCGTTGCGAAACCTTCCATGGAGTCCAGACNNNAGATCGGAAG |
| 39686_1 | TATGAGTGTGGAGTCGTTGCCTTGTAGTCCCAGCTACGTGAAANNNAGATCGGAAG |
| 39686_2 | TATGAGTGTGGAGTCGTTGCCTTGTAGTCCCAGCTACGTGAAANNNAGATCGGAAG |
| 10183_1 | TATGAGTGTGGAGTCGTTGCCCCATCCACTCTCCATACCAGATNNNAGATCGGAAGA |
| 10183_2 | TATGAGTGTGGAGTCGTTGCCCCATCCACTCTCCATACCAGATNNNAGATCGGAAGA |
| 53174_1 | TATGAGTGTGGAGTCGTTGCGCTTTGAAGGATCACGCAGTTTANNNAGATCGGAA |
| 53174_2 | TATGAGTGTGGAGTCGTTGCGCTTTGAAGGATCACGCAGTTTANNNAGATCGGAA |
| 32365_1 | TATGAGTGTGGAGTCGTTGCTTATGCTTTTTGAAGGCATCCGTNNNAGATCGGAAG |
| 32365_2 | TATGAGTGTGGAGTCGTTGCTTATGCTTTTTGAAGGCATCCGTNNNAGATCGGAAG |
| 41299_1 | TATGAGTGTGGAGTCGTTGCTCAAGGTATATAAACACATGCAGGANNNAGATCGG |
| 41299_2 | TATGAGTGTGGAGTCGTTGCTCAAGGTATATAAACACATGCAGGANNNAGATCGG |
| 49325_1 | TATGAGTGTGGAGTCGTTGCAAGTATTGCTGGGATTACAGGCANNNAGATCGGAA |
| 49325_2 | TATGAGTGTGGAGTCGTTGCAAGTATTGCTGGGATTACAGGCANNNAGATCGGAA |

| | |
|---|---|
| 34378_1 | TATGAGTGTGGAGTCGTTGCGGGAGAGGGAGAGCTCTTTTTTTNNNAGATCGGAA |
| 34378_2 | TATGAGTGTGGAGTCGTTGCGGGAGAGGGAGAGCTCTTTTTTTNNNAGATCGGAA |
| 36391_1 | TATGAGTGTGGAGTCGTTGCCTCCAATGGCTCCTTCTCTTTCTNNNAGATCGGAAGA |
| 36391_2 | TATGAGTGTGGAGTCGTTGCCTCCAATGGCTCCTTCTCTTTCTNNNAGATCGGAAGA |
| 50212_1 | TATGAGTGTGGAGTCGTTGCTGTGTGCCAGATACCATTCTAGGNNNAGATCGGAAG |
| 50212_2 | TATGAGTGTGGAGTCGTTGCTGTGTGCCAGATACCATTCTAGGNNNAGATCGGAAG |
| 44602_1 | TATGAGTGTGGAGTCGTTGCGAAACTGTGGAAACGTTGGTGTGNNNAGATCGGAA |
| 44602_2 | TATGAGTGTGGAGTCGTTGCGAAACTGTGGAAACGTTGGTGTGNNNAGATCGGAA |
| 44066_1 | TATGAGTGTGGAGTCGTTGCATGAGTTTTGGGAAGGGATGTCANNNAGATCGGAA |
| 44066_2 | TATGAGTGTGGAGTCGTTGCATGAGTTTTGGGAAGGGATGTCANNNAGATCGGAA |
| 33537_1 | TATGAGTGTGGAGTCGTTGCGATGGACTTACTCTATCGCCCAGNNNAGATCGGAAG |
| 33537_2 | TATGAGTGTGGAGTCGTTGCGATGGACTTACTCTATCGCCCAGNNNAGATCGGAAG |
| 39278_1 | TATGAGTGTGGAGTCGTTGCTGGCTACACAACAAAAAGCAACANNNAGATCGGAAG |
| 39278_2 | TATGAGTGTGGAGTCGTTGCTGGCTACACAACAAAAAGCAACANNNAGATCGGAAG |
| 49471_1 | TATGAGTGTGGAGTCGTTGCAACTGACCATAAATGTAAGGGCTNNNAGATCGGAA |
| 49471_2 | TATGAGTGTGGAGTCGTTGCAACTGACCATAAATGTAAGGGCTNNNAGATCGGAA |
| 47387_1 | TATGAGTGTGGAGTCGTTGCGGGCTAAGGAAAACGGAAATCACNNNAGATCGGAA |
| 47387_2 | TATGAGTGTGGAGTCGTTGCGGGCTAAGGAAAACGGAAATCACNNNAGATCGGAA |
| 44390_1 | TATGAGTGTGGAGTCGTTGCGGGCTTCTTTGGGTGATTTCAAANNNAGATCGGAAG |
| 44390_2 | TATGAGTGTGGAGTCGTTGCGGGCTTCTTTGGGTGATTTCAAANNNAGATCGGAAG |
| 43319_1 | TATGAGTGTGGAGTCGTTGCTAGAGGGGTTTCCATTCTGTTGGNNNAGATCGGAA |
| 43319_2 | TATGAGTGTGGAGTCGTTGCTAGAGGGGTTTCCATTCTGTTGGNNNAGATCGGAA |
| 39472_1 | TATGAGTGTGGAGTCGTTGCATGCCTGTAATCCCACCAGTTTNNNAGATCGGAAGA |
| 39472_2 | TATGAGTGTGGAGTCGTTGCATGCCTGTAATCCCACCAGTTTNNNAGATCGGAAGA |
| 32873_1 | TATGAGTGTGGAGTCGTTGCTCTGGCTTATATGGTTTCTGCTGANNNAGATCGGAA |
| 32873_2 | TATGAGTGTGGAGTCGTTGCTCTGGCTTATATGGTTTCTGCTGANNNAGATCGGAA |
| 39136_1 | TATGAGTGTGGAGTCGTTGCAGGCTGAATAGTTGTCTTGGATCANNNAGATCGGA |
| 39136_2 | TATGAGTGTGGAGTCGTTGCAGGCTGAATAGTTGTCTTGGATCANNNAGATCGGA |
| 45626_1 | TATGAGTGTGGAGTCGTTGCCTCAGCCTTTCAAGTAGTTGGGANNNAGATCGGAAG |
| 45626_2 | TATGAGTGTGGAGTCGTTGCCTCAGCCTTTCAAGTAGTTGGGANNNAGATCGGAAG |
| 52489_1 | TATGAGTGTGGAGTCGTTGCTGCTAGGGTAAGTTTCTCACCACNNNAGATCGGAAG |
| 52489_2 | TATGAGTGTGGAGTCGTTGCTGCTAGGGTAAGTTTCTCACCACNNNAGATCGGAAG |
| 44605_1 | TATGAGTGTGGAGTCGTTGCGTCCCTGTGCTTCTCGGTAGNNNAGATCGGAAGAG |
| 44605_2 | TATGAGTGTGGAGTCGTTGCGTCCCTGTGCTTCTCGGTAGNNNAGATCGGAAGAG |
| 37113_1 | TATGAGTGTGGAGTCGTTGCAATGAACAGGCAACCTACAGGATNNNAGATCGGAA |
| 37113_2 | TATGAGTGTGGAGTCGTTGCAATGAACAGGCAACCTACAGGATNNNAGATCGGAA |
| 48192_1 | TATGAGTGTGGAGTCGTTGCCAGGAAGCTGAGAAGCCAAGGNNNAGATCGGAAG |
| 48192_2 | TATGAGTGTGGAGTCGTTGCCAGGAAGCTGAGAAGCCAAGGNNNAGATCGGAAG |
| 9192_1 | TATGAGTGTGGAGTCGTTGCAGGCAGCAGATGTGAACTGAAATNNNAGATCGGAA |
| 9192_2 | TATGAGTGTGGAGTCGTTGCAGGCAGCAGATGTGAACTGAAATNNNAGATCGGAA |
| 36427_1 | TATGAGTGTGGAGTCGTTGCTGAGGTCTTACCATGTTGTCCAGNNNAGATCGGAAG |
| 36427_2 | TATGAGTGTGGAGTCGTTGCTGAGGTCTTACCATGTTGTCCAGNNNAGATCGGAAG |
| 13270_1 | TATGAGTGTGGAGTCGTTGCGTGTTATAGCAGTTTTGCTGNNNAGATCGGAAGAG |

| | |
|---|---|
| 13270_2 | TATGAGTGTGGAGTCGTTGCGTGTTATAGCAGTTTTGCTGNNNAGATCGGAAGAG |
| 44699_1 | TATGAGTGTGGAGTCGTTGCCTGCTTCTTAGTCTGCACTACCANNNAGATCGGAAGA |
| 44699_2 | TATGAGTGTGGAGTCGTTGCCTGCTTCTTAGTCTGCACTACCANNNAGATCGGAAGA |
| 51194_1 | TATGAGTGTGGAGTCGTTGCAGTAGGGAGACTTCTGAATTGGCNNNAGATCGGAA |
| 51194_2 | TATGAGTGTGGAGTCGTTGCAGTAGGGAGACTTCTGAATTGGCNNNAGATCGGAA |
| 43933_1 | TATGAGTGTGGAGTCGTTGCAACCTGCTCATGTGTATGTCTGTNNNAGATCGGAAG |
| 43933_2 | TATGAGTGTGGAGTCGTTGCAACCTGCTCATGTGTATGTCTGTNNNAGATCGGAAG |
| 11567_1 | TATGAGTGTGGAGTCGTTGCGGGTCCTACAGAACAAAGACAAGGNNNAGATCGGA |
| 11567_2 | TATGAGTGTGGAGTCGTTGCGGGTCCTACAGAACAAAGACAAGGNNNAGATCGGA |
| 3158_1 | TATGAGTGTGGAGTCGTTGCTCAACACCTACGAAGGAAACATCANNNAGATCGGAA |
| 3158_2 | TATGAGTGTGGAGTCGTTGCTCAACACCTACGAAGGAAACATCANNNAGATCGGAA |
| 38868_1 | TATGAGTGTGGAGTCGTTGCAATATGCCCACCTGAGGAACAATNNNAGATCGGAAG |
| 38868_2 | TATGAGTGTGGAGTCGTTGCAATATGCCCACCTGAGGAACAATNNNAGATCGGAAG |
| 49765_1 | TATGAGTGTGGAGTCGTTGCTGCTATCCAGGACTGAAGAACACNNNAGATCGGAAG |
| 49765_2 | TATGAGTGTGGAGTCGTTGCTGCTATCCAGGACTGAAGAACACNNNAGATCGGAAG |
| 44721_1 | TATGAGTGTGGAGTCGTTGCGTGGTTCTGTGATTTTCTCCAGCNNNAGATCGGAAG |
| 44721_2 | TATGAGTGTGGAGTCGTTGCGTGGTTCTGTGATTTTCTCCAGCNNNAGATCGGAAG |
| 8824_1 | TATGAGTGTGGAGTCGTTGCAAGAGAGGAAGCCAGAAAGAAATTNNNAGATCGG |
| 8824_2 | TATGAGTGTGGAGTCGTTGCAAGAGAGGAAGCCAGAAAGAAATTNNNAGATCGG |
| 47004_1 | TATGAGTGTGGAGTCGTTGCGCAAGTAGCTTTGGTTTTTGACTNNNAGATCGGAAG |
| 47004_2 | TATGAGTGTGGAGTCGTTGCGCAAGTAGCTTTGGTTTTTGACTNNNAGATCGGAAG |
| 50338_1 | TATGAGTGTGGAGTCGTTGCGCTGAAACTGGAGTAAGTTCTGCNNNAGATCGGAA |
| 50338_2 | TATGAGTGTGGAGTCGTTGCGCTGAAACTGGAGTAAGTTCTGCNNNAGATCGGAA |
| 44575_1 | TATGAGTGTGGAGTCGTTGCAAGAATCACTTGAACCTGGGAGGNNNAGATCGGAA |
| 44575_2 | TATGAGTGTGGAGTCGTTGCAAGAATCACTTGAACCTGGGAGGNNNAGATCGGAA |
| 46669_1 | TATGAGTGTGGAGTCGTTGCCATGATCTCAGCTCATTGCAACCNNNAGATCGGAAG |
| 46669_2 | TATGAGTGTGGAGTCGTTGCCATGATCTCAGCTCATTGCAACCNNNAGATCGGAAG |
| 32825_1 | TATGAGTGTGGAGTCGTTGCTTCCTGGTTTATTCTTGGGAGGGNNNAGATCGGAAG |
| 32825_2 | TATGAGTGTGGAGTCGTTGCTTCCTGGTTTATTCTTGGGAGGGNNNAGATCGGAAG |
| 33219_1 | TATGAGTGTGGAGTCGTTGCCTCAAGAGCAGAGGTAGGCATGNNNAGATCGGAAG |
| 33219_2 | TATGAGTGTGGAGTCGTTGCCTCAAGAGCAGAGGTAGGCATGNNNAGATCGGAAG |
| 52744_1 | TATGAGTGTGGAGTCGTTGCTTTTAGTAGAGACGGGGCTTCACNNNAGATCGGAA |
| 52744_2 | TATGAGTGTGGAGTCGTTGCTTTTAGTAGAGACGGGGCTTCACNNNAGATCGGAA |
| 36553_1 | TATGAGTGTGGAGTCGTTGCGGAACAAAGTTCCATCAAAGCCTNNNAGATCGGAAG |
| 36553_2 | TATGAGTGTGGAGTCGTTGCGGAACAAAGTTCCATCAAAGCCTNNNAGATCGGAAG |
| 44797_1 | TATGAGTGTGGAGTCGTTGCTAAAAGAGCCAGCTAGCATGGAANNNAGATCGGAA |
| 44797_2 | TATGAGTGTGGAGTCGTTGCTAAAAGAGCCAGCTAGCATGGAANNNAGATCGGAA |
| 32470_1 | TATGAGTGTGGAGTCGTTGCCAGATCACGAGGTCAGGAGATCNNNAGATCGGAAG |
| 32470_2 | TATGAGTGTGGAGTCGTTGCCAGATCACGAGGTCAGGAGATCNNNAGATCGGAAG |
| 51885_1 | TATGAGTGTGGAGTCGTTGCCTTCTGCAGGAGAGGGTAAGAAGNNNAGATCGGAA |
| 51885_2 | TATGAGTGTGGAGTCGTTGCCTTCTGCAGGAGAGGGTAAGAAGNNNAGATCGGAA |
| 42731_1 | TATGAGTGTGGAGTCGTTGCTCTGTAGCAACATGGATGGAACTNNNAGATCGGAA |
| 42731_2 | TATGAGTGTGGAGTCGTTGCTCTGTAGCAACATGGATGGAACTNNNAGATCGGAA |

| | |
|---|---|
| 13410_1 | TATGAGTGTGGAGTCGTTGCTGCCTTGTTTTTAAGTAATACAANNNAGATCGGAAG |
| 13410_2 | TATGAGTGTGGAGTCGTTGCTGCCTTGTTTTTAAGTAATACAANNNAGATCGGAAG |
| 47739_1 | TATGAGTGTGGAGTCGTTGCCAAGTAGCTGGGACTACAGGCNNNAGATCGGAAGA |
| 47739_2 | TATGAGTGTGGAGTCGTTGCCAAGTAGCTGGGACTACAGGCNNNAGATCGGAAGA |
| 12903_1 | TATGAGTGTGGAGTCGTTGCTTGGTCACTTTCATTATAAGAAANNNAGATCGGAAG |
| 12903_2 | TATGAGTGTGGAGTCGTTGCTTGGTCACTTTCATTATAAGAAANNNAGATCGGAAG |
| 33097_1 | TATGAGTGTGGAGTCGTTGCCTCCTGAGTAGCTGGGATTACAGNNNAGATCGGAA |
| 33097_2 | TATGAGTGTGGAGTCGTTGCCTCCTGAGTAGCTGGGATTACAGNNNAGATCGGAA |
| 37889_1 | TATGAGTGTGGAGTCGTTGCCATGATGGTGTGTGCCTGTAATCNNNAGATCGGAA |
| 37889_2 | TATGAGTGTGGAGTCGTTGCCATGATGGTGTGTGCCTGTAATCNNNAGATCGGAA |
| 10158_1 | TATGAGTGTGGAGTCGTTGCACTTTCCCCTTTTGGATTTGACCANNNAGATCGGAAG |
| 10158_2 | TATGAGTGTGGAGTCGTTGCACTTTCCCCTTTTGGATTTGACCANNNAGATCGGAAG |
| 42201_1 | TATGAGTGTGGAGTCGTTGCGGTAAATAGTCTCAGCTCCTCGGNNNAGATCGGAAG |
| 42201_2 | TATGAGTGTGGAGTCGTTGCGGTAAATAGTCTCAGCTCCTCGGNNNAGATCGGAAG |
| 31799_1 | TATGAGTGTGGAGTCGTTGCGTTCAAGCAATTCTCATGCCTCANNNAGATCGGAAG |
| 31799_2 | TATGAGTGTGGAGTCGTTGCGTTCAAGCAATTCTCATGCCTCANNNAGATCGGAAG |
| 51443_1 | TATGAGTGTGGAGTCGTTGCAGCAAAAGGGGACTTTGTTTACGNNNAGATCGGAA |
| 51443_2 | TATGAGTGTGGAGTCGTTGCAGCAAAAGGGGACTTTGTTTACGNNNAGATCGGAA |
| 32042_1 | TATGAGTGTGGAGTCGTTGCTTCAAGTGATTCTCCTGCCTCAGNNNAGATCGGAAG |
| 32042_2 | TATGAGTGTGGAGTCGTTGCTTCAAGTGATTCTCCTGCCTCAGNNNAGATCGGAAG |
| 12538_1 | TATGAGTGTGGAGTCGTTGCGTAAAATGACAAATACTGAAGGTNNNAGATCGGAA |
| 12538_2 | TATGAGTGTGGAGTCGTTGCGTAAAATGACAAATACTGAAGGTNNNAGATCGGAA |
| 51513_1 | TATGAGTGTGGAGTCGTTGCCTCCCTAGGGAATGTTTTAGGCANNNAGATCGGAAG |
| 51513_2 | TATGAGTGTGGAGTCGTTGCCTCCCTAGGGAATGTTTTAGGCANNNAGATCGGAAG |
| 35838_1 | TATGAGTGTGGAGTCGTTGCCTGAGTAGCTGGGATTACAGGTGNNNAGATCGGAA |
| 35838_2 | TATGAGTGTGGAGTCGTTGCCTGAGTAGCTGGGATTACAGGTGNNNAGATCGGAA |
| 46115_1 | TATGAGTGTGGAGTCGTTGCTTCTACCTCCTAGGCTCAAGTGANNNAGATCGGAAG |
| 46115_2 | TATGAGTGTGGAGTCGTTGCTTCTACCTCCTAGGCTCAAGTGANNNAGATCGGAAG |
| 5627_1 | TATGAGTGTGGAGTCGTTGCTCGCATCCCCACACTATTTACTTNNNAGATCGGAAGA |
| 5627_2 | TATGAGTGTGGAGTCGTTGCTCGCATCCCCACACTATTTACTTNNNAGATCGGAAGA |
| 10364_1 | TATGAGTGTGGAGTCGTTGCAGCCAGGGTTAGTTTTGCAATCANNNAGATCGGAA |
| 10364_2 | TATGAGTGTGGAGTCGTTGCAGCCAGGGTTAGTTTTGCAATCANNNAGATCGGAA |
| 40386_1 | TATGAGTGTGGAGTCGTTGCAGATGGTGTTTCAACATGTTGGCNNNAGATCGGAA |
| 40386_2 | TATGAGTGTGGAGTCGTTGCAGATGGTGTTTCAACATGTTGGCNNNAGATCGGAA |
| 30421_1 | TATGAGTGTGGAGTCGTTGCTCCTGTGTGTGCGTAGGTGTATANNNAGATCGGAA |
| 30421_2 | TATGAGTGTGGAGTCGTTGCTCCTGTGTGTGCGTAGGTGTATANNNAGATCGGAA |
| 49056_1 | TATGAGTGTGGAGTCGTTGCCCTTAGCCTCTTGAGTATGGGACNNNAGATCGGAAG |
| 49056_2 | TATGAGTGTGGAGTCGTTGCCCTTAGCCTCTTGAGTATGGGACNNNAGATCGGAAG |
| 52733_1 | TATGAGTGTGGAGTCGTTGCCTGCACCCACTGAACTCCAGNNNAGATCGGAAGAGC |
| 52733_2 | TATGAGTGTGGAGTCGTTGCCTGCACCCACTGAACTCCAGNNNAGATCGGAAGAGC |
| 38498_1 | TATGAGTGTGGAGTCGTTGCAGATAGGATCTTGCTATGCTGTCCNNNAGATCGGAA |
| 38498_2 | TATGAGTGTGGAGTCGTTGCAGATAGGATCTTGCTATGCTGTCCNNNAGATCGGAA |
| 28997_1 | TATGAGTGTGGAGTCGTTGCTCAGAACAGGAGGTGATAGTCCTNNNAGATCGGAA |

| | |
|---|---|
| 28997_2 | TATGAGTGTGGAGTCGTTGCTCAGAACAGGAGGTGATAGTCCTNNNAGATCGGAA |
| 30514_1 | TATGAGTGTGGAGTCGTTGCGTGGGGAAAAGTGAGGAAAGGAGNNNAGATCGG |
| 30514_2 | TATGAGTGTGGAGTCGTTGCGTGGGGAAAAGTGAGGAAAGGAGNNNAGATCGG |
| 44057_1 | TATGAGTGTGGAGTCGTTGCAAAAGGCATCTAGGAGGGGAAAGNNNAGATCGGA |
| 44057_2 | TATGAGTGTGGAGTCGTTGCAAAAGGCATCTAGGAGGGGAAAGNNNAGATCGGA |
| 13565_1 | TATGAGTGTGGAGTCGTTGCGCTCCTTATGAGAATCTGATGNNNAGATCGGAAGA |
| 13565_2 | TATGAGTGTGGAGTCGTTGCGCTCCTTATGAGAATCTGATGNNNAGATCGGAAGA |
| 48508_1 | TATGAGTGTGGAGTCGTTGCGAGCCATCTCATCACCCACTATTNNNAGATCGGAAG |
| 48508_2 | TATGAGTGTGGAGTCGTTGCGAGCCATCTCATCACCCACTATTNNNAGATCGGAAG |
| 30055_1 | TATGAGTGTGGAGTCGTTGCGACTTGTCCCAGGTTAATGGCTANNNAGATCGGAAG |
| 30055_2 | TATGAGTGTGGAGTCGTTGCGACTTGTCCCAGGTTAATGGCTANNNAGATCGGAAG |
| 35938_1 | TATGAGTGTGGAGTCGTTGCGTGTGTGTCTTCTTGTGCTTCTCNNNAGATCGGAAG |
| 35938_2 | TATGAGTGTGGAGTCGTTGCGTGTGTGTCTTCTTGTGCTTCTCNNNAGATCGGAAG |
| 52993_1 | TATGAGTGTGGAGTCGTTGCGCTCTCTCTCTTTGCCTCTTTCTNNNAGATCGGAAGA |
| 52993_2 | TATGAGTGTGGAGTCGTTGCGCTCTCTCTCTTTGCCTCTTTCTNNNAGATCGGAAGA |
| 35250_1 | TATGAGTGTGGAGTCGTTGCTGAACTACTGACCTCAGGCAATCNNNAGATCGGAAG |
| 35250_2 | TATGAGTGTGGAGTCGTTGCTGAACTACTGACCTCAGGCAATCNNNAGATCGGAAG |
| 35665_1 | TATGAGTGTGGAGTCGTTGCTGCCCAGCTACAAATATCAGTGGNNNAGATCGGAAG |
| 35665_2 | TATGAGTGTGGAGTCGTTGCTGCCCAGCTACAAATATCAGTGGNNNAGATCGGAAG |
| 52048_1 | TATGAGTGTGGAGTCGTTGCCTTGGGAGAAAAGTGCATCTTGGNNNAGATCGGAA |
| 52048_2 | TATGAGTGTGGAGTCGTTGCCTTGGGAGAAAAGTGCATCTTGGNNNAGATCGGAA |
| 45928_1 | TATGAGTGTGGAGTCGTTGCGCCTCAAACTCCAAAGCTCAAATNNNAGATCGGAAG |
| 45928_2 | TATGAGTGTGGAGTCGTTGCGCCTCAAACTCCAAAGCTCAAATNNNAGATCGGAAG |
| 50147_1 | TATGAGTGTGGAGTCGTTGCCCTTCTCATTGACCTCAAGGGAANNNAGATCGGAAG |
| 50147_2 | TATGAGTGTGGAGTCGTTGCCCTTCTCATTGACCTCAAGGGAANNNAGATCGGAAG |
| 30370_1 | TATGAGTGTGGAGTCGTTGCATAAAGCTCACCCATCGCCAAATNNNAGATCGGAAG |
| 30370_2 | TATGAGTGTGGAGTCGTTGCATAAAGCTCACCCATCGCCAAATNNNAGATCGGAAG |
| 48073_1 | TATGAGTGTGGAGTCGTTGCCTCCTGAGTAGCTGGGATTACAGNNNAGATCGGAA |
| 48073_2 | TATGAGTGTGGAGTCGTTGCCTCCTGAGTAGCTGGGATTACAGNNNAGATCGGAA |
| 45255_1 | TATGAGTGTGGAGTCGTTGCGTGGGGGGAAGAAAGGAGAATAANNNAGATCGG |
| 45255_2 | TATGAGTGTGGAGTCGTTGCGTGGGGGGAAGAAAGGAGAATAANNNAGATCGG |
| 36178_1 | TATGAGTGTGGAGTCGTTGCTTCTCAAGAGCCTGCAATCATCTNNNAGATCGGAAG |
| 36178_2 | TATGAGTGTGGAGTCGTTGCTTCTCAAGAGCCTGCAATCATCTNNNAGATCGGAAG |
| 42942_1 | TATGAGTGTGGAGTCGTTGCTGGGGGTGCAGATATTTCTTCAANNNAGATCGGAA |
| 42942_2 | TATGAGTGTGGAGTCGTTGCTGGGGGTGCAGATATTTCTTCAANNNAGATCGGAA |
| 37913_1 | TATGAGTGTGGAGTCGTTGCGTATCTCTGCCCTCCATCTGAAGNNNAGATCGGAAG |
| 37913_2 | TATGAGTGTGGAGTCGTTGCGTATCTCTGCCCTCCATCTGAAGNNNAGATCGGAAG |
| 36756_1 | TATGAGTGTGGAGTCGTTGCATAAAGAGATTGTGCTCCCTCCCNNNAGATCGGAAG |
| 36756_2 | TATGAGTGTGGAGTCGTTGCATAAAGAGATTGTGCTCCCTCCCNNNAGATCGGAAG |
| 51087_1 | TATGAGTGTGGAGTCGTTGCAGCTGCCAACAATATACTGGTGTNNNAGATCGGAAG |
| 51087_2 | TATGAGTGTGGAGTCGTTGCAGCTGCCAACAATATACTGGTGTNNNAGATCGGAAG |
| 38960_1 | TATGAGTGTGGAGTCGTTGCATGTCTGGAATGAACCCCACTTTNNNAGATCGGAAG |
| 38960_2 | TATGAGTGTGGAGTCGTTGCATGTCTGGAATGAACCCCACTTTNNNAGATCGGAAG |

| | |
|---|---|
| 41961_1 | TATGAGTGTGGAGTCGTTGCCAATTCTCCTATCTCAGCCTCCCNNNAGATCGGAAGA |
| 41961_2 | TATGAGTGTGGAGTCGTTGCCAATTCTCCTATCTCAGCCTCCCNNNAGATCGGAAGA |
| 12869_1 | TATGAGTGTGGAGTCGTTGCAATAAGTGCTTAAATGTCTTTGGNNNAGATCGGAAG |
| 12869_2 | TATGAGTGTGGAGTCGTTGCAATAAGTGCTTAAATGTCTTTGGNNNAGATCGGAAG |
| 52721_1 | TATGAGTGTGGAGTCGTTGCGTCTTTCTGAGCCACAGGTATCCNNNAGATCGGAAG |
| 52721_2 | TATGAGTGTGGAGTCGTTGCGTCTTTCTGAGCCACAGGTATCCNNNAGATCGGAAG |
| 34913_1 | TATGAGTGTGGAGTCGTTGCCTCCTGAGTAGCTGGGATTACAGNNNAGATCGGAA |
| 34913_2 | TATGAGTGTGGAGTCGTTGCCTCCTGAGTAGCTGGGATTACAGNNNAGATCGGAA |
| 36811_1 | TATGAGTGTGGAGTCGTTGCTATGTGGGACTATTGCATGATGCNNNAGATCGGAA |
| 36811_2 | TATGAGTGTGGAGTCGTTGCTATGTGGGACTATTGCATGATGCNNNAGATCGGAA |
| 2188_1 | TATGAGTGTGGAGTCGTTGCAGATGTTACTGTGTTTTGGGGGANNNAGATCGGAA |
| 2188_2 | TATGAGTGTGGAGTCGTTGCAGATGTTACTGTGTTTTGGGGGANNNAGATCGGAA |
| 51670_1 | TATGAGTGTGGAGTCGTTGCATCTGGTTACCACAGAAAGTCCCNNNAGATCGGAAG |
| 51670_2 | TATGAGTGTGGAGTCGTTGCATCTGGTTACCACAGAAAGTCCCNNNAGATCGGAAG |
| 34780_1 | TATGAGTGTGGAGTCGTTGCTTCAGAGGCGTTTAGTACATGGGNNNAGATCGGAA |
| 34780_2 | TATGAGTGTGGAGTCGTTGCTTCAGAGGCGTTTAGTACATGGGNNNAGATCGGAA |
| 50078_1 | TATGAGTGTGGAGTCGTTGCAGAGACCATGGCTCAAAAGTACTNNNAGATCGGAA |
| 50078_2 | TATGAGTGTGGAGTCGTTGCAGAGACCATGGCTCAAAAGTACTNNNAGATCGGAA |
| 42252_1 | TATGAGTGTGGAGTCGTTGCGAGACACAGACAGACAGACAAGANNNAGATCGGAA |
| 42252_2 | TATGAGTGTGGAGTCGTTGCGAGACACAGACAGACAGACAAGANNNAGATCGGAA |
| 33270_1 | TATGAGTGTGGAGTCGTTGCCTCCCGAGTTCAAGCAGTTCTCNNNAGATCGGAAGA |
| 33270_2 | TATGAGTGTGGAGTCGTTGCCTCCCGAGTTCAAGCAGTTCTCNNNAGATCGGAAGA |
| 42893_1 | TATGAGTGTGGAGTCGTTGCTTGGAGTGCAGTGACATGATCTCNNNAGATCGGAA |
| 42893_2 | TATGAGTGTGGAGTCGTTGCTTGGAGTGCAGTGACATGATCTCNNNAGATCGGAA |
| 44161_1 | TATGAGTGTGGAGTCGTTGCGGGAAGCGACTAATACAAGAGGANNNAGATCGGA |
| 44161_2 | TATGAGTGTGGAGTCGTTGCGGGAAGCGACTAATACAAGAGGANNNAGATCGGA |
| 30828_1 | TATGAGTGTGGAGTCGTTGCGTTGGGAAAGGAGAGATTGGGATNNNAGATCGGA |
| 30828_2 | TATGAGTGTGGAGTCGTTGCGTTGGGAAAGGAGAGATTGGGATNNNAGATCGGA |
| 31866_1 | TATGAGTGTGGAGTCGTTGCAAAAGGAAACAGGACGGTCTGATNNNAGATCGGAA |
| 31866_2 | TATGAGTGTGGAGTCGTTGCAAAAGGAAACAGGACGGTCTGATNNNAGATCGGAA |
| 44273_1 | TATGAGTGTGGAGTCGTTGCCAGGGCAACTGGTTACTAGACATNNNAGATCGGAA |
| 44273_2 | TATGAGTGTGGAGTCGTTGCCAGGGCAACTGGTTACTAGACATNNNAGATCGGAA |
| 43645_1 | TATGAGTGTGGAGTCGTTGCCGAAGAGGAAGGAACTCTGAGACNNNAGATCGGA |
| 43645_2 | TATGAGTGTGGAGTCGTTGCCGAAGAGGAAGGAACTCTGAGACNNNAGATCGGA |
| 49984_1 | TATGAGTGTGGAGTCGTTGCCCACAACTTCTCCCTCCGAAAAGNNNAGATCGGAAG |
| 49984_2 | TATGAGTGTGGAGTCGTTGCCCACAACTTCTCCCTCCGAAAAGNNNAGATCGGAAG |
| 31767_1 | TATGAGTGTGGAGTCGTTGCTCTCTCTCTCTCTCTGTGTGTGTNNNAGATCGGAAGA |
| 31767_2 | TATGAGTGTGGAGTCGTTGCTCTCTCTCTCTCTCTGTGTGTGTNNNAGATCGGAAGA |
| 40726_1 | TATGAGTGTGGAGTCGTTGCATCAAAAGACCTGGGTCAGAGTCNNNAGATCGGAA |
| 40726_2 | TATGAGTGTGGAGTCGTTGCATCAAAAGACCTGGGTCAGAGTCNNNAGATCGGAA |
| 29345_1 | TATGAGTGTGGAGTCGTTGCCTTGGGAGATAAGGGCAGAGAAANNNAGATCGGA |
| 29345_2 | TATGAGTGTGGAGTCGTTGCCTTGGGAGATAAGGGCAGAGAAANNNAGATCGGA |
| 48429_1 | TATGAGTGTGGAGTCGTTGCGGGAGTGAGCAGGTTTTTGTTTTNNNAGATCGGAA |

| | |
|---|---|
| 48429_2 | TATGAGTGTGGAGTCGTTGCGGGAGTGAGCAGGTTTTTGTTTTNNNAGATCGGAA |
| 48615_1 | TATGAGTGTGGAGTCGTTGCCAAGTGATTCTCCTACCTCAGCCNNNAGATCGGAAGA |
| 48615_2 | TATGAGTGTGGAGTCGTTGCCAAGTGATTCTCCTACCTCAGCCNNNAGATCGGAAGA |
| 44327_1 | TATGAGTGTGGAGTCGTTGCAAACTTTCATCTGGAATCGGGGTNNNAGATCGGAAG |
| 44327_2 | TATGAGTGTGGAGTCGTTGCAAACTTTCATCTGGAATCGGGGTNNNAGATCGGAAG |
| 30752_1 | TATGAGTGTGGAGTCGTTGCGGCCATCAAGGTGGTACTTCTATNNNAGATCGGAAG |
| 30752_2 | TATGAGTGTGGAGTCGTTGCGGCCATCAAGGTGGTACTTCTATNNNAGATCGGAAG |
| 49533_1 | TATGAGTGTGGAGTCGTTGCGATGGAGTCTCGCTGTGTCACNNNAGATCGGAAGA |
| 49533_2 | TATGAGTGTGGAGTCGTTGCGATGGAGTCTCGCTGTGTCACNNNAGATCGGAAGA |
| 48141_1 | TATGAGTGTGGAGTCGTTGCCCAAAGTGCTGGAATTACAGGTGNNNAGATCGGAA |
| 48141_2 | TATGAGTGTGGAGTCGTTGCCCAAAGTGCTGGAATTACAGGTGNNNAGATCGGAA |
| 33467_1 | TATGAGTGTGGAGTCGTTGCAAAGAAGAGCTGGTCCCAATTCTNNNAGATCGGAAG |
| 33467_2 | TATGAGTGTGGAGTCGTTGCAAAGAAGAGCTGGTCCCAATTCTNNNAGATCGGAAG |
| 38863_1 | TATGAGTGTGGAGTCGTTGCCACGATGTTGAGCCCAGTTTTACNNNAGATCGGAAG |
| 38863_2 | TATGAGTGTGGAGTCGTTGCCACGATGTTGAGCCCAGTTTTACNNNAGATCGGAAG |
| 35045_1 | TATGAGTGTGGAGTCGTTGCTCTTGAACTTCTGACCTCTGGTGNNNAGATCGGAAG |
| 35045_2 | TATGAGTGTGGAGTCGTTGCTCTTGAACTTCTGACCTCTGGTGNNNAGATCGGAAG |
| 31792_1 | TATGAGTGTGGAGTCGTTGCGCAAAGCACCAAAGACAGAGATTNNNAGATCGGAA |
| 31792_2 | TATGAGTGTGGAGTCGTTGCGCAAAGCACCAAAGACAGAGATTNNNAGATCGGAA |
| 42367_1 | TATGAGTGTGGAGTCGTTGCCGCCCTCATTCTGTCTCTATGAANNNAGATCGGAAGA |
| 42367_2 | TATGAGTGTGGAGTCGTTGCCGCCCTCATTCTGTCTCTATGAANNNAGATCGGAAGA |
| 52644_1 | TATGAGTGTGGAGTCGTTGCTCTCCCTTTCATTGCTCCTTTCANNNAGATCGGAAGA |
| 52644_2 | TATGAGTGTGGAGTCGTTGCTCTCCCTTTCATTGCTCCTTTCANNNAGATCGGAAGA |
| 50105_1 | TATGAGTGTGGAGTCGTTGCGGACTTCTCGCTACATGGGGNNNAGATCGGAAGAG |
| 50105_2 | TATGAGTGTGGAGTCGTTGCGGACTTCTCGCTACATGGGGNNNAGATCGGAAGAG |
| 36559_1 | TATGAGTGTGGAGTCGTTGCACACCTGTGAATGGAGAAGCTAGNNNAGATCGGAA |
| 36559_2 | TATGAGTGTGGAGTCGTTGCACACCTGTGAATGGAGAAGCTAGNNNAGATCGGAA |
| 37825_1 | TATGAGTGTGGAGTCGTTGCCTGAGGCAGGAGAATCACTTGAANNNAGATCGGAA |
| 37825_2 | TATGAGTGTGGAGTCGTTGCCTGAGGCAGGAGAATCACTTGAANNNAGATCGGAA |
| 46563_1 | TATGAGTGTGGAGTCGTTGCTCAACTCCTGGACTCAAGTGATCNNNAGATCGGAAG |
| 46563_2 | TATGAGTGTGGAGTCGTTGCTCAACTCCTGGACTCAAGTGATCNNNAGATCGGAAG |
| 10803_1 | TATGAGTGTGGAGTCGTTGCTGTTCCAGCTTGCTCTCAGAATCNNNAGATCGGAAG |
| 10803_2 | TATGAGTGTGGAGTCGTTGCTGTTCCAGCTTGCTCTCAGAATCNNNAGATCGGAAG |
| 44898_1 | TATGAGTGTGGAGTCGTTGCCAGCCCTTCCTTCCCAGAAAGNNNAGATCGGAAGAG |
| 44898_2 | TATGAGTGTGGAGTCGTTGCCAGCCCTTCCTTCCCAGAAAGNNNAGATCGGAAGAG |
| 40504_1 | TATGAGTGTGGAGTCGTTGCCGTGACTACTTCCTTGCTGTACTNNNAGATCGGAAG |
| 40504_2 | TATGAGTGTGGAGTCGTTGCCGTGACTACTTCCTTGCTGTACTNNNAGATCGGAAG |
| 32022_1 | TATGAGTGTGGAGTCGTTGCTGGGGAAAAGTTGAAAGCATTCCNNNAGATCGGAA |
| 32022_2 | TATGAGTGTGGAGTCGTTGCTGGGGAAAAGTTGAAAGCATTCCNNNAGATCGGAA |
| 51618_1 | TATGAGTGTGGAGTCGTTGCCACAGTTTATTCCTCCAGTCCCTNNNAGATCGGAAGA |
| 51618_2 | TATGAGTGTGGAGTCGTTGCCACAGTTTATTCCTCCAGTCCCTNNNAGATCGGAAGA |
| 10916_1 | TATGAGTGTGGAGTCGTTGCAGTGCCTTTGGAGGAGAGTATGATGTGNNNAGATC |
| 10916_2 | TATGAGTGTGGAGTCGTTGCAGTGCCTTTGGAGGAGAGTATGATGTGNNNAGATC |

| | |
|---|---|
| 46320_1 | TATGAGTGTGGAGTCGTTGCAGTCAGTCTCAGTCTGTCACCTANNNAGATCGGAAG |
| 46320_2 | TATGAGTGTGGAGTCGTTGCAGTCAGTCTCAGTCTGTCACCTANNNAGATCGGAAG |
| 48878_1 | TATGAGTGTGGAGTCGTTGCCCCGAGTAGCTGAGATTATAGGCNNNAGATCGGAA |
| 48878_2 | TATGAGTGTGGAGTCGTTGCCCCGAGTAGCTGAGATTATAGGCNNNAGATCGGAA |
| 44279_1 | TATGAGTGTGGAGTCGTTGCCATGGCCCAATTGAAGAACAACTNNNAGATCGGAAG |
| 44279_2 | TATGAGTGTGGAGTCGTTGCCATGGCCCAATTGAAGAACAACTNNNAGATCGGAAG |
| 37310_1 | TATGAGTGTGGAGTCGTTGCATGACCCTGACTCATCCAATACCNNNAGATCGGAAG |
| 37310_2 | TATGAGTGTGGAGTCGTTGCATGACCCTGACTCATCCAATACCNNNAGATCGGAAG |
| 37758_1 | TATGAGTGTGGAGTCGTTGCACGATCATGGGTCACTGCAGNNNAGATCGGAAGAG |
| 37758_2 | TATGAGTGTGGAGTCGTTGCACGATCATGGGTCACTGCAGNNNAGATCGGAAGAG |
| 50047_1 | TATGAGTGTGGAGTCGTTGCTGGTGTCTCTTCTACTTCCCGTANNNAGATCGGAAG |
| 50047_2 | TATGAGTGTGGAGTCGTTGCTGGTGTCTCTTCTACTTCCCGTANNNAGATCGGAAG |
| 50109_1 | TATGAGTGTGGAGTCGTTGCCTCACCCACATCCAGCTGTTNNNAGATCGGAAGAGC |
| 50109_2 | TATGAGTGTGGAGTCGTTGCCTCACCCACATCCAGCTGTTNNNAGATCGGAAGAGC |
| 43438_1 | TATGAGTGTGGAGTCGTTGCTAGGATTGCCTTGGCTATTCAGGNNNAGATCGGAA |
| 43438_2 | TATGAGTGTGGAGTCGTTGCTAGGATTGCCTTGGCTATTCAGGNNNAGATCGGAA |
| 2449_1 | TATGAGTGTGGAGTCGTTGCGGCACACTCTGGTCTTTCTCTTCNNNAGATCGGAAG |
| 2449_2 | TATGAGTGTGGAGTCGTTGCGGCACACTCTGGTCTTTCTCTTCNNNAGATCGGAAG |
| 12842_1 | TATGAGTGTGGAGTCGTTGCGAATCTCATCCCCTCTCAACNNNAGATCGGAAGAGC |
| 12842_2 | TATGAGTGTGGAGTCGTTGCGAATCTCATCCCCTCTCAACNNNAGATCGGAAGAGC |
| 34754_1 | TATGAGTGTGGAGTCGTTGCTTGAAATCTGGGCAACTGTCTCTNNNAGATCGGAAG |
| 34754_2 | TATGAGTGTGGAGTCGTTGCTTGAAATCTGGGCAACTGTCTCTNNNAGATCGGAAG |
| 39852_1 | TATGAGTGTGGAGTCGTTGCAGGAAAGACCATTGTGATGAGGTNNNAGATCGGAA |
| 39852_2 | TATGAGTGTGGAGTCGTTGCAGGAAAGACCATTGTGATGAGGTNNNAGATCGGAA |
| 4205_1 | TATGAGTGTGGAGTCGTTGCAGGAGCAGGAGTTGAAGTGGATTNNNAGATCGGA |
| 4205_2 | TATGAGTGTGGAGTCGTTGCAGGAGCAGGAGTTGAAGTGGATTNNNAGATCGGA |
| 38008_1 | TATGAGTGTGGAGTCGTTGCGAAGAAGCAGCCAGTCTGAAAAGNNNAGATCGGAA |
| 38008_2 | TATGAGTGTGGAGTCGTTGCGAAGAAGCAGCCAGTCTGAAAAGNNNAGATCGGAA |
| 29078_1 | TATGAGTGTGGAGTCGTTGCGTTCACGGGTTCAAGCTATTCTCNNNAGATCGGAAG |
| 29078_2 | TATGAGTGTGGAGTCGTTGCGTTCACGGGTTCAAGCTATTCTCNNNAGATCGGAAG |
| 36659_1 | TATGAGTGTGGAGTCGTTGCGGCCTTCCTCACCCATGTATATTNNNAGATCGGAAG |
| 36659_2 | TATGAGTGTGGAGTCGTTGCGGCCTTCCTCACCCATGTATATTNNNAGATCGGAAG |
| 30479_1 | TATGAGTGTGGAGTCGTTGCCTGAGGACTCTCAAAGAACAGCTCNNNAGATCGGAA |
| 30479_2 | TATGAGTGTGGAGTCGTTGCCTGAGGACTCTCAAAGAACAGCTCNNNAGATCGGAA |
| 38256_1 | TATGAGTGTGGAGTCGTTGCTGATCTATCGGTTGCAAAGACCANNNAGATCGGAAG |
| 38256_2 | TATGAGTGTGGAGTCGTTGCTGATCTATCGGTTGCAAAGACCANNNAGATCGGAAG |
| 49652_1 | TATGAGTGTGGAGTCGTTGCAGTGCTAGGATTACAGGCATGAGNNNAGATCGGAA |
| 49652_2 | TATGAGTGTGGAGTCGTTGCAGTGCTAGGATTACAGGCATGAGNNNAGATCGGAA |
| 45816_1 | TATGAGTGTGGAGTCGTTGCAGTAGCTGGAATTACAGGCATGTNNNAGATCGGAA |
| 45816_2 | TATGAGTGTGGAGTCGTTGCAGTAGCTGGAATTACAGGCATGTNNNAGATCGGAA |
| 34518_1 | TATGAGTGTGGAGTCGTTGCTTGTGCACTGTAGCAGATAGTGTNNNAGATCGGAA |
| 34518_2 | TATGAGTGTGGAGTCGTTGCTTGTGCACTGTAGCAGATAGTGTNNNAGATCGGAA |
| 3068_1 | TATGAGTGTGGAGTCGTTGCAATCCAGCAGAGAGCTATGAGGTNNNAGATCGGAA |

| | |
|---|---|
| 3068_2 | TATGAGTGTGGAGTCGTTGCAATCCAGCAGAGAGCTATGAGGTNNNAGATCGGAA |
| 43019_1 | TATGAGTGTGGAGTCGTTGCAGCCTACAAACACAGGTAACCAANNNAGATCGGAAG |
| 43019_2 | TATGAGTGTGGAGTCGTTGCAGCCTACAAACACAGGTAACCAANNNAGATCGGAAG |
| 39206_1 | TATGAGTGTGGAGTCGTTGCTTGCTGAGCTGAGAAGTCCTAAGNNNAGATCGGAA |
| 39206_2 | TATGAGTGTGGAGTCGTTGCTTGCTGAGCTGAGAAGTCCTAAGNNNAGATCGGAA |
| 29758_1 | TATGAGTGTGGAGTCGTTGCAAAATGACTGGGTGTTGAGTAGANNNAGATCGGAA |
| 29758_2 | TATGAGTGTGGAGTCGTTGCAAAATGACTGGGTGTTGAGTAGANNNAGATCGGAA |
| 42321_1 | TATGAGTGTGGAGTCGTTGCCACAGAGTCACACATCCTTGAGANNNAGATCGGAAG |
| 42321_2 | TATGAGTGTGGAGTCGTTGCCACAGAGTCACACATCCTTGAGANNNAGATCGGAAG |
| 13459_1 | TATGAGTGTGGAGTCGTTGCAAGGCATATGGAGATGTAGTNNNAGATCGGAAGAG |
| 13459_2 | TATGAGTGTGGAGTCGTTGCAAGGCATATGGAGATGTAGTNNNAGATCGGAAGAG |
| 37316_1 | TATGAGTGTGGAGTCGTTGCAGAGTCCAAACACTGAATAGCACTNNNAGATCGGAA |
| 37316_2 | TATGAGTGTGGAGTCGTTGCAGAGTCCAAACACTGAATAGCACTNNNAGATCGGAA |
| 2845_1 | TATGAGTGTGGAGTCGTTGCGAGAAAGCCACTTATCAGCGAGTNNNAGATCGGAA |
| 2845_2 | TATGAGTGTGGAGTCGTTGCGAGAAAGCCACTTATCAGCGAGTNNNAGATCGGAA |
| 41201_1 | TATGAGTGTGGAGTCGTTGCCCCTCTCTTTCTCGGTACCTTTCNNNAGATCGGAAGA |
| 41201_2 | TATGAGTGTGGAGTCGTTGCCCCTCTCTTTCTCGGTACCTTTCNNNAGATCGGAAGA |
| 53315_1 | TATGAGTGTGGAGTCGTTGCGCCAAGATCCAACTTCAGTGTTTNNNAGATCGGAAG |
| 53315_2 | TATGAGTGTGGAGTCGTTGCGCCAAGATCCAACTTCAGTGTTTNNNAGATCGGAAG |
| 31978_1 | TATGAGTGTGGAGTCGTTGCAATGCGTGTAATCCCAGAACTCTNNNAGATCGGAAG |
| 31978_2 | TATGAGTGTGGAGTCGTTGCAATGCGTGTAATCCCAGAACTCTNNNAGATCGGAAG |
| 40314_1 | TATGAGTGTGGAGTCGTTGCTGAACCGATGAGGAAATAGGCTTNNNAGATCGGAA |
| 40314_2 | TATGAGTGTGGAGTCGTTGCTGAACCGATGAGGAAATAGGCTTNNNAGATCGGAA |
| 51223_1 | TATGAGTGTGGAGTCGTTGCGAAGTTCACAGTCTGCAGCAAATNNNAGATCGGAAG |
| 51223_2 | TATGAGTGTGGAGTCGTTGCGAAGTTCACAGTCTGCAGCAAATNNNAGATCGGAAG |
| 43569_1 | TATGAGTGTGGAGTCGTTGCATCCCAGGTTCAAGCTATTCTCCNNNAGATCGGAAGA |
| 43569_2 | TATGAGTGTGGAGTCGTTGCATCCCAGGTTCAAGCTATTCTCCNNNAGATCGGAAGA |
| 41452_1 | TATGAGTGTGGAGTCGTTGCGAGTTTTGCAGGCCTCAAAGATANNNAGATCGGAAG |
| 41452_2 | TATGAGTGTGGAGTCGTTGCGAGTTTTGCAGGCCTCAAAGATANNNAGATCGGAAG |
| 33180_1 | TATGAGTGTGGAGTCGTTGCGAGGATGCAGCATGAAATCGTACNNNAGATCGGAA |
| 33180_2 | TATGAGTGTGGAGTCGTTGCGAGGATGCAGCATGAAATCGTACNNNAGATCGGAA |
| 48286_1 | TATGAGTGTGGAGTCGTTGCCTCCACTTTCCAGGGTCAAGTAANNNAGATCGGAAG |
| 48286_2 | TATGAGTGTGGAGTCGTTGCCTCCACTTTCCAGGGTCAAGTAANNNAGATCGGAAG |
| 34044_1 | TATGAGTGTGGAGTCGTTGCGAGTAGCTGGGAATATAGGCACCNNNAGATCGGAA |
| 34044_2 | TATGAGTGTGGAGTCGTTGCGAGTAGCTGGGAATATAGGCACCNNNAGATCGGAA |
| 12899_1 | TATGAGTGTGGAGTCGTTGCCACCATTTTTATAGTCCATGTTTNNNAGATCGGAAGA |
| 12899_2 | TATGAGTGTGGAGTCGTTGCCACCATTTTTATAGTCCATGTTTNNNAGATCGGAAGA |
| 50953_1 | TATGAGTGTGGAGTCGTTGCCATGTTCATGATGTGCCCTTGTTNNNAGATCGGAAG |
| 50953_2 | TATGAGTGTGGAGTCGTTGCCATGTTCATGATGTGCCCTTGTTNNNAGATCGGAAG |
| 45265_1 | TATGAGTGTGGAGTCGTTGCTGAAGGGTGGGTCTAAAAAGTCCNNNAGATCGGAA |
| 45265_2 | TATGAGTGTGGAGTCGTTGCTGAAGGGTGGGTCTAAAAAGTCCNNNAGATCGGAA |
| 13760_1 | TATGAGTGTGGAGTCGTTGCACATTTCAACTTTTACTTGCTTNNNAGATCGGAAGAG |
| 13760_2 | TATGAGTGTGGAGTCGTTGCACATTTCAACTTTTACTTGCTTNNNAGATCGGAAGAG |

| | |
|---|---|
| 35307_1 | TATGAGTGTGGAGTCGTTGCCTCCAGGGTTCACACCATTCTCNNNAGATCGGAAGA |
| 35307_2 | TATGAGTGTGGAGTCGTTGCCTCCAGGGTTCACACCATTCTCNNNAGATCGGAAGA |
| 46905_1 | TATGAGTGTGGAGTCGTTGCGAGTGCAGTAGTGCGATCTCAGNNNAGATCGGAAG |
| 46905_2 | TATGAGTGTGGAGTCGTTGCGAGTGCAGTAGTGCGATCTCAGNNNAGATCGGAAG |
| 35061_1 | TATGAGTGTGGAGTCGTTGCGTAGAGACAGAGTTTCACCGTGTNNNAGATCGGAA |
| 35061_2 | TATGAGTGTGGAGTCGTTGCGTAGAGACAGAGTTTCACCGTGTNNNAGATCGGAA |
| 3059_1 | TATGAGTGTGGAGTCGTTGCCTTGCGACGTTCTATGAAGCATGNNNAGATCGGAAG |
| 3059_2 | TATGAGTGTGGAGTCGTTGCCTTGCGACGTTCTATGAAGCATGNNNAGATCGGAAG |
| 51106_1 | TATGAGTGTGGAGTCGTTGCTGGCTTCTCTGAGGAAATGATGTNNNAGATCGGAA |
| 51106_2 | TATGAGTGTGGAGTCGTTGCTGGCTTCTCTGAGGAAATGATGTNNNAGATCGGAA |
| 43490_1 | TATGAGTGTGGAGTCGTTGCAGGTCCTGTGAGATTTATGCTTTNNNAGATCGGAAG |
| 43490_2 | TATGAGTGTGGAGTCGTTGCAGGTCCTGTGAGATTTATGCTTTNNNAGATCGGAAG |
| 37266_1 | TATGAGTGTGGAGTCGTTGCTACTTTGGCATTAGGTGTGTCCCNNNAGATCGGAAG |
| 37266_2 | TATGAGTGTGGAGTCGTTGCTACTTTGGCATTAGGTGTGTCCCNNNAGATCGGAAG |
| 28993_1 | TATGAGTGTGGAGTCGTTGCCCCCAAGGATTGTAGAAATCCCANNNAGATCGGAAG |
| 28993_2 | TATGAGTGTGGAGTCGTTGCCCCCAAGGATTGTAGAAATCCCANNNAGATCGGAAG |
| 43489_1 | TATGAGTGTGGAGTCGTTGCCCATTCTTCTGCCTCAGTCTCTCNNNAGATCGGAAGA |
| 43489_2 | TATGAGTGTGGAGTCGTTGCCCATTCTTCTGCCTCAGTCTCTCNNNAGATCGGAAGA |
| 47898_1 | TATGAGTGTGGAGTCGTTGCGTGCGATCTTGGTTTACTGCAATNNNAGATCGGAAG |
| 47898_2 | TATGAGTGTGGAGTCGTTGCGTGCGATCTTGGTTTACTGCAATNNNAGATCGGAAG |
| 48082_1 | TATGAGTGTGGAGTCGTTGCATCCTTGTCACACTGTGATCTCCNNNAGATCGGAAGA |
| 48082_2 | TATGAGTGTGGAGTCGTTGCATCCTTGTCACACTGTGATCTCCNNNAGATCGGAAGA |
| 47389_1 | TATGAGTGTGGAGTCGTTGCCGATTCTCTTACCTTAGCCTCCCNNNAGATCGGAAGA |
| 47389_2 | TATGAGTGTGGAGTCGTTGCCGATTCTCTTACCTTAGCCTCCCNNNAGATCGGAAGA |
| 51035_1 | TATGAGTGTGGAGTCGTTGCTAGATCGTTTGTCTTCATGGCCANNNAGATCGGAAG |
| 51035_2 | TATGAGTGTGGAGTCGTTGCTAGATCGTTTGTCTTCATGGCCANNNAGATCGGAAG |
| 41243_1 | TATGAGTGTGGAGTCGTTGCAAAACCGCTTGAATGAGAACAGGNNNAGATCGGAA |
| 41243_2 | TATGAGTGTGGAGTCGTTGCAAAACCGCTTGAATGAGAACAGGNNNAGATCGGAA |
| 43954_1 | TATGAGTGTGGAGTCGTTGCCTTAGGATGGGGTCAATCTACCGNNNAGATCGGAA |
| 43954_2 | TATGAGTGTGGAGTCGTTGCCTTAGGATGGGGTCAATCTACCGNNNAGATCGGAA |
| 51056_1 | TATGAGTGTGGAGTCGTTGCAAAGAGGCGAGAGGAGGAGTTTTNNNAGATCGGA |
| 51056_2 | TATGAGTGTGGAGTCGTTGCAAAGAGGCGAGAGGAGGAGTTTTNNNAGATCGGA |
| 41368_1 | TATGAGTGTGGAGTCGTTGCTGTAACTCTTGTCTAGGCTCTGCNNNAGATCGGAAG |
| 41368_2 | TATGAGTGTGGAGTCGTTGCTGTAACTCTTGTCTAGGCTCTGCNNNAGATCGGAAG |
| 30610_1 | TATGAGTGTGGAGTCGTTGCAGAAGCATGTGTGTGTGTCAGTTNNNAGATCGGAA |
| 30610_2 | TATGAGTGTGGAGTCGTTGCAGAAGCATGTGTGTGTGTCAGTTNNNAGATCGGAA |
| 52478_1 | TATGAGTGTGGAGTCGTTGCTCCTTCTGAACTTTTCCCCATGTNNNAGATCGGAAGA |
| 52478_2 | TATGAGTGTGGAGTCGTTGCTCCTTCTGAACTTTTCCCCATGTNNNAGATCGGAAGA |
| 46363_1 | TATGAGTGTGGAGTCGTTGCCTGCTATTTCTGACCTTTGCCACNNNAGATCGGAAGA |
| 46363_2 | TATGAGTGTGGAGTCGTTGCCTGCTATTTCTGACCTTTGCCACNNNAGATCGGAAGA |
| 51202_1 | TATGAGTGTGGAGTCGTTGCGCCAACTTGCAGAAAAGTGGTATNNNAGATCGGAA |
| 51202_2 | TATGAGTGTGGAGTCGTTGCGCCAACTTGCAGAAAAGTGGTATNNNAGATCGGAA |
| 37075_1 | TATGAGTGTGGAGTCGTTGCTCTTTGGTGATGGCATCTCTCATNNNAGATCGGAAG |

| | |
|---|---|
| 37075_2 | TATGAGTGTGGAGTCGTTGCTCTTTGGTGATGGCATCTCTCATNNNAGATCGGAAG |
| 35142_1 | TATGAGTGTGGAGTCGTTGCCAGTAGCTGGGATTACAGGCATGNNNAGATCGGAA |
| 35142_2 | TATGAGTGTGGAGTCGTTGCCAGTAGCTGGGATTACAGGCATGNNNAGATCGGAA |
| 12618_1 | TATGAGTGTGGAGTCGTTGCGCATCATTCACTTTTATCAAATGNNNAGATCGGAAG |
| 12618_2 | TATGAGTGTGGAGTCGTTGCGCATCATTCACTTTTATCAAATGNNNAGATCGGAAG |
| 49211_1 | TATGAGTGTGGAGTCGTTGCCTCTCGAGTAGCTGGGACTACAGNNNAGATCGGAA |
| 49211_2 | TATGAGTGTGGAGTCGTTGCCTCTCGAGTAGCTGGGACTACAGNNNAGATCGGAA |
| 39649_1 | TATGAGTGTGGAGTCGTTGCAAGAAGAAAGCAAGCAAGCAAGTNNNAGATCGGAA |
| 39649_2 | TATGAGTGTGGAGTCGTTGCAAGAAGAAAGCAAGCAAGCAAGTNNNAGATCGGAA |
| 34693_1 | TATGAGTGTGGAGTCGTTGCCACCACCATGCCACACTAAAATTNNNAGATCGGAAG |
| 34693_2 | TATGAGTGTGGAGTCGTTGCCACCACCATGCCACACTAAAATTNNNAGATCGGAAG |
| 42504_1 | TATGAGTGTGGAGTCGTTGCCTGGAGTACAATGGCACGATCTTNNNAGATCGGAAG |
| 42504_2 | TATGAGTGTGGAGTCGTTGCCTGGAGTACAATGGCACGATCTTNNNAGATCGGAAG |
| 49026_1 | TATGAGTGTGGAGTCGTTGCACGCCCGGCTAATTTTTTGTATTNNNAGATCGGAAG |
| 49026_2 | TATGAGTGTGGAGTCGTTGCACGCCCGGCTAATTTTTTGTATTNNNAGATCGGAAG |
| 51325_1 | TATGAGTGTGGAGTCGTTGCCCAGGCCAGAGAACATCATTACTNNNAGATCGGAAG |
| 51325_2 | TATGAGTGTGGAGTCGTTGCCCAGGCCAGAGAACATCATTACTNNNAGATCGGAAG |
| 29797_1 | TATGAGTGTGGAGTCGTTGCCACTGCAGGTACTAGGAAACCTANNNAGATCGGAAG |
| 29797_2 | TATGAGTGTGGAGTCGTTGCCACTGCAGGTACTAGGAAACCTANNNAGATCGGAAG |
| 33123_1 | TATGAGTGTGGAGTCGTTGCGAACGAAAAGTATCTTCCTCGCGNNNAGATCGGAAG |
| 33123_2 | TATGAGTGTGGAGTCGTTGCGAACGAAAAGTATCTTCCTCGCGNNNAGATCGGAAG |
| 51788_1 | TATGAGTGTGGAGTCGTTGCAAACCAAAACTGTGACTGAAGGCNNNAGATCGGAA |
| 51788_2 | TATGAGTGTGGAGTCGTTGCAAACCAAAACTGTGACTGAAGGCNNNAGATCGGAA |
| 51970_1 | TATGAGTGTGGAGTCGTTGCTACACCCTAAAGCCTGAAACCTGNNNAGATCGGAAG |
| 51970_2 | TATGAGTGTGGAGTCGTTGCTACACCCTAAAGCCTGAAACCTGNNNAGATCGGAAG |
| 31990_1 | TATGAGTGTGGAGTCGTTGCGCAGCACTATTCACAATAGCCAGNNNAGATCGGAAG |
| 31990_2 | TATGAGTGTGGAGTCGTTGCGCAGCACTATTCACAATAGCCAGNNNAGATCGGAAG |
| 46477_1 | TATGAGTGTGGAGTCGTTGCTTCAAGCAATTCCCATGCCTCAANNNAGATCGGAAG |
| 46477_2 | TATGAGTGTGGAGTCGTTGCTTCAAGCAATTCCCATGCCTCAANNNAGATCGGAAG |
| 47134_1 | TATGAGTGTGGAGTCGTTGCGCATGTGTCATGTTACAGATCCGNNNAGATCGGAAG |
| 47134_2 | TATGAGTGTGGAGTCGTTGCGCATGTGTCATGTTACAGATCCGNNNAGATCGGAAG |
| 35643_1 | TATGAGTGTGGAGTCGTTGCTGGGATTATAGGAATGTGCCACCNNNAGATCGGAA |
| 35643_2 | TATGAGTGTGGAGTCGTTGCTGGGATTATAGGAATGTGCCACCNNNAGATCGGAA |
| 46806_1 | TATGAGTGTGGAGTCGTTGCCCTCAAGTGATATTCCCTCCTCGNNNAGATCGGAAG |
| 46806_2 | TATGAGTGTGGAGTCGTTGCCCTCAAGTGATATTCCCTCCTCGNNNAGATCGGAAG |
| 13670_1 | TATGAGTGTGGAGTCGTTGCCTGAACTTGATGAGTCCTCTNNNAGATCGGAAGAGC |
| 13670_2 | TATGAGTGTGGAGTCGTTGCCTGAACTTGATGAGTCCTCTNNNAGATCGGAAGAGC |
| 12790_1 | TATGAGTGTGGAGTCGTTGCCATCTCTTGCCTTCATAGTACNNNAGATCGGAAGAG |
| 12790_2 | TATGAGTGTGGAGTCGTTGCCATCTCTTGCCTTCATAGTACNNNAGATCGGAAGAG |
| 40206_1 | TATGAGTGTGGAGTCGTTGCGAGGCAAGAGAGTCGCTTGAANNNAGATCGGAAG |
| 40206_2 | TATGAGTGTGGAGTCGTTGCGAGGCAAGAGAGTCGCTTGAANNNAGATCGGAAG |
| 40830_1 | TATGAGTGTGGAGTCGTTGCGGCTATGCAGTGGTACAAATGACNNNAGATCGGAA |
| 40830_2 | TATGAGTGTGGAGTCGTTGCGGCTATGCAGTGGTACAAATGACNNNAGATCGGAA |

| | |
|---|---|
| 30302_1 | TATGAGTGTGGAGTCGTTGCTCATGATGCATACAGGCAGACCNNNAGATCGGAAGA |
| 30302_2 | TATGAGTGTGGAGTCGTTGCTCATGATGCATACAGGCAGACCNNNAGATCGGAAGA |
| 13506_1 | TATGAGTGTGGAGTCGTTGCACATTTCAACTTTTACTTGCTTNNNAGATCGGAAGAG |
| 13506_2 | TATGAGTGTGGAGTCGTTGCACATTTCAACTTTTACTTGCTTNNNAGATCGGAAGAG |
| 10230_1 | TATGAGTGTGGAGTCGTTGCCTCTCCCTGCAGCTCAGACTATTNNNAGATCGGAAGA |
| 10230_2 | TATGAGTGTGGAGTCGTTGCCTCTCCCTGCAGCTCAGACTATTNNNAGATCGGAAGA |
| 38425_1 | TATGAGTGTGGAGTCGTTGCACAGTGTTTGTCCTCTTTTTGCCNNNAGATCGGAAGA |
| 38425_2 | TATGAGTGTGGAGTCGTTGCACAGTGTTTGTCCTCTTTTTGCCNNNAGATCGGAAGA |
| 39814_1 | TATGAGTGTGGAGTCGTTGCACAGCCAGCCAAGAATCATTAGANNNAGATCGGAAG |
| 39814_2 | TATGAGTGTGGAGTCGTTGCACAGCCAGCCAAGAATCATTAGANNNAGATCGGAAG |
| 35054_1 | TATGAGTGTGGAGTCGTTGCTCAAGTGATTCTCCTGTCTCAGCNNNAGATCGGAAGA |
| 35054_2 | TATGAGTGTGGAGTCGTTGCTCAAGTGATTCTCCTGTCTCAGCNNNAGATCGGAAGA |
| 43174_1 | TATGAGTGTGGAGTCGTTGCGTTCAAGCGATTCTCTTGCCTCNNNAGATCGGAAGA |
| 43174_2 | TATGAGTGTGGAGTCGTTGCGTTCAAGCGATTCTCTTGCCTCNNNAGATCGGAAGA |
| 44646_1 | TATGAGTGTGGAGTCGTTGCCAAGGAAGATATGGGGAGGATGGNNNAGATCGGA |
| 44646_2 | TATGAGTGTGGAGTCGTTGCCAAGGAAGATATGGGGAGGATGGNNNAGATCGGA |
| 41410_1 | TATGAGTGTGGAGTCGTTGCAAAGGACTTGAAAGACAGAGGGGNNNAGATCGGA |
| 41410_2 | TATGAGTGTGGAGTCGTTGCAAAGGACTTGAAAGACAGAGGGGNNNAGATCGGA |
| 41762_1 | TATGAGTGTGGAGTCGTTGCAGTAACTGGGATTACATGCACCCNNNAGATCGGAAG |
| 41762_2 | TATGAGTGTGGAGTCGTTGCAGTAACTGGGATTACATGCACCCNNNAGATCGGAAG |
| 43241_1 | TATGAGTGTGGAGTCGTTGCAGACAGCCCCTGGTACAAAATAGNNNAGATCGGAA |
| 43241_2 | TATGAGTGTGGAGTCGTTGCAGACAGCCCCTGGTACAAAATAGNNNAGATCGGAA |
| 48953_1 | TATGAGTGTGGAGTCGTTGCCCGAGTATCTGGGATTACAGGTGNNNAGATCGGAA |
| 48953_2 | TATGAGTGTGGAGTCGTTGCCCGAGTATCTGGGATTACAGGTGNNNAGATCGGAA |
| 41008_1 | TATGAGTGTGGAGTCGTTGCCCTGTAGTCCCAGCTACTTGGNNNAGATCGGAAGAG |
| 41008_2 | TATGAGTGTGGAGTCGTTGCCCTGTAGTCCCAGCTACTTGGNNNAGATCGGAAGAG |
| 40920_1 | TATGAGTGTGGAGTCGTTGCAGGGCTGAGGAGGAGATCTAAAANNNAGATCGGA |
| 40920_2 | TATGAGTGTGGAGTCGTTGCAGGGCTGAGGAGGAGATCTAAAANNNAGATCGGA |
| 10496_1 | TATGAGTGTGGAGTCGTTGCAATCTTCAGGCTTCCCTGCATTTNNNAGATCGGAAGA |
| 10496_2 | TATGAGTGTGGAGTCGTTGCAATCTTCAGGCTTCCCTGCATTTNNNAGATCGGAAGA |
| 45604_1 | TATGAGTGTGGAGTCGTTGCGAGGTTTTGTCATGTTGCTCAGGNNNAGATCGGAA |
| 45604_2 | TATGAGTGTGGAGTCGTTGCGAGGTTTTGTCATGTTGCTCAGGNNNAGATCGGAA |
| 42510_1 | TATGAGTGTGGAGTCGTTGCGAAAATACCTGCAACGTCACCATNNNAGATCGGAAG |
| 42510_2 | TATGAGTGTGGAGTCGTTGCGAAAATACCTGCAACGTCACCATNNNAGATCGGAAG |
| 36117_1 | TATGAGTGTGGAGTCGTTGCCTGAGGCAGAAGAATTGCTTGAANNNAGATCGGAA |
| 36117_2 | TATGAGTGTGGAGTCGTTGCCTGAGGCAGAAGAATTGCTTGAANNNAGATCGGAA |
| 50231_1 | TATGAGTGTGGAGTCGTTGCAAGGTGGTTGGAGATGACGTTTANNNAGATCGGAA |
| 50231_2 | TATGAGTGTGGAGTCGTTGCAAGGTGGTTGGAGATGACGTTTANNNAGATCGGAA |
| 34092_1 | TATGAGTGTGGAGTCGTTGCGCTTGGTCTCCGTCTCAAAAAATNNNAGATCGGAAGA |
| 34092_2 | TATGAGTGTGGAGTCGTTGCGCTTGGTCTCCGTCTCAAAAAATNNNAGATCGGAAGA |
| 10655_1 | TATGAGTGTGGAGTCGTTGCCACTGTAGGGAGACAGAGGAGAGNNNAGATCGGA |
| 10655_2 | TATGAGTGTGGAGTCGTTGCCACTGTAGGGAGACAGAGGAGAGNNNAGATCGGA |
| 47341_1 | TATGAGTGTGGAGTCGTTGCGGGTTCTTAGCTGTAACTGGCTANNNAGATCGGAAG |

| | |
|---|---|
| 47341_2 | TATGAGTGTGGAGTCGTTGCGGGTTCTTAGCTGTAACTGGCTANNNAGATCGGAAG |
| 47717_1 | TATGAGTGTGGAGTCGTTGCATCTCCTGGGTTCAAGCAATTCTNNNAGATCGGAAG |
| 47717_2 | TATGAGTGTGGAGTCGTTGCATCTCCTGGGTTCAAGCAATTCTNNNAGATCGGAAG |
| 48540_1 | TATGAGTGTGGAGTCGTTGCCTCCCAGGTTCAAGCGATTCTCNNNAGATCGGAAGA |
| 48540_2 | TATGAGTGTGGAGTCGTTGCCTCCCAGGTTCAAGCGATTCTCNNNAGATCGGAAGA |
| 9615_1 | TATGAGTGTGGAGTCGTTGCTTTCTTGACCTCAGATGCAGAGCNNNAGATCGGAAG |
| 9615_2 | TATGAGTGTGGAGTCGTTGCTTTCTTGACCTCAGATGCAGAGCNNNAGATCGGAAG |
| 41377_1 | TATGAGTGTGGAGTCGTTGCGGATTACCTGAGTTCGAGACCAGNNNAGATCGGAA |
| 41377_2 | TATGAGTGTGGAGTCGTTGCGGATTACCTGAGTTCGAGACCAGNNNAGATCGGAA |
| 50120_1 | TATGAGTGTGGAGTCGTTGCAGAATCCTTGGAATCCCAGAAGCNNNAGATCGGAAG |
| 50120_2 | TATGAGTGTGGAGTCGTTGCAGAATCCTTGGAATCCCAGAAGCNNNAGATCGGAAG |
| 48190_1 | TATGAGTGTGGAGTCGTTGCCCACCAAGCCTGGCTAATTTTTTNNNAGATCGGAAGA |
| 48190_2 | TATGAGTGTGGAGTCGTTGCCCACCAAGCCTGGCTAATTTTTTNNNAGATCGGAAGA |
| 46123_1 | TATGAGTGTGGAGTCGTTGCTCAGAAACTATTTGTCAGGCCCANNNAGATCGGAAG |
| 46123_2 | TATGAGTGTGGAGTCGTTGCTCAGAAACTATTTGTCAGGCCCANNNAGATCGGAAG |
| 40508_1 | TATGAGTGTGGAGTCGTTGCGTCTCCTGGGTTCAAGTGATTCTNNNAGATCGGAAG |
| 40508_2 | TATGAGTGTGGAGTCGTTGCGTCTCCTGGGTTCAAGTGATTCTNNNAGATCGGAAG |
| 38007_1 | TATGAGTGTGGAGTCGTTGCGGGGTTTGAAGTCAGACAGATCTNNNAGATCGGAA |
| 38007_2 | TATGAGTGTGGAGTCGTTGCGGGGTTTGAAGTCAGACAGATCTNNNAGATCGGAA |
| 44447_1 | TATGAGTGTGGAGTCGTTGCAGATCCATGTCCTAATCCCCAGANNNAGATCGGAAG |
| 44447_2 | TATGAGTGTGGAGTCGTTGCAGATCCATGTCCTAATCCCCAGANNNAGATCGGAAG |
| 48735_1 | TATGAGTGTGGAGTCGTTGCCAATCTCGGCTCACTGCAATCNNNAGATCGGAAGAG |
| 48735_2 | TATGAGTGTGGAGTCGTTGCCAATCTCGGCTCACTGCAATCNNNAGATCGGAAGAG |
| 6178_1 | TATGAGTGTGGAGTCGTTGCGGAATCAAAGTCTGACAACAGCCANNNAGATCGGA |
| 6178_2 | TATGAGTGTGGAGTCGTTGCGGAATCAAAGTCTGACAACAGCCANNNAGATCGGA |
| 34879_1 | TATGAGTGTGGAGTCGTTGCTTCAAGTGATTCTCCTGCCTCAGNNNAGATCGGAAG |
| 34879_2 | TATGAGTGTGGAGTCGTTGCTTCAAGTGATTCTCCTGCCTCAGNNNAGATCGGAAG |
| 31121_1 | TATGAGTGTGGAGTCGTTGCATACATACACACACACACACACANNNAGATCGGAAG |
| 31121_2 | TATGAGTGTGGAGTCGTTGCATACATACACACACACACACACANNNAGATCGGAAG |
| 47172_1 | TATGAGTGTGGAGTCGTTGCCTGAGTAGGTGGGATTACAGGTGNNNAGATCGGAA |
| 47172_2 | TATGAGTGTGGAGTCGTTGCCTGAGTAGGTGGGATTACAGGTGNNNAGATCGGAA |
| 51305_1 | TATGAGTGTGGAGTCGTTGCGGAGGTGAGGGGCATCTCTGNNNAGATCGGAAGA |
| 51305_2 | TATGAGTGTGGAGTCGTTGCGGAGGTGAGGGGCATCTCTGNNNAGATCGGAAGA |
| 33157_1 | TATGAGTGTGGAGTCGTTGCAATGCAGCTCTCTTCTACCTCTGNNNAGATCGGAAGA |
| 33157_2 | TATGAGTGTGGAGTCGTTGCAATGCAGCTCTCTTCTACCTCTGNNNAGATCGGAAGA |
| 48948_1 | TATGAGTGTGGAGTCGTTGCACGCCCGGCTAATTTTGTATTTTNNNAGATCGGAAG |
| 48948_2 | TATGAGTGTGGAGTCGTTGCACGCCCGGCTAATTTTGTATTTTNNNAGATCGGAAG |
| 35464_1 | TATGAGTGTGGAGTCGTTGCCGCTTGGAATGGTTTCAGAAGAANNNAGATCGGAA |
| 35464_2 | TATGAGTGTGGAGTCGTTGCCGCTTGGAATGGTTTCAGAAGAANNNAGATCGGAA |
| 52315_1 | TATGAGTGTGGAGTCGTTGCCTTGATCTGGGCAGAGCTTCCNNNAGATCGGAAGAG |
| 52315_2 | TATGAGTGTGGAGTCGTTGCCTTGATCTGGGCAGAGCTTCCNNNAGATCGGAAGAG |
| 47798_1 | TATGAGTGTGGAGTCGTTGCGAGACAGTCTTTCTCTGTTGCCCNNNAGATCGGAAG |
| 47798_2 | TATGAGTGTGGAGTCGTTGCGAGACAGTCTTTCTCTGTTGCCCNNNAGATCGGAAG |

| | |
|---|---|
| 44660_1 | TATGAGTGTGGAGTCGTTGCGGCTCAGGTAAGAATAGAGGGTGNNNAGATCGGA |
| 44660_2 | TATGAGTGTGGAGTCGTTGCGGCTCAGGTAAGAATAGAGGGTGNNNAGATCGGA |
| 9981_1 | TATGAGTGTGGAGTCGTTGCGGCTTACCATACCAATCAGCTCCNNNAGATCGGAAG |
| 9981_2 | TATGAGTGTGGAGTCGTTGCGGCTTACCATACCAATCAGCTCCNNNAGATCGGAAG |
| 29122_1 | TATGAGTGTGGAGTCGTTGCATTGGCTGTGACCAAAACAAACANNNAGATCGGAA |
| 29122_2 | TATGAGTGTGGAGTCGTTGCATTGGCTGTGACCAAAACAAACANNNAGATCGGAA |
| 37806_1 | TATGAGTGTGGAGTCGTTGCACTACATCTGCAAGACATCAGCANNNAGATCGGAAG |
| 37806_2 | TATGAGTGTGGAGTCGTTGCACTACATCTGCAAGACATCAGCANNNAGATCGGAAG |
| 52698_1 | TATGAGTGTGGAGTCGTTGCAGGCTACCAGCAAATCCTAAACANNNAGATCGGAAG |
| 52698_2 | TATGAGTGTGGAGTCGTTGCAGGCTACCAGCAAATCCTAAACANNNAGATCGGAAG |
| 29496_1 | ATGAGTGTGGAGTCGTTGCACTGCCTCCTATGGATATAATTGTCTANNNAGATCGG |
| 29496_2 | ATGAGTGTGGAGTCGTTGCACTGCCTCCTATGGATATAATTGTCTANNNAGATCGG |
| 37865_1 | TATGAGTGTGGAGTCGTTGCAAAAAAATAGTGTGGGCAGGGTGNNNAGATCGGA |
| 37865_2 | TATGAGTGTGGAGTCGTTGCAAAAAAATAGTGTGGGCAGGGTGNNNAGATCGGA |
| 41713_1 | TATGAGTGTGGAGTCGTTGCCTATCTCGGCTCACTGCAAACTCNNNAGATCGGAAG |
| 41713_2 | TATGAGTGTGGAGTCGTTGCCTATCTCGGCTCACTGCAAACTCNNNAGATCGGAAG |
| 46598_1 | TATGAGTGTGGAGTCGTTGCGGCACAAGCAATTATCTTCCCTCNNNAGATCGGAAG |
| 46598_2 | TATGAGTGTGGAGTCGTTGCGGCACAAGCAATTATCTTCCCTCNNNAGATCGGAAG |
| 38472_1 | TATGAGTGTGGAGTCGTTGCTCTAATGGTGTTTTCTCCAGCCANNNAGATCGGAAG |
| 38472_2 | TATGAGTGTGGAGTCGTTGCTCTAATGGTGTTTTCTCCAGCCANNNAGATCGGAAG |
| 36766_1 | TATGAGTGTGGAGTCGTTGCCCATCTCACACCAGTCAGAAAGANNNAGATCGGAAG |
| 36766_2 | TATGAGTGTGGAGTCGTTGCCCATCTCACACCAGTCAGAAAGANNNAGATCGGAAG |
| 38724_1 | TATGAGTGTGGAGTCGTTGCCTGCCAAGTAGCTGGGATTACANNNAGATCGGAAG |
| 38724_2 | TATGAGTGTGGAGTCGTTGCCTGCCAAGTAGCTGGGATTACANNNAGATCGGAAG |
| 46743_1 | TATGAGTGTGGAGTCGTTGCCAATGGCATGATCTCAGCTCACTNNNAGATCGGAAG |
| 46743_2 | TATGAGTGTGGAGTCGTTGCCAATGGCATGATCTCAGCTCACTNNNAGATCGGAAG |
| 44060_1 | TATGAGTGTGGAGTCGTTGCGATCACCGTCGTGGATGTGATTNNNAGATCGGAAG |
| 44060_2 | TATGAGTGTGGAGTCGTTGCGATCACCGTCGTGGATGTGATTNNNAGATCGGAAG |
| 37850_1 | TATGAGTGTGGAGTCGTTGCGACCACTTCAGAGAAGAAGGAGANNNAGATCGGAA |
| 37850_2 | TATGAGTGTGGAGTCGTTGCGACCACTTCAGAGAAGAAGGAGANNNAGATCGGAA |
| 32917_1 | TATGAGTGTGGAGTCGTTGCTCTCTTCTGGCTTGTAGGGTTTCNNNAGATCGGAAG |
| 32917_2 | TATGAGTGTGGAGTCGTTGCTCTCTTCTGGCTTGTAGGGTTTCNNNAGATCGGAAG |
| 38779_1 | TATGAGTGTGGAGTCGTTGCAACTGATGGCCTCATGACTGAATNNNAGATCGGAAG |
| 38779_2 | TATGAGTGTGGAGTCGTTGCAACTGATGGCCTCATGACTGAATNNNAGATCGGAAG |
| 41702_1 | TATGAGTGTGGAGTCGTTGCAGAGACAGTCTTGCTCTATTGCCNNNAGATCGGAAG |
| 41702_2 | TATGAGTGTGGAGTCGTTGCAGAGACAGTCTTGCTCTATTGCCNNNAGATCGGAAG |
| 8894_1 | TATGAGTGTGGAGTCGTTGCGGCTATGAGATACTTCAGCTGGTNNNAGATCGGAA |
| 8894_2 | TATGAGTGTGGAGTCGTTGCGGCTATGAGATACTTCAGCTGGTNNNAGATCGGAA |
| 48013_1 | TATGAGTGTGGAGTCGTTGCTCCTGAGTAGCTGGGATTACAGANNNAGATCGGAA |
| 48013_2 | TATGAGTGTGGAGTCGTTGCTCCTGAGTAGCTGGGATTACAGANNNAGATCGGAA |
| 43412_1 | TATGAGTGTGGAGTCGTTGCAGATGGAGTCTCACTCTGTCACCNNNAGATCGGAAG |
| 43412_2 | TATGAGTGTGGAGTCGTTGCAGATGGAGTCTCACTCTGTCACCNNNAGATCGGAAG |
| 3151_1 | TATGAGTGTGGAGTCGTTGCAGATATTCACCATTATAGGTGGGTTTNNNAGATCGG |

| | |
|---|---|
| 3151_2 | TATGAGTGTGGAGTCGTTGCAGATATTCACCATTATAGGTGGGTTTNNNAGATCGG |
| 40510_1 | TATGAGTGTGGAGTCGTTGCCTGAGGCAGGAGAATTGCTTGAANNNAGATCGGAA |
| 40510_2 | TATGAGTGTGGAGTCGTTGCCTGAGGCAGGAGAATTGCTTGAANNNAGATCGGAA |
| 32223_1 | TATGAGTGTGGAGTCGTTGCCCTTGCCTCAATTTGGGACAATTNNNAGATCGGAAG |
| 32223_2 | TATGAGTGTGGAGTCGTTGCCCTTGCCTCAATTTGGGACAATTNNNAGATCGGAAG |
| 53056_1 | TATGAGTGTGGAGTCGTTGCGCTGGTTCCAATCTGACTGAAACNNNAGATCGGAAG |
| 53056_2 | TATGAGTGTGGAGTCGTTGCGCTGGTTCCAATCTGACTGAAACNNNAGATCGGAAG |
| 29520_1 | TATGAGTGTGGAGTCGTTGCGCAAGCACAAGGAACAACTTGAGNNNAGATCGGAA |
| 29520_2 | TATGAGTGTGGAGTCGTTGCGCAAGCACAAGGAACAACTTGAGNNNAGATCGGAA |
| 43388_1 | TATGAGTGTGGAGTCGTTGCTTGAGTAGCTGGGATTACAAGCANNNAGATCGGAA |
| 43388_2 | TATGAGTGTGGAGTCGTTGCTTGAGTAGCTGGGATTACAAGCANNNAGATCGGAA |
| 36682_1 | TATGAGTGTGGAGTCGTTGCTGCACCTGAAGGAAAATACCTGANNNAGATCGGAA |
| 36682_2 | TATGAGTGTGGAGTCGTTGCTGCACCTGAAGGAAAATACCTGANNNAGATCGGAA |
| 43924_1 | TATGAGTGTGGAGTCGTTGCGATAGAGTTTTGCTCCTGTTGCCNNNAGATCGGAAG |
| 43924_2 | TATGAGTGTGGAGTCGTTGCGATAGAGTTTTGCTCCTGTTGCCNNNAGATCGGAAG |
| 30997_1 | TATGAGTGTGGAGTCGTTGCTGCTAGGATATTACAGGTGCCACNNNAGATCGGAA |
| 30997_2 | TATGAGTGTGGAGTCGTTGCTGCTAGGATATTACAGGTGCCACNNNAGATCGGAA |
| 30466_1 | TATGAGTGTGGAGTCGTTGCGTTGCCTTGCCCATTCTTAATAAAAGNNNAGATCGG |
| 30466_2 | TATGAGTGTGGAGTCGTTGCGTTGCCTTGCCCATTCTTAATAAAAGNNNAGATCGG |
| 43818_1 | TATGAGTGTGGAGTCGTTGCGTGCAGAAGAGGACAGCATAGTANNNAGATCGGA |
| 43818_2 | TATGAGTGTGGAGTCGTTGCGTGCAGAAGAGGACAGCATAGTANNNAGATCGGA |
| 1833_1 | TATGAGTGTGGAGTCGTTGCTCATCAGGGGTTGTCACAGTCACNNNAGATCGGAAG |
| 1833_2 | TATGAGTGTGGAGTCGTTGCTCATCAGGGGTTGTCACAGTCACNNNAGATCGGAAG |
| 49374_1 | TATGAGTGTGGAGTCGTTGCCACCTCGGCCTCCAAAAATATTTNNNAGATCGGAAG |
| 49374_2 | TATGAGTGTGGAGTCGTTGCCACCTCGGCCTCCAAAAATATTTNNNAGATCGGAAG |
| 6072_1 | TATGAGTGTGGAGTCGTTGCAGTCAGATTGTCCCTCTTAGTTGNNNAGATCGGAAG |
| 6072_2 | TATGAGTGTGGAGTCGTTGCAGTCAGATTGTCCCTCTTAGTTGNNNAGATCGGAAG |
| 36740_1 | TATGAGTGTGGAGTCGTTGCTTGTTTGATCTGGCCCTGACATANNNAGATCGGAAG |
| 36740_2 | TATGAGTGTGGAGTCGTTGCTTGTTTGATCTGGCCCTGACATANNNAGATCGGAAG |
| 48265_1 | TATGAGTGTGGAGTCGTTGCTGGAGTTCAATGGCATGATCTCGNNNAGATCGGAA |
| 48265_2 | TATGAGTGTGGAGTCGTTGCTGGAGTTCAATGGCATGATCTCGNNNAGATCGGAA |
| 20753_1 | TATGAGTGTGGAGTCGTTGCTCTTTCTCCTGTCTCTCTACNNNAGATCGGAAGAGCA |
| 20753_2 | TATGAGTGTGGAGTCGTTGCTCTTTCTCCTGTCTCTCTACNNNAGATCGGAAGAGCA |
| 32297_1 | TATGAGTGTGGAGTCGTTGCCTCAAGTGATCCTCCCACCTCTNNNAGATCGGAAGAG |
| 32297_2 | TATGAGTGTGGAGTCGTTGCCTCAAGTGATCCTCCCACCTCTNNNAGATCGGAAGAG |
| 48337_1 | TATGAGTGTGGAGTCGTTGCTTCAAGCGATTCTCCTGTCTCTGNNNAGATCGGAAG |
| 48337_2 | TATGAGTGTGGAGTCGTTGCTTCAAGCGATTCTCCTGTCTCTGNNNAGATCGGAAG |
| 40464_1 | TATGAGTGTGGAGTCGTTGCGGTGGTAAGGAAATGTTACCTTGCNNNAGATCGG |
| 40464_2 | TATGAGTGTGGAGTCGTTGCGGTGGTAAGGAAATGTTACCTTGCNNNAGATCGG |
| 32543_1 | TATGAGTGTGGAGTCGTTGCGGGTTGCAATCCTAGTCTCTGATNNNAGATCGGAAG |
| 32543_2 | TATGAGTGTGGAGTCGTTGCGGGTTGCAATCCTAGTCTCTGATNNNAGATCGGAAG |
| 37242_1 | TATGAGTGTGGAGTCGTTGCTTTCAACTGAGGCCTCTGAGATCNNNAGATCGGAAG |
| 37242_2 | TATGAGTGTGGAGTCGTTGCTTTCAACTGAGGCCTCTGAGATCNNNAGATCGGAAG |

| | |
|---|---|
| 41815_1 | TATGAGTGTGGAGTCGTTGCCTCAGCCACCTGAATAGTTGAGANNNAGATCGGAAG |
| 41815_2 | TATGAGTGTGGAGTCGTTGCCTCAGCCACCTGAATAGTTGAGANNNAGATCGGAAG |
| 43528_1 | TATGAGTGTGGAGTCGTTGCCAATGCATTCAGCCATCCTATGTNNNAGATCGGAAG |
| 43528_2 | TATGAGTGTGGAGTCGTTGCCAATGCATTCAGCCATCCTATGTNNNAGATCGGAAG |
| 45548_1 | TATGAGTGTGGAGTCGTTGCAGTAAGGGGGAAGTGCAGTAAAGNNNAGATCGGA |
| 45548_2 | TATGAGTGTGGAGTCGTTGCAGTAAGGGGGAAGTGCAGTAAAGNNNAGATCGGA |
| 48027_1 | TATGAGTGTGGAGTCGTTGCAGACAAGGTCTCATTCTGTCACCNNNAGATCGGAAG |
| 48027_2 | TATGAGTGTGGAGTCGTTGCAGACAAGGTCTCATTCTGTCACCNNNAGATCGGAAG |
| 50806_1 | TATGAGTGTGGAGTCGTTGCCATTTGGACAATCAAGGTGCACANNNAGATCGGAAG |
| 50806_2 | TATGAGTGTGGAGTCGTTGCCATTTGGACAATCAAGGTGCACANNNAGATCGGAAG |
| 50796_1 | TATGAGTGTGGAGTCGTTGCTAAACTAGCCCCTACCTCACAGANNNAGATCGGAAG |
| 50796_2 | TATGAGTGTGGAGTCGTTGCTAAACTAGCCCCTACCTCACAGANNNAGATCGGAAG |
| 13683_1 | TATGAGTGTGGAGTCGTTGCGTTTCTAAGGACTTTTTCTTAGGNNNAGATCGGAAG |
| 13683_2 | TATGAGTGTGGAGTCGTTGCGTTTCTAAGGACTTTTTCTTAGGNNNAGATCGGAAG |
| 30311_1 | TATGAGTGTGGAGTCGTTGCCTCTTAACTGAGGCAAGGAACCCNNNAGATCGGAAG |
| 30311_2 | TATGAGTGTGGAGTCGTTGCCTCTTAACTGAGGCAAGGAACCCNNNAGATCGGAAG |
| 42901_1 | TATGAGTGTGGAGTCGTTGCAACATGGGGGTGCAGATATCTTTNNNAGATCGGAA |
| 42901_2 | TATGAGTGTGGAGTCGTTGCAACATGGGGGTGCAGATATCTTTNNNAGATCGGAA |
| 38132_1 | TATGAGTGTGGAGTCGTTGCACATGGTGAAACCCCGTATCTACNNNAGATCGGAAG |
| 38132_2 | TATGAGTGTGGAGTCGTTGCACATGGTGAAACCCCGTATCTACNNNAGATCGGAAG |
| 30977_1 | TATGAGTGTGGAGTCGTTGCCAGCAGCAGCATTATTCCTTTGANNNAGATCGGAAG |
| 30977_2 | TATGAGTGTGGAGTCGTTGCCAGCAGCAGCATTATTCCTTTGANNNAGATCGGAAG |
| 43037_1 | TATGAGTGTGGAGTCGTTGCTGCATACTCTTTTGACCCAGGAANNNAGATCGGAAG |
| 43037_2 | TATGAGTGTGGAGTCGTTGCTGCATACTCTTTTGACCCAGGAANNNAGATCGGAAG |
| 13398_1 | TATGAGTGTGGAGTCGTTGCATTTACTTCTGGATGAGCTCTNNNAGATCGGAAGAG |
| 13398_2 | TATGAGTGTGGAGTCGTTGCATTTACTTCTGGATGAGCTCTNNNAGATCGGAAGAG |
| 41710_1 | TATGAGTGTGGAGTCGTTGCTGATCTCAGCTCATCTCAACCTCNNNAGATCGGAAGA |
| 41710_2 | TATGAGTGTGGAGTCGTTGCTGATCTCAGCTCATCTCAACCTCNNNAGATCGGAAGA |
| 37407_1 | TATGAGTGTGGAGTCGTTGCGCTTGCAGTGAGCCAAGATTGNNNAGATCGGAAGA |
| 37407_2 | TATGAGTGTGGAGTCGTTGCGCTTGCAGTGAGCCAAGATTGNNNAGATCGGAAGA |
| 50199_1 | TATGAGTGTGGAGTCGTTGCGAGAAGTTGGAGGTGTCACAGATNNNAGATCGGAA |
| 50199_2 | TATGAGTGTGGAGTCGTTGCGAGAAGTTGGAGGTGTCACAGATNNNAGATCGGAA |
| 10598_1 | TATGAGTGTGGAGTCGTTGCCCATCTTTCCCACCTTGAGCTTTNNNAGATCGGAAGA |
| 10598_2 | TATGAGTGTGGAGTCGTTGCCCATCTTTCCCACCTTGAGCTTTNNNAGATCGGAAGA |
| 31915_1 | TATGAGTGTGGAGTCGTTGCACTAGCCTTAGCAACATGGTGAANNNAGATCGGAAG |
| 31915_2 | TATGAGTGTGGAGTCGTTGCACTAGCCTTAGCAACATGGTGAANNNAGATCGGAAG |
| 10583_1 | TATGAGTGTGGAGTCGTTGCCAGCTAAGGAACTGAAGCCTCCTNNNAGATCGGAAG |
| 10583_2 | TATGAGTGTGGAGTCGTTGCCAGCTAAGGAACTGAAGCCTCCTNNNAGATCGGAAG |
| 38916_1 | TATGAGTGTGGAGTCGTTGCAGACACAAGACACTTGTCATCGANNNAGATCGGAAG |
| 38916_2 | TATGAGTGTGGAGTCGTTGCAGACACAAGACACTTGTCATCGANNNAGATCGGAAG |
| 47220_1 | TATGAGTGTGGAGTCGTTGCTCTAGGCACCAGGGATGTAGTAANNNAGATCGGAA |
| 47220_2 | TATGAGTGTGGAGTCGTTGCTCTAGGCACCAGGGATGTAGTAANNNAGATCGGAA |
| 33828_1 | TATGAGTGTGGAGTCGTTGCAAAGGTGGGAGAAGTAGCATCAGNNNAGATCGGA |

| | |
|---|---|
| 33828_2 | TATGAGTGTGGAGTCGTTGCAAAGGTGGGAGAAGTAGCATCAGNNNAGATCGGA |
| 52307_1 | TATGAGTGTGGAGTCGTTGCTTAGCTATACTTGCTGGAGTGGGNNNAGATCGGAA |
| 52307_2 | TATGAGTGTGGAGTCGTTGCTTAGCTATACTTGCTGGAGTGGGNNNAGATCGGAA |
| 44191_1 | TATGAGTGTGGAGTCGTTGCGTGTGGGGTCAACTTCTAAGCTANNNAGATCGGAA |
| 44191_2 | TATGAGTGTGGAGTCGTTGCGTGTGGGGTCAACTTCTAAGCTANNNAGATCGGAA |
| 3944_1 | TATGAGTGTGGAGTCGTTGCGCCTTTGGAAATGTGGTCATGTGNNNAGATCGGAA |
| 3944_2 | TATGAGTGTGGAGTCGTTGCGCCTTTGGAAATGTGGTCATGTGNNNAGATCGGAA |
| 32524_1 | TATGAGTGTGGAGTCGTTGCAACCTGAAAGAACTCCTGGTCAGNNNAGATCGGAA |
| 32524_2 | TATGAGTGTGGAGTCGTTGCAACCTGAAAGAACTCCTGGTCAGNNNAGATCGGAA |
| 36889_1 | TATGAGTGTGGAGTCGTTGCGAAAAGAAAAGCAATGGCAGCATNNNAGATCGGAA |
| 36889_2 | TATGAGTGTGGAGTCGTTGCGAAAAGAAAAGCAATGGCAGCATNNNAGATCGGAA |
| 10007_1 | TATGAGTGTGGAGTCGTTGCACGCCTCAAGTCTTTGTCCTTTTNNNAGATCGGAAGA |
| 10007_2 | TATGAGTGTGGAGTCGTTGCACGCCTCAAGTCTTTGTCCTTTTNNNAGATCGGAAGA |
| 41948_1 | TATGAGTGTGGAGTCGTTGCTTCAAGTGATTCTCCTGCCTCAGNNNAGATCGGAAG |
| 41948_2 | TATGAGTGTGGAGTCGTTGCTTCAAGTGATTCTCCTGCCTCAGNNNAGATCGGAAG |
| 34405_1 | TATGAGTGTGGAGTCGTTGCGCTTTGTCATGGTGAGCTACTTCNNNAGATCGGAAG |
| 34405_2 | TATGAGTGTGGAGTCGTTGCGCTTTGTCATGGTGAGCTACTTCNNNAGATCGGAAG |
| 2319_1 | TATGAGTGTGGAGTCGTTGCTGGCTTTATGGGCATGTTCCAATNNNAGATCGGAAG |
| 2319_2 | TATGAGTGTGGAGTCGTTGCTGGCTTTATGGGCATGTTCCAATNNNAGATCGGAAG |
| 30623_1 | TATGAGTGTGGAGTCGTTGCGCAAAAAGCTCAAAGTTGAAGGCNNNAGATCGGAA |
| 30623_2 | TATGAGTGTGGAGTCGTTGCGCAAAAAGCTCAAAGTTGAAGGCNNNAGATCGGAA |
| 44564_1 | TATGAGTGTGGAGTCGTTGCATTTCTGAGACTACAAGAGCCCGNNNAGATCGGAAG |
| 44564_2 | TATGAGTGTGGAGTCGTTGCATTTCTGAGACTACAAGAGCCCGNNNAGATCGGAAG |
| 48089_1 | TATGAGTGTGGAGTCGTTGCCTCATGCCATGCAGACCAGAANNNAGATCGGAAGA |
| 48089_2 | TATGAGTGTGGAGTCGTTGCCTCATGCCATGCAGACCAGAANNNAGATCGGAAGA |
| 39231_1 | TATGAGTGTGGAGTCGTTGCATCTCTGGTCCTAGCTACTCAGGNNNAGATCGGAAG |
| 39231_2 | TATGAGTGTGGAGTCGTTGCATCTCTGGTCCTAGCTACTCAGGNNNAGATCGGAAG |
| 39198_1 | TATGAGTGTGGAGTCGTTGCAATTTGCACACATGACTCTGGTGNNNAGATCGGAAG |
| 39198_2 | TATGAGTGTGGAGTCGTTGCAATTTGCACACATGACTCTGGTGNNNAGATCGGAAG |
| 40089_1 | TATGAGTGTGGAGTCGTTGCGAAGCGGAATTTGTAGTGAGCTCNNNAGATCGGAA |
| 40089_2 | TATGAGTGTGGAGTCGTTGCGAAGCGGAATTTGTAGTGAGCTCNNNAGATCGGAA |
| 46444_1 | TATGAGTGTGGAGTCGTTGCCTACGCCCGGCTAATTTTTTGTANNNAGATCGGAAG |
| 46444_2 | TATGAGTGTGGAGTCGTTGCCTACGCCCGGCTAATTTTTTGTANNNAGATCGGAAG |
| 39365_1 | TATGAGTGTGGAGTCGTTGCGTTTCAGTGAGTCAAGATGGCACNNNAGATCGGAA |
| 39365_2 | TATGAGTGTGGAGTCGTTGCGTTTCAGTGAGTCAAGATGGCACNNNAGATCGGAA |
| 29207_1 | TATGAGTGTGGAGTCGTTGCGAAAGGGAGATGGGGTAGGTGAGNNNAGATCGG |
| 29207_2 | TATGAGTGTGGAGTCGTTGCGAAAGGGAGATGGGGTAGGTGAGNNNAGATCGG |
| 44117_1 | TATGAGTGTGGAGTCGTTGCAAACGTGAGAATTTGAGTGGCTGNNNAGATCGGAA |
| 44117_2 | TATGAGTGTGGAGTCGTTGCAAACGTGAGAATTTGAGTGGCTGNNNAGATCGGAA |
| 37935_1 | TATGAGTGTGGAGTCGTTGCTGGTCAATATGGTGAAACCCTGTNNNAGATCGGAA |
| 37935_2 | TATGAGTGTGGAGTCGTTGCTGGTCAATATGGTGAAACCCTGTNNNAGATCGGAA |
| 30274_1 | TATGAGTGTGGAGTCGTTGCACAGGAATAAAGCAGTGAAATGACTNNNAGATCGG |
| 30274_2 | TATGAGTGTGGAGTCGTTGCACAGGAATAAAGCAGTGAAATGACTNNNAGATCGG |

| | |
|---|---|
| 46128_1 | TATGAGTGTGGAGTCGTTGCAGACTGAGGCCATTAGTAGTCCTNNNAGATCGGAAG |
| 46128_2 | TATGAGTGTGGAGTCGTTGCAGACTGAGGCCATTAGTAGTCCTNNNAGATCGGAAG |
| 44578_1 | TATGAGTGTGGAGTCGTTGCGTTCTGGCACTCCCTTCTAATCANNNAGATCGGAAGA |
| 44578_2 | TATGAGTGTGGAGTCGTTGCGTTCTGGCACTCCCTTCTAATCANNNAGATCGGAAGA |
| 35558_1 | TATGAGTGTGGAGTCGTTGCCTTATGGTGGGCAGTGTTGTTTTNNNAGATCGGAAG |
| 35558_2 | TATGAGTGTGGAGTCGTTGCCTTATGGTGGGCAGTGTTGTTTTNNNAGATCGGAAG |
| 34520_1 | TATGAGTGTGGAGTCGTTGCTGCAAAATGTGAACCTTGTTGGANNNAGATCGGAAG |
| 34520_2 | TATGAGTGTGGAGTCGTTGCTGCAAAATGTGAACCTTGTTGGANNNAGATCGGAAG |
| 10128_1 | TATGAGTGTGGAGTCGTTGCCTTCCACTGTTTCTCTCCCCAACNNNAGATCGGAAGAG |
| 10128_2 | TATGAGTGTGGAGTCGTTGCCTTCCACTGTTTCTCTCCCCAACNNNAGATCGGAAGAG |
| 38314_1 | TATGAGTGTGGAGTCGTTGCTCCTATGCTCATGGATGGGTAGANNNAGATCGGAAG |
| 38314_2 | TATGAGTGTGGAGTCGTTGCTCCTATGCTCATGGATGGGTAGANNNAGATCGGAAG |
| 39382_1 | TATGAGTGTGGAGTCGTTGCGCCCAAAGCAATTTACAGATTCANNNAGATCGGAAG |
| 39382_2 | TATGAGTGTGGAGTCGTTGCGCCCAAAGCAATTTACAGATTCANNNAGATCGGAAG |
| 38683_1 | TATGAGTGTGGAGTCGTTGCCTGAGGTGGGAGAATCACTTGAGNNNAGATCGGAA |
| 38683_2 | TATGAGTGTGGAGTCGTTGCCTGAGGTGGGAGAATCACTTGAGNNNAGATCGGAA |
| 12719_1 | TATGAGTGTGGAGTCGTTGCCTTTGAGAAGAAAGGTGAGTCNNNAGATCGGAAGA |
| 12719_2 | TATGAGTGTGGAGTCGTTGCCTTTGAGAAGAAAGGTGAGTCNNNAGATCGGAAGA |
| 45032_1 | TATGAGTGTGGAGTCGTTGCGGGTCTAGGTTTCTGTGTAACGTNNNAGATCGGAAG |
| 45032_2 | TATGAGTGTGGAGTCGTTGCGGGTCTAGGTTTCTGTGTAACGTNNNAGATCGGAAG |
| 32612_1 | TATGAGTGTGGAGTCGTTGCTGCTTCACGAGAAAACATACAGTNNNAGATCGGAAG |
| 32612_2 | TATGAGTGTGGAGTCGTTGCTGCTTCACGAGAAAACATACAGTNNNAGATCGGAAG |
| 47513_1 | TATGAGTGTGGAGTCGTTGCACGCCCGGCTAATTTGTTCTAGNNNAGATCGGAAGA |
| 47513_2 | TATGAGTGTGGAGTCGTTGCACGCCCGGCTAATTTGTTCTAGNNNAGATCGGAAGA |
| 49048_1 | TATGAGTGTGGAGTCGTTGCCAGTGGCATGATCTCAGCTTGNNNAGATCGGAAGAG |
| 49048_2 | TATGAGTGTGGAGTCGTTGCCAGTGGCATGATCTCAGCTTGNNNAGATCGGAAGAG |
| 12050_1 | TATGAGTGTGGAGTCGTTGCAGAGAAGTAGCCAGATGGCATGAGNNNAGATCGGA |
| 12050_2 | TATGAGTGTGGAGTCGTTGCAGAGAAGTAGCCAGATGGCATGAGNNNAGATCGGA |
| 33958_1 | TATGAGTGTGGAGTCGTTGCAGAGTACAGTGGTGTGATCATGGNNNAGATCGGAA |
| 33958_2 | TATGAGTGTGGAGTCGTTGCAGAGTACAGTGGTGTGATCATGGNNNAGATCGGAA |
| 34753_1 | TATGAGTGTGGAGTCGTTGCTCTCAGTTCACTGCAACCTCTACNNNAGATCGGAAGA |
| 34753_2 | TATGAGTGTGGAGTCGTTGCTCTCAGTTCACTGCAACCTCTACNNNAGATCGGAAGA |
| 2165_1 | TATGAGTGTGGAGTCGTTGCATCCCAGAAACCTCATCAGGGAANNNAGATCGGAAG |
| 2165_2 | TATGAGTGTGGAGTCGTTGCATCCCAGAAACCTCATCAGGGAANNNAGATCGGAAG |
| 50203_1 | TATGAGTGTGGAGTCGTTGCTTGTCCATTTCCCAGAGTCTCTGNNNAGATCGGAAGA |
| 50203_2 | TATGAGTGTGGAGTCGTTGCTTGTCCATTTCCCAGAGTCTCTGNNNAGATCGGAAGA |
| 49980_1 | TATGAGTGTGGAGTCGTTGCGCTTCTGGCTTGGATGATGTAACNNNAGATCGGAAG |
| 49980_2 | TATGAGTGTGGAGTCGTTGCGCTTCTGGCTTGGATGATGTAACNNNAGATCGGAAG |
| 41246_1 | TATGAGTGTGGAGTCGTTGCCATCCTGGCTAACATGGTGAAACNNNAGATCGGAAG |
| 41246_2 | TATGAGTGTGGAGTCGTTGCCATCCTGGCTAACATGGTGAAACNNNAGATCGGAAG |
| 39784_1 | TATGAGTGTGGAGTCGTTGCCTGTAATCCCAGCTACTCAGGAGNNNAGATCGGAAG |
| 39784_2 | TATGAGTGTGGAGTCGTTGCCTGTAATCCCAGCTACTCAGGAGNNNAGATCGGAAG |
| 51633_1 | TATGAGTGTGGAGTCGTTGCCAGGGCTCTCCTTACTTTCGTTTNNNAGATCGGAAGA |

| | |
|---|---|
| 51633_2 | TATGAGTGTGGAGTCGTTGCCAGGGCTCTCCTTACTTTCGTTTNNNAGATCGGAAGA |
| 50036_1 | TATGAGTGTGGAGTCGTTGCAGGTGATATGAGAGCCCAAATCCNNNAGATCGGAAG |
| 50036_2 | TATGAGTGTGGAGTCGTTGCAGGTGATATGAGAGCCCAAATCCNNNAGATCGGAAG |
| 51580_1 | TATGAGTGTGGAGTCGTTGCCTTCTCATTCTGCAAGTGTTGGGNNNAGATCGGAAG |
| 51580_2 | TATGAGTGTGGAGTCGTTGCCTTCTCATTCTGCAAGTGTTGGGNNNAGATCGGAAG |
| 10639_1 | TATGAGTGTGGAGTCGTTGCTTGGTGTTTTCTTCAGGGACCTGNNNAGATCGGAAG |
| 10639_2 | TATGAGTGTGGAGTCGTTGCTTGGTGTTTTCTTCAGGGACCTGNNNAGATCGGAAG |
| 44569_1 | TATGAGTGTGGAGTCGTTGCGAATGTAAATTGGTGCACCTACTNNNAGATCGGAAG |
| 44569_2 | TATGAGTGTGGAGTCGTTGCGAATGTAAATTGGTGCACCTACTNNNAGATCGGAAG |
| 2532_1 | TATGAGTGTGGAGTCGTTGCTATGCCTCTGTATGTTTGCCAGGNNNAGATCGGAAG |
| 2532_2 | TATGAGTGTGGAGTCGTTGCTATGCCTCTGTATGTTTGCCAGGNNNAGATCGGAAG |
| 34615_1 | TATGAGTGTGGAGTCGTTGCATGCATGCTCAGGAGTGTATGTANNNAGATCGGAAG |
| 34615_2 | TATGAGTGTGGAGTCGTTGCATGCATGCTCAGGAGTGTATGTANNNAGATCGGAAG |
| 47821_1 | TATGAGTGTGGAGTCGTTGCCCTTACCCTCATCACGACTGTAGNNNAGATCGGAAGA |
| 47821_2 | TATGAGTGTGGAGTCGTTGCCCTTACCCTCATCACGACTGTAGNNNAGATCGGAAGA |
| 37947_1 | TATGAGTGTGGAGTCGTTGCCTGAGGCAGGATAATTGCTTGAGNNNAGATCGGAAG |
| 37947_2 | TATGAGTGTGGAGTCGTTGCCTGAGGCAGGATAATTGCTTGAGNNNAGATCGGAAG |
| 33552_1 | TATGAGTGTGGAGTCGTTGCTTCTTCTGGTTCGGTCTTGAGAGNNNAGATCGGAAG |
| 33552_2 | TATGAGTGTGGAGTCGTTGCTTCTTCTGGTTCGGTCTTGAGAGNNNAGATCGGAAG |
| 36281_1 | TATGAGTGTGGAGTCGTTGCACAGGAGATGACAGTCTTCATGTNNNAGATCGGAAG |
| 36281_2 | TATGAGTGTGGAGTCGTTGCACAGGAGATGACAGTCTTCATGTNNNAGATCGGAAG |
| 50148_1 | TATGAGTGTGGAGTCGTTGCGTGAATTTAGGCCAACACAGCTTNNNAGATCGGAAG |
| 50148_2 | TATGAGTGTGGAGTCGTTGCGTGAATTTAGGCCAACACAGCTTNNNAGATCGGAAG |
| 51504_1 | TATGAGTGTGGAGTCGTTGCCTCCACCCCAAATCAACAGGATANNNAGATCGGAAGA |
| 51504_2 | TATGAGTGTGGAGTCGTTGCCTCCACCCCAAATCAACAGGATANNNAGATCGGAAGA |
| 49160_1 | TATGAGTGTGGAGTCGTTGCAGGCTGAAGATGCAGAAAAAACANNNAGATCGGAA |
| 49160_2 | TATGAGTGTGGAGTCGTTGCAGGCTGAAGATGCAGAAAAAACANNNAGATCGGAA |
| 30305_1 | TATGAGTGTGGAGTCGTTGCGTCCTCTCAGTCTTGGTCAGAGANNNAGATCGGAAG |
| 30305_2 | TATGAGTGTGGAGTCGTTGCGTCCTCTCAGTCTTGGTCAGAGANNNAGATCGGAAG |
| 2913_1 | TATGAGTGTGGAGTCGTTGCACATGCTTCCCTCACCATTTTGTNNNAGATCGGAAGA |
| 2913_2 | TATGAGTGTGGAGTCGTTGCACATGCTTCCCTCACCATTTTGTNNNAGATCGGAAGA |
| 29921_1 | TATGAGTGTGGAGTCGTTGCAGAAGCTAGGAATGTCCGAACTGNNNAGATCGGAAG |
| 29921_2 | TATGAGTGTGGAGTCGTTGCAGAAGCTAGGAATGTCCGAACTGNNNAGATCGGAAG |
| 49062_1 | TATGAGTGTGGAGTCGTTGCGTTGTCCAAGTTGGCTTCAAACTNNNAGATCGGAAG |
| 49062_2 | TATGAGTGTGGAGTCGTTGCGTTGTCCAAGTTGGCTTCAAACTNNNAGATCGGAAG |
| 50797_1 | TATGAGTGTGGAGTCGTTGCGCCTGATCTCTCATGAGTAGGACNNNAGATCGGAAG |
| 50797_2 | TATGAGTGTGGAGTCGTTGCGCCTGATCTCTCATGAGTAGGACNNNAGATCGGAAG |
| 40232_1 | TATGAGTGTGGAGTCGTTGCGGGAGACAGGAATTTCACTGACTNNNAGATCGGAAG |
| 40232_2 | TATGAGTGTGGAGTCGTTGCGGGAGACAGGAATTTCACTGACTNNNAGATCGGAAG |
| 45297_1 | TATGAGTGTGGAGTCGTTGCTGATGTTGCCTCCCTTTCATGTANNNAGATCGGAAGA |
| 45297_2 | TATGAGTGTGGAGTCGTTGCTGATGTTGCCTCCCTTTCATGTANNNAGATCGGAAGA |
| 45060_1 | TATGAGTGTGGAGTCGTTGCCGAACGGACGTGAAGAATGTGNNNAGATCGGAAGA |
| 45060_2 | TATGAGTGTGGAGTCGTTGCCGAACGGACGTGAAGAATGTGNNNAGATCGGAAGA |

| | |
|---|---|
| 39367_1 | TATGAGTGTGGAGTCGTTGCATAGTGTGGAGGATGAATTGGGGNNNAGATCGGA |
| 39367_2 | TATGAGTGTGGAGTCGTTGCATAGTGTGGAGGATGAATTGGGGNNNAGATCGGA |
| 50162_1 | TATGAGTGTGGAGTCGTTGCGTATGCCCTAAGTGCTTCTGCTANNNAGATCGGAAG |
| 50162_2 | TATGAGTGTGGAGTCGTTGCGTATGCCCTAAGTGCTTCTGCTANNNAGATCGGAAG |
| 49015_1 | TATGAGTGTGGAGTCGTTGCGCGCCCGGCCTATTTATTTATTNNNAGATCGGAAG |
| 49015_2 | TATGAGTGTGGAGTCGTTGCGCGCCCGGCCTATTTATTTATTNNNAGATCGGAAG |
| 52419_1 | TATGAGTGTGGAGTCGTTGCCTCCACACAAAGAGAACGTCCANNNAGATCGGAAGA |
| 52419_2 | TATGAGTGTGGAGTCGTTGCCTCCACACAAAGAGAACGTCCANNNAGATCGGAAGA |
| 13356_1 | TATGAGTGTGGAGTCGTTGCGGGGGAAAAAAACATCTAATAAANNNAGATCGGAA |
| 13356_2 | TATGAGTGTGGAGTCGTTGCGGGGGAAAAAAACATCTAATAAANNNAGATCGGAA |
| 49451_1 | TATGAGTGTGGAGTCGTTGCACTTTGTCTACAGGCATCTCTCCNNNAGATCGGAAGA |
| 49451_2 | TATGAGTGTGGAGTCGTTGCACTTTGTCTACAGGCATCTCTCCNNNAGATCGGAAGA |
| 39976_1 | TATGAGTGTGGAGTCGTTGCTGGTGTCTCACGCCTGTAATCNNNAGATCGGAAGAG |
| 39976_2 | TATGAGTGTGGAGTCGTTGCTGGTGTCTCACGCCTGTAATCNNNAGATCGGAAGAG |
| 38980_1 | TATGAGTGTGGAGTCGTTGCTGTATCTTTCCTCTGGCATACCANNNAGATCGGAAGA |
| 38980_2 | TATGAGTGTGGAGTCGTTGCTGTATCTTTCCTCTGGCATACCANNNAGATCGGAAGA |
| 35679_1 | TATGAGTGTGGAGTCGTTGCGATTTTGGCTCACTGCAACCTCNNNAGATCGGAAGA |
| 35679_2 | TATGAGTGTGGAGTCGTTGCGATTTTGGCTCACTGCAACCTCNNNAGATCGGAAGA |
| 30713_1 | TATGAGTGTGGAGTCGTTGCTTTGATGGGCCAGCTTTTTCATTNNNAGATCGGAAG |
| 30713_2 | TATGAGTGTGGAGTCGTTGCTTTGATGGGCCAGCTTTTTCATTNNNAGATCGGAAG |
| 39508_1 | TATGAGTGTGGAGTCGTTGCGAGAGAGACAGAAAGACGAAGGANNNAGATCGGA |
| 39508_2 | TATGAGTGTGGAGTCGTTGCGAGAGAGACAGAAAGACGAAGGANNNAGATCGGA |
| 41533_1 | TATGAGTGTGGAGTCGTTGCCGCCCGGCTTTTCTTTTTTTTTTNNNAGATCGGAAGA |
| 41533_2 | TATGAGTGTGGAGTCGTTGCCGCCCGGCTTTTCTTTTTTTTTTNNNAGATCGGAAGA |
| 39603_1 | TATGAGTGTGGAGTCGTTGCACTCCTCAAACCTTTAGGGATGGNNNAGATCGGAAG |
| 39603_2 | TATGAGTGTGGAGTCGTTGCACTCCTCAAACCTTTAGGGATGGNNNAGATCGGAAG |
| 33782_1 | TATGAGTGTGGAGTCGTTGCGAAGACACAAATGGCCAAGAAGTNNNAGATCGGAA |
| 33782_2 | TATGAGTGTGGAGTCGTTGCGAAGACACAAATGGCCAAGAAGTNNNAGATCGGAA |
| 34336_1 | TATGAGTGTGGAGTCGTTGCCCCAAGTCCCCAAAGTCCATTATNNNAGATCGGAAGA |
| 34336_2 | TATGAGTGTGGAGTCGTTGCCCCAAGTCCCCAAAGTCCATTATNNNAGATCGGAAGA |
| 35241_1 | TATGAGTGTGGAGTCGTTGCTACTCAGGCTGGTCTTGAATTCCNNNAGATCGGAAG |
| 35241_2 | TATGAGTGTGGAGTCGTTGCTACTCAGGCTGGTCTTGAATTCCNNNAGATCGGAAG |
| 29317_1 | TATGAGTGTGGAGTCGTTGCAGAGAGAGAGAGGGAGAGAGAGANNNAGATCGG |
| 29317_2 | TATGAGTGTGGAGTCGTTGCAGAGAGAGAGAGGGAGAGAGAGANNNAGATCGG |
| 45094_1 | TATGAGTGTGGAGTCGTTGCAGTGAGATGCAGAGAACCTTGTTNNNAGATCGGAA |
| 45094_2 | TATGAGTGTGGAGTCGTTGCAGTGAGATGCAGAGAACCTTGTTNNNAGATCGGAA |
| 39134_1 | TATGAGTGTGGAGTCGTTGCACTCTACCTTGGACTTTATGCTGTNNNAGATCGGAAG |
| 39134_2 | TATGAGTGTGGAGTCGTTGCACTCTACCTTGGACTTTATGCTGTNNNAGATCGGAAG |
| 41486_1 | TATGAGTGTGGAGTCGTTGCAATCCAACCCTCACAATACTCCCNNNAGATCGGAAGA |
| 41486_2 | TATGAGTGTGGAGTCGTTGCAATCCAACCCTCACAATACTCCCNNNAGATCGGAAGA |
| 31472_1 | TATGAGTGTGGAGTCGTTGCATGACTCCCAAACATCCAGTCAANNNAGATCGGAAG |
| 31472_2 | TATGAGTGTGGAGTCGTTGCATGACTCCCAAACATCCAGTCAANNNAGATCGGAAG |
| 46816_1 | TATGAGTGTGGAGTCGTTGCCGAGTAGCTGGGACTACAGTTGNNNAGATCGGAAG |

| | |
|---|---|
| 46816_2 | TATGAGTGTGGAGTCGTTGCCGAGTAGCTGGGACTACAGTTGNNNAGATCGGAAG |
| 52881_1 | TATGAGTGTGGAGTCGTTGCTTTCTCGTGCTGGGTTGTAAATGNNNAGATCGGAAG |
| 52881_2 | TATGAGTGTGGAGTCGTTGCTTTCTCGTGCTGGGTTGTAAATGNNNAGATCGGAAG |
| 34676_1 | TATGAGTGTGGAGTCGTTGCTTTCCTCTTTGCTTTCTCCTGGANNNAGATCGGAAGA |
| 34676_2 | TATGAGTGTGGAGTCGTTGCTTTCCTCTTTGCTTTCTCCTGGANNNAGATCGGAAGA |
| 32819_1 | TATGAGTGTGGAGTCGTTGCTTCAAGCAATTCTCCTGCCTCAGNNNAGATCGGAAGA |
| 32819_2 | TATGAGTGTGGAGTCGTTGCTTCAAGCAATTCTCCTGCCTCAGNNNAGATCGGAAGA |
| 28884_1 | TATGAGTGTGGAGTCGTTGCTGTCGTCCTAAGCGTTGTTCTTANNNAGATCGGAAG |
| 28884_2 | TATGAGTGTGGAGTCGTTGCTGTCGTCCTAAGCGTTGTTCTTANNNAGATCGGAAG |
| 42215_1 | TATGAGTGTGGAGTCGTTGCAATGATGATGTGTCCATGTGCATNNNAGATCGGAAG |
| 42215_2 | TATGAGTGTGGAGTCGTTGCAATGATGATGTGTCCATGTGCATNNNAGATCGGAAG |
| 39107_1 | TATGAGTGTGGAGTCGTTGCGATATGACCAGATGGCCTGAAGTNNNAGATCGGAA |
| 39107_2 | TATGAGTGTGGAGTCGTTGCGATATGACCAGATGGCCTGAAGTNNNAGATCGGAA |
| 51637_1 | TATGAGTGTGGAGTCGTTGCCATTTATGTGAGCTTCCCTGCAGNNNAGATCGGAAG |
| 51637_2 | TATGAGTGTGGAGTCGTTGCCATTTATGTGAGCTTCCCTGCAGNNNAGATCGGAAG |
| 39964_1 | TATGAGTGTGGAGTCGTTGCGGGAAAAGAAAGACTGGAGATGGNNNAGATCGGA |
| 39964_2 | TATGAGTGTGGAGTCGTTGCGGGAAAAGAAAGACTGGAGATGGNNNAGATCGGA |
| 13549_1 | TATGAGTGTGGAGTCGTTGCTTTAATAAATGAAATTGCTGGGANNNAGATCGGAAG |
| 13549_2 | TATGAGTGTGGAGTCGTTGCTTTAATAAATGAAATTGCTGGGANNNAGATCGGAAG |
| 40143_1 | TATGAGTGTGGAGTCGTTGCTTAGGTTGGAGATACCAGGGGATNNNAGATCGGAA |
| 40143_2 | TATGAGTGTGGAGTCGTTGCTTAGGTTGGAGATACCAGGGGATNNNAGATCGGAA |
| 37997_1 | TATGAGTGTGGAGTCGTTGCATCACTTCTTGCTCCAACACAGANNNAGATCGGAAGA |
| 37997_2 | TATGAGTGTGGAGTCGTTGCATCACTTCTTGCTCCAACACAGANNNAGATCGGAAGA |
| 45330_1 | TATGAGTGTGGAGTCGTTGCAACCTGCCTCCCACTTTATTCAANNNAGATCGGAAGA |
| 45330_2 | TATGAGTGTGGAGTCGTTGCAACCTGCCTCCCACTTTATTCAANNNAGATCGGAAGA |
| 50257_1 | TATGAGTGTGGAGTCGTTGCACTGGGGCAGATATAGATGGAGTNNNAGATCGGAA |
| 50257_2 | TATGAGTGTGGAGTCGTTGCACTGGGGCAGATATAGATGGAGTNNNAGATCGGAA |
| 46730_1 | TATGAGTGTGGAGTCGTTGCTTCAAGCGATTCTCCTGCTTCAANNNAGATCGGAAGA |
| 46730_2 | TATGAGTGTGGAGTCGTTGCTTCAAGCGATTCTCCTGCTTCAANNNAGATCGGAAGA |
| 46622_1 | TATGAGTGTGGAGTCGTTGCCTCCTGAGTAGCTGGGATTACAGNNNAGATCGGAAG |
| 46622_2 | TATGAGTGTGGAGTCGTTGCCTCCTGAGTAGCTGGGATTACAGNNNAGATCGGAAG |
| 44420_1 | TATGAGTGTGGAGTCGTTGCATGCACAGAGTGGATTCATGAGTNNNAGATCGGAA |
| 44420_2 | TATGAGTGTGGAGTCGTTGCATGCACAGAGTGGATTCATGAGTNNNAGATCGGAA |
| 31330_1 | TATGAGTGTGGAGTCGTTGCGTTTCTCCTTCTCTGAGGAACCCNNNAGATCGGAAGA |
| 31330_2 | TATGAGTGTGGAGTCGTTGCGTTTCTCCTTCTCTGAGGAACCCNNNAGATCGGAAGA |
| 36804_1 | TATGAGTGTGGAGTCGTTGCAATACAAAAAATTAGCCGGGCGTNNNAGATCGGAA |
| 36804_2 | TATGAGTGTGGAGTCGTTGCAATACAAAAAATTAGCCGGGCGTNNNAGATCGGAA |
| 35198_1 | TATGAGTGTGGAGTCGTTGCCAATCTTGACTCACTGCAAGCTCNNNAGATCGGAAGA |
| 35198_2 | TATGAGTGTGGAGTCGTTGCCAATCTTGACTCACTGCAAGCTCNNNAGATCGGAAGA |
| 33021_1 | TATGAGTGTGGAGTCGTTGCGGTTCAAACAATTCTCCTGCCTCNNNAGATCGGAAGA |
| 33021_2 | TATGAGTGTGGAGTCGTTGCGGTTCAAACAATTCTCCTGCCTCNNNAGATCGGAAGA |
| 33276_1 | TATGAGTGTGGAGTCGTTGCAGATGGAGTTTCTCTCTGTTGCCNNNAGATCGGAAG |
| 33276_2 | TATGAGTGTGGAGTCGTTGCAGATGGAGTTTCTCTCTGTTGCCNNNAGATCGGAAG |

| | |
|---|---|
| 36475_1 | TATGAGTGTGGAGTCGTTGCTGCTAACTGTCCGAAATTCTCCANNNAGATCGGAAG |
| 36475_2 | TATGAGTGTGGAGTCGTTGCTGCTAACTGTCCGAAATTCTCCANNNAGATCGGAAG |
| 29272_1 | TATGAGTGTGGAGTCGTTGCCAGAAATTTTCCACTCGGGCTTTNNNAGATCGGAAG |
| 29272_2 | TATGAGTGTGGAGTCGTTGCCAGAAATTTTCCACTCGGGCTTTNNNAGATCGGAAG |
| 2702_1 | TATGAGTGTGGAGTCGTTGCTCATCGTGCAGTGTTTTGAGGATNNNAGATCGGAA |
| 2702_2 | TATGAGTGTGGAGTCGTTGCTCATCGTGCAGTGTTTTGAGGATNNNAGATCGGAA |
| 47378_1 | TATGAGTGTGGAGTCGTTGCACGTGGCACATCTAACAGAAGATNNNAGATCGGAA |
| 47378_2 | TATGAGTGTGGAGTCGTTGCACGTGGCACATCTAACAGAAGATNNNAGATCGGAA |
| 2135_1 | TATGAGTGTGGAGTCGTTGCATAATCCAGGAAGAGCACCCTCANNNAGATCGGAAG |
| 2135_2 | TATGAGTGTGGAGTCGTTGCATAATCCAGGAAGAGCACCCTCANNNAGATCGGAAG |
| 50631_1 | TATGAGTGTGGAGTCGTTGCGTTTGTCTGGATCTCCCCCTGNNNAGATCGGAAGAG |
| 50631_2 | TATGAGTGTGGAGTCGTTGCGTTTGTCTGGATCTCCCCCTGNNNAGATCGGAAGAG |
| 42007_1 | TATGAGTGTGGAGTCGTTGCTTTCACGCCATTCTCCTGTCTCNNNAGATCGGAAGAG |
| 42007_2 | TATGAGTGTGGAGTCGTTGCTTTCACGCCATTCTCCTGTCTCNNNAGATCGGAAGAG |
| 45634_1 | TATGAGTGTGGAGTCGTTGCTTCTGAGTTGCTGGGATTACAGGNNNAGATCGGAA |
| 45634_2 | TATGAGTGTGGAGTCGTTGCTTCTGAGTTGCTGGGATTACAGGNNNAGATCGGAA |
| 51424_1 | TATGAGTGTGGAGTCGTTGCCTGGAAGAGGTCAGAGTTCTGTCNNNAGATCGGAA |
| 51424_2 | TATGAGTGTGGAGTCGTTGCCTGGAAGAGGTCAGAGTTCTGTCNNNAGATCGGAA |
| 49579_1 | TATGAGTGTGGAGTCGTTGCTGGCTAGGGCACTGTGTTTATTANNNAGATCGGAA |
| 49579_2 | TATGAGTGTGGAGTCGTTGCTGGCTAGGGCACTGTGTTTATTANNNAGATCGGAA |
| 38513_1 | TATGAGTGTGGAGTCGTTGCCTGAGGCACGAGAATCACTTGAANNNAGATCGGAA |
| 38513_2 | TATGAGTGTGGAGTCGTTGCCTGAGGCACGAGAATCACTTGAANNNAGATCGGAA |
| 3061_1 | TATGAGTGTGGAGTCGTTGCAGAAAGGTTGAGGGGAAATCAGCNNNAGATCGGA |
| 3061_2 | TATGAGTGTGGAGTCGTTGCAGAAAGGTTGAGGGGAAATCAGCNNNAGATCGGA |
| 51862_1 | TATGAGTGTGGAGTCGTTGCAAGGAAATGGGATGTTGGAAGGANNNAGATCGGA |
| 51862_2 | TATGAGTGTGGAGTCGTTGCAAGGAAATGGGATGTTGGAAGGANNNAGATCGGA |
| 10801_1 | TATGAGTGTGGAGTCGTTGCCCCAGCATCTCCCTAACCAATTCNNNAGATCGGAAGA |
| 10801_2 | TATGAGTGTGGAGTCGTTGCCCCAGCATCTCCCTAACCAATTCNNNAGATCGGAAGA |
| 38205_1 | TATGAGTGTGGAGTCGTTGCCAGTAAACAGGATGGAGGACCAANNNAGATCGGAA |
| 38205_2 | TATGAGTGTGGAGTCGTTGCCAGTAAACAGGATGGAGGACCAANNNAGATCGGAA |
| 40989_1 | TATGAGTGTGGAGTCGTTGCAAAGTGTGTAGCATTTCCCCCTTNNNAGATCGGAAG |
| 40989_2 | TATGAGTGTGGAGTCGTTGCAAAGTGTGTAGCATTTCCCCCTTNNNAGATCGGAAG |
| 38311_1 | TATGAGTGTGGAGTCGTTGCCTCCACAGGCAATTTCTTTCAGTNNNAGATCGGAAG |
| 38311_2 | TATGAGTGTGGAGTCGTTGCCTCCACAGGCAATTTCTTTCAGTNNNAGATCGGAAG |
| 33261_1 | TATGAGTGTGGAGTCGTTGCCGATCTCGGTTCACTGCAACCNNNAGATCGGAAGAG |
| 33261_2 | TATGAGTGTGGAGTCGTTGCCGATCTCGGTTCACTGCAACCNNNAGATCGGAAGAG |
| 44742_1 | TATGAGTGTGGAGTCGTTGCGAGGTCAGAAGATCGAGACCATCNNNAGATCGGAA |
| 44742_2 | TATGAGTGTGGAGTCGTTGCGAGGTCAGAAGATCGAGACCATCNNNAGATCGGAA |
| 39097_1 | TATGAGTGTGGAGTCGTTGCGTAGAGAAAACCTGCAGGCCTATNNNAGATCGGAA |
| 39097_2 | TATGAGTGTGGAGTCGTTGCGTAGAGAAAACCTGCAGGCCTATNNNAGATCGGAA |
| 48168_1 | TATGAGTGTGGAGTCGTTGCGAGTAGCTGGGATTATAGGCACCNNNAGATCGGAA |
| 48168_2 | TATGAGTGTGGAGTCGTTGCGAGTAGCTGGGATTATAGGCACCNNNAGATCGGAA |
| 33098_1 | TATGAGTGTGGAGTCGTTGCCAAGTAACTGGGACTACAGGCATNNNAGATCGGAA |

| | |
|---|---|
| 33098_2 | TATGAGTGTGGAGTCGTTGCCAAGTAACTGGGACTACAGGCATNNNAGATCGGAA |
| 2132_1 | TATGAGTGTGGAGTCGTTGCCTGGCTGGTTGGTATTTGGATCANNNAGATCGGAA |
| 2132_2 | TATGAGTGTGGAGTCGTTGCCTGGCTGGTTGGTATTTGGATCANNNAGATCGGAA |
| 49124_1 | TATGAGTGTGGAGTCGTTGCTGGGTTCAATTGATTTTCCTGCCNNNAGATCGGAAG |
| 49124_2 | TATGAGTGTGGAGTCGTTGCTGGGTTCAATTGATTTTCCTGCCNNNAGATCGGAAG |
| 36373_1 | TATGAGTGTGGAGTCGTTGCCCCCAAATGTCAATAGTGCCAAGNNNAGATCGGAAG |
| 36373_2 | TATGAGTGTGGAGTCGTTGCCCCCAAATGTCAATAGTGCCAAGNNNAGATCGGAAG |
| 51661_1 | TATGAGTGTGGAGTCGTTGCGTCATGAGGATGAAGCCCTGTAANNNAGATCGGAA |
| 51661_2 | TATGAGTGTGGAGTCGTTGCGTCATGAGGATGAAGCCCTGTAANNNAGATCGGAA |
| 40984_1 | TATGAGTGTGGAGTCGTTGCAGAAGGCAAGACTCCAAGAACTTNNNAGATCGGAA |
| 40984_2 | TATGAGTGTGGAGTCGTTGCAGAAGGCAAGACTCCAAGAACTTNNNAGATCGGAA |
| 52512_1 | TATGAGTGTGGAGTCGTTGCGGTCAGCTATGCCTAATAGGACCNNNAGATCGGAA |
| 52512_2 | TATGAGTGTGGAGTCGTTGCGGTCAGCTATGCCTAATAGGACCNNNAGATCGGAA |
| 13779_1 | TATGAGTGTGGAGTCGTTGCGTTGTGAAACCTTCCTTGAANNNAGATCGGAAGAG |
| 13779_2 | TATGAGTGTGGAGTCGTTGCGTTGTGAAACCTTCCTTGAANNNAGATCGGAAGAG |
| 35906_1 | TATGAGTGTGGAGTCGTTGCGTGAGAGGCTCAAATAAGGGGAANNNAGATCGGA |
| 35906_2 | TATGAGTGTGGAGTCGTTGCGTGAGAGGCTCAAATAAGGGGAANNNAGATCGGA |
| 35157_1 | TATGAGTGTGGAGTCGTTGCTTAGTCTGTTCTCACACTGCTGTNNNAGATCGGAAG |
| 35157_2 | TATGAGTGTGGAGTCGTTGCTTAGTCTGTTCTCACACTGCTGTNNNAGATCGGAAG |
| 44913_1 | TATGAGTGTGGAGTCGTTGCGGTTTCCAGATGGTGGTTTTCAGNNNAGATCGGAA |
| 44913_2 | TATGAGTGTGGAGTCGTTGCGGTTTCCAGATGGTGGTTTTCAGNNNAGATCGGAA |
| 50371_1 | TATGAGTGTGGAGTCGTTGCAATGCCAGAAAAGGAATGTTCGGNNNAGATCGGAA |
| 50371_2 | TATGAGTGTGGAGTCGTTGCAATGCCAGAAAAGGAATGTTCGGNNNAGATCGGAA |
| 31488_1 | TATGAGTGTGGAGTCGTTGCAACAGGAGTGATGAAAGTGGACANNNAGATCGGA |
| 31488_2 | TATGAGTGTGGAGTCGTTGCAACAGGAGTGATGAAAGTGGACANNNAGATCGGA |
| 32565_1 | TATGAGTGTGGAGTCGTTGCCTCCTGAGTAGCTGGGACTACAGNNNAGATCGGAA |
| 32565_2 | TATGAGTGTGGAGTCGTTGCCTCCTGAGTAGCTGGGACTACAGNNNAGATCGGAA |
| 32224_1 | TATGAGTGTGGAGTCGTTGCCTCAAGCAATCCTCCTACCTCAGNNNAGATCGGAAGA |
| 32224_2 | TATGAGTGTGGAGTCGTTGCCTCAAGCAATCCTCCTACCTCAGNNNAGATCGGAAGA |
| 2941_1 | TATGAGTGTGGAGTCGTTGCACATAATTTGAGGCCCACAGCTTNNNAGATCGGAAG |
| 2941_2 | TATGAGTGTGGAGTCGTTGCACATAATTTGAGGCCCACAGCTTNNNAGATCGGAAG |
| 29593_1 | TATGAGTGTGGAGTCGTTGCTGCTACCAATCAACACACATCTGNNNAGATCGGAAG |
| 29593_2 | TATGAGTGTGGAGTCGTTGCTGCTACCAATCAACACACATCTGNNNAGATCGGAAG |
| 2648_1 | TATGAGTGTGGAGTCGTTGCCAAAAAGGAAAGGGGGCAAGTTGNNNAGATCGGA |
| 2648_2 | TATGAGTGTGGAGTCGTTGCCAAAAAGGAAAGGGGGCAAGTTGNNNAGATCGGA |
| 50149_1 | TATGAGTGTGGAGTCGTTGCAAAAGGCTGAGAAGTGATCCCTTNNNAGATCGGAA |
| 50149_2 | TATGAGTGTGGAGTCGTTGCAAAAGGCTGAGAAGTGATCCCTTNNNAGATCGGAA |
| 10433_1 | TATGAGTGTGGAGTCGTTGCGAAAAATGCAATACAGGCTGGGCNNNAGATCGGAA |
| 10433_2 | TATGAGTGTGGAGTCGTTGCGAAAAATGCAATACAGGCTGGGCNNNAGATCGGAA |
| 6592_1 | TATGAGTGTGGAGTCGTTGCTCTCAGTCCCTCACATTCACCTTNNNAGATCGGAAGA |
| 6592_2 | TATGAGTGTGGAGTCGTTGCTCTCAGTCCCTCACATTCACCTTNNNAGATCGGAAGA |
| 31848_1 | TATGAGTGTGGAGTCGTTGCTGAAAAGCCTCACATAATGCGTCNNNAGATCGGAAG |
| 31848_2 | TATGAGTGTGGAGTCGTTGCTGAAAAGCCTCACATAATGCGTCNNNAGATCGGAAG |

| | |
|---|---|
| 40496_1 | TATGAGTGTGGAGTCGTTGCATTTTTCCAGTCTTTCACCCCCTNNNAGATCGGAAGA |
| 40496_2 | TATGAGTGTGGAGTCGTTGCATTTTTCCAGTCTTTCACCCCCTNNNAGATCGGAAGA |
| 43471_1 | TATGAGTGTGGAGTCGTTGCACACAAGCAGCAATATCACCAACNNNAGATCGGAAG |
| 43471_2 | TATGAGTGTGGAGTCGTTGCACACAAGCAGCAATATCACCAACNNNAGATCGGAAG |
| 35881_1 | TATGAGTGTGGAGTCGTTGCCATGCCTGTAATCCCAGCATTTTNNNAGATCGGAAGA |
| 35881_2 | TATGAGTGTGGAGTCGTTGCCATGCCTGTAATCCCAGCATTTTNNNAGATCGGAAGA |
| 40088_1 | TATGAGTGTGGAGTCGTTGCTGCAAAAGACTTGACCAGATACTNNNAGATCGGAAG |
| 40088_2 | TATGAGTGTGGAGTCGTTGCTGCAAAAGACTTGACCAGATACTNNNAGATCGGAAG |
| 44543_1 | TATGAGTGTGGAGTCGTTGCATAGCAAAATCAGCTTCTGGGGANNNAGATCGGAA |
| 44543_2 | TATGAGTGTGGAGTCGTTGCATAGCAAAATCAGCTTCTGGGGANNNAGATCGGAA |
| 53147_1 | TATGAGTGTGGAGTCGTTGCGCCACCACTGAGAAAACAAGAAGNNNAGATCGGAA |
| 53147_2 | TATGAGTGTGGAGTCGTTGCGCCACCACTGAGAAAACAAGAAGNNNAGATCGGAA |
| 52303_1 | TATGAGTGTGGAGTCGTTGCCTTCAAACGCCATCCAGAACCTNNNAGATCGGAAGA |
| 52303_2 | TATGAGTGTGGAGTCGTTGCCTTCAAACGCCATCCAGAACCTNNNAGATCGGAAGA |
| 46688_1 | TATGAGTGTGGAGTCGTTGCTGGAGTACAGTGGTGCAATATCGNNNAGATCGGAA |
| 46688_2 | TATGAGTGTGGAGTCGTTGCTGGAGTACAGTGGTGCAATATCGNNNAGATCGGAA |
| 46269_1 | TATGAGTGTGGAGTCGTTGCAGGTTCAAGCAATTCTTTTGCCANNNAGATCGGAAG |
| 46269_2 | TATGAGTGTGGAGTCGTTGCAGGTTCAAGCAATTCTTTTGCCANNNAGATCGGAAG |
| 9363_1 | TATGAGTGTGGAGTCGTTGCCCTGAAGCCAGCACCATATTAGGNNNAGATCGGAAG |
| 9363_2 | TATGAGTGTGGAGTCGTTGCCCTGAAGCCAGCACCATATTAGGNNNAGATCGGAAG |
| 34509_1 | TATGAGTGTGGAGTCGTTGCGGGCAATGGCTCCTTATTTTGAGNNNAGATCGGAAG |
| 34509_2 | TATGAGTGTGGAGTCGTTGCGGGCAATGGCTCCTTATTTTGAGNNNAGATCGGAAG |
| 37697_1 | TATGAGTGTGGAGTCGTTGCCTGAAGTAGGAGGATCCCTTGAGNNNAGATCGGAA |
| 37697_2 | TATGAGTGTGGAGTCGTTGCCTGAAGTAGGAGGATCCCTTGAGNNNAGATCGGAA |
| 37624_1 | TATGAGTGTGGAGTCGTTGCGTGGACACCTATAATCCCAGCTANNNAGATCGGAAG |
| 37624_2 | TATGAGTGTGGAGTCGTTGCGTGGACACCTATAATCCCAGCTANNNAGATCGGAAG |
| 45332_1 | TATGAGTGTGGAGTCGTTGCACTTAGCATTGTGTTACAGTTGCCNNNAGATCGGAA |
| 45332_2 | TATGAGTGTGGAGTCGTTGCACTTAGCATTGTGTTACAGTTGCCNNNAGATCGGAA |
| 13165_1 | TATGAGTGTGGAGTCGTTGCGTTGCTTTTCAAAGTGAAGATNNNAGATCGGAAGA |
| 13165_2 | TATGAGTGTGGAGTCGTTGCGTTGCTTTTCAAAGTGAAGATNNNAGATCGGAAGA |
| 29918_1 | TATGAGTGTGGAGTCGTTGCGCCTGAGAACCTGGTAGAAAAGANNNAGATCGGAA |
| 29918_2 | TATGAGTGTGGAGTCGTTGCGCCTGAGAACCTGGTAGAAAAGANNNAGATCGGAA |
| 51827_1 | TATGAGTGTGGAGTCGTTGCGGTCAGGTTCCAAAATGAAGACCNNNAGATCGGAA |
| 51827_2 | TATGAGTGTGGAGTCGTTGCGGTCAGGTTCCAAAATGAAGACCNNNAGATCGGAA |
| 33859_1 | TATGAGTGTGGAGTCGTTGCTGGCATGGATAATCACAAAGCTGNNNAGATCGGAA |
| 33859_2 | TATGAGTGTGGAGTCGTTGCTGGCATGGATAATCACAAAGCTGNNNAGATCGGAA |
| 41888_1 | TATGAGTGTGGAGTCGTTGCGCACAACCATAGCTCACCATAACNNNAGATCGGAAG |
| 41888_2 | TATGAGTGTGGAGTCGTTGCGCACAACCATAGCTCACCATAACNNNAGATCGGAAG |
| 29006_1 | TATGAGTGTGGAGTCGTTGCGCAGAACCTAGGGTGAGTGAAATNNNAGATCGGAA |
| 29006_2 | TATGAGTGTGGAGTCGTTGCGCAGAACCTAGGGTGAGTGAAATNNNAGATCGGAA |
| 41112_1 | TATGAGTGTGGAGTCGTTGCGACCTATTTCTGTCTTCCCTGCTNNNAGATCGGAAGA |
| 41112_2 | TATGAGTGTGGAGTCGTTGCGACCTATTTCTGTCTTCCCTGCTNNNAGATCGGAAGA |
| 36807_1 | TATGAGTGTGGAGTCGTTGCTGGTTATTGACTAAGGGTGTGACNNNAGATCGGAA |

| | |
|---|---|
| 36807_2 | TATGAGTGTGGAGTCGTTGCTGGTTATTGACTAAGGGTGTGACNNNAGATCGGAA |
| 40624_1 | TATGAGTGTGGAGTCGTTGCCTGCATGGCAAAAACCACCTAAANNNAGATCGGAAG |
| 40624_2 | TATGAGTGTGGAGTCGTTGCCTGCATGGCAAAAACCACCTAAANNNAGATCGGAAG |
| 43134_1 | TATGAGTGTGGAGTCGTTGCTTGCCTGTAATCCCAGCAATTTGNNNAGATCGGAAG |
| 43134_2 | TATGAGTGTGGAGTCGTTGCTTGCCTGTAATCCCAGCAATTTGNNNAGATCGGAAG |
| 53305_1 | TATGAGTGTGGAGTCGTTGCCACACACACACACACACACATATNNNAGATCGGAAG |
| 53305_2 | TATGAGTGTGGAGTCGTTGCCACACACACACACACACACATATNNNAGATCGGAAG |
| 2785_1 | TATGAGTGTGGAGTCGTTGCTTCTCATTGGTACTGCCCTAGCTNNNAGATCGGAAG |
| 2785_2 | TATGAGTGTGGAGTCGTTGCTTCTCATTGGTACTGCCCTAGCTNNNAGATCGGAAG |
| 38616_1 | TATGAGTGTGGAGTCGTTGCAGACAAAGTTTCGCTCTTTTCCCNNNAGATCGGAAG |
| 38616_2 | TATGAGTGTGGAGTCGTTGCAGACAAAGTTTCGCTCTTTTCCCNNNAGATCGGAAG |
| 50679_1 | TATGAGTGTGGAGTCGTTGCAGACCTGGAGAAAGGGAGACATANNNAGATCGGA |
| 50679_2 | TATGAGTGTGGAGTCGTTGCAGACCTGGAGAAAGGGAGACATANNNAGATCGGA |
| 44153_1 | TATGAGTGTGGAGTCGTTGCAGACAGGAGAGAAAGTCAGCTTGNNNAGATCGGA |
| 44153_2 | TATGAGTGTGGAGTCGTTGCAGACAGGAGAGAAAGTCAGCTTGNNNAGATCGGA |
| 31572_1 | TATGAGTGTGGAGTCGTTGCAGAGGCACTGTGATCAAGAACACNNNAGATCGGAA |
| 31572_2 | TATGAGTGTGGAGTCGTTGCAGAGGCACTGTGATCAAGAACACNNNAGATCGGAA |
| 47819_1 | TATGAGTGTGGAGTCGTTGCTACTGCCGAGTCTGGATGTCTTCNNNAGATCGGAAG |
| 47819_2 | TATGAGTGTGGAGTCGTTGCTACTGCCGAGTCTGGATGTCTTCNNNAGATCGGAAG |
| 30323_1 | TATGAGTGTGGAGTCGTTGCGACATCATGCGGAAGCAGGTNNNAGATCGGAAGAG |
| 30323_2 | TATGAGTGTGGAGTCGTTGCGACATCATGCGGAAGCAGGTNNNAGATCGGAAGAG |
| 37939_1 | TATGAGTGTGGAGTCGTTGCAAAGAAAAAAGAAGTCCGGGTGCNNNAGATCGGA |
| 37939_2 | TATGAGTGTGGAGTCGTTGCAAAGAAAAAAGAAGTCCGGGTGCNNNAGATCGGA |
| 13795_1 | TATGAGTGTGGAGTCGTTGCGAAAATGGAAAAAAATCTGAACANNNAGATCGGAA |
| 13795_2 | TATGAGTGTGGAGTCGTTGCGAAAATGGAAAAAAATCTGAACANNNAGATCGGAA |
| 12103_1 | TATGAGTGTGGAGTCGTTGCGCCAATTACTAGAGAGAGCTGAAANNNAGATCGGA |
| 12103_2 | TATGAGTGTGGAGTCGTTGCGCCAATTACTAGAGAGAGCTGAAANNNAGATCGGA |
| 219_1 | TATGAGTGTGGAGTCGTTGCTTATCGAGGCTGAGGATTCCTGTNNNAGATCGGAA |
| 219_2 | TATGAGTGTGGAGTCGTTGCTTATCGAGGCTGAGGATTCCTGTNNNAGATCGGAA |
| 35494_1 | TATGAGTGTGGAGTCGTTGCTGATGTTACTCTTGGCAAGGTGTNNNAGATCGGAA |
| 35494_2 | TATGAGTGTGGAGTCGTTGCTGATGTTACTCTTGGCAAGGTGTNNNAGATCGGAA |
| 10165_1 | TATGAGTGTGGAGTCGTTGCCTCACCTTGCATGGCACTATGTTNNNAGATCGGAAG |
| 10165_2 | TATGAGTGTGGAGTCGTTGCCTCACCTTGCATGGCACTATGTTNNNAGATCGGAAG |
| 42783_1 | TATGAGTGTGGAGTCGTTGCGAAACACACATACACACACACCCNNNAGATCGGAAG |
| 42783_2 | TATGAGTGTGGAGTCGTTGCGAAACACACATACACACACACCCNNNAGATCGGAAG |
| 53363_1 | TATGAGTGTGGAGTCGTTGCTTGTGAAGACTTGCCTTGCTTTCNNNAGATCGGAAG |
| 53363_2 | TATGAGTGTGGAGTCGTTGCTTGTGAAGACTTGCCTTGCTTTCNNNAGATCGGAAG |
| 47967_1 | TATGAGTGTGGAGTCGTTGCAAAGCCCCCATGAAGAAGGAAAANNNAGATCGGAA |
| 47967_2 | TATGAGTGTGGAGTCGTTGCAAAGCCCCCATGAAGAAGGAAAANNNAGATCGGAA |
| 33808_1 | TATGAGTGTGGAGTCGTTGCATGGCCAATAAGCACAGGAAAAGNNNAGATCGGAA |
| 33808_2 | TATGAGTGTGGAGTCGTTGCATGGCCAATAAGCACAGGAAAAGNNNAGATCGGAA |
| 49890_1 | TATGAGTGTGGAGTCGTTGCCCCCCAAAAAAGCCTTACATGAANNNAGATCGGAAG |
| 49890_2 | TATGAGTGTGGAGTCGTTGCCCCCCAAAAAAGCCTTACATGAANNNAGATCGGAAG |

| | |
|---|---|
| 36932_1 | TATGAGTGTGGAGTCGTTGCTGAGCTGGTATCAAGCTAACTGGNNNAGATCGGAA |
| 36932_2 | TATGAGTGTGGAGTCGTTGCTGAGCTGGTATCAAGCTAACTGGNNNAGATCGGAA |
| 30695_1 | TATGAGTGTGGAGTCGTTGCTGGGATTTGACTGTTGCTATACAATNNNAGATCGGA |
| 30695_2 | TATGAGTGTGGAGTCGTTGCTGGGATTTGACTGTTGCTATACAATNNNAGATCGGA |
| 50863_1 | TATGAGTGTGGAGTCGTTGCGGTGATGGATGATCGCACCCNNNAGATCGGAAGAG |
| 50863_2 | TATGAGTGTGGAGTCGTTGCGGTGATGGATGATCGCACCCNNNAGATCGGAAGAG |
| 31902_1 | TATGAGTGTGGAGTCGTTGCTCGACTTTATCTTTGGGAGGCAANNNAGATCGGAAG |
| 31902_2 | TATGAGTGTGGAGTCGTTGCTCGACTTTATCTTTGGGAGGCAANNNAGATCGGAAG |
| 49666_1 | TATGAGTGTGGAGTCGTTGCATTCAGCACTTGCCATCTACTGANNNAGATCGGAAG |
| 49666_2 | TATGAGTGTGGAGTCGTTGCATTCAGCACTTGCCATCTACTGANNNAGATCGGAAG |
| 52047_1 | TATGAGTGTGGAGTCGTTGCACAGAGATTTGAGAACTCCCAGGNNNAGATCGGAA |
| 52047_2 | TATGAGTGTGGAGTCGTTGCACAGAGATTTGAGAACTCCCAGGNNNAGATCGGAA |
| 30555_1 | TATGAGTGTGGAGTCGTTGCGCAAACTCAGAACCAGGCAACTNNNAGATCGGAAG |
| 30555_2 | TATGAGTGTGGAGTCGTTGCGCAAACTCAGAACCAGGCAACTNNNAGATCGGAAG |
| 37937_1 | TATGAGTGTGGAGTCGTTGCCCCTGATGTGGAAGATGGTACAANNNAGATCGGAA |
| 37937_2 | TATGAGTGTGGAGTCGTTGCCCCTGATGTGGAAGATGGTACAANNNAGATCGGAA |
| 33100_1 | TATGAGTGTGGAGTCGTTGCCCTAGTAGCTGGGACTACAGGTNNNAGATCGGAAG |
| 33100_2 | TATGAGTGTGGAGTCGTTGCCCTAGTAGCTGGGACTACAGGTNNNAGATCGGAAG |
| 52477_1 | TATGAGTGTGGAGTCGTTGCAGCACCTCTTTCCACCTTAGTTTNNNAGATCGGAAGA |
| 52477_2 | TATGAGTGTGGAGTCGTTGCAGCACCTCTTTCCACCTTAGTTTNNNAGATCGGAAGA |
| 36531_1 | TATGAGTGTGGAGTCGTTGCAAAAGACCCCCAAAGCATTTCAGNNNAGATCGGAAG |
| 36531_2 | TATGAGTGTGGAGTCGTTGCAAAAGACCCCCAAAGCATTTCAGNNNAGATCGGAAG |
| 12480_1 | TATGAGTGTGGAGTCGTTGCTAGAATCCATTTACAAGGACAGNNNAGATCGGAAG |
| 12480_2 | TATGAGTGTGGAGTCGTTGCTAGAATCCATTTACAAGGACAGNNNAGATCGGAAG |
| 38731_1 | TATGAGTGTGGAGTCGTTGCCAGAGCTTGGAAACAAGAAGAGCNNNAGATCGGAA |
| 38731_2 | TATGAGTGTGGAGTCGTTGCCAGAGCTTGGAAACAAGAAGAGCNNNAGATCGGAA |
| 31836_1 | TATGAGTGTGGAGTCGTTGCATAAGAGGAGAGGTTAGTGCCCTNNNAGATCGGAA |
| 31836_2 | TATGAGTGTGGAGTCGTTGCATAAGAGGAGAGGTTAGTGCCCTNNNAGATCGGAA |
| 44028_1 | TATGAGTGTGGAGTCGTTGCTGCTTTCCTATGGGAGAGGGTATNNNAGATCGGAA |
| 44028_2 | TATGAGTGTGGAGTCGTTGCTGCTTTCCTATGGGAGAGGGTATNNNAGATCGGAA |
| 42081_1 | TATGAGTGTGGAGTCGTTGCATTCTGCCAAGAAGCTACCTGAANNNAGATCGGAAG |
| 42081_2 | TATGAGTGTGGAGTCGTTGCATTCTGCCAAGAAGCTACCTGAANNNAGATCGGAAG |
| 34115_1 | TATGAGTGTGGAGTCGTTGCCACCTTGAGGACCATTCACTTCTNNNAGATCGGAAG |
| 34115_2 | TATGAGTGTGGAGTCGTTGCCACCTTGAGGACCATTCACTTCTNNNAGATCGGAAG |
| 43_1 | TATGAGTGTGGAGTCGTTGCGGAGCCTGCCAAAAAGATATGAANNNAGATCGGAA |
| 43_2 | TATGAGTGTGGAGTCGTTGCGGAGCCTGCCAAAAAGATATGAANNNAGATCGGAA |
| 38161_1 | TATGAGTGTGGAGTCGTTGCAAGAAAAAGAAAGCTCAGGCCAGNNNAGATCGGAA |
| 38161_2 | TATGAGTGTGGAGTCGTTGCAAGAAAAAGAAAGCTCAGGCCAGNNNAGATCGGAA |
| 51349_1 | TATGAGTGTGGAGTCGTTGCTCTCACTGGCAGAGTAGAAACACNNNAGATCGGAAG |
| 51349_2 | TATGAGTGTGGAGTCGTTGCTCTCACTGGCAGAGTAGAAACACNNNAGATCGGAAG |
| 34835_1 | TATGAGTGTGGAGTCGTTGCCCACGCCCGGCTAATTTTTATATNNNAGATCGGAAG |
| 34835_2 | TATGAGTGTGGAGTCGTTGCCCACGCCCGGCTAATTTTTATATNNNAGATCGGAAG |
| 37257_1 | TATGAGTGTGGAGTCGTTGCATCACGGTATTTGGATAGGTGGGNNNAGATCGGAA |

| | |
|---|---|
| 37257_2 | TATGAGTGTGGAGTCGTTGCATCACGGTATTTGGATAGGTGGGNNNAGATCGGAA |
| 29319_1 | TATGAGTGTGGAGTCGTTGCCTGCAGTGATATGATCTGGCTCTNNNAGATCGGAAG |
| 29319_2 | TATGAGTGTGGAGTCGTTGCCTGCAGTGATATGATCTGGCTCTNNNAGATCGGAAG |
| 40776_1 | TATGAGTGTGGAGTCGTTGCTCAAAGGAATCTTGCTGCTGAGANNNAGATCGGAA |
| 40776_2 | TATGAGTGTGGAGTCGTTGCTCAAAGGAATCTTGCTGCTGAGANNNAGATCGGAA |
| 38422_1 | TATGAGTGTGGAGTCGTTGCGGGATGCAGAATCTCTCTAGAGCNNNAGATCGGAA |
| 38422_2 | TATGAGTGTGGAGTCGTTGCGGGATGCAGAATCTCTCTAGAGCNNNAGATCGGAA |
| 52799_1 | TATGAGTGTGGAGTCGTTGCTCCTCCATCTTCACTCCTCCCNNNAGATCGGAAGAGC |
| 52799_2 | TATGAGTGTGGAGTCGTTGCTCCTCCATCTTCACTCCTCCCNNNAGATCGGAAGAGC |
| 42881_1 | TATGAGTGTGGAGTCGTTGCCCGTTCTAGGAAGACCAACTAGTNNNAGATCGGAAG |
| 42881_2 | TATGAGTGTGGAGTCGTTGCCCGTTCTAGGAAGACCAACTAGTNNNAGATCGGAAG |
| 37509_1 | TATGAGTGTGGAGTCGTTGCGGGCAGATTACAAGGTCAAGAGANNNAGATCGGA |
| 37509_2 | TATGAGTGTGGAGTCGTTGCGGGCAGATTACAAGGTCAAGAGANNNAGATCGGA |
| 43073_1 | TATGAGTGTGGAGTCGTTGCAGAGACCCTTCTTACCCTGATCANNNAGATCGGAAG |
| 43073_2 | TATGAGTGTGGAGTCGTTGCAGAGACCCTTCTTACCCTGATCANNNAGATCGGAAG |
| 12615_1 | TATGAGTGTGGAGTCGTTGCCTTGGTATTTGTATTCAAGCATCNNNAGATCGGAAG |
| 12615_2 | TATGAGTGTGGAGTCGTTGCCTTGGTATTTGTATTCAAGCATCNNNAGATCGGAAG |
| 42023_1 | TATGAGTGTGGAGTCGTTGCGGTGGACTACTTGATGTCAGGAGNNNAGATCGGAA |
| 42023_2 | TATGAGTGTGGAGTCGTTGCGGTGGACTACTTGATGTCAGGAGNNNAGATCGGAA |
| 42918_1 | TATGAGTGTGGAGTCGTTGCCAGAGGCTTGGATTTCCAGTACANNNAGATCGGAA |
| 42918_2 | TATGAGTGTGGAGTCGTTGCCAGAGGCTTGGATTTCCAGTACANNNAGATCGGAA |
| 32171_1 | TATGAGTGTGGAGTCGTTGCCTCCTGCGTTCAAGCAATTCTGNNNAGATCGGAAGA |
| 32171_2 | TATGAGTGTGGAGTCGTTGCCTCCTGCGTTCAAGCAATTCTGNNNAGATCGGAAGA |
| 10094_1 | TATGAGTGTGGAGTCGTTGCACCGACATTTTACCACCTCTAGTNNNAGATCGGAAGA |
| 10094_2 | TATGAGTGTGGAGTCGTTGCACCGACATTTTACCACCTCTAGTNNNAGATCGGAAGA |
| 53030_1 | TATGAGTGTGGAGTCGTTGCGCCATTTCTTCTCCTTATTGCCCNNNAGATCGGAAGA |
| 53030_2 | TATGAGTGTGGAGTCGTTGCGCCATTTCTTCTCCTTATTGCCCNNNAGATCGGAAGA |
| 48883_1 | TATGAGTGTGGAGTCGTTGCGCGCCCGGCTAATTTTTTTGTANNNAGATCGGAAGA |
| 48883_2 | TATGAGTGTGGAGTCGTTGCGCGCCCGGCTAATTTTTTTGTANNNAGATCGGAAGA |
| 43181_1 | TATGAGTGTGGAGTCGTTGCCAGGTTCACGCCATTCTCCTNNNAGATCGGAAGAGC |
| 43181_2 | TATGAGTGTGGAGTCGTTGCCAGGTTCACGCCATTCTCCTNNNAGATCGGAAGAGC |
| 30454_1 | TATGAGTGTGGAGTCGTTGCAAAGCACCCTGAAGTGAGAGGACANNNAGATCGGA |
| 30454_2 | TATGAGTGTGGAGTCGTTGCAAAGCACCCTGAAGTGAGAGGACANNNAGATCGGA |
| 48421_1 | TATGAGTGTGGAGTCGTTGCAGCTAGAACTAAAGGTGCATGCTNNNAGATCGGAA |
| 48421_2 | TATGAGTGTGGAGTCGTTGCAGCTAGAACTAAAGGTGCATGCTNNNAGATCGGAA |
| 45391_1 | TATGAGTGTGGAGTCGTTGCACACACACACACGAGAAAACAAANNNAGATCGGAA |
| 45391_2 | TATGAGTGTGGAGTCGTTGCACACACACACACGAGAAAACAAANNNAGATCGGAA |
| 42613_1 | TATGAGTGTGGAGTCGTTGCGCCTCCTGAGTAGCTGGAATTACNNNAGATCGGAAG |
| 42613_2 | TATGAGTGTGGAGTCGTTGCGCCTCCTGAGTAGCTGGAATTACNNNAGATCGGAAG |
| 33145_1 | TATGAGTGTGGAGTCGTTGCGGCCAGTTTTGCAGAACATCTTTNNNAGATCGGAAG |
| 33145_2 | TATGAGTGTGGAGTCGTTGCGGCCAGTTTTGCAGAACATCTTTNNNAGATCGGAAG |
| 40195_1 | TATGAGTGTGGAGTCGTTGCTGTCTCGATCTTGGCTTATTCCANNNAGATCGGAAGA |
| 40195_2 | TATGAGTGTGGAGTCGTTGCTGTCTCGATCTTGGCTTATTCCANNNAGATCGGAAGA |

| | |
|---|---|
| 20258_1 | TATGAGTGTGGAGTCGTTGCACACATGTTTTCTACCTTTATTTNNNAGATCGGAAGA |
| 20258_2 | TATGAGTGTGGAGTCGTTGCACACATGTTTTCTACCTTTATTTNNNAGATCGGAAGA |
| 45841_1 | TATGAGTGTGGAGTCGTTGCTAGAGTGCAGTGTTGTGATCTCGNNNAGATCGGAA |
| 45841_2 | TATGAGTGTGGAGTCGTTGCTAGAGTGCAGTGTTGTGATCTCGNNNAGATCGGAA |
| 10982_1 | TATGAGTGTGGAGTCGTTGCAAGATTGGGTTATTTCACGCATCNNNAGATCGGAAG |
| 10982_2 | TATGAGTGTGGAGTCGTTGCAAGATTGGGTTATTTCACGCATCNNNAGATCGGAAG |
| 47101_1 | TATGAGTGTGGAGTCGTTGCGAGATACCCCACAGATTAGGCAGNNNAGATCGGAA |
| 47101_2 | TATGAGTGTGGAGTCGTTGCGAGATACCCCACAGATTAGGCAGNNNAGATCGGAA |
| 31705_1 | TATGAGTGTGGAGTCGTTGCCGTCTTGCTCCTCGATGTGAGNNNAGATCGGAAGAG |
| 31705_2 | TATGAGTGTGGAGTCGTTGCCGTCTTGCTCCTCGATGTGAGNNNAGATCGGAAGAG |
| 52170_1 | TATGAGTGTGGAGTCGTTGCCCCTCCACCATCCATCTTATGTTNNNAGATCGGAAGA |
| 52170_2 | TATGAGTGTGGAGTCGTTGCCCCTCCACCATCCATCTTATGTTNNNAGATCGGAAGA |
| 11012_1 | TATGAGTGTGGAGTCGTTGCGTTTTCTCCCCCACCACCTTGNNNAGATCGGAAGAGC |
| 11012_2 | TATGAGTGTGGAGTCGTTGCGTTTTCTCCCCCACCACCTTGNNNAGATCGGAAGAGC |
| 52522_1 | TATGAGTGTGGAGTCGTTGCCCCTGAAAATGAGTCCTAGCCTTNNNAGATCGGAAG |
| 52522_2 | TATGAGTGTGGAGTCGTTGCCCCTGAAAATGAGTCCTAGCCTTNNNAGATCGGAAG |
| 40497_1 | TATGAGTGTGGAGTCGTTGCACCACCCTGCCATTTTTGTAATGNNNAGATCGGAAG |
| 40497_2 | TATGAGTGTGGAGTCGTTGCACCACCCTGCCATTTTTGTAATGNNNAGATCGGAAG |
| 45106_1 | TATGAGTGTGGAGTCGTTGCTGGCCATGTAACTATCTCTAGTGCNNNAGATCGGAA |
| 45106_2 | TATGAGTGTGGAGTCGTTGCTGGCCATGTAACTATCTCTAGTGCNNNAGATCGGAA |
| 32067_1 | TATGAGTGTGGAGTCGTTGCAAAACAGACACAAAGACCAACGGNNNAGATCGGAA |
| 32067_2 | TATGAGTGTGGAGTCGTTGCAAAACAGACACAAAGACCAACGGNNNAGATCGGAA |
| 10656_1 | TATGAGTGTGGAGTCGTTGCTCCTGGTCTAAAATGTGGAGCAGNNNAGATCGGAA |
| 10656_2 | TATGAGTGTGGAGTCGTTGCTCCTGGTCTAAAATGTGGAGCAGNNNAGATCGGAA |
| 30012_1 | TATGAGTGTGGAGTCGTTGCTCAATGGCACCTAAGTGGAACACNNNAGATCGGAAG |
| 30012_2 | TATGAGTGTGGAGTCGTTGCTCAATGGCACCTAAGTGGAACACNNNAGATCGGAAG |
| 33892_1 | TATGAGTGTGGAGTCGTTGCTCCTGGGTTCAAACGATTCTCATNNNAGATCGGAAG |
| 33892_2 | TATGAGTGTGGAGTCGTTGCTCCTGGGTTCAAACGATTCTCATNNNAGATCGGAAG |
| 41470_1 | TATGAGTGTGGAGTCGTTGCTAAGCCCAAAGAGACACACATCANNNAGATCGGAAG |
| 41470_2 | TATGAGTGTGGAGTCGTTGCTAAGCCCAAAGAGACACACATCANNNAGATCGGAAG |
| 52501_1 | TATGAGTGTGGAGTCGTTGCGTCTACAGCGGGTCCATAGTTAGNNNAGATCGGAA |
| 52501_2 | TATGAGTGTGGAGTCGTTGCGTCTACAGCGGGTCCATAGTTAGNNNAGATCGGAA |
| 38241_1 | TATGAGTGTGGAGTCGTTGCCTTCTCCCTCCAAAGGATCACAANNNAGATCGGAAG |
| 38241_2 | TATGAGTGTGGAGTCGTTGCCTTCTCCCTCCAAAGGATCACAANNNAGATCGGAAG |
| 36016_1 | TATGAGTGTGGAGTCGTTGCAGAGAGAGAGAGAGGCAAAGAGANNNAGATCGG |
| 36016_2 | TATGAGTGTGGAGTCGTTGCAGAGAGAGAGAGAGGCAAAGAGANNNAGATCGG |
| 13365_1 | TATGAGTGTGGAGTCGTTGCAACCTTTAAACATCAAACTCATTNNNAGATCGGAAG |
| 13365_2 | TATGAGTGTGGAGTCGTTGCAACCTTTAAACATCAAACTCATTNNNAGATCGGAAG |
| 10322_1 | TATGAGTGTGGAGTCGTTGCACACCATGTGCTTGCAAGATACTNNNAGATCGGAAG |
| 10322_2 | TATGAGTGTGGAGTCGTTGCACACCATGTGCTTGCAAGATACTNNNAGATCGGAAG |
| 49262_1 | TATGAGTGTGGAGTCGTTGCGGCATAATCTTGGCTCATTGCAANNNAGATCGGAAG |
| 49262_2 | TATGAGTGTGGAGTCGTTGCGGCATAATCTTGGCTCATTGCAANNNAGATCGGAAG |
| 51564_1 | TATGAGTGTGGAGTCGTTGCTTACCTGCACAAAGAGGGGTAAANNNAGATCGGAA |

| | |
|---|---|
| 51564_2 | TATGAGTGTGGAGTCGTTGCTTACCTGCACAAAGAGGGGTAAANNNAGATCGGAA |
| 32660_1 | TATGAGTGTGGAGTCGTTGCCACAATCTCAGCTCACTGCAAGNNNAGATCGGAAGA |
| 32660_2 | TATGAGTGTGGAGTCGTTGCCACAATCTCAGCTCACTGCAAGNNNAGATCGGAAGA |
| 44494_1 | TATGAGTGTGGAGTCGTTGCCATCTGTAAGCTGGAGACCCNNNAGATCGGAAGAG |
| 44494_2 | TATGAGTGTGGAGTCGTTGCCATCTGTAAGCTGGAGACCCNNNAGATCGGAAGAG |
| 43908_1 | TATGAGTGTGGAGTCGTTGCGGCTGACTCAGAATGTGAATAGCNNNAGATCGGAA |
| 43908_2 | TATGAGTGTGGAGTCGTTGCGGCTGACTCAGAATGTGAATAGCNNNAGATCGGAA |
| 52775_1 | TATGAGTGTGGAGTCGTTGCCTGAGAATACCATGGGCTTCTCANNNAGATCGGAAG |
| 52775_2 | TATGAGTGTGGAGTCGTTGCCTGAGAATACCATGGGCTTCTCANNNAGATCGGAAG |
| 38077_1 | TATGAGTGTGGAGTCGTTGCCTCAATGGGAGGGGATTACACAANNNAGATCGGAA |
| 38077_2 | TATGAGTGTGGAGTCGTTGCCTCAATGGGAGGGGATTACACAANNNAGATCGGAA |
| 43868_1 | TATGAGTGTGGAGTCGTTGCAAGGAAGCCACTTGTATAGGTGGNNNAGATCGGAA |
| 43868_2 | TATGAGTGTGGAGTCGTTGCAAGGAAGCCACTTGTATAGGTGGNNNAGATCGGAA |
| 52174_1 | TATGAGTGTGGAGTCGTTGCCTGTTTTCCTCTATCTGGGCTGGNNNAGATCGGAAG |
| 52174_2 | TATGAGTGTGGAGTCGTTGCCTGTTTTCCTCTATCTGGGCTGGNNNAGATCGGAAG |
| 36189_1 | TATGAGTGTGGAGTCGTTGCGCAACAGAGTGAGACTGTGTTTCNNNAGATCGGAA |
| 36189_2 | TATGAGTGTGGAGTCGTTGCGCAACAGAGTGAGACTGTGTTTCNNNAGATCGGAA |
| 36880_1 | TATGAGTGTGGAGTCGTTGCCTCCAATCCCAGAGGACTATCCTNNNAGATCGGAAG |
| 36880_2 | TATGAGTGTGGAGTCGTTGCCTCCAATCCCAGAGGACTATCCTNNNAGATCGGAAG |
| 34366_1 | TATGAGTGTGGAGTCGTTGCCCTACTGGGAAGTGAGGAGCCNNNAGATCGGAAGA |
| 34366_2 | TATGAGTGTGGAGTCGTTGCCCTACTGGGAAGTGAGGAGCCNNNAGATCGGAAGA |
| 46552_1 | TATGAGTGTGGAGTCGTTGCAACCTCTGACTCCCTGGTTCAAGNNNAGATCGGAAG |
| 46552_2 | TATGAGTGTGGAGTCGTTGCAACCTCTGACTCCCTGGTTCAAGNNNAGATCGGAAG |
| 10362_1 | TATGAGTGTGGAGTCGTTGCGGGCATGGCTTCTAAACAACAGANNNAGATCGGAA |
| 10362_2 | TATGAGTGTGGAGTCGTTGCGGGCATGGCTTCTAAACAACAGANNNAGATCGGAA |
| 4587_1 | TATGAGTGTGGAGTCGTTGCCTGTAATGAACGTGAGAGTGCATNNNAGATCGGAA |
| 4587_2 | TATGAGTGTGGAGTCGTTGCCTGTAATGAACGTGAGAGTGCATNNNAGATCGGAA |
| 37852_1 | TATGAGTGTGGAGTCGTTGCGTAGCTTATGTTCATGGCAAGCANNNAGATCGGAAG |
| 37852_2 | TATGAGTGTGGAGTCGTTGCGTAGCTTATGTTCATGGCAAGCANNNAGATCGGAAG |
| 11688_1 | TATGAGTGTGGAGTCGTTGCACTTGGGTTGATTGCGTTTGTGNNNAGATCGGAAG |
| 11688_2 | TATGAGTGTGGAGTCGTTGCACTTGGGTTGATTGCGTTTGTGNNNAGATCGGAAG |
| 33254_1 | TATGAGTGTGGAGTCGTTGCGAGACGGAGTCTTGCTCTGTCNNNAGATCGGAAGA |
| 33254_2 | TATGAGTGTGGAGTCGTTGCGAGACGGAGTCTTGCTCTGTCNNNAGATCGGAAGA |
| 41081_1 | TATGAGTGTGGAGTCGTTGCGTGCACGCAAGTGTGTAGATATTNNNAGATCGGAA |
| 41081_2 | TATGAGTGTGGAGTCGTTGCGTGCACGCAAGTGTGTAGATATTNNNAGATCGGAA |
| 42350_1 | TATGAGTGTGGAGTCGTTGCTGTTGACTGTGGACATACAACCANNNAGATCGGAAG |
| 42350_2 | TATGAGTGTGGAGTCGTTGCTGTTGACTGTGGACATACAACCANNNAGATCGGAAG |
| 12788_1 | TATGAGTGTGGAGTCGTTGCCCATGATTCCCTTTCCTTTGNNNAGATCGGAAGAGCA |
| 12788_2 | TATGAGTGTGGAGTCGTTGCCCATGATTCCCTTTCCTTTGNNNAGATCGGAAGAGCA |
| 36935_1 | TATGAGTGTGGAGTCGTTGCGGAAGGAATAGTAGAGTCAGGCCNNNAGATCGGA |
| 36935_2 | TATGAGTGTGGAGTCGTTGCGGAAGGAATAGTAGAGTCAGGCCNNNAGATCGGA |
| 49434_1 | TATGAGTGTGGAGTCGTTGCTCTGGGGGCTGTCAAATTCATTANNNAGATCGGAAG |
| 49434_2 | TATGAGTGTGGAGTCGTTGCTCTGGGGGCTGTCAAATTCATTANNNAGATCGGAAG |

| | |
|---|---|
| 31204_1 | TATGAGTGTGGAGTCGTTGCTCTGGTGTCTCTTCTCACAAACCNNNAGATCGGAAGA |
| 31204_2 | TATGAGTGTGGAGTCGTTGCTCTGGTGTCTCTTCTCACAAACCNNNAGATCGGAAGA |
| 10081_1 | TATGAGTGTGGAGTCGTTGCGCAGCCATGTACAAACAAGTCTGNNNAGATCGGAAG |
| 10081_2 | TATGAGTGTGGAGTCGTTGCGCAGCCATGTACAAACAAGTCTGNNNAGATCGGAAG |
| 32089_1 | TATGAGTGTGGAGTCGTTGCCCTTCCTGGGAAATTCAGCCTTANNNAGATCGGAAGA |
| 32089_2 | TATGAGTGTGGAGTCGTTGCCCTTCCTGGGAAATTCAGCCTTANNNAGATCGGAAGA |
| 34063_1 | TATGAGTGTGGAGTCGTTGCCAATTGCATGATCTCGGCTCACNNNAGATCGGAAGA |
| 34063_2 | TATGAGTGTGGAGTCGTTGCCAATTGCATGATCTCGGCTCACNNNAGATCGGAAGA |
| 43564_1 | TATGAGTGTGGAGTCGTTGCCTTCTGGGCTCAAGTGATTCTCTNNNAGATCGGAAGA |
| 43564_2 | TATGAGTGTGGAGTCGTTGCCTTCTGGGCTCAAGTGATTCTCTNNNAGATCGGAAGA |
| 48703_1 | TATGAGTGTGGAGTCGTTGCCCCTAGGAGCGAAGCGTTCGNNNAGATCGGAAGAG |
| 48703_2 | TATGAGTGTGGAGTCGTTGCCCCTAGGAGCGAAGCGTTCGNNNAGATCGGAAGAG |
| 52782_1 | TATGAGTGTGGAGTCGTTGCTGAAACCTCACTGCCATGAAAACNNNAGATCGGAAG |
| 52782_2 | TATGAGTGTGGAGTCGTTGCTGAAACCTCACTGCCATGAAAACNNNAGATCGGAAG |
| 51415_1 | TATGAGTGTGGAGTCGTTGCAGTGAAACCCCGTCTCTACTAGANNNAGATCGGAAG |
| 51415_2 | TATGAGTGTGGAGTCGTTGCAGTGAAACCCCGTCTCTACTAGANNNAGATCGGAAG |
| 30858_1 | TATGAGTGTGGAGTCGTTGCCTCATACACACACACACACACACNNNAGATCGGAAGA |
| 30858_2 | TATGAGTGTGGAGTCGTTGCCTCATACACACACACACACACACNNNAGATCGGAAGA |
| 10922_1 | TATGAGTGTGGAGTCGTTGCCTCTGAATGGTCTTTCCCCCACNNNAGATCGGAAGAG |
| 10922_2 | TATGAGTGTGGAGTCGTTGCCTCTGAATGGTCTTTCCCCCACNNNAGATCGGAAGAG |
| 13488_1 | TATGAGTGTGGAGTCGTTGCTTTATTTGTAATTTTTGCTGCTGNNNAGATCGGAAGA |
| 13488_2 | TATGAGTGTGGAGTCGTTGCTTTATTTGTAATTTTTGCTGCTGNNNAGATCGGAAGA |
| 46_1 | TATGAGTGTGGAGTCGTTGCACCAGGGGTGGGATGATACAGGNNNAGATCGGAA |
| 46_2 | TATGAGTGTGGAGTCGTTGCACCAGGGGTGGGATGATACAGGNNNAGATCGGAA |
| 10725_1 | TATGAGTGTGGAGTCGTTGCGGAAACCATGTAGTTTGGGAGACANNNAGATCGGA |
| 10725_2 | TATGAGTGTGGAGTCGTTGCGGAAACCATGTAGTTTGGGAGACANNNAGATCGGA |
| 43603_1 | TATGAGTGTGGAGTCGTTGCTGCAGTTGGTAAAATTCATGCCANNNAGATCGGAAG |
| 43603_2 | TATGAGTGTGGAGTCGTTGCTGCAGTTGGTAAAATTCATGCCANNNAGATCGGAAG |
| 41873_1 | TATGAGTGTGGAGTCGTTGCTCTCCTGCACTTTTGAAGAAACTNNNAGATCGGAAGA |
| 41873_2 | TATGAGTGTGGAGTCGTTGCTCTCCTGCACTTTTGAAGAAACTNNNAGATCGGAAGA |
| 30798_1 | TATGAGTGTGGAGTCGTTGCGGGTGAAAATACAAGGGAAAGGGNNNAGATCGGA |
| 30798_2 | TATGAGTGTGGAGTCGTTGCGGGTGAAAATACAAGGGAAAGGGNNNAGATCGGA |
| 35343_1 | TATGAGTGTGGAGTCGTTGCCACCATGCCCAGCTAATTTTTGTNNNAGATCGGAAGA |
| 35343_2 | TATGAGTGTGGAGTCGTTGCCACCATGCCCAGCTAATTTTTGTNNNAGATCGGAAGA |
| 44976_1 | TATGAGTGTGGAGTCGTTGCTAAAAGGTAAGGGGGGAAGGGAANNNAGATCGGA |
| 44976_2 | TATGAGTGTGGAGTCGTTGCTAAAAGGTAAGGGGGGAAGGGAANNNAGATCGGA |
| 29273_1 | TATGAGTGTGGAGTCGTTGCTCTCAGTGCTCGCTAGTTCTTTTNNNAGATCGGAAGA |
| 29273_2 | TATGAGTGTGGAGTCGTTGCTCTCAGTGCTCGCTAGTTCTTTTNNNAGATCGGAAGA |
| 36711_1 | TATGAGTGTGGAGTCGTTGCCAGAGGTAGAAGGATCGCTTGAGNNNAGATCGGAA |
| 36711_2 | TATGAGTGTGGAGTCGTTGCCAGAGGTAGAAGGATCGCTTGAGNNNAGATCGGAA |
| 36387_1 | TATGAGTGTGGAGTCGTTGCATTGAGTACCGGAAAGAGACAGGNNNAGATCGGAA |
| 36387_2 | TATGAGTGTGGAGTCGTTGCATTGAGTACCGGAAAGAGACAGGNNNAGATCGGAA |
| 42070_1 | TATGAGTGTGGAGTCGTTGCAGCCAAATTTTATGGGAATGAACANNNAGATCGGAA |

| | |
|---|---|
| 42070_2 | TATGAGTGTGGAGTCGTTGCAGCCAAATTTTATGGGAATGAACANNNAGATCGGA |
| 30703_1 | TATGAGTGTGGAGTCGTTGCTTCTCCCTTCTGCTGACTGTAGNNNAGATCGGAAGA |
| 30703_2 | TATGAGTGTGGAGTCGTTGCTTCTCCCTTCTGCTGACTGTAGNNNAGATCGGAAGA |
| 10822_1 | TATGAGTGTGGAGTCGTTGCTTCTTAGGCAGTAGAGGACTGCGNNNAGATCGGAA |
| 10822_2 | TATGAGTGTGGAGTCGTTGCTTCTTAGGCAGTAGAGGACTGCGNNNAGATCGGAA |
| 40591_1 | TATGAGTGTGGAGTCGTTGCACCTCTGGTCTAAAGCCTTGTACNNNAGATCGGAAG |
| 40591_2 | TATGAGTGTGGAGTCGTTGCACCTCTGGTCTAAAGCCTTGTACNNNAGATCGGAAG |
| 30564_1 | TATGAGTGTGGAGTCGTTGCAGGAAAGGATGGGCTCAGAGTNNNAGATCGGAAG |
| 30564_2 | TATGAGTGTGGAGTCGTTGCAGGAAAGGATGGGCTCAGAGTNNNAGATCGGAAG |
| 11527_1 | TATGAGTGTGGAGTCGTTGCCGTCAGAAAGAGGAGGAAGCCNNNAGATCGGAAG |
| 11527_2 | TATGAGTGTGGAGTCGTTGCCGTCAGAAAGAGGAGGAAGCCNNNAGATCGGAAG |
| 42003_1 | TATGAGTGTGGAGTCGTTGCGCTGGTCTTGAACTCCTAACCTTNNNAGATCGGAAG |
| 42003_2 | TATGAGTGTGGAGTCGTTGCGCTGGTCTTGAACTCCTAACCTTNNNAGATCGGAAG |
| 45187_1 | TATGAGTGTGGAGTCGTTGCCTGAGCCCAAGCAGAATCTTTTCNNNAGATCGGAAG |
| 45187_2 | TATGAGTGTGGAGTCGTTGCCTGAGCCCAAGCAGAATCTTTTCNNNAGATCGGAAG |
| 51173_1 | TATGAGTGTGGAGTCGTTGCAATCCTGTTTGTGGTACAGCTCTNNNAGATCGGAAG |
| 51173_2 | TATGAGTGTGGAGTCGTTGCAATCCTGTTTGTGGTACAGCTCTNNNAGATCGGAAG |
| 34604_1 | TATGAGTGTGGAGTCGTTGCGGAGATGGCATATGTCGAGGATTNNNAGATCGGAA |
| 34604_2 | TATGAGTGTGGAGTCGTTGCGGAGATGGCATATGTCGAGGATTNNNAGATCGGAA |
| 12286_1 | TATGAGTGTGGAGTCGTTGCCAACGTGCTACTCATTCTCTGCTANNNAGATCGGAAG |
| 12286_2 | TATGAGTGTGGAGTCGTTGCCAACGTGCTACTCATTCTCTGCTANNNAGATCGGAAG |
| 51688_1 | TATGAGTGTGGAGTCGTTGCCAGTTACCCACACCTCTTTGAGANNNAGATCGGAAG |
| 51688_2 | TATGAGTGTGGAGTCGTTGCCAGTTACCCACACCTCTTTGAGANNNAGATCGGAAG |
| 35714_1 | TATGAGTGTGGAGTCGTTGCCATGCTCTTTCCATCCTCCATCTNNNAGATCGGAAGA |
| 35714_2 | TATGAGTGTGGAGTCGTTGCCATGCTCTTTCCATCCTCCATCTNNNAGATCGGAAGA |
| 46232_1 | TATGAGTGTGGAGTCGTTGCCAAAGTGCTGGGATTACAGGTGTNNNAGATCGGAA |
| 46232_2 | TATGAGTGTGGAGTCGTTGCCAAAGTGCTGGGATTACAGGTGTNNNAGATCGGAA |
| 39953_1 | TATGAGTGTGGAGTCGTTGCTGCATCTAGGACAGAAAGAGAAANNNAGATCGGAA |
| 39953_2 | TATGAGTGTGGAGTCGTTGCTGCATCTAGGACAGAAAGAGAAANNNAGATCGGAA |
| 29902_1 | TATGAGTGTGGAGTCGTTGCTGGCAGTATACAGGTTGAGGTGANNNAGATCGGAA |
| 29902_2 | TATGAGTGTGGAGTCGTTGCTGGCAGTATACAGGTTGAGGTGANNNAGATCGGAA |
| 35717_1 | TATGAGTGTGGAGTCGTTGCTGGAGGTGGAGTCTCATTTTGTCNNNAGATCGGAA |
| 35717_2 | TATGAGTGTGGAGTCGTTGCTGGAGGTGGAGTCTCATTTTGTCNNNAGATCGGAA |
| 2538_1 | TATGAGTGTGGAGTCGTTGCGAAGGGAAAAGGAAGAGCTCAGGNNNAGATCGGA |
| 2538_2 | TATGAGTGTGGAGTCGTTGCGAAGGGAAAAGGAAGAGCTCAGGNNNAGATCGGA |
| 31309_1 | TATGAGTGTGGAGTCGTTGCAGAGAGAGAGAGAGTTTGGAGCTNNNAGATCGGA |
| 31309_2 | TATGAGTGTGGAGTCGTTGCAGAGAGAGAGAGAGTTTGGAGCTNNNAGATCGGA |
| 36191_1 | TATGAGTGTGGAGTCGTTGCTGCTCATTGGTTTCTATTGGGGANNNAGATCGGAAG |
| 36191_2 | TATGAGTGTGGAGTCGTTGCTGCTCATTGGTTTCTATTGGGGANNNAGATCGGAAG |
| 45044_1 | TATGAGTGTGGAGTCGTTGCGAGTAGGCCTGAAACAGTCAAGANNNAGATCGGAA |
| 45044_2 | TATGAGTGTGGAGTCGTTGCGAGTAGGCCTGAAACAGTCAAGANNNAGATCGGAA |
| 32283_1 | TATGAGTGTGGAGTCGTTGCCGGATGTAAGGCCTCTGACTTGNNNAGATCGGAAG |
| 32283_2 | TATGAGTGTGGAGTCGTTGCCGGATGTAAGGCCTCTGACTTGNNNAGATCGGAAG |

45441_1 TATGAGTGTGGAGTCGTTGCCGAAGAAAGCAGAGGGGAAATTCNNNAGATCGGAA
45441_2 TATGAGTGTGGAGTCGTTGCCGAAGAAAGCAGAGGGGAAATTCNNNAGATCGGAA
29405_1 TATGAGTGTGGAGTCGTTGCCTCTCCCCAGCCAAAGAAGAAANNNAGATCGGAAGA
29405_2 TATGAGTGTGGAGTCGTTGCCTCTCCCCAGCCAAAGAAGAAANNNAGATCGGAAGA
35329_1 TATGAGTGTGGAGTCGTTGCAACAATCTGGGTAAGCCTAGCATNNNAGATCGGAAG
35329_2 TATGAGTGTGGAGTCGTTGCAACAATCTGGGTAAGCCTAGCATNNNAGATCGGAAG
38444_1 TATGAGTGTGGAGTCGTTGCCCTCTTCATCTTCCCCTTCCAAANNNAGATCGGAAGAG
38444_2 TATGAGTGTGGAGTCGTTGCCCTCTTCATCTTCCCCTTCCAAANNNAGATCGGAAGAG
46750_1 TATGAGTGTGGAGTCGTTGCAGCTCTTTCTCAGATTTCAGCCANNNAGATCGGAAGA
46750_2 TATGAGTGTGGAGTCGTTGCAGCTCTTTCTCAGATTTCAGCCANNNAGATCGGAAGA
31921_1 TATGAGTGTGGAGTCGTTGCTTTTATAAGGCCAGCATCACCCTNNNAGATCGGAAGA
31921_2 TATGAGTGTGGAGTCGTTGCTTTTATAAGGCCAGCATCACCCTNNNAGATCGGAAGA
40292_1 TATGAGTGTGGAGTCGTTGCGGAAAAGGGGGAAATGTGAAAGGNNNAGATCGGA
40292_2 TATGAGTGTGGAGTCGTTGCGGAAAAGGGGGAAATGTGAAAGGNNNAGATCGGA
42209_1 TATGAGTGTGGAGTCGTTGCTATGATCAAATCACCTGCCACCANNNAGATCGGAAGA
42209_2 TATGAGTGTGGAGTCGTTGCTATGATCAAATCACCTGCCACCANNNAGATCGGAAGA
36115_1 TATGAGTGTGGAGTCGTTGCTCATGTTCCGTGAATGGAAAACCNNNAGATCGGAAG
36115_2 TATGAGTGTGGAGTCGTTGCTCATGTTCCGTGAATGGAAAACCNNNAGATCGGAAG
41316_1 TATGAGTGTGGAGTCGTTGCTGGCTGCTTCTGATGTGATACTTNNNAGATCGGAAG
41316_2 TATGAGTGTGGAGTCGTTGCTGGCTGCTTCTGATGTGATACTTNNNAGATCGGAAG
35519_1 TATGAGTGTGGAGTCGTTGCGCCAGGATTATAGGCATGAGTCANNNAGATCGGAAG
35519_2 TATGAGTGTGGAGTCGTTGCGCCAGGATTATAGGCATGAGTCANNNAGATCGGAAG
33881_1 TATGAGTGTGGAGTCGTTGCCAGTGGTTCAATCTCGGCTCANNNAGATCGGAAGAG
33881_2 TATGAGTGTGGAGTCGTTGCCAGTGGTTCAATCTCGGCTCANNNAGATCGGAAGAG
31485_1 TATGAGTGTGGAGTCGTTGCAAACTTCACGCCAATATTCTGCANNNAGATCGGAAGA
31485_2 TATGAGTGTGGAGTCGTTGCAAACTTCACGCCAATATTCTGCANNNAGATCGGAAGA
49997_1 TATGAGTGTGGAGTCGTTGCGTCCGAACAAATGGCATTGTACANNNAGATCGGAAG
49997_2 TATGAGTGTGGAGTCGTTGCGTCCGAACAAATGGCATTGTACANNNAGATCGGAAG
42302_1 TATGAGTGTGGAGTCGTTGCCATGCCCAGAAAACTTTAGCCAANNNAGATCGGAAG
42302_2 TATGAGTGTGGAGTCGTTGCCATGCCCAGAAAACTTTAGCCAANNNAGATCGGAAG
47121_1 TATGAGTGTGGAGTCGTTGCGGAAGGACTTGAAACCTGACTCANNNAGATCGGAAG
47121_2 TATGAGTGTGGAGTCGTTGCGGAAGGACTTGAAACCTGACTCANNNAGATCGGAAG
39675_1 TATGAGTGTGGAGTCGTTGCAAAACAAAAACACATGCACACGTNNNAGATCGGAAG
39675_2 TATGAGTGTGGAGTCGTTGCAAAACAAAAACACATGCACACGTNNNAGATCGGAAG
38891_1 TATGAGTGTGGAGTCGTTGCGGCTAGGGCTGGTAAAAATGTTGNNNAGATCGGAA
38891_2 TATGAGTGTGGAGTCGTTGCGGCTAGGGCTGGTAAAAATGTTGNNNAGATCGGAA
52854_1 TATGAGTGTGGAGTCGTTGCGTGCCCTATGAGATCAAAGTCCANNNAGATCGGAAG
52854_2 TATGAGTGTGGAGTCGTTGCGTGCCCTATGAGATCAAAGTCCANNNAGATCGGAAG
34985_1 TATGAGTGTGGAGTCGTTGCGAGACAGTCTCGCTCTTGTCCNNNAGATCGGAAGAG
34985_2 TATGAGTGTGGAGTCGTTGCGAGACAGTCTCGCTCTTGTCCNNNAGATCGGAAGAG
41129_1 TATGAGTGTGGAGTCGTTGCCAAAATGTGGTCTACCCAGATGCNNNAGATCGGAAG
41129_2 TATGAGTGTGGAGTCGTTGCCAAAATGTGGTCTACCCAGATGCNNNAGATCGGAAG
47970_1 TATGAGTGTGGAGTCGTTGCCTGGACTCAAGGGATCCTCTTGNNNAGATCGGAAGA

| | |
|---|---|
| 47970_2 | TATGAGTGTGGAGTCGTTGCCTGGACTCAAGGGATCCTCTTGNNNAGATCGGAAGA |
| 52924_1 | TATGAGTGTGGAGTCGTTGCGACACTCTGCCTTTTATGTCCCANNNAGATCGGAAGA |
| 52924_2 | TATGAGTGTGGAGTCGTTGCGACACTCTGCCTTTTATGTCCCANNNAGATCGGAAGA |
| 47628_1 | TATGAGTGTGGAGTCGTTGCCCCAGGTTCAAGCAATTCTCATGNNNAGATCGGAAGA |
| 47628_2 | TATGAGTGTGGAGTCGTTGCCCCAGGTTCAAGCAATTCTCATGNNNAGATCGGAAGA |
| 52172_1 | TATGAGTGTGGAGTCGTTGCAAATTTCCAACATCTGCACCCTCNNNAGATCGGAAGA |
| 52172_2 | TATGAGTGTGGAGTCGTTGCAAATTTCCAACATCTGCACCCTCNNNAGATCGGAAGA |
| 31850_1 | TATGAGTGTGGAGTCGTTGCCAACAGAGTGAGACCATGTCAAGNNNAGATCGGAAC |
| 31850_2 | TATGAGTGTGGAGTCGTTGCCAACAGAGTGAGACCATGTCAAGNNNAGATCGGAAC |
| 30964_1 | TATGAGTGTGGAGTCGTTGCGTACTTTGCCCTCTGTTAGCTCTNNNAGATCGGAAGA |
| 30964_2 | TATGAGTGTGGAGTCGTTGCGTACTTTGCCCTCTGTTAGCTCTNNNAGATCGGAAGA |
| 41408_1 | TATGAGTGTGGAGTCGTTGCGCCATCAATATCACCTACAAGGCNNNAGATCGGAAGA |
| 41408_2 | TATGAGTGTGGAGTCGTTGCGCCATCAATATCACCTACAAGGCNNNAGATCGGAAGA |
| 10131_1 | TATGAGTGTGGAGTCGTTGCTGTGATTCAGGAGAGTGCATTCANNNAGATCGGAAC |
| 10131_2 | TATGAGTGTGGAGTCGTTGCTGTGATTCAGGAGAGTGCATTCANNNAGATCGGAAC |
| 2239_1 | TATGAGTGTGGAGTCGTTGCCACTCCCCACAACGACTTTCATANNNAGATCGGAAGA |
| 2239_2 | TATGAGTGTGGAGTCGTTGCCACTCCCCACAACGACTTTCATANNNAGATCGGAAGA |
| 35993_1 | TATGAGTGTGGAGTCGTTGCCTGAGGAAGGAGGATTGCTTGAGNNNAGATCGGAA |
| 35993_2 | TATGAGTGTGGAGTCGTTGCCTGAGGAAGGAGGATTGCTTGAGNNNAGATCGGAA |
| 47500_1 | TATGAGTGTGGAGTCGTTGCCTGAAGATGTCTGCCCCCAACNNNAGATCGGAAGAG |
| 47500_2 | TATGAGTGTGGAGTCGTTGCCTGAAGATGTCTGCCCCCAACNNNAGATCGGAAGAG |
| 47610_1 | TATGAGTGTGGAGTCGTTGCAATTCAAGCAATTCTCCTGCCTCNNNAGATCGGAAGA |
| 47610_2 | TATGAGTGTGGAGTCGTTGCAATTCAAGCAATTCTCCTGCCTCNNNAGATCGGAAGA |
| 30169_1 | TATGAGTGTGGAGTCGTTGCCTTTGGGGCTTACATCTACTGGTNNNAGATCGGAAG |
| 30169_2 | TATGAGTGTGGAGTCGTTGCCTTTGGGGCTTACATCTACTGGTNNNAGATCGGAAG |
| 45338_1 | TATGAGTGTGGAGTCGTTGCTCTCATGTGCATGGTACTGAACTNNNAGATCGGAAG |
| 45338_2 | TATGAGTGTGGAGTCGTTGCTCTCATGTGCATGGTACTGAACTNNNAGATCGGAAG |
| 39037_1 | TATGAGTGTGGAGTCGTTGCCTCCTTCAACTTCAGCTTCCTGANNNAGATCGGAAGA |
| 39037_2 | TATGAGTGTGGAGTCGTTGCCTCCTTCAACTTCAGCTTCCTGANNNAGATCGGAAGA |
| 30816_1 | TATGAGTGTGGAGTCGTTGCGTAACCACACATCAGAACAGTGCNNNAGATCGGAAG |
| 30816_2 | TATGAGTGTGGAGTCGTTGCGTAACCACACATCAGAACAGTGCNNNAGATCGGAAG |
| 47361_1 | TATGAGTGTGGAGTCGTTGCGTGGAGAGATAGGATTGGTGCATNNNAGATCGGAA |
| 47361_2 | TATGAGTGTGGAGTCGTTGCGTGGAGAGATAGGATTGGTGCATNNNAGATCGGAA |
| 31414_1 | TATGAGTGTGGAGTCGTTGCTCAACTCTTCCCCTCGATAGACTNNNAGATCGGAAGA |
| 31414_2 | TATGAGTGTGGAGTCGTTGCTCAACTCTTCCCCTCGATAGACTNNNAGATCGGAAGA |
| 41608_1 | TATGAGTGTGGAGTCGTTGCCAAGTAGCTGGGACTACAGGCNNNAGATCGGAAGA |
| 41608_2 | TATGAGTGTGGAGTCGTTGCCAAGTAGCTGGGACTACAGGCNNNAGATCGGAAGA |
| 38842_1 | TATGAGTGTGGAGTCGTTGCCAGGAGAATCACTTGAACCCAGGNNNAGATCGGAAC |
| 38842_2 | TATGAGTGTGGAGTCGTTGCCAGGAGAATCACTTGAACCCAGGNNNAGATCGGAAC |
| 39465_1 | TATGAGTGTGGAGTCGTTGCAGTTGTATACTGGAACTGGCCTGNNNAGATCGGAAC |
| 39465_2 | TATGAGTGTGGAGTCGTTGCAGTTGTATACTGGAACTGGCCTGNNNAGATCGGAAC |
| 32952_1 | TATGAGTGTGGAGTCGTTGCAGACAGTGAAAGACATCTGACCTNNNAGATCGGAAC |
| 32952_2 | TATGAGTGTGGAGTCGTTGCAGACAGTGAAAGACATCTGACCTNNNAGATCGGAAC |

| | |
|---|---|
| 41707_1 | TATGAGTGTGGAGTCGTTGCGCTTACTATATCCCCCTCCTCCTNNNAGATCGGAAGA |
| 41707_2 | TATGAGTGTGGAGTCGTTGCGCTTACTATATCCCCCTCCTCCTNNNAGATCGGAAGA |
| 35197_1 | TATGAGTGTGGAGTCGTTGCAACTCCTGGGTTCAAGCCATTATNNNAGATCGGAAG |
| 35197_2 | TATGAGTGTGGAGTCGTTGCAACTCCTGGGTTCAAGCCATTATNNNAGATCGGAAG |
| 2075_1 | TATGAGTGTGGAGTCGTTGCAGTCTCACAACAGCAGAATGACANNNAGATCGGAAG |
| 2075_2 | TATGAGTGTGGAGTCGTTGCAGTCTCACAACAGCAGAATGACANNNAGATCGGAAG |
| 35466_1 | TATGAGTGTGGAGTCGTTGCGGATTACAGGTATGAGCCACCATNNNAGATCGGAA |
| 35466_2 | TATGAGTGTGGAGTCGTTGCGGATTACAGGTATGAGCCACCATNNNAGATCGGAA |
| 47886_1 | TATGAGTGTGGAGTCGTTGCAGTAGAGATGAGGTTTCGCCATGNNNAGATCGGAA |
| 47886_2 | TATGAGTGTGGAGTCGTTGCAGTAGAGATGAGGTTTCGCCATGNNNAGATCGGAA |
| 40900_1 | TATGAGTGTGGAGTCGTTGCGCTCAAATTCACTGAATCACCCCNNNAGATCGGAAG |
| 40900_2 | TATGAGTGTGGAGTCGTTGCGCTCAAATTCACTGAATCACCCCNNNAGATCGGAAG |
| 32841_1 | TATGAGTGTGGAGTCGTTGCGATCTCCTCTCACTGCAAGCTCNNNAGATCGGAAGA |
| 32841_2 | TATGAGTGTGGAGTCGTTGCGATCTCCTCTCACTGCAAGCTCNNNAGATCGGAAGA |
| 40016_1 | TATGAGTGTGGAGTCGTTGCAAAAATTAGCTGGGCATAGTGGCNNNAGATCGGAA |
| 40016_2 | TATGAGTGTGGAGTCGTTGCAAAAATTAGCTGGGCATAGTGGCNNNAGATCGGAA |
| 46823_1 | TATGAGTGTGGAGTCGTTGCAGGGTCTGAGCTGAATTATGTGGNNNAGATCGGAA |
| 46823_2 | TATGAGTGTGGAGTCGTTGCAGGGTCTGAGCTGAATTATGTGGNNNAGATCGGAA |
| 53250_1 | TATGAGTGTGGAGTCGTTGCCTCAGCGGTTACACAATCCCGNNNAGATCGGAAGAG |
| 53250_2 | TATGAGTGTGGAGTCGTTGCCTCAGCGGTTACACAATCCCGNNNAGATCGGAAGAG |
| 47024_1 | TATGAGTGTGGAGTCGTTGCCCACAAAGCATCTGCAGAGTTGNNNAGATCGGAAG |
| 47024_2 | TATGAGTGTGGAGTCGTTGCCCACAAAGCATCTGCAGAGTTGNNNAGATCGGAAG |
| 39761_1 | TATGAGTGTGGAGTCGTTGCGGGAATTGGTTTTGATGGTAGCCNNNAGATCGGAA |
| 39761_2 | TATGAGTGTGGAGTCGTTGCGGGAATTGGTTTTGATGGTAGCCNNNAGATCGGAA |
| 44305_1 | TATGAGTGTGGAGTCGTTGCAGCATGGGAAGGAAACTCTTCATNNNAGATCGGAA |
| 44305_2 | TATGAGTGTGGAGTCGTTGCAGCATGGGAAGGAAACTCTTCATNNNAGATCGGAA |
| 51255_1 | TATGAGTGTGGAGTCGTTGCTGTTTGGAGCTTGCTAAGGCTAGNNNAGATCGGAA |
| 51255_2 | TATGAGTGTGGAGTCGTTGCTGTTTGGAGCTTGCTAAGGCTAGNNNAGATCGGAA |
| 40735_1 | TATGAGTGTGGAGTCGTTGCTGGAGTGCAGTGGTACTATCTTGNNNAGATCGGAA |
| 40735_2 | TATGAGTGTGGAGTCGTTGCTGGAGTGCAGTGGTACTATCTTGNNNAGATCGGAA |
| 33710_1 | TATGAGTGTGGAGTCGTTGCATGTAAATGAATGGGCATGGCTGNNNAGATCGGAA |
| 33710_2 | TATGAGTGTGGAGTCGTTGCATGTAAATGAATGGGCATGGCTGNNNAGATCGGAA |
| 47137_1 | TATGAGTGTGGAGTCGTTGCTCCTTGTTTCCAGGCTAAACAGANNNAGATCGGAAG |
| 47137_2 | TATGAGTGTGGAGTCGTTGCTCCTTGTTTCCAGGCTAAACAGANNNAGATCGGAAG |
| 43904_1 | TATGAGTGTGGAGTCGTTGCCCTTCTGCTTCCCCTTTTTGATGNNNAGATCGGAAGA |
| 43904_2 | TATGAGTGTGGAGTCGTTGCCCTTCTGCTTCCCCTTTTTGATGNNNAGATCGGAAGA |
| 49592_1 | TATGAGTGTGGAGTCGTTGCCAAGTAGCTGGGACTATAGGTGCNNNAGATCGGAA |
| 49592_2 | TATGAGTGTGGAGTCGTTGCCAAGTAGCTGGGACTATAGGTGCNNNAGATCGGAA |
| 43682_1 | TATGAGTGTGGAGTCGTTGCTATTTTCTCGGCGAGATTGATGCNNNAGATCGGAAG |
| 43682_2 | TATGAGTGTGGAGTCGTTGCTATTTTCTCGGCGAGATTGATGCNNNAGATCGGAAG |
| 48091_1 | TATGAGTGTGGAGTCGTTGCGGGCAGGATCTCTCGGAAGANNNAGATCGGAAGAG |
| 48091_2 | TATGAGTGTGGAGTCGTTGCGGGCAGGATCTCTCGGAAGANNNAGATCGGAAGAG |
| 45428_1 | TATGAGTGTGGAGTCGTTGCTGATCTCGGCTCACTGTAAACTCNNNAGATCGGAAG |

| | |
|---|---|
| 45428_2 | TATGAGTGTGGAGTCGTTGCTGATCTCGGCTCACTGTAAACTCNNNAGATCGGAAG |
| 34507_1 | TATGAGTGTGGAGTCGTTGCGATGGTTTCCAGCTTCATCCATGNNNAGATCGGAAG |
| 34507_2 | TATGAGTGTGGAGTCGTTGCGATGGTTTCCAGCTTCATCCATGNNNAGATCGGAAG |
| 30578_1 | TATGAGTGTGGAGTCGTTGCCTTGTTGGGTACCATGTGCTAAAAATNNNAGATCGG |
| 30578_2 | TATGAGTGTGGAGTCGTTGCCTTGTTGGGTACCATGTGCTAAAAATNNNAGATCGG |
| 51075_1 | TATGAGTGTGGAGTCGTTGCGCTGGCTATCTGATGACTCATCANNNAGATCGGAAG |
| 51075_2 | TATGAGTGTGGAGTCGTTGCGCTGGCTATCTGATGACTCATCANNNAGATCGGAAG |
| 32315_1 | TATGAGTGTGGAGTCGTTGCTTTACGATGAATCTTGCTGCTGCNNNAGATCGGAAG |
| 32315_2 | TATGAGTGTGGAGTCGTTGCTTTACGATGAATCTTGCTGCTGCNNNAGATCGGAAG |
| 10024_1 | TATGAGTGTGGAGTCGTTGCGGACAAAGAGAGCAGAGGTGATCNNNAGATCGGA |
| 10024_2 | TATGAGTGTGGAGTCGTTGCGGACAAAGAGAGCAGAGGTGATCNNNAGATCGGA |
| 49165_1 | TATGAGTGTGGAGTCGTTGCTGGGGGGAGATGTTTTACTTTCANNNAGATCGGAA |
| 49165_2 | TATGAGTGTGGAGTCGTTGCTGGGGGGAGATGTTTTACTTTCANNNAGATCGGAA |
| 37194_1 | TATGAGTGTGGAGTCGTTGCTGCTAATACAATGGACGGTATCCNNNAGATCGGAAG |
| 37194_2 | TATGAGTGTGGAGTCGTTGCTGCTAATACAATGGACGGTATCCNNNAGATCGGAAG |
| 49913_1 | TATGAGTGTGGAGTCGTTGCAGAGATGGGGGTAAGGAAAGGTANNNAGATCGGA |
| 49913_2 | TATGAGTGTGGAGTCGTTGCAGAGATGGGGGTAAGGAAAGGTANNNAGATCGGA |
| 53275_1 | TATGAGTGTGGAGTCGTTGCGAATACTGCTGCCCAGGTTCCTTNNNAGATCGGAAG |
| 53275_2 | TATGAGTGTGGAGTCGTTGCGAATACTGCTGCCCAGGTTCCTTNNNAGATCGGAAG |
| 36571_1 | TATGAGTGTGGAGTCGTTGCGCCCAAGATTATGCAACGGTTAGNNNAGATCGGAA |
| 36571_2 | TATGAGTGTGGAGTCGTTGCGCCCAAGATTATGCAACGGTTAGNNNAGATCGGAA |
| 39973_1 | TATGAGTGTGGAGTCGTTGCCATAGGACCAGAGGGGATTTGTGNNNAGATCGGAA |
| 39973_2 | TATGAGTGTGGAGTCGTTGCCATAGGACCAGAGGGGATTTGTGNNNAGATCGGAA |
| 47039_1 | TATGAGTGTGGAGTCGTTGCCCATCATGCTCGGCTAATTTTTGNNNAGATCGGAAG |
| 47039_2 | TATGAGTGTGGAGTCGTTGCCCATCATGCTCGGCTAATTTTTGNNNAGATCGGAAG |
| 48828_1 | TATGAGTGTGGAGTCGTTGCGGAAGGCAAGATTTCCTGTTTCCNNNAGATCGGAAG |
| 48828_2 | TATGAGTGTGGAGTCGTTGCGGAAGGCAAGATTTCCTGTTTCCNNNAGATCGGAAG |
| 46697_1 | TATGAGTGTGGAGTCGTTGCTCTCAAACTCCTAGGTTCAAGCCNNNAGATCGGAAG |
| 46697_2 | TATGAGTGTGGAGTCGTTGCTCTCAAACTCCTAGGTTCAAGCCNNNAGATCGGAAG |
| 35085_1 | TATGAGTGTGGAGTCGTTGCTTTTGAGCCTATGTGTGCCTTTGNNNAGATCGGAAG |
| 35085_2 | TATGAGTGTGGAGTCGTTGCTTTTGAGCCTATGTGTGCCTTTGNNNAGATCGGAAG |
| 52612_1 | TATGAGTGTGGAGTCGTTGCGTTTACTTTACGGTGCTCTGCAGNNNAGATCGGAAG |
| 52612_2 | TATGAGTGTGGAGTCGTTGCGTTTACTTTACGGTGCTCTGCAGNNNAGATCGGAAG |
| 46099_1 | TATGAGTGTGGAGTCGTTGCAGCTTATAGCCTATTATCCCAGGTNNNAGATCGGAA |
| 46099_2 | TATGAGTGTGGAGTCGTTGCAGCTTATAGCCTATTATCCCAGGTNNNAGATCGGAA |
| 44518_1 | TATGAGTGTGGAGTCGTTGCAAGAACCACAACAACACAAAACANNNAGATCGGAAG |
| 44518_2 | TATGAGTGTGGAGTCGTTGCAAGAACCACAACAACACAAAACANNNAGATCGGAAG |
| 51772_1 | TATGAGTGTGGAGTCGTTGCCTACATCAAGGATGCCACCTTCTNNNAGATCGGAAG |
| 51772_2 | TATGAGTGTGGAGTCGTTGCCTACATCAAGGATGCCACCTTCTNNNAGATCGGAAG |
| 33105_1 | TATGAGTGTGGAGTCGTTGCTACATGACCCAAATGTCGTCCTTNNNAGATCGGAAG |
| 33105_2 | TATGAGTGTGGAGTCGTTGCTACATGACCCAAATGTCGTCCTTNNNAGATCGGAAG |
| 44179_1 | TATGAGTGTGGAGTCGTTGCATGTGTGTGTGAGCTTGTCATTGNNNAGATCGGAA |
| 44179_2 | TATGAGTGTGGAGTCGTTGCATGTGTGTGTGAGCTTGTCATTGNNNAGATCGGAA |

| | |
|---|---|
| 41833_1 | TATGAGTGTGGAGTCGTTGCGGAGTCTCACTCTGTCACCCANNNAGATCGGAAGAG |
| 41833_2 | TATGAGTGTGGAGTCGTTGCGGAGTCTCACTCTGTCACCCANNNAGATCGGAAGAG |
| 47901_1 | TATGAGTGTGGAGTCGTTGCGTAACTGGGAATACAGGCAAACGNNNAGATCGGAA |
| 47901_2 | TATGAGTGTGGAGTCGTTGCGTAACTGGGAATACAGGCAAACGNNNAGATCGGAA |
| 43842_1 | TATGAGTGTGGAGTCGTTGCCATGTTGAGCTACTGACAAGCACNNNAGATCGGAAG |
| 43842_2 | TATGAGTGTGGAGTCGTTGCCATGTTGAGCTACTGACAAGCACNNNAGATCGGAAG |
| 39409_1 | TATGAGTGTGGAGTCGTTGCAGGAAACCTAAGTACTCTTGGGCNNNAGATCGGAA |
| 39409_2 | TATGAGTGTGGAGTCGTTGCAGGAAACCTAAGTACTCTTGGGCNNNAGATCGGAA |
| 46886_1 | TATGAGTGTGGAGTCGTTGCAGCCTGTAATCCCAACACTATGGNNNAGATCGGAAG |
| 46886_2 | TATGAGTGTGGAGTCGTTGCAGCCTGTAATCCCAACACTATGGNNNAGATCGGAAG |
| 1999_1 | TATGAGTGTGGAGTCGTTGCCTGTTCCTGCTCTCAGGTCAAAANNNAGATCGGAAG |
| 1999_2 | TATGAGTGTGGAGTCGTTGCCTGTTCCTGCTCTCAGGTCAAAANNNAGATCGGAAG |
| 33656_1 | TATGAGTGTGGAGTCGTTGCCAATGCCAATTGTGGACCTGAAANNNAGATCGGAA |
| 33656_2 | TATGAGTGTGGAGTCGTTGCCAATGCCAATTGTGGACCTGAAANNNAGATCGGAA |
| 43192_1 | TATGAGTGTGGAGTCGTTGCATCTATCTCCCAGGTTCAAGGGANNNAGATCGGAAG |
| 43192_2 | TATGAGTGTGGAGTCGTTGCATCTATCTCCCAGGTTCAAGGGANNNAGATCGGAAG |
| 47171_1 | TATGAGTGTGGAGTCGTTGCCTCTCTGAGTAGCTGGGACTACANNNAGATCGGAAG |
| 47171_2 | TATGAGTGTGGAGTCGTTGCCTCTCTGAGTAGCTGGGACTACANNNAGATCGGAAG |
| 34650_1 | TATGAGTGTGGAGTCGTTGCATTTCAAACTCAGCAACGGAGTCNNNAGATCGGAAG |
| 34650_2 | TATGAGTGTGGAGTCGTTGCATTTCAAACTCAGCAACGGAGTCNNNAGATCGGAAG |
| 7928_1 | TATGAGTGTGGAGTCGTTGCAGGCTGTCTCTCTCTCCCAGTNNNAGATCGGAAGAG |
| 7928_2 | TATGAGTGTGGAGTCGTTGCAGGCTGTCTCTCTCTCCCAGTNNNAGATCGGAAGAG |
| 28928_1 | TATGAGTGTGGAGTCGTTGCATGAGCTCTGTCATTTTTGCTGGNNNAGATCGGAAG |
| 28928_2 | TATGAGTGTGGAGTCGTTGCATGAGCTCTGTCATTTTTGCTGGNNNAGATCGGAAG |
| 34049_1 | TATGAGTGTGGAGTCGTTGCAGATGGATTCTTGCTCTGTCACCNNNAGATCGGAAG |
| 34049_2 | TATGAGTGTGGAGTCGTTGCAGATGGATTCTTGCTCTGTCACCNNNAGATCGGAAG |
| 47732_1 | TATGAGTGTGGAGTCGTTGCAGCAATTCTCCTATCTCAGCCACNNNAGATCGGAAG |
| 47732_2 | TATGAGTGTGGAGTCGTTGCAGCAATTCTCCTATCTCAGCCACNNNAGATCGGAAG |
| 36873_1 | TATGAGTGTGGAGTCGTTGCAAACCCAAGAAATTCCAAGCAGGNNNAGATCGGAA |
| 36873_2 | TATGAGTGTGGAGTCGTTGCAAACCCAAGAAATTCCAAGCAGGNNNAGATCGGAA |
| 32332_1 | TATGAGTGTGGAGTCGTTGCATTTGTTTTTCTGTGCCTGGCTTNNNAGATCGGAAG |
| 32332_2 | TATGAGTGTGGAGTCGTTGCATTTGTTTTTCTGTGCCTGGCTTNNNAGATCGGAAG |
| 46304_1 | TATGAGTGTGGAGTCGTTGCTGTCTGTCTCCCCTCACTAGAATNNNAGATCGGAAG |
| 46304_2 | TATGAGTGTGGAGTCGTTGCTGTCTGTCTCCCCTCACTAGAATNNNAGATCGGAAG |
| 44459_1 | TATGAGTGTGGAGTCGTTGCTGGTAGATCTTGCCTGTAATCCCNNNAGATCGGAAG |
| 44459_2 | TATGAGTGTGGAGTCGTTGCTGGTAGATCTTGCCTGTAATCCCNNNAGATCGGAAG |
| 29807_1 | TATGAGTGTGGAGTCGTTGCGCACCACATTTGCCTTCGTATAANNNAGATCGGAAG |
| 29807_2 | TATGAGTGTGGAGTCGTTGCGCACCACATTTGCCTTCGTATAANNNAGATCGGAAG |
| 30084_1 | TATGAGTGTGGAGTCGTTGCGACTGAATCCTGGCACTTTGAACNNNAGATCGGAAG |
| 30084_2 | TATGAGTGTGGAGTCGTTGCGACTGAATCCTGGCACTTTGAACNNNAGATCGGAAG |
| 49911_1 | TATGAGTGTGGAGTCGTTGCGAGCATTCAGCAACAACTGACTTNNNAGATCGGAAG |
| 49911_2 | TATGAGTGTGGAGTCGTTGCGAGCATTCAGCAACAACTGACTTNNNAGATCGGAAG |
| 10380_1 | TATGAGTGTGGAGTCGTTGCGCCAGTTTTGCTTCTACTTGCAANNNAGATCGGAAG |

| | |
|---|---|
| 10380_2 | TATGAGTGTGGAGTCGTTGCGCCAGTTTTGCTTCTACTTGCAANNNAGATCGGAAG |
| 32685_1 | TATGAGTGTGGAGTCGTTGCCTCAGCCTCCCAAAGTGTGATATNNNAGATCGGAAG |
| 32685_2 | TATGAGTGTGGAGTCGTTGCCTCAGCCTCCCAAAGTGTGATATNNNAGATCGGAAG |
| 50659_1 | TATGAGTGTGGAGTCGTTGCTCCTTATCTGTCAAGTGGCAACANNNAGATCGGAAG |
| 50659_2 | TATGAGTGTGGAGTCGTTGCTCCTTATCTGTCAAGTGGCAACANNNAGATCGGAAG |
| 50197_1 | TATGAGTGTGGAGTCGTTGCGATGTGTCCTTGGATAAGAGCCANNNAGATCGGAA |
| 50197_2 | TATGAGTGTGGAGTCGTTGCGATGTGTCCTTGGATAAGAGCCANNNAGATCGGAA |
| 41853_1 | TATGAGTGTGGAGTCGTTGCCGAGTAGCTGGGATTACAAGCATNNNAGATCGGAA |
| 41853_2 | TATGAGTGTGGAGTCGTTGCCGAGTAGCTGGGATTACAAGCATNNNAGATCGGAA |
| 29748_1 | TATGAGTGTGGAGTCGTTGCAAATGGAATACCCCTTGAGTCCANNNAGATCGGAAG |
| 29748_2 | TATGAGTGTGGAGTCGTTGCAAATGGAATACCCCTTGAGTCCANNNAGATCGGAAG |
| 38680_1 | TATGAGTGTGGAGTCGTTGCCACAGTGGTTCAAGCCTGTAATCNNNAGATCGGAAG |
| 38680_2 | TATGAGTGTGGAGTCGTTGCCACAGTGGTTCAAGCCTGTAATCNNNAGATCGGAAG |
| 29331_1 | TATGAGTGTGGAGTCGTTGCCCTCAGCGTGGAAAATATGGAACNNNAGATCGGAA |
| 29331_2 | TATGAGTGTGGAGTCGTTGCCCTCAGCGTGGAAAATATGGAACNNNAGATCGGAA |
| 44083_1 | TATGAGTGTGGAGTCGTTGCTGTGTGTGTGTCTCTTCTCCATGNNNAGATCGGAAG |
| 44083_2 | TATGAGTGTGGAGTCGTTGCTGTGTGTGTGTCTCTTCTCCATGNNNAGATCGGAAG |
| 35612_1 | TATGAGTGTGGAGTCGTTGCCACGATCTCAGCTCATTGCAAGNNNAGATCGGAAGA |
| 35612_2 | TATGAGTGTGGAGTCGTTGCCACGATCTCAGCTCATTGCAAGNNNAGATCGGAAGA |
| 38864_1 | TATGAGTGTGGAGTCGTTGCTGAAGCTGTGGGGAAAAGAGAATNNNAGATCGGA |
| 38864_2 | TATGAGTGTGGAGTCGTTGCTGAAGCTGTGGGGAAAAGAGAATNNNAGATCGGA |
| 49079_1 | TATGAGTGTGGAGTCGTTGCCACTACGCCCGGCTAATTTTTTANNNAGATCGGAAG |
| 49079_2 | TATGAGTGTGGAGTCGTTGCCACTACGCCCGGCTAATTTTTTANNNAGATCGGAAG |
| 12400_1 | TATGAGTGTGGAGTCGTTGCCTCCACGGCTCTGTCTTGATCNNNAGATCGGAAGAG |
| 12400_2 | TATGAGTGTGGAGTCGTTGCCTCCACGGCTCTGTCTTGATCNNNAGATCGGAAGAG |
| 52370_1 | TATGAGTGTGGAGTCGTTGCGTCTAGGGTGTCCAAAAGCCTATNNNAGATCGGAA |
| 52370_2 | TATGAGTGTGGAGTCGTTGCGTCTAGGGTGTCCAAAAGCCTATNNNAGATCGGAA |
| 50716_1 | TATGAGTGTGGAGTCGTTGCAGACCGTACTCACTTCTACCTGANNNAGATCGGAAG |
| 50716_2 | TATGAGTGTGGAGTCGTTGCAGACCGTACTCACTTCTACCTGANNNAGATCGGAAG |
| 47984_1 | TATGAGTGTGGAGTCGTTGCCTCGGCTTTGATCTGCTAATGTGNNNAGATCGGAAG |
| 47984_2 | TATGAGTGTGGAGTCGTTGCCTCGGCTTTGATCTGCTAATGTGNNNAGATCGGAAG |
| 46483_1 | TATGAGTGTGGAGTCGTTGCGTTAGCAGTGTGGAAAGGGGATANNNAGATCGGA |
| 46483_2 | TATGAGTGTGGAGTCGTTGCGTTAGCAGTGTGGAAAGGGGATANNNAGATCGGA |
| 35693_1 | TATGAGTGTGGAGTCGTTGCCAGAACACAGAATGGAATGCCTGNNNAGATCGGAA |
| 35693_2 | TATGAGTGTGGAGTCGTTGCCAGAACACAGAATGGAATGCCTGNNNAGATCGGAA |
| 41655_1 | TATGAGTGTGGAGTCGTTGCCTAGCAGCTGGGATTACAGATGTNNNAGATCGGAA |
| 41655_2 | TATGAGTGTGGAGTCGTTGCCTAGCAGCTGGGATTACAGATGTNNNAGATCGGAA |
| 42376_1 | TATGAGTGTGGAGTCGTTGCTTGGAGCTCACAGGAAGTTATCCNNNAGATCGGAA |
| 42376_2 | TATGAGTGTGGAGTCGTTGCTTGGAGCTCACAGGAAGTTATCCNNNAGATCGGAA |
| 7251_1 | TATGAGTGTGGAGTCGTTGCTGCTTGTGTCCCCCTAAAATGTTNNNAGATCGGAAG |
| 7251_2 | TATGAGTGTGGAGTCGTTGCTGCTTGTGTCCCCCTAAAATGTTNNNAGATCGGAAG |
| 30944_1 | TATGAGTGTGGAGTCGTTGCTGGAATGAGCCATTCATACAGGTNNNAGATCGGAA |
| 30944_2 | TATGAGTGTGGAGTCGTTGCTGGAATGAGCCATTCATACAGGTNNNAGATCGGAA |

| | |
|---|---|
| 13347_1 | TATGAGTGTGGAGTCGTTGCTTCTGAGAATGACATGAAAATTTNNNAGATCGGAAG |
| 13347_2 | TATGAGTGTGGAGTCGTTGCTTCTGAGAATGACATGAAAATTTNNNAGATCGGAAG |
| 46266_1 | TATGAGTGTGGAGTCGTTGCCTGAGTTCAGGCAATCCACCTACNNNAGATCGGAAG |
| 46266_2 | TATGAGTGTGGAGTCGTTGCCTGAGTTCAGGCAATCCACCTACNNNAGATCGGAAG |
| 28996_1 | TATGAGTGTGGAGTCGTTGCACAAGAGTAGCTTTTGGTGGTCTNNNAGATCGGAA |
| 28996_2 | TATGAGTGTGGAGTCGTTGCACAAGAGTAGCTTTTGGTGGTCTNNNAGATCGGAA |
| 5530_1 | TATGAGTGTGGAGTCGTTGCTTGCTGGGAGAAAGGAAAGTGACNNNAGATCGGA |
| 5530_2 | TATGAGTGTGGAGTCGTTGCTTGCTGGGAGAAAGGAAAGTGACNNNAGATCGGA |
| 51224_1 | TATGAGTGTGGAGTCGTTGCGAGGTTGCAGTGAGCTGAGATCNNNAGATCGGAAG |
| 51224_2 | TATGAGTGTGGAGTCGTTGCGAGGTTGCAGTGAGCTGAGATCNNNAGATCGGAAG |
| 35181_1 | TATGAGTGTGGAGTCGTTGCTACTGAGGACCTGGTACTACAGGNNNAGATCGGAA |
| 35181_2 | TATGAGTGTGGAGTCGTTGCTACTGAGGACCTGGTACTACAGGNNNAGATCGGAA |
| 52857_1 | TATGAGTGTGGAGTCGTTGCCCTCTTCTCACTCTGTCTGTCACNNNAGATCGGAAGA |
| 52857_2 | TATGAGTGTGGAGTCGTTGCCCTCTTCTCACTCTGTCTGTCACNNNAGATCGGAAGA |
| 51355_1 | TATGAGTGTGGAGTCGTTGCCTCTTTGGGTCGAGGATAAGCTTNNNAGATCGGAA |
| 51355_2 | TATGAGTGTGGAGTCGTTGCCTCTTTGGGTCGAGGATAAGCTTNNNAGATCGGAA |
| 49414_1 | TATGAGTGTGGAGTCGTTGCGCTGGAGTACAGTGGTGTAATCTNNNAGATCGGAA |
| 49414_2 | TATGAGTGTGGAGTCGTTGCGCTGGAGTACAGTGGTGTAATCTNNNAGATCGGAA |
| 42164_1 | TATGAGTGTGGAGTCGTTGCGAGTAGCTGGGATTATAGGTGCCNNNAGATCGGAA |
| 42164_2 | TATGAGTGTGGAGTCGTTGCGAGTAGCTGGGATTATAGGTGCCNNNAGATCGGAA |
| 51027_1 | TATGAGTGTGGAGTCGTTGCAACATCTGGGCAATGTAAAGCACNNNAGATCGGAA |
| 51027_2 | TATGAGTGTGGAGTCGTTGCAACATCTGGGCAATGTAAAGCACNNNAGATCGGAA |
| 44298_1 | TATGAGTGTGGAGTCGTTGCGAAGCCTGACCTTGAGAGTGTTANNNAGATCGGAA |
| 44298_2 | TATGAGTGTGGAGTCGTTGCGAAGCCTGACCTTGAGAGTGTTANNNAGATCGGAA |
| 34271_1 | TATGAGTGTGGAGTCGTTGCCTGGACAGCACCATCGAAGTAAGNNNAGATCGGAA |
| 34271_2 | TATGAGTGTGGAGTCGTTGCCTGGACAGCACCATCGAAGTAAGNNNAGATCGGAA |
| 30898_1 | TATGAGTGTGGAGTCGTTGCACTCTCTGTCTCTCTCTCTGTCCNNNAGATCGGAAGA |
| 30898_2 | TATGAGTGTGGAGTCGTTGCACTCTCTGTCTCTCTCTCTGTCCNNNAGATCGGAAGA |
| 12875_1 | TATGAGTGTGGAGTCGTTGCCCCTCTTTTATCTATTCCCTAAANNNAGATCGGAAGA |
| 12875_2 | TATGAGTGTGGAGTCGTTGCCCCTCTTTTATCTATTCCCTAAANNNAGATCGGAAGA |
| 42736_1 | TATGAGTGTGGAGTCGTTGCAGCAATGCTAGAGGAGGAACAAANNNAGATCGGA |
| 42736_2 | TATGAGTGTGGAGTCGTTGCAGCAATGCTAGAGGAGGAACAAANNNAGATCGGA |
| 38685_1 | TATGAGTGTGGAGTCGTTGCGCCAGACCCCATCTCTAAGAAAANNNAGATCGGAAG |
| 38685_2 | TATGAGTGTGGAGTCGTTGCGCCAGACCCCATCTCTAAGAAAANNNAGATCGGAAG |
| 39944_1 | TATGAGTGTGGAGTCGTTGCGATGGTTTGAGGGAGTAATGTGCNNNAGATCGGA |
| 39944_2 | TATGAGTGTGGAGTCGTTGCGATGGTTTGAGGGAGTAATGTGCNNNAGATCGGA |
| 49233_1 | TATGAGTGTGGAGTCGTTGCGAAAAGCAGTTCGTAGAGGGAGANNNAGATCGGA |
| 49233_2 | TATGAGTGTGGAGTCGTTGCGAAAAGCAGTTCGTAGAGGGAGANNNAGATCGGA |
| 47932_1 | TATGAGTGTGGAGTCGTTGCCTGGGTTCAAGTAATTCTCGTGCNNNAGATCGGAAG |
| 47932_2 | TATGAGTGTGGAGTCGTTGCCTGGGTTCAAGTAATTCTCGTGCNNNAGATCGGAAG |
| 33188_1 | TATGAGTGTGGAGTCGTTGCTTTGGTAGCTAGGCTCTTTCCTGNNNAGATCGGAAG |
| 33188_2 | TATGAGTGTGGAGTCGTTGCTTTGGTAGCTAGGCTCTTTCCTGNNNAGATCGGAAG |
| 47774_1 | TATGAGTGTGGAGTCGTTGCCGATCTTGGCTCATTGCATCTTCNNNAGATCGGAAG |

| | |
|---|---|
| 47774_2 | TATGAGTGTGGAGTCGTTGCCGATCTTGGCTCATTGCATCTTCNNNAGATCGGAAG |
| 33173_1 | TATGAGTGTGGAGTCGTTGCCATGCCTGTAATTCCAGCACTTTNNNAGATCGGAAG |
| 33173_2 | TATGAGTGTGGAGTCGTTGCCATGCCTGTAATTCCAGCACTTTNNNAGATCGGAAG |
| 46380_1 | TATGAGTGTGGAGTCGTTGCCCACCATGTGCAGCTTATTTTTGNNNAGATCGGAAG |
| 46380_2 | TATGAGTGTGGAGTCGTTGCCCACCATGTGCAGCTTATTTTTGNNNAGATCGGAAG |
| 11436_1 | TATGAGTGTGGAGTCGTTGCCAGAGCGGACACCTTTGTCCTNNNAGATCGGAAGAG |
| 11436_2 | TATGAGTGTGGAGTCGTTGCCAGAGCGGACACCTTTGTCCTNNNAGATCGGAAGAG |
| 34940_1 | TATGAGTGTGGAGTCGTTGCTTTTGGTAGACACGAGGTTTCACNNNAGATCGGAAG |
| 34940_2 | TATGAGTGTGGAGTCGTTGCTTTTGGTAGACACGAGGTTTCACNNNAGATCGGAAG |
| 29808_1 | TATGAGTGTGGAGTCGTTGCTGCCTTATTGATCTGCAGACAGANNNAGATCGGAAG |
| 29808_2 | TATGAGTGTGGAGTCGTTGCTGCCTTATTGATCTGCAGACAGANNNAGATCGGAAG |
| 2393_1 | TATGAGTGTGGAGTCGTTGCTTACTGGTTTCTACTTCCCAGGGNNNAGATCGGAAG |
| 2393_2 | TATGAGTGTGGAGTCGTTGCTTACTGGTTTCTACTTCCCAGGGNNNAGATCGGAAG |
| 42997_1 | TATGAGTGTGGAGTCGTTGCAGCAGACGTTCTTCTTCCCTTTCNNNAGATCGGAAGA |
| 42997_2 | TATGAGTGTGGAGTCGTTGCAGCAGACGTTCTTCTTCCCTTTCNNNAGATCGGAAGA |
| 32488_1 | TATGAGTGTGGAGTCGTTGCCAAGTAGCTGGGATTACAGGCATNNNAGATCGGAA |
| 32488_2 | TATGAGTGTGGAGTCGTTGCCAAGTAGCTGGGATTACAGGCATNNNAGATCGGAA |
| 41472_1 | TATGAGTGTGGAGTCGTTGCTCAAAAAATGGTGTTGGAATTGCANNNAGATCGGA |
| 41472_2 | TATGAGTGTGGAGTCGTTGCTCAAAAAATGGTGTTGGAATTGCANNNAGATCGGA |
| 46144_1 | TATGAGTGTGGAGTCGTTGCCAAGTAGCTGGGACTACAGGCNNNAGATCGGAAGA |
| 46144_2 | TATGAGTGTGGAGTCGTTGCCAAGTAGCTGGGACTACAGGCNNNAGATCGGAAGA |
| 35292_1 | TATGAGTGTGGAGTCGTTGCCATGATCTCAGCTCAATGCAACCNNNAGATCGGAAG |
| 35292_2 | TATGAGTGTGGAGTCGTTGCCATGATCTCAGCTCAATGCAACCNNNAGATCGGAAG |
| 33284_1 | TATGAGTGTGGAGTCGTTGCCCTCCCAGGTTCAACTGATTCTTNNNAGATCGGAAGA |
| 33284_2 | TATGAGTGTGGAGTCGTTGCCCTCCCAGGTTCAACTGATTCTTNNNAGATCGGAAGA |
| 49616_1 | TATGAGTGTGGAGTCGTTGCAATACACACACACACACACACACNNNAGATCGGAAG |
| 49616_2 | TATGAGTGTGGAGTCGTTGCAATACACACACACACACACACACNNNAGATCGGAAG |
| 36790_1 | TATGAGTGTGGAGTCGTTGCCAGTGGTACAATCTCTGCTCACTNNNAGATCGGAAG |
| 36790_2 | TATGAGTGTGGAGTCGTTGCCAGTGGTACAATCTCTGCTCACTNNNAGATCGGAAG |
| 40172_1 | TATGAGTGTGGAGTCGTTGCCCAAGGCTACCTGATAGGCTAATNNNAGATCGGAAG |
| 40172_2 | TATGAGTGTGGAGTCGTTGCCCAAGGCTACCTGATAGGCTAATNNNAGATCGGAAG |
| 30991_1 | TATGAGTGTGGAGTCGTTGCCCTCCCTCCCTTCCTTTTTCTTTNNNAGATCGGAAGAG |
| 30991_2 | TATGAGTGTGGAGTCGTTGCCCTCCCTCCCTTCCTTTTTCTTTNNNAGATCGGAAGAG |
| 32898_1 | TATGAGTGTGGAGTCGTTGCTTCACCTGGGCTAGTTTCTCATCNNNAGATCGGAAG |
| 32898_2 | TATGAGTGTGGAGTCGTTGCTTCACCTGGGCTAGTTTCTCATCNNNAGATCGGAAG |
| 53073_1 | TATGAGTGTGGAGTCGTTGCAGATCAATTTTCCAACCACAGCGNNNAGATCGGAAG |
| 53073_2 | TATGAGTGTGGAGTCGTTGCAGATCAATTTTCCAACCACAGCGNNNAGATCGGAAG |
| 48740_1 | TATGAGTGTGGAGTCGTTGCCACTGCAATCTTGAACTTCTGGGNNNAGATCGGAAG |
| 48740_2 | TATGAGTGTGGAGTCGTTGCCACTGCAATCTTGAACTTCTGGGNNNAGATCGGAAG |
| 32389_1 | TATGAGTGTGGAGTCGTTGCCCAAGTAGCTGGGACTACAGGNNNAGATCGGAAGA |
| 32389_2 | TATGAGTGTGGAGTCGTTGCCCAAGTAGCTGGGACTACAGGNNNAGATCGGAAGA |
| 38847_1 | TATGAGTGTGGAGTCGTTGCTCAACACAGAACAGAATATGCCTNNNAGATCGGAAG |
| 38847_2 | TATGAGTGTGGAGTCGTTGCTCAACACAGAACAGAATATGCCTNNNAGATCGGAAG |

| | |
|---|---|
| 51791_1 | TATGAGTGTGGAGTCGTTGCTTCTGTCCCTGGCATTTCTCCNNNAGATCGGAAGAGC |
| 51791_2 | TATGAGTGTGGAGTCGTTGCTTCTGTCCCTGGCATTTCTCCNNNAGATCGGAAGAGC |
| 46430_1 | TATGAGTGTGGAGTCGTTGCTCCCAGGTTCAAATGATTCTCGTNNNAGATCGGAAG |
| 46430_2 | TATGAGTGTGGAGTCGTTGCTCCCAGGTTCAAATGATTCTCGTNNNAGATCGGAAG |
| 48465_1 | TATGAGTGTGGAGTCGTTGCCTCCTGGGTTCAAGCGATTCTCNNNAGATCGGAAGA |
| 48465_2 | TATGAGTGTGGAGTCGTTGCCTCCTGGGTTCAAGCGATTCTCNNNAGATCGGAAGA |
| 35541_1 | TATGAGTGTGGAGTCGTTGCAATGCCCAACTTTCAGCATCTTCNNNAGATCGGAAGA |
| 35541_2 | TATGAGTGTGGAGTCGTTGCAATGCCCAACTTTCAGCATCTTCNNNAGATCGGAAGA |
| 51304_1 | TATGAGTGTGGAGTCGTTGCGAGGTGTACCCAACAGCTCATTNNNAGATCGGAAGA |
| 51304_2 | TATGAGTGTGGAGTCGTTGCGAGGTGTACCCAACAGCTCATTNNNAGATCGGAAGA |
| 34018_1 | TATGAGTGTGGAGTCGTTGCTTCTAGCAATTCTCCTGCCTCAGNNNAGATCGGAAGA |
| 34018_2 | TATGAGTGTGGAGTCGTTGCTTCTAGCAATTCTCCTGCCTCAGNNNAGATCGGAAGA |
| 47990_1 | TATGAGTGTGGAGTCGTTGCCCTCTCACATTGCTGCTAGAGATNNNAGATCGGAAG |
| 47990_2 | TATGAGTGTGGAGTCGTTGCCCTCTCACATTGCTGCTAGAGATNNNAGATCGGAAG |
| 41727_1 | TATGAGTGTGGAGTCGTTGCCTCCCAGGTTCATGCCATTCTCNNNAGATCGGAAGAG |
| 41727_2 | TATGAGTGTGGAGTCGTTGCCTCCCAGGTTCATGCCATTCTCNNNAGATCGGAAGAG |
| 50200_1 | TATGAGTGTGGAGTCGTTGCCCGGTGCATTCTGGGAAATTCNNNAGATCGGAAGAG |
| 50200_2 | TATGAGTGTGGAGTCGTTGCCCGGTGCATTCTGGGAAATTCNNNAGATCGGAAGAG |
| 34519_1 | TATGAGTGTGGAGTCGTTGCGAGATGGGGTTTTACAATGCTGCNNNAGATCGGAA |
| 34519_2 | TATGAGTGTGGAGTCGTTGCGAGATGGGGTTTTACAATGCTGCNNNAGATCGGAA |
| 39762_1 | TATGAGTGTGGAGTCGTTGCAAATGGCAAACAGGCATACGAAANNNAGATCGGAA |
| 39762_2 | TATGAGTGTGGAGTCGTTGCAAATGGCAAACAGGCATACGAAANNNAGATCGGAA |
| 30883_1 | TATGAGTGTGGAGTCGTTGCGATGAGAAAGAGCGAGGAGAGAGNNNAGATCGGA |
| 30883_2 | TATGAGTGTGGAGTCGTTGCGATGAGAAAGAGCGAGGAGAGAGNNNAGATCGGA |
| 32446_1 | TATGAGTGTGGAGTCGTTGCCATTCTCCTGCCTCAGCCTCNNNAGATCGGAAGAGCA |
| 32446_2 | TATGAGTGTGGAGTCGTTGCCATTCTCCTGCCTCAGCCTCNNNAGATCGGAAGAGCA |
| 39045_1 | TATGAGTGTGGAGTCGTTGCGGATGTGCTTAGGGGAATCTGATNNNAGATCGGAA |
| 39045_2 | TATGAGTGTGGAGTCGTTGCGGATGTGCTTAGGGGAATCTGATNNNAGATCGGAA |
| 36808_1 | TATGAGTGTGGAGTCGTTGCTGGAAGTTCCAAGCCTGAGTAAANNNAGATCGGAA |
| 36808_2 | TATGAGTGTGGAGTCGTTGCTGGAAGTTCCAAGCCTGAGTAAANNNAGATCGGAA |
| 52097_1 | TATGAGTGTGGAGTCGTTGCAGCATTCAGAGGGAACATAGTCCNNNAGATCGGAA |
| 52097_2 | TATGAGTGTGGAGTCGTTGCAGCATTCAGAGGGAACATAGTCCNNNAGATCGGAA |
| 52583_1 | TATGAGTGTGGAGTCGTTGCCTCCAAGAAGCTAGTTTTTCGGCNNNAGATCGGAAG |
| 52583_2 | TATGAGTGTGGAGTCGTTGCCTCCAAGAAGCTAGTTTTTCGGCNNNAGATCGGAAG |
| 38053_1 | TATGAGTGTGGAGTCGTTGCCAACAACAACAACACACACACACNNNAGATCGGAAG |
| 38053_2 | TATGAGTGTGGAGTCGTTGCCAACAACAACAACACACACACACNNNAGATCGGAAG |
| 35385_1 | TATGAGTGTGGAGTCGTTGCAAGTGTTGGGATTACAGGCATGANNNAGATCGGAA |
| 35385_2 | TATGAGTGTGGAGTCGTTGCAAGTGTTGGGATTACAGGCATGANNNAGATCGGAA |
| 12813_1 | TATGAGTGTGGAGTCGTTGCCAATCCTTTTGAACATACTCATANNNAGATCGGAAGA |
| 12813_2 | TATGAGTGTGGAGTCGTTGCCAATCCTTTTGAACATACTCATANNNAGATCGGAAGA |
| 45458_1 | TATGAGTGTGGAGTCGTTGCGGCCCTCCAATAGTGAATGAGATNNNAGATCGGAA |
| 45458_2 | TATGAGTGTGGAGTCGTTGCGGCCCTCCAATAGTGAATGAGATNNNAGATCGGAA |
| 37101_1 | TATGAGTGTGGAGTCGTTGCTGAAGGTACAAAACTCTCACTGGTNNNAGATCGGAA |

| | |
|---|---|
| 37101_2 | TATGAGTGTGGAGTCGTTGCTGAAGGTACAAAACTCTCACTGGTNNNAGATCGGAA |
| 50398_1 | TATGAGTGTGGAGTCGTTGCCAGCCTGTGAAGGGAAGGAGNNNAGATCGGAAGA |
| 50398_2 | TATGAGTGTGGAGTCGTTGCCAGCCTGTGAAGGGAAGGAGNNNAGATCGGAAGA |
| 35803_1 | TATGAGTGTGGAGTCGTTGCCAGAAGGCCTAATGCACAAGAACNNNAGATCGGAA |
| 35803_2 | TATGAGTGTGGAGTCGTTGCCAGAAGGCCTAATGCACAAGAACNNNAGATCGGAA |
| 32143_1 | TATGAGTGTGGAGTCGTTGCGGGTTCAAACTATTCTCCTGCCTNNNAGATCGGAAG |
| 32143_2 | TATGAGTGTGGAGTCGTTGCGGGTTCAAACTATTCTCCTGCCTNNNAGATCGGAAG |
| 40531_1 | TATGAGTGTGGAGTCGTTGCCAATGTCTTGCTTTGTCACCCAGNNNAGATCGGAAG |
| 40531_2 | TATGAGTGTGGAGTCGTTGCCAATGTCTTGCTTTGTCACCCAGNNNAGATCGGAAG |
| 53081_1 | TATGAGTGTGGAGTCGTTGCCACACACACACACACACACATATNNNAGATCGGAAG |
| 53081_2 | TATGAGTGTGGAGTCGTTGCCACACACACACACACACACATATNNNAGATCGGAAG |
| 40748_1 | TATGAGTGTGGAGTCGTTGCGGCTTCTCCAGTGTTCTATGAGANNNAGATCGGAAG |
| 40748_2 | TATGAGTGTGGAGTCGTTGCGGCTTCTCCAGTGTTCTATGAGANNNAGATCGGAAG |
| 13777_1 | TATGAGTGTGGAGTCGTTGCATAATATAAACCATGTCAGCAACNNNAGATCGGAAG |
| 13777_2 | TATGAGTGTGGAGTCGTTGCATAATATAAACCATGTCAGCAACNNNAGATCGGAAG |
| 41049_1 | TATGAGTGTGGAGTCGTTGCGAACTCACAGAAGCAGAGGGTAGNNNAGATCGGAA |
| 41049_2 | TATGAGTGTGGAGTCGTTGCGAACTCACAGAAGCAGAGGGTAGNNNAGATCGGAA |
| 51662_1 | TATGAGTGTGGAGTCGTTGCCAGGGAACTCTCCTCCTTCAAAANNNAGATCGGAAG |
| 51662_2 | TATGAGTGTGGAGTCGTTGCCAGGGAACTCTCCTCCTTCAAAANNNAGATCGGAAG |
| 32257_1 | TATGAGTGTGGAGTCGTTGCCTACTTAGGCACAAGATGGCTTGNNNAGATCGGAAG |
| 32257_2 | TATGAGTGTGGAGTCGTTGCCTACTTAGGCACAAGATGGCTTGNNNAGATCGGAAG |
| 31752_1 | TATGAGTGTGGAGTCGTTGCTGGAAGTTCCCATTGTGTACACTNNNAGATCGGAAG |
| 31752_2 | TATGAGTGTGGAGTCGTTGCTGGAAGTTCCCATTGTGTACACTNNNAGATCGGAAG |
| 49668_1 | TATGAGTGTGGAGTCGTTGCCACAATCTTAGCTCACTGCAACCNNNAGATCGGAAG |
| 49668_2 | TATGAGTGTGGAGTCGTTGCCACAATCTTAGCTCACTGCAACCNNNAGATCGGAAG |
| 50466_1 | TATGAGTGTGGAGTCGTTGCCTCACACTCACTCACACTTCACTNNNAGATCGGAAGA |
| 50466_2 | TATGAGTGTGGAGTCGTTGCCTCACACTCACTCACACTTCACTNNNAGATCGGAAGA |
| 41282_1 | TATGAGTGTGGAGTCGTTGCGCTGTGTGATAGTTCCAGGAATTNNNAGATCGGAA |
| 41282_2 | TATGAGTGTGGAGTCGTTGCGCTGTGTGATAGTTCCAGGAATTNNNAGATCGGAA |
| 35137_1 | TATGAGTGTGGAGTCGTTGCCCAAGTAGCTGGGACTTCAGGNNNAGATCGGAAGA |
| 35137_2 | TATGAGTGTGGAGTCGTTGCCCAAGTAGCTGGGACTTCAGGNNNAGATCGGAAGA |
| 31269_1 | TATGAGTGTGGAGTCGTTGCACTTGTATGTTTCAGTGCCTTGCNNNAGATCGGAAG |
| 31269_2 | TATGAGTGTGGAGTCGTTGCACTTGTATGTTTCAGTGCCTTGCNNNAGATCGGAAG |
| 52235_1 | TATGAGTGTGGAGTCGTTGCCCTCAGCTGCAAGGAATGTAAAGNNNAGATCGGAA |
| 52235_2 | TATGAGTGTGGAGTCGTTGCCCTCAGCTGCAAGGAATGTAAAGNNNAGATCGGAA |
| 29034_1 | TATGAGTGTGGAGTCGTTGCTGTGTGTGTATATATAGTGTGCGTNNNAGATCGGA |
| 29034_2 | TATGAGTGTGGAGTCGTTGCTGTGTGTGTATATATAGTGTGCGTNNNAGATCGGA |
| 48288_1 | TATGAGTGTGGAGTCGTTGCAGCATGCAGTCATCCCTATTTGTNNNAGATCGGAAG |
| 48288_2 | TATGAGTGTGGAGTCGTTGCAGCATGCAGTCATCCCTATTTGTNNNAGATCGGAAG |
| 48152_1 | TATGAGTGTGGAGTCGTTGCCTCCTTCTGTTCAGCTCTCACTTNNNAGATCGGAAGA |
| 48152_2 | TATGAGTGTGGAGTCGTTGCCTCCTTCTGTTCAGCTCTCACTTNNNAGATCGGAAGA |
| 49214_1 | TATGAGTGTGGAGTCGTTGCAAAGCAGCAGAATTGAGAACCACNNNAGATCGGAA |
| 49214_2 | TATGAGTGTGGAGTCGTTGCAAAGCAGCAGAATTGAGAACCACNNNAGATCGGAA |

| | |
|---|---|
| 52919_1 | TATGAGTGTGGAGTCGTTGCGGTGATCACTGTGGACACTAAGGNNNAGATCGGAA |
| 52919_2 | TATGAGTGTGGAGTCGTTGCGGTGATCACTGTGGACACTAAGGNNNAGATCGGAA |
| 34736_1 | TATGAGTGTGGAGTCGTTGCTTTTTTGAGACGGAGTCTTGCTCNNNAGATCGGAAG |
| 34736_2 | TATGAGTGTGGAGTCGTTGCTTTTTTGAGACGGAGTCTTGCTCNNNAGATCGGAAG |
| 3003_1 | TATGAGTGTGGAGTCGTTGCCCTCTCCCCCTATGCCCTCTTNNNAGATCGGAAGAGC |
| 3003_2 | TATGAGTGTGGAGTCGTTGCCCTCTCCCCCTATGCCCTCTTNNNAGATCGGAAGAGC |
| 44353_1 | TATGAGTGTGGAGTCGTTGCTGGGTCAGTTCTATGCTTGTACCNNNAGATCGGAAG |
| 44353_2 | TATGAGTGTGGAGTCGTTGCTGGGTCAGTTCTATGCTTGTACCNNNAGATCGGAAG |
| 32821_1 | TATGAGTGTGGAGTCGTTGCCCTCCCAAGTAGCTGGAATTACANNNAGATCGGAAG |
| 32821_2 | TATGAGTGTGGAGTCGTTGCCCTCCCAAGTAGCTGGAATTACANNNAGATCGGAAG |
| 37715_1 | TATGAGTGTGGAGTCGTTGCGTGGTGGTTCACACATGTAGTTCNNNAGATCGGAA |
| 37715_2 | TATGAGTGTGGAGTCGTTGCGTGGTGGTTCACACATGTAGTTCNNNAGATCGGAA |
| 32540_1 | TATGAGTGTGGAGTCGTTGCAGAAAGAATGGCAAAGGGTCTGANNNAGATCGGA |
| 32540_2 | TATGAGTGTGGAGTCGTTGCAGAAAGAATGGCAAAGGGTCTGANNNAGATCGGA |
| 10481_1 | TATGAGTGTGGAGTCGTTGCAAACCTTCATGGGTACTGGGGAANNNAGATCGGAA |
| 10481_2 | TATGAGTGTGGAGTCGTTGCAAACCTTCATGGGTACTGGGGAANNNAGATCGGAA |
| 38532_1 | TATGAGTGTGGAGTCGTTGCGTACCACTCCTTTCTCCCAAGAANNNAGATCGGAAGA |
| 38532_2 | TATGAGTGTGGAGTCGTTGCGTACCACTCCTTTCTCCCAAGAANNNAGATCGGAAGA |
| 36986_1 | TATGAGTGTGGAGTCGTTGCGGTAAAGTTCATGCAGTGGGATANNNAGATCGGAA |
| 36986_2 | TATGAGTGTGGAGTCGTTGCGGTAAAGTTCATGCAGTGGGATANNNAGATCGGAA |
| 48197_1 | TATGAGTGTGGAGTCGTTGCCAAGTATCTGGGACTACAGGCACNNNAGATCGGAA |
| 48197_2 | TATGAGTGTGGAGTCGTTGCCAAGTATCTGGGACTACAGGCACNNNAGATCGGAA |
| 41719_1 | TATGAGTGTGGAGTCGTTGCCACTCTGTTTCCCAAGCTAGAGTNNNAGATCGGAAG |
| 41719_2 | TATGAGTGTGGAGTCGTTGCCACTCTGTTTCCCAAGCTAGAGTNNNAGATCGGAAG |
| 36611_1 | TATGAGTGTGGAGTCGTTGCAAGGGGAATGCTCAAGATCTCAGNNNAGATCGGAA |
| 36611_2 | TATGAGTGTGGAGTCGTTGCAAGGGGAATGCTCAAGATCTCAGNNNAGATCGGAA |
| 35805_1 | TATGAGTGTGGAGTCGTTGCCAGGAATTCATCCCACTGAAACANNNAGATCGGAAG |
| 35805_2 | TATGAGTGTGGAGTCGTTGCCAGGAATTCATCCCACTGAAACANNNAGATCGGAAG |
| 47679_1 | TATGAGTGTGGAGTCGTTGCGGTTCATCATGGAGGCAGGTAANNNAGATCGGAAG |
| 47679_2 | TATGAGTGTGGAGTCGTTGCGGTTCATCATGGAGGCAGGTAANNNAGATCGGAAG |
| 51379_1 | TATGAGTGTGGAGTCGTTGCAGATGATGAGTTAGTGGGTGCAGNNNAGATCGGA |
| 51379_2 | TATGAGTGTGGAGTCGTTGCAGATGATGAGTTAGTGGGTGCAGNNNAGATCGGA |
| 30598_1 | TATGAGTGTGGAGTCGTTGCTTGGGTTGGTCAAATGTCTTGTGTNNNAGATCGGA |
| 30598_2 | TATGAGTGTGGAGTCGTTGCTTGGGTTGGTCAAATGTCTTGTGTNNNAGATCGGA |
| 33858_1 | TATGAGTGTGGAGTCGTTGCCATTATCTTGGGGAGTGCCTTCTNNNAGATCGGAAG |
| 33858_2 | TATGAGTGTGGAGTCGTTGCCATTATCTTGGGGAGTGCCTTCTNNNAGATCGGAAG |
| 51418_1 | TATGAGTGTGGAGTCGTTGCTGTGTCTCTTCCAGTCTCACAAGNNNAGATCGGAAG |
| 51418_2 | TATGAGTGTGGAGTCGTTGCTGTGTCTCTTCCAGTCTCACAAGNNNAGATCGGAAG |
| 29876_1 | TATGAGTGTGGAGTCGTTGCGCCTTCAGATAAAATTCCCGAAGCNNNAGATCGGAA |
| 29876_2 | TATGAGTGTGGAGTCGTTGCGCCTTCAGATAAAATTCCCGAAGCNNNAGATCGGAA |
| 36347_1 | TATGAGTGTGGAGTCGTTGCTCATATGCCAGATGTAGAGATGGTNNNAGATCGGA |
| 36347_2 | TATGAGTGTGGAGTCGTTGCTCATATGCCAGATGTAGAGATGGTNNNAGATCGGA |
| 40238_1 | TATGAGTGTGGAGTCGTTGCGGGTTTTTTGTGTCTCTGTCTCCNNNAGATCGGAAG |

| | |
|---|---|
| 40238_2 | TATGAGTGTGGAGTCGTTGCGGGTTTTTTGTGTCTCTGTCTCCNNNAGATCGGAAG |
| 33595_1 | TATGAGTGTGGAGTCGTTGCGTTCAAGCAATTCTCTGCCTCAGNNNAGATCGGAAG |
| 33595_2 | TATGAGTGTGGAGTCGTTGCGTTCAAGCAATTCTCTGCCTCAGNNNAGATCGGAAG |
| 30369_1 | TATGAGTGTGGAGTCGTTGCACATGCTCCTGACTCTATCTCCCNNNAGATCGGAAGA |
| 30369_2 | TATGAGTGTGGAGTCGTTGCACATGCTCCTGACTCTATCTCCCNNNAGATCGGAAGA |
| 47635_1 | TATGAGTGTGGAGTCGTTGCAGAGACAAGAAATAGGGGGTTGGNNNAGATCGGA |
| 47635_2 | TATGAGTGTGGAGTCGTTGCAGAGACAAGAAATAGGGGGTTGGNNNAGATCGGA |
| 47584_1 | TATGAGTGTGGAGTCGTTGCCACAGACCCCCTTCCTGTTCNNNAGATCGGAAGAGC |
| 47584_2 | TATGAGTGTGGAGTCGTTGCCACAGACCCCCTTCCTGTTCNNNAGATCGGAAGAGC |
| 47859_1 | TATGAGTGTGGAGTCGTTGCACGCCCGGCTAATTTTGTATTTTNNNAGATCGGAAG |
| 47859_2 | TATGAGTGTGGAGTCGTTGCACGCCCGGCTAATTTTGTATTTTNNNAGATCGGAAG |
| 34594_1 | TATGAGTGTGGAGTCGTTGCGTGATGCACACCTGTAATTCCAGNNNAGATCGGAAG |
| 34594_2 | TATGAGTGTGGAGTCGTTGCGTGATGCACACCTGTAATTCCAGNNNAGATCGGAAG |
| 35457_1 | TATGAGTGTGGAGTCGTTGCAACCTCTGACTCCCAAGTTGAAGNNNAGATCGGAAG |
| 35457_2 | TATGAGTGTGGAGTCGTTGCAACCTCTGACTCCCAAGTTGAAGNNNAGATCGGAAG |
| 35631_1 | TATGAGTGTGGAGTCGTTGCAGAAGGTCAGCAGATAAACACGTNNNAGATCGGAA |
| 35631_2 | TATGAGTGTGGAGTCGTTGCAGAAGGTCAGCAGATAAACACGTNNNAGATCGGAA |
| 40797_1 | TATGAGTGTGGAGTCGTTGCAGGAGGTAGTCCAGAAGAAGGAANNNAGATCGGA |
| 40797_2 | TATGAGTGTGGAGTCGTTGCAGGAGGTAGTCCAGAAGAAGGAANNNAGATCGGA |
| 47459_1 | TATGAGTGTGGAGTCGTTGCATTGCATGACACTGAGTTTTGGGNNNAGATCGGAA |
| 47459_2 | TATGAGTGTGGAGTCGTTGCATTGCATGACACTGAGTTTTGGGNNNAGATCGGAA |
| 36926_1 | TATGAGTGTGGAGTCGTTGCAGAAAGAAATGCTTGGGACCAGANNNAGATCGGAA |
| 36926_2 | TATGAGTGTGGAGTCGTTGCAGAAAGAAATGCTTGGGACCAGANNNAGATCGGAA |
| 35145_1 | TATGAGTGTGGAGTCGTTGCGACGGAGCCTCACTCTGTTTCNNNAGATCGGAAGAG |
| 35145_2 | TATGAGTGTGGAGTCGTTGCGACGGAGCCTCACTCTGTTTCNNNAGATCGGAAGAG |
| 36421_1 | TATGAGTGTGGAGTCGTTGCGTGTCTGCATTCTTTGACTCAGCNNNAGATCGGAAG |
| 36421_2 | TATGAGTGTGGAGTCGTTGCGTGTCTGCATTCTTTGACTCAGCNNNAGATCGGAAG |
| 13597_1 | TATGAGTGTGGAGTCGTTGCAAGAAAAAAAAGAGGTTATCACANNNAGATCGGAA |
| 13597_2 | TATGAGTGTGGAGTCGTTGCAAGAAAAAAAAGAGGTTATCACANNNAGATCGGAA |
| 52781_1 | TATGAGTGTGGAGTCGTTGCACCACCTGATTTATAGGCCAAGGNNNAGATCGGAAG |
| 52781_2 | TATGAGTGTGGAGTCGTTGCACCACCTGATTTATAGGCCAAGGNNNAGATCGGAAG |
| 36107_1 | TATGAGTGTGGAGTCGTTGCCATCCACAAGAACAGCCAATTGTNNNAGATCGGAAG |
| 36107_2 | TATGAGTGTGGAGTCGTTGCCATCCACAAGAACAGCCAATTGTNNNAGATCGGAAG |
| 38466_1 | TATGAGTGTGGAGTCGTTGCTACCAGTTCCTCCTTGTACCTCTNNNAGATCGGAAGA |
| 38466_2 | TATGAGTGTGGAGTCGTTGCTACCAGTTCCTCCTTGTACCTCTNNNAGATCGGAAGA |
| 34009_1 | TATGAGTGTGGAGTCGTTGCACAATCTCGGCTCACTGCAANNNAGATCGGAAGAGC |
| 34009_2 | TATGAGTGTGGAGTCGTTGCACAATCTCGGCTCACTGCAANNNAGATCGGAAGAGC |
| 29537_1 | TATGAGTGTGGAGTCGTTGCGCAAGTATGTGTGTGTGTGTGTGNNNAGATCGGAA |
| 29537_2 | TATGAGTGTGGAGTCGTTGCGCAAGTATGTGTGTGTGTGTGTGNNNAGATCGGAA |
| 29139_1 | TATGAGTGTGGAGTCGTTGCATAATGAAGTGGGCTAGGTGAGCNNNAGATCGGAA |
| 29139_2 | TATGAGTGTGGAGTCGTTGCATAATGAAGTGGGCTAGGTGAGCNNNAGATCGGAA |
| 33081_1 | TATGAGTGTGGAGTCGTTGCGTGTAGTGAGTGGTGTGATCTCANNNAGATCGGAA |
| 33081_2 | TATGAGTGTGGAGTCGTTGCGTGTAGTGAGTGGTGTGATCTCANNNAGATCGGAA |

| | |
|---|---|
| 42962_1 | TATGAGTGTGGAGTCGTTGCAGACTGTAGGAAACGTGAACACANNNAGATCGGAA |
| 42962_2 | TATGAGTGTGGAGTCGTTGCAGACTGTAGGAAACGTGAACACANNNAGATCGGAA |
| 45355_1 | TATGAGTGTGGAGTCGTTGCAGATTAGAAGTGCAAGCCACTCANNNAGATCGGAA |
| 45355_2 | TATGAGTGTGGAGTCGTTGCAGATTAGAAGTGCAAGCCACTCANNNAGATCGGAA |
| 5400_1 | TATGAGTGTGGAGTCGTTGCAGGTGGGATACACTTCTCCTTCCNNNAGATCGGAAG |
| 5400_2 | TATGAGTGTGGAGTCGTTGCAGGTGGGATACACTTCTCCTTCCNNNAGATCGGAAG |
| 38975_1 | TATGAGTGTGGAGTCGTTGCACTGCTCCTTTGAATCAATGCAANNNAGATCGGAAG |
| 38975_2 | TATGAGTGTGGAGTCGTTGCACTGCTCCTTTGAATCAATGCAANNNAGATCGGAAG |
| 36233_1 | TATGAGTGTGGAGTCGTTGCCCTGACTTCAAGAGCAGGTACTCNNNAGATCGGAAG |
| 36233_2 | TATGAGTGTGGAGTCGTTGCCCTGACTTCAAGAGCAGGTACTCNNNAGATCGGAAG |
| 36654_1 | TATGAGTGTGGAGTCGTTGCCTATACTCCAGCCTAGGCAACAGNNNAGATCGGAAG |
| 36654_2 | TATGAGTGTGGAGTCGTTGCCTATACTCCAGCCTAGGCAACAGNNNAGATCGGAAG |
| 41534_1 | TATGAGTGTGGAGTCGTTGCAGGGTCTTACCATATTGCCCATGNNNAGATCGGAAG |
| 41534_2 | TATGAGTGTGGAGTCGTTGCAGGGTCTTACCATATTGCCCATGNNNAGATCGGAAG |
| 51250_1 | TATGAGTGTGGAGTCGTTGCAGCTCTCCAATAGTCAGGTGTTGNNNAGATCGGAAG |
| 51250_2 | TATGAGTGTGGAGTCGTTGCAGCTCTCCAATAGTCAGGTGTTGNNNAGATCGGAAG |
| 44824_1 | TATGAGTGTGGAGTCGTTGCTCGTTCCTTCAGTGTACTTCTGGNNNAGATCGGAAG |
| 44824_2 | TATGAGTGTGGAGTCGTTGCTCGTTCCTTCAGTGTACTTCTGGNNNAGATCGGAAG |
| 35225_1 | TATGAGTGTGGAGTCGTTGCTCCAGGCTAAAGTGATTCACCTGNNNAGATCGGAAG |
| 35225_2 | TATGAGTGTGGAGTCGTTGCTCCAGGCTAAAGTGATTCACCTGNNNAGATCGGAAG |
| 30889_1 | TATGAGTGTGGAGTCGTTGCGAGCACATCAGCTTCTAGACAGTNNNAGATCGGAAG |
| 30889_2 | TATGAGTGTGGAGTCGTTGCGAGCACATCAGCTTCTAGACAGTNNNAGATCGGAAG |
| 38468_1 | TATGAGTGTGGAGTCGTTGCTTCATCACCCCTCCATTTATCCGNNNAGATCGGAAGA |
| 38468_2 | TATGAGTGTGGAGTCGTTGCTTCATCACCCCTCCATTTATCCGNNNAGATCGGAAGA |
| 47493_1 | TATGAGTGTGGAGTCGTTGCATCTCAGTTCACTGCAACTTCCANNNAGATCGGAAG |
| 47493_2 | TATGAGTGTGGAGTCGTTGCATCTCAGTTCACTGCAACTTCCANNNAGATCGGAAG |
| 39997_1 | TATGAGTGTGGAGTCGTTGCAAAAAGGAAGGAAGGAAGGGAGGNNNAGATCGG |
| 39997_2 | TATGAGTGTGGAGTCGTTGCAAAAAGGAAGGAAGGAAGGGAGGNNNAGATCGG |
| 30014_1 | TATGAGTGTGGAGTCGTTGCATCTGTCTTTGGAAATGAGCCACNNNAGATCGGAAG |
| 30014_2 | TATGAGTGTGGAGTCGTTGCATCTGTCTTTGGAAATGAGCCACNNNAGATCGGAAG |
| 41321_1 | TATGAGTGTGGAGTCGTTGCAGATACTGACTAACCGACCTCCTNNNAGATCGGAAG |
| 41321_2 | TATGAGTGTGGAGTCGTTGCAGATACTGACTAACCGACCTCCTNNNAGATCGGAAG |
| 30134_1 | TATGAGTGTGGAGTCGTTGCAGCAGCAATATGCAGAAAACAGTNNNAGATCGGAA |
| 30134_2 | TATGAGTGTGGAGTCGTTGCAGCAGCAATATGCAGAAAACAGTNNNAGATCGGAA |
| 30288_1 | TATGAGTGTGGAGTCGTTGCTCCTCTTCATGTGTTTGGGGATTNNNAGATCGGAAG |
| 30288_2 | TATGAGTGTGGAGTCGTTGCTCCTCTTCATGTGTTTGGGGATTNNNAGATCGGAAG |
| 37352_1 | TATGAGTGTGGAGTCGTTGCACAGTGGCTCACACCTATAATCCNNNAGATCGGAAG |
| 37352_2 | TATGAGTGTGGAGTCGTTGCACAGTGGCTCACACCTATAATCCNNNAGATCGGAAG |
| 53063_1 | TATGAGTGTGGAGTCGTTGCCATCCACAAGAACAGCCAATTGTNNNAGATCGGAAG |
| 53063_2 | TATGAGTGTGGAGTCGTTGCCATCCACAAGAACAGCCAATTGTNNNAGATCGGAAG |
| 49065_1 | TATGAGTGTGGAGTCGTTGCCCTCCCAGGTTCAAACAGTTCTCNNNAGATCGGAAG |
| 49065_2 | TATGAGTGTGGAGTCGTTGCCCTCCCAGGTTCAAACAGTTCTCNNNAGATCGGAAG |
| 52345_1 | TATGAGTGTGGAGTCGTTGCGGGACCACAGTAACTCAGGAATTNNNAGATCGGAA |

| | |
|---|---|
| 52345_2 | TATGAGTGTGGAGTCGTTGCGGGACCACAGTAACTCAGGAATTNNNAGATCGGAA |
| 50358_1 | TATGAGTGTGGAGTCGTTGCCCAGATCAGACACATAGGAGCATNNNAGATCGGAAG |
| 50358_2 | TATGAGTGTGGAGTCGTTGCCCAGATCAGACACATAGGAGCATNNNAGATCGGAAG |
| 45387_1 | TATGAGTGTGGAGTCGTTGCTAAGTTCCTGTTGCTGTAGCTGTNNNAGATCGGAAG |
| 45387_2 | TATGAGTGTGGAGTCGTTGCTAAGTTCCTGTTGCTGTAGCTGTNNNAGATCGGAAG |
| 38046_1 | TATGAGTGTGGAGTCGTTGCCAGCCAAGGGTGAACACTAAATGNNNAGATCGGAA |
| 38046_2 | TATGAGTGTGGAGTCGTTGCCAGCCAAGGGTGAACACTAAATGNNNAGATCGGAA |
| 33331_1 | TATGAGTGTGGAGTCGTTGCCCAGACCTCAAATAATCCACCCANNNAGATCGGAAG |
| 33331_2 | TATGAGTGTGGAGTCGTTGCCCAGACCTCAAATAATCCACCCANNNAGATCGGAAG |
| 40455_1 | TATGAGTGTGGAGTCGTTGCATGCCTGTAATCTCAGCTACTCGNNNAGATCGGAAG |
| 40455_2 | TATGAGTGTGGAGTCGTTGCATGCCTGTAATCTCAGCTACTCGNNNAGATCGGAAG |
| 13781_1 | TATGAGTGTGGAGTCGTTGCTACTCATCTTTTCCCAGGTATNNNAGATCGGAAGAG |
| 13781_2 | TATGAGTGTGGAGTCGTTGCTACTCATCTTTTCCCAGGTATNNNAGATCGGAAGAG |
| 34282_1 | TATGAGTGTGGAGTCGTTGCAACTAAGGCCCTGCGAGAAATCNNNAGATCGGAAG |
| 34282_2 | TATGAGTGTGGAGTCGTTGCAACTAAGGCCCTGCGAGAAATCNNNAGATCGGAAG |
| 44302_1 | TATGAGTGTGGAGTCGTTGCATCCAGGAGTGGAAAGTTCTCACNNNAGATCGGAA |
| 44302_2 | TATGAGTGTGGAGTCGTTGCATCCAGGAGTGGAAAGTTCTCACNNNAGATCGGAA |
| 32936_1 | TATGAGTGTGGAGTCGTTGCCTTATAGCCAGCAGGGGGATAGNNNAGATCGGAAG |
| 32936_2 | TATGAGTGTGGAGTCGTTGCCTTATAGCCAGCAGGGGGATAGNNNAGATCGGAAG |
| 41827_1 | TATGAGTGTGGAGTCGTTGCAAATGGAGTCTCACTCTGTCACCNNNAGATCGGAAG |
| 41827_2 | TATGAGTGTGGAGTCGTTGCAAATGGAGTCTCACTCTGTCACCNNNAGATCGGAAG |
| 36831_1 | TATGAGTGTGGAGTCGTTGCTCAAATACTGAGCTCCTTGAGGGNNNAGATCGGAAG |
| 36831_2 | TATGAGTGTGGAGTCGTTGCTCAAATACTGAGCTCCTTGAGGGNNNAGATCGGAAG |
| 48199_1 | TATGAGTGTGGAGTCGTTGCACATTGAAATGGAGGGAGAAGCTNNNAGATCGGAA |
| 48199_2 | TATGAGTGTGGAGTCGTTGCACATTGAAATGGAGGGAGAAGCTNNNAGATCGGAA |
| 34722_1 | TATGAGTGTGGAGTCGTTGCCAATGACACGATCTTGGCTCTCTNNNAGATCGGAAG |
| 34722_2 | TATGAGTGTGGAGTCGTTGCCAATGACACGATCTTGGCTCTCTNNNAGATCGGAAG |
| 47496_1 | TATGAGTGTGGAGTCGTTGCGGGGAGTTTTTGTAGGTGACTCTNNNAGATCGGAA |
| 47496_2 | TATGAGTGTGGAGTCGTTGCGGGGAGTTTTTGTAGGTGACTCTNNNAGATCGGAA |
| 45617_1 | TATGAGTGTGGAGTCGTTGCCCCTAGTAGCTGGGACTACAGACNNNAGATCGGAAG |
| 45617_2 | TATGAGTGTGGAGTCGTTGCCCCTAGTAGCTGGGACTACAGACNNNAGATCGGAAG |
| 45240_1 | TATGAGTGTGGAGTCGTTGCGAGCCCCTGGTAAAAATTACTGCNNNAGATCGGAAG |
| 45240_2 | TATGAGTGTGGAGTCGTTGCGAGCCCCTGGTAAAAATTACTGCNNNAGATCGGAAG |
| 41938_1 | TATGAGTGTGGAGTCGTTGCTGCCCAGACTAGAGTACAGTAGTNNNAGATCGGAAG |
| 41938_2 | TATGAGTGTGGAGTCGTTGCTGCCCAGACTAGAGTACAGTAGTNNNAGATCGGAAG |
| 7879_1 | TATGAGTGTGGAGTCGTTGCAACAAAGGAAACACAGAAGACGGNNNAGATCGGAA |
| 7879_2 | TATGAGTGTGGAGTCGTTGCAACAAAGGAAACACAGAAGACGGNNNAGATCGGAA |
| 48648_1 | TATGAGTGTGGAGTCGTTGCCAATCTCAGCTCACTGCAACATCNNNAGATCGGAAG |
| 48648_2 | TATGAGTGTGGAGTCGTTGCCAATCTCAGCTCACTGCAACATCNNNAGATCGGAAG |
| 49754_1 | TATGAGTGTGGAGTCGTTGCTTCCTAGAGCAGCAGGTTTGATTNNNAGATCGGAAG |
| 49754_2 | TATGAGTGTGGAGTCGTTGCTTCCTAGAGCAGCAGGTTTGATTNNNAGATCGGAAG |
| 40679_1 | TATGAGTGTGGAGTCGTTGCTAGCTTGATGGGAATAGCGTTGANNNAGATCGGAA |
| 40679_2 | TATGAGTGTGGAGTCGTTGCTAGCTTGATGGGAATAGCGTTGANNNAGATCGGAA |

| | |
|---|---|
| 37655_1 | TATGAGTGTGGAGTCGTTGCTACCTTAGGTCAGGAGTTCGAGANNNAGATCGGAA |
| 37655_2 | TATGAGTGTGGAGTCGTTGCTACCTTAGGTCAGGAGTTCGAGANNNAGATCGGAA |
| 13752_1 | TATGAGTGTGGAGTCGTTGCGGTCTAAATGATGAAAGTGAAATNNNAGATCGGAA |
| 13752_2 | TATGAGTGTGGAGTCGTTGCGGTCTAAATGATGAAAGTGAAATNNNAGATCGGAA |
| 33025_1 | TATGAGTGTGGAGTCGTTGCTTCACACCATTCTCCTGCCTCNNNAGATCGGAAGAGC |
| 33025_2 | TATGAGTGTGGAGTCGTTGCTTCACACCATTCTCCTGCCTCNNNAGATCGGAAGAGC |
| 43684_1 | TATGAGTGTGGAGTCGTTGCCCTCAGGGCATATTCTCACACCNNNAGATCGGAAGA |
| 43684_2 | TATGAGTGTGGAGTCGTTGCCCTCAGGGCATATTCTCACACCNNNAGATCGGAAGA |
| 33146_1 | TATGAGTGTGGAGTCGTTGCAACAACATGGATGGAACTGGAGANNNAGATCGGAA |
| 33146_2 | TATGAGTGTGGAGTCGTTGCAACAACATGGATGGAACTGGAGANNNAGATCGGAA |
| 40295_1 | TATGAGTGTGGAGTCGTTGCAAACTCTGACTCCCAGGTTCAAGNNNAGATCGGAAG |
| 40295_2 | TATGAGTGTGGAGTCGTTGCAAACTCTGACTCCCAGGTTCAAGNNNAGATCGGAAG |
| 32232_1 | TATGAGTGTGGAGTCGTTGCCCCTCTGCACAGTAATGAGTGAANNNAGATCGGAAG |
| 32232_2 | TATGAGTGTGGAGTCGTTGCCCCTCTGCACAGTAATGAGTGAANNNAGATCGGAAG |
| 51098_1 | TATGAGTGTGGAGTCGTTGCTGTACAGAACCAGGAGGAATTGGNNNAGATCGGAA |
| 51098_2 | TATGAGTGTGGAGTCGTTGCTGTACAGAACCAGGAGGAATTGGNNNAGATCGGAA |
| 37415_1 | TATGAGTGTGGAGTCGTTGCCCAGGAGGATCAAACTTCAGTGANNNAGATCGGAA |
| 37415_2 | TATGAGTGTGGAGTCGTTGCCCAGGAGGATCAAACTTCAGTGANNNAGATCGGAA |
| 38664_1 | TATGAGTGTGGAGTCGTTGCTATTGGGTTTGTGGCTTTTTGCANNNAGATCGGAAG |
| 38664_2 | TATGAGTGTGGAGTCGTTGCTATTGGGTTTGTGGCTTTTTGCANNNAGATCGGAAG |
| 8757_1 | TATGAGTGTGGAGTCGTTGCTTGGCTTCAGTAAGGTGGCATTTNNNAGATCGGAAG |
| 8757_2 | TATGAGTGTGGAGTCGTTGCTTGGCTTCAGTAAGGTGGCATTTNNNAGATCGGAAG |
| 47867_1 | TATGAGTGTGGAGTCGTTGCCTCCCTGGTTCACACCATTCTCNNNAGATCGGAAGAG |
| 47867_2 | TATGAGTGTGGAGTCGTTGCCTCCCTGGTTCACACCATTCTCNNNAGATCGGAAGAG |
| 43533_1 | TATGAGTGTGGAGTCGTTGCTTTCTGTGCCTGGCTTATTTCACNNNAGATCGGAAGA |
| 43533_2 | TATGAGTGTGGAGTCGTTGCTTTCTGTGCCTGGCTTATTTCACNNNAGATCGGAAGA |
| 32238_1 | TATGAGTGTGGAGTCGTTGCGAAGGACAGAAGGACCTGGGAGNNNAGATCGGAA |
| 32238_2 | TATGAGTGTGGAGTCGTTGCGAAGGACAGAAGGACCTGGGAGNNNAGATCGGAA |
| 47887_1 | TATGAGTGTGGAGTCGTTGCGGTTCAAACGATTCTCCTGCATCNNNAGATCGGAAG |
| 47887_2 | TATGAGTGTGGAGTCGTTGCGGTTCAAACGATTCTCCTGCATCNNNAGATCGGAAG |
| 38753_1 | TATGAGTGTGGAGTCGTTGCGTAGTGCAATGGTGGAATCTTGGNNNAGATCGGAA |
| 38753_2 | TATGAGTGTGGAGTCGTTGCGTAGTGCAATGGTGGAATCTTGGNNNAGATCGGAA |
| 49646_1 | TATGAGTGTGGAGTCGTTGCCAATGGCACAATCTCTGCTTACTNNNAGATCGGAAG |
| 49646_2 | TATGAGTGTGGAGTCGTTGCCAATGGCACAATCTCTGCTTACTNNNAGATCGGAAG |
| 29548_1 | TATGAGTGTGGAGTCGTTGCCCGATTGTGAACATGGACTACGTNNNAGATCGGAAG |
| 29548_2 | TATGAGTGTGGAGTCGTTGCCCGATTGTGAACATGGACTACGTNNNAGATCGGAAG |
| 45040_1 | TATGAGTGTGGAGTCGTTGCGTGACAGGGGAAGTCATCTGATTNNNAGATCGGAA |
| 45040_2 | TATGAGTGTGGAGTCGTTGCGTGACAGGGGAAGTCATCTGATTNNNAGATCGGAA |
| 51883_1 | TATGAGTGTGGAGTCGTTGCAAACGATGGGAATTCGAAAAGCCNNNAGATCGGAA |
| 51883_2 | TATGAGTGTGGAGTCGTTGCAAACGATGGGAATTCGAAAAGCCNNNAGATCGGAA |
| 41987_1 | TATGAGTGTGGAGTCGTTGCAGGATGGTCTCGATCTCCTGACNNNAGATCGGAAGA |
| 41987_2 | TATGAGTGTGGAGTCGTTGCAGGATGGTCTCGATCTCCTGACNNNAGATCGGAAGA |
| 38778_1 | TATGAGTGTGGAGTCGTTGCTGGTCCCTAACATCACTTTCCAANNNAGATCGGAAGA |

| | |
|---|---|
| 38778_2 | TATGAGTGTGGAGTCGTTGCTGGTCCCTAACATCACTTTCCAANNNAGATCGGAAGA |
| 36951_1 | TATGAGTGTGGAGTCGTTGCGGGTATGATAGCGCATGTCTGTANNNAGATCGGAA |
| 36951_2 | TATGAGTGTGGAGTCGTTGCGGGTATGATAGCGCATGTCTGTANNNAGATCGGAA |
| 47446_1 | TATGAGTGTGGAGTCGTTGCGCCCTTTTCTTCCCCTTTTTCTCNNNAGATCGGAAGAG |
| 47446_2 | TATGAGTGTGGAGTCGTTGCGCCCTTTTCTTCCCCTTTTTCTCNNNAGATCGGAAGAG |
| 45791_1 | TATGAGTGTGGAGTCGTTGCAGTTAGTAGGGATGGGGTTTCACNNNAGATCGGAA |
| 45791_2 | TATGAGTGTGGAGTCGTTGCAGTTAGTAGGGATGGGGTTTCACNNNAGATCGGAA |
| 43989_1 | TATGAGTGTGGAGTCGTTGCGATATCTCGGCTCACTTCAACCTNNNAGATCGGAAG |
| 43989_2 | TATGAGTGTGGAGTCGTTGCGATATCTCGGCTCACTTCAACCTNNNAGATCGGAAG |
| 42178_1 | TATGAGTGTGGAGTCGTTGCCCTCCAAGGTTCAAGCAATTCTCNNNAGATCGGAAG |
| 42178_2 | TATGAGTGTGGAGTCGTTGCCCTCCAAGGTTCAAGCAATTCTCNNNAGATCGGAAG |
| 39501_1 | TATGAGTGTGGAGTCGTTGCGGCACCTATAATCCCAGCTACTCNNNAGATCGGAAG |
| 39501_2 | TATGAGTGTGGAGTCGTTGCGGCACCTATAATCCCAGCTACTCNNNAGATCGGAAG |
| 36686_1 | TATGAGTGTGGAGTCGTTGCTTCTGACATTATTGGTCTGGGGGNNNAGATCGGAA |
| 36686_2 | TATGAGTGTGGAGTCGTTGCTTCTGACATTATTGGTCTGGGGGNNNAGATCGGAA |
| 12820_1 | TATGAGTGTGGAGTCGTTGCAGATGTTGGTAAGACTTAGTATCNNNAGATCGGAA |
| 12820_2 | TATGAGTGTGGAGTCGTTGCAGATGTTGGTAAGACTTAGTATCNNNAGATCGGAA |
| 46149_1 | TATGAGTGTGGAGTCGTTGCTGGTTAGAGACTAGAATTGGGAAGANNNAGATCGG |
| 46149_2 | TATGAGTGTGGAGTCGTTGCTGGTTAGAGACTAGAATTGGGAAGANNNAGATCGG |
| 10519_1 | TATGAGTGTGGAGTCGTTGCGCTTCTTTCTTTACCCTTGGGCTNNNAGATCGGAAGA |
| 10519_2 | TATGAGTGTGGAGTCGTTGCGCTTCTTTCTTTACCCTTGGGCTNNNAGATCGGAAGA |
| 41570_1 | TATGAGTGTGGAGTCGTTGCCCACTACACCCGGCTAATTTTTTNNNAGATCGGAAGA |
| 41570_2 | TATGAGTGTGGAGTCGTTGCCCACTACACCCGGCTAATTTTTTNNNAGATCGGAAGA |
| 37015_1 | TATGAGTGTGGAGTCGTTGCTTATAGCTCACTGCAGCCTCAAANNNAGATCGGAAG |
| 37015_2 | TATGAGTGTGGAGTCGTTGCTTATAGCTCACTGCAGCCTCAAANNNAGATCGGAAG |
| 42880_1 | TATGAGTGTGGAGTCGTTGCGTCCATGTGCATTTCATCTTCCCNNNAGATCGGAAGA |
| 42880_2 | TATGAGTGTGGAGTCGTTGCGTCCATGTGCATTTCATCTTCCCNNNAGATCGGAAGA |
| 35266_1 | TATGAGTGTGGAGTCGTTGCGTGGTGTGCTCCTCTTCATTCTANNNAGATCGGAAG |
| 35266_2 | TATGAGTGTGGAGTCGTTGCGTGGTGTGCTCCTCTTCATTCTANNNAGATCGGAAG |
| 45964_1 | TATGAGTGTGGAGTCGTTGCCTTCCTGAGTAGCTGGGACTACANNNAGATCGGAAG |
| 45964_2 | TATGAGTGTGGAGTCGTTGCCTTCCTGAGTAGCTGGGACTACANNNAGATCGGAAG |
| 35782_1 | TATGAGTGTGGAGTCGTTGCCAGAGTCTTGCTGTGTCACCTAGNNNAGATCGGAAG |
| 35782_2 | TATGAGTGTGGAGTCGTTGCCAGAGTCTTGCTGTGTCACCTAGNNNAGATCGGAAG |
| 46717_1 | TATGAGTGTGGAGTCGTTGCGCCAAGCCATAAATTACAGTCCCNNNAGATCGGAAG |
| 46717_2 | TATGAGTGTGGAGTCGTTGCGCCAAGCCATAAATTACAGTCCCNNNAGATCGGAAG |
| 51860_1 | TATGAGTGTGGAGTCGTTGCCGTAGATGGGACCTTCCACAAATNNNAGATCGGAAG |
| 51860_2 | TATGAGTGTGGAGTCGTTGCCGTAGATGGGACCTTCCACAAATNNNAGATCGGAAG |
| 7855_1 | TATGAGTGTGGAGTCGTTGCGCAGCATAACAACTCACACCAAANNNAGATCGGAAG |
| 7855_2 | TATGAGTGTGGAGTCGTTGCGCAGCATAACAACTCACACCAAANNNAGATCGGAAG |
| 29737_1 | TATGAGTGTGGAGTCGTTGCAATGTGAGGAGCACGTATTTACGNNNAGATCGGAA |
| 29737_2 | TATGAGTGTGGAGTCGTTGCAATGTGAGGAGCACGTATTTACGNNNAGATCGGAA |
| 34180_1 | TATGAGTGTGGAGTCGTTGCGATCACAAGGTCAGGAGTTCGAGNNNAGATCGGAA |
| 34180_2 | TATGAGTGTGGAGTCGTTGCGATCACAAGGTCAGGAGTTCGAGNNNAGATCGGAA |

| | |
|---|---|
| 39943_1 | TATGAGTGTGGAGTCGTTGCCGCCTTTTTTTCACCCCTTTACTNNNAGATCGGAAGA |
| 39943_2 | TATGAGTGTGGAGTCGTTGCCGCCTTTTTTTCACCCCTTTACTNNNAGATCGGAAGA |
| 41454_1 | TATGAGTGTGGAGTCGTTGCAATGCACAGGAAGCTCAAAAGCNNNAGATCGGAAG |
| 41454_2 | TATGAGTGTGGAGTCGTTGCAATGCACAGGAAGCTCAAAAGCNNNAGATCGGAAG |
| 31710_1 | TATGAGTGTGGAGTCGTTGCGTGTGTGTGTGAGAGAGAGAGAGNNNAGATCGGA |
| 31710_2 | TATGAGTGTGGAGTCGTTGCGTGTGTGTGTGAGAGAGAGAGAGNNNAGATCGGA |
| 9598_1 | TATGAGTGTGGAGTCGTTGCTGTTGCTTGGCTCAGTTCTGTTANNNAGATCGGAAG |
| 9598_2 | TATGAGTGTGGAGTCGTTGCTGTTGCTTGGCTCAGTTCTGTTANNNAGATCGGAAG |
| 47008_1 | TATGAGTGTGGAGTCGTTGCGATAATGACGTGATGGCTGGAGTNNNAGATCGGAA |
| 47008_2 | TATGAGTGTGGAGTCGTTGCGATAATGACGTGATGGCTGGAGTNNNAGATCGGAA |
| 35598_1 | TATGAGTGTGGAGTCGTTGCCAAGCTCTTTTATGTGCCCACAANNNAGATCGGAAG |
| 35598_2 | TATGAGTGTGGAGTCGTTGCCAAGCTCTTTTATGTGCCCACAANNNAGATCGGAAG |
| 46827_1 | TATGAGTGTGGAGTCGTTGCCTCGGCTCACTACAAGCTCTATTNNNAGATCGGAAG |
| 46827_2 | TATGAGTGTGGAGTCGTTGCCTCGGCTCACTACAAGCTCTATTNNNAGATCGGAAG |
| 45836_1 | TATGAGTGTGGAGTCGTTGCCACCACCACAACTGGCTAATTTTNNNAGATCGGAAG |
| 45836_2 | TATGAGTGTGGAGTCGTTGCCACCACCACAACTGGCTAATTTTNNNAGATCGGAAG |
| 36793_1 | TATGAGTGTGGAGTCGTTGCTTTGGAGTCCTCTACCCATCTCTNNNAGATCGGAAGA |
| 36793_2 | TATGAGTGTGGAGTCGTTGCTTTGGAGTCCTCTACCCATCTCTNNNAGATCGGAAGA |
| 48958_1 | TATGAGTGTGGAGTCGTTGCAGGCAAAAATCTTGTTGGGTTCCNNNAGATCGGAA |
| 48958_2 | TATGAGTGTGGAGTCGTTGCAGGCAAAAATCTTGTTGGGTTCCNNNAGATCGGAA |
| 51904_1 | TATGAGTGTGGAGTCGTTGCGGAGTACCCCTTCACCTGTTTACNNNAGATCGGAAG |
| 51904_2 | TATGAGTGTGGAGTCGTTGCGGAGTACCCCTTCACCTGTTTACNNNAGATCGGAAG |
| 29792_1 | TATGAGTGTGGAGTCGTTGCCCATGGTTCCTGATACACCCTTTNNNAGATCGGAAG |
| 29792_2 | TATGAGTGTGGAGTCGTTGCCCATGGTTCCTGATACACCCTTTNNNAGATCGGAAG |
| 44481_1 | TATGAGTGTGGAGTCGTTGCCCAGAGGAACTAGGGAACAAGTCNNNAGATCGGAA |
| 44481_2 | TATGAGTGTGGAGTCGTTGCCCAGAGGAACTAGGGAACAAGTCNNNAGATCGGAA |
| 42267_1 | TATGAGTGTGGAGTCGTTGCAGTTCTATGCCTGTAGCCTATGCNNNAGATCGGAAG |
| 42267_2 | TATGAGTGTGGAGTCGTTGCAGTTCTATGCCTGTAGCCTATGCNNNAGATCGGAAG |
| 42345_1 | TATGAGTGTGGAGTCGTTGCGTTGATGTTGACTGCGGACTTACNNNAGATCGGAA |
| 42345_2 | TATGAGTGTGGAGTCGTTGCGTTGATGTTGACTGCGGACTTACNNNAGATCGGAA |
| 32381_1 | TATGAGTGTGGAGTCGTTGCTGGAGCTGAAATTTAGGGGCATANNNAGATCGGAA |
| 32381_2 | TATGAGTGTGGAGTCGTTGCTGGAGCTGAAATTTAGGGGCATANNNAGATCGGAA |
| 42432_1 | TATGAGTGTGGAGTCGTTGCATACGGCACGTCTTCATGTATCANNNAGATCGGAAG |
| 42432_2 | TATGAGTGTGGAGTCGTTGCATACGGCACGTCTTCATGTATCANNNAGATCGGAAG |
| 40659_1 | TATGAGTGTGGAGTCGTTGCAGGAAAGGAAGGAAAGGAACTCCNNNAGATCGGA |
| 40659_2 | TATGAGTGTGGAGTCGTTGCAGGAAAGGAAGGAAAGGAACTCCNNNAGATCGGA |
| 35223_1 | TATGAGTGTGGAGTCGTTGCCCTCACGAGTAGCTGGGACTATANNNAGATCGGAA |
| 35223_2 | TATGAGTGTGGAGTCGTTGCCCTCACGAGTAGCTGGGACTATANNNAGATCGGAA |
| 51532_1 | TATGAGTGTGGAGTCGTTGCAAACACTGCTGTCTATCTCCAGGNNNAGATCGGAAG |
| 51532_2 | TATGAGTGTGGAGTCGTTGCAAACACTGCTGTCTATCTCCAGGNNNAGATCGGAAG |
| 43180_1 | TATGAGTGTGGAGTCGTTGCGATCTCAGCTCACTGCAACCTNNNAGATCGGAAGAG |
| 43180_2 | TATGAGTGTGGAGTCGTTGCGATCTCAGCTCACTGCAACCTNNNAGATCGGAAGAG |
| 40578_1 | TATGAGTGTGGAGTCGTTGCTAAAGGGACTAGTAGGAGCCAGGNNNAGATCGGA |

| | |
|---|---|
| 40578_2 | TATGAGTGTGGAGTCGTTGCTAAAGGGACTAGTAGGAGCCAGGNNNAGATCGGA |
| 41354_1 | TATGAGTGTGGAGTCGTTGCTCGTTTCGTTTCATTTCGTTTCANNNAGATCGGAAGA |
| 41354_2 | TATGAGTGTGGAGTCGTTGCTCGTTTCGTTTCATTTCGTTTCANNNAGATCGGAAGA |
| 36979_1 | TATGAGTGTGGAGTCGTTGCTGCAGAGTGAATCAACAGAGTGANNNAGATCGGAA |
| 36979_2 | TATGAGTGTGGAGTCGTTGCTGCAGAGTGAATCAACAGAGTGANNNAGATCGGAA |
| 40630_1 | TATGAGTGTGGAGTCGTTGCGAGGTTAGCTAAAGTGGAGAGGANNNAGATCGGA |
| 40630_2 | TATGAGTGTGGAGTCGTTGCGAGGTTAGCTAAAGTGGAGAGGANNNAGATCGGA |
| 30961_1 | TATGAGTGTGGAGTCGTTGCGGCTAAGGAGAGTGACCATTTGANNNAGATCGGAA |
| 30961_2 | TATGAGTGTGGAGTCGTTGCGGCTAAGGAGAGTGACCATTTGANNNAGATCGGAA |
| 45120_1 | TATGAGTGTGGAGTCGTTGCGAACAAAGAGCTCAGGCATCTCNNNAGATCGGAAG |
| 45120_2 | TATGAGTGTGGAGTCGTTGCGAACAAAGAGCTCAGGCATCTCNNNAGATCGGAAG |
| 30150_1 | TATGAGTGTGGAGTCGTTGCACATTAGAATTGGAGGTTAAGAAAGCTNNNAGATC |
| 30150_2 | TATGAGTGTGGAGTCGTTGCACATTAGAATTGGAGGTTAAGAAAGCTNNNAGATC |
| 46373_1 | TATGAGTGTGGAGTCGTTGCGATATCAGGTAATCCGCCCACATNNNAGATCGGAAG |
| 46373_2 | TATGAGTGTGGAGTCGTTGCGATATCAGGTAATCCGCCCACATNNNAGATCGGAAG |
| 31845_1 | TATGAGTGTGGAGTCGTTGCTGGACCTCCTTTCTTTGCAAAACNNNAGATCGGAAG |
| 31845_2 | TATGAGTGTGGAGTCGTTGCTGGACCTCCTTTCTTTGCAAAACNNNAGATCGGAAG |
| 9784_1 | TATGAGTGTGGAGTCGTTGCTTTGGGCATTTGGGGAACACTTANNNAGATCGGAA |
| 9784_2 | TATGAGTGTGGAGTCGTTGCTTTGGGCATTTGGGGAACACTTANNNAGATCGGAA |
| 11130_1 | TATGAGTGTGGAGTCGTTGCATAGGTGTGTACCACCGCAGACNNNAGATCGGAAG |
| 11130_2 | TATGAGTGTGGAGTCGTTGCATAGGTGTGTACCACCGCAGACNNNAGATCGGAAG |
| 43451_1 | TATGAGTGTGGAGTCGTTGCCCCAAGTTCAAGCGATTCTCTTGNNNAGATCGGAAG |
| 43451_2 | TATGAGTGTGGAGTCGTTGCCCCAAGTTCAAGCGATTCTCTTGNNNAGATCGGAAG |
| 51429_1 | TATGAGTGTGGAGTCGTTGCGCAGCCATAATCCTCCTAGGTACNNNAGATCGGAAG |
| 51429_2 | TATGAGTGTGGAGTCGTTGCGCAGCCATAATCCTCCTAGGTACNNNAGATCGGAAG |
| 52188_1 | TATGAGTGTGGAGTCGTTGCCAACTGAGAAATGTTGGGCCTTTNNNAGATCGGAA |
| 52188_2 | TATGAGTGTGGAGTCGTTGCCAACTGAGAAATGTTGGGCCTTTNNNAGATCGGAA |
| 51872_1 | TATGAGTGTGGAGTCGTTGCCACTGTTTTCGGACCTTCCATGNNNAGATCGGAAGA |
| 51872_2 | TATGAGTGTGGAGTCGTTGCCACTGTTTTCGGACCTTCCATGNNNAGATCGGAAGA |
| 47761_1 | TATGAGTGTGGAGTCGTTGCCCACGCCCAGCTAATTTTTGTATNNNAGATCGGAAG |
| 47761_2 | TATGAGTGTGGAGTCGTTGCCCACGCCCAGCTAATTTTTGTATNNNAGATCGGAAG |
| 53193_1 | TATGAGTGTGGAGTCGTTGCTTTGTCATGAACCATCTGTTGCCNNNAGATCGGAAG |
| 53193_2 | TATGAGTGTGGAGTCGTTGCTTTGTCATGAACCATCTGTTGCCNNNAGATCGGAAG |
| 32136_1 | TATGAGTGTGGAGTCGTTGCGAGTTGTCAAAACTCCAATGGGGNNNAGATCGGAA |
| 32136_2 | TATGAGTGTGGAGTCGTTGCGAGTTGTCAAAACTCCAATGGGGNNNAGATCGGAA |
| 10544_1 | TATGAGTGTGGAGTCGTTGCGGGTTCTGTGAAGTTCAGGGTTTNNNAGATCGGAA |
| 10544_2 | TATGAGTGTGGAGTCGTTGCGGGTTCTGTGAAGTTCAGGGTTTNNNAGATCGGAA |
| 12940_1 | TATGAGTGTGGAGTCGTTGCTTCAAAGCGTTCTTCACATTNNNAGATCGGAAGAGC |
| 12940_2 | TATGAGTGTGGAGTCGTTGCTTCAAAGCGTTCTTCACATTNNNAGATCGGAAGAGC |
| 40904_1 | TATGAGTGTGGAGTCGTTGCTATTGTTGGAAGCTGCTGAGACTNNNAGATCGGAA |
| 40904_2 | TATGAGTGTGGAGTCGTTGCTATTGTTGGAAGCTGCTGAGACTNNNAGATCGGAA |
| 29969_1 | TATGAGTGTGGAGTCGTTGCCCCCTCTGTGAGTGTGTGTGTATANNNAGATCGGAA |
| 29969_2 | TATGAGTGTGGAGTCGTTGCCCCCTCTGTGAGTGTGTGTGTATANNNAGATCGGAA |

| | |
|---|---|
| 44222_1 | TATGAGTGTGGAGTCGTTGCCTGTGTAGTGTGGTGGTGAAGTCNNNAGATCGGAA |
| 44222_2 | TATGAGTGTGGAGTCGTTGCCTGTGTAGTGTGGTGGTGAAGTCNNNAGATCGGAA |
| 41993_1 | TATGAGTGTGGAGTCGTTGCCTCAACCTCCTGGGCTCAAGNNNAGATCGGAAGAGC |
| 41993_2 | TATGAGTGTGGAGTCGTTGCCTCAACCTCCTGGGCTCAAGNNNAGATCGGAAGAGC |
| 44886_1 | TATGAGTGTGGAGTCGTTGCCAACAGACCTTGGGTTCAAACTCNNNAGATCGGAAG |
| 44886_2 | TATGAGTGTGGAGTCGTTGCCAACAGACCTTGGGTTCAAACTCNNNAGATCGGAAG |
| 50957_1 | TATGAGTGTGGAGTCGTTGCCTTCAGGATTTCCGTTTCCACTCNNNAGATCGGAAGA |
| 50957_2 | TATGAGTGTGGAGTCGTTGCCTTCAGGATTTCCGTTTCCACTCNNNAGATCGGAAGA |
| 2977_1 | TATGAGTGTGGAGTCGTTGCTGAAGCAATTTAGGGCAAACTATCCNNNAGATCGGA |
| 2977_2 | TATGAGTGTGGAGTCGTTGCTGAAGCAATTTAGGGCAAACTATCCNNNAGATCGGA |
| 47190_1 | TATGAGTGTGGAGTCGTTGCGTAGAGACAGGGTTTCGCTATGTNNNAGATCGGAA |
| 47190_2 | TATGAGTGTGGAGTCGTTGCGTAGAGACAGGGTTTCGCTATGTNNNAGATCGGAA |
| 35642_1 | TATGAGTGTGGAGTCGTTGCGGACTCAAGTGAACCTCATGTCTNNNAGATCGGAAG |
| 35642_2 | TATGAGTGTGGAGTCGTTGCGGACTCAAGTGAACCTCATGTCTNNNAGATCGGAAG |
| 41104_1 | TATGAGTGTGGAGTCGTTGCCCTGCCCCAAACATTCATTTCTTNNNAGATCGGAAGA |
| 41104_2 | TATGAGTGTGGAGTCGTTGCCCTGCCCCAAACATTCATTTCTTNNNAGATCGGAAGA |
| 52241_1 | TATGAGTGTGGAGTCGTTGCCCCCTGGGAAATGCATTCTTTTCNNNAGATCGGAAG |
| 52241_2 | TATGAGTGTGGAGTCGTTGCCCCCTGGGAAATGCATTCTTTTCNNNAGATCGGAAG |
| 40023_1 | TATGAGTGTGGAGTCGTTGCTCGTAAGATCCACCAAGGTCAAGNNNAGATCGGAAG |
| 40023_2 | TATGAGTGTGGAGTCGTTGCTCGTAAGATCCACCAAGGTCAAGNNNAGATCGGAAG |
| 37656_1 | TATGAGTGTGGAGTCGTTGCAATCCAGGTTTGTGCTGTTTACCNNNAGATCGGAAG |
| 37656_2 | TATGAGTGTGGAGTCGTTGCAATCCAGGTTTGTGCTGTTTACCNNNAGATCGGAAG |
| 51104_1 | TATGAGTGTGGAGTCGTTGCCAAAGTGGTACATGCTATGCCTGNNNAGATCGGAAG |
| 51104_2 | TATGAGTGTGGAGTCGTTGCCAAAGTGGTACATGCTATGCCTGNNNAGATCGGAAG |
| 32883_1 | TATGAGTGTGGAGTCGTTGCGTGGGATTCCATTTACAGGAGGTNNNAGATCGGAA |
| 32883_2 | TATGAGTGTGGAGTCGTTGCGTGGGATTCCATTTACAGGAGGTNNNAGATCGGAA |
| 12791_1 | TATGAGTGTGGAGTCGTTGCTTGCAAATCAGTCTCTACAGNNNAGATCGGAAGAGC |
| 12791_2 | TATGAGTGTGGAGTCGTTGCTTGCAAATCAGTCTCTACAGNNNAGATCGGAAGAGC |
| 8999_1 | TATGAGTGTGGAGTCGTTGCTGACTACCCCATTGCCAGGATAANNNAGATCGGAAG |
| 8999_2 | TATGAGTGTGGAGTCGTTGCTGACTACCCCATTGCCAGGATAANNNAGATCGGAAG |
| 45404_1 | TATGAGTGTGGAGTCGTTGCCGTGTGTGTGTATAGGAGATGGANNNAGATCGGAA |
| 45404_2 | TATGAGTGTGGAGTCGTTGCCGTGTGTGTGTATAGGAGATGGANNNAGATCGGAA |
| 751_1 | TATGAGTGTGGAGTCGTTGCTGTCCTGTTTTGCTCACCCTCNNNAGATCGGAAGAGC |
| 751_2 | TATGAGTGTGGAGTCGTTGCTGTCCTGTTTTGCTCACCCTCNNNAGATCGGAAGAGC |
| 43727_1 | TATGAGTGTGGAGTCGTTGCAAGTGTGACTCTTCTCTGTGGTCNNNAGATCGGAAG |
| 43727_2 | TATGAGTGTGGAGTCGTTGCAAGTGTGACTCTTCTCTGTGGTCNNNAGATCGGAAG |
| 40770_1 | TATGAGTGTGGAGTCGTTGCAACAAAAGGCCTTGGTTCTTTGANNNAGATCGGAAG |
| 40770_2 | TATGAGTGTGGAGTCGTTGCAACAAAAGGCCTTGGTTCTTTGANNNAGATCGGAAG |
| 28784_1 | TATGAGTGTGGAGTCGTTGCCACATCATGGAGAATGCGGTTTCNNNAGATCGGAAG |
| 28784_2 | TATGAGTGTGGAGTCGTTGCCACATCATGGAGAATGCGGTTTCNNNAGATCGGAAG |
| 44012_1 | TATGAGTGTGGAGTCGTTGCACTGGCACACTTAGTGTTCTTCTNNNAGATCGGAAG |
| 44012_2 | TATGAGTGTGGAGTCGTTGCACTGGCACACTTAGTGTTCTTCTNNNAGATCGGAAG |
| 43837_1 | TATGAGTGTGGAGTCGTTGCCAACCACAACCTACCAAATGGACNNNAGATCGGAAG |

| | |
|---|---|
| 43837_2 | TATGAGTGTGGAGTCGTTGCCAACCACAACCTACCAAATGGACNNNAGATCGGAAG |
| 53173_1 | TATGAGTGTGGAGTCGTTGCCAACCCAGTGCTACCCATTTTTTNNNAGATCGGAAGA |
| 53173_2 | TATGAGTGTGGAGTCGTTGCCAACCCAGTGCTACCCATTTTTTNNNAGATCGGAAGA |
| 46536_1 | TATGAGTGTGGAGTCGTTGCGTTCAGATGATTCTCCTGCCTCANNNAGATCGGAAG |
| 46536_2 | TATGAGTGTGGAGTCGTTGCGTTCAGATGATTCTCCTGCCTCANNNAGATCGGAAG |
| 30833_1 | TATGAGTGTGGAGTCGTTGCGCTGCAATTGCTGAAACGATTTCNNNAGATCGGAAG |
| 30833_2 | TATGAGTGTGGAGTCGTTGCGCTGCAATTGCTGAAACGATTTCNNNAGATCGGAAG |
| 37006_1 | TATGAGTGTGGAGTCGTTGCACAGAATAAACAGGCAGCCTACANNNAGATCGGAA |
| 37006_2 | TATGAGTGTGGAGTCGTTGCACAGAATAAACAGGCAGCCTACANNNAGATCGGAA |
| 12712_1 | TATGAGTGTGGAGTCGTTGCGAAGAAGAAAACAAAGGTAGAACNNNAGATCGGA |
| 12712_2 | TATGAGTGTGGAGTCGTTGCGAAGAAGAAAACAAAGGTAGAACNNNAGATCGGA |
| 42268_1 | TATGAGTGTGGAGTCGTTGCCTCTCAGCTACTCGGGACTTTACNNNAGATCGGAAG |
| 42268_2 | TATGAGTGTGGAGTCGTTGCCTCTCAGCTACTCGGGACTTTACNNNAGATCGGAAG |
| 49769_1 | TATGAGTGTGGAGTCGTTGCAGCACTAGAACGTCAACTTCCATNNNAGATCGGAAG |
| 49769_2 | TATGAGTGTGGAGTCGTTGCAGCACTAGAACGTCAACTTCCATNNNAGATCGGAAG |
| 52069_1 | TATGAGTGTGGAGTCGTTGCCAAGGAGCTCTCTCTTGTGTCTTNNNAGATCGGAAG |
| 52069_2 | TATGAGTGTGGAGTCGTTGCCAAGGAGCTCTCTCTTGTGTCTTNNNAGATCGGAAG |
| 40644_1 | TATGAGTGTGGAGTCGTTGCGAGACAGATTCTTGCCCCGTCNNNAGATCGGAAGAG |
| 40644_2 | TATGAGTGTGGAGTCGTTGCGAGACAGATTCTTGCCCCGTCNNNAGATCGGAAGAG |
| 45913_1 | TATGAGTGTGGAGTCGTTGCTGATCTTGACTCACTGCAAGCTNNNAGATCGGAAGA |
| 45913_2 | TATGAGTGTGGAGTCGTTGCTGATCTTGACTCACTGCAAGCTNNNAGATCGGAAGA |
| 31054_1 | TATGAGTGTGGAGTCGTTGCTTTATGGAGAGGCATACATGGCANNNAGATCGGAA |
| 31054_2 | TATGAGTGTGGAGTCGTTGCTTTATGGAGAGGCATACATGGCANNNAGATCGGAA |
| 32026_1 | TATGAGTGTGGAGTCGTTGCCCAAGTAGCTGGGACTACAGGNNNAGATCGGAAGA |
| 32026_2 | TATGAGTGTGGAGTCGTTGCCCAAGTAGCTGGGACTACAGGNNNAGATCGGAAGA |
| 52387_1 | TATGAGTGTGGAGTCGTTGCATTTCCCTAGATGGTAATGGGGCNNNAGATCGGAA |
| 52387_2 | TATGAGTGTGGAGTCGTTGCATTTCCCTAGATGGTAATGGGGCNNNAGATCGGAA |
| 49609_1 | TATGAGTGTGGAGTCGTTGCTGTGGCATGAGAGAAATGTTCCTNNNAGATCGGAA |
| 49609_2 | TATGAGTGTGGAGTCGTTGCTGTGGCATGAGAGAAATGTTCCTNNNAGATCGGAA |
| 52162_1 | TATGAGTGTGGAGTCGTTGCTGGACAAATGGTAGCAGCAATTGNNNAGATCGGAA |
| 52162_2 | TATGAGTGTGGAGTCGTTGCTGGACAAATGGTAGCAGCAATTGNNNAGATCGGAA |
| 35432_1 | TATGAGTGTGGAGTCGTTGCCAAGTAGCTGGGACTACAGGCNNNAGATCGGAAGA |
| 35432_2 | TATGAGTGTGGAGTCGTTGCCAAGTAGCTGGGACTACAGGCNNNAGATCGGAAGA |
| 44764_1 | TATGAGTGTGGAGTCGTTGCTAGGCAGGAAAAGAGCAGTAAGGNNNAGATCGGA |
| 44764_2 | TATGAGTGTGGAGTCGTTGCTAGGCAGGAAAAGAGCAGTAAGGNNNAGATCGGA |
| 44269_1 | TATGAGTGTGGAGTCGTTGCTGGAATCACAAGGGGGAAGATAANNNAGATCGGA |
| 44269_2 | TATGAGTGTGGAGTCGTTGCTGGAATCACAAGGGGGAAGATAANNNAGATCGGA |
| 12857_1 | TATGAGTGTGGAGTCGTTGCAGAATATGTACTGTAATAAGGAGANNNAGATCGGA |
| 12857_2 | TATGAGTGTGGAGTCGTTGCAGAATATGTACTGTAATAAGGAGANNNAGATCGGA |
| 37261_1 | TATGAGTGTGGAGTCGTTGCTTGTGAAAGCAAATCAGTAATCCANNNAGATCGGAA |
| 37261_2 | TATGAGTGTGGAGTCGTTGCTTGTGAAAGCAAATCAGTAATCCANNNAGATCGGAA |
| 45976_1 | TATGAGTGTGGAGTCGTTGCCCTTCTGGGTTCAAGCAATTCTCNNNAGATCGGAAG |
| 45976_2 | TATGAGTGTGGAGTCGTTGCCCTTCTGGGTTCAAGCAATTCTCNNNAGATCGGAAG |

| | |
|---|---|
| 36148_1 | TATGAGTGTGGAGTCGTTGCAGCCCATTCTTCTCATAGCTCACNNNAGATCGGAAGA |
| 36148_2 | TATGAGTGTGGAGTCGTTGCAGCCCATTCTTCTCATAGCTCACNNNAGATCGGAAGA |
| 40011_1 | TATGAGTGTGGAGTCGTTGCCCACGAAGAGGGGTAAAGATCTTNNNAGATCGGAA |
| 40011_2 | TATGAGTGTGGAGTCGTTGCCCACGAAGAGGGGTAAAGATCTTNNNAGATCGGAA |
| 47582_1 | TATGAGTGTGGAGTCGTTGCCCTCCTGAGTAGCTGGAACTACANNNAGATCGGAAG |
| 47582_2 | TATGAGTGTGGAGTCGTTGCCCTCCTGAGTAGCTGGAACTACANNNAGATCGGAAG |
| 43003_1 | TATGAGTGTGGAGTCGTTGCTATGTCTCCTGGTTTGTGGTCTGNNNAGATCGGAAG |
| 43003_2 | TATGAGTGTGGAGTCGTTGCTATGTCTCCTGGTTTGTGGTCTGNNNAGATCGGAAG |
| 34235_1 | TATGAGTGTGGAGTCGTTGCAAGTCAACCTACTTCTGTTGGCTNNNAGATCGGAAG |
| 34235_2 | TATGAGTGTGGAGTCGTTGCAAGTCAACCTACTTCTGTTGGCTNNNAGATCGGAAG |
| 40275_1 | TATGAGTGTGGAGTCGTTGCCACTATTGGGATCACAGCTCACTNNNAGATCGGAAG |
| 40275_2 | TATGAGTGTGGAGTCGTTGCCACTATTGGGATCACAGCTCACTNNNAGATCGGAAG |
| 32762_1 | TATGAGTGTGGAGTCGTTGCCGAGAATACTGTTGGTGGTGAGANNNAGATCGGAA |
| 32762_2 | TATGAGTGTGGAGTCGTTGCCGAGAATACTGTTGGTGGTGAGANNNAGATCGGAA |
| 43559_1 | TATGAGTGTGGAGTCGTTGCAGCTGATAACAACTTAGCACTGTNNNAGATCGGAAG |
| 43559_2 | TATGAGTGTGGAGTCGTTGCAGCTGATAACAACTTAGCACTGTNNNAGATCGGAAG |
| 32941_1 | TATGAGTGTGGAGTCGTTGCCATAAATGGGACTTGGGCACTTTNNNAGATCGGAAG |
| 32941_2 | TATGAGTGTGGAGTCGTTGCCATAAATGGGACTTGGGCACTTTNNNAGATCGGAAG |
| 34652_1 | TATGAGTGTGGAGTCGTTGCAAGACAGTAAGACATGGGCAGAGNNNAGATCGGAA |
| 34652_2 | TATGAGTGTGGAGTCGTTGCAAGACAGTAAGACATGGGCAGAGNNNAGATCGGAA |
| 34536_1 | TATGAGTGTGGAGTCGTTGCCAAGGATCCACTGGAGGCTTGNNNAGATCGGAAGA |
| 34536_2 | TATGAGTGTGGAGTCGTTGCCAAGGATCCACTGGAGGCTTGNNNAGATCGGAAGA |
| 13246_1 | TATGAGTGTGGAGTCGTTGCATCCAAACTCTATGAACCTTATTNNNAGATCGGAAGA |
| 13246_2 | TATGAGTGTGGAGTCGTTGCATCCAAACTCTATGAACCTTATTNNNAGATCGGAAGA |
| 42503_1 | TATGAGTGTGGAGTCGTTGCGAGAGAGACAGAGAGAGAGAGGGNNNAGATCGGA |
| 42503_2 | TATGAGTGTGGAGTCGTTGCGAGAGAGACAGAGAGAGAGAGGGNNNAGATCGGA |
| 44594_1 | TATGAGTGTGGAGTCGTTGCTGACTGGAAAGAGTTACTCCGTGNNNAGATCGGAA |
| 44594_2 | TATGAGTGTGGAGTCGTTGCTGACTGGAAAGAGTTACTCCGTGNNNAGATCGGAA |
| 50496_1 | TATGAGTGTGGAGTCGTTGCCTGTGTTATCTGTGGTGCCTTTGNNNAGATCGGAAG |
| 50496_2 | TATGAGTGTGGAGTCGTTGCCTGTGTTATCTGTGGTGCCTTTGNNNAGATCGGAAG |
| 47527_1 | TATGAGTGTGGAGTCGTTGCAAGTTCAAGTGATTCTCCTGCCTNNNAGATCGGAAG |
| 47527_2 | TATGAGTGTGGAGTCGTTGCAAGTTCAAGTGATTCTCCTGCCTNNNAGATCGGAAG |
| 32588_1 | TATGAGTGTGGAGTCGTTGCCTAACTTGAAAGAATGGGGGGGANNNAGATCGGAA |
| 32588_2 | TATGAGTGTGGAGTCGTTGCCTAACTTGAAAGAATGGGGGGGANNNAGATCGGAA |
| 40003_1 | TATGAGTGTGGAGTCGTTGCAGGTTGCAGTGAGTAGAGATCACNNNAGATCGGAA |
| 40003_2 | TATGAGTGTGGAGTCGTTGCAGGTTGCAGTGAGTAGAGATCACNNNAGATCGGAA |
| 32729_1 | TATGAGTGTGGAGTCGTTGCTCCTTGTTCATTCCTACAGCCTGNNNAGATCGGAAGA |
| 32729_2 | TATGAGTGTGGAGTCGTTGCTCCTTGTTCATTCCTACAGCCTGNNNAGATCGGAAGA |
| 53379_1 | TATGAGTGTGGAGTCGTTGCGAGTTTTGCTCTTATTGCCCAGGNNNAGATCGGAAG |
| 53379_2 | TATGAGTGTGGAGTCGTTGCGAGTTTTGCTCTTATTGCCCAGGNNNAGATCGGAAG |
| 12566_1 | TATGAGTGTGGAGTCGTTGCGAAGAAGGAATGAGGCTTAAATANNNAGATCGGAA |
| 12566_2 | TATGAGTGTGGAGTCGTTGCGAAGAAGGAATGAGGCTTAAATANNNAGATCGGAA |
| 10484_1 | TATGAGTGTGGAGTCGTTGCGGTTTGTCTGTGTGTGAGCTCAANNNAGATCGGAA |

| | |
|---|---|
| 10484_2 | TATGAGTGTGGAGTCGTTGCGGTTTGTCTGTGTGTGAGCTCAANNNAGATCGGAA |
| 43194_1 | TATGAGTGTGGAGTCGTTGCGCTCCATACCTGCATAACGACTANNNAGATCGGAAG |
| 43194_2 | TATGAGTGTGGAGTCGTTGCGCTCCATACCTGCATAACGACTANNNAGATCGGAAG |
| 53367_1 | TATGAGTGTGGAGTCGTTGCCACACACATACACACACATACACANNNAGATCGGAAG |
| 53367_2 | TATGAGTGTGGAGTCGTTGCCACACACATACACACACATACACANNNAGATCGGAAG |
| 42240_1 | TATGAGTGTGGAGTCGTTGCTGTTACCCAGGCTAGTCTCAAACNNNAGATCGGAAG |
| 42240_2 | TATGAGTGTGGAGTCGTTGCTGTTACCCAGGCTAGTCTCAAACNNNAGATCGGAAG |
| 35875_1 | TATGAGTGTGGAGTCGTTGCGTGTCCTGCTTTCAAGTTAGCTGNNNAGATCGGAAG |
| 35875_2 | TATGAGTGTGGAGTCGTTGCGTGTCCTGCTTTCAAGTTAGCTGNNNAGATCGGAAG |
| 29869_1 | TATGAGTGTGGAGTCGTTGCAAGACGCAGATTAGTGGCATCTANNNAGATCGGAA |
| 29869_2 | TATGAGTGTGGAGTCGTTGCAAGACGCAGATTAGTGGCATCTANNNAGATCGGAA |
| 36113_1 | TATGAGTGTGGAGTCGTTGCTCTCTAATTACTTGCCGCAGTGTNNNAGATCGGAAG |
| 36113_2 | TATGAGTGTGGAGTCGTTGCTCTCTAATTACTTGCCGCAGTGTNNNAGATCGGAAG |
| 41690_1 | TATGAGTGTGGAGTCGTTGCGAGACAGGGTTTTACCATGTTGCNNNAGATCGGAA |
| 41690_2 | TATGAGTGTGGAGTCGTTGCGAGACAGGGTTTTACCATGTTGCNNNAGATCGGAA |
| 30348_1 | TATGAGTGTGGAGTCGTTGCAATGCTTAATCCAGCACAGTCCGNNNAGATCGGAAG |
| 30348_2 | TATGAGTGTGGAGTCGTTGCAATGCTTAATCCAGCACAGTCCGNNNAGATCGGAAG |
| 43390_1 | TATGAGTGTGGAGTCGTTGCTAGCTGAGATTACAGGGCAAGTGNNNAGATCGGAA |
| 43390_2 | TATGAGTGTGGAGTCGTTGCTAGCTGAGATTACAGGGCAAGTGNNNAGATCGGAA |
| 30184_1 | TATGAGTGTGGAGTCGTTGCGGACAGACGCCAGGTTTAAAAAANNNAGATCGGAA |
| 30184_2 | TATGAGTGTGGAGTCGTTGCGGACAGACGCCAGGTTTAAAAAANNNAGATCGGAA |
| 38470_1 | TATGAGTGTGGAGTCGTTGCACCCTTTCTGAGTGCAGGATTAGNNNAGATCGGAAG |
| 38470_2 | TATGAGTGTGGAGTCGTTGCACCCTTTCTGAGTGCAGGATTAGNNNAGATCGGAAG |
| 36554_1 | TATGAGTGTGGAGTCGTTGCCACTAACCTGCTCTTCAACGAAANNNAGATCGGAAG |
| 36554_2 | TATGAGTGTGGAGTCGTTGCCACTAACCTGCTCTTCAACGAAANNNAGATCGGAAG |
| 35158_1 | TATGAGTGTGGAGTCGTTGCGCCTCCAGAGTAGCTGAGATTACNNNAGATCGGAA |
| 35158_2 | TATGAGTGTGGAGTCGTTGCGCCTCCAGAGTAGCTGAGATTACNNNAGATCGGAA |
| 44463_1 | TATGAGTGTGGAGTCGTTGCTGTGTGTGTGAGTGTGTGTGTATNNNAGATCGGAA |
| 44463_2 | TATGAGTGTGGAGTCGTTGCTGTGTGTGTGAGTGTGTGTGTATNNNAGATCGGAA |
| 33562_1 | TATGAGTGTGGAGTCGTTGCCCTCCCAAAATGCTGAAATCACTNNNAGATCGGAAG |
| 33562_2 | TATGAGTGTGGAGTCGTTGCCCTCCCAAAATGCTGAAATCACTNNNAGATCGGAAG |
| 51270_1 | TATGAGTGTGGAGTCGTTGCACATCCTCTCTGTATCCTTCCCANNNAGATCGGAAGA |
| 51270_2 | TATGAGTGTGGAGTCGTTGCACATCCTCTCTGTATCCTTCCCANNNAGATCGGAAGA |
| 46953_1 | TATGAGTGTGGAGTCGTTGCACTCCTGACCTCAAGTGATTCACNNNAGATCGGAAG |
| 46953_2 | TATGAGTGTGGAGTCGTTGCACTCCTGACCTCAAGTGATTCACNNNAGATCGGAAG |
| 42946_1 | TATGAGTGTGGAGTCGTTGCGTGTGTGTGTGTGTGTGTGTATGNNNAGATCGGAA |
| 42946_2 | TATGAGTGTGGAGTCGTTGCGTGTGTGTGTGTGTGTGTGTATGNNNAGATCGGAA |
| 50510_1 | TATGAGTGTGGAGTCGTTGCGACACTGGGATGAACAAGACACANNNAGATCGGAA |
| 50510_2 | TATGAGTGTGGAGTCGTTGCGACACTGGGATGAACAAGACACANNNAGATCGGAA |
| 49712_1 | TATGAGTGTGGAGTCGTTGCTGTACCGTACACAGTTGCATACANNNAGATCGGAAG |
| 49712_2 | TATGAGTGTGGAGTCGTTGCTGTACCGTACACAGTTGCATACANNNAGATCGGAAG |
| 3005_1 | TATGAGTGTGGAGTCGTTGCCACAGCAGAGAGCAACTTCCTAGNNNAGATCGGAA |
| 3005_2 | TATGAGTGTGGAGTCGTTGCCACAGCAGAGAGCAACTTCCTAGNNNAGATCGGAA |

| | |
|---|---|
| 36209_1 | TATGAGTGTGGAGTCGTTGCAACCTCGAAAAGTCATCTCTGGANNNAGATCGGAAG |
| 36209_2 | TATGAGTGTGGAGTCGTTGCAACCTCGAAAAGTCATCTCTGGANNNAGATCGGAAG |
| 51896_1 | TATGAGTGTGGAGTCGTTGCTCCTCGAGAATGACATAGGGAGGNNNAGATCGGAA |
| 51896_2 | TATGAGTGTGGAGTCGTTGCTCCTCGAGAATGACATAGGGAGGNNNAGATCGGAA |
| 42455_1 | TATGAGTGTGGAGTCGTTGCGGGATTCTCAGTTGAGGGAGATCNNNAGATCGGAA |
| 42455_2 | TATGAGTGTGGAGTCGTTGCGGGATTCTCAGTTGAGGGAGATCNNNAGATCGGAA |
| 46500_1 | TATGAGTGTGGAGTCGTTGCGATCTCAGCTTACTGCAACCTCTNNNAGATCGGAAGA |
| 46500_2 | TATGAGTGTGGAGTCGTTGCGATCTCAGCTTACTGCAACCTCTNNNAGATCGGAAGA |
| 42615_1 | TATGAGTGTGGAGTCGTTGCAGAGAGATGCTAACCTTGTAGCANNNAGATCGGAA |
| 42615_2 | TATGAGTGTGGAGTCGTTGCAGAGAGATGCTAACCTTGTAGCANNNAGATCGGAA |
| 45155_1 | TATGAGTGTGGAGTCGTTGCAGTGGAAACACACAGTAGAGAAAGTNNNAGATCGG |
| 45155_2 | TATGAGTGTGGAGTCGTTGCAGTGGAAACACACAGTAGAGAAAGTNNNAGATCGG |
| 43574_1 | TATGAGTGTGGAGTCGTTGCGCGGGGAATATCTGGGTATCATTNNNAGATCGGAA |
| 43574_2 | TATGAGTGTGGAGTCGTTGCGCGGGGAATATCTGGGTATCATTNNNAGATCGGAA |
| 36906_1 | TATGAGTGTGGAGTCGTTGCTGTAATCCCAGGTACTCAGGAGANNNAGATCGGAA |
| 36906_2 | TATGAGTGTGGAGTCGTTGCTGTAATCCCAGGTACTCAGGAGANNNAGATCGGAA |
| 44003_1 | TATGAGTGTGGAGTCGTTGCGGGTTGCCACTGACCTTACATAGNNNAGATCGGAAG |
| 44003_2 | TATGAGTGTGGAGTCGTTGCGGGTTGCCACTGACCTTACATAGNNNAGATCGGAAG |
| 32059_1 | TATGAGTGTGGAGTCGTTGCCTCTTGAGTAGCTGGGACTACAGNNNAGATCGGAA |
| 32059_2 | TATGAGTGTGGAGTCGTTGCCTCTTGAGTAGCTGGGACTACAGNNNAGATCGGAA |
| 53089_1 | TATGAGTGTGGAGTCGTTGCCTGGAAGTGTTGGACCTAGTACANNNAGATCGGAA |
| 53089_2 | TATGAGTGTGGAGTCGTTGCCTGGAAGTGTTGGACCTAGTACANNNAGATCGGAA |
| 51676_1 | TATGAGTGTGGAGTCGTTGCCCAGCAGGGTCAACTATTTCAAGNNNAGATCGGAAG |
| 51676_2 | TATGAGTGTGGAGTCGTTGCCCAGCAGGGTCAACTATTTCAAGNNNAGATCGGAAG |
| 45596_1 | TATGAGTGTGGAGTCGTTGCCCTGTATGGGTAAAAGGCAGTCTNNNAGATCGGAA |
| 45596_2 | TATGAGTGTGGAGTCGTTGCCCTGTATGGGTAAAAGGCAGTCTNNNAGATCGGAA |
| 40750_1 | TATGAGTGTGGAGTCGTTGCCAGGTTCACGCCATTCTCCTNNNAGATCGGAAGAGCA |
| 40750_2 | TATGAGTGTGGAGTCGTTGCCAGGTTCACGCCATTCTCCTNNNAGATCGGAAGAGCA |
| 50319_1 | TATGAGTGTGGAGTCGTTGCCAGCAAGCACTGAGAACAAATGANNNAGATCGGAA |
| 50319_2 | TATGAGTGTGGAGTCGTTGCCAGCAAGCACTGAGAACAAATGANNNAGATCGGAA |
| 34933_1 | TATGAGTGTGGAGTCGTTGCCTGAGTAGCTGGAACTACAGGACNNNAGATCGGAA |
| 34933_2 | TATGAGTGTGGAGTCGTTGCCTGAGTAGCTGGAACTACAGGACNNNAGATCGGAA |
| 45432_1 | TATGAGTGTGGAGTCGTTGCCTTTCAGAATGGAGCCCACTTTGNNNAGATCGGAAG |
| 45432_2 | TATGAGTGTGGAGTCGTTGCCTTTCAGAATGGAGCCCACTTTGNNNAGATCGGAAG |
| 38191_1 | TATGAGTGTGGAGTCGTTGCGTTGCAATGAGCCGAGATCACNNNAGATCGGAAGA |
| 38191_2 | TATGAGTGTGGAGTCGTTGCGTTGCAATGAGCCGAGATCACNNNAGATCGGAAGA |
| 45621_1 | TATGAGTGTGGAGTCGTTGCGAGTAGCTGAGACTACAGGTGCNNNAGATCGGAAG |
| 45621_2 | TATGAGTGTGGAGTCGTTGCGAGTAGCTGAGACTACAGGTGCNNNAGATCGGAAG |
| 52082_1 | TATGAGTGTGGAGTCGTTGCTGGCATGTGGATATTCAGTTTTCCNNNAGATCGGAA |
| 52082_2 | TATGAGTGTGGAGTCGTTGCTGGCATGTGGATATTCAGTTTTCCNNNAGATCGGAA |
| 40917_1 | TATGAGTGTGGAGTCGTTGCTACCAGTACCATGCTGTTTTGGTNNNAGATCGGAAG |
| 40917_2 | TATGAGTGTGGAGTCGTTGCTACCAGTACCATGCTGTTTTGGTNNNAGATCGGAAG |
| 50561_1 | TATGAGTGTGGAGTCGTTGCAGCACAGATGAAGACAGGAAACANNNAGATCGGAA |

| | |
|---|---|
| 50561_2 | TATGAGTGTGGAGTCGTTGCAGCACAGATGAAGACAGGAAACANNNAGATCGGAA |
| 12913_1 | TATGAGTGTGGAGTCGTTGCATTATGTGAAAAATAAGATGCGGNNNAGATCGGAA |
| 12913_2 | TATGAGTGTGGAGTCGTTGCATTATGTGAAAAATAAGATGCGGNNNAGATCGGAA |
| 33362_1 | TATGAGTGTGGAGTCGTTGCCCTCGCTGGTTCAAGTGATTCTCNNNAGATCGGAAG |
| 33362_2 | TATGAGTGTGGAGTCGTTGCCCTCGCTGGTTCAAGTGATTCTCNNNAGATCGGAAG |
| 40157_1 | TATGAGTGTGGAGTCGTTGCGAGTTGCATGTGTCATGGTCAAGNNNAGATCGGAA |
| 40157_2 | TATGAGTGTGGAGTCGTTGCGAGTTGCATGTGTCATGGTCAAGNNNAGATCGGAA |
| 49023_1 | TATGAGTGTGGAGTCGTTGCCTCCTGAGTAGCTGGGATTACAGNNNAGATCGGAA |
| 49023_2 | TATGAGTGTGGAGTCGTTGCCTCCTGAGTAGCTGGGATTACAGNNNAGATCGGAA |
| 43419_1 | TATGAGTGTGGAGTCGTTGCAGGTGATCTGCCTGAAATGGATTNNNAGATCGGAA |
| 43419_2 | TATGAGTGTGGAGTCGTTGCAGGTGATCTGCCTGAAATGGATTNNNAGATCGGAA |
| 10520_1 | TATGAGTGTGGAGTCGTTGCCTTTGGTCATTGTGTCACTAGGCTNNNAGATCGGAA |
| 10520_2 | TATGAGTGTGGAGTCGTTGCCTTTGGTCATTGTGTCACTAGGCTNNNAGATCGGAA |
| 52624_1 | TATGAGTGTGGAGTCGTTGCCTCTTGGAATTGTGTGCCTGATGNNNAGATCGGAAG |
| 52624_2 | TATGAGTGTGGAGTCGTTGCCTCTTGGAATTGTGTGCCTGATGNNNAGATCGGAAG |
| 38336_1 | TATGAGTGTGGAGTCGTTGCGAACTGGAACAAGACAAGGATGCNNNAGATCGGAA |
| 38336_2 | TATGAGTGTGGAGTCGTTGCGAACTGGAACAAGACAAGGATGCNNNAGATCGGAA |
| 37105_1 | TATGAGTGTGGAGTCGTTGCCCAACGAATCAACAACCTCAGTTNNNAGATCGGAAG |
| 37105_2 | TATGAGTGTGGAGTCGTTGCCCAACGAATCAACAACCTCAGTTNNNAGATCGGAAG |
| 50377_1 | TATGAGTGTGGAGTCGTTGCGTGTCCTCGATGGTCTGGATGNNNAGATCGGAAGA |
| 50377_2 | TATGAGTGTGGAGTCGTTGCGTGTCCTCGATGGTCTGGATGNNNAGATCGGAAGA |
| 36438_1 | TATGAGTGTGGAGTCGTTGCTGTCATTCTTCCTCTGTCCCTTCNNNAGATCGGAAGA |
| 36438_2 | TATGAGTGTGGAGTCGTTGCTGTCATTCTTCCTCTGTCCCTTCNNNAGATCGGAAGA |
| 44936_1 | TATGAGTGTGGAGTCGTTGCTGGACTGTTGGTTTGATGAGGATNNNAGATCGGAA |
| 44936_2 | TATGAGTGTGGAGTCGTTGCTGGACTGTTGGTTTGATGAGGATNNNAGATCGGAA |
| 41271_1 | TATGAGTGTGGAGTCGTTGCGACAGTGTTTGTAGGCATTCAGCNNNAGATCGGAA |
| 41271_2 | TATGAGTGTGGAGTCGTTGCGACAGTGTTTGTAGGCATTCAGCNNNAGATCGGAA |
| 49971_1 | TATGAGTGTGGAGTCGTTGCCACCTAGCCATTCCTCTCCTTTTNNNAGATCGGAAGA |
| 49971_2 | TATGAGTGTGGAGTCGTTGCCACCTAGCCATTCCTCTCCTTTTNNNAGATCGGAAGA |
| 32021_1 | TATGAGTGTGGAGTCGTTGCTACCTACCTCCCTCACAATTTGCNNNAGATCGGAAGA |
| 32021_2 | TATGAGTGTGGAGTCGTTGCTACCTACCTCCCTCACAATTTGCNNNAGATCGGAAGA |
| 43611_1 | TATGAGTGTGGAGTCGTTGCACAGCACTGATTTTGCAGAAGTCNNNAGATCGGAAG |
| 43611_2 | TATGAGTGTGGAGTCGTTGCACAGCACTGATTTTGCAGAAGTCNNNAGATCGGAAG |
| 48623_1 | TATGAGTGTGGAGTCGTTGCGGCATACAGTGGCACAATCATGNNNAGATCGGAAG |
| 48623_2 | TATGAGTGTGGAGTCGTTGCGGCATACAGTGGCACAATCATGNNNAGATCGGAAG |
| 47579_1 | TATGAGTGTGGAGTCGTTGCCTGTCACTCACACATCGAGGTTANNNAGATCGGAAG |
| 47579_2 | TATGAGTGTGGAGTCGTTGCCTGTCACTCACACATCGAGGTTANNNAGATCGGAAG |
| 49703_1 | TATGAGTGTGGAGTCGTTGCCTGGGAGTAGCTGAGACTACAGGNNNAGATCGGAA |
| 49703_2 | TATGAGTGTGGAGTCGTTGCCTGGGAGTAGCTGAGACTACAGGNNNAGATCGGAA |
| 52352_1 | TATGAGTGTGGAGTCGTTGCCCAACAACAAGGATGCAGAAGATNNNAGATCGGAA |
| 52352_2 | TATGAGTGTGGAGTCGTTGCCCAACAACAAGGATGCAGAAGATNNNAGATCGGAA |
| 53058_1 | TATGAGTGTGGAGTCGTTGCCAACTTAGGGATACCAGGAGCTGNNNAGATCGGAA |
| 53058_2 | TATGAGTGTGGAGTCGTTGCCAACTTAGGGATACCAGGAGCTGNNNAGATCGGAA |

| | |
|---|---|
| 34213_1 | TATGAGTGTGGAGTCGTTGCAAACACAAACCTTCTTGGAAGGCNNNAGATCGGAAG |
| 34213_2 | TATGAGTGTGGAGTCGTTGCAAACACAAACCTTCTTGGAAGGCNNNAGATCGGAAG |
| 40224_1 | TATGAGTGTGGAGTCGTTGCACAAGTTCTCTCATCTAGCTGTGTNNNAGATCGGAAG |
| 40224_2 | TATGAGTGTGGAGTCGTTGCACAAGTTCTCTCATCTAGCTGTGTNNNAGATCGGAAG |
| 32474_1 | TATGAGTGTGGAGTCGTTGCGGAGTACAGTGGCACAATCACAGNNNAGATCGGAAG |
| 32474_2 | TATGAGTGTGGAGTCGTTGCGGAGTACAGTGGCACAATCACAGNNNAGATCGGAAG |
| 48558_1 | TATGAGTGTGGAGTCGTTGCCTCCTGAGTAGCTGGGATTACAGNNNAGATCGGAAG |
| 48558_2 | TATGAGTGTGGAGTCGTTGCCTCCTGAGTAGCTGGGATTACAGNNNAGATCGGAAG |
| 42400_1 | TATGAGTGTGGAGTCGTTGCAGCCAGAACATCCAACATCAGATNNNAGATCGGAAG |
| 42400_2 | TATGAGTGTGGAGTCGTTGCAGCCAGAACATCCAACATCAGATNNNAGATCGGAAG |
| 38579_1 | TATGAGTGTGGAGTCGTTGCAGGTTGTAATGAGCCGAGATTGTNNNAGATCGGAAG |
| 38579_2 | TATGAGTGTGGAGTCGTTGCAGGTTGTAATGAGCCGAGATTGTNNNAGATCGGAAG |
| 48542_1 | TATGAGTGTGGAGTCGTTGCCCTTCCAGTTTCAAGCGATTCTCNNNAGATCGGAAGA |
| 48542_2 | TATGAGTGTGGAGTCGTTGCCCTTCCAGTTTCAAGCGATTCTCNNNAGATCGGAAGA |
| 13319_1 | TATGAGTGTGGAGTCGTTGCAGAAAGAGAAAGAAAGAAAGAGANNNAGATCGGA |
| 13319_2 | TATGAGTGTGGAGTCGTTGCAGAAAGAGAAAGAAAGAAAGAGANNNAGATCGGA |
| 10851_1 | TATGAGTGTGGAGTCGTTGCTTATCAGTTCTGTTCTCTGGAGAANNNAGATCGGAAG |
| 10851_2 | TATGAGTGTGGAGTCGTTGCTTATCAGTTCTGTTCTCTGGAGAANNNAGATCGGAAG |
| 12643_1 | TATGAGTGTGGAGTCGTTGCTAGCCTCAATAACAAATAAATGCNNNAGATCGGAAG |
| 12643_2 | TATGAGTGTGGAGTCGTTGCTAGCCTCAATAACAAATAAATGCNNNAGATCGGAAG |
| 35594_1 | TATGAGTGTGGAGTCGTTGCTCACTAGAGGCAAGAACAATCCANNNAGATCGGAAG |
| 35594_2 | TATGAGTGTGGAGTCGTTGCTCACTAGAGGCAAGAACAATCCANNNAGATCGGAAG |
| 33837_1 | TATGAGTGTGGAGTCGTTGCCTGTAGTCCCAGCTACTCAGGAGNNNAGATCGGAAG |
| 33837_2 | TATGAGTGTGGAGTCGTTGCCTGTAGTCCCAGCTACTCAGGAGNNNAGATCGGAAG |
| 12172_1 | TATGAGTGTGGAGTCGTTGCACCAATCTGCATTCTCTGTGGNNNAGATCGGAAGAG |
| 12172_2 | TATGAGTGTGGAGTCGTTGCACCAATCTGCATTCTCTGTGGNNNAGATCGGAAGAG |
| 34011_1 | TATGAGTGTGGAGTCGTTGCTGAGTAACTGGGATTACAGGTGCNNNAGATCGGAAG |
| 34011_2 | TATGAGTGTGGAGTCGTTGCTGAGTAACTGGGATTACAGGTGCNNNAGATCGGAAG |
| 35985_1 | TATGAGTGTGGAGTCGTTGCAATGTCCATCCACAGGAGAATGANNNAGATCGGAAG |
| 35985_2 | TATGAGTGTGGAGTCGTTGCAATGTCCATCCACAGGAGAATGANNNAGATCGGAAG |
| 34034_1 | TATGAGTGTGGAGTCGTTGCAGATGGAATTTCGCTCTTGTTGCNNNAGATCGGAAG |
| 34034_2 | TATGAGTGTGGAGTCGTTGCAGATGGAATTTCGCTCTTGTTGCNNNAGATCGGAAG |
| 9704_1 | TATGAGTGTGGAGTCGTTGCCTGAGGTGGCAAGGAATATCACCNNNAGATCGGAAG |
| 9704_2 | TATGAGTGTGGAGTCGTTGCCTGAGGTGGCAAGGAATATCACCNNNAGATCGGAAG |
| 31568_1 | TATGAGTGTGGAGTCGTTGCTAGATGGCACTCAATCTGTGGTGNNNAGATCGGAAG |
| 31568_2 | TATGAGTGTGGAGTCGTTGCTAGATGGCACTCAATCTGTGGTGNNNAGATCGGAAG |
| 52462_1 | TATGAGTGTGGAGTCGTTGCCCTAGATGTTGTCTGCTCCTGTTNNNAGATCGGAAGA |
| 52462_2 | TATGAGTGTGGAGTCGTTGCCCTAGATGTTGTCTGCTCCTGTTNNNAGATCGGAAGA |
| 29164_1 | TATGAGTGTGGAGTCGTTGCTATAGAACACCCAGCACTGATCCNNNAGATCGGAAGA |
| 29164_2 | TATGAGTGTGGAGTCGTTGCTATAGAACACCCAGCACTGATCCNNNAGATCGGAAGA |
| 47755_1 | TATGAGTGTGGAGTCGTTGCGTTCCCTTCTCTAGCATCCTGTCNNNAGATCGGAAGA |
| 47755_2 | TATGAGTGTGGAGTCGTTGCGTTCCCTTCTCTAGCATCCTGTCNNNAGATCGGAAGA |
| 45036_1 | TATGAGTGTGGAGTCGTTGCCCAAATCTTTCCAACAGAGCTGGNNNAGATCGGAAG |

| | |
|---|---|
| 45036_2 | TATGAGTGTGGAGTCGTTGCCCAAATCTTTCCAACAGAGCTGGNNNAGATCGGAAG |
| 34224_1 | TATGAGTGTGGAGTCGTTGCTTTCTCATGCCCCTTAGTATCCGNNNAGATCGGAAGA |
| 34224_2 | TATGAGTGTGGAGTCGTTGCTTTCTCATGCCCCTTAGTATCCGNNNAGATCGGAAGA |
| 53201_1 | TATGAGTGTGGAGTCGTTGCGTATGAGGTAAGGGGCATATGCANNNAGATCGGA |
| 53201_2 | TATGAGTGTGGAGTCGTTGCGTATGAGGTAAGGGGCATATGCANNNAGATCGGA |
| 52800_1 | TATGAGTGTGGAGTCGTTGCGAGGAATCAGCAGTCTCCAGGNNNAGATCGGAAGA |
| 52800_2 | TATGAGTGTGGAGTCGTTGCGAGGAATCAGCAGTCTCCAGGNNNAGATCGGAAGA |
| 34724_1 | TATGAGTGTGGAGTCGTTGCCAAGTAGCTGGGATTACAGGCATNNNAGATCGGAA |
| 34724_2 | TATGAGTGTGGAGTCGTTGCCAAGTAGCTGGGATTACAGGCATNNNAGATCGGAA |
| 31505_1 | TATGAGTGTGGAGTCGTTGCCTCCTCCAGGTTTCTAATCTGCANNNAGATCGGAAG |
| 31505_2 | TATGAGTGTGGAGTCGTTGCCTCCTCCAGGTTTCTAATCTGCANNNAGATCGGAAG |
| 53404_1 | TATGAGTGTGGAGTCGTTGCGGGTGAGTAAATGTCAATGGCACNNNAGATCGGAA |
| 53404_2 | TATGAGTGTGGAGTCGTTGCGGGTGAGTAAATGTCAATGGCACNNNAGATCGGAA |
| 31229_1 | TATGAGTGTGGAGTCGTTGCACTTCTCTCCTTCTCTCCCTCTCNNNAGATCGGAAGAG |
| 31229_2 | TATGAGTGTGGAGTCGTTGCACTTCTCTCCTTCTCTCCCTCTCNNNAGATCGGAAGAG |
| 42684_1 | TATGAGTGTGGAGTCGTTGCGCAATGGCATGATCTTGGCAGNNNAGATCGGAAGA |
| 42684_2 | TATGAGTGTGGAGTCGTTGCGCAATGGCATGATCTTGGCAGNNNAGATCGGAAGA |
| 32200_1 | TATGAGTGTGGAGTCGTTGCGCAATCTCAGCTCACTGCAAGNNNAGATCGGAAGAG |
| 32200_2 | TATGAGTGTGGAGTCGTTGCGCAATCTCAGCTCACTGCAAGNNNAGATCGGAAGAG |
| 37009_1 | TATGAGTGTGGAGTCGTTGCCCAGCATAACTGAAGCCCAAAAGNNNAGATCGGAAG |
| 37009_2 | TATGAGTGTGGAGTCGTTGCCCAGCATAACTGAAGCCCAAAAGNNNAGATCGGAAG |
| 10411_1 | TATGAGTGTGGAGTCGTTGCTACCTTGAGTGTGGTGATGGGATNNNAGATCGGAA |
| 10411_2 | TATGAGTGTGGAGTCGTTGCTACCTTGAGTGTGGTGATGGGATNNNAGATCGGAA |
| 2751_1 | TATGAGTGTGGAGTCGTTGCGTCTTGCCACGTTATCTTCACCTNNNAGATCGGAAGA |
| 2751_2 | TATGAGTGTGGAGTCGTTGCGTCTTGCCACGTTATCTTCACCTNNNAGATCGGAAGA |
| 31619_1 | TATGAGTGTGGAGTCGTTGCATCAGACTTCAGACCTCAGGGTANNNAGATCGGAAG |
| 31619_2 | TATGAGTGTGGAGTCGTTGCATCAGACTTCAGACCTCAGGGTANNNAGATCGGAAG |
| 40805_1 | TATGAGTGTGGAGTCGTTGCTTAGAGAAGAAGCTTCCAGGCAGNNNAGATCGGAA |
| 40805_2 | TATGAGTGTGGAGTCGTTGCTTAGAGAAGAAGCTTCCAGGCAGNNNAGATCGGAA |
| 31588_1 | TATGAGTGTGGAGTCGTTGCTTCCTGCTACGACACTGAATTCANNNAGATCGGAAG |
| 31588_2 | TATGAGTGTGGAGTCGTTGCTTCCTGCTACGACACTGAATTCANNNAGATCGGAAG |
| 48181_1 | TATGAGTGTGGAGTCGTTGCGGGTTCATGCCATTCTCCTACTTNNNAGATCGGAAG |
| 48181_2 | TATGAGTGTGGAGTCGTTGCGGGTTCATGCCATTCTCCTACTTNNNAGATCGGAAG |
| 51493_1 | TATGAGTGTGGAGTCGTTGCAACTTCAAGCTTCCTGAGTACCTNNNAGATCGGAAG |
| 51493_2 | TATGAGTGTGGAGTCGTTGCAACTTCAAGCTTCCTGAGTACCTNNNAGATCGGAAG |
| 49632_1 | TATGAGTGTGGAGTCGTTGCAAGTAGCTGAGATTACAGGCACCNNNAGATCGGAA |
| 49632_2 | TATGAGTGTGGAGTCGTTGCAAGTAGCTGAGATTACAGGCACCNNNAGATCGGAA |
| 29905_1 | TATGAGTGTGGAGTCGTTGCGGAAAGGAGACAGACCCAAGGNNNAGATCGGAAG |
| 29905_2 | TATGAGTGTGGAGTCGTTGCGGAAAGGAGACAGACCCAAGGNNNAGATCGGAAG |
| 31369_1 | TATGAGTGTGGAGTCGTTGCAACTGAAATTTCACCTCCCCCAANNNAGATCGGAAGA |
| 31369_2 | TATGAGTGTGGAGTCGTTGCAACTGAAATTTCACCTCCCCCAANNNAGATCGGAAGA |
| 625_1 | TATGAGTGTGGAGTCGTTGCCCAAAAGGCGCTATCTCCAATCTNNNAGATCGGAAG |
| 625_2 | TATGAGTGTGGAGTCGTTGCCCAAAAGGCGCTATCTCCAATCTNNNAGATCGGAAG |

| | |
|---|---|
| 46721_1 | TATGAGTGTGGAGTCGTTGCTTCAAGTGATTCTCCTGCCTCAGNNNAGATCGGAAG |
| 46721_2 | TATGAGTGTGGAGTCGTTGCTTCAAGTGATTCTCCTGCCTCAGNNNAGATCGGAAG |
| 35423_1 | TATGAGTGTGGAGTCGTTGCTAGCAGGGTCTCACTCTGTTACTNNNAGATCGGAAG |
| 35423_2 | TATGAGTGTGGAGTCGTTGCTAGCAGGGTCTCACTCTGTTACTNNNAGATCGGAAG |
| 34394_1 | TATGAGTGTGGAGTCGTTGCCGGTGAAGATTCCCTTGATCACANNNAGATCGGAAG |
| 34394_2 | TATGAGTGTGGAGTCGTTGCCGGTGAAGATTCCCTTGATCACANNNAGATCGGAAG |
| 40460_1 | TATGAGTGTGGAGTCGTTGCGAAAGGCAGCTCTAAAGAGGTCANNNAGATCGGAA |
| 40460_2 | TATGAGTGTGGAGTCGTTGCGAAAGGCAGCTCTAAAGAGGTCANNNAGATCGGAA |
| 36948_1 | TATGAGTGTGGAGTCGTTGCCTTCATTGTGATGGCAACCATGTNNNAGATCGGAAG |
| 36948_2 | TATGAGTGTGGAGTCGTTGCCTTCATTGTGATGGCAACCATGTNNNAGATCGGAAG |
| 49248_1 | TATGAGTGTGGAGTCGTTGCAGAGTTTCGTTCTTGTCTCCCAGNNNAGATCGGAAG |
| 49248_2 | TATGAGTGTGGAGTCGTTGCAGAGTTTCGTTCTTGTCTCCCAGNNNAGATCGGAAG |
| 13158_1 | TATGAGTGTGGAGTCGTTGCTGCTGGTTTGTATTTTAGTTTTANNNAGATCGGAAG |
| 13158_2 | TATGAGTGTGGAGTCGTTGCTGCTGGTTTGTATTTTAGTTTTANNNAGATCGGAAG |
| 41241_1 | TATGAGTGTGGAGTCGTTGCCTGAGGCAGGAGAATCACTTGANNNAGATCGGAAG |
| 41241_2 | TATGAGTGTGGAGTCGTTGCCTGAGGCAGGAGAATCACTTGANNNAGATCGGAAG |
| 49546_1 | TATGAGTGTGGAGTCGTTGCACGTGAAGTCTGAGACATTTGGANNNAGATCGGAA |
| 49546_2 | TATGAGTGTGGAGTCGTTGCACGTGAAGTCTGAGACATTTGGANNNAGATCGGAA |
| 46170_1 | TATGAGTGTGGAGTCGTTGCCACCACACCTGGCTAATTTTTGTNNNAGATCGGAAG |
| 46170_2 | TATGAGTGTGGAGTCGTTGCCACCACACCTGGCTAATTTTTGTNNNAGATCGGAAG |
| 47955_1 | TATGAGTGTGGAGTCGTTGCGTGGTCTGTGTATGCATGCAAAANNNAGATCGGAA |
| 47955_2 | TATGAGTGTGGAGTCGTTGCGTGGTCTGTGTATGCATGCAAAANNNAGATCGGAA |
| 37829_1 | TATGAGTGTGGAGTCGTTGCAGAGTCAGGCTATCAGTCCATTGNNNAGATCGGAAG |
| 37829_2 | TATGAGTGTGGAGTCGTTGCAGAGTCAGGCTATCAGTCCATTGNNNAGATCGGAAG |
| 47458_1 | TATGAGTGTGGAGTCGTTGCGAGAAAAGACTGGGGAGGACTTTNNNAGATCGGA |
| 47458_2 | TATGAGTGTGGAGTCGTTGCGAGAAAAGACTGGGGAGGACTTTNNNAGATCGGA |
| 39966_1 | TATGAGTGTGGAGTCGTTGCGTCTCTCTCCCATGTGATGATCCNNNAGATCGGAAG |
| 39966_2 | TATGAGTGTGGAGTCGTTGCGTCTCTCTCCCATGTGATGATCCNNNAGATCGGAAG |
| 30979_1 | TATGAGTGTGGAGTCGTTGCCTCTCTCTCTCTCACACACACACNNNAGATCGGAAGA |
| 30979_2 | TATGAGTGTGGAGTCGTTGCCTCTCTCTCTCTCACACACACACNNNAGATCGGAAGA |
| 36183_1 | TATGAGTGTGGAGTCGTTGCGTAGAGACTAAACCACCAGCCTCNNNAGATCGGAAG |
| 36183_2 | TATGAGTGTGGAGTCGTTGCGTAGAGACTAAACCACCAGCCTCNNNAGATCGGAAG |
| 29649_1 | TATGAGTGTGGAGTCGTTGCTGGGTGGTTTAAATGACAGACACNNNAGATCGGAA |
| 29649_2 | TATGAGTGTGGAGTCGTTGCTGGGTGGTTTAAATGACAGACACNNNAGATCGGAA |
| 2291_1 | TATGAGTGTGGAGTCGTTGCTTCCCAAGTTCTTCACCTTTGGCNNNAGATCGGAAGA |
| 2291_2 | TATGAGTGTGGAGTCGTTGCTTCCCAAGTTCTTCACCTTTGGCNNNAGATCGGAAGA |
| 33179_1 | TATGAGTGTGGAGTCGTTGCTCCACAATCTCACAAGCATCTGANNNAGATCGGAAG |
| 33179_2 | TATGAGTGTGGAGTCGTTGCTCCACAATCTCACAAGCATCTGANNNAGATCGGAAG |
| 29420_1 | TATGAGTGTGGAGTCGTTGCTTTAATGTCTGCGAGGGTCCTGNNNAGATCGGAAG |
| 29420_2 | TATGAGTGTGGAGTCGTTGCTTTAATGTCTGCGAGGGTCCTGNNNAGATCGGAAG |
| 35914_1 | TATGAGTGTGGAGTCGTTGCACTCAAACAGGTACTTGTCCACANNNAGATCGGAAG |
| 35914_2 | TATGAGTGTGGAGTCGTTGCACTCAAACAGGTACTTGTCCACANNNAGATCGGAAG |
| 37957_1 | TATGAGTGTGGAGTCGTTGCGGAAATGGATACTATGGGGTGGGNNNAGATCGGA |

| | |
|---|---|
| 37957_2 | TATGAGTGTGGAGTCGTTGCGGAAATGGATACTATGGGGTGGGNNNAGATCGGA |
| 11548_1 | TATGAGTGTGGAGTCGTTGCCGTCCACAAGTTGTATGTCTTCTGNNNAGATCGGAA |
| 11548_2 | TATGAGTGTGGAGTCGTTGCCGTCCACAAGTTGTATGTCTTCTGNNNAGATCGGAA |
| 50778_1 | TATGAGTGTGGAGTCGTTGCTGCTCTGCATTTTCTTCAAGGAANNNAGATCGGAAG |
| 50778_2 | TATGAGTGTGGAGTCGTTGCTGCTCTGCATTTTCTTCAAGGAANNNAGATCGGAAG |
| 47514_1 | TATGAGTGTGGAGTCGTTGCTTCAAGTGATTCTCCTGCCTCAGNNNAGATCGGAAG |
| 47514_2 | TATGAGTGTGGAGTCGTTGCTTCAAGTGATTCTCCTGCCTCAGNNNAGATCGGAAG |
| 10830_1 | TATGAGTGTGGAGTCGTTGCAGCAACCTGAATATCAGCTTTGANNNAGATCGGAAG |
| 10830_2 | TATGAGTGTGGAGTCGTTGCAGCAACCTGAATATCAGCTTTGANNNAGATCGGAAG |
| 8822_1 | TATGAGTGTGGAGTCGTTGCTGTAAAGTGTTAGCGGGGCAATTNNNAGATCGGAA |
| 8822_2 | TATGAGTGTGGAGTCGTTGCTGTAAAGTGTTAGCGGGGCAATTNNNAGATCGGAA |
| 40490_1 | TATGAGTGTGGAGTCGTTGCATGTACACTATTCGGGTGACAGGNNNAGATCGGAA |
| 40490_2 | TATGAGTGTGGAGTCGTTGCATGTACACTATTCGGGTGACAGGNNNAGATCGGAA |
| 50907_1 | TATGAGTGTGGAGTCGTTGCACTGGTGAGAAAAGGTACAATGANNNAGATCGGAA |
| 50907_2 | TATGAGTGTGGAGTCGTTGCACTGGTGAGAAAAGGTACAATGANNNAGATCGGAA |
| 48594_1 | TATGAGTGTGGAGTCGTTGCAAGCCCTTATCTTCCCTTACTGCNNNAGATCGGAAGA |
| 48594_2 | TATGAGTGTGGAGTCGTTGCAAGCCCTTATCTTCCCTTACTGCNNNAGATCGGAAGA |
| 43658_1 | TATGAGTGTGGAGTCGTTGCCCAGACAAGGAGGTATGCATCTTNNNAGATCGGAA |
| 43658_2 | TATGAGTGTGGAGTCGTTGCCCAGACAAGGAGGTATGCATCTTNNNAGATCGGAA |
| 357_1 | TATGAGTGTGGAGTCGTTGCGCCAGGTTCTCTTTGTTATGGCNNNAGATCGGAAGA |
| 357_2 | TATGAGTGTGGAGTCGTTGCGCCAGGTTCTCTTTGTTATGGCNNNAGATCGGAAGA |
| 40064_1 | TATGAGTGTGGAGTCGTTGCCAACAGAGCAAGACTCCATCTCANNNAGATCGGAAG |
| 40064_2 | TATGAGTGTGGAGTCGTTGCCAACAGAGCAAGACTCCATCTCANNNAGATCGGAAG |
| 30368_1 | TATGAGTGTGGAGTCGTTGCACATGCTCCTGACTCTATCTCCCNNNAGATCGGAAGA |
| 30368_2 | TATGAGTGTGGAGTCGTTGCACATGCTCCTGACTCTATCTCCCNNNAGATCGGAAGA |
| 31119_1 | TATGAGTGTGGAGTCGTTGCGCCAGCCTTCAATGTTCCTTATTNNNAGATCGGAAG |
| 31119_2 | TATGAGTGTGGAGTCGTTGCGCCAGCCTTCAATGTTCCTTATTNNNAGATCGGAAG |
| 48761_1 | TATGAGTGTGGAGTCGTTGCCTTGACGACACTCCTAGCATCATNNNAGATCGGAAG |
| 48761_2 | TATGAGTGTGGAGTCGTTGCCTTGACGACACTCCTAGCATCATNNNAGATCGGAAG |
| 53070_1 | TATGAGTGTGGAGTCGTTGCTGAGATGATATCTGGGAAGCAGCNNNAGATCGGAA |
| 53070_2 | TATGAGTGTGGAGTCGTTGCTGAGATGATATCTGGGAAGCAGCNNNAGATCGGAA |
| 31176_1 | TATGAGTGTGGAGTCGTTGCACAGGCTGAGATTCCTAGAGTCTNNNAGATCGGAA |
| 31176_2 | TATGAGTGTGGAGTCGTTGCACAGGCTGAGATTCCTAGAGTCTNNNAGATCGGAA |
| 50364_1 | TATGAGTGTGGAGTCGTTGCACAGAAGATGCTTCCAACTCCATNNNAGATCGGAAG |
| 50364_2 | TATGAGTGTGGAGTCGTTGCACAGAAGATGCTTCCAACTCCATNNNAGATCGGAAG |
| 9025_1 | TATGAGTGTGGAGTCGTTGCAGGAAGCAATGCAGAATATGGCTNNNAGATCGGAA |
| 9025_2 | TATGAGTGTGGAGTCGTTGCAGGAAGCAATGCAGAATATGGCTNNNAGATCGGAA |
| 48257_1 | TATGAGTGTGGAGTCGTTGCCTGGGGTGAGATGATACAGCATTNNNAGATCGGAA |
| 48257_2 | TATGAGTGTGGAGTCGTTGCCTGGGGTGAGATGATACAGCATTNNNAGATCGGAA |
| 46400_1 | TATGAGTGTGGAGTCGTTGCCTCCTGAGTAGCTGGGACTACAGNNNAGATCGGAA |
| 46400_2 | TATGAGTGTGGAGTCGTTGCCTCCTGAGTAGCTGGGACTACAGNNNAGATCGGAA |
| 49887_1 | TATGAGTGTGGAGTCGTTGCGATAATGACGGTCAGTCGATGGANNNAGATCGGAA |
| 49887_2 | TATGAGTGTGGAGTCGTTGCGATAATGACGGTCAGTCGATGGANNNAGATCGGAA |

41272_1 TATGAGTGTGGAGTCGTTGCTAATCCCATCTACTCAGGAGGCTNNNAGATCGGAAG
41272_2 TATGAGTGTGGAGTCGTTGCTAATCCCATCTACTCAGGAGGCTNNNAGATCGGAAG
39195_1 TATGAGTGTGGAGTCGTTGCTTTCCATCTTCCCACTGACTCTCNNNAGATCGGAAGA
39195_2 TATGAGTGTGGAGTCGTTGCTTTCCATCTTCCCACTGACTCTCNNNAGATCGGAAGA
34167_1 TATGAGTGTGGAGTCGTTGCATTGCCAGGACAGTGAAGGTAATNNNAGATCGGAA
34167_2 TATGAGTGTGGAGTCGTTGCATTGCCAGGACAGTGAAGGTAATNNNAGATCGGAA
41826_1 TATGAGTGTGGAGTCGTTGCCAGGTTCACGCCATTCTCCTNNNAGATCGGAAGAGC
41826_2 TATGAGTGTGGAGTCGTTGCCAGGTTCACGCCATTCTCCTNNNAGATCGGAAGAGC
3025_1 TATGAGTGTGGAGTCGTTGCAGTTCTCTAGCATTCTGTCTTTCGTNNNAGATCGGAA
3025_2 TATGAGTGTGGAGTCGTTGCAGTTCTCTAGCATTCTGTCTTTCGTNNNAGATCGGAA
34424_1 TATGAGTGTGGAGTCGTTGCCATGGATGAAGCTGGAAACCATCNNNAGATCGGAA
34424_2 TATGAGTGTGGAGTCGTTGCCATGGATGAAGCTGGAAACCATCNNNAGATCGGAA
52428_1 TATGAGTGTGGAGTCGTTGCATCCCCTGCTTTATTGCCTTGTANNNAGATCGGAAGA
52428_2 TATGAGTGTGGAGTCGTTGCATCCCCTGCTTTATTGCCTTGTANNNAGATCGGAAGA
47725_1 TATGAGTGTGGAGTCGTTGCACCTTGAGGGCTTACACTTTATTNNNAGATCGGAAG
47725_2 TATGAGTGTGGAGTCGTTGCACCTTGAGGGCTTACACTTTATTNNNAGATCGGAAG
37176_1 TATGAGTGTGGAGTCGTTGCGTGCCAGAGAAGACTATAGGTACGNNNAGATCGGA
37176_2 TATGAGTGTGGAGTCGTTGCGTGCCAGAGAAGACTATAGGTACGNNNAGATCGGA
45540_1 TATGAGTGTGGAGTCGTTGCGACTTTGGAAGATGAGGGTGCTANNNAGATCGGAA
45540_2 TATGAGTGTGGAGTCGTTGCGACTTTGGAAGATGAGGGTGCTANNNAGATCGGAA
47509_1 TATGAGTGTGGAGTCGTTGCAGAAGAGCTACTGATTTGTGTACANNNAGATCGGA
47509_2 TATGAGTGTGGAGTCGTTGCAGAAGAGCTACTGATTTGTGTACANNNAGATCGGA
46867_1 TATGAGTGTGGAGTCGTTGCGCCAACGAATCCACTGATATGTGNNNAGATCGGAAG
46867_2 TATGAGTGTGGAGTCGTTGCGCCAACGAATCCACTGATATGTGNNNAGATCGGAAG
10402_1 TATGAGTGTGGAGTCGTTGCTTGATTTGAAACACACCTCAAGCTNNNAGATCGGAA
10402_2 TATGAGTGTGGAGTCGTTGCTTGATTTGAAACACACCTCAAGCTNNNAGATCGGAA
12892_1 TATGAGTGTGGAGTCGTTGCCTAAGGTATTCAGTCATCATGGNNNAGATCGGAAGA
12892_2 TATGAGTGTGGAGTCGTTGCCTAAGGTATTCAGTCATCATGGNNNAGATCGGAAGA
33419_1 TATGAGTGTGGAGTCGTTGCGAGAGGGAGTTTTGAGACAGAGGNNNAGATCGGA
33419_2 TATGAGTGTGGAGTCGTTGCGAGAGGGAGTTTTGAGACAGAGGNNNAGATCGGA
39637_1 TATGAGTGTGGAGTCGTTGCTAACCAAAAGAGAGCTGGAGTGGNNNAGATCGGA
39637_2 TATGAGTGTGGAGTCGTTGCTAACCAAAAGAGAGCTGGAGTGGNNNAGATCGGA
45676_1 TATGAGTGTGGAGTCGTTGCCAGACACTAGGATGGGATGGATGNNNAGATCGGA
45676_2 TATGAGTGTGGAGTCGTTGCCAGACACTAGGATGGGATGGATGNNNAGATCGGA
35380_1 TATGAGTGTGGAGTCGTTGCCATTCTCCTGCCTCAGCCTCNNNAGATCGGAAGAGCA
35380_2 TATGAGTGTGGAGTCGTTGCCATTCTCCTGCCTCAGCCTCNNNAGATCGGAAGAGCA
33659_1 TATGAGTGTGGAGTCGTTGCCAAGTAGCTGGGACTACAGGCNNNAGATCGGAAGA
33659_2 TATGAGTGTGGAGTCGTTGCCAAGTAGCTGGGACTACAGGCNNNAGATCGGAAGA
42002_1 TATGAGTGTGGAGTCGTTGCCCGGGTTCTAAGAGATTCTCCTGNNNAGATCGGAAG
42002_2 TATGAGTGTGGAGTCGTTGCCCGGGTTCTAAGAGATTCTCCTGNNNAGATCGGAAG
36119_1 TATGAGTGTGGAGTCGTTGCGAGAAAACACATTGGTGCACAGANNNAGATCGGAA
36119_2 TATGAGTGTGGAGTCGTTGCGAGAAAACACATTGGTGCACAGANNNAGATCGGAA
11501_1 TATGAGTGTGGAGTCGTTGCGCGTCTTCTCTGTCCAACTACTGNNNAGATCGGAAG

| | |
|---|---|
| 11501_2 | TATGAGTGTGGAGTCGTTGCGCGTCTTCTCTGTCCAACTACTGNNNAGATCGGAAGA |
| 32275_1 | TATGAGTGTGGAGTCGTTGCCCACCGACATTTGACTCTTTGTTNNNAGATCGGAAGA |
| 32275_2 | TATGAGTGTGGAGTCGTTGCCCACCGACATTTGACTCTTTGTTNNNAGATCGGAAGA |
| 37143_1 | TATGAGTGTGGAGTCGTTGCGCTTTTGCCTCTTTGCTGATCATNNNAGATCGGAAGA |
| 37143_2 | TATGAGTGTGGAGTCGTTGCGCTTTTGCCTCTTTGCTGATCATNNNAGATCGGAAGA |
| 45271_1 | TATGAGTGTGGAGTCGTTGCCTCTTATAGTGACTCAGCCAGCCNNNAGATCGGAAGA |
| 45271_2 | TATGAGTGTGGAGTCGTTGCCTCTTATAGTGACTCAGCCAGCCNNNAGATCGGAAGA |
| 29481_1 | TGAGTGTGGAGTCGTTGCTTCTTCACATTTGTAGGGTTTTTGTACNNNAGATCGGAA |
| 29481_2 | TGAGTGTGGAGTCGTTGCTTCTTCACATTTGTAGGGTTTTTGTACNNNAGATCGGAA |
| 39732_1 | TATGAGTGTGGAGTCGTTGCATTGCTGATGCTACCCTCAGTTTNNNAGATCGGAAGA |
| 39732_2 | TATGAGTGTGGAGTCGTTGCATTGCTGATGCTACCCTCAGTTTNNNAGATCGGAAGA |
| 37581_1 | TATGAGTGTGGAGTCGTTGCTCCCAGTCCATCAGACAAAGTTTNNNAGATCGGAAGA |
| 37581_2 | TATGAGTGTGGAGTCGTTGCTCCCAGTCCATCAGACAAAGTTTNNNAGATCGGAAGA |
| 29157_1 | TATGAGTGTGGAGTCGTTGCCACACAGAGTTCCTAGCACATCTNNNAGATCGGAAGA |
| 29157_2 | TATGAGTGTGGAGTCGTTGCCACACAGAGTTCCTAGCACATCTNNNAGATCGGAAGA |
| 44874_1 | TATGAGTGTGGAGTCGTTGCCTCTAGCCCAGACTTCTCCCTATNNNAGATCGGAAGA |
| 44874_2 | TATGAGTGTGGAGTCGTTGCCTCTAGCCCAGACTTCTCCCTATNNNAGATCGGAAGA |
| 28854_1 | TATGAGTGTGGAGTCGTTGCGCACACACTTTTTTACCGTAAGCNNNAGATCGGAAGA |
| 28854_2 | TATGAGTGTGGAGTCGTTGCGCACACACTTTTTTACCGTAAGCNNNAGATCGGAAGA |
| 47191_1 | TATGAGTGTGGAGTCGTTGCATCATTTCTGAGCAGTCAACCCTNNNAGATCGGAAGA |
| 47191_2 | TATGAGTGTGGAGTCGTTGCATCATTTCTGAGCAGTCAACCCTNNNAGATCGGAAGA |
| 30763_1 | TATGAGTGTGGAGTCGTTGCCTATTTGGCCCCTCAAACAACTGNNNAGATCGGAAGA |
| 30763_2 | TATGAGTGTGGAGTCGTTGCCTATTTGGCCCCTCAAACAACTGNNNAGATCGGAAGA |
| 3440_1 | TATGAGTGTGGAGTCGTTGCGATTCCCTGGTCTTACTCCCTGANNNAGATCGGAAGA |
| 3440_2 | TATGAGTGTGGAGTCGTTGCGATTCCCTGGTCTTACTCCCTGANNNAGATCGGAAGA |
| 43792_1 | TATGAGTGTGGAGTCGTTGCCTGTTGATTCAGACACCTGGAGANNNAGATCGGAAG |
| 43792_2 | TATGAGTGTGGAGTCGTTGCCTGTTGATTCAGACACCTGGAGANNNAGATCGGAAG |
| 51189_1 | TATGAGTGTGGAGTCGTTGCCTCCCTGGGTACGTGTTTTTTTCNNNAGATCGGAAGA |
| 51189_2 | TATGAGTGTGGAGTCGTTGCCTCCCTGGGTACGTGTTTTTTTCNNNAGATCGGAAGA |
| 37202_1 | TATGAGTGTGGAGTCGTTGCCACAGTGGATCACGCATGTAATCNNNAGATCGGAAG |
| 37202_2 | TATGAGTGTGGAGTCGTTGCCACAGTGGATCACGCATGTAATCNNNAGATCGGAAG |
| 34598_1 | TATGAGTGTGGAGTCGTTGCACACAGGATGTTGTGAAAAGCAANNNAGATCGGAAG |
| 34598_2 | TATGAGTGTGGAGTCGTTGCACACAGGATGTTGTGAAAAGCAANNNAGATCGGAAG |
| 47214_1 | TATGAGTGTGGAGTCGTTGCCTCATCCTCTCAAAGTGTTGGGANNNAGATCGGAAG |
| 47214_2 | TATGAGTGTGGAGTCGTTGCCTCATCCTCTCAAAGTGTTGGGANNNAGATCGGAAG |
| 48389_1 | TATGAGTGTGGAGTCGTTGCTGGAAGAACCTCTAAGCCTTTCTNNNAGATCGGAAGA |
| 48389_2 | TATGAGTGTGGAGTCGTTGCTGGAAGAACCTCTAAGCCTTTCTNNNAGATCGGAAGA |
| 43894_1 | TATGAGTGTGGAGTCGTTGCTGAAAGAAGAGTCTGCAGTTGGANNNAGATCGGAA |
| 43894_2 | TATGAGTGTGGAGTCGTTGCTGAAAGAAGAGTCTGCAGTTGGANNNAGATCGGAA |
| 30192_1 | TATGAGTGTGGAGTCGTTGCAGGTTGCAATGGTACCATACGATNNNAGATCGGAAG |
| 30192_2 | TATGAGTGTGGAGTCGTTGCAGGTTGCAATGGTACCATACGATNNNAGATCGGAAG |
| 48391_1 | TATGAGTGTGGAGTCGTTGCAAATGGAGTCTGAGGCTGACGATNNNAGATCGGAA |
| 48391_2 | TATGAGTGTGGAGTCGTTGCAAATGGAGTCTGAGGCTGACGATNNNAGATCGGAA |

| | |
|---|---|
| 35082_1 | TATGAGTGTGGAGTCGTTGCGGCCAGGGTGGTTTATTTCTAGANNNAGATCGGAA |
| 35082_2 | TATGAGTGTGGAGTCGTTGCGGCCAGGGTGGTTTATTTCTAGANNNAGATCGGAA |
| 50649_1 | TATGAGTGTGGAGTCGTTGCCCACACTCTTTGACTTGAGCAAGNNNAGATCGGAAG |
| 50649_2 | TATGAGTGTGGAGTCGTTGCCCACACTCTTTGACTTGAGCAAGNNNAGATCGGAAG |
| 46410_1 | TATGAGTGTGGAGTCGTTGCCTGAAGGGTCTGGGACATTTGTANNNAGATCGGAA |
| 46410_2 | TATGAGTGTGGAGTCGTTGCCTGAAGGGTCTGGGACATTTGTANNNAGATCGGAA |
| 47300_1 | TATGAGTGTGGAGTCGTTGCGTCTCCCTGGTTCAAGTGATTCTNNNAGATCGGAAG |
| 47300_2 | TATGAGTGTGGAGTCGTTGCGTCTCCCTGGTTCAAGTGATTCTNNNAGATCGGAAG |
| 29294_1 | TATGAGTGTGGAGTCGTTGCACGAGTGTGTAAGTATGTGTGCTNNNAGATCGGAA |
| 29294_2 | TATGAGTGTGGAGTCGTTGCACGAGTGTGTAAGTATGTGTGCTNNNAGATCGGAA |
| 43636_1 | TATGAGTGTGGAGTCGTTGCGTGGTGCAATCCTAGCTCAGTACNNNAGATCGGAAG |
| 43636_2 | TATGAGTGTGGAGTCGTTGCGTGGTGCAATCCTAGCTCAGTACNNNAGATCGGAAG |
| 42763_1 | TATGAGTGTGGAGTCGTTGCACACTGGTATTTGTGGGTTTTGGNNNAGATCGGAA |
| 42763_2 | TATGAGTGTGGAGTCGTTGCACACTGGTATTTGTGGGTTTTGGNNNAGATCGGAA |
| 40996_1 | TATGAGTGTGGAGTCGTTGCAATGGGGGTGGGACAAAATCTTANNNAGATCGGAA |
| 40996_2 | TATGAGTGTGGAGTCGTTGCAATGGGGGTGGGACAAAATCTTANNNAGATCGGAA |
| 29782_1 | TATGAGTGTGGAGTCGTTGCACCAAAAACCAAAATCACTCATCTNNNAGATCGGAA |
| 29782_2 | TATGAGTGTGGAGTCGTTGCACCAAAAACCAAAATCACTCATCTNNNAGATCGGAA |
| 2567_1 | TATGAGTGTGGAGTCGTTGCTCTATTCCCAGGCATCCTATGGGNNNAGATCGGAAG |
| 2567_2 | TATGAGTGTGGAGTCGTTGCTCTATTCCCAGGCATCCTATGGGNNNAGATCGGAAG |
| 45988_1 | TATGAGTGTGGAGTCGTTGCCTCAACTGGGATTACAGGTGTGANNNAGATCGGAA |
| 45988_2 | TATGAGTGTGGAGTCGTTGCCTCAACTGGGATTACAGGTGTGANNNAGATCGGAA |
| 40162_1 | TATGAGTGTGGAGTCGTTGCGAAACTTCTTCTGGGAGAAAGCGNNNAGATCGGAA |
| 40162_2 | TATGAGTGTGGAGTCGTTGCGAAACTTCTTCTGGGAGAAAGCGNNNAGATCGGAA |
| 52147_1 | TATGAGTGTGGAGTCGTTGCAAATTGCTGGGATTACAGGCATGNNNAGATCGGAA |
| 52147_2 | TATGAGTGTGGAGTCGTTGCAAATTGCTGGGATTACAGGCATGNNNAGATCGGAA |
| 40994_1 | TATGAGTGTGGAGTCGTTGCATTCAATAGGCCAGCCCAAAATGNNNAGATCGGAAG |
| 40994_2 | TATGAGTGTGGAGTCGTTGCATTCAATAGGCCAGCCCAAAATGNNNAGATCGGAAG |
| 50409_1 | TATGAGTGTGGAGTCGTTGCGAATGGAGATTGCCTCACACAACNNNAGATCGGAA |
| 50409_2 | TATGAGTGTGGAGTCGTTGCGAATGGAGATTGCCTCACACAACNNNAGATCGGAA |
| 30965_1 | TATGAGTGTGGAGTCGTTGCGCTTTCTCTTCATCTGGGTTCCTNNNAGATCGGAAGA |
| 30965_2 | TATGAGTGTGGAGTCGTTGCGCTTTCTCTTCATCTGGGTTCCTNNNAGATCGGAAGA |
| 46815_1 | TATGAGTGTGGAGTCGTTGCTTGGGAAGAATGCCACAGAAGTANNNAGATCGGAA |
| 46815_2 | TATGAGTGTGGAGTCGTTGCTTGGGAAGAATGCCACAGAAGTANNNAGATCGGAA |
| 33673_1 | TATGAGTGTGGAGTCGTTGCGAGATGGAGCCTTGCTTTGTCNNNAGATCGGAAGA |
| 33673_2 | TATGAGTGTGGAGTCGTTGCGAGATGGAGCCTTGCTTTGTCNNNAGATCGGAAGA |
| 40607_1 | TATGAGTGTGGAGTCGTTGCCGGTTGTAATCCCAGCACTTTGNNNAGATCGGAAGA |
| 40607_2 | TATGAGTGTGGAGTCGTTGCCGGTTGTAATCCCAGCACTTTGNNNAGATCGGAAGA |
| 47131_1 | TATGAGTGTGGAGTCGTTGCAACAAGTGCGAGGACAAGAAAAGNNNAGATCGGA |
| 47131_2 | TATGAGTGTGGAGTCGTTGCAACAAGTGCGAGGACAAGAAAAGNNNAGATCGGA |
| 2115_1 | TATGAGTGTGGAGTCGTTGCAGGAGACACAAATTTCTAGGGCANNNAGATCGGAA |
| 2115_2 | TATGAGTGTGGAGTCGTTGCAGGAGACACAAATTTCTAGGGCANNNAGATCGGAA |
| 36441_1 | TATGAGTGTGGAGTCGTTGCGGTACCAGAGTAAGAGAGGCAAGNNNAGATCGGA |

| | |
|---|---|
| 36441_2 | TATGAGTGTGGAGTCGTTGCGGTACCAGAGTAAGAGAGGCAAGNNNAGATCGGA |
| 39128_1 | TATGAGTGTGGAGTCGTTGCGGAGAGGAAGAAAGAGGAAGTGGNNNAGATCGG |
| 39128_2 | TATGAGTGTGGAGTCGTTGCGGAGAGGAAGAAAGAGGAAGTGGNNNAGATCGG |
| 5008_1 | TATGAGTGTGGAGTCGTTGCCTCTTTCTTGATTTGGCTCTGAGCNNNAGATCGGAAG |
| 5008_2 | TATGAGTGTGGAGTCGTTGCCTCTTTCTTGATTTGGCTCTGAGCNNNAGATCGGAAG |
| 51279_1 | TATGAGTGTGGAGTCGTTGCCAAAGAGCTTGTGTAGGGAGTCTNNNAGATCGGAA |
| 51279_2 | TATGAGTGTGGAGTCGTTGCCAAAGAGCTTGTGTAGGGAGTCTNNNAGATCGGAA |
| 49881_1 | TATGAGTGTGGAGTCGTTGCACCAACACCACATTATCTGCTCANNNAGATCGGAAG |
| 49881_2 | TATGAGTGTGGAGTCGTTGCACCAACACCACATTATCTGCTCANNNAGATCGGAAG |
| 36009_1 | TATGAGTGTGGAGTCGTTGCGGGTTGCTTTTTCCCCATTCATGNNNAGATCGGAAG |
| 36009_2 | TATGAGTGTGGAGTCGTTGCGGGTTGCTTTTTCCCCATTCATGNNNAGATCGGAAG |
| 38060_1 | TATGAGTGTGGAGTCGTTGCAGTGCACAATTACACCTGACAGANNNAGATCGGAAG |
| 38060_2 | TATGAGTGTGGAGTCGTTGCAGTGCACAATTACACCTGACAGANNNAGATCGGAAG |
| 48484_1 | TATGAGTGTGGAGTCGTTGCGGATCTCAGCTCACTACAACCTCNNNAGATCGGAAG |
| 48484_2 | TATGAGTGTGGAGTCGTTGCGGATCTCAGCTCACTACAACCTCNNNAGATCGGAAG |
| 32592_1 | TATGAGTGTGGAGTCGTTGCTCCTAGCTACTCTTAGGCAGGAGNNNAGATCGGAAG |
| 32592_2 | TATGAGTGTGGAGTCGTTGCTCCTAGCTACTCTTAGGCAGGAGNNNAGATCGGAAG |
| 35397_1 | TATGAGTGTGGAGTCGTTGCCCTGAGTAGCTAGGACTACAGGTNNNAGATCGGAA |
| 35397_2 | TATGAGTGTGGAGTCGTTGCCCTGAGTAGCTAGGACTACAGGTNNNAGATCGGAA |
| 30958_1 | TATGAGTGTGGAGTCGTTGCCTGCTCCCCTTTCTTTTCCTTTGNNNAGATCGGAAGA |
| 30958_2 | TATGAGTGTGGAGTCGTTGCCTGCTCCCCTTTCTTTTCCTTTGNNNAGATCGGAAGA |
| 29127_1 | TATGAGTGTGGAGTCGTTGCCACCATTCCAAACCAGTGCATTANNNAGATCGGAAG |
| 29127_2 | TATGAGTGTGGAGTCGTTGCCACCATTCCAAACCAGTGCATTANNNAGATCGGAAG |
| 42025_1 | TATGAGTGTGGAGTCGTTGCAGATGACCAGTCTTGGAATTGCTNNNAGATCGGAA |
| 42025_2 | TATGAGTGTGGAGTCGTTGCAGATGACCAGTCTTGGAATTGCTNNNAGATCGGAA |
| 45451_1 | TATGAGTGTGGAGTCGTTGCGTGTCATGAAGCGTTCTAGATGCNNNAGATCGGAA |
| 45451_2 | TATGAGTGTGGAGTCGTTGCGTGTCATGAAGCGTTCTAGATGCNNNAGATCGGAA |
| 47205_1 | TATGAGTGTGGAGTCGTTGCACATTCAGGAGATTCTTGTGCCTNNNAGATCGGAAG |
| 47205_2 | TATGAGTGTGGAGTCGTTGCACATTCAGGAGATTCTTGTGCCTNNNAGATCGGAAG |
| 32318_1 | TATGAGTGTGGAGTCGTTGCTGCTGTAATGAACATGGGAATGCNNNAGATCGGAA |
| 32318_2 | TATGAGTGTGGAGTCGTTGCTGCTGTAATGAACATGGGAATGCNNNAGATCGGAA |
| 52425_1 | TATGAGTGTGGAGTCGTTGCGTCATCAACAGGATTCCAGAGGGNNNAGATCGGAA |
| 52425_2 | TATGAGTGTGGAGTCGTTGCGTCATCAACAGGATTCCAGAGGGNNNAGATCGGAA |
| 5532_1 | TATGAGTGTGGAGTCGTTGCCTCTTGTGCATAAATTGCCCCACNNNAGATCGGAAG |
| 5532_2 | TATGAGTGTGGAGTCGTTGCCTCTTGTGCATAAATTGCCCCACNNNAGATCGGAAG |
| 40289_1 | TATGAGTGTGGAGTCGTTGCCCCTCTCCTGGATGACCTATACANNNAGATCGGAAG |
| 40289_2 | TATGAGTGTGGAGTCGTTGCCCCTCTCCTGGATGACCTATACANNNAGATCGGAAG |
| 47308_1 | TATGAGTGTGGAGTCGTTGCGATCTCGGCTCACTACAAGCTCNNNAGATCGGAAGA |
| 47308_2 | TATGAGTGTGGAGTCGTTGCGATCTCGGCTCACTACAAGCTCNNNAGATCGGAAGA |
| 10816_1 | TATGAGTGTGGAGTCGTTGCTTGAACGCATCCTACTCATTCCCNNNAGATCGGAAGA |
| 10816_2 | TATGAGTGTGGAGTCGTTGCTTGAACGCATCCTACTCATTCCCNNNAGATCGGAAGA |
| 46573_1 | TATGAGTGTGGAGTCGTTGCTTCAAGCTATTCTCCTGCCTCAGNNNAGATCGGAAGA |
| 46573_2 | TATGAGTGTGGAGTCGTTGCTTCAAGCTATTCTCCTGCCTCAGNNNAGATCGGAAGA |

| | |
|---|---|
| 45017_1 | TATGAGTGTGGAGTCGTTGCGTAGCTATCCAGGGGTTATTGCANNNAGATCGGAA |
| 45017_2 | TATGAGTGTGGAGTCGTTGCGTAGCTATCCAGGGGTTATTGCANNNAGATCGGAA |
| 9475_1 | TATGAGTGTGGAGTCGTTGCGATTCTCACCTGACTGAAGGCAGNNNAGATCGGAAG |
| 9475_2 | TATGAGTGTGGAGTCGTTGCGATTCTCACCTGACTGAAGGCAGNNNAGATCGGAAG |
| 12846_1 | TATGAGTGTGGAGTCGTTGCGTGGATTCACAAACACAATCNNNAGATCGGAAGAG |
| 12846_2 | TATGAGTGTGGAGTCGTTGCGTGGATTCACAAACACAATCNNNAGATCGGAAGAG |
| 41406_1 | TATGAGTGTGGAGTCGTTGCATTAGCATGTTCCCAGCCACAATNNNAGATCGGAAG |
| 41406_2 | TATGAGTGTGGAGTCGTTGCATTAGCATGTTCCCAGCCACAATNNNAGATCGGAAG |
| 36558_1 | TATGAGTGTGGAGTCGTTGCGGGGAGGGCTCTAGAAATTTACCNNNAGATCGGAA |
| 36558_2 | TATGAGTGTGGAGTCGTTGCGGGGAGGGCTCTAGAAATTTACCNNNAGATCGGAA |
| 30722_1 | TATGAGTGTGGAGTCGTTGCTTGTTGTTAGAATTGGGAAAACGTNNNAGATCGGA |
| 30722_2 | TATGAGTGTGGAGTCGTTGCTTGTTGTTAGAATTGGGAAAACGTNNNAGATCGGA |
| 42062_1 | TATGAGTGTGGAGTCGTTGCGCACACACTTGATGTCAAACACTNNNAGATCGGAAG |
| 42062_2 | TATGAGTGTGGAGTCGTTGCGCACACACTTGATGTCAAACACTNNNAGATCGGAAG |
| 39433_1 | TATGAGTGTGGAGTCGTTGCATCATCAGTACACAGAAGCCCAGNNNAGATCGGAAG |
| 39433_2 | TATGAGTGTGGAGTCGTTGCATCATCAGTACACAGAAGCCCAGNNNAGATCGGAAG |
| 43925_1 | TATGAGTGTGGAGTCGTTGCCAGGGTCTCACTATGTTACCCAGNNNAGATCGGAAG |
| 43925_2 | TATGAGTGTGGAGTCGTTGCCAGGGTCTCACTATGTTACCCAGNNNAGATCGGAAG |
| 10493_1 | TATGAGTGTGGAGTCGTTGCGATGAGGATAGCTGCTGTGCTTTNNNAGATCGGAA |
| 10493_2 | TATGAGTGTGGAGTCGTTGCGATGAGGATAGCTGCTGTGCTTTNNNAGATCGGAA |
| 52358_1 | TATGAGTGTGGAGTCGTTGCATGTTTGTCAAAAGCATCCGTGTNNNAGATCGGAAG |
| 52358_2 | TATGAGTGTGGAGTCGTTGCATGTTTGTCAAAAGCATCCGTGTNNNAGATCGGAAG |
| 30432_1 | TATGAGTGTGGAGTCGTTGCGCAAAGACTGTTGGTAAGGGTATGNNNAGATCGGA |
| 30432_2 | TATGAGTGTGGAGTCGTTGCGCAAAGACTGTTGGTAAGGGTATGNNNAGATCGGA |
| 29471_1 | TATGAGTGTGGAGTCGTTGCTTGCGGATATTAACCTCTACAGGNNNAGATCGGAAG |
| 29471_2 | TATGAGTGTGGAGTCGTTGCTTGCGGATATTAACCTCTACAGGNNNAGATCGGAAG |
| 46147_1 | TATGAGTGTGGAGTCGTTGCCATGATCTCAGCTCACTGCAATGNNNAGATCGGAAG |
| 46147_2 | TATGAGTGTGGAGTCGTTGCCATGATCTCAGCTCACTGCAATGNNNAGATCGGAAG |
| 47690_1 | TATGAGTGTGGAGTCGTTGCTCTCCTGAGTAGCTGGGATTACANNNAGATCGGAAG |
| 47690_2 | TATGAGTGTGGAGTCGTTGCTCTCCTGAGTAGCTGGGATTACANNNAGATCGGAAG |
| 49480_1 | TATGAGTGTGGAGTCGTTGCGGGTCAGGTTTTAGAGGGGAAAANNNAGATCGGA |
| 49480_2 | TATGAGTGTGGAGTCGTTGCGGGTCAGGTTTTAGAGGGGAAAANNNAGATCGGA |
| 47796_1 | TATGAGTGTGGAGTCGTTGCCAAGAGCGCTAACAGAGGGACNNNAGATCGGAAGA |
| 47796_2 | TATGAGTGTGGAGTCGTTGCCAAGAGCGCTAACAGAGGGACNNNAGATCGGAAGA |
| 30142_1 | TATGAGTGTGGAGTCGTTGCTCTTTCCATGATTTCCATAAATGCANNNAGATCGGAA |
| 30142_2 | TATGAGTGTGGAGTCGTTGCTCTTTCCATGATTTCCATAAATGCANNNAGATCGGAA |
| 29522_1 | TATGAGTGTGGAGTCGTTGCTTTCCAGGACCCTACCAAACAGANNNAGATCGGAAG |
| 29522_2 | TATGAGTGTGGAGTCGTTGCTTTCCAGGACCCTACCAAACAGANNNAGATCGGAAG |
| 45284_1 | TATGAGTGTGGAGTCGTTGCAAACGAGTCTAGGGTTTGACCAANNNAGATCGGAA |
| 45284_2 | TATGAGTGTGGAGTCGTTGCAAACGAGTCTAGGGTTTGACCAANNNAGATCGGAA |
| 51483_1 | TATGAGTGTGGAGTCGTTGCGAGGGAGCAGAAAAAGAAGAGGANNNAGATCGGA |
| 51483_2 | TATGAGTGTGGAGTCGTTGCGAGGGAGCAGAAAAAGAAGAGGANNNAGATCGGA |
| 40408_1 | TATGAGTGTGGAGTCGTTGCGAACCGGCAAAGAAAACTGAGAANNNAGATCGGAA |

| | |
|---|---|
| 40408_2 | TATGAGTGTGGAGTCGTTGCGAACCGGCAAAGAAAACTGAGAANNNAGATCGGAA |
| 30708_1 | TATGAGTGTGGAGTCGTTGCCTCCTCCCACACTAATCACCNNNAGATCGGAAGAGCA |
| 30708_2 | TATGAGTGTGGAGTCGTTGCCTCCTCCCACACTAATCACCNNNAGATCGGAAGAGCA |
| 51492_1 | TATGAGTGTGGAGTCGTTGCCTGAATTCCTGTGCTGCGACTNNNAGATCGGAAGAG |
| 51492_2 | TATGAGTGTGGAGTCGTTGCCTGAATTCCTGTGCTGCGACTNNNAGATCGGAAGAG |
| 51020_1 | TATGAGTGTGGAGTCGTTGCTCTGGACAAGGAAGATGATGAGGNNNAGATCGGA |
| 51020_2 | TATGAGTGTGGAGTCGTTGCTCTGGACAAGGAAGATGATGAGGNNNAGATCGGA |
| 34816_1 | TATGAGTGTGGAGTCGTTGCGATGGACTTACTCTATCGCCCAGNNNAGATCGGAAG |
| 34816_2 | TATGAGTGTGGAGTCGTTGCGATGGACTTACTCTATCGCCCAGNNNAGATCGGAAG |
| 42687_1 | TATGAGTGTGGAGTCGTTGCAGGAGTACAGCTAACCAAGAAGGNNNAGATCGGAA |
| 42687_2 | TATGAGTGTGGAGTCGTTGCAGGAGTACAGCTAACCAAGAAGGNNNAGATCGGAA |
| 48321_1 | TATGAGTGTGGAGTCGTTGCGTATCTAGAACAGTGTCTGGCGTNNNAGATCGGAA |
| 48321_2 | TATGAGTGTGGAGTCGTTGCGTATCTAGAACAGTGTCTGGCGTNNNAGATCGGAA |
| 5213_1 | TATGAGTGTGGAGTCGTTGCTCAGCATATTGCCAGAAAAGGCTNNNAGATCGGAA |
| 5213_2 | TATGAGTGTGGAGTCGTTGCTCAGCATATTGCCAGAAAAGGCTNNNAGATCGGAA |
| 47192_1 | TATGAGTGTGGAGTCGTTGCTTCAAGCAATTCTCCTGCCTCAGNNNAGATCGGAAG |
| 47192_2 | TATGAGTGTGGAGTCGTTGCTTCAAGCAATTCTCCTGCCTCAGNNNAGATCGGAAG |
| 36045_1 | TATGAGTGTGGAGTCGTTGCTAATCCCATCTACTCAGGAGGCTNNNAGATCGGAAG |
| 36045_2 | TATGAGTGTGGAGTCGTTGCTAATCCCATCTACTCAGGAGGCTNNNAGATCGGAAG |
| 46766_1 | TATGAGTGTGGAGTCGTTGCCTACGCCCAGCTAATTTGTGTTTNNNAGATCGGAAG |
| 46766_2 | TATGAGTGTGGAGTCGTTGCCTACGCCCAGCTAATTTGTGTTTNNNAGATCGGAAG |
| 42205_1 | TATGAGTGTGGAGTCGTTGCGAAAATACCTGCAACGTCACCATNNNAGATCGGAAG |
| 42205_2 | TATGAGTGTGGAGTCGTTGCGAAAATACCTGCAACGTCACCATNNNAGATCGGAAG |
| 45716_1 | TATGAGTGTGGAGTCGTTGCTCAGGTCCTAGTCATGTGTACCTNNNAGATCGGAAG |
| 45716_2 | TATGAGTGTGGAGTCGTTGCTCAGGTCCTAGTCATGTGTACCTNNNAGATCGGAAG |
| 36894_1 | TATGAGTGTGGAGTCGTTGCAGCCCTGATGAAGATGTACAAGGNNNAGATCGGAA |
| 36894_2 | TATGAGTGTGGAGTCGTTGCAGCCCTGATGAAGATGTACAAGGNNNAGATCGGAA |
| 33943_1 | TATGAGTGTGGAGTCGTTGCAATCTCGGCTCACTGTAAGCTCNNNAGATCGGAAGA |
| 33943_2 | TATGAGTGTGGAGTCGTTGCAATCTCGGCTCACTGTAAGCTCNNNAGATCGGAAGA |
| 34613_1 | TATGAGTGTGGAGTCGTTGCTGTGCACCAAGGAAATGGATAGANNNAGATCGGAA |
| 34613_2 | TATGAGTGTGGAGTCGTTGCTGTGCACCAAGGAAATGGATAGANNNAGATCGGAA |
| 29864_1 | TATGAGTGTGGAGTCGTTGCGCCACCAAGGATATCACCATCTGNNNAGATCGGAAG |
| 29864_2 | TATGAGTGTGGAGTCGTTGCGCCACCAAGGATATCACCATCTGNNNAGATCGGAAG |
| 39431_1 | TATGAGTGTGGAGTCGTTGCAAGGTCCCAAAATAGCCAAAACGNNNAGATCGGAA |
| 39431_2 | TATGAGTGTGGAGTCGTTGCAAGGTCCCAAAATAGCCAAAACGNNNAGATCGGAA |
| 10854_1 | TATGAGTGTGGAGTCGTTGCATGAGCACAGGTGTTGTTGACAANNNAGATCGGAA |
| 10854_2 | TATGAGTGTGGAGTCGTTGCATGAGCACAGGTGTTGTTGACAANNNAGATCGGAA |
| 35700_1 | TATGAGTGTGGAGTCGTTGCGTACTAGGGTTACAGGTGTGAGCNNNAGATCGGAA |
| 35700_2 | TATGAGTGTGGAGTCGTTGCGTACTAGGGTTACAGGTGTGAGCNNNAGATCGGAA |
| 37279_1 | TATGAGTGTGGAGTCGTTGCCGTATGGAAGGGGTTTTTCACATNNNAGATCGGAA |
| 37279_2 | TATGAGTGTGGAGTCGTTGCCGTATGGAAGGGGTTTTTCACATNNNAGATCGGAA |
| 45425_1 | TATGAGTGTGGAGTCGTTGCAGCACATCTACTTTTGGGGATGTNNNAGATCGGAAG |
| 45425_2 | TATGAGTGTGGAGTCGTTGCAGCACATCTACTTTTGGGGATGTNNNAGATCGGAAG |

| | |
|---|---|
| 8188_1 | TATGAGTGTGGAGTCGTTGCTGCTCAATGGTGCGATCATAGCTNNNAGATCGGAAG |
| 8188_2 | TATGAGTGTGGAGTCGTTGCTGCTCAATGGTGCGATCATAGCTNNNAGATCGGAAG |
| 29735_1 | TATGAGTGTGGAGTCGTTGCGGTGAGTAACTGTGGGTGGTTTNNNAGATCGGAAG |
| 29735_2 | TATGAGTGTGGAGTCGTTGCGGTGAGTAACTGTGGGTGGTTTNNNAGATCGGAAG |
| 45606_1 | TATGAGTGTGGAGTCGTTGCCAGTGGTGTGATCTAGGCTTACTNNNAGATCGGAAG |
| 45606_2 | TATGAGTGTGGAGTCGTTGCCAGTGGTGTGATCTAGGCTTACTNNNAGATCGGAAG |
| 10988_1 | TATGAGTGTGGAGTCGTTGCTAAAGCCTCTGGGAGGTAATGCGNNNAGATCGGAA |
| 10988_2 | TATGAGTGTGGAGTCGTTGCTAAAGCCTCTGGGAGGTAATGCGNNNAGATCGGAA |
| 38770_1 | TATGAGTGTGGAGTCGTTGCATTGAATGTCCATTATGTGCCCGNNNAGATCGGAAG |
| 38770_2 | TATGAGTGTGGAGTCGTTGCATTGAATGTCCATTATGTGCCCGNNNAGATCGGAAG |
| 35837_1 | TATGAGTGTGGAGTCGTTGCCAATAGCAGAATCTCGGCTCACTNNNAGATCGGAAG |
| 35837_2 | TATGAGTGTGGAGTCGTTGCCAATAGCAGAATCTCGGCTCACTNNNAGATCGGAAG |
| 30349_1 | TATGAGTGTGGAGTCGTTGCTGTTCAGCCTGTCCCATTAAGGTNNNAGATCGGAAG |
| 30349_2 | TATGAGTGTGGAGTCGTTGCTGTTCAGCCTGTCCCATTAAGGTNNNAGATCGGAAG |
| 35371_1 | TATGAGTGTGGAGTCGTTGCGCTGGAGTACAATGGTGTGATCTNNNAGATCGGAA |
| 35371_2 | TATGAGTGTGGAGTCGTTGCGCTGGAGTACAATGGTGTGATCTNNNAGATCGGAA |
| 29858_1 | TATGAGTGTGGAGTCGTTGCGGATAAAAATACTTCCAAGGGTTCCANNNAGATCGG |
| 29858_2 | TATGAGTGTGGAGTCGTTGCGGATAAAAATACTTCCAAGGGTTCCANNNAGATCGG |
| 52230_1 | TATGAGTGTGGAGTCGTTGCCTGTGTCCATGTGCTTTCATTGTNNNAGATCGGAAG |
| 52230_2 | TATGAGTGTGGAGTCGTTGCCTGTGTCCATGTGCTTTCATTGTNNNAGATCGGAAG |
| 47701_1 | TATGAGTGTGGAGTCGTTGCCTGTAGCTTTCAACTCCTGGGATNNNAGATCGGAAG |
| 47701_2 | TATGAGTGTGGAGTCGTTGCCTGTAGCTTTCAACTCCTGGGATNNNAGATCGGAAG |
| 32336_1 | TATGAGTGTGGAGTCGTTGCCAATCTCCACCTCCTGAGTTCAANNNAGATCGGAAGA |
| 32336_2 | TATGAGTGTGGAGTCGTTGCCAATCTCCACCTCCTGAGTTCAANNNAGATCGGAAGA |
| 9588_1 | TATGAGTGTGGAGTCGTTGCTAATCCCATCATGAAGCTTGGGCNNNAGATCGGAAG |
| 9588_2 | TATGAGTGTGGAGTCGTTGCTAATCCCATCATGAAGCTTGGGCNNNAGATCGGAAG |
| 43491_1 | TATGAGTGTGGAGTCGTTGCAGTCTCCTGAGTAGTTGGGACTANNNAGATCGGAA |
| 43491_2 | TATGAGTGTGGAGTCGTTGCAGTCTCCTGAGTAGTTGGGACTANNNAGATCGGAA |
| 49305_1 | TATGAGTGTGGAGTCGTTGCTGGAGTGCAGTGGTGTGATCNNNAGATCGGAAGA |
| 49305_2 | TATGAGTGTGGAGTCGTTGCTGGAGTGCAGTGGTGTGATCNNNAGATCGGAAGA |
| 41062_1 | TATGAGTGTGGAGTCGTTGCCCCCATCTCTCCACTTTGCTATTNNNAGATCGGAAGA |
| 41062_2 | TATGAGTGTGGAGTCGTTGCCCCCATCTCTCCACTTTGCTATTNNNAGATCGGAAGA |
| 32392_1 | TATGAGTGTGGAGTCGTTGCTGGAGTGCAGTGGTGTTATCTCNNNAGATCGGAAG |
| 32392_2 | TATGAGTGTGGAGTCGTTGCTGGAGTGCAGTGGTGTTATCTCNNNAGATCGGAAG |
| 32740_1 | TATGAGTGTGGAGTCGTTGCTAAACTCCTATAGGCTTCCCTGCNNNAGATCGGAAG |
| 32740_2 | TATGAGTGTGGAGTCGTTGCTAAACTCCTATAGGCTTCCCTGCNNNAGATCGGAAG |
| 40466_1 | TATGAGTGTGGAGTCGTTGCGCATAGGATGTGTTTGGAGCATGNNNAGATCGGAA |
| 40466_2 | TATGAGTGTGGAGTCGTTGCGCATAGGATGTGTTTGGAGCATGNNNAGATCGGAA |
| 37212_1 | TATGAGTGTGGAGTCGTTGCCTGTAGTCCCAGCTACTCAGGAGNNNAGATCGGAAG |
| 37212_2 | TATGAGTGTGGAGTCGTTGCCTGTAGTCCCAGCTACTCAGGAGNNNAGATCGGAAG |
| 32851_1 | TATGAGTGTGGAGTCGTTGCCAAGTAGCTGGGACTACAGGCNNNAGATCGGAAGA |
| 32851_2 | TATGAGTGTGGAGTCGTTGCCAAGTAGCTGGGACTACAGGCNNNAGATCGGAAGA |
| 36404_1 | TATGAGTGTGGAGTCGTTGCGTTTTGAGTGGGACCAACTTGAGNNNAGATCGGAA |

| | |
|---|---|
| 36404_2 | TATGAGTGTGGAGTCGTTGCGTTTTGAGTGGGACCAACTTGAGNNNAGATCGGAA |
| 10051_1 | TATGAGTGTGGAGTCGTTGCGATATTACCCTGATCCCCTCGCGNNNAGATCGGAAG |
| 10051_2 | TATGAGTGTGGAGTCGTTGCGATATTACCCTGATCCCCTCGCGNNNAGATCGGAAG |
| 38902_1 | TATGAGTGTGGAGTCGTTGCATGGGGAAAAGCGTTCCTATTCANNNAGATCGGAA |
| 38902_2 | TATGAGTGTGGAGTCGTTGCATGGGGAAAAGCGTTCCTATTCANNNAGATCGGAA |
| 51722_1 | TATGAGTGTGGAGTCGTTGCACCTCTGTTGTGATTACCTTGGGNNNAGATCGGAAG |
| 51722_2 | TATGAGTGTGGAGTCGTTGCACCTCTGTTGTGATTACCTTGGGNNNAGATCGGAAG |
| 6415_1 | TATGAGTGTGGAGTCGTTGCGTGAGGTGATGAAGACTGGTGTCNNNAGATCGGAA |
| 6415_2 | TATGAGTGTGGAGTCGTTGCGTGAGGTGATGAAGACTGGTGTCNNNAGATCGGAA |
| 44092_1 | TATGAGTGTGGAGTCGTTGCCCTGCTGCTGAGTTTCCTAATTGNNNAGATCGGAAG |
| 44092_2 | TATGAGTGTGGAGTCGTTGCCCTGCTGCTGAGTTTCCTAATTGNNNAGATCGGAAG |
| 41032_1 | TATGAGTGTGGAGTCGTTGCCTGAGGCAGGAGAATCACTTGAANNNAGATCGGAA |
| 41032_2 | TATGAGTGTGGAGTCGTTGCCTGAGGCAGGAGAATCACTTGAANNNAGATCGGAA |
| 49484_1 | TATGAGTGTGGAGTCGTTGCCAGTCTGGACTTGAACTCCTGACNNNAGATCGGAAG |
| 49484_2 | TATGAGTGTGGAGTCGTTGCCAGTCTGGACTTGAACTCCTGACNNNAGATCGGAAG |
| 47958_1 | TATGAGTGTGGAGTCGTTGCTTTTTTGAGACGGAGTCTGGAGTNNNAGATCGGAA |
| 47958_2 | TATGAGTGTGGAGTCGTTGCTTTTTTGAGACGGAGTCTGGAGTNNNAGATCGGAA |
| 10608_1 | TATGAGTGTGGAGTCGTTGCTGTGTAGGAGGATGGGTAGCTTTNNNAGATCGGAA |
| 10608_2 | TATGAGTGTGGAGTCGTTGCTGTGTAGGAGGATGGGTAGCTTTNNNAGATCGGAA |
| 2916_1 | TATGAGTGTGGAGTCGTTGCTCGGCCTTGTGCATTGTAGAATTNNNAGATCGGAAG |
| 2916_2 | TATGAGTGTGGAGTCGTTGCTCGGCCTTGTGCATTGTAGAATTNNNAGATCGGAAG |
| 44960_1 | TATGAGTGTGGAGTCGTTGCAGACTGACTAGGGCATTCTTTGGNNNAGATCGGAA |
| 44960_2 | TATGAGTGTGGAGTCGTTGCAGACTGACTAGGGCATTCTTTGGNNNAGATCGGAA |
| 36064_1 | TATGAGTGTGGAGTCGTTGCCTGAGGTCAGGAGTTTGAGACCNNNAGATCGGAAG |
| 36064_2 | TATGAGTGTGGAGTCGTTGCCTGAGGTCAGGAGTTTGAGACCNNNAGATCGGAAG |
| 9287_1 | TATGAGTGTGGAGTCGTTGCACCGTGTAGTGAACAATTGCATGNNNAGATCGGAA |
| 9287_2 | TATGAGTGTGGAGTCGTTGCACCGTGTAGTGAACAATTGCATGNNNAGATCGGAA |
| 46379_1 | TATGAGTGTGGAGTCGTTGCATCAAGCTGTTTTGGTTCCTGTGNNNAGATCGGAAG |
| 46379_2 | TATGAGTGTGGAGTCGTTGCATCAAGCTGTTTTGGTTCCTGTGNNNAGATCGGAAG |
| 44792_1 | TATGAGTGTGGAGTCGTTGCGAGTTGGTATGCTGAAGGGAAGANNNAGATCGGA |
| 44792_2 | TATGAGTGTGGAGTCGTTGCGAGTTGGTATGCTGAAGGGAAGANNNAGATCGGA |
| 10229_1 | TATGAGTGTGGAGTCGTTGCAGGTTCTCAAATGCAGGACTATTNNNAGATCGGAAG |
| 10229_2 | TATGAGTGTGGAGTCGTTGCAGGTTCTCAAATGCAGGACTATTNNNAGATCGGAAG |
| 44263_1 | TATGAGTGTGGAGTCGTTGCTCTGTGATCCATGGCTTTGAAGANNNAGATCGGAAG |
| 44263_2 | TATGAGTGTGGAGTCGTTGCTCTGTGATCCATGGCTTTGAAGANNNAGATCGGAAG |
| 50862_1 | TATGAGTGTGGAGTCGTTGCCCTAGGTCGAAGGTTCTTTTTGCNNNAGATCGGAAG |
| 50862_2 | TATGAGTGTGGAGTCGTTGCCCTAGGTCGAAGGTTCTTTTTGCNNNAGATCGGAAG |
| 33237_1 | TATGAGTGTGGAGTCGTTGCATAGGGTCTTACTCAGTCACCCANNNAGATCGGAAG |
| 33237_2 | TATGAGTGTGGAGTCGTTGCATAGGGTCTTACTCAGTCACCCANNNAGATCGGAAG |
| 37798_1 | TATGAGTGTGGAGTCGTTGCGGTGAAACCCCGTCTCTACTAAANNNAGATCGGAAG |
| 37798_2 | TATGAGTGTGGAGTCGTTGCGGTGAAACCCCGTCTCTACTAAANNNAGATCGGAAG |
| 48690_1 | TATGAGTGTGGAGTCGTTGCCTTTCCTTCCCTCATTCCCTCTCNNNAGATCGGAAGAG |
| 48690_2 | TATGAGTGTGGAGTCGTTGCCTTTCCTTCCCTCATTCCCTCTCNNNAGATCGGAAGAG |

| | |
|---|---|
| 43879_1 | TATGAGTGTGGAGTCGTTGCGGGACTGGGTGGAGCTTATAGAGNNNAGATCGGA |
| 43879_2 | TATGAGTGTGGAGTCGTTGCGGGACTGGGTGGAGCTTATAGAGNNNAGATCGGA |
| 42998_1 | TATGAGTGTGGAGTCGTTGCAGCAGACGTTCTTCTTCCCTTTCNNNAGATCGGAAGA |
| 42998_2 | TATGAGTGTGGAGTCGTTGCAGCAGACGTTCTTCTTCCCTTTCNNNAGATCGGAAGA |
| 32735_1 | TATGAGTGTGGAGTCGTTGCATATGCACACAATAACATGCCCTNNNAGATCGGAAG |
| 32735_2 | TATGAGTGTGGAGTCGTTGCATATGCACACAATAACATGCCCTNNNAGATCGGAAG |
| 46774_1 | TATGAGTGTGGAGTCGTTGCCTCCCCAGTAGCTGAGACTACAGNNNAGATCGGAAG |
| 46774_2 | TATGAGTGTGGAGTCGTTGCCTCCCCAGTAGCTGAGACTACAGNNNAGATCGGAAG |
| 46707_1 | TATGAGTGTGGAGTCGTTGCATCTCAGCTCACTGCAGCCTTTANNNAGATCGGAAG |
| 46707_2 | TATGAGTGTGGAGTCGTTGCATCTCAGCTCACTGCAGCCTTTANNNAGATCGGAAG |
| 41284_1 | TATGAGTGTGGAGTCGTTGCCCCGAAACATTTCACATCAGCAANNNAGATCGGAAG |
| 41284_2 | TATGAGTGTGGAGTCGTTGCCCCGAAACATTTCACATCAGCAANNNAGATCGGAAG |
| 43565_1 | TATGAGTGTGGAGTCGTTGCACACTTCTGGTCTCAAAGCTTGANNNAGATCGGAAG |
| 43565_2 | TATGAGTGTGGAGTCGTTGCACACTTCTGGTCTCAAAGCTTGANNNAGATCGGAAG |
| 32304_1 | TATGAGTGTGGAGTCGTTGCCACTGAAAGCGGAGTTTAGGATGNNNAGATCGGAA |
| 32304_2 | TATGAGTGTGGAGTCGTTGCCACTGAAAGCGGAGTTTAGGATGNNNAGATCGGAA |
| 39331_1 | TATGAGTGTGGAGTCGTTGCGACATAAATAACAGATGCCGGCCNNNAGATCGGAA |
| 39331_2 | TATGAGTGTGGAGTCGTTGCGACATAAATAACAGATGCCGGCCNNNAGATCGGAA |
| 39332_1 | TATGAGTGTGGAGTCGTTGCTCAGTAAACACTACTCTTGGGGCNNNAGATCGGAAG |
| 39332_2 | TATGAGTGTGGAGTCGTTGCTCAGTAAACACTACTCTTGGGGCNNNAGATCGGAAG |
| 3295_1 | TATGAGTGTGGAGTCGTTGCAAGGCCAATAGTAACTCCTGGTGNNNAGATCGGAA |
| 3295_2 | TATGAGTGTGGAGTCGTTGCAAGGCCAATAGTAACTCCTGGTGNNNAGATCGGAA |
| 30140_1 | TATGAGTGTGGAGTCGTTGCAGTTCCTTATATCCCAGACCTGGNNNAGATCGGAAG |
| 30140_2 | TATGAGTGTGGAGTCGTTGCAGTTCCTTATATCCCAGACCTGGNNNAGATCGGAAG |
| 42537_1 | TATGAGTGTGGAGTCGTTGCGTGTGTGTGTGTGTTTTTCTGGANNNAGATCGGAA |
| 42537_2 | TATGAGTGTGGAGTCGTTGCGTGTGTGTGTGTGTTTTTCTGGANNNAGATCGGAA |
| 13741_1 | TATGAGTGTGGAGTCGTTGCAGCTGTAGTGTAATCCAATTATTNNNAGATCGGAAG |
| 13741_2 | TATGAGTGTGGAGTCGTTGCAGCTGTAGTGTAATCCAATTATTNNNAGATCGGAAG |
| 44473_1 | TATGAGTGTGGAGTCGTTGCTGTGAAGGGTTGGATGGCTTATTNNNAGATCGGAA |
| 44473_2 | TATGAGTGTGGAGTCGTTGCTGTGAAGGGTTGGATGGCTTATTNNNAGATCGGAA |
| 12761_1 | TATGAGTGTGGAGTCGTTGCGATGATGGCTTCTTCCTAGGNNNAGATCGGAAGAG |
| 12761_2 | TATGAGTGTGGAGTCGTTGCGATGATGGCTTCTTCCTAGGNNNAGATCGGAAGAG |
| 29630_1 | TATGAGTGTGGAGTCGTTGCGCGAGTCCTGGCAAAGAAGATATNNNAGATCGGAA |
| 29630_2 | TATGAGTGTGGAGTCGTTGCGCGAGTCCTGGCAAAGAAGATATNNNAGATCGGAA |
| 50621_1 | TATGAGTGTGGAGTCGTTGCATAGACCTGAAGGACTGGAAAGCNNNAGATCGGAA |
| 50621_2 | TATGAGTGTGGAGTCGTTGCATAGACCTGAAGGACTGGAAAGCNNNAGATCGGAA |
| 30128_1 | TATGAGTGTGGAGTCGTTGCGTGAGATCACAGGGGATTGACTTNNNAGATCGGAA |
| 30128_2 | TATGAGTGTGGAGTCGTTGCGTGAGATCACAGGGGATTGACTTNNNAGATCGGAA |
| 44608_1 | TATGAGTGTGGAGTCGTTGCCAGCTCCCTTTCTAAGTCCTTGTNNNAGATCGGAAGA |
| 44608_2 | TATGAGTGTGGAGTCGTTGCCAGCTCCCTTTCTAAGTCCTTGTNNNAGATCGGAAGA |
| 39425_1 | TATGAGTGTGGAGTCGTTGCAAAACAAAGTTGTGGAGAAGGCCNNNAGATCGGAA |
| 39425_2 | TATGAGTGTGGAGTCGTTGCAAAACAAAGTTGTGGAGAAGGCCNNNAGATCGGAA |
| 34824_1 | TATGAGTGTGGAGTCGTTGCGATGGAGTCTCGCTATGTCATCCNNNAGATCGGAAG |

| | |
|---|---|
| 34824_2 | TATGAGTGTGGAGTCGTTGCGATGGAGTCTCGCTATGTCATCCNNNAGATCGGAAG |
| 47067_1 | TATGAGTGTGGAGTCGTTGCTGAGATCTTGGCTCACTGGAAGNNNAGATCGGAAG |
| 47067_2 | TATGAGTGTGGAGTCGTTGCTGAGATCTTGGCTCACTGGAAGNNNAGATCGGAAG |
| 52084_1 | TATGAGTGTGGAGTCGTTGCTACCCCCCACTCTATCCATATCCNNNAGATCGGAAGA |
| 52084_2 | TATGAGTGTGGAGTCGTTGCTACCCCCCACTCTATCCATATCCNNNAGATCGGAAGA |
| 10570_1 | TATGAGTGTGGAGTCGTTGCGCGAGGCAACATAGGGTACATAANNNAGATCGGAA |
| 10570_2 | TATGAGTGTGGAGTCGTTGCGCGAGGCAACATAGGGTACATAANNNAGATCGGAA |
| 50410_1 | TATGAGTGTGGAGTCGTTGCAACACAGGCAGAGAAACAGGTTANNNAGATCGGAA |
| 50410_2 | TATGAGTGTGGAGTCGTTGCAACACAGGCAGAGAAACAGGTTANNNAGATCGGAA |
| 44495_1 | TATGAGTGTGGAGTCGTTGCAAGGCAGACAAGAATTCAGAGGTNNNAGATCGGAA |
| 44495_2 | TATGAGTGTGGAGTCGTTGCAAGGCAGACAAGAATTCAGAGGTNNNAGATCGGAA |
| 40617_1 | TATGAGTGTGGAGTCGTTGCAAACTAGGAGCTGTGAGTTTGCTNNNAGATCGGAA |
| 40617_2 | TATGAGTGTGGAGTCGTTGCAAACTAGGAGCTGTGAGTTTGCTNNNAGATCGGAA |
| 48534_1 | TATGAGTGTGGAGTCGTTGCCTGAGTAGCTGGAATTACAGGCANNNAGATCGGAA |
| 48534_2 | TATGAGTGTGGAGTCGTTGCCTGAGTAGCTGGAATTACAGGCANNNAGATCGGAA |
| 47337_1 | TATGAGTGTGGAGTCGTTGCCACAAACCAGCCTTTTCTGAAGTNNNAGATCGGAAG |
| 47337_2 | TATGAGTGTGGAGTCGTTGCCACAAACCAGCCTTTTCTGAAGTNNNAGATCGGAAG |
| 12669_1 | TATGAGTGTGGAGTCGTTGCAATCTCCTTGTGGTAAATAAGATNNNAGATCGGAAG |
| 12669_2 | TATGAGTGTGGAGTCGTTGCAATCTCCTTGTGGTAAATAAGATNNNAGATCGGAAG |
| 9987_1 | TATGAGTGTGGAGTCGTTGCGCCTCCCGAGTAGAGAAGTATGANNNAGATCGGAA |
| 9987_2 | TATGAGTGTGGAGTCGTTGCGCCTCCCGAGTAGAGAAGTATGANNNAGATCGGAA |
| 30304_1 | TATGAGTGTGGAGTCGTTGCTTCCTTCTGGCACTCCGGTCNNNAGATCGGAAGAGC |
| 30304_2 | TATGAGTGTGGAGTCGTTGCTTCCTTCTGGCACTCCGGTCNNNAGATCGGAAGAGC |
| 35822_1 | TATGAGTGTGGAGTCGTTGCAGTAGGTGCAATCTTGGCTAAGANNNAGATCGGAA |
| 35822_2 | TATGAGTGTGGAGTCGTTGCAGTAGGTGCAATCTTGGCTAAGANNNAGATCGGAA |
| 10401_1 | TATGAGTGTGGAGTCGTTGCAATAACTAGAACAGGGCCAGGCTNNNAGATCGGAA |
| 10401_2 | TATGAGTGTGGAGTCGTTGCAATAACTAGAACAGGGCCAGGCTNNNAGATCGGAA |
| 35686_1 | TATGAGTGTGGAGTCGTTGCCAGTTCACTGCGGATTCATGATCNNNAGATCGGAAG |
| 35686_2 | TATGAGTGTGGAGTCGTTGCCAGTTCACTGCGGATTCATGATCNNNAGATCGGAAG |
| 48708_1 | TATGAGTGTGGAGTCGTTGCGGTTCAAGCGATTCTTATGCCTCNNNAGATCGGAAG |
| 48708_2 | TATGAGTGTGGAGTCGTTGCGGTTCAAGCGATTCTTATGCCTCNNNAGATCGGAAG |
| 42530_1 | TATGAGTGTGGAGTCGTTGCATGTGTGTGTGTGTGTGTTTTGTNNNAGATCGGAA |
| 42530_2 | TATGAGTGTGGAGTCGTTGCATGTGTGTGTGTGTGTGTTTTGTNNNAGATCGGAA |
| 32165_1 | TATGAGTGTGGAGTCGTTGCCATGATCTCGGCTCACTGCAANNNAGATCGGAAGAG |
| 32165_2 | TATGAGTGTGGAGTCGTTGCCATGATCTCGGCTCACTGCAANNNAGATCGGAAGAG |
| 38871_1 | TATGAGTGTGGAGTCGTTGCTGCAGCCATTTCAGAAACTGTTTNNNAGATCGGAAG |
| 38871_2 | TATGAGTGTGGAGTCGTTGCTGCAGCCATTTCAGAAACTGTTTNNNAGATCGGAAG |
| 53189_1 | TATGAGTGTGGAGTCGTTGCGCTCCCCTCACTCCTCACTTCNNNAGATCGGAAGAGC |
| 53189_2 | TATGAGTGTGGAGTCGTTGCGCTCCCCTCACTCCTCACTTCNNNAGATCGGAAGAGC |
| 13160_1 | TATGAGTGTGGAGTCGTTGCGAATTGAGAGAGGAAGAAGAATTNNNAGATCGGA |
| 13160_2 | TATGAGTGTGGAGTCGTTGCGAATTGAGAGAGGAAGAAGAATTNNNAGATCGGA |
| 31035_1 | TATGAGTGTGGAGTCGTTGCTTTCCCAGGATCCTTGGCTAATTNNNAGATCGGAAG |
| 31035_2 | TATGAGTGTGGAGTCGTTGCTTTCCCAGGATCCTTGGCTAATTNNNAGATCGGAAG |

| | |
|---|---|
| 42422_1 | TATGAGTGTGGAGTCGTTGCTTTTTGCTGAGGTTGGTCTCGAANNNAGATCGGAAG |
| 42422_2 | TATGAGTGTGGAGTCGTTGCTTTTTGCTGAGGTTGGTCTCGAANNNAGATCGGAAG |
| 35002_1 | TATGAGTGTGGAGTCGTTGCGCCTCCCAAGTAGCTAGGATTACNNNAGATCGGAAG |
| 35002_2 | TATGAGTGTGGAGTCGTTGCGCCTCCCAAGTAGCTAGGATTACNNNAGATCGGAAG |
| 50815_1 | TATGAGTGTGGAGTCGTTGCTCCCTCATGTACACACTTCCTTCNNNAGATCGGAAGA |
| 50815_2 | TATGAGTGTGGAGTCGTTGCTCCCTCATGTACACACTTCCTTCNNNAGATCGGAAGA |
| 13637_1 | TATGAGTGTGGAGTCGTTGCGAGAACTGTGTGATATCAGAAANNNAGATCGGAAG |
| 13637_2 | TATGAGTGTGGAGTCGTTGCGAGAACTGTGTGATATCAGAAANNNAGATCGGAAG |
| 29296_1 | TATGAGTGTGGAGTCGTTGCTCCCTGGCAATAGTTCTTTGACANNNAGATCGGAAG |
| 29296_2 | TATGAGTGTGGAGTCGTTGCTCCCTGGCAATAGTTCTTTGACANNNAGATCGGAAG |
| 7454_1 | TATGAGTGTGGAGTCGTTGCGGGTCCTCTCTCATCTTGGTTAGTNNNAGATCGGAA |
| 7454_2 | TATGAGTGTGGAGTCGTTGCGGGTCCTCTCTCATCTTGGTTAGTNNNAGATCGGAA |
| 31109_1 | TATGAGTGTGGAGTCGTTGCCTCTCTCTCTCTCTCTCTCCGNNNAGATCGGAAGAG |
| 31109_2 | TATGAGTGTGGAGTCGTTGCCTCTCTCTCTCTCTCTCTCCGNNNAGATCGGAAGAG |
| 32739_1 | TATGAGTGTGGAGTCGTTGCCATCCCCACTCTCAACACATGTNNNAGATCGGAAGAG |
| 32739_2 | TATGAGTGTGGAGTCGTTGCCATCCCCACTCTCAACACATGTNNNAGATCGGAAGAG |
| 9362_1 | TATGAGTGTGGAGTCGTTGCGGGGCCTTCGAAAGGTTATTAGGNNNAGATCGGAA |
| 9362_2 | TATGAGTGTGGAGTCGTTGCGGGGCCTTCGAAAGGTTATTAGGNNNAGATCGGAA |
| 43201_1 | TATGAGTGTGGAGTCGTTGCCCCTTCCTTCCTTCCTCTCTTTCNNNAGATCGGAAGAG |
| 43201_2 | TATGAGTGTGGAGTCGTTGCCCCTTCCTTCCTTCCTCTCTTTCNNNAGATCGGAAGAG |
| 30929_1 | TATGAGTGTGGAGTCGTTGCTCATTTGCAACAACTAAGCACGANNNAGATCGGAAG |
| 30929_2 | TATGAGTGTGGAGTCGTTGCTCATTTGCAACAACTAAGCACGANNNAGATCGGAAG |
| 42166_1 | TATGAGTGTGGAGTCGTTGCCTGAGTAGCTGGGATTACAGGTGNNNAGATCGGAA |
| 42166_2 | TATGAGTGTGGAGTCGTTGCCTGAGTAGCTGGGATTACAGGTGNNNAGATCGGAA |
| 45292_1 | TATGAGTGTGGAGTCGTTGCGCAAACAACAAACTGCAGTTAGCNNNAGATCGGAAG |
| 45292_2 | TATGAGTGTGGAGTCGTTGCGCAAACAACAAACTGCAGTTAGCNNNAGATCGGAAG |
| 44280_1 | TATGAGTGTGGAGTCGTTGCGCTTGATTATGGGCGTAAGGAACNNNAGATCGGAA |
| 44280_2 | TATGAGTGTGGAGTCGTTGCGCTTGATTATGGGCGTAAGGAACNNNAGATCGGAA |
| 29043_1 | TATGAGTGTGGAGTCGTTGCGAGGTGGGCCTGATAGTACTCTANNNAGATCGGAA |
| 29043_2 | TATGAGTGTGGAGTCGTTGCGAGGTGGGCCTGATAGTACTCTANNNAGATCGGAA |
| 10268_1 | TATGAGTGTGGAGTCGTTGCCTCATTGCTTCTGCTCAACAACCNNNAGATCGGAAGA |
| 10268_2 | TATGAGTGTGGAGTCGTTGCCTCATTGCTTCTGCTCAACAACCNNNAGATCGGAAGA |
| 31895_1 | TATGAGTGTGGAGTCGTTGCGAAAGGAATCGTGGTGTATTGGCNNNAGATCGGAA |
| 31895_2 | TATGAGTGTGGAGTCGTTGCGAAAGGAATCGTGGTGTATTGGCNNNAGATCGGAA |
| 29241_1 | TATGAGTGTGGAGTCGTTGCGAGAAGGCAGGGATGAGATACAGNNNAGATCGGA |
| 29241_2 | TATGAGTGTGGAGTCGTTGCGAGAAGGCAGGGATGAGATACAGNNNAGATCGGA |
| 43911_1 | TATGAGTGTGGAGTCGTTGCTCTCTCTCTCCCTCTTTCTTTCTNNNAGATCGGAAGAG |
| 43911_2 | TATGAGTGTGGAGTCGTTGCTCTCTCTCTCCCTCTTTCTTTCTNNNAGATCGGAAGAG |
| 35563_1 | TATGAGTGTGGAGTCGTTGCCTTCTCTGAGTGTCTGGGATCACNNNAGATCGGAAG |
| 35563_2 | TATGAGTGTGGAGTCGTTGCCTTCTCTGAGTGTCTGGGATCACNNNAGATCGGAAG |
| 45080_1 | TATGAGTGTGGAGTCGTTGCCATCAGTTTGACTTTGACCCACCNNNAGATCGGAAG |
| 45080_2 | TATGAGTGTGGAGTCGTTGCCATCAGTTTGACTTTGACCCACCNNNAGATCGGAAG |
| 31487_1 | TATGAGTGTGGAGTCGTTGCGCCATGAGCCAAGAAATGAAGTTNNNAGATCGGAA |

| | |
|---|---|
| 31487_2 | TATGAGTGTGGAGTCGTTGCGCCATGAGCCAAGAAATGAAGTTNNNAGATCGGAA |
| 31375_1 | TATGAGTGTGGAGTCGTTGCTGATCCAGAAAAACTTAGGCCCANNNAGATCGGAAG |
| 31375_2 | TATGAGTGTGGAGTCGTTGCTGATCCAGAAAAACTTAGGCCCANNNAGATCGGAAG |
| 49779_1 | TATGAGTGTGGAGTCGTTGCAAGACGTCCTGATTCTAGACAGCNNNAGATCGGAAG |
| 49779_2 | TATGAGTGTGGAGTCGTTGCAAGACGTCCTGATTCTAGACAGCNNNAGATCGGAAG |
| 31632_1 | TATGAGTGTGGAGTCGTTGCAGAGAGAAGAAGAAGCCAGAAGCNNNAGATCGGA |
| 31632_2 | TATGAGTGTGGAGTCGTTGCAGAGAGAAGAAGAAGCCAGAAGCNNNAGATCGGA |
| 43251_1 | TATGAGTGTGGAGTCGTTGCCGGCTTCCTAACGACAAAATCAANNNAGATCGGAAG |
| 43251_2 | TATGAGTGTGGAGTCGTTGCCGGCTTCCTAACGACAAAATCAANNNAGATCGGAAG |
| 41997_1 | TATGAGTGTGGAGTCGTTGCAGGTTCAAGTGATTCTTCTGCCTNNNAGATCGGAAG |
| 41997_2 | TATGAGTGTGGAGTCGTTGCAGGTTCAAGTGATTCTTCTGCCTNNNAGATCGGAAG |
| 48127_1 | TATGAGTGTGGAGTCGTTGCCTCAATTTCTCCCTGTCTCTCCCNNNAGATCGGAAGA |
| 48127_2 | TATGAGTGTGGAGTCGTTGCCTCAATTTCTCCCTGTCTCTCCCNNNAGATCGGAAGA |
| 7933_1 | TATGAGTGTGGAGTCGTTGCGGGGAAGCAGCATGATCTCTGCNNNAGATCGGAAG |
| 7933_2 | TATGAGTGTGGAGTCGTTGCGGGGAAGCAGCATGATCTCTGCNNNAGATCGGAAG |
| 37037_1 | TATGAGTGTGGAGTCGTTGCAAGAAATTAGCCAGGCGTAGAGGNNNAGATCGGAA |
| 37037_2 | TATGAGTGTGGAGTCGTTGCAAGAAATTAGCCAGGCGTAGAGGNNNAGATCGGAA |
| 45719_1 | TATGAGTGTGGAGTCGTTGCAGTGCTAGGATTACAGGCATGAGNNNAGATCGGAA |
| 45719_2 | TATGAGTGTGGAGTCGTTGCAGTGCTAGGATTACAGGCATGAGNNNAGATCGGAA |
| 37925_1 | TATGAGTGTGGAGTCGTTGCAAAGTTTCCTACCTTCAGACCCCNNNAGATCGGAAGA |
| 37925_2 | TATGAGTGTGGAGTCGTTGCAAAGTTTCCTACCTTCAGACCCCNNNAGATCGGAAGA |
| 40611_1 | TATGAGTGTGGAGTCGTTGCCTAGCAAGAAAAATGGAGCAGCANNNAGATCGGAA |
| 40611_2 | TATGAGTGTGGAGTCGTTGCCTAGCAAGAAAAATGGAGCAGCANNNAGATCGGAA |
| 33322_1 | TATGAGTGTGGAGTCGTTGCCCTGGGTTCAACCAATTCTCCTANNNAGATCGGAAGA |
| 33322_2 | TATGAGTGTGGAGTCGTTGCCCTGGGTTCAACCAATTCTCCTANNNAGATCGGAAGA |
| 47565_1 | TATGAGTGTGGAGTCGTTGCAAGTATCCCCAGTAAAGGCTGTGNNNAGATCGGAA |
| 47565_2 | TATGAGTGTGGAGTCGTTGCAAGTATCCCCAGTAAAGGCTGTGNNNAGATCGGAA |
| 33251_1 | TATGAGTGTGGAGTCGTTGCCCCCTAAGTAGCTGGGACTACAGNNNAGATCGGAA |
| 33251_2 | TATGAGTGTGGAGTCGTTGCCCCCTAAGTAGCTGGGACTACAGNNNAGATCGGAA |
| 7828_1 | TATGAGTGTGGAGTCGTTGCGCTTGTCTGGAGCTCTTGGATNNNAGATCGGAAGA |
| 7828_2 | TATGAGTGTGGAGTCGTTGCGCTTGTCTGGAGCTCTTGGATNNNAGATCGGAAGA |
| 42327_1 | TATGAGTGTGGAGTCGTTGCGAACATCAAGTCAGGAGTCAGGANNNAGATCGGAA |
| 42327_2 | TATGAGTGTGGAGTCGTTGCGAACATCAAGTCAGGAGTCAGGANNNAGATCGGAA |
| 35323_1 | TATGAGTGTGGAGTCGTTGCCATCACGATGCCCAGCTAATTTCNNNAGATCGGAAG |
| 35323_2 | TATGAGTGTGGAGTCGTTGCCATCACGATGCCCAGCTAATTTCNNNAGATCGGAAG |
| 41217_1 | TATGAGTGTGGAGTCGTTGCTGTGTAGAACAGGCAAACCCATANNNAGATCGGAA |
| 41217_2 | TATGAGTGTGGAGTCGTTGCTGTGTAGAACAGGCAAACCCATANNNAGATCGGAA |
| 48163_1 | TATGAGTGTGGAGTCGTTGCAGGGCATGGTAAAACAATCTGTTNNNAGATCGGAA |
| 48163_2 | TATGAGTGTGGAGTCGTTGCAGGGCATGGTAAAACAATCTGTTNNNAGATCGGAA |
| 51822_1 | TATGAGTGTGGAGTCGTTGCGACCAGCACACCAACTTTATCGNNNAGATCGGAAGA |
| 51822_2 | TATGAGTGTGGAGTCGTTGCGACCAGCACACCAACTTTATCGNNNAGATCGGAAGA |
| 28901_1 | TATGAGTGTGGAGTCGTTGCTGCACGTTTATGTGTGTAGGTCTNNNAGATCGGAAG |
| 28901_2 | TATGAGTGTGGAGTCGTTGCTGCACGTTTATGTGTGTAGGTCTNNNAGATCGGAAG |

| | |
|---|---|
| 30465_1 | TATGAGTGTGGAGTCGTTGCTCTGATATCCCTGGTAGGCCTTGNNNAGATCGGAAG |
| 30465_2 | TATGAGTGTGGAGTCGTTGCTCTGATATCCCTGGTAGGCCTTGNNNAGATCGGAAG |
| 3052_1 | TATGAGTGTGGAGTCGTTGCGGGGCTGTGTAGCGCTGATCNNNAGATCGGAAGAG |
| 3052_2 | TATGAGTGTGGAGTCGTTGCGGGGCTGTGTAGCGCTGATCNNNAGATCGGAAGAG |
| 50998_1 | TATGAGTGTGGAGTCGTTGCACAATGAGACTCTCCAATCCTTCTNNNAGATCGGAAG |
| 50998_2 | TATGAGTGTGGAGTCGTTGCACAATGAGACTCTCCAATCCTTCTNNNAGATCGGAAG |
| 3856_1 | TATGAGTGTGGAGTCGTTGCGCATCACTCCATTGCACTACTCANNNAGATCGGAAGA |
| 3856_2 | TATGAGTGTGGAGTCGTTGCGCATCACTCCATTGCACTACTCANNNAGATCGGAAGA |
| 7982_1 | TATGAGTGTGGAGTCGTTGCCAGACTTTCGTCCAGGCTGGNNNAGATCGGAAGAG |
| 7982_2 | TATGAGTGTGGAGTCGTTGCCAGACTTTCGTCCAGGCTGGNNNAGATCGGAAGAG |
| 43080_1 | TATGAGTGTGGAGTCGTTGCCCCACATATCTGTGAGTCTGTGTNNNAGATCGGAAG |
| 43080_2 | TATGAGTGTGGAGTCGTTGCCCCACATATCTGTGAGTCTGTGTNNNAGATCGGAAG |
| 47601_1 | TATGAGTGTGGAGTCGTTGCTTATGCATTTGGGTAAAAGGGGCNNNAGATCGGAA |
| 47601_2 | TATGAGTGTGGAGTCGTTGCTTATGCATTTGGGTAAAAGGGGCNNNAGATCGGAA |
| 45660_1 | TATGAGTGTGGAGTCGTTGCGGTTCAAGCGATTCTCCTGTTTCNNNAGATCGGAAG |
| 45660_2 | TATGAGTGTGGAGTCGTTGCGGTTCAAGCGATTCTCCTGTTTCNNNAGATCGGAAG |
| 37980_1 | TATGAGTGTGGAGTCGTTGCCAGTTTGGGGTAAAACCACCTTCNNNAGATCGGAAG |
| 37980_2 | TATGAGTGTGGAGTCGTTGCCAGTTTGGGGTAAAACCACCTTCNNNAGATCGGAAG |
| 45194_1 | TATGAGTGTGGAGTCGTTGCTTCGTACCTGTTGGTGTGGAAGNNNAGATCGGAAG |
| 45194_2 | TATGAGTGTGGAGTCGTTGCTTCGTACCTGTTGGTGTGGAAGNNNAGATCGGAAG |
| 13626_1 | TATGAGTGTGGAGTCGTTGCCTCTGAATTCATCTTTTCTTCAANNNAGATCGGAAGA |
| 13626_2 | TATGAGTGTGGAGTCGTTGCCTCTGAATTCATCTTTTCTTCAANNNAGATCGGAAGA |
| 10422_1 | TATGAGTGTGGAGTCGTTGCTGGTAAGTGGGTAGATTCCAGGGNNNAGATCGGAA |
| 10422_2 | TATGAGTGTGGAGTCGTTGCTGGTAAGTGGGTAGATTCCAGGGNNNAGATCGGAA |
| 30484_1 | TATGAGTGTGGAGTCGTTGCAATGAACAGAAGGATAAACTGAGCACNNNAGATCG |
| 30484_2 | TATGAGTGTGGAGTCGTTGCAATGAACAGAAGGATAAACTGAGCACNNNAGATCG |
| 51149_1 | TATGAGTGTGGAGTCGTTGCCTCTCCCCTTCTTACCAAAGTCCNNNAGATCGGAAGA |
| 51149_2 | TATGAGTGTGGAGTCGTTGCCTCTCCCCTTCTTACCAAAGTCCNNNAGATCGGAAGA |
| 47663_1 | TATGAGTGTGGAGTCGTTGCGAGAAAGGGTCTTGTTCTGTTGCNNNAGATCGGAA |
| 47663_2 | TATGAGTGTGGAGTCGTTGCGAGAAAGGGTCTTGTTCTGTTGCNNNAGATCGGAA |
| 32816_1 | TATGAGTGTGGAGTCGTTGCCATTCTCCTGCCTCAGCCTCNNNAGATCGGAAGAGCA |
| 32816_2 | TATGAGTGTGGAGTCGTTGCCATTCTCCTGCCTCAGCCTCNNNAGATCGGAAGAGCA |
| 38334_1 | TATGAGTGTGGAGTCGTTGCAGAGCAAACTAACCACCTTTCCTNNNAGATCGGAAG |
| 38334_2 | TATGAGTGTGGAGTCGTTGCAGAGCAAACTAACCACCTTTCCTNNNAGATCGGAAG |
| 50259_1 | TATGAGTGTGGAGTCGTTGCATCTCCAGCAGTGGTCACAAGNNNAGATCGGAAGA |
| 50259_2 | TATGAGTGTGGAGTCGTTGCATCTCCAGCAGTGGTCACAAGNNNAGATCGGAAGA |
| 10108_1 | TATGAGTGTGGAGTCGTTGCCACACGGGAACCCTAACCAATAGNNNAGATCGGAAG |
| 10108_2 | TATGAGTGTGGAGTCGTTGCCACACGGGAACCCTAACCAATAGNNNAGATCGGAAG |
| 12564_1 | TATGAGTGTGGAGTCGTTGCATTAGGTCCCTTCATTAGAAATCNNNAGATCGGAAG |
| 12564_2 | TATGAGTGTGGAGTCGTTGCATTAGGTCCCTTCATTAGAAATCNNNAGATCGGAAG |
| 53239_1 | TATGAGTGTGGAGTCGTTGCGTCACCTGAGAATATGCCATTGCNNNAGATCGGAAG |
| 53239_2 | TATGAGTGTGGAGTCGTTGCGTCACCTGAGAATATGCCATTGCNNNAGATCGGAAG |
| 40194_1 | TATGAGTGTGGAGTCGTTGCCCTGTAATCCCAGCACTTAGGGNNNAGATCGGAAGA |

| | |
|---|---|
| 40194_2 | TATGAGTGTGGAGTCGTTGCCCTGTAATCCCAGCACTTAGGGNNNAGATCGGAAGA |
| 29625_1 | TATGAGTGTGGAGTCGTTGCTTATAGGCAGCTGACCTAGGAGANNNAGATCGGAA |
| 29625_2 | TATGAGTGTGGAGTCGTTGCTTATAGGCAGCTGACCTAGGAGANNNAGATCGGAA |
| 52288_1 | TATGAGTGTGGAGTCGTTGCCATAGAGGGACCACAGAATCCAGNNNAGATCGGAA |
| 52288_2 | TATGAGTGTGGAGTCGTTGCCATAGAGGGACCACAGAATCCAGNNNAGATCGGAA |
| 8544_1 | TATGAGTGTGGAGTCGTTGCTGTGAAAACTCAGGGTACTGCCCNNNAGATCGGAAG |
| 8544_2 | TATGAGTGTGGAGTCGTTGCTGTGAAAACTCAGGGTACTGCCCNNNAGATCGGAAG |
| 30615_1 | TATGAGTGTGGAGTCGTTGCTGTGTGTGTCTTTCCTAGGACCCNNNAGATCGGAAG |
| 30615_2 | TATGAGTGTGGAGTCGTTGCTGTGTGTGTCTTTCCTAGGACCCNNNAGATCGGAAG |
| 50412_1 | TATGAGTGTGGAGTCGTTGCAATTGAAGCTGAAACACCCATGGNNNAGATCGGAA |
| 50412_2 | TATGAGTGTGGAGTCGTTGCAATTGAAGCTGAAACACCCATGGNNNAGATCGGAA |
| 32121_1 | TATGAGTGTGGAGTCGTTGCTTGATCTTGAACTTCTGGGCTCANNNAGATCGGAAG |
| 32121_2 | TATGAGTGTGGAGTCGTTGCTTGATCTTGAACTTCTGGGCTCANNNAGATCGGAAG |
| 32929_1 | TATGAGTGTGGAGTCGTTGCTAAGCTCACTCACTGCAGAAACANNNAGATCGGAAG |
| 32929_2 | TATGAGTGTGGAGTCGTTGCTAAGCTCACTCACTGCAGAAACANNNAGATCGGAAG |
| 41362_1 | TATGAGTGTGGAGTCGTTGCCTTTTTGTAACTTGCCCCACTGTNNNAGATCGGAAGA |
| 41362_2 | TATGAGTGTGGAGTCGTTGCCTTTTTGTAACTTGCCCCACTGTNNNAGATCGGAAGA |
| 41614_1 | TATGAGTGTGGAGTCGTTGCAGGAGCTGGGACTATAGGCATATNNNAGATCGGAA |
| 41614_2 | TATGAGTGTGGAGTCGTTGCAGGAGCTGGGACTATAGGCATATNNNAGATCGGAA |
| 43612_1 | TATGAGTGTGGAGTCGTTGCCATTTGGAAAACCTCATCTCCGGNNNAGATCGGAAG |
| 43612_2 | TATGAGTGTGGAGTCGTTGCCATTTGGAAAACCTCATCTCCGGNNNAGATCGGAAG |
| 42102_1 | TATGAGTGTGGAGTCGTTGCCACCATTGGACCTATTTGATGGCNNNAGATCGGAAG |
| 42102_2 | TATGAGTGTGGAGTCGTTGCCACCATTGGACCTATTTGATGGCNNNAGATCGGAAG |
| 40264_1 | TATGAGTGTGGAGTCGTTGCACCTAGACTGAGGCCTTATACCTNNNAGATCGGAAG |
| 40264_2 | TATGAGTGTGGAGTCGTTGCACCTAGACTGAGGCCTTATACCTNNNAGATCGGAAG |
| 37057_1 | TATGAGTGTGGAGTCGTTGCTGAAAATGAACTCAAAATGGGTCANNNAGATCGGA |
| 37057_2 | TATGAGTGTGGAGTCGTTGCTGAAAATGAACTCAAAATGGGTCANNNAGATCGGA |
| 47236_1 | TATGAGTGTGGAGTCGTTGCCAGAATCTCAGCTCACTGCAACNNNAGATCGGAAGA |
| 47236_2 | TATGAGTGTGGAGTCGTTGCCAGAATCTCAGCTCACTGCAACNNNAGATCGGAAGA |
| 30949_1 | TATGAGTGTGGAGTCGTTGCTCAAGTCACAAGAAGAAGCTGAANNNAGATCGGAA |
| 30949_2 | TATGAGTGTGGAGTCGTTGCTCAAGTCACAAGAAGAAGCTGAANNNAGATCGGAA |
| 40863_1 | TATGAGTGTGGAGTCGTTGCGCCCTTTATCAGCATAACACCACNNNAGATCGGAAG |
| 40863_2 | TATGAGTGTGGAGTCGTTGCGCCCTTTATCAGCATAACACCACNNNAGATCGGAAG |
| 31264_1 | TATGAGTGTGGAGTCGTTGCTCCATTATTACCTCCAACTCCAGTNNNAGATCGGAAG |
| 31264_2 | TATGAGTGTGGAGTCGTTGCTCCATTATTACCTCCAACTCCAGTNNNAGATCGGAAG |
| 41391_1 | TATGAGTGTGGAGTCGTTGCGCAAAAGCTTCTTTACGCTGAGANNNAGATCGGAAG |
| 41391_2 | TATGAGTGTGGAGTCGTTGCGCAAAAGCTTCTTTACGCTGAGANNNAGATCGGAAG |
| 53255_1 | TATGAGTGTGGAGTCGTTGCTGTTTCATTTTTACCAAGGGCCCNNNAGATCGGAAG |
| 53255_2 | TATGAGTGTGGAGTCGTTGCTGTTTCATTTTTACCAAGGGCCCNNNAGATCGGAAG |
| 38215_1 | TATGAGTGTGGAGTCGTTGCTAGACTGGGTGACAGAGAGAGACNNNAGATCGGA |
| 38215_2 | TATGAGTGTGGAGTCGTTGCTAGACTGGGTGACAGAGAGAGACNNNAGATCGGA |
| 46996_1 | TATGAGTGTGGAGTCGTTGCGGCAAAGTCTGTTTCTGACTCTGNNNAGATCGGAAG |
| 46996_2 | TATGAGTGTGGAGTCGTTGCGGCAAAGTCTGTTTCTGACTCTGNNNAGATCGGAAG |

| | |
|---|---|
| 52011_1 | TATGAGTGTGGAGTCGTTGCCTTGGAAGCCAGTATCACCCTAANNNAGATCGGAAG |
| 52011_2 | TATGAGTGTGGAGTCGTTGCCTTGGAAGCCAGTATCACCCTAANNNAGATCGGAAG |
| 45520_1 | TATGAGTGTGGAGTCGTTGCCCATCCAAGGGTAAGGGAATAGGNNNAGATCGGAA |
| 45520_2 | TATGAGTGTGGAGTCGTTGCCCATCCAAGGGTAAGGGAATAGGNNNAGATCGGAA |
| 37536_1 | TATGAGTGTGGAGTCGTTGCAAAAACGTATGTACCCCTTCACANNNAGATCGGAAG |
| 37536_2 | TATGAGTGTGGAGTCGTTGCAAAAACGTATGTACCCCTTCACANNNAGATCGGAAG |
| 50388_1 | TATGAGTGTGGAGTCGTTGCCAGCCTGTGAAGGGAAGGAGNNNAGATCGGAAGA |
| 50388_2 | TATGAGTGTGGAGTCGTTGCCAGCCTGTGAAGGGAAGGAGNNNAGATCGGAAGA |
| 52473_1 | TATGAGTGTGGAGTCGTTGCAATTTGAGACCCAGGCAGAAGATNNNAGATCGGAA |
| 52473_2 | TATGAGTGTGGAGTCGTTGCAATTTGAGACCCAGGCAGAAGATNNNAGATCGGAA |
| 33803_1 | TATGAGTGTGGAGTCGTTGCATCATAGATTTTGCCAGCACGTCNNNAGATCGGAAG |
| 33803_2 | TATGAGTGTGGAGTCGTTGCATCATAGATTTTGCCAGCACGTCNNNAGATCGGAAG |
| 41103_1 | TATGAGTGTGGAGTCGTTGCTAGAATAGCAGTGACTGTCACCANNNAGATCGGAAG |
| 41103_2 | TATGAGTGTGGAGTCGTTGCTAGAATAGCAGTGACTGTCACCANNNAGATCGGAAG |
| 42431_1 | TATGAGTGTGGAGTCGTTGCGAGAGCAGAGAAAAGAGAGGAGANNNAGATCGGA |
| 42431_2 | TATGAGTGTGGAGTCGTTGCGAGAGCAGAGAAAAGAGAGGAGANNNAGATCGGA |
| 39162_1 | TATGAGTGTGGAGTCGTTGCGAGGACAGAGAACGGCTTGAANNNAGATCGGAAG |
| 39162_2 | TATGAGTGTGGAGTCGTTGCGAGGACAGAGAACGGCTTGAANNNAGATCGGAAG |
| 39475_1 | TATGAGTGTGGAGTCGTTGCAGATGGGGTGACAGAGTATCTCTNNNAGATCGGAA |
| 39475_2 | TATGAGTGTGGAGTCGTTGCAGATGGGGTGACAGAGTATCTCTNNNAGATCGGAA |
| 39939_1 | TATGAGTGTGGAGTCGTTGCATGATAGTGCATCTTTCCCCACANNNAGATCGGAAG |
| 39939_2 | TATGAGTGTGGAGTCGTTGCATGATAGTGCATCTTTCCCCACANNNAGATCGGAAG |
| 38433_1 | TATGAGTGTGGAGTCGTTGCAATATCCGCCCTAAACACCTTGTNNNAGATCGGAAGA |
| 38433_2 | TATGAGTGTGGAGTCGTTGCAATATCCGCCCTAAACACCTTGTNNNAGATCGGAAGA |
| 10592_1 | TATGAGTGTGGAGTCGTTGCGATGACACGTGTAGGTTCACATCTNNNAGATCGGAA |
| 10592_2 | TATGAGTGTGGAGTCGTTGCGATGACACGTGTAGGTTCACATCTNNNAGATCGGAA |
| 39524_1 | TATGAGTGTGGAGTCGTTGCTGGTTCCAAAACATAGAGGAGGGNNNAGATCGGAA |
| 39524_2 | TATGAGTGTGGAGTCGTTGCTGGTTCCAAAACATAGAGGAGGGNNNAGATCGGAA |
| 50089_1 | TATGAGTGTGGAGTCGTTGCTGGAAACTTCTCTCCAGGAACTGNNNAGATCGGAAG |
| 50089_2 | TATGAGTGTGGAGTCGTTGCTGGAAACTTCTCTCCAGGAACTGNNNAGATCGGAAG |
| 32356_1 | TATGAGTGTGGAGTCGTTGCGCATTCCCTAACTCGTTCCAAAGNNNAGATCGGAAG |
| 32356_2 | TATGAGTGTGGAGTCGTTGCGCATTCCCTAACTCGTTCCAAAGNNNAGATCGGAAG |
| 33908_1 | TATGAGTGTGGAGTCGTTGCCAAGATCTTGGCTCACTGCAAGNNNAGATCGGAAGA |
| 33908_2 | TATGAGTGTGGAGTCGTTGCCAAGATCTTGGCTCACTGCAAGNNNAGATCGGAAGA |
| 32452_1 | TATGAGTGTGGAGTCGTTGCCTTGAGGCCAGGAGTTTGAGACNNNAGATCGGAAG |
| 32452_2 | TATGAGTGTGGAGTCGTTGCCTTGAGGCCAGGAGTTTGAGACNNNAGATCGGAAG |
| 36974_1 | TATGAGTGTGGAGTCGTTGCAATACAAAATTAGCCGGGCATGGNNNAGATCGGAA |
| 36974_2 | TATGAGTGTGGAGTCGTTGCAATACAAAATTAGCCGGGCATGGNNNAGATCGGAA |
| 45110_1 | TATGAGTGTGGAGTCGTTGCGGGGGTGTTAGTAAGAGAGAGGANNNAGATCGGA |
| 45110_2 | TATGAGTGTGGAGTCGTTGCGGGGGTGTTAGTAAGAGAGAGGANNNAGATCGGA |
| 33546_1 | TATGAGTGTGGAGTCGTTGCCAGGTTCACGCCATTTTCCTGNNNAGATCGGAAGAG |
| 33546_2 | TATGAGTGTGGAGTCGTTGCCAGGTTCACGCCATTTTCCTGNNNAGATCGGAAGAG |
| 48680_1 | TATGAGTGTGGAGTCGTTGCATCATTCATTCATTCCCCCCACANNNAGATCGGAAGA |

| | |
|---|---|
| 48680_2 | TATGAGTGTGGAGTCGTTGCATCATTCATTCATTCCCCCACANNNAGATCGGAAGA |
| 41889_1 | TATGAGTGTGGAGTCGTTGCCTTATACCACTGCCTCCCAAAGTNNNAGATCGGAAGA |
| 41889_2 | TATGAGTGTGGAGTCGTTGCCTTATACCACTGCCTCCCAAAGTNNNAGATCGGAAGA |
| 34864_1 | TATGAGTGTGGAGTCGTTGCCCACAGTACCCGAGATTACATGTNNNAGATCGGAAG |
| 34864_2 | TATGAGTGTGGAGTCGTTGCCCACAGTACCCGAGATTACATGTNNNAGATCGGAAG |
| 8528_1 | TATGAGTGTGGAGTCGTTGCCCCTGCCACTAGGAGACAGTANNNAGATCGGAAGA |
| 8528_2 | TATGAGTGTGGAGTCGTTGCCCCTGCCACTAGGAGACAGTANNNAGATCGGAAGA |
| 45093_1 | TATGAGTGTGGAGTCGTTGCGGTTGGAGTCAGGTGCTGATATANNNAGATCGGAA |
| 45093_2 | TATGAGTGTGGAGTCGTTGCGGTTGGAGTCAGGTGCTGATATANNNAGATCGGAA |
| 37773_1 | TATGAGTGTGGAGTCGTTGCAAGAAGAAAGGTGCCAGTCAGATNNNAGATCGGAA |
| 37773_2 | TATGAGTGTGGAGTCGTTGCAAGAAGAAAGGTGCCAGTCAGATNNNAGATCGGAA |
| 2011_1 | TATGAGTGTGGAGTCGTTGCAGGGAGATGTAACAGGCGTCACTNNNAGATCGGAA |
| 2011_2 | TATGAGTGTGGAGTCGTTGCAGGGAGATGTAACAGGCGTCACTNNNAGATCGGAA |
| 13474_1 | TATGAGTGTGGAGTCGTTGCCTCTTTGTTTCTCATCAACTGNNNAGATCGGAAGAG |
| 13474_2 | TATGAGTGTGGAGTCGTTGCCTCTTTGTTTCTCATCAACTGNNNAGATCGGAAGAG |
| 50668_1 | TATGAGTGTGGAGTCGTTGCAAACATTTTTCTGGAGGGCCAAGNNNAGATCGGAA |
| 50668_2 | TATGAGTGTGGAGTCGTTGCAAACATTTTTCTGGAGGGCCAAGNNNAGATCGGAA |
| 35135_1 | TATGAGTGTGGAGTCGTTGCTTTTGTGGTGAAATTTGGGTTGTNNNAGATCGGAA |
| 35135_2 | TATGAGTGTGGAGTCGTTGCTTTTGTGGTGAAATTTGGGTTGTNNNAGATCGGAA |
| 36406_1 | TATGAGTGTGGAGTCGTTGCAGGAAAAACAGTCCAGCAAAAGGNNNAGATCGGAA |
| 36406_2 | TATGAGTGTGGAGTCGTTGCAGGAAAAACAGTCCAGCAAAAGGNNNAGATCGGAA |
| 41027_1 | TATGAGTGTGGAGTCGTTGCAGCCTTCTCATATGCCAAGACTTNNNAGATCGGAAG |
| 41027_2 | TATGAGTGTGGAGTCGTTGCAGCCTTCTCATATGCCAAGACTTNNNAGATCGGAAG |
| 45402_1 | TATGAGTGTGGAGTCGTTGCGAGGGTGGTACATGCAGATTTTGNNNAGATCGGAA |
| 45402_2 | TATGAGTGTGGAGTCGTTGCGAGGGTGGTACATGCAGATTTTGNNNAGATCGGAA |
| 48863_1 | TATGAGTGTGGAGTCGTTGCCCAGTGGTGGGGATTACAAGTAGNNNAGATCGGAA |
| 48863_2 | TATGAGTGTGGAGTCGTTGCCCAGTGGTGGGGATTACAAGTAGNNNAGATCGGAA |
| 38446_1 | TATGAGTGTGGAGTCGTTGCCCTAAAAGGACCCGCTCACATAGNNNAGATCGGAAG |
| 38446_2 | TATGAGTGTGGAGTCGTTGCCCTAAAAGGACCCGCTCACATAGNNNAGATCGGAAG |
| 32335_1 | TATGAGTGTGGAGTCGTTGCGAGAAACCATGGTGTCGTCTTTGNNNAGATCGGAA |
| 32335_2 | TATGAGTGTGGAGTCGTTGCGAGAAACCATGGTGTCGTCTTTGNNNAGATCGGAA |
| 35715_1 | TATGAGTGTGGAGTCGTTGCCTTTGGCTGAGAGAGAACAGGATNNNAGATCGGAA |
| 35715_2 | TATGAGTGTGGAGTCGTTGCCTTTGGCTGAGAGAGAACAGGATNNNAGATCGGAA |
| 33663_1 | TATGAGTGTGGAGTCGTTGCCCAAGAGTGCCAGGATTAGAGTTNNNAGATCGGAA |
| 33663_2 | TATGAGTGTGGAGTCGTTGCCCAAGAGTGCCAGGATTAGAGTTNNNAGATCGGAA |
| 31659_1 | TATGAGTGTGGAGTCGTTGCTTTTTGTAATTGCCAGCCCCAATNNNAGATCGGAAG |
| 31659_2 | TATGAGTGTGGAGTCGTTGCTTTTTGTAATTGCCAGCCCCAATNNNAGATCGGAAG |
| 47600_1 | TATGAGTGTGGAGTCGTTGCCCAGTGAGACAGAACTGTTCACTNNNAGATCGGAAG |
| 47600_2 | TATGAGTGTGGAGTCGTTGCCCAGTGAGACAGAACTGTTCACTNNNAGATCGGAAG |
| 49660_1 | TATGAGTGTGGAGTCGTTGCGTTCAAGTGATTCTCCTGCCTCTNNNAGATCGGAAG |
| 49660_2 | TATGAGTGTGGAGTCGTTGCGTTCAAGTGATTCTCCTGCCTCTNNNAGATCGGAAG |
| 48259_1 | TATGAGTGTGGAGTCGTTGCCCTCTGCCTCATGAGTAGCTAAGNNNAGATCGGAAG |
| 48259_2 | TATGAGTGTGGAGTCGTTGCCCTCTGCCTCATGAGTAGCTAAGNNNAGATCGGAAG |

| | |
|---|---|
| 10366_1 | TATGAGTGTGGAGTCGTTGCGACTTATCCATGCTTCCACTGCANNNAGATCGGAAG |
| 10366_2 | TATGAGTGTGGAGTCGTTGCGACTTATCCATGCTTCCACTGCANNNAGATCGGAAG |
| 2038_1 | TATGAGTGTGGAGTCGTTGCCTGAGAAGATATGCGCTGTGGAANNNAGATCGGAA |
| 2038_2 | TATGAGTGTGGAGTCGTTGCCTGAGAAGATATGCGCTGTGGAANNNAGATCGGAA |
| 13012_1 | TATGAGTGTGGAGTCGTTGCTCTCCTTTTATTAATAGGAAGGCNNNAGATCGGAAG |
| 13012_2 | TATGAGTGTGGAGTCGTTGCTCTCCTTTTATTAATAGGAAGGCNNNAGATCGGAAG |
| 44016_1 | TATGAGTGTGGAGTCGTTGCCTTGAAAGCTCACACAGGTTCTGNNNAGATCGGAAG |
| 44016_2 | TATGAGTGTGGAGTCGTTGCCTTGAAAGCTCACACAGGTTCTGNNNAGATCGGAAG |
| 30045_1 | TATGAGTGTGGAGTCGTTGCGCCAAGTGAGTAATTTACAGCTGCNNNAGATCGGAA |
| 30045_2 | TATGAGTGTGGAGTCGTTGCGCCAAGTGAGTAATTTACAGCTGCNNNAGATCGGAA |
| 35872_1 | TATGAGTGTGGAGTCGTTGCATGATAAAAGCAACCACACAGCCNNNAGATCGGAAG |
| 35872_2 | TATGAGTGTGGAGTCGTTGCATGATAAAAGCAACCACACAGCCNNNAGATCGGAAG |
| 36388_1 | TATGAGTGTGGAGTCGTTGCTAACTCAGATCTCTGCTGTTGGGNNNAGATCGGAAG |
| 36388_2 | TATGAGTGTGGAGTCGTTGCTAACTCAGATCTCTGCTGTTGGGNNNAGATCGGAAG |
| 37149_1 | TATGAGTGTGGAGTCGTTGCAGAAGAGGAATGAAGGGAGAGGANNNAGATCGG |
| 37149_2 | TATGAGTGTGGAGTCGTTGCAGAAGAGGAATGAAGGGAGAGGANNNAGATCGG |
| 46284_1 | TATGAGTGTGGAGTCGTTGCCCTCCTGGGTTCAAGTGATTCTCNNNAGATCGGAAG |
| 46284_2 | TATGAGTGTGGAGTCGTTGCCCTCCTGGGTTCAAGTGATTCTCNNNAGATCGGAAG |
| 48430_1 | TATGAGTGTGGAGTCGTTGCAAAGTTCTGGGATTACAGGCGTNNNAGATCGGAAG |
| 48430_2 | TATGAGTGTGGAGTCGTTGCAAAGTTCTGGGATTACAGGCGTNNNAGATCGGAAG |
| 31778_1 | TATGAGTGTGGAGTCGTTGCAGAGAGAGAGAGAGAGAGAGGGANNNAGATCGG |
| 31778_2 | TATGAGTGTGGAGTCGTTGCAGAGAGAGAGAGAGAGAGAGGGANNNAGATCGG |
| 50107_1 | TATGAGTGTGGAGTCGTTGCACTCTAAGTCCAACTCTGCCAAGNNNAGATCGGAAG |
| 50107_2 | TATGAGTGTGGAGTCGTTGCACTCTAAGTCCAACTCTGCCAAGNNNAGATCGGAAG |
| 41926_1 | TATGAGTGTGGAGTCGTTGCTGTTAGCCAGGATGGTCTTGATCNNNAGATCGGAA |
| 41926_2 | TATGAGTGTGGAGTCGTTGCTGTTAGCCAGGATGGTCTTGATCNNNAGATCGGAA |
| 31829_1 | TATGAGTGTGGAGTCGTTGCCACACACAAACACACACACACATNNNAGATCGGAAG |
| 31829_2 | TATGAGTGTGGAGTCGTTGCCACACACAAACACACACACACATNNNAGATCGGAAG |
| 51641_1 | TATGAGTGTGGAGTCGTTGCGGAGAGCAGGAAAGATCTTCCAANNNAGATCGGAA |
| 51641_2 | TATGAGTGTGGAGTCGTTGCGGAGAGCAGGAAAGATCTTCCAANNNAGATCGGAA |
| 50392_1 | TATGAGTGTGGAGTCGTTGCCAGCCTGTGAAGGGAAGGAGNNNAGATCGGAAGA |
| 50392_2 | TATGAGTGTGGAGTCGTTGCCAGCCTGTGAAGGGAAGGAGNNNAGATCGGAAGA |
| 33531_1 | TATGAGTGTGGAGTCGTTGCGAATATCATTGCCCTGAAAGCCCNNNAGATCGGAAG |
| 33531_2 | TATGAGTGTGGAGTCGTTGCGAATATCATTGCCCTGAAAGCCCNNNAGATCGGAAG |
| 44638_1 | TATGAGTGTGGAGTCGTTGCGGTCTTCGATTTTTCTTGCCCTGNNNAGATCGGAAG |
| 44638_2 | TATGAGTGTGGAGTCGTTGCGGTCTTCGATTTTTCTTGCCCTGNNNAGATCGGAAG |
| 29733_1 | TATGAGTGTGGAGTCGTTGCGCTTGGCCCGGGATATTTATAAANNNAGATCGGAA |
| 29733_2 | TATGAGTGTGGAGTCGTTGCGCTTGGCCCGGGATATTTATAAANNNAGATCGGAA |
| 42668_1 | TATGAGTGTGGAGTCGTTGCTATGGTGGTTCACGCCTGTAATCNNNAGATCGGAAG |
| 42668_2 | TATGAGTGTGGAGTCGTTGCTATGGTGGTTCACGCCTGTAATCNNNAGATCGGAAG |
| 42397_1 | TATGAGTGTGGAGTCGTTGCCACACATACACACACACACTCACNNNAGATCGGAAGA |
| 42397_2 | TATGAGTGTGGAGTCGTTGCCACACATACACACACACACTCACNNNAGATCGGAAGA |
| 30917_1 | TATGAGTGTGGAGTCGTTGCAGTTCTCCAAAGTAAATGGCTAGTNNNAGATCGGAA |

30917_2 TATGAGTGTGGAGTCGTTGCAGTTCTCCAAAGTAAATGGCTAGTNNNAGATCGGAA
34232_1 TATGAGTGTGGAGTCGTTGCAGAATACACTTTTCGGACTGCTTNNNAGATCGGAAG
34232_2 TATGAGTGTGGAGTCGTTGCAGAATACACTTTTCGGACTGCTTNNNAGATCGGAAG
37694_1 TATGAGTGTGGAGTCGTTGCAGTGAGTGTGTAAAGGAGTGCAANNNAGATCGGAA
37694_2 TATGAGTGTGGAGTCGTTGCAGTGAGTGTGTAAAGGAGTGCAANNNAGATCGGAA
29211_1 TATGAGTGTGGAGTCGTTGCCTCCACCCATCCCATCTGTTATCNNNAGATCGGAAGA
29211_2 TATGAGTGTGGAGTCGTTGCCTCCACCCATCCCATCTGTTATCNNNAGATCGGAAGA
53278_1 TATGAGTGTGGAGTCGTTGCTTATCAGCTTTGGTGTGATTGCGNNNAGATCGGAAG
53278_2 TATGAGTGTGGAGTCGTTGCTTATCAGCTTTGGTGTGATTGCGNNNAGATCGGAAG
30684_1 TATGAGTGTGGAGTCGTTGCACTGTAGAGGTCAGTGTCTCACNNNAGATCGGAAG
30684_2 TATGAGTGTGGAGTCGTTGCACTGTAGAGGTCAGTGTCTCACNNNAGATCGGAAG
51082_1 TATGAGTGTGGAGTCGTTGCTTTCCCCAAGGCCTCCAAATAGNNNAGATCGGAAGA
51082_2 TATGAGTGTGGAGTCGTTGCTTTCCCCAAGGCCTCCAAATAGNNNAGATCGGAAGA
44320_1 TATGAGTGTGGAGTCGTTGCGAAGCAAGATGTGTGTGCATACANNNAGATCGGAA
44320_2 TATGAGTGTGGAGTCGTTGCGAAGCAAGATGTGTGTGCATACANNNAGATCGGAA
48022_1 TATGAGTGTGGAGTCGTTGCGACCTCAGCTCACTGCAACANNNAGATCGGAAGAGC
48022_2 TATGAGTGTGGAGTCGTTGCGACCTCAGCTCACTGCAACANNNAGATCGGAAGAGC
39233_1 TATGAGTGTGGAGTCGTTGCACCCTTACTTCTGGTAAATGGTATCTNNNAGATCGG
39233_2 TATGAGTGTGGAGTCGTTGCACCCTTACTTCTGGTAAATGGTATCTNNNAGATCGG
47927_1 TATGAGTGTGGAGTCGTTGCGGTCTCGAACTCCTAACCTCAAGNNNAGATCGGAAG
47927_2 TATGAGTGTGGAGTCGTTGCGGTCTCGAACTCCTAACCTCAAGNNNAGATCGGAAG
52770_1 TATGAGTGTGGAGTCGTTGCAATGGATAGGATGAGGGGTAGCTNNNAGATCGGA
52770_2 TATGAGTGTGGAGTCGTTGCAATGGATAGGATGAGGGGTAGCTNNNAGATCGGA
45830_1 TATGAGTGTGGAGTCGTTGCGAGACGGAGTCTTGCTCTGTCNNNAGATCGGAAGA
45830_2 TATGAGTGTGGAGTCGTTGCGAGACGGAGTCTTGCTCTGTCNNNAGATCGGAAGA
50573_1 TATGAGTGTGGAGTCGTTGCACCCCCATTGTAAACTACAGACCNNNAGATCGGAAG
50573_2 TATGAGTGTGGAGTCGTTGCACCCCCATTGTAAACTACAGACCNNNAGATCGGAAG
34429_1 TATGAGTGTGGAGTCGTTGCGGTAGTTTAGGCCCTGGTTTGATNNNAGATCGGAA
34429_2 TATGAGTGTGGAGTCGTTGCGGTAGTTTAGGCCCTGGTTTGATNNNAGATCGGAA
38912_1 TATGAGTGTGGAGTCGTTGCAGCACAGGACCAGATGATTTCAANNNAGATCGGAA
38912_2 TATGAGTGTGGAGTCGTTGCAGCACAGGACCAGATGATTTCAANNNAGATCGGAA
32437_1 TATGAGTGTGGAGTCGTTGCCTGCAAACCGCTCATTCTGTAAANNNAGATCGGAAG
32437_2 TATGAGTGTGGAGTCGTTGCCTGCAAACCGCTCATTCTGTAAANNNAGATCGGAAG
44970_1 TATGAGTGTGGAGTCGTTGCCATTCACCTGAGCTGTCACTACTNNNAGATCGGAAG
44970_2 TATGAGTGTGGAGTCGTTGCCATTCACCTGAGCTGTCACTACTNNNAGATCGGAAG
48760_1 TATGAGTGTGGAGTCGTTGCCCTCCCTGAGTAGCTGGAATTACNNNAGATCGGAAG
48760_2 TATGAGTGTGGAGTCGTTGCCCTCCCTGAGTAGCTGGAATTACNNNAGATCGGAAG
46274_1 TATGAGTGTGGAGTCGTTGCCTCCTGAGTAGCTGGGACTACAGNNNAGATCGGAA
46274_2 TATGAGTGTGGAGTCGTTGCCTCCTGAGTAGCTGGGACTACAGNNNAGATCGGAA
44734_1 TATGAGTGTGGAGTCGTTGCTTTGCTGAAAGTTGGCTGGAAAANNNAGATCGGAA
44734_2 TATGAGTGTGGAGTCGTTGCTTTGCTGAAAGTTGGCTGGAAAANNNAGATCGGAA
53151_1 TATGAGTGTGGAGTCGTTGCGCCTTTGTCTGAACTTTCCCAAANNNAGATCGGAAG
53151_2 TATGAGTGTGGAGTCGTTGCGCCTTTGTCTGAACTTTCCCAAANNNAGATCGGAAG

| | |
|---|---|
| 50610_1 | TATGAGTGTGGAGTCGTTGCGGACAGGAATAAGAGGGCCTATGNNNAGATCGGA |
| 50610_2 | TATGAGTGTGGAGTCGTTGCGGACAGGAATAAGAGGGCCTATGNNNAGATCGGA |
| 36073_1 | TATGAGTGTGGAGTCGTTGCCTAATCCCCAGAACCTGTGAACANNNAGATCGGAAG |
| 36073_2 | TATGAGTGTGGAGTCGTTGCCTAATCCCCAGAACCTGTGAACANNNAGATCGGAAG |
| 45933_1 | TATGAGTGTGGAGTCGTTGCTTGTGGCAGCTCTATCTCAATCANNNAGATCGGAAG |
| 45933_2 | TATGAGTGTGGAGTCGTTGCTTGTGGCAGCTCTATCTCAATCANNNAGATCGGAAG |
| 36484_1 | TATGAGTGTGGAGTCGTTGCTGGAAAACAATGTGGAGATCCCTNNNAGATCGGAA |
| 36484_2 | TATGAGTGTGGAGTCGTTGCTGGAAAACAATGTGGAGATCCCTNNNAGATCGGAA |
| 45792_1 | TATGAGTGTGGAGTCGTTGCAAGAGCTGTTCTTGTCCAGGAATNNNAGATCGGAAG |
| 45792_2 | TATGAGTGTGGAGTCGTTGCAAGAGCTGTTCTTGTCCAGGAATNNNAGATCGGAAG |
| 43741_1 | TATGAGTGTGGAGTCGTTGCTCTCTCACTCAAACACTAAGCCCNNNAGATCGGAAGA |
| 43741_2 | TATGAGTGTGGAGTCGTTGCTCTCTCACTCAAACACTAAGCCCNNNAGATCGGAAGA |
| 51411_1 | TATGAGTGTGGAGTCGTTGCACAAAGTTCTGACCAAAACACCGNNNAGATCGGAAG |
| 51411_2 | TATGAGTGTGGAGTCGTTGCACAAAGTTCTGACCAAAACACCGNNNAGATCGGAAG |
| 49245_1 | TATGAGTGTGGAGTCGTTGCTAGTAGAGACGGGGTTTCTCCATNNNAGATCGGAA |
| 49245_2 | TATGAGTGTGGAGTCGTTGCTAGTAGAGACGGGGTTTCTCCATNNNAGATCGGAA |
| 13243_1 | TATGAGTGTGGAGTCGTTGCTACTGTTTCTTTGAAACTTAAGGNNNAGATCGGAAG |
| 13243_2 | TATGAGTGTGGAGTCGTTGCTACTGTTTCTTTGAAACTTAAGGNNNAGATCGGAAG |
| 39586_1 | TATGAGTGTGGAGTCGTTGCATGTGGGAAAAGAGGTGAAAGGANNNAGATCGGA |
| 39586_2 | TATGAGTGTGGAGTCGTTGCATGTGGGAAAAGAGGTGAAAGGANNNAGATCGGA |
| 33455_1 | TATGAGTGTGGAGTCGTTGCACCAGGGGCTCAGAGATAAAAAGNNNAGATCGGAA |
| 33455_2 | TATGAGTGTGGAGTCGTTGCACCAGGGGCTCAGAGATAAAAAGNNNAGATCGGAA |
| 51114_1 | TATGAGTGTGGAGTCGTTGCGGAAAAATGGGATTGTGGACCAGNNNAGATCGGA |
| 51114_2 | TATGAGTGTGGAGTCGTTGCGGAAAAATGGGATTGTGGACCAGNNNAGATCGGA |
| 30885_1 | TATGAGTGTGGAGTCGTTGCTGGTATGGAGTCAGAAGGTAGGANNNAGATCGGA |
| 30885_2 | TATGAGTGTGGAGTCGTTGCTGGTATGGAGTCAGAAGGTAGGANNNAGATCGGA |
| 35022_1 | TATGAGTGTGGAGTCGTTGCAAGCTATACTGCATGTAGGCACANNNAGATCGGAAG |
| 35022_2 | TATGAGTGTGGAGTCGTTGCAAGCTATACTGCATGTAGGCACANNNAGATCGGAAG |
| 31063_1 | TATGAGTGTGGAGTCGTTGCCAATGGAGCCAGATTGCAGAAAANNNAGATCGGAA |
| 31063_2 | TATGAGTGTGGAGTCGTTGCCAATGGAGCCAGATTGCAGAAAANNNAGATCGGAA |
| 32278_1 | TATGAGTGTGGAGTCGTTGCCTGGAATTACAAGTGTGCACCACNNNAGATCGGAAG |
| 32278_2 | TATGAGTGTGGAGTCGTTGCCTGGAATTACAAGTGTGCACCACNNNAGATCGGAAG |
| 31429_1 | TATGAGTGTGGAGTCGTTGCATAGCAGAACCAATTGTGCCCTANNNAGATCGGAAG |
| 31429_2 | TATGAGTGTGGAGTCGTTGCATAGCAGAACCAATTGTGCCCTANNNAGATCGGAAG |
| 29370_1 | TATGAGTGTGGAGTCGTTGCTAGAGACCATTGTGAGTGTGGTGNNNAGATCGGAA |
| 29370_2 | TATGAGTGTGGAGTCGTTGCTAGAGACCATTGTGAGTGTGGTGNNNAGATCGGAA |
| 50030_1 | TATGAGTGTGGAGTCGTTGCAGACACAGAGAGAGAGATGGGAANNNAGATCGGA |
| 50030_2 | TATGAGTGTGGAGTCGTTGCAGACACAGAGAGAGAGATGGGAANNNAGATCGGA |
| 43589_1 | TATGAGTGTGGAGTCGTTGCCTGTGGAAATGAAGGGAAAGCTGNNNAGATCGGA |
| 43589_2 | TATGAGTGTGGAGTCGTTGCCTGTGGAAATGAAGGGAAAGCTGNNNAGATCGGA |
| 12577_1 | TATGAGTGTGGAGTCGTTGCTCAAAACAAATAAGTTTTCATCAGNNNAGATCGGAA |
| 12577_2 | TATGAGTGTGGAGTCGTTGCTCAAAACAAATAAGTTTTCATCAGNNNAGATCGGAA |
| 34379_1 | TATGAGTGTGGAGTCGTTGCGAGGACATTCTGATGGAGGAGGANNNAGATCGGA |

| | |
|---|---|
| 34379_2 | TATGAGTGTGGAGTCGTTGCGAGGACATTCTGATGGAGGAGGANNNAGATCGGA |
| 50798_1 | TATGAGTGTGGAGTCGTTGCACAAAAAATGGGCTTACTCTGGCNNNAGATCGGAAG |
| 50798_2 | TATGAGTGTGGAGTCGTTGCACAAAAAATGGGCTTACTCTGGCNNNAGATCGGAAG |
| 28814_1 | TATGAGTGTGGAGTCGTTGCAATTGTGCCACTGTAAACATGCANNNAGATCGGAAG |
| 28814_2 | TATGAGTGTGGAGTCGTTGCAATTGTGCCACTGTAAACATGCANNNAGATCGGAAG |
| 49831_1 | TATGAGTGTGGAGTCGTTGCCTTCCCACCATCCTACCTCCTAANNNAGATCGGAAGA |
| 49831_2 | TATGAGTGTGGAGTCGTTGCCTTCCCACCATCCTACCTCCTAANNNAGATCGGAAGA |
| 31128_1 | TATGAGTGTGGAGTCGTTGCATTGCTTCAATCTGCTTCCAAAGNNNAGATCGGAAG |
| 31128_2 | TATGAGTGTGGAGTCGTTGCATTGCTTCAATCTGCTTCCAAAGNNNAGATCGGAAG |
| 36662_1 | TATGAGTGTGGAGTCGTTGCCACCTGTGATCCTAGCTACTCAGNNNAGATCGGAAG |
| 36662_2 | TATGAGTGTGGAGTCGTTGCCACCTGTGATCCTAGCTACTCAGNNNAGATCGGAAG |
| 49208_1 | TATGAGTGTGGAGTCGTTGCTCAAGTCTTTCCCTCAGATGCAANNNAGATCGGAAG |
| 49208_2 | TATGAGTGTGGAGTCGTTGCTCAAGTCTTTCCCTCAGATGCAANNNAGATCGGAAG |
| 44803_1 | TATGAGTGTGGAGTCGTTGCCTGTAGAAGCTGCCCTTACCTTANNNAGATCGGAAG |
| 44803_2 | TATGAGTGTGGAGTCGTTGCCTGTAGAAGCTGCCCTTACCTTANNNAGATCGGAAG |
| 2121_1 | TATGAGTGTGGAGTCGTTGCACACAGTCTGAGGTACTACTGCANNNAGATCGGAAG |
| 2121_2 | TATGAGTGTGGAGTCGTTGCACACAGTCTGAGGTACTACTGCANNNAGATCGGAAG |
| 10750_1 | TATGAGTGTGGAGTCGTTGCCAACATGGATAACCCTCAAGGGTNNNAGATCGGAAG |
| 10750_2 | TATGAGTGTGGAGTCGTTGCCAACATGGATAACCCTCAAGGGTNNNAGATCGGAAG |
| 12414_1 | TATGAGTGTGGAGTCGTTGCGCAGTGTTTCTAATTTCAGGGAGCNNNAGATCGGAA |
| 12414_2 | TATGAGTGTGGAGTCGTTGCGCAGTGTTTCTAATTTCAGGGAGCNNNAGATCGGAA |
| 47909_1 | TATGAGTGTGGAGTCGTTGCAGTAGCTGAGAATACAGATGCCGNNNAGATCGGAA |
| 47909_2 | TATGAGTGTGGAGTCGTTGCAGTAGCTGAGAATACAGATGCCGNNNAGATCGGAA |
| 31224_1 | TATGAGTGTGGAGTCGTTGCTTCTGAGCCACCTAAATTTGGGANNNAGATCGGAAG |
| 31224_2 | TATGAGTGTGGAGTCGTTGCTTCTGAGCCACCTAAATTTGGGANNNAGATCGGAAG |
| 49834_1 | TATGAGTGTGGAGTCGTTGCGTTCAGTCCCAAGGTGTACAGAGNNNAGATCGGAA |
| 49834_2 | TATGAGTGTGGAGTCGTTGCGTTCAGTCCCAAGGTGTACAGAGNNNAGATCGGAA |
| 13797_1 | TATGAGTGTGGAGTCGTTGCCACACTCTCCTATCTTTACTTACNNNAGATCGGAAGA |
| 13797_2 | TATGAGTGTGGAGTCGTTGCCACACTCTCCTATCTTTACTTACNNNAGATCGGAAGA |
| 45778_1 | TATGAGTGTGGAGTCGTTGCGAGTAGCTGGGATTACAGGCATGNNNAGATCGGAA |
| 45778_2 | TATGAGTGTGGAGTCGTTGCGAGTAGCTGGGATTACAGGCATGNNNAGATCGGAA |
| 39242_1 | TATGAGTGTGGAGTCGTTGCCCTATAACTATAGCTGGCCAGGCNNNAGATCGGAAG |
| 39242_2 | TATGAGTGTGGAGTCGTTGCCCTATAACTATAGCTGGCCAGGCNNNAGATCGGAAG |
| 41957_1 | TATGAGTGTGGAGTCGTTGCCCTGGTTTCAAGCAATTCTCCTGNNNAGATCGGAAG |
| 41957_2 | TATGAGTGTGGAGTCGTTGCCCTGGTTTCAAGCAATTCTCCTGNNNAGATCGGAAG |
| 44619_1 | TATGAGTGTGGAGTCGTTGCGTGAGGTGCAAAGCTCGGAGNNNAGATCGGAAGA |
| 44619_2 | TATGAGTGTGGAGTCGTTGCGTGAGGTGCAAAGCTCGGAGNNNAGATCGGAAGA |
| 28992_1 | TATGAGTGTGGAGTCGTTGCTTCCGAAGAGGCAAGAGTGTTTNNNAGATCGGAAG |
| 28992_2 | TATGAGTGTGGAGTCGTTGCTTCCGAAGAGGCAAGAGTGTTTNNNAGATCGGAAG |
| 46041_1 | TATGAGTGTGGAGTCGTTGCTGTGAATACAACAGCTGACCACANNNAGATCGGAAG |
| 46041_2 | TATGAGTGTGGAGTCGTTGCTGTGAATACAACAGCTGACCACANNNAGATCGGAAG |
| 45238_1 | TATGAGTGTGGAGTCGTTGCTGACCAAACTGTGCTGTTAGAGANNNAGATCGGAA |
| 45238_2 | TATGAGTGTGGAGTCGTTGCTGACCAAACTGTGCTGTTAGAGANNNAGATCGGAA |

| | |
|---|---|
| 41072_1 | TATGAGTGTGGAGTCGTTGCGAGGCAGGAGAATGGTGTGAANNNAGATCGGAAG |
| 41072_2 | TATGAGTGTGGAGTCGTTGCGAGGCAGGAGAATGGTGTGAANNNAGATCGGAAG |
| 52606_1 | TATGAGTGTGGAGTCGTTGCCATGTTGACCACCAGGAACTTTCNNNAGATCGGAAG |
| 52606_2 | TATGAGTGTGGAGTCGTTGCCATGTTGACCACCAGGAACTTTCNNNAGATCGGAAG |
| 52246_1 | TATGAGTGTGGAGTCGTTGCGCCTACATTTCCTCTTGGCACTANNNAGATCGGAAGA |
| 52246_2 | TATGAGTGTGGAGTCGTTGCGCCTACATTTCCTCTTGGCACTANNNAGATCGGAAGA |
| 40543_1 | TATGAGTGTGGAGTCGTTGCGGATCCTGCTTTTGCAGAAACTTNNNAGATCGGAAG |
| 40543_2 | TATGAGTGTGGAGTCGTTGCGGATCCTGCTTTTGCAGAAACTTNNNAGATCGGAAG |
| 48478_1 | TATGAGTGTGGAGTCGTTGCCTGGGTTCAAATGATTCTCCTGCNNNAGATCGGAAG |
| 48478_2 | TATGAGTGTGGAGTCGTTGCCTGGGTTCAAATGATTCTCCTGCNNNAGATCGGAAG |
| 36649_1 | TATGAGTGTGGAGTCGTTGCTGTAGCACATTTGACCTAGGCAANNNAGATCGGAAG |
| 36649_2 | TATGAGTGTGGAGTCGTTGCTGTAGCACATTTGACCTAGGCAANNNAGATCGGAAG |
| 49198_1 | TATGAGTGTGGAGTCGTTGCCTCAATCTTGGCTCACTGCAAGNNNAGATCGGAAGA |
| 49198_2 | TATGAGTGTGGAGTCGTTGCCTCAATCTTGGCTCACTGCAAGNNNAGATCGGAAGA |
| 36331_1 | TATGAGTGTGGAGTCGTTGCAACTGAGTGAATAAGCACCACCANNNAGATCGGAAG |
| 36331_2 | TATGAGTGTGGAGTCGTTGCAACTGAGTGAATAAGCACCACCANNNAGATCGGAAG |
| 38788_1 | TATGAGTGTGGAGTCGTTGCACACATTACACACTAGAAGAAGAGANNNAGATCGG |
| 38788_2 | TATGAGTGTGGAGTCGTTGCACACATTACACACTAGAAGAAGAGANNNAGATCGG |
| 33308_1 | TATGAGTGTGGAGTCGTTGCCCTACCAGGTTCACACCATTCTCNNNAGATCGGAAGA |
| 33308_2 | TATGAGTGTGGAGTCGTTGCCCTACCAGGTTCACACCATTCTCNNNAGATCGGAAGA |
| 33071_1 | TATGAGTGTGGAGTCGTTGCCACAATCTTGGCTCACTGCAACNNNAGATCGGAAGA |
| 33071_2 | TATGAGTGTGGAGTCGTTGCCACAATCTTGGCTCACTGCAACNNNAGATCGGAAGA |
| 36055_1 | TATGAGTGTGGAGTCGTTGCTCTTTTTGGGAGACCACAGTTCANNNAGATCGGAAG |
| 36055_2 | TATGAGTGTGGAGTCGTTGCTCTTTTTGGGAGACCACAGTTCANNNAGATCGGAAG |
| 44231_1 | TATGAGTGTGGAGTCGTTGCAAAATGACACCCCAGAGAACAGTNNNAGATCGGAAG |
| 44231_2 | TATGAGTGTGGAGTCGTTGCAAAATGACACCCCAGAGAACAGTNNNAGATCGGAAG |
| 34270_1 | TATGAGTGTGGAGTCGTTGCGGAATCTGCCCTGTGTCGTCNNNAGATCGGAAGAG |
| 34270_2 | TATGAGTGTGGAGTCGTTGCGGAATCTGCCCTGTGTCGTCNNNAGATCGGAAGAG |
| 41748_1 | TATGAGTGTGGAGTCGTTGCTGCCTAGGCCTTCCAAATTTCTANNNAGATCGGAAG |
| 41748_2 | TATGAGTGTGGAGTCGTTGCTGCCTAGGCCTTCCAAATTTCTANNNAGATCGGAAG |
| 44728_1 | TATGAGTGTGGAGTCGTTGCACCACCACAAATAATTTCCCCCTNNNAGATCGGAAGA |
| 44728_2 | TATGAGTGTGGAGTCGTTGCACCACCACAAATAATTTCCCCCTNNNAGATCGGAAGA |
| 34272_1 | TATGAGTGTGGAGTCGTTGCGAGTGTGTGTGTGTGTGTAAAGGNNNAGATCGGA |
| 34272_2 | TATGAGTGTGGAGTCGTTGCGAGTGTGTGTGTGTGTGTAAAGGNNNAGATCGGA |
| 31366_1 | TATGAGTGTGGAGTCGTTGCAGAGAGAGAGAGAGGGAAAGAGGNNNAGATCGG |
| 31366_2 | TATGAGTGTGGAGTCGTTGCAGAGAGAGAGAGAGGGAAAGAGGNNNAGATCGG |
| 10133_1 | TATGAGTGTGGAGTCGTTGCGGTAAGTCCCTGAAACTAGGTAANNNAGATCGGAA |
| 10133_2 | TATGAGTGTGGAGTCGTTGCGGTAAGTCCCTGAAACTAGGTAANNNAGATCGGAA |
| 42905_1 | TATGAGTGTGGAGTCGTTGCATTGTCCAGGTTGGTCTTGAACTNNNAGATCGGAAG |
| 42905_2 | TATGAGTGTGGAGTCGTTGCATTGTCCAGGTTGGTCTTGAACTNNNAGATCGGAAG |
| 53204_1 | TATGAGTGTGGAGTCGTTGCTTTTTTTTGAGACGGAGTCTGGCNNNAGATCGGAAG |
| 53204_2 | TATGAGTGTGGAGTCGTTGCTTTTTTTTGAGACGGAGTCTGGCNNNAGATCGGAAG |
| 50498_1 | TATGAGTGTGGAGTCGTTGCGAGAACCAGAATGGCTTGGAACNNNAGATCGGAAG |

| | |
|---|---|
| 50498_2 | TATGAGTGTGGAGTCGTTGCGAGAACCAGAATGGCTTGGAACNNNAGATCGGAAG |
| 30010_1 | TATGAGTGTGGAGTCGTTGCTGATCGTGAACTGAGAGTTTCCANNNAGATCGGAAG |
| 30010_2 | TATGAGTGTGGAGTCGTTGCTGATCGTGAACTGAGAGTTTCCANNNAGATCGGAAG |
| 46114_1 | TATGAGTGTGGAGTCGTTGCAAGTGCTGGCTTTACAGGTATGANNNAGATCGGAA |
| 46114_2 | TATGAGTGTGGAGTCGTTGCAAGTGCTGGCTTTACAGGTATGANNNAGATCGGAA |
| 48513_1 | TATGAGTGTGGAGTCGTTGCCTGTCAGTTTCCACAGGAGATGANNNAGATCGGAAG |
| 48513_2 | TATGAGTGTGGAGTCGTTGCCTGTCAGTTTCCACAGGAGATGANNNAGATCGGAAG |
| 38553_1 | TATGAGTGTGGAGTCGTTGCATTGTGGCCCGAACCTATAGTTCNNNAGATCGGAAG |
| 38553_2 | TATGAGTGTGGAGTCGTTGCATTGTGGCCCGAACCTATAGTTCNNNAGATCGGAAG |
| 53257_1 | TATGAGTGTGGAGTCGTTGCATGAAACAGCAATGGCAGCATAGNNNAGATCGGAA |
| 53257_2 | TATGAGTGTGGAGTCGTTGCATGAAACAGCAATGGCAGCATAGNNNAGATCGGAA |
| 44946_1 | TATGAGTGTGGAGTCGTTGCGGAGTCCAGATCTAGTGCTGAGANNNAGATCGGAA |
| 44946_2 | TATGAGTGTGGAGTCGTTGCGGAGTCCAGATCTAGTGCTGAGANNNAGATCGGAA |
| 35312_1 | TATGAGTGTGGAGTCGTTGCGAGTAGCTAGGATTACAGGTGCCNNNAGATCGGAA |
| 35312_2 | TATGAGTGTGGAGTCGTTGCGAGTAGCTAGGATTACAGGTGCCNNNAGATCGGAA |
| 42933_1 | TATGAGTGTGGAGTCGTTGCAAAAGAGAGAGAGATCCACACGCNNNAGATCGGAA |
| 42933_2 | TATGAGTGTGGAGTCGTTGCAAAAGAGAGAGAGATCCACACGCNNNAGATCGGAA |
| 37755_1 | TATGAGTGTGGAGTCGTTGCTCCCATTTTCCTTCCTTCCTTCCNNNAGATCGGAAGAG |
| 37755_2 | TATGAGTGTGGAGTCGTTGCTCCCATTTTCCTTCCTTCCTTCCNNNAGATCGGAAGAG |
| 40741_1 | TATGAGTGTGGAGTCGTTGCCTGTGTTCAGGTTTCTGTGTAGANNNAGATCGGAAG |
| 40741_2 | TATGAGTGTGGAGTCGTTGCCTGTGTTCAGGTTTCTGTGTAGANNNAGATCGGAAG |
| 40777_1 | TATGAGTGTGGAGTCGTTGCTAGGTGCAGTCAATATGGGATGGNNNAGATCGGAA |
| 40777_2 | TATGAGTGTGGAGTCGTTGCTAGGTGCAGTCAATATGGGATGGNNNAGATCGGAA |
| 51700_1 | TATGAGTGTGGAGTCGTTGCACTGAGATGTGTGGATAAAGGCTNNNAGATCGGAA |
| 51700_2 | TATGAGTGTGGAGTCGTTGCACTGAGATGTGTGGATAAAGGCTNNNAGATCGGAA |
| 46642_1 | TATGAGTGTGGAGTCGTTGCCTCCTGAGTAGCTGGGACTACAGNNNAGATCGGAA |
| 46642_2 | TATGAGTGTGGAGTCGTTGCCTCCTGAGTAGCTGGGACTACAGNNNAGATCGGAA |
| 49741_1 | TATGAGTGTGGAGTCGTTGCCCTCCTGGGTTCAAGAGATTCTCNNNAGATCGGAAG |
| 49741_2 | TATGAGTGTGGAGTCGTTGCCCTCCTGGGTTCAAGAGATTCTCNNNAGATCGGAAG |
| 39228_1 | TATGAGTGTGGAGTCGTTGCCTGAGGCAGGAGAATCCCTTGNNNAGATCGGAAGA |
| 39228_2 | TATGAGTGTGGAGTCGTTGCCTGAGGCAGGAGAATCCCTTGNNNAGATCGGAAGA |
| 38716_1 | TATGAGTGTGGAGTCGTTGCGCTACTCTGTCTGGTTTTCCCTTNNNAGATCGGAAGA |
| 38716_2 | TATGAGTGTGGAGTCGTTGCGCTACTCTGTCTGGTTTTCCCTTNNNAGATCGGAAGA |
| 53394_1 | TATGAGTGTGGAGTCGTTGCAATGGCATGATCTCAGCATCTCTNNNAGATCGGAAG |
| 53394_2 | TATGAGTGTGGAGTCGTTGCAATGGCATGATCTCAGCATCTCTNNNAGATCGGAAG |
| 12743_1 | TATGAGTGTGGAGTCGTTGCTAAAACCCAGGTAGTTAGATAAGNNNAGATCGGAA |
| 12743_2 | TATGAGTGTGGAGTCGTTGCTAAAACCCAGGTAGTTAGATAAGNNNAGATCGGAA |
| 33774_1 | TATGAGTGTGGAGTCGTTGCGGAGTCTCTGAGGACCTTCAAAGNNNAGATCGGAA |
| 33774_2 | TATGAGTGTGGAGTCGTTGCGGAGTCTCTGAGGACCTTCAAAGNNNAGATCGGAA |
| 38559_1 | TATGAGTGTGGAGTCGTTGCGAAAATTGCTTGAACCCAGGAGGNNNAGATCGGAA |
| 38559_2 | TATGAGTGTGGAGTCGTTGCGAAAATTGCTTGAACCCAGGAGGNNNAGATCGGAA |
| 31333_1 | TATGAGTGTGGAGTCGTTGCTCACCAAAAAACTGTGCATGAGANNNAGATCGGAAG |
| 31333_2 | TATGAGTGTGGAGTCGTTGCTCACCAAAAAACTGTGCATGAGANNNAGATCGGAAG |

| | |
|---|---|
| 50090_1 | TATGAGTGTGGAGTCGTTGCATGTGTAACAGATTGGGCTCACTNNNAGATCGGAA |
| 50090_2 | TATGAGTGTGGAGTCGTTGCATGTGTAACAGATTGGGCTCACTNNNAGATCGGAA |
| 48026_1 | TATGAGTGTGGAGTCGTTGCAAGAAAGCCCCAAGGAAAGAGGNNNAGATCGGAA |
| 48026_2 | TATGAGTGTGGAGTCGTTGCAAGAAAGCCCCAAGGAAAGAGGNNNAGATCGGAA |
| 35110_1 | TATGAGTGTGGAGTCGTTGCAAGCAATCCTTCCATCTCAACCTNNNAGATCGGAAGA |
| 35110_2 | TATGAGTGTGGAGTCGTTGCAAGCAATCCTTCCATCTCAACCTNNNAGATCGGAAGA |
| 380_1 | TATGAGTGTGGAGTCGTTGCCCCTCGTCCGTTTAGAAAGCTCNNNAGATCGGAAGA |
| 380_2 | TATGAGTGTGGAGTCGTTGCCCCTCGTCCGTTTAGAAAGCTCNNNAGATCGGAAGA |
| 30004_1 | TATGAGTGTGGAGTCGTTGCAGACCATCATCAGCACGAACTAGNNNAGATCGGAAG |
| 30004_2 | TATGAGTGTGGAGTCGTTGCAGACCATCATCAGCACGAACTAGNNNAGATCGGAAG |
| 50884_1 | TATGAGTGTGGAGTCGTTGCAGCAACAAAGAATGAAGGCCATGNNNAGATCGGAA |
| 50884_2 | TATGAGTGTGGAGTCGTTGCAGCAACAAAGAATGAAGGCCATGNNNAGATCGGAA |
| 34717_1 | TATGAGTGTGGAGTCGTTGCGTTCAAGCGTTTCTCCTGTTTCANNNAGATCGGAAG |
| 34717_2 | TATGAGTGTGGAGTCGTTGCGTTCAAGCGTTTCTCCTGTTTCANNNAGATCGGAAG |
| 42019_1 | TATGAGTGTGGAGTCGTTGCTGAAGATCACCTTGAGAGAACCCNNNAGATCGGAA |
| 42019_2 | TATGAGTGTGGAGTCGTTGCTGAAGATCACCTTGAGAGAACCCNNNAGATCGGAA |
| 35256_1 | TATGAGTGTGGAGTCGTTGCATACAATCTCTGCCACCCATGTTNNNAGATCGGAAG |
| 35256_2 | TATGAGTGTGGAGTCGTTGCATACAATCTCTGCCACCCATGTTNNNAGATCGGAAG |
| 42124_1 | TATGAGTGTGGAGTCGTTGCACCAACAACCATCAGAGAATAGTNNNAGATCGGAAG |
| 42124_2 | TATGAGTGTGGAGTCGTTGCACCAACAACCATCAGAGAATAGTNNNAGATCGGAAG |
| 48256_1 | TATGAGTGTGGAGTCGTTGCTGGTGCCAAATTTACAACAGTGTNNNAGATCGGAAG |
| 48256_2 | TATGAGTGTGGAGTCGTTGCTGGTGCCAAATTTACAACAGTGTNNNAGATCGGAAG |
| 10560_1 | TATGAGTGTGGAGTCGTTGCTGTCATTGCATTCTTGTTTGGCTNNNAGATCGGAAG |
| 10560_2 | TATGAGTGTGGAGTCGTTGCTGTCATTGCATTCTTGTTTGGCTNNNAGATCGGAAG |
| 10405_1 | TATGAGTGTGGAGTCGTTGCGTGCAAACATACACACAGAGCCTNNNAGATCGGAAG |
| 10405_2 | TATGAGTGTGGAGTCGTTGCGTGCAAACATACACACAGAGCCTNNNAGATCGGAAG |
| 37786_1 | TATGAGTGTGGAGTCGTTGCCTCCATGGAACAGTTTTGGCATTNNNAGATCGGAAG |
| 37786_2 | TATGAGTGTGGAGTCGTTGCCTCCATGGAACAGTTTTGGCATTNNNAGATCGGAAG |
| 52491_1 | TATGAGTGTGGAGTCGTTGCCTCCTGAGTAGCTGGGATTACAGNNNAGATCGGAA |
| 52491_2 | TATGAGTGTGGAGTCGTTGCCTCCTGAGTAGCTGGGATTACAGNNNAGATCGGAA |
| 43214_1 | TATGAGTGTGGAGTCGTTGCACCATCACTGGGATTATTCTGGGNNNAGATCGGAAG |
| 43214_2 | TATGAGTGTGGAGTCGTTGCACCATCACTGGGATTATTCTGGGNNNAGATCGGAAG |
| 31803_1 | TATGAGTGTGGAGTCGTTGCACACTGTTTCTACCAATTGCTGANNNAGATCGGAAG |
| 31803_2 | TATGAGTGTGGAGTCGTTGCACACTGTTTCTACCAATTGCTGANNNAGATCGGAAG |
| 34349_1 | TATGAGTGTGGAGTCGTTGCGGACCTATGATGGGGGATCCTATNNNAGATCGGAA |
| 34349_2 | TATGAGTGTGGAGTCGTTGCGGACCTATGATGGGGGATCCTATNNNAGATCGGAA |
| 37487_1 | TATGAGTGTGGAGTCGTTGCAAGAAAATTAGCCGAGCATGGTGNNNAGATCGGAA |
| 37487_2 | TATGAGTGTGGAGTCGTTGCAAGAAAATTAGCCGAGCATGGTGNNNAGATCGGAA |
| 39714_1 | TATGAGTGTGGAGTCGTTGCAACTAGCCAAAGAACTAGCCCTCNNNAGATCGGAAG |
| 39714_2 | TATGAGTGTGGAGTCGTTGCAACTAGCCAAAGAACTAGCCCTCNNNAGATCGGAAG |
| 12530_1 | TATGAGTGTGGAGTCGTTGCTTTGTGAAGAAATTCAGAATGCNNNAGATCGGAAG |
| 12530_2 | TATGAGTGTGGAGTCGTTGCTTTGTGAAGAAATTCAGAATGCNNNAGATCGGAAG |
| 41750_1 | TATGAGTGTGGAGTCGTTGCAAGAGATTCTCCTACCTCAGCCTNNNAGATCGGAAG |

| | |
|---|---|
| 41750_2 | TATGAGTGTGGAGTCGTTGCAAGAGATTCTCCTACCTCAGCCTNNNAGATCGGAAG |
| 45685_1 | TATGAGTGTGGAGTCGTTGCTAGAGTGCAATGGTGTGATCTCGNNNAGATCGGAA |
| 45685_2 | TATGAGTGTGGAGTCGTTGCTAGAGTGCAATGGTGTGATCTCGNNNAGATCGGAA |
| 32702_1 | TATGAGTGTGGAGTCGTTGCTCAAACTTTTGGGCTCAAGCAATNNNAGATCGGAAG |
| 32702_2 | TATGAGTGTGGAGTCGTTGCTCAAACTTTTGGGCTCAAGCAATNNNAGATCGGAAG |
| 46539_1 | TATGAGTGTGGAGTCGTTGCTTCAAGTGATTCTCCTGCCTCAGNNNAGATCGGAAG |
| 46539_2 | TATGAGTGTGGAGTCGTTGCTTCAAGTGATTCTCCTGCCTCAGNNNAGATCGGAAG |
| 39696_1 | TATGAGTGTGGAGTCGTTGCAGTCTACCAATGAATGCCACTGTNNNAGATCGGAAG |
| 39696_2 | TATGAGTGTGGAGTCGTTGCAGTCTACCAATGAATGCCACTGTNNNAGATCGGAAG |
| 35644_1 | TATGAGTGTGGAGTCGTTGCGTCATTGCTTTTGACCTGCCTAGNNNAGATCGGAAG |
| 35644_2 | TATGAGTGTGGAGTCGTTGCGTCATTGCTTTTGACCTGCCTAGNNNAGATCGGAAG |
| 52098_1 | TATGAGTGTGGAGTCGTTGCCACCATTCCCGGCTAATTTTTGTNNNAGATCGGAAGA |
| 52098_2 | TATGAGTGTGGAGTCGTTGCCACCATTCCCGGCTAATTTTTGTNNNAGATCGGAAGA |
| 52313_1 | TATGAGTGTGGAGTCGTTGCCCAGGCCACAACTGTTTTCTTAGNNNAGATCGGAAG |
| 52313_2 | TATGAGTGTGGAGTCGTTGCCCAGGCCACAACTGTTTTCTTAGNNNAGATCGGAAG |
| 33752_1 | TATGAGTGTGGAGTCGTTGCACATGAGGGGAAATGTTTTCCAANNNAGATCGGAA |
| 33752_2 | TATGAGTGTGGAGTCGTTGCACATGAGGGGAAATGTTTTCCAANNNAGATCGGAA |
| 31173_1 | TATGAGTGTGGAGTCGTTGCCTTTCTTCTTTCTGATTCCCGGCNNNAGATCGGAAGA |
| 31173_2 | TATGAGTGTGGAGTCGTTGCCTTTCTTCTTTCTGATTCCCGGCNNNAGATCGGAAGA |
| 37099_1 | TATGAGTGTGGAGTCGTTGCTAAGGTCATTGTTTTGTGAGGGGNNNAGATCGGAA |
| 37099_2 | TATGAGTGTGGAGTCGTTGCTAAGGTCATTGTTTTGTGAGGGGNNNAGATCGGAA |
| 49582_1 | TATGAGTGTGGAGTCGTTGCCTCCTGTGTAGCTGGGATTACAGNNNAGATCGGAAG |
| 49582_2 | TATGAGTGTGGAGTCGTTGCCTCCTGTGTAGCTGGGATTACAGNNNAGATCGGAAG |
| 42927_1 | TATGAGTGTGGAGTCGTTGCTTCTTCCCATCTGACACTGAGTCNNNAGATCGGAAGA |
| 42927_2 | TATGAGTGTGGAGTCGTTGCTTCTTCCCATCTGACACTGAGTCNNNAGATCGGAAGA |
| 50302_1 | TATGAGTGTGGAGTCGTTGCAGTTCACTAAGACCCTGAGTGCNNNAGATCGGAAGA |
| 50302_2 | TATGAGTGTGGAGTCGTTGCAGTTCACTAAGACCCTGAGTGCNNNAGATCGGAAGA |
| 50701_1 | TATGAGTGTGGAGTCGTTGCCTTAGTCCCAGAAGGCTTCTCAGNNNAGATCGGAAG |
| 50701_2 | TATGAGTGTGGAGTCGTTGCCTTAGTCCCAGAAGGCTTCTCAGNNNAGATCGGAAG |
| 31207_1 | TATGAGTGTGGAGTCGTTGCCCTCATGACCAAATTACCTCCCANNNAGATCGGAAGA |
| 31207_2 | TATGAGTGTGGAGTCGTTGCCCTCATGACCAAATTACCTCCCANNNAGATCGGAAGA |
| 36592_1 | TATGAGTGTGGAGTCGTTGCAAGCACTTCAGACAAAGGGTACTNNNAGATCGGAA |
| 36592_2 | TATGAGTGTGGAGTCGTTGCAAGCACTTCAGACAAAGGGTACTNNNAGATCGGAA |
| 41632_1 | TATGAGTGTGGAGTCGTTGCTTCAAGTGATTCTCCTGCCTCAGNNNAGATCGGAAG |
| 41632_2 | TATGAGTGTGGAGTCGTTGCTTCAAGTGATTCTCCTGCCTCAGNNNAGATCGGAAG |
| 12288_1 | TATGAGTGTGGAGTCGTTGCAGGTCACTTCCAACAGGGAAAATGNNNAGATCGGA |
| 12288_2 | TATGAGTGTGGAGTCGTTGCAGGTCACTTCCAACAGGGAAAATGNNNAGATCGGA |
| 43423_1 | TATGAGTGTGGAGTCGTTGCTGTGGTCTTTCTAAAACACCCCANNNAGATCGGAAG |
| 43423_2 | TATGAGTGTGGAGTCGTTGCTGTGGTCTTTCTAAAACACCCCANNNAGATCGGAAG |
| 46501_1 | TATGAGTGTGGAGTCGTTGCTCTCCCGAGTTCTAAGCGATTTTNNNAGATCGGAAG |
| 46501_2 | TATGAGTGTGGAGTCGTTGCTCTCCCGAGTTCTAAGCGATTTTNNNAGATCGGAAG |
| 50504_1 | TATGAGTGTGGAGTCGTTGCCAGGGCAGTTTGAATCACTTGAGNNNAGATCGGAA |
| 50504_2 | TATGAGTGTGGAGTCGTTGCCAGGGCAGTTTGAATCACTTGAGNNNAGATCGGAA |

| | |
|---|---|
| 37123_1 | TATGAGTGTGGAGTCGTTGCGAGGTCAGGAGATCAAGACCATCNNNAGATCGGAA |
| 37123_2 | TATGAGTGTGGAGTCGTTGCGAGGTCAGGAGATCAAGACCATCNNNAGATCGGAA |
| 41942_1 | TATGAGTGTGGAGTCGTTGCCAGGGGTCAATTTCTCCAAGGTANNNAGATCGGAAG |
| 41942_2 | TATGAGTGTGGAGTCGTTGCCAGGGGTCAATTTCTCCAAGGTANNNAGATCGGAAG |
| 29348_1 | TATGAGTGTGGAGTCGTTGCCTACATTTTTCCCACAGAGCAGCNNNAGATCGGAAG |
| 29348_2 | TATGAGTGTGGAGTCGTTGCCTACATTTTTCCCACAGAGCAGCNNNAGATCGGAAG |
| 33215_1 | TATGAGTGTGGAGTCGTTGCTGACAAATCATCTCACCCCTGTTNNNAGATCGGAAGA |
| 33215_2 | TATGAGTGTGGAGTCGTTGCTGACAAATCATCTCACCCCTGTTNNNAGATCGGAAGA |
| 7372_1 | TATGAGTGTGGAGTCGTTGCCTTCCTTTTCTGCTCTCCATGTCNNNAGATCGGAAGA |
| 7372_2 | TATGAGTGTGGAGTCGTTGCCTTCCTTTTCTGCTCTCCATGTCNNNAGATCGGAAGA |
| 33952_1 | TATGAGTGTGGAGTCGTTGCCCTCCTGGGTTCAAGAGATTCTGNNNAGATCGGAAG |
| 33952_2 | TATGAGTGTGGAGTCGTTGCCCTCCTGGGTTCAAGAGATTCTGNNNAGATCGGAAG |
| 49022_1 | TATGAGTGTGGAGTCGTTGCTTGGGCAACTTCCTGACATAGTANNNAGATCGGAAG |
| 49022_2 | TATGAGTGTGGAGTCGTTGCTTGGGCAACTTCCTGACATAGTANNNAGATCGGAAG |
| 34048_1 | TATGAGTGTGGAGTCGTTGCCTGAGTAGCTGGGATTACAGGTGNNNAGATCGGAA |
| 34048_2 | TATGAGTGTGGAGTCGTTGCCTGAGTAGCTGGGATTACAGGTGNNNAGATCGGAA |
| 33632_1 | TATGAGTGTGGAGTCGTTGCTTCAAGCGATTCTCCTGTCTCAGNNNAGATCGGAAG |
| 33632_2 | TATGAGTGTGGAGTCGTTGCTTCAAGCGATTCTCCTGTCTCAGNNNAGATCGGAAG |
| 46644_1 | TATGAGTGTGGAGTCGTTGCTGTGACTCCGGGAATACTCCTATNNNAGATCGGAAG |
| 46644_2 | TATGAGTGTGGAGTCGTTGCTGTGACTCCGGGAATACTCCTATNNNAGATCGGAAG |
| 49684_1 | TATGAGTGTGGAGTCGTTGCCGAGTAGCTGGGATTACAGGTGNNNAGATCGGAAG |
| 49684_2 | TATGAGTGTGGAGTCGTTGCCGAGTAGCTGGGATTACAGGTGNNNAGATCGGAAG |
| 49074_1 | TATGAGTGTGGAGTCGTTGCCAATCTCAGCTCACTGCAAACTCNNNAGATCGGAAG |
| 49074_2 | TATGAGTGTGGAGTCGTTGCCAATCTCAGCTCACTGCAAACTCNNNAGATCGGAAG |
| 52033_1 | TATGAGTGTGGAGTCGTTGCTGCATGGAATTTGGACAGACCTANNNAGATCGGAA |
| 52033_2 | TATGAGTGTGGAGTCGTTGCTGCATGGAATTTGGACAGACCTANNNAGATCGGAA |
| 38885_1 | TATGAGTGTGGAGTCGTTGCAATATTCACTGCAGCAATCGCATNNNAGATCGGAAG |
| 38885_2 | TATGAGTGTGGAGTCGTTGCAATATTCACTGCAGCAATCGCATNNNAGATCGGAAG |
| 47907_1 | TATGAGTGTGGAGTCGTTGCCCTTGCCGGACTCCTCTCTCNNNAGATCGGAAGAGC |
| 47907_2 | TATGAGTGTGGAGTCGTTGCCCTTGCCGGACTCCTCTCTCNNNAGATCGGAAGAGC |
| 13285_1 | TATGAGTGTGGAGTCGTTGCATAACCAAAGTCACCAATACTTANNNAGATCGGAAG |
| 13285_2 | TATGAGTGTGGAGTCGTTGCATAACCAAAGTCACCAATACTTANNNAGATCGGAAG |
| 53318_1 | TATGAGTGTGGAGTCGTTGCTAGGGCAAAGGGGAATCATTCAANNNAGATCGGAA |
| 53318_2 | TATGAGTGTGGAGTCGTTGCTAGGGCAAAGGGGAATCATTCAANNNAGATCGGAA |
| 31294_1 | TATGAGTGTGGAGTCGTTGCGTCTTTGCCTCAGCTTCCTCATANNNAGATCGGAAGA |
| 31294_2 | TATGAGTGTGGAGTCGTTGCGTCTTTGCCTCAGCTTCCTCATANNNAGATCGGAAGA |
| 7056_1 | TATGAGTGTGGAGTCGTTGCTAAGTACCCTCTTCTCCCTCCCTNNNAGATCGGAAGA |
| 7056_2 | TATGAGTGTGGAGTCGTTGCTAAGTACCCTCTTCTCCCTCCCTNNNAGATCGGAAGA |
| 13000_1 | TATGAGTGTGGAGTCGTTGCGGTTTATCTGGTGGGAATAGNNNAGATCGGAAGAG |
| 13000_2 | TATGAGTGTGGAGTCGTTGCGGTTTATCTGGTGGGAATAGNNNAGATCGGAAGAG |
| 41717_1 | TATGAGTGTGGAGTCGTTGCCACGCCCGGCTAATTTTTTGTATNNNAGATCGGAAG |
| 41717_2 | TATGAGTGTGGAGTCGTTGCCACGCCCGGCTAATTTTTTGTATNNNAGATCGGAAG |
| 52090_1 | TATGAGTGTGGAGTCGTTGCCTGTTCAAGGAAACCAGAGAGGANNNAGATCGGAA |

| | |
|---|---|
| 52090_2 | TATGAGTGTGGAGTCGTTGCCTGTTCAAGGAAACCAGAGAGGANNNAGATCGGAA |
| 38518_1 | TATGAGTGTGGAGTCGTTGCTCCCATATGCCATTCCAGTGATGNNNAGATCGGAAG |
| 38518_2 | TATGAGTGTGGAGTCGTTGCTCCCATATGCCATTCCAGTGATGNNNAGATCGGAAG |
| 43244_1 | TATGAGTGTGGAGTCGTTGCCTGCAGTCCTAGCTACTTGAGAGNNNAGATCGGAAG |
| 43244_2 | TATGAGTGTGGAGTCGTTGCCTGCAGTCCTAGCTACTTGAGAGNNNAGATCGGAAG |
| 53308_1 | TATGAGTGTGGAGTCGTTGCTGAACAGGGGTGTGATATTTGGTNNNAGATCGGAA |
| 53308_2 | TATGAGTGTGGAGTCGTTGCTGAACAGGGGTGTGATATTTGGTNNNAGATCGGAA |
| 12995_1 | TATGAGTGTGGAGTCGTTGCAACTGTTGCGGATATACTTTCNNNAGATCGGAAGAG |
| 12995_2 | TATGAGTGTGGAGTCGTTGCAACTGTTGCGGATATACTTTCNNNAGATCGGAAGAG |
| 35387_1 | TATGAGTGTGGAGTCGTTGCCACCACCATGTCCAGCTAATTTTNNNAGATCGGAAG |
| 35387_2 | TATGAGTGTGGAGTCGTTGCCACCACCATGTCCAGCTAATTTTNNNAGATCGGAAG |
| 45327_1 | TATGAGTGTGGAGTCGTTGCTCCTATTTTCTCCCTCCACCTCTNNNAGATCGGAAGAG |
| 45327_2 | TATGAGTGTGGAGTCGTTGCTCCTATTTTCTCCCTCCACCTCTNNNAGATCGGAAGAG |
| 321_1 | TATGAGTGTGGAGTCGTTGCCCTTGCTATTTCATCAGTAGGCANNNAGATCGGAAG |
| 321_2 | TATGAGTGTGGAGTCGTTGCCCTTGCTATTTCATCAGTAGGCANNNAGATCGGAAG |
| 41066_1 | TATGAGTGTGGAGTCGTTGCCTAAGTGTGGTCAGCCCTCTATGNNNAGATCGGAAG |
| 41066_2 | TATGAGTGTGGAGTCGTTGCCTAAGTGTGGTCAGCCCTCTATGNNNAGATCGGAAG |
| 35945_1 | TATGAGTGTGGAGTCGTTGCGGTGGATCATGAGGTCAAGAGATNNNAGATCGGAA |
| 35945_2 | TATGAGTGTGGAGTCGTTGCGGTGGATCATGAGGTCAAGAGATNNNAGATCGGAA |
| 49267_1 | TATGAGTGTGGAGTCGTTGCAGAAAAGCACAGAGGCAGTTCTANNNAGATCGGAA |
| 49267_2 | TATGAGTGTGGAGTCGTTGCAGAAAAGCACAGAGGCAGTTCTANNNAGATCGGAA |
| 33829_1 | TATGAGTGTGGAGTCGTTGCTACCTCCAATGTTCAGCAACTCANNNAGATCGGAAG |
| 33829_2 | TATGAGTGTGGAGTCGTTGCTACCTCCAATGTTCAGCAACTCANNNAGATCGGAAG |
| 45697_1 | TATGAGTGTGGAGTCGTTGCTCGTGCCAATGTTTTTGGATGAANNNAGATCGGAAG |
| 45697_2 | TATGAGTGTGGAGTCGTTGCTCGTGCCAATGTTTTTGGATGAANNNAGATCGGAAG |
| 40690_1 | TATGAGTGTGGAGTCGTTGCATAGAAGGTGTCAATTTCCCCCCNNNAGATCGGAAG |
| 40690_2 | TATGAGTGTGGAGTCGTTGCATAGAAGGTGTCAATTTCCCCCCNNNAGATCGGAAG |
| 32957_1 | TATGAGTGTGGAGTCGTTGCTGGAGGTGAGATCAATAGTTTCCANNNAGATCGGA |
| 32957_2 | TATGAGTGTGGAGTCGTTGCTGGAGGTGAGATCAATAGTTTCCANNNAGATCGGA |
| 10861_1 | TATGAGTGTGGAGTCGTTGCAGGGGCTGGTGATAATCTCTGTTNNNAGATCGGAA |
| 10861_2 | TATGAGTGTGGAGTCGTTGCAGGGGCTGGTGATAATCTCTGTTNNNAGATCGGAA |
| 34856_1 | TATGAGTGTGGAGTCGTTGCACGCCCGGCTAATTTATTTTTGTNNNAGATCGGAAG |
| 34856_2 | TATGAGTGTGGAGTCGTTGCACGCCCGGCTAATTTATTTTTGTNNNAGATCGGAAG |
| 32720_1 | TATGAGTGTGGAGTCGTTGCTTAGCCATCAGAGAAATGCCAGTNNNAGATCGGAA |
| 32720_2 | TATGAGTGTGGAGTCGTTGCTTAGCCATCAGAGAAATGCCAGTNNNAGATCGGAA |
| 49655_1 | TATGAGTGTGGAGTCGTTGCTTGATCTCCTGACCTTGTGATCCNNNAGATCGGAAG |
| 49655_2 | TATGAGTGTGGAGTCGTTGCTTGATCTCCTGACCTTGTGATCCNNNAGATCGGAAG |
| 45035_1 | TATGAGTGTGGAGTCGTTGCCTCAGATGGGTAAGGGTTTGGAANNNAGATCGGAA |
| 45035_2 | TATGAGTGTGGAGTCGTTGCCTCAGATGGGTAAGGGTTTGGAANNNAGATCGGAA |
| 34410_1 | TATGAGTGTGGAGTCGTTGCTGACATCATTGCCAGAAGGAGGNNNAGATCGGAAG |
| 34410_2 | TATGAGTGTGGAGTCGTTGCTGACATCATTGCCAGAAGGAGGNNNAGATCGGAAG |
| 41556_1 | TATGAGTGTGGAGTCGTTGCCTCCCAAGTAGCTGTGACTACAGNNNAGATCGGAAG |
| 41556_2 | TATGAGTGTGGAGTCGTTGCCTCCCAAGTAGCTGTGACTACAGNNNAGATCGGAAG |

| | |
|---|---|
| 31529_1 | TATGAGTGTGGAGTCGTTGCGGATTTTGAGAGAACAAGGCCACNNNAGATCGGAA |
| 31529_2 | TATGAGTGTGGAGTCGTTGCGGATTTTGAGAGAACAAGGCCACNNNAGATCGGAA |
| 31131_1 | TATGAGTGTGGAGTCGTTGCTTTGAGACGGTCACACTCTTTCCNNNAGATCGGAAG |
| 31131_2 | TATGAGTGTGGAGTCGTTGCTTTGAGACGGTCACACTCTTTCCNNNAGATCGGAAG |
| 40906_1 | TATGAGTGTGGAGTCGTTGCGGTCATTGTCTCTGCCATCTCTTNNNAGATCGGAAGA |
| 40906_2 | TATGAGTGTGGAGTCGTTGCGGTCATTGTCTCTGCCATCTCTTNNNAGATCGGAAGA |
| 35807_1 | TATGAGTGTGGAGTCGTTGCGCGATGGTATGCAATCTCAGTTCNNNAGATCGGAAG |
| 35807_2 | TATGAGTGTGGAGTCGTTGCGCGATGGTATGCAATCTCAGTTCNNNAGATCGGAAG |
| 52517_1 | TATGAGTGTGGAGTCGTTGCCTACCACACACCAGACAGAGAACNNNAGATCGGAAG |
| 52517_2 | TATGAGTGTGGAGTCGTTGCCTACCACACACCAGACAGAGAACNNNAGATCGGAAG |
| 42468_1 | TATGAGTGTGGAGTCGTTGCATCATGACACACTGCAACCTCAANNNAGATCGGAAG |
| 42468_2 | TATGAGTGTGGAGTCGTTGCATCATGACACACTGCAACCTCAANNNAGATCGGAAG |
| 41019_1 | TATGAGTGTGGAGTCGTTGCTTGCTGCTGTTCTCTTAGGCTAANNNAGATCGGAAG |
| 41019_2 | TATGAGTGTGGAGTCGTTGCTTGCTGCTGTTCTCTTAGGCTAANNNAGATCGGAAG |
| 48261_1 | TATGAGTGTGGAGTCGTTGCCCCTTCCCAGTCCCCAGAAGNNNAGATCGGAAGAGC |
| 48261_2 | TATGAGTGTGGAGTCGTTGCCCCTTCCCAGTCCCCAGAAGNNNAGATCGGAAGAGC |
| 29473_1 | TATGAGTGTGGAGTCGTTGCCCACATTCACCAAAGCTGTCATCNNNAGATCGGAAG |
| 29473_2 | TATGAGTGTGGAGTCGTTGCCCACATTCACCAAAGCTGTCATCNNNAGATCGGAAG |
| 42450_1 | TATGAGTGTGGAGTCGTTGCGCCCAGCCTACCATTTTCAGATANNNAGATCGGAAG |
| 42450_2 | TATGAGTGTGGAGTCGTTGCGCCCAGCCTACCATTTTCAGATANNNAGATCGGAAG |
| 3279_1 | TATGAGTGTGGAGTCGTTGCTCAGCACCATCTACTTTCCTGGANNNAGATCGGAAG |
| 3279_2 | TATGAGTGTGGAGTCGTTGCTCAGCACCATCTACTTTCCTGGANNNAGATCGGAAG |
| 50703_1 | TATGAGTGTGGAGTCGTTGCCGCTGGCTCCAAGTATACTGTATNNNAGATCGGAAG |
| 50703_2 | TATGAGTGTGGAGTCGTTGCCGCTGGCTCCAAGTATACTGTATNNNAGATCGGAAG |
| 44018_1 | TATGAGTGTGGAGTCGTTGCGTTTTGATCACACAGATGGCCAANNNAGATCGGAAG |
| 44018_2 | TATGAGTGTGGAGTCGTTGCGTTTTGATCACACAGATGGCCAANNNAGATCGGAAG |
| 47188_1 | TATGAGTGTGGAGTCGTTGCCCCATCATCACACCTGGCTAATTNNNAGATCGGAAGA |
| 47188_2 | TATGAGTGTGGAGTCGTTGCCCCATCATCACACCTGGCTAATTNNNAGATCGGAAGA |
| 44977_1 | TATGAGTGTGGAGTCGTTGCCAGACAGGGTGGGTAAAGACTGNNNAGATCGGAA |
| 44977_2 | TATGAGTGTGGAGTCGTTGCCAGACAGGGTGGGTAAAGACTGNNNAGATCGGAA |
| 30204_1 | TATGAGTGTGGAGTCGTTGCCTGTGGCCAAACGGTTTAACTCNNNAGATCGGAAGA |
| 30204_2 | TATGAGTGTGGAGTCGTTGCCTGTGGCCAAACGGTTTAACTCNNNAGATCGGAAGA |
| 13500_1 | TATGAGTGTGGAGTCGTTGCCCAGGTATATTCATCTCTAAGAANNNAGATCGGAAG |
| 13500_2 | TATGAGTGTGGAGTCGTTGCCCAGGTATATTCATCTCTAAGAANNNAGATCGGAAG |
| 52824_1 | TATGAGTGTGGAGTCGTTGCCCACAGAGATCTCCCTTTGGAAANNNAGATCGGAAG |
| 52824_2 | TATGAGTGTGGAGTCGTTGCCCACAGAGATCTCCCTTTGGAAANNNAGATCGGAAG |
| 40997_1 | TATGAGTGTGGAGTCGTTGCCTCCCTGACTCAAGCAAGATTCTNNNAGATCGGAAG |
| 40997_2 | TATGAGTGTGGAGTCGTTGCCTCCCTGACTCAAGCAAGATTCTNNNAGATCGGAAG |
| 31998_1 | TATGAGTGTGGAGTCGTTGCAATAATGGCATTTGCAGCAACCTNNNAGATCGGAAG |
| 31998_2 | TATGAGTGTGGAGTCGTTGCAATAATGGCATTTGCAGCAACCTNNNAGATCGGAAG |
| 46781_1 | TATGAGTGTGGAGTCGTTGCGGCAGTCAGCTAAGCCATTTGNNNAGATCGGAAGA |
| 46781_2 | TATGAGTGTGGAGTCGTTGCGGCAGTCAGCTAAGCCATTTGNNNAGATCGGAAGA |
| 49625_1 | TATGAGTGTGGAGTCGTTGCCAAGTAGCTGGGATTACAGGCATNNNAGATCGGAA |

| | |
|---|---|
| 49625_2 | TATGAGTGTGGAGTCGTTGCCAAGTAGCTGGGATTACAGGCATNNNAGATCGGAA |
| 39636_1 | TATGAGTGTGGAGTCGTTGCCCTCGTGATTTCTTCTTTGCCTCNNNAGATCGGAAGA |
| 39636_2 | TATGAGTGTGGAGTCGTTGCCCTCGTGATTTCTTCTTTGCCTCNNNAGATCGGAAGA |
| 51888_1 | TATGAGTGTGGAGTCGTTGCCTCCCTTAGCAGCTAGAGAGTTGNNNAGATCGGAAG |
| 51888_2 | TATGAGTGTGGAGTCGTTGCCTCCCTTAGCAGCTAGAGAGTTGNNNAGATCGGAAG |
| 43751_1 | TATGAGTGTGGAGTCGTTGCTGCTCAAAGCTTTTCAGCCTTACNNNAGATCGGAAGA |
| 43751_2 | TATGAGTGTGGAGTCGTTGCTGCTCAAAGCTTTTCAGCCTTACNNNAGATCGGAAGA |
| 29924_1 | TATGAGTGTGGAGTCGTTGCTGTTACTCTGAACCTCCTTGAGGNNNAGATCGGAAG |
| 29924_2 | TATGAGTGTGGAGTCGTTGCTGTTACTCTGAACCTCCTTGAGGNNNAGATCGGAAG |
| 42071_1 | TATGAGTGTGGAGTCGTTGCTCCAAAATGCTTCATTCAAACCCANNNAGATCGGAAG |
| 42071_2 | TATGAGTGTGGAGTCGTTGCTCCAAAATGCTTCATTCAAACCCANNNAGATCGGAAG |
| 39438_1 | TATGAGTGTGGAGTCGTTGCAGAATGAAAAAGCATGCAAGGGTNNNAGATCGGAA |
| 39438_2 | TATGAGTGTGGAGTCGTTGCAGAATGAAAAAGCATGCAAGGGTNNNAGATCGGAA |
| 46666_1 | TATGAGTGTGGAGTCGTTGCGATTACAGAGGTAACCTGCCACANNNAGATCGGAA |
| 46666_2 | TATGAGTGTGGAGTCGTTGCGATTACAGAGGTAACCTGCCACANNNAGATCGGAA |
| 47808_1 | TATGAGTGTGGAGTCGTTGCTTGGGTTCAAGAGATTCTCCTGCNNNAGATCGGAAG |
| 47808_2 | TATGAGTGTGGAGTCGTTGCTTGGGTTCAAGAGATTCTCCTGCNNNAGATCGGAAG |
| 31547_1 | TATGAGTGTGGAGTCGTTGCGATGTCCTTTGTCCTTTTGCTCCNNNAGATCGGAAGA |
| 31547_2 | TATGAGTGTGGAGTCGTTGCGATGTCCTTTGTCCTTTTGCTCCNNNAGATCGGAAGA |
| 33686_1 | TATGAGTGTGGAGTCGTTGCTGTGTAGAACAGGCAAACCCATANNNAGATCGGAA |
| 33686_2 | TATGAGTGTGGAGTCGTTGCTGTGTAGAACAGGCAAACCCATANNNAGATCGGAA |
| 44240_1 | TATGAGTGTGGAGTCGTTGCCTTCCCTGAGACTCTCCACAAAANNNAGATCGGAAGA |
| 44240_2 | TATGAGTGTGGAGTCGTTGCCTTCCCTGAGACTCTCCACAAAANNNAGATCGGAAGA |
| 40825_1 | TATGAGTGTGGAGTCGTTGCAACAAATGGAAACACATCCCAGGNNNAGATCGGAA |
| 40825_2 | TATGAGTGTGGAGTCGTTGCAACAAATGGAAACACATCCCAGGNNNAGATCGGAA |
| 39439_1 | TATGAGTGTGGAGTCGTTGCTGTCCTGTAAGTCCAAGCAATGANNNAGATCGGAAG |
| 39439_2 | TATGAGTGTGGAGTCGTTGCTGTCCTGTAAGTCCAAGCAATGANNNAGATCGGAAG |
| 42203_1 | TATGAGTGTGGAGTCGTTGCGGTGAGTCTGTGGTCATTTTGTGNNNAGATCGGAA |
| 42203_2 | TATGAGTGTGGAGTCGTTGCGGTGAGTCTGTGGTCATTTTGTGNNNAGATCGGAA |
| 36239_1 | TATGAGTGTGGAGTCGTTGCAGGATGAAGTTAGACCTCACATCANNNAGATCGGAA |
| 36239_2 | TATGAGTGTGGAGTCGTTGCAGGATGAAGTTAGACCTCACATCANNNAGATCGGAA |
| 37418_1 | TATGAGTGTGGAGTCGTTGCAAACCCCATCATGTTGTACACCTNNNAGATCGGAAGA |
| 37418_2 | TATGAGTGTGGAGTCGTTGCAAACCCCATCATGTTGTACACCTNNNAGATCGGAAGA |
| 10772_1 | TATGAGTGTGGAGTCGTTGCCAGGAACCTTCAACTAGAGACCTGNNNAGATCGGAA |
| 10772_2 | TATGAGTGTGGAGTCGTTGCCAGGAACCTTCAACTAGAGACCTGNNNAGATCGGAA |
| 52916_1 | TATGAGTGTGGAGTCGTTGCCTTGGGGTCGGCAGGAAATATCNNNAGATCGGAAG |
| 52916_2 | TATGAGTGTGGAGTCGTTGCCTTGGGGTCGGCAGGAAATATCNNNAGATCGGAAG |
| 37443_1 | TATGAGTGTGGAGTCGTTGCCACAAAAAAATTAGCCGGGCATGNNNAGATCGGAA |
| 37443_2 | TATGAGTGTGGAGTCGTTGCCACAAAAAAATTAGCCGGGCATGNNNAGATCGGAA |
| 46948_1 | TATGAGTGTGGAGTCGTTGCACAGAAAGAGAGAAGTGGGGATGNNNAGATCGGA |
| 46948_2 | TATGAGTGTGGAGTCGTTGCACAGAAAGAGAGAAGTGGGGATGNNNAGATCGGA |
| 46007_1 | TATGAGTGTGGAGTCGTTGCGATCTCTGTTCACTGTGACCCTGNNNAGATCGGAAG |
| 46007_2 | TATGAGTGTGGAGTCGTTGCGATCTCTGTTCACTGTGACCCTGNNNAGATCGGAAG |

| | |
|---|---|
| 32077_1 | TATGAGTGTGGAGTCGTTGCCATTCTCCTGCCTCAGCCTCNNNAGATCGGAAGAGCA |
| 32077_2 | TATGAGTGTGGAGTCGTTGCCATTCTCCTGCCTCAGCCTCNNNAGATCGGAAGAGCA |
| 41172_1 | TATGAGTGTGGAGTCGTTGCACTTTTACACTGTTGGTGGGAGTNNNAGATCGGAAG |
| 41172_2 | TATGAGTGTGGAGTCGTTGCACTTTTACACTGTTGGTGGGAGTNNNAGATCGGAAG |
| 53398_1 | TATGAGTGTGGAGTCGTTGCGAGAGGAGAGACAGAGACAGAGANNNAGATCGGA |
| 53398_2 | TATGAGTGTGGAGTCGTTGCGAGAGGAGAGACAGAGACAGAGANNNAGATCGGA |
| 45196_1 | TATGAGTGTGGAGTCGTTGCGGGAAGGAAAGAATGTTTGGCTCNNNAGATCGGAA |
| 45196_2 | TATGAGTGTGGAGTCGTTGCGGGAAGGAAAGAATGTTTGGCTCNNNAGATCGGAA |
| 51351_1 | TATGAGTGTGGAGTCGTTGCCAGGGTCCTGGTGTTAGAATCAANNNAGATCGGAA |
| 51351_2 | TATGAGTGTGGAGTCGTTGCCAGGGTCCTGGTGTTAGAATCAANNNAGATCGGAA |
| 52236_1 | TATGAGTGTGGAGTCGTTGCCGGTGCAGAAGTCAATAGAGAGANNNAGATCGGAA |
| 52236_2 | TATGAGTGTGGAGTCGTTGCCGGTGCAGAAGTCAATAGAGAGANNNAGATCGGAA |
| 53407_1 | TATGAGTGTGGAGTCGTTGCAGAAAGAGAGAGACAAAGGAAAGGANNNAGATCG |
| 53407_2 | TATGAGTGTGGAGTCGTTGCAGAAAGAGAGAGACAAAGGAAAGGANNNAGATCG |
| 28823_1 | TATGAGTGTGGAGTCGTTGCACATGTATGTGTGTGCATATGTGTNNNAGATCGGAA |
| 28823_2 | TATGAGTGTGGAGTCGTTGCACATGTATGTGTGTGCATATGTGTNNNAGATCGGAA |
| 29478_1 | TATGAGTGTGGAGTCGTTGCATTAACTCATAGTTGTGCGAGCANNNAGATCGGAAG |
| 29478_2 | TATGAGTGTGGAGTCGTTGCATTAACTCATAGTTGTGCGAGCANNNAGATCGGAAG |
| 33222_1 | TATGAGTGTGGAGTCGTTGCCGCCCGGCTTTTCTTTTTTTTTTNNNAGATCGGAAGA |
| 33222_2 | TATGAGTGTGGAGTCGTTGCCGCCCGGCTTTTCTTTTTTTTTTNNNAGATCGGAAGA |
| 44842_1 | TATGAGTGTGGAGTCGTTGCGCAGGGGATAGAGATTGGAGAAGNNNAGATCGGA |
| 44842_2 | TATGAGTGTGGAGTCGTTGCGCAGGGGATAGAGATTGGAGAAGNNNAGATCGGA |
| 43259_1 | TATGAGTGTGGAGTCGTTGCTGTAGCAAGTTGGATGGAGCTAGNNNAGATCGGAA |
| 43259_2 | TATGAGTGTGGAGTCGTTGCTGTAGCAAGTTGGATGGAGCTAGNNNAGATCGGAA |
| 37430_1 | TATGAGTGTGGAGTCGTTGCCGATTCCAGAACCAGAATTCACGNNNAGATCGGAAG |
| 37430_2 | TATGAGTGTGGAGTCGTTGCCGATTCCAGAACCAGAATTCACGNNNAGATCGGAAG |
| 30098_1 | TATGAGTGTGGAGTCGTTGCACCAGCGTTGTAGTACAGAAGTTNNNAGATCGGAA |
| 30098_2 | TATGAGTGTGGAGTCGTTGCACCAGCGTTGTAGTACAGAAGTTNNNAGATCGGAA |
| 45206_1 | TATGAGTGTGGAGTCGTTGCTATGGGAGCATGTCTTTGTGTGTNNNAGATCGGAA |
| 45206_2 | TATGAGTGTGGAGTCGTTGCTATGGGAGCATGTCTTTGTGTGTNNNAGATCGGAA |
| 33413_1 | TATGAGTGTGGAGTCGTTGCGGCGAAACCCTGTCTCTACTAAANNNAGATCGGAAG |
| 33413_2 | TATGAGTGTGGAGTCGTTGCGGCGAAACCCTGTCTCTACTAAANNNAGATCGGAAG |
| 10756_1 | TATGAGTGTGGAGTCGTTGCTTCAACATTTCCTTCCTGGAGCTNNNAGATCGGAAGA |
| 10756_2 | TATGAGTGTGGAGTCGTTGCTTCAACATTTCCTTCCTGGAGCTNNNAGATCGGAAGA |
| 30167_1 | TATGAGTGTGGAGTCGTTGCAAGATGGAGTTACTTGCCTGAGGNNNAGATCGGAA |
| 30167_2 | TATGAGTGTGGAGTCGTTGCAAGATGGAGTTACTTGCCTGAGGNNNAGATCGGAA |
| 46101_1 | TATGAGTGTGGAGTCGTTGCTGGCTATTTGGGCTCTTTTTTGGNNNAGATCGGAAG |
| 46101_2 | TATGAGTGTGGAGTCGTTGCTGGCTATTTGGGCTCTTTTTTGGNNNAGATCGGAAG |
| 40013_1 | TATGAGTGTGGAGTCGTTGCATCATTTGAGGTCGGGAGTTTGANNNAGATCGGAA |
| 40013_2 | TATGAGTGTGGAGTCGTTGCATCATTTGAGGTCGGGAGTTTGANNNAGATCGGAA |
| 33211_1 | TATGAGTGTGGAGTCGTTGCACACAAAGGATCAGGAATCAGAANNNAGATCGGAA |
| 33211_2 | TATGAGTGTGGAGTCGTTGCACACAAAGGATCAGGAATCAGAANNNAGATCGGAA |
| 43391_1 | TATGAGTGTGGAGTCGTTGCCGATTCTCCTGCCTCAGTCTCNNNAGATCGGAAGAG |

| | |
|---|---|
| 43391_2 | TATGAGTGTGGAGTCGTTGCCGATTCTCCTGCCTCAGTCTCNNNAGATCGGAAGAG |
| 48492_1 | TATGAGTGTGGAGTCGTTGCAAAGGAACTTACCTTCTGCCACTNNNAGATCGGAAG |
| 48492_2 | TATGAGTGTGGAGTCGTTGCAAAGGAACTTACCTTCTGCCACTNNNAGATCGGAAG |
| 12388_1 | TATGAGTGTGGAGTCGTTGCTATGTGCTAATTTCCATGGCCAGTNNNAGATCGGAA |
| 12388_2 | TATGAGTGTGGAGTCGTTGCTATGTGCTAATTTCCATGGCCAGTNNNAGATCGGAA |
| 43370_1 | TATGAGTGTGGAGTCGTTGCGATCTCCTGACCTCGTGATCTGNNNAGATCGGAAGA |
| 43370_2 | TATGAGTGTGGAGTCGTTGCGATCTCCTGACCTCGTGATCTGNNNAGATCGGAAGA |
| 38797_1 | TATGAGTGTGGAGTCGTTGCTCTGTCTTTCTTATTCCCACCCCNNNAGATCGGAAGA |
| 38797_2 | TATGAGTGTGGAGTCGTTGCTCTGTCTTTCTTATTCCCACCCCNNNAGATCGGAAGA |
| 35650_1 | TATGAGTGTGGAGTCGTTGCTTTAGTAGAGGCGGGGTTTTACCNNNAGATCGGAA |
| 35650_2 | TATGAGTGTGGAGTCGTTGCTTTAGTAGAGGCGGGGTTTTACCNNNAGATCGGAA |
| 49383_1 | TATGAGTGTGGAGTCGTTGCGCTCACTGCAAACTCTGTGTATTNNNAGATCGGAAG |
| 49383_2 | TATGAGTGTGGAGTCGTTGCGCTCACTGCAAACTCTGTGTATTNNNAGATCGGAAG |
| 2286_1 | TATGAGTGTGGAGTCGTTGCTGGGACTAGAGGTACACACACAGNNNAGATCGGAA |
| 2286_2 | TATGAGTGTGGAGTCGTTGCTGGGACTAGAGGTACACACACAGNNNAGATCGGAA |
| 13567_1 | TATGAGTGTGGAGTCGTTGCCTTCATCTGTATTTATAGCCACTNNNAGATCGGAAGA |
| 13567_2 | TATGAGTGTGGAGTCGTTGCCTTCATCTGTATTTATAGCCACTNNNAGATCGGAAGA |
| 50454_1 | TATGAGTGTGGAGTCGTTGCGAGAAAAGGGAGGGAGTCAGTTCNNNAGATCGGA |
| 50454_2 | TATGAGTGTGGAGTCGTTGCGAGAAAAGGGAGGGAGTCAGTTCNNNAGATCGGA |
| 38396_1 | TATGAGTGTGGAGTCGTTGCATGCACTGAAGGTAACCATAGCANNNAGATCGGAA |
| 38396_2 | TATGAGTGTGGAGTCGTTGCATGCACTGAAGGTAACCATAGCANNNAGATCGGAA |
| 40552_1 | TATGAGTGTGGAGTCGTTGCGCACAGGCAACAAAAGCAAAAATNNNAGATCGGAA |
| 40552_2 | TATGAGTGTGGAGTCGTTGCGCACAGGCAACAAAAGCAAAAATNNNAGATCGGAA |
| 33239_1 | TATGAGTGTGGAGTCGTTGCCTCCTGGGTTCTAAGGGATTCTCNNNAGATCGGAAG |
| 33239_2 | TATGAGTGTGGAGTCGTTGCCTCCTGGGTTCTAAGGGATTCTCNNNAGATCGGAAG |
| 44316_1 | TATGAGTGTGGAGTCGTTGCTGTAAAACTGCCCAACTACCTGTNNNAGATCGGAAG |
| 44316_2 | TATGAGTGTGGAGTCGTTGCTGTAAAACTGCCCAACTACCTGTNNNAGATCGGAAG |
| 52182_1 | TATGAGTGTGGAGTCGTTGCTAAAGCAACTTCCCTGTCCAGTTNNNAGATCGGAAG |
| 52182_2 | TATGAGTGTGGAGTCGTTGCTAAAGCAACTTCCCTGTCCAGTTNNNAGATCGGAAG |
| 34880_1 | TATGAGTGTGGAGTCGTTGCCTCCTGGGTTCATGCCATTCTCNNNAGATCGGAAGA |
| 34880_2 | TATGAGTGTGGAGTCGTTGCCTCCTGGGTTCATGCCATTCTCNNNAGATCGGAAGA |
| 47323_1 | TATGAGTGTGGAGTCGTTGCGATTCTTCCGTTTTCCTTTGGCANNNAGATCGGAAGA |
| 47323_2 | TATGAGTGTGGAGTCGTTGCGATTCTTCCGTTTTCCTTTGGCANNNAGATCGGAAGA |
| 52332_1 | TATGAGTGTGGAGTCGTTGCTCTGGGTCAAAGAAATAGTGGGGNNNAGATCGGAA |
| 52332_2 | TATGAGTGTGGAGTCGTTGCTCTGGGTCAAAGAAATAGTGGGGNNNAGATCGGAA |
| 32553_1 | TATGAGTGTGGAGTCGTTGCCGAATAGCTGGGACTACAGGCNNNAGATCGGAAGA |
| 32553_2 | TATGAGTGTGGAGTCGTTGCCGAATAGCTGGGACTACAGGCNNNAGATCGGAAGA |
| 39932_1 | TATGAGTGTGGAGTCGTTGCAGGATTGTCTTGGCTATAAGGGCNNNAGATCGGAA |
| 39932_2 | TATGAGTGTGGAGTCGTTGCAGGATTGTCTTGGCTATAAGGGCNNNAGATCGGAA |
| 28991_1 | TATGAGTGTGGAGTCGTTGCAGTGGTTTGGTTTGTCAAGAAGCNNNAGATCGGAA |
| 28991_2 | TATGAGTGTGGAGTCGTTGCAGTGGTTTGGTTTGTCAAGAAGCNNNAGATCGGAA |
| 44076_1 | TATGAGTGTGGAGTCGTTGCCCTATTTCCTCTCTGGCTCTGTCNNNAGATCGGAAGA |
| 44076_2 | TATGAGTGTGGAGTCGTTGCCCTATTTCCTCTCTGGCTCTGTCNNNAGATCGGAAGA |

| | |
|---|---|
| 51971_1 | TATGAGTGTGGAGTCGTTGCATGGCAATCAAAATCAGTGTGCANNNAGATCGGAA |
| 51971_2 | TATGAGTGTGGAGTCGTTGCATGGCAATCAAAATCAGTGTGCANNNAGATCGGAA |
| 49067_1 | TATGAGTGTGGAGTCGTTGCCTGGGTTCAAGCAATTCTCCTTCNNNAGATCGGAAG |
| 49067_2 | TATGAGTGTGGAGTCGTTGCCTGGGTTCAAGCAATTCTCCTTCNNNAGATCGGAAG |
| 37668_1 | TATGAGTGTGGAGTCGTTGCCCCCACAAAAAAATGACAACTTGTNNNAGATCGGAA |
| 37668_2 | TATGAGTGTGGAGTCGTTGCCCCCACAAAAAAATGACAACTTGTNNNAGATCGGAA |
| 43678_1 | TATGAGTGTGGAGTCGTTGCCCAAGAGAGAGAGTGTGGACACNNNAGATCGGAAG |
| 43678_2 | TATGAGTGTGGAGTCGTTGCCCAAGAGAGAGAGTGTGGACACNNNAGATCGGAAG |
| 48227_1 | TATGAGTGTGGAGTCGTTGCCAGGTTCAAGCAATTCTCCTGCNNNAGATCGGAAGA |
| 48227_2 | TATGAGTGTGGAGTCGTTGCCAGGTTCAAGCAATTCTCCTGCNNNAGATCGGAAGA |
| 37864_1 | TATGAGTGTGGAGTCGTTGCTCACTCAAATCTTTCCTCCTGGGNNNAGATCGGAAGA |
| 37864_2 | TATGAGTGTGGAGTCGTTGCTCACTCAAATCTTTCCTCCTGGGNNNAGATCGGAAGA |
| 10490_1 | TATGAGTGTGGAGTCGTTGCGCTCCTCCCGTAAGTATGCAAAANNNAGATCGGAAG |
| 10490_2 | TATGAGTGTGGAGTCGTTGCGCTCCTCCCGTAAGTATGCAAAANNNAGATCGGAAG |
| 33234_1 | TATGAGTGTGGAGTCGTTGCCTATCTCAGCTCACTGCAAGCNNNAGATCGGAAGAG |
| 33234_2 | TATGAGTGTGGAGTCGTTGCCTATCTCAGCTCACTGCAAGCNNNAGATCGGAAGAG |
| 6129_1 | TATGAGTGTGGAGTCGTTGCCGTTTTTGGTAGTGAGTGTTGCCNNNAGATCGGAAG |
| 6129_2 | TATGAGTGTGGAGTCGTTGCCGTTTTTGGTAGTGAGTGTTGCCNNNAGATCGGAAG |
| 46858_1 | TATGAGTGTGGAGTCGTTGCGTGAATGCAGCAACCTATGTCTCNNNAGATCGGAAG |
| 46858_2 | TATGAGTGTGGAGTCGTTGCGTGAATGCAGCAACCTATGTCTCNNNAGATCGGAAG |
| 36132_1 | TATGAGTGTGGAGTCGTTGCAATCAACCTAGGCACCCATCAATNNNAGATCGGAAG |
| 36132_2 | TATGAGTGTGGAGTCGTTGCAATCAACCTAGGCACCCATCAATNNNAGATCGGAAG |
| 47960_1 | TATGAGTGTGGAGTCGTTGCGATGGTGACGTCTGAAGCCCNNNAGATCGGAAGAG |
| 47960_2 | TATGAGTGTGGAGTCGTTGCGATGGTGACGTCTGAAGCCCNNNAGATCGGAAGAG |
| 2326_1 | TATGAGTGTGGAGTCGTTGCTGCCCTGTTTCTCTAAAGAACCCNNNAGATCGGAAGA |
| 2326_2 | TATGAGTGTGGAGTCGTTGCTGCCCTGTTTCTCTAAAGAACCCNNNAGATCGGAAGA |
| 41329_1 | TATGAGTGTGGAGTCGTTGCAGTCCAGATGAAAAAGGATGATAGCNNNAGATCGG |
| 41329_2 | TATGAGTGTGGAGTCGTTGCAGTCCAGATGAAAAAGGATGATAGCNNNAGATCGG |
| 42380_1 | TATGAGTGTGGAGTCGTTGCGCTTGTCAAAAAAAGCAAAATTTGGANNNAGATCGG |
| 42380_2 | TATGAGTGTGGAGTCGTTGCGCTTGTCAAAAAAAGCAAAATTTGGANNNAGATCGG |
| 44643_1 | TATGAGTGTGGAGTCGTTGCTTAGTTGGAAGGATGGGGTGAAGNNNAGATCGGA |
| 44643_2 | TATGAGTGTGGAGTCGTTGCTTAGTTGGAAGGATGGGGTGAAGNNNAGATCGGA |
| 40910_1 | TATGAGTGTGGAGTCGTTGCAGAGCTGGTAACATTCCTACTGANNNAGATCGGAAG |
| 40910_2 | TATGAGTGTGGAGTCGTTGCAGAGCTGGTAACATTCCTACTGANNNAGATCGGAAG |
| 48209_1 | TATGAGTGTGGAGTCGTTGCGAGTTTTGCTCTTGTTTCCCAGGNNNAGATCGGAAG |
| 48209_2 | TATGAGTGTGGAGTCGTTGCGAGTTTTGCTCTTGTTTCCCAGGNNNAGATCGGAAG |
| 43939_1 | TATGAGTGTGGAGTCGTTGCCACACACACACGGACACACTTANNNAGATCGGAAGA |
| 43939_2 | TATGAGTGTGGAGTCGTTGCCACACACACACGGACACACTTANNNAGATCGGAAGA |
| 30845_1 | TATGAGTGTGGAGTCGTTGCAGAGAGAGAGGCTCCCATTATCANNNAGATCGGAA |
| 30845_2 | TATGAGTGTGGAGTCGTTGCAGAGAGAGAGGCTCCCATTATCANNNAGATCGGAA |
| 33472_1 | TATGAGTGTGGAGTCGTTGCTGTACATTGGGCTTTCTATCAGGANNNAGATCGGAA |
| 33472_2 | TATGAGTGTGGAGTCGTTGCTGTACATTGGGCTTTCTATCAGGANNNAGATCGGAA |
| 46634_1 | TATGAGTGTGGAGTCGTTGCGAGTCTTGCTCTGTTGCCCANNNAGATCGGAAGAGC |

| | |
|---|---|
| 46634_2 | TATGAGTGTGGAGTCGTTGCGAGTCTTGCTCTGTTGCCCANNNAGATCGGAAGAGC |
| 37897_1 | TATGAGTGTGGAGTCGTTGCCAAGAAATGCAAAGGAACGCAGTNNNAGATCGGAA |
| 37897_2 | TATGAGTGTGGAGTCGTTGCCAAGAAATGCAAAGGAACGCAGTNNNAGATCGGAA |
| 10317_1 | TATGAGTGTGGAGTCGTTGCAAAATGCAGGGGACAGTCAAAGTNNNAGATCGGAA |
| 10317_2 | TATGAGTGTGGAGTCGTTGCAAAATGCAGGGGACAGTCAAAGTNNNAGATCGGAA |
| 1924_1 | TATGAGTGTGGAGTCGTTGCAGGCATCAAAGTCTGGAAGAGAGNNNAGATCGGAA |
| 1924_2 | TATGAGTGTGGAGTCGTTGCAGGCATCAAAGTCTGGAAGAGAGNNNAGATCGGAA |
| 34112_1 | TATGAGTGTGGAGTCGTTGCATACAAGCAGAGCGTAGAATGGTNNNAGATCGGAA |
| 34112_2 | TATGAGTGTGGAGTCGTTGCATACAAGCAGAGCGTAGAATGGTNNNAGATCGGAA |
| 41874_1 | TATGAGTGTGGAGTCGTTGCAAAGACAGGGTATTGCCATGTTGNNNAGATCGGAA |
| 41874_2 | TATGAGTGTGGAGTCGTTGCAAAGACAGGGTATTGCCATGTTGNNNAGATCGGAA |
| 41786_1 | TATGAGTGTGGAGTCGTTGCGGGGGAATGCTTTCATCTTTTCCNNNAGATCGGAAG |
| 41786_2 | TATGAGTGTGGAGTCGTTGCGGGGGAATGCTTTCATCTTTTCCNNNAGATCGGAAG |
| 42851_1 | TATGAGTGTGGAGTCGTTGCGAGAACCGGACCATGAACTGTATNNNAGATCGGAA |
| 42851_2 | TATGAGTGTGGAGTCGTTGCGAGAACCGGACCATGAACTGTATNNNAGATCGGAA |
| 51982_1 | TATGAGTGTGGAGTCGTTGCGGAAACAAACCCTGGTTTCAGAGNNNAGATCGGAA |
| 51982_2 | TATGAGTGTGGAGTCGTTGCGGAAACAAACCCTGGTTTCAGAGNNNAGATCGGAA |
| 44052_1 | TATGAGTGTGGAGTCGTTGCGTGTTTTGAGTCCCTGGAGGANNNAGATCGGAAGA |
| 44052_2 | TATGAGTGTGGAGTCGTTGCGTGTTTTGAGTCCCTGGAGGANNNAGATCGGAAGA |
| 41541_1 | TATGAGTGTGGAGTCGTTGCATAGCTAAAATTACAGGCGCCCANNNAGATCGGAAG |
| 41541_2 | TATGAGTGTGGAGTCGTTGCATAGCTAAAATTACAGGCGCCCANNNAGATCGGAAG |
| 43282_1 | TATGAGTGTGGAGTCGTTGCGAAGGAGTCTCACTCTGTTGTCCNNNAGATCGGAAG |
| 43282_2 | TATGAGTGTGGAGTCGTTGCGAAGGAGTCTCACTCTGTTGTCCNNNAGATCGGAAG |
| 50099_1 | TATGAGTGTGGAGTCGTTGCAAGGTTGCAAAATTGGCTCCAATNNNAGATCGGAA |
| 50099_2 | TATGAGTGTGGAGTCGTTGCAAGGTTGCAAAATTGGCTCCAATNNNAGATCGGAA |
| 47952_1 | TATGAGTGTGGAGTCGTTGCCATTGGTATGTAAGGTCCACCCANNNAGATCGGAAG |
| 47952_2 | TATGAGTGTGGAGTCGTTGCCATTGGTATGTAAGGTCCACCCANNNAGATCGGAAG |
| 46439_1 | TATGAGTGTGGAGTCGTTGCAAACTCCTGACCTCATGATCCACNNNAGATCGGAAG |
| 46439_2 | TATGAGTGTGGAGTCGTTGCAAACTCCTGACCTCATGATCCACNNNAGATCGGAAG |
| 33197_1 | TATGAGTGTGGAGTCGTTGCATTTATTTTTTGAGACGGAGTTTCACNNNAGATCGG |
| 33197_2 | TATGAGTGTGGAGTCGTTGCATTTATTTTTTGAGACGGAGTTTCACNNNAGATCGG |
| 48700_1 | TATGAGTGTGGAGTCGTTGCCACTCTGTCTGTCCTCTTTTCCANNNAGATCGGAAGA |
| 48700_2 | TATGAGTGTGGAGTCGTTGCCACTCTGTCTGTCCTCTTTTCCANNNAGATCGGAAGA |
| 50427_1 | TATGAGTGTGGAGTCGTTGCTCATAAAAAGTGCCTGATGTCAGNNNAGATCGGAA |
| 50427_2 | TATGAGTGTGGAGTCGTTGCTCATAAAAAGTGCCTGATGTCAGNNNAGATCGGAA |
| 2082_1 | TATGAGTGTGGAGTCGTTGCCTGTGGAACCTGAATGTGGTACANNNAGATCGGAA |
| 2082_2 | TATGAGTGTGGAGTCGTTGCCTGTGGAACCTGAATGTGGTACANNNAGATCGGAA |
| 46993_1 | TATGAGTGTGGAGTCGTTGCACGCCCGGCTAATTTATTTTTGTNNNAGATCGGAAG |
| 46993_2 | TATGAGTGTGGAGTCGTTGCACGCCCGGCTAATTTATTTTTGTNNNAGATCGGAAG |
| 42819_1 | TATGAGTGTGGAGTCGTTGCCTCTCAGCCCATTGTGCTTACTANNNAGATCGGAAGA |
| 42819_2 | TATGAGTGTGGAGTCGTTGCCTCTCAGCCCATTGTGCTTACTANNNAGATCGGAAGA |
| 49606_1 | TATGAGTGTGGAGTCGTTGCCTGAGTAGCTGGGATTACAGGTGNNNAGATCGGAA |
| 49606_2 | TATGAGTGTGGAGTCGTTGCCTGAGTAGCTGGGATTACAGGTGNNNAGATCGGAA |

| | |
|---|---|
| 49327_1 | TATGAGTGTGGAGTCGTTGCCCCAGAACATTTTCCTCATCCCTNNNAGATCGGAAGA |
| 49327_2 | TATGAGTGTGGAGTCGTTGCCCCAGAACATTTTCCTCATCCCTNNNAGATCGGAAGA |
| 32116_1 | TATGAGTGTGGAGTCGTTGCAAATCTGCTACCTTGAAAGCCCANNNAGATCGGAAG |
| 32116_2 | TATGAGTGTGGAGTCGTTGCAAATCTGCTACCTTGAAAGCCCANNNAGATCGGAAG |
| 45731_1 | TATGAGTGTGGAGTCGTTGCACATCACCACTCCTGGCTAATTTNNNAGATCGGAAGA |
| 45731_2 | TATGAGTGTGGAGTCGTTGCACATCACCACTCCTGGCTAATTTNNNAGATCGGAAGA |
| 45599_1 | TATGAGTGTGGAGTCGTTGCGTATAGTGGCCCAGTCATAGCTCNNNAGATCGGAAG |
| 45599_2 | TATGAGTGTGGAGTCGTTGCGTATAGTGGCCCAGTCATAGCTCNNNAGATCGGAAG |
| 29581_1 | TATGAGTGTGGAGTCGTTGCCCCGAAAAGGTTTGGTGTATGAANNNAGATCGGAA |
| 29581_2 | TATGAGTGTGGAGTCGTTGCCCCGAAAAGGTTTGGTGTATGAANNNAGATCGGAA |
| 34988_1 | TATGAGTGTGGAGTCGTTGCCTAAACCTTGGCCTCCGAAAATGNNNAGATCGGAAG |
| 34988_2 | TATGAGTGTGGAGTCGTTGCCTAAACCTTGGCCTCCGAAAATGNNNAGATCGGAAG |
| 50825_1 | TATGAGTGTGGAGTCGTTGCTCACAATCCCAGATCTTCCTTGGNNNAGATCGGAAG |
| 50825_2 | TATGAGTGTGGAGTCGTTGCTCACAATCCCAGATCTTCCTTGGNNNAGATCGGAAG |
| 31194_1 | TATGAGTGTGGAGTCGTTGCCCCCACTGGATTCTCTCTTCTTCNNNAGATCGGAAGA |
| 31194_2 | TATGAGTGTGGAGTCGTTGCCCCCACTGGATTCTCTCTTCTTCNNNAGATCGGAAGA |
| 38707_1 | TATGAGTGTGGAGTCGTTGCTTCAAGCGATTCTCCTGTCTCAGNNNAGATCGGAAG |
| 38707_2 | TATGAGTGTGGAGTCGTTGCTTCAAGCGATTCTCCTGTCTCAGNNNAGATCGGAAG |
| 29378_1 | TATGAGTGTGGAGTCGTTGCACTGAAAATGAGCCCCACAATTGNNNAGATCGGAAG |
| 29378_2 | TATGAGTGTGGAGTCGTTGCACTGAAAATGAGCCCCACAATTGNNNAGATCGGAAG |
| 13776_1 | TATGAGTGTGGAGTCGTTGCCTCATTCTAGTTCTCTAACAGAGNNNAGATCGGAAG |
| 13776_2 | TATGAGTGTGGAGTCGTTGCCTCATTCTAGTTCTCTAACAGAGNNNAGATCGGAAG |
| 47772_1 | TATGAGTGTGGAGTCGTTGCTGATGTCTCAGAGCCTGATAGGANNNAGATCGGAA |
| 47772_2 | TATGAGTGTGGAGTCGTTGCTGATGTCTCAGAGCCTGATAGGANNNAGATCGGAA |
| 51559_1 | TATGAGTGTGGAGTCGTTGCCCCAAAAAGTCTCCTGAGATCCANNNAGATCGGAAG |
| 51559_2 | TATGAGTGTGGAGTCGTTGCCCCAAAAAGTCTCCTGAGATCCANNNAGATCGGAAG |
| 39272_1 | TATGAGTGTGGAGTCGTTGCTCTGTCTCTGTGTCCTCTCTCTTNNNAGATCGGAAGA |
| 39272_2 | TATGAGTGTGGAGTCGTTGCTCTGTCTCTGTGTCCTCTCTCTTNNNAGATCGGAAGA |
| 44800_1 | TATGAGTGTGGAGTCGTTGCTCCTCCCCTCCATACCTACTTTCNNNAGATCGGAAGA |
| 44800_2 | TATGAGTGTGGAGTCGTTGCTCCTCCCCTCCATACCTACTTTCNNNAGATCGGAAGA |
| 31652_1 | TATGAGTGTGGAGTCGTTGCCCTGAAGTGACTGAGACAATGGANNNAGATCGGAA |
| 31652_2 | TATGAGTGTGGAGTCGTTGCCCTGAAGTGACTGAGACAATGGANNNAGATCGGAA |
| 38801_1 | TATGAGTGTGGAGTCGTTGCAGAAGAAGAAGGAGGAGAAACAGTNNNAGATCGG |
| 38801_2 | TATGAGTGTGGAGTCGTTGCAGAAGAAGAAGGAGGAGAAACAGTNNNAGATCGG |
| 31935_1 | TATGAGTGTGGAGTCGTTGCAACCATCAAAATGAAGCAGACGGNNNAGATCGGAA |
| 31935_2 | TATGAGTGTGGAGTCGTTGCAACCATCAAAATGAAGCAGACGGNNNAGATCGGAA |
| 32023_1 | TATGAGTGTGGAGTCGTTGCGAGTGATTCACCCGGCTAATTTTNNNAGATCGGAAG |
| 32023_2 | TATGAGTGTGGAGTCGTTGCGAGTGATTCACCCGGCTAATTTTNNNAGATCGGAAG |
| 36857_1 | TATGAGTGTGGAGTCGTTGCGGAATTAGTCCTTCTCGAGCCTANNNAGATCGGAAG |
| 36857_2 | TATGAGTGTGGAGTCGTTGCGGAATTAGTCCTTCTCGAGCCTANNNAGATCGGAAG |
| 47817_1 | TATGAGTGTGGAGTCGTTGCCTCGGTTCCGGGGTCATAAGNNNAGATCGGAAGAG |
| 47817_2 | TATGAGTGTGGAGTCGTTGCCTCGGTTCCGGGGTCATAAGNNNAGATCGGAAGAG |
| 50965_1 | TATGAGTGTGGAGTCGTTGCGGGTGTACATTCTACCTAACCCCNNNAGATCGGAAG |

| | |
|---|---|
| 50965_2 | TATGAGTGTGGAGTCGTTGCGGGTGTACATTCTACCTAACCCCNNNAGATCGGAAG |
| 36299_1 | TATGAGTGTGGAGTCGTTGCTAGTCCATAGGAATTCACTGGCCNNNAGATCGGAAG |
| 36299_2 | TATGAGTGTGGAGTCGTTGCTAGTCCATAGGAATTCACTGGCCNNNAGATCGGAAG |
| 9292_1 | TATGAGTGTGGAGTCGTTGCTGGAGTTCACAGCAGGAGAAATGNNNAGATCGGAA |
| 9292_2 | TATGAGTGTGGAGTCGTTGCTGGAGTTCACAGCAGGAGAAATGNNNAGATCGGAA |
| 48302_1 | TATGAGTGTGGAGTCGTTGCCTTCCAGACCCAAAATCTGAGGANNNAGATCGGAAG |
| 48302_2 | TATGAGTGTGGAGTCGTTGCCTTCCAGACCCAAAATCTGAGGANNNAGATCGGAAG |
| 52378_1 | TATGAGTGTGGAGTCGTTGCTTGAAAGCCATCTTTTTGGTGCANNNAGATCGGAAG |
| 52378_2 | TATGAGTGTGGAGTCGTTGCTTGAAAGCCATCTTTTTGGTGCANNNAGATCGGAAG |
| 32382_1 | TATGAGTGTGGAGTCGTTGCGCTAGGAAAGACCAGACCAAAGANNNAGATCGGAA |
| 32382_2 | TATGAGTGTGGAGTCGTTGCGCTAGGAAAGACCAGACCAAAGANNNAGATCGGAA |
| 35971_1 | TATGAGTGTGGAGTCGTTGCAAGAAATCTCAGAGACTCAGCCCNNNAGATCGGAAG |
| 35971_2 | TATGAGTGTGGAGTCGTTGCAAGAAATCTCAGAGACTCAGCCCNNNAGATCGGAAG |
| 40050_1 | TATGAGTGTGGAGTCGTTGCGAGTGCAGTAGCGCAATCTTGNNNAGATCGGAAGA |
| 40050_2 | TATGAGTGTGGAGTCGTTGCGAGTGCAGTAGCGCAATCTTGNNNAGATCGGAAGA |
| 50961_1 | TATGAGTGTGGAGTCGTTGCGACCCACCTCACCATCCAACNNNAGATCGGAAGAGC |
| 50961_2 | TATGAGTGTGGAGTCGTTGCGACCCACCTCACCATCCAACNNNAGATCGGAAGAGC |
| 41502_1 | TATGAGTGTGGAGTCGTTGCAGACAAGGGAAGGGAAGGAAAAANNNAGATCGGA |
| 41502_2 | TATGAGTGTGGAGTCGTTGCAGACAAGGGAAGGGAAGGAAAAANNNAGATCGGA |
| 47946_1 | TATGAGTGTGGAGTCGTTGCTGATCAAATAGGCCACTGAAGCTNNNAGATCGGAA |
| 47946_2 | TATGAGTGTGGAGTCGTTGCTGATCAAATAGGCCACTGAAGCTNNNAGATCGGAA |
| 50876_1 | TATGAGTGTGGAGTCGTTGCCAGTGTCAAGGCTTTGTTTAGCANNNAGATCGGAAG |
| 50876_2 | TATGAGTGTGGAGTCGTTGCCAGTGTCAAGGCTTTGTTTAGCANNNAGATCGGAAG |
| 53241_1 | TATGAGTGTGGAGTCGTTGCAATTGTTCATTATGTGCCCCTGCNNNAGATCGGAAG |
| 53241_2 | TATGAGTGTGGAGTCGTTGCAATTGTTCATTATGTGCCCCTGCNNNAGATCGGAAG |
| 51935_1 | TATGAGTGTGGAGTCGTTGCGCCTTCATCTCTCTATGCTTGGANNNAGATCGGAAG |
| 51935_2 | TATGAGTGTGGAGTCGTTGCGCCTTCATCTCTCTATGCTTGGANNNAGATCGGAAG |
| 9991_1 | TATGAGTGTGGAGTCGTTGCATTTCAGACACAGTCCCACTCTGNNNAGATCGGAAG |
| 9991_2 | TATGAGTGTGGAGTCGTTGCATTTCAGACACAGTCCCACTCTGNNNAGATCGGAAG |
| 44981_1 | TATGAGTGTGGAGTCGTTGCCTGCATTCTTCCCTTGTAGTCCTNNNAGATCGGAAGA |
| 44981_2 | TATGAGTGTGGAGTCGTTGCCTGCATTCTTCCCTTGTAGTCCTNNNAGATCGGAAGA |
| 10709_1 | TATGAGTGTGGAGTCGTTGCATCAACGCCAGGACCACTTTCATNNNAGATCGGAAG |
| 10709_2 | TATGAGTGTGGAGTCGTTGCATCAACGCCAGGACCACTTTCATNNNAGATCGGAAG |
| 39871_1 | TATGAGTGTGGAGTCGTTGCTTGAAGCCAGGAGTTTGAGATCANNNAGATCGGAA |
| 39871_2 | TATGAGTGTGGAGTCGTTGCTTGAAGCCAGGAGTTTGAGATCANNNAGATCGGAA |
| 46489_1 | TATGAGTGTGGAGTCGTTGCAGATGGAGTCTCCCACTGTCANNNAGATCGGAAGA |
| 46489_2 | TATGAGTGTGGAGTCGTTGCAGATGGAGTCTCCCACTGTCANNNAGATCGGAAGA |
| 45052_1 | TATGAGTGTGGAGTCGTTGCCTTGAAAGCAATGGGAAAGTGGTNNNAGATCGGAA |
| 45052_2 | TATGAGTGTGGAGTCGTTGCCTTGAAAGCAATGGGAAAGTGGTNNNAGATCGGAA |
| 43210_1 | TATGAGTGTGGAGTCGTTGCCTTTCTGGCTTTCCCTTCTCTCTNNNAGATCGGAAGA |
| 43210_2 | TATGAGTGTGGAGTCGTTGCCTTTCTGGCTTTCCCTTCTCTCTNNNAGATCGGAAGA |
| 13603_1 | TATGAGTGTGGAGTCGTTGCGAACTACATGAGATGTAAGAATGNNNAGATCGGAA |
| 13603_2 | TATGAGTGTGGAGTCGTTGCGAACTACATGAGATGTAAGAATGNNNAGATCGGAA |

| | |
|---|---|
| 29845_1 | TATGAGTGTGGAGTCGTTGCTGAAGAATCAGGTGGTCGGGTGANNNAGATCGGA |
| 29845_2 | TATGAGTGTGGAGTCGTTGCTGAAGAATCAGGTGGTCGGGTGANNNAGATCGGA |
| 49313_1 | TATGAGTGTGGAGTCGTTGCTTCAAGTGATTCTCCTGCCTCAGNNNAGATCGGAAG |
| 49313_2 | TATGAGTGTGGAGTCGTTGCTTCAAGTGATTCTCCTGCCTCAGNNNAGATCGGAAG |
| 50739_1 | TATGAGTGTGGAGTCGTTGCAATCAGAATCACCTGTGAAGCCTNNNAGATCGGAAG |
| 50739_2 | TATGAGTGTGGAGTCGTTGCAATCAGAATCACCTGTGAAGCCTNNNAGATCGGAAG |
| 47375_1 | TATGAGTGTGGAGTCGTTGCCCTGGTTGTGTGATAATCAGTGCNNNAGATCGGAA |
| 47375_2 | TATGAGTGTGGAGTCGTTGCCCTGGTTGTGTGATAATCAGTGCNNNAGATCGGAA |
| 31049_1 | TATGAGTGTGGAGTCGTTGCCCAAGGAGTGAAACAGGGTAGTTNNNAGATCGGAA |
| 31049_2 | TATGAGTGTGGAGTCGTTGCCCAAGGAGTGAAACAGGGTAGTTNNNAGATCGGAA |
| 40430_1 | TATGAGTGTGGAGTCGTTGCAGAGAGAGAGAGAGAGAAAGGTCANNNAGATCGG |
| 40430_2 | TATGAGTGTGGAGTCGTTGCAGAGAGAGAGAGAGAGAAAGGTCANNNAGATCGG |
| 38054_1 | TATGAGTGTGGAGTCGTTGCAAGGCCTTTGACAAAATTCAGCANNNAGATCGGAAG |
| 38054_2 | TATGAGTGTGGAGTCGTTGCAAGGCCTTTGACAAAATTCAGCANNNAGATCGGAAG |
| 9084_1 | TATGAGTGTGGAGTCGTTGCTGCTGAAGATAGCAAGAGTGACTNNNAGATCGGAA |
| 9084_2 | TATGAGTGTGGAGTCGTTGCTGCTGAAGATAGCAAGAGTGACTNNNAGATCGGAA |
| 36501_1 | TATGAGTGTGGAGTCGTTGCAGAAACCCTTCTCTAAGCCTGTGNNNAGATCGGAAG |
| 36501_2 | TATGAGTGTGGAGTCGTTGCAGAAACCCTTCTCTAAGCCTGTGNNNAGATCGGAAG |
| 34488_1 | TATGAGTGTGGAGTCGTTGCTGGGCATTGATAGGCTCTACTTCNNNAGATCGGAAG |
| 34488_2 | TATGAGTGTGGAGTCGTTGCTGGGCATTGATAGGCTCTACTTCNNNAGATCGGAAG |
| 50243_1 | TATGAGTGTGGAGTCGTTGCCAAAAGTCGGGCATTTTGCTCTANNNAGATCGGAAG |
| 50243_2 | TATGAGTGTGGAGTCGTTGCCAAAAGTCGGGCATTTTGCTCTANNNAGATCGGAAG |
| 45282_1 | TATGAGTGTGGAGTCGTTGCCACAAGGCCTATGAAGCTTTCACNNNAGATCGGAAG |
| 45282_2 | TATGAGTGTGGAGTCGTTGCCACAAGGCCTATGAAGCTTTCACNNNAGATCGGAAG |
| 40260_1 | TATGAGTGTGGAGTCGTTGCTCAATGCATCAAAGAACAAAACCANNNAGATCGGAA |
| 40260_2 | TATGAGTGTGGAGTCGTTGCTCAATGCATCAAAGAACAAAACCANNNAGATCGGAA |
| 29929_1 | TATGAGTGTGGAGTCGTTGCTCCTCTTGTGTTTTTTTGTGCCANNNAGATCGGAAGA |
| 29929_2 | TATGAGTGTGGAGTCGTTGCTCCTCTTGTGTTTTTTTGTGCCANNNAGATCGGAAGA |
| 52625_1 | TATGAGTGTGGAGTCGTTGCCCTCACTTATCAATCACCCCACANNNAGATCGGAAGA |
| 52625_2 | TATGAGTGTGGAGTCGTTGCCCTCACTTATCAATCACCCCACANNNAGATCGGAAGA |
| 43621_1 | TATGAGTGTGGAGTCGTTGCCTCGGCAACATAGTGAGACTTCANNNAGATCGGAAG |
| 43621_2 | TATGAGTGTGGAGTCGTTGCCTCGGCAACATAGTGAGACTTCANNNAGATCGGAAG |
| 53048_1 | TATGAGTGTGGAGTCGTTGCACAAAAGAATGAGGCTGCATTGTNNNAGATCGGAA |
| 53048_2 | TATGAGTGTGGAGTCGTTGCACAAAAGAATGAGGCTGCATTGTNNNAGATCGGAA |
| 43001_1 | TATGAGTGTGGAGTCGTTGCTCTTCACGTTGATACCATTGTGANNNAGATCGGAAG |
| 43001_2 | TATGAGTGTGGAGTCGTTGCTCTTCACGTTGATACCATTGTGANNNAGATCGGAAG |
| 46822_1 | TATGAGTGTGGAGTCGTTGCATGTTCTGGGGAAAGCAGGGNNNAGATCGGAAGA |
| 46822_2 | TATGAGTGTGGAGTCGTTGCATGTTCTGGGGAAAGCAGGGNNNAGATCGGAAGA |
| 53155_1 | TATGAGTGTGGAGTCGTTGCAACTATATTTGCGTGGTCAGCCTNNNAGATCGGAAG |
| 53155_2 | TATGAGTGTGGAGTCGTTGCAACTATATTTGCGTGGTCAGCCTNNNAGATCGGAAG |
| 44507_1 | TATGAGTGTGGAGTCGTTGCCTGGTCTTGAACTTCTGAGCTCANNNAGATCGGAAG |
| 44507_2 | TATGAGTGTGGAGTCGTTGCCTGGTCTTGAACTTCTGAGCTCANNNAGATCGGAAG |
| 51578_1 | TATGAGTGTGGAGTCGTTGCCTTGGAAGGAAGTACCTGGATGGNNNAGATCGGAA |

| | |
|---|---|
| 51578_2 | TATGAGTGTGGAGTCGTTGCCTTGGAAGGAAGTACCTGGATGGNNNAGATCGGAA |
| 34632_1 | TATGAGTGTGGAGTCGTTGCTTTGCTGTTTTTCTCATGTGGCTNNNAGATCGGAAGA |
| 34632_2 | TATGAGTGTGGAGTCGTTGCTTTGCTGTTTTTCTCATGTGGCTNNNAGATCGGAAGA |
| 33561_1 | TATGAGTGTGGAGTCGTTGCCTCCTGGGTTCAAGCGATTCTCNNNAGATCGGAAGA |
| 33561_2 | TATGAGTGTGGAGTCGTTGCCTCCTGGGTTCAAGCGATTCTCNNNAGATCGGAAGA |
| 31867_1 | TATGAGTGTGGAGTCGTTGCGAAGAAAAGCACCGTGTGGGNNNAGATCGGAAGA |
| 31867_2 | TATGAGTGTGGAGTCGTTGCGAAGAAAAGCACCGTGTGGGNNNAGATCGGAAGA |
| 52304_1 | TATGAGTGTGGAGTCGTTGCTGCGGCTAAGATGTCAAAGTACTNNNAGATCGGAA |
| 52304_2 | TATGAGTGTGGAGTCGTTGCTGCGGCTAAGATGTCAAAGTACTNNNAGATCGGAA |
| 34342_1 | TATGAGTGTGGAGTCGTTGCTTCACAATACATCTTGCTGCTGCNNNAGATCGGAAGA |
| 34342_2 | TATGAGTGTGGAGTCGTTGCTTCACAATACATCTTGCTGCTGCNNNAGATCGGAAGA |
| 40262_1 | TATGAGTGTGGAGTCGTTGCTCATGGAGATAGAGAGTAGAAGGANNNAGATCGGA |
| 40262_2 | TATGAGTGTGGAGTCGTTGCTCATGGAGATAGAGAGTAGAAGGANNNAGATCGGA |
| 29454_1 | ATGAGTGTGGAGTCGTTGCTTCACTTAAAGTAGTGGTTCTCAACCTNNNAGATCGGA |
| 29454_2 | ATGAGTGTGGAGTCGTTGCTTCACTTAAAGTAGTGGTTCTCAACCTNNNAGATCGGA |
| 45474_1 | TATGAGTGTGGAGTCGTTGCCCTCTCTCCTTTGGAAATTCCCANNNAGATCGGAAGA |
| 45474_2 | TATGAGTGTGGAGTCGTTGCCCTCTCTCCTTTGGAAATTCCCANNNAGATCGGAAGA |
| 39797_1 | TATGAGTGTGGAGTCGTTGCCACTAGTAATGAGTGGAAAGTGGTNNNAGATCGGA |
| 39797_2 | TATGAGTGTGGAGTCGTTGCCACTAGTAATGAGTGGAAAGTGGTNNNAGATCGGA |
| 39916_1 | TATGAGTGTGGAGTCGTTGCAAAAGAAAAGAAAGTGGGCGTGGNNNAGATCGGA |
| 39916_2 | TATGAGTGTGGAGTCGTTGCAAAAGAAAAGAAAGTGGGCGTGGNNNAGATCGGA |
| 13289_1 | TATGAGTGTGGAGTCGTTGCCTTTAGTAGCCTTCTCTTCATATNNNAGATCGGAAGA |
| 13289_2 | TATGAGTGTGGAGTCGTTGCCTTTAGTAGCCTTCTCTTCATATNNNAGATCGGAAGA |
| 46674_1 | TATGAGTGTGGAGTCGTTGCAGATGGAGTCTTGCTGTGTTGCNNNAGATCGGAAGA |
| 46674_2 | TATGAGTGTGGAGTCGTTGCAGATGGAGTCTTGCTGTGTTGCNNNAGATCGGAAGA |
| 38821_1 | TATGAGTGTGGAGTCGTTGCCTGTAGTCCCAGCTACTCGGAAGNNNAGATCGGAAG |
| 38821_2 | TATGAGTGTGGAGTCGTTGCCTGTAGTCCCAGCTACTCGGAAGNNNAGATCGGAAG |
| 39454_1 | TATGAGTGTGGAGTCGTTGCTAATTCCAGCTACTCGGGAGGTANNNAGATCGGAA |
| 39454_2 | TATGAGTGTGGAGTCGTTGCTAATTCCAGCTACTCGGGAGGTANNNAGATCGGAA |
| 38214_1 | TATGAGTGTGGAGTCGTTGCTATTCCCAGTGCCCAGAAGAATCNNNAGATCGGAAG |
| 38214_2 | TATGAGTGTGGAGTCGTTGCTATTCCCAGTGCCCAGAAGAATCNNNAGATCGGAAG |
| 51774_1 | TATGAGTGTGGAGTCGTTGCCAGGCTGCCCTGGTAGTCATNNNAGATCGGAAGAG |
| 51774_2 | TATGAGTGTGGAGTCGTTGCCAGGCTGCCCTGGTAGTCATNNNAGATCGGAAGAG |
| 38174_1 | TATGAGTGTGGAGTCGTTGCGACATGGACCAAGATGAGGATCANNNAGATCGGAA |
| 38174_2 | TATGAGTGTGGAGTCGTTGCGACATGGACCAAGATGAGGATCANNNAGATCGGAA |
| 36403_1 | TATGAGTGTGGAGTCGTTGCAAGAAACTTGGCATCATAGCAGGNNNAGATCGGAA |
| 36403_2 | TATGAGTGTGGAGTCGTTGCAAGAAACTTGGCATCATAGCAGGNNNAGATCGGAA |
| 18_1 | TATGAGTGTGGAGTCGTTGCGCCCCTAGCCTGTAGTATTTCTTNNNAGATCGGAAGA |
| 18_2 | TATGAGTGTGGAGTCGTTGCGCCCCTAGCCTGTAGTATTTCTTNNNAGATCGGAAGA |
| 48186_1 | TATGAGTGTGGAGTCGTTGCGTTCAAGCAATTCTTCTGCCTCANNNAGATCGGAAGA |
| 48186_2 | TATGAGTGTGGAGTCGTTGCGTTCAAGCAATTCTTCTGCCTCANNNAGATCGGAAGA |
| 51725_1 | TATGAGTGTGGAGTCGTTGCAATAACCACCATCTTGGCCTCTTNNNAGATCGGAAGA |
| 51725_2 | TATGAGTGTGGAGTCGTTGCAATAACCACCATCTTGGCCTCTTNNNAGATCGGAAGA |

| | |
|---|---|
| 42723_1 | TATGAGTGTGGAGTCGTTGCACCTTGGAATACAGCTAACCAGGNNNAGATCGGAA |
| 42723_2 | TATGAGTGTGGAGTCGTTGCACCTTGGAATACAGCTAACCAGGNNNAGATCGGAA |
| 28943_1 | TATGAGTGTGGAGTCGTTGCGGTTATCACATCCTCTCACCCTCNNNAGATCGGAAGA |
| 28943_2 | TATGAGTGTGGAGTCGTTGCGGTTATCACATCCTCTCACCCTCNNNAGATCGGAAGA |
| 50654_1 | TATGAGTGTGGAGTCGTTGCTCAGAGCCTTTCAAAAGCTCTCTNNNAGATCGGAAG |
| 50654_2 | TATGAGTGTGGAGTCGTTGCTCAGAGCCTTTCAAAAGCTCTCTNNNAGATCGGAAG |
| 42271_1 | TATGAGTGTGGAGTCGTTGCATTTTGTGTGTGTGTGTGTGTNNNAGATCGGAA |
| 42271_2 | TATGAGTGTGGAGTCGTTGCATTTTGTGTGTGTGTGTGTGTNNNAGATCGGAA |
| 30028_1 | TATGAGTGTGGAGTCGTTGCAAAAATGGGTGGGTCATTTGCATNNNAGATCGGAA |
| 30028_2 | TATGAGTGTGGAGTCGTTGCAAAAATGGGTGGGTCATTTGCATNNNAGATCGGAA |
| 38736_1 | TATGAGTGTGGAGTCGTTGCAATGGATGAATTGTGTGGCATGTNNNAGATCGGAA |
| 38736_2 | TATGAGTGTGGAGTCGTTGCAATGGATGAATTGTGTGGCATGTNNNAGATCGGAA |
| 50640_1 | TATGAGTGTGGAGTCGTTGCGGAGGGTAGAAGAAGTGGGAATCNNNAGATCGGA |
| 50640_2 | TATGAGTGTGGAGTCGTTGCGGAGGGTAGAAGAAGTGGGAATCNNNAGATCGGA |
| 47349_1 | TATGAGTGTGGAGTCGTTGCAAGCCTCATTCCATTTGCTCCTANNNAGATCGGAAGA |
| 47349_2 | TATGAGTGTGGAGTCGTTGCAAGCCTCATTCCATTTGCTCCTANNNAGATCGGAAGA |
| 53108_1 | TATGAGTGTGGAGTCGTTGCAAAGGGAACAGCAGATGTAGGAGNNNAGATCGGA |
| 53108_2 | TATGAGTGTGGAGTCGTTGCAAAGGGAACAGCAGATGTAGGAGNNNAGATCGGA |
| 45993_1 | TATGAGTGTGGAGTCGTTGCCCAAGTAGCTGGGATTACAGGTGNNNAGATCGGAA |
| 45993_2 | TATGAGTGTGGAGTCGTTGCCCAAGTAGCTGGGATTACAGGTGNNNAGATCGGAA |
| 37953_1 | TATGAGTGTGGAGTCGTTGCCATGGAGAAACCCCGTCTCTATTNNNAGATCGGAAG |
| 37953_2 | TATGAGTGTGGAGTCGTTGCCATGGAGAAACCCCGTCTCTATTNNNAGATCGGAAG |
| 43000_1 | TATGAGTGTGGAGTCGTTGCACACACAAAATTAGCCTGGCATCNNNAGATCGGAAG |
| 43000_2 | TATGAGTGTGGAGTCGTTGCACACACAAAATTAGCCTGGCATCNNNAGATCGGAAG |
| 51381_1 | TATGAGTGTGGAGTCGTTGCCTCAGACTAGCAGGCCTAACAAANNNAGATCGGAAG |
| 51381_2 | TATGAGTGTGGAGTCGTTGCCTCAGACTAGCAGGCCTAACAAANNNAGATCGGAAG |
| 53110_1 | TATGAGTGTGGAGTCGTTGCGGGTCTTGGAGGAATAGGTGAAANNNAGATCGGA |
| 53110_2 | TATGAGTGTGGAGTCGTTGCGGGTCTTGGAGGAATAGGTGAAANNNAGATCGGA |
| 48511_1 | TATGAGTGTGGAGTCGTTGCGATCTCAGAGTTCACTGCAACCTNNNAGATCGGAAG |
| 48511_2 | TATGAGTGTGGAGTCGTTGCGATCTCAGAGTTCACTGCAACCTNNNAGATCGGAAG |
| 40152_1 | TATGAGTGTGGAGTCGTTGCACGTGAACCCTCATTTAGTGTCANNNAGATCGGAAG |
| 40152_2 | TATGAGTGTGGAGTCGTTGCACGTGAACCCTCATTTAGTGTCANNNAGATCGGAAG |
| 49002_1 | TATGAGTGTGGAGTCGTTGCAGGCTAGCATAAACAGTGTAGAGAANNNAGATCGG |
| 49002_2 | TATGAGTGTGGAGTCGTTGCAGGCTAGCATAAACAGTGTAGAGAANNNAGATCGG |
| 33327_1 | TATGAGTGTGGAGTCGTTGCCGAGTAGGTGGGATTACAGGTGNNNAGATCGGAA |
| 33327_2 | TATGAGTGTGGAGTCGTTGCCGAGTAGGTGGGATTACAGGTGNNNAGATCGGAA |
| 47075_1 | TATGAGTGTGGAGTCGTTGCCAAGTAGCTGGGACTACAGGCNNNAGATCGGAAGA |
| 47075_2 | TATGAGTGTGGAGTCGTTGCCAAGTAGCTGGGACTACAGGCNNNAGATCGGAAGA |
| 30112_1 | TATGAGTGTGGAGTCGTTGCGCCATCCTCTATTTTTCAGGTTCANNNAGATCGGAAG |
| 30112_2 | TATGAGTGTGGAGTCGTTGCGCCATCCTCTATTTTTCAGGTTCANNNAGATCGGAAG |
| 40520_1 | TATGAGTGTGGAGTCGTTGCTTGCATTCTGTTCCCCTGATGTANNNAGATCGGAAG |
| 40520_2 | TATGAGTGTGGAGTCGTTGCTTGCATTCTGTTCCCCTGATGTANNNAGATCGGAAG |
| 30298_1 | TATGAGTGTGGAGTCGTTGCTGGGTCAGTATTAAATTTTTGTGCANNNAGATCGGA |

| | |
|---|---|
| 30298_2 | TATGAGTGTGGAGTCGTTGCTGGGTCAGTATTAAATTTTTGTGCANNNAGATCGGA |
| 38700_1 | TATGAGTGTGGAGTCGTTGCGGCATCCTTGTCTTTTTCCAGTTNNNAGATCGGAAGA |
| 38700_2 | TATGAGTGTGGAGTCGTTGCGGCATCCTTGTCTTTTTCCAGTTNNNAGATCGGAAGA |
| 43313_1 | TATGAGTGTGGAGTCGTTGCCAGCCTCTGGTAACTATCCTTCTNNNAGATCGGAAGA |
| 43313_2 | TATGAGTGTGGAGTCGTTGCCAGCCTCTGGTAACTATCCTTCTNNNAGATCGGAAGA |
| 41023_1 | TATGAGTGTGGAGTCGTTGCCACCCATCTTAGACACCCTGAAANNNAGATCGGAAG |
| 41023_2 | TATGAGTGTGGAGTCGTTGCCACCCATCTTAGACACCCTGAAANNNAGATCGGAAG |
| 10018_1 | TATGAGTGTGGAGTCGTTGCAGTTGTATGGCAGGTTGAATTCTNNNAGATCGGAA |
| 10018_2 | TATGAGTGTGGAGTCGTTGCAGTTGTATGGCAGGTTGAATTCTNNNAGATCGGAA |
| 51739_1 | TATGAGTGTGGAGTCGTTGCATGAGTGAGAACAGAAAGCACCTNNNAGATCGGAA |
| 51739_2 | TATGAGTGTGGAGTCGTTGCATGAGTGAGAACAGAAAGCACCTNNNAGATCGGAA |
| 45440_1 | TATGAGTGTGGAGTCGTTGCTCTTATTTGAAGAGCCATGGGGTNNNAGATCGGAA |
| 45440_2 | TATGAGTGTGGAGTCGTTGCTCTTATTTGAAGAGCCATGGGGTNNNAGATCGGAA |
| 47585_1 | TATGAGTGTGGAGTCGTTGCTCTGTCAGGGAAAATTGACTTCCTNNNAGATCGGAA |
| 47585_2 | TATGAGTGTGGAGTCGTTGCTCTGTCAGGGAAAATTGACTTCCTNNNAGATCGGAA |
| 43150_1 | TATGAGTGTGGAGTCGTTGCACCATGCCCGGCTATTTTTTTTTNNNAGATCGGAAGA |
| 43150_2 | TATGAGTGTGGAGTCGTTGCACCATGCCCGGCTATTTTTTTTTNNNAGATCGGAAGA |
| 43298_1 | TATGAGTGTGGAGTCGTTGCGATGGAATCTCACTCTTGTTGCCNNNAGATCGGAAG |
| 43298_2 | TATGAGTGTGGAGTCGTTGCGATGGAATCTCACTCTTGTTGCCNNNAGATCGGAAG |
| 41501_1 | TATGAGTGTGGAGTCGTTGCTCACTTCTAATTCCAAGGCCTACANNNAGATCGGAAG |
| 41501_2 | TATGAGTGTGGAGTCGTTGCTCACTTCTAATTCCAAGGCCTACANNNAGATCGGAAG |
| 36703_1 | TATGAGTGTGGAGTCGTTGCCAGCATAGCATGGTTTTGTCTGTNNNAGATCGGAAG |
| 36703_2 | TATGAGTGTGGAGTCGTTGCCAGCATAGCATGGTTTTGTCTGTNNNAGATCGGAAG |
| 46356_1 | TATGAGTGTGGAGTCGTTGCCTCCCAGGTTCCAGTGATTCTCNNNAGATCGGAAGA |
| 46356_2 | TATGAGTGTGGAGTCGTTGCCTCCCAGGTTCCAGTGATTCTCNNNAGATCGGAAGA |
| 47869_1 | TATGAGTGTGGAGTCGTTGCCACAGCCCAGGTTGTTTTTCTACNNNAGATCGGAAG |
| 47869_2 | TATGAGTGTGGAGTCGTTGCCACAGCCCAGGTTGTTTTTCTACNNNAGATCGGAAG |
| 45996_1 | TATGAGTGTGGAGTCGTTGCCAAGCGAGACAAAACTGTTTGCTNNNAGATCGGAA |
| 45996_2 | TATGAGTGTGGAGTCGTTGCCAAGCGAGACAAAACTGTTTGCTNNNAGATCGGAA |
| 42854_1 | TATGAGTGTGGAGTCGTTGCGTGTGTGTGTGTGTTTTGAGACANNNAGATCGGAA |
| 42854_2 | TATGAGTGTGGAGTCGTTGCGTGTGTGTGTGTGTTTTGAGACANNNAGATCGGAA |
| 39021_1 | TATGAGTGTGGAGTCGTTGCTCGACTTTATCTTTGGGAGGCAANNNAGATCGGAAG |
| 39021_2 | TATGAGTGTGGAGTCGTTGCTCGACTTTATCTTTGGGAGGCAANNNAGATCGGAAG |
| 49901_1 | TATGAGTGTGGAGTCGTTGCTTCTTTCCACCTCCTCCGCGNNNAGATCGGAAGAGCA |
| 49901_2 | TATGAGTGTGGAGTCGTTGCTTCTTTCCACCTCCTCCGCGNNNAGATCGGAAGAGCA |
| 43172_1 | TATGAGTGTGGAGTCGTTGCCAAAGTGCTGGGATTATAGGCGNNNAGATCGGAAG |
| 43172_2 | TATGAGTGTGGAGTCGTTGCCAAAGTGCTGGGATTATAGGCGNNNAGATCGGAAG |
| 42309_1 | TATGAGTGTGGAGTCGTTGCGATCAGAAGAGAGTGGCATGACANNNAGATCGGAA |
| 42309_2 | TATGAGTGTGGAGTCGTTGCGATCAGAAGAGAGTGGCATGACANNNAGATCGGAA |
| 46244_1 | TATGAGTGTGGAGTCGTTGCAAGTTCAAGTGATTCTCCTGCCTNNNAGATCGGAAG |
| 46244_2 | TATGAGTGTGGAGTCGTTGCAAGTTCAAGTGATTCTCCTGCCTNNNAGATCGGAAG |
| 43942_1 | TATGAGTGTGGAGTCGTTGCCAGCCGGTTAGTTGGAAAAGAAGNNNAGATCGGAA |
| 43942_2 | TATGAGTGTGGAGTCGTTGCCAGCCGGTTAGTTGGAAAAGAAGNNNAGATCGGAA |

| | |
|---|---|
| 33366_1 | TATGAGTGTGGAGTCGTTGCTCCAGGTTCAAGTGATTCTCCTGNNNAGATCGGAAG |
| 33366_2 | TATGAGTGTGGAGTCGTTGCTCCAGGTTCAAGTGATTCTCCTGNNNAGATCGGAAG |
| 39092_1 | TATGAGTGTGGAGTCGTTGCTTGTGTGTATTTGGAAGTAGCCANNNAGATCGGAA |
| 39092_2 | TATGAGTGTGGAGTCGTTGCTTGTGTGTATTTGGAAGTAGCCANNNAGATCGGAA |
| 36006_1 | TATGAGTGTGGAGTCGTTGCGAGGCAGGAGAATGGTGTGAANNNAGATCGGAAG |
| 36006_2 | TATGAGTGTGGAGTCGTTGCGAGGCAGGAGAATGGTGTGAANNNAGATCGGAAG |
| 47177_1 | TATGAGTGTGGAGTCGTTGCGTAGAGACGGAGTTTCACCATGTNNNAGATCGGAA |
| 47177_2 | TATGAGTGTGGAGTCGTTGCGTAGAGACGGAGTTTCACCATGTNNNAGATCGGAA |
| 49176_1 | TATGAGTGTGGAGTCGTTGCTCAAGTGATTCTCCTGTCTCAGCNNNAGATCGGAAG |
| 49176_2 | TATGAGTGTGGAGTCGTTGCTCAAGTGATTCTCCTGTCTCAGCNNNAGATCGGAAG |
| 2359_1 | TATGAGTGTGGAGTCGTTGCGAGAAGATTGCTAAGACCCCTGCNNNAGATCGGAA |
| 2359_2 | TATGAGTGTGGAGTCGTTGCGAGAAGATTGCTAAGACCCCTGCNNNAGATCGGAA |
| 47833_1 | TATGAGTGTGGAGTCGTTGCCACGGGATGTGGGCTCATATGNNNAGATCGGAAGA |
| 47833_2 | TATGAGTGTGGAGTCGTTGCCACGGGATGTGGGCTCATATGNNNAGATCGGAAGA |
| 46341_1 | TATGAGTGTGGAGTCGTTGCTTCCATGTTTAGTGCCTCCTTCANNNAGATCGGAAGA |
| 46341_2 | TATGAGTGTGGAGTCGTTGCTTCCATGTTTAGTGCCTCCTTCANNNAGATCGGAAGA |
| 13223_1 | TATGAGTGTGGAGTCGTTGCCTCCTTAAGCCATTCTAAAGTTNNNAGATCGGAAGA |
| 13223_2 | TATGAGTGTGGAGTCGTTGCCTCCTTAAGCCATTCTAAAGTTNNNAGATCGGAAGA |
| 50752_1 | TATGAGTGTGGAGTCGTTGCTTGAGGAGCTGAAGGTGAAGATCNNNAGATCGGAA |
| 50752_2 | TATGAGTGTGGAGTCGTTGCTTGAGGAGCTGAAGGTGAAGATCNNNAGATCGGAA |
| 10423_1 | TATGAGTGTGGAGTCGTTGCCAGAGGGACAGAGACAGAGGTAGNNNAGATCGGA |
| 10423_2 | TATGAGTGTGGAGTCGTTGCCAGAGGGACAGAGACAGAGGTAGNNNAGATCGGA |
| 32876_1 | TATGAGTGTGGAGTCGTTGCCCTCCCGAGTAGCTAGGATTACANNNAGATCGGAAG |
| 32876_2 | TATGAGTGTGGAGTCGTTGCCCTCCCGAGTAGCTAGGATTACANNNAGATCGGAAG |
| 49384_1 | TATGAGTGTGGAGTCGTTGCCAAGCAATTCTCCTGCCTAGGTANNNAGATCGGAAG |
| 49384_2 | TATGAGTGTGGAGTCGTTGCCAAGCAATTCTCCTGCCTAGGTANNNAGATCGGAAG |
| 30337_1 | TATGAGTGTGGAGTCGTTGCAACCGTGAAGGTGAGGAGGAAATNNNAGATCGGA |
| 30337_2 | TATGAGTGTGGAGTCGTTGCAACCGTGAAGGTGAGGAGGAAATNNNAGATCGGA |
| 45997_1 | TATGAGTGTGGAGTCGTTGCCAGTGACATGATCTTGGCTCACTNNNAGATCGGAAG |
| 45997_2 | TATGAGTGTGGAGTCGTTGCCAGTGACATGATCTTGGCTCACTNNNAGATCGGAAG |
| 46989_1 | TATGAGTGTGGAGTCGTTGCCTTCTGGGTTCAGGCGATTCTCNNNAGATCGGAAGA |
| 46989_2 | TATGAGTGTGGAGTCGTTGCCTTCTGGGTTCAGGCGATTCTCNNNAGATCGGAAGA |
| 40390_1 | TATGAGTGTGGAGTCGTTGCGTAGTCCCATCTACTCAGGAGGTNNNAGATCGGAAG |
| 40390_2 | TATGAGTGTGGAGTCGTTGCGTAGTCCCATCTACTCAGGAGGTNNNAGATCGGAAG |
| 47405_1 | TATGAGTGTGGAGTCGTTGCCAGTGGTATGATCTCGGTTCACTNNNAGATCGGAAG |
| 47405_2 | TATGAGTGTGGAGTCGTTGCCAGTGGTATGATCTCGGTTCACTNNNAGATCGGAAG |
| 52747_1 | TATGAGTGTGGAGTCGTTGCACAGCCCCTGATCACAAAACTTANNNAGATCGGAAG |
| 52747_2 | TATGAGTGTGGAGTCGTTGCACAGCCCCTGATCACAAAACTTANNNAGATCGGAAG |
| 47652_1 | TATGAGTGTGGAGTCGTTGCGCCTTCTGAGTAGCTGGGATTACNNNAGATCGGAA |
| 47652_2 | TATGAGTGTGGAGTCGTTGCGCCTTCTGAGTAGCTGGGATTACNNNAGATCGGAA |
| 30340_1 | TATGAGTGTGGAGTCGTTGCCGGAACTGAATAGCGTATCTGGGNNNAGATCGGAA |
| 30340_2 | TATGAGTGTGGAGTCGTTGCCGGAACTGAATAGCGTATCTGGGNNNAGATCGGAA |
| 6162_1 | TATGAGTGTGGAGTCGTTGCATTTAGGCCCCCAGAACTCTTTANNNAGATCGGAAG |

| | |
|---|---|
| 6162_2 | TATGAGTGTGGAGTCGTTGCATTTAGGCCCCAGAACTCTTTANNNAGATCGGAAG |
| 39017_1 | TATGAGTGTGGAGTCGTTGCAATGACTTCAGCAACGTTTCAGGNNNAGATCGGAAG |
| 39017_2 | TATGAGTGTGGAGTCGTTGCAATGACTTCAGCAACGTTTCAGGNNNAGATCGGAAG |
| 13335_1 | TATGAGTGTGGAGTCGTTGCGTGGAGTTTTTGAGATAAATCAANNNAGATCGGAA |
| 13335_2 | TATGAGTGTGGAGTCGTTGCGTGGAGTTTTTGAGATAAATCAANNNAGATCGGAA |
| 40082_1 | TATGAGTGTGGAGTCGTTGCCCCTTACATGAGAGCAGAAACCTNNNAGATCGGAAG |
| 40082_2 | TATGAGTGTGGAGTCGTTGCCCCTTACATGAGAGCAGAAACCTNNNAGATCGGAAG |
| 52350_1 | TATGAGTGTGGAGTCGTTGCTTCTGTTCCACTTCAGATAGGCANNNAGATCGGAAG |
| 52350_2 | TATGAGTGTGGAGTCGTTGCTTCTGTTCCACTTCAGATAGGCANNNAGATCGGAAG |
| 49203_1 | TATGAGTGTGGAGTCGTTGCCAGGCAAAATCATGAAGAGGCAANNNAGATCGGAA |
| 49203_2 | TATGAGTGTGGAGTCGTTGCCAGGCAAAATCATGAAGAGGCAANNNAGATCGGAA |
| 46093_1 | TATGAGTGTGGAGTCGTTGCTCCTGAGTAGCTGCAATTACAGGNNNAGATCGGAA |
| 46093_2 | TATGAGTGTGGAGTCGTTGCTCCTGAGTAGCTGCAATTACAGGNNNAGATCGGAA |
| 12630_1 | TATGAGTGTGGAGTCGTTGCGAAAAGCTCCTATCAAACAAATANNNAGATCGGAAG |
| 12630_2 | TATGAGTGTGGAGTCGTTGCGAAAAGCTCCTATCAAACAAATANNNAGATCGGAAG |
| 12045_1 | TATGAGTGTGGAGTCGTTGCGTTGGCTCAGATGGGAGAGGAGNNNAGATCGGAA |
| 12045_2 | TATGAGTGTGGAGTCGTTGCGTTGGCTCAGATGGGAGAGGAGNNNAGATCGGAA |
| 42195_1 | TATGAGTGTGGAGTCGTTGCAAACAAGGTCTTGCTCTGTTGTCNNNAGATCGGAAG |
| 42195_2 | TATGAGTGTGGAGTCGTTGCAAACAAGGTCTTGCTCTGTTGTCNNNAGATCGGAAG |
| 33992_1 | TATGAGTGTGGAGTCGTTGCGATCTCCATCTCCCAGGTTCAAGNNNAGATCGGAAG |
| 33992_2 | TATGAGTGTGGAGTCGTTGCGATCTCCATCTCCCAGGTTCAAGNNNAGATCGGAAG |
| 52376_1 | TATGAGTGTGGAGTCGTTGCCCCAAAAAAGCCAAGCAATTAGCNNNAGATCGGAAG |
| 52376_2 | TATGAGTGTGGAGTCGTTGCCCCAAAAAAGCCAAGCAATTAGCNNNAGATCGGAAG |
| 30719_1 | TATGAGTGTGGAGTCGTTGCGAGGGAATGACAACTCAGGTCAATNNNAGATCGGA |
| 30719_2 | TATGAGTGTGGAGTCGTTGCGAGGGAATGACAACTCAGGTCAATNNNAGATCGGA |
| 8274_1 | TATGAGTGTGGAGTCGTTGCGACACTTTGGAAGACAATTGGTTNNNAGATCGGAA |
| 8274_2 | TATGAGTGTGGAGTCGTTGCGACACTTTGGAAGACAATTGGTTNNNAGATCGGAA |
| 32013_1 | TATGAGTGTGGAGTCGTTGCACAGGATCTACAAACTTCCCACTNNNAGATCGGAAG |
| 32013_2 | TATGAGTGTGGAGTCGTTGCACAGGATCTACAAACTTCCCACTNNNAGATCGGAAG |
| 35120_1 | TATGAGTGTGGAGTCGTTGCATCTCGTCTTACTGCAACCTCTGNNNAGATCGGAAGA |
| 35120_2 | TATGAGTGTGGAGTCGTTGCATCTCGTCTTACTGCAACCTCTGNNNAGATCGGAAGA |
| 2555_1 | TATGAGTGTGGAGTCGTTGCGCAAGGAATGGAGTATGGAAGGGNNNAGATCGGA |
| 2555_2 | TATGAGTGTGGAGTCGTTGCGCAAGGAATGGAGTATGGAAGGGNNNAGATCGGA |
| 42108_1 | TATGAGTGTGGAGTCGTTGCCCCTGACTGAGATTCCAACTCAANNNAGATCGGAAG |
| 42108_2 | TATGAGTGTGGAGTCGTTGCCCCTGACTGAGATTCCAACTCAANNNAGATCGGAAG |
| 45343_1 | TATGAGTGTGGAGTCGTTGCTCTTAGATGTTTTCCTGCCCTGANNNAGATCGGAAG |
| 45343_2 | TATGAGTGTGGAGTCGTTGCTCTTAGATGTTTTCCTGCCCTGANNNAGATCGGAAG |
| 40168_1 | TATGAGTGTGGAGTCGTTGCTCCCCCATCATTTCATTATGAGTGTNNNAGATCGGAA |
| 40168_2 | TATGAGTGTGGAGTCGTTGCTCCCCCATCATTTCATTATGAGTGTNNNAGATCGGAA |
| 51449_1 | TATGAGTGTGGAGTCGTTGCAGCTGAAGAAACTGGTAAGAGCTNNNAGATCGGAA |
| 51449_2 | TATGAGTGTGGAGTCGTTGCAGCTGAAGAAACTGGTAAGAGCTNNNAGATCGGAA |
| 29056_1 | TATGAGTGTGGAGTCGTTGCAAGAGAGACAGACAGAGTAGGCANNNAGATCGGA |
| 29056_2 | TATGAGTGTGGAGTCGTTGCAAGAGAGACAGACAGAGTAGGCANNNAGATCGGA |

| | |
|---|---|
| 43779_1 | TATGAGTGTGGAGTCGTTGCTTAGGGGTATCAATCCATGGTGTNNNAGATCGGAA |
| 43779_2 | TATGAGTGTGGAGTCGTTGCTTAGGGGTATCAATCCATGGTGTNNNAGATCGGAA |
| 33753_1 | TATGAGTGTGGAGTCGTTGCAGTGCTCTCCTTTCTCAAAGTGANNNAGATCGGAAG |
| 33753_2 | TATGAGTGTGGAGTCGTTGCAGTGCTCTCCTTTCTCAAAGTGANNNAGATCGGAAG |
| 43165_1 | TATGAGTGTGGAGTCGTTGCCCTCCCGAGTAGCTGATTACAGNNNAGATCGGAAGA |
| 43165_2 | TATGAGTGTGGAGTCGTTGCCCTCCCGAGTAGCTGATTACAGNNNAGATCGGAAGA |
| 34230_1 | TATGAGTGTGGAGTCGTTGCTCCACTACAGCTTTCTGAATCCCNNNAGATCGGAAGA |
| 34230_2 | TATGAGTGTGGAGTCGTTGCTCCACTACAGCTTTCTGAATCCCNNNAGATCGGAAGA |
| 13425_1 | TATGAGTGTGGAGTCGTTGCAAAAATTGGTGGGTTATGCTTNNNAGATCGGAAGA |
| 13425_2 | TATGAGTGTGGAGTCGTTGCAAAAATTGGTGGGTTATGCTTNNNAGATCGGAAGA |
| 45563_1 | TATGAGTGTGGAGTCGTTGCGGGTATTTGTGAAAGCCCCTCTCNNNAGATCGGAAG |
| 45563_2 | TATGAGTGTGGAGTCGTTGCGGGTATTTGTGAAAGCCCCTCTCNNNAGATCGGAAG |
| 29984_1 | TATGAGTGTGGAGTCGTTGCACTGTTCCGTTGTCACCCAGNNNAGATCGGAAGAGC |
| 29984_2 | TATGAGTGTGGAGTCGTTGCACTGTTCCGTTGTCACCCAGNNNAGATCGGAAGAGC |
| 39791_1 | TATGAGTGTGGAGTCGTTGCTTAGTTCAAAGCACTCAGCATGCNNNAGATCGGAAG |
| 39791_2 | TATGAGTGTGGAGTCGTTGCTTAGTTCAAAGCACTCAGCATGCNNNAGATCGGAAG |
| 31643_1 | TATGAGTGTGGAGTCGTTGCGGTGCCAGGAAATTGATTTTGGANNNAGATCGGAA |
| 31643_2 | TATGAGTGTGGAGTCGTTGCGGTGCCAGGAAATTGATTTTGGANNNAGATCGGAA |
| 40599_1 | TATGAGTGTGGAGTCGTTGCAGGCAGGAGAATCTCTTGAACTGNNNAGATCGGAA |
| 40599_2 | TATGAGTGTGGAGTCGTTGCAGGCAGGAGAATCTCTTGAACTGNNNAGATCGGAA |
| 36518_1 | TATGAGTGTGGAGTCGTTGCAAAAAATTAGCCAGGTGTGGTGGNNNAGATCGGAA |
| 36518_2 | TATGAGTGTGGAGTCGTTGCAAAAAATTAGCCAGGTGTGGTGGNNNAGATCGGAA |
| 39213_1 | TATGAGTGTGGAGTCGTTGCCATTAGGCTCTGTGTAGGTCCTCNNNAGATCGGAAG |
| 39213_2 | TATGAGTGTGGAGTCGTTGCCATTAGGCTCTGTGTAGGTCCTCNNNAGATCGGAAG |
| 9502_1 | TATGAGTGTGGAGTCGTTGCACACACACATCCCCAACAGAAAGNNNAGATCGGAAG |
| 9502_2 | TATGAGTGTGGAGTCGTTGCACACACACATCCCCAACAGAAAGNNNAGATCGGAAG |
| 34980_1 | TATGAGTGTGGAGTCGTTGCTCAAGTGATTCTTTTGCCTCAGCNNNAGATCGGAAG |
| 34980_2 | TATGAGTGTGGAGTCGTTGCTCAAGTGATTCTTTTGCCTCAGCNNNAGATCGGAAG |
| 46872_1 | TATGAGTGTGGAGTCGTTGCTCAAACTCCTGACCTCGTGATTCNNNAGATCGGAAG |
| 46872_2 | TATGAGTGTGGAGTCGTTGCTCAAACTCCTGACCTCGTGATTCNNNAGATCGGAAG |
| 35622_1 | TATGAGTGTGGAGTCGTTGCTTGTTCCTTAGGTTGGTGTGTCTNNNAGATCGGAAG |
| 35622_2 | TATGAGTGTGGAGTCGTTGCTTGTTCCTTAGGTTGGTGTGTCTNNNAGATCGGAAG |
| 45613_1 | TATGAGTGTGGAGTCGTTGCATAGTGTTTCACTGTCACCCAGGNNNAGATCGGAAG |
| 45613_2 | TATGAGTGTGGAGTCGTTGCATAGTGTTTCACTGTCACCCAGGNNNAGATCGGAAG |
| 37844_1 | TATGAGTGTGGAGTCGTTGCATTGATTGATGGGGTCAGCTGATNNNAGATCGGAA |
| 37844_2 | TATGAGTGTGGAGTCGTTGCATTGATTGATGGGGTCAGCTGATNNNAGATCGGAA |
| 36525_1 | TATGAGTGTGGAGTCGTTGCTTCTCTGTATCTCATGACTGGCTNNNAGATCGGAAG |
| 36525_2 | TATGAGTGTGGAGTCGTTGCTTCTCTGTATCTCATGACTGGCTNNNAGATCGGAAG |
| 39851_1 | TATGAGTGTGGAGTCGTTGCAAAGATGTACTGTGACATGCCCANNNAGATCGGAA |
| 39851_2 | TATGAGTGTGGAGTCGTTGCAAAGATGTACTGTGACATGCCCANNNAGATCGGAA |
| 38366_1 | TATGAGTGTGGAGTCGTTGCACAGATCAACAAGAGCTGAAGGANNNAGATCGGAA |
| 38366_2 | TATGAGTGTGGAGTCGTTGCACAGATCAACAAGAGCTGAAGGANNNAGATCGGAA |
| 30431_1 | TATGAGTGTGGAGTCGTTGCCTCCTGAGTAGCTGGGATTACAGNNNAGATCGGAA |

| | |
|---|---|
| 30431_2 | TATGAGTGTGGAGTCGTTGCCTCCTGAGTAGCTGGGATTACAGNNNAGATCGGAA |
| 13740_1 | TATGAGTGTGGAGTCGTTGCTAACATGCGAATGTAGTTCCNNNAGATCGGAAGAG |
| 13740_2 | TATGAGTGTGGAGTCGTTGCTAACATGCGAATGTAGTTCCNNNAGATCGGAAGAG |
| 46160_1 | TATGAGTGTGGAGTCGTTGCCTCCTGAGTAGCTGGGACTACANNNAGATCGGAAG |
| 46160_2 | TATGAGTGTGGAGTCGTTGCCTCCTGAGTAGCTGGGACTACANNNAGATCGGAAG |
| 34589_1 | TATGAGTGTGGAGTCGTTGCTCATCTCGGGTGAAAGTCCAAATNNNAGATCGGAAG |
| 34589_2 | TATGAGTGTGGAGTCGTTGCTCATCTCGGGTGAAAGTCCAAATNNNAGATCGGAAG |
| 45431_1 | TATGAGTGTGGAGTCGTTGCACCACCATCTCTCTCTAGTTCCANNNAGATCGGAAGA |
| 45431_2 | TATGAGTGTGGAGTCGTTGCACCACCATCTCTCTCTAGTTCCANNNAGATCGGAAGA |
| 46113_1 | TATGAGTGTGGAGTCGTTGCCAGTGATGCAATCTTGGCTCATTNNNAGATCGGAAG |
| 46113_2 | TATGAGTGTGGAGTCGTTGCCAGTGATGCAATCTTGGCTCATTNNNAGATCGGAAG |
| 36270_1 | TATGAGTGTGGAGTCGTTGCTTCTGGTGGGGGATATGGATAGTNNNAGATCGGAA |
| 36270_2 | TATGAGTGTGGAGTCGTTGCTTCTGGTGGGGGATATGGATAGTNNNAGATCGGAA |
| 11532_1 | TATGAGTGTGGAGTCGTTGCTCCACAGACAGTGACTTCTCCCNNNAGATCGGAAGA |
| 11532_2 | TATGAGTGTGGAGTCGTTGCTCCACAGACAGTGACTTCTCCCNNNAGATCGGAAGA |
| 29717_1 | TATGAGTGTGGAGTCGTTGCCAATGAGCTGAGTTGCATTTCCCNNNAGATCGGAAG |
| 29717_2 | TATGAGTGTGGAGTCGTTGCCAATGAGCTGAGTTGCATTTCCCNNNAGATCGGAAG |
| 45345_1 | TATGAGTGTGGAGTCGTTGCAAAAGGAAGAGGGGATAGCAAGGNNNAGATCGGA |
| 45345_2 | TATGAGTGTGGAGTCGTTGCAAAAGGAAGAGGGGATAGCAAGGNNNAGATCGGA |
| 33370_1 | TATGAGTGTGGAGTCGTTGCCGAGTAGCTGGGACTACAGGTNNNAGATCGGAAGA |
| 33370_2 | TATGAGTGTGGAGTCGTTGCCGAGTAGCTGGGACTACAGGTNNNAGATCGGAAGA |
| 45140_1 | TATGAGTGTGGAGTCGTTGCGGCCTGATCAAGACACTGAGAGNNNAGATCGGAAG |
| 45140_2 | TATGAGTGTGGAGTCGTTGCGGCCTGATCAAGACACTGAGAGNNNAGATCGGAAG |
| 48142_1 | TATGAGTGTGGAGTCGTTGCTAACAAAGTCTTCCCCGTCCCTTNNNAGATCGGAAGA |
| 48142_2 | TATGAGTGTGGAGTCGTTGCTAACAAAGTCTTCCCCGTCCCTTNNNAGATCGGAAGA |
| 41338_1 | TATGAGTGTGGAGTCGTTGCATGCTGAGGAAGGAGAATTGCTTNNNAGATCGGAA |
| 41338_2 | TATGAGTGTGGAGTCGTTGCATGCTGAGGAAGGAGAATTGCTTNNNAGATCGGAA |
| 30291_1 | TATGAGTGTGGAGTCGTTGCTGCTGATTTTAAGAGAAATACTGAGAGNNNAGATC |
| 30291_2 | TATGAGTGTGGAGTCGTTGCTGCTGATTTTAAGAGAAATACTGAGAGNNNAGATC |
| 40061_1 | TATGAGTGTGGAGTCGTTGCGTTGGAGGGACTTTGAGAGACATNNNAGATCGGAA |
| 40061_2 | TATGAGTGTGGAGTCGTTGCGTTGGAGGGACTTTGAGAGACATNNNAGATCGGAA |
| 47421_1 | TATGAGTGTGGAGTCGTTGCAAGTAGCTGGGATTACAGGCATCNNNAGATCGGAA |
| 47421_2 | TATGAGTGTGGAGTCGTTGCAAGTAGCTGGGATTACAGGCATCNNNAGATCGGAA |
| 29024_1 | TATGAGTGTGGAGTCGTTGCTTGGACATTTATCTTCCCCCAGGNNNAGATCGGAAG |
| 29024_2 | TATGAGTGTGGAGTCGTTGCTTGGACATTTATCTTCCCCCAGGNNNAGATCGGAAG |
| 31096_1 | TATGAGTGTGGAGTCGTTGCACATTCAGATAAAGCAGCTCCCANNNAGATCGGAAG |
| 31096_2 | TATGAGTGTGGAGTCGTTGCACATTCAGATAAAGCAGCTCCCANNNAGATCGGAAG |
| 47535_1 | TATGAGTGTGGAGTCGTTGCCAGTGGTGCTATCTTGGTTCACTNNNAGATCGGAAG |
| 47535_2 | TATGAGTGTGGAGTCGTTGCCAGTGGTGCTATCTTGGTTCACTNNNAGATCGGAAG |
| 31760_1 | TATGAGTGTGGAGTCGTTGCCTCTCTCTCACCACCTAAAGCTGNNNAGATCGGAAGA |
| 31760_2 | TATGAGTGTGGAGTCGTTGCCTCTCTCTCACCACCTAAAGCTGNNNAGATCGGAAGA |
| 48628_1 | TATGAGTGTGGAGTCGTTGCCTCCTGAGTAGCTGGGACTACAGNNNAGATCGGAA |
| 48628_2 | TATGAGTGTGGAGTCGTTGCCTCCTGAGTAGCTGGGACTACAGNNNAGATCGGAA |

| | |
|---|---|
| 46386_1 | TATGAGTGTGGAGTCGTTGCCATTCTCCTGCCTCAGCCTCNNNAGATCGGAAGAGCA |
| 46386_2 | TATGAGTGTGGAGTCGTTGCCATTCTCCTGCCTCAGCCTCNNNAGATCGGAAGAGCA |
| 47741_1 | TATGAGTGTGGAGTCGTTGCGAGAAGCAAACTGTTCTCTGGGGNNNAGATCGGAA |
| 47741_2 | TATGAGTGTGGAGTCGTTGCGAGAAGCAAACTGTTCTCTGGGGNNNAGATCGGAA |
| 35926_1 | TATGAGTGTGGAGTCGTTGCGGAAGGAAGGAAGAAAGGAAGGANNNAGATCGG |
| 35926_2 | TATGAGTGTGGAGTCGTTGCGGAAGGAAGGAAGAAAGGAAGGANNNAGATCGG |
| 30811_1 | TATGAGTGTGGAGTCGTTGCCTCTGCTAGCATTCACCTCTAGGNNNAGATCGGAAG |
| 30811_2 | TATGAGTGTGGAGTCGTTGCCTCTGCTAGCATTCACCTCTAGGNNNAGATCGGAAG |
| 40405_1 | TATGAGTGTGGAGTCGTTGCTCCCCCCCTTTGATAGGATTAGANNNAGATCGGAAG |
| 40405_2 | TATGAGTGTGGAGTCGTTGCTCCCCCCCTTTGATAGGATTAGANNNAGATCGGAAG |
| 48077_1 | TATGAGTGTGGAGTCGTTGCCTCGTGGTAACACCAGAGTACAANNNAGATCGGAAG |
| 48077_2 | TATGAGTGTGGAGTCGTTGCCTCGTGGTAACACCAGAGTACAANNNAGATCGGAAG |
| 33113_1 | TATGAGTGTGGAGTCGTTGCCCTGTTGATTCCTAGACTGCACTNNNAGATCGGAAG |
| 33113_2 | TATGAGTGTGGAGTCGTTGCCCTGTTGATTCCTAGACTGCACTNNNAGATCGGAAG |
| 53086_1 | TATGAGTGTGGAGTCGTTGCCTAGGTGTACCCAAAAGCGGGNNNAGATCGGAAGA |
| 53086_2 | TATGAGTGTGGAGTCGTTGCCTAGGTGTACCCAAAAGCGGGNNNAGATCGGAAGA |
| 41703_1 | TATGAGTGTGGAGTCGTTGCAGACGGAGTCTCGCTCTGTCNNNAGATCGGAAGAG |
| 41703_2 | TATGAGTGTGGAGTCGTTGCAGACGGAGTCTCGCTCTGTCNNNAGATCGGAAGAG |
| 34764_1 | TATGAGTGTGGAGTCGTTGCCTTCCAGGGTTCAACCGATTCTCNNNAGATCGGAAG |
| 34764_2 | TATGAGTGTGGAGTCGTTGCCTTCCAGGGTTCAACCGATTCTCNNNAGATCGGAAG |
| 12636_1 | TATGAGTGTGGAGTCGTTGCTGAGATTAAGTCCAATCAATCATNNNAGATCGGAAG |
| 12636_2 | TATGAGTGTGGAGTCGTTGCTGAGATTAAGTCCAATCAATCATNNNAGATCGGAAG |
| 35036_1 | TATGAGTGTGGAGTCGTTGCCGATCTCCAATTACTGCAAGCTCNNNAGATCGGAAG |
| 35036_2 | TATGAGTGTGGAGTCGTTGCCGATCTCCAATTACTGCAAGCTCNNNAGATCGGAAG |
| 35560_1 | TATGAGTGTGGAGTCGTTGCTGTATGTTGCCTGTTCACTCTGANNNAGATCGGAAG |
| 35560_2 | TATGAGTGTGGAGTCGTTGCTGTATGTTGCCTGTTCACTCTGANNNAGATCGGAAG |
| 34454_1 | TATGAGTGTGGAGTCGTTGCCCCATCTCCTCCCCTCTCACNNNAGATCGGAAGAGCA |
| 34454_2 | TATGAGTGTGGAGTCGTTGCCCCATCTCCTCCCCTCTCACNNNAGATCGGAAGAGCA |
| 37552_1 | TATGAGTGTGGAGTCGTTGCGCAAACATTGAACTGATCCAGCTNNNAGATCGGAAG |
| 37552_2 | TATGAGTGTGGAGTCGTTGCGCAAACATTGAACTGATCCAGCTNNNAGATCGGAAG |
| 45853_1 | TATGAGTGTGGAGTCGTTGCAGGAGATGATGACACAGACACACNNNAGATCGGAA |
| 45853_2 | TATGAGTGTGGAGTCGTTGCAGGAGATGATGACACAGACACACNNNAGATCGGAA |
| 34887_1 | TATGAGTGTGGAGTCGTTGCCACCACACTCGGCTAATTTTGTTNNNAGATCGGAAG |
| 34887_2 | TATGAGTGTGGAGTCGTTGCCACCACACTCGGCTAATTTTGTTNNNAGATCGGAAG |
| 41064_1 | TATGAGTGTGGAGTCGTTGCAGAATGAACAGATGCAGGGACTTNNNAGATCGGAA |
| 41064_2 | TATGAGTGTGGAGTCGTTGCAGAATGAACAGATGCAGGGACTTNNNAGATCGGAA |
| 37478_1 | TATGAGTGTGGAGTCGTTGCAGAAAAAGAAAACTTGCAGGCCGNNNAGATCGGAA |
| 37478_2 | TATGAGTGTGGAGTCGTTGCAGAAAAAGAAAACTTGCAGGCCGNNNAGATCGGAA |
| 46978_1 | TATGAGTGTGGAGTCGTTGCCAATCTGGGCTCACTGCAAGNNNAGATCGGAAGAG |
| 46978_2 | TATGAGTGTGGAGTCGTTGCCAATCTGGGCTCACTGCAAGNNNAGATCGGAAGAG |
| 42566_1 | TATGAGTGTGGAGTCGTTGCGTGCAGTTTTCTGTGTTTACCGANNNAGATCGGAAG |
| 42566_2 | TATGAGTGTGGAGTCGTTGCGTGCAGTTTTCTGTGTTTACCGANNNAGATCGGAAG |
| 45615_1 | TATGAGTGTGGAGTCGTTGCGCTGACATATCATCAAAGCACCCNNNAGATCGGAAG |

| | |
|---|---|
| 45615_2 | TATGAGTGTGGAGTCGTTGCGCTGACATATCATCAAAGCACCCNNNAGATCGGAAG |
| 32183_1 | TATGAGTGTGGAGTCGTTGCGTTTTACCATGTTGGCCAGTCTGNNNAGATCGGAAG |
| 32183_2 | TATGAGTGTGGAGTCGTTGCGTTTTACCATGTTGGCCAGTCTGNNNAGATCGGAAG |
| 46382_1 | TATGAGTGTGGAGTCGTTGCAACTACAGGAGTGCACCACTATGNNNAGATCGGAA |
| 46382_2 | TATGAGTGTGGAGTCGTTGCAACTACAGGAGTGCACCACTATGNNNAGATCGGAA |
| 37859_1 | TATGAGTGTGGAGTCGTTGCCGTCTGCCAGAAACAGAATTCTGNNNAGATCGGAAG |
| 37859_2 | TATGAGTGTGGAGTCGTTGCCGTCTGCCAGAAACAGAATTCTGNNNAGATCGGAAG |
| 48442_1 | TATGAGTGTGGAGTCGTTGCATGAAGCTGTTCAATCTGCACACNNNAGATCGGAAG |
| 48442_2 | TATGAGTGTGGAGTCGTTGCATGAAGCTGTTCAATCTGCACACNNNAGATCGGAAG |
| 42735_1 | TATGAGTGTGGAGTCGTTGCCACAGGATTGCTCACCTTTGTTTNNNAGATCGGAAG |
| 42735_2 | TATGAGTGTGGAGTCGTTGCCACAGGATTGCTCACCTTTGTTTNNNAGATCGGAAG |
| 32068_1 | TATGAGTGTGGAGTCGTTGCTTCCAGGAGCCTATTGAAGAAGCNNNAGATCGGAA |
| 32068_2 | TATGAGTGTGGAGTCGTTGCTTCCAGGAGCCTATTGAAGAAGCNNNAGATCGGAA |
| 43776_1 | TATGAGTGTGGAGTCGTTGCCTTTCTGTCTTGCACCTGTGAAGNNNAGATCGGAAG |
| 43776_2 | TATGAGTGTGGAGTCGTTGCCTTTCTGTCTTGCACCTGTGAAGNNNAGATCGGAAG |
| 38312_1 | TATGAGTGTGGAGTCGTTGCGGAGACCCTGGTATTGTCACATANNNAGATCGGAA |
| 38312_2 | TATGAGTGTGGAGTCGTTGCGGAGACCCTGGTATTGTCACATANNNAGATCGGAA |
| 41213_1 | TATGAGTGTGGAGTCGTTGCGCTTGGTCTCCGTCTCAAAAAATNNNAGATCGGAAG |
| 41213_2 | TATGAGTGTGGAGTCGTTGCGCTTGGTCTCCGTCTCAAAAAATNNNAGATCGGAAG |
| 10190_1 | TATGAGTGTGGAGTCGTTGCGGTCCTGGAGATGCATTCTAATGGNNNAGATCGGA |
| 10190_2 | TATGAGTGTGGAGTCGTTGCGGTCCTGGAGATGCATTCTAATGGNNNAGATCGGA |
| 47552_1 | TATGAGTGTGGAGTCGTTGCCTCCAAGGAGAACAGAGAGATGGNNNAGATCGGA |
| 47552_2 | TATGAGTGTGGAGTCGTTGCCTCCAAGGAGAACAGAGAGATGGNNNAGATCGGA |
| 47697_1 | TATGAGTGTGGAGTCGTTGCTGGTTCAAGAGATTTTCCTGCCTNNNAGATCGGAAG |
| 47697_2 | TATGAGTGTGGAGTCGTTGCTGGTTCAAGAGATTTTCCTGCCTNNNAGATCGGAAG |
| 40822_1 | TATGAGTGTGGAGTCGTTGCTGAATTCCAAGATGCAACCCCTANNNAGATCGGAAG |
| 40822_2 | TATGAGTGTGGAGTCGTTGCTGAATTCCAAGATGCAACCCCTANNNAGATCGGAAG |
| 50605_1 | TATGAGTGTGGAGTCGTTGCTTGATGTTCTTTCCCTTCTCCCCNNNAGATCGGAAGA |
| 50605_2 | TATGAGTGTGGAGTCGTTGCTTGATGTTCTTTCCCTTCTCCCCNNNAGATCGGAAGA |
| 37172_1 | TATGAGTGTGGAGTCGTTGCACCATCACCCTTTTGATCAGTGTNNNAGATCGGAAG |
| 37172_2 | TATGAGTGTGGAGTCGTTGCACCATCACCCTTTTGATCAGTGTNNNAGATCGGAAG |
| 42515_1 | TATGAGTGTGGAGTCGTTGCGCTCCCTGAGTCTCAAGAAAAGANNNAGATCGGAA |
| 42515_2 | TATGAGTGTGGAGTCGTTGCGCTCCCTGAGTCTCAAGAAAAGANNNAGATCGGAA |
| 13608_1 | TATGAGTGTGGAGTCGTTGCAGACAAGGAAACTAATGATAGAANNNAGATCGGAA |
| 13608_2 | TATGAGTGTGGAGTCGTTGCAGACAAGGAAACTAATGATAGAANNNAGATCGGAA |
| 12646_1 | TATGAGTGTGGAGTCGTTGCCATCAAAACACAGAAACAGGNNNAGATCGGAAGAG |
| 12646_2 | TATGAGTGTGGAGTCGTTGCCATCAAAACACAGAAACAGGNNNAGATCGGAAGAG |
| 35935_1 | TATGAGTGTGGAGTCGTTGCCTGAAAGCGTCTCCAAAAATCCANNNAGATCGGAAG |
| 35935_2 | TATGAGTGTGGAGTCGTTGCCTGAAAGCGTCTCCAAAAATCCANNNAGATCGGAAG |
| 9295_1 | TATGAGTGTGGAGTCGTTGCACAGGGTTTCACTCTCACTTTTGGNNNAGATCGGAA |
| 9295_2 | TATGAGTGTGGAGTCGTTGCACAGGGTTTCACTCTCACTTTTGGNNNAGATCGGAA |
| 50927_1 | TATGAGTGTGGAGTCGTTGCAACCTGGGTTTCCTACGTCAAGNNNAGATCGGAAGA |
| 50927_2 | TATGAGTGTGGAGTCGTTGCAACCTGGGTTTCCTACGTCAAGNNNAGATCGGAAGA |

| | |
|---|---|
| 46538_1 | TATGAGTGTGGAGTCGTTGCGCTCAGGACTAAGCACTGAAGATNNNAGATCGGAA |
| 46538_2 | TATGAGTGTGGAGTCGTTGCGCTCAGGACTAAGCACTGAAGATNNNAGATCGGAA |
| 11044_1 | TATGAGTGTGGAGTCGTTGCACGGATTTTCACTCTCTCAACCCCNNNAGATCGGAAG |
| 11044_2 | TATGAGTGTGGAGTCGTTGCACGGATTTTCACTCTCTCAACCCCNNNAGATCGGAAG |
| 39626_1 | TATGAGTGTGGAGTCGTTGCAGATTTCACCATTTGTAGGCCCTNNNAGATCGGAAG |
| 39626_2 | TATGAGTGTGGAGTCGTTGCAGATTTCACCATTTGTAGGCCCTNNNAGATCGGAAG |
| 49081_1 | TATGAGTGTGGAGTCGTTGCAGGAGGTAGATTACAGAGCCAGTNNNAGATCGGAA |
| 49081_2 | TATGAGTGTGGAGTCGTTGCAGGAGGTAGATTACAGAGCCAGTNNNAGATCGGAA |
| 34923_1 | TATGAGTGTGGAGTCGTTGCTTCAAGTGATTCTCCTGCCTCAGNNNAGATCGGAAG |
| 34923_2 | TATGAGTGTGGAGTCGTTGCTTCAAGTGATTCTCCTGCCTCAGNNNAGATCGGAAG |
| 30096_1 | TATGAGTGTGGAGTCGTTGCGCATACCACTGGATTGAGGATGGNNNAGATCGGAA |
| 30096_2 | TATGAGTGTGGAGTCGTTGCGCATACCACTGGATTGAGGATGGNNNAGATCGGAA |
| 43789_1 | TATGAGTGTGGAGTCGTTGCAACAAAGGGAAAAACCGACAGAGNNNAGATCGGAA |
| 43789_2 | TATGAGTGTGGAGTCGTTGCAACAAAGGGAAAAACCGACAGAGNNNAGATCGGAA |
| 49950_1 | TATGAGTGTGGAGTCGTTGCCACATGGTCCCTGTTCTTCAATGNNNAGATCGGAAG |
| 49950_2 | TATGAGTGTGGAGTCGTTGCCACATGGTCCCTGTTCTTCAATGNNNAGATCGGAAG |
| 40712_1 | TATGAGTGTGGAGTCGTTGCCACGACCTTGGTTTACTGCAACNNNAGATCGGAAGA |
| 40712_2 | TATGAGTGTGGAGTCGTTGCCACGACCTTGGTTTACTGCAACNNNAGATCGGAAGA |
| 50229_1 | TATGAGTGTGGAGTCGTTGCCGTCGAATCCTCCTGACAACTATNNNAGATCGGAAG |
| 50229_2 | TATGAGTGTGGAGTCGTTGCCGTCGAATCCTCCTGACAACTATNNNAGATCGGAAG |
| 44228_1 | TATGAGTGTGGAGTCGTTGCCACGGAGTTTACTCTTCTGACCTNNNAGATCGGAAG |
| 44228_2 | TATGAGTGTGGAGTCGTTGCCACGGAGTTTACTCTTCTGACCTNNNAGATCGGAAG |
| 3078_1 | TATGAGTGTGGAGTCGTTGCGCTGCCTCATCCTCATCGTTATTNNNAGATCGGAAGA |
| 3078_2 | TATGAGTGTGGAGTCGTTGCGCTGCCTCATCCTCATCGTTATTNNNAGATCGGAAGA |
| 47043_1 | TATGAGTGTGGAGTCGTTGCCTCAAGCAATCCTCTCACCTCAGNNNAGATCGGAAGA |
| 47043_2 | TATGAGTGTGGAGTCGTTGCCTCAAGCAATCCTCTCACCTCAGNNNAGATCGGAAGA |
| 29360_1 | TATGAGTGTGGAGTCGTTGCTACGAAAGTGACTCTCCTGAAGGNNNAGATCGGAA |
| 29360_2 | TATGAGTGTGGAGTCGTTGCTACGAAAGTGACTCTCCTGAAGGNNNAGATCGGAA |
| 39839_1 | TATGAGTGTGGAGTCGTTGCGTTTCACCATGTTAGCCAGGATGNNNAGATCGGAAG |
| 39839_2 | TATGAGTGTGGAGTCGTTGCGTTTCACCATGTTAGCCAGGATGNNNAGATCGGAAG |
| 10050_1 | TATGAGTGTGGAGTCGTTGCCGAGCAATTTCCATGCAACAGTTNNNAGATCGGAAG |
| 10050_2 | TATGAGTGTGGAGTCGTTGCCGAGCAATTTCCATGCAACAGTTNNNAGATCGGAAG |
| 41508_1 | TATGAGTGTGGAGTCGTTGCTGCTAATGATACGAGGAGATGAACANNNAGATCGG |
| 41508_2 | TATGAGTGTGGAGTCGTTGCTGCTAATGATACGAGGAGATGAACANNNAGATCGG |
| 49161_1 | TATGAGTGTGGAGTCGTTGCGAAGTCCTGACCTCAAGTGACCNNNAGATCGGAAGA |
| 49161_2 | TATGAGTGTGGAGTCGTTGCGAAGTCCTGACCTCAAGTGACCNNNAGATCGGAAGA |
| 13106_1 | TATGAGTGTGGAGTCGTTGCACAAATTAATATCCAGTAGAGAGANNNAGATCGGA |
| 13106_2 | TATGAGTGTGGAGTCGTTGCACAAATTAATATCCAGTAGAGAGANNNAGATCGGA |
| 36081_1 | TATGAGTGTGGAGTCGTTGCTGTGCATCCATTCTCCAGTAGTTNNNAGATCGGAAG |
| 36081_2 | TATGAGTGTGGAGTCGTTGCTGTGCATCCATTCTCCAGTAGTTNNNAGATCGGAAG |
| 47278_1 | TATGAGTGTGGAGTCGTTGCTGGGAGACTTGTTGTTATGGCTTNNNAGATCGGAA |
| 47278_2 | TATGAGTGTGGAGTCGTTGCTGGGAGACTTGTTGTTATGGCTTNNNAGATCGGAA |
| 34554_1 | TATGAGTGTGGAGTCGTTGCGTTCAAGCGATTCTCCTACCTCANNNAGATCGGAAG |

| | |
|---|---|
| 34554_2 | TATGAGTGTGGAGTCGTTGCGTTCAAGCGATTCTCCTACCTCANNNAGATCGGAAG |
| 44623_1 | TATGAGTGTGGAGTCGTTGCGAGATGGTCTTTTCTCCCAGGTCNNNAGATCGGAAG |
| 44623_2 | TATGAGTGTGGAGTCGTTGCGAGATGGTCTTTTCTCCCAGGTCNNNAGATCGGAAG |
| 5604_1 | TATGAGTGTGGAGTCGTTGCCAACCCATAAGCCTGAACTGGAANNNAGATCGGAAG |
| 5604_2 | TATGAGTGTGGAGTCGTTGCCAACCCATAAGCCTGAACTGGAANNNAGATCGGAAG |
| 53136_1 | TATGAGTGTGGAGTCGTTGCTGGTTTTAGGGTTCAACAGGAGTNNNAGATCGGAA |
| 53136_2 | TATGAGTGTGGAGTCGTTGCTGGTTTTAGGGTTCAACAGGAGTNNNAGATCGGAA |
| 45573_1 | TATGAGTGTGGAGTCGTTGCAAGTGATTTTCCTGCCTCAGCTTNNNAGATCGGAAG |
| 45573_2 | TATGAGTGTGGAGTCGTTGCAAGTGATTTTCCTGCCTCAGCTTNNNAGATCGGAAG |
| 2313_1 | TATGAGTGTGGAGTCGTTGCTGGTCTGTTTTCCGGGTCAATACNNNAGATCGGAAG |
| 2313_2 | TATGAGTGTGGAGTCGTTGCTGGTCTGTTTTCCGGGTCAATACNNNAGATCGGAAG |
| 10557_1 | TATGAGTGTGGAGTCGTTGCGTGTGTGTGTGTGCTAAGCTTTTNNNAGATCGGAAG |
| 10557_2 | TATGAGTGTGGAGTCGTTGCGTGTGTGTGTGTGCTAAGCTTTTNNNAGATCGGAAG |
| 32533_1 | TATGAGTGTGGAGTCGTTGCATTATCCAAGTGGGCCCAATTCANNNAGATCGGAAG |
| 32533_2 | TATGAGTGTGGAGTCGTTGCATTATCCAAGTGGGCCCAATTCANNNAGATCGGAAG |
| 52385_1 | TATGAGTGTGGAGTCGTTGCAGGAAATGAAAGCCCAAAGATGCNNNAGATCGGAA |
| 52385_2 | TATGAGTGTGGAGTCGTTGCAGGAAATGAAAGCCCAAAGATGCNNNAGATCGGAA |
| 34351_1 | TATGAGTGTGGAGTCGTTGCCCTGGCTGGAGTCTGTGGTCNNNAGATCGGAAGAG |
| 34351_2 | TATGAGTGTGGAGTCGTTGCCCTGGCTGGAGTCTGTGGTCNNNAGATCGGAAGAG |
| 30101_1 | TATGAGTGTGGAGTCGTTGCCCCAAGTCACCACAAAAACCTATNNNAGATCGGAAG |
| 30101_2 | TATGAGTGTGGAGTCGTTGCCCCAAGTCACCACAAAAACCTATNNNAGATCGGAAG |
| 12621_1 | TATGAGTGTGGAGTCGTTGCATTAATGCATTCTTCTCCAAATCNNNAGATCGGAAGA |
| 12621_2 | TATGAGTGTGGAGTCGTTGCATTAATGCATTCTTCTCCAAATCNNNAGATCGGAAGA |
| 48033_1 | TATGAGTGTGGAGTCGTTGCCATGATGATAGCGAGTGGGATGANNNAGATCGGAA |
| 48033_2 | TATGAGTGTGGAGTCGTTGCCATGATGATAGCGAGTGGGATGANNNAGATCGGAA |
| 40876_1 | TATGAGTGTGGAGTCGTTGCCAGGCCACAGCACAATCAAATTANNNAGATCGGAAG |
| 40876_2 | TATGAGTGTGGAGTCGTTGCCAGGCCACAGCACAATCAAATTANNNAGATCGGAAG |
| 29504_1 | TATGAGTGTGGAGTCGTTGCTTGAAACTTGGAACTTGAATTCTNNNAGATCGGAAG |
| 29504_2 | TATGAGTGTGGAGTCGTTGCTTGAAACTTGGAACTTGAATTCTNNNAGATCGGAAG |
| 33233_1 | TATGAGTGTGGAGTCGTTGCCTCACTTGTAGCCTCGAACTCTTNNNAGATCGGAAGA |
| 33233_2 | TATGAGTGTGGAGTCGTTGCCTCACTTGTAGCCTCGAACTCTTNNNAGATCGGAAGA |
| 43809_1 | TATGAGTGTGGAGTCGTTGCTCTTTGTCTCATCAGCCTCAGTCNNNAGATCGGAAGA |
| 43809_2 | TATGAGTGTGGAGTCGTTGCTCTTTGTCTCATCAGCCTCAGTCNNNAGATCGGAAGA |
| 48880_1 | TATGAGTGTGGAGTCGTTGCCCGGATTCAAGTGATTCTCCTGTNNNAGATCGGAAG |
| 48880_2 | TATGAGTGTGGAGTCGTTGCCCGGATTCAAGTGATTCTCCTGTNNNAGATCGGAAG |
| 34904_1 | TATGAGTGTGGAGTCGTTGCCTCCCAGGTTCAAGTGATTCTCANNNAGATCGGAAG |
| 34904_2 | TATGAGTGTGGAGTCGTTGCCTCCCAGGTTCAAGTGATTCTCANNNAGATCGGAAG |
| 45744_1 | TATGAGTGTGGAGTCGTTGCTTCAAGTGATTCTCCTGCCTCAGNNNAGATCGGAAG |
| 45744_2 | TATGAGTGTGGAGTCGTTGCTTCAAGTGATTCTCCTGCCTCAGNNNAGATCGGAAG |
| 50193_1 | TATGAGTGTGGAGTCGTTGCGTCAAACAGAAGGCTGCACTTAGNNNAGATCGGAA |
| 50193_2 | TATGAGTGTGGAGTCGTTGCGTCAAACAGAAGGCTGCACTTAGNNNAGATCGGAA |
| 12997_1 | TATGAGTGTGGAGTCGTTGCGAAATATGTGAAAACCAGTAAGTNNNAGATCGGAA |
| 12997_2 | TATGAGTGTGGAGTCGTTGCGAAATATGTGAAAACCAGTAAGTNNNAGATCGGAA |

| | |
|---|---|
| 50150_1 | TATGAGTGTGGAGTCGTTGCCAGCCCAACCTTTCCTTAGTTTGNNNAGATCGGAAGA |
| 50150_2 | TATGAGTGTGGAGTCGTTGCCAGCCCAACCTTTCCTTAGTTTGNNNAGATCGGAAGA |
| 13201_1 | TATGAGTGTGGAGTCGTTGCTGCCTCATATTCATGTTAGATAANNNAGATCGGAAG |
| 13201_2 | TATGAGTGTGGAGTCGTTGCTGCCTCATATTCATGTTAGATAANNNAGATCGGAAG |
| 8242_1 | TATGAGTGTGGAGTCGTTGCGAACAGGGGCAAGAAGACGGACTNNNAGATCGGA |
| 8242_2 | TATGAGTGTGGAGTCGTTGCGAACAGGGGCAAGAAGACGGACTNNNAGATCGGA |
| 50700_1 | TATGAGTGTGGAGTCGTTGCCAGGGACAAAGGCTTCAGATGANNNAGATCGGAAG |
| 50700_2 | TATGAGTGTGGAGTCGTTGCCAGGGACAAAGGCTTCAGATGANNNAGATCGGAAG |
| 45388_1 | TATGAGTGTGGAGTCGTTGCGAGCTGGAGTGCATTTTCCAACNNNAGATCGGAAG |
| 45388_2 | TATGAGTGTGGAGTCGTTGCGAGCTGGAGTGCATTTTCCAACNNNAGATCGGAAG |
| 9168_1 | TATGAGTGTGGAGTCGTTGCTCCTTTAGACTGGAACGTCACATNNNAGATCGGAAG |
| 9168_2 | TATGAGTGTGGAGTCGTTGCTCCTTTAGACTGGAACGTCACATNNNAGATCGGAAG |
| 37024_1 | TATGAGTGTGGAGTCGTTGCTAGCAGGCTTTATGTTGGGAACANNNAGATCGGAA |
| 37024_2 | TATGAGTGTGGAGTCGTTGCTAGCAGGCTTTATGTTGGGAACANNNAGATCGGAA |
| 48728_1 | TATGAGTGTGGAGTCGTTGCCCTGTTGGGTTCAAGCAATTCTTNNNAGATCGGAAG |
| 48728_2 | TATGAGTGTGGAGTCGTTGCCCTGTTGGGTTCAAGCAATTCTTNNNAGATCGGAAG |
| 33520_1 | TATGAGTGTGGAGTCGTTGCGAGTTGTGGTGATAGGGTCTCTGNNNAGATCGGAA |
| 33520_2 | TATGAGTGTGGAGTCGTTGCGAGTTGTGGTGATAGGGTCTCTGNNNAGATCGGAA |
| 47241_1 | TATGAGTGTGGAGTCGTTGCCTCCCAAGTAGCTGGAACTACAGNNNAGATCGGAAG |
| 47241_2 | TATGAGTGTGGAGTCGTTGCCTCCCAAGTAGCTGGAACTACAGNNNAGATCGGAAG |
| 41413_1 | TATGAGTGTGGAGTCGTTGCTGCAACCCTAGTCTCTGATAAAACANNNAGATCGGA |
| 41413_2 | TATGAGTGTGGAGTCGTTGCTGCAACCCTAGTCTCTGATAAAACANNNAGATCGGA |
| 42483_1 | TATGAGTGTGGAGTCGTTGCGAAGAAGAATCCCTGCCTGGTAANNNAGATCGGAA |
| 42483_2 | TATGAGTGTGGAGTCGTTGCGAAGAAGAATCCCTGCCTGGTAANNNAGATCGGAA |
| 31285_1 | TATGAGTGTGGAGTCGTTGCAGATGAAGTCTCACTCTGTCGCNNNAGATCGGAAGA |
| 31285_2 | TATGAGTGTGGAGTCGTTGCAGATGAAGTCTCACTCTGTCGCNNNAGATCGGAAGA |
| 34047_1 | TATGAGTGTGGAGTCGTTGCCGAGTTCACGCCATTCTCCTNNNAGATCGGAAGAGC |
| 34047_2 | TATGAGTGTGGAGTCGTTGCCGAGTTCACGCCATTCTCCTNNNAGATCGGAAGAGC |
| 29507_1 | TATGAGTGTGGAGTCGTTGCCCCATGCACATTGGTTTTCATGANNNAGATCGGAAG |
| 29507_2 | TATGAGTGTGGAGTCGTTGCCCCATGCACATTGGTTTTCATGANNNAGATCGGAAG |
| 37107_1 | TATGAGTGTGGAGTCGTTGCAGAGTTTTCAATTCACTTGGCCCNNNAGATCGGAAG |
| 37107_2 | TATGAGTGTGGAGTCGTTGCAGAGTTTTCAATTCACTTGGCCCNNNAGATCGGAAG |
| 13173_1 | TATGAGTGTGGAGTCGTTGCCTTCAGACCTTGACTTATCTACNNNAGATCGGAAGA |
| 13173_2 | TATGAGTGTGGAGTCGTTGCCTTCAGACCTTGACTTATCTACNNNAGATCGGAAGA |
| 32076_1 | TATGAGTGTGGAGTCGTTGCCTCCTGGGTTCATGCCATTCTCNNNAGATCGGAAGA |
| 32076_2 | TATGAGTGTGGAGTCGTTGCCTCCTGGGTTCATGCCATTCTCNNNAGATCGGAAGA |
| 52249_1 | TATGAGTGTGGAGTCGTTGCCCACCAGGGTAGCAATGTAATTGNNNAGATCGGAA |
| 52249_2 | TATGAGTGTGGAGTCGTTGCCCACCAGGGTAGCAATGTAATTGNNNAGATCGGAA |
| 13297_1 | TATGAGTGTGGAGTCGTTGCGGCAAGGAAAGAGTATTTAAAAANNNAGATCGGAA |
| 13297_2 | TATGAGTGTGGAGTCGTTGCGGCAAGGAAAGAGTATTTAAAAANNNAGATCGGAA |
| 46957_1 | TATGAGTGTGGAGTCGTTGCACAGAGTCTTGCTTTGTTACCCANNNAGATCGGAAG |
| 46957_2 | TATGAGTGTGGAGTCGTTGCACAGAGTCTTGCTTTGTTACCCANNNAGATCGGAAG |
| 44037_1 | TATGAGTGTGGAGTCGTTGCTGCTAAGCACCGTGTAAAGTACTNNNAGATCGGAAG |

| | |
|---|---|
| 44037_2 | TATGAGTGTGGAGTCGTTGCTGCTAAGCACCGTGTAAAGTACTNNNAGATCGGAAG |
| 29132_1 | TATGAGTGTGGAGTCGTTGCTTTCCATCACTCTCTTCCTCTGCNNNAGATCGGAAGA |
| 29132_2 | TATGAGTGTGGAGTCGTTGCTTTCCATCACTCTCTTCCTCTGCNNNAGATCGGAAGA |
| 36251_1 | TATGAGTGTGGAGTCGTTGCACATGGTACTTCAATGGCAGGAANNNAGATCGGAA |
| 36251_2 | TATGAGTGTGGAGTCGTTGCACATGGTACTTCAATGGCAGGAANNNAGATCGGAA |
| 51432_1 | TATGAGTGTGGAGTCGTTGCGCAAGGTCATTCTGGAGATCTGANNNAGATCGGAA |
| 51432_2 | TATGAGTGTGGAGTCGTTGCGCAAGGTCATTCTGGAGATCTGANNNAGATCGGAA |
| 30459_1 | TATGAGTGTGGAGTCGTTGCCAGTTACAGTCCCACATCATGCANNNAGATCGGAAG |
| 30459_2 | TATGAGTGTGGAGTCGTTGCCAGTTACAGTCCCACATCATGCANNNAGATCGGAAG |
| 52068_1 | TATGAGTGTGGAGTCGTTGCAGCCCAACAGCTCATTGAGAANNNAGATCGGAAGA |
| 52068_2 | TATGAGTGTGGAGTCGTTGCAGCCCAACAGCTCATTGAGAANNNAGATCGGAAGA |
| 47746_1 | TATGAGTGTGGAGTCGTTGCCAAGTGTGAGGTCTAAGGCAGAGNNNAGATCGGAA |
| 47746_2 | TATGAGTGTGGAGTCGTTGCCAAGTGTGAGGTCTAAGGCAGAGNNNAGATCGGAA |
| 34436_1 | TATGAGTGTGGAGTCGTTGCCAATGGGAGAACTGAGCGAGGNNNAGATCGGAAG |
| 34436_2 | TATGAGTGTGGAGTCGTTGCCAATGGGAGAACTGAGCGAGGNNNAGATCGGAAG |
| 48439_1 | TATGAGTGTGGAGTCGTTGCTACCACGCACGGCTAATTTTTTTNNNAGATCGGAAG |
| 48439_2 | TATGAGTGTGGAGTCGTTGCTACCACGCACGGCTAATTTTTTTNNNAGATCGGAAG |
| 46938_1 | TATGAGTGTGGAGTCGTTGCTTTTCGTAGAGATGGGGTTTCACNNNAGATCGGAA |
| 46938_2 | TATGAGTGTGGAGTCGTTGCTTTTCGTAGAGATGGGGTTTCACNNNAGATCGGAA |
| 35891_1 | TATGAGTGTGGAGTCGTTGCGAGACAGAGGGAAGGAAGGAAAGNNNAGATCGG |
| 35891_2 | TATGAGTGTGGAGTCGTTGCGAGACAGAGGGAAGGAAGGAAAGNNNAGATCGG |
| 10009_1 | TATGAGTGTGGAGTCGTTGCTGAGTGTTTACTAGGCACTGTGCNNNAGATCGGAA |
| 10009_2 | TATGAGTGTGGAGTCGTTGCTGAGTGTTTACTAGGCACTGTGCNNNAGATCGGAA |
| 32423_1 | TATGAGTGTGGAGTCGTTGCCAGGTTCACGCCATTCTCCTNNNAGATCGGAAGAGC |
| 32423_2 | TATGAGTGTGGAGTCGTTGCCAGGTTCACGCCATTCTCCTNNNAGATCGGAAGAGC |
| 33585_1 | TATGAGTGTGGAGTCGTTGCCCAAGTAGCTGGGATTACAGGTGNNNAGATCGGAA |
| 33585_2 | TATGAGTGTGGAGTCGTTGCCCAAGTAGCTGGGATTACAGGTGNNNAGATCGGAA |
| 43230_1 | TATGAGTGTGGAGTCGTTGCTTGAGAAAAGGTTGCCTTCTGTCNNNAGATCGGAAG |
| 43230_2 | TATGAGTGTGGAGTCGTTGCTTGAGAAAAGGTTGCCTTCTGTCNNNAGATCGGAAG |
| 2879_1 | TATGAGTGTGGAGTCGTTGCTCGCATACTGGAATCTTGCATGTNNNAGATCGGAAG |
| 2879_2 | TATGAGTGTGGAGTCGTTGCTCGCATACTGGAATCTTGCATGTNNNAGATCGGAAG |
| 43242_1 | TATGAGTGTGGAGTCGTTGCAGCTACTTCTCAATTCTGCCTCCNNNAGATCGGAAGA |
| 43242_2 | TATGAGTGTGGAGTCGTTGCAGCTACTTCTCAATTCTGCCTCCNNNAGATCGGAAGA |
| 29569_1 | TATGAGTGTGGAGTCGTTGCAGAGATTCAGGTGCCTCTCATTGNNNAGATCGGAAG |
| 29569_2 | TATGAGTGTGGAGTCGTTGCAGAGATTCAGGTGCCTCTCATTGNNNAGATCGGAAG |
| 30629_1 | TATGAGTGTGGAGTCGTTGCCTGAGGCTTGGAATCTGCTGAGTNNNAGATCGGAA |
| 30629_2 | TATGAGTGTGGAGTCGTTGCCTGAGGCTTGGAATCTGCTGAGTNNNAGATCGGAA |
| 37540_1 | TATGAGTGTGGAGTCGTTGCAACAAAAAACCTGCCAACCATGTNNNAGATCGGAAG |
| 37540_2 | TATGAGTGTGGAGTCGTTGCAACAAAAAACCTGCCAACCATGTNNNAGATCGGAAG |
| 47872_1 | TATGAGTGTGGAGTCGTTGCCCAAGTAGCTGGGATTACAGGTTNNNAGATCGGAA |
| 47872_2 | TATGAGTGTGGAGTCGTTGCCCAAGTAGCTGGGATTACAGGTTNNNAGATCGGAA |
| 34991_1 | TATGAGTGTGGAGTCGTTGCCTCCCAGGTTCAAGCGATTCTCNNNAGATCGGAAGA |
| 34991_2 | TATGAGTGTGGAGTCGTTGCCTCCCAGGTTCAAGCGATTCTCNNNAGATCGGAAGA |

| | |
|---|---|
| 29861_1 | TATGAGTGTGGAGTCGTTGCTTCTTCCTTCCCACCTCATGTGANNNAGATCGGAAGA |
| 29861_2 | TATGAGTGTGGAGTCGTTGCTTCTTCCTTCCCACCTCATGTGANNNAGATCGGAAGA |
| 40642_1 | TATGAGTGTGGAGTCGTTGCCGCCTGTAATCTTGACACTTTGGNNNAGATCGGAAG |
| 40642_2 | TATGAGTGTGGAGTCGTTGCCGCCTGTAATCTTGACACTTTGGNNNAGATCGGAAG |
| 41040_1 | TATGAGTGTGGAGTCGTTGCAGGTCAGAATTCTGGAAAGGCATNNNAGATCGGAA |
| 41040_2 | TATGAGTGTGGAGTCGTTGCAGGTCAGAATTCTGGAAAGGCATNNNAGATCGGAA |
| 36126_1 | TATGAGTGTGGAGTCGTTGCGTTGTGGATACAGCATTCACACCNNNAGATCGGAAG |
| 36126_2 | TATGAGTGTGGAGTCGTTGCGTTGTGGATACAGCATTCACACCNNNAGATCGGAAG |
| 50137_1 | TATGAGTGTGGAGTCGTTGCTACTACCCAGGTGTCTCATTCCTNNNAGATCGGAAGA |
| 50137_2 | TATGAGTGTGGAGTCGTTGCTACTACCCAGGTGTCTCATTCCTNNNAGATCGGAAGA |
| 49402_1 | TATGAGTGTGGAGTCGTTGCTGTATTTTCAGTCAAGGCAGGGTNNNAGATCGGAAG |
| 49402_2 | TATGAGTGTGGAGTCGTTGCTGTATTTTCAGTCAAGGCAGGGTNNNAGATCGGAAG |
| 42661_1 | TATGAGTGTGGAGTCGTTGCGACATTGAAGCAGTTTCCCTACANNNAGATCGGAAG |
| 42661_2 | TATGAGTGTGGAGTCGTTGCGACATTGAAGCAGTTTCCCTACANNNAGATCGGAAG |
| 48146_1 | TATGAGTGTGGAGTCGTTGCTTGACTCACTACAAACTCCGCTTNNNAGATCGGAAGA |
| 48146_2 | TATGAGTGTGGAGTCGTTGCTTGACTCACTACAAACTCCGCTTNNNAGATCGGAAGA |
| 33833_1 | TATGAGTGTGGAGTCGTTGCGAAATCGGCCCACATGATTATGGNNNAGATCGGAAG |
| 33833_2 | TATGAGTGTGGAGTCGTTGCGAAATCGGCCCACATGATTATGGNNNAGATCGGAAG |
| 53296_1 | TATGAGTGTGGAGTCGTTGCCTTGGATGGGGAGGGTTTTTTCNNNAGATCGGAA |
| 53296_2 | TATGAGTGTGGAGTCGTTGCCTTGGATGGGGAGGGTTTTTTCNNNAGATCGGAA |
| 48218_1 | TATGAGTGTGGAGTCGTTGCGATCTGTCTCCTAGGCTGAAGTGNNNAGATCGGAAG |
| 48218_2 | TATGAGTGTGGAGTCGTTGCGATCTGTCTCCTAGGCTGAAGTGNNNAGATCGGAAG |
| 41561_1 | TATGAGTGTGGAGTCGTTGCCAAGAGATCTTCCCACCTCCATCNNNAGATCGGAAGA |
| 41561_2 | TATGAGTGTGGAGTCGTTGCCAAGAGATCTTCCCACCTCCATCNNNAGATCGGAAGA |
| 47751_1 | TATGAGTGTGGAGTCGTTGCCATGATCTTGGCTCATTGCAACCNNNAGATCGGAAG |
| 47751_2 | TATGAGTGTGGAGTCGTTGCCATGATCTTGGCTCATTGCAACCNNNAGATCGGAAG |
| 37215_1 | TATGAGTGTGGAGTCGTTGCATCGCTTGTCATTGCTTACATTTNNNAGATCGGAAGA |
| 37215_2 | TATGAGTGTGGAGTCGTTGCATCGCTTGTCATTGCTTACATTTNNNAGATCGGAAGA |
| 37186_1 | TATGAGTGTGGAGTCGTTGCATAGGAGCAGAAAGGCTGTGATCNNNAGATCGGAA |
| 37186_2 | TATGAGTGTGGAGTCGTTGCATAGGAGCAGAAAGGCTGTGATCNNNAGATCGGAA |
| 48776_1 | TATGAGTGTGGAGTCGTTGCTTGTATTGTTCCAGATCCAGGGGNNNAGATCGGAAG |
| 48776_2 | TATGAGTGTGGAGTCGTTGCTTGTATTGTTCCAGATCCAGGGGNNNAGATCGGAAG |
| 30401_1 | TATGAGTGTGGAGTCGTTGCAAAAATACAGAATTAGCCGGGCGNNNAGATCGGAA |
| 30401_2 | TATGAGTGTGGAGTCGTTGCAAAAATACAGAATTAGCCGGGCGNNNAGATCGGAA |
| 51997_1 | TATGAGTGTGGAGTCGTTGCCCACGCCCTGCTAATTTTTTGTANNNAGATCGGAAGA |
| 51997_2 | TATGAGTGTGGAGTCGTTGCCCACGCCCTGCTAATTTTTTGTANNNAGATCGGAAGA |
| 49073_1 | TATGAGTGTGGAGTCGTTGCGTCGCTGTCATCTGAACAGAATCNNNAGATCGGAAG |
| 49073_2 | TATGAGTGTGGAGTCGTTGCGTCGCTGTCATCTGAACAGAATCNNNAGATCGGAAG |
| 49583_1 | TATGAGTGTGGAGTCGTTGCCCACCACACCTGGCTAATTTTTTNNNAGATCGGAAGA |
| 49583_2 | TATGAGTGTGGAGTCGTTGCCCACCACACCTGGCTAATTTTTTNNNAGATCGGAAGA |
| 33921_1 | TATGAGTGTGGAGTCGTTGCCAAGTAGCTGGGACTACAGGCNNNAGATCGGAAGA |
| 33921_2 | TATGAGTGTGGAGTCGTTGCCAAGTAGCTGGGACTACAGGCNNNAGATCGGAAGA |
| 36614_1 | TATGAGTGTGGAGTCGTTGCTCTTCTGGGCTGAATCAGGATTCNNNAGATCGGAAG |

| | |
|---|---|
| 36614_2 | TATGAGTGTGGAGTCGTTGCTCTTCTGGGCTGAATCAGGATTCNNNAGATCGGAAG |
| 45930_1 | TATGAGTGTGGAGTCGTTGCTTCAAGTGATTCTCCTGCCTCAGNNNAGATCGGAAG |
| 45930_2 | TATGAGTGTGGAGTCGTTGCTTCAAGTGATTCTCCTGCCTCAGNNNAGATCGGAAG |
| 47768_1 | TATGAGTGTGGAGTCGTTGCGTCTTGCTTTATCACACAGGCTGNNNAGATCGGAAG |
| 47768_2 | TATGAGTGTGGAGTCGTTGCGTCTTGCTTTATCACACAGGCTGNNNAGATCGGAAG |
| 45377_1 | TATGAGTGTGGAGTCGTTGCGACTTGGGCATTCATGAAGCTTTNNNAGATCGGAAG |
| 45377_2 | TATGAGTGTGGAGTCGTTGCGACTTGGGCATTCATGAAGCTTTNNNAGATCGGAAG |
| 8517_1 | TATGAGTGTGGAGTCGTTGCTTCCCCTCAGGATGTGTTTGTATAGNNNAGATCGGA |
| 8517_2 | TATGAGTGTGGAGTCGTTGCTTCCCCTCAGGATGTGTTTGTATAGNNNAGATCGGA |
| 3182_1 | TATGAGTGTGGAGTCGTTGCCAAGGCTGCCCTCTAGTTGAATTNNNAGATCGGAAG |
| 3182_2 | TATGAGTGTGGAGTCGTTGCCAAGGCTGCCCTCTAGTTGAATTNNNAGATCGGAAG |
| 2405_1 | TATGAGTGTGGAGTCGTTGCAAGAGAAATCAAGGTCCACCTGCNNNAGATCGGAA |
| 2405_2 | TATGAGTGTGGAGTCGTTGCAAGAGAAATCAAGGTCCACCTGCNNNAGATCGGAA |
| 52344_1 | TATGAGTGTGGAGTCGTTGCCAATACAGGGGGTTGGTTCAGTANNNAGATCGGAA |
| 52344_2 | TATGAGTGTGGAGTCGTTGCCAATACAGGGGGTTGGTTCAGTANNNAGATCGGAA |
| 43275_1 | TATGAGTGTGGAGTCGTTGCCTCGAGTAGCTGGGATTATAGGCNNNAGATCGGAA |
| 43275_2 | TATGAGTGTGGAGTCGTTGCCTCGAGTAGCTGGGATTATAGGCNNNAGATCGGAA |
| 36130_1 | TATGAGTGTGGAGTCGTTGCAATTTCACTCCTGGGGAATGTCANNNAGATCGGAAG |
| 36130_2 | TATGAGTGTGGAGTCGTTGCAATTTCACTCCTGGGGAATGTCANNNAGATCGGAAG |
| 10144_1 | TATGAGTGTGGAGTCGTTGCCAGACAGGAGTGAGTGCTACTTCNNNAGATCGGAA |
| 10144_2 | TATGAGTGTGGAGTCGTTGCCAGACAGGAGTGAGTGCTACTTCNNNAGATCGGAA |
| 49102_1 | TATGAGTGTGGAGTCGTTGCTTCCCTAGCCCTCAAATATTGGGNNNAGATCGGAAG |
| 49102_2 | TATGAGTGTGGAGTCGTTGCTTCCCTAGCCCTCAAATATTGGGNNNAGATCGGAAG |
| 52238_1 | TATGAGTGTGGAGTCGTTGCAGGATGTGTTTTGTGCAAGATCGNNNAGATCGGAA |
| 52238_2 | TATGAGTGTGGAGTCGTTGCAGGATGTGTTTTGTGCAAGATCGNNNAGATCGGAA |
| 29489_1 | TATGAGTGTGGAGTCGTTGCTGCCTTTAACTGTCAAACTGTCAAAANNNAGATCGG |
| 29489_2 | TATGAGTGTGGAGTCGTTGCTGCCTTTAACTGTCAAACTGTCAAAANNNAGATCGG |
| 47764_1 | TATGAGTGTGGAGTCGTTGCAGATCTCAGCTAACTGCAACCTCNNNAGATCGGAAG |
| 47764_2 | TATGAGTGTGGAGTCGTTGCAGATCTCAGCTAACTGCAACCTCNNNAGATCGGAAG |
| 50211_1 | TATGAGTGTGGAGTCGTTGCGGTTGGTACTCATGTCAACACAGNNNAGATCGGAA |
| 50211_2 | TATGAGTGTGGAGTCGTTGCGGTTGGTACTCATGTCAACACAGNNNAGATCGGAA |
| 28921_1 | TATGAGTGTGGAGTCGTTGCCAGAATGGACTGGGTTTTCCCTANNNAGATCGGAAG |
| 28921_2 | TATGAGTGTGGAGTCGTTGCCAGAATGGACTGGGTTTTCCCTANNNAGATCGGAAG |
| 13541_1 | TATGAGTGTGGAGTCGTTGCATGCAAATACATATTCGAAGTTTNNNAGATCGGAAG |
| 13541_2 | TATGAGTGTGGAGTCGTTGCATGCAAATACATATTCGAAGTTTNNNAGATCGGAAG |
| 47069_1 | TATGAGTGTGGAGTCGTTGCGAGTAGCTGGGATTATAGGTGCCNNNAGATCGGAA |
| 47069_2 | TATGAGTGTGGAGTCGTTGCGAGTAGCTGGGATTATAGGTGCCNNNAGATCGGAA |
| 35184_1 | TATGAGTGTGGAGTCGTTGCCTCCTGAGAAGCTGGGATTACAANNNAGATCGGAA |
| 35184_2 | TATGAGTGTGGAGTCGTTGCCTCCTGAGAAGCTGGGATTACAANNNAGATCGGAA |
| 44979_1 | TATGAGTGTGGAGTCGTTGCCAGACTCTTGAGCCAGGTAAGGNNNAGATCGGAAG |
| 44979_2 | TATGAGTGTGGAGTCGTTGCCAGACTCTTGAGCCAGGTAAGGNNNAGATCGGAAG |
| 39789_1 | TATGAGTGTGGAGTCGTTGCATAATTGTTTGAACCTGGCAGGCNNNAGATCGGAA |
| 39789_2 | TATGAGTGTGGAGTCGTTGCATAATTGTTTGAACCTGGCAGGCNNNAGATCGGAA |

| | |
|---|---|
| 51155_1 | TATGAGTGTGGAGTCGTTGCAGTGATATAGATGTGGAGCGCAGNNNAGATCGGAA |
| 51155_2 | TATGAGTGTGGAGTCGTTGCAGTGATATAGATGTGGAGCGCAGNNNAGATCGGAA |
| 31349_1 | TATGAGTGTGGAGTCGTTGCTGATTCCTGTGGCTTGATTCAGANNNAGATCGGAAG |
| 31349_2 | TATGAGTGTGGAGTCGTTGCTGATTCCTGTGGCTTGATTCAGANNNAGATCGGAAG |
| 28929_1 | TATGAGTGTGGAGTCGTTGCTGAAGGAAGTGGAAATAGGTGGGNNNAGATCGGA |
| 28929_2 | TATGAGTGTGGAGTCGTTGCTGAAGGAAGTGGAAATAGGTGGGNNNAGATCGGA |
| 51981_1 | TATGAGTGTGGAGTCGTTGCGACTCTGTCTTCTGTCTGTTGGTNNNAGATCGGAAG |
| 51981_2 | TATGAGTGTGGAGTCGTTGCGACTCTGTCTTCTGTCTGTTGGTNNNAGATCGGAAG |
| 31167_1 | TATGAGTGTGGAGTCGTTGCTGTGTGTATGTTTATCCCACAGGANNNAGATCGGAA |
| 31167_2 | TATGAGTGTGGAGTCGTTGCTGTGTGTATGTTTATCCCACAGGANNNAGATCGGAA |
| 13727_1 | TATGAGTGTGGAGTCGTTGCATGATAAAAGTGGCCTTGTTANNNAGATCGGAAGA |
| 13727_2 | TATGAGTGTGGAGTCGTTGCATGATAAAAGTGGCCTTGTTANNNAGATCGGAAGA |
| 30685_1 | ATGAGTGTGGAGTCGTTGCGCTGTTGACTGTTTATATATTCTGGATNNNAGATCGG |
| 30685_2 | ATGAGTGTGGAGTCGTTGCGCTGTTGACTGTTTATATATTCTGGATNNNAGATCGG |
| 46187_1 | TATGAGTGTGGAGTCGTTGCAGACAGGGTCTCCTTATGTTGCNNNAGATCGGAAGA |
| 46187_2 | TATGAGTGTGGAGTCGTTGCAGACAGGGTCTCCTTATGTTGCNNNAGATCGGAAGA |
| 31307_1 | TATGAGTGTGGAGTCGTTGCCCCAAGTTTATCCCAGTTTGTGGNNNAGATCGGAAG |
| 31307_2 | TATGAGTGTGGAGTCGTTGCCCCAAGTTTATCCCAGTTTGTGGNNNAGATCGGAAG |
| 29825_1 | TATGAGTGTGGAGTCGTTGCATGGTTCTCCAAAATGGGACAGTNNNAGATCGGAA |
| 29825_2 | TATGAGTGTGGAGTCGTTGCATGGTTCTCCAAAATGGGACAGTNNNAGATCGGAA |
| 9850_1 | TATGAGTGTGGAGTCGTTGCGTGGCAGCAGTGGATATTGATGANNNAGATCGGAA |
| 9850_2 | TATGAGTGTGGAGTCGTTGCGTGGCAGCAGTGGATATTGATGANNNAGATCGGAA |
| 7305_1 | TATGAGTGTGGAGTCGTTGCCTGGTTGTGACAGTGTCCTGTAGNNNAGATCGGAA |
| 7305_2 | TATGAGTGTGGAGTCGTTGCCTGGTTGTGACAGTGTCCTGTAGNNNAGATCGGAA |
| 30355_1 | TATGAGTGTGGAGTCGTTGCTCCTTTCTGGACCTTCTTCCTCCNNNAGATCGGAAGA |
| 30355_2 | TATGAGTGTGGAGTCGTTGCTCCTTTCTGGACCTTCTTCCTCCNNNAGATCGGAAGA |
| 33217_1 | TATGAGTGTGGAGTCGTTGCAGACCTTACACCCCTCACAAAAANNNAGATCGGAAG |
| 33217_2 | TATGAGTGTGGAGTCGTTGCAGACCTTACACCCCTCACAAAAANNNAGATCGGAAG |
| 39189_1 | TATGAGTGTGGAGTCGTTGCACTGTGGGCAATTGTAACACAACNNNAGATCGGAAG |
| 39189_2 | TATGAGTGTGGAGTCGTTGCACTGTGGGCAATTGTAACACAACNNNAGATCGGAAG |
| 44582_1 | TATGAGTGTGGAGTCGTTGCCACACACCTACCATGTACCCATANNNAGATCGGAAGA |
| 44582_2 | TATGAGTGTGGAGTCGTTGCCACACACCTACCATGTACCCATANNNAGATCGGAAGA |
| 30146_1 | TATGAGTGTGGAGTCGTTGCGGGCCGATGAAAATCACATTGAANNNAGATCGGAA |
| 30146_2 | TATGAGTGTGGAGTCGTTGCGGGCCGATGAAAATCACATTGAANNNAGATCGGAA |
| 41261_1 | TATGAGTGTGGAGTCGTTGCGACAATGGAAGTGTCTCATCCCTNNNAGATCGGAAG |
| 41261_2 | TATGAGTGTGGAGTCGTTGCGACAATGGAAGTGTCTCATCCCTNNNAGATCGGAAG |
| 49134_1 | TATGAGTGTGGAGTCGTTGCCCAAGTAGCTGGGATTACAGGTGNNNAGATCGGAA |
| 49134_2 | TATGAGTGTGGAGTCGTTGCCCAAGTAGCTGGGATTACAGGTGNNNAGATCGGAA |
| 48160_1 | TATGAGTGTGGAGTCGTTGCGGCGGGATCTAAGTAGGTAGAGANNNAGATCGGA |
| 48160_2 | TATGAGTGTGGAGTCGTTGCGGCGGGATCTAAGTAGGTAGAGANNNAGATCGGA |
| 10232_1 | TATGAGTGTGGAGTCGTTGCTTATTTTGATCACCGTGGGGTGTNNNAGATCGGAAG |
| 10232_2 | TATGAGTGTGGAGTCGTTGCTTATTTTGATCACCGTGGGGTGTNNNAGATCGGAAG |
| 40333_1 | TATGAGTGTGGAGTCGTTGCAAACAAACAAAAAACTGGTGGGGNNNAGATCGGAA |

| | |
|---|---|
| 40333_2 | TATGAGTGTGGAGTCGTTGCAAACAAACAAAAAACTGGTGGGGNNNAGATCGGAA |
| 30344_1 | TATGAGTGTGGAGTCGTTGCAGTTTCTTTATGAAAGCGTATTGTAGCNNNAGATCG |
| 30344_2 | TATGAGTGTGGAGTCGTTGCAGTTTCTTTATGAAAGCGTATTGTAGCNNNAGATCG |
| 43143_1 | TATGAGTGTGGAGTCGTTGCCCAAGCTTGAAGTGCAGTTTTCTNNNAGATCGGAAG |
| 43143_2 | TATGAGTGTGGAGTCGTTGCCCAAGCTTGAAGTGCAGTTTTCTNNNAGATCGGAAG |
| 53352_1 | TATGAGTGTGGAGTCGTTGCTCTTTCTCCCCCGTCTATCTCTTNNNAGATCGGAAGA |
| 53352_2 | TATGAGTGTGGAGTCGTTGCTCTTTCTCCCCCGTCTATCTCTTNNNAGATCGGAAGA |
| 34199_1 | TATGAGTGTGGAGTCGTTGCGTTCCTTTTCTCAAGGCACTGTCNNNAGATCGGAAG |
| 34199_2 | TATGAGTGTGGAGTCGTTGCGTTCCTTTTCTCAAGGCACTGTCNNNAGATCGGAAG |
| 47468_1 | TATGAGTGTGGAGTCGTTGCCCACGCCCAGCTAATTTTTGTATNNNAGATCGGAAG |
| 47468_2 | TATGAGTGTGGAGTCGTTGCCCACGCCCAGCTAATTTTTGTATNNNAGATCGGAAG |
| 40478_1 | TATGAGTGTGGAGTCGTTGCCTATGTGCCCTATCATGCCAGATNNNAGATCGGAAG |
| 40478_2 | TATGAGTGTGGAGTCGTTGCCTATGTGCCCTATCATGCCAGATNNNAGATCGGAAG |
| 45709_1 | TATGAGTGTGGAGTCGTTGCCGAGTAGCTGGGACTACAGGTNNNAGATCGGAAGA |
| 45709_2 | TATGAGTGTGGAGTCGTTGCCGAGTAGCTGGGACTACAGGTNNNAGATCGGAAGA |
| 12972_1 | TATGAGTGTGGAGTCGTTGCCATCAGGTGACCATTAAGTTANNNAGATCGGAAGA |
| 12972_2 | TATGAGTGTGGAGTCGTTGCCATCAGGTGACCATTAAGTTANNNAGATCGGAAGA |
| 38722_1 | TATGAGTGTGGAGTCGTTGCCCACATACCCAACAGCTTCCTTANNNAGATCGGAAGA |
| 38722_2 | TATGAGTGTGGAGTCGTTGCCCACATACCCAACAGCTTCCTTANNNAGATCGGAAGA |
| 35690_1 | TATGAGTGTGGAGTCGTTGCATGCTAAGTAGGGCACTCTGTTCNNNAGATCGGAAG |
| 35690_2 | TATGAGTGTGGAGTCGTTGCATGCTAAGTAGGGCACTCTGTTCNNNAGATCGGAAG |
| 34982_1 | TATGAGTGTGGAGTCGTTGCTTCAAGTGATTCTCCTGCCTCAGNNNAGATCGGAAG |
| 34982_2 | TATGAGTGTGGAGTCGTTGCTTCAAGTGATTCTCCTGCCTCAGNNNAGATCGGAAG |
| 13249_1 | TATGAGTGTGGAGTCGTTGCAAATCGAAGATTTCTATACCACTNNNAGATCGGAAG |
| 13249_2 | TATGAGTGTGGAGTCGTTGCAAATCGAAGATTTCTATACCACTNNNAGATCGGAAG |
| 45564_1 | TATGAGTGTGGAGTCGTTGCCCCCCTAGGAAGTGGATTTGGNNNAGATCGGAAGA |
| 45564_2 | TATGAGTGTGGAGTCGTTGCCCCCCTAGGAAGTGGATTTGGNNNAGATCGGAAGA |
| 51358_1 | TATGAGTGTGGAGTCGTTGCGAAAAAGCAAGGCAAGGTCATCTNNNAGATCGGAA |
| 51358_2 | TATGAGTGTGGAGTCGTTGCGAAAAAGCAAGGCAAGGTCATCTNNNAGATCGGAA |
| 40842_1 | TATGAGTGTGGAGTCGTTGCGAACCAGCATAATCCCATAGCCTNNNAGATCGGAAG |
| 40842_2 | TATGAGTGTGGAGTCGTTGCGAACCAGCATAATCCCATAGCCTNNNAGATCGGAAG |
| 37308_1 | TATGAGTGTGGAGTCGTTGCAGGATAAATGAATATGGGACCACANNNAGATCGGA |
| 37308_2 | TATGAGTGTGGAGTCGTTGCAGGATAAATGAATATGGGACCACANNNAGATCGGA |
| 37382_1 | TATGAGTGTGGAGTCGTTGCTCCTAGCTACTCTGGAGGATGAGNNNAGATCGGAA |
| 37382_2 | TATGAGTGTGGAGTCGTTGCTCCTAGCTACTCTGGAGGATGAGNNNAGATCGGAA |
| 37317_1 | TATGAGTGTGGAGTCGTTGCCTTGGGGTATCAGTACTGCACTTNNNAGATCGGAAG |
| 37317_2 | TATGAGTGTGGAGTCGTTGCCTTGGGGTATCAGTACTGCACTTNNNAGATCGGAAG |
| 49409_1 | TATGAGTGTGGAGTCGTTGCTATGACACGGAACGTTTACCCTTNNNAGATCGGAAG |
| 49409_2 | TATGAGTGTGGAGTCGTTGCTATGACACGGAACGTTTACCCTTNNNAGATCGGAAG |
| 51394_1 | TATGAGTGTGGAGTCGTTGCAAAGCAATTGAATCTCCCTGCTGNNNAGATCGGAAG |
| 51394_2 | TATGAGTGTGGAGTCGTTGCAAAGCAATTGAATCTCCCTGCTGNNNAGATCGGAAG |
| 44451_1 | TATGAGTGTGGAGTCGTTGCCCTGGCACAAGAGATCTAGATCCNNNAGATCGGAAG |
| 44451_2 | TATGAGTGTGGAGTCGTTGCCCTGGCACAAGAGATCTAGATCCNNNAGATCGGAAG |

| | |
|---|---|
| 48366_1 | TATGAGTGTGGAGTCGTTGCAATCTCGGCTCACTACAACCTTTNNNAGATCGGAAG |
| 48366_2 | TATGAGTGTGGAGTCGTTGCAATCTCGGCTCACTACAACCTTTNNNAGATCGGAAG |
| 35140_1 | TATGAGTGTGGAGTCGTTGCTCTGGTTCAGTCTTGGAAGGATGNNNAGATCGGAA |
| 35140_2 | TATGAGTGTGGAGTCGTTGCTCTGGTTCAGTCTTGGAAGGATGNNNAGATCGGAA |
| 12543_1 | TATGAGTGTGGAGTCGTTGCTTCAGGAGTTTTTTATGTTCTTNNNAGATCGGAAG |
| 12543_2 | TATGAGTGTGGAGTCGTTGCTTCAGGAGTTTTTTATGTTCTTNNNAGATCGGAAG |
| 49241_1 | TATGAGTGTGGAGTCGTTGCCCGAGTAGCTGGAATTACAGGTTNNNAGATCGGAA |
| 49241_2 | TATGAGTGTGGAGTCGTTGCCCGAGTAGCTGGAATTACAGGTTNNNAGATCGGAA |
| 47820_1 | TATGAGTGTGGAGTCGTTGCCCACCAAGCCCACCTAATTTTTTNNNAGATCGGAAGA |
| 47820_2 | TATGAGTGTGGAGTCGTTGCCCACCAAGCCCACCTAATTTTTTNNNAGATCGGAAGA |
| 48544_1 | TATGAGTGTGGAGTCGTTGCCCCAGGTTCAAGCAATTCTCTTGNNNAGATCGGAAG |
| 48544_2 | TATGAGTGTGGAGTCGTTGCCCCAGGTTCAAGCAATTCTCTTGNNNAGATCGGAAG |
| 48836_1 | TATGAGTGTGGAGTCGTTGCCATGTTGTGAGTGAGTGTGCATTNNNAGATCGGAA |
| 48836_2 | TATGAGTGTGGAGTCGTTGCCATGTTGTGAGTGAGTGTGCATTNNNAGATCGGAA |
| 31553_1 | TATGAGTGTGGAGTCGTTGCTCTGCTCCCTAAGAGTATGAGGTNNNAGATCGGAAG |
| 31553_2 | TATGAGTGTGGAGTCGTTGCTCTGCTCCCTAAGAGTATGAGGTNNNAGATCGGAAG |
| 39863_1 | TATGAGTGTGGAGTCGTTGCATACTGTGGTCCTTTTTGCCTCTNNNAGATCGGAAG |
| 39863_2 | TATGAGTGTGGAGTCGTTGCATACTGTGGTCCTTTTTGCCTCTNNNAGATCGGAAG |
| 3013_1 | TATGAGTGTGGAGTCGTTGCATCGAACTGTTCTCCCTCTTCCTNNNAGATCGGAAGA |
| 3013_2 | TATGAGTGTGGAGTCGTTGCATCGAACTGTTCTCCCTCTTCCTNNNAGATCGGAAGA |
| 10225_1 | TATGAGTGTGGAGTCGTTGCATGGAAGCCCTAGTGACCTCTTTNNNAGATCGGAAG |
| 10225_2 | TATGAGTGTGGAGTCGTTGCATGGAAGCCCTAGTGACCTCTTTNNNAGATCGGAAG |
| 13150_1 | TATGAGTGTGGAGTCGTTGCTCTTTAATGATTTGGTCATACTGNNNAGATCGGAAG |
| 13150_2 | TATGAGTGTGGAGTCGTTGCTCTTTAATGATTTGGTCATACTGNNNAGATCGGAAG |
| 46845_1 | TATGAGTGTGGAGTCGTTGCATTACAGGCAAGAACCACTTTGCNNNAGATCGGAAG |
| 46845_2 | TATGAGTGTGGAGTCGTTGCATTACAGGCAAGAACCACTTTGCNNNAGATCGGAAG |
| 41963_1 | TATGAGTGTGGAGTCGTTGCAGCTCTGTCTCTAGGGGAAGAATNNNAGATCGGAA |
| 41963_2 | TATGAGTGTGGAGTCGTTGCAGCTCTGTCTCTAGGGGAAGAATNNNAGATCGGAA |
| 37883_1 | TATGAGTGTGGAGTCGTTGCAGGCAGAAGACTTGAATAGGTTCTNNNAGATCGGA |
| 37883_2 | TATGAGTGTGGAGTCGTTGCAGGCAGAAGACTTGAATAGGTTCTNNNAGATCGGA |
| 10621_1 | TATGAGTGTGGAGTCGTTGCGCGAAGGCTTCCTCTGTGTANNNAGATCGGAAGAG |
| 10621_2 | TATGAGTGTGGAGTCGTTGCGCGAAGGCTTCCTCTGTGTANNNAGATCGGAAGAG |
| 32225_1 | TATGAGTGTGGAGTCGTTGCGGCCCAAATTTCACCCATTCAATNNNAGATCGGAAG |
| 32225_2 | TATGAGTGTGGAGTCGTTGCGGCCCAAATTTCACCCATTCAATNNNAGATCGGAAG |
| 32864_1 | TATGAGTGTGGAGTCGTTGCTTCAAGCAATTCTCCTGCCTCAGNNNAGATCGGAAG |
| 32864_2 | TATGAGTGTGGAGTCGTTGCTTCAAGCAATTCTCCTGCCTCAGNNNAGATCGGAAG |
| 52726_1 | TATGAGTGTGGAGTCGTTGCACAAGCTGAGTATGTGCTCTCTCNNNAGATCGGAAG |
| 52726_2 | TATGAGTGTGGAGTCGTTGCACAAGCTGAGTATGTGCTCTCTCNNNAGATCGGAAG |
| 29216_1 | TATGAGTGTGGAGTCGTTGCTGCGTGCATATCTCTGTATCCTCNNNAGATCGGAAG |
| 29216_2 | TATGAGTGTGGAGTCGTTGCTGCGTGCATATCTCTGTATCCTCNNNAGATCGGAAG |
| 31498_1 | TATGAGTGTGGAGTCGTTGCGGGGCAATTTAGTCAGAAGTTGTNNNAGATCGGAA |
| 31498_2 | TATGAGTGTGGAGTCGTTGCGGGGCAATTTAGTCAGAAGTTGTNNNAGATCGGAA |
| 30633_1 | ATGAGTGTGGAGTCGTTGCTTGTTACATCACTGGAAGACAATATAANNNAGATCGG |

| | |
|---|---|
| 30633_2 | ATGAGTGTGGAGTCGTTGCTTGTTACATCACTGGAAGACAATATAANNNAGATCGG |
| 29386_1 | TATGAGTGTGGAGTCGTTGCGTGTCATCCCCAGAACACATGTANNNAGATCGGAAG |
| 29386_2 | TATGAGTGTGGAGTCGTTGCGTGTCATCCCCAGAACACATGTANNNAGATCGGAAG |
| 44021_1 | TATGAGTGTGGAGTCGTTGCATAAAAAGTGACTCTCCCCCACCNNNAGATCGGAAG |
| 44021_2 | TATGAGTGTGGAGTCGTTGCATAAAAAGTGACTCTCCCCCACCNNNAGATCGGAAG |
| 33648_1 | TATGAGTGTGGAGTCGTTGCCTCCCAGGTTCAAGCGATTCTCNNNAGATCGGAAGA |
| 33648_2 | TATGAGTGTGGAGTCGTTGCCTCCCAGGTTCAAGCGATTCTCNNNAGATCGGAAGA |
| 48422_1 | TATGAGTGTGGAGTCGTTGCTTTTAGCAGAGACAGGGTTTCACNNNAGATCGGAA |
| 48422_2 | TATGAGTGTGGAGTCGTTGCTTTTAGCAGAGACAGGGTTTCACNNNAGATCGGAA |
| 5794_1 | TATGAGTGTGGAGTCGTTGCAATACACACACTGCAATGGCAACNNNAGATCGGAAG |
| 5794_2 | TATGAGTGTGGAGTCGTTGCAATACACACACTGCAATGGCAACNNNAGATCGGAAG |
| 13550_1 | TATGAGTGTGGAGTCGTTGCTTTAATAAATGAAATTGCTGGGANNNAGATCGGAA |
| 13550_2 | TATGAGTGTGGAGTCGTTGCTTTAATAAATGAAATTGCTGGGANNNAGATCGGAA |
| 31865_1 | TATGAGTGTGGAGTCGTTGCGCAGTCTATAAGTGGCTGAAGGANNNAGATCGGAA |
| 31865_2 | TATGAGTGTGGAGTCGTTGCGCAGTCTATAAGTGGCTGAAGGANNNAGATCGGAA |
| 48291_1 | TATGAGTGTGGAGTCGTTGCAAGTGTTGGGATTACAGGCTTGANNNAGATCGGAA |
| 48291_2 | TATGAGTGTGGAGTCGTTGCAAGTGTTGGGATTACAGGCTTGANNNAGATCGGAA |
| 42429_1 | TATGAGTGTGGAGTCGTTGCGGAAAACAGCTTTTGCAGGTGTANNNAGATCGGAA |
| 42429_2 | TATGAGTGTGGAGTCGTTGCGGAAAACAGCTTTTGCAGGTGTANNNAGATCGGAA |
| 52812_1 | TATGAGTGTGGAGTCGTTGCCCTGCCCAGACATTTGAAACAAANNNAGATCGGAAG |
| 52812_2 | TATGAGTGTGGAGTCGTTGCCCTGCCCAGACATTTGAAACAAANNNAGATCGGAAG |
| 36366_1 | TATGAGTGTGGAGTCGTTGCAAACGAGAAAGTGGAATGTTTTCTNNNAGATCGGA |
| 36366_2 | TATGAGTGTGGAGTCGTTGCAAACGAGAAAGTGGAATGTTTTCTNNNAGATCGGA |
| 31899_1 | TATGAGTGTGGAGTCGTTGCCAATTCCTGGGCTCAAGCAATCNNNAGATCGGAAGA |
| 31899_2 | TATGAGTGTGGAGTCGTTGCCAATTCCTGGGCTCAAGCAATCNNNAGATCGGAAGA |
| 29801_1 | TATGAGTGTGGAGTCGTTGCCCACTCCTCCATGTTCCTGTTATNNNAGATCGGAAGA |
| 29801_2 | TATGAGTGTGGAGTCGTTGCCCACTCCTCCATGTTCCTGTTATNNNAGATCGGAAGA |
| 40411_1 | TATGAGTGTGGAGTCGTTGCAGCCTCAGGACAAATTGTACCTTNNNAGATCGGAAG |
| 40411_2 | TATGAGTGTGGAGTCGTTGCAGCCTCAGGACAAATTGTACCTTNNNAGATCGGAAG |
| 39163_1 | TATGAGTGTGGAGTCGTTGCGCCCACAACCCATAACTTTCTTTNNNAGATCGGAAGA |
| 39163_2 | TATGAGTGTGGAGTCGTTGCGCCCACAACCCATAACTTTCTTTNNNAGATCGGAAGA |
| 37229_1 | TATGAGTGTGGAGTCGTTGCCTAAGTGATTCTGCCCTCATCCANNNAGATCGGAAG |
| 37229_2 | TATGAGTGTGGAGTCGTTGCCTAAGTGATTCTGCCCTCATCCANNNAGATCGGAAG |
| 50145_1 | TATGAGTGTGGAGTCGTTGCGCTAAGCTGTTACAGAAATCCCCNNNAGATCGGAAG |
| 50145_2 | TATGAGTGTGGAGTCGTTGCGCTAAGCTGTTACAGAAATCCCCNNNAGATCGGAAG |
| 53208_1 | TATGAGTGTGGAGTCGTTGCGAGAGGGAGAGAGAGAGACCAATNNNAGATCGGA |
| 53208_2 | TATGAGTGTGGAGTCGTTGCGAGAGGGAGAGAGAGAGACCAATNNNAGATCGGA |
| 12722_1 | TATGAGTGTGGAGTCGTTGCTGGTTTCAGAACAGTTTAAAAATNNNAGATCGGAAG |
| 12722_2 | TATGAGTGTGGAGTCGTTGCTGGTTTCAGAACAGTTTAAAAATNNNAGATCGGAAG |
| 44492_1 | TATGAGTGTGGAGTCGTTGCAATCTCGACTCACTGCAACTTCTNNNAGATCGGAAG |
| 44492_2 | TATGAGTGTGGAGTCGTTGCAATCTCGACTCACTGCAACTTCTNNNAGATCGGAAG |
| 43106_1 | TATGAGTGTGGAGTCGTTGCCAGTGCACACTTTGAAATGGGTANNNAGATCGGAA |
| 43106_2 | TATGAGTGTGGAGTCGTTGCCAGTGCACACTTTGAAATGGGTANNNAGATCGGAA |

| | |
|---|---|
| 8093_1 | TATGAGTGTGGAGTCGTTGCCCTTACCCAGCCTAGCATTGGTTNNNAGATCGGAAG |
| 8093_2 | TATGAGTGTGGAGTCGTTGCCCTTACCCAGCCTAGCATTGGTTNNNAGATCGGAAG |
| 40886_1 | TATGAGTGTGGAGTCGTTGCCAGATGGAAAAAAGGCTGGATGGNNNAGATCGGA |
| 40886_2 | TATGAGTGTGGAGTCGTTGCCAGATGGAAAAAAGGCTGGATGGNNNAGATCGGA |
| 31105_1 | TATGAGTGTGGAGTCGTTGCTGGGTAAACAGAGTGAGAAGACGNNNAGATCGGA |
| 31105_2 | TATGAGTGTGGAGTCGTTGCTGGGTAAACAGAGTGAGAAGACGNNNAGATCGGA |
| 38548_1 | TATGAGTGTGGAGTCGTTGCGACAGAGCAAGATCCTGTGTCTANNNAGATCGGAA |
| 38548_2 | TATGAGTGTGGAGTCGTTGCGACAGAGCAAGATCCTGTGTCTANNNAGATCGGAA |
| 46271_1 | TATGAGTGTGGAGTCGTTGCGTTCATACCATTCTCCTGCCTCANNNAGATCGGAAGA |
| 46271_2 | TATGAGTGTGGAGTCGTTGCGTTCATACCATTCTCCTGCCTCANNNAGATCGGAAGA |
| 33922_1 | TATGAGTGTGGAGTCGTTGCCACTGTTCACAGGTGGGATCATANNNAGATCGGAA |
| 33922_2 | TATGAGTGTGGAGTCGTTGCCACTGTTCACAGGTGGGATCATANNNAGATCGGAA |
| 41306_1 | TATGAGTGTGGAGTCGTTGCCGGTTCTCTCACCCCTTTGTATANNNAGATCGGAAGA |
| 41306_2 | TATGAGTGTGGAGTCGTTGCCGGTTCTCTCACCCCTTTGTATANNNAGATCGGAAGA |
| 34274_1 | TATGAGTGTGGAGTCGTTGCGGAGGAGGGATAGCATTAGGAGANNNAGATCGGA |
| 34274_2 | TATGAGTGTGGAGTCGTTGCGGAGGAGGGATAGCATTAGGAGANNNAGATCGGA |
| 38936_1 | TATGAGTGTGGAGTCGTTGCAATTGCTTGAAATTGGGAGGTGGNNNAGATCGGAA |
| 38936_2 | TATGAGTGTGGAGTCGTTGCAATTGCTTGAAATTGGGAGGTGGNNNAGATCGGAA |
| 41148_1 | TATGAGTGTGGAGTCGTTGCCCTCCTCCTTCAGCTTATCGATCNNNAGATCGGAAGA |
| 41148_2 | TATGAGTGTGGAGTCGTTGCCCTCCTCCTTCAGCTTATCGATCNNNAGATCGGAAGA |
| 41892_1 | TATGAGTGTGGAGTCGTTGCGTGATCCTTCTGTATCAGCCTCCNNNAGATCGGAAG |
| 41892_2 | TATGAGTGTGGAGTCGTTGCGTGATCCTTCTGTATCAGCCTCCNNNAGATCGGAAG |
| 33815_1 | TATGAGTGTGGAGTCGTTGCAGATCTAGGATGGGCCCAAGTATNNNAGATCGGAA |
| 33815_2 | TATGAGTGTGGAGTCGTTGCAGATCTAGGATGGGCCCAAGTATNNNAGATCGGAA |
| 46842_1 | TATGAGTGTGGAGTCGTTGCCAGGCTAGAGTGCAATGGTATGANNNAGATCGGAA |
| 46842_2 | TATGAGTGTGGAGTCGTTGCCAGGCTAGAGTGCAATGGTATGANNNAGATCGGAA |
| 51286_1 | TATGAGTGTGGAGTCGTTGCTCTTACATATGGGGAAACAGGCCNNNAGATCGGAA |
| 51286_2 | TATGAGTGTGGAGTCGTTGCTCTTACATATGGGGAAACAGGCCNNNAGATCGGAA |
| 37600_1 | TATGAGTGTGGAGTCGTTGCTGTTGAGGACTTAGTATTTGCCTNNNAGATCGGAAG |
| 37600_2 | TATGAGTGTGGAGTCGTTGCTGTTGAGGACTTAGTATTTGCCTNNNAGATCGGAAG |
| 2502_1 | TATGAGTGTGGAGTCGTTGCAGATTGCTTCTTGAGCTTGGGAGNNNAGATCGGAA |
| 2502_2 | TATGAGTGTGGAGTCGTTGCAGATTGCTTCTTGAGCTTGGGAGNNNAGATCGGAA |
| 10527_1 | TATGAGTGTGGAGTCGTTGCTGTGGTTTATGGGATCAGAGTTTNNNAGATCGGAA |
| 10527_2 | TATGAGTGTGGAGTCGTTGCTGTGGTTTATGGGATCAGAGTTTNNNAGATCGGAA |
| 38374_1 | TATGAGTGTGGAGTCGTTGCGTGGATCATGAGGTCAGGAGATCNNNAGATCGGAA |
| 38374_2 | TATGAGTGTGGAGTCGTTGCGTGGATCATGAGGTCAGGAGATCNNNAGATCGGAA |
| 50963_1 | TATGAGTGTGGAGTCGTTGCCCTAGCTCTCGGGTTCTCTTTTCNNNAGATCGGAAGA |
| 50963_2 | TATGAGTGTGGAGTCGTTGCCCTAGCTCTCGGGTTCTCTTTTCNNNAGATCGGAAGA |
| 46316_1 | TATGAGTGTGGAGTCGTTGCATCTCGGCTCACTGTAACCTCTANNNAGATCGGAAG |
| 46316_2 | TATGAGTGTGGAGTCGTTGCATCTCGGCTCACTGTAACCTCTANNNAGATCGGAAG |
| 49594_1 | TATGAGTGTGGAGTCGTTGCTTCTCCTGCCTCAGTCTTACAAGNNNAGATCGGAAG |
| 49594_2 | TATGAGTGTGGAGTCGTTGCTTCTCCTGCCTCAGTCTTACAAGNNNAGATCGGAAG |
| 48404_1 | TATGAGTGTGGAGTCGTTGCTGCTGGGATTACAGATATGAGCCNNNAGATCGGAA |

| | |
|---|---|
| 48404_2 | TATGAGTGTGGAGTCGTTGCTGCTGGGATTACAGATATGAGCCNNNAGATCGGAA |
| 44084_1 | TATGAGTGTGGAGTCGTTGCGAAGAGCTCGGAGTAGCACTGNNNAGATCGGAAGA |
| 44084_2 | TATGAGTGTGGAGTCGTTGCGAAGAGCTCGGAGTAGCACTGNNNAGATCGGAAGA |
| 41069_1 | TATGAGTGTGGAGTCGTTGCCTCAGATTACACACCCAGCTGATNNNAGATCGGAAG |
| 41069_2 | TATGAGTGTGGAGTCGTTGCCTCAGATTACACACCCAGCTGATNNNAGATCGGAAG |
| 49819_1 | TATGAGTGTGGAGTCGTTGCCCGAAAAGGAAGGTGTGTTTGACNNNAGATCGGAA |
| 49819_2 | TATGAGTGTGGAGTCGTTGCCCGAAAAGGAAGGTGTGTTTGACNNNAGATCGGAA |
| 44926_1 | TATGAGTGTGGAGTCGTTGCGAGCGTCGCCAACTTTTCTCNNNAGATCGGAAGAGC |
| 44926_2 | TATGAGTGTGGAGTCGTTGCGAGCGTCGCCAACTTTTCTCNNNAGATCGGAAGAGC |
| 12885_1 | TATGAGTGTGGAGTCGTTGCCCAGGTTTTCAGAAAATGAAGNNNAGATCGGAAGA |
| 12885_2 | TATGAGTGTGGAGTCGTTGCCCAGGTTTTCAGAAAATGAAGNNNAGATCGGAAGA |
| 35365_1 | TATGAGTGTGGAGTCGTTGCCTGAGTAGCTGGGATTACAGACCNNNAGATCGGAA |
| 35365_2 | TATGAGTGTGGAGTCGTTGCCTGAGTAGCTGGGATTACAGACCNNNAGATCGGAA |
| 43064_1 | TATGAGTGTGGAGTCGTTGCGAATACATCACACTGGTGGGTCTNNNAGATCGGAA |
| 43064_2 | TATGAGTGTGGAGTCGTTGCGAATACATCACACTGGTGGGTCTNNNAGATCGGAA |
| 45188_1 | TATGAGTGTGGAGTCGTTGCTCTTAGGCTTGGATCACTGGTTCNNNAGATCGGAAG |
| 45188_2 | TATGAGTGTGGAGTCGTTGCTCTTAGGCTTGGATCACTGGTTCNNNAGATCGGAAG |
| 42185_1 | TATGAGTGTGGAGTCGTTGCGTAGCTGGGACTACAGGCATAAANNNAGATCGGAA |
| 42185_2 | TATGAGTGTGGAGTCGTTGCGTAGCTGGGACTACAGGCATAAANNNAGATCGGAA |
| 33564_1 | TATGAGTGTGGAGTCGTTGCCTCCTGGGTTCAAGCAGTTCTCNNNAGATCGGAAGA |
| 33564_2 | TATGAGTGTGGAGTCGTTGCCTCCTGGGTTCAAGCAGTTCTCNNNAGATCGGAAGA |
| 30605_1 | TATGAGTGTGGAGTCGTTGCTCACATCACATGACATAGGGGAANNNAGATCGGAA |
| 30605_2 | TATGAGTGTGGAGTCGTTGCTCACATCACATGACATAGGGGAANNNAGATCGGAA |
| 32568_1 | TATGAGTGTGGAGTCGTTGCCTATCTCACCTGACTGCAACCTCNNNAGATCGGAAGA |
| 32568_2 | TATGAGTGTGGAGTCGTTGCCTATCTCACCTGACTGCAACCTCNNNAGATCGGAAGA |
| 33496_1 | TATGAGTGTGGAGTCGTTGCAAGGGGAAGGGGAAGCTTTTATTNNNAGATCGGAA |
| 33496_2 | TATGAGTGTGGAGTCGTTGCAAGGGGAAGGGGAAGCTTTTATTNNNAGATCGGAA |
| 39605_1 | TATGAGTGTGGAGTCGTTGCCCCAGGGGAAGGAAATCTCTAACNNNAGATCGGAA |
| 39605_2 | TATGAGTGTGGAGTCGTTGCCCCAGGGGAAGGAAATCTCTAACNNNAGATCGGAA |
| 50103_1 | TATGAGTGTGGAGTCGTTGCTCAAATACCTGCCCTTGCCCNNNAGATCGGAAGAGC |
| 50103_2 | TATGAGTGTGGAGTCGTTGCTCAAATACCTGCCCTTGCCCNNNAGATCGGAAGAGC |
| 52316_1 | TATGAGTGTGGAGTCGTTGCCTGAAAGGCCAAGACAACGATGNNNAGATCGGAAG |
| 52316_2 | TATGAGTGTGGAGTCGTTGCCTGAAAGGCCAAGACAACGATGNNNAGATCGGAAG |
| 47920_1 | TATGAGTGTGGAGTCGTTGCGGCTAAACCAAGGAATTGCTGTCNNNAGATCGGAA |
| 47920_2 | TATGAGTGTGGAGTCGTTGCGGCTAAACCAAGGAATTGCTGTCNNNAGATCGGAA |
| 39577_1 | TATGAGTGTGGAGTCGTTGCTTCAGGCAGTGATGACCTGTTTANNNAGATCGGAAG |
| 39577_2 | TATGAGTGTGGAGTCGTTGCTTCAGGCAGTGATGACCTGTTTANNNAGATCGGAAG |
| 50611_1 | TATGAGTGTGGAGTCGTTGCGACACTGCCTTACACAACTCCTTNNNAGATCGGAAGA |
| 50611_2 | TATGAGTGTGGAGTCGTTGCGACACTGCCTTACACAACTCCTTNNNAGATCGGAAGA |
| 2303_1 | TATGAGTGTGGAGTCGTTGCGATAGTGTTCCCTTGCCAGTCACNNNAGATCGGAAG |
| 2303_2 | TATGAGTGTGGAGTCGTTGCGATAGTGTTCCCTTGCCAGTCACNNNAGATCGGAAG |
| 44097_1 | TATGAGTGTGGAGTCGTTGCAATAGCAAGATCATGTGTCGGGANNNAGATCGGAA |
| 44097_2 | TATGAGTGTGGAGTCGTTGCAATAGCAAGATCATGTGTCGGGANNNAGATCGGAA |

| | |
|---|---|
| 45756_1 | TATGAGTGTGGAGTCGTTGCAGAGACAGGGTTTCACTACGTTGNNNAGATCGGAA |
| 45756_2 | TATGAGTGTGGAGTCGTTGCAGAGACAGGGTTTCACTACGTTGNNNAGATCGGAA |
| 30363_1 | TATGAGTGTGGAGTCGTTGCAGAGTGCAATAGCAGAAACTCGGNNNAGATCGGAA |
| 30363_2 | TATGAGTGTGGAGTCGTTGCAGAGTGCAATAGCAGAAACTCGGNNNAGATCGGAA |
| 44426_1 | TATGAGTGTGGAGTCGTTGCGCCAGCGAGTGACTAATTGATTGNNNAGATCGGAA |
| 44426_2 | TATGAGTGTGGAGTCGTTGCGCCAGCGAGTGACTAATTGATTGNNNAGATCGGAA |
| 31221_1 | TATGAGTGTGGAGTCGTTGCAGAAAGACACCCAAGACAAGACANNNAGATCGGAA |
| 31221_2 | TATGAGTGTGGAGTCGTTGCAGAAAGACACCCAAGACAAGACANNNAGATCGGAA |
| 51526_1 | TATGAGTGTGGAGTCGTTGCGAGTTCTCATGAGGTCTGGTTGTNNNAGATCGGAA |
| 51526_2 | TATGAGTGTGGAGTCGTTGCGAGTTCTCATGAGGTCTGGTTGTNNNAGATCGGAA |
| 34283_1 | TATGAGTGTGGAGTCGTTGCTGTTGTTCAGATTCTTTTGCCCCNNNAGATCGGAAG |
| 34283_2 | TATGAGTGTGGAGTCGTTGCTGTTGTTCAGATTCTTTTGCCCCNNNAGATCGGAAG |
| 10489_1 | TATGAGTGTGGAGTCGTTGCTGGGGGAGATTGATGTCCAGAAANNNAGATCGGA |
| 10489_2 | TATGAGTGTGGAGTCGTTGCTGGGGGAGATTGATGTCCAGAAANNNAGATCGGA |
| 47996_1 | TATGAGTGTGGAGTCGTTGCGTCCTATCAATGTGGGGATCAGGNNNAGATCGGAA |
| 47996_2 | TATGAGTGTGGAGTCGTTGCGTCCTATCAATGTGGGGATCAGGNNNAGATCGGAA |
| 48846_1 | TATGAGTGTGGAGTCGTTGCAAGACTTCTCTGTCCCGTCCCNNNAGATCGGAAGAG |
| 48846_2 | TATGAGTGTGGAGTCGTTGCAAGACTTCTCTGTCCCGTCCCNNNAGATCGGAAGAG |
| 37916_1 | TATGAGTGTGGAGTCGTTGCAGATTCTGGAACCAGATTGACGTNNNAGATCGGAA |
| 37916_2 | TATGAGTGTGGAGTCGTTGCAGATTCTGGAACCAGATTGACGTNNNAGATCGGAA |
| 272_1 | TATGAGTGTGGAGTCGTTGCTCCTCCCACCCCAGACATTTTTCNNNAGATCGGAAGA |
| 272_2 | TATGAGTGTGGAGTCGTTGCTCCTCCCACCCCAGACATTTTTCNNNAGATCGGAAGA |
| 46246_1 | TATGAGTGTGGAGTCGTTGCGTCAACAAGAGAGGCACAATTGTNNNAGATCGGAA |
| 46246_2 | TATGAGTGTGGAGTCGTTGCGTCAACAAGAGAGGCACAATTGTNNNAGATCGGAA |
| 29434_1 | TATGAGTGTGGAGTCGTTGCTTCAAGTGATTCTCCTGCCTCAGNNNAGATCGGAAG |
| 29434_2 | TATGAGTGTGGAGTCGTTGCTTCAAGTGATTCTCCTGCCTCAGNNNAGATCGGAAG |
| 42925_1 | TATGAGTGTGGAGTCGTTGCTCTGTCTCTCTCTGTCTCTGTGTNNNAGATCGGAAGA |
| 42925_2 | TATGAGTGTGGAGTCGTTGCTCTGTCTCTCTCTGTCTCTGTGTNNNAGATCGGAAGA |
| 45871_1 | TATGAGTGTGGAGTCGTTGCCACAATCTTGGCTCACTGCAACNNNAGATCGGAAGA |
| 45871_2 | TATGAGTGTGGAGTCGTTGCCACAATCTTGGCTCACTGCAACNNNAGATCGGAAGA |
| 38326_1 | TATGAGTGTGGAGTCGTTGCAATTATCACAATCTCTCGGGCCANNNAGATCGGAAG |
| 38326_2 | TATGAGTGTGGAGTCGTTGCAATTATCACAATCTCTCGGGCCANNNAGATCGGAAG |
| 33363_1 | TATGAGTGTGGAGTCGTTGCTCCTGAGTAGCTAGGACTACAGGNNNAGATCGGAA |
| 33363_2 | TATGAGTGTGGAGTCGTTGCTCCTGAGTAGCTAGGACTACAGGNNNAGATCGGAA |
| 40550_1 | TATGAGTGTGGAGTCGTTGCAGAGGGAAAGGAATTGTCTGAGTNNNAGATCGGA |
| 40550_2 | TATGAGTGTGGAGTCGTTGCAGAGGGAAAGGAATTGTCTGAGTNNNAGATCGGA |
| 38844_1 | TATGAGTGTGGAGTCGTTGCTCATCTCTAAAACTGAAAGCACCANNNAGATCGGAA |
| 38844_2 | TATGAGTGTGGAGTCGTTGCTCATCTCTAAAACTGAAAGCACCANNNAGATCGGAA |
| 46085_1 | TATGAGTGTGGAGTCGTTGCGCCTCCTGAGTAGCTGAGATTACNNNAGATCGGAAG |
| 46085_2 | TATGAGTGTGGAGTCGTTGCGCCTCCTGAGTAGCTGAGATTACNNNAGATCGGAAG |
| 50604_1 | TATGAGTGTGGAGTCGTTGCCCTCCTATCTCCCCATCACCTATNNNAGATCGGAAGA |
| 50604_2 | TATGAGTGTGGAGTCGTTGCCCTCCTATCTCCCCATCACCTATNNNAGATCGGAAGA |
| 43630_1 | TATGAGTGTGGAGTCGTTGCTAGGTCCTCTGAGAAGTCTCCAGNNNAGATCGGAAG |

| | |
|---|---|
| 43630_2 | TATGAGTGTGGAGTCGTTGCTAGGTCCTCTGAGAAGTCTCCAGNNNAGATCGGAAG |
| 30015_1 | TATGAGTGTGGAGTCGTTGCCCCTGTTAACTGATCAATCTGGANNNAGATCGGAAG |
| 30015_2 | TATGAGTGTGGAGTCGTTGCCCCTGTTAACTGATCAATCTGGANNNAGATCGGAAG |
| 34338_1 | TATGAGTGTGGAGTCGTTGCGGGTACCTCCATTCCTCATGAAGNNNAGATCGGAAG |
| 34338_2 | TATGAGTGTGGAGTCGTTGCGGGTACCTCCATTCCTCATGAAGNNNAGATCGGAAG |
| 36538_1 | TATGAGTGTGGAGTCGTTGCCATAATGCCAGCAAAGCTCACTTNNNAGATCGGAAG |
| 36538_2 | TATGAGTGTGGAGTCGTTGCCATAATGCCAGCAAAGCTCACTTNNNAGATCGGAAG |
| 35282_1 | TATGAGTGTGGAGTCGTTGCGAGATAGGGTCTTGCTCTGTCACNNNAGATCGGAA |
| 35282_2 | TATGAGTGTGGAGTCGTTGCGAGATAGGGTCTTGCTCTGTCACNNNAGATCGGAA |
| 51076_1 | TATGAGTGTGGAGTCGTTGCGATCCAGAGAGGCCAAAACGTNNNAGATCGGAAGA |
| 51076_2 | TATGAGTGTGGAGTCGTTGCGATCCAGAGAGGCCAAAACGTNNNAGATCGGAAGA |
| 3028_1 | TATGAGTGTGGAGTCGTTGCCCAATGAAGCCATTACCTCCCAANNNAGATCGGAAG |
| 3028_2 | TATGAGTGTGGAGTCGTTGCCCAATGAAGCCATTACCTCCCAANNNAGATCGGAAG |
| 39569_1 | TATGAGTGTGGAGTCGTTGCTAGGAGGTAGTCAAGAGTCAGGCNNNAGATCGGAA |
| 39569_2 | TATGAGTGTGGAGTCGTTGCTAGGAGGTAGTCAAGAGTCAGGCNNNAGATCGGAA |
| 47396_1 | TATGAGTGTGGAGTCGTTGCGATCTTGGCTTACTGCAACCTCTNNNAGATCGGAAG |
| 47396_2 | TATGAGTGTGGAGTCGTTGCGATCTTGGCTTACTGCAACCTCTNNNAGATCGGAAG |
| 42052_1 | TATGAGTGTGGAGTCGTTGCTCTGAAATCTAGGGAGAGGTTCCTNNNAGATCGGA |
| 42052_2 | TATGAGTGTGGAGTCGTTGCTCTGAAATCTAGGGAGAGGTTCCTNNNAGATCGGA |
| 49788_1 | TATGAGTGTGGAGTCGTTGCCTGCCGCTACAAGCAACCACNNNAGATCGGAAGAGC |
| 49788_2 | TATGAGTGTGGAGTCGTTGCCTGCCGCTACAAGCAACCACNNNAGATCGGAAGAGC |
| 41517_1 | TATGAGTGTGGAGTCGTTGCGAGACAGGCCTTATTCCATCCCNNNAGATCGGAAGA |
| 41517_2 | TATGAGTGTGGAGTCGTTGCGAGACAGGCCTTATTCCATCCCNNNAGATCGGAAGA |
| 49292_1 | TATGAGTGTGGAGTCGTTGCCTAATGCTATCCCTCCCCCAATCNNNAGATCGGAAGA |
| 49292_2 | TATGAGTGTGGAGTCGTTGCCTAATGCTATCCCTCCCCCAATCNNNAGATCGGAAGA |
| 47698_1 | TATGAGTGTGGAGTCGTTGCCCTTGGTTACTGAGGAAACCGAANNNAGATCGGAA |
| 47698_2 | TATGAGTGTGGAGTCGTTGCCCTTGGTTACTGAGGAAACCGAANNNAGATCGGAA |
| 52611_1 | TATGAGTGTGGAGTCGTTGCCATTCAGACGAGTTTCTGCGATCNNNAGATCGGAAG |
| 52611_2 | TATGAGTGTGGAGTCGTTGCCATTCAGACGAGTTTCTGCGATCNNNAGATCGGAAG |
| 30680_1 | TATGAGTGTGGAGTCGTTGCCGAGTCTCCAAAGTCCAAGGTATTANNNAGATCGGA |
| 30680_2 | TATGAGTGTGGAGTCGTTGCCGAGTCTCCAAAGTCCAAGGTATTANNNAGATCGGA |
| 9056_1 | TATGAGTGTGGAGTCGTTGCTGTCCTCATATACGAACCTACACANNNAGATCGGAA |
| 9056_2 | TATGAGTGTGGAGTCGTTGCTGTCCTCATATACGAACCTACACANNNAGATCGGAA |
| 33469_1 | TATGAGTGTGGAGTCGTTGCCACGCACACACATAAAGGAGAAANNNAGATCGGAA |
| 33469_2 | TATGAGTGTGGAGTCGTTGCCACGCACACACATAAAGGAGAAANNNAGATCGGAA |
| 42379_1 | TATGAGTGTGGAGTCGTTGCACAATAAAAGGGCACATGTCTGCNNNAGATCGGAA |
| 42379_2 | TATGAGTGTGGAGTCGTTGCACAATAAAAGGGCACATGTCTGCNNNAGATCGGAA |
| 38387_1 | TATGAGTGTGGAGTCGTTGCAGAGAACCCAACACAGTACCAAANNNAGATCGGAA |
| 38387_2 | TATGAGTGTGGAGTCGTTGCAGAGAACCCAACACAGTACCAAANNNAGATCGGAA |
| 40053_1 | TATGAGTGTGGAGTCGTTGCAAAGCAGCAAGAGAGAAGTGACTNNNAGATCGGAA |
| 40053_2 | TATGAGTGTGGAGTCGTTGCAAAGCAGCAAGAGAGAAGTGACTNNNAGATCGGAA |
| 45186_1 | TATGAGTGTGGAGTCGTTGCTCAGCAATCTTCCCTCCATTGAANNNAGATCGGAAGA |
| 45186_2 | TATGAGTGTGGAGTCGTTGCTCAGCAATCTTCCCTCCATTGAANNNAGATCGGAAGA |

| | |
|---|---|
| 33153_1 | TATGAGTGTGGAGTCGTTGCATGGAAAGGAACAACTGGTACGANNNAGATCGGAA |
| 33153_2 | TATGAGTGTGGAGTCGTTGCATGGAAAGGAACAACTGGTACGANNNAGATCGGAA |
| 44739_1 | TATGAGTGTGGAGTCGTTGCTTTGCTGAAAGTTGGCTGGAAAANNNAGATCGGAA |
| 44739_2 | TATGAGTGTGGAGTCGTTGCTTTGCTGAAAGTTGGCTGGAAAANNNAGATCGGAA |
| 46625_1 | TATGAGTGTGGAGTCGTTGCGAGCAATTCTCCTGTCTTAGCCTNNNAGATCGGAAG |
| 46625_2 | TATGAGTGTGGAGTCGTTGCGAGCAATTCTCCTGTCTTAGCCTNNNAGATCGGAAG |
| 43597_1 | TATGAGTGTGGAGTCGTTGCCAGACACCAGGGAAAAGAGGTAANNNAGATCGGAA |
| 43597_2 | TATGAGTGTGGAGTCGTTGCCAGACACCAGGGAAAAGAGGTAANNNAGATCGGAA |
| 30928_1 | TATGAGTGTGGAGTCGTTGCTCTCTCCCACTCTTCTTCTCTCCNNNAGATCGGAAGAG |
| 30928_2 | TATGAGTGTGGAGTCGTTGCTCTCTCCCACTCTTCTTCTCTCCNNNAGATCGGAAGAG |
| 13310_1 | TATGAGTGTGGAGTCGTTGCTTGCCTTTCATGAATACCAGNNNAGATCGGAAGAGC |
| 13310_2 | TATGAGTGTGGAGTCGTTGCTTGCCTTTCATGAATACCAGNNNAGATCGGAAGAGC |
| 49922_1 | TATGAGTGTGGAGTCGTTGCTGTCACCCAAGACAAGAGCAATANNNAGATCGGAAG |
| 49922_2 | TATGAGTGTGGAGTCGTTGCTGTCACCCAAGACAAGAGCAATANNNAGATCGGAAG |
| 45583_1 | TATGAGTGTGGAGTCGTTGCCAAGTAGCTGGGACTACAGGCNNNAGATCGGAAGA |
| 45583_2 | TATGAGTGTGGAGTCGTTGCCAAGTAGCTGGGACTACAGGCNNNAGATCGGAAGA |
| 43979_1 | TATGAGTGTGGAGTCGTTGCGGAGAACCAGGGTAGCTGATAAGNNNAGATCGGA |
| 43979_2 | TATGAGTGTGGAGTCGTTGCGGAGAACCAGGGTAGCTGATAAGNNNAGATCGGA |
| 37950_1 | TATGAGTGTGGAGTCGTTGCAAGATGGAGGAAATGGCCACATANNNAGATCGGAA |
| 37950_2 | TATGAGTGTGGAGTCGTTGCAAGATGGAGGAAATGGCCACATANNNAGATCGGAA |
| 37701_1 | TATGAGTGTGGAGTCGTTGCAACACCCAACCCTACAATGGATTNNNAGATCGGAAG |
| 37701_2 | TATGAGTGTGGAGTCGTTGCAACACCCAACCCTACAATGGATTNNNAGATCGGAAG |
| 46720_1 | TATGAGTGTGGAGTCGTTGCTGTAGACAGCATGTAGTTGGGACNNNAGATCGGAA |
| 46720_2 | TATGAGTGTGGAGTCGTTGCTGTAGACAGCATGTAGTTGGGACNNNAGATCGGAA |
| 34005_1 | TATGAGTGTGGAGTCGTTGCTTTCTCTGTGTTGGCTAGGCTGNNNAGATCGGAAGA |
| 34005_2 | TATGAGTGTGGAGTCGTTGCTTTCTCTGTGTTGGCTAGGCTGNNNAGATCGGAAGA |
| 32648_1 | TATGAGTGTGGAGTCGTTGCGAGTAGCTGGGACTACAGGCNNNAGATCGGAAGAG |
| 32648_2 | TATGAGTGTGGAGTCGTTGCGAGTAGCTGGGACTACAGGCNNNAGATCGGAAGAG |
| 41908_1 | TATGAGTGTGGAGTCGTTGCGAGACAAGGCCTCACTCTGTTGNNNAGATCGGAAG |
| 41908_2 | TATGAGTGTGGAGTCGTTGCGAGACAAGGCCTCACTCTGTTGNNNAGATCGGAAG |
| 46312_1 | TATGAGTGTGGAGTCGTTGCTTTCAAGTGATTCTCCTGCCTCANNNAGATCGGAAGA |
| 46312_2 | TATGAGTGTGGAGTCGTTGCTTTCAAGTGATTCTCCTGCCTCANNNAGATCGGAAGA |
| 49472_1 | TATGAGTGTGGAGTCGTTGCCTGGTTCACGCCATTCTCCTNNNAGATCGGAAGAGC |
| 49472_2 | TATGAGTGTGGAGTCGTTGCCTGGTTCACGCCATTCTCCTNNNAGATCGGAAGAGC |
| 44757_1 | TATGAGTGTGGAGTCGTTGCAAGCAGGAGGAGGAGGAGATAAANNNAGATCGGA |
| 44757_2 | TATGAGTGTGGAGTCGTTGCAAGCAGGAGGAGGAGGAGATAAANNNAGATCGGA |
| 48518_1 | TATGAGTGTGGAGTCGTTGCTTCTCTTGGGTATTCCTGGGAGANNNAGATCGGAAG |
| 48518_2 | TATGAGTGTGGAGTCGTTGCTTCTCTTGGGTATTCCTGGGAGANNNAGATCGGAAG |
| 48047_1 | TATGAGTGTGGAGTCGTTGCCAGGTGACCATGAGGACAATCATNNNAGATCGGAA |
| 48047_2 | TATGAGTGTGGAGTCGTTGCCAGGTGACCATGAGGACAATCATNNNAGATCGGAA |
| 33529_1 | TATGAGTGTGGAGTCGTTGCTGGTGGCTCACGTCTGTAATCNNNAGATCGGAAGAG |
| 33529_2 | TATGAGTGTGGAGTCGTTGCTGGTGGCTCACGTCTGTAATCNNNAGATCGGAAGAG |
| 38549_1 | TATGAGTGTGGAGTCGTTGCACCACCTGAAGGGATTAGAGAGANNNAGATCGGAA |

| | |
|---|---|
| 38549_2 | TATGAGTGTGGAGTCGTTGCACCACCTGAAGGGATTAGAGAGANNNAGATCGGAA |
| 42269_1 | TATGAGTGTGGAGTCGTTGCGGTAAATAGTCTCAGCTCCTCGGNNNAGATCGGAAG |
| 42269_2 | TATGAGTGTGGAGTCGTTGCGGTAAATAGTCTCAGCTCCTCGGNNNAGATCGGAAG |
| 47765_1 | TATGAGTGTGGAGTCGTTGCCCACCACACTCGGCTAATTTTTTNNNAGATCGGAAGA |
| 47765_2 | TATGAGTGTGGAGTCGTTGCCCACCACACTCGGCTAATTTTTTNNNAGATCGGAAGA |
| 980_1 | TATGAGTGTGGAGTCGTTGCTGAGTGTCTACTACGTGTCAGGTNNNAGATCGGAAG |
| 980_2 | TATGAGTGTGGAGTCGTTGCTGAGTGTCTACTACGTGTCAGGTNNNAGATCGGAAG |
| 34759_1 | TATGAGTGTGGAGTCGTTGCCAATCTCGCTTCACTGCAACCNNNAGATCGGAAGAGC |
| 34759_2 | TATGAGTGTGGAGTCGTTGCCAATCTCGCTTCACTGCAACCNNNAGATCGGAAGAGC |
| 49483_1 | TATGAGTGTGGAGTCGTTGCTCTTACTCTTTTCTGCCAAAAATCTNNNAGATCGGAAG |
| 49483_2 | TATGAGTGTGGAGTCGTTGCTCTTACTCTTTTCTGCCAAAAATCTNNNAGATCGGAAG |
| 52183_1 | TATGAGTGTGGAGTCGTTGCGGCCATTTCCATATTGACCATGGNNNAGATCGGAAG |
| 52183_2 | TATGAGTGTGGAGTCGTTGCGGCCATTTCCATATTGACCATGGNNNAGATCGGAAG |
| 38777_1 | TATGAGTGTGGAGTCGTTGCGAAAAGCTCCCTCAAATCATGCANNNAGATCGGAAG |
| 38777_2 | TATGAGTGTGGAGTCGTTGCGAAAAGCTCCCTCAAATCATGCANNNAGATCGGAAG |
| 29463_1 | TATGAGTGTGGAGTCGTTGCCTGCTCATCCTCAGGTTTTCCTCNNNAGATCGGAAGA |
| 29463_2 | TATGAGTGTGGAGTCGTTGCCTGCTCATCCTCAGGTTTTCCTCNNNAGATCGGAAGA |
| 41732_1 | TATGAGTGTGGAGTCGTTGCGAGATGGAATCTTGCTTTGTCGCNNNAGATCGGAAG |
| 41732_2 | TATGAGTGTGGAGTCGTTGCGAGATGGAATCTTGCTTTGTCGCNNNAGATCGGAAG |
| 33460_1 | TATGAGTGTGGAGTCGTTGCCTCCAGGTAGCAAACTTCAGAGANNNAGATCGGAAG |
| 33460_2 | TATGAGTGTGGAGTCGTTGCCTCCAGGTAGCAAACTTCAGAGANNNAGATCGGAAG |
| 39110_1 | TATGAGTGTGGAGTCGTTGCATACCATGTCTCCTGTGTTTTCANNNAGATCGGAAGA |
| 39110_2 | TATGAGTGTGGAGTCGTTGCATACCATGTCTCCTGTGTTTTCANNNAGATCGGAAGA |
| 36190_1 | TATGAGTGTGGAGTCGTTGCGGCAGCTCAAAACCAAGAGAATTNNNAGATCGGAAG |
| 36190_2 | TATGAGTGTGGAGTCGTTGCGGCAGCTCAAAACCAAGAGAATTNNNAGATCGGAAG |
| 37709_1 | TATGAGTGTGGAGTCGTTGCGTAGTCCCAGCTACTCGGAGAGNNNAGATCGGAAGA |
| 37709_2 | TATGAGTGTGGAGTCGTTGCGTAGTCCCAGCTACTCGGAGAGNNNAGATCGGAAGA |
| 34645_1 | TATGAGTGTGGAGTCGTTGCAGAGGGGAGATTTCCTCAGGAATNNNAGATCGGAA |
| 34645_2 | TATGAGTGTGGAGTCGTTGCAGAGGGGAGATTTCCTCAGGAATNNNAGATCGGAA |
| 32982_1 | TATGAGTGTGGAGTCGTTGCTTTTTTGGCAAAAGCGGTTTCACNNNAGATCGGAAG |
| 32982_2 | TATGAGTGTGGAGTCGTTGCTTTTTTGGCAAAAGCGGTTTCACNNNAGATCGGAAG |
| 41983_1 | TATGAGTGTGGAGTCGTTGCGGCTAAAGCAATTCTCCCACTTCNNNAGATCGGAAGA |
| 41983_2 | TATGAGTGTGGAGTCGTTGCGGCTAAAGCAATTCTCCCACTTCNNNAGATCGGAAGA |
| 31048_1 | TATGAGTGTGGAGTCGTTGCTTTGAATAGAGTCCCCTAAGGCTNNNAGATCGGAAG |
| 31048_2 | TATGAGTGTGGAGTCGTTGCTTTGAATAGAGTCCCCTAAGGCTNNNAGATCGGAAG |
| 43014_1 | TATGAGTGTGGAGTCGTTGCGATCAGAAGAGAGTGGCATGACANNNAGATCGGAA |
| 43014_2 | TATGAGTGTGGAGTCGTTGCGATCAGAAGAGAGTGGCATGACANNNAGATCGGAA |
| 3103_1 | TATGAGTGTGGAGTCGTTGCTTGTGGATGAGCAGCAGAAAGTTNNNAGATCGGAA |
| 3103_2 | TATGAGTGTGGAGTCGTTGCTTGTGGATGAGCAGCAGAAAGTTNNNAGATCGGAA |
| 31191_1 | TATGAGTGTGGAGTCGTTGCACACCTTTTCATCACTAGGTTCANNNAGATCGGAAGA |
| 31191_2 | TATGAGTGTGGAGTCGTTGCACACCTTTTCATCACTAGGTTCANNNAGATCGGAAGA |
| 34803_1 | TATGAGTGTGGAGTCGTTGCCTTCTGGGTTCAGGCGATTCTCNNNAGATCGGAAGA |
| 34803_2 | TATGAGTGTGGAGTCGTTGCCTTCTGGGTTCAGGCGATTCTCNNNAGATCGGAAGA |

| | |
|---|---|
| 33709_1 | TATGAGTGTGGAGTCGTTGCTACAAAAGTTAGCCAGTCGTGGTNNNAGATCGGAA |
| 33709_2 | TATGAGTGTGGAGTCGTTGCTACAAAAGTTAGCCAGTCGTGGTNNNAGATCGGAA |
| 48094_1 | TATGAGTGTGGAGTCGTTGCGTCGTGTCACAAGAACTTGGTTTNNNAGATCGGAAG |
| 48094_2 | TATGAGTGTGGAGTCGTTGCGTCGTGTCACAAGAACTTGGTTTNNNAGATCGGAAG |
| 44039_1 | TATGAGTGTGGAGTCGTTGCATAGCAAGGGTCTGCAAACTACANNNAGATCGGAA |
| 44039_2 | TATGAGTGTGGAGTCGTTGCATAGCAAGGGTCTGCAAACTACANNNAGATCGGAA |
| 35325_1 | TATGAGTGTGGAGTCGTTGCCCAGGGGTGACTATTTACCAACANNNAGATCGGAAG |
| 35325_2 | TATGAGTGTGGAGTCGTTGCCCAGGGGTGACTATTTACCAACANNNAGATCGGAAG |
| 29878_1 | TATGAGTGTGGAGTCGTTGCCCTGTCTGCCATAAAGACTGTCANNNAGATCGGAAG |
| 29878_2 | TATGAGTGTGGAGTCGTTGCCCTGTCTGCCATAAAGACTGTCANNNAGATCGGAAG |
| 38790_1 | TATGAGTGTGGAGTCGTTGCAGCCACTGCAAAAACAATCCAAANNNAGATCGGAAG |
| 38790_2 | TATGAGTGTGGAGTCGTTGCAGCCACTGCAAAAACAATCCAAANNNAGATCGGAAG |
| 29255_1 | TATGAGTGTGGAGTCGTTGCTAGAGTCACGTCTCCATCTCCATNNNAGATCGGAAG |
| 29255_2 | TATGAGTGTGGAGTCGTTGCTAGAGTCACGTCTCCATCTCCATNNNAGATCGGAAG |
| 45966_1 | TATGAGTGTGGAGTCGTTGCGCAAAGGGAGCAAATCATTCCTTNNNAGATCGGAA |
| 45966_2 | TATGAGTGTGGAGTCGTTGCGCAAAGGGAGCAAATCATTCCTTNNNAGATCGGAA |
| 39245_1 | TATGAGTGTGGAGTCGTTGCATCATTACTGCCACCCATCAGAGNNNAGATCGGAAG |
| 39245_2 | TATGAGTGTGGAGTCGTTGCATCATTACTGCCACCCATCAGAGNNNAGATCGGAAG |
| 3072_1 | TATGAGTGTGGAGTCGTTGCAAGAGGCCAAATGAGTGTAAGGCNNNAGATCGGAA |
| 3072_2 | TATGAGTGTGGAGTCGTTGCAAGAGGCCAAATGAGTGTAAGGCNNNAGATCGGAA |
| 40376_1 | TATGAGTGTGGAGTCGTTGCAAACCATTGTAACTCGAGGACCANNNAGATCGGAAG |
| 40376_2 | TATGAGTGTGGAGTCGTTGCAAACCATTGTAACTCGAGGACCANNNAGATCGGAAG |
| 41489_1 | TATGAGTGTGGAGTCGTTGCATTTTTTCCTGAAGTGGCCACTGNNNAGATCGGAAG |
| 41489_2 | TATGAGTGTGGAGTCGTTGCATTTTTTCCTGAAGTGGCCACTGNNNAGATCGGAAG |
| 34966_1 | TATGAGTGTGGAGTCGTTGCGAGTGCAGGGTGTGATCTCAGNNNAGATCGGAAG |
| 34966_2 | TATGAGTGTGGAGTCGTTGCGAGTGCAGGGTGTGATCTCAGNNNAGATCGGAAG |
| 44668_1 | TATGAGTGTGGAGTCGTTGCCTGCTTTCTGAAAAGAACAGGGTNNNAGATCGGAA |
| 44668_2 | TATGAGTGTGGAGTCGTTGCCTGCTTTCTGAAAAGAACAGGGTNNNAGATCGGAA |
| 33568_1 | TATGAGTGTGGAGTCGTTGCCCAAGTAGCTGGGATTACAGGTGNNNAGATCGGAA |
| 33568_2 | TATGAGTGTGGAGTCGTTGCCCAAGTAGCTGGGATTACAGGTGNNNAGATCGGAA |
| 33987_1 | TATGAGTGTGGAGTCGTTGCGACGGATTCTCGCTCTTTTGCNNNAGATCGGAAGAG |
| 33987_2 | TATGAGTGTGGAGTCGTTGCGACGGATTCTCGCTCTTTTGCNNNAGATCGGAAGAG |
| 30563_1 | TATGAGTGTGGAGTCGTTGCCTAAGTGTGCCATCTGCCATTTCCNNNAGATCGGAA |
| 30563_2 | TATGAGTGTGGAGTCGTTGCCTAAGTGTGCCATCTGCCATTTCCNNNAGATCGGAA |
| 10113_1 | TATGAGTGTGGAGTCGTTGCCAGGAGAGGAGAGTTGCATGAACNNNAGATCGGA |
| 10113_2 | TATGAGTGTGGAGTCGTTGCCAGGAGAGGAGAGTTGCATGAACNNNAGATCGGA |
| 48796_1 | TATGAGTGTGGAGTCGTTGCTTTTCCTTCCTCCTCTTTGCTGTNNNAGATCGGAAGA |
| 48796_2 | TATGAGTGTGGAGTCGTTGCTTTTCCTTCCTCCTCTTTGCTGTNNNAGATCGGAAGA |
| 38719_1 | TATGAGTGTGGAGTCGTTGCGAAGAAGCAACTGCAATATTTGGANNNAGATCGGA |
| 38719_2 | TATGAGTGTGGAGTCGTTGCGAAGAAGCAACTGCAATATTTGGANNNAGATCGGA |
| 51621_1 | TATGAGTGTGGAGTCGTTGCTGCAGGACAGAAATTCCTCATGANNNAGATCGGAA |
| 51621_2 | TATGAGTGTGGAGTCGTTGCTGCAGGACAGAAATTCCTCATGANNNAGATCGGAA |
| 33796_1 | TATGAGTGTGGAGTCGTTGCAACACTCTGGCAACTCAAAAAGCNNNAGATCGGAAG |

| | |
|---|---|
| 33796_2 | TATGAGTGTGGAGTCGTTGCAACACTCTGGCAACTCAAAAAGCNNNAGATCGGAAG |
| 49970_1 | TATGAGTGTGGAGTCGTTGCTGCCTACAAAACAGTCATAGCCTNNNAGATCGGAAG |
| 49970_2 | TATGAGTGTGGAGTCGTTGCTGCCTACAAAACAGTCATAGCCTNNNAGATCGGAAG |
| 47879_1 | TATGAGTGTGGAGTCGTTGCCTGGGTTCAAGCGATTCTCTTTCNNNAGATCGGAAG |
| 47879_2 | TATGAGTGTGGAGTCGTTGCCTGGGTTCAAGCGATTCTCTTTCNNNAGATCGGAAG |
| 41359_1 | TATGAGTGTGGAGTCGTTGCACTTGCTCTTTCCTCTATCTGGGNNNAGATCGGAAGA |
| 41359_2 | TATGAGTGTGGAGTCGTTGCACTTGCTCTTTCCTCTATCTGGGNNNAGATCGGAAGA |
| 33672_1 | TATGAGTGTGGAGTCGTTGCCATTCTCCTGCCTCAGCCTCNNNAGATCGGAAGAGCA |
| 33672_2 | TATGAGTGTGGAGTCGTTGCCATTCTCCTGCCTCAGCCTCNNNAGATCGGAAGAGCA |
| 42039_1 | TATGAGTGTGGAGTCGTTGCTATCACTTGAACCTAGGAGGCAGNNNAGATCGGAA |
| 42039_2 | TATGAGTGTGGAGTCGTTGCTATCACTTGAACCTAGGAGGCAGNNNAGATCGGAA |
| 46575_1 | TATGAGTGTGGAGTCGTTGCATTGTGCTCCATAAACTGCTTGGNNNAGATCGGAAG |
| 46575_2 | TATGAGTGTGGAGTCGTTGCATTGTGCTCCATAAACTGCTTGGNNNAGATCGGAAG |
| 43402_1 | TATGAGTGTGGAGTCGTTGCGGCCTTCGATGATATTGCCAAATNNNAGATCGGAAG |
| 43402_2 | TATGAGTGTGGAGTCGTTGCGGCCTTCGATGATATTGCCAAATNNNAGATCGGAAG |
| 38793_1 | TATGAGTGTGGAGTCGTTGCAAAGTCATCCATGTTGTAGCATGNNNAGATCGGAAG |
| 38793_2 | TATGAGTGTGGAGTCGTTGCAAAGTCATCCATGTTGTAGCATGNNNAGATCGGAAG |
| 51228_1 | TATGAGTGTGGAGTCGTTGCGAGGGAGGTAGGGGGGTCAGNNNAGATCGGAAG |
| 51228_2 | TATGAGTGTGGAGTCGTTGCGAGGGAGGTAGGGGGGTCAGNNNAGATCGGAAG |
| 48867_1 | TATGAGTGTGGAGTCGTTGCTTCTCTTTCGGCACAAACAGTTCNNNAGATCGGAAG |
| 48867_2 | TATGAGTGTGGAGTCGTTGCTTCTCTTTCGGCACAAACAGTTCNNNAGATCGGAAG |
| 47018_1 | TATGAGTGTGGAGTCGTTGCCCGCCATGAACTCTTTCTACAACNNNAGATCGGAAGA |
| 47018_2 | TATGAGTGTGGAGTCGTTGCCCGCCATGAACTCTTTCTACAACNNNAGATCGGAAGA |
| 39880_1 | TATGAGTGTGGAGTCGTTGCAACGCAATCCCCATCAAAATACCNNNAGATCGGAAG |
| 39880_2 | TATGAGTGTGGAGTCGTTGCAACGCAATCCCCATCAAAATACCNNNAGATCGGAAG |
| 29390_1 | TATGAGTGTGGAGTCGTTGCTTTTTTTTTCTTTGAGATGAAGTNNNAGATCGGAAG |
| 29390_2 | TATGAGTGTGGAGTCGTTGCTTTTTTTTTCTTTGAGATGAAGTNNNAGATCGGAAG |
| 44768_1 | TATGAGTGTGGAGTCGTTGCAGGTAGAGAGGAACCAGATTGGANNNAGATCGGAA |
| 44768_2 | TATGAGTGTGGAGTCGTTGCAGGTAGAGAGGAACCAGATTGGANNNAGATCGGAA |
| 12688_1 | TATGAGTGTGGAGTCGTTGCTCCTATTTTATCAGGTATTGTCCNNNAGATCGGAAGA |
| 12688_2 | TATGAGTGTGGAGTCGTTGCTCCTATTTTATCAGGTATTGTCCNNNAGATCGGAAGA |
| 51084_1 | TATGAGTGTGGAGTCGTTGCCAGGTTCAGGTTTTCCACACTTCNNNAGATCGGAAG |
| 51084_2 | TATGAGTGTGGAGTCGTTGCCAGGTTCAGGTTTTCCACACTTCNNNAGATCGGAAG |
| 44505_1 | TATGAGTGTGGAGTCGTTGCCCTAGGATTACAGGCGTGATTCANNNAGATCGGAA |
| 44505_2 | TATGAGTGTGGAGTCGTTGCCCTAGGATTACAGGCGTGATTCANNNAGATCGGAA |
| 35067_1 | TATGAGTGTGGAGTCGTTGCATTGGCATTTTGGTAGGGATTGCNNNAGATCGGAA |
| 35067_2 | TATGAGTGTGGAGTCGTTGCATTGGCATTTTGGTAGGGATTGCNNNAGATCGGAA |
| 35775_1 | TATGAGTGTGGAGTCGTTGCCTCCCAATTCCAAGTCTCACGTANNNAGATCGGAAGA |
| 35775_2 | TATGAGTGTGGAGTCGTTGCCTCCCAATTCCAAGTCTCACGTANNNAGATCGGAAGA |
| 39466_1 | TATGAGTGTGGAGTCGTTGCTTTTTTGAGACGGAGTCTTGCTGNNNAGATCGGAAG |
| 39466_2 | TATGAGTGTGGAGTCGTTGCTTTTTTGAGACGGAGTCTTGCTGNNNAGATCGGAAG |
| 37344_1 | TATGAGTGTGGAGTCGTTGCCTGCTGGGTGGAGATCACTTTATNNNAGATCGGAA |
| 37344_2 | TATGAGTGTGGAGTCGTTGCCTGCTGGGTGGAGATCACTTTATNNNAGATCGGAA |

| | |
|---|---|
| 409_1 | TATGAGTGTGGAGTCGTTGCTTGGAAATTGGCTAGCTGATTCCNNNAGATCGGAAG |
| 409_2 | TATGAGTGTGGAGTCGTTGCTTGGAAATTGGCTAGCTGATTCCNNNAGATCGGAAG |
| 44484_1 | TATGAGTGTGGAGTCGTTGCCCAGACATAATGGCTGACTCCTANNNAGATCGGAAG |
| 44484_2 | TATGAGTGTGGAGTCGTTGCCCAGACATAATGGCTGACTCCTANNNAGATCGGAAG |
| 47528_1 | TATGAGTGTGGAGTCGTTGCGACAAAGTCTCATTCTGTTGCCCNNNAGATCGGAAG |
| 47528_2 | TATGAGTGTGGAGTCGTTGCGACAAAGTCTCATTCTGTTGCCCNNNAGATCGGAAG |
| 40916_1 | TATGAGTGTGGAGTCGTTGCAGATAGCATGACCACACTCTAGCNNNAGATCGGAAG |
| 40916_2 | TATGAGTGTGGAGTCGTTGCAGATAGCATGACCACACTCTAGCNNNAGATCGGAAG |
| 35428_1 | TATGAGTGTGGAGTCGTTGCCTGCAAACTCAGACCCCTAGATTNNNAGATCGGAAG |
| 35428_2 | TATGAGTGTGGAGTCGTTGCCTGCAAACTCAGACCCCTAGATTNNNAGATCGGAAG |
| 39923_1 | TATGAGTGTGGAGTCGTTGCTCCTGGTCAGTTACTATCCCCAANNNAGATCGGAAG |
| 39923_2 | TATGAGTGTGGAGTCGTTGCTCCTGGTCAGTTACTATCCCCAANNNAGATCGGAAG |
| 42037_1 | TATGAGTGTGGAGTCGTTGCGGCAAAATGAAACAGCAGACTCTNNNAGATCGGAA |
| 42037_2 | TATGAGTGTGGAGTCGTTGCGGCAAAATGAAACAGCAGACTCTNNNAGATCGGAA |
| 49488_1 | TATGAGTGTGGAGTCGTTGCTCTACACAGAACTCAATGGCCTCNNNAGATCGGAAG |
| 49488_2 | TATGAGTGTGGAGTCGTTGCTCTACACAGAACTCAATGGCCTCNNNAGATCGGAAG |
| 46813_1 | TATGAGTGTGGAGTCGTTGCATTCACACCATTCTCCTGCCTCNNNAGATCGGAAGAG |
| 46813_2 | TATGAGTGTGGAGTCGTTGCATTCACACCATTCTCCTGCCTCNNNAGATCGGAAGAG |
| 48448_1 | TATGAGTGTGGAGTCGTTGCGGTGACGCTTTTCCATGGATCTANNNAGATCGGAAG |
| 48448_2 | TATGAGTGTGGAGTCGTTGCGGTGACGCTTTTCCATGGATCTANNNAGATCGGAAG |
| 33005_1 | TATGAGTGTGGAGTCGTTGCTTCAAGCAATTCTCCTGCCTCAGNNNAGATCGGAAG |
| 33005_2 | TATGAGTGTGGAGTCGTTGCTTCAAGCAATTCTCCTGCCTCAGNNNAGATCGGAAG |
| 38746_1 | TATGAGTGTGGAGTCGTTGCCATTGGCACAATCTCAGCTCACNNNAGATCGGAAGA |
| 38746_2 | TATGAGTGTGGAGTCGTTGCCATTGGCACAATCTCAGCTCACNNNAGATCGGAAGA |
| 52681_1 | TATGAGTGTGGAGTCGTTGCCAACAGCTAGAAACAAGAAGCCCNNNAGATCGGAA |
| 52681_2 | TATGAGTGTGGAGTCGTTGCCAACAGCTAGAAACAAGAAGCCCNNNAGATCGGAA |
| 44776_1 | TATGAGTGTGGAGTCGTTGCATGGGAACATAGAGCTGCAGAAANNNAGATCGGAA |
| 44776_2 | TATGAGTGTGGAGTCGTTGCATGGGAACATAGAGCTGCAGAAANNNAGATCGGAA |
| 2137_1 | TATGAGTGTGGAGTCGTTGCGTGACCCTTATGAAAGCAGCTCANNNAGATCGGAAG |
| 2137_2 | TATGAGTGTGGAGTCGTTGCGTGACCCTTATGAAAGCAGCTCANNNAGATCGGAAG |
| 48156_1 | TATGAGTGTGGAGTCGTTGCTCTTGAACTCCTGACCTCAAGTGNNNAGATCGGAAG |
| 48156_2 | TATGAGTGTGGAGTCGTTGCTCTTGAACTCCTGACCTCAAGTGNNNAGATCGGAAG |
| 48678_1 | TATGAGTGTGGAGTCGTTGCACATTTGCAGGGAATATCTGGGTNNNAGATCGGAA |
| 48678_2 | TATGAGTGTGGAGTCGTTGCACATTTGCAGGGAATATCTGGGTNNNAGATCGGAA |
| 39858_1 | TATGAGTGTGGAGTCGTTGCTGCAGACCCAATGTGTAAAATGGNNNAGATCGGAA |
| 39858_2 | TATGAGTGTGGAGTCGTTGCTGCAGACCCAATGTGTAAAATGGNNNAGATCGGAA |
| 41199_1 | TATGAGTGTGGAGTCGTTGCCACTCCCGACCCCCAAATTAGNNNAGATCGGAAGAG |
| 41199_2 | TATGAGTGTGGAGTCGTTGCCACTCCCGACCCCCAAATTAGNNNAGATCGGAAGAG |
| 51169_1 | TATGAGTGTGGAGTCGTTGCCCTTTTCACCTCTGGGAGTACAGNNNAGATCGGAAG |
| 51169_2 | TATGAGTGTGGAGTCGTTGCCCTTTTCACCTCTGGGAGTACAGNNNAGATCGGAAG |
| 45306_1 | TATGAGTGTGGAGTCGTTGCACCACAGTAGCAGCAATAACTGANNNAGATCGGAA |
| 45306_2 | TATGAGTGTGGAGTCGTTGCACCACAGTAGCAGCAATAACTGANNNAGATCGGAA |
| 37328_1 | TATGAGTGTGGAGTCGTTGCGACTGAGACAGGAGGATCACTTGNNNAGATCGGAA |

| | |
|---|---|
| 37328_2 | TATGAGTGTGGAGTCGTTGCGACTGAGACAGGAGGATCACTTGNNNAGATCGGAA |
| 36012_1 | TATGAGTGTGGAGTCGTTGCTGAGGTAGCAAACTGCCTAGAAGNNNAGATCGGAA |
| 36012_2 | TATGAGTGTGGAGTCGTTGCTGAGGTAGCAAACTGCCTAGAAGNNNAGATCGGAA |
| 53160_1 | TATGAGTGTGGAGTCGTTGCACTGAATTCATCTTGCCAGTTGGNNNAGATCGGAAG |
| 53160_2 | TATGAGTGTGGAGTCGTTGCACTGAATTCATCTTGCCAGTTGGNNNAGATCGGAAG |
| 32598_1 | TATGAGTGTGGAGTCGTTGCTGCTGGTAAGGATGTGGAGAAAANNNAGATCGGA |
| 32598_2 | TATGAGTGTGGAGTCGTTGCTGCTGGTAAGGATGTGGAGAAAANNNAGATCGGA |
| 36661_1 | TATGAGTGTGGAGTCGTTGCGACTGCTCAAAGGGAAATGCAAANNNAGATCGGAA |
| 36661_2 | TATGAGTGTGGAGTCGTTGCGACTGCTCAAAGGGAAATGCAAANNNAGATCGGAA |
| 46546_1 | TATGAGTGTGGAGTCGTTGCGCTTTGCTTGCTCTGTCATCCNNNAGATCGGAAGAG |
| 46546_2 | TATGAGTGTGGAGTCGTTGCGCTTTGCTTGCTCTGTCATCCNNNAGATCGGAAGAG |
| 46912_1 | TATGAGTGTGGAGTCGTTGCATTACAGACGTGAGCTTACCCTGNNNAGATCGGAAG |
| 46912_2 | TATGAGTGTGGAGTCGTTGCATTACAGACGTGAGCTTACCCTGNNNAGATCGGAAG |
| 37022_1 | TATGAGTGTGGAGTCGTTGCCCAGAGCACTGTAACATGGGATNNNAGATCGGAAG |
| 37022_2 | TATGAGTGTGGAGTCGTTGCCCAGAGCACTGTAACATGGGATNNNAGATCGGAAG |
| 52760_1 | TATGAGTGTGGAGTCGTTGCGAGCAGAATCAAAGAGGGCATTGNNNAGATCGGA |
| 52760_2 | TATGAGTGTGGAGTCGTTGCGAGCAGAATCAAAGAGGGCATTGNNNAGATCGGA |
| 44079_1 | TATGAGTGTGGAGTCGTTGCACTGGAGGATAAAGAGGTTGCTGNNNAGATCGGA |
| 44079_2 | TATGAGTGTGGAGTCGTTGCACTGGAGGATAAAGAGGTTGCTGNNNAGATCGGA |
| 45620_1 | TATGAGTGTGGAGTCGTTGCGATCTCGGCTCACTGTAAGCTCNNNAGATCGGAAGA |
| 45620_2 | TATGAGTGTGGAGTCGTTGCGATCTCGGCTCACTGTAAGCTCNNNAGATCGGAAGA |
| 10606_1 | TATGAGTGTGGAGTCGTTGCAGATCCTGTTGTGTGAAGGTGTTNNNAGATCGGAA |
| 10606_2 | TATGAGTGTGGAGTCGTTGCAGATCCTGTTGTGTGAAGGTGTTNNNAGATCGGAA |
| 46591_1 | TATGAGTGTGGAGTCGTTGCCTCAATTTGGGTTTGCCTGATGTNNNAGATCGGAAG |
| 46591_2 | TATGAGTGTGGAGTCGTTGCCTCAATTTGGGTTTGCCTGATGTNNNAGATCGGAAG |
| 12662_1 | TATGAGTGTGGAGTCGTTGCGTATTAGTGGAAGACTTTAAGAANNNAGATCGGAA |
| 12662_2 | TATGAGTGTGGAGTCGTTGCGTATTAGTGGAAGACTTTAAGAANNNAGATCGGAA |
| 43546_1 | TATGAGTGTGGAGTCGTTGCGGATTACAGATGTGAGCCACAGTNNNAGATCGGAA |
| 43546_2 | TATGAGTGTGGAGTCGTTGCGGATTACAGATGTGAGCCACAGTNNNAGATCGGAA |
| 12758_1 | TATGAGTGTGGAGTCGTTGCAAATCTGTAGGATCAACATAGACNNNAGATCGGAAG |
| 12758_2 | TATGAGTGTGGAGTCGTTGCAAATCTGTAGGATCAACATAGACNNNAGATCGGAAG |
| 32109_1 | TATGAGTGTGGAGTCGTTGCCTCCCAGGTTCAGGTGATTCTCNNNAGATCGGAAGA |
| 32109_2 | TATGAGTGTGGAGTCGTTGCCTCCCAGGTTCAGGTGATTCTCNNNAGATCGGAAGA |
| 33279_1 | TATGAGTGTGGAGTCGTTGCGATCTCAGCTCACTACAACCTCCNNNAGATCGGAAG |
| 33279_2 | TATGAGTGTGGAGTCGTTGCGATCTCAGCTCACTACAACCTCCNNNAGATCGGAAG |
| 31058_1 | TATGAGTGTGGAGTCGTTGCTACCTTTAGACAAGGGTGCTTCCNNNAGATCGGAAG |
| 31058_2 | TATGAGTGTGGAGTCGTTGCTACCTTTAGACAAGGGTGCTTCCNNNAGATCGGAAG |
| 52828_1 | TATGAGTGTGGAGTCGTTGCTTTGATTCCTCCTCAGTCTGTGGNNNAGATCGGAAG |
| 52828_2 | TATGAGTGTGGAGTCGTTGCTTTGATTCCTCCTCAGTCTGTGGNNNAGATCGGAAG |
| 12664_1 | TATGAGTGTGGAGTCGTTGCTGAGATAATAAATTATGGTGTAAGTTNNNAGATCGG |
| 12664_2 | TATGAGTGTGGAGTCGTTGCTGAGATAATAAATTATGGTGTAAGTTNNNAGATCGG |
| 36713_1 | TATGAGTGTGGAGTCGTTGCCACATAATGAGTGTCTGGGTGCTNNNAGATCGGAA |
| 36713_2 | TATGAGTGTGGAGTCGTTGCCACATAATGAGTGTCTGGGTGCTNNNAGATCGGAA |

| | |
|---|---|
| 36131_1 | TATGAGTGTGGAGTCGTTGCGTCTCCTCTCATGTTCGCACTAANNNAGATCGGAAG |
| 36131_2 | TATGAGTGTGGAGTCGTTGCGTCTCCTCTCATGTTCGCACTAANNNAGATCGGAAG |
| 51704_1 | TATGAGTGTGGAGTCGTTGCCTAAGCCAAGTGATTTGAGCGTCNNNAGATCGGAAG |
| 51704_2 | TATGAGTGTGGAGTCGTTGCCTAAGCCAAGTGATTTGAGCGTCNNNAGATCGGAAG |
| 35403_1 | TATGAGTGTGGAGTCGTTGCCTAAAAGCACAAACCACCATCCCNNNAGATCGGAAG |
| 35403_2 | TATGAGTGTGGAGTCGTTGCCTAAAAGCACAAACCACCATCCCNNNAGATCGGAAG |
| 49561_1 | TATGAGTGTGGAGTCGTTGCCAATTCTTGTGCCTCAGCCTTAGNNNAGATCGGAAG |
| 49561_2 | TATGAGTGTGGAGTCGTTGCCAATTCTTGTGCCTCAGCCTTAGNNNAGATCGGAAG |
| 43561_1 | TATGAGTGTGGAGTCGTTGCCTGGAGTACAATGGCACGATCTCNNNAGATCGGAAG |
| 43561_2 | TATGAGTGTGGAGTCGTTGCCTGGAGTACAATGGCACGATCTCNNNAGATCGGAAG |
| 41385_1 | TATGAGTGTGGAGTCGTTGCCAAGAAATGCATATAAAGGCTGATCANNNAGATCGG |
| 41385_2 | TATGAGTGTGGAGTCGTTGCCAAGAAATGCATATAAAGGCTGATCANNNAGATCGG |
| 38883_1 | TATGAGTGTGGAGTCGTTGCGACAGAGCAAGACTCCATCTCAANNNAGATCGGAAG |
| 38883_2 | TATGAGTGTGGAGTCGTTGCGACAGAGCAAGACTCCATCTCAANNNAGATCGGAAG |
| 32831_1 | TATGAGTGTGGAGTCGTTGCAAACTCCTGACCTCAAGTGATCCNNNAGATCGGAAG |
| 32831_2 | TATGAGTGTGGAGTCGTTGCAAACTCCTGACCTCAAGTGATCCNNNAGATCGGAAG |
| 34678_1 | TATGAGTGTGGAGTCGTTGCCCTCAGCTGGGATTACAGGTGNNNAGATCGGAAGA |
| 34678_2 | TATGAGTGTGGAGTCGTTGCCCTCAGCTGGGATTACAGGTGNNNAGATCGGAAGA |
| 33738_1 | TATGAGTGTGGAGTCGTTGCCTGAGGCAGGAGAATAGCTTGAANNNAGATCGGAA |
| 33738_2 | TATGAGTGTGGAGTCGTTGCCTGAGGCAGGAGAATAGCTTGAANNNAGATCGGAA |
| 45822_1 | TATGAGTGTGGAGTCGTTGCGTAGCTGGGAATACAGGCATGTANNNAGATCGGAA |
| 45822_2 | TATGAGTGTGGAGTCGTTGCGTAGCTGGGAATACAGGCATGTANNNAGATCGGAA |
| 34125_1 | TATGAGTGTGGAGTCGTTGCAGCCTCACAAATCCTTTTTACCANNNAGATCGGAAGA |
| 34125_2 | TATGAGTGTGGAGTCGTTGCAGCCTCACAAATCCTTTTTACCANNNAGATCGGAAGA |
| 42021_1 | TATGAGTGTGGAGTCGTTGCGGGGGACCCTAAATGCATCTTTANNNAGATCGGAAG |
| 42021_2 | TATGAGTGTGGAGTCGTTGCGGGGGACCCTAAATGCATCTTTANNNAGATCGGAAG |
| 38319_1 | TATGAGTGTGGAGTCGTTGCAGACACTGCATGTTCCCACTTATNNNAGATCGGAAG |
| 38319_2 | TATGAGTGTGGAGTCGTTGCAGACACTGCATGTTCCCACTTATNNNAGATCGGAAG |
| 35773_1 | TATGAGTGTGGAGTCGTTGCAGTGTCATAAGAGAACCCAGCAGNNNAGATCGGAA |
| 35773_2 | TATGAGTGTGGAGTCGTTGCAGTGTCATAAGAGAACCCAGCAGNNNAGATCGGAA |
| 38201_1 | TATGAGTGTGGAGTCGTTGCCTTGCAGTGAGCCAAGATCACNNNAGATCGGAAGA |
| 38201_2 | TATGAGTGTGGAGTCGTTGCCTTGCAGTGAGCCAAGATCACNNNAGATCGGAAGA |
| 31159_1 | TATGAGTGTGGAGTCGTTGCTTACAAAATCCCGAGACTGTTGCNNNAGATCGGAAG |
| 31159_2 | TATGAGTGTGGAGTCGTTGCTTACAAAATCCCGAGACTGTTGCNNNAGATCGGAAG |
| 39959_1 | TATGAGTGTGGAGTCGTTGCAAAGAGGTTTGTTTGGCTCATGGNNNAGATCGGAA |
| 39959_2 | TATGAGTGTGGAGTCGTTGCAAAGAGGTTTGTTTGGCTCATGGNNNAGATCGGAA |
| 51113_1 | TATGAGTGTGGAGTCGTTGCGCACTTTCTTTCTCTGGGACTTCNNNAGATCGGAAGA |
| 51113_2 | TATGAGTGTGGAGTCGTTGCGCACTTTCTTTCTCTGGGACTTCNNNAGATCGGAAGA |
| 45764_1 | TATGAGTGTGGAGTCGTTGCCAAGTAGCTGGGATTACAGGTGTNNNAGATCGGAA |
| 45764_2 | TATGAGTGTGGAGTCGTTGCCAAGTAGCTGGGATTACAGGTGTNNNAGATCGGAA |
| 51292_1 | TATGAGTGTGGAGTCGTTGCTCCTACTGAACTGAAGGCTGAACNNNAGATCGGAAG |
| 51292_2 | TATGAGTGTGGAGTCGTTGCTCCTACTGAACTGAAGGCTGAACNNNAGATCGGAAG |
| 13415_1 | TATGAGTGTGGAGTCGTTGCGGCATATCTCAAAAACTGATTTNNNAGATCGGAAGA |

| | |
|---|---|
| 13415_2 | TATGAGTGTGGAGTCGTTGCGGCATATCTCAAAAACTGATTTNNNAGATCGGAAGA |
| 45340_1 | TATGAGTGTGGAGTCGTTGCCAGTTCATGTTCCCGCTTACTTCNNNAGATCGGAAGA |
| 45340_2 | TATGAGTGTGGAGTCGTTGCCAGTTCATGTTCCCGCTTACTTCNNNAGATCGGAAGA |
| 12270_1 | TATGAGTGTGGAGTCGTTGCAAGAAAGACAACTCAGAATGGGAGNNNAGATCGGA |
| 12270_2 | TATGAGTGTGGAGTCGTTGCAAGAAAGACAACTCAGAATGGGAGNNNAGATCGGA |
| 35991_1 | TATGAGTGTGGAGTCGTTGCATCAGAGAGGATATTGGGGAGCANNNAGATCGGAA |
| 35991_2 | TATGAGTGTGGAGTCGTTGCATCAGAGAGGATATTGGGGAGCANNNAGATCGGAA |
| 43369_1 | TATGAGTGTGGAGTCGTTGCCAATGGCAAGATCTCAGCTCACTNNNAGATCGGAAG |
| 43369_2 | TATGAGTGTGGAGTCGTTGCCAATGGCAAGATCTCAGCTCACTNNNAGATCGGAAG |
| 52696_1 | TATGAGTGTGGAGTCGTTGCGGCCTCTGTAGCATTTGACTTTGNNNAGATCGGAAG |
| 52696_2 | TATGAGTGTGGAGTCGTTGCGGCCTCTGTAGCATTTGACTTTGNNNAGATCGGAAG |
| 45552_1 | TATGAGTGTGGAGTCGTTGCCCCTTTCCCTGAGAATGTCTTCTNNNAGATCGGAAGA |
| 45552_2 | TATGAGTGTGGAGTCGTTGCCCCTTTCCCTGAGAATGTCTTCTNNNAGATCGGAAGA |
| 37910_1 | TATGAGTGTGGAGTCGTTGCCTGTTGTCCCCAGCTACTCAGNNNAGATCGGAAGAG |
| 37910_2 | TATGAGTGTGGAGTCGTTGCCTGTTGTCCCCAGCTACTCAGNNNAGATCGGAAGAG |
| 42770_1 | TATGAGTGTGGAGTCGTTGCCGTTAGGTGTTGACTATTGCCCANNNAGATCGGAAG |
| 42770_2 | TATGAGTGTGGAGTCGTTGCCGTTAGGTGTTGACTATTGCCCANNNAGATCGGAAG |
| 36201_1 | TATGAGTGTGGAGTCGTTGCTCCCAGCTACTTGAAAGACTGAGNNNAGATCGGAAG |
| 36201_2 | TATGAGTGTGGAGTCGTTGCTCCCAGCTACTTGAAAGACTGAGNNNAGATCGGAAG |
| 51552_1 | TATGAGTGTGGAGTCGTTGCGGATTAGCAGAAGTGCCTGTTTCNNNAGATCGGAAG |
| 51552_2 | TATGAGTGTGGAGTCGTTGCGGATTAGCAGAAGTGCCTGTTTCNNNAGATCGGAAG |
| 50836_1 | TATGAGTGTGGAGTCGTTGCCACCGCTTTGATCATATGCTTGTNNNAGATCGGAAGA |
| 50836_2 | TATGAGTGTGGAGTCGTTGCCACCGCTTTGATCATATGCTTGTNNNAGATCGGAAGA |
| 41792_1 | TATGAGTGTGGAGTCGTTGCCCACAACATCTGGCTGACTTTTTNNNAGATCGGAAGA |
| 41792_2 | TATGAGTGTGGAGTCGTTGCCCACAACATCTGGCTGACTTTTTNNNAGATCGGAAGA |
| 45109_1 | TATGAGTGTGGAGTCGTTGCAGAGCTACGTGGAAATTGAACCTNNNAGATCGGAAG |
| 45109_2 | TATGAGTGTGGAGTCGTTGCAGAGCTACGTGGAAATTGAACCTNNNAGATCGGAAG |
| 44251_1 | TATGAGTGTGGAGTCGTTGCCTCCCAAAAATTCTCAAACCCCCNNNAGATCGGAAGA |
| 44251_2 | TATGAGTGTGGAGTCGTTGCCTCCCAAAAATTCTCAAACCCCCNNNAGATCGGAAGA |
| 52132_1 | TATGAGTGTGGAGTCGTTGCGACCCAGTAGCATGTTGTCTACANNNAGATCGGAAG |
| 52132_2 | TATGAGTGTGGAGTCGTTGCGACCCAGTAGCATGTTGTCTACANNNAGATCGGAAG |
| 50633_1 | TATGAGTGTGGAGTCGTTGCGATGCTGTCAGTCACTAGCTCTTNNNAGATCGGAAGA |
| 50633_2 | TATGAGTGTGGAGTCGTTGCGATGCTGTCAGTCACTAGCTCTTNNNAGATCGGAAGA |
| 38180_1 | TATGAGTGTGGAGTCGTTGCAGAAAAAAACCAGTGACTGCCAGNNNAGATCGGAAG |
| 38180_2 | TATGAGTGTGGAGTCGTTGCAGAAAAAAACCAGTGACTGCCAGNNNAGATCGGAAG |
| 47062_1 | TATGAGTGTGGAGTCGTTGCAGACAGGGTCTCATTTTGTCACTNNNAGATCGGAAG |
| 47062_2 | TATGAGTGTGGAGTCGTTGCAGACAGGGTCTCATTTTGTCACTNNNAGATCGGAAG |
| 36555_1 | TATGAGTGTGGAGTCGTTGCCTCAAGAGTTTTTTGCCTAGGCCNNNAGATCGGAAGA |
| 36555_2 | TATGAGTGTGGAGTCGTTGCCTCAAGAGTTTTTTGCCTAGGCCNNNAGATCGGAAGA |
| 49750_1 | TATGAGTGTGGAGTCGTTGCAGGAAAAAGCTCACTACCCTCTGNNNAGATCGGAAG |
| 49750_2 | TATGAGTGTGGAGTCGTTGCAGGAAAAAGCTCACTACCCTCTGNNNAGATCGGAAG |
| 39421_1 | TATGAGTGTGGAGTCGTTGCAGCCTTTCTAAGAGTGAGCTCAGNNNAGATCGGAAG |
| 39421_2 | TATGAGTGTGGAGTCGTTGCAGCCTTTCTAAGAGTGAGCTCAGNNNAGATCGGAAG |

| | |
|---|---|
| 40469_1 | TATGAGTGTGGAGTCGTTGCTGTGGTATATCCTCACAATGCAANNNAGATCGGAAG |
| 40469_2 | TATGAGTGTGGAGTCGTTGCTGTGGTATATCCTCACAATGCAANNNAGATCGGAAG |
| 46928_1 | TATGAGTGTGGAGTCGTTGCCTTCCGAGTAGCTGGGATTACAGNNNAGATCGGAA |
| 46928_2 | TATGAGTGTGGAGTCGTTGCCTTCCGAGTAGCTGGGATTACAGNNNAGATCGGAA |
| 34945_1 | TATGAGTGTGGAGTCGTTGCGATGGAGTCCTACTCTGTTGTCCNNNAGATCGGAAG |
| 34945_2 | TATGAGTGTGGAGTCGTTGCGATGGAGTCCTACTCTGTTGTCCNNNAGATCGGAAG |
| 33518_1 | TATGAGTGTGGAGTCGTTGCACACACACACATATATCAGCTGANNNAGATCGGAAG |
| 33518_2 | TATGAGTGTGGAGTCGTTGCACACACACACATATATCAGCTGANNNAGATCGGAAG |
| 10378_1 | TATGAGTGTGGAGTCGTTGCCAGTGTTGAGTGGACTTGGTGAANNNAGATCGGAA |
| 10378_2 | TATGAGTGTGGAGTCGTTGCCAGTGTTGAGTGGACTTGGTGAANNNAGATCGGAA |
| 51246_1 | TATGAGTGTGGAGTCGTTGCAAGTCCTTTCAGACACCACATGTNNNAGATCGGAAG |
| 51246_2 | TATGAGTGTGGAGTCGTTGCAAGTCCTTTCAGACACCACATGTNNNAGATCGGAAG |
| 40884_1 | TATGAGTGTGGAGTCGTTGCAACAAGATCTCACTCTGTCACCCNNNAGATCGGAAG |
| 40884_2 | TATGAGTGTGGAGTCGTTGCAACAAGATCTCACTCTGTCACCCNNNAGATCGGAAG |
| 34344_1 | TATGAGTGTGGAGTCGTTGCCTCTCGGGGACCTGTTGATGNNNAGATCGGAAGAG |
| 34344_2 | TATGAGTGTGGAGTCGTTGCCTCTCGGGGACCTGTTGATGNNNAGATCGGAAGAG |
| 50582_1 | TATGAGTGTGGAGTCGTTGCTAGAAGCTGAGGATGGTGTTTCCNNNAGATCGGAA |
| 50582_2 | TATGAGTGTGGAGTCGTTGCTAGAAGCTGAGGATGGTGTTTCCNNNAGATCGGAA |
| 43991_1 | TATGAGTGTGGAGTCGTTGCAGAGAGAGTGTTTGTGTGTGTGTNNNAGATCGGAA |
| 43991_2 | TATGAGTGTGGAGTCGTTGCAGAGAGAGTGTTTGTGTGTGTGTNNNAGATCGGAA |
| 33806_1 | TATGAGTGTGGAGTCGTTGCGAAACTGGAAGACTGGTGATCCTNNNAGATCGGAA |
| 33806_2 | TATGAGTGTGGAGTCGTTGCGAAACTGGAAGACTGGTGATCCTNNNAGATCGGAA |
| 42580_1 | TATGAGTGTGGAGTCGTTGCTCCTTCTTTACTGCTTTCTTTTGCANNNAGATCGGAAG |
| 42580_2 | TATGAGTGTGGAGTCGTTGCTCCTTCTTTACTGCTTTCTTTTGCANNNAGATCGGAAG |
| 32073_1 | TATGAGTGTGGAGTCGTTGCGATTTTGAACTCCTGGCCTCAAGNNNAGATCGGAAG |
| 32073_2 | TATGAGTGTGGAGTCGTTGCGATTTTGAACTCCTGGCCTCAAGNNNAGATCGGAAG |
| 38786_1 | TATGAGTGTGGAGTCGTTGCCCTCCTAGGTTCAAGCGATTCTCNNNAGATCGGAAG |
| 38786_2 | TATGAGTGTGGAGTCGTTGCCCTCCTAGGTTCAAGCGATTCTCNNNAGATCGGAAG |
| 2811_1 | TATGAGTGTGGAGTCGTTGCAAAGTCTGAAGTTTGGCCAGTCTNNNAGATCGGAAG |
| 2811_2 | TATGAGTGTGGAGTCGTTGCAAAGTCTGAAGTTTGGCCAGTCTNNNAGATCGGAAG |
| 51681_1 | TATGAGTGTGGAGTCGTTGCTACTTAGCCAATGGGTAAGGCAGNNNAGATCGGAA |
| 51681_2 | TATGAGTGTGGAGTCGTTGCTACTTAGCCAATGGGTAAGGCAGNNNAGATCGGAA |
| 50919_1 | TATGAGTGTGGAGTCGTTGCCCTGGAATCCAAAAGAAAGTGCANNNAGATCGGAA |
| 50919_2 | TATGAGTGTGGAGTCGTTGCCCTGGAATCCAAAAGAAAGTGCANNNAGATCGGAA |
| 46362_1 | TATGAGTGTGGAGTCGTTGCGAGTAGCTGAGACTACAGGTGTGNNNAGATCGGAA |
| 46362_2 | TATGAGTGTGGAGTCGTTGCGAGTAGCTGAGACTACAGGTGTGNNNAGATCGGAA |
| 31441_1 | TATGAGTGTGGAGTCGTTGCAGAACAGAGACTGTACCTCTCCANNNAGATCGGAAG |
| 31441_2 | TATGAGTGTGGAGTCGTTGCAGAACAGAGACTGTACCTCTCCANNNAGATCGGAAG |
| 39995_1 | TATGAGTGTGGAGTCGTTGCTTAGTCAGGTATGGTGGCATGTGNNNAGATCGGAA |
| 39995_2 | TATGAGTGTGGAGTCGTTGCTTAGTCAGGTATGGTGGCATGTGNNNAGATCGGAA |
| 30390_1 | TATGAGTGTGGAGTCGTTGCCTCAGATGGAGTTTCGCTCTTCCNNNAGATCGGAAG |
| 30390_2 | TATGAGTGTGGAGTCGTTGCCTCAGATGGAGTTTCGCTCTTCCNNNAGATCGGAAG |
| 40190_1 | TATGAGTGTGGAGTCGTTGCTATAAGATGATCATGGGGCCAGGNNNAGATCGGAA |

| | |
|---|---|
| 40190_2 | TATGAGTGTGGAGTCGTTGCTATAAGATGATCATGGGGCCAGGNNNAGATCGGAA |
| 47558_1 | TATGAGTGTGGAGTCGTTGCCCATCACATCCAGCTAGCACTTANNNAGATCGGAAG |
| 47558_2 | TATGAGTGTGGAGTCGTTGCCCATCACATCCAGCTAGCACTTANNNAGATCGGAAG |
| 41001_1 | TATGAGTGTGGAGTCGTTGCAAAGAAAAACCCAGGAATTCCAGNNNAGATCGGAA |
| 41001_2 | TATGAGTGTGGAGTCGTTGCAAAGAAAAACCCAGGAATTCCAGNNNAGATCGGAA |
| 41795_1 | TATGAGTGTGGAGTCGTTGCTTCACTCCATTCTGCTGTCTCAGNNNAGATCGGAAGA |
| 41795_2 | TATGAGTGTGGAGTCGTTGCTTCACTCCATTCTGCTGTCTCAGNNNAGATCGGAAGA |
| 49808_1 | TATGAGTGTGGAGTCGTTGCGGGTTCACTTGAATGGCAAATCANNNAGATCGGAA |
| 49808_2 | TATGAGTGTGGAGTCGTTGCGGGTTCACTTGAATGGCAAATCANNNAGATCGGAA |
| 53027_1 | TATGAGTGTGGAGTCGTTGCGACCCAGCACAGAAAAGTGTAACNNNAGATCGGAA |
| 53027_2 | TATGAGTGTGGAGTCGTTGCGACCCAGCACAGAAAAGTGTAACNNNAGATCGGAA |
| 46233_1 | TATGAGTGTGGAGTCGTTGCATTCCTTGCCATATGGACCTCTCNNNAGATCGGAAG |
| 46233_2 | TATGAGTGTGGAGTCGTTGCATTCCTTGCCATATGGACCTCTCNNNAGATCGGAAG |
| 13540_1 | TATGAGTGTGGAGTCGTTGCGAATATAGACATTTCCCACTCTANNNAGATCGGAAG |
| 13540_2 | TATGAGTGTGGAGTCGTTGCGAATATAGACATTTCCCACTCTANNNAGATCGGAAG |
| 32113_1 | TATGAGTGTGGAGTCGTTGCGAGAATCTCTTGAACCTGGGAGGNNNAGATCGGAA |
| 32113_2 | TATGAGTGTGGAGTCGTTGCGAGAATCTCTTGAACCTGGGAGGNNNAGATCGGAA |
| 42389_1 | TATGAGTGTGGAGTCGTTGCATTTTGTGTGTGTGTGTGTGTGTNNNAGATCGGAA |
| 42389_2 | TATGAGTGTGGAGTCGTTGCATTTTGTGTGTGTGTGTGTGTGTNNNAGATCGGAA |
| 39155_1 | TATGAGTGTGGAGTCGTTGCGGCCCATCTAACAAGAATTGCAANNNAGATCGGAAG |
| 39155_2 | TATGAGTGTGGAGTCGTTGCGGCCCATCTAACAAGAATTGCAANNNAGATCGGAAG |
| 49444_1 | TATGAGTGTGGAGTCGTTGCCAGGTTCAAGCAATTCTCCTGCNNNAGATCGGAAGA |
| 49444_2 | TATGAGTGTGGAGTCGTTGCCAGGTTCAAGCAATTCTCCTGCNNNAGATCGGAAGA |
| 10533_1 | TATGAGTGTGGAGTCGTTGCGCTTTCTCTCTTCCATGGGGTTTNNNAGATCGGAAG |
| 10533_2 | TATGAGTGTGGAGTCGTTGCGCTTTCTCTCTTCCATGGGGTTTNNNAGATCGGAAG |
| 51320_1 | TATGAGTGTGGAGTCGTTGCACCAGCATGGCACATGTATACATNNNAGATCGGAAG |
| 51320_2 | TATGAGTGTGGAGTCGTTGCACCAGCATGGCACATGTATACATNNNAGATCGGAAG |
| 35206_1 | TATGAGTGTGGAGTCGTTGCGACATTGCCAGATCATCATGGTGNNNAGATCGGAA |
| 35206_2 | TATGAGTGTGGAGTCGTTGCGACATTGCCAGATCATCATGGTGNNNAGATCGGAA |
| 38212_1 | TATGAGTGTGGAGTCGTTGCGGGTTTGGAAATAAGAGTCTGCGNNNAGATCGGAA |
| 38212_2 | TATGAGTGTGGAGTCGTTGCGGGTTTGGAAATAAGAGTCTGCGNNNAGATCGGAA |
| 44557_1 | TATGAGTGTGGAGTCGTTGCTCACACTGCTTTCGTAGGATCAANNNAGATCGGAAG |
| 44557_2 | TATGAGTGTGGAGTCGTTGCTCACACTGCTTTCGTAGGATCAANNNAGATCGGAAG |
| 34653_1 | TATGAGTGTGGAGTCGTTGCGCCTCAGTCTTTTCTGGGAAGTANNNAGATCGGAAG |
| 34653_2 | TATGAGTGTGGAGTCGTTGCGCCTCAGTCTTTTCTGGGAAGTANNNAGATCGGAAG |
| 42121_1 | TATGAGTGTGGAGTCGTTGCTCCACTTCACAGGTTCAAGTGATNNNAGATCGGAAG |
| 42121_2 | TATGAGTGTGGAGTCGTTGCTCCACTTCACAGGTTCAAGTGATNNNAGATCGGAAG |
| 45473_1 | TATGAGTGTGGAGTCGTTGCATTTGTGAGGGTAAGGGTATGGGNNNAGATCGGA |
| 45473_2 | TATGAGTGTGGAGTCGTTGCATTTGTGAGGGTAAGGGTATGGGNNNAGATCGGA |
| 46434_1 | TATGAGTGTGGAGTCGTTGCTCAAACTCCTAGGCTCAAGCAATNNNAGATCGGAAG |
| 46434_2 | TATGAGTGTGGAGTCGTTGCTCAAACTCCTAGGCTCAAGCAATNNNAGATCGGAAG |
| 40103_1 | TATGAGTGTGGAGTCGTTGCAAGGATCAGAGATTGGACCATGTNNNAGATCGGAA |
| 40103_2 | TATGAGTGTGGAGTCGTTGCAAGGATCAGAGATTGGACCATGTNNNAGATCGGAA |

| | |
|---|---|
| 36181_1 | TATGAGTGTGGAGTCGTTGCTGTAATCCCAGCTACTCAGGAGANNNAGATCGGAAG |
| 36181_2 | TATGAGTGTGGAGTCGTTGCTGTAATCCCAGCTACTCAGGAGANNNAGATCGGAAG |
| 10388_1 | TATGAGTGTGGAGTCGTTGCATCCGACAATCTAGCAGACACTTNNNAGATCGGAAG |
| 10388_2 | TATGAGTGTGGAGTCGTTGCATCCGACAATCTAGCAGACACTTNNNAGATCGGAAG |
| 45779_1 | TATGAGTGTGGAGTCGTTGCAACGAGCAGGTATGGTTGGTTATNNNAGATCGGAA |
| 45779_2 | TATGAGTGTGGAGTCGTTGCAACGAGCAGGTATGGTTGGTTATNNNAGATCGGAA |
| 52180_1 | TATGAGTGTGGAGTCGTTGCGACTCTCTTTGGCCTCCGTTCNNNAGATCGGAAGAG |
| 52180_2 | TATGAGTGTGGAGTCGTTGCGACTCTCTTTGGCCTCCGTTCNNNAGATCGGAAGAG |
| 31748_1 | TATGAGTGTGGAGTCGTTGCAGTATACCAACACCTGTGCTCAGNNNAGATCGGAAG |
| 31748_2 | TATGAGTGTGGAGTCGTTGCAGTATACCAACACCTGTGCTCAGNNNAGATCGGAAG |
| 35509_1 | TATGAGTGTGGAGTCGTTGCCTCACTGCAAGCTCCTCTTCCNNNAGATCGGAAGAG |
| 35509_2 | TATGAGTGTGGAGTCGTTGCCTCACTGCAAGCTCCTCTTCCNNNAGATCGGAAGAG |
| 34134_1 | TATGAGTGTGGAGTCGTTGCAAGGATAGGAGTTTGAGACCAGCNNNAGATCGGAA |
| 34134_2 | TATGAGTGTGGAGTCGTTGCAAGGATAGGAGTTTGAGACCAGCNNNAGATCGGAA |
| 39419_1 | TATGAGTGTGGAGTCGTTGCACGTCTGTAATCCCAGCACTTAGNNNAGATCGGAAG |
| 39419_2 | TATGAGTGTGGAGTCGTTGCACGTCTGTAATCCCAGCACTTAGNNNAGATCGGAAG |
| 34703_1 | TATGAGTGTGGAGTCGTTGCTTCAAGCGATTCTTCTCCCTCAGNNNAGATCGGAAGA |
| 34703_2 | TATGAGTGTGGAGTCGTTGCTTCAAGCGATTCTTCTCCCTCAGNNNAGATCGGAAGA |
| 29922_1 | TATGAGTGTGGAGTCGTTGCCTCAGCTCCACCGTCAGAGTNNNAGATCGGAAGAGC |
| 29922_2 | TATGAGTGTGGAGTCGTTGCCTCAGCTCCACCGTCAGAGTNNNAGATCGGAAGAGC |
| 36715_1 | TATGAGTGTGGAGTCGTTGCAAAACAGAACATCTGGCCATTCANNNAGATCGGAAG |
| 36715_2 | TATGAGTGTGGAGTCGTTGCAAAACAGAACATCTGGCCATTCANNNAGATCGGAAG |
| 41778_1 | TATGAGTGTGGAGTCGTTGCTTCAAGCAATTCTCCTGCCTCAGNNNAGATCGGAAG |
| 41778_2 | TATGAGTGTGGAGTCGTTGCTTCAAGCAATTCTCCTGCCTCAGNNNAGATCGGAAG |
| 41067_1 | TATGAGTGTGGAGTCGTTGCATCCCTCTTCCCTTCTTTTCACCNNNAGATCGGAAGAG |
| 41067_2 | TATGAGTGTGGAGTCGTTGCATCCCTCTTCCCTTCTTTTCACCNNNAGATCGGAAGAG |
| 29790_1 | TATGAGTGTGGAGTCGTTGCGAACATTAGAAGGTTGGGGTTGCNNNAGATCGGAA |
| 29790_2 | TATGAGTGTGGAGTCGTTGCGAACATTAGAAGGTTGGGGTTGCNNNAGATCGGAA |
| 50858_1 | TATGAGTGTGGAGTCGTTGCGGCATCTGGACTAGATTCAGGAGNNNAGATCGGAA |
| 50858_2 | TATGAGTGTGGAGTCGTTGCGGCATCTGGACTAGATTCAGGAGNNNAGATCGGAA |
| 10730_1 | TATGAGTGTGGAGTCGTTGCTCCTGTCTATCCCCATGTTCCTGNNNAGATCGGAAGA |
| 10730_2 | TATGAGTGTGGAGTCGTTGCTCCTGTCTATCCCCATGTTCCTGNNNAGATCGGAAGA |
| 12847_1 | TATGAGTGTGGAGTCGTTGCTTTCTGGAAAGAGAGAGAAATCNNNAGATCGGAAG |
| 12847_2 | TATGAGTGTGGAGTCGTTGCTTTCTGGAAAGAGAGAGAAATCNNNAGATCGGAAG |
| 47910_1 | TATGAGTGTGGAGTCGTTGCATCTTGGCTCACTGTAACCTCTGNNNAGATCGGAAG |
| 47910_2 | TATGAGTGTGGAGTCGTTGCATCTTGGCTCACTGTAACCTCTGNNNAGATCGGAAG |
| 50788_1 | TATGAGTGTGGAGTCGTTGCCAGCCCCAGACTCAATTAGCTAANNNAGATCGGAAG |
| 50788_2 | TATGAGTGTGGAGTCGTTGCCAGCCCCAGACTCAATTAGCTAANNNAGATCGGAAG |
| 30903_1 | TATGAGTGTGGAGTCGTTGCCTACTACTATCACAGGCCCATGGNNNAGATCGGAAG |
| 30903_2 | TATGAGTGTGGAGTCGTTGCCTACTACTATCACAGGCCCATGGNNNAGATCGGAAG |
| 49557_1 | TATGAGTGTGGAGTCGTTGCCTGGATGAAAGGAATCCCTGGAANNNAGATCGGAA |
| 49557_2 | TATGAGTGTGGAGTCGTTGCCTGGATGAAAGGAATCCCTGGAANNNAGATCGGAA |
| 41630_1 | TATGAGTGTGGAGTCGTTGCGCCTCCTGAGTAGCTGAGATTACNNNAGATCGGAAG |

| | |
|---|---|
| 41630_2 | TATGAGTGTGGAGTCGTTGCGCCTCCTGAGTAGCTGAGATTACNNNAGATCGGAAG |
| 48676_1 | TATGAGTGTGGAGTCGTTGCTTCACACCTTTTCTCTGGGTACCNNNAGATCGGAAGA |
| 48676_2 | TATGAGTGTGGAGTCGTTGCTTCACACCTTTTCTCTGGGTACCNNNAGATCGGAAGA |
| 45050_1 | TATGAGTGTGGAGTCGTTGCTCTTTGAACTCTGCCTGGGTAAGNNNAGATCGGAAG |
| 45050_2 | TATGAGTGTGGAGTCGTTGCTCTTTGAACTCTGCCTGGGTAAGNNNAGATCGGAAG |
| 12575_1 | TATGAGTGTGGAGTCGTTGCAAAAGTTTTTAATACACTGCTGANNNAGATCGGAAG |
| 12575_2 | TATGAGTGTGGAGTCGTTGCAAAAGTTTTTAATACACTGCTGANNNAGATCGGAAG |
| 38344_1 | TATGAGTGTGGAGTCGTTGCTTGTCTGTCAACTCCTGCATCATNNNAGATCGGAAGA |
| 38344_2 | TATGAGTGTGGAGTCGTTGCTTGTCTGTCAACTCCTGCATCATNNNAGATCGGAAGA |
| 46491_1 | TATGAGTGTGGAGTCGTTGCGTGTGTGTGAGTGAGAGAGAGAGNNNAGATCGGA |
| 46491_2 | TATGAGTGTGGAGTCGTTGCGTGTGTGTGAGTGAGAGAGAGAGNNNAGATCGGA |
| 33065_1 | TATGAGTGTGGAGTCGTTGCCTCAACCAATGATCCTGCCTTAGNNNAGATCGGAAGA |
| 33065_2 | TATGAGTGTGGAGTCGTTGCCTCAACCAATGATCCTGCCTTAGNNNAGATCGGAAGA |
| 41819_1 | TATGAGTGTGGAGTCGTTGCCTCTGCTTCAGCCTCCTGAATAGNNNAGATCGGAAGA |
| 41819_2 | TATGAGTGTGGAGTCGTTGCCTCTGCTTCAGCCTCCTGAATAGNNNAGATCGGAAGA |
| 44861_1 | TATGAGTGTGGAGTCGTTGCTGCCAAATGGAACACAGAGGATANNNAGATCGGAAG |
| 44861_2 | TATGAGTGTGGAGTCGTTGCTGCCAAATGGAACACAGAGGATANNNAGATCGGAAG |
| 37433_1 | TATGAGTGTGGAGTCGTTGCCCCAGTAAAGTGAGATGGAGTCCNNNAGATCGGAAG |
| 37433_2 | TATGAGTGTGGAGTCGTTGCCCCAGTAAAGTGAGATGGAGTCCNNNAGATCGGAAG |
| 40816_1 | TATGAGTGTGGAGTCGTTGCGAGATCACGCCATTACACTCCAANNNAGATCGGAAGA |
| 40816_2 | TATGAGTGTGGAGTCGTTGCGAGATCACGCCATTACACTCCAANNNAGATCGGAAGA |
| 34163_1 | TATGAGTGTGGAGTCGTTGCGAGACAAGAGAATCGCTTGAACCNNNAGATCGGAAG |
| 34163_2 | TATGAGTGTGGAGTCGTTGCGAGACAAGAGAATCGCTTGAACCNNNAGATCGGAAG |
| 44121_1 | TATGAGTGTGGAGTCGTTGCTCCACTCTCAGGAGAAGCTGGNNNAGATCGGAAGAG |
| 44121_2 | TATGAGTGTGGAGTCGTTGCTCCACTCTCAGGAGAAGCTGGNNNAGATCGGAAGAG |
| 34402_1 | TATGAGTGTGGAGTCGTTGCCACCTTACCTGGAAAGCACAAAGNNNAGATCGGAAG |
| 34402_2 | TATGAGTGTGGAGTCGTTGCCACCTTACCTGGAAAGCACAAAGNNNAGATCGGAAG |
| 50189_1 | TATGAGTGTGGAGTCGTTGCAAAGGGCAGGAACGGAGGAGNNNAGATCGGAAGA |
| 50189_2 | TATGAGTGTGGAGTCGTTGCAAAGGGCAGGAACGGAGGAGNNNAGATCGGAAGA |
| 42577_1 | TATGAGTGTGGAGTCGTTGCAATCTCTTGACCTCGTGATCCACNNNAGATCGGAAGA |
| 42577_2 | TATGAGTGTGGAGTCGTTGCAATCTCTTGACCTCGTGATCCACNNNAGATCGGAAGA |
| 32794_1 | TATGAGTGTGGAGTCGTTGCATCCTCCTATCTCAACCTCCCAANNNAGATCGGAAGAG |
| 32794_2 | TATGAGTGTGGAGTCGTTGCATCCTCCTATCTCAACCTCCCAANNNAGATCGGAAGAG |
| 43428_1 | TATGAGTGTGGAGTCGTTGCGGCCTTCCAAAGTACTGGGATTANNNAGATCGGAAG |
| 43428_2 | TATGAGTGTGGAGTCGTTGCGGCCTTCCAAAGTACTGGGATTANNNAGATCGGAAG |
| 40072_1 | TATGAGTGTGGAGTCGTTGCACCTGCAAATCCCGTCTGAATTANNNAGATCGGAAGA |
| 40072_2 | TATGAGTGTGGAGTCGTTGCACCTGCAAATCCCGTCTGAATTANNNAGATCGGAAGA |
| 42886_1 | TATGAGTGTGGAGTCGTTGCCCCTAATAGCTGGGATTACAGGCNNNAGATCGGAAG |
| 42886_2 | TATGAGTGTGGAGTCGTTGCCCCTAATAGCTGGGATTACAGGCNNNAGATCGGAAG |
| 38119_1 | TATGAGTGTGGAGTCGTTGCAGGAGACGTTACAACTGATACCANNNAGATCGGAAG |
| 38119_2 | TATGAGTGTGGAGTCGTTGCAGGAGACGTTACAACTGATACCANNNAGATCGGAAG |
| 45911_1 | TATGAGTGTGGAGTCGTTGCTTCCTGGGTTCAAGTGATTCTCANNNAGATCGGAAGA |
| 45911_2 | TATGAGTGTGGAGTCGTTGCTTCCTGGGTTCAAGTGATTCTCANNNAGATCGGAAGA |

| | |
|---|---|
| 29923_1 | TATGAGTGTGGAGTCGTTGCCATCTGTGGGGAAACTGCCANNNAGATCGGAAGAG |
| 29923_2 | TATGAGTGTGGAGTCGTTGCCATCTGTGGGGAAACTGCCANNNAGATCGGAAGAG |
| 10727_1 | TATGAGTGTGGAGTCGTTGCCAAAATCAGGTATCACAAGCTGAANNNAGATCGGA |
| 10727_2 | TATGAGTGTGGAGTCGTTGCCAAAATCAGGTATCACAAGCTGAANNNAGATCGGA |
| 52582_1 | TATGAGTGTGGAGTCGTTGCAATTGCCTTTGCTTCAATCACCANNNAGATCGGAAG |
| 52582_2 | TATGAGTGTGGAGTCGTTGCAATTGCCTTTGCTTCAATCACCANNNAGATCGGAAG |
| 44385_1 | TATGAGTGTGGAGTCGTTGCCCTCCCATCCTAACTATCACTGCNNNAGATCGGAAGA |
| 44385_2 | TATGAGTGTGGAGTCGTTGCCCTCCCATCCTAACTATCACTGCNNNAGATCGGAAGA |
| 51011_1 | TATGAGTGTGGAGTCGTTGCGAAGGGAGGGAAAGAGAGAGAGANNNAGATCGG |
| 51011_2 | TATGAGTGTGGAGTCGTTGCGAAGGGAGGGAAAGAGAGAGAGANNNAGATCGG |
| 34886_1 | TATGAGTGTGGAGTCGTTGCTAGAAGTGCAGTTTCGTTACCCTNNNAGATCGGAAG |
| 34886_2 | TATGAGTGTGGAGTCGTTGCTAGAAGTGCAGTTTCGTTACCCTNNNAGATCGGAAG |
| 47508_1 | TATGAGTGTGGAGTCGTTGCGTTCATGCCATTCTCCTGCTTCNNNAGATCGGAAGAG |
| 47508_2 | TATGAGTGTGGAGTCGTTGCGTTCATGCCATTCTCCTGCTTCNNNAGATCGGAAGAG |
| 46482_1 | TATGAGTGTGGAGTCGTTGCGCGGTGACTGGACTCTAATTCTTNNNAGATCGGAAG |
| 46482_2 | TATGAGTGTGGAGTCGTTGCGCGGTGACTGGACTCTAATTCTTNNNAGATCGGAAG |
| 51928_1 | TATGAGTGTGGAGTCGTTGCTAACAGGAGCTGAAATTGAGGCANNNAGATCGGAA |
| 51928_2 | TATGAGTGTGGAGTCGTTGCTAACAGGAGCTGAAATTGAGGCANNNAGATCGGAA |
| 34986_1 | TATGAGTGTGGAGTCGTTGCGTCTCCTGAGTAGCTGGGACTANNNAGATCGGAAG |
| 34986_2 | TATGAGTGTGGAGTCGTTGCGTCTCCTGAGTAGCTGGGACTANNNAGATCGGAAG |
| 33800_1 | TATGAGTGTGGAGTCGTTGCAGAGGAGAGTTCCGGGAATTTTCNNNAGATCGGAA |
| 33800_2 | TATGAGTGTGGAGTCGTTGCAGAGGAGAGTTCCGGGAATTTTCNNNAGATCGGAA |
| 31284_1 | TATGAGTGTGGAGTCGTTGCAAACCCTCATTTGAAACCCTTGGNNNAGATCGGAAG |
| 31284_2 | TATGAGTGTGGAGTCGTTGCAAACCCTCATTTGAAACCCTTGGNNNAGATCGGAAG |
| 33312_1 | TATGAGTGTGGAGTCGTTGCCTTCAACTGCTGGACTTGTGATCNNNAGATCGGAAG |
| 33312_2 | TATGAGTGTGGAGTCGTTGCCTTCAACTGCTGGACTTGTGATCNNNAGATCGGAAG |
| 39009_1 | TATGAGTGTGGAGTCGTTGCCATGCTTGTTGTCCCAGCTATTCNNNAGATCGGAAG |
| 39009_2 | TATGAGTGTGGAGTCGTTGCCATGCTTGTTGTCCCAGCTATTCNNNAGATCGGAAG |
| 51898_1 | TATGAGTGTGGAGTCGTTGCGGAGAGACATGTCAAGGATCTGGNNNAGATCGGA |
| 51898_2 | TATGAGTGTGGAGTCGTTGCGGAGAGACATGTCAAGGATCTGGNNNAGATCGGA |
| 51821_1 | TATGAGTGTGGAGTCGTTGCGAACTCCTTACCTCAGGTGATCCNNNAGATCGGAAG |
| 51821_2 | TATGAGTGTGGAGTCGTTGCGAACTCCTTACCTCAGGTGATCCNNNAGATCGGAAG |
| 40562_1 | TATGAGTGTGGAGTCGTTGCCTGTAGTACCAGCTACTCGGGAGNNNAGATCGGAA |
| 40562_2 | TATGAGTGTGGAGTCGTTGCCTGTAGTACCAGCTACTCGGGAGNNNAGATCGGAA |
| 45670_1 | TATGAGTGTGGAGTCGTTGCGATCTTGACTCACTGCAACCTCTNNNAGATCGGAAG |
| 45670_2 | TATGAGTGTGGAGTCGTTGCGATCTTGACTCACTGCAACCTCTNNNAGATCGGAAG |
| 51179_1 | TATGAGTGTGGAGTCGTTGCTGGGGGTAAAAGGGACTATAGGGNNNAGATCGGA |
| 51179_2 | TATGAGTGTGGAGTCGTTGCTGGGGGTAAAAGGGACTATAGGGNNNAGATCGGA |
| 32292_1 | TATGAGTGTGGAGTCGTTGCACGTGGAGAAGGAACCTATTGACNNNAGATCGGAA |
| 32292_2 | TATGAGTGTGGAGTCGTTGCACGTGGAGAAGGAACCTATTGACNNNAGATCGGAA |
| 39881_1 | TATGAGTGTGGAGTCGTTGCTTGAACCCAGAAGTTTGAGGTCANNNAGATCGGAA |
| 39881_2 | TATGAGTGTGGAGTCGTTGCTTGAACCCAGAAGTTTGAGGTCANNNAGATCGGAA |
| 10799_1 | TATGAGTGTGGAGTCGTTGCTGGGTTTGTAAGCATGCTGTTTCNNNAGATCGGAAG |

| | |
|---|---|
| 10799_2 | TATGAGTGTGGAGTCGTTGCTGGGTTTGTAAGCATGCTGTTTCNNNAGATCGGAAG |
| 49141_1 | TATGAGTGTGGAGTCGTTGCGCAGAAAATCCAAGTCAACCACANNNAGATCGGAAG |
| 49141_2 | TATGAGTGTGGAGTCGTTGCGCAGAAAATCCAAGTCAACCACANNNAGATCGGAAG |
| 42900_1 | TATGAGTGTGGAGTCGTTGCTACAGAATCCTGAAGACCTGCAGNNNAGATCGGAAG |
| 42900_2 | TATGAGTGTGGAGTCGTTGCTACAGAATCCTGAAGACCTGCAGNNNAGATCGGAAG |
| 48277_1 | TATGAGTGTGGAGTCGTTGCTCTCCTGAGTAGCTGGGATTACANNNAGATCGGAAG |
| 48277_2 | TATGAGTGTGGAGTCGTTGCTCTCCTGAGTAGCTGGGATTACANNNAGATCGGAAG |
| 42847_1 | TATGAGTGTGGAGTCGTTGCAACATGCATTAGTAGGCCTGACANNNAGATCGGAAG |
| 42847_2 | TATGAGTGTGGAGTCGTTGCAACATGCATTAGTAGGCCTGACANNNAGATCGGAAG |
| 31202_1 | TATGAGTGTGGAGTCGTTGCAATTCACATCTATGGGCAATGCCNNNAGATCGGAAG |
| 31202_2 | TATGAGTGTGGAGTCGTTGCAATTCACATCTATGGGCAATGCCNNNAGATCGGAAG |
| 13203_1 | TATGAGTGTGGAGTCGTTGCCACCTTTAATTTTGTAATGGTCTNNNAGATCGGAAGA |
| 13203_2 | TATGAGTGTGGAGTCGTTGCCACCTTTAATTTTGTAATGGTCTNNNAGATCGGAAGA |
| 31272_1 | TATGAGTGTGGAGTCGTTGCACACTGAAGGCTGAACAACTACANNNAGATCGGAAG |
| 31272_2 | TATGAGTGTGGAGTCGTTGCACACTGAAGGCTGAACAACTACANNNAGATCGGAAG |
| 37060_1 | TATGAGTGTGGAGTCGTTGCTACCCTTTCTTCTTCCATCTGCCNNNAGATCGGAAGAG |
| 37060_2 | TATGAGTGTGGAGTCGTTGCTACCCTTTCTTCTTCCATCTGCCNNNAGATCGGAAGAG |
| 38927_1 | TATGAGTGTGGAGTCGTTGCACCAAGGGCCAACTATATATTCCCNNNAGATCGGAAG |
| 38927_2 | TATGAGTGTGGAGTCGTTGCACCAAGGGCCAACTATATATTCCCNNNAGATCGGAAG |
| 28950_1 | TATGAGTGTGGAGTCGTTGCTGCTGCCTTGTGTCTTGAAAATTNNNAGATCGGAAGA |
| 28950_2 | TATGAGTGTGGAGTCGTTGCTGCTGCCTTGTGTCTTGAAAATTNNNAGATCGGAAGA |
| 34599_1 | TATGAGTGTGGAGTCGTTGCGGAGAGCGGACATCTTGAGTATTNNNAGATCGGAAG |
| 34599_2 | TATGAGTGTGGAGTCGTTGCGGAGAGCGGACATCTTGAGTATTNNNAGATCGGAAG |
| 52003_1 | TATGAGTGTGGAGTCGTTGCCATAAAGGTTCTCCAAGTCCCCANNNAGATCGGAAGA |
| 52003_2 | TATGAGTGTGGAGTCGTTGCCATAAAGGTTCTCCAAGTCCCCANNNAGATCGGAAGA |
| 2325_1 | TATGAGTGTGGAGTCGTTGCCCCATTCCTTGGCTGATTCTACCNNNAGATCGGAAGA |
| 2325_2 | TATGAGTGTGGAGTCGTTGCCCCATTCCTTGGCTGATTCTACCNNNAGATCGGAAGA |
| 29045_1 | TATGAGTGTGGAGTCGTTGCAGTTTCTCACTTGAAGCCTCACTNNNAGATCGGAAGA |
| 29045_2 | TATGAGTGTGGAGTCGTTGCAGTTTCTCACTTGAAGCCTCACTNNNAGATCGGAAGA |
| 29566_1 | TATGAGTGTGGAGTCGTTGCCGGTTATGGAATTGCCCTTCAGANNNAGATCGGAAG |
| 29566_2 | TATGAGTGTGGAGTCGTTGCCGGTTATGGAATTGCCCTTCAGANNNAGATCGGAAG |
| 2207_1 | TATGAGTGTGGAGTCGTTGCTGACCAAAATTAGCTGTGGTTGGANNNAGATCGGAA |
| 2207_2 | TATGAGTGTGGAGTCGTTGCTGACCAAAATTAGCTGTGGTTGGANNNAGATCGGAA |
| 36977_1 | TATGAGTGTGGAGTCGTTGCAGGCTAACCAAAGACAGAAGACCNNNAGATCGGAAG |
| 36977_2 | TATGAGTGTGGAGTCGTTGCAGGCTAACCAAAGACAGAAGACCNNNAGATCGGAAG |
| 37611_1 | TATGAGTGTGGAGTCGTTGCGTGCAACTTCCCGGATGTAAAAANNNAGATCGGAAG |
| 37611_2 | TATGAGTGTGGAGTCGTTGCGTGCAACTTCCCGGATGTAAAAANNNAGATCGGAAG |
| 39531_1 | TATGAGTGTGGAGTCGTTGCCTTACACCTGTAATCCCAGCACTNNNAGATCGGAAGA |
| 39531_2 | TATGAGTGTGGAGTCGTTGCCTTACACCTGTAATCCCAGCACTNNNAGATCGGAAGA |
| 31540_1 | TATGAGTGTGGAGTCGTTGCGTGTGTGAGACAGACTTTTGCTCNNNAGATCGGAAG |
| 31540_2 | TATGAGTGTGGAGTCGTTGCGTGTGTGAGACAGACTTTTGCTCNNNAGATCGGAAG |
| 32632_1 | TATGAGTGTGGAGTCGTTGCAGTAGCTGGGATTACAGGTACCTNNNAGATCGGAAG |
| 32632_2 | TATGAGTGTGGAGTCGTTGCAGTAGCTGGGATTACAGGTACCTNNNAGATCGGAAG |

| | |
|---|---|
| 49293_1 | TATGAGTGTGGAGTCGTTGCGCAACTGAAGCTGATTATGAGCCNNNAGATCGGAA |
| 49293_2 | TATGAGTGTGGAGTCGTTGCGCAACTGAAGCTGATTATGAGCCNNNAGATCGGAA |
| 45720_1 | TATGAGTGTGGAGTCGTTGCCCGGCCCATAGCTTCAATCTTTANNNAGATCGGAAG |
| 45720_2 | TATGAGTGTGGAGTCGTTGCCCGGCCCATAGCTTCAATCTTTANNNAGATCGGAAG |
| 39320_1 | TATGAGTGTGGAGTCGTTGCCCTTCCCATGTGTCAATGTCAACNNNAGATCGGAAG |
| 39320_2 | TATGAGTGTGGAGTCGTTGCCCTTCCCATGTGTCAATGTCAACNNNAGATCGGAAG |
| 41551_1 | TATGAGTGTGGAGTCGTTGCCAGTGGTATGATCATGGCTGACTNNNAGATCGGAA |
| 41551_2 | TATGAGTGTGGAGTCGTTGCCAGTGGTATGATCATGGCTGACTNNNAGATCGGAA |
| 12571_1 | TATGAGTGTGGAGTCGTTGCGGAAAACATAAGTCAAGTATCAANNNAGATCGGAA |
| 12571_2 | TATGAGTGTGGAGTCGTTGCGGAAAACATAAGTCAAGTATCAANNNAGATCGGAA |
| 52373_1 | TATGAGTGTGGAGTCGTTGCCCTTGGCTCTAGACTGTTACTCCNNNAGATCGGAAG |
| 52373_2 | TATGAGTGTGGAGTCGTTGCCCTTGGCTCTAGACTGTTACTCCNNNAGATCGGAAG |
| 44137_1 | TATGAGTGTGGAGTCGTTGCTAGTAGGTGTGGTAGTCATCGGTNNNAGATCGGAA |
| 44137_2 | TATGAGTGTGGAGTCGTTGCTAGTAGGTGTGGTAGTCATCGGTNNNAGATCGGAA |
| 42644_1 | TATGAGTGTGGAGTCGTTGCCTTGGGGTCCCTGATTCTAGAACNNNAGATCGGAAG |
| 42644_2 | TATGAGTGTGGAGTCGTTGCCTTGGGGTCCCTGATTCTAGAACNNNAGATCGGAAG |
| 51295_1 | TATGAGTGTGGAGTCGTTGCGCAAACTGTCCATCCATCTGAAGNNNAGATCGGAAG |
| 51295_2 | TATGAGTGTGGAGTCGTTGCGCAAACTGTCCATCCATCTGAAGNNNAGATCGGAAG |
| 52299_1 | TATGAGTGTGGAGTCGTTGCTGACGTTCCTACAGCTATTCCAGNNNAGATCGGAAG |
| 52299_2 | TATGAGTGTGGAGTCGTTGCTGACGTTCCTACAGCTATTCCAGNNNAGATCGGAAG |
| 30683_1 | TATGAGTGTGGAGTCGTTGCATTATATGGCCATCTTGGAGCTCCNNNAGATCGGAA |
| 30683_2 | TATGAGTGTGGAGTCGTTGCATTATATGGCCATCTTGGAGCTCCNNNAGATCGGAA |
| 50020_1 | TATGAGTGTGGAGTCGTTGCAGACCAGGATGATGATGACGATGNNNAGATCGGAA |
| 50020_2 | TATGAGTGTGGAGTCGTTGCAGACCAGGATGATGATGACGATGNNNAGATCGGAA |
| 36494_1 | TATGAGTGTGGAGTCGTTGCTGAGTTACCATCTCATGCCAGTTNNNAGATCGGAAG |
| 36494_2 | TATGAGTGTGGAGTCGTTGCTGAGTTACCATCTCATGCCAGTTNNNAGATCGGAAG |
| 36625_1 | TATGAGTGTGGAGTCGTTGCGTTTCAAGGACTATGCTGCAACCNNNAGATCGGAAG |
| 36625_2 | TATGAGTGTGGAGTCGTTGCGTTTCAAGGACTATGCTGCAACCNNNAGATCGGAAG |
| 8341_1 | TATGAGTGTGGAGTCGTTGCGACAACCAGATCCCTTGACATCTGNNNAGATCGGAA |
| 8341_2 | TATGAGTGTGGAGTCGTTGCGACAACCAGATCCCTTGACATCTGNNNAGATCGGAA |
| 3192_1 | TATGAGTGTGGAGTCGTTGCTGTGATGAGAGGGATTGGAAAAGACNNNAGATCG |
| 3192_2 | TATGAGTGTGGAGTCGTTGCTGTGATGAGAGGGATTGGAAAAGACNNNAGATCG |
| 30214_1 | ATGAGTGTGGAGTCGTTGCTGTCTTTCCAGAATTACTTGTTTCCTNNNAGATCGGAA |
| 30214_2 | ATGAGTGTGGAGTCGTTGCTGTCTTTCCAGAATTACTTGTTTCCTNNNAGATCGGAA |
| 50556_1 | TATGAGTGTGGAGTCGTTGCTTTTAGGGTACATGTGCACAACGNNNAGATCGGAAG |
| 50556_2 | TATGAGTGTGGAGTCGTTGCTTTTAGGGTACATGTGCACAACGNNNAGATCGGAAG |
| 44689_1 | TATGAGTGTGGAGTCGTTGCATCCCTGGTTCTGAATTTACCGTNNNAGATCGGAAG |
| 44689_2 | TATGAGTGTGGAGTCGTTGCATCCCTGGTTCTGAATTTACCGTNNNAGATCGGAAG |
| 52273_1 | TATGAGTGTGGAGTCGTTGCTCATCCCTCTCCTCTCCTTTTCTNNNAGATCGGAAGAG |
| 52273_2 | TATGAGTGTGGAGTCGTTGCTCATCCCTCTCCTCTCCTTTTCTNNNAGATCGGAAGAG |
| 48960_1 | TATGAGTGTGGAGTCGTTGCTTGGAATGGGCTACTTCATGACANNNAGATCGGAAG |
| 48960_2 | TATGAGTGTGGAGTCGTTGCTTGGAATGGGCTACTTCATGACANNNAGATCGGAAG |
| 51128_1 | TATGAGTGTGGAGTCGTTGCACTCTATCACCAGTGGAGCATTGNNNAGATCGGAAG |

| | |
|---|---|
| 51128_2 | TATGAGTGTGGAGTCGTTGCACTCTATCACCAGTGGAGCATTGNNNAGATCGGAAG |
| 39361_1 | TATGAGTGTGGAGTCGTTGCCTTTGCTCTGATGGAGGAGACTTNNNAGATCGGAAG |
| 39361_2 | TATGAGTGTGGAGTCGTTGCCTTTGCTCTGATGGAGGAGACTTNNNAGATCGGAAG |
| 40024_1 | TATGAGTGTGGAGTCGTTGCAGATCCACCAAGGTCAAGAAACTNNNAGATCGGAAG |
| 40024_2 | TATGAGTGTGGAGTCGTTGCAGATCCACCAAGGTCAAGAAACTNNNAGATCGGAAG |
| 42469_1 | TATGAGTGTGGAGTCGTTGCTTTTTGCTGAGGTTGGTCTCGAANNNAGATCGGAAG |
| 42469_2 | TATGAGTGTGGAGTCGTTGCTTTTTGCTGAGGTTGGTCTCGAANNNAGATCGGAAG |
| 44652_1 | TATGAGTGTGGAGTCGTTGCTGAGACTTCCACACCTGAATCTGNNNAGATCGGAAGA |
| 44652_2 | TATGAGTGTGGAGTCGTTGCTGAGACTTCCACACCTGAATCTGNNNAGATCGGAAGA |
| 45561_1 | TATGAGTGTGGAGTCGTTGCGGGCAGGAAGAGAGAACTTCAATNNNAGATCGGAA |
| 45561_2 | TATGAGTGTGGAGTCGTTGCGGGCAGGAAGAGAGAACTTCAATNNNAGATCGGAA |
| 51067_1 | TATGAGTGTGGAGTCGTTGCGAATCAGGACTGAGTGACCTCTGNNNAGATCGGAA |
| 51067_2 | TATGAGTGTGGAGTCGTTGCGAATCAGGACTGAGTGACCTCTGNNNAGATCGGAA |
| 50213_1 | TATGAGTGTGGAGTCGTTGCCCACTGTGTGATGATGGCAAATCNNNAGATCGGAAG |
| 50213_2 | TATGAGTGTGGAGTCGTTGCCCACTGTGTGATGATGGCAAATCNNNAGATCGGAAG |
| 33700_1 | TATGAGTGTGGAGTCGTTGCTGAAGCTCAGAGGTTCAAAGTGGNNNAGATCGGAA |
| 33700_2 | TATGAGTGTGGAGTCGTTGCTGAAGCTCAGAGGTTCAAAGTGGNNNAGATCGGAA |
| 40002_1 | TATGAGTGTGGAGTCGTTGCAGGAAAGAAGGAAAGCCATGGATNNNAGATCGGAA |
| 40002_2 | TATGAGTGTGGAGTCGTTGCAGGAAAGAAGGAAAGCCATGGATNNNAGATCGGAA |
| 42387_1 | TATGAGTGTGGAGTCGTTGCATCTATCTCAGGCTTTGGGGTTGNNNAGATCGGAAG |
| 42387_2 | TATGAGTGTGGAGTCGTTGCATCTATCTCAGGCTTTGGGGTTGNNNAGATCGGAAG |
| 46367_1 | TATGAGTGTGGAGTCGTTGCTCCAGGTTCAAGTGATTCTCCTGNNNAGATCGGAAGA |
| 46367_2 | TATGAGTGTGGAGTCGTTGCTCCAGGTTCAAGTGATTCTCCTGNNNAGATCGGAAGA |
| 40553_1 | TATGAGTGTGGAGTCGTTGCTTAGGTCAACGGACAGAATGGCNNNAGATCGGAAG |
| 40553_2 | TATGAGTGTGGAGTCGTTGCTTAGGTCAACGGACAGAATGGCNNNAGATCGGAAG |
| 49814_1 | TATGAGTGTGGAGTCGTTGCCTTGAGAAGCCTCTGCACCTCNNNAGATCGGAAGAG |
| 49814_2 | TATGAGTGTGGAGTCGTTGCCTTGAGAAGCCTCTGCACCTCNNNAGATCGGAAGAG |
| 52680_1 | TATGAGTGTGGAGTCGTTGCGAAGCTCTTCTGGACACTGTTCTNNNAGATCGGAAGA |
| 52680_2 | TATGAGTGTGGAGTCGTTGCGAAGCTCTTCTGGACACTGTTCTNNNAGATCGGAAGA |
| 52354_1 | TATGAGTGTGGAGTCGTTGCTCCGCCATTTTTCAACTTACCACNNNAGATCGGAAGA |
| 52354_2 | TATGAGTGTGGAGTCGTTGCTCCGCCATTTTTCAACTTACCACNNNAGATCGGAAGA |
| 30904_1 | TATGAGTGTGGAGTCGTTGCCAATATCGTAAACTGCAGCAGGGNNNAGATCGGAA |
| 30904_2 | TATGAGTGTGGAGTCGTTGCCAATATCGTAAACTGCAGCAGGGNNNAGATCGGAA |
| 51363_1 | TATGAGTGTGGAGTCGTTGCGTGCTCAGTGAGAGAGACTTTGANNNAGATCGGAA |
| 51363_2 | TATGAGTGTGGAGTCGTTGCGTGCTCAGTGAGAGAGACTTTGANNNAGATCGGAA |
| 49271_1 | TATGAGTGTGGAGTCGTTGCAAGATGAGGTCTCACTATGTGCCNNNAGATCGGAAG |
| 49271_2 | TATGAGTGTGGAGTCGTTGCAAGATGAGGTCTCACTATGTGCCNNNAGATCGGAAG |
| 10755_1 | TATGAGTGTGGAGTCGTTGCGTGTAACCCTCGAAAACATATGCTNNNAGATCGGAA |
| 10755_2 | TATGAGTGTGGAGTCGTTGCGTGTAACCCTCGAAAACATATGCTNNNAGATCGGAA |
| 43909_1 | TATGAGTGTGGAGTCGTTGCTCTTTCTTTCCTTCCTTCCTTCCTNNNAGATCGGAAGA |
| 43909_2 | TATGAGTGTGGAGTCGTTGCTCTTTCTTTCCTTCCTTCCTTCCTNNNAGATCGGAAGA |
| 52557_1 | TATGAGTGTGGAGTCGTTGCAGGCTGGTAGTTCTAACCAACAGNNNAGATCGGAAG |
| 52557_2 | TATGAGTGTGGAGTCGTTGCAGGCTGGTAGTTCTAACCAACAGNNNAGATCGGAAG |

| | |
|---|---|
| 49945_1 | TATGAGTGTGGAGTCGTTGCCTGGCCAATCAGCAAACGCANNNAGATCGGAAGAG |
| 49945_2 | TATGAGTGTGGAGTCGTTGCCTGGCCAATCAGCAAACGCANNNAGATCGGAAGAG |
| 46873_1 | TATGAGTGTGGAGTCGTTGCACGCCCGGCTAATTTTTTGTATTNNNAGATCGGAAG |
| 46873_2 | TATGAGTGTGGAGTCGTTGCACGCCCGGCTAATTTTTTGTATTNNNAGATCGGAAG |
| 42774_1 | TATGAGTGTGGAGTCGTTGCTGAGAGAGAGAGAGAGAAAGCGANNNAGATCGGA |
| 42774_2 | TATGAGTGTGGAGTCGTTGCTGAGAGAGAGAGAGAGAAAGCGANNNAGATCGGA |
| 10936_1 | TATGAGTGTGGAGTCGTTGCGGAAGTTAGTGCCACATGTAAGGNNNAGATCGGAA |
| 10936_2 | TATGAGTGTGGAGTCGTTGCGGAAGTTAGTGCCACATGTAAGGNNNAGATCGGAA |
| 43082_1 | TATGAGTGTGGAGTCGTTGCGGTATCTATGGGGGATTGGTTCCNNNAGATCGGAA |
| 43082_2 | TATGAGTGTGGAGTCGTTGCGGTATCTATGGGGGATTGGTTCCNNNAGATCGGAA |
| 32790_1 | TATGAGTGTGGAGTCGTTGCGAGGTCAGGAGATTGAGACCATCNNNAGATCGGAA |
| 32790_2 | TATGAGTGTGGAGTCGTTGCGAGGTCAGGAGATTGAGACCATCNNNAGATCGGAA |
| 48385_1 | TATGAGTGTGGAGTCGTTGCACAAGGTCTCATTATGTTGCCCANNNAGATCGGAAG |
| 48385_2 | TATGAGTGTGGAGTCGTTGCACAAGGTCTCATTATGTTGCCCANNNAGATCGGAAG |
| 33733_1 | TATGAGTGTGGAGTCGTTGCCCTGACTTTGTATCCTCCAGCTTNNNAGATCGGAAGA |
| 33733_2 | TATGAGTGTGGAGTCGTTGCCCTGACTTTGTATCCTCCAGCTTNNNAGATCGGAAGA |
| 51430_1 | TATGAGTGTGGAGTCGTTGCCCTGGAATCCTTGAAGGAGGAAGNNNAGATCGGAA |
| 51430_2 | TATGAGTGTGGAGTCGTTGCCCTGGAATCCTTGAAGGAGGAAGNNNAGATCGGAA |
| 31273_1 | TATGAGTGTGGAGTCGTTGCAAGAGTGGGGTGTATTTGGATGTNNNAGATCGGAA |
| 31273_2 | TATGAGTGTGGAGTCGTTGCAAGAGTGGGGTGTATTTGGATGTNNNAGATCGGAA |
| 13252_1 | TATGAGTGTGGAGTCGTTGCAAGTCAGTAAGCAACACTTCNNNAGATCGGAAGAG |
| 13252_2 | TATGAGTGTGGAGTCGTTGCAAGTCAGTAAGCAACACTTCNNNAGATCGGAAGAG |
| 41736_1 | TATGAGTGTGGAGTCGTTGCGATCTTGGCTCACTGCATCCTCNNNAGATCGGAAGA |
| 41736_2 | TATGAGTGTGGAGTCGTTGCGATCTTGGCTCACTGCATCCTCNNNAGATCGGAAGA |
| 9964_1 | TATGAGTGTGGAGTCGTTGCACTTCTCAAGGAAGAAACAAGGANNNAGATCGGAA |
| 9964_2 | TATGAGTGTGGAGTCGTTGCACTTCTCAAGGAAGAAACAAGGANNNAGATCGGAA |
| 32772_1 | TATGAGTGTGGAGTCGTTGCATGTAACAAAATCCAACACCCCCNNNAGATCGGAAG |
| 32772_2 | TATGAGTGTGGAGTCGTTGCATGTAACAAAATCCAACACCCCCNNNAGATCGGAAG |
| 32259_1 | TATGAGTGTGGAGTCGTTGCAGATCCACTGGCTTCACATTCTTNNNAGATCGGAAG |
| 32259_2 | TATGAGTGTGGAGTCGTTGCAGATCCACTGGCTTCACATTCTTNNNAGATCGGAAG |
| 31883_1 | TATGAGTGTGGAGTCGTTGCGCGACAAAGCAAGACTCTGTATCNNNAGATCGGAA |
| 31883_2 | TATGAGTGTGGAGTCGTTGCGCGACAAAGCAAGACTCTGTATCNNNAGATCGGAA |
| 51570_1 | TATGAGTGTGGAGTCGTTGCCTCTGCAAGGTAATCTGGCTGTANNNAGATCGGAAG |
| 51570_2 | TATGAGTGTGGAGTCGTTGCCTCTGCAAGGTAATCTGGCTGTANNNAGATCGGAAG |
| 10980_1 | TATGAGTGTGGAGTCGTTGCTCAGAGCATGCAAATACCCCTCTANNNAGATCGGAA |
| 10980_2 | TATGAGTGTGGAGTCGTTGCTCAGAGCATGCAAATACCCCTCTANNNAGATCGGAA |
| 43250_1 | TATGAGTGTGGAGTCGTTGCAACAACAAAAAACCCAAATGGCCNNNAGATCGGAAG |
| 43250_2 | TATGAGTGTGGAGTCGTTGCAACAACAAAAAACCCAAATGGCCNNNAGATCGGAAG |
| 39140_1 | TATGAGTGTGGAGTCGTTGCAGAGAGATTTTGAGTGTTCTCACCANNNAGATCGGA |
| 39140_2 | TATGAGTGTGGAGTCGTTGCAGAGAGATTTTGAGTGTTCTCACCANNNAGATCGGA |
| 41737_1 | TATGAGTGTGGAGTCGTTGCTTCAAGTGATTCTCCTGCCTCAGNNNAGATCGGAAG |
| 41737_2 | TATGAGTGTGGAGTCGTTGCTTCAAGTGATTCTCCTGCCTCAGNNNAGATCGGAAG |
| 45531_1 | TATGAGTGTGGAGTCGTTGCTCCTAGGACTAAGTGCTCTGGAANNNAGATCGGAA |

| | |
|---|---|
| 45531_2 | TATGAGTGTGGAGTCGTTGCTCCTAGGACTAAGTGCTCTGGAANNNAGATCGGAAG |
| 43777_1 | TATGAGTGTGGAGTCGTTGCAGAAAAAGAAGAAGGAGGAGAGGGNNNAGATCGG |
| 43777_2 | TATGAGTGTGGAGTCGTTGCAGAAAAAGAAGAAGGAGGAGAGGGNNNAGATCGG |
| 52818_1 | TATGAGTGTGGAGTCGTTGCTCTCTGTTGTTCACACTGGACTCNNNAGATCGGAAGA |
| 52818_2 | TATGAGTGTGGAGTCGTTGCTCTCTGTTGTTCACACTGGACTCNNNAGATCGGAAGA |
| 39297_1 | TATGAGTGTGGAGTCGTTGCAGAAAGCCTCTCTTCAGTTTGTANNNAGATCGGAAGA |
| 39297_2 | TATGAGTGTGGAGTCGTTGCAGAAAGCCTCTCTTCAGTTTGTANNNAGATCGGAAGA |
| 48861_1 | TATGAGTGTGGAGTCGTTGCTGGTAGGTTGTATGTTTCCAGGANNNAGATCGGAAG |
| 48861_2 | TATGAGTGTGGAGTCGTTGCTGGTAGGTTGTATGTTTCCAGGANNNAGATCGGAAG |
| 39254_1 | TATGAGTGTGGAGTCGTTGCCAGCCAGGGCAATTCTTTACAAANNNAGATCGGAAG |
| 39254_2 | TATGAGTGTGGAGTCGTTGCCAGCCAGGGCAATTCTTTACAAANNNAGATCGGAAG |
| 34332_1 | TATGAGTGTGGAGTCGTTGCTGAAGGGAAAGGTTGGGAAGAAANNNAGATCGGA |
| 34332_2 | TATGAGTGTGGAGTCGTTGCTGAAGGGAAAGGTTGGGAAGAAANNNAGATCGGA |
| 42035_1 | TATGAGTGTGGAGTCGTTGCCAGGTCTCTACCCCATTTCTGAGNNNAGATCGGAAGA |
| 42035_2 | TATGAGTGTGGAGTCGTTGCCAGGTCTCTACCCCATTTCTGAGNNNAGATCGGAAGA |
| 41878_1 | TATGAGTGTGGAGTCGTTGCAAAGCAATGGTTGACTTCAGCAANNNAGATCGGAAG |
| 41878_2 | TATGAGTGTGGAGTCGTTGCAAAGCAATGGTTGACTTCAGCAANNNAGATCGGAAG |
| 44011_1 | TATGAGTGTGGAGTCGTTGCGCCTGATAATGTTTGGGGAGGTANNNAGATCGGAA |
| 44011_2 | TATGAGTGTGGAGTCGTTGCGCCTGATAATGTTTGGGGAGGTANNNAGATCGGAA |
| 37079_1 | TATGAGTGTGGAGTCGTTGCGAAACACACACACACACACACANNNAGATCGGAAGA |
| 37079_2 | TATGAGTGTGGAGTCGTTGCGAAACACACACACACACACACANNNAGATCGGAAGA |
| 13685_1 | TATGAGTGTGGAGTCGTTGCCTGATCATTTGGCTCTTTTTTTNNNAGATCGGAAGAG |
| 13685_2 | TATGAGTGTGGAGTCGTTGCCTGATCATTTGGCTCTTTTTTTNNNAGATCGGAAGAG |
| 12701_1 | TATGAGTGTGGAGTCGTTGCAACTGAGGAGGATTGCATTTNNNAGATCGGAAGAG |
| 12701_2 | TATGAGTGTGGAGTCGTTGCAACTGAGGAGGATTGCATTTNNNAGATCGGAAGAG |
| 30351_1 | TATGAGTGTGGAGTCGTTGCAAAGTGGGCACATCTCGAAGCNNNAGATCGGAAGAG |
| 30351_2 | TATGAGTGTGGAGTCGTTGCAAAGTGGGCACATCTCGAAGCNNNAGATCGGAAGAG |
| 38972_1 | TATGAGTGTGGAGTCGTTGCCTTCACTGCTGAATTGCACTAAANNNAGATCGGAAGA |
| 38972_2 | TATGAGTGTGGAGTCGTTGCCTTCACTGCTGAATTGCACTAAANNNAGATCGGAAGA |
| 48806_1 | TATGAGTGTGGAGTCGTTGCCCAGCATTTTTTCCCAGTCTTGTNNNAGATCGGAAGA |
| 48806_2 | TATGAGTGTGGAGTCGTTGCCCAGCATTTTTTCCCAGTCTTGTNNNAGATCGGAAGA |
| 52514_1 | TATGAGTGTGGAGTCGTTGCTAAGAAAGCAGGGAGGATATGCCNNNAGATCGGAA |
| 52514_2 | TATGAGTGTGGAGTCGTTGCTAAGAAAGCAGGGAGGATATGCCNNNAGATCGGAA |
| 46862_1 | TATGAGTGTGGAGTCGTTGCCTCCTGGGTTCACACCATTCTCNNNAGATCGGAAGAG |
| 46862_2 | TATGAGTGTGGAGTCGTTGCCTCCTGGGTTCACACCATTCTCNNNAGATCGGAAGAG |
| 33169_1 | TATGAGTGTGGAGTCGTTGCCAGTAAGCCGAGATCATGACACTNNNAGATCGGAAG |
| 33169_2 | TATGAGTGTGGAGTCGTTGCCAGTAAGCCGAGATCATGACACTNNNAGATCGGAAG |
| 49256_1 | TATGAGTGTGGAGTCGTTGCCTCCTGAGTAGCTGGGACTACAGNNNAGATCGGAAG |
| 49256_2 | TATGAGTGTGGAGTCGTTGCCTCCTGAGTAGCTGGGACTACAGNNNAGATCGGAAG |
| 38181_1 | TATGAGTGTGGAGTCGTTGCCTCACTCTTGTAATCCCAGCACTNNNAGATCGGAAGA |
| 38181_2 | TATGAGTGTGGAGTCGTTGCCTCACTCTTGTAATCCCAGCACTNNNAGATCGGAAGA |
| 10855_1 | TATGAGTGTGGAGTCGTTGCAGGAAGTGTGTAGGAGGTGACAANNNAGATCGGAA |
| 10855_2 | TATGAGTGTGGAGTCGTTGCAGGAAGTGTGTAGGAGGTGACAANNNAGATCGGAA |

| | |
|---|---|
| 42313_1 | TATGAGTGTGGAGTCGTTGCTAAAAGCAGCAATGGAAATGCCANNNAGATCGGAA |
| 42313_2 | TATGAGTGTGGAGTCGTTGCTAAAAGCAGCAATGGAAATGCCANNNAGATCGGAA |
| 38991_1 | TATGAGTGTGGAGTCGTTGCTTACAAGGGACATGAAGGACGTCNNNAGATCGGAA |
| 38991_2 | TATGAGTGTGGAGTCGTTGCTTACAAGGGACATGAAGGACGTCNNNAGATCGGAA |
| 1138_1 | TATGAGTGTGGAGTCGTTGCGTTTCCAGGACTACTTGGTGCTTNNNAGATCGGAAG |
| 1138_2 | TATGAGTGTGGAGTCGTTGCGTTTCCAGGACTACTTGGTGCTTNNNAGATCGGAAG |
| 10795_1 | TATGAGTGTGGAGTCGTTGCTTTGTAGCTTCACCAACTGCCTTNNNAGATCGGAAGA |
| 10795_2 | TATGAGTGTGGAGTCGTTGCTTTGTAGCTTCACCAACTGCCTTNNNAGATCGGAAGA |
| 48490_1 | TATGAGTGTGGAGTCGTTGCAAAGGAAAAGCGAATCTCCGGATNNNAGATCGGAA |
| 48490_2 | TATGAGTGTGGAGTCGTTGCAAAGGAAAAGCGAATCTCCGGATNNNAGATCGGAA |
| 13460_1 | TATGAGTGTGGAGTCGTTGCTTTTCAAGTGAGGTAACCAACNNNAGATCGGAAGAG |
| 13460_2 | TATGAGTGTGGAGTCGTTGCTTTTCAAGTGAGGTAACCAACNNNAGATCGGAAGAG |
| 34814_1 | TATGAGTGTGGAGTCGTTGCTGTGGTGTACTAGCTTCTGAACCNNNAGATCGGAAG |
| 34814_2 | TATGAGTGTGGAGTCGTTGCTGTGGTGTACTAGCTTCTGAACCNNNAGATCGGAAG |
| 53187_1 | TATGAGTGTGGAGTCGTTGCACAAACGATCTGCTCTCCTCTTTNNNAGATCGGAAGA |
| 53187_2 | TATGAGTGTGGAGTCGTTGCACAAACGATCTGCTCTCCTCTTTNNNAGATCGGAAGA |
| 50844_1 | TATGAGTGTGGAGTCGTTGCTCTTTGGCCAGTCACAGAAGAATNNNAGATCGGAAG |
| 50844_2 | TATGAGTGTGGAGTCGTTGCTCTTTGGCCAGTCACAGAAGAATNNNAGATCGGAAG |
| 29731_1 | TATGAGTGTGGAGTCGTTGCCCTGTGTTCTCTTTCTTTGCTCGNNNAGATCGGAAGA |
| 29731_2 | TATGAGTGTGGAGTCGTTGCCCTGTGTTCTCTTTCTTTGCTCGNNNAGATCGGAAGA |
| 33396_1 | TATGAGTGTGGAGTCGTTGCGAGCTTGCAGTGAGCCAAGATCNNNAGATCGGAAG |
| 33396_2 | TATGAGTGTGGAGTCGTTGCGAGCTTGCAGTGAGCCAAGATCNNNAGATCGGAAG |
| 34482_1 | TATGAGTGTGGAGTCGTTGCGTGGTGGTGGTGGTGATTTTTATNNNAGATCGGAA |
| 34482_2 | TATGAGTGTGGAGTCGTTGCGTGGTGGTGGTGGTGATTTTTATNNNAGATCGGAA |
| 32239_1 | TATGAGTGTGGAGTCGTTGCGACAAGGTCTCACTCTGTCACCNNNAGATCGGAAGA |
| 32239_2 | TATGAGTGTGGAGTCGTTGCGACAAGGTCTCACTCTGTCACCNNNAGATCGGAAGA |
| 36141_1 | TATGAGTGTGGAGTCGTTGCAGCTATTCCATAGGTACAGCAGCNNNAGATCGGAAG |
| 36141_2 | TATGAGTGTGGAGTCGTTGCAGCTATTCCATAGGTACAGCAGCNNNAGATCGGAAG |
| 40563_1 | TATGAGTGTGGAGTCGTTGCATTACAGTGAAACGCTCTTGCACNNNAGATCGGAAG |
| 40563_2 | TATGAGTGTGGAGTCGTTGCATTACAGTGAAACGCTCTTGCACNNNAGATCGGAAG |
| 39594_1 | TATGAGTGTGGAGTCGTTGCGTCATGCTCTATCCCCTTTCCTTNNNAGATCGGAAGA |
| 39594_2 | TATGAGTGTGGAGTCGTTGCGTCATGCTCTATCCCCTTTCCTTNNNAGATCGGAAGA |
| 44837_1 | TATGAGTGTGGAGTCGTTGCGCTTAGAATCCTCTGGGCCATAGNNNAGATCGGAAG |
| 44837_2 | TATGAGTGTGGAGTCGTTGCGCTTAGAATCCTCTGGGCCATAGNNNAGATCGGAAG |
| 43467_1 | TATGAGTGTGGAGTCGTTGCCCTAGTAGTGGGATTGCTGGATCNNNAGATCGGAA |
| 43467_2 | TATGAGTGTGGAGTCGTTGCCCTAGTAGTGGGATTGCTGGATCNNNAGATCGGAA |
| 39833_1 | TATGAGTGTGGAGTCGTTGCCTGTAGTCCCAGCTACTCAGGAGNNNAGATCGGAAG |
| 39833_2 | TATGAGTGTGGAGTCGTTGCCTGTAGTCCCAGCTACTCAGGAGNNNAGATCGGAAG |
| 47926_1 | TATGAGTGTGGAGTCGTTGCGGTTCAAGTGATTTTCCTGCCTCNNNAGATCGGAAG |
| 47926_2 | TATGAGTGTGGAGTCGTTGCGGTTCAAGTGATTTTCCTGCCTCNNNAGATCGGAAG |
| 41897_1 | TATGAGTGTGGAGTCGTTGCTCCCCTGGTTCAAGTAATTCTCCNNNAGATCGGAAGA |
| 41897_2 | TATGAGTGTGGAGTCGTTGCTCCCCTGGTTCAAGTAATTCTCCNNNAGATCGGAAGA |
| 53103_1 | TATGAGTGTGGAGTCGTTGCAGCCAATATAGAGAGATGCAGGANNNAGATCGGAA |

| | |
|---|---|
| 53103_2 | TATGAGTGTGGAGTCGTTGCAGCCAATATAGAGAGATGCAGGANNNAGATCGGAA |
| 47840_1 | TATGAGTGTGGAGTCGTTGCAAGTAGCTGGGATTACAGGCATCNNNAGATCGGAA |
| 47840_2 | TATGAGTGTGGAGTCGTTGCAAGTAGCTGGGATTACAGGCATCNNNAGATCGGAA |
| 5442_1 | TATGAGTGTGGAGTCGTTGCGGTTCCTGATGGTACTAACTGGGNNNAGATCGGAA |
| 5442_2 | TATGAGTGTGGAGTCGTTGCGGTTCCTGATGGTACTAACTGGGNNNAGATCGGAA |
| 50401_1 | TATGAGTGTGGAGTCGTTGCCAGCCTGTGAAGGGAAGGAGNNNAGATCGGAAGA |
| 50401_2 | TATGAGTGTGGAGTCGTTGCCAGCCTGTGAAGGGAAGGAGNNNAGATCGGAAGA |
| 37196_1 | TATGAGTGTGGAGTCGTTGCTCAGTAGTTTAGGCGTGGTCTTGNNNAGATCGGAA |
| 37196_2 | TATGAGTGTGGAGTCGTTGCTCAGTAGTTTAGGCGTGGTCTTGNNNAGATCGGAA |
| 2440_1 | TATGAGTGTGGAGTCGTTGCTCTTCCAGAACCTTCCCTCTCTCNNNAGATCGGAAGA |
| 2440_2 | TATGAGTGTGGAGTCGTTGCTCTTCCAGAACCTTCCCTCTCTCNNNAGATCGGAAGA |
| 32432_1 | TATGAGTGTGGAGTCGTTGCTTTTTTTCCCCCACAGCTTTGAGNNNAGATCGGAAGA |
| 32432_2 | TATGAGTGTGGAGTCGTTGCTTTTTTTCCCCCACAGCTTTGAGNNNAGATCGGAAGA |
| 31480_1 | TATGAGTGTGGAGTCGTTGCGGCTAAGATGACACGTAGCTACANNNAGATCGGAA |
| 31480_2 | TATGAGTGTGGAGTCGTTGCGGCTAAGATGACACGTAGCTACANNNAGATCGGAA |
| 2003_1 | TATGAGTGTGGAGTCGTTGCTCATGGAGGAGACAGATGACAAANNNAGATCGGAA |
| 2003_2 | TATGAGTGTGGAGTCGTTGCTCATGGAGGAGACAGATGACAAANNNAGATCGGAA |
| 31951_1 | TATGAGTGTGGAGTCGTTGCTGATTGGACAAGACGCTTCCATANNNAGATCGGAAG |
| 31951_2 | TATGAGTGTGGAGTCGTTGCTGATTGGACAAGACGCTTCCATANNNAGATCGGAAG |
| 39549_1 | TATGAGTGTGGAGTCGTTGCAAAAACACACATTGCTTTGCAGTNNNAGATCGGAAG |
| 39549_2 | TATGAGTGTGGAGTCGTTGCAAAAACACACATTGCTTTGCAGTNNNAGATCGGAAG |
| 40985_1 | TATGAGTGTGGAGTCGTTGCGCAGAAGAATCACTTGAACCCAGNNNAGATCGGAA |
| 40985_2 | TATGAGTGTGGAGTCGTTGCGCAGAAGAATCACTTGAACCCAGNNNAGATCGGAA |
| 47311_1 | TATGAGTGTGGAGTCGTTGCGTCTCCCAGGTTCAAGTGATTCTNNNAGATCGGAAG |
| 47311_2 | TATGAGTGTGGAGTCGTTGCGTCTCCCAGGTTCAAGTGATTCTNNNAGATCGGAAG |
| 30531_1 | TATGAGTGTGGAGTCGTTGCATGATTGTGACAGAGGGAGACGTAGNNNAGATCG |
| 30531_2 | TATGAGTGTGGAGTCGTTGCATGATTGTGACAGAGGGAGACGTAGNNNAGATCG |
| 42447_1 | TATGAGTGTGGAGTCGTTGCTCTCTCTCTCTCTCTCTCTCTCTNNNAGATCGGAAG |
| 42447_2 | TATGAGTGTGGAGTCGTTGCTCTCTCTCTCTCTCTCTCTCTCTNNNAGATCGGAAG |
| 38106_1 | TATGAGTGTGGAGTCGTTGCACCACAATACCATTATCACGCCTNNNAGATCGGAAG |
| 38106_2 | TATGAGTGTGGAGTCGTTGCACCACAATACCATTATCACGCCTNNNAGATCGGAAG |
| 50748_1 | TATGAGTGTGGAGTCGTTGCCCGATCATTGCCCAAGGATCTTANNNAGATCGGAAG |
| 50748_2 | TATGAGTGTGGAGTCGTTGCCCGATCATTGCCCAAGGATCTTANNNAGATCGGAAG |
| 33328_1 | TATGAGTGTGGAGTCGTTGCCATGTAGCTGGGACTACAGGCNNNAGATCGGAAGA |
| 33328_2 | TATGAGTGTGGAGTCGTTGCCATGTAGCTGGGACTACAGGCNNNAGATCGGAAGA |
| 49273_1 | TATGAGTGTGGAGTCGTTGCCATTCTCCTGCCTCAGACTACTGNNNAGATCGGAAG |
| 49273_2 | TATGAGTGTGGAGTCGTTGCCATTCTCCTGCCTCAGACTACTGNNNAGATCGGAAG |
| 45865_1 | TATGAGTGTGGAGTCGTTGCCACCATGCCCAGCTAATTTTTGTNNNAGATCGGAAG |
| 45865_2 | TATGAGTGTGGAGTCGTTGCCACCATGCCCAGCTAATTTTTGTNNNAGATCGGAAG |
| 47861_1 | TATGAGTGTGGAGTCGTTGCCACGCCTGGCTAATCTTTTGTATNNNAGATCGGAAG |
| 47861_2 | TATGAGTGTGGAGTCGTTGCCACGCCTGGCTAATCTTTTGTATNNNAGATCGGAAG |
| 41628_1 | TATGAGTGTGGAGTCGTTGCGAAAATGGAACCTCGCTCTGTCNNNAGATCGGAAG |
| 41628_2 | TATGAGTGTGGAGTCGTTGCGAAAATGGAACCTCGCTCTGTCNNNAGATCGGAAG |

| | |
|---|---|
| 45200_1 | TATGAGTGTGGAGTCGTTGCCCTTGGTGTAACGCTGTGAATTTNNNAGATCGGAAG |
| 45200_2 | TATGAGTGTGGAGTCGTTGCCCTTGGTGTAACGCTGTGAATTTNNNAGATCGGAAG |
| 49822_1 | TATGAGTGTGGAGTCGTTGCAATGGAGTGGAGTGGAAACAAGTNNNAGATCGGA |
| 49822_2 | TATGAGTGTGGAGTCGTTGCAATGGAGTGGAGTGGAAACAAGTNNNAGATCGGA |
| 32146_1 | TATGAGTGTGGAGTCGTTGCTCTCCCAAAGTGCTGGAATTACANNNAGATCGGAAG |
| 32146_2 | TATGAGTGTGGAGTCGTTGCTCTCCCAAAGTGCTGGAATTACANNNAGATCGGAAG |
| 31669_1 | TATGAGTGTGGAGTCGTTGCGCAGCGTCTTGCATTTTGACTNNNAGATCGGAAGAG |
| 31669_2 | TATGAGTGTGGAGTCGTTGCGCAGCGTCTTGCATTTTGACTNNNAGATCGGAAGAG |
| 29609_1 | TATGAGTGTGGAGTCGTTGCTGTAGCTCTTGTCTGGAGATCCTNNNAGATCGGAAG |
| 29609_2 | TATGAGTGTGGAGTCGTTGCTGTAGCTCTTGTCTGGAGATCCTNNNAGATCGGAAG |
| 43689_1 | TATGAGTGTGGAGTCGTTGCATGACCCATGATGAAACCCATGANNNAGATCGGAAG |
| 43689_2 | TATGAGTGTGGAGTCGTTGCATGACCCATGATGAAACCCATGANNNAGATCGGAAG |
| 43351_1 | TATGAGTGTGGAGTCGTTGCTCAAGGAATTCCACTGCCTTAGCNNNAGATCGGAAG |
| 43351_2 | TATGAGTGTGGAGTCGTTGCTCAAGGAATTCCACTGCCTTAGCNNNAGATCGGAAG |
| 50773_1 | TATGAGTGTGGAGTCGTTGCGGCCATCTTCCTCCTCGAAAGNNNAGATCGGAAGAG |
| 50773_2 | TATGAGTGTGGAGTCGTTGCGGCCATCTTCCTCCTCGAAAGNNNAGATCGGAAGAG |
| 11200_1 | TATGAGTGTGGAGTCGTTGCGTTACCAAGGGCAGAAACCACNNNAGATCGGAAGA |
| 11200_2 | TATGAGTGTGGAGTCGTTGCGTTACCAAGGGCAGAAACCACNNNAGATCGGAAGA |
| 32674_1 | TATGAGTGTGGAGTCGTTGCGAGATGGAGTGTCGTTCTGTTGCNNNAGATCGGAA |
| 32674_2 | TATGAGTGTGGAGTCGTTGCGAGATGGAGTGTCGTTCTGTTGCNNNAGATCGGAA |
| 31474_1 | TATGAGTGTGGAGTCGTTGCCCCCTTCTCCTGTCAGAATCATTNNNAGATCGGAAGA |
| 31474_2 | TATGAGTGTGGAGTCGTTGCCCCCTTCTCCTGTCAGAATCATTNNNAGATCGGAAGA |
| 34356_1 | TATGAGTGTGGAGTCGTTGCTCCCGGGCTAGTGCTTTATTTANNNAGATCGGAAG |
| 34356_2 | TATGAGTGTGGAGTCGTTGCTCCCGGGCTAGTGCTTTATTTANNNAGATCGGAAG |
| 12797_1 | TATGAGTGTGGAGTCGTTGCCCAATTCAACTTTTTTCCCAGNNNAGATCGGAAGAGC |
| 12797_2 | TATGAGTGTGGAGTCGTTGCCCAATTCAACTTTTTTCCCAGNNNAGATCGGAAGAGC |
| 39604_1 | TATGAGTGTGGAGTCGTTGCTAAAACATTGAAGCCATGCGTGTNNNAGATCGGAA |
| 39604_2 | TATGAGTGTGGAGTCGTTGCTAAAACATTGAAGCCATGCGTGTNNNAGATCGGAA |
| 47206_1 | TATGAGTGTGGAGTCGTTGCCAGTAGTGCAGTCATAGCTCACTNNNAGATCGGAAG |
| 47206_2 | TATGAGTGTGGAGTCGTTGCCAGTAGTGCAGTCATAGCTCACTNNNAGATCGGAAG |
| 47357_1 | TATGAGTGTGGAGTCGTTGCTCTATTCCGACTGGTGTGAGATGNNNAGATCGGAA |
| 47357_2 | TATGAGTGTGGAGTCGTTGCTCTATTCCGACTGGTGTGAGATGNNNAGATCGGAA |
| 8902_1 | TATGAGTGTGGAGTCGTTGCAAAGTACAGAGCAGTTCCTGGGTNNNAGATCGGAA |
| 8902_2 | TATGAGTGTGGAGTCGTTGCAAAGTACAGAGCAGTTCCTGGGTNNNAGATCGGAA |
| 36582_1 | TATGAGTGTGGAGTCGTTGCCAGTAGCATGATGTTGGCTCACTNNNAGATCGGAAG |
| 36582_2 | TATGAGTGTGGAGTCGTTGCCAGTAGCATGATGTTGGCTCACTNNNAGATCGGAAG |
| 31390_1 | TATGAGTGTGGAGTCGTTGCTTAGTGTGAAACTAGCCAGGACCNNNAGATCGGAA |
| 31390_2 | TATGAGTGTGGAGTCGTTGCTTAGTGTGAAACTAGCCAGGACCNNNAGATCGGAA |
| 13077_1 | TATGAGTGTGGAGTCGTTGCGTCATTCTTATTGGTGACGTTNNNAGATCGGAAGAG |
| 13077_2 | TATGAGTGTGGAGTCGTTGCGTCATTCTTATTGGTGACGTTNNNAGATCGGAAGAG |
| 35136_1 | TATGAGTGTGGAGTCGTTGCCAGCTTAGCCATTTGCATGTCTTNNNAGATCGGAAG |
| 35136_2 | TATGAGTGTGGAGTCGTTGCCAGCTTAGCCATTTGCATGTCTTNNNAGATCGGAAG |
| 13334_1 | TATGAGTGTGGAGTCGTTGCCCTTTTGTGTGTTCTTTTATGANNNAGATCGGAAGA |

| | |
|---|---|
| 13334_2 | TATGAGTGTGGAGTCGTTGCCCTTTTGTGTGTTCTTTTATGANNNAGATCGGAAGAG |
| 44616_1 | TATGAGTGTGGAGTCGTTGCCTTGCAGGGCATCAAATCCAGNNNAGATCGGAAGAG |
| 44616_2 | TATGAGTGTGGAGTCGTTGCCTTGCAGGGCATCAAATCCAGNNNAGATCGGAAGAG |
| 13350_1 | TATGAGTGTGGAGTCGTTGCTGTAGTTTATTTCAGGAAGACTGNNNAGATCGGAAG |
| 13350_2 | TATGAGTGTGGAGTCGTTGCTGTAGTTTATTTCAGGAAGACTGNNNAGATCGGAAG |
| 38651_1 | TATGAGTGTGGAGTCGTTGCTAACTTCTCCCTTTTCAGCCATTNNNAGATCGGAAGA |
| 38651_2 | TATGAGTGTGGAGTCGTTGCTAACTTCTCCCTTTTCAGCCATTNNNAGATCGGAAGA |
| 47529_1 | TATGAGTGTGGAGTCGTTGCCAATCTCAGCTCAATGCAACCTCNNNAGATCGGAAGA |
| 47529_2 | TATGAGTGTGGAGTCGTTGCCAATCTCAGCTCAATGCAACCTCNNNAGATCGGAAGA |
| 35074_1 | TATGAGTGTGGAGTCGTTGCCACCACGCCTGGCTAATTTTTTTNNNAGATCGGAAGA |
| 35074_2 | TATGAGTGTGGAGTCGTTGCCACCACGCCTGGCTAATTTTTTTNNNAGATCGGAAGA |
| 47979_1 | TATGAGTGTGGAGTCGTTGCCAAAGTGCTGGGATTACAGGTCTNNNAGATCGGAA |
| 47979_2 | TATGAGTGTGGAGTCGTTGCCAAAGTGCTGGGATTACAGGTCTNNNAGATCGGAA |
| 28977_1 | TATGAGTGTGGAGTCGTTGCTGTCTTCTGGCCTCTGTAGTTTCNNNAGATCGGAAGA |
| 28977_2 | TATGAGTGTGGAGTCGTTGCTGTCTTCTGGCCTCTGTAGTTTCNNNAGATCGGAAGA |
| 5011_1 | TATGAGTGTGGAGTCGTTGCGGGGCATGCTTTTTATACTCAGCNNNAGATCGGAAG |
| 5011_2 | TATGAGTGTGGAGTCGTTGCGGGGCATGCTTTTTATACTCAGCNNNAGATCGGAAG |
| 13632_1 | TATGAGTGTGGAGTCGTTGCTGGGAAGAATGGTTTATGTCNNNAGATCGGAAGAG |
| 13632_2 | TATGAGTGTGGAGTCGTTGCTGGGAAGAATGGTTTATGTCNNNAGATCGGAAGAG |
| 30908_1 | TATGAGTGTGGAGTCGTTGCAAACTACTAGCAAACTGAGGCCANNNAGATCGGAAG |
| 30908_2 | TATGAGTGTGGAGTCGTTGCAAACTACTAGCAAACTGAGGCCANNNAGATCGGAAG |
| 41615_1 | TATGAGTGTGGAGTCGTTGCGATTAGAGCTATGAGCCACCACANNNAGATCGGAAG |
| 41615_2 | TATGAGTGTGGAGTCGTTGCGATTAGAGCTATGAGCCACCACANNNAGATCGGAAG |
| 13644_1 | TATGAGTGTGGAGTCGTTGCCTAGTTTGAGAGAAAGTCCTATGNNNAGATCGGAAG |
| 13644_2 | TATGAGTGTGGAGTCGTTGCCTAGTTTGAGAGAAAGTCCTATGNNNAGATCGGAAG |
| 43020_1 | TATGAGTGTGGAGTCGTTGCCCTGCACCCTAACTAGTATGCAANNNAGATCGGAAGA |
| 43020_2 | TATGAGTGTGGAGTCGTTGCCCTGCACCCTAACTAGTATGCAANNNAGATCGGAAGA |
| 52875_1 | TATGAGTGTGGAGTCGTTGCAGAAAAGGAGAGCTTTGGTTCCTNNNAGATCGGAA |
| 52875_2 | TATGAGTGTGGAGTCGTTGCAGAAAAGGAGAGCTTTGGTTCCTNNNAGATCGGAA |
| 39544_1 | TATGAGTGTGGAGTCGTTGCCCATAGCAAGACTCTGTCTCACANNNAGATCGGAAGA |
| 39544_2 | TATGAGTGTGGAGTCGTTGCCCATAGCAAGACTCTGTCTCACANNNAGATCGGAAGA |
| 37972_1 | TATGAGTGTGGAGTCGTTGCCACTTGAAAATTTGGCACTTGGCNNNAGATCGGAAG |
| 37972_2 | TATGAGTGTGGAGTCGTTGCCACTTGAAAATTTGGCACTTGGCNNNAGATCGGAAG |
| 10236_1 | TATGAGTGTGGAGTCGTTGCGGCTGTCCTCAGATGCCTATTCTNNNAGATCGGAAGA |
| 10236_2 | TATGAGTGTGGAGTCGTTGCGGCTGTCCTCAGATGCCTATTCTNNNAGATCGGAAGA |
| 38034_1 | TATGAGTGTGGAGTCGTTGCTGGCAGTATCGCTGAGTTTTAGTNNNAGATCGGAAG |
| 38034_2 | TATGAGTGTGGAGTCGTTGCTGGCAGTATCGCTGAGTTTTAGTNNNAGATCGGAAG |
| 49932_1 | TATGAGTGTGGAGTCGTTGCCATGATCCTGACTCTGTTCTGGTNNNAGATCGGAAGA |
| 49932_2 | TATGAGTGTGGAGTCGTTGCCATGATCCTGACTCTGTTCTGGTNNNAGATCGGAAGA |
| 30152_1 | TATGAGTGTGGAGTCGTTGCACCCAAAGTGTAAAAGTGCAGGTNNNAGATCGGAA |
| 30152_2 | TATGAGTGTGGAGTCGTTGCACCCAAAGTGTAAAAGTGCAGGTNNNAGATCGGAA |
| 47318_1 | TATGAGTGTGGAGTCGTTGCCTCCCGAGTAGCTGTGACTACAGNNNAGATCGGAAG |
| 47318_2 | TATGAGTGTGGAGTCGTTGCCTCCCGAGTAGCTGTGACTACAGNNNAGATCGGAAG |

| | |
|---|---|
| 7831_1 | TATGAGTGTGGAGTCGTTGCGGCAGACACAATAAAGACACAAANNNAGATCGGAA |
| 7831_2 | TATGAGTGTGGAGTCGTTGCGGCAGACACAATAAAGACACAAANNNAGATCGGAA |
| 40742_1 | TATGAGTGTGGAGTCGTTGCAGATTGCTGGTGGACTGAGAAATNNNAGATCGGAA |
| 40742_2 | TATGAGTGTGGAGTCGTTGCAGATTGCTGGTGGACTGAGAAATNNNAGATCGGAA |
| 47683_1 | TATGAGTGTGGAGTCGTTGCCTGGAGTGTAATGGTGCAGTCTCNNNAGATCGGAA |
| 47683_2 | TATGAGTGTGGAGTCGTTGCCTGGAGTGTAATGGTGCAGTCTCNNNAGATCGGAA |
| 50639_1 | TATGAGTGTGGAGTCGTTGCGTGAGCAGCACAGAAAATCAGACNNNAGATCGGAA |
| 50639_2 | TATGAGTGTGGAGTCGTTGCGTGAGCAGCACAGAAAATCAGACNNNAGATCGGAA |
| 40457_1 | TATGAGTGTGGAGTCGTTGCGGAATGGGAAGCAGGAACTATGANNNAGATCGGA |
| 40457_2 | TATGAGTGTGGAGTCGTTGCGGAATGGGAAGCAGGAACTATGANNNAGATCGGA |
| 46212_1 | TATGAGTGTGGAGTCGTTGCGGTTCAAGTATTCTCCTGCCTCANNNAGATCGGAAG |
| 46212_2 | TATGAGTGTGGAGTCGTTGCGGTTCAAGTATTCTCCTGCCTCANNNAGATCGGAAG |
| 29636_1 | TATGAGTGTGGAGTCGTTGCACACCTGGATAATCACTTCACCTNNNAGATCGGAAG |
| 29636_2 | TATGAGTGTGGAGTCGTTGCACACCTGGATAATCACTTCACCTNNNAGATCGGAAG |
| 49205_1 | TATGAGTGTGGAGTCGTTGCCGATCTCAGCTCACTGTAACCTTNNNAGATCGGAAG |
| 49205_2 | TATGAGTGTGGAGTCGTTGCCGATCTCAGCTCACTGTAACCTTNNNAGATCGGAAG |
| 47894_1 | TATGAGTGTGGAGTCGTTGCTCTGCGTCTGTGTTCATGACTTANNNAGATCGGAAG |
| 47894_2 | TATGAGTGTGGAGTCGTTGCTCTGCGTCTGTGTTCATGACTTANNNAGATCGGAAG |
| 48598_1 | TATGAGTGTGGAGTCGTTGCAAGTGTTGGGATTACAGGTGTGANNNAGATCGGAA |
| 48598_2 | TATGAGTGTGGAGTCGTTGCAAGTGTTGGGATTACAGGTGTGANNNAGATCGGAA |
| 50167_1 | TATGAGTGTGGAGTCGTTGCGCCCCAAAAATCAAACTTACCAANNNAGATCGGAAG |
| 50167_2 | TATGAGTGTGGAGTCGTTGCGCCCCAAAAATCAAACTTACCAANNNAGATCGGAAG |
| 29352_1 | TATGAGTGTGGAGTCGTTGCAATGGCAATATCTTGGTTGTGGCNNNAGATCGGAA |
| 29352_2 | TATGAGTGTGGAGTCGTTGCAATGGCAATATCTTGGTTGTGGCNNNAGATCGGAA |
| 49987_1 | TATGAGTGTGGAGTCGTTGCCTGAAGACATAGCCAAGCTTGTGNNNAGATCGGAA |
| 49987_2 | TATGAGTGTGGAGTCGTTGCCTGAAGACATAGCCAAGCTTGTGNNNAGATCGGAA |
| 39024_1 | TATGAGTGTGGAGTCGTTGCGTTCTGTGTGGGAAATGCATGAANNNAGATCGGAA |
| 39024_2 | TATGAGTGTGGAGTCGTTGCGTTCTGTGTGGGAAATGCATGAANNNAGATCGGAA |
| 13075_1 | TATGAGTGTGGAGTCGTTGCCCTCTCTTTTAATAAACGTTTCTNNNAGATCGGAAGA |
| 13075_2 | TATGAGTGTGGAGTCGTTGCCCTCTCTTTTAATAAACGTTTCTNNNAGATCGGAAGA |
| 47942_1 | TATGAGTGTGGAGTCGTTGCCGCCCAGCCTCATTCTAATTACTNNNAGATCGGAAGA |
| 47942_2 | TATGAGTGTGGAGTCGTTGCCGCCCAGCCTCATTCTAATTACTNNNAGATCGGAAGA |
| 50042_1 | TATGAGTGTGGAGTCGTTGCCTATGGGTCTGCAAAAACCACAGNNNAGATCGGAAG |
| 50042_2 | TATGAGTGTGGAGTCGTTGCCTATGGGTCTGCAAAAACCACAGNNNAGATCGGAAG |
| 33334_1 | TATGAGTGTGGAGTCGTTGCTGCATGATCTCAGCTCACTGTAGNNNAGATCGGAAG |
| 33334_2 | TATGAGTGTGGAGTCGTTGCTGCATGATCTCAGCTCACTGTAGNNNAGATCGGAAG |
| 49255_1 | TATGAGTGTGGAGTCGTTGCTCAGTAGGGATCAGCCATACAGANNNAGATCGGAA |
| 49255_2 | TATGAGTGTGGAGTCGTTGCTCAGTAGGGATCAGCCATACAGANNNAGATCGGAA |
| 29843_1 | TATGAGTGTGGAGTCGTTGCCAGGGGGTTCAAATTCAAGAAAGTNNNAGATCGGA |
| 29843_2 | TATGAGTGTGGAGTCGTTGCCAGGGGGTTCAAATTCAAGAAAGTNNNAGATCGGA |
| 45130_1 | TATGAGTGTGGAGTCGTTGCCACTCTCACTACAACAAACGCTGNNNAGATCGGAAG |
| 45130_2 | TATGAGTGTGGAGTCGTTGCCACTCTCACTACAACAAACGCTGNNNAGATCGGAAG |
| 34976_1 | TATGAGTGTGGAGTCGTTGCTCTCCTGAGTAGCTGGGAATACANNNAGATCGGAA |

| | |
|---|---|
| 34976_2 | TATGAGTGTGGAGTCGTTGCTCTCCTGAGTAGCTGGGAATACANNNAGATCGGAAG |
| 40950_1 | TATGAGTGTGGAGTCGTTGCTGCCTGTAATCTCAGCTACTCAGNNNAGATCGGAAGA |
| 40950_2 | TATGAGTGTGGAGTCGTTGCTGCCTGTAATCTCAGCTACTCAGNNNAGATCGGAAGA |
| 31544_1 | TATGAGTGTGGAGTCGTTGCGAGAGAAAGAGGGGGAGAGAGAGNNNAGATCGGA |
| 31544_2 | TATGAGTGTGGAGTCGTTGCGAGAGAAAGAGGGGGAGAGAGAGNNNAGATCGGA |
| 38085_1 | TATGAGTGTGGAGTCGTTGCTGGCAGGGAGTTGGTTGTTATAANNNAGATCGGAA |
| 38085_2 | TATGAGTGTGGAGTCGTTGCTGGCAGGGAGTTGGTTGTTATAANNNAGATCGGAA |
| 39803_1 | TATGAGTGTGGAGTCGTTGCCGTTATGGGACTGTAAATGGTGCNNNAGATCGGAAG |
| 39803_2 | TATGAGTGTGGAGTCGTTGCCGTTATGGGACTGTAAATGGTGCNNNAGATCGGAAG |
| 48388_1 | TATGAGTGTGGAGTCGTTGCTGCATTCACATTATGTTGCCTCCNNNAGATCGGAAGA |
| 48388_2 | TATGAGTGTGGAGTCGTTGCTGCATTCACATTATGTTGCCTCCNNNAGATCGGAAGA |
| 48509_1 | TATGAGTGTGGAGTCGTTGCTTCAAGTGATTCTCCTGCCTCAGNNNAGATCGGAAGA |
| 48509_2 | TATGAGTGTGGAGTCGTTGCTTCAAGTGATTCTCCTGCCTCAGNNNAGATCGGAAGA |
| 39076_1 | TATGAGTGTGGAGTCGTTGCTCTACCCAGAGGAAAAGAAGTCANNNAGATCGGAAG |
| 39076_2 | TATGAGTGTGGAGTCGTTGCTCTACCCAGAGGAAAAGAAGTCANNNAGATCGGAAG |
| 33210_1 | TATGAGTGTGGAGTCGTTGCACAGAAGTTGAAGAGAGGAAGGANNNAGATCGGA |
| 33210_2 | TATGAGTGTGGAGTCGTTGCACAGAAGTTGAAGAGAGGAAGGANNNAGATCGGA |
| 45037_1 | TATGAGTGTGGAGTCGTTGCAGGAAAGCAAGAGATGGCAGTTANNNAGATCGGAA |
| 45037_2 | TATGAGTGTGGAGTCGTTGCAGGAAAGCAAGAGATGGCAGTTANNNAGATCGGAA |
| 43516_1 | TATGAGTGTGGAGTCGTTGCTTCAAGTGATTCTCCTGTCTCGGNNNAGATCGGAAGA |
| 43516_2 | TATGAGTGTGGAGTCGTTGCTTCAAGTGATTCTCCTGTCTCGGNNNAGATCGGAAGA |
| 34099_1 | TATGAGTGTGGAGTCGTTGCCTATGCGTCTTCTGCAAAGCAATNNNAGATCGGAAGA |
| 34099_2 | TATGAGTGTGGAGTCGTTGCCTATGCGTCTTCTGCAAAGCAATNNNAGATCGGAAGA |
| 46496_1 | TATGAGTGTGGAGTCGTTGCAGTAGCTGGAATTACAGACCTGCNNNAGATCGGAAG |
| 46496_2 | TATGAGTGTGGAGTCGTTGCAGTAGCTGGAATTACAGACCTGCNNNAGATCGGAAG |
| 41361_1 | TATGAGTGTGGAGTCGTTGCACATCTTCGACTGTTTCATTGGCNNNAGATCGGAAGA |
| 41361_2 | TATGAGTGTGGAGTCGTTGCACATCTTCGACTGTTTCATTGGCNNNAGATCGGAAGA |
| 40302_1 | TATGAGTGTGGAGTCGTTGCTTCCTGCAAGATCTCTGGTTTCANNNAGATCGGAAGA |
| 40302_2 | TATGAGTGTGGAGTCGTTGCTTCCTGCAAGATCTCTGGTTTCANNNAGATCGGAAGA |
| 51198_1 | TATGAGTGTGGAGTCGTTGCAAGCAGAAAGAAGAGGTATGGCANNNAGATCGGAA |
| 51198_2 | TATGAGTGTGGAGTCGTTGCAAGCAGAAAGAAGAGGTATGGCANNNAGATCGGAA |
| 48327_1 | TATGAGTGTGGAGTCGTTGCCAGTGACATGATCTCAGCTCACTNNNAGATCGGAAGA |
| 48327_2 | TATGAGTGTGGAGTCGTTGCCAGTGACATGATCTCAGCTCACTNNNAGATCGGAAGA |
| 39343_1 | TATGAGTGTGGAGTCGTTGCTTGAGGGAGGAATGCTTTACCTCNNNAGATCGGAAG |
| 39343_2 | TATGAGTGTGGAGTCGTTGCTTGAGGGAGGAATGCTTTACCTCNNNAGATCGGAAG |
| 35591_1 | TATGAGTGTGGAGTCGTTGCTCCAAGTGATTCTTCTGCCTCAGNNNAGATCGGAAGA |
| 35591_2 | TATGAGTGTGGAGTCGTTGCTCCAAGTGATTCTTCTGCCTCAGNNNAGATCGGAAGA |
| 35746_1 | TATGAGTGTGGAGTCGTTGCTTCAAGTGATTCTCCTGCCTCAGNNNAGATCGGAAGA |
| 35746_2 | TATGAGTGTGGAGTCGTTGCTTCAAGTGATTCTCCTGCCTCAGNNNAGATCGGAAGA |
| 44714_1 | TATGAGTGTGGAGTCGTTGCCCCATAGTCTCATCTTCTGGGTGNNNAGATCGGAAGA |
| 44714_2 | TATGAGTGTGGAGTCGTTGCCCCATAGTCTCATCTTCTGGGTGNNNAGATCGGAAGA |
| 34257_1 | TATGAGTGTGGAGTCGTTGCATGCTAGATGACGAGTTAGTGGGNNNAGATCGGAA |
| 34257_2 | TATGAGTGTGGAGTCGTTGCATGCTAGATGACGAGTTAGTGGGNNNAGATCGGAA |

| | |
|---|---|
| 44411_1 | TATGAGTGTGGAGTCGTTGCCTGCCCTTGAGCTTTCAATTGAANNNAGATCGGAAG |
| 44411_2 | TATGAGTGTGGAGTCGTTGCCTGCCCTTGAGCTTTCAATTGAANNNAGATCGGAAG |
| 35740_1 | TATGAGTGTGGAGTCGTTGCAGAGTAGCTGGGACTACAGATGTNNNAGATCGGAA |
| 35740_2 | TATGAGTGTGGAGTCGTTGCAGAGTAGCTGGGACTACAGATGTNNNAGATCGGAA |
| 47787_1 | TATGAGTGTGGAGTCGTTGCCAGACCTATGGGCTTCCTATGACNNNAGATCGGAAG |
| 47787_2 | TATGAGTGTGGAGTCGTTGCCAGACCTATGGGCTTCCTATGACNNNAGATCGGAAG |
| 9110_1 | TATGAGTGTGGAGTCGTTGCAAGCTCCAACTCAAACGACAACTNNNAGATCGGAAG |
| 9110_2 | TATGAGTGTGGAGTCGTTGCAAGCTCCAACTCAAACGACAACTNNNAGATCGGAAG |
| 30638_1 | TATGAGTGTGGAGTCGTTGCAATCCCAACACCCCCACCTTCTNNNAGATCGGAAGAG |
| 30638_2 | TATGAGTGTGGAGTCGTTGCAATCCCAACACCCCCACCTTCTNNNAGATCGGAAGAG |
| 34776_1 | TATGAGTGTGGAGTCGTTGCCCTCCTGAGTAGCTGAGACTACANNNAGATCGGAAG |
| 34776_2 | TATGAGTGTGGAGTCGTTGCCCTCCTGAGTAGCTGAGACTACANNNAGATCGGAAG |
| 52271_1 | TATGAGTGTGGAGTCGTTGCCCTTAGAGGCTGATGATTGAGCTNNNAGATCGGAA |
| 52271_2 | TATGAGTGTGGAGTCGTTGCCCTTAGAGGCTGATGATTGAGCTNNNAGATCGGAA |
| 34441_1 | TATGAGTGTGGAGTCGTTGCGGATTTCCCTCGGCTAGAAAGAANNNAGATCGGAA |
| 34441_2 | TATGAGTGTGGAGTCGTTGCGGATTTCCCTCGGCTAGAAAGAANNNAGATCGGAA |
| 48529_1 | TATGAGTGTGGAGTCGTTGCTCACTGACCGATATTTGCTGGAANNNAGATCGGAAG |
| 48529_2 | TATGAGTGTGGAGTCGTTGCTCACTGACCGATATTTGCTGGAANNNAGATCGGAAG |
| 33053_1 | TATGAGTGTGGAGTCGTTGCTTCAATAGGTCCCACTCCCAAAGNNNAGATCGGAAG |
| 33053_2 | TATGAGTGTGGAGTCGTTGCTTCAATAGGTCCCACTCCCAAAGNNNAGATCGGAAG |
| 40967_1 | TATGAGTGTGGAGTCGTTGCGGTCCTGGCATAAAAACAGACACNNNAGATCGGAA |
| 40967_2 | TATGAGTGTGGAGTCGTTGCGGTCCTGGCATAAAAACAGACACNNNAGATCGGAA |
| 30718_1 | TATGAGTGTGGAGTCGTTGCCGTGCCCAGCCTATTGTAAGTATNNNAGATCGGAAG |
| 30718_2 | TATGAGTGTGGAGTCGTTGCCGTGCCCAGCCTATTGTAAGTATNNNAGATCGGAAG |
| 47245_1 | TATGAGTGTGGAGTCGTTGCGAGCTGAGATCATGCCATTGAACNNNAGATCGGAA |
| 47245_2 | TATGAGTGTGGAGTCGTTGCGAGCTGAGATCATGCCATTGAACNNNAGATCGGAA |
| 33454_1 | TATGAGTGTGGAGTCGTTGCCCAATCCCCTCAGCATAGTAAGGNNNAGATCGGAAG |
| 33454_2 | TATGAGTGTGGAGTCGTTGCCCAATCCCCTCAGCATAGTAAGGNNNAGATCGGAAG |
| 46484_1 | TATGAGTGTGGAGTCGTTGCCATTTCCCGGGTTCAAGTGATTCNNNAGATCGGAAG |
| 46484_2 | TATGAGTGTGGAGTCGTTGCCATTTCCCGGGTTCAAGTGATTCNNNAGATCGGAAG |
| 42028_1 | TATGAGTGTGGAGTCGTTGCCTGGGTAACAGAGTGAGACACTGNNNAGATCGGAA |
| 42028_2 | TATGAGTGTGGAGTCGTTGCCTGGGTAACAGAGTGAGACACTGNNNAGATCGGAA |
| 34647_1 | TATGAGTGTGGAGTCGTTGCCCTACTGGGAAGTGAGGAGCCNNNAGATCGGAAGA |
| 34647_2 | TATGAGTGTGGAGTCGTTGCCCTACTGGGAAGTGAGGAGCCNNNAGATCGGAAGA |
| 51656_1 | TATGAGTGTGGAGTCGTTGCCCTGCACTAGATTTTGCTTGGAANNNAGATCGGAAG |
| 51656_2 | TATGAGTGTGGAGTCGTTGCCCTGCACTAGATTTTGCTTGGAANNNAGATCGGAAG |
| 39672_1 | TATGAGTGTGGAGTCGTTGCTCCAGTTCATTTCCAAGCCAAACNNNAGATCGGAAG |
| 39672_2 | TATGAGTGTGGAGTCGTTGCTCCAGTTCATTTCCAAGCCAAACNNNAGATCGGAAG |
| 42078_1 | TATGAGTGTGGAGTCGTTGCTTATTGGGCCATGGCTAACTGTANNNAGATCGGAAG |
| 42078_2 | TATGAGTGTGGAGTCGTTGCTTATTGGGCCATGGCTAACTGTANNNAGATCGGAAG |
| 30505_1 | TATGAGTGTGGAGTCGTTGCGGAGGGAAAGTCTAGGGCAAAANNNAGATCGGAA |
| 30505_2 | TATGAGTGTGGAGTCGTTGCGGAGGGAAAGTCTAGGGCAAAANNNAGATCGGAA |
| 34168_1 | TATGAGTGTGGAGTCGTTGCTCAATATGGTGAAACCCCGTCTCNNNAGATCGGAAG |

| | |
|---|---|
| 34168_2 | TATGAGTGTGGAGTCGTTGCTCAATATGGTGAAACCCCGTCTCNNNAGATCGGAAG |
| 43103_1 | TATGAGTGTGGAGTCGTTGCAGGTTCAAGCAATTCTCCTGTCTNNNAGATCGGAAG |
| 43103_2 | TATGAGTGTGGAGTCGTTGCAGGTTCAAGCAATTCTCCTGTCTNNNAGATCGGAAG |
| 33333_1 | TATGAGTGTGGAGTCGTTGCACAATTCTCCTACTTCAGCCTCCNNNAGATCGGAAGA |
| 33333_2 | TATGAGTGTGGAGTCGTTGCACAATTCTCCTACTTCAGCCTCCNNNAGATCGGAAGA |
| 30770_1 | TATGAGTGTGGAGTCGTTGCAGAGAGAGAGAGAGAGAATGGCCNNNAGATCGGA |
| 30770_2 | TATGAGTGTGGAGTCGTTGCAGAGAGAGAGAGAGAGAATGGCCNNNAGATCGGA |
| 35401_1 | TATGAGTGTGGAGTCGTTGCCTCTCGAGCAGCTCAGATTACAGNNNAGATCGGAAG |
| 35401_2 | TATGAGTGTGGAGTCGTTGCCTCTCGAGCAGCTCAGATTACAGNNNAGATCGGAAG |
| 43212_1 | TATGAGTGTGGAGTCGTTGCCAAGTAGCTCAGACTACAGGCATNNNAGATCGGAAG |
| 43212_2 | TATGAGTGTGGAGTCGTTGCCAAGTAGCTCAGACTACAGGCATNNNAGATCGGAAG |
| 9914_1 | TATGAGTGTGGAGTCGTTGCAAACGAAGGGAAGGGAGATCAGANNNAGATCGGA |
| 9914_2 | TATGAGTGTGGAGTCGTTGCAAACGAAGGGAAGGGAGATCAGANNNAGATCGGA |
| 34568_1 | TATGAGTGTGGAGTCGTTGCCTCCCAGGTATTTATGCACCCACNNNAGATCGGAAGA |
| 34568_2 | TATGAGTGTGGAGTCGTTGCCTCCCAGGTATTTATGCACCCACNNNAGATCGGAAGA |
| 50250_1 | TATGAGTGTGGAGTCGTTGCTCCCCACAAACGTGAATGAAATGNNNAGATCGGAAG |
| 50250_2 | TATGAGTGTGGAGTCGTTGCTCCCCACAAACGTGAATGAAATGNNNAGATCGGAAG |
| 51667_1 | TATGAGTGTGGAGTCGTTGCCCCCCGACACACAAATTTCTCTANNNAGATCGGAAGA |
| 51667_2 | TATGAGTGTGGAGTCGTTGCCCCCCGACACACAAATTTCTCTANNNAGATCGGAAGA |
| 2144_1 | TATGAGTGTGGAGTCGTTGCTACAATGTTACCAGTCCTGGTCANNNAGATCGGAAG |
| 2144_2 | TATGAGTGTGGAGTCGTTGCTACAATGTTACCAGTCCTGGTCANNNAGATCGGAAG |
| 42708_1 | TATGAGTGTGGAGTCGTTGCAGTCGGTGGTTGAATACTGATGTNNNAGATCGGAA |
| 42708_2 | TATGAGTGTGGAGTCGTTGCAGTCGGTGGTTGAATACTGATGTNNNAGATCGGAA |
| 42391_1 | TATGAGTGTGGAGTCGTTGCCTCTCTCTCTCTCTCTCTCCCNNNAGATCGGAAGAG |
| 42391_2 | TATGAGTGTGGAGTCGTTGCCTCTCTCTCTCTCTCTCTCCCNNNAGATCGGAAGAG |
| 31382_1 | TATGAGTGTGGAGTCGTTGCACACACACACACACACACAAAATNNNAGATCGGAAG |
| 31382_2 | TATGAGTGTGGAGTCGTTGCACACACACACACACACACAAAATNNNAGATCGGAAG |
| 49107_1 | TATGAGTGTGGAGTCGTTGCGTTTCACCATGTTAGCCAGGATGNNNAGATCGGAAG |
| 49107_2 | TATGAGTGTGGAGTCGTTGCGTTTCACCATGTTAGCCAGGATGNNNAGATCGGAAG |
| 32625_1 | TATGAGTGTGGAGTCGTTGCATCATTTGCAACATGGGTGGAAGNNNAGATCGGAA |
| 32625_2 | TATGAGTGTGGAGTCGTTGCATCATTTGCAACATGGGTGGAAGNNNAGATCGGAA |
| 45921_1 | TATGAGTGTGGAGTCGTTGCACAGCAAAATAAAAAGCCAGCAANNNAGATCGGAA |
| 45921_2 | TATGAGTGTGGAGTCGTTGCACAGCAAAATAAAAAGCCAGCAANNNAGATCGGAA |
| 53120_1 | TATGAGTGTGGAGTCGTTGCTCCCAGTTTGTGGAGAATGATCANNNAGATCGGAAG |
| 53120_2 | TATGAGTGTGGAGTCGTTGCTCCCAGTTTGTGGAGAATGATCANNNAGATCGGAAG |
| 46789_1 | TATGAGTGTGGAGTCGTTGCCTTCCGGTTCAAGCAATTCTCCTNNNAGATCGGAAGA |
| 46789_2 | TATGAGTGTGGAGTCGTTGCCTTCCGGTTCAAGCAATTCTCCTNNNAGATCGGAAGA |
| 52877_1 | TATGAGTGTGGAGTCGTTGCCGATCAGGGAGAGGTGGAGAGNNNAGATCGGAAG |
| 52877_2 | TATGAGTGTGGAGTCGTTGCCGATCAGGGAGAGGTGGAGAGNNNAGATCGGAAG |
| 34189_1 | TATGAGTGTGGAGTCGTTGCCTCAAAACTCCAGCCAATCACAGNNNAGATCGGAAG |
| 34189_2 | TATGAGTGTGGAGTCGTTGCCTCAAAACTCCAGCCAATCACAGNNNAGATCGGAAG |
| 45150_1 | TATGAGTGTGGAGTCGTTGCTCCATGTTCTTGAAGGGGATCAGNNNAGATCGGAA |
| 45150_2 | TATGAGTGTGGAGTCGTTGCTCCATGTTCTTGAAGGGGATCAGNNNAGATCGGAA |

| | |
|---|---|
| 39819_1 | TATGAGTGTGGAGTCGTTGCTTGGAACATAGCCACAGTCATTCNNNAGATCGGAAG |
| 39819_2 | TATGAGTGTGGAGTCGTTGCTTGGAACATAGCCACAGTCATTCNNNAGATCGGAAG |
| 31733_1 | TATGAGTGTGGAGTCGTTGCCCCCTGAACTGGAATACATGGATNNNAGATCGGAAG |
| 31733_2 | TATGAGTGTGGAGTCGTTGCCCCCTGAACTGGAATACATGGATNNNAGATCGGAAG |
| 49132_1 | TATGAGTGTGGAGTCGTTGCGGTTCAAGTGATTTTCCTGCCTCNNNAGATCGGAAG |
| 49132_2 | TATGAGTGTGGAGTCGTTGCGGTTCAAGTGATTTTCCTGCCTCNNNAGATCGGAAG |
| 53192_1 | TATGAGTGTGGAGTCGTTGCGCCTCCCAGATTCAAGTGATTCTNNNAGATCGGAAG |
| 53192_2 | TATGAGTGTGGAGTCGTTGCGCCTCCCAGATTCAAGTGATTCTNNNAGATCGGAAG |
| 52577_1 | TATGAGTGTGGAGTCGTTGCTTGGTGTGCTGGTATAAATCCCANNNAGATCGGAAG |
| 52577_2 | TATGAGTGTGGAGTCGTTGCTTGGTGTGCTGGTATAAATCCCANNNAGATCGGAAG |
| 38838_1 | TATGAGTGTGGAGTCGTTGCGTGCAATCTCAGCTCAATGCAAGNNNAGATCGGAAG |
| 38838_2 | TATGAGTGTGGAGTCGTTGCGTGCAATCTCAGCTCAATGCAAGNNNAGATCGGAAG |
| 215_1 | TATGAGTGTGGAGTCGTTGCACCCAATCTCAGGTATTAGCAGACNNNAGATCGGAA |
| 215_2 | TATGAGTGTGGAGTCGTTGCACCCAATCTCAGGTATTAGCAGACNNNAGATCGGAA |
| 29304_1 | TATGAGTGTGGAGTCGTTGCAAATGGCTTGGGGTGTTTATGTGNNNAGATCGGAA |
| 29304_2 | TATGAGTGTGGAGTCGTTGCAAATGGCTTGGGGTGTTTATGTGNNNAGATCGGAA |
| 53432_1 | TATGAGTGTGGAGTCGTTGCCCTAATGTAAGCACAAGGGTCCTNNNAGATCGGAAG |
| 53432_2 | TATGAGTGTGGAGTCGTTGCCCTAATGTAAGCACAAGGGTCCTNNNAGATCGGAAG |
| 51427_1 | TATGAGTGTGGAGTCGTTGCGTGGGGATGGTTACTTGGTACAANNNAGATCGGAA |
| 51427_2 | TATGAGTGTGGAGTCGTTGCGTGGGGATGGTTACTTGGTACAANNNAGATCGGAA |
| 29208_1 | TATGAGTGTGGAGTCGTTGCTCATTGCTGGATCTTACGGTAGGNNNAGATCGGAAG |
| 29208_2 | TATGAGTGTGGAGTCGTTGCTCATTGCTGGATCTTACGGTAGGNNNAGATCGGAAG |
| 304_1 | TATGAGTGTGGAGTCGTTGCGGACTTATCCCTGGCCTTAGGANNNAGATCGGAAGA |
| 304_2 | TATGAGTGTGGAGTCGTTGCGGACTTATCCCTGGCCTTAGGANNNAGATCGGAAGA |
| 39217_1 | TATGAGTGTGGAGTCGTTGCGAGAATCACTTGAACCTGGGAGTNNNAGATCGGAA |
| 39217_2 | TATGAGTGTGGAGTCGTTGCGAGAATCACTTGAACCTGGGAGTNNNAGATCGGAA |
| 32079_1 | TATGAGTGTGGAGTCGTTGCCATTCTCCTGCCTCAGCCTCNNNAGATCGGAAGAGCA |
| 32079_2 | TATGAGTGTGGAGTCGTTGCCATTCTCCTGCCTCAGCCTCNNNAGATCGGAAGAGCA |
| 51795_1 | TATGAGTGTGGAGTCGTTGCCTTCAGTCTCCCAAGTAGCTGAGNNNAGATCGGAAG |
| 51795_2 | TATGAGTGTGGAGTCGTTGCCTTCAGTCTCCCAAGTAGCTGAGNNNAGATCGGAAG |
| 51998_1 | TATGAGTGTGGAGTCGTTGCTCTGGTTGAAGAAGCAGTCTGGNNNAGATCGGAAG |
| 51998_2 | TATGAGTGTGGAGTCGTTGCTCTGGTTGAAGAAGCAGTCTGGNNNAGATCGGAAG |
| 50323_1 | TATGAGTGTGGAGTCGTTGCGAACTGATCGTGTTCATAGAACTGNNNAGATCGGAA |
| 50323_2 | TATGAGTGTGGAGTCGTTGCGAACTGATCGTGTTCATAGAACTGNNNAGATCGGAA |
| 28946_1 | TATGAGTGTGGAGTCGTTGCGAGTGCAGTGACATGATCATAGCNNNAGATCGGAA |
| 28946_2 | TATGAGTGTGGAGTCGTTGCGAGTGCAGTGACATGATCATAGCNNNAGATCGGAA |
| 42695_1 | TATGAGTGTGGAGTCGTTGCACAAGTGTTGACAAGAATGAGGANNNAGATCGGAA |
| 42695_2 | TATGAGTGTGGAGTCGTTGCACAAGTGTTGACAAGAATGAGGANNNAGATCGGAA |
| 31380_1 | TATGAGTGTGGAGTCGTTGCGTTGTCTTTACCTGGGGCTTCTANNNAGATCGGAAG |
| 31380_2 | TATGAGTGTGGAGTCGTTGCGTTGTCTTTACCTGGGGCTTCTANNNAGATCGGAAG |
| 48095_1 | TATGAGTGTGGAGTCGTTGCCCTCCTGAGTAGCTGCAATTACANNNAGATCGGAAG |
| 48095_2 | TATGAGTGTGGAGTCGTTGCCCTCCTGAGTAGCTGCAATTACANNNAGATCGGAAG |
| 51760_1 | TATGAGTGTGGAGTCGTTGCCCAACAACAAAGGATGAATGCCANNNAGATCGGAA |

| | |
|---|---|
| 51760_2 | TATGAGTGTGGAGTCGTTGCCCAACAACAAAGGATGAATGCCANNNAGATCGGAA |
| 29350_1 | TATGAGTGTGGAGTCGTTGCTTTTCTCTCTCTCTGTCTCCCCANNNAGATCGGAAGA |
| 29350_2 | TATGAGTGTGGAGTCGTTGCTTTTCTCTCTCTCTGTCTCCCCANNNAGATCGGAAGA |
| 45082_1 | TATGAGTGTGGAGTCGTTGCCTGAGGAGTTTCTGTGGGAAGATNNNAGATCGGAA |
| 45082_2 | TATGAGTGTGGAGTCGTTGCCTGAGGAGTTTCTGTGGGAAGATNNNAGATCGGAA |
| 42727_1 | TATGAGTGTGGAGTCGTTGCATAACGGAAGACAACAAGAGCCTNNNAGATCGGAA |
| 42727_2 | TATGAGTGTGGAGTCGTTGCATAACGGAAGACAACAAGAGCCTNNNAGATCGGAA |
| 42696_1 | TATGAGTGTGGAGTCGTTGCGCATGTGCATATCCCTGATTCTTNNNAGATCGGAAG |
| 42696_2 | TATGAGTGTGGAGTCGTTGCGCATGTGCATATCCCTGATTCTTNNNAGATCGGAAG |
| 12359_1 | TATGAGTGTGGAGTCGTTGCCTGCTCTTGGATACTGCACATGTNNNAGATCGGAAG |
| 12359_2 | TATGAGTGTGGAGTCGTTGCCTGCTCTTGGATACTGCACATGTNNNAGATCGGAAG |
| 44132_1 | TATGAGTGTGGAGTCGTTGCGAGGGGACCTACAGTTTTTGTCTNNNAGATCGGAA |
| 44132_2 | TATGAGTGTGGAGTCGTTGCGAGGGGACCTACAGTTTTTGTCTNNNAGATCGGAA |
| 36537_1 | TATGAGTGTGGAGTCGTTGCAGCTGGGACATCACCATTTCATANNNAGATCGGAAG |
| 36537_2 | TATGAGTGTGGAGTCGTTGCAGCTGGGACATCACCATTTCATANNNAGATCGGAAG |
| 35514_1 | TATGAGTGTGGAGTCGTTGCTTGCTTAGGCTAGATGACTACTTTNNNAGATCGGAA |
| 35514_2 | TATGAGTGTGGAGTCGTTGCTTGCTTAGGCTAGATGACTACTTTNNNAGATCGGAA |
| 40598_1 | TATGAGTGTGGAGTCGTTGCACAGGCTTTCCCGATTTAGTAGANNNAGATCGGAAG |
| 40598_2 | TATGAGTGTGGAGTCGTTGCACAGGCTTTCCCGATTTAGTAGANNNAGATCGGAAG |
| 35193_1 | TATGAGTGTGGAGTCGTTGCGGCCACCTGAAAGTACAAATCACNNNAGATCGGAAG |
| 35193_2 | TATGAGTGTGGAGTCGTTGCGGCCACCTGAAAGTACAAATCACNNNAGATCGGAAG |
| 42635_1 | TATGAGTGTGGAGTCGTTGCATGTTTTCAATTCCCTTGGACGTNNNAGATCGGAAG |
| 42635_2 | TATGAGTGTGGAGTCGTTGCATGTTTTCAATTCCCTTGGACGTNNNAGATCGGAAG |
| 49879_1 | TATGAGTGTGGAGTCGTTGCCCTCAGGATTTGGAAGCTCTAGGNNNAGATCGGAA |
| 49879_2 | TATGAGTGTGGAGTCGTTGCCCTCAGGATTTGGAAGCTCTAGGNNNAGATCGGAA |
| 50749_1 | TATGAGTGTGGAGTCGTTGCCCCTAGGCTTTTGGAAGAAGTCTNNNAGATCGGAAG |
| 50749_2 | TATGAGTGTGGAGTCGTTGCCCCTAGGCTTTTGGAAGAAGTCTNNNAGATCGGAAG |
| 52486_1 | TATGAGTGTGGAGTCGTTGCGTATCTCTTTTTGTGGTGCTGGCNNNAGATCGGAAG |
| 52486_2 | TATGAGTGTGGAGTCGTTGCGTATCTCTTTTTGTGGTGCTGGCNNNAGATCGGAAG |
| 36749_1 | TATGAGTGTGGAGTCGTTGCTAATGGCAGTTGAAAAGGCACAGNNNAGATCGGAA |
| 36749_2 | TATGAGTGTGGAGTCGTTGCTAATGGCAGTTGAAAAGGCACAGNNNAGATCGGAA |
| 34948_1 | TATGAGTGTGGAGTCGTTGCCCTCCTGGGTTCAAGTGATTCTCNNNAGATCGGAAG |
| 34948_2 | TATGAGTGTGGAGTCGTTGCCCTCCTGGGTTCAAGTGATTCTCNNNAGATCGGAAG |
| 29821_1 | TATGAGTGTGGAGTCGTTGCCTCAAGTTGACCCACTAGTCCAGNNNAGATCGGAAG |
| 29821_2 | TATGAGTGTGGAGTCGTTGCCTCAAGTTGACCCACTAGTCCAGNNNAGATCGGAAG |
| 3032_1 | TATGAGTGTGGAGTCGTTGCGACAGGAAAGCCTCAAAGGGTTTNNNAGATCGGAA |
| 3032_2 | TATGAGTGTGGAGTCGTTGCGACAGGAAAGCCTCAAAGGGTTTNNNAGATCGGAA |
| 31282_1 | TATGAGTGTGGAGTCGTTGCGGCTGAAGTGTGTTTATGCTTGTNNNAGATCGGAA |
| 31282_2 | TATGAGTGTGGAGTCGTTGCGGCTGAAGTGTGTTTATGCTTGTNNNAGATCGGAA |
| 37951_1 | TATGAGTGTGGAGTCGTTGCCCTTGGAGAGTGGATGAGAGATGNNNAGATCGGA |
| 37951_2 | TATGAGTGTGGAGTCGTTGCCCTTGGAGAGTGGATGAGAGATGNNNAGATCGGA |
| 49310_1 | TATGAGTGTGGAGTCGTTGCTCAAGTGATTCTCCTGTCTCAGCNNNAGATCGGAAG |
| 49310_2 | TATGAGTGTGGAGTCGTTGCTCAAGTGATTCTCCTGTCTCAGCNNNAGATCGGAAG |

| | |
|---|---|
| 13066_1 | TATGAGTGTGGAGTCGTTGCTAGCTTGACATCCTTGGAAANNNAGATCGGAAGAG |
| 13066_2 | TATGAGTGTGGAGTCGTTGCTAGCTTGACATCCTTGGAAANNNAGATCGGAAGAG |
| 45495_1 | TATGAGTGTGGAGTCGTTGCACAGCCCCTCTCCAACTAGTATANNNAGATCGGAAG |
| 45495_2 | TATGAGTGTGGAGTCGTTGCACAGCCCCTCTCCAACTAGTATANNNAGATCGGAAG |
| 35584_1 | TATGAGTGTGGAGTCGTTGCAGGTTCATGCCATTCTCTTGCATNNNAGATCGGAAG |
| 35584_2 | TATGAGTGTGGAGTCGTTGCAGGTTCATGCCATTCTCTTGCATNNNAGATCGGAAG |
| 35484_1 | TATGAGTGTGGAGTCGTTGCTGATCTTAGCTCACTGCAACCTTNNNAGATCGGAAG |
| 35484_2 | TATGAGTGTGGAGTCGTTGCTGATCTTAGCTCACTGCAACCTTNNNAGATCGGAAG |
| 50557_1 | TATGAGTGTGGAGTCGTTGCTCTTTTTTTGAGACAGTCTGGCTGNNNAGATCGGAA |
| 50557_2 | TATGAGTGTGGAGTCGTTGCTCTTTTTTTGAGACAGTCTGGCTGNNNAGATCGGAA |
| 45635_1 | TATGAGTGTGGAGTCGTTGCTTGAGGCTGGGCTAAGTTAGAAGNNNAGATCGGAA |
| 45635_2 | TATGAGTGTGGAGTCGTTGCTTGAGGCTGGGCTAAGTTAGAAGNNNAGATCGGAA |
| 30186_1 | TATGAGTGTGGAGTCGTTGCAGTAACTATTTTGTGTGTGTTGCTNNNAGATCGGAA |
| 30186_2 | TATGAGTGTGGAGTCGTTGCAGTAACTATTTTGTGTGTGTTGCTNNNAGATCGGAA |
| 12729_1 | TATGAGTGTGGAGTCGTTGCTTACATTCTTGTCACAGGTCNNNAGATCGGAAGAGC |
| 12729_2 | TATGAGTGTGGAGTCGTTGCTTACATTCTTGTCACAGGTCNNNAGATCGGAAGAGC |
| 4303_1 | TATGAGTGTGGAGTCGTTGCCAGCTAGGCAGCTTCACTACATCNNNAGATCGGAAG |
| 4303_2 | TATGAGTGTGGAGTCGTTGCCAGCTAGGCAGCTTCACTACATCNNNAGATCGGAAG |
| 38660_1 | TATGAGTGTGGAGTCGTTGCCTCATGCCTGTAGTCCCAGTTACNNNAGATCGGAAG |
| 38660_2 | TATGAGTGTGGAGTCGTTGCCTCATGCCTGTAGTCCCAGTTACNNNAGATCGGAAG |
| 40239_1 | TATGAGTGTGGAGTCGTTGCTTCTCCAAACTTTTGACTTGGGCNNNAGATCGGAAG |
| 40239_2 | TATGAGTGTGGAGTCGTTGCTTCTCCAAACTTTTGACTTGGGCNNNAGATCGGAAG |
| 51599_1 | TATGAGTGTGGAGTCGTTGCCAATAGGACACATCGGGAACTGANNNAGATCGGAA |
| 51599_2 | TATGAGTGTGGAGTCGTTGCCAATAGGACACATCGGGAACTGANNNAGATCGGAA |
| 32688_1 | TATGAGTGTGGAGTCGTTGCGGCATGATATTGGGTCATTGCAANNNAGATCGGAA |
| 32688_2 | TATGAGTGTGGAGTCGTTGCGGCATGATATTGGGTCATTGCAANNNAGATCGGAA |
| 34975_1 | TATGAGTGTGGAGTCGTTGCAGACAGTTTCCTTTTTGTTGCCCNNNAGATCGGAAG |
| 34975_2 | TATGAGTGTGGAGTCGTTGCAGACAGTTTCCTTTTTGTTGCCCNNNAGATCGGAAG |
| 51634_1 | TATGAGTGTGGAGTCGTTGCTATGGCCTGGCTTTTCCTACATTNNNAGATCGGAAG |
| 51634_2 | TATGAGTGTGGAGTCGTTGCTATGGCCTGGCTTTTCCTACATTNNNAGATCGGAAG |
| 52449_1 | TATGAGTGTGGAGTCGTTGCCACACTCAGGGTGAGCTCAGNNNAGATCGGAAGAG |
| 52449_2 | TATGAGTGTGGAGTCGTTGCCACACTCAGGGTGAGCTCAGNNNAGATCGGAAGAG |
| 33534_1 | TATGAGTGTGGAGTCGTTGCTCCTGAGGAGCTGAGACTACATGNNNAGATCGGAA |
| 33534_2 | TATGAGTGTGGAGTCGTTGCTCCTGAGGAGCTGAGACTACATGNNNAGATCGGAA |
| 48245_1 | TATGAGTGTGGAGTCGTTGCACGCCCGGCTAATTTTTTGTATTNNNAGATCGGAAG |
| 48245_2 | TATGAGTGTGGAGTCGTTGCACGCCCGGCTAATTTTTTGTATTNNNAGATCGGAAG |
| 35075_1 | TATGAGTGTGGAGTCGTTGCATTCTTCCTGGGTCAGTCTTGAGNNNAGATCGGAAG |
| 35075_2 | TATGAGTGTGGAGTCGTTGCATTCTTCCTGGGTCAGTCTTGAGNNNAGATCGGAAG |
| 29755_1 | TATGAGTGTGGAGTCGTTGCAAGTTCCTGCTTCTGTCTCTCCTNNNAGATCGGAAGA |
| 29755_2 | TATGAGTGTGGAGTCGTTGCAAGTTCCTGCTTCTGTCTCTCCTNNNAGATCGGAAGA |
| 37513_1 | TATGAGTGTGGAGTCGTTGCAGAAGAAGCTGAAAAGGCAAACCNNNAGATCGGAA |
| 37513_2 | TATGAGTGTGGAGTCGTTGCAGAAGAAGCTGAAAAGGCAAACCNNNAGATCGGAA |
| 36280_1 | TATGAGTGTGGAGTCGTTGCTTTGCGAAATGCTTACTCCACTTNNNAGATCGGAAG |

| | |
|---|---|
| 36280_2 | TATGAGTGTGGAGTCGTTGCTTTGCGAAATGCTTACTCCACTTNNNAGATCGGAAGA |
| 10860_1 | TATGAGTGTGGAGTCGTTGCTCAGTGTTGGTTTCCAGTGTGTTNNNAGATCGGAAG |
| 10860_2 | TATGAGTGTGGAGTCGTTGCTCAGTGTTGGTTTCCAGTGTGTTNNNAGATCGGAAG |
| 3104_1 | TATGAGTGTGGAGTCGTTGCACAGGCTATGGTTCATGTTTTGCNNNAGATCGGAAG |
| 3104_2 | TATGAGTGTGGAGTCGTTGCACAGGCTATGGTTCATGTTTTGCNNNAGATCGGAAG |
| 12610_1 | TATGAGTGTGGAGTCGTTGCCTGAGAGCTTTTGTCTCAAANNNAGATCGGAAGAGC |
| 12610_2 | TATGAGTGTGGAGTCGTTGCCTGAGAGCTTTTGTCTCAAANNNAGATCGGAAGAGC |
| 53125_1 | TATGAGTGTGGAGTCGTTGCGCATGCTTATTTTGCTGACATCCNNNAGATCGGAAGA |
| 53125_2 | TATGAGTGTGGAGTCGTTGCGCATGCTTATTTTGCTGACATCCNNNAGATCGGAAGA |
| 46950_1 | TATGAGTGTGGAGTCGTTGCCTGGAAAAGGAGAAGCTGACAGTNNNAGATCGGAA |
| 46950_2 | TATGAGTGTGGAGTCGTTGCCTGGAAAAGGAGAAGCTGACAGTNNNAGATCGGAA |
| 10595_1 | TATGAGTGTGGAGTCGTTGCTCCTCCCTCCCTTCTTCTCAATTNNNAGATCGGAAGAG |
| 10595_2 | TATGAGTGTGGAGTCGTTGCTCCTCCCTCCCTTCTTCTCAATTNNNAGATCGGAAGAG |
| 37298_1 | TATGAGTGTGGAGTCGTTGCCATTGTTCACCATGATGGCCAGNNNAGATCGGAAGA |
| 37298_2 | TATGAGTGTGGAGTCGTTGCCATTGTTCACCATGATGGCCAGNNNAGATCGGAAGA |
| 36610_1 | TATGAGTGTGGAGTCGTTGCGACTGATGCTACAGTGAGCCATANNNAGATCGGAAG |
| 36610_2 | TATGAGTGTGGAGTCGTTGCGACTGATGCTACAGTGAGCCATANNNAGATCGGAAG |
| 44806_1 | TATGAGTGTGGAGTCGTTGCACTGTTGCAGTAAGGAATCACCANNNAGATCGGAAG |
| 44806_2 | TATGAGTGTGGAGTCGTTGCACTGTTGCAGTAAGGAATCACCANNNAGATCGGAAG |
| 53293_1 | TATGAGTGTGGAGTCGTTGCCTGATCCAGGAGGCCGCATCNNNAGATCGGAAGAGC |
| 53293_2 | TATGAGTGTGGAGTCGTTGCCTGATCCAGGAGGCCGCATCNNNAGATCGGAAGAGC |
| 51071_1 | TATGAGTGTGGAGTCGTTGCGGAGTGTCACAACGGTACAGAATNNNAGATCGGAAG |
| 51071_2 | TATGAGTGTGGAGTCGTTGCGGAGTGTCACAACGGTACAGAATNNNAGATCGGAAG |
| 48736_1 | TATGAGTGTGGAGTCGTTGCTTCACTGCATTCTAACCTGGTGANNNAGATCGGAAGA |
| 48736_2 | TATGAGTGTGGAGTCGTTGCTTCACTGCATTCTAACCTGGTGANNNAGATCGGAAGA |
| 46879_1 | TATGAGTGTGGAGTCGTTGCTTCTCTCAGAGTCCTGGCTTTTCNNNAGATCGGAAGA |
| 46879_2 | TATGAGTGTGGAGTCGTTGCTTCTCTCAGAGTCCTGGCTTTTCNNNAGATCGGAAGA |
| 2461_1 | TATGAGTGTGGAGTCGTTGCGCTGCCACTATCATGCCAAAAAGNNNAGATCGGAAG |
| 2461_2 | TATGAGTGTGGAGTCGTTGCGCTGCCACTATCATGCCAAAAAGNNNAGATCGGAAG |
| 29429_1 | TATGAGTGTGGAGTCGTTGCAGGGGATTGGGCTGTTTTTGGTNNNAGATCGGAAG |
| 29429_2 | TATGAGTGTGGAGTCGTTGCAGGGGATTGGGCTGTTTTTGGTNNNAGATCGGAAG |
| 53323_1 | TATGAGTGTGGAGTCGTTGCGTGCAATCCAGAGTCCTTGGATANNNAGATCGGAAG |
| 53323_2 | TATGAGTGTGGAGTCGTTGCGTGCAATCCAGAGTCCTTGGATANNNAGATCGGAAG |
| 48048_1 | TATGAGTGTGGAGTCGTTGCATGAAAAGGACAAGGGTACTCCGNNNAGATCGGAA |
| 48048_2 | TATGAGTGTGGAGTCGTTGCATGAAAAGGACAAGGGTACTCCGNNNAGATCGGAA |
| 43651_1 | TATGAGTGTGGAGTCGTTGCAACATCAGGTAGGGCGCGTGNNNAGATCGGAAGAG |
| 43651_2 | TATGAGTGTGGAGTCGTTGCAACATCAGGTAGGGCGCGTGNNNAGATCGGAAGAG |
| 40150_1 | TATGAGTGTGGAGTCGTTGCTCCTTCCCACAAAAATGCTTCACNNNAGATCGGAAGA |
| 40150_2 | TATGAGTGTGGAGTCGTTGCTCCTTCCCACAAAAATGCTTCACNNNAGATCGGAAGA |
| 46280_1 | TATGAGTGTGGAGTCGTTGCTTCAAGCAATTCTCCTCCCTCAGNNNAGATCGGAAGA |
| 46280_2 | TATGAGTGTGGAGTCGTTGCTTCAAGCAATTCTCCTCCCTCAGNNNAGATCGGAAGA |
| 45683_1 | TATGAGTGTGGAGTCGTTGCAGCTCTTGAAACTGTGTCATTGCNNNAGATCGGAAG |
| 45683_2 | TATGAGTGTGGAGTCGTTGCAGCTCTTGAAACTGTGTCATTGCNNNAGATCGGAAG |

| | |
|---|---|
| 44075_1 | TATGAGTGTGGAGTCGTTGCGCTGGGTATGTTTCAGTGCATTTNNNAGATCGGAAG |
| 44075_2 | TATGAGTGTGGAGTCGTTGCGCTGGGTATGTTTCAGTGCATTTNNNAGATCGGAAG |
| 38405_1 | TATGAGTGTGGAGTCGTTGCCTGTAATTCCAGCTACTCGGGAGNNNAGATCGGAAG |
| 38405_2 | TATGAGTGTGGAGTCGTTGCCTGTAATTCCAGCTACTCGGGAGNNNAGATCGGAAG |
| 33028_1 | TATGAGTGTGGAGTCGTTGCGGTTCAAATGATTCTCCTGCCTCNNNAGATCGGAAG |
| 33028_2 | TATGAGTGTGGAGTCGTTGCGGTTCAAATGATTCTCCTGCCTCNNNAGATCGGAAG |
| 10223_1 | TATGAGTGTGGAGTCGTTGCAGAGGATGAGAGGTTGTGGGAAANNNAGATCGGA |
| 10223_2 | TATGAGTGTGGAGTCGTTGCAGAGGATGAGAGGTTGTGGGAAANNNAGATCGGA |
| 35144_1 | TATGAGTGTGGAGTCGTTGCCCAAACATGCTTCTCTTCCATCCNNNAGATCGGAAGA |
| 35144_2 | TATGAGTGTGGAGTCGTTGCCCAAACATGCTTCTCTTCCATCCNNNAGATCGGAAGA |
| 39085_1 | TATGAGTGTGGAGTCGTTGCCTTGCCCCTCTATAGTTCCTGTCNNNAGATCGGAAGA |
| 39085_2 | TATGAGTGTGGAGTCGTTGCCTTGCCCCTCTATAGTTCCTGTCNNNAGATCGGAAGA |
| 46506_1 | TATGAGTGTGGAGTCGTTGCCTGGTCTTGAACTCCTGACTTCANNNAGATCGGAAG |
| 46506_2 | TATGAGTGTGGAGTCGTTGCCTGGTCTTGAACTCCTGACTTCANNNAGATCGGAAG |
| 51571_1 | TATGAGTGTGGAGTCGTTGCGTTCCCTGCTGTGTTTTCAGTGNNNAGATCGGAAGA |
| 51571_2 | TATGAGTGTGGAGTCGTTGCGTTCCCTGCTGTGTTTTCAGTGNNNAGATCGGAAGA |
| 32627_1 | TATGAGTGTGGAGTCGTTGCAAAAATCTTGGACACACGGACACNNNAGATCGGAAG |
| 32627_2 | TATGAGTGTGGAGTCGTTGCAAAAATCTTGGACACACGGACACNNNAGATCGGAAG |
| 38554_1 | TATGAGTGTGGAGTCGTTGCAAAAAATTAGCTGGGTGTGGTGGNNNAGATCGGA |
| 38554_2 | TATGAGTGTGGAGTCGTTGCAAAAAATTAGCTGGGTGTGGTGGNNNAGATCGGA |
| 40437_1 | TATGAGTGTGGAGTCGTTGCCTCTAGCCAGGTGGTTCCATATGNNNAGATCGGAAG |
| 40437_2 | TATGAGTGTGGAGTCGTTGCCTCTAGCCAGGTGGTTCCATATGNNNAGATCGGAAG |
| 34341_1 | TATGAGTGTGGAGTCGTTGCAGGACCCAGAGAGTCAAACTTTCNNNAGATCGGAAG |
| 34341_2 | TATGAGTGTGGAGTCGTTGCAGGACCCAGAGAGTCAAACTTTCNNNAGATCGGAAG |
| 31445_1 | TATGAGTGTGGAGTCGTTGCACTCTAGCAAGCATCCTAAGTCCNNNAGATCGGAAG |
| 31445_2 | TATGAGTGTGGAGTCGTTGCACTCTAGCAAGCATCCTAAGTCCNNNAGATCGGAAG |
| 2315_1 | TATGAGTGTGGAGTCGTTGCCCTGGCCTCATTTTACGTTTTGANNNAGATCGGAAG |
| 2315_2 | TATGAGTGTGGAGTCGTTGCCCTGGCCTCATTTTACGTTTTGANNNAGATCGGAAG |
| 42518_1 | TATGAGTGTGGAGTCGTTGCTGTGAGATCAATGGTGATATCCCCNNNAGATCGGAA |
| 42518_2 | TATGAGTGTGGAGTCGTTGCTGTGAGATCAATGGTGATATCCCCNNNAGATCGGAA |
| 52703_1 | TATGAGTGTGGAGTCGTTGCGATGCTTCACTGACACAGTTTGTNNNAGATCGGAAG |
| 52703_2 | TATGAGTGTGGAGTCGTTGCGATGCTTCACTGACACAGTTTGTNNNAGATCGGAAG |
| 41619_1 | TATGAGTGTGGAGTCGTTGCCACATCTTTCTGCTTCCCCAAAGNNNAGATCGGAAGA |
| 41619_2 | TATGAGTGTGGAGTCGTTGCCACATCTTTCTGCTTCCCCAAAGNNNAGATCGGAAGA |
| 39616_1 | TATGAGTGTGGAGTCGTTGCAACAAAAAAAGCCACCCAACTGANNNAGATCGGAAG |
| 39616_2 | TATGAGTGTGGAGTCGTTGCAACAAAAAAAGCCACCCAACTGANNNAGATCGGAAG |
| 48613_1 | TATGAGTGTGGAGTCGTTGCTACATTGATGAAGCCAGTGGAGTNNNAGATCGGAA |
| 48613_2 | TATGAGTGTGGAGTCGTTGCTACATTGATGAAGCCAGTGGAGTNNNAGATCGGAA |
| 38874_1 | TATGAGTGTGGAGTCGTTGCTTCCAGTGATTCTTCTGCCTCAGNNNAGATCGGAAG |
| 38874_2 | TATGAGTGTGGAGTCGTTGCTTCCAGTGATTCTTCTGCCTCAGNNNAGATCGGAAG |
| 32420_1 | TATGAGTGTGGAGTCGTTGCGCACTAGCATTACAGGCATTAGCNNNAGATCGGAAG |
| 32420_2 | TATGAGTGTGGAGTCGTTGCGCACTAGCATTACAGGCATTAGCNNNAGATCGGAAG |
| 49335_1 | TATGAGTGTGGAGTCGTTGCTTGAACTCTGCAAGATGAATGCCNNNAGATCGGAAG |

| | |
|---|---|
| 49335_2 | TATGAGTGTGGAGTCGTTGCTTGAACTCTGCAAGATGAATGCCNNNAGATCGGAAG |
| 29838_1 | TATGAGTGTGGAGTCGTTGCGCTACCTCTACAACCTCACTGTTNNNAGATCGGAAGA |
| 29838_2 | TATGAGTGTGGAGTCGTTGCGCTACCTCTACAACCTCACTGTTNNNAGATCGGAAGA |
| 31189_1 | TATGAGTGTGGAGTCGTTGCTGCTCACAAAAGAAAACCCCATCNNNAGATCGGAAG |
| 31189_2 | TATGAGTGTGGAGTCGTTGCTGCTCACAAAAGAAAACCCCATCNNNAGATCGGAAG |
| 39667_1 | TATGAGTGTGGAGTCGTTGCAAGCAGCTTTTGAAAAAGGGGAGNNNAGATCGGA |
| 39667_2 | TATGAGTGTGGAGTCGTTGCAAGCAGCTTTTGAAAAAGGGGAGNNNAGATCGGA |
| 32651_1 | TATGAGTGTGGAGTCGTTGCAAGATGGTCAGGGAATTGGTAGTNNNAGATCGGA |
| 32651_2 | TATGAGTGTGGAGTCGTTGCAAGATGGTCAGGGAATTGGTAGTNNNAGATCGGA |
| 39907_1 | TATGAGTGTGGAGTCGTTGCAAGTTTTGTTCTTGTTGCCCAGGNNNAGATCGGAAG |
| 39907_2 | TATGAGTGTGGAGTCGTTGCAAGTTTTGTTCTTGTTGCCCAGGNNNAGATCGGAAG |
| 48005_1 | TATGAGTGTGGAGTCGTTGCTAGAGGGAGGATGAAGAGAGGTGNNNAGATCGG |
| 48005_2 | TATGAGTGTGGAGTCGTTGCTAGAGGGAGGATGAAGAGAGGTGNNNAGATCGG |
| 3046_1 | TATGAGTGTGGAGTCGTTGCCTGTCGGCCATCTACAAGTGGATNNNAGATCGGAAG |
| 3046_2 | TATGAGTGTGGAGTCGTTGCCTGTCGGCCATCTACAAGTGGATNNNAGATCGGAAG |
| 40604_1 | TATGAGTGTGGAGTCGTTGCGAGGCATGAGAATGGCTTGAACNNNAGATCGGAAG |
| 40604_2 | TATGAGTGTGGAGTCGTTGCGAGGCATGAGAATGGCTTGAACNNNAGATCGGAAG |
| 42451_1 | TATGAGTGTGGAGTCGTTGCTTCTTTCTGAGACGGAGTCTTGCNNNAGATCGGAAG |
| 42451_2 | TATGAGTGTGGAGTCGTTGCTTCTTTCTGAGACGGAGTCTTGCNNNAGATCGGAAG |
| 34587_1 | TATGAGTGTGGAGTCGTTGCTCAAGAGTCTCACAGTTCCCTTGNNNAGATCGGAAG |
| 34587_2 | TATGAGTGTGGAGTCGTTGCTCAAGAGTCTCACAGTTCCCTTGNNNAGATCGGAAG |
| 48989_1 | TATGAGTGTGGAGTCGTTGCTCTCCCAGGTTCAAGTGATTCTCNNNAGATCGGAAG |
| 48989_2 | TATGAGTGTGGAGTCGTTGCTCTCCCAGGTTCAAGTGATTCTCNNNAGATCGGAAG |
| 36534_1 | TATGAGTGTGGAGTCGTTGCAAAATAACTTGAGGCTGGGAGGTNNNAGATCGGAA |
| 36534_2 | TATGAGTGTGGAGTCGTTGCAAAATAACTTGAGGCTGGGAGGTNNNAGATCGGAA |
| 47790_1 | TATGAGTGTGGAGTCGTTGCCTGAGCTGGGAACATGGTGTCNNNAGATCGGAAGA |
| 47790_2 | TATGAGTGTGGAGTCGTTGCCTGAGCTGGGAACATGGTGTCNNNAGATCGGAAGA |
| 52297_1 | TATGAGTGTGGAGTCGTTGCTCAAAAGACCAGAGAAAATGGCCNNNAGATCGGAA |
| 52297_2 | TATGAGTGTGGAGTCGTTGCTCAAAAGACCAGAGAAAATGGCCNNNAGATCGGAA |
| 49006_1 | TATGAGTGTGGAGTCGTTGCCTTCCAGGTTCACGTGATTCTCCNNNAGATCGGAAG |
| 49006_2 | TATGAGTGTGGAGTCGTTGCCTTCCAGGTTCACGTGATTCTCCNNNAGATCGGAAG |
| 44236_1 | TATGAGTGTGGAGTCGTTGCTTCAGAGCTATGCATTAGGCCATNNNAGATCGGAAG |
| 44236_2 | TATGAGTGTGGAGTCGTTGCTTCAGAGCTATGCATTAGGCCATNNNAGATCGGAAG |
| 37108_1 | TATGAGTGTGGAGTCGTTGCGGTCATTGTAAGGACTTTGGCACNNNAGATCGGAA |
| 37108_2 | TATGAGTGTGGAGTCGTTGCGGTCATTGTAAGGACTTTGGCACNNNAGATCGGAA |
| 33734_1 | TATGAGTGTGGAGTCGTTGCTGTCCTCCCACCATTCCTTTATGNNNAGATCGGAAGA |
| 33734_2 | TATGAGTGTGGAGTCGTTGCTGTCCTCCCACCATTCCTTTATGNNNAGATCGGAAGA |
| 50734_1 | TATGAGTGTGGAGTCGTTGCATGTAGATGCTGTTGTCTCCTGGNNNAGATCGGAAG |
| 50734_2 | TATGAGTGTGGAGTCGTTGCATGTAGATGCTGTTGTCTCCTGGNNNAGATCGGAAG |
| 7981_1 | TATGAGTGTGGAGTCGTTGCTGGGAAGCTCACTTTTGGCATGNNNAGATCGGAAG |
| 7981_2 | TATGAGTGTGGAGTCGTTGCTGGGAAGCTCACTTTTGGCATGNNNAGATCGGAAG |
| 9070_1 | TATGAGTGTGGAGTCGTTGCCTGGAAAGGAGGGGCTTGAATTNNNAGATCGGAA |
| 9070_2 | TATGAGTGTGGAGTCGTTGCCTGGAAAGGAGGGGCTTGAATTNNNAGATCGGAA |

| | |
|---|---|
| 43671_1 | TATGAGTGTGGAGTCGTTGCGATCTGCACCCTGTAAACCCTGNNNAGATCGGAAGA |
| 43671_2 | TATGAGTGTGGAGTCGTTGCGATCTGCACCCTGTAAACCCTGNNNAGATCGGAAGA |
| 52976_1 | TATGAGTGTGGAGTCGTTGCCAAACAATTCTACTGCCTCAGCCNNNAGATCGGAAGA |
| 52976_2 | TATGAGTGTGGAGTCGTTGCCAAACAATTCTACTGCCTCAGCCNNNAGATCGGAAGA |
| 49167_1 | TATGAGTGTGGAGTCGTTGCCAATCTCGGCTCACACTCACTNNNAGATCGGAAGAG |
| 49167_2 | TATGAGTGTGGAGTCGTTGCCAATCTCGGCTCACACTCACTNNNAGATCGGAAGAG |
| 41789_1 | TATGAGTGTGGAGTCGTTGCTGCCTCAATTTCAGGACTTGCTANNNAGATCGGAAG |
| 41789_2 | TATGAGTGTGGAGTCGTTGCTGCCTCAATTTCAGGACTTGCTANNNAGATCGGAAG |
| 32675_1 | TATGAGTGTGGAGTCGTTGCACCTCCCAGATTCAAACGATTCTNNNAGATCGGAAGA |
| 32675_2 | TATGAGTGTGGAGTCGTTGCACCTCCCAGATTCAAACGATTCTNNNAGATCGGAAGA |
| 34472_1 | TATGAGTGTGGAGTCGTTGCTAGTCCCAGCTACTCAAGAGGTTNNNAGATCGGAAG |
| 34472_2 | TATGAGTGTGGAGTCGTTGCTAGTCCCAGCTACTCAAGAGGTTNNNAGATCGGAAG |
| 31860_1 | TATGAGTGTGGAGTCGTTGCCTTTTCCAACACAGGGCTCAAAGNNNAGATCGGAAG |
| 31860_2 | TATGAGTGTGGAGTCGTTGCCTTTTCCAACACAGGGCTCAAAGNNNAGATCGGAAG |
| 38504_1 | TATGAGTGTGGAGTCGTTGCCATCAGGGTGGAAGTATTGGAGTNNNAGATCGGAA |
| 38504_2 | TATGAGTGTGGAGTCGTTGCCATCAGGGTGGAAGTATTGGAGTNNNAGATCGGAA |
| 46885_1 | TATGAGTGTGGAGTCGTTGCCAGGCACTTGTCAAGAAATAGCCNNNAGATCGGAAG |
| 46885_2 | TATGAGTGTGGAGTCGTTGCCAGGCACTTGTCAAGAAATAGCCNNNAGATCGGAAG |
| 40818_1 | TATGAGTGTGGAGTCGTTGCTGAGACAAAAACCATGAACACCTNNNAGATCGGAAG |
| 40818_2 | TATGAGTGTGGAGTCGTTGCTGAGACAAAAACCATGAACACCTNNNAGATCGGAAG |
| 32733_1 | TATGAGTGTGGAGTCGTTGCCTATGTGGTAGGCAGAAGAATGCNNNAGATCGGAA |
| 32733_2 | TATGAGTGTGGAGTCGTTGCCTATGTGGTAGGCAGAAGAATGCNNNAGATCGGAA |
| 34912_1 | TATGAGTGTGGAGTCGTTGCGAGTAGCTGGGATGACAGATACCNNNAGATCGGAA |
| 34912_2 | TATGAGTGTGGAGTCGTTGCGAGTAGCTGGGATGACAGATACCNNNAGATCGGAA |
| 47470_1 | TATGAGTGTGGAGTCGTTGCCAAGTGATTCCCCTGCCTCAGNNNAGATCGGAAGAG |
| 47470_2 | TATGAGTGTGGAGTCGTTGCCAAGTGATTCCCCTGCCTCAGNNNAGATCGGAAGAG |
| 47382_1 | TATGAGTGTGGAGTCGTTGCCTCCTGGGTTCACACCATTCTCNNNAGATCGGAAGAG |
| 47382_2 | TATGAGTGTGGAGTCGTTGCCTCCTGGGTTCACACCATTCTCNNNAGATCGGAAGAG |
| 35226_1 | TATGAGTGTGGAGTCGTTGCACCCAATCAATTGTCTCTTGTGCNNNAGATCGGAAGA |
| 35226_2 | TATGAGTGTGGAGTCGTTGCACCCAATCAATTGTCTCTTGTGCNNNAGATCGGAAGA |
| 34570_1 | TATGAGTGTGGAGTCGTTGCAGGATCCTAGAACCAGCTTTTGTNNNAGATCGGAAG |
| 34570_2 | TATGAGTGTGGAGTCGTTGCAGGATCCTAGAACCAGCTTTTGTNNNAGATCGGAAG |
| 29448_1 | TATGAGTGTGGAGTCGTTGCCTGTAGTCCCAGCTACTCAGGAGNNNAGATCGGAAG |
| 29448_2 | TATGAGTGTGGAGTCGTTGCCTGTAGTCCCAGCTACTCAGGAGNNNAGATCGGAAG |
| 38429_1 | TATGAGTGTGGAGTCGTTGCGTCTCAATCTTCTGACCTCGTGANNNAGATCGGAAG |
| 38429_2 | TATGAGTGTGGAGTCGTTGCGTCTCAATCTTCTGACCTCGTGANNNAGATCGGAAG |
| 53361_1 | TATGAGTGTGGAGTCGTTGCTTTAGACGGAGTCTTGCTCTGTCNNNAGATCGGAAG |
| 53361_2 | TATGAGTGTGGAGTCGTTGCTTTAGACGGAGTCTTGCTCTGTCNNNAGATCGGAAG |
| 29148_1 | TATGAGTGTGGAGTCGTTGCGGGCAAGCTTCTTCTGGATAGATNNNAGATCGGAA |
| 29148_2 | TATGAGTGTGGAGTCGTTGCGGGCAAGCTTCTTCTGGATAGATNNNAGATCGGAA |
| 38584_1 | TATGAGTGTGGAGTCGTTGCGTCCATATTCTCTGGGGCCTAACNNNAGATCGGAAG |
| 38584_2 | TATGAGTGTGGAGTCGTTGCGTCCATATTCTCTGGGGCCTAACNNNAGATCGGAAG |
| 832_1 | TATGAGTGTGGAGTCGTTGCTAGCGAAAATCGGCCCGGCCNNNAGATCGGAAGAG |

| | |
|---|---|
| 832_2 | TATGAGTGTGGAGTCGTTGCTAGCGAAAATCGGCCCGGCCNNNAGATCGGAAGAG |
| 41685_1 | TATGAGTGTGGAGTCGTTGCTTCAAGTGATTCTCCTGCCTCAGNNNAGATCGGAAG |
| 41685_2 | TATGAGTGTGGAGTCGTTGCTTCAAGTGATTCTCCTGCCTCAGNNNAGATCGGAAG |
| 53143_1 | TATGAGTGTGGAGTCGTTGCTTGAAGACCATGTCAGTCTCTGGNNNAGATCGGAAG |
| 53143_2 | TATGAGTGTGGAGTCGTTGCTTGAAGACCATGTCAGTCTCTGGNNNAGATCGGAAG |
| 33347_1 | TATGAGTGTGGAGTCGTTGCTCTGCCTCCTGAGTTCAAACAATNNNAGATCGGAAG |
| 33347_2 | TATGAGTGTGGAGTCGTTGCTCTGCCTCCTGAGTTCAAACAATNNNAGATCGGAAG |
| 49473_1 | TATGAGTGTGGAGTCGTTGCAACCATCCTTCTGACTCAACTGCNNNAGATCGGAAG |
| 49473_2 | TATGAGTGTGGAGTCGTTGCAACCATCCTTCTGACTCAACTGCNNNAGATCGGAAG |
| 39222_1 | TATGAGTGTGGAGTCGTTGCTGATCAAGATTGGAAAGAGGCCANNNAGATCGGAA |
| 39222_2 | TATGAGTGTGGAGTCGTTGCTGATCAAGATTGGAAAGAGGCCANNNAGATCGGAA |
| 2172_1 | TATGAGTGTGGAGTCGTTGCTGGCAAAAGCAAGAGATGAGAGANNNAGATCGGA |
| 2172_2 | TATGAGTGTGGAGTCGTTGCTGGCAAAAGCAAGAGATGAGAGANNNAGATCGGA |
| 41034_1 | TATGAGTGTGGAGTCGTTGCATTGGGAATATTGCCCCTGAACANNNAGATCGGAAG |
| 41034_2 | TATGAGTGTGGAGTCGTTGCATTGGGAATATTGCCCCTGAACANNNAGATCGGAAG |
| 30893_1 | TATGAGTGTGGAGTCGTTGCTAGAGACAGGAGAGAGAAAGCGANNNAGATCGGA |
| 30893_2 | TATGAGTGTGGAGTCGTTGCTAGAGACAGGAGAGAGAAAGCGANNNAGATCGGA |
| 42622_1 | TATGAGTGTGGAGTCGTTGCGCATGATATCGACTCACTGCAACNNNAGATCGGAAG |
| 42622_2 | TATGAGTGTGGAGTCGTTGCGCATGATATCGACTCACTGCAACNNNAGATCGGAAG |
| 5540_1 | TATGAGTGTGGAGTCGTTGCTCCCTATCCAGGCTTTTTGGGTANNNAGATCGGAAG |
| 5540_2 | TATGAGTGTGGAGTCGTTGCTCCCTATCCAGGCTTTTTGGGTANNNAGATCGGAAG |
| 38749_1 | TATGAGTGTGGAGTCGTTGCAACATGAATGGAACTGGAGGTCANNNAGATCGGAA |
| 38749_2 | TATGAGTGTGGAGTCGTTGCAACATGAATGGAACTGGAGGTCANNNAGATCGGAA |
| 38066_1 | TATGAGTGTGGAGTCGTTGCCAAACTCCTGAGCTCGAACAATCNNNAGATCGGAAG |
| 38066_2 | TATGAGTGTGGAGTCGTTGCCAAACTCCTGAGCTCGAACAATCNNNAGATCGGAAG |
| 42048_1 | TATGAGTGTGGAGTCGTTGCAAACTCCCAGGCTCAAGTGATCNNNAGATCGGAAGA |
| 42048_2 | TATGAGTGTGGAGTCGTTGCAAACTCCCAGGCTCAAGTGATCNNNAGATCGGAAGA |
| 43481_1 | TATGAGTGTGGAGTCGTTGCCAAGATCTCGGCTTACTGCAACNNNAGATCGGAAGA |
| 43481_2 | TATGAGTGTGGAGTCGTTGCCAAGATCTCGGCTTACTGCAACNNNAGATCGGAAGA |
| 2388_1 | TATGAGTGTGGAGTCGTTGCTGCTTTCTTATTCAGGTCTGGGCNNNAGATCGGAAG |
| 2388_2 | TATGAGTGTGGAGTCGTTGCTGCTTTCTTATTCAGGTCTGGGCNNNAGATCGGAAG |
| 37256_1 | TATGAGTGTGGAGTCGTTGCATGGAACATAGCAAAGGCAGTTCNNNAGATCGGAA |
| 37256_2 | TATGAGTGTGGAGTCGTTGCATGGAACATAGCAAAGGCAGTTCNNNAGATCGGAA |
| 28957_1 | TATGAGTGTGGAGTCGTTGCATAATGCTGTCTGGTGTTTGACGNNNAGATCGGAA |
| 28957_2 | TATGAGTGTGGAGTCGTTGCATAATGCTGTCTGGTGTTTGACGNNNAGATCGGAA |
| 35944_1 | TATGAGTGTGGAGTCGTTGCATGCAGCTTCCAGAAGAATCAGANNNAGATCGGAA |
| 35944_2 | TATGAGTGTGGAGTCGTTGCATGCAGCTTCCAGAAGAATCAGANNNAGATCGGAA |
| 36202_1 | TATGAGTGTGGAGTCGTTGCCGCCTTCACTGACTGGTAAATTGNNNAGATCGGAAG |
| 36202_2 | TATGAGTGTGGAGTCGTTGCCGCCTTCACTGACTGGTAAATTGNNNAGATCGGAAG |
| 12356_1 | TATGAGTGTGGAGTCGTTGCTTGGTTATTGCTTGTATCCCCAANNNAGATCGGAAG |
| 12356_2 | TATGAGTGTGGAGTCGTTGCTTGGTTATTGCTTGTATCCCCAANNNAGATCGGAAG |
| 9764_1 | TATGAGTGTGGAGTCGTTGCCTTTCAGCAGCCCATCCAAAAACNNNAGATCGGAAG |
| 9764_2 | TATGAGTGTGGAGTCGTTGCCTTTCAGCAGCCCATCCAAAAACNNNAGATCGGAAG |

| | |
|---|---|
| 37767_1 | TATGAGTGTGGAGTCGTTGCTTGGAGCCATAGTTCCACCAAATNNNAGATCGGAAG |
| 37767_2 | TATGAGTGTGGAGTCGTTGCTTGGAGCCATAGTTCCACCAAATNNNAGATCGGAAG |
| 9697_1 | TATGAGTGTGGAGTCGTTGCTGTTGGTGAATAGTCACTGGACCNNNAGATCGGAA |
| 9697_2 | TATGAGTGTGGAGTCGTTGCTGTTGGTGAATAGTCACTGGACCNNNAGATCGGAA |
| 2563_1 | TATGAGTGTGGAGTCGTTGCAACGAAGACGATTAGATTGGGGGNNNAGATCGGA |
| 2563_2 | TATGAGTGTGGAGTCGTTGCAACGAAGACGATTAGATTGGGGGNNNAGATCGGA |
| 50783_1 | TATGAGTGTGGAGTCGTTGCCAGCGTTCTTGAATTGTGTTCCTNNNAGATCGGAAG |
| 50783_2 | TATGAGTGTGGAGTCGTTGCCAGCGTTCTTGAATTGTGTTCCTNNNAGATCGGAAG |
| 12677_1 | TATGAGTGTGGAGTCGTTGCGATAGTGACTATCTCAATAGAGCNNNAGATCGGAA |
| 12677_2 | TATGAGTGTGGAGTCGTTGCGATAGTGACTATCTCAATAGAGCNNNAGATCGGAA |
| 45713_1 | TATGAGTGTGGAGTCGTTGCAGAATGATGTAGTCGTCTGGCAGNNNAGATCGGAA |
| 45713_2 | TATGAGTGTGGAGTCGTTGCAGAATGATGTAGTCGTCTGGCAGNNNAGATCGGAA |
| 2167_1 | TATGAGTGTGGAGTCGTTGCTCTACACACAGAAGCCTTCCAGANNNAGATCGGAAG |
| 2167_2 | TATGAGTGTGGAGTCGTTGCTCTACACACAGAAGCCTTCCAGANNNAGATCGGAAG |
| 38677_1 | TATGAGTGTGGAGTCGTTGCTGATATCAATTTGTCCCACTGGCNNNAGATCGGAAG |
| 38677_2 | TATGAGTGTGGAGTCGTTGCTGATATCAATTTGTCCCACTGGCNNNAGATCGGAAG |
| 11977_1 | TATGAGTGTGGAGTCGTTGCTTAGTTCAGTGCCTGGTGTGATGNNNAGATCGGAA |
| 11977_2 | TATGAGTGTGGAGTCGTTGCTTAGTTCAGTGCCTGGTGTGATGNNNAGATCGGAA |
| 2215_1 | TATGAGTGTGGAGTCGTTGCTGCTACCTTGTTTGATGCTGATGANNNAGATCGGAA |
| 2215_2 | TATGAGTGTGGAGTCGTTGCTGCTACCTTGTTTGATGCTGATGANNNAGATCGGAA |
| 50849_1 | TATGAGTGTGGAGTCGTTGCCCTACAAGCCTACAGTTCCAACANNNAGATCGGAAG |
| 50849_2 | TATGAGTGTGGAGTCGTTGCCCTACAAGCCTACAGTTCCAACANNNAGATCGGAAG |
| 35363_1 | TATGAGTGTGGAGTCGTTGCTGCATAGTATTCCATTGTGCACANNNAGATCGGAAG |
| 35363_2 | TATGAGTGTGGAGTCGTTGCTGCATAGTATTCCATTGTGCACANNNAGATCGGAAG |
| 51606_1 | TATGAGTGTGGAGTCGTTGCGTGTGACTAGAACCCAGTAAGCANNNAGATCGGAA |
| 51606_2 | TATGAGTGTGGAGTCGTTGCGTGTGACTAGAACCCAGTAAGCANNNAGATCGGAA |
| 30130_1 | TATGAGTGTGGAGTCGTTGCCACTGTGGTGGTAAATGCATTTANNNAGATCGGAA |
| 30130_2 | TATGAGTGTGGAGTCGTTGCCACTGTGGTGGTAAATGCATTTANNNAGATCGGAA |
| 33836_1 | TATGAGTGTGGAGTCGTTGCCTAGAGGTTTCTCATTGGCCACTNNNAGATCGGAAG |
| 33836_2 | TATGAGTGTGGAGTCGTTGCCTAGAGGTTTCTCATTGGCCACTNNNAGATCGGAAG |
| 12725_1 | TATGAGTGTGGAGTCGTTGCCTCAATTCTCAGGGACTTTAATNNNAGATCGGAAGA |
| 12725_2 | TATGAGTGTGGAGTCGTTGCCTCAATTCTCAGGGACTTTAATNNNAGATCGGAAGA |
| 29822_1 | TATGAGTGTGGAGTCGTTGCGTTGCTTCTAGTGGGAGTGAAGANNNAGATCGGAA |
| 29822_2 | TATGAGTGTGGAGTCGTTGCGTTGCTTCTAGTGGGAGTGAAGANNNAGATCGGAA |
| 33675_1 | TATGAGTGTGGAGTCGTTGCCTCCTGAGTAGCTGGGACTACAGNNNAGATCGGAA |
| 33675_2 | TATGAGTGTGGAGTCGTTGCCTCCTGAGTAGCTGGGACTACAGNNNAGATCGGAA |
| 31002_1 | TATGAGTGTGGAGTCGTTGCGCTGGTAGTGTAACTCAGTCCAANNNAGATCGGAA |
| 31002_2 | TATGAGTGTGGAGTCGTTGCGCTGGTAGTGTAACTCAGTCCAANNNAGATCGGAA |
| 50811_1 | TATGAGTGTGGAGTCGTTGCGCAGTCAATCAGGGCTTTGTTTNNNAGATCGGAAG |
| 50811_2 | TATGAGTGTGGAGTCGTTGCGCAGTCAATCAGGGCTTTGTTTNNNAGATCGGAAG |
| 43196_1 | TATGAGTGTGGAGTCGTTGCCTCCAGGGTTCAAGCAATTCTCTNNNAGATCGGAAG |
| 43196_2 | TATGAGTGTGGAGTCGTTGCCTCCAGGGTTCAAGCAATTCTCTNNNAGATCGGAAG |
| 5321_1 | TATGAGTGTGGAGTCGTTGCGGTGTAATCATGGGTGCATGGTTNNNAGATCGGAA |

| | |
|---|---|
| 5321_2 | TATGAGTGTGGAGTCGTTGCGGTGTAATCATGGGTGCATGGTTNNNAGATCGGAA |
| 52391_1 | TATGAGTGTGGAGTCGTTGCTGTTATCCCCATTTTCCAGGGAGNNNAGATCGGAAG |
| 52391_2 | TATGAGTGTGGAGTCGTTGCTGTTATCCCCATTTTCCAGGGAGNNNAGATCGGAAG |
| 35032_1 | TATGAGTGTGGAGTCGTTGCGGTACGATATCGACTCACTGCAANNNAGATCGGAAG |
| 35032_2 | TATGAGTGTGGAGTCGTTGCGGTACGATATCGACTCACTGCAANNNAGATCGGAAG |
| 41113_1 | TATGAGTGTGGAGTCGTTGCCCGACATTTGAAAAGAGTGCACTNNNAGATCGGAA |
| 41113_2 | TATGAGTGTGGAGTCGTTGCCCGACATTTGAAAAGAGTGCACTNNNAGATCGGAA |
| 33114_1 | TATGAGTGTGGAGTCGTTGCGGCTGTGTTGGAAGAAAATGTGTNNNAGATCGGAA |
| 33114_2 | TATGAGTGTGGAGTCGTTGCGGCTGTGTTGGAAGAAAATGTGTNNNAGATCGGAA |
| 41257_1 | TATGAGTGTGGAGTCGTTGCGCAAAAAAATCTCCTCCCGATCCNNNAGATCGGAAG |
| 41257_2 | TATGAGTGTGGAGTCGTTGCGCAAAAAAATCTCCTCCCGATCCNNNAGATCGGAAG |
| 33319_1 | TATGAGTGTGGAGTCGTTGCCAGCCTCGACCTTCTAAGCTAAANNNAGATCGGAAG |
| 33319_2 | TATGAGTGTGGAGTCGTTGCCAGCCTCGACCTTCTAAGCTAAANNNAGATCGGAAG |
| 35126_1 | TATGAGTGTGGAGTCGTTGCTTTTTGAGGAACCTCCAAAGCACNNNAGATCGGAAG |
| 35126_2 | TATGAGTGTGGAGTCGTTGCTTTTTGAGGAACCTCCAAAGCACNNNAGATCGGAAG |
| 46990_1 | TATGAGTGTGGAGTCGTTGCTTCACACCATTCTCCTGCCTCNNNAGATCGGAAGAGC |
| 46990_2 | TATGAGTGTGGAGTCGTTGCTTCACACCATTCTCCTGCCTCNNNAGATCGGAAGAGC |
| 50733_1 | TATGAGTGTGGAGTCGTTGCTCTCACCAGAGACAACACATACANNNAGATCGGAAG |
| 50733_2 | TATGAGTGTGGAGTCGTTGCTCTCACCAGAGACAACACATACANNNAGATCGGAAG |
| 47380_1 | TATGAGTGTGGAGTCGTTGCACACAAGGAAGGCAAGTTTTTCCNNNAGATCGGAA |
| 47380_2 | TATGAGTGTGGAGTCGTTGCACACAAGGAAGGCAAGTTTTTCCNNNAGATCGGAA |
| 51036_1 | TATGAGTGTGGAGTCGTTGCGACCACCCAATTGCCAAATACAANNNAGATCGGAAG |
| 51036_2 | TATGAGTGTGGAGTCGTTGCGACCACCCAATTGCCAAATACAANNNAGATCGGAAG |
| 39349_1 | TATGAGTGTGGAGTCGTTGCAAATGGCAACATGTTGTCTCAGGNNNAGATCGGAA |
| 39349_2 | TATGAGTGTGGAGTCGTTGCAAATGGCAACATGTTGTCTCAGGNNNAGATCGGAA |
| 50801_1 | TATGAGTGTGGAGTCGTTGCAATCAAAAGCAATCCCATGAGCCNNNAGATCGGAAG |
| 50801_2 | TATGAGTGTGGAGTCGTTGCAATCAAAAGCAATCCCATGAGCCNNNAGATCGGAAG |
| 32657_1 | TATGAGTGTGGAGTCGTTGCGCTCCCTGGTTCAAACAATTCTCNNNAGATCGGAAG |
| 32657_2 | TATGAGTGTGGAGTCGTTGCGCTCCCTGGTTCAAACAATTCTCNNNAGATCGGAAG |
| 42006_1 | TATGAGTGTGGAGTCGTTGCCCAAGTAGCTGGTACTACAGGTGNNNAGATCGGAA |
| 42006_2 | TATGAGTGTGGAGTCGTTGCCCAAGTAGCTGGTACTACAGGTGNNNAGATCGGAA |
| 1697_1 | TATGAGTGTGGAGTCGTTGCTCACATGGTTGTGGCTGCTATTTNNNAGATCGGAAG |
| 1697_2 | TATGAGTGTGGAGTCGTTGCTCACATGGTTGTGGCTGCTATTTNNNAGATCGGAAG |
| 52956_1 | TATGAGTGTGGAGTCGTTGCGCTTTAGTGATCTATTGCACAGCANNNAGATCGGAA |
| 52956_2 | TATGAGTGTGGAGTCGTTGCGCTTTAGTGATCTATTGCACAGCANNNAGATCGGAA |
| 29201_1 | TATGAGTGTGGAGTCGTTGCAACCTGTAATCTGAACAGCGACTNNNAGATCGGAAG |
| 29201_2 | TATGAGTGTGGAGTCGTTGCAACCTGTAATCTGAACAGCGACTNNNAGATCGGAAG |
| 51870_1 | TATGAGTGTGGAGTCGTTGCCACTGTTTTCGGACCTTCCATGNNNAGATCGGAAGA |
| 51870_2 | TATGAGTGTGGAGTCGTTGCCACTGTTTTCGGACCTTCCATGNNNAGATCGGAAGA |
| 30269_1 | TATGAGTGTGGAGTCGTTGCACGGATAGACTTTTAAACTTCAACCANNNAGATCGG |
| 30269_2 | TATGAGTGTGGAGTCGTTGCACGGATAGACTTTTAAACTTCAACCANNNAGATCGG |
| 32160_1 | TATGAGTGTGGAGTCGTTGCGGATTCAAGAGATTCTCCTGCCTNNNAGATCGGAAG |
| 32160_2 | TATGAGTGTGGAGTCGTTGCGGATTCAAGAGATTCTCCTGCCTNNNAGATCGGAAG |

| | |
|---|---|
| 38493_1 | TATGAGTGTGGAGTCGTTGCGAGAGGGAGGACAAAGGGAAAATNNNAGATCGGA |
| 38493_2 | TATGAGTGTGGAGTCGTTGCGAGAGGGAGGACAAAGGGAAAATNNNAGATCGGA |
| 52412_1 | TATGAGTGTGGAGTCGTTGCCTGGGCTCCCTCTTGATTACTTTNNNAGATCGGAAG |
| 52412_2 | TATGAGTGTGGAGTCGTTGCCTGGGCTCCCTCTTGATTACTTTNNNAGATCGGAAG |
| 42311_1 | TATGAGTGTGGAGTCGTTGCCACAGAGTCACACATCCTTGAGANNNAGATCGGAAG |
| 42311_2 | TATGAGTGTGGAGTCGTTGCCACAGAGTCACACATCCTTGAGANNNAGATCGGAAG |
| 30120_1 | TATGAGTGTGGAGTCGTTGCACAGTTATTTCTAGTGTTGTTGGCANNNAGATCGGA |
| 30120_2 | TATGAGTGTGGAGTCGTTGCACAGTTATTTCTAGTGTTGTTGGCANNNAGATCGGA |
| 44491_1 | TATGAGTGTGGAGTCGTTGCCTAGGTGGGAGGATCACTTGAACNNNAGATCGGAA |
| 44491_2 | TATGAGTGTGGAGTCGTTGCCTAGGTGGGAGGATCACTTGAACNNNAGATCGGAA |
| 38784_1 | TATGAGTGTGGAGTCGTTGCAAAAATAAAAAGTAGGGCCGGGCNNNAGATCGGAA |
| 38784_2 | TATGAGTGTGGAGTCGTTGCAAAAATAAAAAGTAGGGCCGGGCNNNAGATCGGAA |
| 48919_1 | TATGAGTGTGGAGTCGTTGCGGTTCAAACAATTCTCCTGCCTCNNNAGATCGGAAG |
| 48919_2 | TATGAGTGTGGAGTCGTTGCGGTTCAAACAATTCTCCTGCCTCNNNAGATCGGAAG |
| 36365_1 | TATGAGTGTGGAGTCGTTGCACAACTTGTAAGAGGAGTTGGCANNNAGATCGGAA |
| 36365_2 | TATGAGTGTGGAGTCGTTGCACAACTTGTAAGAGGAGTTGGCANNNAGATCGGAA |
| 33305_1 | TATGAGTGTGGAGTCGTTGCCAAGTAGCTGGGACTACAGGCNNNAGATCGGAAGA |
| 33305_2 | TATGAGTGTGGAGTCGTTGCCAAGTAGCTGGGACTACAGGCNNNAGATCGGAAGA |
| 13303_1 | TATGAGTGTGGAGTCGTTGCGTATTCTTGGTTTCTGATGAAATNNNAGATCGGAAG |
| 13303_2 | TATGAGTGTGGAGTCGTTGCGTATTCTTGGTTTCTGATGAAATNNNAGATCGGAAG |
| 34767_1 | TATGAGTGTGGAGTCGTTGCCTCCTGAGTAGCTGGGACTACAGNNNAGATCGGAA |
| 34767_2 | TATGAGTGTGGAGTCGTTGCCTCCTGAGTAGCTGGGACTACAGNNNAGATCGGAA |
| 49832_1 | TATGAGTGTGGAGTCGTTGCTGCTCTCTCATCCTCCTATTCCTNNNAGATCGGAAGA |
| 49832_2 | TATGAGTGTGGAGTCGTTGCTGCTCTCTCATCCTCCTATTCCTNNNAGATCGGAAGA |
| 48899_1 | TATGAGTGTGGAGTCGTTGCGAGTCTCGCTCTTTCGCCAAGNNNAGATCGGAAGAG |
| 48899_2 | TATGAGTGTGGAGTCGTTGCGAGTCTCGCTCTTTCGCCAAGNNNAGATCGGAAGAG |
| 10382_1 | TATGAGTGTGGAGTCGTTGCGCAGGGGCTCAACATACGATAAANNNAGATCGGAA |
| 10382_2 | TATGAGTGTGGAGTCGTTGCGCAGGGGCTCAACATACGATAAANNNAGATCGGAA |
| 40482_1 | TATGAGTGTGGAGTCGTTGCAAAAAATTAGTTGGGCGTGGTGGNNNAGATCGGA |
| 40482_2 | TATGAGTGTGGAGTCGTTGCAAAAAATTAGTTGGGCGTGGTGGNNNAGATCGGA |
| 37690_1 | TATGAGTGTGGAGTCGTTGCCAAGAAACCATCACTGGCACTTTNNNAGATCGGAAG |
| 37690_2 | TATGAGTGTGGAGTCGTTGCCAAGAAACCATCACTGGCACTTTNNNAGATCGGAAG |
| 52868_1 | TATGAGTGTGGAGTCGTTGCCAGGAGCTGGCAGAGTTCATGNNNAGATCGGAAGA |
| 52868_2 | TATGAGTGTGGAGTCGTTGCCAGGAGCTGGCAGAGTTCATGNNNAGATCGGAAGA |
| 47862_1 | TATGAGTGTGGAGTCGTTGCAGGCTGGAGTATAGTGGAGTGATNNNAGATCGGA |
| 47862_2 | TATGAGTGTGGAGTCGTTGCAGGCTGGAGTATAGTGGAGTGATNNNAGATCGGA |
| 34130_1 | TATGAGTGTGGAGTCGTTGCCACTCCTGTAATCCCAGCACTTTNNNAGATCGGAAGA |
| 34130_2 | TATGAGTGTGGAGTCGTTGCCACTCCTGTAATCCCAGCACTTTNNNAGATCGGAAGA |
| 49670_1 | TATGAGTGTGGAGTCGTTGCCTCAAGGGATCCTTCCACCTTACNNNAGATCGGAAG |
| 49670_2 | TATGAGTGTGGAGTCGTTGCCTCAAGGGATCCTTCCACCTTACNNNAGATCGGAAG |
| 34385_1 | TATGAGTGTGGAGTCGTTGCTTTGAGCCAGGATCTCTCTTAGCNNNAGATCGGAAG |
| 34385_2 | TATGAGTGTGGAGTCGTTGCTTTGAGCCAGGATCTCTCTTAGCNNNAGATCGGAAG |
| 44880_1 | TATGAGTGTGGAGTCGTTGCTTAGTTCATTTGCATGGCAGCTGNNNAGATCGGAAG |

| | |
|---|---|
| 44880_2 | TATGAGTGTGGAGTCGTTGCTTAGTTCATTTGCATGGCAGCTGNNNAGATCGGAAG |
| 46120_1 | TATGAGTGTGGAGTCGTTGCGTTCAAGCAGTTCTCCTACCTCANNNAGATCGGAAG |
| 46120_2 | TATGAGTGTGGAGTCGTTGCGTTCAAGCAGTTCTCCTACCTCANNNAGATCGGAAG |
| 28958_1 | TATGAGTGTGGAGTCGTTGCGATGAGGTCTCACTCTGTCTGTCNNNAGATCGGAAG |
| 28958_2 | TATGAGTGTGGAGTCGTTGCGATGAGGTCTCACTCTGTCTGTCNNNAGATCGGAAG |
| 2489_1 | TATGAGTGTGGAGTCGTTGCTCTGGCAATGGGTTCATTCTCAGNNNAGATCGGAAG |
| 2489_2 | TATGAGTGTGGAGTCGTTGCTCTGGCAATGGGTTCATTCTCAGNNNAGATCGGAAG |
| 51038_1 | TATGAGTGTGGAGTCGTTGCGAGGACAAGGAAATGGAGGAACTNNNAGATCGGA |
| 51038_2 | TATGAGTGTGGAGTCGTTGCGAGGACAAGGAAATGGAGGAACTNNNAGATCGGA |
| 47644_1 | TATGAGTGTGGAGTCGTTGCAGACAGTTCCCAAATGAAGAGGTNNNAGATCGGAA |
| 47644_2 | TATGAGTGTGGAGTCGTTGCAGACAGTTCCCAAATGAAGAGGTNNNAGATCGGAA |
| 12487_1 | TATGAGTGTGGAGTCGTTGCGAATGATGATCACACCACTGNNNAGATCGGAAGAG |
| 12487_2 | TATGAGTGTGGAGTCGTTGCGAATGATGATCACACCACTGNNNAGATCGGAAGAG |
| 29333_1 | TATGAGTGTGGAGTCGTTGCGGATGAGAGCATTCACAGGAGATNNNAGATCGGAA |
| 29333_2 | TATGAGTGTGGAGTCGTTGCGGATGAGAGCATTCACAGGAGATNNNAGATCGGAA |
| 52193_1 | TATGAGTGTGGAGTCGTTGCGCTGTCCTCTACCCAGATCTTTCNNNAGATCGGAAGA |
| 52193_2 | TATGAGTGTGGAGTCGTTGCGCTGTCCTCTACCCAGATCTTTCNNNAGATCGGAAGA |
| 40826_1 | TATGAGTGTGGAGTCGTTGCACAGAAACCTTAGACATCCATTCGANNNAGATCGGA |
| 40826_2 | TATGAGTGTGGAGTCGTTGCACAGAAACCTTAGACATCCATTCGANNNAGATCGGA |
| 29410_1 | TATGAGTGTGGAGTCGTTGCCATTCTGGGACAGCCAAGTCTNNNAGATCGGAAGAG |
| 29410_2 | TATGAGTGTGGAGTCGTTGCCATTCTGGGACAGCCAAGTCTNNNAGATCGGAAGAG |
| 39948_1 | TATGAGTGTGGAGTCGTTGCCTGCCTCCTGGTTCAAATGATTCNNNAGATCGGAAG |
| 39948_2 | TATGAGTGTGGAGTCGTTGCCTGCCTCCTGGTTCAAATGATTCNNNAGATCGGAAG |
| 42277_1 | TATGAGTGTGGAGTCGTTGCTGTGTGTGTGTGTGTATATGTGTGNNNAGATCGGA |
| 42277_2 | TATGAGTGTGGAGTCGTTGCTGTGTGTGTGTGTGTATATGTGTGNNNAGATCGGA |
| 50093_1 | TATGAGTGTGGAGTCGTTGCTACAGAAGAACGACCACAACCTTNNNAGATCGGAAG |
| 50093_2 | TATGAGTGTGGAGTCGTTGCTACAGAAGAACGACCACAACCTTNNNAGATCGGAAG |
| 45183_1 | TATGAGTGTGGAGTCGTTGCTTGTGTTCAGAAGATGGGAGAGGNNNAGATCGGA |
| 45183_2 | TATGAGTGTGGAGTCGTTGCTTGTGTTCAGAAGATGGGAGAGGNNNAGATCGGA |
| 10707_1 | TATGAGTGTGGAGTCGTTGCCCAGAAGTGACAACTGTCCAAAANNNAGATCGGAA |
| 10707_2 | TATGAGTGTGGAGTCGTTGCCCAGAAGTGACAACTGTCCAAAANNNAGATCGGAA |
| 12763_1 | TATGAGTGTGGAGTCGTTGCTAGAAAAGGAGACACAAAAGAAANNNAGATCGGA |
| 12763_2 | TATGAGTGTGGAGTCGTTGCTAGAAAAGGAGACACAAAAGAAANNNAGATCGGA |
| 39989_1 | TATGAGTGTGGAGTCGTTGCAGTATCAGTGTTTCAGGGGTTCCNNNAGATCGGAA |
| 39989_2 | TATGAGTGTGGAGTCGTTGCAGTATCAGTGTTTCAGGGGTTCCNNNAGATCGGAA |
| 31570_1 | TATGAGTGTGGAGTCGTTGCTTGATGACCTTGACAGTGATGCTNNNAGATCGGAAG |
| 31570_2 | TATGAGTGTGGAGTCGTTGCTTGATGACCTTGACAGTGATGCTNNNAGATCGGAAG |
| 2501_1 | TATGAGTGTGGAGTCGTTGCTGGAAAGTACCCAGAGTGAGGAGNNNAGATCGGA |
| 2501_2 | TATGAGTGTGGAGTCGTTGCTGGAAAGTACCCAGAGTGAGGAGNNNAGATCGGA |
| 45122_1 | TATGAGTGTGGAGTCGTTGCAAGTATTAGGTCTTCCTGCTCCGNNNAGATCGGAAG |
| 45122_2 | TATGAGTGTGGAGTCGTTGCAAGTATTAGGTCTTCCTGCTCCGNNNAGATCGGAAG |
| 9102_1 | TATGAGTGTGGAGTCGTTGCTGAATGTGTCAGCTTGCTTGACANNNAGATCGGAAG |
| 9102_2 | TATGAGTGTGGAGTCGTTGCTGAATGTGTCAGCTTGCTTGACANNNAGATCGGAAG |

| | |
|---|---|
| 48375_1 | TATGAGTGTGGAGTCGTTGCGGTCACGATGTCCCCAACACNNNAGATCGGAAGAGC |
| 48375_2 | TATGAGTGTGGAGTCGTTGCGGTCACGATGTCCCCAACACNNNAGATCGGAAGAGC |
| 41060_1 | TATGAGTGTGGAGTCGTTGCAAAACCTCAGTGGGTGTGATACANNNAGATCGGAA |
| 41060_2 | TATGAGTGTGGAGTCGTTGCAAAACCTCAGTGGGTGTGATACANNNAGATCGGAA |
| 31686_1 | TATGAGTGTGGAGTCGTTGCTAACCAAATCCAGCAGCACATTGNNNAGATCGGAAG |
| 31686_2 | TATGAGTGTGGAGTCGTTGCTAACCAAATCCAGCAGCACATTGNNNAGATCGGAAG |
| 37288_1 | TATGAGTGTGGAGTCGTTGCATTCTTTCTCTGCCCCTTTGAGANNNAGATCGGAAGA |
| 37288_2 | TATGAGTGTGGAGTCGTTGCATTCTTTCTCTGCCCCTTTGAGANNNAGATCGGAAGA |
| 51354_1 | TATGAGTGTGGAGTCGTTGCAATGAGGACCAGTTGGCTACATTNNNAGATCGGAA |
| 51354_2 | TATGAGTGTGGAGTCGTTGCAATGAGGACCAGTTGGCTACATTNNNAGATCGGAA |
| 38698_1 | TATGAGTGTGGAGTCGTTGCTTGAAAGAAGCATCTTGCCACTGNNNAGATCGGAAG |
| 38698_2 | TATGAGTGTGGAGTCGTTGCTTGAAAGAAGCATCTTGCCACTGNNNAGATCGGAAG |
| 37874_1 | TATGAGTGTGGAGTCGTTGCATCACCAAAGTGCTATCCCTTGTNNNAGATCGGAAG |
| 37874_2 | TATGAGTGTGGAGTCGTTGCATCACCAAAGTGCTATCCCTTGTNNNAGATCGGAAG |
| 38591_1 | TATGAGTGTGGAGTCGTTGCGGAACTGCAAATCAAAATCGCAANNNAGATCGGAA |
| 38591_2 | TATGAGTGTGGAGTCGTTGCGGAACTGCAAATCAAAATCGCAANNNAGATCGGAA |
| 35300_1 | TATGAGTGTGGAGTCGTTGCCCAAGCCCGGCTAATTTTTTGTANNNAGATCGGAAG |
| 35300_2 | TATGAGTGTGGAGTCGTTGCCCAAGCCCGGCTAATTTTTTGTANNNAGATCGGAAG |
| 29683_1 | TATGAGTGTGGAGTCGTTGCCTCCATTATGAGAGGCTGTGCTANNNAGATCGGAAG |
| 29683_2 | TATGAGTGTGGAGTCGTTGCCTCCATTATGAGAGGCTGTGCTANNNAGATCGGAAG |
| 46243_1 | TATGAGTGTGGAGTCGTTGCCGAGTAAAAGCAGAAGGCAAAGGNNNAGATCGGA |
| 46243_2 | TATGAGTGTGGAGTCGTTGCCGAGTAAAAGCAGAAGGCAAAGGNNNAGATCGGA |
| 46628_1 | TATGAGTGTGGAGTCGTTGCCATATCAGCTCTCTGCAACCTCTNNNAGATCGGAAGA |
| 46628_2 | TATGAGTGTGGAGTCGTTGCCATATCAGCTCTCTGCAACCTCTNNNAGATCGGAAGA |
| 29512_1 | TATGAGTGTGGAGTCGTTGCCTCCTTGCTTTAATCCTTAAACANNNAGATCGGAAGA |
| 29512_2 | TATGAGTGTGGAGTCGTTGCCTCCTTGCTTTAATCCTTAAACANNNAGATCGGAAGA |
| 51692_1 | TATGAGTGTGGAGTCGTTGCTTCAGAACTGCTTAGCATGGTCANNNAGATCGGAAG |
| 51692_2 | TATGAGTGTGGAGTCGTTGCTTCAGAACTGCTTAGCATGGTCANNNAGATCGGAAG |
| 47016_1 | TATGAGTGTGGAGTCGTTGCGCGTATGACTACACCTGGACATTNNNAGATCGGAAG |
| 47016_2 | TATGAGTGTGGAGTCGTTGCGCGTATGACTACACCTGGACATTNNNAGATCGGAAG |
| 52895_1 | TATGAGTGTGGAGTCGTTGCGCCATCACAGCCAATGACGGNNNAGATCGGAAGAG |
| 52895_2 | TATGAGTGTGGAGTCGTTGCGCCATCACAGCCAATGACGGNNNAGATCGGAAGAG |
| 35786_1 | TATGAGTGTGGAGTCGTTGCACAGACACCCAGCTAATTGTCATNNNAGATCGGAAG |
| 35786_2 | TATGAGTGTGGAGTCGTTGCACAGACACCCAGCTAATTGTCATNNNAGATCGGAAG |
| 13581_1 | TATGAGTGTGGAGTCGTTGCTTTATTTTTTTGAGACAGAGTCTNNNAGATCGGAAG |
| 13581_2 | TATGAGTGTGGAGTCGTTGCTTTATTTTTTTGAGACAGAGTCTNNNAGATCGGAAG |
| 39408_1 | TATGAGTGTGGAGTCGTTGCGACCAGCCTGGCTAACATAGTAANNNAGATCGGAA |
| 39408_2 | TATGAGTGTGGAGTCGTTGCGACCAGCCTGGCTAACATAGTAANNNAGATCGGAA |
| 52945_1 | TATGAGTGTGGAGTCGTTGCGGCAAAAAACACATCTCTGCTGANNNAGATCGGAAG |
| 52945_2 | TATGAGTGTGGAGTCGTTGCGGCAAAAAACACATCTCTGCTGANNNAGATCGGAAG |
| 50331_1 | TATGAGTGTGGAGTCGTTGCTAAAAATGCAGTGGCCCAATCATNNNAGATCGGAAG |
| 50331_2 | TATGAGTGTGGAGTCGTTGCTAAAAATGCAGTGGCCCAATCATNNNAGATCGGAAG |
| 39692_1 | TATGAGTGTGGAGTCGTTGCAATATGAGGACCTGCTATGGGTCNNNAGATCGGAA |

| | |
|---|---|
| 39692_2 | TATGAGTGTGGAGTCGTTGCAATATGAGGACCTGCTATGGGTCNNNAGATCGGAA |
| 38928_1 | TATGAGTGTGGAGTCGTTGCGAAAACAGGAGGTGGAGCTTGNNNAGATCGGAAG |
| 38928_2 | TATGAGTGTGGAGTCGTTGCGAAAACAGGAGGTGGAGCTTGNNNAGATCGGAAG |
| 29491_1 | ATGAGTGTGGAGTCGTTGCAAATATCGACTGTCCAAGATCTACAAANNNAGATCGG |
| 29491_2 | ATGAGTGTGGAGTCGTTGCAAATATCGACTGTCCAAGATCTACAAANNNAGATCGG |
| 41487_1 | TATGAGTGTGGAGTCGTTGCTCTCAAAAAGGGAGGGGTGTATGNNNAGATCGGAA |
| 41487_2 | TATGAGTGTGGAGTCGTTGCTCTCAAAAAGGGAGGGGTGTATGNNNAGATCGGAA |
| 43389_1 | TATGAGTGTGGAGTCGTTGCCTCCCAGGTTCATGCCATTCTCNNNAGATCGGAAGAG |
| 43389_2 | TATGAGTGTGGAGTCGTTGCCTCCCAGGTTCATGCCATTCTCNNNAGATCGGAAGAG |
| 390_1 | TATGAGTGTGGAGTCGTTGCTAAGTTTCTGACTGTCCAGCCCTNNNAGATCGGAAG |
| 390_2 | TATGAGTGTGGAGTCGTTGCTAAGTTTCTGACTGTCCAGCCCTNNNAGATCGGAAG |
| 33566_1 | TATGAGTGTGGAGTCGTTGCCTGGGTTCAATTGATTCTCCTGCNNNAGATCGGAAG |
| 33566_2 | TATGAGTGTGGAGTCGTTGCCTGGGTTCAATTGATTCTCCTGCNNNAGATCGGAAG |
| 37439_1 | TATGAGTGTGGAGTCGTTGCGGATAAAGTATCTGCCAGCCTGANNNAGATCGGAA |
| 37439_2 | TATGAGTGTGGAGTCGTTGCGGATAAAGTATCTGCCAGCCTGANNNAGATCGGAA |
| 40971_1 | TATGAGTGTGGAGTCGTTGCCTGTAATCCCAGCTACTCAGGAGNNNAGATCGGAAG |
| 40971_2 | TATGAGTGTGGAGTCGTTGCCTGTAATCCCAGCTACTCAGGAGNNNAGATCGGAAG |
| 36338_1 | TATGAGTGTGGAGTCGTTGCGGTATCTAGTCATGTTCTCGTCACANNNAGATCGGA |
| 36338_2 | TATGAGTGTGGAGTCGTTGCGGTATCTAGTCATGTTCTCGTCACANNNAGATCGGA |
| 35779_1 | TATGAGTGTGGAGTCGTTGCATTCCCTGGTTCGAGTGATTCTCNNNAGATCGGAAG |
| 35779_2 | TATGAGTGTGGAGTCGTTGCATTCCCTGGTTCGAGTGATTCTCNNNAGATCGGAAG |
| 32532_1 | TATGAGTGTGGAGTCGTTGCAGCAACATCAAACCTGAGAGTCANNNAGATCGGAAG |
| 32532_2 | TATGAGTGTGGAGTCGTTGCAGCAACATCAAACCTGAGAGTCANNNAGATCGGAAG |
| 31011_1 | TATGAGTGTGGAGTCGTTGCTGGACCAGGCTCTTTTTATCACANNNAGATCGGAAG |
| 31011_2 | TATGAGTGTGGAGTCGTTGCTGGACCAGGCTCTTTTTATCACANNNAGATCGGAAG |
| 46153_1 | TATGAGTGTGGAGTCGTTGCCGAGTAGCTGGGACTACAGGTNNNAGATCGGAAGA |
| 46153_2 | TATGAGTGTGGAGTCGTTGCCGAGTAGCTGGGACTACAGGTNNNAGATCGGAAGA |
| 34713_1 | TATGAGTGTGGAGTCGTTGCCCATGCCCGGCTAATTTTTTGTANNNAGATCGGAAG |
| 34713_2 | TATGAGTGTGGAGTCGTTGCCCATGCCCGGCTAATTTTTTGTANNNAGATCGGAAG |
| 7966_1 | TATGAGTGTGGAGTCGTTGCAGGTCTCACTGTCACTCCAGCTANNNAGATCGGAAG |
| 7966_2 | TATGAGTGTGGAGTCGTTGCAGGTCTCACTGTCACTCCAGCTANNNAGATCGGAAG |
| 53266_1 | TATGAGTGTGGAGTCGTTGCAAGTAGATGTCAGCTGATGGGTGNNNAGATCGGAA |
| 53266_2 | TATGAGTGTGGAGTCGTTGCAAGTAGATGTCAGCTGATGGGTGNNNAGATCGGAA |
| 28948_1 | TATGAGTGTGGAGTCGTTGCGCTGAAATTCTGTAGCAGTGTTGTNNNAGATCGGAA |
| 28948_2 | TATGAGTGTGGAGTCGTTGCGCTGAAATTCTGTAGCAGTGTTGTNNNAGATCGGAA |
| 37247_1 | TATGAGTGTGGAGTCGTTGCCCCACTGTACACTTAGGCTACACNNNAGATCGGAAG |
| 37247_2 | TATGAGTGTGGAGTCGTTGCCCCACTGTACACTTAGGCTACACNNNAGATCGGAAG |
| 35501_1 | TATGAGTGTGGAGTCGTTGCGCCCGGCCTCTAAAAGTATTTCTNNNAGATCGGAAG |
| 35501_2 | TATGAGTGTGGAGTCGTTGCGCCCGGCCTCTAAAAGTATTTCTNNNAGATCGGAAG |
| 41992_1 | TATGAGTGTGGAGTCGTTGCGGGTTCAAGTGATTCTCCTGTCTNNNAGATCGGAAG |
| 41992_2 | TATGAGTGTGGAGTCGTTGCGGGTTCAAGTGATTCTCCTGTCTNNNAGATCGGAAG |
| 38261_1 | TATGAGTGTGGAGTCGTTGCATAAATCCGCACATCACGAGGTANNNAGATCGGAAG |
| 38261_2 | TATGAGTGTGGAGTCGTTGCATAAATCCGCACATCACGAGGTANNNAGATCGGAAG |

| | |
|---|---|
| 51649_1 | TATGAGTGTGGAGTCGTTGCGCAGTAGCATTCCAGGTTCCTAANNNAGATCGGAAG |
| 51649_2 | TATGAGTGTGGAGTCGTTGCGCAGTAGCATTCCAGGTTCCTAANNNAGATCGGAAG |
| 50074_1 | TATGAGTGTGGAGTCGTTGCCTGAGGATCAAAGAGAGCCAGTTNNNAGATCGGAA |
| 50074_2 | TATGAGTGTGGAGTCGTTGCCTGAGGATCAAAGAGAGCCAGTTNNNAGATCGGAA |
| 48248_1 | TATGAGTGTGGAGTCGTTGCTCTTCTTGATTCAACCTTGGTAGGNNNAGATCGGAA |
| 48248_2 | TATGAGTGTGGAGTCGTTGCTCTTCTTGATTCAACCTTGGTAGGNNNAGATCGGAA |
| 10338_1 | TATGAGTGTGGAGTCGTTGCTTGTCTCCTTCCTTCCAGAGGAGNNNAGATCGGAAG |
| 10338_2 | TATGAGTGTGGAGTCGTTGCTTGTCTCCTTCCTTCCAGAGGAGNNNAGATCGGAAG |
| 51699_1 | TATGAGTGTGGAGTCGTTGCTCAGGAACGCATTCTCTTTCTCANNNAGATCGGAAGA |
| 51699_2 | TATGAGTGTGGAGTCGTTGCTCAGGAACGCATTCTCTTTCTCANNNAGATCGGAAGA |
| 12889_1 | TATGAGTGTGGAGTCGTTGCCGGCTTTTAAGTGTGATTTTANNNAGATCGGAAGAG |
| 12889_2 | TATGAGTGTGGAGTCGTTGCCGGCTTTTAAGTGTGATTTTANNNAGATCGGAAGAG |
| 52338_1 | TATGAGTGTGGAGTCGTTGCAAAAGCAAGCCCAAGACACAAATNNNAGATCGGAA |
| 52338_2 | TATGAGTGTGGAGTCGTTGCAAAAGCAAGCCCAAGACACAAATNNNAGATCGGAA |
| 49996_1 | TATGAGTGTGGAGTCGTTGCCAGGTTGGTAGGATAATCGACCCNNNAGATCGGAA |
| 49996_2 | TATGAGTGTGGAGTCGTTGCCAGGTTGGTAGGATAATCGACCCNNNAGATCGGAA |
| 38693_1 | TATGAGTGTGGAGTCGTTGCCAAAGTTCTCCAAGCCCGAATTTNNNAGATCGGAAG |
| 38693_2 | TATGAGTGTGGAGTCGTTGCCAAAGTTCTCCAAGCCCGAATTTNNNAGATCGGAAG |
| 29536_1 | TATGAGTGTGGAGTCGTTGCGTGTTAGTGTGTCAGGAGACTGANNNAGATCGGAA |
| 29536_2 | TATGAGTGTGGAGTCGTTGCGTGTTAGTGTGTCAGGAGACTGANNNAGATCGGAA |
| 52420_1 | TATGAGTGTGGAGTCGTTGCAGAAGTTGCCCTTGTTATGTCCTNNNAGATCGGAAG |
| 52420_2 | TATGAGTGTGGAGTCGTTGCAGAAGTTGCCCTTGTTATGTCCTNNNAGATCGGAAG |
| 40870_1 | TATGAGTGTGGAGTCGTTGCCCAGACAAATAAAACCCCACAACANNNAGATCGGAA |
| 40870_2 | TATGAGTGTGGAGTCGTTGCCCAGACAAATAAAACCCCACAACANNNAGATCGGAA |
| 32550_1 | TATGAGTGTGGAGTCGTTGCAAAGGGGACAAAGGAGAGACAAGNNNAGATCGGA |
| 32550_2 | TATGAGTGTGGAGTCGTTGCAAAGGGGACAAAGGAGAGACAAGNNNAGATCGGA |
| 33783_1 | TATGAGTGTGGAGTCGTTGCGCCTGTAATCCCAGATACTCCAGNNNAGATCGGAAG |
| 33783_2 | TATGAGTGTGGAGTCGTTGCGCCTGTAATCCCAGATACTCCAGNNNAGATCGGAAG |
| 41544_1 | TATGAGTGTGGAGTCGTTGCTTTCACTCTATCACCAAGGCTGGNNNAGATCGGAAG |
| 41544_2 | TATGAGTGTGGAGTCGTTGCTTTCACTCTATCACCAAGGCTGGNNNAGATCGGAAG |
| 45700_1 | TATGAGTGTGGAGTCGTTGCAAAGACAAGACTGCTGTTTGTGGNNNAGATCGGAA |
| 45700_2 | TATGAGTGTGGAGTCGTTGCAAAGACAAGACTGCTGTTTGTGGNNNAGATCGGAA |
| 38939_1 | TATGAGTGTGGAGTCGTTGCGGGGAAAGAGAAGGGATAGCAAANNNAGATCGGA |
| 38939_2 | TATGAGTGTGGAGTCGTTGCGGGGAAAGAGAAGGGATAGCAAANNNAGATCGGA |
| 33022_1 | TATGAGTGTGGAGTCGTTGCTCAAGTGATTCTCCCACCTTAGCNNNAGATCGGAAGA |
| 33022_2 | TATGAGTGTGGAGTCGTTGCTCAAGTGATTCTCCCACCTTAGCNNNAGATCGGAAGA |
| 50970_1 | TATGAGTGTGGAGTCGTTGCCAGCAAACTCACAGATGAAAGGGNNNAGATCGGAA |
| 50970_2 | TATGAGTGTGGAGTCGTTGCCAGCAAACTCACAGATGAAAGGGNNNAGATCGGAA |
| 763_1 | TATGAGTGTGGAGTCGTTGCATCGTACCTGAGTTCCTCTCTGGNNNAGATCGGAAG |
| 763_2 | TATGAGTGTGGAGTCGTTGCATCGTACCTGAGTTCCTCTCTGGNNNAGATCGGAAG |
| 29474_1 | TATGAGTGTGGAGTCGTTGCTTTGAGTCTTACTTCACCTTGCTNNNAGATCGGAAGA |
| 29474_2 | TATGAGTGTGGAGTCGTTGCTTTGAGTCTTACTTCACCTTGCTNNNAGATCGGAAGA |
| 30394_1 | TATGAGTGTGGAGTCGTTGCTGTAATCCTAGCACTTTGGGAGGNNNAGATCGGAA |

| | |
|---|---|
| 30394_2 | TATGAGTGTGGAGTCGTTGCTGTAATCCTAGCACTTTGGGAGGNNNAGATCGGAA |
| 30675_1 | TATGAGTGTGGAGTCGTTGCTAGACCAACACCCACAGCACTTTNNNAGATCGGAAG |
| 30675_2 | TATGAGTGTGGAGTCGTTGCTAGACCAACACCCACAGCACTTTNNNAGATCGGAAG |
| 40888_1 | TATGAGTGTGGAGTCGTTGCACTTCATGTCTTTAGCCCCAGTTNNNAGATCGGAAGA |
| 40888_2 | TATGAGTGTGGAGTCGTTGCACTTCATGTCTTTAGCCCCAGTTNNNAGATCGGAAGA |
| 37292_1 | TATGAGTGTGGAGTCGTTGCTGACAAAAGTGGCTGCAGAAAAANNNAGATCGGAA |
| 37292_2 | TATGAGTGTGGAGTCGTTGCTGACAAAAGTGGCTGCAGAAAAANNNAGATCGGAA |
| 46146_1 | TATGAGTGTGGAGTCGTTGCCCAGTAAGCCCTGAGATCTCTTCNNNAGATCGGAAG |
| 46146_2 | TATGAGTGTGGAGTCGTTGCCCAGTAAGCCCTGAGATCTCTTCNNNAGATCGGAAG |
| 33724_1 | TATGAGTGTGGAGTCGTTGCAAGGGATGGAGGAAGATCTACCANNNAGATCGGAA |
| 33724_2 | TATGAGTGTGGAGTCGTTGCAAGGGATGGAGGAAGATCTACCANNNAGATCGGAA |
| 41528_1 | TATGAGTGTGGAGTCGTTGCTGGGCAGTGTGACTCATGTTTATNNNAGATCGGAA |
| 41528_2 | TATGAGTGTGGAGTCGTTGCTGGGCAGTGTGACTCATGTTTATNNNAGATCGGAA |
| 13162_1 | TATGAGTGTGGAGTCGTTGCAATTCCAGAAAAGGCTGTCTNNNAGATCGGAAGAG |
| 13162_2 | TATGAGTGTGGAGTCGTTGCAATTCCAGAAAAGGCTGTCTNNNAGATCGGAAGAG |
| 45916_1 | TATGAGTGTGGAGTCGTTGCACCACACCCAGCTAATTTTGGTANNNAGATCGGAAG |
| 45916_2 | TATGAGTGTGGAGTCGTTGCACCACACCCAGCTAATTTTGGTANNNAGATCGGAAG |
| 31901_1 | TATGAGTGTGGAGTCGTTGCAAAATCTGGACTTTGTGCTGTGGNNNAGATCGGAA |
| 31901_2 | TATGAGTGTGGAGTCGTTGCAAAATCTGGACTTTGTGCTGTGGNNNAGATCGGAA |
| 37557_1 | TATGAGTGTGGAGTCGTTGCGGTGTTTTTCTCAGCCATCGTAGNNNAGATCGGAAG |
| 37557_2 | TATGAGTGTGGAGTCGTTGCGGTGTTTTTCTCAGCCATCGTAGNNNAGATCGGAAG |
| 34147_1 | TATGAGTGTGGAGTCGTTGCTGTTAGTCTTTCAAGAATGCCCTNNNAGATCGGAAG |
| 34147_2 | TATGAGTGTGGAGTCGTTGCTGTTAGTCTTTCAAGAATGCCCTNNNAGATCGGAAG |
| 9575_1 | TATGAGTGTGGAGTCGTTGCATCACCTCCAAAATCCCTCATGCNNNAGATCGGAAGA |
| 9575_2 | TATGAGTGTGGAGTCGTTGCATCACCTCCAAAATCCCTCATGCNNNAGATCGGAAGA |
| 50192_1 | TATGAGTGTGGAGTCGTTGCTCCCTGGTAGTTATTGGAAAGCCNNNAGATCGGAAG |
| 50192_2 | TATGAGTGTGGAGTCGTTGCTCCCTGGTAGTTATTGGAAAGCCNNNAGATCGGAAG |
| 31687_1 | TATGAGTGTGGAGTCGTTGCACTCAGTTTCTGCAGATTGTTGCNNNAGATCGGAAG |
| 31687_2 | TATGAGTGTGGAGTCGTTGCACTCAGTTTCTGCAGATTGTTGCNNNAGATCGGAAG |
| 48876_1 | TATGAGTGTGGAGTCGTTGCGACTATTTTTGCAGGCTTCAGGGNNNAGATCGGAA |
| 48876_2 | TATGAGTGTGGAGTCGTTGCGACTATTTTTGCAGGCTTCAGGGNNNAGATCGGAA |
| 2270_1 | TATGAGTGTGGAGTCGTTGCCATCACGGCTCACTGAAGCTTTANNNAGATCGGAAG |
| 2270_2 | TATGAGTGTGGAGTCGTTGCCATCACGGCTCACTGAAGCTTTANNNAGATCGGAAG |
| 34052_1 | TATGAGTGTGGAGTCGTTGCTTCAAGCGATTCTCCTGTCTCAGNNNAGATCGGAAG |
| 34052_2 | TATGAGTGTGGAGTCGTTGCTTCAAGCGATTCTCCTGTCTCAGNNNAGATCGGAAG |
| 40443_1 | TATGAGTGTGGAGTCGTTGCCTGGGTGACAGAACAAGACTCTGNNNAGATCGGAA |
| 40443_2 | TATGAGTGTGGAGTCGTTGCCTGGGTGACAGAACAAGACTCTGNNNAGATCGGAA |
| 41760_1 | TATGAGTGTGGAGTCGTTGCACATGGTGAAGCCCTAACTCTACNNNAGATCGGAAG |
| 41760_2 | TATGAGTGTGGAGTCGTTGCACATGGTGAAGCCCTAACTCTACNNNAGATCGGAAG |
| 49776_1 | TATGAGTGTGGAGTCGTTGCAAAAATGTCCTGGAGCTGTCTCTNNNAGATCGGAAG |
| 49776_2 | TATGAGTGTGGAGTCGTTGCAAAAATGTCCTGGAGCTGTCTCTNNNAGATCGGAAG |
| 51540_1 | TATGAGTGTGGAGTCGTTGCGAGAGTGGGAGCAGGTTTCCTAGNNNAGATCGGAA |
| 51540_2 | TATGAGTGTGGAGTCGTTGCGAGAGTGGGAGCAGGTTTCCTAGNNNAGATCGGAA |

| | |
|---|---|
| 50551_1 | TATGAGTGTGGAGTCGTTGCGCTTGAAAGAGAGTAGCCAGACTNNNAGATCGGAA |
| 50551_2 | TATGAGTGTGGAGTCGTTGCGCTTGAAAGAGAGTAGCCAGACTNNNAGATCGGAA |
| 33411_1 | TATGAGTGTGGAGTCGTTGCACATAGGAGGTAGAGACTGCAGTNNNAGATCGGAA |
| 33411_2 | TATGAGTGTGGAGTCGTTGCACATAGGAGGTAGAGACTGCAGTNNNAGATCGGAA |
| 37863_1 | TATGAGTGTGGAGTCGTTGCGAAAGCTGAAGGTTGTTGGTCTTNNNAGATCGGAA |
| 37863_2 | TATGAGTGTGGAGTCGTTGCGAAAGCTGAAGGTTGTTGGTCTTNNNAGATCGGAA |
| 40905_1 | TATGAGTGTGGAGTCGTTGCACCTCTATGACCCAAACACCTTCNNNAGATCGGAAGA |
| 40905_2 | TATGAGTGTGGAGTCGTTGCACCTCTATGACCCAAACACCTTCNNNAGATCGGAAGA |
| 28824_1 | TATGAGTGTGGAGTCGTTGCACACACACACACACACAGAGATANNNAGATCGGAAG |
| 28824_2 | TATGAGTGTGGAGTCGTTGCACACACACACACACACAGAGATANNNAGATCGGAAG |
| 50288_1 | TATGAGTGTGGAGTCGTTGCACAAGACCCTGAATGCACTAAGTNNNAGATCGGAAG |
| 50288_2 | TATGAGTGTGGAGTCGTTGCACAAGACCCTGAATGCACTAAGTNNNAGATCGGAAG |
| 49580_1 | TATGAGTGTGGAGTCGTTGCTCAGCCCTGACGTAACACAATTANNNAGATCGGAAG |
| 49580_2 | TATGAGTGTGGAGTCGTTGCTCAGCCCTGACGTAACACAATTANNNAGATCGGAAG |
| 45459_1 | TATGAGTGTGGAGTCGTTGCGGGTTGAGCCATTTGAGGAAAAANNNAGATCGGAA |
| 45459_2 | TATGAGTGTGGAGTCGTTGCGGGTTGAGCCATTTGAGGAAAAANNNAGATCGGAA |
| 36370_1 | TATGAGTGTGGAGTCGTTGCAAACAGGGATTCTCTTCACACCANNNAGATCGGAAG |
| 36370_2 | TATGAGTGTGGAGTCGTTGCAAACAGGGATTCTCTTCACACCANNNAGATCGGAAG |
| 51742_1 | TATGAGTGTGGAGTCGTTGCTGAGTCTCAGGGTCTTCTTTCAANNNAGATCGGAAG |
| 51742_2 | TATGAGTGTGGAGTCGTTGCTGAGTCTCAGGGTCTTCTTTCAANNNAGATCGGAAG |
| 30729_1 | TATGAGTGTGGAGTCGTTGCGCCTTTCTGAGCTTGAGTTTTCTTNNNAGATCGGAAG |
| 30729_2 | TATGAGTGTGGAGTCGTTGCGCCTTTCTGAGCTTGAGTTTTCTTNNNAGATCGGAAG |
| 44667_1 | TATGAGTGTGGAGTCGTTGCCACAGGAAGTGGTTTCTGGGTCNNNAGATCGGAAG |
| 44667_2 | TATGAGTGTGGAGTCGTTGCCACAGGAAGTGGTTTCTGGGTCNNNAGATCGGAAG |
| 29276_1 | TATGAGTGTGGAGTCGTTGCACAGAGTTGGGGACATAACTTGGNNNAGATCGGAA |
| 29276_2 | TATGAGTGTGGAGTCGTTGCACAGAGTTGGGGACATAACTTGGNNNAGATCGGAA |
| 46437_1 | TATGAGTGTGGAGTCGTTGCACACGTATTTCTGCTTGGTTGTGNNNAGATCGGAAG |
| 46437_2 | TATGAGTGTGGAGTCGTTGCACACGTATTTCTGCTTGGTTGTGNNNAGATCGGAAG |
| 49466_1 | TATGAGTGTGGAGTCGTTGCCCAGATGTCTTGGCTATGACAGTNNNAGATCGGAAG |
| 49466_2 | TATGAGTGTGGAGTCGTTGCCCAGATGTCTTGGCTATGACAGTNNNAGATCGGAAG |
| 45813_1 | TATGAGTGTGGAGTCGTTGCCGCCAAGGACTACAGGAACTTNNNAGATCGGAAGA |
| 45813_2 | TATGAGTGTGGAGTCGTTGCCGCCAAGGACTACAGGAACTTNNNAGATCGGAAGA |
| 51698_1 | TATGAGTGTGGAGTCGTTGCATCTCCTCCTCAAACAATCAGCANNNAGATCGGAAGA |
| 51698_2 | TATGAGTGTGGAGTCGTTGCATCTCCTCCTCAAACAATCAGCANNNAGATCGGAAGA |
| 30636_1 | TATGAGTGTGGAGTCGTTGCCAATTTTACAGAGTGCTGCCTGCNNNAGATCGGAAG |
| 30636_2 | TATGAGTGTGGAGTCGTTGCCAATTTTACAGAGTGCTGCCTGCNNNAGATCGGAAG |
| 40286_1 | TATGAGTGTGGAGTCGTTGCAGAATCCAGTCCAATTGATAGCTNNNAGATCGGAAG |
| 40286_2 | TATGAGTGTGGAGTCGTTGCAGAATCCAGTCCAATTGATAGCTNNNAGATCGGAAG |
| 50450_1 | TATGAGTGTGGAGTCGTTGCCTGGGAGTGTCCTTGGTTCTAAANNNAGATCGGAAG |
| 50450_2 | TATGAGTGTGGAGTCGTTGCCTGGGAGTGTCCTTGGTTCTAAANNNAGATCGGAAG |
| 43838_1 | TATGAGTGTGGAGTCGTTGCGCACTCAGAACCAATTGTCTGAGNNNAGATCGGAAG |
| 43838_2 | TATGAGTGTGGAGTCGTTGCGCACTCAGAACCAATTGTCTGAGNNNAGATCGGAAG |
| 10078_1 | TATGAGTGTGGAGTCGTTGCATGCCTGAGAATCAGTGAGGACTNNNAGATCGGAA |

| | |
|---|---|
| 10078_2 | TATGAGTGTGGAGTCGTTGCATGCCTGAGAATCAGTGAGGACTNNNAGATCGGAA |
| 29914_1 | TATGAGTGTGGAGTCGTTGCGCTGCAGAGAAGGCTTTGTTTGNNNAGATCGGAAG |
| 29914_2 | TATGAGTGTGGAGTCGTTGCGCTGCAGAGAAGGCTTTGTTTGNNNAGATCGGAAG |
| 52883_1 | TATGAGTGTGGAGTCGTTGCCACATTGAAAAGCTGGGAGAAGGNNNAGATCGGAA |
| 52883_2 | TATGAGTGTGGAGTCGTTGCCACATTGAAAAGCTGGGAGAAGGNNNAGATCGGAA |
| 51892_1 | TATGAGTGTGGAGTCGTTGCGGTTCTCCAGAAAGACCTGTCTTNNNAGATCGGAAG |
| 51892_2 | TATGAGTGTGGAGTCGTTGCGGTTCTCCAGAAAGACCTGTCTTNNNAGATCGGAAG |
| 29701_1 | TATGAGTGTGGAGTCGTTGCTTCTGCTTTGCAAAGATTTCCCANNNAGATCGGAAG |
| 29701_2 | TATGAGTGTGGAGTCGTTGCTTCTGCTTTGCAAAGATTTCCCANNNAGATCGGAAG |
| 2429_1 | TATGAGTGTGGAGTCGTTGCTACCTGGCTGACTACAGGAACTCNNNAGATCGGAAG |
| 2429_2 | TATGAGTGTGGAGTCGTTGCTACCTGGCTGACTACAGGAACTCNNNAGATCGGAAG |
| 48124_1 | TATGAGTGTGGAGTCGTTGCGGCAGCCAATTCATAGGTAACACNNNAGATCGGAA |
| 48124_2 | TATGAGTGTGGAGTCGTTGCGGCAGCCAATTCATAGGTAACACNNNAGATCGGAA |
| 39882_1 | TATGAGTGTGGAGTCGTTGCCCTCATGGAGCTTACTTCATGGTNNNAGATCGGAAG |
| 39882_2 | TATGAGTGTGGAGTCGTTGCCCTCATGGAGCTTACTTCATGGTNNNAGATCGGAAG |
| 10263_1 | TATGAGTGTGGAGTCGTTGCATTTTGCCTTTGCCTGCTTCATCNNNAGATCGGAAGA |
| 10263_2 | TATGAGTGTGGAGTCGTTGCATTTTGCCTTTGCCTGCTTCATCNNNAGATCGGAAGA |
| 12907_1 | TATGAGTGTGGAGTCGTTGCTTTGCTGATTTTCATCTACTTCANNNAGATCGGAAGA |
| 12907_2 | TATGAGTGTGGAGTCGTTGCTTTGCTGATTTTCATCTACTTCANNNAGATCGGAAGA |
| 46526_1 | TATGAGTGTGGAGTCGTTGCCTTCCTGAGTAGGTGGGATCACANNNAGATCGGAA |
| 46526_2 | TATGAGTGTGGAGTCGTTGCCTTCCTGAGTAGGTGGGATCACANNNAGATCGGAA |
| 49470_1 | TATGAGTGTGGAGTCGTTGCGAGTAGCTGGGATTATAGGCACCNNNAGATCGGAA |
| 49470_2 | TATGAGTGTGGAGTCGTTGCGAGTAGCTGGGATTATAGGCACCNNNAGATCGGAA |
| 32281_1 | TATGAGTGTGGAGTCGTTGCTCTAATCCAAACTAGAGCTGGGCNNNAGATCGGAAG |
| 32281_2 | TATGAGTGTGGAGTCGTTGCTCTAATCCAAACTAGAGCTGGGCNNNAGATCGGAAG |
| 42636_1 | TATGAGTGTGGAGTCGTTGCTGCTAGGATATACATGGGAGTGCNNNAGATCGGAA |
| 42636_2 | TATGAGTGTGGAGTCGTTGCTGCTAGGATATACATGGGAGTGCNNNAGATCGGAA |
| 40793_1 | TATGAGTGTGGAGTCGTTGCATTGCTTGAATCTGAGAGGCAGANNNAGATCGGAA |
| 40793_2 | TATGAGTGTGGAGTCGTTGCATTGCTTGAATCTGAGAGGCAGANNNAGATCGGAA |
| 38506_1 | TATGAGTGTGGAGTCGTTGCAGCAAAGCTGCAGGATATACAAANNNAGATCGGAA |
| 38506_2 | TATGAGTGTGGAGTCGTTGCAGCAAAGCTGCAGGATATACAAANNNAGATCGGAA |
| 28878_1 | TATGAGTGTGGAGTCGTTGCTCTGCCCAAGAGACCCTCATATANNNAGATCGGAAG |
| 28878_2 | TATGAGTGTGGAGTCGTTGCTCTGCCCAAGAGACCCTCATATANNNAGATCGGAAG |
| 52820_1 | TATGAGTGTGGAGTCGTTGCCTCCTTGGAGAGCTTAGAGGGTNNNAGATCGGAAG |
| 52820_2 | TATGAGTGTGGAGTCGTTGCCTCCTTGGAGAGCTTAGAGGGTNNNAGATCGGAAG |
| 31594_1 | TATGAGTGTGGAGTCGTTGCAGAGAGAGAGAGAGAGAGAGTTTTNNNAGATCGG |
| 31594_2 | TATGAGTGTGGAGTCGTTGCAGAGAGAGAGAGAGAGAGAGTTTTNNNAGATCGG |
| 12774_1 | TATGAGTGTGGAGTCGTTGCTAAGTGATTCTTCCTTGTGTTTANNNAGATCGGAAG |
| 12774_2 | TATGAGTGTGGAGTCGTTGCTAAGTGATTCTTCCTTGTGTTTANNNAGATCGGAAG |
| 38281_1 | TATGAGTGTGGAGTCGTTGCTCTCAGAATGCACAAAGCCAATGNNNAGATCGGAAG |
| 38281_2 | TATGAGTGTGGAGTCGTTGCTCTCAGAATGCACAAAGCCAATGNNNAGATCGGAAG |
| 29368_1 | TATGAGTGTGGAGTCGTTGCAGGAAGTTGCCAGCAAAAAGTTTNNNAGATCGGAA |
| 29368_2 | TATGAGTGTGGAGTCGTTGCAGGAAGTTGCCAGCAAAAAGTTTNNNAGATCGGAA |

| | |
|---|---|
| 43673_1 | TATGAGTGTGGAGTCGTTGCCAGGTTTAGAAGGGGTGTCTGGNNNAGATCGGAAG |
| 43673_2 | TATGAGTGTGGAGTCGTTGCCAGGTTTAGAAGGGGTGTCTGGNNNAGATCGGAAG |
| 36339_1 | TATGAGTGTGGAGTCGTTGCGAAGACACCCCAATACAGCTGTANNNAGATCGGAAG |
| 36339_2 | TATGAGTGTGGAGTCGTTGCGAAGACACCCCAATACAGCTGTANNNAGATCGGAAG |
| 47042_1 | TATGAGTGTGGAGTCGTTGCTCAGCCTTCCGAGTAGCTAATTTNNNAGATCGGAAG |
| 47042_2 | TATGAGTGTGGAGTCGTTGCTCAGCCTTCCGAGTAGCTAATTTNNNAGATCGGAAG |
| 10149_1 | TATGAGTGTGGAGTCGTTGCATTCTGTGAGGTGGTAGTGATGCNNNAGATCGGAA |
| 10149_2 | TATGAGTGTGGAGTCGTTGCATTCTGTGAGGTGGTAGTGATGCNNNAGATCGGAA |
| 49299_1 | TATGAGTGTGGAGTCGTTGCGGAGATGCAGTTACTTGGAAACGNNNAGATCGGAA |
| 49299_2 | TATGAGTGTGGAGTCGTTGCGGAGATGCAGTTACTTGGAAACGNNNAGATCGGAA |
| 48189_1 | TATGAGTGTGGAGTCGTTGCGCTCACATCCAGCTCCTTGAGNNNAGATCGGAAGAG |
| 48189_2 | TATGAGTGTGGAGTCGTTGCGCTCACATCCAGCTCCTTGAGNNNAGATCGGAAGAG |
| 47028_1 | TATGAGTGTGGAGTCGTTGCTCCTTGGTCAACTGTATGTTCCANNNAGATCGGAAG |
| 47028_2 | TATGAGTGTGGAGTCGTTGCTCCTTGGTCAACTGTATGTTCCANNNAGATCGGAAG |
| 26290_1 | TATGAGTGTGGAGTCGTTGCGGTGTTGAGTATGTTAATTTGATNNNAGATCGGAA |
| 26290_2 | TATGAGTGTGGAGTCGTTGCGGTGTTGAGTATGTTAATTTGATNNNAGATCGGAA |
| 35473_1 | TATGAGTGTGGAGTCGTTGCTTGGCTCTCTCTTTGTCTACTGTNNNAGATCGGAAGA |
| 35473_2 | TATGAGTGTGGAGTCGTTGCTTGGCTCTCTCTTTGTCTACTGTNNNAGATCGGAAGA |
| 42284_1 | TATGAGTGTGGAGTCGTTGCGTGTGTGTGTGTGTGTATACGTANNNAGATCGGAA |
| 42284_2 | TATGAGTGTGGAGTCGTTGCGTGTGTGTGTGTGTGTATACGTANNNAGATCGGAA |
| 51026_1 | TATGAGTGTGGAGTCGTTGCAAGTGCTGGGATTATAAGCGTGANNNAGATCGGAA |
| 51026_2 | TATGAGTGTGGAGTCGTTGCAAGTGCTGGGATTATAAGCGTGANNNAGATCGGAA |
| 38670_1 | TATGAGTGTGGAGTCGTTGCATAAGGGAGTAAGGGAGCTAGGGNNNAGATCGGA |
| 38670_2 | TATGAGTGTGGAGTCGTTGCATAAGGGAGTAAGGGAGCTAGGGNNNAGATCGGA |
| 35499_1 | TATGAGTGTGGAGTCGTTGCAAATCTGGGAGCCAAGAAATCCTNNNAGATCGGAA |
| 35499_2 | TATGAGTGTGGAGTCGTTGCAAATCTGGGAGCCAAGAAATCCTNNNAGATCGGAA |
| 42423_1 | TATGAGTGTGGAGTCGTTGCGAGGGAAGGAGAGAGAGAGAGAGNNNAGATCGG |
| 42423_2 | TATGAGTGTGGAGTCGTTGCGAGGGAAGGAGAGAGAGAGAGAGNNNAGATCGG |
| 39138_1 | TATGAGTGTGGAGTCGTTGCACCTATTGAGTGCAGTGTACACTNNNAGATCGGAAG |
| 39138_2 | TATGAGTGTGGAGTCGTTGCACCTATTGAGTGCAGTGTACACTNNNAGATCGGAAG |
| 44430_1 | TATGAGTGTGGAGTCGTTGCCATTGCAGGTGATTGACTCCAAGNNNAGATCGGAAG |
| 44430_2 | TATGAGTGTGGAGTCGTTGCCATTGCAGGTGATTGACTCCAAGNNNAGATCGGAAG |
| 29627_1 | TATGAGTGTGGAGTCGTTGCTCATTGGAACAGCCTAATCAATATGANNNAGATCGG |
| 29627_2 | TATGAGTGTGGAGTCGTTGCTCATTGGAACAGCCTAATCAATATGANNNAGATCGG |
| 49315_1 | TATGAGTGTGGAGTCGTTGCCTCCCGAGTAGCAGAGATTACAGNNNAGATCGGAA |
| 49315_2 | TATGAGTGTGGAGTCGTTGCCTCCCGAGTAGCAGAGATTACAGNNNAGATCGGAA |
| 39341_1 | TATGAGTGTGGAGTCGTTGCAGAGCGAAAAATGTGACTGTGTTNNNAGATCGGAA |
| 39341_2 | TATGAGTGTGGAGTCGTTGCAGAGCGAAAAATGTGACTGTGTTNNNAGATCGGAA |
| 8523_1 | TATGAGTGTGGAGTCGTTGCAGTCATGGACGGTAAAAACTAGGNNNAGATCGGAA |
| 8523_2 | TATGAGTGTGGAGTCGTTGCAGTCATGGACGGTAAAAACTAGGNNNAGATCGGAA |
| 42353_1 | TATGAGTGTGGAGTCGTTGCTGGTCATGCTCAATACATGGTCANNNAGATCGGAAG |
| 42353_2 | TATGAGTGTGGAGTCGTTGCTGGTCATGCTCAATACATGGTCANNNAGATCGGAAG |
| 38695_1 | TATGAGTGTGGAGTCGTTGCTTCCTGAAGGCAAAGGGAATACANNNAGATCGGAA |

| | |
|---|---|
| 38695_2 | TATGAGTGTGGAGTCGTTGCTTCCTGAAGGCAAAGGGAATACANNNAGATCGGAA |
| 51482_1 | TATGAGTGTGGAGTCGTTGCGAACCAAGTCAGGGGAAGTATGTNNNAGATCGGAA |
| 51482_2 | TATGAGTGTGGAGTCGTTGCGAACCAAGTCAGGGGAAGTATGTNNNAGATCGGAA |
| 50935_1 | TATGAGTGTGGAGTCGTTGCCTGAGACGACTCCTCGAAAAGTGNNNAGATCGGAA |
| 50935_2 | TATGAGTGTGGAGTCGTTGCCTGAGACGACTCCTCGAAAAGTGNNNAGATCGGAA |
| 31555_1 | TATGAGTGTGGAGTCGTTGCACATCAAAATTCGTGGTCCCTCTNNNAGATCGGAAG |
| 31555_2 | TATGAGTGTGGAGTCGTTGCACATCAAAATTCGTGGTCCCTCTNNNAGATCGGAAG |
| 34355_1 | TATGAGTGTGGAGTCGTTGCGGGAACGTGACCTTATTTGCAAANNNAGATCGGAA |
| 34355_2 | TATGAGTGTGGAGTCGTTGCGGGAACGTGACCTTATTTGCAAANNNAGATCGGAA |
| 2703_1 | TATGAGTGTGGAGTCGTTGCACTTTGCTCCATAACATCTGGATNNNAGATCGGAAG |
| 2703_2 | TATGAGTGTGGAGTCGTTGCACTTTGCTCCATAACATCTGGATNNNAGATCGGAAG |
| 53130_1 | TATGAGTGTGGAGTCGTTGCACAAAAACTTTCTGAGCATGCCANNNAGATCGGAAG |
| 53130_2 | TATGAGTGTGGAGTCGTTGCACAAAAACTTTCTGAGCATGCCANNNAGATCGGAAG |
| 13630_1 | TATGAGTGTGGAGTCGTTGCTTGTACTAAAGACCTGACAACNNNAGATCGGAAGAG |
| 13630_2 | TATGAGTGTGGAGTCGTTGCTTGTACTAAAGACCTGACAACNNNAGATCGGAAGAG |
| 51585_1 | TATGAGTGTGGAGTCGTTGCTTCCAATGCTGATGAAGAGGACANNNAGATCGGAA |
| 51585_2 | TATGAGTGTGGAGTCGTTGCTTCCAATGCTGATGAAGAGGACANNNAGATCGGAA |
| 42742_1 | TATGAGTGTGGAGTCGTTGCAAGCTGGTTCTTTGAACAGATGANNNAGATCGGAA |
| 42742_2 | TATGAGTGTGGAGTCGTTGCAAGCTGGTTCTTTGAACAGATGANNNAGATCGGAA |
| 41020_1 | TATGAGTGTGGAGTCGTTGCTGACAGCTTCACCTACTTCCTACNNNAGATCGGAAGA |
| 41020_2 | TATGAGTGTGGAGTCGTTGCTGACAGCTTCACCTACTTCCTACNNNAGATCGGAAGA |
| 44180_1 | TATGAGTGTGGAGTCGTTGCGGGGTGCTTGACAAAATACCTTCNNNAGATCGGAA |
| 44180_2 | TATGAGTGTGGAGTCGTTGCGGGGTGCTTGACAAAATACCTTCNNNAGATCGGAA |
| 971_1 | TATGAGTGTGGAGTCGTTGCAAGCCATCCCTGACACTCTGTCNNNAGATCGGAAGA |
| 971_2 | TATGAGTGTGGAGTCGTTGCAAGCCATCCCTGACACTCTGTCNNNAGATCGGAAGA |
| 39843_1 | TATGAGTGTGGAGTCGTTGCAGAAAATCCCTGAGTTGAGCAGANNNAGATCGGAA |
| 39843_2 | TATGAGTGTGGAGTCGTTGCAGAAAATCCCTGAGTTGAGCAGANNNAGATCGGAA |
| 9980_1 | TATGAGTGTGGAGTCGTTGCAACTGTGCTCGCTTCAGAGAAATNNNAGATCGGAAG |
| 9980_2 | TATGAGTGTGGAGTCGTTGCAACTGTGCTCGCTTCAGAGAAATNNNAGATCGGAAG |
| 29379_1 | TATGAGTGTGGAGTCGTTGCCCTCCAGTTCCTATCTGTTGAGCNNNAGATCGGAAG |
| 29379_2 | TATGAGTGTGGAGTCGTTGCCCTCCAGTTCCTATCTGTTGAGCNNNAGATCGGAAG |
| 32306_1 | TATGAGTGTGGAGTCGTTGCAGAGAGAGAGAGAGGGAGAGAGANNNAGATCGG |
| 32306_2 | TATGAGTGTGGAGTCGTTGCAGAGAGAGAGAGAGGGAGAGAGANNNAGATCGG |
| 53047_1 | TATGAGTGTGGAGTCGTTGCACGTATGTTTTTGCAGCACTGTTNNNAGATCGGAAG |
| 53047_2 | TATGAGTGTGGAGTCGTTGCACGTATGTTTTTGCAGCACTGTTNNNAGATCGGAAG |
| 30502_1 | TATGAGTGTGGAGTCGTTGCAAAATTAGGTCTTTGGCTGGGCANNNAGATCGGAA |
| 30502_2 | TATGAGTGTGGAGTCGTTGCAAAATTAGGTCTTTGGCTGGGCANNNAGATCGGAA |
| 28864_1 | TATGAGTGTGGAGTCGTTGCTCTGTGTGATGAACTTCTCTCCCNNNAGATCGGAAG |
| 28864_2 | TATGAGTGTGGAGTCGTTGCTCTGTGTGATGAACTTCTCTCCCNNNAGATCGGAAG |
| 38631_1 | TATGAGTGTGGAGTCGTTGCAGCAAACTAGCCGAAAAGCAAATNNNAGATCGGAA |
| 38631_2 | TATGAGTGTGGAGTCGTTGCAGCAAACTAGCCGAAAAGCAAATNNNAGATCGGAA |
| 50013_1 | TATGAGTGTGGAGTCGTTGCGAGCCTCTTCCTCATATCTCCCTNNNAGATCGGAAGA |
| 50013_2 | TATGAGTGTGGAGTCGTTGCGAGCCTCTTCCTCATATCTCCCTNNNAGATCGGAAGA |

| | |
|---|---|
| 50780_1 | TATGAGTGTGGAGTCGTTGCACTCTTGCCAGAAGAAAATGCAGNNNAGATCGGAAG |
| 50780_2 | TATGAGTGTGGAGTCGTTGCACTCTTGCCAGAAGAAAATGCAGNNNAGATCGGAAG |
| 8967_1 | TATGAGTGTGGAGTCGTTGCACACACACACTCAAAAAGGTTTCTNNNAGATCGGAAG |
| 8967_2 | TATGAGTGTGGAGTCGTTGCACACACACACTCAAAAAGGTTTCTNNNAGATCGGAAG |
| 30975_1 | TATGAGTGTGGAGTCGTTGCGTCACTATGGGACTACGGAACTCNNNAGATCGGAAG |
| 30975_2 | TATGAGTGTGGAGTCGTTGCGTCACTATGGGACTACGGAACTCNNNAGATCGGAAG |
| 47868_1 | TATGAGTGTGGAGTCGTTGCGTGTGTCTTGGGGATGATCTTCTNNNAGATCGGAAG |
| 47868_2 | TATGAGTGTGGAGTCGTTGCGTGTGTCTTGGGGATGATCTTCTNNNAGATCGGAAG |
| 13282_1 | TATGAGTGTGGAGTCGTTGCGTACTAGGTAATGGTCCATAACNNNAGATCGGAAGA |
| 13282_2 | TATGAGTGTGGAGTCGTTGCGTACTAGGTAATGGTCCATAACNNNAGATCGGAAGA |
| 47423_1 | TATGAGTGTGGAGTCGTTGCTGGAGTCCAGCAAACTAAGATCCNNNAGATCGGAAG |
| 47423_2 | TATGAGTGTGGAGTCGTTGCTGGAGTCCAGCAAACTAAGATCCNNNAGATCGGAAG |
| 34954_1 | TATGAGTGTGGAGTCGTTGCGAGTTTTTGCTCTTGTCACCCAGNNNAGATCGGAAGA |
| 34954_2 | TATGAGTGTGGAGTCGTTGCGAGTTTTTGCTCTTGTCACCCAGNNNAGATCGGAAGA |
| 46936_1 | TATGAGTGTGGAGTCGTTGCAACAGTTCTCCATGGTGGTTGTANNNAGATCGGAAG |
| 46936_2 | TATGAGTGTGGAGTCGTTGCAACAGTTCTCCATGGTGGTTGTANNNAGATCGGAAG |
| 34075_1 | TATGAGTGTGGAGTCGTTGCCCAGGGTTCAAGCAATTCTCCTANNNAGATCGGAAGA |
| 34075_2 | TATGAGTGTGGAGTCGTTGCCCAGGGTTCAAGCAATTCTCCTANNNAGATCGGAAGA |
| 43977_1 | TATGAGTGTGGAGTCGTTGCAAGAAAGGAAAGAAGGAAGGAGANNNAGATCGGA |
| 43977_2 | TATGAGTGTGGAGTCGTTGCAAGAAAGGAAAGAAGGAAGGAGANNNAGATCGGA |
| 29093_1 | TATGAGTGTGGAGTCGTTGCCCTCTCTCCCTCTGTCTCTGTTANNNAGATCGGAAGAG |
| 29093_2 | TATGAGTGTGGAGTCGTTGCCCTCTCTCCCTCTGTCTCTGTTANNNAGATCGGAAGAG |
| 28777_1 | TATGAGTGTGGAGTCGTTGCGTGTGTGTGTTCCCAAGGAAAAANNNAGATCGGAAG |
| 28777_2 | TATGAGTGTGGAGTCGTTGCGTGTGTGTGTTCCCAAGGAAAAANNNAGATCGGAAG |
| 34551_1 | TATGAGTGTGGAGTCGTTGCTGCCTTTCCAATCACATTCCCTANNNAGATCGGAAGAG |
| 34551_2 | TATGAGTGTGGAGTCGTTGCTGCCTTTCCAATCACATTCCCTANNNAGATCGGAAGAG |
| 13536_1 | TATGAGTGTGGAGTCGTTGCATACACTCAGGAGTATGTATGTANNNAGATCGGAAG |
| 13536_2 | TATGAGTGTGGAGTCGTTGCATACACTCAGGAGTATGTATGTANNNAGATCGGAAG |
| 42248_1 | TATGAGTGTGGAGTCGTTGCTGGTCCCATGCGTATGTTGATTANNNAGATCGGAAG |
| 42248_2 | TATGAGTGTGGAGTCGTTGCTGGTCCCATGCGTATGTTGATTANNNAGATCGGAAG |
| 46148_1 | TATGAGTGTGGAGTCGTTGCACCACCACACTGGGATTTTGTATNNNAGATCGGAAGA |
| 46148_2 | TATGAGTGTGGAGTCGTTGCACCACCACACTGGGATTTTGTATNNNAGATCGGAAGA |
| 47921_1 | TATGAGTGTGGAGTCGTTGCACTGGATTCAAACGATTCTCCCANNNAGATCGGAAGA |
| 47921_2 | TATGAGTGTGGAGTCGTTGCACTGGATTCAAACGATTCTCCCANNNAGATCGGAAGA |
| 38747_1 | TATGAGTGTGGAGTCGTTGCCCTAAATACTTGGGAGGCTGAGGNNNAGATCGGAAG |
| 38747_2 | TATGAGTGTGGAGTCGTTGCCCTAAATACTTGGGAGGCTGAGGNNNAGATCGGAAG |
| 48816_1 | TATGAGTGTGGAGTCGTTGCCTGTCTCAGCTTCCCAAGTAGTTNNNAGATCGGAAGA |
| 48816_2 | TATGAGTGTGGAGTCGTTGCCTGTCTCAGCTTCCCAAGTAGTTNNNAGATCGGAAGA |
| 35215_1 | TATGAGTGTGGAGTCGTTGCATGCCCACAAACACAAGTTTTCANNNAGATCGGAAGA |
| 35215_2 | TATGAGTGTGGAGTCGTTGCATGCCCACAAACACAAGTTTTCANNNAGATCGGAAGA |
| 40600_1 | TATGAGTGTGGAGTCGTTGCAGAAAAATGGGGAGGGTATGGTGNNNAGATCGGA |
| 40600_2 | TATGAGTGTGGAGTCGTTGCAGAAAAATGGGGAGGGTATGGTGNNNAGATCGGA |
| 49816_1 | TATGAGTGTGGAGTCGTTGCAGTGGTGATGCCCATTCACATNNNAGATCGGAAGAG |

| | |
|---|---|
| 49816_2 | TATGAGTGTGGAGTCGTTGCAGTGGTGATGCCCATTCACATNNNAGATCGGAAGAG |
| 48298_1 | TATGAGTGTGGAGTCGTTGCAATACAGGAGAATGGGAACCCTGNNNAGATCGGAA |
| 48298_2 | TATGAGTGTGGAGTCGTTGCAATACAGGAGAATGGGAACCCTGNNNAGATCGGAA |
| 34229_1 | TATGAGTGTGGAGTCGTTGCGCTATGTATTCCAATGGGGGGATNNNAGATCGGAA |
| 34229_2 | TATGAGTGTGGAGTCGTTGCGCTATGTATTCCAATGGGGGGATNNNAGATCGGAA |
| 29665_1 | TATGAGTGTGGAGTCGTTGCATCACGCAGATGAAGAGAGGCNNNAGATCGGAAGA |
| 29665_2 | TATGAGTGTGGAGTCGTTGCATCACGCAGATGAAGAGAGGCNNNAGATCGGAAGA |
| 34925_1 | TATGAGTGTGGAGTCGTTGCAAAGTGCTGGGATTACAGACGTCNNNAGATCGGAA |
| 34925_2 | TATGAGTGTGGAGTCGTTGCAAAGTGCTGGGATTACAGACGTCNNNAGATCGGAA |
| 9897_1 | TATGAGTGTGGAGTCGTTGCCAAACTCTGACCTGATTCAGCGANNNAGATCGGAAG |
| 9897_2 | TATGAGTGTGGAGTCGTTGCCAAACTCTGACCTGATTCAGCGANNNAGATCGGAAG |
| 13678_1 | TATGAGTGTGGAGTCGTTGCTACAAACGTATGGAATGCTTNNNAGATCGGAAGAG |
| 13678_2 | TATGAGTGTGGAGTCGTTGCTACAAACGTATGGAATGCTTNNNAGATCGGAAGAG |
| 50348_1 | TATGAGTGTGGAGTCGTTGCAGGTGTCTCTCTACAGGTCAACTNNNAGATCGGAAG |
| 50348_2 | TATGAGTGTGGAGTCGTTGCAGGTGTCTCTCTACAGGTCAACTNNNAGATCGGAAG |
| 32813_1 | TATGAGTGTGGAGTCGTTGCTTCAAGTGATTCTCCTGCCTCAGNNNAGATCGGAAG |
| 32813_2 | TATGAGTGTGGAGTCGTTGCTTCAAGTGATTCTCCTGCCTCAGNNNAGATCGGAAG |
| 34685_1 | TATGAGTGTGGAGTCGTTGCCCACGCCCAGCTAATTTTTGTATNNNAGATCGGAAG |
| 34685_2 | TATGAGTGTGGAGTCGTTGCCCACGCCCAGCTAATTTTTGTATNNNAGATCGGAAG |
| 37623_1 | TATGAGTGTGGAGTCGTTGCCTGAGGTCAGGAGTTTGAGACCNNNAGATCGGAAG |
| 37623_2 | TATGAGTGTGGAGTCGTTGCCTGAGGTCAGGAGTTTGAGACCNNNAGATCGGAAG |
| 32496_1 | TATGAGTGTGGAGTCGTTGCCTCCTGAGAAGCTGGGATTACAGNNNAGATCGGAA |
| 32496_2 | TATGAGTGTGGAGTCGTTGCCTCCTGAGAAGCTGGGATTACAGNNNAGATCGGAA |
| 52574_1 | TATGAGTGTGGAGTCGTTGCTCTGGCCAAGTAGCAAATACCATNNNAGATCGGAAG |
| 52574_2 | TATGAGTGTGGAGTCGTTGCTCTGGCCAAGTAGCAAATACCATNNNAGATCGGAAG |
| 39827_1 | TATGAGTGTGGAGTCGTTGCTGATTCACTACACAAGAATTTGCANNNAGATCGGAA |
| 39827_2 | TATGAGTGTGGAGTCGTTGCTGATTCACTACACAAGAATTTGCANNNAGATCGGAA |
| 33350_1 | TATGAGTGTGGAGTCGTTGCGCCTCCTGAGTAGTTGGGATTACNNNAGATCGGAAG |
| 33350_2 | TATGAGTGTGGAGTCGTTGCGCCTCCTGAGTAGTTGGGATTACNNNAGATCGGAAG |
| 7832_1 | TATGAGTGTGGAGTCGTTGCACAAAATCCAAAAAGCTAGAAGAAGANNNAGATCG |
| 7832_2 | TATGAGTGTGGAGTCGTTGCACAAAATCCAAAAAGCTAGAAGAAGANNNAGATCG |
| 10065_1 | TATGAGTGTGGAGTCGTTGCCGAGACCCTGAGCTCTCCCGNNNAGATCGGAAGAG |
| 10065_2 | TATGAGTGTGGAGTCGTTGCCGAGACCCTGAGCTCTCCCGNNNAGATCGGAAGAG |
| 47281_1 | TATGAGTGTGGAGTCGTTGCTGTAGAGATAGGGTTTCGCCATGNNNAGATCGGAA |
| 47281_2 | TATGAGTGTGGAGTCGTTGCTGTAGAGATAGGGTTTCGCCATGNNNAGATCGGAA |
| 35014_1 | TATGAGTGTGGAGTCGTTGCCCATCTCCTGACCTCATGATCTGNNNAGATCGGAAGA |
| 35014_2 | TATGAGTGTGGAGTCGTTGCCCATCTCCTGACCTCATGATCTGNNNAGATCGGAAGA |
| 44303_1 | TATGAGTGTGGAGTCGTTGCTACTCCTCCTCCTCATCCTGTACNNNAGATCGGAAGA |
| 44303_2 | TATGAGTGTGGAGTCGTTGCTACTCCTCCTCCTCATCCTGTACNNNAGATCGGAAGA |
| 47885_1 | TATGAGTGTGGAGTCGTTGCCAAGTAGCTGGGATTACAGGTGTNNNAGATCGGAA |
| 47885_2 | TATGAGTGTGGAGTCGTTGCCAAGTAGCTGGGATTACAGGTGTNNNAGATCGGAA |
| 45658_1 | TATGAGTGTGGAGTCGTTGCGATCTCGGCTCACCACAACTTNNNAGATCGGAAGAG |
| 45658_2 | TATGAGTGTGGAGTCGTTGCGATCTCGGCTCACCACAACTTNNNAGATCGGAAGAG |

| | |
|---|---|
| 38718_1 | TATGAGTGTGGAGTCGTTGCTACATGCTTTTCTCACTGGTTGGNNNAGATCGGAAG |
| 38718_2 | TATGAGTGTGGAGTCGTTGCTACATGCTTTTCTCACTGGTTGGNNNAGATCGGAAG |
| 30985_1 | TATGAGTGTGGAGTCGTTGCGAAACCTGGATGGGATAGGAGACNNNAGATCGGA |
| 30985_2 | TATGAGTGTGGAGTCGTTGCGAAACCTGGATGGGATAGGAGACNNNAGATCGGA |
| 47556_1 | TATGAGTGTGGAGTCGTTGCAGAGGTCCAAATACCCTGCAAATNNNAGATCGGAAG |
| 47556_2 | TATGAGTGTGGAGTCGTTGCAGAGGTCCAAATACCCTGCAAATNNNAGATCGGAAG |
| 12789_1 | TATGAGTGTGGAGTCGTTGCTACAAAGGTAAGAGGAGACTGNNNAGATCGGAAG |
| 12789_2 | TATGAGTGTGGAGTCGTTGCTACAAAGGTAAGAGGAGACTGNNNAGATCGGAAG |
| 50034_1 | TATGAGTGTGGAGTCGTTGCGGACACCAAAAACCTCTTGCATTNNNAGATCGGAAG |
| 50034_2 | TATGAGTGTGGAGTCGTTGCGGACACCAAAAACCTCTTGCATTNNNAGATCGGAAG |
| 44743_1 | TATGAGTGTGGAGTCGTTGCAGCAATCTCTCTTGGTCATGGAGNNNAGATCGGAAG |
| 44743_2 | TATGAGTGTGGAGTCGTTGCAGCAATCTCTCTTGGTCATGGAGNNNAGATCGGAAG |
| 48766_1 | TATGAGTGTGGAGTCGTTGCTTCACGACATTCTCCTGCCTCNNNAGATCGGAAGAGC |
| 48766_2 | TATGAGTGTGGAGTCGTTGCTTCACGACATTCTCCTGCCTCNNNAGATCGGAAGAGC |
| 32679_1 | TATGAGTGTGGAGTCGTTGCCTCCCGAGTAGCTGGAATTACAGNNNAGATCGGAAG |
| 32679_2 | TATGAGTGTGGAGTCGTTGCCTCCCGAGTAGCTGGAATTACAGNNNAGATCGGAAG |
| 40581_1 | TATGAGTGTGGAGTCGTTGCGATGTCTGTGACAGTGAAGGACTNNNAGATCGGAA |
| 40581_2 | TATGAGTGTGGAGTCGTTGCGATGTCTGTGACAGTGAAGGACTNNNAGATCGGAA |
| 28803_1 | TATGAGTGTGGAGTCGTTGCACTACACAACCTATCTCTCACCANNNAGATCGGAAGA |
| 28803_2 | TATGAGTGTGGAGTCGTTGCACTACACAACCTATCTCTCACCANNNAGATCGGAAGA |
| 12652_1 | TATGAGTGTGGAGTCGTTGCGTAGTGGAAAGTAATGATCTCTTNNNAGATCGGAAG |
| 12652_2 | TATGAGTGTGGAGTCGTTGCGTAGTGGAAAGTAATGATCTCTTNNNAGATCGGAAG |
| 39487_1 | TATGAGTGTGGAGTCGTTGCACTGACATAGACACGGATAGCAGNNNAGATCGGAA |
| 39487_2 | TATGAGTGTGGAGTCGTTGCACTGACATAGACACGGATAGCAGNNNAGATCGGAA |
| 31738_1 | TATGAGTGTGGAGTCGTTGCGCAGCATAATGCAATTCTCTTGCNNNAGATCGGAAG |
| 31738_2 | TATGAGTGTGGAGTCGTTGCGCAGCATAATGCAATTCTCTTGCNNNAGATCGGAAG |
| 29115_1 | TATGAGTGTGGAGTCGTTGCTATTCAGTTGGGAGAATGGAGCANNNAGATCGGAA |
| 29115_2 | TATGAGTGTGGAGTCGTTGCTATTCAGTTGGGAGAATGGAGCANNNAGATCGGAA |
| 8515_1 | TATGAGTGTGGAGTCGTTGCATGTGCCTGGAGAGGAGCCTGAANNNAGATCGGAA |
| 8515_2 | TATGAGTGTGGAGTCGTTGCATGTGCCTGGAGAGGAGCCTGAANNNAGATCGGAA |
| 51646_1 | TATGAGTGTGGAGTCGTTGCCTGAATCCCTTTGCCTACCCATANNNAGATCGGAAGA |
| 51646_2 | TATGAGTGTGGAGTCGTTGCCTGAATCCCTTTGCCTACCCATANNNAGATCGGAAGA |
| 50383_1 | TATGAGTGTGGAGTCGTTGCGCCATGTATTTGGGAACGTTGTTNNNAGATCGGAAG |
| 50383_2 | TATGAGTGTGGAGTCGTTGCGCCATGTATTTGGGAACGTTGTTNNNAGATCGGAAG |
| 51136_1 | TATGAGTGTGGAGTCGTTGCAGATGGAGCCTCATTTTCTCTCCNNNAGATCGGAAG |
| 51136_2 | TATGAGTGTGGAGTCGTTGCAGATGGAGCCTCATTTTCTCTCCNNNAGATCGGAAG |
| 47993_1 | TATGAGTGTGGAGTCGTTGCTTTCACCATGTTAGCCAGGATGANNNAGATCGGAAG |
| 47993_2 | TATGAGTGTGGAGTCGTTGCTTTCACCATGTTAGCCAGGATGANNNAGATCGGAAG |
| 34277_1 | TATGAGTGTGGAGTCGTTGCGTGGCTTGTTTCTCAGACTGTTCNNNAGATCGGAAG |
| 34277_2 | TATGAGTGTGGAGTCGTTGCGTGGCTTGTTTCTCAGACTGTTCNNNAGATCGGAAG |
| 45850_1 | TATGAGTGTGGAGTCGTTGCCGCGCCCAGCTAATTTTTTGTATNNNAGATCGGAAG |
| 45850_2 | TATGAGTGTGGAGTCGTTGCCGCGCCCAGCTAATTTTTTGTATNNNAGATCGGAAG |
| 43531_1 | TATGAGTGTGGAGTCGTTGCATTTCTTTTGCTGTGCAGAAGCTNNNAGATCGGAAG |

| | |
|---|---|
| 43531_2 | TATGAGTGTGGAGTCGTTGCATTTCTTTTGCTGTGCAGAAGCTNNNAGATCGGAAG |
| 41661_1 | TATGAGTGTGGAGTCGTTGCTTCACACCATTCTTCTGCCTCAGNNNAGATCGGAAGA |
| 41661_2 | TATGAGTGTGGAGTCGTTGCTTCACACCATTCTTCTGCCTCAGNNNAGATCGGAAGA |
| 41430_1 | TATGAGTGTGGAGTCGTTGCAGAGCTCAGTGACTTCACAAGAGNNNAGATCGGAA |
| 41430_2 | TATGAGTGTGGAGTCGTTGCAGAGCTCAGTGACTTCACAAGAGNNNAGATCGGAA |
| 12718_1 | TATGAGTGTGGAGTCGTTGCCAAGAAGGAAGGGAAATTAAAGNNNAGATCGGAA |
| 12718_2 | TATGAGTGTGGAGTCGTTGCCAAGAAGGAAGGGAAATTAAAGNNNAGATCGGAA |
| 28972_1 | TATGAGTGTGGAGTCGTTGCGTCACGTGCCAGAGTTCAGTATANNNAGATCGGAA |
| 28972_2 | TATGAGTGTGGAGTCGTTGCGTCACGTGCCAGAGTTCAGTATANNNAGATCGGAA |
| 43877_1 | TATGAGTGTGGAGTCGTTGCTTTACAACTTCCATGCATGAGGCNNNAGATCGGAAG |
| 43877_2 | TATGAGTGTGGAGTCGTTGCTTTACAACTTCCATGCATGAGGCNNNAGATCGGAAG |
| 35883_1 | TATGAGTGTGGAGTCGTTGCGCATCCGTATTTCTGCAAAGGACNNNAGATCGGAAG |
| 35883_2 | TATGAGTGTGGAGTCGTTGCGCATCCGTATTTCTGCAAAGGACNNNAGATCGGAAG |
| 4765_1 | TATGAGTGTGGAGTCGTTGCTTACAGGTGCACAACATCATGTCCNNNAGATCGGAA |
| 4765_2 | TATGAGTGTGGAGTCGTTGCTTACAGGTGCACAACATCATGTCCNNNAGATCGGAA |
| 30376_1 | TATGAGTGTGGAGTCGTTGCCCGGTATGTAATTTAACATTCAANNNAGATCGGAAG |
| 30376_2 | TATGAGTGTGGAGTCGTTGCCCGGTATGTAATTTAACATTCAANNNAGATCGGAAG |
| 38971_1 | TATGAGTGTGGAGTCGTTGCTCATGGAGGAAGCTGACATTGAANNNAGATCGGAA |
| 38971_2 | TATGAGTGTGGAGTCGTTGCTCATGGAGGAAGCTGACATTGAANNNAGATCGGAA |
| 44498_1 | TATGAGTGTGGAGTCGTTGCTGAAATCACTGCCAGGAATCAGANNNAGATCGGAA |
| 44498_2 | TATGAGTGTGGAGTCGTTGCTGAAATCACTGCCAGGAATCAGANNNAGATCGGAA |
| 44204_1 | TATGAGTGTGGAGTCGTTGCTTGTCTTCAGTGTGGCAGTTTTGNNNAGATCGGAAG |
| 44204_2 | TATGAGTGTGGAGTCGTTGCTTGTCTTCAGTGTGGCAGTTTTGNNNAGATCGGAAG |
| 39838_1 | TATGAGTGTGGAGTCGTTGCTGAGTGAAGAAGAGAATGACCGGNNNAGATCGGA |
| 39838_2 | TATGAGTGTGGAGTCGTTGCTGAGTGAAGAAGAGAATGACCGGNNNAGATCGGA |
| 32295_1 | TATGAGTGTGGAGTCGTTGCTTGGTGGGCACTTAGGTTGATTANNNAGATCGGAA |
| 32295_2 | TATGAGTGTGGAGTCGTTGCTTGGTGGGCACTTAGGTTGATTANNNAGATCGGAA |
| 2577_1 | TATGAGTGTGGAGTCGTTGCTGGCTCTCTGGCACTAAATATGGNNNAGATCGGAAG |
| 2577_2 | TATGAGTGTGGAGTCGTTGCTGGCTCTCTGGCACTAAATATGGNNNAGATCGGAAG |
| 40234_1 | TATGAGTGTGGAGTCGTTGCTGCAAATCACGATAGTAACGATCTNNNAGATCGGAA |
| 40234_2 | TATGAGTGTGGAGTCGTTGCTGCAAATCACGATAGTAACGATCTNNNAGATCGGAA |
| 30886_1 | TATGAGTGTGGAGTCGTTGCGATAACTTGAGCCCAGGAGTTCANNNAGATCGGAA |
| 30886_2 | TATGAGTGTGGAGTCGTTGCGATAACTTGAGCCCAGGAGTTCANNNAGATCGGAA |
| 38742_1 | TATGAGTGTGGAGTCGTTGCTTCAAGTGATTCCCCTGCTTCAGNNNAGATCGGAAG |
| 38742_2 | TATGAGTGTGGAGTCGTTGCTTCAAGTGATTCCCCTGCTTCAGNNNAGATCGGAAG |
| 42300_1 | TATGAGTGTGGAGTCGTTGCACAATCTTGGCTCACTACAACCTNNNAGATCGGAAG |
| 42300_2 | TATGAGTGTGGAGTCGTTGCACAATCTTGGCTCACTACAACCTNNNAGATCGGAAG |
| 33670_1 | TATGAGTGTGGAGTCGTTGCCTGGAGTATAGTGGTGCAACCTTNNNAGATCGGAA |
| 33670_2 | TATGAGTGTGGAGTCGTTGCCTGGAGTATAGTGGTGCAACCTTNNNAGATCGGAA |
| 47442_1 | TATGAGTGTGGAGTCGTTGCTTCAAGTGATTCTCCTGCCTCAANNNAGATCGGAAG |
| 47442_2 | TATGAGTGTGGAGTCGTTGCTTCAAGTGATTCTCCTGCCTCAANNNAGATCGGAAG |
| 13653_1 | TATGAGTGTGGAGTCGTTGCAAAATTGCAGAACGAATATAAGTNNNAGATCGGAA |
| 13653_2 | TATGAGTGTGGAGTCGTTGCAAAATTGCAGAACGAATATAAGTNNNAGATCGGAA |

| | |
|---|---|
| 49572_1 | TATGAGTGTGGAGTCGTTGCGAAACGGAGTCTTGCTCTGTTGNNNAGATCGGAAG |
| 49572_2 | TATGAGTGTGGAGTCGTTGCGAAACGGAGTCTTGCTCTGTTGNNNAGATCGGAAG |
| 9953_1 | TATGAGTGTGGAGTCGTTGCTCCCACCCCTAGCCAAGATTTTTNNNAGATCGGAAGA |
| 9953_2 | TATGAGTGTGGAGTCGTTGCTCCCACCCCTAGCCAAGATTTTTNNNAGATCGGAAGA |
| 38254_1 | TATGAGTGTGGAGTCGTTGCATGCTAAAGGGGGTTTTTCAAGCNNNAGATCGGAA |
| 38254_2 | TATGAGTGTGGAGTCGTTGCATGCTAAAGGGGGTTTTTCAAGCNNNAGATCGGAA |
| 30230_1 | TATGAGTGTGGAGTCGTTGCTGCTGCTTTTATCTTTGTTATCACTNNNAGATCGGAA |
| 30230_2 | TATGAGTGTGGAGTCGTTGCTGCTGCTTTTATCTTTGTTATCACTNNNAGATCGGAA |
| 37834_1 | TATGAGTGTGGAGTCGTTGCACAAAAAAAGGCCATGTGAGGACNNNAGATCGGAA |
| 37834_2 | TATGAGTGTGGAGTCGTTGCACAAAAAAAGGCCATGTGAGGACNNNAGATCGGAA |
| 44288_1 | TATGAGTGTGGAGTCGTTGCATTTTCATGGTGACTGATGCCAGNNNAGATCGGAAG |
| 44288_2 | TATGAGTGTGGAGTCGTTGCATTTTCATGGTGACTGATGCCAGNNNAGATCGGAAG |
| 13131_1 | TATGAGTGTGGAGTCGTTGCGTAACTCTTCATCCGTTGACNNNAGATCGGAAGAGC |
| 13131_2 | TATGAGTGTGGAGTCGTTGCGTAACTCTTCATCCGTTGACNNNAGATCGGAAGAGC |
| 30336_1 | TATGAGTGTGGAGTCGTTGCGTGGTCAAGCGTTTCCATTAACTNNNAGATCGGAAG |
| 30336_2 | TATGAGTGTGGAGTCGTTGCGTGGTCAAGCGTTTCCATTAACTNNNAGATCGGAAG |
| 35909_1 | TATGAGTGTGGAGTCGTTGCCCCACAAACGACAAACAAACAAANNNAGATCGGAAG |
| 35909_2 | TATGAGTGTGGAGTCGTTGCCCCACAAACGACAAACAAACAAANNNAGATCGGAAG |
| 30611_1 | TATGAGTGTGGAGTCGTTGCAGGTACAGAGTCTCACACCTGTANNNAGATCGGAAG |
| 30611_2 | TATGAGTGTGGAGTCGTTGCAGGTACAGAGTCTCACACCTGTANNNAGATCGGAAG |
| 32186_1 | TATGAGTGTGGAGTCGTTGCTCCTGGAGGAAAAAGTCAGTAGCNNNAGATCGGAA |
| 32186_2 | TATGAGTGTGGAGTCGTTGCTCCTGGAGGAAAAAGTCAGTAGCNNNAGATCGGAA |
| 36900_1 | TATGAGTGTGGAGTCGTTGCAAGAAAACAAGATTTGGCTGGGCNNNAGATCGGAA |
| 36900_2 | TATGAGTGTGGAGTCGTTGCAAGAAAACAAGATTTGGCTGGGCNNNAGATCGGAA |
| 50519_1 | TATGAGTGTGGAGTCGTTGCGAGGGATTGGAACACTCAGACTTNNNAGATCGGAA |
| 50519_2 | TATGAGTGTGGAGTCGTTGCGAGGGATTGGAACACTCAGACTTNNNAGATCGGAA |
| 458_1 | TATGAGTGTGGAGTCGTTGCAGGAAGTGTGACCTGGGAGTTCNNNAGATCGGAAG |
| 458_2 | TATGAGTGTGGAGTCGTTGCAGGAAGTGTGACCTGGGAGTTCNNNAGATCGGAAG |
| 30329_1 | TATGAGTGTGGAGTCGTTGCTGGCCTTCAATCCTTCCATTTCCNNNAGATCGGAAGA |
| 30329_2 | TATGAGTGTGGAGTCGTTGCTGGCCTTCAATCCTTCCATTTCCNNNAGATCGGAAGA |
| 29736_1 | TATGAGTGTGGAGTCGTTGCGTTGCAAGGCTTGTATCAGTAGCNNNAGATCGGAA |
| 29736_2 | TATGAGTGTGGAGTCGTTGCGTTGCAAGGCTTGTATCAGTAGCNNNAGATCGGAA |
| 46455_1 | TATGAGTGTGGAGTCGTTGCCCACCACGCCTGACTAATTTTTTNNNAGATCGGAAGA |
| 46455_2 | TATGAGTGTGGAGTCGTTGCCCACCACGCCTGACTAATTTTTTNNNAGATCGGAAGA |
| 43480_1 | TATGAGTGTGGAGTCGTTGCCACTACGCCCGGCTAATTTTTTNNNAGATCGGAAGA |
| 43480_2 | TATGAGTGTGGAGTCGTTGCCACTACGCCCGGCTAATTTTTTNNNAGATCGGAAGA |
| 29900_1 | TATGAGTGTGGAGTCGTTGCACCACGAAAAACAGAGTGAAAGCNNNAGATCGGAA |
| 29900_2 | TATGAGTGTGGAGTCGTTGCACCACGAAAAACAGAGTGAAAGCNNNAGATCGGAA |
| 28825_1 | TATGAGTGTGGAGTCGTTGCAATCAGACAGACCTGGGTTTGAANNNAGATCGGAA |
| 28825_2 | TATGAGTGTGGAGTCGTTGCAATCAGACAGACCTGGGTTTGAANNNAGATCGGAA |
| 50086_1 | TATGAGTGTGGAGTCGTTGCCTCAGGTCTGTGAACACAGTCTTNNNAGATCGGAAG |
| 50086_2 | TATGAGTGTGGAGTCGTTGCCTCAGGTCTGTGAACACAGTCTTNNNAGATCGGAAG |
| 40987_1 | TATGAGTGTGGAGTCGTTGCATTCCCTTCCTCTCTCCCTTCTTNNNAGATCGGAAGAG |

| | |
|---|---|
| 40987_2 | TATGAGTGTGGAGTCGTTGCATTCCCTTCCTCTCTCCCTTCTTNNNAGATCGGAAGAG |
| 30711_1 | TATGAGTGTGGAGTCGTTGCGAAGCTATGACTCTGAGGTCAAGNNNAGATCGGAA |
| 30711_2 | TATGAGTGTGGAGTCGTTGCGAAGCTATGACTCTGAGGTCAAGNNNAGATCGGAA |
| 45435_1 | TATGAGTGTGGAGTCGTTGCTACCAGCTGTCTTGCAAACAATGNNNAGATCGGAAG |
| 45435_2 | TATGAGTGTGGAGTCGTTGCTACCAGCTGTCTTGCAAACAATGNNNAGATCGGAAG |
| 32711_1 | TATGAGTGTGGAGTCGTTGCCATGCTTGTAATCCCAGCACTTTNNNAGATCGGAAG |
| 32711_2 | TATGAGTGTGGAGTCGTTGCCATGCTTGTAATCCCAGCACTTTNNNAGATCGGAAG |
| 50552_1 | TATGAGTGTGGAGTCGTTGCAATTCTGGCCTTCTGGAGGAAATNNNAGATCGGAA |
| 50552_2 | TATGAGTGTGGAGTCGTTGCAATTCTGGCCTTCTGGAGGAAATNNNAGATCGGAA |
| 31147_1 | TATGAGTGTGGAGTCGTTGCATACAGGCATTTTACGTGTGTGCNNNAGATCGGAAG |
| 31147_2 | TATGAGTGTGGAGTCGTTGCATACAGGCATTTTACGTGTGTGCNNNAGATCGGAAG |
| 40663_1 | TATGAGTGTGGAGTCGTTGCGGACAACACTGCAAGACTCTTTCNNNAGATCGGAAG |
| 40663_2 | TATGAGTGTGGAGTCGTTGCGGACAACACTGCAAGACTCTTTCNNNAGATCGGAAG |
| 30141_1 | TATGAGTGTGGAGTCGTTGCGACTTGCTTATCCAACTGCACAGNNNAGATCGGAAG |
| 30141_2 | TATGAGTGTGGAGTCGTTGCGACTTGCTTATCCAACTGCACAGNNNAGATCGGAAG |
| 29337_1 | TATGAGTGTGGAGTCGTTGCCCTGTAATCCTAGCAGTTTGGGANNNAGATCGGAAG |
| 29337_2 | TATGAGTGTGGAGTCGTTGCCCTGTAATCCTAGCAGTTTGGGANNNAGATCGGAAG |
| 41269_1 | TATGAGTGTGGAGTCGTTGCTCTAACAAGTAATGGAAGGCGGGNNNAGATCGGAA |
| 41269_2 | TATGAGTGTGGAGTCGTTGCTCTAACAAGTAATGGAAGGCGGGNNNAGATCGGAA |
| 39182_1 | TATGAGTGTGGAGTCGTTGCGGATTTTTGGTCAAGAGCCAGACNNNAGATCGGAA |
| 39182_2 | TATGAGTGTGGAGTCGTTGCGGATTTTTGGTCAAGAGCCAGACNNNAGATCGGAA |
| 37007_1 | TATGAGTGTGGAGTCGTTGCCTTGCTTTGGGTGGCTGAAATAANNNAGATCGGAA |
| 37007_2 | TATGAGTGTGGAGTCGTTGCCTTGCTTTGGGTGGCTGAAATAANNNAGATCGGAA |
| 35354_1 | TATGAGTGTGGAGTCGTTGCCTGAATAGCTGGGATTACAGGCANNNAGATCGGAA |
| 35354_2 | TATGAGTGTGGAGTCGTTGCCTGAATAGCTGGGATTACAGGCANNNAGATCGGAA |
| 36315_1 | TATGAGTGTGGAGTCGTTGCAACAGTTCAGTGAGTTCCTGACANNNAGATCGGAAG |
| 36315_2 | TATGAGTGTGGAGTCGTTGCAACAGTTCAGTGAGTTCCTGACANNNAGATCGGAAG |
| 50765_1 | TATGAGTGTGGAGTCGTTGCCTAGGGACACCACCTGTTCCTANNNAGATCGGAAGA |
| 50765_2 | TATGAGTGTGGAGTCGTTGCCTAGGGACACCACCTGTTCCTANNNAGATCGGAAGA |
| 34729_1 | TATGAGTGTGGAGTCGTTGCCATGATCTTGGCTCACTGCAAACNNNAGATCGGAAG |
| 34729_2 | TATGAGTGTGGAGTCGTTGCCATGATCTTGGCTCACTGCAAACNNNAGATCGGAAG |
| 38016_1 | TATGAGTGTGGAGTCGTTGCAACCAACCAAGAATTGATGCAGANNNAGATCGGAA |
| 38016_2 | TATGAGTGTGGAGTCGTTGCAACCAACCAAGAATTGATGCAGANNNAGATCGGAA |
| 48220_1 | TATGAGTGTGGAGTCGTTGCTCCACCAGGGAAAGATCAATTCANNNAGATCGGAAG |
| 48220_2 | TATGAGTGTGGAGTCGTTGCTCCACCAGGGAAAGATCAATTCANNNAGATCGGAAG |
| 37137_1 | TATGAGTGTGGAGTCGTTGCAACCGAAGGCTTTATTCCTCCATNNNAGATCGGAAG |
| 37137_2 | TATGAGTGTGGAGTCGTTGCAACCGAAGGCTTTATTCCTCCATNNNAGATCGGAAG |
| 38593_1 | TATGAGTGTGGAGTCGTTGCGCCTCTCAAGTACCTGGGATTACNNNAGATCGGAAG |
| 38593_2 | TATGAGTGTGGAGTCGTTGCGCCTCTCAAGTACCTGGGATTACNNNAGATCGGAAG |
| 44986_1 | TATGAGTGTGGAGTCGTTGCCCCACACAATACTGAGGCTGTAANNNAGATCGGAAG |
| 44986_2 | TATGAGTGTGGAGTCGTTGCCCCACACAATACTGAGGCTGTAANNNAGATCGGAAG |
| 51070_1 | TATGAGTGTGGAGTCGTTGCTTGAATCTGGGAGAAGGAGGTTGNNNAGATCGGA |
| 51070_2 | TATGAGTGTGGAGTCGTTGCTTGAATCTGGGAGAAGGAGGTTGNNNAGATCGGA |

34731_1 TATGAGTGTGGAGTCGTTGCCCTCCCAGGTTCAAGTGATTCTCNNNAGATCGGAAG
34731_2 TATGAGTGTGGAGTCGTTGCCCTCCCAGGTTCAAGTGATTCTCNNNAGATCGGAAG
13005_1 TATGAGTGTGGAGTCGTTGCCAAAGAAGTCCTGGATGATAATNNNAGATCGGAAG
13005_2 TATGAGTGTGGAGTCGTTGCCAAAGAAGTCCTGGATGATAATNNNAGATCGGAAG
35208_1 TATGAGTGTGGAGTCGTTGCAGACGGAGTCTCGCTCTGTCNNNAGATCGGAAGAG
35208_2 TATGAGTGTGGAGTCGTTGCAGACGGAGTCTCGCTCTGTCNNNAGATCGGAAGAG
33699_1 TATGAGTGTGGAGTCGTTGCTTGGACCCTAACCGCTGTTATTTNNNAGATCGGAAG
33699_2 TATGAGTGTGGAGTCGTTGCTTGGACCCTAACCGCTGTTATTTNNNAGATCGGAAG
41767_1 TATGAGTGTGGAGTCGTTGCGACCTCGGCTTACTGCAAACTCNNNAGATCGGAAGA
41767_2 TATGAGTGTGGAGTCGTTGCGACCTCGGCTTACTGCAAACTCNNNAGATCGGAAGA
48693_1 TATGAGTGTGGAGTCGTTGCCACAATCTTGGCTCACTGCAAGNNNAGATCGGAAGA
48693_2 TATGAGTGTGGAGTCGTTGCCACAATCTTGGCTCACTGCAAGNNNAGATCGGAAGA
46228_1 TATGAGTGTGGAGTCGTTGCTGAAGTGCAGTGGTGTGATCTCNNNAGATCGGAAG
46228_2 TATGAGTGTGGAGTCGTTGCTGAAGTGCAGTGGTGTGATCTCNNNAGATCGGAAG
41809_1 TATGAGTGTGGAGTCGTTGCCCCACCACTATGCTCAGCTAATTNNNAGATCGGAAGA
41809_2 TATGAGTGTGGAGTCGTTGCCCCACCACTATGCTCAGCTAATTNNNAGATCGGAAGA
36448_1 TATGAGTGTGGAGTCGTTGCCTCATCACAAGCAGCAGTTAGATNNNAGATCGGAAG
36448_2 TATGAGTGTGGAGTCGTTGCCTCATCACAAGCAGCAGTTAGATNNNAGATCGGAAG
37788_1 TATGAGTGTGGAGTCGTTGCAAAAGGAGGAAAGTGAGACGACANNNAGATCGGA
37788_2 TATGAGTGTGGAGTCGTTGCAAAAGGAGGAAAGTGAGACGACANNNAGATCGGA
29992_1 TATGAGTGTGGAGTCGTTGCTACCTTCATCAACATCTGAGGCANNNAGATCGGAAG
29992_2 TATGAGTGTGGAGTCGTTGCTACCTTCATCAACATCTGAGGCANNNAGATCGGAAG
34083_1 TATGAGTGTGGAGTCGTTGCCACCTTAGCCTCCTGATTAGCTGNNNAGATCGGAAG
34083_2 TATGAGTGTGGAGTCGTTGCCACCTTAGCCTCCTGATTAGCTGNNNAGATCGGAAG
40321_1 TATGAGTGTGGAGTCGTTGCGCCAAGCAAAGATTTGAGAGTGTNNNAGATCGGAA
40321_2 TATGAGTGTGGAGTCGTTGCGCCAAGCAAAGATTTGAGAGTGTNNNAGATCGGAA
29623_1 TATGAGTGTGGAGTCGTTGCCGGGGTGAAGTGGAGAAGAATTANNNAGATCGGA
29623_2 TATGAGTGTGGAGTCGTTGCCGGGGTGAAGTGGAGAAGAATTANNNAGATCGGA
12773_1 TATGAGTGTGGAGTCGTTGCTAGTTCTATCCAAATGAGGAAAANNNAGATCGGAA
12773_2 TATGAGTGTGGAGTCGTTGCTAGTTCTATCCAAATGAGGAAAANNNAGATCGGAA
34712_1 TATGAGTGTGGAGTCGTTGCAGCAATTCTCCTGCCTTACTCTCNNNAGATCGGAAGA
34712_2 TATGAGTGTGGAGTCGTTGCAGCAATTCTCCTGCCTTACTCTCNNNAGATCGGAAGA
32754_1 TATGAGTGTGGAGTCGTTGCAAAAAGAAGGGCAGGGAGAAGAANNNAGATCGGA
32754_2 TATGAGTGTGGAGTCGTTGCAAAAAGAAGGGCAGGGAGAAGAANNNAGATCGGA
13463_1 TATGAGTGTGGAGTCGTTGCTGAAAATTAAGTGCTATGGTAAGNNNAGATCGGAA
13463_2 TATGAGTGTGGAGTCGTTGCTGAAAATTAAGTGCTATGGTAAGNNNAGATCGGAA
42461_1 TATGAGTGTGGAGTCGTTGCCTGAGTAGCTGGGAATATAGGCGNNNAGATCGGAA
42461_2 TATGAGTGTGGAGTCGTTGCCTGAGTAGCTGGGAATATAGGCGNNNAGATCGGAA
37495_1 TATGAGTGTGGAGTCGTTGCATTTTTCCTTCCCCTCTAGCTCCNNNAGATCGGAAGA
37495_2 TATGAGTGTGGAGTCGTTGCATTTTTCCTTCCCCTCTAGCTCCNNNAGATCGGAAGA
43668_1 TATGAGTGTGGAGTCGTTGCACGTCCATTTGCTTCAGTAGGATNNNAGATCGGAAG
43668_2 TATGAGTGTGGAGTCGTTGCACGTCCATTTGCTTCAGTAGGATNNNAGATCGGAAG
49322_1 TATGAGTGTGGAGTCGTTGCCCTCCCAGGTACAAGCTATTCTCNNNAGATCGGAAG

| | |
|---|---|
| 49322_2 | TATGAGTGTGGAGTCGTTGCCCTCCCAGGTACAAGCTATTCTCNNNAGATCGGAAG |
| 33517_1 | TATGAGTGTGGAGTCGTTGCATAACAGATGCTCGTGAGGTTGTNNNAGATCGGAA |
| 33517_2 | TATGAGTGTGGAGTCGTTGCATAACAGATGCTCGTGAGGTTGTNNNAGATCGGAA |
| 38109_1 | TATGAGTGTGGAGTCGTTGCCAATTGAGTGATGGGCATGATGANNNAGATCGGAA |
| 38109_2 | TATGAGTGTGGAGTCGTTGCCAATTGAGTGATGGGCATGATGANNNAGATCGGAA |
| 52708_1 | TATGAGTGTGGAGTCGTTGCCCCATTTCTAGGTATCCGTGAGGNNNAGATCGGAAG |
| 52708_2 | TATGAGTGTGGAGTCGTTGCCCCATTTCTAGGTATCCGTGAGGNNNAGATCGGAAG |
| 31538_1 | TATGAGTGTGGAGTCGTTGCTTCCATCCATAAGCTTGAGACCCNNNAGATCGGAAG |
| 31538_2 | TATGAGTGTGGAGTCGTTGCTTCCATCCATAAGCTTGAGACCCNNNAGATCGGAAG |
| 44607_1 | TATGAGTGTGGAGTCGTTGCAGTAACTCAGCCACTCAGTCATCNNNAGATCGGAAG |
| 44607_2 | TATGAGTGTGGAGTCGTTGCAGTAACTCAGCCACTCAGTCATCNNNAGATCGGAAG |
| 34340_1 | TATGAGTGTGGAGTCGTTGCTGTGAATGTCTGAAAAACTGCCCNNNAGATCGGAAG |
| 34340_2 | TATGAGTGTGGAGTCGTTGCTGTGAATGTCTGAAAAACTGCCCNNNAGATCGGAAG |
| 46381_1 | TATGAGTGTGGAGTCGTTGCCAATTCTCCTGTCTCAGTCTCCCNNNAGATCGGAAGA |
| 46381_2 | TATGAGTGTGGAGTCGTTGCCAATTCTCCTGTCTCAGTCTCCCNNNAGATCGGAAGA |
| 50934_1 | TATGAGTGTGGAGTCGTTGCACGTGTGTTCCTACTGGTACTTCNNNAGATCGGAAG |
| 50934_2 | TATGAGTGTGGAGTCGTTGCACGTGTGTTCCTACTGGTACTTCNNNAGATCGGAAG |
| 45421_1 | TATGAGTGTGGAGTCGTTGCGCTGGACATCAGATTGAGCACTANNNAGATCGGAA |
| 45421_2 | TATGAGTGTGGAGTCGTTGCGCTGGACATCAGATTGAGCACTANNNAGATCGGAA |
| 28989_1 | TATGAGTGTGGAGTCGTTGCACTGGTTCTGATCCTTTGGTTGANNNAGATCGGAAG |
| 28989_2 | TATGAGTGTGGAGTCGTTGCACTGGTTCTGATCCTTTGGTTGANNNAGATCGGAAG |
| 48210_1 | TATGAGTGTGGAGTCGTTGCACGCCTGGATGATTTTTTGGTACNNNAGATCGGAAG |
| 48210_2 | TATGAGTGTGGAGTCGTTGCACGCCTGGATGATTTTTTGGTACNNNAGATCGGAAG |
| 32818_1 | TATGAGTGTGGAGTCGTTGCCATCTCACTACAACCTCCCACAANNNAGATCGGAAGA |
| 32818_2 | TATGAGTGTGGAGTCGTTGCCATCTCACTACAACCTCCCACAANNNAGATCGGAAGA |
| 47888_1 | TATGAGTGTGGAGTCGTTGCTGTCACAGGGTATCATGTGTGAGNNNAGATCGGAA |
| 47888_2 | TATGAGTGTGGAGTCGTTGCTGTCACAGGGTATCATGTGTGAGNNNAGATCGGAA |
| 43069_1 | TATGAGTGTGGAGTCGTTGCAAGAGGAACCACCAACTGAAAGTNNNAGATCGGAA |
| 43069_2 | TATGAGTGTGGAGTCGTTGCAAGAGGAACCACCAACTGAAAGTNNNAGATCGGAA |
| 41024_1 | TATGAGTGTGGAGTCGTTGCACACTAAAGTCCATTCCCCCAAANNNAGATCGGAAG |
| 41024_2 | TATGAGTGTGGAGTCGTTGCACACTAAAGTCCATTCCCCCAAANNNAGATCGGAAG |
| 49677_1 | TATGAGTGTGGAGTCGTTGCTTTGCCTTTCAAGTGGTGTTCTTNNNAGATCGGAAG |
| 49677_2 | TATGAGTGTGGAGTCGTTGCTTTGCCTTTCAAGTGGTGTTCTTNNNAGATCGGAAG |
| 51478_1 | TATGAGTGTGGAGTCGTTGCAGTCCCTGTAAGGCTGTTGTTTCNNNAGATCGGAAG |
| 51478_2 | TATGAGTGTGGAGTCGTTGCAGTCCCTGTAAGGCTGTTGTTTCNNNAGATCGGAAG |
| 36024_1 | TATGAGTGTGGAGTCGTTGCTATCACTTGAACCTAGGAGGCAGNNNAGATCGGAA |
| 36024_2 | TATGAGTGTGGAGTCGTTGCTATCACTTGAACCTAGGAGGCAGNNNAGATCGGAA |
| 50487_1 | TATGAGTGTGGAGTCGTTGCCTGTCTCACCCAGGTAAATCACANNNAGATCGGAAG |
| 50487_2 | TATGAGTGTGGAGTCGTTGCCTGTCTCACCCAGGTAAATCACANNNAGATCGGAAG |
| 50097_1 | TATGAGTGTGGAGTCGTTGCGTGTGGACATCAGAGAATCCCACNNNAGATCGGAA |
| 50097_2 | TATGAGTGTGGAGTCGTTGCGTGTGGACATCAGAGAATCCCACNNNAGATCGGAA |
| 13198_1 | TATGAGTGTGGAGTCGTTGCCTACTACCTTGTATCAACCTCTNNNAGATCGGAAGAG |
| 13198_2 | TATGAGTGTGGAGTCGTTGCCTACTACCTTGTATCAACCTCTNNNAGATCGGAAGAG |

| | |
|---|---|
| 38856_1 | TATGAGTGTGGAGTCGTTGCTACCTCTATGCACAAAAGCCAGANNNAGATCGGAAG |
| 38856_2 | TATGAGTGTGGAGTCGTTGCTACCTCTATGCACAAAAGCCAGANNNAGATCGGAAG |
| 52829_1 | TATGAGTGTGGAGTCGTTGCGCAGCTTAGACATCCCTTCTCTTNNNAGATCGGAAGA |
| 52829_2 | TATGAGTGTGGAGTCGTTGCGCAGCTTAGACATCCCTTCTCTTNNNAGATCGGAAGA |
| 47353_1 | TATGAGTGTGGAGTCGTTGCGCTCCTACCATGCATTGTGTTTTNNNAGATCGGAAGA |
| 47353_2 | TATGAGTGTGGAGTCGTTGCGCTCCTACCATGCATTGTGTTTTNNNAGATCGGAAGA |
| 42094_1 | TATGAGTGTGGAGTCGTTGCGCATCACCCTGATACCATAGCTTNNNAGATCGGAAGA |
| 42094_2 | TATGAGTGTGGAGTCGTTGCGCATCACCCTGATACCATAGCTTNNNAGATCGGAAGA |
| 50191_1 | TATGAGTGTGGAGTCGTTGCAATAGAGATGCCTGACTGAAGCCNNNAGATCGGAAG |
| 50191_2 | TATGAGTGTGGAGTCGTTGCAATAGAGATGCCTGACTGAAGCCNNNAGATCGGAAG |
| 31140_1 | TATGAGTGTGGAGTCGTTGCATCCACCTTGAAAGTGTGTCTGTNNNAGATCGGAAG |
| 31140_2 | TATGAGTGTGGAGTCGTTGCATCCACCTTGAAAGTGTGTCTGTNNNAGATCGGAAG |
| 3034_1 | TATGAGTGTGGAGTCGTTGCGCTGCCTCTTCTATCATCCAGACNNNAGATCGGAAGA |
| 3034_2 | TATGAGTGTGGAGTCGTTGCGCTGCCTCTTCTATCATCCAGACNNNAGATCGGAAGA |
| 47225_1 | TATGAGTGTGGAGTCGTTGCGAGACAGGGTTTCACTCTGTTGTNNNAGATCGGAAG |
| 47225_2 | TATGAGTGTGGAGTCGTTGCGAGACAGGGTTTCACTCTGTTGTNNNAGATCGGAAG |
| 46202_1 | TATGAGTGTGGAGTCGTTGCCCCAGCCAAAACAATGCTTTCTANNNAGATCGGAAGA |
| 46202_2 | TATGAGTGTGGAGTCGTTGCCCCAGCCAAAACAATGCTTTCTANNNAGATCGGAAGA |
| 50434_1 | TATGAGTGTGGAGTCGTTGCGATGGAGGGATTTAGCTTCTCCANNNAGATCGGAAG |
| 50434_2 | TATGAGTGTGGAGTCGTTGCGATGGAGGGATTTAGCTTCTCCANNNAGATCGGAAG |
| 42786_1 | TATGAGTGTGGAGTCGTTGCCATGTACTCCCCTCATCCGATTCNNNAGATCGGAAGA |
| 42786_2 | TATGAGTGTGGAGTCGTTGCCATGTACTCCCCTCATCCGATTCNNNAGATCGGAAGA |
| 31237_1 | TATGAGTGTGGAGTCGTTGCAAGCAGTGGTTAGGGTTAGATGGNNNAGATCGGAA |
| 31237_2 | TATGAGTGTGGAGTCGTTGCAAGCAGTGGTTAGGGTTAGATGGNNNAGATCGGAA |
| 3169_1 | TATGAGTGTGGAGTCGTTGCGGGAGGTGGCTAGACTAGTTGATNNNAGATCGGAA |
| 3169_2 | TATGAGTGTGGAGTCGTTGCGGGAGGTGGCTAGACTAGTTGATNNNAGATCGGAA |
| 45936_1 | TATGAGTGTGGAGTCGTTGCGATCAGTGTGTCGATCTCTTCCANNNAGATCGGAAGA |
| 45936_2 | TATGAGTGTGGAGTCGTTGCGATCAGTGTGTCGATCTCTTCCANNNAGATCGGAAGA |
| 11838_1 | TATGAGTGTGGAGTCGTTGCCCATTTTTCCCTAAAAGGCCTCTNNNAGATCGGAAGA |
| 11838_2 | TATGAGTGTGGAGTCGTTGCCCATTTTTCCCTAAAAGGCCTCTNNNAGATCGGAAGA |
| 37171_1 | TATGAGTGTGGAGTCGTTGCCAGTATCTGCAGAGCCAATAAGGNNNAGATCGGAAG |
| 37171_2 | TATGAGTGTGGAGTCGTTGCCAGTATCTGCAGAGCCAATAAGGNNNAGATCGGAAG |
| 29028_1 | TATGAGTGTGGAGTCGTTGCAAGTCTGGATCACATAGGGCTTGNNNAGATCGGAAG |
| 29028_2 | TATGAGTGTGGAGTCGTTGCAAGTCTGGATCACATAGGGCTTGNNNAGATCGGAAG |
| 46340_1 | TATGAGTGTGGAGTCGTTGCACCACGCCCAGCTAATTTTTTTTNNNAGATCGGAAGA |
| 46340_2 | TATGAGTGTGGAGTCGTTGCACCACGCCCAGCTAATTTTTTTTNNNAGATCGGAAGA |
| 45826_1 | TATGAGTGTGGAGTCGTTGCATCCATAGCCTTTGAGTGCATCANNNAGATCGGAAGA |
| 45826_2 | TATGAGTGTGGAGTCGTTGCATCCATAGCCTTTGAGTGCATCANNNAGATCGGAAGA |
| 34327_1 | TATGAGTGTGGAGTCGTTGCAAAATCCAGCAGGGCAGTAACTANNNAGATCGGAAG |
| 34327_2 | TATGAGTGTGGAGTCGTTGCAAAATCCAGCAGGGCAGTAACTANNNAGATCGGAAG |
| 43280_1 | TATGAGTGTGGAGTCGTTGCGGTTGCATGAGCCAAGATCAAGNNNAGATCGGAAG |
| 43280_2 | TATGAGTGTGGAGTCGTTGCGGTTGCATGAGCCAAGATCAAGNNNAGATCGGAAG |
| 30648_1 | TATGAGTGTGGAGTCGTTGCCTGTTAGATGGTCTTGCCCTGATNNNAGATCGGAAG |

| | |
|---|---|
| 30648_2 | TATGAGTGTGGAGTCGTTGCCTGTTAGATGGTCTTGCCCTGATNNNAGATCGGAAG |
| 33423_1 | TATGAGTGTGGAGTCGTTGCTGGGCTCAGTGAAAACAAGAAACNNNAGATCGGAA |
| 33423_2 | TATGAGTGTGGAGTCGTTGCTGGGCTCAGTGAAAACAAGAAACNNNAGATCGGAA |
| 10514_1 | TATGAGTGTGGAGTCGTTGCAATCAGCTGTCAAGTTCCACAGTNNNAGATCGGAAG |
| 10514_2 | TATGAGTGTGGAGTCGTTGCAATCAGCTGTCAAGTTCCACAGTNNNAGATCGGAAG |
| 39617_1 | TATGAGTGTGGAGTCGTTGCTGAGGCAGGAGAATCTCTTGAACNNNAGATCGGAA |
| 39617_2 | TATGAGTGTGGAGTCGTTGCTGAGGCAGGAGAATCTCTTGAACNNNAGATCGGAA |
| 34417_1 | TATGAGTGTGGAGTCGTTGCCCTGTAGTCCCAGTTACTAGGGANNNAGATCGGAAG |
| 34417_2 | TATGAGTGTGGAGTCGTTGCCCTGTAGTCCCAGTTACTAGGGANNNAGATCGGAAG |
| 45618_1 | TATGAGTGTGGAGTCGTTGCCTGGCCAAGTTCTTTCTTGAAGGNNNAGATCGGAAG |
| 45618_2 | TATGAGTGTGGAGTCGTTGCCTGGCCAAGTTCTTTCTTGAAGGNNNAGATCGGAAG |
| 41711_1 | TATGAGTGTGGAGTCGTTGCATGCACCCAGACAGGTAACAAAANNNAGATCGGAA |
| 41711_2 | TATGAGTGTGGAGTCGTTGCATGCACCCAGACAGGTAACAAAANNNAGATCGGAA |
| 13010_1 | TATGAGTGTGGAGTCGTTGCGGCTATTTTTCCACTGATTAAATNNNAGATCGGAAG |
| 13010_2 | TATGAGTGTGGAGTCGTTGCGGCTATTTTTCCACTGATTAAATNNNAGATCGGAAG |
| 30469_1 | TATGAGTGTGGAGTCGTTGCTGAGGATGAAGCCAAATCTTGCTNNNAGATCGGAA |
| 30469_2 | TATGAGTGTGGAGTCGTTGCTGAGGATGAAGCCAAATCTTGCTNNNAGATCGGAA |
| 36207_1 | TATGAGTGTGGAGTCGTTGCGATCACTTGAGCCCAGGAGTTTANNNAGATCGGAA |
| 36207_2 | TATGAGTGTGGAGTCGTTGCGATCACTTGAGCCCAGGAGTTTANNNAGATCGGAA |
| 41764_1 | TATGAGTGTGGAGTCGTTGCCTCCTGAGTAGCTGGGATTACAGNNNAGATCGGAA |
| 41764_2 | TATGAGTGTGGAGTCGTTGCCTCCTGAGTAGCTGGGATTACAGNNNAGATCGGAA |
| 34365_1 | TATGAGTGTGGAGTCGTTGCACTCCAGTTCTGCTCCTTAGGATNNNAGATCGGAAG |
| 34365_2 | TATGAGTGTGGAGTCGTTGCACTCCAGTTCTGCTCCTTAGGATNNNAGATCGGAAG |
| 41858_1 | TATGAGTGTGGAGTCGTTGCGAGTAGCTGGGACTATAGGCACNNNAGATCGGAAG |
| 41858_2 | TATGAGTGTGGAGTCGTTGCGAGTAGCTGGGACTATAGGCACNNNAGATCGGAAG |
| 31444_1 | TATGAGTGTGGAGTCGTTGCTCTGTCTCTCTTCCTTCCTGTCTNNNAGATCGGAAGA |
| 31444_2 | TATGAGTGTGGAGTCGTTGCTCTGTCTCTCTTCCTTCCTGTCTNNNAGATCGGAAGA |
| 36151_1 | TATGAGTGTGGAGTCGTTGCACCCTTCCCAGCCTCTAGTATCNNNAGATCGGAAGAG |
| 36151_2 | TATGAGTGTGGAGTCGTTGCACCCTTCCCAGCCTCTAGTATCNNNAGATCGGAAGAG |
| 39013_1 | TATGAGTGTGGAGTCGTTGCCTGAGAATTCCTTGAACCCAGGANNNAGATCGGAAG |
| 39013_2 | TATGAGTGTGGAGTCGTTGCCTGAGAATTCCTTGAACCCAGGANNNAGATCGGAAG |
| 49012_1 | TATGAGTGTGGAGTCGTTGCTAGAACCATCACGCTATTAGGGCNNNAGATCGGAAG |
| 49012_2 | TATGAGTGTGGAGTCGTTGCTAGAACCATCACGCTATTAGGGCNNNAGATCGGAAG |
| 38003_1 | TATGAGTGTGGAGTCGTTGCTCTGCTACAATCAGAGTAGACTCANNNAGATCGGAA |
| 38003_2 | TATGAGTGTGGAGTCGTTGCTCTGCTACAATCAGAGTAGACTCANNNAGATCGGAA |
| 35406_1 | TATGAGTGTGGAGTCGTTGCCCGTTCTCTTGCTTCAGCCTNNNAGATCGGAAGAGC |
| 35406_2 | TATGAGTGTGGAGTCGTTGCCCGTTCTCTTGCTTCAGCCTNNNAGATCGGAAGAGC |
| 41346_1 | TATGAGTGTGGAGTCGTTGCACAAAAACAGTTTCTGGTGCCAANNNAGATCGGAAG |
| 41346_2 | TATGAGTGTGGAGTCGTTGCACAAAAACAGTTTCTGGTGCCAANNNAGATCGGAAG |
| 50367_1 | TATGAGTGTGGAGTCGTTGCTTAGGTAGGTGATCTCAGCCTGTNNNAGATCGGAA |
| 50367_2 | TATGAGTGTGGAGTCGTTGCTTAGGTAGGTGATCTCAGCCTGTNNNAGATCGGAA |
| 38537_1 | TATGAGTGTGGAGTCGTTGCATAATTCTCAGGGCAAGGCAACTNNNAGATCGGAA |
| 38537_2 | TATGAGTGTGGAGTCGTTGCATAATTCTCAGGGCAAGGCAACTNNNAGATCGGAA |

| | |
|---|---|
| 10649_1 | TATGAGTGTGGAGTCGTTGCTATGGGTGTGCTTGTGTGTAAGGNNNAGATCGGAA |
| 10649_2 | TATGAGTGTGGAGTCGTTGCTATGGGTGTGCTTGTGTGTAAGGNNNAGATCGGAA |
| 30448_1 | TATGAGTGTGGAGTCGTTGCTAACAGAGAAACAGAGGTCGGGTNNNAGATCGGAA |
| 30448_2 | TATGAGTGTGGAGTCGTTGCTAACAGAGAAACAGAGGTCGGGTNNNAGATCGGAA |
| 45824_1 | TATGAGTGTGGAGTCGTTGCGACAGGGTATCACTCTATCACCCNNNAGATCGGAAGA |
| 45824_2 | TATGAGTGTGGAGTCGTTGCGACAGGGTATCACTCTATCACCCNNNAGATCGGAAGA |
| 50771_1 | TATGAGTGTGGAGTCGTTGCGTTAGATTCGGATGGGGTAGGTCNNNAGATCGGAA |
| 50771_2 | TATGAGTGTGGAGTCGTTGCGTTAGATTCGGATGGGGTAGGTCNNNAGATCGGAA |
| 48244_1 | TATGAGTGTGGAGTCGTTGCGAAATACAGGTGTCTGCCAACACNNNAGATCGGAAG |
| 48244_2 | TATGAGTGTGGAGTCGTTGCGAAATACAGGTGTCTGCCAACACNNNAGATCGGAAG |
| 36123_1 | TATGAGTGTGGAGTCGTTGCCCCTCTTCATACACGGTACTGTGNNNAGATCGGAAGA |
| 36123_2 | TATGAGTGTGGAGTCGTTGCCCCTCTTCATACACGGTACTGTGNNNAGATCGGAAGA |
| 39578_1 | TATGAGTGTGGAGTCGTTGCCAAGATGCAGCTAAAGCAGTGTTNNNAGATCGGAA |
| 39578_2 | TATGAGTGTGGAGTCGTTGCCAAGATGCAGCTAAAGCAGTGTTNNNAGATCGGAA |
| 10587_1 | TATGAGTGTGGAGTCGTTGCCTACACCATTCGCATCCCTTCTTNNNAGATCGGAAGA |
| 10587_2 | TATGAGTGTGGAGTCGTTGCCTACACCATTCGCATCCCTTCTTNNNAGATCGGAAGA |
| 52195_1 | TATGAGTGTGGAGTCGTTGCCCAGTCAATCAGCGGCAGAGNNNAGATCGGAAGAG |
| 52195_2 | TATGAGTGTGGAGTCGTTGCCCAGTCAATCAGCGGCAGAGNNNAGATCGGAAGAG |
| 52192_1 | TATGAGTGTGGAGTCGTTGCCATTCACCTCTGCTGGTTTGAAGNNNAGATCGGAAGA |
| 52192_2 | TATGAGTGTGGAGTCGTTGCCATTCACCTCTGCTGGTTTGAAGNNNAGATCGGAAGA |
| 32329_1 | TATGAGTGTGGAGTCGTTGCCCCTCTCTTTCTCGGTACCTTTCNNNAGATCGGAAGA |
| 32329_2 | TATGAGTGTGGAGTCGTTGCCCCTCTCTTTCTCGGTACCTTTCNNNAGATCGGAAGA |
| 50391_1 | TATGAGTGTGGAGTCGTTGCCAGCCTGTGAAGGGAAGGAGNNNAGATCGGAAGA |
| 50391_2 | TATGAGTGTGGAGTCGTTGCCAGCCTGTGAAGGGAAGGAGNNNAGATCGGAAGA |
| 39697_1 | TATGAGTGTGGAGTCGTTGCAGAAAGGTGGTTACCAGAGACTGNNNAGATCGGAA |
| 39697_2 | TATGAGTGTGGAGTCGTTGCAGAAAGGTGGTTACCAGAGACTGNNNAGATCGGAA |
| 45678_1 | TATGAGTGTGGAGTCGTTGCCTGAAGTGCAGTGGTGCTATCTCNNNAGATCGGAAG |
| 45678_2 | TATGAGTGTGGAGTCGTTGCCTGAAGTGCAGTGGTGCTATCTCNNNAGATCGGAAG |
| 46098_1 | TATGAGTGTGGAGTCGTTGCCTTGAAAGAGAGTGGGATGGGAANNNAGATCGGAA |
| 46098_2 | TATGAGTGTGGAGTCGTTGCCTTGAAAGAGAGTGGGATGGGAANNNAGATCGGAA |
| 49287_1 | TATGAGTGTGGAGTCGTTGCCTGGAATTACAGGCACACACAACNNNAGATCGGAAG |
| 49287_2 | TATGAGTGTGGAGTCGTTGCCTGGAATTACAGGCACACACAACNNNAGATCGGAAG |
| 45076_1 | TATGAGTGTGGAGTCGTTGCATCATTGCAGGCAAGAAGCATTCNNNAGATCGGAAG |
| 45076_2 | TATGAGTGTGGAGTCGTTGCATCATTGCAGGCAAGAAGCATTCNNNAGATCGGAAG |
| 31706_1 | TATGAGTGTGGAGTCGTTGCGAACAAGAAAGAGGAGGAGGAGGNNNAGATCGGA |
| 31706_2 | TATGAGTGTGGAGTCGTTGCGAACAAGAAAGAGGAGGAGGAGGNNNAGATCGGA |
| 29567_1 | TATGAGTGTGGAGTCGTTGCATTTCTGGAGAGTGACTGTGGTCNNNAGATCGGAA |
| 29567_2 | TATGAGTGTGGAGTCGTTGCATTTCTGGAGAGTGACTGTGGTCNNNAGATCGGAA |
| 46006_1 | TATGAGTGTGGAGTCGTTGCCACCACACACGGCTAATTTTTGTNNNAGATCGGAAGA |
| 46006_2 | TATGAGTGTGGAGTCGTTGCCACCACACACGGCTAATTTTTGTNNNAGATCGGAAGA |
| 53232_1 | TATGAGTGTGGAGTCGTTGCAAAGCATTCCTCCAAGATCCACANNNAGATCGGAAGA |
| 53232_2 | TATGAGTGTGGAGTCGTTGCAAAGCATTCCTCCAAGATCCACANNNAGATCGGAAGA |
| 38675_1 | TATGAGTGTGGAGTCGTTGCGAAACCAGGGACTCTCAGAATCTNNNAGATCGGAAG |

| | |
|---|---|
| 38675_2 | TATGAGTGTGGAGTCGTTGCGAAACCAGGGACTCTCAGAATCTNNNAGATCGGAAG |
| 53254_1 | TATGAGTGTGGAGTCGTTGCAGAGAATTGGGCTTCGACTCATTNNNAGATCGGAAG |
| 53254_2 | TATGAGTGTGGAGTCGTTGCAGAGAATTGGGCTTCGACTCATTNNNAGATCGGAAG |
| 11201_1 | TATGAGTGTGGAGTCGTTGCCCCAATGCTGCCTATTGTCAGANNNAGATCGGAAGA |
| 11201_2 | TATGAGTGTGGAGTCGTTGCCCCAATGCTGCCTATTGTCAGANNNAGATCGGAAGA |
| 41068_1 | TATGAGTGTGGAGTCGTTGCCCTAGGCGACAGAACAAGACTCNNNAGATCGGAAGA |
| 41068_2 | TATGAGTGTGGAGTCGTTGCCCTAGGCGACAGAACAAGACTCNNNAGATCGGAAGA |
| 41155_1 | TATGAGTGTGGAGTCGTTGCCGGAGACTGGGTAATTTGGGTAANNNAGATCGGAA |
| 41155_2 | TATGAGTGTGGAGTCGTTGCCGGAGACTGGGTAATTTGGGTAANNNAGATCGGAA |
| 12800_1 | TATGAGTGTGGAGTCGTTGCAAAACCCCTATGTACCACTGNNNAGATCGGAAGAGC |
| 12800_2 | TATGAGTGTGGAGTCGTTGCAAAACCCCTATGTACCACTGNNNAGATCGGAAGAGC |
| 33843_1 | TATGAGTGTGGAGTCGTTGCTTTTCTCTTGCTCCCTCTCTCACNNNAGATCGGAAGAG |
| 33843_2 | TATGAGTGTGGAGTCGTTGCTTTTCTCTTGCTCCCTCTCTCACNNNAGATCGGAAGAG |
| 41393_1 | TATGAGTGTGGAGTCGTTGCAGTATTGGATTCTGGACTTCCCCNNNAGATCGGAAG |
| 41393_2 | TATGAGTGTGGAGTCGTTGCAGTATTGGATTCTGGACTTCCCCNNNAGATCGGAAG |
| 39390_1 | TATGAGTGTGGAGTCGTTGCTGAGAGGGGAAGGGAACTAACATNNNAGATCGGA |
| 39390_2 | TATGAGTGTGGAGTCGTTGCTGAGAGGGGAAGGGAACTAACATNNNAGATCGGA |
| 48881_1 | TATGAGTGTGGAGTCGTTGCGTACAGTGGCATGTTTTCAGCTCNNNAGATCGGAAG |
| 48881_2 | TATGAGTGTGGAGTCGTTGCGTACAGTGGCATGTTTTCAGCTCNNNAGATCGGAAG |
| 37841_1 | TATGAGTGTGGAGTCGTTGCACCATATGACTATTCACACTGCCTNNNAGATCGGAAG |
| 37841_2 | TATGAGTGTGGAGTCGTTGCACCATATGACTATTCACACTGCCTNNNAGATCGGAAG |
| 43888_1 | TATGAGTGTGGAGTCGTTGCTTTTTGAGACAGGGTCTTGCTCTNNNAGATCGGAAG |
| 43888_2 | TATGAGTGTGGAGTCGTTGCTTTTTGAGACAGGGTCTTGCTCTNNNAGATCGGAAG |
| 43270_1 | TATGAGTGTGGAGTCGTTGCCCACCACACCTGGCTAATTTTTTNNNAGATCGGAAGA |
| 43270_2 | TATGAGTGTGGAGTCGTTGCCCACCACACCTGGCTAATTTTTTNNNAGATCGGAAGA |
| 35283_1 | TATGAGTGTGGAGTCGTTGCACGATCTAAGCTCACTGCAACTTNNNAGATCGGAAG |
| 35283_2 | TATGAGTGTGGAGTCGTTGCACGATCTAAGCTCACTGCAACTTNNNAGATCGGAAG |
| 41310_1 | TATGAGTGTGGAGTCGTTGCGATCACAAGGTCAGGAGATCGAGNNNAGATCGGAA |
| 41310_2 | TATGAGTGTGGAGTCGTTGCGATCACAAGGTCAGGAGATCGAGNNNAGATCGGAA |
| 33975_1 | TATGAGTGTGGAGTCGTTGCCTCCCATCTCAGCCTCTCTAGTANNNAGATCGGAAGA |
| 33975_2 | TATGAGTGTGGAGTCGTTGCCTCCCATCTCAGCCTCTCTAGTANNNAGATCGGAAGA |
| 34420_1 | TATGAGTGTGGAGTCGTTGCCTGAGGCATGAGAATTGCTTGAGNNNAGATCGGAA |
| 34420_2 | TATGAGTGTGGAGTCGTTGCCTGAGGCATGAGAATTGCTTGAGNNNAGATCGGAA |
| 43557_1 | TATGAGTGTGGAGTCGTTGCCAAGTAGCTGGGACTACAGGTGNNNAGATCGGAAG |
| 43557_2 | TATGAGTGTGGAGTCGTTGCCAAGTAGCTGGGACTACAGGTGNNNAGATCGGAAG |
| 43980_1 | TATGAGTGTGGAGTCGTTGCCTGGAGTGTAGTGGTGTGATCATNNNAGATCGGAA |
| 43980_2 | TATGAGTGTGGAGTCGTTGCCTGGAGTGTAGTGGTGTGATCATNNNAGATCGGAA |
| 12915_1 | TATGAGTGTGGAGTCGTTGCTTTGGAATTAAGCTCTTTGTTATNNNAGATCGGAAG |
| 12915_2 | TATGAGTGTGGAGTCGTTGCTTTGGAATTAAGCTCTTTGTTATNNNAGATCGGAAG |
| 50063_1 | TATGAGTGTGGAGTCGTTGCCACACACACACATACACACACATNNNAGATCGGAAGA |
| 50063_2 | TATGAGTGTGGAGTCGTTGCCACACACACACATACACACACATNNNAGATCGGAAGA |
| 31138_1 | TATGAGTGTGGAGTCGTTGCCCATGACTTCCAGGAACTCTCTTNNNAGATCGGAAGA |
| 31138_2 | TATGAGTGTGGAGTCGTTGCCCATGACTTCCAGGAACTCTCTTNNNAGATCGGAAGA |

| | |
|---|---|
| 49736_1 | TATGAGTGTGGAGTCGTTGCTCTGAATGCACGTCTCATACCTTNNNAGATCGGAAG |
| 49736_2 | TATGAGTGTGGAGTCGTTGCTCTGAATGCACGTCTCATACCTTNNNAGATCGGAAG |
| 35666_1 | TATGAGTGTGGAGTCGTTGCCATGCTCTTTCCATCCTCCATCTNNNAGATCGGAAGA |
| 35666_2 | TATGAGTGTGGAGTCGTTGCCATGCTCTTTCCATCCTCCATCTNNNAGATCGGAAGA |
| 51794_1 | TATGAGTGTGGAGTCGTTGCACCACTAAATAGGGAAGGAGGGANNNAGATCGGA |
| 51794_2 | TATGAGTGTGGAGTCGTTGCACCACTAAATAGGGAAGGAGGGANNNAGATCGGA |
| 32359_1 | TATGAGTGTGGAGTCGTTGCGCTCAGCCTAGATGTTGTTCTTGNNNAGATCGGAAG |
| 32359_2 | TATGAGTGTGGAGTCGTTGCGCTCAGCCTAGATGTTGTTCTTGNNNAGATCGGAAG |
| 35335_1 | TATGAGTGTGGAGTCGTTGCTTCGAGTAGCTGAGATTACAGGCNNNAGATCGGAA |
| 35335_2 | TATGAGTGTGGAGTCGTTGCTTCGAGTAGCTGAGATTACAGGCNNNAGATCGGAA |
| 13606_1 | TATGAGTGTGGAGTCGTTGCGTAATAGGAGCTAGTATCAGAAANNNAGATCGGAA |
| 13606_2 | TATGAGTGTGGAGTCGTTGCGTAATAGGAGCTAGTATCAGAAANNNAGATCGGAA |
| 44148_1 | TATGAGTGTGGAGTCGTTGCTCCTTCCTCCCACACATAAAGTGNNNAGATCGGAAG |
| 44148_2 | TATGAGTGTGGAGTCGTTGCTCCTTCCTCCCACACATAAAGTGNNNAGATCGGAAG |
| 29817_1 | TATGAGTGTGGAGTCGTTGCGCACTTACTGTCTTCAACCCCTANNNAGATCGGAAGA |
| 29817_2 | TATGAGTGTGGAGTCGTTGCGCACTTACTGTCTTCAACCCCTANNNAGATCGGAAGA |
| 52936_1 | TATGAGTGTGGAGTCGTTGCCGGTCTGGTTTCCTTTACGTTTANNNAGATCGGAAG |
| 52936_2 | TATGAGTGTGGAGTCGTTGCCGGTCTGGTTTCCTTTACGTTTANNNAGATCGGAAG |
| 33268_1 | TATGAGTGTGGAGTCGTTGCCTCCTGACCTCAAGTGATCCATCNNNAGATCGGAAG |
| 33268_2 | TATGAGTGTGGAGTCGTTGCCTCCTGACCTCAAGTGATCCATCNNNAGATCGGAAG |
| 38351_1 | TATGAGTGTGGAGTCGTTGCTGCATTCATATGTTCATCACAGCANNNAGATCGGAA |
| 38351_2 | TATGAGTGTGGAGTCGTTGCTGCATTCATATGTTCATCACAGCANNNAGATCGGAA |
| 51986_1 | TATGAGTGTGGAGTCGTTGCGCCTAGTTCTCCAGAATGTAGGGNNNAGATCGGAA |
| 51986_2 | TATGAGTGTGGAGTCGTTGCGCCTAGTTCTCCAGAATGTAGGGNNNAGATCGGAA |
| 44866_1 | TATGAGTGTGGAGTCGTTGCAGTTTGTTCTGCTTTGTCTCTGCNNNAGATCGGAAG |
| 44866_2 | TATGAGTGTGGAGTCGTTGCAGTTTGTTCTGCTTTGTCTCTGCNNNAGATCGGAAG |
| 51518_1 | TATGAGTGTGGAGTCGTTGCTGCCCTGATTGTAAGTTTCCTGANNNAGATCGGAAG |
| 51518_2 | TATGAGTGTGGAGTCGTTGCTGCCCTGATTGTAAGTTTCCTGANNNAGATCGGAAG |
| 30174_1 | TATGAGTGTGGAGTCGTTGCCCATGCAGGTAAATAGCTTTTTAGTNNNAGATCGGA |
| 30174_2 | TATGAGTGTGGAGTCGTTGCCCATGCAGGTAAATAGCTTTTTAGTNNNAGATCGGA |
| 38557_1 | TATGAGTGTGGAGTCGTTGCTGGCAGAGCTGAGATTCAAACTCNNNAGATCGGAA |
| 38557_2 | TATGAGTGTGGAGTCGTTGCTGGCAGAGCTGAGATTCAAACTCNNNAGATCGGAA |
| 2168_1 | TATGAGTGTGGAGTCGTTGCCAGAAGGGAGAATAAGGGGCAGANNNAGATCGGA |
| 2168_2 | TATGAGTGTGGAGTCGTTGCCAGAAGGGAGAATAAGGGGCAGANNNAGATCGGA |
| 41962_1 | TATGAGTGTGGAGTCGTTGCGCCTCTTGAGTAGCTGGGATTACNNNAGATCGGAA |
| 41962_2 | TATGAGTGTGGAGTCGTTGCGCCTCTTGAGTAGCTGGGATTACNNNAGATCGGAA |
| 33309_1 | TATGAGTGTGGAGTCGTTGCCTGGTCTTGAACTCATGGTCTCANNNAGATCGGAAG |
| 33309_2 | TATGAGTGTGGAGTCGTTGCCTGGTCTTGAACTCATGGTCTCANNNAGATCGGAAG |
| 39624_1 | TATGAGTGTGGAGTCGTTGCAAATCTGAACCCTCGAACATTGCNNNAGATCGGAAG |
| 39624_2 | TATGAGTGTGGAGTCGTTGCAAATCTGAACCCTCGAACATTGCNNNAGATCGGAAG |
| 35079_1 | TATGAGTGTGGAGTCGTTGCCAGTGGTGCAATCTCAGCTCANNNAGATCGGAAGA |
| 35079_2 | TATGAGTGTGGAGTCGTTGCCAGTGGTGCAATCTCAGCTCANNNAGATCGGAAGA |
| 32857_1 | TATGAGTGTGGAGTCGTTGCGTTTTTGCAGAGACAGGGTCTTGNNNAGATCGGAA |

| | |
|---|---|
| 32857_2 | TATGAGTGTGGAGTCGTTGCGTTTTTGCAGAGACAGGGTCTTGNNNAGATCGGAAG |
| 40629_1 | TATGAGTGTGGAGTCGTTGCCCAAGAGCTCCAACCTTGTAAGANNNAGATCGGAAG |
| 40629_2 | TATGAGTGTGGAGTCGTTGCCCAAGAGCTCCAACCTTGTAAGANNNAGATCGGAAG |
| 49442_1 | TATGAGTGTGGAGTCGTTGCGAACCTTGAGCCCTGAATCAACTNNNAGATCGGAAG |
| 49442_2 | TATGAGTGTGGAGTCGTTGCGAACCTTGAGCCCTGAATCAACTNNNAGATCGGAAG |
| 31931_1 | TATGAGTGTGGAGTCGTTGCTCCTTGAATTGCTAACCTCTGCTNNNAGATCGGAAGA |
| 31931_2 | TATGAGTGTGGAGTCGTTGCTCCTTGAATTGCTAACCTCTGCTNNNAGATCGGAAGA |
| 49339_1 | TATGAGTGTGGAGTCGTTGCCTGGAAGTGTAGTGGTGTGATCANNNAGATCGGAA |
| 49339_2 | TATGAGTGTGGAGTCGTTGCCTGGAAGTGTAGTGGTGTGATCANNNAGATCGGAA |
| 40052_1 | TATGAGTGTGGAGTCGTTGCCTAGGAGATGGAAGTTGCAGTGANNNAGATCGGAA |
| 40052_2 | TATGAGTGTGGAGTCGTTGCCTAGGAGATGGAAGTTGCAGTGANNNAGATCGGAA |
| 42654_1 | TATGAGTGTGGAGTCGTTGCGCTCTGCAATAAACAGATCCACGNNNAGATCGGAAG |
| 42654_2 | TATGAGTGTGGAGTCGTTGCGCTCTGCAATAAACAGATCCACGNNNAGATCGGAAG |
| 47026_1 | TATGAGTGTGGAGTCGTTGCGGACTACACGTATTGACCAGTGTNNNAGATCGGAAG |
| 47026_2 | TATGAGTGTGGAGTCGTTGCGGACTACACGTATTGACCAGTGTNNNAGATCGGAAG |
| 44020_1 | TATGAGTGTGGAGTCGTTGCTGCAGAGAGTGAAGGCATATTGANNNAGATCGGAA |
| 44020_2 | TATGAGTGTGGAGTCGTTGCTGCAGAGAGTGAAGGCATATTGANNNAGATCGGAA |
| 44964_1 | TATGAGTGTGGAGTCGTTGCAAGCAGAGTGATGGTTAGAAGCANNNAGATCGGAA |
| 44964_2 | TATGAGTGTGGAGTCGTTGCAAGCAGAGTGATGGTTAGAAGCANNNAGATCGGAA |
| 41313_1 | TATGAGTGTGGAGTCGTTGCACAAACAACCCCATCAAAAAGGGNNNAGATCGGAAG |
| 41313_2 | TATGAGTGTGGAGTCGTTGCACAAACAACCCCATCAAAAAGGGNNNAGATCGGAAG |
| 37533_1 | TATGAGTGTGGAGTCGTTGCCTTGTCAGAAATGGCATCTAGGCNNNAGATCGGAAG |
| 37533_2 | TATGAGTGTGGAGTCGTTGCCTTGTCAGAAATGGCATCTAGGCNNNAGATCGGAAG |
| 41672_1 | TATGAGTGTGGAGTCGTTGCTTTCAAGTGATTCTCCTGCCTCANNNAGATCGGAAGA |
| 41672_2 | TATGAGTGTGGAGTCGTTGCTTTCAAGTGATTCTCCTGCCTCANNNAGATCGGAAGA |
| 42996_1 | TATGAGTGTGGAGTCGTTGCGCCTCCAAAAGTGCTGGAATTACNNNAGATCGGAAG |
| 42996_2 | TATGAGTGTGGAGTCGTTGCGCCTCCAAAAGTGCTGGAATTACNNNAGATCGGAAG |
| 10459_1 | TATGAGTGTGGAGTCGTTGCATTGAAGGAAACCCGGTAGGAGANNNAGATCGGAA |
| 10459_2 | TATGAGTGTGGAGTCGTTGCATTGAAGGAAACCCGGTAGGAGANNNAGATCGGAA |
| 44588_1 | TATGAGTGTGGAGTCGTTGCTCCTGCTTGTTTTGGTCAATGTCNNNAGATCGGAAGA |
| 44588_2 | TATGAGTGTGGAGTCGTTGCTCCTGCTTGTTTTGGTCAATGTCNNNAGATCGGAAGA |
| 36666_1 | TATGAGTGTGGAGTCGTTGCAAACTGAACCCAACATAAGCAGANNNAGATCGGAAG |
| 36666_2 | TATGAGTGTGGAGTCGTTGCAAACTGAACCCAACATAAGCAGANNNAGATCGGAAG |
| 40847_1 | TATGAGTGTGGAGTCGTTGCAGGGGAACTCTCTTAGTACCTGGNNNAGATCGGAAG |
| 40847_2 | TATGAGTGTGGAGTCGTTGCAGGGGAACTCTCTTAGTACCTGGNNNAGATCGGAAG |
| 35062_1 | TATGAGTGTGGAGTCGTTGCTTACTTCACATGTCACGTGACCTNNNAGATCGGAAGA |
| 35062_2 | TATGAGTGTGGAGTCGTTGCTTACTTCACATGTCACGTGACCTNNNAGATCGGAAGA |
| 2645_1 | TATGAGTGTGGAGTCGTTGCCCCCAGACGAAAATACCAAATGCNNNAGATCGGAAG |
| 2645_2 | TATGAGTGTGGAGTCGTTGCCCCCAGACGAAAATACCAAATGCNNNAGATCGGAAG |
| 42244_1 | TATGAGTGTGGAGTCGTTGCGATGAGAAGCACCTCCTAGAAGGNNNAGATCGGAAG |
| 42244_2 | TATGAGTGTGGAGTCGTTGCGATGAGAAGCACCTCCTAGAAGGNNNAGATCGGAAG |
| 30856_1 | TATGAGTGTGGAGTCGTTGCTGGTCGATATCAATTTTGCGTCGNNNAGATCGGAAG |
| 30856_2 | TATGAGTGTGGAGTCGTTGCTGGTCGATATCAATTTTGCGTCGNNNAGATCGGAAG |

| | |
|---|---|
| 13063_1 | TATGAGTGTGGAGTCGTTGCTTCTGCAAAATCATAACTGTTATNNNAGATCGGAAGA |
| 13063_2 | TATGAGTGTGGAGTCGTTGCTTCTGCAAAATCATAACTGTTATNNNAGATCGGAAGA |
| 43068_1 | TATGAGTGTGGAGTCGTTGCCATACACACACACACACACACACNNNAGATCGGAAGA |
| 43068_2 | TATGAGTGTGGAGTCGTTGCCATACACACACACACACACACACNNNAGATCGGAAGA |
| 48987_1 | TATGAGTGTGGAGTCGTTGCAATAACAAGCCCCTAAGGACCACNNNAGATCGGAAG |
| 48987_2 | TATGAGTGTGGAGTCGTTGCAATAACAAGCCCCTAAGGACCACNNNAGATCGGAAG |
| 42904_1 | TATGAGTGTGGAGTCGTTGCAATTGAATCATGGGGGCAGTTTCNNNAGATCGGAA |
| 42904_2 | TATGAGTGTGGAGTCGTTGCAATTGAATCATGGGGGCAGTTTCNNNAGATCGGAA |
| 12783_1 | TATGAGTGTGGAGTCGTTGCTGAAGATAAAGTCAGCCATGNNNAGATCGGAAGAG |
| 12783_2 | TATGAGTGTGGAGTCGTTGCTGAAGATAAAGTCAGCCATGNNNAGATCGGAAGAG |
| 51686_1 | TATGAGTGTGGAGTCGTTGCTGTCCTTTCTCTTTACTGTGGGGNNNAGATCGGAAGA |
| 51686_2 | TATGAGTGTGGAGTCGTTGCTGTCCTTTCTCTTTACTGTGGGGNNNAGATCGGAAGA |
| 34868_1 | TATGAGTGTGGAGTCGTTGCATCTTGGCTCACTGTAACCTCTGNNNAGATCGGAAGA |
| 34868_2 | TATGAGTGTGGAGTCGTTGCATCTTGGCTCACTGTAACCTCTGNNNAGATCGGAAGA |
| 39540_1 | TATGAGTGTGGAGTCGTTGCAAGGTGGAAGAATCTGAGCAAGANNNAGATCGGAA |
| 39540_2 | TATGAGTGTGGAGTCGTTGCAAGGTGGAAGAATCTGAGCAAGANNNAGATCGGAA |
| 35188_1 | TATGAGTGTGGAGTCGTTGCCAAGTAGCTGGGACCACAGTAGNNNAGATCGGAAG |
| 35188_2 | TATGAGTGTGGAGTCGTTGCCAAGTAGCTGGGACCACAGTAGNNNAGATCGGAAG |
| 49663_1 | TATGAGTGTGGAGTCGTTGCCAAGCAATTCTCCTTTCTCAGCCNNNAGATCGGAAGA |
| 49663_2 | TATGAGTGTGGAGTCGTTGCCAAGCAATTCTCCTTTCTCAGCCNNNAGATCGGAAGA |
| 52830_1 | TATGAGTGTGGAGTCGTTGCAAGAAAATAAATCAGGCTGGGCGNNNAGATCGGAA |
| 52830_2 | TATGAGTGTGGAGTCGTTGCAAGAAAATAAATCAGGCTGGGCGNNNAGATCGGAA |
| 34177_1 | TATGAGTGTGGAGTCGTTGCGCCAAGTTTGAGAAACCCTGATCNNNAGATCGGAAG |
| 34177_2 | TATGAGTGTGGAGTCGTTGCGCCAAGTTTGAGAAACCCTGATCNNNAGATCGGAAG |
| 50416_1 | TATGAGTGTGGAGTCGTTGCCTGCAGGATCCATGTAAGACTGANNNAGATCGGAA |
| 50416_2 | TATGAGTGTGGAGTCGTTGCCTGCAGGATCCATGTAAGACTGANNNAGATCGGAA |
| 35956_1 | TATGAGTGTGGAGTCGTTGCACCAAAAAATGAGGGAGACTGGTNNNAGATCGGAA |
| 35956_2 | TATGAGTGTGGAGTCGTTGCACCAAAAAATGAGGGAGACTGGTNNNAGATCGGAA |
| 44349_1 | TATGAGTGTGGAGTCGTTGCGTCCATTGGCCCTTCCCTAGNNNAGATCGGAAGAGCA |
| 44349_2 | TATGAGTGTGGAGTCGTTGCGTCCATTGGCCCTTCCCTAGNNNAGATCGGAAGAGCA |
| 32560_1 | TATGAGTGTGGAGTCGTTGCCCAAGTAGCTGGGATTATAGGCANNNAGATCGGAA |
| 32560_2 | TATGAGTGTGGAGTCGTTGCCCAAGTAGCTGGGATTATAGGCANNNAGATCGGAA |
| 43260_1 | TATGAGTGTGGAGTCGTTGCGAGTGCAATGGTGATCTCAGCTCNNNAGATCGGAA |
| 43260_2 | TATGAGTGTGGAGTCGTTGCGAGTGCAATGGTGATCTCAGCTCNNNAGATCGGAA |
| 46074_1 | TATGAGTGTGGAGTCGTTGCGCTTTCCAGGTTCAAGCAATTCTNNNAGATCGGAAGA |
| 46074_2 | TATGAGTGTGGAGTCGTTGCGCTTTCCAGGTTCAAGCAATTCTNNNAGATCGGAAGA |
| 41500_1 | TATGAGTGTGGAGTCGTTGCCTCTTCAAGCTAGCTCAGTGTCTNNNAGATCGGAAGA |
| 41500_2 | TATGAGTGTGGAGTCGTTGCCTCTTCAAGCTAGCTCAGTGTCTNNNAGATCGGAAGA |
| 31170_1 | TATGAGTGTGGAGTCGTTGCTGTGTGATTAGTGTAGTGGCCAANNNAGATCGGAA |
| 31170_2 | TATGAGTGTGGAGTCGTTGCTGTGTGATTAGTGTAGTGGCCAANNNAGATCGGAA |
| 41686_1 | TATGAGTGTGGAGTCGTTGCCCATCCGATGAGCATAAGGTGATNNNAGATCGGAA |
| 41686_2 | TATGAGTGTGGAGTCGTTGCCCATCCGATGAGCATAAGGTGATNNNAGATCGGAA |
| 46795_1 | TATGAGTGTGGAGTCGTTGCTTCTGGACCTTGATTGAGAAGGGNNNAGATCGGAA |

| | |
|---|---|
| 46795_2 | TATGAGTGTGGAGTCGTTGCTTCTGGACCTTGATTGAGAAGGGNNNAGATCGGAA |
| 29521_1 | TATGAGTGTGGAGTCGTTGCCTGTGCTGAGAGAGAGAGACTCTNNNAGATCGGAA |
| 29521_2 | TATGAGTGTGGAGTCGTTGCCTGTGCTGAGAGAGAGAGACTCTNNNAGATCGGAA |
| 44472_1 | TATGAGTGTGGAGTCGTTGCGTGTGTGTGTGTGTGTGTGTATANNNAGATCGGAA |
| 44472_2 | TATGAGTGTGGAGTCGTTGCGTGTGTGTGTGTGTGTGTGTATANNNAGATCGGAA |
| 35708_1 | TATGAGTGTGGAGTCGTTGCAATCTCGGCTCACTGCAATCTCNNNAGATCGGAAGA |
| 35708_2 | TATGAGTGTGGAGTCGTTGCAATCTCGGCTCACTGCAATCTCNNNAGATCGGAAGA |
| 45097_1 | TATGAGTGTGGAGTCGTTGCAACTGCTCTTGTTCTCAGATGCTNNNAGATCGGAAG |
| 45097_2 | TATGAGTGTGGAGTCGTTGCAACTGCTCTTGTTCTCAGATGCTNNNAGATCGGAAG |
| 43659_1 | TATGAGTGTGGAGTCGTTGCAAGGAAAGATTTGGGGTTCCAGANNNAGATCGGAA |
| 43659_2 | TATGAGTGTGGAGTCGTTGCAAGGAAAGATTTGGGGTTCCAGANNNAGATCGGAA |
| 32047_1 | TATGAGTGTGGAGTCGTTGCTGGAGTGCAGTGGTACAATCTCNNNAGATCGGAAG |
| 32047_2 | TATGAGTGTGGAGTCGTTGCTGGAGTGCAGTGGTACAATCTCNNNAGATCGGAAG |
| 2673_1 | TATGAGTGTGGAGTCGTTGCTGCGAAAATGTTCATTGCACCATNNNAGATCGGAAG |
| 2673_2 | TATGAGTGTGGAGTCGTTGCTGCGAAAATGTTCATTGCACCATNNNAGATCGGAAG |
| 42624_1 | TATGAGTGTGGAGTCGTTGCCTGCTTGACTGTTGTTGGCATATNNNAGATCGGAAG |
| 42624_2 | TATGAGTGTGGAGTCGTTGCCTGCTTGACTGTTGTTGGCATATNNNAGATCGGAAG |
| 41547_1 | TATGAGTGTGGAGTCGTTGCGCCTACAGAGTAGCTGGGATTACNNNAGATCGGAA |
| 41547_2 | TATGAGTGTGGAGTCGTTGCGCCTACAGAGTAGCTGGGATTACNNNAGATCGGAA |
| 48175_1 | TATGAGTGTGGAGTCGTTGCATTACAGGCATGAACCACAATGCNNNAGATCGGAAG |
| 48175_2 | TATGAGTGTGGAGTCGTTGCATTACAGGCATGAACCACAATGCNNNAGATCGGAAG |
| 13353_1 | TATGAGTGTGGAGTCGTTGCAATGGAATACTAAAAAAACCTGCNNNAGATCGGAAG |
| 13353_2 | TATGAGTGTGGAGTCGTTGCAATGGAATACTAAAAAAACCTGCNNNAGATCGGAAG |
| 40228_1 | TATGAGTGTGGAGTCGTTGCAGAGCCAAAGTAGTCTGTTTTTGTNNNAGATCGGAA |
| 40228_2 | TATGAGTGTGGAGTCGTTGCAGAGCCAAAGTAGTCTGTTTTTGTNNNAGATCGGAA |
| 49224_1 | TATGAGTGTGGAGTCGTTGCTTCAAGTGATTCTCCTGCCTCAGNNNAGATCGGAAG |
| 49224_2 | TATGAGTGTGGAGTCGTTGCTTCAAGTGATTCTCCTGCCTCAGNNNAGATCGGAAG |
| 40472_1 | TATGAGTGTGGAGTCGTTGCAATACAATGGGTTGGTGCAAAGTNNNAGATCGGAA |
| 40472_2 | TATGAGTGTGGAGTCGTTGCAATACAATGGGTTGGTGCAAAGTNNNAGATCGGAA |
| 47906_1 | TATGAGTGTGGAGTCGTTGCCTCTCAAGGTGCTGGGATTACAGNNNAGATCGGAA |
| 47906_2 | TATGAGTGTGGAGTCGTTGCCTCTCAAGGTGCTGGGATTACAGNNNAGATCGGAA |
| 47532_1 | TATGAGTGTGGAGTCGTTGCAGTGTGGTGGTCTTGTTTCTTCANNNAGATCGGAAG |
| 47532_2 | TATGAGTGTGGAGTCGTTGCAGTGTGGTGGTCTTGTTTCTTCANNNAGATCGGAAG |
| 52579_1 | TATGAGTGTGGAGTCGTTGCGGTTGGCTGTACCCAGGCTCNNNAGATCGGAAGAG |
| 52579_2 | TATGAGTGTGGAGTCGTTGCGGTTGGCTGTACCCAGGCTCNNNAGATCGGAAGAG |
| 48007_1 | TATGAGTGTGGAGTCGTTGCGAGTAGGTGGGATTACAGGCACNNNAGATCGGAA |
| 48007_2 | TATGAGTGTGGAGTCGTTGCGAGTAGGTGGGATTACAGGCACNNNAGATCGGAA |
| 44639_1 | TATGAGTGTGGAGTCGTTGCGCCAGGAAAGGAAATTGGTGTTTNNNAGATCGGAA |
| 44639_2 | TATGAGTGTGGAGTCGTTGCGCCAGGAAAGGAAATTGGTGTTTNNNAGATCGGAA |
| 30529_1 | TATGAGTGTGGAGTCGTTGCTGAAACTAGGAAGCAGGGACTCANNNAGATCGGAA |
| 30529_2 | TATGAGTGTGGAGTCGTTGCTGAAACTAGGAAGCAGGGACTCANNNAGATCGGAA |
| 42728_1 | TATGAGTGTGGAGTCGTTGCAAAGAGAAAGACGGAAGGGACTTNNNAGATCGGA |
| 42728_2 | TATGAGTGTGGAGTCGTTGCAAAGAGAAAGACGGAAGGGACTTNNNAGATCGGA |

| | |
|---|---|
| 46981_1 | TATGAGTGTGGAGTCGTTGCCAGGCACACTGATTTTAGCAACANNNAGATCGGAAG |
| 46981_2 | TATGAGTGTGGAGTCGTTGCCAGGCACACTGATTTTAGCAACANNNAGATCGGAAG |
| 39190_1 | TATGAGTGTGGAGTCGTTGCCTGGTCTCAAACTCCTGATCTCANNNAGATCGGAAG |
| 39190_2 | TATGAGTGTGGAGTCGTTGCCTGGTCTCAAACTCCTGATCTCANNNAGATCGGAAG |
| 37573_1 | TATGAGTGTGGAGTCGTTGCGGCTGTAGTGGAGAAAAAGCTTGNNNAGATCGGAA |
| 37573_2 | TATGAGTGTGGAGTCGTTGCGGCTGTAGTGGAGAAAAAGCTTGNNNAGATCGGAA |
| 46467_1 | TATGAGTGTGGAGTCGTTGCTCCTGAGTAGCTGGGATTACAGANNNAGATCGGAA |
| 46467_2 | TATGAGTGTGGAGTCGTTGCTCCTGAGTAGCTGGGATTACAGANNNAGATCGGAA |
| 38014_1 | TATGAGTGTGGAGTCGTTGCCTCGTACAATCACCTTGGAAAGCNNNAGATCGGAAG |
| 38014_2 | TATGAGTGTGGAGTCGTTGCCTCGTACAATCACCTTGGAAAGCNNNAGATCGGAAG |
| 51154_1 | TATGAGTGTGGAGTCGTTGCGAAGAGGTGCAGGTCCACAGNNNAGATCGGAAGA |
| 51154_2 | TATGAGTGTGGAGTCGTTGCGAAGAGGTGCAGGTCCACAGNNNAGATCGGAAGA |
| 51170_1 | TATGAGTGTGGAGTCGTTGCTATAATTCTCTGCAAGCCTGGCTNNNAGATCGGAAG |
| 51170_2 | TATGAGTGTGGAGTCGTTGCTATAATTCTCTGCAAGCCTGGCTNNNAGATCGGAAG |
| 2284_1 | TATGAGTGTGGAGTCGTTGCTTTCCCACATTTCCTGGAGGAACNNNAGATCGGAAG |
| 2284_2 | TATGAGTGTGGAGTCGTTGCTTTCCCACATTTCCTGGAGGAACNNNAGATCGGAAG |
| 46004_1 | TATGAGTGTGGAGTCGTTGCGAGTTTTGCTCTTGTTGTCCAGGNNNAGATCGGAAG |
| 46004_2 | TATGAGTGTGGAGTCGTTGCGAGTTTTGCTCTTGTTGTCCAGGNNNAGATCGGAAG |
| 30547_1 | TATGAGTGTGGAGTCGTTGCTTCCCACGTGCCTTCATGGANNNAGATCGGAAGAGC |
| 30547_2 | TATGAGTGTGGAGTCGTTGCTTCCCACGTGCCTTCATGGANNNAGATCGGAAGAGC |
| 40348_1 | TATGAGTGTGGAGTCGTTGCCTGAGGCAGGAGAATTGCTTGAANNNAGATCGGAA |
| 40348_2 | TATGAGTGTGGAGTCGTTGCCTGAGGCAGGAGAATTGCTTGAANNNAGATCGGAA |
| 37692_1 | TATGAGTGTGGAGTCGTTGCAAGGCAAGTGACTGCTATGTAGTNNNAGATCGGAA |
| 37692_2 | TATGAGTGTGGAGTCGTTGCAAGGCAAGTGACTGCTATGTAGTNNNAGATCGGAA |
| 44111_1 | TATGAGTGTGGAGTCGTTGCATAGAAGCCTCCCAGTGTTTTGTNNNAGATCGGAAG |
| 44111_2 | TATGAGTGTGGAGTCGTTGCATAGAAGCCTCCCAGTGTTTTGTNNNAGATCGGAAG |
| 50628_1 | TATGAGTGTGGAGTCGTTGCCTAAGCAGGAGTTCTGATTTGCCNNNAGATCGGAAG |
| 50628_2 | TATGAGTGTGGAGTCGTTGCCTAAGCAGGAGTTCTGATTTGCCNNNAGATCGGAAG |
| 30292_1 | TATGAGTGTGGAGTCGTTGCACATCTGTGGTGTGTTGGCTTATNNNAGATCGGAAG |
| 30292_2 | TATGAGTGTGGAGTCGTTGCACATCTGTGGTGTGTTGGCTTATNNNAGATCGGAAG |
| 34285_1 | TATGAGTGTGGAGTCGTTGCCACTGTGCCCTCATTTCTTTTGTNNNAGATCGGAAGA |
| 34285_2 | TATGAGTGTGGAGTCGTTGCCACTGTGCCCTCATTTCTTTTGTNNNAGATCGGAAGA |
| 52010_1 | TATGAGTGTGGAGTCGTTGCCTAAACCTCCGCAGAGCTGTCNNNAGATCGGAAGAG |
| 52010_2 | TATGAGTGTGGAGTCGTTGCCTAAACCTCCGCAGAGCTGTCNNNAGATCGGAAGAG |
| 36157_1 | TATGAGTGTGGAGTCGTTGCAATCCTCAGTAGCCACTTGTGAGNNNAGATCGGAAG |
| 36157_2 | TATGAGTGTGGAGTCGTTGCAATCCTCAGTAGCCACTTGTGAGNNNAGATCGGAAG |
| 42642_1 | TATGAGTGTGGAGTCGTTGCATTGGAAACACGACCGCTAAAAGNNNAGATCGGAA |
| 42642_2 | TATGAGTGTGGAGTCGTTGCATTGGAAACACGACCGCTAAAAGNNNAGATCGGAA |
| 38124_1 | TATGAGTGTGGAGTCGTTGCTGAGGCAGGAGAATCTCTTGAACNNNAGATCGGAA |
| 38124_2 | TATGAGTGTGGAGTCGTTGCTGAGGCAGGAGAATCTCTTGAACNNNAGATCGGAA |
| 36005_1 | TATGAGTGTGGAGTCGTTGCGCGACAAAGCAAGACTCTGTATCNNNAGATCGGAA |
| 36005_2 | TATGAGTGTGGAGTCGTTGCGCGACAAAGCAAGACTCTGTATCNNNAGATCGGAA |
| 13004_1 | TATGAGTGTGGAGTCGTTGCAATTTCTGAGGAGTGGTTTTATANNNAGATCGGAA |

| | |
|---|---|
| 13004_2 | TATGAGTGTGGAGTCGTTGCAATTTCTGAGGAGTGGTTTTATANNNAGATCGGAAG |
| 38208_1 | TATGAGTGTGGAGTCGTTGCGAGGCAGAAGAATCACTTGAACCNNNAGATCGGAA |
| 38208_2 | TATGAGTGTGGAGTCGTTGCGAGGCAGAAGAATCACTTGAACCNNNAGATCGGAA |
| 33845_1 | TATGAGTGTGGAGTCGTTGCAGAGATTGAATGCTACAGGAAGCANNNAGATCGGA |
| 33845_2 | TATGAGTGTGGAGTCGTTGCAGAGATTGAATGCTACAGGAAGCANNNAGATCGGA |
| 29718_1 | TATGAGTGTGGAGTCGTTGCATGATGATTGTCCAGCAGGAGATNNNAGATCGGAA |
| 29718_2 | TATGAGTGTGGAGTCGTTGCATGATGATTGTCCAGCAGGAGATNNNAGATCGGAA |
| 49128_1 | TATGAGTGTGGAGTCGTTGCTGCTCTTTGGTTGGCTTTATAGCNNNAGATCGGAAG |
| 49128_2 | TATGAGTGTGGAGTCGTTGCTGCTCTTTGGTTGGCTTTATAGCNNNAGATCGGAAG |
| 32345_1 | TATGAGTGTGGAGTCGTTGCAGACAGGGTCTCATTTTGTCACCNNNAGATCGGAAG |
| 32345_2 | TATGAGTGTGGAGTCGTTGCAGACAGGGTCTCATTTTGTCACCNNNAGATCGGAAG |
| 46850_1 | TATGAGTGTGGAGTCGTTGCGAGAGGGGGAACCTAGGAGATTANNNAGATCGGA |
| 46850_2 | TATGAGTGTGGAGTCGTTGCGAGAGGGGGAACCTAGGAGATTANNNAGATCGGA |
| 42287_1 | TATGAGTGTGGAGTCGTTGCTTCCATGTGCTTTCTGGGCATTTNNNAGATCGGAAG |
| 42287_2 | TATGAGTGTGGAGTCGTTGCTTCCATGTGCTTTCTGGGCATTTNNNAGATCGGAAG |
| 32260_1 | TATGAGTGTGGAGTCGTTGCCCCCTTCCTGCTACATTCTGAAANNNAGATCGGAAGA |
| 32260_2 | TATGAGTGTGGAGTCGTTGCCCCCTTCCTGCTACATTCTGAAANNNAGATCGGAAGA |
| 37554_1 | TATGAGTGTGGAGTCGTTGCTCAACAAGCCTTAGACAGCATCANNNAGATCGGAAG |
| 37554_2 | TATGAGTGTGGAGTCGTTGCTCAACAAGCCTTAGACAGCATCANNNAGATCGGAAG |
| 49554_1 | TATGAGTGTGGAGTCGTTGCAGGTTCAAGCAATTCTCCTGTCTNNNAGATCGGAAG |
| 49554_2 | TATGAGTGTGGAGTCGTTGCAGGTTCAAGCAATTCTCCTGTCTNNNAGATCGGAAG |
| 10238_1 | TATGAGTGTGGAGTCGTTGCCTGCGGGAAATACATTTAGCCCTNNNAGATCGGAAG |
| 10238_2 | TATGAGTGTGGAGTCGTTGCCTGCGGGAAATACATTTAGCCCTNNNAGATCGGAAG |
| 8360_1 | TATGAGTGTGGAGTCGTTGCTTTGTGTTGATTGCTTTCCAGACANNNAGATCGGAA |
| 8360_2 | TATGAGTGTGGAGTCGTTGCTTTGTGTTGATTGCTTTCCAGACANNNAGATCGGAA |
| 38858_1 | TATGAGTGTGGAGTCGTTGCTACAAAAATTAGCTGGGCGTGGTNNNAGATCGGAA |
| 38858_2 | TATGAGTGTGGAGTCGTTGCTACAAAAATTAGCTGGGCGTGGTNNNAGATCGGAA |
| 52672_1 | TATGAGTGTGGAGTCGTTGCCTCTGCACCATGGTTTCAAAACANNNAGATCGGAAG |
| 52672_2 | TATGAGTGTGGAGTCGTTGCCTCTGCACCATGGTTTCAAAACANNNAGATCGGAAG |
| 53172_1 | TATGAGTGTGGAGTCGTTGCGCCTCAATTGTAGCACCACTTTTNNNAGATCGGAAG |
| 53172_2 | TATGAGTGTGGAGTCGTTGCGCCTCAATTGTAGCACCACTTTTNNNAGATCGGAAG |
| 36621_1 | TATGAGTGTGGAGTCGTTGCTTTCATCACCCTAGAAAGTGGCCNNNAGATCGGAAG |
| 36621_2 | TATGAGTGTGGAGTCGTTGCTTTCATCACCCTAGAAAGTGGCCNNNAGATCGGAAG |
| 38038_1 | TATGAGTGTGGAGTCGTTGCTTCTACTACAAACACCACCCTGGNNNAGATCGGAAG |
| 38038_2 | TATGAGTGTGGAGTCGTTGCTTCTACTACAAACACCACCCTGGNNNAGATCGGAAG |
| 33480_1 | TATGAGTGTGGAGTCGTTGCACACACACAACCAGAGTACAAGANNNAGATCGGAA |
| 33480_2 | TATGAGTGTGGAGTCGTTGCACACACACAACCAGAGTACAAGANNNAGATCGGAA |
| 30064_1 | TATGAGTGTGGAGTCGTTGCTGAGAAGCAACATGTAAGACACTNNNAGATCGGAA |
| 30064_2 | TATGAGTGTGGAGTCGTTGCTGAGAAGCAACATGTAAGACACTNNNAGATCGGAA |
| 45234_1 | TATGAGTGTGGAGTCGTTGCGTCACCCTGTACAGTTTCAGCTANNNAGATCGGAAG |
| 45234_2 | TATGAGTGTGGAGTCGTTGCGTCACCCTGTACAGTTTCAGCTANNNAGATCGGAAG |
| 37076_1 | TATGAGTGTGGAGTCGTTGCCAGAGGAGTCACTTGAACCTGGNNNAGATCGGAAG |
| 37076_2 | TATGAGTGTGGAGTCGTTGCCAGAGGAGTCACTTGAACCTGGNNNAGATCGGAAG |

49071_1 TATGAGTGTGGAGTCGTTGCCCTGGATGAGAGATCCTTACTGCNNNAGATCGGAAG
49071_2 TATGAGTGTGGAGTCGTTGCCCTGGATGAGAGATCCTTACTGCNNNAGATCGGAAG
43207_1 TATGAGTGTGGAGTCGTTGCCCCTTCATCAGTTTCCAGCAATCNNNAGATCGGAAGA
43207_2 TATGAGTGTGGAGTCGTTGCCCCTTCATCAGTTTCCAGCAATCNNNAGATCGGAAGA
42154_1 TATGAGTGTGGAGTCGTTGCCTGAGTAGCTGGGATTACAGGTGNNNAGATCGGAA
42154_2 TATGAGTGTGGAGTCGTTGCCTGAGTAGCTGGGATTACAGGTGNNNAGATCGGAA
47853_1 TATGAGTGTGGAGTCGTTGCATAGATCCTGGCTGCAATAAGGGNNNAGATCGGAA
47853_2 TATGAGTGTGGAGTCGTTGCATAGATCCTGGCTGCAATAAGGGNNNAGATCGGAA
39210_1 TATGAGTGTGGAGTCGTTGCAAACACTGAAGTCTAGTTGACCANNNAGATCGGAAG
39210_2 TATGAGTGTGGAGTCGTTGCAAACACTGAAGTCTAGTTGACCANNNAGATCGGAAG
40267_1 TATGAGTGTGGAGTCGTTGCTGAGCCAAAAGTGTTAGTTGTGCNNNAGATCGGAA
40267_2 TATGAGTGTGGAGTCGTTGCTGAGCCAAAAGTGTTAGTTGTGCNNNAGATCGGAA
37039_1 TATGAGTGTGGAGTCGTTGCAGGGGTGGTGTTAGGATTCAATTNNNAGATCGGAA
37039_2 TATGAGTGTGGAGTCGTTGCAGGGGTGGTGTTAGGATTCAATTNNNAGATCGGAA
37590_1 TATGAGTGTGGAGTCGTTGCTCCAGGTCAAGACTATGTTGGTGNNNAGATCGGAA
37590_2 TATGAGTGTGGAGTCGTTGCTCCAGGTCAAGACTATGTTGGTGNNNAGATCGGAA
32138_1 TATGAGTGTGGAGTCGTTGCCATGATCTTGGCTCACTGCAAACNNNAGATCGGAAG
32138_2 TATGAGTGTGGAGTCGTTGCCATGATCTTGGCTCACTGCAAACNNNAGATCGGAAG
37402_1 TATGAGTGTGGAGTCGTTGCCAGTCAGACGGCTCATATGGTAGNNNAGATCGGAA
37402_2 TATGAGTGTGGAGTCGTTGCCAGTCAGACGGCTCATATGGTAGNNNAGATCGGAA
51607_1 TATGAGTGTGGAGTCGTTGCCTCAGCTGTGGTGCCATCATNNNAGATCGGAAGAG
51607_2 TATGAGTGTGGAGTCGTTGCCTCAGCTGTGGTGCCATCATNNNAGATCGGAAGAG
35287_1 TATGAGTGTGGAGTCGTTGCCTCGGGTTTATGCCATTCTCGAGNNNAGATCGGAAG
35287_2 TATGAGTGTGGAGTCGTTGCCTCGGGTTTATGCCATTCTCGAGNNNAGATCGGAAG
44134_1 TATGAGTGTGGAGTCGTTGCAACCCAAAGTATGCGAGTCATCTNNNAGATCGGAAG
44134_2 TATGAGTGTGGAGTCGTTGCAACCCAAAGTATGCGAGTCATCTNNNAGATCGGAAG
31346_1 TATGAGTGTGGAGTCGTTGCGTGCTGAGGTTGGGAAAATTTGANNNAGATCGGAA
31346_2 TATGAGTGTGGAGTCGTTGCGTGCTGAGGTTGGGAAAATTTGANNNAGATCGGAA
12558_1 TATGAGTGTGGAGTCGTTGCTGTGTAAAAAAAATATGTCCATGNNNAGATCGGAA
12558_2 TATGAGTGTGGAGTCGTTGCTGTGTAAAAAAAATATGTCCATGNNNAGATCGGAA
34798_1 TATGAGTGTGGAGTCGTTGCCAGCTGGGATAACAGGTGCATNNNAGATCGGAAGA
34798_2 TATGAGTGTGGAGTCGTTGCCAGCTGGGATAACAGGTGCATNNNAGATCGGAAGA
40749_1 TATGAGTGTGGAGTCGTTGCAAGCCTCCAGTACAGCAGTATTCNNNAGATCGGAAG
40749_2 TATGAGTGTGGAGTCGTTGCAAGCCTCCAGTACAGCAGTATTCNNNAGATCGGAAG
35778_1 TATGAGTGTGGAGTCGTTGCGCTGGAGTGTAGTGATGTGATCTNNNAGATCGGAA
35778_2 TATGAGTGTGGAGTCGTTGCGCTGGAGTGTAGTGATGTGATCTNNNAGATCGGAA
34993_1 TATGAGTGTGGAGTCGTTGCTCAGAGTTCACTGTAGCCTCAACNNNAGATCGGAAG
34993_2 TATGAGTGTGGAGTCGTTGCTCAGAGTTCACTGTAGCCTCAACNNNAGATCGGAAG
53102_1 TATGAGTGTGGAGTCGTTGCGTGTAATCCCAGCTACTCGAGAGNNNAGATCGGAA
53102_2 TATGAGTGTGGAGTCGTTGCGTGTAATCCCAGCTACTCGAGAGNNNAGATCGGAA
32960_1 TATGAGTGTGGAGTCGTTGCGCCAATATGCCTGATGAACATCGNNNAGATCGGAAG
32960_2 TATGAGTGTGGAGTCGTTGCGCCAATATGCCTGATGAACATCGNNNAGATCGGAAG
29809_1 TATGAGTGTGGAGTCGTTGCTGATGGTGGGGGGAACATTTATTNNNAGATCGGAA

| | |
|---|---|
| 29809_2 | TATGAGTGTGGAGTCGTTGCTGATGGTGGGGGGAACATTTATTNNNAGATCGGAA |
| 51809_1 | TATGAGTGTGGAGTCGTTGCATCCTAAGTTGTTGACCTGACCCNNNAGATCGGAAG |
| 51809_2 | TATGAGTGTGGAGTCGTTGCATCCTAAGTTGTTGACCTGACCCNNNAGATCGGAAG |
| 52805_1 | TATGAGTGTGGAGTCGTTGCGTCATCACATCTGCAGATACCCTNNNAGATCGGAAG |
| 52805_2 | TATGAGTGTGGAGTCGTTGCGTCATCACATCTGCAGATACCCTNNNAGATCGGAAG |
| 33482_1 | TATGAGTGTGGAGTCGTTGCTAATTCCAGCATTTTGCAAGGCTNNNAGATCGGAAG |
| 33482_2 | TATGAGTGTGGAGTCGTTGCTAATTCCAGCATTTTGCAAGGCTNNNAGATCGGAAG |
| 36248_1 | TATGAGTGTGGAGTCGTTGCCAGGAGAAGGTCAGTGAGAAAGANNNAGATCGGA |
| 36248_2 | TATGAGTGTGGAGTCGTTGCCAGGAGAAGGTCAGTGAGAAAGANNNAGATCGGA |
| 10170_1 | TATGAGTGTGGAGTCGTTGCCGCTTCTGTGTGTCTAGAAATGTNNNAGATCGGAAG |
| 10170_2 | TATGAGTGTGGAGTCGTTGCCGCTTCTGTGTGTCTAGAAATGTNNNAGATCGGAAG |
| 29577_1 | TATGAGTGTGGAGTCGTTGCGGGAAGAAAGCATGGGCAATAAANNNAGATCGGA |
| 29577_2 | TATGAGTGTGGAGTCGTTGCGGGAAGAAAGCATGGGCAATAAANNNAGATCGGA |
| 12983_1 | TATGAGTGTGGAGTCGTTGCACCATTGTCACAATGTATATGANNNAGATCGGAAGA |
| 12983_2 | TATGAGTGTGGAGTCGTTGCACCATTGTCACAATGTATATGANNNAGATCGGAAGA |
| 37720_1 | TATGAGTGTGGAGTCGTTGCTTGTCCTCAACTGGGTGATAAGCNNNAGATCGGAAG |
| 37720_2 | TATGAGTGTGGAGTCGTTGCTTGTCCTCAACTGGGTGATAAGCNNNAGATCGGAAG |
| 13736_1 | TATGAGTGTGGAGTCGTTGCTCCACATTTTAAAAACTAAGCTTNNNAGATCGGAAG |
| 13736_2 | TATGAGTGTGGAGTCGTTGCTCCACATTTTAAAAACTAAGCTTNNNAGATCGGAAG |
| 53004_1 | TATGAGTGTGGAGTCGTTGCATGAAGTTTTGCTCTTCTTGCCCNNNAGATCGGAAG |
| 53004_2 | TATGAGTGTGGAGTCGTTGCATGAAGTTTTGCTCTTCTTGCCCNNNAGATCGGAAG |
| 47964_1 | TATGAGTGTGGAGTCGTTGCCCAGACAAGGTCTCATTCTGTCANNNAGATCGGAAG |
| 47964_2 | TATGAGTGTGGAGTCGTTGCCCAGACAAGGTCTCATTCTGTCANNNAGATCGGAAG |
| 43827_1 | TATGAGTGTGGAGTCGTTGCTGGGTTGCAGGAGAGAATTCTTTNNNAGATCGGAA |
| 43827_2 | TATGAGTGTGGAGTCGTTGCTGGGTTGCAGGAGAGAATTCTTTNNNAGATCGGAA |
| 31302_1 | TATGAGTGTGGAGTCGTTGCCCCCGAGATGAAACACTGAGTAANNNAGATCGGAA |
| 31302_2 | TATGAGTGTGGAGTCGTTGCCCCCGAGATGAAACACTGAGTAANNNAGATCGGAA |
| 52058_1 | TATGAGTGTGGAGTCGTTGCGAGTGAGAATATGCGGTGTTTGGNNNAGATCGGAA |
| 52058_2 | TATGAGTGTGGAGTCGTTGCGAGTGAGAATATGCGGTGTTTGGNNNAGATCGGAA |
| 49920_1 | TATGAGTGTGGAGTCGTTGCGCTTACTCTACCTTAGCTGGACCNNNAGATCGGAAG |
| 49920_2 | TATGAGTGTGGAGTCGTTGCGCTTACTCTACCTTAGCTGGACCNNNAGATCGGAAG |
| 40434_1 | TATGAGTGTGGAGTCGTTGCCAGAATCACGTCCTTTGCAACAANNNAGATCGGAAG |
| 40434_2 | TATGAGTGTGGAGTCGTTGCCAGAATCACGTCCTTTGCAACAANNNAGATCGGAAG |
| 52251_1 | TATGAGTGTGGAGTCGTTGCTGCAAAGGTGGTTCAACATATGCNNNAGATCGGAA |
| 52251_2 | TATGAGTGTGGAGTCGTTGCTGCAAAGGTGGTTCAACATATGCNNNAGATCGGAA |
| 29572_1 | TATGAGTGTGGAGTCGTTGCACCTTCTGCTTTTCTAGGTGTGTNNNAGATCGGAAG |
| 29572_2 | TATGAGTGTGGAGTCGTTGCACCTTCTGCTTTTCTAGGTGTGTNNNAGATCGGAAG |
| 44164_1 | TATGAGTGTGGAGTCGTTGCTATCACAGGAGGGTTCCAATTGGNNNAGATCGGAA |
| 44164_2 | TATGAGTGTGGAGTCGTTGCTATCACAGGAGGGTTCCAATTGGNNNAGATCGGAA |
| 10381_1 | TATGAGTGTGGAGTCGTTGCACAGTTCTGGCAATTAGTGACACANNNAGATCGGAA |
| 10381_2 | TATGAGTGTGGAGTCGTTGCACAGTTCTGGCAATTAGTGACACANNNAGATCGGAA |
| 42628_1 | TATGAGTGTGGAGTCGTTGCCGCGTGTGTTGGAATACTACTTANNNAGATCGGAAG |
| 42628_2 | TATGAGTGTGGAGTCGTTGCCGCGTGTGTTGGAATACTACTTANNNAGATCGGAAG |

| | |
|---|---|
| 5290_1 | TATGAGTGTGGAGTCGTTGCGTGGGTGGACATGATCGGTTTTTNNNAGATCGGAA |
| 5290_2 | TATGAGTGTGGAGTCGTTGCGTGGGTGGACATGATCGGTTTTTNNNAGATCGGAA |
| 49337_1 | TATGAGTGTGGAGTCGTTGCAGGATTGTTTTGGCTTTCTAGGTNNNAGATCGGAAG |
| 49337_2 | TATGAGTGTGGAGTCGTTGCAGGATTGTTTTGGCTTTCTAGGTNNNAGATCGGAAG |
| 49897_1 | TATGAGTGTGGAGTCGTTGCCCCCACCCCATCCAAAAAAAAAANNNAGATCGGAAG |
| 49897_2 | TATGAGTGTGGAGTCGTTGCCCCCACCCCATCCAAAAAAAAAANNNAGATCGGAAG |
| 35718_1 | TATGAGTGTGGAGTCGTTGCCCTGGACTGACTACAACCTCAAANNNAGATCGGAAG |
| 35718_2 | TATGAGTGTGGAGTCGTTGCCCTGGACTGACTACAACCTCAAANNNAGATCGGAAG |
| 38560_1 | TATGAGTGTGGAGTCGTTGCTCTGTACTCCTATGGCATCATGCNNNAGATCGGAAG |
| 38560_2 | TATGAGTGTGGAGTCGTTGCTCTGTACTCCTATGGCATCATGCNNNAGATCGGAAG |
| 49623_1 | TATGAGTGTGGAGTCGTTGCCCATGTAGCTGGTACTACAGGTGNNNAGATCGGAA |
| 49623_2 | TATGAGTGTGGAGTCGTTGCCCATGTAGCTGGTACTACAGGTGNNNAGATCGGAA |
| 13265_1 | TATGAGTGTGGAGTCGTTGCGTCAGCATAACTTGATATTTTGANNNAGATCGGAAG |
| 13265_2 | TATGAGTGTGGAGTCGTTGCGTCAGCATAACTTGATATTTTGANNNAGATCGGAAG |
| 38704_1 | TATGAGTGTGGAGTCGTTGCACAAATGGAAACACATCCCATGCNNNAGATCGGAAG |
| 38704_2 | TATGAGTGTGGAGTCGTTGCACAAATGGAAACACATCCCATGCNNNAGATCGGAAG |
| 50910_1 | TATGAGTGTGGAGTCGTTGCTTCTGTGGCCTACCTTCTGATTCNNNAGATCGGAAGA |
| 50910_2 | TATGAGTGTGGAGTCGTTGCTTCTGTGGCCTACCTTCTGATTCNNNAGATCGGAAGA |
| 36972_1 | TATGAGTGTGGAGTCGTTGCTCGCATAAAATTGGCCCTTGTTTNNNAGATCGGAAG |
| 36972_2 | TATGAGTGTGGAGTCGTTGCTCGCATAAAATTGGCCCTTGTTTNNNAGATCGGAAG |
| 35448_1 | TATGAGTGTGGAGTCGTTGCCAGGGTATAGTGCAGTGACATGANNNAGATCGGAA |
| 35448_2 | TATGAGTGTGGAGTCGTTGCCAGGGTATAGTGCAGTGACATGANNNAGATCGGAA |
| 45248_1 | TATGAGTGTGGAGTCGTTGCAAGGACATGCTACCCCCTAAAAGNNNAGATCGGAAG |
| 45248_2 | TATGAGTGTGGAGTCGTTGCAAGGACATGCTACCCCCTAAAAGNNNAGATCGGAAG |
| 35583_1 | TATGAGTGTGGAGTCGTTGCCAGTCTCAGCTCACTACATCCTCNNNAGATCGGAAGA |
| 35583_2 | TATGAGTGTGGAGTCGTTGCCAGTCTCAGCTCACTACATCCTCNNNAGATCGGAAGA |
| 6824_1 | TATGAGTGTGGAGTCGTTGCTGCTTGTTGCCAATTGAATTCCTNNNAGATCGGAAG |
| 6824_2 | TATGAGTGTGGAGTCGTTGCTGCTTGTTGCCAATTGAATTCCTNNNAGATCGGAAG |
| 31595_1 | TATGAGTGTGGAGTCGTTGCCGACTTGCCTACTGGAAAACAAGNNNAGATCGGAA |
| 31595_2 | TATGAGTGTGGAGTCGTTGCCGACTTGCCTACTGGAAAACAAGNNNAGATCGGAA |
| 13625_1 | TATGAGTGTGGAGTCGTTGCGCAAAGTTTATTTTCTCTTTCTTNNNAGATCGGAAGA |
| 13625_2 | TATGAGTGTGGAGTCGTTGCGCAAAGTTTATTTTCTCTTTCTTNNNAGATCGGAAGA |
| 34398_1 | TATGAGTGTGGAGTCGTTGCGGGTGAATAATCAAGCAGGCATCNNNAGATCGGAA |
| 34398_2 | TATGAGTGTGGAGTCGTTGCGGGTGAATAATCAAGCAGGCATCNNNAGATCGGAA |
| 32270_1 | TATGAGTGTGGAGTCGTTGCGTCAGTTTGCAGACAACAGAGTCNNNAGATCGGAA |
| 32270_2 | TATGAGTGTGGAGTCGTTGCGTCAGTTTGCAGACAACAGAGTCNNNAGATCGGAA |
| 13405_1 | TATGAGTGTGGAGTCGTTGCAGAACTCACAGATTCTTTCTTATNNNAGATCGGAAG |
| 13405_2 | TATGAGTGTGGAGTCGTTGCAGAACTCACAGATTCTTTCTTATNNNAGATCGGAAG |
| 47471_1 | TATGAGTGTGGAGTCGTTGCTTCAAGTGATTCTCCTGCCTCAGNNNAGATCGGAAG |
| 47471_2 | TATGAGTGTGGAGTCGTTGCTTCAAGTGATTCTCCTGCCTCAGNNNAGATCGGAAG |
| 41218_1 | TATGAGTGTGGAGTCGTTGCAAGAAAATTAGCTGGGCATGGTGNNNAGATCGGAA |
| 41218_2 | TATGAGTGTGGAGTCGTTGCAAGAAAATTAGCTGGGCATGGTGNNNAGATCGGAA |
| 48231_1 | TATGAGTGTGGAGTCGTTGCCGTGCCCAGCTGATTTTTTTTTNNNAGATCGGAAGA |

| | |
|---|---|
| 48231_2 | TATGAGTGTGGAGTCGTTGCCGTGCCCAGCTGATTTTTTTTTTNNNAGATCGGAAGA |
| 12626_1 | TATGAGTGTGGAGTCGTTGCACATAGGAGAAGACGATCAGNNNAGATCGGAAGAG |
| 12626_2 | TATGAGTGTGGAGTCGTTGCACATAGGAGAAGACGATCAGNNNAGATCGGAAGAG |
| 12970_1 | TATGAGTGTGGAGTCGTTGCCAAAGTCCCATAGAACACTATGNNNAGATCGGAAGA |
| 12970_2 | TATGAGTGTGGAGTCGTTGCCAAAGTCCCATAGAACACTATGNNNAGATCGGAAGA |
| 41844_1 | TATGAGTGTGGAGTCGTTGCATCCCTGGGTTCAAGTGATTCTCNNNAGATCGGAAGA |
| 41844_2 | TATGAGTGTGGAGTCGTTGCATCCCTGGGTTCAAGTGATTCTCNNNAGATCGGAAGA |
| 46544_1 | TATGAGTGTGGAGTCGTTGCGTAGAGACAGGGTTTCACTGTGTNNNAGATCGGAA |
| 46544_2 | TATGAGTGTGGAGTCGTTGCGTAGAGACAGGGTTTCACTGTGTNNNAGATCGGAA |
| 50187_1 | TATGAGTGTGGAGTCGTTGCTCTCATTACCATCACACCTGCAANNNAGATCGGAAGA |
| 50187_2 | TATGAGTGTGGAGTCGTTGCTCTCATTACCATCACACCTGCAANNNAGATCGGAAGA |
| 31714_1 | TATGAGTGTGGAGTCGTTGCTCAGAGGAAATGAGGAAATGGGGNNNAGATCGGA |
| 31714_2 | TATGAGTGTGGAGTCGTTGCTCAGAGGAAATGAGGAAATGGGGNNNAGATCGGA |
| 32190_1 | TATGAGTGTGGAGTCGTTGCCTCCCAGGTTCATGCCATTCTCNNNAGATCGGAAGAG |
| 32190_2 | TATGAGTGTGGAGTCGTTGCCTCCCAGGTTCATGCCATTCTCNNNAGATCGGAAGAG |
| 50253_1 | TATGAGTGTGGAGTCGTTGCGGAGGGGGTATAACACATAGCTGNNNAGATCGGAA |
| 50253_2 | TATGAGTGTGGAGTCGTTGCGGAGGGGGTATAACACATAGCTGNNNAGATCGGAA |
| 44527_1 | TATGAGTGTGGAGTCGTTGCAAAGTGAGGTGCTAAGAGTCAGGNNNAGATCGGAA |
| 44527_2 | TATGAGTGTGGAGTCGTTGCAAAGTGAGGTGCTAAGAGTCAGGNNNAGATCGGAA |
| 49360_1 | TATGAGTGTGGAGTCGTTGCTGCTGGGATTATAGAAGTGAGCCNNNAGATCGGAA |
| 49360_2 | TATGAGTGTGGAGTCGTTGCTGCTGGGATTATAGAAGTGAGCCNNNAGATCGGAA |
| 7839_1 | TATGAGTGTGGAGTCGTTGCCTGGCTGGAGAGCACGATTTTNNNAGATCGGAAGA |
| 7839_2 | TATGAGTGTGGAGTCGTTGCCTGGCTGGAGAGCACGATTTTNNNAGATCGGAAGA |
| 41340_1 | TATGAGTGTGGAGTCGTTGCTTTTGAGAGCTTAGAATTGCCCGNNNAGATCGGAAG |
| 41340_2 | TATGAGTGTGGAGTCGTTGCTTTTGAGAGCTTAGAATTGCCCGNNNAGATCGGAAG |
| 39051_1 | TATGAGTGTGGAGTCGTTGCTTCACAGAACTCAGGAAAGTGCTNNNAGATCGGAAG |
| 39051_2 | TATGAGTGTGGAGTCGTTGCTTCACAGAACTCAGGAAAGTGCTNNNAGATCGGAAG |
| 51074_1 | TATGAGTGTGGAGTCGTTGCCGTGTATAGGCTTATGGGGACAANNNAGATCGGAA |
| 51074_2 | TATGAGTGTGGAGTCGTTGCCGTGTATAGGCTTATGGGGACAANNNAGATCGGAA |
| 41966_1 | TATGAGTGTGGAGTCGTTGCCGAGTAGCTGGGACTACAGGTNNNAGATCGGAAGA |
| 41966_2 | TATGAGTGTGGAGTCGTTGCCGAGTAGCTGGGACTACAGGTNNNAGATCGGAAGA |
| 32305_1 | TATGAGTGTGGAGTCGTTGCGTTCAAGAGATTCTCCTGCCTCANNNAGATCGGAAGA |
| 32305_2 | TATGAGTGTGGAGTCGTTGCGTTCAAGAGATTCTCCTGCCTCANNNAGATCGGAAGA |
| 32037_1 | TATGAGTGTGGAGTCGTTGCTCCAAATGACACTTTGTTGCTGTNNNAGATCGGAAGA |
| 32037_2 | TATGAGTGTGGAGTCGTTGCTCCAAATGACACTTTGTTGCTGTNNNAGATCGGAAGA |
| 48330_1 | TATGAGTGTGGAGTCGTTGCTTCAAGTGATTCTCCTGCCTCAANNNAGATCGGAAGA |
| 48330_2 | TATGAGTGTGGAGTCGTTGCTTCAAGTGATTCTCCTGCCTCAANNNAGATCGGAAGA |
| 48826_1 | TATGAGTGTGGAGTCGTTGCCAATCTTGGCTCACTTCAAGCTCNNNAGATCGGAAGA |
| 48826_2 | TATGAGTGTGGAGTCGTTGCCAATCTTGGCTCACTTCAAGCTCNNNAGATCGGAAGA |
| 2399_1 | TATGAGTGTGGAGTCGTTGCCCATCTCTCTCCTGCTGTAATGCNNNAGATCGGAAGA |
| 2399_2 | TATGAGTGTGGAGTCGTTGCCCATCTCTCTCCTGCTGTAATGCNNNAGATCGGAAGA |
| 48980_1 | TATGAGTGTGGAGTCGTTGCCACTTCAGCCTCTTTCTGGGATANNNAGATCGGAAGA |
| 48980_2 | TATGAGTGTGGAGTCGTTGCCACTTCAGCCTCTTTCTGGGATANNNAGATCGGAAGA |

| | |
|---|---|
| 41866_1 | TATGAGTGTGGAGTCGTTGCAACCTCCGCCTTCTAGATTCAAGNNNAGATCGGAAG |
| 41866_2 | TATGAGTGTGGAGTCGTTGCAACCTCCGCCTTCTAGATTCAAGNNNAGATCGGAAG |
| 44421_1 | TATGAGTGTGGAGTCGTTGCATTGTGGAACGATGATTCTGGGANNNAGATCGGAA |
| 44421_2 | TATGAGTGTGGAGTCGTTGCATTGTGGAACGATGATTCTGGGANNNAGATCGGAA |
| 41995_1 | TATGAGTGTGGAGTCGTTGCACGATCTCAGCTCACTGCAAGNNNAGATCGGAAGAG |
| 41995_2 | TATGAGTGTGGAGTCGTTGCACGATCTCAGCTCACTGCAAGNNNAGATCGGAAGAG |
| 35951_1 | TATGAGTGTGGAGTCGTTGCGAAAGTGTGCACATTCTTTGGCNNNAGATCGGAAG |
| 35951_2 | TATGAGTGTGGAGTCGTTGCGAAAGTGTGCACATTCTTTGGCNNNAGATCGGAAG |
| 28917_1 | TATGAGTGTGGAGTCGTTGCGCTCTCAGGAGGGAACAATACAANNNAGATCGGAA |
| 28917_2 | TATGAGTGTGGAGTCGTTGCGCTCTCAGGAGGGAACAATACAANNNAGATCGGAA |
| 35268_1 | TATGAGTGTGGAGTCGTTGCAAGATGATATTTGAGAGCCCCCCNNNAGATCGGAAG |
| 35268_2 | TATGAGTGTGGAGTCGTTGCAAGATGATATTTGAGAGCCCCCCNNNAGATCGGAAG |
| 36955_1 | TATGAGTGTGGAGTCGTTGCGCATCTATAGTGTAGCACAGTGCNNNAGATCGGAAG |
| 36955_2 | TATGAGTGTGGAGTCGTTGCGCATCTATAGTGTAGCACAGTGCNNNAGATCGGAAG |
| 35231_1 | TATGAGTGTGGAGTCGTTGCCCCTCACCAGATTGAATCCTCAANNNAGATCGGAAG |
| 35231_2 | TATGAGTGTGGAGTCGTTGCCCCTCACCAGATTGAATCCTCAANNNAGATCGGAAG |
| 53405_1 | TATGAGTGTGGAGTCGTTGCCCCTGACTTTCTCATTATGGGCTNNNAGATCGGAAG |
| 53405_2 | TATGAGTGTGGAGTCGTTGCCCCTGACTTTCTCATTATGGGCTNNNAGATCGGAAG |
| 45479_1 | TATGAGTGTGGAGTCGTTGCATTGTGCCTCTAGGACCTAGTCTNNNAGATCGGAAG |
| 45479_2 | TATGAGTGTGGAGTCGTTGCATTGTGCCTCTAGGACCTAGTCTNNNAGATCGGAAG |
| 37999_1 | TATGAGTGTGGAGTCGTTGCCTGTTCAGTTCCCAGGTAAAGGANNNAGATCGGAAG |
| 37999_2 | TATGAGTGTGGAGTCGTTGCCTGTTCAGTTCCCAGGTAAAGGANNNAGATCGGAAG |
| 38033_1 | TATGAGTGTGGAGTCGTTGCACAAAACAAAAAACAGGTTGCGGNNNAGATCGGAA |
| 38033_2 | TATGAGTGTGGAGTCGTTGCACAAAACAAAAAACAGGTTGCGGNNNAGATCGGAA |
| 45800_1 | TATGAGTGTGGAGTCGTTGCGATCTCGGCTCACTGCAATCTCNNNAGATCGGAAGA |
| 45800_2 | TATGAGTGTGGAGTCGTTGCGATCTCGGCTCACTGCAATCTCNNNAGATCGGAAGA |
| 40104_1 | TATGAGTGTGGAGTCGTTGCTGTTTGGTATACCTCCTAGGGGTNNNAGATCGGAAG |
| 40104_2 | TATGAGTGTGGAGTCGTTGCTGTTTGGTATACCTCCTAGGGGTNNNAGATCGGAAG |
| 33776_1 | TATGAGTGTGGAGTCGTTGCAAACCTGGCACAGAAAGACAAACNNNAGATCGGAA |
| 33776_2 | TATGAGTGTGGAGTCGTTGCAAACCTGGCACAGAAAGACAAACNNNAGATCGGAA |
| 29406_1 | TATGAGTGTGGAGTCGTTGCCCACCTCTTACCGATTTCTTCCANNNAGATCGGAAGA |
| 29406_2 | TATGAGTGTGGAGTCGTTGCCCACCTCTTACCGATTTCTTCCANNNAGATCGGAAGA |
| 38330_1 | TATGAGTGTGGAGTCGTTGCGTGCATACTTGTAGACCCAGTCANNNAGATCGGAAG |
| 38330_2 | TATGAGTGTGGAGTCGTTGCGTGCATACTTGTAGACCCAGTCANNNAGATCGGAAG |
| 42573_1 | TATGAGTGTGGAGTCGTTGCCCACTTGTTGATTGATGGGCATTNNNAGATCGGAAG |
| 42573_2 | TATGAGTGTGGAGTCGTTGCCCACTTGTTGATTGATGGGCATTNNNAGATCGGAAG |
| 50235_1 | TATGAGTGTGGAGTCGTTGCGGGCCTTCTTCATCCAGTTTTTCNNNAGATCGGAAGA |
| 50235_2 | TATGAGTGTGGAGTCGTTGCGGGCCTTCTTCATCCAGTTTTTCNNNAGATCGGAAGA |
| 36572_1 | TATGAGTGTGGAGTCGTTGCGATCACGAGGTCAGGAGTTCAAGNNNAGATCGGAA |
| 36572_2 | TATGAGTGTGGAGTCGTTGCGATCACGAGGTCAGGAGTTCAAGNNNAGATCGGAA |
| 34688_1 | TATGAGTGTGGAGTCGTTGCCAGGTTCAAGCAATTCTCCTGCNNNAGATCGGAAGA |
| 34688_2 | TATGAGTGTGGAGTCGTTGCCAGGTTCAAGCAATTCTCCTGCNNNAGATCGGAAGA |
| 43797_1 | TATGAGTGTGGAGTCGTTGCACTTGCTGTACAAATGGCATCTTNNNAGATCGGAAG |

| | |
|---|---|
| 43797_2 | TATGAGTGTGGAGTCGTTGCACTTGCTGTACAAATGGCATCTTNNNAGATCGGAAG |
| 38486_1 | TATGAGTGTGGAGTCGTTGCCAGAATAACCTGGGGCCTCTATCNNNAGATCGGAAG |
| 38486_2 | TATGAGTGTGGAGTCGTTGCCAGAATAACCTGGGGCCTCTATCNNNAGATCGGAAG |
| 51364_1 | TATGAGTGTGGAGTCGTTGCCCCTAGGTGCACATTCTCTTTCTNNNAGATCGGAAGA |
| 51364_2 | TATGAGTGTGGAGTCGTTGCCCCTAGGTGCACATTCTCTTTCTNNNAGATCGGAAGA |
| 29836_1 | TATGAGTGTGGAGTCGTTGCAGTAATTTGTTCCCTCTCCCAGGNNNAGATCGGAAG |
| 29836_2 | TATGAGTGTGGAGTCGTTGCAGTAATTTGTTCCCTCTCCCAGGNNNAGATCGGAAG |
| 35992_1 | TATGAGTGTGGAGTCGTTGCTCCCACATTGGAAAAGCAGTACTNNNAGATCGGAAG |
| 35992_2 | TATGAGTGTGGAGTCGTTGCTCCCACATTGGAAAAGCAGTACTNNNAGATCGGAAG |
| 33868_1 | TATGAGTGTGGAGTCGTTGCTTCCATGACAGCACATACAGACTNNNAGATCGGAAG |
| 33868_2 | TATGAGTGTGGAGTCGTTGCTTCCATGACAGCACATACAGACTNNNAGATCGGAAG |
| 30645_1 | TATGAGTGTGGAGTCGTTGCGCAGTCACTCTCTCTTTCTGCTTNNNAGATCGGAAGA |
| 30645_2 | TATGAGTGTGGAGTCGTTGCGCAGTCACTCTCTCTTTCTGCTTNNNAGATCGGAAGA |
| 10082_1 | TATGAGTGTGGAGTCGTTGCCCTCTCACCCCCTTATCTCTCTGNNNAGATCGGAAGA |
| 10082_2 | TATGAGTGTGGAGTCGTTGCCCTCTCACCCCCTTATCTCTCTGNNNAGATCGGAAGA |
| 36243_1 | TATGAGTGTGGAGTCGTTGCATGTAGAAGGGAAGTCAGCACAGNNNAGATCGGA |
| 36243_2 | TATGAGTGTGGAGTCGTTGCATGTAGAAGGGAAGTCAGCACAGNNNAGATCGGA |
| 33976_1 | TATGAGTGTGGAGTCGTTGCCCAAGTAACTGGGACTACAGGCNNNAGATCGGAAG |
| 33976_2 | TATGAGTGTGGAGTCGTTGCCCAAGTAACTGGGACTACAGGCNNNAGATCGGAAG |
| 46543_1 | TATGAGTGTGGAGTCGTTGCAGGAGGATGTGTACTGTCTCAGANNNAGATCGGAA |
| 46543_2 | TATGAGTGTGGAGTCGTTGCAGGAGGATGTGTACTGTCTCAGANNNAGATCGGAA |
| 32887_1 | TATGAGTGTGGAGTCGTTGCCAGTGAGCTGAGATCATGTCACTNNNAGATCGGAA |
| 32887_2 | TATGAGTGTGGAGTCGTTGCCAGTGAGCTGAGATCATGTCACTNNNAGATCGGAA |
| 12922_1 | TATGAGTGTGGAGTCGTTGCCAAGCTTGTTGAAGAAGACANNNAGATCGGAAGAG |
| 12922_2 | TATGAGTGTGGAGTCGTTGCCAAGCTTGTTGAAGAAGACANNNAGATCGGAAGAG |
| 44736_1 | TATGAGTGTGGAGTCGTTGCGCCATAGTTCCAGAGGTCATTGANNNAGATCGGAA |
| 44736_2 | TATGAGTGTGGAGTCGTTGCGCCATAGTTCCAGAGGTCATTGANNNAGATCGGAA |
| 52507_1 | TATGAGTGTGGAGTCGTTGCTGATACAGAGCTGGAGGAGAAGANNNAGATCGGA |
| 52507_2 | TATGAGTGTGGAGTCGTTGCTGATACAGAGCTGGAGGAGAAGANNNAGATCGGA |
| 53053_1 | TATGAGTGTGGAGTCGTTGCACAAGGAATACATTTGGGGTTCCNNNAGATCGGAA |
| 53053_2 | TATGAGTGTGGAGTCGTTGCACAAGGAATACATTTGGGGTTCCNNNAGATCGGAA |
| 46647_1 | TATGAGTGTGGAGTCGTTGCCCGAGTAGCTGTATTACAGGCATNNNAGATCGGAA |
| 46647_2 | TATGAGTGTGGAGTCGTTGCCCGAGTAGCTGTATTACAGGCATNNNAGATCGGAA |
| 43060_1 | TATGAGTGTGGAGTCGTTGCCAAACCTGCTCTCTTGTGAATCCNNNAGATCGGAAG |
| 43060_2 | TATGAGTGTGGAGTCGTTGCCAAACCTGCTCTCTTGTGAATCCNNNAGATCGGAAG |
| 37790_1 | TATGAGTGTGGAGTCGTTGCTAGTTCCATCCAATGAAACCCCCNNNAGATCGGAAG |
| 37790_2 | TATGAGTGTGGAGTCGTTGCTAGTTCCATCCAATGAAACCCCCNNNAGATCGGAAG |
| 48164_1 | TATGAGTGTGGAGTCGTTGCTATTAGCTGGGATTACAGGTGCCNNNAGATCGGAA |
| 48164_2 | TATGAGTGTGGAGTCGTTGCTATTAGCTGGGATTACAGGTGCCNNNAGATCGGAA |
| 4622_1 | TATGAGTGTGGAGTCGTTGCGTGATTGCCACAAAATTCACCCANNNAGATCGGAAG |
| 4622_2 | TATGAGTGTGGAGTCGTTGCGTGATTGCCACAAAATTCACCCANNNAGATCGGAAG |
| 33083_1 | TATGAGTGTGGAGTCGTTGCCTCAAGTGATCCTTCCACCTCAGNNNAGATCGGAAG |
| 33083_2 | TATGAGTGTGGAGTCGTTGCCTCAAGTGATCCTTCCACCTCAGNNNAGATCGGAAG |

| | |
|---|---|
| 38403_1 | TATGAGTGTGGAGTCGTTGCGGCTGAGGTAGAAGAATCGCTTANNNAGATCGGAA |
| 38403_2 | TATGAGTGTGGAGTCGTTGCGGCTGAGGTAGAAGAATCGCTTANNNAGATCGGAA |
| 50650_1 | TATGAGTGTGGAGTCGTTGCCCTTGATGTCCCCAGATCTTGACNNNAGATCGGAAG |
| 50650_2 | TATGAGTGTGGAGTCGTTGCCCTTGATGTCCCCAGATCTTGACNNNAGATCGGAAG |
| 13763_1 | TATGAGTGTGGAGTCGTTGCCTTCCTATTGGTTCGTTTTAGNNNAGATCGGAAGA |
| 13763_2 | TATGAGTGTGGAGTCGTTGCCTTCCTATTGGTTCGTTTTAGNNNAGATCGGAAGA |
| 47578_1 | TATGAGTGTGGAGTCGTTGCCCTCCCAGGTACAAACGATTCTCNNNAGATCGGAAG |
| 47578_2 | TATGAGTGTGGAGTCGTTGCCCTCCCAGGTACAAACGATTCTCNNNAGATCGGAAG |
| 44068_1 | TATGAGTGTGGAGTCGTTGCTGAAGGATGTGGGTCATTGTTCTNNNAGATCGGAA |
| 44068_2 | TATGAGTGTGGAGTCGTTGCTGAAGGATGTGGGTCATTGTTCTNNNAGATCGGAA |
| 45213_1 | TATGAGTGTGGAGTCGTTGCCTGGGCATTTGTGTGAAAGCATANNNAGATCGGAA |
| 45213_2 | TATGAGTGTGGAGTCGTTGCCTGGGCATTTGTGTGAAAGCATANNNAGATCGGAA |
| 52005_1 | TATGAGTGTGGAGTCGTTGCTGTTTGGTTTTCTGTCCTTGCAANNNAGATCGGAAG |
| 52005_2 | TATGAGTGTGGAGTCGTTGCTGTTTGGTTTTCTGTCCTTGCAANNNAGATCGGAAG |
| 30624_1 | TATGAGTGTGGAGTCGTTGCCGCCAATTTGTAATTACGTGTAAATTNNNAGATCGG |
| 30624_2 | TATGAGTGTGGAGTCGTTGCCGCCAATTTGTAATTACGTGTAAATTNNNAGATCGG |
| 2267_1 | TATGAGTGTGGAGTCGTTGCTCACCACACCCATGCTTTTGTTANNNAGATCGGAAGA |
| 2267_2 | TATGAGTGTGGAGTCGTTGCTCACCACACCCATGCTTTTGTTANNNAGATCGGAAGA |
| 44098_1 | TATGAGTGTGGAGTCGTTGCCTCTTTCACTGCCCACACATTCNNNAGATCGGAAGAG |
| 44098_2 | TATGAGTGTGGAGTCGTTGCCTCTTTCACTGCCCACACATTCNNNAGATCGGAAGAG |
| 38615_1 | TATGAGTGTGGAGTCGTTGCAGAAATAACAAAGATCAGGGCAGANNNAGATCGGA |
| 38615_2 | TATGAGTGTGGAGTCGTTGCAGAAATAACAAAGATCAGGGCAGANNNAGATCGGA |
| 49752_1 | TATGAGTGTGGAGTCGTTGCGGACCCCGTAGCAGCTAAGCNNNAGATCGGAAGAG |
| 49752_2 | TATGAGTGTGGAGTCGTTGCGGACCCCGTAGCAGCTAAGCNNNAGATCGGAAGAG |
| 46869_1 | TATGAGTGTGGAGTCGTTGCGAGTAGCTGGGATTACAGGCATNNNAGATCGGAAG |
| 46869_2 | TATGAGTGTGGAGTCGTTGCGAGTAGCTGGGATTACAGGCATNNNAGATCGGAAG |
| 39670_1 | TATGAGTGTGGAGTCGTTGCCCTGGATGACAGAGTGAGCAAAGNNNAGATCGGAA |
| 39670_2 | TATGAGTGTGGAGTCGTTGCCCTGGATGACAGAGTGAGCAAAGNNNAGATCGGAA |
| 42606_1 | TATGAGTGTGGAGTCGTTGCCTCTCAGCTACTCGGGACTTTACNNNAGATCGGAAG |
| 42606_2 | TATGAGTGTGGAGTCGTTGCCTCTCAGCTACTCGGGACTTTACNNNAGATCGGAAG |
| 36946_1 | TATGAGTGTGGAGTCGTTGCAAACTAAGCTGGCACTAGAAGGANNNAGATCGGAA |
| 36946_2 | TATGAGTGTGGAGTCGTTGCAAACTAAGCTGGCACTAGAAGGANNNAGATCGGAA |
| 43898_1 | TATGAGTGTGGAGTCGTTGCGACCTCCCAGGCTAACTTTTGTANNNAGATCGGAAG |
| 43898_2 | TATGAGTGTGGAGTCGTTGCGACCTCCCAGGCTAACTTTTGTANNNAGATCGGAAG |
| 50488_1 | TATGAGTGTGGAGTCGTTGCATCACATCTCCATGCCCTGAAATNNNAGATCGGAAG |
| 50488_2 | TATGAGTGTGGAGTCGTTGCATCACATCTCCATGCCCTGAAATNNNAGATCGGAAG |
| 42548_1 | TATGAGTGTGGAGTCGTTGCTCTATCAGAATGATGCTGGCCTCNNNAGATCGGAAG |
| 42548_2 | TATGAGTGTGGAGTCGTTGCTCTATCAGAATGATGCTGGCCTCNNNAGATCGGAAG |
| 34196_1 | TATGAGTGTGGAGTCGTTGCGCTACTCCAGAGACAGAGCAGNNNAGATCGGAAGA |
| 34196_2 | TATGAGTGTGGAGTCGTTGCGCTACTCCAGAGACAGAGCAGNNNAGATCGGAAGA |
| 39119_1 | TATGAGTGTGGAGTCGTTGCGGTAAATGTGTCATGGGGGTTTGNNNAGATCGGAA |
| 39119_2 | TATGAGTGTGGAGTCGTTGCGGTAAATGTGTCATGGGGGTTTGNNNAGATCGGAA |
| 29288_1 | TATGAGTGTGGAGTCGTTGCACATGCGTCTTTGCTTCTCTCTANNNAGATCGGAAGA |

| | |
|---|---|
| 29288_2 | TATGAGTGTGGAGTCGTTGCACATGCGTCTTTGCTTCTCTCTANNNAGATCGGAAGA |
| 49448_1 | TATGAGTGTGGAGTCGTTGCTTGTACTGTCTGCCATTGATTGGNNNAGATCGGAAG |
| 49448_2 | TATGAGTGTGGAGTCGTTGCTTGTACTGTCTGCCATTGATTGGNNNAGATCGGAAG |
| 3196_1 | TATGAGTGTGGAGTCGTTGCTGAGGCAGGAATCAAAGACAGATNNNAGATCGGAA |
| 3196_2 | TATGAGTGTGGAGTCGTTGCTGAGGCAGGAATCAAAGACAGATNNNAGATCGGAA |
| 47285_1 | TATGAGTGTGGAGTCGTTGCGTGATCCCAGCCTTGTAATGGTANNNAGATCGGAAG |
| 47285_2 | TATGAGTGTGGAGTCGTTGCGTGATCCCAGCCTTGTAATGGTANNNAGATCGGAAG |
| 42846_1 | TATGAGTGTGGAGTCGTTGCTGAACTCCTGAGCTCAAGCAATCNNNAGATCGGAAG |
| 42846_2 | TATGAGTGTGGAGTCGTTGCTGAACTCCTGAGCTCAAGCAATCNNNAGATCGGAAG |
| 48092_1 | TATGAGTGTGGAGTCGTTGCAGTAGCTTTCCCAGAGAGAGTCTNNNAGATCGGAAG |
| 48092_2 | TATGAGTGTGGAGTCGTTGCAGTAGCTTTCCCAGAGAGAGTCTNNNAGATCGGAAG |
| 39003_1 | TATGAGTGTGGAGTCGTTGCATGCTTATGTACAGCTTGTGGGANNNAGATCGGAAG |
| 39003_2 | TATGAGTGTGGAGTCGTTGCATGCTTATGTACAGCTTGTGGGANNNAGATCGGAAG |
| 52008_1 | TATGAGTGTGGAGTCGTTGCCAGGCCAGGAGCATAATCTCTAGNNNAGATCGGAAG |
| 52008_2 | TATGAGTGTGGAGTCGTTGCCAGGCCAGGAGCATAATCTCTAGNNNAGATCGGAAG |
| 37667_1 | TATGAGTGTGGAGTCGTTGCGACAGAGCAAGACTCCATCTCAANNNAGATCGGAAG |
| 37667_2 | TATGAGTGTGGAGTCGTTGCGACAGAGCAAGACTCCATCTCAANNNAGATCGGAAG |
| 45254_1 | TATGAGTGTGGAGTCGTTGCAATGGAAATGTTTAGTGGCGACGNNNAGATCGGAA |
| 45254_2 | TATGAGTGTGGAGTCGTTGCAATGGAAATGTTTAGTGGCGACGNNNAGATCGGAA |
| 46753_1 | TATGAGTGTGGAGTCGTTGCGCCTCCCAAGTACCTGAGATTACNNNAGATCGGAAGA |
| 46753_2 | TATGAGTGTGGAGTCGTTGCGCCTCCCAAGTACCTGAGATTACNNNAGATCGGAAGA |
| 46559_1 | TATGAGTGTGGAGTCGTTGCGATCTCAACTCACAGCAGCCTCNNNAGATCGGAAGAG |
| 46559_2 | TATGAGTGTGGAGTCGTTGCGATCTCAACTCACAGCAGCCTCNNNAGATCGGAAGAG |
| 33425_1 | TATGAGTGTGGAGTCGTTGCTCACAAAAGACCCAGAACATCCANNNAGATCGGAAG |
| 33425_2 | TATGAGTGTGGAGTCGTTGCTCACAAAAGACCCAGAACATCCANNNAGATCGGAAG |
| 41384_1 | TATGAGTGTGGAGTCGTTGCAAAAGAAGCTAGCTGAAATGCCTNNNAGATCGGAAG |
| 41384_2 | TATGAGTGTGGAGTCGTTGCAAAAGAAGCTAGCTGAAATGCCTNNNAGATCGGAAG |
| 30758_1 | TATGAGTGTGGAGTCGTTGCCATGTATGTGTTTTTGGTGGGGGNNNAGATCGGAAG |
| 30758_2 | TATGAGTGTGGAGTCGTTGCCATGTATGTGTTTTTGGTGGGGGNNNAGATCGGAAG |
| 32653_1 | TATGAGTGTGGAGTCGTTGCACTTCAGGTCTCAAGTGATCCACNNNAGATCGGAAGA |
| 32653_2 | TATGAGTGTGGAGTCGTTGCACTTCAGGTCTCAAGTGATCCACNNNAGATCGGAAGA |
| 48577_1 | TATGAGTGTGGAGTCGTTGCCCACGCCCAGCTAATTTTGTATTNNNAGATCGGAAGA |
| 48577_2 | TATGAGTGTGGAGTCGTTGCCCACGCCCAGCTAATTTTGTATTNNNAGATCGGAAGA |
| 49138_1 | TATGAGTGTGGAGTCGTTGCCTCCTGGGTTCAAGTGATTCTCANNNAGATCGGAAGA |
| 49138_2 | TATGAGTGTGGAGTCGTTGCCTCCTGGGTTCAAGTGATTCTCANNNAGATCGGAAGA |
| 34921_1 | TATGAGTGTGGAGTCGTTGCAATCTCGGCTCACTACAACCTTTNNNAGATCGGAAGA |
| 34921_2 | TATGAGTGTGGAGTCGTTGCAATCTCGGCTCACTACAACCTTTNNNAGATCGGAAGA |
| 52002_1 | TATGAGTGTGGAGTCGTTGCAATTGCTTCCAGTTCAAACCACCNNNAGATCGGAAGA |
| 52002_2 | TATGAGTGTGGAGTCGTTGCAATTGCTTCCAGTTCAAACCACCNNNAGATCGGAAGA |
| 45427_1 | TATGAGTGTGGAGTCGTTGCCTGGGATTACAGGCATGAGCTATNNNAGATCGGAAG |
| 45427_2 | TATGAGTGTGGAGTCGTTGCCTGGGATTACAGGCATGAGCTATNNNAGATCGGAAG |
| 53237_1 | TATGAGTGTGGAGTCGTTGCTGGCAAAACCCCACATCTACTAANNNAGATCGGAAGA |
| 53237_2 | TATGAGTGTGGAGTCGTTGCTGGCAAAACCCCACATCTACTAANNNAGATCGGAAGA |

| | |
|---|---|
| 48289_1 | TATGAGTGTGGAGTCGTTGCAACCACATGAGACTCTGATCCACNNNAGATCGGAAG |
| 48289_2 | TATGAGTGTGGAGTCGTTGCAACCACATGAGACTCTGATCCACNNNAGATCGGAAG |
| 41399_1 | TATGAGTGTGGAGTCGTTGCCCTAAGCCGTTTGAAGACAGTTGNNNAGATCGGAA |
| 41399_2 | TATGAGTGTGGAGTCGTTGCCCTAAGCCGTTTGAAGACAGTTGNNNAGATCGGAA |
| 52878_1 | TATGAGTGTGGAGTCGTTGCCAATTTACCAGTGGGGAGAGCTNNNAGATCGGAAG |
| 52878_2 | TATGAGTGTGGAGTCGTTGCCAATTTACCAGTGGGGAGAGCTNNNAGATCGGAAG |
| 42969_1 | TATGAGTGTGGAGTCGTTGCGCAGCCCAATTTGTATTCCAGAGNNNAGATCGGAAG |
| 42969_2 | TATGAGTGTGGAGTCGTTGCGCAGCCCAATTTGTATTCCAGAGNNNAGATCGGAAG |
| 51031_1 | TATGAGTGTGGAGTCGTTGCTGACTCTAGTTAGGGCCAAGAGANNNAGATCGGAA |
| 51031_2 | TATGAGTGTGGAGTCGTTGCTGACTCTAGTTAGGGCCAAGAGANNNAGATCGGAA |
| 43959_1 | TATGAGTGTGGAGTCGTTGCGAGGGTCGATGGCCAGAAACNNNAGATCGGAAGA |
| 43959_2 | TATGAGTGTGGAGTCGTTGCGAGGGTCGATGGCCAGAAACNNNAGATCGGAAGA |
| 45698_1 | TATGAGTGTGGAGTCGTTGCCTCCTGAGTAGCTGGGACTACAGNNNAGATCGGAA |
| 45698_2 | TATGAGTGTGGAGTCGTTGCCTCCTGAGTAGCTGGGACTACAGNNNAGATCGGAA |
| 43509_1 | TATGAGTGTGGAGTCGTTGCGAATTTCACTCTTGTTGCCCAGGNNNAGATCGGAAG |
| 43509_2 | TATGAGTGTGGAGTCGTTGCGAATTTCACTCTTGTTGCCCAGGNNNAGATCGGAAG |
| 37934_1 | TATGAGTGTGGAGTCGTTGCGGGAACATAGTAAGGTCCCTGTCNNNAGATCGGAA |
| 37934_2 | TATGAGTGTGGAGTCGTTGCGGGAACATAGTAAGGTCCCTGTCNNNAGATCGGAA |
| 44466_1 | TATGAGTGTGGAGTCGTTGCGTTCTCATTATGGAGTGTGTGCGNNNAGATCGGAA |
| 44466_2 | TATGAGTGTGGAGTCGTTGCGTTCTCATTATGGAGTGTGTGCGNNNAGATCGGAA |
| 51323_1 | TATGAGTGTGGAGTCGTTGCTGCTGTTACTGATCAGGTACCACNNNAGATCGGAAG |
| 51323_2 | TATGAGTGTGGAGTCGTTGCTGCTGTTACTGATCAGGTACCACNNNAGATCGGAAG |
| 43149_1 | TATGAGTGTGGAGTCGTTGCCATGTGAAGGATCATGTCAAAGGTNNNAGATCGGA |
| 43149_2 | TATGAGTGTGGAGTCGTTGCCATGTGAAGGATCATGTCAAAGGTNNNAGATCGGA |
| 38978_1 | TATGAGTGTGGAGTCGTTGCTGGTAGTGGCTTTGGGAAAAGATNNNAGATCGGAA |
| 38978_2 | TATGAGTGTGGAGTCGTTGCTGGTAGTGGCTTTGGGAAAAGATNNNAGATCGGAA |
| 32325_1 | TATGAGTGTGGAGTCGTTGCGTGTGTGTGCATGTGTGTATGAANNNAGATCGGAA |
| 32325_2 | TATGAGTGTGGAGTCGTTGCGTGTGTGTGCATGTGTGTATGAANNNAGATCGGAA |
| 29349_1 | TATGAGTGTGGAGTCGTTGCAGGATTACTGCAAGGGTAAACAGANNNAGATCGGA |
| 29349_2 | TATGAGTGTGGAGTCGTTGCAGGATTACTGCAAGGGTAAACAGANNNAGATCGGA |
| 29892_1 | TATGAGTGTGGAGTCGTTGCGAGCATGGTGGTGTTTTGTTTTGNNNAGATCGGAA |
| 29892_2 | TATGAGTGTGGAGTCGTTGCGAGCATGGTGGTGTTTTGTTTTGNNNAGATCGGAA |
| 49707_1 | TATGAGTGTGGAGTCGTTGCTACCCGCATCATAGCTACAAAGTNNNAGATCGGAAG |
| 49707_2 | TATGAGTGTGGAGTCGTTGCTACCCGCATCATAGCTACAAAGTNNNAGATCGGAAG |
| 44873_1 | TATGAGTGTGGAGTCGTTGCGAGTCACAGAGGAACACAGAGAGNNNAGATCGGA |
| 44873_2 | TATGAGTGTGGAGTCGTTGCGAGTCACAGAGGAACACAGAGAGNNNAGATCGGA |
| 30051_1 | TATGAGTGTGGAGTCGTTGCCCCGGTGAGTCACAGAATCTTAGNNNAGATCGGAA |
| 30051_2 | TATGAGTGTGGAGTCGTTGCCCCGGTGAGTCACAGAATCTTAGNNNAGATCGGAA |
| 29323_1 | TATGAGTGTGGAGTCGTTGCATTGGCTAGACTGTGAGCTGAAANNNAGATCGGAA |
| 29323_2 | TATGAGTGTGGAGTCGTTGCATTGGCTAGACTGTGAGCTGAAANNNAGATCGGAA |
| 40471_1 | TATGAGTGTGGAGTCGTTGCAGGATTACAAGGCTACAGTGAGCNNNAGATCGGAA |
| 40471_2 | TATGAGTGTGGAGTCGTTGCAGGATTACAAGGCTACAGTGAGCNNNAGATCGGAA |
| 39157_1 | TATGAGTGTGGAGTCGTTGCGGGTTCAAGTGATTCTCCTGTCTNNNAGATCGGAAG |

| | |
|---|---|
| 39157_2 | TATGAGTGTGGAGTCGTTGCGGGTTCAAGTGATTCTCCTGTCTNNNAGATCGGAAG |
| 41090_1 | TATGAGTGTGGAGTCGTTGCCACCAAGTCCAGCCTATACTCAANNNAGATCGGAAG |
| 41090_2 | TATGAGTGTGGAGTCGTTGCCACCAAGTCCAGCCTATACTCAANNNAGATCGGAAG |
| 10097_1 | TATGAGTGTGGAGTCGTTGCCCCTAAAGCATGGGGAGCTTTAGNNNAGATCGGAA |
| 10097_2 | TATGAGTGTGGAGTCGTTGCCCCTAAAGCATGGGGAGCTTTAGNNNAGATCGGAA |
| 11102_1 | TATGAGTGTGGAGTCGTTGCGAAATATGGGCTGCACTGAGCANNNAGATCGGAAG |
| 11102_2 | TATGAGTGTGGAGTCGTTGCGAAATATGGGCTGCACTGAGCANNNAGATCGGAAG |
| 35846_1 | TATGAGTGTGGAGTCGTTGCGATTCAAGCAATTCTCCTGCCTCNNNAGATCGGAAG |
| 35846_2 | TATGAGTGTGGAGTCGTTGCGATTCAAGCAATTCTCCTGCCTCNNNAGATCGGAAG |
| 37746_1 | TATGAGTGTGGAGTCGTTGCAAGAAGGTGAGGTTGGAAAGGAGNNNAGATCGGA |
| 37746_2 | TATGAGTGTGGAGTCGTTGCAAGAAGGTGAGGTTGGAAAGGAGNNNAGATCGGA |
| 46696_1 | TATGAGTGTGGAGTCGTTGCATGGGGTTTCTCCATGTTCATCANNNAGATCGGAAG |
| 46696_2 | TATGAGTGTGGAGTCGTTGCATGGGGTTTCTCCATGTTCATCANNNAGATCGGAAG |
| 41412_1 | TATGAGTGTGGAGTCGTTGCACCGGATGAACACCTCTTTCTTTNNNAGATCGGAAG |
| 41412_2 | TATGAGTGTGGAGTCGTTGCACCGGATGAACACCTCTTTCTTTNNNAGATCGGAAG |
| 2077_1 | TATGAGTGTGGAGTCGTTGCTGGAGCAAAGGAATTGTCTTGACANNNAGATCGGA |
| 2077_2 | TATGAGTGTGGAGTCGTTGCTGGAGCAAAGGAATTGTCTTGACANNNAGATCGGA |
| 45543_1 | TATGAGTGTGGAGTCGTTGCAGGGTGCAGATGAAACATCTCATNNNAGATCGGAA |
| 45543_2 | TATGAGTGTGGAGTCGTTGCAGGGTGCAGATGAAACATCTCATNNNAGATCGGAA |
| 28844_1 | TATGAGTGTGGAGTCGTTGCGGCAAGCAACAAGATGGTAACTTNNNAGATCGGAA |
| 28844_2 | TATGAGTGTGGAGTCGTTGCGGCAAGCAACAAGATGGTAACTTNNNAGATCGGAA |
| 36020_1 | TATGAGTGTGGAGTCGTTGCACACAAAACTCTCCCTCCCAAATNNNAGATCGGAAGA |
| 36020_2 | TATGAGTGTGGAGTCGTTGCACACAAAACTCTCCCTCCCAAATNNNAGATCGGAAGA |
| 38219_1 | TATGAGTGTGGAGTCGTTGCTAATCCCAGCTACTAGTGAGGCTNNNAGATCGGAAG |
| 38219_2 | TATGAGTGTGGAGTCGTTGCTAATCCCAGCTACTAGTGAGGCTNNNAGATCGGAAG |
| 8237_1 | TATGAGTGTGGAGTCGTTGCTCTCAGCTCACTTCAGCTCACTCNNNAGATCGGAAGA |
| 8237_2 | TATGAGTGTGGAGTCGTTGCTCTCAGCTCACTTCAGCTCACTCNNNAGATCGGAAGA |
| 49667_1 | TATGAGTGTGGAGTCGTTGCCCATTCTTCCTGTGTCAGCCTCNNNAGATCGGAAGAG |
| 49667_2 | TATGAGTGTGGAGTCGTTGCCCATTCTTCCTGTGTCAGCCTCNNNAGATCGGAAGAG |
| 8340_1 | TATGAGTGTGGAGTCGTTGCTCAGAGCCGTAACAGCTTCCAGNNNAGATCGGAAG |
| 8340_2 | TATGAGTGTGGAGTCGTTGCTCAGAGCCGTAACAGCTTCCAGNNNAGATCGGAAG |
| 38049_1 | TATGAGTGTGGAGTCGTTGCTTTTGGGGAAAATGAAGCATCCCNNNAGATCGGAA |
| 38049_2 | TATGAGTGTGGAGTCGTTGCTTTTGGGGAAAATGAAGCATCCCNNNAGATCGGAA |
| 34113_1 | TATGAGTGTGGAGTCGTTGCGTGTCCATTAGAGGTCATGGCTANNNAGATCGGAA |
| 34113_2 | TATGAGTGTGGAGTCGTTGCGTGTCCATTAGAGGTCATGGCTANNNAGATCGGAA |
| 33438_1 | TATGAGTGTGGAGTCGTTGCAAATATTGGTTGGACACAGTGGCNNNAGATCGGAA |
| 33438_2 | TATGAGTGTGGAGTCGTTGCAAATATTGGTTGGACACAGTGGCNNNAGATCGGAA |
| 30750_1 | TATGAGTGTGGAGTCGTTGCCTGTGGTCTCAGGACTTCTTTGTNNNAGATCGGAAG |
| 30750_2 | TATGAGTGTGGAGTCGTTGCCTGTGGTCTCAGGACTTCTTTGTNNNAGATCGGAAG |
| 3089_1 | TATGAGTGTGGAGTCGTTGCTGACTGGGAAGAGACTATGATGGAANNNAGATCGG |
| 3089_2 | TATGAGTGTGGAGTCGTTGCTGACTGGGAAGAGACTATGATGGAANNNAGATCGG |
| 9206_1 | TATGAGTGTGGAGTCGTTGCTACCCTGTACCCTGGTTCTCTTGNNNAGATCGGAAG |
| 9206_2 | TATGAGTGTGGAGTCGTTGCTACCCTGTACCCTGGTTCTCTTGNNNAGATCGGAAG |

| | |
|---|---|
| 13242_1 | TATGAGTGTGGAGTCGTTGCGGTCTTCACATTCATTTTCTACNNNAGATCGGAAGAG |
| 13242_2 | TATGAGTGTGGAGTCGTTGCGGTCTTCACATTCATTTTCTACNNNAGATCGGAAGAG |
| 34745_1 | TATGAGTGTGGAGTCGTTGCCTTCCGAGTAGCTGGGATTACAGNNNAGATCGGAAG |
| 34745_2 | TATGAGTGTGGAGTCGTTGCCTTCCGAGTAGCTGGGATTACAGNNNAGATCGGAAG |
| 39688_1 | TATGAGTGTGGAGTCGTTGCCACAGGGCCAATTAGACATTTGGNNNAGATCGGAAG |
| 39688_2 | TATGAGTGTGGAGTCGTTGCCACAGGGCCAATTAGACATTTGGNNNAGATCGGAAG |
| 10887_1 | TATGAGTGTGGAGTCGTTGCCGCTCTGTAGCCCTGTAGCCNNNAGATCGGAAGAGC |
| 10887_2 | TATGAGTGTGGAGTCGTTGCCGCTCTGTAGCCCTGTAGCCNNNAGATCGGAAGAGC |
| 44128_1 | TATGAGTGTGGAGTCGTTGCGGGGGATTACATGGCAGATAACTNNNAGATCGGAA |
| 44128_2 | TATGAGTGTGGAGTCGTTGCGGGGGATTACATGGCAGATAACTNNNAGATCGGAA |
| 46311_1 | TATGAGTGTGGAGTCGTTGCGATCTCGGCTCATTGCAAGCTCNNNAGATCGGAAGA |
| 46311_2 | TATGAGTGTGGAGTCGTTGCGATCTCGGCTCATTGCAAGCTCNNNAGATCGGAAGA |
| 962_1 | TATGAGTGTGGAGTCGTTGCGTAAAATGTGCATTCCCTTTGCAGNNNAGATCGGAA |
| 962_2 | TATGAGTGTGGAGTCGTTGCGTAAAATGTGCATTCCCTTTGCAGNNNAGATCGGAA |
| 48258_1 | TATGAGTGTGGAGTCGTTGCCAGAGGAAAGTGGGGAATCAAGANNNAGATCGGA |
| 48258_2 | TATGAGTGTGGAGTCGTTGCCAGAGGAAAGTGGGGAATCAAGANNNAGATCGGA |
| 38859_1 | TATGAGTGTGGAGTCGTTGCAGAAAACCCAATCCAAGAATGCANNNAGATCGGAAG |
| 38859_2 | TATGAGTGTGGAGTCGTTGCAGAAAACCCAATCCAAGAATGCANNNAGATCGGAAG |
| 37833_1 | TATGAGTGTGGAGTCGTTGCGTGTCACTGTGGAAGGCTACTAANNNAGATCGGAA |
| 37833_2 | TATGAGTGTGGAGTCGTTGCGTGTCACTGTGGAAGGCTACTAANNNAGATCGGAA |
| 36445_1 | TATGAGTGTGGAGTCGTTGCGAACTATGAATTGCACTCCAGCCNNNAGATCGGAAG |
| 36445_2 | TATGAGTGTGGAGTCGTTGCGAACTATGAATTGCACTCCAGCCNNNAGATCGGAAG |
| 44149_1 | TATGAGTGTGGAGTCGTTGCATGTTTATTTCCCCTTCCTGCCTNNNAGATCGGAAGA |
| 44149_2 | TATGAGTGTGGAGTCGTTGCATGTTTATTTCCCCTTCCTGCCTNNNAGATCGGAAGA |
| 36504_1 | TATGAGTGTGGAGTCGTTGCCCCACTTTGAACTGAGAGGAAGANNNAGATCGGAA |
| 36504_2 | TATGAGTGTGGAGTCGTTGCCCCACTTTGAACTGAGAGGAAGANNNAGATCGGAA |
| 30425_1 | TATGAGTGTGGAGTCGTTGCTCTCACTCACTCCATCACCCGAANNNAGATCGGAAGA |
| 30425_2 | TATGAGTGTGGAGTCGTTGCTCTCACTCACTCCATCACCCGAANNNAGATCGGAAGA |
| 43433_1 | TATGAGTGTGGAGTCGTTGCTTCTGGGTTCAAGCAATTCTCCTNNNAGATCGGAAG |
| 43433_2 | TATGAGTGTGGAGTCGTTGCTTCTGGGTTCAAGCAATTCTCCTNNNAGATCGGAAG |
| 43563_1 | TATGAGTGTGGAGTCGTTGCTCACTGTAGCCTTGAATTCCTGGNNNAGATCGGAAG |
| 43563_2 | TATGAGTGTGGAGTCGTTGCTCACTGTAGCCTTGAATTCCTGGNNNAGATCGGAAG |
| 43525_1 | TATGAGTGTGGAGTCGTTGCTAGTGGTGCAATCACAGCTTACTNNNAGATCGGAAG |
| 43525_2 | TATGAGTGTGGAGTCGTTGCTAGTGGTGCAATCACAGCTTACTNNNAGATCGGAAG |
| 51639_1 | TATGAGTGTGGAGTCGTTGCTTGGGTCCAAGGCGCATAGGNNNAGATCGGAAGAG |
| 51639_2 | TATGAGTGTGGAGTCGTTGCTTGGGTCCAAGGCGCATAGGNNNAGATCGGAAGAG |
| 8539_1 | TATGAGTGTGGAGTCGTTGCGAAGGGCAAGAAAATTGAACCCTGNNNAGATCGGA |
| 8539_2 | TATGAGTGTGGAGTCGTTGCGAAGGGCAAGAAAATTGAACCCTGNNNAGATCGGA |
| 30830_1 | TATGAGTGTGGAGTCGTTGCTGTTTTTGACGATTCTTGAGGCCNNNAGATCGGAAG |
| 30830_2 | TATGAGTGTGGAGTCGTTGCTGTTTTTGACGATTCTTGAGGCCNNNAGATCGGAAG |
| 33186_1 | TATGAGTGTGGAGTCGTTGCGAATCAACCGTTCTGGACCCAGNNNAGATCGGAAGA |
| 33186_2 | TATGAGTGTGGAGTCGTTGCGAATCAACCGTTCTGGACCCAGNNNAGATCGGAAGA |
| 45876_1 | TATGAGTGTGGAGTCGTTGCTTCAAGCAATTCTTCTGCCTTGGNNNAGATCGGAAG |

| | |
|---|---|
| 45876_2 | TATGAGTGTGGAGTCGTTGCTTCAAGCAATTCTTCTGCCTTGGNNNAGATCGGAAG |
| 44289_1 | TATGAGTGTGGAGTCGTTGCCCAATTGAGGGTCTTCAGGAGAANNNAGATCGGAA |
| 44289_2 | TATGAGTGTGGAGTCGTTGCCCAATTGAGGGTCTTCAGGAGAANNNAGATCGGAA |
| 39197_1 | TATGAGTGTGGAGTCGTTGCGGGGACTTCGGTTCTATAGTCACNNNAGATCGGAAG |
| 39197_2 | TATGAGTGTGGAGTCGTTGCGGGGACTTCGGTTCTATAGTCACNNNAGATCGGAAG |
| 32654_1 | TATGAGTGTGGAGTCGTTGCGTGCATGAGATGATCAGCTGTTCNNNAGATCGGAAG |
| 32654_2 | TATGAGTGTGGAGTCGTTGCGTGCATGAGATGATCAGCTGTTCNNNAGATCGGAAG |
| 47022_1 | TATGAGTGTGGAGTCGTTGCACCACGCCTGGCTAATTTTTTTTNNNAGATCGGAAGA |
| 47022_2 | TATGAGTGTGGAGTCGTTGCACCACGCCTGGCTAATTTTTTTTNNNAGATCGGAAGA |
| 44784_1 | TATGAGTGTGGAGTCGTTGCATGAATCCCTGAAGGAGAGAGGANNNAGATCGGAA |
| 44784_2 | TATGAGTGTGGAGTCGTTGCATGAATCCCTGAAGGAGAGAGGANNNAGATCGGAA |
| 43976_1 | TATGAGTGTGGAGTCGTTGCTCTGGAGACGTTTTTGGAGTCATNNNAGATCGGAAG |
| 43976_2 | TATGAGTGTGGAGTCGTTGCTCTGGAGACGTTTTTGGAGTCATNNNAGATCGGAAG |
| 41851_1 | TATGAGTGTGGAGTCGTTGCGATGGAGTCTCATTCTGTCTCCCNNNAGATCGGAAG |
| 41851_2 | TATGAGTGTGGAGTCGTTGCGATGGAGTCTCATTCTGTCTCCCNNNAGATCGGAAG |
| 41526_1 | TATGAGTGTGGAGTCGTTGCTAGTCTTATGCGACCACCATTGTNNNAGATCGGAAG |
| 41526_2 | TATGAGTGTGGAGTCGTTGCTAGTCTTATGCGACCACCATTGTNNNAGATCGGAAG |
| 34813_1 | TATGAGTGTGGAGTCGTTGCTTAGTAGAGATGGGGGAAGAGGGNNNAGATCGGA |
| 34813_2 | TATGAGTGTGGAGTCGTTGCTTAGTAGAGATGGGGGAAGAGGGNNNAGATCGGA |
| 37035_1 | TATGAGTGTGGAGTCGTTGCGAGGGGAAGAAAGGAGAAGACAGNNNAGATCGGA |
| 37035_2 | TATGAGTGTGGAGTCGTTGCGAGGGGAAGAAAGGAGAAGACAGNNNAGATCGGA |
| 49290_1 | TATGAGTGTGGAGTCGTTGCCTCACTGCAACCTTTGTCTTCTGNNNAGATCGGAAGA |
| 49290_2 | TATGAGTGTGGAGTCGTTGCCTCACTGCAACCTTTGTCTTCTGNNNAGATCGGAAGA |
| 42938_1 | TATGAGTGTGGAGTCGTTGCTGAAGTCCAGGAATCCAAAGAGGNNNAGATCGGAA |
| 42938_2 | TATGAGTGTGGAGTCGTTGCTGAAGTCCAGGAATCCAAAGAGGNNNAGATCGGAA |
| 32700_1 | TATGAGTGTGGAGTCGTTGCGACGGAGTCTCACTCTGTTGCNNNAGATCGGAAGAG |
| 32700_2 | TATGAGTGTGGAGTCGTTGCGACGGAGTCTCACTCTGTTGCNNNAGATCGGAAGAG |
| 29003_1 | TATGAGTGTGGAGTCGTTGCTGTGTGTATCCGTAATCTCAGAANNNAGATCGGAAG |
| 29003_2 | TATGAGTGTGGAGTCGTTGCTGTGTGTATCCGTAATCTCAGAANNNAGATCGGAAG |
| 39886_1 | TATGAGTGTGGAGTCGTTGCTGAGATACCACTTCACATCCACTNNNAGATCGGAAG |
| 39886_2 | TATGAGTGTGGAGTCGTTGCTGAGATACCACTTCACATCCACTNNNAGATCGGAAG |
| 4304_1 | TATGAGTGTGGAGTCGTTGCGATGGCAAATGGGATTACAGATGTNNNAGATCGGA |
| 4304_2 | TATGAGTGTGGAGTCGTTGCGATGGCAAATGGGATTACAGATGTNNNAGATCGGA |
| 50366_1 | TATGAGTGTGGAGTCGTTGCAGACTTTGTGCTCAGTGGTATGTNNNAGATCGGAAG |
| 50366_2 | TATGAGTGTGGAGTCGTTGCAGACTTTGTGCTCAGTGGTATGTNNNAGATCGGAAG |
| 13039_1 | TATGAGTGTGGAGTCGTTGCCATCAAAATCACAGTGGATGNNNAGATCGGAAGAG |
| 13039_2 | TATGAGTGTGGAGTCGTTGCCATCAAAATCACAGTGGATGNNNAGATCGGAAGAG |
| 39344_1 | TATGAGTGTGGAGTCGTTGCCTGTAATCCCAGCTACTCAGGAGNNNAGATCGGAAG |
| 39344_2 | TATGAGTGTGGAGTCGTTGCCTGTAATCCCAGCTACTCAGGAGNNNAGATCGGAAG |
| 47571_1 | TATGAGTGTGGAGTCGTTGCCTGGGTTCAAATGATTCTCCTGCNNNAGATCGGAAG |
| 47571_2 | TATGAGTGTGGAGTCGTTGCCTGGGTTCAAATGATTCTCCTGCNNNAGATCGGAAG |
| 39237_1 | TATGAGTGTGGAGTCGTTGCATACCTGGGGGAAGACTGTCATANNNAGATCGGAA |
| 39237_2 | TATGAGTGTGGAGTCGTTGCATACCTGGGGGAAGACTGTCATANNNAGATCGGAA |

| | |
|---|---|
| 39591_1 | TATGAGTGTGGAGTCGTTGCTGGGTCTAGGTTAGAAGTGGGATNNNAGATCGGAA |
| 39591_2 | TATGAGTGTGGAGTCGTTGCTGGGTCTAGGTTAGAAGTGGGATNNNAGATCGGAA |
| 34315_1 | TATGAGTGTGGAGTCGTTGCTGGAAGTATGCTTTCTCTTGAGTNNNAGATCGGAAG |
| 34315_2 | TATGAGTGTGGAGTCGTTGCTGGAAGTATGCTTTCTCTTGAGTNNNAGATCGGAAG |
| 45644_1 | TATGAGTGTGGAGTCGTTGCAAACTCCTGAGGTCAAGTGATCCNNNAGATCGGAAG |
| 45644_2 | TATGAGTGTGGAGTCGTTGCAAACTCCTGAGGTCAAGTGATCCNNNAGATCGGAAG |
| 13447_1 | TATGAGTGTGGAGTCGTTGCATTATGTGAGAAGGTGAATGATNNNAGATCGGAAG |
| 13447_2 | TATGAGTGTGGAGTCGTTGCATTATGTGAGAAGGTGAATGATNNNAGATCGGAAG |
| 11083_1 | TATGAGTGTGGAGTCGTTGCTGCTTGAAAAGACACAAAGCAGTNNNAGATCGGAA |
| 11083_2 | TATGAGTGTGGAGTCGTTGCTGCTTGAAAAGACACAAAGCAGTNNNAGATCGGAA |
| 35894_1 | TATGAGTGTGGAGTCGTTGCGTGTGGAATGCATGGAAATGTCANNNAGATCGGAA |
| 35894_2 | TATGAGTGTGGAGTCGTTGCGTGTGGAATGCATGGAAATGTCANNNAGATCGGAA |
| 41914_1 | TATGAGTGTGGAGTCGTTGCTATCTCCCAATGCTATCCCTCCCNNNAGATCGGAAGA |
| 41914_2 | TATGAGTGTGGAGTCGTTGCTATCTCCCAATGCTATCCCTCCCNNNAGATCGGAAGA |
| 49924_1 | TATGAGTGTGGAGTCGTTGCGGGCCAAAAGATCTGTTCATTCCNNNAGATCGGAAG |
| 49924_2 | TATGAGTGTGGAGTCGTTGCGGGCCAAAAGATCTGTTCATTCCNNNAGATCGGAAG |
| 45648_1 | TATGAGTGTGGAGTCGTTGCGTTTCAAGCGATTCTTCTGCCTCNNNAGATCGGAAG |
| 45648_2 | TATGAGTGTGGAGTCGTTGCGTTTCAAGCGATTCTTCTGCCTCNNNAGATCGGAAG |
| 33040_1 | TATGAGTGTGGAGTCGTTGCCTCTCGAGTAGCTGGGATTACAGNNNAGATCGGAA |
| 33040_2 | TATGAGTGTGGAGTCGTTGCCTCTCGAGTAGCTGGGATTACAGNNNAGATCGGAA |
| 48838_1 | TATGAGTGTGGAGTCGTTGCACAACCTCTTTGGAAGACAGGTTNNNAGATCGGAAG |
| 48838_2 | TATGAGTGTGGAGTCGTTGCACAACCTCTTTGGAAGACAGGTTNNNAGATCGGAAG |
| 13003_1 | TATGAGTGTGGAGTCGTTGCATAGTGACTATCTCAATAGAACCNNNAGATCGGAAG |
| 13003_2 | TATGAGTGTGGAGTCGTTGCATAGTGACTATCTCAATAGAACCNNNAGATCGGAAG |
| 12491_1 | TATGAGTGTGGAGTCGTTGCAATAGCTAAATATTGGAAACCACNNNAGATCGGAAG |
| 12491_2 | TATGAGTGTGGAGTCGTTGCAATAGCTAAATATTGGAAACCACNNNAGATCGGAAG |
| 38184_1 | TATGAGTGTGGAGTCGTTGCAGCCTCTAGAAGCTGAAAAAGGCNNNAGATCGGAA |
| 38184_2 | TATGAGTGTGGAGTCGTTGCAGCCTCTAGAAGCTGAAAAAGGCNNNAGATCGGAA |
| 29912_1 | TATGAGTGTGGAGTCGTTGCAGCTTCCTGGAGTTCGATGATTTNNNAGATCGGAAG |
| 29912_2 | TATGAGTGTGGAGTCGTTGCAGCTTCCTGGAGTTCGATGATTTNNNAGATCGGAAG |
| 52601_1 | TATGAGTGTGGAGTCGTTGCTGTGTCCTTCTGTTTCTTGGGAGNNNAGATCGGAAG |
| 52601_2 | TATGAGTGTGGAGTCGTTGCTGTGTCCTTCTGTTTCTTGGGAGNNNAGATCGGAAG |
| 43253_1 | TATGAGTGTGGAGTCGTTGCTTGGTTCTTTGAGGAATTGCCACNNNAGATCGGAAG |
| 43253_2 | TATGAGTGTGGAGTCGTTGCTTGGTTCTTTGAGGAATTGCCACNNNAGATCGGAAG |
| 42291_1 | TATGAGTGTGGAGTCGTTGCAAGTTTTCAGCAATGGTCACACCNNNAGATCGGAAG |
| 42291_2 | TATGAGTGTGGAGTCGTTGCAAGTTTTCAGCAATGGTCACACCNNNAGATCGGAAG |
| 50837_1 | TATGAGTGTGGAGTCGTTGCAAGGGACTTTTCACATAGGCCATNNNAGATCGGAAG |
| 50837_2 | TATGAGTGTGGAGTCGTTGCAAGGGACTTTTCACATAGGCCATNNNAGATCGGAAG |
| 42421_1 | TATGAGTGTGGAGTCGTTGCCCTGTCTTTTCCCTCAGCTCTGNNNAGATCGGAAGAG |
| 42421_2 | TATGAGTGTGGAGTCGTTGCCCTGTCTTTTCCCTCAGCTCTGNNNAGATCGGAAGAG |
| 31930_1 | TATGAGTGTGGAGTCGTTGCGGGTTCAAAAGCAGATTTGAGCANNNAGATCGGAA |
| 31930_2 | TATGAGTGTGGAGTCGTTGCGGGTTCAAAAGCAGATTTGAGCANNNAGATCGGAA |
| 47253_1 | TATGAGTGTGGAGTCGTTGCTATCTCTCGGGTTCAAGGGATTCNNNAGATCGGAAG |

| | |
|---|---|
| 47253_2 | TATGAGTGTGGAGTCGTTGCTATCTCTCGGGTTCAAGGGATTCNNNAGATCGGAAG |
| 44360_1 | TATGAGTGTGGAGTCGTTGCCTTGCAGGAAACCATCAAGCTTTNNNAGATCGGAAG |
| 44360_2 | TATGAGTGTGGAGTCGTTGCCTTGCAGGAAACCATCAAGCTTTNNNAGATCGGAAG |
| 52867_1 | TATGAGTGTGGAGTCGTTGCTTCCTTCTGACTTGGACTTCCTGNNNAGATCGGAAGA |
| 52867_2 | TATGAGTGTGGAGTCGTTGCTTCCTTCTGACTTGGACTTCCTGNNNAGATCGGAAGA |
| 50920_1 | TATGAGTGTGGAGTCGTTGCCAAATGGACAGGAGGGTTTTGACNNNAGATCGGAA |
| 50920_2 | TATGAGTGTGGAGTCGTTGCCAAATGGACAGGAGGGTTTTGACNNNAGATCGGAA |
| 43903_1 | TATGAGTGTGGAGTCGTTGCTAGAGAGAGAGAGCGTGATGATTNNNAGATCGGAA |
| 43903_2 | TATGAGTGTGGAGTCGTTGCTAGAGAGAGAGAGCGTGATGATTNNNAGATCGGAA |
| 2054_1 | TATGAGTGTGGAGTCGTTGCAGCATTCCATTTCACAGGAAACANNNAGATCGGAAG |
| 2054_2 | TATGAGTGTGGAGTCGTTGCAGCATTCCATTTCACAGGAAACANNNAGATCGGAAG |
| 39044_1 | TATGAGTGTGGAGTCGTTGCCAGTGACATGATCTCAGCTCACTNNNAGATCGGAAGA |
| 39044_2 | TATGAGTGTGGAGTCGTTGCCAGTGACATGATCTCAGCTCACTNNNAGATCGGAAGA |
| 32544_1 | TATGAGTGTGGAGTCGTTGCTTTGCAACTTGTCATGGGAAACANNNAGATCGGAAG |
| 32544_2 | TATGAGTGTGGAGTCGTTGCTTTGCAACTTGTCATGGGAAACANNNAGATCGGAAG |
| 38914_1 | TATGAGTGTGGAGTCGTTGCCCAGAATGAGCAACAGTGGTTTTNNNAGATCGGAAG |
| 38914_2 | TATGAGTGTGGAGTCGTTGCCCAGAATGAGCAACAGTGGTTTTNNNAGATCGGAAG |
| 10458_1 | TATGAGTGTGGAGTCGTTGCACTGTCTAAACGGATGGATGTAGANNNAGATCGGAA |
| 10458_2 | TATGAGTGTGGAGTCGTTGCACTGTCTAAACGGATGGATGTAGANNNAGATCGGAA |
| 30536_1 | TATGAGTGTGGAGTCGTTGCAATGAATATGTGGGAGGCCTGTTNNNAGATCGGAAG |
| 30536_2 | TATGAGTGTGGAGTCGTTGCAATGAATATGTGGGAGGCCTGTTNNNAGATCGGAAG |
| 38406_1 | TATGAGTGTGGAGTCGTTGCTGGGAGAATCAACTAAACCCAGGNNNAGATCGGAAG |
| 38406_2 | TATGAGTGTGGAGTCGTTGCTGGGAGAATCAACTAAACCCAGGNNNAGATCGGAAG |
| 35735_1 | TATGAGTGTGGAGTCGTTGCTGGTAGTTTCTTTTGCTGTGCAGNNNAGATCGGAAG |
| 35735_2 | TATGAGTGTGGAGTCGTTGCTGGTAGTTTCTTTTGCTGTGCAGNNNAGATCGGAAG |
| 52864_1 | TATGAGTGTGGAGTCGTTGCCTCGCTCTGCCCTGAACCTCNNNAGATCGGAAGAGCA |
| 52864_2 | TATGAGTGTGGAGTCGTTGCCTCGCTCTGCCCTGAACCTCNNNAGATCGGAAGAGCA |
| 42746_1 | TATGAGTGTGGAGTCGTTGCGCTGATTCCATGACTTTGCTGTTNNNAGATCGGAAGA |
| 42746_2 | TATGAGTGTGGAGTCGTTGCGCTGATTCCATGACTTTGCTGTTNNNAGATCGGAAGA |
| 32371_1 | TATGAGTGTGGAGTCGTTGCTGCATAGCTGTTTCTGCAGGTATNNNAGATCGGAAG |
| 32371_2 | TATGAGTGTGGAGTCGTTGCTGCATAGCTGTTTCTGCAGGTATNNNAGATCGGAAG |
| 52179_1 | TATGAGTGTGGAGTCGTTGCGTCTCACCTGGCTAAGAGAGAAGNNNAGATCGGAAG |
| 52179_2 | TATGAGTGTGGAGTCGTTGCGTCTCACCTGGCTAAGAGAGAAGNNNAGATCGGAAG |
| 13671_1 | TATGAGTGTGGAGTCGTTGCTTTTTTTTTTTTGGAGACAGAGTNNNAGATCGGAAGA |
| 13671_2 | TATGAGTGTGGAGTCGTTGCTTTTTTTTTTTTGGAGACAGAGTNNNAGATCGGAAGA |
| 45545_1 | TATGAGTGTGGAGTCGTTGCTGTGACCCAAGTTAGAATCCCAGNNNAGATCGGAAG |
| 45545_2 | TATGAGTGTGGAGTCGTTGCTGTGACCCAAGTTAGAATCCCAGNNNAGATCGGAAG |
| 3045_1 | TATGAGTGTGGAGTCGTTGCCTGGAGGAAAAGTCTGGTAGGGANNNAGATCGGAA |
| 3045_2 | TATGAGTGTGGAGTCGTTGCCTGGAGGAAAAGTCTGGTAGGGANNNAGATCGGAA |
| 50684_1 | TATGAGTGTGGAGTCGTTGCTGCATTTGCTCACACTGACATTTNNNAGATCGGAAGA |
| 50684_2 | TATGAGTGTGGAGTCGTTGCTGCATTTGCTCACACTGACATTTNNNAGATCGGAAGA |
| 45754_1 | TATGAGTGTGGAGTCGTTGCTTCAAGCGATTCTCCTTCCTCAGNNNAGATCGGAAGA |
| 45754_2 | TATGAGTGTGGAGTCGTTGCTTCAAGCGATTCTCCTTCCTCAGNNNAGATCGGAAGA |

| | |
|---|---|
| 43076_1 | TATGAGTGTGGAGTCGTTGCTTATGGAGACCAAGAAGTCCTGCNNNAGATCGGAA |
| 43076_2 | TATGAGTGTGGAGTCGTTGCTTATGGAGACCAAGAAGTCCTGCNNNAGATCGGAA |
| 39496_1 | TATGAGTGTGGAGTCGTTGCACACCACTTCGTACCTACTAGGANNNAGATCGGAAG |
| 39496_2 | TATGAGTGTGGAGTCGTTGCACACCACTTCGTACCTACTAGGANNNAGATCGGAAG |
| 42394_1 | TATGAGTGTGGAGTCGTTGCGCACACAGAAACACACAAAACTANNNAGATCGGAAG |
| 42394_2 | TATGAGTGTGGAGTCGTTGCGCACACAGAAACACACAAAACTANNNAGATCGGAAG |
| 37217_1 | TATGAGTGTGGAGTCGTTGCTTCTGGGTTCTCTTTTCTGGTCTNNNAGATCGGAAGA |
| 37217_2 | TATGAGTGTGGAGTCGTTGCTTCTGGGTTCTCTTTTCTGGTCTNNNAGATCGGAAGA |
| 13359_1 | TATGAGTGTGGAGTCGTTGCAAAGAATGTTTGTATACAAAGCTNNNAGATCGGAA |
| 13359_2 | TATGAGTGTGGAGTCGTTGCAAAGAATGTTTGTATACAAAGCTNNNAGATCGGAA |
| 44830_1 | TATGAGTGTGGAGTCGTTGCCGGGTAGGAAGGCAGGCTAGNNNAGATCGGAAGA |
| 44830_2 | TATGAGTGTGGAGTCGTTGCCGGGTAGGAAGGCAGGCTAGNNNAGATCGGAAGA |
| 46952_1 | TATGAGTGTGGAGTCGTTGCAGACAGCTCACTGACTCTATTGCNNNAGATCGGAAG |
| 46952_2 | TATGAGTGTGGAGTCGTTGCAGACAGCTCACTGACTCTATTGCNNNAGATCGGAAG |
| 30037_1 | TATGAGTGTGGAGTCGTTGCAAGGCATGACTTTTAATCGTGGCNNNAGATCGGAA |
| 30037_2 | TATGAGTGTGGAGTCGTTGCAAGGCATGACTTTTAATCGTGGCNNNAGATCGGAA |
| 37989_1 | TATGAGTGTGGAGTCGTTGCGTGGCAAGGTAGTGGTACAGTAGNNNAGATCGGA |
| 37989_2 | TATGAGTGTGGAGTCGTTGCGTGGCAAGGTAGTGGTACAGTAGNNNAGATCGGA |
| 51838_1 | TATGAGTGTGGAGTCGTTGCTTTCACCCCATTGTCACCCATTTNNNAGATCGGAAGA |
| 51838_2 | TATGAGTGTGGAGTCGTTGCTTTCACCCCATTGTCACCCATTTNNNAGATCGGAAGA |
| 44563_1 | TATGAGTGTGGAGTCGTTGCGGAGAACCAGGGTAGCTGATAAGNNNAGATCGGA |
| 44563_2 | TATGAGTGTGGAGTCGTTGCGGAGAACCAGGGTAGCTGATAAGNNNAGATCGGA |
| 43881_1 | TATGAGTGTGGAGTCGTTGCGCCAGAGGATCAGGTAGGAAAAANNNAGATCGGA |
| 43881_2 | TATGAGTGTGGAGTCGTTGCGCCAGAGGATCAGGTAGGAAAAANNNAGATCGGA |
| 45712_1 | TATGAGTGTGGAGTCGTTGCCCACCACACCTGGCTAATTTTTTNNNAGATCGGAAGA |
| 45712_2 | TATGAGTGTGGAGTCGTTGCCCACCACACCTGGCTAATTTTTTNNNAGATCGGAAGA |
| 43955_1 | TATGAGTGTGGAGTCGTTGCCAGAAAGATGCTTCCAAAGGCAANNNAGATCGGAA |
| 43955_2 | TATGAGTGTGGAGTCGTTGCCAGAAAGATGCTTCCAAAGGCAANNNAGATCGGAA |
| 47033_1 | TATGAGTGTGGAGTCGTTGCGCTGGAATACAATGGTGTGATCGNNNAGATCGGAA |
| 47033_2 | TATGAGTGTGGAGTCGTTGCGCTGGAATACAATGGTGTGATCGNNNAGATCGGAA |
| 9609_1 | TATGAGTGTGGAGTCGTTGCTTCTCTCTCTCTTCCCTGCTAGCNNNAGATCGGAAGA |
| 9609_2 | TATGAGTGTGGAGTCGTTGCTTCTCTCTCTCTTCCCTGCTAGCNNNAGATCGGAAGA |
| 29728_1 | TATGAGTGTGGAGTCGTTGCACAAAGGATGAAACACTAGCTAGANNNAGATCGGA |
| 29728_2 | TATGAGTGTGGAGTCGTTGCACAAAGGATGAAACACTAGCTAGANNNAGATCGGA |
| 51391_1 | TATGAGTGTGGAGTCGTTGCGAACAGCCACGCATGTAATAGTCNNNAGATCGGAAG |
| 51391_2 | TATGAGTGTGGAGTCGTTGCGAACAGCCACGCATGTAATAGTCNNNAGATCGGAAG |
| 48119_1 | TATGAGTGTGGAGTCGTTGCCCCAAGTGACTTAGGGAAGAGAGNNNAGATCGGAA |
| 48119_2 | TATGAGTGTGGAGTCGTTGCCCCAAGTGACTTAGGGAAGAGAGNNNAGATCGGAA |
| 45681_1 | TATGAGTGTGGAGTCGTTGCGAAACCTGGGATTTGCTCAGTTGNNNAGATCGGAA |
| 45681_2 | TATGAGTGTGGAGTCGTTGCGAAACCTGGGATTTGCTCAGTTGNNNAGATCGGAA |
| 42662_1 | TATGAGTGTGGAGTCGTTGCTCAAGTGGAATGTGACCTGATGANNNAGATCGGAA |
| 42662_2 | TATGAGTGTGGAGTCGTTGCTCAAGTGGAATGTGACCTGATGANNNAGATCGGAA |
| 34877_1 | TATGAGTGTGGAGTCGTTGCCCTCCTGGGTTCAAGCAATTCTCNNNAGATCGGAAG |

| | |
|---|---|
| 34877_2 | TATGAGTGTGGAGTCGTTGCCCTCCTGGGTTCAAGCAATTCTCNNNAGATCGGAAG |
| 43184_1 | TATGAGTGTGGAGTCGTTGCCCACGCTCGGCTAATTTTTTGTANNNAGATCGGAAG |
| 43184_2 | TATGAGTGTGGAGTCGTTGCCCACGCTCGGCTAATTTTTTGTANNNAGATCGGAAG |
| 48174_1 | TATGAGTGTGGAGTCGTTGCCTGGGTTCAAGCAATTCCTATGCNNNAGATCGGAAG |
| 48174_2 | TATGAGTGTGGAGTCGTTGCCTGGGTTCAAGCAATTCCTATGCNNNAGATCGGAAG |
| 30467_1 | TATGAGTGTGGAGTCGTTGCCCTTCTACTTTAGTTGCCTTGCCNNNAGATCGGAAGA |
| 30467_2 | TATGAGTGTGGAGTCGTTGCCCTTCTACTTTAGTTGCCTTGCCNNNAGATCGGAAGA |
| 34750_1 | TATGAGTGTGGAGTCGTTGCCACAATCTCAGCTCACTGCAATCNNNAGATCGGAAG |
| 34750_2 | TATGAGTGTGGAGTCGTTGCCACAATCTCAGCTCACTGCAATCNNNAGATCGGAAG |
| 42869_1 | TATGAGTGTGGAGTCGTTGCATGACTGTGCTGTTAGAGGGATCNNNAGATCGGAA |
| 42869_2 | TATGAGTGTGGAGTCGTTGCATGACTGTGCTGTTAGAGGGATCNNNAGATCGGAA |
| 44048_1 | TATGAGTGTGGAGTCGTTGCATTGCAAGACCATGACAGAGGATNNNAGATCGGAA |
| 44048_2 | TATGAGTGTGGAGTCGTTGCATTGCAAGACCATGACAGAGGATNNNAGATCGGAA |
| 41549_1 | TATGAGTGTGGAGTCGTTGCGCACGCCCAGCTAATTTTTTTTTNNNAGATCGGAAG |
| 41549_2 | TATGAGTGTGGAGTCGTTGCGCACGCCCAGCTAATTTTTTTTTNNNAGATCGGAAG |
| 13180_1 | TATGAGTGTGGAGTCGTTGCGAGTTCTTTGTCTTCAGAAAAGNNNAGATCGGAAG |
| 13180_2 | TATGAGTGTGGAGTCGTTGCGAGTTCTTTGTCTTCAGAAAAGNNNAGATCGGAAG |
| 29289_1 | TATGAGTGTGGAGTCGTTGCCACAGAGACACACACACATGAACNNNAGATCGGAAG |
| 29289_2 | TATGAGTGTGGAGTCGTTGCCACAGAGACACACACACATGAACNNNAGATCGGAAG |
| 43027_1 | TATGAGTGTGGAGTCGTTGCAGAGAGAGACAGAGAAAGAGGGANNNAGATCGGA |
| 43027_2 | TATGAGTGTGGAGTCGTTGCAGAGAGAGACAGAGAAAGAGGGANNNAGATCGGA |
| 9950_1 | TATGAGTGTGGAGTCGTTGCAGCAGCAGCATGGCATAGTAAATNNNAGATCGGAA |
| 9950_2 | TATGAGTGTGGAGTCGTTGCAGCAGCAGCATGGCATAGTAAATNNNAGATCGGAA |
| 34450_1 | TATGAGTGTGGAGTCGTTGCGAGCCAACCTGGAGCTGACCNNNAGATCGGAAGAG |
| 34450_2 | TATGAGTGTGGAGTCGTTGCGAGCCAACCTGGAGCTGACCNNNAGATCGGAAGAG |
| 41400_1 | TATGAGTGTGGAGTCGTTGCCGCCTGTAATCCCAACACTTTGNNNAGATCGGAAGA |
| 41400_2 | TATGAGTGTGGAGTCGTTGCCGCCTGTAATCCCAACACTTTGNNNAGATCGGAAGA |
| 43739_1 | TATGAGTGTGGAGTCGTTGCGCTAGAACTACGGCTCAACCATANNNAGATCGGAAG |
| 43739_2 | TATGAGTGTGGAGTCGTTGCGCTAGAACTACGGCTCAACCATANNNAGATCGGAAG |
| 52943_1 | TATGAGTGTGGAGTCGTTGCGGTCATCTATCCCATGTGAAGCTNNNAGATCGGAAG |
| 52943_2 | TATGAGTGTGGAGTCGTTGCGGTCATCTATCCCATGTGAAGCTNNNAGATCGGAAG |
| 29891_1 | TATGAGTGTGGAGTCGTTGCTTGTTGTTGTCATGGATGTGGTGNNNAGATCGGAA |
| 29891_2 | TATGAGTGTGGAGTCGTTGCTTGTTGTTGTCATGGATGTGGTGNNNAGATCGGAA |
| 29500_1 | TATGAGTGTGGAGTCGTTGCGATTGTTTTTCTTCCTCGCATTNNNAGATCGGAAGAG |
| 29500_2 | TATGAGTGTGGAGTCGTTGCGATTGTTTTTCTTCCTCGCATTNNNAGATCGGAAGAG |
| 46089_1 | TATGAGTGTGGAGTCGTTGCAGGAAAGGAAATTGCTTGAGTGCNNNAGATCGGAA |
| 46089_2 | TATGAGTGTGGAGTCGTTGCAGGAAAGGAAATTGCTTGAGTGCNNNAGATCGGAA |
| 29455_1 | TATGAGTGTGGAGTCGTTGCGTGACAACTGACAGCTACTCAGGNNNAGATCGGAA |
| 29455_2 | TATGAGTGTGGAGTCGTTGCGTGACAACTGACAGCTACTCAGGNNNAGATCGGAA |
| 44185_1 | TATGAGTGTGGAGTCGTTGCCCTAAAGTGCTGTCAGTTCCTCANNNAGATCGGAAG |
| 44185_2 | TATGAGTGTGGAGTCGTTGCCCTAAAGTGCTGTCAGTTCCTCANNNAGATCGGAAG |
| 2360_1 | TATGAGTGTGGAGTCGTTGCGAGAGGGAGAGAAATCTGCTTCCNNNAGATCGGAA |
| 2360_2 | TATGAGTGTGGAGTCGTTGCGAGAGGGAGAGAAATCTGCTTCCNNNAGATCGGAA |

| | |
|---|---|
| 52810_1 | TATGAGTGTGGAGTCGTTGCAAGATAAGACACGAGACAGGCAGNNNAGATCGGAA |
| 52810_2 | TATGAGTGTGGAGTCGTTGCAAGATAAGACACGAGACAGGCAGNNNAGATCGGAA |
| 41162_1 | TATGAGTGTGGAGTCGTTGCTTCATTGCAACCTGATCTGCTTTNNNAGATCGGAAGA |
| 41162_2 | TATGAGTGTGGAGTCGTTGCTTCATTGCAACCTGATCTGCTTTNNNAGATCGGAAGA |
| 43677_1 | TATGAGTGTGGAGTCGTTGCCCAAGGCTGAACTCAACTTTGTTNNNAGATCGGAAG |
| 43677_2 | TATGAGTGTGGAGTCGTTGCCCAAGGCTGAACTCAACTTTGTTNNNAGATCGGAAG |
| 49823_1 | TATGAGTGTGGAGTCGTTGCGTAACACAGAAATGGTGAGGTGCNNNAGATCGGAA |
| 49823_2 | TATGAGTGTGGAGTCGTTGCGTAACACAGAAATGGTGAGGTGCNNNAGATCGGAA |
| 40396_1 | TATGAGTGTGGAGTCGTTGCCCTAGTGAGAGTGAGACCACAACNNNAGATCGGAA |
| 40396_2 | TATGAGTGTGGAGTCGTTGCCCTAGTGAGAGTGAGACCACAACNNNAGATCGGAA |
| 36798_1 | TATGAGTGTGGAGTCGTTGCCCCCAGAACCATGAGAGAGAAAANNNAGATCGGAA |
| 36798_2 | TATGAGTGTGGAGTCGTTGCCCCCAGAACCATGAGAGAGAAAANNNAGATCGGAA |
| 48949_1 | TATGAGTGTGGAGTCGTTGCCAACCCCTTTTCATGCATCCATGNNNAGATCGGAAGA |
| 48949_2 | TATGAGTGTGGAGTCGTTGCCAACCCCTTTTCATGCATCCATGNNNAGATCGGAAGA |
| 45770_1 | TATGAGTGTGGAGTCGTTGCAAGAAACTTGGGACAAGGGGGNNNAGATCGGAAG |
| 45770_2 | TATGAGTGTGGAGTCGTTGCAAGAAACTTGGGACAAGGGGGNNNAGATCGGAAG |
| 45501_1 | TATGAGTGTGGAGTCGTTGCTTGGACTTGGAGGACAGATCAAGNNNAGATCGGAA |
| 45501_2 | TATGAGTGTGGAGTCGTTGCTTGGACTTGGAGGACAGATCAAGNNNAGATCGGAA |
| 48886_1 | TATGAGTGTGGAGTCGTTGCATGACAGAGTGAAGTGAGTCACCNNNAGATCGGAA |
| 48886_2 | TATGAGTGTGGAGTCGTTGCATGACAGAGTGAAGTGAGTCACCNNNAGATCGGAA |
| 10398_1 | TATGAGTGTGGAGTCGTTGCTGTTTTGGAAACCGGAGTGAAGTNNNAGATCGGAA |
| 10398_2 | TATGAGTGTGGAGTCGTTGCTGTTTTGGAAACCGGAGTGAAGTNNNAGATCGGAA |
| 50046_1 | TATGAGTGTGGAGTCGTTGCCCCTGGGTTTTATCCTGCTTTTGNNNAGATCGGAAGA |
| 50046_2 | TATGAGTGTGGAGTCGTTGCCCCTGGGTTTTATCCTGCTTTTGNNNAGATCGGAAGA |
| 35344_1 | TATGAGTGTGGAGTCGTTGCGAGTCTCACTCTATTGCCTAGGCNNNAGATCGGAAG |
| 35344_2 | TATGAGTGTGGAGTCGTTGCGAGTCTCACTCTATTGCCTAGGCNNNAGATCGGAAG |
| 42045_1 | TATGAGTGTGGAGTCGTTGCGCTTTGGGCATACTTCAAAATGCNNNAGATCGGAAG |
| 42045_2 | TATGAGTGTGGAGTCGTTGCGCTTTGGGCATACTTCAAAATGCNNNAGATCGGAAG |
| 33832_1 | TATGAGTGTGGAGTCGTTGCAACCCCCCAAATTATACCAACCANNNAGATCGGAAGA |
| 33832_2 | TATGAGTGTGGAGTCGTTGCAACCCCCCAAATTATACCAACCANNNAGATCGGAAGA |
| 45209_1 | TATGAGTGTGGAGTCGTTGCATGAGATGAGCTGTGGGCAATATNNNAGATCGGAA |
| 45209_2 | TATGAGTGTGGAGTCGTTGCATGAGATGAGCTGTGGGCAATATNNNAGATCGGAA |
| 13057_1 | TATGAGTGTGGAGTCGTTGCGGAAACATCATGAAGAATAAAGANNNAGATCGGAA |
| 13057_2 | TATGAGTGTGGAGTCGTTGCGGAAACATCATGAAGAATAAAGANNNAGATCGGAA |
| 2002_1 | TATGAGTGTGGAGTCGTTGCAACAGCTGCAGACAGAGAACAAANNNAGATCGGAA |
| 2002_2 | TATGAGTGTGGAGTCGTTGCAACAGCTGCAGACAGAGAACAAANNNAGATCGGAA |
| 12799_1 | TATGAGTGTGGAGTCGTTGCTAGAAACAAGACAGATGTTAAACNNNAGATCGGAA |
| 12799_2 | TATGAGTGTGGAGTCGTTGCTAGAAACAAGACAGATGTTAAACNNNAGATCGGAA |
| 52972_1 | TATGAGTGTGGAGTCGTTGCGGGATTGCGAGAGTTTGTTTCAANNNAGATCGGAA |
| 52972_2 | TATGAGTGTGGAGTCGTTGCGGGATTGCGAGAGTTTGTTTCAANNNAGATCGGAA |
| 51882_1 | TATGAGTGTGGAGTCGTTGCGCCTTCTGCCATGATTGTAAGTTNNNAGATCGGAAG |
| 51882_2 | TATGAGTGTGGAGTCGTTGCGCCTTCTGCCATGATTGTAAGTTNNNAGATCGGAAG |
| 38638_1 | TATGAGTGTGGAGTCGTTGCTGACTCTTGGGACTTTGTCTGTTNNNAGATCGGAAG |

| | |
|---|---|
| 38638_2 | TATGAGTGTGGAGTCGTTGCTGACTCTTGGGACTTTGTCTGTTNNNAGATCGGAAG |
| 37647_1 | TATGAGTGTGGAGTCGTTGCAGATGAGTGTGGTAGTAAAAGCCANNNAGATCGGA |
| 37647_2 | TATGAGTGTGGAGTCGTTGCAGATGAGTGTGGTAGTAAAAGCCANNNAGATCGGA |
| 34098_1 | TATGAGTGTGGAGTCGTTGCGTGCCTGTAATCCCAGCTACTCNNNAGATCGGAAGA |
| 34098_2 | TATGAGTGTGGAGTCGTTGCGTGCCTGTAATCCCAGCTACTCNNNAGATCGGAAGA |
| 34996_1 | TATGAGTGTGGAGTCGTTGCGATAATGACGTGATGGCTGGAGTNNNAGATCGGAA |
| 34996_2 | TATGAGTGTGGAGTCGTTGCGATAATGACGTGATGGCTGGAGTNNNAGATCGGAA |
| 29928_1 | TATGAGTGTGGAGTCGTTGCAAGTAGGCAACATTCAGTTTGCCNNNAGATCGGAAG |
| 29928_2 | TATGAGTGTGGAGTCGTTGCAAGTAGGCAACATTCAGTTTGCCNNNAGATCGGAAG |
| 33811_1 | TATGAGTGTGGAGTCGTTGCAAAAACAAGAAGCTGCCTCTTGGNNNAGATCGGAA |
| 33811_2 | TATGAGTGTGGAGTCGTTGCAAAAACAAGAAGCTGCCTCTTGGNNNAGATCGGAA |
| 52970_1 | TATGAGTGTGGAGTCGTTGCGTCCCCCACCTAAATCTCATGTTNNNAGATCGGAAGA |
| 52970_2 | TATGAGTGTGGAGTCGTTGCGTCCCCCACCTAAATCTCATGTTNNNAGATCGGAAGA |
| 46615_1 | TATGAGTGTGGAGTCGTTGCGATTAGCTGGGACTACAGGCATGNNNAGATCGGAA |
| 46615_2 | TATGAGTGTGGAGTCGTTGCGATTAGCTGGGACTACAGGCATGNNNAGATCGGAA |
| 44732_1 | TATGAGTGTGGAGTCGTTGCAGGTGAGCATGGAGTGAGTTTTTNNNAGATCGGAA |
| 44732_2 | TATGAGTGTGGAGTCGTTGCAGGTGAGCATGGAGTGAGTTTTTNNNAGATCGGAA |
| 48405_1 | TATGAGTGTGGAGTCGTTGCAACACACACACACCTATGAGCCNNNAGATCGGAAGA |
| 48405_2 | TATGAGTGTGGAGTCGTTGCAACACACACACACCTATGAGCCNNNAGATCGGAAGA |
| 50960_1 | TATGAGTGTGGAGTCGTTGCTGAGGGGCTTGATGAAAACCTTANNNAGATCGGAA |
| 50960_2 | TATGAGTGTGGAGTCGTTGCTGAGGGGCTTGATGAAAACCTTANNNAGATCGGAA |
| 50735_1 | TATGAGTGTGGAGTCGTTGCCATAGTATCACCTTTTCAGCGCGNNNAGATCGGAAG |
| 50735_2 | TATGAGTGTGGAGTCGTTGCCATAGTATCACCTTTTCAGCGCGNNNAGATCGGAAG |
| 43145_1 | TATGAGTGTGGAGTCGTTGCAGTGAGATGGAGGTCCTGAACTANNNAGATCGGAA |
| 43145_2 | TATGAGTGTGGAGTCGTTGCAGTGAGATGGAGGTCCTGAACTANNNAGATCGGAA |
| 44415_1 | TATGAGTGTGGAGTCGTTGCAGAGCAGAGGCAGGAAATAGTTTNNNAGATCGGAA |
| 44415_2 | TATGAGTGTGGAGTCGTTGCAGAGCAGAGGCAGGAAATAGTTTNNNAGATCGGAA |
| 50747_1 | TATGAGTGTGGAGTCGTTGCACCATCTCCTTGTTTCTGTCACTNNNAGATCGGAAGA |
| 50747_2 | TATGAGTGTGGAGTCGTTGCACCATCTCCTTGTTTCTGTCACTNNNAGATCGGAAGA |
| 50490_1 | TATGAGTGTGGAGTCGTTGCACAAACAAGTGAGGAGAGGGTTTNNNAGATCGGAA |
| 50490_2 | TATGAGTGTGGAGTCGTTGCACAAACAAGTGAGGAGAGGGTTTNNNAGATCGGAA |
| 53001_1 | TATGAGTGTGGAGTCGTTGCCAAGAGAGAAATCCACCCCATGANNNAGATCGGAAG |
| 53001_2 | TATGAGTGTGGAGTCGTTGCCAAGAGAGAAATCCACCCCATGANNNAGATCGGAAG |
| 1246_1 | TATGAGTGTGGAGTCGTTGCTACAAGTCTGCACTTACGTGTGTNNNAGATCGGAAG |
| 1246_2 | TATGAGTGTGGAGTCGTTGCTACAAGTCTGCACTTACGTGTGTNNNAGATCGGAAG |
| 40969_1 | TATGAGTGTGGAGTCGTTGCGAAGAGGACGCCAAAAAATGGAANNNAGATCGGAA |
| 40969_2 | TATGAGTGTGGAGTCGTTGCGAAGAGGACGCCAAAAAATGGAANNNAGATCGGAA |
| 40732_1 | TATGAGTGTGGAGTCGTTGCAGTGGGAGAGAAAGAGGACTGTANNNAGATCGGA |
| 40732_2 | TATGAGTGTGGAGTCGTTGCAGTGGGAGAGAAAGAGGACTGTANNNAGATCGGA |
| 42466_1 | TATGAGTGTGGAGTCGTTGCTCTGGCTAGAGAGGTCATTCCNNNAGATCGGAAGAG |
| 42466_2 | TATGAGTGTGGAGTCGTTGCTCTGGCTAGAGAGGTCATTCCNNNAGATCGGAAGAG |
| 46589_1 | TATGAGTGTGGAGTCGTTGCTGAACTAAAATCCAAAGGCTGCCNNNAGATCGGAAG |
| 46589_2 | TATGAGTGTGGAGTCGTTGCTGAACTAAAATCCAAAGGCTGCCNNNAGATCGGAAG |

| | |
|---|---|
| 47195_1 | TATGAGTGTGGAGTCGTTGCGCTAGGATCACAGATTCACACCANNNAGATCGGAAG |
| 47195_2 | TATGAGTGTGGAGTCGTTGCGCTAGGATCACAGATTCACACCANNNAGATCGGAAG |
| 37151_1 | TATGAGTGTGGAGTCGTTGCCCCTTGCAACAATCTCACCTTTTNNNAGATCGGAAGA |
| 37151_2 | TATGAGTGTGGAGTCGTTGCCCCTTGCAACAATCTCACCTTTTNNNAGATCGGAAGA |
| 40473_1 | TATGAGTGTGGAGTCGTTGCCAGAGTTGAAGGGAAAGGGGATTNNNAGATCGGA |
| 40473_2 | TATGAGTGTGGAGTCGTTGCCAGAGTTGAAGGGAAAGGGGATTNNNAGATCGGA |
| 47343_1 | TATGAGTGTGGAGTCGTTGCTTCACCACATTCTCCTGCCTTAGNNNAGATCGGAAGA |
| 47343_2 | TATGAGTGTGGAGTCGTTGCTTCACCACATTCTCCTGCCTTAGNNNAGATCGGAAGA |
| 40044_1 | TATGAGTGTGGAGTCGTTGCCAGAGTGGGATTCTGTCTCAAAANNNAGATCGGAA |
| 40044_2 | TATGAGTGTGGAGTCGTTGCCAGAGTGGGATTCTGTCTCAAAANNNAGATCGGAA |
| 33115_1 | TATGAGTGTGGAGTCGTTGCCGGGTTTCACTTTCTTGGTTCTCNNNAGATCGGAAG |
| 33115_2 | TATGAGTGTGGAGTCGTTGCCGGGTTTCACTTTCTTGGTTCTCNNNAGATCGGAAG |
| 35148_1 | TATGAGTGTGGAGTCGTTGCGTAGGTGGGACTACAGGAATGTGNNNAGATCGGA |
| 35148_2 | TATGAGTGTGGAGTCGTTGCGTAGGTGGGACTACAGGAATGTGNNNAGATCGGA |
| 39945_1 | TATGAGTGTGGAGTCGTTGCTTTTTTTGAGAGGGAGTCTCGCTNNNAGATCGGAAG |
| 39945_2 | TATGAGTGTGGAGTCGTTGCTTTTTTTGAGAGGGAGTCTCGCTNNNAGATCGGAAG |
| 44124_1 | TATGAGTGTGGAGTCGTTGCTAGAAATGCCTGAACAGAGACCCNNNAGATCGGAA |
| 44124_2 | TATGAGTGTGGAGTCGTTGCTAGAAATGCCTGAACAGAGACCCNNNAGATCGGAA |
| 28778_1 | TATGAGTGTGGAGTCGTTGCAGATTGCTGGGTCAAGTGGTATTNNNAGATCGGAA |
| 28778_2 | TATGAGTGTGGAGTCGTTGCAGATTGCTGGGTCAAGTGGTATTNNNAGATCGGAA |
| 39744_1 | TATGAGTGTGGAGTCGTTGCGAAGAAACTGAGGCACAGAGAGANNNAGATCGGA |
| 39744_2 | TATGAGTGTGGAGTCGTTGCGAAGAAACTGAGGCACAGAGAGANNNAGATCGGA |
| 12992_1 | TATGAGTGTGGAGTCGTTGCCATGATCGTACTTGTCCTATTATNNNAGATCGGAAG |
| 12992_2 | TATGAGTGTGGAGTCGTTGCCATGATCGTACTTGTCCTATTATNNNAGATCGGAAG |
| 41304_1 | TATGAGTGTGGAGTCGTTGCCACAAACGTAGTGATCTTGCCAGNNNAGATCGGAAG |
| 41304_2 | TATGAGTGTGGAGTCGTTGCCACAAACGTAGTGATCTTGCCAGNNNAGATCGGAAG |
| 49469_1 | TATGAGTGTGGAGTCGTTGCTTGCTATGGATAAGGGCTTCCAGNNNAGATCGGAA |
| 49469_2 | TATGAGTGTGGAGTCGTTGCTTGCTATGGATAAGGGCTTCCAGNNNAGATCGGAA |
| 38179_1 | TATGAGTGTGGAGTCGTTGCGGCTCAAACCTGTAATCCCAGTANNNAGATCGGAAG |
| 38179_2 | TATGAGTGTGGAGTCGTTGCGGCTCAAACCTGTAATCCCAGTANNNAGATCGGAAG |
| 37669_1 | TATGAGTGTGGAGTCGTTGCTGGTTTTCCATTCCTGAGTTACTNNNAGATCGGAAG |
| 37669_2 | TATGAGTGTGGAGTCGTTGCTGGTTTTCCATTCCTGAGTTACTNNNAGATCGGAAG |
| 37355_1 | TATGAGTGTGGAGTCGTTGCTTAGTTGGGAGTAGTGGCTCATGNNNAGATCGGAA |
| 37355_2 | TATGAGTGTGGAGTCGTTGCTTAGTTGGGAGTAGTGGCTCATGNNNAGATCGGAA |
| 53354_1 | TATGAGTGTGGAGTCGTTGCTCTGTCTCTCTCTCTCCTTCCNNNAGATCGGAAGAG |
| 53354_2 | TATGAGTGTGGAGTCGTTGCTCTGTCTCTCTCTCTCCTTCCNNNAGATCGGAAGAG |
| 37021_1 | TATGAGTGTGGAGTCGTTGCAAAAAAAACAATGCTCTCCTGCANNNAGATCGGAAG |
| 37021_2 | TATGAGTGTGGAGTCGTTGCAAAAAAAACAATGCTCTCCTGCANNNAGATCGGAAG |
| 48561_1 | TATGAGTGTGGAGTCGTTGCTAGGTTCAAGTGATTCTCCTGCCNNNAGATCGGAAG |
| 48561_2 | TATGAGTGTGGAGTCGTTGCTAGGTTCAAGTGATTCTCCTGCCNNNAGATCGGAAG |
| 38564_1 | TATGAGTGTGGAGTCGTTGCCATACACACACACACACACAACANNNAGATCGGAAG |
| 38564_2 | TATGAGTGTGGAGTCGTTGCCATACACACACACACACACAACANNNAGATCGGAAG |
| 43807_1 | TATGAGTGTGGAGTCGTTGCACAGAACAGCCAAAATCATGGTACNNNAGATCGGA |

| | |
|---|---|
| 43807_2 | TATGAGTGTGGAGTCGTTGCACAGAACAGCCAAAATCATGGTACNNNAGATCGGAA |
| 48783_1 | TATGAGTGTGGAGTCGTTGCTCTGCCTCCTAGATTCCAGTGATNNNAGATCGGAAGA |
| 48783_2 | TATGAGTGTGGAGTCGTTGCTCTGCCTCCTAGATTCCAGTGATNNNAGATCGGAAGA |
| 10944_1 | TATGAGTGTGGAGTCGTTGCTCACAGTGGTCACAGCTACTTGNNNAGATCGGAAGA |
| 10944_2 | TATGAGTGTGGAGTCGTTGCTCACAGTGGTCACAGCTACTTGNNNAGATCGGAAGA |
| 35817_1 | TATGAGTGTGGAGTCGTTGCTCCTGGGTTCAAAGTGATTCTCCNNNAGATCGGAAGA |
| 35817_2 | TATGAGTGTGGAGTCGTTGCTCCTGGGTTCAAAGTGATTCTCCNNNAGATCGGAAGA |
| 49574_1 | TATGAGTGTGGAGTCGTTGCAATCTCGGCTCACATCAACTTCTNNNAGATCGGAAGA |
| 49574_2 | TATGAGTGTGGAGTCGTTGCAATCTCGGCTCACATCAACTTCTNNNAGATCGGAAGA |
| 38672_1 | TATGAGTGTGGAGTCGTTGCCTTCTGGGAAAATGGAGGGAGAANNNAGATCGGAA |
| 38672_2 | TATGAGTGTGGAGTCGTTGCCTTCTGGGAAAATGGAGGGAGAANNNAGATCGGAA |
| 50833_1 | TATGAGTGTGGAGTCGTTGCAACCAATGTTCCCTGAAGACTCANNNAGATCGGAAGA |
| 50833_2 | TATGAGTGTGGAGTCGTTGCAACCAATGTTCCCTGAAGACTCANNNAGATCGGAAGA |
| 49888_1 | TATGAGTGTGGAGTCGTTGCGTCCATTCTCCTGGAGAGTAAGCNNNAGATCGGAAG |
| 49888_2 | TATGAGTGTGGAGTCGTTGCGTCCATTCTCCTGGAGAGTAAGCNNNAGATCGGAAG |
| 41880_1 | TATGAGTGTGGAGTCGTTGCCTCCTGGGTTCACACCATTCTCNNNAGATCGGAAGAG |
| 41880_2 | TATGAGTGTGGAGTCGTTGCCTCCTGGGTTCACACCATTCTCNNNAGATCGGAAGAG |
| 39728_1 | TATGAGTGTGGAGTCGTTGCGAGGCAGGTGAATCGTTTGAACNNNAGATCGGAAG |
| 39728_2 | TATGAGTGTGGAGTCGTTGCGAGGCAGGTGAATCGTTTGAACNNNAGATCGGAAG |
| 49541_1 | TATGAGTGTGGAGTCGTTGCGCCCAGCCATGAATGTTCTTTTTNNNAGATCGGAAGA |
| 49541_2 | TATGAGTGTGGAGTCGTTGCGCCCAGCCATGAATGTTCTTTTTNNNAGATCGGAAGA |
| 31068_1 | TATGAGTGTGGAGTCGTTGCGGTCAGTATTATTGGCCTCAGCTNNNAGATCGGAAG |
| 31068_2 | TATGAGTGTGGAGTCGTTGCGGTCAGTATTATTGGCCTCAGCTNNNAGATCGGAAG |
| 8139_1 | TATGAGTGTGGAGTCGTTGCAATCCTCCAGTCCTAAGGGTGGNNNAGATCGGAAGA |
| 8139_2 | TATGAGTGTGGAGTCGTTGCAATCCTCCAGTCCTAAGGGTGGNNNAGATCGGAAGA |
| 33867_1 | TATGAGTGTGGAGTCGTTGCACACATACTCGTTGAGAGCTCTGNNNAGATCGGAAG |
| 33867_2 | TATGAGTGTGGAGTCGTTGCACACATACTCGTTGAGAGCTCTGNNNAGATCGGAAG |
| 9091_1 | TATGAGTGTGGAGTCGTTGCGTAAGCTTTTTGTGAGCTGTCACTNNNAGATCGGAA |
| 9091_2 | TATGAGTGTGGAGTCGTTGCGTAAGCTTTTTGTGAGCTGTCACTNNNAGATCGGAA |
| 41934_1 | TATGAGTGTGGAGTCGTTGCCTCAGTGCAACCTCCACCTCNNNAGATCGGAAGAGCA |
| 41934_2 | TATGAGTGTGGAGTCGTTGCCTCAGTGCAACCTCCACCTCNNNAGATCGGAAGAGCA |
| 34513_1 | TATGAGTGTGGAGTCGTTGCGATAGTAGAGATGTCCAGCACCCNNNAGATCGGAAG |
| 34513_2 | TATGAGTGTGGAGTCGTTGCGATAGTAGAGATGTCCAGCACCCNNNAGATCGGAAG |
| 49559_1 | TATGAGTGTGGAGTCGTTGCGTCTACATCAAAGGCCAGGGTTANNNAGATCGGAAG |
| 49559_2 | TATGAGTGTGGAGTCGTTGCGTCTACATCAAAGGCCAGGGTTANNNAGATCGGAAG |
| 13272_1 | TATGAGTGTGGAGTCGTTGCGAGTGACACAATGGGAAAAGNNNAGATCGGAAGAC |
| 13272_2 | TATGAGTGTGGAGTCGTTGCGAGTGACACAATGGGAAAAGNNNAGATCGGAAGAC |
| 48255_1 | TATGAGTGTGGAGTCGTTGCAGCCTCACTCTGTTACTCTGTTGNNNAGATCGGAAGA |
| 48255_2 | TATGAGTGTGGAGTCGTTGCAGCCTCACTCTGTTACTCTGTTGNNNAGATCGGAAGA |
| 34140_1 | TATGAGTGTGGAGTCGTTGCAGGGGAGTAGGGGAAAGAAGATANNNAGATCGGA |
| 34140_2 | TATGAGTGTGGAGTCGTTGCAGGGGAGTAGGGGAAAGAAGATANNNAGATCGGA |
| 47762_1 | TATGAGTGTGGAGTCGTTGCCACCAAGCCTGGCTAATTTTTGTNNNAGATCGGAAGA |
| 47762_2 | TATGAGTGTGGAGTCGTTGCCACCAAGCCTGGCTAATTTTTGTNNNAGATCGGAAGA |

| | |
|---|---|
| 52616_1 | TATGAGTGTGGAGTCGTTGCCTGGCCTCTCCCTGTTTTCTTATNNNAGATCGGAAGA |
| 52616_2 | TATGAGTGTGGAGTCGTTGCCTGGCCTCTCCCTGTTTTCTTATNNNAGATCGGAAGA |
| 39499_1 | TATGAGTGTGGAGTCGTTGCGGAGGTTGCAGTGAGTCATGATANNNAGATCGGAA |
| 39499_2 | TATGAGTGTGGAGTCGTTGCGGAGGTTGCAGTGAGTCATGATANNNAGATCGGAA |
| 39328_1 | TATGAGTGTGGAGTCGTTGCCAGAATGGACTTTGGTGACTTGGNNNAGATCGGAA |
| 39328_2 | TATGAGTGTGGAGTCGTTGCCAGAATGGACTTTGGTGACTTGGNNNAGATCGGAA |
| 39010_1 | TATGAGTGTGGAGTCGTTGCAGTCTACCCAAATGAGAAGAAACAGANNNAGATCG |
| 39010_2 | TATGAGTGTGGAGTCGTTGCAGTCTACCCAAATGAGAAGAAACAGANNNAGATCG |
| 47642_1 | TATGAGTGTGGAGTCGTTGCAAGTGATCCTCCTGCCTTGATTTNNNAGATCGGAAG |
| 47642_2 | TATGAGTGTGGAGTCGTTGCAAGTGATCCTCCTGCCTTGATTTNNNAGATCGGAAG |
| 41537_1 | TATGAGTGTGGAGTCGTTGCGAGTAGCTGGGATTACAGGAGTGNNNAGATCGGA |
| 41537_2 | TATGAGTGTGGAGTCGTTGCGAGTAGCTGGGATTACAGGAGTGNNNAGATCGGA |
| 32394_1 | TATGAGTGTGGAGTCGTTGCCAAGTAGCTGGGATTACAGGCATNNNAGATCGGAA |
| 32394_2 | TATGAGTGTGGAGTCGTTGCCAAGTAGCTGGGATTACAGGCATNNNAGATCGGAA |
| 36351_1 | TATGAGTGTGGAGTCGTTGCACTGCCAAGCTCACTCAATAAGANNNAGATCGGAAG |
| 36351_2 | TATGAGTGTGGAGTCGTTGCACTGCCAAGCTCACTCAATAAGANNNAGATCGGAAG |
| 30849_1 | TATGAGTGTGGAGTCGTTGCGCACACCAAAAGGGGAGAGATTANNNAGATCGGAA |
| 30849_2 | TATGAGTGTGGAGTCGTTGCGCACACCAAAAGGGGAGAGATTANNNAGATCGGAA |
| 43829_1 | TATGAGTGTGGAGTCGTTGCCTGAAACCACAGACATCCGTTTCNNNAGATCGGAAG |
| 43829_2 | TATGAGTGTGGAGTCGTTGCCTGAAACCACAGACATCCGTTTCNNNAGATCGGAAG |
| 49303_1 | TATGAGTGTGGAGTCGTTGCTGATCTCACCTCAATGCAACCTCNNNAGATCGGAAGA |
| 49303_2 | TATGAGTGTGGAGTCGTTGCTGATCTCACCTCAATGCAACCTCNNNAGATCGGAAGA |
| 53258_1 | TATGAGTGTGGAGTCGTTGCTTCATAGGATCACAACTTGGCCANNNAGATCGGAAG |
| 53258_2 | TATGAGTGTGGAGTCGTTGCTTCATAGGATCACAACTTGGCCANNNAGATCGGAAG |
| 32311_1 | TATGAGTGTGGAGTCGTTGCTTGATGCTGTTGTTGCTTTCTGTNNNAGATCGGAAG |
| 32311_2 | TATGAGTGTGGAGTCGTTGCTTGATGCTGTTGTTGCTTTCTGTNNNAGATCGGAAG |
| 43965_1 | TATGAGTGTGGAGTCGTTGCGGGAGGGAGGGAAAGAAAAGAAANNNAGATCGG |
| 43965_2 | TATGAGTGTGGAGTCGTTGCGGGAGGGAGGGAAAGAAAAGAAANNNAGATCGG |
| 10142_1 | TATGAGTGTGGAGTCGTTGCGCCAAGAGGAGATAACAAGGTGGNNNAGATCGGA |
| 10142_2 | TATGAGTGTGGAGTCGTTGCGCCAAGAGGAGATAACAAGGTGGNNNAGATCGGA |
| 45038_1 | TATGAGTGTGGAGTCGTTGCCACTGGCTGAACTCTTAGGTAGGNNNAGATCGGAA |
| 45038_2 | TATGAGTGTGGAGTCGTTGCCACTGGCTGAACTCTTAGGTAGGNNNAGATCGGAA |
| 42424_1 | TATGAGTGTGGAGTCGTTGCTGCGAGAGAGAGAGAGAGAAAGANNNAGATCGGA |
| 42424_2 | TATGAGTGTGGAGTCGTTGCTGCGAGAGAGAGAGAGAGAAAGANNNAGATCGGA |
| 42899_1 | TATGAGTGTGGAGTCGTTGCTGTGTGTGAGAGATGGAGTCTTGNNNAGATCGGAA |
| 42899_2 | TATGAGTGTGGAGTCGTTGCTGTGTGTGAGAGATGGAGTCTTGNNNAGATCGGAA |
| 10836_1 | TATGAGTGTGGAGTCGTTGCAAAGAAAACAGTATGTGCTCCTCANNNAGATCGGAA |
| 10836_2 | TATGAGTGTGGAGTCGTTGCAAAGAAAACAGTATGTGCTCCTCANNNAGATCGGAA |
| 44456_1 | TATGAGTGTGGAGTCGTTGCGCCTCTTTCACAAACACATAGCANNNAGATCGGAAG |
| 44456_2 | TATGAGTGTGGAGTCGTTGCGCCTCTTTCACAAACACATAGCANNNAGATCGGAAG |
| 42798_1 | TATGAGTGTGGAGTCGTTGCTCACAAAAACACAGCACATCTGGNNNAGATCGGAAG |
| 42798_2 | TATGAGTGTGGAGTCGTTGCTCACAAAAACACAGCACATCTGGNNNAGATCGGAAG |
| 13647_1 | TATGAGTGTGGAGTCGTTGCGATACAGAAGTAGAGGAAAGTAANNNAGATCGGA |

| | |
|---|---|
| 13647_2 | TATGAGTGTGGAGTCGTTGCGATACAGAAGTAGAGGAAAGTAANNNAGATCGGA |
| 34960_1 | TATGAGTGTGGAGTCGTTGCGTGTGATCTCAGTTCATGGCAACNNNAGATCGGAAG |
| 34960_2 | TATGAGTGTGGAGTCGTTGCGTGTGATCTCAGTTCATGGCAACNNNAGATCGGAAG |
| 13507_1 | TATGAGTGTGGAGTCGTTGCTTTGTCTAGTCAAAACTGTGANNNAGATCGGAAGAG |
| 13507_2 | TATGAGTGTGGAGTCGTTGCTTTGTCTAGTCAAAACTGTGANNNAGATCGGAAGAG |
| 30541_1 | TATGAGTGTGGAGTCGTTGCTCCAGCCTCCCCATTTATTCAGANNNAGATCGGAAGA |
| 30541_2 | TATGAGTGTGGAGTCGTTGCTCCAGCCTCCCCATTTATTCAGANNNAGATCGGAAGA |
| 50118_1 | TATGAGTGTGGAGTCGTTGCTACATTTGCCCCTAGTTTCCTCCNNNAGATCGGAAGA |
| 50118_2 | TATGAGTGTGGAGTCGTTGCTACATTTGCCCCTAGTTTCCTCCNNNAGATCGGAAGA |
| 41037_1 | TATGAGTGTGGAGTCGTTGCCTCTCATTTGGACTGCTGGGATANNNAGATCGGAAG |
| 41037_2 | TATGAGTGTGGAGTCGTTGCCTCTCATTTGGACTGCTGGGATANNNAGATCGGAAG |
| 29052_1 | TATGAGTGTGGAGTCGTTGCTTTTTATGAGGGCACTGATCCCANNNAGATCGGAAG |
| 29052_2 | TATGAGTGTGGAGTCGTTGCTTTTTATGAGGGCACTGATCCCANNNAGATCGGAAG |
| 28875_1 | TATGAGTGTGGAGTCGTTGCAGAAGGGAAAGAGGAGGAGAAGANNNAGATCGG |
| 28875_2 | TATGAGTGTGGAGTCGTTGCAGAAGGGAAAGAGGAGGAGAAGANNNAGATCGG |
| 33853_1 | TATGAGTGTGGAGTCGTTGCCCCCAAAAGTCCCCAAATAGCTANNNAGATCGGAAG |
| 33853_2 | TATGAGTGTGGAGTCGTTGCCCCCAAAAGTCCCCAAATAGCTANNNAGATCGGAAG |
| 30869_1 | TATGAGTGTGGAGTCGTTGCAGGAGTCAATGGGAAATCTGGTTNNNAGATCGGAA |
| 30869_2 | TATGAGTGTGGAGTCGTTGCAGGAGTCAATGGGAAATCTGGTTNNNAGATCGGAA |
| 52731_1 | TATGAGTGTGGAGTCGTTGCATATTCTCAAATGGTGGGGAGGGNNNAGATCGGAA |
| 52731_2 | TATGAGTGTGGAGTCGTTGCATATTCTCAAATGGTGGGGAGGGNNNAGATCGGAA |
| 29611_1 | TATGAGTGTGGAGTCGTTGCAACAGAGTAACCAGAACTCGAGCNNNAGATCGGAA |
| 29611_2 | TATGAGTGTGGAGTCGTTGCAACAGAGTAACCAGAACTCGAGCNNNAGATCGGAA |
| 52109_1 | TATGAGTGTGGAGTCGTTGCCCAATGGAAGGCACTGTTAGGATNNNAGATCGGAA |
| 52109_2 | TATGAGTGTGGAGTCGTTGCCCAATGGAAGGCACTGTTAGGATNNNAGATCGGAA |
| 13231_1 | TATGAGTGTGGAGTCGTTGCCCAAATAGCTAGGACTCCAGNNNAGATCGGAAGAG |
| 13231_2 | TATGAGTGTGGAGTCGTTGCCCAAATAGCTAGGACTCCAGNNNAGATCGGAAGAG |
| 50245_1 | TATGAGTGTGGAGTCGTTGCGGCCTCTGTCTTCCTTCTAAGTCNNNAGATCGGAAGA |
| 50245_2 | TATGAGTGTGGAGTCGTTGCGGCCTCTGTCTTCCTTCTAAGTCNNNAGATCGGAAGA |
| 47570_1 | TATGAGTGTGGAGTCGTTGCCTTGAATTTCCAGGCTCAAGCAANNNAGATCGGAAG |
| 47570_2 | TATGAGTGTGGAGTCGTTGCCTTGAATTTCCAGGCTCAAGCAANNNAGATCGGAAG |
| 53382_1 | TATGAGTGTGGAGTCGTTGCCATCCCCATCACTCTCCTTTCTCNNNAGATCGGAAGA |
| 53382_2 | TATGAGTGTGGAGTCGTTGCCATCCCCATCACTCTCCTTTCTCNNNAGATCGGAAGA |
| 30341_1 | TATGAGTGTGGAGTCGTTGCTTGGGAAAACCAAAGCCAGAGAANNNAGATCGGAA |
| 30341_2 | TATGAGTGTGGAGTCGTTGCTTGGGAAAACCAAAGCCAGAGAANNNAGATCGGAA |
| 51663_1 | TATGAGTGTGGAGTCGTTGCAACAAAGAGTCTCTTCTCCCTGANNNAGATCGGAAG |
| 51663_2 | TATGAGTGTGGAGTCGTTGCAACAAAGAGTCTCTTCTCCCTGANNNAGATCGGAAG |
| 46865_1 | TATGAGTGTGGAGTCGTTGCCCTTCTGGCACATGTTCATTGATNNNAGATCGGAAG |
| 46865_2 | TATGAGTGTGGAGTCGTTGCCCTTCTGGCACATGTTCATTGATNNNAGATCGGAAG |
| 41234_1 | TATGAGTGTGGAGTCGTTGCCAAGGTCAAGAGATCGAGACCATNNNAGATCGGAA |
| 41234_2 | TATGAGTGTGGAGTCGTTGCCAAGGTCAAGAGATCGAGACCATNNNAGATCGGAA |
| 48037_1 | TATGAGTGTGGAGTCGTTGCTTCAGAAGGAATGGTACCAGCTCNNNAGATCGGAA |
| 48037_2 | TATGAGTGTGGAGTCGTTGCTTCAGAAGGAATGGTACCAGCTCNNNAGATCGGAA |

| | |
|---|---|
| 39664_1 | TATGAGTGTGGAGTCGTTGCTGTCTGGTACATGATAAGTGTGCANNNAGATCGGA |
| 39664_2 | TATGAGTGTGGAGTCGTTGCTGTCTGGTACATGATAAGTGTGCANNNAGATCGGA |
| 34192_1 | TATGAGTGTGGAGTCGTTGCTTTGCCACGACATACTTTGTAGCNNNAGATCGGAAG |
| 34192_2 | TATGAGTGTGGAGTCGTTGCTTTGCCACGACATACTTTGTAGCNNNAGATCGGAAG |
| 44217_1 | TATGAGTGTGGAGTCGTTGCAAACTGAAATGTCAAGAGCAGCCNNNAGATCGGAA |
| 44217_2 | TATGAGTGTGGAGTCGTTGCAAACTGAAATGTCAAGAGCAGCCNNNAGATCGGAA |
| 49858_1 | TATGAGTGTGGAGTCGTTGCAGGGAAGAAAACTGGCTCTTGTTNNNAGATCGGAA |
| 49858_2 | TATGAGTGTGGAGTCGTTGCAGGGAAGAAAACTGGCTCTTGTTNNNAGATCGGAA |
| 30857_1 | TATGAGTGTGGAGTCGTTGCTGTGTGGTGATCTGAAAGACAGTNNNAGATCGGAA |
| 30857_2 | TATGAGTGTGGAGTCGTTGCTGTGTGGTGATCTGAAAGACAGTNNNAGATCGGAA |
| 37341_1 | TATGAGTGTGGAGTCGTTGCAGAAGATTTCTTGAGCCCAGGAGNNNAGATCGGAA |
| 37341_2 | TATGAGTGTGGAGTCGTTGCAGAAGATTTCTTGAGCCCAGGAGNNNAGATCGGAA |
| 52538_1 | TATGAGTGTGGAGTCGTTGCAAGTTTGCTGTCTCCTTTGTTCGNNNAGATCGGAAG |
| 52538_2 | TATGAGTGTGGAGTCGTTGCAAGTTTGCTGTCTCCTTTGTTCGNNNAGATCGGAAG |
| 35564_1 | TATGAGTGTGGAGTCGTTGCTCTGCCTCACAAGTTCAAGTGATNNNAGATCGGAAG |
| 35564_2 | TATGAGTGTGGAGTCGTTGCTCTGCCTCACAAGTTCAAGTGATNNNAGATCGGAAG |
| 34697_1 | TATGAGTGTGGAGTCGTTGCCGCCCGGCTTTTCTTTTTTTTTNNNAGATCGGAAGA |
| 34697_2 | TATGAGTGTGGAGTCGTTGCCGCCCGGCTTTTCTTTTTTTTTNNNAGATCGGAAGA |
| 35876_1 | TATGAGTGTGGAGTCGTTGCATGGAGAAACCCCGTCTCTACTANNNAGATCGGAAG |
| 35876_2 | TATGAGTGTGGAGTCGTTGCATGGAGAAACCCCGTCTCTACTANNNAGATCGGAAG |
| 51242_1 | TATGAGTGTGGAGTCGTTGCTGTTGTGCACATATACCCCAGAANNNAGATCGGAAG |
| 51242_2 | TATGAGTGTGGAGTCGTTGCTGTTGTGCACATATACCCCAGAANNNAGATCGGAAG |
| 30244_1 | TATGAGTGTGGAGTCGTTGCACTCTTGTGGCATAACTGGATCANNNAGATCGGAAG |
| 30244_2 | TATGAGTGTGGAGTCGTTGCACTCTTGTGGCATAACTGGATCANNNAGATCGGAAG |
| 33839_1 | TATGAGTGTGGAGTCGTTGCCTTCCCAGCTCAAGTGATCCTCNNNAGATCGGAAGA |
| 33839_2 | TATGAGTGTGGAGTCGTTGCCTTCCCAGCTCAAGTGATCCTCNNNAGATCGGAAGA |
| 40049_1 | TATGAGTGTGGAGTCGTTGCCTGTAGTCCCAGCTACTCAGGAGNNNAGATCGGAAG |
| 40049_2 | TATGAGTGTGGAGTCGTTGCCTGTAGTCCCAGCTACTCAGGAGNNNAGATCGGAAG |
| 30437_1 | TATGAGTGTGGAGTCGTTGCATCTTTTCCACCAAGACTGTGGTNNNAGATCGGAAG |
| 30437_2 | TATGAGTGTGGAGTCGTTGCATCTTTTCCACCAAGACTGTGGTNNNAGATCGGAAG |
| 34905_1 | TATGAGTGTGGAGTCGTTGCCTGGGTTCAAGCGATTCTCCTNNNAGATCGGAAGAG |
| 34905_2 | TATGAGTGTGGAGTCGTTGCCTGGGTTCAAGCGATTCTCCTNNNAGATCGGAAGAG |
| 32613_1 | TATGAGTGTGGAGTCGTTGCAACCAACTCCTCAATCACCTGAANNNAGATCGGAAG |
| 32613_2 | TATGAGTGTGGAGTCGTTGCAACCAACTCCTCAATCACCTGAANNNAGATCGGAAG |
| 13493_1 | TATGAGTGTGGAGTCGTTGCAAAAAGTTCAGGATGTGTAGTNNNAGATCGGAAGA |
| 13493_2 | TATGAGTGTGGAGTCGTTGCAAAAAGTTCAGGATGTGTAGTNNNAGATCGGAAGA |
| 34334_1 | TATGAGTGTGGAGTCGTTGCTGTCCTTTGCAGGGATCCTATTCNNNAGATCGGAAG |
| 34334_2 | TATGAGTGTGGAGTCGTTGCTGTCCTTTGCAGGGATCCTATTCNNNAGATCGGAAG |
| 8349_1 | TATGAGTGTGGAGTCGTTGCAGTTATGCTCTTTCTCCCAGGCTANNNAGATCGGAAG |
| 8349_2 | TATGAGTGTGGAGTCGTTGCAGTTATGCTCTTTCTCCCAGGCTANNNAGATCGGAAG |
| 10397_1 | TATGAGTGTGGAGTCGTTGCCACCCATCCAATCAGTGTTCAGTNNNAGATCGGAAG |
| 10397_2 | TATGAGTGTGGAGTCGTTGCCACCCATCCAATCAGTGTTCAGTNNNAGATCGGAAG |
| 41298_1 | TATGAGTGTGGAGTCGTTGCGTCATCCTGCAAGTGACTCAAAGNNNAGATCGGAAG |

| | |
|---|---|
| 41298_2 | TATGAGTGTGGAGTCGTTGCGTCATCCTGCAAGTGACTCAAAGNNNAGATCGGAAG |
| 34714_1 | TATGAGTGTGGAGTCGTTGCCTGTTGCTCAGGCTATAGTCTCANNNAGATCGGAAG |
| 34714_2 | TATGAGTGTGGAGTCGTTGCCTGTTGCTCAGGCTATAGTCTCANNNAGATCGGAAG |
| 47207_1 | TATGAGTGTGGAGTCGTTGCCATTCTCCTGCCTCAGACTACTGNNNAGATCGGAAGA |
| 47207_2 | TATGAGTGTGGAGTCGTTGCCATTCTCCTGCCTCAGACTACTGNNNAGATCGGAAGA |
| 44356_1 | TATGAGTGTGGAGTCGTTGCAGAGGCAAGAGACAAAACAAAGCNNNAGATCGGAA |
| 44356_2 | TATGAGTGTGGAGTCGTTGCAGAGGCAAGAGACAAAACAAAGCNNNAGATCGGAA |
| 29098_1 | TATGAGTGTGGAGTCGTTGCCGCCTTTAGTTTTACAGCCTCTGNNNAGATCGGAAGA |
| 29098_2 | TATGAGTGTGGAGTCGTTGCCGCCTTTAGTTTTACAGCCTCTGNNNAGATCGGAAGA |
| 32009_1 | TATGAGTGTGGAGTCGTTGCGAACAGTTTCATCACCCCCAAAGNNNAGATCGGAAG |
| 32009_2 | TATGAGTGTGGAGTCGTTGCGAACAGTTTCATCACCCCCAAAGNNNAGATCGGAAG |
| 13673_1 | TATGAGTGTGGAGTCGTTGCCTTCCTGAAAACAACGTTCTNNNAGATCGGAAGAGC |
| 13673_2 | TATGAGTGTGGAGTCGTTGCCTTCCTGAAAACAACGTTCTNNNAGATCGGAAGAGC |
| 40865_1 | TATGAGTGTGGAGTCGTTGCTTGGCTCCTAATCTCATTTGGCTNNNAGATCGGAAGA |
| 40865_2 | TATGAGTGTGGAGTCGTTGCTTGGCTCCTAATCTCATTTGGCTNNNAGATCGGAAGA |
| 34273_1 | TATGAGTGTGGAGTCGTTGCATGCTAGATGACGAGTTAGTGGGNNNAGATCGGAA |
| 34273_2 | TATGAGTGTGGAGTCGTTGCATGCTAGATGACGAGTTAGTGGGNNNAGATCGGAA |
| 49771_1 | TATGAGTGTGGAGTCGTTGCTCATGTCCCTACTTCCCTTTTCGNNNAGATCGGAAGA |
| 49771_2 | TATGAGTGTGGAGTCGTTGCTCATGTCCCTACTTCCCTTTTCGNNNAGATCGGAAGA |
| 38849_1 | TATGAGTGTGGAGTCGTTGCCCATAGACTTTACCCTCCTTGGCNNNAGATCGGAAGA |
| 38849_2 | TATGAGTGTGGAGTCGTTGCCCATAGACTTTACCCTCCTTGGCNNNAGATCGGAAGA |
| 35959_1 | TATGAGTGTGGAGTCGTTGCCAAGATTATGCCCCTGTACTCCANNNAGATCGGAAG |
| 35959_2 | TATGAGTGTGGAGTCGTTGCCAAGATTATGCCCCTGTACTCCANNNAGATCGGAAG |
| 40008_1 | TATGAGTGTGGAGTCGTTGCGTGAAGCAGAGATGGAAGCATTCNNNAGATCGGAA |
| 40008_2 | TATGAGTGTGGAGTCGTTGCGTGAAGCAGAGATGGAAGCATTCNNNAGATCGGAA |
| 10765_1 | TATGAGTGTGGAGTCGTTGCGAGCAGGAACAAGAGCTGCTTATNNNAGATCGGAA |
| 10765_2 | TATGAGTGTGGAGTCGTTGCGAGCAGGAACAAGAGCTGCTTATNNNAGATCGGAA |
| 39849_1 | TATGAGTGTGGAGTCGTTGCAATGTTACTTTCTGACAGGCCCANNNAGATCGGAAG |
| 39849_2 | TATGAGTGTGGAGTCGTTGCAATGTTACTTTCTGACAGGCCCANNNAGATCGGAAG |
| 968_1 | TATGAGTGTGGAGTCGTTGCGGAAAGGCTGCTGGAGGGGCNNNAGATCGGAAGA |
| 968_2 | TATGAGTGTGGAGTCGTTGCGGAAAGGCTGCTGGAGGGGCNNNAGATCGGAAGA |
| 31688_1 | TATGAGTGTGGAGTCGTTGCGTGTGTGTGTGTGTATTTGGGAGNNNAGATCGGAA |
| 31688_2 | TATGAGTGTGGAGTCGTTGCGTGTGTGTGTGTGTATTTGGGAGNNNAGATCGGAA |
| 45842_1 | TATGAGTGTGGAGTCGTTGCATTTGGCATTTTGGCACTTACCANNNAGATCGGAAG |
| 45842_2 | TATGAGTGTGGAGTCGTTGCATTTGGCATTTTGGCACTTACCANNNAGATCGGAAG |
| 31853_1 | TATGAGTGTGGAGTCGTTGCTCAGGAAAGTGCTGTACTTGTGANNNAGATCGGAA |
| 31853_2 | TATGAGTGTGGAGTCGTTGCTCAGGAAAGTGCTGTACTTGTGANNNAGATCGGAA |
| 50174_1 | TATGAGTGTGGAGTCGTTGCGCTCATGGCTGATTTTAGTCCAANNNAGATCGGAAG |
| 50174_2 | TATGAGTGTGGAGTCGTTGCGCTCATGGCTGATTTTAGTCCAANNNAGATCGGAAG |
| 49363_1 | TATGAGTGTGGAGTCGTTGCTGAGACAGAGTCTTGCTTTGTCANNNAGATCGGAAG |
| 49363_2 | TATGAGTGTGGAGTCGTTGCTGAGACAGAGTCTTGCTTTGTCANNNAGATCGGAAG |
| 43018_1 | TATGAGTGTGGAGTCGTTGCAAATGGCTGATTCTAGGGCTAGGNNNAGATCGGAA |
| 43018_2 | TATGAGTGTGGAGTCGTTGCAAATGGCTGATTCTAGGGCTAGGNNNAGATCGGAA |

| | |
|---|---|
| 52293_1 | TATGAGTGTGGAGTCGTTGCTCAAAAGACCAGAGAAAATGGCCNNNAGATCGGAA |
| 52293_2 | TATGAGTGTGGAGTCGTTGCTCAAAAGACCAGAGAAAATGGCCNNNAGATCGGAA |
| 51086_1 | TATGAGTGTGGAGTCGTTGCTGTTTTCATTAGCAGGGCATGAANNNAGATCGGAAG |
| 51086_2 | TATGAGTGTGGAGTCGTTGCTGTTTTCATTAGCAGGGCATGAANNNAGATCGGAAG |
| 49962_1 | TATGAGTGTGGAGTCGTTGCCGCCCCATACATCCACATAGTTTNNNAGATCGGAAGA |
| 49962_2 | TATGAGTGTGGAGTCGTTGCCGCCCCATACATCCACATAGTTTNNNAGATCGGAAGA |
| 28863_1 | TATGAGTGTGGAGTCGTTGCCCCAATGTTCATGGTGGTTGTTTNNNAGATCGGAAG |
| 28863_2 | TATGAGTGTGGAGTCGTTGCCCCAATGTTCATGGTGGTTGTTTNNNAGATCGGAAG |
| 39265_1 | TATGAGTGTGGAGTCGTTGCGTTAGATGGGCGTGGTGGAGNNNAGATCGGAAGA |
| 39265_2 | TATGAGTGTGGAGTCGTTGCGTTAGATGGGCGTGGTGGAGNNNAGATCGGAAGA |
| 51406_1 | TATGAGTGTGGAGTCGTTGCACAAAGTTCTGACCAAAACACCGNNNAGATCGGAAG |
| 51406_2 | TATGAGTGTGGAGTCGTTGCACAAAGTTCTGACCAAAACACCGNNNAGATCGGAAG |
| 29152_1 | TATGAGTGTGGAGTCGTTGCTTTACCCTGTCACACTTTCTCCCNNNAGATCGGAAGA |
| 29152_2 | TATGAGTGTGGAGTCGTTGCTTTACCCTGTCACACTTTCTCCCNNNAGATCGGAAGA |
| 29705_1 | TATGAGTGTGGAGTCGTTGCGGAATCCTCCACTCTCTTCACTGNNNAGATCGGAAGA |
| 29705_2 | TATGAGTGTGGAGTCGTTGCGGAATCCTCCACTCTCTTCACTGNNNAGATCGGAAGA |
| 29253_1 | TATGAGTGTGGAGTCGTTGCTTGGCCTTCTACCGTTGTCATAANNNAGATCGGAAG |
| 29253_2 | TATGAGTGTGGAGTCGTTGCTTGGCCTTCTACCGTTGTCATAANNNAGATCGGAAG |
| 38989_1 | TATGAGTGTGGAGTCGTTGCTGCCTAAAGAGTTTGAATGCATGANNNAGATCGGA |
| 38989_2 | TATGAGTGTGGAGTCGTTGCTGCCTAAAGAGTTTGAATGCATGANNNAGATCGGA |
| 43320_1 | TATGAGTGTGGAGTCGTTGCCCACCTAAGCTCAAGTGATCCTTNNNAGATCGGAAG |
| 43320_2 | TATGAGTGTGGAGTCGTTGCCCACCTAAGCTCAAGTGATCCTTNNNAGATCGGAAG |
| 41937_1 | TATGAGTGTGGAGTCGTTGCCTCACTGTAACCTCCACCTTCTGNNNAGATCGGAAGA |
| 41937_2 | TATGAGTGTGGAGTCGTTGCCTCACTGTAACCTCCACCTTCTGNNNAGATCGGAAGA |
| 43330_1 | TATGAGTGTGGAGTCGTTGCTGCTAGGATTACAGGATTGAGCCNNNAGATCGGAA |
| 43330_2 | TATGAGTGTGGAGTCGTTGCTGCTAGGATTACAGGATTGAGCCNNNAGATCGGAA |
| 2705_1 | TATGAGTGTGGAGTCGTTGCGAGCAACTCTTGTGCTAGCTGATNNNAGATCGGAAG |
| 2705_2 | TATGAGTGTGGAGTCGTTGCGAGCAACTCTTGTGCTAGCTGATNNNAGATCGGAAG |
| 44956_1 | TATGAGTGTGGAGTCGTTGCATTCCTGCCAGTCTTCAGAAAGTNNNAGATCGGAAG |
| 44956_2 | TATGAGTGTGGAGTCGTTGCATTCCTGCCAGTCTTCAGAAAGTNNNAGATCGGAAG |
| 52117_1 | TATGAGTGTGGAGTCGTTGCTTGGTCTGGATTTCTGCCTCTACNNNAGATCGGAAG |
| 52117_2 | TATGAGTGTGGAGTCGTTGCTTGGTCTGGATTTCTGCCTCTACNNNAGATCGGAAG |
| 42885_1 | TATGAGTGTGGAGTCGTTGCTCAAGTCTGAAGGCATTTTGCTGNNNAGATCGGAAG |
| 42885_2 | TATGAGTGTGGAGTCGTTGCTCAAGTCTGAAGGCATTTTGCTGNNNAGATCGGAAG |
| 42022_1 | TATGAGTGTGGAGTCGTTGCGTGTCCTGCTTTCAAGTTAGCTGNNNAGATCGGAAG |
| 42022_2 | TATGAGTGTGGAGTCGTTGCGTGTCCTGCTTTCAAGTTAGCTGNNNAGATCGGAAG |
| 35050_1 | TATGAGTGTGGAGTCGTTGCTTCAAGCGATTCTACTGCCTCAGNNNAGATCGGAAG |
| 35050_2 | TATGAGTGTGGAGTCGTTGCTTCAAGCGATTCTACTGCCTCAGNNNAGATCGGAAG |
| 12727_1 | TATGAGTGTGGAGTCGTTGCGGTCTGGCTATATATAACACTTTNNNAGATCGGAAG |
| 12727_2 | TATGAGTGTGGAGTCGTTGCGGTCTGGCTATATATAACACTTTNNNAGATCGGAAG |
| 40677_1 | TATGAGTGTGGAGTCGTTGCACTATCTGTATCTGGTCACTCCCANNNAGATCGGAAG |
| 40677_2 | TATGAGTGTGGAGTCGTTGCACTATCTGTATCTGGTCACTCCCANNNAGATCGGAAG |
| 52843_1 | TATGAGTGTGGAGTCGTTGCAGAGGGCTCCCTGCTTATATAGANNNAGATCGGAA |

| | |
|---|---|
| 52843_2 | TATGAGTGTGGAGTCGTTGCAGAGGGCTCCCTGCTTATATAGANNNAGATCGGAAG |
| 42075_1 | TATGAGTGTGGAGTCGTTGCTATCCTGACTTTGCAGTAAGCTGNNNAGATCGGAAG |
| 42075_2 | TATGAGTGTGGAGTCGTTGCTATCCTGACTTTGCAGTAAGCTGNNNAGATCGGAAG |
| 42913_1 | TATGAGTGTGGAGTCGTTGCACACTATGTACTAGGCTCTGTGCNNNAGATCGGAAG |
| 42913_2 | TATGAGTGTGGAGTCGTTGCACACTATGTACTAGGCTCTGTGCNNNAGATCGGAAG |
| 48630_1 | TATGAGTGTGGAGTCGTTGCAGTAGCTAGGATTACAGGCATGCNNNAGATCGGAA |
| 48630_2 | TATGAGTGTGGAGTCGTTGCAGTAGCTAGGATTACAGGCATGCNNNAGATCGGAA |
| 36878_1 | TATGAGTGTGGAGTCGTTGCCGCGAAACAGTTGAGAGATACAGNNNAGATCGGAA |
| 36878_2 | TATGAGTGTGGAGTCGTTGCCGCGAAACAGTTGAGAGATACAGNNNAGATCGGAA |
| 53071_1 | TATGAGTGTGGAGTCGTTGCCGAGGCTACAGTGAGCTATGATTNNNAGATCGGAA |
| 53071_2 | TATGAGTGTGGAGTCGTTGCCGAGGCTACAGTGAGCTATGATTNNNAGATCGGAA |
| 12486_1 | TATGAGTGTGGAGTCGTTGCTGTTCTGGATTTAAAGAAATACCNNNAGATCGGAAG |
| 12486_2 | TATGAGTGTGGAGTCGTTGCTGTTCTGGATTTAAAGAAATACCNNNAGATCGGAAG |
| 29968_1 | TATGAGTGTGGAGTCGTTGCCCCCTCTGTGAGTGTGTGTGTATANNNAGATCGGAA |
| 29968_2 | TATGAGTGTGGAGTCGTTGCCCCCTCTGTGAGTGTGTGTGTATANNNAGATCGGAA |
| 10638_1 | TATGAGTGTGGAGTCGTTGCGTTTTCTCGTTTCCCTTTGCCTGNNNAGATCGGAAGA |
| 10638_2 | TATGAGTGTGGAGTCGTTGCGTTTTCTCGTTTCCCTTTGCCTGNNNAGATCGGAAGA |
| 12924_1 | TATGAGTGTGGAGTCGTTGCTCTAACTTCTCCGAGAACACNNNAGATCGGAAGAGC |
| 12924_2 | TATGAGTGTGGAGTCGTTGCTCTAACTTCTCCGAGAACACNNNAGATCGGAAGAGC |
| 51399_1 | TATGAGTGTGGAGTCGTTGCAAAATACAAAAACCAGCCAGGCANNNAGATCGGAA |
| 51399_2 | TATGAGTGTGGAGTCGTTGCAAAATACAAAAACCAGCCAGGCANNNAGATCGGAA |
| 48329_1 | TATGAGTGTGGAGTCGTTGCCTCAAGTGGTCCTCATACCTCAGNNNAGATCGGAAG |
| 48329_2 | TATGAGTGTGGAGTCGTTGCCTCAAGTGGTCCTCATACCTCAGNNNAGATCGGAAG |
| 10540_1 | TATGAGTGTGGAGTCGTTGCACACAGTGGTAGTTGCTATTGTTNNNAGATCGGAAG |
| 10540_2 | TATGAGTGTGGAGTCGTTGCACACAGTGGTAGTTGCTATTGTTNNNAGATCGGAAG |
| 36918_1 | TATGAGTGTGGAGTCGTTGCAAACAGAAAGGGTTTGCTTCTCGNNNAGATCGGAA |
| 36918_2 | TATGAGTGTGGAGTCGTTGCAAACAGAAAGGGTTTGCTTCTCGNNNAGATCGGAA |
| 3689_1 | TATGAGTGTGGAGTCGTTGCGCTTCTTTTTCGTTCCGCGAATANNNAGATCGGAAG |
| 3689_2 | TATGAGTGTGGAGTCGTTGCGCTTCTTTTTCGTTCCGCGAATANNNAGATCGGAAG |
| 13299_1 | TATGAGTGTGGAGTCGTTGCTTAGTGTCTGGCTTAAATTCCNNNAGATCGGAAGAG |
| 13299_2 | TATGAGTGTGGAGTCGTTGCTTAGTGTCTGGCTTAAATTCCNNNAGATCGGAAGAG |
| 47882_1 | TATGAGTGTGGAGTCGTTGCCAGGTTCAAGCAATTCTCCTGCNNNAGATCGGAAGA |
| 47882_2 | TATGAGTGTGGAGTCGTTGCCAGGTTCAAGCAATTCTCCTGCNNNAGATCGGAAGA |
| 50702_1 | TATGAGTGTGGAGTCGTTGCCACTACAGGGAGAGATTGCTGTTNNNAGATCGGAA |
| 50702_2 | TATGAGTGTGGAGTCGTTGCCACTACAGGGAGAGATTGCTGTTNNNAGATCGGAA |
| 13594_1 | TATGAGTGTGGAGTCGTTGCAAGTCGGTAAGTAAAGAAAGCNNNAGATCGGAAGA |
| 13594_2 | TATGAGTGTGGAGTCGTTGCAAGTCGGTAAGTAAAGAAAGCNNNAGATCGGAAGA |
| 51175_1 | TATGAGTGTGGAGTCGTTGCAATCAGATACCACCTCCCCAAACNNNAGATCGGAAG |
| 51175_2 | TATGAGTGTGGAGTCGTTGCAATCAGATACCACCTCCCCAAACNNNAGATCGGAAG |
| 35251_1 | TATGAGTGTGGAGTCGTTGCAGGTTCTTCTGTCTGTGTAGCTGNNNAGATCGGAAG |
| 35251_2 | TATGAGTGTGGAGTCGTTGCAGGTTCTTCTGTCTGTGTAGCTGNNNAGATCGGAAG |
| 30462_1 | TATGAGTGTGGAGTCGTTGCTGCGGGACATAAAATCTGGCTNNNAGATCGGAAGA |
| 30462_2 | TATGAGTGTGGAGTCGTTGCTGCGGGACATAAAATCTGGCTNNNAGATCGGAAGA |

| | |
|---|---|
| 45932_1 | TATGAGTGTGGAGTCGTTGCCACTACCTGGACTCAAGTGATCCNNNAGATCGGAAG |
| 45932_2 | TATGAGTGTGGAGTCGTTGCCACTACCTGGACTCAAGTGATCCNNNAGATCGGAAG |
| 10948_1 | TATGAGTGTGGAGTCGTTGCAGAACCTACGGATGCAGAGAAGAGNNNAGATCGG |
| 10948_2 | TATGAGTGTGGAGTCGTTGCAGAACCTACGGATGCAGAGAAGAGNNNAGATCGG |
| 40803_1 | TATGAGTGTGGAGTCGTTGCAAATTCAAGGCAGCTGCAGAAATNNNAGATCGGAA |
| 40803_2 | TATGAGTGTGGAGTCGTTGCAAATTCAAGGCAGCTGCAGAAATNNNAGATCGGAA |
| 31014_1 | TATGAGTGTGGAGTCGTTGCTCAAGTAGGTAGGACTACAGGCANNNAGATCGGAA |
| 31014_2 | TATGAGTGTGGAGTCGTTGCTCAAGTAGGTAGGACTACAGGCANNNAGATCGGAA |
| 37169_1 | TATGAGTGTGGAGTCGTTGCTACAAATCTGTACAGCCTAGGGCNNNAGATCGGAAG |
| 37169_2 | TATGAGTGTGGAGTCGTTGCTACAAATCTGTACAGCCTAGGGCNNNAGATCGGAAG |
| 46174_1 | TATGAGTGTGGAGTCGTTGCTTTTGCAGTGCTATTGGGAGAGTNNNAGATCGGAA |
| 46174_2 | TATGAGTGTGGAGTCGTTGCTTTTGCAGTGCTATTGGGAGAGTNNNAGATCGGAA |
| 13528_1 | TATGAGTGTGGAGTCGTTGCTTTAAACCTTACAACTGAATGACNNNAGATCGGAAG |
| 13528_2 | TATGAGTGTGGAGTCGTTGCTTTAAACCTTACAACTGAATGACNNNAGATCGGAAG |
| 30306_1 | TATGAGTGTGGAGTCGTTGCCTGGGTTGAGTGGACACATGATGNNNAGATCGGAA |
| 30306_2 | TATGAGTGTGGAGTCGTTGCCTGGGTTGAGTGGACACATGATGNNNAGATCGGAA |
| 32844_1 | TATGAGTGTGGAGTCGTTGCCCTCCCAGGTTCAAGCAATTTTTNNNAGATCGGAAG |
| 32844_2 | TATGAGTGTGGAGTCGTTGCCCTCCCAGGTTCAAGCAATTTTTNNNAGATCGGAAG |
| 46087_1 | TATGAGTGTGGAGTCGTTGCCACCATGACCGGCTAATTTTTGTNNNAGATCGGAAG |
| 46087_2 | TATGAGTGTGGAGTCGTTGCCACCATGACCGGCTAATTTTTGTNNNAGATCGGAAG |
| 37245_1 | TATGAGTGTGGAGTCGTTGCCACCTGAATTCTACCAGGGGTACNNNAGATCGGAAG |
| 37245_2 | TATGAGTGTGGAGTCGTTGCCACCTGAATTCTACCAGGGGTACNNNAGATCGGAAG |
| 35667_1 | TATGAGTGTGGAGTCGTTGCCTTTGGCTGAGAGAGAACAGGATNNNAGATCGGAA |
| 35667_2 | TATGAGTGTGGAGTCGTTGCCTTTGGCTGAGAGAGAACAGGATNNNAGATCGGAA |
| 45029_1 | TATGAGTGTGGAGTCGTTGCTAGCAAGACTAGGCAACAAGGAGNNNAGATCGGAA |
| 45029_2 | TATGAGTGTGGAGTCGTTGCTAGCAAGACTAGGCAACAAGGAGNNNAGATCGGAA |
| 47260_1 | TATGAGTGTGGAGTCGTTGCTCTTTAGTAGAGGTGGGGTTTCACNNNAGATCGGA |
| 47260_2 | TATGAGTGTGGAGTCGTTGCTCTTTAGTAGAGGTGGGGTTTCACNNNAGATCGGA |
| 45925_1 | TATGAGTGTGGAGTCGTTGCGCTTCCCATGTTCAAGCAATTCTNNNAGATCGGAAG |
| 45925_2 | TATGAGTGTGGAGTCGTTGCGCTTCCCATGTTCAAGCAATTCTNNNAGATCGGAAG |
| 39661_1 | TATGAGTGTGGAGTCGTTGCCTCCTGAGTAGCTGGGACTACANNNAGATCGGAAG |
| 39661_2 | TATGAGTGTGGAGTCGTTGCCTCCTGAGTAGCTGGGACTACANNNAGATCGGAAG |
| 38632_1 | TATGAGTGTGGAGTCGTTGCCTCTGGAGTTATAGGAGGTGGGANNNAGATCGGA |
| 38632_2 | TATGAGTGTGGAGTCGTTGCCTCTGGAGTTATAGGAGGTGGGANNNAGATCGGA |
| 9046_1 | TATGAGTGTGGAGTCGTTGCGGTACTCACAGCCAAACTCACATNNNAGATCGGAAG |
| 9046_2 | TATGAGTGTGGAGTCGTTGCGGTACTCACAGCCAAACTCACATNNNAGATCGGAAG |
| 13804_1 | TATGAGTGTGGAGTCGTTGCGTTTTTTTATGTAAGCATCTCCTNNNAGATCGGAAG |
| 13804_2 | TATGAGTGTGGAGTCGTTGCGTTTTTTTATGTAAGCATCTCCTNNNAGATCGGAAG |
| 12757_1 | TATGAGTGTGGAGTCGTTGCGGAACACATTAATTTTCTTCTCTNNNAGATCGGAAG |
| 12757_2 | TATGAGTGTGGAGTCGTTGCGGAACACATTAATTTTCTTCTCTNNNAGATCGGAAG |
| 41848_1 | TATGAGTGTGGAGTCGTTGCTAACAACCTCAAATTCCTGGGCTNNNAGATCGGAAG |
| 41848_2 | TATGAGTGTGGAGTCGTTGCTAACAACCTCAAATTCCTGGGCTNNNAGATCGGAAG |
| 10832_1 | TATGAGTGTGGAGTCGTTGCGGAAAATAATAGCAGGCATGGTGGNNNAGATCGG |

| | |
|---|---|
| 10832_2 | TATGAGTGTGGAGTCGTTGCGGAAAATAATAGCAGGCATGGTGGNNNAGATCGGA |
| 30071_1 | TATGAGTGTGGAGTCGTTGCTCTCACCCCCTTTCTAAAATGGANNNAGATCGGAAG |
| 30071_2 | TATGAGTGTGGAGTCGTTGCTCTCACCCCCTTTCTAAAATGGANNNAGATCGGAAG |
| 34375_1 | TATGAGTGTGGAGTCGTTGCAGGAAGGTGAGGGAAAATTTGGANNNAGATCGGA |
| 34375_2 | TATGAGTGTGGAGTCGTTGCAGGAAGGTGAGGGAAAATTTGGANNNAGATCGGA |
| 52181_1 | TATGAGTGTGGAGTCGTTGCTCAAATGCATCTCCCCCTGTTTANNNAGATCGGAAGA |
| 52181_2 | TATGAGTGTGGAGTCGTTGCTCAAATGCATCTCCCCCTGTTTANNNAGATCGGAAGA |
| 37082_1 | TATGAGTGTGGAGTCGTTGCTTGCTCTGGAGAAAATAGGTTGTNNNAGATCGGAA |
| 37082_2 | TATGAGTGTGGAGTCGTTGCTTGCTCTGGAGAAAATAGGTTGTNNNAGATCGGAA |
| 48125_1 | TATGAGTGTGGAGTCGTTGCACTTATGCACTCGAAAGAGGAGGNNNAGATCGGAA |
| 48125_2 | TATGAGTGTGGAGTCGTTGCACTTATGCACTCGAAAGAGGAGGNNNAGATCGGAA |
| 10457_1 | TATGAGTGTGGAGTCGTTGCAGGCTGTGTGGTATACCTAAGGANNNAGATCGGAA |
| 10457_2 | TATGAGTGTGGAGTCGTTGCAGGCTGTGTGGTATACCTAAGGANNNAGATCGGAA |
| 37556_1 | TATGAGTGTGGAGTCGTTGCGTTGTGAAGTCCATGGGAAAAGGNNNAGATCGGA |
| 37556_2 | TATGAGTGTGGAGTCGTTGCGTTGTGAAGTCCATGGGAAAAGGNNNAGATCGGA |
| 49394_1 | TATGAGTGTGGAGTCGTTGCGAGAACGGGGTCTCGCTATATTGNNNAGATCGGAA |
| 49394_2 | TATGAGTGTGGAGTCGTTGCGAGAACGGGGTCTCGCTATATTGNNNAGATCGGAA |
| 32866_1 | TATGAGTGTGGAGTCGTTGCCTCCCGAGTACCTGAGACTACAGNNNAGATCGGAAG |
| 32866_2 | TATGAGTGTGGAGTCGTTGCCTCCCGAGTACCTGAGACTACAGNNNAGATCGGAAG |
| 45223_1 | TATGAGTGTGGAGTCGTTGCATTCTGCTCCATCTTCCTACAGCNNNAGATCGGAAGA |
| 45223_2 | TATGAGTGTGGAGTCGTTGCATTCTGCTCCATCTTCCTACAGCNNNAGATCGGAAGA |
| 46937_1 | TATGAGTGTGGAGTCGTTGCTTCAAGCAATTTTCCTGCCTCAGNNNAGATCGGAAG |
| 46937_2 | TATGAGTGTGGAGTCGTTGCTTCAAGCAATTTTCCTGCCTCAGNNNAGATCGGAAG |
| 47519_1 | TATGAGTGTGGAGTCGTTGCGGAAAGGAAAACGATGCTAAGGGNNNAGATCGGA |
| 47519_2 | TATGAGTGTGGAGTCGTTGCGGAAAGGAAAACGATGCTAAGGGNNNAGATCGGA |
| 49859_1 | TATGAGTGTGGAGTCGTTGCCTACCGTAACCCACCCAATACTTNNNAGATCGGAAGA |
| 49859_2 | TATGAGTGTGGAGTCGTTGCCTACCGTAACCCACCCAATACTTNNNAGATCGGAAGA |
| 34042_1 | TATGAGTGTGGAGTCGTTGCCCTCCCAAGTAGCTGGAATTACANNNAGATCGGAAG |
| 34042_2 | TATGAGTGTGGAGTCGTTGCCCTCCCAAGTAGCTGGAATTACANNNAGATCGGAAG |
| 29000_1 | TATGAGTGTGGAGTCGTTGCGTATGGAGCCTTCACAAAATGGCNNNAGATCGGAA |
| 29000_2 | TATGAGTGTGGAGTCGTTGCGTATGGAGCCTTCACAAAATGGCNNNAGATCGGAA |
| 13483_1 | TATGAGTGTGGAGTCGTTGCATAAGTTTTTCAGTTTGGGAATCNNNAGATCGGAAG |
| 13483_2 | TATGAGTGTGGAGTCGTTGCATAAGTTTTTCAGTTTGGGAATCNNNAGATCGGAAG |
| 45190_1 | TATGAGTGTGGAGTCGTTGCTTATGGGGATGTGGATGAAGCAANNNAGATCGGA |
| 45190_2 | TATGAGTGTGGAGTCGTTGCTTATGGGGATGTGGATGAAGCAANNNAGATCGGA |
| 40635_1 | TATGAGTGTGGAGTCGTTGCCATGCCTGTAATCCCAGCATTTTNNNAGATCGGAAG |
| 40635_2 | TATGAGTGTGGAGTCGTTGCCATGCCTGTAATCCCAGCATTTTNNNAGATCGGAAG |
| 32322_1 | TATGAGTGTGGAGTCGTTGCGCCTCAGGATACAAACAAACCACNNNAGATCGGAAG |
| 32322_2 | TATGAGTGTGGAGTCGTTGCGCCTCAGGATACAAACAAACCACNNNAGATCGGAAG |
| 39262_1 | TATGAGTGTGGAGTCGTTGCGATCATAGCAAGCCTCAGAGCTNNNAGATCGGAAG |
| 39262_2 | TATGAGTGTGGAGTCGTTGCGATCATAGCAAGCCTCAGAGCTNNNAGATCGGAAG |
| 38959_1 | TATGAGTGTGGAGTCGTTGCAGCAGCACATCAAAAAGCTTCAANNNAGATCGGAAG |
| 38959_2 | TATGAGTGTGGAGTCGTTGCAGCAGCACATCAAAAAGCTTCAANNNAGATCGGAAG |

| | |
|---|---|
| 49689_1 | TATGAGTGTGGAGTCGTTGCCACCACGCTCGACTAATTTTGTANNNAGATCGGAAG |
| 49689_2 | TATGAGTGTGGAGTCGTTGCCACCACGCTCGACTAATTTTGTANNNAGATCGGAAG |
| 50188_1 | TATGAGTGTGGAGTCGTTGCTTCCGTTTCTTTTACGTTTCCCGNNNAGATCGGAAGA |
| 50188_2 | TATGAGTGTGGAGTCGTTGCTTCCGTTTCTTTTACGTTTCCCGNNNAGATCGGAAGA |
| 45786_1 | TATGAGTGTGGAGTCGTTGCAGTGCTGAGATTATAGGCGTGAGNNNAGATCGGAA |
| 45786_2 | TATGAGTGTGGAGTCGTTGCAGTGCTGAGATTATAGGCGTGAGNNNAGATCGGAA |
| 9651_1 | TATGAGTGTGGAGTCGTTGCCTGACAAATTGGGATTCACCACANNNAGATCGGAAG |
| 9651_2 | TATGAGTGTGGAGTCGTTGCCTGACAAATTGGGATTCACCACANNNAGATCGGAAG |
| 37524_1 | TATGAGTGTGGAGTCGTTGCCCTTGGTGGTTCTTTTTTGAGACANNNAGATCGGAA |
| 37524_2 | TATGAGTGTGGAGTCGTTGCCCTTGGTGGTTCTTTTTTGAGACANNNAGATCGGAA |
| 41728_1 | TATGAGTGTGGAGTCGTTGCGTGCCTTTCTACACTCAGACTCANNNAGATCGGAAG |
| 41728_2 | TATGAGTGTGGAGTCGTTGCGTGCCTTTCTACACTCAGACTCANNNAGATCGGAAG |
| 41658_1 | TATGAGTGTGGAGTCGTTGCAGTGATTCTTCTTCCTCAGCCTCNNNAGATCGGAAGA |
| 41658_2 | TATGAGTGTGGAGTCGTTGCAGTGATTCTTCTTCCTCAGCCTCNNNAGATCGGAAGA |
| 29116_1 | TATGAGTGTGGAGTCGTTGCCTAGATGGGCCTCCAAAAGAACTNNNAGATCGGAA |
| 29116_2 | TATGAGTGTGGAGTCGTTGCCTAGATGGGCCTCCAAAAGAACTNNNAGATCGGAA |
| 28879_1 | TATGAGTGTGGAGTCGTTGCTGTGTGTGGATGTGTGTGTCTAANNNAGATCGGAA |
| 28879_2 | TATGAGTGTGGAGTCGTTGCTGTGTGTGGATGTGTGTGTCTAANNNAGATCGGAA |
| 34275_1 | TATGAGTGTGGAGTCGTTGCAGTAGATAGGCATCTCTCTCCCCNNNAGATCGGAAG |
| 34275_2 | TATGAGTGTGGAGTCGTTGCAGTAGATAGGCATCTCTCTCCCCNNNAGATCGGAAG |
| 29704_1 | TATGAGTGTGGAGTCGTTGCATGTTTACATGAACAACAGCAGTNNNAGATCGGAAG |
| 29704_2 | TATGAGTGTGGAGTCGTTGCATGTTTACATGAACAACAGCAGTNNNAGATCGGAAG |
| 34263_1 | TATGAGTGTGGAGTCGTTGCCGTACACCTGTAGTCTCAGCTACNNNAGATCGGAAG |
| 34263_2 | TATGAGTGTGGAGTCGTTGCCGTACACCTGTAGTCTCAGCTACNNNAGATCGGAAG |
| 32950_1 | TATGAGTGTGGAGTCGTTGCAGGTTGAGGCAACAGGATTACTTNNNAGATCGGAA |
| 32950_2 | TATGAGTGTGGAGTCGTTGCAGGTTGAGGCAACAGGATTACTTNNNAGATCGGAA |
| 34422_1 | TATGAGTGTGGAGTCGTTGCGGTAAGTGGTGTAGTAGGGTGTGNNNAGATCGGA |
| 34422_2 | TATGAGTGTGGAGTCGTTGCGGTAAGTGGTGTAGTAGGGTGTGNNNAGATCGGA |
| 43655_1 | TATGAGTGTGGAGTCGTTGCCCAGACAAGGAGGTATGCATCTTNNNAGATCGGAA |
| 43655_2 | TATGAGTGTGGAGTCGTTGCCCAGACAAGGAGGTATGCATCTTNNNAGATCGGAA |
| 43400_1 | TATGAGTGTGGAGTCGTTGCAATCTCGTCTCACTGCAAACTCTNNNAGATCGGAAGA |
| 43400_2 | TATGAGTGTGGAGTCGTTGCAATCTCGTCTCACTGCAAACTCTNNNAGATCGGAAGA |
| 47057_1 | TATGAGTGTGGAGTCGTTGCCTGAGTAGCTGGGATTACAGGTGNNNAGATCGGAA |
| 47057_2 | TATGAGTGTGGAGTCGTTGCCTGAGTAGCTGGGATTACAGGTGNNNAGATCGGAA |
| 12660_1 | TATGAGTGTGGAGTCGTTGCTTCTAAAAGTCATGTTAGACAGTNNNAGATCGGAAG |
| 12660_2 | TATGAGTGTGGAGTCGTTGCTTCTAAAAGTCATGTTAGACAGTNNNAGATCGGAAG |
| 39889_1 | TATGAGTGTGGAGTCGTTGCTTCAAAATGACCCTTTCCCTTGCNNNAGATCGGAAGA |
| 39889_2 | TATGAGTGTGGAGTCGTTGCTTCAAAATGACCCTTTCCCTTGCNNNAGATCGGAAGA |
| 38097_1 | TATGAGTGTGGAGTCGTTGCAACACACATGAATGATTGCCAGGNNNAGATCGGAA |
| 38097_2 | TATGAGTGTGGAGTCGTTGCAACACACATGAATGATTGCCAGGNNNAGATCGGAA |
| 30773_1 | TATGAGTGTGGAGTCGTTGCGCAGTGTGATGTGTTTGTCTTCANNNAGATCGGAAG |
| 30773_2 | TATGAGTGTGGAGTCGTTGCGCAGTGTGATGTGTTTGTCTTCANNNAGATCGGAAG |
| 30389_1 | TATGAGTGTGGAGTCGTTGCACACACACACACACGTATCTATGNNNAGATCGGAAG |

| | |
|---|---|
| 30389_2 | TATGAGTGTGGAGTCGTTGCACACACACACACACGTATCTATGNNNAGATCGGAAG |
| 40421_1 | TATGAGTGTGGAGTCGTTGCATTGTTGCCTATGCTTTCTCACGNNNAGATCGGAAG |
| 40421_2 | TATGAGTGTGGAGTCGTTGCATTGTTGCCTATGCTTTCTCACGNNNAGATCGGAAG |
| 34216_1 | TATGAGTGTGGAGTCGTTGCTTCCATGAGAGGAGAAAACCTGGNNNAGATCGGAA |
| 34216_2 | TATGAGTGTGGAGTCGTTGCTTCCATGAGAGGAGAAAACCTGGNNNAGATCGGAA |
| 32236_1 | TATGAGTGTGGAGTCGTTGCCTCAAGCAATCTTCCTGCCTTGNNNAGATCGGAAGA |
| 32236_2 | TATGAGTGTGGAGTCGTTGCCTCAAGCAATCTTCCTGCCTTGNNNAGATCGGAAGA |
| 12951_1 | TATGAGTGTGGAGTCGTTGCGTTCTAGAACTTACCCTTTTTGNNNAGATCGGAAGA |
| 12951_2 | TATGAGTGTGGAGTCGTTGCGTTCTAGAACTTACCCTTTTTGNNNAGATCGGAAGA |
| 38999_1 | TATGAGTGTGGAGTCGTTGCGAGAACGGTGTGAATCTGGGAGNNNAGATCGGAA |
| 38999_2 | TATGAGTGTGGAGTCGTTGCGAGAACGGTGTGAATCTGGGAGNNNAGATCGGAA |
| 52481_1 | TATGAGTGTGGAGTCGTTGCTGGTCTCTGATTTGTAACCTCGTNNNAGATCGGAAG |
| 52481_2 | TATGAGTGTGGAGTCGTTGCTGGTCTCTGATTTGTAACCTCGTNNNAGATCGGAAG |
| 38042_1 | TATGAGTGTGGAGTCGTTGCGGATGATAATCCGTCCTGGTTGANNNAGATCGGAA |
| 38042_2 | TATGAGTGTGGAGTCGTTGCGGATGATAATCCGTCCTGGTTGANNNAGATCGGAA |
| 13241_1 | TATGAGTGTGGAGTCGTTGCAAAACAATTTGCACATACCATATNNNAGATCGGAAG |
| 13241_2 | TATGAGTGTGGAGTCGTTGCAAAACAATTTGCACATACCATATNNNAGATCGGAAG |
| 49601_1 | TATGAGTGTGGAGTCGTTGCCCTAGTAGCTGGGACTACAGGTANNNAGATCGGAA |
| 49601_2 | TATGAGTGTGGAGTCGTTGCCCTAGTAGCTGGGACTACAGGTANNNAGATCGGAA |
| 40223_1 | TATGAGTGTGGAGTCGTTGCTGCAGTCAGAAAAACAGAAAGTACANNNAGATCGG |
| 40223_2 | TATGAGTGTGGAGTCGTTGCTGCAGTCAGAAAAACAGAAAGTACANNNAGATCGG |
| 47866_1 | TATGAGTGTGGAGTCGTTGCCCCAACCAATCAGTAGCACCTATNNNAGATCGGAAG |
| 47866_2 | TATGAGTGTGGAGTCGTTGCCCCAACCAATCAGTAGCACCTATNNNAGATCGGAAG |
| 34802_1 | TATGAGTGTGGAGTCGTTGCGCTGGTCTTGAACTCCTAACCTTNNNAGATCGGAAG |
| 34802_2 | TATGAGTGTGGAGTCGTTGCGCTGGTCTTGAACTCCTAACCTTNNNAGATCGGAAG |
| 36072_1 | TATGAGTGTGGAGTCGTTGCTCCCTTTCAAACCTAAAACCCCANNNAGATCGGAAGA |
| 36072_2 | TATGAGTGTGGAGTCGTTGCTCCCTTTCAAACCTAAAACCCCANNNAGATCGGAAGA |
| 52725_1 | TATGAGTGTGGAGTCGTTGCTGTCCTCTAAACATTTCCAGGGGNNNAGATCGGAAG |
| 52725_2 | TATGAGTGTGGAGTCGTTGCTGTCCTCTAAACATTTCCAGGGGNNNAGATCGGAAG |
| 30601_1 | TATGAGTGTGGAGTCGTTGCACCTTGCTGAAGAAGTGCAAATNNNAGATCGGAAG |
| 30601_2 | TATGAGTGTGGAGTCGTTGCACCTTGCTGAAGAAGTGCAAATNNNAGATCGGAAG |
| 30240_1 | TATGAGTGTGGAGTCGTTGCCCAGCAGCACACAACAGTTATTTNNNAGATCGGAAG |
| 30240_2 | TATGAGTGTGGAGTCGTTGCCCAGCAGCACACAACAGTTATTTNNNAGATCGGAAG |
| 36774_1 | TATGAGTGTGGAGTCGTTGCGAGTTCCCCTCAGTAACCTCAACNNNAGATCGGAAG |
| 36774_2 | TATGAGTGTGGAGTCGTTGCGAGTTCCCCTCAGTAACCTCAACNNNAGATCGGAAG |
| 34214_1 | TATGAGTGTGGAGTCGTTGCGTTGCAGTGAGCTGAGATCATGNNNAGATCGGAAG |
| 34214_2 | TATGAGTGTGGAGTCGTTGCGTTGCAGTGAGCTGAGATCATGNNNAGATCGGAAG |
| 36318_1 | TATGAGTGTGGAGTCGTTGCGAGAGAGAATGCTCCATGGCTTANNNAGATCGGAA |
| 36318_2 | TATGAGTGTGGAGTCGTTGCGAGAGAGAATGCTCCATGGCTTANNNAGATCGGAA |
| 49362_1 | TATGAGTGTGGAGTCGTTGCGAGTAGCTGGGACTATAGGCAACNNNAGATCGGAA |
| 49362_2 | TATGAGTGTGGAGTCGTTGCGAGTAGCTGGGACTATAGGCAACNNNAGATCGGAA |
| 39551_1 | TATGAGTGTGGAGTCGTTGCAAGAAAGAAGACGTAAAGGCCCTNNNAGATCGGAA |
| 39551_2 | TATGAGTGTGGAGTCGTTGCAAGAAAGAAGACGTAAAGGCCCTNNNAGATCGGAA |

| | |
|---|---|
| 48969_1 | TATGAGTGTGGAGTCGTTGCACCCAGTTTTTGGAAAATTAGCCTNNNAGATCGGAA |
| 48969_2 | TATGAGTGTGGAGTCGTTGCACCCAGTTTTTGGAAAATTAGCCTNNNAGATCGGAA |
| 50455_1 | TATGAGTGTGGAGTCGTTGCTTCTTGCTTGAAAAAAATGCCGCNNNAGATCGGAAG |
| 50455_2 | TATGAGTGTGGAGTCGTTGCTTCTTGCTTGAAAAAAATGCCGCNNNAGATCGGAAG |
| 44086_1 | TATGAGTGTGGAGTCGTTGCGGAAAATGTTGAAAGGAGCCTCCNNNAGATCGGAA |
| 44086_2 | TATGAGTGTGGAGTCGTTGCGGAAAATGTTGAAAGGAGCCTCCNNNAGATCGGAA |
| 38024_1 | TATGAGTGTGGAGTCGTTGCGTTTTTGTGCCAGTACCAGTACCNNNAGATCGGAAG |
| 38024_2 | TATGAGTGTGGAGTCGTTGCGTTTTTGTGCCAGTACCAGTACCNNNAGATCGGAAG |
| 34133_1 | TATGAGTGTGGAGTCGTTGCGAATCACTGGGAGTCTGCCTAAANNNAGATCGGAA |
| 34133_2 | TATGAGTGTGGAGTCGTTGCGAATCACTGGGAGTCTGCCTAAANNNAGATCGGAA |
| 13364_1 | TATGAGTGTGGAGTCGTTGCATTGAGACATGAAGAACAGACNNNAGATCGGAAGA |
| 13364_2 | TATGAGTGTGGAGTCGTTGCATTGAGACATGAAGAACAGACNNNAGATCGGAAGA |
| 49562_1 | TATGAGTGTGGAGTCGTTGCTTGACAGGGTCTTGCTCTATCACNNNAGATCGGAAG |
| 49562_2 | TATGAGTGTGGAGTCGTTGCTTGACAGGGTCTTGCTCTATCACNNNAGATCGGAAG |
| 40804_1 | TATGAGTGTGGAGTCGTTGCTTTAGCATGAGGCCACAACAAACNNNAGATCGGAAG |
| 40804_2 | TATGAGTGTGGAGTCGTTGCTTTAGCATGAGGCCACAACAAACNNNAGATCGGAAG |
| 35301_1 | TATGAGTGTGGAGTCGTTGCAGTATCACTCTATCCCTCAGGCTNNNAGATCGGAAG |
| 35301_2 | TATGAGTGTGGAGTCGTTGCAGTATCACTCTATCCCTCAGGCTNNNAGATCGGAAG |
| 43074_1 | TATGAGTGTGGAGTCGTTGCTTCAGGCCAATATCCTTGGTGAANNNAGATCGGAAG |
| 43074_2 | TATGAGTGTGGAGTCGTTGCTTCAGGCCAATATCCTTGGTGAANNNAGATCGGAAG |
| 31199_1 | TATGAGTGTGGAGTCGTTGCACACGAACTCACACATTTTTACCCNNNAGATCGGAAG |
| 31199_2 | TATGAGTGTGGAGTCGTTGCACACGAACTCACACATTTTTACCCNNNAGATCGGAAG |
| 5761_1 | TATGAGTGTGGAGTCGTTGCGGCACAAGCACACACATTCTTTTNNNAGATCGGAAG |
| 5761_2 | TATGAGTGTGGAGTCGTTGCGGCACAAGCACACACATTCTTTTNNNAGATCGGAAG |
| 37093_1 | TATGAGTGTGGAGTCGTTGCCCAAAGCTACGTCTCTCAGAGTTNNNAGATCGGAAG |
| 37093_2 | TATGAGTGTGGAGTCGTTGCCCAAAGCTACGTCTCTCAGAGTTNNNAGATCGGAAG |
| 29230_1 | TATGAGTGTGGAGTCGTTGCACACACACACAAGAGCACATCTANNNAGATCGGAAG |
| 29230_2 | TATGAGTGTGGAGTCGTTGCACACACACACAAGAGCACATCTANNNAGATCGGAAG |
| 35784_1 | TATGAGTGTGGAGTCGTTGCAGTAGCTGGGACTACAGGTATGTNNNAGATCGGAA |
| 35784_2 | TATGAGTGTGGAGTCGTTGCAGTAGCTGGGACTACAGGTATGTNNNAGATCGGAA |
| 47593_1 | TATGAGTGTGGAGTCGTTGCAAGTAGCTGGGATTACAGGCATCNNNAGATCGGAA |
| 47593_2 | TATGAGTGTGGAGTCGTTGCAAGTAGCTGGGATTACAGGCATCNNNAGATCGGAA |
| 46412_1 | TATGAGTGTGGAGTCGTTGCTTCAAGTGATTCTCCTGCCTCAGNNNAGATCGGAAG |
| 46412_2 | TATGAGTGTGGAGTCGTTGCTTCAAGTGATTCTCCTGCCTCAGNNNAGATCGGAAG |
| 48634_1 | TATGAGTGTGGAGTCGTTGCGAGAGTGTTGAAAGGGTCGGAGNNNAGATCGGAA |
| 48634_2 | TATGAGTGTGGAGTCGTTGCGAGAGTGTTGAAAGGGTCGGAGNNNAGATCGGAA |
| 28971_1 | TATGAGTGTGGAGTCGTTGCCCCACCATAGCTGTGTACCTATCNNNAGATCGGAAG |
| 28971_2 | TATGAGTGTGGAGTCGTTGCCCCACCATAGCTGTGTACCTATCNNNAGATCGGAAG |
| 41456_1 | TATGAGTGTGGAGTCGTTGCTGAGAGCATCACTGACATGTTGANNNAGATCGGAA |
| 41456_2 | TATGAGTGTGGAGTCGTTGCTGAGAGCATCACTGACATGTTGANNNAGATCGGAA |
| 36755_1 | TATGAGTGTGGAGTCGTTGCAGTGTTTTTGCTCCATTGCTGTTNNNAGATCGGAAG |
| 36755_2 | TATGAGTGTGGAGTCGTTGCAGTGTTTTTGCTCCATTGCTGTTNNNAGATCGGAAG |
| 53287_1 | TATGAGTGTGGAGTCGTTGCCTGATCCAGGAGGCCGCATCNNNAGATCGGAAGAG |

| | |
|---|---|
| 53287_2 | TATGAGTGTGGAGTCGTTGCCTGATCCAGGAGGCCGCATCNNNAGATCGGAAGAG |
| 12603_1 | TATGAGTGTGGAGTCGTTGCCTACAAAAGCTTCCACTAAATTCNNNAGATCGGAAGA |
| 12603_2 | TATGAGTGTGGAGTCGTTGCCTACAAAAGCTTCCACTAAATTCNNNAGATCGGAAGA |
| 50738_1 | TATGAGTGTGGAGTCGTTGCGGGTTGTAGAAGACACACAGGATNNNAGATCGGAA |
| 50738_2 | TATGAGTGTGGAGTCGTTGCGGGTTGTAGAAGACACACAGGATNNNAGATCGGAA |
| 47636_1 | TATGAGTGTGGAGTCGTTGCCCTCTGGAGTATCTGGGACTACANNNAGATCGGAAG |
| 47636_2 | TATGAGTGTGGAGTCGTTGCCCTCTGGAGTATCTGGGACTACANNNAGATCGGAAG |
| 13442_1 | TATGAGTGTGGAGTCGTTGCTTTGGTTTCCATAACCTAAGTATNNNAGATCGGAAG |
| 13442_2 | TATGAGTGTGGAGTCGTTGCTTTGGTTTCCATAACCTAAGTATNNNAGATCGGAAG |
| 40584_1 | TATGAGTGTGGAGTCGTTGCACTGCAATTTTCATTTGAGGGGGNNNAGATCGGAA |
| 40584_2 | TATGAGTGTGGAGTCGTTGCACTGCAATTTTCATTTGAGGGGGNNNAGATCGGAA |
| 40365_1 | TATGAGTGTGGAGTCGTTGCCAATGGCACAATCTTGGCACANNNAGATCGGAAGA |
| 40365_2 | TATGAGTGTGGAGTCGTTGCCAATGGCACAATCTTGGCACANNNAGATCGGAAGA |
| 35328_1 | TATGAGTGTGGAGTCGTTGCGACTTCAAGAATAGAGCCACGGANNNAGATCGGAA |
| 35328_2 | TATGAGTGTGGAGTCGTTGCGACTTCAAGAATAGAGCCACGGANNNAGATCGGAA |
| 51537_1 | TATGAGTGTGGAGTCGTTGCCCAGAGGTTGAATTTGCAAACCTNNNAGATCGGAAG |
| 51537_2 | TATGAGTGTGGAGTCGTTGCCCAGAGGTTGAATTTGCAAACCTNNNAGATCGGAAG |
| 31771_1 | TATGAGTGTGGAGTCGTTGCCTGAGCTCAGGAGTTCAAAACCANNNAGATCGGAA |
| 31771_2 | TATGAGTGTGGAGTCGTTGCCTGAGCTCAGGAGTTCAAAACCANNNAGATCGGAA |
| 49775_1 | TATGAGTGTGGAGTCGTTGCTTGATTTGGTTCAACATGCAGCTNNNAGATCGGAAG |
| 49775_2 | TATGAGTGTGGAGTCGTTGCTTGATTTGGTTCAACATGCAGCTNNNAGATCGGAAG |
| 39600_1 | TATGAGTGTGGAGTCGTTGCAGTGCTGATCATGCAGAACCTTANNNAGATCGGAAG |
| 39600_2 | TATGAGTGTGGAGTCGTTGCAGTGCTGATCATGCAGAACCTTANNNAGATCGGAAG |
| 36217_1 | TATGAGTGTGGAGTCGTTGCCCACTGTTTCTGGAATATGCCACNNNAGATCGGAAG |
| 36217_2 | TATGAGTGTGGAGTCGTTGCCCACTGTTTCTGGAATATGCCACNNNAGATCGGAAG |
| 29222_1 | TATGAGTGTGGAGTCGTTGCAGAGAACAAGAGTCTCTGCTTGGNNNAGATCGGAA |
| 29222_2 | TATGAGTGTGGAGTCGTTGCAGAGAACAAGAGTCTCTGCTTGGNNNAGATCGGAA |
| 38879_1 | TATGAGTGTGGAGTCGTTGCTTGTCTTCAGTGAGGAGGTACACNNNAGATCGGAA |
| 38879_2 | TATGAGTGTGGAGTCGTTGCTTGTCTTCAGTGAGGAGGTACACNNNAGATCGGAA |
| 48591_1 | TATGAGTGTGGAGTCGTTGCAGACAGAGTCTCACTTTGTCACCNNNAGATCGGAAG |
| 48591_2 | TATGAGTGTGGAGTCGTTGCAGACAGAGTCTCACTTTGTCACCNNNAGATCGGAAG |
| 30009_1 | TATGAGTGTGGAGTCGTTGCGCATCTCATGAAATAAAATGCCTCCNNNAGATCGGA |
| 30009_2 | TATGAGTGTGGAGTCGTTGCGCATCTCATGAAATAAAATGCCTCCNNNAGATCGGA |
| 13479_1 | TATGAGTGTGGAGTCGTTGCGGAATTTCCAGTGTATTGTTTGNNNAGATCGGAAGA |
| 13479_2 | TATGAGTGTGGAGTCGTTGCGGAATTTCCAGTGTATTGTTTGNNNAGATCGGAAGA |
| 40958_1 | TATGAGTGTGGAGTCGTTGCTATGAGGGTTGACATTGGACCTGNNNAGATCGGAA |
| 40958_2 | TATGAGTGTGGAGTCGTTGCTATGAGGGTTGACATTGGACCTGNNNAGATCGGAA |
| 31023_1 | TATGAGTGTGGAGTCGTTGCCTCCTGAGTAGCTGGGATTACAGNNNAGATCGGAA |
| 31023_2 | TATGAGTGTGGAGTCGTTGCCTCCTGAGTAGCTGGGATTACAGNNNAGATCGGAA |
| 53357_1 | TATGAGTGTGGAGTCGTTGCTCTCTCTCCTTCCTTCCTTCCTTNNNAGATCGGAAGAG |
| 53357_2 | TATGAGTGTGGAGTCGTTGCTCTCTCTCCTTCCTTCCTTCCTTNNNAGATCGGAAGAG |
| 47147_1 | TATGAGTGTGGAGTCGTTGCGGGATCAAGCAGTGAAATTCTGGNNNAGATCGGAA |
| 47147_2 | TATGAGTGTGGAGTCGTTGCGGGATCAAGCAGTGAAATTCTGGNNNAGATCGGAA |

| | |
|---|---|
| 45227_1 | TATGAGTGTGGAGTCGTTGCGACTAGCTGTGCCTGTAACCTAGNNNAGATCGGAAG |
| 45227_2 | TATGAGTGTGGAGTCGTTGCGACTAGCTGTGCCTGTAACCTAGNNNAGATCGGAAG |
| 39156_1 | TATGAGTGTGGAGTCGTTGCGGGAAAGGACACCCTCTTCAATANNNAGATCGGAAG |
| 39156_2 | TATGAGTGTGGAGTCGTTGCGGGAAAGGACACCCTCTTCAATANNNAGATCGGAAG |
| 41970_1 | TATGAGTGTGGAGTCGTTGCGATCTCGAACTCCTGAGCTCAAGNNNAGATCGGAAG |
| 41970_2 | TATGAGTGTGGAGTCGTTGCGATCTCGAACTCCTGAGCTCAAGNNNAGATCGGAAG |
| 34166_1 | TATGAGTGTGGAGTCGTTGCGGCTCAAAGTCATCAGGAAGAGTNNNAGATCGGAA |
| 34166_2 | TATGAGTGTGGAGTCGTTGCGGCTCAAAGTCATCAGGAAGAGTNNNAGATCGGAA |
| 43322_1 | TATGAGTGTGGAGTCGTTGCTTCAAGTGATTCTCCTGCCTCAGNNNAGATCGGAAG |
| 43322_2 | TATGAGTGTGGAGTCGTTGCTTCAAGTGATTCTCCTGCCTCAGNNNAGATCGGAAG |
| 50864_1 | TATGAGTGTGGAGTCGTTGCCGATTTCCTCTGCCTCTCTCTTTNNNAGATCGGAAGA |
| 50864_2 | TATGAGTGTGGAGTCGTTGCCGATTTCCTCTGCCTCTCTCTTTNNNAGATCGGAAGA |
| 40442_1 | TATGAGTGTGGAGTCGTTGCAAAGAAAGTATGTCACCTGCCCANNNAGATCGGAAG |
| 40442_2 | TATGAGTGTGGAGTCGTTGCAAAGAAAGTATGTCACCTGCCCANNNAGATCGGAAG |
| 44907_1 | TATGAGTGTGGAGTCGTTGCCCAGGTGCCTTGTTTTTGATGAANNNAGATCGGAAG |
| 44907_2 | TATGAGTGTGGAGTCGTTGCCCAGGTGCCTTGTTTTTGATGAANNNAGATCGGAAG |
| 44788_1 | TATGAGTGTGGAGTCGTTGCAAGAGTCTGTTAGGAGTAAGCGCNNNAGATCGGAA |
| 44788_2 | TATGAGTGTGGAGTCGTTGCAAGAGTCTGTTAGGAGTAAGCGCNNNAGATCGGAA |
| 40249_1 | TATGAGTGTGGAGTCGTTGCTGAGAATTACTGTGGCTCTTCCCNNNAGATCGGAAG |
| 40249_2 | TATGAGTGTGGAGTCGTTGCTGAGAATTACTGTGGCTCTTCCCNNNAGATCGGAAG |
| 43915_1 | TATGAGTGTGGAGTCGTTGCACGTATAATTTTTTGCACTGCTACCANNNAGATCGGA |
| 43915_2 | TATGAGTGTGGAGTCGTTGCACGTATAATTTTTTGCACTGCTACCANNNAGATCGGA |
| 41542_1 | TATGAGTGTGGAGTCGTTGCACCACGTCCGGCTAATTTTTTTTNNNAGATCGGAAGA |
| 41542_2 | TATGAGTGTGGAGTCGTTGCACCACGTCCGGCTAATTTTTTTTNNNAGATCGGAAGA |
| 37597_1 | TATGAGTGTGGAGTCGTTGCACCTTCTCCAAAAATGTCCTGGANNNAGATCGGAAG |
| 37597_2 | TATGAGTGTGGAGTCGTTGCACCTTCTCCAAAAATGTCCTGGANNNAGATCGGAAG |
| 35728_1 | TATGAGTGTGGAGTCGTTGCCATTCTAACAGTCAGGCCCTTCTNNNAGATCGGAAGA |
| 35728_2 | TATGAGTGTGGAGTCGTTGCCATTCTAACAGTCAGGCCCTTCTNNNAGATCGGAAGA |
| 49025_1 | TATGAGTGTGGAGTCGTTGCGCTACCATGCCCAGCTAATTTTTNNNAGATCGGAAGA |
| 49025_2 | TATGAGTGTGGAGTCGTTGCGCTACCATGCCCAGCTAATTTTTNNNAGATCGGAAGA |
| 48917_1 | TATGAGTGTGGAGTCGTTGCCCTCTGTCTCCTAAGTTCAAGCANNNAGATCGGAAGA |
| 48917_2 | TATGAGTGTGGAGTCGTTGCCCTCTGTCTCCTAAGTTCAAGCANNNAGATCGGAAGA |
| 50141_1 | TATGAGTGTGGAGTCGTTGCCTGTGTCACATTCCCGGAAGATANNNAGATCGGAAG |
| 50141_2 | TATGAGTGTGGAGTCGTTGCCTGTGTCACATTCCCGGAAGATANNNAGATCGGAAG |
| 33359_1 | TATGAGTGTGGAGTCGTTGCGATCTCACCTCACCACAACCTCNNNAGATCGGAAGAG |
| 33359_2 | TATGAGTGTGGAGTCGTTGCGATCTCACCTCACCACAACCTCNNNAGATCGGAAGAG |
| 45019_1 | TATGAGTGTGGAGTCGTTGCAGTGGACGGAAAAACTCAGGTATNNNAGATCGGAA |
| 45019_2 | TATGAGTGTGGAGTCGTTGCAGTGGACGGAAAAACTCAGGTATNNNAGATCGGAA |
| 31008_1 | TATGAGTGTGGAGTCGTTGCTTGAGACGGAGTCTTACTCTGTCNNNAGATCGGAAG |
| 31008_2 | TATGAGTGTGGAGTCGTTGCTTGAGACGGAGTCTTACTCTGTCNNNAGATCGGAAG |
| 53314_1 | TATGAGTGTGGAGTCGTTGCTTTTTTTTGAGGCGGAGTCTCACNNNAGATCGGAAG |
| 53314_2 | TATGAGTGTGGAGTCGTTGCTTTTTTTTGAGGCGGAGTCTCACNNNAGATCGGAAG |
| 13224_1 | TATGAGTGTGGAGTCGTTGCCTCCTTAAGCCATTCTAAAGTTNNNAGATCGGAAGA |

| | |
|---|---|
| 13224_2 | TATGAGTGTGGAGTCGTTGCCTCCTTAAGCCATTCTAAAGTTNNNAGATCGGAAGA |
| 32475_1 | TATGAGTGTGGAGTCGTTGCATAATTCTCATGCCTCAGCCTCCNNNAGATCGGAAGA |
| 32475_2 | TATGAGTGTGGAGTCGTTGCATAATTCTCATGCCTCAGCCTCCNNNAGATCGGAAGA |
| 33313_1 | TATGAGTGTGGAGTCGTTGCCCTCCCAGGTTCAAGTAGTTCTTNNNAGATCGGAAG |
| 33313_2 | TATGAGTGTGGAGTCGTTGCCCTCCCAGGTTCAAGTAGTTCTTNNNAGATCGGAAG |
| 36449_1 | TATGAGTGTGGAGTCGTTGCGAGACAGAGGGAAGGAAGGAAAGNNNAGATCGG |
| 36449_2 | TATGAGTGTGGAGTCGTTGCGAGACAGAGGGAAGGAAGGAAAGNNNAGATCGG |
| 29366_1 | TATGAGTGTGGAGTCGTTGCACTTTCTCCCAATCTGTAGGTTGNNNAGATCGGAAG |
| 29366_2 | TATGAGTGTGGAGTCGTTGCACTTTCTCCCAATCTGTAGGTTGNNNAGATCGGAAG |
| 45975_1 | TATGAGTGTGGAGTCGTTGCCACTGTAACCTCTGCCTCATAGGNNNAGATCGGAAG |
| 45975_2 | TATGAGTGTGGAGTCGTTGCCACTGTAACCTCTGCCTCATAGGNNNAGATCGGAAG |
| 32915_1 | TATGAGTGTGGAGTCGTTGCTCATGCTGCTGACAGTCTAATGANNNAGATCGGAAG |
| 32915_2 | TATGAGTGTGGAGTCGTTGCTCATGCTGCTGACAGTCTAATGANNNAGATCGGAAG |
| 30654_1 | TATGAGTGTGGAGTCGTTGCCCCCTCACCTCCTGGATAGGNNNAGATCGGAAGAGC |
| 30654_2 | TATGAGTGTGGAGTCGTTGCCCCCTCACCTCCTGGATAGGNNNAGATCGGAAGAGC |
| 12735_1 | TATGAGTGTGGAGTCGTTGCGAAGAATTGCAAAAGAAGTTAGANNNAGATCGGAA |
| 12735_2 | TATGAGTGTGGAGTCGTTGCGAAGAATTGCAAAAGAAGTTAGANNNAGATCGGAA |
| 41704_1 | TATGAGTGTGGAGTCGTTGCTGGAGTACAGTGGCAAGATCTTGNNNAGATCGGAA |
| 41704_2 | TATGAGTGTGGAGTCGTTGCTGGAGTACAGTGGCAAGATCTTGNNNAGATCGGAA |
| 42405_1 | TATGAGTGTGGAGTCGTTGCTCTTTCTTTCTTTCTTTTCTCACTTGTNNNAGATCGGA |
| 42405_2 | TATGAGTGTGGAGTCGTTGCTCTTTCTTTCTTTCTTTTCTCACTTGTNNNAGATCGGA |
| 30616_1 | TATGAGTGTGGAGTCGTTGCTGGTTTCTTACCTAGTCAGCATAACNNNAGATCGGA |
| 30616_2 | TATGAGTGTGGAGTCGTTGCTGGTTTCTTACCTAGTCAGCATAACNNNAGATCGGA |
| 43122_1 | TATGAGTGTGGAGTCGTTGCACCATGCAAACCAAGAGCAAAAANNNAGATCGGAA |
| 43122_2 | TATGAGTGTGGAGTCGTTGCACCATGCAAACCAAGAGCAAAAANNNAGATCGGAA |
| 43199_1 | TATGAGTGTGGAGTCGTTGCCTCTCCTCTCTCCTCTCTCTCTGNNNAGATCGGAAGAG |
| 43199_2 | TATGAGTGTGGAGTCGTTGCCTCTCCTCTCTCCTCTCTCTCTGNNNAGATCGGAAGAG |
| 51650_1 | TATGAGTGTGGAGTCGTTGCAGGATCTTGCTAAGTTTCCCAGGNNNAGATCGGAAG |
| 51650_2 | TATGAGTGTGGAGTCGTTGCAGGATCTTGCTAAGTTTCCCAGGNNNAGATCGGAAG |
| 38619_1 | TATGAGTGTGGAGTCGTTGCTACAAAAAAATTAGCCAGGCGTGGNNNAGATCGGAA |
| 38619_2 | TATGAGTGTGGAGTCGTTGCTACAAAAAAATTAGCCAGGCGTGGNNNAGATCGGAA |
| 32635_1 | TATGAGTGTGGAGTCGTTGCTCTCTCAAAGTCACCCTCACTTGNNNAGATCGGAAGA |
| 32635_2 | TATGAGTGTGGAGTCGTTGCTCTCTCAAAGTCACCCTCACTTGNNNAGATCGGAAGA |
| 43070_1 | TATGAGTGTGGAGTCGTTGCTCCAAATACAGCCCCACTCTTACNNNAGATCGGAAGA |
| 43070_2 | TATGAGTGTGGAGTCGTTGCTCCAAATACAGCCCCACTCTTACNNNAGATCGGAAGA |
| 40173_1 | TATGAGTGTGGAGTCGTTGCTGCAAAAGAGAGTGAGACTTCGTNNNAGATCGGAA |
| 40173_2 | TATGAGTGTGGAGTCGTTGCTGCAAAAGAGAGTGAGACTTCGTNNNAGATCGGAA |
| 30482_1 | TATGAGTGTGGAGTCGTTGCCACACTTTAACTCTTCCAAAAGCCCNNNAGATCGGAA |
| 30482_2 | TATGAGTGTGGAGTCGTTGCCACACTTTAACTCTTCCAAAAGCCCNNNAGATCGGAA |
| 10680_1 | TATGAGTGTGGAGTCGTTGCGCAAATATCAACGACCTGTCAGCNNNAGATCGGAAG |
| 10680_2 | TATGAGTGTGGAGTCGTTGCGCAAATATCAACGACCTGTCAGCNNNAGATCGGAAG |
| 33184_1 | TATGAGTGTGGAGTCGTTGCGAAAGGACTTGGTTGCTGTGATCNNNAGATCGGAA |
| 33184_2 | TATGAGTGTGGAGTCGTTGCGAAAGGACTTGGTTGCTGTGATCNNNAGATCGGAA |

| | |
|---|---|
| 42264_1 | TATGAGTGTGGAGTCGTTGCTTGGTTGAGAATTATGTCCCCCCNNNAGATCGGAAG |
| 42264_2 | TATGAGTGTGGAGTCGTTGCTTGGTTGAGAATTATGTCCCCCCNNNAGATCGGAAG |
| 39984_1 | TATGAGTGTGGAGTCGTTGCCATCCATCGGTAGCAGACTGTTCNNNAGATCGGAAG |
| 39984_2 | TATGAGTGTGGAGTCGTTGCCATCCATCGGTAGCAGACTGTTCNNNAGATCGGAAG |
| 32817_1 | TATGAGTGTGGAGTCGTTGCTTCAAGCAATTCTCCTGCCTCAGNNNAGATCGGAAG |
| 32817_2 | TATGAGTGTGGAGTCGTTGCTTCAAGCAATTCTCCTGCCTCAGNNNAGATCGGAAG |
| 12737_1 | TATGAGTGTGGAGTCGTTGCAAGGAAATTCAGTTACAGGAGNNNAGATCGGAAGA |
| 12737_2 | TATGAGTGTGGAGTCGTTGCAAGGAAATTCAGTTACAGGAGNNNAGATCGGAAGA |
| 49250_1 | TATGAGTGTGGAGTCGTTGCAGATAGGGTTTCACTCTGTTGCCNNNAGATCGGAAG |
| 49250_2 | TATGAGTGTGGAGTCGTTGCAGATAGGGTTTCACTCTGTTGCCNNNAGATCGGAAG |
| 48269_1 | TATGAGTGTGGAGTCGTTGCTGCATTCTTGCCAACATCTGTTTNNNAGATCGGAAGA |
| 48269_2 | TATGAGTGTGGAGTCGTTGCTGCATTCTTGCCAACATCTGTTTNNNAGATCGGAAGA |
| 12314_1 | TATGAGTGTGGAGTCGTTGCATAATGTTAGGTGTGCTAGGCCTNNNAGATCGGAA |
| 12314_2 | TATGAGTGTGGAGTCGTTGCATAATGTTAGGTGTGCTAGGCCTNNNAGATCGGAA |
| 38860_1 | TATGAGTGTGGAGTCGTTGCGCACAGAAGACACAAACAAATGGNNNAGATCGGAA |
| 38860_2 | TATGAGTGTGGAGTCGTTGCGCACAGAAGACACAAACAAATGGNNNAGATCGGAA |
| 41123_1 | TATGAGTGTGGAGTCGTTGCTAGCAATGGAGAAACCCACTTCANNNAGATCGGAAG |
| 41123_2 | TATGAGTGTGGAGTCGTTGCTAGCAATGGAGAAACCCACTTCANNNAGATCGGAAG |
| 37638_1 | TATGAGTGTGGAGTCGTTGCTCGAAATGAGATTTGGGCTGGTANNNAGATCGGAA |
| 37638_2 | TATGAGTGTGGAGTCGTTGCTCGAAATGAGATTTGGGCTGGTANNNAGATCGGAA |
| 13092_1 | TATGAGTGTGGAGTCGTTGCCATCTGAATCAACAATGTGATAANNNAGATCGGAAG |
| 13092_2 | TATGAGTGTGGAGTCGTTGCCATCTGAATCAACAATGTGATAANNNAGATCGGAAG |
| 38855_1 | TATGAGTGTGGAGTCGTTGCCCCATGTAAATCTGAAACCCAGCNNNAGATCGGAAG |
| 38855_2 | TATGAGTGTGGAGTCGTTGCCCCATGTAAATCTGAAACCCAGCNNNAGATCGGAAG |
| 45367_1 | TATGAGTGTGGAGTCGTTGCGTCATGGTTTGCTTCCTTCTAGCNNNAGATCGGAAG |
| 45367_2 | TATGAGTGTGGAGTCGTTGCGTCATGGTTTGCTTCCTTCTAGCNNNAGATCGGAAG |
| 30649_1 | TATGAGTGTGGAGTCGTTGCTGTTCTTTTATGCTTTGGTCCTTCTNNNAGATCGGAA |
| 30649_2 | TATGAGTGTGGAGTCGTTGCTGTTCTTTTATGCTTTGGTCCTTCTNNNAGATCGGAA |
| 31160_1 | TATGAGTGTGGAGTCGTTGCGCAAGAAGGCAACTGTCTATGTGNNNAGATCGGAA |
| 31160_2 | TATGAGTGTGGAGTCGTTGCGCAAGAAGGCAACTGTCTATGTGNNNAGATCGGAA |
| 37630_1 | TATGAGTGTGGAGTCGTTGCAATCACTTTACCCCTGGAGATGGNNNAGATCGGAAG |
| 37630_2 | TATGAGTGTGGAGTCGTTGCAATCACTTTACCCCTGGAGATGGNNNAGATCGGAAG |
| 44230_1 | TATGAGTGTGGAGTCGTTGCGGTTTCAAATGTCCAACTGCCATNNNAGATCGGAAG |
| 44230_2 | TATGAGTGTGGAGTCGTTGCGGTTTCAAATGTCCAACTGCCATNNNAGATCGGAAG |
| 38711_1 | TATGAGTGTGGAGTCGTTGCAGACATTTATGTGGCCAACAAACANNNAGATCGGAA |
| 38711_2 | TATGAGTGTGGAGTCGTTGCAGACATTTATGTGGCCAACAAACANNNAGATCGGAA |
| 33341_1 | TATGAGTGTGGAGTCGTTGCCGCCTCCTGGATTGAAATGATTCNNNAGATCGGAAG |
| 33341_2 | TATGAGTGTGGAGTCGTTGCCGCCTCCTGGATTGAAATGATTCNNNAGATCGGAAG |
| 48819_1 | TATGAGTGTGGAGTCGTTGCCTCACTGTAACCTCTGCCTCTTGNNNAGATCGGAAGA |
| 48819_2 | TATGAGTGTGGAGTCGTTGCCTCACTGTAACCTCTGCCTCTTGNNNAGATCGGAAGA |
| 35554_1 | TATGAGTGTGGAGTCGTTGCATCTCCACCTCCTGAATTCAAGGNNNAGATCGGAAG |
| 35554_2 | TATGAGTGTGGAGTCGTTGCATCTCCACCTCCTGAATTCAAGGNNNAGATCGGAAG |
| 52594_1 | TATGAGTGTGGAGTCGTTGCATCTTCCTTACGTTCCTGTTCCCNNNAGATCGGAAGA |

| | |
|---|---|
| 52594_2 | TATGAGTGTGGAGTCGTTGCATCTTCCTTACGTTCCTGTTCCCNNNAGATCGGAAGA |
| 45153_1 | TATGAGTGTGGAGTCGTTGCTGGTATCAGAACAGGAATCTGCANNNAGATCGGAA |
| 45153_2 | TATGAGTGTGGAGTCGTTGCTGGTATCAGAACAGGAATCTGCANNNAGATCGGAA |
| 49717_1 | TATGAGTGTGGAGTCGTTGCCAATCTTCCTACCTCAACCTCCCNNNAGATCGGAAGA |
| 49717_2 | TATGAGTGTGGAGTCGTTGCCAATCTTCCTACCTCAACCTCCCNNNAGATCGGAAGA |
| 47027_1 | TATGAGTGTGGAGTCGTTGCATGGATATTGAGGGGAGGTGGTANNNAGATCGGA |
| 47027_2 | TATGAGTGTGGAGTCGTTGCATGGATATTGAGGGGAGGTGGTANNNAGATCGGA |
| 47497_1 | TATGAGTGTGGAGTCGTTGCTCTGTGAAAAATGGCATTGGTGTNNNAGATCGGAA |
| 47497_2 | TATGAGTGTGGAGTCGTTGCTCTGTGAAAAATGGCATTGGTGTNNNAGATCGGAA |
| 46452_1 | TATGAGTGTGGAGTCGTTGCGAGATAAAGTCTGGCTCTGTCGCNNNAGATCGGAA |
| 46452_2 | TATGAGTGTGGAGTCGTTGCGAGATAAAGTCTGGCTCTGTCGCNNNAGATCGGAA |
| 43417_1 | TATGAGTGTGGAGTCGTTGCCAGTGGCATGATCTTGGCTCANNNAGATCGGAAGA |
| 43417_2 | TATGAGTGTGGAGTCGTTGCCAGTGGCATGATCTTGGCTCANNNAGATCGGAAGA |
| 43994_1 | TATGAGTGTGGAGTCGTTGCGCTTATGATGGCACAAAAACAGCNNNAGATCGGAA |
| 43994_2 | TATGAGTGTGGAGTCGTTGCGCTTATGATGGCACAAAAACAGCNNNAGATCGGAA |
| 32064_1 | TATGAGTGTGGAGTCGTTGCTGCAATTTACATGACAACCCCACNNNAGATCGGAAG |
| 32064_2 | TATGAGTGTGGAGTCGTTGCTGCAATTTACATGACAACCCCACNNNAGATCGGAAG |
| 36464_1 | TATGAGTGTGGAGTCGTTGCGTTCCAGATACTGAAGAGGCTGANNNAGATCGGAA |
| 36464_2 | TATGAGTGTGGAGTCGTTGCGTTCCAGATACTGAAGAGGCTGANNNAGATCGGAA |
| 37444_1 | TATGAGTGTGGAGTCGTTGCGGAGAGTAAGAGGCAGCTATCACNNNAGATCGGAA |
| 37444_2 | TATGAGTGTGGAGTCGTTGCGGAGAGTAAGAGGCAGCTATCACNNNAGATCGGAA |
| 52587_1 | TATGAGTGTGGAGTCGTTGCTTTTTACCGTGTTTGGCCTTCAANNNAGATCGGAAG |
| 52587_2 | TATGAGTGTGGAGTCGTTGCTTTTTACCGTGTTTGGCCTTCAANNNAGATCGGAAG |
| 30590_1 | TATGAGTGTGGAGTCGTTGCTCGTGTCTTTCACCGATTGAGATACNNNAGATCGGA |
| 30590_2 | TATGAGTGTGGAGTCGTTGCTCGTGTCTTTCACCGATTGAGATACNNNAGATCGGA |
| 46976_1 | TATGAGTGTGGAGTCGTTGCCAGGTTCAAGCAATTCTCCTGCNNNAGATCGGAAGA |
| 46976_2 | TATGAGTGTGGAGTCGTTGCCAGGTTCAAGCAATTCTCCTGCNNNAGATCGGAAGA |
| 51675_1 | TATGAGTGTGGAGTCGTTGCACACACTCACACCCGTAAACATANNNAGATCGGAAG |
| 51675_2 | TATGAGTGTGGAGTCGTTGCACACACTCACACCCGTAAACATANNNAGATCGGAAG |
| 40811_1 | TATGAGTGTGGAGTCGTTGCTGCTGCTGTCATTTTTGTGGTAGNNNAGATCGGAAG |
| 40811_2 | TATGAGTGTGGAGTCGTTGCTGCTGCTGTCATTTTTGTGGTAGNNNAGATCGGAAG |
| 12928_1 | TATGAGTGTGGAGTCGTTGCCATCATTTCTGGAGTCTCAGNNNAGATCGGAAGAGC |
| 12928_2 | TATGAGTGTGGAGTCGTTGCCATCATTTCTGGAGTCTCAGNNNAGATCGGAAGAGC |
| 30956_1 | TATGAGTGTGGAGTCGTTGCACAGAGAGACAGAGAGAGAAGCTNNNAGATCGGA |
| 30956_2 | TATGAGTGTGGAGTCGTTGCACAGAGAGACAGAGAGAGAAGCTNNNAGATCGGA |
| 29430_1 | TATGAGTGTGGAGTCGTTGCTCTCTTTAAATGCATGCTGGCCANNNAGATCGGAAG |
| 29430_2 | TATGAGTGTGGAGTCGTTGCTCTCTTTAAATGCATGCTGGCCANNNAGATCGGAAG |
| 40597_1 | TATGAGTGTGGAGTCGTTGCGGTTCTTTCACTGGGCAGAATTGNNNAGATCGGAA |
| 40597_2 | TATGAGTGTGGAGTCGTTGCGGTTCTTTCACTGGGCAGAATTGNNNAGATCGGAA |
| 37708_1 | TATGAGTGTGGAGTCGTTGCAGGATCATTCTCGACTGCTTTGANNNAGATCGGAAG |
| 37708_2 | TATGAGTGTGGAGTCGTTGCAGGATCATTCTCGACTGCTTTGANNNAGATCGGAAG |
| 40924_1 | TATGAGTGTGGAGTCGTTGCCTGGGCAACAGAGCAATATTCTGNNNAGATCGGAA |
| 40924_2 | TATGAGTGTGGAGTCGTTGCCTGGGCAACAGAGCAATATTCTGNNNAGATCGGAA |

| | |
|---|---|
| 13088_1 | TATGAGTGTGGAGTCGTTGCAGTATTTTGGGGAATTTTGAATTNNNAGATCGGAAG |
| 13088_2 | TATGAGTGTGGAGTCGTTGCAGTATTTTGGGGAATTTTGAATTNNNAGATCGGAAG |
| 44641_1 | TATGAGTGTGGAGTCGTTGCGAGTATGAGTCATGGGGTCACTGNNNAGATCGGAA |
| 44641_2 | TATGAGTGTGGAGTCGTTGCGAGTATGAGTCATGGGGTCACTGNNNAGATCGGAA |
| 37608_1 | TATGAGTGTGGAGTCGTTGCCACATACCTCACATACCTGCAGTNNNAGATCGGAAG |
| 37608_2 | TATGAGTGTGGAGTCGTTGCCACATACCTCACATACCTGCAGTNNNAGATCGGAAG |
| 36200_1 | TATGAGTGTGGAGTCGTTGCACATGGCATTCAAAAGACACCAGNNNAGATCGGAAG |
| 36200_2 | TATGAGTGTGGAGTCGTTGCACATGGCATTCAAAAGACACCAGNNNAGATCGGAAG |
| 45914_1 | TATGAGTGTGGAGTCGTTGCTAGTTGGGACTACAGATGTGTGCNNNAGATCGGAA |
| 45914_2 | TATGAGTGTGGAGTCGTTGCTAGTTGGGACTACAGATGTGTGCNNNAGATCGGAA |
| 9580_1 | TATGAGTGTGGAGTCGTTGCACAGCAAACATAGTAAATGGTGGCNNNAGATCGGA |
| 9580_2 | TATGAGTGTGGAGTCGTTGCACAGCAAACATAGTAAATGGTGGCNNNAGATCGGA |
| 13596_1 | TATGAGTGTGGAGTCGTTGCTTGCATTGATATGATGGATATTGNNNAGATCGGAAG |
| 13596_2 | TATGAGTGTGGAGTCGTTGCTTGCATTGATATGATGGATATTGNNNAGATCGGAAG |
| 12296_1 | TATGAGTGTGGAGTCGTTGCCCCTCCTCAGGACACACCTAAACNNNAGATCGGAAG |
| 12296_2 | TATGAGTGTGGAGTCGTTGCCCCTCCTCAGGACACACCTAAACNNNAGATCGGAAG |
| 45748_1 | TATGAGTGTGGAGTCGTTGCACTCAATGTGACCTTGAAGTGGGNNNAGATCGGAA |
| 45748_2 | TATGAGTGTGGAGTCGTTGCACTCAATGTGACCTTGAAGTGGGNNNAGATCGGAA |
| 52665_1 | TATGAGTGTGGAGTCGTTGCGTGTCCTACCTCTGGGAAAGAAGNNNAGATCGGAA |
| 52665_2 | TATGAGTGTGGAGTCGTTGCGTGTCCTACCTCTGGGAAAGAAGNNNAGATCGGAA |
| 39141_1 | TATGAGTGTGGAGTCGTTGCTCAACCTAAATGTCCATCAACAGANNNAGATCGGAA |
| 39141_2 | TATGAGTGTGGAGTCGTTGCTCAACCTAAATGTCCATCAACAGANNNAGATCGGAA |
| 40766_1 | TATGAGTGTGGAGTCGTTGCATCACGGCTGTTCAAAGTTGAAGNNNAGATCGGAAG |
| 40766_2 | TATGAGTGTGGAGTCGTTGCATCACGGCTGTTCAAAGTTGAAGNNNAGATCGGAAG |
| 29016_1 | TATGAGTGTGGAGTCGTTGCTCAAATCCCACAAATCAGCTTCGNNNAGATCGGAAG |
| 29016_2 | TATGAGTGTGGAGTCGTTGCTCAAATCCCACAAATCAGCTTCGNNNAGATCGGAAG |
| 29032_1 | TATGAGTGTGGAGTCGTTGCTCATGCAGACTCACACATACACANNNAGATCGGAAG |
| 29032_2 | TATGAGTGTGGAGTCGTTGCTCATGCAGACTCACACATACACANNNAGATCGGAAG |
| 30238_1 | TATGAGTGTGGAGTCGTTGCACTTTCATGCAGACTAATGAGCGNNNAGATCGGAAG |
| 30238_2 | TATGAGTGTGGAGTCGTTGCACTTTCATGCAGACTAATGAGCGNNNAGATCGGAAG |
| 48593_1 | TATGAGTGTGGAGTCGTTGCAACTTAGCTGTATGTGGTGGTGTNNNAGATCGGAA |
| 48593_2 | TATGAGTGTGGAGTCGTTGCAACTTAGCTGTATGTGGTGGTGTNNNAGATCGGAA |
| 49514_1 | TATGAGTGTGGAGTCGTTGCGAGGCCATTACTCTCTCTGCATTNNNAGATCGGAAG |
| 49514_2 | TATGAGTGTGGAGTCGTTGCGAGGCCATTACTCTCTCTGCATTNNNAGATCGGAAG |
| 45056_1 | TATGAGTGTGGAGTCGTTGCTGCTTCCTTCCTGAGTTTTGAGTNNNAGATCGGAAG |
| 45056_2 | TATGAGTGTGGAGTCGTTGCTGCTTCCTTCCTGAGTTTTGAGTNNNAGATCGGAAG |
| 39878_1 | TATGAGTGTGGAGTCGTTGCAGAAAAGCAAGGAAAAGAAACCCANNNAGATCGGA |
| 39878_2 | TATGAGTGTGGAGTCGTTGCAGAAAAGCAAGGAAAAGAAACCCANNNAGATCGGA |
| 46658_1 | TATGAGTGTGGAGTCGTTGCCCTCCTGAGTTCACACCATTCTCNNNAGATCGGAAGA |
| 46658_2 | TATGAGTGTGGAGTCGTTGCCCTCCTGAGTTCACACCATTCTCNNNAGATCGGAAGA |
| 51933_1 | TATGAGTGTGGAGTCGTTGCTCAAAGAGTTCAGATCCCCATGTNNNAGATCGGAAG |
| 51933_2 | TATGAGTGTGGAGTCGTTGCTCAAAGAGTTCAGATCCCCATGTNNNAGATCGGAAG |
| 46140_1 | TATGAGTGTGGAGTCGTTGCCTCCCAGGTTCAAGCGATTCTNNNAGATCGGAAGAG |

| | |
|---|---|
| 46140_2 | TATGAGTGTGGAGTCGTTGCCTCCCAGGTTCAAGCGATTCTNNNAGATCGGAAGAG |
| 33981_1 | TATGAGTGTGGAGTCGTTGCGATCTCCACTCACTGCAACCTCNNNAGATCGGAAGAG |
| 33981_2 | TATGAGTGTGGAGTCGTTGCGATCTCCACTCACTGCAACCTCNNNAGATCGGAAGAG |
| 50314_1 | TATGAGTGTGGAGTCGTTGCAGAGCTCAATCCCAAGATAGCTCNNNAGATCGGAAG |
| 50314_2 | TATGAGTGTGGAGTCGTTGCAGAGCTCAATCCCAAGATAGCTCNNNAGATCGGAAG |
| 30914_1 | TATGAGTGTGGAGTCGTTGCGTCACCACTATTGGGAATGGGATNNNAGATCGGAA |
| 30914_2 | TATGAGTGTGGAGTCGTTGCGTCACCACTATTGGGAATGGGATNNNAGATCGGAA |
| 10419_1 | TATGAGTGTGGAGTCGTTGCTAGCATTGTGCTGAGGAATGTGGNNNAGATCGGAA |
| 10419_2 | TATGAGTGTGGAGTCGTTGCTAGCATTGTGCTGAGGAATGTGGNNNAGATCGGAA |
| 37445_1 | TATGAGTGTGGAGTCGTTGCTTGACCTACTCACATTCCCTTGGNNNAGATCGGAAGA |
| 37445_2 | TATGAGTGTGGAGTCGTTGCTTGACCTACTCACATTCCCTTGGNNNAGATCGGAAGA |
| 33300_1 | TATGAGTGTGGAGTCGTTGCCAAGTAGCTGGGACTACAGGCNNNAGATCGGAAGA |
| 33300_2 | TATGAGTGTGGAGTCGTTGCCAAGTAGCTGGGACTACAGGCNNNAGATCGGAAGA |
| 12557_1 | TATGAGTGTGGAGTCGTTGCAAAACTGAGTAGTGTTGTTAAAGNNNAGATCGGAA |
| 12557_2 | TATGAGTGTGGAGTCGTTGCAAAACTGAGTAGTGTTGTTAAAGNNNAGATCGGAA |
| 33064_1 | TATGAGTGTGGAGTCGTTGCCGAAGCAGGAAGAGATAGCAGATNNNAGATCGGAA |
| 33064_2 | TATGAGTGTGGAGTCGTTGCCGAAGCAGGAAGAGATAGCAGATNNNAGATCGGAA |
| 46613_1 | TATGAGTGTGGAGTCGTTGCCTATTCTTCTGCCTCAGTCTCCCNNNAGATCGGAAGA |
| 46613_2 | TATGAGTGTGGAGTCGTTGCCTATTCTTCTGCCTCAGTCTCCCNNNAGATCGGAAGA |
| 51186_1 | TATGAGTGTGGAGTCGTTGCTTCACTCAACTACTGGTGCTACCNNNAGATCGGAAGA |
| 51186_2 | TATGAGTGTGGAGTCGTTGCTTCACTCAACTACTGGTGCTACCNNNAGATCGGAAGA |
| 44324_1 | TATGAGTGTGGAGTCGTTGCTGGTGCTGATTGGGTTGTCTTATNNNAGATCGGAAG |
| 44324_2 | TATGAGTGTGGAGTCGTTGCTGGTGCTGATTGGGTTGTCTTATNNNAGATCGGAAG |
| 36633_1 | TATGAGTGTGGAGTCGTTGCAAAGTATTCAAGGTTTGGCACGGNNNAGATCGGAA |
| 36633_2 | TATGAGTGTGGAGTCGTTGCAAAGTATTCAAGGTTTGGCACGGNNNAGATCGGAA |
| 10646_1 | TATGAGTGTGGAGTCGTTGCGGATGTTCAGAGCAGTGTATGGGNNNAGATCGGAA |
| 10646_2 | TATGAGTGTGGAGTCGTTGCGGATGTTCAGAGCAGTGTATGGGNNNAGATCGGAA |
| 29381_1 | TATGAGTGTGGAGTCGTTGCATGTAGGGGTGTGTGAATGTGAGNNNAGATCGGAA |
| 29381_2 | TATGAGTGTGGAGTCGTTGCATGTAGGGGTGTGTGAATGTGAGNNNAGATCGGAA |
| 42443_1 | TATGAGTGTGGAGTCGTTGCGAGAGAGAGAGGAGAGAGAGAGANNNAGATCGGA |
| 42443_2 | TATGAGTGTGGAGTCGTTGCGAGAGAGAGAGGAGAGAGAGAGANNNAGATCGGA |
| 38255_1 | TATGAGTGTGGAGTCGTTGCTTACGGCAGCAGTTTCTCAAAAGNNNAGATCGGAAG |
| 38255_2 | TATGAGTGTGGAGTCGTTGCTTACGGCAGCAGTTTCTCAAAAGNNNAGATCGGAAG |
| 37965_1 | TATGAGTGTGGAGTCGTTGCCAGTGAGCTCTGATAACACCACTNNNAGATCGGAAG |
| 37965_2 | TATGAGTGTGGAGTCGTTGCCAGTGAGCTCTGATAACACCACTNNNAGATCGGAAG |
| 12667_1 | TATGAGTGTGGAGTCGTTGCAATTTTATCACACAACAGTCATCNNNAGATCGGAAGA |
| 12667_2 | TATGAGTGTGGAGTCGTTGCAATTTTATCACACAACAGTCATCNNNAGATCGGAAGA |
| 43685_1 | TATGAGTGTGGAGTCGTTGCGCATGACTCAACAGGATGAGAGANNNAGATCGGAA |
| 43685_2 | TATGAGTGTGGAGTCGTTGCGCATGACTCAACAGGATGAGAGANNNAGATCGGAA |
| 45244_1 | TATGAGTGTGGAGTCGTTGCAGAGGGTAAGGGATTCCAAGAGANNNAGATCGGAA |
| 45244_2 | TATGAGTGTGGAGTCGTTGCAGAGGGTAAGGGATTCCAAGAGANNNAGATCGGAA |
| 41824_1 | TATGAGTGTGGAGTCGTTGCCAAGTAGCTGGGACTACAGGCNNNAGATCGGAAGA |
| 41824_2 | TATGAGTGTGGAGTCGTTGCCAAGTAGCTGGGACTACAGGCNNNAGATCGGAAGA |

| | |
|---|---|
| 39080_1 | TATGAGTGTGGAGTCGTTGCACTGATCGTGTTCATAGAACTGANNNAGATCGGAAG |
| 39080_2 | TATGAGTGTGGAGTCGTTGCACTGATCGTGTTCATAGAACTGANNNAGATCGGAAG |
| 12894_1 | TATGAGTGTGGAGTCGTTGCTTTCCTTGAGCTGTTTTCTGNNNAGATCGGAAGAGC |
| 12894_2 | TATGAGTGTGGAGTCGTTGCTTTCCTTGAGCTGTTTTCTGNNNAGATCGGAAGAGC |
| 51964_1 | TATGAGTGTGGAGTCGTTGCTTCCTGTGTTAGTTTGCTGAGGANNNAGATCGGAAG |
| 51964_2 | TATGAGTGTGGAGTCGTTGCTTCCTGTGTTAGTTTGCTGAGGANNNAGATCGGAAG |
| 38940_1 | TATGAGTGTGGAGTCGTTGCTTGAGCTCAGGAGTTTGAAACCTNNNAGATCGGAA |
| 38940_2 | TATGAGTGTGGAGTCGTTGCTTGAGCTCAGGAGTTTGAAACCTNNNAGATCGGAA |
| 37368_1 | TATGAGTGTGGAGTCGTTGCTGAGACCGAGTTTTCACTCTTGTNNNAGATCGGAAG |
| 37368_2 | TATGAGTGTGGAGTCGTTGCTGAGACCGAGTTTTCACTCTTGTNNNAGATCGGAAG |
| 40038_1 | TATGAGTGTGGAGTCGTTGCCATTGCCTGCCAAACAAAATGAANNNAGATCGGAAG |
| 40038_2 | TATGAGTGTGGAGTCGTTGCCATTGCCTGCCAAACAAAATGAANNNAGATCGGAAG |
| 13080_1 | TATGAGTGTGGAGTCGTTGCGATAAGGAAGGGTGAGAGTTNNNAGATCGGAAGA |
| 13080_2 | TATGAGTGTGGAGTCGTTGCGATAAGGAAGGGTGAGAGTTNNNAGATCGGAAGA |
| 50947_1 | TATGAGTGTGGAGTCGTTGCCTTTCTGGAAGCTGATGGGAAAGNNNAGATCGGAA |
| 50947_2 | TATGAGTGTGGAGTCGTTGCCTTTCTGGAAGCTGATGGGAAAGNNNAGATCGGAA |
| 38137_1 | TATGAGTGTGGAGTCGTTGCTGGTTGACTCACACTAACCAAGANNNAGATCGGAAG |
| 38137_2 | TATGAGTGTGGAGTCGTTGCTGGTTGACTCACACTAACCAAGANNNAGATCGGAAG |
| 41288_1 | TATGAGTGTGGAGTCGTTGCGAAAGAAACCCTCTGTCCCTGATNNNAGATCGGAAG |
| 41288_2 | TATGAGTGTGGAGTCGTTGCGAAAGAAACCCTCTGTCCCTGATNNNAGATCGGAAG |
| 3147_1 | TATGAGTGTGGAGTCGTTGCCGGCTCCTCAGATAATGAGTACTNNNAGATCGGAAG |
| 3147_2 | TATGAGTGTGGAGTCGTTGCCGGCTCCTCAGATAATGAGTACTNNNAGATCGGAAG |
| 48468_1 | TATGAGTGTGGAGTCGTTGCTCAAGTGATTCTCCTGCCTTAGCNNNAGATCGGAAG |
| 48468_2 | TATGAGTGTGGAGTCGTTGCTCAAGTGATTCTCCTGCCTTAGCNNNAGATCGGAAG |
| 52283_1 | TATGAGTGTGGAGTCGTTGCTCCTGAGTAGCTGGGATTACAGANNNAGATCGGAA |
| 52283_2 | TATGAGTGTGGAGTCGTTGCTCCTGAGTAGCTGGGATTACAGANNNAGATCGGAA |
| 46698_1 | TATGAGTGTGGAGTCGTTGCCTCCTGAGTAGCTGGGATTACAGNNNAGATCGGAA |
| 46698_2 | TATGAGTGTGGAGTCGTTGCCTCCTGAGTAGCTGGGATTACAGNNNAGATCGGAA |
| 29277_1 | TATGAGTGTGGAGTCGTTGCCGTGAAATGTGGCCCATAGAATCNNNAGATCGGAA |
| 29277_2 | TATGAGTGTGGAGTCGTTGCCGTGAAATGTGGCCCATAGAATCNNNAGATCGGAA |
| 12686_1 | TATGAGTGTGGAGTCGTTGCTTTTAGCTTTTTGGAAGACTAAANNNAGATCGGAAG |
| 12686_2 | TATGAGTGTGGAGTCGTTGCTTTTAGCTTTTTGGAAGACTAAANNNAGATCGGAAG |
| 28904_1 | TATGAGTGTGGAGTCGTTGCATTTCTTTGCTCCCCTTTGATGCNNNAGATCGGAAGA |
| 28904_2 | TATGAGTGTGGAGTCGTTGCATTTCTTTGCTCCCCTTTGATGCNNNAGATCGGAAGA |
| 31567_1 | TATGAGTGTGGAGTCGTTGCCAGGACTAAGACACCTATCACCCNNNAGATCGGAAG |
| 31567_2 | TATGAGTGTGGAGTCGTTGCCAGGACTAAGACACCTATCACCCNNNAGATCGGAAG |
| 12481_1 | TATGAGTGTGGAGTCGTTGCCTTTGTTAGAGCAATACTCAATNNNAGATCGGAAGA |
| 12481_2 | TATGAGTGTGGAGTCGTTGCCTTTGTTAGAGCAATACTCAATNNNAGATCGGAAGA |
| 41758_1 | TATGAGTGTGGAGTCGTTGCAAGTGCAATGGTACAATCTTGGCNNNAGATCGGAA |
| 41758_2 | TATGAGTGTGGAGTCGTTGCAAGTGCAATGGTACAATCTTGGCNNNAGATCGGAA |
| 40001_1 | TATGAGTGTGGAGTCGTTGCATGACATAATACAGCGGCATCCTNNNAGATCGGAAG |
| 40001_2 | TATGAGTGTGGAGTCGTTGCATGACATAATACAGCGGCATCCTNNNAGATCGGAAG |
| 43200_1 | TATGAGTGTGGAGTCGTTGCGGTCACTCAAAACCATCTAAGGTNNNAGATCGGAAG |

| | |
|---|---|
| 43200_2 | TATGAGTGTGGAGTCGTTGCGGTCACTCAAAACCATCTAAGGTNNNAGATCGGAAG |
| 48882_1 | TATGAGTGTGGAGTCGTTGCTCTTTGCAGGCACTTACTTTCTGNNNAGATCGGAAG |
| 48882_2 | TATGAGTGTGGAGTCGTTGCTCTTTGCAGGCACTTACTTTCTGNNNAGATCGGAAG |
| 33443_1 | TATGAGTGTGGAGTCGTTGCAAGGAGGCCTGTGGAGTCTATATNNNAGATCGGAA |
| 33443_2 | TATGAGTGTGGAGTCGTTGCAAGGAGGCCTGTGGAGTCTATATNNNAGATCGGAA |
| 47312_1 | TATGAGTGTGGAGTCGTTGCTTCAAGTGATTCTCCTGCCTCAGNNNAGATCGGAAG |
| 47312_2 | TATGAGTGTGGAGTCGTTGCTTCAAGTGATTCTCCTGCCTCAGNNNAGATCGGAAG |
| 41151_1 | TATGAGTGTGGAGTCGTTGCAAAAAATTAGCAGGGTGTGGTGGNNNAGATCGGA |
| 41151_2 | TATGAGTGTGGAGTCGTTGCAAAAAATTAGCAGGGTGTGGTGGNNNAGATCGGA |
| 49649_1 | TATGAGTGTGGAGTCGTTGCCAGGAAAACTCAATGGAGAGGGANNNAGATCGGA |
| 49649_2 | TATGAGTGTGGAGTCGTTGCCAGGAAAACTCAATGGAGAGGGANNNAGATCGGA |
| 7801_1 | TATGAGTGTGGAGTCGTTGCGATGAAATCCTGCTGCTCTGTNNNAGATCGGAAGAG |
| 7801_2 | TATGAGTGTGGAGTCGTTGCGATGAAATCCTGCTGCTCTGTNNNAGATCGGAAGAG |
| 52392_1 | TATGAGTGTGGAGTCGTTGCTTTCTCTTCCAACAGATTGCCCTNNNAGATCGGAAGA |
| 52392_2 | TATGAGTGTGGAGTCGTTGCTTTCTCTTCCAACAGATTGCCCTNNNAGATCGGAAGA |
| 42111_1 | TATGAGTGTGGAGTCGTTGCAGTAGCTGGGACTACAGGTATGTNNNAGATCGGAA |
| 42111_2 | TATGAGTGTGGAGTCGTTGCAGTAGCTGGGACTACAGGTATGTNNNAGATCGGAA |
| 37886_1 | TATGAGTGTGGAGTCGTTGCCCAACATGGTGAAACCTTGTCTCNNNAGATCGGAAG |
| 37886_2 | TATGAGTGTGGAGTCGTTGCCCAACATGGTGAAACCTTGTCTCNNNAGATCGGAAG |
| 47505_1 | TATGAGTGTGGAGTCGTTGCATAGAAGCAGTGATTGTGCTCCTNNNAGATCGGAAG |
| 47505_2 | TATGAGTGTGGAGTCGTTGCATAGAAGCAGTGATTGTGCTCCTNNNAGATCGGAAG |
| 43919_1 | TATGAGTGTGGAGTCGTTGCTCTTGAGCAGTTTTCCTTCTAGTNNNAGATCGGAAG |
| 43919_2 | TATGAGTGTGGAGTCGTTGCTCTTGAGCAGTTTTCCTTCTAGTNNNAGATCGGAAG |
| 12552_1 | TATGAGTGTGGAGTCGTTGCGTGTGAGTGCAAAAAAGAAGNNNAGATCGGAAGA |
| 12552_2 | TATGAGTGTGGAGTCGTTGCGTGTGAGTGCAAAAAAGAAGNNNAGATCGGAAGA |
| 53206_1 | TATGAGTGTGGAGTCGTTGCCAAAAACGAGTTTCTGCAGAGGTNNNAGATCGGAA |
| 53206_2 | TATGAGTGTGGAGTCGTTGCCAAAAACGAGTTTCTGCAGAGGTNNNAGATCGGAA |
| 52732_1 | TATGAGTGTGGAGTCGTTGCCAGATATGTCCCTTGCCACTAGGNNNAGATCGGAAG |
| 52732_2 | TATGAGTGTGGAGTCGTTGCCAGATATGTCCCTTGCCACTAGGNNNAGATCGGAAG |
| 52615_1 | TATGAGTGTGGAGTCGTTGCTCCAGTGTTTCTTTGGCTTTTCCNNNAGATCGGAAGA |
| 52615_2 | TATGAGTGTGGAGTCGTTGCTCCAGTGTTTCTTTGGCTTTTCCNNNAGATCGGAAGA |
| 37515_1 | TATGAGTGTGGAGTCGTTGCCAGGCACCTATAATCCCAGCTACNNNAGATCGGAAG |
| 37515_2 | TATGAGTGTGGAGTCGTTGCCAGGCACCTATAATCCCAGCTACNNNAGATCGGAAG |
| 10706_1 | TATGAGTGTGGAGTCGTTGCGAGTTCCTCCATCAGCCAAGATGNNNAGATCGGAAG |
| 10706_2 | TATGAGTGTGGAGTCGTTGCGAGTTCCTCCATCAGCCAAGATGNNNAGATCGGAAG |
| 47053_1 | TATGAGTGTGGAGTCGTTGCCCAAGTAGCTGGGATTACAGGTGNNNAGATCGGAA |
| 47053_2 | TATGAGTGTGGAGTCGTTGCCCAAGTAGCTGGGATTACAGGTGNNNAGATCGGAA |
| 34434_1 | TATGAGTGTGGAGTCGTTGCGGAGGGGGGAGAGAGAATGAATANNNAGATCGG |
| 34434_2 | TATGAGTGTGGAGTCGTTGCGGAGGGGGGAGAGAGAATGAATANNNAGATCGG |
| 36853_1 | TATGAGTGTGGAGTCGTTGCAAGCAATCTGGAGAAGACTCAGTNNNAGATCGGAA |
| 36853_2 | TATGAGTGTGGAGTCGTTGCAAGCAATCTGGAGAAGACTCAGTNNNAGATCGGAA |
| 2327_1 | TATGAGTGTGGAGTCGTTGCAGCAAAACCCTGTGGGTCTAGAANNNAGATCGGAA |
| 2327_2 | TATGAGTGTGGAGTCGTTGCAGCAAAACCCTGTGGGTCTAGAANNNAGATCGGAA |

| | |
|---|---|
| 52472_1 | TATGAGTGTGGAGTCGTTGCCATAAGACTAGGAATGGAGGGGCNNNAGATCGGAA |
| 52472_2 | TATGAGTGTGGAGTCGTTGCCATAAGACTAGGAATGGAGGGGCNNNAGATCGGAA |
| 53161_1 | TATGAGTGTGGAGTCGTTGCCCTGCCATTTGAGCCAACATAATNNNAGATCGGAAGA |
| 53161_2 | TATGAGTGTGGAGTCGTTGCCCTGCCATTTGAGCCAACATAATNNNAGATCGGAAGA |
| 32180_1 | TATGAGTGTGGAGTCGTTGCCCTTCACTATCCCCCATTTCCAANNNAGATCGGAAGA |
| 32180_2 | TATGAGTGTGGAGTCGTTGCCCTTCACTATCCCCCATTTCCAANNNAGATCGGAAGA |
| 39926_1 | TATGAGTGTGGAGTCGTTGCTTCCACAAAACAACCACAGCAATNNNAGATCGGAAGA |
| 39926_2 | TATGAGTGTGGAGTCGTTGCTTCCACAAAACAACCACAGCAATNNNAGATCGGAAGA |
| 53131_1 | TATGAGTGTGGAGTCGTTGCTGAGAGAATGGGTATCAGTGGTTNNNAGATCGGAA |
| 53131_2 | TATGAGTGTGGAGTCGTTGCTGAGAGAATGGGTATCAGTGGTTNNNAGATCGGAA |
| 33226_1 | TATGAGTGTGGAGTCGTTGCGAGTAGCTGGGATTACAGGCATGNNNAGATCGGAA |
| 33226_2 | TATGAGTGTGGAGTCGTTGCGAGTAGCTGGGATTACAGGCATGNNNAGATCGGAA |
| 46521_1 | TATGAGTGTGGAGTCGTTGCCGTTTACCCTCTAAACCAATGCCNNNAGATCGGAAGA |
| 46521_2 | TATGAGTGTGGAGTCGTTGCCGTTTACCCTCTAAACCAATGCCNNNAGATCGGAAGA |
| 50955_1 | TATGAGTGTGGAGTCGTTGCTGGTTTTACAGAGGTCGCCATAANNNAGATCGGAAG |
| 50955_2 | TATGAGTGTGGAGTCGTTGCTGGTTTTACAGAGGTCGCCATAANNNAGATCGGAAG |
| 33785_1 | TATGAGTGTGGAGTCGTTGCCACGCCTGTAATTCCAACACTTTNNNAGATCGGAAGA |
| 33785_2 | TATGAGTGTGGAGTCGTTGCCACGCCTGTAATTCCAACACTTTNNNAGATCGGAAGA |
| 6180_1 | TATGAGTGTGGAGTCGTTGCTATTTGCACACTTCAGATGGGTAANNNAGATCGGAA |
| 6180_2 | TATGAGTGTGGAGTCGTTGCTATTTGCACACTTCAGATGGGTAANNNAGATCGGAA |
| 45436_1 | TATGAGTGTGGAGTCGTTGCCACACACACATGCACACACTATTNNNAGATCGGAAGA |
| 45436_2 | TATGAGTGTGGAGTCGTTGCCACACACACATGCACACACTATTNNNAGATCGGAAGA |
| 37230_1 | TATGAGTGTGGAGTCGTTGCCTCTGGAGGAGGGAAAGAAATTTNNNAGATCGGAA |
| 37230_2 | TATGAGTGTGGAGTCGTTGCCTCTGGAGGAGGGAAAGAAATTTNNNAGATCGGAA |
| 2575_1 | TATGAGTGTGGAGTCGTTGCGAGTTTGAAGACCTAGCCCATGGNNNAGATCGGAA |
| 2575_2 | TATGAGTGTGGAGTCGTTGCGAGTTTGAAGACCTAGCCCATGGNNNAGATCGGAA |
| 35326_1 | TATGAGTGTGGAGTCGTTGCATGGAATCTCAGTCTGTTGTCCANNNAGATCGGAAG |
| 35326_2 | TATGAGTGTGGAGTCGTTGCATGGAATCTCAGTCTGTTGTCCANNNAGATCGGAAG |
| 43124_1 | TATGAGTGTGGAGTCGTTGCTGACTCATCTAATGGTCTGACACANNNAGATCGGAA |
| 43124_2 | TATGAGTGTGGAGTCGTTGCTGACTCATCTAATGGTCTGACACANNNAGATCGGAA |
| 13015_1 | TATGAGTGTGGAGTCGTTGCGAACAACCCCTTATACTTAATTGNNNAGATCGGAAGA |
| 13015_2 | TATGAGTGTGGAGTCGTTGCGAACAACCCCTTATACTTAATTGNNNAGATCGGAAGA |
| 31630_1 | TATGAGTGTGGAGTCGTTGCAGAGAGAGAGAGAGAGTGAAGCCNNNAGATCGGA |
| 31630_2 | TATGAGTGTGGAGTCGTTGCAGAGAGAGAGAGAGAGTGAAGCCNNNAGATCGGA |
| 53362_1 | TATGAGTGTGGAGTCGTTGCCCTCCCATTTCTCCCTTCTTCTCNNNAGATCGGAAGAG |
| 53362_2 | TATGAGTGTGGAGTCGTTGCCCTCCCATTTCTCCCTTCTTCTCNNNAGATCGGAAGAG |
| 50629_1 | TATGAGTGTGGAGTCGTTGCCATGGGATGTGTTTGGTTGGTTGNNNAGATCGGAA |
| 50629_2 | TATGAGTGTGGAGTCGTTGCCATGGGATGTGTTTGGTTGGTTGNNNAGATCGGAA |
| 51383_1 | TATGAGTGTGGAGTCGTTGCAGCTCAAACTGGACCTGATCATTNNNAGATCGGAAG |
| 51383_2 | TATGAGTGTGGAGTCGTTGCAGCTCAAACTGGACCTGATCATTNNNAGATCGGAAG |
| 29118_1 | TATGAGTGTGGAGTCGTTGCCAAGGACTGAGGAATACTGCAGTNNNAGATCGGAA |
| 29118_2 | TATGAGTGTGGAGTCGTTGCCAAGGACTGAGGAATACTGCAGTNNNAGATCGGAA |
| 33074_1 | TATGAGTGTGGAGTCGTTGCGTAGAGATAGGGTTTCGCCATGTNNNAGATCGGAA |

| | |
|---|---|
| 33074_2 | TATGAGTGTGGAGTCGTTGCGTAGAGATAGGGTTTCGCCATGTNNNAGATCGGAA |
| 43866_1 | TATGAGTGTGGAGTCGTTGCCGGGGGGAATCTCACAACTTTATNNNAGATCGGAAG |
| 43866_2 | TATGAGTGTGGAGTCGTTGCCGGGGGGAATCTCACAACTTTATNNNAGATCGGAAG |
| 11344_1 | TATGAGTGTGGAGTCGTTGCGTGCTAAAAGTGGAACAGTCCAANNNAGATCGGAA |
| 11344_2 | TATGAGTGTGGAGTCGTTGCGTGCTAAAAGTGGAACAGTCCAANNNAGATCGGAA |
| 32602_1 | TATGAGTGTGGAGTCGTTGCGGCATGGTATTGTGTGCTTGTATNNNAGATCGGAAG |
| 32602_2 | TATGAGTGTGGAGTCGTTGCGGCATGGTATTGTGTGCTTGTATNNNAGATCGGAAG |
| 33904_1 | TATGAGTGTGGAGTCGTTGCTTCACACCATTCTCCTGCCTCNNNAGATCGGAAGAGC |
| 33904_2 | TATGAGTGTGGAGTCGTTGCTTCACACCATTCTCCTGCCTCNNNAGATCGGAAGAGC |
| 38811_1 | TATGAGTGTGGAGTCGTTGCTGCAGCACTATTTACAACTGTGGNNNAGATCGGAAG |
| 38811_2 | TATGAGTGTGGAGTCGTTGCTGCAGCACTATTTACAACTGTGGNNNAGATCGGAAG |
| 40975_1 | TATGAGTGTGGAGTCGTTGCCATTTACACTCTTGCTGGTGGTGNNNAGATCGGAAG |
| 40975_2 | TATGAGTGTGGAGTCGTTGCCATTTACACTCTTGCTGGTGGTGNNNAGATCGGAAG |
| 37548_1 | TATGAGTGTGGAGTCGTTGCCAAAAACAAAAAACCTCGTGGCANNNAGATCGGAAG |
| 37548_2 | TATGAGTGTGGAGTCGTTGCCAAAAACAAAAAACCTCGTGGCANNNAGATCGGAAG |
| 5798_1 | TATGAGTGTGGAGTCGTTGCGGGTTCCACACTTGGTCAACANNNAGATCGGAAGAG |
| 5798_2 | TATGAGTGTGGAGTCGTTGCGGGTTCCACACTTGGTCAACANNNAGATCGGAAGAG |
| 13742_1 | TATGAGTGTGGAGTCGTTGCCATCATAAACTAGTTTTTCAGGTNNNAGATCGGAAG |
| 13742_2 | TATGAGTGTGGAGTCGTTGCCATCATAAACTAGTTTTTCAGGTNNNAGATCGGAAG |
| 31830_1 | TATGAGTGTGGAGTCGTTGCCTCAAGCAATCCTCCTACCTCAGNNNAGATCGGAAGA |
| 31830_2 | TATGAGTGTGGAGTCGTTGCCTCAAGCAATCCTCCTACCTCAGNNNAGATCGGAAGA |
| 50339_1 | TATGAGTGTGGAGTCGTTGCCTAGGACTATGTGGTGAGCAAGGNNNAGATCGGAA |
| 50339_2 | TATGAGTGTGGAGTCGTTGCCTAGGACTATGTGGTGAGCAAGGNNNAGATCGGAA |
| 30764_1 | TATGAGTGTGGAGTCGTTGCCTTCAGATTCCGGAGTTCCTTGANNNAGATCGGAAG |
| 30764_2 | TATGAGTGTGGAGTCGTTGCCTTCAGATTCCGGAGTTCCTTGANNNAGATCGGAAG |
| 51978_1 | TATGAGTGTGGAGTCGTTGCTTCTGTTAGGAAGGAGGAAAGCCNNNAGATCGGAA |
| 51978_2 | TATGAGTGTGGAGTCGTTGCTTCTGTTAGGAAGGAGGAAAGCCNNNAGATCGGAA |
| 30574_1 | TATGAGTGTGGAGTCGTTGCGCATTTCAGACAGTTTGAGTCAGACNNNAGATCGGA |
| 30574_2 | TATGAGTGTGGAGTCGTTGCGCATTTCAGACAGTTTGAGTCAGACNNNAGATCGGA |
| 40612_1 | TATGAGTGTGGAGTCGTTGCACAGGAGTGAGGGCTCAATATTGNNNAGATCGGAA |
| 40612_2 | TATGAGTGTGGAGTCGTTGCACAGGAGTGAGGGCTCAATATTGNNNAGATCGGAA |
| 13127_1 | TATGAGTGTGGAGTCGTTGCAAAAATGACATTGAATTCTGGACNNNAGATCGGAAG |
| 13127_2 | TATGAGTGTGGAGTCGTTGCAAAAATGACATTGAATTCTGGACNNNAGATCGGAAG |
| 35579_1 | TATGAGTGTGGAGTCGTTGCTTCATGATGCCAGGCCAATTTTTNNNAGATCGGAAG |
| 35579_2 | TATGAGTGTGGAGTCGTTGCTTCATGATGCCAGGCCAATTTTTNNNAGATCGGAAG |
| 50311_1 | TATGAGTGTGGAGTCGTTGCCCACTGAACCCAACTCCTCTCNNNAGATCGGAAGAG |
| 50311_2 | TATGAGTGTGGAGTCGTTGCCCACTGAACCCAACTCCTCTCNNNAGATCGGAAGAG |
| 48943_1 | TATGAGTGTGGAGTCGTTGCTGATCTCAACTCACTGCAACCTCNNNAGATCGGAAGA |
| 48943_2 | TATGAGTGTGGAGTCGTTGCTGATCTCAACTCACTGCAACCTCNNNAGATCGGAAGA |
| 39199_1 | TATGAGTGTGGAGTCGTTGCAAAAAGCAACCTATGGGGGAGATNNNAGATCGGAA |
| 39199_2 | TATGAGTGTGGAGTCGTTGCAAAAAGCAACCTATGGGGGAGATNNNAGATCGGAA |
| 35205_1 | TATGAGTGTGGAGTCGTTGCTACAGTTCAAAGGAACGGTTTGCNNNAGATCGGAA |
| 35205_2 | TATGAGTGTGGAGTCGTTGCTACAGTTCAAAGGAACGGTTTGCNNNAGATCGGAA |

| | |
|---|---|
| 50044_1 | TATGAGTGTGGAGTCGTTGCGAGACCCAAGGAAGAGTTGATGTNNNAGATCGGAA |
| 50044_2 | TATGAGTGTGGAGTCGTTGCGAGACCCAAGGAAGAGTTGATGTNNNAGATCGGAA |
| 40767_1 | TATGAGTGTGGAGTCGTTGCAATGAGAAAACTGGGGATGAGCANNNAGATCGGAA |
| 40767_2 | TATGAGTGTGGAGTCGTTGCAATGAGAAAACTGGGGATGAGCANNNAGATCGGAA |
| 42177_1 | TATGAGTGTGGAGTCGTTGCGACTCACTGTAATCTCTGCCTCCNNNAGATCGGAAGA |
| 42177_2 | TATGAGTGTGGAGTCGTTGCGACTCACTGTAATCTCTGCCTCCNNNAGATCGGAAGA |
| 47673_1 | TATGAGTGTGGAGTCGTTGCAGATGATTTTCCTGCCTCAACCTNNNAGATCGGAAGA |
| 47673_2 | TATGAGTGTGGAGTCGTTGCAGATGATTTTCCTGCCTCAACCTNNNAGATCGGAAGA |
| 2766_1 | TATGAGTGTGGAGTCGTTGCGTTTTGTTTCTGGCATGTGCCTNNNAGATCGGAAGA |
| 2766_2 | TATGAGTGTGGAGTCGTTGCGTTTTGTTTCTGGCATGTGCCTNNNAGATCGGAAGA |
| 38626_1 | TATGAGTGTGGAGTCGTTGCGTTGCAGGTAGCTGAGTTCATGNNNAGATCGGAAGA |
| 38626_2 | TATGAGTGTGGAGTCGTTGCGTTGCAGGTAGCTGAGTTCATGNNNAGATCGGAAGA |
| 33928_1 | TATGAGTGTGGAGTCGTTGCGAGACAGGTTTTTCCCATGTTGCNNNAGATCGGAAG |
| 33928_2 | TATGAGTGTGGAGTCGTTGCGAGACAGGTTTTTCCCATGTTGCNNNAGATCGGAAG |
| 13748_1 | TATGAGTGTGGAGTCGTTGCAGTCATTATTTGTAATCTGGATGNNNAGATCGGAAG |
| 13748_2 | TATGAGTGTGGAGTCGTTGCAGTCATTATTTGTAATCTGGATGNNNAGATCGGAAG |
| 38086_1 | TATGAGTGTGGAGTCGTTGCCTTGTACCCTGTGACCTTGCTATNNNAGATCGGAAGA |
| 38086_2 | TATGAGTGTGGAGTCGTTGCCTTGTACCCTGTGACCTTGCTATNNNAGATCGGAAGA |
| 42197_1 | TATGAGTGTGGAGTCGTTGCGGGGTGCTGGAACATGGTAAATANNNAGATCGGAA |
| 42197_2 | TATGAGTGTGGAGTCGTTGCGGGGTGCTGGAACATGGTAAATANNNAGATCGGAA |
| 52596_1 | TATGAGTGTGGAGTCGTTGCAGAGATGATGCAGTGGGATGATCNNNAGATCGGAA |
| 52596_2 | TATGAGTGTGGAGTCGTTGCAGAGATGATGCAGTGGGATGATCNNNAGATCGGAA |
| 37471_1 | TATGAGTGTGGAGTCGTTGCTTTCTGATCTGTGTGGTGATGGTNNNAGATCGGAAG |
| 37471_2 | TATGAGTGTGGAGTCGTTGCTTTCTGATCTGTGTGGTGATGGTNNNAGATCGGAAG |
| 35097_1 | TATGAGTGTGGAGTCGTTGCCAAGTAGCTGGGACTACAGGCNNNAGATCGGAAGA |
| 35097_2 | TATGAGTGTGGAGTCGTTGCCAAGTAGCTGGGACTACAGGCNNNAGATCGGAAGA |
| 28760_1 | TATGAGTGTGGAGTCGTTGCAACTCCCCCTTTCATAGTCAACCNNNAGATCGGAAGA |
| 28760_2 | TATGAGTGTGGAGTCGTTGCAACTCCCCCTTTCATAGTCAACCNNNAGATCGGAAGA |
| 52994_1 | TATGAGTGTGGAGTCGTTGCCCCAATCTCAGCTCACTACAACTNNNAGATCGGAAGA |
| 52994_2 | TATGAGTGTGGAGTCGTTGCCCCAATCTCAGCTCACTACAACTNNNAGATCGGAAGA |
| 35517_1 | TATGAGTGTGGAGTCGTTGCAGACGGAGTCTCTCTCTGTCGNNNAGATCGGAAGAG |
| 35517_2 | TATGAGTGTGGAGTCGTTGCAGACGGAGTCTCTCTCTGTCGNNNAGATCGGAAGAG |
| 8933_1 | TATGAGTGTGGAGTCGTTGCTACTAGCCAAGCCACTTTTGCAANNNAGATCGGAAGA |
| 8933_2 | TATGAGTGTGGAGTCGTTGCTACTAGCCAAGCCACTTTTGCAANNNAGATCGGAAGA |
| 34979_1 | TATGAGTGTGGAGTCGTTGCCATGCGCCATTATTTCCAGCTAANNNAGATCGGAAGA |
| 34979_2 | TATGAGTGTGGAGTCGTTGCCATGCGCCATTATTTCCAGCTAANNNAGATCGGAAGA |
| 38525_1 | TATGAGTGTGGAGTCGTTGCCAGGATCTAGTTGCAGGCACTAANNNAGATCGGAAG |
| 38525_2 | TATGAGTGTGGAGTCGTTGCCAGGATCTAGTTGCAGGCACTAANNNAGATCGGAAG |
| 47048_1 | TATGAGTGTGGAGTCGTTGCCTGCTGTTTCTCAATTGCCTTCANNNAGATCGGAAGA |
| 47048_2 | TATGAGTGTGGAGTCGTTGCCTGCTGTTTCTCAATTGCCTTCANNNAGATCGGAAGA |
| 48620_1 | TATGAGTGTGGAGTCGTTGCAGACAGGGTCTCACTCTCTCACNNNAGATCGGAAGA |
| 48620_2 | TATGAGTGTGGAGTCGTTGCAGACAGGGTCTCACTCTCTCACNNNAGATCGGAAGA |
| 28903_1 | TATGAGTGTGGAGTCGTTGCGCAATGTGACTTTTGGCAAGTTCNNNAGATCGGAAG |

| | |
|---|---|
| 28903_2 | TATGAGTGTGGAGTCGTTGCGCAATGTGACTTTTGGCAAGTTCNNNAGATCGGAAG |
| 32666_1 | TATGAGTGTGGAGTCGTTGCGTTCAGGCAATTCTCCTACCTCANNNAGATCGGAAG |
| 32666_2 | TATGAGTGTGGAGTCGTTGCGTTCAGGCAATTCTCCTACCTCANNNAGATCGGAAG |
| 9562_1 | TATGAGTGTGGAGTCGTTGCTGGTCCTAAGGTTTTCTGGTTGCNNNAGATCGGAAG |
| 9562_2 | TATGAGTGTGGAGTCGTTGCTGGTCCTAAGGTTTTCTGGTTGCNNNAGATCGGAAG |
| 50705_1 | TATGAGTGTGGAGTCGTTGCAGACAAGGTCAGTTAGCCATCATNNNAGATCGGAA |
| 50705_2 | TATGAGTGTGGAGTCGTTGCAGACAAGGTCAGTTAGCCATCATNNNAGATCGGAA |
| 49524_1 | TATGAGTGTGGAGTCGTTGCCCAGGACTACAGGCATATGCTACNNNAGATCGGAAG |
| 49524_2 | TATGAGTGTGGAGTCGTTGCCCAGGACTACAGGCATATGCTACNNNAGATCGGAAG |
| 2006_1 | TATGAGTGTGGAGTCGTTGCTCTCTGGTTATCAGACTGGCAAANNNAGATCGGAAG |
| 2006_2 | TATGAGTGTGGAGTCGTTGCTCTCTGGTTATCAGACTGGCAAANNNAGATCGGAAG |
| 44441_1 | TATGAGTGTGGAGTCGTTGCTGATCACAACTCACTGTAGCCTCNNNAGATCGGAAG |
| 44441_2 | TATGAGTGTGGAGTCGTTGCTGATCACAACTCACTGTAGCCTCNNNAGATCGGAAG |
| 48504_1 | TATGAGTGTGGAGTCGTTGCTTGATTGTGTTCCTCTCCTCTCCNNNAGATCGGAAGA |
| 48504_2 | TATGAGTGTGGAGTCGTTGCTTGATTGTGTTCCTCTCCTCTCCNNNAGATCGGAAGA |
| 35348_1 | TATGAGTGTGGAGTCGTTGCCTCAGACTAGCAGGCCTAACAAANNNAGATCGGAAG |
| 35348_2 | TATGAGTGTGGAGTCGTTGCCTCAGACTAGCAGGCCTAACAAANNNAGATCGGAAG |
| 47290_1 | TATGAGTGTGGAGTCGTTGCGTGATCTCGGCTCAACTCACTGNNNAGATCGGAAGA |
| 47290_2 | TATGAGTGTGGAGTCGTTGCGTGATCTCGGCTCAACTCACTGNNNAGATCGGAAGA |
| 52395_1 | TATGAGTGTGGAGTCGTTGCCATGCTCCTCTTGCTCAGAAAAGNNNAGATCGGAAG |
| 52395_2 | TATGAGTGTGGAGTCGTTGCCATGCTCCTCTTGCTCAGAAAAGNNNAGATCGGAAG |
| 48982_1 | TATGAGTGTGGAGTCGTTGCACCAATGATCAGGTTTTCACAAANNNAGATCGGAAG |
| 48982_2 | TATGAGTGTGGAGTCGTTGCACCAATGATCAGGTTTTCACAAANNNAGATCGGAAG |
| 41388_1 | TATGAGTGTGGAGTCGTTGCACATTGTTGCTGAGTGACACTTGNNNAGATCGGAAG |
| 41388_2 | TATGAGTGTGGAGTCGTTGCACATTGTTGCTGAGTGACACTTGNNNAGATCGGAAG |
| 33775_1 | TATGAGTGTGGAGTCGTTGCTCATTCAAGGCAACTACGAACACNNNAGATCGGAAG |
| 33775_2 | TATGAGTGTGGAGTCGTTGCTCATTCAAGGCAACTACGAACACNNNAGATCGGAAG |
| 38592_1 | TATGAGTGTGGAGTCGTTGCCCAAGGAGGCAAAAAACTTGTACANNNAGATCGGA |
| 38592_2 | TATGAGTGTGGAGTCGTTGCCCAAGGAGGCAAAAAACTTGTACANNNAGATCGGA |
| 34085_1 | TATGAGTGTGGAGTCGTTGCTTCAAGCAATTCTCCTGCCTCAGNNNAGATCGGAAG |
| 34085_2 | TATGAGTGTGGAGTCGTTGCTTCAAGCAATTCTCCTGCCTCAGNNNAGATCGGAAG |
| 8543_1 | TATGAGTGTGGAGTCGTTGCTGAAAGTGAAATTGCTGGGTATCANNNAGATCGGA |
| 8543_2 | TATGAGTGTGGAGTCGTTGCTGAAAGTGAAATTGCTGGGTATCANNNAGATCGGA |
| 38393_1 | TATGAGTGTGGAGTCGTTGCCAGACTGCAGCCATTATTACAGCNNNAGATCGGAAG |
| 38393_2 | TATGAGTGTGGAGTCGTTGCCAGACTGCAGCCATTATTACAGCNNNAGATCGGAAG |
| 45208_1 | TATGAGTGTGGAGTCGTTGCCCCACAGTGAATCTGATGAGGAANNNAGATCGGAA |
| 45208_2 | TATGAGTGTGGAGTCGTTGCCCCACAGTGAATCTGATGAGGAANNNAGATCGGAA |
| 30739_1 | TATGAGTGTGGAGTCGTTGCGAGGGGGCAGGTAGCTTATTATTNNNAGATCGGAA |
| 30739_2 | TATGAGTGTGGAGTCGTTGCGAGGGGGCAGGTAGCTTATTATTNNNAGATCGGAA |
| 50110_1 | TATGAGTGTGGAGTCGTTGCACCTTCACTGCCCAAAAGACTTANNNAGATCGGAAG |
| 50110_2 | TATGAGTGTGGAGTCGTTGCACCTTCACTGCCCAAAAGACTTANNNAGATCGGAAG |
| 38687_1 | TATGAGTGTGGAGTCGTTGCACATTTATGCAGCCAACAGACACNNNAGATCGGAAG |
| 38687_2 | TATGAGTGTGGAGTCGTTGCACATTTATGCAGCCAACAGACACNNNAGATCGGAAG |

| | |
|---|---|
| 45334_1 | TATGAGTGTGGAGTCGTTGCCTCAAGCAATTCTCCCATGTCAGNNNAGATCGGAAG |
| 45334_2 | TATGAGTGTGGAGTCGTTGCCTCAAGCAATTCTCCCATGTCAGNNNAGATCGGAAG |
| 47467_1 | TATGAGTGTGGAGTCGTTGCGTGTCTGGATTCTTTCATTCAACGTNNNAGATCGGA |
| 47467_2 | TATGAGTGTGGAGTCGTTGCGTGTCTGGATTCTTTCATTCAACGTNNNAGATCGGA |
| 47365_1 | TATGAGTGTGGAGTCGTTGCCCTATTTGCCTGGGGAGGATTTNNNAGATCGGAAG |
| 47365_2 | TATGAGTGTGGAGTCGTTGCCCTATTTGCCTGGGGAGGATTTNNNAGATCGGAAG |
| 13775_1 | TATGAGTGTGGAGTCGTTGCCTCATTTGACTATAGAGCTAAGTNNNAGATCGGAAG |
| 13775_2 | TATGAGTGTGGAGTCGTTGCCTCATTTGACTATAGAGCTAAGTNNNAGATCGGAAG |
| 30880_1 | TATGAGTGTGGAGTCGTTGCAATATTGTTTGCTCTGTGGCTGGNNNAGATCGGAAG |
| 30880_2 | TATGAGTGTGGAGTCGTTGCAATATTGTTTGCTCTGTGGCTGGNNNAGATCGGAAG |
| 51166_1 | TATGAGTGTGGAGTCGTTGCGGTGCCTAGAGTATGCTTTCCTANNNAGATCGGAAG |
| 51166_2 | TATGAGTGTGGAGTCGTTGCGGTGCCTAGAGTATGCTTTCCTANNNAGATCGGAAG |
| 34260_1 | TATGAGTGTGGAGTCGTTGCATCACCTGAGTTTGGGAGTTTGANNNAGATCGGAA |
| 34260_2 | TATGAGTGTGGAGTCGTTGCATCACCTGAGTTTGGGAGTTTGANNNAGATCGGAA |
| 13577_1 | TATGAGTGTGGAGTCGTTGCAAGACAGCTGTGTCCTAATCNNNAGATCGGAAGAG |
| 13577_2 | TATGAGTGTGGAGTCGTTGCAAGACAGCTGTGTCCTAATCNNNAGATCGGAAGAG |
| 45878_1 | TATGAGTGTGGAGTCGTTGCTTCGAGCAATTCTCCTGACTCAGNNNAGATCGGAAG |
| 45878_2 | TATGAGTGTGGAGTCGTTGCTTCGAGCAATTCTCCTGACTCAGNNNAGATCGGAAG |
| 47721_1 | TATGAGTGTGGAGTCGTTGCCTGAGTAGCTGGGATTACAGGTGNNNAGATCGGAA |
| 47721_2 | TATGAGTGTGGAGTCGTTGCCTGAGTAGCTGGGATTACAGGTGNNNAGATCGGAA |
| 32615_1 | TATGAGTGTGGAGTCGTTGCAAAAGGGAGAGAAGAGAACAGGTNNNAGATCGGA |
| 32615_2 | TATGAGTGTGGAGTCGTTGCAAAAGGGAGAGAAGAGAACAGGTNNNAGATCGGA |
| 13375_1 | TATGAGTGTGGAGTCGTTGCTTCTTTCCCTCCACTATGTTATNNNAGATCGGAAGAG |
| 13375_2 | TATGAGTGTGGAGTCGTTGCTTCTTTCCCTCCACTATGTTATNNNAGATCGGAAGAG |
| 48970_1 | TATGAGTGTGGAGTCGTTGCTGAAACGCAAGATGAGCTGTTACNNNAGATCGGAA |
| 48970_2 | TATGAGTGTGGAGTCGTTGCTGAAACGCAAGATGAGCTGTTACNNNAGATCGGAA |
| 39414_1 | TATGAGTGTGGAGTCGTTGCTTGAGCTCAGCAGTTCAAGATCANNNAGATCGGAAG |
| 39414_2 | TATGAGTGTGGAGTCGTTGCTTGAGCTCAGCAGTTCAAGATCANNNAGATCGGAAG |
| 40546_1 | TATGAGTGTGGAGTCGTTGCCCACAGGAGAATCACTTGAACCTNNNAGATCGGAAG |
| 40546_2 | TATGAGTGTGGAGTCGTTGCCCACAGGAGAATCACTTGAACCTNNNAGATCGGAAG |
| 37633_1 | TATGAGTGTGGAGTCGTTGCAGGCTAAGGTAGGAGGATCACTTNNNAGATCGGAA |
| 37633_2 | TATGAGTGTGGAGTCGTTGCAGGCTAAGGTAGGAGGATCACTTNNNAGATCGGAA |
| 44296_1 | TATGAGTGTGGAGTCGTTGCGGAACTAGGCAAGCAGATGTCTANNNAGATCGGAA |
| 44296_2 | TATGAGTGTGGAGTCGTTGCGGAACTAGGCAAGCAGATGTCTANNNAGATCGGAA |
| 38536_1 | TATGAGTGTGGAGTCGTTGCGCCTCCAGAGTAGTTGGGACTANNNAGATCGGAAG |
| 38536_2 | TATGAGTGTGGAGTCGTTGCGCCTCCAGAGTAGTTGGGACTANNNAGATCGGAAG |
| 13211_1 | TATGAGTGTGGAGTCGTTGCAAACCCCATCTCTATCTTAAAATNNNAGATCGGAAGA |
| 13211_2 | TATGAGTGTGGAGTCGTTGCAAACCCCATCTCTATCTTAAAATNNNAGATCGGAAGA |
| 34782_1 | TATGAGTGTGGAGTCGTTGCGATCTCAGCTCACTGCAACCTCNNNAGATCGGAAGA |
| 34782_2 | TATGAGTGTGGAGTCGTTGCGATCTCAGCTCACTGCAACCTCNNNAGATCGGAAGA |
| 43575_1 | TATGAGTGTGGAGTCGTTGCTTTGCAGAAGTGAATCTGGGGTANNNAGATCGGAA |
| 43575_2 | TATGAGTGTGGAGTCGTTGCTTTGCAGAAGTGAATCTGGGGTANNNAGATCGGAA |
| 40200_1 | TATGAGTGTGGAGTCGTTGCTTCCCTAACCTACACTGCCAAAANNNAGATCGGAAGA |

| | |
|---|---|
| 40200_2 | TATGAGTGTGGAGTCGTTGCTTCCCTAACCTACACTGCCAAAANNNAGATCGGAAGA |
| 49158_1 | TATGAGTGTGGAGTCGTTGCGTGACTCAGCAGAAAAAGGGAACNNNAGATCGGAA |
| 49158_2 | TATGAGTGTGGAGTCGTTGCGTGACTCAGCAGAAAAAGGGAACNNNAGATCGGAA |
| 38246_1 | TATGAGTGTGGAGTCGTTGCCTGGGTGACAGAGCAAGACTCNNNAGATCGGAAGA |
| 38246_2 | TATGAGTGTGGAGTCGTTGCCTGGGTGACAGAGCAAGACTCNNNAGATCGGAAGA |
| 50766_1 | TATGAGTGTGGAGTCGTTGCCTACTATGTGCTCCTATGCCACCNNNAGATCGGAAGA |
| 50766_2 | TATGAGTGTGGAGTCGTTGCCTACTATGTGCTCCTATGCCACCNNNAGATCGGAAGA |
| 35046_1 | TATGAGTGTGGAGTCGTTGCTCAGTCTCCAGAGTAACTCGGATNNNAGATCGGAAG |
| 35046_2 | TATGAGTGTGGAGTCGTTGCTCAGTCTCCAGAGTAACTCGGATNNNAGATCGGAAG |
| 51421_1 | TATGAGTGTGGAGTCGTTGCTGTACCCAACAGCTCATTGAGAANNNAGATCGGAAG |
| 51421_2 | TATGAGTGTGGAGTCGTTGCTGTACCCAACAGCTCATTGAGAANNNAGATCGGAAG |
| 31999_1 | TATGAGTGTGGAGTCGTTGCCATTAGACTGGTGGTTCTAGGCANNNAGATCGGAAG |
| 31999_2 | TATGAGTGTGGAGTCGTTGCCATTAGACTGGTGGTTCTAGGCANNNAGATCGGAAG |
| 31838_1 | TATGAGTGTGGAGTCGTTGCTCTCTCTCTGGGGAAGTTTTGTGNNNAGATCGGAAG |
| 31838_2 | TATGAGTGTGGAGTCGTTGCTCTCTCTCTGGGGAAGTTTTGTGNNNAGATCGGAAG |
| 37400_1 | TATGAGTGTGGAGTCGTTGCGGAGAATCACTTGAGTCTGGGAGNNNAGATCGGAA |
| 37400_2 | TATGAGTGTGGAGTCGTTGCGGAGAATCACTTGAGTCTGGGAGNNNAGATCGGAA |
| 42281_1 | TATGAGTGTGGAGTCGTTGCTGCTATACCAACAGCCATGAGTTNNNAGATCGGAAG |
| 42281_2 | TATGAGTGTGGAGTCGTTGCTGCTATACCAACAGCCATGAGTTNNNAGATCGGAAG |
| 53099_1 | TATGAGTGTGGAGTCGTTGCAGCCATCTAACAGTCAGTCTTCCNNNAGATCGGAAGA |
| 53099_2 | TATGAGTGTGGAGTCGTTGCAGCCATCTAACAGTCAGTCTTCCNNNAGATCGGAAGA |
| 45761_1 | TATGAGTGTGGAGTCGTTGCCCTTCAGGGCAAAGAGTGACTAANNNAGATCGGAAG |
| 45761_2 | TATGAGTGTGGAGTCGTTGCCCTTCAGGGCAAAGAGTGACTAANNNAGATCGGAAG |
| 10340_1 | TATGAGTGTGGAGTCGTTGCGACGGGGAAAGAGAGAACAGAAGNNNAGATCGGA |
| 10340_2 | TATGAGTGTGGAGTCGTTGCGACGGGGAAAGAGAGAACAGAAGNNNAGATCGGA |
| 29935_1 | TATGAGTGTGGAGTCGTTGCACTATAGAAAGAGAGGGGAAGGCNNNAGATCGGAA |
| 29935_2 | TATGAGTGTGGAGTCGTTGCACTATAGAAAGAGAGGGGAAGGCNNNAGATCGGAA |
| 30628_1 | TATGAGTGTGGAGTCGTTGCGGACAAAAGTTTGAATCCTGGCTCNNNAGATCGGAA |
| 30628_2 | TATGAGTGTGGAGTCGTTGCGGACAAAAGTTTGAATCCTGGCTCNNNAGATCGGAA |
| 50082_1 | TATGAGTGTGGAGTCGTTGCGAACTCCTTATCGCCCAGTGTCNNNAGATCGGAAGAG |
| 50082_2 | TATGAGTGTGGAGTCGTTGCGAACTCCTTATCGCCCAGTGTCNNNAGATCGGAAGAG |
| 39211_1 | TATGAGTGTGGAGTCGTTGCTATTATTGTGCTGCACCTCAGGTNNNAGATCGGAAGA |
| 39211_2 | TATGAGTGTGGAGTCGTTGCTATTATTGTGCTGCACCTCAGGTNNNAGATCGGAAGA |
| 29400_1 | TATGAGTGTGGAGTCGTTGCACTTTTCTTTGCTTGTCAGATGCCNNNAGATCGGAAG |
| 29400_2 | TATGAGTGTGGAGTCGTTGCACTTTTCTTTGCTTGTCAGATGCCNNNAGATCGGAAG |
| 44780_1 | TATGAGTGTGGAGTCGTTGCCTTTTATCTCCCCTTTCCAGGCTNNNAGATCGGAAGAG |
| 44780_2 | TATGAGTGTGGAGTCGTTGCCTTTTATCTCCCCTTTCCAGGCTNNNAGATCGGAAGAG |
| 44990_1 | TATGAGTGTGGAGTCGTTGCGGCTTGTTAGACCACAAACATCCNNNAGATCGGAAG |
| 44990_2 | TATGAGTGTGGAGTCGTTGCGGCTTGTTAGACCACAAACATCCNNNAGATCGGAAG |
| 35806_1 | TATGAGTGTGGAGTCGTTGCGAAAGACCCCTAGCAAGGCTAATNNNAGATCGGAAG |
| 35806_2 | TATGAGTGTGGAGTCGTTGCGAAAGACCCCTAGCAAGGCTAATNNNAGATCGGAAG |
| 50106_1 | TATGAGTGTGGAGTCGTTGCGAGAGAACAGACACAGACCTGTCNNNAGATCGGAAG |
| 50106_2 | TATGAGTGTGGAGTCGTTGCGAGAGAACAGACACAGACCTGTCNNNAGATCGGAAG |

| | |
|---|---|
| 33385_1 | TATGAGTGTGGAGTCGTTGCATTCTGTAGCAGCAAAGAGCCTCNNNAGATCGGAAG |
| 33385_2 | TATGAGTGTGGAGTCGTTGCATTCTGTAGCAGCAAAGAGCCTCNNNAGATCGGAAG |
| 5606_1 | TATGAGTGTGGAGTCGTTGCTGGTTATTGCTGGCCTTGTAGAANNNAGATCGGAAG |
| 5606_2 | TATGAGTGTGGAGTCGTTGCTGGTTATTGCTGGCCTTGTAGAANNNAGATCGGAAG |
| 40487_1 | TATGAGTGTGGAGTCGTTGCCTATAAAGTACCTGGGCATGGGGNNNAGATCGGAAG |
| 40487_2 | TATGAGTGTGGAGTCGTTGCCTATAAAGTACCTGGGCATGGGGNNNAGATCGGAAG |
| 49357_1 | TATGAGTGTGGAGTCGTTGCGACTCTGATCCTGAGCCTCTCTCNNNAGATCGGAAGA |
| 49357_2 | TATGAGTGTGGAGTCGTTGCGACTCTGATCCTGAGCCTCTCTCNNNAGATCGGAAGA |
| 45558_1 | TATGAGTGTGGAGTCGTTGCTAGAGAAGGGCGTAAAGTTCCAGNNNAGATCGGAA |
| 45558_2 | TATGAGTGTGGAGTCGTTGCTAGAGAAGGGCGTAAAGTTCCAGNNNAGATCGGAA |
| 37900_1 | TATGAGTGTGGAGTCGTTGCCTCACATCTGTAATCCCAGCACTNNNAGATCGGAAGA |
| 37900_2 | TATGAGTGTGGAGTCGTTGCCTCACATCTGTAATCCCAGCACTNNNAGATCGGAAGA |
| 32404_1 | TATGAGTGTGGAGTCGTTGCGTGTGGTGTCCTCAAAAACTGACNNNAGATCGGAAG |
| 32404_2 | TATGAGTGTGGAGTCGTTGCGTGTGGTGTCCTCAAAAACTGACNNNAGATCGGAAG |
| 29147_1 | TATGAGTGTGGAGTCGTTGCAGAACATGGCCTGAAGCTTATGANNNAGATCGGAAG |
| 29147_2 | TATGAGTGTGGAGTCGTTGCAGAACATGGCCTGAAGCTTATGANNNAGATCGGAAG |
| 37067_1 | TATGAGTGTGGAGTCGTTGCGGAGATAGTGGCAGGCCTAATTTNNNAGATCGGAAG |
| 37067_2 | TATGAGTGTGGAGTCGTTGCGGAGATAGTGGCAGGCCTAATTTNNNAGATCGGAAG |
| 36999_1 | TATGAGTGTGGAGTCGTTGCATGCTCAATTGCAGTGTGTATGGNNNAGATCGGAAG |
| 36999_2 | TATGAGTGTGGAGTCGTTGCATGCTCAATTGCAGTGTGTATGGNNNAGATCGGAAG |
| 30210_1 | TATGAGTGTGGAGTCGTTGCATTTCTTGTAAAGTCGAGCTGCANNNAGATCGGAAG |
| 30210_2 | TATGAGTGTGGAGTCGTTGCATTTCTTGTAAAGTCGAGCTGCANNNAGATCGGAAG |
| 48519_1 | TATGAGTGTGGAGTCGTTGCCCACCACACCCAGGTAATTTTTGNNNAGATCGGAAGA |
| 48519_2 | TATGAGTGTGGAGTCGTTGCCCACCACACCCAGGTAATTTTTGNNNAGATCGGAAGA |
| 38634_1 | TATGAGTGTGGAGTCGTTGCAAGAAACCTCCTCTTCCTGTGTCNNNAGATCGGAAGA |
| 38634_2 | TATGAGTGTGGAGTCGTTGCAAGAAACCTCCTCTTCCTGTGTCNNNAGATCGGAAGA |
| 50542_1 | TATGAGTGTGGAGTCGTTGCCCACCTCTATCCCAAAAGAACCANNNAGATCGGAAGA |
| 50542_2 | TATGAGTGTGGAGTCGTTGCCCACCTCTATCCCAAAAGAACCANNNAGATCGGAAGA |
| 48105_1 | TATGAGTGTGGAGTCGTTGCTAGGTTCAAGTGATTCTCCTGCCNNNAGATCGGAAGA |
| 48105_2 | TATGAGTGTGGAGTCGTTGCTAGGTTCAAGTGATTCTCCTGCCNNNAGATCGGAAGA |
| 42435_1 | TATGAGTGTGGAGTCGTTGCCTTCAGGTCAGTTGTCCTTCTCANNNAGATCGGAAGA |
| 42435_2 | TATGAGTGTGGAGTCGTTGCCTTCAGGTCAGTTGTCCTTCTCANNNAGATCGGAAGA |
| 53065_1 | TATGAGTGTGGAGTCGTTGCCTCTCCCACAAAAAAAGCCACATNNNAGATCGGAAGA |
| 53065_2 | TATGAGTGTGGAGTCGTTGCCTCTCCCACAAAAAAAGCCACATNNNAGATCGGAAGA |
| 30018_1 | TATGAGTGTGGAGTCGTTGCTACTGAAGTACTCTGAGACGGGGNNNAGATCGGAAG |
| 30018_2 | TATGAGTGTGGAGTCGTTGCTACTGAAGTACTCTGAGACGGGGNNNAGATCGGAAG |
| 48921_1 | TATGAGTGTGGAGTCGTTGCGACAGAATCTCACTCTCTCACCCNNNAGATCGGAAGA |
| 48921_2 | TATGAGTGTGGAGTCGTTGCGACAGAATCTCACTCTCTCACCCNNNAGATCGGAAGA |
| 37161_1 | TATGAGTGTGGAGTCGTTGCTTTGCTCTCTCTTAGGGGACCTANNNAGATCGGAAGA |
| 37161_2 | TATGAGTGTGGAGTCGTTGCTTTGCTCTCTCTTAGGGGACCTANNNAGATCGGAAGA |
| 12592_1 | TATGAGTGTGGAGTCGTTGCCCTTCTTATCTAGTTAGATGCTTNNNAGATCGGAAGA |
| 12592_2 | TATGAGTGTGGAGTCGTTGCCCTTCTTATCTAGTTAGATGCTTNNNAGATCGGAAGA |
| 43578_1 | TATGAGTGTGGAGTCGTTGCACATTTGCAGGGAATATCTGGGTNNNAGATCGGAAG |

| | |
|---|---|
| 43578_2 | TATGAGTGTGGAGTCGTTGCACATTTGCAGGGAATATCTGGGTNNNAGATCGGAA |
| 2989_1 | TATGAGTGTGGAGTCGTTGCATTACAAGATGAGCAGTAAGGCCNNNAGATCGGAA |
| 2989_2 | TATGAGTGTGGAGTCGTTGCATTACAAGATGAGCAGTAAGGCCNNNAGATCGGAA |
| 43399_1 | TATGAGTGTGGAGTCGTTGCCTGGGTTCAAATGATTCTCCTGCNNNAGATCGGAAG |
| 43399_2 | TATGAGTGTGGAGTCGTTGCCTGGGTTCAAATGATTCTCCTGCNNNAGATCGGAAG |
| 48113_1 | TATGAGTGTGGAGTCGTTGCTCTGTTCATGCCCTTTACCTACTNNNAGATCGGAAGA |
| 48113_2 | TATGAGTGTGGAGTCGTTGCTCTGTTCATGCCCTTTACCTACTNNNAGATCGGAAGA |
| 40658_1 | TATGAGTGTGGAGTCGTTGCTCTAACAGTCATCATGGTAACACCANNNAGATCGGA |
| 40658_2 | TATGAGTGTGGAGTCGTTGCTCTAACAGTCATCATGGTAACACCANNNAGATCGGA |
| 32370_1 | TATGAGTGTGGAGTCGTTGCACTATCTGATCATATGCAGTGCTNNNAGATCGGAAG |
| 32370_2 | TATGAGTGTGGAGTCGTTGCACTATCTGATCATATGCAGTGCTNNNAGATCGGAAG |
| 41314_1 | TATGAGTGTGGAGTCGTTGCTTGTGTGTATTTGGAAGTAGCCANNNAGATCGGAA |
| 41314_2 | TATGAGTGTGGAGTCGTTGCTTGTGTGTATTTGGAAGTAGCCANNNAGATCGGAA |
| 11666_1 | TATGAGTGTGGAGTCGTTGCTTTTTGGGCACACATGCTCTGGNNNAGATCGGAAGA |
| 11666_2 | TATGAGTGTGGAGTCGTTGCTTTTTGGGCACACATGCTCTGGNNNAGATCGGAAGA |
| 52789_1 | TATGAGTGTGGAGTCGTTGCGTGCGGAAGACCAAAAGAGAAAANNNAGATCGGA |
| 52789_2 | TATGAGTGTGGAGTCGTTGCGTGCGGAAGACCAAAAGAGAAAANNNAGATCGGA |
| 44893_1 | TATGAGTGTGGAGTCGTTGCGAGAAGGAGTTTGGTTTGGTGTGNNNAGATCGGA |
| 44893_2 | TATGAGTGTGGAGTCGTTGCGAGAAGGAGTTTGGTTTGGTGTGNNNAGATCGGA |
| 31448_1 | TATGAGTGTGGAGTCGTTGCTGCTGTACTTCAACCCGTAGATCNNNAGATCGGAAG |
| 31448_2 | TATGAGTGTGGAGTCGTTGCTGCTGTACTTCAACCCGTAGATCNNNAGATCGGAAG |
| 37870_1 | TATGAGTGTGGAGTCGTTGCTAAATAAATAAAGCGTGGGCCGGNNNAGATCGGAA |
| 37870_2 | TATGAGTGTGGAGTCGTTGCTAAATAAATAAAGCGTGGGCCGGNNNAGATCGGAA |
| 34470_1 | TATGAGTGTGGAGTCGTTGCGAGTCCATTTCAAGGTCAGGAGANNNAGATCGGAA |
| 34470_2 | TATGAGTGTGGAGTCGTTGCGAGTCCATTTCAAGGTCAGGAGANNNAGATCGGAA |
| 29008_1 | TATGAGTGTGGAGTCGTTGCAGAGATCTGAAGGTGGAGAAGGANNNAGATCGGA |
| 29008_2 | TATGAGTGTGGAGTCGTTGCAGAGATCTGAAGGTGGAGAAGGANNNAGATCGGA |
| 51461_1 | TATGAGTGTGGAGTCGTTGCCAGTGGTTCGCACCTATAATCCTNNNAGATCGGAAG |
| 51461_2 | TATGAGTGTGGAGTCGTTGCCAGTGGTTCGCACCTATAATCCTNNNAGATCGGAAG |
| 36904_1 | TATGAGTGTGGAGTCGTTGCTCCAAGTTGAACTCCTTTTAGGTNNNAGATCGGAAG |
| 36904_2 | TATGAGTGTGGAGTCGTTGCTCCAAGTTGAACTCCTTTTAGGTNNNAGATCGGAAG |
| 45460_1 | TATGAGTGTGGAGTCGTTGCAGAGTGGATAGAGGGAAGACACANNNAGATCGGA |
| 45460_2 | TATGAGTGTGGAGTCGTTGCAGAGTGGATAGAGGGAAGACACANNNAGATCGGA |
| 50534_1 | TATGAGTGTGGAGTCGTTGCGGTGTAACCAATCCCCTCTGAATNNNAGATCGGAAG |
| 50534_2 | TATGAGTGTGGAGTCGTTGCGGTGTAACCAATCCCCTCTGAATNNNAGATCGGAAG |
| 52205_1 | TATGAGTGTGGAGTCGTTGCGTGGTGTGAAAGCAGAGGAAAAANNNAGATCGGA |
| 52205_2 | TATGAGTGTGGAGTCGTTGCGTGGTGTGAAAGCAGAGGAAAAANNNAGATCGGA |
| 50949_1 | TATGAGTGTGGAGTCGTTGCTAAAGTGAAATGAGGTGGTGGCTNNNAGATCGGAA |
| 50949_2 | TATGAGTGTGGAGTCGTTGCTAAAGTGAAATGAGGTGGTGGCTNNNAGATCGGAA |
| 36316_1 | TATGAGTGTGGAGTCGTTGCGCTGCAGTGAACTATGATTGCATNNNAGATCGGAA |
| 36316_2 | TATGAGTGTGGAGTCGTTGCGCTGCAGTGAACTATGATTGCATNNNAGATCGGAA |
| 40243_1 | TATGAGTGTGGAGTCGTTGCAATAGAGAGGGAAGGACACTCCANNNAGATCGGAA |
| 40243_2 | TATGAGTGTGGAGTCGTTGCAATAGAGAGGGAAGGACACTCCANNNAGATCGGAA |

| | |
|---|---|
| 34821_1 | TATGAGTGTGGAGTCGTTGCGGTTCAAATGATTCTCCTGCCTCNNNAGATCGGAAG |
| 34821_2 | TATGAGTGTGGAGTCGTTGCGGTTCAAATGATTCTCCTGCCTCNNNAGATCGGAAG |
| 51423_1 | TATGAGTGTGGAGTCGTTGCGTCAGACCTGAGCTAAAACTCCANNNAGATCGGAAG |
| 51423_2 | TATGAGTGTGGAGTCGTTGCGTCAGACCTGAGCTAAAACTCCANNNAGATCGGAAG |
| 45547_1 | TATGAGTGTGGAGTCGTTGCTCCCATTATTGTGCTCCAGCTTANNNAGATCGGAAGA |
| 45547_2 | TATGAGTGTGGAGTCGTTGCTCCCATTATTGTGCTCCAGCTTANNNAGATCGGAAGA |
| 10246_1 | TATGAGTGTGGAGTCGTTGCGGGTTCAAGTGATTCAGTGGCATNNNAGATCGGAA |
| 10246_2 | TATGAGTGTGGAGTCGTTGCGGGTTCAAGTGATTCAGTGGCATNNNAGATCGGAA |
| 42803_1 | TATGAGTGTGGAGTCGTTGCCTGAGGGACTCCAAGTCAGTAAGNNNAGATCGGAA |
| 42803_2 | TATGAGTGTGGAGTCGTTGCCTGAGGGACTCCAAGTCAGTAAGNNNAGATCGGAA |
| 48402_1 | TATGAGTGTGGAGTCGTTGCGTAGCTGTCACCCTCTTTTCTGANNNAGATCGGAAG |
| 48402_2 | TATGAGTGTGGAGTCGTTGCGTAGCTGTCACCCTCTTTTCTGANNNAGATCGGAAG |
| 51584_1 | TATGAGTGTGGAGTCGTTGCGAAAGGTGGAAAGAAATGGGTGGNNNAGATCGGA |
| 51584_2 | TATGAGTGTGGAGTCGTTGCGAAAGGTGGAAAGAAATGGGTGGNNNAGATCGGA |
| 50891_1 | TATGAGTGTGGAGTCGTTGCTGAATTTTGCCAAAGGGAAGGTCNNNAGATCGGAA |
| 50891_2 | TATGAGTGTGGAGTCGTTGCTGAATTTTGCCAAAGGGAAGGTCNNNAGATCGGAA |
| 8852_1 | TATGAGTGTGGAGTCGTTGCCTCCTAGCTCAGATGCTCTCTGTNNNAGATCGGAAG |
| 8852_2 | TATGAGTGTGGAGTCGTTGCCTCCTAGCTCAGATGCTCTCTGTNNNAGATCGGAAG |
| 13045_1 | TATGAGTGTGGAGTCGTTGCACGTCAGAGACTTTGTATAAAGNNNAGATCGGAAG |
| 13045_2 | TATGAGTGTGGAGTCGTTGCACGTCAGAGACTTTGTATAAAGNNNAGATCGGAAG |
| 49688_1 | TATGAGTGTGGAGTCGTTGCGGTCAGTTGTGCCTACTTCTGAGNNNAGATCGGAAG |
| 49688_2 | TATGAGTGTGGAGTCGTTGCGGTCAGTTGTGCCTACTTCTGAGNNNAGATCGGAAG |
| 30111_1 | TATGAGTGTGGAGTCGTTGCTGGGGGGCACTGCATTATAAATANNNAGATCGGAA |
| 30111_2 | TATGAGTGTGGAGTCGTTGCTGGGGGGCACTGCATTATAAATANNNAGATCGGAA |
| 46969_1 | TATGAGTGTGGAGTCGTTGCCACCACACCTGGCCTATATTGAANNNAGATCGGAAG |
| 46969_2 | TATGAGTGTGGAGTCGTTGCCACCACACCTGGCCTATATTGAANNNAGATCGGAAG |
| 36186_1 | TATGAGTGTGGAGTCGTTGCAGCATGGTGGCATATACCTGTAGNNNAGATCGGAA |
| 36186_2 | TATGAGTGTGGAGTCGTTGCAGCATGGTGGCATATACCTGTAGNNNAGATCGGAA |
| 41462_1 | TATGAGTGTGGAGTCGTTGCAGAACAAATCTGGAGGCATTACANNNAGATCGGAA |
| 41462_2 | TATGAGTGTGGAGTCGTTGCAGAACAAATCTGGAGGCATTACANNNAGATCGGAA |
| 37835_1 | TATGAGTGTGGAGTCGTTGCTGGAAGAATCACTTGAGACCAGGNNNAGATCGGAA |
| 37835_2 | TATGAGTGTGGAGTCGTTGCTGGAAGAATCACTTGAGACCAGGNNNAGATCGGAA |
| 49797_1 | TATGAGTGTGGAGTCGTTGCCACAACAACCTTGGTAGGGGATANNNAGATCGGAA |
| 49797_2 | TATGAGTGTGGAGTCGTTGCCACAACAACCTTGGTAGGGGATANNNAGATCGGAA |
| 52941_1 | TATGAGTGTGGAGTCGTTGCGAGGCTAGAGGAGTTTGGATCAGNNNAGATCGGA |
| 52941_2 | TATGAGTGTGGAGTCGTTGCGAGGCTAGAGGAGTTTGGATCAGNNNAGATCGGA |
| 30471_1 | TATGAGTGTGGAGTCGTTGCAGCACGAAGTCCTCACACATCANNNAGATCGGAAGA |
| 30471_2 | TATGAGTGTGGAGTCGTTGCAGCACGAAGTCCTCACACATCANNNAGATCGGAAGA |
| 31780_1 | TATGAGTGTGGAGTCGTTGCAGAGAGAGAGAGAGAGAGAGGGANNNAGATCGG |
| 31780_2 | TATGAGTGTGGAGTCGTTGCAGAGAGAGAGAGAGAGAGAGGGANNNAGATCGG |
| 11768_1 | TATGAGTGTGGAGTCGTTGCCCCCAGGTTGTGTAGATTCCCANNNAGATCGGAAGA |
| 11768_2 | TATGAGTGTGGAGTCGTTGCCCCCAGGTTGTGTAGATTCCCANNNAGATCGGAAGA |
| 41426_1 | TATGAGTGTGGAGTCGTTGCAATGTTTCATGCTCTCCAACTGCNNNAGATCGGAAG |

| | |
|---|---|
| 41426_2 | TATGAGTGTGGAGTCGTTGCAATGTTTCATGCTCTCCAACTGCNNNAGATCGGAAGA |
| 32563_1 | TATGAGTGTGGAGTCGTTGCGTTTCAAGCAATTCTCTGCCTCANNNAGATCGGAAGA |
| 32563_2 | TATGAGTGTGGAGTCGTTGCGTTTCAAGCAATTCTCTGCCTCANNNAGATCGGAAGA |
| 36815_1 | TATGAGTGTGGAGTCGTTGCCGGCTGTTTGTGATAGCAAAAACNNNAGATCGGAA |
| 36815_2 | TATGAGTGTGGAGTCGTTGCCGGCTGTTTGTGATAGCAAAAACNNNAGATCGGAA |
| 43950_1 | TATGAGTGTGGAGTCGTTGCCCTGTCTTTTCCCTCAGCTCTGNNNAGATCGGAAGAG |
| 43950_2 | TATGAGTGTGGAGTCGTTGCCCTGTCTTTTCCCTCAGCTCTGNNNAGATCGGAAGAG |
| 31707_1 | TATGAGTGTGGAGTCGTTGCCCAGCAGATTTCCTCTCACATCTNNNAGATCGGAAGA |
| 31707_2 | TATGAGTGTGGAGTCGTTGCCCAGCAGATTTCCTCTCACATCTNNNAGATCGGAAGA |
| 45971_1 | TATGAGTGTGGAGTCGTTGCTAGGACAATAGTGGAGGGAAGGTNNNAGATCGGA |
| 45971_2 | TATGAGTGTGGAGTCGTTGCTAGGACAATAGTGGAGGGAAGGTNNNAGATCGGA |
| 8923_1 | TATGAGTGTGGAGTCGTTGCAAGGAGTATTATAGGCCAGGCGTNNNAGATCGGAA |
| 8923_2 | TATGAGTGTGGAGTCGTTGCAAGGAGTATTATAGGCCAGGCGTNNNAGATCGGAA |
| 47871_1 | TATGAGTGTGGAGTCGTTGCGAACTTCGGACACACAACCTGANNNAGATCGGAAGA |
| 47871_2 | TATGAGTGTGGAGTCGTTGCGAACTTCGGACACACAACCTGANNNAGATCGGAAGA |
| 10063_1 | TATGAGTGTGGAGTCGTTGCGGTCCATACAACTGGCCTTCTTCNNNAGATCGGAAGA |
| 10063_2 | TATGAGTGTGGAGTCGTTGCGGTCCATACAACTGGCCTTCTTCNNNAGATCGGAAGA |
| 35532_1 | TATGAGTGTGGAGTCGTTGCGATCTCGGCTCACTGTAAGCTCNNNAGATCGGAAGA |
| 35532_2 | TATGAGTGTGGAGTCGTTGCGATCTCGGCTCACTGTAAGCTCNNNAGATCGGAAGA |
| 47259_1 | TATGAGTGTGGAGTCGTTGCAGCTGAGCACAAACTGAAAAACANNNAGATCGGAA |
| 47259_2 | TATGAGTGTGGAGTCGTTGCAGCTGAGCACAAACTGAAAAACANNNAGATCGGAA |
| 2024_1 | TATGAGTGTGGAGTCGTTGCGCACACATAAAACCTTGTGGCAANNNAGATCGGAAG |
| 2024_2 | TATGAGTGTGGAGTCGTTGCGCACACATAAAACCTTGTGGCAANNNAGATCGGAAG |
| 33493_1 | TATGAGTGTGGAGTCGTTGCGTGGTATTTGTTGCCTGCTTTCTNNNAGATCGGAAGA |
| 33493_2 | TATGAGTGTGGAGTCGTTGCGTGGTATTTGTTGCCTGCTTTCTNNNAGATCGGAAGA |
| 41039_1 | TATGAGTGTGGAGTCGTTGCCTCCTCCTTGTACCTCTGGTAGANNNAGATCGGAAGA |
| 41039_2 | TATGAGTGTGGAGTCGTTGCCTCCTCCTTGTACCTCTGGTAGANNNAGATCGGAAGA |
| 50948_1 | TATGAGTGTGGAGTCGTTGCCTTAGGGCCATGGTTTTCAATCCNNNAGATCGGAAGA |
| 50948_2 | TATGAGTGTGGAGTCGTTGCCTTAGGGCCATGGTTTTCAATCCNNNAGATCGGAAGA |
| 51374_1 | TATGAGTGTGGAGTCGTTGCAAAAGGTCTAGGGCTGAAGTCTGNNNAGATCGGAA |
| 51374_2 | TATGAGTGTGGAGTCGTTGCAAAAGGTCTAGGGCTGAAGTCTGNNNAGATCGGAA |
| 50008_1 | TATGAGTGTGGAGTCGTTGCTCTCTGCCTTTGATGAGAGACACNNNAGATCGGAAG |
| 50008_2 | TATGAGTGTGGAGTCGTTGCTCTCTGCCTTTGATGAGAGACACNNNAGATCGGAAG |
| 46761_1 | TATGAGTGTGGAGTCGTTGCCTCCCAGGTTCAAGCGATTCTCNNNAGATCGGAAGA |
| 46761_2 | TATGAGTGTGGAGTCGTTGCCTCCCAGGTTCAAGCGATTCTCNNNAGATCGGAAGA |
| 43265_1 | TATGAGTGTGGAGTCGTTGCTTCAAGCAAATCTCCTGCCTCAGNNNAGATCGGAAGA |
| 43265_2 | TATGAGTGTGGAGTCGTTGCTTCAAGCAAATCTCCTGCCTCAGNNNAGATCGGAAGA |
| 45783_1 | TATGAGTGTGGAGTCGTTGCGGGCGATACAAACGTTACAGAAGNNNAGATCGGAA |
| 45783_2 | TATGAGTGTGGAGTCGTTGCGGGCGATACAAACGTTACAGAAGNNNAGATCGGAA |
| 36230_1 | TATGAGTGTGGAGTCGTTGCTGGTATCTTTCTTTCCCCCACAGNNNAGATCGGAAGA |
| 36230_2 | TATGAGTGTGGAGTCGTTGCTGGTATCTTTCTTTCCCCCACAGNNNAGATCGGAAGA |
| 39218_1 | TATGAGTGTGGAGTCGTTGCACAAAGAAATGTATTGCTCGGCCNNNAGATCGGAA |
| 39218_2 | TATGAGTGTGGAGTCGTTGCACAAAGAAATGTATTGCTCGGCCNNNAGATCGGAA |

| | |
|---|---|
| 40956_1 | TATGAGTGTGGAGTCGTTGCTAGTATGTGTGGAAATGTCAGGTNNNAGATCGGAA |
| 40956_2 | TATGAGTGTGGAGTCGTTGCTAGTATGTGTGGAAATGTCAGGTNNNAGATCGGAA |
| 30676_1 | TATGAGTGTGGAGTCGTTGCTCAGCTAGGTTTCCCAGGTATATTCNNNAGATCGGA |
| 30676_2 | TATGAGTGTGGAGTCGTTGCTCAGCTAGGTTTCCCAGGTATATTCNNNAGATCGGA |
| 13301_1 | TATGAGTGTGGAGTCGTTGCATACCATGTTTTGTTTATCCATTNNNAGATCGGAAGA |
| 13301_2 | TATGAGTGTGGAGTCGTTGCATACCATGTTTTGTTTATCCATTNNNAGATCGGAAGA |
| 52651_1 | TATGAGTGTGGAGTCGTTGCACAGGCAAAGATACCAAGTCCTTNNNAGATCGGAAG |
| 52651_2 | TATGAGTGTGGAGTCGTTGCACAGGCAAAGATACCAAGTCCTTNNNAGATCGGAAG |
| 28770_1 | TATGAGTGTGGAGTCGTTGCTTGAGGCAGGAGAATCACTTGAANNNAGATCGGAA |
| 28770_2 | TATGAGTGTGGAGTCGTTGCTTGAGGCAGGAGAATCACTTGAANNNAGATCGGAA |
| 38100_1 | TATGAGTGTGGAGTCGTTGCCAGGTCAGGAGATTGAGACCATCNNNAGATCGGAA |
| 38100_2 | TATGAGTGTGGAGTCGTTGCCAGGTCAGGAGATTGAGACCATCNNNAGATCGGAA |
| 49494_1 | TATGAGTGTGGAGTCGTTGCTGTTAGCCAGGATGGTCTCAATCNNNAGATCGGAAG |
| 49494_2 | TATGAGTGTGGAGTCGTTGCTGTTAGCCAGGATGGTCTCAATCNNNAGATCGGAAG |
| 49529_1 | TATGAGTGTGGAGTCGTTGCCCATCTTGGCTCACTACAACCTNNNAGATCGGAAGA |
| 49529_2 | TATGAGTGTGGAGTCGTTGCCCATCTTGGCTCACTACAACCTNNNAGATCGGAAGA |
| 12633_1 | TATGAGTGTGGAGTCGTTGCGCATCGTCATGAGTATGTACNNNAGATCGGAAGAG |
| 12633_2 | TATGAGTGTGGAGTCGTTGCGCATCGTCATGAGTATGTACNNNAGATCGGAAGAG |
| 13207_1 | TATGAGTGTGGAGTCGTTGCTTAGACAGTACCATGTTGATATGNNNAGATCGGAAG |
| 13207_2 | TATGAGTGTGGAGTCGTTGCTTAGACAGTACCATGTTGATATGNNNAGATCGGAAG |
| 53164_1 | TATGAGTGTGGAGTCGTTGCCCGGAGCTACTCAACTGAACTAANNNAGATCGGAAG |
| 53164_2 | TATGAGTGTGGAGTCGTTGCCCGGAGCTACTCAACTGAACTAANNNAGATCGGAAG |
| 29651_1 | TATGAGTGTGGAGTCGTTGCAAAAAATTAGCTGGGCATGGTGGNNNAGATCGGAA |
| 29651_2 | TATGAGTGTGGAGTCGTTGCAAAAAATTAGCTGGGCATGGTGGNNNAGATCGGAA |
| 34452_1 | TATGAGTGTGGAGTCGTTGCATCTCCTCCCCTCTCACCCCNNNAGATCGGAAGAGCA |
| 34452_2 | TATGAGTGTGGAGTCGTTGCATCTCCTCCCCTCTCACCCCNNNAGATCGGAAGAGCA |
| 40739_1 | TATGAGTGTGGAGTCGTTGCGTGAAACCCCATCTCTACCAGAANNNAGATCGGAAG |
| 40739_2 | TATGAGTGTGGAGTCGTTGCGTGAAACCCCATCTCTACCAGAANNNAGATCGGAAG |
| 29125_1 | TATGAGTGTGGAGTCGTTGCTAGATGCATCCACATTCTGCTGTNNNAGATCGGAAG |
| 29125_2 | TATGAGTGTGGAGTCGTTGCTAGATGCATCCACATTCTGCTGTNNNAGATCGGAAG |
| 42151_1 | TATGAGTGTGGAGTCGTTGCGGAGTCTCACTCTGTCACCCANNNAGATCGGAAGAG |
| 42151_2 | TATGAGTGTGGAGTCGTTGCGGAGTCTCACTCTGTCACCCANNNAGATCGGAAGAG |
| 42088_1 | TATGAGTGTGGAGTCGTTGCTTCAAAGAGCTAGGTGGGAAACCNNNAGATCGGAA |
| 42088_2 | TATGAGTGTGGAGTCGTTGCTTCAAAGAGCTAGGTGGGAAACCNNNAGATCGGAA |
| 40615_1 | TATGAGTGTGGAGTCGTTGCAGTTGCTTTTCTATGCCTGGAGANNNAGATCGGAAG |
| 40615_2 | TATGAGTGTGGAGTCGTTGCAGTTGCTTTTCTATGCCTGGAGANNNAGATCGGAAG |
| 10178_1 | TATGAGTGTGGAGTCGTTGCACAGGATGTACACTAGGAGACCTNNNAGATCGGAA |
| 10178_2 | TATGAGTGTGGAGTCGTTGCACAGGATGTACACTAGGAGACCTNNNAGATCGGAA |
| 37390_1 | TATGAGTGTGGAGTCGTTGCTGTCCATGCAAAGATGAAAAGGCNNNAGATCGGAA |
| 37390_2 | TATGAGTGTGGAGTCGTTGCTGTCCATGCAAAGATGAAAAGGCNNNAGATCGGAA |
| 51433_1 | TATGAGTGTGGAGTCGTTGCGGGGAGAGTAAACGAGCACTATGNNNAGATCGGA |
| 51433_2 | TATGAGTGTGGAGTCGTTGCGGGGAGAGTAAACGAGCACTATGNNNAGATCGGA |
| 44345_1 | TATGAGTGTGGAGTCGTTGCGGGGAGGACTGGAAACCCTGNNNAGATCGGAAGA |

| | |
|---|---|
| 44345_2 | TATGAGTGTGGAGTCGTTGCGGGGAGGACTGGAAACCCTGNNNAGATCGGAAGA |
| 36767_1 | TATGAGTGTGGAGTCGTTGCCCTGGCTCTCTTCCTTTCATCAANNNAGATCGGAAGA |
| 36767_2 | TATGAGTGTGGAGTCGTTGCCCTGGCTCTCTTCCTTTCATCAANNNAGATCGGAAGA |
| 52463_1 | TATGAGTGTGGAGTCGTTGCAGGCACTTTCTTGTGTTCCAATGNNNAGATCGGAAG |
| 52463_2 | TATGAGTGTGGAGTCGTTGCAGGCACTTTCTTGTGTTCCAATGNNNAGATCGGAAG |
| 38347_1 | TATGAGTGTGGAGTCGTTGCAAAGAATGCAGAAGCTGACAACCNNNAGATCGGAA |
| 38347_2 | TATGAGTGTGGAGTCGTTGCAAAGAATGCAGAAGCTGACAACCNNNAGATCGGAA |
| 44237_1 | TATGAGTGTGGAGTCGTTGCAGGAATTATTGGTGGGCACTGATNNNAGATCGGAA |
| 44237_2 | TATGAGTGTGGAGTCGTTGCAGGAATTATTGGTGGGCACTGATNNNAGATCGGAA |
| 29517_1 | TATGAGTGTGGAGTCGTTGCAGAAATACCAAGCTCTCCAATCNNNAGATCGGAAGA |
| 29517_2 | TATGAGTGTGGAGTCGTTGCAGAAATACCAAGCTCTCCAATCNNNAGATCGGAAGA |
| 39459_1 | TATGAGTGTGGAGTCGTTGCCAAGCAAGCAAGCAAGCAATAATNNNAGATCGGAA |
| 39459_2 | TATGAGTGTGGAGTCGTTGCCAAGCAAGCAAGCAAGCAATAATNNNAGATCGGAA |
| 35471_1 | TATGAGTGTGGAGTCGTTGCACAATCTCGGCTCACTGCAAGNNNAGATCGGAAGAG |
| 35471_2 | TATGAGTGTGGAGTCGTTGCACAATCTCGGCTCACTGCAAGNNNAGATCGGAAGAG |
| 31148_1 | TATGAGTGTGGAGTCGTTGCTCCATCCTTCCACTACCAAACTCNNNAGATCGGAAGA |
| 31148_2 | TATGAGTGTGGAGTCGTTGCTCCATCCTTCCACTACCAAACTCNNNAGATCGGAAGA |
| 49227_1 | TATGAGTGTGGAGTCGTTGCCAATGCCTGCTCTCTTGGAAAAANNNAGATCGGAAG |
| 49227_2 | TATGAGTGTGGAGTCGTTGCCAATGCCTGCTCTCTTGGAAAAANNNAGATCGGAAG |
| 40073_1 | TATGAGTGTGGAGTCGTTGCGCCAAAAGCAGGATTGGTGATAGNNNAGATCGGAA |
| 40073_2 | TATGAGTGTGGAGTCGTTGCGCCAAAAGCAGGATTGGTGATAGNNNAGATCGGAA |
| 50941_1 | TATGAGTGTGGAGTCGTTGCCCCTGTACCTTTACACCAACCTTNNNAGATCGGAAGA |
| 50941_2 | TATGAGTGTGGAGTCGTTGCCCCTGTACCTTTACACCAACCTTNNNAGATCGGAAGA |
| 47832_1 | TATGAGTGTGGAGTCGTTGCCCTCCAGAGCTCAAGCAATTCTCNNNAGATCGGAAG |
| 47832_2 | TATGAGTGTGGAGTCGTTGCCCTCCAGAGCTCAAGCAATTCTCNNNAGATCGGAAG |
| 29041_1 | TATGAGTGTGGAGTCGTTGCTGTGTCTGGTGTGTGTGTATTCANNNAGATCGGAAG |
| 29041_2 | TATGAGTGTGGAGTCGTTGCTGTGTCTGGTGTGTGTGTATTCANNNAGATCGGAAG |
| 36862_1 | TATGAGTGTGGAGTCGTTGCTGGACACTTGCTATGTGAATAGTNNNAGATCGGAAG |
| 36862_2 | TATGAGTGTGGAGTCGTTGCTGGACACTTGCTATGTGAATAGTNNNAGATCGGAAG |
| 47971_1 | TATGAGTGTGGAGTCGTTGCCGGAGTTTTGCTTTTATGTTGCCNNNAGATCGGAAG |
| 47971_2 | TATGAGTGTGGAGTCGTTGCCGGAGTTTTGCTTTTATGTTGCCNNNAGATCGGAAG |
| 49711_1 | TATGAGTGTGGAGTCGTTGCGTTGAACTAGGTGACAGTGAGGTNNNAGATCGGAA |
| 49711_2 | TATGAGTGTGGAGTCGTTGCGTTGAACTAGGTGACAGTGAGGTNNNAGATCGGAA |
| 30517_1 | TATGAGTGTGGAGTCGTTGCTTTGTTTAGTAGGATGAGGGCTGTNNNAGATCGGA |
| 30517_2 | TATGAGTGTGGAGTCGTTGCTTTGTTTAGTAGGATGAGGGCTGTNNNAGATCGGA |
| 49154_1 | TATGAGTGTGGAGTCGTTGCGTGCCCACCAGCTAATTATTGTANNNAGATCGGAAG |
| 49154_2 | TATGAGTGTGGAGTCGTTGCGTGCCCACCAGCTAATTATTGTANNNAGATCGGAAG |
| 40675_1 | TATGAGTGTGGAGTCGTTGCCCTGTTACTCAGCTACTCAGGACNNNAGATCGGAAG |
| 40675_2 | TATGAGTGTGGAGTCGTTGCCCTGTTACTCAGCTACTCAGGACNNNAGATCGGAAG |
| 42623_1 | TATGAGTGTGGAGTCGTTGCTGTCAGGGATAGGAGAATGTTGCNNNAGATCGGAA |
| 42623_2 | TATGAGTGTGGAGTCGTTGCTGTCAGGGATAGGAGAATGTTGCNNNAGATCGGAA |
| 52728_1 | TATGAGTGTGGAGTCGTTGCACTCAGAGTCAGAGACTTGGTCTNNNAGATCGGAAG |
| 52728_2 | TATGAGTGTGGAGTCGTTGCACTCAGAGTCAGAGACTTGGTCTNNNAGATCGGAAG |

| | |
|---|---|
| 34045_1 | TATGAGTGTGGAGTCGTTGCGACTGAGTCTCTCACTGTCGCNNNAGATCGGAAGAG |
| 34045_2 | TATGAGTGTGGAGTCGTTGCGACTGAGTCTCTCACTGTCGCNNNAGATCGGAAGAG |
| 2265_1 | TATGAGTGTGGAGTCGTTGCAGCATCAAACGGGGCAAACATTANNNAGATCGGAA |
| 2265_2 | TATGAGTGTGGAGTCGTTGCAGCATCAAACGGGGCAAACATTANNNAGATCGGAA |
| 31296_1 | TATGAGTGTGGAGTCGTTGCATCTTGCTCTTTTGTCCAGGCTANNNAGATCGGAAG |
| 31296_2 | TATGAGTGTGGAGTCGTTGCATCTTGCTCTTTTGTCCAGGCTANNNAGATCGGAAG |
| 43332_1 | TATGAGTGTGGAGTCGTTGCCAATCTCGGTTCACTGCAAAGTCNNNAGATCGGAAG |
| 43332_2 | TATGAGTGTGGAGTCGTTGCCAATCTCGGTTCACTGCAAAGTCNNNAGATCGGAAG |
| 12766_1 | TATGAGTGTGGAGTCGTTGCGAGATGGTCCATTTCATTCATNNNAGATCGGAAGAG |
| 12766_2 | TATGAGTGTGGAGTCGTTGCGAGATGGTCCATTTCATTCATNNNAGATCGGAAGAG |
| 32877_1 | TATGAGTGTGGAGTCGTTGCGAGTGCAATGGCGTGATCTTGNNNAGATCGGAAGA |
| 32877_2 | TATGAGTGTGGAGTCGTTGCGAGTGCAATGGCGTGATCTTGNNNAGATCGGAAGA |
| 31629_1 | TATGAGTGTGGAGTCGTTGCACACATCCCTCTTCTCTCTCTCANNNAGATCGGAAGA |
| 31629_2 | TATGAGTGTGGAGTCGTTGCACACATCCCTCTTCTCTCTCTCANNNAGATCGGAAGA |
| 36634_1 | TATGAGTGTGGAGTCGTTGCAAAGCACAATCCAGCAACTTCTCNNNAGATCGGAAG |
| 36634_2 | TATGAGTGTGGAGTCGTTGCAAAGCACAATCCAGCAACTTCTCNNNAGATCGGAAG |
| 32731_1 | TATGAGTGTGGAGTCGTTGCACAGGAAACTACAGTTGTCAGCANNNAGATCGGAA |
| 32731_2 | TATGAGTGTGGAGTCGTTGCACAGGAAACTACAGTTGTCAGCANNNAGATCGGAA |
| 2971_1 | TATGAGTGTGGAGTCGTTGCATAAACTGCATTGAGCTGCGTTTNNNAGATCGGAAG |
| 2971_2 | TATGAGTGTGGAGTCGTTGCATAAACTGCATTGAGCTGCGTTTNNNAGATCGGAAG |
| 30981_1 | TATGAGTGTGGAGTCGTTGCTGTATATGTGTGCGTGCAAATCCNNNAGATCGGAAG |
| 30981_2 | TATGAGTGTGGAGTCGTTGCTGTATATGTGTGCGTGCAAATCCNNNAGATCGGAAG |
| 2030_1 | TATGAGTGTGGAGTCGTTGCGCATCTTCACATCACTGTCGGATNNNAGATCGGAAG |
| 2030_2 | TATGAGTGTGGAGTCGTTGCGCATCTTCACATCACTGTCGGATNNNAGATCGGAAG |
| 38454_1 | TATGAGTGTGGAGTCGTTGCGTAGCTCACACCTGTAATCCCAANNNAGATCGGAAG |
| 38454_2 | TATGAGTGTGGAGTCGTTGCGTAGCTCACACCTGTAATCCCAANNNAGATCGGAAG |
| 50326_1 | TATGAGTGTGGAGTCGTTGCGTGTGATGCAGACAAAAGAGGTCNNNAGATCGGAA |
| 50326_2 | TATGAGTGTGGAGTCGTTGCGTGTGATGCAGACAAAAGAGGTCNNNAGATCGGAA |
| 37353_1 | TATGAGTGTGGAGTCGTTGCTGCCCATCTGTAAAATGGCAATGNNNAGATCGGAAG |
| 37353_2 | TATGAGTGTGGAGTCGTTGCTGCCCATCTGTAAAATGGCAATGNNNAGATCGGAAG |
| 31961_1 | TATGAGTGTGGAGTCGTTGCGCCATTTGTGTGGAGCCATATACNNNAGATCGGAAG |
| 31961_2 | TATGAGTGTGGAGTCGTTGCGCCATTTGTGTGGAGCCATATACNNNAGATCGGAAG |
| 40115_1 | TATGAGTGTGGAGTCGTTGCTGGAGACATTACAACTGACCACANNNAGATCGGAAG |
| 40115_2 | TATGAGTGTGGAGTCGTTGCTGGAGACATTACAACTGACCACANNNAGATCGGAAG |
| 37302_1 | TATGAGTGTGGAGTCGTTGCTGTGTGTGTGTGTGTGTAAGAGANNNAGATCGGAA |
| 37302_2 | TATGAGTGTGGAGTCGTTGCTGTGTGTGTGTGTGTGTAAGAGANNNAGATCGGAA |
| 49215_1 | TATGAGTGTGGAGTCGTTGCTTTGCCTGAGCTGAGGTTGTATANNNAGATCGGAA |
| 49215_2 | TATGAGTGTGGAGTCGTTGCTTTGCCTGAGCTGAGGTTGTATANNNAGATCGGAA |
| 42494_1 | TATGAGTGTGGAGTCGTTGCAACTGCACACTTTCAAACAACCANNNAGATCGGAAG |
| 42494_2 | TATGAGTGTGGAGTCGTTGCAACTGCACACTTTCAAACAACCANNNAGATCGGAAG |
| 33542_1 | TATGAGTGTGGAGTCGTTGCTTCACACCATTCTCCTGCCTCNNNAGATCGGAAGAGC |
| 33542_2 | TATGAGTGTGGAGTCGTTGCTTCACACCATTCTCCTGCCTCNNNAGATCGGAAGAGC |
| 13495_1 | TATGAGTGTGGAGTCGTTGCTTGCTTTGGGATCACATTATTTNNNAGATCGGAAGA |

| | |
|---|---|
| 13495_2 | TATGAGTGTGGAGTCGTTGCTTGCTTTGGGATCACATTATTTNNNAGATCGGAAGA |
| 7293_1 | TATGAGTGTGGAGTCGTTGCAGAGAAGCTATGGGAAAACAGCANNNAGATCGGAA |
| 7293_2 | TATGAGTGTGGAGTCGTTGCAGAGAAGCTATGGGAAAACAGCANNNAGATCGGAA |
| 52266_1 | TATGAGTGTGGAGTCGTTGCATTGGAGAAATGGGCTCATTCCTNNNAGATCGGAAG |
| 52266_2 | TATGAGTGTGGAGTCGTTGCATTGGAGAAATGGGCTCATTCCTNNNAGATCGGAAG |
| 47270_1 | TATGAGTGTGGAGTCGTTGCTTAGGGTACATGTGCACATTGTGNNNAGATCGGAAG |
| 47270_2 | TATGAGTGTGGAGTCGTTGCTTAGGGTACATGTGCACATTGTGNNNAGATCGGAAG |
| 52724_1 | TATGAGTGTGGAGTCGTTGCGCTTACCAGTTGAAGAGGTTCAGNNNAGATCGGAAG |
| 52724_2 | TATGAGTGTGGAGTCGTTGCGCTTACCAGTTGAAGAGGTTCAGNNNAGATCGGAAG |
| 28920_1 | TATGAGTGTGGAGTCGTTGCGGGATTGGATGGTTAGGGAGAAGNNNAGATCGGA |
| 28920_2 | TATGAGTGTGGAGTCGTTGCGGGATTGGATGGTTAGGGAGAAGNNNAGATCGGA |
| 48962_1 | TATGAGTGTGGAGTCGTTGCCAAGAGATTCTTCACCTCAGCCTNNNAGATCGGAAGA |
| 48962_2 | TATGAGTGTGGAGTCGTTGCCAAGAGATTCTTCACCTCAGCCTNNNAGATCGGAAGA |
| 37928_1 | TATGAGTGTGGAGTCGTTGCTTACAGAGTTCCCATATGCCTCGNNNAGATCGGAAGA |
| 37928_2 | TATGAGTGTGGAGTCGTTGCTTACAGAGTTCCCATATGCCTCGNNNAGATCGGAAGA |
| 50942_1 | TATGAGTGTGGAGTCGTTGCAACCTGGCTCATTTGAATCCTCTNNNAGATCGGAAGA |
| 50942_2 | TATGAGTGTGGAGTCGTTGCAACCTGGCTCATTTGAATCCTCTNNNAGATCGGAAGA |
| 45062_1 | TATGAGTGTGGAGTCGTTGCACATGTGCAGGTTTGTTACATGGNNNAGATCGGAAG |
| 45062_2 | TATGAGTGTGGAGTCGTTGCACATGTGCAGGTTTGTTACATGGNNNAGATCGGAAG |
| 45656_1 | TATGAGTGTGGAGTCGTTGCGAGTAGCTGGGACTTCAGGCNNNAGATCGGAAGAG |
| 45656_2 | TATGAGTGTGGAGTCGTTGCGAGTAGCTGGGACTTCAGGCNNNAGATCGGAAGAG |
| 48726_1 | TATGAGTGTGGAGTCGTTGCCTCCTGAGTAGCTGGGATTACAGNNNAGATCGGAAG |
| 48726_2 | TATGAGTGTGGAGTCGTTGCCTCCTGAGTAGCTGGGATTACAGNNNAGATCGGAAG |
| 35368_1 | TATGAGTGTGGAGTCGTTGCTGACACAGCTCAGAGCAAATTTGNNNAGATCGGAAG |
| 35368_2 | TATGAGTGTGGAGTCGTTGCTGACACAGCTCAGAGCAAATTTGNNNAGATCGGAAG |
| 50180_1 | TATGAGTGTGGAGTCGTTGCGTCAGAAGGACTCCTACATGAGCNNNAGATCGGAAG |
| 50180_2 | TATGAGTGTGGAGTCGTTGCGTCAGAAGGACTCCTACATGAGCNNNAGATCGGAAG |
| 42752_1 | TATGAGTGTGGAGTCGTTGCAAGGTGGTCTGCAGGTATACAAGNNNAGATCGGAA |
| 42752_2 | TATGAGTGTGGAGTCGTTGCAAGGTGGTCTGCAGGTATACAAGNNNAGATCGGAA |
| 34480_1 | TATGAGTGTGGAGTCGTTGCAAAAACAGTTACGGGACAGAAGANNNAGATCGGAA |
| 34480_2 | TATGAGTGTGGAGTCGTTGCAAAAACAGTTACGGGACAGAAGANNNAGATCGGAA |
| 46071_1 | TATGAGTGTGGAGTCGTTGCCTCTGGAATAGCTGGGACTACAGNNNAGATCGGAAG |
| 46071_2 | TATGAGTGTGGAGTCGTTGCCTCTGGAATAGCTGGGACTACAGNNNAGATCGGAAG |
| 33653_1 | TATGAGTGTGGAGTCGTTGCGAGTTTTGCTCTTGTTGCCCAGNNNAGATCGGAAGA |
| 33653_2 | TATGAGTGTGGAGTCGTTGCGAGTTTTGCTCTTGTTGCCCAGNNNAGATCGGAAGA |
| 45177_1 | TATGAGTGTGGAGTCGTTGCGGTGGAGGAAGAAAAGCAAACAANNNAGATCGGA |
| 45177_2 | TATGAGTGTGGAGTCGTTGCGGTGGAGGAAGAAAAGCAAACAANNNAGATCGGA |
| 36034_1 | TATGAGTGTGGAGTCGTTGCTTTCACAGGCAATGTCCATCCTANNNAGATCGGAAGA |
| 36034_2 | TATGAGTGTGGAGTCGTTGCTTTCACAGGCAATGTCCATCCTANNNAGATCGGAAGA |
| 48857_1 | TATGAGTGTGGAGTCGTTGCCATTCTCCTGCCTCAGCCTCNNNAGATCGGAAGAGCA |
| 48857_2 | TATGAGTGTGGAGTCGTTGCCATTCTCCTGCCTCAGCCTCNNNAGATCGGAAGAGCA |
| 40317_1 | TATGAGTGTGGAGTCGTTGCTCTCCAGCTGTGGCAGATATTAGNNNAGATCGGAAG |
| 40317_2 | TATGAGTGTGGAGTCGTTGCTCTCCAGCTGTGGCAGATATTAGNNNAGATCGGAAG |

| | |
|---|---|
| 43358_1 | TATGAGTGTGGAGTCGTTGCGTTTCACCATGTTAGCCAGGATGNNNAGATCGGAAG |
| 43358_2 | TATGAGTGTGGAGTCGTTGCGTTTCACCATGTTAGCCAGGATGNNNAGATCGGAAG |
| 36543_1 | TATGAGTGTGGAGTCGTTGCGCAAACAAGAATGGGTCTCTTCCNNNAGATCGGAAG |
| 36543_2 | TATGAGTGTGGAGTCGTTGCGCAAACAAGAATGGGTCTCTTCCNNNAGATCGGAAG |
| 29308_1 | TATGAGTGTGGAGTCGTTGCATCCACAGAACACACAGAGCTTANNNAGATCGGAAG |
| 29308_2 | TATGAGTGTGGAGTCGTTGCATCCACAGAACACACAGAGCTTANNNAGATCGGAAG |
| 39657_1 | TATGAGTGTGGAGTCGTTGCTGCCCAATTTCTGCTGAATATGCNNNAGATCGGAAG |
| 39657_2 | TATGAGTGTGGAGTCGTTGCTGCCCAATTTCTGCTGAATATGCNNNAGATCGGAAG |
| 35389_1 | TATGAGTGTGGAGTCGTTGCCAGAGTAGCTGAGATTACAGGCANNNAGATCGGAA |
| 35389_2 | TATGAGTGTGGAGTCGTTGCCAGAGTAGCTGAGATTACAGGCANNNAGATCGGAA |
| 49584_1 | TATGAGTGTGGAGTCGTTGCGAGGGAATCTCTCACTCTGTCACNNNAGATCGGAAG |
| 49584_2 | TATGAGTGTGGAGTCGTTGCGAGGGAATCTCTCACTCTGTCACNNNAGATCGGAAG |
| 39750_1 | TATGAGTGTGGAGTCGTTGCTGACCGACACCACATTCAAGATANNNAGATCGGAAG |
| 39750_2 | TATGAGTGTGGAGTCGTTGCTGACCGACACCACATTCAAGATANNNAGATCGGAAG |
| 30290_1 | TATGAGTGTGGAGTCGTTGCTGTCGCTCAAATCCTTTCCTAGTNNNAGATCGGAAG |
| 30290_2 | TATGAGTGTGGAGTCGTTGCTGTCGCTCAAATCCTTTCCTAGTNNNAGATCGGAAG |
| 47995_1 | TATGAGTGTGGAGTCGTTGCTGGTAGACTCTTCCTCAGTCACANNNAGATCGGAAG |
| 47995_2 | TATGAGTGTGGAGTCGTTGCTGGTAGACTCTTCCTCAGTCACANNNAGATCGGAAG |
| 45407_1 | TATGAGTGTGGAGTCGTTGCACAAAGTTCTGACCAAAACACCGNNNAGATCGGAAG |
| 45407_2 | TATGAGTGTGGAGTCGTTGCACAAAGTTCTGACCAAAACACCGNNNAGATCGGAAG |
| 45834_1 | TATGAGTGTGGAGTCGTTGCCATTCTCCTGCCTCAGCCTCNNNAGATCGGAAGAGC |
| 45834_2 | TATGAGTGTGGAGTCGTTGCCATTCTCCTGCCTCAGCCTCNNNAGATCGGAAGAGC |
| 52086_1 | TATGAGTGTGGAGTCGTTGCTCATCGACATAGAAGTCCACGAGNNNAGATCGGAA |
| 52086_2 | TATGAGTGTGGAGTCGTTGCTCATCGACATAGAAGTCCACGAGNNNAGATCGGAA |
| 13069_1 | TATGAGTGTGGAGTCGTTGCGAATACATCACCACTAGCAANNNAGATCGGAAGAG |
| 13069_2 | TATGAGTGTGGAGTCGTTGCGAATACATCACCACTAGCAANNNAGATCGGAAGAG |
| 33394_1 | TATGAGTGTGGAGTCGTTGCCATTTGGTTGATCTATGGGCAGTNNNAGATCGGAA |
| 33394_2 | TATGAGTGTGGAGTCGTTGCCATTTGGTTGATCTATGGGCAGTNNNAGATCGGAA |
| 1009_1 | TATGAGTGTGGAGTCGTTGCATGTGTGTATGTACAGTGGCTTGTNNNAGATCGGA |
| 1009_2 | TATGAGTGTGGAGTCGTTGCATGTGTGTATGTACAGTGGCTTGTNNNAGATCGGA |
| 29746_1 | TATGAGTGTGGAGTCGTTGCTATTGGCAGTTGGAGGAAGAGTGNNNAGATCGGA |
| 29746_2 | TATGAGTGTGGAGTCGTTGCTATTGGCAGTTGGAGGAAGAGTGNNNAGATCGGA |
| 42667_1 | TATGAGTGTGGAGTCGTTGCCCCCCCTCCTCAAGTTTGATAAANNNAGATCGGAAG |
| 42667_2 | TATGAGTGTGGAGTCGTTGCCCCCCCTCCTCAAGTTTGATAAANNNAGATCGGAAG |
| 39735_1 | TATGAGTGTGGAGTCGTTGCATATCACTGCAGCCTTTACCTCCNNNAGATCGGAAGA |
| 39735_2 | TATGAGTGTGGAGTCGTTGCATATCACTGCAGCCTTTACCTCCNNNAGATCGGAAGA |
| 10184_1 | TATGAGTGTGGAGTCGTTGCGAAGCCCCTCTCTCTAACTCCATNNNAGATCGGAAGA |
| 10184_2 | TATGAGTGTGGAGTCGTTGCGAAGCCCCTCTCTCTAACTCCATNNNAGATCGGAAGA |
| 32830_1 | TATGAGTGTGGAGTCGTTGCGTCAAGTGATTCTCCTGCCTCANNNAGATCGGAAGA |
| 32830_2 | TATGAGTGTGGAGTCGTTGCGTCAAGTGATTCTCCTGCCTCANNNAGATCGGAAGA |
| 35172_1 | TATGAGTGTGGAGTCGTTGCCCTAGTGAGTAGCTGGGACTACANNNAGATCGGAA |
| 35172_2 | TATGAGTGTGGAGTCGTTGCCCTAGTGAGTAGCTGGGACTACANNNAGATCGGAA |
| 50244_1 | TATGAGTGTGGAGTCGTTGCCACACTCTACTGTACTGACGCCNNNAGATCGGAAGA |

| | |
|---|---|
| 50244_2 | TATGAGTGTGGAGTCGTTGCCACACTCTACTGTACTGACGCCNNNAGATCGGAAGA |
| 50071_1 | TATGAGTGTGGAGTCGTTGCATATATTAGCTGGGCATGGTGGCNNNAGATCGGAA |
| 50071_2 | TATGAGTGTGGAGTCGTTGCATATATTAGCTGGGCATGGTGGCNNNAGATCGGAA |
| 44553_1 | TATGAGTGTGGAGTCGTTGCAGAGAGAGAGAGAGAGAGAGAGAGANNNAGATC |
| 44553_2 | TATGAGTGTGGAGTCGTTGCAGAGAGAGAGAGAGAGAGAGAGAGANNNAGATC |
| 10447_1 | TATGAGTGTGGAGTCGTTGCCTGAGTTTCTCCTGGGTGTACACNNNAGATCGGAAG |
| 10447_2 | TATGAGTGTGGAGTCGTTGCCTGAGTTTCTCCTGGGTGTACACNNNAGATCGGAAG |
| 39736_1 | TATGAGTGTGGAGTCGTTGCGGTAATGTGACTTGCCTGAGTTCNNNAGATCGGAAG |
| 39736_2 | TATGAGTGTGGAGTCGTTGCGGTAATGTGACTTGCCTGAGTTCNNNAGATCGGAAG |
| 35284_1 | TATGAGTGTGGAGTCGTTGCTGGTGCAATCTCAGTCACTGTAANNNAGATCGGAAG |
| 35284_2 | TATGAGTGTGGAGTCGTTGCTGGTGCAATCTCAGTCACTGTAANNNAGATCGGAAG |
| 3054_1 | TATGAGTGTGGAGTCGTTGCTGGCTTTGTCACAGGTTGTTGATNNNAGATCGGAAG |
| 3054_2 | TATGAGTGTGGAGTCGTTGCTGGCTTTGTCACAGGTTGTTGATNNNAGATCGGAAG |
| 30395_1 | TATGAGTGTGGAGTCGTTGCTTTTTTCCAAGATAGAGTCAAACNNNAGATCGGAAG |
| 30395_2 | TATGAGTGTGGAGTCGTTGCTTTTTTCCAAGATAGAGTCAAACNNNAGATCGGAAG |
| 34179_1 | TATGAGTGTGGAGTCGTTGCGGACATTGTGTGCGTCAAGAAAANNNAGATCGGAA |
| 34179_2 | TATGAGTGTGGAGTCGTTGCGGACATTGTGTGCGTCAAGAAAANNNAGATCGGAA |
| 51535_1 | TATGAGTGTGGAGTCGTTGCCGGCACACGTATACCTATGTAACNNNAGATCGGAAG |
| 51535_2 | TATGAGTGTGGAGTCGTTGCCGGCACACGTATACCTATGTAACNNNAGATCGGAAG |
| 43213_1 | TATGAGTGTGGAGTCGTTGCGGCTTTAGAACTGGATAGCAGGTNNNAGATCGGAA |
| 43213_2 | TATGAGTGTGGAGTCGTTGCGGCTTTAGAACTGGATAGCAGGTNNNAGATCGGAA |
| 39193_1 | TATGAGTGTGGAGTCGTTGCGCAAATGATTTCTGCCCATGGAANNNAGATCGGAAG |
| 39193_2 | TATGAGTGTGGAGTCGTTGCGCAAATGATTTCTGCCCATGGAANNNAGATCGGAAG |
| 51273_1 | TATGAGTGTGGAGTCGTTGCAACCGGTCACTTGGACATATTCANNNAGATCGGAAG |
| 51273_2 | TATGAGTGTGGAGTCGTTGCAACCGGTCACTTGGACATATTCANNNAGATCGGAAG |
| 29571_1 | TATGAGTGTGGAGTCGTTGCTGAGCATGATTCAATGTTGTTGCNNNAGATCGGAAG |
| 29571_2 | TATGAGTGTGGAGTCGTTGCTGAGCATGATTCAATGTTGTTGCNNNAGATCGGAAG |
| 34403_1 | TATGAGTGTGGAGTCGTTGCGAACGTCCTTCTGCTACTGCTATNNNAGATCGGAAG |
| 34403_2 | TATGAGTGTGGAGTCGTTGCGAACGTCCTTCTGCTACTGCTATNNNAGATCGGAAG |
| 34977_1 | TATGAGTGTGGAGTCGTTGCGAGACGGAGTCTTGCTCTGTCNNNAGATCGGAAGA |
| 34977_2 | TATGAGTGTGGAGTCGTTGCGAGACGGAGTCTTGCTCTGTCNNNAGATCGGAAGA |
| 34498_1 | TATGAGTGTGGAGTCGTTGCGGGCTCCTCACTTCCCAGTANNNAGATCGGAAGAGC |
| 34498_2 | TATGAGTGTGGAGTCGTTGCGGGCTCCTCACTTCCCAGTANNNAGATCGGAAGAGC |
| 39451_1 | TATGAGTGTGGAGTCGTTGCGACTGTCTTTCCGTGTATGTTGGNNNAGATCGGAAG |
| 39451_2 | TATGAGTGTGGAGTCGTTGCGACTGTCTTTCCGTGTATGTTGGNNNAGATCGGAAG |
| 12402_1 | TATGAGTGTGGAGTCGTTGCACTCTGGTAACAGCCCAAAAAGTGNNNAGATCGGAA |
| 12402_2 | TATGAGTGTGGAGTCGTTGCACTCTGGTAACAGCCCAAAAAGTGNNNAGATCGGAA |
| 13643_1 | TATGAGTGTGGAGTCGTTGCTCTTTAATAGAAAACCTTTGGACNNNAGATCGGAAG |
| 13643_2 | TATGAGTGTGGAGTCGTTGCTCTTTAATAGAAAACCTTTGGACNNNAGATCGGAAG |
| 13058_1 | TATGAGTGTGGAGTCGTTGCAGGAGAAAAGAAAACCATTTAAANNNAGATCGGAA |
| 13058_2 | TATGAGTGTGGAGTCGTTGCAGGAGAAAAGAAAACCATTTAAANNNAGATCGGAA |
| 31397_1 | TATGAGTGTGGAGTCGTTGCACCCTTTAGTCGTAGTTGTGAGGNNNAGATCGGAAG |
| 31397_2 | TATGAGTGTGGAGTCGTTGCACCCTTTAGTCGTAGTTGTGAGGNNNAGATCGGAAG |

| | |
|---|---|
| 50400_1 | TATGAGTGTGGAGTCGTTGCCAGCCTGTGAAGGGAAGGAGNNNAGATCGGAAGA |
| 50400_2 | TATGAGTGTGGAGTCGTTGCCAGCCTGTGAAGGGAAGGAGNNNAGATCGGAAGA |
| 43010_1 | TATGAGTGTGGAGTCGTTGCAGGGGACTAGTGAAGAGATGTCANNNAGATCGGA |
| 43010_2 | TATGAGTGTGGAGTCGTTGCAGGGGACTAGTGAAGAGATGTCANNNAGATCGGA |
| 28941_1 | TATGAGTGTGGAGTCGTTGCCATCCTAACCACTGCATCTAGCTNNNAGATCGGAAG |
| 28941_2 | TATGAGTGTGGAGTCGTTGCCATCCTAACCACTGCATCTAGCTNNNAGATCGGAAG |
| 53331_1 | TATGAGTGTGGAGTCGTTGCCCCAGGCTGGTTTCAAATTTCATNNNAGATCGGAAG |
| 53331_2 | TATGAGTGTGGAGTCGTTGCCCCAGGCTGGTTTCAAATTTCATNNNAGATCGGAAG |
| 47249_1 | TATGAGTGTGGAGTCGTTGCGACTGAGAAGTACTCCCCTAGGTNNNAGATCGGAA |
| 47249_2 | TATGAGTGTGGAGTCGTTGCGACTGAGAAGTACTCCCCTAGGTNNNAGATCGGAA |
| 46086_1 | TATGAGTGTGGAGTCGTTGCCCTCTCAAAGTGCTGGAACTACANNNAGATCGGAAG |
| 46086_2 | TATGAGTGTGGAGTCGTTGCCCTCTCAAAGTGCTGGAACTACANNNAGATCGGAAG |
| 10374_1 | TATGAGTGTGGAGTCGTTGCTGGTTTTCAATGAGAGGTGGACANNNAGATCGGAA |
| 10374_2 | TATGAGTGTGGAGTCGTTGCTGGTTTTCAATGAGAGGTGGACANNNAGATCGGAA |
| 13533_1 | TATGAGTGTGGAGTCGTTGCCAAGGTATACATTTAGGATTTGTNNNAGATCGGAAG |
| 13533_2 | TATGAGTGTGGAGTCGTTGCCAAGGTATACATTTAGGATTTGTNNNAGATCGGAAG |
| 48618_1 | TATGAGTGTGGAGTCGTTGCCACCACGCCCAGTTGTATTTTTANNNAGATCGGAAG |
| 48618_2 | TATGAGTGTGGAGTCGTTGCCACCACGCCCAGTTGTATTTTTANNNAGATCGGAAG |
| 43620_1 | TATGAGTGTGGAGTCGTTGCCTTCTCGTGGCCCAAGCTGANNNAGATCGGAAGAG |
| 43620_2 | TATGAGTGTGGAGTCGTTGCCTTCTCGTGGCCCAAGCTGANNNAGATCGGAAGAG |
| 12672_1 | TATGAGTGTGGAGTCGTTGCAACCTAACGGAGTTCTCTATNNNAGATCGGAAGAGC |
| 12672_2 | TATGAGTGTGGAGTCGTTGCAACCTAACGGAGTTCTCTATNNNAGATCGGAAGAGC |
| 44590_1 | TATGAGTGTGGAGTCGTTGCCACACACACACACACACACATATNNNAGATCGGAAG |
| 44590_2 | TATGAGTGTGGAGTCGTTGCCACACACACACACACACACATATNNNAGATCGGAAG |
| 37409_1 | TATGAGTGTGGAGTCGTTGCAAAGAAAAAGGACTTTGGCAGCCNNNAGATCGGAA |
| 37409_2 | TATGAGTGTGGAGTCGTTGCAAAGAAAAAGGACTTTGGCAGCCNNNAGATCGGAA |
| 44242_1 | TATGAGTGTGGAGTCGTTGCATTTGAGAGCAAACACATCCTGCNNNAGATCGGAAG |
| 44242_2 | TATGAGTGTGGAGTCGTTGCATTTGAGAGCAAACACATCCTGCNNNAGATCGGAAG |
| 36812_1 | TATGAGTGTGGAGTCGTTGCTTCACCGATCTCCTTGTCACATTNNNAGATCGGAAGA |
| 36812_2 | TATGAGTGTGGAGTCGTTGCTTCACCGATCTCCTTGTCACATTNNNAGATCGGAAGA |
| 51127_1 | TATGAGTGTGGAGTCGTTGCTGTAACCAAGTGTCTAGCTACGGNNNAGATCGGAA |
| 51127_2 | TATGAGTGTGGAGTCGTTGCTGTAACCAAGTGTCTAGCTACGGNNNAGATCGGAA |
| 42920_1 | TATGAGTGTGGAGTCGTTGCGAGATGGGGTTTCAACATGTTGGNNNAGATCGGAA |
| 42920_2 | TATGAGTGTGGAGTCGTTGCGAGATGGGGTTTCAACATGTTGGNNNAGATCGGAA |
| 45633_1 | TATGAGTGTGGAGTCGTTGCGGAGTACAGTGGCCCAATCTCNNNAGATCGGAAGA |
| 45633_2 | TATGAGTGTGGAGTCGTTGCGGAGTACAGTGGCCCAATCTCNNNAGATCGGAAGA |
| 12152_1 | TATGAGTGTGGAGTCGTTGCTTCTTGCAATCTCCCCTAAACGGNNNAGATCGGAAG |
| 12152_2 | TATGAGTGTGGAGTCGTTGCTTCTTGCAATCTCCCCTAAACGGNNNAGATCGGAAG |
| 43110_1 | TATGAGTGTGGAGTCGTTGCATCAGAGAGGATATTGGGGAGCANNNAGATCGGA |
| 43110_2 | TATGAGTGTGGAGTCGTTGCATCAGAGAGGATATTGGGGAGCANNNAGATCGGA |
| 2013_1 | TATGAGTGTGGAGTCGTTGCCTGACCCCCTCCCCAGTTATAAANNNAGATCGGAAG |
| 2013_2 | TATGAGTGTGGAGTCGTTGCCTGACCCCCTCCCCAGTTATAAANNNAGATCGGAAG |
| 9880_1 | TATGAGTGTGGAGTCGTTGCGTCAAGTAGCGGAAAATGTCTTCANNNAGATCGGA |

| | |
|---|---|
| 9880_2 | TATGAGTGTGGAGTCGTTGCGTCAAGTAGCGGAAAATGTCTTCANNNAGATCGGAA |
| 50219_1 | TATGAGTGTGGAGTCGTTGCTCCAGTCTTTTCTTCCAGGCAATNNNAGATCGGAAGA |
| 50219_2 | TATGAGTGTGGAGTCGTTGCTCCAGTCTTTTCTTCCAGGCAATNNNAGATCGGAAGA |
| 49512_1 | TATGAGTGTGGAGTCGTTGCGCTTTGAACTCCATCCACCTCTANNNAGATCGGAAGA |
| 49512_2 | TATGAGTGTGGAGTCGTTGCGCTTTGAACTCCATCCACCTCTANNNAGATCGGAAGA |
| 32429_1 | TATGAGTGTGGAGTCGTTGCGATGGAGTCTCTCGTTCTGTCACNNNAGATCGGAAG |
| 32429_2 | TATGAGTGTGGAGTCGTTGCGATGGAGTCTCTCGTTCTGTCACNNNAGATCGGAAG |
| 38390_1 | TATGAGTGTGGAGTCGTTGCGTGATAGAGCAAGACCCTGTCTCNNNAGATCGGAAG |
| 38390_2 | TATGAGTGTGGAGTCGTTGCGTGATAGAGCAAGACCCTGTCTCNNNAGATCGGAAG |
| 33748_1 | TATGAGTGTGGAGTCGTTGCGAACACTGCCCAAAATATCCAGGNNNAGATCGGAAG |
| 33748_2 | TATGAGTGTGGAGTCGTTGCGAACACTGCCCAAAATATCCAGGNNNAGATCGGAAG |
| 50570_1 | TATGAGTGTGGAGTCGTTGCAGACAGTTGCCCTTTTACCGTATNNNAGATCGGAAGA |
| 50570_2 | TATGAGTGTGGAGTCGTTGCAGACAGTTGCCCTTTTACCGTATNNNAGATCGGAAGA |
| 47510_1 | TATGAGTGTGGAGTCGTTGCATTCTCCTTTCTCAGCCTTCCAGNNNAGATCGGAAGA |
| 47510_2 | TATGAGTGTGGAGTCGTTGCATTCTCCTTTCTCAGCCTTCCAGNNNAGATCGGAAGA |
| 34711_1 | TATGAGTGTGGAGTCGTTGCAAAGTGCTGGGATTACAGGCAAGNNNAGATCGGAA |
| 34711_2 | TATGAGTGTGGAGTCGTTGCAAAGTGCTGGGATTACAGGCAAGNNNAGATCGGAA |
| 31361_1 | TATGAGTGTGGAGTCGTTGCCATCCCACTCCAAAAACATTGCTNNNAGATCGGAAGA |
| 31361_2 | TATGAGTGTGGAGTCGTTGCCATCCCACTCCAAAAACATTGCTNNNAGATCGGAAGA |
| 32551_1 | TATGAGTGTGGAGTCGTTGCTCTCACTCTGTTCTTTCACCCAGNNNAGATCGGAAGA |
| 32551_2 | TATGAGTGTGGAGTCGTTGCTCTCACTCTGTTCTTTCACCCAGNNNAGATCGGAAGA |
| 42734_1 | TATGAGTGTGGAGTCGTTGCGTCCATTCAGGCTGTTATACCCANNNAGATCGGAAGA |
| 42734_2 | TATGAGTGTGGAGTCGTTGCGTCCATTCAGGCTGTTATACCCANNNAGATCGGAAGA |
| 41158_1 | TATGAGTGTGGAGTCGTTGCTACGCTAAGTGAATAGGCCAGACNNNAGATCGGAA |
| 41158_2 | TATGAGTGTGGAGTCGTTGCTACGCTAAGTGAATAGGCCAGACNNNAGATCGGAA |
| 50138_1 | TATGAGTGTGGAGTCGTTGCGTCATCTTCCACACTTCCTGTCTNNNAGATCGGAAGA |
| 50138_2 | TATGAGTGTGGAGTCGTTGCGTCATCTTCCACACTTCCTGTCTNNNAGATCGGAAGA |
| 51212_1 | TATGAGTGTGGAGTCGTTGCCAGGAGTTCGAGACTATACTGGCNNNAGATCGGAA |
| 51212_2 | TATGAGTGTGGAGTCGTTGCCAGGAGTTCGAGACTATACTGGCNNNAGATCGGAA |
| 32367_1 | TATGAGTGTGGAGTCGTTGCCCATGTCTCGAACTCCTGATCTCNNNAGATCGGAAGA |
| 32367_2 | TATGAGTGTGGAGTCGTTGCCCATGTCTCGAACTCCTGATCTCNNNAGATCGGAAGA |
| 51812_1 | TATGAGTGTGGAGTCGTTGCTTCTTCCTTTTCCAGAGTGCCATNNNAGATCGGAAGA |
| 51812_2 | TATGAGTGTGGAGTCGTTGCTTCTTCCTTTTCCAGAGTGCCATNNNAGATCGGAAGA |
| 41111_1 | TATGAGTGTGGAGTCGTTGCTGTTTTCCGACATGGCTATACCANNNAGATCGGAAGA |
| 41111_2 | TATGAGTGTGGAGTCGTTGCTGTTTTCCGACATGGCTATACCANNNAGATCGGAAGA |
| 41875_1 | TATGAGTGTGGAGTCGTTGCTGTATTTTTCACAGAGACGGGGTNNNAGATCGGAAG |
| 41875_2 | TATGAGTGTGGAGTCGTTGCTGTATTTTTCACAGAGACGGGGTNNNAGATCGGAAG |
| 36716_1 | TATGAGTGTGGAGTCGTTGCTGTGACAGCAAGGATGATGACAANNNAGATCGGAA |
| 36716_2 | TATGAGTGTGGAGTCGTTGCTGTGACAGCAAGGATGATGACAANNNAGATCGGAA |
| 46798_1 | TATGAGTGTGGAGTCGTTGCCAGCTTAGCCATTTGCATGTCTTNNNAGATCGGAAGA |
| 46798_2 | TATGAGTGTGGAGTCGTTGCCAGCTTAGCCATTTGCATGTCTTNNNAGATCGGAAGA |
| 42283_1 | TATGAGTGTGGAGTCGTTGCACTTCAACTTCCTGAGTAGCTGGNNNAGATCGGAAG |
| 42283_2 | TATGAGTGTGGAGTCGTTGCACTTCAACTTCCTGAGTAGCTGGNNNAGATCGGAAG |

| | |
|---|---|
| 51778_1 | TATGAGTGTGGAGTCGTTGCAAGACTGCATCCCAGAAGTTTCTNNNAGATCGGAAG |
| 51778_2 | TATGAGTGTGGAGTCGTTGCAAGACTGCATCCCAGAAGTTTCTNNNAGATCGGAAG |
| 31500_1 | TATGAGTGTGGAGTCGTTGCCTCTCCCTCTCTTCAACAGGTTCNNNAGATCGGAAGA |
| 31500_2 | TATGAGTGTGGAGTCGTTGCCTCTCCCTCTCTTCAACAGGTTCNNNAGATCGGAAGA |
| 46982_1 | TATGAGTGTGGAGTCGTTGCTTAGAGTGCAATGGTGGGATCTCNNNAGATCGGAA |
| 46982_2 | TATGAGTGTGGAGTCGTTGCTTAGAGTGCAATGGTGGGATCTCNNNAGATCGGAA |
| 46794_1 | TATGAGTGTGGAGTCGTTGCGTGAAGAATGTCAATGGCAGCTTNNNAGATCGGAA |
| 46794_2 | TATGAGTGTGGAGTCGTTGCGTGAAGAATGTCAATGGCAGCTTNNNAGATCGGAA |
| 44598_1 | TATGAGTGTGGAGTCGTTGCATCCTTCTAGACTCCCACTGACTNNNAGATCGGAAGA |
| 44598_2 | TATGAGTGTGGAGTCGTTGCATCCTTCTAGACTCCCACTGACTNNNAGATCGGAAGA |
| 45611_1 | TATGAGTGTGGAGTCGTTGCTCAAGAGATTCTACTGCCTCAGCNNNAGATCGGAAG |
| 45611_2 | TATGAGTGTGGAGTCGTTGCTCAAGAGATTCTACTGCCTCAGCNNNAGATCGGAAG |
| 30884_1 | TATGAGTGTGGAGTCGTTGCGAGTCCAATACACCTCGAGACAGNNNAGATCGGAAG |
| 30884_2 | TATGAGTGTGGAGTCGTTGCGAGTCCAATACACCTCGAGACAGNNNAGATCGGAAG |
| 47240_1 | TATGAGTGTGGAGTCGTTGCTACACTGGCCCTTTTACTGTCAANNNAGATCGGAAG |
| 47240_2 | TATGAGTGTGGAGTCGTTGCTACACTGGCCCTTTTACTGTCAANNNAGATCGGAAG |
| 29298_1 | TATGAGTGTGGAGTCGTTGCACATTTCTTCTGTATCCCCCACCNNNAGATCGGAAGA |
| 29298_2 | TATGAGTGTGGAGTCGTTGCACATTTCTTCTGTATCCCCCACCNNNAGATCGGAAGA |
| 52049_1 | TATGAGTGTGGAGTCGTTGCCCAATGCTGACATGTCATAGCAGNNNAGATCGGAAG |
| 52049_2 | TATGAGTGTGGAGTCGTTGCCCAATGCTGACATGTCATAGCAGNNNAGATCGGAAG |
| 43163_1 | TATGAGTGTGGAGTCGTTGCCTGGGATTACAAGCATGTGACACNNNAGATCGGAAG |
| 43163_2 | TATGAGTGTGGAGTCGTTGCCTGGGATTACAAGCATGTGACACNNNAGATCGGAAG |
| 30332_1 | TATGAGTGTGGAGTCGTTGCGGCAACTTAGACTCAAGCTGGTTNNNAGATCGGAAG |
| 30332_2 | TATGAGTGTGGAGTCGTTGCGGCAACTTAGACTCAAGCTGGTTNNNAGATCGGAAG |
| 49149_1 | TATGAGTGTGGAGTCGTTGCCCTCCTGAGTTCAAGCGATTCTCNNNAGATCGGAAG |
| 49149_2 | TATGAGTGTGGAGTCGTTGCCCTCCTGAGTTCAAGCGATTCTCNNNAGATCGGAAG |
| 31869_1 | TATGAGTGTGGAGTCGTTGCATGGAGAAACCCCGTCTCTACTANNNAGATCGGAAG |
| 31869_2 | TATGAGTGTGGAGTCGTTGCATGGAGAAACCCCGTCTCTACTANNNAGATCGGAAG |
| 49419_1 | TATGAGTGTGGAGTCGTTGCCCGGATTCAAGCAATTCACCTACNNNAGATCGGAAG |
| 49419_2 | TATGAGTGTGGAGTCGTTGCCCGGATTCAAGCAATTCACCTACNNNAGATCGGAAG |
| 36348_1 | TATGAGTGTGGAGTCGTTGCCTCTTACACAGGGTAGTCTTGGGNNNAGATCGGAAG |
| 36348_2 | TATGAGTGTGGAGTCGTTGCCTCTTACACAGGGTAGTCTTGGGNNNAGATCGGAAG |
| 42915_1 | TATGAGTGTGGAGTCGTTGCTGCGTGTGTGTGTGTGTGTATATNNNAGATCGGAA |
| 42915_2 | TATGAGTGTGGAGTCGTTGCTGCGTGTGTGTGTGTGTGTATATNNNAGATCGGAA |
| 32506_1 | TATGAGTGTGGAGTCGTTGCGACAATGGAAGTGTCTCATCCCTNNNAGATCGGAAG |
| 32506_2 | TATGAGTGTGGAGTCGTTGCGACAATGGAAGTGTCTCATCCCTNNNAGATCGGAAG |
| 52148_1 | TATGAGTGTGGAGTCGTTGCGCAGGAATTTACCCAATGCAGAGNNNAGATCGGAA |
| 52148_2 | TATGAGTGTGGAGTCGTTGCGCAGGAATTTACCCAATGCAGAGNNNAGATCGGAA |
| 39500_1 | TATGAGTGTGGAGTCGTTGCTGCAACATCATCAGCCATTAGGANNNAGATCGGAAG |
| 39500_2 | TATGAGTGTGGAGTCGTTGCTGCAACATCATCAGCCATTAGGANNNAGATCGGAAG |
| 36473_1 | TATGAGTGTGGAGTCGTTGCGGGAGATTGAGGCAAGAGAATCANNNAGATCGGA |
| 36473_2 | TATGAGTGTGGAGTCGTTGCGGGAGATTGAGGCAAGAGAATCANNNAGATCGGA |
| 44971_1 | TATGAGTGTGGAGTCGTTGCCAGCGATTCTTGCTTCTCCAAATNNNAGATCGGAAG |

| | |
|---|---|
| 44971_2 | TATGAGTGTGGAGTCGTTGCCAGCGATTCTTGCTTCTCCAAATNNNAGATCGGAAGA |
| 43581_1 | TATGAGTGTGGAGTCGTTGCTTTAGACACAGGATGACTGGAGCNNNAGATCGGAA |
| 43581_2 | TATGAGTGTGGAGTCGTTGCTTTAGACACAGGATGACTGGAGCNNNAGATCGGAA |
| 48911_1 | TATGAGTGTGGAGTCGTTGCCAAAACAAAGGCGTTTGCAGTTTNNNAGATCGGAAG |
| 48911_2 | TATGAGTGTGGAGTCGTTGCCAAAACAAAGGCGTTTGCAGTTTNNNAGATCGGAAG |
| 32528_1 | TATGAGTGTGGAGTCGTTGCTTTTCCTTCAAGTCAGAGGTTCCNNNAGATCGGAAGA |
| 32528_2 | TATGAGTGTGGAGTCGTTGCTTTTCCTTCAAGTCAGAGGTTCCNNNAGATCGGAAGA |
| 43193_1 | TATGAGTGTGGAGTCGTTGCAGTGGTTCACTGTAGGTTTTCCANNNAGATCGGAAG |
| 43193_2 | TATGAGTGTGGAGTCGTTGCAGTGGTTCACTGTAGGTTTTCCANNNAGATCGGAAG |
| 45859_1 | TATGAGTGTGGAGTCGTTGCGGTGTGATCTCAACACATTGCAANNNAGATCGGAAG |
| 45859_2 | TATGAGTGTGGAGTCGTTGCGGTGTGATCTCAACACATTGCAANNNAGATCGGAAG |
| 7777_1 | TATGAGTGTGGAGTCGTTGCATAGAAATAACAGAGTGGAGGCCNNNAGATCGGAA |
| 7777_2 | TATGAGTGTGGAGTCGTTGCATAGAAATAACAGAGTGGAGGCCNNNAGATCGGAA |
| 52292_1 | TATGAGTGTGGAGTCGTTGCTACAGGCTCTAGAGAGGGAGAAGNNNAGATCGGAA |
| 52292_2 | TATGAGTGTGGAGTCGTTGCTACAGGCTCTAGAGAGGGAGAAGNNNAGATCGGAA |
| 33541_1 | TATGAGTGTGGAGTCGTTGCCTTGGCTCACTGCAACCTCTANNNAGATCGGAAGAG |
| 33541_2 | TATGAGTGTGGAGTCGTTGCCTTGGCTCACTGCAACCTCTANNNAGATCGGAAGAG |
| 47616_1 | TATGAGTGTGGAGTCGTTGCAGAGATGAGACTAGGTGGGAGACNNNAGATCGGA |
| 47616_2 | TATGAGTGTGGAGTCGTTGCAGAGATGAGACTAGGTGGGAGACNNNAGATCGGA |
| 28970_1 | TATGAGTGTGGAGTCGTTGCTAACATGTTTCAAGGAGCCAGGANNNAGATCGGAA |
| 28970_2 | TATGAGTGTGGAGTCGTTGCTAACATGTTTCAAGGAGCCAGGANNNAGATCGGAA |
| 31608_1 | TATGAGTGTGGAGTCGTTGCAGGTGGCAGTGATTCCATAATGTNNNAGATCGGAA |
| 31608_2 | TATGAGTGTGGAGTCGTTGCAGGTGGCAGTGATTCCATAATGTNNNAGATCGGAA |
| 41153_1 | TATGAGTGTGGAGTCGTTGCTGGTCAAAGCCATTCAACAAGTCNNNAGATCGGAAG |
| 41153_2 | TATGAGTGTGGAGTCGTTGCTGGTCAAAGCCATTCAACAAGTCNNNAGATCGGAAG |
| 47695_1 | TATGAGTGTGGAGTCGTTGCTGCCTGCCACATTTTTGTACTTTNNNAGATCGGAAGA |
| 47695_2 | TATGAGTGTGGAGTCGTTGCTGCCTGCCACATTTTTGTACTTTNNNAGATCGGAAGA |
| 44987_1 | TATGAGTGTGGAGTCGTTGCGACTCTCCAGCTTGGGTTTTTGNNNAGATCGGAAGA |
| 44987_2 | TATGAGTGTGGAGTCGTTGCGACTCTCCAGCTTGGGTTTTTGNNNAGATCGGAAGA |
| 44758_1 | TATGAGTGTGGAGTCGTTGCCCTGGCTCATCTTATCTCATCCCNNNAGATCGGAAGA |
| 44758_2 | TATGAGTGTGGAGTCGTTGCCCTGGCTCATCTTATCTCATCCCNNNAGATCGGAAGA |
| 47976_1 | TATGAGTGTGGAGTCGTTGCAGGAAGGTCTCTATCTCCTGACCNNNAGATCGGAAG |
| 47976_2 | TATGAGTGTGGAGTCGTTGCAGGAAGGTCTCTATCTCCTGACCNNNAGATCGGAAG |
| 53190_1 | TATGAGTGTGGAGTCGTTGCCTGGGATTACAGGTGTGAGCTACNNNAGATCGGAA |
| 53190_2 | TATGAGTGTGGAGTCGTTGCCTGGGATTACAGGTGTGAGCTACNNNAGATCGGAA |
| 39221_1 | TATGAGTGTGGAGTCGTTGCTGTTCAATAACACTGGATGCTGCNNNAGATCGGAAG |
| 39221_2 | TATGAGTGTGGAGTCGTTGCTGTTCAATAACACTGGATGCTGCNNNAGATCGGAAG |
| 43297_1 | TATGAGTGTGGAGTCGTTGCACACACACACACACAAATACACTNNNAGATCGGAAGA |
| 43297_2 | TATGAGTGTGGAGTCGTTGCACACACACACACACAAATACACTNNNAGATCGGAAGA |
| 32468_1 | TATGAGTGTGGAGTCGTTGCATCCTGAATCTTGCAAAGGCTTTNNNAGATCGGAAG |
| 32468_2 | TATGAGTGTGGAGTCGTTGCATCCTGAATCTTGCAAAGGCTTTNNNAGATCGGAAG |
| 35207_1 | TATGAGTGTGGAGTCGTTGCGGATCTCGGCTCACTGCAAGNNNAGATCGGAAGAG |
| 35207_2 | TATGAGTGTGGAGTCGTTGCGGATCTCGGCTCACTGCAAGNNNAGATCGGAAGAG |

| | |
|---|---|
| 40241_1 | TATGAGTGTGGAGTCGTTGCGCTGGGTTCTGCAAATAAGACAGNNNAGATCGGAA |
| 40241_2 | TATGAGTGTGGAGTCGTTGCGCTGGGTTCTGCAAATAAGACAGNNNAGATCGGAA |
| 48365_1 | TATGAGTGTGGAGTCGTTGCGTTCAAGAGATTCTCCTGCCTCANNNAGATCGGAAG |
| 48365_2 | TATGAGTGTGGAGTCGTTGCGTTCAAGAGATTCTCCTGCCTCANNNAGATCGGAAG |
| 49502_1 | TATGAGTGTGGAGTCGTTGCACCACGCCCAGCTAATTTTTTTTNNNAGATCGGAAGA |
| 49502_2 | TATGAGTGTGGAGTCGTTGCACCACGCCCAGCTAATTTTTTTTNNNAGATCGGAAGA |
| 43424_1 | TATGAGTGTGGAGTCGTTGCCCTTCCTGAGTAGCTGGGATTACNNNAGATCGGAAG |
| 43424_2 | TATGAGTGTGGAGTCGTTGCCCTTCCTGAGTAGCTGGGATTACNNNAGATCGGAAG |
| 48039_1 | TATGAGTGTGGAGTCGTTGCGAATACAGGCGTGTGCTACCATNNNAGATCGGAAG |
| 48039_2 | TATGAGTGTGGAGTCGTTGCGAATACAGGCGTGTGCTACCATNNNAGATCGGAAG |
| 40908_1 | TATGAGTGTGGAGTCGTTGCCTGAGACAGGAGAATGGCATGAANNNAGATCGGAA |
| 40908_2 | TATGAGTGTGGAGTCGTTGCCTGAGACAGGAGAATGGCATGAANNNAGATCGGAA |
| 39120_1 | TATGAGTGTGGAGTCGTTGCTCATAAAAAGTGCCTGATGTCAGNNNAGATCGGAA |
| 39120_2 | TATGAGTGTGGAGTCGTTGCTCATAAAAAGTGCCTGATGTCAGNNNAGATCGGAA |
| 10883_1 | TATGAGTGTGGAGTCGTTGCGTGCCTGGAGACAGAACCATTTNNNAGATCGGAAG |
| 10883_2 | TATGAGTGTGGAGTCGTTGCGTGCCTGGAGACAGAACCATTTNNNAGATCGGAAG |
| 37970_1 | TATGAGTGTGGAGTCGTTGCTGTCTGGGGAAACTGATTCAAGTNNNAGATCGGAA |
| 37970_2 | TATGAGTGTGGAGTCGTTGCTGTCTGGGGAAACTGATTCAAGTNNNAGATCGGAA |
| 40720_1 | TATGAGTGTGGAGTCGTTGCGTTCAAAAGGGAGAGGTTGGGTANNNAGATCGGA |
| 40720_2 | TATGAGTGTGGAGTCGTTGCGTTCAAAAGGGAGAGGTTGGGTANNNAGATCGGA |
| 13747_1 | TATGAGTGTGGAGTCGTTGCGATTCAAGTGCTGTTTCAACNNNAGATCGGAAGAG |
| 13747_2 | TATGAGTGTGGAGTCGTTGCGATTCAAGTGCTGTTTCAACNNNAGATCGGAAGAG |
| 48525_1 | TATGAGTGTGGAGTCGTTGCTGGTACAATCTTAGCTCACTGCANNNAGATCGGAAG |
| 48525_2 | TATGAGTGTGGAGTCGTTGCTGGTACAATCTTAGCTCACTGCANNNAGATCGGAAG |
| 39102_1 | TATGAGTGTGGAGTCGTTGCTCAAGGAGAAGACATTTTCCCACTNNNAGATCGGAA |
| 39102_2 | TATGAGTGTGGAGTCGTTGCTCAAGGAGAAGACATTTTCCCACTNNNAGATCGGAA |
| 38098_1 | TATGAGTGTGGAGTCGTTGCGCGCCTGGCCAATTTTATACAATNNNAGATCGGAAG |
| 38098_2 | TATGAGTGTGGAGTCGTTGCGCGCCTGGCCAATTTTATACAATNNNAGATCGGAAG |
| 49662_1 | TATGAGTGTGGAGTCGTTGCGAGTAGCTGGGATTATAGGTGCCNNNAGATCGGAA |
| 49662_2 | TATGAGTGTGGAGTCGTTGCGAGTAGCTGGGATTATAGGTGCCNNNAGATCGGAA |
| 45242_1 | TATGAGTGTGGAGTCGTTGCATGCTTCTCATTGTACAGAGGCANNNAGATCGGAAG |
| 45242_2 | TATGAGTGTGGAGTCGTTGCATGCTTCTCATTGTACAGAGGCANNNAGATCGGAAG |
| 2302_1 | TATGAGTGTGGAGTCGTTGCAAAGACATGCATCTGTCCCTCACNNNAGATCGGAAG |
| 2302_2 | TATGAGTGTGGAGTCGTTGCAAAGACATGCATCTGTCCCTCACNNNAGATCGGAAG |
| 47485_1 | TATGAGTGTGGAGTCGTTGCTCCTGAATAGCTGGGACTACAGANNNAGATCGGAA |
| 47485_2 | TATGAGTGTGGAGTCGTTGCTCCTGAATAGCTGGGACTACAGANNNAGATCGGAA |
| 37879_1 | TATGAGTGTGGAGTCGTTGCGATTGATTCCTTAGCACACGAGCNNNAGATCGGAAG |
| 37879_2 | TATGAGTGTGGAGTCGTTGCGATTGATTCCTTAGCACACGAGCNNNAGATCGGAAG |
| 52604_1 | TATGAGTGTGGAGTCGTTGCTTTCTACACGCTAAGACGTCAGGNNNAGATCGGAAG |
| 52604_2 | TATGAGTGTGGAGTCGTTGCTTTCTACACGCTAAGACGTCAGGNNNAGATCGGAAG |
| 10615_1 | TATGAGTGTGGAGTCGTTGCGAGAGAGAGACGCTGTGTGTTTTNNNAGATCGGAA |
| 10615_2 | TATGAGTGTGGAGTCGTTGCGAGAGAGAGACGCTGTGTGTTTTNNNAGATCGGAA |
| 42193_1 | TATGAGTGTGGAGTCGTTGCTTTGGCCATTTGGGTGTTTCTTTNNNAGATCGGAAG |

| | |
|---|---|
| 42193_2 | TATGAGTGTGGAGTCGTTGCTTTGGCCATTTGGGTGTTTCTTTNNNAGATCGGAAG |
| 51685_1 | TATGAGTGTGGAGTCGTTGCAATGAAGCTGCCTTTGAGTTGTCNNNAGATCGGAAG |
| 51685_2 | TATGAGTGTGGAGTCGTTGCAATGAAGCTGCCTTTGAGTTGTCNNNAGATCGGAAG |
| 33965_1 | TATGAGTGTGGAGTCGTTGCACCACACCCGGCTAATTTTTTTTNNNAGATCGGAAGA |
| 33965_2 | TATGAGTGTGGAGTCGTTGCACCACACCCGGCTAATTTTTTTTNNNAGATCGGAAGA |
| 39729_1 | TATGAGTGTGGAGTCGTTGCAGATCGAGACCATCTTGGCTAACNNNAGATCGGAAG |
| 39729_2 | TATGAGTGTGGAGTCGTTGCAGATCGAGACCATCTTGGCTAACNNNAGATCGGAAG |
| 42750_1 | TATGAGTGTGGAGTCGTTGCAAGTCCTAAGATCCGTAGTTGGCNNNAGATCGGAA |
| 42750_2 | TATGAGTGTGGAGTCGTTGCAAGTCCTAAGATCCGTAGTTGGCNNNAGATCGGAA |
| 52861_1 | TATGAGTGTGGAGTCGTTGCTTGTCTGTCTTCTCTGTTGCCTGNNNAGATCGGAAGA |
| 52861_2 | TATGAGTGTGGAGTCGTTGCTTGTCTGTCTTCTCTGTTGCCTGNNNAGATCGGAAGA |
| 31650_1 | TATGAGTGTGGAGTCGTTGCCACTTAGAATTGCCTTTCCAGCANNNAGATCGGAAG |
| 31650_2 | TATGAGTGTGGAGTCGTTGCCACTTAGAATTGCCTTTCCAGCANNNAGATCGGAAG |
| 43870_1 | TATGAGTGTGGAGTCGTTGCAGTTAGGTAAAGAAGGCAGGCAANNNAGATCGGA |
| 43870_2 | TATGAGTGTGGAGTCGTTGCAGTTAGGTAAAGAAGGCAGGCAANNNAGATCGGA |
| 50468_1 | TATGAGTGTGGAGTCGTTGCACTATCCTGACTTTCAGCTGGTGNNNAGATCGGAAG |
| 50468_2 | TATGAGTGTGGAGTCGTTGCACTATCCTGACTTTCAGCTGGTGNNNAGATCGGAAG |
| 36688_1 | TATGAGTGTGGAGTCGTTGCAACCAGACAAAAAGCAGAACACCNNNAGATCGGAA |
| 36688_2 | TATGAGTGTGGAGTCGTTGCAACCAGACAAAAAGCAGAACACCNNNAGATCGGAA |
| 50178_1 | TATGAGTGTGGAGTCGTTGCAATAAAAGGGGGAGCATAGACGGNNNAGATCGGA |
| 50178_2 | TATGAGTGTGGAGTCGTTGCAATAAAAGGGGGAGCATAGACGGNNNAGATCGGA |
| 37059_1 | TATGAGTGTGGAGTCGTTGCCACTCACACCATTCAAAGACTCANNNAGATCGGAAGA |
| 37059_2 | TATGAGTGTGGAGTCGTTGCCACTCACACCATTCAAAGACTCANNNAGATCGGAAGA |
| 37961_1 | TATGAGTGTGGAGTCGTTGCCTGAGGTCAGGAATTCAAGACCANNNAGATCGGAA |
| 37961_2 | TATGAGTGTGGAGTCGTTGCCTGAGGTCAGGAATTCAAGACCANNNAGATCGGAA |
| 3012_1 | TATGAGTGTGGAGTCGTTGCGGCTGCTTCCTTCTCTAAAGCTTNNNAGATCGGAAGA |
| 3012_2 | TATGAGTGTGGAGTCGTTGCGGCTGCTTCCTTCTCTAAAGCTTNNNAGATCGGAAGA |
| 46724_1 | TATGAGTGTGGAGTCGTTGCGAGGTGGCAGATGTAGGTGTTATNNNAGATCGGAA |
| 46724_2 | TATGAGTGTGGAGTCGTTGCGAGGTGGCAGATGTAGGTGTTATNNNAGATCGGAA |
| 34453_1 | TATGAGTGTGGAGTCGTTGCCCCATCTCCTCCCCTCTCACNNNAGATCGGAAGAGCA |
| 34453_2 | TATGAGTGTGGAGTCGTTGCCCCATCTCCTCCCCTCTCACNNNAGATCGGAAGAGCA |
| 52903_1 | TATGAGTGTGGAGTCGTTGCGTGCTTAGAGACAGAGGATGTGGNNNAGATCGGAA |
| 52903_2 | TATGAGTGTGGAGTCGTTGCGTGCTTAGAGACAGAGGATGTGGNNNAGATCGGAA |
| 13744_1 | TATGAGTGTGGAGTCGTTGCTCTTTTACTGAGAATCTTTTCTGNNNAGATCGGAAGA |
| 13744_2 | TATGAGTGTGGAGTCGTTGCTCTTTTACTGAGAATCTTTTCTGNNNAGATCGGAAGA |
| 44896_1 | TATGAGTGTGGAGTCGTTGCGAGAAGGGAGGGATTGGAACTTCNNNAGATCGGA |
| 44896_2 | TATGAGTGTGGAGTCGTTGCGAGAAGGGAGGGATTGGAACTTCNNNAGATCGGA |
| 52333_1 | TATGAGTGTGGAGTCGTTGCCTCCTTGGACTGCTCCTTCTTGNNNAGATCGGAAGA |
| 52333_2 | TATGAGTGTGGAGTCGTTGCCTCCTTGGACTGCTCCTTCTTGNNNAGATCGGAAGA |
| 39084_1 | TATGAGTGTGGAGTCGTTGCAGTCTGAGCCAGGAGATCATTTGNNNAGATCGGAA |
| 39084_2 | TATGAGTGTGGAGTCGTTGCAGTCTGAGCCAGGAGATCATTTGNNNAGATCGGAA |
| 43506_1 | TATGAGTGTGGAGTCGTTGCCTCCTGAGTAGCTGGGATTACAGNNNAGATCGGAA |
| 43506_2 | TATGAGTGTGGAGTCGTTGCCTCCTGAGTAGCTGGGATTACAGNNNAGATCGGAA |

| | |
|---|---|
| 38310_1 | TATGAGTGTGGAGTCGTTGCACACAGCCTTGATTACTGTAGCTNNNAGATCGGAAG |
| 38310_2 | TATGAGTGTGGAGTCGTTGCACACAGCCTTGATTACTGTAGCTNNNAGATCGGAAG |
| 29416_1 | TATGAGTGTGGAGTCGTTGCGCTTCATCCTGTCTCATCTGCATNNNAGATCGGAAGA |
| 29416_2 | TATGAGTGTGGAGTCGTTGCGCTTCATCCTGTCTCATCTGCATNNNAGATCGGAAGA |
| 33415_1 | TATGAGTGTGGAGTCGTTGCAAAGAACAGACCAAAGCCAAGTCNNNAGATCGGAA |
| 33415_2 | TATGAGTGTGGAGTCGTTGCAAAGAACAGACCAAAGCCAAGTCNNNAGATCGGAA |
| 34918_1 | TATGAGTGTGGAGTCGTTGCTGTGTGTCTCTCTAGGACTTTGCNNNAGATCGGAAG |
| 34918_2 | TATGAGTGTGGAGTCGTTGCTGTGTGTCTCTCTAGGACTTTGCNNNAGATCGGAAG |
| 39921_1 | TATGAGTGTGGAGTCGTTGCGCTAGAGAATCTCTCCAACCTGGNNNAGATCGGAAG |
| 39921_2 | TATGAGTGTGGAGTCGTTGCGCTAGAGAATCTCTCCAACCTGGNNNAGATCGGAAG |
| 32397_1 | TATGAGTGTGGAGTCGTTGCTGGATTCCAATTGAGAATGTGTGACNNNAGATCGG |
| 32397_2 | TATGAGTGTGGAGTCGTTGCTGGATTCCAATTGAGAATGTGTGACNNNAGATCGG |
| 50529_1 | TATGAGTGTGGAGTCGTTGCAGGAGGAAGAGAAGGACAGAGAANNNAGATCGG |
| 50529_2 | TATGAGTGTGGAGTCGTTGCAGGAGGAAGAGAAGGACAGAGAANNNAGATCGG |
| 12909_1 | TATGAGTGTGGAGTCGTTGCAGACACCGAAGAGAAAAAATTNNNAGATCGGAAGA |
| 12909_2 | TATGAGTGTGGAGTCGTTGCAGACACCGAAGAGAAAAAATTNNNAGATCGGAAGA |
| 31070_1 | TATGAGTGTGGAGTCGTTGCGGATGGGGATATGTAAACCTGCTNNNAGATCGGAA |
| 31070_2 | TATGAGTGTGGAGTCGTTGCGGATGGGGATATGTAAACCTGCTNNNAGATCGGAA |
| 32453_1 | TATGAGTGTGGAGTCGTTGCGCTCATGCCTATAATTCCAGCACNNNAGATCGGAAG |
| 32453_2 | TATGAGTGTGGAGTCGTTGCGCTCATGCCTATAATTCCAGCACNNNAGATCGGAAG |
| 51622_1 | TATGAGTGTGGAGTCGTTGCACATGCACATACAGTCACACTCTNNNAGATCGGAAG |
| 51622_2 | TATGAGTGTGGAGTCGTTGCACATGCACATACAGTCACACTCTNNNAGATCGGAAG |
| 39354_1 | TATGAGTGTGGAGTCGTTGCGTAAACTGAAATCACGCCACTGTNNNAGATCGGAAG |
| 39354_2 | TATGAGTGTGGAGTCGTTGCGTAAACTGAAATCACGCCACTGTNNNAGATCGGAAG |
| 50730_1 | TATGAGTGTGGAGTCGTTGCACACTGCTCGGACCAAGAAGNNNAGATCGGAAGAG |
| 50730_2 | TATGAGTGTGGAGTCGTTGCACACTGCTCGGACCAAGAAGNNNAGATCGGAAGAG |
| 41002_1 | TATGAGTGTGGAGTCGTTGCAAAAAATTGCCGGACATGGTGATNNNAGATCGGAA |
| 41002_2 | TATGAGTGTGGAGTCGTTGCAAAAAATTGCCGGACATGGTGATNNNAGATCGGAA |
| 12580_1 | TATGAGTGTGGAGTCGTTGCTTTAATTCCTGCAAGAGAAATACNNNAGATCGGAAG |
| 12580_2 | TATGAGTGTGGAGTCGTTGCTTTAATTCCTGCAAGAGAAATACNNNAGATCGGAAG |
| 45151_1 | TATGAGTGTGGAGTCGTTGCGGGTGAGAAGTACTAGCTGCTTTNNNAGATCGGAA |
| 45151_2 | TATGAGTGTGGAGTCGTTGCGGGTGAGAAGTACTAGCTGCTTTNNNAGATCGGAA |
| 43394_1 | TATGAGTGTGGAGTCGTTGCTCCAGGTTCAAGTGATTCTCCTGNNNAGATCGGAAG |
| 43394_2 | TATGAGTGTGGAGTCGTTGCTCCAGGTTCAAGTGATTCTCCTGNNNAGATCGGAAG |
| 34309_1 | TATGAGTGTGGAGTCGTTGCAAGTCTCCATAAATGTGGGGACCNNNAGATCGGAA |
| 34309_2 | TATGAGTGTGGAGTCGTTGCAAGTCTCCATAAATGTGGGGACCNNNAGATCGGAA |
| 34149_1 | TATGAGTGTGGAGTCGTTGCAAAGCTGCACACCTACAACAATCNNNAGATCGGAAG |
| 34149_2 | TATGAGTGTGGAGTCGTTGCAAAGCTGCACACCTACAACAATCNNNAGATCGGAAG |
| 52409_1 | TATGAGTGTGGAGTCGTTGCTTCAATTGCAGAGAAGTGAAGGCNNNAGATCGGAA |
| 52409_2 | TATGAGTGTGGAGTCGTTGCTTCAATTGCAGAGAAGTGAAGGCNNNAGATCGGAA |
| 44436_1 | TATGAGTGTGGAGTCGTTGCTAGAGTGCAGTGGTACAATCTCGNNNAGATCGGAA |
| 44436_2 | TATGAGTGTGGAGTCGTTGCTAGAGTGCAGTGGTACAATCTCGNNNAGATCGGAA |
| 5760_1 | TATGAGTGTGGAGTCGTTGCGTTGGAAGGAGGGCTGAGGGGANNNAGATCGGA |

| | |
|---|---|
| 5760_2 | TATGAGTGTGGAGTCGTTGCGTTGGAAGGAGGGCTGAGGGGANNNAGATCGGAA |
| 29930_1 | TATGAGTGTGGAGTCGTTGCAACTCTGGTAGAGATGTGATCCANNNAGATCGGAA |
| 29930_2 | TATGAGTGTGGAGTCGTTGCAACTCTGGTAGAGATGTGATCCANNNAGATCGGAA |
| 45439_1 | TATGAGTGTGGAGTCGTTGCAGATTCTATTTGCCTGCTCAGCANNNAGATCGGAAG |
| 45439_2 | TATGAGTGTGGAGTCGTTGCAGATTCTATTTGCCTGCTCAGCANNNAGATCGGAAG |
| 32221_1 | TATGAGTGTGGAGTCGTTGCCCTCCTGGGTTCAAGTGATTCTCNNNAGATCGGAAG |
| 32221_2 | TATGAGTGTGGAGTCGTTGCCCTCCTGGGTTCAAGTGATTCTCNNNAGATCGGAAG |
| 37981_1 | TATGAGTGTGGAGTCGTTGCGTAATCCCAATCCCAGCACTTTGNNNAGATCGGAAG |
| 37981_2 | TATGAGTGTGGAGTCGTTGCGTAATCCCAATCCCAGCACTTTGNNNAGATCGGAAG |
| 30382_1 | TATGAGTGTGGAGTCGTTGCCCGGTATGTAATTTAACATTCAANNNAGATCGGAAG |
| 30382_2 | TATGAGTGTGGAGTCGTTGCCCGGTATGTAATTTAACATTCAANNNAGATCGGAAG |
| 47404_1 | TATGAGTGTGGAGTCGTTGCAAACAGGAAATTAGTTGCGAGCCNNNAGATCGGAA |
| 47404_2 | TATGAGTGTGGAGTCGTTGCAAACAGGAAATTAGTTGCGAGCCNNNAGATCGGAA |
| 32669_1 | TATGAGTGTGGAGTCGTTGCCCTCCTGAGTAGCTGGGACTATANNNAGATCGGAAG |
| 32669_2 | TATGAGTGTGGAGTCGTTGCCCTCCTGAGTAGCTGGGACTATANNNAGATCGGAAG |
| 30195_1 | TATGAGTGTGGAGTCGTTGCTAGGATACCCTACCACTACACCCNNNAGATCGGAAG |
| 30195_2 | TATGAGTGTGGAGTCGTTGCTAGGATACCCTACCACTACACCCNNNAGATCGGAAG |
| 13344_1 | TATGAGTGTGGAGTCGTTGCACCACACAAGACATTTTTCTANNNAGATCGGAAGAG |
| 13344_2 | TATGAGTGTGGAGTCGTTGCACCACACAAGACATTTTTCTANNNAGATCGGAAGAG |
| 52364_1 | TATGAGTGTGGAGTCGTTGCACGCTGACTTTGCTCTCACTGNNNAGATCGGAAGAG |
| 52364_2 | TATGAGTGTGGAGTCGTTGCACGCTGACTTTGCTCTCACTGNNNAGATCGGAAGAG |
| 35430_1 | TATGAGTGTGGAGTCGTTGCTAGAGTGCAGTGAGTGATTTCGGNNNAGATCGGAA |
| 35430_2 | TATGAGTGTGGAGTCGTTGCTAGAGTGCAGTGAGTGATTTCGGNNNAGATCGGAA |
| 47076_1 | TATGAGTGTGGAGTCGTTGCAGTAGCTAGGAGTACAGGCTCATNNNAGATCGGAA |
| 47076_2 | TATGAGTGTGGAGTCGTTGCAGTAGCTAGGAGTACAGGCTCATNNNAGATCGGAA |
| 33052_1 | TATGAGTGTGGAGTCGTTGCGGCCATTCTCCTATCTTAGCCTCNNNAGATCGGAAGA |
| 33052_2 | TATGAGTGTGGAGTCGTTGCGGCCATTCTCCTATCTTAGCCTCNNNAGATCGGAAGA |
| 34103_1 | TATGAGTGTGGAGTCGTTGCAACCTACTTTTGAGTGGAAGCCTNNNAGATCGGAAG |
| 34103_2 | TATGAGTGTGGAGTCGTTGCAACCTACTTTTGAGTGGAAGCCTNNNAGATCGGAAG |
| 43459_1 | TATGAGTGTGGAGTCGTTGCCGTGGTAACCAGGATGATCTTGANNNAGATCGGAA |
| 43459_2 | TATGAGTGTGGAGTCGTTGCCGTGGTAACCAGGATGATCTTGANNNAGATCGGAA |
| 45294_1 | TATGAGTGTGGAGTCGTTGCAGTTTCAGGTGTATGGCTCACATNNNAGATCGGAAG |
| 45294_2 | TATGAGTGTGGAGTCGTTGCAGTTTCAGGTGTATGGCTCACATNNNAGATCGGAAG |
| 31329_1 | TATGAGTGTGGAGTCGTTGCACACAGGGCTTTGCAGGATATATNNNAGATCGGAA |
| 31329_2 | TATGAGTGTGGAGTCGTTGCACACAGGGCTTTGCAGGATATATNNNAGATCGGAA |
| 42605_1 | TATGAGTGTGGAGTCGTTGCGAACTCCTGGCTTCAAGCAATCNNNAGATCGGAAGA |
| 42605_2 | TATGAGTGTGGAGTCGTTGCGAACTCCTGGCTTCAAGCAATCNNNAGATCGGAAGA |
| 40841_1 | TATGAGTGTGGAGTCGTTGCGTAGGTAGGATTACAGGCTCGTGNNNAGATCGGAA |
| 40841_2 | TATGAGTGTGGAGTCGTTGCGTAGGTAGGATTACAGGCTCGTGNNNAGATCGGAA |
| 36557_1 | TATGAGTGTGGAGTCGTTGCAATGCTGGGAATGCCAAGATCTANNNAGATCGGAA |
| 36557_2 | TATGAGTGTGGAGTCGTTGCAATGCTGGGAATGCCAAGATCTANNNAGATCGGAA |
| 50793_1 | TATGAGTGTGGAGTCGTTGCTACAGACATCAGCACACTCCATCNNNAGATCGGAAG |
| 50793_2 | TATGAGTGTGGAGTCGTTGCTACAGACATCAGCACACTCCATCNNNAGATCGGAAG |

| | |
|---|---|
| 42333_1 | TATGAGTGTGGAGTCGTTGCCACACACTCTTCATGAAGCTGTCNNNAGATCGGAAG |
| 42333_2 | TATGAGTGTGGAGTCGTTGCCACACACTCTTCATGAAGCTGTCNNNAGATCGGAAG |
| 38399_1 | TATGAGTGTGGAGTCGTTGCAAAGGTGGAACAACTCAAAGCACNNNAGATCGGAA |
| 38399_2 | TATGAGTGTGGAGTCGTTGCAAAGGTGGAACAACTCAAAGCACNNNAGATCGGAA |
| 47902_1 | TATGAGTGTGGAGTCGTTGCTTCTTCTTTCCAAATCCCAGCCTNNNAGATCGGAAGA |
| 47902_2 | TATGAGTGTGGAGTCGTTGCTTCTTCTTTCCAAATCCCAGCCTNNNAGATCGGAAGA |
| 35322_1 | TATGAGTGTGGAGTCGTTGCCTCCTGAGTAGCTGGGACTACAGNNNAGATCGGAA |
| 35322_2 | TATGAGTGTGGAGTCGTTGCCTCCTGAGTAGCTGGGACTACAGNNNAGATCGGAA |
| 36677_1 | TATGAGTGTGGAGTCGTTGCAAACTTTTGGAGTTGAAGCCCACNNNAGATCGGAAG |
| 36677_2 | TATGAGTGTGGAGTCGTTGCAAACTTTTGGAGTTGAAGCCCACNNNAGATCGGAAG |
| 32433_1 | TATGAGTGTGGAGTCGTTGCATTCTCCTGCTTTAGCCTCCCTANNNAGATCGGAAGA |
| 32433_2 | TATGAGTGTGGAGTCGTTGCATTCTCCTGCTTTAGCCTCCCTANNNAGATCGGAAGA |
| 45862_1 | TATGAGTGTGGAGTCGTTGCCCTCCTAGGTTCACATGCTTCTCNNNAGATCGGAAGA |
| 45862_2 | TATGAGTGTGGAGTCGTTGCCCTCCTAGGTTCACATGCTTCTCNNNAGATCGGAAGA |
| 33669_1 | TATGAGTGTGGAGTCGTTGCACGATCTCAGCTCACTGCAAGNNNAGATCGGAAGAG |
| 33669_2 | TATGAGTGTGGAGTCGTTGCACGATCTCAGCTCACTGCAAGNNNAGATCGGAAGAG |
| 44692_1 | TATGAGTGTGGAGTCGTTGCGAAGGGAAGGGGAAAAGGACTAGNNNAGATCGG |
| 44692_2 | TATGAGTGTGGAGTCGTTGCGAAGGGAAGGGGAAAAGGACTAGNNNAGATCGG |
| 35212_1 | TATGAGTGTGGAGTCGTTGCCCTCCCAAGTAGCTGGAATTACANNNAGATCGGAAG |
| 35212_2 | TATGAGTGTGGAGTCGTTGCCCTCCCAAGTAGCTGGAATTACANNNAGATCGGAAG |
| 39704_1 | TATGAGTGTGGAGTCGTTGCAGGCACATAACACATGAAAACCANNNAGATCGGAA |
| 39704_2 | TATGAGTGTGGAGTCGTTGCAGGCACATAACACATGAAAACCANNNAGATCGGAA |
| 49794_1 | TATGAGTGTGGAGTCGTTGCTTATTTGAAGTCCACTGGGGTCCNNNAGATCGGAAG |
| 49794_2 | TATGAGTGTGGAGTCGTTGCTTATTTGAAGTCCACTGGGGTCCNNNAGATCGGAAG |
| 41395_1 | TATGAGTGTGGAGTCGTTGCGCACTTCCATTTGCACATTTGTTNNNAGATCGGAAG |
| 41395_2 | TATGAGTGTGGAGTCGTTGCGCACTTCCATTTGCACATTTGTTNNNAGATCGGAAG |
| 31003_1 | TATGAGTGTGGAGTCGTTGCACTATACCTGGGGCTGAAGTCATNNNAGATCGGAA |
| 31003_2 | TATGAGTGTGGAGTCGTTGCACTATACCTGGGGCTGAAGTCATNNNAGATCGGAA |
| 49643_1 | TATGAGTGTGGAGTCGTTGCGAGACAAAGTCTTGCTCTGTTGCNNNAGATCGGAAG |
| 49643_2 | TATGAGTGTGGAGTCGTTGCGAGACAAAGTCTTGCTCTGTTGCNNNAGATCGGAAG |
| 45394_1 | TATGAGTGTGGAGTCGTTGCCCAGAGTTCTTTGTGTCATGACCNNNAGATCGGAAG |
| 45394_2 | TATGAGTGTGGAGTCGTTGCCCAGAGTTCTTTGTGTCATGACCNNNAGATCGGAAG |
| 41211_1 | TATGAGTGTGGAGTCGTTGCACACACACACATAGCTCATAGGGNNNAGATCGGAAG |
| 41211_2 | TATGAGTGTGGAGTCGTTGCACACACACACATAGCTCATAGGGNNNAGATCGGAAG |
| 47700_1 | TATGAGTGTGGAGTCGTTGCGAGTAGCTGGGATTACAGGCATGNNNAGATCGGAA |
| 47700_2 | TATGAGTGTGGAGTCGTTGCGAGTAGCTGGGATTACAGGCATGNNNAGATCGGAA |
| 6435_1 | TATGAGTGTGGAGTCGTTGCTGCTCCATGAATAACAGTGAGGTNNNAGATCGGAA |
| 6435_2 | TATGAGTGTGGAGTCGTTGCTGCTCCATGAATAACAGTGAGGTNNNAGATCGGAA |
| 50073_1 | TATGAGTGTGGAGTCGTTGCTTCAAGTGATTCTCCTGCCTCAGNNNAGATCGGAAG |
| 50073_2 | TATGAGTGTGGAGTCGTTGCTTCAAGTGATTCTCCTGCCTCAGNNNAGATCGGAAG |
| 46262_1 | TATGAGTGTGGAGTCGTTGCAGGCTTAGTTGTATGTGGATGGGNNNAGATCGGAA |
| 46262_2 | TATGAGTGTGGAGTCGTTGCAGGCTTAGTTGTATGTGGATGGGNNNAGATCGGAA |
| 34086_1 | TATGAGTGTGGAGTCGTTGCGTTTCACCATGTTAGCCAGGATGNNNAGATCGGAAG |

| | |
|---|---|
| 34086_2 | TATGAGTGTGGAGTCGTTGCGTTTCACCATGTTAGCCAGGATGNNNAGATCGGAAG |
| 31145_1 | TATGAGTGTGGAGTCGTTGCAGCTCCCTGAGGATTTGTTTCATNNNAGATCGGAAG |
| 31145_2 | TATGAGTGTGGAGTCGTTGCAGCTCCCTGAGGATTTGTTTCATNNNAGATCGGAAG |
| 29047_1 | TATGAGTGTGGAGTCGTTGCAACTGTTGGCAGAGGTGGTATAGNNNAGATCGGAA |
| 29047_2 | TATGAGTGTGGAGTCGTTGCAACTGTTGGCAGAGGTGGTATAGNNNAGATCGGAA |
| 53249_1 | TATGAGTGTGGAGTCGTTGCGACGCCTAAGACTCTCCCTCTTNNNAGATCGGAAGA |
| 53249_2 | TATGAGTGTGGAGTCGTTGCGACGCCTAAGACTCTCCCTCTTNNNAGATCGGAAGA |
| 51954_1 | TATGAGTGTGGAGTCGTTGCTGATCTCAGGCACATTCTGACTCNNNAGATCGGAAG |
| 51954_2 | TATGAGTGTGGAGTCGTTGCTGATCTCAGGCACATTCTGACTCNNNAGATCGGAAG |
| 47213_1 | TATGAGTGTGGAGTCGTTGCGCGCCCAGCTAGTATCAACTAATNNNAGATCGGAAG |
| 47213_2 | TATGAGTGTGGAGTCGTTGCGCGCCCAGCTAGTATCAACTAATNNNAGATCGGAAG |
| 52786_1 | TATGAGTGTGGAGTCGTTGCAGAAAGACAGATTGCCCTTTCCTNNNAGATCGGAAG |
| 52786_2 | TATGAGTGTGGAGTCGTTGCAGAAAGACAGATTGCCCTTTCCTNNNAGATCGGAAG |
| 30397_1 | TATGAGTGTGGAGTCGTTGCATCCAAATCAAAGTGTTGGCAGGNNNAGATCGGAA |
| 30397_2 | TATGAGTGTGGAGTCGTTGCATCCAAATCAAAGTGTTGGCAGGNNNAGATCGGAA |
| 34926_1 | TATGAGTGTGGAGTCGTTGCCTCCTGAGTAGCTGGGATTACAGNNNAGATCGGAA |
| 34926_2 | TATGAGTGTGGAGTCGTTGCCTCCTGAGTAGCTGGGATTACAGNNNAGATCGGAA |
| 13694_1 | TATGAGTGTGGAGTCGTTGCAGTAAATAGACAGCCTTTCTGNNNAGATCGGAAGA |
| 13694_2 | TATGAGTGTGGAGTCGTTGCAGTAAATAGACAGCCTTTCTGNNNAGATCGGAAGA |
| 34150_1 | TATGAGTGTGGAGTCGTTGCCTTGTTCATGTCCTTTGCCCATTNNNAGATCGGAAGA |
| 34150_2 | TATGAGTGTGGAGTCGTTGCCTTGTTCATGTCCTTTGCCCATTNNNAGATCGGAAGA |
| 36584_1 | TATGAGTGTGGAGTCGTTGCAGGGATAAACCTTTCTCAGAGGAANNNAGATCGGA |
| 36584_2 | TATGAGTGTGGAGTCGTTGCAGGGATAAACCTTTCTCAGAGGAANNNAGATCGGA |
| 40753_1 | TATGAGTGTGGAGTCGTTGCCTTTGAGAAGGAGGAAGTGGTCANNNAGATCGGAA |
| 40753_2 | TATGAGTGTGGAGTCGTTGCCTTTGAGAAGGAGGAAGTGGTCANNNAGATCGGAA |
| 45507_1 | TATGAGTGTGGAGTCGTTGCTTCTTCTTGCAGTCAGTACCTCCNNNAGATCGGAAGA |
| 45507_2 | TATGAGTGTGGAGTCGTTGCTTCTTCTTGCAGTCAGTACCTCCNNNAGATCGGAAGA |
| 47795_1 | TATGAGTGTGGAGTCGTTGCGCCTGGCCCTCCTTATTTTTCTANNNAGATCGGAAGA |
| 47795_2 | TATGAGTGTGGAGTCGTTGCGCCTGGCCCTCCTTATTTTTCTANNNAGATCGGAAGA |
| 49475_1 | TATGAGTGTGGAGTCGTTGCGACAGTTTTCACGAGCCTTCTGANNNAGATCGGAAG |
| 49475_2 | TATGAGTGTGGAGTCGTTGCGACAGTTTTCACGAGCCTTCTGANNNAGATCGGAAG |
| 48635_1 | TATGAGTGTGGAGTCGTTGCCAATGACACAGTCTCGGATCACTNNNAGATCGGAAG |
| 48635_2 | TATGAGTGTGGAGTCGTTGCCAATGACACAGTCTCGGATCACTNNNAGATCGGAAG |
| 36788_1 | TATGAGTGTGGAGTCGTTGCTGCCTCATGACAGAATGTGTACANNNAGATCGGAAG |
| 36788_2 | TATGAGTGTGGAGTCGTTGCTGCCTCATGACAGAATGTGTACANNNAGATCGGAAG |
| 34359_1 | TATGAGTGTGGAGTCGTTGCGTGTACCCAACAGCTCATTGAGANNNAGATCGGAAG |
| 34359_2 | TATGAGTGTGGAGTCGTTGCGTGTACCCAACAGCTCATTGAGANNNAGATCGGAAG |
| 12794_1 | TATGAGTGTGGAGTCGTTGCTTAGGAAGTTATCAGAAACCAACNNNAGATCGGAA |
| 12794_2 | TATGAGTGTGGAGTCGTTGCTTAGGAAGTTATCAGAAACCAACNNNAGATCGGAA |
| 37561_1 | TATGAGTGTGGAGTCGTTGCCTGGACTCTGCTGCAGGTAATAANNNAGATCGGAA |
| 37561_2 | TATGAGTGTGGAGTCGTTGCCTGGACTCTGCTGCAGGTAATAANNNAGATCGGAA |
| 29194_1 | TATGAGTGTGGAGTCGTTGCCCATCTATGCGTTGGTAGGGTTANNNAGATCGGAA |
| 29194_2 | TATGAGTGTGGAGTCGTTGCCCATCTATGCGTTGGTAGGGTTANNNAGATCGGAA |

| | |
|---|---|
| 40934_1 | TATGAGTGTGGAGTCGTTGCTTTCACAGGTGTATATGTGCCGANNNAGATCGGAAG |
| 40934_2 | TATGAGTGTGGAGTCGTTGCTTTCACAGGTGTATATGTGCCGANNNAGATCGGAAG |
| 41569_1 | TATGAGTGTGGAGTCGTTGCCTCCTGAGTAGCCGAGATTACAGNNNAGATCGGAAG |
| 41569_2 | TATGAGTGTGGAGTCGTTGCCTCCTGAGTAGCCGAGATTACAGNNNAGATCGGAAG |
| 45496_1 | TATGAGTGTGGAGTCGTTGCGGTAAGCGCTAAGTTCATGCTTCNNNAGATCGGAAG |
| 45496_2 | TATGAGTGTGGAGTCGTTGCGGTAAGCGCTAAGTTCATGCTTCNNNAGATCGGAAG |
| 49751_1 | TATGAGTGTGGAGTCGTTGCGAAAGAGAAGGGCAGGACTCTAGNNNAGATCGGA |
| 49751_2 | TATGAGTGTGGAGTCGTTGCGAAAGAGAAGGGCAGGACTCTAGNNNAGATCGGA |
| 29603_1 | TATGAGTGTGGAGTCGTTGCTAAACTGTGTGTGTGTGTGTGTGNNNAGATCGGAA |
| 29603_2 | TATGAGTGTGGAGTCGTTGCTAAACTGTGTGTGTGTGTGTGTGNNNAGATCGGAA |
| 42660_1 | TATGAGTGTGGAGTCGTTGCCTGTAGTCCCAGCTACTCAGGAGNNNAGATCGGAAG |
| 42660_2 | TATGAGTGTGGAGTCGTTGCCTGTAGTCCCAGCTACTCAGGAGNNNAGATCGGAAG |
| 30957_1 | TATGAGTGTGGAGTCGTTGCAATGCTCAGTCCTTATCCATCCCNNNAGATCGGAAGA |
| 30957_2 | TATGAGTGTGGAGTCGTTGCAATGCTCAGTCCTTATCCATCCCNNNAGATCGGAAGA |
| 33120_1 | TATGAGTGTGGAGTCGTTGCCCACACAAAAATTAGCTGGGCATNNNAGATCGGAAG |
| 33120_2 | TATGAGTGTGGAGTCGTTGCCCACACAAAAATTAGCTGGGCATNNNAGATCGGAAG |
| 36944_1 | TATGAGTGTGGAGTCGTTGCCTCATGCCTGTAATCCCAGAACTNNNAGATCGGAAG |
| 36944_2 | TATGAGTGTGGAGTCGTTGCCTCATGCCTGTAATCCCAGAACTNNNAGATCGGAAG |
| 42560_1 | TATGAGTGTGGAGTCGTTGCTCAAACAGTTCTCTACCTCAGCCNNNAGATCGGAAGA |
| 42560_2 | TATGAGTGTGGAGTCGTTGCTCAAACAGTTCTCTACCTCAGCCNNNAGATCGGAAGA |
| 31773_1 | TATGAGTGTGGAGTCGTTGCCCAGGGGCTTGGAAATATGAGATNNNAGATCGGAA |
| 31773_2 | TATGAGTGTGGAGTCGTTGCCCAGGGGCTTGGAAATATGAGATNNNAGATCGGAA |
| 10733_1 | TATGAGTGTGGAGTCGTTGCGTTTCCTGGCAGAAGGCATTTACNNNAGATCGGAAG |
| 10733_2 | TATGAGTGTGGAGTCGTTGCGTTTCCTGGCAGAAGGCATTTACNNNAGATCGGAAG |
| 10546_1 | TATGAGTGTGGAGTCGTTGCCCTCTGGGCTCATGTTGAGTTTTNNNAGATCGGAAG |
| 10546_2 | TATGAGTGTGGAGTCGTTGCCCTCTGGGCTCATGTTGAGTTTTNNNAGATCGGAAG |
| 34870_1 | TATGAGTGTGGAGTCGTTGCCATTCTCTCACCTCAGCTTCTCANNNAGATCGGAAGA |
| 34870_2 | TATGAGTGTGGAGTCGTTGCCATTCTCTCACCTCAGCTTCTCANNNAGATCGGAAGA |
| 40698_1 | TATGAGTGTGGAGTCGTTGCATGACAAAAGACCCCGAATAGCTNNNAGATCGGAAG |
| 40698_2 | TATGAGTGTGGAGTCGTTGCATGACAAAAGACCCCGAATAGCTNNNAGATCGGAAG |
| 28797_1 | TATGAGTGTGGAGTCGTTGCGTTTCACCATGTTGCTCAGTCTGNNNAGATCGGAAG |
| 28797_2 | TATGAGTGTGGAGTCGTTGCGTTTCACCATGTTGCTCAGTCTGNNNAGATCGGAAG |
| 29248_1 | TATGAGTGTGGAGTCGTTGCAAGATGATATGCTGTCACTGGCTNNNAGATCGGAAG |
| 29248_2 | TATGAGTGTGGAGTCGTTGCAAGATGATATGCTGTCACTGGCTNNNAGATCGGAAG |
| 38306_1 | TATGAGTGTGGAGTCGTTGCACCAAATTGATAGACTGCTAGCTNNNAGATCGGAAG |
| 38306_2 | TATGAGTGTGGAGTCGTTGCACCAAATTGATAGACTGCTAGCTNNNAGATCGGAAG |
| 29029_1 | TATGAGTGTGGAGTCGTTGCAGACAGCTGGGAAGACCTATAGTNNNAGATCGGAA |
| 29029_2 | TATGAGTGTGGAGTCGTTGCAGACAGCTGGGAAGACCTATAGTNNNAGATCGGAA |
| 33381_1 | TATGAGTGTGGAGTCGTTGCTCCAGACCTCAACAAATTGTCCTNNNAGATCGGAAG |
| 33381_2 | TATGAGTGTGGAGTCGTTGCTCCAGACCTCAACAAATTGTCCTNNNAGATCGGAAG |
| 47670_1 | TATGAGTGTGGAGTCGTTGCTTCAAGCAATTCTCTTGCCTCAGNNNAGATCGGAAG |
| 47670_2 | TATGAGTGTGGAGTCGTTGCTTCAAGCAATTCTCTTGCCTCAGNNNAGATCGGAAG |
| 44542_1 | TATGAGTGTGGAGTCGTTGCCACACACACACTACACACATACACNNNAGATCGGAAG |

| | |
|---|---|
| 44542_2 | TATGAGTGTGGAGTCGTTGCCACACACACACTACACACATACACNNNAGATCGGAAG |
| 31150_1 | TATGAGTGTGGAGTCGTTGCCACATACACAGACACACACACACNNNAGATCGGAAG |
| 31150_2 | TATGAGTGTGGAGTCGTTGCCACATACACAGACACACACACACNNNAGATCGGAAG |
| 45533_1 | TATGAGTGTGGAGTCGTTGCTGTCAGTGGGAGAGGAAGAAATGNNNAGATCGGA |
| 45533_2 | TATGAGTGTGGAGTCGTTGCTGTCAGTGGGAGAGGAAGAAATGNNNAGATCGGA |
| 31708_1 | TATGAGTGTGGAGTCGTTGCATAGGCTGCTTACATCTGGGATGNNNAGATCGGAA |
| 31708_2 | TATGAGTGTGGAGTCGTTGCATAGGCTGCTTACATCTGGGATGNNNAGATCGGAA |
| 46030_1 | TATGAGTGTGGAGTCGTTGCCTCAAGCAATCCTCCTACCTCAGNNNAGATCGGAAG |
| 46030_2 | TATGAGTGTGGAGTCGTTGCCTCAAGCAATCCTCCTACCTCAGNNNAGATCGGAAG |
| 36261_1 | TATGAGTGTGGAGTCGTTGCCTGCTGCTGCTTTACCAACTAAGNNNAGATCGGAAG |
| 36261_2 | TATGAGTGTGGAGTCGTTGCCTGCTGCTGCTTTACCAACTAAGNNNAGATCGGAAG |
| 42190_1 | TATGAGTGTGGAGTCGTTGCCTCCTGAGTAGCTGGGACTACAGNNNAGATCGGAA |
| 42190_2 | TATGAGTGTGGAGTCGTTGCCTCCTGAGTAGCTGGGACTACAGNNNAGATCGGAA |
| 41146_1 | TATGAGTGTGGAGTCGTTGCTGCCTGTAATCCCAGCTATTTGGNNNAGATCGGAAG |
| 41146_2 | TATGAGTGTGGAGTCGTTGCTGCCTGTAATCCCAGCTATTTGGNNNAGATCGGAAG |
| 39334_1 | TATGAGTGTGGAGTCGTTGCGATGTTTTGTACCCTGCTACCCANNNAGATCGGAAG |
| 39334_2 | TATGAGTGTGGAGTCGTTGCGATGTTTTGTACCCTGCTACCCANNNAGATCGGAAG |
| 49274_1 | TATGAGTGTGGAGTCGTTGCTACCTCAGCCTCCCAAAAATGTTNNNAGATCGGAAG |
| 49274_2 | TATGAGTGTGGAGTCGTTGCTACCTCAGCCTCCCAAAAATGTTNNNAGATCGGAAG |
| 52693_1 | TATGAGTGTGGAGTCGTTGCTGCTGTATTTGCTTCTTCTGTGCNNNAGATCGGAAG |
| 52693_2 | TATGAGTGTGGAGTCGTTGCTGCTGTATTTGCTTCTTCTGTGCNNNAGATCGGAAG |
| 47633_1 | TATGAGTGTGGAGTCGTTGCCTAAGGCAGGTGACGACACGNNNAGATCGGAAGAG |
| 47633_2 | TATGAGTGTGGAGTCGTTGCCTAAGGCAGGTGACGACACGNNNAGATCGGAAGAG |
| 13548_1 | TATGAGTGTGGAGTCGTTGCGAGGATTAAAAAGGACATGTAAGNNNAGATCGGAA |
| 13548_2 | TATGAGTGTGGAGTCGTTGCGAGGATTAAAAAGGACATGTAAGNNNAGATCGGAA |
| 48774_1 | TATGAGTGTGGAGTCGTTGCTTAGACATATGTATGTGGCGCCANNNAGATCGGAA |
| 48774_2 | TATGAGTGTGGAGTCGTTGCTTAGACATATGTATGTGGCGCCANNNAGATCGGAA |
| 46742_1 | TATGAGTGTGGAGTCGTTGCGCTGGCTCGAGTTTTTCTTCTTTNNNAGATCGGAAG |
| 46742_2 | TATGAGTGTGGAGTCGTTGCGCTGGCTCGAGTTTTTCTTCTTTNNNAGATCGGAAG |
| 38795_1 | TATGAGTGTGGAGTCGTTGCTGAACAGACAACCCACAGAATCTNNNAGATCGGAAG |
| 38795_2 | TATGAGTGTGGAGTCGTTGCTGAACAGACAACCCACAGAATCTNNNAGATCGGAAG |
| 49918_1 | TATGAGTGTGGAGTCGTTGCCTTTGAAAAGTGCCGAGTGAGTCNNNAGATCGGAA |
| 49918_2 | TATGAGTGTGGAGTCGTTGCCTTTGAAAAGTGCCGAGTGAGTCNNNAGATCGGAA |
| 13757_1 | TATGAGTGTGGAGTCGTTGCTTCTTGCTTCTGTAAGAGATGNNNAGATCGGAAGAG |
| 13757_2 | TATGAGTGTGGAGTCGTTGCTTCTTGCTTCTGTAAGAGATGNNNAGATCGGAAGAG |
| 39753_1 | TATGAGTGTGGAGTCGTTGCGACAGACTTCTTCAAGGGGCATANNNAGATCGGAA |
| 39753_2 | TATGAGTGTGGAGTCGTTGCGACAGACTTCTTCAAGGGGCATANNNAGATCGGAA |
| 46517_1 | TATGAGTGTGGAGTCGTTGCCGAGTAGCTGGGATTACAAGTGTNNNAGATCGGAA |
| 46517_2 | TATGAGTGTGGAGTCGTTGCCGAGTAGCTGGGATTACAAGTGTNNNAGATCGGAA |
| 33609_1 | TATGAGTGTGGAGTCGTTGCGAGATGGAGTCTTACTCTGTCGCNNNAGATCGGAAG |
| 33609_2 | TATGAGTGTGGAGTCGTTGCGAGATGGAGTCTTACTCTGTCGCNNNAGATCGGAAG |
| 11950_1 | TATGAGTGTGGAGTCGTTGCATCTCTCCCGCCTCACTCTTGTCNNNAGATCGGAAGA |
| 11950_2 | TATGAGTGTGGAGTCGTTGCATCTCTCCCGCCTCACTCTTGTCNNNAGATCGGAAGA |

| | |
|---|---|
| 37547_1 | TATGAGTGTGGAGTCGTTGCAAGAAAACCCTTCATCCCAACACNNNAGATCGGAAG |
| 37547_2 | TATGAGTGTGGAGTCGTTGCAAGAAAACCCTTCATCCCAACACNNNAGATCGGAAG |
| 39274_1 | TATGAGTGTGGAGTCGTTGCTCTGGCCCTTTATTAGCAAATGCNNNAGATCGGAAG |
| 39274_2 | TATGAGTGTGGAGTCGTTGCTCTGGCCCTTTATTAGCAAATGCNNNAGATCGGAAG |
| 30982_1 | TATGAGTGTGGAGTCGTTGCCTTGGGGGTGGAGGATCTGATTCNNNAGATCGGAA |
| 30982_2 | TATGAGTGTGGAGTCGTTGCCTTGGGGGTGGAGGATCTGATTCNNNAGATCGGAA |
| 31085_1 | TATGAGTGTGGAGTCGTTGCTTTTTTGTGCATGTTGTCCTGGANNNAGATCGGAAG |
| 31085_2 | TATGAGTGTGGAGTCGTTGCTTTTTTGTGCATGTTGTCCTGGANNNAGATCGGAAG |
| 32736_1 | TATGAGTGTGGAGTCGTTGCAAGGACTCCATCATGCAGCTAAANNNAGATCGGAAG |
| 32736_2 | TATGAGTGTGGAGTCGTTGCAAGGACTCCATCATGCAGCTAAANNNAGATCGGAAG |
| 52989_1 | TATGAGTGTGGAGTCGTTGCTTGTCCCTGTCTCTATGTCTCCTNNNAGATCGGAAGA |
| 52989_2 | TATGAGTGTGGAGTCGTTGCTTGTCCCTGTCTCTATGTCTCCTNNNAGATCGGAAGA |
| 34669_1 | TATGAGTGTGGAGTCGTTGCCTTGCACCATGTACCTGGAAAAGNNNAGATCGGAAG |
| 34669_2 | TATGAGTGTGGAGTCGTTGCCTTGCACCATGTACCTGGAAAAGNNNAGATCGGAAG |
| 42952_1 | TATGAGTGTGGAGTCGTTGCCAGGTTCAAGCAATTCTCCTGCNNNAGATCGGAAGA |
| 42952_2 | TATGAGTGTGGAGTCGTTGCCAGGTTCAAGCAATTCTCCTGCNNNAGATCGGAAGA |
| 44756_1 | TATGAGTGTGGAGTCGTTGCGAGATCCCATCTGTTCCTTGGAANNNAGATCGGAAG |
| 44756_2 | TATGAGTGTGGAGTCGTTGCGAGATCCCATCTGTTCCTTGGAANNNAGATCGGAAG |
| 2090_1 | TATGAGTGTGGAGTCGTTGCATAGTTGTAATTCACCTGAGGCCANNNAGATCGGAA |
| 2090_2 | TATGAGTGTGGAGTCGTTGCATAGTTGTAATTCACCTGAGGCCANNNAGATCGGAA |
| 42157_1 | TATGAGTGTGGAGTCGTTGCCATGATCTCGGCTCACTGCAATNNNAGATCGGAAGA |
| 42157_2 | TATGAGTGTGGAGTCGTTGCCATGATCTCGGCTCACTGCAATNNNAGATCGGAAGA |
| 52050_1 | TATGAGTGTGGAGTCGTTGCTTGAGATGAGATTTGGGTGGAGGNNNAGATCGGA |
| 52050_2 | TATGAGTGTGGAGTCGTTGCTTGAGATGAGATTTGGGTGGAGGNNNAGATCGGA |
| 47620_1 | TATGAGTGTGGAGTCGTTGCCAGAAGGGAGATGATAGAGCCACNNNAGATCGGAA |
| 47620_2 | TATGAGTGTGGAGTCGTTGCCAGAAGGGAGATGATAGAGCCACNNNAGATCGGAA |
| 50930_1 | TATGAGTGTGGAGTCGTTGCTCCAGCACTAAAATGGATGGAGGNNNAGATCGGAA |
| 50930_2 | TATGAGTGTGGAGTCGTTGCTCCAGCACTAAAATGGATGGAGGNNNAGATCGGAA |
| 39946_1 | TATGAGTGTGGAGTCGTTGCGCAACAGAGTGAAACCCTGATTCNNNAGATCGGAA |
| 39946_2 | TATGAGTGTGGAGTCGTTGCGCAACAGAGTGAAACCCTGATTCNNNAGATCGGAA |
| 34586_1 | TATGAGTGTGGAGTCGTTGCGGATGGAGGTGTTTGGATCTCATNNNAGATCGGAA |
| 34586_2 | TATGAGTGTGGAGTCGTTGCGGATGGAGGTGTTTGGATCTCATNNNAGATCGGAA |
| 45104_1 | TATGAGTGTGGAGTCGTTGCAGGTACAGAAAGGGATTTGGAGCNNNAGATCGGA |
| 45104_2 | TATGAGTGTGGAGTCGTTGCAGGTACAGAAAGGGATTTGGAGCNNNAGATCGGA |
| 42992_1 | TATGAGTGTGGAGTCGTTGCAGCAGTGTTACTGAATGCAAAGANNNAGATCGGAA |
| 42992_2 | TATGAGTGTGGAGTCGTTGCAGCAGTGTTACTGAATGCAAAGANNNAGATCGGAA |
| 39125_1 | TATGAGTGTGGAGTCGTTGCGAGTCAAGACCTGTTAGTGTGCTNNNAGATCGGAA |
| 39125_2 | TATGAGTGTGGAGTCGTTGCGAGTCAAGACCTGTTAGTGTGCTNNNAGATCGGAA |
| 37165_1 | TATGAGTGTGGAGTCGTTGCTCCTGCAGTGGAGGATAGTAATGNNNAGATCGGAA |
| 37165_2 | TATGAGTGTGGAGTCGTTGCTCCTGCAGTGGAGGATAGTAATGNNNAGATCGGAA |
| 45607_1 | TATGAGTGTGGAGTCGTTGCAGACGGAGTCTCACTTTTGTCACNNNAGATCGGAAG |
| 45607_2 | TATGAGTGTGGAGTCGTTGCAGACGGAGTCTCACTTTTGTCACNNNAGATCGGAAG |
| 43078_1 | TATGAGTGTGGAGTCGTTGCCTTGCTGGTTCTGTTTCTTGGTGNNNAGATCGGAAG |

| | |
|---|---|
| 43078_2 | TATGAGTGTGGAGTCGTTGCCTTGCTGGTTCTGTTTCTTGGTGNNNAGATCGGAAG |
| 32814_1 | TATGAGTGTGGAGTCGTTGCTGTCTAGGCTGGTCTTGAACTTCNNNAGATCGGAAG |
| 32814_2 | TATGAGTGTGGAGTCGTTGCTGTCTAGGCTGGTCTTGAACTTCNNNAGATCGGAAG |
| 43236_1 | TATGAGTGTGGAGTCGTTGCATGGATTTTTCCCACTCACTCCANNNAGATCGGAAGA |
| 43236_2 | TATGAGTGTGGAGTCGTTGCATGGATTTTTCCCACTCACTCCANNNAGATCGGAAGA |
| 10182_1 | TATGAGTGTGGAGTCGTTGCATGGTTAGACAGCAGGGAGTGATNNNAGATCGGAA |
| 10182_2 | TATGAGTGTGGAGTCGTTGCATGGTTAGACAGCAGGGAGTGATNNNAGATCGGAA |
| 8170_1 | TATGAGTGTGGAGTCGTTGCCTGGGGTAAGTTTTCAAGGCTGTCNNNAGATCGGAA |
| 8170_2 | TATGAGTGTGGAGTCGTTGCCTGGGGTAAGTTTTCAAGGCTGTCNNNAGATCGGAA |
| 44249_1 | TATGAGTGTGGAGTCGTTGCTGAAGCGACAAAGAAAGATGAGCNNNAGATCGGAA |
| 44249_2 | TATGAGTGTGGAGTCGTTGCTGAAGCGACAAAGAAAGATGAGCNNNAGATCGGAA |
| 46979_1 | TATGAGTGTGGAGTCGTTGCAATGGGGTCTTACCATATTGCCCNNNAGATCGGAAG |
| 46979_2 | TATGAGTGTGGAGTCGTTGCAATGGGGTCTTACCATATTGCCCNNNAGATCGGAAG |
| 10085_1 | TATGAGTGTGGAGTCGTTGCGGGCTGGTTTAGAGATCCTCTGTNNNAGATCGGAA |
| 10085_2 | TATGAGTGTGGAGTCGTTGCGGGCTGGTTTAGAGATCCTCTGTNNNAGATCGGAA |
| 45247_1 | TATGAGTGTGGAGTCGTTGCGAACAGAGTCTAGAGGACCTTGCNNNAGATCGGAA |
| 45247_2 | TATGAGTGTGGAGTCGTTGCGAACAGAGTCTAGAGGACCTTGCNNNAGATCGGAA |
| 48188_1 | TATGAGTGTGGAGTCGTTGCGACTCCAGGTTGTCCATCTGCNNNAGATCGGAAGAG |
| 48188_2 | TATGAGTGTGGAGTCGTTGCGACTCCAGGTTGTCCATCTGCNNNAGATCGGAAGAG |
| 34227_1 | TATGAGTGTGGAGTCGTTGCTGTTTGGTCCTTAGGATAAGGCANNNAGATCGGAA |
| 34227_2 | TATGAGTGTGGAGTCGTTGCTGTTTGGTCCTTAGGATAAGGCANNNAGATCGGAA |
| 31052_1 | TATGAGTGTGGAGTCGTTGCTGTTTCATGAGGTAGTGGAAGCANNNAGATCGGAA |
| 31052_2 | TATGAGTGTGGAGTCGTTGCTGTTTCATGAGGTAGTGGAAGCANNNAGATCGGAA |
| 38833_1 | TATGAGTGTGGAGTCGTTGCACCTCAGCTACAACTGACAAACANNNAGATCGGAAG |
| 38833_2 | TATGAGTGTGGAGTCGTTGCACCTCAGCTACAACTGACAAACANNNAGATCGGAAG |
| 43657_1 | TATGAGTGTGGAGTCGTTGCCAAGGAGGCATACATCTTGCAAGNNNAGATCGGAA |
| 43657_2 | TATGAGTGTGGAGTCGTTGCCAAGGAGGCATACATCTTGCAAGNNNAGATCGGAA |
| 38271_1 | TATGAGTGTGGAGTCGTTGCAGCCAACAGATCTTTGACAAAGCNNNAGATCGGAAG |
| 38271_2 | TATGAGTGTGGAGTCGTTGCAGCCAACAGATCTTTGACAAAGCNNNAGATCGGAAG |
| 39077_1 | TATGAGTGTGGAGTCGTTGCGTGAGCAGAGATAGCGCAGATNNNAGATCGGAAG |
| 39077_2 | TATGAGTGTGGAGTCGTTGCGTGAGCAGAGATAGCGCAGATNNNAGATCGGAAG |
| 35633_1 | TATGAGTGTGGAGTCGTTGCACACTGCAAGTTCAGCTTTGATCNNNAGATCGGAAG |
| 35633_2 | TATGAGTGTGGAGTCGTTGCACACTGCAAGTTCAGCTTTGATCNNNAGATCGGAAG |
| 30814_1 | TATGAGTGTGGAGTCGTTGCTCCTTTCCATTCCAACCCCATAGNNNAGATCGGAAGA |
| 30814_2 | TATGAGTGTGGAGTCGTTGCTCCTTTCCATTCCAACCCCATAGNNNAGATCGGAAGA |
| 39340_1 | TATGAGTGTGGAGTCGTTGCCCACGCACGGCTAATTTTTGTATNNNAGATCGGAAG |
| 39340_2 | TATGAGTGTGGAGTCGTTGCCCACGCACGGCTAATTTTTGTATNNNAGATCGGAAG |
| 10643_1 | TATGAGTGTGGAGTCGTTGCAAAGTGGCATACTACTAGCGTGGNNNAGATCGGAA |
| 10643_2 | TATGAGTGTGGAGTCGTTGCAAAGTGGCATACTACTAGCGTGGNNNAGATCGGAA |
| 48345_1 | TATGAGTGTGGAGTCGTTGCCCATGCCCGGCTAATTTTTTGTANNNAGATCGGAAG |
| 48345_2 | TATGAGTGTGGAGTCGTTGCCCATGCCCGGCTAATTTTTTGTANNNAGATCGGAAG |
| 29518_1 | TATGAGTGTGGAGTCGTTGCTAAAGGACACAACATACAGGTATNNNAGATCGGAA |
| 29518_2 | TATGAGTGTGGAGTCGTTGCTAAAGGACACAACATACAGGTATNNNAGATCGGAA |

| | |
|---|---|
| 35713_1 | TATGAGTGTGGAGTCGTTGCTGCCCAGCTACAAATATCAGTGGNNNAGATCGGAAG |
| 35713_2 | TATGAGTGTGGAGTCGTTGCTGCCCAGCTACAAATATCAGTGGNNNAGATCGGAAG |
| 43784_1 | TATGAGTGTGGAGTCGTTGCAGCTGAATGAGGTATGTGTTCTCTNNNAGATCGGAA |
| 43784_2 | TATGAGTGTGGAGTCGTTGCAGCTGAATGAGGTATGTGTTCTCTNNNAGATCGGAA |
| 40643_1 | TATGAGTGTGGAGTCGTTGCTTGATGCAGATGACGGTGGATATNNNAGATCGGAA |
| 40643_2 | TATGAGTGTGGAGTCGTTGCTTGATGCAGATGACGGTGGATATNNNAGATCGGAA |
| 31975_1 | TATGAGTGTGGAGTCGTTGCGGTGAAGAGAGCAGGAATAGAGGNNNAGATCGGA |
| 31975_2 | TATGAGTGTGGAGTCGTTGCGGTGAAGAGAGCAGGAATAGAGGNNNAGATCGGA |
| 38308_1 | TATGAGTGTGGAGTCGTTGCAAGGAGAAGAAGGAGAAGGAGGANNNAGATCGG |
| 38308_2 | TATGAGTGTGGAGTCGTTGCAAGGAGAAGAAGGAGAAGGAGGANNNAGATCGG |
| 53050_1 | TATGAGTGTGGAGTCGTTGCGAGGCAGGAGAATGGTGTGAANNNAGATCGGAAG |
| 53050_2 | TATGAGTGTGGAGTCGTTGCGAGGCAGGAGAATGGTGTGAANNNAGATCGGAAG |
| 35275_1 | TATGAGTGTGGAGTCGTTGCAAACATGCAAAAGTGAGGGGAAGNNNAGATCGGA |
| 35275_2 | TATGAGTGTGGAGTCGTTGCAAACATGCAAAAGTGAGGGGAAGNNNAGATCGGA |
| 46067_1 | TATGAGTGTGGAGTCGTTGCCTTCCAGGTTCACGCCATTCTCNNNAGATCGGAAGA |
| 46067_2 | TATGAGTGTGGAGTCGTTGCCTTCCAGGTTCACGCCATTCTCNNNAGATCGGAAGA |
| 53388_1 | TATGAGTGTGGAGTCGTTGCAAATTGAGGGGACTAAGGCTGAGNNNAGATCGGA |
| 53388_2 | TATGAGTGTGGAGTCGTTGCAAATTGAGGGGACTAAGGCTGAGNNNAGATCGGA |
| 49645_1 | TATGAGTGTGGAGTCGTTGCTTTATGGATTCTCAGTGCGTGGTNNNAGATCGGAAG |
| 49645_2 | TATGAGTGTGGAGTCGTTGCTTTATGGATTCTCAGTGCGTGGTNNNAGATCGGAAG |
| 30780_1 | TATGAGTGTGGAGTCGTTGCTGCATGGGTAAATTGTGTGTCACNNNAGATCGGAA |
| 30780_2 | TATGAGTGTGGAGTCGTTGCTGCATGGGTAAATTGTGTGTCACNNNAGATCGGAA |
| 31410_1 | TATGAGTGTGGAGTCGTTGCCCTCTTTGAGCAGGTGACATTTGNNNAGATCGGAAG |
| 31410_2 | TATGAGTGTGGAGTCGTTGCCCTCTTTGAGCAGGTGACATTTGNNNAGATCGGAAG |
| 53378_1 | TATGAGTGTGGAGTCGTTGCTCTCCTCTCGTTTCTCTCTGTCTNNNAGATCGGAAGA |
| 53378_2 | TATGAGTGTGGAGTCGTTGCTCTCCTCTCGTTTCTCTCTGTCTNNNAGATCGGAAGA |
| 49678_1 | TATGAGTGTGGAGTCGTTGCTCCCGTATTCAAGCAATTCTCCTNNNAGATCGGAAGA |
| 49678_2 | TATGAGTGTGGAGTCGTTGCTCCCGTATTCAAGCAATTCTCCTNNNAGATCGGAAGA |
| 50443_1 | TATGAGTGTGGAGTCGTTGCTTCTGCAGGGAACTGATATCACCNNNAGATCGGAAG |
| 50443_2 | TATGAGTGTGGAGTCGTTGCTTCTGCAGGGAACTGATATCACCNNNAGATCGGAAG |
| 40962_1 | TATGAGTGTGGAGTCGTTGCTCAAACTCAGTGTGAACTACTCANNNAGATCGGAAG |
| 40962_2 | TATGAGTGTGGAGTCGTTGCTCAAACTCAGTGTGAACTACTCANNNAGATCGGAAG |
| 28808_1 | TATGAGTGTGGAGTCGTTGCCCCAGTTTCAAGCAATTCTCCTGNNNAGATCGGAAG |
| 28808_2 | TATGAGTGTGGAGTCGTTGCCCCAGTTTCAAGCAATTCTCCTGNNNAGATCGGAAG |
| 2200_1 | TATGAGTGTGGAGTCGTTGCCTTTGCATGCCTACTCCATTGGANNNAGATCGGAAG |
| 2200_2 | TATGAGTGTGGAGTCGTTGCCTTTGCATGCCTACTCCATTGGANNNAGATCGGAAG |
| 48090_1 | TATGAGTGTGGAGTCGTTGCCCACTTGCCAGGAACACATTCNNNAGATCGGAAGAG |
| 48090_2 | TATGAGTGTGGAGTCGTTGCCCACTTGCCAGGAACACATTCNNNAGATCGGAAGAG |
| 28988_1 | TATGAGTGTGGAGTCGTTGCACATGTGGCTGTGTCTTTCTTTGNNNAGATCGGAAG |
| 28988_2 | TATGAGTGTGGAGTCGTTGCACATGTGGCTGTGTCTTTCTTTGNNNAGATCGGAAG |
| 48552_1 | TATGAGTGTGGAGTCGTTGCTCATTCTCACAGGCCTGATTTCANNNAGATCGGAAG |
| 48552_2 | TATGAGTGTGGAGTCGTTGCTCATTCTCACAGGCCTGATTTCANNNAGATCGGAAG |
| 45418_1 | TATGAGTGTGGAGTCGTTGCATGGCATTAGCAGCAATTTGGAGNNNAGATCGGAA |

| | |
|---|---|
| 45418_2 | TATGAGTGTGGAGTCGTTGCATGGCATTAGCAGCAATTTGGAGNNNAGATCGGAA |
| 32948_1 | TATGAGTGTGGAGTCGTTGCCAGAGATGACGGGAAAACATTCCNNNAGATCGGAA |
| 32948_2 | TATGAGTGTGGAGTCGTTGCCAGAGATGACGGGAAAACATTCCNNNAGATCGGAA |
| 42120_1 | TATGAGTGTGGAGTCGTTGCGAGTAGCTGGGATTACAGGCATGNNNAGATCGGAA |
| 42120_2 | TATGAGTGTGGAGTCGTTGCGAGTAGCTGGGATTACAGGCATGNNNAGATCGGAA |
| 2618_1 | TATGAGTGTGGAGTCGTTGCGTTTACATGAGCAGAGCACATCTGNNNAGATCGGA |
| 2618_2 | TATGAGTGTGGAGTCGTTGCGTTTACATGAGCAGAGCACATCTGNNNAGATCGGA |
| 44883_1 | TATGAGTGTGGAGTCGTTGCCCATCGATCCACCACTGAGTATCNNNAGATCGGAAG |
| 44883_2 | TATGAGTGTGGAGTCGTTGCCCATCGATCCACCACTGAGTATCNNNAGATCGGAAG |
| 47440_1 | TATGAGTGTGGAGTCGTTGCGAGACAGAGTCTTGCTTTGTCACNNNAGATCGGAAG |
| 47440_2 | TATGAGTGTGGAGTCGTTGCGAGACAGAGTCTTGCTTTGTCACNNNAGATCGGAAG |
| 40130_1 | TATGAGTGTGGAGTCGTTGCACCTTTGCAAATACACTAGCACTNNNAGATCGGAAG |
| 40130_2 | TATGAGTGTGGAGTCGTTGCACCTTTGCAAATACACTAGCACTNNNAGATCGGAAG |
| 32938_1 | TATGAGTGTGGAGTCGTTGCAATCTGTCCTCATTGCTTTTGCCNNNAGATCGGAAGA |
| 32938_2 | TATGAGTGTGGAGTCGTTGCAATCTGTCCTCATTGCTTTTGCCNNNAGATCGGAAGA |
| 32363_1 | TATGAGTGTGGAGTCGTTGCGCATATATGTAGGCCGGGTTGTANNNAGATCGGAA |
| 32363_2 | TATGAGTGTGGAGTCGTTGCGCATATATGTAGGCCGGGTTGTANNNAGATCGGAA |
| 52237_1 | TATGAGTGTGGAGTCGTTGCGTCACCGTCATCAAAGACCAAAGNNNAGATCGGAAG |
| 52237_2 | TATGAGTGTGGAGTCGTTGCGTCACCGTCATCAAAGACCAAAGNNNAGATCGGAAG |
| 47502_1 | TATGAGTGTGGAGTCGTTGCAAGTTCAAGTGATTCTCCTGCCTNNNAGATCGGAAG |
| 47502_2 | TATGAGTGTGGAGTCGTTGCAAGTTCAAGTGATTCTCCTGCCTNNNAGATCGGAAG |
| 39095_1 | TATGAGTGTGGAGTCGTTGCTGGCTGCTTCTGATGTGATACTTNNNAGATCGGAAG |
| 39095_2 | TATGAGTGTGGAGTCGTTGCTGGCTGCTTCTGATGTGATACTTNNNAGATCGGAAG |
| 33615_1 | TATGAGTGTGGAGTCGTTGCCTTCTGAGTAGCTGGGACTACAGNNNAGATCGGAA |
| 33615_2 | TATGAGTGTGGAGTCGTTGCCTTCTGAGTAGCTGGGACTACAGNNNAGATCGGAA |
| 43453_1 | TATGAGTGTGGAGTCGTTGCCCTTCCAAGTAACTGGGACTACANNNAGATCGGAAG |
| 43453_2 | TATGAGTGTGGAGTCGTTGCCCTTCCAAGTAACTGGGACTACANNNAGATCGGAAG |
| 52414_1 | TATGAGTGTGGAGTCGTTGCCTTGGCATCTCTCCTGGATTTCTNNNAGATCGGAAG |
| 52414_2 | TATGAGTGTGGAGTCGTTGCCTTGGCATCTCTCCTGGATTTCTNNNAGATCGGAAG |
| 34408_1 | TATGAGTGTGGAGTCGTTGCACCTTTGCTGAGGATTGAGTAGGNNNAGATCGGAA |
| 34408_2 | TATGAGTGTGGAGTCGTTGCACCTTTGCTGAGGATTGAGTAGGNNNAGATCGGAA |
| 50472_1 | TATGAGTGTGGAGTCGTTGCCACTACCTTCCTTCTCTTGGCTTNNNAGATCGGAAGA |
| 50472_2 | TATGAGTGTGGAGTCGTTGCCACTACCTTCCTTCTCTTGGCTTNNNAGATCGGAAGA |
| 37473_1 | TATGAGTGTGGAGTCGTTGCCTTTGGCAAATAGACTTGGAGCCNNNAGATCGGAA |
| 37473_2 | TATGAGTGTGGAGTCGTTGCCTTTGGCAAATAGACTTGGAGCCNNNAGATCGGAA |
| 34210_1 | TATGAGTGTGGAGTCGTTGCTCCTTCCTTCCTTTCATTCGTCCNNNAGATCGGAAGA |
| 34210_2 | TATGAGTGTGGAGTCGTTGCTCCTTCCTTCCTTTCATTCGTCCNNNAGATCGGAAGA |
| 35843_1 | TATGAGTGTGGAGTCGTTGCCCACGCCCAGCTAATTTTTTGTANNNAGATCGGAAG |
| 35843_2 | TATGAGTGTGGAGTCGTTGCCCACGCCCAGCTAATTTTTTGTANNNAGATCGGAAG |
| 2621_1 | TATGAGTGTGGAGTCGTTGCTCTTTTTCTGAACCTGGCCTCTGNNNAGATCGGAAG |
| 2621_2 | TATGAGTGTGGAGTCGTTGCTCTTTTTCTGAACCTGGCCTCTGNNNAGATCGGAAG |
| 37239_1 | TATGAGTGTGGAGTCGTTGCGGCTAGGTCAGTTCTTTGGAGAANNNAGATCGGAA |
| 37239_2 | TATGAGTGTGGAGTCGTTGCGGCTAGGTCAGTTCTTTGGAGAANNNAGATCGGAA |

| | |
|---|---|
| 52242_1 | TATGAGTGTGGAGTCGTTGCAAACAAGGCTAGAAAAAGCTGGGNNNAGATCGGAA |
| 52242_2 | TATGAGTGTGGAGTCGTTGCAAACAAGGCTAGAAAAAGCTGGGNNNAGATCGGAA |
| 50265_1 | TATGAGTGTGGAGTCGTTGCCTCCATGTTGACCAAGCCTCTACNNNAGATCGGAAG |
| 50265_2 | TATGAGTGTGGAGTCGTTGCCTCCATGTTGACCAAGCCTCTACNNNAGATCGGAAG |
| 36075_1 | TATGAGTGTGGAGTCGTTGCTGCTTCCACCCAGTTATTAGACCNNNAGATCGGAAG |
| 36075_2 | TATGAGTGTGGAGTCGTTGCTGCTTCCACCCAGTTATTAGACCNNNAGATCGGAAG |
| 29314_1 | TATGAGTGTGGAGTCGTTGCTCCCCACTTAGTCCACTCATACTNNNAGATCGGAAGA |
| 29314_2 | TATGAGTGTGGAGTCGTTGCTCCCCACTTAGTCCACTCATACTNNNAGATCGGAAGA |
| 34706_1 | TATGAGTGTGGAGTCGTTGCCTGGAGTGCAATGGTGTGATCTCNNNAGATCGGAA |
| 34706_2 | TATGAGTGTGGAGTCGTTGCCTGGAGTGCAATGGTGTGATCTCNNNAGATCGGAA |
| 42848_1 | TATGAGTGTGGAGTCGTTGCATGTGTGTGTATATGTGTGGGTATNNNAGATCGGA |
| 42848_2 | TATGAGTGTGGAGTCGTTGCATGTGTGTGTATATGTGTGGGTATNNNAGATCGGA |
| 39192_1 | TATGAGTGTGGAGTCGTTGCACTGAGGTAGGAGGATCACTTGANNNAGATCGGAA |
| 39192_2 | TATGAGTGTGGAGTCGTTGCACTGAGGTAGGAGGATCACTTGANNNAGATCGGAA |
| 47480_1 | TATGAGTGTGGAGTCGTTGCTAGGAGAATCGACAGCTGTGATGNNNAGATCGGAA |
| 47480_2 | TATGAGTGTGGAGTCGTTGCTAGGAGAATCGACAGCTGTGATGNNNAGATCGGAA |
| 40544_1 | TATGAGTGTGGAGTCGTTGCGAATTTCGTTCTTGTTGCCCAGANNNAGATCGGAAG |
| 40544_2 | TATGAGTGTGGAGTCGTTGCGAATTTCGTTCTTGTTGCCCAGANNNAGATCGGAAG |
| 49495_1 | TATGAGTGTGGAGTCGTTGCGAAGCAAACTCTGGTAGTCTTGCNNNAGATCGGAA |
| 49495_2 | TATGAGTGTGGAGTCGTTGCGAAGCAAACTCTGGTAGTCTTGCNNNAGATCGGAA |
| 52105_1 | TATGAGTGTGGAGTCGTTGCCGGGTCAGTGAAAAAACGATCAGNNNAGATCGGAA |
| 52105_2 | TATGAGTGTGGAGTCGTTGCCGGGTCAGTGAAAAAACGATCAGNNNAGATCGGAA |
| 31946_1 | TATGAGTGTGGAGTCGTTGCACCTGATAGAGCTGAATGACACANNNAGATCGGAA |
| 31946_2 | TATGAGTGTGGAGTCGTTGCACCTGATAGAGCTGAATGACACANNNAGATCGGAA |
| 38528_1 | TATGAGTGTGGAGTCGTTGCGTTTGAAGCAATTCTCCTGCCTCNNNAGATCGGAAG |
| 38528_2 | TATGAGTGTGGAGTCGTTGCGTTTGAAGCAATTCTCCTGCCTCNNNAGATCGGAAG |
| 28898_1 | TATGAGTGTGGAGTCGTTGCCTTTGAGCATCCAGGCAAAAAGANNNAGATCGGAA |
| 28898_2 | TATGAGTGTGGAGTCGTTGCCTTTGAGCATCCAGGCAAAAAGANNNAGATCGGAA |
| 52852_1 | TATGAGTGTGGAGTCGTTGCCTTTCAGGAGCTCCATACACACGNNNAGATCGGAAG |
| 52852_2 | TATGAGTGTGGAGTCGTTGCCTTTCAGGAGCTCCATACACACGNNNAGATCGGAAG |
| 37531_1 | TATGAGTGTGGAGTCGTTGCTCTTTGGTAACGTAGGCAAAACTNNNAGATCGGAAG |
| 37531_2 | TATGAGTGTGGAGTCGTTGCTCTTTGGTAACGTAGGCAAAACTNNNAGATCGGAAG |
| 43566_1 | TATGAGTGTGGAGTCGTTGCCACCTCCTAAAGTGCTGGGATTANNNAGATCGGAAG |
| 43566_2 | TATGAGTGTGGAGTCGTTGCCACCTCCTAAAGTGCTGGGATTANNNAGATCGGAAG |
| 42343_1 | TATGAGTGTGGAGTCGTTGCGTGTGTGTGTGTGTGTGTGTATGNNNAGATCGGAA |
| 42343_2 | TATGAGTGTGGAGTCGTTGCGTGTGTGTGTGTGTGTGTGTATGNNNAGATCGGAA |
| 52832_1 | TATGAGTGTGGAGTCGTTGCTCATCTCTGGACTTCACCTCCTNNNAGATCGGAAGAG |
| 52832_2 | TATGAGTGTGGAGTCGTTGCTCATCTCTGGACTTCACCTCCTNNNAGATCGGAAGAG |
| 32646_1 | TATGAGTGTGGAGTCGTTGCCACTGGGTGGAAAAAAGAGCAAGNNNAGATCGGA |
| 32646_2 | TATGAGTGTGGAGTCGTTGCCACTGGGTGGAAAAAAGAGCAAGNNNAGATCGGA |
| 41651_1 | TATGAGTGTGGAGTCGTTGCATCTCAGCTGACTGTAACCTCTGNNNAGATCGGAAG |
| 41651_2 | TATGAGTGTGGAGTCGTTGCATCTCAGCTGACTGTAACCTCTGNNNAGATCGGAAG |
| 38516_1 | TATGAGTGTGGAGTCGTTGCAATCGCCTGTAATTCTAGCACCTNNNAGATCGGAAG |

| | |
|---|---|
| 38516_2 | TATGAGTGTGGAGTCGTTGCAATCGCCTGTAATTCTAGCACCTNNNAGATCGGAAG |
| 51500_1 | TATGAGTGTGGAGTCGTTGCCGTAGTGCTTGTCTTTGTTCCAGNNNAGATCGGAAG |
| 51500_2 | TATGAGTGTGGAGTCGTTGCCGTAGTGCTTGTCTTTGTTCCAGNNNAGATCGGAAG |
| 51225_1 | TATGAGTGTGGAGTCGTTGCAGTCATAGAACTGCCTCAAGGTCNNNAGATCGGAAG |
| 51225_2 | TATGAGTGTGGAGTCGTTGCAGTCATAGAACTGCCTCAAGGTCNNNAGATCGGAAG |
| 35705_1 | TATGAGTGTGGAGTCGTTGCTGCAACTTCTGTCTCCTAGGTTCNNNAGATCGGAAG |
| 35705_2 | TATGAGTGTGGAGTCGTTGCTGCAACTTCTGTCTCCTAGGTTCNNNAGATCGGAAG |
| 53244_1 | TATGAGTGTGGAGTCGTTGCAAGTACAGGTGACCACTTCTTCCNNNAGATCGGAAG |
| 53244_2 | TATGAGTGTGGAGTCGTTGCAAGTACAGGTGACCACTTCTTCCNNNAGATCGGAAG |
| 13307_1 | TATGAGTGTGGAGTCGTTGCCTCCTTAAGCCATTCTAAAGTTNNNAGATCGGAAGA |
| 13307_2 | TATGAGTGTGGAGTCGTTGCCTCCTTAAGCCATTCTAAAGTTNNNAGATCGGAAGA |
| 50292_1 | TATGAGTGTGGAGTCGTTGCGATGTCCTAAGCCCAGATACCTGNNNAGATCGGAAG |
| 50292_2 | TATGAGTGTGGAGTCGTTGCGATGTCCTAAGCCCAGATACCTGNNNAGATCGGAAG |
| 47104_1 | TATGAGTGTGGAGTCGTTGCAAAAGTCACCAGTCTCAGCCAATNNNAGATCGGAAG |
| 47104_2 | TATGAGTGTGGAGTCGTTGCAAAAGTCACCAGTCTCAGCCAATNNNAGATCGGAAG |
| 29084_1 | TATGAGTGTGGAGTCGTTGCACTTGTGTGTGTGTGCTGAAAAANNNAGATCGGAA |
| 29084_2 | TATGAGTGTGGAGTCGTTGCACTTGTGTGTGTGTGCTGAAAAANNNAGATCGGAA |
| 48487_1 | TATGAGTGTGGAGTCGTTGCTCCCCAACTTTTTGTTTTGATCTGTNNNAGATCGGAA |
| 48487_2 | TATGAGTGTGGAGTCGTTGCTCCCCAACTTTTTGTTTTGATCTGTNNNAGATCGGAA |
| 10623_1 | TATGAGTGTGGAGTCGTTGCTCAGCAAATGAGATGGTTTGCCTNNNAGATCGGAAG |
| 10623_2 | TATGAGTGTGGAGTCGTTGCTCAGCAAATGAGATGGTTTGCCTNNNAGATCGGAAG |
| 48370_1 | TATGAGTGTGGAGTCGTTGCCATCTCGCCCGGCTAATTTTTTANNNAGATCGGAAG |
| 48370_2 | TATGAGTGTGGAGTCGTTGCCATCTCGCCCGGCTAATTTTTTANNNAGATCGGAAG |
| 50458_1 | TATGAGTGTGGAGTCGTTGCCTCACACTCACTCACACTTCACTNNNAGATCGGAAGA |
| 50458_2 | TATGAGTGTGGAGTCGTTGCCTCACACTCACTCACACTTCACTNNNAGATCGGAAGA |
| 35744_1 | TATGAGTGTGGAGTCGTTGCCTGGAGTGTAATGGTGAGCTCTCNNNAGATCGGAA |
| 35744_2 | TATGAGTGTGGAGTCGTTGCCTGGAGTGTAATGGTGAGCTCTCNNNAGATCGGAA |
| 52629_1 | TATGAGTGTGGAGTCGTTGCGACACAGAGGACTGCTATTGCTANNNAGATCGGAA |
| 52629_2 | TATGAGTGTGGAGTCGTTGCGACACAGAGGACTGCTATTGCTANNNAGATCGGAA |
| 35588_1 | TATGAGTGTGGAGTCGTTGCTTGCTGCAATTATAGGCATCTGCNNNAGATCGGAAG |
| 35588_2 | TATGAGTGTGGAGTCGTTGCTTGCTGCAATTATAGGCATCTGCNNNAGATCGGAAG |
| 45886_1 | TATGAGTGTGGAGTCGTTGCCTCAGGCAATTCTCCTACCTCAGNNNAGATCGGAAG |
| 45886_2 | TATGAGTGTGGAGTCGTTGCCTCAGGCAATTCTCCTACCTCAGNNNAGATCGGAAG |
| 46505_1 | TATGAGTGTGGAGTCGTTGCTTCAAGTGATTCTCCTGCCTCAGNNNAGATCGGAAG |
| 46505_2 | TATGAGTGTGGAGTCGTTGCTTCAAGTGATTCTCCTGCCTCAGNNNAGATCGGAAG |
| 38803_1 | TATGAGTGTGGAGTCGTTGCATCCTGGCTAACACAGTGAAACTNNNAGATCGGAAG |
| 38803_2 | TATGAGTGTGGAGTCGTTGCATCCTGGCTAACACAGTGAAACTNNNAGATCGGAAG |
| 45810_1 | TATGAGTGTGGAGTCGTTGCAGAGTGCAATGGTGTAATCTCGGNNNAGATCGGAA |
| 45810_2 | TATGAGTGTGGAGTCGTTGCAGAGTGCAATGGTGTAATCTCGGNNNAGATCGGAA |
| 38474_1 | TATGAGTGTGGAGTCGTTGCAACTGCACTAGAGTGGAGACAACNNNAGATCGGAA |
| 38474_2 | TATGAGTGTGGAGTCGTTGCAACTGCACTAGAGTGGAGACAACNNNAGATCGGAA |
| 10032_1 | TATGAGTGTGGAGTCGTTGCTGAAGTTGGTGGGGAATGTTTATNNNAGATCGGAA |
| 10032_2 | TATGAGTGTGGAGTCGTTGCTGAAGTTGGTGGGGAATGTTTATNNNAGATCGGAA |

| | |
|---|---|
| 5207_1 | TATGAGTGTGGAGTCGTTGCACTGGAATCTGATGGGACATTCTNNNAGATCGGAAG |
| 5207_2 | TATGAGTGTGGAGTCGTTGCACTGGAATCTGATGGGACATTCTNNNAGATCGGAAG |
| 44082_1 | TATGAGTGTGGAGTCGTTGCGTCACCACTGCCCAAATTTGTAGNNNAGATCGGAAG |
| 44082_2 | TATGAGTGTGGAGTCGTTGCGTCACCACTGCCCAAATTTGTAGNNNAGATCGGAAG |
| 30436_1 | TATGAGTGTGGAGTCGTTGCATCTTTTCCACCAAGACTGTGGTNNNAGATCGGAAG |
| 30436_2 | TATGAGTGTGGAGTCGTTGCATCTTTTCCACCAAGACTGTGGTNNNAGATCGGAAG |
| 50332_1 | TATGAGTGTGGAGTCGTTGCATGCCTTCACTGTGTCACATTTGNNNAGATCGGAAG |
| 50332_2 | TATGAGTGTGGAGTCGTTGCATGCCTTCACTGTGTCACATTTGNNNAGATCGGAAG |
| 43699_1 | TATGAGTGTGGAGTCGTTGCAGCCTTTGGGAAAGAAGGAGGNNNAGATCGGAAG |
| 43699_2 | TATGAGTGTGGAGTCGTTGCAGCCTTTGGGAAAGAAGGAGGNNNAGATCGGAAG |
| 47987_1 | TATGAGTGTGGAGTCGTTGCCTAGGATGGTCTCGATCTCCTGANNNAGATCGGAAG |
| 47987_2 | TATGAGTGTGGAGTCGTTGCCTAGGATGGTCTCGATCTCCTGANNNAGATCGGAAG |
| 50525_1 | TATGAGTGTGGAGTCGTTGCAGAGAGGTAAGCAGGCATTGAAANNNAGATCGGA |
| 50525_2 | TATGAGTGTGGAGTCGTTGCAGAGAGGTAAGCAGGCATTGAAANNNAGATCGGA |
| 33850_1 | TATGAGTGTGGAGTCGTTGCAGGCTTTCTCTAACTGTGAAGGTNNNAGATCGGAAG |
| 33850_2 | TATGAGTGTGGAGTCGTTGCAGGCTTTCTCTAACTGTGAAGGTNNNAGATCGGAAG |
| 30566_1 | TATGAGTGTGGAGTCGTTGCCCCATGCCATCATCCCTACCCNNNAGATCGGAAGAGC |
| 30566_2 | TATGAGTGTGGAGTCGTTGCCCCATGCCATCATCCCTACCCNNNAGATCGGAAGAGC |
| 42167_1 | TATGAGTGTGGAGTCGTTGCGAGATCTCGGCTCACTACGACNNNAGATCGGAAGA |
| 42167_2 | TATGAGTGTGGAGTCGTTGCGAGATCTCGGCTCACTACGACNNNAGATCGGAAGA |
| 47779_1 | TATGAGTGTGGAGTCGTTGCTCTTGACTCACTGTAACCTCTGCNNNAGATCGGAAG |
| 47779_2 | TATGAGTGTGGAGTCGTTGCTCTTGACTCACTGTAACCTCTGCNNNAGATCGGAAG |
| 28949_1 | TATGAGTGTGGAGTCGTTGCAGCCACATTTACCAAGTTGTAAAANNNAGATCGGAA |
| 28949_2 | TATGAGTGTGGAGTCGTTGCAGCCACATTTACCAAGTTGTAAAANNNAGATCGGAA |
| 6231_1 | TATGAGTGTGGAGTCGTTGCGCCACCAGGCACCAATATTTTTNNNAGATCGGAAGA |
| 6231_2 | TATGAGTGTGGAGTCGTTGCGCCACCAGGCACCAATATTTTTNNNAGATCGGAAGA |
| 43466_1 | TATGAGTGTGGAGTCGTTGCTTCAAGTGATTCTCCTGCCTCAGNNNAGATCGGAAG |
| 43466_2 | TATGAGTGTGGAGTCGTTGCTTCAAGTGATTCTCCTGCCTCAGNNNAGATCGGAAG |
| 46657_1 | TATGAGTGTGGAGTCGTTGCCAGTGCTGAGCTTTTCTTTCAGGNNNAGATCGGAAG |
| 46657_2 | TATGAGTGTGGAGTCGTTGCCAGTGCTGAGCTTTTCTTTCAGGNNNAGATCGGAAG |
| 30246_1 | TATGAGTGTGGAGTCGTTGCTCTGCTCACTCAGACCTAAATGANNNAGATCGGAAG |
| 30246_2 | TATGAGTGTGGAGTCGTTGCTCTGCTCACTCAGACCTAAATGANNNAGATCGGAAG |
| 33694_1 | TATGAGTGTGGAGTCGTTGCATTGCTTGAATCTGAGAGACGGANNNAGATCGGAA |
| 33694_2 | TATGAGTGTGGAGTCGTTGCATTGCTTGAATCTGAGAGACGGANNNAGATCGGAA |
| 33295_1 | TATGAGTGTGGAGTCGTTGCTTAGTTCAAGTGAGTCTCCTGCCNNNAGATCGGAAG |
| 33295_2 | TATGAGTGTGGAGTCGTTGCTTAGTTCAAGTGAGTCTCCTGCCNNNAGATCGGAAG |
| 52289_1 | TATGAGTGTGGAGTCGTTGCCTAGGTGCTCAGCTATCTTGAGGNNNAGATCGGAAG |
| 52289_2 | TATGAGTGTGGAGTCGTTGCCTAGGTGCTCAGCTATCTTGAGGNNNAGATCGGAAG |
| 43733_1 | TATGAGTGTGGAGTCGTTGCCCCTCTCATCCTTCACTTCTGACNNNAGATCGGAAGA |
| 43733_2 | TATGAGTGTGGAGTCGTTGCCCCTCTCATCCTTCACTTCTGACNNNAGATCGGAAGA |
| 49424_1 | TATGAGTGTGGAGTCGTTGCACAGGGAACACTGCTTTTTCACNNNAGATCGGAAG |
| 49424_2 | TATGAGTGTGGAGTCGTTGCACAGGGAACACTGCTTTTTCACNNNAGATCGGAAG |
| 29977_1 | TATGAGTGTGGAGTCGTTGCAAAACACTCGAAGGAGCTGAGATNNNAGATCGGAA |

| | |
|---|---|
| 29977_2 | TATGAGTGTGGAGTCGTTGCAAAACACTCGAAGGAGCTGAGATNNNAGATCGGAA |
| 44567_1 | TATGAGTGTGGAGTCGTTGCATGGCATTCACAGCAACCTAGATNNNAGATCGGAAG |
| 44567_2 | TATGAGTGTGGAGTCGTTGCATGGCATTCACAGCAACCTAGATNNNAGATCGGAAG |
| 53216_1 | TATGAGTGTGGAGTCGTTGCTTTGGCCTCAAACTTCAGGTGANNNAGATCGGAAGA |
| 53216_2 | TATGAGTGTGGAGTCGTTGCTTTGGCCTCAAACTTCAGGTGANNNAGATCGGAAGA |
| 52019_1 | TATGAGTGTGGAGTCGTTGCTCCTAACTCTGTGGGTTGAACTGNNNAGATCGGAAG |
| 52019_2 | TATGAGTGTGGAGTCGTTGCTCCTAACTCTGTGGGTTGAACTGNNNAGATCGGAAG |
| 40856_1 | TATGAGTGTGGAGTCGTTGCTGTACCTGTTTACAATGTACAAACTGNNNAGATCGG |
| 40856_2 | TATGAGTGTGGAGTCGTTGCTGTACCTGTTTACAATGTACAAACTGNNNAGATCGG |
| 50452_1 | TATGAGTGTGGAGTCGTTGCTAACTAGGGAGGCTTAGGTGGAANNNAGATCGGAA |
| 50452_2 | TATGAGTGTGGAGTCGTTGCTAACTAGGGAGGCTTAGGTGGAANNNAGATCGGAA |
| 52844_1 | TATGAGTGTGGAGTCGTTGCCTACTGATGCCCTGGAATAGCAANNNAGATCGGAAG |
| 52844_2 | TATGAGTGTGGAGTCGTTGCCTACTGATGCCCTGGAATAGCAANNNAGATCGGAAG |
| 31015_1 | TATGAGTGTGGAGTCGTTGCGAAAGTGTTGAACAGGACAGAGCNNNAGATCGGAA |
| 31015_2 | TATGAGTGTGGAGTCGTTGCGAAAGTGTTGAACAGGACAGAGCNNNAGATCGGAA |
| 50012_1 | TATGAGTGTGGAGTCGTTGCGCTGTTTGCTTCACACATGCAGNNNAGATCGGAAGA |
| 50012_2 | TATGAGTGTGGAGTCGTTGCGCTGTTTGCTTCACACATGCAGNNNAGATCGGAAGA |
| 31225_1 | TATGAGTGTGGAGTCGTTGCAGGTTTCTGCTGTGCTAATGGTANNNAGATCGGAAG |
| 31225_2 | TATGAGTGTGGAGTCGTTGCAGGTTTCTGCTGTGCTAATGGTANNNAGATCGGAAG |
| 36568_1 | TATGAGTGTGGAGTCGTTGCTAGGGACGTTTTGGAAATGCAACNNNAGATCGGAA |
| 36568_2 | TATGAGTGTGGAGTCGTTGCTAGGGACGTTTTGGAAATGCAACNNNAGATCGGAA |
| 39151_1 | TATGAGTGTGGAGTCGTTGCGCCGAGATCGTGCTCAAAAAATANNNAGATCGGAAG |
| 39151_2 | TATGAGTGTGGAGTCGTTGCGCCGAGATCGTGCTCAAAAAATANNNAGATCGGAAG |
| 33203_1 | TATGAGTGTGGAGTCGTTGCATTCATCCTCTCTTTCTTCGGCANNNAGATCGGAAGA |
| 33203_2 | TATGAGTGTGGAGTCGTTGCATTCATCCTCTCTTTCTTCGGCANNNAGATCGGAAGA |
| 35656_1 | TATGAGTGTGGAGTCGTTGCGGCCTGGATTCTGTCTAATCTGANNNAGATCGGAAG |
| 35656_2 | TATGAGTGTGGAGTCGTTGCGGCCTGGATTCTGTCTAATCTGANNNAGATCGGAAG |
| 46219_1 | TATGAGTGTGGAGTCGTTGCCTGGAATGCAATGGTGTGATCACNNNAGATCGGAA |
| 46219_2 | TATGAGTGTGGAGTCGTTGCCTGGAATGCAATGGTGTGATCACNNNAGATCGGAA |
| 49509_1 | TATGAGTGTGGAGTCGTTGCATTCAAGCGATTATCCTGCCTCANNNAGATCGGAAG |
| 49509_2 | TATGAGTGTGGAGTCGTTGCATTCAAGCGATTATCCTGCCTCANNNAGATCGGAAG |
| 34626_1 | TATGAGTGTGGAGTCGTTGCGGTGTGTTCCTCAGGGCTATTTANNNAGATCGGAAG |
| 34626_2 | TATGAGTGTGGAGTCGTTGCGGTGTGTTCCTCAGGGCTATTTANNNAGATCGGAAG |
| 45449_1 | TATGAGTGTGGAGTCGTTGCTTTCCCAGGTTGGTAACACTGTCNNNAGATCGGAAG |
| 45449_2 | TATGAGTGTGGAGTCGTTGCTTTCCCAGGTTGGTAACACTGTCNNNAGATCGGAAG |
| 30942_1 | TATGAGTGTGGAGTCGTTGCAGGTTGAAGAGAGCTGTCCTTTGNNNAGATCGGAA |
| 30942_2 | TATGAGTGTGGAGTCGTTGCAGGTTGAAGAGAGCTGTCCTTTGNNNAGATCGGAA |
| 47740_1 | TATGAGTGTGGAGTCGTTGCTTGGAGTAGGAGCTGAGGAGGNNNAGATCGGAAG |
| 47740_2 | TATGAGTGTGGAGTCGTTGCTTGGAGTAGGAGCTGAGGAGGNNNAGATCGGAAG |
| 47116_1 | TATGAGTGTGGAGTCGTTGCCTGAGGCAGGTAGATCACTTGAGNNNAGATCGGAA |
| 47116_2 | TATGAGTGTGGAGTCGTTGCCTGAGGCAGGTAGATCACTTGAGNNNAGATCGGAA |
| 45123_1 | TATGAGTGTGGAGTCGTTGCATCAGCATATGCACACACACAACNNNAGATCGGAAG |
| 45123_2 | TATGAGTGTGGAGTCGTTGCATCAGCATATGCACACACACAACNNNAGATCGGAAG |

| | |
|---|---|
| 30182_1 | TATGAGTGTGGAGTCGTTGCGTGTTGGGATTACAGGTTTGAGCNNNAGATCGGAA |
| 30182_2 | TATGAGTGTGGAGTCGTTGCGTGTTGGGATTACAGGTTTGAGCNNNAGATCGGAA |
| 43700_1 | TATGAGTGTGGAGTCGTTGCTAAGCACCATGTTGGCCTCTGNNNAGATCGGAAGAG |
| 43700_2 | TATGAGTGTGGAGTCGTTGCTAAGCACCATGTTGGCCTCTGNNNAGATCGGAAGAG |
| 37314_1 | TATGAGTGTGGAGTCGTTGCCCTCCCAGGTTCAAGTGATTCTCNNNAGATCGGAAG |
| 37314_2 | TATGAGTGTGGAGTCGTTGCCCTCCCAGGTTCAAGTGATTCTCNNNAGATCGGAAG |
| 48526_1 | TATGAGTGTGGAGTCGTTGCCCTCACAAATAGCTGGGACTACANNNAGATCGGAAG |
| 48526_2 | TATGAGTGTGGAGTCGTTGCCCTCACAAATAGCTGGGACTACANNNAGATCGGAAG |
| 53274_1 | TATGAGTGTGGAGTCGTTGCGTTCCCCAAACTCTCTCCCTCNNNAGATCGGAAGAGC |
| 53274_2 | TATGAGTGTGGAGTCGTTGCGTTCCCCAAACTCTCTCCCTCNNNAGATCGGAAGAGC |
| 41442_1 | TATGAGTGTGGAGTCGTTGCCACTCCTCTAATCCCAGCACTTTNNNAGATCGGAAGA |
| 41442_2 | TATGAGTGTGGAGTCGTTGCCACTCCTCTAATCCCAGCACTTTNNNAGATCGGAAGA |
| 3015_1 | TATGAGTGTGGAGTCGTTGCTCTGCATCCCTTGTTTCCTGATGNNNAGATCGGAAGA |
| 3015_2 | TATGAGTGTGGAGTCGTTGCTCTGCATCCCTTGTTTCCTGATGNNNAGATCGGAAGA |
| 40823_1 | TATGAGTGTGGAGTCGTTGCGAATTACAAGGCATGCCCTTGACNNNAGATCGGAAG |
| 40823_2 | TATGAGTGTGGAGTCGTTGCGAATTACAAGGCATGCCCTTGACNNNAGATCGGAAG |
| 46206_1 | TATGAGTGTGGAGTCGTTGCTTCAAGTGATTCTCCTGCCTCAGNNNAGATCGGAAG |
| 46206_2 | TATGAGTGTGGAGTCGTTGCTTCAAGTGATTCTCCTGCCTCAGNNNAGATCGGAAG |
| 37941_1 | TATGAGTGTGGAGTCGTTGCTGGCCAATATGGTGAAACCTCATNNNAGATCGGAAG |
| 37941_2 | TATGAGTGTGGAGTCGTTGCTGGCCAATATGGTGAAACCTCATNNNAGATCGGAAG |
| 41646_1 | TATGAGTGTGGAGTCGTTGCTTTGCCAATTGTCCCTCTACAGTNNNAGATCGGAAGA |
| 41646_2 | TATGAGTGTGGAGTCGTTGCTTTGCCAATTGTCCCTCTACAGTNNNAGATCGGAAGA |
| 41221_1 | TATGAGTGTGGAGTCGTTGCTACAAAAAATTAGCCGGGTGCAGNNNAGATCGGAA |
| 41221_2 | TATGAGTGTGGAGTCGTTGCTACAAAAAATTAGCCGGGTGCAGNNNAGATCGGAA |
| 32971_1 | TATGAGTGTGGAGTCGTTGCGAATGACACAATGGACTTTGGGGNNNAGATCGGAA |
| 32971_2 | TATGAGTGTGGAGTCGTTGCGAATGACACAATGGACTTTGGGGNNNAGATCGGAA |
| 51244_1 | TATGAGTGTGGAGTCGTTGCTGGGATGGGGAGCGAAATTTATANNNAGATCGGAA |
| 51244_2 | TATGAGTGTGGAGTCGTTGCTGGGATGGGGAGCGAAATTTATANNNAGATCGGAA |
| 45590_1 | TATGAGTGTGGAGTCGTTGCAGGTTCATAGCTGGACTTGGAAGNNNAGATCGGAA |
| 45590_2 | TATGAGTGTGGAGTCGTTGCAGGTTCATAGCTGGACTTGGAAGNNNAGATCGGAA |
| 29738_1 | TATGAGTGTGGAGTCGTTGCATTGTGTTTACCAGGCCATCTCTNNNAGATCGGAAG |
| 29738_2 | TATGAGTGTGGAGTCGTTGCATTGTGTTTACCAGGCCATCTCTNNNAGATCGGAAG |
| 10305_1 | TATGAGTGTGGAGTCGTTGCCCCATACAATCTTGGCTCTGCTTNNNAGATCGGAAGA |
| 10305_2 | TATGAGTGTGGAGTCGTTGCCCCATACAATCTTGGCTCTGCTTNNNAGATCGGAAGA |
| 2316_1 | TATGAGTGTGGAGTCGTTGCAAATAAGTGTGCCACCATCACGANNNAGATCGGAAG |
| 2316_2 | TATGAGTGTGGAGTCGTTGCAAATAAGTGTGCCACCATCACGANNNAGATCGGAAG |
| 30388_1 | TATGAGTGTGGAGTCGTTGCTGTATACACACACACACGTATCTGTNNNAGATCGGA |
| 30388_2 | TATGAGTGTGGAGTCGTTGCTGTATACACACACACACGTATCTGTNNNAGATCGGA |
| 43568_1 | TATGAGTGTGGAGTCGTTGCTCCCGGGTTCAAATGATTCTCTTNNNAGATCGGAAG |
| 43568_2 | TATGAGTGTGGAGTCGTTGCTCCCGGGTTCAAATGATTCTCTTNNNAGATCGGAAG |
| 10253_1 | TATGAGTGTGGAGTCGTTGCAGAGATTCCTAAGGGAGTGTGGGNNNAGATCGGA |
| 10253_2 | TATGAGTGTGGAGTCGTTGCAGAGATTCCTAAGGGAGTGTGGGNNNAGATCGGA |
| 51010_1 | TATGAGTGTGGAGTCGTTGCGCTCCTGTAGTCTCAGCTACTTGNNNAGATCGGAAG |

| | |
|---|---|
| 51010_2 | TATGAGTGTGGAGTCGTTGCGCTCCTGTAGTCTCAGCTACTTGNNNAGATCGGAAG |
| 34305_1 | TATGAGTGTGGAGTCGTTGCGGTAGTGGAGTGTTGACCAATCANNNAGATCGGAA |
| 34305_2 | TATGAGTGTGGAGTCGTTGCGGTAGTGGAGTGTTGACCAATCANNNAGATCGGAA |
| 31645_1 | TATGAGTGTGGAGTCGTTGCCTGGTGGTGTTTTTCTCCATGACNNNAGATCGGAAG |
| 31645_2 | TATGAGTGTGGAGTCGTTGCCTGGTGGTGTTTTTCTCCATGACNNNAGATCGGAAG |
| 12960_1 | TATGAGTGTGGAGTCGTTGCATCTACAACTCTTCCTTCTACTNNNAGATCGGAAGAG |
| 12960_2 | TATGAGTGTGGAGTCGTTGCATCTACAACTCTTCCTTCTACTNNNAGATCGGAAGAG |
| 35505_1 | TATGAGTGTGGAGTCGTTGCCAATCTCCACCTCCTGAGTTCAANNNAGATCGGAAG |
| 35505_2 | TATGAGTGTGGAGTCGTTGCCAATCTCCACCTCCTGAGTTCAANNNAGATCGGAAG |
| 44818_1 | TATGAGTGTGGAGTCGTTGCGGAAAACCAGCTGCTACATTCAANNNAGATCGGAAG |
| 44818_2 | TATGAGTGTGGAGTCGTTGCGGAAAACCAGCTGCTACATTCAANNNAGATCGGAAG |
| 12482_1 | TATGAGTGTGGAGTCGTTGCCTTGGGATTCAGAATTTTGAGNNNAGATCGGAAGA |
| 12482_2 | TATGAGTGTGGAGTCGTTGCCTTGGGATTCAGAATTTTGAGNNNAGATCGGAAGA |
| 44339_1 | TATGAGTGTGGAGTCGTTGCACCCAGATGGCCAAAGATGATTANNNAGATCGGAA |
| 44339_2 | TATGAGTGTGGAGTCGTTGCACCCAGATGGCCAAAGATGATTANNNAGATCGGAA |
| 9183_1 | TATGAGTGTGGAGTCGTTGCAGCATAGTGAGTACACTGTTCCANNNAGATCGGAAG |
| 9183_2 | TATGAGTGTGGAGTCGTTGCAGCATAGTGAGTACACTGTTCCANNNAGATCGGAAG |
| 30414_1 | TATGAGTGTGGAGTCGTTGCAGAAACACATACACAAGGATGCACTNNNAGATCGG |
| 30414_2 | TATGAGTGTGGAGTCGTTGCAGAAACACATACACAAGGATGCACTNNNAGATCGG |
| 13501_1 | TATGAGTGTGGAGTCGTTGCACTATGGCAAGACTCAGTTTNNNAGATCGGAAGAG |
| 13501_2 | TATGAGTGTGGAGTCGTTGCACTATGGCAAGACTCAGTTTNNNAGATCGGAAGAG |
| 10580_1 | TATGAGTGTGGAGTCGTTGCACGATTCCTTTGATTTGGAAACANNNAGATCGGAAG |
| 10580_2 | TATGAGTGTGGAGTCGTTGCACGATTCCTTTGATTTGGAAACANNNAGATCGGAAG |
| 29316_1 | TATGAGTGTGGAGTCGTTGCAGTCAATTTGAGGAGAGTTGCCANNNAGATCGGAA |
| 29316_2 | TATGAGTGTGGAGTCGTTGCAGTCAATTTGAGGAGAGTTGCCANNNAGATCGGAA |
| 10175_1 | TATGAGTGTGGAGTCGTTGCATACTCCCAAAAGTGTGGGTCCTNNNAGATCGGAAG |
| 10175_2 | TATGAGTGTGGAGTCGTTGCATACTCCCAAAAGTGTGGGTCCTNNNAGATCGGAAG |
| 44058_1 | TATGAGTGTGGAGTCGTTGCAGGAACATTCAGGAACTCCACATNNNAGATCGGAAG |
| 44058_2 | TATGAGTGTGGAGTCGTTGCAGGAACATTCAGGAACTCCACATNNNAGATCGGAAG |
| 31734_1 | TATGAGTGTGGAGTCGTTGCGGAAGTAGAAGGTGGAGGGAAAGNNNAGATCGG |
| 31734_2 | TATGAGTGTGGAGTCGTTGCGGAAGTAGAAGGTGGAGGGAAAGNNNAGATCGG |
| 46088_1 | TATGAGTGTGGAGTCGTTGCTCAAGTCAAGGTCCAATGATCCANNNAGATCGGAAG |
| 46088_2 | TATGAGTGTGGAGTCGTTGCTCAAGTCAAGGTCCAATGATCCANNNAGATCGGAAG |
| 47539_1 | TATGAGTGTGGAGTCGTTGCCTCAAACTCCTGACCGTGTGATCNNNAGATCGGAAG |
| 47539_2 | TATGAGTGTGGAGTCGTTGCCTCAAACTCCTGACCGTGTGATCNNNAGATCGGAAG |
| 37228_1 | TATGAGTGTGGAGTCGTTGCATTCAGAGAGGAGCAACAGGAAGNNNAGATCGGA |
| 37228_2 | TATGAGTGTGGAGTCGTTGCATTCAGAGAGGAGCAACAGGAAGNNNAGATCGGA |
| 11221_1 | TATGAGTGTGGAGTCGTTGCGAAGCATGTGAGAACACAGCAAGNNNAGATCGGAA |
| 11221_2 | TATGAGTGTGGAGTCGTTGCGAAGCATGTGAGAACACAGCAAGNNNAGATCGGAA |
| 42976_1 | TATGAGTGTGGAGTCGTTGCCACAGGCTAGATAGAGTGCAGTGNNNAGATCGGAA |
| 42976_2 | TATGAGTGTGGAGTCGTTGCCACAGGCTAGATAGAGTGCAGTGNNNAGATCGGAA |
| 47299_1 | TATGAGTGTGGAGTCGTTGCGTTCAAGCAATTCTCCTGCCTCNNNAGATCGGAAGA |
| 47299_2 | TATGAGTGTGGAGTCGTTGCGTTCAAGCAATTCTCCTGCCTCNNNAGATCGGAAGA |

| | |
|---|---|
| 38258_1 | TATGAGTGTGGAGTCGTTGCACCACATGTACAAAGAAGAGCTCANNNAGATCGGAA |
| 38258_2 | TATGAGTGTGGAGTCGTTGCACCACATGTACAAAGAAGAGCTCANNNAGATCGGAA |
| 29447_1 | TATGAGTGTGGAGTCGTTGCCCTGATTCATCTCATTGGGACACGNNNAGATCGGAA |
| 29447_2 | TATGAGTGTGGAGTCGTTGCCCTGATTCATCTCATTGGGACACGNNNAGATCGGAA |
| 29425_1 | TATGAGTGTGGAGTCGTTGCCACGAGACGTCCATCGACATCNNNAGATCGGAAGAG |
| 29425_2 | TATGAGTGTGGAGTCGTTGCCACGAGACGTCCATCGACATCNNNAGATCGGAAGAG |
| 32119_1 | TATGAGTGTGGAGTCGTTGCAGGGAAACTCACACATTCAGGAANNNAGATCGGAA |
| 32119_2 | TATGAGTGTGGAGTCGTTGCAGGGAAACTCACACATTCAGGAANNNAGATCGGAA |
| 31877_1 | TATGAGTGTGGAGTCGTTGCTTGTGTACATTTTTTGGCTGGGCNNNAGATCGGAAG |
| 31877_2 | TATGAGTGTGGAGTCGTTGCTTGTGTACATTTTTTGGCTGGGCNNNAGATCGGAAG |
| 48562_1 | TATGAGTGTGGAGTCGTTGCCACCATGCCCAGCTAATTTTTGTNNNAGATCGGAAGA |
| 48562_2 | TATGAGTGTGGAGTCGTTGCCACCATGCCCAGCTAATTTTTGTNNNAGATCGGAAGA |
| 34580_1 | TATGAGTGTGGAGTCGTTGCGAGAGGATGGAGGCTGCATTCNNNAGATCGGAAGA |
| 34580_2 | TATGAGTGTGGAGTCGTTGCGAGAGGATGGAGGCTGCATTCNNNAGATCGGAAGA |
| 50954_1 | TATGAGTGTGGAGTCGTTGCACACAGTGATTCCAACTCTCCTTNNNAGATCGGAAGA |
| 50954_2 | TATGAGTGTGGAGTCGTTGCACACAGTGATTCCAACTCTCCTTNNNAGATCGGAAGA |
| 46218_1 | TATGAGTGTGGAGTCGTTGCAGAGTACAGTGGTTCAATCACGGNNNAGATCGGAA |
| 46218_2 | TATGAGTGTGGAGTCGTTGCAGAGTACAGTGGTTCAATCACGGNNNAGATCGGAA |
| 42134_1 | TATGAGTGTGGAGTCGTTGCTCCAGGTTCAAGTGATTCTCCTGNNNAGATCGGAAG |
| 42134_2 | TATGAGTGTGGAGTCGTTGCTCCAGGTTCAAGTGATTCTCCTGNNNAGATCGGAAG |
| 43529_1 | TATGAGTGTGGAGTCGTTGCAGGTAGTTTTTCAACCCACACCTNNNAGATCGGAAGA |
| 43529_2 | TATGAGTGTGGAGTCGTTGCAGGTAGTTTTTCAACCCACACCTNNNAGATCGGAAGA |
| 45814_1 | TATGAGTGTGGAGTCGTTGCGCACTGCAATTGGAGAAGAATGTNNNAGATCGGAA |
| 45814_2 | TATGAGTGTGGAGTCGTTGCGCACTGCAATTGGAGAAGAATGTNNNAGATCGGAA |
| 31510_1 | TATGAGTGTGGAGTCGTTGCACACACACACACACAAACACATTNNNAGATCGGAAGA |
| 31510_2 | TATGAGTGTGGAGTCGTTGCACACACACACACACAAACACATTNNNAGATCGGAAGA |
| 46417_1 | TATGAGTGTGGAGTCGTTGCGTTCACACCATTCTCCTCCCTCNNNAGATCGGAAGAG |
| 46417_2 | TATGAGTGTGGAGTCGTTGCGTTCACACCATTCTCCTCCCTCNNNAGATCGGAAGAG |
| 53028_1 | TATGAGTGTGGAGTCGTTGCTTGCAATGAACCAAGATCATGCCNNNAGATCGGAAG |
| 53028_2 | TATGAGTGTGGAGTCGTTGCTTGCAATGAACCAAGATCATGCCNNNAGATCGGAAG |
| 34141_1 | TATGAGTGTGGAGTCGTTGCGGTCACGGTGTTTTGTAACCTAANNNAGATCGGAAG |
| 34141_2 | TATGAGTGTGGAGTCGTTGCGGTCACGGTGTTTTGTAACCTAANNNAGATCGGAAG |
| 52819_1 | TATGAGTGTGGAGTCGTTGCCCAGAGGTGTAACTGCCCAGNNNAGATCGGAAGAG |
| 52819_2 | TATGAGTGTGGAGTCGTTGCCCAGAGGTGTAACTGCCCAGNNNAGATCGGAAGAG |
| 38426_1 | TATGAGTGTGGAGTCGTTGCGTAGAAGGGAGCTGCAGTTTCTANNNAGATCGGAA |
| 38426_2 | TATGAGTGTGGAGTCGTTGCGTAGAAGGGAGCTGCAGTTTCTANNNAGATCGGAA |
| 51190_1 | TATGAGTGTGGAGTCGTTGCCAAAGTGGCAAGATTTCCCCTGNNNAGATCGGAAGA |
| 51190_2 | TATGAGTGTGGAGTCGTTGCCAAAGTGGCAAGATTTCCCCTGNNNAGATCGGAAGA |
| 47931_1 | TATGAGTGTGGAGTCGTTGCAGCTCCCAGTATAACCAGAGACTNNNAGATCGGAAG |
| 47931_2 | TATGAGTGTGGAGTCGTTGCAGCTCCCAGTATAACCAGAGACTNNNAGATCGGAAG |
| 36901_1 | TATGAGTGTGGAGTCGTTGCTTTCGTCAAATAACCCACAGCACNNNAGATCGGAAGA |
| 36901_2 | TATGAGTGTGGAGTCGTTGCTTTCGTCAAATAACCCACAGCACNNNAGATCGGAAGA |
| 50869_1 | TATGAGTGTGGAGTCGTTGCATGAGAGAAAAGGAGTGCAGGAGNNNAGATCGGA |

| | |
|---|---|
| 50869_2 | TATGAGTGTGGAGTCGTTGCATGAGAGAAAAGGAGTGCAGGAGNNNAGATCGGA |
| 35712_1 | TATGAGTGTGGAGTCGTTGCTACAGTTCAAAGGAACGGTTTGCNNNAGATCGGAA |
| 35712_2 | TATGAGTGTGGAGTCGTTGCTACAGTTCAAAGGAACGGTTTGCNNNAGATCGGAA |
| 47855_1 | TATGAGTGTGGAGTCGTTGCTAGCATAATCTTGGCTCACTGCANNNAGATCGGAAG |
| 47855_2 | TATGAGTGTGGAGTCGTTGCTAGCATAATCTTGGCTCACTGCANNNAGATCGGAAG |
| 41503_1 | TATGAGTGTGGAGTCGTTGCTGGAACACAGTAAGAGGGGAATGNNNAGATCGGA |
| 41503_2 | TATGAGTGTGGAGTCGTTGCTGGAACACAGTAAGAGGGGAATGNNNAGATCGGA |
| 38263_1 | TATGAGTGTGGAGTCGTTGCTCCTTTTTGACAATAGGCCTCCANNNAGATCGGAAG |
| 38263_2 | TATGAGTGTGGAGTCGTTGCTCCTTTTTGACAATAGGCCTCCANNNAGATCGGAAG |
| 51856_1 | TATGAGTGTGGAGTCGTTGCCATACTACTTGCTGTCCCTGACANNNAGATCGGAAG |
| 51856_2 | TATGAGTGTGGAGTCGTTGCCATACTACTTGCTGTCCCTGACANNNAGATCGGAAG |
| 45821_1 | TATGAGTGTGGAGTCGTTGCCCTCCCAGGCTCAAGTAATTCTTNNNAGATCGGAAG |
| 45821_2 | TATGAGTGTGGAGTCGTTGCCCTCCCAGGCTCAAGTAATTCTTNNNAGATCGGAAG |
| 49725_1 | TATGAGTGTGGAGTCGTTGCTGGAGTACAGTGGTGCAATCATGNNNAGATCGGAA |
| 49725_2 | TATGAGTGTGGAGTCGTTGCTGGAGTACAGTGGTGCAATCATGNNNAGATCGGAA |
| 12826_1 | TATGAGTGTGGAGTCGTTGCCATTGATGAGTAGCTTTTTCAGNNNAGATCGGAAGA |
| 12826_2 | TATGAGTGTGGAGTCGTTGCCATTGATGAGTAGCTTTTTCAGNNNAGATCGGAAGA |
| 52653_1 | TATGAGTGTGGAGTCGTTGCTGAAGACAAGACGATGAGACAGGNNNAGATCGGA |
| 52653_2 | TATGAGTGTGGAGTCGTTGCTGAAGACAAGACGATGAGACAGGNNNAGATCGGA |
| 8277_1 | TATGAGTGTGGAGTCGTTGCGGACAACCGTCATCTCTAGCTCNNNAGATCGGAAGA |
| 8277_2 | TATGAGTGTGGAGTCGTTGCGGACAACCGTCATCTCTAGCTCNNNAGATCGGAAGA |
| 36903_1 | TATGAGTGTGGAGTCGTTGCACAAAAGTTAGCCAGACATTGTGGNNNAGATCGGA |
| 36903_2 | TATGAGTGTGGAGTCGTTGCACAAAAGTTAGCCAGACATTGTGGNNNAGATCGGA |
| 42053_1 | TATGAGTGTGGAGTCGTTGCTCCTGTAAATGGGTAGTTGGACCNNNAGATCGGAA |
| 42053_2 | TATGAGTGTGGAGTCGTTGCTCCTGTAAATGGGTAGTTGGACCNNNAGATCGGAA |
| 42444_1 | TATGAGTGTGGAGTCGTTGCTCACATTAGCCTCAAACTCCCAANNNAGATCGGAAGA |
| 42444_2 | TATGAGTGTGGAGTCGTTGCTCACATTAGCCTCAAACTCCCAANNNAGATCGGAAGA |
| 42771_1 | TATGAGTGTGGAGTCGTTGCATGCAATCGTGTCTTTTGAAGCANNNAGATCGGAAG |
| 42771_2 | TATGAGTGTGGAGTCGTTGCATGCAATCGTGTCTTTTGAAGCANNNAGATCGGAAG |
| 46902_1 | TATGAGTGTGGAGTCGTTGCCTAGTCTTGTGGAGGTGTCACTCNNNAGATCGGAAG |
| 46902_2 | TATGAGTGTGGAGTCGTTGCCTAGTCTTGTGGAGGTGTCACTCNNNAGATCGGAAG |
| 50913_1 | TATGAGTGTGGAGTCGTTGCTCAGGGCAGATTGATCCATGTAGNNNAGATCGGAA |
| 50913_2 | TATGAGTGTGGAGTCGTTGCTCAGGGCAGATTGATCCATGTAGNNNAGATCGGAA |
| 52207_1 | TATGAGTGTGGAGTCGTTGCTGCTAACTGGTGTACAATGCTCANNNAGATCGGAAG |
| 52207_2 | TATGAGTGTGGAGTCGTTGCTGCTAACTGGTGTACAATGCTCANNNAGATCGGAAG |
| 11316_1 | TATGAGTGTGGAGTCGTTGCATTTGATGGCTGAGGAAACTGGGANNNAGATCGGA |
| 11316_2 | TATGAGTGTGGAGTCGTTGCATTTGATGGCTGAGGAAACTGGGANNNAGATCGGA |
| 31291_1 | TATGAGTGTGGAGTCGTTGCCAAACTGCTCAGCTCTCATTGACNNNAGATCGGAAG |
| 31291_2 | TATGAGTGTGGAGTCGTTGCCAAACTGCTCAGCTCTCATTGACNNNAGATCGGAAG |
| 36915_1 | TATGAGTGTGGAGTCGTTGCAATGGCCGATTTCTCCCACTTATNNNAGATCGGAAG |
| 36915_2 | TATGAGTGTGGAGTCGTTGCAATGGCCGATTTCTCCCACTTATNNNAGATCGGAAG |
| 30492_1 | TATGAGTGTGGAGTCGTTGCCATACTCCCAGGGTGGGTACTCNNNAGATCGGAAGA |
| 30492_2 | TATGAGTGTGGAGTCGTTGCCATACTCCCAGGGTGGGTACTCNNNAGATCGGAAGA |

| | |
|---|---|
| 52907_1 | TATGAGTGTGGAGTCGTTGCGACAAGGTCTCACTCTGTCACCNNNAGATCGGAAGA |
| 52907_2 | TATGAGTGTGGAGTCGTTGCGACAAGGTCTCACTCTGTCACCNNNAGATCGGAAGA |
| 33281_1 | TATGAGTGTGGAGTCGTTGCGGTTCAAGTGATTCTCCTGCTTCNNNAGATCGGAAG |
| 33281_2 | TATGAGTGTGGAGTCGTTGCGGTTCAAGTGATTCTCCTGCTTCNNNAGATCGGAAG |
| 51738_1 | TATGAGTGTGGAGTCGTTGCCACCACTTTTGGACAGGAGTCTANNNAGATCGGAAG |
| 51738_2 | TATGAGTGTGGAGTCGTTGCCACCACTTTTGGACAGGAGTCTANNNAGATCGGAAG |
| 32510_1 | TATGAGTGTGGAGTCGTTGCGCTTTTCTTTCATTTGGCTTGCANNNAGATCGGAAGA |
| 32510_2 | TATGAGTGTGGAGTCGTTGCGCTTTTCTTTCATTTGGCTTGCANNNAGATCGGAAGA |
| 39660_1 | TATGAGTGTGGAGTCGTTGCCTGAGACAGGAGGATTGCTTGAGNNNAGATCGGAA |
| 39660_2 | TATGAGTGTGGAGTCGTTGCCTGAGACAGGAGGATTGCTTGAGNNNAGATCGGAA |
| 34494_1 | TATGAGTGTGGAGTCGTTGCCAGAGGCACCACTCACCTCCNNNAGATCGGAAGAGC |
| 34494_2 | TATGAGTGTGGAGTCGTTGCCAGAGGCACCACTCACCTCCNNNAGATCGGAAGAGC |
| 40786_1 | TATGAGTGTGGAGTCGTTGCTTCTTCAGTCATCCAGACAAGGGNNNAGATCGGAAG |
| 40786_2 | TATGAGTGTGGAGTCGTTGCTTCTTCAGTCATCCAGACAAGGGNNNAGATCGGAAG |
| 5553_1 | TATGAGTGTGGAGTCGTTGCTGCTGTGGCATTTCTGACACATANNNAGATCGGAAG |
| 5553_2 | TATGAGTGTGGAGTCGTTGCTGCTGTGGCATTTCTGACACATANNNAGATCGGAAG |
| 40201_1 | TATGAGTGTGGAGTCGTTGCGGCACCTATAATCCCAGCTACTCNNNAGATCGGAAG |
| 40201_2 | TATGAGTGTGGAGTCGTTGCGGCACCTATAATCCCAGCTACTCNNNAGATCGGAAG |
| 42452_1 | TATGAGTGTGGAGTCGTTGCTTTCACTCTTGTTGCCTAGGCTGNNNAGATCGGAAG |
| 42452_2 | TATGAGTGTGGAGTCGTTGCTTTCACTCTTGTTGCCTAGGCTGNNNAGATCGGAAG |
| 32054_1 | TATGAGTGTGGAGTCGTTGCGGAGTGCAGTGGTACGATCTCNNNAGATCGGAAGA |
| 32054_2 | TATGAGTGTGGAGTCGTTGCGGAGTGCAGTGGTACGATCTCNNNAGATCGGAAGA |
| 7894_1 | TATGAGTGTGGAGTCGTTGCGCAGTTGACGGACAGATCATCATNNNAGATCGGAA |
| 7894_2 | TATGAGTGTGGAGTCGTTGCGCAGTTGACGGACAGATCATCATNNNAGATCGGAA |
| 36095_1 | TATGAGTGTGGAGTCGTTGCGCATTGACTTCTGGCTCTGATTCNNNAGATCGGAAG |
| 36095_2 | TATGAGTGTGGAGTCGTTGCGCATTGACTTCTGGCTCTGATTCNNNAGATCGGAAG |
| 43305_1 | TATGAGTGTGGAGTCGTTGCTTCAAGTGATTCTCCTGCCTCAGNNNAGATCGGAAG |
| 43305_2 | TATGAGTGTGGAGTCGTTGCTTCAAGTGATTCTCCTGCCTCAGNNNAGATCGGAAG |
| 32518_1 | TATGAGTGTGGAGTCGTTGCGGATGAGTAGGGAGGCAAAGAAGNNNAGATCGGA |
| 32518_2 | TATGAGTGTGGAGTCGTTGCGGATGAGTAGGGAGGCAAAGAAGNNNAGATCGGA |
| 40873_1 | TATGAGTGTGGAGTCGTTGCCTTCCAGTAACAAGACTTGCTGCNNNAGATCGGAAG |
| 40873_2 | TATGAGTGTGGAGTCGTTGCCTTCCAGTAACAAGACTTGCTGCNNNAGATCGGAAG |
| 44606_1 | TATGAGTGTGGAGTCGTTGCTCCATTTGGGACAAGAAAGGGATNNNAGATCGGAA |
| 44606_2 | TATGAGTGTGGAGTCGTTGCTCCATTTGGGACAAGAAAGGGATNNNAGATCGGAA |
| 8003_1 | TATGAGTGTGGAGTCGTTGCGGACCTGGGAAATACTTCCAAACTNNNAGATCGGAA |
| 8003_2 | TATGAGTGTGGAGTCGTTGCGGACCTGGGAAATACTTCCAAACTNNNAGATCGGAA |
| 44090_1 | TATGAGTGTGGAGTCGTTGCATTATGTTCTGGGACATGGCACTNNNAGATCGGAAG |
| 44090_2 | TATGAGTGTGGAGTCGTTGCATTATGTTCTGGGACATGGCACTNNNAGATCGGAAG |
| 9771_1 | TATGAGTGTGGAGTCGTTGCGTGTGGAGAAAGAGGCCAGTTTANNNAGATCGGAA |
| 9771_2 | TATGAGTGTGGAGTCGTTGCGTGTGGAGAAAGAGGCCAGTTTANNNAGATCGGAA |
| 36179_1 | TATGAGTGTGGAGTCGTTGCAGAACACCCAACAGAGACTAGTCNNNAGATCGGAAG |
| 36179_2 | TATGAGTGTGGAGTCGTTGCAGAACACCCAACAGAGACTAGTCNNNAGATCGGAAG |
| 52255_1 | TATGAGTGTGGAGTCGTTGCAGCTTTCTCTGGAAACTTGAGAANNNAGATCGGAAG |

| | |
|---|---|
| 52255_2 | TATGAGTGTGGAGTCGTTGCAGCTTTCTCTGGAAACTTGAGAANNNAGATCGGAAG |
| 53262_1 | TATGAGTGTGGAGTCGTTGCTTCCCACCAGAAGCCATTCTATCNNNAGATCGGAAGA |
| 53262_2 | TATGAGTGTGGAGTCGTTGCTTCCCACCAGAAGCCATTCTATCNNNAGATCGGAAGA |
| 40654_1 | TATGAGTGTGGAGTCGTTGCAGGTAGCCCAGGAATTTCAAGTTNNNAGATCGGAAG |
| 40654_2 | TATGAGTGTGGAGTCGTTGCAGGTAGCCCAGGAATTTCAAGTTNNNAGATCGGAAG |
| 12013_1 | TATGAGTGTGGAGTCGTTGCCTGCATCTACACTGCCTGCAGNNNAGATCGGAAGAG |
| 12013_2 | TATGAGTGTGGAGTCGTTGCCTGCATCTACACTGCCTGCAGNNNAGATCGGAAGAG |
| 51015_1 | TATGAGTGTGGAGTCGTTGCGATCACGAGGTCAGGAGTTTGAGNNNAGATCGGAA |
| 51015_2 | TATGAGTGTGGAGTCGTTGCGATCACGAGGTCAGGAGTTTGAGNNNAGATCGGAA |
| 35674_1 | TATGAGTGTGGAGTCGTTGCACTTTTAGTAGAGACGGGGTTTCCNNNAGATCGGAA |
| 35674_2 | TATGAGTGTGGAGTCGTTGCACTTTTAGTAGAGACGGGGTTTCCNNNAGATCGGAA |
| 51024_1 | TATGAGTGTGGAGTCGTTGCAATGTTTGGGCTGAAATGTCTGGNNNAGATCGGAAG |
| 51024_2 | TATGAGTGTGGAGTCGTTGCAATGTTTGGGCTGAAATGTCTGGNNNAGATCGGAAG |
| 52134_1 | TATGAGTGTGGAGTCGTTGCAAAAATTAGTTGGGTGTGGTGGCNNNAGATCGGAA |
| 52134_2 | TATGAGTGTGGAGTCGTTGCAAAAATTAGTTGGGTGTGGTGGCNNNAGATCGGAA |
| 10279_1 | TATGAGTGTGGAGTCGTTGCTCTGGTAATGGCGGACTAGATGANNNAGATCGGAAG |
| 10279_2 | TATGAGTGTGGAGTCGTTGCTCTGGTAATGGCGGACTAGATGANNNAGATCGGAAG |
| 30522_1 | TATGAGTGTGGAGTCGTTGCCCACCTCATTTCCTATGCTAGGCNNNAGATCGGAAGA |
| 30522_2 | TATGAGTGTGGAGTCGTTGCCCACCTCATTTCCTATGCTAGGCNNNAGATCGGAAGA |
| 36877_1 | TATGAGTGTGGAGTCGTTGCTGGTCTACAAAACAGGACTTCCTNNNAGATCGGAAG |
| 36877_2 | TATGAGTGTGGAGTCGTTGCTGGTCTACAAAACAGGACTTCCTNNNAGATCGGAAG |
| 51613_1 | TATGAGTGTGGAGTCGTTGCCCACGCCCGGCTAATTTTTATTNNNAGATCGGAAGA |
| 51613_2 | TATGAGTGTGGAGTCGTTGCCCACGCCCGGCTAATTTTTATTNNNAGATCGGAAGA |
| 41515_1 | TATGAGTGTGGAGTCGTTGCGCACTAAGGCTCTTTACTTGCACNNNAGATCGGAAGA |
| 41515_2 | TATGAGTGTGGAGTCGTTGCGCACTAAGGCTCTTTACTTGCACNNNAGATCGGAAGA |
| 39098_1 | TATGAGTGTGGAGTCGTTGCTCAAAACATTCCAAATGGTATCTGTNNNAGATCGGAA |
| 39098_2 | TATGAGTGTGGAGTCGTTGCTCAAAACATTCCAAATGGTATCTGTNNNAGATCGGAA |
| 41635_1 | TATGAGTGTGGAGTCGTTGCCTCAAGTCATCCTCCCACCATAGNNNAGATCGGAAGA |
| 41635_2 | TATGAGTGTGGAGTCGTTGCCTCAAGTCATCCTCCCACCATAGNNNAGATCGGAAGA |
| 31677_1 | TATGAGTGTGGAGTCGTTGCTACCTGAGGATAGGGAACCTTGTNNNAGATCGGAAG |
| 31677_2 | TATGAGTGTGGAGTCGTTGCTACCTGAGGATAGGGAACCTTGTNNNAGATCGGAAG |
| 42683_1 | TATGAGTGTGGAGTCGTTGCTTTATGGGCATTTGGGTTGGTTCNNNAGATCGGAAG |
| 42683_2 | TATGAGTGTGGAGTCGTTGCTTTATGGGCATTTGGGTTGGTTCNNNAGATCGGAAG |
| 2562_1 | TATGAGTGTGGAGTCGTTGCTTAGTCTCTGAGAAACCTGGGGGNNNAGATCGGAAG |
| 2562_2 | TATGAGTGTGGAGTCGTTGCTTAGTCTCTGAGAAACCTGGGGGNNNAGATCGGAAG |
| 41494_1 | TATGAGTGTGGAGTCGTTGCTTTACTTCATGAGGTCAGCACCANNNAGATCGGAAGA |
| 41494_2 | TATGAGTGTGGAGTCGTTGCTTTACTTCATGAGGTCAGCACCANNNAGATCGGAAGA |
| 39369_1 | TATGAGTGTGGAGTCGTTGCAAAAAATTAGCCAGGTGTGGTGGNNNAGATCGGAA |
| 39369_2 | TATGAGTGTGGAGTCGTTGCAAAAAATTAGCCAGGTGTGGTGGNNNAGATCGGAA |
| 51319_1 | TATGAGTGTGGAGTCGTTGCGGGGAGATTTGACAAGTCAGAGANNNAGATCGGAA |
| 51319_2 | TATGAGTGTGGAGTCGTTGCGGGGAGATTTGACAAGTCAGAGANNNAGATCGGAA |
| 33208_1 | TATGAGTGTGGAGTCGTTGCAAGTTCTTGCATTCAAAGGGTCGNNNAGATCGGAAG |
| 33208_2 | TATGAGTGTGGAGTCGTTGCAAGTTCTTGCATTCAAAGGGTCGNNNAGATCGGAAG |

| | |
|---|---|
| 52226_1 | TATGAGTGTGGAGTCGTTGCGCAGATGTTCTTTGTCTTGCTGTNNNAGATCGGAAG |
| 52226_2 | TATGAGTGTGGAGTCGTTGCGCAGATGTTCTTTGTCTTGCTGTNNNAGATCGGAAG |
| 36579_1 | TATGAGTGTGGAGTCGTTGCAGAAAAGTCAAACACACCATTGTNNNAGATCGGAAG |
| 36579_2 | TATGAGTGTGGAGTCGTTGCAGAAAAGTCAAACACACCATTGTNNNAGATCGGAAG |
| 46714_1 | TATGAGTGTGGAGTCGTTGCCCACCACGCCTGACTAATTTTTTNNNAGATCGGAAGA |
| 46714_2 | TATGAGTGTGGAGTCGTTGCCCACCACGCCTGACTAATTTTTTNNNAGATCGGAAGA |
| 37995_1 | TATGAGTGTGGAGTCGTTGCGATGTACCCACACACAATGCATCNNNAGATCGGAAG |
| 37995_2 | TATGAGTGTGGAGTCGTTGCGATGTACCCACACACAATGCATCNNNAGATCGGAAG |
| 37483_1 | TATGAGTGTGGAGTCGTTGCGAGACACAAGACACATGCACAAANNNAGATCGGAA |
| 37483_2 | TATGAGTGTGGAGTCGTTGCGAGACACAAGACACATGCACAAANNNAGATCGGAA |
| 13136_1 | TATGAGTGTGGAGTCGTTGCCGCCAGCGAATAAATAATAAATANNNAGATCGGAAG |
| 13136_2 | TATGAGTGTGGAGTCGTTGCCGCCAGCGAATAAATAATAAATANNNAGATCGGAAG |
| 10661_1 | TATGAGTGTGGAGTCGTTGCACCTTTGAAAGCCACTACCACAGNNNAGATCGGAAG |
| 10661_2 | TATGAGTGTGGAGTCGTTGCACCTTTGAAAGCCACTACCACAGNNNAGATCGGAAG |
| 10991_1 | TATGAGTGTGGAGTCGTTGCCTCAGCCCAACCTTGCTCCAAGANNNAGATCGGAAG |
| 10991_2 | TATGAGTGTGGAGTCGTTGCCTCAGCCCAACCTTGCTCCAAGANNNAGATCGGAAG |
| 32130_1 | TATGAGTGTGGAGTCGTTGCCGCCATCTTGAAACATGAACTGANNNAGATCGGAAG |
| 32130_2 | TATGAGTGTGGAGTCGTTGCCGCCATCTTGAAACATGAACTGANNNAGATCGGAAG |
| 38103_1 | TATGAGTGTGGAGTCGTTGCTTCTGACTCTAATGCTACAAACCTNNNAGATCGGAAG |
| 38103_2 | TATGAGTGTGGAGTCGTTGCTTCTGACTCTAATGCTACAAACCTNNNAGATCGGAAG |
| 40773_1 | TATGAGTGTGGAGTCGTTGCAACAGAGATGGTAGCTTCCATGTNNNAGATCGGAA |
| 40773_2 | TATGAGTGTGGAGTCGTTGCAACAGAGATGGTAGCTTCCATGTNNNAGATCGGAA |
| 50029_1 | TATGAGTGTGGAGTCGTTGCTTTTGAGACAGGATCCCACTCTGNNNAGATCGGAAG |
| 50029_2 | TATGAGTGTGGAGTCGTTGCTTTTGAGACAGGATCCCACTCTGNNNAGATCGGAAG |
| 13546_1 | TATGAGTGTGGAGTCGTTGCGAAGCCGAACATTATTTTATGAANNNAGATCGGAAG |
| 13546_2 | TATGAGTGTGGAGTCGTTGCGAAGCCGAACATTATTTTATGAANNNAGATCGGAAG |
| 13155_1 | TATGAGTGTGGAGTCGTTGCTCTTCAGTATCAGGACAAGGNNNAGATCGGAAGAG |
| 13155_2 | TATGAGTGTGGAGTCGTTGCTCTTCAGTATCAGGACAAGGNNNAGATCGGAAGAG |
| 32038_1 | TATGAGTGTGGAGTCGTTGCGTTTCCTTGTTTGCCATCATCCANNNAGATCGGAAGA |
| 32038_2 | TATGAGTGTGGAGTCGTTGCGTTTCCTTGTTTGCCATCATCCANNNAGATCGGAAGA |
| 43231_1 | TATGAGTGTGGAGTCGTTGCCGTCTCAACAGAAGAAAGGAGGANNNAGATCGGAA |
| 43231_2 | TATGAGTGTGGAGTCGTTGCCGTCTCAACAGAAGAAAGGAGGANNNAGATCGGAA |
| 53181_1 | TATGAGTGTGGAGTCGTTGCGGCCAACCTGGAGATTACTCTTTNNNAGATCGGAAG |
| 53181_2 | TATGAGTGTGGAGTCGTTGCGGCCAACCTGGAGATTACTCTTTNNNAGATCGGAAG |
| 34622_1 | TATGAGTGTGGAGTCGTTGCCAATGGCAACCCGAAAATCAATGNNNAGATCGGAAG |
| 34622_2 | TATGAGTGTGGAGTCGTTGCCAATGGCAACCCGAAAATCAATGNNNAGATCGGAAG |
| 35274_1 | TATGAGTGTGGAGTCGTTGCGGTTTTGAACACCTGTCCTCAAGNNNAGATCGGAAG |
| 35274_2 | TATGAGTGTGGAGTCGTTGCGGTTTTGAACACCTGTCCTCAAGNNNAGATCGGAAG |
| 53157_1 | TATGAGTGTGGAGTCGTTGCGGTAAAGTGCTGGCTCTAACTTGNNNAGATCGGAA |
| 53157_2 | TATGAGTGTGGAGTCGTTGCGGTAAAGTGCTGGCTCTAACTTGNNNAGATCGGAA |
| 42538_1 | TATGAGTGTGGAGTCGTTGCCTGGGTTCAAGAAATTCTCCTGCNNNAGATCGGAAG |
| 42538_2 | TATGAGTGTGGAGTCGTTGCCTGGGTTCAAGAAATTCTCCTGCNNNAGATCGGAAG |
| 29555_1 | TATGAGTGTGGAGTCGTTGCCAACGCTTGGTTATTGGGAGATGNNNAGATCGGAA |

| | |
|---|---|
| 29555_2 | TATGAGTGTGGAGTCGTTGCCAACGCTTGGTTATTGGGAGATGNNNAGATCGGAA |
| 41133_1 | TATGAGTGTGGAGTCGTTGCGATGCTCAAGGGCAGAGATTCTANNNAGATCGGAA |
| 41133_2 | TATGAGTGTGGAGTCGTTGCGATGCTCAAGGGCAGAGATTCTANNNAGATCGGAA |
| 35658_1 | TATGAGTGTGGAGTCGTTGCTGTGTCAGATCAGAGTTTCACAANNNAGATCGGAAG |
| 35658_2 | TATGAGTGTGGAGTCGTTGCTGTGTCAGATCAGAGTTTCACAANNNAGATCGGAAG |
| 48017_1 | TATGAGTGTGGAGTCGTTGCCACCTCCAACGTTCAAGTGATTCNNNAGATCGGAAG |
| 48017_2 | TATGAGTGTGGAGTCGTTGCCACCTCCAACGTTCAAGTGATTCNNNAGATCGGAAG |
| 46300_1 | TATGAGTGTGGAGTCGTTGCTCAACCTCCTGAGTATCTGGGATNNNAGATCGGAAG |
| 46300_2 | TATGAGTGTGGAGTCGTTGCTCAACCTCCTGAGTATCTGGGATNNNAGATCGGAAG |
| 46619_1 | TATGAGTGTGGAGTCGTTGCGTTCAAGTGATTCTCCTGCCTCANNNAGATCGGAAG |
| 46619_2 | TATGAGTGTGGAGTCGTTGCGTTCAAGTGATTCTCCTGCCTCANNNAGATCGGAAG |
| 35217_1 | TATGAGTGTGGAGTCGTTGCATCCTAAGCTAAGCCAACCAGTCNNNAGATCGGAAG |
| 35217_2 | TATGAGTGTGGAGTCGTTGCATCCTAAGCTAAGCCAACCAGTCNNNAGATCGGAAG |
| 42780_1 | TATGAGTGTGGAGTCGTTGCCCAGGACCACCTAAAACAAACACNNNAGATCGGAAG |
| 42780_2 | TATGAGTGTGGAGTCGTTGCCCAGGACCACCTAAAACAAACACNNNAGATCGGAAG |
| 1451_1 | TATGAGTGTGGAGTCGTTGCATTGGGAAGCTGAGGTAGGACTTNNNAGATCGGAA |
| 1451_2 | TATGAGTGTGGAGTCGTTGCATTGGGAAGCTGAGGTAGGACTTNNNAGATCGGAA |
| 43218_1 | TATGAGTGTGGAGTCGTTGCCAGTCATGTGGAACTGAGAGTCANNNAGATCGGAA |
| 43218_2 | TATGAGTGTGGAGTCGTTGCCAGTCATGTGGAACTGAGAGTCANNNAGATCGGAA |
| 12594_1 | TATGAGTGTGGAGTCGTTGCCTCTTAAGGACCCATCTAGATNNNAGATCGGAAGAG |
| 12594_2 | TATGAGTGTGGAGTCGTTGCCTCTTAAGGACCCATCTAGATNNNAGATCGGAAGAG |
| 46860_1 | TATGAGTGTGGAGTCGTTGCAAAGGCCTTTCATGTGCAGAAAGNNNAGATCGGAA |
| 46860_2 | TATGAGTGTGGAGTCGTTGCAAAGGCCTTTCATGTGCAGAAAGNNNAGATCGGAA |
| 42587_1 | TATGAGTGTGGAGTCGTTGCGTTGTTGGACAGAATGACAGGTGNNNAGATCGGAA |
| 42587_2 | TATGAGTGTGGAGTCGTTGCGTTGTTGGACAGAATGACAGGTGNNNAGATCGGAA |
| 39256_1 | TATGAGTGTGGAGTCGTTGCGATGAATCCCCCCAATTATCCCANNNAGATCGGAAG |
| 39256_2 | TATGAGTGTGGAGTCGTTGCGATGAATCCCCCCAATTATCCCANNNAGATCGGAAG |
| 37289_1 | TATGAGTGTGGAGTCGTTGCTAATCCCAAGCTATTCAGGAGGCNNNAGATCGGAAG |
| 37289_2 | TATGAGTGTGGAGTCGTTGCTAATCCCAAGCTATTCAGGAGGCNNNAGATCGGAAG |
| 46958_1 | TATGAGTGTGGAGTCGTTGCTTGCTTTGGCTACAACTTCAAGTNNNAGATCGGAAG |
| 46958_2 | TATGAGTGTGGAGTCGTTGCTTGCTTTGGCTACAACTTCAAGTNNNAGATCGGAAG |
| 48792_1 | TATGAGTGTGGAGTCGTTGCAAGGAACTGGTTTGGGATTCTGTNNNAGATCGGAA |
| 48792_2 | TATGAGTGTGGAGTCGTTGCAAGGAACTGGTTTGGGATTCTGTNNNAGATCGGAA |
| 34241_1 | TATGAGTGTGGAGTCGTTGCTGAATGTATGTGAACGTACATGCANNNAGATCGGA |
| 34241_2 | TATGAGTGTGGAGTCGTTGCTGAATGTATGTGAACGTACATGCANNNAGATCGGA |
| 31974_1 | TATGAGTGTGGAGTCGTTGCCCAGCCACAGCCATAGATAGAAANNNAGATCGGAAG |
| 31974_2 | TATGAGTGTGGAGTCGTTGCCCAGCCACAGCCATAGATAGAAANNNAGATCGGAAG |
| 39214_1 | TATGAGTGTGGAGTCGTTGCCTCAAACTCCTGACCTCGTGATCNNNAGATCGGAAG |
| 39214_2 | TATGAGTGTGGAGTCGTTGCCTCAAACTCCTGACCTCGTGATCNNNAGATCGGAAG |
| 44673_1 | TATGAGTGTGGAGTCGTTGCGCTGTCAGAGTGCCTTGTAAATGNNNAGATCGGAAG |
| 44673_2 | TATGAGTGTGGAGTCGTTGCGCTGTCAGAGTGCCTTGTAAATGNNNAGATCGGAAG |
| 10572_1 | TATGAGTGTGGAGTCGTTGCTACTTCTCTGACAATCCCAGCCANNNAGATCGGAAGA |
| 10572_2 | TATGAGTGTGGAGTCGTTGCTACTTCTCTGACAATCCCAGCCANNNAGATCGGAAGA |

| | |
|---|---|
| 47091_1 | TATGAGTGTGGAGTCGTTGCCATCTTCTGGGCTCAAGTGATCCNNNAGATCGGAAG |
| 47091_2 | TATGAGTGTGGAGTCGTTGCCATCTTCTGGGCTCAAGTGATCCNNNAGATCGGAAG |
| 522_1 | TATGAGTGTGGAGTCGTTGCTGTCTGTTTTGTCTACTGCAGCTNNNAGATCGGAAG |
| 522_2 | TATGAGTGTGGAGTCGTTGCTGTCTGTTTTGTCTACTGCAGCTNNNAGATCGGAAG |
| 46695_1 | TATGAGTGTGGAGTCGTTGCGTTTCACCATGTTAGCCAGGATGNNNAGATCGGAAG |
| 46695_2 | TATGAGTGTGGAGTCGTTGCGTTTCACCATGTTAGCCAGGATGNNNAGATCGGAAG |
| 30801_1 | TATGAGTGTGGAGTCGTTGCGGGAGAAAGGAGAGAGGAGGTNNNAGATCGGAA |
| 30801_2 | TATGAGTGTGGAGTCGTTGCGGGAGAAAGGAGAGAGGAGGTNNNAGATCGGAA |
| 49433_1 | TATGAGTGTGGAGTCGTTGCTGATAAGAATCTGGGCCATCAGGNNNAGATCGGAA |
| 49433_2 | TATGAGTGTGGAGTCGTTGCTGATAAGAATCTGGGCCATCAGGNNNAGATCGGAA |
| 43652_1 | TATGAGTGTGGAGTCGTTGCCCATAAGAAGTTCCCTCTGGAGGNNNAGATCGGAAG |
| 43652_2 | TATGAGTGTGGAGTCGTTGCCCATAAGAAGTTCCCTCTGGAGGNNNAGATCGGAAG |
| 48136_1 | TATGAGTGTGGAGTCGTTGCCTGAGTAGCTGGGATGGGATTACNNNAGATCGGAA |
| 48136_2 | TATGAGTGTGGAGTCGTTGCCTGAGTAGCTGGGATGGGATTACNNNAGATCGGAA |
| 29794_1 | TATGAGTGTGGAGTCGTTGCTACAGAGGTGCCAGCTATTTCTGNNNAGATCGGAAG |
| 29794_2 | TATGAGTGTGGAGTCGTTGCTACAGAGGTGCCAGCTATTTCTGNNNAGATCGGAAG |
| 48944_1 | TATGAGTGTGGAGTCGTTGCCAAACTTCCTGTGGGTCACCTCNNNAGATCGGAAGA |
| 48944_2 | TATGAGTGTGGAGTCGTTGCCAAACTTCCTGTGGGTCACCTCNNNAGATCGGAAGA |
| 29421_1 | TATGAGTGTGGAGTCGTTGCACAGGCCTCACCACCATCACNNNAGATCGGAAGAGC |
| 29421_2 | TATGAGTGTGGAGTCGTTGCACAGGCCTCACCACCATCACNNNAGATCGGAAGAGC |
| 43759_1 | TATGAGTGTGGAGTCGTTGCATTTATAGCAGGAGGGATGTGCCNNNAGATCGGAA |
| 43759_2 | TATGAGTGTGGAGTCGTTGCATTTATAGCAGGAGGGATGTGCCNNNAGATCGGAA |
| 46908_1 | TATGAGTGTGGAGTCGTTGCAGTGATTCTTGTACCTCAGCCTCNNNAGATCGGAAG |
| 46908_2 | TATGAGTGTGGAGTCGTTGCAGTGATTCTTGTACCTCAGCCTCNNNAGATCGGAAG |
| 53230_1 | TATGAGTGTGGAGTCGTTGCGAGTGAGAGAGAGAGACAAGTGCNNNAGATCGGA |
| 53230_2 | TATGAGTGTGGAGTCGTTGCGAGTGAGAGAGAGAGACAAGTGCNNNAGATCGGA |
| 32044_1 | TATGAGTGTGGAGTCGTTGCTGTTGAATGAAAGTGGACGTGTGNNNAGATCGGAA |
| 32044_2 | TATGAGTGTGGAGTCGTTGCTGTTGAATGAAAGTGGACGTGTGNNNAGATCGGAA |
| 35178_1 | TATGAGTGTGGAGTCGTTGCCACGCCGAGCTAATTTCTGTATTNNNAGATCGGAAG |
| 35178_2 | TATGAGTGTGGAGTCGTTGCCACGCCGAGCTAATTTCTGTATTNNNAGATCGGAAG |
| 52853_1 | TATGAGTGTGGAGTCGTTGCTCAGATTTTGGCAACATCTCAGCNNNAGATCGGAAG |
| 52853_2 | TATGAGTGTGGAGTCGTTGCTCAGATTTTGGCAACATCTCAGCNNNAGATCGGAAG |
| 40713_1 | TATGAGTGTGGAGTCGTTGCTGGAAAAACCCCTTAGACCTAGCNNNAGATCGGAAG |
| 40713_2 | TATGAGTGTGGAGTCGTTGCTGGAAAAACCCCTTAGACCTAGCNNNAGATCGGAAG |
| 36806_1 | TATGAGTGTGGAGTCGTTGCCAGTTACTTGGGAAGGTGAGGAANNNAGATCGGAA |
| 36806_2 | TATGAGTGTGGAGTCGTTGCCAGTTACTTGGGAAGGTGAGGAANNNAGATCGGAA |
| 44019_1 | TATGAGTGTGGAGTCGTTGCATTTTGCTTGTGGAACTCTGCTCNNNAGATCGGAAG |
| 44019_2 | TATGAGTGTGGAGTCGTTGCATTTTGCTTGTGGAACTCTGCTCNNNAGATCGGAAG |
| 38021_1 | TATGAGTGTGGAGTCGTTGCTGCAAGAATATCAGGTTGCACACNNNAGATCGGAAG |
| 38021_2 | TATGAGTGTGGAGTCGTTGCTGCAAGAATATCAGGTTGCACACNNNAGATCGGAAG |
| 41806_1 | TATGAGTGTGGAGTCGTTGCGCACGCCAATGGTACAATCTTTANNNAGATCGGAAG |
| 41806_2 | TATGAGTGTGGAGTCGTTGCGCACGCCAATGGTACAATCTTTANNNAGATCGGAAG |
| 33614_1 | TATGAGTGTGGAGTCGTTGCCTCCCAGGTTCAGGTGATTCTCNNNAGATCGGAAGA |

| | |
|---|---|
| 33614_2 | TATGAGTGTGGAGTCGTTGCCTCCCAGGTTCAGGTGATTCTCNNNAGATCGGAAGA |
| 40879_1 | TATGAGTGTGGAGTCGTTGCGAGAATGGCATGAACCCAGGAGNNNAGATCGGAAG |
| 40879_2 | TATGAGTGTGGAGTCGTTGCGAGAATGGCATGAACCCAGGAGNNNAGATCGGAAG |
| 36349_1 | TATGAGTGTGGAGTCGTTGCCCACGTCCTTGAACTTCTCAGTANNNAGATCGGAAGA |
| 36349_2 | TATGAGTGTGGAGTCGTTGCCCACGTCCTTGAACTTCTCAGTANNNAGATCGGAAGA |
| 35394_1 | TATGAGTGTGGAGTCGTTGCCTAGAGTGAAGTGGCAGGATCTCNNNAGATCGGAA |
| 35394_2 | TATGAGTGTGGAGTCGTTGCCTAGAGTGAAGTGGCAGGATCTCNNNAGATCGGAA |
| 36526_1 | TATGAGTGTGGAGTCGTTGCTGCCAAGAACATACACTGGGTAANNNAGATCGGAAG |
| 36526_2 | TATGAGTGTGGAGTCGTTGCTGCCAAGAACATACACTGGGTAANNNAGATCGGAAG |
| 44454_1 | TATGAGTGTGGAGTCGTTGCTCTCAAACTCCTGACCTCAAGTGNNNAGATCGGAAGA |
| 44454_2 | TATGAGTGTGGAGTCGTTGCTCTCAAACTCCTGACCTCAAGTGNNNAGATCGGAAGA |
| 30132_1 | TATGAGTGTGGAGTCGTTGCTGTGTGTGCCTGTAAATGAGAGANNNAGATCGGAA |
| 30132_2 | TATGAGTGTGGAGTCGTTGCTGTGTGTGCCTGTAAATGAGAGANNNAGATCGGAA |
| 38169_1 | TATGAGTGTGGAGTCGTTGCGAATGGGAAGTGGACAGCTGTNNNAGATCGGAAGA |
| 38169_2 | TATGAGTGTGGAGTCGTTGCGAATGGGAAGTGGACAGCTGTNNNAGATCGGAAGA |
| 29220_1 | TATGAGTGTGGAGTCGTTGCTGCATGTATGTATGTATGCCACANNNAGATCGGAAG |
| 29220_2 | TATGAGTGTGGAGTCGTTGCTGCATGTATGTATGTATGCCACANNNAGATCGGAAG |
| 53370_1 | TATGAGTGTGGAGTCGTTGCCCCATGTCGAGAGCTGTATACTTNNNAGATCGGAAG |
| 53370_2 | TATGAGTGTGGAGTCGTTGCCCCATGTCGAGAGCTGTATACTTNNNAGATCGGAAG |
| 38355_1 | TATGAGTGTGGAGTCGTTGCACAAACAACCCCATCAACAAGTGNNNAGATCGGAAG |
| 38355_2 | TATGAGTGTGGAGTCGTTGCACAAACAACCCCATCAACAAGTGNNNAGATCGGAAG |
| 49949_1 | TATGAGTGTGGAGTCGTTGCACCAACCTATTCCATTTCTGCCTNNNAGATCGGAAGA |
| 49949_2 | TATGAGTGTGGAGTCGTTGCACCAACCTATTCCATTTCTGCCTNNNAGATCGGAAGA |
| 49036_1 | TATGAGTGTGGAGTCGTTGCGAGTAGCTGGGATGACAGATACCNNNAGATCGGAA |
| 49036_2 | TATGAGTGTGGAGTCGTTGCGAGTAGCTGGGATGACAGATACCNNNAGATCGGAA |
| 2693_1 | TATGAGTGTGGAGTCGTTGCATGTAGGTCCACAGACACCAGATNNNAGATCGGAAG |
| 2693_2 | TATGAGTGTGGAGTCGTTGCATGTAGGTCCACAGACACCAGATNNNAGATCGGAAG |
| 30538_1 | TATGAGTGTGGAGTCGTTGCACATATAGCCAGGAAAGGGTCCCNNNAGATCGGAA |
| 30538_2 | TATGAGTGTGGAGTCGTTGCACATATAGCCAGGAAAGGGTCCCNNNAGATCGGAA |
| 29686_1 | TATGAGTGTGGAGTCGTTGCTAAAGCCCCTTTTCTCATCTGCTNNNAGATCGGAAGA |
| 29686_2 | TATGAGTGTGGAGTCGTTGCTAAAGCCCCTTTTCTCATCTGCTNNNAGATCGGAAGA |
| 32249_1 | TATGAGTGTGGAGTCGTTGCCTTTCTCTTTCTGCTCCCCCTACNNNAGATCGGAAGAG |
| 32249_2 | TATGAGTGTGGAGTCGTTGCCTTTCTCTTTCTGCTCCCCCTACNNNAGATCGGAAGAG |
| 42445_1 | TATGAGTGTGGAGTCGTTGCTCACATTAGCCTCAAACTCCCAANNNAGATCGGAAGA |
| 42445_2 | TATGAGTGTGGAGTCGTTGCTCACATTAGCCTCAAACTCCCAANNNAGATCGGAAGA |
| 48332_1 | TATGAGTGTGGAGTCGTTGCCCTCCTGGTTCAAGTGATCCTCNNNAGATCGGAAGA |
| 48332_2 | TATGAGTGTGGAGTCGTTGCCCTCCTGGTTCAAGTGATCCTCNNNAGATCGGAAGA |
| 53052_1 | TATGAGTGTGGAGTCGTTGCGGGCACTGAGAAACAGAAACAAANNNAGATCGGAA |
| 53052_2 | TATGAGTGTGGAGTCGTTGCGGGCACTGAGAAACAGAAACAAANNNAGATCGGAA |
| 41715_1 | TATGAGTGTGGAGTCGTTGCGATGGAGTCTTACTCTGTCACCCNNNAGATCGGAAG |
| 41715_2 | TATGAGTGTGGAGTCGTTGCGATGGAGTCTTACTCTGTCACCCNNNAGATCGGAAG |
| 43477_1 | TATGAGTGTGGAGTCGTTGCCAGGTTCAAGCAATTCTCCTGCNNNAGATCGGAAGA |
| 43477_2 | TATGAGTGTGGAGTCGTTGCCAGGTTCAAGCAATTCTCCTGCNNNAGATCGGAAGA |

| | |
|---|---|
| 48767_1 | TATGAGTGTGGAGTCGTTGCCAGACGGTTAGAGCTGATTCTCANNNAGATCGGAAG |
| 48767_2 | TATGAGTGTGGAGTCGTTGCCAGACGGTTAGAGCTGATTCTCANNNAGATCGGAAG |
| 39873_1 | TATGAGTGTGGAGTCGTTGCAAAAATGAGAAAACCTGGGCCTGNNNAGATCGGAA |
| 39873_2 | TATGAGTGTGGAGTCGTTGCAAAAATGAGAAAACCTGGGCCTGNNNAGATCGGAA |
| 44636_1 | TATGAGTGTGGAGTCGTTGCCATGGAAGAGGAGTCCTGATCAGNNNAGATCGGAA |
| 44636_2 | TATGAGTGTGGAGTCGTTGCCATGGAAGAGGAGTCCTGATCAGNNNAGATCGGAA |
| 29039_1 | TATGAGTGTGGAGTCGTTGCCCCACTTACAACTTGATGATCCGNNNAGATCGGAAG |
| 29039_2 | TATGAGTGTGGAGTCGTTGCCCCACTTACAACTTGATGATCCGNNNAGATCGGAAG |
| 49593_1 | TATGAGTGTGGAGTCGTTGCTAGTAGCTGGGATTACAGGCATGNNNAGATCGGAA |
| 49593_2 | TATGAGTGTGGAGTCGTTGCTAGTAGCTGGGATTACAGGCATGNNNAGATCGGAA |
| 49631_1 | TATGAGTGTGGAGTCGTTGCAGATGGGTCAGGTTGCCAAATATNNNAGATCGGAA |
| 49631_2 | TATGAGTGTGGAGTCGTTGCAGATGGGTCAGGTTGCCAAATATNNNAGATCGGAA |
| 31110_1 | TATGAGTGTGGAGTCGTTGCTCCTGGAGCTGTATATTGCTTCCNNNAGATCGGAAG |
| 31110_2 | TATGAGTGTGGAGTCGTTGCTCCTGGAGCTGTATATTGCTTCCNNNAGATCGGAAG |
| 36681_1 | TATGAGTGTGGAGTCGTTGCTAGGTGTGACCAATGTTTCCCATNNNAGATCGGAAG |
| 36681_2 | TATGAGTGTGGAGTCGTTGCTAGGTGTGACCAATGTTTCCCATNNNAGATCGGAAG |
| 43034_1 | TATGAGTGTGGAGTCGTTGCATGGACACTTTGATTGTTTCCGTNNNAGATCGGAAG |
| 43034_2 | TATGAGTGTGGAGTCGTTGCATGGACACTTTGATTGTTTCCGTNNNAGATCGGAAG |
| 8_1 | TATGAGTGTGGAGTCGTTGCGCAAACACCAAGGGAAAACAACANNNAGATCGGAA |
| 8_2 | TATGAGTGTGGAGTCGTTGCGCAAACACCAAGGGAAAACAACANNNAGATCGGAA |
| 10119_1 | TATGAGTGTGGAGTCGTTGCTCAATAAGGTCCCAGAAGAGGCANNNAGATCGGAA |
| 10119_2 | TATGAGTGTGGAGTCGTTGCTCAATAAGGTCCCAGAAGAGGCANNNAGATCGGAA |
| 40324_1 | TATGAGTGTGGAGTCGTTGCGAACAGTGAAGGGAGAATGGTCTNNNAGATCGGAA |
| 40324_2 | TATGAGTGTGGAGTCGTTGCGAACAGTGAAGGGAGAATGGTCTNNNAGATCGGAA |
| 50822_1 | TATGAGTGTGGAGTCGTTGCCCTCCAAGGAAGACCAATACTGGNNNAGATCGGAAG |
| 50822_2 | TATGAGTGTGGAGTCGTTGCCCTCCAAGGAAGACCAATACTGGNNNAGATCGGAAG |
| 38104_1 | TATGAGTGTGGAGTCGTTGCAAAGAATTGGAAAGCTTGTGGGGNNNAGATCGGAA |
| 38104_2 | TATGAGTGTGGAGTCGTTGCAAAGAATTGGAAAGCTTGTGGGGNNNAGATCGGAA |
| 44116_1 | TATGAGTGTGGAGTCGTTGCCCACACCAAGTCAGAAATTCCAGNNNAGATCGGAAG |
| 44116_2 | TATGAGTGTGGAGTCGTTGCCCACACCAAGTCAGAAATTCCAGNNNAGATCGGAAG |
| 39775_1 | TATGAGTGTGGAGTCGTTGCTATATTCCTTGTAGCCCCCAGGANNNAGATCGGAAG |
| 39775_2 | TATGAGTGTGGAGTCGTTGCTATATTCCTTGTAGCCCCCAGGANNNAGATCGGAAG |
| 41308_1 | TATGAGTGTGGAGTCGTTGCTGGTGTGCAATATGGTTTTTCTGTNNNAGATCGGAA |
| 41308_2 | TATGAGTGTGGAGTCGTTGCTGGTGTGCAATATGGTTTTTCTGTNNNAGATCGGAA |
| 41696_1 | TATGAGTGTGGAGTCGTTGCCTGAGTAGCTAGGATTACAGGCGNNNAGATCGGAA |
| 41696_2 | TATGAGTGTGGAGTCGTTGCCTGAGTAGCTAGGATTACAGGCGNNNAGATCGGAA |
| 13641_1 | TATGAGTGTGGAGTCGTTGCTGAATCACTAGCCTCTAAATTATNNNAGATCGGAAG |
| 13641_2 | TATGAGTGTGGAGTCGTTGCTGAATCACTAGCCTCTAAATTATNNNAGATCGGAAG |
| 32878_1 | TATGAGTGTGGAGTCGTTGCCTCTGATTCCCAGGTTCAAGTGANNNAGATCGGAAG |
| 32878_2 | TATGAGTGTGGAGTCGTTGCCTCTGATTCCCAGGTTCAAGTGANNNAGATCGGAAG |
| 11595_1 | TATGAGTGTGGAGTCGTTGCCTGGCTTCACTGGGATATCCGANNNAGATCGGAAGA |
| 11595_2 | TATGAGTGTGGAGTCGTTGCCTGGCTTCACTGGGATATCCGANNNAGATCGGAAGA |
| 51132_1 | TATGAGTGTGGAGTCGTTGCGCAGGAGAATTGCTTGAACCTGNNNAGATCGGAAG |

| | |
|---|---|
| 51132_2 | TATGAGTGTGGAGTCGTTGCGCAGGAGAATTGCTTGAACCTGNNNAGATCGGAAG |
| 52107_1 | TATGAGTGTGGAGTCGTTGCCAATGAGAGGGAGAGAAGAGCTGNNNAGATCGGA |
| 52107_2 | TATGAGTGTGGAGTCGTTGCCAATGAGAGGGAGAGAAGAGCTGNNNAGATCGGA |
| 40506_1 | TATGAGTGTGGAGTCGTTGCCGTGGTCTTCTCTAGATGGTGAGNNNAGATCGGAA |
| 40506_2 | TATGAGTGTGGAGTCGTTGCCGTGGTCTTCTCTAGATGGTGAGNNNAGATCGGAA |
| 45084_1 | TATGAGTGTGGAGTCGTTGCATGGAAACTCTACGTCCAGTGACNNNAGATCGGAAG |
| 45084_2 | TATGAGTGTGGAGTCGTTGCATGGAAACTCTACGTCCAGTGACNNNAGATCGGAAG |
| 33162_1 | TATGAGTGTGGAGTCGTTGCGAAAAGTTTTTGGGCCGAGATGANNNAGATCGGAA |
| 33162_2 | TATGAGTGTGGAGTCGTTGCGAAAAGTTTTTGGGCCGAGATGANNNAGATCGGAA |
| 46606_1 | TATGAGTGTGGAGTCGTTGCCCTCCCAGCTATTCATCATGGTTNNNAGATCGGAAGA |
| 46606_2 | TATGAGTGTGGAGTCGTTGCCCTCCCAGCTATTCATCATGGTTNNNAGATCGGAAGA |
| 32070_1 | TATGAGTGTGGAGTCGTTGCTGCACAGCAAAAGAAATAACCGANNNAGATCGGAA |
| 32070_2 | TATGAGTGTGGAGTCGTTGCTGCACAGCAAAAGAAATAACCGANNNAGATCGGAA |
| 29312_1 | TATGAGTGTGGAGTCGTTGCTCTGCAGCTGCTTCAAAATTAGGNNNAGATCGGAAG |
| 29312_2 | TATGAGTGTGGAGTCGTTGCTCTGCAGCTGCTTCAAAATTAGGNNNAGATCGGAAG |
| 10743_1 | TATGAGTGTGGAGTCGTTGCAGATTTGCGGGGAGTTCATTTGTNNNAGATCGGAA |
| 10743_2 | TATGAGTGTGGAGTCGTTGCAGATTTGCGGGGAGTTCATTTGTNNNAGATCGGAA |
| 43057_1 | TATGAGTGTGGAGTCGTTGCCTCTGTGAGGTGATGGATACGTTNNNAGATCGGAA |
| 43057_2 | TATGAGTGTGGAGTCGTTGCCTCTGTGAGGTGATGGATACGTTNNNAGATCGGAA |
| 47982_1 | TATGAGTGTGGAGTCGTTGCTTCCTTGTTTCTGCAGATGAGGANNNAGATCGGAAG |
| 47982_2 | TATGAGTGTGGAGTCGTTGCTTCCTTGTTTCTGCAGATGAGGANNNAGATCGGAAG |
| 48045_1 | TATGAGTGTGGAGTCGTTGCCTTGGCTCCCCTTCTACACTATCNNNAGATCGGAAGA |
| 48045_2 | TATGAGTGTGGAGTCGTTGCCTTGGCTCCCCTTCTACACTATCNNNAGATCGGAAGA |
| 46595_1 | TATGAGTGTGGAGTCGTTGCCTCAAGTGATCCACCCATCTCAGNNNAGATCGGAAG |
| 46595_2 | TATGAGTGTGGAGTCGTTGCCTCAAGTGATCCACCCATCTCAGNNNAGATCGGAAG |
| 52461_1 | TATGAGTGTGGAGTCGTTGCTCTATTTTAGGCGACTCCAGCTCNNNAGATCGGAAG |
| 52461_2 | TATGAGTGTGGAGTCGTTGCTCTATTTTAGGCGACTCCAGCTCNNNAGATCGGAAG |
| 41465_1 | TATGAGTGTGGAGTCGTTGCAAAACCTAACTTTGCTCAAGCGTNNNAGATCGGAAG |
| 41465_2 | TATGAGTGTGGAGTCGTTGCAAAACCTAACTTTGCTCAAGCGTNNNAGATCGGAAG |
| 10996_1 | TATGAGTGTGGAGTCGTTGCCGAGATTTCAAGCCAGTGGATCTNNNAGATCGGAA |
| 10996_2 | TATGAGTGTGGAGTCGTTGCCGAGATTTCAAGCCAGTGGATCTNNNAGATCGGAA |
| 52450_1 | TATGAGTGTGGAGTCGTTGCGGTAGAAAAGGCTTCTTGGGAGANNNAGATCGGAA |
| 52450_2 | TATGAGTGTGGAGTCGTTGCGGTAGAAAAGGCTTCTTGGGAGANNNAGATCGGAA |
| 36256_1 | TATGAGTGTGGAGTCGTTGCAGAGAAGGCTAGGTTATGTCACANNNAGATCGGAA |
| 36256_2 | TATGAGTGTGGAGTCGTTGCAGAGAAGGCTAGGTTATGTCACANNNAGATCGGAA |
| 44429_1 | TATGAGTGTGGAGTCGTTGCATTCTGGAAACTGTTGGTCCCTTNNNAGATCGGAAG |
| 44429_2 | TATGAGTGTGGAGTCGTTGCATTCTGGAAACTGTTGGTCCCTTNNNAGATCGGAAG |
| 51672_1 | TATGAGTGTGGAGTCGTTGCAGCTGGAAGTATTTACTGGTGCANNNAGATCGGAA |
| 51672_2 | TATGAGTGTGGAGTCGTTGCAGCTGGAAGTATTTACTGGTGCANNNAGATCGGAA |
| 42865_1 | TATGAGTGTGGAGTCGTTGCTTTTTTTAGACGGAGTCTCGCTCNNNAGATCGGAAG |
| 42865_2 | TATGAGTGTGGAGTCGTTGCTTTTTTTAGACGGAGTCTCGCTCNNNAGATCGGAAG |
| 46532_1 | TATGAGTGTGGAGTCGTTGCAGGTTCAAGTGATTCTCTTGCCTNNNAGATCGGAAG |
| 46532_2 | TATGAGTGTGGAGTCGTTGCAGGTTCAAGTGATTCTCTTGCCTNNNAGATCGGAAG |

| | |
|---|---|
| 35203_1 | TATGAGTGTGGAGTCGTTGCCACCAACAAGCCTGGCTAATTTTNNNAGATCGGAAG |
| 35203_2 | TATGAGTGTGGAGTCGTTGCCACCAACAAGCCTGGCTAATTTTNNNAGATCGGAAG |
| 41366_1 | TATGAGTGTGGAGTCGTTGCCACAATAAGCTGCATTTCCTTTGCNNNAGATCGGAA |
| 41366_2 | TATGAGTGTGGAGTCGTTGCCACAATAAGCTGCATTTCCTTTGCNNNAGATCGGAA |
| 53364_1 | TATGAGTGTGGAGTCGTTGCCCCTCTCTACCTCTCTCCTTCTCNNNAGATCGGAAGAG |
| 53364_2 | TATGAGTGTGGAGTCGTTGCCCCTCTCTACCTCTCTCCTTCTCNNNAGATCGGAAGAG |
| 38951_1 | TATGAGTGTGGAGTCGTTGCGAGTGGATATCTTACAGGCCAGGNNNAGATCGGAA |
| 38951_2 | TATGAGTGTGGAGTCGTTGCGAGTGGATATCTTACAGGCCAGGNNNAGATCGGAA |
| 30650_1 | TATGAGTGTGGAGTCGTTGCTGTTCTTTTATGCTTTGGTCCTTCTNNNAGATCGGAA |
| 30650_2 | TATGAGTGTGGAGTCGTTGCTGTTCTTTTATGCTTTGGTCCTTCTNNNAGATCGGAA |
| 32348_1 | TATGAGTGTGGAGTCGTTGCAGAGAGAGAGACAGAGATGGCTCNNNAGATCGGA |
| 32348_2 | TATGAGTGTGGAGTCGTTGCAGAGAGAGAGACAGAGATGGCTCNNNAGATCGGA |
| 50951_1 | TATGAGTGTGGAGTCGTTGCAGCATGACCATTCTCCACATCTTNNNAGATCGGAAG |
| 50951_2 | TATGAGTGTGGAGTCGTTGCAGCATGACCATTCTCCACATCTTNNNAGATCGGAAG |
| 35218_1 | TATGAGTGTGGAGTCGTTGCGGCTTCAGTTCACAGTCATGATGNNNAGATCGGAAG |
| 35218_2 | TATGAGTGTGGAGTCGTTGCGGCTTCAGTTCACAGTCATGATGNNNAGATCGGAAG |
| 826_1 | TATGAGTGTGGAGTCGTTGCTCTCAACAAACCAGTCCCGAAGNNNAGATCGGAAGA |
| 826_2 | TATGAGTGTGGAGTCGTTGCTCTCAACAAACCAGTCCCGAAGNNNAGATCGGAAGA |
| 51238_1 | TATGAGTGTGGAGTCGTTGCTCGTAGCTCCAAAATGTCTTCCTNNNAGATCGGAAG |
| 51238_2 | TATGAGTGTGGAGTCGTTGCTCGTAGCTCCAAAATGTCTTCCTNNNAGATCGGAAG |
| 9992_1 | TATGAGTGTGGAGTCGTTGCCTAATTGCGAACTCCTGAGGTCCNNNAGATCGGAAG |
| 9992_2 | TATGAGTGTGGAGTCGTTGCCTAATTGCGAACTCCTGAGGTCCNNNAGATCGGAAG |
| 29616_1 | TATGAGTGTGGAGTCGTTGCTTCAACACCAGATACACCCAGACNNNAGATCGGAAG |
| 29616_2 | TATGAGTGTGGAGTCGTTGCTTCAACACCAGATACACCCAGACNNNAGATCGGAAG |
| 29533_1 | TATGAGTGTGGAGTCGTTGCGCCTTAAAACACCTAGGTTAGTTTGTGNNNAGATCG |
| 29533_2 | TATGAGTGTGGAGTCGTTGCGCCTTAAAACACCTAGGTTAGTTTGTGNNNAGATCG |
| 31371_1 | TATGAGTGTGGAGTCGTTGCATGAGGCAGTTGAACTTGTGAGANNNAGATCGGAA |
| 31371_2 | TATGAGTGTGGAGTCGTTGCATGAGGCAGTTGAACTTGTGAGANNNAGATCGGAA |
| 39695_1 | TATGAGTGTGGAGTCGTTGCAAAAGTCTCTCCATCCAGTGTCANNNAGATCGGAAG |
| 39695_2 | TATGAGTGTGGAGTCGTTGCAAAAGTCTCTCCATCCAGTGTCANNNAGATCGGAAG |
| 35450_1 | TATGAGTGTGGAGTCGTTGCTGGTAGAGATGCAGTTTCACCATNNNAGATCGGAA |
| 35450_2 | TATGAGTGTGGAGTCGTTGCTGGTAGAGATGCAGTTTCACCATNNNAGATCGGAA |
| 13076_1 | TATGAGTGTGGAGTCGTTGCTTCCTCTAGAGTATGTAATCTCTNNNAGATCGGAAG |
| 13076_2 | TATGAGTGTGGAGTCGTTGCTTCCTCTAGAGTATGTAATCTCTNNNAGATCGGAAG |
| 31789_1 | TATGAGTGTGGAGTCGTTGCAACTTCCCAACTATGAAGCCAGTNNNAGATCGGAAG |
| 31789_2 | TATGAGTGTGGAGTCGTTGCAACTTCCCAACTATGAAGCCAGTNNNAGATCGGAAG |
| 39428_1 | TATGAGTGTGGAGTCGTTGCTCCAGAAATGAAGCCAATTGACTNNNAGATCGGAAG |
| 39428_2 | TATGAGTGTGGAGTCGTTGCTCCAGAAATGAAGCCAATTGACTNNNAGATCGGAAG |
| 46396_1 | TATGAGTGTGGAGTCGTTGCGTCTCGAACTTCTGACCTTGTGANNNAGATCGGAAG |
| 46396_2 | TATGAGTGTGGAGTCGTTGCGTCTCGAACTTCTGACCTTGTGANNNAGATCGGAAG |
| 53114_1 | TATGAGTGTGGAGTCGTTGCGGTTTACACAGATCGGAAGTTGCNNNAGATCGGAA |
| 53114_2 | TATGAGTGTGGAGTCGTTGCGGTTTACACAGATCGGAAGTTGCNNNAGATCGGAA |
| 12751_1 | TATGAGTGTGGAGTCGTTGCAATGTACCACTGAAATTTTTTCTNNNAGATCGGAAG |

| | |
|---|---|
| 12751_2 | TATGAGTGTGGAGTCGTTGCAATGTACCACTGAAATTTTTCTNNNAGATCGGAAG |
| 35418_1 | TATGAGTGTGGAGTCGTTGCTCCAAGAGTGTTGTGAGCAAAACNNNAGATCGGAA |
| 35418_2 | TATGAGTGTGGAGTCGTTGCTCCAAGAGTGTTGTGAGCAAAACNNNAGATCGGAA |
| 35556_1 | TATGAGTGTGGAGTCGTTGCCATTCTCCTGCCTCAGCCTCNNNAGATCGGAAGAGCA |
| 35556_2 | TATGAGTGTGGAGTCGTTGCCATTCTCCTGCCTCAGCCTCNNNAGATCGGAAGAGCA |
| 8329_1 | TATGAGTGTGGAGTCGTTGCCTCCTGGTTCATCACAACCTCTNNNAGATCGGAAGAG |
| 8329_2 | TATGAGTGTGGAGTCGTTGCCTCCTGGTTCATCACAACCTCTNNNAGATCGGAAGAG |
| 37801_1 | TATGAGTGTGGAGTCGTTGCCATGCCTGTAGTCCCATCTACTCNNNAGATCGGAAG |
| 37801_2 | TATGAGTGTGGAGTCGTTGCCATGCCTGTAGTCCCATCTACTCNNNAGATCGGAAG |
| 35878_1 | TATGAGTGTGGAGTCGTTGCAATACAAAATTAGCCGGGCATGGNNNAGATCGGAA |
| 35878_2 | TATGAGTGTGGAGTCGTTGCAATACAAAATTAGCCGGGCATGGNNNAGATCGGAA |
| 34857_1 | TATGAGTGTGGAGTCGTTGCAAAGTGCTGGGATTACAGGCATNNNAGATCGGAAG |
| 34857_2 | TATGAGTGTGGAGTCGTTGCAAAGTGCTGGGATTACAGGCATNNNAGATCGGAAG |
| 29804_1 | TATGAGTGTGGAGTCGTTGCTCCTTCCTAATGTTTACTGTGTCANNNAGATCGGAAG |
| 29804_2 | TATGAGTGTGGAGTCGTTGCTCCTTCCTAATGTTTACTGTGTCANNNAGATCGGAAG |
| 47727_1 | TATGAGTGTGGAGTCGTTGCAGATTTCCTTCCCTTGGATGCTTNNNAGATCGGAAG |
| 47727_2 | TATGAGTGTGGAGTCGTTGCAGATTTCCTTCCCTTGGATGCTTNNNAGATCGGAAG |
| 39484_1 | TATGAGTGTGGAGTCGTTGCGATTTCGGCTCACTGCAAACTTCNNNAGATCGGAAG |
| 39484_2 | TATGAGTGTGGAGTCGTTGCGATTTCGGCTCACTGCAAACTTCNNNAGATCGGAAG |
| 51595_1 | TATGAGTGTGGAGTCGTTGCCGAAACGTGAAGTCGTCTGTTACNNNAGATCGGAA |
| 51595_2 | TATGAGTGTGGAGTCGTTGCCGAAACGTGAAGTCGTCTGTTACNNNAGATCGGAA |
| 34201_1 | TATGAGTGTGGAGTCGTTGCAGGGCACCATAATTTTGTTCATGTNNNAGATCGGAA |
| 34201_2 | TATGAGTGTGGAGTCGTTGCAGGGCACCATAATTTTGTTCATGTNNNAGATCGGAA |
| 37931_1 | TATGAGTGTGGAGTCGTTGCCTCATGCCTGTAATCCTAGCACTNNNAGATCGGAAG |
| 37931_2 | TATGAGTGTGGAGTCGTTGCCTCATGCCTGTAATCCTAGCACTNNNAGATCGGAAG |
| 36199_1 | TATGAGTGTGGAGTCGTTGCTGAGTAAGAAGTCCCAAAGACCGNNNAGATCGGAA |
| 36199_2 | TATGAGTGTGGAGTCGTTGCTGAGTAAGAAGTCCCAAAGACCGNNNAGATCGGAA |
| 52901_1 | TATGAGTGTGGAGTCGTTGCTCAGGGTATGTTTGACTCAAGGGNNNAGATCGGAA |
| 52901_2 | TATGAGTGTGGAGTCGTTGCTCAGGGTATGTTTGACTCAAGGGNNNAGATCGGAA |
| 53259_1 | TATGAGTGTGGAGTCGTTGCCTTTGGTCACATCGAACACCTTGNNNAGATCGGAAG |
| 53259_2 | TATGAGTGTGGAGTCGTTGCCTTTGGTCACATCGAACACCTTGNNNAGATCGGAAG |
| 46898_1 | TATGAGTGTGGAGTCGTTGCCTAAGCACCCAGCCAGAAATAGANNNAGATCGGAA |
| 46898_2 | TATGAGTGTGGAGTCGTTGCCTAAGCACCCAGCCAGAAATAGANNNAGATCGGAA |
| 44408_1 | TATGAGTGTGGAGTCGTTGCAATCAGCCAGAAACAAAAGAGGCNNNAGATCGGAA |
| 44408_2 | TATGAGTGTGGAGTCGTTGCAATCAGCCAGAAACAAAAGAGGCNNNAGATCGGAA |
| 31580_1 | TATGAGTGTGGAGTCGTTGCCATCTAACCAAGTGGGCAAAAGGNNNAGATCGGAA |
| 31580_2 | TATGAGTGTGGAGTCGTTGCCATCTAACCAAGTGGGCAAAAGGNNNAGATCGGAA |
| 39248_1 | TATGAGTGTGGAGTCGTTGCCAGGTTCAAGCGATGCTCTTGNNNAGATCGGAAGA |
| 39248_2 | TATGAGTGTGGAGTCGTTGCCAGGTTCAAGCGATGCTCTTGNNNAGATCGGAAGA |
| 45317_1 | TATGAGTGTGGAGTCGTTGCTCTAATGGCTAATAAACTTGGAGCANNNAGATCGGA |
| 45317_2 | TATGAGTGTGGAGTCGTTGCTCTAATGGCTAATAAACTTGGAGCANNNAGATCGGA |
| 41849_1 | TATGAGTGTGGAGTCGTTGCCTCCAAACTCTGAGTAGCACTGTNNNAGATCGGAAG |
| 41849_2 | TATGAGTGTGGAGTCGTTGCCTCCAAACTCTGAGTAGCACTGTNNNAGATCGGAAG |

| | |
|---|---|
| 36912_1 | TATGAGTGTGGAGTCGTTGCGGAAACATACCAAGACCCTGTCTNNNAGATCGGAAG |
| 36912_2 | TATGAGTGTGGAGTCGTTGCGGAAACATACCAAGACCCTGTCTNNNAGATCGGAAG |
| 37320_1 | TATGAGTGTGGAGTCGTTGCAACACAAGGCCTGAAAATCATGGNNNAGATCGGAA |
| 37320_2 | TATGAGTGTGGAGTCGTTGCAACACAAGGCCTGAAAATCATGGNNNAGATCGGAA |
| 44346_1 | TATGAGTGTGGAGTCGTTGCGTTGGGAGGGAGGGAGGTAGNNNAGATCGGAAG |
| 44346_2 | TATGAGTGTGGAGTCGTTGCGTTGGGAGGGAGGGAGGTAGNNNAGATCGGAAG |
| 36872_1 | TATGAGTGTGGAGTCGTTGCGTAATCCCAAGACTTTGTGAGGCNNNAGATCGGAAG |
| 36872_2 | TATGAGTGTGGAGTCGTTGCGTAATCCCAAGACTTTGTGAGGCNNNAGATCGGAAG |
| 29727_1 | TATGAGTGTGGAGTCGTTGCTGGAAGGAACTTAGGACTTTCAGNNNAGATCGGAA |
| 29727_2 | TATGAGTGTGGAGTCGTTGCTGGAAGGAACTTAGGACTTTCAGNNNAGATCGGAA |
| 35711_1 | TATGAGTGTGGAGTCGTTGCGGTTCAAGCAATTCTCCTGTCTCNNNAGATCGGAAG |
| 35711_2 | TATGAGTGTGGAGTCGTTGCGGTTCAAGCAATTCTCCTGTCTCNNNAGATCGGAAG |
| 43990_1 | TATGAGTGTGGAGTCGTTGCAGACATAACCAAGACCAGGCAATNNNAGATCGGAAG |
| 43990_2 | TATGAGTGTGGAGTCGTTGCAGACATAACCAAGACCAGGCAATNNNAGATCGGAAG |
| 39970_1 | TATGAGTGTGGAGTCGTTGCTAGGACAGAATTGAGGCTTCAGCNNNAGATCGGAA |
| 39970_2 | TATGAGTGTGGAGTCGTTGCTAGGACAGAATTGAGGCTTCAGCNNNAGATCGGAA |
| 32868_1 | TATGAGTGTGGAGTCGTTGCCCTCCCAAGTAGCTGGAATTACANNNAGATCGGAAG |
| 32868_2 | TATGAGTGTGGAGTCGTTGCCCTCCCAAGTAGCTGGAATTACANNNAGATCGGAAG |
| 30253_1 | TATGAGTGTGGAGTCGTTGCGCCATCCCCTGTCAGAGATTTATNNNAGATCGGAAG |
| 30253_2 | TATGAGTGTGGAGTCGTTGCGCCATCCCCTGTCAGAGATTTATNNNAGATCGGAAG |
| 29486_1 | TATGAGTGTGGAGTCGTTGCTCCATGTCACTCAGGTAAGAGCCATNNNAGATCGGA |
| 29486_2 | TATGAGTGTGGAGTCGTTGCTCCATGTCACTCAGGTAAGAGCCATNNNAGATCGGA |
| 37807_1 | TATGAGTGTGGAGTCGTTGCCATCATGGCAAAACCCAGTTTCTNNNAGATCGGAAG |
| 37807_2 | TATGAGTGTGGAGTCGTTGCCATCATGGCAAAACCCAGTTTCTNNNAGATCGGAAG |
| 39321_1 | TATGAGTGTGGAGTCGTTGCTGGGCATGAAAGATCAGACTTCANNNAGATCGGAA |
| 39321_2 | TATGAGTGTGGAGTCGTTGCTGGGCATGAAAGATCAGACTTCANNNAGATCGGAA |
| 12950_1 | TATGAGTGTGGAGTCGTTGCCTAACTCCAGATAACTCCCTNNNAGATCGGAAGAGC |
| 12950_2 | TATGAGTGTGGAGTCGTTGCCTAACTCCAGATAACTCCCTNNNAGATCGGAAGAGC |
| 12816_1 | TATGAGTGTGGAGTCGTTGCGAACTCAAAGACAATATTCTTGTNNNAGATCGGAAG |
| 12816_2 | TATGAGTGTGGAGTCGTTGCGAACTCAAAGACAATATTCTTGTNNNAGATCGGAAG |
| 42217_1 | TATGAGTGTGGAGTCGTTGCATGAAAAGGGGAGACAGAAAGCTNNNAGATCGGA |
| 42217_2 | TATGAGTGTGGAGTCGTTGCATGAAAAGGGGAGACAGAAAGCTNNNAGATCGGA |
| 2955_1 | TATGAGTGTGGAGTCGTTGCGCATACCTTCTCTGGAAAGGGTTNNNAGATCGGAAG |
| 2955_2 | TATGAGTGTGGAGTCGTTGCGCATACCTTCTCTGGAAAGGGTTNNNAGATCGGAAG |
| 48041_1 | TATGAGTGTGGAGTCGTTGCATGTCATGGAGGACTGTAAGTCANNNAGATCGGAA |
| 48041_2 | TATGAGTGTGGAGTCGTTGCATGTCATGGAGGACTGTAAGTCANNNAGATCGGAA |
| 50006_1 | TATGAGTGTGGAGTCGTTGCTTTCTTAGATGTGGCTGTGGACANNNAGATCGGAAG |
| 50006_2 | TATGAGTGTGGAGTCGTTGCTTTCTTAGATGTGGCTGTGGACANNNAGATCGGAAG |
| 13638_1 | TATGAGTGTGGAGTCGTTGCTGTATGTTATTTAACAACAAGCANNNAGATCGGAAG |
| 13638_2 | TATGAGTGTGGAGTCGTTGCTGTATGTTATTTAACAACAAGCANNNAGATCGGAAG |
| 42323_1 | TATGAGTGTGGAGTCGTTGCCACAGAGTCACACATCCTTGAGANNNAGATCGGAAG |
| 42323_2 | TATGAGTGTGGAGTCGTTGCCACAGAGTCACACATCCTTGAGANNNAGATCGGAAG |
| 35156_1 | TATGAGTGTGGAGTCGTTGCTTTCTACTACACGTCCAAGCTGANNNAGATCGGAAG |

| | |
|---|---|
| 35156_2 | TATGAGTGTGGAGTCGTTGCTTTCTACTACACGTCCAAGCTGANNNAGATCGGAAG |
| 5797_1 | TATGAGTGTGGAGTCGTTGCTGGGGCAAAGGACAGAGAGATTANNNAGATCGGA |
| 5797_2 | TATGAGTGTGGAGTCGTTGCTGGGGCAAAGGACAGAGAGATTANNNAGATCGGA |
| 28840_1 | TATGAGTGTGGAGTCGTTGCAAATCAATAAATTCGGCTGGGCGNNNAGATCGGAA |
| 28840_2 | TATGAGTGTGGAGTCGTTGCAAATCAATAAATTCGGCTGGGCGNNNAGATCGGAA |
| 49840_1 | TATGAGTGTGGAGTCGTTGCTTCTGAAGAATAGACCATGGGCCNNNAGATCGGAA |
| 49840_2 | TATGAGTGTGGAGTCGTTGCTTCTGAAGAATAGACCATGGGCCNNNAGATCGGAA |
| 34019_1 | TATGAGTGTGGAGTCGTTGCTTTACGCCATTCTCCAGCCTCNNNAGATCGGAAGAG |
| 34019_2 | TATGAGTGTGGAGTCGTTGCTTTACGCCATTCTCCAGCCTCNNNAGATCGGAAGAG |
| 29325_1 | TATGAGTGTGGAGTCGTTGCAGCTGTATGTGGTTACATGTGGTNNNAGATCGGAA |
| 29325_2 | TATGAGTGTGGAGTCGTTGCAGCTGTATGTGGTTACATGTGGTNNNAGATCGGAA |
| 50581_1 | TATGAGTGTGGAGTCGTTGCAGGCCAAATATGCAGGTCAAAACNNNAGATCGGAA |
| 50581_2 | TATGAGTGTGGAGTCGTTGCAGGCCAAATATGCAGGTCAAAACNNNAGATCGGAA |
| 44135_1 | TATGAGTGTGGAGTCGTTGCACACTACCACGCAGGAAAGCNNNAGATCGGAAGAG |
| 44135_2 | TATGAGTGTGGAGTCGTTGCACACTACCACGCAGGAAAGCNNNAGATCGGAAGAG |
| 31312_1 | TATGAGTGTGGAGTCGTTGCGAGAGAGAGAACCAAAGCCTGAANNNAGATCGGA |
| 31312_2 | TATGAGTGTGGAGTCGTTGCGAGAGAGAGAACCAAAGCCTGAANNNAGATCGGA |
| 41401_1 | TATGAGTGTGGAGTCGTTGCTCTCAAAGCGGGAAGAAATGAGANNNAGATCGGAA |
| 41401_2 | TATGAGTGTGGAGTCGTTGCTCTCAAAGCGGGAAGAAATGAGANNNAGATCGGAA |
| 33935_1 | TATGAGTGTGGAGTCGTTGCAAGTGTTGGGATTACAGGCTTGANNNAGATCGGAA |
| 33935_2 | TATGAGTGTGGAGTCGTTGCAAGTGTTGGGATTACAGGCTTGANNNAGATCGGAA |
| 52298_1 | TATGAGTGTGGAGTCGTTGCAATATGGGTGTCCAGATTCTGCCNNNAGATCGGAAG |
| 52298_2 | TATGAGTGTGGAGTCGTTGCAATATGGGTGTCCAGATTCTGCCNNNAGATCGGAAG |
| 32204_1 | TATGAGTGTGGAGTCGTTGCTAAAGTCGGTCCTCATTCAGGTCNNNAGATCGGAAG |
| 32204_2 | TATGAGTGTGGAGTCGTTGCTAAAGTCGGTCCTCATTCAGGTCNNNAGATCGGAAG |
| 51811_1 | TATGAGTGTGGAGTCGTTGCTTTCCAGTGGCTGCTACACAATANNNAGATCGGAAG |
| 51811_2 | TATGAGTGTGGAGTCGTTGCTTTCCAGTGGCTGCTACACAATANNNAGATCGGAAG |
| 39161_1 | TATGAGTGTGGAGTCGTTGCAGAAAAGAAAAAGCCACATGCCANNNAGATCGGAA |
| 39161_2 | TATGAGTGTGGAGTCGTTGCAGAAAAGAAAAAGCCACATGCCANNNAGATCGGAA |
| 42235_1 | TATGAGTGTGGAGTCGTTGCAGGGTTTCATCTCTCAATGTCCTNNNAGATCGGAAG |
| 42235_2 | TATGAGTGTGGAGTCGTTGCAGGGTTTCATCTCTCAATGTCCTNNNAGATCGGAAG |
| 42342_1 | TATGAGTGTGGAGTCGTTGCAGTGTTCTCGCAATTGAATGCATNNNAGATCGGAAG |
| 42342_2 | TATGAGTGTGGAGTCGTTGCAGTGTTCTCGCAATTGAATGCATNNNAGATCGGAAG |
| 32027_1 | TATGAGTGTGGAGTCGTTGCCTCCCAGGTTCAGGCAATTCTCNNNAGATCGGAAGA |
| 32027_2 | TATGAGTGTGGAGTCGTTGCCTCCCAGGTTCAGGCAATTCTCNNNAGATCGGAAGA |
| 38767_1 | TATGAGTGTGGAGTCGTTGCTCTAAATATTGACTGGGGCTGGGNNNAGATCGGAA |
| 38767_2 | TATGAGTGTGGAGTCGTTGCTCTAAATATTGACTGGGGCTGGGNNNAGATCGGAA |
| 39673_1 | TATGAGTGTGGAGTCGTTGCATGATTGCACCACTGTACTCCAGNNNAGATCGGAAG |
| 39673_2 | TATGAGTGTGGAGTCGTTGCATGATTGCACCACTGTACTCCAGNNNAGATCGGAAG |
| 43258_1 | TATGAGTGTGGAGTCGTTGCGTGTGTGTGTGTGTGTGTGTATANNNAGATCGGAA |
| 43258_2 | TATGAGTGTGGAGTCGTTGCGTGTGTGTGTGTGTGTGTGTATANNNAGATCGGAA |
| 37917_1 | TATGAGTGTGGAGTCGTTGCCAATGGGAATGCACAGTTTCCTTNNNAGATCGGAAG |
| 37917_2 | TATGAGTGTGGAGTCGTTGCCAATGGGAATGCACAGTTTCCTTNNNAGATCGGAAG |

| | |
|---|---|
| 50306_1 | TATGAGTGTGGAGTCGTTGCTGAGGTAGCAAACTGCCTAGAAGNNNAGATCGGAAG |
| 50306_2 | TATGAGTGTGGAGTCGTTGCTGAGGTAGCAAACTGCCTAGAAGNNNAGATCGGAAG |
| 52133_1 | TATGAGTGTGGAGTCGTTGCATGGTGCTCATTTTTGCTTGGAANNNAGATCGGAAG |
| 52133_2 | TATGAGTGTGGAGTCGTTGCATGGTGCTCATTTTTGCTTGGAANNNAGATCGGAAG |
| 45293_1 | TATGAGTGTGGAGTCGTTGCTCCTCCCTCTTTCCCTCTCGNNNAGATCGGAAGAGCAC |
| 45293_2 | TATGAGTGTGGAGTCGTTGCTCCTCCCTCTTTCCCTCTCGNNNAGATCGGAAGAGCAC |
| 51886_1 | TATGAGTGTGGAGTCGTTGCCTCCCTTAGCAGCTAGAGAGTTGNNNAGATCGGAAG |
| 51886_2 | TATGAGTGTGGAGTCGTTGCCTCCCTTAGCAGCTAGAGAGTTGNNNAGATCGGAAG |
| 7783_1 | TATGAGTGTGGAGTCGTTGCGTTACTCTCCAGATCTGCCTTTCCNNNAGATCGGAAG |
| 7783_2 | TATGAGTGTGGAGTCGTTGCGTTACTCTCCAGATCTGCCTTTCCNNNAGATCGGAAG |
| 41367_1 | TATGAGTGTGGAGTCGTTGCCAGTGAGCTATGATCATGCCACTNNNAGATCGGAAG |
| 41367_2 | TATGAGTGTGGAGTCGTTGCCAGTGAGCTATGATCATGCCACTNNNAGATCGGAAG |
| 49851_1 | TATGAGTGTGGAGTCGTTGCGCATGAGAAGGCTAGGTTTCTCTNNNAGATCGGAAG |
| 49851_2 | TATGAGTGTGGAGTCGTTGCGCATGAGAAGGCTAGGTTTCTCTNNNAGATCGGAAG |
| 12917_1 | TATGAGTGTGGAGTCGTTGCGATGAGGAGGAAGAGATGATNNNAGATCGGAAGA |
| 12917_2 | TATGAGTGTGGAGTCGTTGCGATGAGGAGGAAGAGATGATNNNAGATCGGAAGA |
| 42852_1 | TATGAGTGTGGAGTCGTTGCATGGGCATTTGGACTGGTTCTATNNNAGATCGGAAG |
| 42852_2 | TATGAGTGTGGAGTCGTTGCATGGGCATTTGGACTGGTTCTATNNNAGATCGGAAG |
| 29054_1 | TATGAGTGTGGAGTCGTTGCAGATCAAATTGGAGTGACTGGGGNNNAGATCGGAA |
| 29054_2 | TATGAGTGTGGAGTCGTTGCAGATCAAATTGGAGTGACTGGGGNNNAGATCGGAA |
| 42428_1 | TATGAGTGTGGAGTCGTTGCGAGGGAGGGTTAGGGAACAAAATNNNAGATCGGA |
| 42428_2 | TATGAGTGTGGAGTCGTTGCGAGGGAGGGTTAGGGAACAAAATNNNAGATCGGA |
| 36588_1 | TATGAGTGTGGAGTCGTTGCGAGGCTTTATGACAAAAGGAGGCNNNAGATCGGAA |
| 36588_2 | TATGAGTGTGGAGTCGTTGCGAGGCTTTATGACAAAAGGAGGCNNNAGATCGGAA |
| 30587_1 | TATGAGTGTGGAGTCGTTGCAGAATTTGTCAGCATGTTCCCTCTNNNAGATCGGAAG |
| 30587_2 | TATGAGTGTGGAGTCGTTGCAGAATTTGTCAGCATGTTCCCTCTNNNAGATCGGAAG |
| 34768_1 | TATGAGTGTGGAGTCGTTGCCGTGCCTGGCTAATTTGTGTATTNNNAGATCGGAAG |
| 34768_2 | TATGAGTGTGGAGTCGTTGCCGTGCCTGGCTAATTTGTGTATTNNNAGATCGGAAG |
| 43756_1 | TATGAGTGTGGAGTCGTTGCCTCGGAGAAGTAGAGCGCACNNNAGATCGGAAGAG |
| 43756_2 | TATGAGTGTGGAGTCGTTGCCTCGGAGAAGTAGAGCGCACNNNAGATCGGAAGAG |
| 37427_1 | TATGAGTGTGGAGTCGTTGCGGAGAATGGTGACTCGAGTGACNNNAGATCGGAAG |
| 37427_2 | TATGAGTGTGGAGTCGTTGCGGAGAATGGTGACTCGAGTGACNNNAGATCGGAAG |
| 38083_1 | TATGAGTGTGGAGTCGTTGCGTGAGCCAAGATCTCAGTCTCAANNNAGATCGGAAG |
| 38083_2 | TATGAGTGTGGAGTCGTTGCGTGAGCCAAGATCTCAGTCTCAANNNAGATCGGAAG |
| 47666_1 | TATGAGTGTGGAGTCGTTGCAATTCTTGTGCCTCCCGAGTAGNNNAGATCGGAAGA |
| 47666_2 | TATGAGTGTGGAGTCGTTGCAATTCTTGTGCCTCCCGAGTAGNNNAGATCGGAAGA |
| 35294_1 | TATGAGTGTGGAGTCGTTGCTCTTGGGAGGTTGTATGTTTCCANNNAGATCGGAAG |
| 35294_2 | TATGAGTGTGGAGTCGTTGCTCTTGGGAGGTTGTATGTTTCCANNNAGATCGGAAG |
| 44348_1 | TATGAGTGTGGAGTCGTTGCGAGATGAGTGTGTCTGGTTTGGANNNAGATCGGAA |
| 44348_2 | TATGAGTGTGGAGTCGTTGCGAGATGAGTGTGTCTGGTTTGGANNNAGATCGGAA |
| 36032_1 | TATGAGTGTGGAGTCGTTGCGAAAGAAAGGAAAACGCTGTCCANNNAGATCGGAA |
| 36032_2 | TATGAGTGTGGAGTCGTTGCGAAAGAAAGGAAAACGCTGTCCANNNAGATCGGAA |
| 40338_1 | TATGAGTGTGGAGTCGTTGCCTGAGGCAGGAGAATTGCTTGAANNNAGATCGGAA |

| | |
|---|---|
| 40338_2 | TATGAGTGTGGAGTCGTTGCCTGAGGCAGGAGAATTGCTTGAANNNAGATCGGAA |
| 34625_1 | TATGAGTGTGGAGTCGTTGCAAGTGGGGACAGCAAAAAATGTCNNNAGATCGGAA |
| 34625_2 | TATGAGTGTGGAGTCGTTGCAAGTGGGGACAGCAAAAAATGTCNNNAGATCGGAA |
| 47688_1 | TATGAGTGTGGAGTCGTTGCTTCAAGTGTTTCTCCTGCCTCAGNNNAGATCGGAAG |
| 47688_2 | TATGAGTGTGGAGTCGTTGCTTCAAGTGTTTCTCCTGCCTCAGNNNAGATCGGAAG |
| 52505_1 | TATGAGTGTGGAGTCGTTGCGACAGACTGCTGCTTCTAGTGATNNNAGATCGGAAG |
| 52505_2 | TATGAGTGTGGAGTCGTTGCGACAGACTGCTGCTTCTAGTGATNNNAGATCGGAAG |
| 43945_1 | TATGAGTGTGGAGTCGTTGCATTCATCACAGACCACAACCAGTNNNAGATCGGAAG |
| 43945_2 | TATGAGTGTGGAGTCGTTGCATTCATCACAGACCACAACCAGTNNNAGATCGGAAG |
| 43386_1 | TATGAGTGTGGAGTCGTTGCAACACGCCCAGCTAATTTTTGTNNNAGATCGGAAG |
| 43386_2 | TATGAGTGTGGAGTCGTTGCAACACGCCCAGCTAATTTTTGTNNNAGATCGGAAG |
| 46686_1 | TATGAGTGTGGAGTCGTTGCGAGTAGCTGGGATTACAGGCATGNNNAGATCGGAA |
| 46686_2 | TATGAGTGTGGAGTCGTTGCGAGTAGCTGGGATTACAGGCATGNNNAGATCGGAA |
| 49933_1 | TATGAGTGTGGAGTCGTTGCGGGATCTCAGGGAAAGCAGATTANNNAGATCGGAA |
| 49933_2 | TATGAGTGTGGAGTCGTTGCGGGATCTCAGGGAAAGCAGATTANNNAGATCGGAA |
| 44709_1 | TATGAGTGTGGAGTCGTTGCGTCTTTGTGTCTACCTACCCAGGNNNAGATCGGAAG |
| 44709_2 | TATGAGTGTGGAGTCGTTGCGTCTTTGTGTCTACCTACCCAGGNNNAGATCGGAAG |
| 31162_1 | TATGAGTGTGGAGTCGTTGCTGGGAAAGTTCTGCCTTCAACTANNNAGATCGGAAG |
| 31162_2 | TATGAGTGTGGAGTCGTTGCTGGGAAAGTTCTGCCTTCAACTANNNAGATCGGAAG |
| 34448_1 | TATGAGTGTGGAGTCGTTGCCTCTCTCTAGCTATCCCCCTCTCNNNAGATCGGAAGA |
| 34448_2 | TATGAGTGTGGAGTCGTTGCCTCTCTCTAGCTATCCCCCTCTCNNNAGATCGGAAGA |
| 44315_1 | TATGAGTGTGGAGTCGTTGCGGCTAGACTATGCAGACCCTTTTNNNAGATCGGAAG |
| 44315_2 | TATGAGTGTGGAGTCGTTGCGGCTAGACTATGCAGACCCTTTTNNNAGATCGGAAG |
| 40151_1 | TATGAGTGTGGAGTCGTTGCTGTTTGATGAAATCCAGTTGGCANNNAGATCGGAAG |
| 40151_2 | TATGAGTGTGGAGTCGTTGCTGTTTGATGAAATCCAGTTGGCANNNAGATCGGAAG |
| 37115_1 | TATGAGTGTGGAGTCGTTGCAATGTCTGGTCACAATGATTGCANNNAGATCGGAAG |
| 37115_2 | TATGAGTGTGGAGTCGTTGCAATGTCTGGTCACAATGATTGCANNNAGATCGGAAG |
| 38242_1 | TATGAGTGTGGAGTCGTTGCAACTCTAAGCAACATCACGTCCTNNNAGATCGGAAG |
| 38242_2 | TATGAGTGTGGAGTCGTTGCAACTCTAAGCAACATCACGTCCTNNNAGATCGGAAG |
| 8500_1 | TATGAGTGTGGAGTCGTTGCATGGGCAAAAAGAGGAACTGAANNNAGATCGGAA |
| 8500_2 | TATGAGTGTGGAGTCGTTGCATGGGCAAAAAGAGGAACTGAANNNAGATCGGAA |
| 37170_1 | TATGAGTGTGGAGTCGTTGCAAACAAGTAGGGGAGAGTGTTCCNNNAGATCGGAA |
| 37170_2 | TATGAGTGTGGAGTCGTTGCAAACAAGTAGGGGAGAGTGTTCCNNNAGATCGGAA |
| 13491_1 | TATGAGTGTGGAGTCGTTGCTGATTTCATGGTCTGACTATAAGNNNAGATCGGAAG |
| 13491_2 | TATGAGTGTGGAGTCGTTGCTGATTTCATGGTCTGACTATAAGNNNAGATCGGAAG |
| 45643_1 | TATGAGTGTGGAGTCGTTGCCCTCTCAGGTTCAAGGGATTCTCNNNAGATCGGAAG |
| 45643_2 | TATGAGTGTGGAGTCGTTGCCCTCTCAGGTTCAAGGGATTCTCNNNAGATCGGAAG |
| 53226_1 | TATGAGTGTGGAGTCGTTGCTTTGGCCTCAAACTTCAGGTGANNNAGATCGGAAGA |
| 53226_2 | TATGAGTGTGGAGTCGTTGCTTTGGCCTCAAACTTCAGGTGANNNAGATCGGAAGA |
| 40907_1 | TATGAGTGTGGAGTCGTTGCGGCAGGTGTAGAGGAGAGATAGANNNAGATCGGA |
| 40907_2 | TATGAGTGTGGAGTCGTTGCGGCAGGTGTAGAGGAGAGATAGANNNAGATCGGA |
| 8060_1 | TATGAGTGTGGAGTCGTTGCACGGTCTTGCTCTGTCGCCATANNNAGATCGGAAGA |
| 8060_2 | TATGAGTGTGGAGTCGTTGCACGGTCTTGCTCTGTCGCCATANNNAGATCGGAAGA |

| | |
|---|---|
| 2311_1 | TATGAGTGTGGAGTCGTTGCGCAAGCAATTTGGTAAGTTTGGACNNNAGATCGGAA |
| 2311_2 | TATGAGTGTGGAGTCGTTGCGCAAGCAATTTGGTAAGTTTGGACNNNAGATCGGAA |
| 43984_1 | TATGAGTGTGGAGTCGTTGCAGGCTGAGGGGTTTTCATAGATGNNNAGATCGGAA |
| 43984_2 | TATGAGTGTGGAGTCGTTGCAGGCTGAGGGGTTTTCATAGATGNNNAGATCGGAA |
| 39640_1 | TATGAGTGTGGAGTCGTTGCAATGCCAGCATCACTTATCTTGCNNNAGATCGGAAGA |
| 39640_2 | TATGAGTGTGGAGTCGTTGCAATGCCAGCATCACTTATCTTGCNNNAGATCGGAAGA |
| 47082_1 | TATGAGTGTGGAGTCGTTGCTCTCATGACTCAACCTCCCTAGTNNNAGATCGGAAGA |
| 47082_2 | TATGAGTGTGGAGTCGTTGCTCTCATGACTCAACCTCCCTAGTNNNAGATCGGAAGA |
| 33250_1 | TATGAGTGTGGAGTCGTTGCCTCCCGAGTAGCTGAGACTACAGNNNAGATCGGAAG |
| 33250_2 | TATGAGTGTGGAGTCGTTGCCTCCCGAGTAGCTGAGACTACAGNNNAGATCGGAAG |
| 12503_1 | TATGAGTGTGGAGTCGTTGCCAGTAAGTAACGATGCTCTTTANNNAGATCGGAAGA |
| 12503_2 | TATGAGTGTGGAGTCGTTGCCAGTAAGTAACGATGCTCTTTANNNAGATCGGAAGA |
| 39613_1 | TATGAGTGTGGAGTCGTTGCCCTGTGATGTGCTAGGAACCATANNNAGATCGGAAG |
| 39613_2 | TATGAGTGTGGAGTCGTTGCCCTGTGATGTGCTAGGAACCATANNNAGATCGGAAG |
| 33814_1 | TATGAGTGTGGAGTCGTTGCTGGACACAGTTACCCATACCTTCNNNAGATCGGAAGA |
| 33814_2 | TATGAGTGTGGAGTCGTTGCTGGACACAGTTACCCATACCTTCNNNAGATCGGAAGA |
| 30540_1 | TATGAGTGTGGAGTCGTTGCCCGTGCTTAGAGCAACAACAATTNNNAGATCGGAAG |
| 30540_2 | TATGAGTGTGGAGTCGTTGCCCGTGCTTAGAGCAACAACAATTNNNAGATCGGAAG |
| 47445_1 | TATGAGTGTGGAGTCGTTGCGCAGCTATCGTAAAATGGGTTGANNNAGATCGGAA[illegible] |
| 47445_2 | TATGAGTGTGGAGTCGTTGCGCAGCTATCGTAAAATGGGTTGANNNAGATCGGAA[illegible] |
| 43177_1 | TATGAGTGTGGAGTCGTTGCCATTCTCCTGCCTCAGCCTCNNNAGATCGGAAGAGCA |
| 43177_2 | TATGAGTGTGGAGTCGTTGCCATTCTCCTGCCTCAGCCTCNNNAGATCGGAAGAGCA |
| 29227_1 | TATGAGTGTGGAGTCGTTGCGAGGGGATAAGGGCTGATATCAGNNNAGATCGGAA |
| 29227_2 | TATGAGTGTGGAGTCGTTGCGAGGGGATAAGGGCTGATATCAGNNNAGATCGGAA |
| 44223_1 | TATGAGTGTGGAGTCGTTGCCCAAGTAGGAGAAGCAAGTAGGGNNNAGATCGGAA |
| 44223_2 | TATGAGTGTGGAGTCGTTGCCCAAGTAGGAGAAGCAAGTAGGGNNNAGATCGGAA |
| 50855_1 | TATGAGTGTGGAGTCGTTGCCGGTTGCATTCACACTGAACATTNNNAGATCGGAAGA |
| 50855_2 | TATGAGTGTGGAGTCGTTGCCGGTTGCATTCACACTGAACATTNNNAGATCGGAAGA |
| 51703_1 | TATGAGTGTGGAGTCGTTGCAGAGGTGGCTTCAATCATCATGTNNNAGATCGGAAG |
| 51703_2 | TATGAGTGTGGAGTCGTTGCAGAGGTGGCTTCAATCATCATGTNNNAGATCGGAAG |
| 50220_1 | TATGAGTGTGGAGTCGTTGCATGAGTGCAAGTGGGTTTTAGGANNNAGATCGGAA |
| 50220_2 | TATGAGTGTGGAGTCGTTGCATGAGTGCAAGTGGGTTTTAGGANNNAGATCGGAA |
| 33547_1 | TATGAGTGTGGAGTCGTTGCGCTCAAGTGATCCTTCCATCTCANNNAGATCGGAAGA |
| 33547_2 | TATGAGTGTGGAGTCGTTGCGCTCAAGTGATCCTTCCATCTCANNNAGATCGGAAGA |
| 34884_1 | TATGAGTGTGGAGTCGTTGCACCACGTCCGGCTAATTTTTTTTNNNAGATCGGAAGA |
| 34884_2 | TATGAGTGTGGAGTCGTTGCACCACGTCCGGCTAATTTTTTTTNNNAGATCGGAAGA |
| 28925_1 | TATGAGTGTGGAGTCGTTGCCAGTCAAGGCATCACAAGTCATTNNNAGATCGGAAG |
| 28925_2 | TATGAGTGTGGAGTCGTTGCCAGTCAAGGCATCACAAGTCATTNNNAGATCGGAAG |
| 49571_1 | TATGAGTGTGGAGTCGTTGCTGTGCCAATATCAGCTAGGTGTTNNNAGATCGGAAG |
| 49571_2 | TATGAGTGTGGAGTCGTTGCTGTGCCAATATCAGCTAGGTGTTNNNAGATCGGAAG |
| 41602_1 | TATGAGTGTGGAGTCGTTGCTTCCTACGTCAGTGTTTTAGGGGNNNAGATCGGAAG |
| 41602_2 | TATGAGTGTGGAGTCGTTGCTTCCTACGTCAGTGTTTTAGGGGNNNAGATCGGAAG |
| 34738_1 | TATGAGTGTGGAGTCGTTGCAGTGCAGTGGTGTGATCTCGNNNAGATCGGAAGAG |

| | |
|---|---|
| 34738_2 | TATGAGTGTGGAGTCGTTGCAGTGCAGTGGTGTGATCTCGNNNAGATCGGAAGAG |
| 30671_1 | TATGAGTGTGGAGTCGTTGCTAATACTGGCTGAGTGCAAGGGTNNNAGATCGGAA |
| 30671_2 | TATGAGTGTGGAGTCGTTGCTAATACTGGCTGAGTGCAAGGGTNNNAGATCGGAA |
| 10035_1 | TATGAGTGTGGAGTCGTTGCGCGGGAGGAGATGCTGTATATTTNNNAGATCGGAA |
| 10035_2 | TATGAGTGTGGAGTCGTTGCGCGGGAGGAGATGCTGTATATTTNNNAGATCGGAA |
| 49010_1 | TATGAGTGTGGAGTCGTTGCGCATGATCTTGGCTCACTACAACNNNAGATCGGAAG |
| 49010_2 | TATGAGTGTGGAGTCGTTGCGCATGATCTTGGCTCACTACAACNNNAGATCGGAAG |
| 37480_1 | TATGAGTGTGGAGTCGTTGCTTCAGTCAGCTCTGGATCATTGGNNNAGATCGGAAG |
| 37480_2 | TATGAGTGTGGAGTCGTTGCTTCAGTCAGCTCTGGATCATTGGNNNAGATCGGAAG |
| 45270_1 | TATGAGTGTGGAGTCGTTGCGGAGTGGGGACCCTACTTTCTANNNAGATCGGAAG |
| 45270_2 | TATGAGTGTGGAGTCGTTGCGGAGTGGGGACCCTACTTTCTANNNAGATCGGAAG |
| 35030_1 | TATGAGTGTGGAGTCGTTGCCCCAAAGTGCTAGGATTACAGGTNNNAGATCGGAA |
| 35030_2 | TATGAGTGTGGAGTCGTTGCCCCAAAGTGCTAGGATTACAGGTNNNAGATCGGAA |
| 33036_1 | TATGAGTGTGGAGTCGTTGCTGGAGTATAATGGCACAATCCCGNNNAGATCGGAA |
| 33036_2 | TATGAGTGTGGAGTCGTTGCTGGAGTATAATGGCACAATCCCGNNNAGATCGGAA |
| 46601_1 | TATGAGTGTGGAGTCGTTGCCTTCCCGAGTAGCTGGAAATACANNNAGATCGGAAG |
| 46601_2 | TATGAGTGTGGAGTCGTTGCCTTCCCGAGTAGCTGGAAATACANNNAGATCGGAAG |
| 32114_1 | TATGAGTGTGGAGTCGTTGCCTGAAAATTAGCTGGGTGTGGTGNNNAGATCGGAA |
| 32114_2 | TATGAGTGTGGAGTCGTTGCCTGAAAATTAGCTGGGTGTGGTGNNNAGATCGGAA |
| 32752_1 | TATGAGTGTGGAGTCGTTGCGGGGAACTGTCAGACTTGATCTTNNNAGATCGGAA |
| 32752_2 | TATGAGTGTGGAGTCGTTGCGGGGAACTGTCAGACTTGATCTTNNNAGATCGGAA |
| 48705_1 | TATGAGTGTGGAGTCGTTGCCAATACCCCAACCTACACAGCANNNAGATCGGAAGA |
| 48705_2 | TATGAGTGTGGAGTCGTTGCCAATACCCCAACCTACACAGCANNNAGATCGGAAGA |
| 42943_1 | TATGAGTGTGGAGTCGTTGCGCACCTGCTTGGGAATCAAATTANNNAGATCGGAAG |
| 42943_2 | TATGAGTGTGGAGTCGTTGCGCACCTGCTTGGGAATCAAATTANNNAGATCGGAAG |
| 40431_1 | TATGAGTGTGGAGTCGTTGCATGACTACTGTAAGTTGAGTGCANNNAGATCGGAA |
| 40431_2 | TATGAGTGTGGAGTCGTTGCATGACTACTGTAAGTTGAGTGCANNNAGATCGGAA |
| 44952_1 | TATGAGTGTGGAGTCGTTGCTTCCTAGCTCTGAAACAGGAACCNNNAGATCGGAAG |
| 44952_2 | TATGAGTGTGGAGTCGTTGCTTCCTAGCTCTGAAACAGGAACCNNNAGATCGGAAG |
| 47803_1 | TATGAGTGTGGAGTCGTTGCCAAGTAGCTGGGATTACAGGCATNNNAGATCGGAA |
| 47803_2 | TATGAGTGTGGAGTCGTTGCCAAGTAGCTGGGATTACAGGCATNNNAGATCGGAA |
| 52245_1 | TATGAGTGTGGAGTCGTTGCCTCCATGGGCTCCTGTGCTANNNAGATCGGAAGAGC |
| 52245_2 | TATGAGTGTGGAGTCGTTGCCTCCATGGGCTCCTGTGCTANNNAGATCGGAAGAGC |
| 37768_1 | TATGAGTGTGGAGTCGTTGCCACACTCTTTTTCGTCTCTGTGTNNNAGATCGGAAGA |
| 37768_2 | TATGAGTGTGGAGTCGTTGCCACACTCTTTTTCGTCTCTGTGTNNNAGATCGGAAGA |
| 45525_1 | TATGAGTGTGGAGTCGTTGCCTTCCTGGACAACCAAACAAGACNNNAGATCGGAAG |
| 45525_2 | TATGAGTGTGGAGTCGTTGCCTTCCTGGACAACCAAACAAGACNNNAGATCGGAAG |
| 52569_1 | TATGAGTGTGGAGTCGTTGCGTTTTAGTCTGTGGTTGAAGGCCNNNAGATCGGAA |
| 52569_2 | TATGAGTGTGGAGTCGTTGCGTTTTAGTCTGTGGTTGAAGGCCNNNAGATCGGAA |
| 2067_1 | TATGAGTGTGGAGTCGTTGCTTTTCTGTAAGCTATCCCTCACANNNAGATCGGAAGA |
| 2067_2 | TATGAGTGTGGAGTCGTTGCTTTTCTGTAAGCTATCCCTCACANNNAGATCGGAAGA |
| 49845_1 | TATGAGTGTGGAGTCGTTGCAGGAATGTGACAGAGAAGAGCTGNNNAGATCGGA |
| 49845_2 | TATGAGTGTGGAGTCGTTGCAGGAATGTGACAGAGAAGAGCTGNNNAGATCGGA |

| | |
|---|---|
| 47135_1 | TATGAGTGTGGAGTCGTTGCTTTTTTGAGTTGCAGTCTCGCTCNNNAGATCGGAAG |
| 47135_2 | TATGAGTGTGGAGTCGTTGCTTTTTTGAGTTGCAGTCTCGCTCNNNAGATCGGAAG |
| 48975_1 | TATGAGTGTGGAGTCGTTGCTTGTGGGTTGTCTGTTTACTCCANNNAGATCGGAAG |
| 48975_2 | TATGAGTGTGGAGTCGTTGCTTGTGGGTTGTCTGTTTACTCCANNNAGATCGGAAG |
| 45252_1 | TATGAGTGTGGAGTCGTTGCTGCATGGAACTATCTCCTGAGTGNNNAGATCGGAAG |
| 45252_2 | TATGAGTGTGGAGTCGTTGCTGCATGGAACTATCTCCTGAGTGNNNAGATCGGAAG |
| 36860_1 | TATGAGTGTGGAGTCGTTGCGGGCAGGAAGTGACCATATACAANNNAGATCGGAA |
| 36860_2 | TATGAGTGTGGAGTCGTTGCGGGCAGGAAGTGACCATATACAANNNAGATCGGAA |
| 5456_1 | TATGAGTGTGGAGTCGTTGCGGACATTACAGGCCCCAAGAATTNNNAGATCGGAA |
| 5456_2 | TATGAGTGTGGAGTCGTTGCGGACATTACAGGCCCCAAGAATTNNNAGATCGGAA |
| 38004_1 | TATGAGTGTGGAGTCGTTGCAACAGAACAAAAAAGGAGGTCAGNNNAGATCGGA |
| 38004_2 | TATGAGTGTGGAGTCGTTGCAACAGAACAAAAAAGGAGGTCAGNNNAGATCGGA |
| 8797_1 | TATGAGTGTGGAGTCGTTGCAGACAAACAAGTGTAGGTGCCTTNNNAGATCGGAA |
| 8797_2 | TATGAGTGTGGAGTCGTTGCAGACAAACAAGTGTAGGTGCCTTNNNAGATCGGAA |
| 40779_1 | TATGAGTGTGGAGTCGTTGCTTCTTCGTGGTTAGGTGTCTCTGNNNAGATCGGAAG |
| 40779_2 | TATGAGTGTGGAGTCGTTGCTTCTTCGTGGTTAGGTGTCTCTGNNNAGATCGGAAG |
| 29639_1 | TATGAGTGTGGAGTCGTTGCAGTCTGGAATTTGTGTTTGCTGTNNNAGATCGGAAG |
| 29639_2 | TATGAGTGTGGAGTCGTTGCAGTCTGGAATTTGTGTTTGCTGTNNNAGATCGGAAG |
| 53104_1 | TATGAGTGTGGAGTCGTTGCACAAATCCTAAGATGCACCACCTNNNAGATCGGAAG |
| 53104_2 | TATGAGTGTGGAGTCGTTGCACAAATCCTAAGATGCACCACCTNNNAGATCGGAAG |
| 30301_1 | TATGAGTGTGGAGTCGTTGCTGTGTCAACAAATGGTAGTTCAGGNNNAGATCGGA |
| 30301_2 | TATGAGTGTGGAGTCGTTGCTGTGTCAACAAATGGTAGTTCAGGNNNAGATCGGA |
| 29269_1 | TATGAGTGTGGAGTCGTTGCAGAGACAAGAGAGAGAACGGAGANNNAGATCGGA |
| 29269_2 | TATGAGTGTGGAGTCGTTGCAGAGACAAGAGAGAGAACGGAGANNNAGATCGGA |
| 31968_1 | TATGAGTGTGGAGTCGTTGCTAATGCTACTTGGGAGGCTGAAGNNNAGATCGGAA |
| 31968_2 | TATGAGTGTGGAGTCGTTGCTAATGCTACTTGGGAGGCTGAAGNNNAGATCGGAA |
| 52078_1 | TATGAGTGTGGAGTCGTTGCTTTCCAGAGGAACGATCAGACAGNNNAGATCGGAA |
| 52078_2 | TATGAGTGTGGAGTCGTTGCTTTCCAGAGGAACGATCAGACAGNNNAGATCGGAA |
| 43012_1 | TATGAGTGTGGAGTCGTTGCTCTCTTCAGTGCCTCTTTCAGTGNNNAGATCGGAAGA |
| 43012_2 | TATGAGTGTGGAGTCGTTGCTCTCTTCAGTGCCTCTTTCAGTGNNNAGATCGGAAGA |
| 42403_1 | TATGAGTGTGGAGTCGTTGCCAATCTCGGCTTGCTGTAAACTCNNNAGATCGGAAG |
| 42403_2 | TATGAGTGTGGAGTCGTTGCCAATCTCGGCTTGCTGTAAACTCNNNAGATCGGAAG |
| 31742_1 | TATGAGTGTGGAGTCGTTGCTGGCCATTGTCAGCAATAAACTTNNNAGATCGGAAG |
| 31742_2 | TATGAGTGTGGAGTCGTTGCTGGCCATTGTCAGCAATAAACTTNNNAGATCGGAAG |
| 48624_1 | TATGAGTGTGGAGTCGTTGCCTGGTTCTGCTACTGCTCAGAGNNNAGATCGGAAGA |
| 48624_2 | TATGAGTGTGGAGTCGTTGCCTGGTTCTGCTACTGCTCAGAGNNNAGATCGGAAGA |
| 47342_1 | TATGAGTGTGGAGTCGTTGCGCTACCACACCCAGCTAATTTTTNNNAGATCGGAAGA |
| 47342_2 | TATGAGTGTGGAGTCGTTGCGCTACCACACCCAGCTAATTTTTNNNAGATCGGAAGA |
| 43056_1 | TATGAGTGTGGAGTCGTTGCGGACACAAACATTCAGTCCATGGNNNAGATCGGAA |
| 43056_2 | TATGAGTGTGGAGTCGTTGCGGACACAAACATTCAGTCCATGGNNNAGATCGGAA |
| 30002_1 | TATGAGTGTGGAGTCGTTGCACCAAGCATCTGAGCTATTGGAANNNAGATCGGAA |
| 30002_2 | TATGAGTGTGGAGTCGTTGCACCAAGCATCTGAGCTATTGGAANNNAGATCGGAA |
| 41999_1 | TATGAGTGTGGAGTCGTTGCGATCTGGGCTCACTGAAACTTCTNNNAGATCGGAAG |

| | |
|---|---|
| 41999_2 | TATGAGTGTGGAGTCGTTGCGATCTGGGCTCACTGAAACTTCTNNNAGATCGGAAG |
| 43285_1 | TATGAGTGTGGAGTCGTTGCCCCTTTCAGTAGATTGGCTGTCTNNNAGATCGGAAG |
| 43285_2 | TATGAGTGTGGAGTCGTTGCCCCTTTCAGTAGATTGGCTGTCTNNNAGATCGGAAG |
| 45961_1 | TATGAGTGTGGAGTCGTTGCGTTCAAGGGATTATCCTGCCTCANNNAGATCGGAAG |
| 45961_2 | TATGAGTGTGGAGTCGTTGCGTTCAAGGGATTATCCTGCCTCANNNAGATCGGAAG |
| 44591_1 | TATGAGTGTGGAGTCGTTGCCTGCCTCTGTGTGTTTCATGAACNNNAGATCGGAAG |
| 44591_2 | TATGAGTGTGGAGTCGTTGCCTGCCTCTGTGTGTTTCATGAACNNNAGATCGGAAG |
| 10257_1 | TATGAGTGTGGAGTCGTTGCTTCATCACATGTGCAGGTGTAGGNNNAGATCGGAA |
| 10257_2 | TATGAGTGTGGAGTCGTTGCTTCATCACATGTGCAGGTGTAGGNNNAGATCGGAA |
| 31598_1 | TATGAGTGTGGAGTCGTTGCACAACTCGTGTACCTAGAGGACTNNNAGATCGGAAG |
| 31598_2 | TATGAGTGTGGAGTCGTTGCACAACTCGTGTACCTAGAGGACTNNNAGATCGGAAG |
| 35263_1 | TATGAGTGTGGAGTCGTTGCTGAGACTCAAGAAAAGTAGCGCTNNNAGATCGGAA |
| 35263_2 | TATGAGTGTGGAGTCGTTGCTGAGACTCAAGAAAAGTAGCGCTNNNAGATCGGAA |
| 29444_1 | TATGAGTGTGGAGTCGTTGCATTTCTGTTGAGTGTTTTGGGTTCNNNAGATCGGAA |
| 29444_2 | TATGAGTGTGGAGTCGTTGCATTTCTGTTGAGTGTTTTGGGTTCNNNAGATCGGAA |
| 49178_1 | TATGAGTGTGGAGTCGTTGCGTTAGCCAGGATGGTCTCGATCNNNAGATCGGAAG |
| 49178_2 | TATGAGTGTGGAGTCGTTGCGTTAGCCAGGATGGTCTCGATCNNNAGATCGGAAG |
| 29997_1 | TATGAGTGTGGAGTCGTTGCGTCAGCAGCCATAATTGTTGTCANNNAGATCGGAAG |
| 29997_2 | TATGAGTGTGGAGTCGTTGCGTCAGCAGCCATAATTGTTGTCANNNAGATCGGAAG |
| 38430_1 | TATGAGTGTGGAGTCGTTGCGTTAGAATGTGAGTGGGGAGGAANNNAGATCGGA |
| 38430_2 | TATGAGTGTGGAGTCGTTGCGTTAGAATGTGAGTGGGGAGGAANNNAGATCGGA |
| 37952_1 | TATGAGTGTGGAGTCGTTGCGAAAAAAATTAGCCCAGGCATGGNNNAGATCGGAA |
| 37952_2 | TATGAGTGTGGAGTCGTTGCGAAAAAAATTAGCCCAGGCATGGNNNAGATCGGAA |
| 41206_1 | TATGAGTGTGGAGTCGTTGCAAAAGACTCACCCCTACAAGGTCNNNAGATCGGAAG |
| 41206_2 | TATGAGTGTGGAGTCGTTGCAAAAGACTCACCCCTACAAGGTCNNNAGATCGGAAG |
| 30521_1 | TATGAGTGTGGAGTCGTTGCAGTTTGCTAATGCTGAGAGACCCNNNAGATCGGAA |
| 30521_2 | TATGAGTGTGGAGTCGTTGCAGTTTGCTAATGCTGAGAGACCCNNNAGATCGGAA |
| 38608_1 | TATGAGTGTGGAGTCGTTGCTTTTTGCTCCACACCAGGTAGTANNNAGATCGGAAG |
| 38608_2 | TATGAGTGTGGAGTCGTTGCTTTTTGCTCCACACCAGGTAGTANNNAGATCGGAAG |
| 44888_1 | TATGAGTGTGGAGTCGTTGCGAAAATCTCCTGGTTGGGCTCTNNNAGATCGGAAGA |
| 44888_2 | TATGAGTGTGGAGTCGTTGCGAAAATCTCCTGGTTGGGCTCTNNNAGATCGGAAGA |
| 33241_1 | TATGAGTGTGGAGTCGTTGCCAGCTCACTGCAGTTTTGACCNNNAGATCGGAAGAG |
| 33241_2 | TATGAGTGTGGAGTCGTTGCCAGCTCACTGCAGTTTTGACCNNNAGATCGGAAGAG |
| 43235_1 | TATGAGTGTGGAGTCGTTGCTTCAAGCGATTCTCCTGTCTCAGNNNAGATCGGAAG |
| 43235_2 | TATGAGTGTGGAGTCGTTGCTTCAAGCGATTCTCCTGTCTCAGNNNAGATCGGAAG |
| 47100_1 | TATGAGTGTGGAGTCGTTGCCATCTCCCAGGCTCAAGAGATCNNNAGATCGGAAGA |
| 47100_2 | TATGAGTGTGGAGTCGTTGCCATCTCCCAGGCTCAAGAGATCNNNAGATCGGAAGA |
| 43692_1 | TATGAGTGTGGAGTCGTTGCACCAAGAGAACCACGTAGAATCCNNNAGATCGGAA |
| 43692_2 | TATGAGTGTGGAGTCGTTGCACCAAGAGAACCACGTAGAATCCNNNAGATCGGAA |
| 45233_1 | TATGAGTGTGGAGTCGTTGCCAAAGTGCTGGGATTACAGGTGTNNNAGATCGGAA |
| 45233_2 | TATGAGTGTGGAGTCGTTGCCAAAGTGCTGGGATTACAGGTGTNNNAGATCGGAA |
| 42686_1 | TATGAGTGTGGAGTCGTTGCGTTGTAGGACAAGAGCCTGTGTANNNAGATCGGAA |
| 42686_2 | TATGAGTGTGGAGTCGTTGCGTTGTAGGACAAGAGCCTGTGTANNNAGATCGGAA |

| | |
|---|---|
| 33274_1 | TATGAGTGTGGAGTCGTTGCGATAGGGTCTTGCTCCGTCACNNNAGATCGGAAGAG |
| 33274_2 | TATGAGTGTGGAGTCGTTGCGATAGGGTCTTGCTCCGTCACNNNAGATCGGAAGAG |
| 43387_1 | TATGAGTGTGGAGTCGTTGCGCTTCTCCCTTGCATGTTCTTTTNNNAGATCGGAAGA |
| 43387_2 | TATGAGTGTGGAGTCGTTGCGCTTCTCCCTTGCATGTTCTTTTNNNAGATCGGAAGA |
| 2237_1 | TATGAGTGTGGAGTCGTTGCAAAGAGATGCACAGGCACACATANNNAGATCGGAA |
| 2237_2 | TATGAGTGTGGAGTCGTTGCAAAGAGATGCACAGGCACACATANNNAGATCGGAA |
| 38688_1 | TATGAGTGTGGAGTCGTTGCCTCCTGGGTTCAAGCGATTCTCNNNAGATCGGAAGA |
| 38688_2 | TATGAGTGTGGAGTCGTTGCCTCCTGGGTTCAAGCGATTCTCNNNAGATCGGAAGA |
| 49083_1 | TATGAGTGTGGAGTCGTTGCTGAACTCCTGAGCTCAAGTGATCNNNAGATCGGAAG |
| 49083_2 | TATGAGTGTGGAGTCGTTGCTGAACTCCTGAGCTCAAGTGATCNNNAGATCGGAAG |
| 45518_1 | TATGAGTGTGGAGTCGTTGCCCAGAAGAGTGAAAGACAGGGAANNNAGATCGGA |
| 45518_2 | TATGAGTGTGGAGTCGTTGCCCAGAAGAGTGAAAGACAGGGAANNNAGATCGGA |
| 10010_1 | TATGAGTGTGGAGTCGTTGCTCCCAAAGAGGCATTTCCTGAACNNNAGATCGGAAG |
| 10010_2 | TATGAGTGTGGAGTCGTTGCTCCCAAAGAGGCATTTCCTGAACNNNAGATCGGAAG |
| 42743_1 | TATGAGTGTGGAGTCGTTGCTGAGTCCAGTGAGTTACGATCACNNNAGATCGGAAG |
| 42743_2 | TATGAGTGTGGAGTCGTTGCTGAGTCCAGTGAGTTACGATCACNNNAGATCGGAAG |
| 38423_1 | TATGAGTGTGGAGTCGTTGCTGCAAGGCTCCTGTTCAAAAATCNNNAGATCGGAAG |
| 38423_2 | TATGAGTGTGGAGTCGTTGCTGCAAGGCTCCTGTTCAAAAATCNNNAGATCGGAAG |
| 42489_1 | TATGAGTGTGGAGTCGTTGCGGAGGGAGGGAGTGAAGGAANNNAGATCGGAAG |
| 42489_2 | TATGAGTGTGGAGTCGTTGCGGAGGGAGGGAGTGAAGGAANNNAGATCGGAAG |
| 43663_1 | TATGAGTGTGGAGTCGTTGCAACTGCCTCTTGAGAAAGGTGAANNNAGATCGGAA |
| 43663_2 | TATGAGTGTGGAGTCGTTGCAACTGCCTCTTGAGAAAGGTGAANNNAGATCGGAA |
| 29770_1 | TATGAGTGTGGAGTCGTTGCAATCGTCGTGTACAGCTTTACACNNNAGATCGGAAG |
| 29770_2 | TATGAGTGTGGAGTCGTTGCAATCGTCGTGTACAGCTTTACACNNNAGATCGGAAG |
| 49237_1 | TATGAGTGTGGAGTCGTTGCCAAGTAGCTGGGACTACAGGTGNNNAGATCGGAAG |
| 49237_2 | TATGAGTGTGGAGTCGTTGCCAAGTAGCTGGGACTACAGGTGNNNAGATCGGAAG |
| 46495_1 | TATGAGTGTGGAGTCGTTGCGCTCCTGGGTTCAAAGGATTCTNNNAGATCGGAAGA |
| 46495_2 | TATGAGTGTGGAGTCGTTGCGCTCCTGGGTTCAAAGGATTCTNNNAGATCGGAAGA |
| 13427_1 | TATGAGTGTGGAGTCGTTGCACATAAAATGTGGTAACATACACNNNAGATCGGAAG |
| 13427_2 | TATGAGTGTGGAGTCGTTGCACATAAAATGTGGTAACATACACNNNAGATCGGAAG |
| 40454_1 | TATGAGTGTGGAGTCGTTGCTGCAGTATGGAAAGGGAAAGACANNNAGATCGGAA |
| 40454_2 | TATGAGTGTGGAGTCGTTGCTGCAGTATGGAAAGGGAAAGACANNNAGATCGGAA |
| 47229_1 | TATGAGTGTGGAGTCGTTGCCAATCTCGGCTCACTGTAACCTCNNNAGATCGGAAGA |
| 47229_2 | TATGAGTGTGGAGTCGTTGCCAATCTCGGCTCACTGTAACCTCNNNAGATCGGAAGA |
| 38892_1 | TATGAGTGTGGAGTCGTTGCACGGTCAGGAAGAGTCAAAAAGANNNAGATCGGAA |
| 38892_2 | TATGAGTGTGGAGTCGTTGCACGGTCAGGAAGAGTCAAAAAGANNNAGATCGGAA |
| 29480_1 | TATGAGTGTGGAGTCGTTGCCATTCAGTGGATTGTGGACCCTNNNAGATCGGAAGA |
| 29480_2 | TATGAGTGTGGAGTCGTTGCCATTCAGTGGATTGTGGACCCTNNNAGATCGGAAGA |
| 32440_1 | TATGAGTGTGGAGTCGTTGCCTCTCGAGTTGCTGGAACTACAGNNNAGATCGGAAG |
| 32440_2 | TATGAGTGTGGAGTCGTTGCCTCTCGAGTTGCTGGAACTACAGNNNAGATCGGAAG |
| 36545_1 | TATGAGTGTGGAGTCGTTGCTTGCTGCTGGTGATTTTGGTTTTNNNAGATCGGAAG |
| 36545_2 | TATGAGTGTGGAGTCGTTGCTTGCTGCTGGTGATTTTGGTTTTNNNAGATCGGAAG |
| 43008_1 | TATGAGTGTGGAGTCGTTGCAGCTACTTCTGCTGGTAGGAATGNNNAGATCGGAA |

| | |
|---|---|
| 43008_2 | TATGAGTGTGGAGTCGTTGCAGCTACTTCTGCTGGTAGGAATGNNNAGATCGGAA |
| 52900_1 | TATGAGTGTGGAGTCGTTGCAGAAAAGCTATGGGAGAAAGCCTNNNAGATCGGAA |
| 52900_2 | TATGAGTGTGGAGTCGTTGCAGAAAAGCTATGGGAGAAAGCCTNNNAGATCGGAA |
| 47136_1 | TATGAGTGTGGAGTCGTTGCGAACTTCTGACCTCAGGTGATCCNNNAGATCGGAAG |
| 47136_2 | TATGAGTGTGGAGTCGTTGCGAACTTCTGACCTCAGGTGATCCNNNAGATCGGAAG |
| 43743_1 | TATGAGTGTGGAGTCGTTGCCACGCTCCACAGTATGATTTCACNNNAGATCGGAAG |
| 43743_2 | TATGAGTGTGGAGTCGTTGCCACGCTCCACAGTATGATTTCACNNNAGATCGGAAG |
| 50485_1 | TATGAGTGTGGAGTCGTTGCCTATTCTGCATTACCCAGGGAGGNNNAGATCGGAAG |
| 50485_2 | TATGAGTGTGGAGTCGTTGCCTATTCTGCATTACCCAGGGAGGNNNAGATCGGAAG |
| 4867_1 | TATGAGTGTGGAGTCGTTGCACTTTCTCACAGAGCTCTCCAGTNNNAGATCGGAAG |
| 4867_2 | TATGAGTGTGGAGTCGTTGCACTTTCTCACAGAGCTCTCCAGTNNNAGATCGGAAG |
| 42639_1 | TATGAGTGTGGAGTCGTTGCTCATAGCTCACTGTAGCCTTGACNNNAGATCGGAAG |
| 42639_2 | TATGAGTGTGGAGTCGTTGCTCATAGCTCACTGTAGCCTTGACNNNAGATCGGAAG |
| 28754_1 | TATGAGTGTGGAGTCGTTGCCTGCAATACTGGCCAAGAATGTCNNNAGATCGGAAG |
| 28754_2 | TATGAGTGTGGAGTCGTTGCCTGCAATACTGGCCAAGAATGTCNNNAGATCGGAAG |
| 35133_1 | TATGAGTGTGGAGTCGTTGCTTCAAACCATTCTCCTGCCTCAGNNNAGATCGGAAGA |
| 35133_2 | TATGAGTGTGGAGTCGTTGCTTCAAACCATTCTCCTGCCTCAGNNNAGATCGGAAGA |
| 40266_1 | TATGAGTGTGGAGTCGTTGCCACCCCCAACAGAATTCACTTTCNNNAGATCGGAAGA |
| 40266_2 | TATGAGTGTGGAGTCGTTGCCACCCCCAACAGAATTCACTTTCNNNAGATCGGAAGA |
| 31417_1 | TATGAGTGTGGAGTCGTTGCTGAGTCGTTCTGTTTTGTTGGAGNNNAGATCGGAA |
| 31417_2 | TATGAGTGTGGAGTCGTTGCTGAGTCGTTCTGTTTTGTTGGAGNNNAGATCGGAA |
| 49674_1 | TATGAGTGTGGAGTCGTTGCGCCTCCTGAGTAGCTGGTATTACNNNAGATCGGAAG |
| 49674_2 | TATGAGTGTGGAGTCGTTGCGCCTCCTGAGTAGCTGGTATTACNNNAGATCGGAAG |
| 28899_1 | TATGAGTGTGGAGTCGTTGCCTACTCTCACTTCCCATACACCGNNNAGATCGGAAGA |
| 28899_2 | TATGAGTGTGGAGTCGTTGCCTACTCTCACTTCCCATACACCGNNNAGATCGGAAGA |
| 39411_1 | TATGAGTGTGGAGTCGTTGCCTTCCGCCTAATCCCTGGACNNNAGATCGGAAGAGC |
| 39411_2 | TATGAGTGTGGAGTCGTTGCCTTCCGCCTAATCCCTGGACNNNAGATCGGAAGAGC |
| 47858_1 | TATGAGTGTGGAGTCGTTGCTAATCTGAGCACTTTGGGAGTCCNNNAGATCGGAAG |
| 47858_2 | TATGAGTGTGGAGTCGTTGCTAATCTGAGCACTTTGGGAGTCCNNNAGATCGGAAG |
| 41891_1 | TATGAGTGTGGAGTCGTTGCCAGACTCTTGCTCTATCACCCAGNNNAGATCGGAAG |
| 41891_2 | TATGAGTGTGGAGTCGTTGCCAGACTCTTGCTCTATCACCCAGNNNAGATCGGAAG |
| 51766_1 | TATGAGTGTGGAGTCGTTGCGAGTGAGAATATGCGGTGTTTGGNNNAGATCGGAA |
| 51766_2 | TATGAGTGTGGAGTCGTTGCGAGTGAGAATATGCGGTGTTTGGNNNAGATCGGAA |
| 33373_1 | TATGAGTGTGGAGTCGTTGCTGAGGTAGGAGAATCGCTTGAACNNNAGATCGGAA |
| 33373_2 | TATGAGTGTGGAGTCGTTGCTGAGGTAGGAGAATCGCTTGAACNNNAGATCGGAA |
| 48051_1 | TATGAGTGTGGAGTCGTTGCCCCCGTCTGGCCTAAAATACATTNNNAGATCGGAAG |
| 48051_2 | TATGAGTGTGGAGTCGTTGCCCCCGTCTGGCCTAAAATACATTNNNAGATCGGAAG |
| 8532_1 | TATGAGTGTGGAGTCGTTGCTGGAGTATGTCAAGAATGGCCAGAGNNNAGATCGG |
| 8532_2 | TATGAGTGTGGAGTCGTTGCTGGAGTATGTCAAGAATGGCCAGAGNNNAGATCGG |
| 10072_1 | TATGAGTGTGGAGTCGTTGCTCATGCATTGTCTGTCCTTTCCCNNNAGATCGGAAGA |
| 10072_2 | TATGAGTGTGGAGTCGTTGCTCATGCATTGTCTGTCCTTTCCCNNNAGATCGGAAGA |
| 51612_1 | TATGAGTGTGGAGTCGTTGCCCACGCCCGGCTAATTTTTTATTNNNAGATCGGAAGA |
| 51612_2 | TATGAGTGTGGAGTCGTTGCCCACGCCCGGCTAATTTTTTATTNNNAGATCGGAAGA |

| | |
|---|---|
| 43353_1 | TATGAGTGTGGAGTCGTTGCTTGGCTAAAGGCGACGTAAAATGNNNAGATCGGAA |
| 43353_2 | TATGAGTGTGGAGTCGTTGCTTGGCTAAAGGCGACGTAAAATGNNNAGATCGGAA |
| 47677_1 | TATGAGTGTGGAGTCGTTGCTCCTGCTTTCCAAATCATGAATCCNNNAGATCGGAAG |
| 47677_2 | TATGAGTGTGGAGTCGTTGCTCCTGCTTTCCAAATCATGAATCCNNNAGATCGGAAG |
| 52667_1 | TATGAGTGTGGAGTCGTTGCTCCTAATGATTGGTTGCATGTGCNNNAGATCGGAAG |
| 52667_2 | TATGAGTGTGGAGTCGTTGCTCCTAATGATTGGTTGCATGTGCNNNAGATCGGAAG |
| 44683_1 | TATGAGTGTGGAGTCGTTGCGACACACATTTGGGGCTTGATAGNNNAGATCGGAA |
| 44683_2 | TATGAGTGTGGAGTCGTTGCGACACACATTTGGGGCTTGATAGNNNAGATCGGAA |
| 36564_1 | TATGAGTGTGGAGTCGTTGCGGCCAACATGGTGAAATCTCATCNNNAGATCGGAAG |
| 36564_2 | TATGAGTGTGGAGTCGTTGCGGCCAACATGGTGAAATCTCATCNNNAGATCGGAAG |
| 36386_1 | TATGAGTGTGGAGTCGTTGCCTGATGGATGCTCAAATCCCTGANNNAGATCGGAAG |
| 36386_2 | TATGAGTGTGGAGTCGTTGCCTGATGGATGCTCAAATCCCTGANNNAGATCGGAAG |
| 34236_1 | TATGAGTGTGGAGTCGTTGCAAAGCTTTAGAAAAGGTCCACGCNNNAGATCGGAA |
| 34236_2 | TATGAGTGTGGAGTCGTTGCAAAGCTTTAGAAAAGGTCCACGCNNNAGATCGGAA |
| 10033_1 | TATGAGTGTGGAGTCGTTGCAGAACCTACTTTGTCCCTGAGCTNNNAGATCGGAAG |
| 10033_2 | TATGAGTGTGGAGTCGTTGCAGAACCTACTTTGTCCCTGAGCTNNNAGATCGGAAG |
| 42516_1 | TATGAGTGTGGAGTCGTTGCATTTCAGACAGTCTTGCTCTGCTNNNAGATCGGAAG |
| 42516_2 | TATGAGTGTGGAGTCGTTGCATTTCAGACAGTCTTGCTCTGCTNNNAGATCGGAAG |
| 29851_1 | TATGAGTGTGGAGTCGTTGCTCATTACACCAGTTCGTCCCTTTNNNAGATCGGAAGA |
| 29851_2 | TATGAGTGTGGAGTCGTTGCTCATTACACCAGTTCGTCCCTTTNNNAGATCGGAAGA |
| 47011_1 | TATGAGTGTGGAGTCGTTGCCCACCATGCCCAGCTAATTTTTTNNNAGATCGGAAGA |
| 47011_2 | TATGAGTGTGGAGTCGTTGCCCACCATGCCCAGCTAATTTTTTNNNAGATCGGAAGA |
| 49868_1 | TATGAGTGTGGAGTCGTTGCGAGCCAGATTATAGAGGTCGCAANNNAGATCGGAA |
| 49868_2 | TATGAGTGTGGAGTCGTTGCGAGCCAGATTATAGAGGTCGCAANNNAGATCGGAA |
| 37826_1 | TATGAGTGTGGAGTCGTTGCCTCATGCCTATGATCGTAGCACTNNNAGATCGGAAG |
| 37826_2 | TATGAGTGTGGAGTCGTTGCCTCATGCCTATGATCGTAGCACTNNNAGATCGGAAG |
| 29422_1 | TATGAGTGTGGAGTCGTTGCCAGCAGTAGGGAATCTGGGATCTNNNAGATCGGAA |
| 29422_2 | TATGAGTGTGGAGTCGTTGCCAGCAGTAGGGAATCTGGGATCTNNNAGATCGGAA |
| 47672_1 | TATGAGTGTGGAGTCGTTGCGAGACGGAGTTTCGTTCTTGTTGNNNAGATCGGAA |
| 47672_2 | TATGAGTGTGGAGTCGTTGCGAGACGGAGTTTCGTTCTTGTTGNNNAGATCGGAA |
| 40436_1 | TATGAGTGTGGAGTCGTTGCCTGTAGTTCTAGCTGCTCTGGAGNNNAGATCGGAAG |
| 40436_2 | TATGAGTGTGGAGTCGTTGCCTGTAGTTCTAGCTGCTCTGGAGNNNAGATCGGAAG |
| 10062_1 | TATGAGTGTGGAGTCGTTGCTGCGAGACGTACCTGTATTGTTCNNNAGATCGGAAG |
| 10062_2 | TATGAGTGTGGAGTCGTTGCTGCGAGACGTACCTGTATTGTTCNNNAGATCGGAAG |
| 48817_1 | TATGAGTGTGGAGTCGTTGCCAAGTAGCTGGGACTACAGGTACNNNAGATCGGAA |
| 48817_2 | TATGAGTGTGGAGTCGTTGCCAAGTAGCTGGGACTACAGGTACNNNAGATCGGAA |
| 32578_1 | TATGAGTGTGGAGTCGTTGCCATTCTCCTGCCTCAGCCTCNNNAGATCGGAAGAGCA |
| 32578_2 | TATGAGTGTGGAGTCGTTGCCATTCTCCTGCCTCAGCCTCNNNAGATCGGAAGAGCA |
| 42273_1 | TATGAGTGTGGAGTCGTTGCTTATAGCTCACTGCAGCCTCAAANNNAGATCGGAAG |
| 42273_2 | TATGAGTGTGGAGTCGTTGCTTATAGCTCACTGCAGCCTCAAANNNAGATCGGAAG |
| 40226_1 | TATGAGTGTGGAGTCGTTGCCGGCCACAGTGCTTATACTTTTCNNNAGATCGGAAG |
| 40226_2 | TATGAGTGTGGAGTCGTTGCCGGCCACAGTGCTTATACTTTTCNNNAGATCGGAAG |
| 13592_1 | TATGAGTGTGGAGTCGTTGCAAACCTCTCTGAAAATTAATTGGNNNAGATCGGAAG |

| | |
|---|---|
| 13592_2 | TATGAGTGTGGAGTCGTTGCAAACCTCTCTGAAAATTAATTGGNNNAGATCGGAAG |
| 42528_1 | TATGAGTGTGGAGTCGTTGCGCATCCAGCCTTGAAATGTTTTCNNNAGATCGGAAG |
| 42528_2 | TATGAGTGTGGAGTCGTTGCGCATCCAGCCTTGAAATGTTTTCNNNAGATCGGAAG |
| 51456_1 | TATGAGTGTGGAGTCGTTGCACATAGCTCACTACAGCCTTGACNNNAGATCGGAAG |
| 51456_2 | TATGAGTGTGGAGTCGTTGCACATAGCTCACTACAGCCTTGACNNNAGATCGGAAG |
| 46773_1 | TATGAGTGTGGAGTCGTTGCCCAGGATCAAACGATTCTCATGCNNNAGATCGGAAG |
| 46773_2 | TATGAGTGTGGAGTCGTTGCCCAGGATCAAACGATTCTCATGCNNNAGATCGGAAG |
| 43042_1 | TATGAGTGTGGAGTCGTTGCAGCAATTCTTCTACCTCAGCCTCNNNAGATCGGAAGA |
| 43042_2 | TATGAGTGTGGAGTCGTTGCAGCAATTCTTCTACCTCAGCCTCNNNAGATCGGAAGA |
| 49319_1 | TATGAGTGTGGAGTCGTTGCGTCTTGAACTCCTGACTTCGTGANNNAGATCGGAAG |
| 49319_2 | TATGAGTGTGGAGTCGTTGCGTCTTGAACTCCTGACTTCGTGANNNAGATCGGAAG |
| 49694_1 | TATGAGTGTGGAGTCGTTGCGACACAGCAGAGTCCTAGAACAANNNAGATCGGAA |
| 49694_2 | TATGAGTGTGGAGTCGTTGCGACACAGCAGAGTCCTAGAACAANNNAGATCGGAA |
| 32958_1 | TATGAGTGTGGAGTCGTTGCACATAGCATAGTTTTGGGGGGAGNNNAGATCGGAA |
| 32958_2 | TATGAGTGTGGAGTCGTTGCACATAGCATAGTTTTGGGGGGAGNNNAGATCGGAA |
| 12548_1 | TATGAGTGTGGAGTCGTTGCGTTATGAAGGTAAATGGATTCACNNNAGATCGGAA |
| 12548_2 | TATGAGTGTGGAGTCGTTGCGTTATGAAGGTAAATGGATTCACNNNAGATCGGAA |
| 44258_1 | TATGAGTGTGGAGTCGTTGCATCTCCCATCCCAGAAGCTTTTTNNNAGATCGGAAGA |
| 44258_2 | TATGAGTGTGGAGTCGTTGCATCTCCCATCCCAGAAGCTTTTTNNNAGATCGGAAGA |
| 37736_1 | TATGAGTGTGGAGTCGTTGCATATGCACCCAATACAGGAGGACNNNAGATCGGAA |
| 37736_2 | TATGAGTGTGGAGTCGTTGCATATGCACCCAATACAGGAGGACNNNAGATCGGAA |
| 37598_1 | TATGAGTGTGGAGTCGTTGCACATACATCATGGTGCTGCTTTTNNNAGATCGGAAG |
| 37598_2 | TATGAGTGTGGAGTCGTTGCACATACATCATGGTGCTGCTTTTNNNAGATCGGAAG |
| 40380_1 | TATGAGTGTGGAGTCGTTGCATAGAAACTGCAGTGAGATGCCANNNAGATCGGAA |
| 40380_2 | TATGAGTGTGGAGTCGTTGCATAGAAACTGCAGTGAGATGCCANNNAGATCGGAA |
| 38350_1 | TATGAGTGTGGAGTCGTTGCACAAATGGAAACACATCCCATGTNNNAGATCGGAAG |
| 38350_2 | TATGAGTGTGGAGTCGTTGCACAAATGGAAACACATCCCATGTNNNAGATCGGAAG |
| 35661_1 | TATGAGTGTGGAGTCGTTGCCAGTGGCCAGATCTAAACTCACTNNNAGATCGGAAG |
| 35661_2 | TATGAGTGTGGAGTCGTTGCCAGTGGCCAGATCTAAACTCACTNNNAGATCGGAAG |
| 31612_1 | TATGAGTGTGGAGTCGTTGCCAAGTCCCATGCTGAAAAGATGTNNNAGATCGGAA |
| 31612_2 | TATGAGTGTGGAGTCGTTGCCAAGTCCCATGCTGAAAAGATGTNNNAGATCGGAA |
| 37489_1 | TATGAGTGTGGAGTCGTTGCCGCACAGCATTCTTGGGAATTTANNNAGATCGGAAG |
| 37489_2 | TATGAGTGTGGAGTCGTTGCCGCACAGCATTCTTGGGAATTTANNNAGATCGGAAG |
| 42462_1 | TATGAGTGTGGAGTCGTTGCACACACACACACACACACAAATTNNNAGATCGGAAG |
| 42462_2 | TATGAGTGTGGAGTCGTTGCACACACACACACACACACAAATTNNNAGATCGGAAG |
| 51785_1 | TATGAGTGTGGAGTCGTTGCCTCACACTTTTACCAGCCACAAGNNNAGATCGGAAG |
| 51785_2 | TATGAGTGTGGAGTCGTTGCCTCACACTTTTACCAGCCACAAGNNNAGATCGGAAG |
| 44452_1 | TATGAGTGTGGAGTCGTTGCCGTTAGGGACCTTGAAACGATCTNNNAGATCGGAA |
| 44452_2 | TATGAGTGTGGAGTCGTTGCCGTTAGGGACCTTGAAACGATCTNNNAGATCGGAA |
| 45984_1 | TATGAGTGTGGAGTCGTTGCAGGCTTCCACATAGTTGGGATTTNNNAGATCGGAAG |
| 45984_2 | TATGAGTGTGGAGTCGTTGCAGGCTTCCACATAGTTGGGATTTNNNAGATCGGAAG |
| 5803_1 | TATGAGTGTGGAGTCGTTGCCTTACCTGCCTCTATACAACCCGANNNAGATCGGAAG |
| 5803_2 | TATGAGTGTGGAGTCGTTGCCTTACCTGCCTCTATACAACCCGANNNAGATCGGAAG |

| | |
|---|---|
| 49163_1 | TATGAGTGTGGAGTCGTTGCGAGAGAACTGAGCCGATTTGAGANNNAGATCGGAA |
| 49163_2 | TATGAGTGTGGAGTCGTTGCGAGAGAACTGAGCCGATTTGAGANNNAGATCGGAA |
| 46638_1 | TATGAGTGTGGAGTCGTTGCCCTCCTGGGTTCAAGTAATCCTCNNNAGATCGGAAG |
| 46638_2 | TATGAGTGTGGAGTCGTTGCCCTCCTGGGTTCAAGTAATCCTCNNNAGATCGGAAG |
| 53288_1 | TATGAGTGTGGAGTCGTTGCCTGATCCAGGAGGCCGCATCNNNAGATCGGAAGAG |
| 53288_2 | TATGAGTGTGGAGTCGTTGCCTGATCCAGGAGGCCGCATCNNNAGATCGGAAGAG |
| 30510_1 | TATGAGTGTGGAGTCGTTGCAAGAGCCTGATACTTGGGTTGGNNNAGATCGGAAG |
| 30510_2 | TATGAGTGTGGAGTCGTTGCAAGAGCCTGATACTTGGGTTGGNNNAGATCGGAAG |
| 36506_1 | TATGAGTGTGGAGTCGTTGCATGCCTGTACGTCTTTCTTTCCTNNNAGATCGGAAGA |
| 36506_2 | TATGAGTGTGGAGTCGTTGCATGCCTGTACGTCTTTCTTTCCTNNNAGATCGGAAGA |
| 51460_1 | TATGAGTGTGGAGTCGTTGCCCATGGAATTGTGGGACTGTTTCNNNAGATCGGAAG |
| 51460_2 | TATGAGTGTGGAGTCGTTGCCCATGGAATTGTGGGACTGTTTCNNNAGATCGGAAG |
| 31463_1 | TATGAGTGTGGAGTCGTTGCGGGGAGAGAGAGAGAAAGAGAGANNNAGATCGG |
| 31463_2 | TATGAGTGTGGAGTCGTTGCGGGGAGAGAGAGAGAAAGAGAGANNNAGATCGG |
| 46740_1 | TATGAGTGTGGAGTCGTTGCTTTAGTGTAGCCCTACTTCGCAGNNNAGATCGGAAG |
| 46740_2 | TATGAGTGTGGAGTCGTTGCTTTAGTGTAGCCCTACTTCGCAGNNNAGATCGGAAG |
| 30013_1 | TATGAGTGTGGAGTCGTTGCACTACTGCCTTCAATGGCACCTNNNAGATCGGAAGA |
| 30013_2 | TATGAGTGTGGAGTCGTTGCACTACTGCCTTCAATGGCACCTNNNAGATCGGAAGA |
| 30361_1 | TATGAGTGTGGAGTCGTTGCCTAAAGGAAGGGGGAAATATTAANNNAGATCGGAA |
| 30361_2 | TATGAGTGTGGAGTCGTTGCCTAAAGGAAGGGGGAAATATTAANNNAGATCGGAA |
| 12835_1 | TATGAGTGTGGAGTCGTTGCGGATGAAGTAGAATTCCAAAAACNNNAGATCGGAA |
| 12835_2 | TATGAGTGTGGAGTCGTTGCGGATGAAGTAGAATTCCAAAAACNNNAGATCGGAA |
| 42130_1 | TATGAGTGTGGAGTCGTTGCCTGGTCTCAAACTCTTGGACTCANNNAGATCGGAAG |
| 42130_2 | TATGAGTGTGGAGTCGTTGCCTGGTCTCAAACTCTTGGACTCANNNAGATCGGAAG |
| 33018_1 | TATGAGTGTGGAGTCGTTGCGAGTAGCTGGGATTACAGGCATGNNNAGATCGGAA |
| 33018_2 | TATGAGTGTGGAGTCGTTGCGAGTAGCTGGGATTACAGGCATGNNNAGATCGGAA |
| 45348_1 | TATGAGTGTGGAGTCGTTGCGAAAACACACACACATTTGCCACNNNAGATCGGAAG |
| 45348_2 | TATGAGTGTGGAGTCGTTGCGAAAACACACACACATTTGCCACNNNAGATCGGAAG |
| 35114_1 | TATGAGTGTGGAGTCGTTGCGTTCACTCCATTCTCCTGCCTCNNNAGATCGGAAGAG |
| 35114_2 | TATGAGTGTGGAGTCGTTGCGTTCACTCCATTCTCCTGCCTCNNNAGATCGGAAGAG |
| 52902_1 | TATGAGTGTGGAGTCGTTGCAAGGGCTGCTGATGATTTTTCACNNNAGATCGGAAG |
| 52902_2 | TATGAGTGTGGAGTCGTTGCAAGGGCTGCTGATGATTTTTCACNNNAGATCGGAAG |
| 35412_1 | TATGAGTGTGGAGTCGTTGCCAAGTAGCTGGGACTACAGGCNNNAGATCGGAAGA |
| 35412_2 | TATGAGTGTGGAGTCGTTGCCAAGTAGCTGGGACTACAGGCNNNAGATCGGAAGA |
| 47289_1 | TATGAGTGTGGAGTCGTTGCGGAGTGCAATGGTGTGATCTCAGNNNAGATCGGAA |
| 47289_2 | TATGAGTGTGGAGTCGTTGCGGAGTGCAATGGTGTGATCTCAGNNNAGATCGGAA |
| 30211_1 | TATGAGTGTGGAGTCGTTGCAGTCAACAAATTGGTTTACTGGGTNNNAGATCGGAA |
| 30211_2 | TATGAGTGTGGAGTCGTTGCAGTCAACAAATTGGTTTACTGGGTNNNAGATCGGAA |
| 42599_1 | TATGAGTGTGGAGTCGTTGCACTATCTCAGCTCACTGCAAGCNNNAGATCGGAAGA |
| 42599_2 | TATGAGTGTGGAGTCGTTGCACTATCTCAGCTCACTGCAAGCNNNAGATCGGAAGA |
| 52219_1 | TATGAGTGTGGAGTCGTTGCTTGTCTAGAAAGGGATCAGGCAGNNNAGATCGGAA |
| 52219_2 | TATGAGTGTGGAGTCGTTGCTTGTCTAGAAAGGGATCAGGCAGNNNAGATCGGAA |
| 35102_1 | TATGAGTGTGGAGTCGTTGCCTGGGATTACAGGTGTGAGCTACNNNAGATCGGAA |

| | |
|---|---|
| 35102_2 | TATGAGTGTGGAGTCGTTGCCTGGGATTACAGGTGTGAGCTACNNNAGATCGGAA |
| 34649_1 | TATGAGTGTGGAGTCGTTGCTCTTGATGAACCTCCTGAGTTGTNNNAGATCGGAAG |
| 34649_2 | TATGAGTGTGGAGTCGTTGCTCTTGATGAACCTCCTGAGTTGTNNNAGATCGGAAG |
| 48918_1 | TATGAGTGTGGAGTCGTTGCCTGTGCCCGGCTAATTTTTGTANNNAGATCGGAAG |
| 48918_2 | TATGAGTGTGGAGTCGTTGCCTGTGCCCGGCTAATTTTTGTANNNAGATCGGAAG |
| 35108_1 | TATGAGTGTGGAGTCGTTGCCAGTCTGGACTTGAACTCCTGACNNNAGATCGGAAG |
| 35108_2 | TATGAGTGTGGAGTCGTTGCCAGTCTGGACTTGAACTCCTGACNNNAGATCGGAAG |
| 32630_1 | TATGAGTGTGGAGTCGTTGCGAGACTTGCTCTATTACCCAGGCNNNAGATCGGAAG |
| 32630_2 | TATGAGTGTGGAGTCGTTGCGAGACTTGCTCTATTACCCAGGCNNNAGATCGGAAG |
| 33155_1 | TATGAGTGTGGAGTCGTTGCTCATTTGGTGAGTTACTGGCAGANNNAGATCGGAA |
| 33155_2 | TATGAGTGTGGAGTCGTTGCTCATTTGGTGAGTTACTGGCAGANNNAGATCGGAA |
| 1083_1 | TATGAGTGTGGAGTCGTTGCTGTCTAGGTCTTGTAGGTGTGGTNNNAGATCGGAA |
| 1083_2 | TATGAGTGTGGAGTCGTTGCTGTCTAGGTCTTGTAGGTGTGGTNNNAGATCGGAA |
| 48823_1 | TATGAGTGTGGAGTCGTTGCCTCCCGAGTAGCTGGAATTACAGNNNAGATCGGAA |
| 48823_2 | TATGAGTGTGGAGTCGTTGCCTCCCGAGTAGCTGGAATTACAGNNNAGATCGGAA |
| 31158_1 | TATGAGTGTGGAGTCGTTGCACTACTAATCAACCAGAGGCTGCNNNAGATCGGAAG |
| 31158_2 | TATGAGTGTGGAGTCGTTGCACTACTAATCAACCAGAGGCTGCNNNAGATCGGAAG |
| 36535_1 | TATGAGTGTGGAGTCGTTGCAAAGATCAGTGCAAGGGAAAAGCNNNAGATCGGA |
| 36535_2 | TATGAGTGTGGAGTCGTTGCAAAGATCAGTGCAAGGGAAAAGCNNNAGATCGGA |
| 5496_1 | TATGAGTGTGGAGTCGTTGCAGCCTCCCGAGTATTAGTAGTTGNNNAGATCGGAAG |
| 5496_2 | TATGAGTGTGGAGTCGTTGCAGCCTCCCGAGTATTAGTAGTTGNNNAGATCGGAAG |
| 43502_1 | TATGAGTGTGGAGTCGTTGCGTTCAAGTGATTCTCCTGCCTTGNNNAGATCGGAAG |
| 43502_2 | TATGAGTGTGGAGTCGTTGCGTTCAAGTGATTCTCCTGCCTTGNNNAGATCGGAAG |
| 47929_1 | TATGAGTGTGGAGTCGTTGCCCTCTCAGGTTCAAGCGATTCTCNNNAGATCGGAAG |
| 47929_2 | TATGAGTGTGGAGTCGTTGCCCTCTCAGGTTCAAGCGATTCTCNNNAGATCGGAAG |
| 52840_1 | TATGAGTGTGGAGTCGTTGCACCTTCGAGAACATGATTGTGGANNNAGATCGGAA |
| 52840_2 | TATGAGTGTGGAGTCGTTGCACCTTCGAGAACATGATTGTGGANNNAGATCGGAA |
| 29040_1 | TATGAGTGTGGAGTCGTTGCTTTAGACAGAAAGCGTGACCTCANNNAGATCGGAA |
| 29040_2 | TATGAGTGTGGAGTCGTTGCTTTAGACAGAAAGCGTGACCTCANNNAGATCGGAA |
| 49603_1 | TATGAGTGTGGAGTCGTTGCCCTGCAAGTAACTGGGACTACAGNNNAGATCGGAA |
| 49603_2 | TATGAGTGTGGAGTCGTTGCCCTGCAAGTAACTGGGACTACAGNNNAGATCGGAA |
| 38076_1 | TATGAGTGTGGAGTCGTTGCAAAAAATTAGCCAGGTGTGGTGGNNNAGATCGGAA |
| 38076_2 | TATGAGTGTGGAGTCGTTGCAAAAAATTAGCCAGGTGTGGTGGNNNAGATCGGAA |
| 31163_1 | TATGAGTGTGGAGTCGTTGCAGACATAGTCTCTGCCAATGGTGNNNAGATCGGAA |
| 31163_2 | TATGAGTGTGGAGTCGTTGCAGACATAGTCTCTGCCAATGGTGNNNAGATCGGAA |
| 37771_1 | TATGAGTGTGGAGTCGTTGCAACAAATGGCGTATCCCAATACANNNAGATCGGAAG |
| 37771_2 | TATGAGTGTGGAGTCGTTGCAACAAATGGCGTATCCCAATACANNNAGATCGGAAG |
| 36155_1 | TATGAGTGTGGAGTCGTTGCGATTGCAGTGAGCTGAGATTGTGNNNAGATCGGAA |
| 36155_2 | TATGAGTGTGGAGTCGTTGCGATTGCAGTGAGCTGAGATTGTGNNNAGATCGGAA |
| 49898_1 | TATGAGTGTGGAGTCGTTGCACCCGAGTCAAACATTTGGTAGANNNAGATCGGAA |
| 49898_2 | TATGAGTGTGGAGTCGTTGCACCCGAGTCAAACATTTGGTAGANNNAGATCGGAA |
| 9851_1 | TATGAGTGTGGAGTCGTTGCATGTAGTGCTGGATTTGATGGGGNNNAGATCGGAA |
| 9851_2 | TATGAGTGTGGAGTCGTTGCATGTAGTGCTGGATTTGATGGGGNNNAGATCGGAA |

| | |
|---|---|
| 41202_1 | TATGAGTGTGGAGTCGTTGCTTCTGAAGCAACCAGATACTGCTNNNAGATCGGAAG |
| 41202_2 | TATGAGTGTGGAGTCGTTGCTTCTGAAGCAACCAGATACTGCTNNNAGATCGGAAG |
| 51280_1 | TATGAGTGTGGAGTCGTTGCAGCTGCCTCTACCTAGGAAAGTANNNAGATCGGAAG |
| 51280_2 | TATGAGTGTGGAGTCGTTGCAGCTGCCTCTACCTAGGAAAGTANNNAGATCGGAAG |
| 36165_1 | TATGAGTGTGGAGTCGTTGCAACACGAATTCACTGTTGCGAATNNNAGATCGGAAG |
| 36165_2 | TATGAGTGTGGAGTCGTTGCAACACGAATTCACTGTTGCGAATNNNAGATCGGAAG |
| 33499_1 | TATGAGTGTGGAGTCGTTGCAACCAAACCAAACTTCTGCCTTCNNNAGATCGGAAGA |
| 33499_2 | TATGAGTGTGGAGTCGTTGCAACCAAACCAAACTTCTGCCTTCNNNAGATCGGAAGA |
| 37330_1 | TATGAGTGTGGAGTCGTTGCGTTGACATTTCCATGCCTTCTTANNNAGATCGGAAGA |
| 37330_2 | TATGAGTGTGGAGTCGTTGCGTTGACATTTCCATGCCTTCTTANNNAGATCGGAAGA |
| 29026_1 | TATGAGTGTGGAGTCGTTGCATGAGACAGAGAGACAGACAGGTNNNAGATCGGAA |
| 29026_2 | TATGAGTGTGGAGTCGTTGCATGAGACAGAGAGACAGACAGGTNNNAGATCGGAA |
| 48581_1 | TATGAGTGTGGAGTCGTTGCCTCCCGAGTAGCTGGAATTACAGNNNAGATCGGAAG |
| 48581_2 | TATGAGTGTGGAGTCGTTGCCTCCCGAGTAGCTGGAATTACAGNNNAGATCGGAAG |
| 31306_1 | TATGAGTGTGGAGTCGTTGCGCGTGTGTGTGTGTTGTAAGTTANNNAGATCGGAAG |
| 31306_2 | TATGAGTGTGGAGTCGTTGCGCGTGTGTGTGTGTTGTAAGTTANNNAGATCGGAAG |
| 40456_1 | TATGAGTGTGGAGTCGTTGCTTCAAGCGATTCTCCTGAGTGGNNNAGATCGGAAGA |
| 40456_2 | TATGAGTGTGGAGTCGTTGCTTCAAGCGATTCTCCTGAGTGGNNNAGATCGGAAGA |
| 32970_1 | TATGAGTGTGGAGTCGTTGCGGAACAGGGCGATTTTACATCATNNNAGATCGGAAG |
| 32970_2 | TATGAGTGTGGAGTCGTTGCGGAACAGGGCGATTTTACATCATNNNAGATCGGAAG |
| 36450_1 | TATGAGTGTGGAGTCGTTGCAGTCAAAGGCAACGAGGATTCTANNNAGATCGGAAG |
| 36450_2 | TATGAGTGTGGAGTCGTTGCAGTCAAAGGCAACGAGGATTCTANNNAGATCGGAAG |
| 35696_1 | TATGAGTGTGGAGTCGTTGCAAACAAGAGCTTTGCATTCAGGCNNNAGATCGGAAG |
| 35696_2 | TATGAGTGTGGAGTCGTTGCAAACAAGAGCTTTGCATTCAGGCNNNAGATCGGAAG |
| 36357_1 | TATGAGTGTGGAGTCGTTGCATGCCAGGTCTAGTACAACGATGNNNAGATCGGAAG |
| 36357_2 | TATGAGTGTGGAGTCGTTGCATGCCAGGTCTAGTACAACGATGNNNAGATCGGAAG |
| 40054_1 | TATGAGTGTGGAGTCGTTGCGGGAGGAAATCAGTAGGCAATCANNNAGATCGGAA |
| 40054_2 | TATGAGTGTGGAGTCGTTGCGGGAGGAAATCAGTAGGCAATCANNNAGATCGGAA |
| 7930_1 | TATGAGTGTGGAGTCGTTGCTTGTATTGGGCCATGAGGTAGTTGNNNAGATCGGAA |
| 7930_2 | TATGAGTGTGGAGTCGTTGCTTGTATTGGGCCATGAGGTAGTTGNNNAGATCGGAA |
| 31152_1 | TATGAGTGTGGAGTCGTTGCTCTTGGTGCAAAAATCTTCAACANNNAGATCGGAAGA |
| 31152_2 | TATGAGTGTGGAGTCGTTGCTCTTGGTGCAAAAATCTTCAACANNNAGATCGGAAGA |
| 43254_1 | TATGAGTGTGGAGTCGTTGCGAGATCTCAGCTCACTGCAACCNNNAGATCGGAAGA |
| 43254_2 | TATGAGTGTGGAGTCGTTGCGAGATCTCAGCTCACTGCAACCNNNAGATCGGAAGA |
| 38544_1 | TATGAGTGTGGAGTCGTTGCGTCTCTTCCAGGTAAAACGTCTTNNNAGATCGGAAGA |
| 38544_2 | TATGAGTGTGGAGTCGTTGCGTCTCTTCCAGGTAAAACGTCTTNNNAGATCGGAAGA |
| 2023_1 | TATGAGTGTGGAGTCGTTGCTGTGCCTCTTTCTTTCTGAGCAANNNAGATCGGAAGA |
| 2023_2 | TATGAGTGTGGAGTCGTTGCTGTGCCTCTTTCTTTCTGAGCAANNNAGATCGGAAGA |
| 43301_1 | TATGAGTGTGGAGTCGTTGCTTCAAACGATCCTCCTGCCTTAGNNNAGATCGGAAGA |
| 43301_2 | TATGAGTGTGGAGTCGTTGCTTCAAACGATCCTCCTGCCTTAGNNNAGATCGGAAGA |
| 31933_1 | TATGAGTGTGGAGTCGTTGCTTTCAAGAATGAAGCTGTGGACCNNNAGATCGGAAG |
| 31933_2 | TATGAGTGTGGAGTCGTTGCTTTCAAGAATGAAGCTGTGGACCNNNAGATCGGAAG |
| 9215_1 | TATGAGTGTGGAGTCGTTGCGGTCCTCTATCTAGGCTGTTGTGNNNAGATCGGAAG |

| | |
|---|---|
| 9215_2 | TATGAGTGTGGAGTCGTTGCGGTCCTCTATCTAGGCTGTTGTGNNNAGATCGGAAG |
| 10224_1 | TATGAGTGTGGAGTCGTTGCAGGGGTCATGGGTGATACAGAAANNNAGATCGGAA |
| 10224_2 | TATGAGTGTGGAGTCGTTGCAGGGGTCATGGGTGATACAGAAANNNAGATCGGAA |
| 10604_1 | TATGAGTGTGGAGTCGTTGCGAATCTATTGCCTTCCCGGGTTTNNNAGATCGGAAGA |
| 10604_2 | TATGAGTGTGGAGTCGTTGCGAATCTATTGCCTTCCCGGGTTTNNNAGATCGGAAGA |
| 31758_1 | TATGAGTGTGGAGTCGTTGCCTGAGCCACCTAAAGCTGAGAATNNNAGATCGGAAG |
| 31758_2 | TATGAGTGTGGAGTCGTTGCCTGAGCCACCTAAAGCTGAGAATNNNAGATCGGAAG |
| 29172_1 | TATGAGTGTGGAGTCGTTGCCAGGCCTATGTAATGAAGCTTGCNNNAGATCGGAAG |
| 29172_2 | TATGAGTGTGGAGTCGTTGCCAGGCCTATGTAATGAAGCTTGCNNNAGATCGGAAG |
| 42849_1 | TATGAGTGTGGAGTCGTTGCGCAGCATGATCATGGTTCATTGANNNAGATCGGAAG |
| 42849_2 | TATGAGTGTGGAGTCGTTGCGCAGCATGATCATGGTTCATTGANNNAGATCGGAAG |
| 31196_1 | TATGAGTGTGGAGTCGTTGCTCATAGCATATTCCACTCAGCCCNNNAGATCGGAAGA |
| 31196_2 | TATGAGTGTGGAGTCGTTGCTCATAGCATATTCCACTCAGCCCNNNAGATCGGAAGA |
| 46256_1 | TATGAGTGTGGAGTCGTTGCCCAGGCTGATCTTGAACTCCTAGNNNAGATCGGAAG |
| 46256_2 | TATGAGTGTGGAGTCGTTGCCCAGGCTGATCTTGAACTCCTAGNNNAGATCGGAAG |
| 46962_1 | TATGAGTGTGGAGTCGTTGCGCCTGCCCTGCTAATTTTTGTTANNNAGATCGGAAGA |
| 46962_2 | TATGAGTGTGGAGTCGTTGCGCCTGCCCTGCTAATTTTTGTTANNNAGATCGGAAGA |
| 12777_1 | TATGAGTGTGGAGTCGTTGCCTTTATGAAGTGTTTCACCAAANNNAGATCGGAAGA |
| 12777_2 | TATGAGTGTGGAGTCGTTGCCTTTATGAAGTGTTTCACCAAANNNAGATCGGAAGA |
| 49824_1 | TATGAGTGTGGAGTCGTTGCTTTCCCTCATATTGCCATTCCCANNNAGATCGGAAGA |
| 49824_2 | TATGAGTGTGGAGTCGTTGCTTTCCCTCATATTGCCATTCCCANNNAGATCGGAAGA |
| 42680_1 | TATGAGTGTGGAGTCGTTGCCAGAACTTGGTGTAACTTCAGCCNNNAGATCGGAAG |
| 42680_2 | TATGAGTGTGGAGTCGTTGCCAGAACTTGGTGTAACTTCAGCCNNNAGATCGGAAG |
| 50597_1 | TATGAGTGTGGAGTCGTTGCGGCGCATGTATAGTTCATGGTTCNNNAGATCGGAAG |
| 50597_2 | TATGAGTGTGGAGTCGTTGCGGCGCATGTATAGTTCATGGTTCNNNAGATCGGAAG |
| 32698_1 | TATGAGTGTGGAGTCGTTGCCTCCTGGGTTCACACCATTCTCNNNAGATCGGAAGAG |
| 32698_2 | TATGAGTGTGGAGTCGTTGCCTCCTGGGTTCACACCATTCTCNNNAGATCGGAAGAG |
| 43413_1 | TATGAGTGTGGAGTCGTTGCCCTCCTACGCTAAAGTGATCCTCNNNAGATCGGAAGA |
| 43413_2 | TATGAGTGTGGAGTCGTTGCCCTCCTACGCTAAAGTGATCCTCNNNAGATCGGAAGA |
| 43757_1 | TATGAGTGTGGAGTCGTTGCCTCGGAGAAGTAGAGCGCACNNNAGATCGGAAGAG |
| 43757_2 | TATGAGTGTGGAGTCGTTGCCTCGGAGAAGTAGAGCGCACNNNAGATCGGAAGAG |
| 31057_1 | TATGAGTGTGGAGTCGTTGCTGTGTAACCTCCACCACCATAAGNNNAGATCGGAAGA |
| 31057_2 | TATGAGTGTGGAGTCGTTGCTGTGTAACCTCCACCACCATAAGNNNAGATCGGAAGA |
| 33412_1 | TATGAGTGTGGAGTCGTTGCCTTACACCTGTAATCCCAGCACTNNNAGATCGGAAGA |
| 33412_2 | TATGAGTGTGGAGTCGTTGCCTTACACCTGTAATCCCAGCACTNNNAGATCGGAAGA |
| 48285_1 | TATGAGTGTGGAGTCGTTGCAATCAGCCACTTCTCTTAGGAGCNNNAGATCGGAAGA |
| 48285_2 | TATGAGTGTGGAGTCGTTGCAATCAGCCACTTCTCTTAGGAGCNNNAGATCGGAAGA |
| 44820_1 | TATGAGTGTGGAGTCGTTGCCATCCGGAAGGCTTAGATCTTGTNNNAGATCGGAAG |
| 44820_2 | TATGAGTGTGGAGTCGTTGCCATCCGGAAGGCTTAGATCTTGTNNNAGATCGGAAG |
| 34863_1 | TATGAGTGTGGAGTCGTTGCCAAGTAGCTGGGACTACAGGTGNNNAGATCGGAAG |
| 34863_2 | TATGAGTGTGGAGTCGTTGCCAAGTAGCTGGGACTACAGGTGNNNAGATCGGAAG |
| 39123_1 | TATGAGTGTGGAGTCGTTGCAACTGCCATGGAGACTGCTTTATNNNAGATCGGAAG |
| 39123_2 | TATGAGTGTGGAGTCGTTGCAACTGCCATGGAGACTGCTTTATNNNAGATCGGAAG |

| | |
|---|---|
| 50566_1 | TATGAGTGTGGAGTCGTTGCACCAAGATGCCAAGGCAAATAACNNNAGATCGGAAG |
| 50566_2 | TATGAGTGTGGAGTCGTTGCACCAAGATGCCAAGGCAAATAACNNNAGATCGGAAG |
| 38292_1 | TATGAGTGTGGAGTCGTTGCGGTTGCCATCCTAGTCTCTGATTNNNAGATCGGAAGA |
| 38292_2 | TATGAGTGTGGAGTCGTTGCGGTTGCCATCCTAGTCTCTGATTNNNAGATCGGAAGA |
| 52635_1 | TATGAGTGTGGAGTCGTTGCTCATCAACAGTCCTTCCAACCATNNNAGATCGGAAGA |
| 52635_2 | TATGAGTGTGGAGTCGTTGCTCATCAACAGTCCTTCCAACCATNNNAGATCGGAAGA |
| 49148_1 | TATGAGTGTGGAGTCGTTGCAACTTCAAGTGATTCTCCTGCCTNNNAGATCGGAAGA |
| 49148_2 | TATGAGTGTGGAGTCGTTGCAACTTCAAGTGATTCTCCTGCCTNNNAGATCGGAAGA |
| 31604_1 | TATGAGTGTGGAGTCGTTGCGAACTCAAGGAACTGAAACTGGCNNNAGATCGGAAG |
| 31604_2 | TATGAGTGTGGAGTCGTTGCGAACTCAAGGAACTGAAACTGGCNNNAGATCGGAAG |
| 48651_1 | TATGAGTGTGGAGTCGTTGCCAAGTAGCTGGGACTACAGGTGNNNAGATCGGAAG |
| 48651_2 | TATGAGTGTGGAGTCGTTGCCAAGTAGCTGGGACTACAGGTGNNNAGATCGGAAG |
| 42341_1 | TATGAGTGTGGAGTCGTTGCATCCCAACTACTCAGGAGACTGANNNAGATCGGAAG |
| 42341_2 | TATGAGTGTGGAGTCGTTGCATCCCAACTACTCAGGAGACTGANNNAGATCGGAAG |
| 43771_1 | TATGAGTGTGGAGTCGTTGCACGTTCTGGAGAGAATCACCTTCNNNAGATCGGAAG |
| 43771_2 | TATGAGTGTGGAGTCGTTGCACGTTCTGGAGAGAATCACCTTCNNNAGATCGGAAG |
| 51138_1 | TATGAGTGTGGAGTCGTTGCTCAGCTTATTCTCCCCAGTTCACNNNAGATCGGAAGA |
| 51138_2 | TATGAGTGTGGAGTCGTTGCTCAGCTTATTCTCCCCAGTTCACNNNAGATCGGAAGA |
| 37047_1 | TATGAGTGTGGAGTCGTTGCCACGTCACAGTTATGCAGCTATGNNNAGATCGGAAG |
| 37047_2 | TATGAGTGTGGAGTCGTTGCCACGTCACAGTTATGCAGCTATGNNNAGATCGGAAG |
| 3105_1 | TATGAGTGTGGAGTCGTTGCTTGGGCTACTGATTTGGGGAGATNNNAGATCGGAAG |
| 3105_2 | TATGAGTGTGGAGTCGTTGCTTGGGCTACTGATTTGGGGAGATNNNAGATCGGAAG |
| 33266_1 | TATGAGTGTGGAGTCGTTGCCTCGATCTCTTGACCTCGTGATCNNNAGATCGGAAGA |
| 33266_2 | TATGAGTGTGGAGTCGTTGCCTCGATCTCTTGACCTCGTGATCNNNAGATCGGAAGA |
| 47991_1 | TATGAGTGTGGAGTCGTTGCAGATGGAGGTCTCACTATGTTGCNNNAGATCGGAAG |
| 47991_2 | TATGAGTGTGGAGTCGTTGCAGATGGAGGTCTCACTATGTTGCNNNAGATCGGAAG |
| 49311_1 | TATGAGTGTGGAGTCGTTGCATGCTGGAATTACAAGCATGAGCNNNAGATCGGAAG |
| 49311_2 | TATGAGTGTGGAGTCGTTGCATGCTGGAATTACAAGCATGAGCNNNAGATCGGAAG |
| 39681_1 | TATGAGTGTGGAGTCGTTGCCACTCCTATATTCCTTGCACCGTNNNAGATCGGAAGA |
| 39681_2 | TATGAGTGTGGAGTCGTTGCCACTCCTATATTCCTTGCACCGTNNNAGATCGGAAGA |
| 39840_1 | TATGAGTGTGGAGTCGTTGCAGTCCTAACCATCATGAGAAACANNNAGATCGGAAG |
| 39840_2 | TATGAGTGTGGAGTCGTTGCAGTCCTAACCATCATGAGAAACANNNAGATCGGAAG |
| 33900_1 | TATGAGTGTGGAGTCGTTGCCTCACTGGAAGCTTCATCTCCTGNNNAGATCGGAAGA |
| 33900_2 | TATGAGTGTGGAGTCGTTGCCTCACTGGAAGCTTCATCTCCTGNNNAGATCGGAAGA |
| 47139_1 | TATGAGTGTGGAGTCGTTGCAGTTCCTGGTTGAGAGCTTCAATNNNAGATCGGAAG |
| 47139_2 | TATGAGTGTGGAGTCGTTGCAGTTCCTGGTTGAGAGCTTCAATNNNAGATCGGAAG |
| 13738_1 | TATGAGTGTGGAGTCGTTGCCATCATTTCTGGAGTCTCAGNNNAGATCGGAAGAGC |
| 13738_2 | TATGAGTGTGGAGTCGTTGCCATCATTTCTGGAGTCTCAGNNNAGATCGGAAGAGC |
| 7897_1 | TATGAGTGTGGAGTCGTTGCGTGGAAATGAGGCCTCGGTATGTNNNAGATCGGAA |
| 7897_2 | TATGAGTGTGGAGTCGTTGCGTGGAAATGAGGCCTCGGTATGTNNNAGATCGGAA |
| 50669_1 | TATGAGTGTGGAGTCGTTGCGGTCATGGCTTAGCAGAGAGTATNNNAGATCGGAAG |
| 50669_2 | TATGAGTGTGGAGTCGTTGCGGTCATGGCTTAGCAGAGAGTATNNNAGATCGGAAG |
| 47776_1 | TATGAGTGTGGAGTCGTTGCTTCAAGCAGTTTTCATGCCTCAGNNNAGATCGGAAGA |

| | |
|---|---|
| 47776_2 | TATGAGTGTGGAGTCGTTGCTTCAAGCAGTTTTCATGCCTCAGNNNAGATCGGAAG |
| 51126_1 | TATGAGTGTGGAGTCGTTGCAAAATGATGAATTGGCACCCCTGNNNAGATCGGAAG |
| 51126_2 | TATGAGTGTGGAGTCGTTGCAAAATGATGAATTGGCACCCCTGNNNAGATCGGAAG |
| 10686_1 | TATGAGTGTGGAGTCGTTGCTAAGCACACTTTGTTGCTAGGCCNNNAGATCGGAAG |
| 10686_2 | TATGAGTGTGGAGTCGTTGCTAAGCACACTTTGTTGCTAGGCCNNNAGATCGGAAG |
| 32836_1 | TATGAGTGTGGAGTCGTTGCCTCCTGAGTAGCTGGGACTACAGNNNAGATCGGAAG |
| 32836_2 | TATGAGTGTGGAGTCGTTGCCTCCTGAGTAGCTGGGACTACAGNNNAGATCGGAAG |
| 52024_1 | TATGAGTGTGGAGTCGTTGCTGAAACACACCCTCTCTACCAAGNNNAGATCGGAAG |
| 52024_2 | TATGAGTGTGGAGTCGTTGCTGAAACACACCCTCTCTACCAAGNNNAGATCGGAAG |
| 37329_1 | TATGAGTGTGGAGTCGTTGCAAAAGAAACAGGGCATCTTGTCANNNAGATCGGAA |
| 37329_2 | TATGAGTGTGGAGTCGTTGCAAAAGAAACAGGGCATCTTGTCANNNAGATCGGAA |
| 50269_1 | TATGAGTGTGGAGTCGTTGCTAGCCTTTCTCACAGAACACTCCNNNAGATCGGAAGA |
| 50269_2 | TATGAGTGTGGAGTCGTTGCTAGCCTTTCTCACAGAACACTCCNNNAGATCGGAAGA |
| 51239_1 | TATGAGTGTGGAGTCGTTGCGGAAAGACCGGCTATCCTAAGAGNNNAGATCGGAA |
| 51239_2 | TATGAGTGTGGAGTCGTTGCGGAAAGACCGGCTATCCTAAGAGNNNAGATCGGAA |
| 41093_1 | TATGAGTGTGGAGTCGTTGCGGCTCTGAGGCAATATGAAGAGGNNNAGATCGGAA |
| 41093_2 | TATGAGTGTGGAGTCGTTGCGGCTCTGAGGCAATATGAAGAGGNNNAGATCGGAA |
| 32715_1 | TATGAGTGTGGAGTCGTTGCAGGTGAACAATAAGTAACCAAAGAAANNNAGATCG |
| 32715_2 | TATGAGTGTGGAGTCGTTGCAGGTGAACAATAAGTAACCAAAGAAANNNAGATCG |
| 39894_1 | TATGAGTGTGGAGTCGTTGCCATCTCCTGGGATCAAGGGATTCNNNAGATCGGAAG |
| 39894_2 | TATGAGTGTGGAGTCGTTGCCATCTCCTGGGATCAAGGGATTCNNNAGATCGGAAG |
| 51638_1 | TATGAGTGTGGAGTCGTTGCAAGGATGATGAGCTCAGAGTGAGNNNAGATCGGAA |
| 51638_2 | TATGAGTGTGGAGTCGTTGCAAGGATGATGAGCTCAGAGTGAGNNNAGATCGGAA |
| 37299_1 | TATGAGTGTGGAGTCGTTGCGCCTTACAAGAAATGCTAGACGGNNNAGATCGGAA |
| 37299_2 | TATGAGTGTGGAGTCGTTGCGCCTTACAAGAAATGCTAGACGGNNNAGATCGGAA |
| 45260_1 | TATGAGTGTGGAGTCGTTGCTCCATAAAATGTGGACAAGGGGTNNNAGATCGGAA |
| 45260_2 | TATGAGTGTGGAGTCGTTGCTCCATAAAATGTGGACAAGGGGTNNNAGATCGGAA |
| 702_1 | TATGAGTGTGGAGTCGTTGCCACTCATGTCTCAGTCTCACAGANNNAGATCGGAAG |
| 702_2 | TATGAGTGTGGAGTCGTTGCCACTCATGTCTCAGTCTCACAGANNNAGATCGGAAG |
| 12931_1 | TATGAGTGTGGAGTCGTTGCCTCTCTGAAGGACATCTACAANNNAGATCGGAAGAG |
| 12931_2 | TATGAGTGTGGAGTCGTTGCCTCTCTGAAGGACATCTACAANNNAGATCGGAAGAG |
| 49710_1 | TATGAGTGTGGAGTCGTTGCGCCAACGGTACATGTTCATAGTGNNNAGATCGGAAG |
| 49710_2 | TATGAGTGTGGAGTCGTTGCGCCAACGGTACATGTTCATAGTGNNNAGATCGGAAG |
| 48920_1 | TATGAGTGTGGAGTCGTTGCACCTCCATAGACAGCCATCAAAGNNNAGATCGGAAG |
| 48920_2 | TATGAGTGTGGAGTCGTTGCACCTCCATAGACAGCCATCAAAGNNNAGATCGGAAG |
| 49157_1 | TATGAGTGTGGAGTCGTTGCCTGGAGTGCAATGGTGTGATTTCNNNAGATCGGAA |
| 49157_2 | TATGAGTGTGGAGTCGTTGCCTGGAGTGCAATGGTGTGATTTCNNNAGATCGGAA |
| 52026_1 | TATGAGTGTGGAGTCGTTGCGATGGTTTCCAGCTTCATCCATGNNNAGATCGGAAG |
| 52026_2 | TATGAGTGTGGAGTCGTTGCGATGGTTTCCAGCTTCATCCATGNNNAGATCGGAAG |
| 38182_1 | TATGAGTGTGGAGTCGTTGCGATGTGAACACCTTGCCAAAGAANNNAGATCGGAAG |
| 38182_2 | TATGAGTGTGGAGTCGTTGCGATGTGAACACCTTGCCAAAGAANNNAGATCGGAAG |
| 5690_1 | TATGAGTGTGGAGTCGTTGCTGAACCGTTACTGTATCCCAAGANNNAGATCGGAAG |
| 5690_2 | TATGAGTGTGGAGTCGTTGCTGAACCGTTACTGTATCCCAAGANNNAGATCGGAAG |

| | |
|---|---|
| 49581_1 | TATGAGTGTGGAGTCGTTGCCTCCTGAGTAGCTGGCAATAGAGNNNAGATCGGAA |
| 49581_2 | TATGAGTGTGGAGTCGTTGCCTCCTGAGTAGCTGGCAATAGAGNNNAGATCGGAA |
| 52163_1 | TATGAGTGTGGAGTCGTTGCGCGCATATATCGGCAACATAGTCNNNAGATCGGAAG |
| 52163_2 | TATGAGTGTGGAGTCGTTGCGCGCATATATCGGCAACATAGTCNNNAGATCGGAAG |
| 50116_1 | TATGAGTGTGGAGTCGTTGCCTTTGCACAGTTACCAACCTGAGNNNAGATCGGAAG |
| 50116_2 | TATGAGTGTGGAGTCGTTGCCTTTGCACAGTTACCAACCTGAGNNNAGATCGGAAG |
| 48894_1 | TATGAGTGTGGAGTCGTTGCAGTTTCTCCATGTTGGTTAGGCTNNNAGATCGGAAG |
| 48894_2 | TATGAGTGTGGAGTCGTTGCAGTTTCTCCATGTTGGTTAGGCTNNNAGATCGGAAG |
| 51814_1 | TATGAGTGTGGAGTCGTTGCCAAATGTTGGTGTCAAAGGCTCTNNNAGATCGGAA |
| 51814_2 | TATGAGTGTGGAGTCGTTGCCAAATGTTGGTGTCAAAGGCTCTNNNAGATCGGAA |
| 39019_1 | TATGAGTGTGGAGTCGTTGCAAAATCTGGACTTTGTGCTGTGGNNNAGATCGGAA |
| 39019_2 | TATGAGTGTGGAGTCGTTGCAAAATCTGGACTTTGTGCTGTGGNNNAGATCGGAA |
| 30106_1 | TATGAGTGTGGAGTCGTTGCGAAGTCTAAGTCCTTATCCCCTTTNNNAGATCGGAAG |
| 30106_2 | TATGAGTGTGGAGTCGTTGCGAAGTCTAAGTCCTTATCCCCTTTNNNAGATCGGAAG |
| 45409_1 | TATGAGTGTGGAGTCGTTGCACAAAGTTCTGACCAAAACACCGNNNAGATCGGAAG |
| 45409_2 | TATGAGTGTGGAGTCGTTGCACAAAGTTCTGACCAAAACACCGNNNAGATCGGAAG |
| 38881_1 | TATGAGTGTGGAGTCGTTGCTCATCACAGGCCAAAAAAATGGTNNNAGATCGGAA |
| 38881_2 | TATGAGTGTGGAGTCGTTGCTCATCACAGGCCAAAAAAATGGTNNNAGATCGGAA |
| 52769_1 | TATGAGTGTGGAGTCGTTGCCTAAGGCTCTTTTGGCATCCCTANNNAGATCGGAAG |
| 52769_2 | TATGAGTGTGGAGTCGTTGCCTAAGGCTCTTTTGGCATCCCTANNNAGATCGGAAG |
| 36936_1 | TATGAGTGTGGAGTCGTTGCTTGGAAGCCATTATCCTAAGCGANNNAGATCGGAAG |
| 36936_2 | TATGAGTGTGGAGTCGTTGCTTGGAAGCCATTATCCTAAGCGANNNAGATCGGAAG |
| 48633_1 | TATGAGTGTGGAGTCGTTGCGAAGTTATTCCGTTGGTGCATCCNNNAGATCGGAAG |
| 48633_2 | TATGAGTGTGGAGTCGTTGCGAAGTTATTCCGTTGGTGCATCCNNNAGATCGGAAG |
| 40897_1 | TATGAGTGTGGAGTCGTTGCAGAGTCAGCAGACACAGCAAATANNNAGATCGGAA |
| 40897_2 | TATGAGTGTGGAGTCGTTGCAGAGTCAGCAGACACAGCAAATANNNAGATCGGAA |
| 51533_1 | TATGAGTGTGGAGTCGTTGCCAAATGTATATGCGGCTCCCAAGNNNAGATCGGAAG |
| 51533_2 | TATGAGTGTGGAGTCGTTGCCAAATGTATATGCGGCTCCCAAGNNNAGATCGGAAG |
| 35858_1 | TATGAGTGTGGAGTCGTTGCGTTCAAGCGATTCTCCTACCTCANNNAGATCGGAAG |
| 35858_2 | TATGAGTGTGGAGTCGTTGCGTTCAAGCGATTCTCCTACCTCANNNAGATCGGAAG |
| 31437_1 | TATGAGTGTGGAGTCGTTGCGGGGGTTGAGGATAAGAGTAAGCNNNAGATCGGA |
| 31437_2 | TATGAGTGTGGAGTCGTTGCGGGGGTTGAGGATAAGAGTAAGCNNNAGATCGGA |
| 52052_1 | TATGAGTGTGGAGTCGTTGCATAATGTGTGTGGCTCTCATGGANNNAGATCGGAA |
| 52052_2 | TATGAGTGTGGAGTCGTTGCATAATGTGTGTGGCTCTCATGGANNNAGATCGGAA |
| 40555_1 | TATGAGTGTGGAGTCGTTGCGGAAGGATGAAGGGTTGGGAATANNNAGATCGGA |
| 40555_2 | TATGAGTGTGGAGTCGTTGCGGAAGGATGAAGGGTTGGGAATANNNAGATCGGA |
| 32976_1 | TATGAGTGTGGAGTCGTTGCGAAGGGAACTCAAGGCAAACAAANNNAGATCGGAA |
| 32976_2 | TATGAGTGTGGAGTCGTTGCGAAGGGAACTCAAGGCAAACAAANNNAGATCGGAA |
| 30310_1 | TATGAGTGTGGAGTCGTTGCTATGAAAGACTCCCCCTCTCAGCNNNAGATCGGAAG |
| 30310_2 | TATGAGTGTGGAGTCGTTGCTATGAAAGACTCCCCCTCTCAGCNNNAGATCGGAAG |
| 52836_1 | TATGAGTGTGGAGTCGTTGCCAGTTTCTCCACCATGGTCTCTGNNNAGATCGGAAG |
| 52836_2 | TATGAGTGTGGAGTCGTTGCCAGTTTCTCCACCATGGTCTCTGNNNAGATCGGAAG |
| 12505_1 | TATGAGTGTGGAGTCGTTGCTTTATCAAAAAAGAACTCAACCANNNAGATCGGAAG |

| | |
|---|---|
| 12505_2 | TATGAGTGTGGAGTCGTTGCTTTATCAAAAAAGAACTCAACCANNNAGATCGGAAG |
| 50549_1 | TATGAGTGTGGAGTCGTTGCCTACGACCTTGAGGCACTTCTACNNNAGATCGGAAG |
| 50549_2 | TATGAGTGTGGAGTCGTTGCCTACGACCTTGAGGCACTTCTACNNNAGATCGGAAG |
| 34146_1 | TATGAGTGTGGAGTCGTTGCTCATGGCTATTGTTTGAAGGCACNNNAGATCGGAAG |
| 34146_2 | TATGAGTGTGGAGTCGTTGCTCATGGCTATTGTTTGAAGGCACNNNAGATCGGAAG |
| 35056_1 | TATGAGTGTGGAGTCGTTGCACAGATATCTCAAGTCCCTGCTTNNNAGATCGGAAG |
| 35056_2 | TATGAGTGTGGAGTCGTTGCACAGATATCTCAAGTCCCTGCTTNNNAGATCGGAAG |
| 29252_1 | TATGAGTGTGGAGTCGTTGCAGTGACTTTCTCCTTCTCCCCTANNNAGATCGGAAGA |
| 29252_2 | TATGAGTGTGGAGTCGTTGCAGTGACTTTCTCCTTCTCCCCTANNNAGATCGGAAGA |
| 48809_1 | TATGAGTGTGGAGTCGTTGCCCAAGTAGCTGGGATTATAGGGGNNNAGATCGGAA |
| 48809_2 | TATGAGTGTGGAGTCGTTGCCCAAGTAGCTGGGATTATAGGGGNNNAGATCGGAA |
| 47333_1 | TATGAGTGTGGAGTCGTTGCGAGACGGATTCTTGCTGTGTTGNNNAGATCGGAAG |
| 47333_2 | TATGAGTGTGGAGTCGTTGCGAGACGGATTCTTGCTGTGTTGNNNAGATCGGAAG |
| 9074_1 | TATGAGTGTGGAGTCGTTGCGGTGTGTGAAGGACAGAGTACCTNNNAGATCGGAA |
| 9074_2 | TATGAGTGTGGAGTCGTTGCGGTGTGTGAAGGACAGAGTACCTNNNAGATCGGAA |
| 36307_1 | TATGAGTGTGGAGTCGTTGCGCAGTGGGTATCATTTTCTGCTCNNNAGATCGGAAG |
| 36307_2 | TATGAGTGTGGAGTCGTTGCGCAGTGGGTATCATTTTCTGCTCNNNAGATCGGAAG |
| 13438_1 | TATGAGTGTGGAGTCGTTGCTTCATGGGATTAGAAGAATTCAANNNAGATCGGAA |
| 13438_2 | TATGAGTGTGGAGTCGTTGCTTCATGGGATTAGAAGAATTCAANNNAGATCGGAA |
| 43514_1 | TATGAGTGTGGAGTCGTTGCGTTCAAGCAATTCTCCTGCATCANNNAGATCGGAAG |
| 43514_2 | TATGAGTGTGGAGTCGTTGCGTTCAAGCAATTCTCCTGCATCANNNAGATCGGAAG |
| 29513_1 | TATGAGTGTGGAGTCGTTGCCCATCCTCCAGTGGGTGAAGNNNAGATCGGAAGAG |
| 29513_2 | TATGAGTGTGGAGTCGTTGCCCATCCTCCAGTGGGTGAAGNNNAGATCGGAAGAG |
| 12107_1 | TATGAGTGTGGAGTCGTTGCCTTGTCAACTGCATTTCCTCTTGCNNNAGATCGGAAG |
| 12107_2 | TATGAGTGTGGAGTCGTTGCCTTGTCAACTGCATTTCCTCTTGCNNNAGATCGGAAG |
| 46852_1 | TATGAGTGTGGAGTCGTTGCGATCTTGACTCACTGCAACCTCTNNNAGATCGGAAG |
| 46852_2 | TATGAGTGTGGAGTCGTTGCGATCTTGACTCACTGCAACCTCTNNNAGATCGGAAG |
| 36335_1 | TATGAGTGTGGAGTCGTTGCGGACCACTACAATAAGCACAAGCNNNAGATCGGAA |
| 36335_2 | TATGAGTGTGGAGTCGTTGCGGACCACTACAATAAGCACAAGCNNNAGATCGGAA |
| 45490_1 | TATGAGTGTGGAGTCGTTGCCTGACACTGCCCATCTTTGTTTTNNNAGATCGGAAGA |
| 45490_2 | TATGAGTGTGGAGTCGTTGCCTGACACTGCCCATCTTTGTTTTNNNAGATCGGAAGA |
| 40988_1 | TATGAGTGTGGAGTCGTTGCGTAGGGTGATTAGAGGGCTTGAGNNNAGATCGGA |
| 40988_2 | TATGAGTGTGGAGTCGTTGCGTAGGGTGATTAGAGGGCTTGAGNNNAGATCGGA |
| 41722_1 | TATGAGTGTGGAGTCGTTGCCGGCTAATTTGTGACTTGTGTGTNNNAGATCGGAAG |
| 41722_2 | TATGAGTGTGGAGTCGTTGCCGGCTAATTTGTGACTTGTGTGTNNNAGATCGGAAG |
| 51111_1 | TATGAGTGTGGAGTCGTTGCAATTTCCATTTCCCAACTCCCCTNNNAGATCGGAAGA |
| 51111_2 | TATGAGTGTGGAGTCGTTGCAATTTCCATTTCCCAACTCCCCTNNNAGATCGGAAGA |
| 38752_1 | TATGAGTGTGGAGTCGTTGCAATCCTAGCTACTTTGGAGGCTGNNNAGATCGGAAG |
| 38752_2 | TATGAGTGTGGAGTCGTTGCAATCCTAGCTACTTTGGAGGCTGNNNAGATCGGAAG |
| 33926_1 | TATGAGTGTGGAGTCGTTGCATCCCAGGTTCAAGTGATTCTCCNNNAGATCGGAAG |
| 33926_2 | TATGAGTGTGGAGTCGTTGCATCCCAGGTTCAAGTGATTCTCCNNNAGATCGGAAG |
| 37160_1 | TATGAGTGTGGAGTCGTTGCGAAGGAAAGGAAGGAAGGAAGGANNNAGATCGG |
| 37160_2 | TATGAGTGTGGAGTCGTTGCGAAGGAAAGGAAGGAAGGAAGGANNNAGATCGG |

| | |
|---|---|
| 33326_1 | TATGAGTGTGGAGTCGTTGCTCCTGGGTTCAAATGATTCTCGTNNNAGATCGGAAG |
| 33326_2 | TATGAGTGTGGAGTCGTTGCTCCTGGGTTCAAATGATTCTCGTNNNAGATCGGAAG |
| 48592_1 | TATGAGTGTGGAGTCGTTGCCTCCTGGACTCAAGCAACCTTCNNNAGATCGGAAGA |
| 48592_2 | TATGAGTGTGGAGTCGTTGCCTCCTGGACTCAAGCAACCTTCNNNAGATCGGAAGA |
| 30125_1 | TATGAGTGTGGAGTCGTTGCGGATCCATGGCAGAGAAAAAAGGNNNAGATCGGA |
| 30125_2 | TATGAGTGTGGAGTCGTTGCGGATCCATGGCAGAGAAAAAAGGNNNAGATCGGA |
| 32489_1 | TATGAGTGTGGAGTCGTTGCGTTCAAGCGATTCTCTTGCCTCNNNAGATCGGAAGA |
| 32489_2 | TATGAGTGTGGAGTCGTTGCGTTCAAGCGATTCTCTTGCCTCNNNAGATCGGAAGA |
| 47111_1 | TATGAGTGTGGAGTCGTTGCCCACACCCGGCTAATTTTGTATTNNNAGATCGGAAGA |
| 47111_2 | TATGAGTGTGGAGTCGTTGCCCACACCCGGCTAATTTTGTATTNNNAGATCGGAAGA |
| 34987_1 | TATGAGTGTGGAGTCGTTGCGAGAAAGGGTCTTGTTCTGTTGCNNNAGATCGGAA |
| 34987_2 | TATGAGTGTGGAGTCGTTGCGAGAAAGGGTCTTGTTCTGTTGCNNNAGATCGGAA |
| 5563_1 | TATGAGTGTGGAGTCGTTGCTTGCCAGAAGGTATAACTGAAGANNNAGATCGGAA |
| 5563_2 | TATGAGTGTGGAGTCGTTGCTTGCCAGAAGGTATAACTGAAGANNNAGATCGGAA |
| 52542_1 | TATGAGTGTGGAGTCGTTGCAAGATGGGTAATGAATGAGGGGGNNNAGATCGGA |
| 52542_2 | TATGAGTGTGGAGTCGTTGCAAGATGGGTAATGAATGAGGGGGNNNAGATCGGA |
| 43168_1 | TATGAGTGTGGAGTCGTTGCCTCCTGAGTAGCTGAGCTTACAGNNNAGATCGGAAG |
| 43168_2 | TATGAGTGTGGAGTCGTTGCCTCCTGAGTAGCTGAGCTTACAGNNNAGATCGGAAG |
| 39196_1 | TATGAGTGTGGAGTCGTTGCCACCCATGCCTTATAATGCCTTGNNNAGATCGGAAGA |
| 39196_2 | TATGAGTGTGGAGTCGTTGCCACCCATGCCTTATAATGCCTTGNNNAGATCGGAAGA |
| 33606_1 | TATGAGTGTGGAGTCGTTGCAATTTCCCAGGCTCAAGCAATTCNNNAGATCGGAAG |
| 33606_2 | TATGAGTGTGGAGTCGTTGCAATTTCCCAGGCTCAAGCAATTCNNNAGATCGGAAG |
| 45433_1 | TATGAGTGTGGAGTCGTTGCCACGTGTATGGAATGTGTGTGTGNNNAGATCGGAA |
| 45433_2 | TATGAGTGTGGAGTCGTTGCCACGTGTATGGAATGTGTGTGTGNNNAGATCGGAA |
| 35943_1 | TATGAGTGTGGAGTCGTTGCGGTGGATCATGAGGTCAAGAGATNNNAGATCGGAA |
| 35943_2 | TATGAGTGTGGAGTCGTTGCGGTGGATCATGAGGTCAAGAGATNNNAGATCGGAA |
| 33822_1 | TATGAGTGTGGAGTCGTTGCGAACAGGGAAGGTTTAGCCAGTANNNAGATCGGAA |
| 33822_2 | TATGAGTGTGGAGTCGTTGCGAACAGGGAAGGTTTAGCCAGTANNNAGATCGGAA |
| 43511_1 | TATGAGTGTGGAGTCGTTGCTCTGGGTTCAAGTGATTCTCCTGNNNAGATCGGAAG |
| 43511_2 | TATGAGTGTGGAGTCGTTGCTCTGGGTTCAAGTGATTCTCCTGNNNAGATCGGAAG |
| 42835_1 | TATGAGTGTGGAGTCGTTGCGGTGAGTCTGTGGTCATTTTGTGNNNAGATCGGAA |
| 42835_2 | TATGAGTGTGGAGTCGTTGCGGTGAGTCTGTGGTCATTTTGTGNNNAGATCGGAA |
| 34187_1 | TATGAGTGTGGAGTCGTTGCAAGGAGAGATGAGGGAGTCAGAANNNAGATCGGA |
| 34187_2 | TATGAGTGTGGAGTCGTTGCAAGGAGAGATGAGGGAGTCAGAANNNAGATCGGA |
| 41159_1 | TATGAGTGTGGAGTCGTTGCAATCCCAGGTGTTCTATGAGTGANNNAGATCGGAA |
| 41159_2 | TATGAGTGTGGAGTCGTTGCAATCCCAGGTGTTCTATGAGTGANNNAGATCGGAA |
| 52650_1 | TATGAGTGTGGAGTCGTTGCCCTTACCAGTTGCTATTGTTGGCNNNAGATCGGAAG |
| 52650_2 | TATGAGTGTGGAGTCGTTGCCCTTACCAGTTGCTATTGTTGGCNNNAGATCGGAAG |
| 43041_1 | TATGAGTGTGGAGTCGTTGCATGAAATAATGGCATTCGCAGCANNNAGATCGGAA |
| 43041_2 | TATGAGTGTGGAGTCGTTGCATGAAATAATGGCATTCGCAGCANNNAGATCGGAA |
| 10301_1 | TATGAGTGTGGAGTCGTTGCGCTCAGACCTTCAGAACAATGTTNNNAGATCGGAAG |
| 10301_2 | TATGAGTGTGGAGTCGTTGCGCTCAGACCTTCAGAACAATGTTNNNAGATCGGAAG |
| 42196_1 | TATGAGTGTGGAGTCGTTGCCTGGAGTACAATGGCACGATCTTNNNAGATCGGAA |

| | |
|---|---|
| 42196_2 | TATGAGTGTGGAGTCGTTGCCTGGAGTACAATGGCACGATCTTNNNAGATCGGAAG |
| 46458_1 | TATGAGTGTGGAGTCGTTGCCTCTCGAGTAGCTGGGATTACAGNNNAGATCGGAAG |
| 46458_2 | TATGAGTGTGGAGTCGTTGCCTCTCGAGTAGCTGGGATTACAGNNNAGATCGGAAG |
| 40255_1 | TATGAGTGTGGAGTCGTTGCGGATAAGCAAAACAGGGTCCAACNNNAGATCGGAA |
| 40255_2 | TATGAGTGTGGAGTCGTTGCGGATAAGCAAAACAGGGTCCAACNNNAGATCGGAA |
| 50750_1 | TATGAGTGTGGAGTCGTTGCCATGAGATACAGCATGCAAGAGCNNNAGATCGGAA |
| 50750_2 | TATGAGTGTGGAGTCGTTGCCATGAGATACAGCATGCAAGAGCNNNAGATCGGAA |
| 30158_1 | TATGAGTGTGGAGTCGTTGCGCTCTACTGTGGTAAGTGTCCAANNNAGATCGGAAG |
| 30158_2 | TATGAGTGTGGAGTCGTTGCGCTCTACTGTGGTAAGTGTCCAANNNAGATCGGAAG |
| 39823_1 | TATGAGTGTGGAGTCGTTGCTTACAAGCATGAGCCACTGTACTNNNAGATCGGAAG |
| 39823_2 | TATGAGTGTGGAGTCGTTGCTTACAAGCATGAGCCACTGTACTNNNAGATCGGAAG |
| 30516_1 | TATGAGTGTGGAGTCGTTGCGACTGAAATAGCTGGGTGTTTTAGGNNNAGATCGG |
| 30516_2 | TATGAGTGTGGAGTCGTTGCGACTGAAATAGCTGGGTGTTTTAGGNNNAGATCGG |
| 52813_1 | TATGAGTGTGGAGTCGTTGCAGTCCTAATTCCTGCATCCAAGGNNNAGATCGGAAG |
| 52813_2 | TATGAGTGTGGAGTCGTTGCAGTCCTAATTCCTGCATCCAAGGNNNAGATCGGAAG |
| 10186_1 | TATGAGTGTGGAGTCGTTGCGTTCAAGAGCAGTGGTAGCCAATNNNAGATCGGAA |
| 10186_2 | TATGAGTGTGGAGTCGTTGCGTTCAAGAGCAGTGGTAGCCAATNNNAGATCGGAA |
| 51615_1 | TATGAGTGTGGAGTCGTTGCGCTGAGACATGAGAGGCCTCNNNAGATCGGAAGAG |
| 51615_2 | TATGAGTGTGGAGTCGTTGCGCTGAGACATGAGAGGCCTCNNNAGATCGGAAGAG |
| 53261_1 | TATGAGTGTGGAGTCGTTGCTTCCTGAAATGCCATCACTACCANNNAGATCGGAAGA |
| 53261_2 | TATGAGTGTGGAGTCGTTGCTTCCTGAAATGCCATCACTACCANNNAGATCGGAAGA |
| 2000_1 | TATGAGTGTGGAGTCGTTGCGGCTTTGCTAGGGCATTTCAAAANNNAGATCGGAAG |
| 2000_2 | TATGAGTGTGGAGTCGTTGCGGCTTTGCTAGGGCATTTCAAAANNNAGATCGGAAG |
| 36150_1 | TATGAGTGTGGAGTCGTTGCCCCATTCCTCCAACTTCTGAACTNNNAGATCGGAAGA |
| 36150_2 | TATGAGTGTGGAGTCGTTGCCCCATTCCTCCAACTTCTGAACTNNNAGATCGGAAGA |
| 42290_1 | TATGAGTGTGGAGTCGTTGCTATTTATAGAACTGGGCAGGGGCNNNAGATCGGAA |
| 42290_2 | TATGAGTGTGGAGTCGTTGCTATTTATAGAACTGGGCAGGGGCNNNAGATCGGAA |
| 52562_1 | TATGAGTGTGGAGTCGTTGCACTTCTGCCCAGTGTAAGATTGTNNNAGATCGGAAG |
| 52562_2 | TATGAGTGTGGAGTCGTTGCACTTCTGCCCAGTGTAAGATTGTNNNAGATCGGAAG |
| 45393_1 | TATGAGTGTGGAGTCGTTGCTTTGGGAGAGATGGAAGGAAAGGNNNAGATCGGA |
| 45393_2 | TATGAGTGTGGAGTCGTTGCTTTGGGAGAGATGGAAGGAAAGGNNNAGATCGGA |
| 41383_1 | TATGAGTGTGGAGTCGTTGCCCTGCTCTGATTTACGTCTTCCTNNNAGATCGGAAGA |
| 41383_2 | TATGAGTGTGGAGTCGTTGCCCTGCTCTGATTTACGTCTTCCTNNNAGATCGGAAGA |
| 13729_1 | TATGAGTGTGGAGTCGTTGCATTGTTGTATCACCAGTTAGTTNNNAGATCGGAAGA |
| 13729_2 | TATGAGTGTGGAGTCGTTGCATTGTTGTATCACCAGTTAGTTNNNAGATCGGAAGA |
| 43512_1 | TATGAGTGTGGAGTCGTTGCGGGTTCAAGTGATTCTCCTGTCTNNNAGATCGGAAG |
| 43512_2 | TATGAGTGTGGAGTCGTTGCGGGTTCAAGTGATTCTCCTGTCTNNNAGATCGGAAG |
| 35651_1 | TATGAGTGTGGAGTCGTTGCTTTAGTAGAGGCGGGGTTTTACCNNNAGATCGGAA |
| 35651_2 | TATGAGTGTGGAGTCGTTGCTTTAGTAGAGGCGGGGTTTTACCNNNAGATCGGAA |
| 53335_1 | TATGAGTGTGGAGTCGTTGCTGCTCACGGATAGAAAGCATCAANNNAGATCGGAAG |
| 53335_2 | TATGAGTGTGGAGTCGTTGCTGCTCACGGATAGAAAGCATCAANNNAGATCGGAAG |
| 10112_1 | TATGAGTGTGGAGTCGTTGCTTCTCAACATCCCTTTCCTGCACNNNAGATCGGAAGA |
| 10112_2 | TATGAGTGTGGAGTCGTTGCTTCTCAACATCCCTTTCCTGCACNNNAGATCGGAAGA |

| | |
|---|---|
| 30706_1 | TATGAGTGTGGAGTCGTTGCTCACAAATCGTTCTTTGGGCAGNNNAGATCGGAAGA |
| 30706_2 | TATGAGTGTGGAGTCGTTGCTCACAAATCGTTCTTTGGGCAGNNNAGATCGGAAGA |
| 37503_1 | TATGAGTGTGGAGTCGTTGCGAAAAGAAAAAACCACCGTACGGNNNAGATCGGAA |
| 37503_2 | TATGAGTGTGGAGTCGTTGCGAAAAGAAAAAACCACCGTACGGNNNAGATCGGAA |
| 37644_1 | TATGAGTGTGGAGTCGTTGCTGAGGCAGAAAATTGCTTGAACCNNNAGATCGGAA |
| 37644_2 | TATGAGTGTGGAGTCGTTGCTGAGGCAGAAAATTGCTTGAACCNNNAGATCGGAA |
| 46645_1 | TATGAGTGTGGAGTCGTTGCGACAGAGTCTCACCCTTTCACCNNNAGATCGGAAGA |
| 46645_2 | TATGAGTGTGGAGTCGTTGCGACAGAGTCTCACCCTTTCACCNNNAGATCGGAAGA |
| 42330_1 | TATGAGTGTGGAGTCGTTGCTCCTTTTCCTGGAGGTACTAGCTNNNAGATCGGAAG |
| 42330_2 | TATGAGTGTGGAGTCGTTGCTCCTTTTCCTGGAGGTACTAGCTNNNAGATCGGAAG |
| 40692_1 | TATGAGTGTGGAGTCGTTGCAGCCCATTCCTATTGGTTCCATTNNNAGATCGGAAGA |
| 40692_2 | TATGAGTGTGGAGTCGTTGCAGCCCATTCCTATTGGTTCCATTNNNAGATCGGAAGA |
| 41436_1 | TATGAGTGTGGAGTCGTTGCTTGAGCAGACCTTGGAAGTTTCTNNNAGATCGGAAG |
| 41436_2 | TATGAGTGTGGAGTCGTTGCTTGAGCAGACCTTGGAAGTTTCTNNNAGATCGGAAG |
| 45437_1 | TATGAGTGTGGAGTCGTTGCTATGCAAACACACACACACAGACNNNAGATCGGAAG |
| 45437_2 | TATGAGTGTGGAGTCGTTGCTATGCAAACACACACACACAGACNNNAGATCGGAAG |
| 38663_1 | TATGAGTGTGGAGTCGTTGCTATGTCCTTCCTCCAGCAAACTTNNNAGATCGGAAGA |
| 38663_2 | TATGAGTGTGGAGTCGTTGCTATGTCCTTCCTCCAGCAAACTTNNNAGATCGGAAGA |
| 53006_1 | TATGAGTGTGGAGTCGTTGCACTCTTGAACAATGTGGGGGTTANNNAGATCGGAA |
| 53006_2 | TATGAGTGTGGAGTCGTTGCACTCTTGAACAATGTGGGGGTTANNNAGATCGGAA |
| 34382_1 | TATGAGTGTGGAGTCGTTGCGGAATCTGCCCTGTGTCGTCNNNAGATCGGAAGAG |
| 34382_2 | TATGAGTGTGGAGTCGTTGCGGAATCTGCCCTGTGTCGTCNNNAGATCGGAAGAG |
| 46691_1 | TATGAGTGTGGAGTCGTTGCGATGGGAGGATCACTTGATGTCANNNAGATCGGAA |
| 46691_2 | TATGAGTGTGGAGTCGTTGCGATGGGAGGATCACTTGATGTCANNNAGATCGGAA |
| 42558_1 | TATGAGTGTGGAGTCGTTGCGTCACCCCCTCTCAATTTTTTGGNNNAGATCGGAAGA |
| 42558_2 | TATGAGTGTGGAGTCGTTGCGTCACCCCCTCTCAATTTTTTGGNNNAGATCGGAAGA |
| 30511_1 | TATGAGTGTGGAGTCGTTGCTCTTAAAGCCCCCAGATTCATGCNNNAGATCGGAAG |
| 30511_2 | TATGAGTGTGGAGTCGTTGCTCTTAAAGCCCCCAGATTCATGCNNNAGATCGGAAG |
| 12509_1 | TATGAGTGTGGAGTCGTTGCGCCTTGGTAACTTTTTCTAATTANNNAGATCGGAAG |
| 12509_2 | TATGAGTGTGGAGTCGTTGCGCCTTGGTAACTTTTTCTAATTANNNAGATCGGAAG |
| 10095_1 | TATGAGTGTGGAGTCGTTGCTGGTTCCTGGCTCATAACTCTTGNNNAGATCGGAAG |
| 10095_2 | TATGAGTGTGGAGTCGTTGCTGGTTCCTGGCTCATAACTCTTGNNNAGATCGGAAG |
| 52593_1 | TATGAGTGTGGAGTCGTTGCGAGGATGTAGGGCAAAAATGAGCNNNAGATCGGA |
| 52593_2 | TATGAGTGTGGAGTCGTTGCGAGGATGTAGGGCAAAAATGAGCNNNAGATCGGA |
| 33549_1 | TATGAGTGTGGAGTCGTTGCCTCCTAAGTAGCTGGGATTGCAGNNNAGATCGGAA |
| 33549_2 | TATGAGTGTGGAGTCGTTGCCTCCTAAGTAGCTGGGATTGCAGNNNAGATCGGAA |
| 44966_1 | TATGAGTGTGGAGTCGTTGCGTTTCTTAGCCATGTGCCCATTTNNNAGATCGGAAG |
| 44966_2 | TATGAGTGTGGAGTCGTTGCGTTTCTTAGCCATGTGCCCATTTNNNAGATCGGAAG |
| 13040_1 | TATGAGTGTGGAGTCGTTGCCTCTTTGTTTCTCATCAACTGNNNAGATCGGAAGAG |
| 13040_2 | TATGAGTGTGGAGTCGTTGCCTCTTTGTTTCTCATCAACTGNNNAGATCGGAAGAG |
| 42225_1 | TATGAGTGTGGAGTCGTTGCAGTTGGAGTCTCACTGTGTCATCNNNAGATCGGAAG |
| 42225_2 | TATGAGTGTGGAGTCGTTGCAGTTGGAGTCTCACTGTGTCATCNNNAGATCGGAAG |
| 38641_1 | TATGAGTGTGGAGTCGTTGCAAGAACCGTGCAACCATAAAAGGNNNAGATCGGAA |

| | |
|---|---|
| 38641_2 | TATGAGTGTGGAGTCGTTGCAAGAACCGTGCAACCATAAAAGGNNNAGATCGGAA |
| 10136_1 | TATGAGTGTGGAGTCGTTGCTGGTTTCCTGAATTCTGTCACAGTNNNAGATCGGAA |
| 10136_2 | TATGAGTGTGGAGTCGTTGCTGGTTTCCTGAATTCTGTCACAGTNNNAGATCGGAA |
| 34215_1 | TATGAGTGTGGAGTCGTTGCGGTTGCAATCCTAGTCTCTGACANNNAGATCGGAAG |
| 34215_2 | TATGAGTGTGGAGTCGTTGCGGTTGCAATCCTAGTCTCTGACANNNAGATCGGAAG |
| 33962_1 | TATGAGTGTGGAGTCGTTGCCTTCACAAGCAGCTGGAATTACANNNAGATCGGAAG |
| 33962_2 | TATGAGTGTGGAGTCGTTGCCTTCACAAGCAGCTGGAATTACANNNAGATCGGAAG |
| 33507_1 | TATGAGTGTGGAGTCGTTGCTGAGAAACTCATTCCTTGTCCCCNNNAGATCGGAAG |
| 33507_2 | TATGAGTGTGGAGTCGTTGCTGAGAAACTCATTCCTTGTCCCCNNNAGATCGGAAG |
| 36966_1 | TATGAGTGTGGAGTCGTTGCAAGACTCCAGAGAAAGAAAGCCCNNNAGATCGGAA |
| 36966_2 | TATGAGTGTGGAGTCGTTGCAAGACTCCAGAGAAAGAAAGCCCNNNAGATCGGAA |
| 48787_1 | TATGAGTGTGGAGTCGTTGCTGACACCAGAACGGACCAATATTNNNAGATCGGAAG |
| 48787_2 | TATGAGTGTGGAGTCGTTGCTGACACCAGAACGGACCAATATTNNNAGATCGGAAG |
| 45690_1 | TATGAGTGTGGAGTCGTTGCGGAGGGGAGAAGCACCGATCNNNAGATCGGAAGA |
| 45690_2 | TATGAGTGTGGAGTCGTTGCGGAGGGGAGAAGCACCGATCNNNAGATCGGAAGA |
| 33481_1 | TATGAGTGTGGAGTCGTTGCTGCAGGTAAAGTGCTTTTTTGTTNNNAGATCGGAAG |
| 33481_2 | TATGAGTGTGGAGTCGTTGCTGCAGGTAAAGTGCTTTTTTGTTNNNAGATCGGAAG |
| 38210_1 | TATGAGTGTGGAGTCGTTGCTTGAAAGGAGGCAAAAAGGTCACNNNAGATCGGAA |
| 38210_2 | TATGAGTGTGGAGTCGTTGCTTGAAAGGAGGCAAAAAGGTCACNNNAGATCGGAA |
| 48476_1 | TATGAGTGTGGAGTCGTTGCTGCAGGTTATTTTGCGATGAAGGNNNAGATCGGAA |
| 48476_2 | TATGAGTGTGGAGTCGTTGCTGCAGGTTATTTTGCGATGAAGGNNNAGATCGGAA |
| 8804_1 | TATGAGTGTGGAGTCGTTGCCCCCACCCTTCCTTTCTCTACTTNNNAGATCGGAAGAG |
| 8804_2 | TATGAGTGTGGAGTCGTTGCCCCCACCCTTCCTTTCTCTACTTNNNAGATCGGAAGAG |
| 53289_1 | TATGAGTGTGGAGTCGTTGCCTGATCCAGGAGGCCGCATCNNNAGATCGGAAGAG |
| 53289_2 | TATGAGTGTGGAGTCGTTGCCTGATCCAGGAGGCCGCATCNNNAGATCGGAAGAG |
| 10787_1 | TATGAGTGTGGAGTCGTTGCATGAAGTTCGGGAGCTATTGCAGNNNAGATCGGAA |
| 10787_2 | TATGAGTGTGGAGTCGTTGCATGAAGTTCGGGAGCTATTGCAGNNNAGATCGGAA |
| 49800_1 | TATGAGTGTGGAGTCGTTGCGCTGAAATTCACTGAGGTCCCTANNNAGATCGGAAG |
| 49800_2 | TATGAGTGTGGAGTCGTTGCGCTGAAATTCACTGAGGTCCCTANNNAGATCGGAAG |
| 47328_1 | TATGAGTGTGGAGTCGTTGCTTATGAGACAGGGTCTTGCTCTGNNNAGATCGGAA |
| 47328_2 | TATGAGTGTGGAGTCGTTGCTTATGAGACAGGGTCTTGCTCTGNNNAGATCGGAA |
| 30321_1 | TATGAGTGTGGAGTCGTTGCATTGTGTGACCGTGTCCTTTGAANNNAGATCGGAAG |
| 30321_2 | TATGAGTGTGGAGTCGTTGCATTGTGTGACCGTGTCCTTTGAANNNAGATCGGAAG |
| 39787_1 | TATGAGTGTGGAGTCGTTGCGTTCTGGTTCAGGAAAGCTTGACNNNAGATCGGAA |
| 39787_2 | TATGAGTGTGGAGTCGTTGCGTTCTGGTTCAGGAAAGCTTGACNNNAGATCGGAA |
| 10669_1 | TATGAGTGTGGAGTCGTTGCACGAAGCCAAGATCCTGAGATTCNNNAGATCGGAA |
| 10669_2 | TATGAGTGTGGAGTCGTTGCACGAAGCCAAGATCCTGAGATTCNNNAGATCGGAA |
| 40060_1 | TATGAGTGTGGAGTCGTTGCAAATGGACTGCCCTTTACCTAGCNNNAGATCGGAAG |
| 40060_2 | TATGAGTGTGGAGTCGTTGCAAATGGACTGCCCTTTACCTAGCNNNAGATCGGAAG |
| 36958_1 | TATGAGTGTGGAGTCGTTGCACAAACAGGCTATGCAATGTAACANNNAGATCGGA |
| 36958_2 | TATGAGTGTGGAGTCGTTGCACAAACAGGCTATGCAATGTAACANNNAGATCGGA |
| 43788_1 | TATGAGTGTGGAGTCGTTGCATATATCAGCGCCTTGGAGAAGCNNNAGATCGGAA |
| 43788_2 | TATGAGTGTGGAGTCGTTGCATATATCAGCGCCTTGGAGAAGCNNNAGATCGGAA |

| | |
|---|---|
| 38001_1 | TATGAGTGTGGAGTCGTTGCATGGGCAAAACAGGGTTATACCTNNNAGATCGGAA |
| 38001_2 | TATGAGTGTGGAGTCGTTGCATGGGCAAAACAGGGTTATACCTNNNAGATCGGAA |
| 43863_1 | TATGAGTGTGGAGTCGTTGCTCCATCCTTGCAATAATCCCACANNNAGATCGGAAGA |
| 43863_2 | TATGAGTGTGGAGTCGTTGCTCCATCCTTGCAATAATCCCACANNNAGATCGGAAGA |
| 50234_1 | TATGAGTGTGGAGTCGTTGCTTCCTGGAGCTCTGTCCCTATTANNNAGATCGGAAG |
| 50234_2 | TATGAGTGTGGAGTCGTTGCTTCCTGGAGCTCTGTCCCTATTANNNAGATCGGAAG |
| 42012_1 | TATGAGTGTGGAGTCGTTGCTTCAAGTGATTCTCCTGCCTCAGNNNAGATCGGAAG |
| 42012_2 | TATGAGTGTGGAGTCGTTGCTTCAAGTGATTCTCCTGCCTCAGNNNAGATCGGAAG |
| 37743_1 | TATGAGTGTGGAGTCGTTGCAGAAAATTGCCAAACCCCTCATGNNNAGATCGGAAG |
| 37743_2 | TATGAGTGTGGAGTCGTTGCAGAAAATTGCCAAACCCCTCATGNNNAGATCGGAAG |
| 46398_1 | TATGAGTGTGGAGTCGTTGCGATGGTCTCGATCTCCTGACTTCNNNAGATCGGAAG |
| 46398_2 | TATGAGTGTGGAGTCGTTGCGATGGTCTCGATCTCCTGACTTCNNNAGATCGGAAG |
| 44568_1 | TATGAGTGTGGAGTCGTTGCGGCTTCTCAGGCATGATCTGTATNNNAGATCGGAAG |
| 44568_2 | TATGAGTGTGGAGTCGTTGCGGCTTCTCAGGCATGATCTGTATNNNAGATCGGAAG |
| 39312_1 | TATGAGTGTGGAGTCGTTGCCAGTGAGCCAAGATCACACTACTNNNAGATCGGAAG |
| 39312_2 | TATGAGTGTGGAGTCGTTGCCAGTGAGCCAAGATCACACTACTNNNAGATCGGAAG |
| 43500_1 | TATGAGTGTGGAGTCGTTGCCCTCCCAAGTAGCTGGATTACAGNNNAGATCGGAAG |
| 43500_2 | TATGAGTGTGGAGTCGTTGCCCTCCCAAGTAGCTGGATTACAGNNNAGATCGGAAG |
| 52131_1 | TATGAGTGTGGAGTCGTTGCCTGTGCCTCATAATATGCCATGCNNNAGATCGGAAG |
| 52131_2 | TATGAGTGTGGAGTCGTTGCCTGTGCCTCATAATATGCCATGCNNNAGATCGGAAG |
| 34961_1 | TATGAGTGTGGAGTCGTTGCCTTTCCATGGCCTCAACTCTACTNNNAGATCGGAAGA |
| 34961_2 | TATGAGTGTGGAGTCGTTGCCTTTCCATGGCCTCAACTCTACTNNNAGATCGGAAGA |
| 36896_1 | TATGAGTGTGGAGTCGTTGCAAAATAATATGCCAGGCACGGTGNNNAGATCGGAA |
| 36896_2 | TATGAGTGTGGAGTCGTTGCAAAATAATATGCCAGGCACGGTGNNNAGATCGGAA |
| 12689_1 | TATGAGTGTGGAGTCGTTGCAAATTGGATTGAGGTAAATTGTTNNNAGATCGGAA |
| 12689_2 | TATGAGTGTGGAGTCGTTGCAAATTGGATTGAGGTAAATTGTTNNNAGATCGGAA |
| 48984_1 | TATGAGTGTGGAGTCGTTGCGCTGGAGTGTAGTGGTACAATCTNNNAGATCGGAA |
| 48984_2 | TATGAGTGTGGAGTCGTTGCGCTGGAGTGTAGTGGTACAATCTNNNAGATCGGAA |
| 36284_1 | TATGAGTGTGGAGTCGTTGCGTTACTTTTGATGTCCAGCCTGCNNNAGATCGGAAG |
| 36284_2 | TATGAGTGTGGAGTCGTTGCGTTACTTTTGATGTCCAGCCTGCNNNAGATCGGAAG |
| 40179_1 | TATGAGTGTGGAGTCGTTGCAGGTACTGGGATTGTCATTGGACNNNAGATCGGAA |
| 40179_2 | TATGAGTGTGGAGTCGTTGCAGGTACTGGGATTGTCATTGGACNNNAGATCGGAA |
| 52841_1 | TATGAGTGTGGAGTCGTTGCTCAGAGGCTCGCAAGAAAGAAGNNNAGATCGGAAG |
| 52841_2 | TATGAGTGTGGAGTCGTTGCTCAGAGGCTCGCAAGAAAGAAGNNNAGATCGGAAG |
| 31995_1 | TATGAGTGTGGAGTCGTTGCTACCAGATGGATTCACAGCCAAANNNAGATCGGAAG |
| 31995_2 | TATGAGTGTGGAGTCGTTGCTACCAGATGGATTCACAGCCAAANNNAGATCGGAAG |
| 48187_1 | TATGAGTGTGGAGTCGTTGCTTCAAGTGATTCTCCTGCCTCAGNNNAGATCGGAAG |
| 48187_2 | TATGAGTGTGGAGTCGTTGCTTCAAGTGATTCTCCTGCCTCAGNNNAGATCGGAAG |
| 41977_1 | TATGAGTGTGGAGTCGTTGCTCCGGATTAGCTGGAATTACAGGNNNAGATCGGAA |
| 41977_2 | TATGAGTGTGGAGTCGTTGCTCCGGATTAGCTGGAATTACAGGNNNAGATCGGAA |
| 53252_1 | TATGAGTGTGGAGTCGTTGCCTGGCAGGCTGACTGACTATATTNNNAGATCGGAAG |
| 53252_2 | TATGAGTGTGGAGTCGTTGCCTGGCAGGCTGACTGACTATATTNNNAGATCGGAAG |
| 52423_1 | TATGAGTGTGGAGTCGTTGCCTTTTTCACCTTCCTTCTAGCGCNNNAGATCGGAAGA |

| | |
|---|---|
| 52423_2 | TATGAGTGTGGAGTCGTTGCCTTTTTCACCTTCCTTCTAGCGCNNNAGATCGGAAGA |
| 45325_1 | TATGAGTGTGGAGTCGTTGCAGGTGAAATCCACATAAAAGCCTNNNAGATCGGAA |
| 45325_2 | TATGAGTGTGGAGTCGTTGCAGGTGAAATCCACATAAAAGCCTNNNAGATCGGAA |
| 36638_1 | TATGAGTGTGGAGTCGTTGCGTGGATCATTTGAGGTCAGGAGTNNNAGATCGGAA |
| 36638_2 | TATGAGTGTGGAGTCGTTGCGTGGATCATTTGAGGTCAGGAGTNNNAGATCGGAA |
| 40835_1 | TATGAGTGTGGAGTCGTTGCCAGGATGGTGGTTGCTATAGGTTNNNAGATCGGAA |
| 40835_2 | TATGAGTGTGGAGTCGTTGCCAGGATGGTGGTTGCTATAGGTTNNNAGATCGGAA |
| 47338_1 | TATGAGTGTGGAGTCGTTGCTACTCTGCCATCTTGCTCTTGTTNNNAGATCGGAAGA |
| 47338_2 | TATGAGTGTGGAGTCGTTGCTACTCTGCCATCTTGCTCTTGTTNNNAGATCGGAAGA |
| 31489_1 | TATGAGTGTGGAGTCGTTGCGTGACGTACATCAATTTGCCGATNNNAGATCGGAAG |
| 31489_2 | TATGAGTGTGGAGTCGTTGCGTGACGTACATCAATTTGCCGATNNNAGATCGGAAG |
| 51617_1 | TATGAGTGTGGAGTCGTTGCTGGCTGATATGACAACACTTTGCNNNAGATCGGAAG |
| 51617_2 | TATGAGTGTGGAGTCGTTGCTGGCTGATATGACAACACTTTGCNNNAGATCGGAAG |
| 39464_1 | TATGAGTGTGGAGTCGTTGCTTGCATAGCTATCAGAGGCATCANNNAGATCGGAAG |
| 39464_2 | TATGAGTGTGGAGTCGTTGCTTGCATAGCTATCAGAGGCATCANNNAGATCGGAAG |
| 44620_1 | TATGAGTGTGGAGTCGTTGCCATCACCCAGGCTCACCAGGNNNAGATCGGAAGAG |
| 44620_2 | TATGAGTGTGGAGTCGTTGCCATCACCCAGGCTCACCAGGNNNAGATCGGAAGAG |
| 29036_1 | TATGAGTGTGGAGTCGTTGCCCAATAGAAAAGGCATCATGCGTNNNAGATCGGAA |
| 29036_2 | TATGAGTGTGGAGTCGTTGCCCAATAGAAAAGGCATCATGCGTNNNAGATCGGAA |
| 42621_1 | TATGAGTGTGGAGTCGTTGCATGTTGTGTGGGTGGGTGTATATNNNAGATCGGAA |
| 42621_2 | TATGAGTGTGGAGTCGTTGCATGTTGTGTGGGTGGGTGTATATNNNAGATCGGAA |
| 48805_1 | TATGAGTGTGGAGTCGTTGCTTCAAGTGATTCTCCTGCCTCAGNNNAGATCGGAAG |
| 48805_2 | TATGAGTGTGGAGTCGTTGCTTCAAGTGATTCTCCTGCCTCAGNNNAGATCGGAAG |
| 38307_1 | TATGAGTGTGGAGTCGTTGCGACTTGAAGCCAACAAAGGTCAANNNAGATCGGAA |
| 38307_2 | TATGAGTGTGGAGTCGTTGCGACTTGAAGCCAACAAAGGTCAANNNAGATCGGAA |
| 38120_1 | TATGAGTGTGGAGTCGTTGCAGCAAAGGAAACAATCAACAGAGTNNNAGATCGGA |
| 38120_2 | TATGAGTGTGGAGTCGTTGCAGCAAAGGAAACAATCAACAGAGTNNNAGATCGGA |
| 48769_1 | TATGAGTGTGGAGTCGTTGCAGACTCTGCCTCCAAATTCCAAGNNNAGATCGGAAG |
| 48769_2 | TATGAGTGTGGAGTCGTTGCAGACTCTGCCTCCAAATTCCAAGNNNAGATCGGAAG |
| 10894_1 | TATGAGTGTGGAGTCGTTGCGAGGAAGAAGCCGGCCCAGCNNNAGATCGGAAGA |
| 10894_2 | TATGAGTGTGGAGTCGTTGCGAGGAAGAAGCCGGCCCAGCNNNAGATCGGAAGA |
| 13264_1 | TATGAGTGTGGAGTCGTTGCGATTGGAGTGGTATGATCATAGNNNAGATCGGAAG |
| 13264_2 | TATGAGTGTGGAGTCGTTGCGATTGGAGTGGTATGATCATAGNNNAGATCGGAAG |
| 51263_1 | TATGAGTGTGGAGTCGTTGCCTCCTTGAGTCTGACAGGTCTTCNNNAGATCGGAAG |
| 51263_2 | TATGAGTGTGGAGTCGTTGCCTCCTTGAGTCTGACAGGTCTTCNNNAGATCGGAAG |
| 46289_1 | TATGAGTGTGGAGTCGTTGCAGACGGGATTTCTCCATATTGGCNNNAGATCGGAAG |
| 46289_2 | TATGAGTGTGGAGTCGTTGCAGACGGGATTTCTCCATATTGGCNNNAGATCGGAAG |
| 52269_1 | TATGAGTGTGGAGTCGTTGCTGGGGATTTCTCACCTCTTTCTGNNNAGATCGGAAG |
| 52269_2 | TATGAGTGTGGAGTCGTTGCTGGGGATTTCTCACCTCTTTCTGNNNAGATCGGAAG |
| 34907_1 | TATGAGTGTGGAGTCGTTGCGCCACCATGCCAGTTTACTTTTTNNNAGATCGGAAGA |
| 34907_2 | TATGAGTGTGGAGTCGTTGCGCCACCATGCCAGTTTACTTTTTNNNAGATCGGAAGA |
| 36167_1 | TATGAGTGTGGAGTCGTTGCATACATCCGTGGAGATTCAGAGCNNNAGATCGGAA |
| 36167_2 | TATGAGTGTGGAGTCGTTGCATACATCCGTGGAGATTCAGAGCNNNAGATCGGAA |

| | |
|---|---|
| 31562_1 | TATGAGTGTGGAGTCGTTGCGAGTCATACAGTGAGAAGCAGCTNNNAGATCGGAA |
| 31562_2 | TATGAGTGTGGAGTCGTTGCGAGTCATACAGTGAGAAGCAGCTNNNAGATCGGAA |
| 38915_1 | TATGAGTGTGGAGTCGTTGCCAAAGCAGTAGTTTTCAGGCACANNNAGATCGGAAG |
| 38915_2 | TATGAGTGTGGAGTCGTTGCCAAAGCAGTAGTTTTCAGGCACANNNAGATCGGAAG |
| 42930_1 | TATGAGTGTGGAGTCGTTGCGTAGTAGCACCATCTCGGCTCNNNAGATCGGAAGAG |
| 42930_2 | TATGAGTGTGGAGTCGTTGCGTAGTAGCACCATCTCGGCTCNNNAGATCGGAAGAG |
| 52488_1 | TATGAGTGTGGAGTCGTTGCCTTTTTCTAGGGCGGCTCCGNNNAGATCGGAAGAGC |
| 52488_2 | TATGAGTGTGGAGTCGTTGCCTTTTTCTAGGGCGGCTCCGNNNAGATCGGAAGAGC |
| 44482_1 | TATGAGTGTGGAGTCGTTGCGATTGGGACCTATGGACATTGGTNNNAGATCGGAA |
| 44482_2 | TATGAGTGTGGAGTCGTTGCGATTGGGACCTATGGACATTGGTNNNAGATCGGAA |
| 44645_1 | TATGAGTGTGGAGTCGTTGCGAGAAGGCAGGGGTTGGTTGNNNAGATCGGAAGA |
| 44645_2 | TATGAGTGTGGAGTCGTTGCGAGAAGGCAGGGGTTGGTTGNNNAGATCGGAAGA |
| 2953_1 | TATGAGTGTGGAGTCGTTGCAAGACTGAGTTGTGGTCTTCGTTNNNAGATCGGAAG |
| 2953_2 | TATGAGTGTGGAGTCGTTGCAAGACTGAGTTGTGGTCTTCGTTNNNAGATCGGAAG |
| 43830_1 | TATGAGTGTGGAGTCGTTGCAGCCTTTCAGTGAGAGGTCTTTTNNNAGATCGGAAG |
| 43830_2 | TATGAGTGTGGAGTCGTTGCAGCCTTTCAGTGAGAGGTCTTTTNNNAGATCGGAAG |
| 46494_1 | TATGAGTGTGGAGTCGTTGCTTCCAGTGATTTTCCTGCCTCAGNNNAGATCGGAAG |
| 46494_2 | TATGAGTGTGGAGTCGTTGCTTCCAGTGATTTTCCTGCCTCAGNNNAGATCGGAAG |
| 45041_1 | TATGAGTGTGGAGTCGTTGCTTTGTGGCTGGTACTTGTTTCTGNNNAGATCGGAAG |
| 45041_2 | TATGAGTGTGGAGTCGTTGCTTTGTGGCTGGTACTTGTTTCTGNNNAGATCGGAAG |
| 43937_1 | TATGAGTGTGGAGTCGTTGCACTTTGATGGGTCATACCTGTGGNNNAGATCGGAAG |
| 43937_2 | TATGAGTGTGGAGTCGTTGCACTTTGATGGGTCATACCTGTGGNNNAGATCGGAAG |
| 40166_1 | TATGAGTGTGGAGTCGTTGCCACTTGTAATCCCAGCACTTTGGNNNAGATCGGAAG |
| 40166_2 | TATGAGTGTGGAGTCGTTGCCACTTGTAATCCCAGCACTTTGGNNNAGATCGGAAG |
| 31242_1 | TATGAGTGTGGAGTCGTTGCAAGCAAGGGAAGAGAACAAGAGTNNNAGATCGGA |
| 31242_2 | TATGAGTGTGGAGTCGTTGCAAGCAAGGGAAGAGAACAAGAGTNNNAGATCGGA |
| 52445_1 | TATGAGTGTGGAGTCGTTGCCTAATCATTTCGGTGGGCATTGGNNNAGATCGGAAG |
| 52445_2 | TATGAGTGTGGAGTCGTTGCCTAATCATTTCGGTGGGCATTGGNNNAGATCGGAAG |
| 49515_1 | TATGAGTGTGGAGTCGTTGCCCAAGTAGCTTGATAGTCCCTCANNNAGATCGGAAG |
| 49515_2 | TATGAGTGTGGAGTCGTTGCCCAAGTAGCTTGATAGTCCCTCANNNAGATCGGAAG |
| 39415_1 | TATGAGTGTGGAGTCGTTGCAGAAAAGAAAAAAACGGCCAGGCNNNAGATCGGAA |
| 39415_2 | TATGAGTGTGGAGTCGTTGCAGAAAAGAAAAAAACGGCCAGGCNNNAGATCGGAA |
| 36224_1 | TATGAGTGTGGAGTCGTTGCTGGAAACAACTATCCTCAAGGCANNNAGATCGGAAG |
| 36224_2 | TATGAGTGTGGAGTCGTTGCTGGAAACAACTATCCTCAAGGCANNNAGATCGGAAG |
| 34561_1 | TATGAGTGTGGAGTCGTTGCTCTGGGGATGGTCCTAAGAGGNNNAGATCGGAAGA |
| 34561_2 | TATGAGTGTGGAGTCGTTGCTCTGGGGATGGTCCTAAGAGGNNNAGATCGGAAGA |
| 10556_1 | TATGAGTGTGGAGTCGTTGCCTGAGGCTGGGTTACAGGTTTTTNNNAGATCGGAA |
| 10556_2 | TATGAGTGTGGAGTCGTTGCCTGAGGCTGGGTTACAGGTTTTTNNNAGATCGGAA |
| 13761_1 | TATGAGTGTGGAGTCGTTGCCAATATAATCTCTGTGGCAATCNNNAGATCGGAAGA |
| 13761_2 | TATGAGTGTGGAGTCGTTGCCAATATAATCTCTGTGGCAATCNNNAGATCGGAAGA |
| 41013_1 | TATGAGTGTGGAGTCGTTGCATGAGATTCTGACGCAGGTAGTCNNNAGATCGGAA |
| 41013_2 | TATGAGTGTGGAGTCGTTGCATGAGATTCTGACGCAGGTAGTCNNNAGATCGGAA |
| 44284_1 | TATGAGTGTGGAGTCGTTGCCTTTGCGATGTGGAACACACTAANNNAGATCGGAAG |

| | |
|---|---|
| 44284_2 | TATGAGTGTGGAGTCGTTGCCTTTGCGATGTGGAACACACTAANNNAGATCGGAAG |
| 41613_1 | TATGAGTGTGGAGTCGTTGCTTCAAGTGATTCTCCTGCCTCAGNNNAGATCGGAAG |
| 41613_2 | TATGAGTGTGGAGTCGTTGCTTCAAGTGATTCTCCTGCCTCAGNNNAGATCGGAAG |
| 42954_1 | TATGAGTGTGGAGTCGTTGCATGTGTGTGTGACAGAGTCTCTCNNNAGATCGGAAG |
| 42954_2 | TATGAGTGTGGAGTCGTTGCATGTGTGTGTGACAGAGTCTCTCNNNAGATCGGAAG |
| 51737_1 | TATGAGTGTGGAGTCGTTGCTCATGGTGACAATAACATGGAAAANNNAGATCGGA |
| 51737_2 | TATGAGTGTGGAGTCGTTGCTCATGGTGACAATAACATGGAAAANNNAGATCGGA |
| 35316_1 | TATGAGTGTGGAGTCGTTGCGGGATCTCGGCTTACTGCAAGNNNAGATCGGAAGA |
| 35316_2 | TATGAGTGTGGAGTCGTTGCGGGATCTCGGCTTACTGCAAGNNNAGATCGGAAGA |
| 33897_1 | TATGAGTGTGGAGTCGTTGCCAAGTAGCTGGGACTACAGGCNNNAGATCGGAAGA |
| 33897_2 | TATGAGTGTGGAGTCGTTGCCAAGTAGCTGGGACTACAGGCNNNAGATCGGAAGA |
| 34898_1 | TATGAGTGTGGAGTCGTTGCGCCTTCTGAGTAGCTGGGATTACNNNAGATCGGAAG |
| 34898_2 | TATGAGTGTGGAGTCGTTGCGCCTTCTGAGTAGCTGGGATTACNNNAGATCGGAAG |
| 51881_1 | TATGAGTGTGGAGTCGTTGCATTTCAGACTCCACCTCCAACTCNNNAGATCGGAAGA |
| 51881_2 | TATGAGTGTGGAGTCGTTGCATTTCAGACTCCACCTCCAACTCNNNAGATCGGAAGA |
| 34683_1 | TATGAGTGTGGAGTCGTTGCAGGCACTGTTACGAAGACTGAATNNNAGATCGGAA |
| 34683_2 | TATGAGTGTGGAGTCGTTGCAGGCACTGTTACGAAGACTGAATNNNAGATCGGAA |
| 44446_1 | TATGAGTGTGGAGTCGTTGCCAGTGGCACAATCATGGCTCNNNAGATCGGAAGAG |
| 44446_2 | TATGAGTGTGGAGTCGTTGCCAGTGGCACAATCATGGCTCNNNAGATCGGAAGAG |
| 51019_1 | TATGAGTGTGGAGTCGTTGCTTAGAGATGCCACCTGAAGCATTNNNAGATCGGAAG |
| 51019_2 | TATGAGTGTGGAGTCGTTGCTTAGAGATGCCACCTGAAGCATTNNNAGATCGGAAG |
| 52400_1 | TATGAGTGTGGAGTCGTTGCTCCCCTCCAATACTACAAACAGCNNNAGATCGGAAGA |
| 52400_2 | TATGAGTGTGGAGTCGTTGCTCCCCTCCAATACTACAAACAGCNNNAGATCGGAAGA |
| 38621_1 | TATGAGTGTGGAGTCGTTGCTGTCAAGAAATTGGATAGCCAGGANNNAGATCGGA |
| 38621_2 | TATGAGTGTGGAGTCGTTGCTGTCAAGAAATTGGATAGCCAGGANNNAGATCGGA |
| 33380_1 | TATGAGTGTGGAGTCGTTGCTCTACCCACTCTCTCTCTTGGTCNNNAGATCGGAAGA |
| 33380_2 | TATGAGTGTGGAGTCGTTGCTCTACCCACTCTCTCTCTTGGTCNNNAGATCGGAAGA |
| 35199_1 | TATGAGTGTGGAGTCGTTGCGGTATGATCTCAGCTCACCACAANNNAGATCGGAAG |
| 35199_2 | TATGAGTGTGGAGTCGTTGCGGTATGATCTCAGCTCACCACAANNNAGATCGGAAG |
| 4427_1 | TATGAGTGTGGAGTCGTTGCTTTCCTGGTCTTGTCAATGCACTNNNAGATCGGAAGA |
| 4427_2 | TATGAGTGTGGAGTCGTTGCTTTCCTGGTCTTGTCAATGCACTNNNAGATCGGAAGA |
| 33645_1 | TATGAGTGTGGAGTCGTTGCGATGAGATTTCACCATGTTGGCCNNNAGATCGGAAG |
| 33645_2 | TATGAGTGTGGAGTCGTTGCGATGAGATTTCACCATGTTGGCCNNNAGATCGGAAG |
| 42664_1 | TATGAGTGTGGAGTCGTTGCCCACGGATACCAAATTCTGAGGANNNAGATCGGAAG |
| 42664_2 | TATGAGTGTGGAGTCGTTGCCCACGGATACCAAATTCTGAGGANNNAGATCGGAAG |
| 45551_1 | TATGAGTGTGGAGTCGTTGCCCTTTTTCACGTTAGCCTTCTGGNNNAGATCGGAAGA |
| 45551_2 | TATGAGTGTGGAGTCGTTGCCCTTTTTCACGTTAGCCTTCTGGNNNAGATCGGAAGA |
| 39740_1 | TATGAGTGTGGAGTCGTTGCTAGTGAGCTGAGATTGCACAACTNNNAGATCGGAA |
| 39740_2 | TATGAGTGTGGAGTCGTTGCTAGTGAGCTGAGATTGCACAACTNNNAGATCGGAA |
| 38870_1 | TATGAGTGTGGAGTCGTTGCGTACAGGGACAACAGCAAAAACANNNAGATCGGAA |
| 38870_2 | TATGAGTGTGGAGTCGTTGCGTACAGGGACAACAGCAAAAACANNNAGATCGGAA |
| 44493_1 | TATGAGTGTGGAGTCGTTGCGTGATCTCAGCTCACTGTAACCTNNNAGATCGGAAG |
| 44493_2 | TATGAGTGTGGAGTCGTTGCGTGATCTCAGCTCACTGTAACCTNNNAGATCGGAAG |

| | |
|---|---|
| 51508_1 | TATGAGTGTGGAGTCGTTGCATTCCTCTAATTGCTGCAGCCAGNNNAGATCGGAAG |
| 51508_2 | TATGAGTGTGGAGTCGTTGCATTCCTCTAATTGCTGCAGCCAGNNNAGATCGGAAG |
| 49799_1 | TATGAGTGTGGAGTCGTTGCGTGTGCGTCTTCGTTATCAACAANNNAGATCGGAAG |
| 49799_2 | TATGAGTGTGGAGTCGTTGCGTGTGCGTCTTCGTTATCAACAANNNAGATCGGAAG |
| 10703_1 | TATGAGTGTGGAGTCGTTGCATCAAAGGGGGAAATGGCATGAANNNAGATCGGA |
| 10703_2 | TATGAGTGTGGAGTCGTTGCATCAAAGGGGGAAATGGCATGAANNNAGATCGGA |
| 41095_1 | TATGAGTGTGGAGTCGTTGCCAGTACAGATGCTGCTGAGATCTNNNAGATCGGAA |
| 41095_2 | TATGAGTGTGGAGTCGTTGCCAGTACAGATGCTGCTGAGATCTNNNAGATCGGAA |
| 40947_1 | TATGAGTGTGGAGTCGTTGCCGCCTATCTTTATCACCCACTCANNNAGATCGGAAGA |
| 40947_2 | TATGAGTGTGGAGTCGTTGCCGCCTATCTTTATCACCCACTCANNNAGATCGGAAGA |
| 47144_1 | TATGAGTGTGGAGTCGTTGCCCAAGTATCTGGGACTACAGGTGNNNAGATCGGAA |
| 47144_2 | TATGAGTGTGGAGTCGTTGCCCAAGTATCTGGGACTACAGGTGNNNAGATCGGAA |
| 35691_1 | TATGAGTGTGGAGTCGTTGCATTATCAGGTCACCACAACCACCNNNAGATCGGAAG |
| 35691_2 | TATGAGTGTGGAGTCGTTGCATTATCAGGTCACCACAACCACCNNNAGATCGGAAG |
| 34592_1 | TATGAGTGTGGAGTCGTTGCATTCCTGGCTGGGGTCTGTGNNNAGATCGGAAGAG |
| 34592_2 | TATGAGTGTGGAGTCGTTGCATTCCTGGCTGGGGTCTGTGNNNAGATCGGAAGAG |
| 43725_1 | TATGAGTGTGGAGTCGTTGCGGGGATGATAGGGTTGGTGATTTNNNAGATCGGA |
| 43725_2 | TATGAGTGTGGAGTCGTTGCGGGGATGATAGGGTTGGTGATTTNNNAGATCGGA |
| 29460_1 | TATGAGTGTGGAGTCGTTGCGGTATGCCTGTATCATATTTTTTNNNAGATCGGAAG |
| 29460_2 | TATGAGTGTGGAGTCGTTGCGGTATGCCTGTATCATATTTTTTNNNAGATCGGAAG |
| 51187_1 | TATGAGTGTGGAGTCGTTGCCTGCATGTAATTTGTCCTGGTGGNNNAGATCGGAAG |
| 51187_2 | TATGAGTGTGGAGTCGTTGCCTGCATGTAATTTGTCCTGGTGGNNNAGATCGGAAG |
| 48367_1 | TATGAGTGTGGAGTCGTTGCCTGAGTAGCTGGAACTACAGGACNNNAGATCGGAA |
| 48367_2 | TATGAGTGTGGAGTCGTTGCCTGAGTAGCTGGAACTACAGGACNNNAGATCGGAA |
| 41122_1 | TATGAGTGTGGAGTCGTTGCTTGCAAAGATAAGGAACCAACCTNNNAGATCGGAA |
| 41122_2 | TATGAGTGTGGAGTCGTTGCTTGCAAAGATAAGGAACCAACCTNNNAGATCGGAA |
| 45524_1 | TATGAGTGTGGAGTCGTTGCCAGTGGCAGGCACATAATAGGTANNNAGATCGGAA |
| 45524_2 | TATGAGTGTGGAGTCGTTGCCAGTGGCAGGCACATAATAGGTANNNAGATCGGAA |
| 36847_1 | TATGAGTGTGGAGTCGTTGCGGTGCAAAGAATGTGATTTCTGGNNNAGATCGGAA |
| 36847_2 | TATGAGTGTGGAGTCGTTGCGGTGCAAAGAATGTGATTTCTGGNNNAGATCGGAA |
| 51047_1 | TATGAGTGTGGAGTCGTTGCCCCGGCTGGAAACTCTTTATACTNNNAGATCGGAAG |
| 51047_2 | TATGAGTGTGGAGTCGTTGCCCCGGCTGGAAACTCTTTATACTNNNAGATCGGAAG |
| 51605_1 | TATGAGTGTGGAGTCGTTGCGGAGGTGTACCCAACAGCTCNNNAGATCGGAAGAG |
| 51605_2 | TATGAGTGTGGAGTCGTTGCGGAGGTGTACCCAACAGCTCNNNAGATCGGAAGAG |
| 47347_1 | TATGAGTGTGGAGTCGTTGCAATAGCTGGGATTACAGGTGCATNNNAGATCGGAA |
| 47347_2 | TATGAGTGTGGAGTCGTTGCAATAGCTGGGATTACAGGTGCATNNNAGATCGGAA |
| 38087_1 | TATGAGTGTGGAGTCGTTGCTTGGGGTTTTTGTGTTTTGTGGGNNNAGATCGGAA |
| 38087_2 | TATGAGTGTGGAGTCGTTGCTTGGGGTTTTTGTGTTTTGTGGGNNNAGATCGGAA |
| 42908_1 | TATGAGTGTGGAGTCGTTGCCACACACGTACACATATACACACANNNAGATCGGAA |
| 42908_2 | TATGAGTGTGGAGTCGTTGCCACACACGTACACATATACACACANNNAGATCGGAA |
| 42557_1 | TATGAGTGTGGAGTCGTTGCATATGTGTGTGTGTGTGTGTGTGNNNAGATCGGAA |
| 42557_2 | TATGAGTGTGGAGTCGTTGCATATGTGTGTGTGTGTGTGTGTGNNNAGATCGGAA |
| 31620_1 | TATGAGTGTGGAGTCGTTGCGAGAGCTGCCTGAGATGGTAAGNNNAGATCGGAA |

| | |
|---|---|
| 31620_2 | TATGAGTGTGGAGTCGTTGCGAGAGCTGCCTGAGATGGTAAGNNNAGATCGGAAG |
| 32787_1 | TATGAGTGTGGAGTCGTTGCGCTTGAGTGCACAAGTTTGAGATNNNAGATCGGAA |
| 32787_2 | TATGAGTGTGGAGTCGTTGCGCTTGAGTGCACAAGTTTGAGATNNNAGATCGGAA |
| 48275_1 | TATGAGTGTGGAGTCGTTGCATTCAAGCGATTCTCCTGCCTTANNNAGATCGGAAG |
| 48275_2 | TATGAGTGTGGAGTCGTTGCATTCAAGCGATTCTCCTGCCTTANNNAGATCGGAAG |
| 34555_1 | TATGAGTGTGGAGTCGTTGCTCTGGGGATGGTCCTAAGAGGNNNAGATCGGAAGA |
| 34555_2 | TATGAGTGTGGAGTCGTTGCTCTGGGGATGGTCCTAAGAGGNNNAGATCGGAAGA |
| 41045_1 | TATGAGTGTGGAGTCGTTGCCCAGCTTTATTCTGCCTTTCCAGNNNAGATCGGAAGA |
| 41045_2 | TATGAGTGTGGAGTCGTTGCCCAGCTTTATTCTGCCTTTCCAGNNNAGATCGGAAGA |
| 29695_1 | TATGAGTGTGGAGTCGTTGCAAATTGAGCCTTTCCCCTTCCTGNNNAGATCGGAAGA |
| 29695_2 | TATGAGTGTGGAGTCGTTGCAAATTGAGCCTTTCCCCTTCCTGNNNAGATCGGAAGA |
| 46770_1 | TATGAGTGTGGAGTCGTTGCTTCAAGACATTCTCCTGCCTCAGNNNAGATCGGAAG |
| 46770_2 | TATGAGTGTGGAGTCGTTGCTTCAAGACATTCTCCTGCCTCAGNNNAGATCGGAAG |
| 46783_1 | TATGAGTGTGGAGTCGTTGCGCCAGAATAGTCTCGATCTCCTGNNNAGATCGGAAG |
| 46783_2 | TATGAGTGTGGAGTCGTTGCGCCAGAATAGTCTCGATCTCCTGNNNAGATCGGAAG |
| 49207_1 | TATGAGTGTGGAGTCGTTGCCATTCCCAAGGTTCTTGCCAAGNNNAGATCGGAAGA |
| 49207_2 | TATGAGTGTGGAGTCGTTGCCATTCCCAAGGTTCTTGCCAAGNNNAGATCGGAAGA |
| 50708_1 | TATGAGTGTGGAGTCGTTGCCACAGATGGTCAGCGAGCAGNNNAGATCGGAAGAG |
| 50708_2 | TATGAGTGTGGAGTCGTTGCCACAGATGGTCAGCGAGCAGNNNAGATCGGAAGAG |
| 41953_1 | TATGAGTGTGGAGTCGTTGCCAAGTAGCTGGGATTACAGGCATNNNAGATCGGAA |
| 41953_2 | TATGAGTGTGGAGTCGTTGCCAAGTAGCTGGGATTACAGGCATNNNAGATCGGAA |
| 44462_1 | TATGAGTGTGGAGTCGTTGCGAAGTTGGGGGGAATACACAGATNNNAGATCGGA |
| 44462_2 | TATGAGTGTGGAGTCGTTGCGAAGTTGGGGGGAATACACAGATNNNAGATCGGA |
| 38618_1 | TATGAGTGTGGAGTCGTTGCTTAGTAACTGAAAGAGGGCCAGGNNNAGATCGGAA |
| 38618_2 | TATGAGTGTGGAGTCGTTGCTTAGTAACTGAAAGAGGGCCAGGNNNAGATCGGAA |
| 41766_1 | TATGAGTGTGGAGTCGTTGCCTTTCACCTCAGCCTTCCAGATANNNAGATCGGAAGA |
| 41766_2 | TATGAGTGTGGAGTCGTTGCCTTTCACCTCAGCCTTCCAGATANNNAGATCGGAAGA |
| 51991_1 | TATGAGTGTGGAGTCGTTGCAGGCCTATAAATTGTCCAAGCCTNNNAGATCGGAAG |
| 51991_2 | TATGAGTGTGGAGTCGTTGCAGGCCTATAAATTGTCCAAGCCTNNNAGATCGGAAG |
| 37185_1 | TATGAGTGTGGAGTCGTTGCTAACAGCAAGGTGATCGTCTTGANNNAGATCGGAA |
| 37185_2 | TATGAGTGTGGAGTCGTTGCTAACAGCAAGGTGATCGTCTTGANNNAGATCGGAA |
| 48570_1 | TATGAGTGTGGAGTCGTTGCGGGAACTCCTTGCAGAAAACATTNNNAGATCGGAA |
| 48570_2 | TATGAGTGTGGAGTCGTTGCGGGAACTCCTTGCAGAAAACATTNNNAGATCGGAA |
| 35492_1 | TATGAGTGTGGAGTCGTTGCCTCAAGAGGATCACTTTGTCCCANNNAGATCGGAAG |
| 35492_2 | TATGAGTGTGGAGTCGTTGCCTCAAGAGGATCACTTTGTCCCANNNAGATCGGAAG |
| 50683_1 | TATGAGTGTGGAGTCGTTGCGGTGTCTCTAGGGTCCCATTTTTNNNAGATCGGAAG |
| 50683_2 | TATGAGTGTGGAGTCGTTGCGGTGTCTCTAGGGTCCCATTTTTNNNAGATCGGAAG |
| 31491_1 | TATGAGTGTGGAGTCGTTGCTGTCAGTAATCTCTGAGTGCTGANNNAGATCGGAAG |
| 31491_2 | TATGAGTGTGGAGTCGTTGCTGTCAGTAATCTCTGAGTGCTGANNNAGATCGGAAG |
| 29963_1 | TATGAGTGTGGAGTCGTTGCTGGGTTTGGTAACTTGGTCATCTNNNAGATCGGAAG |
| 29963_2 | TATGAGTGTGGAGTCGTTGCTGGGTTTGGTAACTTGGTCATCTNNNAGATCGGAAG |
| 44576_1 | TATGAGTGTGGAGTCGTTGCATCACCTAAGCTGGAGTGCAGNNNAGATCGGAAGA |
| 44576_2 | TATGAGTGTGGAGTCGTTGCATCACCTAAGCTGGAGTGCAGNNNAGATCGGAAGA |

| | |
|---|---|
| 32523_1 | TATGAGTGTGGAGTCGTTGCATCCAGGTTGCCAAAGCTACTAANNNAGATCGGAAG |
| 32523_2 | TATGAGTGTGGAGTCGTTGCATCCAGGTTGCCAAAGCTACTAANNNAGATCGGAAG |
| 45405_1 | TATGAGTGTGGAGTCGTTGCGTAATTGGCTCACACCTTCACACNNNAGATCGGAAG |
| 45405_2 | TATGAGTGTGGAGTCGTTGCGTAATTGGCTCACACCTTCACACNNNAGATCGGAAG |
| 13225_1 | TATGAGTGTGGAGTCGTTGCCCATTCTCAACATCATCATAAATNNNAGATCGGAAGA |
| 13225_2 | TATGAGTGTGGAGTCGTTGCCCATTCTCAACATCATCATAAATNNNAGATCGGAAGA |
| 43227_1 | TATGAGTGTGGAGTCGTTGCGTGCAATGGTGTTATCTTGGCTTNNNAGATCGGAAG |
| 43227_2 | TATGAGTGTGGAGTCGTTGCGTGCAATGGTGTTATCTTGGCTTNNNAGATCGGAAG |
| 30049_1 | TATGAGTGTGGAGTCGTTGCGTTCAAAAGGTTGTCCCAGTTGANNNAGATCGGAA |
| 30049_2 | TATGAGTGTGGAGTCGTTGCGTTCAAAAGGTTGTCCCAGTTGANNNAGATCGGAA |
| 29915_1 | TATGAGTGTGGAGTCGTTGCACCTGAGACAGAAGGAGAAATTCTNNNAGATCGGA |
| 29915_2 | TATGAGTGTGGAGTCGTTGCACCTGAGACAGAAGGAGAAATTCTNNNAGATCGGA |
| 39928_1 | TATGAGTGTGGAGTCGTTGCCTGTAGTCCCAGCTACTCAGGAGNNNAGATCGGAAG |
| 39928_2 | TATGAGTGTGGAGTCGTTGCCTGTAGTCCCAGCTACTCAGGAGNNNAGATCGGAAG |
| 45522_1 | TATGAGTGTGGAGTCGTTGCTCAGGAGAGAGAGCACAGATACANNNAGATCGGAA |
| 45522_2 | TATGAGTGTGGAGTCGTTGCTCAGGAGAGAGAGCACAGATACANNNAGATCGGAA |
| 30000_1 | TATGAGTGTGGAGTCGTTGCTATAATACTCCACTGGGGCCTCANNNAGATCGGAAG |
| 30000_2 | TATGAGTGTGGAGTCGTTGCTATAATACTCCACTGGGGCCTCANNNAGATCGGAAG |
| 3063_1 | TATGAGTGTGGAGTCGTTGCCTGACCGCCTGATGAGAATGAAANNNAGATCGGAA |
| 3063_2 | TATGAGTGTGGAGTCGTTGCCTGACCGCCTGATGAGAATGAAANNNAGATCGGAA |
| 33193_1 | TATGAGTGTGGAGTCGTTGCCAGGAGAATTGCTTGAACTTGGGNNNAGATCGGAA |
| 33193_2 | TATGAGTGTGGAGTCGTTGCCAGGAGAATTGCTTGAACTTGGGNNNAGATCGGAA |
| 39705_1 | TATGAGTGTGGAGTCGTTGCGACATGTGACTCATACCCAACCTNNNAGATCGGAAG |
| 39705_2 | TATGAGTGTGGAGTCGTTGCGACATGTGACTCATACCCAACCTNNNAGATCGGAAG |
| 40660_1 | TATGAGTGTGGAGTCGTTGCTACCAATCGAGTTCAACTCCCTCNNNAGATCGGAAGA |
| 40660_2 | TATGAGTGTGGAGTCGTTGCTACCAATCGAGTTCAACTCCCTCNNNAGATCGGAAGA |
| 36238_1 | TATGAGTGTGGAGTCGTTGCTAAATGTGAAAAATCCCAGCCGGNNNAGATCGGAA |
| 36238_2 | TATGAGTGTGGAGTCGTTGCTAAATGTGAAAAATCCCAGCCGGNNNAGATCGGAA |
| 46270_1 | TATGAGTGTGGAGTCGTTGCCTGTATCAGCTCACTGCAACCTCNNNAGATCGGAAGA |
| 46270_2 | TATGAGTGTGGAGTCGTTGCCTGTATCAGCTCACTGCAACCTCNNNAGATCGGAAGA |
| 50655_1 | TATGAGTGTGGAGTCGTTGCTCTGCCACTTATTAGCTGTGTGTNNNAGATCGGAAG |
| 50655_2 | TATGAGTGTGGAGTCGTTGCTCTGCCACTTATTAGCTGTGTGTNNNAGATCGGAAG |
| 10796_1 | TATGAGTGTGGAGTCGTTGCCACACAGTGCAGCATAATGCAATNNNAGATCGGAAG |
| 10796_2 | TATGAGTGTGGAGTCGTTGCCACACAGTGCAGCATAATGCAATNNNAGATCGGAAG |
| 34160_1 | TATGAGTGTGGAGTCGTTGCTATCCATTTGGTGTGTGGGTAGGNNNAGATCGGAA |
| 34160_2 | TATGAGTGTGGAGTCGTTGCTATCCATTTGGTGTGTGGGTAGGNNNAGATCGGAA |
| 50784_1 | TATGAGTGTGGAGTCGTTGCAAATATGTGCTGGGAGAGTGAGGNNNAGATCGGA |
| 50784_2 | TATGAGTGTGGAGTCGTTGCAAATATGTGCTGGGAGAGTGAGGNNNAGATCGGA |
| 41063_1 | TATGAGTGTGGAGTCGTTGCAGGCATGAGAAATCACCCTAAGANNNAGATCGGAA |
| 41063_2 | TATGAGTGTGGAGTCGTTGCAGGCATGAGAAATCACCCTAAGANNNAGATCGGAA |
| 30589_1 | TATGAGTGTGGAGTCGTTGCGGGGTAGTGTAAATCCACTCTGCTNNNAGATCGGAA |
| 30589_2 | TATGAGTGTGGAGTCGTTGCGGGGTAGTGTAAATCCACTCTGCTNNNAGATCGGAA |
| 52720_1 | TATGAGTGTGGAGTCGTTGCAGCATGAGTGGTTCGGGATTATTNNNAGATCGGAA |

| | |
|---|---|
| 52720_2 | TATGAGTGTGGAGTCGTTGCAGCATGAGTGGTTCGGGATTATTNNNAGATCGGAA |
| 34435_1 | TATGAGTGTGGAGTCGTTGCTCTGTATCATGGAGTTTGCCCTCNNNAGATCGGAAG |
| 34435_2 | TATGAGTGTGGAGTCGTTGCTCTGTATCATGGAGTTTGCCCTCNNNAGATCGGAAG |
| 46016_1 | TATGAGTGTGGAGTCGTTGCGAGAACCTCCTCCCCTTCCTTTCNNNAGATCGGAAGA |
| 46016_2 | TATGAGTGTGGAGTCGTTGCGAGAACCTCCTCCCCTTCCTTTCNNNAGATCGGAAGA |
| 47735_1 | TATGAGTGTGGAGTCGTTGCGGCACAATCTCGACTCACTGTATNNNAGATCGGAAG |
| 47735_2 | TATGAGTGTGGAGTCGTTGCGGCACAATCTCGACTCACTGTATNNNAGATCGGAAG |
| 50906_1 | TATGAGTGTGGAGTCGTTGCTGACTTCTGTCCTCAGGAAGTTGNNNAGATCGGAAG |
| 50906_2 | TATGAGTGTGGAGTCGTTGCTGACTTCTGTCCTCAGGAAGTTGNNNAGATCGGAAG |
| 1262_1 | TATGAGTGTGGAGTCGTTGCGTCCACAGACCAATAGCATAGCANNNAGATCGGAAG |
| 1262_2 | TATGAGTGTGGAGTCGTTGCGTCCACAGACCAATAGCATAGCANNNAGATCGGAAG |
| 12769_1 | TATGAGTGTGGAGTCGTTGCTAAATGTCATTTGCTGTGAGTATNNNAGATCGGAAG |
| 12769_2 | TATGAGTGTGGAGTCGTTGCTAAATGTCATTTGCTGTGAGTATNNNAGATCGGAAG |
| 46856_1 | TATGAGTGTGGAGTCGTTGCCAAGTAGCTGGGACTACAGGCNNNAGATCGGAAGA |
| 46856_2 | TATGAGTGTGGAGTCGTTGCCAAGTAGCTGGGACTACAGGCNNNAGATCGGAAGA |
| 32655_1 | TATGAGTGTGGAGTCGTTGCAAGTTCAGGAAGCAAGAGTCTGTNNNAGATCGGAA |
| 32655_2 | TATGAGTGTGGAGTCGTTGCAAGTTCAGGAAGCAAGAGTCTGTNNNAGATCGGAA |
| 48339_1 | TATGAGTGTGGAGTCGTTGCCTGATTAGCTGGGATTACAGGCANNNAGATCGGAA |
| 48339_2 | TATGAGTGTGGAGTCGTTGCCTGATTAGCTGGGATTACAGGCANNNAGATCGGAA |
| 47051_1 | TATGAGTGTGGAGTCGTTGCTTGGAAGGAAGTCACCATGCATANNNAGATCGGAA |
| 47051_2 | TATGAGTGTGGAGTCGTTGCTTGGAAGGAAGTCACCATGCATANNNAGATCGGAA |
| 37315_1 | TATGAGTGTGGAGTCGTTGCTGAGAGAACCTTGAACTTTTGCTNNNAGATCGGAAG |
| 37315_2 | TATGAGTGTGGAGTCGTTGCTGAGAGAACCTTGAACTTTTGCTNNNAGATCGGAAG |
| 41195_1 | TATGAGTGTGGAGTCGTTGCACTCACAAAAAAATTGCACCTGCNNNAGATCGGAAG |
| 41195_2 | TATGAGTGTGGAGTCGTTGCACTCACAAAAAAATTGCACCTGCNNNAGATCGGAAG |
| 49878_1 | TATGAGTGTGGAGTCGTTGCACCTACGGGCATTTGCATAAATGNNNAGATCGGAAG |
| 49878_2 | TATGAGTGTGGAGTCGTTGCACCTACGGGCATTTGCATAAATGNNNAGATCGGAAG |
| 39413_1 | TATGAGTGTGGAGTCGTTGCGCTCTGGATGAGAAAACTGAGGANNNAGATCGGAA |
| 39413_2 | TATGAGTGTGGAGTCGTTGCGCTCTGGATGAGAAAACTGAGGANNNAGATCGGAA |
| 29482_1 | TATGAGTGTGGAGTCGTTGCCTGTTTCTCACAGCTGGAGTTGATNNNAGATCGGAA |
| 29482_2 | TATGAGTGTGGAGTCGTTGCCTGTTTCTCACAGCTGGAGTTGATNNNAGATCGGAA |
| 46119_1 | TATGAGTGTGGAGTCGTTGCCACCACCACGCCTAATTTTTCTTNNNAGATCGGAAGA |
| 46119_2 | TATGAGTGTGGAGTCGTTGCCACCACCACGCCTAATTTTTCTTNNNAGATCGGAAGA |
| 40922_1 | TATGAGTGTGGAGTCGTTGCACAGGAGCTGCTACCACTTATTCNNNAGATCGGAAG |
| 40922_2 | TATGAGTGTGGAGTCGTTGCACAGGAGCTGCTACCACTTATTCNNNAGATCGGAAG |
| 50384_1 | TATGAGTGTGGAGTCGTTGCCCTCTTCCTCCTCCCCTTCATCNNNAGATCGGAAGAG |
| 50384_2 | TATGAGTGTGGAGTCGTTGCCCTCTTCCTCCTCCCCTTCATCNNNAGATCGGAAGAG |
| 34524_1 | TATGAGTGTGGAGTCGTTGCGTTGCTGAGTCTAGACACCAGAANNNAGATCGGAA |
| 34524_2 | TATGAGTGTGGAGTCGTTGCGTTGCTGAGTCTAGACACCAGAANNNAGATCGGAA |
| 50386_1 | TATGAGTGTGGAGTCGTTGCCACCTGCGAAAACAGAAATAGCANNNAGATCGGAA |
| 50386_2 | TATGAGTGTGGAGTCGTTGCCACCTGCGAAAACAGAAATAGCANNNAGATCGGAA |
| 49228_1 | TATGAGTGTGGAGTCGTTGCGGCACAATATCGACTCATTGCAANNNAGATCGGAAG |
| 49228_2 | TATGAGTGTGGAGTCGTTGCGGCACAATATCGACTCATTGCAANNNAGATCGGAAG |

| | |
|---|---|
| 48738_1 | TATGAGTGTGGAGTCGTTGCTGCCATTGGGATTTTGATAGGGANNNAGATCGGAA |
| 48738_2 | TATGAGTGTGGAGTCGTTGCTGCCATTGGGATTTTGATAGGGANNNAGATCGGAA |
| 10636_1 | TATGAGTGTGGAGTCGTTGCGTTATGCTCTGTGTGGATCTGTGNNNAGATCGGAAG |
| 10636_2 | TATGAGTGTGGAGTCGTTGCGTTATGCTCTGTGTGGATCTGTGNNNAGATCGGAAG |
| 39050_1 | TATGAGTGTGGAGTCGTTGCGTGGGCAAAAGACATGATCAGACNNNAGATCGGAA |
| 39050_2 | TATGAGTGTGGAGTCGTTGCGTGGGCAAAAGACATGATCAGACNNNAGATCGGAA |
| 32902_1 | TATGAGTGTGGAGTCGTTGCCCTTTGAGCCCTCATTCACATTCNNNAGATCGGAAGA |
| 32902_2 | TATGAGTGTGGAGTCGTTGCCCTTTGAGCCCTCATTCACATTCNNNAGATCGGAAGA |
| 46759_1 | TATGAGTGTGGAGTCGTTGCATGTCTGGCCATGACCACTATTTNNNAGATCGGAAG |
| 46759_2 | TATGAGTGTGGAGTCGTTGCATGTCTGGCCATGACCACTATTTNNNAGATCGGAAG |
| 37718_1 | TATGAGTGTGGAGTCGTTGCAACAATCGCTTGGGAGGCTATATNNNAGATCGGAA |
| 37718_2 | TATGAGTGTGGAGTCGTTGCAACAATCGCTTGGGAGGCTATATNNNAGATCGGAA |
| 37116_1 | TATGAGTGTGGAGTCGTTGCCTGTAGTTCCAGCTACTTGGGAGNNNAGATCGGAAG |
| 37116_2 | TATGAGTGTGGAGTCGTTGCCTGTAGTTCCAGCTACTTGGGAGNNNAGATCGGAAG |
| 10808_1 | TATGAGTGTGGAGTCGTTGCAGGATTTGGTGCTATGGCAAAGCNNNAGATCGGAA |
| 10808_2 | TATGAGTGTGGAGTCGTTGCAGGATTTGGTGCTATGGCAAAGCNNNAGATCGGAA |
| 47553_1 | TATGAGTGTGGAGTCGTTGCAGGTTCAAGTGATTCTCTTGGCTNNNAGATCGGAAG |
| 47553_2 | TATGAGTGTGGAGTCGTTGCAGGTTCAAGTGATTCTCTTGGCTNNNAGATCGGAAG |
| 41664_1 | TATGAGTGTGGAGTCGTTGCTTTTGAAGAGACAGCGTTTCACCNNNAGATCGGAAG |
| 41664_2 | TATGAGTGTGGAGTCGTTGCTTTTGAAGAGACAGCGTTTCACCNNNAGATCGGAAG |
| 8058_1 | TATGAGTGTGGAGTCGTTGCATCAGAGCCAAGCAGGACTTCTANNNAGATCGGAAG |
| 8058_2 | TATGAGTGTGGAGTCGTTGCATCAGAGCCAAGCAGGACTTCTANNNAGATCGGAAG |
| 29588_1 | TATGAGTGTGGAGTCGTTGCGCCCCTAGAATCTGTCTTGATGTNNNAGATCGGAAG |
| 29588_2 | TATGAGTGTGGAGTCGTTGCGCCCCTAGAATCTGTCTTGATGTNNNAGATCGGAAG |
| 43261_1 | TATGAGTGTGGAGTCGTTGCGACCTCAGCTCACTACAACCTCNNNAGATCGGAAGAG |
| 43261_2 | TATGAGTGTGGAGTCGTTGCGACCTCAGCTCACTACAACCTCNNNAGATCGGAAGAG |
| 42059_1 | TATGAGTGTGGAGTCGTTGCAACACTACCATTGAACACTCCCANNNAGATCGGAAG |
| 42059_2 | TATGAGTGTGGAGTCGTTGCAACACTACCATTGAACACTCCCANNNAGATCGGAAG |
| 13196_1 | TATGAGTGTGGAGTCGTTGCTTTAAGAAAGGGAGATTTTAAGCNNNAGATCGGAA |
| 13196_2 | TATGAGTGTGGAGTCGTTGCTTTAAGAAAGGGAGATTTTAAGCNNNAGATCGGAA |
| 39853_1 | TATGAGTGTGGAGTCGTTGCCCTATCACTTGTTCACTTTGGGCNNNAGATCGGAAG |
| 39853_2 | TATGAGTGTGGAGTCGTTGCCCTATCACTTGTTCACTTTGGGCNNNAGATCGGAAG |
| 2298_1 | TATGAGTGTGGAGTCGTTGCAGTTGGTCTTATGTCGTCTGCACNNNAGATCGGAAG |
| 2298_2 | TATGAGTGTGGAGTCGTTGCAGTTGGTCTTATGTCGTCTGCACNNNAGATCGGAAG |
| 29640_1 | TATGAGTGTGGAGTCGTTGCTAACAACGGGGACACATTCTAGGNNNAGATCGGAA |
| 29640_2 | TATGAGTGTGGAGTCGTTGCTAACAACGGGGACACATTCTAGGNNNAGATCGGAA |
| 52560_1 | TATGAGTGTGGAGTCGTTGCAAAAGCCTTCATTCTCCTGCATGNNNAGATCGGAAG |
| 52560_2 | TATGAGTGTGGAGTCGTTGCAAAAGCCTTCATTCTCCTGCATGNNNAGATCGGAAG |
| 48912_1 | TATGAGTGTGGAGTCGTTGCAGATGGATTCTCGCTCTGTTACCNNNAGATCGGAAG |
| 48912_2 | TATGAGTGTGGAGTCGTTGCAGATGGATTCTCGCTCTGTTACCNNNAGATCGGAAG |
| 33112_1 | TATGAGTGTGGAGTCGTTGCGCAAATACCCCAGTGGATTTAGCNNNAGATCGGAAG |
| 33112_2 | TATGAGTGTGGAGTCGTTGCGCAAATACCCCAGTGGATTTAGCNNNAGATCGGAAG |
| 43499_1 | TATGAGTGTGGAGTCGTTGCAATGCTCTGGGATTGTTTTCATTNNNAGATCGGAAG |

| | |
|---|---|
| 43499_2 | TATGAGTGTGGAGTCGTTGCAATGCTCTGGGATTGTTTTCATTNNNAGATCGGAAG |
| 36870_1 | TATGAGTGTGGAGTCGTTGCAGAGGACAGAGTTACGTGTTAGCNNNAGATCGGAA |
| 36870_2 | TATGAGTGTGGAGTCGTTGCAGAGGACAGAGTTACGTGTTAGCNNNAGATCGGAA |
| 38769_1 | TATGAGTGTGGAGTCGTTGCTCCTTGCAACAGAACAGAAAGTTNNNAGATCGGAAG |
| 38769_2 | TATGAGTGTGGAGTCGTTGCTCCTTGCAACAGAACAGAAAGTTNNNAGATCGGAAG |
| 3116_1 | TATGAGTGTGGAGTCGTTGCACTGTCAGGCAGAAGACACTGGNNNAGATCGGAAG |
| 3116_2 | TATGAGTGTGGAGTCGTTGCACTGTCAGGCAGAAGACACTGGNNNAGATCGGAAG |
| 50508_1 | TATGAGTGTGGAGTCGTTGCCTCTTTCGTCTCTACTGCCCCNNNAGATCGGAAGAGC |
| 50508_2 | TATGAGTGTGGAGTCGTTGCCTCTTTCGTCTCTACTGCCCCNNNAGATCGGAAGAGC |
| 51157_1 | TATGAGTGTGGAGTCGTTGCGCCACCTAAAAAGGAGAGGAGAGNNNAGATCGGA |
| 51157_2 | TATGAGTGTGGAGTCGTTGCGCCACCTAAAAAGGAGAGGAGAGNNNAGATCGGA |
| 5238_1 | TATGAGTGTGGAGTCGTTGCCCTGGGGAAAGGTTTTGGAGTTTNNNAGATCGGAA |
| 5238_2 | TATGAGTGTGGAGTCGTTGCCCTGGGGAAAGGTTTTGGAGTTTNNNAGATCGGAA |
| 52807_1 | TATGAGTGTGGAGTCGTTGCCTAGAGGAATAAGGGGGGACAGGNNNAGATCGGA |
| 52807_2 | TATGAGTGTGGAGTCGTTGCCTAGAGGAATAAGGGGGGACAGGNNNAGATCGGA |
| 34576_1 | TATGAGTGTGGAGTCGTTGCACAGACTTTAGATGGTGGGTTGGNNNAGATCGGAA |
| 34576_2 | TATGAGTGTGGAGTCGTTGCACAGACTTTAGATGGTGGGTTGGNNNAGATCGGAA |
| 33557_1 | TATGAGTGTGGAGTCGTTGCCTGAGTAGCTGGAACTATAGGCGNNNAGATCGGAA |
| 33557_2 | TATGAGTGTGGAGTCGTTGCCTGAGTAGCTGGAACTATAGGCGNNNAGATCGGAA |
| 34761_1 | TATGAGTGTGGAGTCGTTGCGATCTCGGCTCACTGCAACATCNNNAGATCGGAAGA |
| 34761_2 | TATGAGTGTGGAGTCGTTGCGATCTCGGCTCACTGCAACATCNNNAGATCGGAAGA |
| 34209_1 | TATGAGTGTGGAGTCGTTGCCTCACGCCTGTAATCCTAACACTNNNAGATCGGAAGA |
| 34209_2 | TATGAGTGTGGAGTCGTTGCCTCACGCCTGTAATCCTAACACTNNNAGATCGGAAGA |
| 31339_1 | TATGAGTGTGGAGTCGTTGCGCTCAGGCTATCTCGAATAGTGANNNAGATCGGAA |
| 31339_2 | TATGAGTGTGGAGTCGTTGCGCTCAGGCTATCTCGAATAGTGANNNAGATCGGAA |
| 30019_1 | TATGAGTGTGGAGTCGTTGCTATGACAATATCCCCTGTGCTGGNNNAGATCGGAAG |
| 30019_2 | TATGAGTGTGGAGTCGTTGCTATGACAATATCCCCTGTGCTGGNNNAGATCGGAAG |
| 30573_1 | TATGAGTGTGGAGTCGTTGCGACCAGTGGAAGGTTTCAGAGTCNNNAGATCGGAA |
| 30573_2 | TATGAGTGTGGAGTCGTTGCGACCAGTGGAAGGTTTCAGAGTCNNNAGATCGGAA |
| 30236_1 | TATGAGTGTGGAGTCGTTGCCCATCCACTCCCAATATGGCANNNAGATCGGAAGAG |
| 30236_2 | TATGAGTGTGGAGTCGTTGCCCATCCACTCCCAATATGGCANNNAGATCGGAAGAG |
| 32623_1 | TATGAGTGTGGAGTCGTTGCGGCAGGAACAACTAGAACAAAGGNNNAGATCGGA |
| 32623_2 | TATGAGTGTGGAGTCGTTGCGGCAGGAACAACTAGAACAAAGGNNNAGATCGGA |
| 45370_1 | TATGAGTGTGGAGTCGTTGCTGTCCACACACAGAAAACTCTCANNNAGATCGGAAG |
| 45370_2 | TATGAGTGTGGAGTCGTTGCTGTCCACACACAGAAAACTCTCANNNAGATCGGAAG |
| 50266_1 | TATGAGTGTGGAGTCGTTGCTGGGAGGAAGAAGTGGGAAATTTNNNAGATCGGA |
| 50266_2 | TATGAGTGTGGAGTCGTTGCTGGGAGGAAGAAGTGGGAAATTTNNNAGATCGGA |
| 31428_1 | TATGAGTGTGGAGTCGTTGCTTATTGGGCAACAGGCAGATACTNNNAGATCGGAA |
| 31428_2 | TATGAGTGTGGAGTCGTTGCTTATTGGGCAACAGGCAGATACTNNNAGATCGGAA |
| 33971_1 | TATGAGTGTGGAGTCGTTGCCCATGTACCTGGGATTACAGACGNNNAGATCGGAA |
| 33971_2 | TATGAGTGTGGAGTCGTTGCCCATGTACCTGGGATTACAGACGNNNAGATCGGAA |
| 4534_1 | TATGAGTGTGGAGTCGTTGCACTCTCAGGTTAGATGAGGGCTTNNNAGATCGGAA |
| 4534_2 | TATGAGTGTGGAGTCGTTGCACTCTCAGGTTAGATGAGGGCTTNNNAGATCGGAA |

| | |
|---|---|
| 48015_1 | TATGAGTGTGGAGTCGTTGCTTCAAGTGATTCTCCTGCCTCAGNNNAGATCGGAAGA |
| 48015_2 | TATGAGTGTGGAGTCGTTGCTTCAAGTGATTCTCCTGCCTCAGNNNAGATCGGAAGA |
| 37508_1 | TATGAGTGTGGAGTCGTTGCGGAAAAGACAGCATGGTAGATGCNNNAGATCGGAA |
| 37508_2 | TATGAGTGTGGAGTCGTTGCGGAAAAGACAGCATGGTAGATGCNNNAGATCGGAA |
| 52690_1 | TATGAGTGTGGAGTCGTTGCCACATCTGAATGCTCACCATGTCNNNAGATCGGAAGA |
| 52690_2 | TATGAGTGTGGAGTCGTTGCCACATCTGAATGCTCACCATGTCNNNAGATCGGAAGA |
| 43017_1 | TATGAGTGTGGAGTCGTTGCCTGAAGTTCCACCTACTTGGGAANNNAGATCGGAAG |
| 43017_2 | TATGAGTGTGGAGTCGTTGCCTGAAGTTCCACCTACTTGGGAANNNAGATCGGAAG |
| 53381_1 | TATGAGTGTGGAGTCGTTGCTCTCTCTCTTTCTTTCCCTCCCTNNNAGATCGGAAGAG |
| 53381_2 | TATGAGTGTGGAGTCGTTGCTCTCTCTCTTTCTTTCCCTCCCTNNNAGATCGGAAGAG |
| 13624_1 | TATGAGTGTGGAGTCGTTGCAAAAAGTCCTCCATAAATTATGTNNNAGATCGGAAGA |
| 13624_2 | TATGAGTGTGGAGTCGTTGCAAAAAGTCCTCCATAAATTATGTNNNAGATCGGAAGA |
| 33257_1 | TATGAGTGTGGAGTCGTTGCGAGAGAGAGCAGTTTGAAGGTGANNNAGATCGGAA |
| 33257_2 | TATGAGTGTGGAGTCGTTGCGAGAGAGAGCAGTTTGAAGGTGANNNAGATCGGAA |
| 46918_1 | TATGAGTGTGGAGTCGTTGCGTGAAGGGGAGAAAGAAAGCAACNNNAGATCGGAA |
| 46918_2 | TATGAGTGTGGAGTCGTTGCGTGAAGGGGAGAAAGAAAGCAACNNNAGATCGGAA |
| 3162_1 | TATGAGTGTGGAGTCGTTGCTTCCTCTGCTCCGTGCTGCCTANNNAGATCGGAAGAG |
| 3162_2 | TATGAGTGTGGAGTCGTTGCTTCCTCTGCTCCGTGCTGCCTANNNAGATCGGAAGAG |
| 49730_1 | TATGAGTGTGGAGTCGTTGCCCCTAGTATCTGGGACTACAGGTNNNAGATCGGAAG |
| 49730_2 | TATGAGTGTGGAGTCGTTGCCCCTAGTATCTGGGACTACAGGTNNNAGATCGGAAG |
| 28856_1 | TATGAGTGTGGAGTCGTTGCCACAATTGTGGAGGCTGATTCTGNNNAGATCGGAAG |
| 28856_2 | TATGAGTGTGGAGTCGTTGCCACAATTGTGGAGGCTGATTCTGNNNAGATCGGAAG |
| 12980_1 | TATGAGTGTGGAGTCGTTGCATTTTGGCTACATTAAGCTTTTNNNAGATCGGAAGAG |
| 12980_2 | TATGAGTGTGGAGTCGTTGCATTTTGGCTACATTAAGCTTTTNNNAGATCGGAAGAG |
| 29427_1 | TATGAGTGTGGAGTCGTTGCATCTCTACTCTGAGTCCCATGCCNNNAGATCGGAAGA |
| 29427_2 | TATGAGTGTGGAGTCGTTGCATCTCTACTCTGAGTCCCATGCCNNNAGATCGGAAGA |
| 43545_1 | TATGAGTGTGGAGTCGTTGCTCCTCTCTGCTTGTTCTTTTTGCNNNAGATCGGAAGAG |
| 43545_2 | TATGAGTGTGGAGTCGTTGCTCCTCTCTGCTTGTTCTTTTTGCNNNAGATCGGAAGAG |
| 30008_1 | TATGAGTGTGGAGTCGTTGCACTCAATATAGCTGTTTGGGCCANNNAGATCGGAAG |
| 30008_2 | TATGAGTGTGGAGTCGTTGCACTCAATATAGCTGTTTGGGCCANNNAGATCGGAAG |
| 38166_1 | TATGAGTGTGGAGTCGTTGCGTGTAAAGGAGGTGAGAGAAGGGNNNAGATCGGA |
| 38166_2 | TATGAGTGTGGAGTCGTTGCGTGTAAAGGAGGTGAGAGAAGGGNNNAGATCGGA |
| 38286_1 | TATGAGTGTGGAGTCGTTGCAGGCAGAGAATCACTTGAACCTTNNNAGATCGGAAG |
| 38286_2 | TATGAGTGTGGAGTCGTTGCAGGCAGAGAATCACTTGAACCTTNNNAGATCGGAAG |
| 37562_1 | TATGAGTGTGGAGTCGTTGCGGGAGTTTCGTAAGGTTACCACTNNNAGATCGGAAG |
| 37562_2 | TATGAGTGTGGAGTCGTTGCGGGAGTTTCGTAAGGTTACCACTNNNAGATCGGAAG |
| 34223_1 | TATGAGTGTGGAGTCGTTGCGAAGACATACAAATGGCCAACGANNNAGATCGGAAG |
| 34223_2 | TATGAGTGTGGAGTCGTTGCGAAGACATACAAATGGCCAACGANNNAGATCGGAAG |
| 3113_1 | TATGAGTGTGGAGTCGTTGCCTTTCTCGGCATCTGGAAGCTTCNNNAGATCGGAAGA |
| 3113_2 | TATGAGTGTGGAGTCGTTGCCTTTCTCGGCATCTGGAAGCTTCNNNAGATCGGAAGA |
| 39914_1 | TATGAGTGTGGAGTCGTTGCTCCGTATTTTATGGTTGGGGGTTNNNAGATCGGAAG |
| 39914_2 | TATGAGTGTGGAGTCGTTGCTCCGTATTTTATGGTTGGGGGTTNNNAGATCGGAAG |
| 52468_1 | TATGAGTGTGGAGTCGTTGCACCCATGCCAGGCTTCTAAAATANNNAGATCGGAAGA |

| | |
|---|---|
| 52468_2 | TATGAGTGTGGAGTCGTTGCACCCATGCCAGGCTTCTAAAATANNNAGATCGGAAG |
| 33970_1 | TATGAGTGTGGAGTCGTTGCGCCTCCTGAGTAGTTGGGATTACNNNAGATCGGAA |
| 33970_2 | TATGAGTGTGGAGTCGTTGCGCCTCCTGAGTAGTTGGGATTACNNNAGATCGGAA |
| 39238_1 | TATGAGTGTGGAGTCGTTGCACAAGAACGCTGCTTGATCATTTNNNAGATCGGAAG |
| 39238_2 | TATGAGTGTGGAGTCGTTGCACAAGAACGCTGCTTGATCATTTNNNAGATCGGAAG |
| 44470_1 | TATGAGTGTGGAGTCGTTGCCACTCATCCTCACAGCCTGAAGNNNAGATCGGAAGA |
| 44470_2 | TATGAGTGTGGAGTCGTTGCCACTCATCCTCACAGCCTGAAGNNNAGATCGGAAGA |
| 37485_1 | TATGAGTGTGGAGTCGTTGCCAACGATGCTTTAGATTGATCACTNNNAGATCGGAA |
| 37485_2 | TATGAGTGTGGAGTCGTTGCCAACGATGCTTTAGATTGATCACTNNNAGATCGGAA |
| 49762_1 | TATGAGTGTGGAGTCGTTGCTTCCAGGAAGCTTTTTGTTTCCCNNNAGATCGGAAG |
| 49762_2 | TATGAGTGTGGAGTCGTTGCTTCCAGGAAGCTTTTTGTTTCCCNNNAGATCGGAAG |
| 3048_1 | TATGAGTGTGGAGTCGTTGCGCTGTAACCTCCACCACTCAGNNNAGATCGGAAGAG |
| 3048_2 | TATGAGTGTGGAGTCGTTGCGCTGTAACCTCCACCACTCAGNNNAGATCGGAAGAG |
| 39908_1 | TATGAGTGTGGAGTCGTTGCAACATCAATTCCTCTGGACACCANNNAGATCGGAAG |
| 39908_2 | TATGAGTGTGGAGTCGTTGCAACATCAATTCCTCTGGACACCANNNAGATCGGAAG |
| 33017_1 | TATGAGTGTGGAGTCGTTGCAATCTAGAATGCAGTGGTGTGGTNNNAGATCGGAA |
| 33017_2 | TATGAGTGTGGAGTCGTTGCAATCTAGAATGCAGTGGTGTGGTNNNAGATCGGAA |
| 45740_1 | TATGAGTGTGGAGTCGTTGCTGCGAGAAAGTCATTGGCATTTTNNNAGATCGGAA |
| 45740_2 | TATGAGTGTGGAGTCGTTGCTGCGAGAAAGTCATTGGCATTTTNNNAGATCGGAA |
| 44940_1 | TATGAGTGTGGAGTCGTTGCATGAATGAGGCCTCTTGGTCAATNNNAGATCGGAA |
| 44940_2 | TATGAGTGTGGAGTCGTTGCATGAATGAGGCCTCTTGGTCAATNNNAGATCGGAA |
| 42921_1 | TATGAGTGTGGAGTCGTTGCCCAGACAGAAGCTGAGGAAGTACNNNAGATCGGAA |
| 42921_2 | TATGAGTGTGGAGTCGTTGCCCAGACAGAAGCTGAGGAAGTACNNNAGATCGGAA |
| 39658_1 | TATGAGTGTGGAGTCGTTGCATGGCACTGTTTTTCTAACCTGCNNNAGATCGGAAG |
| 39658_2 | TATGAGTGTGGAGTCGTTGCATGGCACTGTTTTTCTAACCTGCNNNAGATCGGAAG |
| 46843_1 | TATGAGTGTGGAGTCGTTGCCTCCTGAGTAGCTGGGATTACAGNNNAGATCGGAA |
| 46843_2 | TATGAGTGTGGAGTCGTTGCCTCCTGAGTAGCTGGGATTACAGNNNAGATCGGAA |
| 37177_1 | TATGAGTGTGGAGTCGTTGCTTTATGGCCAGAGCACAACATTGNNNAGATCGGAAG |
| 37177_2 | TATGAGTGTGGAGTCGTTGCTTTATGGCCAGAGCACAACATTGNNNAGATCGGAAG |
| 52415_1 | TATGAGTGTGGAGTCGTTGCGGTGACCAGGAGAGGATCTGATANNNAGATCGGA |
| 52415_2 | TATGAGTGTGGAGTCGTTGCGGTGACCAGGAGAGGATCTGATANNNAGATCGGA |
| 30737_1 | TATGAGTGTGGAGTCGTTGCCAAAAGGAGGGCAAGAAGAAAGGNNNAGATCGGA |
| 30737_2 | TATGAGTGTGGAGTCGTTGCCAAAAGGAGGGCAAGAAGAAAGGNNNAGATCGGA |
| 52001_1 | TATGAGTGTGGAGTCGTTGCACTCTCTCATCAGGACTTCCAGANNNAGATCGGAAG |
| 52001_2 | TATGAGTGTGGAGTCGTTGCACTCTCTCATCAGGACTTCCAGANNNAGATCGGAAG |
| 36509_1 | TATGAGTGTGGAGTCGTTGCGGTCTCATTTGTGCTACCCTGTANNNAGATCGGAAG |
| 36509_2 | TATGAGTGTGGAGTCGTTGCGGTCTCATTTGTGCTACCCTGTANNNAGATCGGAAG |
| 30427_1 | TATGAGTGTGGAGTCGTTGCGACCTGACGTTTGGAAGAGCAATNNNAGATCGGAA |
| 30427_2 | TATGAGTGTGGAGTCGTTGCGACCTGACGTTTGGAAGAGCAATNNNAGATCGGAA |
| 48516_1 | TATGAGTGTGGAGTCGTTGCGCTCAAGTGATGGTCCTACTTCANNNAGATCGGAAG |
| 48516_2 | TATGAGTGTGGAGTCGTTGCGCTCAAGTGATGGTCCTACTTCANNNAGATCGGAAG |
| 51952_1 | TATGAGTGTGGAGTCGTTGCGGTTTCCTTACTGTTTGGTGCTGNNNAGATCGGAAG |
| 51952_2 | TATGAGTGTGGAGTCGTTGCGGTTTCCTTACTGTTTGGTGCTGNNNAGATCGGAAG |

| | |
|---|---|
| 41971_1 | TATGAGTGTGGAGTCGTTGCGATTATCAGCACCTACCACCACANNNAGATCGGAAG |
| 41971_2 | TATGAGTGTGGAGTCGTTGCGATTATCAGCACCTACCACCACANNNAGATCGGAAG |
| 42104_1 | TATGAGTGTGGAGTCGTTGCCCTCCTGGGTTCAAGTGATTCTCNNNAGATCGGAAG |
| 42104_2 | TATGAGTGTGGAGTCGTTGCCCTCCTGGGTTCAAGTGATTCTCNNNAGATCGGAAG |
| 46438_1 | TATGAGTGTGGAGTCGTTGCAGAAGGAGTCTCACTTTGTCACCNNNAGATCGGAAG |
| 46438_2 | TATGAGTGTGGAGTCGTTGCAGAAGGAGTCTCACTTTGTCACCNNNAGATCGGAAG |
| 33181_1 | TATGAGTGTGGAGTCGTTGCCAAGACCTAAGCATGGAAGACGANNNAGATCGGAA |
| 33181_2 | TATGAGTGTGGAGTCGTTGCCAAGACCTAAGCATGGAAGACGANNNAGATCGGAA |
| 36476_1 | TATGAGTGTGGAGTCGTTGCAGTATAATTGCCTGAACCCAGCANNNAGATCGGAAG |
| 36476_2 | TATGAGTGTGGAGTCGTTGCAGTATAATTGCCTGAACCCAGCANNNAGATCGGAAG |
| 43831_1 | TATGAGTGTGGAGTCGTTGCATCTGCAGCATCCAAGTGAAAACNNNAGATCGGAAG |
| 43831_2 | TATGAGTGTGGAGTCGTTGCATCTGCAGCATCCAAGTGAAAACNNNAGATCGGAAG |
| 31924_1 | TATGAGTGTGGAGTCGTTGCTACAGCAAGTGTCTGTGAGAACANNNAGATCGGAA |
| 31924_2 | TATGAGTGTGGAGTCGTTGCTACAGCAAGTGTCTGTGAGAACANNNAGATCGGAA |
| 33863_1 | TATGAGTGTGGAGTCGTTGCAGGAGAATTGCTTGAACCTGGAANNNAGATCGGAA |
| 33863_2 | TATGAGTGTGGAGTCGTTGCAGGAGAATTGCTTGAACCTGGAANNNAGATCGGAA |
| 39395_1 | TATGAGTGTGGAGTCGTTGCGGAGTGATGTCATGACCCTTTGGNNNAGATCGGAA |
| 39395_2 | TATGAGTGTGGAGTCGTTGCGGAGTGATGTCATGACCCTTTGGNNNAGATCGGAA |
| 34468_1 | TATGAGTGTGGAGTCGTTGCACTTGTACATGCATGTTCACAGCNNNAGATCGGAAG |
| 34468_2 | TATGAGTGTGGAGTCGTTGCACTTGTACATGCATGTTCACAGCNNNAGATCGGAAG |
| 39952_1 | TATGAGTGTGGAGTCGTTGCAAACAGCATGTCCCATTCTAGGANNNAGATCGGAAG |
| 39952_2 | TATGAGTGTGGAGTCGTTGCAAACAGCATGTCCCATTCTAGGANNNAGATCGGAAG |
| 38467_1 | TATGAGTGTGGAGTCGTTGCTGAGCATTACAATTGTGCAGCTTNNNAGATCGGAAG |
| 38467_2 | TATGAGTGTGGAGTCGTTGCTGAGCATTACAATTGTGCAGCTTNNNAGATCGGAAG |
| 32600_1 | TATGAGTGTGGAGTCGTTGCTGTAACAAGCAGGTCCTTAGGTCNNNAGATCGGAA |
| 32600_2 | TATGAGTGTGGAGTCGTTGCTGTAACAAGCAGGTCCTTAGGTCNNNAGATCGGAA |
| 43129_1 | TATGAGTGTGGAGTCGTTGCTGTGAGATGTTTGCAGTTTGTCCNNNAGATCGGAAG |
| 43129_2 | TATGAGTGTGGAGTCGTTGCTGTGAGATGTTTGCAGTTTGTCCNNNAGATCGGAAG |
| 53138_1 | TATGAGTGTGGAGTCGTTGCAAAGGCTAACTACCGAACTTGGANNNAGATCGGAA |
| 53138_2 | TATGAGTGTGGAGTCGTTGCAAAGGCTAACTACCGAACTTGGANNNAGATCGGAA |
| 43097_1 | TATGAGTGTGGAGTCGTTGCGAGTACAATGGCATGATCTTGGCNNNAGATCGGAA |
| 43097_2 | TATGAGTGTGGAGTCGTTGCGAGTACAATGGCATGATCTTGGCNNNAGATCGGAA |
| 52940_1 | TATGAGTGTGGAGTCGTTGCCCACACCCGGCTAATTTTTTGTANNNAGATCGGAAG |
| 52940_2 | TATGAGTGTGGAGTCGTTGCCCACACCCGGCTAATTTTTTGTANNNAGATCGGAAG |
| 35545_1 | TATGAGTGTGGAGTCGTTGCCCTGCCACCAAATTCAGCTAATTNNNAGATCGGAAG |
| 35545_2 | TATGAGTGTGGAGTCGTTGCCCTGCCACCAAATTCAGCTAATTNNNAGATCGGAAG |
| 45838_1 | TATGAGTGTGGAGTCGTTGCGTTCAAGCGATTCTCCTACCTCANNNAGATCGGAAG |
| 45838_2 | TATGAGTGTGGAGTCGTTGCGTTCAAGCGATTCTCCTACCTCANNNAGATCGGAAG |
| 41226_1 | TATGAGTGTGGAGTCGTTGCTAAAGTCGGTCCTCATTCAGGTCNNNAGATCGGAAG |
| 41226_2 | TATGAGTGTGGAGTCGTTGCTAAAGTCGGTCCTCATTCAGGTCNNNAGATCGGAAG |
| 48866_1 | TATGAGTGTGGAGTCGTTGCCCACCATGCCCAGAGAATTTTTTNNNAGATCGGAAG |
| 48866_2 | TATGAGTGTGGAGTCGTTGCCCACCATGCCCAGAGAATTTTTTNNNAGATCGGAAG |
| 13662_1 | TATGAGTGTGGAGTCGTTGCCAACATCTTTTACATTCTGCAANNNAGATCGGAAGA |

| | |
|---|---|
| 13662_2 | TATGAGTGTGGAGTCGTTGCCAACATCTTTTACATTCTGCAANNNAGATCGGAAGAG |
| 40587_1 | TATGAGTGTGGAGTCGTTGCTGCATGGCAGGAATATTTTTCTCCNNNAGATCGGAA |
| 40587_2 | TATGAGTGTGGAGTCGTTGCTGCATGGCAGGAATATTTTTCTCCNNNAGATCGGAA |
| 38605_1 | TATGAGTGTGGAGTCGTTGCAGTCACCTGGAACCTCTATGACTNNNAGATCGGAAG |
| 38605_2 | TATGAGTGTGGAGTCGTTGCAGTCACCTGGAACCTCTATGACTNNNAGATCGGAAG |
| 35623_1 | TATGAGTGTGGAGTCGTTGCCCTCCCAGATTCATGCCATTCTCNNNAGATCGGAAGA |
| 35623_2 | TATGAGTGTGGAGTCGTTGCCCTCCCAGATTCATGCCATTCTCNNNAGATCGGAAGA |
| 52014_1 | TATGAGTGTGGAGTCGTTGCTGTGAACTCAGGTAGAGGGTACANNNAGATCGGAA |
| 52014_2 | TATGAGTGTGGAGTCGTTGCTGTGAACTCAGGTAGAGGGTACANNNAGATCGGAA |
| 49007_1 | TATGAGTGTGGAGTCGTTGCTACTTGTTCCTTTCCGATCTGGANNNAGATCGGAAGA |
| 49007_2 | TATGAGTGTGGAGTCGTTGCTACTTGTTCCTTTCCGATCTGGANNNAGATCGGAAGA |
| 29449_1 | TATGAGTGTGGAGTCGTTGCCCTTTGGTATTGCCCTATAATAANNNAGATCGGAAGA |
| 29449_2 | TATGAGTGTGGAGTCGTTGCCCTTTGGTATTGCCCTATAATAANNNAGATCGGAAGA |
| 47159_1 | TATGAGTGTGGAGTCGTTGCTTCTGGGTTCAAGCAATTCTCCTNNNAGATCGGAAGA |
| 47159_2 | TATGAGTGTGGAGTCGTTGCTTCTGGGTTCAAGCAATTCTCCTNNNAGATCGGAAGA |
| 37990_1 | TATGAGTGTGGAGTCGTTGCGTTGCAGTGAGCCAAGATCACNNNAGATCGGAAGA |
| 37990_2 | TATGAGTGTGGAGTCGTTGCGTTGCAGTGAGCCAAGATCACNNNAGATCGGAAGA |
| 9746_1 | TATGAGTGTGGAGTCGTTGCCACACCTGATCACATCCACACACNNNAGATCGGAAGA |
| 9746_2 | TATGAGTGTGGAGTCGTTGCCACACCTGATCACATCCACACACNNNAGATCGGAAGA |
| 48503_1 | TATGAGTGTGGAGTCGTTGCATGTGTTTTTATCACAGCCGGGNNNAGATCGGAAG |
| 48503_2 | TATGAGTGTGGAGTCGTTGCATGTGTTTTTATCACAGCCGGGNNNAGATCGGAAG |
| 41242_1 | TATGAGTGTGGAGTCGTTGCCTGTAATCCCAGCTACTCAGGAGNNNAGATCGGAAG |
| 41242_2 | TATGAGTGTGGAGTCGTTGCCTGTAATCCCAGCTACTCAGGAGNNNAGATCGGAAG |
| 48440_1 | TATGAGTGTGGAGTCGTTGCCTCAAGTGATCCTTCCACCTCAGNNNAGATCGGAAGA |
| 48440_2 | TATGAGTGTGGAGTCGTTGCCTCAAGTGATCCTTCCACCTCAGNNNAGATCGGAAGA |
| 13467_1 | TATGAGTGTGGAGTCGTTGCGTTTTCCTGCTATTCATGTTTCNNNAGATCGGAAGAG |
| 13467_2 | TATGAGTGTGGAGTCGTTGCGTTTTCCTGCTATTCATGTTTCNNNAGATCGGAAGAG |
| 35529_1 | TATGAGTGTGGAGTCGTTGCATAGGTTTTCACCATGCTTCCCANNNAGATCGGAAGA |
| 35529_2 | TATGAGTGTGGAGTCGTTGCATAGGTTTTCACCATGCTTCCCANNNAGATCGGAAGA |
| 39380_1 | TATGAGTGTGGAGTCGTTGCAAAATTAGCCAGGTGTGATGGTGNNNAGATCGGAA |
| 39380_2 | TATGAGTGTGGAGTCGTTGCAAAATTAGCCAGGTGTGATGGTGNNNAGATCGGAA |
| 43833_1 | TATGAGTGTGGAGTCGTTGCGGAAGGAGAAACACCTGCTAAGANNNAGATCGGAA |
| 43833_2 | TATGAGTGTGGAGTCGTTGCGGAAGGAGAAACACCTGCTAAGANNNAGATCGGAA |
| 48458_1 | TATGAGTGTGGAGTCGTTGCGATGTGTCATAATTTGGCCTCCGNNNAGATCGGAAG |
| 48458_2 | TATGAGTGTGGAGTCGTTGCGATGTGTCATAATTTGGCCTCCGNNNAGATCGGAAG |
| 29993_1 | TATGAGTGTGGAGTCGTTGCTACACGTTGAATTGAGGGGGATTNNNAGATCGGAA |
| 29993_2 | TATGAGTGTGGAGTCGTTGCTACACGTTGAATTGAGGGGGATTNNNAGATCGGAA |
| 40398_1 | TATGAGTGTGGAGTCGTTGCTACACTCACATAGCCACTATCCCNNNAGATCGGAAGA |
| 40398_2 | TATGAGTGTGGAGTCGTTGCTACACTCACATAGCCACTATCCCNNNAGATCGGAAGA |
| 43198_1 | TATGAGTGTGGAGTCGTTGCCCTCCTGGGTTCAAGTGATTCTCNNNAGATCGGAAG |
| 43198_2 | TATGAGTGTGGAGTCGTTGCCCTCCTGGGTTCAAGTGATTCTCNNNAGATCGGAAG |
| 30296_1 | TATGAGTGTGGAGTCGTTGCGGAGCATTGTGTGTGCATTAAGTNNNAGATCGGAA |
| 30296_2 | TATGAGTGTGGAGTCGTTGCGGAGCATTGTGTGTGCATTAAGTNNNAGATCGGAA |

| | |
|---|---|
| 13706_1 | TATGAGTGTGGAGTCGTTGCTGTTTCAAAAAAAATCAAACTGTNNNAGATCGGAAG |
| 13706_2 | TATGAGTGTGGAGTCGTTGCTGTTTCAAAAAAAATCAAACTGTNNNAGATCGGAAG |
| 38806_1 | TATGAGTGTGGAGTCGTTGCAGAGAGAATAACCTCACAAGAGCANNNAGATCGGA |
| 38806_2 | TATGAGTGTGGAGTCGTTGCAGAGAGAATAACCTCACAAGAGCANNNAGATCGGA |
| 29060_1 | TATGAGTGTGGAGTCGTTGCATCAAGCAGATTTTTAGCAGCCCNNNAGATCGGAAG |
| 29060_2 | TATGAGTGTGGAGTCGTTGCATCAAGCAGATTTTTAGCAGCCCNNNAGATCGGAAG |
| 39359_1 | TATGAGTGTGGAGTCGTTGCGTAATCCCAGGACTTTTGGAAGCNNNAGATCGGAAG |
| 39359_2 | TATGAGTGTGGAGTCGTTGCGTAATCCCAGGACTTTTGGAAGCNNNAGATCGGAAG |
| 40097_1 | TATGAGTGTGGAGTCGTTGCGAGGCTGAGACAGAAGAATCACTNNNAGATCGGAA |
| 40097_2 | TATGAGTGTGGAGTCGTTGCGAGGCTGAGACAGAAGAATCACTNNNAGATCGGAA |
| 48427_1 | TATGAGTGTGGAGTCGTTGCCCTCTCGAGTAGCTGGGACTATANNNAGATCGGAAG |
| 48427_2 | TATGAGTGTGGAGTCGTTGCCCTCTCGAGTAGCTGGGACTATANNNAGATCGGAAG |
| 40792_1 | TATGAGTGTGGAGTCGTTGCAGAAAGAAAGAAAGAAGCTCCAGCNNNAGATCGGA |
| 40792_2 | TATGAGTGTGGAGTCGTTGCAGAAAGAAAGAAAGAAGCTCCAGCNNNAGATCGGA |
| 50258_1 | TATGAGTGTGGAGTCGTTGCGGACTTGGATGGTCTCAGTGATTNNNAGATCGGAA |
| 50258_2 | TATGAGTGTGGAGTCGTTGCGGACTTGGATGGTCTCAGTGATTNNNAGATCGGAA |
| 36252_1 | TATGAGTGTGGAGTCGTTGCATGATAGCAGATGCAGATCACCCNNNAGATCGGAAG |
| 36252_2 | TATGAGTGTGGAGTCGTTGCATGATAGCAGATGCAGATCACCCNNNAGATCGGAAG |
| 38283_1 | TATGAGTGTGGAGTCGTTGCGAGCAATAACCATCACTGCAGTCNNNAGATCGGAAG |
| 38283_2 | TATGAGTGTGGAGTCGTTGCGAGCAATAACCATCACTGCAGTCNNNAGATCGGAAG |
| 13014_1 | TATGAGTGTGGAGTCGTTGCATTTAACTCCTTGGGTTTTTTAGNNNAGATCGGAAG |
| 13014_2 | TATGAGTGTGGAGTCGTTGCATTTAACTCCTTGGGTTTTTTAGNNNAGATCGGAAG |
| 38713_1 | TATGAGTGTGGAGTCGTTGCTATTTCAGGCTTCTTGGAGAGCANNNAGATCGGAAG |
| 38713_2 | TATGAGTGTGGAGTCGTTGCTATTTCAGGCTTCTTGGAGAGCANNNAGATCGGAAG |
| 40695_1 | TATGAGTGTGGAGTCGTTGCGGCAGGAGAATCACTTGAACCTANNNAGATCGGAA |
| 40695_2 | TATGAGTGTGGAGTCGTTGCGGCAGGAGAATCACTTGAACCTANNNAGATCGGAA |
| 47280_1 | TATGAGTGTGGAGTCGTTGCCTCCTGGGTAGCTGGGACTATAGNNNAGATCGGAA |
| 47280_2 | TATGAGTGTGGAGTCGTTGCCTCCTGGGTAGCTGGGACTATAGNNNAGATCGGAA |
| 44547_1 | TATGAGTGTGGAGTCGTTGCCTTAGCCGCTGTGAATAAAGCTCNNNAGATCGGAAG |
| 44547_2 | TATGAGTGTGGAGTCGTTGCCTTAGCCGCTGTGAATAAAGCTCNNNAGATCGGAAG |
| 50492_1 | TATGAGTGTGGAGTCGTTGCCATATCGGCCAGCACCTGTGNNNAGATCGGAAGAG |
| 50492_2 | TATGAGTGTGGAGTCGTTGCCATATCGGCCAGCACCTGTGNNNAGATCGGAAGAG |
| 46183_1 | TATGAGTGTGGAGTCGTTGCCCCGAGTAGCTGGGACTATAGGNNNAGATCGGAAG |
| 46183_2 | TATGAGTGTGGAGTCGTTGCCCCGAGTAGCTGGGACTATAGGNNNAGATCGGAAG |
| 48563_1 | TATGAGTGTGGAGTCGTTGCGCCTCCCAGATTCAAGCAATTTTNNNAGATCGGAAG |
| 48563_2 | TATGAGTGTGGAGTCGTTGCGCCTCCCAGATTCAAGCAATTTTNNNAGATCGGAAG |
| 28984_1 | TATGAGTGTGGAGTCGTTGCGGTTGTGTCCGTATATGTGCATGNNNAGATCGGAA |
| 28984_2 | TATGAGTGTGGAGTCGTTGCGGTTGTGTCCGTATATGTGCATGNNNAGATCGGAA |
| 39778_1 | TATGAGTGTGGAGTCGTTGCCTATAGCAAGACAGCCAAGCCTANNNAGATCGGAAG |
| 39778_2 | TATGAGTGTGGAGTCGTTGCCTATAGCAAGACAGCCAAGCCTANNNAGATCGGAAG |
| 35443_1 | TATGAGTGTGGAGTCGTTGCCCTAAAGTGCTGGCATACTAGGTNNNAGATCGGAA |
| 35443_2 | TATGAGTGTGGAGTCGTTGCCCTAAAGTGCTGGCATACTAGGTNNNAGATCGGAA |
| 30717_1 | TATGAGTGTGGAGTCGTTGCAACGGCCCTGGTGTTTTAAGAANNNAGATCGGAAG |

| | |
|---|---|
| 30717_2 | TATGAGTGTGGAGTCGTTGCAACGGCCCTGGTGTTTTAAGAANNNAGATCGGAAGA |
| 35086_1 | TATGAGTGTGGAGTCGTTGCTTTTGTGCAGTGGGTTGAAATGTNNNAGATCGGAA |
| 35086_2 | TATGAGTGTGGAGTCGTTGCTTTTGTGCAGTGGGTTGAAATGTNNNAGATCGGAA |
| 38515_1 | TATGAGTGTGGAGTCGTTGCATTCCAACCAGGACAAACATCCTNNNAGATCGGAAGA |
| 38515_2 | TATGAGTGTGGAGTCGTTGCATTCCAACCAGGACAAACATCCTNNNAGATCGGAAGA |
| 31779_1 | TATGAGTGTGGAGTCGTTGCAGAGAGAGAGAGAGAGAGAGGGANNNAGATCGGA |
| 31779_2 | TATGAGTGTGGAGTCGTTGCAGAGAGAGAGAGAGAGAGAGGGANNNAGATCGGA |
| 50698_1 | TATGAGTGTGGAGTCGTTGCGGAACTGAGTTTTGACCAAGCTCNNNAGATCGGAAG |
| 50698_2 | TATGAGTGTGGAGTCGTTGCGGAACTGAGTTTTGACCAAGCTCNNNAGATCGGAAG |
| 52549_1 | TATGAGTGTGGAGTCGTTGCATTGCTTTCTATCAGGGCCTCTCNNNAGATCGGAAGA |
| 52549_2 | TATGAGTGTGGAGTCGTTGCATTGCTTTCTATCAGGGCCTCTCNNNAGATCGGAAGA |
| 44965_1 | TATGAGTGTGGAGTCGTTGCGTCACTCACACATGGGAGCTCNNNAGATCGGAAGAG |
| 44965_2 | TATGAGTGTGGAGTCGTTGCGTCACTCACACATGGGAGCTCNNNAGATCGGAAGAG |
| 47722_1 | TATGAGTGTGGAGTCGTTGCTCTTTTTTGCAGCCACAGTCTTCNNNAGATCGGAAGA |
| 47722_2 | TATGAGTGTGGAGTCGTTGCTCTTTTTTGCAGCCACAGTCTTCNNNAGATCGGAAGA |
| 29818_1 | TATGAGTGTGGAGTCGTTGCATCTCAGCAATAAAGTCACAGCGNNNAGATCGGAAG |
| 29818_2 | TATGAGTGTGGAGTCGTTGCATCTCAGCAATAAAGTCACAGCGNNNAGATCGGAAG |
| 45781_1 | TATGAGTGTGGAGTCGTTGCCCACCACGTCTGGCTAATTTTTTNNNAGATCGGAAGA |
| 45781_2 | TATGAGTGTGGAGTCGTTGCCCACCACGTCTGGCTAATTTTTTNNNAGATCGGAAGA |
| 33185_1 | TATGAGTGTGGAGTCGTTGCTGCCAATCAAAGCCAAAGACAATNNNAGATCGGAAG |
| 33185_2 | TATGAGTGTGGAGTCGTTGCTGCCAATCAAAGCCAAAGACAATNNNAGATCGGAAG |
| 46012_1 | TATGAGTGTGGAGTCGTTGCCACCTTCCTGAGTAGTTGGGATTNNNAGATCGGAAG |
| 46012_2 | TATGAGTGTGGAGTCGTTGCCACCTTCCTGAGTAGTTGGGATTNNNAGATCGGAAG |
| 48340_1 | TATGAGTGTGGAGTCGTTGCGATCTCAGTTCACTGCAACCTCCNNNAGATCGGAAGA |
| 48340_2 | TATGAGTGTGGAGTCGTTGCGATCTCAGTTCACTGCAACCTCCNNNAGATCGGAAGA |
| 46992_1 | TATGAGTGTGGAGTCGTTGCCCAAGTTCAAGTGATCCTTCTGCNNNAGATCGGAAGA |
| 46992_2 | TATGAGTGTGGAGTCGTTGCCCAAGTTCAAGTGATCCTTCTGCNNNAGATCGGAAGA |
| 49190_1 | TATGAGTGTGGAGTCGTTGCTTCTGCACTGTGTTGTCCTATGANNNAGATCGGAAGA |
| 49190_2 | TATGAGTGTGGAGTCGTTGCTTCTGCACTGTGTTGTCCTATGANNNAGATCGGAAGA |
| 39122_1 | TATGAGTGTGGAGTCGTTGCACTGTACATATTTTTGTGGTACATGTGNNNAGATCG |
| 39122_2 | TATGAGTGTGGAGTCGTTGCACTGTACATATTTTTGTGGTACATGTGNNNAGATCG |
| 39776_1 | TATGAGTGTGGAGTCGTTGCTTTTGGCTAAGATCAAGTGTGAAANNNAGATCGGAA |
| 39776_2 | TATGAGTGTGGAGTCGTTGCTTTTGGCTAAGATCAAGTGTGAAANNNAGATCGGAA |
| 45095_1 | TATGAGTGTGGAGTCGTTGCAGATAGAGACAGACAAGACAGACTNNNAGATCGGA |
| 45095_2 | TATGAGTGTGGAGTCGTTGCAGATAGAGACAGACAAGACAGACTNNNAGATCGGA |
| 44143_1 | TATGAGTGTGGAGTCGTTGCGTTTTACAACTGAGCAAACCTGCNNNAGATCGGAAG |
| 44143_2 | TATGAGTGTGGAGTCGTTGCGTTTTACAACTGAGCAAACCTGCNNNAGATCGGAAG |
| 33589_1 | TATGAGTGTGGAGTCGTTGCCGAATAGCTAGGATTACAGGCGTNNNAGATCGGAA |
| 33589_2 | TATGAGTGTGGAGTCGTTGCCGAATAGCTAGGATTACAGGCGTNNNAGATCGGAA |
| 41349_1 | TATGAGTGTGGAGTCGTTGCACCCCATGGAGATAGAGAGCAATNNNAGATCGGAA |
| 41349_2 | TATGAGTGTGGAGTCGTTGCACCCCATGGAGATAGAGAGCAATNNNAGATCGGAA |
| 51802_1 | TATGAGTGTGGAGTCGTTGCCCTATCCTGCCCCTGTATAAACCNNNAGATCGGAAGA |
| 51802_2 | TATGAGTGTGGAGTCGTTGCCCTATCCTGCCCCTGTATAAACCNNNAGATCGGAAGA |

| | |
|---|---|
| 38163_1 | TATGAGTGTGGAGTCGTTGCATAAGACCACACATGATCTGGCCNNNAGATCGGAAG |
| 38163_2 | TATGAGTGTGGAGTCGTTGCATAAGACCACACATGATCTGGCCNNNAGATCGGAAG |
| 47540_1 | TATGAGTGTGGAGTCGTTGCATGCACTACCATGCCTAGTTACANNNAGATCGGAAG |
| 47540_2 | TATGAGTGTGGAGTCGTTGCATGCACTACCATGCCTAGTTACANNNAGATCGGAAG |
| 29023_1 | TATGAGTGTGGAGTCGTTGCAGGGTGACAAAATCAGTCAGGTTNNNAGATCGGAA |
| 29023_2 | TATGAGTGTGGAGTCGTTGCAGGGTGACAAAATCAGTCAGGTTNNNAGATCGGAA |
| 45847_1 | TATGAGTGTGGAGTCGTTGCGAGTTTTGCTCTTGTTGCCCAGNNNAGATCGGAAGA |
| 45847_2 | TATGAGTGTGGAGTCGTTGCGAGTTTTGCTCTTGTTGCCCAGNNNAGATCGGAAGA |
| 38543_1 | TATGAGTGTGGAGTCGTTGCCTGCAAGCCAGCATTATGGAAATNNNAGATCGGAAG |
| 38543_2 | TATGAGTGTGGAGTCGTTGCCTGCAAGCCAGCATTATGGAAATNNNAGATCGGAAG |
| 45550_1 | TATGAGTGTGGAGTCGTTGCTCTAAACTGACCCTGGAAATGCANNNAGATCGGAAG |
| 45550_2 | TATGAGTGTGGAGTCGTTGCTCTAAACTGACCCTGGAAATGCANNNAGATCGGAAG |
| 51049_1 | TATGAGTGTGGAGTCGTTGCGGCCTTGTGGGAGATATTAGTGTNNNAGATCGGAA |
| 51049_2 | TATGAGTGTGGAGTCGTTGCGGCCTTGTGGGAGATATTAGTGTNNNAGATCGGAA |
| 52268_1 | TATGAGTGTGGAGTCGTTGCCATCCTCAGCTCCACTCTACACNNNAGATCGGAAGAG |
| 52268_2 | TATGAGTGTGGAGTCGTTGCCATCCTCAGCTCCACTCTACACNNNAGATCGGAAGAG |
| 50861_1 | TATGAGTGTGGAGTCGTTGCTTCTCATTTCCTGTACAGACATGANNNAGATCGGAAG |
| 50861_2 | TATGAGTGTGGAGTCGTTGCTTCTCATTTCCTGTACAGACATGANNNAGATCGGAAG |
| 41741_1 | TATGAGTGTGGAGTCGTTGCAATACAGCACACTGATGGGTCTTNNNAGATCGGAAG |
| 41741_2 | TATGAGTGTGGAGTCGTTGCAATACAGCACACTGATGGGTCTTNNNAGATCGGAAG |
| 41491_1 | TATGAGTGTGGAGTCGTTGCAGACTGGCCTTACAAGAAATGGTNNNAGATCGGAA |
| 41491_2 | TATGAGTGTGGAGTCGTTGCAGACTGGCCTTACAAGAAATGGTNNNAGATCGGAA |
| 44947_1 | TATGAGTGTGGAGTCGTTGCCGGTCCCTTCCACCTTAGTAATCNNNAGATCGGAAGA |
| 44947_2 | TATGAGTGTGGAGTCGTTGCCGGTCCCTTCCACCTTAGTAATCNNNAGATCGGAAGA |
| 41730_1 | TATGAGTGTGGAGTCGTTGCCATTCTCCTGCCTCAGCCTCNNNAGATCGGAAGAGCA |
| 41730_2 | TATGAGTGTGGAGTCGTTGCCATTCTCCTGCCTCAGCCTCNNNAGATCGGAAGAGCA |
| 38023_1 | TATGAGTGTGGAGTCGTTGCACCAGATACACAAGGAAGAGCTGNNNAGATCGGAA |
| 38023_2 | TATGAGTGTGGAGTCGTTGCACCAGATACACAAGGAAGAGCTGNNNAGATCGGAA |
| 47030_1 | TATGAGTGTGGAGTCGTTGCCTCAAGCAATCCTCCCATCTCAGNNNAGATCGGAAGA |
| 47030_2 | TATGAGTGTGGAGTCGTTGCCTCAAGCAATCCTCCCATCTCAGNNNAGATCGGAAGA |
| 37162_1 | TATGAGTGTGGAGTCGTTGCCAGAACAAATCAATCCACGTGGGNNNAGATCGGAAG |
| 37162_2 | TATGAGTGTGGAGTCGTTGCCAGAACAAATCAATCCACGTGGGNNNAGATCGGAAG |
| 41344_1 | TATGAGTGTGGAGTCGTTGCGTGACTGTGGGTAAGCAAACAAANNNAGATCGGAA |
| 41344_2 | TATGAGTGTGGAGTCGTTGCGTGACTGTGGGTAAGCAAACAAANNNAGATCGGAA |
| 33163_1 | TATGAGTGTGGAGTCGTTGCATGCAGCCCCCAAAAATATTGAGNNNAGATCGGAAG |
| 33163_2 | TATGAGTGTGGAGTCGTTGCATGCAGCCCCCAAAAATATTGAGNNNAGATCGGAAG |
| 31385_1 | TATGAGTGTGGAGTCGTTGCCCTGTGGAGGTGAAAACATTTGANNNAGATCGGAA |
| 31385_2 | TATGAGTGTGGAGTCGTTGCCCTGTGGAGGTGAAAACATTTGANNNAGATCGGAA |
| 51781_1 | TATGAGTGTGGAGTCGTTGCGGTCTTTTCTGACTGGCTTCTCTNNNAGATCGGAAGA |
| 51781_2 | TATGAGTGTGGAGTCGTTGCGGTCTTTTCTGACTGGCTTCTCTNNNAGATCGGAAGA |
| 38673_1 | TATGAGTGTGGAGTCGTTGCAGGAAGGAGAATCACTTGAACCCNNNAGATCGGAA |
| 38673_2 | TATGAGTGTGGAGTCGTTGCAGGAAGGAGAATCACTTGAACCCNNNAGATCGGAA |
| 47001_1 | TATGAGTGTGGAGTCGTTGCGATCTCTGCTCAGTACAGCCTACNNNAGATCGGAAG |

| | |
|---|---|
| 47001_2 | TATGAGTGTGGAGTCGTTGCGATCTCTGCTCAGTACAGCCTACNNNAGATCGGAAG |
| 31200_1 | TATGAGTGTGGAGTCGTTGCATATGCCAAGTCTGCAGTCTACCNNNAGATCGGAAG |
| 31200_2 | TATGAGTGTGGAGTCGTTGCATATGCCAAGTCTGCAGTCTACCNNNAGATCGGAAG |
| 44922_1 | TATGAGTGTGGAGTCGTTGCTAAACCTGCTAGGGGGTTAGAGANNNAGATCGGAA |
| 44922_2 | TATGAGTGTGGAGTCGTTGCTAAACCTGCTAGGGGGTTAGAGANNNAGATCGGAA |
| 47780_1 | TATGAGTGTGGAGTCGTTGCTTCAAGTGATTCTCCTGCCTCAGNNNAGATCGGAAG |
| 47780_2 | TATGAGTGTGGAGTCGTTGCTTCAAGTGATTCTCCTGCCTCAGNNNAGATCGGAAG |
| 48165_1 | TATGAGTGTGGAGTCGTTGCGAGTAGCTGGGATTATAGGCACCNNNAGATCGGAA |
| 48165_2 | TATGAGTGTGGAGTCGTTGCGAGTAGCTGGGATTATAGGCACCNNNAGATCGGAA |
| 31703_1 | TATGAGTGTGGAGTCGTTGCTTTGCTGTACTCCCTCTGCAGNNNAGATCGGAAGAG |
| 31703_2 | TATGAGTGTGGAGTCGTTGCTTTGCTGTACTCCCTCTGCAGNNNAGATCGGAAGAG |
| 41882_1 | TATGAGTGTGGAGTCGTTGCCCTGAGTAGCTGGGATTATAGGCNNNAGATCGGAA |
| 41882_2 | TATGAGTGTGGAGTCGTTGCCCTGAGTAGCTGGGATTATAGGCNNNAGATCGGAA |
| 40754_1 | TATGAGTGTGGAGTCGTTGCACCTTGTAAGCTTTGTGTTGTGGNNNAGATCGGAAG |
| 40754_2 | TATGAGTGTGGAGTCGTTGCACCTTGTAAGCTTTGTGTTGTGGNNNAGATCGGAAG |
| 13705_1 | TATGAGTGTGGAGTCGTTGCTGTTTCAAAAAAAATCAAACTGTNNNAGATCGGAAG |
| 13705_2 | TATGAGTGTGGAGTCGTTGCTGTTTCAAAAAAAATCAAACTGTNNNAGATCGGAAG |
| 42809_1 | TATGAGTGTGGAGTCGTTGCCAGGTACCTAGATGCATAGCCTGNNNAGATCGGAAG |
| 42809_2 | TATGAGTGTGGAGTCGTTGCCAGGTACCTAGATGCATAGCCTGNNNAGATCGGAAG |
| 4009_1 | TATGAGTGTGGAGTCGTTGCGGATTTTACCCAAGGTCACAGAACNNNAGATCGGAA |
| 4009_2 | TATGAGTGTGGAGTCGTTGCGGATTTTACCCAAGGTCACAGAACNNNAGATCGGAA |
| 37614_1 | TATGAGTGTGGAGTCGTTGCTTATAAGGCCGAGATGGGATTGCNNNAGATCGGAA |
| 37614_2 | TATGAGTGTGGAGTCGTTGCTTATAAGGCCGAGATGGGATTGCNNNAGATCGGAA |
| 51702_1 | TATGAGTGTGGAGTCGTTGCCTTCCAAGTAGCTAGGACCACAGNNNAGATCGGAAG |
| 51702_2 | TATGAGTGTGGAGTCGTTGCCTTCCAAGTAGCTAGGACCACAGNNNAGATCGGAAG |
| 47543_1 | TATGAGTGTGGAGTCGTTGCGATGCTGATCTGACAATGGGTTGNNNAGATCGGAA |
| 47543_2 | TATGAGTGTGGAGTCGTTGCGATGCTGATCTGACAATGGGTTGNNNAGATCGGAA |
| 31658_1 | TATGAGTGTGGAGTCGTTGCCTACCAGCTCCTACCTTTGCTACNNNAGATCGGAAGA |
| 31658_2 | TATGAGTGTGGAGTCGTTGCCTACCAGCTCCTACCTTTGCTACNNNAGATCGGAAGA |
| 13451_1 | TATGAGTGTGGAGTCGTTGCCTTTTCCTGACCATTAGCATNNNAGATCGGAAGAGC |
| 13451_2 | TATGAGTGTGGAGTCGTTGCCTTTTCCTGACCATTAGCATNNNAGATCGGAAGAGC |
| 40827_1 | TATGAGTGTGGAGTCGTTGCATTTATGCACCCATGTCAGAGCANNNAGATCGGAAG |
| 40827_2 | TATGAGTGTGGAGTCGTTGCATTTATGCACCCATGTCAGAGCANNNAGATCGGAAG |
| 12918_1 | TATGAGTGTGGAGTCGTTGCAAGATCAAGAAAATTTAGCTGTTNNNAGATCGGAAG |
| 12918_2 | TATGAGTGTGGAGTCGTTGCAAGATCAAGAAAATTTAGCTGTTNNNAGATCGGAAG |
| 40808_1 | TATGAGTGTGGAGTCGTTGCAGAAAAATTGCTTGAACCCCAGGNNNAGATCGGAAG |
| 40808_2 | TATGAGTGTGGAGTCGTTGCAGAAAAATTGCTTGAACCCCAGGNNNAGATCGGAAG |
| 37698_1 | TATGAGTGTGGAGTCGTTGCAGTCCTGACACAGACAAGTTGAGNNNAGATCGGAA |
| 37698_2 | TATGAGTGTGGAGTCGTTGCAGTCCTGACACAGACAAGTTGAGNNNAGATCGGAA |
| 50051_1 | TATGAGTGTGGAGTCGTTGCGAAGGAAGGAAGGAAGGAAGGAANNNAGATCGG |
| 50051_2 | TATGAGTGTGGAGTCGTTGCGAAGGAAGGAAGGAAGGAAGGAANNNAGATCGG |
| 31403_1 | TATGAGTGTGGAGTCGTTGCGGAAATGCAATCACAGATGCAGANNNAGATCGGAA |
| 31403_2 | TATGAGTGTGGAGTCGTTGCGGAAATGCAATCACAGATGCAGANNNAGATCGGAA |

| | |
|---|---|
| 34090_1 | TATGAGTGTGGAGTCGTTGCCATTCTCCTGCCTCAGCCTCNNNAGATCGGAAGAGCA |
| 34090_2 | TATGAGTGTGGAGTCGTTGCCATTCTCCTGCCTCAGCCTCNNNAGATCGGAAGAGCA |
| 12661_1 | TATGAGTGTGGAGTCGTTGCTCACACAAAAAAATGTTAAGGATNNNAGATCGGAAG |
| 12661_2 | TATGAGTGTGGAGTCGTTGCTCACACAAAAAAATGTTAAGGATNNNAGATCGGAAG |
| 38508_1 | TATGAGTGTGGAGTCGTTGCAAAGGGCATACTTTAGCTCCAGTNNNAGATCGGAAG |
| 38508_2 | TATGAGTGTGGAGTCGTTGCAAAGGGCATACTTTAGCTCCAGTNNNAGATCGGAAG |
| 35720_1 | TATGAGTGTGGAGTCGTTGCTGAGCCAAGCCAACTAGTACTTTNNNAGATCGGAAG |
| 35720_2 | TATGAGTGTGGAGTCGTTGCTGAGCCAAGCCAACTAGTACTTTNNNAGATCGGAAG |
| 31939_1 | TATGAGTGTGGAGTCGTTGCAAAAAATTAGCTGGGTGTGGTGGNNNAGATCGGAA |
| 31939_2 | TATGAGTGTGGAGTCGTTGCAAAAAATTAGCTGGGTGTGGTGGNNNAGATCGGAA |
| 51441_1 | TATGAGTGTGGAGTCGTTGCAAACTTTCTGGACTGGACGGTTANNNAGATCGGAAG |
| 51441_2 | TATGAGTGTGGAGTCGTTGCAAACTTTCTGGACTGGACGGTTANNNAGATCGGAAG |
| 52140_1 | TATGAGTGTGGAGTCGTTGCCCAGTACCCAACAAGAGACATCANNNAGATCGGAAG |
| 52140_2 | TATGAGTGTGGAGTCGTTGCCCAGTACCCAACAAGAGACATCANNNAGATCGGAAG |
| 49830_1 | TATGAGTGTGGAGTCGTTGCTTCTCTTCCCGGTCACATTGATTNNNAGATCGGAAGA |
| 49830_2 | TATGAGTGTGGAGTCGTTGCTTCTCTTCCCGGTCACATTGATTNNNAGATCGGAAGA |
| 34937_1 | TATGAGTGTGGAGTCGTTGCGAGTATCTGGGATTACAGGCGTGNNNAGATCGGAA |
| 34937_2 | TATGAGTGTGGAGTCGTTGCGAGTATCTGGGATTACAGGCGTGNNNAGATCGGAA |
| 30594_1 | TATGAGTGTGGAGTCGTTGCTCACAAGGTCAGGAGTTTGAGACNNNAGATCGGAA |
| 30594_2 | TATGAGTGTGGAGTCGTTGCTCACAAGGTCAGGAGTTTGAGACNNNAGATCGGAA |
| 44253_1 | TATGAGTGTGGAGTCGTTGCGAGTTCCATTTCCTCCCCACATTNNNAGATCGGAAGA |
| 44253_2 | TATGAGTGTGGAGTCGTTGCGAGTTCCATTTCCTCCCCACATTNNNAGATCGGAAGA |
| 38068_1 | TATGAGTGTGGAGTCGTTGCTGCAGTGAAGTATGATCACACCANNNAGATCGGAAG |
| 38068_2 | TATGAGTGTGGAGTCGTTGCTGCAGTGAAGTATGATCACACCANNNAGATCGGAAG |
| 31879_1 | TATGAGTGTGGAGTCGTTGCTTGGCAACATAGCAAGATCTCCTNNNAGATCGGAAG |
| 31879_2 | TATGAGTGTGGAGTCGTTGCTTGGCAACATAGCAAGATCTCCTNNNAGATCGGAAG |
| 48377_1 | TATGAGTGTGGAGTCGTTGCGACAGGGTCTCACTATATTGCCCNNNAGATCGGAAG |
| 48377_2 | TATGAGTGTGGAGTCGTTGCGACAGGGTCTCACTATATTGCCCNNNAGATCGGAAG |
| 37683_1 | TATGAGTGTGGAGTCGTTGCAGTTCTGCTTACTAAGGAAGAGTTNNNAGATCGGAA |
| 37683_2 | TATGAGTGTGGAGTCGTTGCAGTTCTGCTTACTAAGGAAGAGTTNNNAGATCGGAA |
| 29204_1 | TATGAGTGTGGAGTCGTTGCATTTCTCCAAGGAGCCCTCATTTNNNAGATCGGAAGA |
| 29204_2 | TATGAGTGTGGAGTCGTTGCATTTCTCCAAGGAGCCCTCATTTNNNAGATCGGAAGA |
| 45998_1 | TATGAGTGTGGAGTCGTTGCTCCCCGGACTAAAACTAGAAGGANNNAGATCGGAAG |
| 45998_2 | TATGAGTGTGGAGTCGTTGCTCCCCGGACTAAAACTAGAAGGANNNAGATCGGAAG |
| 37259_1 | TATGAGTGTGGAGTCGTTGCTGGAATTCAGCAACACGTCAAAANNNAGATCGGAAG |
| 37259_2 | TATGAGTGTGGAGTCGTTGCTGGAATTCAGCAACACGTCAAAANNNAGATCGGAAG |
| 29608_1 | TATGAGTGTGGAGTCGTTGCTGACCTTTACATAGTGAAGGCCGNNNAGATCGGAAG |
| 29608_2 | TATGAGTGTGGAGTCGTTGCTGACCTTTACATAGTGAAGGCCGNNNAGATCGGAAG |
| 38036_1 | TATGAGTGTGGAGTCGTTGCACAAAACAAAAAACAGGTTGCGGNNNAGATCGGAA |
| 38036_2 | TATGAGTGTGGAGTCGTTGCACAAAACAAAAAACAGGTTGCGGNNNAGATCGGAA |
| 43610_1 | TATGAGTGTGGAGTCGTTGCCTCAGCCAGTGAAGGGGTAAACNNNAGATCGGAAG |
| 43610_2 | TATGAGTGTGGAGTCGTTGCCTCAGCCAGTGAAGGGGTAAACNNNAGATCGGAAG |
| 13473_1 | TATGAGTGTGGAGTCGTTGCCCCACCTTCTGTTTACTTTAGNNNAGATCGGAAGAGC |

| | |
|---|---|
| 13473_2 | TATGAGTGTGGAGTCGTTGCCCCACCTTCTGTTTACTTTAGNNNAGATCGGAAGAGC |
| 2592_1 | TATGAGTGTGGAGTCGTTGCCTCTCCTTCCTTGGTTTCTGTGGNNNAGATCGGAAGA |
| 2592_2 | TATGAGTGTGGAGTCGTTGCCTCTCCTTCCTTGGTTTCTGTGGNNNAGATCGGAAGA |
| 42895_1 | TATGAGTGTGGAGTCGTTGCTGGAAGATTAGTTCTAGAACCCACTNNNAGATCGGA |
| 42895_2 | TATGAGTGTGGAGTCGTTGCTGGAAGATTAGTTCTAGAACCCACTNNNAGATCGGA |
| 50132_1 | TATGAGTGTGGAGTCGTTGCGTTTGAGAATGGCATGTTCCTCCNNNAGATCGGAAG |
| 50132_2 | TATGAGTGTGGAGTCGTTGCGTTTGAGAATGGCATGTTCCTCCNNNAGATCGGAAG |
| 46344_1 | TATGAGTGTGGAGTCGTTGCCTCCTGAGTAGCTGGGATTACAGNNNAGATCGGAAG |
| 46344_2 | TATGAGTGTGGAGTCGTTGCCTCCTGAGTAGCTGGGATTACAGNNNAGATCGGAAG |
| 40101_1 | TATGAGTGTGGAGTCGTTGCTCTATGGGATACAGCACAAGCAANNNAGATCGGAAG |
| 40101_2 | TATGAGTGTGGAGTCGTTGCTCTATGGGATACAGCACAAGCAANNNAGATCGGAAG |
| 52581_1 | TATGAGTGTGGAGTCGTTGCCAAACAACAGGGTCTTGGTTTGTNNNAGATCGGAAG |
| 52581_2 | TATGAGTGTGGAGTCGTTGCCAAACAACAGGGTCTTGGTTTGTNNNAGATCGGAAG |
| 12704_1 | TATGAGTGTGGAGTCGTTGCGAGAAGGGAGAAAAAGGAAGNNNAGATCGGAAGA |
| 12704_2 | TATGAGTGTGGAGTCGTTGCGAGAAGGGAGAAAAAGGAAGNNNAGATCGGAAGA |
| 52202_1 | TATGAGTGTGGAGTCGTTGCGAGGCAGGAGAATAGCTTGAACTNNNAGATCGGAA |
| 52202_2 | TATGAGTGTGGAGTCGTTGCGAGGCAGGAGAATAGCTTGAACTNNNAGATCGGAA |
| 33726_1 | TATGAGTGTGGAGTCGTTGCTCTTTGGGACCAACACCATGTATNNNAGATCGGAAGA |
| 33726_2 | TATGAGTGTGGAGTCGTTGCTCTTTGGGACCAACACCATGTATNNNAGATCGGAAGA |
| 40269_1 | TATGAGTGTGGAGTCGTTGCATCCCTTCCACCTAGGTAACAGANNNAGATCGGAAGA |
| 40269_2 | TATGAGTGTGGAGTCGTTGCATCCCTTCCACCTAGGTAACAGANNNAGATCGGAAGA |
| 47870_1 | TATGAGTGTGGAGTCGTTGCGAGATGGACTCTTGCTCTGTCGNNNAGATCGGAAGA |
| 47870_2 | TATGAGTGTGGAGTCGTTGCGAGATGGACTCTTGCTCTGTCGNNNAGATCGGAAGA |
| 29010_1 | TATGAGTGTGGAGTCGTTGCAACTTTGTAATGTTGGCAGCTGGNNNAGATCGGAAG |
| 29010_2 | TATGAGTGTGGAGTCGTTGCAACTTTGTAATGTTGGCAGCTGGNNNAGATCGGAAG |
| 36360_1 | TATGAGTGTGGAGTCGTTGCAACTGCACACCGTAAAAAGATCTNNNAGATCGGAAG |
| 36360_2 | TATGAGTGTGGAGTCGTTGCAACTGCACACCGTAAAAAGATCTNNNAGATCGGAAG |
| 32907_1 | TATGAGTGTGGAGTCGTTGCCAACTGGATTGTGGTGATGGTTGNNNAGATCGGAAG |
| 32907_2 | TATGAGTGTGGAGTCGTTGCCAACTGGATTGTGGTGATGGTTGNNNAGATCGGAAG |
| 35026_1 | TATGAGTGTGGAGTCGTTGCCAATCTCAGCTCACTGCAACTTCNNNAGATCGGAAGA |
| 35026_2 | TATGAGTGTGGAGTCGTTGCCAATCTCAGCTCACTGCAACTTCNNNAGATCGGAAGA |
| 42472_1 | TATGAGTGTGGAGTCGTTGCAGCCAAGAGTGCTAGAGGAAAAANNNAGATCGGAA |
| 42472_2 | TATGAGTGTGGAGTCGTTGCAGCCAAGAGTGCTAGAGGAAAAANNNAGATCGGAA |
| 52794_1 | TATGAGTGTGGAGTCGTTGCGTGGTGAGGCTGTGGCTATAGNNNAGATCGGAAGA |
| 52794_2 | TATGAGTGTGGAGTCGTTGCGTGGTGAGGCTGTGGCTATAGNNNAGATCGGAAGA |
| 42592_1 | TATGAGTGTGGAGTCGTTGCCTGGTCTTGAACTCCTGACTTCANNNAGATCGGAAGA |
| 42592_2 | TATGAGTGTGGAGTCGTTGCCTGGTCTTGAACTCCTGACTTCANNNAGATCGGAAGA |
| 42249_1 | TATGAGTGTGGAGTCGTTGCCCTTTTTGTTTCACCTTTGCAGCNNNAGATCGGAAGA |
| 42249_2 | TATGAGTGTGGAGTCGTTGCCCTTTTTGTTTCACCTTTGCAGCNNNAGATCGGAAGA |
| 36137_1 | TATGAGTGTGGAGTCGTTGCGAAAACATTCTAAGCCTCCCACCNNNAGATCGGAAGA |
| 36137_2 | TATGAGTGTGGAGTCGTTGCGAAAACATTCTAAGCCTCCCACCNNNAGATCGGAAGA |
| 40153_1 | TATGAGTGTGGAGTCGTTGCGATCTCGGCTCACTGCAACTTCNNNAGATCGGAAGAC |
| 40153_2 | TATGAGTGTGGAGTCGTTGCGATCTCGGCTCACTGCAACTTCNNNAGATCGGAAGAC |

| | |
|---|---|
| 49479_1 | TATGAGTGTGGAGTCGTTGCTATCTCAGTGAGCAGAGGGATGANNNAGATCGGAA |
| 49479_2 | TATGAGTGTGGAGTCGTTGCTATCTCAGTGAGCAGAGGGATGANNNAGATCGGAA |
| 34081_1 | TATGAGTGTGGAGTCGTTGCCTCACTGCAACTTCCATCTACCANNNAGATCGGAAGA |
| 34081_2 | TATGAGTGTGGAGTCGTTGCCTCACTGCAACTTCCATCTACCANNNAGATCGGAAGA |
| 36168_1 | TATGAGTGTGGAGTCGTTGCACAATCAGCAGGCATTTACATAGGNNNAGATCGGAA |
| 36168_2 | TATGAGTGTGGAGTCGTTGCACAATCAGCAGGCATTTACATAGGNNNAGATCGGAA |
| 2026_1 | TATGAGTGTGGAGTCGTTGCCACGCACAAAGAATCAACTGCAANNNAGATCGGAAG |
| 2026_2 | TATGAGTGTGGAGTCGTTGCCACGCACAAAGAATCAACTGCAANNNAGATCGGAAG |
| 39515_1 | TATGAGTGTGGAGTCGTTGCATCTCAGCTCACTGCAACCTCTANNNAGATCGGAAGA |
| 39515_2 | TATGAGTGTGGAGTCGTTGCATCTCAGCTCACTGCAACCTCTANNNAGATCGGAAGA |
| 44924_1 | TATGAGTGTGGAGTCGTTGCAGAAGGGATCAGGAGATAGAGCCNNNAGATCGGAA |
| 44924_2 | TATGAGTGTGGAGTCGTTGCAGAAGGGATCAGGAGATAGAGCCNNNAGATCGGAA |
| 47516_1 | TATGAGTGTGGAGTCGTTGCTGAAAGCATTTCCTTCTCCTCCANNNAGATCGGAAGA |
| 47516_2 | TATGAGTGTGGAGTCGTTGCTGAAAGCATTTCCTTCTCCTCCANNNAGATCGGAAGA |
| 35493_1 | TATGAGTGTGGAGTCGTTGCCAGCCTCTTGAGTAGTTGGGATTNNNAGATCGGAAG |
| 35493_2 | TATGAGTGTGGAGTCGTTGCCAGCCTCTTGAGTAGTTGGGATTNNNAGATCGGAAG |
| 31560_1 | TATGAGTGTGGAGTCGTTGCTGTCCACTTGAATGGTGACTACANNNAGATCGGAAG |
| 31560_2 | TATGAGTGTGGAGTCGTTGCTGTCCACTTGAATGGTGACTACANNNAGATCGGAAG |
| 47202_1 | TATGAGTGTGGAGTCGTTGCTTCAAGCAATTCTCCTGCCTCAGNNNAGATCGGAAGA |
| 47202_2 | TATGAGTGTGGAGTCGTTGCTTCAAGCAATTCTCCTGCCTCAGNNNAGATCGGAAGA |
| 37312_1 | TATGAGTGTGGAGTCGTTGCAGTTGCATGTAACAAACAGTGCTNNNAGATCGGAAG |
| 37312_2 | TATGAGTGTGGAGTCGTTGCAGTTGCATGTAACAAACAGTGCTNNNAGATCGGAAG |
| 31842_1 | TATGAGTGTGGAGTCGTTGCACAACATGGTCCTCAGTTAGTCGNNNAGATCGGAAG |
| 31842_2 | TATGAGTGTGGAGTCGTTGCACAACATGGTCCTCAGTTAGTCGNNNAGATCGGAAG |
| 33852_1 | TATGAGTGTGGAGTCGTTGCGACTGCTGGTGACAAGGATATGANNNAGATCGGAA |
| 33852_2 | TATGAGTGTGGAGTCGTTGCGACTGCTGGTGACAAGGATATGANNNAGATCGGAA |
| 30831_1 | TATGAGTGTGGAGTCGTTGCTGAGTATAACCAGATAGCCGTGCNNNAGATCGGAAG |
| 30831_2 | TATGAGTGTGGAGTCGTTGCTGAGTATAACCAGATAGCCGTGCNNNAGATCGGAAG |
| 49532_1 | TATGAGTGTGGAGTCGTTGCCAAGTAGCTGGGACTACAGGTACNNNAGATCGGAAG |
| 49532_2 | TATGAGTGTGGAGTCGTTGCCAAGTAGCTGGGACTACAGGTACNNNAGATCGGAAG |
| 37847_1 | TATGAGTGTGGAGTCGTTGCTCTGTAGAATGCTACCAAGAGGTNNNAGATCGGAAG |
| 37847_2 | TATGAGTGTGGAGTCGTTGCTCTGTAGAATGCTACCAAGAGGTNNNAGATCGGAAG |
| 35345_1 | TATGAGTGTGGAGTCGTTGCTCTCAGCCTCCTAAAGTACTGGANNNAGATCGGAAGA |
| 35345_2 | TATGAGTGTGGAGTCGTTGCTCTCAGCCTCCTAAAGTACTGGANNNAGATCGGAAGA |
| 47438_1 | TATGAGTGTGGAGTCGTTGCACCGTGTTAGTCAGGAATGTCTCNNNAGATCGGAAG |
| 47438_2 | TATGAGTGTGGAGTCGTTGCACCGTGTTAGTCAGGAATGTCTCNNNAGATCGGAAG |
| 41516_1 | TATGAGTGTGGAGTCGTTGCCCCAGCACTTAGGACATAGACAANNNAGATCGGAAG |
| 41516_2 | TATGAGTGTGGAGTCGTTGCCCCAGCACTTAGGACATAGACAANNNAGATCGGAAG |
| 46325_1 | TATGAGTGTGGAGTCGTTGCTTCAAGTGATTCTCCTGCCTCAGNNNAGATCGGAAGA |
| 46325_2 | TATGAGTGTGGAGTCGTTGCTTCAAGTGATTCTCCTGCCTCAGNNNAGATCGGAAGA |
| 36753_1 | TATGAGTGTGGAGTCGTTGCAATCCCAACTATTCAGGAGGCTGNNNAGATCGGAAG |
| 36753_2 | TATGAGTGTGGAGTCGTTGCAATCCCAACTATTCAGGAGGCTGNNNAGATCGGAAG |
| 31134_1 | TATGAGTGTGGAGTCGTTGCGGAGGAGCTCAAAGGAAAAGAGANNNAGATCGGAA |

| | |
|---|---|
| 31134_2 | TATGAGTGTGGAGTCGTTGCGGAGGAGCTCAAAGGAAAAGAGANNNAGATCGGA |
| 35216_1 | TATGAGTGTGGAGTCGTTGCGCCCCACATCTAGACACTGAAATNNNAGATCGGAAG |
| 35216_2 | TATGAGTGTGGAGTCGTTGCGCCCCACATCTAGACACTGAAATNNNAGATCGGAAG |
| 31642_1 | TATGAGTGTGGAGTCGTTGCCCTCACCCAACACAGACAAAATCNNNAGATCGGAAG |
| 31642_2 | TATGAGTGTGGAGTCGTTGCCCTCACCCAACACAGACAAAATCNNNAGATCGGAAG |
| 33680_1 | TATGAGTGTGGAGTCGTTGCTTTCCGAGGTAAACTTCCTGGAANNNAGATCGGAAG |
| 33680_2 | TATGAGTGTGGAGTCGTTGCTTTCCGAGGTAAACTTCCTGGAANNNAGATCGGAAG |
| 46446_1 | TATGAGTGTGGAGTCGTTGCCTCTCGAGTAGCTGGGACTACAANNNAGATCGGAA |
| 46446_2 | TATGAGTGTGGAGTCGTTGCCTCTCGAGTAGCTGGGACTACAANNNAGATCGGAA |
| 8961_1 | TATGAGTGTGGAGTCGTTGCTGAGGTTCAAATGGCTAGCAAGTNNNAGATCGGAA |
| 8961_2 | TATGAGTGTGGAGTCGTTGCTGAGGTTCAAATGGCTAGCAAGTNNNAGATCGGAA |
| 40242_1 | TATGAGTGTGGAGTCGTTGCCTTTGTTCCCAGGATGGAGTGTANNNAGATCGGAA |
| 40242_2 | TATGAGTGTGGAGTCGTTGCCTTTGTTCCCAGGATGGAGTGTANNNAGATCGGAA |
| 50254_1 | TATGAGTGTGGAGTCGTTGCGTATCCTGACTTAGACGGGACACNNNAGATCGGAA |
| 50254_2 | TATGAGTGTGGAGTCGTTGCGTATCCTGACTTAGACGGGACACNNNAGATCGGAA |
| 36082_1 | TATGAGTGTGGAGTCGTTGCTAGCTGAGCATTATGTTGTGTGCNNNAGATCGGAA |
| 36082_2 | TATGAGTGTGGAGTCGTTGCTAGCTGAGCATTATGTTGTGTGCNNNAGATCGGAA |
| 35098_1 | TATGAGTGTGGAGTCGTTGCTTCAAGTGATTCTCCTGCCTCAGNNNAGATCGGAAG |
| 35098_2 | TATGAGTGTGGAGTCGTTGCTTCAAGTGATTCTCCTGCCTCAGNNNAGATCGGAAG |
| 40507_1 | TATGAGTGTGGAGTCGTTGCGGTCACAGAGTAGTAGAGAGAGTNNNAGATCGGA |
| 40507_2 | TATGAGTGTGGAGTCGTTGCGGTCACAGAGTAGTAGAGAGAGTNNNAGATCGGA |
| 49955_1 | TATGAGTGTGGAGTCGTTGCCAGCTCCTTGTTTCTGTTTCCTGNNNAGATCGGAAGA |
| 49955_2 | TATGAGTGTGGAGTCGTTGCCAGCTCCTTGTTTCTGTTTCCTGNNNAGATCGGAAGA |
| 658_1 | TATGAGTGTGGAGTCGTTGCAGTACAGCTCACATGGTTCCTCTNNNAGATCGGAAG |
| 658_2 | TATGAGTGTGGAGTCGTTGCAGTACAGCTCACATGGTTCCTCTNNNAGATCGGAAG |
| 47287_1 | TATGAGTGTGGAGTCGTTGCCTCCCAAGTAGCTGTGACTACAGNNNAGATCGGAAG |
| 47287_2 | TATGAGTGTGGAGTCGTTGCCTCCCAAGTAGCTGTGACTACAGNNNAGATCGGAAG |
| 34930_1 | TATGAGTGTGGAGTCGTTGCTTTCACTCTATCACCAAGGCTGGNNNAGATCGGAAG |
| 34930_2 | TATGAGTGTGGAGTCGTTGCTTTCACTCTATCACCAAGGCTGGNNNAGATCGGAAG |
| 42066_1 | TATGAGTGTGGAGTCGTTGCCTGAAGTGCATTAGCCACATTCCNNNAGATCGGAAG |
| 42066_2 | TATGAGTGTGGAGTCGTTGCCTGAAGTGCATTAGCCACATTCCNNNAGATCGGAAG |
| 34015_1 | TATGAGTGTGGAGTCGTTGCGGGTTTCACACTGTTAGTCAGGANNNAGATCGGAA |
| 34015_2 | TATGAGTGTGGAGTCGTTGCGGGTTTCACACTGTTAGTCAGGANNNAGATCGGAA |
| 48781_1 | TATGAGTGTGGAGTCGTTGCTTACAGCAACCTTTATTCGTGGCNNNAGATCGGAAG |
| 48781_2 | TATGAGTGTGGAGTCGTTGCTTACAGCAACCTTTATTCGTGGCNNNAGATCGGAAG |
| 40992_1 | TATGAGTGTGGAGTCGTTGCCATGTGGAGAAATTGGAAGCCTCNNNAGATCGGAA |
| 40992_2 | TATGAGTGTGGAGTCGTTGCCATGTGGAGAAATTGGAAGCCTCNNNAGATCGGAA |
| 46738_1 | TATGAGTGTGGAGTCGTTGCAGAGTGTAGTGGTGCAATCTCTGNNNAGATCGGAA |
| 46738_2 | TATGAGTGTGGAGTCGTTGCAGAGTGTAGTGGTGCAATCTCTGNNNAGATCGGAA |
| 51435_1 | TATGAGTGTGGAGTCGTTGCGACCTCAGTGCGTGGTTCTGNNNAGATCGGAAGAG |
| 51435_2 | TATGAGTGTGGAGTCGTTGCGACCTCAGTGCGTGGTTCTGNNNAGATCGGAAGAG |
| 37458_1 | TATGAGTGTGGAGTCGTTGCTTTCCTGGATTTTGCGCTGTTTANNNAGATCGGAAG |
| 37458_2 | TATGAGTGTGGAGTCGTTGCTTTCCTGGATTTTGCGCTGTTTANNNAGATCGGAAG |

| | |
|---|---|
| 13602_1 | TATGAGTGTGGAGTCGTTGCAGAGGTCAGGTACAAATGAANNNAGATCGGAAGAG |
| 13602_2 | TATGAGTGTGGAGTCGTTGCAGAGGTCAGGTACAAATGAANNNAGATCGGAAGAG |
| 13530_1 | TATGAGTGTGGAGTCGTTGCGTAATTTTGGTCTCATTATGTCANNNAGATCGGAAG |
| 13530_2 | TATGAGTGTGGAGTCGTTGCGTAATTTTGGTCTCATTATGTCANNNAGATCGGAAG |
| 47200_1 | TATGAGTGTGGAGTCGTTGCGATGCAATCATAGCTCACTGCAGNNNAGATCGGAAG |
| 47200_2 | TATGAGTGTGGAGTCGTTGCGATGCAATCATAGCTCACTGCAGNNNAGATCGGAAG |
| 53284_1 | TATGAGTGTGGAGTCGTTGCATCTGAAAGGAGGTATCTCACGCNNNAGATCGGAAG |
| 53284_2 | TATGAGTGTGGAGTCGTTGCATCTGAAAGGAGGTATCTCACGCNNNAGATCGGAAG |
| 30809_1 | TATGAGTGTGGAGTCGTTGCCATTCATGCCTCTGTCTTAGGGTNNNAGATCGGAAG |
| 30809_2 | TATGAGTGTGGAGTCGTTGCCATTCATGCCTCTGTCTTAGGGTNNNAGATCGGAAG |
| 47157_1 | TATGAGTGTGGAGTCGTTGCGCGTGATAATTACAGGTGTGAGCNNNAGATCGGAA |
| 47157_2 | TATGAGTGTGGAGTCGTTGCGCGTGATAATTACAGGTGTGAGCNNNAGATCGGAA |
| 50327_1 | TATGAGTGTGGAGTCGTTGCCAAGTGACAGAGGAGAACCTGAANNNAGATCGGAA |
| 50327_2 | TATGAGTGTGGAGTCGTTGCCAAGTGACAGAGGAGAACCTGAANNNAGATCGGAA |
| 37062_1 | TATGAGTGTGGAGTCGTTGCTGGCCAGTCTCATCAATTAGTTGTNNNAGATCGGAA |
| 37062_2 | TATGAGTGTGGAGTCGTTGCTGGCCAGTCTCATCAATTAGTTGTNNNAGATCGGAA |
| 40020_1 | TATGAGTGTGGAGTCGTTGCGTGAGCTCCAGAATACCAAGGAANNNAGATCGGAA |
| 40020_2 | TATGAGTGTGGAGTCGTTGCGTGAGCTCCAGAATACCAAGGAANNNAGATCGGAA |
| 47966_1 | TATGAGTGTGGAGTCGTTGCGTTCAAGAGATTCTCCTGCCTCANNNAGATCGGAAG |
| 47966_2 | TATGAGTGTGGAGTCGTTGCGTTCAAGAGATTCTCCTGCCTCANNNAGATCGGAAG |
| 40671_1 | TATGAGTGTGGAGTCGTTGCAGGATCTTTTGAACCCAGGAGACNNNAGATCGGAAG |
| 40671_2 | TATGAGTGTGGAGTCGTTGCAGGATCTTTTGAACCCAGGAGACNNNAGATCGGAAG |
| 34169_1 | TATGAGTGTGGAGTCGTTGCATCCCTCAGATCACATTTCCCACNNNAGATCGGAAGA |
| 34169_2 | TATGAGTGTGGAGTCGTTGCATCCCTCAGATCACATTTCCCACNNNAGATCGGAAGA |
| 41600_1 | TATGAGTGTGGAGTCGTTGCCACATGCCTTTGACTCTGGATTGNNNAGATCGGAAG |
| 41600_2 | TATGAGTGTGGAGTCGTTGCCACATGCCTTTGACTCTGGATTGNNNAGATCGGAAG |
| 28861_1 | TATGAGTGTGGAGTCGTTGCATCCCCTGGAAATCATGATTGGTNNNAGATCGGAAG |
| 28861_2 | TATGAGTGTGGAGTCGTTGCATCCCCTGGAAATCATGATTGGTNNNAGATCGGAAG |
| 2120_1 | TATGAGTGTGGAGTCGTTGCCTGCAAGGTTTGGAAATACTGCANNNAGATCGGAAG |
| 2120_2 | TATGAGTGTGGAGTCGTTGCCTGCAAGGTTTGGAAATACTGCANNNAGATCGGAAG |
| 52755_1 | TATGAGTGTGGAGTCGTTGCCCTCTCAAGGGTGTCAGCTGNNNAGATCGGAAGAG |
| 52755_2 | TATGAGTGTGGAGTCGTTGCCCTCTCAAGGGTGTCAGCTGNNNAGATCGGAAGAG |
| 12208_1 | TATGAGTGTGGAGTCGTTGCGGCTCTCACAACAGAGATTCACANNNAGATCGGAAG |
| 12208_2 | TATGAGTGTGGAGTCGTTGCGGCTCTCACAACAGAGATTCACANNNAGATCGGAAG |
| 38717_1 | TATGAGTGTGGAGTCGTTGCTTGTGAAAATGACCATACTGCCANNNAGATCGGAAG |
| 38717_2 | TATGAGTGTGGAGTCGTTGCTTGTGAAAATGACCATACTGCCANNNAGATCGGAAG |
| 5561_1 | TATGAGTGTGGAGTCGTTGCGAGATGCTGAGAAAGCACACAGANNNAGATCGGAA |
| 5561_2 | TATGAGTGTGGAGTCGTTGCGAGATGCTGAGAAAGCACACAGANNNAGATCGGAA |
| 43595_1 | TATGAGTGTGGAGTCGTTGCGGCAGTAACCTGAGAGGATCTTCNNNAGATCGGAA |
| 43595_2 | TATGAGTGTGGAGTCGTTGCGGCAGTAACCTGAGAGGATCTTCNNNAGATCGGAA |
| 52982_1 | TATGAGTGTGGAGTCGTTGCCCTCCCTCTCTGTCTCTCTTTCTNNNAGATCGGAAGAG |
| 52982_2 | TATGAGTGTGGAGTCGTTGCCCTCCCTCTCTGTCTCTCTTTCTNNNAGATCGGAAGAG |
| 45009_1 | TATGAGTGTGGAGTCGTTGCCAGGAGAGGTGTAGGTGTACATGNNNAGATCGGAA |

| | |
|---|---|
| 45009_2 | TATGAGTGTGGAGTCGTTGCCAGGAGAGGTGTAGGTGTACATGNNNAGATCGGA |
| 29519_1 | TATGAGTGTGGAGTCGTTGCTGCTGGAGAGAAAGGAGCAAAAGNNNAGATCGGA |
| 29519_2 | TATGAGTGTGGAGTCGTTGCTGCTGGAGAGAAAGGAGCAAAAGNNNAGATCGGA |
| 43722_1 | TATGAGTGTGGAGTCGTTGCGTAGTCCTGGTGGGGTTCTTGNNNAGATCGGAAGA |
| 43722_2 | TATGAGTGTGGAGTCGTTGCGTAGTCCTGGTGGGGTTCTTGNNNAGATCGGAAGA |
| 34968_1 | TATGAGTGTGGAGTCGTTGCGTTCCAGTGATTCTCCTGCCTTANNNAGATCGGAAG |
| 34968_2 | TATGAGTGTGGAGTCGTTGCGTTCCAGTGATTCTCCTGCCTTANNNAGATCGGAAG |
| 48715_1 | TATGAGTGTGGAGTCGTTGCCCAGCTGACCACACACAGAGNNNAGATCGGAAGAG |
| 48715_2 | TATGAGTGTGGAGTCGTTGCCCAGCTGACCACACACAGAGNNNAGATCGGAAGAG |
| 31806_1 | TATGAGTGTGGAGTCGTTGCATTGGGGATTAGGGTTTCAGCATNNNAGATCGGAA |
| 31806_2 | TATGAGTGTGGAGTCGTTGCATTGGGGATTAGGGTTTCAGCATNNNAGATCGGAA |
| 48372_1 | TATGAGTGTGGAGTCGTTGCGAGACGGAGTCTTGCTCTGTCNNNAGATCGGAAGA |
| 48372_2 | TATGAGTGTGGAGTCGTTGCGAGACGGAGTCTTGCTCTGTCNNNAGATCGGAAGA |
| 35669_1 | TATGAGTGTGGAGTCGTTGCCAAGTGTTGGATGTCTGCATGTTNNNAGATCGGAA |
| 35669_2 | TATGAGTGTGGAGTCGTTGCCAAGTGTTGGATGTCTGCATGTTNNNAGATCGGAA |
| 29007_1 | TATGAGTGTGGAGTCGTTGCACAATTCCACTTCACTTCTGGGTNNNAGATCGGAAG |
| 29007_2 | TATGAGTGTGGAGTCGTTGCACAATTCCACTTCACTTCTGGGTNNNAGATCGGAAG |
| 46654_1 | TATGAGTGTGGAGTCGTTGCTTCAAGCAATTCTCCTGCCTCAGNNNAGATCGGAAG |
| 46654_2 | TATGAGTGTGGAGTCGTTGCTTCAAGCAATTCTCCTGCCTCAGNNNAGATCGGAAG |
| 34533_1 | TATGAGTGTGGAGTCGTTGCCGTGTTCTCATTGTTCAACTCCCNNNAGATCGGAAGA |
| 34533_2 | TATGAGTGTGGAGTCGTTGCCGTGTTCTCATTGTTCAACTCCCNNNAGATCGGAAGA |
| 45487_1 | TATGAGTGTGGAGTCGTTGCTTGTTGAGACGATGGTGATCAGTNNNAGATCGGAA |
| 45487_2 | TATGAGTGTGGAGTCGTTGCTTGTTGAGACGATGGTGATCAGTNNNAGATCGGAA |
| 49180_1 | TATGAGTGTGGAGTCGTTGCTCTTGTGTGAATCTCTCTATGCANNNAGATCGGAAG |
| 49180_2 | TATGAGTGTGGAGTCGTTGCTCTTGTGTGAATCTCTCTATGCANNNAGATCGGAAG |
| 46941_1 | TATGAGTGTGGAGTCGTTGCCAAACTCCTCAGCTCAAGCAATCNNNAGATCGGAAG |
| 46941_2 | TATGAGTGTGGAGTCGTTGCCAAACTCCTCAGCTCAAGCAATCNNNAGATCGGAAG |
| 32319_1 | TATGAGTGTGGAGTCGTTGCCTGTAGTCCCAGCTACTTGAGAGNNNAGATCGGAAG |
| 32319_2 | TATGAGTGTGGAGTCGTTGCCTGTAGTCCCAGCTACTTGAGAGNNNAGATCGGAAG |
| 51226_1 | TATGAGTGTGGAGTCGTTGCGGGGTAGGAGGTTGATTCTGATGNNNAGATCGGA |
| 51226_2 | TATGAGTGTGGAGTCGTTGCGGGGTAGGAGGTTGATTCTGATGNNNAGATCGGA |
| 49943_1 | TATGAGTGTGGAGTCGTTGCACTGTTATGGGATGAGTGGTTCCNNNAGATCGGAA |
| 49943_2 | TATGAGTGTGGAGTCGTTGCACTGTTATGGGATGAGTGGTTCCNNNAGATCGGAA |
| 37391_1 | TATGAGTGTGGAGTCGTTGCGATTGGAGCTACTCAGAAGCCTTNNNAGATCGGAA |
| 37391_2 | TATGAGTGTGGAGTCGTTGCGATTGGAGCTACTCAGAAGCCTTNNNAGATCGGAA |
| 51547_1 | TATGAGTGTGGAGTCGTTGCCCTAAACAACTGGGGATCCTCAANNNAGATCGGAAG |
| 51547_2 | TATGAGTGTGGAGTCGTTGCCCTAAACAACTGGGGATCCTCAANNNAGATCGGAAG |
| 32217_1 | TATGAGTGTGGAGTCGTTGCCAACACTGCAGTCATGGTCTTGNNNAGATCGGAAGA |
| 32217_2 | TATGAGTGTGGAGTCGTTGCCAACACTGCAGTCATGGTCTTGNNNAGATCGGAAGA |
| 38293_1 | TATGAGTGTGGAGTCGTTGCGAGAATGGCGTGAGACTCTGTCNNNAGATCGGAAG |
| 38293_2 | TATGAGTGTGGAGTCGTTGCGAGAATGGCGTGAGACTCTGTCNNNAGATCGGAAG |
| 48149_1 | TATGAGTGTGGAGTCGTTGCCTGCTTTGCATGCTTTCTCTTCTNNNAGATCGGAAGA |
| 48149_2 | TATGAGTGTGGAGTCGTTGCCTGCTTTGCATGCTTTCTCTTCTNNNAGATCGGAAGA |

| | |
|---|---|
| 40429_1 | TATGAGTGTGGAGTCGTTGCGAGGTAGTGAACAAGATCCAGCANNNAGATCGGAA |
| 40429_2 | TATGAGTGTGGAGTCGTTGCGAGGTAGTGAACAAGATCCAGCANNNAGATCGGAA |
| 42934_1 | TATGAGTGTGGAGTCGTTGCATCTTGGCTCACTACAACCTCTGNNNAGATCGGAAG |
| 42934_2 | TATGAGTGTGGAGTCGTTGCATCTTGGCTCACTACAACCTCTGNNNAGATCGGAAG |
| 46473_1 | TATGAGTGTGGAGTCGTTGCTTCAAGTGATTCTCCTGCCTCAGNNNAGATCGGAAG |
| 46473_2 | TATGAGTGTGGAGTCGTTGCTTCAAGTGATTCTCCTGCCTCAGNNNAGATCGGAAG |
| 45243_1 | TATGAGTGTGGAGTCGTTGCTTTGTCCAGCTGTTTCTATTCGCNNNAGATCGGAAG |
| 45243_2 | TATGAGTGTGGAGTCGTTGCTTTGTCCAGCTGTTTCTATTCGCNNNAGATCGGAAG |
| 46883_1 | TATGAGTGTGGAGTCGTTGCCTCCCAGGTTCACGCCATTCNNNAGATCGGAAGAGC |
| 46883_2 | TATGAGTGTGGAGTCGTTGCCTCCCAGGTTCACGCCATTCNNNAGATCGGAAGAGC |
| 39706_1 | TATGAGTGTGGAGTCGTTGCAATAGCCATTGTGACTGGTGTGANNNAGATCGGAA |
| 39706_2 | TATGAGTGTGGAGTCGTTGCAATAGCCATTGTGACTGGTGTGANNNAGATCGGAA |
| 13047_1 | TATGAGTGTGGAGTCGTTGCCACAGAGGAAGAGAATCTCCNNNAGATCGGAAGAG |
| 13047_2 | TATGAGTGTGGAGTCGTTGCCACAGAGGAAGAGAATCTCCNNNAGATCGGAAGAG |
| 33619_1 | TATGAGTGTGGAGTCGTTGCCTCGACCTCCTAACCTCGTGATCNNNAGATCGGAAG |
| 33619_2 | TATGAGTGTGGAGTCGTTGCCTCGACCTCCTAACCTCGTGATCNNNAGATCGGAAG |
| 38808_1 | TATGAGTGTGGAGTCGTTGCTCATTAGGATGTCTTGCACACCTNNNAGATCGGAAG |
| 38808_2 | TATGAGTGTGGAGTCGTTGCTCATTAGGATGTCTTGCACACCTNNNAGATCGGAAG |
| 49340_1 | TATGAGTGTGGAGTCGTTGCCTGGAGTGCAATGACGTGATCTCNNNAGATCGGAA |
| 49340_2 | TATGAGTGTGGAGTCGTTGCCTGGAGTGCAATGACGTGATCTCNNNAGATCGGAA |
| 40809_1 | TATGAGTGTGGAGTCGTTGCTGCTCAACAGTACCTAACACAGANNNAGATCGGAAG |
| 40809_2 | TATGAGTGTGGAGTCGTTGCTGCTCAACAGTACCTAACACAGANNNAGATCGGAAG |
| 38924_1 | TATGAGTGTGGAGTCGTTGCGTTATGTACTGCGTGTTGGTGTCNNNAGATCGGAAG |
| 38924_2 | TATGAGTGTGGAGTCGTTGCGTTATGTACTGCGTGTTGGTGTCNNNAGATCGGAAG |
| 37601_1 | TATGAGTGTGGAGTCGTTGCTGTACTTCTCACTTTCTGCTCCCNNNAGATCGGAAGA |
| 37601_2 | TATGAGTGTGGAGTCGTTGCTGTACTTCTCACTTTCTGCTCCCNNNAGATCGGAAGA |
| 39954_1 | TATGAGTGTGGAGTCGTTGCAGGGGTTTGTGTGTAGAAGACAGNNNAGATCGGAA |
| 39954_2 | TATGAGTGTGGAGTCGTTGCAGGGGTTTGTGTGTAGAAGACAGNNNAGATCGGAA |
| 41192_1 | TATGAGTGTGGAGTCGTTGCCTGGCTGAAGTCTATGGTCCTTGNNNAGATCGGAAG |
| 41192_2 | TATGAGTGTGGAGTCGTTGCCTGGCTGAAGTCTATGGTCCTTGNNNAGATCGGAAG |
| 44760_1 | TATGAGTGTGGAGTCGTTGCAAGTAACCTGCTATTGCCTGTGANNNAGATCGGAAG |
| 44760_2 | TATGAGTGTGGAGTCGTTGCAAGTAACCTGCTATTGCCTGTGANNNAGATCGGAAG |
| 45628_1 | TATGAGTGTGGAGTCGTTGCAGGTTCAAGCGATTCTCATGTCTNNNAGATCGGAAG |
| 45628_2 | TATGAGTGTGGAGTCGTTGCAGGTTCAAGCGATTCTCATGTCTNNNAGATCGGAAG |
| 29732_1 | TATGAGTGTGGAGTCGTTGCTTTTGTCCTTCCATCCAGACTCCNNNAGATCGGAAGA |
| 29732_2 | TATGAGTGTGGAGTCGTTGCTTTTGTCCTTCCATCCAGACTCCNNNAGATCGGAAGA |
| 49481_1 | TATGAGTGTGGAGTCGTTGCTTTTGAGCCTATGTGTGCCTTTGNNNAGATCGGAAG |
| 49481_2 | TATGAGTGTGGAGTCGTTGCTTTTGAGCCTATGTGTGCCTTTGNNNAGATCGGAAG |
| 50328_1 | TATGAGTGTGGAGTCGTTGCAAACCGGATCTGTAAAACTGGGTNNNAGATCGGAA |
| 50328_2 | TATGAGTGTGGAGTCGTTGCAAACCGGATCTGTAAAACTGGGTNNNAGATCGGAA |
| 39737_1 | TATGAGTGTGGAGTCGTTGCGGAAAACCGGGGGAGTTATTTANNNAGATCGGAA |
| 39737_2 | TATGAGTGTGGAGTCGTTGCGGAAAACCGGGGGAGTTATTTANNNAGATCGGAA |
| 52764_1 | TATGAGTGTGGAGTCGTTGCCTCTGAAGGAAGAGACACAGGACNNNAGATCGGAA |

| | |
|---|---|
| 52764_2 | TATGAGTGTGGAGTCGTTGCCTCTGAAGGAAGAGACACAGGACNNNAGATCGGAA |
| 42546_1 | TATGAGTGTGGAGTCGTTGCAAATGAACATCCAAGGGCAAAGGNNNAGATCGGAA |
| 42546_2 | TATGAGTGTGGAGTCGTTGCAAATGAACATCCAAGGGCAAAGGNNNAGATCGGAA |
| 49802_1 | TATGAGTGTGGAGTCGTTGCCTAGGGGTCTGGCGTTAGATAACNNNAGATCGGAA |
| 49802_2 | TATGAGTGTGGAGTCGTTGCCTAGGGGTCTGGCGTTAGATAACNNNAGATCGGAA |
| 34304_1 | TATGAGTGTGGAGTCGTTGCCCTTAGAAACGCAGGTGGTTTTTNNNAGATCGGAAG |
| 34304_2 | TATGAGTGTGGAGTCGTTGCCCTTAGAAACGCAGGTGGTTTTTNNNAGATCGGAAG |
| 47261_1 | TATGAGTGTGGAGTCGTTGCTTCGTTCTTTGGTTTCCTAGGGTNNNAGATCGGAAG |
| 47261_2 | TATGAGTGTGGAGTCGTTGCTTCGTTCTTTGGTTTCCTAGGGTNNNAGATCGGAAG |
| 36440_1 | TATGAGTGTGGAGTCGTTGCTTTTGTCAGGAGAGAAAGGGTCANNNAGATCGGAA |
| 36440_2 | TATGAGTGTGGAGTCGTTGCTTTTGTCAGGAGAGAAAGGGTCANNNAGATCGGAA |
| 44368_1 | TATGAGTGTGGAGTCGTTGCCTGGGCAACATGGTAAAACTCTGNNNAGATCGGAA |
| 44368_2 | TATGAGTGTGGAGTCGTTGCCTGGGCAACATGGTAAAACTCTGNNNAGATCGGAA |
| 53210_1 | TATGAGTGTGGAGTCGTTGCGTAGCTGGGATTACAAGTGTCCANNNAGATCGGAA |
| 53210_2 | TATGAGTGTGGAGTCGTTGCGTAGCTGGGATTACAAGTGTCCANNNAGATCGGAA |
| 52043_1 | TATGAGTGTGGAGTCGTTGCAGGCAAGTTCTCTGATGACTCTGNNNAGATCGGAAG |
| 52043_2 | TATGAGTGTGGAGTCGTTGCAGGCAAGTTCTCTGATGACTCTGNNNAGATCGGAAG |
| 46365_1 | TATGAGTGTGGAGTCGTTGCAATCTGACGGATCGAAGGCATTTNNNAGATCGGAA |
| 46365_2 | TATGAGTGTGGAGTCGTTGCAATCTGACGGATCGAAGGCATTTNNNAGATCGGAA |
| 44740_1 | TATGAGTGTGGAGTCGTTGCCTGTGTTGCAACTTCTCAACTCCNNNAGATCGGAAGA |
| 44740_2 | TATGAGTGTGGAGTCGTTGCCTGTGTTGCAACTTCTCAACTCCNNNAGATCGGAAGA |
| 37911_1 | TATGAGTGTGGAGTCGTTGCAGGTATAAGTGGTTGGAATGGGGNNNAGATCGGA |
| 37911_2 | TATGAGTGTGGAGTCGTTGCAGGTATAAGTGGTTGGAATGGGGNNNAGATCGGA |
| 31389_1 | TATGAGTGTGGAGTCGTTGCTCCCATAGGTACTTCTCTTGGGANNNAGATCGGAAG |
| 31389_2 | TATGAGTGTGGAGTCGTTGCTCCCATAGGTACTTCTCTTGGGANNNAGATCGGAAG |
| 52701_1 | TATGAGTGTGGAGTCGTTGCAATGCTTCTCCTCTTGGCTCTTTNNNAGATCGGAAGA |
| 52701_2 | TATGAGTGTGGAGTCGTTGCAATGCTTCTCCTCTTGGCTCTTTNNNAGATCGGAAGA |
| 35870_1 | TATGAGTGTGGAGTCGTTGCATGAGATGTTCAGGTCACGTTGANNNAGATCGGAA |
| 35870_2 | TATGAGTGTGGAGTCGTTGCATGAGATGTTCAGGTCACGTTGANNNAGATCGGAA |
| 51900_1 | TATGAGTGTGGAGTCGTTGCAGAGGGTTGTGTAAAGCTCAGAGNNNAGATCGGAA |
| 51900_2 | TATGAGTGTGGAGTCGTTGCAGAGGGTTGTGTAAAGCTCAGAGNNNAGATCGGAA |
| 46285_1 | TATGAGTGTGGAGTCGTTGCTCTCCTGAGTAGCTGGGATTACANNNAGATCGGAAG |
| 46285_2 | TATGAGTGTGGAGTCGTTGCTCTCCTGAGTAGCTGGGATTACANNNAGATCGGAAG |
| 45809_1 | TATGAGTGTGGAGTCGTTGCGGATTCACGCCATTCTTCTGCNNNAGATCGGAAGAG |
| 45809_2 | TATGAGTGTGGAGTCGTTGCGGATTCACGCCATTCTTCTGCNNNAGATCGGAAGAG |
| 43085_1 | TATGAGTGTGGAGTCGTTGCAGAGAGAGACAGAGAGAGGGAACNNNAGATCGGA |
| 43085_2 | TATGAGTGTGGAGTCGTTGCAGAGAGAGACAGAGAGAGGGAACNNNAGATCGGA |
| 8235_1 | TATGAGTGTGGAGTCGTTGCAGTGGTGCATTCTAGGCTCACCANNNAGATCGGAAG |
| 8235_2 | TATGAGTGTGGAGTCGTTGCAGTGGTGCATTCTAGGCTCACCANNNAGATCGGAAG |
| 39227_1 | TATGAGTGTGGAGTCGTTGCCTGCTCGAAATCATTAGCCACTGNNNAGATCGGAAG |
| 39227_2 | TATGAGTGTGGAGTCGTTGCCTGCTCGAAATCATTAGCCACTGNNNAGATCGGAAG |
| 52063_1 | TATGAGTGTGGAGTCGTTGCGGGCTCCTCACTTCCCAGTANNNAGATCGGAAGAGC |
| 52063_2 | TATGAGTGTGGAGTCGTTGCGGGCTCCTCACTTCCCAGTANNNAGATCGGAAGAGC |

| | |
|---|---|
| 30560_1 | TATGAGTGTGGAGTCGTTGCAGACTTTGTCACACACACACGTNNNAGATCGGAAGA |
| 30560_2 | TATGAGTGTGGAGTCGTTGCAGACTTTGTCACACACACACGTNNNAGATCGGAAGA |
| 39635_1 | TATGAGTGTGGAGTCGTTGCAGGAAGGAAGGAGAGAGAGAAAGANNNAGATCG |
| 39635_2 | TATGAGTGTGGAGTCGTTGCAGGAAGGAAGGAGAGAGAGAAAGANNNAGATCG |
| 50758_1 | TATGAGTGTGGAGTCGTTGCTTACCTATTCTTGCAGCCCATCANNNAGATCGGAAGA |
| 50758_2 | TATGAGTGTGGAGTCGTTGCTTACCTATTCTTGCAGCCCATCANNNAGATCGGAAGA |
| 29097_1 | TATGAGTGTGGAGTCGTTGCACAATGGGGCTGTTCCTCTAAATNNNAGATCGGAAG |
| 29097_2 | TATGAGTGTGGAGTCGTTGCACAATGGGGCTGTTCCTCTAAATNNNAGATCGGAAG |
| 49774_1 | TATGAGTGTGGAGTCGTTGCGGGAAGAGTTATTTGTGCAGTGGNNNAGATCGGA |
| 49774_2 | TATGAGTGTGGAGTCGTTGCGGGAAGAGTTATTTGTGCAGTGGNNNAGATCGGA |
| 41571_1 | TATGAGTGTGGAGTCGTTGCATCATTCTTGGGTGTTTCTCGGANNNAGATCGGAAG |
| 41571_2 | TATGAGTGTGGAGTCGTTGCATCATTCTTGGGTGTTTCTCGGANNNAGATCGGAAG |
| 10308_1 | TATGAGTGTGGAGTCGTTGCGGAGACAACCTCATGTGCTCATTNNNAGATCGGAAG |
| 10308_2 | TATGAGTGTGGAGTCGTTGCGGAGACAACCTCATGTGCTCATTNNNAGATCGGAAG |
| 51538_1 | TATGAGTGTGGAGTCGTTGCCACCAAGCTCTCACTCCAGAGNNNAGATCGGAAGAG |
| 51538_2 | TATGAGTGTGGAGTCGTTGCCACCAAGCTCTCACTCCAGAGNNNAGATCGGAAGAG |
| 31679_1 | TATGAGTGTGGAGTCGTTGCGTAACATCAGTCTGGAGGGAGTGNNNAGATCGGAA |
| 31679_2 | TATGAGTGTGGAGTCGTTGCGTAACATCAGTCTGGAGGGAGTGNNNAGATCGGAA |
| 46039_1 | TATGAGTGTGGAGTCGTTGCCTCCCAGGTTCATGCCATTCTCNNNAGATCGGAAGA |
| 46039_2 | TATGAGTGTGGAGTCGTTGCCTCCCAGGTTCATGCCATTCTCNNNAGATCGGAAGA |
| 50664_1 | TATGAGTGTGGAGTCGTTGCGGAAACAGAGGTGGGAAGGATAANNNAGATCGGA |
| 50664_2 | TATGAGTGTGGAGTCGTTGCGGAAACAGAGGTGGGAAGGATAANNNAGATCGGA |
| 33019_1 | TATGAGTGTGGAGTCGTTGCGTGGTATGATCTTGGCTCATTGCNNNAGATCGGAAG |
| 33019_2 | TATGAGTGTGGAGTCGTTGCGTGGTATGATCTTGGCTCATTGCNNNAGATCGGAAG |
| 44814_1 | TATGAGTGTGGAGTCGTTGCTCTGTCCTATGAAGCTCTGCTTGNNNAGATCGGAAG |
| 44814_2 | TATGAGTGTGGAGTCGTTGCTCTGTCCTATGAAGCTCTGCTTGNNNAGATCGGAAG |
| 32240_1 | TATGAGTGTGGAGTCGTTGCACATGGTGAAACCTCGTCTCTACNNNAGATCGGAAG |
| 32240_2 | TATGAGTGTGGAGTCGTTGCACATGGTGAAACCTCGTCTCTACNNNAGATCGGAAG |
| 32782_1 | TATGAGTGTGGAGTCGTTGCAAACCCACACATCTATGGTCAATNNNAGATCGGAAG |
| 32782_2 | TATGAGTGTGGAGTCGTTGCAAACCCACACATCTATGGTCAATNNNAGATCGGAAG |
| 41089_1 | TATGAGTGTGGAGTCGTTGCAAAGAAAAATTAGCGAGGCGAGGNNNAGATCGGA |
| 41089_2 | TATGAGTGTGGAGTCGTTGCAAAGAAAAATTAGCGAGGCGAGGNNNAGATCGGA |
| 47605_1 | TATGAGTGTGGAGTCGTTGCGATGGAAACTGGGGGACCATGNNNAGATCGGAAG |
| 47605_2 | TATGAGTGTGGAGTCGTTGCGATGGAAACTGGGGGACCATGNNNAGATCGGAAG |
| 51393_1 | TATGAGTGTGGAGTCGTTGCACTTCCTAAGGATGTGCCAACATNNNAGATCGGAAG |
| 51393_2 | TATGAGTGTGGAGTCGTTGCACTTCCTAAGGATGTGCCAACATNNNAGATCGGAAG |
| 51487_1 | TATGAGTGTGGAGTCGTTGCAGTGTATACTGCTTGGGTGATGGNNNAGATCGGAA |
| 51487_2 | TATGAGTGTGGAGTCGTTGCAGTGTATACTGCTTGGGTGATGGNNNAGATCGGAA |
| 11117_1 | TATGAGTGTGGAGTCGTTGCTCAGCTCCCAAAGCAGTAGAATTANNNAGATCGGAA |
| 11117_2 | TATGAGTGTGGAGTCGTTGCTCAGCTCCCAAAGCAGTAGAATTANNNAGATCGGAA |
| 51083_1 | TATGAGTGTGGAGTCGTTGCACCCAGTGGGAGGTAATTGAATCNNNAGATCGGAA |
| 51083_2 | TATGAGTGTGGAGTCGTTGCACCCAGTGGGAGGTAATTGAATCNNNAGATCGGAA |
| 48471_1 | TATGAGTGTGGAGTCGTTGCGGTGAAGGCTTCAGAGTTGAGTANNNAGATCGGAA |

| | |
|---|---|
| 48471_2 | TATGAGTGTGGAGTCGTTGCGGTGAAGGCTTCAGAGTTGAGTANNNAGATCGGAA |
| 42115_1 | TATGAGTGTGGAGTCGTTGCGGGATTGCAGGATCAAATGGTAGNNNAGATCGGAA |
| 42115_2 | TATGAGTGTGGAGTCGTTGCGGGATTGCAGGATCAAATGGTAGNNNAGATCGGAA |
| 49784_1 | TATGAGTGTGGAGTCGTTGCGATTGCTCCCTTCTTTGTTTCCCNNNAGATCGGAAGA |
| 49784_2 | TATGAGTGTGGAGTCGTTGCGATTGCTCCCTTCTTTGTTTCCCNNNAGATCGGAAGA |
| 4971_1 | TATGAGTGTGGAGTCGTTGCGAGGGCTATGAGCAGAAATCCTGNNNAGATCGGAA |
| 4971_2 | TATGAGTGTGGAGTCGTTGCGAGGGCTATGAGCAGAAATCCTGNNNAGATCGGAA |
| 41847_1 | TATGAGTGTGGAGTCGTTGCCAATGGCATGATCTCAGCTCACNNNAGATCGGAAGA |
| 41847_2 | TATGAGTGTGGAGTCGTTGCCAATGGCATGATCTCAGCTCACNNNAGATCGGAAGA |
| 48053_1 | TATGAGTGTGGAGTCGTTGCGATCTCGGCTCACTACAAGCTCNNNAGATCGGAAGA |
| 48053_2 | TATGAGTGTGGAGTCGTTGCGATCTCGGCTCACTACAAGCTCNNNAGATCGGAAGA |
| 13256_1 | TATGAGTGTGGAGTCGTTGCAATTCAAAACCCATTTCCTAATTNNNAGATCGGAAGA |
| 13256_2 | TATGAGTGTGGAGTCGTTGCAATTCAAAACCCATTTCCTAATTNNNAGATCGGAAGA |
| 619_1 | TATGAGTGTGGAGTCGTTGCCCCTAAAGTCACCCATGGACATCNNNAGATCGGAAG |
| 619_2 | TATGAGTGTGGAGTCGTTGCCCCTAAAGTCACCCATGGACATCNNNAGATCGGAAG |
| 32710_1 | TATGAGTGTGGAGTCGTTGCCAAGGTCAAGAGATCGAGACCATNNNAGATCGGAA |
| 32710_2 | TATGAGTGTGGAGTCGTTGCCAAGGTCAAGAGATCGAGACCATNNNAGATCGGAA |
| 43744_1 | TATGAGTGTGGAGTCGTTGCTTTGTTCAGCTCTGCTCATTTCCNNNAGATCGGAAGA |
| 43744_2 | TATGAGTGTGGAGTCGTTGCTTTGTTCAGCTCTGCTCATTTCCNNNAGATCGGAAGA |
| 51930_1 | TATGAGTGTGGAGTCGTTGCCAATACCTCTCCACTGACACCATNNNAGATCGGAAGA |
| 51930_2 | TATGAGTGTGGAGTCGTTGCCAATACCTCTCCACTGACACCATNNNAGATCGGAAGA |
| 28771_1 | TATGAGTGTGGAGTCGTTGCGTCGTCACAGACATTCCTGTTTTNNNAGATCGGAAG |
| 28771_2 | TATGAGTGTGGAGTCGTTGCGTCGTCACAGACATTCCTGTTTTNNNAGATCGGAAG |
| 53396_1 | TATGAGTGTGGAGTCGTTGCGTGTGTGTGTGAGAGAGAGAGAGNNNAGATCGGA |
| 53396_2 | TATGAGTGTGGAGTCGTTGCGTGTGTGTGTGAGAGAGAGAGAGNNNAGATCGGA |
| 52683_1 | TATGAGTGTGGAGTCGTTGCAGTTTTCATGAGGGTCTGGCTAGNNNAGATCGGAA |
| 52683_2 | TATGAGTGTGGAGTCGTTGCAGTTTTCATGAGGGTCTGGCTAGNNNAGATCGGAA |
| 44719_1 | TATGAGTGTGGAGTCGTTGCCGGTTCGAGTAATCTAGGAGTGGNNNAGATCGGAA |
| 44719_2 | TATGAGTGTGGAGTCGTTGCCGGTTCGAGTAATCTAGGAGTGGNNNAGATCGGAA |
| 51125_1 | TATGAGTGTGGAGTCGTTGCTCTGTCCCCCTTATCTTCCCTCNNNAGATCGGAAGAG |
| 51125_2 | TATGAGTGTGGAGTCGTTGCTCTGTCCCCCTTATCTTCCCTCNNNAGATCGGAAGAG |
| 34057_1 | TATGAGTGTGGAGTCGTTGCGCTTCCCAGTTTCAAGCAAATCTNNNAGATCGGAAG |
| 34057_2 | TATGAGTGTGGAGTCGTTGCGCTTCCCAGTTTCAAGCAAATCTNNNAGATCGGAAG |
| 34577_1 | TATGAGTGTGGAGTCGTTGCTACCCATGATGGAACCCATGATGNNNAGATCGGAAG |
| 34577_2 | TATGAGTGTGGAGTCGTTGCTACCCATGATGGAACCCATGATGNNNAGATCGGAAG |
| 48721_1 | TATGAGTGTGGAGTCGTTGCCACAATCTTGGCTCACTGCAAGNNNAGATCGGAAGA |
| 48721_2 | TATGAGTGTGGAGTCGTTGCCACAATCTTGGCTCACTGCAAGNNNAGATCGGAAGA |
| 10970_1 | TATGAGTGTGGAGTCGTTGCACCATGTCACTGGGCTTCCATTTNNNAGATCGGAAG |
| 10970_2 | TATGAGTGTGGAGTCGTTGCACCATGTCACTGGGCTTCCATTTNNNAGATCGGAAG |
| 46534_1 | TATGAGTGTGGAGTCGTTGCGCCTGGATCTCTATACCCTGATGNNNAGATCGGAAG |
| 46534_2 | TATGAGTGTGGAGTCGTTGCGCCTGGATCTCTATACCCTGATGNNNAGATCGGAAG |
| 47122_1 | TATGAGTGTGGAGTCGTTGCCTGGACAGTCTGGAGAGAGAGAGNNNAGATCGGA |
| 47122_2 | TATGAGTGTGGAGTCGTTGCCTGGACAGTCTGGAGAGAGAGAGNNNAGATCGGA |

| | |
|---|---|
| 43632_1 | TATGAGTGTGGAGTCGTTGCTAGAATGGCTCAGGTGGGTTTTCNNNAGATCGGAA |
| 43632_2 | TATGAGTGTGGAGTCGTTGCTAGAATGGCTCAGGTGGGTTTTCNNNAGATCGGAA |
| 31969_1 | TATGAGTGTGGAGTCGTTGCCTGAGGCAGGAGAATGGAATGAANNNAGATCGGA |
| 31969_2 | TATGAGTGTGGAGTCGTTGCCTGAGGCAGGAGAATGGAATGAANNNAGATCGGA |
| 40486_1 | TATGAGTGTGGAGTCGTTGCCGTAATCCCAGCTACTCAGGAGNNNAGATCGGAAGA |
| 40486_2 | TATGAGTGTGGAGTCGTTGCCGTAATCCCAGCTACTCAGGAGNNNAGATCGGAAGA |
| 51398_1 | TATGAGTGTGGAGTCGTTGCGGGCGACTCTGTCTCAAAAAAAANNNAGATCGGAA |
| 51398_2 | TATGAGTGTGGAGTCGTTGCGGGCGACTCTGTCTCAAAAAAAANNNAGATCGGAA |
| 41769_1 | TATGAGTGTGGAGTCGTTGCCAGAGTCTCAAGAAACTGGCTCTNNNAGATCGGAAG |
| 41769_2 | TATGAGTGTGGAGTCGTTGCCAGAGTCTCAAGAAACTGGCTCTNNNAGATCGGAAG |
| 47623_1 | TATGAGTGTGGAGTCGTTGCGTCTCCCTATGTTGTCCAGACTGNNNAGATCGGAAG |
| 47623_2 | TATGAGTGTGGAGTCGTTGCGTCTCCCTATGTTGTCCAGACTGNNNAGATCGGAAG |
| 45159_1 | TATGAGTGTGGAGTCGTTGCGATACGGCGCGAAAGTGCTTNNNAGATCGGAAGAG |
| 45159_2 | TATGAGTGTGGAGTCGTTGCGATACGGCGCGAAAGTGCTTNNNAGATCGGAAGAG |
| 52817_1 | TATGAGTGTGGAGTCGTTGCATCAGCACACACCCTCTTCTTTANNNAGATCGGAAGA |
| 52817_2 | TATGAGTGTGGAGTCGTTGCATCAGCACACACCCTCTTCTTTANNNAGATCGGAAGA |
| 44931_1 | TATGAGTGTGGAGTCGTTGCTCTGGTCTTTTTCATGTGAGCCTNNNAGATCGGAAG |
| 44931_2 | TATGAGTGTGGAGTCGTTGCTCTGGTCTTTTTCATGTGAGCCTNNNAGATCGGAAG |
| 36834_1 | TATGAGTGTGGAGTCGTTGCGCCCAACAGTGTATCTTGTTCTGNNNAGATCGGAAG |
| 36834_2 | TATGAGTGTGGAGTCGTTGCGCCCAACAGTGTATCTTGTTCTGNNNAGATCGGAAG |
| 33771_1 | TATGAGTGTGGAGTCGTTGCTGTCTAAATGAAGCCAGCCACTANNNAGATCGGAAG |
| 33771_2 | TATGAGTGTGGAGTCGTTGCTGTCTAAATGAAGCCAGCCACTANNNAGATCGGAAG |
| 53167_1 | TATGAGTGTGGAGTCGTTGCAATGGATTACAAACACAGTGGCCNNNAGATCGGAA |
| 53167_2 | TATGAGTGTGGAGTCGTTGCAATGGATTACAAACACAGTGGCCNNNAGATCGGAA |
| 44398_1 | TATGAGTGTGGAGTCGTTGCGTTTAGAATGTCCTTGTGTCGGCNNNAGATCGGAAG |
| 44398_2 | TATGAGTGTGGAGTCGTTGCGTTTAGAATGTCCTTGTGTCGGCNNNAGATCGGAAG |
| 48645_1 | TATGAGTGTGGAGTCGTTGCCCAGACTGCAGCATCTCTTTCTANNNAGATCGGAAG |
| 48645_2 | TATGAGTGTGGAGTCGTTGCCCAGACTGCAGCATCTCTTTCTANNNAGATCGGAAG |
| 49535_1 | TATGAGTGTGGAGTCGTTGCGCCTACTGAGTAGCTGGGATTACNNNAGATCGGAA |
| 49535_2 | TATGAGTGTGGAGTCGTTGCGCCTACTGAGTAGCTGGGATTACNNNAGATCGGAA |
| 29462_1 | TATGAGTGTGGAGTCGTTGCCACTCTGGTGCTCTGTGTTCATANNNAGATCGGAAG |
| 29462_2 | TATGAGTGTGGAGTCGTTGCCACTCTGGTGCTCTGTGTTCATANNNAGATCGGAAG |
| 46001_1 | TATGAGTGTGGAGTCGTTGCCATGGTGCAATCTCAGCTAACTGNNNAGATCGGAAG |
| 46001_2 | TATGAGTGTGGAGTCGTTGCCATGGTGCAATCTCAGCTAACTGNNNAGATCGGAAG |
| 8906_1 | TATGAGTGTGGAGTCGTTGCCACAGTCCAGGTTGAGAAACCAANNNAGATCGGAA |
| 8906_2 | TATGAGTGTGGAGTCGTTGCCACAGTCCAGGTTGAGAAACCAANNNAGATCGGAA |
| 29849_1 | TATGAGTGTGGAGTCGTTGCGCTTCTTCTGCAGGATGGAAATGNNNAGATCGGAA |
| 29849_2 | TATGAGTGTGGAGTCGTTGCGCTTCTTCTGCAGGATGGAAATGNNNAGATCGGAA |
| 33404_1 | TATGAGTGTGGAGTCGTTGCACAGAGGAGAAAATTGGACAGCTNNNAGATCGGAA |
| 33404_2 | TATGAGTGTGGAGTCGTTGCACAGAGGAGAAAATTGGACAGCTNNNAGATCGGAA |
| 34539_1 | TATGAGTGTGGAGTCGTTGCCAAACGTCTTCTCTCAGCAAGTGNNNAGATCGGAAG |
| 34539_2 | TATGAGTGTGGAGTCGTTGCCAAACGTCTTCTCTCAGCAAGTGNNNAGATCGGAAG |
| 43335_1 | TATGAGTGTGGAGTCGTTGCGAGTCGGAGTCTCACTCTGTCANNNAGATCGGAAGA |

| | |
|---|---|
| 43335_2 | TATGAGTGTGGAGTCGTTGCGAGTCGGAGTCTCACTCTGTCANNNAGATCGGAAGA |
| 36719_1 | TATGAGTGTGGAGTCGTTGCGCTCATGTCTGTAATCCCAACACNNNAGATCGGAAG |
| 36719_2 | TATGAGTGTGGAGTCGTTGCGCTCATGTCTGTAATCCCAACACNNNAGATCGGAAG |
| 53152_1 | TATGAGTGTGGAGTCGTTGCAAACCAAAACCACACACACACAANNNAGATCGGAAG |
| 53152_2 | TATGAGTGTGGAGTCGTTGCAAACCAAAACCACACACACACAANNNAGATCGGAAG |
| 40083_1 | TATGAGTGTGGAGTCGTTGCAGAAGAAAATGAAGCTGGGGACTNNNAGATCGGA |
| 40083_2 | TATGAGTGTGGAGTCGTTGCAGAAGAAAATGAAGCTGGGGACTNNNAGATCGGA |
| 32025_1 | TATGAGTGTGGAGTCGTTGCGCATCTTGAAACCGCTGGATTTANNNAGATCGGAAG |
| 32025_2 | TATGAGTGTGGAGTCGTTGCGCATCTTGAAACCGCTGGATTTANNNAGATCGGAAG |
| 35604_1 | TATGAGTGTGGAGTCGTTGCGGGTGTAGGTTTTGCTTTGTTGTNNNAGATCGGAA |
| 35604_2 | TATGAGTGTGGAGTCGTTGCGGGTGTAGGTTTTGCTTTGTTGTNNNAGATCGGAA |
| 41219_1 | TATGAGTGTGGAGTCGTTGCGTGGGATTCCATTTACAGGAGGTNNNAGATCGGAA |
| 41219_2 | TATGAGTGTGGAGTCGTTGCGTGGGATTCCATTTACAGGAGGTNNNAGATCGGAA |
| 35798_1 | TATGAGTGTGGAGTCGTTGCTATGACGTATGAGCCACCATGTCNNNAGATCGGAAG |
| 35798_2 | TATGAGTGTGGAGTCGTTGCTATGACGTATGAGCCACCATGTCNNNAGATCGGAAG |
| 48317_1 | TATGAGTGTGGAGTCGTTGCATTATAGGCATGCAACACCATGCNNNAGATCGGAAG |
| 48317_2 | TATGAGTGTGGAGTCGTTGCATTATAGGCATGCAACACCATGCNNNAGATCGGAAG |
| 40328_1 | TATGAGTGTGGAGTCGTTGCATATCCAAGCAGAGGCCACATAGNNNAGATCGGAA |
| 40328_2 | TATGAGTGTGGAGTCGTTGCATATCCAAGCAGAGGCCACATAGNNNAGATCGGAA |
| 33983_1 | TATGAGTGTGGAGTCGTTGCCAGGTAGCTGGGACTATAGGCANNNAGATCGGAAG |
| 33983_2 | TATGAGTGTGGAGTCGTTGCCAGGTAGCTGGGACTATAGGCANNNAGATCGGAAG |
| 32668_1 | TATGAGTGTGGAGTCGTTGCCCTGCCTCAACCTCTCTATTACCNNNAGATCGGAAGA |
| 32668_2 | TATGAGTGTGGAGTCGTTGCCCTGCCTCAACCTCTCTATTACCNNNAGATCGGAAGA |
| 52073_1 | TATGAGTGTGGAGTCGTTGCAGTGATACAGTTTGGCTCTGTGTNNNAGATCGGAA |
| 52073_2 | TATGAGTGTGGAGTCGTTGCAGTGATACAGTTTGGCTCTGTGTNNNAGATCGGAA |
| 33647_1 | TATGAGTGTGGAGTCGTTGCATGATCTCAGCTCACTGCAATCTNNNAGATCGGAAG |
| 33647_2 | TATGAGTGTGGAGTCGTTGCATGATCTCAGCTCACTGCAATCTNNNAGATCGGAAG |
| 43416_1 | TATGAGTGTGGAGTCGTTGCCTCCTGAGTAGCTGGGACTACAGNNNAGATCGGAA |
| 43416_2 | TATGAGTGTGGAGTCGTTGCCTCCTGAGTAGCTGGGACTACAGNNNAGATCGGAA |
| 41765_1 | TATGAGTGTGGAGTCGTTGCCTGAGTAGCTGGGATTACAGGTGNNNAGATCGGAA |
| 41765_2 | TATGAGTGTGGAGTCGTTGCCTGAGTAGCTGGGATTACAGGTGNNNAGATCGGAA |
| 32056_1 | TATGAGTGTGGAGTCGTTGCCATTCTCCTGCCTCAGCCTCNNNAGATCGGAAGAGCA |
| 32056_2 | TATGAGTGTGGAGTCGTTGCCATTCTCCTGCCTCAGCCTCNNNAGATCGGAAGAGCA |
| 36354_1 | TATGAGTGTGGAGTCGTTGCAGCTAACAAGCCACACAAAAACANNNAGATCGGAAG |
| 36354_2 | TATGAGTGTGGAGTCGTTGCAGCTAACAAGCCACACAAAAACANNNAGATCGGAAG |
| 40914_1 | TATGAGTGTGGAGTCGTTGCCACCTTTAGTCCCAGCTACTCAGNNNAGATCGGAAG |
| 40914_2 | TATGAGTGTGGAGTCGTTGCCACCTTTAGTCCCAGCTACTCAGNNNAGATCGGAAG |
| 49556_1 | TATGAGTGTGGAGTCGTTGCTACATTCCTCCTTCCCTGGCTATNNNAGATCGGAAGA |
| 49556_2 | TATGAGTGTGGAGTCGTTGCTACATTCCTCCTTCCCTGGCTATNNNAGATCGGAAGA |
| 31116_1 | TATGAGTGTGGAGTCGTTGCTGTAGTGTTGTGGCAAGATCTCANNNAGATCGGAA |
| 31116_2 | TATGAGTGTGGAGTCGTTGCTGTAGTGTTGTGGCAAGATCTCANNNAGATCGGAA |
| 44804_1 | TATGAGTGTGGAGTCGTTGCCTTTGGTCATGGCCATTGTCTGNNNAGATCGGAAGA |
| 44804_2 | TATGAGTGTGGAGTCGTTGCCTTTGGTCATGGCCATTGTCTGNNNAGATCGGAAGA |

| | |
|---|---|
| 41041_1 | TATGAGTGTGGAGTCGTTGCGCGCAAATTACATGAAGAGCTCTNNNAGATCGGAAG |
| 41041_2 | TATGAGTGTGGAGTCGTTGCGCGCAAATTACATGAAGAGCTCTNNNAGATCGGAAG |
| 10251_1 | TATGAGTGTGGAGTCGTTGCGAGGTTCCACTATTGGCTCCATGNNNAGATCGGAAG |
| 10251_2 | TATGAGTGTGGAGTCGTTGCGAGGTTCCACTATTGGCTCCATGNNNAGATCGGAAG |
| 32998_1 | TATGAGTGTGGAGTCGTTGCCTGGGTTCAAGCAATTCTCGTGNNNAGATCGGAAGA |
| 32998_2 | TATGAGTGTGGAGTCGTTGCCTGGGTTCAAGCAATTCTCGTGNNNAGATCGGAAGA |
| 41698_1 | TATGAGTGTGGAGTCGTTGCCTGAGTAGCTGGAACTACAGGTGNNNAGATCGGAA |
| 41698_2 | TATGAGTGTGGAGTCGTTGCCTGAGTAGCTGGAACTACAGGTGNNNAGATCGGAA |
| 46944_1 | TATGAGTGTGGAGTCGTTGCTCTTGACTCACTGCAACCTCTACNNNAGATCGGAAGA |
| 46944_2 | TATGAGTGTGGAGTCGTTGCTCTTGACTCACTGCAACCTCTACNNNAGATCGGAAGA |
| 11346_1 | TATGAGTGTGGAGTCGTTGCGCTCAGTTAGGACGAACCCTGAGNNNAGATCGGAA |
| 11346_2 | TATGAGTGTGGAGTCGTTGCGCTCAGTTAGGACGAACCCTGAGNNNAGATCGGAA |
| 36779_1 | TATGAGTGTGGAGTCGTTGCAGGGTGCTATTCTAACTTCAGACANNNAGATCGGAA |
| 36779_2 | TATGAGTGTGGAGTCGTTGCAGGGTGCTATTCTAACTTCAGACANNNAGATCGGAA |
| 48733_1 | TATGAGTGTGGAGTCGTTGCTGCAATGGAGTGATCTTGACTCANNNAGATCGGAAG |
| 48733_2 | TATGAGTGTGGAGTCGTTGCTGCAATGGAGTGATCTTGACTCANNNAGATCGGAAG |
| 33095_1 | TATGAGTGTGGAGTCGTTGCTCTGGAATAGCTGGGATTACAGGNNNAGATCGGAA |
| 33095_2 | TATGAGTGTGGAGTCGTTGCTCTGGAATAGCTGGGATTACAGGNNNAGATCGGAA |
| 38781_1 | TATGAGTGTGGAGTCGTTGCTCACGGTTTCAATGGACACTTTTNNNAGATCGGAAG |
| 38781_2 | TATGAGTGTGGAGTCGTTGCTCACGGTTTCAATGGACACTTTTNNNAGATCGGAAG |
| 47215_1 | TATGAGTGTGGAGTCGTTGCCAACCTCCCAGACTCAAGTGATCNNNAGATCGGAAG |
| 47215_2 | TATGAGTGTGGAGTCGTTGCCAACCTCCCAGACTCAAGTGATCNNNAGATCGGAAG |
| 31022_1 | TATGAGTGTGGAGTCGTTGCGGTCTCTGCCTGATAATCCAGAGNNNAGATCGGAAG |
| 31022_2 | TATGAGTGTGGAGTCGTTGCGGTCTCTGCCTGATAATCCAGAGNNNAGATCGGAAG |
| 10020_1 | TATGAGTGTGGAGTCGTTGCTCCTCCTCTCCCACTAAAGGGTANNNAGATCGGAAG |
| 10020_2 | TATGAGTGTGGAGTCGTTGCTCCTCCTCTCCCACTAAAGGGTANNNAGATCGGAAG |
| 33888_1 | TATGAGTGTGGAGTCGTTGCCCTCCCAGGTTCAAGTGATACTCNNNAGATCGGAAG |
| 33888_2 | TATGAGTGTGGAGTCGTTGCCCTCCCAGGTTCAAGTGATACTCNNNAGATCGGAAG |
| 33671_1 | TATGAGTGTGGAGTCGTTGCGCAAGATCATGGCTCACTGTAACNNNAGATCGGAAG |
| 33671_2 | TATGAGTGTGGAGTCGTTGCGCAAGATCATGGCTCACTGTAACNNNAGATCGGAAG |
| 50657_1 | TATGAGTGTGGAGTCGTTGCGCCTCATGTCAGGACTTTGAAACNNNAGATCGGAAG |
| 50657_2 | TATGAGTGTGGAGTCGTTGCGCCTCATGTCAGGACTTTGAAACNNNAGATCGGAAG |
| 50916_1 | TATGAGTGTGGAGTCGTTGCCAGCTTCCACTGACCTGACTCNNNAGATCGGAAGAG |
| 50916_2 | TATGAGTGTGGAGTCGTTGCCAGCTTCCACTGACCTGACTCNNNAGATCGGAAGAG |
| 34462_1 | TATGAGTGTGGAGTCGTTGCTCAAGATAATTCTCCGAGCCCTGNNNAGATCGGAAG |
| 34462_2 | TATGAGTGTGGAGTCGTTGCTCAAGATAATTCTCCGAGCCCTGNNNAGATCGGAAG |
| 45423_1 | TATGAGTGTGGAGTCGTTGCGGAGAGTGAGAGAGAACTTGCTCNNNAGATCGGAA |
| 45423_2 | TATGAGTGTGGAGTCGTTGCGGAGAGTGAGAGAGAACTTGCTCNNNAGATCGGAA |
| 37081_1 | TATGAGTGTGGAGTCGTTGCGGATGAGAGAAGGTGGTGAAAGTNNNAGATCGGA |
| 37081_2 | TATGAGTGTGGAGTCGTTGCGGATGAGAGAAGGTGGTGAAAGTNNNAGATCGGA |
| 11170_1 | TATGAGTGTGGAGTCGTTGCCACTCCCATTAGCCTGTGGTCCCNNNAGATCGGAAG |
| 11170_2 | TATGAGTGTGGAGTCGTTGCCACTCCCATTAGCCTGTGGTCCCNNNAGATCGGAAG |
| 34891_1 | TATGAGTGTGGAGTCGTTGCCCAAGTAGCTGGTACTACAGGTGNNNAGATCGGAA |

| | |
|---|---|
| 34891_2 | TATGAGTGTGGAGTCGTTGCCCAAGTAGCTGGTACTACAGGTGNNNAGATCGGAA |
| 52862_1 | TATGAGTGTGGAGTCGTTGCTTATAAGTGGGGAAACTGAGGCCNNNAGATCGGAA |
| 52862_2 | TATGAGTGTGGAGTCGTTGCTTATAAGTGGGGAAACTGAGGCCNNNAGATCGGAA |
| 30939_1 | TATGAGTGTGGAGTCGTTGCGCATGGAGAAGAGCACTTACAACNNNAGATCGGAA |
| 30939_2 | TATGAGTGTGGAGTCGTTGCGCATGGAGAAGAGCACTTACAACNNNAGATCGGAA |
| 13064_1 | TATGAGTGTGGAGTCGTTGCGTTTGCCTTTTTTTCATGTAAATNNNAGATCGGAAGA |
| 13064_2 | TATGAGTGTGGAGTCGTTGCGTTTGCCTTTTTTTCATGTAAATNNNAGATCGGAAGA |
| 46614_1 | TATGAGTGTGGAGTCGTTGCCAGGTGGGAATACAACTGTCTCANNNAGATCGGAA |
| 46614_2 | TATGAGTGTGGAGTCGTTGCCAGGTGGGAATACAACTGTCTCANNNAGATCGGAA |
| 38094_1 | TATGAGTGTGGAGTCGTTGCAGGAGGATCACTTGAACCTAGGANNNAGATCGGAA |
| 38094_2 | TATGAGTGTGGAGTCGTTGCAGGAGGATCACTTGAACCTAGGANNNAGATCGGAA |
| 45450_1 | TATGAGTGTGGAGTCGTTGCGCTCCACAAAGGCAGGAAATATGNNNAGATCGGAA |
| 45450_2 | TATGAGTGTGGAGTCGTTGCGCTCCACAAAGGCAGGAAATATGNNNAGATCGGAA |
| 36129_1 | TATGAGTGTGGAGTCGTTGCCCTGTAATCCCAACACCTTGAGANNNAGATCGGAAG |
| 36129_2 | TATGAGTGTGGAGTCGTTGCCCTGTAATCCCAACACCTTGAGANNNAGATCGGAAG |
| 42004_1 | TATGAGTGTGGAGTCGTTGCCCAAGTAGCTGGGATGACAGGNNNAGATCGGAAGA |
| 42004_2 | TATGAGTGTGGAGTCGTTGCCCAAGTAGCTGGGATGACAGGNNNAGATCGGAAGA |
| 32195_1 | TATGAGTGTGGAGTCGTTGCTGTGGTTTTGATTTGAAAGCCCANNNAGATCGGAAG |
| 32195_2 | TATGAGTGTGGAGTCGTTGCTGTGGTTTTGATTTGAAAGCCCANNNAGATCGGAAG |
| 37423_1 | TATGAGTGTGGAGTCGTTGCCCCAAGAGTTTGAGACCTCATCTNNNAGATCGGAAG |
| 37423_2 | TATGAGTGTGGAGTCGTTGCCCCAAGAGTTTGAGACCTCATCTNNNAGATCGGAAG |
| 37042_1 | TATGAGTGTGGAGTCGTTGCTAGATTGACTCTAAGGACTGGGTNNNAGATCGGAA |
| 37042_2 | TATGAGTGTGGAGTCGTTGCTAGATTGACTCTAAGGACTGGGTNNNAGATCGGAA |
| 37055_1 | TATGAGTGTGGAGTCGTTGCGTAATCTCAGCTACTCAGGAGGCNNNAGATCGGAAG |
| 37055_2 | TATGAGTGTGGAGTCGTTGCGTAATCTCAGCTACTCAGGAGGCNNNAGATCGGAAG |
| 46512_1 | TATGAGTGTGGAGTCGTTGCGATTCTCCTGCCTCAATCTCCCNNNAGATCGGAAGAG |
| 46512_2 | TATGAGTGTGGAGTCGTTGCGATTCTCCTGCCTCAATCTCCCNNNAGATCGGAAGAG |
| 47412_1 | TATGAGTGTGGAGTCGTTGCTGCTTTTGTTGGAAAATGCACAANNNAGATCGGAAG |
| 47412_2 | TATGAGTGTGGAGTCGTTGCTGCTTTTGTTGGAAAATGCACAANNNAGATCGGAAG |
| 52027_1 | TATGAGTGTGGAGTCGTTGCCTTCCTGTGTCCATGTGATCTCANNNAGATCGGAAGA |
| 52027_2 | TATGAGTGTGGAGTCGTTGCCTTCCTGTGTCCATGTGATCTCANNNAGATCGGAAGA |
| 53369_1 | TATGAGTGTGGAGTCGTTGCCCTCTCTCTCTCTCTCTCTCTCTNNNAGATCGGAAGAG |
| 53369_2 | TATGAGTGTGGAGTCGTTGCCCTCTCTCTCTCTCTCTCTCTCTNNNAGATCGGAAGAG |
| 41303_1 | TATGAGTGTGGAGTCGTTGCAAGAGCCAGACCCCTTTTCATAGNNNAGATCGGAAG |
| 41303_2 | TATGAGTGTGGAGTCGTTGCAAGAGCCAGACCCCTTTTCATAGNNNAGATCGGAAG |
| 52401_1 | TATGAGTGTGGAGTCGTTGCTCGATCCATAAAGCCAGGAGATGNNNAGATCGGAA |
| 52401_2 | TATGAGTGTGGAGTCGTTGCTCGATCCATAAAGCCAGGAGATGNNNAGATCGGAA |
| 13686_1 | TATGAGTGTGGAGTCGTTGCCTGATCATTTGGCTCTTTTTTTNNNAGATCGGAAGAG |
| 13686_2 | TATGAGTGTGGAGTCGTTGCCTGATCATTTGGCTCTTTTTTTNNNAGATCGGAAGAG |
| 34547_1 | TATGAGTGTGGAGTCGTTGCGATACCAGGACCTAGCCTACTGTNNNAGATCGGAAG |
| 34547_2 | TATGAGTGTGGAGTCGTTGCGATACCAGGACCTAGCCTACTGTNNNAGATCGGAAG |
| 46793_1 | TATGAGTGTGGAGTCGTTGCAACAGTTGAGCTTACCTCTCAGCNNNAGATCGGAAG |
| 46793_2 | TATGAGTGTGGAGTCGTTGCAACAGTTGAGCTTACCTCTCAGCNNNAGATCGGAAG |

| | |
|---|---|
| 45949_1 | TATGAGTGTGGAGTCGTTGCCAGAATCTCACTCTGTCACCCANNNAGATCGGAAGA |
| 45949_2 | TATGAGTGTGGAGTCGTTGCCAGAATCTCACTCTGTCACCCANNNAGATCGGAAGA |
| 9967_1 | TATGAGTGTGGAGTCGTTGCTAAGCACAGGAGGGAATCTCTCGNNNAGATCGGAA |
| 9967_2 | TATGAGTGTGGAGTCGTTGCTAAGCACAGGAGGGAATCTCTCGNNNAGATCGGAA |
| 41991_1 | TATGAGTGTGGAGTCGTTGCGTCCTGAGTAGCTGGAATACAGGNNNAGATCGGAA |
| 41991_2 | TATGAGTGTGGAGTCGTTGCGTCCTGAGTAGCTGGAATACAGGNNNAGATCGGAA |
| 32350_1 | TATGAGTGTGGAGTCGTTGCATGTACACTCACACAGACACACANNNAGATCGGAAG |
| 32350_2 | TATGAGTGTGGAGTCGTTGCATGTACACTCACACAGACACACANNNAGATCGGAAG |
| 36816_1 | TATGAGTGTGGAGTCGTTGCATAAAAGCAGGTGGCAAATAGGCNNNAGATCGGAA |
| 36816_2 | TATGAGTGTGGAGTCGTTGCATAAAAGCAGGTGGCAAATAGGCNNNAGATCGGAA |
| 30373_1 | TATGAGTGTGGAGTCGTTGCTCCTGAGCTCTATGTTGTTAACAGANNNAGATCGGA |
| 30373_2 | TATGAGTGTGGAGTCGTTGCTCCTGAGCTCTATGTTGTTAACAGANNNAGATCGGA |
| 50355_1 | TATGAGTGTGGAGTCGTTGCTAAGGAGGGAAGGAAGAAAAGGCNNNAGATCGGA |
| 50355_2 | TATGAGTGTGGAGTCGTTGCTAAGGAGGGAAGGAAGAAAAGGCNNNAGATCGGA |
| 47609_1 | TATGAGTGTGGAGTCGTTGCGGAGTAGCTAGGATTACAGGTGCNNNAGATCGGAA |
| 47609_2 | TATGAGTGTGGAGTCGTTGCGGAGTAGCTAGGATTACAGGTGCNNNAGATCGGAA |
| 32265_1 | TATGAGTGTGGAGTCGTTGCGCACCCCTAAAGATAGTTCCCAANNNAGATCGGAAG |
| 32265_2 | TATGAGTGTGGAGTCGTTGCGCACCCCTAAAGATAGTTCCCAANNNAGATCGGAAG |
| 44241_1 | TATGAGTGTGGAGTCGTTGCGTGTCTGTTACTGACCCACATGANNNAGATCGGAAG |
| 44241_2 | TATGAGTGTGGAGTCGTTGCGTGTCTGTTACTGACCCACATGANNNAGATCGGAAG |
| 28821_1 | TATGAGTGTGGAGTCGTTGCTGAAAATGGCATGAACAGGAACANNNAGATCGGAA |
| 28821_2 | TATGAGTGTGGAGTCGTTGCTGAAAATGGCATGAACAGGAACANNNAGATCGGAA |
| 46017_1 | TATGAGTGTGGAGTCGTTGCGTTCAAGCGATTCTCATGCCTCNNNAGATCGGAAGA |
| 46017_2 | TATGAGTGTGGAGTCGTTGCGTTCAAGCGATTCTCATGCCTCNNNAGATCGGAAGA |
| 48681_1 | TATGAGTGTGGAGTCGTTGCTTTGCAGAAGTGAATCTGGGGTANNNAGATCGGAA |
| 48681_2 | TATGAGTGTGGAGTCGTTGCTTTGCAGAAGTGAATCTGGGGTANNNAGATCGGAA |
| 46131_1 | TATGAGTGTGGAGTCGTTGCGAGTAACTGGGACTACAGGTGTGNNNAGATCGGAA |
| 46131_2 | TATGAGTGTGGAGTCGTTGCGAGTAACTGGGACTACAGGTGTGNNNAGATCGGAA |
| 43090_1 | TATGAGTGTGGAGTCGTTGCGCCTCACCACTCCTTATTGCTATNNNAGATCGGAAGA |
| 43090_2 | TATGAGTGTGGAGTCGTTGCGCCTCACCACTCCTTATTGCTATNNNAGATCGGAAGA |
| 35732_1 | TATGAGTGTGGAGTCGTTGCTCCTGAGTAGCTGGGATTACAGANNNAGATCGGAA |
| 35732_2 | TATGAGTGTGGAGTCGTTGCTCCTGAGTAGCTGGGATTACAGANNNAGATCGGAA |
| 10552_1 | TATGAGTGTGGAGTCGTTGCATTTGCCCAGGACTGATGGTTTTNNNAGATCGGAAG |
| 10552_2 | TATGAGTGTGGAGTCGTTGCATTTGCCCAGGACTGATGGTTTTNNNAGATCGGAAG |
| 49517_1 | TATGAGTGTGGAGTCGTTGCGCATCTTGTCTATCACCACATGCNNNAGATCGGAAG |
| 49517_2 | TATGAGTGTGGAGTCGTTGCGCATCTTGTCTATCACCACATGCNNNAGATCGGAAG |
| 45010_1 | TATGAGTGTGGAGTCGTTGCGGGCAGTGCATGACTTTTATCTCNNNAGATCGGAAG |
| 45010_2 | TATGAGTGTGGAGTCGTTGCGGGCAGTGCATGACTTTTATCTCNNNAGATCGGAAG |
| 46257_1 | TATGAGTGTGGAGTCGTTGCCACGCCCGGCTAATTTTTTTTTTNNNAGATCGGAAGA |
| 46257_2 | TATGAGTGTGGAGTCGTTGCCACGCCCGGCTAATTTTTTTTTTNNNAGATCGGAAGA |
| 44144_1 | TATGAGTGTGGAGTCGTTGCAAGACAAAGTACGATTCCCAGGGNNNAGATCGGAA |
| 44144_2 | TATGAGTGTGGAGTCGTTGCAAGACAAAGTACGATTCCCAGGGNNNAGATCGGAA |
| 41307_1 | TATGAGTGTGGAGTCGTTGCCTATTTCCAGGAAGCAGGTAGCANNNAGATCGGAA |

| | |
|---|---|
| 41307_2 | TATGAGTGTGGAGTCGTTGCCTATTTCCAGGAAGCAGGTAGCANNNAGATCGGAA |
| 32012_1 | TATGAGTGTGGAGTCGTTGCCTTCAACACCACACTTTCAGCATNNNAGATCGGAAGA |
| 32012_2 | TATGAGTGTGGAGTCGTTGCCTTCAACACCACACTTTCAGCATNNNAGATCGGAAGA |
| 39011_1 | TATGAGTGTGGAGTCGTTGCCAAGATTCAGTCCAAATTGCAGGTNNNAGATCGGAA |
| 39011_2 | TATGAGTGTGGAGTCGTTGCCAAGATTCAGTCCAAATTGCAGGTNNNAGATCGGAA |
| 12872_1 | TATGAGTGTGGAGTCGTTGCATCCCATGAATAGAAAGAGATGNNNAGATCGGAAG |
| 12872_2 | TATGAGTGTGGAGTCGTTGCATCCCATGAATAGAAAGAGATGNNNAGATCGGAAG |
| 31318_1 | TATGAGTGTGGAGTCGTTGCCAAACTCTATCACGCCAAACACANNNAGATCGGAAG |
| 31318_2 | TATGAGTGTGGAGTCGTTGCCAAACTCTATCACGCCAAACACANNNAGATCGGAAG |
| 13205_1 | TATGAGTGTGGAGTCGTTGCAAGTTGTGATCATGTATCTGTATNNNAGATCGGAAG |
| 13205_2 | TATGAGTGTGGAGTCGTTGCAAGTTGTGATCATGTATCTGTATNNNAGATCGGAAG |
| 13054_1 | TATGAGTGTGGAGTCGTTGCTATTGTGGAATGAAATTGTGATGNNNAGATCGGAA |
| 13054_2 | TATGAGTGTGGAGTCGTTGCTATTGTGGAATGAAATTGTGATGNNNAGATCGGAA |
| 28967_1 | TATGAGTGTGGAGTCGTTGCTCTCTCTGTCTCTGTCTGTGTCTNNNAGATCGGAAGA |
| 28967_2 | TATGAGTGTGGAGTCGTTGCTCTCTCTGTCTCTGTCTGTGTCTNNNAGATCGGAAGA |
| 31024_1 | TATGAGTGTGGAGTCGTTGCTTGATGGGGAGAACTAGCTATGCNNNAGATCGGAA |
| 31024_2 | TATGAGTGTGGAGTCGTTGCTTGATGGGGAGAACTAGCTATGCNNNAGATCGGAA |
| 52282_1 | TATGAGTGTGGAGTCGTTGCGTGGGAGAAAGACTGAAACAACGNNNAGATCGGA |
| 52282_2 | TATGAGTGTGGAGTCGTTGCGTGGGAGAAAGACTGAAACAACGNNNAGATCGGA |
| 36689_1 | TATGAGTGTGGAGTCGTTGCCTGTAATACCAGCTGCTCAGGAGNNNAGATCGGAAG |
| 36689_2 | TATGAGTGTGGAGTCGTTGCCTGTAATACCAGCTGCTCAGGAGNNNAGATCGGAAG |
| 43032_1 | TATGAGTGTGGAGTCGTTGCGGATGACTTGAGCCCAGGAATTANNNAGATCGGAA |
| 43032_2 | TATGAGTGTGGAGTCGTTGCGGATGACTTGAGCCCAGGAATTANNNAGATCGGAA |
| 40636_1 | TATGAGTGTGGAGTCGTTGCAAGAGGCAGCAGACAAGTTTCTANNNAGATCGGAA |
| 40636_2 | TATGAGTGTGGAGTCGTTGCAAGAGGCAGCAGACAAGTTTCTANNNAGATCGGAA |
| 10121_1 | TATGAGTGTGGAGTCGTTGCGGAAGAGAAGAAAAGGGCTCGAGNNNAGATCGGA |
| 10121_2 | TATGAGTGTGGAGTCGTTGCGGAAGAGAAGAAAAGGGCTCGAGNNNAGATCGGA |
| 47786_1 | TATGAGTGTGGAGTCGTTGCGAAATGGGGTTTTGCCATGTTGNNNAGATCGGAAG |
| 47786_2 | TATGAGTGTGGAGTCGTTGCGAAATGGGGTTTTGCCATGTTGNNNAGATCGGAAG |
| 12744_1 | TATGAGTGTGGAGTCGTTGCATTTGATAACCAACACTTAGAACNNNAGATCGGAAG |
| 12744_2 | TATGAGTGTGGAGTCGTTGCATTTGATAACCAACACTTAGAACNNNAGATCGGAAG |
| 40312_1 | TATGAGTGTGGAGTCGTTGCAGAACATGACATGACCTGTGTGANNNAGATCGGAA |
| 40312_2 | TATGAGTGTGGAGTCGTTGCAGAACATGACATGACCTGTGTGANNNAGATCGGAA |
| 29235_1 | TATGAGTGTGGAGTCGTTGCAACCTGGATGGATTTGGAGACAANNNAGATCGGAA |
| 29235_2 | TATGAGTGTGGAGTCGTTGCAACCTGGATGGATTTGGAGACAANNNAGATCGGAA |
| 126_1 | TATGAGTGTGGAGTCGTTGCTTTCTGAGCCTTCTTGCCCCAGNNNAGATCGGAAGA |
| 126_2 | TATGAGTGTGGAGTCGTTGCTTTCTGAGCCTTCTTGCCCCAGNNNAGATCGGAAGA |
| 47046_1 | TATGAGTGTGGAGTCGTTGCAAACTCCTGGATCAAGCAATCCTNNNAGATCGGAAG |
| 47046_2 | TATGAGTGTGGAGTCGTTGCAAACTCCTGGATCAAGCAATCCTNNNAGATCGGAAG |
| 12628_1 | TATGAGTGTGGAGTCGTTGCAAAGTTGTATTTTGGGTAAAAGANNNAGATCGGAA |
| 12628_2 | TATGAGTGTGGAGTCGTTGCAAAGTTGTATTTTGGGTAAAAGANNNAGATCGGAA |
| 39493_1 | TATGAGTGTGGAGTCGTTGCTCTGGTTCACTCATTTCTGCTCCNNNAGATCGGAAGA |
| 39493_2 | TATGAGTGTGGAGTCGTTGCTCTGGTTCACTCATTTCTGCTCCNNNAGATCGGAAGA |

| | |
|---|---|
| 42486_1 | TATGAGTGTGGAGTCGTTGCTTTGTTGGGGGGAAAGTAAGGAANNNAGATCGGA |
| 42486_2 | TATGAGTGTGGAGTCGTTGCTTTGTTGGGGGGAAAGTAAGGAANNNAGATCGGA |
| 29899_1 | TATGAGTGTGGAGTCGTTGCCCACACTGGAAGTACCTACAACANNNAGATCGGAAG |
| 29899_2 | TATGAGTGTGGAGTCGTTGCCCACACTGGAAGTACCTACAACANNNAGATCGGAAG |
| 41563_1 | TATGAGTGTGGAGTCGTTGCGCTAATCTCGGCTCACTGCAAGNNNAGATCGGAAGA |
| 41563_2 | TATGAGTGTGGAGTCGTTGCGCTAATCTCGGCTCACTGCAAGNNNAGATCGGAAGA |
| 41096_1 | TATGAGTGTGGAGTCGTTGCAAACAGTACAGATGCTGCTGAGANNNAGATCGGAA |
| 41096_2 | TATGAGTGTGGAGTCGTTGCAAACAGTACAGATGCTGCTGAGANNNAGATCGGAA |
| 32781_1 | TATGAGTGTGGAGTCGTTGCATGTACACAGTCTTAGCCTCCAGNNNAGATCGGAAG |
| 32781_2 | TATGAGTGTGGAGTCGTTGCATGTACACAGTCTTAGCCTCCAGNNNAGATCGGAAG |
| 49724_1 | TATGAGTGTGGAGTCGTTGCGCCTGGGATGGTGGAACTATATTNNNAGATCGGAA |
| 49724_2 | TATGAGTGTGGAGTCGTTGCGCCTGGGATGGTGGAACTATATTNNNAGATCGGAA |
| 44999_1 | TATGAGTGTGGAGTCGTTGCCAGCAGAAAAGGGAACGTCATTTNNNAGATCGGAA |
| 44999_2 | TATGAGTGTGGAGTCGTTGCCAGCAGAAAAGGGAACGTCATTTNNNAGATCGGAA |
| 42464_1 | TATGAGTGTGGAGTCGTTGCTTTTCTTGGTCTTGCTCTGTTGCNNNAGATCGGAAGA |
| 42464_2 | TATGAGTGTGGAGTCGTTGCTTTTCTTGGTCTTGCTCTGTTGCNNNAGATCGGAAGA |
| 9670_1 | TATGAGTGTGGAGTCGTTGCAAAGCACTATGGCACTAGAAGCCNNNAGATCGGAA |
| 9670_2 | TATGAGTGTGGAGTCGTTGCAAAGCACTATGGCACTAGAAGCCNNNAGATCGGAA |
| 33448_1 | TATGAGTGTGGAGTCGTTGCCTTCTAGGGCTGTTTTGAGCAACNNNAGATCGGAAG |
| 33448_2 | TATGAGTGTGGAGTCGTTGCCTTCTAGGGCTGTTTTGAGCAACNNNAGATCGGAAG |
| 32103_1 | TATGAGTGTGGAGTCGTTGCGAACAAAAGGAAGTCAGGCAAGGNNNAGATCGGA |
| 32103_2 | TATGAGTGTGGAGTCGTTGCGAACAAAAGGAAGTCAGGCAAGGNNNAGATCGGA |
| 13453_1 | TATGAGTGTGGAGTCGTTGCGGATTCTTTAAAGGGGTTGTNNNAGATCGGAAGAG |
| 13453_2 | TATGAGTGTGGAGTCGTTGCGGATTCTTTAAAGGGGTTGTNNNAGATCGGAAGAG |
| 41945_1 | TATGAGTGTGGAGTCGTTGCGAACTCCTGACCTAGTGATCCACNNNAGATCGGAAG |
| 41945_2 | TATGAGTGTGGAGTCGTTGCGAACTCCTGACCTAGTGATCCACNNNAGATCGGAAG |
| 12690_1 | TATGAGTGTGGAGTCGTTGCACAGCATTATATGGAATACAGATNNNAGATCGGAAG |
| 12690_2 | TATGAGTGTGGAGTCGTTGCACAGCATTATATGGAATACAGATNNNAGATCGGAAG |
| 52229_1 | TATGAGTGTGGAGTCGTTGCACCCCCATTTATTGCCTAGTGAANNNAGATCGGAAG |
| 52229_2 | TATGAGTGTGGAGTCGTTGCACCCCCATTTATTGCCTAGTGAANNNAGATCGGAAG |
| 39836_1 | TATGAGTGTGGAGTCGTTGCAGGAGAATCACTGGAACCCAAAANNNAGATCGGAA |
| 39836_2 | TATGAGTGTGGAGTCGTTGCAGGAGAATCACTGGAACCCAAAANNNAGATCGGAA |
| 35033_1 | TATGAGTGTGGAGTCGTTGCAGTAGAGATGGGGTTTCAGCATGNNNAGATCGGAA |
| 35033_2 | TATGAGTGTGGAGTCGTTGCAGTAGAGATGGGGTTTCAGCATGNNNAGATCGGAA |
| 34850_1 | TATGAGTGTGGAGTCGTTGCCTGACTCAGTCTGTTCACTGGTTNNNAGATCGGAAG |
| 34850_2 | TATGAGTGTGGAGTCGTTGCCTGACTCAGTCTGTTCACTGGTTNNNAGATCGGAAG |
| 37646_1 | TATGAGTGTGGAGTCGTTGCTGCCAAGAAAATTCCAGAACTGCNNNAGATCGGAAG |
| 37646_2 | TATGAGTGTGGAGTCGTTGCTGCCAAGAAAATTCCAGAACTGCNNNAGATCGGAAG |
| 35808_1 | TATGAGTGTGGAGTCGTTGCTTTTTTGAGATGGAGTCTGCCCTNNNAGATCGGAAG |
| 35808_2 | TATGAGTGTGGAGTCGTTGCTTTTTTGAGATGGAGTCTGCCCTNNNAGATCGGAAG |
| 49247_1 | TATGAGTGTGGAGTCGTTGCCTCCCAGGTTCAAGCGATTCTCNNNAGATCGGAAGA |
| 49247_2 | TATGAGTGTGGAGTCGTTGCCTCCCAGGTTCAAGCGATTCTCNNNAGATCGGAAGA |
| 41331_1 | TATGAGTGTGGAGTCGTTGCCTGGGTTCAAGAGACTCTTTTGCNNNAGATCGGAAG |

| | |
|---|---|
| 41331_2 | TATGAGTGTGGAGTCGTTGCCTGGGTTCAAGAGACTCTTTGCNNNAGATCGGAAG |
| 13756_1 | TATGAGTGTGGAGTCGTTGCAAAAACTTTCCCAGTGAAGTATNNNAGATCGGAAGA |
| 13756_2 | TATGAGTGTGGAGTCGTTGCAAAAACTTTCCCAGTGAAGTATNNNAGATCGGAAGA |
| 42208_1 | TATGAGTGTGGAGTCGTTGCAACTCCAATTCCACTCAGCAGAANNNAGATCGGAAG |
| 42208_2 | TATGAGTGTGGAGTCGTTGCAACTCCAATTCCACTCAGCAGAANNNAGATCGGAAG |
| 42956_1 | TATGAGTGTGGAGTCGTTGCCTCCTTCTACCTCCTCTCCTCTCNNNAGATCGGAAGAG |
| 42956_2 | TATGAGTGTGGAGTCGTTGCCTCCTTCTACCTCCTCTCCTCTCNNNAGATCGGAAGAG |
| 52765_1 | TATGAGTGTGGAGTCGTTGCTCTTGCCGTCTTTGCAGAATTTTNNNAGATCGGAAG |
| 52765_2 | TATGAGTGTGGAGTCGTTGCTCTTGCCGTCTTTGCAGAATTTTNNNAGATCGGAAG |
| 36456_1 | TATGAGTGTGGAGTCGTTGCAGTCTTCAAGTCTATACCCCCANNNAGATCGGAAG |
| 36456_2 | TATGAGTGTGGAGTCGTTGCAGTCTTCAAGTCTATACCCCCANNNAGATCGGAAG |
| 41605_1 | TATGAGTGTGGAGTCGTTGCGTTTCACCATGTTAGCCAGGATGNNNAGATCGGAAG |
| 41605_2 | TATGAGTGTGGAGTCGTTGCGTTTCACCATGTTAGCCAGGATGNNNAGATCGGAAG |
| 38485_1 | TATGAGTGTGGAGTCGTTGCGGTGAAACCCCGTCTCTACTAAANNNAGATCGGAAG |
| 38485_2 | TATGAGTGTGGAGTCGTTGCGGTGAAACCCCGTCTCTACTAAANNNAGATCGGAAG |
| 34353_1 | TATGAGTGTGGAGTCGTTGCCTCTCTCCTGATGATGATGGTGGNNNAGATCGGAAG |
| 34353_2 | TATGAGTGTGGAGTCGTTGCCTCTCTCCTGATGATGATGGTGGNNNAGATCGGAAG |
| 51894_1 | TATGAGTGTGGAGTCGTTGCTTCTGCCTCTTTTCCTACCTTCCNNNAGATCGGAAGA |
| 51894_2 | TATGAGTGTGGAGTCGTTGCTTCTGCCTCTTTTCCTACCTTCCNNNAGATCGGAAGA |
| 46003_1 | TATGAGTGTGGAGTCGTTGCAAACGTCCCAAGCTACTAAGAGGNNNAGATCGGAA |
| 46003_2 | TATGAGTGTGGAGTCGTTGCAAACGTCCCAAGCTACTAAGAGGNNNAGATCGGAA |
| 35970_1 | TATGAGTGTGGAGTCGTTGCAGAGCGACACTCCATCTCAAAAANNNAGATCGGAAG |
| 35970_2 | TATGAGTGTGGAGTCGTTGCAGAGCGACACTCCATCTCAAAAANNNAGATCGGAAG |
| 50616_1 | TATGAGTGTGGAGTCGTTGCTTATGAGCCTAACTAGCAGCAGGNNNAGATCGGAA |
| 50616_2 | TATGAGTGTGGAGTCGTTGCTTATGAGCCTAACTAGCAGCAGGNNNAGATCGGAA |
| 7983_1 | TATGAGTGTGGAGTCGTTGCATTTCCTGCCTCCAGAGTAGCTNNNAGATCGGAAGA |
| 7983_2 | TATGAGTGTGGAGTCGTTGCATTTCCTGCCTCCAGAGTAGCTNNNAGATCGGAAGA |
| 28954_1 | TATGAGTGTGGAGTCGTTGCAGAGCGTGTGTATATGTGTGTGTNNNAGATCGGAA |
| 28954_2 | TATGAGTGTGGAGTCGTTGCAGAGCGTGTGTATATGTGTGTGTNNNAGATCGGAA |
| 13266_1 | TATGAGTGTGGAGTCGTTGCATTTTTTTTTTTGCTGTTTTGACNNNAGATCGGAAGA |
| 13266_2 | TATGAGTGTGGAGTCGTTGCATTTTTTTTTTTGCTGTTTTGACNNNAGATCGGAAGA |
| 45782_1 | TATGAGTGTGGAGTCGTTGCCAATTTCAGCTCACTGCAACCTCNNNAGATCGGAAG |
| 45782_2 | TATGAGTGTGGAGTCGTTGCCAATTTCAGCTCACTGCAACCTCNNNAGATCGGAAG |
| 36197_1 | TATGAGTGTGGAGTCGTTGCCTGGGTGACAGAGTGAAACACTANNNAGATCGGAA |
| 36197_2 | TATGAGTGTGGAGTCGTTGCCTGGGTGACAGAGTGAAACACTANNNAGATCGGAA |
| 34251_1 | TATGAGTGTGGAGTCGTTGCAAAAAACACACATGGCCTCATCCNNNAGATCGGAAG |
| 34251_2 | TATGAGTGTGGAGTCGTTGCAAAAAACACACATGGCCTCATCCNNNAGATCGGAAG |
| 34256_1 | TATGAGTGTGGAGTCGTTGCGAGAGAGAGAGAGAGAGAGATGANNNAGATCGG |
| 34256_2 | TATGAGTGTGGAGTCGTTGCGAGAGAGAGAGAGAGAGAGATGANNNAGATCGG |
| 46057_1 | TATGAGTGTGGAGTCGTTGCTTCAAGTGATTCTCCTGCCTCAGNNNAGATCGGAAG |
| 46057_2 | TATGAGTGTGGAGTCGTTGCTTCAAGTGATTCTCCTGCCTCAGNNNAGATCGGAAG |
| 44855_1 | TATGAGTGTGGAGTCGTTGCAAAAAAATGGGCATCCTGGGATGNNNAGATCGGAA |
| 44855_2 | TATGAGTGTGGAGTCGTTGCAAAAAAATGGGCATCCTGGGATGNNNAGATCGGAA |

| | |
|---|---|
| 49119_1 | TATGAGTGTGGAGTCGTTGCCACTCAACTGTTGCCTTCTTGAGNNNAGATCGGAAG |
| 49119_2 | TATGAGTGTGGAGTCGTTGCCACTCAACTGTTGCCTTCTTGAGNNNAGATCGGAAG |
| 38824_1 | TATGAGTGTGGAGTCGTTGCGATGAAACTGGACCCTTACCTGTNNNAGATCGGAAG |
| 38824_2 | TATGAGTGTGGAGTCGTTGCGATGAAACTGGACCCTTACCTGTNNNAGATCGGAAG |
| 33224_1 | TATGAGTGTGGAGTCGTTGCCTCCTGAGTAGCTGGGACTACAGNNNAGATCGGAA |
| 33224_2 | TATGAGTGTGGAGTCGTTGCCTCCTGAGTAGCTGGGACTACAGNNNAGATCGGAA |
| 10424_1 | TATGAGTGTGGAGTCGTTGCCACACACCTATCTGACAGCTGAGNNNAGATCGGAAG |
| 10424_2 | TATGAGTGTGGAGTCGTTGCCACACACCTATCTGACAGCTGAGNNNAGATCGGAAG |
| 46537_1 | TATGAGTGTGGAGTCGTTGCGTGCCCAGCCTATTTTTGACTTTNNNAGATCGGAAG |
| 46537_2 | TATGAGTGTGGAGTCGTTGCGTGCCCAGCCTATTTTTGACTTTNNNAGATCGGAAG |
| 35595_1 | TATGAGTGTGGAGTCGTTGCCACAAGCAATTCACCTTCCTCAGNNNAGATCGGAAG |
| 35595_2 | TATGAGTGTGGAGTCGTTGCCACAAGCAATTCACCTTCCTCAGNNNAGATCGGAAG |
| 32104_1 | TATGAGTGTGGAGTCGTTGCAAGTAGCTGGGATTACAGGCACNNNAGATCGGAAG |
| 32104_2 | TATGAGTGTGGAGTCGTTGCAAGTAGCTGGGATTACAGGCACNNNAGATCGGAAG |
| 3019_1 | TATGAGTGTGGAGTCGTTGCATCATCTGGTGCTACCTGAGGGNNNAGATCGGAAG |
| 3019_2 | TATGAGTGTGGAGTCGTTGCATCATCTGGTGCTACCTGAGGGNNNAGATCGGAAG |
| 52798_1 | TATGAGTGTGGAGTCGTTGCTGGAGCTCCCAGATCTGCTCNNNAGATCGGAAGAG |
| 52798_2 | TATGAGTGTGGAGTCGTTGCTGGAGCTCCCAGATCTGCTCNNNAGATCGGAAGAG |
| 37068_1 | TATGAGTGTGGAGTCGTTGCGAATGCAGGAGAATGGTGTGAACNNNAGATCGGA |
| 37068_2 | TATGAGTGTGGAGTCGTTGCGAATGCAGGAGAATGGTGTGAACNNNAGATCGGA |
| 42363_1 | TATGAGTGTGGAGTCGTTGCTAATATTTCACAACGCACCCGTGNNNAGATCGGAAG |
| 42363_2 | TATGAGTGTGGAGTCGTTGCTAATATTTCACAACGCACCCGTGNNNAGATCGGAAG |
| 37665_1 | TATGAGTGTGGAGTCGTTGCAGTATCTCCTTCTTCCTTGCACCNNNAGATCGGAAGA |
| 37665_2 | TATGAGTGTGGAGTCGTTGCAGTATCTCCTTCTTCCTTGCACCNNNAGATCGGAAGA |
| 40601_1 | TATGAGTGTGGAGTCGTTGCCTGGGCATAGCAGAACAAACAAANNNAGATCGGAA |
| 40601_2 | TATGAGTGTGGAGTCGTTGCCTGGGCATAGCAGAACAAACAAANNNAGATCGGAA |
| 31431_1 | TATGAGTGTGGAGTCGTTGCCAAGGCCAGAGATGTTGCTAAACNNNAGATCGGAA |
| 31431_2 | TATGAGTGTGGAGTCGTTGCCAAGGCCAGAGATGTTGCTAAACNNNAGATCGGAA |
| 37284_1 | TATGAGTGTGGAGTCGTTGCCGCTTGCAGTTCTCAGATCAAATNNNAGATCGGAAG |
| 37284_2 | TATGAGTGTGGAGTCGTTGCCGCTTGCAGTTCTCAGATCAAATNNNAGATCGGAAG |
| 38890_1 | TATGAGTGTGGAGTCGTTGCGAAAAACAGACACGAGCGCTTATNNNAGATCGGAA |
| 38890_2 | TATGAGTGTGGAGTCGTTGCGAAAAACAGACACGAGCGCTTATNNNAGATCGGAA |
| 52115_1 | TATGAGTGTGGAGTCGTTGCGGCAAATTTCAGGACAGTGAGAGNNNAGATCGGAA |
| 52115_2 | TATGAGTGTGGAGTCGTTGCGGCAAATTTCAGGACAGTGAGAGNNNAGATCGGAA |
| 33189_1 | TATGAGTGTGGAGTCGTTGCTCAATGAATGGGAAACCAAGTGCNNNAGATCGGAA |
| 33189_2 | TATGAGTGTGGAGTCGTTGCTCAATGAATGGGAAACCAAGTGCNNNAGATCGGAA |
| 37717_1 | TATGAGTGTGGAGTCGTTGCCAGCTTGTTAGAACTGGTCCAGANNNAGATCGGAA |
| 37717_2 | TATGAGTGTGGAGTCGTTGCCAGCTTGTTAGAACTGGTCCAGANNNAGATCGGAA |
| 42505_1 | TATGAGTGTGGAGTCGTTGCGGGGTGCTGGAACATGGTAAATANNNAGATCGGA |
| 42505_2 | TATGAGTGTGGAGTCGTTGCGGGGTGCTGGAACATGGTAAATANNNAGATCGGA |
| 37550_1 | TATGAGTGTGGAGTCGTTGCGTCAGACAGATTCACAGCCAAATNNNAGATCGGAAG |
| 37550_2 | TATGAGTGTGGAGTCGTTGCGTCAGACAGATTCACAGCCAAATNNNAGATCGGAAG |
| 52922_1 | TATGAGTGTGGAGTCGTTGCTGAACTGGCATGAAGGGTTAGGNNNAGATCGGAA |

| | |
|---|---|
| 52922_2 | TATGAGTGTGGAGTCGTTGCTGAACTGGCATGAAGGGTTAGGNNNAGATCGGAAG |
| 47624_1 | TATGAGTGTGGAGTCGTTGCGAAGGAGTGAAAGCTGAGGAGAGNNNAGATCGGA |
| 47624_2 | TATGAGTGTGGAGTCGTTGCGAAGGAGTGAAAGCTGAGGAGAGNNNAGATCGGA |
| 33915_1 | TATGAGTGTGGAGTCGTTGCACCTTAGCCTCCCAAAGTGTTAGNNNAGATCGGAAG |
| 33915_2 | TATGAGTGTGGAGTCGTTGCACCTTAGCCTCCCAAAGTGTTAGNNNAGATCGGAAG |
| 40307_1 | TATGAGTGTGGAGTCGTTGCTGCCCATAGAAGTTCCATCCAAANNNAGATCGGAAG |
| 40307_2 | TATGAGTGTGGAGTCGTTGCTGCCCATAGAAGTTCCATCCAAANNNAGATCGGAAG |
| 50999_1 | TATGAGTGTGGAGTCGTTGCCCATGCCCAGCTGATTTTTGTATNNNAGATCGGAAG |
| 50999_2 | TATGAGTGTGGAGTCGTTGCCCATGCCCAGCTGATTTTTGTATNNNAGATCGGAAG |
| 13390_1 | TATGAGTGTGGAGTCGTTGCCGTTAAGTCGTAGCCCTTAANNNAGATCGGAAGAG |
| 13390_2 | TATGAGTGTGGAGTCGTTGCCGTTAAGTCGTAGCCCTTAANNNAGATCGGAAGAG |
| 34631_1 | TATGAGTGTGGAGTCGTTGCAGAGATGGGGCACTATGGAGATANNNAGATCGGA |
| 34631_2 | TATGAGTGTGGAGTCGTTGCAGAGATGGGGCACTATGGAGATANNNAGATCGGA |
| 33344_1 | TATGAGTGTGGAGTCGTTGCGATTACAGGCATGCACCACAATTNNNAGATCGGAAG |
| 33344_2 | TATGAGTGTGGAGTCGTTGCGATTACAGGCATGCACCACAATTNNNAGATCGGAAG |
| 2984_1 | TATGAGTGTGGAGTCGTTGCTCTAAGCTTCTCGTCACTGGTNNNAGATCGGAAGAG |
| 2984_2 | TATGAGTGTGGAGTCGTTGCTCTAAGCTTCTCGTCACTGGTNNNAGATCGGAAGAG |
| 30250_1 | TATGAGTGTGGAGTCGTTGCCTCAATGACAAGACCAGTCCTCANNNAGATCGGAAG |
| 30250_2 | TATGAGTGTGGAGTCGTTGCCTCAATGACAAGACCAGTCCTCANNNAGATCGGAAG |
| 12651_1 | TATGAGTGTGGAGTCGTTGCTGAAAAAATAGGTAGCTTTTTGTNNNAGATCGGAAG |
| 12651_2 | TATGAGTGTGGAGTCGTTGCTGAAAAAATAGGTAGCTTTTTGTNNNAGATCGGAAG |
| 13185_1 | TATGAGTGTGGAGTCGTTGCCATCCAAGTTCTTCTTACTACTTNNNAGATCGGAAGA |
| 13185_2 | TATGAGTGTGGAGTCGTTGCCATCCAAGTTCTTCTTACTACTTNNNAGATCGGAAGA |
| 29720_1 | TATGAGTGTGGAGTCGTTGCCCCTGAGTTTGAAGGAGGTAGACNNNAGATCGGAA |
| 29720_2 | TATGAGTGTGGAGTCGTTGCCCCTGAGTTTGAAGGAGGTAGACNNNAGATCGGAA |
| 46052_1 | TATGAGTGTGGAGTCGTTGCCTTCCGAGTAGCTGGGATTACAGNNNAGATCGGAAG |
| 46052_2 | TATGAGTGTGGAGTCGTTGCCTTCCGAGTAGCTGGGATTACAGNNNAGATCGGAAG |
| 34646_1 | TATGAGTGTGGAGTCGTTGCCCTACTGGGAAGTGAGGAGCCNNNAGATCGGAAGA |
| 34646_2 | TATGAGTGTGGAGTCGTTGCCCTACTGGGAAGTGAGGAGCCNNNAGATCGGAAGA |
| 13321_1 | TATGAGTGTGGAGTCGTTGCAGAAAGAGAAAGAAAGAAAGAGANNNAGATCGGA |
| 13321_2 | TATGAGTGTGGAGTCGTTGCAGAAAGAGAAAGAAAGAAAGAGANNNAGATCGGA |
| 12956_1 | TATGAGTGTGGAGTCGTTGCGATGTGACAAGATTGTGGAGNNNAGATCGGAAGAG |
| 12956_2 | TATGAGTGTGGAGTCGTTGCGATGTGACAAGATTGTGGAGNNNAGATCGGAAGAG |
| 36225_1 | TATGAGTGTGGAGTCGTTGCTCTGACAGCTGCAAAGAAAAGTGNNNAGATCGGAA |
| 36225_2 | TATGAGTGTGGAGTCGTTGCTCTGACAGCTGCAAAGAAAAGTGNNNAGATCGGAA |
| 44190_1 | TATGAGTGTGGAGTCGTTGCAATGTCTTCTCTCGTTCTGCTCCNNNAGATCGGAAGA |
| 44190_2 | TATGAGTGTGGAGTCGTTGCAATGTCTTCTCTCGTTCTGCTCCNNNAGATCGGAAGA |
| 35364_1 | TATGAGTGTGGAGTCGTTGCGCTTACTAAAACCTCTGCCTCCTNNNAGATCGGAAGA |
| 35364_2 | TATGAGTGTGGAGTCGTTGCGCTTACTAAAACCTCTGCCTCCTNNNAGATCGGAAGA |
| 46821_1 | TATGAGTGTGGAGTCGTTGCCAAGTAGCTGGGACTACAGGCNNNAGATCGGAAGA |
| 46821_2 | TATGAGTGTGGAGTCGTTGCCAAGTAGCTGGGACTACAGGCNNNAGATCGGAAGA |
| 49803_1 | TATGAGTGTGGAGTCGTTGCTTGCCCCTTTCTCAAGTGAAAACNNNAGATCGGAAG |
| 49803_2 | TATGAGTGTGGAGTCGTTGCTTGCCCCTTTCTCAAGTGAAAACNNNAGATCGGAAG |

| | |
|---|---|
| 32055_1 | TATGAGTGTGGAGTCGTTGCCATTCTCCTGCCTCAGCCTCNNNAGATCGGAAGAGCA |
| 32055_2 | TATGAGTGTGGAGTCGTTGCCATTCTCCTGCCTCAGCCTCNNNAGATCGGAAGAGCA |
| 30915_1 | TATGAGTGTGGAGTCGTTGCCCATGTGGTCTCTCATCTCCATTNNNAGATCGGAAGA |
| 30915_2 | TATGAGTGTGGAGTCGTTGCCCATGTGGTCTCTCATCTCCATTNNNAGATCGGAAGA |
| 39132_1 | TATGAGTGTGGAGTCGTTGCTAAAGCGGTGCTGTGTAAAAGTGNNNAGATCGGAA |
| 39132_2 | TATGAGTGTGGAGTCGTTGCTAAAGCGGTGCTGTGTAAAAGTGNNNAGATCGGAA |
| 41134_1 | TATGAGTGTGGAGTCGTTGCTAGAGTGCACTAATACACCCAGCNNNAGATCGGAAG |
| 41134_2 | TATGAGTGTGGAGTCGTTGCTAGAGTGCACTAATACACCCAGCNNNAGATCGGAAG |
| 31470_1 | TATGAGTGTGGAGTCGTTGCCTCCCTATTTCCTTTCCCTGCTTNNNAGATCGGAAGAG |
| 31470_2 | TATGAGTGTGGAGTCGTTGCCTCCCTATTTCCTTTCCCTGCTTNNNAGATCGGAAGAG |
| 37065_1 | TATGAGTGTGGAGTCGTTGCGAGCTTGTAGTGAGCCGAGATCNNNAGATCGGAAG |
| 37065_2 | TATGAGTGTGGAGTCGTTGCGAGCTTGTAGTGAGCCGAGATCNNNAGATCGGAAG |
| 32243_1 | TATGAGTGTGGAGTCGTTGCTGGGAGGAGTGAGTATACCAAGTNNNAGATCGGAA |
| 32243_2 | TATGAGTGTGGAGTCGTTGCTGGGAGGAGTGAGTATACCAAGTNNNAGATCGGAA |
| 37706_1 | TATGAGTGTGGAGTCGTTGCCTCCTGTAGTCCCAGCTACTCGNNNAGATCGGAAGA |
| 37706_2 | TATGAGTGTGGAGTCGTTGCCTCCTGTAGTCCCAGCTACTCGNNNAGATCGGAAGA |
| 2776_1 | TATGAGTGTGGAGTCGTTGCGGACACACACACTGACATAAGCTNNNAGATCGGAAG |
| 2776_2 | TATGAGTGTGGAGTCGTTGCGGACACACACACTGACATAAGCTNNNAGATCGGAAG |
| 44665_1 | TATGAGTGTGGAGTCGTTGCCAAAGGAAACTGCCTGAGGAAAGNNNAGATCGGAA |
| 44665_2 | TATGAGTGTGGAGTCGTTGCCAAAGGAAACTGCCTGAGGAAAGNNNAGATCGGAA |
| 31241_1 | TATGAGTGTGGAGTCGTTGCACGGTCATTTCCACAGAGATTTTNNNAGATCGGAAG |
| 31241_2 | TATGAGTGTGGAGTCGTTGCACGGTCATTTCCACAGAGATTTTNNNAGATCGGAAG |
| 47689_1 | TATGAGTGTGGAGTCGTTGCCAGGGTTCAAGTGATTCTTCTGCNNNAGATCGGAAG |
| 47689_2 | TATGAGTGTGGAGTCGTTGCCAGGGTTCAAGTGATTCTTCTGCNNNAGATCGGAAG |
| 39370_1 | TATGAGTGTGGAGTCGTTGCGTTGCAGTGAGCCAAGATCATGNNNAGATCGGAAG |
| 39370_2 | TATGAGTGTGGAGTCGTTGCGTTGCAGTGAGCCAAGATCATGNNNAGATCGGAAG |
| 51728_1 | TATGAGTGTGGAGTCGTTGCCCCGGAGAGGTCAGAGTCCTNNNAGATCGGAAGAG |
| 51728_2 | TATGAGTGTGGAGTCGTTGCCCCGGAGAGGTCAGAGTCCTNNNAGATCGGAAGAG |
| 52689_1 | TATGAGTGTGGAGTCGTTGCCCCTTCTAGGTCACATGCTTTCTNNNAGATCGGAAGA |
| 52689_2 | TATGAGTGTGGAGTCGTTGCCCCTTCTAGGTCACATGCTTTCTNNNAGATCGGAAGA |
| 13232_1 | TATGAGTGTGGAGTCGTTGCGATACTTACTGTAGTTCTCAGAGNNNAGATCGGAAG |
| 13232_2 | TATGAGTGTGGAGTCGTTGCGATACTTACTGTAGTTCTCAGAGNNNAGATCGGAAG |
| 44026_1 | TATGAGTGTGGAGTCGTTGCTCGGAATGAAAAATGGTTCAGCCNNNAGATCGGAA |
| 44026_2 | TATGAGTGTGGAGTCGTTGCTCGGAATGAAAAATGGTTCAGCCNNNAGATCGGAA |
| 50430_1 | TATGAGTGTGGAGTCGTTGCCTCAGGGTATAGGCCAACAATGTNNNAGATCGGAA |
| 50430_2 | TATGAGTGTGGAGTCGTTGCCTCAGGGTATAGGCCAACAATGTNNNAGATCGGAA |
| 41955_1 | TATGAGTGTGGAGTCGTTGCCCATCTCAGTTCACTGCAAACTCNNNAGATCGGAAGA |
| 41955_2 | TATGAGTGTGGAGTCGTTGCCCATCTCAGTTCACTGCAAACTCNNNAGATCGGAAGA |
| 51519_1 | TATGAGTGTGGAGTCGTTGCTCTACCCTCTTACGTGTCTCAGTNNNAGATCGGAAGA |
| 51519_2 | TATGAGTGTGGAGTCGTTGCTCTACCCTCTTACGTGTCTCAGTNNNAGATCGGAAGA |
| 36394_1 | TATGAGTGTGGAGTCGTTGCGCCAACATGCGAAACCATATCTCNNNAGATCGGAAG |
| 36394_2 | TATGAGTGTGGAGTCGTTGCGCCAACATGCGAAACCATATCTCNNNAGATCGGAAG |
| 34740_1 | TATGAGTGTGGAGTCGTTGCTTTAGTGCAGACAGGGTTTCACTNNNAGATCGGAAG |

| | |
|---|---|
| 34740_2 | TATGAGTGTGGAGTCGTTGCTTTAGTGCAGACAGGGTTTCACTNNNAGATCGGAAG |
| 51450_1 | TATGAGTGTGGAGTCGTTGCGAATTCTGCCAACAACCACATGANNNAGATCGGAAG |
| 51450_2 | TATGAGTGTGGAGTCGTTGCGAATTCTGCCAACAACCACATGANNNAGATCGGAAG |
| 39994_1 | TATGAGTGTGGAGTCGTTGCAACATCTTCACCAATTAGTCCAGANNNAGATCGGAA |
| 39994_2 | TATGAGTGTGGAGTCGTTGCAACATCTTCACCAATTAGTCCAGANNNAGATCGGAA |
| 30559_1 | TATGAGTGTGGAGTCGTTGCACATCCTGGAGAAAAGGGAAGTGNNNAGATCGGA |
| 30559_2 | TATGAGTGTGGAGTCGTTGCACATCCTGGAGAAAAGGGAAGTGNNNAGATCGGA |
| 46732_1 | TATGAGTGTGGAGTCGTTGCGACTCCTGGGTTCAAGTGATTCTNNNAGATCGGAAG |
| 46732_2 | TATGAGTGTGGAGTCGTTGCGACTCCTGGGTTCAAGTGATTCTNNNAGATCGGAAG |
| 50632_1 | TATGAGTGTGGAGTCGTTGCGTGAAGAATGGGTGCAACATTCANNNAGATCGGAA |
| 50632_2 | TATGAGTGTGGAGTCGTTGCGTGAAGAATGGGTGCAACATTCANNNAGATCGGAA |
| 31183_1 | TATGAGTGTGGAGTCGTTGCTTCCTTTTGTCATCTGCTCCCATNNNAGATCGGAAGA |
| 31183_2 | TATGAGTGTGGAGTCGTTGCTTCCTTTTGTCATCTGCTCCCATNNNAGATCGGAAGA |
| 37234_1 | TATGAGTGTGGAGTCGTTGCAATCAAACTTTGCCCTTCCTTCCNNNAGATCGGAAGA |
| 37234_2 | TATGAGTGTGGAGTCGTTGCAATCAAACTTTGCCCTTCCTTCCNNNAGATCGGAAGA |
| 50912_1 | TATGAGTGTGGAGTCGTTGCTTTGTCTCTGGAGAATTCCGGTTNNNAGATCGGAAG |
| 50912_2 | TATGAGTGTGGAGTCGTTGCTTTGTCTCTGGAGAATTCCGGTTNNNAGATCGGAAG |
| 30281_1 | TATGAGTGTGGAGTCGTTGCTTCCAGAGCAAGAATTGACTCGANNNAGATCGGAA |
| 30281_2 | TATGAGTGTGGAGTCGTTGCTTCCAGAGCAAGAATTGACTCGANNNAGATCGGAA |
| 10702_1 | TATGAGTGTGGAGTCGTTGCGGGCAGGAAGAGTGGTGTACATANNNAGATCGGA |
| 10702_2 | TATGAGTGTGGAGTCGTTGCGGGCAGGAAGAGTGGTGTACATANNNAGATCGGA |
| 39748_1 | TATGAGTGTGGAGTCGTTGCGACTTCCCAACTGATGTCTCCAGNNNAGATCGGAAG |
| 39748_2 | TATGAGTGTGGAGTCGTTGCGACTTCCCAACTGATGTCTCCAGNNNAGATCGGAAG |
| 48734_1 | TATGAGTGTGGAGTCGTTGCGAACTCTGGACCTCTGGTTATCCNNNAGATCGGAAG |
| 48734_2 | TATGAGTGTGGAGTCGTTGCGAACTCTGGACCTCTGGTTATCCNNNAGATCGGAAG |
| 38878_1 | TATGAGTGTGGAGTCGTTGCTAGAATAAGCCACCAAGCAGAGGNNNAGATCGGAA |
| 38878_2 | TATGAGTGTGGAGTCGTTGCTAGAATAAGCCACCAAGCAGAGGNNNAGATCGGAA |
| 37713_1 | TATGAGTGTGGAGTCGTTGCGAGCTTGATGTCATAAGCAACCANNNAGATCGGAA |
| 37713_2 | TATGAGTGTGGAGTCGTTGCGAGCTTGATGTCATAAGCAACCANNNAGATCGGAA |
| 37425_1 | TATGAGTGTGGAGTCGTTGCTCATCAACAGAATGGAATTGTGGTNNNAGATCGGA |
| 37425_2 | TATGAGTGTGGAGTCGTTGCTCATCAACAGAATGGAATTGTGGTNNNAGATCGGA |
| 13032_1 | TATGAGTGTGGAGTCGTTGCATTTTAGCTAAAGATTTACCAGCNNNAGATCGGAAG |
| 13032_2 | TATGAGTGTGGAGTCGTTGCATTTTAGCTAAAGATTTACCAGCNNNAGATCGGAAG |
| 29190_1 | TATGAGTGTGGAGTCGTTGCCCTAATCCCATGAGCCTGACTACNNNAGATCGGAAG |
| 29190_2 | TATGAGTGTGGAGTCGTTGCCCTAATCCCATGAGCCTGACTACNNNAGATCGGAAG |
| 49610_1 | TATGAGTGTGGAGTCGTTGCCGGCAGCTTTCACTTTGACTATGNNNAGATCGGAAG |
| 49610_2 | TATGAGTGTGGAGTCGTTGCCGGCAGCTTTCACTTTGACTATGNNNAGATCGGAAG |
| 30494_1 | TATGAGTGTGGAGTCGTTGCAGAAAACGGGGAGCCAAGGAAAAANNNAGATCGG |
| 30494_2 | TATGAGTGTGGAGTCGTTGCAGAAAACGGGGAGCCAAGGAAAAANNNAGATCGG |
| 51990_1 | TATGAGTGTGGAGTCGTTGCCAAAACCCTTCAGCATCTTCCTGNNNAGATCGGAAGA |
| 51990_2 | TATGAGTGTGGAGTCGTTGCCAAAACCCTTCAGCATCTTCCTGNNNAGATCGGAAGA |
| 42498_1 | TATGAGTGTGGAGTCGTTGCAGGGAGAGAGAGAGAGAGAGAGANNNAGATCGG |
| 42498_2 | TATGAGTGTGGAGTCGTTGCAGGGAGAGAGAGAGAGAGAGAGANNNAGATCGG |

| | |
|---|---|
| 48179_1 | TATGAGTGTGGAGTCGTTGCCTGGAGCAGGACTCTATTTACCCNNNAGATCGGAAG |
| 48179_2 | TATGAGTGTGGAGTCGTTGCCTGGAGCAGGACTCTATTTACCCNNNAGATCGGAAG |
| 40092_1 | TATGAGTGTGGAGTCGTTGCCTGAGGCAGGAGAATCACTTGAANNNAGATCGGAA |
| 40092_2 | TATGAGTGTGGAGTCGTTGCCTGAGGCAGGAGAATCACTTGAANNNAGATCGGAA |
| 48514_1 | TATGAGTGTGGAGTCGTTGCAAGTGCTGAGATTACAGGCATGANNNAGATCGGAA |
| 48514_2 | TATGAGTGTGGAGTCGTTGCAAGTGCTGAGATTACAGGCATGANNNAGATCGGAA |
| 41294_1 | TATGAGTGTGGAGTCGTTGCTCTGCTCTGAGAAAATAACGGAANNNAGATCGGAA |
| 41294_2 | TATGAGTGTGGAGTCGTTGCTCTGCTCTGAGAAAATAACGGAANNNAGATCGGAA |
| 45131_1 | TATGAGTGTGGAGTCGTTGCAGCATAGCCCACAAAAGAAATCGNNNAGATCGGAA |
| 45131_2 | TATGAGTGTGGAGTCGTTGCAGCATAGCCCACAAAAGAAATCGNNNAGATCGGAA |
| 31862_1 | TATGAGTGTGGAGTCGTTGCCCTGGGCTTTCAGAGACTATGTTNNNAGATCGGAAG |
| 31862_2 | TATGAGTGTGGAGTCGTTGCCCTGGGCTTTCAGAGACTATGTTNNNAGATCGGAAG |
| 39309_1 | TATGAGTGTGGAGTCGTTGCTTGGGCTTGGTTTGATCTTACCANNNAGATCGGAAG |
| 39309_2 | TATGAGTGTGGAGTCGTTGCTTGGGCTTGGTTTGATCTTACCANNNAGATCGGAAG |
| 45787_1 | TATGAGTGTGGAGTCGTTGCGTAGCTGATATTACAGGTCGGCANNNAGATCGGAA |
| 45787_2 | TATGAGTGTGGAGTCGTTGCGTAGCTGATATTACAGGTCGGCANNNAGATCGGAA |
| 39957_1 | TATGAGTGTGGAGTCGTTGCTTCCCTTTTACCCTTGGTGTAGGNNNAGATCGGAAG |
| 39957_2 | TATGAGTGTGGAGTCGTTGCTTCCCTTTTACCCTTGGTGTAGGNNNAGATCGGAAG |
| 35189_1 | TATGAGTGTGGAGTCGTTGCACCTCCTGGAATCAAGCAATTCTNNNAGATCGGAAG |
| 35189_2 | TATGAGTGTGGAGTCGTTGCACCTCCTGGAATCAAGCAATTCTNNNAGATCGGAAG |
| 32312_1 | TATGAGTGTGGAGTCGTTGCCCACCTGGTCTCCACAAAAAATCNNNAGATCGGAAG |
| 32312_2 | TATGAGTGTGGAGTCGTTGCCCACCTGGTCTCCACAAAAAATCNNNAGATCGGAAG |
| 36953_1 | TATGAGTGTGGAGTCGTTGCTCAGCCCCAGAACAACTAAATGANNNAGATCGGAAG |
| 36953_2 | TATGAGTGTGGAGTCGTTGCTCAGCCCCAGAACAACTAAATGANNNAGATCGGAAG |
| 39079_1 | TATGAGTGTGGAGTCGTTGCGAAAGCAGAATTTGGAGGCTGAGNNNAGATCGGAA |
| 39079_2 | TATGAGTGTGGAGTCGTTGCGAAAGCAGAATTTGGAGGCTGAGNNNAGATCGGAA |
| 35793_1 | TATGAGTGTGGAGTCGTTGCCTTCCCGGATTCATGCAATTCTCNNNAGATCGGAAGA |
| 35793_2 | TATGAGTGTGGAGTCGTTGCCTTCCCGGATTCATGCAATTCTCNNNAGATCGGAAGA |
| 50697_1 | TATGAGTGTGGAGTCGTTGCATCCAAGCATGAGGATTTAGGCTNNNAGATCGGAA |
| 50697_2 | TATGAGTGTGGAGTCGTTGCATCCAAGCATGAGGATTTAGGCTNNNAGATCGGAA |
| 30381_1 | TATGAGTGTGGAGTCGTTGCCCGGTATGTAATTTAACATTCAANNNAGATCGGAAG |
| 30381_2 | TATGAGTGTGGAGTCGTTGCCCGGTATGTAATTTAACATTCAANNNAGATCGGAAG |
| 33817_1 | TATGAGTGTGGAGTCGTTGCAGAACCCTTTCTTGTGGAGTGTTNNNAGATCGGAAG |
| 33817_2 | TATGAGTGTGGAGTCGTTGCAGAACCCTTTCTTGTGGAGTGTTNNNAGATCGGAAG |
| 37832_1 | TATGAGTGTGGAGTCGTTGCTCCTTCACCTCCAAAAGTTTCCTNNNAGATCGGAAGA |
| 37832_2 | TATGAGTGTGGAGTCGTTGCTCCTTCACCTCCAAAAGTTTCCTNNNAGATCGGAAGA |
| 31402_1 | TATGAGTGTGGAGTCGTTGCTTGCTTATGGTGCTAGGTACAGGNNNAGATCGGAA |
| 31402_2 | TATGAGTGTGGAGTCGTTGCTTGCTTATGGTGCTAGGTACAGGNNNAGATCGGAA |
| 9827_1 | TATGAGTGTGGAGTCGTTGCATTTCGGGGGACTAAGGAGTTGANNNAGATCGGAA |
| 9827_2 | TATGAGTGTGGAGTCGTTGCATTTCGGGGGACTAAGGAGTTGANNNAGATCGGAA |
| 49558_1 | TATGAGTGTGGAGTCGTTGCGAGTAGCTGGGATTACAGGCATGNNNAGATCGGAA |
| 49558_2 | TATGAGTGTGGAGTCGTTGCGAGTAGCTGGGATTACAGGCATGNNNAGATCGGAA |
| 144_1 | TATGAGTGTGGAGTCGTTGCGTTTGTTGCTGTTGACTTTGAGGTNNNAGATCGGAA |

| | |
|---|---|
| 144_2 | TATGAGTGTGGAGTCGTTGCGTTTGTTGCTGTTGACTTTGAGGTNNNAGATCGGAA |
| 45729_1 | TATGAGTGTGGAGTCGTTGCGTGATGTGGAGCGGATGGAGNNNAGATCGGAAGA |
| 45729_2 | TATGAGTGTGGAGTCGTTGCGTGATGTGGAGCGGATGGAGNNNAGATCGGAAGA |
| 10573_1 | TATGAGTGTGGAGTCGTTGCTGGGGATAGGTGTGGAGAATGTTNNNAGATCGGA |
| 10573_2 | TATGAGTGTGGAGTCGTTGCTGGGGATAGGTGTGGAGAATGTTNNNAGATCGGA |
| 33038_1 | TATGAGTGTGGAGTCGTTGCTTCAAGTGATTCTCCTGCCTCAGNNNAGATCGGAAG |
| 33038_2 | TATGAGTGTGGAGTCGTTGCTTCAAGTGATTCTCCTGCCTCAGNNNAGATCGGAAG |
| 43957_1 | TATGAGTGTGGAGTCGTTGCGACTAGGGAAAGTTTGCTGCAATNNNAGATCGGAA |
| 43957_2 | TATGAGTGTGGAGTCGTTGCGACTAGGGAAAGTTTGCTGCAATNNNAGATCGGAA |
| 52185_1 | TATGAGTGTGGAGTCGTTGCATCAAGCAGACACTTCCCAGAATNNNAGATCGGAAG |
| 52185_2 | TATGAGTGTGGAGTCGTTGCATCAAGCAGACACTTCCCAGAATNNNAGATCGGAAG |
| 45199_1 | TATGAGTGTGGAGTCGTTGCCTCTTAGCTTTACAGCCCTTCCANNNAGATCGGAAGA |
| 45199_2 | TATGAGTGTGGAGTCGTTGCCTCTTAGCTTTACAGCCCTTCCANNNAGATCGGAAGA |
| 30972_1 | TATGAGTGTGGAGTCGTTGCAACGTCTTCTCTGCCACTCTAAGNNNAGATCGGAAG |
| 30972_2 | TATGAGTGTGGAGTCGTTGCAACGTCTTCTCTGCCACTCTAAGNNNAGATCGGAAG |
| 37553_1 | TATGAGTGTGGAGTCGTTGCTTCAAGGGAGGGAAAATGGAGTCNNNAGATCGGA |
| 37553_2 | TATGAGTGTGGAGTCGTTGCTTCAAGGGAGGGAAAATGGAGTCNNNAGATCGGA |
| 35603_1 | TATGAGTGTGGAGTCGTTGCTGCAATTGTTTTCAGAGTCTGCANNNAGATCGGAAG |
| 35603_2 | TATGAGTGTGGAGTCGTTGCTGCAATTGTTTTCAGAGTCTGCANNNAGATCGGAAG |
| 40290_1 | TATGAGTGTGGAGTCGTTGCTGGAGTGCAGTGGTACAATCTCNNNAGATCGGAAG |
| 40290_2 | TATGAGTGTGGAGTCGTTGCTGGAGTGCAGTGGTACAATCTCNNNAGATCGGAAG |
| 33794_1 | TATGAGTGTGGAGTCGTTGCTCTTGTGCCGGTTTTCAAAGAAANNNAGATCGGAAG |
| 33794_2 | TATGAGTGTGGAGTCGTTGCTCTTGTGCCGGTTTTCAAAGAAANNNAGATCGGAAG |
| 42812_1 | TATGAGTGTGGAGTCGTTGCTTTTGCAGCAATTGGATGGAACTNNNAGATCGGAA |
| 42812_2 | TATGAGTGTGGAGTCGTTGCTTTTGCAGCAATTGGATGGAACTNNNAGATCGGAA |
| 35930_1 | TATGAGTGTGGAGTCGTTGCCTTCCCTCATTGCAGTGAGAAAGNNNAGATCGGAAG |
| 35930_2 | TATGAGTGTGGAGTCGTTGCCTTCCCTCATTGCAGTGAGAAAGNNNAGATCGGAAG |
| 4272_1 | TATGAGTGTGGAGTCGTTGCCCTTTCTTTCTTGGTCGTGGGTANNNAGATCGGAAG |
| 4272_2 | TATGAGTGTGGAGTCGTTGCCCTTTCTTTCTTGGTCGTGGGTANNNAGATCGGAAG |
| 38958_1 | TATGAGTGTGGAGTCGTTGCGAACATGCTTCCATCACCTGAACNNNAGATCGGAAG |
| 38958_2 | TATGAGTGTGGAGTCGTTGCGAACATGCTTCCATCACCTGAACNNNAGATCGGAAG |
| 51557_1 | TATGAGTGTGGAGTCGTTGCGTGCATTTTGAGAAGGGGGTGNNNAGATCGGAAG |
| 51557_2 | TATGAGTGTGGAGTCGTTGCGTGCATTTTGAGAAGGGGGTGNNNAGATCGGAAG |
| 44735_1 | TATGAGTGTGGAGTCGTTGCTTATGAAGAGATCAGCGACCCTGNNNAGATCGGAA |
| 44735_2 | TATGAGTGTGGAGTCGTTGCTTATGAAGAGATCAGCGACCCTGNNNAGATCGGAA |
| 48978_1 | TATGAGTGTGGAGTCGTTGCGGATCAAGTAATTTGCCTGCCTCNNNAGATCGGAAG |
| 48978_2 | TATGAGTGTGGAGTCGTTGCGGATCAAGTAATTTGCCTGCCTCNNNAGATCGGAAG |
| 12494_1 | TATGAGTGTGGAGTCGTTGCGAACTTCATCTGCAAGTGAGNNNAGATCGGAAGAG |
| 12494_2 | TATGAGTGTGGAGTCGTTGCGAACTTCATCTGCAAGTGAGNNNAGATCGGAAGAG |
| 32080_1 | TATGAGTGTGGAGTCGTTGCAAAAGTTTCCAGGCATGGCTTTANNNAGATCGGAAG |
| 32080_2 | TATGAGTGTGGAGTCGTTGCAAAAGTTTCCAGGCATGGCTTTANNNAGATCGGAAG |
| 41743_1 | TATGAGTGTGGAGTCGTTGCTCAGGTCCATTTGATCCAGTGTTNNNAGATCGGAAG |
| 41743_2 | TATGAGTGTGGAGTCGTTGCTCAGGTCCATTTGATCCAGTGTTNNNAGATCGGAAG |

| | |
|---|---|
| 31358_1 | TATGAGTGTGGAGTCGTTGCTGTAAGACCAACTGTCCTAACGTNNNAGATCGGAAG |
| 31358_2 | TATGAGTGTGGAGTCGTTGCTGTAAGACCAACTGTCCTAACGTNNNAGATCGGAAG |
| 50648_1 | TATGAGTGTGGAGTCGTTGCGGCAGGGAAAAAGGGGGATTATANNNAGATCGGA |
| 50648_2 | TATGAGTGTGGAGTCGTTGCGGCAGGGAAAAAGGGGGATTATANNNAGATCGGA |
| 37440_1 | TATGAGTGTGGAGTCGTTGCATCAGCGAAACAATTGCAACTCANNNAGATCGGAAG |
| 37440_2 | TATGAGTGTGGAGTCGTTGCATCAGCGAAACAATTGCAACTCANNNAGATCGGAAG |
| 9483_1 | TATGAGTGTGGAGTCGTTGCAGGTCAGTCAGACAATCAGACAGNNNAGATCGGAA |
| 9483_2 | TATGAGTGTGGAGTCGTTGCAGGTCAGTCAGACAATCAGACAGNNNAGATCGGAA |
| 38574_1 | TATGAGTGTGGAGTCGTTGCTTTTGACAGAGTCTTGCTGTGTCNNNAGATCGGAAG |
| 38574_2 | TATGAGTGTGGAGTCGTTGCTTTTGACAGAGTCTTGCTGTGTCNNNAGATCGGAAG |
| 36140_1 | TATGAGTGTGGAGTCGTTGCACAACAGCAGAAGTTTCTTTGCANNNAGATCGGAAG |
| 36140_2 | TATGAGTGTGGAGTCGTTGCACAACAGCAGAAGTTTCTTTGCANNNAGATCGGAAG |
| 2731_1 | TATGAGTGTGGAGTCGTTGCCTACTCACTCTGCCTTTTGCACTNNNAGATCGGAAGA |
| 2731_2 | TATGAGTGTGGAGTCGTTGCCTACTCACTCTGCCTTTTGCACTNNNAGATCGGAAGA |
| 53080_1 | TATGAGTGTGGAGTCGTTGCTTTGATATGAAAGTGGCTGCAGANNNAGATCGGAA |
| 53080_2 | TATGAGTGTGGAGTCGTTGCTTTGATATGAAAGTGGCTGCAGANNNAGATCGGAA |
| 13558_1 | TATGAGTGTGGAGTCGTTGCGGCAGCAATGGAAAACTAATNNNAGATCGGAAGAG |
| 13558_2 | TATGAGTGTGGAGTCGTTGCGGCAGCAATGGAAAACTAATNNNAGATCGGAAGAG |
| 263_1 | TATGAGTGTGGAGTCGTTGCCAGTGTCAGCATTTTCCTGCTCNNNAGATCGGAAGA |
| 263_2 | TATGAGTGTGGAGTCGTTGCCAGTGTCAGCATTTTCCTGCTCNNNAGATCGGAAGA |
| 33773_1 | TATGAGTGTGGAGTCGTTGCTCATCAGAGTGAACAGACAACCTNNNAGATCGGAAG |
| 33773_2 | TATGAGTGTGGAGTCGTTGCTCATCAGAGTGAACAGACAACCTNNNAGATCGGAAG |
| 45269_1 | TATGAGTGTGGAGTCGTTGCCAAAGGTCCTAGCTAGATTGGGTNNNAGATCGGAA |
| 45269_2 | TATGAGTGTGGAGTCGTTGCCAAAGGTCCTAGCTAGATTGGGTNNNAGATCGGAA |
| 38041_1 | TATGAGTGTGGAGTCGTTGCTAGCATCTCAGCACTTTGGTCAANNNAGATCGGAAG |
| 38041_2 | TATGAGTGTGGAGTCGTTGCTAGCATCTCAGCACTTTGGTCAANNNAGATCGGAAG |
| 38040_1 | TATGAGTGTGGAGTCGTTGCTTGTTATGGGAACCACCTCACTTNNNAGATCGGAAG |
| 38040_2 | TATGAGTGTGGAGTCGTTGCTTGTTATGGGAACCACCTCACTTNNNAGATCGGAAG |
| 48954_1 | TATGAGTGTGGAGTCGTTGCGAGTAGCTGGGATTACAGGTGTGNNNAGATCGGAA |
| 48954_2 | TATGAGTGTGGAGTCGTTGCGAGTAGCTGGGATTACAGGTGTGNNNAGATCGGAA |
| 51153_1 | TATGAGTGTGGAGTCGTTGCCCCCCAAAAGAGGACAGATTGTANNNAGATCGGAA |
| 51153_2 | TATGAGTGTGGAGTCGTTGCCCCCCAAAAGAGGACAGATTGTANNNAGATCGGAA |
| 37648_1 | TATGAGTGTGGAGTCGTTGCTGATGGTCACAGGACTGGAAATTNNNAGATCGGAA |
| 37648_2 | TATGAGTGTGGAGTCGTTGCTGATGGTCACAGGACTGGAAATTNNNAGATCGGAA |
| 35507_1 | TATGAGTGTGGAGTCGTTGCTGGCCTGATTCAAAGAGATGTCANNNAGATCGGAA |
| 35507_2 | TATGAGTGTGGAGTCGTTGCTGGCCTGATTCAAAGAGATGTCANNNAGATCGGAA |
| 40846_1 | TATGAGTGTGGAGTCGTTGCCTCCAACTGGGTAACAGAGTGAGNNNAGATCGGAA |
| 40846_2 | TATGAGTGTGGAGTCGTTGCCTCCAACTGGGTAACAGAGTGAGNNNAGATCGGAA |
| 32909_1 | TATGAGTGTGGAGTCGTTGCTAGGCAAAGGACATGAACAGACANNNAGATCGGAA |
| 32909_2 | TATGAGTGTGGAGTCGTTGCTAGGCAAAGGACATGAACAGACANNNAGATCGGAA |
| 43355_1 | TATGAGTGTGGAGTCGTTGCCATTCTCTCACCTCAGCTTCTCANNNAGATCGGAAGA |
| 43355_2 | TATGAGTGTGGAGTCGTTGCCATTCTCTCACCTCAGCTTCTCANNNAGATCGGAAGA |
| 29497_1 | TATGAGTGTGGAGTCGTTGCAAAACTTTAGAATTCCATGTGGGNNNAGATCGGAAG |

| | |
|---|---|
| 29497_2 | TATGAGTGTGGAGTCGTTGCAAAACTTTAGAATTCCATGTGGGNNNAGATCGGAAG |
| 41612_1 | TATGAGTGTGGAGTCGTTGCCACGGAGTTAGGACTAAGGCATGNNNAGATCGGAA |
| 41612_2 | TATGAGTGTGGAGTCGTTGCCACGGAGTTAGGACTAAGGCATGNNNAGATCGGAA |
| 2529_1 | TATGAGTGTGGAGTCGTTGCGACAGCTAGGAGCAAGATGAAGGNNNAGATCGGAA |
| 2529_2 | TATGAGTGTGGAGTCGTTGCGACAGCTAGGAGCAAGATGAAGGNNNAGATCGGAA |
| 47914_1 | TATGAGTGTGGAGTCGTTGCTTTGTCCCTCTCTATCTACCGGTNNNAGATCGGAAGA |
| 47914_2 | TATGAGTGTGGAGTCGTTGCTTTGTCCCTCTCTATCTACCGGTNNNAGATCGGAAGA |
| 12948_1 | TATGAGTGTGGAGTCGTTGCTGGAAGAATAGATACAAGCATAANNNAGATCGGAA |
| 12948_2 | TATGAGTGTGGAGTCGTTGCTGGAAGAATAGATACAAGCATAANNNAGATCGGAA |
| 47617_1 | TATGAGTGTGGAGTCGTTGCTGTCAAGGTCAGAGGCCAATAAANNNAGATCGGAA |
| 47617_2 | TATGAGTGTGGAGTCGTTGCTGTCAAGGTCAGAGGCCAATAAANNNAGATCGGAA |
| 30698_1 | TATGAGTGTGGAGTCGTTGCAGAGACACACACAAGAGGAAGGCNNNAGATCGGAA |
| 30698_2 | TATGAGTGTGGAGTCGTTGCAGAGACACACACAAGAGGAAGGCNNNAGATCGGAA |
| 34254_1 | TATGAGTGTGGAGTCGTTGCCTGAGTGAGCAGTAACGAAGGTNNNAGATCGGAAG |
| 34254_2 | TATGAGTGTGGAGTCGTTGCCTGAGTGAGCAGTAACGAAGGTNNNAGATCGGAAG |
| 6146_1 | TATGAGTGTGGAGTCGTTGCTGTAGCCATTCATTTTCCACCCTNNNAGATCGGAAGA |
| 6146_2 | TATGAGTGTGGAGTCGTTGCTGTAGCCATTCATTTTCCACCCTNNNAGATCGGAAGA |
| 39268_1 | TATGAGTGTGGAGTCGTTGCCTAGGCAACAAAGCAAGACTCCANNNAGATCGGAAG |
| 39268_2 | TATGAGTGTGGAGTCGTTGCCTAGGCAACAAAGCAAGACTCCANNNAGATCGGAAG |
| 42061_1 | TATGAGTGTGGAGTCGTTGCCCTGAAGTGAAGGATACCCTGTCNNNAGATCGGAAG |
| 42061_2 | TATGAGTGTGGAGTCGTTGCCCTGAAGTGAAGGATACCCTGTCNNNAGATCGGAAG |
| 42408_1 | TATGAGTGTGGAGTCGTTGCCAGGGTCTCACTATGTTACCCAGNNNAGATCGGAAG |
| 42408_2 | TATGAGTGTGGAGTCGTTGCCAGGGTCTCACTATGTTACCCAGNNNAGATCGGAAG |
| 40180_1 | TATGAGTGTGGAGTCGTTGCGAAAGTAGATTTTGGAGCTGGGCNNNAGATCGGAA |
| 40180_2 | TATGAGTGTGGAGTCGTTGCGAAAGTAGATTTTGGAGCTGGGCNNNAGATCGGAA |
| 45711_1 | TATGAGTGTGGAGTCGTTGCGAGTGCAGTGGTGTGATCTCAGNNNAGATCGGAAG |
| 45711_2 | TATGAGTGTGGAGTCGTTGCGAGTGCAGTGGTGTGATCTCAGNNNAGATCGGAAG |
| 52835_1 | TATGAGTGTGGAGTCGTTGCCATGAGCTTGGGGTTCCTGTNNNAGATCGGAAGAG |
| 52835_2 | TATGAGTGTGGAGTCGTTGCCATGAGCTTGGGGTTCCTGTNNNAGATCGGAAGAG |
| 40208_1 | TATGAGTGTGGAGTCGTTGCACAGCCCTTCCCATTATCAACATNNNAGATCGGAAGA |
| 40208_2 | TATGAGTGTGGAGTCGTTGCACAGCCCTTCCCATTATCAACATNNNAGATCGGAAGA |
| 43159_1 | TATGAGTGTGGAGTCGTTGCCCACGCCCAGCCAATTTTTATATNNNAGATCGGAAGA |
| 43159_2 | TATGAGTGTGGAGTCGTTGCCCACGCCCAGCCAATTTTTATATNNNAGATCGGAAGA |
| 50737_1 | TATGAGTGTGGAGTCGTTGCAAGGACATGTTCCCAGGAGTATGNNNAGATCGGAA |
| 50737_2 | TATGAGTGTGGAGTCGTTGCAAGGACATGTTCCCAGGAGTATGNNNAGATCGGAA |
| 41309_1 | TATGAGTGTGGAGTCGTTGCCCCCATGTGCTTAGATTGGAAGANNNAGATCGGAAG |
| 41309_2 | TATGAGTGTGGAGTCGTTGCCCCCATGTGCTTAGATTGGAAGANNNAGATCGGAAG |
| 40252_1 | TATGAGTGTGGAGTCGTTGCGGGAAGGAGACAAGCAAATTGTGNNNAGATCGGAA |
| 40252_2 | TATGAGTGTGGAGTCGTTGCGGGAAGGAGACAAGCAAATTGTGNNNAGATCGGAA |
| 42412_1 | TATGAGTGTGGAGTCGTTGCTTCTCCCCTCTCCTTCCACTTATNNNAGATCGGAAGAG |
| 42412_2 | TATGAGTGTGGAGTCGTTGCTTCTCCCCTCTCCTTCCACTTATNNNAGATCGGAAGAG |
| 32101_1 | TATGAGTGTGGAGTCGTTGCGTCCTGGCTTTTTTGGGGTTATTNNNAGATCGGAAG |
| 32101_2 | TATGAGTGTGGAGTCGTTGCGTCCTGGCTTTTTTGGGGTTATTNNNAGATCGGAAG |

| | |
|---|---|
| 34681_1 | TATGAGTGTGGAGTCGTTGCGCGAGACTCTGCCATGTAAAAAANNNAGATCGGAA |
| 34681_2 | TATGAGTGTGGAGTCGTTGCGCGAGACTCTGCCATGTAAAAAANNNAGATCGGAA |
| 48723_1 | TATGAGTGTGGAGTCGTTGCCAGATCACTTGAGGTCAGGAGTTNNNAGATCGGAA |
| 48723_2 | TATGAGTGTGGAGTCGTTGCCAGATCACTTGAGGTCAGGAGTTNNNAGATCGGAA |
| 37331_1 | TATGAGTGTGGAGTCGTTGCGCCTTCATTTGTCTGTCTTCACCNNNAGATCGGAAGA |
| 37331_2 | TATGAGTGTGGAGTCGTTGCGCCTTCATTTGTCTGTCTTCACCNNNAGATCGGAAGA |
| 39877_1 | TATGAGTGTGGAGTCGTTGCTGAAACCCCGTCTCTACCAAAAANNNAGATCGGAAG |
| 39877_2 | TATGAGTGTGGAGTCGTTGCTGAAACCCCGTCTCTACCAAAAANNNAGATCGGAAG |
| 49896_1 | TATGAGTGTGGAGTCGTTGCCTCATGCCTATAATCCCAGCACTNNNAGATCGGAAG |
| 49896_2 | TATGAGTGTGGAGTCGTTGCCTCATGCCTATAATCCCAGCACTNNNAGATCGGAAG |
| 39315_1 | TATGAGTGTGGAGTCGTTGCGAGCAAGACTCTGTCTCAAAAAANNNAGATCGGAA |
| 39315_2 | TATGAGTGTGGAGTCGTTGCGAGCAAGACTCTGTCTCAAAAAANNNAGATCGGAA |
| 46387_1 | TATGAGTGTGGAGTCGTTGCCCACCATGCCCAGCTAATTTTTTNNNAGATCGGAAGA |
| 46387_2 | TATGAGTGTGGAGTCGTTGCCCACCATGCCCAGCTAATTTTTTNNNAGATCGGAAGA |
| 32932_1 | TATGAGTGTGGAGTCGTTGCCACGAGGTCCAGAGATTGAGACNNNAGATCGGAAG |
| 32932_2 | TATGAGTGTGGAGTCGTTGCCACGAGGTCCAGAGATTGAGACNNNAGATCGGAAG |
| 49994_1 | TATGAGTGTGGAGTCGTTGCTATCATTTCAGGTGCATGGAGGTNNNAGATCGGAA |
| 49994_2 | TATGAGTGTGGAGTCGTTGCTATCATTTCAGGTGCATGGAGGTNNNAGATCGGAA |
| 43407_1 | TATGAGTGTGGAGTCGTTGCAGAGTGGGAAAAGATGAAAGCCTNNNAGATCGGA |
| 43407_2 | TATGAGTGTGGAGTCGTTGCAGAGTGGGAAAAGATGAAAGCCTNNNAGATCGGA |
| 39255_1 | TATGAGTGTGGAGTCGTTGCTGTAATCCCAGCTACTCAGGAGANNNAGATCGGAAG |
| 39255_2 | TATGAGTGTGGAGTCGTTGCTGTAATCCCAGCTACTCAGGAGANNNAGATCGGAAG |
| 35512_1 | TATGAGTGTGGAGTCGTTGCATGATCTCAGCTCACTGCAATCTNNNAGATCGGAAG |
| 35512_2 | TATGAGTGTGGAGTCGTTGCATGATCTCAGCTCACTGCAATCTNNNAGATCGGAAG |
| 33463_1 | TATGAGTGTGGAGTCGTTGCAATTCAGCTGTTTTGCTTTACGCNNNAGATCGGAAG |
| 33463_2 | TATGAGTGTGGAGTCGTTGCAATTCAGCTGTTTTGCTTTACGCNNNAGATCGGAAG |
| 34059_1 | TATGAGTGTGGAGTCGTTGCAGTGCTGGGATTATAGGCATGAGNNNAGATCGGAA |
| 34059_2 | TATGAGTGTGGAGTCGTTGCAGTGCTGGGATTATAGGCATGAGNNNAGATCGGAA |
| 13672_1 | TATGAGTGTGGAGTCGTTGCTAAAAGACGAGGGAGATCCTNNNAGATCGGAAGAG |
| 13672_2 | TATGAGTGTGGAGTCGTTGCTAAAAGACGAGGGAGATCCTNNNAGATCGGAAGAG |
| 32584_1 | TATGAGTGTGGAGTCGTTGCGAGACGGAGTCTTGCTCTGTCNNNAGATCGGAAGA |
| 32584_2 | TATGAGTGTGGAGTCGTTGCGAGACGGAGTCTTGCTCTGTCNNNAGATCGGAAGA |
| 46578_1 | TATGAGTGTGGAGTCGTTGCAAGTAGCTGGGATTACAAGTGCANNNAGATCGGAA |
| 46578_2 | TATGAGTGTGGAGTCGTTGCAAGTAGCTGGGATTACAAGTGCANNNAGATCGGAA |
| 12817_1 | TATGAGTGTGGAGTCGTTGCGAAACCTAGTCACATTGGAGNNNAGATCGGAAGAG |
| 12817_2 | TATGAGTGTGGAGTCGTTGCGAAACCTAGTCACATTGGAGNNNAGATCGGAAGAG |
| 38370_1 | TATGAGTGTGGAGTCGTTGCGCTAACCATCCAAGTTCCTGGTANNNAGATCGGAAG |
| 38370_2 | TATGAGTGTGGAGTCGTTGCGCTAACCATCCAAGTTCCTGGTANNNAGATCGGAAG |
| 36563_1 | TATGAGTGTGGAGTCGTTGCCCTCCCGAGTCTTTTCTATGCATNNNAGATCGGAAGA |
| 36563_2 | TATGAGTGTGGAGTCGTTGCCCTCCCGAGTCTTTTCTATGCATNNNAGATCGGAAGA |
| 42262_1 | TATGAGTGTGGAGTCGTTGCGCACTCTGCTCTTCTTGGATCTANNNAGATCGGAAG |
| 42262_2 | TATGAGTGTGGAGTCGTTGCGCACTCTGCTCTTCTTGGATCTANNNAGATCGGAAG |
| 41317_1 | TATGAGTGTGGAGTCGTTGCTCAAAACATTCCAAATGGTATCTGTNNNAGATCGGA |

| | |
|---|---|
| 41317_2 | TATGAGTGTGGAGTCGTTGCTCAAAACATTCCAAATGGTATCTGTNNNAGATCGGA |
| 35420_1 | TATGAGTGTGGAGTCGTTGCAGATGAGGGTCACACTATGTTGCNNNAGATCGGAA |
| 35420_2 | TATGAGTGTGGAGTCGTTGCAGATGAGGGTCACACTATGTTGCNNNAGATCGGAA |
| 13345_1 | TATGAGTGTGGAGTCGTTGCACATTGAAGATGCTTTTTTTTTNNNAGATCGGAAG |
| 13345_2 | TATGAGTGTGGAGTCGTTGCACATTGAAGATGCTTTTTTTTTNNNAGATCGGAAG |
| 32159_1 | TATGAGTGTGGAGTCGTTGCCCTCCCAAAGTGCTGAGATTACANNNAGATCGGAAG |
| 32159_2 | TATGAGTGTGGAGTCGTTGCCCTCCCAAAGTGCTGAGATTACANNNAGATCGGAAG |
| 10442_1 | TATGAGTGTGGAGTCGTTGCATACTCCCTTGGGTAACTCCACCNNNAGATCGGAAG |
| 10442_2 | TATGAGTGTGGAGTCGTTGCATACTCCCTTGGGTAACTCCACCNNNAGATCGGAAG |
| 44834_1 | TATGAGTGTGGAGTCGTTGCGGTGCAAGGGAGACAGCTTGNNNAGATCGGAAGA |
| 44834_2 | TATGAGTGTGGAGTCGTTGCGGTGCAAGGGAGACAGCTTGNNNAGATCGGAAGA |
| 39980_1 | TATGAGTGTGGAGTCGTTGCACATGGGAGGGAGACGTTATAGANNNAGATCGGA |
| 39980_2 | TATGAGTGTGGAGTCGTTGCACATGGGAGGGAGACGTTATAGANNNAGATCGGA |
| 53194_1 | TATGAGTGTGGAGTCGTTGCCTCTCTCTCTGACAGGATCTTGCNNNAGATCGGAAG |
| 53194_2 | TATGAGTGTGGAGTCGTTGCCTCTCTCTCTGACAGGATCTTGCNNNAGATCGGAAG |
| 29605_1 | TATGAGTGTGGAGTCGTTGCGGGACATTCGCCTCTTAACAATGNNNAGATCGGAAG |
| 29605_2 | TATGAGTGTGGAGTCGTTGCGGGACATTCGCCTCTTAACAATGNNNAGATCGGAAG |
| 46301_1 | TATGAGTGTGGAGTCGTTGCCTCCTGAGTAGCTGGAACTACAGNNNAGATCGGAA |
| 46301_2 | TATGAGTGTGGAGTCGTTGCCTCCTGAGTAGCTGGAACTACAGNNNAGATCGGAA |
| 31193_1 | TATGAGTGTGGAGTCGTTGCAAGGTAGGTTTCTGAATGGTCCCNNNAGATCGGAA |
| 31193_2 | TATGAGTGTGGAGTCGTTGCAAGGTAGGTTTCTGAATGGTCCCNNNAGATCGGAA |
| 30070_1 | TATGAGTGTGGAGTCGTTGCATTGTCCTGTCCCACTCACTTACNNNAGATCGGAAGA |
| 30070_2 | TATGAGTGTGGAGTCGTTGCATTGTCCTGTCCCACTCACTTACNNNAGATCGGAAGA |
| 43101_1 | TATGAGTGTGGAGTCGTTGCTCAACAAAGCCAAACAAAGACGTNNNAGATCGGAA |
| 43101_2 | TATGAGTGTGGAGTCGTTGCTCAACAAAGCCAAACAAAGACGTNNNAGATCGGAA |
| 38901_1 | TATGAGTGTGGAGTCGTTGCGCATTGGTAGCTCACACATCAAANNNAGATCGGAAG |
| 38901_2 | TATGAGTGTGGAGTCGTTGCGCATTGGTAGCTCACACATCAAANNNAGATCGGAAG |
| 51390_1 | TATGAGTGTGGAGTCGTTGCTGGCCATTTGAAGCAATAACGTTNNNAGATCGGAAG |
| 51390_2 | TATGAGTGTGGAGTCGTTGCTGGCCATTTGAAGCAATAACGTTNNNAGATCGGAAG |
| 33592_1 | TATGAGTGTGGAGTCGTTGCGGGTTCAAGTGATTCTCCTGTCTNNNAGATCGGAAG |
| 33592_2 | TATGAGTGTGGAGTCGTTGCGGGTTCAAGTGATTCTCCTGTCTNNNAGATCGGAAG |
| 12133_1 | TATGAGTGTGGAGTCGTTGCAGTGGGATGGATGTCTCGATTCCNNNAGATCGGAA |
| 12133_2 | TATGAGTGTGGAGTCGTTGCAGTGGGATGGATGTCTCGATTCCNNNAGATCGGAA |
| 49046_1 | TATGAGTGTGGAGTCGTTGCGATCGAATGTGTCTGATGCAAGGNNNAGATCGGAA |
| 49046_2 | TATGAGTGTGGAGTCGTTGCGATCGAATGTGTCTGATGCAAGGNNNAGATCGGAA |
| 40693_1 | TATGAGTGTGGAGTCGTTGCTATGTGCCAATGTTCATAGCAGCNNNAGATCGGAAG |
| 40693_2 | TATGAGTGTGGAGTCGTTGCTATGTGCCAATGTTCATAGCAGCNNNAGATCGGAAG |
| 33026_1 | TATGAGTGTGGAGTCGTTGCTCTCCCGGGTTCAAATGATTCTCNNNAGATCGGAAG |
| 33026_2 | TATGAGTGTGGAGTCGTTGCTCTCCCGGGTTCAAATGATTCTCNNNAGATCGGAAG |
| 50555_1 | TATGAGTGTGGAGTCGTTGCTATTGGTTGGTCTCTCAGAGCTCNNNAGATCGGAAG |
| 50555_2 | TATGAGTGTGGAGTCGTTGCTATTGGTTGGTCTCTCAGAGCTCNNNAGATCGGAAG |
| 50952_1 | TATGAGTGTGGAGTCGTTGCTACATTCTGCCTCAGTGCTTTCTNNNAGATCGGAAGA |
| 50952_2 | TATGAGTGTGGAGTCGTTGCTACATTCTGCCTCAGTGCTTTCTNNNAGATCGGAAGA |

| | |
|---|---|
| 50645_1 | TATGAGTGTGGAGTCGTTGCGCACACACAGAGGTTCCTAAAAANNNAGATCGGAA |
| 50645_2 | TATGAGTGTGGAGTCGTTGCGCACACACAGAGGTTCCTAAAAANNNAGATCGGAA |
| 52913_1 | TATGAGTGTGGAGTCGTTGCCACTCAAGATCCCAAACTCCAGANNNAGATCGGAAG |
| 52913_2 | TATGAGTGTGGAGTCGTTGCCACTCAAGATCCCAAACTCCAGANNNAGATCGGAAG |
| 48104_1 | TATGAGTGTGGAGTCGTTGCCCTGTCTTTGGTTTGCTAAGGTTNNNAGATCGGAAG |
| 48104_2 | TATGAGTGTGGAGTCGTTGCCCTGTCTTTGGTTTGCTAAGGTTNNNAGATCGGAAG |
| 52951_1 | TATGAGTGTGGAGTCGTTGCCAAGCACTGAGATGGAGGTCTCNNNAGATCGGAAG |
| 52951_2 | TATGAGTGTGGAGTCGTTGCCAAGCACTGAGATGGAGGTCTCNNNAGATCGGAAG |
| 33246_1 | TATGAGTGTGGAGTCGTTGCGATTCTCCTACCTCAGCCTTCTGNNNAGATCGGAAGA |
| 33246_2 | TATGAGTGTGGAGTCGTTGCGATTCTCCTACCTCAGCCTTCTGNNNAGATCGGAAGA |
| 46427_1 | TATGAGTGTGGAGTCGTTGCATGATCTCAGTTCACTGCAACCTNNNAGATCGGAAG |
| 46427_2 | TATGAGTGTGGAGTCGTTGCATGATCTCAGTTCACTGCAACCTNNNAGATCGGAAG |
| 9329_1 | TATGAGTGTGGAGTCGTTGCATCCTTGTGATTTGGGGTTTGGGNNNAGATCGGAA |
| 9329_2 | TATGAGTGTGGAGTCGTTGCATCCTTGTGATTTGGGGTTTGGGNNNAGATCGGAA |
| 41821_1 | TATGAGTGTGGAGTCGTTGCCAGTAGCTGGGATTACAGGCATGNNNAGATCGGAA |
| 41821_2 | TATGAGTGTGGAGTCGTTGCCAGTAGCTGGGATTACAGGCATGNNNAGATCGGAA |
| 49349_1 | TATGAGTGTGGAGTCGTTGCTGGGGGCATATACACAGTAATGGNNNAGATCGGAA |
| 49349_2 | TATGAGTGTGGAGTCGTTGCTGGGGGCATATACACAGTAATGGNNNAGATCGGAA |
| 31790_1 | TATGAGTGTGGAGTCGTTGCAGGAGAGACAATGGTGTAAGTTCCNNNAGATCGGA |
| 31790_2 | TATGAGTGTGGAGTCGTTGCAGGAGAGACAATGGTGTAAGTTCCNNNAGATCGGA |
| 30499_1 | TATGAGTGTGGAGTCGTTGCAGCGGTCAATGAGCATCTGAAATNNNAGATCGGAA |
| 30499_2 | TATGAGTGTGGAGTCGTTGCAGCGGTCAATGAGCATCTGAAATNNNAGATCGGAA |
| 37147_1 | TATGAGTGTGGAGTCGTTGCCTCAAACCCTCTAAAGCCTCAGTNNNAGATCGGAAG |
| 37147_2 | TATGAGTGTGGAGTCGTTGCCTCAAACCCTCTAAAGCCTCAGTNNNAGATCGGAAG |
| 46408_1 | TATGAGTGTGGAGTCGTTGCGTTCAAGCAATTCTCTGCCTCAGNNNAGATCGGAAG |
| 46408_2 | TATGAGTGTGGAGTCGTTGCGTTCAAGCAATTCTCTGCCTCAGNNNAGATCGGAAG |
| 13176_1 | TATGAGTGTGGAGTCGTTGCTACATTTGACTCGGAGAAAAATNNNAGATCGGAAG |
| 13176_2 | TATGAGTGTGGAGTCGTTGCTACATTTGACTCGGAGAAAAATNNNAGATCGGAAG |
| 44032_1 | TATGAGTGTGGAGTCGTTGCTCTAACAGTCTATGGCACACTGGNNNAGATCGGAAG |
| 44032_2 | TATGAGTGTGGAGTCGTTGCTCTAACAGTCTATGGCACACTGGNNNAGATCGGAAG |
| 34617_1 | TATGAGTGTGGAGTCGTTGCTCTGTCTGAAACAAACACACACGNNNAGATCGGAAG |
| 34617_2 | TATGAGTGTGGAGTCGTTGCTCTGTCTGAAACAAACACACACGNNNAGATCGGAAG |
| 36884_1 | TATGAGTGTGGAGTCGTTGCCAAAAACCAAACCAAACCAAACCANNNAGATCGGAA |
| 36884_2 | TATGAGTGTGGAGTCGTTGCCAAAAACCAAACCAAACCAAACCANNNAGATCGGAA |
| 31262_1 | TATGAGTGTGGAGTCGTTGCCACAAGCACATACACATCAGCTCNNNAGATCGGAAG |
| 31262_2 | TATGAGTGTGGAGTCGTTGCCACAAGCACATACACATCAGCTCNNNAGATCGGAAG |
| 10797_1 | TATGAGTGTGGAGTCGTTGCAAAACACACCACCAAAATGCTGGNNNAGATCGGAAG |
| 10797_2 | TATGAGTGTGGAGTCGTTGCAAAACACACCACCAAAATGCTGGNNNAGATCGGAAG |
| 51938_1 | TATGAGTGTGGAGTCGTTGCGGGCAACAGTGACTTTGATTACCNNNAGATCGGAA |
| 51938_2 | TATGAGTGTGGAGTCGTTGCGGGCAACAGTGACTTTGATTACCNNNAGATCGGAA |
| 41353_1 | TATGAGTGTGGAGTCGTTGCAGGTCAAGAAAGCTATGTGGGAGNNNAGATCGGA |
| 41353_2 | TATGAGTGTGGAGTCGTTGCAGGTCAAGAAAGCTATGTGGGAGNNNAGATCGGA |
| 52933_1 | TATGAGTGTGGAGTCGTTGCGTGTCCAAGGACTTTTTCGTCTGNNNAGATCGGAAG |

| | |
|---|---|
| 52933_2 | TATGAGTGTGGAGTCGTTGCGTGTCCAAGGACTTTTCGTCTGNNNAGATCGGAAG |
| 42947_1 | TATGAGTGTGGAGTCGTTGCTAGGTCTTTCACCAAGTCCACAGNNNAGATCGGAAG |
| 42947_2 | TATGAGTGTGGAGTCGTTGCTAGGTCTTTCACCAAGTCCACAGNNNAGATCGGAAG |
| 36128_1 | TATGAGTGTGGAGTCGTTGCGAGAGCAGTTTCATGCATCCATCNNNAGATCGGAAG |
| 36128_2 | TATGAGTGTGGAGTCGTTGCGAGAGCAGTTTCATGCATCCATCNNNAGATCGGAAG |
| 38372_1 | TATGAGTGTGGAGTCGTTGCGCAAAGGAGAAGGACTACTTTCCNNNAGATCGGAA |
| 38372_2 | TATGAGTGTGGAGTCGTTGCGCAAAGGAGAAGGACTACTTTCCNNNAGATCGGAA |
| 10480_1 | TATGAGTGTGGAGTCGTTGCCAGACAGAGTGAAAGGCCTTGAANNNAGATCGGAA |
| 10480_2 | TATGAGTGTGGAGTCGTTGCCAGACAGAGTGAAAGGCCTTGAANNNAGATCGGAA |
| 2219_1 | TATGAGTGTGGAGTCGTTGCAAATGACTGTGCTAACAGCCTGANNNAGATCGGAA |
| 2219_2 | TATGAGTGTGGAGTCGTTGCAAATGACTGTGCTAACAGCCTGANNNAGATCGGAA |
| 52592_1 | TATGAGTGTGGAGTCGTTGCTTTCTTTCACTCTGAGAGGCTCCNNNAGATCGGAAG |
| 52592_2 | TATGAGTGTGGAGTCGTTGCTTTCTTTCACTCTGAGAGGCTCCNNNAGATCGGAAG |
| 49709_1 | TATGAGTGTGGAGTCGTTGCTGTGACTACAGGTGAACATCCAGNNNAGATCGGAA |
| 49709_2 | TATGAGTGTGGAGTCGTTGCTGTGACTACAGGTGAACATCCAGNNNAGATCGGAA |
| 47850_1 | TATGAGTGTGGAGTCGTTGCTCCGATAATGAGCTCAGTCAACANNNAGATCGGAAG |
| 47850_2 | TATGAGTGTGGAGTCGTTGCTCCGATAATGAGCTCAGTCAACANNNAGATCGGAAG |
| 37663_1 | TATGAGTGTGGAGTCGTTGCTGTCACTTTTCTGCTCTAGGTGANNNAGATCGGAAG |
| 37663_2 | TATGAGTGTGGAGTCGTTGCTGTCACTTTTCTGCTCTAGGTGANNNAGATCGGAAG |
| 29475_1 | TATGAGTGTGGAGTCGTTGCGGATCTCTACGACTGAAGAGCCTNNNAGATCGGAA |
| 29475_2 | TATGAGTGTGGAGTCGTTGCGGATCTCTACGACTGAAGAGCCTNNNAGATCGGAA |
| 51843_1 | TATGAGTGTGGAGTCGTTGCAAGGGAGCTTGAACTTAGAGTGGNNNAGATCGGAA |
| 51843_2 | TATGAGTGTGGAGTCGTTGCAAGGGAGCTTGAACTTAGAGTGGNNNAGATCGGAA |
| 48130_1 | TATGAGTGTGGAGTCGTTGCCTCAGGAATCCAGAGGGAGCTCNNNAGATCGGAAG |
| 48130_2 | TATGAGTGTGGAGTCGTTGCCTCAGGAATCCAGAGGGAGCTCNNNAGATCGGAAG |
| 32722_1 | TATGAGTGTGGAGTCGTTGCCCAGTGGATCATGAGATCAGGAGNNNAGATCGGAA |
| 32722_2 | TATGAGTGTGGAGTCGTTGCCCAGTGGATCATGAGATCAGGAGNNNAGATCGGAA |
| 48737_1 | TATGAGTGTGGAGTCGTTGCTTCAAGAGATTCTCCTGCCTCAGNNNAGATCGGAAG |
| 48737_2 | TATGAGTGTGGAGTCGTTGCTTCAAGAGATTCTCCTGCCTCAGNNNAGATCGGAAG |
| 42894_1 | TATGAGTGTGGAGTCGTTGCGTGTGAAGAGAACACCAAACAGGNNNAGATCGGAA |
| 42894_2 | TATGAGTGTGGAGTCGTTGCGTGTGAAGAGAACACCAAACAGGNNNAGATCGGAA |
| 35544_1 | TATGAGTGTGGAGTCGTTGCGTTTCACCATGTTAGCCAGGATGNNNAGATCGGAAG |
| 35544_2 | TATGAGTGTGGAGTCGTTGCGTTTCACCATGTTAGCCAGGATGNNNAGATCGGAAG |
| 46756_1 | TATGAGTGTGGAGTCGTTGCCTTTCAAATCAAAGGGAAGCCGTNNNAGATCGGAA |
| 46756_2 | TATGAGTGTGGAGTCGTTGCCTTTCAAATCAAAGGGAAGCCGTNNNAGATCGGAA |
| 51419_1 | TATGAGTGTGGAGTCGTTGCGAAGCAAGGTCAAAGGTGAACTGNNNAGATCGGAA |
| 51419_2 | TATGAGTGTGGAGTCGTTGCGAAGCAAGGTCAAAGGTGAACTGNNNAGATCGGAA |
| 34684_1 | TATGAGTGTGGAGTCGTTGCCCTCTTCCCTGGCACATTATCTTNNNAGATCGGAAGA |
| 34684_2 | TATGAGTGTGGAGTCGTTGCCCTCTTCCCTGGCACATTATCTTNNNAGATCGGAAGA |
| 37681_1 | TATGAGTGTGGAGTCGTTGCCACTCAGCATGGGTAACAAAGTGNNNAGATCGGAA |
| 37681_2 | TATGAGTGTGGAGTCGTTGCCACTCAGCATGGGTAACAAAGTGNNNAGATCGGAA |
| 39282_1 | TATGAGTGTGGAGTCGTTGCGGAAGTATTGATGGCATGTGCTGNNNAGATCGGAA |
| 39282_2 | TATGAGTGTGGAGTCGTTGCGGAAGTATTGATGGCATGTGCTGNNNAGATCGGAA |

| | |
|---|---|
| 41052_1 | TATGAGTGTGGAGTCGTTGCCGTTTATACACTGCTAGTGGGCANNNAGATCGGAAG |
| 41052_2 | TATGAGTGTGGAGTCGTTGCCGTTTATACACTGCTAGTGGGCANNNAGATCGGAAG |
| 29192_1 | TATGAGTGTGGAGTCGTTGCAAACCAACCAACCAAGATGGAACNNNAGATCGGAAG |
| 29192_2 | TATGAGTGTGGAGTCGTTGCAAACCAACCAACCAAGATGGAACNNNAGATCGGAAG |
| 43472_1 | TATGAGTGTGGAGTCGTTGCCACCACGCCCACTAATTTTTGTANNNAGATCGGAAGA |
| 43472_2 | TATGAGTGTGGAGTCGTTGCCACCACGCCCACTAATTTTTGTANNNAGATCGGAAGA |
| 37766_1 | TATGAGTGTGGAGTCGTTGCTTTGAAAAGGCCTGCTTACAAGGNNNAGATCGGAAG |
| 37766_2 | TATGAGTGTGGAGTCGTTGCTTTGAAAAGGCCTGCTTACAAGGNNNAGATCGGAAG |
| 9017_1 | TATGAGTGTGGAGTCGTTGCCAACCCAGTAAGATTGCCTTGCTNNNAGATCGGAAGA |
| 9017_2 | TATGAGTGTGGAGTCGTTGCCAACCCAGTAAGATTGCCTTGCTNNNAGATCGGAAGA |
| 34527_1 | TATGAGTGTGGAGTCGTTGCCCTGTAATCCTAGCACTTTGGGANNNAGATCGGAAG |
| 34527_2 | TATGAGTGTGGAGTCGTTGCCCTGTAATCCTAGCACTTTGGGANNNAGATCGGAAG |
| 49485_1 | TATGAGTGTGGAGTCGTTGCGGACTACAGGAGCATGCTACAATNNNAGATCGGAAG |
| 49485_2 | TATGAGTGTGGAGTCGTTGCGGACTACAGGAGCATGCTACAATNNNAGATCGGAAG |
| 10538_1 | TATGAGTGTGGAGTCGTTGCCCCTCAATGGACTACAGCTGTTTNNNAGATCGGAAGA |
| 10538_2 | TATGAGTGTGGAGTCGTTGCCCCTCAATGGACTACAGCTGTTTNNNAGATCGGAAGA |
| 30457_1 | TATGAGTGTGGAGTCGTTGCGAAATGCCTGATGAAGGGAAGTTTTNNNAGATCGGA |
| 30457_2 | TATGAGTGTGGAGTCGTTGCGAAATGCCTGATGAAGGGAAGTTTTNNNAGATCGGA |
| 38766_1 | TATGAGTGTGGAGTCGTTGCATAGATACTGCGGACTACTGGGTNNNAGATCGGAAG |
| 38766_2 | TATGAGTGTGGAGTCGTTGCATAGATACTGCGGACTACTGGGTNNNAGATCGGAAG |
| 52160_1 | TATGAGTGTGGAGTCGTTGCGAGGTGTACCCAACAGCTCATTNNNAGATCGGAAGA |
| 52160_2 | TATGAGTGTGGAGTCGTTGCGAGGTGTACCCAACAGCTCATTNNNAGATCGGAAGA |
| 13719_1 | TATGAGTGTGGAGTCGTTGCTTGTATGTTAATTTCCTAACCAGNNNAGATCGGAAGA |
| 13719_2 | TATGAGTGTGGAGTCGTTGCTTGTATGTTAATTTCCTAACCAGNNNAGATCGGAAGA |
| 51053_1 | TATGAGTGTGGAGTCGTTGCGCCTGGAGATTGAGTTATGTGGANNNAGATCGGAA |
| 51053_2 | TATGAGTGTGGAGTCGTTGCGCCTGGAGATTGAGTTATGTGGANNNAGATCGGAA |
| 33206_1 | TATGAGTGTGGAGTCGTTGCTACTAGTCAGGTCCTGAGGGTTTNNNAGATCGGAAG |
| 33206_2 | TATGAGTGTGGAGTCGTTGCTACTAGTCAGGTCCTGAGGGTTTNNNAGATCGGAAG |
| 32993_1 | TATGAGTGTGGAGTCGTTGCGGTTCAAACATTTCTCCTGCCTCNNNAGATCGGAAGA |
| 32993_2 | TATGAGTGTGGAGTCGTTGCGGTTCAAACATTTCTCCTGCCTCNNNAGATCGGAAGA |
| 298_1 | TATGAGTGTGGAGTCGTTGCTGATAAAAGCCAAGGACATGTGGNNNAGATCGGAA |
| 298_2 | TATGAGTGTGGAGTCGTTGCTGATAAAAGCCAAGGACATGTGGNNNAGATCGGAA |
| 30034_1 | TATGAGTGTGGAGTCGTTGCATAAGAAAGGCTGCAGAGGTTCANNNAGATCGGAA |
| 30034_2 | TATGAGTGTGGAGTCGTTGCATAAGAAAGGCTGCAGAGGTTCANNNAGATCGGAA |
| 52480_1 | TATGAGTGTGGAGTCGTTGCGCAAGCAAAGTTGAGGACAGATTNNNAGATCGGAA |
| 52480_2 | TATGAGTGTGGAGTCGTTGCGCAAGCAAAGTTGAGGACAGATTNNNAGATCGGAA |
| 48449_1 | TATGAGTGTGGAGTCGTTGCCATACACACGAGCCCTTTTTTGTNNNAGATCGGAAGA |
| 48449_2 | TATGAGTGTGGAGTCGTTGCCATACACACGAGCCCTTTTTTGTNNNAGATCGGAAGA |
| 4016_1 | TATGAGTGTGGAGTCGTTGCGGTGAGAAACATGAGTTGGCTCANNNAGATCGGAA |
| 4016_2 | TATGAGTGTGGAGTCGTTGCGGTGAGAAACATGAGTTGGCTCANNNAGATCGGAA |
| 29795_1 | TATGAGTGTGGAGTCGTTGCAAGATATGGTCAATCTGGCCAGGNNNAGATCGGAAG |
| 29795_2 | TATGAGTGTGGAGTCGTTGCAAGATATGGTCAATCTGGCCAGGNNNAGATCGGAAG |
| 44295_1 | TATGAGTGTGGAGTCGTTGCTGAACCTGTCATCCCCTAGATCTNNNAGATCGGAAGA |

| | |
|---|---|
| 44295_2 | TATGAGTGTGGAGTCGTTGCTGAACCTGTCATCCCCTAGATCTNNNAGATCGGAAG |
| 29106_1 | TATGAGTGTGGAGTCGTTGCATACACAGACACACACACACACTNNNAGATCGGAAG |
| 29106_2 | TATGAGTGTGGAGTCGTTGCATACACAGACACACACACACACTNNNAGATCGGAAG |
| 50091_1 | TATGAGTGTGGAGTCGTTGCAATGATGTCCATGTGCTAATGCCNNNAGATCGGAAG |
| 50091_2 | TATGAGTGTGGAGTCGTTGCAATGATGTCCATGTGCTAATGCCNNNAGATCGGAAG |
| 51100_1 | TATGAGTGTGGAGTCGTTGCCTGGCTAACACGGTAAACCTAGTNNNAGATCGGAAG |
| 51100_2 | TATGAGTGTGGAGTCGTTGCCTGGCTAACACGGTAAACCTAGTNNNAGATCGGAAG |
| 36176_1 | TATGAGTGTGGAGTCGTTGCCAAGAAGAGGCAGATGGCAAAATNNNAGATCGGA |
| 36176_2 | TATGAGTGTGGAGTCGTTGCCAAGAAGAGGCAGATGGCAAAATNNNAGATCGGA |
| 31784_1 | TATGAGTGTGGAGTCGTTGCAGATAGATGCTAAGGAGGTAGGTNNNAGATCGGA |
| 31784_2 | TATGAGTGTGGAGTCGTTGCAGATAGATGCTAAGGAGGTAGGTNNNAGATCGGA |
| 49906_1 | TATGAGTGTGGAGTCGTTGCTACAGTGCCTTCTGAGAGTGTTCNNNAGATCGGAAG |
| 49906_2 | TATGAGTGTGGAGTCGTTGCTACAGTGCCTTCTGAGAGTGTTCNNNAGATCGGAAG |
| 51506_1 | TATGAGTGTGGAGTCGTTGCTAGCAACATTCCTGGCCTCTATCNNNAGATCGGAAG |
| 51506_2 | TATGAGTGTGGAGTCGTTGCTAGCAACATTCCTGGCCTCTATCNNNAGATCGGAAG |
| 38866_1 | TATGAGTGTGGAGTCGTTGCACCATGATCAGGTGGGTTTCATANNNAGATCGGAA |
| 38866_2 | TATGAGTGTGGAGTCGTTGCACCATGATCAGGTGGGTTTCATANNNAGATCGGAA |
| 30234_1 | TATGAGTGTGGAGTCGTTGCAAAAACGACAACTTTTTTCTTTTGTTTNNNAGATCGG |
| 30234_2 | TATGAGTGTGGAGTCGTTGCAAAAACGACAACTTTTTTCTTTTGTTTNNNAGATCGG |
| 45127_1 | TATGAGTGTGGAGTCGTTGCAAGCATTCTTGAAGTGGGCTTTCNNNAGATCGGAAG |
| 45127_2 | TATGAGTGTGGAGTCGTTGCAAGCATTCTTGAAGTGGGCTTTCNNNAGATCGGAAG |
| 2847_1 | TATGAGTGTGGAGTCGTTGCGGGATAGTAGGTGCTCAGTGAGTNNNAGATCGGA |
| 2847_2 | TATGAGTGTGGAGTCGTTGCGGGATAGTAGGTGCTCAGTGAGTNNNAGATCGGA |
| 51023_1 | TATGAGTGTGGAGTCGTTGCCAGCTGCACAGATAAGAGGAGTTNNNAGATCGGAA |
| 51023_2 | TATGAGTGTGGAGTCGTTGCCAGCTGCACAGATAAGAGGAGTTNNNAGATCGGAA |
| 48169_1 | TATGAGTGTGGAGTCGTTGCGGAGTACAATGGTGCGATCTCAGNNNAGATCGGAA |
| 48169_2 | TATGAGTGTGGAGTCGTTGCGGAGTACAATGGTGCGATCTCAGNNNAGATCGGAA |
| 33156_1 | TATGAGTGTGGAGTCGTTGCGGCATCCAAATCAGCAAAGAGAANNNAGATCGGAA |
| 33156_2 | TATGAGTGTGGAGTCGTTGCGGCATCCAAATCAGCAAAGAGAANNNAGATCGGAA |
| 50810_1 | TATGAGTGTGGAGTCGTTGCTCCTTCCTACACAGTTCTTTGCANNNAGATCGGAAGA |
| 50810_2 | TATGAGTGTGGAGTCGTTGCTCCTTCCTACACAGTTCTTTGCANNNAGATCGGAAGA |
| 8195_1 | TATGAGTGTGGAGTCGTTGCTGTCCACGATCACTTCTTTTGTTNNNAGATCGGAAGA |
| 8195_2 | TATGAGTGTGGAGTCGTTGCTGTCCACGATCACTTCTTTTGTTNNNAGATCGGAAGA |
| 7837_1 | TATGAGTGTGGAGTCGTTGCGCAACTCTGGCTTAGACTGTGAGNNNAGATCGGAA |
| 7837_2 | TATGAGTGTGGAGTCGTTGCGCAACTCTGGCTTAGACTGTGAGNNNAGATCGGAA |
| 51368_1 | TATGAGTGTGGAGTCGTTGCGTGGGGTAAAGCATAAGAGGGAANNNAGATCGGA |
| 51368_2 | TATGAGTGTGGAGTCGTTGCGTGGGGTAAAGCATAAGAGGGAANNNAGATCGGA |
| 49624_1 | TATGAGTGTGGAGTCGTTGCCTCAAGTGATTCTCCCACCTCAGNNNAGATCGGAAG |
| 49624_2 | TATGAGTGTGGAGTCGTTGCCTCAAGTGATTCTCCCACCTCAGNNNAGATCGGAAG |
| 29646_1 | TATGAGTGTGGAGTCGTTGCCTCCTCCCCAAGACAGATGGAGNNNAGATCGGAAG |
| 29646_2 | TATGAGTGTGGAGTCGTTGCCTCCTCCCCAAGACAGATGGAGNNNAGATCGGAAG |
| 46659_1 | TATGAGTGTGGAGTCGTTGCCCATGCCCGGCTAATTTTTTGTANNNAGATCGGAAG |
| 46659_2 | TATGAGTGTGGAGTCGTTGCCCATGCCCGGCTAATTTTTTGTANNNAGATCGGAAG |

| | |
|---|---|
| 35244_1 | TATGAGTGTGGAGTCGTTGCCCTGAGTATCTGGGATTACAGGCNNNAGATCGGAA |
| 35244_2 | TATGAGTGTGGAGTCGTTGCCCTGAGTATCTGGGATTACAGGCNNNAGATCGGAA |
| 47178_1 | TATGAGTGTGGAGTCGTTGCAGACGAAGTCTCACTCTGTTGCNNNAGATCGGAAGA |
| 47178_2 | TATGAGTGTGGAGTCGTTGCAGACGAAGTCTCACTCTGTTGCNNNAGATCGGAAGA |
| 45147_1 | TATGAGTGTGGAGTCGTTGCACCTTTTTGGAGCAAGAGAAGGANNNAGATCGGAA |
| 45147_2 | TATGAGTGTGGAGTCGTTGCACCTTTTTGGAGCAAGAGAAGGANNNAGATCGGAA |
| 35668_1 | TATGAGTGTGGAGTCGTTGCTGGAGGTGGAGTCTCATTTTGTCNNNAGATCGGAA |
| 35668_2 | TATGAGTGTGGAGTCGTTGCTGGAGGTGGAGTCTCATTTTGTCNNNAGATCGGAA |
| 47843_1 | TATGAGTGTGGAGTCGTTGCAGTGCTGGGATTACAGGTGTAAGNNNAGATCGGAA |
| 47843_2 | TATGAGTGTGGAGTCGTTGCAGTGCTGGGATTACAGGTGTAAGNNNAGATCGGAA |
| 44691_1 | TATGAGTGTGGAGTCGTTGCACTAAAGGGGTTGCCAAGAAGAANNNAGATCGGAA |
| 44691_2 | TATGAGTGTGGAGTCGTTGCACTAAAGGGGTTGCCAAGAAGAANNNAGATCGGAA |
| 51592_1 | TATGAGTGTGGAGTCGTTGCTCTCTGACATTGCTATGGAGAGCNNNAGATCGGAAG |
| 51592_2 | TATGAGTGTGGAGTCGTTGCTCTCTGACATTGCTATGGAGAGCNNNAGATCGGAAG |
| 44917_1 | TATGAGTGTGGAGTCGTTGCGGTGTGTAAGCAGCCTAGGTTTANNNAGATCGGAA |
| 44917_2 | TATGAGTGTGGAGTCGTTGCGGTGTGTAAGCAGCCTAGGTTTANNNAGATCGGAA |
| 41416_1 | TATGAGTGTGGAGTCGTTGCCCCACACATTCATCATTGGGAATNNNAGATCGGAAG |
| 41416_2 | TATGAGTGTGGAGTCGTTGCCCCACACATTCATCATTGGGAATNNNAGATCGGAAG |
| 49183_1 | TATGAGTGTGGAGTCGTTGCAAAAAATATGTGGTGCTGTTTGCNNNAGATCGGAA |
| 49183_2 | TATGAGTGTGGAGTCGTTGCAAAAAATATGTGGTGCTGTTTGCNNNAGATCGGAA |
| 38162_1 | TATGAGTGTGGAGTCGTTGCAGAGAGAGAGAGAGAGAAATCAGANNNAGATCGG |
| 38162_2 | TATGAGTGTGGAGTCGTTGCAGAGAGAGAGAGAGAGAAATCAGANNNAGATCGG |
| 32253_1 | TATGAGTGTGGAGTCGTTGCCCCCCCAAAAAAAAGTGTGTGTANNNAGATCGGAAG |
| 32253_2 | TATGAGTGTGGAGTCGTTGCCCCCCCAAAAAAAAGTGTGTGTANNNAGATCGGAAG |
| 29247_1 | TATGAGTGTGGAGTCGTTGCTCATTGGAGAAGGCCAGATTGAANNNAGATCGGAA |
| 29247_2 | TATGAGTGTGGAGTCGTTGCTCATTGGAGAAGGCCAGATTGAANNNAGATCGGAA |
| 33320_1 | TATGAGTGTGGAGTCGTTGCCTAGTAGCTGGGACTACAGGCANNNAGATCGGAAG |
| 33320_2 | TATGAGTGTGGAGTCGTTGCCTAGTAGCTGGGACTACAGGCANNNAGATCGGAAG |
| 43247_1 | TATGAGTGTGGAGTCGTTGCAGGAAATAGTTAGCCAGGTGGTGNNNAGATCGGAA |
| 43247_2 | TATGAGTGTGGAGTCGTTGCAGGAAATAGTTAGCCAGGTGGTGNNNAGATCGGAA |
| 50799_1 | TATGAGTGTGGAGTCGTTGCACAGAGAGAAGAATACAGCAGCCNNNAGATCGGAA |
| 50799_2 | TATGAGTGTGGAGTCGTTGCACAGAGAGAAGAATACAGCAGCCNNNAGATCGGAA |
| 32845_1 | TATGAGTGTGGAGTCGTTGCGATAAGGTCTCACTCTGTCACCCNNNAGATCGGAAG |
| 32845_2 | TATGAGTGTGGAGTCGTTGCGATAAGGTCTCACTCTGTCACCCNNNAGATCGGAAG |
| 37880_1 | TATGAGTGTGGAGTCGTTGCCTTTTTGGGGTGAAGCTGATCAGNNNAGATCGGAA |
| 37880_2 | TATGAGTGTGGAGTCGTTGCCTTTTTGGGGTGAAGCTGATCAGNNNAGATCGGAA |
| 37144_1 | TATGAGTGTGGAGTCGTTGCGCAGAACTACCATTTGACCAAGCNNNAGATCGGAAG |
| 37144_2 | TATGAGTGTGGAGTCGTTGCGCAGAACTACCATTTGACCAAGCNNNAGATCGGAAG |
| 38187_1 | TATGAGTGTGGAGTCGTTGCACAAAGCTGACTGAGAATTTCCANNNAGATCGGAAG |
| 38187_2 | TATGAGTGTGGAGTCGTTGCACAAAGCTGACTGAGAATTTCCANNNAGATCGGAAG |
| 46447_1 | TATGAGTGTGGAGTCGTTGCATACCAAATGATGTGGGCTCAGTNNNAGATCGGAA |
| 46447_2 | TATGAGTGTGGAGTCGTTGCATACCAAATGATGTGGGCTCAGTNNNAGATCGGAA |
| 33924_1 | TATGAGTGTGGAGTCGTTGCAAAAGCCTCTGCAGGAAGATCATNNNAGATCGGAA |

| | |
|---|---|
| 33924_2 | TATGAGTGTGGAGTCGTTGCAAAAGCCTCTGCAGGAAGATCATNNNAGATCGGAA |
| 43333_1 | TATGAGTGTGGAGTCGTTGCGTAGCTGAGATCACAGGCATACANNNAGATCGGAA |
| 43333_2 | TATGAGTGTGGAGTCGTTGCGTAGCTGAGATCACAGGCATACANNNAGATCGGAA |
| 38287_1 | TATGAGTGTGGAGTCGTTGCAGAAGAAGAAGAAGAGGAGGAGANNNAGATCGG |
| 38287_2 | TATGAGTGTGGAGTCGTTGCAGAAGAAGAAGAAGAGGAGGAGANNNAGATCGG |
| 33060_1 | TATGAGTGTGGAGTCGTTGCGCTTACTGCTGCCTTGACTTCNNNAGATCGGAAGAG |
| 33060_2 | TATGAGTGTGGAGTCGTTGCGCTTACTGCTGCCTTGACTTCNNNAGATCGGAAGAG |
| 44218_1 | TATGAGTGTGGAGTCGTTGCTAGGGAACACTAGCAGAGATGGANNNAGATCGGA |
| 44218_2 | TATGAGTGTGGAGTCGTTGCTAGGGAACACTAGCAGAGATGGANNNAGATCGGA |
| 36562_1 | TATGAGTGTGGAGTCGTTGCAGCATTTCTGATTGGTTGCCTTCNNNAGATCGGAAG |
| 36562_2 | TATGAGTGTGGAGTCGTTGCAGCATTTCTGATTGGTTGCCTTCNNNAGATCGGAAG |
| 43374_1 | TATGAGTGTGGAGTCGTTGCAGGTTCAAGCAATTCTCCTGTCTNNNAGATCGGAAG |
| 43374_2 | TATGAGTGTGGAGTCGTTGCAGGTTCAAGCAATTCTCCTGTCTNNNAGATCGGAAG |
| 51531_1 | TATGAGTGTGGAGTCGTTGCTATGTCCCTTCTGTGCCAAGTTTNNNAGATCGGAAG |
| 51531_2 | TATGAGTGTGGAGTCGTTGCTATGTCCCTTCTGTGCCAAGTTTNNNAGATCGGAAG |
| 38270_1 | TATGAGTGTGGAGTCGTTGCAATGTATGCTCTGCAGTTGTTGGNNNAGATCGGAA |
| 38270_2 | TATGAGTGTGGAGTCGTTGCAATGTATGCTCTGCAGTTGTTGGNNNAGATCGGAA |
| 30732_1 | TATGAGTGTGGAGTCGTTGCGTCTGGGCTTCAAGAGGCTATTGNNNAGATCGGAA |
| 30732_2 | TATGAGTGTGGAGTCGTTGCGTCTGGGCTTCAAGAGGCTATTGNNNAGATCGGAA |
| 13123_1 | TATGAGTGTGGAGTCGTTGCAGATGTTGACATTTTAATAAGCANNNAGATCGGAAG |
| 13123_2 | TATGAGTGTGGAGTCGTTGCAGATGTTGACATTTTAATAAGCANNNAGATCGGAAG |
| 35533_1 | TATGAGTGTGGAGTCGTTGCCACTCCTCTGATTCAGAAGGCTTNNNAGATCGGAAG |
| 35533_2 | TATGAGTGTGGAGTCGTTGCCACTCCTCTGATTCAGAAGGCTTNNNAGATCGGAAG |
| 50569_1 | TATGAGTGTGGAGTCGTTGCTTGCCTTCTTCACTGTAGCTTCTNNNAGATCGGAAGA |
| 50569_2 | TATGAGTGTGGAGTCGTTGCTTGCCTTCTTCACTGTAGCTTCTNNNAGATCGGAAGA |
| 49758_1 | TATGAGTGTGGAGTCGTTGCCTACTCCTGGTCTCTCTCCCTAGNNNAGATCGGAAGA |
| 49758_2 | TATGAGTGTGGAGTCGTTGCCTACTCCTGGTCTCTCTCCCTAGNNNAGATCGGAAGA |
| 48812_1 | TATGAGTGTGGAGTCGTTGCAAGTAGAGATGGGGTTTTGCCATNNNAGATCGGAA |
| 48812_2 | TATGAGTGTGGAGTCGTTGCAAGTAGAGATGGGGTTTTGCCATNNNAGATCGGAA |
| 44998_1 | TATGAGTGTGGAGTCGTTGCGACAGGGTCACATAAGTGGATGTNNNAGATCGGAA |
| 44998_2 | TATGAGTGTGGAGTCGTTGCGACAGGGTCACATAAGTGGATGTNNNAGATCGGAA |
| 30608_1 | TATGAGTGTGGAGTCGTTGCGTCCCAGTGATTAAACTGAGGTCANNNAGATCGGA |
| 30608_2 | TATGAGTGTGGAGTCGTTGCGTCCCAGTGATTAAACTGAGGTCANNNAGATCGGA |
| 29962_1 | TATGAGTGTGGAGTCGTTGCCTCTCACCAGGCTCTTGTTCAATNNNAGATCGGAAG |
| 29962_2 | TATGAGTGTGGAGTCGTTGCCTCTCACCAGGCTCTTGTTCAATNNNAGATCGGAAG |
| 34809_1 | TATGAGTGTGGAGTCGTTGCCACCCCCGGCTAATTTTTTGTATNNNAGATCGGAAG |
| 34809_2 | TATGAGTGTGGAGTCGTTGCCACCCCCGGCTAATTTTTTGTATNNNAGATCGGAAG |
| 53057_1 | TATGAGTGTGGAGTCGTTGCTGCAATCAGCTATAATCCTTCCCTNNNAGATCGGAAG |
| 53057_2 | TATGAGTGTGGAGTCGTTGCTGCAATCAGCTATAATCCTTCCCTNNNAGATCGGAAG |
| 7972_1 | TATGAGTGTGGAGTCGTTGCGCTCATGGACACTATCCTCCATCTNNNAGATCGGAAG |
| 7972_2 | TATGAGTGTGGAGTCGTTGCGCTCATGGACACTATCCTCCATCTNNNAGATCGGAAG |
| 37735_1 | TATGAGTGTGGAGTCGTTGCGAAAGAGCAGGACAGTGCAAAATNNNAGATCGGA |
| 37735_2 | TATGAGTGTGGAGTCGTTGCGAAAGAGCAGGACAGTGCAAAATNNNAGATCGGA |

| | |
|---|---|
| 36091_1 | TATGAGTGTGGAGTCGTTGCGTCACCAAAATCAGAAAGCTCCGNNNAGATCGGAAG |
| 36091_2 | TATGAGTGTGGAGTCGTTGCGTCACCAAAATCAGAAAGCTCCGNNNAGATCGGAAG |
| 31523_1 | TATGAGTGTGGAGTCGTTGCGCTTTCTCCTGGTACCTGAGAAANNNAGATCGGAAG |
| 31523_2 | TATGAGTGTGGAGTCGTTGCGCTTTCTCCTGGTACCTGAGAAANNNAGATCGGAAG |
| 41478_1 | TATGAGTGTGGAGTCGTTGCAAACTCTGAACAATGCCTGAAGCNNNAGATCGGAAG |
| 41478_2 | TATGAGTGTGGAGTCGTTGCAAACTCTGAACAATGCCTGAAGCNNNAGATCGGAAG |
| 39388_1 | TATGAGTGTGGAGTCGTTGCATACAAAAATTAGCCAGGCGTGGNNNAGATCGGAA |
| 39388_2 | TATGAGTGTGGAGTCGTTGCATACAAAAATTAGCCAGGCGTGGNNNAGATCGGAA |
| 53112_1 | TATGAGTGTGGAGTCGTTGCGATGCACCGTGTTTATTGGACATNNNAGATCGGAAG |
| 53112_2 | TATGAGTGTGGAGTCGTTGCGATGCACCGTGTTTATTGGACATNNNAGATCGGAAG |
| 50967_1 | TATGAGTGTGGAGTCGTTGCTTGAAACGTGGCAGACTCTTTTCNNNAGATCGGAAG |
| 50967_2 | TATGAGTGTGGAGTCGTTGCTTGAAACGTGGCAGACTCTTTTCNNNAGATCGGAAG |
| 52165_1 | TATGAGTGTGGAGTCGTTGCAGACCCAAGCATGTGTATGTTCANNNAGATCGGAAG |
| 52165_2 | TATGAGTGTGGAGTCGTTGCAGACCCAAGCATGTGTATGTTCANNNAGATCGGAAG |
| 29642_1 | TATGAGTGTGGAGTCGTTGCCGAGTAGCTGAGATTACAGGCATNNNAGATCGGAA |
| 29642_2 | TATGAGTGTGGAGTCGTTGCCGAGTAGCTGAGATTACAGGCATNNNAGATCGGAA |
| 37658_1 | TATGAGTGTGGAGTCGTTGCAGGTATGTTGATGTGGTGGGAAANNNAGATCGGA |
| 37658_2 | TATGAGTGTGGAGTCGTTGCAGGTATGTTGATGTGGTGGGAAANNNAGATCGGA |
| 35829_1 | TATGAGTGTGGAGTCGTTGCTAGAGTACAATGGCGGGATCTTGNNNAGATCGGAA |
| 35829_2 | TATGAGTGTGGAGTCGTTGCTAGAGTACAATGGCGGGATCTTGNNNAGATCGGAA |
| 44211_1 | TATGAGTGTGGAGTCGTTGCAAAGTGGACACTCCTAGCTTCAGNNNAGATCGGAAG |
| 44211_2 | TATGAGTGTGGAGTCGTTGCAAAGTGGACACTCCTAGCTTCAGNNNAGATCGGAAG |
| 38460_1 | TATGAGTGTGGAGTCGTTGCTGTCTTCTCCAGGCTAAATGTTGTNNNAGATCGGAA |
| 38460_2 | TATGAGTGTGGAGTCGTTGCTGTCTTCTCCAGGCTAAATGTTGTNNNAGATCGGAA |
| 46133_1 | TATGAGTGTGGAGTCGTTGCCAAGTAGGTGAGACTACAGGCACNNNAGATCGGAA |
| 46133_2 | TATGAGTGTGGAGTCGTTGCCAAGTAGGTGAGACTACAGGCACNNNAGATCGGAA |
| 51840_1 | TATGAGTGTGGAGTCGTTGCAAGACGAGGATCTCCCTATGTTGNNNAGATCGGAAG |
| 51840_2 | TATGAGTGTGGAGTCGTTGCAAGACGAGGATCTCCCTATGTTGNNNAGATCGGAAG |
| 44376_1 | TATGAGTGTGGAGTCGTTGCCTAGAAACACACAATGGCCCCTCNNNAGATCGGAAG |
| 44376_2 | TATGAGTGTGGAGTCGTTGCCTAGAAACACACAATGGCCCCTCNNNAGATCGGAAG |
| 44751_1 | TATGAGTGTGGAGTCGTTGCCTCTGTCTTGAGACCTTAGGCAGNNNAGATCGGAAG |
| 44751_2 | TATGAGTGTGGAGTCGTTGCCTCTGTCTTGAGACCTTAGGCAGNNNAGATCGGAAG |
| 48137_1 | TATGAGTGTGGAGTCGTTGCGGGTTTCACTGTGTTAGCTAGGANNNAGATCGGAA |
| 48137_2 | TATGAGTGTGGAGTCGTTGCGGGTTTCACTGTGTTAGCTAGGANNNAGATCGGAA |
| 13446_1 | TATGAGTGTGGAGTCGTTGCCACCATTTTTATAGTCCATGTTTNNNAGATCGGAAGA |
| 13446_2 | TATGAGTGTGGAGTCGTTGCCACCATTTTTATAGTCCATGTTTNNNAGATCGGAAGA |
| 39693_1 | TATGAGTGTGGAGTCGTTGCATGCCTCCAGTCTCACTGAAATTNNNAGATCGGAAG |
| 39693_2 | TATGAGTGTGGAGTCGTTGCATGCCTCCAGTCTCACTGAAATTNNNAGATCGGAAG |
| 43968_1 | TATGAGTGTGGAGTCGTTGCTCACTGTACGTAGGTATCTGGGTNNNAGATCGGAA |
| 43968_2 | TATGAGTGTGGAGTCGTTGCTCACTGTACGTAGGTATCTGGGTNNNAGATCGGAA |
| 41622_1 | TATGAGTGTGGAGTCGTTGCGATTCTCACTCTGTCACCTAGGCNNNAGATCGGAAG |
| 41622_2 | TATGAGTGTGGAGTCGTTGCGATTCTCACTCTGTCACCTAGGCNNNAGATCGGAAG |
| 10104_1 | TATGAGTGTGGAGTCGTTGCGGAGTACGAAAGCATGAAGGTGTNNNAGATCGGA |

| | |
|---|---|
| 10104_2 | TATGAGTGTGGAGTCGTTGCGGAGTACGAAAGCATGAAGGTGTNNNAGATCGGA |
| 31142_1 | TATGAGTGTGGAGTCGTTGCGTACCGAGTCTCACCCTATTGTCNNNAGATCGGAAG |
| 31142_2 | TATGAGTGTGGAGTCGTTGCGTACCGAGTCTCACCCTATTGTCNNNAGATCGGAAG |
| 43693_1 | TATGAGTGTGGAGTCGTTGCCTTCGTGTACCCGCTGGATTTCNNNAGATCGGAAGA |
| 43693_2 | TATGAGTGTGGAGTCGTTGCCTTCGTGTACCCGCTGGATTTCNNNAGATCGGAAGA |
| 43115_1 | TATGAGTGTGGAGTCGTTGCGAGACAATGAGGACAAGGGTAGGNNNAGATCGGA |
| 43115_2 | TATGAGTGTGGAGTCGTTGCGAGACAATGAGGACAAGGGTAGGNNNAGATCGGA |
| 50807_1 | TATGAGTGTGGAGTCGTTGCGCTCAGTGCTTCTTTGGTGATTTNNNAGATCGGAAG |
| 50807_2 | TATGAGTGTGGAGTCGTTGCGCTCAGTGCTTCTTTGGTGATTTNNNAGATCGGAAG |
| 35234_1 | TATGAGTGTGGAGTCGTTGCGGTGATTTGAACTCCAGCTGTTCNNNAGATCGGAAG |
| 35234_2 | TATGAGTGTGGAGTCGTTGCGGTGATTTGAACTCCAGCTGTTCNNNAGATCGGAAG |
| 45464_1 | TATGAGTGTGGAGTCGTTGCGAACACCCGAACCAAATAATCCCNNNAGATCGGAAG |
| 45464_2 | TATGAGTGTGGAGTCGTTGCGAACACCCGAACCAAATAATCCCNNNAGATCGGAAG |
| 40730_1 | TATGAGTGTGGAGTCGTTGCAGCCTTCTGAGTAGCTGGTACTANNNAGATCGGAAG |
| 40730_2 | TATGAGTGTGGAGTCGTTGCAGCCTTCTGAGTAGCTGGTACTANNNAGATCGGAAG |
| 44690_1 | TATGAGTGTGGAGTCGTTGCCCACATAAGTTGGTTGGGTGTTCNNNAGATCGGAAG |
| 44690_2 | TATGAGTGTGGAGTCGTTGCCCACATAAGTTGGTTGGGTGTTCNNNAGATCGGAAG |
| 45225_1 | TATGAGTGTGGAGTCGTTGCTGCTCAGAGCGCAGTACTTACNNNAGATCGGAAGAG |
| 45225_2 | TATGAGTGTGGAGTCGTTGCTGCTCAGAGCGCAGTACTTACNNNAGATCGGAAGAG |
| 44157_1 | TATGAGTGTGGAGTCGTTGCTTTCACATATGGTTTCAGGGGGTNNNAGATCGGAAG |
| 44157_2 | TATGAGTGTGGAGTCGTTGCTTTCACATATGGTTTCAGGGGGTNNNAGATCGGAAG |
| 32018_1 | TATGAGTGTGGAGTCGTTGCGCCTGAGCACCTATCTACCATTTNNNAGATCGGAAG |
| 32018_2 | TATGAGTGTGGAGTCGTTGCGCCTGAGCACCTATCTACCATTTNNNAGATCGGAAG |
| 47572_1 | TATGAGTGTGGAGTCGTTGCAGACGGAGTCTCATTCTGTTGCNNNAGATCGGAAGA |
| 47572_2 | TATGAGTGTGGAGTCGTTGCAGACGGAGTCTCATTCTGTTGCNNNAGATCGGAAGA |
| 39483_1 | TATGAGTGTGGAGTCGTTGCTTGTTGTGGGGCTTTTTCACTTTNNNAGATCGGAAG |
| 39483_2 | TATGAGTGTGGAGTCGTTGCTTGTTGTGGGGCTTTTTCACTTTNNNAGATCGGAAG |
| 43638_1 | TATGAGTGTGGAGTCGTTGCATGCACACACAGGTACACAGATANNNAGATCGGAAG |
| 43638_2 | TATGAGTGTGGAGTCGTTGCATGCACACACAGGTACACAGATANNNAGATCGGAAG |
| 43687_1 | TATGAGTGTGGAGTCGTTGCCTATGACCAATGATGGGACCCATNNNAGATCGGAAG |
| 43687_2 | TATGAGTGTGGAGTCGTTGCCTATGACCAATGATGGGACCCATNNNAGATCGGAAG |
| 51693_1 | TATGAGTGTGGAGTCGTTGCCTTGTGAGTGAGAACATGCAGTGNNNAGATCGGAA |
| 51693_2 | TATGAGTGTGGAGTCGTTGCCTTGTGAGTGAGAACATGCAGTGNNNAGATCGGAA |
| 50554_1 | TATGAGTGTGGAGTCGTTGCCCTTCAGTGTGAACGGAGTAGATNNNAGATCGGAA |
| 50554_2 | TATGAGTGTGGAGTCGTTGCCCTTCAGTGTGAACGGAGTAGATNNNAGATCGGAA |
| 36637_1 | TATGAGTGTGGAGTCGTTGCGTGACCCAAACAGAAGGGAAAACNNNAGATCGGAA |
| 36637_2 | TATGAGTGTGGAGTCGTTGCGTGACCCAAACAGAAGGGAAAACNNNAGATCGGAA |
| 52336_1 | TATGAGTGTGGAGTCGTTGCGACCTGTTGAAGTTTATTCGGCCNNNAGATCGGAAG |
| 52336_2 | TATGAGTGTGGAGTCGTTGCGACCTGTTGAAGTTTATTCGGCCNNNAGATCGGAAG |
| 52129_1 | TATGAGTGTGGAGTCGTTGCACACTTGCCTTTTCTTTGAGGTGNNNAGATCGGAAG |
| 52129_2 | TATGAGTGTGGAGTCGTTGCACACTTGCCTTTTCTTTGAGGTGNNNAGATCGGAAG |
| 43139_1 | TATGAGTGTGGAGTCGTTGCCAGAGAATCAGTGAAGCTCTCCANNNAGATCGGAAG |
| 43139_2 | TATGAGTGTGGAGTCGTTGCCAGAGAATCAGTGAAGCTCTCCANNNAGATCGGAAG |

| | |
|---|---|
| 43763_1 | TATGAGTGTGGAGTCGTTGCCTGTCATCTCAGCACTTTGGGAGNNNAGATCGGAAG |
| 43763_2 | TATGAGTGTGGAGTCGTTGCCTGTCATCTCAGCACTTTGGGAGNNNAGATCGGAAG |
| 12588_1 | TATGAGTGTGGAGTCGTTGCATATCAAGGTTATTGGTAAGTGANNNAGATCGGAA |
| 12588_2 | TATGAGTGTGGAGTCGTTGCATATCAAGGTTATTGGTAAGTGANNNAGATCGGAA |
| 33633_1 | TATGAGTGTGGAGTCGTTGCAAGCCTGAAGTGCCAGAGAATTANNNAGATCGGAA |
| 33633_2 | TATGAGTGTGGAGTCGTTGCAAGCCTGAAGTGCCAGAGAATTANNNAGATCGGAA |
| 10849_1 | TATGAGTGTGGAGTCGTTGCATTTGCAGCAACCTAGGTGGGAANNNAGATCGGAA |
| 10849_2 | TATGAGTGTGGAGTCGTTGCATTTGCAGCAACCTAGGTGGGAANNNAGATCGGAA |
| 44914_1 | TATGAGTGTGGAGTCGTTGCCAGTCCAACCTCTTCATGGTACANNNAGATCGGAAG |
| 44914_2 | TATGAGTGTGGAGTCGTTGCCAGTCCAACCTCTTCATGGTACANNNAGATCGGAAG |
| 50356_1 | TATGAGTGTGGAGTCGTTGCTTAGCAGGATTTGGCAGCAAATGNNNAGATCGGAA |
| 50356_2 | TATGAGTGTGGAGTCGTTGCTTAGCAGGATTTGGCAGCAAATGNNNAGATCGGAA |
| 10187_1 | TATGAGTGTGGAGTCGTTGCCTCACTCACTGCAACCTGTGTGNNNAGATCGGAAGA |
| 10187_2 | TATGAGTGTGGAGTCGTTGCCTCACTCACTGCAACCTGTGTGNNNAGATCGGAAGA |
| 52325_1 | TATGAGTGTGGAGTCGTTGCGTCGATCTCCTGACCTTGTGATCNNNAGATCGGAAG |
| 52325_2 | TATGAGTGTGGAGTCGTTGCGTCGATCTCCTGACCTTGTGATCNNNAGATCGGAAG |
| 9984_1 | TATGAGTGTGGAGTCGTTGCTGATACTGTCGCAGGTGCTAAACNNNAGATCGGAAG |
| 9984_2 | TATGAGTGTGGAGTCGTTGCTGATACTGTCGCAGGTGCTAAACNNNAGATCGGAAG |
| 49145_1 | TATGAGTGTGGAGTCGTTGCGTGAAATGGTGTTGAACTACCCCNNNAGATCGGAAG |
| 49145_2 | TATGAGTGTGGAGTCGTTGCGTGAAATGGTGTTGAACTACCCCNNNAGATCGGAAG |
| 36193_1 | TATGAGTGTGGAGTCGTTGCGGCCTTGGGATTATCCACCTATTNNNAGATCGGAAG |
| 36193_2 | TATGAGTGTGGAGTCGTTGCGGCCTTGGGATTATCCACCTATTNNNAGATCGGAAG |
| 11015_1 | TATGAGTGTGGAGTCGTTGCCAGAGACAATCAGAGGATCACGANNNAGATCGGAA |
| 11015_2 | TATGAGTGTGGAGTCGTTGCCAGAGACAATCAGAGGATCACGANNNAGATCGGAA |
| 49818_1 | TATGAGTGTGGAGTCGTTGCAGATCACCTGTCCCTCCTCCNNNAGATCGGAAGAGC |
| 49818_2 | TATGAGTGTGGAGTCGTTGCAGATCACCTGTCCCTCCTCCNNNAGATCGGAAGAGC |
| 12762_1 | TATGAGTGTGGAGTCGTTGCTAGATTGTCTATGGAATTAGCAANNNAGATCGGAAG |
| 12762_2 | TATGAGTGTGGAGTCGTTGCTAGATTGTCTATGGAATTAGCAANNNAGATCGGAAG |
| 37679_1 | TATGAGTGTGGAGTCGTTGCCATTCACAATGGAACACCACCAANNNAGATCGGAAG |
| 37679_2 | TATGAGTGTGGAGTCGTTGCCATTCACAATGGAACACCACCAANNNAGATCGGAAG |
| 35186_1 | TATGAGTGTGGAGTCGTTGCTATCTCCCGGGTTCAAGTGATTCNNNAGATCGGAAG |
| 35186_2 | TATGAGTGTGGAGTCGTTGCTATCTCCCGGGTTCAAGTGATTCNNNAGATCGGAAG |
| 39782_1 | TATGAGTGTGGAGTCGTTGCAGCCAAAAAGTGGAAACAACACANNNAGATCGGAA |
| 39782_2 | TATGAGTGTGGAGTCGTTGCAGCCAAAAAGTGGAAACAACACANNNAGATCGGAA |
| 39831_1 | TATGAGTGTGGAGTCGTTGCTTTAGGGACCATTACCTTTACCTTTNNNAGATCGGAA |
| 39831_2 | TATGAGTGTGGAGTCGTTGCTTTAGGGACCATTACCTTTACCTTTNNNAGATCGGAA |
| 42844_1 | TATGAGTGTGGAGTCGTTGCTTTGCTATGTTACTGAGGCTGGTNNNAGATCGGAAG |
| 42844_2 | TATGAGTGTGGAGTCGTTGCTTTGCTATGTTACTGAGGCTGGTNNNAGATCGGAAG |
| 9954_1 | TATGAGTGTGGAGTCGTTGCCACGCACATGAATATTGGCCTTCNNNAGATCGGAAG |
| 9954_2 | TATGAGTGTGGAGTCGTTGCCACGCACATGAATATTGGCCTTCNNNAGATCGGAAG |
| 46294_1 | TATGAGTGTGGAGTCGTTGCTCTCAAACTCCTGACCTCAAGTGNNNAGATCGGAAG |
| 46294_2 | TATGAGTGTGGAGTCGTTGCTCTCAAACTCCTGACCTCAAGTGNNNAGATCGGAAG |
| 38492_1 | TATGAGTGTGGAGTCGTTGCATACTCATCACAAGCCTTGGAGGNNNAGATCGGAAG |

| | |
|---|---|
| 38492_2 | TATGAGTGTGGAGTCGTTGCATACTCATCACAAGCCTTGGAGGNNNAGATCGGAAG |
| 46972_1 | TATGAGTGTGGAGTCGTTGCGTTCAACCTATTCTCCTGCCTCANNNAGATCGGAAGA |
| 46972_2 | TATGAGTGTGGAGTCGTTGCGTTCAACCTATTCTCCTGCCTCANNNAGATCGGAAGA |
| 52444_1 | TATGAGTGTGGAGTCGTTGCTCTGGACAGTTTGTCAATGAGCTNNNAGATCGGAAG |
| 52444_2 | TATGAGTGTGGAGTCGTTGCTCTGGACAGTTTGTCAATGAGCTNNNAGATCGGAAG |
| 39099_1 | TATGAGTGTGGAGTCGTTGCTACCTGAAGCACCTAGTCTACCTNNNAGATCGGAAG |
| 39099_2 | TATGAGTGTGGAGTCGTTGCTACCTGAAGCACCTAGTCTACCTNNNAGATCGGAAG |
| 41115_1 | TATGAGTGTGGAGTCGTTGCAGAGATGTTAGGTGTGCTGATGGNNNAGATCGGAA |
| 41115_2 | TATGAGTGTGGAGTCGTTGCAGAGATGTTAGGTGTGCTGATGGNNNAGATCGGAA |
| 32741_1 | TATGAGTGTGGAGTCGTTGCAACAGGAGAATCCCTTGAACTCGNNNAGATCGGAAG |
| 32741_2 | TATGAGTGTGGAGTCGTTGCAACAGGAGAATCCCTTGAACTCGNNNAGATCGGAAG |
| 41842_1 | TATGAGTGTGGAGTCGTTGCCACTACGCCCGGCTAATTTTTTNNNAGATCGGAAGA |
| 41842_2 | TATGAGTGTGGAGTCGTTGCCACTACGCCCGGCTAATTTTTTNNNAGATCGGAAGA |
| 49894_1 | TATGAGTGTGGAGTCGTTGCAAGCAGTCTTCAAACTGACTTGCNNNAGATCGGAAG |
| 49894_2 | TATGAGTGTGGAGTCGTTGCAAGCAGTCTTCAAACTGACTTGCNNNAGATCGGAAG |
| 41960_1 | TATGAGTGTGGAGTCGTTGCCCTGGTTTCAAGCAATTCTCCTGNNNAGATCGGAAG |
| 41960_2 | TATGAGTGTGGAGTCGTTGCCCTGGTTTCAAGCAATTCTCCTGNNNAGATCGGAAG |
| 29061_1 | TATGAGTGTGGAGTCGTTGCGACCCTGTGTCCACTCTAAGTGNNNAGATCGGAAGA |
| 29061_2 | TATGAGTGTGGAGTCGTTGCGACCCTGTGTCCACTCTAAGTGNNNAGATCGGAAGA |
| 34584_1 | TATGAGTGTGGAGTCGTTGCGAGGGGAGGGGAAGGGAAAGNNNAGATCGGAAG |
| 34584_2 | TATGAGTGTGGAGTCGTTGCGAGGGGAGGGGAAGGGAAAGNNNAGATCGGAAG |
| 29492_1 | TATGAGTGTGGAGTCGTTGCCGTGGCTTTTTATGGGGACTTAANNNAGATCGGAAG |
| 29492_2 | TATGAGTGTGGAGTCGTTGCCGTGGCTTTTTATGGGGACTTAANNNAGATCGGAAG |
| 40791_1 | TATGAGTGTGGAGTCGTTGCACATGCATGAACACACACGTTAGNNNAGATCGGAAG |
| 40791_2 | TATGAGTGTGGAGTCGTTGCACATGCATGAACACACACGTTAGNNNAGATCGGAAG |
| 34585_1 | TATGAGTGTGGAGTCGTTGCCTCCCAGGTTCAAGTGATCCTCNNNAGATCGGAAGA |
| 34585_2 | TATGAGTGTGGAGTCGTTGCCTCCCAGGTTCAAGTGATCCTCNNNAGATCGGAAGA |
| 41804_1 | TATGAGTGTGGAGTCGTTGCGATTTCGGTTCACTGCAAGCTCNNNAGATCGGAAGA |
| 41804_2 | TATGAGTGTGGAGTCGTTGCGATTTCGGTTCACTGCAAGCTCNNNAGATCGGAAGA |
| 33762_1 | TATGAGTGTGGAGTCGTTGCGGAGAATCTCTTGAGCCCAGTAGNNNAGATCGGAA |
| 33762_2 | TATGAGTGTGGAGTCGTTGCGGAGAATCTCTTGAGCCCAGTAGNNNAGATCGGAA |
| 38984_1 | TATGAGTGTGGAGTCGTTGCTCTGTTTATGCCTGGCTTCATGTNNNAGATCGGAAG |
| 38984_2 | TATGAGTGTGGAGTCGTTGCTCTGTTTATGCCTGGCTTCATGTNNNAGATCGGAAG |
| 46442_1 | TATGAGTGTGGAGTCGTTGCACGTTCAAGTAATTCTCCTGCCTNNNAGATCGGAAG |
| 46442_2 | TATGAGTGTGGAGTCGTTGCACGTTCAAGTAATTCTCCTGCCTNNNAGATCGGAAG |
| 43274_1 | TATGAGTGTGGAGTCGTTGCACAGGTTTAGTATTCCCCCATTACTNNNAGATCGGAA |
| 43274_2 | TATGAGTGTGGAGTCGTTGCACAGGTTTAGTATTCCCCCATTACTNNNAGATCGGAA |
| 47466_1 | TATGAGTGTGGAGTCGTTGCCTCAATCTCAGCTCAATGCAACCNNNAGATCGGAAG |
| 47466_2 | TATGAGTGTGGAGTCGTTGCCTCAATCTCAGCTCAATGCAACCNNNAGATCGGAAG |
| 51465_1 | TATGAGTGTGGAGTCGTTGCGTTTTCATCAGCTCTTGGCAGTTNNNAGATCGGAAG |
| 51465_2 | TATGAGTGTGGAGTCGTTGCGTTTTCATCAGCTCTTGGCAGTTNNNAGATCGGAAG |
| 31040_1 | TATGAGTGTGGAGTCGTTGCCCAGGGCAGGAACATGTAATCTANNNAGATCGGAA |
| 31040_2 | TATGAGTGTGGAGTCGTTGCCCAGGGCAGGAACATGTAATCTANNNAGATCGGAA |

| | |
|---|---|
| 38048_1 | TATGAGTGTGGAGTCGTTGCCCTGAAAAGAGCCACCTAGTTCTNNNAGATCGGAAG |
| 38048_2 | TATGAGTGTGGAGTCGTTGCCCTGAAAAGAGCCACCTAGTTCTNNNAGATCGGAAG |
| 31541_1 | TATGAGTGTGGAGTCGTTGCCAGGATGCAAAGACTTAGTACCANNNAGATCGGAA |
| 31541_2 | TATGAGTGTGGAGTCGTTGCCAGGATGCAAAGACTTAGTACCANNNAGATCGGAA |
| 38552_1 | TATGAGTGTGGAGTCGTTGCCACACACACTCACTCTGTCTCTCNNNAGATCGGAAGA |
| 38552_2 | TATGAGTGTGGAGTCGTTGCCACACACACTCACTCTGTCTCTCNNNAGATCGGAAGA |
| 41900_1 | TATGAGTGTGGAGTCGTTGCCAACTCCTGACCTCAAGTGATCTNNNAGATCGGAAG |
| 41900_2 | TATGAGTGTGGAGTCGTTGCCAACTCCTGACCTCAAGTGATCTNNNAGATCGGAAG |
| 43719_1 | TATGAGTGTGGAGTCGTTGCGCTAACCAGCGTATACTCCATCANNNAGATCGGAAG |
| 43719_2 | TATGAGTGTGGAGTCGTTGCGCTAACCAGCGTATACTCCATCANNNAGATCGGAAG |
| 40862_1 | TATGAGTGTGGAGTCGTTGCGCCCCTTACAAATTTGAAACCCANNNAGATCGGAAG |
| 40862_2 | TATGAGTGTGGAGTCGTTGCGCCCCTTACAAATTTGAAACCCANNNAGATCGGAAG |
| 50248_1 | TATGAGTGTGGAGTCGTTGCGGTGCATACCTGTCCTCATAGACNNNAGATCGGAAG |
| 50248_2 | TATGAGTGTGGAGTCGTTGCGGTGCATACCTGTCCTCATAGACNNNAGATCGGAAG |
| 40782_1 | TATGAGTGTGGAGTCGTTGCAAATGAAGGGCTTACTCTCCCAGNNNAGATCGGAAG |
| 40782_2 | TATGAGTGTGGAGTCGTTGCAAATGAAGGGCTTACTCTCCCAGNNNAGATCGGAAG |
| 45273_1 | TATGAGTGTGGAGTCGTTGCTTGAGTCCCTGCTCTATTTCAGGNNNAGATCGGAAG |
| 45273_2 | TATGAGTGTGGAGTCGTTGCTTGAGTCCCTGCTCTATTTCAGGNNNAGATCGGAAG |
| 12786_1 | TATGAGTGTGGAGTCGTTGCGGAAAGAGGAAGGAGATCTGNNNAGATCGGAAGA |
| 12786_2 | TATGAGTGTGGAGTCGTTGCGGAAAGAGGAAGGAGATCTGNNNAGATCGGAAGA |
| 29049_1 | TATGAGTGTGGAGTCGTTGCACCTCTTCACAACACACTTGAGTNNNAGATCGGAAG |
| 29049_2 | TATGAGTGTGGAGTCGTTGCACCTCTTCACAACACACTTGAGTNNNAGATCGGAAG |
| 34771_1 | TATGAGTGTGGAGTCGTTGCCCACCACGCCCAGATAATTTTTTNNNAGATCGGAAGA |
| 34771_2 | TATGAGTGTGGAGTCGTTGCCCACCACGCCCAGATAATTTTTTNNNAGATCGGAAGA |
| 48845_1 | TATGAGTGTGGAGTCGTTGCTGATGGAGAGTTTTGGTGAGGAGNNNAGATCGGA |
| 48845_2 | TATGAGTGTGGAGTCGTTGCTGATGGAGAGTTTTGGTGAGGAGNNNAGATCGGA |
| 51798_1 | TATGAGTGTGGAGTCGTTGCATGCCTAGAATGTCTTTCCCAGGNNNAGATCGGAAG |
| 51798_2 | TATGAGTGTGGAGTCGTTGCATGCCTAGAATGTCTTTCCCAGGNNNAGATCGGAAG |
| 47522_1 | TATGAGTGTGGAGTCGTTGCACCACACCTGGCTACTTTTTGTANNNAGATCGGAAG |
| 47522_2 | TATGAGTGTGGAGTCGTTGCACCACACCTGGCTACTTTTTGTANNNAGATCGGAAG |
| 10437_1 | TATGAGTGTGGAGTCGTTGCGCCTTTGTGGTCCTCTAGATTCCNNNAGATCGGAAG |
| 10437_2 | TATGAGTGTGGAGTCGTTGCGCCTTTGTGGTCCTCTAGATTCCNNNAGATCGGAAG |
| 40051_1 | TATGAGTGTGGAGTCGTTGCAAAAGGAGTCAGCAAATGCACTCNNNAGATCGGAA |
| 40051_2 | TATGAGTGTGGAGTCGTTGCAAAAGGAGTCAGCAAATGCACTCNNNAGATCGGAA |
| 50800_1 | TATGAGTGTGGAGTCGTTGCAAGAAGGTGTGGGGTCAGAAATTNNNAGATCGGA |
| 50800_2 | TATGAGTGTGGAGTCGTTGCAAGAAGGTGTGGGGTCAGAAATTNNNAGATCGGA |
| 41140_1 | TATGAGTGTGGAGTCGTTGCCACACCTGTAATCCCAGCACTTTNNNAGATCGGAAGA |
| 41140_2 | TATGAGTGTGGAGTCGTTGCCACACCTGTAATCCCAGCACTTTNNNAGATCGGAAGA |
| 46360_1 | TATGAGTGTGGAGTCGTTGCCAGTGATGCAACTTCAGCTCACNNNAGATCGGAAGA |
| 46360_2 | TATGAGTGTGGAGTCGTTGCCAGTGATGCAACTTCAGCTCACNNNAGATCGGAAGA |
| 33375_1 | TATGAGTGTGGAGTCGTTGCTCCTAAATCTCCACACCCAGAGANNNAGATCGGAAG |
| 33375_2 | TATGAGTGTGGAGTCGTTGCTCCTAAATCTCCACACCCAGAGANNNAGATCGGAAG |
| 51642_1 | TATGAGTGTGGAGTCGTTGCCCCCCAGGGAGAAGATACATTACNNNAGATCGGAA |

| | |
|---|---|
| 51642_2 | TATGAGTGTGGAGTCGTTGCCCCCCAGGGAGAAGATACATTACNNNAGATCGGAAG |
| 44112_1 | TATGAGTGTGGAGTCGTTGCGCTCCTCTCCATTGCTCCTAAGNNNAGATCGGAAGAG |
| 44112_2 | TATGAGTGTGGAGTCGTTGCGCTCCTCTCCATTGCTCCTAAGNNNAGATCGGAAGAG |
| 28922_1 | TATGAGTGTGGAGTCGTTGCAATTGATTCAGTTGGGTCAGGGANNNAGATCGGAA |
| 28922_2 | TATGAGTGTGGAGTCGTTGCAATTGATTCAGTTGGGTCAGGGANNNAGATCGGAA |
| 44402_1 | TATGAGTGTGGAGTCGTTGCCCTCCTTTGGGGGCAATTTTGNNNAGATCGGAAGAG |
| 44402_2 | TATGAGTGTGGAGTCGTTGCCCTCCTTTGGGGGCAATTTTGNNNAGATCGGAAGAG |
| 38895_1 | TATGAGTGTGGAGTCGTTGCCCAAAAAAGAGCCCAAATAGCCANNNAGATCGGAAG |
| 38895_2 | TATGAGTGTGGAGTCGTTGCCCAAAAAAGAGCCCAAATAGCCANNNAGATCGGAAG |
| 46353_1 | TATGAGTGTGGAGTCGTTGCACAGGAGAGTGGATAATCTGGGANNNAGATCGGA |
| 46353_2 | TATGAGTGTGGAGTCGTTGCACAGGAGAGTGGATAATCTGGGANNNAGATCGGA |
| 2_1 | TATGAGTGTGGAGTCGTTGCAGCAGGCATGAAACACCTAAACANNNAGATCGGAA |
| 2_2 | TATGAGTGTGGAGTCGTTGCAGCAGGCATGAAACACCTAAACANNNAGATCGGAA |
| 10485_1 | TATGAGTGTGGAGTCGTTGCACAGTTTGTGGAGGAGTATGCAANNNAGATCGGAA |
| 10485_2 | TATGAGTGTGGAGTCGTTGCACAGTTTGTGGAGGAGTATGCAANNNAGATCGGAA |
| 9767_1 | TATGAGTGTGGAGTCGTTGCTCATGGCCTCTCCCTTCTGTTTANNNAGATCGGAAGA |
| 9767_2 | TATGAGTGTGGAGTCGTTGCTCATGGCCTCTCCCTTCTGTTTANNNAGATCGGAAGA |
| 39760_1 | TATGAGTGTGGAGTCGTTGCCAAGGACAGCTGTATTTTGCCATNNNAGATCGGAAG |
| 39760_2 | TATGAGTGTGGAGTCGTTGCCAAGGACAGCTGTATTTTGCCATNNNAGATCGGAAG |
| 10215_1 | TATGAGTGTGGAGTCGTTGCTTCCAGGATAGTGGCCTTTCAGANNNAGATCGGAAG |
| 10215_2 | TATGAGTGTGGAGTCGTTGCTTCCAGGATAGTGGCCTTTCAGANNNAGATCGGAAG |
| 40878_1 | TATGAGTGTGGAGTCGTTGCTGCGAGCCACAATTGTAAGAATGNNNAGATCGGAA |
| 40878_2 | TATGAGTGTGGAGTCGTTGCTGCGAGCCACAATTGTAAGAATGNNNAGATCGGAA |
| 41228_1 | TATGAGTGTGGAGTCGTTGCACTCAAACAGGTACTTGTCCACANNNAGATCGGAAG |
| 41228_2 | TATGAGTGTGGAGTCGTTGCACTCAAACAGGTACTTGTCCACANNNAGATCGGAAG |
| 32872_1 | TATGAGTGTGGAGTCGTTGCCATTCTCCTGCCTCAGCCTCNNNAGATCGGAAGAGCA |
| 32872_2 | TATGAGTGTGGAGTCGTTGCCATTCTCCTGCCTCAGCCTCNNNAGATCGGAAGAGCA |
| 42202_1 | TATGAGTGTGGAGTCGTTGCGGTGAGTCTGTGGTCATTTTGTGNNNAGATCGGAA |
| 42202_2 | TATGAGTGTGGAGTCGTTGCGGTGAGTCTGTGGTCATTTTGTGNNNAGATCGGAA |
| 43801_1 | TATGAGTGTGGAGTCGTTGCCCACATAAATATGAGGGAGCCCTNNNAGATCGGAAG |
| 43801_2 | TATGAGTGTGGAGTCGTTGCCCACATAAATATGAGGGAGCCCTNNNAGATCGGAAG |
| 40719_1 | TATGAGTGTGGAGTCGTTGCACGGAAATTAGCTGAGGGTAGTGNNNAGATCGGA |
| 40719_2 | TATGAGTGTGGAGTCGTTGCACGGAAATTAGCTGAGGGTAGTGNNNAGATCGGA |
| 47625_1 | TATGAGTGTGGAGTCGTTGCGACGGAGTCTCGTTCTGTCACNNNAGATCGGAAGAG |
| 47625_2 | TATGAGTGTGGAGTCGTTGCGACGGAGTCTCGTTCTGTCACNNNAGATCGGAAGAG |
| 10022_1 | TATGAGTGTGGAGTCGTTGCCTGTTTGTCAGGATTACCCTCGGNNNAGATCGGAAG |
| 10022_2 | TATGAGTGTGGAGTCGTTGCCTGTTTGTCAGGATTACCCTCGGNNNAGATCGGAAG |
| 33458_1 | TATGAGTGTGGAGTCGTTGCCACTCTAGTCTGGGCAACAGAGNNNAGATCGGAAG |
| 33458_2 | TATGAGTGTGGAGTCGTTGCCACTCTAGTCTGGGCAACAGAGNNNAGATCGGAAG |
| 36401_1 | TATGAGTGTGGAGTCGTTGCGGGTTGCAATCCTAGTCTGTGATNNNAGATCGGAAG |
| 36401_2 | TATGAGTGTGGAGTCGTTGCGGGTTGCAATCCTAGTCTGTGATNNNAGATCGGAAG |
| 33735_1 | TATGAGTGTGGAGTCGTTGCTTGCGCATGTTTCATAATGAAGANNNAGATCGGAAG |
| 33735_2 | TATGAGTGTGGAGTCGTTGCTTGCGCATGTTTCATAATGAAGANNNAGATCGGAAG |

| | |
|---|---|
| 45422_1 | TATGAGTGTGGAGTCGTTGCAAACCAAACATTGTTCTTGGGGGNNNAGATCGGAA |
| 45422_2 | TATGAGTGTGGAGTCGTTGCAAACCAAACATTGTTCTTGGGGGNNNAGATCGGAA |
| 49039_1 | TATGAGTGTGGAGTCGTTGCGAGTATCTGGGATTACAGGCGTGNNNAGATCGGAA |
| 49039_2 | TATGAGTGTGGAGTCGTTGCGAGTATCTGGGATTACAGGCGTGNNNAGATCGGAA |
| 31919_1 | TATGAGTGTGGAGTCGTTGCCATTGGCTCACACTTGTAATCCCNNNAGATCGGAAG |
| 31919_2 | TATGAGTGTGGAGTCGTTGCCATTGGCTCACACTTGTAATCCCNNNAGATCGGAAG |
| 45257_1 | TATGAGTGTGGAGTCGTTGCCCTTGAGGGCATTGCAGATATTGNNNAGATCGGAA |
| 45257_2 | TATGAGTGTGGAGTCGTTGCCCTTGAGGGCATTGCAGATATTGNNNAGATCGGAA |
| 45136_1 | TATGAGTGTGGAGTCGTTGCTGTTAGATGTGAGTGTTGGAGCANNNAGATCGGAA |
| 45136_2 | TATGAGTGTGGAGTCGTTGCTGTTAGATGTGAGTGTTGGAGCANNNAGATCGGAA |
| 48488_1 | TATGAGTGTGGAGTCGTTGCAAAGTGCTGGGATTACAGGCATNNNAGATCGGAAG |
| 48488_2 | TATGAGTGTGGAGTCGTTGCAAAGTGCTGGGATTACAGGCATNNNAGATCGGAAG |
| 50226_1 | TATGAGTGTGGAGTCGTTGCCTCCCTTCCTAGAACATCATCCGNNNAGATCGGAAGA |
| 50226_2 | TATGAGTGTGGAGTCGTTGCCTCCCTTCCTAGAACATCATCCGNNNAGATCGGAAGA |
| 43730_1 | TATGAGTGTGGAGTCGTTGCCTCCTGGCCTTGACCTTCAGNNNAGATCGGAAGAGC |
| 43730_2 | TATGAGTGTGGAGTCGTTGCCTCCTGGCCTTGACCTTCAGNNNAGATCGGAAGAGC |
| 48112_1 | TATGAGTGTGGAGTCGTTGCCTGAGTAGCTGGGATTACAGGTGNNNAGATCGGAA |
| 48112_2 | TATGAGTGTGGAGTCGTTGCCTGAGTAGCTGGGATTACAGGTGNNNAGATCGGAA |
| 37313_1 | TATGAGTGTGGAGTCGTTGCGGGTGATAGAGTGAGTCCCTTTCNNNAGATCGGAA |
| 37313_2 | TATGAGTGTGGAGTCGTTGCGGGTGATAGAGTGAGTCCCTTTCNNNAGATCGGAA |
| 37285_1 | TATGAGTGTGGAGTCGTTGCGAGGGTAGTGATGAAGACAGTGGNNNAGATCGGA |
| 37285_2 | TATGAGTGTGGAGTCGTTGCGAGGGTAGTGATGAAGACAGTGGNNNAGATCGGA |
| 48837_1 | TATGAGTGTGGAGTCGTTGCCTCCTGAGTAGCTGGGACTACAGNNNAGATCGGAA |
| 48837_2 | TATGAGTGTGGAGTCGTTGCCTCCTGAGTAGCTGGGACTACAGNNNAGATCGGAA |
| 39955_1 | TATGAGTGTGGAGTCGTTGCGAATAGGAATTACTGGGGGTGGGNNNAGATCGGA |
| 39955_2 | TATGAGTGTGGAGTCGTTGCGAATAGGAATTACTGGGGGTGGGNNNAGATCGGA |
| 32746_1 | TATGAGTGTGGAGTCGTTGCGAGGTCAGGAGATTGAGACCATCNNNAGATCGGAA |
| 32746_2 | TATGAGTGTGGAGTCGTTGCGAGGTCAGGAGATTGAGACCATCNNNAGATCGGAA |
| 43907_1 | TATGAGTGTGGAGTCGTTGCGAGTAGAACTGCTCAGATGTGCTNNNAGATCGGAA |
| 43907_2 | TATGAGTGTGGAGTCGTTGCGAGTAGAACTGCTCAGATGTGCTNNNAGATCGGAA |
| 39001_1 | TATGAGTGTGGAGTCGTTGCGCAAGAAAAAAACCAAACAATCCCANNNAGATCGGA |
| 39001_2 | TATGAGTGTGGAGTCGTTGCGCAAGAAAAAAACCAAACAATCCCANNNAGATCGGA |
| 30613_1 | TATGAGTGTGGAGTCGTTGCCATCAGTGCTACCCTCCCATCTANNNAGATCGGAAGA |
| 30613_2 | TATGAGTGTGGAGTCGTTGCCATCAGTGCTACCCTCCCATCTANNNAGATCGGAAGA |
| 49296_1 | TATGAGTGTGGAGTCGTTGCCCCATCTCTCAGAGGTTTTGCTANNNAGATCGGAAG |
| 49296_2 | TATGAGTGTGGAGTCGTTGCCCCATCTCTCAGAGGTTTTGCTANNNAGATCGGAAG |
| 38789_1 | TATGAGTGTGGAGTCGTTGCCTAGTCCACCATGACCAAGTAGGNNNAGATCGGAAG |
| 38789_2 | TATGAGTGTGGAGTCGTTGCCTAGTCCACCATGACCAAGTAGGNNNAGATCGGAAG |
| 42179_1 | TATGAGTGTGGAGTCGTTGCCCTCCCAAGTAGCTAGGACTACANNNAGATCGGAAG |
| 42179_2 | TATGAGTGTGGAGTCGTTGCCCTCCCAAGTAGCTAGGACTACANNNAGATCGGAAG |
| 48742_1 | TATGAGTGTGGAGTCGTTGCCCCAAAGTGCTAGGATTACAGGTNNNAGATCGGAA |
| 48742_2 | TATGAGTGTGGAGTCGTTGCCCCAAAGTGCTAGGATTACAGGTNNNAGATCGGAA |
| 43873_1 | TATGAGTGTGGAGTCGTTGCGCTGCTGTCTTCAAGTAAGCATGNNNAGATCGGAAG |

| | |
|---|---|
| 43873_2 | TATGAGTGTGGAGTCGTTGCGCTGCTGTCTTCAAGTAAGCATGNNNAGATCGGAAG |
| 48099_1 | TATGAGTGTGGAGTCGTTGCGTTCAGCAGTGTGGGCCATGNNNAGATCGGAAGAG |
| 48099_2 | TATGAGTGTGGAGTCGTTGCGTTCAGCAGTGTGGGCCATGNNNAGATCGGAAGAG |
| 52077_1 | TATGAGTGTGGAGTCGTTGCAAATACCCATACTCCAGACCCCANNNAGATCGGAAG |
| 52077_2 | TATGAGTGTGGAGTCGTTGCAAATACCCATACTCCAGACCCCANNNAGATCGGAAG |
| 30245_1 | TATGAGTGTGGAGTCGTTGCTTGGCTGATTTGGCTGGTTTTAGNNNAGATCGGAA |
| 30245_2 | TATGAGTGTGGAGTCGTTGCTTGGCTGATTTGGCTGGTTTTAGNNNAGATCGGAA |
| 44244_1 | TATGAGTGTGGAGTCGTTGCAGCTGTGAGGGAGGTAAAAAACTNNNAGATCGGA |
| 44244_2 | TATGAGTGTGGAGTCGTTGCAGCTGTGAGGGAGGTAAAAAACTNNNAGATCGGA |
| 31004_1 | TATGAGTGTGGAGTCGTTGCCATTTCCCTCAGTCTCCCATTCANNNAGATCGGAAGA |
| 31004_2 | TATGAGTGTGGAGTCGTTGCCATTTCCCTCAGTCTCCCATTCANNNAGATCGGAAGA |
| 52580_1 | TATGAGTGTGGAGTCGTTGCTTTTGTGTAGGGGGAAGAAGAGGNNNAGATCGGA |
| 52580_2 | TATGAGTGTGGAGTCGTTGCTTTTGTGTAGGGGGAAGAAGAGGNNNAGATCGGA |
| 37122_1 | TATGAGTGTGGAGTCGTTGCGGGTAATCCTTCTACATGCCAGTNNNAGATCGGAAG |
| 37122_2 | TATGAGTGTGGAGTCGTTGCGGGTAATCCTTCTACATGCCAGTNNNAGATCGGAAG |
| 40185_1 | TATGAGTGTGGAGTCGTTGCGATCACAAGGTCAGGAGATCGAGNNNAGATCGGAA |
| 40185_2 | TATGAGTGTGGAGTCGTTGCGATCACAAGGTCAGGAGATCGAGNNNAGATCGGAA |
| 39785_1 | TATGAGTGTGGAGTCGTTGCGGAAAAGCAAATCAGAGCCACAANNNAGATCGGAA |
| 39785_2 | TATGAGTGTGGAGTCGTTGCGGAAAAGCAAATCAGAGCCACAANNNAGATCGGAA |
| 34974_1 | TATGAGTGTGGAGTCGTTGCAGTCTCCTGAGTAGCTGGGATTNNNAGATCGGAAG |
| 34974_2 | TATGAGTGTGGAGTCGTTGCAGTCTCCTGAGTAGCTGGGATTNNNAGATCGGAAG |
| 47495_1 | TATGAGTGTGGAGTCGTTGCTTTCACAGAGCCAATTTGTGCATNNNAGATCGGAAG |
| 47495_2 | TATGAGTGTGGAGTCGTTGCTTTCACAGAGCCAATTTGTGCATNNNAGATCGGAAG |
| 47071_1 | TATGAGTGTGGAGTCGTTGCAGTGCTGGAATTACAGGTGTGATNNNAGATCGGAA |
| 47071_2 | TATGAGTGTGGAGTCGTTGCAGTGCTGGAATTACAGGTGTGATNNNAGATCGGAA |
| 13212_1 | TATGAGTGTGGAGTCGTTGCCCACTGTTTTTCATTTTTAGTACNNNAGATCGGAAGA |
| 13212_2 | TATGAGTGTGGAGTCGTTGCCCACTGTTTTTCATTTTTAGTACNNNAGATCGGAAGA |
| 37969_1 | TATGAGTGTGGAGTCGTTGCGGGGAGACTCTAAATGACCACTTNNNAGATCGGAA |
| 37969_2 | TATGAGTGTGGAGTCGTTGCGGGGAGACTCTAAATGACCACTTNNNAGATCGGAA |
| 51583_1 | TATGAGTGTGGAGTCGTTGCGCTGGCAGCTTTGTCAATAGTGNNNAGATCGGAAG |
| 51583_2 | TATGAGTGTGGAGTCGTTGCGCTGGCAGCTTTGTCAATAGTGNNNAGATCGGAAG |
| 32570_1 | TATGAGTGTGGAGTCGTTGCTTCACACCATTCCCTTGCCTCNNNAGATCGGAAGAGC |
| 32570_2 | TATGAGTGTGGAGTCGTTGCTTCACACCATTCCCTTGCCTCNNNAGATCGGAAGAGC |
| 43354_1 | TATGAGTGTGGAGTCGTTGCCCGGATTCAAACGATTCTCCTCTNNNAGATCGGAAG |
| 43354_2 | TATGAGTGTGGAGTCGTTGCCCGGATTCAAACGATTCTCCTCTNNNAGATCGGAAG |
| 42484_1 | TATGAGTGTGGAGTCGTTGCAGTGCTGGGATTATAGGCATGAGNNNAGATCGGAA |
| 42484_2 | TATGAGTGTGGAGTCGTTGCAGTGCTGGGATTATAGGCATGAGNNNAGATCGGAA |
| 32875_1 | TATGAGTGTGGAGTCGTTGCCAGGTTCAAGCAATTCTCCTGCNNNAGATCGGAAGA |
| 32875_2 | TATGAGTGTGGAGTCGTTGCCAGGTTCAAGCAATTCTCCTGCNNNAGATCGGAAGA |
| 10176_1 | TATGAGTGTGGAGTCGTTGCACCTTTTCCCTTGGATCTCTCCTNNNAGATCGGAAGA |
| 10176_2 | TATGAGTGTGGAGTCGTTGCACCTTTTCCCTTGGATCTCTCCTNNNAGATCGGAAGA |
| 40674_1 | TATGAGTGTGGAGTCGTTGCTCTGCATTTCCATCTGAGGTACCNNNAGATCGGAAG |
| 40674_2 | TATGAGTGTGGAGTCGTTGCTCTGCATTTCCATCTGAGGTACCNNNAGATCGGAAG |

| | |
|---|---|
| 45707_1 | TATGAGTGTGGAGTCGTTGCTCCAGGTTCAAGTGATTCTCCTGNNNAGATCGGAAG |
| 45707_2 | TATGAGTGTGGAGTCGTTGCTCCAGGTTCAAGTGATTCTCCTGNNNAGATCGGAAG |
| 39224_1 | TATGAGTGTGGAGTCGTTGCCTGTGTGTGTGTGTGTGTGTATANNNAGATCGGAA |
| 39224_2 | TATGAGTGTGGAGTCGTTGCCTGTGTGTGTGTGTGTGTGTATANNNAGATCGGAA |
| 43161_1 | TATGAGTGTGGAGTCGTTGCCGATCTCCTGACCTCATGATCTGNNNAGATCGGAAG |
| 43161_2 | TATGAGTGTGGAGTCGTTGCCGATCTCCTGACCTCATGATCTGNNNAGATCGGAAG |
| 52995_1 | TATGAGTGTGGAGTCGTTGCCTTCCTTCCTTGGTTTCCTTCCTNNNAGATCGGAAGA |
| 52995_2 | TATGAGTGTGGAGTCGTTGCCTTCCTTCCTTGGTTTCCTTCCTNNNAGATCGGAAGA |
| 32485_1 | TATGAGTGTGGAGTCGTTGCCTCAAGAGATCCTCCCACCTTAGNNNAGATCGGAAG |
| 32485_2 | TATGAGTGTGGAGTCGTTGCCTCAAGAGATCCTCCCACCTTAGNNNAGATCGGAAG |
| 32601_1 | TATGAGTGTGGAGTCGTTGCGCAGAGGTTGGTTCATGATGTTTNNNAGATCGGAA |
| 32601_2 | TATGAGTGTGGAGTCGTTGCGCAGAGGTTGGTTCATGATGTTTNNNAGATCGGAA |
| 46189_1 | TATGAGTGTGGAGTCGTTGCTAGCTGGGACTATACATGTGTGCNNNAGATCGGAA |
| 46189_2 | TATGAGTGTGGAGTCGTTGCTAGCTGGGACTATACATGTGTGCNNNAGATCGGAA |
| 29615_1 | TATGAGTGTGGAGTCGTTGCTGAAAACAAAGAGTTCAAAGGTCANNNAGATCGGA |
| 29615_2 | TATGAGTGTGGAGTCGTTGCTGAAAACAAAGAGTTCAAAGGTCANNNAGATCGGA |
| 50565_1 | TATGAGTGTGGAGTCGTTGCACAGCCAGAGAATGAAGTGGTTTNNNAGATCGGAA |
| 50565_2 | TATGAGTGTGGAGTCGTTGCACAGCCAGAGAATGAAGTGGTTTNNNAGATCGGAA |
| 44193_1 | TATGAGTGTGGAGTCGTTGCCATACTGTCTGGTGCTTGCTTTTNNNAGATCGGAAG |
| 44193_2 | TATGAGTGTGGAGTCGTTGCCATACTGTCTGGTGCTTGCTTTTNNNAGATCGGAAG |
| 52935_1 | TATGAGTGTGGAGTCGTTGCCTCACGCTGCTATAAAGAACTGCNNNAGATCGGAAG |
| 52935_2 | TATGAGTGTGGAGTCGTTGCCTCACGCTGCTATAAAGAACTGCNNNAGATCGGAAG |
| 35701_1 | TATGAGTGTGGAGTCGTTGCCAAAACTGCTGGGATTACAGGTGNNNAGATCGGAA |
| 35701_2 | TATGAGTGTGGAGTCGTTGCCAAAACTGCTGGGATTACAGGTGNNNAGATCGGAA |
| 28935_1 | TATGAGTGTGGAGTCGTTGCCACATTCAACTCACTAGCGCAAANNNAGATCGGAAG |
| 28935_2 | TATGAGTGTGGAGTCGTTGCCACATTCAACTCACTAGCGCAAANNNAGATCGGAAG |
| 31666_1 | TATGAGTGTGGAGTCGTTGCTGGGCTCAAAGTAGACAAATCCANNNAGATCGGAA |
| 31666_2 | TATGAGTGTGGAGTCGTTGCTGGGCTCAAAGTAGACAAATCCANNNAGATCGGAA |
| 28851_1 | TATGAGTGTGGAGTCGTTGCGTGTCATGCAGGTGTACTTTGACNNNAGATCGGAA |
| 28851_2 | TATGAGTGTGGAGTCGTTGCGTGTCATGCAGGTGTACTTTGACNNNAGATCGGAA |
| 45735_1 | TATGAGTGTGGAGTCGTTGCTTCAAGCAATTGTTCTGCCTCAGNNNAGATCGGAAG |
| 45735_2 | TATGAGTGTGGAGTCGTTGCTTCAAGCAATTGTTCTGCCTCAGNNNAGATCGGAAG |
| 40834_1 | TATGAGTGTGGAGTCGTTGCCTGAGGCAGGAGAATCACTTGAGNNNAGATCGGAA |
| 40834_2 | TATGAGTGTGGAGTCGTTGCCTGAGGCAGGAGAATCACTTGAGNNNAGATCGGAA |
| 41724_1 | TATGAGTGTGGAGTCGTTGCCTGTCACCAAGGCTGGACTATAGNNNAGATCGGAA |
| 41724_2 | TATGAGTGTGGAGTCGTTGCCTGTCACCAAGGCTGGACTATAGNNNAGATCGGAA |
| 42944_1 | TATGAGTGTGGAGTCGTTGCGGAGGACAGTGTGAGGAAGAATTNNNAGATCGGA |
| 42944_2 | TATGAGTGTGGAGTCGTTGCGGAGGACAGTGTGAGGAAGAATTNNNAGATCGGA |
| 34281_1 | TATGAGTGTGGAGTCGTTGCCCTGCCCTCTTGTGATTGTTTTTNNNAGATCGGAAGA |
| 34281_2 | TATGAGTGTGGAGTCGTTGCCCTGCCCTCTTGTGATTGTTTTTNNNAGATCGGAAGA |
| 39539_1 | TATGAGTGTGGAGTCGTTGCGGTAAAACCAAGGGAGTATGGGTNNNAGATCGGA |
| 39539_2 | TATGAGTGTGGAGTCGTTGCGGTAAAACCAAGGGAGTATGGGTNNNAGATCGGA |
| 43334_1 | TATGAGTGTGGAGTCGTTGCGGAGTGCAATGTCGTGATCTTGNNNAGATCGGAAG |

| | |
|---|---|
| 43334_2 | TATGAGTGTGGAGTCGTTGCGGAGTGCAATGTCGTGATCTTGNNNAGATCGGAAG |
| 33527_1 | TATGAGTGTGGAGTCGTTGCTGGGCAAAAGATATGAAGAGGCANNNAGATCGGA |
| 33527_2 | TATGAGTGTGGAGTCGTTGCTGGGCAAAAGATATGAAGAGGCANNNAGATCGGA |
| 52934_1 | TATGAGTGTGGAGTCGTTGCCAGGAAAAATCTGTGGGCACATTNNNAGATCGGAA |
| 52934_2 | TATGAGTGTGGAGTCGTTGCCAGGAAAAATCTGTGGGCACATTNNNAGATCGGAA |
| 10989_1 | TATGAGTGTGGAGTCGTTGCCTCCACCCCAGGTTATGACAGACNNNAGATCGGAAG |
| 10989_2 | TATGAGTGTGGAGTCGTTGCCTCCACCCCAGGTTATGACAGACNNNAGATCGGAAG |
| 43752_1 | TATGAGTGTGGAGTCGTTGCGCCAGCTGAGCAATCCTATTTTCNNNAGATCGGAAG |
| 43752_2 | TATGAGTGTGGAGTCGTTGCGCCAGCTGAGCAATCCTATTTTCNNNAGATCGGAAG |
| 35974_1 | TATGAGTGTGGAGTCGTTGCTTCAAGATCAGCCTGGACAACATNNNAGATCGGAAG |
| 35974_2 | TATGAGTGTGGAGTCGTTGCTTCAAGATCAGCCTGGACAACATNNNAGATCGGAAG |
| 44504_1 | TATGAGTGTGGAGTCGTTGCGAAAGTGGGACAGTGAGAGTGAANNNAGATCGGA |
| 44504_2 | TATGAGTGTGGAGTCGTTGCGAAAGTGGGACAGTGAGAGTGAANNNAGATCGGA |
| 50023_1 | TATGAGTGTGGAGTCGTTGCCATCAGTGACTTCCACGAAACTTNNNAGATCGGAAG |
| 50023_2 | TATGAGTGTGGAGTCGTTGCCATCAGTGACTTCCACGAAACTTNNNAGATCGGAAG |
| 40871_1 | TATGAGTGTGGAGTCGTTGCCTCCTAGGGGAAGGCAGAGTATANNNAGATCGGAA |
| 40871_2 | TATGAGTGTGGAGTCGTTGCCTCCTAGGGGAAGGCAGAGTATANNNAGATCGGAA |
| 39650_1 | TATGAGTGTGGAGTCGTTGCTACAATGGACTTTGGGACTTGGGNNNAGATCGGAA |
| 39650_2 | TATGAGTGTGGAGTCGTTGCTACAATGGACTTTGGGACTTGGGNNNAGATCGGAA |
| 46159_1 | TATGAGTGTGGAGTCGTTGCAAACTCCTGGGCTCAAGCAATCNNNAGATCGGAAGA |
| 46159_2 | TATGAGTGTGGAGTCGTTGCAAACTCCTGGGCTCAAGCAATCNNNAGATCGGAAGA |
| 34612_1 | TATGAGTGTGGAGTCGTTGCCAACATACTGAGAGCCCATCTGTNNNAGATCGGAAG |
| 34612_2 | TATGAGTGTGGAGTCGTTGCCAACATACTGAGAGCCCATCTGTNNNAGATCGGAAG |
| 33070_1 | TATGAGTGTGGAGTCGTTGCGGACTTCCATGGCTCAAGTGATCNNNAGATCGGAAG |
| 33070_2 | TATGAGTGTGGAGTCGTTGCGGACTTCCATGGCTCAAGTGATCNNNAGATCGGAAG |
| 43356_1 | TATGAGTGTGGAGTCGTTGCCGCATCCCATCAACTCATGTCTTNNNAGATCGGAAGA |
| 43356_2 | TATGAGTGTGGAGTCGTTGCCGCATCCCATCAACTCATGTCTTNNNAGATCGGAAGA |
| 37660_1 | TATGAGTGTGGAGTCGTTGCACACTCCTCTTTCTGCTTTCTGGNNNAGATCGGAAGA |
| 37660_2 | TATGAGTGTGGAGTCGTTGCACACTCCTCTTTCTGCTTTCTGGNNNAGATCGGAAGA |
| 40276_1 | TATGAGTGTGGAGTCGTTGCTGGATTATGGATGCTGAACCAGTNNNAGATCGGAA |
| 40276_2 | TATGAGTGTGGAGTCGTTGCTGGATTATGGATGCTGAACCAGTNNNAGATCGGAA |
| 30874_1 | TATGAGTGTGGAGTCGTTGCCCCTTGCCTAATCTAGGACTTCCNNNAGATCGGAAGA |
| 30874_2 | TATGAGTGTGGAGTCGTTGCCCCTTGCCTAATCTAGGACTTCCNNNAGATCGGAAGA |
| 48353_1 | TATGAGTGTGGAGTCGTTGCTTCAGAGGCGTTTAGTACATGGGNNNAGATCGGAA |
| 48353_2 | TATGAGTGTGGAGTCGTTGCTTCAGAGGCGTTTAGTACATGGGNNNAGATCGGAA |
| 48941_1 | TATGAGTGTGGAGTCGTTGCAGTTGAACCCAGGTACCTTTCTCNNNAGATCGGAAG |
| 48941_2 | TATGAGTGTGGAGTCGTTGCAGTTGAACCCAGGTACCTTTCTCNNNAGATCGGAAG |
| 8545_1 | TATGAGTGTGGAGTCGTTGCTTCCAGACCCTATCTCAGAGCCNNNAGATCGGAAGA |
| 8545_2 | TATGAGTGTGGAGTCGTTGCTTCCAGACCCTATCTCAGAGCCNNNAGATCGGAAGA |
| 39639_1 | TATGAGTGTGGAGTCGTTGCTTGCCTTCTAACTCCTGAACTCANNNAGATCGGAAGA |
| 39639_2 | TATGAGTGTGGAGTCGTTGCTTGCCTTCTAACTCCTGAACTCANNNAGATCGGAAGA |
| 39087_1 | TATGAGTGTGGAGTCGTTGCCCCCATGTGCTTAGATTGGAAGANNNAGATCGGAA |
| 39087_2 | TATGAGTGTGGAGTCGTTGCCCCCATGTGCTTAGATTGGAAGANNNAGATCGGAA |

| | |
|---|---|
| 29145_1 | TATGAGTGTGGAGTCGTTGCTGTTGGTTGAACCTAACAGTCGGNNNAGATCGGAAG |
| 29145_2 | TATGAGTGTGGAGTCGTTGCTGTTGGTTGAACCTAACAGTCGGNNNAGATCGGAAG |
| 44544_1 | TATGAGTGTGGAGTCGTTGCTGGAATTGAGAGGGAGACAACTGNNNAGATCGGAA |
| 44544_2 | TATGAGTGTGGAGTCGTTGCTGGAATTGAGAGGGAGACAACTGNNNAGATCGGAA |
| 50215_1 | TATGAGTGTGGAGTCGTTGCCACATCAGGCACAAATTCCAGAGNNNAGATCGGAAG |
| 50215_2 | TATGAGTGTGGAGTCGTTGCCACATCAGGCACAAATTCCAGAGNNNAGATCGGAAG |
| 33013_1 | TATGAGTGTGGAGTCGTTGCCATTCTCCTGCCTCAGCCTCNNNAGATCGGAAGAGCA |
| 33013_2 | TATGAGTGTGGAGTCGTTGCCATTCTCCTGCCTCAGCCTCNNNAGATCGGAAGAGCA |
| 28918_1 | TATGAGTGTGGAGTCGTTGCAGAATCTGATGAGGTAAGCAGGCNNNAGATCGGAA |
| 28918_2 | TATGAGTGTGGAGTCGTTGCAGAATCTGATGAGGTAAGCAGGCNNNAGATCGGAA |
| 34020_1 | TATGAGTGTGGAGTCGTTGCCAAGTAGCTGGGACTACAGGCNNNAGATCGGAAGA |
| 34020_2 | TATGAGTGTGGAGTCGTTGCCAAGTAGCTGGGACTACAGGCNNNAGATCGGAAGA |
| 34123_1 | TATGAGTGTGGAGTCGTTGCATGTCTTACATGGTAGCAGGCAANNNAGATCGGAAG |
| 34123_2 | TATGAGTGTGGAGTCGTTGCATGTCTTACATGGTAGCAGGCAANNNAGATCGGAAG |
| 29231_1 | TATGAGTGTGGAGTCGTTGCTCAGAGATAATGCCCCCAAGATGNNNAGATCGGAAG |
| 29231_2 | TATGAGTGTGGAGTCGTTGCTCAGAGATAATGCCCCCAAGATGNNNAGATCGGAAG |
| 50768_1 | TATGAGTGTGGAGTCGTTGCCTAAACCAGCCCTGCAAACGNNNAGATCGGAAGAGC |
| 50768_2 | TATGAGTGTGGAGTCGTTGCCTAAACCAGCCCTGCAAACGNNNAGATCGGAAGAGC |
| 39387_1 | TATGAGTGTGGAGTCGTTGCGGAGCAGCTTGCAATACCATATGNNNAGATCGGAAG |
| 39387_2 | TATGAGTGTGGAGTCGTTGCGGAGCAGCTTGCAATACCATATGNNNAGATCGGAAG |
| 41588_1 | TATGAGTGTGGAGTCGTTGCGTTCACTATAACCTCTGCTCCCANNNAGATCGGAAGA |
| 41588_2 | TATGAGTGTGGAGTCGTTGCGTTCACTATAACCTCTGCTCCCANNNAGATCGGAAGA |
| 30960_1 | TATGAGTGTGGAGTCGTTGCCAGTTGAGCTTTGAGCAGTGATGNNNAGATCGGAAG |
| 30960_2 | TATGAGTGTGGAGTCGTTGCCAGTTGAGCTTTGAGCAGTGATGNNNAGATCGGAAG |
| 40000_1 | TATGAGTGTGGAGTCGTTGCGCTCTGATGACCCAAATTGTCAGNNNAGATCGGAAG |
| 40000_2 | TATGAGTGTGGAGTCGTTGCGCTCTGATGACCCAAATTGTCAGNNNAGATCGGAAG |
| 47085_1 | TATGAGTGTGGAGTCGTTGCTTCAGTGGTATGATCACGGTTCANNNAGATCGGAAG |
| 47085_2 | TATGAGTGTGGAGTCGTTGCTTCAGTGGTATGATCACGGTTCANNNAGATCGGAAG |
| 42027_1 | TATGAGTGTGGAGTCGTTGCCTGTCATCCCAGCACTTTGAGAGNNNAGATCGGAAG |
| 42027_2 | TATGAGTGTGGAGTCGTTGCCTGTCATCCCAGCACTTTGAGAGNNNAGATCGGAAG |
| 39456_1 | TATGAGTGTGGAGTCGTTGCTCAGGTGATTCTTATGCCTCAGCNNNAGATCGGAAGA |
| 39456_2 | TATGAGTGTGGAGTCGTTGCTCAGGTGATTCTTATGCCTCAGCNNNAGATCGGAAGA |
| 33054_1 | TATGAGTGTGGAGTCGTTGCTGGAGTTTGTTATTGGTCTGTTCANNNAGATCGGAA |
| 33054_2 | TATGAGTGTGGAGTCGTTGCTGGAGTTTGTTATTGGTCTGTTCANNNAGATCGGAA |
| 37728_1 | TATGAGTGTGGAGTCGTTGCGAGGAACTGCCAGACTGTTTTTCNNNAGATCGGAAG |
| 37728_2 | TATGAGTGTGGAGTCGTTGCGAGGAACTGCCAGACTGTTTTTCNNNAGATCGGAAG |
| 29257_1 | TATGAGTGTGGAGTCGTTGCACACAAACACACACACACACATTNNNAGATCGGAAGA |
| 29257_2 | TATGAGTGTGGAGTCGTTGCACACAAACACACACACACACATTNNNAGATCGGAAGA |
| 51463_1 | TATGAGTGTGGAGTCGTTGCAGGAGAGTTTTGGCATCAAAAGCNNNAGATCGGAAG |
| 51463_2 | TATGAGTGTGGAGTCGTTGCAGGAGAGTTTTGGCATCAAAAGCNNNAGATCGGAAG |
| 10477_1 | TATGAGTGTGGAGTCGTTGCATTTAGTGCCCAGCAAGCATACANNNAGATCGGAAG |
| 10477_2 | TATGAGTGTGGAGTCGTTGCATTTAGTGCCCAGCAAGCATACANNNAGATCGGAAG |
| 33824_1 | TATGAGTGTGGAGTCGTTGCAGTCCAAAGGCTGTTCTTTAGGANNNAGATCGGAAG |

| | |
|---|---|
| 33824_2 | TATGAGTGTGGAGTCGTTGCAGTCCAAAGGCTGTTCTTTAGGANNNAGATCGGAAG |
| 52093_1 | TATGAGTGTGGAGTCGTTGCCTGCATGGTAACGATTACATGCANNNAGATCGGAAG |
| 52093_2 | TATGAGTGTGGAGTCGTTGCCTGCATGGTAACGATTACATGCANNNAGATCGGAAG |
| 43816_1 | TATGAGTGTGGAGTCGTTGCATGAGAACTGAAAGTCACCACGTNNNAGATCGGAA |
| 43816_2 | TATGAGTGTGGAGTCGTTGCATGAGAACTGAAAGTCACCACGTNNNAGATCGGAA |
| 47599_1 | TATGAGTGTGGAGTCGTTGCATCAGGATGTCTGGCTGTGTTCNNNAGATCGGAAG |
| 47599_2 | TATGAGTGTGGAGTCGTTGCATCAGGATGTCTGGCTGTGTTCNNNAGATCGGAAG |
| 30909_1 | TATGAGTGTGGAGTCGTTGCGAAGGGTCAGACTTGCTCAAAACNNNAGATCGGAA |
| 30909_2 | TATGAGTGTGGAGTCGTTGCGAAGGGTCAGACTTGCTCAAAACNNNAGATCGGAA |
| 29011_1 | TATGAGTGTGGAGTCGTTGCAGTCAAGATTCCCTAAGCAGCAANNNAGATCGGAAG |
| 29011_2 | TATGAGTGTGGAGTCGTTGCAGTCAAGATTCCCTAAGCAGCAANNNAGATCGGAAG |
| 46800_1 | TATGAGTGTGGAGTCGTTGCCTCCTGAGTAGGTGGGACTACAGNNNAGATCGGAA |
| 46800_2 | TATGAGTGTGGAGTCGTTGCCTCCTGAGTAGGTGGGACTACAGNNNAGATCGGAA |
| 12562_1 | TATGAGTGTGGAGTCGTTGCATTCCTTATTGACATAGCTTAGCNNNAGATCGGAAG |
| 12562_2 | TATGAGTGTGGAGTCGTTGCATTCCTTATTGACATAGCTTAGCNNNAGATCGGAAG |
| 10665_1 | TATGAGTGTGGAGTCGTTGCTAGGATATGGGCCCAAGGAGAAGNNNAGATCGGA |
| 10665_2 | TATGAGTGTGGAGTCGTTGCTAGGATATGGGCCCAAGGAGAAGNNNAGATCGGA |
| 33731_1 | TATGAGTGTGGAGTCGTTGCCATGTGCGTTTCCATGGTATCAANNNAGATCGGAAG |
| 33731_2 | TATGAGTGTGGAGTCGTTGCCATGTGCGTTTCCATGGTATCAANNNAGATCGGAAG |
| 40438_1 | TATGAGTGTGGAGTCGTTGCGGCACCTATAATCCCAGCTACTCNNNAGATCGGAAG |
| 40438_2 | TATGAGTGTGGAGTCGTTGCGGCACCTATAATCCCAGCTACTCNNNAGATCGGAAG |
| 12591_1 | TATGAGTGTGGAGTCGTTGCGTTTAAACTACTATGAGAAAAGGANNNAGATCGGA |
| 12591_2 | TATGAGTGTGGAGTCGTTGCGTTTAAACTACTATGAGAAAAGGANNNAGATCGGA |
| 44693_1 | TATGAGTGTGGAGTCGTTGCTCCTCCATAGCAAGCATGAAACTNNNAGATCGGAAG |
| 44693_2 | TATGAGTGTGGAGTCGTTGCTCCTCCATAGCAAGCATGAAACTNNNAGATCGGAAG |
| 2041_1 | TATGAGTGTGGAGTCGTTGCTTTCCCTGACCTGGAAGAGTGAANNNAGATCGGAAG |
| 2041_2 | TATGAGTGTGGAGTCGTTGCTTTCCCTGACCTGGAAGAGTGAANNNAGATCGGAAG |
| 12796_1 | TATGAGTGTGGAGTCGTTGCATTCTGTAGATAATGTCTTGGATNNNAGATCGGAAG |
| 12796_2 | TATGAGTGTGGAGTCGTTGCATTCTGTAGATAATGTCTTGGATNNNAGATCGGAAG |
| 41810_1 | TATGAGTGTGGAGTCGTTGCCAATCTTGGCTCACTGCAATCTCNNNAGATCGGAAG |
| 41810_2 | TATGAGTGTGGAGTCGTTGCCAATCTTGGCTCACTGCAATCTCNNNAGATCGGAAG |
| 39295_1 | TATGAGTGTGGAGTCGTTGCGGAGGTCTTTTGTCAATATGCCGNNNAGATCGGAAG |
| 39295_2 | TATGAGTGTGGAGTCGTTGCGGAGGTCTTTTGTCAATATGCCGNNNAGATCGGAAG |
| 49692_1 | TATGAGTGTGGAGTCGTTGCCTGTTGCATGAGGGACGTAAAATNNNAGATCGGAA |
| 49692_2 | TATGAGTGTGGAGTCGTTGCCTGTTGCATGAGGGACGTAAAATNNNAGATCGGAA |
| 45031_1 | TATGAGTGTGGAGTCGTTGCAACTCGGTCACGGTCATATTTCANNNAGATCGGAAG |
| 45031_2 | TATGAGTGTGGAGTCGTTGCAACTCGGTCACGGTCATATTTCANNNAGATCGGAAG |
| 32234_1 | TATGAGTGTGGAGTCGTTGCTCTGGGTGCTGAGTCTCTAATTGNNNAGATCGGAAG |
| 32234_2 | TATGAGTGTGGAGTCGTTGCTCTGGGTGCTGAGTCTCTAATTGNNNAGATCGGAAG |
| 32509_1 | TATGAGTGTGGAGTCGTTGCTGGAATTCCTCTTGCTAGATAGGGNNNAGATCGGAA |
| 32509_2 | TATGAGTGTGGAGTCGTTGCTGGAATTCCTCTTGCTAGATAGGGNNNAGATCGGAA |
| 43729_1 | TATGAGTGTGGAGTCGTTGCGGACTGAACCCGAGGATTCACNNNAGATCGGAAGA |
| 43729_2 | TATGAGTGTGGAGTCGTTGCGGACTGAACCCGAGGATTCACNNNAGATCGGAAGA |

| | |
|---|---|
| 47436_1 | TATGAGTGTGGAGTCGTTGCGCAGTGGCACAATCATAACCTTTNNNAGATCGGAAG |
| 47436_2 | TATGAGTGTGGAGTCGTTGCGCAGTGGCACAATCATAACCTTTNNNAGATCGGAAG |
| 34062_1 | TATGAGTGTGGAGTCGTTGCTTCAAGTGATTCTCCTGCCTCAGNNNAGATCGGAAG |
| 34062_2 | TATGAGTGTGGAGTCGTTGCTTCAAGTGATTCTCCTGCCTCAGNNNAGATCGGAAG |
| 41901_1 | TATGAGTGTGGAGTCGTTGCGGTTCAAGTGATTCTCTTGCCTCNNNAGATCGGAAG |
| 41901_2 | TATGAGTGTGGAGTCGTTGCGGTTCAAGTGATTCTCTTGCCTCNNNAGATCGGAAG |
| 39868_1 | TATGAGTGTGGAGTCGTTGCTCTGTTTCAAGGCTCAGACCATTNNNAGATCGGAAG |
| 39868_2 | TATGAGTGTGGAGTCGTTGCTCTGTTTCAAGGCTCAGACCATTNNNAGATCGGAAG |
| 3896_1 | TATGAGTGTGGAGTCGTTGCAGGATGTTTGTACCTGTCCCACTNNNAGATCGGAAG |
| 3896_2 | TATGAGTGTGGAGTCGTTGCAGGATGTTTGTACCTGTCCCACTNNNAGATCGGAAG |
| 955_1 | TATGAGTGTGGAGTCGTTGCTATTATCGGCAAGGCCTGGTAGCNNNAGATCGGAA |
| 955_2 | TATGAGTGTGGAGTCGTTGCTATTATCGGCAAGGCCTGGTAGCNNNAGATCGGAA |
| 50343_1 | TATGAGTGTGGAGTCGTTGCAGTTAGACACTAGCTCTGGCAACNNNAGATCGGAAG |
| 50343_2 | TATGAGTGTGGAGTCGTTGCAGTTAGACACTAGCTCTGGCAACNNNAGATCGGAAG |
| 44777_1 | TATGAGTGTGGAGTCGTTGCAGAGGCTGGGAAAGACATGAAATNNNAGATCGGA |
| 44777_2 | TATGAGTGTGGAGTCGTTGCAGAGGCTGGGAAAGACATGAAATNNNAGATCGGA |
| 30156_1 | TATGAGTGTGGAGTCGTTGCTTTTGTCTGTTTATGCTTGGGCANNNAGATCGGAAG |
| 30156_2 | TATGAGTGTGGAGTCGTTGCTTTTGTCTGTTTATGCTTGGGCANNNAGATCGGAAG |
| 7821_1 | TATGAGTGTGGAGTCGTTGCGCCACTATATACACACACTGCCANNNAGATCGGAAG |
| 7821_2 | TATGAGTGTGGAGTCGTTGCGCCACTATATACACACACTGCCANNNAGATCGGAAG |
| 51121_1 | TATGAGTGTGGAGTCGTTGCGGGACCTGATGTGAACGTACAANNNAGATCGGAAG |
| 51121_2 | TATGAGTGTGGAGTCGTTGCGGGACCTGATGTGAACGTACAANNNAGATCGGAAG |
| 49807_1 | TATGAGTGTGGAGTCGTTGCGAAATTTAGTCTTCCTGGCCGTGNNNAGATCGGAAG |
| 49807_2 | TATGAGTGTGGAGTCGTTGCGAAATTTAGTCTTCCTGGCCGTGNNNAGATCGGAAG |
| 36330_1 | TATGAGTGTGGAGTCGTTGCAAAGCAGAAGTGCCCTAATCTCANNNAGATCGGAAG |
| 36330_2 | TATGAGTGTGGAGTCGTTGCAAAGCAGAAGTGCCCTAATCTCANNNAGATCGGAAG |
| 12778_1 | TATGAGTGTGGAGTCGTTGCGTCATTGGTACAGTCTCAAATANNNAGATCGGAAGA |
| 12778_2 | TATGAGTGTGGAGTCGTTGCGTCATTGGTACAGTCTCAAATANNNAGATCGGAAGA |
| 51416_1 | TATGAGTGTGGAGTCGTTGCTTTCCAGGGATGTTCCTTGTTGANNNAGATCGGAAG |
| 51416_2 | TATGAGTGTGGAGTCGTTGCTTTCCAGGGATGTTCCTTGTTGANNNAGATCGGAAG |
| 47685_1 | TATGAGTGTGGAGTCGTTGCCTCCTCTCCATTCCAGTCCATTCNNNAGATCGGAAGA |
| 47685_2 | TATGAGTGTGGAGTCGTTGCCTCCTCTCCATTCCAGTCCATTCNNNAGATCGGAAGA |
| 595_1 | TATGAGTGTGGAGTCGTTGCACTTGTGACCTGTAGTGGCTGTNNNAGATCGGAAG |
| 595_2 | TATGAGTGTGGAGTCGTTGCACTTGTGACCTGTAGTGGCTGTNNNAGATCGGAAG |
| 44992_1 | TATGAGTGTGGAGTCGTTGCGGATGTCCAAGAAACCCCAAAAGNNNAGATCGGAA |
| 44992_2 | TATGAGTGTGGAGTCGTTGCGGATGTCCAAGAAACCCCAAAAGNNNAGATCGGAA |
| 3044_1 | TATGAGTGTGGAGTCGTTGCCTATGTTGTCAGGGGCAATGTGANNNAGATCGGAA |
| 3044_2 | TATGAGTGTGGAGTCGTTGCCTATGTTGTCAGGGGCAATGTGANNNAGATCGGAA |
| 38581_1 | TATGAGTGTGGAGTCGTTGCTGGGGCTCCTTTGTAATTTCTCTNNNAGATCGGAAG |
| 38581_2 | TATGAGTGTGGAGTCGTTGCTGGGGCTCCTTTGTAATTTCTCTNNNAGATCGGAAG |
| 44418_1 | TATGAGTGTGGAGTCGTTGCGGGCCAGACTTTGTGAGGTTATANNNAGATCGGAA |
| 44418_2 | TATGAGTGTGGAGTCGTTGCGGGCCAGACTTTGTGAGGTTATANNNAGATCGGAA |
| 48730_1 | TATGAGTGTGGAGTCGTTGCCTCCTGGGTTCAAGTGATTCTCANNNAGATCGGAAG |

| | |
|---|---|
| 48730_2 | TATGAGTGTGGAGTCGTTGCCTCCTGGGTTCAAGTGATTCTCANNNAGATCGGAAG |
| 45765_1 | TATGAGTGTGGAGTCGTTGCGAGTAGCTGAGACTACAGGTGTGNNNAGATCGGAA |
| 45765_2 | TATGAGTGTGGAGTCGTTGCGAGTAGCTGAGACTACAGGTGTGNNNAGATCGGAA |
| 38157_1 | TATGAGTGTGGAGTCGTTGCAAATTGAAGGCAAGAAGGCACTCNNNAGATCGGAA |
| 38157_2 | TATGAGTGTGGAGTCGTTGCAAATTGAAGGCAAGAAGGCACTCNNNAGATCGGAA |
| 11002_1 | TATGAGTGTGGAGTCGTTGCAGTCTGTTTTGCCAGGCTGGAGNNNAGATCGGAAG |
| 11002_2 | TATGAGTGTGGAGTCGTTGCAGTCTGTTTTGCCAGGCTGGAGNNNAGATCGGAAG |
| 29074_1 | TATGAGTGTGGAGTCGTTGCGTTAGGCCTCCTCCTTCTTTCAANNNAGATCGGAAGA |
| 29074_2 | TATGAGTGTGGAGTCGTTGCGTTAGGCCTCCTCCTTCTTTCAANNNAGATCGGAAGA |
| 46519_1 | TATGAGTGTGGAGTCGTTGCGGTTCAAACAATTCTCCTGCCTCNNNAGATCGGAAG |
| 46519_2 | TATGAGTGTGGAGTCGTTGCGGTTCAAACAATTCTCCTGCCTCNNNAGATCGGAAG |
| 38381_1 | TATGAGTGTGGAGTCGTTGCCTTGAGCTCAGGAGTTGGAGATTNNNAGATCGGAA |
| 38381_2 | TATGAGTGTGGAGTCGTTGCCTTGAGCTCAGGAGTTGGAGATTNNNAGATCGGAA |
| 49127_1 | TATGAGTGTGGAGTCGTTGCTAGAAACGGGGTTTCGCCATGNNNAGATCGGAAGA |
| 49127_2 | TATGAGTGTGGAGTCGTTGCTAGAAACGGGGTTTCGCCATGNNNAGATCGGAAGA |
| 38600_1 | TATGAGTGTGGAGTCGTTGCAACTTCTTGGATTTGTGGGTTGCNNNAGATCGGAAG |
| 38600_2 | TATGAGTGTGGAGTCGTTGCAACTTCTTGGATTTGTGGGTTGCNNNAGATCGGAAG |
| 37565_1 | TATGAGTGTGGAGTCGTTGCTCTTTGCTGAGAAGCTCTGCTTANNNAGATCGGAAG |
| 37565_2 | TATGAGTGTGGAGTCGTTGCTCTTTGCTGAGAAGCTCTGCTTANNNAGATCGGAAG |
| 35557_1 | TATGAGTGTGGAGTCGTTGCGAGCTAGAAGTCTGGCCTCTTACNNNAGATCGGAAG |
| 35557_2 | TATGAGTGTGGAGTCGTTGCGAGCTAGAAGTCTGGCCTCTTACNNNAGATCGGAAG |
| 31513_1 | TATGAGTGTGGAGTCGTTGCCCTTGTGGTTCATGTTTCAGCAANNNAGATCGGAAG |
| 31513_2 | TATGAGTGTGGAGTCGTTGCCCTTGTGGTTCATGTTTCAGCAANNNAGATCGGAAG |
| 48762_1 | TATGAGTGTGGAGTCGTTGCTGTTTATGTCCTCTGCCCACTTTNNNAGATCGGAAGA |
| 48762_2 | TATGAGTGTGGAGTCGTTGCTGTTTATGTCCTCTGCCCACTTTNNNAGATCGGAAGA |
| 39148_1 | TATGAGTGTGGAGTCGTTGCTCACACTCAGAATTGATTGCTGCNNNAGATCGGAAG |
| 39148_2 | TATGAGTGTGGAGTCGTTGCTCACACTCAGAATTGATTGCTGCNNNAGATCGGAAG |
| 30994_1 | TATGAGTGTGGAGTCGTTGCCATCTCATTACTGCCTGGTCTGTNNNAGATCGGAAG |
| 30994_2 | TATGAGTGTGGAGTCGTTGCCATCTCATTACTGCCTGGTCTGTNNNAGATCGGAAG |
| 45624_1 | TATGAGTGTGGAGTCGTTGCCTGGAGTGCAATGGTGTGATCTCNNNAGATCGGAA |
| 45624_2 | TATGAGTGTGGAGTCGTTGCCTGGAGTGCAATGGTGTGATCTCNNNAGATCGGAA |
| 46790_1 | TATGAGTGTGGAGTCGTTGCCTATCTTGGCTCACTGCAAGCTCNNNAGATCGGAAG |
| 46790_2 | TATGAGTGTGGAGTCGTTGCCTATCTTGGCTCACTGCAAGCTCNNNAGATCGGAAG |
| 50113_1 | TATGAGTGTGGAGTCGTTGCTCTCACTTTCAGACTGCTCACTCNNNAGATCGGAAGA |
| 50113_2 | TATGAGTGTGGAGTCGTTGCTCTCACTTTCAGACTGCTCACTCNNNAGATCGGAAGA |
| 13379_1 | TATGAGTGTGGAGTCGTTGCACAATGCTTAGAAACTAATACACNNNAGATCGGAAG |
| 13379_2 | TATGAGTGTGGAGTCGTTGCACAATGCTTAGAAACTAATACACNNNAGATCGGAAG |
| 37780_1 | TATGAGTGTGGAGTCGTTGCTGAAAGTGACTGAAGTACCTAGGGNNNAGATCGGA |
| 37780_2 | TATGAGTGTGGAGTCGTTGCTGAAAGTGACTGAAGTACCTAGGGNNNAGATCGGA |
| 1882_1 | TATGAGTGTGGAGTCGTTGCGGAAGCTGTGGGGAGAAGCAGNNNAGATCGGAAG |
| 1882_2 | TATGAGTGTGGAGTCGTTGCGGAAGCTGTGGGGAGAAGCAGNNNAGATCGGAAG |
| 52371_1 | TATGAGTGTGGAGTCGTTGCGCAGAGAAGAGTGAGAGGAAGACNNNAGATCGGA |
| 52371_2 | TATGAGTGTGGAGTCGTTGCGCAGAGAAGAGTGAGAGGAAGACNNNAGATCGGA |

| | |
|---|---|
| 53214_1 | TATGAGTGTGGAGTCGTTGCTGTGTACATCAGAATCACCTGGGNNNAGATCGGAAG |
| 53214_2 | TATGAGTGTGGAGTCGTTGCTGTGTACATCAGAATCACCTGGGNNNAGATCGGAAG |
| 5367_1 | TATGAGTGTGGAGTCGTTGCTTGGCCTCAGGATACTCAGGTTTNNNAGATCGGAAG |
| 5367_2 | TATGAGTGTGGAGTCGTTGCTTGGCCTCAGGATACTCAGGTTTNNNAGATCGGAAG |
| 31956_1 | TATGAGTGTGGAGTCGTTGCCTGTAATCCCAGCTACTCAGGAGNNNAGATCGGAAG |
| 31956_2 | TATGAGTGTGGAGTCGTTGCCTGTAATCCCAGCTACTCAGGAGNNNAGATCGGAAG |
| 10502_1 | TATGAGTGTGGAGTCGTTGCGTATGTTGTGCGGCTGTGTGTNNNAGATCGGAAGA |
| 10502_2 | TATGAGTGTGGAGTCGTTGCGTATGTTGTGCGGCTGTGTGTNNNAGATCGGAAGA |
| 45651_1 | TATGAGTGTGGAGTCGTTGCTAGTTGGGACTACAGGTGCCTNNNAGATCGGAAGA |
| 45651_2 | TATGAGTGTGGAGTCGTTGCTAGTTGGGACTACAGGTGCCTNNNAGATCGGAAGA |
| 13739_1 | TATGAGTGTGGAGTCGTTGCCTCTTTGTTTCTCATCAACTGNNNAGATCGGAAGAG |
| 13739_2 | TATGAGTGTGGAGTCGTTGCCTCTTTGTTTCTCATCAACTGNNNAGATCGGAAGAG |
| 32973_1 | TATGAGTGTGGAGTCGTTGCCTATGCAGCAGTACCACAACAAGNNNAGATCGGAAG |
| 32973_2 | TATGAGTGTGGAGTCGTTGCCTATGCAGCAGTACCACAACAAGNNNAGATCGGAAG |
| 32973_3 | TATGAGTGTGGAGTCGTTGCCTATGCAGCAGTACCACAACAAGNNNAGATCGGAAG |
| 32973_4 | TATGAGTGTGGAGTCGTTGCCTATGCAGCAGTACCACAACAAGNNNAGATCGGAAG |
| 32973_5 | TATGAGTGTGGAGTCGTTGCCTATGCAGCAGTACCACAACAAGNNNAGATCGGAAG |
| 47464_1 | TATGAGTGTGGAGTCGTTGCCCTCCCAGATTCATGCCATTCTCNNNAGATCGGAAGA |
| 47464_2 | TATGAGTGTGGAGTCGTTGCCCTCCCAGATTCATGCCATTCTCNNNAGATCGGAAGA |
| 47464_3 | TATGAGTGTGGAGTCGTTGCCCTCCCAGATTCATGCCATTCTCNNNAGATCGGAAGA |
| 47464_4 | TATGAGTGTGGAGTCGTTGCCCTCCCAGATTCATGCCATTCTCNNNAGATCGGAAGA |
| 47464_5 | TATGAGTGTGGAGTCGTTGCCCTCCCAGATTCATGCCATTCTCNNNAGATCGGAAGA |
| 48309_1 | TATGAGTGTGGAGTCGTTGCTTACTATATTGGCCAGGCTGGTCNNNAGATCGGAAG |
| 48309_2 | TATGAGTGTGGAGTCGTTGCTTACTATATTGGCCAGGCTGGTCNNNAGATCGGAAG |
| 48309_3 | TATGAGTGTGGAGTCGTTGCTTACTATATTGGCCAGGCTGGTCNNNAGATCGGAAG |
| 48309_4 | TATGAGTGTGGAGTCGTTGCTTACTATATTGGCCAGGCTGGTCNNNAGATCGGAAG |
| 48309_5 | TATGAGTGTGGAGTCGTTGCTTACTATATTGGCCAGGCTGGTCNNNAGATCGGAAG |
| 45828_1 | TATGAGTGTGGAGTCGTTGCCTCCTGAGTAGCTGGGACTACAGNNNAGATCGGAA |
| 45828_2 | TATGAGTGTGGAGTCGTTGCCTCCTGAGTAGCTGGGACTACAGNNNAGATCGGAA |
| 45828_3 | TATGAGTGTGGAGTCGTTGCCTCCTGAGTAGCTGGGACTACAGNNNAGATCGGAA |
| 45828_4 | TATGAGTGTGGAGTCGTTGCCTCCTGAGTAGCTGGGACTACAGNNNAGATCGGAA |
| 45828_5 | TATGAGTGTGGAGTCGTTGCCTCCTGAGTAGCTGGGACTACAGNNNAGATCGGAA |
| 40163_1 | TATGAGTGTGGAGTCGTTGCCTCCATGCTCTGAACTCTAGTGGNNNAGATCGGAAG |
| 40163_2 | TATGAGTGTGGAGTCGTTGCCTCCATGCTCTGAACTCTAGTGGNNNAGATCGGAAG |
| 40163_3 | TATGAGTGTGGAGTCGTTGCCTCCATGCTCTGAACTCTAGTGGNNNAGATCGGAAG |
| 40163_4 | TATGAGTGTGGAGTCGTTGCCTCCATGCTCTGAACTCTAGTGGNNNAGATCGGAAG |
| 40163_5 | TATGAGTGTGGAGTCGTTGCCTCCATGCTCTGAACTCTAGTGGNNNAGATCGGAAG |
| 37365_1 | TATGAGTGTGGAGTCGTTGCTTTGTGATGCAATCATTGTGGCCNNNAGATCGGAAG |
| 37365_2 | TATGAGTGTGGAGTCGTTGCTTTGTGATGCAATCATTGTGGCCNNNAGATCGGAAG |
| 37365_3 | TATGAGTGTGGAGTCGTTGCTTTGTGATGCAATCATTGTGGCCNNNAGATCGGAAG |
| 37365_4 | TATGAGTGTGGAGTCGTTGCTTTGTGATGCAATCATTGTGGCCNNNAGATCGGAAG |
| 37365_5 | TATGAGTGTGGAGTCGTTGCTTTGTGATGCAATCATTGTGGCCNNNAGATCGGAAG |
| 37612_1 | TATGAGTGTGGAGTCGTTGCAAAGAAAGATGTAGACCGGCAGANNNAGATCGGA |

| | |
|---|---|
| 37612_2 | TATGAGTGTGGAGTCGTTGCAAAGAAAGATGTAGACCGGCAGANNNAGATCGGAA |
| 37612_3 | TATGAGTGTGGAGTCGTTGCAAAGAAAGATGTAGACCGGCAGANNNAGATCGGAA |
| 37612_4 | TATGAGTGTGGAGTCGTTGCAAAGAAAGATGTAGACCGGCAGANNNAGATCGGAA |
| 37612_5 | TATGAGTGTGGAGTCGTTGCAAAGAAAGATGTAGACCGGCAGANNNAGATCGGAA |
| 35887_1 | TATGAGTGTGGAGTCGTTGCGAGTTGGAGTCTCACTGGGTGNNNAGATCGGAAGA |
| 35887_2 | TATGAGTGTGGAGTCGTTGCGAGTTGGAGTCTCACTGGGTGNNNAGATCGGAAGA |
| 35887_3 | TATGAGTGTGGAGTCGTTGCGAGTTGGAGTCTCACTGGGTGNNNAGATCGGAAGA |
| 35887_4 | TATGAGTGTGGAGTCGTTGCGAGTTGGAGTCTCACTGGGTGNNNAGATCGGAAGA |
| 35887_5 | TATGAGTGTGGAGTCGTTGCGAGTTGGAGTCTCACTGGGTGNNNAGATCGGAAGA |
| 36604_1 | TATGAGTGTGGAGTCGTTGCCAGAAAGTCAATTACGGCTCACCNNNAGATCGGAAG |
| 36604_2 | TATGAGTGTGGAGTCGTTGCCAGAAAGTCAATTACGGCTCACCNNNAGATCGGAAG |
| 36604_3 | TATGAGTGTGGAGTCGTTGCCAGAAAGTCAATTACGGCTCACCNNNAGATCGGAAG |
| 36604_4 | TATGAGTGTGGAGTCGTTGCCAGAAAGTCAATTACGGCTCACCNNNAGATCGGAAG |
| 36604_5 | TATGAGTGTGGAGTCGTTGCCAGAAAGTCAATTACGGCTCACCNNNAGATCGGAAG |
| 39404_1 | TATGAGTGTGGAGTCGTTGCAGGGATCTTAGAGGATGGGTCAANNNAGATCGGAA |
| 39404_2 | TATGAGTGTGGAGTCGTTGCAGGGATCTTAGAGGATGGGTCAANNNAGATCGGAA |
| 39404_3 | TATGAGTGTGGAGTCGTTGCAGGGATCTTAGAGGATGGGTCAANNNAGATCGGAA |
| 39404_4 | TATGAGTGTGGAGTCGTTGCAGGGATCTTAGAGGATGGGTCAANNNAGATCGGAA |
| 39404_5 | TATGAGTGTGGAGTCGTTGCAGGGATCTTAGAGGATGGGTCAANNNAGATCGGAA |
| 48895_1 | TATGAGTGTGGAGTCGTTGCCATCCCCTCAAAATGCAGTTCAGNNNAGATCGGAAGA |
| 48895_2 | TATGAGTGTGGAGTCGTTGCCATCCCCTCAAAATGCAGTTCAGNNNAGATCGGAAGA |
| 48895_3 | TATGAGTGTGGAGTCGTTGCCATCCCCTCAAAATGCAGTTCAGNNNAGATCGGAAGA |
| 48895_4 | TATGAGTGTGGAGTCGTTGCCATCCCCTCAAAATGCAGTTCAGNNNAGATCGGAAGA |
| 48895_5 | TATGAGTGTGGAGTCGTTGCCATCCCCTCAAAATGCAGTTCAGNNNAGATCGGAAGA |
| 42092_1 | TATGAGTGTGGAGTCGTTGCACCAAATGCAGAACTTGGAACTGNNNAGATCGGAAG |
| 42092_2 | TATGAGTGTGGAGTCGTTGCACCAAATGCAGAACTTGGAACTGNNNAGATCGGAAG |
| 42092_3 | TATGAGTGTGGAGTCGTTGCACCAAATGCAGAACTTGGAACTGNNNAGATCGGAAG |
| 42092_4 | TATGAGTGTGGAGTCGTTGCACCAAATGCAGAACTTGGAACTGNNNAGATCGGAAG |
| 42092_5 | TATGAGTGTGGAGTCGTTGCACCAAATGCAGAACTTGGAACTGNNNAGATCGGAAG |
| 32867_1 | TATGAGTGTGGAGTCGTTGCTTCAAGTGATTCTCCTGCCTCAGNNNAGATCGGAAGA |
| 32867_2 | TATGAGTGTGGAGTCGTTGCTTCAAGTGATTCTCCTGCCTCAGNNNAGATCGGAAGA |
| 32867_3 | TATGAGTGTGGAGTCGTTGCTTCAAGTGATTCTCCTGCCTCAGNNNAGATCGGAAGA |
| 32867_4 | TATGAGTGTGGAGTCGTTGCTTCAAGTGATTCTCCTGCCTCAGNNNAGATCGGAAGA |
| 32867_5 | TATGAGTGTGGAGTCGTTGCTTCAAGTGATTCTCCTGCCTCAGNNNAGATCGGAAGA |
| 31155_1 | TATGAGTGTGGAGTCGTTGCCCTTCCCCACTGAACCTTTCTAANNNAGATCGGAAGA |
| 31155_2 | TATGAGTGTGGAGTCGTTGCCCTTCCCCACTGAACCTTTCTAANNNAGATCGGAAGA |
| 31155_3 | TATGAGTGTGGAGTCGTTGCCCTTCCCCACTGAACCTTTCTAANNNAGATCGGAAGA |
| 31155_4 | TATGAGTGTGGAGTCGTTGCCCTTCCCCACTGAACCTTTCTAANNNAGATCGGAAGA |
| 31155_5 | TATGAGTGTGGAGTCGTTGCCCTTCCCCACTGAACCTTTCTAANNNAGATCGGAAGA |
| 47239_1 | TATGAGTGTGGAGTCGTTGCGTGTTGCGAACATAGCTTACTCANNNAGATCGGAAG |
| 47239_2 | TATGAGTGTGGAGTCGTTGCGTGTTGCGAACATAGCTTACTCANNNAGATCGGAAG |
| 47239_3 | TATGAGTGTGGAGTCGTTGCGTGTTGCGAACATAGCTTACTCANNNAGATCGGAAG |
| 47239_4 | TATGAGTGTGGAGTCGTTGCGTGTTGCGAACATAGCTTACTCANNNAGATCGGAAG |

| | |
|---|---|
| 47239_5 | TATGAGTGTGGAGTCGTTGCGTGTTGCGAACATAGCTTACTCANNNAGATCGGAAG |
| 9997_1 | TATGAGTGTGGAGTCGTTGCGTGTGCCAATTGCTGTTTCAAGTNNNAGATCGGAAG |
| 9997_2 | TATGAGTGTGGAGTCGTTGCGTGTGCCAATTGCTGTTTCAAGTNNNAGATCGGAAG |
| 9997_3 | TATGAGTGTGGAGTCGTTGCGTGTGCCAATTGCTGTTTCAAGTNNNAGATCGGAAG |
| 9997_4 | TATGAGTGTGGAGTCGTTGCGTGTGCCAATTGCTGTTTCAAGTNNNAGATCGGAAG |
| 9997_5 | TATGAGTGTGGAGTCGTTGCGTGTGCCAATTGCTGTTTCAAGTNNNAGATCGGAAG |
| 32705_1 | TATGAGTGTGGAGTCGTTGCTCTCAAACTGCTGACCTCATGATNNNAGATCGGAAG |
| 32705_2 | TATGAGTGTGGAGTCGTTGCTCTCAAACTGCTGACCTCATGATNNNAGATCGGAAG |
| 32705_3 | TATGAGTGTGGAGTCGTTGCTCTCAAACTGCTGACCTCATGATNNNAGATCGGAAG |
| 32705_4 | TATGAGTGTGGAGTCGTTGCTCTCAAACTGCTGACCTCATGATNNNAGATCGGAAG |
| 32705_5 | TATGAGTGTGGAGTCGTTGCTCTCAAACTGCTGACCTCATGATNNNAGATCGGAAG |
| 30746_1 | TATGAGTGTGGAGTCGTTGCAAACAGACACTCTCAGACATGCTNNNAGATCGGAAG |
| 30746_2 | TATGAGTGTGGAGTCGTTGCAAACAGACACTCTCAGACATGCTNNNAGATCGGAAG |
| 30746_3 | TATGAGTGTGGAGTCGTTGCAAACAGACACTCTCAGACATGCTNNNAGATCGGAAG |
| 30746_4 | TATGAGTGTGGAGTCGTTGCAAACAGACACTCTCAGACATGCTNNNAGATCGGAAG |
| 30746_5 | TATGAGTGTGGAGTCGTTGCAAACAGACACTCTCAGACATGCTNNNAGATCGGAAG |
| 33031_1 | TATGAGTGTGGAGTCGTTGCGAGTAGCTGGTACTACAGGTGTGNNNAGATCGGAA |
| 33031_2 | TATGAGTGTGGAGTCGTTGCGAGTAGCTGGTACTACAGGTGTGNNNAGATCGGAA |
| 33031_3 | TATGAGTGTGGAGTCGTTGCGAGTAGCTGGTACTACAGGTGTGNNNAGATCGGAA |
| 33031_4 | TATGAGTGTGGAGTCGTTGCGAGTAGCTGGTACTACAGGTGTGNNNAGATCGGAA |
| 33031_5 | TATGAGTGTGGAGTCGTTGCGAGTAGCTGGTACTACAGGTGTGNNNAGATCGGAA |
| 50286_1 | TATGAGTGTGGAGTCGTTGCGAGAGAGAGAGAGAGAGACCTCANNNAGATCGGA |
| 50286_2 | TATGAGTGTGGAGTCGTTGCGAGAGAGAGAGAGAGAGACCTCANNNAGATCGGA |
| 50286_3 | TATGAGTGTGGAGTCGTTGCGAGAGAGAGAGAGAGAGACCTCANNNAGATCGGA |
| 50286_4 | TATGAGTGTGGAGTCGTTGCGAGAGAGAGAGAGAGAGACCTCANNNAGATCGGA |
| 50286_5 | TATGAGTGTGGAGTCGTTGCGAGAGAGAGAGAGAGAGACCTCANNNAGATCGGA |
| 37126_1 | TATGAGTGTGGAGTCGTTGCAGTGAGCCGAGATTGTGTCATTNNNAGATCGGAAG |
| 37126_2 | TATGAGTGTGGAGTCGTTGCAGTGAGCCGAGATTGTGTCATTNNNAGATCGGAAG |
| 37126_3 | TATGAGTGTGGAGTCGTTGCAGTGAGCCGAGATTGTGTCATTNNNAGATCGGAAG |
| 37126_4 | TATGAGTGTGGAGTCGTTGCAGTGAGCCGAGATTGTGTCATTNNNAGATCGGAAG |
| 37126_5 | TATGAGTGTGGAGTCGTTGCAGTGAGCCGAGATTGTGTCATTNNNAGATCGGAAG |
| 37977_1 | TATGAGTGTGGAGTCGTTGCGGGACCGTTTTGCAGATTCTATGNNNAGATCGGAA |
| 37977_2 | TATGAGTGTGGAGTCGTTGCGGGACCGTTTTGCAGATTCTATGNNNAGATCGGAA |
| 37977_3 | TATGAGTGTGGAGTCGTTGCGGGACCGTTTTGCAGATTCTATGNNNAGATCGGAA |
| 37977_4 | TATGAGTGTGGAGTCGTTGCGGGACCGTTTTGCAGATTCTATGNNNAGATCGGAA |
| 37977_5 | TATGAGTGTGGAGTCGTTGCGGGACCGTTTTGCAGATTCTATGNNNAGATCGGAA |
| 39562_1 | TATGAGTGTGGAGTCGTTGCCTGTACTCCCAGCTATTCAGGTGNNNAGATCGGAAG |
| 39562_2 | TATGAGTGTGGAGTCGTTGCCTGTACTCCCAGCTATTCAGGTGNNNAGATCGGAAG |
| 39562_3 | TATGAGTGTGGAGTCGTTGCCTGTACTCCCAGCTATTCAGGTGNNNAGATCGGAAG |
| 39562_4 | TATGAGTGTGGAGTCGTTGCCTGTACTCCCAGCTATTCAGGTGNNNAGATCGGAAG |
| 39562_5 | TATGAGTGTGGAGTCGTTGCCTGTACTCCCAGCTATTCAGGTGNNNAGATCGGAAG |
| 42377_1 | TATGAGTGTGGAGTCGTTGCGCAATTTTCTTACCTCAGCCTCCNNNAGATCGGAAGA |
| 42377_2 | TATGAGTGTGGAGTCGTTGCGCAATTTTCTTACCTCAGCCTCCNNNAGATCGGAAGA |

| | |
|---|---|
| 42377_3 | TATGAGTGTGGAGTCGTTGCGCAATTTTCTTACCTCAGCCTCCNNNAGATCGGAAGA |
| 42377_4 | TATGAGTGTGGAGTCGTTGCGCAATTTTCTTACCTCAGCCTCCNNNAGATCGGAAGA |
| 42377_5 | TATGAGTGTGGAGTCGTTGCGCAATTTTCTTACCTCAGCCTCCNNNAGATCGGAAGA |
| 33245_1 | TATGAGTGTGGAGTCGTTGCCTCCTGAGTAGCTGGGACTACAGNNNAGATCGGAAG |
| 33245_2 | TATGAGTGTGGAGTCGTTGCCTCCTGAGTAGCTGGGACTACAGNNNAGATCGGAAG |
| 33245_3 | TATGAGTGTGGAGTCGTTGCCTCCTGAGTAGCTGGGACTACAGNNNAGATCGGAAG |
| 33245_4 | TATGAGTGTGGAGTCGTTGCCTCCTGAGTAGCTGGGACTACAGNNNAGATCGGAAG |
| 33245_5 | TATGAGTGTGGAGTCGTTGCCTCCTGAGTAGCTGGGACTACAGNNNAGATCGGAAG |
| 34892_1 | TATGAGTGTGGAGTCGTTGCTGCACCACTATGGAAAACAGTCANNNAGATCGGAAG |
| 34892_2 | TATGAGTGTGGAGTCGTTGCTGCACCACTATGGAAAACAGTCANNNAGATCGGAAG |
| 34892_3 | TATGAGTGTGGAGTCGTTGCTGCACCACTATGGAAAACAGTCANNNAGATCGGAAG |
| 34892_4 | TATGAGTGTGGAGTCGTTGCTGCACCACTATGGAAAACAGTCANNNAGATCGGAAG |
| 34892_5 | TATGAGTGTGGAGTCGTTGCTGCACCACTATGGAAAACAGTCANNNAGATCGGAAG |
| 49001_1 | TATGAGTGTGGAGTCGTTGCGTTTCACCATGTTAGCCAGGATGNNNAGATCGGAAG |
| 49001_2 | TATGAGTGTGGAGTCGTTGCGTTTCACCATGTTAGCCAGGATGNNNAGATCGGAAG |
| 49001_3 | TATGAGTGTGGAGTCGTTGCGTTTCACCATGTTAGCCAGGATGNNNAGATCGGAAG |
| 49001_4 | TATGAGTGTGGAGTCGTTGCGTTTCACCATGTTAGCCAGGATGNNNAGATCGGAAG |
| 49001_5 | TATGAGTGTGGAGTCGTTGCGTTTCACCATGTTAGCCAGGATGNNNAGATCGGAAG |
| 32978_1 | TATGAGTGTGGAGTCGTTGCAACATGGATGAAACTGGAGGTCANNNAGATCGGAA |
| 32978_2 | TATGAGTGTGGAGTCGTTGCAACATGGATGAAACTGGAGGTCANNNAGATCGGAA |
| 32978_3 | TATGAGTGTGGAGTCGTTGCAACATGGATGAAACTGGAGGTCANNNAGATCGGAA |
| 32978_4 | TATGAGTGTGGAGTCGTTGCAACATGGATGAAACTGGAGGTCANNNAGATCGGAA |
| 32978_5 | TATGAGTGTGGAGTCGTTGCAACATGGATGAAACTGGAGGTCANNNAGATCGGAA |
| 53178_1 | TATGAGTGTGGAGTCGTTGCTATTGAGTCCCCACATTAGGCTGNNNAGATCGGAAG |
| 53178_2 | TATGAGTGTGGAGTCGTTGCTATTGAGTCCCCACATTAGGCTGNNNAGATCGGAAG |
| 53178_3 | TATGAGTGTGGAGTCGTTGCTATTGAGTCCCCACATTAGGCTGNNNAGATCGGAAG |
| 53178_4 | TATGAGTGTGGAGTCGTTGCTATTGAGTCCCCACATTAGGCTGNNNAGATCGGAAG |
| 53178_5 | TATGAGTGTGGAGTCGTTGCTATTGAGTCCCCACATTAGGCTGNNNAGATCGGAAG |
| 13413_1 | TATGAGTGTGGAGTCGTTGCAAACTGATTTTTGTTGTTGTAGTNNNAGATCGGAAG |
| 13413_2 | TATGAGTGTGGAGTCGTTGCAAACTGATTTTTGTTGTTGTAGTNNNAGATCGGAAG |
| 13413_3 | TATGAGTGTGGAGTCGTTGCAAACTGATTTTTGTTGTTGTAGTNNNAGATCGGAAG |
| 13413_4 | TATGAGTGTGGAGTCGTTGCAAACTGATTTTTGTTGTTGTAGTNNNAGATCGGAAG |
| 13413_5 | TATGAGTGTGGAGTCGTTGCAAACTGATTTTTGTTGTTGTAGTNNNAGATCGGAAG |
| 47533_1 | TATGAGTGTGGAGTCGTTGCCCACACCCGGCTAATTTTTTGTANNNAGATCGGAAGA |
| 47533_2 | TATGAGTGTGGAGTCGTTGCCCACACCCGGCTAATTTTTTGTANNNAGATCGGAAGA |
| 47533_3 | TATGAGTGTGGAGTCGTTGCCCACACCCGGCTAATTTTTTGTANNNAGATCGGAAGA |
| 47533_4 | TATGAGTGTGGAGTCGTTGCCCACACCCGGCTAATTTTTTGTANNNAGATCGGAAGA |
| 47533_5 | TATGAGTGTGGAGTCGTTGCCCACACCCGGCTAATTTTTTGTANNNAGATCGGAAGA |
| 13529_1 | TATGAGTGTGGAGTCGTTGCTTTTAAAGAGGTAGATTCCCAATNNNAGATCGGAAG |
| 13529_2 | TATGAGTGTGGAGTCGTTGCTTTTAAAGAGGTAGATTCCCAATNNNAGATCGGAAG |
| 13529_3 | TATGAGTGTGGAGTCGTTGCTTTTAAAGAGGTAGATTCCCAATNNNAGATCGGAAG |
| 13529_4 | TATGAGTGTGGAGTCGTTGCTTTTAAAGAGGTAGATTCCCAATNNNAGATCGGAAG |
| 13529_5 | TATGAGTGTGGAGTCGTTGCTTTTAAAGAGGTAGATTCCCAATNNNAGATCGGAAG |

| | |
|---|---|
| 30866_1 | TATGAGTGTGGAGTCGTTGCCCTATTTGCAGTCACTCCTCACTNNNAGATCGGAAGA |
| 30866_2 | TATGAGTGTGGAGTCGTTGCCCTATTTGCAGTCACTCCTCACTNNNAGATCGGAAGA |
| 30866_3 | TATGAGTGTGGAGTCGTTGCCCTATTTGCAGTCACTCCTCACTNNNAGATCGGAAGA |
| 30866_4 | TATGAGTGTGGAGTCGTTGCCCTATTTGCAGTCACTCCTCACTNNNAGATCGGAAGA |
| 30866_5 | TATGAGTGTGGAGTCGTTGCCCTATTTGCAGTCACTCCTCACTNNNAGATCGGAAGA |
| 36177_1 | TATGAGTGTGGAGTCGTTGCCAAGACATGACTCAGCATGAAAGTNNNAGATCGGA |
| 36177_2 | TATGAGTGTGGAGTCGTTGCCAAGACATGACTCAGCATGAAAGTNNNAGATCGGA |
| 36177_3 | TATGAGTGTGGAGTCGTTGCCAAGACATGACTCAGCATGAAAGTNNNAGATCGGA |
| 36177_4 | TATGAGTGTGGAGTCGTTGCCAAGACATGACTCAGCATGAAAGTNNNAGATCGGA |
| 36177_5 | TATGAGTGTGGAGTCGTTGCCAAGACATGACTCAGCATGAAAGTNNNAGATCGGA |
| 32775_1 | TATGAGTGTGGAGTCGTTGCCTTGAGCCAGGTGAAAATGAAGGNNNAGATCGGAA |
| 32775_2 | TATGAGTGTGGAGTCGTTGCCTTGAGCCAGGTGAAAATGAAGGNNNAGATCGGAA |
| 32775_3 | TATGAGTGTGGAGTCGTTGCCTTGAGCCAGGTGAAAATGAAGGNNNAGATCGGAA |
| 32775_4 | TATGAGTGTGGAGTCGTTGCCTTGAGCCAGGTGAAAATGAAGGNNNAGATCGGAA |
| 32775_5 | TATGAGTGTGGAGTCGTTGCCTTGAGCCAGGTGAAAATGAAGGNNNAGATCGGAA |
| 52053_1 | TATGAGTGTGGAGTCGTTGCCCCCTCTCACTCTTTGGTAACTCNNNAGATCGGAAGA |
| 52053_2 | TATGAGTGTGGAGTCGTTGCCCCCTCTCACTCTTTGGTAACTCNNNAGATCGGAAGA |
| 52053_3 | TATGAGTGTGGAGTCGTTGCCCCCTCTCACTCTTTGGTAACTCNNNAGATCGGAAGA |
| 52053_4 | TATGAGTGTGGAGTCGTTGCCCCCTCTCACTCTTTGGTAACTCNNNAGATCGGAAGA |
| 52053_5 | TATGAGTGTGGAGTCGTTGCCCCCTCTCACTCTTTGGTAACTCNNNAGATCGGAAGA |
| 30173_1 | TATGAGTGTGGAGTCGTTGCGGGGCTCAAACTGCTTCATAAAANNNAGATCGGAA |
| 30173_2 | TATGAGTGTGGAGTCGTTGCGGGGCTCAAACTGCTTCATAAAANNNAGATCGGAA |
| 30173_3 | TATGAGTGTGGAGTCGTTGCGGGGCTCAAACTGCTTCATAAAANNNAGATCGGAA |
| 30173_4 | TATGAGTGTGGAGTCGTTGCGGGGCTCAAACTGCTTCATAAAANNNAGATCGGAA |
| 30173_5 | TATGAGTGTGGAGTCGTTGCGGGGCTCAAACTGCTTCATAAAANNNAGATCGGAA |
| 39509_1 | TATGAGTGTGGAGTCGTTGCCCTAAGGTCAGGAGTTCAAGACCNNNAGATCGGAA |
| 39509_2 | TATGAGTGTGGAGTCGTTGCCCTAAGGTCAGGAGTTCAAGACCNNNAGATCGGAA |
| 39509_3 | TATGAGTGTGGAGTCGTTGCCCTAAGGTCAGGAGTTCAAGACCNNNAGATCGGAA |
| 39509_4 | TATGAGTGTGGAGTCGTTGCCCTAAGGTCAGGAGTTCAAGACCNNNAGATCGGAA |
| 39509_5 | TATGAGTGTGGAGTCGTTGCCCTAAGGTCAGGAGTTCAAGACCNNNAGATCGGAA |
| 36036_1 | TATGAGTGTGGAGTCGTTGCGGCAGAGACTGGAGTGATGTAGNNNAGATCGGAA |
| 36036_2 | TATGAGTGTGGAGTCGTTGCGGCAGAGACTGGAGTGATGTAGNNNAGATCGGAA |
| 36036_3 | TATGAGTGTGGAGTCGTTGCGGCAGAGACTGGAGTGATGTAGNNNAGATCGGAA |
| 36036_4 | TATGAGTGTGGAGTCGTTGCGGCAGAGACTGGAGTGATGTAGNNNAGATCGGAA |
| 36036_5 | TATGAGTGTGGAGTCGTTGCGGCAGAGACTGGAGTGATGTAGNNNAGATCGGAA |
| 32788_1 | TATGAGTGTGGAGTCGTTGCCAATGAGATACCACTGCACACCANNNAGATCGGAAG |
| 32788_2 | TATGAGTGTGGAGTCGTTGCCAATGAGATACCACTGCACACCANNNAGATCGGAAG |
| 32788_3 | TATGAGTGTGGAGTCGTTGCCAATGAGATACCACTGCACACCANNNAGATCGGAAG |
| 32788_4 | TATGAGTGTGGAGTCGTTGCCAATGAGATACCACTGCACACCANNNAGATCGGAAG |
| 32788_5 | TATGAGTGTGGAGTCGTTGCCAATGAGATACCACTGCACACCANNNAGATCGGAAG |
| 52318_1 | TATGAGTGTGGAGTCGTTGCGTTGGGGTTGCATCCTATACAGANNNAGATCGGAA |
| 52318_2 | TATGAGTGTGGAGTCGTTGCGTTGGGGTTGCATCCTATACAGANNNAGATCGGAA |
| 52318_3 | TATGAGTGTGGAGTCGTTGCGTTGGGGTTGCATCCTATACAGANNNAGATCGGAA |

| | |
|---|---|
| 52318_4 | TATGAGTGTGGAGTCGTTGCGTTGGGGTTGCATCCTATACAGANNNAGATCGGAAG |
| 52318_5 | TATGAGTGTGGAGTCGTTGCGTTGGGGTTGCATCCTATACAGANNNAGATCGGAAG |
| 31589_1 | TATGAGTGTGGAGTCGTTGCAATGGATCAGTGGATGGATGCATNNNAGATCGGAA |
| 31589_2 | TATGAGTGTGGAGTCGTTGCAATGGATCAGTGGATGGATGCATNNNAGATCGGAA |
| 31589_3 | TATGAGTGTGGAGTCGTTGCAATGGATCAGTGGATGGATGCATNNNAGATCGGAA |
| 31589_4 | TATGAGTGTGGAGTCGTTGCAATGGATCAGTGGATGGATGCATNNNAGATCGGAA |
| 31589_5 | TATGAGTGTGGAGTCGTTGCAATGGATCAGTGGATGGATGCATNNNAGATCGGAA |
| 50802_1 | TATGAGTGTGGAGTCGTTGCTGAAGGGAAAGTGGAAGTAGGTGNNNAGATCGGA |
| 50802_2 | TATGAGTGTGGAGTCGTTGCTGAAGGGAAAGTGGAAGTAGGTGNNNAGATCGGA |
| 50802_3 | TATGAGTGTGGAGTCGTTGCTGAAGGGAAAGTGGAAGTAGGTGNNNAGATCGGA |
| 50802_4 | TATGAGTGTGGAGTCGTTGCTGAAGGGAAAGTGGAAGTAGGTGNNNAGATCGGA |
| 50802_5 | TATGAGTGTGGAGTCGTTGCTGAAGGGAAAGTGGAAGTAGGTGNNNAGATCGGA |
| 44342_1 | TATGAGTGTGGAGTCGTTGCAAGACTTCTGTGCTCGCTAATGANNNAGATCGGAAG |
| 44342_2 | TATGAGTGTGGAGTCGTTGCAAGACTTCTGTGCTCGCTAATGANNNAGATCGGAAG |
| 44342_3 | TATGAGTGTGGAGTCGTTGCAAGACTTCTGTGCTCGCTAATGANNNAGATCGGAAG |
| 44342_4 | TATGAGTGTGGAGTCGTTGCAAGACTTCTGTGCTCGCTAATGANNNAGATCGGAAG |
| 44342_5 | TATGAGTGTGGAGTCGTTGCAAGACTTCTGTGCTCGCTAATGANNNAGATCGGAAG |
| 13547_1 | TATGAGTGTGGAGTCGTTGCTTCCACTTTCTTTTGAAAAATTTNNNAGATCGGAAGA |
| 13547_2 | TATGAGTGTGGAGTCGTTGCTTCCACTTTCTTTTGAAAAATTTNNNAGATCGGAAGA |
| 13547_3 | TATGAGTGTGGAGTCGTTGCTTCCACTTTCTTTTGAAAAATTTNNNAGATCGGAAGA |
| 13547_4 | TATGAGTGTGGAGTCGTTGCTTCCACTTTCTTTTGAAAAATTTNNNAGATCGGAAGA |
| 13547_5 | TATGAGTGTGGAGTCGTTGCTTCCACTTTCTTTTGAAAAATTTNNNAGATCGGAAGA |
| 42874_1 | TATGAGTGTGGAGTCGTTGCTATGGTATGGTGTGTGTGTGTGTNNNAGATCGGAA |
| 42874_2 | TATGAGTGTGGAGTCGTTGCTATGGTATGGTGTGTGTGTGTGTNNNAGATCGGAA |
| 42874_3 | TATGAGTGTGGAGTCGTTGCTATGGTATGGTGTGTGTGTGTGTNNNAGATCGGAA |
| 42874_4 | TATGAGTGTGGAGTCGTTGCTATGGTATGGTGTGTGTGTGTGTNNNAGATCGGAA |
| 42874_5 | TATGAGTGTGGAGTCGTTGCTATGGTATGGTGTGTGTGTGTGTNNNAGATCGGAA |
| 51462_1 | TATGAGTGTGGAGTCGTTGCATTTGAGACCAACCTAGGCAACANNNAGATCGGAAG |
| 51462_2 | TATGAGTGTGGAGTCGTTGCATTTGAGACCAACCTAGGCAACANNNAGATCGGAAG |
| 51462_3 | TATGAGTGTGGAGTCGTTGCATTTGAGACCAACCTAGGCAACANNNAGATCGGAAG |
| 51462_4 | TATGAGTGTGGAGTCGTTGCATTTGAGACCAACCTAGGCAACANNNAGATCGGAAG |
| 51462_5 | TATGAGTGTGGAGTCGTTGCATTTGAGACCAACCTAGGCAACANNNAGATCGGAAG |
| 36478_1 | TATGAGTGTGGAGTCGTTGCAGTGCCAAAAACTTACAGCCAATNNNAGATCGGAAG |
| 36478_2 | TATGAGTGTGGAGTCGTTGCAGTGCCAAAAACTTACAGCCAATNNNAGATCGGAAG |
| 36478_3 | TATGAGTGTGGAGTCGTTGCAGTGCCAAAAACTTACAGCCAATNNNAGATCGGAAG |
| 36478_4 | TATGAGTGTGGAGTCGTTGCAGTGCCAAAAACTTACAGCCAATNNNAGATCGGAAG |
| 36478_5 | TATGAGTGTGGAGTCGTTGCAGTGCCAAAAACTTACAGCCAATNNNAGATCGGAAG |
| 31239_1 | TATGAGTGTGGAGTCGTTGCAGTATAGTGTGCAGGGACCAAACNNNAGATCGGAA |
| 31239_2 | TATGAGTGTGGAGTCGTTGCAGTATAGTGTGCAGGGACCAAACNNNAGATCGGAA |
| 31239_3 | TATGAGTGTGGAGTCGTTGCAGTATAGTGTGCAGGGACCAAACNNNAGATCGGAA |
| 31239_4 | TATGAGTGTGGAGTCGTTGCAGTATAGTGTGCAGGGACCAAACNNNAGATCGGAA |
| 31239_5 | TATGAGTGTGGAGTCGTTGCAGTATAGTGTGCAGGGACCAAACNNNAGATCGGAA |
| 33200_1 | TATGAGTGTGGAGTCGTTGCGGCTGAGGTAGGAGAATAGCTTGNNNAGATCGGAA |

| | |
|---|---|
| 33200_2 | TATGAGTGTGGAGTCGTTGCGGCTGAGGTAGGAGAATAGCTTGNNNAGATCGGAA |
| 33200_3 | TATGAGTGTGGAGTCGTTGCGGCTGAGGTAGGAGAATAGCTTGNNNAGATCGGAA |
| 33200_4 | TATGAGTGTGGAGTCGTTGCGGCTGAGGTAGGAGAATAGCTTGNNNAGATCGGAA |
| 33200_5 | TATGAGTGTGGAGTCGTTGCGGCTGAGGTAGGAGAATAGCTTGNNNAGATCGGAA |
| 31763_1 | TATGAGTGTGGAGTCGTTGCTTGTGCTACAAATGGCTCCAAAGNNNAGATCGGAAG |
| 31763_2 | TATGAGTGTGGAGTCGTTGCTTGTGCTACAAATGGCTCCAAAGNNNAGATCGGAAG |
| 31763_3 | TATGAGTGTGGAGTCGTTGCTTGTGCTACAAATGGCTCCAAAGNNNAGATCGGAAG |
| 31763_4 | TATGAGTGTGGAGTCGTTGCTTGTGCTACAAATGGCTCCAAAGNNNAGATCGGAAG |
| 31763_5 | TATGAGTGTGGAGTCGTTGCTTGTGCTACAAATGGCTCCAAAGNNNAGATCGGAAG |
| 42133_1 | TATGAGTGTGGAGTCGTTGCTTCAAGTGATTCTCCTGCCTCAGNNNAGATCGGAAGA |
| 42133_2 | TATGAGTGTGGAGTCGTTGCTTCAAGTGATTCTCCTGCCTCAGNNNAGATCGGAAGA |
| 42133_3 | TATGAGTGTGGAGTCGTTGCTTCAAGTGATTCTCCTGCCTCAGNNNAGATCGGAAGA |
| 42133_4 | TATGAGTGTGGAGTCGTTGCTTCAAGTGATTCTCCTGCCTCAGNNNAGATCGGAAGA |
| 42133_5 | TATGAGTGTGGAGTCGTTGCTTCAAGTGATTCTCCTGCCTCAGNNNAGATCGGAAGA |
| 31433_1 | TATGAGTGTGGAGTCGTTGCTAGAGATCTAGGAATCGCCAGGANNNAGATCGGAA |
| 31433_2 | TATGAGTGTGGAGTCGTTGCTAGAGATCTAGGAATCGCCAGGANNNAGATCGGAA |
| 31433_3 | TATGAGTGTGGAGTCGTTGCTAGAGATCTAGGAATCGCCAGGANNNAGATCGGAA |
| 31433_4 | TATGAGTGTGGAGTCGTTGCTAGAGATCTAGGAATCGCCAGGANNNAGATCGGAA |
| 31433_5 | TATGAGTGTGGAGTCGTTGCTAGAGATCTAGGAATCGCCAGGANNNAGATCGGAA |
| 29436_1 | TATGAGTGTGGAGTCGTTGCGCAAGGAAAGAGTATTTAAAAACNNNAGATCGGAA |
| 29436_2 | TATGAGTGTGGAGTCGTTGCGCAAGGAAAGAGTATTTAAAAACNNNAGATCGGAA |
| 29436_3 | TATGAGTGTGGAGTCGTTGCGCAAGGAAAGAGTATTTAAAAACNNNAGATCGGAA |
| 29436_4 | TATGAGTGTGGAGTCGTTGCGCAAGGAAAGAGTATTTAAAAACNNNAGATCGGAA |
| 29436_5 | TATGAGTGTGGAGTCGTTGCGCAAGGAAAGAGTATTTAAAAACNNNAGATCGGAA |
| 30303_1 | TATGAGTGTGGAGTCGTTGCGATAACCAGCATTGAGCACAACCNNNAGATCGGAAG |
| 30303_2 | TATGAGTGTGGAGTCGTTGCGATAACCAGCATTGAGCACAACCNNNAGATCGGAAG |
| 30303_3 | TATGAGTGTGGAGTCGTTGCGATAACCAGCATTGAGCACAACCNNNAGATCGGAAG |
| 30303_4 | TATGAGTGTGGAGTCGTTGCGATAACCAGCATTGAGCACAACCNNNAGATCGGAAG |
| 30303_5 | TATGAGTGTGGAGTCGTTGCGATAACCAGCATTGAGCACAACCNNNAGATCGGAAG |
| 44181_1 | TATGAGTGTGGAGTCGTTGCTTCCTCTACATATCCTGAGCCCANNNAGATCGGAAGA |
| 44181_2 | TATGAGTGTGGAGTCGTTGCTTCCTCTACATATCCTGAGCCCANNNAGATCGGAAGA |
| 44181_3 | TATGAGTGTGGAGTCGTTGCTTCCTCTACATATCCTGAGCCCANNNAGATCGGAAGA |
| 44181_4 | TATGAGTGTGGAGTCGTTGCTTCCTCTACATATCCTGAGCCCANNNAGATCGGAAGA |
| 44181_5 | TATGAGTGTGGAGTCGTTGCTTCCTCTACATATCCTGAGCCCANNNAGATCGGAAGA |
| 48072_1 | TATGAGTGTGGAGTCGTTGCGCATAATCTCAGCTCACTGCAAGNNNAGATCGGAAG |
| 48072_2 | TATGAGTGTGGAGTCGTTGCGCATAATCTCAGCTCACTGCAAGNNNAGATCGGAAG |
| 48072_3 | TATGAGTGTGGAGTCGTTGCGCATAATCTCAGCTCACTGCAAGNNNAGATCGGAAG |
| 48072_4 | TATGAGTGTGGAGTCGTTGCGCATAATCTCAGCTCACTGCAAGNNNAGATCGGAAG |
| 48072_5 | TATGAGTGTGGAGTCGTTGCGCATAATCTCAGCTCACTGCAAGNNNAGATCGGAAG |
| 49951_1 | TATGAGTGTGGAGTCGTTGCCGCTCTACTCTCACTACCAAGTCNNNAGATCGGAAGA |
| 49951_2 | TATGAGTGTGGAGTCGTTGCCGCTCTACTCTCACTACCAAGTCNNNAGATCGGAAGA |
| 49951_3 | TATGAGTGTGGAGTCGTTGCCGCTCTACTCTCACTACCAAGTCNNNAGATCGGAAGA |
| 49951_4 | TATGAGTGTGGAGTCGTTGCCGCTCTACTCTCACTACCAAGTCNNNAGATCGGAAGA |

| | |
|---|---|
| 49951_5 | TATGAGTGTGGAGTCGTTGCCGCTCTACTCTCACTACCAAGTCNNNAGATCGGAAGA |
| 6472_1 | TATGAGTGTGGAGTCGTTGCTGGAAGTCTAAGCAGCATGCTTCNNNAGATCGGAAG |
| 6472_2 | TATGAGTGTGGAGTCGTTGCTGGAAGTCTAAGCAGCATGCTTCNNNAGATCGGAAG |
| 6472_3 | TATGAGTGTGGAGTCGTTGCTGGAAGTCTAAGCAGCATGCTTCNNNAGATCGGAAG |
| 6472_4 | TATGAGTGTGGAGTCGTTGCTGGAAGTCTAAGCAGCATGCTTCNNNAGATCGGAAG |
| 6472_5 | TATGAGTGTGGAGTCGTTGCTGGAAGTCTAAGCAGCATGCTTCNNNAGATCGGAAG |
| 33132_1 | TATGAGTGTGGAGTCGTTGCTCTCCGTGGTTTTACAAGGATGANNNAGATCGGAAG |
| 33132_2 | TATGAGTGTGGAGTCGTTGCTCTCCGTGGTTTTACAAGGATGANNNAGATCGGAAG |
| 33132_3 | TATGAGTGTGGAGTCGTTGCTCTCCGTGGTTTTACAAGGATGANNNAGATCGGAAG |
| 33132_4 | TATGAGTGTGGAGTCGTTGCTCTCCGTGGTTTTACAAGGATGANNNAGATCGGAAG |
| 33132_5 | TATGAGTGTGGAGTCGTTGCTCTCCGTGGTTTTACAAGGATGANNNAGATCGGAAG |
| 34595_1 | TATGAGTGTGGAGTCGTTGCACCCTGTCTCAAAAAAGAATCCGNNNAGATCGGAAG |
| 34595_2 | TATGAGTGTGGAGTCGTTGCACCCTGTCTCAAAAAAGAATCCGNNNAGATCGGAAG |
| 34595_3 | TATGAGTGTGGAGTCGTTGCACCCTGTCTCAAAAAAGAATCCGNNNAGATCGGAAG |
| 34595_4 | TATGAGTGTGGAGTCGTTGCACCCTGTCTCAAAAAAGAATCCGNNNAGATCGGAAG |
| 34595_5 | TATGAGTGTGGAGTCGTTGCACCCTGTCTCAAAAAAGAATCCGNNNAGATCGGAAG |
| 49493_1 | TATGAGTGTGGAGTCGTTGCTCAGCTTCCCAAAGTGCTAGAATNNNAGATCGGAAG |
| 49493_2 | TATGAGTGTGGAGTCGTTGCTCAGCTTCCCAAAGTGCTAGAATNNNAGATCGGAAG |
| 49493_3 | TATGAGTGTGGAGTCGTTGCTCAGCTTCCCAAAGTGCTAGAATNNNAGATCGGAAG |
| 49493_4 | TATGAGTGTGGAGTCGTTGCTCAGCTTCCCAAAGTGCTAGAATNNNAGATCGGAAG |
| 49493_5 | TATGAGTGTGGAGTCGTTGCTCAGCTTCCCAAAGTGCTAGAATNNNAGATCGGAAG |
| 46418_1 | TATGAGTGTGGAGTCGTTGCCAGGTTCACGCCATTCTCCTNNNAGATCGGAAGAGCA |
| 46418_2 | TATGAGTGTGGAGTCGTTGCCAGGTTCACGCCATTCTCCTNNNAGATCGGAAGAGCA |
| 46418_3 | TATGAGTGTGGAGTCGTTGCCAGGTTCACGCCATTCTCCTNNNAGATCGGAAGAGCA |
| 46418_4 | TATGAGTGTGGAGTCGTTGCCAGGTTCACGCCATTCTCCTNNNAGATCGGAAGAGCA |
| 46418_5 | TATGAGTGTGGAGTCGTTGCCAGGTTCACGCCATTCTCCTNNNAGATCGGAAGAGCA |
| 32268_1 | TATGAGTGTGGAGTCGTTGCCAGTGAAGAGAAGAGGAAGGCATNNNAGATCGGAA |
| 32268_2 | TATGAGTGTGGAGTCGTTGCCAGTGAAGAGAAGAGGAAGGCATNNNAGATCGGAA |
| 32268_3 | TATGAGTGTGGAGTCGTTGCCAGTGAAGAGAAGAGGAAGGCATNNNAGATCGGAA |
| 32268_4 | TATGAGTGTGGAGTCGTTGCCAGTGAAGAGAAGAGGAAGGCATNNNAGATCGGAA |
| 32268_5 | TATGAGTGTGGAGTCGTTGCCAGTGAAGAGAAGAGGAAGGCATNNNAGATCGGAA |
| 30602_1 | TATGAGTGTGGAGTCGTTGCAAGGGTATAAAGAGGCAGGTCTGANNNAGATCGGA |
| 30602_2 | TATGAGTGTGGAGTCGTTGCAAGGGTATAAAGAGGCAGGTCTGANNNAGATCGGA |
| 30602_3 | TATGAGTGTGGAGTCGTTGCAAGGGTATAAAGAGGCAGGTCTGANNNAGATCGGA |
| 30602_4 | TATGAGTGTGGAGTCGTTGCAAGGGTATAAAGAGGCAGGTCTGANNNAGATCGGA |
| 30602_5 | TATGAGTGTGGAGTCGTTGCAAGGGTATAAAGAGGCAGGTCTGANNNAGATCGGA |
| 47549_1 | TATGAGTGTGGAGTCGTTGCTCCCAGGTTCAATGATTCTCCTGNNNAGATCGGAAGA |
| 47549_2 | TATGAGTGTGGAGTCGTTGCTCCCAGGTTCAATGATTCTCCTGNNNAGATCGGAAGA |
| 47549_3 | TATGAGTGTGGAGTCGTTGCTCCCAGGTTCAATGATTCTCCTGNNNAGATCGGAAGA |
| 47549_4 | TATGAGTGTGGAGTCGTTGCTCCCAGGTTCAATGATTCTCCTGNNNAGATCGGAAGA |
| 47549_5 | TATGAGTGTGGAGTCGTTGCTCCCAGGTTCAATGATTCTCCTGNNNAGATCGGAAGA |
| 39445_1 | TATGAGTGTGGAGTCGTTGCAAAAGAAGAGAGTGAGGGAAGGGNNNAGATCGGA |
| 39445_2 | TATGAGTGTGGAGTCGTTGCAAAAGAAGAGAGTGAGGGAAGGGNNNAGATCGGA |

| | |
|---|---|
| 39445_3 | TATGAGTGTGGAGTCGTTGCAAAAGAAGAGAGTGAGGGAAGGGNNNAGATCGG |
| 39445_4 | TATGAGTGTGGAGTCGTTGCAAAAGAAGAGAGTGAGGGAAGGGNNNAGATCGG |
| 39445_5 | TATGAGTGTGGAGTCGTTGCAAAAGAAGAGAGTGAGGGAAGGGNNNAGATCGG |
| 45860_1 | TATGAGTGTGGAGTCGTTGCCCAAGTAGCTGGGATTATAGGCGNNNAGATCGGAA |
| 45860_2 | TATGAGTGTGGAGTCGTTGCCCAAGTAGCTGGGATTATAGGCGNNNAGATCGGAA |
| 45860_3 | TATGAGTGTGGAGTCGTTGCCCAAGTAGCTGGGATTATAGGCGNNNAGATCGGAA |
| 45860_4 | TATGAGTGTGGAGTCGTTGCCCAAGTAGCTGGGATTATAGGCGNNNAGATCGGAA |
| 45860_5 | TATGAGTGTGGAGTCGTTGCCCAAGTAGCTGGGATTATAGGCGNNNAGATCGGAA |
| 44717_1 | TATGAGTGTGGAGTCGTTGCAACTTAGGCCTTCTACAGCACAANNNAGATCGGAAG |
| 44717_2 | TATGAGTGTGGAGTCGTTGCAACTTAGGCCTTCTACAGCACAANNNAGATCGGAAG |
| 44717_3 | TATGAGTGTGGAGTCGTTGCAACTTAGGCCTTCTACAGCACAANNNAGATCGGAAG |
| 44717_4 | TATGAGTGTGGAGTCGTTGCAACTTAGGCCTTCTACAGCACAANNNAGATCGGAAG |
| 44717_5 | TATGAGTGTGGAGTCGTTGCAACTTAGGCCTTCTACAGCACAANNNAGATCGGAAG |
| 2617_1 | TATGAGTGTGGAGTCGTTGCCACTGATTGCTCTTAGGTCTGGCNNNAGATCGGAAG |
| 2617_2 | TATGAGTGTGGAGTCGTTGCCACTGATTGCTCTTAGGTCTGGCNNNAGATCGGAAG |
| 2617_3 | TATGAGTGTGGAGTCGTTGCCACTGATTGCTCTTAGGTCTGGCNNNAGATCGGAAG |
| 2617_4 | TATGAGTGTGGAGTCGTTGCCACTGATTGCTCTTAGGTCTGGCNNNAGATCGGAAG |
| 2617_5 | TATGAGTGTGGAGTCGTTGCCACTGATTGCTCTTAGGTCTGGCNNNAGATCGGAAG |
| 12724_1 | TATGAGTGTGGAGTCGTTGCTTTGTGGACCTAATTATAAAGGTNNNAGATCGGAAG |
| 12724_2 | TATGAGTGTGGAGTCGTTGCTTTGTGGACCTAATTATAAAGGTNNNAGATCGGAAG |
| 12724_3 | TATGAGTGTGGAGTCGTTGCTTTGTGGACCTAATTATAAAGGTNNNAGATCGGAAG |
| 12724_4 | TATGAGTGTGGAGTCGTTGCTTTGTGGACCTAATTATAAAGGTNNNAGATCGGAAG |
| 12724_5 | TATGAGTGTGGAGTCGTTGCTTTGTGGACCTAATTATAAAGGTNNNAGATCGGAAG |
| 44912_1 | TATGAGTGTGGAGTCGTTGCCTGTTTCTGTGGCTAATGCAACANNNAGATCGGAAG |
| 44912_2 | TATGAGTGTGGAGTCGTTGCCTGTTTCTGTGGCTAATGCAACANNNAGATCGGAAG |
| 44912_3 | TATGAGTGTGGAGTCGTTGCCTGTTTCTGTGGCTAATGCAACANNNAGATCGGAAG |
| 44912_4 | TATGAGTGTGGAGTCGTTGCCTGTTTCTGTGGCTAATGCAACANNNAGATCGGAAG |
| 44912_5 | TATGAGTGTGGAGTCGTTGCCTGTTTCTGTGGCTAATGCAACANNNAGATCGGAAG |
| 39053_1 | TATGAGTGTGGAGTCGTTGCAGCAAAAAGGTGGCCTTGTTATGNNNAGATCGGAA |
| 39053_2 | TATGAGTGTGGAGTCGTTGCAGCAAAAAGGTGGCCTTGTTATGNNNAGATCGGAA |
| 39053_3 | TATGAGTGTGGAGTCGTTGCAGCAAAAAGGTGGCCTTGTTATGNNNAGATCGGAA |
| 39053_4 | TATGAGTGTGGAGTCGTTGCAGCAAAAAGGTGGCCTTGTTATGNNNAGATCGGAA |
| 39053_5 | TATGAGTGTGGAGTCGTTGCAGCAAAAAGGTGGCCTTGTTATGNNNAGATCGGAA |
| 38335_1 | TATGAGTGTGGAGTCGTTGCAAATGATTGCATAGAGCCAGGTGNNNAGATCGGAA |
| 38335_2 | TATGAGTGTGGAGTCGTTGCAAATGATTGCATAGAGCCAGGTGNNNAGATCGGAA |
| 38335_3 | TATGAGTGTGGAGTCGTTGCAAATGATTGCATAGAGCCAGGTGNNNAGATCGGAA |
| 38335_4 | TATGAGTGTGGAGTCGTTGCAAATGATTGCATAGAGCCAGGTGNNNAGATCGGAA |
| 38335_5 | TATGAGTGTGGAGTCGTTGCAAATGATTGCATAGAGCCAGGTGNNNAGATCGGAA |
| 49863_1 | TATGAGTGTGGAGTCGTTGCCAAGAACAGGGAAAGGACGTCTANNNAGATCGGAA |
| 49863_2 | TATGAGTGTGGAGTCGTTGCCAAGAACAGGGAAAGGACGTCTANNNAGATCGGAA |
| 49863_3 | TATGAGTGTGGAGTCGTTGCCAAGAACAGGGAAAGGACGTCTANNNAGATCGGAA |
| 49863_4 | TATGAGTGTGGAGTCGTTGCCAAGAACAGGGAAAGGACGTCTANNNAGATCGGAA |
| 49863_5 | TATGAGTGTGGAGTCGTTGCCAAGAACAGGGAAAGGACGTCTANNNAGATCGGAA |

| | |
|---|---|
| 38427_1 | TATGAGTGTGGAGTCGTTGCCATGCCTGTAATCCCAGCTACTCNNNAGATCGGAAGA |
| 38427_2 | TATGAGTGTGGAGTCGTTGCCATGCCTGTAATCCCAGCTACTCNNNAGATCGGAAGA |
| 38427_3 | TATGAGTGTGGAGTCGTTGCCATGCCTGTAATCCCAGCTACTCNNNAGATCGGAAGA |
| 38427_4 | TATGAGTGTGGAGTCGTTGCCATGCCTGTAATCCCAGCTACTCNNNAGATCGGAAGA |
| 38427_5 | TATGAGTGTGGAGTCGTTGCCATGCCTGTAATCCCAGCTACTCNNNAGATCGGAAGA |
| 52081_1 | TATGAGTGTGGAGTCGTTGCCATACTTGCATATAACCAGCGCANNNAGATCGGAAGA |
| 52081_2 | TATGAGTGTGGAGTCGTTGCCATACTTGCATATAACCAGCGCANNNAGATCGGAAGA |
| 52081_3 | TATGAGTGTGGAGTCGTTGCCATACTTGCATATAACCAGCGCANNNAGATCGGAAGA |
| 52081_4 | TATGAGTGTGGAGTCGTTGCCATACTTGCATATAACCAGCGCANNNAGATCGGAAGA |
| 52081_5 | TATGAGTGTGGAGTCGTTGCCATACTTGCATATAACCAGCGCANNNAGATCGGAAGA |
| 46107_1 | TATGAGTGTGGAGTCGTTGCGTCTCCTGGGTTCAAGTGATTCTNNNAGATCGGAAG |
| 46107_2 | TATGAGTGTGGAGTCGTTGCGTCTCCTGGGTTCAAGTGATTCTNNNAGATCGGAAG |
| 46107_3 | TATGAGTGTGGAGTCGTTGCGTCTCCTGGGTTCAAGTGATTCTNNNAGATCGGAAG |
| 46107_4 | TATGAGTGTGGAGTCGTTGCGTCTCCTGGGTTCAAGTGATTCTNNNAGATCGGAAG |
| 46107_5 | TATGAGTGTGGAGTCGTTGCGTCTCCTGGGTTCAAGTGATTCTNNNAGATCGGAAG |
| 36717_1 | TATGAGTGTGGAGTCGTTGCCTTCTCAAAGGACCACCTACGAANNNAGATCGGAAGA |
| 36717_2 | TATGAGTGTGGAGTCGTTGCCTTCTCAAAGGACCACCTACGAANNNAGATCGGAAGA |
| 36717_3 | TATGAGTGTGGAGTCGTTGCCTTCTCAAAGGACCACCTACGAANNNAGATCGGAAGA |
| 36717_4 | TATGAGTGTGGAGTCGTTGCCTTCTCAAAGGACCACCTACGAANNNAGATCGGAAGA |
| 36717_5 | TATGAGTGTGGAGTCGTTGCCTTCTCAAAGGACCACCTACGAANNNAGATCGGAAGA |
| 39793_1 | TATGAGTGTGGAGTCGTTGCGAGGGTTGGATGGATCAGTACAGNNNAGATCGGAA |
| 39793_2 | TATGAGTGTGGAGTCGTTGCGAGGGTTGGATGGATCAGTACAGNNNAGATCGGAA |
| 39793_3 | TATGAGTGTGGAGTCGTTGCGAGGGTTGGATGGATCAGTACAGNNNAGATCGGAA |
| 39793_4 | TATGAGTGTGGAGTCGTTGCGAGGGTTGGATGGATCAGTACAGNNNAGATCGGAA |
| 39793_5 | TATGAGTGTGGAGTCGTTGCGAGGGTTGGATGGATCAGTACAGNNNAGATCGGAA |
| 53068_1 | TATGAGTGTGGAGTCGTTGCAACCAAACCAATAATATGCATGCTNNNAGATCGGAA |
| 53068_2 | TATGAGTGTGGAGTCGTTGCAACCAAACCAATAATATGCATGCTNNNAGATCGGAA |
| 53068_3 | TATGAGTGTGGAGTCGTTGCAACCAAACCAATAATATGCATGCTNNNAGATCGGAA |
| 53068_4 | TATGAGTGTGGAGTCGTTGCAACCAAACCAATAATATGCATGCTNNNAGATCGGAA |
| 53068_5 | TATGAGTGTGGAGTCGTTGCAACCAAACCAATAATATGCATGCTNNNAGATCGGAA |
| 45020_1 | TATGAGTGTGGAGTCGTTGCTAGTTATGACCTGCTTCATCCGGNNNAGATCGGAAGA |
| 45020_2 | TATGAGTGTGGAGTCGTTGCTAGTTATGACCTGCTTCATCCGGNNNAGATCGGAAGA |
| 45020_3 | TATGAGTGTGGAGTCGTTGCTAGTTATGACCTGCTTCATCCGGNNNAGATCGGAAGA |
| 45020_4 | TATGAGTGTGGAGTCGTTGCTAGTTATGACCTGCTTCATCCGGNNNAGATCGGAAGA |
| 45020_5 | TATGAGTGTGGAGTCGTTGCTAGTTATGACCTGCTTCATCCGGNNNAGATCGGAAGA |
| 10967_1 | TATGAGTGTGGAGTCGTTGCCCAGAGGCTATTCCAGTGATTGANNNAGATCGGAAG |
| 10967_2 | TATGAGTGTGGAGTCGTTGCCCAGAGGCTATTCCAGTGATTGANNNAGATCGGAAG |
| 10967_3 | TATGAGTGTGGAGTCGTTGCCCAGAGGCTATTCCAGTGATTGANNNAGATCGGAAG |
| 10967_4 | TATGAGTGTGGAGTCGTTGCCCAGAGGCTATTCCAGTGATTGANNNAGATCGGAAG |
| 10967_5 | TATGAGTGTGGAGTCGTTGCCCAGAGGCTATTCCAGTGATTGANNNAGATCGGAAG |
| 10116_1 | TATGAGTGTGGAGTCGTTGCGCGGTTCTTCCCAGTGATTGTAANNNAGATCGGAAG |
| 10116_2 | TATGAGTGTGGAGTCGTTGCGCGGTTCTTCCCAGTGATTGTAANNNAGATCGGAAG |
| 10116_3 | TATGAGTGTGGAGTCGTTGCGCGGTTCTTCCCAGTGATTGTAANNNAGATCGGAAG |

| | |
|---|---|
| 10116_4 | TATGAGTGTGGAGTCGTTGCGCGGTTCTTCCCAGTGATTGTAANNNAGATCGGAAG |
| 10116_5 | TATGAGTGTGGAGTCGTTGCGCGGTTCTTCCCAGTGATTGTAANNNAGATCGGAAG |
| 50255_1 | TATGAGTGTGGAGTCGTTGCTACTGGTAGAGTCTTACTGCCCANNNAGATCGGAAG |
| 50255_2 | TATGAGTGTGGAGTCGTTGCTACTGGTAGAGTCTTACTGCCCANNNAGATCGGAAG |
| 50255_3 | TATGAGTGTGGAGTCGTTGCTACTGGTAGAGTCTTACTGCCCANNNAGATCGGAAG |
| 50255_4 | TATGAGTGTGGAGTCGTTGCTACTGGTAGAGTCTTACTGCCCANNNAGATCGGAAG |
| 50255_5 | TATGAGTGTGGAGTCGTTGCTACTGGTAGAGTCTTACTGCCCANNNAGATCGGAAG |
| 35520_1 | TATGAGTGTGGAGTCGTTGCATACTGCCATGCCCAGCTAATTTNNNAGATCGGAAGA |
| 35520_2 | TATGAGTGTGGAGTCGTTGCATACTGCCATGCCCAGCTAATTTNNNAGATCGGAAGA |
| 35520_3 | TATGAGTGTGGAGTCGTTGCATACTGCCATGCCCAGCTAATTTNNNAGATCGGAAGA |
| 35520_4 | TATGAGTGTGGAGTCGTTGCATACTGCCATGCCCAGCTAATTTNNNAGATCGGAAGA |
| 35520_5 | TATGAGTGTGGAGTCGTTGCATACTGCCATGCCCAGCTAATTTNNNAGATCGGAAGA |
| 47489_1 | TATGAGTGTGGAGTCGTTGCTGTGTGGTGTATTTATGTGGGGTNNNAGATCGGAA |
| 47489_2 | TATGAGTGTGGAGTCGTTGCTGTGTGGTGTATTTATGTGGGGTNNNAGATCGGAA |
| 47489_3 | TATGAGTGTGGAGTCGTTGCTGTGTGGTGTATTTATGTGGGGTNNNAGATCGGAA |
| 47489_4 | TATGAGTGTGGAGTCGTTGCTGTGTGGTGTATTTATGTGGGGTNNNAGATCGGAA |
| 47489_5 | TATGAGTGTGGAGTCGTTGCTGTGTGGTGTATTTATGTGGGGTNNNAGATCGGAA |
| 32058_1 | TATGAGTGTGGAGTCGTTGCTTCTGGGTTCAAGCAATTCTCCTNNNAGATCGGAAGA |
| 32058_2 | TATGAGTGTGGAGTCGTTGCTTCTGGGTTCAAGCAATTCTCCTNNNAGATCGGAAGA |
| 32058_3 | TATGAGTGTGGAGTCGTTGCTTCTGGGTTCAAGCAATTCTCCTNNNAGATCGGAAGA |
| 32058_4 | TATGAGTGTGGAGTCGTTGCTTCTGGGTTCAAGCAATTCTCCTNNNAGATCGGAAGA |
| 32058_5 | TATGAGTGTGGAGTCGTTGCTTCTGGGTTCAAGCAATTCTCCTNNNAGATCGGAAGA |
| 34497_1 | TATGAGTGTGGAGTCGTTGCGATCTCGACTCACTGTAACCTCCNNNAGATCGGAAGA |
| 34497_2 | TATGAGTGTGGAGTCGTTGCGATCTCGACTCACTGTAACCTCCNNNAGATCGGAAGA |
| 34497_3 | TATGAGTGTGGAGTCGTTGCGATCTCGACTCACTGTAACCTCCNNNAGATCGGAAGA |
| 34497_4 | TATGAGTGTGGAGTCGTTGCGATCTCGACTCACTGTAACCTCCNNNAGATCGGAAGA |
| 34497_5 | TATGAGTGTGGAGTCGTTGCGATCTCGACTCACTGTAACCTCCNNNAGATCGGAAGA |
| 13480_1 | TATGAGTGTGGAGTCGTTGCATTCATTTCAACTTGGTACATTTNNNAGATCGGAAGA |
| 13480_2 | TATGAGTGTGGAGTCGTTGCATTCATTTCAACTTGGTACATTTNNNAGATCGGAAGA |
| 13480_3 | TATGAGTGTGGAGTCGTTGCATTCATTTCAACTTGGTACATTTNNNAGATCGGAAGA |
| 13480_4 | TATGAGTGTGGAGTCGTTGCATTCATTTCAACTTGGTACATTTNNNAGATCGGAAGA |
| 13480_5 | TATGAGTGTGGAGTCGTTGCATTCATTTCAACTTGGTACATTTNNNAGATCGGAAGA |
| 2173_1 | TATGAGTGTGGAGTCGTTGCCCAATGACCCAGGAAAAACGAGANNNAGATCGGAA |
| 2173_2 | TATGAGTGTGGAGTCGTTGCCCAATGACCCAGGAAAAACGAGANNNAGATCGGAA |
| 2173_3 | TATGAGTGTGGAGTCGTTGCCCAATGACCCAGGAAAAACGAGANNNAGATCGGAA |
| 2173_4 | TATGAGTGTGGAGTCGTTGCCCAATGACCCAGGAAAAACGAGANNNAGATCGGAA |
| 2173_5 | TATGAGTGTGGAGTCGTTGCCCAATGACCCAGGAAAAACGAGANNNAGATCGGAA |
| 36678_1 | TATGAGTGTGGAGTCGTTGCGCTACAGAACGAGACTCCATCTCNNNAGATCGGAAG |
| 36678_2 | TATGAGTGTGGAGTCGTTGCGCTACAGAACGAGACTCCATCTCNNNAGATCGGAAG |
| 36678_3 | TATGAGTGTGGAGTCGTTGCGCTACAGAACGAGACTCCATCTCNNNAGATCGGAAG |
| 36678_4 | TATGAGTGTGGAGTCGTTGCGCTACAGAACGAGACTCCATCTCNNNAGATCGGAAG |
| 36678_5 | TATGAGTGTGGAGTCGTTGCGCTACAGAACGAGACTCCATCTCNNNAGATCGGAAG |
| 30180_1 | TATGAGTGTGGAGTCGTTGCCAGACAAGAAGACCTGCTATGGTNNNAGATCGGAA |

| | |
|---|---|
| 30180_2 | TATGAGTGTGGAGTCGTTGCCAGACAAGAAGACCTGCTATGGTNNNAGATCGGAA |
| 30180_3 | TATGAGTGTGGAGTCGTTGCCAGACAAGAAGACCTGCTATGGTNNNAGATCGGAA |
| 30180_4 | TATGAGTGTGGAGTCGTTGCCAGACAAGAAGACCTGCTATGGTNNNAGATCGGAA |
| 30180_5 | TATGAGTGTGGAGTCGTTGCCAGACAAGAAGACCTGCTATGGTNNNAGATCGGAA |
| 46998_1 | TATGAGTGTGGAGTCGTTGCCAGATGGAGTCTCACTCTGTCACNNNAGATCGGAAG |
| 46998_2 | TATGAGTGTGGAGTCGTTGCCAGATGGAGTCTCACTCTGTCACNNNAGATCGGAAG |
| 46998_3 | TATGAGTGTGGAGTCGTTGCCAGATGGAGTCTCACTCTGTCACNNNAGATCGGAAG |
| 46998_4 | TATGAGTGTGGAGTCGTTGCCAGATGGAGTCTCACTCTGTCACNNNAGATCGGAAG |
| 46998_5 | TATGAGTGTGGAGTCGTTGCCAGATGGAGTCTCACTCTGTCACNNNAGATCGGAAG |
| 44465_1 | TATGAGTGTGGAGTCGTTGCAAAGTCACCTAGTGGAACCCTTCNNNAGATCGGAAG |
| 44465_2 | TATGAGTGTGGAGTCGTTGCAAAGTCACCTAGTGGAACCCTTCNNNAGATCGGAAG |
| 44465_3 | TATGAGTGTGGAGTCGTTGCAAAGTCACCTAGTGGAACCCTTCNNNAGATCGGAAG |
| 44465_4 | TATGAGTGTGGAGTCGTTGCAAAGTCACCTAGTGGAACCCTTCNNNAGATCGGAAG |
| 44465_5 | TATGAGTGTGGAGTCGTTGCAAAGTCACCTAGTGGAACCCTTCNNNAGATCGGAAG |
| 5336_1 | TATGAGTGTGGAGTCGTTGCAGACACTAGGCAGTCATACGGATNNNAGATCGGAA |
| 5336_2 | TATGAGTGTGGAGTCGTTGCAGACACTAGGCAGTCATACGGATNNNAGATCGGAA |
| 5336_3 | TATGAGTGTGGAGTCGTTGCAGACACTAGGCAGTCATACGGATNNNAGATCGGAA |
| 5336_4 | TATGAGTGTGGAGTCGTTGCAGACACTAGGCAGTCATACGGATNNNAGATCGGAA |
| 5336_5 | TATGAGTGTGGAGTCGTTGCAGACACTAGGCAGTCATACGGATNNNAGATCGGAA |
| 50417_1 | TATGAGTGTGGAGTCGTTGCAATGTCCATCCACAGGAGAATGANNNAGATCGGAA |
| 50417_2 | TATGAGTGTGGAGTCGTTGCAATGTCCATCCACAGGAGAATGANNNAGATCGGAA |
| 50417_3 | TATGAGTGTGGAGTCGTTGCAATGTCCATCCACAGGAGAATGANNNAGATCGGAA |
| 50417_4 | TATGAGTGTGGAGTCGTTGCAATGTCCATCCACAGGAGAATGANNNAGATCGGAA |
| 50417_5 | TATGAGTGTGGAGTCGTTGCAATGTCCATCCACAGGAGAATGANNNAGATCGGAA |
| 29215_1 | TATGAGTGTGGAGTCGTTGCTAAAGTCTATCTCCCCTCTCCCCNNNAGATCGGAAGA |
| 29215_2 | TATGAGTGTGGAGTCGTTGCTAAAGTCTATCTCCCCTCTCCCCNNNAGATCGGAAGA |
| 29215_3 | TATGAGTGTGGAGTCGTTGCTAAAGTCTATCTCCCCTCTCCCCNNNAGATCGGAAGA |
| 29215_4 | TATGAGTGTGGAGTCGTTGCTAAAGTCTATCTCCCCTCTCCCCNNNAGATCGGAAGA |
| 29215_5 | TATGAGTGTGGAGTCGTTGCTAAAGTCTATCTCCCCTCTCCCCNNNAGATCGGAAGA |
| 30219_1 | TATGAGTGTGGAGTCGTTGCATCTGCCATGTTGTTCAGATCCTNNNAGATCGGAAG |
| 30219_2 | TATGAGTGTGGAGTCGTTGCATCTGCCATGTTGTTCAGATCCTNNNAGATCGGAAG |
| 30219_3 | TATGAGTGTGGAGTCGTTGCATCTGCCATGTTGTTCAGATCCTNNNAGATCGGAAG |
| 30219_4 | TATGAGTGTGGAGTCGTTGCATCTGCCATGTTGTTCAGATCCTNNNAGATCGGAAG |
| 30219_5 | TATGAGTGTGGAGTCGTTGCATCTGCCATGTTGTTCAGATCCTNNNAGATCGGAAG |
| 12682_1 | TATGAGTGTGGAGTCGTTGCATTTCCACAGAATCTTTTAACAGNNNAGATCGGAAG |
| 12682_2 | TATGAGTGTGGAGTCGTTGCATTTCCACAGAATCTTTTAACAGNNNAGATCGGAAG |
| 12682_3 | TATGAGTGTGGAGTCGTTGCATTTCCACAGAATCTTTTAACAGNNNAGATCGGAAG |
| 12682_4 | TATGAGTGTGGAGTCGTTGCATTTCCACAGAATCTTTTAACAGNNNAGATCGGAAG |
| 12682_5 | TATGAGTGTGGAGTCGTTGCATTTCCACAGAATCTTTTAACAGNNNAGATCGGAAG |
| 38131_1 | TATGAGTGTGGAGTCGTTGCTATTTTCTCCCAGTCTGTGGCTTNNNAGATCGGAAGA |
| 38131_2 | TATGAGTGTGGAGTCGTTGCTATTTTCTCCCAGTCTGTGGCTTNNNAGATCGGAAGA |
| 38131_3 | TATGAGTGTGGAGTCGTTGCTATTTTCTCCCAGTCTGTGGCTTNNNAGATCGGAAGA |
| 38131_4 | TATGAGTGTGGAGTCGTTGCTATTTTCTCCCAGTCTGTGGCTTNNNAGATCGGAAGA |

| | |
|---|---|
| 38131_5 | TATGAGTGTGGAGTCGTTGCTATTTTCTCCCAGTCTGTGGCTTNNNAGATCGGAAG |
| 45364_1 | TATGAGTGTGGAGTCGTTGCGCGCTATAGCCACGAAAGTGANNNAGATCGGAAGA |
| 45364_2 | TATGAGTGTGGAGTCGTTGCGCGCTATAGCCACGAAAGTGANNNAGATCGGAAGA |
| 45364_3 | TATGAGTGTGGAGTCGTTGCGCGCTATAGCCACGAAAGTGANNNAGATCGGAAGA |
| 45364_4 | TATGAGTGTGGAGTCGTTGCGCGCTATAGCCACGAAAGTGANNNAGATCGGAAGA |
| 45364_5 | TATGAGTGTGGAGTCGTTGCGCGCTATAGCCACGAAAGTGANNNAGATCGGAAGA |
| 39529_1 | TATGAGTGTGGAGTCGTTGCGATGCCCTCTCTCACTACTCCTANNNAGATCGGAAGA |
| 39529_2 | TATGAGTGTGGAGTCGTTGCGATGCCCTCTCTCACTACTCCTANNNAGATCGGAAGA |
| 39529_3 | TATGAGTGTGGAGTCGTTGCGATGCCCTCTCTCACTACTCCTANNNAGATCGGAAGA |
| 39529_4 | TATGAGTGTGGAGTCGTTGCGATGCCCTCTCTCACTACTCCTANNNAGATCGGAAGA |
| 39529_5 | TATGAGTGTGGAGTCGTTGCGATGCCCTCTCTCACTACTCCTANNNAGATCGGAAGA |
| 40281_1 | TATGAGTGTGGAGTCGTTGCACCATTCAACCTAGCAACCTCATNNNAGATCGGAAG |
| 40281_2 | TATGAGTGTGGAGTCGTTGCACCATTCAACCTAGCAACCTCATNNNAGATCGGAAG |
| 40281_3 | TATGAGTGTGGAGTCGTTGCACCATTCAACCTAGCAACCTCATNNNAGATCGGAAG |
| 40281_4 | TATGAGTGTGGAGTCGTTGCACCATTCAACCTAGCAACCTCATNNNAGATCGGAAG |
| 40281_5 | TATGAGTGTGGAGTCGTTGCACCATTCAACCTAGCAACCTCATNNNAGATCGGAAG |
| 13421_1 | TATGAGTGTGGAGTCGTTGCACACAATCTGGATAATATTGAGANNNAGATCGGAA |
| 13421_2 | TATGAGTGTGGAGTCGTTGCACACAATCTGGATAATATTGAGANNNAGATCGGAA |
| 13421_3 | TATGAGTGTGGAGTCGTTGCACACAATCTGGATAATATTGAGANNNAGATCGGAA |
| 13421_4 | TATGAGTGTGGAGTCGTTGCACACAATCTGGATAATATTGAGANNNAGATCGGAA |
| 13421_5 | TATGAGTGTGGAGTCGTTGCACACAATCTGGATAATATTGAGANNNAGATCGGAA |
| 37049_1 | TATGAGTGTGGAGTCGTTGCCTGATGTAGGAGGATGGCTTGAGNNNAGATCGGA |
| 37049_2 | TATGAGTGTGGAGTCGTTGCCTGATGTAGGAGGATGGCTTGAGNNNAGATCGGA |
| 37049_3 | TATGAGTGTGGAGTCGTTGCCTGATGTAGGAGGATGGCTTGAGNNNAGATCGGA |
| 37049_4 | TATGAGTGTGGAGTCGTTGCCTGATGTAGGAGGATGGCTTGAGNNNAGATCGGA |
| 37049_5 | TATGAGTGTGGAGTCGTTGCCTGATGTAGGAGGATGGCTTGAGNNNAGATCGGA |
| 47316_1 | TATGAGTGTGGAGTCGTTGCTGGTCTGTTCAAGGATCCAACTTNNNAGATCGGAAG |
| 47316_2 | TATGAGTGTGGAGTCGTTGCTGGTCTGTTCAAGGATCCAACTTNNNAGATCGGAAG |
| 47316_3 | TATGAGTGTGGAGTCGTTGCTGGTCTGTTCAAGGATCCAACTTNNNAGATCGGAAG |
| 47316_4 | TATGAGTGTGGAGTCGTTGCTGGTCTGTTCAAGGATCCAACTTNNNAGATCGGAAG |
| 47316_5 | TATGAGTGTGGAGTCGTTGCTGGTCTGTTCAAGGATCCAACTTNNNAGATCGGAAG |
| 36287_1 | TATGAGTGTGGAGTCGTTGCAATACGAGCATCCCCAAGGAAATNNNAGATCGGAA |
| 36287_2 | TATGAGTGTGGAGTCGTTGCAATACGAGCATCCCCAAGGAAATNNNAGATCGGAA |
| 36287_3 | TATGAGTGTGGAGTCGTTGCAATACGAGCATCCCCAAGGAAATNNNAGATCGGAA |
| 36287_4 | TATGAGTGTGGAGTCGTTGCAATACGAGCATCCCCAAGGAAATNNNAGATCGGAA |
| 36287_5 | TATGAGTGTGGAGTCGTTGCAATACGAGCATCCCCAAGGAAATNNNAGATCGGAA |
| 30447_1 | TATGAGTGTGGAGTCGTTGCCCCTATACACCCATATCGGCCNNNAGATCGGAAGAG |
| 30447_2 | TATGAGTGTGGAGTCGTTGCCCCTATACACCCATATCGGCCNNNAGATCGGAAGAG |
| 30447_3 | TATGAGTGTGGAGTCGTTGCCCCTATACACCCATATCGGCCNNNAGATCGGAAGAG |
| 30447_4 | TATGAGTGTGGAGTCGTTGCCCCTATACACCCATATCGGCCNNNAGATCGGAAGAG |
| 30447_5 | TATGAGTGTGGAGTCGTTGCCCCTATACACCCATATCGGCCNNNAGATCGGAAGAG |
| 44833_1 | TATGAGTGTGGAGTCGTTGCTTACCAGTCCCCTCAGCGATNNNAGATCGGAAGAGC |
| 44833_2 | TATGAGTGTGGAGTCGTTGCTTACCAGTCCCCTCAGCGATNNNAGATCGGAAGAGC |

| | |
|---|---|
| 44833_3 | TATGAGTGTGGAGTCGTTGCTTACCAGTCCCCTCAGCGATNNNAGATCGGAAGAGC |
| 44833_4 | TATGAGTGTGGAGTCGTTGCTTACCAGTCCCCTCAGCGATNNNAGATCGGAAGAGC |
| 44833_5 | TATGAGTGTGGAGTCGTTGCTTACCAGTCCCCTCAGCGATNNNAGATCGGAAGAGC |
| 12878_1 | TATGAGTGTGGAGTCGTTGCTTTGTCAATTCCAGAATACAAANNNAGATCGGAAG |
| 12878_2 | TATGAGTGTGGAGTCGTTGCTTTGTCAATTCCAGAATACAAANNNAGATCGGAAG |
| 12878_3 | TATGAGTGTGGAGTCGTTGCTTTGTCAATTCCAGAATACAAANNNAGATCGGAAG |
| 12878_4 | TATGAGTGTGGAGTCGTTGCTTTGTCAATTCCAGAATACAAANNNAGATCGGAAG |
| 12878_5 | TATGAGTGTGGAGTCGTTGCTTTGTCAATTCCAGAATACAAANNNAGATCGGAAG |
| 34935_1 | TATGAGTGTGGAGTCGTTGCTTGAATACCCATTGCCCACTCTTNNNAGATCGGAAGA |
| 34935_2 | TATGAGTGTGGAGTCGTTGCTTGAATACCCATTGCCCACTCTTNNNAGATCGGAAGA |
| 34935_3 | TATGAGTGTGGAGTCGTTGCTTGAATACCCATTGCCCACTCTTNNNAGATCGGAAGA |
| 34935_4 | TATGAGTGTGGAGTCGTTGCTTGAATACCCATTGCCCACTCTTNNNAGATCGGAAGA |
| 34935_5 | TATGAGTGTGGAGTCGTTGCTTGAATACCCATTGCCCACTCTTNNNAGATCGGAAGA |
| 37898_1 | TATGAGTGTGGAGTCGTTGCGTATAAAAAAGTGGGCCAGGAGCNNNAGATCGGAA |
| 37898_2 | TATGAGTGTGGAGTCGTTGCGTATAAAAAAGTGGGCCAGGAGCNNNAGATCGGAA |
| 37898_3 | TATGAGTGTGGAGTCGTTGCGTATAAAAAAGTGGGCCAGGAGCNNNAGATCGGAA |
| 37898_4 | TATGAGTGTGGAGTCGTTGCGTATAAAAAAGTGGGCCAGGAGCNNNAGATCGGAA |
| 37898_5 | TATGAGTGTGGAGTCGTTGCGTATAAAAAAGTGGGCCAGGAGCNNNAGATCGGAA |
| 49235_1 | TATGAGTGTGGAGTCGTTGCGGTTCAAGCAATTATCCTGCCTCNNNAGATCGGAAG |
| 49235_2 | TATGAGTGTGGAGTCGTTGCGGTTCAAGCAATTATCCTGCCTCNNNAGATCGGAAG |
| 49235_3 | TATGAGTGTGGAGTCGTTGCGGTTCAAGCAATTATCCTGCCTCNNNAGATCGGAAG |
| 49235_4 | TATGAGTGTGGAGTCGTTGCGGTTCAAGCAATTATCCTGCCTCNNNAGATCGGAAG |
| 49235_5 | TATGAGTGTGGAGTCGTTGCGGTTCAAGCAATTATCCTGCCTCNNNAGATCGGAAG |
| 964_1 | TATGAGTGTGGAGTCGTTGCGCCCTCTTCTTCCACTGATTCGNNNAGATCGGAAGAG |
| 964_2 | TATGAGTGTGGAGTCGTTGCGCCCTCTTCTTCCACTGATTCGNNNAGATCGGAAGAG |
| 964_3 | TATGAGTGTGGAGTCGTTGCGCCCTCTTCTTCCACTGATTCGNNNAGATCGGAAGAG |
| 964_4 | TATGAGTGTGGAGTCGTTGCGCCCTCTTCTTCCACTGATTCGNNNAGATCGGAAGAG |
| 964_5 | TATGAGTGTGGAGTCGTTGCGCCCTCTTCTTCCACTGATTCGNNNAGATCGGAAGAG |
| 8315_1 | TATGAGTGTGGAGTCGTTGCTCTTGTTCAGCAGATTCCACCCTNNNAGATCGGAAGA |
| 8315_2 | TATGAGTGTGGAGTCGTTGCTCTTGTTCAGCAGATTCCACCCTNNNAGATCGGAAGA |
| 8315_3 | TATGAGTGTGGAGTCGTTGCTCTTGTTCAGCAGATTCCACCCTNNNAGATCGGAAGA |
| 8315_4 | TATGAGTGTGGAGTCGTTGCTCTTGTTCAGCAGATTCCACCCTNNNAGATCGGAAGA |
| 8315_5 | TATGAGTGTGGAGTCGTTGCTCTTGTTCAGCAGATTCCACCCTNNNAGATCGGAAGA |
| 43170_1 | TATGAGTGTGGAGTCGTTGCTCTCGAACTTCTGGTCTCAAGTGNNNAGATCGGAAG |
| 43170_2 | TATGAGTGTGGAGTCGTTGCTCTCGAACTTCTGGTCTCAAGTGNNNAGATCGGAAG |
| 43170_3 | TATGAGTGTGGAGTCGTTGCTCTCGAACTTCTGGTCTCAAGTGNNNAGATCGGAAG |
| 43170_4 | TATGAGTGTGGAGTCGTTGCTCTCGAACTTCTGGTCTCAAGTGNNNAGATCGGAAG |
| 43170_5 | TATGAGTGTGGAGTCGTTGCTCTCGAACTTCTGGTCTCAAGTGNNNAGATCGGAAG |
| 46011_1 | TATGAGTGTGGAGTCGTTGCCTAGAGTGCAATGGCAGGATCTTNNNAGATCGGAA |
| 46011_2 | TATGAGTGTGGAGTCGTTGCCTAGAGTGCAATGGCAGGATCTTNNNAGATCGGAA |
| 46011_3 | TATGAGTGTGGAGTCGTTGCCTAGAGTGCAATGGCAGGATCTTNNNAGATCGGAA |
| 46011_4 | TATGAGTGTGGAGTCGTTGCCTAGAGTGCAATGGCAGGATCTTNNNAGATCGGAA |
| 46011_5 | TATGAGTGTGGAGTCGTTGCCTAGAGTGCAATGGCAGGATCTTNNNAGATCGGAA |

| | |
|---|---|
| 45149_1 | TATGAGTGTGGAGTCGTTGCCACTCTTCCCCATCACACACTTANNNAGATCGGAAGA |
| 45149_2 | TATGAGTGTGGAGTCGTTGCCACTCTTCCCCATCACACACTTANNNAGATCGGAAGA |
| 45149_3 | TATGAGTGTGGAGTCGTTGCCACTCTTCCCCATCACACACTTANNNAGATCGGAAGA |
| 45149_4 | TATGAGTGTGGAGTCGTTGCCACTCTTCCCCATCACACACTTANNNAGATCGGAAGA |
| 45149_5 | TATGAGTGTGGAGTCGTTGCCACTCTTCCCCATCACACACTTANNNAGATCGGAAGA |
| 1547_1 | TATGAGTGTGGAGTCGTTGCACACTGCTGGAGAGGTAAGTCTTNNNAGATCGGAA |
| 1547_2 | TATGAGTGTGGAGTCGTTGCACACTGCTGGAGAGGTAAGTCTTNNNAGATCGGAA |
| 1547_3 | TATGAGTGTGGAGTCGTTGCACACTGCTGGAGAGGTAAGTCTTNNNAGATCGGAA |
| 1547_4 | TATGAGTGTGGAGTCGTTGCACACTGCTGGAGAGGTAAGTCTTNNNAGATCGGAA |
| 1547_5 | TATGAGTGTGGAGTCGTTGCACACTGCTGGAGAGGTAAGTCTTNNNAGATCGGAA |
| 44059_1 | TATGAGTGTGGAGTCGTTGCTCTTTCCATTAGGATGTTGGGGGNNNAGATCGGAA |
| 44059_2 | TATGAGTGTGGAGTCGTTGCTCTTTCCATTAGGATGTTGGGGGNNNAGATCGGAA |
| 44059_3 | TATGAGTGTGGAGTCGTTGCTCTTTCCATTAGGATGTTGGGGGNNNAGATCGGAA |
| 44059_4 | TATGAGTGTGGAGTCGTTGCTCTTTCCATTAGGATGTTGGGGGNNNAGATCGGAA |
| 44059_5 | TATGAGTGTGGAGTCGTTGCTCTTTCCATTAGGATGTTGGGGGNNNAGATCGGAA |
| 48387_1 | TATGAGTGTGGAGTCGTTGCTTTGTTGGGGCCAAGAAAAAACTNNNAGATCGGAA |
| 48387_2 | TATGAGTGTGGAGTCGTTGCTTTGTTGGGGCCAAGAAAAAACTNNNAGATCGGAA |
| 48387_3 | TATGAGTGTGGAGTCGTTGCTTTGTTGGGGCCAAGAAAAAACTNNNAGATCGGAA |
| 48387_4 | TATGAGTGTGGAGTCGTTGCTTTGTTGGGGCCAAGAAAAAACTNNNAGATCGGAA |
| 48387_5 | TATGAGTGTGGAGTCGTTGCTTTGTTGGGGCCAAGAAAAAACTNNNAGATCGGAA |
| 31020_1 | TATGAGTGTGGAGTCGTTGCACCCCTCAGTCCTATTCTAAGCTNNNAGATCGGAAGA |
| 31020_2 | TATGAGTGTGGAGTCGTTGCACCCCTCAGTCCTATTCTAAGCTNNNAGATCGGAAGA |
| 31020_3 | TATGAGTGTGGAGTCGTTGCACCCCTCAGTCCTATTCTAAGCTNNNAGATCGGAAGA |
| 31020_4 | TATGAGTGTGGAGTCGTTGCACCCCTCAGTCCTATTCTAAGCTNNNAGATCGGAAGA |
| 31020_5 | TATGAGTGTGGAGTCGTTGCACCCCTCAGTCCTATTCTAAGCTNNNAGATCGGAAGA |
| 10684_1 | TATGAGTGTGGAGTCGTTGCCGTGGTAAGCACAATTTCTTGCCNNNAGATCGGAAG |
| 10684_2 | TATGAGTGTGGAGTCGTTGCCGTGGTAAGCACAATTTCTTGCCNNNAGATCGGAAG |
| 10684_3 | TATGAGTGTGGAGTCGTTGCCGTGGTAAGCACAATTTCTTGCCNNNAGATCGGAAG |
| 10684_4 | TATGAGTGTGGAGTCGTTGCCGTGGTAAGCACAATTTCTTGCCNNNAGATCGGAAG |
| 10684_5 | TATGAGTGTGGAGTCGTTGCCGTGGTAAGCACAATTTCTTGCCNNNAGATCGGAAG |
| 42322_1 | TATGAGTGTGGAGTCGTTGCCACAGAGTCACACATCCTTGAGANNNAGATCGGAAG |
| 42322_2 | TATGAGTGTGGAGTCGTTGCCACAGAGTCACACATCCTTGAGANNNAGATCGGAAG |
| 42322_3 | TATGAGTGTGGAGTCGTTGCCACAGAGTCACACATCCTTGAGANNNAGATCGGAAG |
| 42322_4 | TATGAGTGTGGAGTCGTTGCCACAGAGTCACACATCCTTGAGANNNAGATCGGAAG |
| 42322_5 | TATGAGTGTGGAGTCGTTGCCACAGAGTCACACATCCTTGAGANNNAGATCGGAAG |
| 47939_1 | TATGAGTGTGGAGTCGTTGCCTGGGTTCAAGTGATTCTTCTGCNNNAGATCGGAAG |
| 47939_2 | TATGAGTGTGGAGTCGTTGCCTGGGTTCAAGTGATTCTTCTGCNNNAGATCGGAAG |
| 47939_3 | TATGAGTGTGGAGTCGTTGCCTGGGTTCAAGTGATTCTTCTGCNNNAGATCGGAAG |
| 47939_4 | TATGAGTGTGGAGTCGTTGCCTGGGTTCAAGTGATTCTTCTGCNNNAGATCGGAAG |
| 47939_5 | TATGAGTGTGGAGTCGTTGCCTGGGTTCAAGTGATTCTTCTGCNNNAGATCGGAAG |
| 33555_1 | TATGAGTGTGGAGTCGTTGCCTCTCGGGTTCACACCATTCTCNNNAGATCGGAAGAG |
| 33555_2 | TATGAGTGTGGAGTCGTTGCCTCTCGGGTTCACACCATTCTCNNNAGATCGGAAGAG |
| 33555_3 | TATGAGTGTGGAGTCGTTGCCTCTCGGGTTCACACCATTCTCNNNAGATCGGAAGAG |

| | |
|---|---|
| 33555_4 | TATGAGTGTGGAGTCGTTGCCTCTCGGGTTCACACCATTCTCNNNAGATCGGAAGAG |
| 33555_5 | TATGAGTGTGGAGTCGTTGCCTCTCGGGTTCACACCATTCTCNNNAGATCGGAAGAG |
| 40952_1 | TATGAGTGTGGAGTCGTTGCGCAAGACCCTGTCTCCAAAAAAANNNAGATCGGAAG |
| 40952_2 | TATGAGTGTGGAGTCGTTGCGCAAGACCCTGTCTCCAAAAAAANNNAGATCGGAAG |
| 40952_3 | TATGAGTGTGGAGTCGTTGCGCAAGACCCTGTCTCCAAAAAAANNNAGATCGGAAG |
| 40952_4 | TATGAGTGTGGAGTCGTTGCGCAAGACCCTGTCTCCAAAAAAANNNAGATCGGAAG |
| 40952_5 | TATGAGTGTGGAGTCGTTGCGCAAGACCCTGTCTCCAAAAAAANNNAGATCGGAAG |
| 52876_1 | TATGAGTGTGGAGTCGTTGCCCAATGGCATCGAGTCCCTGNNNAGATCGGAAGAG |
| 52876_2 | TATGAGTGTGGAGTCGTTGCCCAATGGCATCGAGTCCCTGNNNAGATCGGAAGAG |
| 52876_3 | TATGAGTGTGGAGTCGTTGCCCAATGGCATCGAGTCCCTGNNNAGATCGGAAGAG |
| 52876_4 | TATGAGTGTGGAGTCGTTGCCCAATGGCATCGAGTCCCTGNNNAGATCGGAAGAG |
| 52876_5 | TATGAGTGTGGAGTCGTTGCCCAATGGCATCGAGTCCCTGNNNAGATCGGAAGAG |
| 29677_1 | TATGAGTGTGGAGTCGTTGCCTATTCCTTCACCATCAAGGCCANNNAGATCGGAAG |
| 29677_2 | TATGAGTGTGGAGTCGTTGCCTATTCCTTCACCATCAAGGCCANNNAGATCGGAAG |
| 29677_3 | TATGAGTGTGGAGTCGTTGCCTATTCCTTCACCATCAAGGCCANNNAGATCGGAAG |
| 29677_4 | TATGAGTGTGGAGTCGTTGCCTATTCCTTCACCATCAAGGCCANNNAGATCGGAAG |
| 29677_5 | TATGAGTGTGGAGTCGTTGCCTATTCCTTCACCATCAAGGCCANNNAGATCGGAAG |
| 45773_1 | TATGAGTGTGGAGTCGTTGCCTGGAGTGTAGTGGCATGATCTCNNNAGATCGGAA |
| 45773_2 | TATGAGTGTGGAGTCGTTGCCTGGAGTGTAGTGGCATGATCTCNNNAGATCGGAA |
| 45773_3 | TATGAGTGTGGAGTCGTTGCCTGGAGTGTAGTGGCATGATCTCNNNAGATCGGAA |
| 45773_4 | TATGAGTGTGGAGTCGTTGCCTGGAGTGTAGTGGCATGATCTCNNNAGATCGGAA |
| 45773_5 | TATGAGTGTGGAGTCGTTGCCTGGAGTGTAGTGGCATGATCTCNNNAGATCGGAA |
| 45116_1 | TATGAGTGTGGAGTCGTTGCAGGCAGTTAGACATTCTGGAAGTNNNAGATCGGAA |
| 45116_2 | TATGAGTGTGGAGTCGTTGCAGGCAGTTAGACATTCTGGAAGTNNNAGATCGGAA |
| 45116_3 | TATGAGTGTGGAGTCGTTGCAGGCAGTTAGACATTCTGGAAGTNNNAGATCGGAA |
| 45116_4 | TATGAGTGTGGAGTCGTTGCAGGCAGTTAGACATTCTGGAAGTNNNAGATCGGAA |
| 45116_5 | TATGAGTGTGGAGTCGTTGCAGGCAGTTAGACATTCTGGAAGTNNNAGATCGGAA |
| 42821_1 | TATGAGTGTGGAGTCGTTGCATGAAACAAGGAGTTTGCATGGGNNNAGATCGGAA |
| 42821_2 | TATGAGTGTGGAGTCGTTGCATGAAACAAGGAGTTTGCATGGGNNNAGATCGGAA |
| 42821_3 | TATGAGTGTGGAGTCGTTGCATGAAACAAGGAGTTTGCATGGGNNNAGATCGGAA |
| 42821_4 | TATGAGTGTGGAGTCGTTGCATGAAACAAGGAGTTTGCATGGGNNNAGATCGGAA |
| 42821_5 | TATGAGTGTGGAGTCGTTGCATGAAACAAGGAGTTTGCATGGGNNNAGATCGGAA |
| 34929_1 | TATGAGTGTGGAGTCGTTGCTTCAAGTGATTCTCCTGCCTCAGNNNAGATCGGAAG |
| 34929_2 | TATGAGTGTGGAGTCGTTGCTTCAAGTGATTCTCCTGCCTCAGNNNAGATCGGAAG |
| 34929_3 | TATGAGTGTGGAGTCGTTGCTTCAAGTGATTCTCCTGCCTCAGNNNAGATCGGAAG |
| 34929_4 | TATGAGTGTGGAGTCGTTGCTTCAAGTGATTCTCCTGCCTCAGNNNAGATCGGAAG |
| 34929_5 | TATGAGTGTGGAGTCGTTGCTTCAAGTGATTCTCCTGCCTCAGNNNAGATCGGAAG |
| 42611_1 | TATGAGTGTGGAGTCGTTGCCTTCTGAGTAGCTGGGACTACAGNNNAGATCGGAA |
| 42611_2 | TATGAGTGTGGAGTCGTTGCCTTCTGAGTAGCTGGGACTACAGNNNAGATCGGAA |
| 42611_3 | TATGAGTGTGGAGTCGTTGCCTTCTGAGTAGCTGGGACTACAGNNNAGATCGGAA |
| 42611_4 | TATGAGTGTGGAGTCGTTGCCTTCTGAGTAGCTGGGACTACAGNNNAGATCGGAA |
| 42611_5 | TATGAGTGTGGAGTCGTTGCCTTCTGAGTAGCTGGGACTACAGNNNAGATCGGAA |
| 29576_1 | TATGAGTGTGGAGTCGTTGCTCGTCATCGTGGAAGACATAGACNNNAGATCGGAA |

| | |
|---|---|
| 29576_2 | TATGAGTGTGGAGTCGTTGCTCGTCATCGTGGAAGACATAGACNNNAGATCGGAA |
| 29576_3 | TATGAGTGTGGAGTCGTTGCTCGTCATCGTGGAAGACATAGACNNNAGATCGGAA |
| 29576_4 | TATGAGTGTGGAGTCGTTGCTCGTCATCGTGGAAGACATAGACNNNAGATCGGAA |
| 29576_5 | TATGAGTGTGGAGTCGTTGCTCGTCATCGTGGAAGACATAGACNNNAGATCGGAA |
| 50680_1 | TATGAGTGTGGAGTCGTTGCCACAGACATGTCCACAAAGCATTNNNAGATCGGAAG |
| 50680_2 | TATGAGTGTGGAGTCGTTGCCACAGACATGTCCACAAAGCATTNNNAGATCGGAAG |
| 50680_3 | TATGAGTGTGGAGTCGTTGCCACAGACATGTCCACAAAGCATTNNNAGATCGGAAG |
| 50680_4 | TATGAGTGTGGAGTCGTTGCCACAGACATGTCCACAAAGCATTNNNAGATCGGAAG |
| 50680_5 | TATGAGTGTGGAGTCGTTGCCACAGACATGTCCACAAAGCATTNNNAGATCGGAAG |
| 31556_1 | TATGAGTGTGGAGTCGTTGCTGGCAGAGACAAGTGTACATCAANNNAGATCGGAA |
| 31556_2 | TATGAGTGTGGAGTCGTTGCTGGCAGAGACAAGTGTACATCAANNNAGATCGGAA |
| 31556_3 | TATGAGTGTGGAGTCGTTGCTGGCAGAGACAAGTGTACATCAANNNAGATCGGAA |
| 31556_4 | TATGAGTGTGGAGTCGTTGCTGGCAGAGACAAGTGTACATCAANNNAGATCGGAA |
| 31556_5 | TATGAGTGTGGAGTCGTTGCTGGCAGAGACAAGTGTACATCAANNNAGATCGGAA |
| 7849_1 | TATGAGTGTGGAGTCGTTGCTAGTGAGAGGAAGTTTCTAAAAAGTTTNNNAGATC |
| 7849_2 | TATGAGTGTGGAGTCGTTGCTAGTGAGAGGAAGTTTCTAAAAAGTTTNNNAGATC |
| 7849_3 | TATGAGTGTGGAGTCGTTGCTAGTGAGAGGAAGTTTCTAAAAAGTTTNNNAGATC |
| 7849_4 | TATGAGTGTGGAGTCGTTGCTAGTGAGAGGAAGTTTCTAAAAAGTTTNNNAGATC |
| 7849_5 | TATGAGTGTGGAGTCGTTGCTAGTGAGAGGAAGTTTCTAAAAAGTTTNNNAGATC |
| 35123_1 | TATGAGTGTGGAGTCGTTGCCCTCCCACCATGCTAAGATTACANNNAGATCGGAAG |
| 35123_2 | TATGAGTGTGGAGTCGTTGCCCTCCCACCATGCTAAGATTACANNNAGATCGGAAG |
| 35123_3 | TATGAGTGTGGAGTCGTTGCCCTCCCACCATGCTAAGATTACANNNAGATCGGAAG |
| 35123_4 | TATGAGTGTGGAGTCGTTGCCCTCCCACCATGCTAAGATTACANNNAGATCGGAAG |
| 35123_5 | TATGAGTGTGGAGTCGTTGCCCTCCCACCATGCTAAGATTACANNNAGATCGGAAG |
| 49032_1 | TATGAGTGTGGAGTCGTTGCAGTAGCTGGGACTACAGGTATGTNNNAGATCGGAA |
| 49032_2 | TATGAGTGTGGAGTCGTTGCAGTAGCTGGGACTACAGGTATGTNNNAGATCGGAA |
| 49032_3 | TATGAGTGTGGAGTCGTTGCAGTAGCTGGGACTACAGGTATGTNNNAGATCGGAA |
| 49032_4 | TATGAGTGTGGAGTCGTTGCAGTAGCTGGGACTACAGGTATGTNNNAGATCGGAA |
| 49032_5 | TATGAGTGTGGAGTCGTTGCAGTAGCTGGGACTACAGGTATGTNNNAGATCGGAA |
| 37820_1 | TATGAGTGTGGAGTCGTTGCATGAGTGGTTGATAGAGGTTCGGNNNAGATCGGA |
| 37820_2 | TATGAGTGTGGAGTCGTTGCATGAGTGGTTGATAGAGGTTCGGNNNAGATCGGA |
| 37820_3 | TATGAGTGTGGAGTCGTTGCATGAGTGGTTGATAGAGGTTCGGNNNAGATCGGA |
| 37820_4 | TATGAGTGTGGAGTCGTTGCATGAGTGGTTGATAGAGGTTCGGNNNAGATCGGA |
| 37820_5 | TATGAGTGTGGAGTCGTTGCATGAGTGGTTGATAGAGGTTCGGNNNAGATCGGA |
| 12608_1 | TATGAGTGTGGAGTCGTTGCCCAGGAATAGTACCATATTCATTNNNAGATCGGAAG |
| 12608_2 | TATGAGTGTGGAGTCGTTGCCCAGGAATAGTACCATATTCATTNNNAGATCGGAAG |
| 12608_3 | TATGAGTGTGGAGTCGTTGCCCAGGAATAGTACCATATTCATTNNNAGATCGGAAG |
| 12608_4 | TATGAGTGTGGAGTCGTTGCCCAGGAATAGTACCATATTCATTNNNAGATCGGAAG |
| 12608_5 | TATGAGTGTGGAGTCGTTGCCCAGGAATAGTACCATATTCATTNNNAGATCGGAAG |
| 30085_1 | TATGAGTGTGGAGTCGTTGCTGTTTAGTCAGTGTTGGGGGAATNNNAGATCGGAA |
| 30085_2 | TATGAGTGTGGAGTCGTTGCTGTTTAGTCAGTGTTGGGGGAATNNNAGATCGGAA |
| 30085_3 | TATGAGTGTGGAGTCGTTGCTGTTTAGTCAGTGTTGGGGGAATNNNAGATCGGAA |
| 30085_4 | TATGAGTGTGGAGTCGTTGCTGTTTAGTCAGTGTTGGGGGAATNNNAGATCGGAA |

| | |
|---|---|
| 30085_5 | TATGAGTGTGGAGTCGTTGCTGTTTAGTCAGTGTTGGGGGAATNNNAGATCGGAA |
| 34404_1 | TATGAGTGTGGAGTCGTTGCCCTACTGGGAAGTGAGGAGCCNNNAGATCGGAAGA |
| 34404_2 | TATGAGTGTGGAGTCGTTGCCCTACTGGGAAGTGAGGAGCCNNNAGATCGGAAGA |
| 34404_3 | TATGAGTGTGGAGTCGTTGCCCTACTGGGAAGTGAGGAGCCNNNAGATCGGAAGA |
| 34404_4 | TATGAGTGTGGAGTCGTTGCCCTACTGGGAAGTGAGGAGCCNNNAGATCGGAAGA |
| 34404_5 | TATGAGTGTGGAGTCGTTGCCCTACTGGGAAGTGAGGAGCCNNNAGATCGGAAGA |
| 45098_1 | TATGAGTGTGGAGTCGTTGCTAAGTAACTTCCCGCTTCCTCTGNNNAGATCGGAAGA |
| 45098_2 | TATGAGTGTGGAGTCGTTGCTAAGTAACTTCCCGCTTCCTCTGNNNAGATCGGAAGA |
| 45098_3 | TATGAGTGTGGAGTCGTTGCTAAGTAACTTCCCGCTTCCTCTGNNNAGATCGGAAGA |
| 45098_4 | TATGAGTGTGGAGTCGTTGCTAAGTAACTTCCCGCTTCCTCTGNNNAGATCGGAAGA |
| 45098_5 | TATGAGTGTGGAGTCGTTGCTAAGTAACTTCCCGCTTCCTCTGNNNAGATCGGAAGA |
| 43195_1 | TATGAGTGTGGAGTCGTTGCCAGTCACACTAGCCATCACTCATNNNAGATCGGAAGA |
| 43195_2 | TATGAGTGTGGAGTCGTTGCCAGTCACACTAGCCATCACTCATNNNAGATCGGAAGA |
| 43195_3 | TATGAGTGTGGAGTCGTTGCCAGTCACACTAGCCATCACTCATNNNAGATCGGAAGA |
| 43195_4 | TATGAGTGTGGAGTCGTTGCCAGTCACACTAGCCATCACTCATNNNAGATCGGAAGA |
| 43195_5 | TATGAGTGTGGAGTCGTTGCCAGTCACACTAGCCATCACTCATNNNAGATCGGAAGA |
| 47394_1 | TATGAGTGTGGAGTCGTTGCGTACTTCCTCAGCTTCTGTCTGGNNNAGATCGGAAGA |
| 47394_2 | TATGAGTGTGGAGTCGTTGCGTACTTCCTCAGCTTCTGTCTGGNNNAGATCGGAAGA |
| 47394_3 | TATGAGTGTGGAGTCGTTGCGTACTTCCTCAGCTTCTGTCTGGNNNAGATCGGAAGA |
| 47394_4 | TATGAGTGTGGAGTCGTTGCGTACTTCCTCAGCTTCTGTCTGGNNNAGATCGGAAGA |
| 47394_5 | TATGAGTGTGGAGTCGTTGCGTACTTCCTCAGCTTCTGTCTGGNNNAGATCGGAAGA |
| 33610_1 | TATGAGTGTGGAGTCGTTGCTCTGGGTTCAAGTGATTCTCCTGNNNAGATCGGAAG |
| 33610_2 | TATGAGTGTGGAGTCGTTGCTCTGGGTTCAAGTGATTCTCCTGNNNAGATCGGAAG |
| 33610_3 | TATGAGTGTGGAGTCGTTGCTCTGGGTTCAAGTGATTCTCCTGNNNAGATCGGAAG |
| 33610_4 | TATGAGTGTGGAGTCGTTGCTCTGGGTTCAAGTGATTCTCCTGNNNAGATCGGAAG |
| 33610_5 | TATGAGTGTGGAGTCGTTGCTCTGGGTTCAAGTGATTCTCCTGNNNAGATCGGAAG |
| 50725_1 | TATGAGTGTGGAGTCGTTGCAAACTATCCCAGTGGCAGTGAATNNNAGATCGGAAG |
| 50725_2 | TATGAGTGTGGAGTCGTTGCAAACTATCCCAGTGGCAGTGAATNNNAGATCGGAAG |
| 50725_3 | TATGAGTGTGGAGTCGTTGCAAACTATCCCAGTGGCAGTGAATNNNAGATCGGAAG |
| 50725_4 | TATGAGTGTGGAGTCGTTGCAAACTATCCCAGTGGCAGTGAATNNNAGATCGGAAG |
| 50725_5 | TATGAGTGTGGAGTCGTTGCAAACTATCCCAGTGGCAGTGAATNNNAGATCGGAAG |
| 10479_1 | TATGAGTGTGGAGTCGTTGCGCAATCGTGTTCAACTTGACAACANNNAGATCGGAA |
| 10479_2 | TATGAGTGTGGAGTCGTTGCGCAATCGTGTTCAACTTGACAACANNNAGATCGGAA |
| 10479_3 | TATGAGTGTGGAGTCGTTGCGCAATCGTGTTCAACTTGACAACANNNAGATCGGAA |
| 10479_4 | TATGAGTGTGGAGTCGTTGCGCAATCGTGTTCAACTTGACAACANNNAGATCGGAA |
| 10479_5 | TATGAGTGTGGAGTCGTTGCGCAATCGTGTTCAACTTGACAACANNNAGATCGGAA |
| 52673_1 | TATGAGTGTGGAGTCGTTGCGGCTCTACCTGGATCCATACAAGNNNAGATCGGAAG |
| 52673_2 | TATGAGTGTGGAGTCGTTGCGGCTCTACCTGGATCCATACAAGNNNAGATCGGAAG |
| 52673_3 | TATGAGTGTGGAGTCGTTGCGGCTCTACCTGGATCCATACAAGNNNAGATCGGAAG |
| 52673_4 | TATGAGTGTGGAGTCGTTGCGGCTCTACCTGGATCCATACAAGNNNAGATCGGAAG |
| 52673_5 | TATGAGTGTGGAGTCGTTGCGGCTCTACCTGGATCCATACAAGNNNAGATCGGAAG |
| 29750_1 | TATGAGTGTGGAGTCGTTGCCTTGAGAGATAGGGGCCTCTTTGNNNAGATCGGAA |
| 29750_2 | TATGAGTGTGGAGTCGTTGCCTTGAGAGATAGGGGCCTCTTTGNNNAGATCGGAA |

| | |
|---|---|
| 29750_3 | TATGAGTGTGGAGTCGTTGCCTTGAGAGATAGGGGCCTCTTTGNNNAGATCGGAA |
| 29750_4 | TATGAGTGTGGAGTCGTTGCCTTGAGAGATAGGGGCCTCTTTGNNNAGATCGGAA |
| 29750_5 | TATGAGTGTGGAGTCGTTGCCTTGAGAGATAGGGGCCTCTTTGNNNAGATCGGAA |
| 39829_1 | TATGAGTGTGGAGTCGTTGCGTCAGGGATTTGGGGGAAAGATNNNAGATCGGAA |
| 39829_2 | TATGAGTGTGGAGTCGTTGCGTCAGGGATTTGGGGGAAAGATNNNAGATCGGAA |
| 39829_3 | TATGAGTGTGGAGTCGTTGCGTCAGGGATTTGGGGGAAAGATNNNAGATCGGAA |
| 39829_4 | TATGAGTGTGGAGTCGTTGCGTCAGGGATTTGGGGGAAAGATNNNAGATCGGAA |
| 39829_5 | TATGAGTGTGGAGTCGTTGCGTCAGGGATTTGGGGGAAAGATNNNAGATCGGAA |
| 31678_1 | TATGAGTGTGGAGTCGTTGCTTCACGGAAAATAAAAGTGGCCGNNNAGATCGGAA |
| 31678_2 | TATGAGTGTGGAGTCGTTGCTTCACGGAAAATAAAAGTGGCCGNNNAGATCGGAA |
| 31678_3 | TATGAGTGTGGAGTCGTTGCTTCACGGAAAATAAAAGTGGCCGNNNAGATCGGAA |
| 31678_4 | TATGAGTGTGGAGTCGTTGCTTCACGGAAAATAAAAGTGGCCGNNNAGATCGGAA |
| 31678_5 | TATGAGTGTGGAGTCGTTGCTTCACGGAAAATAAAAGTGGCCGNNNAGATCGGAA |
| 28871_1 | TATGAGTGTGGAGTCGTTGCGGAGGCAGAGCATGATAAAGAGANNNAGATCGGA |
| 28871_2 | TATGAGTGTGGAGTCGTTGCGGAGGCAGAGCATGATAAAGAGANNNAGATCGGA |
| 28871_3 | TATGAGTGTGGAGTCGTTGCGGAGGCAGAGCATGATAAAGAGANNNAGATCGGA |
| 28871_4 | TATGAGTGTGGAGTCGTTGCGGAGGCAGAGCATGATAAAGAGANNNAGATCGGA |
| 28871_5 | TATGAGTGTGGAGTCGTTGCGGAGGCAGAGCATGATAAAGAGANNNAGATCGGA |
| 47107_1 | TATGAGTGTGGAGTCGTTGCTCAATTGCTGGCCATTTGACTTTNNNAGATCGGAAG |
| 47107_2 | TATGAGTGTGGAGTCGTTGCTCAATTGCTGGCCATTTGACTTTNNNAGATCGGAAG |
| 47107_3 | TATGAGTGTGGAGTCGTTGCTCAATTGCTGGCCATTTGACTTTNNNAGATCGGAAG |
| 47107_4 | TATGAGTGTGGAGTCGTTGCTCAATTGCTGGCCATTTGACTTTNNNAGATCGGAAG |
| 47107_5 | TATGAGTGTGGAGTCGTTGCTCAATTGCTGGCCATTTGACTTTNNNAGATCGGAAG |
| 10673_1 | TATGAGTGTGGAGTCGTTGCATGACAACAAAAACCGTGCCATCNNNAGATCGGAAG |
| 10673_2 | TATGAGTGTGGAGTCGTTGCATGACAACAAAAACCGTGCCATCNNNAGATCGGAAG |
| 10673_3 | TATGAGTGTGGAGTCGTTGCATGACAACAAAAACCGTGCCATCNNNAGATCGGAAG |
| 10673_4 | TATGAGTGTGGAGTCGTTGCATGACAACAAAAACCGTGCCATCNNNAGATCGGAAG |
| 10673_5 | TATGAGTGTGGAGTCGTTGCATGACAACAAAAACCGTGCCATCNNNAGATCGGAAG |
| 51727_1 | TATGAGTGTGGAGTCGTTGCAGCAGCTAAGAAGTCTTGAAGCTNNNAGATCGGAA |
| 51727_2 | TATGAGTGTGGAGTCGTTGCAGCAGCTAAGAAGTCTTGAAGCTNNNAGATCGGAA |
| 51727_3 | TATGAGTGTGGAGTCGTTGCAGCAGCTAAGAAGTCTTGAAGCTNNNAGATCGGAA |
| 51727_4 | TATGAGTGTGGAGTCGTTGCAGCAGCTAAGAAGTCTTGAAGCTNNNAGATCGGAA |
| 51727_5 | TATGAGTGTGGAGTCGTTGCAGCAGCTAAGAAGTCTTGAAGCTNNNAGATCGGAA |
| 3577_1 | TATGAGTGTGGAGTCGTTGCCTGGGTTCAAGAATTTCTCCAGTNNNAGATCGGAAG |
| 3577_2 | TATGAGTGTGGAGTCGTTGCCTGGGTTCAAGAATTTCTCCAGTNNNAGATCGGAAG |
| 3577_3 | TATGAGTGTGGAGTCGTTGCCTGGGTTCAAGAATTTCTCCAGTNNNAGATCGGAAG |
| 3577_4 | TATGAGTGTGGAGTCGTTGCCTGGGTTCAAGAATTTCTCCAGTNNNAGATCGGAAG |
| 3577_5 | TATGAGTGTGGAGTCGTTGCCTGGGTTCAAGAATTTCTCCAGTNNNAGATCGGAAG |
| 33256_1 | TATGAGTGTGGAGTCGTTGCTTCTCAAGTGCCTACTCTCACACNNNAGATCGGAAGA |
| 33256_2 | TATGAGTGTGGAGTCGTTGCTTCTCAAGTGCCTACTCTCACACNNNAGATCGGAAGA |
| 33256_3 | TATGAGTGTGGAGTCGTTGCTTCTCAAGTGCCTACTCTCACACNNNAGATCGGAAGA |
| 33256_4 | TATGAGTGTGGAGTCGTTGCTTCTCAAGTGCCTACTCTCACACNNNAGATCGGAAGA |
| 33256_5 | TATGAGTGTGGAGTCGTTGCTTCTCAAGTGCCTACTCTCACACNNNAGATCGGAAGA |

| | |
|---|---|
| 51920_1 | TATGAGTGTGGAGTCGTTGCGTGGTGTGTAGAAAAGGCCTTTGNNNAGATCGGAA |
| 51920_2 | TATGAGTGTGGAGTCGTTGCGTGGTGTGTAGAAAAGGCCTTTGNNNAGATCGGAA |
| 51920_3 | TATGAGTGTGGAGTCGTTGCGTGGTGTGTAGAAAAGGCCTTTGNNNAGATCGGAA |
| 51920_4 | TATGAGTGTGGAGTCGTTGCGTGGTGTGTAGAAAAGGCCTTTGNNNAGATCGGAA |
| 51920_5 | TATGAGTGTGGAGTCGTTGCGTGGTGTGTAGAAAAGGCCTTTGNNNAGATCGGAA |
| 46464_1 | TATGAGTGTGGAGTCGTTGCTTTCACCATACTAGCCAGGATGGNNNAGATCGGAAG |
| 46464_2 | TATGAGTGTGGAGTCGTTGCTTTCACCATACTAGCCAGGATGGNNNAGATCGGAAG |
| 46464_3 | TATGAGTGTGGAGTCGTTGCTTTCACCATACTAGCCAGGATGGNNNAGATCGGAAG |
| 46464_4 | TATGAGTGTGGAGTCGTTGCTTTCACCATACTAGCCAGGATGGNNNAGATCGGAAG |
| 46464_5 | TATGAGTGTGGAGTCGTTGCTTTCACCATACTAGCCAGGATGGNNNAGATCGGAAG |
| 40568_1 | TATGAGTGTGGAGTCGTTGCGGGGTAGGAAACACAGTCTTGAANNNAGATCGGAA |
| 40568_2 | TATGAGTGTGGAGTCGTTGCGGGGTAGGAAACACAGTCTTGAANNNAGATCGGAA |
| 40568_3 | TATGAGTGTGGAGTCGTTGCGGGGTAGGAAACACAGTCTTGAANNNAGATCGGAA |
| 40568_4 | TATGAGTGTGGAGTCGTTGCGGGGTAGGAAACACAGTCTTGAANNNAGATCGGAA |
| 40568_5 | TATGAGTGTGGAGTCGTTGCGGGGTAGGAAACACAGTCTTGAANNNAGATCGGAA |
| 33504_1 | TATGAGTGTGGAGTCGTTGCCAGTCCTGACTTAGCATGGAAGTNNNAGATCGGAAG |
| 33504_2 | TATGAGTGTGGAGTCGTTGCCAGTCCTGACTTAGCATGGAAGTNNNAGATCGGAAG |
| 33504_3 | TATGAGTGTGGAGTCGTTGCCAGTCCTGACTTAGCATGGAAGTNNNAGATCGGAAG |
| 33504_4 | TATGAGTGTGGAGTCGTTGCCAGTCCTGACTTAGCATGGAAGTNNNAGATCGGAAG |
| 33504_5 | TATGAGTGTGGAGTCGTTGCCAGTCCTGACTTAGCATGGAAGTNNNAGATCGGAAG |
| 35833_1 | TATGAGTGTGGAGTCGTTGCCACTTTGCCCAGCCTACATTTTTNNNAGATCGGAAGA |
| 35833_2 | TATGAGTGTGGAGTCGTTGCCACTTTGCCCAGCCTACATTTTTNNNAGATCGGAAGA |
| 35833_3 | TATGAGTGTGGAGTCGTTGCCACTTTGCCCAGCCTACATTTTTNNNAGATCGGAAGA |
| 35833_4 | TATGAGTGTGGAGTCGTTGCCACTTTGCCCAGCCTACATTTTTNNNAGATCGGAAGA |
| 35833_5 | TATGAGTGTGGAGTCGTTGCCACTTTGCCCAGCCTACATTTTTNNNAGATCGGAAGA |
| 30073_1 | TATGAGTGTGGAGTCGTTGCCACCGTATTACTTTGCTGCAGAANNNAGATCGGAAGA |
| 30073_2 | TATGAGTGTGGAGTCGTTGCCACCGTATTACTTTGCTGCAGAANNNAGATCGGAAGA |
| 30073_3 | TATGAGTGTGGAGTCGTTGCCACCGTATTACTTTGCTGCAGAANNNAGATCGGAAGA |
| 30073_4 | TATGAGTGTGGAGTCGTTGCCACCGTATTACTTTGCTGCAGAANNNAGATCGGAAGA |
| 30073_5 | TATGAGTGTGGAGTCGTTGCCACCGTATTACTTTGCTGCAGAANNNAGATCGGAAGA |
| 2367_1 | TATGAGTGTGGAGTCGTTGCGCCTCAGAGAAGTGTAAGACAGCNNNAGATCGGAA |
| 2367_2 | TATGAGTGTGGAGTCGTTGCGCCTCAGAGAAGTGTAAGACAGCNNNAGATCGGAA |
| 2367_3 | TATGAGTGTGGAGTCGTTGCGCCTCAGAGAAGTGTAAGACAGCNNNAGATCGGAA |
| 2367_4 | TATGAGTGTGGAGTCGTTGCGCCTCAGAGAAGTGTAAGACAGCNNNAGATCGGAA |
| 2367_5 | TATGAGTGTGGAGTCGTTGCGCCTCAGAGAAGTGTAAGACAGCNNNAGATCGGAA |
| 53374_1 | TATGAGTGTGGAGTCGTTGCAAGCAATTCTCCTGCCTCAAGTANNNAGATCGGAAGA |
| 53374_2 | TATGAGTGTGGAGTCGTTGCAAGCAATTCTCCTGCCTCAAGTANNNAGATCGGAAGA |
| 53374_3 | TATGAGTGTGGAGTCGTTGCAAGCAATTCTCCTGCCTCAAGTANNNAGATCGGAAGA |
| 53374_4 | TATGAGTGTGGAGTCGTTGCAAGCAATTCTCCTGCCTCAAGTANNNAGATCGGAAGA |
| 53374_5 | TATGAGTGTGGAGTCGTTGCAAGCAATTCTCCTGCCTCAAGTANNNAGATCGGAAGA |
| 29780_1 | TATGAGTGTGGAGTCGTTGCTGCACATTTTCAAACTGTCTGCTNNNAGATCGGAAGA |
| 29780_2 | TATGAGTGTGGAGTCGTTGCTGCACATTTTCAAACTGTCTGCTNNNAGATCGGAAGA |
| 29780_3 | TATGAGTGTGGAGTCGTTGCTGCACATTTTCAAACTGTCTGCTNNNAGATCGGAAGA |

| | |
|---|---|
| 29780_4 | TATGAGTGTGGAGTCGTTGCTGCACATTTTCAAACTGTCTGCTNNNAGATCGGAAGA |
| 29780_5 | TATGAGTGTGGAGTCGTTGCTGCACATTTTCAAACTGTCTGCTNNNAGATCGGAAGA |
| 13280_1 | TATGAGTGTGGAGTCGTTGCCCGTCTCTACTAAAAATACAAAANNNAGATCGGAAGA |
| 13280_2 | TATGAGTGTGGAGTCGTTGCCCGTCTCTACTAAAAATACAAAANNNAGATCGGAAGA |
| 13280_3 | TATGAGTGTGGAGTCGTTGCCCGTCTCTACTAAAAATACAAAANNNAGATCGGAAGA |
| 13280_4 | TATGAGTGTGGAGTCGTTGCCCGTCTCTACTAAAAATACAAAANNNAGATCGGAAGA |
| 13280_5 | TATGAGTGTGGAGTCGTTGCCCGTCTCTACTAAAAATACAAAANNNAGATCGGAAGA |
| 49146_1 | TATGAGTGTGGAGTCGTTGCTTCAAGTGATTCTCCTGCCTCAGNNNAGATCGGAAGA |
| 49146_2 | TATGAGTGTGGAGTCGTTGCTTCAAGTGATTCTCCTGCCTCAGNNNAGATCGGAAGA |
| 49146_3 | TATGAGTGTGGAGTCGTTGCTTCAAGTGATTCTCCTGCCTCAGNNNAGATCGGAAGA |
| 49146_4 | TATGAGTGTGGAGTCGTTGCTTCAAGTGATTCTCCTGCCTCAGNNNAGATCGGAAGA |
| 49146_5 | TATGAGTGTGGAGTCGTTGCTTCAAGTGATTCTCCTGCCTCAGNNNAGATCGGAAGA |
| 47006_1 | TATGAGTGTGGAGTCGTTGCTTCAAGTGATTCTCCTGCCTCAGNNNAGATCGGAAGA |
| 47006_2 | TATGAGTGTGGAGTCGTTGCTTCAAGTGATTCTCCTGCCTCAGNNNAGATCGGAAGA |
| 47006_3 | TATGAGTGTGGAGTCGTTGCTTCAAGTGATTCTCCTGCCTCAGNNNAGATCGGAAGA |
| 47006_4 | TATGAGTGTGGAGTCGTTGCTTCAAGTGATTCTCCTGCCTCAGNNNAGATCGGAAGA |
| 47006_5 | TATGAGTGTGGAGTCGTTGCTTCAAGTGATTCTCCTGCCTCAGNNNAGATCGGAAGA |
| 32415_1 | TATGAGTGTGGAGTCGTTGCTTCAAGGCCCTTTACAGTCTAGCNNNAGATCGGAAGA |
| 32415_2 | TATGAGTGTGGAGTCGTTGCTTCAAGGCCCTTTACAGTCTAGCNNNAGATCGGAAGA |
| 32415_3 | TATGAGTGTGGAGTCGTTGCTTCAAGGCCCTTTACAGTCTAGCNNNAGATCGGAAGA |
| 32415_4 | TATGAGTGTGGAGTCGTTGCTTCAAGGCCCTTTACAGTCTAGCNNNAGATCGGAAGA |
| 32415_5 | TATGAGTGTGGAGTCGTTGCTTCAAGGCCCTTTACAGTCTAGCNNNAGATCGGAAGA |
| 40556_1 | TATGAGTGTGGAGTCGTTGCTCTCCTTCCTTTTATCTGCACCCNNNAGATCGGAAGAG |
| 40556_2 | TATGAGTGTGGAGTCGTTGCTCTCCTTCCTTTTATCTGCACCCNNNAGATCGGAAGAG |
| 40556_3 | TATGAGTGTGGAGTCGTTGCTCTCCTTCCTTTTATCTGCACCCNNNAGATCGGAAGAG |
| 40556_4 | TATGAGTGTGGAGTCGTTGCTCTCCTTCCTTTTATCTGCACCCNNNAGATCGGAAGAG |
| 40556_5 | TATGAGTGTGGAGTCGTTGCTCTCCTTCCTTTTATCTGCACCCNNNAGATCGGAAGAG |
| 13791_1 | TATGAGTGTGGAGTCGTTGCTCCCCTCTCCTCTAATTAAAANNNAGATCGGAAGAGC |
| 13791_2 | TATGAGTGTGGAGTCGTTGCTCCCCTCTCCTCTAATTAAAANNNAGATCGGAAGAGC |
| 13791_3 | TATGAGTGTGGAGTCGTTGCTCCCCTCTCCTCTAATTAAAANNNAGATCGGAAGAGC |
| 13791_4 | TATGAGTGTGGAGTCGTTGCTCCCCTCTCCTCTAATTAAAANNNAGATCGGAAGAGC |
| 13791_5 | TATGAGTGTGGAGTCGTTGCTCCCCTCTCCTCTAATTAAAANNNAGATCGGAAGAGC |
| 42227_1 | TATGAGTGTGGAGTCGTTGCAGTTGGAGTCTCACTGTGTCATCNNNAGATCGGAAG |
| 42227_2 | TATGAGTGTGGAGTCGTTGCAGTTGGAGTCTCACTGTGTCATCNNNAGATCGGAAG |
| 42227_3 | TATGAGTGTGGAGTCGTTGCAGTTGGAGTCTCACTGTGTCATCNNNAGATCGGAAG |
| 42227_4 | TATGAGTGTGGAGTCGTTGCAGTTGGAGTCTCACTGTGTCATCNNNAGATCGGAAG |
| 42227_5 | TATGAGTGTGGAGTCGTTGCAGTTGGAGTCTCACTGTGTCATCNNNAGATCGGAAG |
| 52196_1 | TATGAGTGTGGAGTCGTTGCGCAGTCCATTCTAACCCAGATGANNNAGATCGGAAG |
| 52196_2 | TATGAGTGTGGAGTCGTTGCGCAGTCCATTCTAACCCAGATGANNNAGATCGGAAG |
| 52196_3 | TATGAGTGTGGAGTCGTTGCGCAGTCCATTCTAACCCAGATGANNNAGATCGGAAG |
| 52196_4 | TATGAGTGTGGAGTCGTTGCGCAGTCCATTCTAACCCAGATGANNNAGATCGGAAG |
| 52196_5 | TATGAGTGTGGAGTCGTTGCGCAGTCCATTCTAACCCAGATGANNNAGATCGGAAG |
| 41677_1 | TATGAGTGTGGAGTCGTTGCCACGCCTGGCTACTTTTTGTATTNNNAGATCGGAAGA |

| | |
|---|---|
| 41677_2 | TATGAGTGTGGAGTCGTTGCCACGCCTGGCTACTTTTTGTATTNNNAGATCGGAAGA |
| 41677_3 | TATGAGTGTGGAGTCGTTGCCACGCCTGGCTACTTTTTGTATTNNNAGATCGGAAGA |
| 41677_4 | TATGAGTGTGGAGTCGTTGCCACGCCTGGCTACTTTTTGTATTNNNAGATCGGAAGA |
| 41677_5 | TATGAGTGTGGAGTCGTTGCCACGCCTGGCTACTTTTTGTATTNNNAGATCGGAAGA |
| 46042_1 | TATGAGTGTGGAGTCGTTGCGTGACATTTCAAATTGGGCCTGANNNAGATCGGAAG |
| 46042_2 | TATGAGTGTGGAGTCGTTGCGTGACATTTCAAATTGGGCCTGANNNAGATCGGAAG |
| 46042_3 | TATGAGTGTGGAGTCGTTGCGTGACATTTCAAATTGGGCCTGANNNAGATCGGAAG |
| 46042_4 | TATGAGTGTGGAGTCGTTGCGTGACATTTCAAATTGGGCCTGANNNAGATCGGAAG |
| 46042_5 | TATGAGTGTGGAGTCGTTGCGTGACATTTCAAATTGGGCCTGANNNAGATCGGAAG |
| 12716_1 | TATGAGTGTGGAGTCGTTGCGAGATTTCAGGTCATTATCAATCNNNAGATCGGAAG |
| 12716_2 | TATGAGTGTGGAGTCGTTGCGAGATTTCAGGTCATTATCAATCNNNAGATCGGAAG |
| 12716_3 | TATGAGTGTGGAGTCGTTGCGAGATTTCAGGTCATTATCAATCNNNAGATCGGAAG |
| 12716_4 | TATGAGTGTGGAGTCGTTGCGAGATTTCAGGTCATTATCAATCNNNAGATCGGAAG |
| 12716_5 | TATGAGTGTGGAGTCGTTGCGAGATTTCAGGTCATTATCAATCNNNAGATCGGAAG |
| 43880_1 | TATGAGTGTGGAGTCGTTGCGTGTCTTTCTGTCCTCTCTCCAGNNNAGATCGGAAGA |
| 43880_2 | TATGAGTGTGGAGTCGTTGCGTGTCTTTCTGTCCTCTCTCCAGNNNAGATCGGAAGA |
| 43880_3 | TATGAGTGTGGAGTCGTTGCGTGTCTTTCTGTCCTCTCTCCAGNNNAGATCGGAAGA |
| 43880_4 | TATGAGTGTGGAGTCGTTGCGTGTCTTTCTGTCCTCTCTCCAGNNNAGATCGGAAGA |
| 43880_5 | TATGAGTGTGGAGTCGTTGCGTGTCTTTCTGTCCTCTCTCCAGNNNAGATCGGAAGA |
| 8501_1 | TATGAGTGTGGAGTCGTTGCATGTACTCAAAGTGGAATTGATGGATNNNAGATCGG |
| 8501_2 | TATGAGTGTGGAGTCGTTGCATGTACTCAAAGTGGAATTGATGGATNNNAGATCGG |
| 8501_3 | TATGAGTGTGGAGTCGTTGCATGTACTCAAAGTGGAATTGATGGATNNNAGATCGG |
| 8501_4 | TATGAGTGTGGAGTCGTTGCATGTACTCAAAGTGGAATTGATGGATNNNAGATCGG |
| 8501_5 | TATGAGTGTGGAGTCGTTGCATGTACTCAAAGTGGAATTGATGGATNNNAGATCGG |
| 34076_1 | TATGAGTGTGGAGTCGTTGCCAATCAGCCTATCTCTGCATCCCNNNAGATCGGAAGA |
| 34076_2 | TATGAGTGTGGAGTCGTTGCCAATCAGCCTATCTCTGCATCCCNNNAGATCGGAAGA |
| 34076_3 | TATGAGTGTGGAGTCGTTGCCAATCAGCCTATCTCTGCATCCCNNNAGATCGGAAGA |
| 34076_4 | TATGAGTGTGGAGTCGTTGCCAATCAGCCTATCTCTGCATCCCNNNAGATCGGAAGA |
| 34076_5 | TATGAGTGTGGAGTCGTTGCCAATCAGCCTATCTCTGCATCCCNNNAGATCGGAAGA |
| 42485_1 | TATGAGTGTGGAGTCGTTGCAGAAAGAAGGAAGGAAGGAAGGANNNAGATCGGA |
| 42485_2 | TATGAGTGTGGAGTCGTTGCAGAAAGAAGGAAGGAAGGAAGGANNNAGATCGGA |
| 42485_3 | TATGAGTGTGGAGTCGTTGCAGAAAGAAGGAAGGAAGGAAGGANNNAGATCGGA |
| 42485_4 | TATGAGTGTGGAGTCGTTGCAGAAAGAAGGAAGGAAGGAAGGANNNAGATCGGA |
| 42485_5 | TATGAGTGTGGAGTCGTTGCAGAAAGAAGGAAGGAAGGAAGGANNNAGATCGGA |
| 50769_1 | TATGAGTGTGGAGTCGTTGCCATTTTCTTCAAGGAAGGCACGTNNNAGATCGGAAG |
| 50769_2 | TATGAGTGTGGAGTCGTTGCCATTTTCTTCAAGGAAGGCACGTNNNAGATCGGAAG |
| 50769_3 | TATGAGTGTGGAGTCGTTGCCATTTTCTTCAAGGAAGGCACGTNNNAGATCGGAAG |
| 50769_4 | TATGAGTGTGGAGTCGTTGCCATTTTCTTCAAGGAAGGCACGTNNNAGATCGGAAG |
| 50769_5 | TATGAGTGTGGAGTCGTTGCCATTTTCTTCAAGGAAGGCACGTNNNAGATCGGAAG |
| 41477_1 | TATGAGTGTGGAGTCGTTGCTATGACCCGTAACCGTCTCCTATNNNAGATCGGAAGA |
| 41477_2 | TATGAGTGTGGAGTCGTTGCTATGACCCGTAACCGTCTCCTATNNNAGATCGGAAGA |
| 41477_3 | TATGAGTGTGGAGTCGTTGCTATGACCCGTAACCGTCTCCTATNNNAGATCGGAAGA |
| 41477_4 | TATGAGTGTGGAGTCGTTGCTATGACCCGTAACCGTCTCCTATNNNAGATCGGAAGA |

| | |
|---|---|
| 41477_5 | TATGAGTGTGGAGTCGTTGCTATGACCCGTAACCGTCTCCTATNNNAGATCGGAAG |
| 43930_1 | TATGAGTGTGGAGTCGTTGCACACACAAAAGGCCTTCATCCTANNNAGATCGGAAG |
| 43930_2 | TATGAGTGTGGAGTCGTTGCACACACAAAAGGCCTTCATCCTANNNAGATCGGAAG |
| 43930_3 | TATGAGTGTGGAGTCGTTGCACACACAAAAGGCCTTCATCCTANNNAGATCGGAAG |
| 43930_4 | TATGAGTGTGGAGTCGTTGCACACACAAAAGGCCTTCATCCTANNNAGATCGGAAG |
| 43930_5 | TATGAGTGTGGAGTCGTTGCACACACAAAAGGCCTTCATCCTANNNAGATCGGAAG |
| 13526_1 | TATGAGTGTGGAGTCGTTGCATTGTTGGCATTTTTTAACAATNNNAGATCGGAAG |
| 13526_2 | TATGAGTGTGGAGTCGTTGCATTGTTGGCATTTTTTAACAATNNNAGATCGGAAG |
| 13526_3 | TATGAGTGTGGAGTCGTTGCATTGTTGGCATTTTTTAACAATNNNAGATCGGAAG |
| 13526_4 | TATGAGTGTGGAGTCGTTGCATTGTTGGCATTTTTTAACAATNNNAGATCGGAAG |
| 13526_5 | TATGAGTGTGGAGTCGTTGCATTGTTGGCATTTTTTAACAATNNNAGATCGGAAG |
| 40661_1 | TATGAGTGTGGAGTCGTTGCCCACCACATCTGGCTAGTTTTTNNNAGATCGGAAGA |
| 40661_2 | TATGAGTGTGGAGTCGTTGCCCACCACATCTGGCTAGTTTTTNNNAGATCGGAAGA |
| 40661_3 | TATGAGTGTGGAGTCGTTGCCCACCACATCTGGCTAGTTTTTNNNAGATCGGAAGA |
| 40661_4 | TATGAGTGTGGAGTCGTTGCCCACCACATCTGGCTAGTTTTTNNNAGATCGGAAGA |
| 40661_5 | TATGAGTGTGGAGTCGTTGCCCACCACATCTGGCTAGTTTTTNNNAGATCGGAAGA |
| 12590_1 | TATGAGTGTGGAGTCGTTGCCCTCTATTTGTAAACCTCAGANNNAGATCGGAAGA |
| 12590_2 | TATGAGTGTGGAGTCGTTGCCCTCTATTTGTAAACCTCAGANNNAGATCGGAAGA |
| 12590_3 | TATGAGTGTGGAGTCGTTGCCCTCTATTTGTAAACCTCAGANNNAGATCGGAAGA |
| 12590_4 | TATGAGTGTGGAGTCGTTGCCCTCTATTTGTAAACCTCAGANNNAGATCGGAAGA |
| 12590_5 | TATGAGTGTGGAGTCGTTGCCCTCTATTTGTAAACCTCAGANNNAGATCGGAAGA |
| 41322_1 | TATGAGTGTGGAGTCGTTGCACATGACATATCACGCGAGATCANNNAGATCGGAAG |
| 41322_2 | TATGAGTGTGGAGTCGTTGCACATGACATATCACGCGAGATCANNNAGATCGGAAG |
| 41322_3 | TATGAGTGTGGAGTCGTTGCACATGACATATCACGCGAGATCANNNAGATCGGAAG |
| 41322_4 | TATGAGTGTGGAGTCGTTGCACATGACATATCACGCGAGATCANNNAGATCGGAAG |
| 41322_5 | TATGAGTGTGGAGTCGTTGCACATGACATATCACGCGAGATCANNNAGATCGGAAG |
| 12522_1 | TATGAGTGTGGAGTCGTTGCCAAAGGTTATTCTTAATCCATCCNNNAGATCGGAAG |
| 12522_2 | TATGAGTGTGGAGTCGTTGCCAAAGGTTATTCTTAATCCATCCNNNAGATCGGAAG |
| 12522_3 | TATGAGTGTGGAGTCGTTGCCAAAGGTTATTCTTAATCCATCCNNNAGATCGGAAG |
| 12522_4 | TATGAGTGTGGAGTCGTTGCCAAAGGTTATTCTTAATCCATCCNNNAGATCGGAAG |
| 12522_5 | TATGAGTGTGGAGTCGTTGCCAAAGGTTATTCTTAATCCATCCNNNAGATCGGAAG |
| 10631_1 | TATGAGTGTGGAGTCGTTGCCTGAGAAGCAGAGGGAGAGGAATNNNAGATCGGA |
| 10631_2 | TATGAGTGTGGAGTCGTTGCCTGAGAAGCAGAGGGAGAGGAATNNNAGATCGGA |
| 10631_3 | TATGAGTGTGGAGTCGTTGCCTGAGAAGCAGAGGGAGAGGAATNNNAGATCGGA |
| 10631_4 | TATGAGTGTGGAGTCGTTGCCTGAGAAGCAGAGGGAGAGGAATNNNAGATCGGA |
| 10631_5 | TATGAGTGTGGAGTCGTTGCCTGAGAAGCAGAGGGAGAGGAATNNNAGATCGGA |
| 45789_1 | TATGAGTGTGGAGTCGTTGCCCCGAGTAGCTGAGATTATAGGCNNNAGATCGGAA |
| 45789_2 | TATGAGTGTGGAGTCGTTGCCCCGAGTAGCTGAGATTATAGGCNNNAGATCGGAA |
| 45789_3 | TATGAGTGTGGAGTCGTTGCCCCGAGTAGCTGAGATTATAGGCNNNAGATCGGAA |
| 45789_4 | TATGAGTGTGGAGTCGTTGCCCCGAGTAGCTGAGATTATAGGCNNNAGATCGGAA |
| 45789_5 | TATGAGTGTGGAGTCGTTGCCCCGAGTAGCTGAGATTATAGGCNNNAGATCGGAA |
| 12726_1 | TATGAGTGTGGAGTCGTTGCTTTTCTGTCCTTGAACTTCTTCNNNAGATCGGAAGAG |
| 12726_2 | TATGAGTGTGGAGTCGTTGCTTTTCTGTCCTTGAACTTCTTCNNNAGATCGGAAGAG |

| | |
|---|---|
| 12726_3 | TATGAGTGTGGAGTCGTTGCTTTTCTGTCCTTGAACTTCTTCNNNAGATCGGAAGAG |
| 12726_4 | TATGAGTGTGGAGTCGTTGCTTTTCTGTCCTTGAACTTCTTCNNNAGATCGGAAGAG |
| 12726_5 | TATGAGTGTGGAGTCGTTGCTTTTCTGTCCTTGAACTTCTTCNNNAGATCGGAAGAG |
| 46077_1 | TATGAGTGTGGAGTCGTTGCTTCACCCCATTCTCCTGCCTAAGNNNAGATCGGAAGA |
| 46077_2 | TATGAGTGTGGAGTCGTTGCTTCACCCCATTCTCCTGCCTAAGNNNAGATCGGAAGA |
| 46077_3 | TATGAGTGTGGAGTCGTTGCTTCACCCCATTCTCCTGCCTAAGNNNAGATCGGAAGA |
| 46077_4 | TATGAGTGTGGAGTCGTTGCTTCACCCCATTCTCCTGCCTAAGNNNAGATCGGAAGA |
| 46077_5 | TATGAGTGTGGAGTCGTTGCTTCACCCCATTCTCCTGCCTAAGNNNAGATCGGAAGA |
| 13681_1 | TATGAGTGTGGAGTCGTTGCTCTATTTGTGTGGCAGTTATTNNNAGATCGGAAGAG |
| 13681_2 | TATGAGTGTGGAGTCGTTGCTCTATTTGTGTGGCAGTTATTNNNAGATCGGAAGAG |
| 13681_3 | TATGAGTGTGGAGTCGTTGCTCTATTTGTGTGGCAGTTATTNNNAGATCGGAAGAG |
| 13681_4 | TATGAGTGTGGAGTCGTTGCTCTATTTGTGTGGCAGTTATTNNNAGATCGGAAGAG |
| 13681_5 | TATGAGTGTGGAGTCGTTGCTCTATTTGTGTGGCAGTTATTNNNAGATCGGAAGAG |
| 31972_1 | TATGAGTGTGGAGTCGTTGCACTAACAGAAGCTTTTCCCAGGTNNNAGATCGGAAG |
| 31972_2 | TATGAGTGTGGAGTCGTTGCACTAACAGAAGCTTTTCCCAGGTNNNAGATCGGAAG |
| 31972_3 | TATGAGTGTGGAGTCGTTGCACTAACAGAAGCTTTTCCCAGGTNNNAGATCGGAAG |
| 31972_4 | TATGAGTGTGGAGTCGTTGCACTAACAGAAGCTTTTCCCAGGTNNNAGATCGGAAG |
| 31972_5 | TATGAGTGTGGAGTCGTTGCACTAACAGAAGCTTTTCCCAGGTNNNAGATCGGAAG |
| 48668_1 | TATGAGTGTGGAGTCGTTGCCACCATGCCCAGCTAAGTTTTTTNNNAGATCGGAAGA |
| 48668_2 | TATGAGTGTGGAGTCGTTGCCACCATGCCCAGCTAAGTTTTTTNNNAGATCGGAAGA |
| 48668_3 | TATGAGTGTGGAGTCGTTGCCACCATGCCCAGCTAAGTTTTTTNNNAGATCGGAAGA |
| 48668_4 | TATGAGTGTGGAGTCGTTGCCACCATGCCCAGCTAAGTTTTTTNNNAGATCGGAAGA |
| 48668_5 | TATGAGTGTGGAGTCGTTGCCACCATGCCCAGCTAAGTTTTTTNNNAGATCGGAAGA |
| 24657_1 | TATGAGTGTGGAGTCGTTGCCTCCAATCTCTACTGTAATAAGANNNAGATCGGAAGA |
| 24657_2 | TATGAGTGTGGAGTCGTTGCCTCCAATCTCTACTGTAATAAGANNNAGATCGGAAGA |
| 24657_3 | TATGAGTGTGGAGTCGTTGCCTCCAATCTCTACTGTAATAAGANNNAGATCGGAAGA |
| 24657_4 | TATGAGTGTGGAGTCGTTGCCTCCAATCTCTACTGTAATAAGANNNAGATCGGAAGA |
| 24657_5 | TATGAGTGTGGAGTCGTTGCCTCCAATCTCTACTGTAATAAGANNNAGATCGGAAGA |
| 33870_1 | TATGAGTGTGGAGTCGTTGCTGAGCATGGCAATAGTAAGAGCANNNAGATCGGAA |
| 33870_2 | TATGAGTGTGGAGTCGTTGCTGAGCATGGCAATAGTAAGAGCANNNAGATCGGAA |
| 33870_3 | TATGAGTGTGGAGTCGTTGCTGAGCATGGCAATAGTAAGAGCANNNAGATCGGAA |
| 33870_4 | TATGAGTGTGGAGTCGTTGCTGAGCATGGCAATAGTAAGAGCANNNAGATCGGAA |
| 33870_5 | TATGAGTGTGGAGTCGTTGCTGAGCATGGCAATAGTAAGAGCANNNAGATCGGAA |
| 34070_1 | TATGAGTGTGGAGTCGTTGCTAGAGTGCAGTGGTGTGATCTTGNNNAGATCGGAA |
| 34070_2 | TATGAGTGTGGAGTCGTTGCTAGAGTGCAGTGGTGTGATCTTGNNNAGATCGGAA |
| 34070_3 | TATGAGTGTGGAGTCGTTGCTAGAGTGCAGTGGTGTGATCTTGNNNAGATCGGAA |
| 34070_4 | TATGAGTGTGGAGTCGTTGCTAGAGTGCAGTGGTGTGATCTTGNNNAGATCGGAA |
| 34070_5 | TATGAGTGTGGAGTCGTTGCTAGAGTGCAGTGGTGTGATCTTGNNNAGATCGGAA |
| 29383_1 | TATGAGTGTGGAGTCGTTGCCAGAAGGAAAATCAATGCCCTCCNNNAGATCGGAAG |
| 29383_2 | TATGAGTGTGGAGTCGTTGCCAGAAGGAAAATCAATGCCCTCCNNNAGATCGGAAG |
| 29383_3 | TATGAGTGTGGAGTCGTTGCCAGAAGGAAAATCAATGCCCTCCNNNAGATCGGAAG |
| 29383_4 | TATGAGTGTGGAGTCGTTGCCAGAAGGAAAATCAATGCCCTCCNNNAGATCGGAAG |
| 29383_5 | TATGAGTGTGGAGTCGTTGCCAGAAGGAAAATCAATGCCCTCCNNNAGATCGGAAG |

| | |
|---|---|
| 47715_1 | TATGAGTGTGGAGTCGTTGCGATCTCAGCTCACTGCACTACAANNNAGATCGGAAG |
| 47715_2 | TATGAGTGTGGAGTCGTTGCGATCTCAGCTCACTGCACTACAANNNAGATCGGAAG |
| 47715_3 | TATGAGTGTGGAGTCGTTGCGATCTCAGCTCACTGCACTACAANNNAGATCGGAAG |
| 47715_4 | TATGAGTGTGGAGTCGTTGCGATCTCAGCTCACTGCACTACAANNNAGATCGGAAG |
| 47715_5 | TATGAGTGTGGAGTCGTTGCGATCTCAGCTCACTGCACTACAANNNAGATCGGAAG |
| 52504_1 | TATGAGTGTGGAGTCGTTGCGGTGGGGTGAGTAGGTACAAAAANNNAGATCGGA |
| 52504_2 | TATGAGTGTGGAGTCGTTGCGGTGGGGTGAGTAGGTACAAAAANNNAGATCGGA |
| 52504_3 | TATGAGTGTGGAGTCGTTGCGGTGGGGTGAGTAGGTACAAAAANNNAGATCGGA |
| 52504_4 | TATGAGTGTGGAGTCGTTGCGGTGGGGTGAGTAGGTACAAAAANNNAGATCGGA |
| 52504_5 | TATGAGTGTGGAGTCGTTGCGGTGGGGTGAGTAGGTACAAAAANNNAGATCGGA |
| 52290_1 | TATGAGTGTGGAGTCGTTGCCCTTTGGCTCTCACCACAGCNNNAGATCGGAAGAGC |
| 52290_2 | TATGAGTGTGGAGTCGTTGCCCTTTGGCTCTCACCACAGCNNNAGATCGGAAGAGC |
| 52290_3 | TATGAGTGTGGAGTCGTTGCCCTTTGGCTCTCACCACAGCNNNAGATCGGAAGAGC |
| 52290_4 | TATGAGTGTGGAGTCGTTGCCCTTTGGCTCTCACCACAGCNNNAGATCGGAAGAGC |
| 52290_5 | TATGAGTGTGGAGTCGTTGCCCTTTGGCTCTCACCACAGCNNNAGATCGGAAGAGC |
| 38708_1 | TATGAGTGTGGAGTCGTTGCTCCATTGTGATTGCAGAAGTTTGTNNNAGATCGGAA |
| 38708_2 | TATGAGTGTGGAGTCGTTGCTCCATTGTGATTGCAGAAGTTTGTNNNAGATCGGAA |
| 38708_3 | TATGAGTGTGGAGTCGTTGCTCCATTGTGATTGCAGAAGTTTGTNNNAGATCGGAA |
| 38708_4 | TATGAGTGTGGAGTCGTTGCTCCATTGTGATTGCAGAAGTTTGTNNNAGATCGGAA |
| 38708_5 | TATGAGTGTGGAGTCGTTGCTCCATTGTGATTGCAGAAGTTTGTNNNAGATCGGAA |
| 32252_1 | TATGAGTGTGGAGTCGTTGCGCCAAAAAAAAGGAGTCTCAGTANNNAGATCGGAA |
| 32252_2 | TATGAGTGTGGAGTCGTTGCGCCAAAAAAAAGGAGTCTCAGTANNNAGATCGGAA |
| 32252_3 | TATGAGTGTGGAGTCGTTGCGCCAAAAAAAAGGAGTCTCAGTANNNAGATCGGAA |
| 32252_4 | TATGAGTGTGGAGTCGTTGCGCCAAAAAAAAGGAGTCTCAGTANNNAGATCGGAA |
| 32252_5 | TATGAGTGTGGAGTCGTTGCGCCAAAAAAAAGGAGTCTCAGTANNNAGATCGGAA |
| 12576_1 | TATGAGTGTGGAGTCGTTGCTGCTTAGGGATTTTCTTATTGATNNNAGATCGGAAG |
| 12576_2 | TATGAGTGTGGAGTCGTTGCTGCTTAGGGATTTTCTTATTGATNNNAGATCGGAAG |
| 12576_3 | TATGAGTGTGGAGTCGTTGCTGCTTAGGGATTTTCTTATTGATNNNAGATCGGAAG |
| 12576_4 | TATGAGTGTGGAGTCGTTGCTGCTTAGGGATTTTCTTATTGATNNNAGATCGGAAG |
| 12576_5 | TATGAGTGTGGAGTCGTTGCTGCTTAGGGATTTTCTTATTGATNNNAGATCGGAAG |
| 47266_1 | TATGAGTGTGGAGTCGTTGCCACCATGCCCAGCTAATTTTTGTNNNAGATCGGAAG |
| 47266_2 | TATGAGTGTGGAGTCGTTGCCACCATGCCCAGCTAATTTTTGTNNNAGATCGGAAG |
| 47266_3 | TATGAGTGTGGAGTCGTTGCCACCATGCCCAGCTAATTTTTGTNNNAGATCGGAAG |
| 47266_4 | TATGAGTGTGGAGTCGTTGCCACCATGCCCAGCTAATTTTTGTNNNAGATCGGAAG |
| 47266_5 | TATGAGTGTGGAGTCGTTGCCACCATGCCCAGCTAATTTTTGTNNNAGATCGGAAG |
| 31324_1 | TATGAGTGTGGAGTCGTTGCTCAAGCAAAAAACACATCTTGCCNNNAGATCGGAAG |
| 31324_2 | TATGAGTGTGGAGTCGTTGCTCAAGCAAAAAACACATCTTGCCNNNAGATCGGAAG |
| 31324_3 | TATGAGTGTGGAGTCGTTGCTCAAGCAAAAAACACATCTTGCCNNNAGATCGGAAG |
| 31324_4 | TATGAGTGTGGAGTCGTTGCTCAAGCAAAAAACACATCTTGCCNNNAGATCGGAAG |
| 31324_5 | TATGAGTGTGGAGTCGTTGCTCAAGCAAAAAACACATCTTGCCNNNAGATCGGAAG |
| 39424_1 | TATGAGTGTGGAGTCGTTGCGATGAAGGCAACAATTAGAGCCCNNNAGATCGGAA |
| 39424_2 | TATGAGTGTGGAGTCGTTGCGATGAAGGCAACAATTAGAGCCCNNNAGATCGGAA |
| 39424_3 | TATGAGTGTGGAGTCGTTGCGATGAAGGCAACAATTAGAGCCCNNNAGATCGGAA |

| | |
|---|---|
| 39424_4 | TATGAGTGTGGAGTCGTTGCGATGAAGGCAACAATTAGAGCCCNNNAGATCGGAA |
| 39424_5 | TATGAGTGTGGAGTCGTTGCGATGAAGGCAACAATTAGAGCCCNNNAGATCGGAA |
| 29910_1 | TATGAGTGTGGAGTCGTTGCGACTGTGCCAAGCCCTTGAATATNNNAGATCGGAAG |
| 29910_2 | TATGAGTGTGGAGTCGTTGCGACTGTGCCAAGCCCTTGAATATNNNAGATCGGAAG |
| 29910_3 | TATGAGTGTGGAGTCGTTGCGACTGTGCCAAGCCCTTGAATATNNNAGATCGGAAG |
| 29910_4 | TATGAGTGTGGAGTCGTTGCGACTGTGCCAAGCCCTTGAATATNNNAGATCGGAAG |
| 29910_5 | TATGAGTGTGGAGTCGTTGCGACTGTGCCAAGCCCTTGAATATNNNAGATCGGAAG |
| 8317_1 | TATGAGTGTGGAGTCGTTGCAGACAATGCAGGTGTGATCCCTNNNAGATCGGAAG |
| 8317_2 | TATGAGTGTGGAGTCGTTGCAGACAATGCAGGTGTGATCCCTNNNAGATCGGAAG |
| 8317_3 | TATGAGTGTGGAGTCGTTGCAGACAATGCAGGTGTGATCCCTNNNAGATCGGAAG |
| 8317_4 | TATGAGTGTGGAGTCGTTGCAGACAATGCAGGTGTGATCCCTNNNAGATCGGAAG |
| 8317_5 | TATGAGTGTGGAGTCGTTGCAGACAATGCAGGTGTGATCCCTNNNAGATCGGAAG |
| 33681_1 | TATGAGTGTGGAGTCGTTGCCCGGCTGAATCACTGTTTCTAAGNNNAGATCGGAAG |
| 33681_2 | TATGAGTGTGGAGTCGTTGCCCGGCTGAATCACTGTTTCTAAGNNNAGATCGGAAG |
| 33681_3 | TATGAGTGTGGAGTCGTTGCCCGGCTGAATCACTGTTTCTAAGNNNAGATCGGAAG |
| 33681_4 | TATGAGTGTGGAGTCGTTGCCCGGCTGAATCACTGTTTCTAAGNNNAGATCGGAAG |
| 33681_5 | TATGAGTGTGGAGTCGTTGCCCGGCTGAATCACTGTTTCTAAGNNNAGATCGGAAG |
| 13593_1 | TATGAGTGTGGAGTCGTTGCTCAAGTGTTGATGTAGTAATAAATTNNNAGATCGGA |
| 13593_2 | TATGAGTGTGGAGTCGTTGCTCAAGTGTTGATGTAGTAATAAATTNNNAGATCGGA |
| 13593_3 | TATGAGTGTGGAGTCGTTGCTCAAGTGTTGATGTAGTAATAAATTNNNAGATCGGA |
| 13593_4 | TATGAGTGTGGAGTCGTTGCTCAAGTGTTGATGTAGTAATAAATTNNNAGATCGGA |
| 13593_5 | TATGAGTGTGGAGTCGTTGCTCAAGTGTTGATGTAGTAATAAATTNNNAGATCGGA |
| 10093_1 | TATGAGTGTGGAGTCGTTGCTGTGTTGGGTTTAGGTTTTGGGTNNNAGATCGGAA |
| 10093_2 | TATGAGTGTGGAGTCGTTGCTGTGTTGGGTTTAGGTTTTGGGTNNNAGATCGGAA |
| 10093_3 | TATGAGTGTGGAGTCGTTGCTGTGTTGGGTTTAGGTTTTGGGTNNNAGATCGGAA |
| 10093_4 | TATGAGTGTGGAGTCGTTGCTGTGTTGGGTTTAGGTTTTGGGTNNNAGATCGGAA |
| 10093_5 | TATGAGTGTGGAGTCGTTGCTGTGTTGGGTTTAGGTTTTGGGTNNNAGATCGGAA |
| 47462_1 | TATGAGTGTGGAGTCGTTGCGCAAAGATCTGTGTAGGGGAAGANNNAGATCGGAA |
| 47462_2 | TATGAGTGTGGAGTCGTTGCGCAAAGATCTGTGTAGGGGAAGANNNAGATCGGAA |
| 47462_3 | TATGAGTGTGGAGTCGTTGCGCAAAGATCTGTGTAGGGGAAGANNNAGATCGGAA |
| 47462_4 | TATGAGTGTGGAGTCGTTGCGCAAAGATCTGTGTAGGGGAAGANNNAGATCGGAA |
| 47462_5 | TATGAGTGTGGAGTCGTTGCGCAAAGATCTGTGTAGGGGAAGANNNAGATCGGAA |
| 32763_1 | TATGAGTGTGGAGTCGTTGCTAGAGATCTGCTGTACGACGTTGNNNAGATCGGAA |
| 32763_2 | TATGAGTGTGGAGTCGTTGCTAGAGATCTGCTGTACGACGTTGNNNAGATCGGAA |
| 32763_3 | TATGAGTGTGGAGTCGTTGCTAGAGATCTGCTGTACGACGTTGNNNAGATCGGAA |
| 32763_4 | TATGAGTGTGGAGTCGTTGCTAGAGATCTGCTGTACGACGTTGNNNAGATCGGAA |
| 32763_5 | TATGAGTGTGGAGTCGTTGCTAGAGATCTGCTGTACGACGTTGNNNAGATCGGAA |
| 37695_1 | TATGAGTGTGGAGTCGTTGCTCATTTGCAAAAGGTGTAAACACANNNAGATCGGAA |
| 37695_2 | TATGAGTGTGGAGTCGTTGCTCATTTGCAAAAGGTGTAAACACANNNAGATCGGAA |
| 37695_3 | TATGAGTGTGGAGTCGTTGCTCATTTGCAAAAGGTGTAAACACANNNAGATCGGAA |
| 37695_4 | TATGAGTGTGGAGTCGTTGCTCATTTGCAAAAGGTGTAAACACANNNAGATCGGAA |
| 37695_5 | TATGAGTGTGGAGTCGTTGCTCATTTGCAAAAGGTGTAAACACANNNAGATCGGAA |
| 41094_1 | TATGAGTGTGGAGTCGTTGCATTGAATGACAACACAGGGCATGNNNAGATCGGAA |

| | |
|---|---|
| 41094_2 | TATGAGTGTGGAGTCGTTGCATTGAATGACAACACAGGGCATGNNNAGATCGGAA |
| 41094_3 | TATGAGTGTGGAGTCGTTGCATTGAATGACAACACAGGGCATGNNNAGATCGGAA |
| 41094_4 | TATGAGTGTGGAGTCGTTGCATTGAATGACAACACAGGGCATGNNNAGATCGGAA |
| 41094_5 | TATGAGTGTGGAGTCGTTGCATTGAATGACAACACAGGGCATGNNNAGATCGGAA |
| 35395_1 | TATGAGTGTGGAGTCGTTGCCAAGTAGCTGGGACTACAGGCNNNAGATCGGAAGA |
| 35395_2 | TATGAGTGTGGAGTCGTTGCCAAGTAGCTGGGACTACAGGCNNNAGATCGGAAGA |
| 35395_3 | TATGAGTGTGGAGTCGTTGCCAAGTAGCTGGGACTACAGGCNNNAGATCGGAAGA |
| 35395_4 | TATGAGTGTGGAGTCGTTGCCAAGTAGCTGGGACTACAGGCNNNAGATCGGAAGA |
| 35395_5 | TATGAGTGTGGAGTCGTTGCCAAGTAGCTGGGACTACAGGCNNNAGATCGGAAGA |
| 10548_1 | TATGAGTGTGGAGTCGTTGCGCATGTTCACACACACACTGTTTNNNAGATCGGAAG |
| 10548_2 | TATGAGTGTGGAGTCGTTGCGCATGTTCACACACACACTGTTTNNNAGATCGGAAG |
| 10548_3 | TATGAGTGTGGAGTCGTTGCGCATGTTCACACACACACTGTTTNNNAGATCGGAAG |
| 10548_4 | TATGAGTGTGGAGTCGTTGCGCATGTTCACACACACACTGTTTNNNAGATCGGAAG |
| 10548_5 | TATGAGTGTGGAGTCGTTGCGCATGTTCACACACACACTGTTTNNNAGATCGGAAG |
| 11007_1 | TATGAGTGTGGAGTCGTTGCCCGCAGACCATGAAACCTAGAANNNAGATCGGAAG |
| 11007_2 | TATGAGTGTGGAGTCGTTGCCCGCAGACCATGAAACCTAGAANNNAGATCGGAAG |
| 11007_3 | TATGAGTGTGGAGTCGTTGCCCGCAGACCATGAAACCTAGAANNNAGATCGGAAG |
| 11007_4 | TATGAGTGTGGAGTCGTTGCCCGCAGACCATGAAACCTAGAANNNAGATCGGAAG |
| 11007_5 | TATGAGTGTGGAGTCGTTGCCCGCAGACCATGAAACCTAGAANNNAGATCGGAAG |
| 37265_1 | TATGAGTGTGGAGTCGTTGCCTGTAGTCCCAGCTACTCAGGAANNNAGATCGGAAG |
| 37265_2 | TATGAGTGTGGAGTCGTTGCCTGTAGTCCCAGCTACTCAGGAANNNAGATCGGAAG |
| 37265_3 | TATGAGTGTGGAGTCGTTGCCTGTAGTCCCAGCTACTCAGGAANNNAGATCGGAAG |
| 37265_4 | TATGAGTGTGGAGTCGTTGCCTGTAGTCCCAGCTACTCAGGAANNNAGATCGGAAG |
| 37265_5 | TATGAGTGTGGAGTCGTTGCCTGTAGTCCCAGCTACTCAGGAANNNAGATCGGAAG |
| 50687_1 | TATGAGTGTGGAGTCGTTGCTTGGCTTCCTCTTCTGCAAAATGNNNAGATCGGAAG |
| 50687_2 | TATGAGTGTGGAGTCGTTGCTTGGCTTCCTCTTCTGCAAAATGNNNAGATCGGAAG |
| 50687_3 | TATGAGTGTGGAGTCGTTGCTTGGCTTCCTCTTCTGCAAAATGNNNAGATCGGAAG |
| 50687_4 | TATGAGTGTGGAGTCGTTGCTTGGCTTCCTCTTCTGCAAAATGNNNAGATCGGAAG |
| 50687_5 | TATGAGTGTGGAGTCGTTGCTTGGCTTCCTCTTCTGCAAAATGNNNAGATCGGAAG |
| 47465_1 | TATGAGTGTGGAGTCGTTGCGAGGACAATTCAGACAGATTGCANNNAGATCGGAA |
| 47465_2 | TATGAGTGTGGAGTCGTTGCGAGGACAATTCAGACAGATTGCANNNAGATCGGAA |
| 47465_3 | TATGAGTGTGGAGTCGTTGCGAGGACAATTCAGACAGATTGCANNNAGATCGGAA |
| 47465_4 | TATGAGTGTGGAGTCGTTGCGAGGACAATTCAGACAGATTGCANNNAGATCGGAA |
| 47465_5 | TATGAGTGTGGAGTCGTTGCGAGGACAATTCAGACAGATTGCANNNAGATCGGAA |
| 9698_1 | TATGAGTGTGGAGTCGTTGCTTCAACTCTGTGTGTGTGAGACCNNNAGATCGGAAG |
| 9698_2 | TATGAGTGTGGAGTCGTTGCTTCAACTCTGTGTGTGTGAGACCNNNAGATCGGAAG |
| 9698_3 | TATGAGTGTGGAGTCGTTGCTTCAACTCTGTGTGTGTGAGACCNNNAGATCGGAAG |
| 9698_4 | TATGAGTGTGGAGTCGTTGCTTCAACTCTGTGTGTGTGAGACCNNNAGATCGGAAG |
| 9698_5 | TATGAGTGTGGAGTCGTTGCTTCAACTCTGTGTGTGTGAGACCNNNAGATCGGAAG |
| 39956_1 | TATGAGTGTGGAGTCGTTGCGCCTTCCCTGATCACTCAAGTTANNNAGATCGGAAG |
| 39956_2 | TATGAGTGTGGAGTCGTTGCGCCTTCCCTGATCACTCAAGTTANNNAGATCGGAAG |
| 39956_3 | TATGAGTGTGGAGTCGTTGCGCCTTCCCTGATCACTCAAGTTANNNAGATCGGAAG |
| 39956_4 | TATGAGTGTGGAGTCGTTGCGCCTTCCCTGATCACTCAAGTTANNNAGATCGGAAG |

| | |
|---|---|
| 39956_5 | TATGAGTGTGGAGTCGTTGCGCCTTCCCTGATCACTCAAGTTANNNAGATCGGAAGA |
| 45662_1 | TATGAGTGTGGAGTCGTTGCCTTTAGCCTCCTGAGTAGCTGAGNNNAGATCGGAAG |
| 45662_2 | TATGAGTGTGGAGTCGTTGCCTTTAGCCTCCTGAGTAGCTGAGNNNAGATCGGAAG |
| 45662_3 | TATGAGTGTGGAGTCGTTGCCTTTAGCCTCCTGAGTAGCTGAGNNNAGATCGGAAG |
| 45662_4 | TATGAGTGTGGAGTCGTTGCCTTTAGCCTCCTGAGTAGCTGAGNNNAGATCGGAAG |
| 45662_5 | TATGAGTGTGGAGTCGTTGCCTTTAGCCTCCTGAGTAGCTGAGNNNAGATCGGAAG |
| 13362_1 | TATGAGTGTGGAGTCGTTGCATTGAGACATGAAGAACAGACNNNAGATCGGAAGA |
| 13362_2 | TATGAGTGTGGAGTCGTTGCATTGAGACATGAAGAACAGACNNNAGATCGGAAGA |
| 13362_3 | TATGAGTGTGGAGTCGTTGCATTGAGACATGAAGAACAGACNNNAGATCGGAAGA |
| 13362_4 | TATGAGTGTGGAGTCGTTGCATTGAGACATGAAGAACAGACNNNAGATCGGAAGA |
| 13362_5 | TATGAGTGTGGAGTCGTTGCATTGAGACATGAAGAACAGACNNNAGATCGGAAGA |
| 29966_1 | TATGAGTGTGGAGTCGTTGCTCATGGATGAACTGGTTAGTCCGNNNAGATCGGAAG |
| 29966_2 | TATGAGTGTGGAGTCGTTGCTCATGGATGAACTGGTTAGTCCGNNNAGATCGGAAG |
| 29966_3 | TATGAGTGTGGAGTCGTTGCTCATGGATGAACTGGTTAGTCCGNNNAGATCGGAAG |
| 29966_4 | TATGAGTGTGGAGTCGTTGCTCATGGATGAACTGGTTAGTCCGNNNAGATCGGAAG |
| 29966_5 | TATGAGTGTGGAGTCGTTGCTCATGGATGAACTGGTTAGTCCGNNNAGATCGGAAG |
| 44103_1 | TATGAGTGTGGAGTCGTTGCTCTTTCCCTCTCCAACCCTTTTCNNNAGATCGGAAGAG |
| 44103_2 | TATGAGTGTGGAGTCGTTGCTCTTTCCCTCTCCAACCCTTTTCNNNAGATCGGAAGAG |
| 44103_3 | TATGAGTGTGGAGTCGTTGCTCTTTCCCTCTCCAACCCTTTTCNNNAGATCGGAAGAG |
| 44103_4 | TATGAGTGTGGAGTCGTTGCTCTTTCCCTCTCCAACCCTTTTCNNNAGATCGGAAGAG |
| 44103_5 | TATGAGTGTGGAGTCGTTGCTCTTTCCCTCTCCAACCCTTTTCNNNAGATCGGAAGAG |
| 37613_1 | TATGAGTGTGGAGTCGTTGCTAGTTGGTTCATGGTTCTGCAGGNNNAGATCGGAAG |
| 37613_2 | TATGAGTGTGGAGTCGTTGCTAGTTGGTTCATGGTTCTGCAGGNNNAGATCGGAAG |
| 37613_3 | TATGAGTGTGGAGTCGTTGCTAGTTGGTTCATGGTTCTGCAGGNNNAGATCGGAAG |
| 37613_4 | TATGAGTGTGGAGTCGTTGCTAGTTGGTTCATGGTTCTGCAGGNNNAGATCGGAAG |
| 37613_5 | TATGAGTGTGGAGTCGTTGCTAGTTGGTTCATGGTTCTGCAGGNNNAGATCGGAAG |
| 46463_1 | TATGAGTGTGGAGTCGTTGCTTCAAGTGATTCTCCTGCCTCAGNNNAGATCGGAAGA |
| 46463_2 | TATGAGTGTGGAGTCGTTGCTTCAAGTGATTCTCCTGCCTCAGNNNAGATCGGAAGA |
| 46463_3 | TATGAGTGTGGAGTCGTTGCTTCAAGTGATTCTCCTGCCTCAGNNNAGATCGGAAGA |
| 46463_4 | TATGAGTGTGGAGTCGTTGCTTCAAGTGATTCTCCTGCCTCAGNNNAGATCGGAAGA |
| 46463_5 | TATGAGTGTGGAGTCGTTGCTTCAAGTGATTCTCCTGCCTCAGNNNAGATCGGAAGA |
| 42898_1 | TATGAGTGTGGAGTCGTTGCTTTCACTCCACTGAAAGGCATCANNNAGATCGGAAGA |
| 42898_2 | TATGAGTGTGGAGTCGTTGCTTTCACTCCACTGAAAGGCATCANNNAGATCGGAAGA |
| 42898_3 | TATGAGTGTGGAGTCGTTGCTTTCACTCCACTGAAAGGCATCANNNAGATCGGAAGA |
| 42898_4 | TATGAGTGTGGAGTCGTTGCTTTCACTCCACTGAAAGGCATCANNNAGATCGGAAGA |
| 42898_5 | TATGAGTGTGGAGTCGTTGCTTTCACTCCACTGAAAGGCATCANNNAGATCGGAAGA |
| 35605_1 | TATGAGTGTGGAGTCGTTGCGATGAACAAGGGCAGACTGATTGNNNAGATCGGAA |
| 35605_2 | TATGAGTGTGGAGTCGTTGCGATGAACAAGGGCAGACTGATTGNNNAGATCGGAA |
| 35605_3 | TATGAGTGTGGAGTCGTTGCGATGAACAAGGGCAGACTGATTGNNNAGATCGGAA |
| 35605_4 | TATGAGTGTGGAGTCGTTGCGATGAACAAGGGCAGACTGATTGNNNAGATCGGAA |
| 35605_5 | TATGAGTGTGGAGTCGTTGCGATGAACAAGGGCAGACTGATTGNNNAGATCGGAA |
| 37034_1 | TATGAGTGTGGAGTCGTTGCTCCAAGATTGCCGAATAGGAACANNNAGATCGGAAG |
| 37034_2 | TATGAGTGTGGAGTCGTTGCTCCAAGATTGCCGAATAGGAACANNNAGATCGGAAG |

| | |
|---|---|
| 37034_3 | TATGAGTGTGGAGTCGTTGCTCCAAGATTGCCGAATAGGAACANNNAGATCGGAA |
| 37034_4 | TATGAGTGTGGAGTCGTTGCTCCAAGATTGCCGAATAGGAACANNNAGATCGGAA |
| 37034_5 | TATGAGTGTGGAGTCGTTGCTCCAAGATTGCCGAATAGGAACANNNAGATCGGAA |
| 45466_1 | TATGAGTGTGGAGTCGTTGCAATTGTGGTCCCTGACTGATACCNNNAGATCGGAAG |
| 45466_2 | TATGAGTGTGGAGTCGTTGCAATTGTGGTCCCTGACTGATACCNNNAGATCGGAAG |
| 45466_3 | TATGAGTGTGGAGTCGTTGCAATTGTGGTCCCTGACTGATACCNNNAGATCGGAAG |
| 45466_4 | TATGAGTGTGGAGTCGTTGCAATTGTGGTCCCTGACTGATACCNNNAGATCGGAAG |
| 45466_5 | TATGAGTGTGGAGTCGTTGCAATTGTGGTCCCTGACTGATACCNNNAGATCGGAAG |
| 30748_1 | TATGAGTGTGGAGTCGTTGCAGTTGTTGTCTGTGGGAGGTTAGNNNAGATCGGAA |
| 30748_2 | TATGAGTGTGGAGTCGTTGCAGTTGTTGTCTGTGGGAGGTTAGNNNAGATCGGAA |
| 30748_3 | TATGAGTGTGGAGTCGTTGCAGTTGTTGTCTGTGGGAGGTTAGNNNAGATCGGAA |
| 30748_4 | TATGAGTGTGGAGTCGTTGCAGTTGTTGTCTGTGGGAGGTTAGNNNAGATCGGAA |
| 30748_5 | TATGAGTGTGGAGTCGTTGCAGTTGTTGTCTGTGGGAGGTTAGNNNAGATCGGAA |
| 48937_1 | TATGAGTGTGGAGTCGTTGCCCTCCTGGGTTCAAACGATTCTNNNAGATCGGAAGA |
| 48937_2 | TATGAGTGTGGAGTCGTTGCCCTCCTGGGTTCAAACGATTCTNNNAGATCGGAAGA |
| 48937_3 | TATGAGTGTGGAGTCGTTGCCCTCCTGGGTTCAAACGATTCTNNNAGATCGGAAGA |
| 48937_4 | TATGAGTGTGGAGTCGTTGCCCTCCTGGGTTCAAACGATTCTNNNAGATCGGAAGA |
| 48937_5 | TATGAGTGTGGAGTCGTTGCCCTCCTGGGTTCAAACGATTCTNNNAGATCGGAAGA |
| 46497_1 | TATGAGTGTGGAGTCGTTGCATCTCAGTTCACTGCAACCTCTGNNNAGATCGGAAG |
| 46497_2 | TATGAGTGTGGAGTCGTTGCATCTCAGTTCACTGCAACCTCTGNNNAGATCGGAAG |
| 46497_3 | TATGAGTGTGGAGTCGTTGCATCTCAGTTCACTGCAACCTCTGNNNAGATCGGAAG |
| 46497_4 | TATGAGTGTGGAGTCGTTGCATCTCAGTTCACTGCAACCTCTGNNNAGATCGGAAG |
| 46497_5 | TATGAGTGTGGAGTCGTTGCATCTCAGTTCACTGCAACCTCTGNNNAGATCGGAAG |
| 13328_1 | TATGAGTGTGGAGTCGTTGCTAAAACCCTGTCTCTACTAAAAGNNNAGATCGGAAG |
| 13328_2 | TATGAGTGTGGAGTCGTTGCTAAAACCCTGTCTCTACTAAAAGNNNAGATCGGAAG |
| 13328_3 | TATGAGTGTGGAGTCGTTGCTAAAACCCTGTCTCTACTAAAAGNNNAGATCGGAAG |
| 13328_4 | TATGAGTGTGGAGTCGTTGCTAAAACCCTGTCTCTACTAAAAGNNNAGATCGGAAG |
| 13328_5 | TATGAGTGTGGAGTCGTTGCTAAAACCCTGTCTCTACTAAAAGNNNAGATCGGAAG |
| 53098_1 | TATGAGTGTGGAGTCGTTGCTCTTGAGCCTGTGGATGCAAATANNNAGATCGGAAG |
| 53098_2 | TATGAGTGTGGAGTCGTTGCTCTTGAGCCTGTGGATGCAAATANNNAGATCGGAAG |
| 53098_3 | TATGAGTGTGGAGTCGTTGCTCTTGAGCCTGTGGATGCAAATANNNAGATCGGAAG |
| 53098_4 | TATGAGTGTGGAGTCGTTGCTCTTGAGCCTGTGGATGCAAATANNNAGATCGGAAG |
| 53098_5 | TATGAGTGTGGAGTCGTTGCTCTTGAGCCTGTGGATGCAAATANNNAGATCGGAAG |
| 41239_1 | TATGAGTGTGGAGTCGTTGCGTGTGTGTCTTCTTGTGCTTCTCNNNAGATCGGAAG |
| 41239_2 | TATGAGTGTGGAGTCGTTGCGTGTGTGTCTTCTTGTGCTTCTCNNNAGATCGGAAG |
| 41239_3 | TATGAGTGTGGAGTCGTTGCGTGTGTGTCTTCTTGTGCTTCTCNNNAGATCGGAAG |
| 41239_4 | TATGAGTGTGGAGTCGTTGCGTGTGTGTCTTCTTGTGCTTCTCNNNAGATCGGAAG |
| 41239_5 | TATGAGTGTGGAGTCGTTGCGTGTGTGTCTTCTTGTGCTTCTCNNNAGATCGGAAG |
| 34190_1 | TATGAGTGTGGAGTCGTTGCTTTGAAGACTGAGGCTCTAGTCCNNNAGATCGGAAG |
| 34190_2 | TATGAGTGTGGAGTCGTTGCTTTGAAGACTGAGGCTCTAGTCCNNNAGATCGGAAG |
| 34190_3 | TATGAGTGTGGAGTCGTTGCTTTGAAGACTGAGGCTCTAGTCCNNNAGATCGGAAG |
| 34190_4 | TATGAGTGTGGAGTCGTTGCTTTGAAGACTGAGGCTCTAGTCCNNNAGATCGGAAG |
| 34190_5 | TATGAGTGTGGAGTCGTTGCTTTGAAGACTGAGGCTCTAGTCCNNNAGATCGGAAG |

| | |
|---|---|
| 43449_1 | TATGAGTGTGGAGTCGTTGCTTCAAGCGATTCTCCTGCCTTAGNNNAGATCGGAAGA |
| 43449_2 | TATGAGTGTGGAGTCGTTGCTTCAAGCGATTCTCCTGCCTTAGNNNAGATCGGAAGA |
| 43449_3 | TATGAGTGTGGAGTCGTTGCTTCAAGCGATTCTCCTGCCTTAGNNNAGATCGGAAGA |
| 43449_4 | TATGAGTGTGGAGTCGTTGCTTCAAGCGATTCTCCTGCCTTAGNNNAGATCGGAAGA |
| 43449_5 | TATGAGTGTGGAGTCGTTGCTTCAAGCGATTCTCCTGCCTTAGNNNAGATCGGAAGA |
| 30677_1 | TATGAGTGTGGAGTCGTTGCCGAATGTAGTTCCTCAGGGTAACNNNAGATCGGAAG |
| 30677_2 | TATGAGTGTGGAGTCGTTGCCGAATGTAGTTCCTCAGGGTAACNNNAGATCGGAAG |
| 30677_3 | TATGAGTGTGGAGTCGTTGCCGAATGTAGTTCCTCAGGGTAACNNNAGATCGGAAG |
| 30677_4 | TATGAGTGTGGAGTCGTTGCCGAATGTAGTTCCTCAGGGTAACNNNAGATCGGAAG |
| 30677_5 | TATGAGTGTGGAGTCGTTGCCGAATGTAGTTCCTCAGGGTAACNNNAGATCGGAAG |
| 49976_1 | TATGAGTGTGGAGTCGTTGCGACCACAAGATCCAGTACACAGTNNNAGATCGGAAG |
| 49976_2 | TATGAGTGTGGAGTCGTTGCGACCACAAGATCCAGTACACAGTNNNAGATCGGAAG |
| 49976_3 | TATGAGTGTGGAGTCGTTGCGACCACAAGATCCAGTACACAGTNNNAGATCGGAAG |
| 49976_4 | TATGAGTGTGGAGTCGTTGCGACCACAAGATCCAGTACACAGTNNNAGATCGGAAG |
| 49976_5 | TATGAGTGTGGAGTCGTTGCGACCACAAGATCCAGTACACAGTNNNAGATCGGAAG |
| 10030_1 | TATGAGTGTGGAGTCGTTGCATCTGTTCCACCACTTTGCTGTTNNNAGATCGGAAGA |
| 10030_2 | TATGAGTGTGGAGTCGTTGCATCTGTTCCACCACTTTGCTGTTNNNAGATCGGAAGA |
| 10030_3 | TATGAGTGTGGAGTCGTTGCATCTGTTCCACCACTTTGCTGTTNNNAGATCGGAAGA |
| 10030_4 | TATGAGTGTGGAGTCGTTGCATCTGTTCCACCACTTTGCTGTTNNNAGATCGGAAGA |
| 10030_5 | TATGAGTGTGGAGTCGTTGCATCTGTTCCACCACTTTGCTGTTNNNAGATCGGAAGA |
| 37558_1 | TATGAGTGTGGAGTCGTTGCTGCTTCCCACATTTTGCATGTAGNNNAGATCGGAAGA |
| 37558_2 | TATGAGTGTGGAGTCGTTGCTGCTTCCCACATTTTGCATGTAGNNNAGATCGGAAGA |
| 37558_3 | TATGAGTGTGGAGTCGTTGCTGCTTCCCACATTTTGCATGTAGNNNAGATCGGAAGA |
| 37558_4 | TATGAGTGTGGAGTCGTTGCTGCTTCCCACATTTTGCATGTAGNNNAGATCGGAAGA |
| 37558_5 | TATGAGTGTGGAGTCGTTGCTGCTTCCCACATTTTGCATGTAGNNNAGATCGGAAGA |
| 29005_1 | TATGAGTGTGGAGTCGTTGCCCAAATGGTCAGAAAGAGAGGGTNNNAGATCGGAA |
| 29005_2 | TATGAGTGTGGAGTCGTTGCCCAAATGGTCAGAAAGAGAGGGTNNNAGATCGGAA |
| 29005_3 | TATGAGTGTGGAGTCGTTGCCCAAATGGTCAGAAAGAGAGGGTNNNAGATCGGAA |
| 29005_4 | TATGAGTGTGGAGTCGTTGCCCAAATGGTCAGAAAGAGAGGGTNNNAGATCGGAA |
| 29005_5 | TATGAGTGTGGAGTCGTTGCCCAAATGGTCAGAAAGAGAGGGTNNNAGATCGGAA |
| 50497_1 | TATGAGTGTGGAGTCGTTGCTCCAAGGTATTGTTCAGACCTCANNNAGATCGGAAGA |
| 50497_2 | TATGAGTGTGGAGTCGTTGCTCCAAGGTATTGTTCAGACCTCANNNAGATCGGAAGA |
| 50497_3 | TATGAGTGTGGAGTCGTTGCTCCAAGGTATTGTTCAGACCTCANNNAGATCGGAAGA |
| 50497_4 | TATGAGTGTGGAGTCGTTGCTCCAAGGTATTGTTCAGACCTCANNNAGATCGGAAGA |
| 50497_5 | TATGAGTGTGGAGTCGTTGCTCCAAGGTATTGTTCAGACCTCANNNAGATCGGAAGA |
| 13539_1 | TATGAGTGTGGAGTCGTTGCAAATGTCATATATACCTAGCACANNNAGATCGGAAGA |
| 13539_2 | TATGAGTGTGGAGTCGTTGCAAATGTCATATATACCTAGCACANNNAGATCGGAAGA |
| 13539_3 | TATGAGTGTGGAGTCGTTGCAAATGTCATATATACCTAGCACANNNAGATCGGAAGA |
| 13539_4 | TATGAGTGTGGAGTCGTTGCAAATGTCATATATACCTAGCACANNNAGATCGGAAGA |
| 13539_5 | TATGAGTGTGGAGTCGTTGCAAATGTCATATATACCTAGCACANNNAGATCGGAAGA |
| 36516_1 | TATGAGTGTGGAGTCGTTGCCACCTGTAATCCCAGCAATTTGGNNNAGATCGGAAGA |
| 36516_2 | TATGAGTGTGGAGTCGTTGCCACCTGTAATCCCAGCAATTTGGNNNAGATCGGAAGA |
| 36516_3 | TATGAGTGTGGAGTCGTTGCCACCTGTAATCCCAGCAATTTGGNNNAGATCGGAAGA |

| | |
|---|---|
| 36516_4 | TATGAGTGTGGAGTCGTTGCCACCTGTAATCCCAGCAATTTGGNNNAGATCGGAAG |
| 36516_5 | TATGAGTGTGGAGTCGTTGCCACCTGTAATCCCAGCAATTTGGNNNAGATCGGAAG |
| 8146_1 | TATGAGTGTGGAGTCGTTGCGGAGACAAGAAGACAAGCTTTTGCNNNAGATCGGA |
| 8146_2 | TATGAGTGTGGAGTCGTTGCGGAGACAAGAAGACAAGCTTTTGCNNNAGATCGGA |
| 8146_3 | TATGAGTGTGGAGTCGTTGCGGAGACAAGAAGACAAGCTTTTGCNNNAGATCGGA |
| 8146_4 | TATGAGTGTGGAGTCGTTGCGGAGACAAGAAGACAAGCTTTTGCNNNAGATCGGA |
| 8146_5 | TATGAGTGTGGAGTCGTTGCGGAGACAAGAAGACAAGCTTTTGCNNNAGATCGGA |
| 43328_1 | TATGAGTGTGGAGTCGTTGCATTTTGGGTCTTTTGTGGTTCCANNNAGATCGGAAG |
| 43328_2 | TATGAGTGTGGAGTCGTTGCATTTTGGGTCTTTTGTGGTTCCANNNAGATCGGAAG |
| 43328_3 | TATGAGTGTGGAGTCGTTGCATTTTGGGTCTTTTGTGGTTCCANNNAGATCGGAAG |
| 43328_4 | TATGAGTGTGGAGTCGTTGCATTTTGGGTCTTTTGTGGTTCCANNNAGATCGGAAG |
| 43328_5 | TATGAGTGTGGAGTCGTTGCATTTTGGGTCTTTTGTGGTTCCANNNAGATCGGAAG |
| 48228_1 | TATGAGTGTGGAGTCGTTGCCTAGGTTCAAGCGATTCTCTTGCNNNAGATCGGAAG |
| 48228_2 | TATGAGTGTGGAGTCGTTGCCTAGGTTCAAGCGATTCTCTTGCNNNAGATCGGAAG |
| 48228_3 | TATGAGTGTGGAGTCGTTGCCTAGGTTCAAGCGATTCTCTTGCNNNAGATCGGAAG |
| 48228_4 | TATGAGTGTGGAGTCGTTGCCTAGGTTCAAGCGATTCTCTTGCNNNAGATCGGAAG |
| 48228_5 | TATGAGTGTGGAGTCGTTGCCTAGGTTCAAGCGATTCTCTTGCNNNAGATCGGAAG |
| 40711_1 | TATGAGTGTGGAGTCGTTGCTGGCCAACATGGTGAAACTACTANNNAGATCGGAAG |
| 40711_2 | TATGAGTGTGGAGTCGTTGCTGGCCAACATGGTGAAACTACTANNNAGATCGGAAG |
| 40711_3 | TATGAGTGTGGAGTCGTTGCTGGCCAACATGGTGAAACTACTANNNAGATCGGAAG |
| 40711_4 | TATGAGTGTGGAGTCGTTGCTGGCCAACATGGTGAAACTACTANNNAGATCGGAAG |
| 40711_5 | TATGAGTGTGGAGTCGTTGCTGGCCAACATGGTGAAACTACTANNNAGATCGGAAG |
| 34400_1 | TATGAGTGTGGAGTCGTTGCATATCTAAGCAGAGACGGAGGGTNNNAGATCGGAA |
| 34400_2 | TATGAGTGTGGAGTCGTTGCATATCTAAGCAGAGACGGAGGGTNNNAGATCGGAA |
| 34400_3 | TATGAGTGTGGAGTCGTTGCATATCTAAGCAGAGACGGAGGGTNNNAGATCGGAA |
| 34400_4 | TATGAGTGTGGAGTCGTTGCATATCTAAGCAGAGACGGAGGGTNNNAGATCGGAA |
| 34400_5 | TATGAGTGTGGAGTCGTTGCATATCTAAGCAGAGACGGAGGGTNNNAGATCGGAA |
| 46991_1 | TATGAGTGTGGAGTCGTTGCGGGAGGCCTAGATGAAACAAGAANNNAGATCGGA |
| 46991_2 | TATGAGTGTGGAGTCGTTGCGGGAGGCCTAGATGAAACAAGAANNNAGATCGGA |
| 46991_3 | TATGAGTGTGGAGTCGTTGCGGGAGGCCTAGATGAAACAAGAANNNAGATCGGA |
| 46991_4 | TATGAGTGTGGAGTCGTTGCGGGAGGCCTAGATGAAACAAGAANNNAGATCGGA |
| 46991_5 | TATGAGTGTGGAGTCGTTGCGGGAGGCCTAGATGAAACAAGAANNNAGATCGGA |
| 31857_1 | TATGAGTGTGGAGTCGTTGCCTTCTGTCACAGTGTGGCTAGATNNNAGATCGGAAG |
| 31857_2 | TATGAGTGTGGAGTCGTTGCCTTCTGTCACAGTGTGGCTAGATNNNAGATCGGAAG |
| 31857_3 | TATGAGTGTGGAGTCGTTGCCTTCTGTCACAGTGTGGCTAGATNNNAGATCGGAAG |
| 31857_4 | TATGAGTGTGGAGTCGTTGCCTTCTGTCACAGTGTGGCTAGATNNNAGATCGGAAG |
| 31857_5 | TATGAGTGTGGAGTCGTTGCCTTCTGTCACAGTGTGGCTAGATNNNAGATCGGAAG |
| 40406_1 | TATGAGTGTGGAGTCGTTGCACTCTTGCTTTTCTGGTTTGTGGNNNAGATCGGAAG |
| 40406_2 | TATGAGTGTGGAGTCGTTGCACTCTTGCTTTTCTGGTTTGTGGNNNAGATCGGAAG |
| 40406_3 | TATGAGTGTGGAGTCGTTGCACTCTTGCTTTTCTGGTTTGTGGNNNAGATCGGAAG |
| 40406_4 | TATGAGTGTGGAGTCGTTGCACTCTTGCTTTTCTGGTTTGTGGNNNAGATCGGAAG |
| 40406_5 | TATGAGTGTGGAGTCGTTGCACTCTTGCTTTTCTGGTTTGTGGNNNAGATCGGAAG |
| 52678_1 | TATGAGTGTGGAGTCGTTGCCCAGCTGGTCTTTTTACCCTTTGNNNAGATCGGAAGA |

| | |
|---|---|
| 52678_2 | TATGAGTGTGGAGTCGTTGCCCAGCTGGTCTTTTTACCCTTTGNNNAGATCGGAAGA |
| 52678_3 | TATGAGTGTGGAGTCGTTGCCCAGCTGGTCTTTTTACCCTTTGNNNAGATCGGAAGA |
| 52678_4 | TATGAGTGTGGAGTCGTTGCCCAGCTGGTCTTTTTACCCTTTGNNNAGATCGGAAGA |
| 52678_5 | TATGAGTGTGGAGTCGTTGCCCAGCTGGTCTTTTTACCCTTTGNNNAGATCGGAAGA |
| 35315_1 | TATGAGTGTGGAGTCGTTGCATTTGTGTTTCAAGAGCTGCCAGNNNAGATCGGAAG |
| 35315_2 | TATGAGTGTGGAGTCGTTGCATTTGTGTTTCAAGAGCTGCCAGNNNAGATCGGAAG |
| 35315_3 | TATGAGTGTGGAGTCGTTGCATTTGTGTTTCAAGAGCTGCCAGNNNAGATCGGAAG |
| 35315_4 | TATGAGTGTGGAGTCGTTGCATTTGTGTTTCAAGAGCTGCCAGNNNAGATCGGAAG |
| 35315_5 | TATGAGTGTGGAGTCGTTGCATTTGTGTTTCAAGAGCTGCCAGNNNAGATCGGAAG |
| 49907_1 | TATGAGTGTGGAGTCGTTGCGGGAAGGATTCTACAACCCAACTNNNAGATCGGAAG |
| 49907_2 | TATGAGTGTGGAGTCGTTGCGGGAAGGATTCTACAACCCAACTNNNAGATCGGAAG |
| 49907_3 | TATGAGTGTGGAGTCGTTGCGGGAAGGATTCTACAACCCAACTNNNAGATCGGAAG |
| 49907_4 | TATGAGTGTGGAGTCGTTGCGGGAAGGATTCTACAACCCAACTNNNAGATCGGAAG |
| 49907_5 | TATGAGTGTGGAGTCGTTGCGGGAAGGATTCTACAACCCAACTNNNAGATCGGAAG |
| 39360_1 | TATGAGTGTGGAGTCGTTGCGAAGTGGAGATATGGGCATCAGANNNAGATCGGAA |
| 39360_2 | TATGAGTGTGGAGTCGTTGCGAAGTGGAGATATGGGCATCAGANNNAGATCGGAA |
| 39360_3 | TATGAGTGTGGAGTCGTTGCGAAGTGGAGATATGGGCATCAGANNNAGATCGGAA |
| 39360_4 | TATGAGTGTGGAGTCGTTGCGAAGTGGAGATATGGGCATCAGANNNAGATCGGAA |
| 39360_5 | TATGAGTGTGGAGTCGTTGCGAAGTGGAGATATGGGCATCAGANNNAGATCGGAA |
| 13320_1 | TATGAGTGTGGAGTCGTTGCTCCTCTAACAATCTTATTCAAGANNNAGATCGGAAGA |
| 13320_2 | TATGAGTGTGGAGTCGTTGCTCCTCTAACAATCTTATTCAAGANNNAGATCGGAAGA |
| 13320_3 | TATGAGTGTGGAGTCGTTGCTCCTCTAACAATCTTATTCAAGANNNAGATCGGAAGA |
| 13320_4 | TATGAGTGTGGAGTCGTTGCTCCTCTAACAATCTTATTCAAGANNNAGATCGGAAGA |
| 13320_5 | TATGAGTGTGGAGTCGTTGCTCCTCTAACAATCTTATTCAAGANNNAGATCGGAAGA |
| 34952_1 | TATGAGTGTGGAGTCGTTGCTTCAAGTGATTCTCCTGCCTCAGNNNAGATCGGAAGA |
| 34952_2 | TATGAGTGTGGAGTCGTTGCTTCAAGTGATTCTCCTGCCTCAGNNNAGATCGGAAGA |
| 34952_3 | TATGAGTGTGGAGTCGTTGCTTCAAGTGATTCTCCTGCCTCAGNNNAGATCGGAAGA |
| 34952_4 | TATGAGTGTGGAGTCGTTGCTTCAAGTGATTCTCCTGCCTCAGNNNAGATCGGAAGA |
| 34952_5 | TATGAGTGTGGAGTCGTTGCTTCAAGTGATTCTCCTGCCTCAGNNNAGATCGGAAGA |
| 35347_1 | TATGAGTGTGGAGTCGTTGCATGATCTCAGCTCACTGCAACAGNNNAGATCGGAAG |
| 35347_2 | TATGAGTGTGGAGTCGTTGCATGATCTCAGCTCACTGCAACAGNNNAGATCGGAAG |
| 35347_3 | TATGAGTGTGGAGTCGTTGCATGATCTCAGCTCACTGCAACAGNNNAGATCGGAAG |
| 35347_4 | TATGAGTGTGGAGTCGTTGCATGATCTCAGCTCACTGCAACAGNNNAGATCGGAAG |
| 35347_5 | TATGAGTGTGGAGTCGTTGCATGATCTCAGCTCACTGCAACAGNNNAGATCGGAAG |
| 34751_1 | TATGAGTGTGGAGTCGTTGCTTTTTCTGCTGGACACTCTACGTNNNAGATCGGAAGA |
| 34751_2 | TATGAGTGTGGAGTCGTTGCTTTTTCTGCTGGACACTCTACGTNNNAGATCGGAAGA |
| 34751_3 | TATGAGTGTGGAGTCGTTGCTTTTTCTGCTGGACACTCTACGTNNNAGATCGGAAGA |
| 34751_4 | TATGAGTGTGGAGTCGTTGCTTTTTCTGCTGGACACTCTACGTNNNAGATCGGAAGA |
| 34751_5 | TATGAGTGTGGAGTCGTTGCTTTTTCTGCTGGACACTCTACGTNNNAGATCGGAAGA |
| 30618_1 | TATGAGTGTGGAGTCGTTGCTTTGCTGTTTTGACAGAGTCTCACNNNAGATCGGAA |
| 30618_2 | TATGAGTGTGGAGTCGTTGCTTTGCTGTTTTGACAGAGTCTCACNNNAGATCGGAA |
| 30618_3 | TATGAGTGTGGAGTCGTTGCTTTGCTGTTTTGACAGAGTCTCACNNNAGATCGGAA |
| 30618_4 | TATGAGTGTGGAGTCGTTGCTTTGCTGTTTTGACAGAGTCTCACNNNAGATCGGAA |

| | |
|---|---|
| 30618_5 | TATGAGTGTGGAGTCGTTGCTTTTGCTGTTTTGACAGAGTCTCACNNNAGATCGGAA |
| 13616_1 | TATGAGTGTGGAGTCGTTGCCCTAGTTGTCACCTTTGTATAATNNNAGATCGGAAGA |
| 13616_2 | TATGAGTGTGGAGTCGTTGCCCTAGTTGTCACCTTTGTATAATNNNAGATCGGAAGA |
| 13616_3 | TATGAGTGTGGAGTCGTTGCCCTAGTTGTCACCTTTGTATAATNNNAGATCGGAAGA |
| 13616_4 | TATGAGTGTGGAGTCGTTGCCCTAGTTGTCACCTTTGTATAATNNNAGATCGGAAGA |
| 13616_5 | TATGAGTGTGGAGTCGTTGCCCTAGTTGTCACCTTTGTATAATNNNAGATCGGAAGA |
| 37759_1 | TATGAGTGTGGAGTCGTTGCTATGAAAACCTACAACTGACGGCNNNAGATCGGAAG |
| 37759_2 | TATGAGTGTGGAGTCGTTGCTATGAAAACCTACAACTGACGGCNNNAGATCGGAAG |
| 37759_3 | TATGAGTGTGGAGTCGTTGCTATGAAAACCTACAACTGACGGCNNNAGATCGGAAG |
| 37759_4 | TATGAGTGTGGAGTCGTTGCTATGAAAACCTACAACTGACGGCNNNAGATCGGAAG |
| 37759_5 | TATGAGTGTGGAGTCGTTGCTATGAAAACCTACAACTGACGGCNNNAGATCGGAAG |
| 51251_1 | TATGAGTGTGGAGTCGTTGCAGAGAGGAGCAAAGTCAGTGTACNNNAGATCGGAA |
| 51251_2 | TATGAGTGTGGAGTCGTTGCAGAGAGGAGCAAAGTCAGTGTACNNNAGATCGGAA |
| 51251_3 | TATGAGTGTGGAGTCGTTGCAGAGAGGAGCAAAGTCAGTGTACNNNAGATCGGAA |
| 51251_4 | TATGAGTGTGGAGTCGTTGCAGAGAGGAGCAAAGTCAGTGTACNNNAGATCGGAA |
| 51251_5 | TATGAGTGTGGAGTCGTTGCAGAGAGGAGCAAAGTCAGTGTACNNNAGATCGGAA |
| 44831_1 | TATGAGTGTGGAGTCGTTGCTTTCTGACTCTGCTTTTGCCTTGNNNAGATCGGAAGA |
| 44831_2 | TATGAGTGTGGAGTCGTTGCTTTCTGACTCTGCTTTTGCCTTGNNNAGATCGGAAGA |
| 44831_3 | TATGAGTGTGGAGTCGTTGCTTTCTGACTCTGCTTTTGCCTTGNNNAGATCGGAAGA |
| 44831_4 | TATGAGTGTGGAGTCGTTGCTTTCTGACTCTGCTTTTGCCTTGNNNAGATCGGAAGA |
| 44831_5 | TATGAGTGTGGAGTCGTTGCTTTCTGACTCTGCTTTTGCCTTGNNNAGATCGGAAGA |
| 48929_1 | TATGAGTGTGGAGTCGTTGCCAACCCCTTTTCATGCATCCATGNNNAGATCGGAAGA |
| 48929_2 | TATGAGTGTGGAGTCGTTGCCAACCCCTTTTCATGCATCCATGNNNAGATCGGAAGA |
| 48929_3 | TATGAGTGTGGAGTCGTTGCCAACCCCTTTTCATGCATCCATGNNNAGATCGGAAGA |
| 48929_4 | TATGAGTGTGGAGTCGTTGCCAACCCCTTTTCATGCATCCATGNNNAGATCGGAAGA |
| 48929_5 | TATGAGTGTGGAGTCGTTGCCAACCCCTTTTCATGCATCCATGNNNAGATCGGAAGA |
| 43061_1 | TATGAGTGTGGAGTCGTTGCACTGTAAGAACAAACAGCCTGAGNNNAGATCGGAA |
| 43061_2 | TATGAGTGTGGAGTCGTTGCACTGTAAGAACAAACAGCCTGAGNNNAGATCGGAA |
| 43061_3 | TATGAGTGTGGAGTCGTTGCACTGTAAGAACAAACAGCCTGAGNNNAGATCGGAA |
| 43061_4 | TATGAGTGTGGAGTCGTTGCACTGTAAGAACAAACAGCCTGAGNNNAGATCGGAA |
| 43061_5 | TATGAGTGTGGAGTCGTTGCACTGTAAGAACAAACAGCCTGAGNNNAGATCGGAA |
| 40613_1 | TATGAGTGTGGAGTCGTTGCAAATGGTCCTCTCACACATGGTTNNNAGATCGGAAG |
| 40613_2 | TATGAGTGTGGAGTCGTTGCAAATGGTCCTCTCACACATGGTTNNNAGATCGGAAG |
| 40613_3 | TATGAGTGTGGAGTCGTTGCAAATGGTCCTCTCACACATGGTTNNNAGATCGGAAG |
| 40613_4 | TATGAGTGTGGAGTCGTTGCAAATGGTCCTCTCACACATGGTTNNNAGATCGGAAG |
| 40613_5 | TATGAGTGTGGAGTCGTTGCAAATGGTCCTCTCACACATGGTTNNNAGATCGGAAG |
| 34737_1 | TATGAGTGTGGAGTCGTTGCCAGTGGTGCAAACACAGCTTGNNNAGATCGGAAGA |
| 34737_2 | TATGAGTGTGGAGTCGTTGCCAGTGGTGCAAACACAGCTTGNNNAGATCGGAAGA |
| 34737_3 | TATGAGTGTGGAGTCGTTGCCAGTGGTGCAAACACAGCTTGNNNAGATCGGAAGA |
| 34737_4 | TATGAGTGTGGAGTCGTTGCCAGTGGTGCAAACACAGCTTGNNNAGATCGGAAGA |
| 34737_5 | TATGAGTGTGGAGTCGTTGCCAGTGGTGCAAACACAGCTTGNNNAGATCGGAAGA |
| 51063_1 | TATGAGTGTGGAGTCGTTGCTTCAGACCATTTGAGAAGAGGGGNNNAGATCGGAA |
| 51063_2 | TATGAGTGTGGAGTCGTTGCTTCAGACCATTTGAGAAGAGGGGNNNAGATCGGAA |

| | |
|---|---|
| 51063_3 | TATGAGTGTGGAGTCGTTGCTTCAGACCATTTGAGAAGAGGGGNNNAGATCGGAA |
| 51063_4 | TATGAGTGTGGAGTCGTTGCTTCAGACCATTTGAGAAGAGGGGNNNAGATCGGAA |
| 51063_5 | TATGAGTGTGGAGTCGTTGCTTCAGACCATTTGAGAAGAGGGGNNNAGATCGGAA |
| 37956_1 | TATGAGTGTGGAGTCGTTGCATCCTAACCTCATCTCCCAGTCTNNNAGATCGGAAGA |
| 37956_2 | TATGAGTGTGGAGTCGTTGCATCCTAACCTCATCTCCCAGTCTNNNAGATCGGAAGA |
| 37956_3 | TATGAGTGTGGAGTCGTTGCATCCTAACCTCATCTCCCAGTCTNNNAGATCGGAAGA |
| 37956_4 | TATGAGTGTGGAGTCGTTGCATCCTAACCTCATCTCCCAGTCTNNNAGATCGGAAGA |
| 37956_5 | TATGAGTGTGGAGTCGTTGCATCCTAACCTCATCTCCCAGTCTNNNAGATCGGAAGA |
| 52305_1 | TATGAGTGTGGAGTCGTTGCCTTCAGCCTTTCCCCAGGATCNNNAGATCGGAAGAG |
| 52305_2 | TATGAGTGTGGAGTCGTTGCCTTCAGCCTTTCCCCAGGATCNNNAGATCGGAAGAG |
| 52305_3 | TATGAGTGTGGAGTCGTTGCCTTCAGCCTTTCCCCAGGATCNNNAGATCGGAAGAG |
| 52305_4 | TATGAGTGTGGAGTCGTTGCCTTCAGCCTTTCCCCAGGATCNNNAGATCGGAAGAG |
| 52305_5 | TATGAGTGTGGAGTCGTTGCCTTCAGCCTTTCCCCAGGATCNNNAGATCGGAAGAG |
| 36272_1 | TATGAGTGTGGAGTCGTTGCTCTGCTGCCTGTGGATAAGATACNNNAGATCGGAAG |
| 36272_2 | TATGAGTGTGGAGTCGTTGCTCTGCTGCCTGTGGATAAGATACNNNAGATCGGAAG |
| 36272_3 | TATGAGTGTGGAGTCGTTGCTCTGCTGCCTGTGGATAAGATACNNNAGATCGGAAG |
| 36272_4 | TATGAGTGTGGAGTCGTTGCTCTGCTGCCTGTGGATAAGATACNNNAGATCGGAAG |
| 36272_5 | TATGAGTGTGGAGTCGTTGCTCTGCTGCCTGTGGATAAGATACNNNAGATCGGAAG |
| 13445_1 | TATGAGTGTGGAGTCGTTGCTTTATACCTGATTTTTCCAACACNNNAGATCGGAAGA |
| 13445_2 | TATGAGTGTGGAGTCGTTGCTTTATACCTGATTTTTCCAACACNNNAGATCGGAAGA |
| 13445_3 | TATGAGTGTGGAGTCGTTGCTTTATACCTGATTTTTCCAACACNNNAGATCGGAAGA |
| 13445_4 | TATGAGTGTGGAGTCGTTGCTTTATACCTGATTTTTCCAACACNNNAGATCGGAAGA |
| 13445_5 | TATGAGTGTGGAGTCGTTGCTTTATACCTGATTTTTCCAACACNNNAGATCGGAAGA |
| 13416_1 | TATGAGTGTGGAGTCGTTGCCACCATTCATTTCATTTTGTTTTNNNAGATCGGAAGA |
| 13416_2 | TATGAGTGTGGAGTCGTTGCCACCATTCATTTCATTTTGTTTTNNNAGATCGGAAGA |
| 13416_3 | TATGAGTGTGGAGTCGTTGCCACCATTCATTTCATTTTGTTTTNNNAGATCGGAAGA |
| 13416_4 | TATGAGTGTGGAGTCGTTGCCACCATTCATTTCATTTTGTTTTNNNAGATCGGAAGA |
| 13416_5 | TATGAGTGTGGAGTCGTTGCCACCATTCATTTCATTTTGTTTTNNNAGATCGGAAGA |
| 39805_1 | TATGAGTGTGGAGTCGTTGCCCACCACCAAAAAAAGAGACAACTNNNAGATCGGAA |
| 39805_2 | TATGAGTGTGGAGTCGTTGCCCACCACCAAAAAAAGAGACAACTNNNAGATCGGAA |
| 39805_3 | TATGAGTGTGGAGTCGTTGCCCACCACCAAAAAAAGAGACAACTNNNAGATCGGAA |
| 39805_4 | TATGAGTGTGGAGTCGTTGCCCACCACCAAAAAAAGAGACAACTNNNAGATCGGAA |
| 39805_5 | TATGAGTGTGGAGTCGTTGCCCACCACCAAAAAAAGAGACAACTNNNAGATCGGAA |
| 52946_1 | TATGAGTGTGGAGTCGTTGCCTCCATGAGTTGTCCTGCTACATNNNAGATCGGAAG |
| 52946_2 | TATGAGTGTGGAGTCGTTGCCTCCATGAGTTGTCCTGCTACATNNNAGATCGGAAG |
| 52946_3 | TATGAGTGTGGAGTCGTTGCCTCCATGAGTTGTCCTGCTACATNNNAGATCGGAAG |
| 52946_4 | TATGAGTGTGGAGTCGTTGCCTCCATGAGTTGTCCTGCTACATNNNAGATCGGAAG |
| 52946_5 | TATGAGTGTGGAGTCGTTGCCTCCATGAGTTGTCCTGCTACATNNNAGATCGGAAG |
| 47806_1 | TATGAGTGTGGAGTCGTTGCCTCCTGAGTAGCTGGGATTACAGNNNAGATCGGAA |
| 47806_2 | TATGAGTGTGGAGTCGTTGCCTCCTGAGTAGCTGGGATTACAGNNNAGATCGGAA |
| 47806_3 | TATGAGTGTGGAGTCGTTGCCTCCTGAGTAGCTGGGATTACAGNNNAGATCGGAA |
| 47806_4 | TATGAGTGTGGAGTCGTTGCCTCCTGAGTAGCTGGGATTACAGNNNAGATCGGAA |
| 47806_5 | TATGAGTGTGGAGTCGTTGCCTCCTGAGTAGCTGGGATTACAGNNNAGATCGGAA |

| | |
|---|---|
| 34172_1 | TATGAGTGTGGAGTCGTTGCAAAGGGATCACTGCTTCTGAGAANNNAGATCGGAA |
| 34172_2 | TATGAGTGTGGAGTCGTTGCAAAGGGATCACTGCTTCTGAGAANNNAGATCGGAA |
| 34172_3 | TATGAGTGTGGAGTCGTTGCAAAGGGATCACTGCTTCTGAGAANNNAGATCGGAA |
| 34172_4 | TATGAGTGTGGAGTCGTTGCAAAGGGATCACTGCTTCTGAGAANNNAGATCGGAA |
| 34172_5 | TATGAGTGTGGAGTCGTTGCAAAGGGATCACTGCTTCTGAGAANNNAGATCGGAA |
| 29338_1 | TATGAGTGTGGAGTCGTTGCTGATGGACACTTAGGTTGATTCCANNNAGATCGGAA |
| 29338_2 | TATGAGTGTGGAGTCGTTGCTGATGGACACTTAGGTTGATTCCANNNAGATCGGAA |
| 29338_3 | TATGAGTGTGGAGTCGTTGCTGATGGACACTTAGGTTGATTCCANNNAGATCGGAA |
| 29338_4 | TATGAGTGTGGAGTCGTTGCTGATGGACACTTAGGTTGATTCCANNNAGATCGGAA |
| 29338_5 | TATGAGTGTGGAGTCGTTGCTGATGGACACTTAGGTTGATTCCANNNAGATCGGAA |
| 9265_1 | TATGAGTGTGGAGTCGTTGCTTGTGACCCTCCTAAGGATCCTGNNNAGATCGGAAG |
| 9265_2 | TATGAGTGTGGAGTCGTTGCTTGTGACCCTCCTAAGGATCCTGNNNAGATCGGAAG |
| 9265_3 | TATGAGTGTGGAGTCGTTGCTTGTGACCCTCCTAAGGATCCTGNNNAGATCGGAAG |
| 9265_4 | TATGAGTGTGGAGTCGTTGCTTGTGACCCTCCTAAGGATCCTGNNNAGATCGGAAG |
| 9265_5 | TATGAGTGTGGAGTCGTTGCTTGTGACCCTCCTAAGGATCCTGNNNAGATCGGAAG |
| 37719_1 | TATGAGTGTGGAGTCGTTGCAAAATGAGACCCAGTCATTTGCANNNAGATCGGAAG |
| 37719_2 | TATGAGTGTGGAGTCGTTGCAAAATGAGACCCAGTCATTTGCANNNAGATCGGAAG |
| 37719_3 | TATGAGTGTGGAGTCGTTGCAAAATGAGACCCAGTCATTTGCANNNAGATCGGAAG |
| 37719_4 | TATGAGTGTGGAGTCGTTGCAAAATGAGACCCAGTCATTTGCANNNAGATCGGAAG |
| 37719_5 | TATGAGTGTGGAGTCGTTGCAAAATGAGACCCAGTCATTTGCANNNAGATCGGAAG |
| 35910_1 | TATGAGTGTGGAGTCGTTGCAGCTACAGAGACAGGGGATAAGTNNNAGATCGGAA |
| 35910_2 | TATGAGTGTGGAGTCGTTGCAGCTACAGAGACAGGGGATAAGTNNNAGATCGGAA |
| 35910_3 | TATGAGTGTGGAGTCGTTGCAGCTACAGAGACAGGGGATAAGTNNNAGATCGGAA |
| 35910_4 | TATGAGTGTGGAGTCGTTGCAGCTACAGAGACAGGGGATAAGTNNNAGATCGGAA |
| 35910_5 | TATGAGTGTGGAGTCGTTGCAGCTACAGAGACAGGGGATAAGTNNNAGATCGGAA |
| 43836_1 | TATGAGTGTGGAGTCGTTGCAGAGAGAGAGAACGCACTAGCTANNNAGATCGGAA |
| 43836_2 | TATGAGTGTGGAGTCGTTGCAGAGAGAGAGAACGCACTAGCTANNNAGATCGGAA |
| 43836_3 | TATGAGTGTGGAGTCGTTGCAGAGAGAGAGAACGCACTAGCTANNNAGATCGGAA |
| 43836_4 | TATGAGTGTGGAGTCGTTGCAGAGAGAGAGAACGCACTAGCTANNNAGATCGGAA |
| 43836_5 | TATGAGTGTGGAGTCGTTGCAGAGAGAGAGAACGCACTAGCTANNNAGATCGGAA |
| 44694_1 | TATGAGTGTGGAGTCGTTGCAGCGAGTTGTGAAGACTGAAGATNNNAGATCGGAA |
| 44694_2 | TATGAGTGTGGAGTCGTTGCAGCGAGTTGTGAAGACTGAAGATNNNAGATCGGAA |
| 44694_3 | TATGAGTGTGGAGTCGTTGCAGCGAGTTGTGAAGACTGAAGATNNNAGATCGGAA |
| 44694_4 | TATGAGTGTGGAGTCGTTGCAGCGAGTTGTGAAGACTGAAGATNNNAGATCGGAA |
| 44694_5 | TATGAGTGTGGAGTCGTTGCAGCGAGTTGTGAAGACTGAAGATNNNAGATCGGAA |
| 37595_1 | TATGAGTGTGGAGTCGTTGCACAAGAAGACCATAAAGGCCCATNNNAGATCGGAA |
| 37595_2 | TATGAGTGTGGAGTCGTTGCACAAGAAGACCATAAAGGCCCATNNNAGATCGGAA |
| 37595_3 | TATGAGTGTGGAGTCGTTGCACAAGAAGACCATAAAGGCCCATNNNAGATCGGAA |
| 37595_4 | TATGAGTGTGGAGTCGTTGCACAAGAAGACCATAAAGGCCCATNNNAGATCGGAA |
| 37595_5 | TATGAGTGTGGAGTCGTTGCACAAGAAGACCATAAAGGCCCATNNNAGATCGGAA |
| 41190_1 | TATGAGTGTGGAGTCGTTGCACATGGTGAAACCTCGTCTCTACNNNAGATCGGAAG |
| 41190_2 | TATGAGTGTGGAGTCGTTGCACATGGTGAAACCTCGTCTCTACNNNAGATCGGAAG |
| 41190_3 | TATGAGTGTGGAGTCGTTGCACATGGTGAAACCTCGTCTCTACNNNAGATCGGAAG |

| | |
|---|---|
| 41190_4 | TATGAGTGTGGAGTCGTTGCACATGGTGAAACCTCGTCTCTACNNNAGATCGGAAG |
| 41190_5 | TATGAGTGTGGAGTCGTTGCACATGGTGAAACCTCGTCTCTACNNNAGATCGGAAG |
| 5031_1 | TATGAGTGTGGAGTCGTTGCAAAGTTACAGCAATCTGGGTGGANNNAGATCGGAA |
| 5031_2 | TATGAGTGTGGAGTCGTTGCAAAGTTACAGCAATCTGGGTGGANNNAGATCGGAA |
| 5031_3 | TATGAGTGTGGAGTCGTTGCAAAGTTACAGCAATCTGGGTGGANNNAGATCGGAA |
| 5031_4 | TATGAGTGTGGAGTCGTTGCAAAGTTACAGCAATCTGGGTGGANNNAGATCGGAA |
| 5031_5 | TATGAGTGTGGAGTCGTTGCAAAGTTACAGCAATCTGGGTGGANNNAGATCGGAA |
| 49438_1 | TATGAGTGTGGAGTCGTTGCCATGCCCGGCTAAGTTTTTGTATNNNAGATCGGAAG |
| 49438_2 | TATGAGTGTGGAGTCGTTGCCATGCCCGGCTAAGTTTTTGTATNNNAGATCGGAAG |
| 49438_3 | TATGAGTGTGGAGTCGTTGCCATGCCCGGCTAAGTTTTTGTATNNNAGATCGGAAG |
| 49438_4 | TATGAGTGTGGAGTCGTTGCCATGCCCGGCTAAGTTTTTGTATNNNAGATCGGAAG |
| 49438_5 | TATGAGTGTGGAGTCGTTGCCATGCCCGGCTAAGTTTTTGTATNNNAGATCGGAAG |
| 5373_1 | TATGAGTGTGGAGTCGTTGCGCCAGTTCAAGCTGTGTTGTTTTNNNAGATCGGAAG |
| 5373_2 | TATGAGTGTGGAGTCGTTGCGCCAGTTCAAGCTGTGTTGTTTTNNNAGATCGGAAG |
| 5373_3 | TATGAGTGTGGAGTCGTTGCGCCAGTTCAAGCTGTGTTGTTTTNNNAGATCGGAAG |
| 5373_4 | TATGAGTGTGGAGTCGTTGCGCCAGTTCAAGCTGTGTTGTTTTNNNAGATCGGAAG |
| 5373_5 | TATGAGTGTGGAGTCGTTGCGCCAGTTCAAGCTGTGTTGTTTTNNNAGATCGGAAG |
| 37388_1 | TATGAGTGTGGAGTCGTTGCGGAAGGAAGACACCATCTTGGAANNNAGATCGGAA |
| 37388_2 | TATGAGTGTGGAGTCGTTGCGGAAGGAAGACACCATCTTGGAANNNAGATCGGAA |
| 37388_3 | TATGAGTGTGGAGTCGTTGCGGAAGGAAGACACCATCTTGGAANNNAGATCGGAA |
| 37388_4 | TATGAGTGTGGAGTCGTTGCGGAAGGAAGACACCATCTTGGAANNNAGATCGGAA |
| 37388_5 | TATGAGTGTGGAGTCGTTGCGGAAGGAAGACACCATCTTGGAANNNAGATCGGAA |
| 51640_1 | TATGAGTGTGGAGTCGTTGCGAAAACAGCTTGGGGAAAGTGAGNNNAGATCGGA |
| 51640_2 | TATGAGTGTGGAGTCGTTGCGAAAACAGCTTGGGGAAAGTGAGNNNAGATCGGA |
| 51640_3 | TATGAGTGTGGAGTCGTTGCGAAAACAGCTTGGGGAAAGTGAGNNNAGATCGGA |
| 51640_4 | TATGAGTGTGGAGTCGTTGCGAAAACAGCTTGGGGAAAGTGAGNNNAGATCGGA |
| 51640_5 | TATGAGTGTGGAGTCGTTGCGAAAACAGCTTGGGGAAAGTGAGNNNAGATCGGA |
| 48306_1 | TATGAGTGTGGAGTCGTTGCCCTCATGGGTTCACACCATTCTCNNNAGATCGGAAGA |
| 48306_2 | TATGAGTGTGGAGTCGTTGCCCTCATGGGTTCACACCATTCTCNNNAGATCGGAAGA |
| 48306_3 | TATGAGTGTGGAGTCGTTGCCCTCATGGGTTCACACCATTCTCNNNAGATCGGAAGA |
| 48306_4 | TATGAGTGTGGAGTCGTTGCCCTCATGGGTTCACACCATTCTCNNNAGATCGGAAGA |
| 48306_5 | TATGAGTGTGGAGTCGTTGCCCTCATGGGTTCACACCATTCTCNNNAGATCGGAAGA |
| 43895_1 | TATGAGTGTGGAGTCGTTGCTATTACATACGTGGGGGGCTTTTNNNAGATCGGAAG |
| 43895_2 | TATGAGTGTGGAGTCGTTGCTATTACATACGTGGGGGGCTTTTNNNAGATCGGAAG |
| 43895_3 | TATGAGTGTGGAGTCGTTGCTATTACATACGTGGGGGGCTTTTNNNAGATCGGAAG |
| 43895_4 | TATGAGTGTGGAGTCGTTGCTATTACATACGTGGGGGGCTTTTNNNAGATCGGAAG |
| 43895_5 | TATGAGTGTGGAGTCGTTGCTATTACATACGTGGGGGGCTTTTNNNAGATCGGAAG |
| 29989_1 | TATGAGTGTGGAGTCGTTGCTGCTCTTCTTCGAAGTAAAATGTGTNNNAGATCGGA |
| 29989_2 | TATGAGTGTGGAGTCGTTGCTGCTCTTCTTCGAAGTAAAATGTGTNNNAGATCGGA |
| 29989_3 | TATGAGTGTGGAGTCGTTGCTGCTCTTCTTCGAAGTAAAATGTGTNNNAGATCGGA |
| 29989_4 | TATGAGTGTGGAGTCGTTGCTGCTCTTCTTCGAAGTAAAATGTGTNNNAGATCGGA |
| 29989_5 | TATGAGTGTGGAGTCGTTGCTGCTCTTCTTCGAAGTAAAATGTGTNNNAGATCGGA |
| 43713_1 | TATGAGTGTGGAGTCGTTGCTCTCCCTGAGAAGTTGAAAACCANNNAGATCGGAAG |

| | |
|---|---|
| 43713_2 | TATGAGTGTGGAGTCGTTGCTCTCCCTGAGAAGTTGAAAACCANNNAGATCGGAAG |
| 43713_3 | TATGAGTGTGGAGTCGTTGCTCTCCCTGAGAAGTTGAAAACCANNNAGATCGGAAG |
| 43713_4 | TATGAGTGTGGAGTCGTTGCTCTCCCTGAGAAGTTGAAAACCANNNAGATCGGAAG |
| 43713_5 | TATGAGTGTGGAGTCGTTGCTCTCCCTGAGAAGTTGAAAACCANNNAGATCGGAAG |
| 47461_1 | TATGAGTGTGGAGTCGTTGCGCAAAGATCTGTGTAGGGGAAGANNNAGATCGGAA |
| 47461_2 | TATGAGTGTGGAGTCGTTGCGCAAAGATCTGTGTAGGGGAAGANNNAGATCGGAA |
| 47461_3 | TATGAGTGTGGAGTCGTTGCGCAAAGATCTGTGTAGGGGAAGANNNAGATCGGAA |
| 47461_4 | TATGAGTGTGGAGTCGTTGCGCAAAGATCTGTGTAGGGGAAGANNNAGATCGGAA |
| 47461_5 | TATGAGTGTGGAGTCGTTGCGCAAAGATCTGTGTAGGGGAAGANNNAGATCGGAA |
| 43517_1 | TATGAGTGTGGAGTCGTTGCCCCAGGTTCAAGCAATTCTCTTGNNNAGATCGGAAGA |
| 43517_2 | TATGAGTGTGGAGTCGTTGCCCCAGGTTCAAGCAATTCTCTTGNNNAGATCGGAAGA |
| 43517_3 | TATGAGTGTGGAGTCGTTGCCCCAGGTTCAAGCAATTCTCTTGNNNAGATCGGAAGA |
| 43517_4 | TATGAGTGTGGAGTCGTTGCCCCAGGTTCAAGCAATTCTCTTGNNNAGATCGGAAGA |
| 43517_5 | TATGAGTGTGGAGTCGTTGCCCCAGGTTCAAGCAATTCTCTTGNNNAGATCGGAAGA |
| 36773_1 | TATGAGTGTGGAGTCGTTGCTCCTGGCCTCTGTTCTAATTGTTNNNAGATCGGAAGA |
| 36773_2 | TATGAGTGTGGAGTCGTTGCTCCTGGCCTCTGTTCTAATTGTTNNNAGATCGGAAGA |
| 36773_3 | TATGAGTGTGGAGTCGTTGCTCCTGGCCTCTGTTCTAATTGTTNNNAGATCGGAAGA |
| 36773_4 | TATGAGTGTGGAGTCGTTGCTCCTGGCCTCTGTTCTAATTGTTNNNAGATCGGAAGA |
| 36773_5 | TATGAGTGTGGAGTCGTTGCTCCTGGCCTCTGTTCTAATTGTTNNNAGATCGGAAGA |
| 13468_1 | TATGAGTGTGGAGTCGTTGCGAGAAAGAAGCAATTGTGATGNNNAGATCGGAAGA |
| 13468_2 | TATGAGTGTGGAGTCGTTGCGAGAAAGAAGCAATTGTGATGNNNAGATCGGAAGA |
| 13468_3 | TATGAGTGTGGAGTCGTTGCGAGAAAGAAGCAATTGTGATGNNNAGATCGGAAGA |
| 13468_4 | TATGAGTGTGGAGTCGTTGCGAGAAAGAAGCAATTGTGATGNNNAGATCGGAAGA |
| 13468_5 | TATGAGTGTGGAGTCGTTGCGAGAAAGAAGCAATTGTGATGNNNAGATCGGAAGA |
| 52337_1 | TATGAGTGTGGAGTCGTTGCTATGGACACCCTGCAGATACATGNNNAGATCGGAAG |
| 52337_2 | TATGAGTGTGGAGTCGTTGCTATGGACACCCTGCAGATACATGNNNAGATCGGAAG |
| 52337_3 | TATGAGTGTGGAGTCGTTGCTATGGACACCCTGCAGATACATGNNNAGATCGGAAG |
| 52337_4 | TATGAGTGTGGAGTCGTTGCTATGGACACCCTGCAGATACATGNNNAGATCGGAAG |
| 52337_5 | TATGAGTGTGGAGTCGTTGCTATGGACACCCTGCAGATACATGNNNAGATCGGAAG |
| 38128_1 | TATGAGTGTGGAGTCGTTGCCAGGCATTGAACACACCAGTTAGNNNAGATCGGAAG |
| 38128_2 | TATGAGTGTGGAGTCGTTGCCAGGCATTGAACACACCAGTTAGNNNAGATCGGAAG |
| 38128_3 | TATGAGTGTGGAGTCGTTGCCAGGCATTGAACACACCAGTTAGNNNAGATCGGAAG |
| 38128_4 | TATGAGTGTGGAGTCGTTGCCAGGCATTGAACACACCAGTTAGNNNAGATCGGAAG |
| 38128_5 | TATGAGTGTGGAGTCGTTGCCAGGCATTGAACACACCAGTTAGNNNAGATCGGAAG |
| 49304_1 | TATGAGTGTGGAGTCGTTGCCAACTGGGACTATAGGCACCAGNNNAGATCGGAAGA |
| 49304_2 | TATGAGTGTGGAGTCGTTGCCAACTGGGACTATAGGCACCAGNNNAGATCGGAAGA |
| 49304_3 | TATGAGTGTGGAGTCGTTGCCAACTGGGACTATAGGCACCAGNNNAGATCGGAAGA |
| 49304_4 | TATGAGTGTGGAGTCGTTGCCAACTGGGACTATAGGCACCAGNNNAGATCGGAAGA |
| 49304_5 | TATGAGTGTGGAGTCGTTGCCAACTGGGACTATAGGCACCAGNNNAGATCGGAAGA |
| 40592_1 | TATGAGTGTGGAGTCGTTGCGGAGGGATCCTTCTGGTCAATTTNNNAGATCGGAAG |
| 40592_2 | TATGAGTGTGGAGTCGTTGCGGAGGGATCCTTCTGGTCAATTTNNNAGATCGGAAG |
| 40592_3 | TATGAGTGTGGAGTCGTTGCGGAGGGATCCTTCTGGTCAATTTNNNAGATCGGAAG |
| 40592_4 | TATGAGTGTGGAGTCGTTGCGGAGGGATCCTTCTGGTCAATTTNNNAGATCGGAAG |

| | |
|---|---|
| 40592_5 | TATGAGTGTGGAGTCGTTGCGGAGGGATCCTTCTGGTCAATTTNNNAGATCGGAA |
| 39412_1 | TATGAGTGTGGAGTCGTTGCCTCTCTGCTAACCACACTCCAATNNNAGATCGGAAGA |
| 39412_2 | TATGAGTGTGGAGTCGTTGCCTCTCTGCTAACCACACTCCAATNNNAGATCGGAAGA |
| 39412_3 | TATGAGTGTGGAGTCGTTGCCTCTCTGCTAACCACACTCCAATNNNAGATCGGAAGA |
| 39412_4 | TATGAGTGTGGAGTCGTTGCCTCTCTGCTAACCACACTCCAATNNNAGATCGGAAGA |
| 39412_5 | TATGAGTGTGGAGTCGTTGCCTCTCTGCTAACCACACTCCAATNNNAGATCGGAAGA |
| 44942_1 | TATGAGTGTGGAGTCGTTGCTCCTTAGTTCTACTTCCCTCCGTNNNAGATCGGAAGA |
| 44942_2 | TATGAGTGTGGAGTCGTTGCTCCTTAGTTCTACTTCCCTCCGTNNNAGATCGGAAGA |
| 44942_3 | TATGAGTGTGGAGTCGTTGCTCCTTAGTTCTACTTCCCTCCGTNNNAGATCGGAAGA |
| 44942_4 | TATGAGTGTGGAGTCGTTGCTCCTTAGTTCTACTTCCCTCCGTNNNAGATCGGAAGA |
| 44942_5 | TATGAGTGTGGAGTCGTTGCTCCTTAGTTCTACTTCCCTCCGTNNNAGATCGGAAGA |
| 36536_1 | TATGAGTGTGGAGTCGTTGCGCATGCACCTGTAGTCTCAACTANNNAGATCGGAAG |
| 36536_2 | TATGAGTGTGGAGTCGTTGCGCATGCACCTGTAGTCTCAACTANNNAGATCGGAAG |
| 36536_3 | TATGAGTGTGGAGTCGTTGCGCATGCACCTGTAGTCTCAACTANNNAGATCGGAAG |
| 36536_4 | TATGAGTGTGGAGTCGTTGCGCATGCACCTGTAGTCTCAACTANNNAGATCGGAAG |
| 36536_5 | TATGAGTGTGGAGTCGTTGCGCATGCACCTGTAGTCTCAACTANNNAGATCGGAAG |
| 31026_1 | TATGAGTGTGGAGTCGTTGCAACCAAAAGCAGCAAAGGACAATNNNAGATCGGAA |
| 31026_2 | TATGAGTGTGGAGTCGTTGCAACCAAAAGCAGCAAAGGACAATNNNAGATCGGAA |
| 31026_3 | TATGAGTGTGGAGTCGTTGCAACCAAAAGCAGCAAAGGACAATNNNAGATCGGAA |
| 31026_4 | TATGAGTGTGGAGTCGTTGCAACCAAAAGCAGCAAAGGACAATNNNAGATCGGAA |
| 31026_5 | TATGAGTGTGGAGTCGTTGCAACCAAAAGCAGCAAAGGACAATNNNAGATCGGAA |
| 46899_1 | TATGAGTGTGGAGTCGTTGCGCTGCCCTACCACCTGATAATTANNNAGATCGGAAG |
| 46899_2 | TATGAGTGTGGAGTCGTTGCGCTGCCCTACCACCTGATAATTANNNAGATCGGAAG |
| 46899_3 | TATGAGTGTGGAGTCGTTGCGCTGCCCTACCACCTGATAATTANNNAGATCGGAAG |
| 46899_4 | TATGAGTGTGGAGTCGTTGCGCTGCCCTACCACCTGATAATTANNNAGATCGGAAG |
| 46899_5 | TATGAGTGTGGAGTCGTTGCGCTGCCCTACCACCTGATAATTANNNAGATCGGAAG |
| 39766_1 | TATGAGTGTGGAGTCGTTGCGGGTTTGAGTGCCAATAGGAGTANNNAGATCGGAA |
| 39766_2 | TATGAGTGTGGAGTCGTTGCGGGTTTGAGTGCCAATAGGAGTANNNAGATCGGAA |
| 39766_3 | TATGAGTGTGGAGTCGTTGCGGGTTTGAGTGCCAATAGGAGTANNNAGATCGGAA |
| 39766_4 | TATGAGTGTGGAGTCGTTGCGGGTTTGAGTGCCAATAGGAGTANNNAGATCGGAA |
| 39766_5 | TATGAGTGTGGAGTCGTTGCGGGTTTGAGTGCCAATAGGAGTANNNAGATCGGAA |
| 12767_1 | TATGAGTGTGGAGTCGTTGCGTTGCTCTATTACTAAAACTGTTNNNAGATCGGAAG |
| 12767_2 | TATGAGTGTGGAGTCGTTGCGTTGCTCTATTACTAAAACTGTTNNNAGATCGGAAG |
| 12767_3 | TATGAGTGTGGAGTCGTTGCGTTGCTCTATTACTAAAACTGTTNNNAGATCGGAAG |
| 12767_4 | TATGAGTGTGGAGTCGTTGCGTTGCTCTATTACTAAAACTGTTNNNAGATCGGAAG |
| 12767_5 | TATGAGTGTGGAGTCGTTGCGTTGCTCTATTACTAAAACTGTTNNNAGATCGGAAG |
| 51059_1 | TATGAGTGTGGAGTCGTTGCTGGCTGAGAATGAAAGGGAAAGANNNAGATCGGA |
| 51059_2 | TATGAGTGTGGAGTCGTTGCTGGCTGAGAATGAAAGGGAAAGANNNAGATCGGA |
| 51059_3 | TATGAGTGTGGAGTCGTTGCTGGCTGAGAATGAAAGGGAAAGANNNAGATCGGA |
| 51059_4 | TATGAGTGTGGAGTCGTTGCTGGCTGAGAATGAAAGGGAAAGANNNAGATCGGA |
| 51059_5 | TATGAGTGTGGAGTCGTTGCTGGCTGAGAATGAAAGGGAAAGANNNAGATCGGA |
| 33204_1 | TATGAGTGTGGAGTCGTTGCATTACAACACACTTGGGGACTGTNNNAGATCGGAAG |
| 33204_2 | TATGAGTGTGGAGTCGTTGCATTACAACACACTTGGGGACTGTNNNAGATCGGAAG |

| | |
|---|---|
| 33204_3 | TATGAGTGTGGAGTCGTTGCATTACAACACACTTGGGGACTGTNNNAGATCGGAAG |
| 33204_4 | TATGAGTGTGGAGTCGTTGCATTACAACACACTTGGGGACTGTNNNAGATCGGAAG |
| 33204_5 | TATGAGTGTGGAGTCGTTGCATTACAACACACTTGGGGACTGTNNNAGATCGGAAG |
| 51558_1 | TATGAGTGTGGAGTCGTTGCGGTTTTTGGGAGTGGGTTGAAAGNNNAGATCGGA |
| 51558_2 | TATGAGTGTGGAGTCGTTGCGGTTTTTGGGAGTGGGTTGAAAGNNNAGATCGGA |
| 51558_3 | TATGAGTGTGGAGTCGTTGCGGTTTTTGGGAGTGGGTTGAAAGNNNAGATCGGA |
| 51558_4 | TATGAGTGTGGAGTCGTTGCGGTTTTTGGGAGTGGGTTGAAAGNNNAGATCGGA |
| 51558_5 | TATGAGTGTGGAGTCGTTGCGGTTTTTGGGAGTGGGTTGAAAGNNNAGATCGGA |
| 8215_1 | TATGAGTGTGGAGTCGTTGCATGTGGAGCTAGTACTTGTGCAGNNNAGATCGGAA |
| 8215_2 | TATGAGTGTGGAGTCGTTGCATGTGGAGCTAGTACTTGTGCAGNNNAGATCGGAA |
| 8215_3 | TATGAGTGTGGAGTCGTTGCATGTGGAGCTAGTACTTGTGCAGNNNAGATCGGAA |
| 8215_4 | TATGAGTGTGGAGTCGTTGCATGTGGAGCTAGTACTTGTGCAGNNNAGATCGGAA |
| 8215_5 | TATGAGTGTGGAGTCGTTGCATGTGGAGCTAGTACTTGTGCAGNNNAGATCGGAA |
| 44093_1 | TATGAGTGTGGAGTCGTTGCCAGGAAAGGAAAGTGTGGGTTTGNNNAGATCGGA |
| 44093_2 | TATGAGTGTGGAGTCGTTGCCAGGAAAGGAAAGTGTGGGTTTGNNNAGATCGGA |
| 44093_3 | TATGAGTGTGGAGTCGTTGCCAGGAAAGGAAAGTGTGGGTTTGNNNAGATCGGA |
| 44093_4 | TATGAGTGTGGAGTCGTTGCCAGGAAAGGAAAGTGTGGGTTTGNNNAGATCGGA |
| 44093_5 | TATGAGTGTGGAGTCGTTGCCAGGAAAGGAAAGTGTGGGTTTGNNNAGATCGGA |
| 35571_1 | TATGAGTGTGGAGTCGTTGCGTTTCACTCTTGTTGCCCAGAATNNNAGATCGGAAG |
| 35571_2 | TATGAGTGTGGAGTCGTTGCGTTTCACTCTTGTTGCCCAGAATNNNAGATCGGAAG |
| 35571_3 | TATGAGTGTGGAGTCGTTGCGTTTCACTCTTGTTGCCCAGAATNNNAGATCGGAAG |
| 35571_4 | TATGAGTGTGGAGTCGTTGCGTTTCACTCTTGTTGCCCAGAATNNNAGATCGGAAG |
| 35571_5 | TATGAGTGTGGAGTCGTTGCGTTTCACTCTTGTTGCCCAGAATNNNAGATCGGAAG |
| 48445_1 | TATGAGTGTGGAGTCGTTGCGGCGGTGTTACATAGCAAAAGAANNNAGATCGGAA |
| 48445_2 | TATGAGTGTGGAGTCGTTGCGGCGGTGTTACATAGCAAAAGAANNNAGATCGGAA |
| 48445_3 | TATGAGTGTGGAGTCGTTGCGGCGGTGTTACATAGCAAAAGAANNNAGATCGGAA |
| 48445_4 | TATGAGTGTGGAGTCGTTGCGGCGGTGTTACATAGCAAAAGAANNNAGATCGGAA |
| 48445_5 | TATGAGTGTGGAGTCGTTGCGGCGGTGTTACATAGCAAAAGAANNNAGATCGGAA |
| 33953_1 | TATGAGTGTGGAGTCGTTGCACTTGCAGAACTGTGAGAATTGCNNNAGATCGGAA |
| 33953_2 | TATGAGTGTGGAGTCGTTGCACTTGCAGAACTGTGAGAATTGCNNNAGATCGGAA |
| 33953_3 | TATGAGTGTGGAGTCGTTGCACTTGCAGAACTGTGAGAATTGCNNNAGATCGGAA |
| 33953_4 | TATGAGTGTGGAGTCGTTGCACTTGCAGAACTGTGAGAATTGCNNNAGATCGGAA |
| 33953_5 | TATGAGTGTGGAGTCGTTGCACTTGCAGAACTGTGAGAATTGCNNNAGATCGGAA |
| 33691_1 | TATGAGTGTGGAGTCGTTGCATTCGTGAAGTTGAAGTGGCTCANNNAGATCGGAA |
| 33691_2 | TATGAGTGTGGAGTCGTTGCATTCGTGAAGTTGAAGTGGCTCANNNAGATCGGAA |
| 33691_3 | TATGAGTGTGGAGTCGTTGCATTCGTGAAGTTGAAGTGGCTCANNNAGATCGGAA |
| 33691_4 | TATGAGTGTGGAGTCGTTGCATTCGTGAAGTTGAAGTGGCTCANNNAGATCGGAA |
| 33691_5 | TATGAGTGTGGAGTCGTTGCATTCGTGAAGTTGAAGTGGCTCANNNAGATCGGAA |
| 12572_1 | TATGAGTGTGGAGTCGTTGCCATAGAAACTGGAAAGTACAGTANNNAGATCGGAA |
| 12572_2 | TATGAGTGTGGAGTCGTTGCCATAGAAACTGGAAAGTACAGTANNNAGATCGGAA |
| 12572_3 | TATGAGTGTGGAGTCGTTGCCATAGAAACTGGAAAGTACAGTANNNAGATCGGAA |
| 12572_4 | TATGAGTGTGGAGTCGTTGCCATAGAAACTGGAAAGTACAGTANNNAGATCGGAA |
| 12572_5 | TATGAGTGTGGAGTCGTTGCCATAGAAACTGGAAAGTACAGTANNNAGATCGGAA |

| | |
|---|---|
| 34466_1 | TATGAGTGTGGAGTCGTTGCTGCCCCTTATGCAAGACATTTTGNNNAGATCGGAAG |
| 34466_2 | TATGAGTGTGGAGTCGTTGCTGCCCCTTATGCAAGACATTTTGNNNAGATCGGAAG |
| 34466_3 | TATGAGTGTGGAGTCGTTGCTGCCCCTTATGCAAGACATTTTGNNNAGATCGGAAG |
| 34466_4 | TATGAGTGTGGAGTCGTTGCTGCCCCTTATGCAAGACATTTTGNNNAGATCGGAAG |
| 34466_5 | TATGAGTGTGGAGTCGTTGCTGCCCCTTATGCAAGACATTTTGNNNAGATCGGAAG |
| 34387_1 | TATGAGTGTGGAGTCGTTGCCTTTGCTTCCTCTTCACCTTTGGNNNAGATCGGAAGA |
| 34387_2 | TATGAGTGTGGAGTCGTTGCCTTTGCTTCCTCTTCACCTTTGGNNNAGATCGGAAGA |
| 34387_3 | TATGAGTGTGGAGTCGTTGCCTTTGCTTCCTCTTCACCTTTGGNNNAGATCGGAAGA |
| 34387_4 | TATGAGTGTGGAGTCGTTGCCTTTGCTTCCTCTTCACCTTTGGNNNAGATCGGAAGA |
| 34387_5 | TATGAGTGTGGAGTCGTTGCCTTTGCTTCCTCTTCACCTTTGGNNNAGATCGGAAGA |
| 52382_1 | TATGAGTGTGGAGTCGTTGCTGTGTATAGGGAAAGGCTGTTGGNNNAGATCGGAA |
| 52382_2 | TATGAGTGTGGAGTCGTTGCTGTGTATAGGGAAAGGCTGTTGGNNNAGATCGGAA |
| 52382_3 | TATGAGTGTGGAGTCGTTGCTGTGTATAGGGAAAGGCTGTTGGNNNAGATCGGAA |
| 52382_4 | TATGAGTGTGGAGTCGTTGCTGTGTATAGGGAAAGGCTGTTGGNNNAGATCGGAA |
| 52382_5 | TATGAGTGTGGAGTCGTTGCTGTGTATAGGGAAAGGCTGTTGGNNNAGATCGGAA |
| 12514_1 | TATGAGTGTGGAGTCGTTGCAATAACAAACCATTTTCATGAGTNNNAGATCGGAAG |
| 12514_2 | TATGAGTGTGGAGTCGTTGCAATAACAAACCATTTTCATGAGTNNNAGATCGGAAG |
| 12514_3 | TATGAGTGTGGAGTCGTTGCAATAACAAACCATTTTCATGAGTNNNAGATCGGAAG |
| 12514_4 | TATGAGTGTGGAGTCGTTGCAATAACAAACCATTTTCATGAGTNNNAGATCGGAAG |
| 12514_5 | TATGAGTGTGGAGTCGTTGCAATAACAAACCATTTTCATGAGTNNNAGATCGGAAG |
| 43703_1 | TATGAGTGTGGAGTCGTTGCGGTGACAGGCGGATAGTTATAGGNNNAGATCGGAA |
| 43703_2 | TATGAGTGTGGAGTCGTTGCGGTGACAGGCGGATAGTTATAGGNNNAGATCGGAA |
| 43703_3 | TATGAGTGTGGAGTCGTTGCGGTGACAGGCGGATAGTTATAGGNNNAGATCGGAA |
| 43703_4 | TATGAGTGTGGAGTCGTTGCGGTGACAGGCGGATAGTTATAGGNNNAGATCGGAA |
| 43703_5 | TATGAGTGTGGAGTCGTTGCGGTGACAGGCGGATAGTTATAGGNNNAGATCGGAA |
| 13292_1 | TATGAGTGTGGAGTCGTTGCATTTTACCATGATATACCCTGAANNNAGATCGGAAG |
| 13292_2 | TATGAGTGTGGAGTCGTTGCATTTTACCATGATATACCCTGAANNNAGATCGGAAG |
| 13292_3 | TATGAGTGTGGAGTCGTTGCATTTTACCATGATATACCCTGAANNNAGATCGGAAG |
| 13292_4 | TATGAGTGTGGAGTCGTTGCATTTTACCATGATATACCCTGAANNNAGATCGGAAG |
| 13292_5 | TATGAGTGTGGAGTCGTTGCATTTTACCATGATATACCCTGAANNNAGATCGGAAG |
| 38570_1 | TATGAGTGTGGAGTCGTTGCGGAGAGAGAGAGTGAGAGAGACANNNAGATCGGA |
| 38570_2 | TATGAGTGTGGAGTCGTTGCGGAGAGAGAGAGTGAGAGAGACANNNAGATCGGA |
| 38570_3 | TATGAGTGTGGAGTCGTTGCGGAGAGAGAGAGTGAGAGAGACANNNAGATCGGA |
| 38570_4 | TATGAGTGTGGAGTCGTTGCGGAGAGAGAGAGTGAGAGAGACANNNAGATCGGA |
| 38570_5 | TATGAGTGTGGAGTCGTTGCGGAGAGAGAGAGTGAGAGAGACANNNAGATCGGA |
| 13255_1 | TATGAGTGTGGAGTCGTTGCGAGTGAAATTAGATACATTCACCNNNAGATCGGAAG |
| 13255_2 | TATGAGTGTGGAGTCGTTGCGAGTGAAATTAGATACATTCACCNNNAGATCGGAAG |
| 13255_3 | TATGAGTGTGGAGTCGTTGCGAGTGAAATTAGATACATTCACCNNNAGATCGGAAG |
| 13255_4 | TATGAGTGTGGAGTCGTTGCGAGTGAAATTAGATACATTCACCNNNAGATCGGAAG |
| 13255_5 | TATGAGTGTGGAGTCGTTGCGAGTGAAATTAGATACATTCACCNNNAGATCGGAAG |
| 46560_1 | TATGAGTGTGGAGTCGTTGCCCTTTCATTTCCAGTCTCTGTGCNNNAGATCGGAAGA |
| 46560_2 | TATGAGTGTGGAGTCGTTGCCCTTTCATTTCCAGTCTCTGTGCNNNAGATCGGAAGA |
| 46560_3 | TATGAGTGTGGAGTCGTTGCCCTTTCATTTCCAGTCTCTGTGCNNNAGATCGGAAGA |

| | |
|---|---|
| 46560_4 | TATGAGTGTGGAGTCGTTGCCCTTTCATTTCCAGTCTCTGTGCNNNAGATCGGAAGA |
| 46560_5 | TATGAGTGTGGAGTCGTTGCCCTTTCATTTCCAGTCTCTGTGCNNNAGATCGGAAGA |
| 41016_1 | TATGAGTGTGGAGTCGTTGCTACCATTCAGGACATAGGCATGGNNNAGATCGGAAG |
| 41016_2 | TATGAGTGTGGAGTCGTTGCTACCATTCAGGACATAGGCATGGNNNAGATCGGAAG |
| 41016_3 | TATGAGTGTGGAGTCGTTGCTACCATTCAGGACATAGGCATGGNNNAGATCGGAAG |
| 41016_4 | TATGAGTGTGGAGTCGTTGCTACCATTCAGGACATAGGCATGGNNNAGATCGGAAG |
| 41016_5 | TATGAGTGTGGAGTCGTTGCTACCATTCAGGACATAGGCATGGNNNAGATCGGAAG |
| 41590_1 | TATGAGTGTGGAGTCGTTGCTTCAAGTGATTCTCCTGCCTCAGNNNAGATCGGAAGA |
| 41590_2 | TATGAGTGTGGAGTCGTTGCTTCAAGTGATTCTCCTGCCTCAGNNNAGATCGGAAGA |
| 41590_3 | TATGAGTGTGGAGTCGTTGCTTCAAGTGATTCTCCTGCCTCAGNNNAGATCGGAAGA |
| 41590_4 | TATGAGTGTGGAGTCGTTGCTTCAAGTGATTCTCCTGCCTCAGNNNAGATCGGAAGA |
| 41590_5 | TATGAGTGTGGAGTCGTTGCTTCAAGTGATTCTCCTGCCTCAGNNNAGATCGGAAGA |
| 46737_1 | TATGAGTGTGGAGTCGTTGCCCCCTGTTCAAGCAATTCTTCTGNNNAGATCGGAAGA |
| 46737_2 | TATGAGTGTGGAGTCGTTGCCCCCTGTTCAAGCAATTCTTCTGNNNAGATCGGAAGA |
| 46737_3 | TATGAGTGTGGAGTCGTTGCCCCCTGTTCAAGCAATTCTTCTGNNNAGATCGGAAGA |
| 46737_4 | TATGAGTGTGGAGTCGTTGCCCCCTGTTCAAGCAATTCTTCTGNNNAGATCGGAAGA |
| 46737_5 | TATGAGTGTGGAGTCGTTGCCCCCTGTTCAAGCAATTCTTCTGNNNAGATCGGAAGA |
| 48129_1 | TATGAGTGTGGAGTCGTTGCGATGGCCCTAATCTGAATCCCATNNNAGATCGGAAGA |
| 48129_2 | TATGAGTGTGGAGTCGTTGCGATGGCCCTAATCTGAATCCCATNNNAGATCGGAAGA |
| 48129_3 | TATGAGTGTGGAGTCGTTGCGATGGCCCTAATCTGAATCCCATNNNAGATCGGAAGA |
| 48129_4 | TATGAGTGTGGAGTCGTTGCGATGGCCCTAATCTGAATCCCATNNNAGATCGGAAGA |
| 48129_5 | TATGAGTGTGGAGTCGTTGCGATGGCCCTAATCTGAATCCCATNNNAGATCGGAAGA |
| 12755_1 | TATGAGTGTGGAGTCGTTGCAAAATGAAAAGAAGGTACTGTACNNNAGATCGGAAG |
| 12755_2 | TATGAGTGTGGAGTCGTTGCAAAATGAAAAGAAGGTACTGTACNNNAGATCGGAAG |
| 12755_3 | TATGAGTGTGGAGTCGTTGCAAAATGAAAAGAAGGTACTGTACNNNAGATCGGAAG |
| 12755_4 | TATGAGTGTGGAGTCGTTGCAAAATGAAAAGAAGGTACTGTACNNNAGATCGGAAG |
| 12755_5 | TATGAGTGTGGAGTCGTTGCAAAATGAAAAGAAGGTACTGTACNNNAGATCGGAAG |
| 34689_1 | TATGAGTGTGGAGTCGTTGCCAATCTGGGCTCACTGCAAGNNNAGATCGGAAGAGC |
| 34689_2 | TATGAGTGTGGAGTCGTTGCCAATCTGGGCTCACTGCAAGNNNAGATCGGAAGAGC |
| 34689_3 | TATGAGTGTGGAGTCGTTGCCAATCTGGGCTCACTGCAAGNNNAGATCGGAAGAGC |
| 34689_4 | TATGAGTGTGGAGTCGTTGCCAATCTGGGCTCACTGCAAGNNNAGATCGGAAGAGC |
| 34689_5 | TATGAGTGTGGAGTCGTTGCCAATCTGGGCTCACTGCAAGNNNAGATCGGAAGAGC |
| 45344_1 | TATGAGTGTGGAGTCGTTGCTGAATCTGCAAGCAAAAGAAGGGNNNAGATCGGAA |
| 45344_2 | TATGAGTGTGGAGTCGTTGCTGAATCTGCAAGCAAAAGAAGGGNNNAGATCGGAA |
| 45344_3 | TATGAGTGTGGAGTCGTTGCTGAATCTGCAAGCAAAAGAAGGGNNNAGATCGGAA |
| 45344_4 | TATGAGTGTGGAGTCGTTGCTGAATCTGCAAGCAAAAGAAGGGNNNAGATCGGAA |
| 45344_5 | TATGAGTGTGGAGTCGTTGCTGAATCTGCAAGCAAAAGAAGGGNNNAGATCGGAA |
| 36722_1 | TATGAGTGTGGAGTCGTTGCCGACCAGGCATCTATGGAGATTANNNAGATCGGAAG |
| 36722_2 | TATGAGTGTGGAGTCGTTGCCGACCAGGCATCTATGGAGATTANNNAGATCGGAAG |
| 36722_3 | TATGAGTGTGGAGTCGTTGCCGACCAGGCATCTATGGAGATTANNNAGATCGGAAG |
| 36722_4 | TATGAGTGTGGAGTCGTTGCCGACCAGGCATCTATGGAGATTANNNAGATCGGAAG |
| 36722_5 | TATGAGTGTGGAGTCGTTGCCGACCAGGCATCTATGGAGATTANNNAGATCGGAAG |
| 52537_1 | TATGAGTGTGGAGTCGTTGCTTTTTCATGAAAAGGAACAGTCTCTNNNAGATCGGA |

| | |
|---|---|
| 52537_2 | TATGAGTGTGGAGTCGTTGCTTTTTCATGAAAAGGAACAGTCTCTNNNAGATCGGA |
| 52537_3 | TATGAGTGTGGAGTCGTTGCTTTTTCATGAAAAGGAACAGTCTCTNNNAGATCGGA |
| 52537_4 | TATGAGTGTGGAGTCGTTGCTTTTTCATGAAAAGGAACAGTCTCTNNNAGATCGGA |
| 52537_5 | TATGAGTGTGGAGTCGTTGCTTTTTCATGAAAAGGAACAGTCTCTNNNAGATCGGA |
| 47662_1 | TATGAGTGTGGAGTCGTTGCGGGCTCTGTGTTTCCTTAGGTAANNNAGATCGGAAG |
| 47662_2 | TATGAGTGTGGAGTCGTTGCGGGCTCTGTGTTTCCTTAGGTAANNNAGATCGGAAG |
| 47662_3 | TATGAGTGTGGAGTCGTTGCGGGCTCTGTGTTTCCTTAGGTAANNNAGATCGGAAG |
| 47662_4 | TATGAGTGTGGAGTCGTTGCGGGCTCTGTGTTTCCTTAGGTAANNNAGATCGGAAG |
| 47662_5 | TATGAGTGTGGAGTCGTTGCGGGCTCTGTGTTTCCTTAGGTAANNNAGATCGGAAG |
| 35904_1 | TATGAGTGTGGAGTCGTTGCCAACGCATCTTCACTGATGGACNNNAGATCGGAAGA |
| 35904_2 | TATGAGTGTGGAGTCGTTGCCAACGCATCTTCACTGATGGACNNNAGATCGGAAGA |
| 35904_3 | TATGAGTGTGGAGTCGTTGCCAACGCATCTTCACTGATGGACNNNAGATCGGAAGA |
| 35904_4 | TATGAGTGTGGAGTCGTTGCCAACGCATCTTCACTGATGGACNNNAGATCGGAAGA |
| 35904_5 | TATGAGTGTGGAGTCGTTGCCAACGCATCTTCACTGATGGACNNNAGATCGGAAGA |
| 36249_1 | TATGAGTGTGGAGTCGTTGCTCTCAAACTCCTGACCTCAAGTGNNNAGATCGGAAG |
| 36249_2 | TATGAGTGTGGAGTCGTTGCTCTCAAACTCCTGACCTCAAGTGNNNAGATCGGAAG |
| 36249_3 | TATGAGTGTGGAGTCGTTGCTCTCAAACTCCTGACCTCAAGTGNNNAGATCGGAAG |
| 36249_4 | TATGAGTGTGGAGTCGTTGCTCTCAAACTCCTGACCTCAAGTGNNNAGATCGGAAG |
| 36249_5 | TATGAGTGTGGAGTCGTTGCTCTCAAACTCCTGACCTCAAGTGNNNAGATCGGAAG |
| 46283_1 | TATGAGTGTGGAGTCGTTGCACCAACCCTAGCTAAAGGACATGNNNAGATCGGAAG |
| 46283_2 | TATGAGTGTGGAGTCGTTGCACCAACCCTAGCTAAAGGACATGNNNAGATCGGAAG |
| 46283_3 | TATGAGTGTGGAGTCGTTGCACCAACCCTAGCTAAAGGACATGNNNAGATCGGAAG |
| 46283_4 | TATGAGTGTGGAGTCGTTGCACCAACCCTAGCTAAAGGACATGNNNAGATCGGAAG |
| 46283_5 | TATGAGTGTGGAGTCGTTGCACCAACCCTAGCTAAAGGACATGNNNAGATCGGAAG |
| 42682_1 | TATGAGTGTGGAGTCGTTGCGGGTGGCTAGAGAACAGGAATAGNNNAGATCGGA |
| 42682_2 | TATGAGTGTGGAGTCGTTGCGGGTGGCTAGAGAACAGGAATAGNNNAGATCGGA |
| 42682_3 | TATGAGTGTGGAGTCGTTGCGGGTGGCTAGAGAACAGGAATAGNNNAGATCGGA |
| 42682_4 | TATGAGTGTGGAGTCGTTGCGGGTGGCTAGAGAACAGGAATAGNNNAGATCGGA |
| 42682_5 | TATGAGTGTGGAGTCGTTGCGGGTGGCTAGAGAACAGGAATAGNNNAGATCGGA |
| 29779_1 | TATGAGTGTGGAGTCGTTGCTGTACACCATAGGGCTCAGTTTCNNNAGATCGGAAG |
| 29779_2 | TATGAGTGTGGAGTCGTTGCTGTACACCATAGGGCTCAGTTTCNNNAGATCGGAAG |
| 29779_3 | TATGAGTGTGGAGTCGTTGCTGTACACCATAGGGCTCAGTTTCNNNAGATCGGAAG |
| 29779_4 | TATGAGTGTGGAGTCGTTGCTGTACACCATAGGGCTCAGTTTCNNNAGATCGGAAG |
| 29779_5 | TATGAGTGTGGAGTCGTTGCTGTACACCATAGGGCTCAGTTTCNNNAGATCGGAAG |
| 52570_1 | TATGAGTGTGGAGTCGTTGCATCTTTAGAGAACGTGTGGGCTTNNNAGATCGGAA |
| 52570_2 | TATGAGTGTGGAGTCGTTGCATCTTTAGAGAACGTGTGGGCTTNNNAGATCGGAA |
| 52570_3 | TATGAGTGTGGAGTCGTTGCATCTTTAGAGAACGTGTGGGCTTNNNAGATCGGAA |
| 52570_4 | TATGAGTGTGGAGTCGTTGCATCTTTAGAGAACGTGTGGGCTTNNNAGATCGGAA |
| 52570_5 | TATGAGTGTGGAGTCGTTGCATCTTTAGAGAACGTGTGGGCTTNNNAGATCGGAA |
| 30666_1 | TATGAGTGTGGAGTCGTTGCGAAGGTTGGGGAAGCTTGAGGATNNNAGATCGGA |
| 30666_2 | TATGAGTGTGGAGTCGTTGCGAAGGTTGGGGAAGCTTGAGGATNNNAGATCGGA |
| 30666_3 | TATGAGTGTGGAGTCGTTGCGAAGGTTGGGGAAGCTTGAGGATNNNAGATCGGA |
| 30666_4 | TATGAGTGTGGAGTCGTTGCGAAGGTTGGGGAAGCTTGAGGATNNNAGATCGGA |

| | |
|---|---|
| 30666_5 | TATGAGTGTGGAGTCGTTGCGAAGGTTGGGGAAGCTTGAGGATNNNAGATCGGA |
| 581_1 | TATGAGTGTGGAGTCGTTGCAAGAATCCTCAAATCACAGTGGCTNNNAGATCGGAA |
| 581_2 | TATGAGTGTGGAGTCGTTGCAAGAATCCTCAAATCACAGTGGCTNNNAGATCGGAA |
| 581_3 | TATGAGTGTGGAGTCGTTGCAAGAATCCTCAAATCACAGTGGCTNNNAGATCGGAA |
| 581_4 | TATGAGTGTGGAGTCGTTGCAAGAATCCTCAAATCACAGTGGCTNNNAGATCGGAA |
| 581_5 | TATGAGTGTGGAGTCGTTGCAAGAATCCTCAAATCACAGTGGCTNNNAGATCGGAA |
| 47997_1 | TATGAGTGTGGAGTCGTTGCCTCCTGAGTAGCTGGGACTACAGNNNAGATCGGAA |
| 47997_2 | TATGAGTGTGGAGTCGTTGCCTCCTGAGTAGCTGGGACTACAGNNNAGATCGGAA |
| 47997_3 | TATGAGTGTGGAGTCGTTGCCTCCTGAGTAGCTGGGACTACAGNNNAGATCGGAA |
| 47997_4 | TATGAGTGTGGAGTCGTTGCCTCCTGAGTAGCTGGGACTACAGNNNAGATCGGAA |
| 47997_5 | TATGAGTGTGGAGTCGTTGCCTCCTGAGTAGCTGGGACTACAGNNNAGATCGGAA |
| 41097_1 | TATGAGTGTGGAGTCGTTGCTGTATATGCAGACCACTTAGCACTNNNAGATCGGAA |
| 41097_2 | TATGAGTGTGGAGTCGTTGCTGTATATGCAGACCACTTAGCACTNNNAGATCGGAA |
| 41097_3 | TATGAGTGTGGAGTCGTTGCTGTATATGCAGACCACTTAGCACTNNNAGATCGGAA |
| 41097_4 | TATGAGTGTGGAGTCGTTGCTGTATATGCAGACCACTTAGCACTNNNAGATCGGAA |
| 41097_5 | TATGAGTGTGGAGTCGTTGCTGTATATGCAGACCACTTAGCACTNNNAGATCGGAA |
| 12684_1 | TATGAGTGTGGAGTCGTTGCATCAGAACCTGGTAAATATCTTTNNNAGATCGGAAG |
| 12684_2 | TATGAGTGTGGAGTCGTTGCATCAGAACCTGGTAAATATCTTTNNNAGATCGGAAG |
| 12684_3 | TATGAGTGTGGAGTCGTTGCATCAGAACCTGGTAAATATCTTTNNNAGATCGGAAG |
| 12684_4 | TATGAGTGTGGAGTCGTTGCATCAGAACCTGGTAAATATCTTTNNNAGATCGGAAG |
| 12684_5 | TATGAGTGTGGAGTCGTTGCATCAGAACCTGGTAAATATCTTTNNNAGATCGGAAG |
| 8201_1 | TATGAGTGTGGAGTCGTTGCCGGTGGCAAGAACTTACTTCNNNAGATCGGAAGAG |
| 8201_2 | TATGAGTGTGGAGTCGTTGCCGGTGGCAAGAACTTACTTCNNNAGATCGGAAGAG |
| 8201_3 | TATGAGTGTGGAGTCGTTGCCGGTGGCAAGAACTTACTTCNNNAGATCGGAAGAG |
| 8201_4 | TATGAGTGTGGAGTCGTTGCCGGTGGCAAGAACTTACTTCNNNAGATCGGAAGAG |
| 8201_5 | TATGAGTGTGGAGTCGTTGCCGGTGGCAAGAACTTACTTCNNNAGATCGGAAGAG |
| 51030_1 | TATGAGTGTGGAGTCGTTGCTGACTCTAGTTAGGGCCAAGAGANNNAGATCGGAA |
| 51030_2 | TATGAGTGTGGAGTCGTTGCTGACTCTAGTTAGGGCCAAGAGANNNAGATCGGAA |
| 51030_3 | TATGAGTGTGGAGTCGTTGCTGACTCTAGTTAGGGCCAAGAGANNNAGATCGGAA |
| 51030_4 | TATGAGTGTGGAGTCGTTGCTGACTCTAGTTAGGGCCAAGAGANNNAGATCGGAA |
| 51030_5 | TATGAGTGTGGAGTCGTTGCTGACTCTAGTTAGGGCCAAGAGANNNAGATCGGAA |
| 51494_1 | TATGAGTGTGGAGTCGTTGCTGGATCTAGGTTTGCACCACTTTNNNAGATCGGAAG |
| 51494_2 | TATGAGTGTGGAGTCGTTGCTGGATCTAGGTTTGCACCACTTTNNNAGATCGGAAG |
| 51494_3 | TATGAGTGTGGAGTCGTTGCTGGATCTAGGTTTGCACCACTTTNNNAGATCGGAAG |
| 51494_4 | TATGAGTGTGGAGTCGTTGCTGGATCTAGGTTTGCACCACTTTNNNAGATCGGAAG |
| 51494_5 | TATGAGTGTGGAGTCGTTGCTGGATCTAGGTTTGCACCACTTTNNNAGATCGGAAG |
| 40781_1 | TATGAGTGTGGAGTCGTTGCTGTGTGTGATAGAATGGCTGTGANNNAGATCGGAA |
| 40781_2 | TATGAGTGTGGAGTCGTTGCTGTGTGTGATAGAATGGCTGTGANNNAGATCGGAA |
| 40781_3 | TATGAGTGTGGAGTCGTTGCTGTGTGTGATAGAATGGCTGTGANNNAGATCGGAA |
| 40781_4 | TATGAGTGTGGAGTCGTTGCTGTGTGTGATAGAATGGCTGTGANNNAGATCGGAA |
| 40781_5 | TATGAGTGTGGAGTCGTTGCTGTGTGTGATAGAATGGCTGTGANNNAGATCGGAA |
| 51716_1 | TATGAGTGTGGAGTCGTTGCACAAGGTATGGCTTCCAATGTCTNNNAGATCGGAAG |
| 51716_2 | TATGAGTGTGGAGTCGTTGCACAAGGTATGGCTTCCAATGTCTNNNAGATCGGAAG |

| | |
|---|---|
| 51716_3 | TATGAGTGTGGAGTCGTTGCACAAGGTATGGCTTCCAATGTCTNNNAGATCGGAAG |
| 51716_4 | TATGAGTGTGGAGTCGTTGCACAAGGTATGGCTTCCAATGTCTNNNAGATCGGAAG |
| 51716_5 | TATGAGTGTGGAGTCGTTGCACAAGGTATGGCTTCCAATGTCTNNNAGATCGGAAG |
| 40618_1 | TATGAGTGTGGAGTCGTTGCCTGAGGCAGGAGAATTGCTTGAANNNAGATCGGAA |
| 40618_2 | TATGAGTGTGGAGTCGTTGCCTGAGGCAGGAGAATTGCTTGAANNNAGATCGGAA |
| 40618_3 | TATGAGTGTGGAGTCGTTGCCTGAGGCAGGAGAATTGCTTGAANNNAGATCGGAA |
| 40618_4 | TATGAGTGTGGAGTCGTTGCCTGAGGCAGGAGAATTGCTTGAANNNAGATCGGAA |
| 40618_5 | TATGAGTGTGGAGTCGTTGCCTGAGGCAGGAGAATTGCTTGAANNNAGATCGGAA |
| 39598_1 | TATGAGTGTGGAGTCGTTGCTGTACACAGTTGGGTAGCTTGTANNNAGATCGGAA |
| 39598_2 | TATGAGTGTGGAGTCGTTGCTGTACACAGTTGGGTAGCTTGTANNNAGATCGGAA |
| 39598_3 | TATGAGTGTGGAGTCGTTGCTGTACACAGTTGGGTAGCTTGTANNNAGATCGGAA |
| 39598_4 | TATGAGTGTGGAGTCGTTGCTGTACACAGTTGGGTAGCTTGTANNNAGATCGGAA |
| 39598_5 | TATGAGTGTGGAGTCGTTGCTGTACACAGTTGGGTAGCTTGTANNNAGATCGGAA |
| 10370_1 | TATGAGTGTGGAGTCGTTGCTGGAGGGAGTAGAGAACAAAGCANNNAGATCGGA |
| 10370_2 | TATGAGTGTGGAGTCGTTGCTGGAGGGAGTAGAGAACAAAGCANNNAGATCGGA |
| 10370_3 | TATGAGTGTGGAGTCGTTGCTGGAGGGAGTAGAGAACAAAGCANNNAGATCGGA |
| 10370_4 | TATGAGTGTGGAGTCGTTGCTGGAGGGAGTAGAGAACAAAGCANNNAGATCGGA |
| 10370_5 | TATGAGTGTGGAGTCGTTGCTGGAGGGAGTAGAGAACAAAGCANNNAGATCGGA |
| 36680_1 | TATGAGTGTGGAGTCGTTGCAAAAAACAAAAGAGCCACCACCANNNAGATCGGAA |
| 36680_2 | TATGAGTGTGGAGTCGTTGCAAAAAACAAAAGAGCCACCACCANNNAGATCGGAA |
| 36680_3 | TATGAGTGTGGAGTCGTTGCAAAAAACAAAAGAGCCACCACCANNNAGATCGGAA |
| 36680_4 | TATGAGTGTGGAGTCGTTGCAAAAAACAAAAGAGCCACCACCANNNAGATCGGAA |
| 36680_5 | TATGAGTGTGGAGTCGTTGCAAAAAACAAAAGAGCCACCACCANNNAGATCGGAA |
| 48965_1 | TATGAGTGTGGAGTCGTTGCACTGCTGTCATGAAAGCAAGATGNNNAGATCGGAA |
| 48965_2 | TATGAGTGTGGAGTCGTTGCACTGCTGTCATGAAAGCAAGATGNNNAGATCGGAA |
| 48965_3 | TATGAGTGTGGAGTCGTTGCACTGCTGTCATGAAAGCAAGATGNNNAGATCGGAA |
| 48965_4 | TATGAGTGTGGAGTCGTTGCACTGCTGTCATGAAAGCAAGATGNNNAGATCGGAA |
| 48965_5 | TATGAGTGTGGAGTCGTTGCACTGCTGTCATGAAAGCAAGATGNNNAGATCGGAA |
| 28882_1 | TATGAGTGTGGAGTCGTTGCGCCAATTACAGGAAGCCAAGTTTNNNAGATCGGAAG |
| 28882_2 | TATGAGTGTGGAGTCGTTGCGCCAATTACAGGAAGCCAAGTTTNNNAGATCGGAAG |
| 28882_3 | TATGAGTGTGGAGTCGTTGCGCCAATTACAGGAAGCCAAGTTTNNNAGATCGGAAG |
| 28882_4 | TATGAGTGTGGAGTCGTTGCGCCAATTACAGGAAGCCAAGTTTNNNAGATCGGAAG |
| 28882_5 | TATGAGTGTGGAGTCGTTGCGCCAATTACAGGAAGCCAAGTTTNNNAGATCGGAAG |
| 36206_1 | TATGAGTGTGGAGTCGTTGCGTGGCAAAACCCTGTCTCTACTANNNAGATCGGAAG |
| 36206_2 | TATGAGTGTGGAGTCGTTGCGTGGCAAAACCCTGTCTCTACTANNNAGATCGGAAG |
| 36206_3 | TATGAGTGTGGAGTCGTTGCGTGGCAAAACCCTGTCTCTACTANNNAGATCGGAAG |
| 36206_4 | TATGAGTGTGGAGTCGTTGCGTGGCAAAACCCTGTCTCTACTANNNAGATCGGAAG |
| 36206_5 | TATGAGTGTGGAGTCGTTGCGTGGCAAAACCCTGTCTCTACTANNNAGATCGGAAG |
| 42801_1 | TATGAGTGTGGAGTCGTTGCTGCATCATTGTTAGGATAAGCCCTNNNAGATCGGAA |
| 42801_2 | TATGAGTGTGGAGTCGTTGCTGCATCATTGTTAGGATAAGCCCTNNNAGATCGGAA |
| 42801_3 | TATGAGTGTGGAGTCGTTGCTGCATCATTGTTAGGATAAGCCCTNNNAGATCGGAA |
| 42801_4 | TATGAGTGTGGAGTCGTTGCTGCATCATTGTTAGGATAAGCCCTNNNAGATCGGAA |
| 42801_5 | TATGAGTGTGGAGTCGTTGCTGCATCATTGTTAGGATAAGCCCTNNNAGATCGGAA |

| | |
|---|---|
| 49283_1 | TATGAGTGTGGAGTCGTTGCCAATCTCAGCTCACTGCAACTTCNNNAGATCGGAAGA |
| 49283_2 | TATGAGTGTGGAGTCGTTGCCAATCTCAGCTCACTGCAACTTCNNNAGATCGGAAGA |
| 49283_3 | TATGAGTGTGGAGTCGTTGCCAATCTCAGCTCACTGCAACTTCNNNAGATCGGAAGA |
| 49283_4 | TATGAGTGTGGAGTCGTTGCCAATCTCAGCTCACTGCAACTTCNNNAGATCGGAAGA |
| 49283_5 | TATGAGTGTGGAGTCGTTGCCAATCTCAGCTCACTGCAACTTCNNNAGATCGGAAGA |
| 39127_1 | TATGAGTGTGGAGTCGTTGCAAAGAAAGAAACAGCAAGAGCCANNNAGATCGGAA |
| 39127_2 | TATGAGTGTGGAGTCGTTGCAAAGAAAGAAACAGCAAGAGCCANNNAGATCGGAA |
| 39127_3 | TATGAGTGTGGAGTCGTTGCAAAGAAAGAAACAGCAAGAGCCANNNAGATCGGAA |
| 39127_4 | TATGAGTGTGGAGTCGTTGCAAAGAAAGAAACAGCAAGAGCCANNNAGATCGGAA |
| 39127_5 | TATGAGTGTGGAGTCGTTGCAAAGAAAGAAACAGCAAGAGCCANNNAGATCGGAA |
| 36628_1 | TATGAGTGTGGAGTCGTTGCTATTCCTGGAAGCCTCAATCGAGNNNAGATCGGAAG |
| 36628_2 | TATGAGTGTGGAGTCGTTGCTATTCCTGGAAGCCTCAATCGAGNNNAGATCGGAAG |
| 36628_3 | TATGAGTGTGGAGTCGTTGCTATTCCTGGAAGCCTCAATCGAGNNNAGATCGGAAG |
| 36628_4 | TATGAGTGTGGAGTCGTTGCTATTCCTGGAAGCCTCAATCGAGNNNAGATCGGAAG |
| 36628_5 | TATGAGTGTGGAGTCGTTGCTATTCCTGGAAGCCTCAATCGAGNNNAGATCGGAAG |
| 29788_1 | TATGAGTGTGGAGTCGTTGCAGGATATGCTAGATGGTACTTTTGTNNNAGATCGGA |
| 29788_2 | TATGAGTGTGGAGTCGTTGCAGGATATGCTAGATGGTACTTTTGTNNNAGATCGGA |
| 29788_3 | TATGAGTGTGGAGTCGTTGCAGGATATGCTAGATGGTACTTTTGTNNNAGATCGGA |
| 29788_4 | TATGAGTGTGGAGTCGTTGCAGGATATGCTAGATGGTACTTTTGTNNNAGATCGGA |
| 29788_5 | TATGAGTGTGGAGTCGTTGCAGGATATGCTAGATGGTACTTTTGTNNNAGATCGGA |
| 31137_1 | TATGAGTGTGGAGTCGTTGCCCATGACTTCCAGGAACTCTCTTNNNAGATCGGAAGA |
| 31137_2 | TATGAGTGTGGAGTCGTTGCCCATGACTTCCAGGAACTCTCTTNNNAGATCGGAAGA |
| 31137_3 | TATGAGTGTGGAGTCGTTGCCCATGACTTCCAGGAACTCTCTTNNNAGATCGGAAGA |
| 31137_4 | TATGAGTGTGGAGTCGTTGCCCATGACTTCCAGGAACTCTCTTNNNAGATCGGAAGA |
| 31137_5 | TATGAGTGTGGAGTCGTTGCCCATGACTTCCAGGAACTCTCTTNNNAGATCGGAAGA |
| 9623_1 | TATGAGTGTGGAGTCGTTGCTGCTGCTAGATGGAAATCACAGCNNNAGATCGGAAG |
| 9623_2 | TATGAGTGTGGAGTCGTTGCTGCTGCTAGATGGAAATCACAGCNNNAGATCGGAAG |
| 9623_3 | TATGAGTGTGGAGTCGTTGCTGCTGCTAGATGGAAATCACAGCNNNAGATCGGAAG |
| 9623_4 | TATGAGTGTGGAGTCGTTGCTGCTGCTAGATGGAAATCACAGCNNNAGATCGGAAG |
| 9623_5 | TATGAGTGTGGAGTCGTTGCTGCTGCTAGATGGAAATCACAGCNNNAGATCGGAAG |
| 36971_1 | TATGAGTGTGGAGTCGTTGCACACACACACACACACAAAAGAGNNNAGATCGGAAG |
| 36971_2 | TATGAGTGTGGAGTCGTTGCACACACACACACACACAAAAGAGNNNAGATCGGAAG |
| 36971_3 | TATGAGTGTGGAGTCGTTGCACACACACACACACACAAAAGAGNNNAGATCGGAAG |
| 36971_4 | TATGAGTGTGGAGTCGTTGCACACACACACACACACAAAAGAGNNNAGATCGGAAG |
| 36971_5 | TATGAGTGTGGAGTCGTTGCACACACACACACACACAAAAGAGNNNAGATCGGAAG |
| 35826_1 | TATGAGTGTGGAGTCGTTGCGCAATCTTGGCTCACTGCAAGNNNAGATCGGAAGAG |
| 35826_2 | TATGAGTGTGGAGTCGTTGCGCAATCTTGGCTCACTGCAAGNNNAGATCGGAAGAG |
| 35826_3 | TATGAGTGTGGAGTCGTTGCGCAATCTTGGCTCACTGCAAGNNNAGATCGGAAGAG |
| 35826_4 | TATGAGTGTGGAGTCGTTGCGCAATCTTGGCTCACTGCAAGNNNAGATCGGAAGAG |
| 35826_5 | TATGAGTGTGGAGTCGTTGCGCAATCTTGGCTCACTGCAAGNNNAGATCGGAAGAG |
| 44326_1 | TATGAGTGTGGAGTCGTTGCAGAACAGCCTGAATGGTCTCTTTNNNAGATCGGAAG |
| 44326_2 | TATGAGTGTGGAGTCGTTGCAGAACAGCCTGAATGGTCTCTTTNNNAGATCGGAAG |
| 44326_3 | TATGAGTGTGGAGTCGTTGCAGAACAGCCTGAATGGTCTCTTTNNNAGATCGGAAG |

| | |
|---|---|
| 44326_4 | TATGAGTGTGGAGTCGTTGCAGAACAGCCTGAATGGTCTCTTTNNNAGATCGGAAG |
| 44326_5 | TATGAGTGTGGAGTCGTTGCAGAACAGCCTGAATGGTCTCTTTNNNAGATCGGAAG |
| 31101_1 | TATGAGTGTGGAGTCGTTGCAGAATGGATAGCTGCCTCTATGCNNNAGATCGGAAG |
| 31101_2 | TATGAGTGTGGAGTCGTTGCAGAATGGATAGCTGCCTCTATGCNNNAGATCGGAAG |
| 31101_3 | TATGAGTGTGGAGTCGTTGCAGAATGGATAGCTGCCTCTATGCNNNAGATCGGAAG |
| 31101_4 | TATGAGTGTGGAGTCGTTGCAGAATGGATAGCTGCCTCTATGCNNNAGATCGGAAG |
| 31101_5 | TATGAGTGTGGAGTCGTTGCAGAATGGATAGCTGCCTCTATGCNNNAGATCGGAAG |
| 44168_1 | TATGAGTGTGGAGTCGTTGCATCATGAACTGCACAGACTGAGTNNNAGATCGGAAG |
| 44168_2 | TATGAGTGTGGAGTCGTTGCATCATGAACTGCACAGACTGAGTNNNAGATCGGAAG |
| 44168_3 | TATGAGTGTGGAGTCGTTGCATCATGAACTGCACAGACTGAGTNNNAGATCGGAAG |
| 44168_4 | TATGAGTGTGGAGTCGTTGCATCATGAACTGCACAGACTGAGTNNNAGATCGGAAG |
| 44168_5 | TATGAGTGTGGAGTCGTTGCATCATGAACTGCACAGACTGAGTNNNAGATCGGAAG |
| 44425_1 | TATGAGTGTGGAGTCGTTGCCAGGAATCAAACTGCACTGTCACNNNAGATCGGAAG |
| 44425_2 | TATGAGTGTGGAGTCGTTGCCAGGAATCAAACTGCACTGTCACNNNAGATCGGAAG |
| 44425_3 | TATGAGTGTGGAGTCGTTGCCAGGAATCAAACTGCACTGTCACNNNAGATCGGAAG |
| 44425_4 | TATGAGTGTGGAGTCGTTGCCAGGAATCAAACTGCACTGTCACNNNAGATCGGAAG |
| 44425_5 | TATGAGTGTGGAGTCGTTGCCAGGAATCAAACTGCACTGTCACNNNAGATCGGAAG |
| 36264_1 | TATGAGTGTGGAGTCGTTGCAGCAGAAGTAAGTGGACTTTGCTNNNAGATCGGAA |
| 36264_2 | TATGAGTGTGGAGTCGTTGCAGCAGAAGTAAGTGGACTTTGCTNNNAGATCGGAA |
| 36264_3 | TATGAGTGTGGAGTCGTTGCAGCAGAAGTAAGTGGACTTTGCTNNNAGATCGGAA |
| 36264_4 | TATGAGTGTGGAGTCGTTGCAGCAGAAGTAAGTGGACTTTGCTNNNAGATCGGAA |
| 36264_5 | TATGAGTGTGGAGTCGTTGCAGCAGAAGTAAGTGGACTTTGCTNNNAGATCGGAA |
| 52209_1 | TATGAGTGTGGAGTCGTTGCTTTTACCTGTCAGACCACTCAGGNNNAGATCGGAAG |
| 52209_2 | TATGAGTGTGGAGTCGTTGCTTTTACCTGTCAGACCACTCAGGNNNAGATCGGAAG |
| 52209_3 | TATGAGTGTGGAGTCGTTGCTTTTACCTGTCAGACCACTCAGGNNNAGATCGGAAG |
| 52209_4 | TATGAGTGTGGAGTCGTTGCTTTTACCTGTCAGACCACTCAGGNNNAGATCGGAAG |
| 52209_5 | TATGAGTGTGGAGTCGTTGCTTTTACCTGTCAGACCACTCAGGNNNAGATCGGAAG |
| 32610_1 | TATGAGTGTGGAGTCGTTGCCCAAGCTTGTGGTACATTCAGTGNNNAGATCGGAAG |
| 32610_2 | TATGAGTGTGGAGTCGTTGCCCAAGCTTGTGGTACATTCAGTGNNNAGATCGGAAG |
| 32610_3 | TATGAGTGTGGAGTCGTTGCCCAAGCTTGTGGTACATTCAGTGNNNAGATCGGAAG |
| 32610_4 | TATGAGTGTGGAGTCGTTGCCCAAGCTTGTGGTACATTCAGTGNNNAGATCGGAAG |
| 32610_5 | TATGAGTGTGGAGTCGTTGCCCAAGCTTGTGGTACATTCAGTGNNNAGATCGGAAG |
| 13680_1 | TATGAGTGTGGAGTCGTTGCTCTATTTGTGTGGCAGTTATTNNNAGATCGGAAGAG |
| 13680_2 | TATGAGTGTGGAGTCGTTGCTCTATTTGTGTGGCAGTTATTNNNAGATCGGAAGAG |
| 13680_3 | TATGAGTGTGGAGTCGTTGCTCTATTTGTGTGGCAGTTATTNNNAGATCGGAAGAG |
| 13680_4 | TATGAGTGTGGAGTCGTTGCTCTATTTGTGTGGCAGTTATTNNNAGATCGGAAGAG |
| 13680_5 | TATGAGTGTGGAGTCGTTGCTCTATTTGTGTGGCAGTTATTNNNAGATCGGAAGAG |
| 53198_1 | TATGAGTGTGGAGTCGTTGCGAAGGAAGGAAGGAAGGAAGGAANNNAGATCGG |
| 53198_2 | TATGAGTGTGGAGTCGTTGCGAAGGAAGGAAGGAAGGAAGGAANNNAGATCGG |
| 53198_3 | TATGAGTGTGGAGTCGTTGCGAAGGAAGGAAGGAAGGAAGGAANNNAGATCGG |
| 53198_4 | TATGAGTGTGGAGTCGTTGCGAAGGAAGGAAGGAAGGAAGGAANNNAGATCGG |
| 53198_5 | TATGAGTGTGGAGTCGTTGCGAAGGAAGGAAGGAAGGAAGGAANNNAGATCGG |
| 40149_1 | TATGAGTGTGGAGTCGTTGCCTTGGGTGACAGCAAGACAAAATNNNAGATCGGAA |

| | |
|---|---|
| 40149_2 | TATGAGTGTGGAGTCGTTGCCTTGGGTGACAGCAAGACAAAATNNNAGATCGGAA |
| 40149_3 | TATGAGTGTGGAGTCGTTGCCTTGGGTGACAGCAAGACAAAATNNNAGATCGGAA |
| 40149_4 | TATGAGTGTGGAGTCGTTGCCTTGGGTGACAGCAAGACAAAATNNNAGATCGGAA |
| 40149_5 | TATGAGTGTGGAGTCGTTGCCTTGGGTGACAGCAAGACAAAATNNNAGATCGGAA |
| 39571_1 | TATGAGTGTGGAGTCGTTGCAAGCAGGTGTAGCAATCCTAGTCNNNAGATCGGAAG |
| 39571_2 | TATGAGTGTGGAGTCGTTGCAAGCAGGTGTAGCAATCCTAGTCNNNAGATCGGAAG |
| 39571_3 | TATGAGTGTGGAGTCGTTGCAAGCAGGTGTAGCAATCCTAGTCNNNAGATCGGAAG |
| 39571_4 | TATGAGTGTGGAGTCGTTGCAAGCAGGTGTAGCAATCCTAGTCNNNAGATCGGAAG |
| 39571_5 | TATGAGTGTGGAGTCGTTGCAAGCAGGTGTAGCAATCCTAGTCNNNAGATCGGAAG |
| 42106_1 | TATGAGTGTGGAGTCGTTGCATCTCCTGGGTTCAAGCAATTCTNNNAGATCGGAAG |
| 42106_2 | TATGAGTGTGGAGTCGTTGCATCTCCTGGGTTCAAGCAATTCTNNNAGATCGGAAG |
| 42106_3 | TATGAGTGTGGAGTCGTTGCATCTCCTGGGTTCAAGCAATTCTNNNAGATCGGAAG |
| 42106_4 | TATGAGTGTGGAGTCGTTGCATCTCCTGGGTTCAAGCAATTCTNNNAGATCGGAAG |
| 42106_5 | TATGAGTGTGGAGTCGTTGCATCTCCTGGGTTCAAGCAATTCTNNNAGATCGGAAG |
| 30534_1 | TATGAGTGTGGAGTCGTTGCCTACGTTCATACAGACAAACAGACATNNNAGATCGG |
| 30534_2 | TATGAGTGTGGAGTCGTTGCCTACGTTCATACAGACAAACAGACATNNNAGATCGG |
| 30534_3 | TATGAGTGTGGAGTCGTTGCCTACGTTCATACAGACAAACAGACATNNNAGATCGG |
| 30534_4 | TATGAGTGTGGAGTCGTTGCCTACGTTCATACAGACAAACAGACATNNNAGATCGG |
| 30534_5 | TATGAGTGTGGAGTCGTTGCCTACGTTCATACAGACAAACAGACATNNNAGATCGG |
| 38869_1 | TATGAGTGTGGAGTCGTTGCAAGGCTTAGAAACGACAACGAGANNNAGATCGGAA |
| 38869_2 | TATGAGTGTGGAGTCGTTGCAAGGCTTAGAAACGACAACGAGANNNAGATCGGAA |
| 38869_3 | TATGAGTGTGGAGTCGTTGCAAGGCTTAGAAACGACAACGAGANNNAGATCGGAA |
| 38869_4 | TATGAGTGTGGAGTCGTTGCAAGGCTTAGAAACGACAACGAGANNNAGATCGGAA |
| 38869_5 | TATGAGTGTGGAGTCGTTGCAAGGCTTAGAAACGACAACGAGANNNAGATCGGAA |
| 36939_1 | TATGAGTGTGGAGTCGTTGCCCTCCCAAAGTGCCGAGATTNNNAGATCGGAAGAGC |
| 36939_2 | TATGAGTGTGGAGTCGTTGCCCTCCCAAAGTGCCGAGATTNNNAGATCGGAAGAGC |
| 36939_3 | TATGAGTGTGGAGTCGTTGCCCTCCCAAAGTGCCGAGATTNNNAGATCGGAAGAGC |
| 36939_4 | TATGAGTGTGGAGTCGTTGCCCTCCCAAAGTGCCGAGATTNNNAGATCGGAAGAGC |
| 36939_5 | TATGAGTGTGGAGTCGTTGCCCTCCCAAAGTGCCGAGATTNNNAGATCGGAAGAGC |
| 50329_1 | TATGAGTGTGGAGTCGTTGCTTGAAATCAGGAGCTTGAGACCANNNAGATCGGAA |
| 50329_2 | TATGAGTGTGGAGTCGTTGCTTGAAATCAGGAGCTTGAGACCANNNAGATCGGAA |
| 50329_3 | TATGAGTGTGGAGTCGTTGCTTGAAATCAGGAGCTTGAGACCANNNAGATCGGAA |
| 50329_4 | TATGAGTGTGGAGTCGTTGCTTGAAATCAGGAGCTTGAGACCANNNAGATCGGAA |
| 50329_5 | TATGAGTGTGGAGTCGTTGCTTGAAATCAGGAGCTTGAGACCANNNAGATCGGAA |
| 28801_1 | TATGAGTGTGGAGTCGTTGCTTAGTGCCTGGGTGACGAAATAANNNAGATCGGAA |
| 28801_2 | TATGAGTGTGGAGTCGTTGCTTAGTGCCTGGGTGACGAAATAANNNAGATCGGAA |
| 28801_3 | TATGAGTGTGGAGTCGTTGCTTAGTGCCTGGGTGACGAAATAANNNAGATCGGAA |
| 28801_4 | TATGAGTGTGGAGTCGTTGCTTAGTGCCTGGGTGACGAAATAANNNAGATCGGAA |
| 28801_5 | TATGAGTGTGGAGTCGTTGCTTAGTGCCTGGGTGACGAAATAANNNAGATCGGAA |
| 52000_1 | TATGAGTGTGGAGTCGTTGCCGCCAAGGCATATGTACTTTCTTNNNAGATCGGAAG |
| 52000_2 | TATGAGTGTGGAGTCGTTGCCGCCAAGGCATATGTACTTTCTTNNNAGATCGGAAG |
| 52000_3 | TATGAGTGTGGAGTCGTTGCCGCCAAGGCATATGTACTTTCTTNNNAGATCGGAAG |
| 52000_4 | TATGAGTGTGGAGTCGTTGCCGCCAAGGCATATGTACTTTCTTNNNAGATCGGAAG |

| | |
|---|---|
| 52000_5 | TATGAGTGTGGAGTCGTTGCCGCCAAGGCATATGTACTTTCTTNNNAGATCGGAAG |
| 35593_1 | TATGAGTGTGGAGTCGTTGCTGAGCTGCCAAAATTTGAGATGGNNNAGATCGGAA |
| 35593_2 | TATGAGTGTGGAGTCGTTGCTGAGCTGCCAAAATTTGAGATGGNNNAGATCGGAA |
| 35593_3 | TATGAGTGTGGAGTCGTTGCTGAGCTGCCAAAATTTGAGATGGNNNAGATCGGAA |
| 35593_4 | TATGAGTGTGGAGTCGTTGCTGAGCTGCCAAAATTTGAGATGGNNNAGATCGGAA |
| 35593_5 | TATGAGTGTGGAGTCGTTGCTGAGCTGCCAAAATTTGAGATGGNNNAGATCGGAA |
| 29933_1 | TATGAGTGTGGAGTCGTTGCCACAGAAAGCATTTTGGGGTAGGNNNAGATCGGAA |
| 29933_2 | TATGAGTGTGGAGTCGTTGCCACAGAAAGCATTTTGGGGTAGGNNNAGATCGGAA |
| 29933_3 | TATGAGTGTGGAGTCGTTGCCACAGAAAGCATTTTGGGGTAGGNNNAGATCGGAA |
| 29933_4 | TATGAGTGTGGAGTCGTTGCCACAGAAAGCATTTTGGGGTAGGNNNAGATCGGAA |
| 29933_5 | TATGAGTGTGGAGTCGTTGCCACAGAAAGCATTTTGGGGTAGGNNNAGATCGGAA |
| 46797_1 | TATGAGTGTGGAGTCGTTGCGAGTAGCTGGGACTACAGGCANNNAGATCGGAAGA |
| 46797_2 | TATGAGTGTGGAGTCGTTGCGAGTAGCTGGGACTACAGGCANNNAGATCGGAAGA |
| 46797_3 | TATGAGTGTGGAGTCGTTGCGAGTAGCTGGGACTACAGGCANNNAGATCGGAAGA |
| 46797_4 | TATGAGTGTGGAGTCGTTGCGAGTAGCTGGGACTACAGGCANNNAGATCGGAAGA |
| 46797_5 | TATGAGTGTGGAGTCGTTGCGAGTAGCTGGGACTACAGGCANNNAGATCGGAAGA |
| 8536_1 | TATGAGTGTGGAGTCGTTGCAAGCAAAAATGCCATGTTCCACATCTNNNAGATCGG |
| 8536_2 | TATGAGTGTGGAGTCGTTGCAAGCAAAAATGCCATGTTCCACATCTNNNAGATCGG |
| 8536_3 | TATGAGTGTGGAGTCGTTGCAAGCAAAAATGCCATGTTCCACATCTNNNAGATCGG |
| 8536_4 | TATGAGTGTGGAGTCGTTGCAAGCAAAAATGCCATGTTCCACATCTNNNAGATCGG |
| 8536_5 | TATGAGTGTGGAGTCGTTGCAAGCAAAAATGCCATGTTCCACATCTNNNAGATCGG |
| 12665_1 | TATGAGTGTGGAGTCGTTGCCCCAGAAAGGTAAGAAAATTAAANNNAGATCGGAA |
| 12665_2 | TATGAGTGTGGAGTCGTTGCCCCAGAAAGGTAAGAAAATTAAANNNAGATCGGAA |
| 12665_3 | TATGAGTGTGGAGTCGTTGCCCCAGAAAGGTAAGAAAATTAAANNNAGATCGGAA |
| 12665_4 | TATGAGTGTGGAGTCGTTGCCCCAGAAAGGTAAGAAAATTAAANNNAGATCGGAA |
| 12665_5 | TATGAGTGTGGAGTCGTTGCCCCAGAAAGGTAAGAAAATTAAANNNAGATCGGAA |
| 29286_1 | TATGAGTGTGGAGTCGTTGCCATATGTGCATGTGTGTGTCTCTNNNAGATCGGAAG |
| 29286_2 | TATGAGTGTGGAGTCGTTGCCATATGTGCATGTGTGTGTCTCTNNNAGATCGGAAG |
| 29286_3 | TATGAGTGTGGAGTCGTTGCCATATGTGCATGTGTGTGTCTCTNNNAGATCGGAAG |
| 29286_4 | TATGAGTGTGGAGTCGTTGCCATATGTGCATGTGTGTGTCTCTNNNAGATCGGAAG |
| 29286_5 | TATGAGTGTGGAGTCGTTGCCATATGTGCATGTGTGTGTCTCTNNNAGATCGGAAG |
| 44336_1 | TATGAGTGTGGAGTCGTTGCGACTTCTGAGGGTGGATTCATCANNNAGATCGGAA |
| 44336_2 | TATGAGTGTGGAGTCGTTGCGACTTCTGAGGGTGGATTCATCANNNAGATCGGAA |
| 44336_3 | TATGAGTGTGGAGTCGTTGCGACTTCTGAGGGTGGATTCATCANNNAGATCGGAA |
| 44336_4 | TATGAGTGTGGAGTCGTTGCGACTTCTGAGGGTGGATTCATCANNNAGATCGGAA |
| 44336_5 | TATGAGTGTGGAGTCGTTGCGACTTCTGAGGGTGGATTCATCANNNAGATCGGAA |
| 12568_1 | TATGAGTGTGGAGTCGTTGCGAAAACACTTTAAACCTCTTTTGNNNAGATCGGAAG |
| 12568_2 | TATGAGTGTGGAGTCGTTGCGAAAACACTTTAAACCTCTTTTGNNNAGATCGGAAG |
| 12568_3 | TATGAGTGTGGAGTCGTTGCGAAAACACTTTAAACCTCTTTTGNNNAGATCGGAAG |
| 12568_4 | TATGAGTGTGGAGTCGTTGCGAAAACACTTTAAACCTCTTTTGNNNAGATCGGAAG |
| 12568_5 | TATGAGTGTGGAGTCGTTGCGAAAACACTTTAAACCTCTTTTGNNNAGATCGGAAG |
| 31647_1 | TATGAGTGTGGAGTCGTTGCTTCACAAGAGCAATGTAGGAGCTNNNAGATCGGAA |
| 31647_2 | TATGAGTGTGGAGTCGTTGCTTCACAAGAGCAATGTAGGAGCTNNNAGATCGGAA |

| | |
|---|---|
| 31647_3 | TATGAGTGTGGAGTCGTTGCTTCACAAGAGCAATGTAGGAGCTNNNAGATCGGAAC |
| 31647_4 | TATGAGTGTGGAGTCGTTGCTTCACAAGAGCAATGTAGGAGCTNNNAGATCGGAAC |
| 31647_5 | TATGAGTGTGGAGTCGTTGCTTCACAAGAGCAATGTAGGAGCTNNNAGATCGGAAC |
| 45008_1 | TATGAGTGTGGAGTCGTTGCGTTCCAAGATTCTGCCTACTGGANNNAGATCGGAAG |
| 45008_2 | TATGAGTGTGGAGTCGTTGCGTTCCAAGATTCTGCCTACTGGANNNAGATCGGAAG |
| 45008_3 | TATGAGTGTGGAGTCGTTGCGTTCCAAGATTCTGCCTACTGGANNNAGATCGGAAG |
| 45008_4 | TATGAGTGTGGAGTCGTTGCGTTCCAAGATTCTGCCTACTGGANNNAGATCGGAAG |
| 45008_5 | TATGAGTGTGGAGTCGTTGCGTTCCAAGATTCTGCCTACTGGANNNAGATCGGAAG |
| 47953_1 | TATGAGTGTGGAGTCGTTGCTTCAAGTGATTCTCCTGCCTCAGNNNAGATCGGAAGA |
| 47953_2 | TATGAGTGTGGAGTCGTTGCTTCAAGTGATTCTCCTGCCTCAGNNNAGATCGGAAGA |
| 47953_3 | TATGAGTGTGGAGTCGTTGCTTCAAGTGATTCTCCTGCCTCAGNNNAGATCGGAAGA |
| 47953_4 | TATGAGTGTGGAGTCGTTGCTTCAAGTGATTCTCCTGCCTCAGNNNAGATCGGAAGA |
| 47953_5 | TATGAGTGTGGAGTCGTTGCTTCAAGTGATTCTCCTGCCTCAGNNNAGATCGGAAGA |
| 39239_1 | TATGAGTGTGGAGTCGTTGCTCTAACCATTTGCCTCACAGACANNNAGATCGGAAGA |
| 39239_2 | TATGAGTGTGGAGTCGTTGCTCTAACCATTTGCCTCACAGACANNNAGATCGGAAGA |
| 39239_3 | TATGAGTGTGGAGTCGTTGCTCTAACCATTTGCCTCACAGACANNNAGATCGGAAGA |
| 39239_4 | TATGAGTGTGGAGTCGTTGCTCTAACCATTTGCCTCACAGACANNNAGATCGGAAGA |
| 39239_5 | TATGAGTGTGGAGTCGTTGCTCTAACCATTTGCCTCACAGACANNNAGATCGGAAGA |
| 51235_1 | TATGAGTGTGGAGTCGTTGCTTAGCCCTCAGAGTAGGTACCTCNNNAGATCGGAAG |
| 51235_2 | TATGAGTGTGGAGTCGTTGCTTAGCCCTCAGAGTAGGTACCTCNNNAGATCGGAAG |
| 51235_3 | TATGAGTGTGGAGTCGTTGCTTAGCCCTCAGAGTAGGTACCTCNNNAGATCGGAAG |
| 51235_4 | TATGAGTGTGGAGTCGTTGCTTAGCCCTCAGAGTAGGTACCTCNNNAGATCGGAAG |
| 51235_5 | TATGAGTGTGGAGTCGTTGCTTAGCCCTCAGAGTAGGTACCTCNNNAGATCGGAAG |
| 31983_1 | TATGAGTGTGGAGTCGTTGCAAAAGCAACCCTAACACTACGGANNNAGATCGGAAG |
| 31983_2 | TATGAGTGTGGAGTCGTTGCAAAAGCAACCCTAACACTACGGANNNAGATCGGAAG |
| 31983_3 | TATGAGTGTGGAGTCGTTGCAAAAGCAACCCTAACACTACGGANNNAGATCGGAAG |
| 31983_4 | TATGAGTGTGGAGTCGTTGCAAAAGCAACCCTAACACTACGGANNNAGATCGGAAG |
| 31983_5 | TATGAGTGTGGAGTCGTTGCAAAAGCAACCCTAACACTACGGANNNAGATCGGAAG |
| 31994_1 | TATGAGTGTGGAGTCGTTGCGACATGATGCCACAAGTGGAAAANNNAGATCGGAAC |
| 31994_2 | TATGAGTGTGGAGTCGTTGCGACATGATGCCACAAGTGGAAAANNNAGATCGGAAC |
| 31994_3 | TATGAGTGTGGAGTCGTTGCGACATGATGCCACAAGTGGAAAANNNAGATCGGAAC |
| 31994_4 | TATGAGTGTGGAGTCGTTGCGACATGATGCCACAAGTGGAAAANNNAGATCGGAAC |
| 31994_5 | TATGAGTGTGGAGTCGTTGCGACATGATGCCACAAGTGGAAAANNNAGATCGGAAC |
| 43303_1 | TATGAGTGTGGAGTCGTTGCGAGTTTTGCTCTTGTTGCCCAGNNNAGATCGGAAGA |
| 43303_2 | TATGAGTGTGGAGTCGTTGCGAGTTTTGCTCTTGTTGCCCAGNNNAGATCGGAAGA |
| 43303_3 | TATGAGTGTGGAGTCGTTGCGAGTTTTGCTCTTGTTGCCCAGNNNAGATCGGAAGA |
| 43303_4 | TATGAGTGTGGAGTCGTTGCGAGTTTTGCTCTTGTTGCCCAGNNNAGATCGGAAGA |
| 43303_5 | TATGAGTGTGGAGTCGTTGCGAGTTTTGCTCTTGTTGCCCAGNNNAGATCGGAAGA |
| 45589_1 | TATGAGTGTGGAGTCGTTGCCTTCTGAGTAGCTGGGACTACAGNNNAGATCGGAAC |
| 45589_2 | TATGAGTGTGGAGTCGTTGCCTTCTGAGTAGCTGGGACTACAGNNNAGATCGGAAC |
| 45589_3 | TATGAGTGTGGAGTCGTTGCCTTCTGAGTAGCTGGGACTACAGNNNAGATCGGAAC |
| 45589_4 | TATGAGTGTGGAGTCGTTGCCTTCTGAGTAGCTGGGACTACAGNNNAGATCGGAAC |
| 45589_5 | TATGAGTGTGGAGTCGTTGCCTTCTGAGTAGCTGGGACTACAGNNNAGATCGGAAC |

| | |
|---|---|
| 42982_1 | TATGAGTGTGGAGTCGTTGCGAGAGAGAGAAAGAGGCAGACAGNNNAGATCGGA |
| 42982_2 | TATGAGTGTGGAGTCGTTGCGAGAGAGAGAAAGAGGCAGACAGNNNAGATCGGA |
| 42982_3 | TATGAGTGTGGAGTCGTTGCGAGAGAGAGAAAGAGGCAGACAGNNNAGATCGGA |
| 42982_4 | TATGAGTGTGGAGTCGTTGCGAGAGAGAGAAAGAGGCAGACAGNNNAGATCGGA |
| 42982_5 | TATGAGTGTGGAGTCGTTGCGAGAGAGAGAAAGAGGCAGACAGNNNAGATCGGA |
| 40614_1 | TATGAGTGTGGAGTCGTTGCATGTAGTCCCAGCTACTGAGGAGNNNAGATCGGAAC |
| 40614_2 | TATGAGTGTGGAGTCGTTGCATGTAGTCCCAGCTACTGAGGAGNNNAGATCGGAAC |
| 40614_3 | TATGAGTGTGGAGTCGTTGCATGTAGTCCCAGCTACTGAGGAGNNNAGATCGGAAC |
| 40614_4 | TATGAGTGTGGAGTCGTTGCATGTAGTCCCAGCTACTGAGGAGNNNAGATCGGAAC |
| 40614_5 | TATGAGTGTGGAGTCGTTGCATGTAGTCCCAGCTACTGAGGAGNNNAGATCGGAAC |
| 48477_1 | TATGAGTGTGGAGTCGTTGCCATCACCATGCCCAGCTAATTTTNNNAGATCGGAAGA |
| 48477_2 | TATGAGTGTGGAGTCGTTGCCATCACCATGCCCAGCTAATTTTNNNAGATCGGAAGA |
| 48477_3 | TATGAGTGTGGAGTCGTTGCCATCACCATGCCCAGCTAATTTTNNNAGATCGGAAGA |
| 48477_4 | TATGAGTGTGGAGTCGTTGCCATCACCATGCCCAGCTAATTTTNNNAGATCGGAAGA |
| 48477_5 | TATGAGTGTGGAGTCGTTGCCATCACCATGCCCAGCTAATTTTNNNAGATCGGAAGA |
| 32946_1 | TATGAGTGTGGAGTCGTTGCAAGACGGGAGTAGAAAAAGGCAANNNAGATCGGAA |
| 32946_2 | TATGAGTGTGGAGTCGTTGCAAGACGGGAGTAGAAAAAGGCAANNNAGATCGGAA |
| 32946_3 | TATGAGTGTGGAGTCGTTGCAAGACGGGAGTAGAAAAAGGCAANNNAGATCGGAA |
| 32946_4 | TATGAGTGTGGAGTCGTTGCAAGACGGGAGTAGAAAAAGGCAANNNAGATCGGAA |
| 32946_5 | TATGAGTGTGGAGTCGTTGCAAGACGGGAGTAGAAAAAGGCAANNNAGATCGGAA |
| 44510_1 | TATGAGTGTGGAGTCGTTGCCACACACACACACACACACATATNNNAGATCGGAAGA |
| 44510_2 | TATGAGTGTGGAGTCGTTGCCACACACACACACACACACATATNNNAGATCGGAAGA |
| 44510_3 | TATGAGTGTGGAGTCGTTGCCACACACACACACACACACATATNNNAGATCGGAAGA |
| 44510_4 | TATGAGTGTGGAGTCGTTGCCACACACACACACACACACATATNNNAGATCGGAAGA |
| 44510_5 | TATGAGTGTGGAGTCGTTGCCACACACACACACACACACATATNNNAGATCGGAAGA |
| 35915_1 | TATGAGTGTGGAGTCGTTGCAACATGGTGAAACCTGGTCTCTANNNAGATCGGAAG |
| 35915_2 | TATGAGTGTGGAGTCGTTGCAACATGGTGAAACCTGGTCTCTANNNAGATCGGAAG |
| 35915_3 | TATGAGTGTGGAGTCGTTGCAACATGGTGAAACCTGGTCTCTANNNAGATCGGAAG |
| 35915_4 | TATGAGTGTGGAGTCGTTGCAACATGGTGAAACCTGGTCTCTANNNAGATCGGAAG |
| 35915_5 | TATGAGTGTGGAGTCGTTGCAACATGGTGAAACCTGGTCTCTANNNAGATCGGAAG |
| 53069_1 | TATGAGTGTGGAGTCGTTGCTGACAGACGACACAGAGATTGTTNNNAGATCGGAAC |
| 53069_2 | TATGAGTGTGGAGTCGTTGCTGACAGACGACACAGAGATTGTTNNNAGATCGGAAC |
| 53069_3 | TATGAGTGTGGAGTCGTTGCTGACAGACGACACAGAGATTGTTNNNAGATCGGAAC |
| 53069_4 | TATGAGTGTGGAGTCGTTGCTGACAGACGACACAGAGATTGTTNNNAGATCGGAAC |
| 53069_5 | TATGAGTGTGGAGTCGTTGCTGACAGACGACACAGAGATTGTTNNNAGATCGGAAC |
| 36930_1 | TATGAGTGTGGAGTCGTTGCTGAGTGACCTGATGTGTAGTGACNNNAGATCGGAAC |
| 36930_2 | TATGAGTGTGGAGTCGTTGCTGAGTGACCTGATGTGTAGTGACNNNAGATCGGAAC |
| 36930_3 | TATGAGTGTGGAGTCGTTGCTGAGTGACCTGATGTGTAGTGACNNNAGATCGGAAC |
| 36930_4 | TATGAGTGTGGAGTCGTTGCTGAGTGACCTGATGTGTAGTGACNNNAGATCGGAAC |
| 36930_5 | TATGAGTGTGGAGTCGTTGCTGAGTGACCTGATGTGTAGTGACNNNAGATCGGAAC |
| 47647_1 | TATGAGTGTGGAGTCGTTGCTATGAAGCAGTCACTCCCCTTTCNNNAGATCGGAAGA |
| 47647_2 | TATGAGTGTGGAGTCGTTGCTATGAAGCAGTCACTCCCCTTTCNNNAGATCGGAAGA |
| 47647_3 | TATGAGTGTGGAGTCGTTGCTATGAAGCAGTCACTCCCCTTTCNNNAGATCGGAAGA |

| | |
|---|---|
| 47647_4 | TATGAGTGTGGAGTCGTTGCTATGAAGCAGTCACTCCCCTTTCNNNAGATCGGAAG |
| 47647_5 | TATGAGTGTGGAGTCGTTGCTATGAAGCAGTCACTCCCCTTTCNNNAGATCGGAAG |
| 35358_1 | TATGAGTGTGGAGTCGTTGCGCTAGAGTACAGTGGTGCAATCTNNNAGATCGGAA |
| 35358_2 | TATGAGTGTGGAGTCGTTGCGCTAGAGTACAGTGGTGCAATCTNNNAGATCGGAA |
| 35358_3 | TATGAGTGTGGAGTCGTTGCGCTAGAGTACAGTGGTGCAATCTNNNAGATCGGAA |
| 35358_4 | TATGAGTGTGGAGTCGTTGCGCTAGAGTACAGTGGTGCAATCTNNNAGATCGGAA |
| 35358_5 | TATGAGTGTGGAGTCGTTGCGCTAGAGTACAGTGGTGCAATCTNNNAGATCGGAA |
| 45033_1 | TATGAGTGTGGAGTCGTTGCCTGGGAAGGCTTCTGTCTCTTTNNNAGATCGGAAGA |
| 45033_2 | TATGAGTGTGGAGTCGTTGCCTGGGAAGGCTTCTGTCTCTTTNNNAGATCGGAAGA |
| 45033_3 | TATGAGTGTGGAGTCGTTGCCTGGGAAGGCTTCTGTCTCTTTNNNAGATCGGAAGA |
| 45033_4 | TATGAGTGTGGAGTCGTTGCCTGGGAAGGCTTCTGTCTCTTTNNNAGATCGGAAGA |
| 45033_5 | TATGAGTGTGGAGTCGTTGCCTGGGAAGGCTTCTGTCTCTTTNNNAGATCGGAAGA |
| 52806_1 | TATGAGTGTGGAGTCGTTGCAAATGACCACAAACCGCTTGGNNNAGATCGGAAGA |
| 52806_2 | TATGAGTGTGGAGTCGTTGCAAATGACCACAAACCGCTTGGNNNAGATCGGAAGA |
| 52806_3 | TATGAGTGTGGAGTCGTTGCAAATGACCACAAACCGCTTGGNNNAGATCGGAAGA |
| 52806_4 | TATGAGTGTGGAGTCGTTGCAAATGACCACAAACCGCTTGGNNNAGATCGGAAGA |
| 52806_5 | TATGAGTGTGGAGTCGTTGCAAATGACCACAAACCGCTTGGNNNAGATCGGAAGA |
| 37938_1 | TATGAGTGTGGAGTCGTTGCGCTGGGAGAACTCTGAATAGGAANNNAGATCGGAA |
| 37938_2 | TATGAGTGTGGAGTCGTTGCGCTGGGAGAACTCTGAATAGGAANNNAGATCGGAA |
| 37938_3 | TATGAGTGTGGAGTCGTTGCGCTGGGAGAACTCTGAATAGGAANNNAGATCGGAA |
| 37938_4 | TATGAGTGTGGAGTCGTTGCGCTGGGAGAACTCTGAATAGGAANNNAGATCGGAA |
| 37938_5 | TATGAGTGTGGAGTCGTTGCGCTGGGAGAACTCTGAATAGGAANNNAGATCGGAA |
| 41680_1 | TATGAGTGTGGAGTCGTTGCCCAGGTTCAAGCGATTCTCTTTGNNNAGATCGGAAG |
| 41680_2 | TATGAGTGTGGAGTCGTTGCCCAGGTTCAAGCGATTCTCTTTGNNNAGATCGGAAG |
| 41680_3 | TATGAGTGTGGAGTCGTTGCCCAGGTTCAAGCGATTCTCTTTGNNNAGATCGGAAG |
| 41680_4 | TATGAGTGTGGAGTCGTTGCCCAGGTTCAAGCGATTCTCTTTGNNNAGATCGGAAG |
| 41680_5 | TATGAGTGTGGAGTCGTTGCCCAGGTTCAAGCGATTCTCTTTGNNNAGATCGGAAG |
| 31220_1 | TATGAGTGTGGAGTCGTTGCCTCCTCAAAAAGCAGAGCAAACANNNAGATCGGAAG |
| 31220_2 | TATGAGTGTGGAGTCGTTGCCTCCTCAAAAAGCAGAGCAAACANNNAGATCGGAAG |
| 31220_3 | TATGAGTGTGGAGTCGTTGCCTCCTCAAAAAGCAGAGCAAACANNNAGATCGGAAG |
| 31220_4 | TATGAGTGTGGAGTCGTTGCCTCCTCAAAAAGCAGAGCAAACANNNAGATCGGAAG |
| 31220_5 | TATGAGTGTGGAGTCGTTGCCTCCTCAAAAAGCAGAGCAAACANNNAGATCGGAAG |
| 33039_1 | TATGAGTGTGGAGTCGTTGCCTCCCTGGTTCTAGCGATTCTGNNNAGATCGGAAGA |
| 33039_2 | TATGAGTGTGGAGTCGTTGCCTCCCTGGTTCTAGCGATTCTGNNNAGATCGGAAGA |
| 33039_3 | TATGAGTGTGGAGTCGTTGCCTCCCTGGTTCTAGCGATTCTGNNNAGATCGGAAGA |
| 33039_4 | TATGAGTGTGGAGTCGTTGCCTCCCTGGTTCTAGCGATTCTGNNNAGATCGGAAGA |
| 33039_5 | TATGAGTGTGGAGTCGTTGCCTCCCTGGTTCTAGCGATTCTGNNNAGATCGGAAGA |
| 42793_1 | TATGAGTGTGGAGTCGTTGCAAATCATGGCATTCTCAGCAACCNNNAGATCGGAAG |
| 42793_2 | TATGAGTGTGGAGTCGTTGCAAATCATGGCATTCTCAGCAACCNNNAGATCGGAAG |
| 42793_3 | TATGAGTGTGGAGTCGTTGCAAATCATGGCATTCTCAGCAACCNNNAGATCGGAAG |
| 42793_4 | TATGAGTGTGGAGTCGTTGCAAATCATGGCATTCTCAGCAACCNNNAGATCGGAAG |
| 42793_5 | TATGAGTGTGGAGTCGTTGCAAATCATGGCATTCTCAGCAACCNNNAGATCGGAAG |
| 30669_1 | TATGAGTGTGGAGTCGTTGCGTAAGCCATTGAGCATCAGAGTCNNNAGATCGGAA |

| | |
|---|---|
| 30669_2 | TATGAGTGTGGAGTCGTTGCGTAAGCCATTGAGCATCAGAGTCNNNAGATCGGAAG |
| 30669_3 | TATGAGTGTGGAGTCGTTGCGTAAGCCATTGAGCATCAGAGTCNNNAGATCGGAAG |
| 30669_4 | TATGAGTGTGGAGTCGTTGCGTAAGCCATTGAGCATCAGAGTCNNNAGATCGGAAG |
| 30669_5 | TATGAGTGTGGAGTCGTTGCGTAAGCCATTGAGCATCAGAGTCNNNAGATCGGAAG |
| 42861_1 | TATGAGTGTGGAGTCGTTGCGCTGAAGTGCAATGGTGTGATTTNNNAGATCGGAA |
| 42861_2 | TATGAGTGTGGAGTCGTTGCGCTGAAGTGCAATGGTGTGATTTNNNAGATCGGAA |
| 42861_3 | TATGAGTGTGGAGTCGTTGCGCTGAAGTGCAATGGTGTGATTTNNNAGATCGGAA |
| 42861_4 | TATGAGTGTGGAGTCGTTGCGCTGAAGTGCAATGGTGTGATTTNNNAGATCGGAA |
| 42861_5 | TATGAGTGTGGAGTCGTTGCGCTGAAGTGCAATGGTGTGATTTNNNAGATCGGAA |
| 52416_1 | TATGAGTGTGGAGTCGTTGCACCCCCCTTTTTAGAGCAATGAGNNNAGATCGGAAG |
| 52416_2 | TATGAGTGTGGAGTCGTTGCACCCCCCTTTTTAGAGCAATGAGNNNAGATCGGAAG |
| 52416_3 | TATGAGTGTGGAGTCGTTGCACCCCCCTTTTTAGAGCAATGAGNNNAGATCGGAAG |
| 52416_4 | TATGAGTGTGGAGTCGTTGCACCCCCCTTTTTAGAGCAATGAGNNNAGATCGGAAG |
| 52416_5 | TATGAGTGTGGAGTCGTTGCACCCCCCTTTTTAGAGCAATGAGNNNAGATCGGAAG |
| 52656_1 | TATGAGTGTGGAGTCGTTGCAGAGGACCCCAAATAATACAGCGNNNAGATCGGAA |
| 52656_2 | TATGAGTGTGGAGTCGTTGCAGAGGACCCCAAATAATACAGCGNNNAGATCGGAA |
| 52656_3 | TATGAGTGTGGAGTCGTTGCAGAGGACCCCAAATAATACAGCGNNNAGATCGGAA |
| 52656_4 | TATGAGTGTGGAGTCGTTGCAGAGGACCCCAAATAATACAGCGNNNAGATCGGAA |
| 52656_5 | TATGAGTGTGGAGTCGTTGCAGAGGACCCCAAATAATACAGCGNNNAGATCGGAA |
| 53184_1 | TATGAGTGTGGAGTCGTTGCTCTATAGTGAGCATCGCCTGAACNNNAGATCGGAAG |
| 53184_2 | TATGAGTGTGGAGTCGTTGCTCTATAGTGAGCATCGCCTGAACNNNAGATCGGAAG |
| 53184_3 | TATGAGTGTGGAGTCGTTGCTCTATAGTGAGCATCGCCTGAACNNNAGATCGGAAG |
| 53184_4 | TATGAGTGTGGAGTCGTTGCTCTATAGTGAGCATCGCCTGAACNNNAGATCGGAAG |
| 53184_5 | TATGAGTGTGGAGTCGTTGCTCTATAGTGAGCATCGCCTGAACNNNAGATCGGAAG |
| 13508_1 | TATGAGTGTGGAGTCGTTGCGAGTATCTAGTAATTTGGAAGGTNNNAGATCGGAA |
| 13508_2 | TATGAGTGTGGAGTCGTTGCGAGTATCTAGTAATTTGGAAGGTNNNAGATCGGAA |
| 13508_3 | TATGAGTGTGGAGTCGTTGCGAGTATCTAGTAATTTGGAAGGTNNNAGATCGGAA |
| 13508_4 | TATGAGTGTGGAGTCGTTGCGAGTATCTAGTAATTTGGAAGGTNNNAGATCGGAA |
| 13508_5 | TATGAGTGTGGAGTCGTTGCGAGTATCTAGTAATTTGGAAGGTNNNAGATCGGAA |
| 35783_1 | TATGAGTGTGGAGTCGTTGCAGCCTTGACCTCATGGACTAAAGNNNAGATCGGAAG |
| 35783_2 | TATGAGTGTGGAGTCGTTGCAGCCTTGACCTCATGGACTAAAGNNNAGATCGGAAG |
| 35783_3 | TATGAGTGTGGAGTCGTTGCAGCCTTGACCTCATGGACTAAAGNNNAGATCGGAAG |
| 35783_4 | TATGAGTGTGGAGTCGTTGCAGCCTTGACCTCATGGACTAAAGNNNAGATCGGAAG |
| 35783_5 | TATGAGTGTGGAGTCGTTGCAGCCTTGACCTCATGGACTAAAGNNNAGATCGGAAG |
| 29332_1 | TATGAGTGTGGAGTCGTTGCCACACATGTTCACACACACACATNNNAGATCGGAAG |
| 29332_2 | TATGAGTGTGGAGTCGTTGCCACACATGTTCACACACACACATNNNAGATCGGAAG |
| 29332_3 | TATGAGTGTGGAGTCGTTGCCACACATGTTCACACACACACATNNNAGATCGGAAG |
| 29332_4 | TATGAGTGTGGAGTCGTTGCCACACATGTTCACACACACACATNNNAGATCGGAAG |
| 29332_5 | TATGAGTGTGGAGTCGTTGCCACACATGTTCACACACACACATNNNAGATCGGAAG |
| 50399_1 | TATGAGTGTGGAGTCGTTGCCAGCCTGTGAAGGGAAGGAGNNNAGATCGGAAGA |
| 50399_2 | TATGAGTGTGGAGTCGTTGCCAGCCTGTGAAGGGAAGGAGNNNAGATCGGAAGA |
| 50399_3 | TATGAGTGTGGAGTCGTTGCCAGCCTGTGAAGGGAAGGAGNNNAGATCGGAAGA |
| 50399_4 | TATGAGTGTGGAGTCGTTGCCAGCCTGTGAAGGGAAGGAGNNNAGATCGGAAGA |

| | |
|---|---|
| 50399_5 | TATGAGTGTGGAGTCGTTGCCAGCCTGTGAAGGGAAGGAGNNNAGATCGGAAGA |
| 46999_1 | TATGAGTGTGGAGTCGTTGCCACCAAGTAACACCACACACACNNNAGATCGGAAGA |
| 46999_2 | TATGAGTGTGGAGTCGTTGCCACCAAGTAACACCACACACACNNNAGATCGGAAGA |
| 46999_3 | TATGAGTGTGGAGTCGTTGCCACCAAGTAACACCACACACACNNNAGATCGGAAGA |
| 46999_4 | TATGAGTGTGGAGTCGTTGCCACCAAGTAACACCACACACACNNNAGATCGGAAGA |
| 46999_5 | TATGAGTGTGGAGTCGTTGCCACCAAGTAACACCACACACACNNNAGATCGGAAGA |
| 38079_1 | TATGAGTGTGGAGTCGTTGCAAAATGGTGGCAGAAGAACACAGNNNAGATCGGA |
| 38079_2 | TATGAGTGTGGAGTCGTTGCAAAATGGTGGCAGAAGAACACAGNNNAGATCGGA |
| 38079_3 | TATGAGTGTGGAGTCGTTGCAAAATGGTGGCAGAAGAACACAGNNNAGATCGGA |
| 38079_4 | TATGAGTGTGGAGTCGTTGCAAAATGGTGGCAGAAGAACACAGNNNAGATCGGA |
| 38079_5 | TATGAGTGTGGAGTCGTTGCAAAATGGTGGCAGAAGAACACAGNNNAGATCGGA |
| 43639_1 | TATGAGTGTGGAGTCGTTGCAAAGCAAAACACTACATGAGGCCNNNAGATCGGAA |
| 43639_2 | TATGAGTGTGGAGTCGTTGCAAAGCAAAACACTACATGAGGCCNNNAGATCGGAA |
| 43639_3 | TATGAGTGTGGAGTCGTTGCAAAGCAAAACACTACATGAGGCCNNNAGATCGGAA |
| 43639_4 | TATGAGTGTGGAGTCGTTGCAAAGCAAAACACTACATGAGGCCNNNAGATCGGAA |
| 43639_5 | TATGAGTGTGGAGTCGTTGCAAAGCAAAACACTACATGAGGCCNNNAGATCGGAA |
| 12637_1 | TATGAGTGTGGAGTCGTTGCCCAGATCTAAGTAATAAGAAGCANNNAGATCGGAA |
| 12637_2 | TATGAGTGTGGAGTCGTTGCCCAGATCTAAGTAATAAGAAGCANNNAGATCGGAA |
| 12637_3 | TATGAGTGTGGAGTCGTTGCCCAGATCTAAGTAATAAGAAGCANNNAGATCGGAA |
| 12637_4 | TATGAGTGTGGAGTCGTTGCCCAGATCTAAGTAATAAGAAGCANNNAGATCGGAA |
| 12637_5 | TATGAGTGTGGAGTCGTTGCCCAGATCTAAGTAATAAGAAGCANNNAGATCGGAA |
| 33360_1 | TATGAGTGTGGAGTCGTTGCTTTCCTCAACCCCTATCCCCTATNNNAGATCGGAAGA |
| 33360_2 | TATGAGTGTGGAGTCGTTGCTTTCCTCAACCCCTATCCCCTATNNNAGATCGGAAGA |
| 33360_3 | TATGAGTGTGGAGTCGTTGCTTTCCTCAACCCCTATCCCCTATNNNAGATCGGAAGA |
| 33360_4 | TATGAGTGTGGAGTCGTTGCTTTCCTCAACCCCTATCCCCTATNNNAGATCGGAAGA |
| 33360_5 | TATGAGTGTGGAGTCGTTGCTTTCCTCAACCCCTATCCCCTATNNNAGATCGGAAGA |
| 12886_1 | TATGAGTGTGGAGTCGTTGCTGCTCTCTTCTTCCAAGATGNNNAGATCGGAAGAGC |
| 12886_2 | TATGAGTGTGGAGTCGTTGCTGCTCTCTTCTTCCAAGATGNNNAGATCGGAAGAGC |
| 12886_3 | TATGAGTGTGGAGTCGTTGCTGCTCTCTTCTTCCAAGATGNNNAGATCGGAAGAGC |
| 12886_4 | TATGAGTGTGGAGTCGTTGCTGCTCTCTTCTTCCAAGATGNNNAGATCGGAAGAGC |
| 12886_5 | TATGAGTGTGGAGTCGTTGCTGCTCTCTTCTTCCAAGATGNNNAGATCGGAAGAGC |
| 37072_1 | TATGAGTGTGGAGTCGTTGCGAAAGGGCTAGGGATTTTGGGTANNNAGATCGGA |
| 37072_2 | TATGAGTGTGGAGTCGTTGCGAAAGGGCTAGGGATTTTGGGTANNNAGATCGGA |
| 37072_3 | TATGAGTGTGGAGTCGTTGCGAAAGGGCTAGGGATTTTGGGTANNNAGATCGGA |
| 37072_4 | TATGAGTGTGGAGTCGTTGCGAAAGGGCTAGGGATTTTGGGTANNNAGATCGGA |
| 37072_5 | TATGAGTGTGGAGTCGTTGCGAAAGGGCTAGGGATTTTGGGTANNNAGATCGGA |
| 29951_1 | TATGAGTGTGGAGTCGTTGCTCCAGGTATGAGAGAGAAAACAGANNNAGATCGGA |
| 29951_2 | TATGAGTGTGGAGTCGTTGCTCCAGGTATGAGAGAGAAAACAGANNNAGATCGGA |
| 29951_3 | TATGAGTGTGGAGTCGTTGCTCCAGGTATGAGAGAGAAAACAGANNNAGATCGGA |
| 29951_4 | TATGAGTGTGGAGTCGTTGCTCCAGGTATGAGAGAGAAAACAGANNNAGATCGGA |
| 29951_5 | TATGAGTGTGGAGTCGTTGCTCCAGGTATGAGAGAGAAAACAGANNNAGATCGGA |
| 44442_1 | TATGAGTGTGGAGTCGTTGCTCCTCAGCATGACGATGTTACTGNNNAGATCGGAAG |
| 44442_2 | TATGAGTGTGGAGTCGTTGCTCCTCAGCATGACGATGTTACTGNNNAGATCGGAAG |

| | |
|---|---|
| 44442_3 | TATGAGTGTGGAGTCGTTGCTCCTCAGCATGACGATGTTACTGNNNAGATCGGAAG |
| 44442_4 | TATGAGTGTGGAGTCGTTGCTCCTCAGCATGACGATGTTACTGNNNAGATCGGAAG |
| 44442_5 | TATGAGTGTGGAGTCGTTGCTCCTCAGCATGACGATGTTACTGNNNAGATCGGAAG |
| 10664_1 | TATGAGTGTGGAGTCGTTGCGCTGGCTTCTTCTGTCCTGTAAGNNNAGATCGGAAG |
| 10664_2 | TATGAGTGTGGAGTCGTTGCGCTGGCTTCTTCTGTCCTGTAAGNNNAGATCGGAAG |
| 10664_3 | TATGAGTGTGGAGTCGTTGCGCTGGCTTCTTCTGTCCTGTAAGNNNAGATCGGAAG |
| 10664_4 | TATGAGTGTGGAGTCGTTGCGCTGGCTTCTTCTGTCCTGTAAGNNNAGATCGGAAG |
| 10664_5 | TATGAGTGTGGAGTCGTTGCGCTGGCTTCTTCTGTCCTGTAAGNNNAGATCGGAAG |
| 52827_1 | TATGAGTGTGGAGTCGTTGCCACTACTGTCTCAGTCCTGTCACNNNAGATCGGAAG |
| 52827_2 | TATGAGTGTGGAGTCGTTGCCACTACTGTCTCAGTCCTGTCACNNNAGATCGGAAG |
| 52827_3 | TATGAGTGTGGAGTCGTTGCCACTACTGTCTCAGTCCTGTCACNNNAGATCGGAAG |
| 52827_4 | TATGAGTGTGGAGTCGTTGCCACTACTGTCTCAGTCCTGTCACNNNAGATCGGAAG |
| 52827_5 | TATGAGTGTGGAGTCGTTGCCACTACTGTCTCAGTCCTGTCACNNNAGATCGGAAG |
| 40678_1 | TATGAGTGTGGAGTCGTTGCAGCAGTGCTCAGAGGGAAATTTANNNAGATCGGAA |
| 40678_2 | TATGAGTGTGGAGTCGTTGCAGCAGTGCTCAGAGGGAAATTTANNNAGATCGGAA |
| 40678_3 | TATGAGTGTGGAGTCGTTGCAGCAGTGCTCAGAGGGAAATTTANNNAGATCGGAA |
| 40678_4 | TATGAGTGTGGAGTCGTTGCAGCAGTGCTCAGAGGGAAATTTANNNAGATCGGAA |
| 40678_5 | TATGAGTGTGGAGTCGTTGCAGCAGTGCTCAGAGGGAAATTTANNNAGATCGGAA |
| 40837_1 | TATGAGTGTGGAGTCGTTGCTGGCCTTACATAGGAGGTAGACANNNAGATCGGAA |
| 40837_2 | TATGAGTGTGGAGTCGTTGCTGGCCTTACATAGGAGGTAGACANNNAGATCGGAA |
| 40837_3 | TATGAGTGTGGAGTCGTTGCTGGCCTTACATAGGAGGTAGACANNNAGATCGGAA |
| 40837_4 | TATGAGTGTGGAGTCGTTGCTGGCCTTACATAGGAGGTAGACANNNAGATCGGAA |
| 40837_5 | TATGAGTGTGGAGTCGTTGCTGGCCTTACATAGGAGGTAGACANNNAGATCGGAA |
| 39052_1 | TATGAGTGTGGAGTCGTTGCGCCTCAAGAGGTACTGATGACATNNNAGATCGGAAG |
| 39052_2 | TATGAGTGTGGAGTCGTTGCGCCTCAAGAGGTACTGATGACATNNNAGATCGGAAG |
| 39052_3 | TATGAGTGTGGAGTCGTTGCGCCTCAAGAGGTACTGATGACATNNNAGATCGGAAG |
| 39052_4 | TATGAGTGTGGAGTCGTTGCGCCTCAAGAGGTACTGATGACATNNNAGATCGGAAG |
| 39052_5 | TATGAGTGTGGAGTCGTTGCGCCTCAAGAGGTACTGATGACATNNNAGATCGGAAG |
| 29361_1 | TATGAGTGTGGAGTCGTTGCATGAGAGAGAGGGAGAAATGGGANNNAGATCGGA |
| 29361_2 | TATGAGTGTGGAGTCGTTGCATGAGAGAGAGGGAGAAATGGGANNNAGATCGGA |
| 29361_3 | TATGAGTGTGGAGTCGTTGCATGAGAGAGAGGGAGAAATGGGANNNAGATCGGA |
| 29361_4 | TATGAGTGTGGAGTCGTTGCATGAGAGAGAGGGAGAAATGGGANNNAGATCGGA |
| 29361_5 | TATGAGTGTGGAGTCGTTGCATGAGAGAGAGGGAGAAATGGGANNNAGATCGGA |
| 34152_1 | TATGAGTGTGGAGTCGTTGCTTCTTTTACTTTGGGCTGAGGGTNNNAGATCGGAAG |
| 34152_2 | TATGAGTGTGGAGTCGTTGCTTCTTTTACTTTGGGCTGAGGGTNNNAGATCGGAAG |
| 34152_3 | TATGAGTGTGGAGTCGTTGCTTCTTTTACTTTGGGCTGAGGGTNNNAGATCGGAAG |
| 34152_4 | TATGAGTGTGGAGTCGTTGCTTCTTTTACTTTGGGCTGAGGGTNNNAGATCGGAAG |
| 34152_5 | TATGAGTGTGGAGTCGTTGCTTCTTTTACTTTGGGCTGAGGGTNNNAGATCGGAAG |
| 10000_1 | TATGAGTGTGGAGTCGTTGCGACCATTTGCCAATAGGAATGCCNNNAGATCGGAAG |
| 10000_2 | TATGAGTGTGGAGTCGTTGCGACCATTTGCCAATAGGAATGCCNNNAGATCGGAAG |
| 10000_3 | TATGAGTGTGGAGTCGTTGCGACCATTTGCCAATAGGAATGCCNNNAGATCGGAAG |
| 10000_4 | TATGAGTGTGGAGTCGTTGCGACCATTTGCCAATAGGAATGCCNNNAGATCGGAAG |
| 10000_5 | TATGAGTGTGGAGTCGTTGCGACCATTTGCCAATAGGAATGCCNNNAGATCGGAAG |

| | |
|---|---|
| 30165_1 | ATGAGTGTGGAGTCGTTGCTGCACACTTTGGAAATATACAGCTNNNAGATCGGAAG |
| 30165_2 | ATGAGTGTGGAGTCGTTGCTGCACACTTTGGAAATATACAGCTNNNAGATCGGAAG |
| 30165_3 | ATGAGTGTGGAGTCGTTGCTGCACACTTTGGAAATATACAGCTNNNAGATCGGAAG |
| 30165_4 | ATGAGTGTGGAGTCGTTGCTGCACACTTTGGAAATATACAGCTNNNAGATCGGAAG |
| 30165_5 | ATGAGTGTGGAGTCGTTGCTGCACACTTTGGAAATATACAGCTNNNAGATCGGAAG |
| 35673_1 | TATGAGTGTGGAGTCGTTGCCTTCAGCAGCACTACTGGACATANNNAGATCGGAAG |
| 35673_2 | TATGAGTGTGGAGTCGTTGCCTTCAGCAGCACTACTGGACATANNNAGATCGGAAG |
| 35673_3 | TATGAGTGTGGAGTCGTTGCCTTCAGCAGCACTACTGGACATANNNAGATCGGAAG |
| 35673_4 | TATGAGTGTGGAGTCGTTGCCTTCAGCAGCACTACTGGACATANNNAGATCGGAAG |
| 35673_5 | TATGAGTGTGGAGTCGTTGCCTTCAGCAGCACTACTGGACATANNNAGATCGGAAG |
| 9813_1 | TATGAGTGTGGAGTCGTTGCTCAGTGGCACAAAGTACACTCACNNNAGATCGGAAG |
| 9813_2 | TATGAGTGTGGAGTCGTTGCTCAGTGGCACAAAGTACACTCACNNNAGATCGGAAG |
| 9813_3 | TATGAGTGTGGAGTCGTTGCTCAGTGGCACAAAGTACACTCACNNNAGATCGGAAG |
| 9813_4 | TATGAGTGTGGAGTCGTTGCTCAGTGGCACAAAGTACACTCACNNNAGATCGGAAG |
| 9813_5 | TATGAGTGTGGAGTCGTTGCTCAGTGGCACAAAGTACACTCACNNNAGATCGGAAG |
| 49530_1 | TATGAGTGTGGAGTCGTTGCGCACAATCTCGACTTACTGCAAGNNNAGATCGGAAG |
| 49530_2 | TATGAGTGTGGAGTCGTTGCGCACAATCTCGACTTACTGCAAGNNNAGATCGGAAG |
| 49530_3 | TATGAGTGTGGAGTCGTTGCGCACAATCTCGACTTACTGCAAGNNNAGATCGGAAG |
| 49530_4 | TATGAGTGTGGAGTCGTTGCGCACAATCTCGACTTACTGCAAGNNNAGATCGGAAG |
| 49530_5 | TATGAGTGTGGAGTCGTTGCGCACAATCTCGACTTACTGCAAGNNNAGATCGGAAG |
| 33880_1 | TATGAGTGTGGAGTCGTTGCCTGAGGTGGGAAGATAGCTTGAGNNNAGATCGGA |
| 33880_2 | TATGAGTGTGGAGTCGTTGCCTGAGGTGGGAAGATAGCTTGAGNNNAGATCGGA |
| 33880_3 | TATGAGTGTGGAGTCGTTGCCTGAGGTGGGAAGATAGCTTGAGNNNAGATCGGA |
| 33880_4 | TATGAGTGTGGAGTCGTTGCCTGAGGTGGGAAGATAGCTTGAGNNNAGATCGGA |
| 33880_5 | TATGAGTGTGGAGTCGTTGCCTGAGGTGGGAAGATAGCTTGAGNNNAGATCGGA |
| 31596_1 | TATGAGTGTGGAGTCGTTGCGGATCCCAACCACTCATTCAGAANNNAGATCGGAAG |
| 31596_2 | TATGAGTGTGGAGTCGTTGCGGATCCCAACCACTCATTCAGAANNNAGATCGGAAG |
| 31596_3 | TATGAGTGTGGAGTCGTTGCGGATCCCAACCACTCATTCAGAANNNAGATCGGAAG |
| 31596_4 | TATGAGTGTGGAGTCGTTGCGGATCCCAACCACTCATTCAGAANNNAGATCGGAAG |
| 31596_5 | TATGAGTGTGGAGTCGTTGCGGATCCCAACCACTCATTCAGAANNNAGATCGGAAG |
| 36933_1 | TATGAGTGTGGAGTCGTTGCTCTATGGTTTCTAGTTGCCATTGCNNNAGATCGGAA |
| 36933_2 | TATGAGTGTGGAGTCGTTGCTCTATGGTTTCTAGTTGCCATTGCNNNAGATCGGAA |
| 36933_3 | TATGAGTGTGGAGTCGTTGCTCTATGGTTTCTAGTTGCCATTGCNNNAGATCGGAA |
| 36933_4 | TATGAGTGTGGAGTCGTTGCTCTATGGTTTCTAGTTGCCATTGCNNNAGATCGGAA |
| 36933_5 | TATGAGTGTGGAGTCGTTGCTCTATGGTTTCTAGTTGCCATTGCNNNAGATCGGAA |
| 9957_1 | TATGAGTGTGGAGTCGTTGCCTTTGGAATGGGGAGAAGCCTTTNNNAGATCGGAA |
| 9957_2 | TATGAGTGTGGAGTCGTTGCCTTTGGAATGGGGAGAAGCCTTTNNNAGATCGGAA |
| 9957_3 | TATGAGTGTGGAGTCGTTGCCTTTGGAATGGGGAGAAGCCTTTNNNAGATCGGAA |
| 9957_4 | TATGAGTGTGGAGTCGTTGCCTTTGGAATGGGGAGAAGCCTTTNNNAGATCGGAA |
| 9957_5 | TATGAGTGTGGAGTCGTTGCCTTTGGAATGGGGAGAAGCCTTTNNNAGATCGGAA |
| 42438_1 | TATGAGTGTGGAGTCGTTGCGGGGAGAGAAGTGAGAGACTAGTNNNAGATCGGA |
| 42438_2 | TATGAGTGTGGAGTCGTTGCGGGGAGAGAAGTGAGAGACTAGTNNNAGATCGGA |
| 42438_3 | TATGAGTGTGGAGTCGTTGCGGGGAGAGAAGTGAGAGACTAGTNNNAGATCGGA |

| | |
|---|---|
| 42438_4 | TATGAGTGTGGAGTCGTTGCGGGGAGAGAAGTGAGAGACTAGTNNNAGATCGGA |
| 42438_5 | TATGAGTGTGGAGTCGTTGCGGGGAGAGAAGTGAGAGACTAGTNNNAGATCGGA |
| 42495_1 | TATGAGTGTGGAGTCGTTGCGCTTGGGAGAAAGTAAGGGAAGANNNAGATCGGA |
| 42495_2 | TATGAGTGTGGAGTCGTTGCGCTTGGGAGAAAGTAAGGGAAGANNNAGATCGGA |
| 42495_3 | TATGAGTGTGGAGTCGTTGCGCTTGGGAGAAAGTAAGGGAAGANNNAGATCGGA |
| 42495_4 | TATGAGTGTGGAGTCGTTGCGCTTGGGAGAAAGTAAGGGAAGANNNAGATCGGA |
| 42495_5 | TATGAGTGTGGAGTCGTTGCGCTTGGGAGAAAGTAAGGGAAGANNNAGATCGGA |
| 41312_1 | TATGAGTGTGGAGTCGTTGCCTGTTTTTATCAAGGGGCCTGGNNNAGATCGGAAG |
| 41312_2 | TATGAGTGTGGAGTCGTTGCCTGTTTTTATCAAGGGGCCTGGNNNAGATCGGAAG |
| 41312_3 | TATGAGTGTGGAGTCGTTGCCTGTTTTTATCAAGGGGCCTGGNNNAGATCGGAAG |
| 41312_4 | TATGAGTGTGGAGTCGTTGCCTGTTTTTATCAAGGGGCCTGGNNNAGATCGGAAG |
| 41312_5 | TATGAGTGTGGAGTCGTTGCCTGTTTTTATCAAGGGGCCTGGNNNAGATCGGAAG |
| 35727_1 | TATGAGTGTGGAGTCGTTGCCAAGTAGCTGGGACTACAGGCNNNAGATCGGAAGA |
| 35727_2 | TATGAGTGTGGAGTCGTTGCCAAGTAGCTGGGACTACAGGCNNNAGATCGGAAGA |
| 35727_3 | TATGAGTGTGGAGTCGTTGCCAAGTAGCTGGGACTACAGGCNNNAGATCGGAAGA |
| 35727_4 | TATGAGTGTGGAGTCGTTGCCAAGTAGCTGGGACTACAGGCNNNAGATCGGAAGA |
| 35727_5 | TATGAGTGTGGAGTCGTTGCCAAGTAGCTGGGACTACAGGCNNNAGATCGGAAGA |
| 52530_1 | TATGAGTGTGGAGTCGTTGCATGCCCATGTATTCAAAGTCCCTNNNAGATCGGAAG |
| 52530_2 | TATGAGTGTGGAGTCGTTGCATGCCCATGTATTCAAAGTCCCTNNNAGATCGGAAG |
| 52530_3 | TATGAGTGTGGAGTCGTTGCATGCCCATGTATTCAAAGTCCCTNNNAGATCGGAAG |
| 52530_4 | TATGAGTGTGGAGTCGTTGCATGCCCATGTATTCAAAGTCCCTNNNAGATCGGAAG |
| 52530_5 | TATGAGTGTGGAGTCGTTGCATGCCCATGTATTCAAAGTCCCTNNNAGATCGGAAG |
| 44593_1 | TATGAGTGTGGAGTCGTTGCGGAATGAAAGACACAGATGGCAANNNAGATCGGA |
| 44593_2 | TATGAGTGTGGAGTCGTTGCGGAATGAAAGACACAGATGGCAANNNAGATCGGA |
| 44593_3 | TATGAGTGTGGAGTCGTTGCGGAATGAAAGACACAGATGGCAANNNAGATCGGA |
| 44593_4 | TATGAGTGTGGAGTCGTTGCGGAATGAAAGACACAGATGGCAANNNAGATCGGA |
| 44593_5 | TATGAGTGTGGAGTCGTTGCGGAATGAAAGACACAGATGGCAANNNAGATCGGA |
| 29938_1 | TATGAGTGTGGAGTCGTTGCAGGAACAGCAGTGGACATCTTAGNNNAGATCGGAA |
| 29938_2 | TATGAGTGTGGAGTCGTTGCAGGAACAGCAGTGGACATCTTAGNNNAGATCGGAA |
| 29938_3 | TATGAGTGTGGAGTCGTTGCAGGAACAGCAGTGGACATCTTAGNNNAGATCGGAA |
| 29938_4 | TATGAGTGTGGAGTCGTTGCAGGAACAGCAGTGGACATCTTAGNNNAGATCGGAA |
| 29938_5 | TATGAGTGTGGAGTCGTTGCAGGAACAGCAGTGGACATCTTAGNNNAGATCGGAA |
| 45848_1 | TATGAGTGTGGAGTCGTTGCCCCAGGTTCAAGCAATTCTCTTGNNNAGATCGGAAG |
| 45848_2 | TATGAGTGTGGAGTCGTTGCCCCAGGTTCAAGCAATTCTCTTGNNNAGATCGGAAG |
| 45848_3 | TATGAGTGTGGAGTCGTTGCCCCAGGTTCAAGCAATTCTCTTGNNNAGATCGGAAG |
| 45848_4 | TATGAGTGTGGAGTCGTTGCCCCAGGTTCAAGCAATTCTCTTGNNNAGATCGGAAG |
| 45848_5 | TATGAGTGTGGAGTCGTTGCCCCAGGTTCAAGCAATTCTCTTGNNNAGATCGGAAG |
| 10003_1 | TATGAGTGTGGAGTCGTTGCGGAAAGCCTCCTCTGGAACATTNNNAGATCGGAAGA |
| 10003_2 | TATGAGTGTGGAGTCGTTGCGGAAAGCCTCCTCTGGAACATTNNNAGATCGGAAGA |
| 10003_3 | TATGAGTGTGGAGTCGTTGCGGAAAGCCTCCTCTGGAACATTNNNAGATCGGAAGA |
| 10003_4 | TATGAGTGTGGAGTCGTTGCGGAAAGCCTCCTCTGGAACATTNNNAGATCGGAAGA |
| 10003_5 | TATGAGTGTGGAGTCGTTGCGGAAAGCCTCCTCTGGAACATTNNNAGATCGGAAGA |
| 47742_1 | TATGAGTGTGGAGTCGTTGCATACCTGACTCCCACGACCTCNNNAGATCGGAAGAG |

| | |
|---|---|
| 47742_2 | TATGAGTGTGGAGTCGTTGCATACCTGACTCCCACGACCTCNNNAGATCGGAAGAG |
| 47742_3 | TATGAGTGTGGAGTCGTTGCATACCTGACTCCCACGACCTCNNNAGATCGGAAGAG |
| 47742_4 | TATGAGTGTGGAGTCGTTGCATACCTGACTCCCACGACCTCNNNAGATCGGAAGAG |
| 47742_5 | TATGAGTGTGGAGTCGTTGCATACCTGACTCCCACGACCTCNNNAGATCGGAAGAG |
| 50879_1 | TATGAGTGTGGAGTCGTTGCTAGAGTGCAGTAACACGATCTCGNNNAGATCGGAA |
| 50879_2 | TATGAGTGTGGAGTCGTTGCTAGAGTGCAGTAACACGATCTCGNNNAGATCGGAA |
| 50879_3 | TATGAGTGTGGAGTCGTTGCTAGAGTGCAGTAACACGATCTCGNNNAGATCGGAA |
| 50879_4 | TATGAGTGTGGAGTCGTTGCTAGAGTGCAGTAACACGATCTCGNNNAGATCGGAA |
| 50879_5 | TATGAGTGTGGAGTCGTTGCTAGAGTGCAGTAACACGATCTCGNNNAGATCGGAA |
| 34699_1 | TATGAGTGTGGAGTCGTTGCAAACTCCTGGGCTCAAGCAATCNNNAGATCGGAAGA |
| 34699_2 | TATGAGTGTGGAGTCGTTGCAAACTCCTGGGCTCAAGCAATCNNNAGATCGGAAGA |
| 34699_3 | TATGAGTGTGGAGTCGTTGCAAACTCCTGGGCTCAAGCAATCNNNAGATCGGAAGA |
| 34699_4 | TATGAGTGTGGAGTCGTTGCAAACTCCTGGGCTCAAGCAATCNNNAGATCGGAAGA |
| 34699_5 | TATGAGTGTGGAGTCGTTGCAAACTCCTGGGCTCAAGCAATCNNNAGATCGGAAGA |
| 40230_1 | TATGAGTGTGGAGTCGTTGCCACAAGAGTTTTGGCACCATGAANNNAGATCGGAA |
| 40230_2 | TATGAGTGTGGAGTCGTTGCCACAAGAGTTTTGGCACCATGAANNNAGATCGGAA |
| 40230_3 | TATGAGTGTGGAGTCGTTGCCACAAGAGTTTTGGCACCATGAANNNAGATCGGAA |
| 40230_4 | TATGAGTGTGGAGTCGTTGCCACAAGAGTTTTGGCACCATGAANNNAGATCGGAA |
| 40230_5 | TATGAGTGTGGAGTCGTTGCCACAAGAGTTTTGGCACCATGAANNNAGATCGGAA |
| 52263_1 | TATGAGTGTGGAGTCGTTGCTTCCTGCCTTGTGTTGGAATTTCNNNAGATCGGAAG |
| 52263_2 | TATGAGTGTGGAGTCGTTGCTTCCTGCCTTGTGTTGGAATTTCNNNAGATCGGAAG |
| 52263_3 | TATGAGTGTGGAGTCGTTGCTTCCTGCCTTGTGTTGGAATTTCNNNAGATCGGAAG |
| 52263_4 | TATGAGTGTGGAGTCGTTGCTTCCTGCCTTGTGTTGGAATTTCNNNAGATCGGAAG |
| 52263_5 | TATGAGTGTGGAGTCGTTGCTTCCTGCCTTGTGTTGGAATTTCNNNAGATCGGAAG |
| 10815_1 | TATGAGTGTGGAGTCGTTGCCCCCGTAGTAATGGAATAGAGCCNNNAGATCGGAA |
| 10815_2 | TATGAGTGTGGAGTCGTTGCCCCCGTAGTAATGGAATAGAGCCNNNAGATCGGAA |
| 10815_3 | TATGAGTGTGGAGTCGTTGCCCCCGTAGTAATGGAATAGAGCCNNNAGATCGGAA |
| 10815_4 | TATGAGTGTGGAGTCGTTGCCCCCGTAGTAATGGAATAGAGCCNNNAGATCGGAA |
| 10815_5 | TATGAGTGTGGAGTCGTTGCCCCCGTAGTAATGGAATAGAGCCNNNAGATCGGAA |
| 41396_1 | TATGAGTGTGGAGTCGTTGCGGGAAGCTTGGGGTTTTGTAAAGNNNAGATCGGA |
| 41396_2 | TATGAGTGTGGAGTCGTTGCGGGAAGCTTGGGGTTTTGTAAAGNNNAGATCGGA |
| 41396_3 | TATGAGTGTGGAGTCGTTGCGGGAAGCTTGGGGTTTTGTAAAGNNNAGATCGGA |
| 41396_4 | TATGAGTGTGGAGTCGTTGCGGGAAGCTTGGGGTTTTGTAAAGNNNAGATCGGA |
| 41396_5 | TATGAGTGTGGAGTCGTTGCGGGAAGCTTGGGGTTTTGTAAAGNNNAGATCGGA |
| 19430_1 | TATGAGTGTGGAGTCGTTGCGGAGACTAGAATTGAATCTCTAANNNAGATCGGAA |
| 19430_2 | TATGAGTGTGGAGTCGTTGCGGAGACTAGAATTGAATCTCTAANNNAGATCGGAA |
| 19430_3 | TATGAGTGTGGAGTCGTTGCGGAGACTAGAATTGAATCTCTAANNNAGATCGGAA |
| 19430_4 | TATGAGTGTGGAGTCGTTGCGGAGACTAGAATTGAATCTCTAANNNAGATCGGAA |
| 19430_5 | TATGAGTGTGGAGTCGTTGCGGAGACTAGAATTGAATCTCTAANNNAGATCGGAA |
| 44644_1 | TATGAGTGTGGAGTCGTTGCTGCGAGCTACGAAGAGGGAGNNNAGATCGGAAGA |
| 44644_2 | TATGAGTGTGGAGTCGTTGCTGCGAGCTACGAAGAGGGAGNNNAGATCGGAAGA |
| 44644_3 | TATGAGTGTGGAGTCGTTGCTGCGAGCTACGAAGAGGGAGNNNAGATCGGAAGA |
| 44644_4 | TATGAGTGTGGAGTCGTTGCTGCGAGCTACGAAGAGGGAGNNNAGATCGGAAGA |

| | |
|---|---|
| 44644_5 | TATGAGTGTGGAGTCGTTGCTGCGAGCTACGAAGAGGGAGNNNAGATCGGAAGA |
| 48107_1 | TATGAGTGTGGAGTCGTTGCCGCTAAGATGTCATTGCGGATGNNNAGATCGGAAG |
| 48107_2 | TATGAGTGTGGAGTCGTTGCCGCTAAGATGTCATTGCGGATGNNNAGATCGGAAG |
| 48107_3 | TATGAGTGTGGAGTCGTTGCCGCTAAGATGTCATTGCGGATGNNNAGATCGGAAG |
| 48107_4 | TATGAGTGTGGAGTCGTTGCCGCTAAGATGTCATTGCGGATGNNNAGATCGGAAG |
| 48107_5 | TATGAGTGTGGAGTCGTTGCCGCTAAGATGTCATTGCGGATGNNNAGATCGGAAG |
| 10640_1 | TATGAGTGTGGAGTCGTTGCTCTTACTGGGTGCTTTTCCACTGNNNAGATCGGAAG |
| 10640_2 | TATGAGTGTGGAGTCGTTGCTCTTACTGGGTGCTTTTCCACTGNNNAGATCGGAAG |
| 10640_3 | TATGAGTGTGGAGTCGTTGCTCTTACTGGGTGCTTTTCCACTGNNNAGATCGGAAG |
| 10640_4 | TATGAGTGTGGAGTCGTTGCTCTTACTGGGTGCTTTTCCACTGNNNAGATCGGAAG |
| 10640_5 | TATGAGTGTGGAGTCGTTGCTCTTACTGGGTGCTTTTCCACTGNNNAGATCGGAAG |
| 48035_1 | TATGAGTGTGGAGTCGTTGCCAGGTTTGTTGGCCATCTGAATTNNNAGATCGGAAG |
| 48035_2 | TATGAGTGTGGAGTCGTTGCCAGGTTTGTTGGCCATCTGAATTNNNAGATCGGAAG |
| 48035_3 | TATGAGTGTGGAGTCGTTGCCAGGTTTGTTGGCCATCTGAATTNNNAGATCGGAAG |
| 48035_4 | TATGAGTGTGGAGTCGTTGCCAGGTTTGTTGGCCATCTGAATTNNNAGATCGGAAG |
| 48035_5 | TATGAGTGTGGAGTCGTTGCCAGGTTTGTTGGCCATCTGAATTNNNAGATCGGAAG |
| 42114_1 | TATGAGTGTGGAGTCGTTGCAGCCTCCTGAGTTCAAGTGATTCNNNAGATCGGAAG |
| 42114_2 | TATGAGTGTGGAGTCGTTGCAGCCTCCTGAGTTCAAGTGATTCNNNAGATCGGAAG |
| 42114_3 | TATGAGTGTGGAGTCGTTGCAGCCTCCTGAGTTCAAGTGATTCNNNAGATCGGAAG |
| 42114_4 | TATGAGTGTGGAGTCGTTGCAGCCTCCTGAGTTCAAGTGATTCNNNAGATCGGAAG |
| 42114_5 | TATGAGTGTGGAGTCGTTGCAGCCTCCTGAGTTCAAGTGATTCNNNAGATCGGAAG |
| 35029_1 | TATGAGTGTGGAGTCGTTGCAGATGGAGTCTCACTCTGTTGCNNNAGATCGGAAGA |
| 35029_2 | TATGAGTGTGGAGTCGTTGCAGATGGAGTCTCACTCTGTTGCNNNAGATCGGAAGA |
| 35029_3 | TATGAGTGTGGAGTCGTTGCAGATGGAGTCTCACTCTGTTGCNNNAGATCGGAAGA |
| 35029_4 | TATGAGTGTGGAGTCGTTGCAGATGGAGTCTCACTCTGTTGCNNNAGATCGGAAGA |
| 35029_5 | TATGAGTGTGGAGTCGTTGCAGATGGAGTCTCACTCTGTTGCNNNAGATCGGAAGA |
| 39208_1 | TATGAGTGTGGAGTCGTTGCGTGTGGTGGGTGTCAAAACATATNNNAGATCGGAA |
| 39208_2 | TATGAGTGTGGAGTCGTTGCGTGTGGTGGGTGTCAAAACATATNNNAGATCGGAA |
| 39208_3 | TATGAGTGTGGAGTCGTTGCGTGTGGTGGGTGTCAAAACATATNNNAGATCGGAA |
| 39208_4 | TATGAGTGTGGAGTCGTTGCGTGTGGTGGGTGTCAAAACATATNNNAGATCGGAA |
| 39208_5 | TATGAGTGTGGAGTCGTTGCGTGTGGTGGGTGTCAAAACATATNNNAGATCGGAA |
| 49343_1 | TATGAGTGTGGAGTCGTTGCCTCTCAAGTAGCTGGGACTACAGNNNAGATCGGAA |
| 49343_2 | TATGAGTGTGGAGTCGTTGCCTCTCAAGTAGCTGGGACTACAGNNNAGATCGGAA |
| 49343_3 | TATGAGTGTGGAGTCGTTGCCTCTCAAGTAGCTGGGACTACAGNNNAGATCGGAA |
| 49343_4 | TATGAGTGTGGAGTCGTTGCCTCTCAAGTAGCTGGGACTACAGNNNAGATCGGAA |
| 49343_5 | TATGAGTGTGGAGTCGTTGCCTCTCAAGTAGCTGGGACTACAGNNNAGATCGGAA |
| 51491_1 | TATGAGTGTGGAGTCGTTGCCACCATTCATGCCTGGCTAATTTNNNAGATCGGAAG |
| 51491_2 | TATGAGTGTGGAGTCGTTGCCACCATTCATGCCTGGCTAATTTNNNAGATCGGAAG |
| 51491_3 | TATGAGTGTGGAGTCGTTGCCACCATTCATGCCTGGCTAATTTNNNAGATCGGAAG |
| 51491_4 | TATGAGTGTGGAGTCGTTGCCACCATTCATGCCTGGCTAATTTNNNAGATCGGAAG |
| 51491_5 | TATGAGTGTGGAGTCGTTGCCACCATTCATGCCTGGCTAATTTNNNAGATCGGAAG |
| 49936_1 | TATGAGTGTGGAGTCGTTGCAGCTCAGCTTCTAGACAGTTTCANNNAGATCGGAAG |
| 49936_2 | TATGAGTGTGGAGTCGTTGCAGCTCAGCTTCTAGACAGTTTCANNNAGATCGGAAG |

| | |
|---|---|
| 49936_3 | TATGAGTGTGGAGTCGTTGCAGCTCAGCTTCTAGACAGTTTCANNNAGATCGGAAG |
| 49936_4 | TATGAGTGTGGAGTCGTTGCAGCTCAGCTTCTAGACAGTTTCANNNAGATCGGAAG |
| 49936_5 | TATGAGTGTGGAGTCGTTGCAGCTCAGCTTCTAGACAGTTTCANNNAGATCGGAAG |
| 30123_1 | TATGAGTGTGGAGTCGTTGCTTAGCTGCCTTGGTAGAGATTGCNNNAGATCGGAAG |
| 30123_2 | TATGAGTGTGGAGTCGTTGCTTAGCTGCCTTGGTAGAGATTGCNNNAGATCGGAAG |
| 30123_3 | TATGAGTGTGGAGTCGTTGCTTAGCTGCCTTGGTAGAGATTGCNNNAGATCGGAAG |
| 30123_4 | TATGAGTGTGGAGTCGTTGCTTAGCTGCCTTGGTAGAGATTGCNNNAGATCGGAAG |
| 30123_5 | TATGAGTGTGGAGTCGTTGCTTAGCTGCCTTGGTAGAGATTGCNNNAGATCGGAAG |
| 28980_1 | TATGAGTGTGGAGTCGTTGCTGACTCATACCGTTTTGTTCCCTNNNAGATCGGAAGA |
| 28980_2 | TATGAGTGTGGAGTCGTTGCTGACTCATACCGTTTTGTTCCCTNNNAGATCGGAAGA |
| 28980_3 | TATGAGTGTGGAGTCGTTGCTGACTCATACCGTTTTGTTCCCTNNNAGATCGGAAGA |
| 28980_4 | TATGAGTGTGGAGTCGTTGCTGACTCATACCGTTTTGTTCCCTNNNAGATCGGAAGA |
| 28980_5 | TATGAGTGTGGAGTCGTTGCTGACTCATACCGTTTTGTTCCCTNNNAGATCGGAAGA |
| 51148_1 | TATGAGTGTGGAGTCGTTGCCATCAATTGAGTGCTGTGGGAAGNNNAGATCGGAA |
| 51148_2 | TATGAGTGTGGAGTCGTTGCCATCAATTGAGTGCTGTGGGAAGNNNAGATCGGAA |
| 51148_3 | TATGAGTGTGGAGTCGTTGCCATCAATTGAGTGCTGTGGGAAGNNNAGATCGGAA |
| 51148_4 | TATGAGTGTGGAGTCGTTGCCATCAATTGAGTGCTGTGGGAAGNNNAGATCGGAA |
| 51148_5 | TATGAGTGTGGAGTCGTTGCCATCAATTGAGTGCTGTGGGAAGNNNAGATCGGAA |
| 31332_1 | TATGAGTGTGGAGTCGTTGCTGACTGAAGGAGCTGAGGGATATNNNAGATCGGAA |
| 31332_2 | TATGAGTGTGGAGTCGTTGCTGACTGAAGGAGCTGAGGGATATNNNAGATCGGAA |
| 31332_3 | TATGAGTGTGGAGTCGTTGCTGACTGAAGGAGCTGAGGGATATNNNAGATCGGAA |
| 31332_4 | TATGAGTGTGGAGTCGTTGCTGACTGAAGGAGCTGAGGGATATNNNAGATCGGAA |
| 31332_5 | TATGAGTGTGGAGTCGTTGCTGACTGAAGGAGCTGAGGGATATNNNAGATCGGAA |
| 50406_1 | TATGAGTGTGGAGTCGTTGCTGTGCAGTCCACTATCTTAGCTCNNNAGATCGGAAG |
| 50406_2 | TATGAGTGTGGAGTCGTTGCTGTGCAGTCCACTATCTTAGCTCNNNAGATCGGAAG |
| 50406_3 | TATGAGTGTGGAGTCGTTGCTGTGCAGTCCACTATCTTAGCTCNNNAGATCGGAAG |
| 50406_4 | TATGAGTGTGGAGTCGTTGCTGTGCAGTCCACTATCTTAGCTCNNNAGATCGGAAG |
| 50406_5 | TATGAGTGTGGAGTCGTTGCTGTGCAGTCCACTATCTTAGCTCNNNAGATCGGAAG |
| 38945_1 | TATGAGTGTGGAGTCGTTGCCTAAGTCAGATAGCAAGCTACCTNNNAGATCGGAAG |
| 38945_2 | TATGAGTGTGGAGTCGTTGCCTAAGTCAGATAGCAAGCTACCTNNNAGATCGGAAG |
| 38945_3 | TATGAGTGTGGAGTCGTTGCCTAAGTCAGATAGCAAGCTACCTNNNAGATCGGAAG |
| 38945_4 | TATGAGTGTGGAGTCGTTGCCTAAGTCAGATAGCAAGCTACCTNNNAGATCGGAAG |
| 38945_5 | TATGAGTGTGGAGTCGTTGCCTAAGTCAGATAGCAAGCTACCTNNNAGATCGGAAG |
| 10613_1 | TATGAGTGTGGAGTCGTTGCTGAATTGCTATGCTGCTGGCTTTNNNAGATCGGAAG |
| 10613_2 | TATGAGTGTGGAGTCGTTGCTGAATTGCTATGCTGCTGGCTTTNNNAGATCGGAAG |
| 10613_3 | TATGAGTGTGGAGTCGTTGCTGAATTGCTATGCTGCTGGCTTTNNNAGATCGGAAG |
| 10613_4 | TATGAGTGTGGAGTCGTTGCTGAATTGCTATGCTGCTGGCTTTNNNAGATCGGAAG |
| 10613_5 | TATGAGTGTGGAGTCGTTGCTGAATTGCTATGCTGCTGGCTTTNNNAGATCGGAAG |
| 42689_1 | TATGAGTGTGGAGTCGTTGCGGACCGAAAATTTTCCAGAGTGGNNNAGATCGGAA |
| 42689_2 | TATGAGTGTGGAGTCGTTGCGGACCGAAAATTTTCCAGAGTGGNNNAGATCGGAA |
| 42689_3 | TATGAGTGTGGAGTCGTTGCGGACCGAAAATTTTCCAGAGTGGNNNAGATCGGAA |
| 42689_4 | TATGAGTGTGGAGTCGTTGCGGACCGAAAATTTTCCAGAGTGGNNNAGATCGGAA |
| 42689_5 | TATGAGTGTGGAGTCGTTGCGGACCGAAAATTTTCCAGAGTGGNNNAGATCGGAA |

| | |
|---|---|
| 2589_1 | TATGAGTGTGGAGTCGTTGCAGGAGGGGGAAGGAAATATGTGGNNNAGATCGGA |
| 2589_2 | TATGAGTGTGGAGTCGTTGCAGGAGGGGGAAGGAAATATGTGGNNNAGATCGGA |
| 2589_3 | TATGAGTGTGGAGTCGTTGCAGGAGGGGGAAGGAAATATGTGGNNNAGATCGGA |
| 2589_4 | TATGAGTGTGGAGTCGTTGCAGGAGGGGGAAGGAAATATGTGGNNNAGATCGGA |
| 2589_5 | TATGAGTGTGGAGTCGTTGCAGGAGGGGGAAGGAAATATGTGGNNNAGATCGGA |
| 45369_1 | TATGAGTGTGGAGTCGTTGCACTCCTGAGCTCAAGGAATCATCNNNAGATCGGAAG |
| 45369_2 | TATGAGTGTGGAGTCGTTGCACTCCTGAGCTCAAGGAATCATCNNNAGATCGGAAG |
| 45369_3 | TATGAGTGTGGAGTCGTTGCACTCCTGAGCTCAAGGAATCATCNNNAGATCGGAAG |
| 45369_4 | TATGAGTGTGGAGTCGTTGCACTCCTGAGCTCAAGGAATCATCNNNAGATCGGAAG |
| 45369_5 | TATGAGTGTGGAGTCGTTGCACTCCTGAGCTCAAGGAATCATCNNNAGATCGGAAG |
| 32676_1 | TATGAGTGTGGAGTCGTTGCTACGATCTCGGCTCACTGGAANNNAGATCGGAAGAG |
| 32676_2 | TATGAGTGTGGAGTCGTTGCTACGATCTCGGCTCACTGGAANNNAGATCGGAAGAG |
| 32676_3 | TATGAGTGTGGAGTCGTTGCTACGATCTCGGCTCACTGGAANNNAGATCGGAAGAG |
| 32676_4 | TATGAGTGTGGAGTCGTTGCTACGATCTCGGCTCACTGGAANNNAGATCGGAAGAG |
| 32676_5 | TATGAGTGTGGAGTCGTTGCTACGATCTCGGCTCACTGGAANNNAGATCGGAAGAG |
| 12863_1 | TATGAGTGTGGAGTCGTTGCAAAACACAGGTACATTATGGTNNNAGATCGGAAGA |
| 12863_2 | TATGAGTGTGGAGTCGTTGCAAAACACAGGTACATTATGGTNNNAGATCGGAAGA |
| 12863_3 | TATGAGTGTGGAGTCGTTGCAAAACACAGGTACATTATGGTNNNAGATCGGAAGA |
| 12863_4 | TATGAGTGTGGAGTCGTTGCAAAACACAGGTACATTATGGTNNNAGATCGGAAGA |
| 12863_5 | TATGAGTGTGGAGTCGTTGCAAAACACAGGTACATTATGGTNNNAGATCGGAAGA |
| 39897_1 | TATGAGTGTGGAGTCGTTGCTAGAGGATCATGACTGTGTCCCTNNNAGATCGGAAG |
| 39897_2 | TATGAGTGTGGAGTCGTTGCTAGAGGATCATGACTGTGTCCCTNNNAGATCGGAAG |
| 39897_3 | TATGAGTGTGGAGTCGTTGCTAGAGGATCATGACTGTGTCCCTNNNAGATCGGAAG |
| 39897_4 | TATGAGTGTGGAGTCGTTGCTAGAGGATCATGACTGTGTCCCTNNNAGATCGGAAG |
| 39897_5 | TATGAGTGTGGAGTCGTTGCTAGAGGATCATGACTGTGTCCCTNNNAGATCGGAAG |
| 30060_1 | TGAGTGTGGAGTCGTTGCTGAAGTTGCTAAGGTAGTAGTTTCCANNNAGATCGGAA |
| 30060_2 | TGAGTGTGGAGTCGTTGCTGAAGTTGCTAAGGTAGTAGTTTCCANNNAGATCGGAA |
| 30060_3 | TGAGTGTGGAGTCGTTGCTGAAGTTGCTAAGGTAGTAGTTTCCANNNAGATCGGAA |
| 30060_4 | TGAGTGTGGAGTCGTTGCTGAAGTTGCTAAGGTAGTAGTTTCCANNNAGATCGGAA |
| 30060_5 | TGAGTGTGGAGTCGTTGCTGAAGTTGCTAAGGTAGTAGTTTCCANNNAGATCGGAA |
| 42795_1 | TATGAGTGTGGAGTCGTTGCTTATCCTATTGGTTCTGCCCCTCNNNAGATCGGAAGA |
| 42795_2 | TATGAGTGTGGAGTCGTTGCTTATCCTATTGGTTCTGCCCCTCNNNAGATCGGAAGA |
| 42795_3 | TATGAGTGTGGAGTCGTTGCTTATCCTATTGGTTCTGCCCCTCNNNAGATCGGAAGA |
| 42795_4 | TATGAGTGTGGAGTCGTTGCTTATCCTATTGGTTCTGCCCCTCNNNAGATCGGAAGA |
| 42795_5 | TATGAGTGTGGAGTCGTTGCTTATCCTATTGGTTCTGCCCCTCNNNAGATCGGAAGA |
| 43206_1 | TATGAGTGTGGAGTCGTTGCTTCTCCCCCTCTTTCTCTCTCTTNNNAGATCGGAAGAG |
| 43206_2 | TATGAGTGTGGAGTCGTTGCTTCTCCCCCTCTTTCTCTCTCTTNNNAGATCGGAAGAG |
| 43206_3 | TATGAGTGTGGAGTCGTTGCTTCTCCCCCTCTTTCTCTCTCTTNNNAGATCGGAAGAG |
| 43206_4 | TATGAGTGTGGAGTCGTTGCTTCTCCCCCTCTTTCTCTCTCTTNNNAGATCGGAAGAG |
| 43206_5 | TATGAGTGTGGAGTCGTTGCTTCTCCCCCTCTTTCTCTCTCTTNNNAGATCGGAAGAG |
| 34388_1 | TATGAGTGTGGAGTCGTTGCTAATGAGACATCTTGGGCATGGGNNNAGATCGGAA |
| 34388_2 | TATGAGTGTGGAGTCGTTGCTAATGAGACATCTTGGGCATGGGNNNAGATCGGAA |
| 34388_3 | TATGAGTGTGGAGTCGTTGCTAATGAGACATCTTGGGCATGGGNNNAGATCGGAA |

| | |
|---|---|
| 34388_4 | TATGAGTGTGGAGTCGTTGCTAATGAGACATCTTGGGCATGGGNNNAGATCGGAA |
| 34388_5 | TATGAGTGTGGAGTCGTTGCTAATGAGACATCTTGGGCATGGGNNNAGATCGGAA |
| 40441_1 | TATGAGTGTGGAGTCGTTGCTAATCTCAACACTTTGGGAGGCANNNAGATCGGAAG |
| 40441_2 | TATGAGTGTGGAGTCGTTGCTAATCTCAACACTTTGGGAGGCANNNAGATCGGAAG |
| 40441_3 | TATGAGTGTGGAGTCGTTGCTAATCTCAACACTTTGGGAGGCANNNAGATCGGAAG |
| 40441_4 | TATGAGTGTGGAGTCGTTGCTAATCTCAACACTTTGGGAGGCANNNAGATCGGAAG |
| 40441_5 | TATGAGTGTGGAGTCGTTGCTAATCTCAACACTTTGGGAGGCANNNAGATCGGAAG |
| 35733_1 | TATGAGTGTGGAGTCGTTGCCCAAGTAGCTGGGACTACAGGNNNAGATCGGAAGA |
| 35733_2 | TATGAGTGTGGAGTCGTTGCCCAAGTAGCTGGGACTACAGGNNNAGATCGGAAGA |
| 35733_3 | TATGAGTGTGGAGTCGTTGCCCAAGTAGCTGGGACTACAGGNNNAGATCGGAAGA |
| 35733_4 | TATGAGTGTGGAGTCGTTGCCCAAGTAGCTGGGACTACAGGNNNAGATCGGAAGA |
| 35733_5 | TATGAGTGTGGAGTCGTTGCCCAAGTAGCTGGGACTACAGGNNNAGATCGGAAGA |
| 48049_1 | TATGAGTGTGGAGTCGTTGCAGTAATTCTCCTGCCTCAGCTTCNNNAGATCGGAAGA |
| 48049_2 | TATGAGTGTGGAGTCGTTGCAGTAATTCTCCTGCCTCAGCTTCNNNAGATCGGAAGA |
| 48049_3 | TATGAGTGTGGAGTCGTTGCAGTAATTCTCCTGCCTCAGCTTCNNNAGATCGGAAGA |
| 48049_4 | TATGAGTGTGGAGTCGTTGCAGTAATTCTCCTGCCTCAGCTTCNNNAGATCGGAAGA |
| 48049_5 | TATGAGTGTGGAGTCGTTGCAGTAATTCTCCTGCCTCAGCTTCNNNAGATCGGAAGA |
| 37891_1 | TATGAGTGTGGAGTCGTTGCTTTCCTCTGGCTGCCTATTTCTTNNNAGATCGGAAGA |
| 37891_2 | TATGAGTGTGGAGTCGTTGCTTTCCTCTGGCTGCCTATTTCTTNNNAGATCGGAAGA |
| 37891_3 | TATGAGTGTGGAGTCGTTGCTTTCCTCTGGCTGCCTATTTCTTNNNAGATCGGAAGA |
| 37891_4 | TATGAGTGTGGAGTCGTTGCTTTCCTCTGGCTGCCTATTTCTTNNNAGATCGGAAGA |
| 37891_5 | TATGAGTGTGGAGTCGTTGCTTTCCTCTGGCTGCCTATTTCTTNNNAGATCGGAAGA |
| 34655_1 | TATGAGTGTGGAGTCGTTGCCGAGGAATGAGGAATGACTCAGANNNAGATCGGAA |
| 34655_2 | TATGAGTGTGGAGTCGTTGCCGAGGAATGAGGAATGACTCAGANNNAGATCGGAA |
| 34655_3 | TATGAGTGTGGAGTCGTTGCCGAGGAATGAGGAATGACTCAGANNNAGATCGGAA |
| 34655_4 | TATGAGTGTGGAGTCGTTGCCGAGGAATGAGGAATGACTCAGANNNAGATCGGAA |
| 34655_5 | TATGAGTGTGGAGTCGTTGCCGAGGAATGAGGAATGACTCAGANNNAGATCGGAA |
| 10486_1 | TATGAGTGTGGAGTCGTTGCATCCATGCTCTGTGTTCTAACAANNNAGATCGGAAGA |
| 10486_2 | TATGAGTGTGGAGTCGTTGCATCCATGCTCTGTGTTCTAACAANNNAGATCGGAAGA |
| 10486_3 | TATGAGTGTGGAGTCGTTGCATCCATGCTCTGTGTTCTAACAANNNAGATCGGAAGA |
| 10486_4 | TATGAGTGTGGAGTCGTTGCATCCATGCTCTGTGTTCTAACAANNNAGATCGGAAGA |
| 10486_5 | TATGAGTGTGGAGTCGTTGCATCCATGCTCTGTGTTCTAACAANNNAGATCGGAAGA |
| 36667_1 | TATGAGTGTGGAGTCGTTGCAGGTTTCTCCCACAGTACTTGATNNNAGATCGGAAGA |
| 36667_2 | TATGAGTGTGGAGTCGTTGCAGGTTTCTCCCACAGTACTTGATNNNAGATCGGAAGA |
| 36667_3 | TATGAGTGTGGAGTCGTTGCAGGTTTCTCCCACAGTACTTGATNNNAGATCGGAAGA |
| 36667_4 | TATGAGTGTGGAGTCGTTGCAGGTTTCTCCCACAGTACTTGATNNNAGATCGGAAGA |
| 36667_5 | TATGAGTGTGGAGTCGTTGCAGGTTTCTCCCACAGTACTTGATNNNAGATCGGAAGA |
| 52855_1 | TATGAGTGTGGAGTCGTTGCCTTCTCCTGCCTAGCACCATCNNNAGATCGGAAGAGC |
| 52855_2 | TATGAGTGTGGAGTCGTTGCCTTCTCCTGCCTAGCACCATCNNNAGATCGGAAGAGC |
| 52855_3 | TATGAGTGTGGAGTCGTTGCCTTCTCCTGCCTAGCACCATCNNNAGATCGGAAGAGC |
| 52855_4 | TATGAGTGTGGAGTCGTTGCCTTCTCCTGCCTAGCACCATCNNNAGATCGGAAGAGC |
| 52855_5 | TATGAGTGTGGAGTCGTTGCCTTCTCCTGCCTAGCACCATCNNNAGATCGGAAGAGC |
| 2269_1 | TATGAGTGTGGAGTCGTTGCGGAGGTGGGGAGATGAAGCTTTANNNAGATCGGA |

| | |
|---|---|
| 2269_2 | TATGAGTGTGGAGTCGTTGCGGAGGTGGGGAGATGAAGCTTTANNNAGATCGGA |
| 2269_3 | TATGAGTGTGGAGTCGTTGCGGAGGTGGGGAGATGAAGCTTTANNNAGATCGGA |
| 2269_4 | TATGAGTGTGGAGTCGTTGCGGAGGTGGGGAGATGAAGCTTTANNNAGATCGGA |
| 2269_5 | TATGAGTGTGGAGTCGTTGCGGAGGTGGGGAGATGAAGCTTTANNNAGATCGGA |
| 40929_1 | TATGAGTGTGGAGTCGTTGCACAACTTTTCCCAAACTCGGTTTNNNAGATCGGAAGA |
| 40929_2 | TATGAGTGTGGAGTCGTTGCACAACTTTTCCCAAACTCGGTTTNNNAGATCGGAAGA |
| 40929_3 | TATGAGTGTGGAGTCGTTGCACAACTTTTCCCAAACTCGGTTTNNNAGATCGGAAGA |
| 40929_4 | TATGAGTGTGGAGTCGTTGCACAACTTTTCCCAAACTCGGTTTNNNAGATCGGAAGA |
| 40929_5 | TATGAGTGTGGAGTCGTTGCACAACTTTTCCCAAACTCGGTTTNNNAGATCGGAAGA |
| 40722_1 | TATGAGTGTGGAGTCGTTGCGCCTGTAATCCCAGCACTTTGNNNAGATCGGAAGAG |
| 40722_2 | TATGAGTGTGGAGTCGTTGCGCCTGTAATCCCAGCACTTTGNNNAGATCGGAAGAG |
| 40722_3 | TATGAGTGTGGAGTCGTTGCGCCTGTAATCCCAGCACTTTGNNNAGATCGGAAGAG |
| 40722_4 | TATGAGTGTGGAGTCGTTGCGCCTGTAATCCCAGCACTTTGNNNAGATCGGAAGAG |
| 40722_5 | TATGAGTGTGGAGTCGTTGCGCCTGTAATCCCAGCACTTTGNNNAGATCGGAAGAG |
| 47094_1 | TATGAGTGTGGAGTCGTTGCAGTTGGAACTACAGAAGTGAGCCNNNAGATCGGAA |
| 47094_2 | TATGAGTGTGGAGTCGTTGCAGTTGGAACTACAGAAGTGAGCCNNNAGATCGGAA |
| 47094_3 | TATGAGTGTGGAGTCGTTGCAGTTGGAACTACAGAAGTGAGCCNNNAGATCGGAA |
| 47094_4 | TATGAGTGTGGAGTCGTTGCAGTTGGAACTACAGAAGTGAGCCNNNAGATCGGAA |
| 47094_5 | TATGAGTGTGGAGTCGTTGCAGTTGGAACTACAGAAGTGAGCCNNNAGATCGGAA |
| 43501_1 | TATGAGTGTGGAGTCGTTGCTCTCTCAGGTTCAAGCGATTCTCNNNAGATCGGAAG |
| 43501_2 | TATGAGTGTGGAGTCGTTGCTCTCTCAGGTTCAAGCGATTCTCNNNAGATCGGAAG |
| 43501_3 | TATGAGTGTGGAGTCGTTGCTCTCTCAGGTTCAAGCGATTCTCNNNAGATCGGAAG |
| 43501_4 | TATGAGTGTGGAGTCGTTGCTCTCTCAGGTTCAAGCGATTCTCNNNAGATCGGAAG |
| 43501_5 | TATGAGTGTGGAGTCGTTGCTCTCTCAGGTTCAAGCGATTCTCNNNAGATCGGAAG |
| 42496_1 | TATGAGTGTGGAGTCGTTGCGCTTGTGGCTGACCTGTATTTTTNNNAGATCGGAAG |
| 42496_2 | TATGAGTGTGGAGTCGTTGCGCTTGTGGCTGACCTGTATTTTTNNNAGATCGGAAG |
| 42496_3 | TATGAGTGTGGAGTCGTTGCGCTTGTGGCTGACCTGTATTTTTNNNAGATCGGAAG |
| 42496_4 | TATGAGTGTGGAGTCGTTGCGCTTGTGGCTGACCTGTATTTTTNNNAGATCGGAAG |
| 42496_5 | TATGAGTGTGGAGTCGTTGCGCTTGTGGCTGACCTGTATTTTTNNNAGATCGGAAG |
| 38495_1 | TATGAGTGTGGAGTCGTTGCTTTTCTTTTAGCCCCTTCTCCCCNNNAGATCGGAAGA |
| 38495_2 | TATGAGTGTGGAGTCGTTGCTTTTCTTTTAGCCCCTTCTCCCCNNNAGATCGGAAGA |
| 38495_3 | TATGAGTGTGGAGTCGTTGCTTTTCTTTTAGCCCCTTCTCCCCNNNAGATCGGAAGA |
| 38495_4 | TATGAGTGTGGAGTCGTTGCTTTTCTTTTAGCCCCTTCTCCCCNNNAGATCGGAAGA |
| 38495_5 | TATGAGTGTGGAGTCGTTGCTTTTCTTTTAGCCCCTTCTCCCCNNNAGATCGGAAGA |
| 42230_1 | TATGAGTGTGGAGTCGTTGCGTTGAACAAAATAGGATCGCCCANNNAGATCGGAA |
| 42230_2 | TATGAGTGTGGAGTCGTTGCGTTGAACAAAATAGGATCGCCCANNNAGATCGGAA |
| 42230_3 | TATGAGTGTGGAGTCGTTGCGTTGAACAAAATAGGATCGCCCANNNAGATCGGAA |
| 42230_4 | TATGAGTGTGGAGTCGTTGCGTTGAACAAAATAGGATCGCCCANNNAGATCGGAA |
| 42230_5 | TATGAGTGTGGAGTCGTTGCGTTGAACAAAATAGGATCGCCCANNNAGATCGGAA |
| 53389_1 | TATGAGTGTGGAGTCGTTGCGCCTCCCAGATTCAAGCAATTTTNNNAGATCGGAAG |
| 53389_2 | TATGAGTGTGGAGTCGTTGCGCCTCCCAGATTCAAGCAATTTTNNNAGATCGGAAG |
| 53389_3 | TATGAGTGTGGAGTCGTTGCGCCTCCCAGATTCAAGCAATTTTNNNAGATCGGAAG |
| 53389_4 | TATGAGTGTGGAGTCGTTGCGCCTCCCAGATTCAAGCAATTTTNNNAGATCGGAAG |

| | |
|---|---|
| 53389_5 | TATGAGTGTGGAGTCGTTGCGCCTCCCAGATTCAAGCAATTTTNNNAGATCGGAAG |
| 32748_1 | TATGAGTGTGGAGTCGTTGCCACACACACACACACTACAAAGGNNNAGATCGGAAG |
| 32748_2 | TATGAGTGTGGAGTCGTTGCCACACACACACACACTACAAAGGNNNAGATCGGAAG |
| 32748_3 | TATGAGTGTGGAGTCGTTGCCACACACACACACACTACAAAGGNNNAGATCGGAAG |
| 32748_4 | TATGAGTGTGGAGTCGTTGCCACACACACACACACTACAAAGGNNNAGATCGGAAG |
| 32748_5 | TATGAGTGTGGAGTCGTTGCCACACACACACACACTACAAAGGNNNAGATCGGAAG |
| 39273_1 | TATGAGTGTGGAGTCGTTGCCCAGTGTTCAGTGCAACCTAATCNNNAGATCGGAAG |
| 39273_2 | TATGAGTGTGGAGTCGTTGCCCAGTGTTCAGTGCAACCTAATCNNNAGATCGGAAG |
| 39273_3 | TATGAGTGTGGAGTCGTTGCCCAGTGTTCAGTGCAACCTAATCNNNAGATCGGAAG |
| 39273_4 | TATGAGTGTGGAGTCGTTGCCCAGTGTTCAGTGCAACCTAATCNNNAGATCGGAAG |
| 39273_5 | TATGAGTGTGGAGTCGTTGCCCAGTGTTCAGTGCAACCTAATCNNNAGATCGGAAG |
| 44101_1 | TATGAGTGTGGAGTCGTTGCGAGTTCAGGGAGGGGACTAGATNNNAGATCGGAA |
| 44101_2 | TATGAGTGTGGAGTCGTTGCGAGTTCAGGGAGGGGACTAGATNNNAGATCGGAA |
| 44101_3 | TATGAGTGTGGAGTCGTTGCGAGTTCAGGGAGGGGACTAGATNNNAGATCGGAA |
| 44101_4 | TATGAGTGTGGAGTCGTTGCGAGTTCAGGGAGGGGACTAGATNNNAGATCGGAA |
| 44101_5 | TATGAGTGTGGAGTCGTTGCGAGTTCAGGGAGGGGACTAGATNNNAGATCGGAA |
| 40637_1 | TATGAGTGTGGAGTCGTTGCGGCAAAACCCCGTCTCTACTAAANNNAGATCGGAAG |
| 40637_2 | TATGAGTGTGGAGTCGTTGCGGCAAAACCCCGTCTCTACTAAANNNAGATCGGAAG |
| 40637_3 | TATGAGTGTGGAGTCGTTGCGGCAAAACCCCGTCTCTACTAAANNNAGATCGGAAG |
| 40637_4 | TATGAGTGTGGAGTCGTTGCGGCAAAACCCCGTCTCTACTAAANNNAGATCGGAAG |
| 40637_5 | TATGAGTGTGGAGTCGTTGCGGCAAAACCCCGTCTCTACTAAANNNAGATCGGAAG |
| 40156_1 | TATGAGTGTGGAGTCGTTGCGGCAAAAAAACGGCAAAATGTTGNNNAGATCGGAA |
| 40156_2 | TATGAGTGTGGAGTCGTTGCGGCAAAAAAACGGCAAAATGTTGNNNAGATCGGAA |
| 40156_3 | TATGAGTGTGGAGTCGTTGCGGCAAAAAAACGGCAAAATGTTGNNNAGATCGGAA |
| 40156_4 | TATGAGTGTGGAGTCGTTGCGGCAAAAAAACGGCAAAATGTTGNNNAGATCGGAA |
| 40156_5 | TATGAGTGTGGAGTCGTTGCGGCAAAAAAACGGCAAAATGTTGNNNAGATCGGAA |
| 39834_1 | TATGAGTGTGGAGTCGTTGCAAAAAGAAAAAAATTGGCCGGCCNNNAGATCGGAA |
| 39834_2 | TATGAGTGTGGAGTCGTTGCAAAAAGAAAAAAATTGGCCGGCCNNNAGATCGGAA |
| 39834_3 | TATGAGTGTGGAGTCGTTGCAAAAAGAAAAAAATTGGCCGGCCNNNAGATCGGAA |
| 39834_4 | TATGAGTGTGGAGTCGTTGCAAAAAGAAAAAAATTGGCCGGCCNNNAGATCGGAA |
| 39834_5 | TATGAGTGTGGAGTCGTTGCAAAAAGAAAAAAATTGGCCGGCCNNNAGATCGGAA |
| 50095_1 | TATGAGTGTGGAGTCGTTGCTTGTCTAGCTTTCCTGGTGACTCNNNAGATCGGAAG |
| 50095_2 | TATGAGTGTGGAGTCGTTGCTTGTCTAGCTTTCCTGGTGACTCNNNAGATCGGAAG |
| 50095_3 | TATGAGTGTGGAGTCGTTGCTTGTCTAGCTTTCCTGGTGACTCNNNAGATCGGAAG |
| 50095_4 | TATGAGTGTGGAGTCGTTGCTTGTCTAGCTTTCCTGGTGACTCNNNAGATCGGAAG |
| 50095_5 | TATGAGTGTGGAGTCGTTGCTTGTCTAGCTTTCCTGGTGACTCNNNAGATCGGAAG |
| 48376_1 | TATGAGTGTGGAGTCGTTGCCTGTGAGGTGAGGTGTGGACNNNAGATCGGAAGA |
| 48376_2 | TATGAGTGTGGAGTCGTTGCCTGTGAGGTGAGGTGTGGACNNNAGATCGGAAGA |
| 48376_3 | TATGAGTGTGGAGTCGTTGCCTGTGAGGTGAGGTGTGGACNNNAGATCGGAAGA |
| 48376_4 | TATGAGTGTGGAGTCGTTGCCTGTGAGGTGAGGTGTGGACNNNAGATCGGAAGA |
| 48376_5 | TATGAGTGTGGAGTCGTTGCCTGTGAGGTGAGGTGTGGACNNNAGATCGGAAGA |
| 32341_1 | TATGAGTGTGGAGTCGTTGCTTTCTTACAGTTCTGGAGGCTGGNNNAGATCGGAAG |
| 32341_2 | TATGAGTGTGGAGTCGTTGCTTTCTTACAGTTCTGGAGGCTGGNNNAGATCGGAAG |

| | |
|---|---|
| 32341_3 | TATGAGTGTGGAGTCGTTGCTTTCTTACAGTTCTGGAGGCTGGNNNAGATCGGAAG |
| 32341_4 | TATGAGTGTGGAGTCGTTGCTTTCTTACAGTTCTGGAGGCTGGNNNAGATCGGAAG |
| 32341_5 | TATGAGTGTGGAGTCGTTGCTTTCTTACAGTTCTGGAGGCTGGNNNAGATCGGAAG |
| 33433_1 | TATGAGTGTGGAGTCGTTGCAAAACGTTTCCTGACTCCAATGCNNNAGATCGGAAG |
| 33433_2 | TATGAGTGTGGAGTCGTTGCAAAACGTTTCCTGACTCCAATGCNNNAGATCGGAAG |
| 33433_3 | TATGAGTGTGGAGTCGTTGCAAAACGTTTCCTGACTCCAATGCNNNAGATCGGAAG |
| 33433_4 | TATGAGTGTGGAGTCGTTGCAAAACGTTTCCTGACTCCAATGCNNNAGATCGGAAG |
| 33433_5 | TATGAGTGTGGAGTCGTTGCAAAACGTTTCCTGACTCCAATGCNNNAGATCGGAAG |
| 57378_1 | TGAGTGTGGAGTCGTTGCAAAGACTAATAATACCTTCCTTAGGGTNNNAGATCGGA |
| 64710_1 | TGAGTGTGGAGTCGTTGCGGTCTTTATTCTTTCTCTCTTCCTGCNNNAGATCGGAAG |
| 67501_1 | TGAGTGTGGAGTCGTTGCTGCCAGAAGAAGTCTATTCCTAAAGANNNAGATCGGAA |
| 59231_1 | ATGAGTGTGGAGTCGTTGCTCTTCCAGTATGTACATACGAACCANNNAGATCGGAA |
| 67481_1 | ATGAGTGTGGAGTCGTTGCTCTCCAGTATTTTTTTCAACCACANNNAGATCGGAAG |
| 63074_1 | ATGAGTGTGGAGTCGTTGCAGCTGAACATCTGAAAACAAAACANNNAGATCGGAA |
| 64415_1 | ATGAGTGTGGAGTCGTTGCTGTTTTGGGAAATATATGTGCTATAGTNNNAGATCGG |
| 66151_1 | ATGAGTGTGGAGTCGTTGCTCCAAGAGAAAAGAAAAAAGGGCANNNAGATCGGA |
| 66185_1 | ATGAGTGTGGAGTCGTTGCAGTGTCTCATCAAAGCAAAAAGCANNNAGATCGGAA |
| 66243_1 | ATGAGTGTGGAGTCGTTGCAGAATGAGGAGAGGAAAGTTAGAAGTNNNAGATCG |
| 69377_1 | ATGAGTGTGGAGTCGTTGCTCCGTGTACATATATATAGACAGATGGNNNAGATCGG |
| 69541_1 | ATGAGTGTGGAGTCGTTGCCCATATTATCAAGCTTTATTTATGGCCNNNAGATCGGA |
| 12_1 | TATGAGTGTGGAGTCGTTGCTTGGCCTCACAATCATATTTGGTNNNAGATCGGAAG |
| 27_1 | TATGAGTGTGGAGTCGTTGCGCAACTCTTTAGACAGAAAGGTAAACANNNAGATCG |
| 40_1 | TATGAGTGTGGAGTCGTTGCTCCAGGTTTGGGTTAGGAAATGTNNNAGATCGGAA |
| 63_1 | TATGAGTGTGGAGTCGTTGCACCTCTCTCTTACAGGAACTCTTGTNNNAGATCGGAA |
| 75_1 | TATGAGTGTGGAGTCGTTGCTCTTTCCCTAGAGGTGCTGAACANNNAGATCGGAAG |
| 128_1 | TATGAGTGTGGAGTCGTTGCACACACATACATAAACATCTACACACANNNAGATCGG |
| 129_1 | TATGAGTGTGGAGTCGTTGCGGGTGTCTCGCTCTGTTATATGTNNNAGATCGGAAG |
| 149_1 | TATGAGTGTGGAGTCGTTGCACTGGATCAGGCTAATTGACACANNNAGATCGGAAG |
| 150_1 | TATGAGTGTGGAGTCGTTGCCACAAACTCATCAGTAGATTTGACACANNNAGATCG |
| 192_1 | TATGAGTGTGGAGTCGTTGCCTTGTGTTTGGGTCTTACTAGTTACAANNNAGATCG |
| 208_1 | TATGAGTGTGGAGTCGTTGCACAAGAGAAGAGCTGCTGTAAAANNNAGATCGGAA |
| 273_1 | TATGAGTGTGGAGTCGTTGCACCAAATAGCCAAAGCAATAGTGAGANNNAGATCG |
| 345_1 | TATGAGTGTGGAGTCGTTGCTTTCTTGGAACTCTAGTCTTGAGANNNAGATCGGAA |
| 367_1 | TATGAGTGTGGAGTCGTTGCGCATGTATAGCATTTACACTATCGTGTNNNAGATCG |
| 369_1 | TATGAGTGTGGAGTCGTTGCACCATAGAAATGACCAGAGGACANNNAGATCGGAA |
| 379_1 | TATGAGTGTGGAGTCGTTGCGTTATCTGTGTGGGATACATTCCAAGNNNAGATCGG |
| 420_1 | TATGAGTGTGGAGTCGTTGCGGTGGATCAGGAAATGTGTGACANNNAGATCGGAA |
| 456_1 | TATGAGTGTGGAGTCGTTGCCGCACAATATATCTGGATACATATACANNNAGATCG |
| 476_1 | TATGAGTGTGGAGTCGTTGCTCTAGGGCAGTCTCATTGCTACANNNAGATCGGAAG |
| 494_1 | TATGAGTGTGGAGTCGTTGCTGTTGTGCATAGATACTTGATAGTACANNNAGATCG |
| 506_1 | TATGAGTGTGGAGTCGTTGCGATGGACACCTAGCTTGCTTACANNNAGATCGGAAG |
| 512_1 | TATGAGTGTGGAGTCGTTGCTTTCAAGAATTCATAGGACATGTTACANNNAGATCG |
| 513_1 | TATGAGTGTGGAGTCGTTGCGCATTTTCTACTCTTTTGAACAGTTGTNNNAGATCGG |

| | |
|---|---|
| 607_1 | TATGAGTGTGGAGTCGTTGCACCAGATAAAAGAAAATTCCTGACCANNNAGATCGG |
| 742_1 | TATGAGTGTGGAGTCGTTGCATTTCTAATGTCTGCAACTTTCATGANNNAGATCGGA |
| 770_1 | TATGAGTGTGGAGTCGTTGCTTGGATCAGGGATCAGATTCCCANNNAGATCGGAAG |
| 784_1 | TATGAGTGTGGAGTCGTTGCGGAATGAGTTGGGTGATTTCCCANNNAGATCGGAA |
| 793_1 | TATGAGTGTGGAGTCGTTGCAGCACATGGATATTTGATAAAAAGCCANNNAGATCG |
| 813_1 | TATGAGTGTGGAGTCGTTGCAACACACACACTACACATAAATCATTTNNNAGATCGG |
| 969_1 | TATGAGTGTGGAGTCGTTGCGCATCAATGAACACTAAGTTGTATCCANNNAGATCG |
| 1048_1 | TATGAGTGTGGAGTCGTTGCTTTCTTTAGGCAAATACCCAGTAGTNNNAGATCGGA |
| 1085_1 | TATGAGTGTGGAGTCGTTGCTGTTGTTTCACACATAGTACTTTCCANNNAGATCGGA |
| 1251_1 | TATGAGTGTGGAGTCGTTGCTTTTCCCAATATCCTACTCTTTTTCTTNNNAGATCGGA |
| 1258_1 | TATGAGTGTGGAGTCGTTGCTATTTACTGTGCTCAGCCAGCAGAGNNNAGATCGGA |
| 1283_1 | TATGAGTGTGGAGTCGTTGCTAATGTGGGGAAGGAGTGTAGCANNNAGATCGGA |
| 1293_1 | TATGAGTGTGGAGTCGTTGCTGGCTTCTACTTCTTATGTCTTGGAAANNNAGATCGG |
| 1300_1 | TATGAGTGTGGAGTCGTTGCCACTTCTTCACTAGATTTCTTCCAACANNNAGATCGG |
| 1337_1 | TATGAGTGTGGAGTCGTTGCGACATTTTAGCTGTACCTGATTTTATCNNNAGATCGG |
| 1341_1 | TATGAGTGTGGAGTCGTTGCTGGTAATTGTGTATTTATTGTCAGGCANNNAGATCG |
| 1344_1 | TATGAGTGTGGAGTCGTTGCACCAGAGACCTAATTTTCAGGCANNNAGATCGGAAG |
| 1419_1 | TATGAGTGTGGAGTCGTTGCGTCACATTGAACTTGATAAGATGGCANNNAGATCGG |
| 1467_1 | TATGAGTGTGGAGTCGTTGCGCTAACTAGAGTTCTTACTGTTGTTTTNNNAGATCGG |
| 1642_1 | TATGAGTGTGGAGTCGTTGCGTGTGTGTGTGTCATTTTCCTTTTTCTNNNAGATCGG |
| 1645_1 | TATGAGTGTGGAGTCGTTGCACCTGGCATCAGCAAAAGTAGAANNNAGATCGGAA |
| 1666_1 | TATGAGTGTGGAGTCGTTGCTCAACTCAAGATGGATAAAATACCTTTNNNAGATCG |
| 1667_1 | TATGAGTGTGGAGTCGTTGCACAAGAAGTAACAAGTTTTGGCAAGGNNNAGATCG |
| 1803_1 | TATGAGTGTGGAGTCGTTGCCACTTTTGATTTTGTCAATTGAACAGTNNNAGATCGG |
| 1804_1 | TATGAGTGTGGAGTCGTTGCGCTTGGGAGGGTATTCAAATATTCAGTNNNAGATCG |
| 1838_1 | TATGAGTGTGGAGTCGTTGCTATTTGCACTGTCATGTTTATTGCAGTNNNAGATCGG |
| 1846_1 | TATGAGTGTGGAGTCGTTGCAGATACAGCAAATTGTGGTAAAATTTTNNNAGATCG |
| 1850_1 | TATGAGTGTGGAGTCGTTGCATAGCTTGTTTCCTTCCACTCTATGNNNAGATCGGAA |
| 1851_1 | TATGAGTGTGGAGTCGTTGCTGAAACCAATAAGGAAAAGTGAATTTTNNNAGATCG |
| 1859_1 | TATGAGTGTGGAGTCGTTGCCGCAAGTACCTTTTTGCTATAATGACTNNNAGATCGG |
| 1879_1 | TATGAGTGTGGAGTCGTTGCTGGTAGAATGTTGCACAACTTTTNNNAGATCGGAAG |
| 1908_1 | TATGAGTGTGGAGTCGTTGCTCTTTCAGTATGTCTTCAAATTCACTANNNAGATCGG |
| 2153_1 | TATGAGTGTGGAGTCGTTGCGAAGAAAAATTGCTAAAAGCATTGAAANNNAGATC |
| 2155_1 | TATGAGTGTGGAGTCGTTGCTTTACCAAGATTTTTGCATTGGCCATANNNAGATCGG |
| 2184_1 | TATGAGTGTGGAGTCGTTGCTGTCACTTTGATTGATATGTCAGTTCANNNAGATCG |
| 2206_1 | TATGAGTGTGGAGTCGTTGCTCTTTTCCCTATGGGATGGTACATACTNNNAGATCGG |
| 2213_1 | TATGAGTGTGGAGTCGTTGCCGTCCATAACAGTAAAGATTATGTTCANNNAGATCG |
| 2323_1 | TATGAGTGTGGAGTCGTTGCGGATTGCTGGACCGAAGGATAGAANNNAGATCGGA |
| 2354_1 | TATGAGTGTGGAGTCGTTGCCCCTTCACTGAAAAACATAAAAGAAGANNNAGATCG |
| 2357_1 | TATGAGTGTGGAGTCGTTGCTCCAAAAAGAGTAGGAAACAAGAAGANNNAGATCG |
| 2376_1 | TATGAGTGTGGAGTCGTTGCTGAAGAATAACATCAAAGCTGGAAGANNNAGATCG |
| 2390_1 | TATGAGTGTGGAGTCGTTGCAAAGTCAATATCTTGGTCTCGTTCCTANNNAGATCGG |
| 2392_1 | TATGAGTGTGGAGTCGTTGCGCAGCTATCTCTTCAACATACCAACCNNNAGATCGGA |

| | |
|---|---|
| 2609_1 | TATGAGTGTGGAGTCGTTGCCAGTCGTTTGCTAGAGCAAGAGANNNAGATCGGAA |
| 2716_1 | TATGAGTGTGGAGTCGTTGCGCATAGTGATATTAGGGTAAAGCAAGTNNNAGATC |
| 2765_1 | TATGAGTGTGGAGTCGTTGCTGCTATTTGTCAAGAGTAAAGACTAGANNNAGATCG |
| 2788_1 | TATGAGTGTGGAGTCGTTGCCTGAACCTATTGAGATGATCACATGATNNNAGATCG |
| 2815_1 | TATGAGTGTGGAGTCGTTGCAGGTATCAGGAACAAAACAAAGGTTTTNNNAGATC |
| 2821_1 | TATGAGTGTGGAGTCGTTGCGCTGCAATAAACATACAAGAGCAAGTANNNAGATCG |
| 2829_1 | TATGAGTGTGGAGTCGTTGCAGACAAGACCTTCCTTTGGTCAANNNAGATCGGAAG |
| 2840_1 | TATGAGTGTGGAGTCGTTGCAATCTTTTCTTCCTCCTTTCCTTCCNNNAGATCGGAAG |
| 2880_1 | TATGAGTGTGGAGTCGTTGCTTCTATTTCACTTCTTGGAAGACTGCCNNNAGATCGG |
| 2894_1 | TATGAGTGTGGAGTCGTTGCTCTTTATCTGTGTCCATAAAGCATTTNNNAGATCGGA |
| 2895_1 | TATGAGTGTGGAGTCGTTGCTTCTCACCCATGAAATTGGCAAAAATTNNNAGATCG |
| 3120_1 | TATGAGTGTGGAGTCGTTGCGGATTTTGATCGGGTTGCATTGTNNNAGATCGGAA |
| 3210_1 | TATGAGTGTGGAGTCGTTGCTGTCCATCCTTGTCTGGTTTTGATATNNNAGATCGGA |
| 3220_1 | TATGAGTGTGGAGTCGTTGCCACTTATCTCCCTTTTTCCTTTACTATNNNAGATCGGA |
| 3224_1 | TATGAGTGTGGAGTCGTTGCGTGAGACTAAGGGATACGTGGGANNNAGATCGGA |
| 3230_1 | TATGAGTGTGGAGTCGTTGCAAGAATCCCTATTTGGCAGATCTGAAANNNAGATCG |
| 3256_1 | TATGAGTGTGGAGTCGTTGCATACGGTGAAACCATGAAGTATAAAGNNNAGATCG |
| 3265_1 | TATGAGTGTGGAGTCGTTGCATGGTGATTTGTGCCTGAAATCTCNNNAGATCGGAA |
| 3274_1 | TATGAGTGTGGAGTCGTTGCCTGTGATTTCCTATGGTGCTTTAGGGANNNAGATCG |
| 3285_1 | TATGAGTGTGGAGTCGTTGCAGTTGTAAAGTCTCACAATATGCCATCNNNAGATCG |
| 3306_1 | TATGAGTGTGGAGTCGTTGCTGTATCTGGCTTCTTCCACATTGTAATNNNAGATCGG |
| 3330_1 | TATGAGTGTGGAGTCGTTGCAAAAAGGCAATTCCGCAATAAGGTTATNNNAGATCG |
| 3349_1 | TATGAGTGTGGAGTCGTTGCTGGATAGCTCTGAAAAACTGATAATGANNNAGATCG |
| 3371_1 | TATGAGTGTGGAGTCGTTGCACGCAAAGGAACACTTCTATCATGANNNAGATCGGA |
| 3372_1 | TATGAGTGTGGAGTCGTTGCTACATCCAAACATATGTACAAAAACACNNNAGATCG |
| 3399_1 | TATGAGTGTGGAGTCGTTGCCAGGACCTCACCAGAAGATATAAATGANNNAGATCG |
| 3436_1 | TATGAGTGTGGAGTCGTTGCGGTATTTTCACATTCTTCATGGTGTGTNNNAGATCG |
| 3496_1 | TATGAGTGTGGAGTCGTTGCGATGGCAACATAATAATGTGGGANNNAGATCGGAA |
| 3528_1 | TATGAGTGTGGAGTCGTTGCTCCTTCCTTCCTGTCTTTCTTTGACTANNNAGATCGGA |
| 3541_1 | TATGAGTGTGGAGTCGTTGCCATAACAAACCTTACATTTTACCTGTTNNNAGATCGG |
| 3561_1 | TATGAGTGTGGAGTCGTTGCCCTGAAATACACACATCCACTCTTCTGNNNAGATCGG |
| 3596_1 | TATGAGTGTGGAGTCGTTGCAACAATGAAAAAGTAGTCAGGCAGCCTNNNAGATC |
| 3637_1 | TATGAGTGTGGAGTCGTTGCTCAAACTGCTACTACAAAATAGTTTCTNNNAGATCGG |
| 3640_1 | TATGAGTGTGGAGTCGTTGCACCTCCACATGTTTATCCAAAATTTGANNNAGATCGG |
| 3650_1 | TATGAGTGTGGAGTCGTTGCCCTGTGCCAAATGTGTTATACAAATGANNNAGATCG |
| 3656_1 | TATGAGTGTGGAGTCGTTGCTCTTCCATTTGGCTAAAGAGATTTTGANNNAGATCG |
| 3682_1 | TATGAGTGTGGAGTCGTTGCAAGAGCCAATTAGTGGGCCAATANNNAGATCGGAA |
| 3684_1 | TATGAGTGTGGAGTCGTTGCCAGCCGAAATATTCATTTGTGTTTGTNNNAGATCGG |
| 3685_1 | TATGAGTGTGGAGTCGTTGCGTGTTTTATAAGTGCCTTTCTAAATGCNNNAGATCG |
| 3694_1 | TATGAGTGTGGAGTCGTTGCCAGCACACCGCATGGATCTAATANNNAGATCGGAAG |
| 3714_1 | TATGAGTGTGGAGTCGTTGCGTGCCCATGGTTTAGTGTCCATANNNAGATCGGAAG |
| 3817_1 | TATGAGTGTGGAGTCGTTGCCTCCAGAAAGTCTTGTCCACCTANNNAGATCGGAAG |
| 3906_1 | TATGAGTGTGGAGTCGTTGCCCTGACAGAAGACAGTGTTTCCAGAANNNAGATCGG |

| | |
|---|---|
| 3932_1 | TATGAGTGTGGAGTCGTTGCAGATAGCCTCTTAGGATACAAAGGGACNNNAGATCG |
| 3941_1 | TATGAGTGTGGAGTCGTTGCCTGTCTTTCACATAAAACATCATCTTTNNNAGATCGG |
| 3951_1 | TATGAGTGTGGAGTCGTTGCAGTTGCATTTTTCCAAAGAAATGATAANNNAGATCG |
| 3993_1 | TATGAGTGTGGAGTCGTTGCCAACACCATAGCACGTACTTACAAANNNAGATCGGA |
| 4004_1 | TATGAGTGTGGAGTCGTTGCCTCTGTTCTAGATCCTGAACTATCAATNNNAGATCGG |
| 4051_1 | TATGAGTGTGGAGTCGTTGCTAGATGTGTGTATGTGGGTATATGTGTNNNAGATC |
| 4084_1 | TATGAGTGTGGAGTCGTTGCCTTTCCCCTTTGTGAGCCACTTANNNAGATCGGAAGA |
| 4094_1 | TATGAGTGTGGAGTCGTTGCAAATTCAGCCTGGGACATCCTTANNNAGATCGGAAG |
| 4163_1 | TATGAGTGTGGAGTCGTTGCAGTTGAGGAATTCAAAATTTATTCCAGNNNAGATCG |
| 4176_1 | TATGAGTGTGGAGTCGTTGCTTCTGCTAAACAACCTCAATGAAAATCNNNAGATCGG |
| 4254_1 | TATGAGTGTGGAGTCGTTGCATCACATCAGTTAGAAACCCCTGTTTCNNNAGATCGG |
| 4255_1 | TATGAGTGTGGAGTCGTTGCATCAACTGAAATCAGGATCTTGAAGACNNNAGATCG |
| 4256_1 | TATGAGTGTGGAGTCGTTGCTAAAATCACAAGCCATAGAGAACATGGNNNAGATCG |
| 4257_1 | TATGAGTGTGGAGTCGTTGCACCTGAATGGAATTGGAGACTATTGTCNNNAGATCG |
| 4301_1 | TATGAGTGTGGAGTCGTTGCGGCGAATACCTGTATCTGGGAACNNNAGATCGGAA |
| 4313_1 | TATGAGTGTGGAGTCGTTGCCAGGAGCAACAAATATCTTCACCAAGANNNAGATCG |
| 4381_1 | TATGAGTGTGGAGTCGTTGCTGTGGTAGTGGAGTAACAACCACNNNAGATCGGAA |
| 4394_1 | TATGAGTGTGGAGTCGTTGCTTAGGGAGTTCTCGATAGAAATACCACNNNAGATCG |
| 4429_1 | TATGAGTGTGGAGTCGTTGCTTTTATAGAGGAAAAGTCTCGCCANNNAGATCGGAA |
| 4436_1 | TATGAGTGTGGAGTCGTTGCACATAGTCTCAATTATACCTCTGGTCTNNNAGATCGG |
| 4440_1 | TATGAGTGTGGAGTCGTTGCACACATATTCCACCCAACAGCACNNNAGATCGGAAG |
| 4441_1 | TATGAGTGTGGAGTCGTTGCTCTGAACTGACAAAGAAATGTTTATTGNNNAGATCG |
| 4453_1 | TATGAGTGTGGAGTCGTTGCTATTAGGATGACTTGATCCAGAGCTGANNNAGATCG |
| 4497_1 | TATGAGTGTGGAGTCGTTGCTGGTAGTACAGTATAGGAAGTCCTCACNNNAGATCG |
| 4526_1 | TATGAGTGTGGAGTCGTTGCGGATATTTTGACACTGCATTTTCCGNNNAGATCGGA |
| 4543_1 | TATGAGTGTGGAGTCGTTGCACACACAAATTCATACAACAGGCCAGNNNAGATCGG |
| 4576_1 | TATGAGTGTGGAGTCGTTGCGACAAAAATACAGTTGTTGAGAGAGACNNNAGATC |
| 4687_1 | TATGAGTGTGGAGTCGTTGCTGCTGTATTCAGTTGCCACCTACNNNAGATCGGAAG |
| 4762_1 | TATGAGTGTGGAGTCGTTGCGTGTGTGTATCATTTATTTTGACACAANNNAGATCG |
| 4780_1 | TATGAGTGTGGAGTCGTTGCTCAGCCAAAAGGTAAACTTGAACCNNNAGATCGGAA |
| 4803_1 | TATGAGTGTGGAGTCGTTGCTATGTGTATGGGTGTGTGTATATATCTNNNAGATCG |
| 4836_1 | TATGAGTGTGGAGTCGTTGCCTCAATAGTCTCTTCTGAATGTTGATANNNAGATCGG |
| 4854_1 | TATGAGTGTGGAGTCGTTGCTCTATGTGTGTATTACTAGAGCCTTCTNNNAGATCGG |
| 4897_1 | TATGAGTGTGGAGTCGTTGCTAACAAGACCAACAAGGTACTATTACCNNNAGATCG |
| 4898_1 | TATGAGTGTGGAGTCGTTGCAACTAGACATGTTGTGTAGCAGAGTAGNNNAGATC |
| 4900_1 | TATGAGTGTGGAGTCGTTGCAAGATACCATCTCACAGCATCAGAATGNNNAGATCG |
| 4912_1 | TATGAGTGTGGAGTCGTTGCCGTCGTAGAATCTGTATGATTGTTGTTNNNAGATCG |
| 4919_1 | TATGAGTGTGGAGTCGTTGCTATTACCTTTCTAGTAGTTGCATGTTTNNNAGATCGG |
| 4937_1 | TATGAGTGTGGAGTCGTTGCAGAGAGAGAGGAGAGACTGAGTGANNNAGATCGG |
| 4951_1 | TATGAGTGTGGAGTCGTTGCATGTATGTATGTATGTATGTTGAAGGCNNNAGATCG |
| 4954_1 | TATGAGTGTGGAGTCGTTGCTAGACCTTTTGGTGCTCTGTTCATCATNNNAGATCGG |
| 4959_1 | TATGAGTGTGGAGTCGTTGCTGGAGGATATTTTTGTTTGGTTAGTGANNNAGATCG |
| 4962_1 | TATGAGTGTGGAGTCGTTGCTACACACACAATCTCTCTCCCCCNNNAGATCGGAAGA |

| | |
|---|---|
| 4969_1 | TATGAGTGTGGAGTCGTTGCTGTTTATAGTGTGAGCGTGTATAATAGNNNAGATCG |
| 5000_1 | TATGAGTGTGGAGTCGTTGCATGTAACAAGATGTGCCCAGATTCTTTNNNAGATCG |
| 5078_1 | TATGAGTGTGGAGTCGTTGCGAAGTCTCCTGTTAGTTCTGTCCCNNNAGATCGGAA |
| 5113_1 | TATGAGTGTGGAGTCGTTGCTCAGCAAGAAGAGATTATACCAAGACANNNAGATCG |
| 5119_1 | TATGAGTGTGGAGTCGTTGCTGAATTGCTCTGGAATCACTGTCAAAANNNAGATCG |
| 5156_1 | TATGAGTGTGGAGTCGTTGCTAGACTTTATGTGGTATGTACACTTGANNNAGATCG |
| 5216_1 | TATGAGTGTGGAGTCGTTGCTACACACCCACACAAATCCTATTGGTNNNAGATCGGA |
| 5223_1 | TATGAGTGTGGAGTCGTTGCGTGTGTTGTTGGAAACTGGACCTTANNNAGATCGG |
| 5234_1 | TATGAGTGTGGAGTCGTTGCTGTTTGCATGCTTTGATATATACTAGCNNNAGATCG |
| 5349_1 | TATGAGTGTGGAGTCGTTGCCTCTCCCCGTCTTTGTCATTCAGNNNAGATCGGAAGA |
| 5407_1 | TATGAGTGTGGAGTCGTTGCTGGCTATGGTTTGAGACCCTTGANNNAGATCGGAAG |
| 5491_1 | TATGAGTGTGGAGTCGTTGCTCTCCTGGTAATTCTGAGCTGCTNNNAGATCGGAAG |
| 5569_1 | TATGAGTGTGGAGTCGTTGCCACACAGACAACGATATCTTATTCAGCNNNAGATCG |
| 5577_1 | TATGAGTGTGGAGTCGTTGCTAGGGGAAGCTGTTTTGAAGAGCNNNAGATCGGAA |
| 5586_1 | TATGAGTGTGGAGTCGTTGCGTCATTGTTATCATTCCTTTTCAGAGCNNNAGATCGG |
| 5616_1 | TATGAGTGTGGAGTCGTTGCCTGGAGTAACTTGGAGTGATCATCTTCNNNAGATCG |
| 5772_1 | TATGAGTGTGGAGTCGTTGCGAGTTGGCTTATCCCTTAGATTTCATCNNNAGATCGG |
| 5836_1 | TATGAGTGTGGAGTCGTTGCAGAAAGGGAGATTTGGAAATCCCTNNNAGATCGGA |
| 5839_1 | TATGAGTGTGGAGTCGTTGCTGTGTGTGTATCTTCAACTTTGGCNNNAGATCGGAA |
| 5908_1 | TATGAGTGTGGAGTCGTTGCAACTTAGTCTGAGGGATTTGGCANNNAGATCGGAA |
| 5935_1 | TATGAGTGTGGAGTCGTTGCCTGGGCCATATTGTATATTGCTGCNNNAGATCGGAA |
| 5939_1 | TATGAGTGTGGAGTCGTTGCTACATACCACCACGCCCATATTTTTANNNAGATCGGA |
| 5946_1 | TATGAGTGTGGAGTCGTTGCGGCTTGTTCTAATAAATTTCAGTCTGCNNNAGATCGG |
| 5951_1 | TATGAGTGTGGAGTCGTTGCCCCTTAGTATTGGCAGTTTCTGCNNNAGATCGGAAG |
| 6015_1 | TATGAGTGTGGAGTCGTTGCTTGAATTCCTACTATTTGTCAGATTGCNNNAGATCGG |
| 6068_1 | TATGAGTGTGGAGTCGTTGCACGCTTTCTAAAGTACTATGTTTTTGCNNNAGATCGG |
| 6115_1 | TATGAGTGTGGAGTCGTTGCCAACAATTCCACTGCTCAGCATCNNNAGATCGGAAG |
| 6187_1 | TATGAGTGTGGAGTCGTTGCCTGGTTCTGTCTCTGGAGAACTCNNNAGATCGGAAG |
| 6226_1 | TATGAGTGTGGAGTCGTTGCGGTCAAATTCAGTCACCTCCCTCNNNAGATCGGAAG |
| 6235_1 | TATGAGTGTGGAGTCGTTGCTTCACACATCCACAGCATATGGANNNAGATCGGAAG |
| 6249_1 | TATGAGTGTGGAGTCGTTGCTCTCAGCATCTCTAGCTGTCCTCNNNAGATCGGAAGA |
| 6404_1 | TATGAGTGTGGAGTCGTTGCACAATAGGCTGCCTGCAATACTANNNAGATCGGAAG |
| 6526_1 | TATGAGTGTGGAGTCGTTGCTCTCTCCCAAATATCCTTCTAGCTCTNNNAGATCGGA |
| 6537_1 | TATGAGTGTGGAGTCGTTGCGCACCACATCCACATAACGTTTCNNNAGATCGGAAG |
| 6625_1 | TATGAGTGTGGAGTCGTTGCTGTATGTGTGTGTTCTTCCTACCCTCNNNAGATCGGA |
| 6626_1 | TATGAGTGTGGAGTCGTTGCCTATCAGCAGCTTTTACATACGGCTNNNAGATCGGA |
| 6652_1 | TATGAGTGTGGAGTCGTTGCTAGGTCAGGAGTGGTAGCTCAAGNNNAGATCGGAA |
| 6668_1 | TATGAGTGTGGAGTCGTTGCTGACTAACAAAAAGAATACCACAGAAGNNNAGATCG |
| 6671_1 | TATGAGTGTGGAGTCGTTGCTTTGATTTTTGACACATAAGAATTGCANNNAGATCG |
| 6680_1 | TATGAGTGTGGAGTCGTTGCATTGAAGGAGTATTCATTCTCCATAGCNNNAGATCG |
| 6685_1 | TATGAGTGTGGAGTCGTTGCGATTTACTTCTCAAGGCTACATTTTGGNNNAGATCG |
| 6728_1 | TATGAGTGTGGAGTCGTTGCAGCCCCTTCTTTTCTTCCAGTAGNNNAGATCGGAAGA |
| 6837_1 | TATGAGTGTGGAGTCGTTGCTTCTGACAAAGTTTTGCTCCTGTGGNNNAGATCGGA |

| | |
|---|---|
| 6842_1 | TATGAGTGTGGAGTCGTTGCTCTGTCTCTGTCTCTCTCTTATCTCTTNNNAGATCGGA |
| 6862_1 | TATGAGTGTGGAGTCGTTGCAAAACAACCTAAGAACATTTACCAAAANNNAGATCG |
| 6869_1 | TATGAGTGTGGAGTCGTTGCATCTATCCATCCTTCCACATATCTATTNNNAGATCGGA |
| 6877_1 | TATGAGTGTGGAGTCGTTGCCTCTCTTTCTTTCTCTGCTGGTCTTTNNNAGATCGGAA |
| 6947_1 | TATGAGTGTGGAGTCGTTGCCTCTCTCTCTCTCTCTGTTTGAAGAGNNNAGATCGGA |
| 6955_1 | TATGAGTGTGGAGTCGTTGCACTAAATCAGCTTCAATCATCACAGCCNNNAGATCGG |
| 7040_1 | TATGAGTGTGGAGTCGTTGCAGGAAACAGGTAGAAGGAGACTATATCNNNAGATC |
| 7041_1 | TATGAGTGTGGAGTCGTTGCTCAATAGTACCCCCTACCTCACCNNNAGATCGGAAGA |
| 7105_1 | TATGAGTGTGGAGTCGTTGCGCACCCACCTTTTTAGGTAGGAGNNNAGATCGGAAG |
| 7165_1 | TATGAGTGTGGAGTCGTTGCACCTATCTACCTATCTGTCTATTCATCNNNAGATCGG |
| 7166_1 | TATGAGTGTGGAGTCGTTGCCTACCTACCTACCTATTCATCTCTATCNNNAGATCGGA |
| 7260_1 | TATGAGTGTGGAGTCGTTGCTGGCTACATGGTCACAGCTAGTANNNAGATCGGAA |
| 7284_1 | TATGAGTGTGGAGTCGTTGCCTATTCCTATTCTATCTTCCTAGAGCTNNNAGATCGG |
| 7294_1 | TATGAGTGTGGAGTCGTTGCTGTGTGTGTTTCTCTTTTCTTTTGGAANNNAGATCGG |
| 7297_1 | TATGAGTGTGGAGTCGTTGCAAAAATGGCACCGGAAAAGAATGAGAANNNAGATC |
| 7307_1 | TATGAGTGTGGAGTCGTTGCAGGAGTTGACATGTGTTTCTATATGCANNNAGATCG |
| 7349_1 | TATGAGTGTGGAGTCGTTGCTACTTCAACCCAGTGACAACACGNNNAGATCGGAAG |
| 7487_1 | TATGAGTGTGGAGTCGTTGCATGGGACAGAAACAATTTGCATTACTANNNAGATCG |
| 7498_1 | TATGAGTGTGGAGTCGTTGCGTGTTTATTTTGGTAAAATGGGCAGGANNNAGATC |
| 7510_1 | TATGAGTGTGGAGTCGTTGCCAGTCTCATGGTAAAAAGTACAACAGGNNNAGATCG |
| 7560_1 | TATGAGTGTGGAGTCGTTGCTGAGGGTGAGAGAAGGAAAGAGGNNNAGATCGG |
| 7573_1 | TATGAGTGTGGAGTCGTTGCCAGTTGGTGATGGACAAGAGAGGNNNAGATCGGA |
| 7692_1 | TATGAGTGTGGAGTCGTTGCAATAACTTACGAGCTTCCCAGGGNNNAGATCGGAAG |
| 7701_1 | TATGAGTGTGGAGTCGTTGCGGTACTGTTCTGAGAGCTTTATGTGNNNAGATCGG |
| 7719_1 | TATGAGTGTGGAGTCGTTGCTGGAAATCTATTTGGTTCCTAAACAANNNAGATCGG |
| 7935_1 | TATGAGTGTGGAGTCGTTGCAGGCTGCGTTTATAGTTTTATATCTTTNNNAGATCGG |
| 7997_1 | TATGAGTGTGGAGTCGTTGCAATCTACATAAAACAAGGCAAAACGGGNNNAGATC |
| 8017_1 | TATGAGTGTGGAGTCGTTGCGCCCTCACCAAGAACCTAATTATACTGNNNAGATCGG |
| 8090_1 | TATGAGTGTGGAGTCGTTGCTTTTAGATCTAGCCATGTTGACATTTANNNAGATCGG |
| 8116_1 | TATGAGTGTGGAGTCGTTGCGCATTTTCTTATCTTCGGTTTTCATTTNNNAGATCGG |
| 8121_1 | TATGAGTGTGGAGTCGTTGCAGCATTTCATTCTCTTTTCTTTGTTGGNNNAGATCGG |
| 8192_1 | TATGAGTGTGGAGTCGTTGCTCCTTTCCATTTTGGGTGTTTTACTTTNNNAGATCGG |
| 8196_1 | TATGAGTGTGGAGTCGTTGCTACTTTTAGATTGAGGTTTGCCCATCANNNAGATCG |
| 8366_1 | TATGAGTGTGGAGTCGTTGCCTCTGACCAAGACAATAATTTTTACTGNNNAGATCGG |
| 8408_1 | TATGAGTGTGGAGTCGTTGCTCTAAACCTCATCCAATATTATGCTCTNNNAGATCGG |
| 8580_1 | TATGAGTGTGGAGTCGTTGCACTTTCAGCCCACTTGTTTCAAACNNNAGATCGGAAG |
| 8593_1 | TATGAGTGTGGAGTCGTTGCTGTATAATTGTTTATAGTGTGAGCGTGNNNAGATCG |
| 8612_1 | TATGAGTGTGGAGTCGTTGCTTTGAGTACTACCCATTCTCGGTGNNNAGATCGGAA |
| 8632_1 | TATGAGTGTGGAGTCGTTGCACCTACCTACCTATTCATCTCTATCGTNNNAGATCGG |
| 8649_1 | TATGAGTGTGGAGTCGTTGCCCCAGAAGAGAAGTATGTAAGACTGTGNNNAGATC |
| 8735_1 | TATGAGTGTGGAGTCGTTGCATGGATTTGCTTCTCCTCCCTTGNNNAGATCGGAAG |
| 8749_1 | TATGAGTGTGGAGTCGTTGCTGAACATCAGATCTCTGGGCTTGNNNAGATCGGAAG |
| 8791_1 | TATGAGTGTGGAGTCGTTGCTGTATTCTGTCAAAGAAGGATATGTTGNNNAGATCG |

| | |
|---|---|
| 8796_1 | TATGAGTGTGGAGTCGTTGCTTTAGATGGAGCCTGCTCTGTTGNNNAGATCGGAAG |
| 8806_1 | TATGAGTGTGGAGTCGTTGCTCCAAATAACTATGGCATCCAACAGANNNAGATCGG |
| 8926_1 | TATGAGTGTGGAGTCGTTGCCTGCAAATGGGTAGGAGTTGAACTANNNAGATCGG |
| 8934_1 | TATGAGTGTGGAGTCGTTGCGCATGTATCCCTTACCCAGCAATNNNAGATCGGAAG |
| 8940_1 | TATGAGTGTGGAGTCGTTGCAAATCAGTAACTGCAATGGCAATNNNAGATCGGAA |
| 8947_1 | TATGAGTGTGGAGTCGTTGCCCCCCGAATACAGTCATTGCAATNNNAGATCGGAAG |
| 8970_1 | TATGAGTGTGGAGTCGTTGCTTGTTTTGAAGGAGAAAGAATAGTCTTNNNAGATCG |
| 8974_1 | TATGAGTGTGGAGTCGTTGCATCACTGAAATTTTGGCAGGAATNNNAGATCGGAA |
| 8996_1 | TATGAGTGTGGAGTCGTTGCCATTGGGACAATTAGATGAAAAGATTTNNNAGATCG |
| 9012_1 | TATGAGTGTGGAGTCGTTGCTTCATATTCTTTGCCCTTTCCCAAATTNNNAGATCGGA |
| 9016_1 | TATGAGTGTGGAGTCGTTGCTGCCACAATCTAGTACCACAAACTAAANNNAGATCG |
| 9223_1 | TATGAGTGTGGAGTCGTTGCCACACAACGAGCAAAGCTTTCATNNNAGATCGGAAG |
| 9231_1 | TATGAGTGTGGAGTCGTTGCAGCCCATTTGAAGAGAGGAAGATNNNAGATCGGAA |
| 9441_1 | TATGAGTGTGGAGTCGTTGCTCAATACCTCATAGTGCTAGTGCTACTNNNAGATCGG |
| 9532_1 | TATGAGTGTGGAGTCGTTGCACCAGTTTCAGTGAAGTATAGAACACTNNNAGATCG |
| 9569_1 | TATGAGTGTGGAGTCGTTGCGTCTCTTCTAATATGCCAGGCACTNNNAGATCGGAA |
| 9602_1 | TATGAGTGTGGAGTCGTTGCTATGTGTGTGATATACTTGCGTGTCTCNNNAGATCG |
| 9628_1 | TATGAGTGTGGAGTCGTTGCACAGAAGAAGGAAGAAAATTTGGGACTNNNAGATC |
| 9635_1 | TATGAGTGTGGAGTCGTTGCATTCTCGGATGCTCAAGTTCCTTATTNNNAGATCGGA |
| 9659_1 | TATGAGTGTGGAGTCGTTGCAAGACCTCTAAGTGTCAATGCTACTNNNAGATCGGA |
| 9718_1 | TATGAGTGTGGAGTCGTTGCTGGAATGACCAAAGAAAGCACCTNNNAGATCGGAA |
| 9761_1 | TATGAGTGTGGAGTCGTTGCGTTACATAGGGCACATTGACCCTNNNAGATCGGAAG |
| 9768_1 | TATGAGTGTGGAGTCGTTGCCACTATTTATAGCCCCACCCCTNNNAGATCGGAAGA |
| 9781_1 | TATGAGTGTGGAGTCGTTGCTAAACACAATCTTCAAGGCCCCTNNNAGATCGGAAG |
| 9899_1 | TATGAGTGTGGAGTCGTTGCAATGCAGGGTCTCACTGTATCCTNNNAGATCGGAAG |
| 9900_1 | TATGAGTGTGGAGTCGTTGCATGTGCTCAGACTTTTACATTATCCTNNNAGATCGGA |
| 9918_1 | TATGAGTGTGGAGTCGTTGCGTCAAATTCTTCTTCCGGTATCTCCTNNNAGATCGGA |
| 9960_1 | TATGAGTGTGGAGTCGTTGCTCTGTCTGTCTGCCTGTATTCCTNNNAGATCGGAAGA |
| 10025_1 | TATGAGTGTGGAGTCGTTGCTGGAAAAACATAATAGAAGAGTTTCCTNNNAGATCG |
| 10056_1 | TATGAGTGTGGAGTCGTTGCTCCTTTGTTCTTTTCAAACTGAAAGCTNNNAGATCGG |
| 10191_1 | TATGAGTGTGGAGTCGTTGCTTACCTCCAACATGGAAAGAGCTNNNAGATCGGAAG |
| 10243_1 | TATGAGTGTGGAGTCGTTGCTTGCAGGAAAACACATTTGAGCTNNNAGATCGGAA |
| 10777_1 | TATGAGTGTGGAGTCGTTGCACAAGAGGGAGTATTATTTTGTGCTNNNAGATCGG |
| 11033_1 | TATGAGTGTGGAGTCGTTGCAAGGCTCAGGTAATTTCTCTGCCNNNAGATCGGAAG |
| 11099_1 | TATGAGTGTGGAGTCGTTGCGCTTCATAGCTGTTCTCCCAAAANNNAGATCGGAAG |
| 11166_1 | TATGAGTGTGGAGTCGTTGCGCCACTCATTCATCTTTTTGTATGGGNNNAGATCGGA |
| 11188_1 | TATGAGTGTGGAGTCGTTGCTCCTATGGGTCTCCTTCCTCTCTNNNAGATCGGAAGA |
| 11229_1 | TATGAGTGTGGAGTCGTTGCTACTCATATTTGGTTTTAGTTGCCAAANNNAGATCGG |
| 11272_1 | TATGAGTGTGGAGTCGTTGCACACATAGACACACACTCATCAAANNNAGATCGGAA |
| 11276_1 | TATGAGTGTGGAGTCGTTGCGAAACATCCTCTTGAAAACACTAGGTANNNAGATCG |
| 11437_1 | TATGAGTGTGGAGTCGTTGCGCACAAAAGTATTGCCAGGTTTCTNNNAGATCGGAA |
| 11444_1 | TATGAGTGTGGAGTCGTTGCTTCAAGGCTGGTAAGAATAATTTTCTNNNAGATCGG |
| 11485_1 | TATGAGTGTGGAGTCGTTGCATTTTCTCTTCCTCTCCCTCTCTCTGNNNAGATCGGAA |

| | |
|---|---|
| 11498_1 | TATGAGTGTGGAGTCGTTGCAGAGGTGGAAAGGCACATAAAGTNNNAGATCGGA |
| 11605_1 | TATGAGTGTGGAGTCGTTGCGGAAAAATATGAATTCTCAGAGACAGTNNNAGATC |
| 11624_1 | TATGAGTGTGGAGTCGTTGCACCATAATAGCAAAGACATCTTACAGTNNNAGATCG |
| 11628_1 | TATGAGTGTGGAGTCGTTGCCAAATGCTGACACCCAAAAGCAANNNAGATCGGAAG |
| 11652_1 | TATGAGTGTGGAGTCGTTGCCTGACCAAGTAAGGTTTATTCCAGTNNNAGATCGGA |
| 11671_1 | TATGAGTGTGGAGTCGTTGCTGTCCATAATATGCTGTTGTTAGCAGTNNNAGATCG |
| 11700_1 | TATGAGTGTGGAGTCGTTGCGATAGAATCTCTGACCAAGACAATAATNNNAGATCG |
| 11714_1 | TATGAGTGTGGAGTCGTTGCGCATAATGTGTGATTTCAAATGTCAGTNNNAGATCG |
| 11741_1 | TATGAGTGTGGAGTCGTTGCAGTCTGGCTAGAACAAATGCTAGAATTNNNAGATCG |
| 11805_1 | TATGAGTGTGGAGTCGTTGCAATACATGTCAGAAAAACGAAATGAGTNNNAGATC |
| 11806_1 | TATGAGTGTGGAGTCGTTGCGGGTAACTACGTGGCAACAAGAANNNAGATCGGAA |
| 11824_1 | TATGAGTGTGGAGTCGTTGCCCTGCTTATTTGTTTTGTTTCCTGAGTNNNAGATCGG |
| 11852_1 | TATGAGTGTGGAGTCGTTGCTGTCAGAAGGCCTGGGAAATAGTNNNAGATCGGAA |
| 11856_1 | TATGAGTGTGGAGTCGTTGCACCCTAATGCCATTCCAATAATAGTNNNAGATCGGA |
| 11996_1 | TATGAGTGTGGAGTCGTTGCTCCAGAAAAATCTCCCAACAAAGGTNNNAGATCGGA |
| 12012_1 | TATGAGTGTGGAGTCGTTGCAAAAGGTGTCCGGTAATCTGGAANNNAGATCGGAA |
| 12032_1 | TATGAGTGTGGAGTCGTTGCCCAACCACCATTAGGAAACAGGTNNNAGATCGGAAG |
| 12140_1 | TATGAGTGTGGAGTCGTTGCGTTGATGCATTAGAAGTTTCTAGGTNNNAGATCGG |
| 12216_1 | TATGAGTGTGGAGTCGTTGCCACACAAATACACTCCTTTATTCCACANNNAGATCGG |
| 12252_1 | TATGAGTGTGGAGTCGTTGCTTGCTGATGAACATTTCGGTTGCTNNNAGATCGGAA |
| 12269_1 | TATGAGTGTGGAGTCGTTGCTGCCATTATTCTTTGGACAAATGGTNNNAGATCGGA |
| 12317_1 | TATGAGTGTGGAGTCGTTGCAGACTGTAGAATGCATTGTTATATGGTNNNAGATCG |
| 12320_1 | TATGAGTGTGGAGTCGTTGCTTATCCCCAAGATACTTTTGGTATGGTNNNAGATCG |
| 12346_1 | TATGAGTGTGGAGTCGTTGCAAGAATGAGTTGATGGCATTTGCAACNNNAGATCG |
| 12393_1 | TATGAGTGTGGAGTCGTTGCAGAAGGATCCTGAATAATTTCAAACANNNAGATCGG |
| 53679_1 | TATGAGTGTGGAGTCGTTGCGCCAGTTCCATATAACTAGTGTTTTCANNNAGATCGG |
| 53990_1 | TATGAGTGTGGAGTCGTTGCTGGTTATTGTGAGCATTAGAATACAAANNNAGATCG |
| 54647_1 | TATGAGTGTGGAGTCGTTGCGGAAGAAGAAAAAGAAAGAGAAGAATCNNNAGAT |
| 54867_1 | TATGAGTGTGGAGTCGTTGCGACGGAGTCTTACTCTATCACCCNNNAGATCGGAAG |
| 55201_1 | TATGAGTGTGGAGTCGTTGCCAATCTCGGCTTACTGCAAACTCNNNAGATCGGAAG |
| 55350_1 | TATGAGTGTGGAGTCGTTGCCGTGTAATTTGAAAACAGAGACAAAGANNNAGATC |
| 55386_1 | TATGAGTGTGGAGTCGTTGCACAGAACAGTGAAATATACACACTAGTNNNAGATCG |
| 55498_1 | TATGAGTGTGGAGTCGTTGCTGGGACCATTATAAATAAGTTTATGCTNNNAGATCG |
| 55543_1 | TATGAGTGTGGAGTCGTTGCTCTGTGAAATAGCTGAAAAGAAATTTGNNNAGATCG |
| 55588_1 | TATGAGTGTGGAGTCGTTGCTTAGGAAGCTTAGTTTGGCCAGANNNAGATCGGAA |
| 55624_1 | TATGAGTGTGGAGTCGTTGCAAAGAAAAGAAAAAAAGGAAATCCAGTNNNAGATC |
| 56864_1 | TATGAGTGTGGAGTCGTTGCCCAGTGGGTTCCAAATAAACCAANNNAGATCGGAAG |
| 57031_1 | TATGAGTGTGGAGTCGTTGCTCAATCAGTAGTGTTTCAATACATTGANNNAGATCG |
| 57036_1 | TATGAGTGTGGAGTCGTTGCACCTCAGAAAATCAGTATCAATATCCANNNAGATCG |
| 57296_1 | TATGAGTGTGGAGTCGTTGCTCCTTACAATCCAAATCAGGTGGTNNNAGATCGGAA |
| 58134_1 | TATGAGTGTGGAGTCGTTGCCTTCAAGCAATTCTCCTGCCTCANNNAGATCGGAAGA |
| 58407_1 | TATGAGTGTGGAGTCGTTGCGTTCAAGAGATTCTCCTGCCTCANNNAGATCGGAAG |
| 58885_1 | TATGAGTGTGGAGTCGTTGCTGTTGGCATAATTGAGAATACTTTCTGNNNAGATCG |

| | |
|---|---|
| 59169_1 | TATGAGTGTGGAGTCGTTGCACAATGTAGAATAGAGTGGAGACATCANNNAGATC |
| 59223_1 | TATGAGTGTGGAGTCGTTGCGGTTCAAGCAATTCTCCTGTCTCNNNAGATCGGAAG |
| 59506_1 | TATGAGTGTGGAGTCGTTGCTCAAGTGATTCTTCCACCTCACCNNNAGATCGGAAGA |
| 60308_1 | TATGAGTGTGGAGTCGTTGCCGAGTTCAAGCAATTCTCCTGTCNNNAGATCGGAAG |
| 61991_1 | TATGAGTGTGGAGTCGTTGCACACAGTAATTCAAGTGTAGATTTCACNNNAGATCG |
| 62032_1 | TATGAGTGTGGAGTCGTTGCCTGTTAGGTCCAGGACTCCTTTGNNNAGATCGGAAG |
| 62043_1 | TATGAGTGTGGAGTCGTTGCAACCAGGCAGAGTTTTACTGGAANNNAGATCGGAA |
| 62233_1 | TATGAGTGTGGAGTCGTTGCATGCACATCCTCCCATATACCTTNNNAGATCGGAAGA |
| 62451_1 | TATGAGTGTGGAGTCGTTGCTTTTAGGCATAGGTGAAGAGGGCNNNAGATCGGAA |
| 62558_1 | TATGAGTGTGGAGTCGTTGCGCAACTTTATGATATGTTGGTAGAGCANNNAGATCG |
| 62734_1 | TATGAGTGTGGAGTCGTTGCGGAGGAGAAAGAAAAAATCTCAATCANNNAGATCG |
| 62976_1 | TATGAGTGTGGAGTCGTTGCTCAGTGTGTCTAGAGATCTAATGTACANNNAGATCG |
| 63020_1 | TATGAGTGTGGAGTCGTTGCGAAAAGAAAAGAAAGAAAGAAGAAGCANNNAGAT |
| 63219_1 | TATGAGTGTGGAGTCGTTGCTGACTATTGATGACTAAGGATTACACANNNAGATCG |
| 63230_1 | TATGAGTGTGGAGTCGTTGCGGTTCAAACAATTCTCCTGCCTCNNNAGATCGGAAG |
| 63285_1 | TATGAGTGTGGAGTCGTTGCTGCTCCTAAAAATACTAAGCAACTACCNNNAGATCGG |
| 63401_1 | TATGAGTGTGGAGTCGTTGCACACACACATGCATAGACAGATCTNNNAGATCGGAA |
| 63983_1 | TATGAGTGTGGAGTCGTTGCACCACTGGAATATTGTATTGTACAGTGNNNAGATCG |
| 64164_1 | TATGAGTGTGGAGTCGTTGCAAAGAGGGTGAGATGATACCTGCNNNAGATCGGAA |
| 64228_1 | TATGAGTGTGGAGTCGTTGCAAACGATGATACCTAAGGCAGCANNNAGATCGGAA |
| 64248_1 | TATGAGTGTGGAGTCGTTGCTGACATAGCATAAATTGTTGAAGACTGNNNAGATCG |
| 65368_1 | TATGAGTGTGGAGTCGTTGCTCATGGATAAAGATTGATTCTTAGCCANNNAGATCG |
| 65394_1 | TATGAGTGTGGAGTCGTTGCCAGTGCTTAGTGTTGTGGACATGNNNAGATCGGAA |
| 65969_1 | TATGAGTGTGGAGTCGTTGCGATTGAAGCAATTCTCCTGCCTCNNNAGATCGGAAG |
| 66182_1 | TATGAGTGTGGAGTCGTTGCAGTCATAGACATTATGAAAAGATGCCANNNAGATCG |
| 66218_1 | TATGAGTGTGGAGTCGTTGCTGGATGATTTTTCTATTTGACCTTGGANNNAGATCG |
| 66272_1 | TATGAGTGTGGAGTCGTTGCGCCCTGTGGAAATATTTTGAAGCNNNAGATCGGAAG |
| 66515_1 | TATGAGTGTGGAGTCGTTGCAATGGCCGTTTCAAAATAACCGTNNNAGATCGGAAG |
| 66568_1 | TATGAGTGTGGAGTCGTTGCTTCAAGTGATTCTCCTGCCTCAGNNNAGATCGGAAG |
| 66597_1 | TATGAGTGTGGAGTCGTTGCATGAAATAATGGCATTCGCAGCANNNAGATCGGAAG |
| 66620_1 | TATGAGTGTGGAGTCGTTGCAGGCCAGCTGAAACTATCTTGAANNNAGATCGGAAG |
| 66688_1 | TATGAGTGTGGAGTCGTTGCCAGTCTGGACGTCTCTGCATATTNNNAGATCGGAAG |
| 66703_1 | TATGAGTGTGGAGTCGTTGCTCTGTGATTTTTCCCCCCATCATNNNAGATCGGAAGA |
| 66717_1 | TATGAGTGTGGAGTCGTTGCCCATGTTTTTTTTTAGACTTCAGCACANNNAGATCGG |
| 66881_1 | TATGAGTGTGGAGTCGTTGCCCATCTATGCCTCTCAAAGTGCTNNNAGATCGGAAGA |
| 66975_1 | TATGAGTGTGGAGTCGTTGCAGACAGGGTCTCATTCTGTTGTCNNNAGATCGGAAG |
| 67072_1 | TATGAGTGTGGAGTCGTTGCTATGCCAGACTTCCTGAATGTCTNNNAGATCGGAAG |
| 67312_1 | TATGAGTGTGGAGTCGTTGCTCCCCAGAGAAAAGTAAAAGGTGANNNAGATCGGA |
| 67313_1 | TATGAGTGTGGAGTCGTTGCAGTTCTGTTTTGATTCAAGGATCTTGANNNAGATCG |
| 67455_1 | TATGAGTGTGGAGTCGTTGCTGCAGAAAAGGTTATATGAAAATGCANNNAGATCG |
| 67696_1 | TATGAGTGTGGAGTCGTTGCAGCAAGGTTTCAAGATACAAGGTNNNAGATCGGAA |
| 67720_1 | TATGAGTGTGGAGTCGTTGCCGCGCATATATGTATATAGAGTGTGCNNNAGATCGG |
| 67764_1 | TATGAGTGTGGAGTCGTTGCGATTTTCCTCCACGGTTCCTAGTNNNAGATCGGAAG |

| | |
|---|---|
| 67796_1 | TATGAGTGTGGAGTCGTTGCGCATGATCTTGGCTCACTACAACNNNAGATCGGAAG |
| 67829_1 | TATGAGTGTGGAGTCGTTGCTACACCCCTTCCTCTATTGGTTCNNNAGATCGGAAGA |
| 68177_1 | TATGAGTGTGGAGTCGTTGCTCTTCAGATTGTATAAATCTTTCCCCTNNNAGATCGG |
| 68256_1 | TATGAGTGTGGAGTCGTTGCTGTAGCTTACAGATGAATGAAAAACANNNAGATCG |
| 68268_1 | TATGAGTGTGGAGTCGTTGCTGTGTGTGTATATATATGTGTGTGTGTNNNAGATCG |
| 68392_1 | TATGAGTGTGGAGTCGTTGCACAAAAACTAAGATGTCGCTGGGNNNAGATCGGAA |
| 68611_1 | TATGAGTGTGGAGTCGTTGCCCAATTCACATAATGTTTATGACAGTGNNNAGATCG |
| 68642_1 | TATGAGTGTGGAGTCGTTGCGCAACCTCTACCTCTCATGTTCANNNAGATCGGAAGA |
| 68652_1 | TATGAGTGTGGAGTCGTTGCAGGGACCGAGGTGCTATTTTATGNNNAGATCGGAA |
| 68803_1 | TATGAGTGTGGAGTCGTTGCGTCTCCCCTATCTATCTCTATCTCTCTNNNAGATCGGA |
| 68820_1 | TATGAGTGTGGAGTCGTTGCGAGAGAGAGAGAGAGACGAGACANNNAGATCGGA |
| 68829_1 | TATGAGTGTGGAGTCGTTGCAGAAAGAAAGAAAGAAAGAAAGAAGGANNNAGAT |
| 68890_1 | TATGAGTGTGGAGTCGTTGCTATCTTGCAAACTCTGGAGCCATNNNAGATCGGAAG |
| 69034_1 | TATGAGTGTGGAGTCGTTGCAGTGCCTACTTTATATATAGAGTGCCTNNNAGATCG |
| 69094_1 | TATGAGTGTGGAGTCGTTGCGAGGGAAGGAAGGAAGGAAAAGANNNAGATCGG |
| 69177_1 | TATGAGTGTGGAGTCGTTGCTGGCTCATAAAACATTATTTATACCCTNNNAGATCGG |
| 69206_1 | TATGAGTGTGGAGTCGTTGCCCACATTCACATAACCTTTATTATGGTNNNAGATCGG |
| 69358_1 | TATGAGTGTGGAGTCGTTGCCCATCTTTCTGGAGAACTGTAATTACANNNAGATCG |
| 69523_1 | TATGAGTGTGGAGTCGTTGCAGATCTCCCTTTGTTTATGTTTCTTCTNNNAGATCGG |
| 69762_1 | TATGAGTGTGGAGTCGTTGCTCTTTTTCTTTCTTTCTCTCTGTTGGTNNNAGATCGGA |
| 69765_1 | TATGAGTGTGGAGTCGTTGCTCTATTTCTCTCTCTCTCTCTCTCTNNNAGATCGGAAG |
| 69869_1 | TATGAGTGTGGAGTCGTTGCTGTACACAGATGATAGATATATGCACTNNNAGATCG |
| 69901_1 | TATGAGTGTGGAGTCGTTGCTGCTCTGGAGAAAGGACTTTTGTNNNAGATCGGAA |
| 70237_1 | TATGAGTGTGGAGTCGTTGCTCTGGATCAAGAATTAGCCCAGGNNNAGATCGGAA |
| 70266_1 | TATGAGTGTGGAGTCGTTGCTGATCTCATCTCACTGCAACCTCNNNAGATCGGAAGA |
| 70267_1 | TATGAGTGTGGAGTCGTTGCTGGACAGAGTCTCACTTTGTTGCNNNAGATCGGAAG |
| 70300_1 | TATGAGTGTGGAGTCGTTGCACTCCCTATCTCTTCTTCAAAATCAGTNNNAGATCGG |
| 70414_1 | TATGAGTGTGGAGTCGTTGCCTGGAGTGTAGTTGTGCAATCTCNNNAGATCGGAA |
| 70450_1 | TATGAGTGTGGAGTCGTTGCAGAGAGAAAGAGAAAGAAGAAGAGAGANNNAGAT |
| 3_1 | TATGAGTGTGGAGTCGTTGCACAGCCTCAGAATAAGTTTCTTGGTNNNAGATCGGA |
| 26_1 | TATGAGTGTGGAGTCGTTGCATGGGTCTGTTTCTGGTTTTGGTNNNAGATCGGAAG |
| 44_1 | TATGAGTGTGGAGTCGTTGCAGGTAACAAAACCCCTCAATAAATGTNNNAGATCGG |
| 65_1 | TATGAGTGTGGAGTCGTTGCGTCAACAACTTGCTAGAAATGGAACANNNAGATCGG |
| 71_1 | TATGAGTGTGGAGTCGTTGCTGTATGTATGTTTTTGACAACCATGTNNNAGATCGG |
| 87_1 | TATGAGTGTGGAGTCGTTGCGCACATTATTTGTACAGGCACTAACANNNAGATCGG |
| 90_1 | TATGAGTGTGGAGTCGTTGCTGGTGTATAATAGCAGACAGGTAACANNNAGATCG |
| 107_1 | TATGAGTGTGGAGTCGTTGCCTCCTCCCAGTGCAAATAACACANNNAGATCGGAAG |
| 112_1 | TATGAGTGTGGAGTCGTTGCTCCAAACATATGTACAAAAACACACANNNAGATCGG |
| 151_1 | TATGAGTGTGGAGTCGTTGCAGGACAAAAACTGCAGAAACTGTNNNAGATCGGAA |
| 233_1 | TATGAGTGTGGAGTCGTTGCAGATGTCAATATGCTAATTAGGCACANNNAGATCGG |
| 285_1 | TATGAGTGTGGAGTCGTTGCCATAGCCTCCAAGCATCTTCACANNNAGATCGGAAG |
| 289_1 | TATGAGTGTGGAGTCGTTGCGCAAAGATGAATACAGTACAGTCTGTNNNAGATCG |
| 305_1 | TATGAGTGTGGAGTCGTTGCGACCTCCAAACTAAAGGAACAGGAATNNNAGATCG |

| | |
|---|---|
| 312_1 | TATGAGTGTGGAGTCGTTGCCGGCTTCAAAATACACAAAGAAGACANNNAGATCGG |
| 323_1 | TATGAGTGTGGAGTCGTTGCAGAAAGAACCACAGAGACAGACANNNAGATCGGAA |
| 343_1 | TATGAGTGTGGAGTCGTTGCTGGTTCTGCTTCTAAAATCTCAGTGTNNNAGATCGG |
| 375_1 | TATGAGTGTGGAGTCGTTGCTGTGTGCCTACCATATTACTAGGACANNNAGATCGG |
| 385_1 | TATGAGTGTGGAGTCGTTGCTCTGTGTCTGTTATATGTAGACATGTNNNAGATCGG |
| 387_1 | TATGAGTGTGGAGTCGTTGCTTATATTGGCCCATACTGGAATGTGTNNNAGATCGG |
| 448_1 | TATGAGTGTGGAGTCGTTGCCGATCTGGTCCTCTAAAGACAGGAACNNNAGATCGG |
| 451_1 | TATGAGTGTGGAGTCGTTGCTGATGAACATACGTAGCTTTTGTGTNNNAGATCGGA |
| 483_1 | TATGAGTGTGGAGTCGTTGCACAGTAAGTCAGGGATAAAGTCTACANNNAGATCG |
| 500_1 | TATGAGTGTGGAGTCGTTGCAGACAGACTAGGAAAAAGTGCTTGTNNNAGATCGG |
| 507_1 | TATGAGTGTGGAGTCGTTGCTCTTGGTTCACGGAGTTTCTTGTNNNAGATCGGAAG |
| 547_1 | TATGAGTGTGGAGTCGTTGCTGTGTGAGTGAGTAGATATGTAACCANNNAGATCG |
| 597_1 | TATGAGTGTGGAGTCGTTGCAGAGAGAGAGTGAATAAAAGGGACCANNNAGATC |
| 614_1 | TATGAGTGTGGAGTCGTTGCCCAGGTTTGAAGCTTCTGAAATTNNNAGATCGGAAG |
| 639_1 | TATGAGTGTGGAGTCGTTGCTCCTTCTGGAATAAAACATGTTACCANNNAGATCGG |
| 642_1 | TATGAGTGTGGAGTCGTTGCAGGATTTGATCAAGGAAGAAAACCCANNNAGATCG |
| 643_1 | TATGAGTGTGGAGTCGTTGCTGTTCCATTATCACTTCAAAGGAATTNNNAGATCGG |
| 650_1 | TATGAGTGTGGAGTCGTTGCTTGCCTATGTTTTCTTTCCGTAGTTNNNAGATCGGAA |
| 651_1 | TATGAGTGTGGAGTCGTTGCAGTTTGTTTACCACATGTCCTTAGANNNAGATCGGA |
| 654_1 | TATGAGTGTGGAGTCGTTGCACCACAAATCTCCTAGTCACCCANNNAGATCGGAAGA |
| 684_1 | TATGAGTGTGGAGTCGTTGCATGAAAAATTGTAGCAGCCCCCANNNAGATCGGAAG |
| 716_1 | TATGAGTGTGGAGTCGTTGCATGATTCTTTTGTACTGGGCCCANNNAGATCGGAAG |
| 758_1 | TATGAGTGTGGAGTCGTTGCGCATATTAGAAGCGTAAAGTCCCANNNAGATCGGAA |
| 792_1 | TATGAGTGTGGAGTCGTTGCATTATCCACACACTCAGCTTTGAAGTNNNAGATCGG |
| 815_1 | TATGAGTGTGGAGTCGTTGCGGGTGGGCCAGAAAGATATGATTNNNAGATCGGAA |
| 820_1 | TATGAGTGTGGAGTCGTTGCGAAACTGACTTGTTTTGCAGCCANNNAGATCGGAAG |
| 833_1 | TATGAGTGTGGAGTCGTTGCCTTCAGGAGTTTGGGATACAAATATTNNNAGATCGG |
| 876_1 | TATGAGTGTGGAGTCGTTGCACACAGTTGATTCCATATATTGGCCANNNAGATCGG |
| 890_1 | TATGAGTGTGGAGTCGTTGCAACAATAGTAGATAGAGGCCAGGTGCNNNAGATCG |
| 913_1 | TATGAGTGTGGAGTCGTTGCACTTGGGTTCCACAGAATGTTAGTTTNNNAGATCGG |
| 934_1 | TATGAGTGTGGAGTCGTTGCGGACAGGGCATCATTCTAGACTTNNNAGATCGGAAG |
| 1016_1 | TATGAGTGTGGAGTCGTTGCAGATGAAATGGCAATAGCTAGTCCANNNAGATCGG |
| 1111_1 | TATGAGTGTGGAGTCGTTGCCCCCAGTACCCTATTCATTTGATGTGNNNAGATCGGA |
| 1114_1 | TATGAGTGTGGAGTCGTTGCATAAAATGTGTGTGTGTGTATACGTATNNNAGATCG |
| 1146_1 | TATGAGTGTGGAGTCGTTGCAGACAGTTTATTGCAGGATCTAAGCANNNAGATCGG |
| 1153_1 | TATGAGTGTGGAGTCGTTGCCTGAAAAACGGAAGGAAATCTTGTCNNNAGATCGG |
| 1192_1 | TATGAGTGTGGAGTCGTTGCTAGATCCTTTCCTCTGCTCAGCANNNAGATCGGAAG |
| 1199_1 | TATGAGTGTGGAGTCGTTGCTCCCTAGACTAACAGCTTCAGCANNNAGATCGGAAG |
| 1200_1 | TATGAGTGTGGAGTCGTTGCCCAAGCTTCTTTCTTCTGAAACATANNNAGATCGGAA |
| 1205_1 | TATGAGTGTGGAGTCGTTGCATTTGCATCTCCTTCCAAGTAACTCTNNNAGATCGGA |
| 1207_1 | TATGAGTGTGGAGTCGTTGCCCTTGTGTGTTCAGGTAGTACTCTACNNNAGATCGG |
| 1230_1 | TATGAGTGTGGAGTCGTTGCTGGGTGGAAAAGAGAATAGATTGGAANNNAGATC |
| 1245_1 | TATGAGTGTGGAGTCGTTGCGAGAGAGAGATAGACCCAGTCTCACTNNNAGATCG |

| | |
|---|---|
| 1281_1 | TATGAGTGTGGAGTCGTTGCTACACAAAGAAGACAGCAAACTAAAANNNAGATCGG |
| 1314_1 | TATGAGTGTGGAGTCGTTGCGTGCTGGACAATTTCAAAAGGCANNNAGATCGGAAC |
| 1321_1 | TATGAGTGTGGAGTCGTTGCGTTGCTTTCATTCAGTTGATCCAGACNNNAGATCGGA |
| 1322_1 | TATGAGTGTGGAGTCGTTGCTGTGTGTGTGTGTATAGTAACTAGTTNNNAGATCGG |
| 1383_1 | TATGAGTGTGGAGTCGTTGCTAAATAGGATTATAGGTGCGTGCCNNNAGATCGGA |
| 1386_1 | TATGAGTGTGGAGTCGTTGCCAGTTGAAGGGGTGGATATGGTTNNNAGATCGGAA |
| 1396_1 | TATGAGTGTGGAGTCGTTGCTATCAAGATCTACTGGCTGGGCANNNAGATCGGAAG |
| 1458_1 | TATGAGTGTGGAGTCGTTGCCACGACATAATGGAAACAACTAATGTNNNAGATCGG |
| 1488_1 | TATGAGTGTGGAGTCGTTGCCCAAACTGCCCCTTTTTATAAGGAGANNNAGATCGGA |
| 1511_1 | TATGAGTGTGGAGTCGTTGCTCCTCTCTCAGGAAATAAAGGCTACANNNAGATCGGA |
| 1528_1 | TATGAGTGTGGAGTCGTTGCTCAGAAACAGAAAGACAAAAGCCACANNNAGATCGG |
| 1535_1 | TATGAGTGTGGAGTCGTTGCAAGCTACATGGCTATATGTATATGCANNNAGATCGG |
| 1556_1 | TATGAGTGTGGAGTCGTTGCTTCTAACTCTGGGCTATGGCTGTNNNAGATCGGAAG |
| 1563_1 | TATGAGTGTGGAGTCGTTGCGGAAGAGATCTATAAACTGCACTGCANNNAGATCGG |
| 1568_1 | TATGAGTGTGGAGTCGTTGCCAACGCTTTCATGATGGCCATTTNNNAGATCGGAAGA |
| 1577_1 | TATGAGTGTGGAGTCGTTGCTGTTATGCATCTGTACTATCTGCACANNNAGATCGGA |
| 1582_1 | TATGAGTGTGGAGTCGTTGCACAATCACATATTCTCAGTAGCTGCANNNAGATCGGA |
| 1654_1 | TATGAGTGTGGAGTCGTTGCTTGAAGGGTTACTATTTTGTGACTTTNNNAGATCGGA |
| 1658_1 | TATGAGTGTGGAGTCGTTGCGCTACCACCAATACATATTTGTAGGGNNNAGATCGGA |
| 1663_1 | TATGAGTGTGGAGTCGTTGCTGGAGATGGTAGTTATGAACTTGCANNNAGATCGGA |
| 1720_1 | TATGAGTGTGGAGTCGTTGCTGGGCTTCTAAACTGATGAAGAATCANNNAGATCGG |
| 1721_1 | TATGAGTGTGGAGTCGTTGCACTCAACTCCCAATAAACAGGAATCANNNAGATCGGA |
| 1731_1 | TATGAGTGTGGAGTCGTTGCACCTTTTACCCCAGAAGACTTCAGTANNNAGATCGGA |
| 1775_1 | TATGAGTGTGGAGTCGTTGCTGGACAAACTGTTACCTATTAGATCANNNAGATCGGA |
| 1789_1 | TATGAGTGTGGAGTCGTTGCATGGGAGTAGACAGAAAGGAAAGAAGNNNAGATCG |
| 1813_1 | TATGAGTGTGGAGTCGTTGCAGTGAAAAGAGCCAAACAGGAAAAGNNNAGATCGG |
| 1822_1 | TATGAGTGTGGAGTCGTTGCTTTTACTTCATCTCCTTTGTTGGTTTNNNAGATCGGAA |
| 1849_1 | TATGAGTGTGGAGTCGTTGCAAGGTTGAAAACGAATGGACGAAGATNNNAGATCG |
| 1865_1 | TATGAGTGTGGAGTCGTTGCAGATTGTGGAAATACAAGGGATTTTNNNAGATCGGA |
| 1899_1 | TATGAGTGTGGAGTCGTTGCTGTCATCTGTCTTTTTCTTCTGCTCANNNAGATCGGAA |
| 1906_1 | TATGAGTGTGGAGTCGTTGCTTTCAGGGAAGAGGGAAACATTCTGTNNNAGATCGG |
| 1909_1 | TATGAGTGTGGAGTCGTTGCCGATCATTCCACCAAGAAAAATCTCANNNAGATCGGA |
| 1925_1 | TATGAGTGTGGAGTCGTTGCAGGGAAAGAAATGCTAACAGTTTTNNNAGATCGGAA |
| 1954_1 | TATGAGTGTGGAGTCGTTGCAGTCCTTTCAGTCCTGAGTATTCTCANNNAGATCGGA |
| 1955_1 | TATGAGTGTGGAGTCGTTGCATAAGATTAGTCAAGGATTCAGGCCANNNAGATCGG |
| 1960_1 | TATGAGTGTGGAGTCGTTGCTCTGCTCTGGGTGTATATGTGTGATANNNAGATCGG |
| 2001_1 | TATGAGTGTGGAGTCGTTGCGAAAGCCAACATTTCTGTGAAATTGTNNNAGATCGG |
| 2092_1 | TATGAGTGTGGAGTCGTTGCCGTATTGTTTTGTGGCTGCTTTTGTNNNAGATCGGAA |
| 2095_1 | TATGAGTGTGGAGTCGTTGCGTTGCTCTGACTAGGACTTCCAAAACNNNAGATCGGA |
| 2157_1 | TATGAGTGTGGAGTCGTTGCTAGCAGCAGACACAAACTGTATGAAANNNAGATCGG |
| 2169_1 | TATGAGTGTGGAGTCGTTGCGTGTGCTTTCAGTTCATCATGTCAATNNNAGATCGGA |
| 2174_1 | TATGAGTGTGGAGTCGTTGCATGAGAAGCTGAATGCAAGAGCTCTANNNAGATCGG |
| 2193_1 | TATGAGTGTGGAGTCGTTGCTCTGCAACATTTTCTTTATCCGTTCANNNAGATCGGAA |

| | |
|---|---|
| 2217_1 | TATGAGTGTGGAGTCGTTGCTGATTGGTGTTACAAATGGTGTTCANNNAGATCGGA |
| 2218_1 | TATGAGTGTGGAGTCGTTGCGGTACTTGGTCACAGATGTGTTCANNNAGATCGGAA |
| 2305_1 | TATGAGTGTGGAGTCGTTGCACCTAATATTTCTGGGGAAAGAAAGANNNAGATCGG |
| 2337_1 | TATGAGTGTGGAGTCGTTGCAGAAACAGCATCTATCATGCCAAGANNNAGATCGGA |
| 2346_1 | TATGAGTGTGGAGTCGTTGCTATTCTTCCAACCCAAATGTCCAACANNNAGATCGGA |
| 2356_1 | TATGAGTGTGGAGTCGTTGCGGGTAAATTCTGATAAGGGACAGTCCNNNAGATCGG |
| 2368_1 | TATGAGTGTGGAGTCGTTGCCTAATGGGTTCACCTTTCTTGCTGNNNAGATCGGAAG |
| 2374_1 | TATGAGTGTGGAGTCGTTGCGAGATAGAATGCTGAATGTTGTCCCNNNAGATCGGA |
| 2384_1 | TATGAGTGTGGAGTCGTTGCACCCCTTTTACTTTACAGTCTGAAGANNNAGATCGGA |
| 2387_1 | TATGAGTGTGGAGTCGTTGCATGTGCTGGCAAGAAAGTGAAGANNNAGATCGGAA |
| 2409_1 | TATGAGTGTGGAGTCGTTGCACAACCCAATAATTGAGAATAGGCAANNNAGATCGG |
| 2410_1 | TATGAGTGTGGAGTCGTTGCTCCTCAAAGGGGAATTACTGTAAGANNNAGATCGGA |
| 2468_1 | TATGAGTGTGGAGTCGTTGCAAATGCAAAAGACTGTAGGTCTCTGANNNAGATCGG |
| 2602_1 | TATGAGTGTGGAGTCGTTGCATAGCATTGCTGGAGATAAAAGAGANNNAGATCGG |
| 2611_1 | TATGAGTGTGGAGTCGTTGCACCTCTCTGTACCAAGATAGAAGAGANNNAGATCGG |
| 2628_1 | TATGAGTGTGGAGTCGTTGCTCCTGGTAAGAAAGCAAAAAGAGAGANNNAGATCG |
| 2636_1 | TATGAGTGTGGAGTCGTTGCCCATTGAGGATCCATAGCTTGAATCNNNAGATCGGA |
| 2670_1 | TATGAGTGTGGAGTCGTTGCACACTAGTCCTTAGTTGAATAGGAGANNNAGATCGG |
| 2713_1 | TATGAGTGTGGAGTCGTTGCCCAAGAGATGAGAGACCTCTATGAGANNNAGATCGG |
| 2725_1 | TATGAGTGTGGAGTCGTTGCTTCTAGCTCATTTTCTCAATCTGAGANNNAGATCGGA |
| 2726_1 | TATGAGTGTGGAGTCGTTGCGCATCGTATATCCCTTTGGAAGTTTGNNNAGATCGGA |
| 2739_1 | TATGAGTGTGGAGTCGTTGCGAGAGATGACTAGCAGGAATTGCAANNNAGATCGG |
| 2748_1 | TATGAGTGTGGAGTCGTTGCCACATTGTATTAGTCCATTCCCACATNNNAGATCGGA |
| 2769_1 | TATGAGTGTGGAGTCGTTGCATAGGGATGATTTCAGCTTAGCATAANNNAGATCGG |
| 2834_1 | TATGAGTGTGGAGTCGTTGCTTGCTCCTTTATCAATGATCTGTTGANNNAGATCGGA |
| 2846_1 | TATGAGTGTGGAGTCGTTGCCCTTTTCTAATCGATCATAGCTGTTCNNNAGATCGGA |
| 2855_1 | TATGAGTGTGGAGTCGTTGCGTAACTTCTAATTGGCCATCTTGCTGNNNAGATCGGA |
| 2886_1 | TATGAGTGTGGAGTCGTTGCAAAGCTCATAACATTGGATTTGGCNNNAGATCGGAA |
| 2921_1 | TATGAGTGTGGAGTCGTTGCAGAAAGTATCAAGTATCTGGACAGGANNNAGATCGG |
| 2927_1 | TATGAGTGTGGAGTCGTTGCAGACAGAAAGAAAGAAGGTGAAGANNNAGATCGGA |
| 3001_1 | TATGAGTGTGGAGTCGTTGCTGCCCATCATTATTATTGCGATCTTTNNNAGATCGGA |
| 3014_1 | TATGAGTGTGGAGTCGTTGCCTCTCTGGTGGTAAAAGTTTGATGTGNNNAGATCGG |
| 3067_1 | TATGAGTGTGGAGTCGTTGCTTCTTTGCATATGTTTTGCTAAGCCTNNNAGATCGGA |
| 3075_1 | TATGAGTGTGGAGTCGTTGCTACCAAAACAATGACCTCAAACTGTCNNNAGATCGGA |
| 3080_1 | TATGAGTGTGGAGTCGTTGCAAACACAAATGAACAATCTCAACGATNNNAGATCGGA |
| 3115_1 | TATGAGTGTGGAGTCGTTGCAATATCTGTGAAGCCTAAAAAGCAATANNNAGATCG |
| 3190_1 | TATGAGTGTGGAGTCGTTGCCTCCCATTATTTACTACCTAGCTTGGTNNNAGATCGGA |
| 3195_1 | TATGAGTGTGGAGTCGTTGCAGGCTTATCTGGTTTCTAGATGGGANNNAGATCGGA |
| 3257_1 | TATGAGTGTGGAGTCGTTGCGAATGAATGCCTAATGTGTGCAAACANNNAGATCGG |
| 3268_1 | TATGAGTGTGGAGTCGTTGCAGAAAGAAAATTAGTGGTGGGAGTGGNNNAGATCG |
| 3284_1 | TATGAGTGTGGAGTCGTTGCTGTATGTCCATTTATCTGTTGCTGGANNNAGATCGGA |
| 3336_1 | TATGAGTGTGGAGTCGTTGCTGGGAAAACAATTGGAAAAGTTTGGANNNAGATCG |
| 3343_1 | TATGAGTGTGGAGTCGTTGCGGTAGCATAAAATACACACTGCTTTGNNNAGATCGG |

| | |
|---|---|
| 3347_1 | TATGAGTGTGGAGTCGTTGCCACACACAAACACATCCAGCAAGNNNAGATCGGAAG |
| 3348_1 | TATGAGTGTGGAGTCGTTGCCTGTAAGGGCACAGGCATAATGANNNAGATCGGAA |
| 3351_1 | TATGAGTGTGGAGTCGTTGCAGGCTGGAGTACAGGAAACATGANNNAGATCGGAA |
| 3355_1 | TATGAGTGTGGAGTCGTTGCACACCATCATTATTCAGAGGACATGANNNAGATCGG |
| 3433_1 | TATGAGTGTGGAGTCGTTGCAGTCTCATTTCCTATTTCCAACCTGANNNAGATCGGA |
| 3439_1 | TATGAGTGTGGAGTCGTTGCGTTTAGTACAGGGTTCCACTCTGTTANNNAGATCGG |
| 3452_1 | TATGAGTGTGGAGTCGTTGCCCCAGGATTGTAATGAGTGTCTCATANNNAGATCGG |
| 3477_1 | TATGAGTGTGGAGTCGTTGCAAATGCAGATGTGTCATGCATACTACNNNAGATCGG |
| 3488_1 | TATGAGTGTGGAGTCGTTGCGGCAGGTATATCTCATTGCTGACTTNNNAGATCGGA |
| 3526_1 | TATGAGTGTGGAGTCGTTGCTTGAAAGAAGAACCTCCCAACTAATANNNAGATCGG |
| 3527_1 | TATGAGTGTGGAGTCGTTGCCACACATACATCCCTACTCTTCACTTNNNAGATCGGAA |
| 3584_1 | TATGAGTGTGGAGTCGTTGCATGTGGCTTTTTCTGTCTACACTTGTNNNAGATCGGA |
| 3601_1 | TATGAGTGTGGAGTCGTTGCGCACTCGTGTTTTTAGTTTAGATTGANNNAGATCGG |
| 3603_1 | TATGAGTGTGGAGTCGTTGCTGGGGGAAAATGATTGATACTATTGANNNAGATCG |
| 3604_1 | TATGAGTGTGGAGTCGTTGCACACACAGAAAACATCTCCCTATTGANNNAGATCGGA |
| 3679_1 | TATGAGTGTGGAGTCGTTGCCTGCCTGCCTGCCTGTATAAATANNNAGATCGGAAG |
| 3735_1 | TATGAGTGTGGAGTCGTTGCAAACTACATGGAAATGGAAAAGTGCANNNAGATCG |
| 3795_1 | TATGAGTGTGGAGTCGTTGCTTTGGCTGTTATTGCCTACGTTATCCNNNAGATCGGA |
| 3806_1 | TATGAGTGTGGAGTCGTTGCTGAAGAAAAATTTTGGAGTAGGACCANNNAGATCG |
| 3845_1 | TATGAGTGTGGAGTCGTTGCCTCCATTTCGTGTACTAACACATGTGNNNAGATCGGA |
| 3909_1 | TATGAGTGTGGAGTCGTTGCACCCACACAGACATTTATCTACATCTNNNAGATCGGA |
| 3942_1 | TATGAGTGTGGAGTCGTTGCAAACAACTTCTCTTTTGAGTTCCCANNNAGATCGGAA |
| 3978_1 | TATGAGTGTGGAGTCGTTGCTATGTGATGATTGATGAGTGCACCAANNNAGATCGG |
| 3986_1 | TATGAGTGTGGAGTCGTTGCTGTCAAAGAATCAGCAGTGAGCAAGANNNAGATCG |
| 3987_1 | TATGAGTGTGGAGTCGTTGCGCAGCCTGGACTAGAAGAGATGANNNAGATCGGAA |
| 4022_1 | TATGAGTGTGGAGTCGTTGCTGGAGAGAATACCTGAATCAGTGTGGNNNAGATCG |
| 4086_1 | TATGAGTGTGGAGTCGTTGCCACACACACAGTTTTCATAAGTGTTTNNNAGATCGGA |
| 4093_1 | TATGAGTGTGGAGTCGTTGCAAACAGTCATTATTCAGAGCAACTCANNNAGATCGG |
| 4169_1 | TATGAGTGTGGAGTCGTTGCCTTCAACACACAATTCCTAATATGCCNNNAGATCGGA |
| 4203_1 | TATGAGTGTGGAGTCGTTGCGTGTATGTGCGTTTATTAGTGAGGGTNNNAGATCG |
| 4204_1 | TATGAGTGTGGAGTCGTTGCACCAAGAGACAGAAGGGTAATAGCANNNAGATCGG |
| 4222_1 | TATGAGTGTGGAGTCGTTGCCTTGGGTGTTTGTAGGACTTACTTCANNNAGATCGG |
| 4225_1 | TATGAGTGTGGAGTCGTTGCTACAGATGTGGCTCTAGGCAAACNNNAGATCGGAA |
| 4260_1 | TATGAGTGTGGAGTCGTTGCAGTTGCATGGATAGTTGTTCAAATCGNNNAGATCGG |
| 4265_1 | TATGAGTGTGGAGTCGTTGCAATGAAGAACATCCATCCATCCATCCNNNAGATCGGA |
| 4270_1 | TATGAGTGTGGAGTCGTTGCAGAACAAATGTTGACCACATATCAACNNNAGATCGG |
| 4273_1 | TATGAGTGTGGAGTCGTTGCACTTCCTGCTCTCATTCTCTCAACNNNAGATCGGAAG |
| 4280_1 | TATGAGTGTGGAGTCGTTGCAACTACTGGCAAGGATGCAGAACNNNAGATCGGAA |
| 4306_1 | TATGAGTGTGGAGTCGTTGCAGGAAAATGAATGGAGCTAGAGGTNNNAGATCGG |
| 4343_1 | TATGAGTGTGGAGTCGTTGCAATCTTTCCACCTAAGCCATCAAGTANNNAGATCGGA |
| 4449_1 | TATGAGTGTGGAGTCGTTGCACAGTAGGTAACAAATGCTAGTTTGTNNNAGATCGG |
| 4455_1 | TATGAGTGTGGAGTCGTTGCGTGTATGAGAGAGAGAAATGCTAACCNNNAGATCG |
| 4458_1 | TATGAGTGTGGAGTCGTTGCCCAATCTCCCTCTGCTTCCTTTTTATNNNAGATCGGAA |

| | |
|---|---|
| 4486_1 | TATGAGTGTGGAGTCGTTGCTGTGTCTGTAAAAGAAGGGAGATCACNNNAGATCG |
| 4496_1 | TATGAGTGTGGAGTCGTTGCGCAGATCCAGTGTGCTTCTCAAATATNNNAGATCGG |
| 4506_1 | TATGAGTGTGGAGTCGTTGCCTGGCACTGTAATCCTATAGTCTCACNNNAGATCGGA |
| 4520_1 | TATGAGTGTGGAGTCGTTGCACTATCAACTTCTGCTGAGGCATAAANNNAGATCGG |
| 4591_1 | TATGAGTGTGGAGTCGTTGCGCTGGTTCACATCCTTGGAGATTNNNAGATCGGAAG |
| 4626_1 | TATGAGTGTGGAGTCGTTGCGAAGCTGGCATATATTCTCTCACAGNNNAGATCGGA |
| 4627_1 | TATGAGTGTGGAGTCGTTGCAGGGTTCCTTACTCTTTGATTTTCTGNNNAGATCGGA |
| 4642_1 | TATGAGTGTGGAGTCGTTGCTTTGGTGATGTGTTTAGCATGAAGCNNNAGATCGGA |
| 4652_1 | TATGAGTGTGGAGTCGTTGCACTTTCCAGGCTATGTCCTTTGACNNNAGATCGGAAG |
| 4661_1 | TATGAGTGTGGAGTCGTTGCTGTATGACTGTGTGTGTGTCTCTATGNNNAGATCGG |
| 4711_1 | TATGAGTGTGGAGTCGTTGCTGTGTGTGTAGCAATCTCTCTCTCTANNNAGATCGGA |
| 4734_1 | TATGAGTGTGGAGTCGTTGCTGTGTTCTTGCTCTCACTCTCTCACCCNNNAGATCGGA |
| 4741_1 | TATGAGTGTGGAGTCGTTGCGTCTCCCATACTTTCCCTATCTCTCTNNNAGATCGGAA |
| 4815_1 | TATGAGTGTGGAGTCGTTGCTGTCAAGGTCATGATTCTTATCCACCNNNAGATCGGA |
| 4835_1 | TATGAGTGTGGAGTCGTTGCTGACAGGGGACAGTGTTTCCAGNNNAGATCGGAAG |
| 4842_1 | TATGAGTGTGGAGTCGTTGCGGCATATGGGATACAATGAAAAGACCNNNAGATCG |
| 4862_1 | TATGAGTGTGGAGTCGTTGCTTTGTTCATCAGTGGACACTTAGCTTNNNAGATCGGA |
| 4880_1 | TATGAGTGTGGAGTCGTTGCTGCGTTTGTGTCACTTTATTCATACCNNNAGATCGGA |
| 4884_1 | TATGAGTGTGGAGTCGTTGCACATACCAGGGAACAGAACAGAGANNNAGATCGGA |
| 4891_1 | TATGAGTGTGGAGTCGTTGCACAGATACAGAGGTGACTGGTACCNNNAGATCGGA |
| 4999_1 | TATGAGTGTGGAGTCGTTGCCTGGAGAATATACATACACACACACANNNAGATCGG |
| 5038_1 | TATGAGTGTGGAGTCGTTGCTGGCTACGTGTATCTCTCCACTTNNNAGATCGGAAGA |
| 5040_1 | TATGAGTGTGGAGTCGTTGCGTATGTATTTTCCTGAGGGATGTCCCNNNAGATCGG |
| 5044_1 | TATGAGTGTGGAGTCGTTGCTTTTGATGTTTAGTTTTGAACTGGCTNNNAGATCGGA |
| 5045_1 | TATGAGTGTGGAGTCGTTGCACCACCTTACCTTACCCTTATCCCNNNAGATCGGAAGA |
| 5103_1 | TATGAGTGTGGAGTCGTTGCGTGTGTGTTTCCACCTAAGTTGATAGNNNAGATCGG |
| 5112_1 | TATGAGTGTGGAGTCGTTGCACATTTGAGGGGTGAAGTAAGCCNNNAGATCGGAA |
| 5126_1 | TATGAGTGTGGAGTCGTTGCTTCAAAACCAACCTATTTGACAGCATNNNAGATCGGA |
| 5137_1 | TATGAGTGTGGAGTCGTTGCATAAAGTATGCCTGCTCTGAGCCNNNAGATCGGAAG |
| 5197_1 | TATGAGTGTGGAGTCGTTGCCTTCTGTTCGTTCTTTCATTTGTTCNNNAGATCGGAAG |
| 5198_1 | TATGAGTGTGGAGTCGTTGCCTGTCCCTCTTCATGTTATGGCCNNNAGATCGGAAGA |
| 5214_1 | TATGAGTGTGGAGTCGTTGCTGAGTTTACCTATCACTAAGTTGGCCNNNAGATCGGA |
| 5258_1 | TATGAGTGTGGAGTCGTTGCAGCACTGTCAGATTTTTAGTGCCNNNAGATCGGAAG |
| 5265_1 | TATGAGTGTGGAGTCGTTGCTGGTTGATATGGGAAATGAGATATCTNNNAGATCG |
| 5284_1 | TATGAGTGTGGAGTCGTTGCGGTGGTGGGCAGAAGATTGTATTANNNAGATCGGA |
| 5302_1 | TATGAGTGTGGAGTCGTTGCTCAGAAGAGAGCATGCACTATCCNNNAGATCGGAAG |
| 5396_1 | TATGAGTGTGGAGTCGTTGCTACTCAATCTCAGGGCAGCATTGNNNAGATCGGAAG |
| 5416_1 | TATGAGTGTGGAGTCGTTGCTCATCTGCCTTTCACACAGTTCCNNNAGATCGGAAGA |
| 5495_1 | TATGAGTGTGGAGTCGTTGCATGTTACCAAAAGACCCAGCAATTCNNNAGATCGGA |
| 5629_1 | TATGAGTGTGGAGTCGTTGCATCTTGACCCAAACTTTCGTGCTNNNAGATCGGAAGA |
| 5642_1 | TATGAGTGTGGAGTCGTTGCGCAGAGGTGCTATTTAGGGTAGCNNNAGATCGGAA |
| 5669_1 | TATGAGTGTGGAGTCGTTGCTCTGTCATTGCTGAGGTGTTTCCNNNAGATCGGAAG |
| 5687_1 | TATGAGTGTGGAGTCGTTGCCCACACCATTTTCTCTTTCAAGGCNNNAGATCGGAAG |

| | |
|---|---|
| 5717_1 | TATGAGTGTGGAGTCGTTGCTGAGACTGGAAAGATAAAAGAGAGGCNNNAGATCG |
| 5734_1 | TATGAGTGTGGAGTCGTTGCAGAGATTCATAGTTTGGATTGGCATGNNNAGATCGG |
| 5748_1 | TATGAGTGTGGAGTCGTTGCATTCCCTACTTCACTGATAGGGCNNNAGATCGGAAG |
| 5783_1 | TATGAGTGTGGAGTCGTTGCCATATTACTCCCAAAGCCATGGCNNNAGATCGGAAG |
| 5851_1 | TATGAGTGTGGAGTCGTTGCAAGAGGAAGCTAATCTGTGAGGATACNNNAGATCG |
| 5944_1 | TATGAGTGTGGAGTCGTTGCGTGTGTAAGTTTGGGCTAGGGAGNNNAGATCGGA |
| 6034_1 | TATGAGTGTGGAGTCGTTGCTCCAGTGAAAAACATATTTTGCTTGCNNNAGATCGG |
| 6074_1 | TATGAGTGTGGAGTCGTTGCTCCCCAAACCAGATCTATAGCATACANNNAGATCGGA |
| 6084_1 | TATGAGTGTGGAGTCGTTGCGACACTCCTAAACAGCAGTGGAGNNNAGATCGGAA |
| 6086_1 | TATGAGTGTGGAGTCGTTGCAGCAACAGGTGGTGAGTTGAATCNNNAGATCGGAA |
| 6127_1 | TATGAGTGTGGAGTCGTTGCGCCGGGTTTTTGTTCCCCTAGATTTAGNNNAGATCG |
| 6225_1 | TATGAGTGTGGAGTCGTTGCCCATTTTTCGCTTTCTCTGAGCANNNAGATCGGAAGA |
| 6355_1 | TATGAGTGTGGAGTCGTTGCAAAACAACAATAACCCACACACCAATNNNAGATCGG |
| 6629_1 | TATGAGTGTGGAGTCGTTGCATCATGGACTGCTGTGTATCTCTNNNAGATCGGAAG |
| 6664_1 | TATGAGTGTGGAGTCGTTGCTGTGTGTGAGATAAAGTCTCACTGTGNNNAGATCG |
| 6796_1 | TATGAGTGTGGAGTCGTTGCCTCTCTCCTCTCTCTCTCATCTCTTTNNNAGATCGGAA |
| 6799_1 | TATGAGTGTGGAGTCGTTGCTGCTCTTGTTTTCATTCTCTTGCTCTNNNAGATCGGA |
| 6856_1 | TATGAGTGTGGAGTCGTTGCTCTTTGTTTTTCTGTGTGCATTTGCTNNNAGATCGGA |
| 6882_1 | TATGAGTGTGGAGTCGTTGCCCAGATAAAGAGGAGGTGGAATGGGNNNAGATCG |
| 6939_1 | TATGAGTGTGGAGTCGTTGCCAGACTTTTATGTGCAAACTCAGTCTNNNAGATCGG |
| 6954_1 | TATGAGTGTGGAGTCGTTGCTGCTGCTCACAGACAAGTGTTATTAGNNNAGATCGG |
| 6977_1 | TATGAGTGTGGAGTCGTTGCTTTCCTCTTTCTTTGTCTTGTTCAGNNNAGATCGGAA |
| 6998_1 | TATGAGTGTGGAGTCGTTGCATGAGAGAGAGAGAGATACAGGGTNNNAGATCGG |
| 7016_1 | TATGAGTGTGGAGTCGTTGCTTCCTTTGCTCAAATCAGAGAAGAGNNNAGATCGGA |
| 7021_1 | TATGAGTGTGGAGTCGTTGCGGTGCATGTAGACCAAATAGGAAAAGNNNAGATCG |
| 7022_1 | TATGAGTGTGGAGTCGTTGCGTTGACCCCATACCTTTGCTGTTNNNAGATCGGAAG |
| 7024_1 | TATGAGTGTGGAGTCGTTGCGAGATATATGCATGCACCCACCCNNNAGATCGGAAG |
| 7047_1 | TATGAGTGTGGAGTCGTTGCAACCAACCCTTTTCCGTATCTATGAGNNNAGATCGGA |
| 7050_1 | TATGAGTGTGGAGTCGTTGCTCTTTTTCATGGTGTCTGTCACCTAGNNNAGATCGGA |
| 7053_1 | TATGAGTGTGGAGTCGTTGCCAGCAATCAATGAGATAGGAGACATNNNAGATCGG |
| 7064_1 | TATGAGTGTGGAGTCGTTGCAATAGAGCGCTGCAAAAGGAGAGNNNAGATCGGA |
| 7136_1 | TATGAGTGTGGAGTCGTTGCTGTGTGTATTTGGTTGGCTAGTTAGTNNNAGATCGG |
| 7184_1 | TATGAGTGTGGAGTCGTTGCATGAAGGAATCCTCCCCTATATGGAGNNNAGATCGG |
| 7227_1 | TATGAGTGTGGAGTCGTTGCTTACCAGCTCTTTGAAGGAATATTCANNNAGATCGG |
| 7274_1 | TATGAGTGTGGAGTCGTTGCACATGCCCAAAAGAAGCCTCTAGNNNAGATCGGAAG |
| 7342_1 | TATGAGTGTGGAGTCGTTGCACATTTTATAGCCTACTTTTCGTCATNNNAGATCGGA |
| 7365_1 | TATGAGTGTGGAGTCGTTGCTCTTTCCCCTCTCTCTCTCTCCTGNNNAGATCGGAAGA |
| 7378_1 | TATGAGTGTGGAGTCGTTGCCTCTGTTTTCAGTGAGGGAATTAGGTNNNAGATCGG |
| 7379_1 | TATGAGTGTGGAGTCGTTGCCCCCAAATCAACAGTGGCTAATAANNNAGATCGGAA |
| 7481_1 | TATGAGTGTGGAGTCGTTGCGTGTTCTTTGGCTTGCATTCTCTTTCNNNAGATCGGA |
| 7501_1 | TATGAGTGTGGAGTCGTTGCACAGTCCAGAGACATCTTTGAAGGNNNAGATCGGA |
| 7528_1 | TATGAGTGTGGAGTCGTTGCTTTCCTGATGCTCAAATCCCAGGNNNAGATCGGAAG |
| 7529_1 | TATGAGTGTGGAGTCGTTGCGTGATTTTGGAAGTTTAGTTCCCAGGNNNAGATCGG |

| | |
|---|---|
| 7558_1 | TATGAGTGTGGAGTCGTTGCAGTGTTTTGTTGTGGGTGGTTCTNNNAGATCGGAA |
| 7643_1 | TATGAGTGTGGAGTCGTTGCAAAGAGTGGTGTGTGTGTGTAGGNNNAGATCGGA |
| 7731_1 | TATGAGTGTGGAGTCGTTGCAGAGCAATGCACATGGATTAGGGNNNAGATCGGAA |
| 7787_1 | TATGAGTGTGGAGTCGTTGCCCAAGATAACAATTTGAGGTGATGGGNNNAGATCG |
| 7857_1 | TATGAGTGTGGAGTCGTTGCCTGCCAAAGAAAGGGAATTTGGGNNNAGATCGGAA |
| 7868_1 | TATGAGTGTGGAGTCGTTGCTTTCCTTTCACAGGACCACAATCANNNAGATCGGAAG |
| 7943_1 | TATGAGTGTGGAGTCGTTGCTGCTACTTTTGCATTGATTTCACTGGNNNAGATCGG |
| 7944_1 | TATGAGTGTGGAGTCGTTGCACTAGAGGAAGCGAGAAGACTGGNNNAGATCGGA |
| 7953_1 | TATGAGTGTGGAGTCGTTGCTCCCAAAGAAAACAAAGAAACCTGGNNNAGATCGG |
| 7959_1 | TATGAGTGTGGAGTCGTTGCACCAGATGATAAATCACAGTACCTGGNNNAGATCGG |
| 8006_1 | TATGAGTGTGGAGTCGTTGCGGTCTGTCCACTAAGAAAGTCTGGNNNAGATCGGA |
| 8074_1 | TATGAGTGTGGAGTCGTTGCTGCTTCTCTAATTCTTTCACTGTGANNNAGATCGGAA |
| 8159_1 | TATGAGTGTGGAGTCGTTGCAGAGTGGTGCAAGATAGCCAATGNNNAGATCGGAA |
| 8228_1 | TATGAGTGTGGAGTCGTTGCATTGCTGTACCTTTTGCTGAATTCAANNNAGATCGG |
| 8268_1 | TATGAGTGTGGAGTCGTTGCGCTCTCGCTCTCAAAAATAGGATGNNNAGATCGGAA |
| 8358_1 | TATGAGTGTGGAGTCGTTGCACATTGATGAGGGGAATATAAAGGGANNNAGATCG |
| 8370_1 | TATGAGTGTGGAGTCGTTGCTCACAGATTGAAAAGGAACCAGTNNNAGATCGGAA |
| 8405_1 | TATGAGTGTGGAGTCGTTGCCGCAAGCCATAGAAGAAAAGCTGNNNAGATCGGAA |
| 8462_1 | TATGAGTGTGGAGTCGTTGCCTTCTGTTGGCTCTGTTTTGCTGNNNAGATCGGAAG |
| 8473_1 | TATGAGTGTGGAGTCGTTGCCGGGTTTGGTTTGATCTCATCTGNNNAGATCGGAAG |
| 8537_1 | TATGAGTGTGGAGTCGTTGCAGCACATACACAGAAACATTCTTCTGNNNAGATCGG |
| 8550_1 | TATGAGTGTGGAGTCGTTGCGTGGGTTTTTCTTTTCTGGTTTTCTGNNNAGATCGGA |
| 8571_1 | TATGAGTGTGGAGTCGTTGCTCAGTGCATTATGTACACTTCAGTGNNNAGATCGGA |
| 8599_1 | TATGAGTGTGGAGTCGTTGCCAGAAGCATTGTCTTATCCAAGGTGNNNAGATCGGA |
| 8630_1 | TATGAGTGTGGAGTCGTTGCTGCGTATGTGTATTATTGCTATGGTGNNNAGATCGG |
| 8646_1 | TATGAGTGTGGAGTCGTTGCCACACACTTCATCTAGCTAACTCAGTNNNAGATCGGA |
| 8683_1 | TATGAGTGTGGAGTCGTTGCGAGAGAGAGAATCACCTTGTCAGTNNNAGATCGGA |
| 8692_1 | TATGAGTGTGGAGTCGTTGCTCAGAAAAATGCAGAGAAATTTTGTGNNNAGATCG |
| 8711_1 | TATGAGTGTGGAGTCGTTGCTTTTTGCTCTCACTGAAACAATTTTTNNNAGATCGGA |
| 8732_1 | TATGAGTGTGGAGTCGTTGCCATCCCCACCATACTTGACCTTGNNNAGATCGGAAGA |
| 8745_1 | TATGAGTGTGGAGTCGTTGCGCTACAGATGATAGATAGAATAGCTTGNNNAGATCG |
| 8827_1 | TATGAGTGTGGAGTCGTTGCACATGCCAATACCACACTGTTTGNNNAGATCGGAAG |
| 8845_1 | TATGAGTGTGGAGTCGTTGCGCATCACATTGAACTCTTCATAGAGTNNNAGATCGG |
| 8887_1 | TATGAGTGTGGAGTCGTTGCCTGGAGAATCCTGACTGAAACACCTANNNAGATCGG |
| 8895_1 | TATGAGTGTGGAGTCGTTGCTGGCTGTGTGTCTTTGGTGAAATNNNAGATCGGAA |
| 8960_1 | TATGAGTGTGGAGTCGTTGCTGGATGGTAACAAATTGTCAGATGGANNNAGATCG |
| 8965_1 | TATGAGTGTGGAGTCGTTGCGGGTGTGGGGTAAGGAGAGAATNNNAGATCGGAA |
| 9003_1 | TATGAGTGTGGAGTCGTTGCTCCCCCTCCATGTATATGCATAANNNAGATCGGAAGA |
| 9031_1 | TATGAGTGTGGAGTCGTTGCCAGAAGGAAGCATTGCTAATATAGGANNNAGATCG |
| 9186_1 | TATGAGTGTGGAGTCGTTGCGCCCCACAAGATCAGAAACTCATNNNAGATCGGAAG |
| 9200_1 | TATGAGTGTGGAGTCGTTGCTTATGAGAAAACAAAAGGCGCACTGANNNAGATCG |
| 9225_1 | TATGAGTGTGGAGTCGTTGCAGCATTTTCTTATCTTCGGTTTTCATNNNAGATCGGA |
| 9249_1 | TATGAGTGTGGAGTCGTTGCACTGCCAGGGCTTTATTTTCAGTNNNAGATCGGAAG |

| | |
|---|---|
| 9466_1 | TATGAGTGTGGAGTCGTTGCCATCACCTCAGGGCTCCCTTATNNNAGATCGGAAGAG |
| 9479_1 | TATGAGTGTGGAGTCGTTGCGTGTCCCTGTAGCAGCGATTTATNNNAGATCGGAAG |
| 9511_1 | TATGAGTGTGGAGTCGTTGCTGCCCCTTACAACTTTGTCAACTNNNAGATCGGAAGA |
| 9536_1 | TATGAGTGTGGAGTCGTTGCCCTCCTCATAGTTTATTCATCTGCCANNNAGATCGGA |
| 9576_1 | TATGAGTGTGGAGTCGTTGCGTGCATATACTAGACTTCGAATCACTNNNAGATCGG |
| 9599_1 | TATGAGTGTGGAGTCGTTGCAGTGGAATGCTGTTATATATGTGCGTNNNAGATCGG |
| 9616_1 | TATGAGTGTGGAGTCGTTGCACATAGGGGACATGAAAAGGACTNNNAGATCGGAA |
| 9625_1 | TATGAGTGTGGAGTCGTTGCACAAAAGAAGAATAATGACAGGGACTNNNAGATCG |
| 9640_1 | TATGAGTGTGGAGTCGTTGCACAGTATTTTGACCTAGGGATTGACTNNNAGATCGG |
| 9725_1 | TATGAGTGTGGAGTCGTTGCTCACTTACACGGTGTTTTCACCTNNNAGATCGGAAGA |
| 9766_1 | TATGAGTGTGGAGTCGTTGCGTTTCCTTGAAGGTTTATTTTACCCTNNNAGATCGGA |
| 9792_1 | TATGAGTGTGGAGTCGTTGCTCTTCCCTGAAAACACTGATAGATCCNNNAGATCGGA |
| 9805_1 | TATGAGTGTGGAGTCGTTGCCCCATACATTCACACACAACTAGAGTNNNAGATCGGA |
| 9838_1 | TATGAGTGTGGAGTCGTTGCTGCCTGTCTATCTATATCTAGCCTNNNAGATCGGAAG |
| 9891_1 | TATGAGTGTGGAGTCGTTGCTGGGGAAGAAAAAGTTTTATCCAGGTNNNAGATCG |
| 9895_1 | TATGAGTGTGGAGTCGTTGCGCATTTTTCTACCTTGCATACATCCTNNNAGATCGGA |
| 9910_1 | TATGAGTGTGGAGTCGTTGCGAACTTGATTGGTTCTATTTCTCTGGNNNAGATCGGA |
| 9919_1 | TATGAGTGTGGAGTCGTTGCTTTTAGCTGTACCTGATTTTATCTCCTNNNAGATCGGA |
| 9974_1 | TATGAGTGTGGAGTCGTTGCTCCCTAACAAGAAATCCCTTCCTNNNAGATCGGAAGA |
| 10014_1 | TATGAGTGTGGAGTCGTTGCACATCTCTCTTGTTTTACTTGTTCCTNNNAGATCGGAA |
| 10125_1 | TATGAGTGTGGAGTCGTTGCTCTCTGAAGAACACTAACATACAGCTNNNAGATCGGA |
| 10259_1 | TATGAGTGTGGAGTCGTTGCACAAGTTTAGACCAAGGAAACTAGCTNNNAGATCGG |
| 10285_1 | TATGAGTGTGGAGTCGTTGCATTACCAGTATGGACGGGTAGCTNNNAGATCGGAAG |
| 10429_1 | TATGAGTGTGGAGTCGTTGCGCTAGAATATGAATGTCATGAGGGCTNNNAGATCGG |
| 10487_1 | TATGAGTGTGGAGTCGTTGCTATGGCTATGTGAGATGATGGCTNNNAGATCGGAAG |
| 10494_1 | TATGAGTGTGGAGTCGTTGCTGGCTTGTGACTATAGAGACTTGGTNNNAGATCGGA |
| 10582_1 | TATGAGTGTGGAGTCGTTGCACCTTTTCCTTCCTTTTCCTAATGCTNNNAGATCGGAA |
| 10804_1 | TATGAGTGTGGAGTCGTTGCTAACAGAGCCAATCATGCTTGCTNNNAGATCGGAAG |
| 11056_1 | TATGAGTGTGGAGTCGTTGCTGGGTATCAAGGAAACCATCCTATCCNNNAGATCGGA |
| 11096_1 | TATGAGTGTGGAGTCGTTGCGAGACATAGAACAGACACTTTCAGGTNNNAGATCGG |
| 11199_1 | TATGAGTGTGGAGTCGTTGCTCGCCAAGTTTAGTTCTACTTGACTTNNNAGATCGGA |
| 11210_1 | TATGAGTGTGGAGTCGTTGCTGAAGAGATGGCCAACAACCAAANNNAGATCGGAAG |
| 11287_1 | TATGAGTGTGGAGTCGTTGCTTCCTGGTTACATTCATCAAAAGAAANNNAGATCGGA |
| 11296_1 | TATGAGTGTGGAGTCGTTGCCACTGTATTTCTAGTTAGCTCTGTCTNNNAGATCGGA |
| 11341_1 | TATGAGTGTGGAGTCGTTGCGCTTTCTCCTTTGTTACTCTCATTCTNNNAGATCGGAA |
| 11357_1 | TATGAGTGTGGAGTCGTTGCACCCAGTAACTACTCTTCTTACTTCTNNNAGATCGGAA |
| 11397_1 | TATGAGTGTGGAGTCGTTGCTGTAAGAAACATTGGTTTGAGTTCTNNNAGATCGGA |
| 11438_1 | TATGAGTGTGGAGTCGTTGCGTGTTTGAAATGTCTGTTCATATGCNNNAGATCGGA |
| 11516_1 | TATGAGTGTGGAGTCGTTGCGTGCCATACACACACATTGACAANNNAGATCGGAAG |
| 11533_1 | TATGAGTGTGGAGTCGTTGCCAGTGTGATGCATTTTCAAAGTTCCTNNNAGATCGGA |
| 11593_1 | TATGAGTGTGGAGTCGTTGCAGCAAACTAAAAGTATACAACACAGTNNNAGATCGG |
| 11684_1 | TATGAGTGTGGAGTCGTTGCGTGTGCTCTTGGGTCAATGCAANNNAGATCGGAAGA |
| 11733_1 | TATGAGTGTGGAGTCGTTGCTGCCCTCCTGATTAGAACCTCAANNNAGATCGGAAGA |

| | |
|---|---|
| 11780_1 | TATGAGTGTGGAGTCGTTGCGGCTGGTGCTCAAATCTCTTCAANNNAGATCGGAAG |
| 11813_1 | TATGAGTGTGGAGTCGTTGCGGTGGTGATTCTAAAGGACATGAGTNNNAGATCGG |
| 11872_1 | TATGAGTGTGGAGTCGTTGCGCAAGAGAGAAGGCAATCCTAGTNNNAGATCGGAA |
| 11902_1 | TATGAGTGTGGAGTCGTTGCGAATACGACCGTAACAGTGAGAANNNAGATCGGAA |
| 11914_1 | TATGAGTGTGGAGTCGTTGCAAGCTCGGAAATAACTGATCCACCATNNNAGATCGG |
| 11923_1 | TATGAGTGTGGAGTCGTTGCCAGCTTCTCTTCTTAGTCATTTAGAANNNAGATCGGA |
| 11924_1 | TATGAGTGTGGAGTCGTTGCAGGAAATCATCTTGCAAAATACACGTNNNAGATCGG |
| 11929_1 | TATGAGTGTGGAGTCGTTGCAGTTAGCATGGGGAGATCTCGAANNNAGATCGGAA |
| 11937_1 | TATGAGTGTGGAGTCGTTGCAAGCTGACATGCAATCACAGGAANNNAGATCGGAA |
| 11943_1 | TATGAGTGTGGAGTCGTTGCGTGCAGTTACCAATATCCACCGTNNNAGATCGGAAG |
| 11951_1 | TATGAGTGTGGAGTCGTTGCCTCAAGTTTTCAGATTTGGTGAAGTGNNNAGATCGG |
| 11953_1 | TATGAGTGTGGAGTCGTTGCTTTCCCCATTCGGTATGATGAGGNNNAGATCGGAAG |
| 12071_1 | TATGAGTGTGGAGTCGTTGCCTTTTTCAGCACCAACATGCTCAAAANNNAGATCGGA |
| 12135_1 | TATGAGTGTGGAGTCGTTGCCCTATGGCATCTAAAGAAGGAGTGTANNNAGATCG |
| 12139_1 | TATGAGTGTGGAGTCGTTGCAAATCTCTTCAGTTTCTGCAACACTGNNNAGATCGGA |
| 12187_1 | TATGAGTGTGGAGTCGTTGCTGGACTTACGAGACATACCTACANNNAGATCGGAAG |
| 12236_1 | TATGAGTGTGGAGTCGTTGCGAAGGCTTGATAACTCCTATATGGGTNNNAGATCGG |
| 12267_1 | TATGAGTGTGGAGTCGTTGCTGGAAAACCTTGATTCTTTTGGGTNNNAGATCGGAA |
| 12300_1 | TATGAGTGTGGAGTCGTTGCTGTGTGCATCTATGTGCCTGTAANNNAGATCGGAAG |
| 12308_1 | TATGAGTGTGGAGTCGTTGCTACACACTCCATGGTCTGATGGTNNNAGATCGGAAG |
| 12372_1 | TATGAGTGTGGAGTCGTTGCATGATTACCTGATGTTGCTAAAACANNNAGATCGGA |
| 12404_1 | TATGAGTGTGGAGTCGTTGCACTTCTGCTATTCCAACAAAGAAACANNNAGATCGG |
| 53436_1 | TATGAGTGTGGAGTCGTTGCAAGAAGAAGATGGCAATGTTCCTNNNAGATCGGAA |
| 53476_1 | TATGAGTGTGGAGTCGTTGCCAACACCTCCCATCCTCTCAATCNNNAGATCGGAAGA |
| 53600_1 | TATGAGTGTGGAGTCGTTGCTCTTGAACCCATACATAGCTACAAGANNNAGATCGG |
| 53636_1 | TATGAGTGTGGAGTCGTTGCCCTCCTGGGTTCAAACAATTCTTNNNAGATCGGAAG |
| 53676_1 | TATGAGTGTGGAGTCGTTGCTGTTGGAATTCTGAATTGATGACACTNNNAGATCGG |
| 53770_1 | TATGAGTGTGGAGTCGTTGCTCACAACATAAGACAAGACAAAGTGANNNAGATCG |
| 54070_1 | TATGAGTGTGGAGTCGTTGCCCTTTGATGCTTAGTTTATTGAGGGTNNNAGATCGG |
| 54267_1 | TATGAGTGTGGAGTCGTTGCTCTGAAGAAAGATAGAAAAAATGAGGGNNNAGAT |
| 54322_1 | TATGAGTGTGGAGTCGTTGCCTCCCATCTCAGCTTCCCAAATANNNAGATCGGAAGA |
| 54346_1 | TATGAGTGTGGAGTCGTTGCCTAGGATTACAGACACCTGCCACNNNAGATCGGAAG |
| 54356_1 | TATGAGTGTGGAGTCGTTGCCTGGAGTGACAGCCTTGTATATANNNAGATCGGAA |
| 54597_1 | TATGAGTGTGGAGTCGTTGCGAGTACTGTGGCCTGATCATCGNNNAGATCGGAAG |
| 54795_1 | TATGAGTGTGGAGTCGTTGCTTCAAAGTAAACTATCTTGACAGTTCANNNAGATCG |
| 54855_1 | TATGAGTGTGGAGTCGTTGCCTCCAGGGTTCATGCCATTCTCNNNAGATCGGAAGA |
| 54876_1 | TATGAGTGTGGAGTCGTTGCGTGATCTTGGTTCACTGCAACCNNNAGATCGGAAGA |
| 55034_1 | TATGAGTGTGGAGTCGTTGCAGCAAAATGTTAGCAAATTGAATTCANNNAGATCGG |
| 55052_1 | TATGAGTGTGGAGTCGTTGCGCCTCCTGAGTAGCTGAGATTACNNNAGATCGGAAG |
| 55085_1 | TATGAGTGTGGAGTCGTTGCCTTCCCAGGTTCAAGCAATTCTCNNNAGATCGGAAG |
| 55492_1 | TATGAGTGTGGAGTCGTTGCGCATTTCAGTGCATAATATTCTCTTGTNNNAGATCGG |
| 55629_1 | TATGAGTGTGGAGTCGTTGCTCAGGTGAAGAAAGAAGAAATATGGTNNNAGATCG |
| 56063_1 | TATGAGTGTGGAGTCGTTGCAGTGCAAAAAAGCAGTAAATTATGGCNNNAGATCG |

| | |
|---|---|
| 56412_1 | TATGAGTGTGGAGTCGTTGCAGGCACTTTAGAAATAATGTAGTCCTNNNAGATCGG |
| 56447_1 | TATGAGTGTGGAGTCGTTGCTGCTTACCATAAATCATGAGCTAAGCNNNAGATCGG |
| 56522_1 | TATGAGTGTGGAGTCGTTGCATGCCCTACATATCAAAGATGCTNNNAGATCGGAAG |
| 56523_1 | TATGAGTGTGGAGTCGTTGCTCTGCTCTCTACTTTGTTACTGATGTNNNAGATCGGA |
| 56584_1 | TATGAGTGTGGAGTCGTTGCTGGCCCATTTTAGGAAGTAGTCCNNNAGATCGGAAG |
| 56662_1 | TATGAGTGTGGAGTCGTTGCAGGGAAGAAAGAAAGAAAAAAATGGANNNAGATC |
| 56764_1 | TATGAGTGTGGAGTCGTTGCCTTCCCAAGTAGCTGGAACTACANNNAGATCGGAAG |
| 56868_1 | TATGAGTGTGGAGTCGTTGCAGAAGAATATGTGAGAGGAGCTGANNNAGATCGG |
| 56944_1 | TATGAGTGTGGAGTCGTTGCAAGAAAAAGAAAAATTGCCTCATTGTNNNAGATCGG |
| 56963_1 | TATGAGTGTGGAGTCGTTGCCAAGAGATTCTCCTGTCTCAGCCNNNAGATCGGAAG |
| 56981_1 | TATGAGTGTGGAGTCGTTGCCTAGGTTCAAGCAATTCACCTGCNNNAGATCGGAAG |
| 57145_1 | TATGAGTGTGGAGTCGTTGCAAAGCAAAACCTTACATTTTTGATCANNNAGATCGG |
| 57486_1 | TATGAGTGTGGAGTCGTTGCTGTGATGTTTTTTGTTGAAATTTGCANNNAGATCGG |
| 57645_1 | TATGAGTGTGGAGTCGTTGCTGACAAAACCACAGTAACTTCTTTCANNNAGATCGG |
| 57758_1 | TATGAGTGTGGAGTCGTTGCACTGTGACTTTTGAATTTTGTGATACANNNAGATCG |
| 57774_1 | TATGAGTGTGGAGTCGTTGCGGAAGAAAGAAGAAGGAAGAAGAAGGNNNAGAT |
| 57849_1 | TATGAGTGTGGAGTCGTTGCCTCCTGAGTAGCTGGAACTACAGNNNAGATCGGAA |
| 58151_1 | TATGAGTGTGGAGTCGTTGCAAACAATTCTCCTTCCTCAGCCTNNNAGATCGGAAGA |
| 58266_1 | TATGAGTGTGGAGTCGTTGCAACTTGCTAATCATGTGAACAAAAATNNNAGATCGG |
| 58298_1 | TATGAGTGTGGAGTCGTTGCACGTTTCAAAAATTAGAACGCTTTGTNNNAGATCGG |
| 58346_1 | TATGAGTGTGGAGTCGTTGCGCCTAATTTCTCTGTCTAGGACTTCTNNNAGATCGGA |
| 58415_1 | TATGAGTGTGGAGTCGTTGCTCAGAAAACAAATCCAAAGTCACTCANNNAGATCGG |
| 58675_1 | TATGAGTGTGGAGTCGTTGCCTCCTGGGTTCACACCATTCTCNNNAGATCGGAAGA |
| 58770_1 | TATGAGTGTGGAGTCGTTGCCAGTGGCTTGATCTCAGCTTACTNNNAGATCGGAAG |
| 58779_1 | TATGAGTGTGGAGTCGTTGCCAGTCTCCCAAGTAGCTGGAATTNNNAGATCGGAAG |
| 59129_1 | TATGAGTGTGGAGTCGTTGCTCTGCTCTGTGAGTTTCTCAGTCNNNAGATCGGAAG |
| 59154_1 | TATGAGTGTGGAGTCGTTGCTGGCAAGAAGAAAGATAAAAGAGACANNNAGATCG |
| 59193_1 | TATGAGTGTGGAGTCGTTGCAGTACGTAGGCAATATTAGTACTTGCNNNAGATCGG |
| 59201_1 | TATGAGTGTGGAGTCGTTGCGCAAATCAGTTAGCAAATCAGTTACCNNNAGATCGG |
| 59355_1 | TATGAGTGTGGAGTCGTTGCGGCTAACAAACAGAAAAAAGAAGAGGNNNAGATCG |
| 59412_1 | TATGAGTGTGGAGTCGTTGCAGGTAGAAAAGATGGAAACATGCTNNNAGATCGGA |
| 59462_1 | TATGAGTGTGGAGTCGTTGCGTTCAAGCGATTCTTCTGCCTCNNNAGATCGGAAGA |
| 59618_1 | TATGAGTGTGGAGTCGTTGCGGTATCTATCACTTGAATTGATGGGTNNNAGATCGG |
| 59755_1 | TATGAGTGTGGAGTCGTTGCACACACGATTTTTTTTAGACCTGTCANNNAGATCGGA |
| 60047_1 | TATGAGTGTGGAGTCGTTGCTGCAGTAGTTTCATAGTTTTAGGTCANNNAGATCGG |
| 60049_1 | TATGAGTGTGGAGTCGTTGCACTTTCTGAACTTACTTCTCTGGAGTNNNAGATCGGA |
| 60134_1 | TATGAGTGTGGAGTCGTTGCACTTGCCCCAACTTGGACTATAGNNNAGATCGGAAG |
| 60319_1 | TATGAGTGTGGAGTCGTTGCTCATTTATCTGTATGTTGTCTTGCCTNNNAGATCGGA |
| 61988_1 | TATGAGTGTGGAGTCGTTGCAGGCTGGAATGTAATGGTGCTATNNNAGATCGGAA |
| 62076_1 | TATGAGTGTGGAGTCGTTGCATCTTGATTCACTGCAACCTCCTNNNAGATCGGAAGA |
| 62232_1 | TATGAGTGTGGAGTCGTTGCAGCCATACAAGTCTATAATGGTGTCTNNNAGATCGG |
| 62790_1 | TATGAGTGTGGAGTCGTTGCCAGTCCCACAAGTTCCTGAGTATNNNAGATCGGAAG |
| 62918_1 | TATGAGTGTGGAGTCGTTGCAGTCACAAAGTAAAGCATGTGGCNNNAGATCGGAA |

| | |
|---|---|
| 62944_1 | TATGAGTGTGGAGTCGTTGCTGCTACCATTCATATAAAAAAGGGGGNNNAGATCGG |
| 63186_1 | TATGAGTGTGGAGTCGTTGCTGGATCCTATGTGTTTTTTTGTGTGTNNNAGATCGG |
| 63301_1 | TATGAGTGTGGAGTCGTTGCCTCCTGAGTAGCTGGGATTACAGNNNAGATCGGAA |
| 63396_1 | TATGAGTGTGGAGTCGTTGCGCCTATTTTTGTATAGTCTCTGAGCTNNNAGATCGG |
| 63851_1 | TATGAGTGTGGAGTCGTTGCCAATGGTGCAATCTCAGCTCACNNNAGATCGGAAGA |
| 63869_1 | TATGAGTGTGGAGTCGTTGCGTTCAAGCGATTCTCTTGCCTCNNNAGATCGGAAGA |
| 64253_1 | TATGAGTGTGGAGTCGTTGCTGTGTGTGTGTGTATGTATAAAGTNNNAGATCGG |
| 64260_1 | TATGAGTGTGGAGTCGTTGCTGGCTATCAAATCAATACCATTAGGGNNNAGATCGG |
| 64296_1 | TATGAGTGTGGAGTCGTTGCGCCTCCAATATGAATTTCTTGTCTGTNNNAGATCGGA |
| 64307_1 | TATGAGTGTGGAGTCGTTGCGCCTATCTTTCCTGTATGTAAACACTNNNAGATCGGA |
| 64522_1 | TATGAGTGTGGAGTCGTTGCTCTTTTTGTTTTGTTTGATCGTTTGTNNNAGATCGGA |
| 64579_1 | TATGAGTGTGGAGTCGTTGCTGAGATACTCATCTGGGTTTTATGGANNNAGATCGG |
| 64677_1 | TATGAGTGTGGAGTCGTTGCAGAGGAAAAACTATAAACACTGAGGANNNAGATCG |
| 64700_1 | TATGAGTGTGGAGTCGTTGCCTTTGGAGAGGGGAATGAGGAATNNNAGATCGGA |
| 64711_1 | TATGAGTGTGGAGTCGTTGCTTTGTTTATGATTTCTCCTTTGACCTNNNAGATCGGA |
| 64734_1 | TATGAGTGTGGAGTCGTTGCTGCAAATATCTTGAGGGACTATCACANNNAGATCGG |
| 64744_1 | TATGAGTGTGGAGTCGTTGCAGATAGATAATCCTTCTGGTTCTGTCNNNAGATCGG |
| 64815_1 | TATGAGTGTGGAGTCGTTGCTTTGTGGAAGAAAGGAAAGGGGANNNAGATCGGA |
| 64911_1 | TATGAGTGTGGAGTCGTTGCTTTCTGTAGCATTTTCCAATGTCANNNAGATCGGAAG |
| 64979_1 | TATGAGTGTGGAGTCGTTGCAGTCTTACTCTGTCACCTAGCCTNNNAGATCGGAAGA |
| 64981_1 | TATGAGTGTGGAGTCGTTGCTCCAAGACGGAGTCTCACTTTTTNNNAGATCGGAAG |
| 64993_1 | TATGAGTGTGGAGTCGTTGCTCACATAGATATCTCTGGCTAGGAGANNNAGATCGG |
| 65010_1 | TATGAGTGTGGAGTCGTTGCTGTGTGTGTATGTGTATATACGTGTGNNNAGATCGG |
| 65046_1 | TATGAGTGTGGAGTCGTTGCTTCTTCATCCAGTCATCCACTCANNNAGATCGGAAGA |
| 65102_1 | TATGAGTGTGGAGTCGTTGCTGTCTATGCCACATTTTCTTTATTCANNNAGATCGGA |
| 65360_1 | TATGAGTGTGGAGTCGTTGCTGGGGGGTATAGTCAATGCTATTNNNAGATCGGAA |
| 65395_1 | TATGAGTGTGGAGTCGTTGCGGCCCACAAAGCTGAAAGTATTTNNNAGATCGGAAG |
| 65533_1 | TATGAGTGTGGAGTCGTTGCACCTGAATTCAAATAGAGTTCTCCCANNNAGATCGG |
| 65563_1 | TATGAGTGTGGAGTCGTTGCAGTTTTGGACTCCTTTTTTGGGCNNNAGATCGGAAG |
| 65621_1 | TATGAGTGTGGAGTCGTTGCAGTCCTGCAAAATGATGAATACAACTNNNAGATCGG |
| 65846_1 | TATGAGTGTGGAGTCGTTGCACTTTTTTTTTTTTGAGATGGGGTCTNNNAGATCGG |
| 65942_1 | TATGAGTGTGGAGTCGTTGCAGAGAACTGGGAAGCTTTGAGAGNNNAGATCGGA |
| 66172_1 | TATGAGTGTGGAGTCGTTGCTGAATGGATATCACAAGGAACCANNNAGATCGGAA |
| 66193_1 | TATGAGTGTGGAGTCGTTGCTGATGATGATAAGGACACATGGCANNNAGATCGGA |
| 66454_1 | TATGAGTGTGGAGTCGTTGCCGGTGTTTTCTTCCTCCTAATAATAANNNAGATCGGA |
| 66651_1 | TATGAGTGTGGAGTCGTTGCTCCTGAATGAAACTGGTCTCCAGNNNAGATCGGAAG |
| 66679_1 | TATGAGTGTGGAGTCGTTGCTGAATATACATGAAGGAGTTAGGAGTNNNAGATCG |
| 66715_1 | TATGAGTGTGGAGTCGTTGCCATTCTCAGCAAACTAACACGGGNNNAGATCGGAAG |
| 66736_1 | TATGAGTGTGGAGTCGTTGCCAATTACAGCATGGGAGTGAAGGNNNAGATCGGAA |
| 66896_1 | TATGAGTGTGGAGTCGTTGCTTGCCTTTCAAAACTACCTTGGGNNNAGATCGGAAG |
| 66937_1 | TATGAGTGTGGAGTCGTTGCGACAAGAGTCTCTCCCTGTTGCNNNAGATCGGAAGA |
| 67044_1 | TATGAGTGTGGAGTCGTTGCTCTGGCCTCATCCGTGATTTTATNNNAGATCGGAAGA |
| 67165_1 | TATGAGTGTGGAGTCGTTGCGATGTGGTGGCTTACAGGCTTANNNAGATCGGAAG |

| | |
|---|---|
| 67351_1 | TATGAGTGTGGAGTCGTTGCTGAAGTCTGAGAACAGGCACTACNNNAGATCGGAA |
| 67541_1 | TATGAGTGTGGAGTCGTTGCTGTGATCCTTAGGGGAAGAGCTANNNAGATCGGAA |
| 67583_1 | TATGAGTGTGGAGTCGTTGCTGAGTGTATATGTACTGTGTGTGTGTNNNAGATCG |
| 67636_1 | TATGAGTGTGGAGTCGTTGCTCTTTTTGAGACAGCGTCTTGCNNNAGATCGGAAGA |
| 67731_1 | TATGAGTGTGGAGTCGTTGCCCCTGAAGACCCTAAGGAACAAANNNAGATCGGAA |
| 67871_1 | TATGAGTGTGGAGTCGTTGCTCTGAAGAACCCTGACTAAAATACCANNNAGATCGG |
| 67925_1 | TATGAGTGTGGAGTCGTTGCAGCAACAGTGTTTACTAAAAGAGGTNNNAGATCGG |
| 67927_1 | TATGAGTGTGGAGTCGTTGCTATTTTGAGACGGAGCAAGACTCNNNAGATCGGAA |
| 67985_1 | TATGAGTGTGGAGTCGTTGCTGTGTGTATGTGTATATATACGCACGNNNAGATCGG |
| 67986_1 | TATGAGTGTGGAGTCGTTGCAGGAACATTGTAATATAATGCCAGGTNNNAGATCG |
| 68054_1 | TATGAGTGTGGAGTCGTTGCACCAAAGTATTCCAAAGTCCATGTNNNAGATCGGAA |
| 68194_1 | TATGAGTGTGGAGTCGTTGCATGTATTCATAAGTGTAAAGCAGCANNNAGATCGG |
| 68336_1 | TATGAGTGTGGAGTCGTTGCGGAGTAGCTAGGATTACAGGCACNNNAGATCGGAA |
| 68779_1 | TATGAGTGTGGAGTCGTTGCGGAACTAACTCATGTGATTATGGAGGNNNAGATCG |
| 69030_1 | TATGAGTGTGGAGTCGTTGCTACAGCATTAGGTCTCACCCAAGNNNAGATCGGAAG |
| 69077_1 | TATGAGTGTGGAGTCGTTGCATACACGTGTATATGTATATGTACGTGNNNAGATCG |
| 69078_1 | TATGAGTGTGGAGTCGTTGCAGCACTATGTCTTGAAAAGTTGTTGTNNNAGATCGG |
| 69152_1 | TATGAGTGTGGAGTCGTTGCGATCTTGGCTCACTGCAACCTCNNNAGATCGGAAGA |
| 69154_1 | TATGAGTGTGGAGTCGTTGCCAAAAGAAGACATACACACGGCCNNNAGATCGGAA |
| 69159_1 | TATGAGTGTGGAGTCGTTGCAACTTGCAGGATTGTAAGTTGCCNNNAGATCGGAA |
| 69382_1 | TATGAGTGTGGAGTCGTTGCGGCTCACTGTAGCTTCAAATTCCNNNAGATCGGAAG |
| 69419_1 | TATGAGTGTGGAGTCGTTGCTCTTGGATCTGTGGATTTATGGTNNNAGATCGGAAG |
| 69458_1 | TATGAGTGTGGAGTCGTTGCGTTGGAGTGCAATGACGTGATCNNNAGATCGGAAG |
| 69773_1 | TATGAGTGTGGAGTCGTTGCAAAAGAAAGAGAAAGAAAAGGAAAGGANNNAGAT |
| 69808_1 | TATGAGTGTGGAGTCGTTGCTCTCTCACAGTTAGAATTTGGTCTGTNNNAGATCGG |
| 69820_1 | TATGAGTGTGGAGTCGTTGCAGAGAGAAAGAAAGAAAGAAAGAAAGANNNAGAT |
| 69827_1 | TATGAGTGTGGAGTCGTTGCTCTCTCTTTCTCTCTTTCTCTCTTTCTNNNAGATCGGAA |
| 70013_1 | TATGAGTGTGGAGTCGTTGCAGAAAGAAAGAAAAATATGTGAGTGGANNNAGAT |
| 70236_1 | TATGAGTGTGGAGTCGTTGCCAAACTGCTGGTCTCAAGTGATCNNNAGATCGGAAG |
| 70280_1 | TATGAGTGTGGAGTCGTTGCAGTTTCTAGAACTTTTTCTTGCAGCANNNAGATCGG |
| 70440_1 | TATGAGTGTGGAGTCGTTGCCAGAGATTAGGTACCGATACATAAGANNNAGATCG |
| 14_1 | TATGAGTGTGGAGTCGTTGCTCTCTAGGCTTTTCCCCTTTGGTNNNAGATCGGAAGA |
| 32_1 | TATGAGTGTGGAGTCGTTGCTCCCAAAGGACATACACTAAAATGTNNNAGATCGGA |
| 47_1 | TATGAGTGTGGAGTCGTTGCAGATTGTGGTGGTCATTTTACAATGTNNNAGATCGG |
| 68_1 | TATGAGTGTGGAGTCGTTGCCCACCCAATGCACATACCTGTTTCNNNAGATCGGAAG |
| 98_1 | TATGAGTGTGGAGTCGTTGCAGGATTCCAAACTACTCCAAACACANNNAGATCGGA |
| 101_1 | TATGAGTGTGGAGTCGTTGCTATCCCCTACCACATCCAACACANNNAGATCGGAAGA |
| 119_1 | TATGAGTGTGGAGTCGTTGCCACACCCTGCTGTAATATCTGATGTNNNAGATCGGA |
| 124_1 | TATGAGTGTGGAGTCGTTGCGGTTCTCATTGCAGGCAAATATGTNNNAGATCGGAA |
| 134_1 | TATGAGTGTGGAGTCGTTGCTTGGTCACTTTGTCACATTCTACTGTNNNAGATCGGA |
| 159_1 | TATGAGTGTGGAGTCGTTGCTCCCACTATTCCATGAAACTACACANNNAGATCGGAA |
| 161_1 | TATGAGTGTGGAGTCGTTGCGTCTACTTAGTTGTGGAGTTCGTGTNNNAGATCGGA |
| 181_1 | TATGAGTGTGGAGTCGTTGCCCTCCAAATTGTGAAATTACCCACANNNAGATCGGA |

| | |
|---|---|
| 198_1 | TATGAGTGTGGAGTCGTTGCTTTCCCCTAAGGATTACCCCTGTNNNAGATCGGAAG |
| 202_1 | TATGAGTGTGGAGTCGTTGCAAGGGAGAGAAATGAGAGCCTGTNNNAGATCGGA |
| 205_1 | TATGAGTGTGGAGTCGTTGCTCCTCCCTTCTATATTTATGCCTGTNNNAGATCGGAA |
| 216_1 | TATGAGTGTGGAGTCGTTGCTGGTTCTCAACAAGATGAAGCACANNNAGATCGGA |
| 251_1 | TATGAGTGTGGAGTCGTTGCTGAAGCAGTTTCCTCTTATCCCATANNNAGATCGGAA |
| 308_1 | TATGAGTGTGGAGTCGTTGCTGAACTGCAGAGTAGCATAAAGACANNNAGATCGG |
| 370_1 | TATGAGTGTGGAGTCGTTGCTTTCCCTGATCTAACCCAGGTGTNNNAGATCGGAAG |
| 373_1 | TATGAGTGTGGAGTCGTTGCGCTTGGTTTCCCTGAATAGGACANNNAGATCGGAAG |
| 382_1 | TATGAGTGTGGAGTCGTTGCTGGGAACTTTATCTTGTTGGGTGTNNNAGATCGGA |
| 397_1 | TATGAGTGTGGAGTCGTTGCACACAGCTACCTGAAAGATGACANNNAGATCGGAA |
| 431_1 | TATGAGTGTGGAGTCGTTGCAGTTCTGACCCATACCCAAAATACANNNAGATCGGA |
| 444_1 | TATGAGTGTGGAGTCGTTGCACAGAATGTAGAGCATTGGTTGTGTNNNAGATCGG |
| 457_1 | TATGAGTGTGGAGTCGTTGCGAGGGTGTATGACCTTTGCGATATTNNNAGATCGG |
| 462_1 | TATGAGTGTGGAGTCGTTGCCCTCTTTTTCATCCTGCCTGATCAGNNNAGATCGGAA |
| 526_1 | TATGAGTGTGGAGTCGTTGCACAAACCAATCCTTCCCCTTTTTCANNNAGATCGGAA |
| 530_1 | TATGAGTGTGGAGTCGTTGCTGTCAAGATGCTACAATGGAAACCANNNAGATCGG |
| 533_1 | TATGAGTGTGGAGTCGTTGCGATGAGAAGGCCTAAAGCAACCANNNAGATCGGAA |
| 546_1 | TATGAGTGTGGAGTCGTTGCATGTTTATGACAAGAGCCTATTTGTNNNAGATCGGA |
| 564_1 | TATGAGTGTGGAGTCGTTGCTCCCAAACCCTGTTTTATTCTTTGTNNNAGATCGGAA |
| 574_1 | TATGAGTGTGGAGTCGTTGCACATCTCTCTCTCTTGAACTCACCANNNAGATCGGAA |
| 587_1 | TATGAGTGTGGAGTCGTTGCTGCTGTGGCTACTATACAGTTTTGTNNNAGATCGGA |
| 589_1 | TATGAGTGTGGAGTCGTTGCAGAGAGAGGAGAGATACTGAGACCANNNAGATCG |
| 591_1 | TATGAGTGTGGAGTCGTTGCACATACACACACACATCATAGACCANNNAGATCGGA |
| 592_1 | TATGAGTGTGGAGTCGTTGCAATGTCCTTTGGTGTAGATTTTTGTNNNAGATCGGA |
| 629_1 | TATGAGTGTGGAGTCGTTGCTGAAAGAATGGTGTTGAAAGTACCANNNAGATCGG |
| 631_1 | TATGAGTGTGGAGTCGTTGCTTGTAGTGAAACTGCAGAGTACCANNNAGATCGGA |
| 652_1 | TATGAGTGTGGAGTCGTTGCAAAGGATTGGAAGAAAATACACCCANNNAGATCGG |
| 655_1 | TATGAGTGTGGAGTCGTTGCTTGCCCCATCTCAGTCTACTATCTGNNNAGATCGGAA |
| 681_1 | TATGAGTGTGGAGTCGTTGCAGAGTAGATTGGAGTAGGAAGAGGTNNNAGATCG |
| 713_1 | TATGAGTGTGGAGTCGTTGCGAGAAGTCTAAGATCAAAGGCCCANNNAGATCGGA |
| 733_1 | TATGAGTGTGGAGTCGTTGCCGCCTACATGTTCTGGCATCATTNNNAGATCGGAAG |
| 740_1 | TATGAGTGTGGAGTCGTTGCACAGACAGTGGTACACATATCCCANNNAGATCGGAA |
| 741_1 | TATGAGTGTGGAGTCGTTGCAATGAGAGATGTTTTGGATATCCCANNNAGATCGG |
| 753_1 | TATGAGTGTGGAGTCGTTGCAGGCTACTTTCAGATCATAGTCATTNNNAGATCGGA |
| 776_1 | TATGAGTGTGGAGTCGTTGCGCCATTGTCCTACAGAATCACGANNNAGATCGGAAG |
| 790_1 | TATGAGTGTGGAGTCGTTGCAATGAAGAAGCTGAGGTATAGCCACNNNAGATCGG |
| 791_1 | TATGAGTGTGGAGTCGTTGCTAAGACCCCTCCTAAAAAGTGCCATNNNAGATCGGA |
| 804_1 | TATGAGTGTGGAGTCGTTGCTTGCGTGAAGGTTTGTTGGGATTNNNAGATCGGAA |
| 807_1 | TATGAGTGTGGAGTCGTTGCAGTCAAATGGTCTGTCAATAAGCCANNNAGATCGGA |
| 824_1 | TATGAGTGTGGAGTCGTTGCGTTTGTTTTCTTTGGTCTTCCCTCCNNNAGATCGGAA |
| 845_1 | TATGAGTGTGGAGTCGTTGCTGAAACAATGCTTAGGCATACTATTNNNAGATCGGA |
| 859_1 | TATGAGTGTGGAGTCGTTGCCCTTGGCTAAAAGGGACCAACATAANNNAGATCGGA |
| 893_1 | TATGAGTGTGGAGTCGTTGCCGTACACACACACGTATATGCCANNNAGATCGGAAG |

| | |
|---|---|
| 897_1 | TATGAGTGTGGAGTCGTTGCTTGAAATGAAGCTGAGACTGCCANNNAGATCGGAA |
| 898_1 | TATGAGTGTGGAGTCGTTGCAAAGCAGGATGCAAAGGGAACTTNNNAGATCGGAA |
| 904_1 | TATGAGTGTGGAGTCGTTGCAAAGTCTCACTCTGTCCCCACTTNNNAGATCGGAAGA |
| 961_1 | TATGAGTGTGGAGTCGTTGCTCCAGGGTGTCTTTCCATATCCANNNAGATCGGAAG |
| 970_1 | TATGAGTGTGGAGTCGTTGCACAACTTCATCGGGTAACTTATCCANNNAGATCGGA |
| 973_1 | TATGAGTGTGGAGTCGTTGCGCCCCTTGAACTGAAAAGCACTATCNNNAGATCGGA |
| 989_1 | TATGAGTGTGGAGTCGTTGCTCACCATGCTTTACATTACCTCCANNNAGATCGGAAG |
| 990_1 | TATGAGTGTGGAGTCGTTGCAAACTGATAAAGCAAACCCTCCANNNAGATCGGAAG |
| 1014_1 | TATGAGTGTGGAGTCGTTGCTCAGGGTCATAAAGTTGTTCTCCANNNAGATCGGAA |
| 1043_1 | TATGAGTGTGGAGTCGTTGCGTCATAAACACAGGAGTTCAGCTATNNNAGATCGGA |
| 1045_1 | TATGAGTGTGGAGTCGTTGCGTATGCTTCCAGAAGAGATCAGCATNNNAGATCGG |
| 1057_1 | TATGAGTGTGGAGTCGTTGCAGTTTTTACCTGTTCTTGACTTCCANNNAGATCGGAA |
| 1079_1 | TATGAGTGTGGAGTCGTTGCTCAGGAGGCTGTACATATAAGTCCGNNNAGATCGG |
| 1088_1 | TATGAGTGTGGAGTCGTTGCGGCTTTCTTTCCTTTTCTGCCATGNNNAGATCGGAAG |
| 1092_1 | TATGAGTGTGGAGTCGTTGCCCCACAGACACATACGTATTTTCCANNNAGATCGGAA |
| 1099_1 | TATGAGTGTGGAGTCGTTGCACCAGAAGAAAAATTGCTAAAAGCANNNAGATCGG |
| 1115_1 | TATGAGTGTGGAGTCGTTGCTGGAAATGTAATTGCTGGACCACANNNAGATCGGA |
| 1119_1 | TATGAGTGTGGAGTCGTTGCTAGACATGTAACTGAAAGACCAGCANNNAGATCGG |
| 1122_1 | TATGAGTGTGGAGTCGTTGCTAACTTCACAAAAGGACCAAGCANNNAGATCGGAAG |
| 1125_1 | TATGAGTGTGGAGTCGTTGCACCAAGTCTGGAATACAGTGATGCANNNAGATCGG |
| 1160_1 | TATGAGTGTGGAGTCGTTGCTCCTCAGTGACATTTCTAACATCTTNNNAGATCGGAA |
| 1163_1 | TATGAGTGTGGAGTCGTTGCTCCAGTGTCCATAATCATTACAGCANNNAGATCGGA |
| 1175_1 | TATGAGTGTGGAGTCGTTGCTTCAAAAATGGCATTGACGGATACANNNAGATCGG |
| 1178_1 | TATGAGTGTGGAGTCGTTGCTCAAATTGCAAAGAAAGGCAGCANNNAGATCGGAA |
| 1196_1 | TATGAGTGTGGAGTCGTTGCGTCAGCAGCCCTATCTATAATGTNNNAGATCGGAAG |
| 1212_1 | TATGAGTGTGGAGTCGTTGCGCCTCTGATCAGTAAAGAGAGCANNNAGATCGGAA |
| 1216_1 | TATGAGTGTGGAGTCGTTGCGTGCTGCAGATTCTTCCTCTCAAAGNNNAGATCGGA |
| 1228_1 | TATGAGTGTGGAGTCGTTGCGTGTATTTGTAGGAGTGCCATGAGANNNAGATCGG |
| 1241_1 | TATGAGTGTGGAGTCGTTGCGAGCTACAGCATAACTGATGAGCANNNAGATCGGA |
| 1261_1 | TATGAGTGTGGAGTCGTTGCCCTAAAAATCCTAGAGTAGGCCAGGNNNAGATCGG |
| 1275_1 | TATGAGTGTGGAGTCGTTGCCATAGCACATACAATACAGACAGTTNNNAGATCGGA |
| 1276_1 | TATGAGTGTGGAGTCGTTGCTGAATGTAAGAGTTACAGGACAGTTNNNAGATCGG |
| 1288_1 | TATGAGTGTGGAGTCGTTGCTTCCTGTTGAATCTGGACAAAGTTANNNAGATCGGA |
| 1301_1 | TATGAGTGTGGAGTCGTTGCGGGAGGTTTTCGTTTTCTAAGACTGNNNAGATCGG |
| 1305_1 | TATGAGTGTGGAGTCGTTGCGAGGGGACGATTATCTTGGCTTTTANNNAGATCGG |
| 1306_1 | TATGAGTGTGGAGTCGTTGCAACCCAAACTTTTTCTCTCCGCANNNAGATCGGAAGA |
| 1318_1 | TATGAGTGTGGAGTCGTTGCTGGGGACTTACTAGATTGTCCAAGCNNNAGATCGG |
| 1319_1 | TATGAGTGTGGAGTCGTTGCAGAGAGAGAGTAGAAAGAAAGGCANNNAGATCGG |
| 1343_1 | TATGAGTGTGGAGTCGTTGCCTTAGTGAGGAGGTAGTTAGCAAGGNNNAGATCGG |
| 1346_1 | TATGAGTGTGGAGTCGTTGCAGGATCAGAGAGGGCAGTCTCTNNNAGATCGGAAG |
| 1347_1 | TATGAGTGTGGAGTCGTTGCAACTCACTGAAGCCTCTCCTGGANNNAGATCGGAAG |
| 1348_1 | TATGAGTGTGGAGTCGTTGCATTCATCTAGAGAGAGAGAGAGGCANNNAGATCGG |
| 1362_1 | TATGAGTGTGGAGTCGTTGCTGTCCATTCCCTCATAAGACTTTCTNNNAGATCGGAA |

| | |
|---|---|
| 1370_1 | TATGAGTGTGGAGTCGTTGCCCTGAAGCTCTGTTCATTTGTTTGGNNNAGATCGGA |
| 1375_1 | TATGAGTGTGGAGTCGTTGCTCTTGATCTGGTCATCTTTTAGGCANNNAGATCGGA |
| 1378_1 | TATGAGTGTGGAGTCGTTGCATGAATTTGAGTTCTGAGAGTGGGTNNNAGATCGG |
| 1388_1 | TATGAGTGTGGAGTCGTTGCGGCTTATCCAGACTGAAAATCTCACNNNAGATCGGA |
| 1392_1 | TATGAGTGTGGAGTCGTTGCGGATGGTTTTGGTACCTGTGTAATGNNNAGATCGG |
| 1399_1 | TATGAGTGTGGAGTCGTTGCCCCAAAGAGCCGATGGTATAAATCCNNNAGATCGGA |
| 1401_1 | TATGAGTGTGGAGTCGTTGCTCTCTTTCTCTTTTCAGTTTGGGCANNNAGATCGGAA |
| 1406_1 | TATGAGTGTGGAGTCGTTGCTGGAATGTAATGGCAAAATCACAACNNNAGATCGG |
| 1412_1 | TATGAGTGTGGAGTCGTTGCTGTGTGTGTTTGGGATGGTAGAGTANNNAGATCGG |
| 1432_1 | TATGAGTGTGGAGTCGTTGCACACGAGAAACCTGTTAGATGTTNNNAGATCGGAA |
| 1443_1 | TATGAGTGTGGAGTCGTTGCAAAGTACAAGACTTGGGGGTCTATCNNNAGATCGG |
| 1453_1 | TATGAGTGTGGAGTCGTTGCAGACAAAACAGAGATAGAGTTGGCANNNAGATCGG |
| 1455_1 | TATGAGTGTGGAGTCGTTGCAGAGGAATGCAAAGCCAAAACATAANNNAGATCGG |
| 1460_1 | TATGAGTGTGGAGTCGTTGCTGCTGAAAAATACAACTCCAGGTGTNNNAGATCGGA |
| 1463_1 | TATGAGTGTGGAGTCGTTGCCTTGCTCTAGGAATTGCAAAATGCANNNAGATCGGA |
| 1477_1 | TATGAGTGTGGAGTCGTTGCACTGTATGGAATTGAACAGAATGCANNNAGATCGG |
| 1483_1 | TATGAGTGTGGAGTCGTTGCCCACAAAAGCTATGTCATGAATGCANNNAGATCGGA |
| 1492_1 | TATGAGTGTGGAGTCGTTGCGCTCACATGCTCTCAGTGAATAGTANNNAGATCGGA |
| 1493_1 | TATGAGTGTGGAGTCGTTGCGTCTGTATAACAATATGCACATGCANNNAGATCGGA |
| 1494_1 | TATGAGTGTGGAGTCGTTGCCAACCCATTTATCTGCACATGCANNNAGATCGGAAG |
| 1506_1 | TATGAGTGTGGAGTCGTTGCAGTTTTGGGTCTTTCTAGCATGCANNNAGATCGGAA |
| 1512_1 | TATGAGTGTGGAGTCGTTGCTACACAGGAGCTCCAAATCAGGANNNAGATCGGAA |
| 1513_1 | TATGAGTGTGGAGTCGTTGCATCAGACTGGCTTTGAGATTTAGCANNNAGATCGGA |
| 1519_1 | TATGAGTGTGGAGTCGTTGCAATGCAGATTGAGATCTTTGAGGAANNNAGATCGG |
| 1523_1 | TATGAGTGTGGAGTCGTTGCTGGAGGTGGCTTTATGCTTTGTTNNNAGATCGGAA |
| 1559_1 | TATGAGTGTGGAGTCGTTGCCTTAGGCACACTGGGAGAATAGATGNNNAGATCGG |
| 1565_1 | TATGAGTGTGGAGTCGTTGCTCCTCATCCTTTTCAACTCTTGGAANNNAGATCGGAA |
| 1588_1 | TATGAGTGTGGAGTCGTTGCCCTCATCTTTCCCTCTTTCTTTCAANNNAGATCGGAAG |
| 1591_1 | TATGAGTGTGGAGTCGTTGCACACTTTTGACTATGGCTTTCATTTNNNAGATCGGAA |
| 1592_1 | TATGAGTGTGGAGTCGTTGCACTGTAGACGTAGTCACTCTGCANNNAGATCGGAAG |
| 1611_1 | TATGAGTGTGGAGTCGTTGCTGCTACTTACAGGAAATGTATGAGANNNAGATCGG |
| 1633_1 | TATGAGTGTGGAGTCGTTGCTGTAAATGTCTGTGAGAGTGTGCANNNAGATCGGA |
| 1637_1 | TATGAGTGTGGAGTCGTTGCATACACTGTCAGAGACACACTTTNNNAGATCGGAAG |
| 1644_1 | TATGAGTGTGGAGTCGTTGCAGTGGGAATTTTGAGGAAATTGCANNNAGATCGGA |
| 1661_1 | TATGAGTGTGGAGTCGTTGCCTGATTTTGAATGGCCAGTAGGTGTNNNAGATCGG |
| 1676_1 | TATGAGTGTGGAGTCGTTGCTGTGGAGGTTTTTCTGAGTTGCANNNAGATCGGAA |
| 1679_1 | TATGAGTGTGGAGTCGTTGCCATTATTTGCCTGCCATGTTGCANNNAGATCGGAAG |
| 1716_1 | TATGAGTGTGGAGTCGTTGCCATCCTTCTGCATTGCATGCTTTNNNAGATCGGAAGA |
| 1718_1 | TATGAGTGTGGAGTCGTTGCGGATTGGTAGATAGGTGAATGCAGGNNNAGATCG |
| 1732_1 | TATGAGTGTGGAGTCGTTGCGCAGCTAGAGGACTGAAAACATCANNNAGATCGGA |
| 1733_1 | TATGAGTGTGGAGTCGTTGCCCCCAACCCCATCCTCTATCTTTNNNAGATCGGAAGA |
| 1742_1 | TATGAGTGTGGAGTCGTTGCAGACAGAGAGGGAAAGAGTACATCANNNAGATCG |
| 1759_1 | TATGAGTGTGGAGTCGTTGCCGGGGTAGTACCTAAGACCATTCTCNNNAGATCGGA |

| | |
|---|---|
| 1760_1 | TATGAGTGTGGAGTCGTTGCACCCAGAGAGAAGGAGGTTCTTTNNNAGATCGGAA |
| 1761_1 | TATGAGTGTGGAGTCGTTGCTGAAAATGTCTTTGATGGCTCATCANNNAGATCGGA |
| 1762_1 | TATGAGTGTGGAGTCGTTGCAATGTTTTGGCACCATGTTCTTTNNNAGATCGGAAG |
| 1778_1 | TATGAGTGTGGAGTCGTTGCTGAGTTCCCTAAATTGGCAGTTTNNNAGATCGGAAG |
| 1798_1 | TATGAGTGTGGAGTCGTTGCTGTGGCAAAACACATAACTTTATCANNNAGATCGGA |
| 1811_1 | TATGAGTGTGGAGTCGTTGCAGCCATAAGGTGAAAGACTACTTTTNNNAGATCGGA |
| 1819_1 | TATGAGTGTGGAGTCGTTGCAGAGAGAGAGAGATCATAGCACTCANNNAGATCGG |
| 1825_1 | TATGAGTGTGGAGTCGTTGCTGTATCCCTCGCCCAAAATGTTTNNNAGATCGGAAG |
| 1853_1 | TATGAGTGTGGAGTCGTTGCAAACCAAAAGTCCAACTCTAGTAACNNNAGATCGGA |
| 1861_1 | TATGAGTGTGGAGTCGTTGCAAGGTGAAAATGGAGAGCAGTGTCANNNAGATCG |
| 1956_1 | TATGAGTGTGGAGTCGTTGCTATCCCAAAGCTGCTAAAACTCAGTNNNAGATCGGA |
| 1985_1 | TATGAGTGTGGAGTCGTTGCCAGTTTCCCCTCTCAGGCTTTTTNNNAGATCGGAAGA |
| 2060_1 | TATGAGTGTGGAGTCGTTGCTCTACCATTCTTTCCTTGCTGTCANNNAGATCGGAAG |
| 2074_1 | TATGAGTGTGGAGTCGTTGCGTGTGTTTGCTTTTAGCCTTTGAANNNAGATCGGAA |
| 2109_1 | TATGAGTGTGGAGTCGTTGCTGAGTGTTCAGAGTTACTTCATTCANNNAGATCGGA |
| 2134_1 | TATGAGTGTGGAGTCGTTGCGCATATTTTCCAGAACTGTAGAGCTNNNAGATCGGA |
| 2156_1 | TATGAGTGTGGAGTCGTTGCCTGAAATAGTCCCAGACAAATGGGGNNNAGATCGG |
| 2164_1 | TATGAGTGTGGAGTCGTTGCAGTGCCTAAGGTTGCAAATCTTCANNNAGATCGGAA |
| 2180_1 | TATGAGTGTGGAGTCGTTGCACAACCCCGAAATGGAATAAGTAGGNNNAGATCGG |
| 2196_1 | TATGAGTGTGGAGTCGTTGCTGGAAGAGATTTTACTCAAGGTTCANNNAGATCGGA |
| 2203_1 | TATGAGTGTGGAGTCGTTGCTTAGTGTGGAAACTTGGGGTTCANNNAGATCGGAA |
| 2205_1 | TATGAGTGTGGAGTCGTTGCATAGAGGGGTCTTTGAGTTTTCTAANNNAGATCGGA |
| 2212_1 | TATGAGTGTGGAGTCGTTGCAGTAAAGTGAAACAATCGGCTTCACNNNAGATCGGA |
| 2241_1 | TATGAGTGTGGAGTCGTTGCGCTGTGAAGTCAAGGTCACTATTCCNNNAGATCGGA |
| 2261_1 | TATGAGTGTGGAGTCGTTGCGCACGATGTGGTAATTAGTTTTCANNNAGATCGGAA |
| 2276_1 | TATGAGTGTGGAGTCGTTGCACGCGCTGCCAAGTATAAAAAGANNNAGATCGGAA |
| 2289_1 | TATGAGTGTGGAGTCGTTGCAGCCAGAGATTAGTAAGCACAAAGANNNAGATCGG |
| 2320_1 | TATGAGTGTGGAGTCGTTGCCTTGGCTCAGTGACAGTCTCTAGTTNNNAGATCGGA |
| 2328_1 | TATGAGTGTGGAGTCGTTGCCCCTCAGCATCTTCTGTATACAAGANNNAGATCGGAA |
| 2381_1 | TATGAGTGTGGAGTCGTTGCGACCAAAATGGTAAGAGTTGGAAGANNNAGATCGG |
| 2389_1 | TATGAGTGTGGAGTCGTTGCCCAAAAAGTGAAGTCAATGTGAAGANNNAGATCGG |
| 2395_1 | TATGAGTGTGGAGTCGTTGCCAGGAAGGGTGGCATAACTAAGANNNAGATCGGAA |
| 2404_1 | TATGAGTGTGGAGTCGTTGCTGGGAAAGAAGAAAGGACAGTAAGANNNAGATCG |
| 2412_1 | TATGAGTGTGGAGTCGTTGCCACTTTTGGAGGGAGGATGTTCATANNNAGATCGG |
| 2414_1 | TATGAGTGTGGAGTCGTTGCTCAGGAAAAGAATGAGAGGCAGGTTNNNAGATCG |
| 2416_1 | TATGAGTGTGGAGTCGTTGCGAAAGTCTGAAAGACTAGGCACTGCNNNAGATCGG |
| 2421_1 | TATGAGTGTGGAGTCGTTGCTGGAGTGGGAGATTAGAAACAGANNNAGATCGGA |
| 2425_1 | TATGAGTGTGGAGTCGTTGCACCAACATACAGACAGTTCAACAGANNNAGATCGGA |
| 2428_1 | TATGAGTGTGGAGTCGTTGCCTCTTACCATGCACTGTGGCTTTATNNNAGATCGGAA |
| 2448_1 | TATGAGTGTGGAGTCGTTGCACAAGATTAGAAGCACAGACAGANNNAGATCGGAA |
| 2485_1 | TATGAGTGTGGAGTCGTTGCTCAGATGGGAACTCACAGAAGTCTGNNNAGATCGG |
| 2499_1 | TATGAGTGTGGAGTCGTTGCTGTACTTTGGAATAAAGTTGCCAGANNNAGATCGGA |
| 2527_1 | TATGAGTGTGGAGTCGTTGCTTGGGGAGTGGTAGTAGAAAACACCNNNAGATCGG |

| | |
|---|---|
| 2542_1 | TATGAGTGTGGAGTCGTTGCTTTCTGTTGTGATAGCCATGCAGANNNAGATCGGAA |
| 2557_1 | TATGAGTGTGGAGTCGTTGCCTTTAGATGGGTCTGCTTGCAGANNNAGATCGGAAG |
| 2572_1 | TATGAGTGTGGAGTCGTTGCGCCACTGAAGAGCAACCAAAATAATNNNAGATCGGA |
| 2612_1 | TATGAGTGTGGAGTCGTTGCCAGGGTGTGATAAATGGCAGACTACNNNAGATCGGA |
| 2627_1 | TATGAGTGTGGAGTCGTTGCAGAGCCTAAATTCTCATCCTCATGCNNNAGATCGGAA |
| 2662_1 | TATGAGTGTGGAGTCGTTGCTGATTTTGGACTTTCTGGTCTCCAGNNNAGATCGGAA |
| 2714_1 | TATGAGTGTGGAGTCGTTGCCCCAACCCCGAAATAATCTCTTGAANNNAGATCGGAA |
| 2730_1 | TATGAGTGTGGAGTCGTTGCGCTAACCCAATCTTACAATCAGCAANNNAGATCGGAA |
| 2754_1 | TATGAGTGTGGAGTCGTTGCTAAGGTCATGCCATCTCTGAATAGANNNAGATCGGA |
| 2781_1 | TATGAGTGTGGAGTCGTTGCAGGTGTGTAGGACGTTTTCTCTAGANNNAGATCGGA |
| 2789_1 | TATGAGTGTGGAGTCGTTGCACAAGATCCACAAGGCAATATAGCANNNAGATCGGA |
| 2791_1 | TATGAGTGTGGAGTCGTTGCGGGCTGGTACACATTTTTGCAATTTNNNAGATCGGA |
| 2799_1 | TATGAGTGTGGAGTCGTTGCACTAACAAGTTGATGACATGGATGTNNNAGATCGGA |
| 2802_1 | TATGAGTGTGGAGTCGTTGCTCTCCATGAAACATGACCTTAGGAANNNAGATCGGA |
| 2841_1 | TATGAGTGTGGAGTCGTTGCTCCTAAAGCAACCAAAGATTTACGANNNAGATCGGA |
| 2856_1 | TATGAGTGTGGAGTCGTTGCTCCACAATATCTTCAGAAATGCGANNNAGATCGGAA |
| 2857_1 | TATGAGTGTGGAGTCGTTGCAGGAGACAAAACAGACCTCCTCTTTNNNAGATCGGA |
| 2859_1 | TATGAGTGTGGAGTCGTTGCCATTTTGTCTAATGAGAGCCCCTCANNNAGATCGGAA |
| 2866_1 | TATGAGTGTGGAGTCGTTGCGATACACATGCCTACGTCTTCGANNNAGATCGGAAG |
| 2884_1 | TATGAGTGTGGAGTCGTTGCCACACACACATACATCTCTCCATGTNNNAGATCGGAA |
| 2898_1 | TATGAGTGTGGAGTCGTTGCAGCCACATCCCTTAGAATAGAAGGANNNAGATCGGA |
| 2904_1 | TATGAGTGTGGAGTCGTTGCACACACACATACATATAGCCATTGTNNNAGATCGGAA |
| 2906_1 | TATGAGTGTGGAGTCGTTGCACAGTCTTTTGTGATAGCTCCTGTTNNNAGATCGGAA |
| 2943_1 | TATGAGTGTGGAGTCGTTGCAGTCATTCAATCCTTCCACTGCAGCNNNAGATCGGAA |
| 2949_1 | TATGAGTGTGGAGTCGTTGCTCATTACCTGGGAAAATGTCAGGANNNAGATCGGAA |
| 2950_1 | TATGAGTGTGGAGTCGTTGCTACTGCAATGAAAATTTTTGGCTTGNNNAGATCGGA |
| 2956_1 | TATGAGTGTGGAGTCGTTGCATCCCTTTGTCATGTAACAGAGGANNNAGATCGGAA |
| 2958_1 | TATGAGTGTGGAGTCGTTGCTCATCTTGTTTGCCTATAGAGAGGGNNNAGATCGGA |
| 2970_1 | TATGAGTGTGGAGTCGTTGCGGAATGGAACCAAGTATCGTTATGTNNNAGATCGGA |
| 2980_1 | TATGAGTGTGGAGTCGTTGCGAAAGTTGATTTTGCCAAGGCTAATNNNAGATCGGA |
| 2991_1 | TATGAGTGTGGAGTCGTTGCAAAACAAAATCCGAGTTTTTGAGGANNNAGATCGGA |
| 3002_1 | TATGAGTGTGGAGTCGTTGCATAATACTGGTCACCTTCTTGCCTNNNAGATCGGAAG |
| 3039_1 | TATGAGTGTGGAGTCGTTGCAGGGGGAAAAATGACATATAACGGANNNAGATCGG |
| 3107_1 | TATGAGTGTGGAGTCGTTGCCTAATCGGGCAGTCTTCTGTGTTTTNNNAGATCGGA |
| 3121_1 | TATGAGTGTGGAGTCGTTGCTCTCTACAGGCACATATGCTTTAGTNNNAGATCGGAA |
| 3126_1 | TATGAGTGTGGAGTCGTTGCAAATTCAAGGCGTTCTCTCAACAANNNAGATCGGAA |
| 3176_1 | TATGAGTGTGGAGTCGTTGCAGTAGGTAAGGGCCAGATATCTTGTNNNAGATCGGA |
| 3181_1 | TATGAGTGTGGAGTCGTTGCATCCCAATACTATTTACCCAAAGGANNNAGATCGGAA |
| 3205_1 | TATGAGTGTGGAGTCGTTGCACCACAGAAATAAAAGGAACTGGGANNNAGATCGG |
| 3208_1 | TATGAGTGTGGAGTCGTTGCTCAAAGATTCCAGAAGAGGAAGAGGNNNAGATCGG |
| 3223_1 | TATGAGTGTGGAGTCGTTGCTGTTCGCTTTCTTGTTTCTGGGANNNAGATCGGAAG |
| 3241_1 | TATGAGTGTGGAGTCGTTGCGAAGGAAGCTTGGACAATGAGTACANNNAGATCGG |
| 3243_1 | TATGAGTGTGGAGTCGTTGCACTGTAGCATATAGGGTGAAATGGANNNAGATCGG |

| | |
|---|---|
| 3245_1 | TATGAGTGTGGAGTCGTTGCTCTGAAAGGAAAGGTAGAGAATGGANNNAGATCG |
| 3267_1 | TATGAGTGTGGAGTCGTTGCTGGCATGAGCTTATTTACAACTGGANNNAGATCGGA |
| 3273_1 | TATGAGTGTGGAGTCGTTGCCCTGTTTTCATTCCCATTTTTCCCANNNAGATCGGAAG |
| 3314_1 | TATGAGTGTGGAGTCGTTGCAGCAAGATCCCCTAACCTGAGANNNAGATCGGAAG |
| 3318_1 | TATGAGTGTGGAGTCGTTGCCATACCCGACTTGCTTGCTTATTTANNNAGATCGGAA |
| 3352_1 | TATGAGTGTGGAGTCGTTGCTCACACCTATACTTTGGCAACATGANNNAGATCGGA |
| 3368_1 | TATGAGTGTGGAGTCGTTGCCCTTGTGCTTCTGGTATGCATGANNNAGATCGGAAG |
| 3378_1 | TATGAGTGTGGAGTCGTTGCATGCTATCAAAACTTATGCAGATGANNNAGATCGGA |
| 3384_1 | TATGAGTGTGGAGTCGTTGCAGAGAAATATACCTACGACAGTTGTNNNAGATCGG |
| 3388_1 | TATGAGTGTGGAGTCGTTGCTCTATCACCTCAAATGACACCACATNNNAGATCGGAA |
| 3413_1 | TATGAGTGTGGAGTCGTTGCACTCTTTACTGGAATACATTGTGGTNNNAGATCGGA |
| 3416_1 | TATGAGTGTGGAGTCGTTGCCTTTCATACCAGGAAATCGAAGAAGNNNAGATCGG |
| 3427_1 | TATGAGTGTGGAGTCGTTGCTCTCTTTTTCCCATTATGTGACTGANNNAGATCGGAA |
| 3428_1 | TATGAGTGTGGAGTCGTTGCATAACTAAAAGGGGAGAAAAGCTTGNNNAGATCGG |
| 3455_1 | TATGAGTGTGGAGTCGTTGCACTGTCAATAGCCATCTTTGTTACCNNNAGATCGGAA |
| 3457_1 | TATGAGTGTGGAGTCGTTGCAGTAATCCTTACAACTCTCAGCTGANNNAGATCGGA |
| 3469_1 | TATGAGTGTGGAGTCGTTGCACTCACTGCAATCTCAAATTCATGGNNNAGATCGGA |
| 3480_1 | TATGAGTGTGGAGTCGTTGCAAGGATATGGAGGAGAAACAGGAGGNNNAGATCG |
| 3489_1 | TATGAGTGTGGAGTCGTTGCCTGTGTGTGTGTATGTGAATCCTTANNNAGATCGGA |
| 3490_1 | TATGAGTGTGGAGTCGTTGCGTCATATCATAGAAGCTGCCTCTGANNNAGATCGGA |
| 3492_1 | TATGAGTGTGGAGTCGTTGCAGTTTGGAGTGTTACATGCTAGAGTNNNAGATCGG |
| 3545_1 | TATGAGTGTGGAGTCGTTGCTCCCACTCTCTTGGCTATTACAAATNNNAGATCGGAA |
| 3549_1 | TATGAGTGTGGAGTCGTTGCTCCTGATTTTCTTCTGTGTTCCTTTNNNAGATCGGAA |
| 3567_1 | TATGAGTGTGGAGTCGTTGCTGCTTACATGATTTAGCTACTGTGANNNAGATCGGA |
| 3571_1 | TATGAGTGTGGAGTCGTTGCTTTAGTTGATGGACATTTAGGCTGGNNNAGATCGG |
| 3579_1 | TATGAGTGTGGAGTCGTTGCACAGAAAGAGTGAGAAATCGTGTGANNNAGATCGG |
| 3581_1 | TATGAGTGTGGAGTCGTTGCCTGTTACCCAGACAATGGTGTGANNNAGATCGGAA |
| 3583_1 | TATGAGTGTGGAGTCGTTGCTCTTCACATAGAACATAGTGTGTGANNNAGATCGGA |
| 3588_1 | TATGAGTGTGGAGTCGTTGCTGTTGGTAGTCTCTGGGTTGTGANNNAGATCGGAA |
| 3593_1 | TATGAGTGTGGAGTCGTTGCGGTGATGGCTTCTGTGTTATTACCTNNNAGATCGGA |
| 3606_1 | TATGAGTGTGGAGTCGTTGCATGACAGCAGCACCATTTATTGANNNAGATCGGAAG |
| 3613_1 | TATGAGTGTGGAGTCGTTGCCTGATTTGGATCAAGTGTCAAAGTGNNNAGATCGG |
| 3617_1 | TATGAGTGTGGAGTCGTTGCCCTGGCCCACAGTAGATTGTTTCNNNAGATCGGAAG |
| 3626_1 | TATGAGTGTGGAGTCGTTGCACAGCTACTTGACGTTAGTTGGANNNAGATCGGAA |
| 3639_1 | TATGAGTGTGGAGTCGTTGCTCCCTCTGCAAATTCTTAGGAATGANNNAGATCGGA |
| 3663_1 | TATGAGTGTGGAGTCGTTGCGCTGCCATTTCTGTGTTTATACAGGNNNAGATCGGA |
| 3669_1 | TATGAGTGTGGAGTCGTTGCGTGCAGATCATGGTTTTGTTGTTGTNNNAGATCGGA |
| 3681_1 | TATGAGTGTGGAGTCGTTGCTGTGTGTGTGTTCACCAAGAGATATNNNAGATCGG |
| 3702_1 | TATGAGTGTGGAGTCGTTGCTAGTATCCTCCTCAGTTCTCCCCCTNNNAGATCGGAA |
| 3704_1 | TATGAGTGTGGAGTCGTTGCTTCTTTGGGGTGTGACCATATGTTCNNNAGATCGGA |
| 3723_1 | TATGAGTGTGGAGTCGTTGCTGTCACTACAGGATGATCTAGGCTCNNNAGATCGGA |
| 3734_1 | TATGAGTGTGGAGTCGTTGCGTGGTGTGTGGCATGAAAAATCTAGNNNAGATCGG |
| 3746_1 | TATGAGTGTGGAGTCGTTGCTGAGAATCAAGCTGTTTTGTTTTGTNNNAGATCGGA |

| | |
|---|---|
| 3763_1 | TATGAGTGTGGAGTCGTTGCGTGGAAAGTTGTGGGGGTTGATANNNAGATCGGA |
| 3767_1 | TATGAGTGTGGAGTCGTTGCTCCAACCAGTGTCTTATCTTTTTGTNNNAGATCGGAA |
| 3770_1 | TATGAGTGTGGAGTCGTTGCGCGGAGAGGGAGGGATAACTATANNNAGATCGGA |
| 3814_1 | TATGAGTGTGGAGTCGTTGCGCATGTAAGTTCGCCAGAAATTTTCNNNAGATCGGA |
| 3820_1 | TATGAGTGTGGAGTCGTTGCATCCAGTAAGTGAGAAAAGATGTGANNNAGATCGG |
| 3835_1 | TATGAGTGTGGAGTCGTTGCGTCTGCACTTTCATCCAGGAGTATTNNNAGATCGGA |
| 3842_1 | TATGAGTGTGGAGTCGTTGCAGTTTCTCATGTTTGTGTGGGTGAGNNNAGATCGG |
| 3921_1 | TATGAGTGTGGAGTCGTTGCTTTTATTTGACAGAGATGCCTCAGGNNNAGATCGGA |
| 3946_1 | TATGAGTGTGGAGTCGTTGCGGCTTTCTAACACTTGGTAGGATATNNNAGATCGGA |
| 3955_1 | TATGAGTGTGGAGTCGTTGCGTGTTCCTCAAATTTTGCTTTGAACNNNAGATCGGA |
| 3957_1 | TATGAGTGTGGAGTCGTTGCATCAGAAACCATTTCACTTCAGCCANNNAGATCGGA |
| 3967_1 | TATGAGTGTGGAGTCGTTGCAATGGACAACACAGGTCAAGTCATCNNNAGATCGG |
| 3968_1 | TATGAGTGTGGAGTCGTTGCACACAGTAGTATATGATTCGGCCATNNNAGATCGGA |
| 3971_1 | TATGAGTGTGGAGTCGTTGCCCCACATGGCAAGTTTACTTTTTCTNNNAGATCGGAA |
| 3980_1 | TATGAGTGTGGAGTCGTTGCGCCTTTGGTGGAGAAGAGAGGTANNNAGATCGGA |
| 3990_1 | TATGAGTGTGGAGTCGTTGCTTGAGCATCCTGGGGCTATGAATACNNNAGATCGG |
| 3998_1 | TATGAGTGTGGAGTCGTTGCACTGTGACAACAACTTTGGGGTANNNAGATCGGAA |
| 4007_1 | TATGAGTGTGGAGTCGTTGCCCATTCAAATCTAGGAACGCTACACNNNAGATCGGA |
| 4027_1 | TATGAGTGTGGAGTCGTTGCTCCCCCAAACCATTTGAAAGTTAGANNNAGATCGGA |
| 4029_1 | TATGAGTGTGGAGTCGTTGCATGTGTGCGTTTTAGACATTAGTCCNNNAGATCGGA |
| 4037_1 | TATGAGTGTGGAGTCGTTGCTCATGGTTTTGTGTTGTAGGTACACNNNAGATCGGA |
| 4041_1 | TATGAGTGTGGAGTCGTTGCGTTAGTGTCAGAAGCCACGTGTANNNAGATCGGAA |
| 4042_1 | TATGAGTGTGGAGTCGTTGCTACAAGTCATGGCTCACTACAGCCNNNAGATCGGAA |
| 4043_1 | TATGAGTGTGGAGTCGTTGCCATCACATTGTACGCCTGGTGTANNNAGATCGGAAG |
| 4067_1 | TATGAGTGTGGAGTCGTTGCGGTGTCATCTAGGTAGTGGATTTGGNNNAGATCGG |
| 4069_1 | TATGAGTGTGGAGTCGTTGCCCTTTTCTGAGCAACTGATGACATTNNNAGATCGGA |
| 4081_1 | TATGAGTGTGGAGTCGTTGCTGTAGTTTGGTGGTATCAGTGATGANNNAGATCGG |
| 4119_1 | TATGAGTGTGGAGTCGTTGCTTTTTGTTTGTTTCCCAGTGCATTTNNNAGATCGGAA |
| 4153_1 | TATGAGTGTGGAGTCGTTGCTGATAGAAACATTTTGGCATTAGCTNNNAGATCGGA |
| 4156_1 | TATGAGTGTGGAGTCGTTGCCCTGAGCCTGGCCTTGTTATTTANNNAGATCGGAAG |
| 4162_1 | TATGAGTGTGGAGTCGTTGCTCCCAACAAAACCTTGGCCTTTANNNAGATCGGAAG |
| 4186_1 | TATGAGTGTGGAGTCGTTGCTACAAAGAGTGGTAAGTGAGCATTTNNNAGATCGG |
| 4224_1 | TATGAGTGTGGAGTCGTTGCGATTTTTCAGGTTTTTCTGGACCCTNNNAGATCGGA |
| 4235_1 | TATGAGTGTGGAGTCGTTGCGTGGAGAGAAAACTGTGTGATTCAGNNNAGATCGG |
| 4237_1 | TATGAGTGTGGAGTCGTTGCGTGTGTGGGTGTGAGTATGTGTACANNNAGATCGG |
| 4253_1 | TATGAGTGTGGAGTCGTTGCCCCTACCTACCTACCTACCTACAACNNNAGATCGGAA |
| 4268_1 | TATGAGTGTGGAGTCGTTGCAAGAGAGCGAGAGAGAGAAAGCATANNNAGATCG |
| 4284_1 | TATGAGTGTGGAGTCGTTGCTGCTAAAAAGTACATTGAGAGCCATNNNAGATCGG |
| 4288_1 | TATGAGTGTGGAGTCGTTGCATCTTATCATCTGCAAGTGGAAACANNNAGATCGGA |
| 4312_1 | TATGAGTGTGGAGTCGTTGCAGTCAATTACATACTGATGAGTGAACNNNAGATCGG |
| 4314_1 | TATGAGTGTGGAGTCGTTGCATGACAACTGGCACAAAACAAACCTNNNAGATCGGA |
| 4324_1 | TATGAGTGTGGAGTCGTTGCTATGTTGCACAGGGTGGTCTAACNNNAGATCGGAA |
| 4326_1 | TATGAGTGTGGAGTCGTTGCTCATCCAGTCAAAATGTGAAGTAACNNNAGATCGGA |

| | |
|---|---|
| 4331_1 | TATGAGTGTGGAGTCGTTGCGTCCAGTCATTCTCTCCAACACAGANNNAGATCGGAA |
| 4335_1 | TATGAGTGTGGAGTCGTTGCGTGTAACCCAAGCAGTAAATTTCAGNNNAGATCGGA |
| 4341_1 | TATGAGTGTGGAGTCGTTGCCCCAAAGCAAAACAAAACCTAACACNNNAGATCGGA |
| 4378_1 | TATGAGTGTGGAGTCGTTGCATCCTAGAAAGAGGTGATTGTCAGGNNNAGATCGG |
| 4382_1 | TATGAGTGTGGAGTCGTTGCCAGACCATGAGTACCCTTCAAAGAANNNAGATCGGA |
| 4384_1 | TATGAGTGTGGAGTCGTTGCGCTTCTCTTTCCTATGGAACCACNNNAGATCGGAAGA |
| 4396_1 | TATGAGTGTGGAGTCGTTGCGGAACTCTCTCTGAAATGTGTGCTTNNNAGATCGGA |
| 4408_1 | TATGAGTGTGGAGTCGTTGCTGATTGATGGCATTTGGAATGGTTCNNNAGATCGGA |
| 4410_1 | TATGAGTGTGGAGTCGTTGCTTGCTTACCCCTAAAAAGCCCACNNNAGATCGGAAGA |
| 4412_1 | TATGAGTGTGGAGTCGTTGCACAAAAATGGCCCATAATCCCACNNNAGATCGGAAG |
| 4414_1 | TATGAGTGTGGAGTCGTTGCGTATCCATGTGTTTGAATTACTGCCNNNAGATCGGA |
| 4422_1 | TATGAGTGTGGAGTCGTTGCGAGAAACCTGGAGGGCATTATGTTTNNNAGATCGG |
| 4472_1 | TATGAGTGTGGAGTCGTTGCGTTATCCAAGCTGAACTCAAACTCCNNNAGATCGGA |
| 4473_1 | TATGAGTGTGGAGTCGTTGCACATGCACAAAAGTGAAATAGCCTTNNNAGATCGGA |
| 4477_1 | TATGAGTGTGGAGTCGTTGCGCTTCTTGGATCTGTGAATTGAGTTNNNAGATCGGA |
| 4527_1 | TATGAGTGTGGAGTCGTTGCGTCTGTGTGTATGCCTATGTGTCTGNNNAGATCGGA |
| 4545_1 | TATGAGTGTGGAGTCGTTGCACACACACACACCAGTTGATAGCACNNNAGATCGGA |
| 4549_1 | TATGAGTGTGGAGTCGTTGCAAGAGAGAAGGAAGAGGGAACTCCANNNAGATCGG |
| 4563_1 | TATGAGTGTGGAGTCGTTGCAAGACCAACAAGGTACTATTACCATNNNAGATCGGA |
| 4565_1 | TATGAGTGTGGAGTCGTTGCTGATCTGTTTTCTACGGTATTCCATNNNAGATCGGAA |
| 4567_1 | TATGAGTGTGGAGTCGTTGCCTGACCACACCACGTAATCATCTNNNAGATCGGAAGA |
| 4570_1 | TATGAGTGTGGAGTCGTTGCCACTGCAAGGAGAGGAAAGAGACNNNAGATCGGAA |
| 4585_1 | TATGAGTGTGGAGTCGTTGCCACACCAGAGAAACAGAACCAATGTNNNAGATCGGA |
| 4592_1 | TATGAGTGTGGAGTCGTTGCTCATAGATAGATGGCAGCAGATATCNNNAGATCGGA |
| 4593_1 | TATGAGTGTGGAGTCGTTGCACTTAGAAAAAGAAACACACACGACNNNAGATCGGA |
| 4594_1 | TATGAGTGTGGAGTCGTTGCTGACTATTACCCAGTGTGTGTTTTTNNNAGATCGGAA |
| 4598_1 | TATGAGTGTGGAGTCGTTGCCAAAACACTCTTCAGAACCTCATCNNNAGATCGGAAG |
| 4617_1 | TATGAGTGTGGAGTCGTTGCGAATAGCCCAAGCAATCCTGGACNNNAGATCGGAAG |
| 4638_1 | TATGAGTGTGGAGTCGTTGCTGCCATACATCCACATATTACCGANNNAGATCGGAAG |
| 4647_1 | TATGAGTGTGGAGTCGTTGCAGGATAGTGGTGGGAACAGACAANNNAGATCGGAA |
| 4668_1 | TATGAGTGTGGAGTCGTTGCCCATCTCTTATCCCTTTGTCATGTNNNAGATCGGAAGA |
| 4678_1 | TATGAGTGTGGAGTCGTTGCAAATGCCACATTCTGGGTGGATTNNNAGATCGGAAG |
| 4680_1 | TATGAGTGTGGAGTCGTTGCTTCCTCTTTCTTGTGGGGATTTCTGNNNAGATCGGAA |
| 4697_1 | TATGAGTGTGGAGTCGTTGCGAGAGAGAGAGAAGAGAGCACTGTCNNNAGATCGG |
| 4705_1 | TATGAGTGTGGAGTCGTTGCAAGAGAAAGAAATCATGAGGAGACTNNNAGATCGG |
| 4706_1 | TATGAGTGTGGAGTCGTTGCTTTCACAACTATGTTCTTCCTCTGANNNAGATCGGAAG |
| 4714_1 | TATGAGTGTGGAGTCGTTGCATGTGTGTTTGGCAGCATAAATTGNNNAGATCGGAA |
| 4726_1 | TATGAGTGTGGAGTCGTTGCAATGCGATCTATGAAAAGAGCCANNNAGATCGGAAG |
| 4743_1 | TATGAGTGTGGAGTCGTTGCTACTGGTGTGTGTGTAGATAGTGGTNNNAGATCGGA |
| 4752_1 | TATGAGTGTGGAGTCGTTGCGTGAGTAGCTGGGATTACAAGCTTANNNAGATCGGA |
| 4763_1 | TATGAGTGTGGAGTCGTTGCGTTCCCATATTATGTCCATAGCTGCNNNAGATCGGAA |
| 4771_1 | TATGAGTGTGGAGTCGTTGCGCTACAACAAAAACCCCAGAACCNNNAGATCGGAAG |
| 4791_1 | TATGAGTGTGGAGTCGTTGCCGAAGAAACATAGCTGGAAAACACCNNNAGATCGGA |

| | |
|---|---|
| 4793_1 | TATGAGTGTGGAGTCGTTGCATTTCTTTCTCTCTCCTGGAGCANNNAGATCGGAA |
| 4798_1 | TATGAGTGTGGAGTCGTTGCATTGTCATCTGGCCCATTACACCNNNAGATCGGAAG |
| 4843_1 | TATGAGTGTGGAGTCGTTGCGTGTCAGTGTTGTAGGACTTTTTCCNNNAGATCGGA |
| 4849_1 | TATGAGTGTGGAGTCGTTGCTAGGCATTTAGACTGGTTCCACACNNNAGATCGGAA |
| 4868_1 | TATGAGTGTGGAGTCGTTGCAGACTTATAAGGGCAAATGCCATGCNNNAGATCGG |
| 4877_1 | TATGAGTGTGGAGTCGTTGCAAGAATGAATTGATGGCACTTGCAGNNNAGATCGG |
| 4888_1 | TATGAGTGTGGAGTCGTTGCGTGTTTGTGTGTTTGTGACTACAGNNNAGATCGGA |
| 4906_1 | TATGAGTGTGGAGTCGTTGCCAGCCATGAAAATGAAGGAAACCCNNNAGATCGGA |
| 4915_1 | TATGAGTGTGGAGTCGTTGCATGTGTGTGTTTGGGAGTAGAGTGTNNNAGATCGG |
| 4922_1 | TATGAGTGTGGAGTCGTTGCTGCCAAAACTTGTTTACATTCACCCNNNAGATCGGAA |
| 4939_1 | TATGAGTGTGGAGTCGTTGCCCTTGATGGACCAGAAATTATGGGGNNNAGATCGG |
| 4957_1 | TATGAGTGTGGAGTCGTTGCATACACACAAAATTAGGCGGGAGTGNNNAGATCGG |
| 4960_1 | TATGAGTGTGGAGTCGTTGCTGGTAGAAAATCAAAGGTGTATGTGNNNAGATCGG |
| 5021_1 | TATGAGTGTGGAGTCGTTGCTTGGAGTGGTCTAATACAGGGTTNNNAGATCGGAA |
| 5050_1 | TATGAGTGTGGAGTCGTTGCGTAAGGATGGGGTAACTATTCTAGCNNNAGATCGG |
| 5068_1 | TATGAGTGTGGAGTCGTTGCTGACTTGAAACAAACATTTTGGGCTNNNAGATCGGA |
| 5073_1 | TATGAGTGTGGAGTCGTTGCAAACCACATGTATGGTAGTAGTCCCNNNAGATCGGA |
| 5074_1 | TATGAGTGTGGAGTCGTTGCTGATTTGCCTCTCACTTCGAATGTNNNAGATCGGAA |
| 5100_1 | TATGAGTGTGGAGTCGTTGCAAGGGATTGTGTTTGTATGTGTGCANNNAGATCGG |
| 5138_1 | TATGAGTGTGGAGTCGTTGCTAAGTGGAAAACACAGTTCTCCTGGNNNAGATCGG |
| 5151_1 | TATGAGTGTGGAGTCGTTGCTTAGCATCAGGAGAAACAGAAACAANNNAGATCGG |
| 5171_1 | TATGAGTGTGGAGTCGTTGCAATAGGAAGCCTACAGTGTTCTTTANNNAGATCGGA |
| 5183_1 | TATGAGTGTGGAGTCGTTGCAGGTAGGTAAGTGGAGAGAGTGAGANNNAGATCG |
| 5209_1 | TATGAGTGTGGAGTCGTTGCATGCAATGATGGAAACTGGGAAGTNNNAGATCGGA |
| 5228_1 | TATGAGTGTGGAGTCGTTGCTGGCTTTGACTAGCAGTCTATACTCNNNAGATCGGA |
| 5250_1 | TATGAGTGTGGAGTCGTTGCTACAAAAGAAGCTGGGATCTGCCNNNAGATCGGAA |
| 5254_1 | TATGAGTGTGGAGTCGTTGCTGTGTAGAAGCAGAGGGATATTTCTNNNAGATCGG |
| 5293_1 | TATGAGTGTGGAGTCGTTGCACGCACACTACATTTTGTTTCATCCNNNAGATCGGAA |
| 5297_1 | TATGAGTGTGGAGTCGTTGCCCTGTGTGTATCAACAATGTGATCCNNNAGATCGGA |
| 5304_1 | TATGAGTGTGGAGTCGTTGCACACCTGACAAAGGACTTCTATCCNNNAGATCGGAA |
| 5362_1 | TATGAGTGTGGAGTCGTTGCAAGGGATGAGGCTAGAATAAAGGGANNNAGATCG |
| 5364_1 | TATGAGTGTGGAGTCGTTGCTTGTCTTCCTAGAGCCTCAGTCCNNNAGATCGGAAG |
| 5371_1 | TATGAGTGTGGAGTCGTTGCTTGTGTGTTTGTGTTTCTGTGAGTGNNNAGATCGGA |
| 5408_1 | TATGAGTGTGGAGTCGTTGCCCACACAGAGCAGTAGATCTTCCNNNAGATCGGAAG |
| 5426_1 | TATGAGTGTGGAGTCGTTGCTGTGAGTTTCTTCCCAAGAATTTCCNNNAGATCGGA |
| 5478_1 | TATGAGTGTGGAGTCGTTGCCTGTGGAACCCCAACAAATACAACTNNNAGATCGGA |
| 5480_1 | TATGAGTGTGGAGTCGTTGCTGAGAATGAACAGGTGGAGAAGCNNNAGATCGGA |
| 5508_1 | TATGAGTGTGGAGTCGTTGCCGCATAACAATGTCTTGGACAGCNNNAGATCGGAAG |
| 5510_1 | TATGAGTGTGGAGTCGTTGCTGGATGCTTTGCATTTCTACAGCNNNAGATCGGAAG |
| 5520_1 | TATGAGTGTGGAGTCGTTGCTGGTGTGTATTTCTTCTCCCAGCNNNAGATCGGAAG |
| 5534_1 | TATGAGTGTGGAGTCGTTGCCCCTAAGTTCTCTGACCGTCCTTNNNAGATCGGAAGA |
| 5550_1 | TATGAGTGTGGAGTCGTTGCAATACACCCCTCCTCTTAGCCAAAGNNNAGATCGGAA |
| 5646_1 | TATGAGTGTGGAGTCGTTGCCTTAGCCCCATGTAGAACTTATAGTNNNAGATCGGA |

| | |
|---|---|
| 5681_1 | TATGAGTGTGGAGTCGTTGCGATTTCAGGGCAAAGAACAAGGCNNNAGATCGGAA |
| 5682_1 | TATGAGTGTGGAGTCGTTGCCTACTGGTTCTGCTCCTTTAGAAANNNAGATCGGAA |
| 5742_1 | TATGAGTGTGGAGTCGTTGCCCCCAACCATGGCCTAAACCTATTANNNAGATCGGAA |
| 5762_1 | TATGAGTGTGGAGTCGTTGCAAATATACCCACAAAGGCTGGGCNNNAGATCGGAA |
| 5784_1 | TATGAGTGTGGAGTCGTTGCATGTTTAGTCTGATTGCCATGGCNNNAGATCGGAAG |
| 5831_1 | TATGAGTGTGGAGTCGTTGCCCAGGTAGTGTACATTTTACCCAGTNNNAGATCGGA |
| 5860_1 | TATGAGTGTGGAGTCGTTGCCTGCATGAATTGGAAGATCATTCTTNNNAGATCGGA |
| 5868_1 | TATGAGTGTGGAGTCGTTGCAGAGGGTACTGAATAACTCACCTCANNNAGATCGGA |
| 5881_1 | TATGAGTGTGGAGTCGTTGCCCCAAAAGACAATGGAGAAAAATCTNNNAGATCGG |
| 5890_1 | TATGAGTGTGGAGTCGTTGCCCAAGATGTTCCCTAGCAAATCAGANNNAGATCGGA |
| 5898_1 | TATGAGTGTGGAGTCGTTGCACTGCAAGGATAAACTCAAAACTGCNNNAGATCGGA |
| 5907_1 | TATGAGTGTGGAGTCGTTGCATACAACAGGCTGCTCTTACTGCNNNAGATCGGAAG |
| 5916_1 | TATGAGTGTGGAGTCGTTGCCACTCTACTCTGCAACTTCCTGCNNNAGATCGGAAGA |
| 5919_1 | TATGAGTGTGGAGTCGTTGCCACACACACATATACACCACATCTCNNNAGATCGGAA |
| 5943_1 | TATGAGTGTGGAGTCGTTGCCTTTGTATTGTATTCAGGAGGCACANNNAGATCGGA |
| 5945_1 | TATGAGTGTGGAGTCGTTGCGCCATTCATTTTGAAAAGTCTCTGCNNNAGATCGGA |
| 5952_1 | TATGAGTGTGGAGTCGTTGCATCTGCAGCCATTCTCACCTTTANNNAGATCGGAAGA |
| 5967_1 | TATGAGTGTGGAGTCGTTGCAGCCATTTGAGAATTACAAACGTGCNNNAGATCGGA |
| 5973_1 | TATGAGTGTGGAGTCGTTGCAAATGATTGTGCTACTGGTTTACTTNNNAGATCGGA |
| 5979_1 | TATGAGTGTGGAGTCGTTGCGGGACCTCCAAATAATCTAAGCAGGNNNAGATCGG |
| 6018_1 | TATGAGTGTGGAGTCGTTGCCCCACAATCCCTTAGGAGTTATTGCNNNAGATCGGA |
| 6020_1 | TATGAGTGTGGAGTCGTTGCCCACACATCTTACTCTACAACTTGCNNNAGATCGGAA |
| 6022_1 | TATGAGTGTGGAGTCGTTGCATGAATCTCCCAACCTCACTTGCNNNAGATCGGAAGA |
| 6023_1 | TATGAGTGTGGAGTCGTTGCCCTTCTCTCTCATTCTCTCACTTGCNNNAGATCGGAAG |
| 6026_1 | TATGAGTGTGGAGTCGTTGCGGGGTTTTCATATAATTTGGGGTGTNNNAGATCGG |
| 6031_1 | TATGAGTGTGGAGTCGTTGCGGCTTCATCTTTCATTGGCTTGCNNNAGATCGGAAG |
| 6032_1 | TATGAGTGTGGAGTCGTTGCGTATATGTGTGCTTGGTGCTTGCNNNAGATCGGAA |
| 6036_1 | TATGAGTGTGGAGTCGTTGCACTGCATGACAACAATGTATCTTGCNNNAGATCGGA |
| 6069_1 | TATGAGTGTGGAGTCGTTGCCAACACTTGGTCTAGTCGGTCTTNNNAGATCGGAAG |
| 6114_1 | TATGAGTGTGGAGTCGTTGCCACCTAACCAGGAAGACAGCATCNNNAGATCGGAAG |
| 6167_1 | TATGAGTGTGGAGTCGTTGCTTGATGTAGATCCTAGTGGTGATCTNNNAGATCGGA |
| 6228_1 | TATGAGTGTGGAGTCGTTGCTAATTCATGCTGCTTTCTCAACTGANNNAGATCGGAA |
| 6242_1 | TATGAGTGTGGAGTCGTTGCCCCCTGATAATCCTCATACCCCTGANNNAGATCGGAA |
| 6283_1 | TATGAGTGTGGAGTCGTTGCACTTGGGAAAATATGGGGGCAAGNNNAGATCGGA |
| 6314_1 | TATGAGTGTGGAGTCGTTGCGAAGGGTCCCTGGTCTATCTCTCNNNAGATCGGAAG |
| 6315_1 | TATGAGTGTGGAGTCGTTGCTGAACCAGGAATAATTTTGCTAGCANNNAGATCGGA |
| 6320_1 | TATGAGTGTGGAGTCGTTGCAGAAGATGAGACTAGACTGTGAGGANNNAGATCG |
| 6363_1 | TATGAGTGTGGAGTCGTTGCTCTGGGAGATTTTCTGCTCTACTCTNNNAGATCGGA |
| 6392_1 | TATGAGTGTGGAGTCGTTGCTGATTTCCAAAACTGAGCTGGTCNNNAGATCGGAAG |
| 6436_1 | TATGAGTGTGGAGTCGTTGCCCAGTGCAATCACTCATGCATTCNNNAGATCGGAAG |
| 6477_1 | TATGAGTGTGGAGTCGTTGCATGTCTACTTCAGGGTTCAGCTTTANNNAGATCGGA |
| 6484_1 | TATGAGTGTGGAGTCGTTGCGCACTTGGGTTCTGTGTTTCTTCNNNAGATCGGAAG |
| 6494_1 | TATGAGTGTGGAGTCGTTGCTCACAAGTGGAGATATTTGCAACTTNNNAGATCGGA |

| | |
|---|---|
| 6548_1 | TATGAGTGTGGAGTCGTTGCCCCTCTAAAGTCTCTGGAAATTTTCNNNAGATCGGAA |
| 6615_1 | TATGAGTGTGGAGTCGTTGCACAGGTTTTGCACCGTTTACAAGNNNAGATCGGAAG |
| 6630_1 | TATGAGTGTGGAGTCGTTGCTTCTTTCTGCATGGTCTTGTTCTGTNNNAGATCGGAA |
| 6654_1 | TATGAGTGTGGAGTCGTTGCTTCTTTCTTAGACAGGGTCTCCTGCNNNAGATCGGA |
| 6661_1 | TATGAGTGTGGAGTCGTTGCACCCTTCTTGCTGTATTTCTCTTCCNNNAGATCGGAA |
| 6667_1 | TATGAGTGTGGAGTCGTTGCAGTGCTGGAAAGGAACACAGAAGNNNAGATCGGA |
| 6687_1 | TATGAGTGTGGAGTCGTTGCTTTTCTTAGAGATAGGTCTCGCTCTNNNAGATCGGA |
| 6691_1 | TATGAGTGTGGAGTCGTTGCCTATCTCTCTCTTTCTTGGAGGCTGNNNAGATCGGAA |
| 6702_1 | TATGAGTGTGGAGTCGTTGCCTTTGGTATCAGGTTAGTGCTTGCTNNNAGATCGGA |
| 6749_1 | TATGAGTGTGGAGTCGTTGCCCTGTGTCCAGACTTTTCTCTGTTTNNNAGATCGGAA |
| 6751_1 | TATGAGTGTGGAGTCGTTGCCTTCTTGCTTTCCTTCAACTAAGTANNNAGATCGGAA |
| 6758_1 | TATGAGTGTGGAGTCGTTGCCCTTCATTCATCTGTAAGTGGGCANNNAGATCGGAA |
| 6813_1 | TATGAGTGTGGAGTCGTTGCCCACCCCCTGTGAGATACTACAGNNNAGATCGGAAG |
| 6816_1 | TATGAGTGTGGAGTCGTTGCCAGAGAGACAGAGGTGTGTATGTATNNNAGATCGG |
| 6846_1 | TATGAGTGTGGAGTCGTTGCCTCCCATAGTTGACAGAATGCCAGANNNAGATCGGA |
| 6851_1 | TATGAGTGTGGAGTCGTTGCCACTCCTTCTCTCTCTCTTCCTANNNAGATCGGAAG |
| 6867_1 | TATGAGTGTGGAGTCGTTGCGGGGCCAATAATATCAAAGACATTGNNNAGATCGG |
| 6870_1 | TATGAGTGTGGAGTCGTTGCATAAAATCACATGAGGCAATTCCTTNNNAGATCGGA |
| 6903_1 | TATGAGTGTGGAGTCGTTGCAGTGTAGCCTTGACCTGTCAATCTCNNNAGATCGGA |
| 6928_1 | TATGAGTGTGGAGTCGTTGCGCAATGTTGTGTTCACAGTCTAATTNNNAGATCGGA |
| 6940_1 | TATGAGTGTGGAGTCGTTGCCTGAGGAACAGTCATGTGAGTACATNNNAGATCGG |
| 6950_1 | TATGAGTGTGGAGTCGTTGCTCTCTCACTTCCCCATCACAGTAATNNNAGATCGGAA |
| 6969_1 | TATGAGTGTGGAGTCGTTGCGTCCTGGGGAAATGGGACATCAANNNAGATCGGAA |
| 6975_1 | TATGAGTGTGGAGTCGTTGCAACCTCGTTCTCTATACACACATACNNNAGATCGGAA |
| 6994_1 | TATGAGTGTGGAGTCGTTGCCTCTCACTCTCGTTGGTAAGGTTTANNNAGATCGGA |
| 6995_1 | TATGAGTGTGGAGTCGTTGCTTCTTTAGGCAGAACTATTGTTCAGNNNAGATCGGA |
| 7018_1 | TATGAGTGTGGAGTCGTTGCACAAAGGCCTGGAATTGGAAGAGNNNAGATCGGA |
| 7019_1 | TATGAGTGTGGAGTCGTTGCTAGCCTTGGTAACCCCATTTTTACCNNNAGATCGGAA |
| 7033_1 | TATGAGTGTGGAGTCGTTGCCCTCTCTGTTTCTCTCCAACTCCCNNNAGATCGGAAG |
| 7090_1 | TATGAGTGTGGAGTCGTTGCTGTATGGGTGTGTGTTTATCCAGTTNNNAGATCGGA |
| 7124_1 | TATGAGTGTGGAGTCGTTGCGTGCAAACTGCTTATAGGTGTGGTANNNAGATCGG |
| 7173_1 | TATGAGTGTGGAGTCGTTGCGTGCTTCCTAACCAGTCACTGAGNNNAGATCGGAAG |
| 7179_1 | TATGAGTGTGGAGTCGTTGCGTGTGTTTCTGTGCTATGAAAAACTNNNAGATCGGA |
| 7212_1 | TATGAGTGTGGAGTCGTTGCGTCTACTGAAGATGGGCCTTGAGNNNAGATCGGAA |
| 7214_1 | TATGAGTGTGGAGTCGTTGCGCTCATGCATATTTTACCCAGGCNNNAGATCGGAAG |
| 7215_1 | TATGAGTGTGGAGTCGTTGCTCTTTCAACGGACTTTTGCTCTGNNNAGATCGGAAG |
| 7226_1 | TATGAGTGTGGAGTCGTTGCCACCATCTTCCCCAAAGATCACATGNNNAGATCGGAA |
| 7236_1 | TATGAGTGTGGAGTCGTTGCTAGGGGGAGAGAGACAGGGATAGNNNAGATCGG |
| 7240_1 | TATGAGTGTGGAGTCGTTGCGTCTCACAGCCAAGTGGCTATAGNNNAGATCGGAA |
| 7276_1 | TATGAGTGTGGAGTCGTTGCTGGTGATTCTTTGTTTTCCTGCCTANNNAGATCGGAA |
| 7279_1 | TATGAGTGTGGAGTCGTTGCTGATTGTATGAACACCTGGGATCTGNNNAGATCGG |
| 7285_1 | TATGAGTGTGGAGTCGTTGCCAAGGAGATAGTTTGCAGACGAAAGNNNAGATCGG |
| 7287_1 | TATGAGTGTGGAGTCGTTGCTTTCAGTTTTGTCAGTTTCTGCTCCNNNAGATCGGAA |

| | |
|---|---|
| 7288_1 | TATGAGTGTGGAGTCGTTGCGACACAGCTCTCTTCTTATGAGGACNNNAGATCGGA |
| 7302_1 | TATGAGTGTGGAGTCGTTGCAGTGCCTCAAACTCCTCTACTAAACNNNAGATCGGAA |
| 7311_1 | TATGAGTGTGGAGTCGTTGCCCTCTACAACCATTGCTTGGGATCTNNNAGATCGGA |
| 7313_1 | TATGAGTGTGGAGTCGTTGCTGCTCTACGCCATTCGTCATTAGNNNAGATCGGAAG |
| 7327_1 | TATGAGTGTGGAGTCGTTGCTGCCCCTGTATAAGTAGTGATGTNNNAGATCGGAAG |
| 7347_1 | TATGAGTGTGGAGTCGTTGCCTCTTCCTCCCTTCTCCTGTTGATANNNAGATCGGAA |
| 7348_1 | TATGAGTGTGGAGTCGTTGCAGCAAAAAGCACAAGCCATAACGNNNAGATCGGAA |
| 7426_1 | TATGAGTGTGGAGTCGTTGCAATCCAAAATCTGCAGGGTGAACAGNNNAGATCGG |
| 7465_1 | TATGAGTGTGGAGTCGTTGCCCCAGGAATGGATTGTTTCAAAGGNNNAGATCGGA |
| 7509_1 | TATGAGTGTGGAGTCGTTGCAGGTCATGTTGCCTGTAAACAGGNNNAGATCGGAA |
| 7551_1 | TATGAGTGTGGAGTCGTTGCGCTCATGAATGTTACAGCTCAGGNNNAGATCGGAA |
| 7579_1 | TATGAGTGTGGAGTCGTTGCTCACTCACATACACACTACTGAGGTNNNAGATCGGA |
| 7581_1 | TATGAGTGTGGAGTCGTTGCTCTCCCAAAGTGTGTAGTGTTCTNNNAGATCGGAAG |
| 7634_1 | TATGAGTGTGGAGTCGTTGCACCCTTCAGACCAATAATGCTAGGNNNAGATCGGAA |
| 7655_1 | TATGAGTGTGGAGTCGTTGCGGTGGCCCAATGTATATCTATACGGNNNAGATCGGA |
| 7668_1 | TATGAGTGTGGAGTCGTTGCGAGAGAGAGAGTAGTGTTTGCGGNNNAGATCGGA |
| 7726_1 | TATGAGTGTGGAGTCGTTGCCCAGCCTGTTAGCATTTCTAGGGNNNAGATCGGAAG |
| 7764_1 | TATGAGTGTGGAGTCGTTGCTCCCTGAGAAAGGCGATATATTTTGNNNAGATCGGA |
| 7800_1 | TATGAGTGTGGAGTCGTTGCAATAGCCTTCCTAAAGGCCTGGGNNNAGATCGGAA |
| 7829_1 | TATGAGTGTGGAGTCGTTGCTGGAAATGTGATGAAGACAGTGGGNNNAGATCGG |
| 7923_1 | TATGAGTGTGGAGTCGTTGCTTCCATCAGCATGTCAGATTTATGGNNNAGATCGGA |
| 7934_1 | TATGAGTGTGGAGTCGTTGCTCATTCCTCATTTGACCCACTGGNNNAGATCGGAAGA |
| 7938_1 | TATGAGTGTGGAGTCGTTGCCAGATTTCAGAACTCATTGCACTGGNNNAGATCGGA |
| 7999_1 | TATGAGTGTGGAGTCGTTGCCCACACGTTCTGTAGCTATCTGGNNNAGATCGGAAG |
| 8028_1 | TATGAGTGTGGAGTCGTTGCTTTTAGATGATGGAGCCATGCATGCNNNAGATCGGA |
| 8040_1 | TATGAGTGTGGAGTCGTTGCAGGTTTTCACACCACATTCTAGTGGNNNAGATCGGA |
| 8056_1 | TATGAGTGTGGAGTCGTTGCAGTGAAATCTGAATAAGGTGGGTGGNNNAGATCG |
| 8072_1 | TATGAGTGTGGAGTCGTTGCCATTGGTGCAAAGCAGTATGTGGNNNAGATCGGAA |
| 8081_1 | TATGAGTGTGGAGTCGTTGCTATCTGTTCTTGGCTGCTTGTGGNNNAGATCGGAAG |
| 8084_1 | TATGAGTGTGGAGTCGTTGCGCATTGTGGGTTCTGACAATTGGNNNAGATCGGAA |
| 8124_1 | TATGAGTGTGGAGTCGTTGCTCACATGCACACACACTTTATTTGGNNNAGATCGGA |
| 8126_1 | TATGAGTGTGGAGTCGTTGCCCACATTTTGTCTGTCTGACTTTGGNNNAGATCGGA |
| 8130_1 | TATGAGTGTGGAGTCGTTGCTCCTTGCTTGCTTGTTTATGTTTGGNNNAGATCGGA |
| 8131_1 | TATGAGTGTGGAGTCGTTGCTGCCTTGTTCTATCACAATTTTGGCNNNAGATCGGAA |
| 8163_1 | TATGAGTGTGGAGTCGTTGCACACTAGCACTAACGGCAGAATGNNNAGATCGGAA |
| 8217_1 | TATGAGTGTGGAGTCGTTGCTACTCTGCAACTTCCTGCTTTTTNNNAGATCGGAAGA |
| 8219_1 | TATGAGTGTGGAGTCGTTGCGGGCAATGCATAAGTGTTCACATTTNNNAGATCGGA |
| 8220_1 | TATGAGTGTGGAGTCGTTGCACCCTTATGATTTGACATCAACAGTNNNAGATCGGA |
| 8234_1 | TATGAGTGTGGAGTCGTTGCTAAGATATGTGTGTGCGCCTATGACNNNAGATCGGA |
| 8265_1 | TATGAGTGTGGAGTCGTTGCGAGGGGCTAATATTGGCAGGATGNNNAGATCGGA |
| 8324_1 | TATGAGTGTGGAGTCGTTGCATCAAATGGGAGAGTGTAAAAAGGCNNNAGATCGG |
| 8345_1 | TATGAGTGTGGAGTCGTTGCAGTATTCATCTGTTGCCCAGACTGNNNAGATCGGAA |
| 8364_1 | TATGAGTGTGGAGTCGTTGCTCACATCAAATCCTGATCAGTACTGNNNAGATCGGA |

| | |
|---|---|
| 8437_1 | TATGAGTGTGGAGTCGTTGCCAACAGGAGAAACAGAACAAAACGANNNAGATCGG |
| 8448_1 | TATGAGTGTGGAGTCGTTGCTCAGATGGAAAGGATTATGTGGCTGNNNAGATCGG |
| 8461_1 | TATGAGTGTGGAGTCGTTGCTTGCTGCTGTTATTGTGTTGCTGNNNAGATCGGAAG |
| 8478_1 | TATGAGTGTGGAGTCGTTGCCATATCCATGGCTTCCACCTCTGNNNAGATCGGAAGA |
| 8484_1 | TATGAGTGTGGAGTCGTTGCATGTCACGCGATCAAGATCTCTGNNNAGATCGGAAG |
| 8547_1 | TATGAGTGTGGAGTCGTTGCGACAAAATTGCAAAGGCATTTCTGNNNAGATCGGAA |
| 8558_1 | TATGAGTGTGGAGTCGTTGCTCAGTCAACTTCTCTGCTTACAGTGNNNAGATCGGAA |
| 8573_1 | TATGAGTGTGGAGTCGTTGCGAAGAGATGTGGGGAAGAGAGTGNNNAGATCGGA |
| 8578_1 | TATGAGTGTGGAGTCGTTGCTCTCTCAATAGTGGTACCATGAGTGNNNAGATCGGA |
| 8579_1 | TATGAGTGTGGAGTCGTTGCACAGAAGGATGGAGAATGATGAGTGNNNAGATCGG |
| 8584_1 | TATGAGTGTGGAGTCGTTGCGGGTCACAAAAACATAGACACTGTCNNNAGATCGGA |
| 8602_1 | TATGAGTGTGGAGTCGTTGCCAGAAAAGAGAGCCAAACAGGTGNNNAGATCGGAA |
| 8603_1 | TATGAGTGTGGAGTCGTTGCAATGAAGCAGAGAACCACAGGTGNNNAGATCGGAA |
| 8626_1 | TATGAGTGTGGAGTCGTTGCCGAGATGTTGAGTTTACTTTGGGTGNNNAGATCGGA |
| 8645_1 | TATGAGTGTGGAGTCGTTGCGGTCGCTAAGTGAATTATGGATGTGNNNAGATCGGA |
| 8655_1 | TATGAGTGTGGAGTCGTTGCACACCTGCTTTATTTGAACTCTGTGNNNAGATCGGAA |
| 8664_1 | TATGAGTGTGGAGTCGTTGCTTCAGCACATCCTCATCAGTGTGNNNAGATCGGAAGA |
| 8783_1 | TATGAGTGTGGAGTCGTTGCTGGAATAGAAGTCAGTGGGGTTGNNNAGATCGGAA |
| 8795_1 | TATGAGTGTGGAGTCGTTGCTTGTCCACAGACACACTCTGTTGNNNAGATCGGAAGA |
| 8837_1 | TATGAGTGTGGAGTCGTTGCCAAGTCCCAGGCTACAGTGGTTCNNNAGATCGGAAG |
| 8838_1 | TATGAGTGTGGAGTCGTTGCCCTTTTGGCATTTTGGGCTTTTGNNNAGATCGGAAGA |
| 8851_1 | TATGAGTGTGGAGTCGTTGCGACCATAGACACATACCACTACACCNNNAGATCGGAA |
| 8868_1 | TATGAGTGTGGAGTCGTTGCACGCAAGGATTGGTCTTCCAAATNNNAGATCGGAAG |
| 8873_1 | TATGAGTGTGGAGTCGTTGCACCCTAAAACTTAGTGGCTTACAAANNNAGATCGGA |
| 8969_1 | TATGAGTGTGGAGTCGTTGCTCAGTGGATTAGAGACTAAGTAGGTNNNAGATCGGA |
| 8977_1 | TATGAGTGTGGAGTCGTTGCTCAGGTATCTCGTGTAAGTGGAGTNNNAGATCGGAA |
| 8997_1 | TATGAGTGTGGAGTCGTTGCGGTGCCAGGAATCGGGAAATAATNNNAGATCGGAA |
| 9022_1 | TATGAGTGTGGAGTCGTTGCACTGGAATATGAGCCCCTATATTTCNNNAGATCGGAA |
| 9086_1 | TATGAGTGTGGAGTCGTTGCGTAGAAACAATCCAAGGGCCCATNNNAGATCGGAAG |
| 9151_1 | TATGAGTGTGGAGTCGTTGCAAGTTGTGGCTTCCTGAATGCATNNNAGATCGGAAG |
| 9172_1 | TATGAGTGTGGAGTCGTTGCCAGGGCCTTTGTTCTTGTTGCATNNNAGATCGGAAGA |
| 9181_1 | TATGAGTGTGGAGTCGTTGCTGAAGCAAACCCTGCAAGATCATNNNAGATCGGAAG |
| 9190_1 | TATGAGTGTGGAGTCGTTGCAGTTCTCCAAAGGGGAATACATTCANNNAGATCGGA |
| 9219_1 | TATGAGTGTGGAGTCGTTGCCTGTGAACCTATTGTCAAGCTCACTNNNAGATCGGAA |
| 9244_1 | TATGAGTGTGGAGTCGTTGCTCCCAGAAGCATCGTATCCAGATNNNAGATCGGAAG |
| 9263_1 | TATGAGTGTGGAGTCGTTGCAAGTGTACTGATTCAGCTTAGAGATNNNAGATCGGA |
| 9303_1 | TATGAGTGTGGAGTCGTTGCACCTACACAATTGCAAAACTTCTCTNNNAGATCGGAA |
| 9328_1 | TATGAGTGTGGAGTCGTTGCAGTTGCATTTTTCCAAAGAAATGATNNNAGATCGGA |
| 9330_1 | TATGAGTGTGGAGTCGTTGCAGGAAAGGAGGAACGGGAATGATNNNAGATCGGA |
| 9393_1 | TATGAGTGTGGAGTCGTTGCGTTCATCATTTGGGGAAGAAAAAGTNNNAGATCGGA |
| 9403_1 | TATGAGTGTGGAGTCGTTGCAGAACCTAAATTGGATCGAGCCTTTNNNAGATCGGA |
| 9416_1 | TATGAGTGTGGAGTCGTTGCTCTGCCAAAAGAGACCTATCTTCANNNAGATCGGAAG |
| 9481_1 | TATGAGTGTGGAGTCGTTGCTTGGGCTGCTTCCATATTATTACAANNNAGATCGGAA |

9492_1 TATGAGTGTGGAGTCGTTGCTGATGGAAGAAGACGGAAGAAACTNNNAGATCGG
9498_1 TATGAGTGTGGAGTCGTTGCTTGCCCAGAAAGCATGGTAAACTNNNAGATCGGAAG
9539_1 TATGAGTGTGGAGTCGTTGCGAGCATGATGATGTGGTATCACACTNNNAGATCGG
9546_1 TATGAGTGTGGAGTCGTTGCTGCCCATTTATCTGTTGATGACACTNNNAGATCGGA
9547_1 TATGAGTGTGGAGTCGTTGCGTCATTTGGCCTATGCAATACACTNNNAGATCGGAA
9552_1 TATGAGTGTGGAGTCGTTGCCTTTGCTGGATAGAAAATAACCACTNNNAGATCGG
9556_1 TATGAGTGTGGAGTCGTTGCGCACAAACAAACCAAACATCCACNNNAGATCGGAAG
9561_1 TATGAGTGTGGAGTCGTTGCACCACATGACTTAGTAACTCCACTNNNAGATCGGAAG
9568_1 TATGAGTGTGGAGTCGTTGCACAAATACTCAGAATCAGGAGCACTNNNAGATCGGA
9571_1 TATGAGTGTGGAGTCGTTGCCTTGGTGATGAAGCTATGGCACTNNNAGATCGGAA
9581_1 TATGAGTGTGGAGTCGTTGCTGTCCCCGATATATGTCCTCACTNNNAGATCGGAAGA
9597_1 TATGAGTGTGGAGTCGTTGCTGTTTTGAATCCTGCTTCTTTCACTNNNAGATCGGAA
9624_1 TATGAGTGTGGAGTCGTTGCTCAGTTTCAGATTTTTCAAGGGACTNNNAGATCGG
9665_1 TATGAGTGTGGAGTCGTTGCCCCACAGAACCTATGAGATTCTACTNNNAGATCGGA
9668_1 TATGAGTGTGGAGTCGTTGCCCCAAACAAAACCTTTGAGAGTACTNNNAGATCGG
9705_1 TATGAGTGTGGAGTCGTTGCTGAGAGAGCAGAGAAAGAATAACCTNNNAGATCGG
9716_1 TATGAGTGTGGAGTCGTTGCGAAGGCTTGTGTGTATACCACCTNNNAGATCGGAAG
9822_1 TATGAGTGTGGAGTCGTTGCTTAGCCTTTCCAATGTGAAGCCTNNNAGATCGGAAG
9890_1 TATGAGTGTGGAGTCGTTGCTTTTTGTCCAAGTCTAGAGAATCCTNNNAGATCGGA
9930_1 TATGAGTGTGGAGTCGTTGCTTTCAACTTCTCTTTTGTTTCTCCTNNNAGATCGGAAG
9958_1 TATGAGTGTGGAGTCGTTGCTCCTTTTAGATACTCAGTAGTGGGANNNAGATCGGA
10226_1 TATGAGTGTGGAGTCGTTGCACCTGAAACTACTGAAGAATGAGCTNNNAGATCGGA
10389_1 TATGAGTGTGGAGTCGTTGCTGTGAAAGGTAAGTAGATTCAGGCTNNNAGATCGG
10455_1 TATGAGTGTGGAGTCGTTGCACTTTCCATCTTGTGTCTTCCCAAANNNAGATCGGAA
10518_1 TATGAGTGTGGAGTCGTTGCTTCCTCTTTTAGGGTCTCTGGCTNNNAGATCGGAAG
10617_1 TATGAGTGTGGAGTCGTTGCTTCATCTGACTTCAATGTTGATGCTNNNAGATCGGA
10648_1 TATGAGTGTGGAGTCGTTGCATCTTATTCCCGGATGGACTGCTNNNAGATCGGAAG
10682_1 TATGAGTGTGGAGTCGTTGCGTTTTCTAATGAGCCTGATGCATGCNNNAGATCGG
10737_1 TATGAGTGTGGAGTCGTTGCGCTGTTCTTATGTGTTTGTAGTGCTNNNAGATCGG
10856_1 TATGAGTGTGGAGTCGTTGCGTCCACCCTTATCTGTACTTTTGCTNNNAGATCGGAA
10859_1 TATGAGTGTGGAGTCGTTGCAGCCCTGGTTACAATCTTTTGCTNNNAGATCGGAAG
10880_1 TATGAGTGTGGAGTCGTTGCATCAGAAACTTCTTGGGAGGCATAGNNNAGATCGG
10937_1 TATGAGTGTGGAGTCGTTGCTGTTCCATCTAAACCACAAGCATCTNNNAGATCGGAA
10968_1 TATGAGTGTGGAGTCGTTGCGCCTGGTGCAAATAAGAGTGAGANNNAGATCGGAA
10983_1 TATGAGTGTGGAGTCGTTGCTCTAAAAATGAACTACCCCTGAACANNNAGATCGGA
10987_1 TATGAGTGTGGAGTCGTTGCCGTACATTCTATTTTGGCTCCCANNNAGATCGGAAGA
11020_1 TATGAGTGTGGAGTCGTTGCTCAAGACTGCTTCTAACCTCTGTNNNAGATCGGAAG
11059_1 TATGAGTGTGGAGTCGTTGCTGCCACAGAACAGAAAAACATTTGTNNNAGATCGGA
11066_1 TATGAGTGTGGAGTCGTTGCGTTTGTATGCTCATCCTCTGAAGACNNNAGATCGGA
11070_1 TATGAGTGTGGAGTCGTTGCACAAATACACTCTAGTGCTCAAAAANNNAGATCGG
11072_1 TATGAGTGTGGAGTCGTTGCGTTTTGTTTAGGTGGCCCACTCTNNNAGATCGGAAG
11086_1 TATGAGTGTGGAGTCGTTGCACATACACATCACACATACCATGANNNAGATCGGAA
11089_1 TATGAGTGTGGAGTCGTTGCATGCCTATAAAAGACCATCACAAAANNNAGATCGG

| | |
|---|---|
| 11114_1 | TATGAGTGTGGAGTCGTTGCACAGACTACACATGAAAAAGCCTCTNNNAGATCGGA |
| 11158_1 | TATGAGTGTGGAGTCGTTGCTGATGGGGAGGAGGAATAACACAAANNNAGATCGG |
| 11165_1 | TATGAGTGTGGAGTCGTTGCCCTAGAGTAACTGTGTGAGGTAAAANNNAGATCGG |
| 11243_1 | TATGAGTGTGGAGTCGTTGCTCTTTGAAGGTTCAGGTCTAAGTCTNNNAGATCGGA |
| 11268_1 | TATGAGTGTGGAGTCGTTGCTCTCTGACACCAAACACATCAAANNNAGATCGGAAG |
| 11284_1 | TATGAGTGTGGAGTCGTTGCTCTGATTACTGGCAACGATGTCTNNNAGATCGGAAG |
| 11286_1 | TATGAGTGTGGAGTCGTTGCTGAAAGGAAAGCAGTAAAACTGTCTNNNAGATCGG |
| 11298_1 | TATGAGTGTGGAGTCGTTGCCACAGGCACAGAAGAGACAGAAANNNAGATCGGAA |
| 11301_1 | TATGAGTGTGGAGTCGTTGCTCTATTTAGTCATGCCACCGCTCAANNNAGATCGGAA |
| 11317_1 | TATGAGTGTGGAGTCGTTGCTTCTTATTCTGCATACTGGGAGAAANNNAGATCGGA |
| 11348_1 | TATGAGTGTGGAGTCGTTGCTGTAGGTTCCGAATATTTCAGGAAANNNAGATCGGA |
| 11371_1 | TATGAGTGTGGAGTCGTTGCGAAAAATCACTGACTAAAGGCAGGTNNNAGATCGG |
| 11385_1 | TATGAGTGTGGAGTCGTTGCAGCAGTAATGGCAGTGGTATGAAANNNAGATCGGA |
| 11388_1 | TATGAGTGTGGAGTCGTTGCACCCGCCAAAATTTATGTCTTTCTCNNNAGATCGGAA |
| 11415_1 | TATGAGTGTGGAGTCGTTGCACCCCACTGAGTTTTAGGAATTTCTNNNAGATCGGA |
| 11430_1 | TATGAGTGTGGAGTCGTTGCGCCTAATCGTCCCCACAAAATCTTANNNAGATCGGAA |
| 11433_1 | TATGAGTGTGGAGTCGTTGCGCGTATACATTTTTCACCTCTTTCTNNNAGATCGGAA |
| 11447_1 | TATGAGTGTGGAGTCGTTGCTGTATGCCTGGAGAAACAAACAANNNAGATCGGAA |
| 11471_1 | TATGAGTGTGGAGTCGTTGCAGCAACAACAATAATGTCCTAACAANNNAGATCGGA |
| 11494_1 | TATGAGTGTGGAGTCGTTGCTGCTCATCTTAGGGACAAGTAAGAANNNAGATCGG |
| 11495_1 | TATGAGTGTGGAGTCGTTGCTGAGCCATAGAAAGCCATAGAAAGTNNNAGATCGG |
| 11508_1 | TATGAGTGTGGAGTCGTTGCCCCGCAAAAGTTTCTCCTACCTTGGNNNAGATCGGA |
| 11520_1 | TATGAGTGTGGAGTCGTTGCTCCCAAACCTCATGCTTTGACAANNNAGATCGGAAG |
| 11523_1 | TATGAGTGTGGAGTCGTTGCTGTTAGAAGGTGTAAAAGGCTACAANNNAGATCGG |
| 11536_1 | TATGAGTGTGGAGTCGTTGCAAGCCATTTACATGCGAGGAAGTGANNNAGATCGG |
| 11541_1 | TATGAGTGTGGAGTCGTTGCGCAATCAAACAGCAGAACACCAANNNAGATCGGAA |
| 11542_1 | TATGAGTGTGGAGTCGTTGCACTCTCATTCCACAAAGCAGAAGTNNNAGATCGGAA |
| 11544_1 | TATGAGTGTGGAGTCGTTGCTCTTCTTTTCATAGCCAGCAGAAGTNNNAGATCGGA |
| 11545_1 | TATGAGTGTGGAGTCGTTGCATGAATGCAGGTTTAGAGAAATGGTNNNAGATCGG |
| 11568_1 | TATGAGTGTGGAGTCGTTGCCCAGAGGTGGGATTTCTTTAGAATGNNNAGATCGG |
| 11574_1 | TATGAGTGTGGAGTCGTTGCCAGGGGAGGTGCTAAGAATATAAGTNNNAGATCGG |
| 11581_1 | TATGAGTGTGGAGTCGTTGCAAAGGAAGATGGAAAGCAGCCAANNNAGATCGGA |
| 11588_1 | TATGAGTGTGGAGTCGTTGCTTGTTGCTGAACCATTCAACAGTNNNAGATCGGAAG |
| 11589_1 | TATGAGTGTGGAGTCGTTGCTTTCTTCCAGATAGGGTACACTTCTNNNAGATCGGA |
| 11594_1 | TATGAGTGTGGAGTCGTTGCCATTGTTGGAAACTGAGTGACCCAANNNAGATCGG |
| 11606_1 | TATGAGTGTGGAGTCGTTGCGGACTTGTATCCAGATTGAGACAGTNNNAGATCGG |
| 11612_1 | TATGAGTGTGGAGTCGTTGCTGTGTGTGCATGTATGTATGACAGTNNNAGATCGG |
| 11613_1 | TATGAGTGTGGAGTCGTTGCTGTATTAGGCCATAGGGGTCCAANNNAGATCGGAA |
| 11617_1 | TATGAGTGTGGAGTCGTTGCACAACTCAGAGGATGAAGATACAGTNNNAGATCGG |
| 11638_1 | TATGAGTGTGGAGTCGTTGCTACTAGACACCATGCTTCCATCTTGNNNAGATCGGAA |
| 11680_1 | TATGAGTGTGGAGTCGTTGCTCTTTTTGATGGGTATTCTGGCAGTNNNAGATCGGA |
| 11706_1 | TATGAGTGTGGAGTCGTTGCTTGGTATGGGCTAAGAGCTCAGTNNNAGATCGGAA |
| 11707_1 | TATGAGTGTGGAGTCGTTGCCCCTTATGCTTGCCAAATTGCAANNNAGATCGGAAG |

| | |
|---|---|
| 11719_1 | TATGAGTGTGGAGTCGTTGCGGATGGGATCAAGAGGCAAAGAATTNNNAGATCGG |
| 11727_1 | TATGAGTGTGGAGTCGTTGCCCCTCTCAATGAATAACTGTTCAGTNNNAGATCGGAA |
| 11740_1 | TATGAGTGTGGAGTCGTTGCTGACACCTGAGTTGGAGAAGAGTNNNAGATCGGAA |
| 11744_1 | TATGAGTGTGGAGTCGTTGCGTATCCATCACCCTAGAAGTTCCTTNNNAGATCGGAA |
| 11764_1 | TATGAGTGTGGAGTCGTTGCAGACACAACCTCTGAATAATAGAGTNNNAGATCGGA |
| 11786_1 | TATGAGTGTGGAGTCGTTGCGTTAGAGTACAGTGGCAAGACCTCGNNNAGATCGGA |
| 11788_1 | TATGAGTGTGGAGTCGTTGCACACATTAGAGGGAAATTTGTTCAANNNAGATCGGA |
| 11811_1 | TATGAGTGTGGAGTCGTTGCTCCATGTCTTATTTCCAGTTTGGGNNNAGATCGGAAG |
| 11842_1 | TATGAGTGTGGAGTCGTTGCTAACAAAAGCCCACGAAGAGTTTTCNNNAGATCGGA |
| 11843_1 | TATGAGTGTGGAGTCGTTGCCCAGACCCTAGCCATCTGATACANNNAGATCGGAAGA |
| 11844_1 | TATGAGTGTGGAGTCGTTGCGGAAAGGCTGAAGGAAACCAGAANNNAGATCGGAA |
| 11846_1 | TATGAGTGTGGAGTCGTTGCCCATTTGTCTACCCACCTATCCACCNNNAGATCGGAAG |
| 11855_1 | TATGAGTGTGGAGTCGTTGCCCTTGGAACCAAACTGTCCAGAANNNAGATCGGAAG |
| 11866_1 | TATGAGTGTGGAGTCGTTGCTGGGAATAAAGACTGGTACAGAGACNNNAGATCGG |
| 11879_1 | TATGAGTGTGGAGTCGTTGCACTCTGGGTATGTAGCTCATCTAGTNNNAGATCGGA |
| 11916_1 | TATGAGTGTGGAGTCGTTGCTGCCATCTATATTTGAAAGAAACGTNNNAGATCGGA |
| 11918_1 | TATGAGTGTGGAGTCGTTGCAACAGTCAGAATTGGATTACAGCCTNNNAGATCGGA |
| 11919_1 | TATGAGTGTGGAGTCGTTGCTGGATCGCTGTAGAGTAGCATTGATNNNAGATCGGA |
| 11928_1 | TATGAGTGTGGAGTCGTTGCTAAGGATTCCAAACTACTCCAAACANNNAGATCGGAA |
| 11944_1 | TATGAGTGTGGAGTCGTTGCCGTATCTTTGAAAACCAATCACCGTNNNAGATCGGAA |
| 11945_1 | TATGAGTGTGGAGTCGTTGCCTATCCTGTGCTTTCGTTACCGTNNNAGATCGGAAGA |
| 11957_1 | TATGAGTGTGGAGTCGTTGCGTGTGTATGTGCATGCCTAGGAANNNAGATCGGAAG |
| 12015_1 | TATGAGTGTGGAGTCGTTGCACAGGAGAGAGTATAACTTCAAGGTNNNAGATCGGA |
| 12018_1 | TATGAGTGTGGAGTCGTTGCAGAATCATCTGGATTTGTAGAAGGTNNNAGATCGGA |
| 12024_1 | TATGAGTGTGGAGTCGTTGCGGACATCATTTTCTTTCTTCAACGCNNNAGATCGGAA |
| 12026_1 | TATGAGTGTGGAGTCGTTGCCCAAAGCCAAATGAAAAGATAAGGTNNNAGATCGGA |
| 12027_1 | TATGAGTGTGGAGTCGTTGCACTGGAGATGTGGCTATTCTAAGGTNNNAGATCGGA |
| 12049_1 | TATGAGTGTGGAGTCGTTGCTTTCTTTGTACGATGCCATGTTCTGNNNAGATCGGAA |
| 12063_1 | TATGAGTGTGGAGTCGTTGCTCTGAAATCACTCGGAATGTACAGTNNNAGATCGGA |
| 12072_1 | TATGAGTGTGGAGTCGTTGCGGTTGCTATACAGGACAAGAGGTNNNAGATCGGAA |
| 12080_1 | TATGAGTGTGGAGTCGTTGCAGAAGAAACAGGAGTTCAGAGAGGTNNNAGATCGG |
| 12106_1 | TATGAGTGTGGAGTCGTTGCAGTGCTCATGACAGAAATAATGAGGTNNNAGATCGG |
| 12109_1 | TATGAGTGTGGAGTCGTTGCGTTTGCTACTGGATTCACTGAGGTNNNAGATCGGAA |
| 12114_1 | TATGAGTGTGGAGTCGTTGCTTTCCAAGTCTCCTTTACCCTGTGANNNAGATCGGAAG |
| 12124_1 | TATGAGTGTGGAGTCGTTGCAGTTATGACCTGGATACACATAGGTNNNAGATCGGA |
| 12184_1 | TATGAGTGTGGAGTCGTTGCGTCTCTCTAGTTTCCAGTTGACACCNNNAGATCGGAA |
| 12282_1 | TATGAGTGTGGAGTCGTTGCCCCGACTTTCCTTTTATCCAAAGAGNNNAGATCGGAA |
| 12310_1 | TATGAGTGTGGAGTCGTTGCTGAACAGCTTATACTTTGTGATGGTNNNAGATCGGA |
| 12311_1 | TATGAGTGTGGAGTCGTTGCGCACCTTGAGCCTTCCTCATTATGANNNAGATCGGAA |
| 12325_1 | TATGAGTGTGGAGTCGTTGCAGACACAAAGAAGGGAAACTGGTNNNAGATCGGAA |
| 12329_1 | TATGAGTGTGGAGTCGTTGCACAGTGAGAGTTACAAAACACTGGTNNNAGATCGGA |
| 12347_1 | TATGAGTGTGGAGTCGTTGCGATGGTCAAATCAAATTACTCTGGTNNNAGATCGGA |
| 12354_1 | TATGAGTGTGGAGTCGTTGCTGGTTGTGTATGTGTGCTAAAAACANNNAGATCGGA |

| | |
|---|---|
| 12398_1 | TATGAGTGTGGAGTCGTTGCGGACAATGTGGTTAGGGGTTGTTNNNAGATCGGAA |
| 12410_1 | TATGAGTGTGGAGTCGTTGCTTTTGAGATTTCTCACCTCCTTGGTNNNAGATCGGAA |
| 12413_1 | TATGAGTGTGGAGTCGTTGCACTGTCTTTGGATCACAGATAAACANNNAGATCGGA |
| 53662_1 | TATGAGTGTGGAGTCGTTGCAGTTGAATTTAGATGATCCTGCATTNNNAGATCGGA |
| 53820_1 | TATGAGTGTGGAGTCGTTGCGGCTCAGGTTATTCTCTCACCTCNNNAGATCGGAAG |
| 53884_1 | TATGAGTGTGGAGTCGTTGCACGTATATGTATGTGTGTGTGTGNNNAGATCGG |
| 53958_1 | TATGAGTGTGGAGTCGTTGCACAAAGGATAACAACCATGTGATCANNNAGATCGG |
| 54118_1 | TATGAGTGTGGAGTCGTTGCGCCTACAATCCACTAGTACTTTTATTNNNAGATCGGA |
| 54171_1 | TATGAGTGTGGAGTCGTTGCAGATTTCAGCAGTTTCATCTTGTGTNNNAGATCGGA |
| 54224_1 | TATGAGTGTGGAGTCGTTGCTTCAAGTGATTCTCCTGCCTCAGNNNAGATCGGAAG |
| 54333_1 | TATGAGTGTGGAGTCGTTGCCCGAGTAGCTGGGATTACAGATGNNNAGATCGGAA |
| 54354_1 | TATGAGTGTGGAGTCGTTGCAGGTTCAAGTGATTCTCTTGCCTNNNAGATCGGAAG |
| 54366_1 | TATGAGTGTGGAGTCGTTGCTCTCCTGGGTTCAAGTGATTCTCNNNAGATCGGAAG |
| 54367_1 | TATGAGTGTGGAGTCGTTGCGAGATAGGACCTTGCTCTACTGCNNNAGATCGGAAG |
| 54464_1 | TATGAGTGTGGAGTCGTTGCAGCTGGACCAAGAAGATGAAAGTNNNAGATCGGAA |
| 54579_1 | TATGAGTGTGGAGTCGTTGCAGTTGCTTGAATTCAGAACAAGTCANNNAGATCGGA |
| 54596_1 | TATGAGTGTGGAGTCGTTGCTTCAAGTGATTCTCCTGCCTCAGNNNAGATCGGAAG |
| 54606_1 | TATGAGTGTGGAGTCGTTGCATGCCTGTAATCCCAGCACTTTNNNAGATCGGAAGA |
| 54687_1 | TATGAGTGTGGAGTCGTTGCTGCAAACACCTAAATCTAACCATCANNNAGATCGGA |
| 54695_1 | TATGAGTGTGGAGTCGTTGCACTATGCTGAAGGAAAAATCCATCTNNNAGATCGGA |
| 54720_1 | TATGAGTGTGGAGTCGTTGCCTCATGTGATCATCCCACCTCAGNNNAGATCGGAAG |
| 54724_1 | TATGAGTGTGGAGTCGTTGCCACAATCTCGGTTGACTGCAACNNNAGATCGGAAGA |
| 54835_1 | TATGAGTGTGGAGTCGTTGCAAGTACTCAAAACTCATCATTTCCTAANNNAGATCGG |
| 54909_1 | TATGAGTGTGGAGTCGTTGCAGCATTTTCTCTGTATTGTCTTTCTNNNAGATCGGAA |
| 55046_1 | TATGAGTGTGGAGTCGTTGCCAATGGCATGATTTCAGCTCACTNNNAGATCGGAAG |
| 55151_1 | TATGAGTGTGGAGTCGTTGCTGATATCAGGGTAATATTGGCCTCANNNAGATCGGA |
| 55152_1 | TATGAGTGTGGAGTCGTTGCGGTTCTGCTACCTCAATTCTTACACNNNAGATCGGAA |
| 55174_1 | TATGAGTGTGGAGTCGTTGCTCCTAATGATTAGTGATGTAGAGCANNNAGATCGG |
| 55232_1 | TATGAGTGTGGAGTCGTTGCTGCTACATATAGGCAAATAAAGCAANNNAGATCGG |
| 55276_1 | TATGAGTGTGGAGTCGTTGCACTTCACCAATGAATAAGAAGGGGANNNAGATCGG |
| 55277_1 | TATGAGTGTGGAGTCGTTGCAGGCAATATGAGATTTTCCCTAAGTNNNAGATCGGA |
| 55433_1 | TATGAGTGTGGAGTCGTTGCAAGATGGGGTGCATTCATGTGATNNNAGATCGGAA |
| 55461_1 | TATGAGTGTGGAGTCGTTGCCTGAGTAGCTGGGACTACAGGTANNNAGATCGGAA |
| 55601_1 | TATGAGTGTGGAGTCGTTGCAACTTCTTTGAGGAACAGTGCACNNNAGATCGGAAG |
| 55640_1 | TATGAGTGTGGAGTCGTTGCTCTGAATAATCAAATCAGGGTCACTNNNAGATCGGA |
| 55681_1 | TATGAGTGTGGAGTCGTTGCAGCCATGAGACAGCAATACAAACNNNAGATCGGAA |
| 55709_1 | TATGAGTGTGGAGTCGTTGCTCAGCAAGAATTCCTCTAGATAGCANNNAGATCGGA |
| 55766_1 | TATGAGTGTGGAGTCGTTGCAGAAAGAAGAAGAAAGGCTGAATGCNNNAGATCG |
| 55803_1 | TATGAGTGTGGAGTCGTTGCCTGGAATTACAGGCATGTGCAACNNNAGATCGGAA |
| 55822_1 | TATGAGTGTGGAGTCGTTGCCCTTCCAAGTAGCTGGGATTACANNNAGATCGGAAG |
| 55835_1 | TATGAGTGTGGAGTCGTTGCTCAACAATGACCCATCTGAAAAAGANNNAGATCGGA |
| 55851_1 | TATGAGTGTGGAGTCGTTGCTTCACCATATTAGCAGAATAAGGAAAANNNAGATCG |
| 55857_1 | TATGAGTGTGGAGTCGTTGCACCTGTTGTGCTTTTATTCCAATGTNNNAGATCGGAA |

| | |
|---|---|
| 56100_1 | TATGAGTGTGGAGTCGTTGCGCAGAATGATTTTACTAATTGTGCCANNNAGATCGG |
| 56260_1 | TATGAGTGTGGAGTCGTTGCTGAATAGCTGGGTTTACAGGCATNNNAGATCGGAA |
| 56422_1 | TATGAGTGTGGAGTCGTTGCAGAATAGTCCAAACAGTACTTACCTNNNAGATCGGA |
| 56520_1 | TATGAGTGTGGAGTCGTTGCTGTTACAGATTCTGGTTTATTGGCTNNNAGATCGGA |
| 56540_1 | TATGAGTGTGGAGTCGTTGCAGCTCCACCTAAATAACTCAAGAGGNNNAGATCGGA |
| 56668_1 | TATGAGTGTGGAGTCGTTGCTGTTTTTAGTAGAGACAGGGTTTCACNNNAGATCGG |
| 56673_1 | TATGAGTGTGGAGTCGTTGCAGCTCAAAAGTTTCTACAGCTCTNNNAGATCGGAAG |
| 56750_1 | TATGAGTGTGGAGTCGTTGCCTCACTGTAGCCTCAAACTCCTTNNNAGATCGGAAGA |
| 56867_1 | TATGAGTGTGGAGTCGTTGCATTCAGGAGTTGTTTGTTGCACGNNNAGATCGGAAG |
| 57008_1 | TATGAGTGTGGAGTCGTTGCATCTCCTGGGTTCAAGCAATTCTNNNAGATCGGAAG |
| 57101_1 | TATGAGTGTGGAGTCGTTGCCTCCCGAGTAGCTGCAATTACAGNNNAGATCGGAAG |
| 57151_1 | TATGAGTGTGGAGTCGTTGCTCAGAAAAAATAGCCTCAAGAAACTNNNAGATCGGA |
| 57214_1 | TATGAGTGTGGAGTCGTTGCCCTCCCGAGTAGCTGAGACTACNNNAGATCGGAAGA |
| 57233_1 | TATGAGTGTGGAGTCGTTGCTAATTCTGCCCTTGGGACAACATNNNAGATCGGAAG |
| 57363_1 | TATGAGTGTGGAGTCGTTGCTGGTTCCTCTAAAATCTTACCTTCTNNNAGATCGGAA |
| 57382_1 | TATGAGTGTGGAGTCGTTGCCCAGATAGGTTCAGGACTTGATAGTNNNAGATCGG |
| 57419_1 | TATGAGTGTGGAGTCGTTGCACAGTGTTTCCTCTTCCAATTTTGANNNAGATCGGAA |
| 57491_1 | TATGAGTGTGGAGTCGTTGCGAAGATGGCTATCTACAGACAATGANNNAGATCGG |
| 57502_1 | TATGAGTGTGGAGTCGTTGCAGAGAGAGAGAGAGAGAGAGAGAGAGANNNAGATC |
| 57506_1 | TATGAGTGTGGAGTCGTTGCTGTTTTCCACCTTTTTCTTTTGTTCTNNNAGATCGGAA |
| 57513_1 | TATGAGTGTGGAGTCGTTGCTGGAGTTGATTTTGAAGAGAAATCANNNAGATCGG |
| 57620_1 | TATGAGTGTGGAGTCGTTGCACAATCTTGAAGAAGCAGATGTCTGNNNAGATCGGA |
| 57880_1 | TATGAGTGTGGAGTCGTTGCTGGGTTCAAACAATTCTCTTGCCNNNAGATCGGAAG |
| 57909_1 | TATGAGTGTGGAGTCGTTGCTTCAAGTGATTCTCCTGCCTCAGNNNAGATCGGAAG |
| 57934_1 | TATGAGTGTGGAGTCGTTGCTCCAATCTGTGATTTCTTGAAACGTNNNAGATCGGA |
| 58017_1 | TATGAGTGTGGAGTCGTTGCATCTTGGCTCACTACAACCTCTGNNNAGATCGGAAGA |
| 58065_1 | TATGAGTGTGGAGTCGTTGCACGTTTCTTATGATAAGGGATGGGTNNNAGATCGG |
| 58121_1 | TATGAGTGTGGAGTCGTTGCAGAGAGAGAGAGAGAGAGAGAGAGAGANNNAGATC |
| 58176_1 | TATGAGTGTGGAGTCGTTGCAAGATGGGGTGCATTCATGTGATNNNAGATCGGAA |
| 58268_1 | TATGAGTGTGGAGTCGTTGCAGGGTGATTGAAGGTAACATTTACTNNNAGATCGG |
| 58291_1 | TATGAGTGTGGAGTCGTTGCACAAATTTTCTGAGTATAGCTGTGTNNNAGATCGGA |
| 58297_1 | TATGAGTGTGGAGTCGTTGCTGCATCTTTGATCCAATTCAACACANNNAGATCGGAA |
| 58354_1 | TATGAGTGTGGAGTCGTTGCTATCTGCTTCCTGGGTTCAAACGNNNAGATCGGAAG |
| 58418_1 | TATGAGTGTGGAGTCGTTGCTCCTTCTATCCCATCTAGATGAGCTNNNAGATCGGAA |
| 58439_1 | TATGAGTGTGGAGTCGTTGCACTGATTACTCTTTCTCTTTCCTCTNNNAGATCGGAAG |
| 58464_1 | TATGAGTGTGGAGTCGTTGCAGTCTATGCCAAGTAAATCGAAGGTNNNAGATCGG |
| 58568_1 | TATGAGTGTGGAGTCGTTGCACAAGTGAACAAAGGTCTCTGGTNNNAGATCGGAA |
| 58707_1 | TATGAGTGTGGAGTCGTTGCGCTGGAGTACAGTAGCATGATCTNNNAGATCGGAA |
| 58846_1 | TATGAGTGTGGAGTCGTTGCAGAGAGAGAGAGAGAGAGAGAGAGAGAGANNNAGATC |
| 58864_1 | TATGAGTGTGGAGTCGTTGCTCCTTTAGGACAATTCCAGATTCCANNNAGATCGGAA |
| 58879_1 | TATGAGTGTGGAGTCGTTGCACCATTTCCAGTGCTTTTTATTCCTNNNAGATCGGAA |
| 58949_1 | TATGAGTGTGGAGTCGTTGCGCTATTCAACTTTGTTGGGAAAAGTNNNAGATCGGA |
| 58973_1 | TATGAGTGTGGAGTCGTTGCTCTAACTGATTTCATACAACTGCGTNNNAGATCGGAA |

| | |
|---|---|
| 59020_1 | TATGAGTGTGGAGTCGTTGCTGCCTAGGCTACTAAGATCTACTGTNNNAGATCGGA |
| 59074_1 | TATGAGTGTGGAGTCGTTGCCCCACAGGTTTTGGTAAGTTGTGNNNAGATCGGAA |
| 59151_1 | TATGAGTGTGGAGTCGTTGCGAGGTAAGAGAGATCAAGTATGGAANNNAGATCG |
| 59203_1 | TATGAGTGTGGAGTCGTTGCAATCACTGGATCATAGAAAGATGGANNNAGATCGG |
| 59405_1 | TATGAGTGTGGAGTCGTTGCTTCAAGTGATTCTCCCACCTCAGNNNAGATCGGAAG |
| 59410_1 | TATGAGTGTGGAGTCGTTGCTGGGATACAGTAAAAACAGTACTACANNNAGATCG |
| 59426_1 | TATGAGTGTGGAGTCGTTGCACACACATACATACACACATACACCNNNAGATCGGAA |
| 59588_1 | TATGAGTGTGGAGTCGTTGCGAAAGACATCCATGTTCATTTCTCTNNNAGATCGGA |
| 59596_1 | TATGAGTGTGGAGTCGTTGCACAAAAAAAACCAGTTATCTCCCCANNNAGATCGGA |
| 59926_1 | TATGAGTGTGGAGTCGTTGCACTACCCTGAAACTAAGAACTTTCANNNAGATCGGA |
| 59934_1 | TATGAGTGTGGAGTCGTTGCACACTTCCTTGTCCTCTATCATTGANNNAGATCGGAA |
| 60166_1 | TATGAGTGTGGAGTCGTTGCTGAAGCTGGATCCTTATCTTACACTNNNAGATCGGA |
| 60173_1 | TATGAGTGTGGAGTCGTTGCAACAACAAAAAAAACTAACCATCACANNNAGATCGG |
| 60200_1 | TATGAGTGTGGAGTCGTTGCCCATTATTTGACTGCTAAATTTGCTCANNNAGATCGG |
| 60204_1 | TATGAGTGTGGAGTCGTTGCAGAGTCGGAACTATATTGCTGAACTNNNAGATCGGA |
| 60295_1 | TATGAGTGTGGAGTCGTTGCCCAGTATCAAGCGATTCTCCTCANNNAGATCGGAAG |
| 61990_1 | TATGAGTGTGGAGTCGTTGCCGGCAGATGGATGGAGTCTTATTNNNAGATCGGAA |
| 62052_1 | TATGAGTGTGGAGTCGTTGCGGCTGATGTAGGAGGATCTCTTGNNNAGATCGGAA |
| 62099_1 | TATGAGTGTGGAGTCGTTGCACAGGAGAGAGGGGTATGACATANNNAGATCGGA |
| 62279_1 | TATGAGTGTGGAGTCGTTGCTGTGCCAGATGTTTGATTCATTACTNNNAGATCGGA |
| 62390_1 | TATGAGTGTGGAGTCGTTGCTGTTCCAACAACACAGAATCTTTGTNNNAGATCGGA |
| 62439_1 | TATGAGTGTGGAGTCGTTGCCTCCTGAGTAGCTGGGACTACAGNNNAGATCGGAAG |
| 62474_1 | TATGAGTGTGGAGTCGTTGCATAGCTAGGACTATAGGCATGCGNNNAGATCGGAA |
| 62497_1 | TATGAGTGTGGAGTCGTTGCCCTCCTGGCTTCAATCAGTTCTCNNNAGATCGGAAGA |
| 62560_1 | TATGAGTGTGGAGTCGTTGCCCCAATACAGTAATAGCTGGAGACANNNAGATCGGA |
| 62562_1 | TATGAGTGTGGAGTCGTTGCTTCAGTGGAAGATTTACATGTGAAANNNAGATCGG |
| 62617_1 | TATGAGTGTGGAGTCGTTGCAGTTCTGAAATGCTACCTCTAAGCANNNAGATCGGA |
| 62633_1 | TATGAGTGTGGAGTCGTTGCCCTCAAGGCACAATCTATCATATGANNNAGATCGGA |
| 62669_1 | TATGAGTGTGGAGTCGTTGCAGTCTTTGGTTTACCTGTGGTGTNNNAGATCGGAAG |
| 62707_1 | TATGAGTGTGGAGTCGTTGCTGTTTTTAGATTCTAGCCTTGTCCANNNAGATCGGAA |
| 62815_1 | TATGAGTGTGGAGTCGTTGCCAAGTAGCTGGGATTACAGGTGTNNNAGATCGGAA |
| 62881_1 | TATGAGTGTGGAGTCGTTGCATTCAAGTGATTCTCCTGCCTCANNNAGATCGGAAG |
| 62899_1 | TATGAGTGTGGAGTCGTTGCTCAGCCTCAGAAATCAGAGACACNNNAGATCGGAAG |
| 63000_1 | TATGAGTGTGGAGTCGTTGCTGGTGCATCTACATTAGGGAATACTNNNAGATCGGA |
| 63030_1 | TATGAGTGTGGAGTCGTTGCTGGGAATACAGGTAGAGATAACCACNNNAGATCGG |
| 63325_1 | TATGAGTGTGGAGTCGTTGCTCAAGCGATTTTCATGTCTCAGCNNNAGATCGGAAG |
| 63420_1 | TATGAGTGTGGAGTCGTTGCATTCTTTTCTGGCAGGCTTTTGGNNNAGATCGGAAG |
| 63422_1 | TATGAGTGTGGAGTCGTTGCTGTATGTATGTCTTTGATTGGGTGTNNNAGATCGGA |
| 63641_1 | TATGAGTGTGGAGTCGTTGCTCAGAATCACAAACATTCCTTACCANNNAGATCGGAA |
| 63856_1 | TATGAGTGTGGAGTCGTTGCTCTTTCTCTCTCTCTCATCCCTNNNAGATCGGAAGA |
| 63932_1 | TATGAGTGTGGAGTCGTTGCTGGGTTCAAAGATTCTTCTGCCTNNNAGATCGGAAG |
| 63973_1 | TATGAGTGTGGAGTCGTTGCGGTCTTACTTTATGAACCAAGTCGTNNNAGATCGGA |
| 63981_1 | TATGAGTGTGGAGTCGTTGCAGAATAACCTAACCAAAAAGCAGGTNNNAGATCGG |

| | |
|---|---|
| 64063_1 | TATGAGTGTGGAGTCGTTGCATCCAATGGAAACGACTGACTGANNNAGATCGGAA |
| 64082_1 | TATGAGTGTGGAGTCGTTGCTCCAATCCATCATTGATGAGTACCTNNNAGATCGGA |
| 64133_1 | TATGAGTGTGGAGTCGTTGCACTCTAGAACTGCTAAGACAACTGGNNNAGATCGGA |
| 64163_1 | TATGAGTGTGGAGTCGTTGCAACTGAGGTAAAGTGATCTGCCTNNNAGATCGGAA |
| 64222_1 | TATGAGTGTGGAGTCGTTGCTGATCAAATTGGCTTCTATCTGTGANNNAGATCGGA |
| 64233_1 | TATGAGTGTGGAGTCGTTGCATTACATGTTCATGGATTGGAAGAANNNAGATCGG |
| 64236_1 | TATGAGTGTGGAGTCGTTGCCAGGAGTTGAACTGAGAACAGAANNNAGATCGGAA |
| 64258_1 | TATGAGTGTGGAGTCGTTGCCCACACACACAAAAAAGAAAAACCANNNAGATCGGA |
| 64456_1 | TATGAGTGTGGAGTCGTTGCTGTAGTGGTGCAATCTTGACTCANNNAGATCGGAAG |
| 64566_1 | TATGAGTGTGGAGTCGTTGCTCATACCATGGAATACAGTACAGCANNNAGATCGGA |
| 64799_1 | TATGAGTGTGGAGTCGTTGCCCATGATGATTAGTGATGTTGAGCANNNAGATCGG |
| 64946_1 | TATGAGTGTGGAGTCGTTGCTGTCATGAAACTTGGCAGAAAATCANNNAGATCGG |
| 65131_1 | TATGAGTGTGGAGTCGTTGCTCCCTATATACCAGTCTCAGATTCTNNNAGATCGGAA |
| 65143_1 | TATGAGTGTGGAGTCGTTGCTCTCTTTCAGCTTTTCTTCTGTCANNNAGATCGGAAG |
| 65267_1 | TATGAGTGTGGAGTCGTTGCTGTATTATTGGTTAGTGAAGTCCCTNNNAGATCGGA |
| 65269_1 | TATGAGTGTGGAGTCGTTGCTCCTTTGGTTGGAATGAAAAATTGTNNNAGATCGGA |
| 65306_1 | TATGAGTGTGGAGTCGTTGCAAACATAAATGTCACATTCCCTTCTNNNAGATCGGAA |
| 65310_1 | TATGAGTGTGGAGTCGTTGCAGCATTTTCTCCACATGTTTGATCANNNAGATCGGAA |
| 65366_1 | TATGAGTGTGGAGTCGTTGCCCCATACCCATCTTGTCTAAACCANNNAGATCGGAAG |
| 65471_1 | TATGAGTGTGGAGTCGTTGCCAAGCCATAAAAAGCCGATGAGANNNAGATCGGAA |
| 65479_1 | TATGAGTGTGGAGTCGTTGCATTAGGAGAGTGAGGTTCTGGGANNNAGATCGGA |
| 65526_1 | TATGAGTGTGGAGTCGTTGCGGCTCCAAAACCTAGACTCTTCANNNAGATCGGAAG |
| 65664_1 | TATGAGTGTGGAGTCGTTGCATGGAGTCTCTCTCTGTCACCANNNAGATCGGAAGA |
| 65665_1 | TATGAGTGTGGAGTCGTTGCAAAGGCAAAGTTCCATGATAGGCNNNAGATCGGAA |
| 65696_1 | TATGAGTGTGGAGTCGTTGCAGTTTGGAGATGTAGAAAAAAGCANNNAGATCGGA |
| 65711_1 | TATGAGTGTGGAGTCGTTGCTGTTTTTCTTTTTGAGACGGAGTCTNNNAGATCGGA |
| 65766_1 | TATGAGTGTGGAGTCGTTGCACTTCCTGGGCTCAGCTACTATANNNAGATCGGAAG |
| 65825_1 | TATGAGTGTGGAGTCGTTGCACTCATTTTGGCTGAATAAAAGGCANNNAGATCGGA |
| 65826_1 | TATGAGTGTGGAGTCGTTGCTCAACCCACAAAAATCAGTTGCANNNAGATCGGAAG |
| 65870_1 | TATGAGTGTGGAGTCGTTGCGGGGTCAGGAGTTTGAGACTAACNNNAGATCGGAA |
| 65880_1 | TATGAGTGTGGAGTCGTTGCTAATACGCCATGATCAAGCAGGANNNAGATCGGAA |
| 65994_1 | TATGAGTGTGGAGTCGTTGCCAGTGGTATGATCTTGGCTCACTNNNAGATCGGAAG |
| 66065_1 | TATGAGTGTGGAGTCGTTGCTCTCTCTCCTTCTTTCTCTCTTTCTNNNAGATCGGAAG |
| 66079_1 | TATGAGTGTGGAGTCGTTGCGAGATGGAGTTTCGTTCTGTTGCNNNAGATCGGAA |
| 66101_1 | TATGAGTGTGGAGTCGTTGCAAGGCTTCATTCTGTCAAGGTGTNNNAGATCGGAAG |
| 66106_1 | TATGAGTGTGGAGTCGTTGCTCTCTCTCTCTCTCTCTCTCTCTCTNNNAGATCGGAAG |
| 66132_1 | TATGAGTGTGGAGTCGTTGCTGACAAAGATGGTATTTCAAGTGAGANNNAGATCG |
| 66187_1 | TATGAGTGTGGAGTCGTTGCTCTTCATTCTCTACGGTCACAGCNNNAGATCGGAAGA |
| 66202_1 | TATGAGTGTGGAGTCGTTGCTGCCTAATTTTGTTTGTGTGATTCANNNAGATCGGA |
| 66214_1 | TATGAGTGTGGAGTCGTTGCCTGCAGCATAGACTTTAGAGGCTNNNAGATCGGAA |
| 66251_1 | TATGAGTGTGGAGTCGTTGCTGGTCATAGTGAAAACAGAAGACCANNNAGATCGG |
| 66392_1 | TATGAGTGTGGAGTCGTTGCGGCACACATGGAATATATATATGCANNNAGATCGGA |
| 66408_1 | TATGAGTGTGGAGTCGTTGCTCCACAGTTAGAATGTAAGCATCANNNAGATCGGAA |

| | |
|---|---|
| 66429_1 | TATGAGTGTGGAGTCGTTGCAGCCAATCTTGAAAAAGTACCAAGTNNNAGATCGGA |
| 66520_1 | TATGAGTGTGGAGTCGTTGCAGAGTTGAATACATTTGGGCCTACTNNNAGATCGGA |
| 66576_1 | TATGAGTGTGGAGTCGTTGCACACCCCAATAAATCCACTGTAAGTNNNAGATCGGA |
| 66627_1 | TATGAGTGTGGAGTCGTTGCTTCTTTGTGGTTACTATGAGGCTNNNAGATCGGAAG |
| 66729_1 | TATGAGTGTGGAGTCGTTGCACACAACAATTGAGTATCTCAGTGGNNNAGATCGGA |
| 66774_1 | TATGAGTGTGGAGTCGTTGCGTGTACGTGTGTGTGTGTGTATGNNNAGATCGGAA |
| 66804_1 | TATGAGTGTGGAGTCGTTGCAGGGTAGACATTGTTTTCTCCACTNNNAGATCGGAA |
| 67071_1 | TATGAGTGTGGAGTCGTTGCTTTGCTTATTGGAGACAGGGTCTNNNAGATCGGAAG |
| 67078_1 | TATGAGTGTGGAGTCGTTGCAGTGGTCTAAAATCTCCAATCCANNNAGATCGGAAG |
| 67145_1 | TATGAGTGTGGAGTCGTTGCCACTACCTCCTGAGTTCAAGTGANNNAGATCGGAAG |
| 67178_1 | TATGAGTGTGGAGTCGTTGCCCAACTTGCCTTCACAGAAATGTNNNAGATCGGAAG |
| 67296_1 | TATGAGTGTGGAGTCGTTGCAGATGCTGGGAATAAATGAGAGCTNNNAGATCGGA |
| 67346_1 | TATGAGTGTGGAGTCGTTGCTTGTGTGCTCATTCTTACCCAAANNNAGATCGGAAG |
| 67466_1 | TATGAGTGTGGAGTCGTTGCAGCCTACAGAAACAACTGGGAAANNNAGATCGGAA |
| 67494_1 | TATGAGTGTGGAGTCGTTGCAGGTGTATCAGAATTTCGGATTCCANNNAGATCGGA |
| 67553_1 | TATGAGTGTGGAGTCGTTGCTTCCAGTACTTTCTCACACCTGGNNNAGATCGGAAG |
| 67673_1 | TATGAGTGTGGAGTCGTTGCAGAGAGAGAGAGAGAGAGAGAGAGAGANNNAGATC |
| 67760_1 | TATGAGTGTGGAGTCGTTGCACACACATACTCCAAAAAACTGAANNNAGATCGGAA |
| 67770_1 | TATGAGTGTGGAGTCGTTGCTCTCTCTCTCTCTCTCTCTCTCTCTNNNAGATCGGAAG |
| 67827_1 | TATGAGTGTGGAGTCGTTGCTCATCTGTTGATGTACACTTGGGTNNNAGATCGGAA |
| 67907_1 | TATGAGTGTGGAGTCGTTGCGTGACAGAGTCTTGCTTTTGTCGNNNAGATCGGAAG |
| 68007_1 | TATGAGTGTGGAGTCGTTGCACATACTGGTGCACTTATGTACCANNNAGATCGGAA |
| 68016_1 | TATGAGTGTGGAGTCGTTGCTCCTTCAGGTTTTCTACATGTAAGTNNNAGATCGGA |
| 68032_1 | TATGAGTGTGGAGTCGTTGCGCCTCTTGAGTAGCTGGGATTATNNNAGATCGGAA |
| 68123_1 | TATGAGTGTGGAGTCGTTGCGGTACCAGCCTTCAAACCTAAGANNNAGATCGGAAG |
| 68153_1 | TATGAGTGTGGAGTCGTTGCGTACCATTACCATCACAAATGTACANNNAGATCGGA |
| 68216_1 | TATGAGTGTGGAGTCGTTGCAGAACTTGAAGCTGTAAGAAAATGTNNNAGATCGG |
| 68222_1 | TATGAGTGTGGAGTCGTTGCACACACACACACAAACATATACACANNNAGATCGGA |
| 68282_1 | TATGAGTGTGGAGTCGTTGCGAGAGAAAAAGGCAACCACAGACNNNAGATCGGAA |
| 68310_1 | TATGAGTGTGGAGTCGTTGCTCATTGCTGTTCAATTGATGACACANNNAGATCGGA |
| 68326_1 | TATGAGTGTGGAGTCGTTGCGACAGACAGACAGAGACAGACAGNNNAGATCGGAA |
| 68364_1 | TATGAGTGTGGAGTCGTTGCAGAGAGAGAGAGAGAGAGAGAGAGAGANNNAGATC |
| 68393_1 | TATGAGTGTGGAGTCGTTGCTGTTTGGTTTGGTTTGGTTTGGTNNNAGATCGGAA |
| 68444_1 | TATGAGTGTGGAGTCGTTGCAGTGGACCAATACTTCAGGTTGANNNAGATCGGAA |
| 68449_1 | TATGAGTGTGGAGTCGTTGCAGAGAGGAAGAGAGGAAGAAAGAGANNNAGATC |
| 68477_1 | TATGAGTGTGGAGTCGTTGCAGTCCTAAAATCAAGGTATCAGCAGANNNAGATCGG |
| 68511_1 | TATGAGTGTGGAGTCGTTGCGAGAAATGGGGAAAGACTGGTCTNNNAGATCGGA |
| 68625_1 | TATGAGTGTGGAGTCGTTGCTTTATCTCACAGTAACACACATGTGNNNAGATCGGA |
| 68634_1 | TATGAGTGTGGAGTCGTTGCTGCTGTTTTCTCAAATGCGAAGATNNNAGATCGGAA |
| 68788_1 | TATGAGTGTGGAGTCGTTGCGAAGGAAGGAAGGAAGGAAGGAANNNAGATCGG |
| 68841_1 | TATGAGTGTGGAGTCGTTGCAGAGAGAGAGAGAAAGAAAGAAAGGANNNAGATC |
| 69029_1 | TATGAGTGTGGAGTCGTTGCACACACACACACACACATATATTCTNNNAGATCGGAA |
| 69050_1 | TATGAGTGTGGAGTCGTTGCTCTCTCTCTCTCTCTCTCTCTCTCTNNNAGATCGGAAG |

| | |
|---|---|
| 69059_1 | TATGAGTGTGGAGTCGTTGCAGACAGAGACAGAGACAGAGACANNNAGATCGGA |
| 69081_1 | TATGAGTGTGGAGTCGTTGCTCCTTGTACTGGTTTTTGTTGTTGTNNNAGATCGGA |
| 69088_1 | TATGAGTGTGGAGTCGTTGCTTGACATTGGTGCTATCCACAGANNNAGATCGGAAG |
| 69144_1 | TATGAGTGTGGAGTCGTTGCGGAAGGAAGGAAGGAAAGAAGGANNNAGATCGG |
| 69182_1 | TATGAGTGTGGAGTCGTTGCAGAGAGAGACAGAAAGAGAGAGCTNNNAGATCGG |
| 69191_1 | TATGAGTGTGGAGTCGTTGCACACCATAAATGATCCAGAATCCANNNAGATCGGAA |
| 69318_1 | TATGAGTGTGGAGTCGTTGCAGGAGTCATTACATAGGTGCAACANNNAGATCGGA |
| 69343_1 | TATGAGTGTGGAGTCGTTGCACCAAAACATTAGGAGTAGTTGAGTNNNAGATCGG |
| 69373_1 | TATGAGTGTGGAGTCGTTGCTCTCCCATGATACAGTTATATCCTGNNNAGATCGGAA |
| 69399_1 | TATGAGTGTGGAGTCGTTGCTCCTTCCTTCCTTTTCTTTCTGTCTNNNAGATCGGAAG |
| 69558_1 | TATGAGTGTGGAGTCGTTGCGTGATGAGCTCAAGACAAGTTTAGANNNAGATCGG |
| 69690_1 | TATGAGTGTGGAGTCGTTGCTGATGGAGTATAACTCTGTCAACCANNNAGATCGGA |
| 69713_1 | TATGAGTGTGGAGTCGTTGCACAAGTATTATCAGATCAAGGCCCTNNNAGATCGGA |
| 69714_1 | TATGAGTGTGGAGTCGTTGCTCCTACAAAAATGCTCACATACACANNNAGATCGGAA |
| 69728_1 | TATGAGTGTGGAGTCGTTGCAGCTCACTGCAATCTTGAACTCTNNNAGATCGGAAG |
| 69743_1 | TATGAGTGTGGAGTCGTTGCGAAGGAAGGAAGGAAGGAAGGAANNNAGATCGG |
| 69754_1 | TATGAGTGTGGAGTCGTTGCTGGTCTCAACCTCATCAAAATGTTGNNNAGATCGGA |
| 69796_1 | TATGAGTGTGGAGTCGTTGCTCAGCCCTGAAATGTAGGATATTCTNNNAGATCGGA |
| 69802_1 | TATGAGTGTGGAGTCGTTGCTGTGTTGCTATTTGTGAACTTGTCANNNAGATCGGA |
| 69850_1 | TATGAGTGTGGAGTCGTTGCTGCAGAAATAGCTTCAAAGGCTTNNNAGATCGGAAG |
| 69865_1 | TATGAGTGTGGAGTCGTTGCATACAAAAATTAGCCGGGCATGGNNNAGATCGGAA |
| 69942_1 | TATGAGTGTGGAGTCGTTGCAAGAAAGAAAGAAAGAAAAACAACCCNNNAGATCG |
| 69963_1 | TATGAGTGTGGAGTCGTTGCCACATTGACCATGACTGAGCATGNNNAGATCGGAAG |
| 69964_1 | TATGAGTGTGGAGTCGTTGCTGGAGTTATCACTCTTGGATGAGACNNNAGATCGGA |
| 70027_1 | TATGAGTGTGGAGTCGTTGCAATCAGCTGACCACATAAATGCGNNNAGATCGGAAG |
| 70192_1 | TATGAGTGTGGAGTCGTTGCAGTGATTCTCATGCTTCAGCCTCNNNAGATCGGAAG |
| 70262_1 | TATGAGTGTGGAGTCGTTGCAGAAAGAAAGAAAGAAAGAAAGAGAGANNNAGAT |
| 70343_1 | TATGAGTGTGGAGTCGTTGCTTAGAGGCTCAGAACTAATGGGANNNAGATCGGAA |
| 70349_1 | TATGAGTGTGGAGTCGTTGCGCACGTGTTTCTATGTGTCTGTCNNNAGATCGGAAG |
| 70425_1 | TATGAGTGTGGAGTCGTTGCCTCCTGAGCTCAAGTGATCCTCNNNAGATCGGAAGA |
| 70509_1 | TATGAGTGTGGAGTCGTTGCTCTCTTTCTCTCTGTCTCTCTCTCTNNNAGATCGGAAG |
| 10_1 | TATGAGTGTGGAGTCGTTGCTCCAAGACTGTGACTGAATTTGGTNNNAGATCGGAA |
| 11_1 | TATGAGTGTGGAGTCGTTGCATCAGACCTGTGAGTTGATTTGGTNNNAGATCGGAA |
| 15_1 | TATGAGTGTGGAGTCGTTGCTACTGTGGAAAGAGAGCTTTGGTNNNAGATCGGAA |
| 24_1 | TATGAGTGTGGAGTCGTTGCCCAAACATTTTCTGAGCACCAACANNNAGATCGGAA |
| 25_1 | TATGAGTGTGGAGTCGTTGCCCTTATCACCAACAGTGACCAACANNNAGATCGGAA |
| 28_1 | TATGAGTGTGGAGTCGTTGCTCCATGCTTATTGGTTACCCAACANNNAGATCGGAA |
| 39_1 | TATGAGTGTGGAGTCGTTGCGCTCTCTCTTTCTCTGGAGAATCCNNNAGATCGGAAG |
| 52_1 | TATGAGTGTGGAGTCGTTGCTGCCAACTTTTGTGGTAGGAATGTNNNAGATCGGAA |
| 55_1 | TATGAGTGTGGAGTCGTTGCCCTTATCTGTGACTGGTCAGAACANNNAGATCGGAA |
| 56_1 | TATGAGTGTGGAGTCGTTGCTATGGAAAACAGCGTGGAGAACANNNAGATCGGAA |
| 57_1 | TATGAGTGTGGAGTCGTTGCCACAGAGTACCTTTCCAAACATGTNNNAGATCGGAA |
| 60_1 | TATGAGTGTGGAGTCGTTGCAACAACTACAACAGGCAGGAACANNNAGATCGGAA |

| | |
|---|---|
| 70_1 | TATGAGTGTGGAGTCGTTGCTACATACCACCTCCCACACAACANNNAGATCGGAAGA |
| 78_1 | TATGAGTGTGGAGTCGTTGCACAAGGCAACATTTGATGTGAACANNNAGATCGGAA |
| 83_1 | TATGAGTGTGGAGTCGTTGCACATGATGGGGAGGAGGAATAACANNNAGATCGGA |
| 86_1 | TATGAGTGTGGAGTCGTTGCGTAAGTGCATGTATAGGGGCATGTNNNAGATCGGA |
| 93_1 | TATGAGTGTGGAGTCGTTGCTCTGCATCTCAATGTTTCTCATGTNNNAGATCGGAAG |
| 95_1 | TATGAGTGTGGAGTCGTTGCACCTCTCTACCTTGGAGTTCATGTNNNAGATCGGAA |
| 96_1 | TATGAGTGTGGAGTCGTTGCATCTTCCTAGAGCTGTAAAACACANNNAGATCGGAA |
| 99_1 | TATGAGTGTGGAGTCGTTGCACCCTGACCAATACAGAAACACANNNAGATCGGAAG |
| 102_1 | TATGAGTGTGGAGTCGTTGCAGGGTTTCTAGATGTTCCAGATGTNNNAGATCGGAA |
| 136_1 | TATGAGTGTGGAGTCGTTGCTTACTCCTAACCCTCAGGAAGCCNNNAGATCGGAAGA |
| 141_1 | TATGAGTGTGGAGTCGTTGCGGGACAGAGAGACAATGCTTATGTNNNAGATCGGA |
| 152_1 | TATGAGTGTGGAGTCGTTGCACCCATGTACATACAACATACACANNNAGATCGGAA |
| 157_1 | TATGAGTGTGGAGTCGTTGCAATCTGGATCATGTTGCATACACANNNAGATCGGAA |
| 174_1 | TATGAGTGTGGAGTCGTTGCTAGTCCTGAACCTTGTGCTTTCTCNNNAGATCGGAAG |
| 180_1 | TATGAGTGTGGAGTCGTTGCATCGTTCTAGACAAAGACTACTGTNNNAGATCGGAA |
| 185_1 | TATGAGTGTGGAGTCGTTGCTTTCCCATCAAGTGCTATTACTGTNNNAGATCGGAAG |
| 188_1 | TATGAGTGTGGAGTCGTTGCACGGAAACAAAACCTCAACCTGTNNNAGATCGGAAG |
| 195_1 | TATGAGTGTGGAGTCGTTGCTGCTGACTTGAGGATATTACCTGTNNNAGATCGGAA |
| 200_1 | TATGAGTGTGGAGTCGTTGCAGGGACAGTTTTTCTTTTCCCTGTNNNAGATCGGAA |
| 201_1 | TATGAGTGTGGAGTCGTTGCAGGCATGATTTCCCTTTATCCACANNNAGATCGGAA |
| 204_1 | TATGAGTGTGGAGTCGTTGCAGTCATGTGTGGCTTTTGATTGANNNAGATCGGAA |
| 206_1 | TATGAGTGTGGAGTCGTTGCACAACCATTACCCAAAAAGGGAGANNNAGATCGGAA |
| 211_1 | TATGAGTGTGGAGTCGTTGCAGGGTCTAGATGGTATTTTCCACANNNAGATCGGAA |
| 214_1 | TATGAGTGTGGAGTCGTTGCAAAACAAAAAGCCTCCCTCCTGTNNNAGATCGGAAGA |
| 218_1 | TATGAGTGTGGAGTCGTTGCGTCCAGAATGCTCACATTCCTGTNNNAGATCGGAAGA |
| 236_1 | TATGAGTGTGGAGTCGTTGCTTTTCATGACTTTGAACTAGCTGTNNNAGATCGGAA |
| 237_1 | TATGAGTGTGGAGTCGTTGCAAATGAAAATGAAGCTGCGCTGTNNNAGATCGGAA |
| 240_1 | TATGAGTGTGGAGTCGTTGCCCATACTGAGCTCCTCTGCTAAANNNAGATCGGAAGA |
| 254_1 | TATGAGTGTGGAGTCGTTGCGAACCACCTTGTGAGCACAGATTCNNNAGATCGGAA |
| 260_1 | TATGAGTGTGGAGTCGTTGCTGTGGTTTTCTTCATGTTTGCTGTNNNAGATCGGAA |
| 278_1 | TATGAGTGTGGAGTCGTTGCGCATCCTTTAGCACTTGGGAAAAGNNNAGATCGGAA |
| 286_1 | TATGAGTGTGGAGTCGTTGCAGTGGGTTTTCATCTGTCTTCACANNNAGATCGGAA |
| 291_1 | TATGAGTGTGGAGTCGTTGCCACACAAATAGAAGCCCTCTGTGANNNAGATCGGAA |
| 297_1 | TATGAGTGTGGAGTCGTTGCAGTAAGAGAAATGCCCTTGTCTGTNNNAGATCGGAA |
| 303_1 | TATGAGTGTGGAGTCGTTGCGGTATTGCTGTCTTGCTGTGAGANNNAGATCGGAA |
| 315_1 | TATGAGTGTGGAGTCGTTGCTGTGTCTCCTCTAAAATTCGGGTGNNNAGATCGGAA |
| 320_1 | TATGAGTGTGGAGTCGTTGCGGTAGCCACAGTTCTATTTTCTGTNNNAGATCGGAA |
| 322_1 | TATGAGTGTGGAGTCGTTGCAGTATCCAATACACACACAGACANNNAGATCGGAAG |
| 332_1 | TATGAGTGTGGAGTCGTTGCGGTGTGATGGTCAGTATGAAGTGTNNNAGATCGGA |
| 337_1 | TATGAGTGTGGAGTCGTTGCCACAGGAATGGAAAGGAAGAGACANNNAGATCGGA |
| 342_1 | TATGAGTGTGGAGTCGTTGCGTGCATTTTTATTGGAGGCAGTGTNNNAGATCGGAA |
| 351_1 | TATGAGTGTGGAGTCGTTGCAGCAGTCACCAAAATACTGAGTGTNNNAGATCGGAA |
| 356_1 | TATGAGTGTGGAGTCGTTGCCCCTTCTTGAGTTGGTGTTAGTGTNNNAGATCGGAA |

| | |
|---|---|
| 359_1 | TATGAGTGTGGAGTCGTTGCAGAAGGTCAGGAAGGTCAGGAAAANNNAGATCGG |
| 374_1 | TATGAGTGTGGAGTCGTTGCCCCTGCTGTGTCAAAATAGGTGTNNNAGATCGGAAG |
| 376_1 | TATGAGTGTGGAGTCGTTGCAGCAGTACCAATGTAATTAGGTGTNNNAGATCGGA |
| 403_1 | TATGAGTGTGGAGTCGTTGCAGATCAGTGTGAGTAGGACTGACANNNAGATCGGA |
| 416_1 | TATGAGTGTGGAGTCGTTGCAAAGAAGGGGAAATAAGAGTGTGTNNNAGATCGG |
| 427_1 | TATGAGTGTGGAGTCGTTGCTTCACAGAATGGCAGATGCCTGAANNNAGATCGGA |
| 430_1 | TATGAGTGTGGAGTCGTTGCGCATAACTACTCCTGGTGTGTGTNNNAGATCGGAAG |
| 435_1 | TATGAGTGTGGAGTCGTTGCCAGTGATGCTGGGCAAATTTACACNNNAGATCGGA |
| 438_1 | TATGAGTGTGGAGTCGTTGCAGGTAAAGACTATATGGCCCATGTNNNAGATCGGA |
| 442_1 | TATGAGTGTGGAGTCGTTGCTTTCGCATTCCTACCAGTTGTGTNNNAGATCGGAAG |
| 454_1 | TATGAGTGTGGAGTCGTTGCTGCTTTGATAACAGGGTGGTTTCCNNNAGATCGGAA |
| 459_1 | TATGAGTGTGGAGTCGTTGCTGTATCTGGCTTCTTCCACATTGTNNNAGATCGGAAG |
| 460_1 | TATGAGTGTGGAGTCGTTGCCCTTGATAGAAGTGCCTGTGAACANNNAGATCGGA |
| 461_1 | TATGAGTGTGGAGTCGTTGCAGCCAGAAGAAAACACACACTACANNNAGATCGGA |
| 463_1 | TATGAGTGTGGAGTCGTTGCGCATGAAACTTGAAGGCCATTGTNNNAGATCGGAA |
| 477_1 | TATGAGTGTGGAGTCGTTGCTCCTATCACTTCCCTGTTGCTACANNNAGATCGGAAG |
| 478_1 | TATGAGTGTGGAGTCGTTGCGTGTGTGTAACTCCCAAATCTACANNNAGATCGGAA |
| 479_1 | TATGAGTGTGGAGTCGTTGCATGCAGCAGAGGTTTACATCTACANNNAGATCGGAA |
| 496_1 | TATGAGTGTGGAGTCGTTGCTCTCAAAAGTCAATGAACCGTACANNNAGATCGGAA |
| 501_1 | TATGAGTGTGGAGTCGTTGCACCTTCTCCAGGTTGTCTGTACANNNAGATCGGAAG |
| 502_1 | TATGAGTGTGGAGTCGTTGCCACTCATTCCTGCTTGGTCTTGTNNNAGATCGGAAGA |
| 504_1 | TATGAGTGTGGAGTCGTTGCTGCTGAACTCTTTCTTGCCTTACANNNAGATCGGAAG |
| 511_1 | TATGAGTGTGGAGTCGTTGCTGATAGCCACAGTGAGAAAGTTGTNNNAGATCGGA |
| 534_1 | TATGAGTGTGGAGTCGTTGCCTTGGTATACTGGAGAAGTCAGCANNNAGATCGGA |
| 539_1 | TATGAGTGTGGAGTCGTTGCAGTCAAGAAAGCTTTGGAGAACCANNNAGATCGGA |
| 552_1 | TATGAGTGTGGAGTCGTTGCGTTCATTGTTGTGTCCAACACCANNNAGATCGGAAG |
| 557_1 | TATGAGTGTGGAGTCGTTGCACATCCTCTTTGACCTTCCTTTGTNNNAGATCGGAAG |
| 558_1 | TATGAGTGTGGAGTCGTTGCTCTCAAAGGGACAAAAAGACACCANNNAGATCGGA |
| 571_1 | TATGAGTGTGGAGTCGTTGCACTCCCTTCGTGAATTTCCCTTTCNNNAGATCGGAAG |
| 582_1 | TATGAGTGTGGAGTCGTTGCCAGTGAGGACATTTGGAAAGACCANNNAGATCGGA |
| 594_1 | TATGAGTGTGGAGTCGTTGCATTGGAAGGAGATGTCAGGACCANNNAGATCGGAA |
| 596_1 | TATGAGTGTGGAGTCGTTGCTCTTGTGTGAAATGCTTAGGACCANNNAGATCGGAA |
| 637_1 | TATGAGTGTGGAGTCGTTGCTCCCCTCAGGTTTTGTTATTACCANNNAGATCGGAAG |
| 657_1 | TATGAGTGTGGAGTCGTTGCGTTCTCAAGTCCTGCCTGAAAGAGNNNAGATCGGAA |
| 661_1 | TATGAGTGTGGAGTCGTTGCTCCTCATAGTAGAAGCACAACATTNNNAGATCGGAA |
| 664_1 | TATGAGTGTGGAGTCGTTGCTTCTGGGTCTCACTTCCCTCATAANNNAGATCGGAAG |
| 665_1 | TATGAGTGTGGAGTCGTTGCACCACATTTTCTTTGTGGAACATTNNNAGATCGGAA |
| 678_1 | TATGAGTGTGGAGTCGTTGCTTTACTCAACTTCCCTCACCCCANNNAGATCGGAAGA |
| 680_1 | TATGAGTGTGGAGTCGTTGCACATGCTCTGCTAACTTTTTCTCTNNNAGATCGGAAG |
| 689_1 | TATGAGTGTGGAGTCGTTGCTTTTATTCACTGGAGTGGCACTGTNNNAGATCGGAA |
| 692_1 | TATGAGTGTGGAGTCGTTGCATTATGTGGTTTCTTTGGCCCCANNNAGATCGGAAG |
| 699_1 | TATGAGTGTGGAGTCGTTGCAGGGGATGACTATAGCTTGGATGTNNNAGATCGGA |
| 706_1 | TATGAGTGTGGAGTCGTTGCCGAGTAGAAATCAGCCCAGCATTNNNAGATCGGAA |

| | |
|---|---|
| 714_1 | TATGAGTGTGGAGTCGTTGCACCAAATCCAAATAGGCATGATGTNNNAGATCGGAA |
| 745_1 | TATGAGTGTGGAGTCGTTGCTGCCAAGTCCAGTTTTTATCCCANNNAGATCGGAAG |
| 761_1 | TATGAGTGTGGAGTCGTTGCAGAAAAACAGTCTGGAGGTTCATTNNNAGATCGGA |
| 768_1 | TATGAGTGTGGAGTCGTTGCCCAAATGTTCAGTCAATTGTCCCANNNAGATCGGAA |
| 783_1 | TATGAGTGTGGAGTCGTTGCTGGCCTTGATATTCAAATTTCCCANNNAGATCGGAA |
| 786_1 | TATGAGTGTGGAGTCGTTGCACACAATTCCAAAGTCCTGAGATTNNNAGATCGGAA |
| 800_1 | TATGAGTGTGGAGTCGTTGCGCTATTTCCCAGTAACCACTGATCNNNAGATCGGAA |
| 802_1 | TATGAGTGTGGAGTCGTTGCCCAGAAGAAATGTAATGCAAGCCANNNAGATCGGA |
| 805_1 | TATGAGTGTGGAGTCGTTGCTTTGTTTTGATTCAGTGAAGCCANNNAGATCGGAAG |
| 810_1 | TATGAGTGTGGAGTCGTTGCTCTAGCTTTGGAAATGGTAAGCCANNNAGATCGGAA |
| 812_1 | TATGAGTGTGGAGTCGTTGCCCTACTTAGCCTTCATAACAGCCANNNAGATCGGAAG |
| 814_1 | TATGAGTGTGGAGTCGTTGCGACTGGAGTACGTTGGCATGATTNNNAGATCGGAA |
| 816_1 | TATGAGTGTGGAGTCGTTGCGCACTCATATGCCTATCCCTATGTNNNAGATCGGAAG |
| 836_1 | TATGAGTGTGGAGTCGTTGCAGAGAGTATGTATCAGGTGAGCCANNNAGATCGGA |
| 843_1 | TATGAGTGTGGAGTCGTTGCTGTGTGGTGGAAGCTTTGTCCTNNNAGATCGGAAG |
| 849_1 | TATGAGTGTGGAGTCGTTGCGTGGGGGAGTAATTATTGGCTATTNNNAGATCGGA |
| 850_1 | TATGAGTGTGGAGTCGTTGCATTTCTTTGGCTCTTTGGGTACTTNNNAGATCGGAA |
| 855_1 | TATGAGTGTGGAGTCGTTGCAATGCCTCCCTGGTCAGTCTATTNNNAGATCGGAAG |
| 866_1 | TATGAGTGTGGAGTCGTTGCATAGTGTCAGTTTGAATGGGCCANNNAGATCGGAA |
| 870_1 | TATGAGTGTGGAGTCGTTGCAATGTAAATCAGGATGTGGGCCANNNAGATCGGAA |
| 887_1 | TATGAGTGTGGAGTCGTTGCTCACCACAGATCCTATTGAAACTTNNNAGATCGGAA |
| 894_1 | TATGAGTGTGGAGTCGTTGCTCCTGTTCCCAACTTTATATGCCANNNAGATCGGAAG |
| 923_1 | TATGAGTGTGGAGTCGTTGCGCAAATACCACAGCACTTCACTTNNNAGATCGGAAG |
| 926_1 | TATGAGTGTGGAGTCGTTGCGCAGACATAAAGACCTTTTTGCCANNNAGATCGGAA |
| 927_1 | TATGAGTGTGGAGTCGTTGCGTTCAGGTTCAGGAGCAAATCCANNNAGATCGGAA |
| 928_1 | TATGAGTGTGGAGTCGTTGCTGAATTGTCTCCCTCTCAAATCCANNNAGATCGGAAG |
| 952_1 | TATGAGTGTGGAGTCGTTGCAGCAGTCTAAGATAGCCAGATCCANNNAGATCGGAA |
| 957_1 | TATGAGTGTGGAGTCGTTGCAGCAAATAATTCACCCGTGATCCANNNAGATCGGAA |
| 976_1 | TATGAGTGTGGAGTCGTTGCGTATTCTGTTAGACTTGGGCCATGNNNAGATCGGAA |
| 981_1 | TATGAGTGTGGAGTCGTTGCTGCCTGGCCCACAAAGATACTTTANNNAGATCGGAA |
| 987_1 | TATGAGTGTGGAGTCGTTGCATTCTCCCTGGGGTAAGTCAACTTNNNAGATCGGAA |
| 995_1 | TATGAGTGTGGAGTCGTTGCTGAGACTGTAAGTGAAGCCTCCANNNAGATCGGAA |
| 997_1 | TATGAGTGTGGAGTCGTTGCTTTAGCTCTTGCCTCCCTACCTTNNNAGATCGGAAGA |
| 1021_1 | TATGAGTGTGGAGTCGTTGCGTTATCTTCTCCAGTCTGCTACCANNNAGATCGGAAG |
| 1022_1 | TATGAGTGTGGAGTCGTTGCATTAGTCCATCCATAGCTGGTCCANNNAGATCGGAA |
| 1026_1 | TATGAGTGTGGAGTCGTTGCGAATGCCACTTTTTCCATGTCCANNNAGATCGGAAG |
| 1033_1 | TATGAGTGTGGAGTCGTTGCAGGGTTCCAGTGTTCTTGTACCTCNNNAGATCGGAA |
| 1038_1 | TATGAGTGTGGAGTCGTTGCAATCCAGACAGGGTAACAGCAACANNNAGATCGGA |
| 1041_1 | TATGAGTGTGGAGTCGTTGCGGCTGGAGTGCTGTAGTAGGATAANNNAGATCGGA |
| 1049_1 | TATGAGTGTGGAGTCGTTGCTTCTGGCTTCCTTCAGTATTCCANNNAGATCGGAAGA |
| 1054_1 | TATGAGTGTGGAGTCGTTGCCACCACACCCAGACATACTTACTTTNNNAGATCGGAA |
| 1069_1 | TATGAGTGTGGAGTCGTTGCACTTACTGCTGCCTTCAAAGCTTNNNAGATCGGAAG |
| 1075_1 | TATGAGTGTGGAGTCGTTGCGTGTTGGGCTAAGAAGGAAGCTTNNNAGATCGGAA |

| | |
|---|---|
| 1076_1 | TATGAGTGTGGAGTCGTTGCTGTCTTCATTCCTGAAATGTTCCANNNAGATCGGAAG |
| 1089_1 | TATGAGTGTGGAGTCGTTGCTCTTCAAGTGTCGACCAAACTCCTNNNAGATCGGAAG |
| 1097_1 | TATGAGTGTGGAGTCGTTGCGAGAAAGATCCTGCCTTGGGAAAANNNAGATCGGA |
| 1129_1 | TATGAGTGTGGAGTCGTTGCTTCATCTCTCCAGTTCAAGTGATCNNNAGATCGGAAG |
| 1130_1 | TATGAGTGTGGAGTCGTTGCAGTTGATTCCATATATTGGCCACTNNNAGATCGGAA |
| 1137_1 | TATGAGTGTGGAGTCGTTGCAGTGGGGAAACTAAGGCCAAAATGNNNAGATCGGA |
| 1142_1 | TATGAGTGTGGAGTCGTTGCATTTGAGTGGAAGTCACGTGGATGNNNAGATCGGA |
| 1143_1 | TATGAGTGTGGAGTCGTTGCCATCTCAGGCTCATCTTGTGCTTNNNAGATCGGAAGA |
| 1144_1 | TATGAGTGTGGAGTCGTTGCAAAACCGATCTCTCTCTCTCTTNNNAGATCGGAAG |
| 1157_1 | TATGAGTGTGGAGTCGTTGCAAACAATTGCACAAAGCAATCTTNNNAGATCGGAAG |
| 1162_1 | TATGAGTGTGGAGTCGTTGCTCAGAGAAAGAGGCAGAGAAACCTNNNAGATCGGA |
| 1166_1 | TATGAGTGTGGAGTCGTTGCTCTTCAATGCTTTCTAACCCAGCANNNAGATCGGAAG |
| 1168_1 | TATGAGTGTGGAGTCGTTGCTCCAATCAGCCATTGATCCATTGGNNNAGATCGGAA |
| 1177_1 | TATGAGTGTGGAGTCGTTGCGGCAGGGATGATTCTCAGGTTTATNNNAGATCGGA |
| 1182_1 | TATGAGTGTGGAGTCGTTGCTGTGTGTGTGTGTCTCCAATAAATNNNAGATCGGAA |
| 1184_1 | TATGAGTGTGGAGTCGTTGCACCCTATTCATTTGATGTGCAGCANNNAGATCGGAA |
| 1191_1 | TATGAGTGTGGAGTCGTTGCACACACAAACGTTTTCAAACCTGTNNNAGATCGGAA |
| 1201_1 | TATGAGTGTGGAGTCGTTGCGGCAGCTAATAAATGTGTTCAGCANNNAGATCGGA |
| 1213_1 | TATGAGTGTGGAGTCGTTGCCACTGACTGTTTCTGCTTATTCTTNNNAGATCGGAAG |
| 1214_1 | TATGAGTGTGGAGTCGTTGCTACCCATCATTTGCCTGAAAGACCNNNAGATCGGAA |
| 1215_1 | TATGAGTGTGGAGTCGTTGCTTAGTCAATGTGCAGTGCTGTGATNNNAGATCGGAA |
| 1223_1 | TATGAGTGTGGAGTCGTTGCTCTGTGTCTGGATAGCTGAGAGTGNNNAGATCGGA |
| 1227_1 | TATGAGTGTGGAGTCGTTGCGCATCGCAGGATTTTAGGCAAAGGNNNAGATCGGA |
| 1232_1 | TATGAGTGTGGAGTCGTTGCAGAGGCTGTTGCCAATTGTTCTTNNNAGATCGGAAG |
| 1233_1 | TATGAGTGTGGAGTCGTTGCTCTATTTTGCACAATTTGCCCCAANNNAGATCGGAAG |
| 1235_1 | TATGAGTGTGGAGTCGTTGCGAGTAGGGTCAGTTCTCGCTCTTTNNNAGATCGGAA |
| 1237_1 | TATGAGTGTGGAGTCGTTGCGGCATCTGAGAGTAAACATGAGCANNNAGATCGGA |
| 1242_1 | TATGAGTGTGGAGTCGTTGCAACTCCACATGAAACACTGAGCANNNAGATCGGAAG |
| 1249_1 | TATGAGTGTGGAGTCGTTGCTGATTCTAGTGTCCCAGTGAGCANNNAGATCGGAAG |
| 1257_1 | TATGAGTGTGGAGTCGTTGCCAACAGCTGCAATGGGCCAAATAANNNAGATCGGA |
| 1259_1 | TATGAGTGTGGAGTCGTTGCGCAAAGGAAAGAAAGACCATAGCANNNAGATCGGA |
| 1260_1 | TATGAGTGTGGAGTCGTTGCAGCGCACATTTCCTTTAGAAGTTNNNAGATCGGAAG |
| 1264_1 | TATGAGTGTGGAGTCGTTGCCACAGTCAAGGACAGGGTTTACTTNNNAGATCGGA |
| 1277_1 | TATGAGTGTGGAGTCGTTGCTGAGAGGTGATCAGGAAAGCTAGANNNAGATCGG |
| 1294_1 | TATGAGTGTGGAGTCGTTGCACTGAGGATGTAGAGGATGAAGGANNNAGATCGG |
| 1303_1 | TATGAGTGTGGAGTCGTTGCTCCTCTGATCTCTGCTATTACGCANNNAGATCGGAAG |
| 1320_1 | TATGAGTGTGGAGTCGTTGCTTGGTTGGACATTGAGAAAGGCANNNAGATCGGAA |
| 1328_1 | TATGAGTGTGGAGTCGTTGCAAGACCATTTGCTGATGAAGGCANNNAGATCGGAA |
| 1330_1 | TATGAGTGTGGAGTCGTTGCCCTGTTTTGTGGTGGTTCGACGTTNNNAGATCGGAA |
| 1332_1 | TATGAGTGTGGAGTCGTTGCTGATTTAGGTATGATGCCAGGAGCNNNAGATCGGA |
| 1342_1 | TATGAGTGTGGAGTCGTTGCTGGTCATCTCTGGTGGTAAGGTTNNNAGATCGGAA |
| 1350_1 | TATGAGTGTGGAGTCGTTGCAGAACTGTTACAAAATAGGAGGCANNNAGATCGGA |
| 1353_1 | TATGAGTGTGGAGTCGTTGCTGGTTTTGGACACCTAGGTTGATCNNNAGATCGGAA |

| | |
|---|---|
| 1354_1 | TATGAGTGTGGAGTCGTTGCACTTCGTCTTTTCAGAATGAGGCANNNAGATCGGAA |
| 1393_1 | TATGAGTGTGGAGTCGTTGCAGTGTTTATGCAAAGGTATGGGCANNNAGATCGGA |
| 1411_1 | TATGAGTGTGGAGTCGTTGCGTGTGTCTGTGTAGACATGGTCTANNNAGATCGGA |
| 1415_1 | TATGAGTGTGGAGTCGTTGCACCAAAAGATGCCTGGAAATGTTNNNAGATCGGAA |
| 1424_1 | TATGAGTGTGGAGTCGTTGCAATCAACCACCAATCTCTGGCCATNNNAGATCGGAA |
| 1433_1 | TATGAGTGTGGAGTCGTTGCGCAATAACACAGATGAAGGATGTTNNNAGATCGGA |
| 1447_1 | TATGAGTGTGGAGTCGTTGCTATGTGTTTTCAGAATGGGCGTGGNNNAGATCGGA |
| 1465_1 | TATGAGTGTGGAGTCGTTGCACACATATCCAATCCCAAATGCANNNAGATCGGAAG |
| 1473_1 | TATGAGTGTGGAGTCGTTGCCTGCATGCCTTTTTCTTTTGTTTNNNAGATCGGAAGA |
| 1491_1 | TATGAGTGTGGAGTCGTTGCTGTGTGTGTAGTCTTGCTGTGTTNNNAGATCGGAAG |
| 1497_1 | TATGAGTGTGGAGTCGTTGCTGATTTCCATGCTGACCAACATTGNNNAGATCGGAA |
| 1498_1 | TATGAGTGTGGAGTCGTTGCTTCATCAGTGCCTTTTTGACGTTANNNAGATCGGAA |
| 1502_1 | TATGAGTGTGGAGTCGTTGCAAGGGAAAGAAGGATTTCCATGCANNNAGATCGGA |
| 1505_1 | TATGAGTGTGGAGTCGTTGCATCACCCTCAGCTAATTTTTGCTTNNNAGATCGGAAG |
| 1520_1 | TATGAGTGTGGAGTCGTTGCAATATGCTGTGTTACAGAGATGCANNNAGATCGGA |
| 1524_1 | TATGAGTGTGGAGTCGTTGCTTTCTGCTTAGCATTTGACAGCTCNNNAGATCGGAA |
| 1538_1 | TATGAGTGTGGAGTCGTTGCGTGGCTCAAAACAACACAAAAGANNNAGATCGGAA |
| 1548_1 | TATGAGTGTGGAGTCGTTGCTCAGTGTGGCCATAAAGTTATGCANNNAGATCGGA |
| 1549_1 | TATGAGTGTGGAGTCGTTGCTGTGCCTGTGGAAGTATTTATGCANNNAGATCGGA |
| 1550_1 | TATGAGTGTGGAGTCGTTGCAGTGAACACGTGAATGTTTATGCANNNAGATCGGA |
| 1553_1 | TATGAGTGTGGAGTCGTTGCAAATCCAAGCTAGGGGAAACACAGNNNAGATCGGA |
| 1555_1 | TATGAGTGTGGAGTCGTTGCTTGGGCTGTAATTTCCAACTGCANNNAGATCGGAAG |
| 1570_1 | TATGAGTGTGGAGTCGTTGCGCTCTCATGGTAAGCTGCCATTTNNNAGATCGGAAG |
| 1573_1 | TATGAGTGTGGAGTCGTTGCACTAGCAAATTGAATCCACCTGCANNNAGATCGGAA |
| 1626_1 | TATGAGTGTGGAGTCGTTGCTGTATATGTGTGCTTGGTGCTTNNNAGATCGGAAG |
| 1629_1 | TATGAGTGTGGAGTCGTTGCTACATAAAGACAGCCCCAGAGGANNNAGATCGGAA |
| 1638_1 | TATGAGTGTGGAGTCGTTGCATTGTAACCTCCATCATCCCCACGNNNAGATCGGAAG |
| 1646_1 | TATGAGTGTGGAGTCGTTGCTCTCTGTGTGTGGACATGAAGCTTNNNAGATCGGAA |
| 1647_1 | TATGAGTGTGGAGTCGTTGCAGAGGTTTTGTTGATTGCCTACCANNNAGATCGGAA |
| 1650_1 | TATGAGTGTGGAGTCGTTGCTGACTTCTCCCATAGCTAATTGCANNNAGATCGGAA |
| 1657_1 | TATGAGTGTGGAGTCGTTGCTCCCTCATCCTCTTCACATCCCTTNNNAGATCGGAAGA |
| 1669_1 | TATGAGTGTGGAGTCGTTGCTGAGAAGCAATACACCATCTTGCANNNAGATCGGAA |
| 1675_1 | TATGAGTGTGGAGTCGTTGCACATAATGAGCCCCCAGTAATTGTNNNAGATCGGAA |
| 1682_1 | TATGAGTGTGGAGTCGTTGCAGGCTCCCATAAGCTACTCCTTTNNNAGATCGGAAG |
| 1685_1 | TATGAGTGTGGAGTCGTTGCGGATCATCAGTCGATGTATTTGCANNNAGATCGGAA |
| 1692_1 | TATGAGTGTGGAGTCGTTGCTCAAATACATGCCATGCAGCTTTNNNAGATCGGAAG |
| 1700_1 | TATGAGTGTGGAGTCGTTGCCTGTATGGGAGAGCAGTAGCTTTNNNAGATCGGAA |
| 1701_1 | TATGAGTGTGGAGTCGTTGCTTACAGGGCCATCCATCTTCTTTGNNNAGATCGGAA |
| 1708_1 | TATGAGTGTGGAGTCGTTGCGCTCATGTTGTATGTCTTTTTGCANNNAGATCGGAA |
| 1717_1 | TATGAGTGTGGAGTCGTTGCCCACCTGAAATGGGAATCATTACCNNNAGATCGGAA |
| 1723_1 | TATGAGTGTGGAGTCGTTGCGTTGTCTTTGTGCACATGAGGAGANNNAGATCGGA |
| 1727_1 | TATGAGTGTGGAGTCGTTGCCTGGTGACTGAATTTAGAGCAGTGNNNAGATCGGA |
| 1750_1 | TATGAGTGTGGAGTCGTTGCTCTTCTCCAAATCCATGCCATCANNNAGATCGGAAGA |

| | |
|---|---|
| 1764_1 | TATGAGTGTGGAGTCGTTGCACACATTTACACCTTGGTCATCANNNAGATCGGAAGA |
| 1767_1 | TATGAGTGTGGAGTCGTTGCTCTTTCCCTCTCTCAGTGGGTTGTNNNAGATCGGAAG |
| 1772_1 | TATGAGTGTGGAGTCGTTGCTCATCTAGCAATTCCATTCCTGGANNNAGATCGGAAG |
| 1786_1 | TATGAGTGTGGAGTCGTTGCACTGGCTGTGTAAAACTGGAGTTTNNNAGATCGGAA |
| 1792_1 | TATGAGTGTGGAGTCGTTGCGTGCATACTAATGCCAGGGTATCANNNAGATCGGAA |
| 1796_1 | TATGAGTGTGGAGTCGTTGCTTTTGTGTTGGGGGAGAATCAGAGNNNAGATCGGA |
| 1808_1 | TATGAGTGTGGAGTCGTTGCGGAAGTAAAAGTCAGTTGACCCCANNNAGATCGGAA |
| 1820_1 | TATGAGTGTGGAGTCGTTGCGCTCCAGTACGATCTCATCTTAGCNNNAGATCGGAAG |
| 1831_1 | TATGAGTGTGGAGTCGTTGCTCTTGGAGGAGATGCAGATACTCANNNAGATCGGAA |
| 1835_1 | TATGAGTGTGGAGTCGTTGCGATGGTGTTGGTGGACATTTAGGTNNNAGATCGGA |
| 1848_1 | TATGAGTGTGGAGTCGTTGCGAGACGAGCTTTTGCTTAGGCTATNNNAGATCGGAA |
| 1857_1 | TATGAGTGTGGAGTCGTTGCAGGCTTACTTATGACAGATCCTCANNNAGATCGGAAG |
| 1860_1 | TATGAGTGTGGAGTCGTTGCCAGAGGAGAGAAGAAGGGAGTGATNNNAGATCGG |
| 1863_1 | TATGAGTGTGGAGTCGTTGCCCAGATAATGAAGATGCTGCTGGNNNAGATCGGAAG |
| 1870_1 | TATGAGTGTGGAGTCGTTGCATCACTTTCTCAAATCATCCACCGNNNAGATCGGAAG |
| 1871_1 | TATGAGTGTGGAGTCGTTGCCTGTGTGAACTCATCTTTTCCTCANNNAGATCGGAAG |
| 1877_1 | TATGAGTGTGGAGTCGTTGCGACCTAGAGCTTGGCAAGAGCATANNNAGATCGGAA |
| 1886_1 | TATGAGTGTGGAGTCGTTGCCCCTTCTGGTGTGTATGCCTTTTNNNAGATCGGAAGA |
| 1888_1 | TATGAGTGTGGAGTCGTTGCCAGTGTTCTATCGCTTGCCTTTTNNNAGATCGGAAGA |
| 1892_1 | TATGAGTGTGGAGTCGTTGCAGCATTGTGGTGAAGGAGCTTTTNNNAGATCGGAAG |
| 1893_1 | TATGAGTGTGGAGTCGTTGCCCAGAATGATTGGCCTATGCTCANNNAGATCGGAAG |
| 1895_1 | TATGAGTGTGGAGTCGTTGCCCCCATGCTCTTGTACATACTTTTNNNAGATCGGAAGA |
| 1947_1 | TATGAGTGTGGAGTCGTTGCCAAGGAGAAGGTTTCTGTGTCTCANNNAGATCGGAA |
| 1964_1 | TATGAGTGTGGAGTCGTTGCCAAACAGATGAACACCACAAGGTGNNNAGATCGGAA |
| 1966_1 | TATGAGTGTGGAGTCGTTGCATGTGGAATGTGCCTAGCAAAACTNNNAGATCGGAA |
| 1971_1 | TATGAGTGTGGAGTCGTTGCTTTCCAACACCACAAGAAGCACTANNNAGATCGGAAG |
| 1979_1 | TATGAGTGTGGAGTCGTTGCAATGACAAGGCAAAATGGCAAGATNNNAGATCGGA |
| 1981_1 | TATGAGTGTGGAGTCGTTGCTTTGGCAACACCATGCACTTTTTNNNAGATCGGAAGA |
| 1984_1 | TATGAGTGTGGAGTCGTTGCGCATGTTCCTTCAGTCACAGTCANNNAGATCGGAAGA |
| 1995_1 | TATGAGTGTGGAGTCGTTGCGAGAGTAGCTGAGGTTCAGAGAGANNNAGATCGGA |
| 2014_1 | TATGAGTGTGGAGTCGTTGCCCTACTTGGGGAAGCCTAAAACTGNNNAGATCGGAA |
| 2020_1 | TATGAGTGTGGAGTCGTTGCGCTTCGGTATGAAGTAACGGTCANNNAGATCGGAAG |
| 2037_1 | TATGAGTGTGGAGTCGTTGCTTGTTTTCATGTGGTTTTGGGTTTNNNAGATCGGAAG |
| 2042_1 | TATGAGTGTGGAGTCGTTGCTTTGTCATGCATGTGTTCATGTCANNNAGATCGGAAG |
| 2045_1 | TATGAGTGTGGAGTCGTTGCTCACTTAGCCTCTGTGGATGTCANNNAGATCGGAAGA |
| 2046_1 | TATGAGTGTGGAGTCGTTGCGTCACTTTCAATGTCATGATGTCANNNAGATCGGAAG |
| 2048_1 | TATGAGTGTGGAGTCGTTGCTTGTAACCATGGGTGATGACGGANNNAGATCGGAAG |
| 2061_1 | TATGAGTGTGGAGTCGTTGCTGAAAACTGTAAAGCACTCTGTCANNNAGATCGGAA |
| 2064_1 | TATGAGTGTGGAGTCGTTGCACATGCCTGATTGAAAGTTTCCTGANNNAGATCGGA |
| 2066_1 | TATGAGTGTGGAGTCGTTGCGCAGCAAGATACAGAAAAGTGTCANNNAGATCGGA |
| 2073_1 | TATGAGTGTGGAGTCGTTGCTTTCTGCCTGTGTGATACTGCATTNNNAGATCGGAAG |
| 2076_1 | TATGAGTGTGGAGTCGTTGCAGAGAGTAAGAGATGGGCAGACTGNNNAGATCGGA |
| 2088_1 | TATGAGTGTGGAGTCGTTGCCATCAGTAGAAATGGCAACTGCCTNNNAGATCGGAA |

| | |
|---|---|
| 2096_1 | TATGAGTGTGGAGTCGTTGCTTCTGCCTAGGTTACGACATTCANNNAGATCGGAAG |
| 2104_1 | TATGAGTGTGGAGTCGTTGCAAAGGACTAGACTTGAGACCAGGCNNNAGATCGGA |
| 2110_1 | TATGAGTGTGGAGTCGTTGCAGATGCCTTTGGCTGTTCATTCANNNAGATCGGAAG |
| 2126_1 | TATGAGTGTGGAGTCGTTGCCAACAGCAACAATAAGAGCTAGCCNNNAGATCGGAA |
| 2128_1 | TATGAGTGTGGAGTCGTTGCATGCACAATTTCCTTGCTTATTCANNNAGATCGGAAG |
| 2131_1 | TATGAGTGTGGAGTCGTTGCAGCAGTGGAATCATGAGCAATTTTNNNAGATCGGA |
| 2161_1 | TATGAGTGTGGAGTCGTTGCGCTTCACCTCTAGGCATTGTATCCNNNAGATCGGAA |
| 2166_1 | TATGAGTGTGGAGTCGTTGCTTCGGATCATGTTGTGGAAGTGTNNNAGATCGGAA |
| 2171_1 | TATGAGTGTGGAGTCGTTGCTCTTCTGGGAGTTTCCAATGGTATNNNAGATCGGAA |
| 2177_1 | TATGAGTGTGGAGTCGTTGCCAGTGTGGCACAGATCTTCTTCANNNAGATCGGAAG |
| 2181_1 | TATGAGTGTGGAGTCGTTGCGTCCCTGTAAGGCTGTTTTCTTCANNNAGATCGGAA |
| 2185_1 | TATGAGTGTGGAGTCGTTGCGAGATATGAGACAGTGCCAGGTGTNNNAGATCGGA |
| 2195_1 | TATGAGTGTGGAGTCGTTGCTGTGTGTGTGTGAAGAAAGGTTCANNNAGATCGGA |
| 2208_1 | TATGAGTGTGGAGTCGTTGCTGCCTTAGAACCTGAAATTCTGCTNNNAGATCGGAA |
| 2216_1 | TATGAGTGTGGAGTCGTTGCAAGCTCCAGGGCTATCCTTGAGAGNNNAGATCGGA |
| 2230_1 | TATGAGTGTGGAGTCGTTGCTGTGAGACTGTGGTTCATGATTTCANNNAGATCGGA |
| 2245_1 | TATGAGTGTGGAGTCGTTGCCACCACCAGAACATGGTCTTTCANNNAGATCGGAAG |
| 2262_1 | TATGAGTGTGGAGTCGTTGCTGGCTGCATCTTTCAATGTTTTCANNNAGATCGGAA |
| 2278_1 | TATGAGTGTGGAGTCGTTGCTGGGAGTAATGACTGAGGAAAGGANNNAGATCGG |
| 2285_1 | TATGAGTGTGGAGTCGTTGCACGCATTCTGAGGACTTGAAAAGANNNAGATCGGA |
| 2309_1 | TATGAGTGTGGAGTCGTTGCGCAGGAAAGCATGGTTTGATATGGNNNAGATCGGA |
| 2330_1 | TATGAGTGTGGAGTCGTTGCTGCTAAATTCCTTTGGAACCAAGANNNAGATCGGAA |
| 2344_1 | TATGAGTGTGGAGTCGTTGCTGGAGTGGAGGTTAGTAGAGGGANNNAGATCGGA |
| 2351_1 | TATGAGTGTGGAGTCGTTGCCTGGTAGGCCATAAAGCCAATGATNNNAGATCGGA |
| 2370_1 | TATGAGTGTGGAGTCGTTGCTCACTGCATTCCTCCTCTTTCAGGNNNAGATCGGAAG |
| 2375_1 | TATGAGTGTGGAGTCGTTGCCCCCAAGGCAACCATTTTCACATCNNNAGATCGGAAG |
| 2401_1 | TATGAGTGTGGAGTCGTTGCTTCTTATAAAGCTCCCCACGATGCNNNAGATCGGAA |
| 2427_1 | TATGAGTGTGGAGTCGTTGCAGAGGGAAAAATGAAGCTAACAGANNNAGATCGG |
| 2438_1 | TATGAGTGTGGAGTCGTTGCGGAACCCTTGCACTTTACACAGANNNAGATCGGAAG |
| 2443_1 | TATGAGTGTGGAGTCGTTGCAACAAATCTGCATTTCTGCACAGANNNAGATCGGAA |
| 2458_1 | TATGAGTGTGGAGTCGTTGCGGCAAGGTCAAAAAGAATGACAGANNNAGATCGG |
| 2459_1 | TATGAGTGTGGAGTCGTTGCGCTCTAACTTGTTTCACACCTGGCGNNNAGATCGGA |
| 2466_1 | TATGAGTGTGGAGTCGTTGCTGGAAAAGTGGTAGAGGCTACAGANNNAGATCGG |
| 2471_1 | TATGAGTGTGGAGTCGTTGCAACTAATAGGCAGGGATTTCCTCANNNAGATCGGAA |
| 2475_1 | TATGAGTGTGGAGTCGTTGCAATGCCATGGTGAAAGAATTGGTNNNAGATCGGAA |
| 2478_1 | TATGAGTGTGGAGTCGTTGCAGTAAGCACATTGACAAACCCAGANNNAGATCGGA |
| 2486_1 | TATGAGTGTGGAGTCGTTGCTGGTTCTCTGTGTCATCCCCATAGNNNAGATCGGAA |
| 2498_1 | TATGAGTGTGGAGTCGTTGCATTCTGATCCCTGATCTGCCAGANNNAGATCGGAAG |
| 2504_1 | TATGAGTGTGGAGTCGTTGCAATCCAACAGGCTTAGGACCCTANNNAGATCGGAAG |
| 2507_1 | TATGAGTGTGGAGTCGTTGCAGGAAGTACATGAAGGTCTCCAGANNNAGATCGGA |
| 2518_1 | TATGAGTGTGGAGTCGTTGCAAAGCAACTAGGACAACAGCAGANNNAGATCGGAA |
| 2526_1 | TATGAGTGTGGAGTCGTTGCTCTGTAGCCTCAACTTCTAGCAGANNNAGATCGGAA |
| 2541_1 | TATGAGTGTGGAGTCGTTGCAATGGTCTAACCAAACATGCAGANNNAGATCGGAA |

| | |
|---|---|
| 2552_1 | TATGAGTGTGGAGTCGTTGCTCAGCAGCATCAAAAATCCAAGGANNNAGATCGGAA |
| 2558_1 | TATGAGTGTGGAGTCGTTGCATCAACAGCACCTGAGGGATTCTTNNNAGATCGGAA |
| 2571_1 | TATGAGTGTGGAGTCGTTGCACCATCTCAAGCTATAGGCTCAGANNNAGATCGGAA |
| 2586_1 | TATGAGTGTGGAGTCGTTGCCAGCAGATTCTATACCCTTCAGANNNAGATCGGAAGA |
| 2596_1 | TATGAGTGTGGAGTCGTTGCGGAGTTCAGTTCAAAGCCATCAGANNNAGATCGGAA |
| 2606_1 | TATGAGTGTGGAGTCGTTGCCACTGAAAGCCCACTGAAAGAGANNNAGATCGGAA |
| 2607_1 | TATGAGTGTGGAGTCGTTGCCTCATTCATTCTCTTGCTGGGTACNNNAGATCGGAAG |
| 2613_1 | TATGAGTGTGGAGTCGTTGCTGACGTGTCTTCTGTAAACAGAGANNNAGATCGGAA |
| 2626_1 | TATGAGTGTGGAGTCGTTGCAAAAACTCTTGGAACCTTCAGAGANNNAGATCGGAA |
| 2631_1 | TATGAGTGTGGAGTCGTTGCCGCTTAGACTGCAGAATCACTTTANNNAGATCGGAA |
| 2639_1 | TATGAGTGTGGAGTCGTTGCGTCCTTGAACTCCTCAGCTCAAACNNNAGATCGGAAG |
| 2647_1 | TATGAGTGTGGAGTCGTTGCTGTGAAAGGAGGTAAGCTGAGAGANNNAGATCGGA |
| 2651_1 | TATGAGTGTGGAGTCGTTGCAGACACACATTCAGAGGCTAGAGANNNAGATCGGA |
| 2653_1 | TATGAGTGTGGAGTCGTTGCGGGGCAGTCCATTGATAAATACTTNNNAGATCGGAA |
| 2655_1 | TATGAGTGTGGAGTCGTTGCGAGGAGAAATGGAAGCAAAGGAGANNNAGATCGG |
| 2675_1 | TATGAGTGTGGAGTCGTTGCGTAAGCATGCATGTGCAAGTATCANNNAGATCGGAA |
| 2677_1 | TATGAGTGTGGAGTCGTTGCTCTGGAATACAGTGATGCAGTCACNNNAGATCGGAA |
| 2681_1 | TATGAGTGTGGAGTCGTTGCACTGTATTGCCAAATATGGGGAGANNNAGATCGGA |
| 2684_1 | TATGAGTGTGGAGTCGTTGCACTACACACACTGTCTTGAATGCCNNNAGATCGGAA |
| 2695_1 | TATGAGTGTGGAGTCGTTGCAACTGGGAGTAAAAGTTAGCAGGCNNNAGATCGGA |
| 2700_1 | TATGAGTGTGGAGTCGTTGCTAATGGCTCTCAGCTCCAAATCCANNNAGATCGGAA |
| 2706_1 | TATGAGTGTGGAGTCGTTGCCCCCAGGAGTAGTGCACAAATATNNNAGATCGGAAG |
| 2733_1 | TATGAGTGTGGAGTCGTTGCAAGCCTCTTCCCATGAAGAGAGAANNNAGATCGGAA |
| 2737_1 | TATGAGTGTGGAGTCGTTGCTCAAAGGAAAAGGTAACTGTGAGANNNAGATCGGA |
| 2741_1 | TATGAGTGTGGAGTCGTTGCGCTAACACAACCATGGCTTGAGANNNAGATCGGAAG |
| 2743_1 | TATGAGTGTGGAGTCGTTGCAGCCACAGAAGACTGGAAGAAGTCNNNAGATCGGA |
| 2750_1 | TATGAGTGTGGAGTCGTTGCCTCTTCCTGCATGCCTTTTGAGANNNAGATCGGAAGA |
| 2762_1 | TATGAGTGTGGAGTCGTTGCCCTCTCAACCCACCTTTTGAAAATNNNAGATCGGAAG |
| 2773_1 | TATGAGTGTGGAGTCGTTGCGGTGGAGTTTCTTTTCCTCAATCTNNNAGATCGGAA |
| 2797_1 | TATGAGTGTGGAGTCGTTGCGTTTTCCTGTGATTGCCTCAGAGCNNNAGATCGGAA |
| 2800_1 | TATGAGTGTGGAGTCGTTGCATGTGAGAGAAAGAGAGAGGTGGGNNNAGATCGG |
| 2814_1 | TATGAGTGTGGAGTCGTTGCTGTCCCAAGAGTTGTTGCTTTAGANNNAGATCGGAA |
| 2817_1 | TATGAGTGTGGAGTCGTTGCCTTAGTCTGGCAAGCTGTTTAGANNNAGATCGGAAG |
| 2828_1 | TATGAGTGTGGAGTCGTTGCCTGAAAATGAGTGTTCCTCCTCCTNNNAGATCGGAA |
| 2837_1 | TATGAGTGTGGAGTCGTTGCTAGGCTGGAGTGGCACAATCATAANNNAGATCGGA |
| 2852_1 | TATGAGTGTGGAGTCGTTGCGTCAACCGGTCATTAGGTATTTCTNNNAGATCGGAA |
| 2865_1 | TATGAGTGTGGAGTCGTTGCTCTTGATTTGCCTCTCACTTCGANNNAGATCGGAAGA |
| 2867_1 | TATGAGTGTGGAGTCGTTGCTTTCTTGCTTTGGTCTTGCAATTCNNNAGATCGGAAG |
| 2869_1 | TATGAGTGTGGAGTCGTTGCAGGTAGTGGCATGGGTTATGTGTGNNNAGATCGGA |
| 2871_1 | TATGAGTGTGGAGTCGTTGCTCTGGGCTTCAGGTTCTGAGAAATNNNAGATCGGAA |
| 2885_1 | TATGAGTGTGGAGTCGTTGCGTAGGCAAGCTGGTAACAAAAGAGNNNAGATCGGA |
| 2902_1 | TATGAGTGTGGAGTCGTTGCTGAGGAGATACCTTGAAGGAAGGANNNAGATCGGA |
| 2907_1 | TATGAGTGTGGAGTCGTTGCCTCTGCATGTCTGAATTTGGTGGGNNNAGATCGGAA |

| | |
|---|---|
| 2911_1 | TATGAGTGTGGAGTCGTTGCTGGTGGTGTTTGTGACAATAAGGANNNAGATCGGA |
| 2946_1 | TATGAGTGTGGAGTCGTTGCGTGTGTGTGTGTTGTGGTGTAGACNNNAGATCGGA |
| 2957_1 | TATGAGTGTGGAGTCGTTGCCTTTGTCATTTTTAGGTGTTGCAANNNAGATCGGAA |
| 2972_1 | TATGAGTGTGGAGTCGTTGCGAGAACATGAATGGATCTGGAGGANNNAGATCGG |
| 2973_1 | TATGAGTGTGGAGTCGTTGCGGGAGCTGGAGATTGTTCTTATGGNNNAGATCGGA |
| 3004_1 | TATGAGTGTGGAGTCGTTGCAAGCCAAGAAAAGAACCACTAGGANNNAGATCGGA |
| 3008_1 | TATGAGTGTGGAGTCGTTGCGTGTGTTTCCTTGTATGGCTATCTNNNAGATCGGAA |
| 3021_1 | TATGAGTGTGGAGTCGTTGCTGTGTGTGTTTGTAGGAGAGAAGANNNAGATCGG |
| 3022_1 | TATGAGTGTGGAGTCGTTGCTGCTCATTATTTTTGAGGGTAGGANNNAGATCGGA |
| 3024_1 | TATGAGTGTGGAGTCGTTGCTGTTTTGCAATATCCGTGGTAGGANNNAGATCGGA |
| 3026_1 | TATGAGTGTGGAGTCGTTGCTTCTACTACCCAGTGGGAATTTGGNNNAGATCGGAA |
| 3056_1 | TATGAGTGTGGAGTCGTTGCAAATCCTGTTGTCTGCAAAGGGANNNAGATCGGAA |
| 3108_1 | TATGAGTGTGGAGTCGTTGCAGAAATGGCGGAAGTTGGAGTATTNNNAGATCGG |
| 3112_1 | TATGAGTGTGGAGTCGTTGCTCTTCCAAGGAGTATTGCAGGGANNNAGATCGGAA |
| 3117_1 | TATGAGTGTGGAGTCGTTGCTGTGCCTAGAATGAGTTCAGGGANNNAGATCGGAA |
| 3118_1 | TATGAGTGTGGAGTCGTTGCGTGGTGGCTAAAAGTTTCAGGGANNNAGATCGGAA |
| 3122_1 | TATGAGTGTGGAGTCGTTGCTGTGCTGCTGTGATTATGTGTGNNNAGATCGGAAG |
| 3172_1 | TATGAGTGTGGAGTCGTTGCTCCAGCCTTTTATTTGAATGGGGANNNAGATCGGAA |
| 3179_1 | TATGAGTGTGGAGTCGTTGCCCATCAGTGGCAGGCTGGATATATNNNAGATCGGA |
| 3194_1 | TATGAGTGTGGAGTCGTTGCACACCTTTCTGTGAAAAGATGGGANNNAGATCGGA |
| 3199_1 | TATGAGTGTGGAGTCGTTGCAAAAACAATTGTCCAGCTAGTGGANNNAGATCGGA |
| 3203_1 | TATGAGTGTGGAGTCGTTGCTGAGGGTAAAAAGAACAACTGGGANNNAGATCGG |
| 3214_1 | TATGAGTGTGGAGTCGTTGCAGTCTCAGTACAAACTTCCTGGGANNNAGATCGGAA |
| 3219_1 | TATGAGTGTGGAGTCGTTGCGCGATTCCCACATTTTCATTCAATNNNAGATCGGAAG |
| 3221_1 | TATGAGTGTGGAGTCGTTGCTGCTGCCATTTGAATGATCTATCANNNAGATCGGAA |
| 3231_1 | TATGAGTGTGGAGTCGTTGCGGGAACAGCAGATAGTGGGGAAAANNNAGATCGG |
| 3233_1 | TATGAGTGTGGAGTCGTTGCTCTCTCTCTCACTCTCAATTGGGANNNAGATCGGAAG |
| 3235_1 | TATGAGTGTGGAGTCGTTGCCAGGTGCTGTGTTTTCAAGGAGAGNNNAGATCGGA |
| 3236_1 | TATGAGTGTGGAGTCGTTGCTCTTGCTACCTAGCTATCTTGGGANNNAGATCGGAA |
| 3242_1 | TATGAGTGTGGAGTCGTTGCTATCTGAAAGTGGGGGCAGTTGTANNNAGATCGGA |
| 3250_1 | TATGAGTGTGGAGTCGTTGCGGACTTGATGCTCTTATCCATGGANNNAGATCGGAA |
| 3252_1 | TATGAGTGTGGAGTCGTTGCGGAGGTTGTCAGAAAGTTCATGGANNNAGATCGGA |
| 3255_1 | TATGAGTGTGGAGTCGTTGCTTGCCAGGATTCCCATAGATGGANNNAGATCGGAA |
| 3261_1 | TATGAGTGTGGAGTCGTTGCTTGGAACCTTCAGAATAGTTACCTNNNAGATCGGAA |
| 3264_1 | TATGAGTGTGGAGTCGTTGCCCCTCCTTGACCATTCAGGTTCAANNNAGATCGGAAG |
| 3286_1 | TATGAGTGTGGAGTCGTTGCATCTCCCAACTGCCTTTAGCATCTNNNAGATCGGAAG |
| 3289_1 | TATGAGTGTGGAGTCGTTGCAGTGGACCTTAGCCAATTCTGGANNNAGATCGGAA |
| 3292_1 | TATGAGTGTGGAGTCGTTGCGATAGGCAGTTTACCACTGGGTAANNNAGATCGGA |
| 3296_1 | TATGAGTGTGGAGTCGTTGCTGGGGGTTCTGGAAAATTCTACTGNNNAGATCGGA |
| 3310_1 | TATGAGTGTGGAGTCGTTGCTCTTGCTCCTTGTATCCTGTGGANNNAGATCGGAAG |
| 3315_1 | TATGAGTGTGGAGTCGTTGCTGTGGATTATGAGCAACTTGTGGANNNAGATCGGA |
| 3319_1 | TATGAGTGTGGAGTCGTTGCGGTGAGGTTTCTCAAAGGATTGGANNNAGATCGGA |
| 3320_1 | TATGAGTGTGGAGTCGTTGCCTACCGCCAATGTTCACTCTAAGCNNNAGATCGGAA |

| | |
|---|---|
| 3321_1 | TATGAGTGTGGAGTCGTTGCTTCAGTTGTTCCAGAACTGCTTGTNNNAGATCGGAA |
| 3325_1 | TATGAGTGTGGAGTCGTTGCCCCTTTGAAAGTAGAGAGAGGTGTNNNAGATCGGA |
| 3326_1 | TATGAGTGTGGAGTCGTTGCCAGAGTAGTTCAGTGAGGCCATCANNNAGATCGGA |
| 3329_1 | TATGAGTGTGGAGTCGTTGCTTTGGCTTATCTTCCCAGTTGGANNNAGATCGGAAG |
| 3339_1 | TATGAGTGTGGAGTCGTTGCATTGTATGTGTCCATGATGAGTGCNNNAGATCGGA |
| 3346_1 | TATGAGTGTGGAGTCGTTGCTTCTAGCTCCGTCTCTTGTGAAGGNNNAGATCGGAA |
| 3361_1 | TATGAGTGTGGAGTCGTTGCCCATGTAACCTCCCACTACCACCTNNNAGATCGGAAG |
| 3363_1 | TATGAGTGTGGAGTCGTTGCGGTGTGTAGGTGTAATGCCATGANNNAGATCGGAA |
| 3369_1 | TATGAGTGTGGAGTCGTTGCCTTGCACATTGGTCTAGTGCTTTCNNNAGATCGGAA |
| 3374_1 | TATGAGTGTGGAGTCGTTGCCATAATTGCTCCGACTAGGACATCCNNNAGATCGGA |
| 3382_1 | TATGAGTGTGGAGTCGTTGCAGGTGACATTATGCTTGGAGATGANNNAGATCGGA |
| 3390_1 | TATGAGTGTGGAGTCGTTGCTCCAGACCAGGCTCGTAATAAACANNNAGATCGGAA |
| 3392_1 | TATGAGTGTGGAGTCGTTGCTGTGTGTGTGTGTAAGCTGATGANNNAGATCGGAA |
| 3394_1 | TATGAGTGTGGAGTCGTTGCGCTAAACCCACCTATACAGAGGACNNNAGATCGGAA |
| 3398_1 | TATGAGTGTGGAGTCGTTGCAAGAGACGTGTCAGGGAACTATGANNNAGATCGGA |
| 3400_1 | TATGAGTGTGGAGTCGTTGCTGGGGTGGTATTTGATTCCTATGANNNAGATCGGA |
| 3408_1 | TATGAGTGTGGAGTCGTTGCTTTTCTTCCTCTGAGCCCTGAAAGNNNAGATCGGAAG |
| 3414_1 | TATGAGTGTGGAGTCGTTGCTCTGTTATCATTGGGGATGGGAAGNNNAGATCGGA |
| 3415_1 | TATGAGTGTGGAGTCGTTGCTGATAAGGCCAGAAAAATCACCATNNNAGATCGGA |
| 3421_1 | TATGAGTGTGGAGTCGTTGCTGGTAGTGAACTGTGTTTCACTGANNNAGATCGGA |
| 3425_1 | TATGAGTGTGGAGTCGTTGCACTTGCCTCTTCTTTACTCAGGACNNNAGATCGGAAG |
| 3442_1 | TATGAGTGTGGAGTCGTTGCCTACTTGTCAGGGCTGGAACTATTNNNAGATCGGAA |
| 3443_1 | TATGAGTGTGGAGTCGTTGCTGTGTACTATGCCAAGTGTTGATANNNAGATCGGAA |
| 3447_1 | TATGAGTGTGGAGTCGTTGCGAGAGGAGTATCACAGGATGGAATNNNAGATCGG |
| 3451_1 | TATGAGTGTGGAGTCGTTGCAGATGCATACCATCATGCCAGCTNNNAGATCGGAAG |
| 3459_1 | TATGAGTGTGGAGTCGTTGCGCAAAGGGAAAAACAACTCCATGTNNNAGATCGGA |
| 3465_1 | TATGAGTGTGGAGTCGTTGCTCTTCTCTCTGTCACATTGCCTCTNNNAGATCGGAAG |
| 3466_1 | TATGAGTGTGGAGTCGTTGCAGATATTGGTGTGTTTCAGGCTGANNNAGATCGGA |
| 3470_1 | TATGAGTGTGGAGTCGTTGCAACAGTAGCAGGAACATGGCTCATNNNAGATCGGA |
| 3472_1 | TATGAGTGTGGAGTCGTTGCTGGGTTCCAATTCTATTTGGCTGANNNAGATCGGAA |
| 3483_1 | TATGAGTGTGGAGTCGTTGCGTTTGAAGACTAAAGGATTTGGGGNNNAGATCGGA |
| 3491_1 | TATGAGTGTGGAGTCGTTGCCTGGCTTATTTCACCGACCTCATANNNAGATCGGAAG |
| 3506_1 | TATGAGTGTGGAGTCGTTGCATCTGGTGTCTCCTCCTTTTCTGANNNAGATCGGAAG |
| 3516_1 | TATGAGTGTGGAGTCGTTGCAAGGAAGGAAGGAAAACCAGTGANNNAGATCGGA |
| 3523_1 | TATGAGTGTGGAGTCGTTGCGAAAACGCAGATTCCATTCAGTGANNNAGATCGGA |
| 3524_1 | TATGAGTGTGGAGTCGTTGCTCCTTGGCTCATAACAAAGAGTGANNNAGATCGGAA |
| 3529_1 | TATGAGTGTGGAGTCGTTGCCACAACCGTGTCCATGTAAGTACNNNAGATCGGAAG |
| 3535_1 | TATGAGTGTGGAGTCGTTGCCCTTCCTTCTTTGTACCGGTTTCTNNNAGATCGGAAG |
| 3552_1 | TATGAGTGTGGAGTCGTTGCAGTGGAGCATGTAGTAGATGGTGANNNAGATCGG |
| 3556_1 | TATGAGTGTGGAGTCGTTGCATGGAATATTGGGCTTGAGGACACNNNAGATCGGA |
| 3559_1 | TATGAGTGTGGAGTCGTTGCGCTGGGCACTTTCCATTATCTAGNNNAGATCGGAAG |
| 3562_1 | TATGAGTGTGGAGTCGTTGCTGCTGAGACAAATTATCCATGTGANNNAGATCGGAA |
| 3563_1 | TATGAGTGTGGAGTCGTTGCTGTTTTCATCCATGGTTCGTGACTNNNAGATCGGAA |

| | |
|---|---|
| 3568_1 | TATGAGTGTGGAGTCGTTGCGGGAATAGCCTTGCATTCATTCATNNNAGATCGGAA |
| 3570_1 | TATGAGTGTGGAGTCGTTGCCAAGGCCAGTTAGTTAGCTGTGANNNAGATCGGAA |
| 3572_1 | TATGAGTGTGGAGTCGTTGCTGGGATATTTAGGACAACCAAGGGNNNAGATCGGA |
| 3573_1 | TATGAGTGTGGAGTCGTTGCGTGCGGGTATGTTGACCAATAACCNNNAGATCGGA |
| 3574_1 | TATGAGTGTGGAGTCGTTGCGGTTCAAGTCCAAAGGCAAGGAAANNNAGATCGGA |
| 3578_1 | TATGAGTGTGGAGTCGTTGCTTCTGGGGAGTATGCAAATTTCTCNNNAGATCGGAA |
| 3580_1 | TATGAGTGTGGAGTCGTTGCGGACCAACCTCTGTTCTAGATCCTNNNAGATCGGAA |
| 3585_1 | TATGAGTGTGGAGTCGTTGCCATTGACCTAAACAGGCTTGTGANNNAGATCGGAAG |
| 3589_1 | TATGAGTGTGGAGTCGTTGCATGCAGACCAAAAATGGTTGTGANNNAGATCGGAA |
| 3590_1 | TATGAGTGTGGAGTCGTTGCGGAGGTGATTATGGCTTGGATCTGNNNAGATCGGA |
| 3600_1 | TATGAGTGTGGAGTCGTTGCAATCTCCATTGTTCACAGTAACCTNNNAGATCGGAAG |
| 3605_1 | TATGAGTGTGGAGTCGTTGCACTGGAGAAAGTAGGGTGTATTGANNNAGATCGG |
| 3619_1 | TATGAGTGTGGAGTCGTTGCTCAGGCTTATACCCATTTCCTTGANNNAGATCGGAAG |
| 3622_1 | TATGAGTGTGGAGTCGTTGCGTTGGTCAAATGATTATGGAGGCCNNNAGATCGGA |
| 3629_1 | TATGAGTGTGGAGTCGTTGCAATGCCACACAATGTTTCCTTCTGNNNAGATCGGAA |
| 3630_1 | TATGAGTGTGGAGTCGTTGCTCCACCTTTCTGCTTCTTCCTTTNNNAGATCGGAAGA |
| 3634_1 | TATGAGTGTGGAGTCGTTGCAGATGTGAAAGACTTATCGGTTGANNNAGATCGGA |
| 3636_1 | TATGAGTGTGGAGTCGTTGCGCACTGGTGTTTCTCAAACTTCTCNNNAGATCGGAA |
| 3648_1 | TATGAGTGTGGAGTCGTTGCATGTGTTTGTTTCTTCCAGCCTTGNNNAGATCGGAA |
| 3649_1 | TATGAGTGTGGAGTCGTTGCGGCTATGATTGAGAGACAGTTTGANNNAGATCGGA |
| 3653_1 | TATGAGTGTGGAGTCGTTGCCCTAGACCTTTTCCTCCCTGTACANNNAGATCGGAAG |
| 3664_1 | TATGAGTGTGGAGTCGTTGCTGTCAGCAATAGTCTGCTTTTTGANNNAGATCGGAA |
| 3665_1 | TATGAGTGTGGAGTCGTTGCTGAAACGGCACAATCATAGGTCACNNNAGATCGGA |
| 3666_1 | TATGAGTGTGGAGTCGTTGCCTGTGTGTAAGGTTTTGGACCCACNNNAGATCGGA |
| 3680_1 | TATGAGTGTGGAGTCGTTGCGGGCAGGGAGAGTAGTTACTAGGTNNNAGATCGG |
| 3688_1 | TATGAGTGTGGAGTCGTTGCTTGGCACTACCCTCTCCTCTTTACNNNAGATCGGAAG |
| 3697_1 | TATGAGTGTGGAGTCGTTGCTTCGACTGAACACTTTGCTTTCCTNNNAGATCGGAAG |
| 3712_1 | TATGAGTGTGGAGTCGTTGCCTGTTCCTGGTTGTCATGCCATANNNAGATCGGAAG |
| 3732_1 | TATGAGTGTGGAGTCGTTGCCCTTCATGGCTTTAGCCTTTCCAGNNNAGATCGGAAG |
| 3739_1 | TATGAGTGTGGAGTCGTTGCTTGTGTGTAGCGGTTTGAATGTTGNNNAGATCGGA |
| 3751_1 | TATGAGTGTGGAGTCGTTGCACCCACTGATGGTGTTCAGGATANNNAGATCGGAA |
| 3756_1 | TATGAGTGTGGAGTCGTTGCAAAATCTCCAGGGCTCTTTACTAANNNAGATCGGAA |
| 3760_1 | TATGAGTGTGGAGTCGTTGCGCTCCTTCTACATTCATTTCCAGTNNNAGATCGGAAG |
| 3773_1 | TATGAGTGTGGAGTCGTTGCGCTGAGGGATGGTACAGCCTATANNNAGATCGGAA |
| 3779_1 | TATGAGTGTGGAGTCGTTGCATTGATTTCTGCAACTGTGGAAACNNNAGATCGGAA |
| 3783_1 | TATGAGTGTGGAGTCGTTGCTACTTATTTTCTCTTGCCTGGCCTNNNAGATCGGAAG |
| 3789_1 | TATGAGTGTGGAGTCGTTGCGAGATTACACAGCAAAGCCTGAGTNNNAGATCGGA |
| 3798_1 | TATGAGTGTGGAGTCGTTGCAACGTCCAAGTGCAGGTATGTNNNAGATCGGAAGA |
| 3799_1 | TATGAGTGTGGAGTCGTTGCCCTGGTGTTGCACTTTGAGACTANNNAGATCGGAAG |
| 3802_1 | TATGAGTGTGGAGTCGTTGCCACAAAAGTGAGGACCCCAGTATANNNAGATCGGA |
| 3803_1 | TATGAGTGTGGAGTCGTTGCATAGGCAAGACCCATACCAGTACTNNNAGATCGGAA |
| 3805_1 | TATGAGTGTGGAGTCGTTGCTGGGTCATTTGGCTTGATTCATTANNNAGATCGGAA |
| 3818_1 | TATGAGTGTGGAGTCGTTGCAGTTGAAGCATTTGACATTGACCANNNAGATCGGAA |

| | |
|---|---|
| 3824_1 | TATGAGTGTGGAGTCGTTGCGTATTGCTCTAAGGAACTCCCTGANNNAGATCGGAA |
| 3825_1 | TATGAGTGTGGAGTCGTTGCACTCTACTCCACACTCTCCCCTANNNAGATCGGAAGA |
| 3831_1 | TATGAGTGTGGAGTCGTTGCGTTTGATTGATGAGGTCGATGGTGNNNAGATCGGA |
| 3833_1 | TATGAGTGTGGAGTCGTTGCATGTGTGTGCATTTTGGGAGGGTANNNAGATCGGA |
| 3840_1 | TATGAGTGTGGAGTCGTTGCCCAGGCCTCTAATTTTGTCTTCCTNNNAGATCGGAAG |
| 3846_1 | TATGAGTGTGGAGTCGTTGCTCTGGGGGTTAGTTTTGCTCCTANNNAGATCGGAAG |
| 3858_1 | TATGAGTGTGGAGTCGTTGCTCAGCTCTGGTAAGGGCTACTAGTNNNAGATCGGAA |
| 3860_1 | TATGAGTGTGGAGTCGTTGCGTGTAGTGGTGCAATCCTGGTTTANNNAGATCGGAA |
| 3862_1 | TATGAGTGTGGAGTCGTTGCTGTCATTCCTCTGTGTGTGGACATNNNAGATCGGAA |
| 3863_1 | TATGAGTGTGGAGTCGTTGCTTGCCCTACTTGCATTTCAGCTANNNAGATCGGAAGA |
| 3865_1 | TATGAGTGTGGAGTCGTTGCGCCCCGTGATTTTGTAGGTTTAGANNNAGATCGGAA |
| 3875_1 | TATGAGTGTGGAGTCGTTGCGAGCGTGAATGAAAATACAGCTGGNNNAGATCGGA |
| 3876_1 | TATGAGTGTGGAGTCGTTGCGTGGATTTTTCCATGAGGATGCCANNNAGATCGGAA |
| 3902_1 | TATGAGTGTGGAGTCGTTGCTTCTTTCTCCCCAGTGGCATCTANNNAGATCGGAAGA |
| 3910_1 | TATGAGTGTGGAGTCGTTGCGACCTTTAGCCTCTAGCTTCTCTCTNNNAGATCGGAA |
| 3927_1 | TATGAGTGTGGAGTCGTTGCTGTTGCACTTGTGAATTTATGGCCNNNAGATCGGAA |
| 3933_1 | TATGAGTGTGGAGTCGTTGCAAAGCAGGTAGCTTGAACAATGAGNNNAGATCGGA |
| 3934_1 | TATGAGTGTGGAGTCGTTGCCCAGCGCTCACGTTTTATTTGTACNNNAGATCGGAAG |
| 3945_1 | TATGAGTGTGGAGTCGTTGCTGTCTGTCTTTTCTCACTCAAGGNNNAGATCGGAAGA |
| 3947_1 | TATGAGTGTGGAGTCGTTGCTCATGTTGCTGGAGCTCAGAGTANNNAGATCGGAAG |
| 3948_1 | TATGAGTGTGGAGTCGTTGCTGTGAAATCTTCTGCCTGTCTCCANNNAGATCGGAAG |
| 3949_1 | TATGAGTGTGGAGTCGTTGCTTAGGTTCTTCCCACTCCACATCTNNNAGATCGGAAG |
| 3963_1 | TATGAGTGTGGAGTCGTTGCAGAAAATGCTCAAGCAAGCAGGAGNNNAGATCGGA |
| 3972_1 | TATGAGTGTGGAGTCGTTGCTGTGTGTGTAGACAGTCTTGCTATGNNNAGATCGGA |
| 3975_1 | TATGAGTGTGGAGTCGTTGCTGAGGCTACATGAAACACAGGTANNNAGATCGGAA |
| 3996_1 | TATGAGTGTGGAGTCGTTGCCATAAGCAAGCAAGCAATGACCAGNNNAGATCGGAA |
| 3997_1 | TATGAGTGTGGAGTCGTTGCATTGGCTCTGTCTTTCCTGGAGAANNNAGATCGGAA |
| 3999_1 | TATGAGTGTGGAGTCGTTGCTTATGCTCCGGGATACATGGGTANNNAGATCGGAAG |
| 4013_1 | TATGAGTGTGGAGTCGTTGCCCTGCTAGCTCTGTTCTCTGGTANNNAGATCGGAAGA |
| 4018_1 | TATGAGTGTGGAGTCGTTGCCCATCCTGGTTTGAGAAGCAGAATNNNAGATCGGAA |
| 4026_1 | TATGAGTGTGGAGTCGTTGCGAACCTGATGACCAATGGATTGATNNNAGATCGGAA |
| 4028_1 | TATGAGTGTGGAGTCGTTGCTCAGATGGTGTCTCATTTTGTCGNNNAGATCGGAAG |
| 4039_1 | TATGAGTGTGGAGTCGTTGCGAGGGACAGAGAGCACTTACTACANNNAGATCGGA |
| 4053_1 | TATGAGTGTGGAGTCGTTGCCACAGGAAAAAGGGTGCTCACTTTNNNAGATCGGAA |
| 4068_1 | TATGAGTGTGGAGTCGTTGCCCTGGACTGATGGAAAGTAAGAGGNNNAGATCGGA |
| 4092_1 | TATGAGTGTGGAGTCGTTGCGGGCATGGCTTCTAAAGATTTCTCNNNAGATCGGAA |
| 4100_1 | TATGAGTGTGGAGTCGTTGCCCCCCCTTGGGATTCCTTTCCTTANNNAGATCGGAAGA |
| 4101_1 | TATGAGTGTGGAGTCGTTGCTTGTCCTTCCCTTGGTATCCATGGNNNAGATCGGAAG |
| 4121_1 | TATGAGTGTGGAGTCGTTGCACTAGCCAATCCCCCATCAGTTANNNAGATCGGAAGA |
| 4124_1 | TATGAGTGTGGAGTCGTTGCTGGCAAGTCCTGAGTGAGAGTTANNNAGATCGGAA |
| 4130_1 | TATGAGTGTGGAGTCGTTGCGCAAGGGAGAGATCCTGAGGTTANNNAGATCGGAA |
| 4149_1 | TATGAGTGTGGAGTCGTTGCGACCCCTGGTGCTTGTGAATTTANNNAGATCGGAAG |
| 4165_1 | TATGAGTGTGGAGTCGTTGCTGTCCAAAAATGTTTCAGCTGGGANNNAGATCGGAA |

| | |
|---|---|
| 4166_1 | TATGAGTGTGGAGTCGTTGCAGGGTCACAATATTTATCAGGGCANNNAGATCGGA |
| 4168_1 | TATGAGTGTGGAGTCGTTGCATCTCTGTGAAGCAGTCAGATTGCNNNAGATCGGAA |
| 4174_1 | TATGAGTGTGGAGTCGTTGCTCTGTAGCCATTGGATGACCATTTNNNAGATCGGAA |
| 4183_1 | TATGAGTGTGGAGTCGTTGCTGCCCATGTTCCAAGGCATTTANNNAGATCGGAAG |
| 4193_1 | TATGAGTGTGGAGTCGTTGCGAGAGCGAGCCAGGTTTCTTTTANNNAGATCGGAA |
| 4198_1 | TATGAGTGTGGAGTCGTTGCTGTTTCCTGGTTTGCTGAACTGANNNAGATCGGAAG |
| 4202_1 | TATGAGTGTGGAGTCGTTGCAATACTGGTACACACCACCACACTNNNAGATCGGAA |
| 4206_1 | TATGAGTGTGGAGTCGTTGCTGGTGTGGTGTGTGTGTGTAAAAANNNAGATCGGA |
| 4210_1 | TATGAGTGTGGAGTCGTTGCCGGGCTATTTCTGGAAGGATTGAANNNAGATCGGA |
| 4211_1 | TATGAGTGTGGAGTCGTTGCCTGAAGTTGGGAGAGGGGAAAACNNNAGATCGGA |
| 4220_1 | TATGAGTGTGGAGTCGTTGCCTCCTGACCTCACTCGATTCTCTCNNNAGATCGGAAG |
| 4228_1 | TATGAGTGTGGAGTCGTTGCCACATCGGAGGTCAGCAAATTTGGNNNAGATCGGA |
| 4231_1 | TATGAGTGTGGAGTCGTTGCAAGGCTGGAGTAAAAGCAATGTCCNNNAGATCGGA |
| 4236_1 | TATGAGTGTGGAGTCGTTGCTTGGCAGAGAGTAATAAGAGCCAGNNNAGATCGGA |
| 4242_1 | TATGAGTGTGGAGTCGTTGCTCTGAGGCCTCTTTCCTTCTAAACNNNAGATCGGAAG |
| 4258_1 | TATGAGTGTGGAGTCGTTGCCACATACATATACACACGGAAGCANNNAGATCGGAA |
| 4261_1 | TATGAGTGTGGAGTCGTTGCTGGCTTGGTAAACTGTTTCTTGGGNNNAGATCGGAA |
| 4300_1 | TATGAGTGTGGAGTCGTTGCAGTACGCAAATGTACCTGGGAACNNNAGATCGGAA |
| 4311_1 | TATGAGTGTGGAGTCGTTGCTAAGGTGCACAGAAGCTCTGAACNNNAGATCGGAA |
| 4320_1 | TATGAGTGTGGAGTCGTTGCGGAGAGAGAGAGAAGCTTTGATGGNNNAGATCGG |
| 4322_1 | TATGAGTGTGGAGTCGTTGCATACACAAATATACACGCACATGCNNNAGATCGGAA |
| 4323_1 | TATGAGTGTGGAGTCGTTGCGTGGGTGCAGTTCTCTCTCTAACNNNAGATCGGAAG |
| 4333_1 | TATGAGTGTGGAGTCGTTGCCTTCCTCCTCCCATCTGAAACACNNNAGATCGGAAGA |
| 4340_1 | TATGAGTGTGGAGTCGTTGCTGCTAAGTGCAGCAACAATAACACNNNAGATCGGAA |
| 4345_1 | TATGAGTGTGGAGTCGTTGCCCTTTCGGTTATACACACACACACNNNAGATCGGAAG |
| 4359_1 | TATGAGTGTGGAGTCGTTGCAGTGAGGCAGAGATTTGCATACAANNNAGATCGGA |
| 4362_1 | TATGAGTGTGGAGTCGTTGCAATACAGTGGGCCAGGCATACAGNNNAGATCGGAA |
| 4363_1 | TATGAGTGTGGAGTCGTTGCGCACGCACTATGCTATTTCACACNNNAGATCGGAAG |
| 4373_1 | TATGAGTGTGGAGTCGTTGCTACCCAGTTTTCAGTGCCTACACNNNAGATCGGAAGA |
| 4374_1 | TATGAGTGTGGAGTCGTTGCTGAACCACCATGCTCAGCAATACTNNNAGATCGGAA |
| 4386_1 | TATGAGTGTGGAGTCGTTGCCTTTACTATTCTGCCTAGTGGTCTNNNAGATCGGAAG |
| 4389_1 | TATGAGTGTGGAGTCGTTGCGGTGCTATGGAATGGAGACAGAATNNNAGATCGGA |
| 4395_1 | TATGAGTGTGGAGTCGTTGCACCTTTTGGTCCATGACATACCACNNNAGATCGGAA |
| 4400_1 | TATGAGTGTGGAGTCGTTGCCCTTTCCTTAGGGTTTGCTGTTCTNNNAGATCGGAAG |
| 4405_1 | TATGAGTGTGGAGTCGTTGCTGAGAGCCCCATAATATGCTGTCTNNNAGATCGGAA |
| 4406_1 | TATGAGTGTGGAGTCGTTGCAGATAAAACGCAAGTCAGGGACTGNNNAGATCGGA |
| 4433_1 | TATGAGTGTGGAGTCGTTGCTGGAAATGCTCTGTTTTCTTCCACNNNAGATCGGAA |
| 4442_1 | TATGAGTGTGGAGTCGTTGCCTGACAACAACTAATCCGTTCTCCNNNAGATCGGAAG |
| 4456_1 | TATGAGTGTGGAGTCGTTGCGTGTGTGTAAACAAGTCCTGTCTCNNNAGATCGGAA |
| 4462_1 | TATGAGTGTGGAGTCGTTGCTTGTACCCATTTTTGAGCGTTCATNNNAGATCGGAA |
| 4481_1 | TATGAGTGTGGAGTCGTTGCTGTGTGGATTAGAGTGACCATCACNNNAGATCGGA |
| 4490_1 | TATGAGTGTGGAGTCGTTGCGGGTGTTCTCTCATTTTCTTGTTGNNNAGATCGGAA |
| 4493_1 | TATGAGTGTGGAGTCGTTGCCTCAAAGGAGGATTTGAAGGAGCTNNNAGATCGGA |

| | |
|---|---|
| 4494_1 | TATGAGTGTGGAGTCGTTGCTAGTAACAGACTGGGACCCTCACNNNAGATCGGAAG |
| 4512_1 | TATGAGTGTGGAGTCGTTGCACCTCCTTACAAATCAGCAGTCACNNNAGATCGGAAG |
| 4513_1 | TATGAGTGTGGAGTCGTTGCAGTTGCTCCTTGAATTGGATACCANNNAGATCGGAA |
| 4516_1 | TATGAGTGTGGAGTCGTTGCACCTATATGCTAGATGGCCCACTGNNNAGATCGGAA |
| 4522_1 | TATGAGTGTGGAGTCGTTGCACACTCGCCATCTTACAAACATAGNNNAGATCGGAAG |
| 4544_1 | TATGAGTGTGGAGTCGTTGCACAAACACCATATTTGGCCAAGACNNNAGATCGGAA |
| 4551_1 | TATGAGTGTGGAGTCGTTGCCTGGAACCATCAGCCTGAATCTAANNNAGATCGGAA |
| 4573_1 | TATGAGTGTGGAGTCGTTGCGTGAGAGGCAAGAAGGAGTAAGGANNNAGATCGG |
| 4574_1 | TATGAGTGTGGAGTCGTTGCGAAACTTTTGCTCTGTTGTCCAGGNNNAGATCGGAA |
| 4579_1 | TATGAGTGTGGAGTCGTTGCCAATGTGGCAAGGATTATGACAGGNNNAGATCGGA |
| 4606_1 | TATGAGTGTGGAGTCGTTGCGAAAGTAACAGATGCTGTGGCAAANNNAGATCGGA |
| 4611_1 | TATGAGTGTGGAGTCGTTGCAGACCCTTGAACAATTTGGGGACNNNAGATCGGAAG |
| 4615_1 | TATGAGTGTGGAGTCGTTGCGCTGTCTATACCATTTCTGGACCTCNNNAGATCGGAA |
| 4618_1 | TATGAGTGTGGAGTCGTTGCATCAGACCCTGCAATATCTGGACNNNAGATCGGAAG |
| 4625_1 | TATGAGTGTGGAGTCGTTGCAAGGATAAGAACATGCCAAATTCANNNAGATCGGA |
| 4635_1 | TATGAGTGTGGAGTCGTTGCTGATTTTGGGGCAGAACTTTCTATNNNAGATCGGAA |
| 4646_1 | TATGAGTGTGGAGTCGTTGCTCACTAGGTACTAGCCCCTTGACNNNAGATCGGAAG |
| 4653_1 | TATGAGTGTGGAGTCGTTGCATTTTCCCACCCGTCAAATTCATTNNNAGATCGGAAG |
| 4663_1 | TATGAGTGTGGAGTCGTTGCAGGGTCTCCTAAAATGTCATCTGANNNAGATCGGAA |
| 4666_1 | TATGAGTGTGGAGTCGTTGCATAAAGACATATCAGTGTGGCCATNNNAGATCGGAA |
| 4693_1 | TATGAGTGTGGAGTCGTTGCGTCACCTCCCTGAGATAGGAGGGANNNAGATCGGA |
| 4715_1 | TATGAGTGTGGAGTCGTTGCGCACACAGGGTACTAAGTGGTACNNNAGATCGGAA |
| 4722_1 | TATGAGTGTGGAGTCGTTGCTATGCACTTGGATGGAATTGTCCANNNAGATCGGAA |
| 4729_1 | TATGAGTGTGGAGTCGTTGCCAGAGAGAGGCTTAGGTCCGATTGNNNAGATCGGA |
| 4731_1 | TATGAGTGTGGAGTCGTTGCTCCTCGACTTCCAATGAGGTTACNNNAGATCGGAAG |
| 4739_1 | TATGAGTGTGGAGTCGTTGCTTTTCCTGCTTCTTAGTGTGGTTTNNNAGATCGGAAG |
| 4744_1 | TATGAGTGTGGAGTCGTTGCCCCTATTTGGCAGATCTGAAAACCNNNAGATCGGAA |
| 4746_1 | TATGAGTGTGGAGTCGTTGCGGAGCAAGAACTGAAAGGATTAGCNNNAGATCGGA |
| 4748_1 | TATGAGTGTGGAGTCGTTGCATCCTTTCAAAGACCAAGCTTTTGNNNAGATCGGAA |
| 4753_1 | TATGAGTGTGGAGTCGTTGCAGGAGAGAAGAGATGGGGAAACCNNNAGATCGGA |
| 4810_1 | TATGAGTGTGGAGTCGTTGCGTAACTGAGGACAGGGAGCTATGCNNNAGATCGGA |
| 4812_1 | TATGAGTGTGGAGTCGTTGCCCAAATCAGATTCAATGGCCACCNNNAGATCGGAAG |
| 4814_1 | TATGAGTGTGGAGTCGTTGCATCTGTGGGAGTGCATTCATTTTTNNNAGATCGGAA |
| 4832_1 | TATGAGTGTGGAGTCGTTGCGGAGCTTTGCTGAATCTTTTGCAGNNNAGATCGGAA |
| 4840_1 | TATGAGTGTGGAGTCGTTGCTGTGTGTCTCTCATTAGTTTCACCNNNAGATCGGAAG |
| 4847_1 | TATGAGTGTGGAGTCGTTGCGTATTCTGGATCCCTTTCTGTTGTNNNAGATCGGAAG |
| 4863_1 | TATGAGTGTGGAGTCGTTGCAATCTGACTGAGACAGGCACACAANNNAGATCGGA |
| 4870_1 | TATGAGTGTGGAGTCGTTGCGGTCACACTTGATTTTCACACTTGNNNAGATCGGAA |
| 4874_1 | TATGAGTGTGGAGTCGTTGCAGTAAAATGTCAAAGGCGACTGACNNNAGATCGGA |
| 4914_1 | TATGAGTGTGGAGTCGTTGCAGTTGTTTCCCACTTGAACTCCTGNNNAGATCGGAAG |
| 4916_1 | TATGAGTGTGGAGTCGTTGCACCTCCCATGAGATACTAACACCCNNNAGATCGGAAG |
| 4924_1 | TATGAGTGTGGAGTCGTTGCAGCATGGCAGTAAATCCTTGATCTNNNAGATCGGAA |
| 4934_1 | TATGAGTGTGGAGTCGTTGCTGAAAGTGGCGTATTGAAATACCCNNNAGATCGGA |

| | |
|---|---|
| 4943_1 | TATGAGTGTGGAGTCGTTGCACAAAGGCAGAAACTGTAACCCCNNNAGATCGGAA |
| 4945_1 | TATGAGTGTGGAGTCGTTGCATACAGGAAATGCAGCAACATAGANNNAGATCGGA |
| 4946_1 | TATGAGTGTGGAGTCGTTGCAGCAAAATGAAAATGCAAGACCCCNNNAGATCGGA |
| 4953_1 | TATGAGTGTGGAGTCGTTGCTTGTGTGTCTTGATGGATTCAGAGNNNAGATCGGA |
| 4967_1 | TATGAGTGTGGAGTCGTTGCAGCAGGCACCGTATATTTGTTCTGNNNAGATCGGAA |
| 4982_1 | TATGAGTGTGGAGTCGTTGCTGTATGTGGAGAGAGAGAATCCGTNNNAGATCGGA |
| 4985_1 | TATGAGTGTGGAGTCGTTGCCAACCTGCTATACATTAGGTCCCCNNNAGATCGGAA |
| 4986_1 | TATGAGTGTGGAGTCGTTGCGGAGTAAGTCAAAGATGTGTCCCCNNNAGATCGGA |
| 4987_1 | TATGAGTGTGGAGTCGTTGCCTGAGACAGAATCTCACTGTGTGTNNNAGATCGGAA |
| 4990_1 | TATGAGTGTGGAGTCGTTGCCCCTTTGGATTTCTCGTGGATAGTNNNAGATCGGAA |
| 4993_1 | TATGAGTGTGGAGTCGTTGCTATGTCTGCTTTCTCAGAAGCCCNNNAGATCGGAAG |
| 5001_1 | TATGAGTGTGGAGTCGTTGCTTCAGAGAAAAAGAACCTGGGACCNNNAGATCGGA |
| 5006_1 | TATGAGTGTGGAGTCGTTGCAGGAACTTGATCATTGTGTAGCCCNNNAGATCGGAA |
| 5009_1 | TATGAGTGTGGAGTCGTTGCAAGCAGTCTGCAAACCAGAAAGANNNAGATCGGAA |
| 5016_1 | TATGAGTGTGGAGTCGTTGCCTGGTTGGGGTTTGTTGTGAATCANNNAGATCGGA |
| 5017_1 | TATGAGTGTGGAGTCGTTGCGTCAGAACTTATGTGTTTGTGCGTNNNAGATCGGAA |
| 5020_1 | TATGAGTGTGGAGTCGTTGCTGTAGCTTTGTTTTGCCTGACTAAANNNAGATCGGA |
| 5047_1 | TATGAGTGTGGAGTCGTTGCGCAGACATCAAGGCAATTTATCCCNNNAGATCGGAA |
| 5056_1 | TATGAGTGTGGAGTCGTTGCAGTGAGTGCATGTAACAATGGTGANNNAGATCGGA |
| 5080_1 | TATGAGTGTGGAGTCGTTGCACAAACAGCTACTGTCTCATTCCCNNNAGATCGGAAG |
| 5081_1 | TATGAGTGTGGAGTCGTTGCAGTGAGGAAGAGGTGACTATTCCCNNNAGATCGGA |
| 5083_1 | TATGAGTGTGGAGTCGTTGCACTGGAAGAGTAAGGAACTTCCCNNNAGATCGGAA |
| 5087_1 | TATGAGTGTGGAGTCGTTGCCTGTCTTCTCTAGGAAACCTTCCCNNNAGATCGGAAG |
| 5095_1 | TATGAGTGTGGAGTCGTTGCTACATGAGCAACACCAGTTTCCCNNNAGATCGGAAG |
| 5104_1 | TATGAGTGTGGAGTCGTTGCACTAGGCTCAAGCCATCTCCCTNNNAGATCGGAAGA |
| 5131_1 | TATGAGTGTGGAGTCGTTGCAACAATGGGCTGAATCAGGAGATANNNAGATCGGA |
| 5133_1 | TATGAGTGTGGAGTCGTTGCACTCCTTGTAAGTTTGCAGAGCCNNNAGATCGGAAG |
| 5141_1 | TATGAGTGTGGAGTCGTTGCCGCCGGACCACTTGTTTGTTTTGTTTTNNNAGATCGG |
| 5158_1 | TATGAGTGTGGAGTCGTTGCAGGAGAACAGAAGAAAGAGCAATCNNNAGATCGG |
| 5163_1 | TATGAGTGTGGAGTCGTTGCCTGCATCTTGCTATAGTTGCCTAANNNAGATCGGAA |
| 5176_1 | TATGAGTGTGGAGTCGTTGCACCCCTCTCTTTTATCTCAGGCCNNNAGATCGGAAGA |
| 5191_1 | TATGAGTGTGGAGTCGTTGCGAGGAACAGTTTGAAACACTTGGCNNNAGATCGGA |
| 5195_1 | TATGAGTGTGGAGTCGTTGCGGTGGCTATTACTGAAATCGCAATNNNAGATCGGA |
| 5204_1 | TATGAGTGTGGAGTCGTTGCCTTTCATCTGTCAGCATCTGGCCNNNAGATCGGAAG |
| 5205_1 | TATGAGTGTGGAGTCGTTGCTGTGACGTGTATGTGAGTGGTGTGNNNAGATCGGA |
| 5212_1 | TATGAGTGTGGAGTCGTTGCGCCAAGTGAAAAGAGCCAAACAGNNNAGATCGGAA |
| 5242_1 | TATGAGTGTGGAGTCGTTGCGATTCTACTTTCCTGACCCTGCCNNNAGATCGGAAGA |
| 5245_1 | TATGAGTGTGGAGTCGTTGCACACTATTCTGCATCCAGTATCCANNNAGATCGGAAG |
| 5247_1 | TATGAGTGTGGAGTCGTTGCTGCCAGAAAAGTTTGAAATTGGCTNNNAGATCGGA |
| 5248_1 | TATGAGTGTGGAGTCGTTGCTATGGCATGAAAAATTATGCTGCCNNNAGATCGGAA |
| 5251_1 | TATGAGTGTGGAGTCGTTGCACTAAGAGAACAAGTGTCTCTGCCNNNAGATCGGAA |
| 5252_1 | TATGAGTGTGGAGTCGTTGCTCACTTGGGACTTTCTCAGTTCAANNNAGATCGGAA |
| 5273_1 | TATGAGTGTGGAGTCGTTGCGAAAACTAGTGCAGTCATTTGGCTNNNAGATCGGA |

| | |
|---|---|
| 5275_1 | TATGAGTGTGGAGTCGTTGCTCCTCAGTTGCTCTTCTCAAATCCNNNAGATCGGAAG |
| 5281_1 | TATGAGTGTGGAGTCGTTGCGCATTTGTGTTGCTTTCAACATCCNNNAGATCGGAA |
| 5307_1 | TATGAGTGTGGAGTCGTTGCGTATGATGCAAAGGTCCCATCTCANNNAGATCGGAA |
| 5317_1 | TATGAGTGTGGAGTCGTTGCACAAACATTGCAGCTCTTTACTCCNNNAGATCGGAAG |
| 5333_1 | TATGAGTGTGGAGTCGTTGCCCCTACTTCCTTCTTCTTTCCTCCNNNAGATCGGAAGA |
| 5339_1 | TATGAGTGTGGAGTCGTTGCTCTAGCATGAAGCTGTTTGCTCCNNNAGATCGGAAG |
| 5370_1 | TATGAGTGTGGAGTCGTTGCCCACTGTCTTCAATCAAAAGGTCCNNNAGATCGGAA |
| 5380_1 | TATGAGTGTGGAGTCGTTGCTTATTACTCGCCTGGCAATGTCCNNNAGATCGGAAG |
| 5382_1 | TATGAGTGTGGAGTCGTTGCTGGTTGATGGTGAGAAACTGTCCNNNAGATCGGAA |
| 5397_1 | TATGAGTGTGGAGTCGTTGCTCCATAGCTCTCTATGCCCTTCCNNNAGATCGGAAGA |
| 5399_1 | TATGAGTGTGGAGTCGTTGCGTTGTGTGAAGTCCTCTCCTTCCNNNAGATCGGAAG |
| 5403_1 | TATGAGTGTGGAGTCGTTGCTCTGATCATCTGGAGAACCCTTGANNNAGATCGGAA |
| 5413_1 | TATGAGTGTGGAGTCGTTGCTCCTTTCATGTAGCAGCTTCTTCCNNNAGATCGGAAG |
| 5414_1 | TATGAGTGTGGAGTCGTTGCCCAATGAGCTGTCCAGTTATGCTNNNAGATCGGAAG |
| 5420_1 | TATGAGTGTGGAGTCGTTGCTGCCTGAAATTTGACTTCTGTTCCNNNAGATCGGAA |
| 5423_1 | TATGAGTGTGGAGTCGTTGCTGGGATTGTTGTATACGGTGTTCCNNNAGATCGGA |
| 5432_1 | TATGAGTGTGGAGTCGTTGCTTTGTGCTCCAGTTAGGCATATCTNNNAGATCGGAA |
| 5444_1 | TATGAGTGTGGAGTCGTTGCGGTCAATTTATGGCTGTGGTTTTGNNNAGATCGGA |
| 5448_1 | TATGAGTGTGGAGTCGTTGCTGTGTGTTTCCTAAGCAGAAAAGCNNNAGATCGGA |
| 5458_1 | TATGAGTGTGGAGTCGTTGCTGCTTCTCTGTGCACTTCTGTTACNNNAGATCGGAAG |
| 5471_1 | TATGAGTGTGGAGTCGTTGCCATTTACAATCTGCGGGTCAAGCNNNAGATCGGAAG |
| 5488_1 | TATGAGTGTGGAGTCGTTGCTTTTCCAGGTACCAATGGGAAGCNNNAGATCGGAA |
| 5494_1 | TATGAGTGTGGAGTCGTTGCACTGCTTTTCCTCTGCAGAAAGTNNNAGATCGGAAG |
| 5504_1 | TATGAGTGTGGAGTCGTTGCACACATGAGGACAAAGAAAACAGANNNAGATCGGA |
| 5515_1 | TATGAGTGTGGAGTCGTTGCGGGAAATCTTTTCAATGGACCAGCNNNAGATCGGA |
| 5529_1 | TATGAGTGTGGAGTCGTTGCTCCAAATGTATCAGGCTATCCAGCNNNAGATCGGAA |
| 5531_1 | TATGAGTGTGGAGTCGTTGCATGTGTGTTTGGGGTTTGTTTGTTNNNAGATCGGA |
| 5565_1 | TATGAGTGTGGAGTCGTTGCAGTCCTGTAAACATTGTCACTGAGNNNAGATCGGAA |
| 5573_1 | TATGAGTGTGGAGTCGTTGCAGCCAGGCAAATTGGAAATGAGTNNNAGATCGGAA |
| 5584_1 | TATGAGTGTGGAGTCGTTGCATTTTTGGCATTTCCTGCAGAGCNNNAGATCGGAAG |
| 5587_1 | TATGAGTGTGGAGTCGTTGCGTAGAGGCAGTTTTCAAAGAGAGCNNNAGATCGGA |
| 5594_1 | TATGAGTGTGGAGTCGTTGCGCACAGGGAAAATATTGGTAGAGCNNNAGATCGGA |
| 5633_1 | TATGAGTGTGGAGTCGTTGCATTTCGGAAGACACTGCTACTAGCNNNAGATCGGAA |
| 5639_1 | TATGAGTGTGGAGTCGTTGCGTAGGTGGTAGGTGGTAGGTTTTCNNNAGATCGGA |
| 5643_1 | TATGAGTGTGGAGTCGTTGCCCAGGCTCTTGAGTTACTGTAGCNNNAGATCGGAAG |
| 5662_1 | TATGAGTGTGGAGTCGTTGCATTTGAAAGTACGGGTAGGACGCNNNAGATCGGAA |
| 5672_1 | TATGAGTGTGGAGTCGTTGCGCCAGGTTTTGGTATCAGGAATGCNNNAGATCGGA |
| 5683_1 | TATGAGTGTGGAGTCGTTGCTCTGAGGAATTGTACTGCAAGGCNNNAGATCGGAA |
| 5695_1 | TATGAGTGTGGAGTCGTTGCAACTGTGAAGGATCAAGACAGGCNNNAGATCGGAA |
| 5740_1 | TATGAGTGTGGAGTCGTTGCTCACAATGTTGGGGTACTTTAGGCNNNAGATCGGA |
| 5755_1 | TATGAGTGTGGAGTCGTTGCCCTGGTTTCTCTTCCTTTACAGCANNNAGATCGGAAG |
| 5756_1 | TATGAGTGTGGAGTCGTTGCGTCCCTGTAGTTGAGGGTGTCTTNNNAGATCGGAA |
| 5757_1 | TATGAGTGTGGAGTCGTTGCATCCCTAAACCATTCCAATGGGCNNNAGATCGGAAG |

| | |
|---|---|
| 5767_1 | TATGAGTGTGGAGTCGTTGCGTTTGGTGATGTTCCTCCAGATTTNNNAGATCGGAA |
| 5790_1 | TATGAGTGTGGAGTCGTTGCCACTTGTAGGTTGATGCGCATTTGNNNAGATCGGAA |
| 5796_1 | TATGAGTGTGGAGTCGTTGCATTAGCTGGCATGAAGCACTGTNNNAGATCGGAAG |
| 5821_1 | TATGAGTGTGGAGTCGTTGCATGGGAGGCTACATATAGGCTAGANNNAGATCGGA |
| 5828_1 | TATGAGTGTGGAGTCGTTGCGCTGGTTTTTCTGGTAAGATTGGCNNNAGATCGGA |
| 5830_1 | TATGAGTGTGGAGTCGTTGCTCGGAATGAACAAACATGATTGGCNNNAGATCGG |
| 5843_1 | TATGAGTGTGGAGTCGTTGCAACAAAAATTGCCTGCTGTGCTTNNNAGATCGGAAG |
| 5879_1 | TATGAGTGTGGAGTCGTTGCTCAAGAGGAGGACAAACAGATGCNNNAGATCGGAA |
| 5894_1 | TATGAGTGTGGAGTCGTTGCCGTTTTCCCAGAGCAGATTTATGCNNNAGATCGGAA |
| 5928_1 | TATGAGTGTGGAGTCGTTGCCTGACCATTACTGCACCAGTCAANNNAGATCGGAAG |
| 5931_1 | TATGAGTGTGGAGTCGTTGCCTTGATCTTGTGGCACCTTGATCTNNNAGATCGGAA |
| 5961_1 | TATGAGTGTGGAGTCGTTGCGAGTGTGGTTCTTCCATGCTCTTNNNAGATCGGAAG |
| 5976_1 | TATGAGTGTGGAGTCGTTGCAATGAATGGAGGGATGATGGTGCNNNAGATCGG |
| 5978_1 | TATGAGTGTGGAGTCGTTGCCAGTGACTTACACAAAGTTGGTGCNNNAGATCGGA |
| 5985_1 | TATGAGTGTGGAGTCGTTGCTGCCAAGGCTTATAAAAACTGTGCNNNAGATCGGAA |
| 5999_1 | TATGAGTGTGGAGTCGTTGCCCAACACCTAGGCTGAAGTATAGTNNNAGATCGGAA |
| 6012_1 | TATGAGTGTGGAGTCGTTGCGGCTGTCATCTCAGAAAAGATTGCNNNAGATCGGA |
| 6014_1 | TATGAGTGTGGAGTCGTTGCACAGCCAGTGTCTTCATAGTGAANNNAGATCGGAAG |
| 6027_1 | TATGAGTGTGGAGTCGTTGCCCTGGGAGTATCTGTCTAGCATCTNNNAGATCGGAA |
| 6028_1 | TATGAGTGTGGAGTCGTTGCGTTAGGGTTGCAATGAAACGTGTGNNNAGATCGGA |
| 6040_1 | TATGAGTGTGGAGTCGTTGCAAACACTAGCCCCATGCTCAGTTCNNNAGATCGGAA |
| 6045_1 | TATGAGTGTGGAGTCGTTGCGTTGGAATGGAAATGAAACGAGGTNNNAGATCGG |
| 6051_1 | TATGAGTGTGGAGTCGTTGCTGTGGTGAAAAGAAATGTTGTTGCNNNAGATCGGA |
| 6062_1 | TATGAGTGTGGAGTCGTTGCGGGGTACTGAAGAGCTTATTCTCCNNNAGATCGGA |
| 6073_1 | TATGAGTGTGGAGTCGTTGCCTGGAGCAGGGAAGAGTGAAATCNNNAGATCGGA |
| 6094_1 | TATGAGTGTGGAGTCGTTGCGTTACAAGAACCCAACGCACATCNNNAGATCGGAAG |
| 6103_1 | TATGAGTGTGGAGTCGTTGCAATGCCCTCTCCTTTTCACCATCNNNAGATCGGAAGA |
| 6109_1 | TATGAGTGTGGAGTCGTTGCTGGGTGGGGAAATATGACTGAGAANNNAGATCGG |
| 6124_1 | TATGAGTGTGGAGTCGTTGCGGAAAGATGGATTTGTGTGGAGGTNNNAGATCGG |
| 6132_1 | TATGAGTGTGGAGTCGTTGCTCTCTTTCTACTCCCATCTCCAGANNNAGATCGGAAG |
| 6152_1 | TATGAGTGTGGAGTCGTTGCAGAGAGAACCCTGTCCAAGGATCNNNAGATCGGAA |
| 6183_1 | TATGAGTGTGGAGTCGTTGCTAAAGGCTGAGATGTCCCAACTCNNNAGATCGGAAG |
| 6220_1 | TATGAGTGTGGAGTCGTTGCTTTGACTTAGAGGTAGCCCCCTCNNNAGATCGGAAG |
| 6263_1 | TATGAGTGTGGAGTCGTTGCCCTGCAAAGGTCAAAATCAGCTCNNNAGATCGGAAG |
| 6265_1 | TATGAGTGTGGAGTCGTTGCACAGTCATTATTCAGAGCAACTCANNNAGATCGGAA |
| 6276_1 | TATGAGTGTGGAGTCGTTGCTAGGGCTTCAGAGATGACTGATCTTNNNAGATCGG |
| 6278_1 | TATGAGTGTGGAGTCGTTGCCACTATTCCAGGCACATAGGCTCNNNAGATCGGAAG |
| 6304_1 | TATGAGTGTGGAGTCGTTGCATGCAAAGTACTGTTGTGGGAGCANNNAGATCGGA |
| 6316_1 | TATGAGTGTGGAGTCGTTGCTGGGTATGCAACTCTCCTCTCTCNNNAGATCGGAAG |
| 6322_1 | TATGAGTGTGGAGTCGTTGCGGATCTCTCCTAACAGGGGTCTCNNNAGATCGGAAG |
| 6332_1 | TATGAGTGTGGAGTCGTTGCCCCATTAGACTGTCCAGGCTCTTCNNNAGATCGGAA |
| 6372_1 | TATGAGTGTGGAGTCGTTGCCCCTCAAACAAAGGGAATCCTGTNNNAGATCGGAAG |
| 6379_1 | TATGAGTGTGGAGTCGTTGCTGCCTTCGAGAGAAATCTGTGAGNNNAGATCGGAA |

| | |
|---|---|
| 6394_1 | TATGAGTGTGGAGTCGTTGCTCTGGGATGCAGTAGAACATGTCNNNAGATCGGAA |
| 6439_1 | TATGAGTGTGGAGTCGTTGCGGATCAGTTCTGCCTGGAGATTCNNNAGATCGGAAG |
| 6460_1 | TATGAGTGTGGAGTCGTTGCACTGACAATTCCTAACCCTCCTCTNNNAGATCGGAAG |
| 6465_1 | TATGAGTGTGGAGTCGTTGCTCAGTAACCAACGCTGGAGAAAANNNAGATCGGAA |
| 6468_1 | TATGAGTGTGGAGTCGTTGCTCAGCTTTAGCCACAGTAGCTTCNNNAGATCGGAAG |
| 6479_1 | TATGAGTGTGGAGTCGTTGCGGCTGAATACCCTGCTTCTCTTCNNNAGATCGGAAG |
| 6482_1 | TATGAGTGTGGAGTCGTTGCCCAGTAAGTAGCCTAGAGCAGTTNNNAGATCGGAAG |
| 6490_1 | TATGAGTGTGGAGTCGTTGCAGGGTAGAATACTTGGGTAAGGCANNNAGATCGGA |
| 6502_1 | TATGAGTGTGGAGTCGTTGCTTGTTACCAGCCCAGATTGGTTCNNNAGATCGGAAG |
| 6516_1 | TATGAGTGTGGAGTCGTTGCCTACACTAGGTCCCGTGCATTTCNNNAGATCGGAAG |
| 6552_1 | TATGAGTGTGGAGTCGTTGCCGTCAGGGATGTAGACGATGTTGNNNAGATCGGAA |
| 6566_1 | TATGAGTGTGGAGTCGTTGCCCAGGCTCAGCTTTAGTACCAGAANNNAGATCGGAA |
| 6586_1 | TATGAGTGTGGAGTCGTTGCGGGGTGCAGAAAGGGTAAGAAAGNNNAGATCGGA |
| 6591_1 | TATGAGTGTGGAGTCGTTGCGAGGAAGGAGAAAGTCCCTTGTCANNNAGATCGGA |
| 6628_1 | TATGAGTGTGGAGTCGTTGCACTTCAGCAGGACATTTCTGTGCTNNNAGATCGGAA |
| 6646_1 | TATGAGTGTGGAGTCGTTGCGACAGCAGGAAGCAGAGATCAAGNNNAGATCGGA |
| 6647_1 | TATGAGTGTGGAGTCGTTGCGCTGGGGAAAGTGGTCTATCAAGNNNAGATCGGAA |
| 6653_1 | TATGAGTGTGGAGTCGTTGCACAACTTCTGGCTTCCTTTCCTCCNNNAGATCGGAAG |
| 6655_1 | TATGAGTGTGGAGTCGTTGCTTGTACCCACTACCACAGTCAAGNNNAGATCGGAAG |
| 6669_1 | TATGAGTGTGGAGTCGTTGCTCTCTCTCTCCCTTGCCACCATTNNNAGATCGGAAGA |
| 6677_1 | TATGAGTGTGGAGTCGTTGCTGGTGTGTGCTTTCGTAGAGAAGNNNAGATCGGAA |
| 6684_1 | TATGAGTGTGGAGTCGTTGCGGGATAGCAATGAATTGCCGAAGNNNAGATCGGAA |
| 6693_1 | TATGAGTGTGGAGTCGTTGCAGCCCTCCTAGCCACCTAAAGAATNNNAGATCGGAA |
| 6696_1 | TATGAGTGTGGAGTCGTTGCTCAGTTGGCCCATTTTTCCACAGCNNNAGATCGGAAG |
| 6698_1 | TATGAGTGTGGAGTCGTTGCAACCCTGGCTAAAACAGTGAACTCNNNAGATCGGAA |
| 6707_1 | TATGAGTGTGGAGTCGTTGCTACCACAATTGTAAGCCTCCTGAGNNNAGATCGGAA |
| 6724_1 | TATGAGTGTGGAGTCGTTGCCTGGAGCTGAGACATTCTTCTCCANNNAGATCGGAA |
| 6725_1 | TATGAGTGTGGAGTCGTTGCGCGTGCGTGTATGTGAATTGAAGNNNAGATCGGAA |
| 6726_1 | TATGAGTGTGGAGTCGTTGCTAGAATCTCGCTCTGTCTCCCATTNNNAGATCGGAAG |
| 6731_1 | TATGAGTGTGGAGTCGTTGCGGGGGAAAATGGTAGTTTGAGCTANNNAGATCGGA |
| 6741_1 | TATGAGTGTGGAGTCGTTGCTCTCCACTTTCCCAAGTCATTCATNNNAGATCGGAAG |
| 6748_1 | TATGAGTGTGGAGTCGTTGCAAGCAATTCAGCATCTGTGGCTANNNAGATCGGAAG |
| 6752_1 | TATGAGTGTGGAGTCGTTGCTTTACTGGGGACGTTCCAAACAGNNNAGATCGGAAG |
| 6764_1 | TATGAGTGTGGAGTCGTTGCAATCTTTTCCAAGTGGGAGAACAGNNNAGATCGGA |
| 6784_1 | TATGAGTGTGGAGTCGTTGCACGCTATCCACTACTTTGCACAGNNNAGATCGGAAG |
| 6791_1 | TATGAGTGTGGAGTCGTTGCCAAAAACCTCAACTATTCCTCCTCNNNAGATCGGAAG |
| 6801_1 | TATGAGTGTGGAGTCGTTGCAGAAGCATGAATGGCAGAGACAGNNNAGATCGGA |
| 6806_1 | TATGAGTGTGGAGTCGTTGCACCAAGCAGTGCCAAAAAGTACTNNNAGATCGGAAG |
| 6807_1 | TATGAGTGTGGAGTCGTTGCAAGGGCAAAAGGACATTAGTGGCANNNAGATCGGA |
| 6811_1 | TATGAGTGTGGAGTCGTTGCCCCAGAATAGCCTGTAACCATCCTNNNAGATCGGAAG |
| 6817_1 | TATGAGTGTGGAGTCGTTGCGTTTGCACCTGTGTCTCTCAGTATNNNAGATCGGAAG |
| 6825_1 | TATGAGTGTGGAGTCGTTGCCTCCTGCACTGATTTTACCTTCCANNNAGATCGGAAG |
| 6834_1 | TATGAGTGTGGAGTCGTTGCCTTTGTAACCTGCTGTCCTACAGCNNNAGATCGGAAG |

| | |
|---|---|
| 6836_1 | TATGAGTGTGGAGTCGTTGCAGGAGCCACATTTTCTTGACCAGNNNAGATCGGAAG |
| 6840_1 | TATGAGTGTGGAGTCGTTGCAGGATACAGCAAGGTGTCTTTCCANNNAGATCGGAA |
| 6841_1 | TATGAGTGTGGAGTCGTTGCGTTTTGTACAGAGGGGAAGGTGAGNNNAGATCGGA |
| 6844_1 | TATGAGTGTGGAGTCGTTGCGAATGTACGCTGATACGTTTCACTNNNAGATCGGAAG |
| 6864_1 | TATGAGTGTGGAGTCGTTGCTGGCTTTTGTCAGTTCTCCATTCANNNAGATCGGAAG |
| 6872_1 | TATGAGTGTGGAGTCGTTGCACTCAAAAAGCTGCAACAATCCAGNNNAGATCGGAA |
| 6874_1 | TATGAGTGTGGAGTCGTTGCCTTGTCGATAGCTTGCTGGTGTACNNNAGATCGGAA |
| 6878_1 | TATGAGTGTGGAGTCGTTGCTGACCATGCAGGTTTTATTCCAGNNNAGATCGGAAG |
| 6879_1 | TATGAGTGTGGAGTCGTTGCCCCTATTCTCTGTGGATGTCACCANNNAGATCGGAAG |
| 6883_1 | TATGAGTGTGGAGTCGTTGCGTTGTGCTCTCTCTCTCTTACTCTNNNAGATCGGAAGA |
| 6894_1 | TATGAGTGTGGAGTCGTTGCTGCTTCTGTTTCTTTTGAGAACCTNNNAGATCGGAAG |
| 6917_1 | TATGAGTGTGGAGTCGTTGCCAGACATACACATCAGCGGATGCTCCNNNAGATCGGA |
| 6921_1 | TATGAGTGTGGAGTCGTTGCAAAGCCATAAAGGCACAATGCAGNNNAGATCGGAAG |
| 6924_1 | TATGAGTGTGGAGTCGTTGCCTCTCTTTCTTCTCCCTCAGTTGCNNNAGATCGGAAGA |
| 6927_1 | TATGAGTGTGGAGTCGTTGCACAGCTACCACCCATTCTTGGATTNNNAGATCGGAAG |
| 6962_1 | TATGAGTGTGGAGTCGTTGCGCTTGTTTGACCTTCAGTAGGCAANNNAGATCGGAA |
| 6965_1 | TATGAGTGTGGAGTCGTTGCAGCGTATCACCTGTCTAGCTCAGNNNAGATCGGAAG |
| 6972_1 | TATGAGTGTGGAGTCGTTGCTTTCTTCTCCTGGGAATTTCAGAGNNNAGATCGGAAG |
| 6976_1 | TATGAGTGTGGAGTCGTTGCTGAGAGTGTGTGTGTGTTTGTAGTNNNAGATCGGAA |
| 6984_1 | TATGAGTGTGGAGTCGTTGCGATGGATGTGCGGTTGGATACGTANNNAGATCGGA |
| 6987_1 | TATGAGTGTGGAGTCGTTGCACTGGGACATTGAGAAGAAGGATGNNNAGATCGGA |
| 7009_1 | TATGAGTGTGGAGTCGTTGCGCAGACTGGACATTTCAGGGGAAANNNAGATCGGA |
| 7015_1 | TATGAGTGTGGAGTCGTTGCTAAGGAATAGGGTGAGAGGGCACANNNAGATCGGA |
| 7034_1 | TATGAGTGTGGAGTCGTTGCATTCAGATGTTCCTAGGTGTGTGGNNNAGATCGGAA |
| 7042_1 | TATGAGTGTGGAGTCGTTGCCCCACATGATTACTGAGGCAGAGNNNAGATCGGAAG |
| 7045_1 | TATGAGTGTGGAGTCGTTGCAACACAATCTCAGCTTACGGCAGNNNAGATCGGAAG |
| 7052_1 | TATGAGTGTGGAGTCGTTGCCTGGCTAAATTCTGGCGGCTTCTTNNNAGATCGGAAG |
| 7061_1 | TATGAGTGTGGAGTCGTTGCTCCTGCATTCTACTGGTTCTGTTNNNAGATCGGAAGA |
| 7067_1 | TATGAGTGTGGAGTCGTTGCCATCAACCATGCAGGTAAAGACANNNAGATCGGAAG |
| 7069_1 | TATGAGTGTGGAGTCGTTGCTAATGGTTTCCGCAGGATGAGAGNNNAGATCGGAAG |
| 7074_1 | TATGAGTGTGGAGTCGTTGCCCCTTCTTTCCTGCCTTGGATATTNNNAGATCGGAAGA |
| 7080_1 | TATGAGTGTGGAGTCGTTGCGTGACATCGAGGACCTTAGGAAAGNNNAGATCGGA |
| 7095_1 | TATGAGTGTGGAGTCGTTGCTGTGAGTGTGTGTTTGAAGAGGAGNNNAGATCGGA |
| 7102_1 | TATGAGTGTGGAGTCGTTGCCAAATGGTTGAATGAATGGTGAAAAAGNNNAGATCG |
| 7120_1 | TATGAGTGTGGAGTCGTTGCTAAAGCAGGTAAAGCCATGGGAGNNNAGATCGGAA |
| 7123_1 | TATGAGTGTGGAGTCGTTGCTTGTTCTTGCTCTCTCTTCCCTCTNNNAGATCGGAAGA |
| 7134_1 | TATGAGTGTGGAGTCGTTGCCCCACCCTCCAGAATGGTTCTATTNNNAGATCGGAAG |
| 7135_1 | TATGAGTGTGGAGTCGTTGCTAGCAGGGAGATCTAGACTTCACGNNNAGATCGGAA |
| 7156_1 | TATGAGTGTGGAGTCGTTGCTCCACAGTTTTGAGCTTACTCCATNNNAGATCGGAAG |
| 7163_1 | TATGAGTGTGGAGTCGTTGCAAGAAACCAACCTAAGAAGGAGGCNNNAGATCGGA |
| 7171_1 | TATGAGTGTGGAGTCGTTGCGACTGATAGGATCGGTGTGTACGGNNNAGATCGGA |
| 7200_1 | TATGAGTGTGGAGTCGTTGCGCATAAACAACCACACATGGTTACNNNAGATCGGAAG |
| 7203_1 | TATGAGTGTGGAGTCGTTGCAGTGTAGCCTTGACCTGTCAATCTNNNAGATCGGAAG |

| | |
|---|---|
| 7233_1 | TATGAGTGTGGAGTCGTTGCCACCCAACAATGGTTACAACTGGTNNNAGATCGGAA |
| 7237_1 | TATGAGTGTGGAGTCGTTGCAGGATTGTGGGTTGACTGTTGTTNNNAGATCGGAA |
| 7252_1 | TATGAGTGTGGAGTCGTTGCGGTCTTTCGTTCATGTACTCAGCTNNNAGATCGGAA |
| 7254_1 | TATGAGTGTGGAGTCGTTGCTTGAGATGGAGTACAGTGTTGCAGNNNAGATCGGA |
| 7266_1 | TATGAGTGTGGAGTCGTTGCAAAGCGTAATGACACAAACACCGNNNAGATCGGAA |
| 7268_1 | TATGAGTGTGGAGTCGTTGCCCTGCTCACTTCAGAACTGCTAGNNNAGATCGGAAG |
| 7269_1 | TATGAGTGTGGAGTCGTTGCGGAATAGAGAAGTCCAAGAAGCCTNNNAGATCGGA |
| 7292_1 | TATGAGTGTGGAGTCGTTGCCACATGCACGCCTATATTAGTCTGNNNAGATCGGAA |
| 7295_1 | TATGAGTGTGGAGTCGTTGCTCTTATTGGCCAAAGCTGGGTAGNNNAGATCGGAA |
| 7299_1 | TATGAGTGTGGAGTCGTTGCAAAGTGCAGAGGTGCAATGATAGCNNNAGATCGGA |
| 7317_1 | TATGAGTGTGGAGTCGTTGCATGTCCTCATGAATAGCCATTCCANNNAGATCGGAA |
| 7326_1 | TATGAGTGTGGAGTCGTTGCCTGAGGTGAAGAAATAGGGAGGGANNNAGATCGG |
| 7344_1 | TATGAGTGTGGAGTCGTTGCCTTTCGTCTCTCTCTCTTTCTGTGNNNAGATCGGAAG |
| 7352_1 | TATGAGTGTGGAGTCGTTGCTTGACCATTATTTTGGTGCTTTCCNNNAGATCGGAAG |
| 7360_1 | TATGAGTGTGGAGTCGTTGCTAGCCTTGTCTCTCTGTGTCCATTNNNAGATCGGAAG |
| 7368_1 | TATGAGTGTGGAGTCGTTGCACATATAGGCTCACCATTGAGACGNNNAGATCGGAA |
| 7370_1 | TATGAGTGTGGAGTCGTTGCTCAGAGAACAAACACAATTGGACGNNNAGATCGGA |
| 7373_1 | TATGAGTGTGGAGTCGTTGCCTCAAAGATGTAGGAACTGTGCTGNNNAGATCGGA |
| 7384_1 | TATGAGTGTGGAGTCGTTGCAGCGTCTTTCATTGCTCATTTACGNNNAGATCGGAA |
| 7386_1 | TATGAGTGTGGAGTCGTTGCTGGTGTTTCACTCTTGTGATCTTGNNNAGATCGGAA |
| 7389_1 | TATGAGTGTGGAGTCGTTGCTCTCGCTCTCTCTTGTATGTGGGTNNNAGATCGGAA |
| 7412_1 | TATGAGTGTGGAGTCGTTGCATCGCAAATCTGATGAGGTTCCGNNNAGATCGGAA |
| 7427_1 | TATGAGTGTGGAGTCGTTGCTACTAAGAAAAACTGGTGGGTGCGNNNAGATCGGA |
| 7445_1 | TATGAGTGTGGAGTCGTTGCACTCTTACTTTGCCAGCTCCAATNNNAGATCGGAAGA |
| 7457_1 | TATGAGTGTGGAGTCGTTGCGCCAGGGGAGTATTACTGAAAAGGNNNAGATCGGA |
| 7458_1 | TATGAGTGTGGAGTCGTTGCGCACGCAAAAGAGAGTGAAAAGGNNNAGATCGGA |
| 7459_1 | TATGAGTGTGGAGTCGTTGCGATAGGAGATGTTCCTGGTTGTCCNNNAGATCGGA |
| 7483_1 | TATGAGTGTGGAGTCGTTGCTGTGTTTCCCTTTATTGTGCTTCTNNNAGATCGGAAG |
| 7505_1 | TATGAGTGTGGAGTCGTTGCGCCACACATGAAACATTCCTAAGGNNNAGATCGGAA |
| 7511_1 | TATGAGTGTGGAGTCGTTGCAAGGAAGCACCAAGTTTCACAGGNNNAGATCGGAA |
| 7533_1 | TATGAGTGTGGAGTCGTTGCATAGTTGTTGAAATGGGGCCAGGNNNAGATCGGAA |
| 7540_1 | TATGAGTGTGGAGTCGTTGCCAGAAGCCAAATGTATGAGCAGGNNNAGATCGGAA |
| 7556_1 | TATGAGTGTGGAGTCGTTGCTGTGTGAAGAATTATGCTGACCTTNNNAGATCGGAA |
| 7592_1 | TATGAGTGTGGAGTCGTTGCTGGGATGTGTTTTATGTAGGGAGGNNNAGATCGGA |
| 7625_1 | TATGAGTGTGGAGTCGTTGCTTTGCGATCCCATCACTACATAGGNNNAGATCGGAA |
| 7628_1 | TATGAGTGTGGAGTCGTTGCTCTCAGGGTGCTTGAGGTATAGGNNNAGATCGGAA |
| 7629_1 | TATGAGTGTGGAGTCGTTGCGCACATCTTTATGGGTCAATAGGANNNAGATCGGAA |
| 7632_1 | TATGAGTGTGGAGTCGTTGCCCCCTTTATGAAATCTGCCCTAGGNNNAGATCGGAA |
| 7642_1 | TATGAGTGTGGAGTCGTTGCTCCTGTTCCAGTTGCTTACTTTCTNNNAGATCGGAAG |
| 7656_1 | TATGAGTGTGGAGTCGTTGCAGGTGTCTCTGATATACTCACCGGNNNAGATCGGAA |
| 7673_1 | TATGAGTGTGGAGTCGTTGCTCTGCTCACTTACATTCAGTTTCTNNNAGATCGGAAG |
| 7721_1 | TATGAGTGTGGAGTCGTTGCTTTAGCTCACAGGACTAGTGAGGGNNNAGATCGGA |
| 7730_1 | TATGAGTGTGGAGTCGTTGCAGAGACAGAGAGTGCATAAGGGANNNAGATCGGA |

| | |
|---|---|
| 7732_1 | TATGAGTGTGGAGTCGTTGCCAAGGGGATAAGGGGACTTAGGGNNNAGATCGGA |
| 7744_1 | TATGAGTGTGGAGTCGTTGCTTTCTTTTCCTCTGGGCACATACANNNAGATCGGAAG |
| 7750_1 | TATGAGTGTGGAGTCGTTGCTGCTGACCTCTGAATACAAAACAANNNAGATCGGAA |
| 7759_1 | TATGAGTGTGGAGTCGTTGCTCTATATTTGGGGCAGAGTGGGGNNNAGATCGGAA |
| 7769_1 | TATGAGTGTGGAGTCGTTGCTGCCCATTTTGAAATTGTTGGGGNNNAGATCGGAAG |
| 7770_1 | TATGAGTGTGGAGTCGTTGCTTGTGGGCTGTGATCAGAAAAGTNNNAGATCGGAA |
| 7774_1 | TATGAGTGTGGAGTCGTTGCCCACATACCCCAGTAACAATGGGNNNAGATCGGAAG |
| 7778_1 | TATGAGTGTGGAGTCGTTGCTTTAGGCATGTTTACTGGAAAAGTNNNAGATCGGAA |
| 7815_1 | TATGAGTGTGGAGTCGTTGCGCCCTAGAGGAAGTCATTGTTTCCNNNAGATCGGAA |
| 7816_1 | TATGAGTGTGGAGTCGTTGCGATATTTTTGAGCCTTGGCTGGGNNNAGATCGGAA |
| 7833_1 | TATGAGTGTGGAGTCGTTGCACCCAAGGAGTATCTATCAGTGGGNNNAGATCGGA |
| 7871_1 | TATGAGTGTGGAGTCGTTGCAACTTATGTGGCAGTCACAATGGNNNAGATCGGAA |
| 7878_1 | TATGAGTGTGGAGTCGTTGCAGGTCACCAGGCTAAATGAATGGNNNAGATCGGAA |
| 7890_1 | TATGAGTGTGGAGTCGTTGCGCAACATGCTATACCTTTCCATGGNNNAGATCGGAA |
| 7915_1 | TATGAGTGTGGAGTCGTTGCCCTTGAAGATGCCTCTCCCTTCACNNNAGATCGGAAG |
| 7921_1 | TATGAGTGTGGAGTCGTTGCAAAGCCATGGTGACTTCTGTATGGNNNAGATCGGA |
| 7946_1 | TATGAGTGTGGAGTCGTTGCTGACTGAAACATAGGTGAGACTGGNNNAGATCGGA |
| 7958_1 | TATGAGTGTGGAGTCGTTGCACTTCCTGTTCTCTCTCTACCTGGNNNAGATCGGAAG |
| 7968_1 | TATGAGTGTGGAGTCGTTGCTAATGCTTTCCTTGTACAGCCTGGNNNAGATCGGAA |
| 7988_1 | TATGAGTGTGGAGTCGTTGCAACTGAAGAACCTTTCTGGCTGGNNNAGATCGGAAG |
| 8027_1 | TATGAGTGTGGAGTCGTTGCATTACCCAGTTTGGATGCAGTGGNNNAGATCGGAA |
| 8033_1 | TATGAGTGTGGAGTCGTTGCAATACCTCTTCTAGGGAGGAAGGCNNNAGATCGGA |
| 8046_1 | TATGAGTGTGGAGTCGTTGCTCCCCAAGTTGTTTAGAAGGTGGNNNAGATCGGAA |
| 8064_1 | TATGAGTGTGGAGTCGTTGCAAATAACATCGATGGGGTGGTGGNNNAGATCGGAA |
| 8071_1 | TATGAGTGTGGAGTCGTTGCACCAGATGGCCACTTTGAGAAGTNNNAGATCGGAA |
| 8103_1 | TATGAGTGTGGAGTCGTTGCTCCTCTTTCACATACACTCCTTGGNNNAGATCGGAAG |
| 8133_1 | TATGAGTGTGGAGTCGTTGCCAGATGCAGATACGTGTGTTTGGNNNAGATCGGAA |
| 8137_1 | TATGAGTGTGGAGTCGTTGCTCAGCCAAGGAACAGACTTTTGGNNNAGATCGGAA |
| 8144_1 | TATGAGTGTGGAGTCGTTGCACTGCATCTGGGCATTCAAAATGNNNAGATCGGAAG |
| 8158_1 | TATGAGTGTGGAGTCGTTGCCTGATGGTGAGAGGTTCCCAATGNNNAGATCGGAA |
| 8168_1 | TATGAGTGTGGAGTCGTTGCCCAGGAAGACTATAGGCCGAATGNNNAGATCGGAA |
| 8175_1 | TATGAGTGTGGAGTCGTTGCTCCCCTGAGTTCATAAGCTGAATGNNNAGATCGGAA |
| 8177_1 | TATGAGTGTGGAGTCGTTGCGCAGTGGGTACATCCACCTAATGNNNAGATCGGAAG |
| 8179_1 | TATGAGTGTGGAGTCGTTGCGCAGCAGGAATAGCAAGGTAATGNNNAGATCGGAA |
| 8180_1 | TATGAGTGTGGAGTCGTTGCGGGGGAAATGCTTTGTGCATTTANNNAGATCGGAA |
| 8184_1 | TATGAGTGTGGAGTCGTTGCGGGGTTTGGTAGAACCCAACATGNNNAGATCGGAA |
| 8222_1 | TATGAGTGTGGAGTCGTTGCCTGTCATACACACGTACTTGCATGNNNAGATCGGAA |
| 8226_1 | TATGAGTGTGGAGTCGTTGCATTCAGGCCAAACTGTCAACAGTNNNAGATCGGAAG |
| 8231_1 | TATGAGTGTGGAGTCGTTGCGGAAATTGGCTCATTGGCTCATGNNNAGATCGGAA |
| 8238_1 | TATGAGTGTGGAGTCGTTGCACTGTTCTTCCTACGAGCTTCATGNNNAGATCGGAAG |
| 8262_1 | TATGAGTGTGGAGTCGTTGCAGAGGTTGTTCGGGGGAATATAGANNNAGATCGGA |
| 8272_1 | TATGAGTGTGGAGTCGTTGCCTGGTGACAAAAAGGGTAAGGACTNNNAGATCGGA |
| 8288_1 | TATGAGTGTGGAGTCGTTGCGTTAGTGGGAGATACAATTCCCCCNNNAGATCGGAA |

| | |
|---|---|
| 8309_1 | TATGAGTGTGGAGTCGTTGCGGTGGGAGATGACAGTGTGTATGNNNAGATCGGA |
| 8327_1 | TATGAGTGTGGAGTCGTTGCATGCTATCTAGTAACTGGCCACTGNNNAGATCGGAA |
| 8335_1 | TATGAGTGTGGAGTCGTTGCACTAAACACTTTGCCTTTGCACTGNNNAGATCGGAA |
| 8342_1 | TATGAGTGTGGAGTCGTTGCGCCAAGACTGTTTTGTGAAGACTGNNNAGATCGGA |
| 8344_1 | TATGAGTGTGGAGTCGTTGCCACTCTGAAACCTCACCAGACTGNNNAGATCGGAAG |
| 8348_1 | TATGAGTGTGGAGTCGTTGCAGGCTTAGATAACCACAAGGACTGNNNAGATCGGA |
| 8351_1 | TATGAGTGTGGAGTCGTTGCTTTTTCCCCAAGATGGTGGACTGNNNAGATCGGAAG |
| 8355_1 | TATGAGTGTGGAGTCGTTGCTTCACAACTGTCTCTTGGTGACTGNNNAGATCGGAA |
| 8363_1 | TATGAGTGTGGAGTCGTTGCGGGAATGAACTGAGGCAGTACTGNNNAGATCGGA |
| 8365_1 | TATGAGTGTGGAGTCGTTGCTCAGTGGAGAATCCCTGTGGATTNNNAGATCGGAA |
| 8367_1 | TATGAGTGTGGAGTCGTTGCTGGTCAGTCATGAGAATTTTCCAANNNAGATCGGAA |
| 8375_1 | TATGAGTGTGGAGTCGTTGCTCCCTGTTGAGGTTGCTTTTCCATNNNAGATCGGAA |
| 8410_1 | TATGAGTGTGGAGTCGTTGCCTGAGAAGCTACTGGACAAGCTGNNNAGATCGGAA |
| 8414_1 | TATGAGTGTGGAGTCGTTGCGGATGTCAGTATCACCCGTATCAANNNAGATCGGAA |
| 8418_1 | TATGAGTGTGGAGTCGTTGCGGGTGGGTGTTATAAGTCAGCTGNNNAGATCGGAA |
| 8419_1 | TATGAGTGTGGAGTCGTTGCGAGTGAGGGTTGAAGAAGAGCTGNNNAGATCGGA |
| 8420_1 | TATGAGTGTGGAGTCGTTGCGCCACACCCAGCTAGTATTGTTNNNAGATCGGAAGA |
| 8436_1 | TATGAGTGTGGAGTCGTTGCAGCATACATACATACAAGCAACCTNNNAGATCGGAA |
| 8443_1 | TATGAGTGTGGAGTCGTTGCGTTCCTCTATAGGCGAGATGTCCTNNNAGATCGGAA |
| 8466_1 | TATGAGTGTGGAGTCGTTGCTTTATTAGCGTCACTCAGAATCTGNNNAGATCGGAA |
| 8475_1 | TATGAGTGTGGAGTCGTTGCTGCCTCCTTACCTCAAAAACTCTGNNNAGATCGGAAG |
| 8529_1 | TATGAGTGTGGAGTCGTTGCCAGGGTGTTATTCTGTCATCCAAGNNNAGATCGGAA |
| 8554_1 | TATGAGTGTGGAGTCGTTGCGCATTTGTTGTCCTCCATGAAGTGNNNAGATCGGAA |
| 8555_1 | TATGAGTGTGGAGTCGTTGCGGGAGGAGACACAACAACAAAAAGNNNAGATCGG |
| 8560_1 | TATGAGTGTGGAGTCGTTGCGTTCAAAGTGACAGCTTCCAGTGNNNAGATCGGAA |
| 8562_1 | TATGAGTGTGGAGTCGTTGCCACCTCCCAGGAATATTTTTGTCTNNNAGATCGGAAG |
| 8576_1 | TATGAGTGTGGAGTCGTTGCCTCACTCTTATCACCCTGGAGTGNNNAGATCGGAAG |
| 8582_1 | TATGAGTGTGGAGTCGTTGCAGCCTGAGTGTTACGTGAGTGNNNAGATCGGAAGA |
| 8586_1 | TATGAGTGTGGAGTCGTTGCACACCCCAAAGTTCACTCATAGTGNNNAGATCGGAA |
| 8598_1 | TATGAGTGTGGAGTCGTTGCGAAGGAGCTTTGTGTGTGTTTTCTNNNAGATCGGA |
| 8601_1 | TATGAGTGTGGAGTCGTTGCCCCACAAAGGAACAATGTAAGGTGNNNAGATCGGA |
| 8617_1 | TATGAGTGTGGAGTCGTTGCTAACATTGGTTGCCTATGGGGTGNNNAGATCGGAA |
| 8629_1 | TATGAGTGTGGAGTCGTTGCGTCAGCTCTTATGTTACACCTGGGNNNAGATCGGAA |
| 8640_1 | TATGAGTGTGGAGTCGTTGCGTCCACTAGGGCAAGGAAATGTGNNNAGATCGGAA |
| 8642_1 | TATGAGTGTGGAGTCGTTGCCCATTTTCTGTGTTGGGGTTTCCTNNNAGATCGGAA |
| 8647_1 | TATGAGTGTGGAGTCGTTGCTGGTGTGACTACTTCAGGTATGTGNNNAGATCGGA |
| 8652_1 | TATGAGTGTGGAGTCGTTGCCTTGCTTGAATGGAGTTGCTGTGNNNAGATCGGAA |
| 8675_1 | TATGAGTGTGGAGTCGTTGCACAAAGCACAAGACATGTGTGTGNNNAGATCGGAA |
| 8680_1 | TATGAGTGTGGAGTCGTTGCTGGAGAACCCTAACACACATTGTGNNNAGATCGGAA |
| 8687_1 | TATGAGTGTGGAGTCGTTGCGGCAGCATAGTACACATTGTTGTGNNNAGATCGGA |
| 8700_1 | TATGAGTGTGGAGTCGTTGCACACAAGACGTTGACTTGACATTGNNNAGATCGGAA |
| 8704_1 | TATGAGTGTGGAGTCGTTGCTTCCAGCTACCTTCTTGCCATTGNNNAGATCGGAAGA |
| 8718_1 | TATGAGTGTGGAGTCGTTGCAGGGTGTCTGTGTGAGATGATTGNNNAGATCGGAA |

| | |
|---|---|
| 8721_1 | TATGAGTGTGGAGTCGTTGCACACACACACTTGGTACATTTTTNNNAGATCGGAAGA |
| 8731_1 | TATGAGTGTGGAGTCGTTGCAAGAGGCAACCAAAGTCCTTCTGNNNAGATCGGAAG |
| 8741_1 | TATGAGTGTGGAGTCGTTGCGTAGAAAAGTAGGTGAGAACACGTNNNAGATCGGA |
| 8760_1 | TATGAGTGTGGAGTCGTTGCGCACCACACTGTCTTGATTCTTGNNNAGATCGGAAGA |
| 8765_1 | TATGAGTGTGGAGTCGTTGCGCAATCTTCGCTCTGCTTTCTTGNNNAGATCGGAAGA |
| 8874_1 | TATGAGTGTGGAGTCGTTGCAGGGATGGTATGAGATCAAGGTCANNNAGATCGGA |
| 8897_1 | TATGAGTGTGGAGTCGTTGCACACACATACATCATGGTTGAAATNNNAGATCGGAA |
| 8907_1 | TATGAGTGTGGAGTCGTTGCTGCTATACGTGTCAGTGAGGAGTNNNAGATCGGAA |
| 8914_1 | TATGAGTGTGGAGTCGTTGCTCTTCCTTCATTGCACTGTCCACTCNNNAGATCGGAAG |
| 8917_1 | TATGAGTGTGGAGTCGTTGCAGCAATGACTATTTTGCAGGGTCANNNAGATCGGAA |
| 8928_1 | TATGAGTGTGGAGTCGTTGCCCCTCCCTCCTAAAACCAAATGGNNNAGATCGGAAGA |
| 8939_1 | TATGAGTGTGGAGTCGTTGCGGACCAAGAATAGCTGAGGCAATNNNAGATCGGAA |
| 8941_1 | TATGAGTGTGGAGTCGTTGCTTCTCTAGCTCCCTCATGGCAATNNNAGATCGGAAGA |
| 8946_1 | TATGAGTGTGGAGTCGTTGCAGTTCTCCAAGTGGTTGTGCAATNNNAGATCGGAAG |
| 8950_1 | TATGAGTGTGGAGTCGTTGCAGCAATCGTCCTTTCACCTCAATNNNAGATCGGAAGA |
| 8955_1 | TATGAGTGTGGAGTCGTTGCGCTTTTTGAGATGGAGGTCTCTCTNNNAGATCGGAA |
| 9004_1 | TATGAGTGTGGAGTCGTTGCATGGATTCCTCCTGTCTCTCTCTCNNNAGATCGGAAG |
| 9020_1 | TATGAGTGTGGAGTCGTTGCTCTTTGTCATGTCCTGTGGCTTTCNNNAGATCGGAAG |
| 9037_1 | TATGAGTGTGGAGTCGTTGCTTAGGTAGCCATCAGCACTGAGTNNNAGATCGGAAG |
| 9039_1 | TATGAGTGTGGAGTCGTTGCACATACAAGGGTTGTGTCCACATNNNAGATCGGAAG |
| 9050_1 | TATGAGTGTGGAGTCGTTGCACACACACAGACATAACAGACATNNNAGATCGGAAG |
| 9071_1 | TATGAGTGTGGAGTCGTTGCATGCCAACATACGGTGAAACCATNNNAGATCGGAAG |
| 9077_1 | TATGAGTGTGGAGTCGTTGCACAGTTGGCAATAACCTGACCATNNNAGATCGGAAG |
| 9085_1 | TATGAGTGTGGAGTCGTTGCAGCAAAGCTGAAAAATAGCCCATNNNAGATCGGAA |
| 9117_1 | TATGAGTGTGGAGTCGTTGCAGTAGAGATGGTGTGTTGTGTGTGNNNAGATCGGA |
| 9130_1 | TATGAGTGTGGAGTCGTTGCTCTTCTACCGTATGAGCCAGCATNNNAGATCGGAAG |
| 9138_1 | TATGAGTGTGGAGTCGTTGCTGTGCTGGTATTGACCCTGTTTCNNNAGATCGGAAG |
| 9210_1 | TATGAGTGTGGAGTCGTTGCAGTGCAGAAAAGGTGGCATTTATTNNNAGATCGGA |
| 9212_1 | TATGAGTGTGGAGTCGTTGCCGCACTAGGCATTGTACCTTCATNNNAGATCGGAAG |
| 9228_1 | TATGAGTGTGGAGTCGTTGCGTAAAAGCCATCAGGATGACACACNNNAGATCGGA |
| 9261_1 | TATGAGTGTGGAGTCGTTGCAAGCTCTGAGAGAGAGAGAGAGATNNNAGATCGG |
| 9273_1 | TATGAGTGTGGAGTCGTTGCAGGGCAGAGAAAAGAGGTGAGATNNNAGATCGGA |
| 9296_1 | TATGAGTGTGGAGTCGTTGCTGGCACTCAGTTATCGAAACTTCANNNAGATCGGAA |
| 9320_1 | TATGAGTGTGGAGTCGTTGCAGAAGGGTGACTTGAGCTCTAGNNNAGATCGGAAG |
| 9321_1 | TATGAGTGTGGAGTCGTTGCTTGCTGATACACACAAGCTGGATNNNAGATCGGAAG |
| 9323_1 | TATGAGTGTGGAGTCGTTGCTTTTTGTTGGAGTGGGTGTGGATNNNAGATCGGAA |
| 9344_1 | TATGAGTGTGGAGTCGTTGCGCCTGAAGTATGTGTGTGCTGATNNNAGATCGGAA |
| 9364_1 | TATGAGTGTGGAGTCGTTGCACACTCTTTGGGTGCTTGTTGATNNNAGATCGGAAG |
| 9367_1 | TATGAGTGTGGAGTCGTTGCCCATCACACCTCATTTCTCCTTCANNNAGATCGGAAGA |
| 9421_1 | TATGAGTGTGGAGTCGTTGCCAGGCATCAGAGGTGTAATCTATNNNAGATCGGAA |
| 9449_1 | TATGAGTGTGGAGTCGTTGCAACTCCTCAGGAAGGGATGGTATNNNAGATCGGAA |
| 9464_1 | TATGAGTGTGGAGTCGTTGCGTGCATGCTGTAGTCATTTCTTCANNNAGATCGGAA |
| 9496_1 | TATGAGTGTGGAGTCGTTGCTGCCTGAAAAACAGAGGGAAACTNNNAGATCGGAA |

| | |
|---|---|
| 9513_1 | TATGAGTGTGGAGTCGTTGCACACACTATCCAACCTCAAGAACTNNNAGATCGGAAG |
| 9544_1 | TATGAGTGTGGAGTCGTTGCTCCACTCATCATCCATATCTTGGTNNNAGATCGGAAG |
| 9555_1 | TATGAGTGTGGAGTCGTTGCACTTCTGTTCAATTTGCTACCACTNNNAGATCGGAAG |
| 9573_1 | TATGAGTGTGGAGTCGTTGCACACTTGGGTATCTTTCATGCACTNNNAGATCGGAAG |
| 9582_1 | TATGAGTGTGGAGTCGTTGCTGTGAACCTATTGTCAAGCTCACTNNNAGATCGGAA |
| 9587_1 | TATGAGTGTGGAGTCGTTGCTTTTGTATCAGCATATTCCTTGCCNNNAGATCGGAAG |
| 9596_1 | TATGAGTGTGGAGTCGTTGCACCACTTTCTTGCATTCTTTCACTNNNAGATCGGAAG |
| 9600_1 | TATGAGTGTGGAGTCGTTGCGGAGCATGCATAAGGTCCAACTTNNNAGATCGGAAG |
| 9601_1 | TATGAGTGTGGAGTCGTTGCAGTACAGCCATAAGGTGAAAGACTNNNAGATCGGA |
| 9607_1 | TATGAGTGTGGAGTCGTTGCCAAGACTGTGAATCCATAGGGCAGNNNAGATCGGA |
| 9617_1 | TATGAGTGTGGAGTCGTTGCCCTGATTCCTTCCCCACAGTCTANNNAGATCGGAAGA |
| 9619_1 | TATGAGTGTGGAGTCGTTGCATGTTCTTTGAGTACAGCAGGACTNNNAGATCGGAA |
| 9630_1 | TATGAGTGTGGAGTCGTTGCGGAAGAGTACCAGACCATGGACTNNNAGATCGGAA |
| 9632_1 | TATGAGTGTGGAGTCGTTGCACCTTGACCTAAACTTCATGACTNNNAGATCGGAAG |
| 9660_1 | TATGAGTGTGGAGTCGTTGCCCTTGTGTTGTAGAGCTGCTACTNNNAGATCGGAAG |
| 9666_1 | TATGAGTGTGGAGTCGTTGCTGGGTTGCAGGAGAGAAAGTACTNNNAGATCGGAA |
| 9726_1 | TATGAGTGTGGAGTCGTTGCTCATTGCCCTAAACAGAAAGACCTNNNAGATCGGAA |
| 9734_1 | TATGAGTGTGGAGTCGTTGCGCCCATCATACACAAGTATGACCTNNNAGATCGGAA |
| 9762_1 | TATGAGTGTGGAGTCGTTGCTGTGTAAAAGTTTCCCTTGACCCTNNNAGATCGGAAG |
| 9785_1 | TATGAGTGTGGAGTCGTTGCTCAGCTCTTCTGTCATTTCCCCTNNNAGATCGGAAGA |
| 9806_1 | TATGAGTGTGGAGTCGTTGCACAAAGAAGTATCCCAATCTCCCTNNNAGATCGGAAG |
| 9812_1 | TATGAGTGTGGAGTCGTTGCACAGTGCCACAACAAACATGAGATNNNAGATCGGAA |
| 9816_1 | TATGAGTGTGGAGTCGTTGCGCTGACTTCCCAGTATTTTTCCCTNNNAGATCGGAAG |
| 9832_1 | TATGAGTGTGGAGTCGTTGCTAGGCAGGGAGAAGTAGATGAGGANNNAGATCGG |
| 9842_1 | TATGAGTGTGGAGTCGTTGCGAACTAGATATGCACTTGCGCCTNNNAGATCGGAAG |
| 9854_1 | TATGAGTGTGGAGTCGTTGCATGGAAGAAACAAAGTATGGGCCTNNNAGATCGGA |
| 9864_1 | TATGAGTGTGGAGTCGTTGCAGTAGTGCTTAGAGGAAAATGCCTNNNAGATCGGA |
| 9883_1 | TATGAGTGTGGAGTCGTTGCACTATGTTCGTCTCCAATTGCCTNNNAGATCGGAAGA |
| 9888_1 | TATGAGTGTGGAGTCGTTGCATGGTTGTCAACTGCCAAATCCTNNNAGATCGGAAG |
| 9903_1 | TATGAGTGTGGAGTCGTTGCCCCCTACAAGAGTTCACACTCCTNNNAGATCGGAAGA |
| 9926_1 | TATGAGTGTGGAGTCGTTGCAGACACCAGTGAATAGATTCTCCTNNNAGATCGGAA |
| 9933_1 | TATGAGTGTGGAGTCGTTGCGTTCCCCTGCACCATCTCAGANNNAGATCGGAAGAG |
| 9947_1 | TATGAGTGTGGAGTCGTTGCTCCAGTCAACCACTTCAAATTCCTNNNAGATCGGAAG |
| 9948_1 | TATGAGTGTGGAGTCGTTGCTTGGCTTGAGAAAGAGAAATTCCTNNNAGATCGGAA |
| 10174_1 | TATGAGTGTGGAGTCGTTGCTGGTCCTAATAAGATCTCTCAGCTNNNAGATCGGAA |
| 10180_1 | TATGAGTGTGGAGTCGTTGCTGGGACTGCATTATTGATTCAGCTNNNAGATCGGAA |
| 10250_1 | TATGAGTGTGGAGTCGTTGCTCACTCTCTCTCTTATCCATAGCTNNNAGATCGGAAG |
| 10290_1 | TATGAGTGTGGAGTCGTTGCACTTTTTGTGTGTCCTGGTAGCTNNNAGATCGGAAG |
| 10400_1 | TATGAGTGTGGAGTCGTTGCTGGACCTAATTTTGTCTGAGGCTNNNAGATCGGAAG |
| 10417_1 | TATGAGTGTGGAGTCGTTGCTCTGAATAACTCTTATGCACGGCTNNNAGATCGGAA |
| 10434_1 | TATGAGTGTGGAGTCGTTGCTGGCTGTCTAAAATAAATGGGGCTNNNAGATCGGA |
| 10436_1 | TATGAGTGTGGAGTCGTTGCGAGAAATCTGTAGGTTTGGGGCTNNNAGATCGGAA |
| 10578_1 | TATGAGTGTGGAGTCGTTGCTGTATGGATTTGAATCCCAATGCTNNNAGATCGGAA |

| | |
|---|---|
| 10645_1 | TATGAGTGTGGAGTCGTTGCTGAAAGGAAAAGGTAAGACTGCTNNNAGATCGGAA |
| 10753_1 | TATGAGTGTGGAGTCGTTGCTGCCAGAAATTGTACACTGTGCTNNNAGATCGGAAG |
| 10781_1 | TATGAGTGTGGAGTCGTTGCATGTCACAACACATGACAATTGCTNNNAGATCGGAA |
| 10833_1 | TATGAGTGTGGAGTCGTTGCAAATCATCAACTTCTCCATTTGCTNNNAGATCGGAAG |
| 10876_1 | TATGAGTGTGGAGTCGTTGCAGGCTGAGTGAAAATAGCCAATCTNNNAGATCGGA |
| 10955_1 | TATGAGTGTGGAGTCGTTGCTCTGGCTCCATTTCTTGAAGAGACNNNAGATCGGAA |
| 10958_1 | TATGAGTGTGGAGTCGTTGCTCTGTTAGAAGGGGGAAGAGGTGNNNAGATCGGA |
| 10997_1 | TATGAGTGTGGAGTCGTTGCTGGCTTTACGTAGCTCAATCCTGNNNAGATCGGAAG |
| 11003_1 | TATGAGTGTGGAGTCGTTGCGGAGGAGAGGACCAAACAGATGANNNAGATCGGA |
| 11030_1 | TATGAGTGTGGAGTCGTTGCTCCAAGATCCAGTGGTACATCTGTNNNAGATCGGAA |
| 11051_1 | TATGAGTGTGGAGTCGTTGCATAAACCCTGGAATGCAGAAGGCNNNAGATCGGAA |
| 11054_1 | TATGAGTGTGGAGTCGTTGCCCAAGACCCATTGAGAAACTTCCTNNNAGATCGGAA |
| 11060_1 | TATGAGTGTGGAGTCGTTGCCGCACACACATGAACAACAAAAANNNAGATCGGAAG |
| 11067_1 | TATGAGTGTGGAGTCGTTGCTGGGGTCTATAGTAGGACACTCTNNNAGATCGGAA |
| 11073_1 | TATGAGTGTGGAGTCGTTGCACCAGTTACTTTGTATCCCACTCTNNNAGATCGGAAG |
| 11092_1 | TATGAGTGTGGAGTCGTTGCCAAGGAGCAACAGTCTATGTGTCTNNNAGATCGGA |
| 11095_1 | TATGAGTGTGGAGTCGTTGCAAGTCTCCTGTAATCCCACCTCTNNNAGATCGGAAGA |
| 11129_1 | TATGAGTGTGGAGTCGTTGCGTGTACCTAAAACCCCTCTCCTCTNNNAGATCGGAAG |
| 11131_1 | TATGAGTGTGGAGTCGTTGCTGAGCAGTGGTTGTGGTAGAAAANNNAGATCGGA |
| 11147_1 | TATGAGTGTGGAGTCGTTGCGCCTGTCTGTGCACAAATGAAAANNNAGATCGGAA |
| 11154_1 | TATGAGTGTGGAGTCGTTGCTCGCCAATAGAAAACAGTTGAAAANNNAGATCGGA |
| 11174_1 | TATGAGTGTGGAGTCGTTGCGGTGGAAGGTAGGTGTGATCTCTNNNAGATCGGAA |
| 11185_1 | TATGAGTGTGGAGTCGTTGCCACACTTATGCCATGTCCACAAANNNAGATCGGAAG |
| 11195_1 | TATGAGTGTGGAGTCGTTGCTCTGGTGTTTCCTTCTCTCTCTCTNNNAGATCGGAAG |
| 11198_1 | TATGAGTGTGGAGTCGTTGCTGCTTCCAGAGGTTTCTCTCTCTNNNAGATCGGAAG |
| 11207_1 | TATGAGTGTGGAGTCGTTGCTAGAGATACCGAGATATGGTCTCTNNNAGATCGGA |
| 11208_1 | TATGAGTGTGGAGTCGTTGCACTCTTAGATGCAGCAAACCAAANNNAGATCGGAAG |
| 11244_1 | TATGAGTGTGGAGTCGTTGCAAAAATGAAACGTTGGAAAGCAAANNNAGATCGGA |
| 11249_1 | TATGAGTGTGGAGTCGTTGCTTCAGAGGGGAGAAAAAGGCAAANNNAGATCGGA |
| 11250_1 | TATGAGTGTGGAGTCGTTGCACACACACATACACCAAGCAACACNNNAGATCGGAA |
| 11251_1 | TATGAGTGTGGAGTCGTTGCATGTTCTTCATCCCCTTCTCCCTGNNNAGATCGGAAG |
| 11257_1 | TATGAGTGTGGAGTCGTTGCGTTCTTCAACTGTGATTGGCAAANNNAGATCGGAAG |
| 11265_1 | TATGAGTGTGGAGTCGTTGCAAAACAACCCAACAGCTCAAAATNNNAGATCGGAAG |
| 11281_1 | TATGAGTGTGGAGTCGTTGCATCCAATGACAGCATGTATTCAAANNNAGATCGGAA |
| 11304_1 | TATGAGTGTGGAGTCGTTGCTTCCTAATGACCGTATAGCAGAAANNNAGATCGGAA |
| 11322_1 | TATGAGTGTGGAGTCGTTGCGCCTACACAGGTTCGTTTTCTGGTTGANNNAGATCG |
| 11363_1 | TATGAGTGTGGAGTCGTTGCCATACTCTGTCTACCCTACCAGGCNNNAGATCGGAAG |
| 11364_1 | TATGAGTGTGGAGTCGTTGCCTCTCTCTAGGACAAGTGTGTGAGNNNAGATCGGA |
| 11368_1 | TATGAGTGTGGAGTCGTTGCGCTTTATGGTAAGGGGCTGGAAANNNAGATCGGAA |
| 11401_1 | TATGAGTGTGGAGTCGTTGCTGTTATCTTTTGGTGCCAGGTTCTNNNAGATCGGAA |
| 11406_1 | TATGAGTGTGGAGTCGTTGCTTGTGGAGAAGGAAGAGGGAAGGANNNAGATCGG |
| 11411_1 | TATGAGTGTGGAGTCGTTGCTGTTGCTTTGTTCCTGTTTGTTCTNNNAGATCGGAAG |
| 11413_1 | TATGAGTGTGGAGTCGTTGCATGGGGAGGGGCAGTATGATAAANNNAGATCGGA |

| | |
|---|---|
| 11421_1 | TATGAGTGTGGAGTCGTTGCATGTATTGAGACGGAGCTTGCTCTNNNAGATCGGAA |
| 11422_1 | TATGAGTGTGGAGTCGTTGCTCTCACACAAACAAATCACCTAAANNNAGATCGGAAG |
| 11424_1 | TATGAGTGTGGAGTCGTTGCTTCATCATTGGTGGGCACTTAGGTNNNAGATCGGAA |
| 11428_1 | TATGAGTGTGGAGTCGTTGCCGATGCCATGTGTCTTCCTTTCTNNNAGATCGGAAGA |
| 11431_1 | TATGAGTGTGGAGTCGTTGCACTGGTGCGTGTGTTTCTCTAAANNNAGATCGGAAG |
| 11449_1 | TATGAGTGTGGAGTCGTTGCAACACAGTTTCTGAGACAAACAANNNAGATCGGAAG |
| 11453_1 | TATGAGTGTGGAGTCGTTGCTTTCTGGCCATTGTATTGAAACAANNNAGATCGGAA |
| 11470_1 | TATGAGTGTGGAGTCGTTGCTGGAGGAAGTGGGGAAATAACAANNNAGATCGGA |
| 11481_1 | TATGAGTGTGGAGTCGTTGCGCTCCTGATGCCAAGATACACAANNNAGATCGGAAG |
| 11505_1 | TATGAGTGTGGAGTCGTTGCTGCTGCCTCTCAAACACACATATCNNNAGATCGGAAG |
| 11506_1 | TATGAGTGTGGAGTCGTTGCAAGCCTGGAACTAAGTGAAAACAGNNNAGATCGGA |
| 11509_1 | TATGAGTGTGGAGTCGTTGCCCAAGGATCAGGAAGATCCAGAGCNNNAGATCGGA |
| 11528_1 | TATGAGTGTGGAGTCGTTGCATTGAGCTTCTTGAAATGGTACAANNNAGATCGGAA |
| 11539_1 | TATGAGTGTGGAGTCGTTGCGCAATTCAGGCACAAGAAAGAAGTNNNAGATCGGA |
| 11543_1 | TATGAGTGTGGAGTCGTTGCTCTCTACTGTGTGGTCTCCTCTTTNNNAGATCGGAAG |
| 11556_1 | TATGAGTGTGGAGTCGTTGCAGAAACTTGAATTTACCTTGGGGGNNNAGATCGGA |
| 11563_1 | TATGAGTGTGGAGTCGTTGCACCTTTCCTACAGTCCCTGAAGTNNNAGATCGGAAG |
| 11575_1 | TATGAGTGTGGAGTCGTTGCCACTTGCTTCTGTACCCCTAAGTNNNAGATCGGAAGA |
| 11591_1 | TATGAGTGTGGAGTCGTTGCAAGTAGAGCCCAGTGGTAACAGTNNNAGATCGGAA |
| 11604_1 | TATGAGTGTGGAGTCGTTGCTCACCCTCATTCTTCAAAGACAGTNNNAGATCGGAAG |
| 11618_1 | TATGAGTGTGGAGTCGTTGCACATGCCAATGATATCAGGGGTCANNNAGATCGGA |
| 11626_1 | TATGAGTGTGGAGTCGTTGCAAGCCAGTTGAACAGATACAAAAANNNAGATCGGA |
| 11644_1 | TATGAGTGTGGAGTCGTTGCTCCTCAAAGATCTTGCAATCCAGTNNNAGATCGGAA |
| 11645_1 | TATGAGTGTGGAGTCGTTGCACTTCATCTTCCCCACCACTAAGANNNAGATCGGAAG |
| 11650_1 | TATGAGTGTGGAGTCGTTGCGAGAGAGGAGTGTATGTTGTCAGCNNNAGATCGGA |
| 11655_1 | TATGAGTGTGGAGTCGTTGCTCCAAACGTGCATACAGTAGCAANNNAGATCGGAAG |
| 11656_1 | TATGAGTGTGGAGTCGTTGCCATCAGAGACACACCAAAGCAGTNNNAGATCGGAAG |
| 11661_1 | TATGAGTGTGGAGTCGTTGCGCAAGGAAAGATCCATCAGCAGTNNNAGATCGGAA |
| 11662_1 | TATGAGTGTGGAGTCGTTGCATAAAAAGTAGAAGTTGGGGCTGGNNNAGATCGGA |
| 11679_1 | TATGAGTGTGGAGTCGTTGCACAGAGAGAAGCAAATGTGGCAANNNAGATCGGAA |
| 11683_1 | TATGAGTGTGGAGTCGTTGCCCTGATTTCAAAGTCTCCCCTCCANNNAGATCGGAAG |
| 11685_1 | TATGAGTGTGGAGTCGTTGCGGTTTTGCATACTCTTACATGCAANNNAGATCGGAA |
| 11692_1 | TATGAGTGTGGAGTCGTTGCGTCACTCCTTGTATGTTTTGCAGTNNNAGATCGGAA |
| 11702_1 | TATGAGTGTGGAGTCGTTGCCTAATAAACTCTGTTGCCAGGGGANNNAGATCGGAA |
| 11703_1 | TATGAGTGTGGAGTCGTTGCACAGAGGGGAAAGAGATACTCAGTNNNAGATCGGA |
| 11711_1 | TATGAGTGTGGAGTCGTTGCACTTAGCAGCAAAAGCAGTCAGTNNNAGATCGGAA |
| 11721_1 | TATGAGTGTGGAGTCGTTGCGCAAGTGAAGATGAGATTTGGGAGNNNAGATCGG |
| 11725_1 | TATGAGTGTGGAGTCGTTGCGTCCATTCACAAACTCAGGATCCTNNNAGATCGGAA |
| 11728_1 | TATGAGTGTGGAGTCGTTGCTGCTCTTTCTGTCTGTCTTTCAGTNNNAGATCGGAAG |
| 11731_1 | TATGAGTGTGGAGTCGTTGCCCATCAGAGGAGAAATTTTTCAGTNNNAGATCGGAA |
| 11736_1 | TATGAGTGTGGAGTCGTTGCTGGGATGCTGAGATACAGAAGAGTNNNAGATCGGA |
| 11763_1 | TATGAGTGTGGAGTCGTTGCGTTTTCACTGGGCCAACATATTACNNNAGATCGGAA |
| 11767_1 | TATGAGTGTGGAGTCGTTGCTGCCAAGAAACAAGTCTGGTCAANNNAGATCGGAA |

| | |
|---|---|
| 11773_1 | TATGAGTGTGGAGTCGTTGCGACCTCCTGCCAGTCAAATTCAANNNAGATCGGAAG |
| 11778_1 | TATGAGTGTGGAGTCGTTGCAGCCTAGTCAACGTAGAGAAAACCNNNAGATCGGA |
| 11836_1 | TATGAGTGTGGAGTCGTTGCTCCTTTCATTTTTGGCCTTGCCTANNNAGATCGGAAG |
| 11860_1 | TATGAGTGTGGAGTCGTTGCCACACACAGAGACATACGCAGAANNNAGATCGGAA |
| 11883_1 | TATGAGTGTGGAGTCGTTGCGTAAGCCAGACATTCTATGCAGCTNNNAGATCGGAA |
| 11891_1 | TATGAGTGTGGAGTCGTTGCGAGGAGCTAGGTATGCAGGAGAANNNAGATCGGA |
| 11893_1 | TATGAGTGTGGAGTCGTTGCCAGGAAGAAGGGTACCTGGAGAANNNAGATCGGA |
| 11898_1 | TATGAGTGTGGAGTCGTTGCTCCAGATTGAAAAACACCTGAGAANNNAGATCGGA |
| 11904_1 | TATGAGTGTGGAGTCGTTGCAACTGCAGGGGTAATGGTAGGCTANNNAGATCGGA |
| 11905_1 | TATGAGTGTGGAGTCGTTGCAGTACATGGTGGGGCTCATTAGTNNNAGATCGGAA |
| 11925_1 | TATGAGTGTGGAGTCGTTGCTAATTTCACATTTCAGTGAGGCTGNNNAGATCGGAA |
| 11927_1 | TATGAGTGTGGAGTCGTTGCTGAATACCCAAAGCAATCACGAANNNAGATCGGAAG |
| 11932_1 | TATGAGTGTGGAGTCGTTGCCATATAGAGTGGTGTGTGCACAGTANNNAGATCGG |
| 11954_1 | TATGAGTGTGGAGTCGTTGCGGCACAAAAGTTCTCCTGAGGAANNNAGATCGGAA |
| 11960_1 | TATGAGTGTGGAGTCGTTGCTGTCTTTCTCTGCCACCCAGGATANNNAGATCGGAA |
| 11965_1 | TATGAGTGTGGAGTCGTTGCTGGACTTCCTCTCAACTACGGAANNNAGATCGGAAG |
| 12000_1 | TATGAGTGTGGAGTCGTTGCTCCAGTAGGTTCTGTTTCTCTGGTNNNAGATCGGAA |
| 12006_1 | TATGAGTGTGGAGTCGTTGCTACCTTTTCCCTTGGACCTGGAANNNAGATCGGAAG |
| 12035_1 | TATGAGTGTGGAGTCGTTGCACAGGTGGTACGATCATAGCTCATNNNAGATCGGA |
| 12048_1 | TATGAGTGTGGAGTCGTTGCAGTGAACAGGAATCACAGCATACANNNAGATCGGA |
| 12068_1 | TATGAGTGTGGAGTCGTTGCTCTGGAAAACTGCATACTTCAGGTNNNAGATCGGAA |
| 12079_1 | TATGAGTGTGGAGTCGTTGCCCTTCAGTAAAGCAACAGAGAGGTNNNAGATCGGA |
| 12090_1 | TATGAGTGTGGAGTCGTTGCTAAAGGCCAAGAGCGAATCTGAANNNAGATCGGAA |
| 12096_1 | TATGAGTGTGGAGTCGTTGCCGGATGTCGCCGGATTTTATGATCNNNAGATCGGAA |
| 12115_1 | TATGAGTGTGGAGTCGTTGCTCTCATCTTCATCACAAGTGAGGTNNNAGATCGGAA |
| 12123_1 | TATGAGTGTGGAGTCGTTGCTGTATATGAACCTGGTCTCCTTGGNNNAGATCGGAA |
| 12126_1 | TATGAGTGTGGAGTCGTTGCCCACTCCTTTTCCACTGTGTGAANNNAGATCGGAAG |
| 12128_1 | TATGAGTGTGGAGTCGTTGCTACACTCACACACAATCACATGCANNNAGATCGGAA |
| 12137_1 | TATGAGTGTGGAGTCGTTGCGCAATGGTAAAGAAAGGGCTAGGTNNNAGATCGG |
| 12146_1 | TATGAGTGTGGAGTCGTTGCACCAAAAAGTGCAGAAAGGTAGGTNNNAGATCGGA |
| 12161_1 | TATGAGTGTGGAGTCGTTGCGGAAACAAAGAGAGAAGAGGTGGTNNNAGATCGG |
| 12174_1 | TATGAGTGTGGAGTCGTTGCGCTGGGCTTGGGAATTCAGATAANNNAGATCGGAA |
| 12180_1 | TATGAGTGTGGAGTCGTTGCCTGCATTCATGATATTCCCAGCCTNNNAGATCGGAAG |
| 12192_1 | TATGAGTGTGGAGTCGTTGCGCACACTGACCTCACCCATATAANNNAGATCGGAAG |
| 12215_1 | TATGAGTGTGGAGTCGTTGCAAAGAGGAAGTGGGGAACCCTAANNNAGATCGGA |
| 12219_1 | TATGAGTGTGGAGTCGTTGCACATATCACAGGTTGAAAAAGCGANNNAGATCGGA |
| 12220_1 | TATGAGTGTGGAGTCGTTGCTGGTTGGATGTAATGATTGTGAGTNNNAGATCGGA |
| 12226_1 | TATGAGTGTGGAGTCGTTGCTTGCTACAGTTCTGTGCCTTGTTNNNAGATCGGAAG |
| 12227_1 | TATGAGTGTGGAGTCGTTGCGAGAGAGCGCGCTAAAAAGCTAANNNAGATCGGAA |
| 12231_1 | TATGAGTGTGGAGTCGTTGCCTTAGTGCCATCTTGCCAGCTAANNNAGATCGGAAG |
| 12238_1 | TATGAGTGTGGAGTCGTTGCCAAACCTCGACCACTGGTAAAAATNNNAGATCGGAA |
| 12256_1 | TATGAGTGTGGAGTCGTTGCACACACACACAAAAGAACTCTCCANNNAGATCGGAA |
| 12275_1 | TATGAGTGTGGAGTCGTTGCGCAGGGAGGCTATTGTAAATGGTNNNAGATCGGAA |

| | |
|---|---|
| 12291_1 | TATGAGTGTGGAGTCGTTGCTTGAAATGGAAATATGCTCATGGTNNNAGATCGGAA |
| 12301_1 | TATGAGTGTGGAGTCGTTGCCCAATCTTTTCCCTATGGGATGGTNNNAGATCGGAAC |
| 12351_1 | TATGAGTGTGGAGTCGTTGCATAGTAATAAAGAGGTGCGGAGAGNNNAGATCGGA |
| 12352_1 | TATGAGTGTGGAGTCGTTGCAGGTAGCATATCCGTGGAAAAACANNNAGATCGGA |
| 12360_1 | TATGAGTGTGGAGTCGTTGCTCCCTGAACTTGTCTATCAGTGGTNNNAGATCGGAAC |
| 12364_1 | TATGAGTGTGGAGTCGTTGCGATGTCTCTAGGAACCATAGTGGTNNNAGATCGGAA |
| 12368_1 | TATGAGTGTGGAGTCGTTGCTGAGCACTGACTTCTGGAAAACANNNAGATCGGAAG |
| 12370_1 | TATGAGTGTGGAGTCGTTGCTGAATGAAACAGCTCTGAAAACANNNAGATCGGAAG |
| 12376_1 | TATGAGTGTGGAGTCGTTGCGAGAGAGAGAGACAGAGCATCTATNNNAGATCGGA |
| 12382_1 | TATGAGTGTGGAGTCGTTGCACTGATTGGTGCCAGTTTACAATCNNNAGATCGGAA |
| 12384_1 | TATGAGTGTGGAGTCGTTGCGGAGATATCATGTGCATGCAAACANNNAGATCGGA |
| 12386_1 | TATGAGTGTGGAGTCGTTGCGCATTTCTCTTTAGATGGGGTTGGNNNAGATCGGAA |
| 12406_1 | TATGAGTGTGGAGTCGTTGCTCAACGTGTCCCCCTAAATTCATGNNNAGATCGGAAG |
| 12411_1 | TATGAGTGTGGAGTCGTTGCTCCTTCCCACTTTGCAAATAAACANNNAGATCGGAAG |
| 53482_1 | TATGAGTGTGGAGTCGTTGCGTGGAGGGAAGTTTCAGAACCTANNNAGATCGGAA |
| 53488_1 | TATGAGTGTGGAGTCGTTGCTCTCTTCCAAAATCAACCCTGAGTNNNAGATCGGAAG |
| 53515_1 | TATGAGTGTGGAGTCGTTGCTGAGAACCAAACTAGCCATCAAGTNNNAGATCGGAA |
| 53616_1 | TATGAGTGTGGAGTCGTTGCTCTGCCAAAATCTTAGTGTCCTCANNNAGATCGGAAG |
| 53659_1 | TATGAGTGTGGAGTCGTTGCTGCTGTATTTATTGCCTCAATGTTNNNAGATCGGAAG |
| 53666_1 | TATGAGTGTGGAGTCGTTGCGTGAAACACTATGTCATTCAAGGCTNNNAGATCGGA |
| 53674_1 | TATGAGTGTGGAGTCGTTGCTTTGGCAAAGTGTAGATTTCTCTTNNNAGATCGGAA |
| 53697_1 | TATGAGTGTGGAGTCGTTGCCAGAATGTGGTCCCAGAAAAGAACNNNAGATCGGA |
| 53699_1 | TATGAGTGTGGAGTCGTTGCACAAGTTTCAAAAGGACATGTACANNNAGATCGGAA |
| 53714_1 | TATGAGTGTGGAGTCGTTGCCAAGTAGCTGGGACTACAGGCNNNAGATCGGAAGA |
| 53718_1 | TATGAGTGTGGAGTCGTTGCATATGGTGGTTTCAGCGACAGATNNNAGATCGGAA |
| 53719_1 | TATGAGTGTGGAGTCGTTGCCTTTATCTCCTGGGCTCAAGTGANNNAGATCGGAAG |
| 53781_1 | TATGAGTGTGGAGTCGTTGCTGGGGAGCTAAAATTTTCTCTTTGNNNAGATCGGAA |
| 53799_1 | TATGAGTGTGGAGTCGTTGCCAAGTAGCTGGGAATATAGGCGTNNNAGATCGGAA |
| 53838_1 | TATGAGTGTGGAGTCGTTGCGTGGGGAGATGGTAGATGAAATGANNNAGATCGG |
| 53900_1 | TATGAGTGTGGAGTCGTTGCAACATTCAAGCAATTCTCCTGCCNNNAGATCGGAAGA |
| 54001_1 | TATGAGTGTGGAGTCGTTGCCCTCCCAGGTTCAAGAGATTCTCNNNAGATCGGAAG |
| 54079_1 | TATGAGTGTGGAGTCGTTGCTGTCCCTGTAAACCAAAGAGTTCANNNAGATCGGAA |
| 54103_1 | TATGAGTGTGGAGTCGTTGCAGGTCTTGTTTTTGCAACGTTACANNNAGATCGGAA |
| 54112_1 | TATGAGTGTGGAGTCGTTGCCAAGCCTGTAATTCCAGCACTTTNNNAGATCGGAAG |
| 54240_1 | TATGAGTGTGGAGTCGTTGCAGTTTTCTTTGCTGTGCAGAAACTNNNAGATCGGAA |
| 54268_1 | TATGAGTGTGGAGTCGTTGCAGGTACAACTCTATGGCATTGACANNNAGATCGGAA |
| 54309_1 | TATGAGTGTGGAGTCGTTGCAGAGTAGCTGGAATTATAGGTGCCNNNAGATCGGA |
| 54348_1 | TATGAGTGTGGAGTCGTTGCCAATCTCCACTCACTGCAAACTCNNNAGATCGGAAGA |
| 54349_1 | TATGAGTGTGGAGTCGTTGCGCTGAGGCCATTTTTAGACATGGNNNAGATCGGAAG |
| 54394_1 | TATGAGTGTGGAGTCGTTGCCCACCTCCTCATTTATCTAGAGCCNNNAGATCGGAAG |
| 54427_1 | TATGAGTGTGGAGTCGTTGCTGGAGGATGGAGGATAAAATGACTNNNAGATCGGA |
| 54435_1 | TATGAGTGTGGAGTCGTTGCAGTATAACAACTTTCCTGAGTGCCTNNNAGATCGGA |
| 54478_1 | TATGAGTGTGGAGTCGTTGCAGATATGCAGATGGAAAACTAGCANNNAGATCGGA |

| | |
|---|---|
| 54529_1 | TATGAGTGTGGAGTCGTTGCACCTCCCAGGTTCAAGTAATTCTTNNNAGATCGGAAG |
| 54544_1 | TATGAGTGTGGAGTCGTTGCTGACTGATTTCATTGTAGTTGGCCNNNAGATCGGAA |
| 54545_1 | TATGAGTGTGGAGTCGTTGCACACATACCTACTTACAACCCTGGNNNAGATCGGAAG |
| 54558_1 | TATGAGTGTGGAGTCGTTGCTCACAATGAAATATGGACTTCAGGTNNNAGATCGGA |
| 54582_1 | TATGAGTGTGGAGTCGTTGCACTGAGACTAAACACCAAGACACANNNAGATCGGAA |
| 54627_1 | TATGAGTGTGGAGTCGTTGCTTGCATATGGCTATAGTTTTTCCTNNNAGATCGGAA |
| 54656_1 | TATGAGTGTGGAGTCGTTGCTGGATGGTAGCTATTCCCAGTACNNNAGATCGGAAG |
| 54662_1 | TATGAGTGTGGAGTCGTTGCTTCAAGTGATTCTCCTGCCTCAGNNNAGATCGGAAGA |
| 54719_1 | TATGAGTGTGGAGTCGTTGCGTGCGATCTTGTATCACTGCAAGNNNAGATCGGAAG |
| 54744_1 | TATGAGTGTGGAGTCGTTGCGCCTTCCAGTTTCAAGGGATTCTNNNAGATCGGAAGA |
| 54815_1 | TATGAGTGTGGAGTCGTTGCTGAAATTCAAATTCTACTGCCCACANNNAGATCGGAA |
| 54860_1 | TATGAGTGTGGAGTCGTTGCCCATTCTCCTGCCTCAACCTTNNNAGATCGGAAGAGCA |
| 54866_1 | TATGAGTGTGGAGTCGTTGCGATCTCTGCTCACTACAAGCTCCNNNAGATCGGAAGA |
| 54913_1 | TATGAGTGTGGAGTCGTTGCTGTATCACAAAGAGCTATCCCTCANNNAGATCGGAA |
| 54925_1 | TATGAGTGTGGAGTCGTTGCACCTTGGGAATTGTAGTTTATGGTNNNAGATCGGAA |
| 54928_1 | TATGAGTGTGGAGTCGTTGCGTTGGTAATTCACAGTGAAACGGTNNNAGATCGGAA |
| 54940_1 | TATGAGTGTGGAGTCGTTGCTCTCAGTAAATAATAGAAAAGCAGACCNNNAGATCG |
| 54950_1 | TATGAGTGTGGAGTCGTTGCTGGAGAAGGGAAATCTTTTTCTCANNNAGATCGGAA |
| 54957_1 | TATGAGTGTGGAGTCGTTGCAATGGAGTTGACTTTCACATTTGTNNNAGATCGGAA |
| 55008_1 | TATGAGTGTGGAGTCGTTGCGTTGTTTCTGTTCTCCTTTGTTTTCTNNNAGATCGGAA |
| 55025_1 | TATGAGTGTGGAGTCGTTGCAGGAATGGATTTTGCATTTTATCAAGTNNNAGATCG |
| 55032_1 | TATGAGTGTGGAGTCGTTGCACCTATGAATGTACCAGTTGGAGGNNNAGATCGGAA |
| 55067_1 | TATGAGTGTGGAGTCGTTGCGTGAGTAGCTGGGACTACAGATGNNNAGATCGGAA |
| 55077_1 | TATGAGTGTGGAGTCGTTGCGGTTCAAGTGATTCTTCTGCCTCNNNAGATCGGAAGA |
| 55133_1 | TATGAGTGTGGAGTCGTTGCAGCTGCAAATTATAAACTGAGTAAGANNNAGATCGG |
| 55142_1 | TATGAGTGTGGAGTCGTTGCCACGTTTTCACTCCTTTCTTGCTANNNAGATCGGAAGA |
| 55218_1 | TATGAGTGTGGAGTCGTTGCGATCTCGGCTTTGGTATCAGGATNNNAGATCGGAAG |
| 55255_1 | TATGAGTGTGGAGTCGTTGCGACATAAACAGAGTGATTTGGGAAANNNAGATCGG |
| 55261_1 | TATGAGTGTGGAGTCGTTGCCATTCTGTGGGTTGTCTTTTCACTNNNAGATCGGAAG |
| 55298_1 | TATGAGTGTGGAGTCGTTGCCTCCTGAGTAGCTGGGACTACAGNNNAGATCGGAAG |
| 55303_1 | TATGAGTGTGGAGTCGTTGCTTCAAGTGATTCTCCTGCCTCAGNNNAGATCGGAAGA |
| 55342_1 | TATGAGTGTGGAGTCGTTGCTGTTTTGTTTTTGAGACGGAGTCTNNNAGATCGGAA |
| 55458_1 | TATGAGTGTGGAGTCGTTGCGGCCAGTGGAATGTGGATAAAAGNNNAGATCGGAA |
| 55502_1 | TATGAGTGTGGAGTCGTTGCTGCTTTCATGTGAAAAAATTAGAGCTNNNAGATCGG |
| 55574_1 | TATGAGTGTGGAGTCGTTGCCTCCTGAGTAGCTGGGACTACAGNNNAGATCGGAAG |
| 55589_1 | TATGAGTGTGGAGTCGTTGCAATAATTCTGTCTCAGCCTCCCGNNNAGATCGGAAGA |
| 55602_1 | TATGAGTGTGGAGTCGTTGCCAATCTCAGCTCACTGCAACTTCNNNAGATCGGAAGA |
| 55628_1 | TATGAGTGTGGAGTCGTTGCAGCTATGCCTGTATAGTTCTGGACNNNAGATCGGAA |
| 55637_1 | TATGAGTGTGGAGTCGTTGCTCCCATTCAAAAATGTCTCTGTGTGNNNAGATCGGAA |
| 55686_1 | TATGAGTGTGGAGTCGTTGCTGAGCATCCCTAATCTGAAATCCANNNAGATCGGAA |
| 55723_1 | TATGAGTGTGGAGTCGTTGCGATTCTCCTGCCTCAGCCTCNNNAGATCGGAAGAGCA |
| 55801_1 | TATGAGTGTGGAGTCGTTGCCTCAGACAATCCTCCTACCTCAGNNNAGATCGGAAGA |
| 55805_1 | TATGAGTGTGGAGTCGTTGCTCCCAGGTTCAAGTGATCTTCTGNNNAGATCGGAAGA |

| | |
|---|---|
| 55821_1 | TATGAGTGTGGAGTCGTTGCACCATCTCAGCTCACTGCAAGNNNAGATCGGAAGAG |
| 55961_1 | TATGAGTGTGGAGTCGTTGCTCCCAAATCCATGTCTTAGTGTGTNNNAGATCGGAA |
| 55966_1 | TATGAGTGTGGAGTCGTTGCAGCAGAAGTAGCTAAGCTTACACANNNAGATCGGA |
| 56016_1 | TATGAGTGTGGAGTCGTTGCCATTCTCCTGCCTCAGCCTCNNNAGATCGGAAGAGCA |
| 56065_1 | TATGAGTGTGGAGTCGTTGCAGAATGACTGTCGGGTAAAGTACANNNAGATCGGA |
| 56069_1 | TATGAGTGTGGAGTCGTTGCGGAAACAATTGTTCATGTCACAGTNNNAGATCGGAA |
| 56076_1 | TATGAGTGTGGAGTCGTTGCGCAAGAAATCTACCTTCCAAGGACNNNAGATCGGAA |
| 56098_1 | TATGAGTGTGGAGTCGTTGCTCCCAAAGAATCCTGACACTACTGNNNAGATCGGAA |
| 56110_1 | TATGAGTGTGGAGTCGTTGCTGAGCTCTGGATCACATTTTTGANNNAGATCGGAA |
| 56203_1 | TATGAGTGTGGAGTCGTTGCACCCTTTCCTAGAGTTTCTTCACCNNNAGATCGGAAG |
| 56207_1 | TATGAGTGTGGAGTCGTTGCACGATTATGGTAAGTGAAAGAAAGCANNNAGATCG |
| 56224_1 | TATGAGTGTGGAGTCGTTGCTGCAACAACAACAAAAACATTTGANNNAGATCGGAA |
| 56227_1 | TATGAGTGTGGAGTCGTTGCACCTATCACAAAGAGGTTCAAACCNNNAGATCGGAA |
| 56311_1 | TATGAGTGTGGAGTCGTTGCTCTTTGTTGTCATCATTAGTGCCANNNAGATCGGAAG |
| 56337_1 | TATGAGTGTGGAGTCGTTGCGGAATGCGATGGTGTGATCTTGNNNAGATCGGAAG |
| 56365_1 | TATGAGTGTGGAGTCGTTGCCCATCTCCTTGGTAGTAAGTGAGTNNNAGATCGGAA |
| 56415_1 | TATGAGTGTGGAGTCGTTGCCCTTGAGTCACATCCCAAAGTCANNNAGATCGGAAG |
| 56438_1 | TATGAGTGTGGAGTCGTTGCCAGGGAACTAGATGGATGTAAGCANNNAGATCGGA |
| 56488_1 | TATGAGTGTGGAGTCGTTGCTCTCCTACATTAGCCTCCCAAGANNNAGATCGGAAGA |
| 56569_1 | TATGAGTGTGGAGTCGTTGCTTCAAGTGATTCTCCTGCCTCAGNNNAGATCGGAAG |
| 56634_1 | TATGAGTGTGGAGTCGTTGCACCAGAAACATAATCCCAAAGAGTNNNAGATCGGAA |
| 56661_1 | TATGAGTGTGGAGTCGTTGCACGTGCAACAACATAACAATGAGANNNAGATCGGA |
| 56814_1 | TATGAGTGTGGAGTCGTTGCCTCACTGTAACCTCCACCTCTCANNNAGATCGGAAGA |
| 56881_1 | TATGAGTGTGGAGTCGTTGCTGATATGAACCTTTTGTCTTCTGCTNNNAGATCGGAA |
| 56882_1 | TATGAGTGTGGAGTCGTTGCCCAACAGATTTTGTCTATGGCACANNNAGATCGGAA |
| 56939_1 | TATGAGTGTGGAGTCGTTGCACCAAACAATACCATGAAGCAAGTNNNAGATCGGAA |
| 56947_1 | TATGAGTGTGGAGTCGTTGCACACAAGGCAGACAAGAAAATAGCNNNAGATCGGA |
| 56972_1 | TATGAGTGTGGAGTCGTTGCGAGGGGAAAGAGAGAGAACACAGNNNAGATCGGA |
| 56992_1 | TATGAGTGTGGAGTCGTTGCCATTCTCCTGCCTCAGCCTCNNNAGATCGGAAGAGCA |
| 57051_1 | TATGAGTGTGGAGTCGTTGCAGAATTTCTAAGGCCTCTTCAGCTNNNAGATCGGAA |
| 57066_1 | TATGAGTGTGGAGTCGTTGCAGAAGTCACAGTAAAAGGATGGGTNNNAGATCGGA |
| 57076_1 | TATGAGTGTGGAGTCGTTGCTCAGGATCCAGAGTATCAAACAGTNNNAGATCGGAA |
| 57099_1 | TATGAGTGTGGAGTCGTTGCCCATCTCAGCTCACTGTAACCTCNNNAGATCGGAAGA |
| 57131_1 | TATGAGTGTGGAGTCGTTGCACTGGACGGTTATTCATTCTCTCTNNNAGATCGGAAG |
| 57175_1 | TATGAGTGTGGAGTCGTTGCTTTGCAACTTAGATGAAATGGACANNNAGATCGGAA |
| 57232_1 | TATGAGTGTGGAGTCGTTGCTGTGCACTAAAATTTTAGGTTGCANNNAGATCGGAA |
| 57278_1 | TATGAGTGTGGAGTCGTTGCTTCAAGCGATTCTCCTGCCTTAGNNNAGATCGGAAG |
| 57279_1 | TATGAGTGTGGAGTCGTTGCCCAGGCGGAGACTTTTATTCTCTNNNAGATCGGAAG |
| 57289_1 | TATGAGTGTGGAGTCGTTGCTTCAAGCAATTCTCCTGCCTCAGNNNAGATCGGAAG |
| 57353_1 | TATGAGTGTGGAGTCGTTGCACCAGTGTTTGATCATCTTCATGGNNNAGATCGGAA |
| 57365_1 | TATGAGTGTGGAGTCGTTGCCCAAGGCTTTTCATAATTCCCTCANNNAGATCGGAAG |
| 57386_1 | TATGAGTGTGGAGTCGTTGCACACACAACTCTCTTTCATCAAGTNNNAGATCGGAAG |
| 57389_1 | TATGAGTGTGGAGTCGTTGCAGCCTTGGAGAAATAAGGGATTCTNNNAGATCGGA |

| | |
|---|---|
| 57392_1 | TATGAGTGTGGAGTCGTTGCAGCAGGTAAAACATGGTAGAAACANNNAGATCGGA |
| 57448_1 | TATGAGTGTGGAGTCGTTGCTTCACACCATTCTCCTGCCTCNNNAGATCGGAAGAGC |
| 57485_1 | TATGAGTGTGGAGTCGTTGCGGGAAAAACCTACAGCTAACATCANNNAGATCGGA |
| 57488_1 | TATGAGTGTGGAGTCGTTGCCGGTAGATATGTAGACCATGGAGTNNNAGATCGGA |
| 57497_1 | TATGAGTGTGGAGTCGTTGCACAAGGATTTTGAAGAAACCATCANNNAGATCGGA |
| 57674_1 | TATGAGTGTGGAGTCGTTGCCGAGTAGCTGGGATTACAGGTACNNNAGATCGGAA |
| 57689_1 | TATGAGTGTGGAGTCGTTGCGCCTCCCAAGTAGCTAGGATTACNNNAGATCGGAAG |
| 57690_1 | TATGAGTGTGGAGTCGTTGCCCTCAGCTCTTGGACAAAACTTCNNNAGATCGGAAG |
| 57691_1 | TATGAGTGTGGAGTCGTTGCAAGTAGCTGGAATTACAGACGCANNNAGATCGGAA |
| 57714_1 | TATGAGTGTGGAGTCGTTGCGCCTCCCAAGTTCAAGTGATTCTNNNAGATCGGAAG |
| 57759_1 | TATGAGTGTGGAGTCGTTGCAGAGGTAGATGGTAAGGACTGACANNNAGATCGG |
| 57768_1 | TATGAGTGTGGAGTCGTTGCTGGGTAATTAGGTTTTGCACAAGTNNNAGATCGGA |
| 57779_1 | TATGAGTGTGGAGTCGTTGCCCCAGGTTTGGTCTGATATGTTCANNNAGATCGGAA |
| 57783_1 | TATGAGTGTGGAGTCGTTGCAAAACTATGGTAGTCAGATGACCANNNAGATCGGA |
| 57790_1 | TATGAGTGTGGAGTCGTTGCTGCTCAACTTTTTATCCAGCTGTGNNNAGATCGGAA |
| 57805_1 | TATGAGTGTGGAGTCGTTGCCAGGATGGAAGACAAAATGTGACANNNAGATCGGA |
| 57807_1 | TATGAGTGTGGAGTCGTTGCACTTGAATGTAAGATCTGGAGCCANNNAGATCGGA |
| 57835_1 | TATGAGTGTGGAGTCGTTGCCTCCCAAGTAGCTGAGACTACAGNNNAGATCGGAAG |
| 57878_1 | TATGAGTGTGGAGTCGTTGCTTCAAGCGATTCTCCTGTCTCAGNNNAGATCGGAAG |
| 57900_1 | TATGAGTGTGGAGTCGTTGCTGCACTAGACAACAATGGATGGANNNAGATCGGAA |
| 57906_1 | TATGAGTGTGGAGTCGTTGCCAACCTTCAGGGCTCAAATGATCNNNAGATCGGAAG |
| 57935_1 | TATGAGTGTGGAGTCGTTGCGCTGAGGATAAGAGAGAAGAGACANNNAGATCGG |
| 57970_1 | TATGAGTGTGGAGTCGTTGCGGTATGTTGACCCTGAATTTCAGCNNNAGATCGGAA |
| 57989_1 | TATGAGTGTGGAGTCGTTGCAGTTCAGGTAAGGTGGAAAGAAAGTNNNAGATCG |
| 58058_1 | TATGAGTGTGGAGTCGTTGCAGGACATGTAGCAGAAAGATGTCANNNAGATCGGA |
| 58108_1 | TATGAGTGTGGAGTCGTTGCGCTTCTCCTTTCTCCTTCTTATTGCNNNAGATCGGAAG |
| 58141_1 | TATGAGTGTGGAGTCGTTGCCTCCTGAGTAGCTGGGACTACANNNAGATCGGAAG |
| 58169_1 | TATGAGTGTGGAGTCGTTGCAATACAGTGGCATGATCTCAGCTNNNAGATCGGAAG |
| 58296_1 | TATGAGTGTGGAGTCGTTGCTGGTAAATGAAAGTTTGCATAGCTNNNAGATCGGA |
| 58308_1 | TATGAGTGTGGAGTCGTTGCTGCTTTTACACAAAAAAGGCAATTNNNAGATCGGAA |
| 58324_1 | TATGAGTGTGGAGTCGTTGCTAGATCCCCCAGTTTTGTTCAGGNNNAGATCGGAAG |
| 58422_1 | TATGAGTGTGGAGTCGTTGCTGATGGATACCTTCATGAGCTTGANNNAGATCGGAA |
| 58427_1 | TATGAGTGTGGAGTCGTTGCACAAAGAAATAGAGAGACAAGGAGCNNNAGATCG |
| 58433_1 | TATGAGTGTGGAGTCGTTGCCCCCTACCTCACACCAATACAAANNNAGATCGGAAGA |
| 58510_1 | TATGAGTGTGGAGTCGTTGCCCATTCTGTTTGAGTATGCTGAGANNNAGATCGGAA |
| 58526_1 | TATGAGTGTGGAGTCGTTGCGTACAGTGGCACAAACACAGTTTNNNAGATCGGAA |
| 58584_1 | TATGAGTGTGGAGTCGTTGCTCTTCACTTCTAGACCATCTTGTTGANNNAGATCGGA |
| 58608_1 | TATGAGTGTGGAGTCGTTGCTGCATACGTGTTACAAACCTCACNNNAGATCGGAAG |
| 58616_1 | TATGAGTGTGGAGTCGTTGCGAGTTTCAGTCTTGTTTCCCAGGNNNAGATCGGAAG |
| 58622_1 | TATGAGTGTGGAGTCGTTGCATATTGGCTCACTGAAACCTCCANNNAGATCGGAAG |
| 58655_1 | TATGAGTGTGGAGTCGTTGCCAATGGCGCAATCTCAGCTCNNNAGATCGGAAGAG |
| 58672_1 | TATGAGTGTGGAGTCGTTGCAGTAGCTGAGATTACAGGTGTGCNNNAGATCGGAA |
| 58691_1 | TATGAGTGTGGAGTCGTTGCCATTCCGAGTAGCTGGAACTACANNNAGATCGGAAG |

| | |
|---|---|
| 58695_1 | TATGAGTGTGGAGTCGTTGCGGGTAGCTGGGAATATAGGTGCNNNAGATCGGAA |
| 58719_1 | TATGAGTGTGGAGTCGTTGCCACACTTATACTGTCCTGCTCACTNNNAGATCGGAAG |
| 58737_1 | TATGAGTGTGGAGTCGTTGCGTGCTATGGGCTATTGATGCAATTNNNAGATCGGA |
| 58738_1 | TATGAGTGTGGAGTCGTTGCTGGAAAGAACTGATCCACATAGGTNNNAGATCGGA |
| 58788_1 | TATGAGTGTGGAGTCGTTGCCTAGAGTGCAGTGGTGTGATCTCNNNAGATCGGAA |
| 58871_1 | TATGAGTGTGGAGTCGTTGCTGCCATTCGTACTGAAACTATTCCNNNAGATCGGAA |
| 58878_1 | TATGAGTGTGGAGTCGTTGCGGTGTCTGTTTGTTGTCTTTTGGANNNAGATCGGAA |
| 58944_1 | TATGAGTGTGGAGTCGTTGCTGATCTTCCACCTCTCTGTATCCTNNNAGATCGGAAG |
| 58953_1 | TATGAGTGTGGAGTCGTTGCTCAAGGCAGAAAATACAAAAAAGCCNNNAGATCGG |
| 59021_1 | TATGAGTGTGGAGTCGTTGCTGTTGATCTCATAGAAGTGGAGAGTNNNAGATCGG |
| 59050_1 | TATGAGTGTGGAGTCGTTGCTGAAACTTACATGCTACTTTGCAGANNNAGATCGGA |
| 59116_1 | TATGAGTGTGGAGTCGTTGCTGTTTGGGATTTTTTGTTTGTGCTNNNAGATCGGAA |
| 59150_1 | TATGAGTGTGGAGTCGTTGCACACACACACACACATACATATTCANNNAGATCGGAA |
| 59159_1 | TATGAGTGTGGAGTCGTTGCAGTCACATTTTCATAGTTCCTTCCTNNNAGATCGGAA |
| 59160_1 | TATGAGTGTGGAGTCGTTGCTGACAAAATGGAAAGTCAGTTTTCTGNNNAGATCGG |
| 59171_1 | TATGAGTGTGGAGTCGTTGCAGAAAAGAAGGGGATAATCCAATCTNNNAGATCGG |
| 59175_1 | TATGAGTGTGGAGTCGTTGCTGCTCTGTAGGATTTCTCAACTCTNNNAGATCGGAA |
| 59245_1 | TATGAGTGTGGAGTCGTTGCTGGGTTGGCCTTATTCTCCTTTTNNNAGATCGGAAG |
| 59276_1 | TATGAGTGTGGAGTCGTTGCATCTTGACTCACTCCAACCTCTGNNNAGATCGGAAGA |
| 59284_1 | TATGAGTGTGGAGTCGTTGCAACTTCTGCCTACTGAGCTCAAGNNNAGATCGGAAG |
| 59357_1 | TATGAGTGTGGAGTCGTTGCTGCAACAACAACAAAAACATTTGANNNAGATCGGAA |
| 59388_1 | TATGAGTGTGGAGTCGTTGCTAGTTCACTGTAGCCTTGACCTGNNNAGATCGGAAG |
| 59416_1 | TATGAGTGTGGAGTCGTTGCTGCAGCCAACAAACATATGAAAGANNNAGATCGGA |
| 59423_1 | TATGAGTGTGGAGTCGTTGCGAACGTTACCAGCAGGTGATTTCNNNAGATCGGAA |
| 59425_1 | TATGAGTGTGGAGTCGTTGCACACACACATACATAAAACATCACANNNAGATCGGA |
| 59469_1 | TATGAGTGTGGAGTCGTTGCGTAGCTGGGATTACAGGTACACANNNAGATCGGAA |
| 59503_1 | TATGAGTGTGGAGTCGTTGCCAGAGGGCTGGTCTTTTATTCCTNNNAGATCGGAAG |
| 59530_1 | TATGAGTGTGGAGTCGTTGCGCAATAGCATGATCTTGGCTCACNNNAGATCGGAAG |
| 59545_1 | TATGAGTGTGGAGTCGTTGCTGCCACATGATTCAAAACCAAAGANNNAGATCGGAA |
| 59579_1 | TATGAGTGTGGAGTCGTTGCGAATTGTTGGCAGCCATGACTAGNNNAGATCGGAA |
| 59583_1 | TATGAGTGTGGAGTCGTTGCTCTCTGCATTTTTGGAAGTGAAGANNNAGATCGGAA |
| 59584_1 | TATGAGTGTGGAGTCGTTGCTGTTGTAAAAGCTTTGGTAAGGCTNNNAGATCGGA |
| 59586_1 | TATGAGTGTGGAGTCGTTGCTGAGCAAAGGTATGACTAGACAGTNNNAGATCGGA |
| 59615_1 | TATGAGTGTGGAGTCGTTGCAGTTGGTTGTTTGAGAAGATAAACANNNAGATCGG |
| 59716_1 | TATGAGTGTGGAGTCGTTGCAGTAACTGGGACTATAAGTGCACANNNAGATCGGA |
| 59940_1 | TATGAGTGTGGAGTCGTTGCGGGACCACCATTAGCAATAAAACANNNAGATCGGAA |
| 59948_1 | TATGAGTGTGGAGTCGTTGCAGTCTGTCTGTCTGTCTACACTACNNNAGATCGGAA |
| 59950_1 | TATGAGTGTGGAGTCGTTGCACTTGCTAATGGGAAGACATTTCTNNNAGATCGGAA |
| 60065_1 | TATGAGTGTGGAGTCGTTGCCTCCTGAGTAGCTGGGACTACAGNNNAGATCGGAA |
| 60101_1 | TATGAGTGTGGAGTCGTTGCCTGGATTCAAGCCATTCTTCTGCNNNAGATCGGAAG |
| 60167_1 | TATGAGTGTGGAGTCGTTGCGAATCCCATGATGAACATACTGGANNNAGATCGGAA |
| 60240_1 | TATGAGTGTGGAGTCGTTGCCATCCCAGAGTTGCAGAGATAGTNNNAGATCGGAA |
| 60268_1 | TATGAGTGTGGAGTCGTTGCGGTTCTAAGTAGGAAGTAGCTGAGGNNNAGATCGG |

| | |
|---|---|
| 62025_1 | TATGAGTGTGGAGTCGTTGCAGAGAGAATTTGAGACAAGGGACCNNNAGATCGG |
| 62026_1 | TATGAGTGTGGAGTCGTTGCTCAGTACAAGGCATAATAGTGTGGTNNNAGATCGG |
| 62049_1 | TATGAGTGTGGAGTCGTTGCACCAAGATTGAACCAGGAAGAAANNNAGATCGGAA |
| 62056_1 | TATGAGTGTGGAGTCGTTGCGATTCAAGCGATTCTTCTGGCTCNNNAGATCGGAAG |
| 62114_1 | TATGAGTGTGGAGTCGTTGCTTTGCAAACTATCCATCTGGCAANNNAGATCGGAAG |
| 62131_1 | TATGAGTGTGGAGTCGTTGCCGTGCAGATAGGAAGTTTTGGAANNNAGATCGGAA |
| 62240_1 | TATGAGTGTGGAGTCGTTGCACACAGAAACTTGTTCATAGCAGCNNNAGATCGGAA |
| 62267_1 | TATGAGTGTGGAGTCGTTGCTGAGATGACAGGAAATTTGGTTTTNNNAGATCGGA |
| 62268_1 | TATGAGTGTGGAGTCGTTGCAACACACCACAAACTTTTAGAACTNNNAGATCGGAA |
| 62308_1 | TATGAGTGTGGAGTCGTTGCTCACATGTGGTGGAAGATTTTCTTNNNAGATCGGAA |
| 62335_1 | TATGAGTGTGGAGTCGTTGCTGGAGTGTAATGGCACAATCTCANNNAGATCGGAA |
| 62369_1 | TATGAGTGTGGAGTCGTTGCTGGAACCTGTGAATGTTACTTTGTNNNAGATCGGAA |
| 62370_1 | TATGAGTGTGGAGTCGTTGCCTCAAGTGATCCTCCCAACTCAGNNNAGATCGGAAG |
| 62379_1 | TATGAGTGTGGAGTCGTTGCGTTCCAAGAATTACTCCTTTCTCTACANNNAGATCGG |
| 62383_1 | TATGAGTGTGGAGTCGTTGCTGCTCCTGATTTTGATATGAGGGTNNNAGATCGGAA |
| 62392_1 | TATGAGTGTGGAGTCGTTGCAGCAAACTGAATTCATCTGAACATNNNAGATCGGAA |
| 62412_1 | TATGAGTGTGGAGTCGTTGCTTCAAGCAATTCTCCTGCCTCAGNNNAGATCGGAAG |
| 62533_1 | TATGAGTGTGGAGTCGTTGCCTCTTGGGTTCACGCCATTCTCNNNAGATCGGAAGA |
| 62559_1 | TATGAGTGTGGAGTCGTTGCAGAGAACCCAGACATAAATCCACANNNAGATCGGAA |
| 62565_1 | TATGAGTGTGGAGTCGTTGCCCAAAGTGCTGGGATTACAAGTGNNNAGATCGGAA |
| 62597_1 | TATGAGTGTGGAGTCGTTGCTCTTCAATTGCATCTAGTTTGGAACANNNAGATCGG |
| 62608_1 | TATGAGTGTGGAGTCGTTGCAGTAGCAGGAGTACACTGTAAAATNNNAGATCGGA |
| 62635_1 | TATGAGTGTGGAGTCGTTGCTCCCATATGAATGTGTGTGTATGTNNNAGATCGGAA |
| 62676_1 | TATGAGTGTGGAGTCGTTGCTCATGAGGAGTTTTTTGGTGAGGANNNAGATCGGA |
| 62837_1 | TATGAGTGTGGAGTCGTTGCTTCATGAAGTTCTCCTGCCTCAGNNNAGATCGGAAG |
| 62889_1 | TATGAGTGTGGAGTCGTTGCCAATCTCGGCTCACTGCAAGNNNAGATCGGAAGAG |
| 62926_1 | TATGAGTGTGGAGTCGTTGCTGCTTAGATGTAGTTACGCACTGANNNAGATCGGAA |
| 62932_1 | TATGAGTGTGGAGTCGTTGCTGTTGTGTCGTATGTTTTACCACANNNAGATCGGAA |
| 62937_1 | TATGAGTGTGGAGTCGTTGCTGCTAAAGAAAAGCAGCAATGACTNNNAGATCGGA |
| 62942_1 | TATGAGTGTGGAGTCGTTGCTGGTATTCTTTGGTATCACTGGGGNNNAGATCGGA |
| 62983_1 | TATGAGTGTGGAGTCGTTGCACCATTTCACATTTTTTGAGTCCANNNAGATCGGAAG |
| 63034_1 | TATGAGTGTGGAGTCGTTGCAGATTTATGGGTGTTAGGGAAGTTNNNAGATCGGA |
| 63045_1 | TATGAGTGTGGAGTCGTTGCAGTCTAAAGTTATCAGCATAGGCCANNNAGATCGGA |
| 63134_1 | TATGAGTGTGGAGTCGTTGCGGAAACTTTGTCCATTCCTAGTGCNNNAGATCGGAA |
| 63142_1 | TATGAGTGTGGAGTCGTTGCTTCAAGTGATTCTCCTGCCTCAGNNNAGATCGGAAG |
| 63188_1 | TATGAGTGTGGAGTCGTTGCAGCAATCTCAAAAGGGTACGTACTNNNAGATCGGA |
| 63225_1 | TATGAGTGTGGAGTCGTTGCGGCATGGAATTCTTCACACTAACCNNNAGATCGGAA |
| 63243_1 | TATGAGTGTGGAGTCGTTGCTGTGTTTTTAGTAGACACGGGGTNNNAGATCGGAA |
| 63256_1 | TATGAGTGTGGAGTCGTTGCGGTTCAAGTGATTCTCCTGCTTCNNNAGATCGGAAG |
| 63258_1 | TATGAGTGTGGAGTCGTTGCTTCAAGTGATTCTCCTGCCTCAGNNNAGATCGGAAG |
| 63271_1 | TATGAGTGTGGAGTCGTTGCAATCTCAGCTCACTGCAACCTCNNNAGATCGGAAGA |
| 63277_1 | TATGAGTGTGGAGTCGTTGCGCGCACGTGTATGAAATATTTCCTNNNAGATCGGAA |
| 63288_1 | TATGAGTGTGGAGTCGTTGCAGAGAGAGAGAGAGAGAGAAAGCANNNAGATCGG |

| | |
|---|---|
| 63303_1 | TATGAGTGTGGAGTCGTTGCTTGAAGCGATTCTCCTGCCTTAGNNNAGATCGGAAG |
| 63364_1 | TATGAGTGTGGAGTCGTTGCTCCTTGGTACTCACTTCATTAGCCNNNAGATCGGAAG |
| 63391_1 | TATGAGTGTGGAGTCGTTGCCTCAAGTGATCCTACCACCTCAGNNNAGATCGGAAG |
| 63400_1 | TATGAGTGTGGAGTCGTTGCTGGTCGTCAATTTACAAGACATGCNNNAGATCGGAA |
| 63436_1 | TATGAGTGTGGAGTCGTTGCTCATGCTGCTTGTGAATGTATTGANNNAGATCGGAA |
| 63477_1 | TATGAGTGTGGAGTCGTTGCTTGTGAAAATTACCACACTGCCANNNAGATCGGAAG |
| 63488_1 | TATGAGTGTGGAGTCGTTGCGGTAGGTCTCTTCATTTCAGGGTNNNAGATCGGAAG |
| 63493_1 | TATGAGTGTGGAGTCGTTGCTGGAGTGCAGTGGTGTAATCTCNNNAGATCGGAAG |
| 63510_1 | TATGAGTGTGGAGTCGTTGCCATAGAATCTCACTCTGTCACCCANNNAGATCGGAAG |
| 63560_1 | TATGAGTGTGGAGTCGTTGCACTTCGGTATCCTCTCTTGATTCCNNNAGATCGGAAG |
| 63568_1 | TATGAGTGTGGAGTCGTTGCTGTAATACCAGCTACTCAGAAGGCNNNAGATCGGAA |
| 63576_1 | TATGAGTGTGGAGTCGTTGCGTCTCCTGGGTTCAAGTGATTCTNNNAGATCGGAAG |
| 63616_1 | TATGAGTGTGGAGTCGTTGCTGTTTAGTACCCTGCTTTTCTCANNNAGATCGGAAGA |
| 63666_1 | TATGAGTGTGGAGTCGTTGCACTTGACATGTTTACTGTTGCTTTGNNNAGATCGGA |
| 63671_1 | TATGAGTGTGGAGTCGTTGCAGTTTTGCTTACCTGTTCTGTTGTNNNAGATCGGAA |
| 63673_1 | TATGAGTGTGGAGTCGTTGCGTCCTATCTTCTGACCACAACTCANNNAGATCGGAAG |
| 63757_1 | TATGAGTGTGGAGTCGTTGCGGACAAAGGAAGGAAAGAGGAGANNNAGATCGG |
| 63766_1 | TATGAGTGTGGAGTCGTTGCTAACCTGGATGGAATTGGAGACCNNNAGATCGGAA |
| 63771_1 | TATGAGTGTGGAGTCGTTGCACTACATCTCGAGAGTGTAGAACANNNAGATCGGA |
| 63780_1 | TATGAGTGTGGAGTCGTTGCTCTTTTTTGAGACAGAGTCTTGCTNNNAGATCGGAA |
| 63857_1 | TATGAGTGTGGAGTCGTTGCAGGGTCTCAATCTATCTTCCAAGCNNNAGATCGGAA |
| 63859_1 | TATGAGTGTGGAGTCGTTGCTCCATATCCATGAGGAAAACAGCANNNAGATCGGAA |
| 63885_1 | TATGAGTGTGGAGTCGTTGCTCTGGCCTCAAAAATATTTCAGCCNNNAGATCGGAA |
| 63967_1 | TATGAGTGTGGAGTCGTTGCTGAATACCTGCATGGAGTTACAACNNNAGATCGGAA |
| 64029_1 | TATGAGTGTGGAGTCGTTGCAGTAAACCCATCCAGGCTGAAATNNNAGATCGGAAG |
| 64038_1 | TATGAGTGTGGAGTCGTTGCTCTTTGTCTAGTCATCTGTCAGCANNNAGATCGGAA |
| 64080_1 | TATGAGTGTGGAGTCGTTGCAGAACTGTATGTGTTGAAGGCCANNNAGATCGGAA |
| 64101_1 | TATGAGTGTGGAGTCGTTGCTGAAACATTTCAGAACATTTGGGCNNNAGATCGGAA |
| 64143_1 | TATGAGTGTGGAGTCGTTGCTATGGAAGACATGATACAGCCCCNNNAGATCGGAA |
| 64191_1 | TATGAGTGTGGAGTCGTTGCAGGTTCAAGCAATTCTCCCTCAGNNNAGATCGGAAG |
| 64229_1 | TATGAGTGTGGAGTCGTTGCATTTCCTGGCTTCAGACATTCCTNNNAGATCGGAAG |
| 64235_1 | TATGAGTGTGGAGTCGTTGCAGCACATCAAAAAGCTAATCCACCNNNAGATCGGAA |
| 64247_1 | TATGAGTGTGGAGTCGTTGCAGCAAGAGGAATTTTGCAAGTTCANNNAGATCGGA |
| 64264_1 | TATGAGTGTGGAGTCGTTGCAGCTCTATGTCTCTGGAGATCTCTNNNAGATCGGAA |
| 64275_1 | TATGAGTGTGGAGTCGTTGCTCCTCAACAAAATACCAGCAAACCNNNAGATCGGAA |
| 64282_1 | TATGAGTGTGGAGTCGTTGCTGGTTCCATAAGAATTTCAGGACTNNNAGATCGGAA |
| 64301_1 | TATGAGTGTGGAGTCGTTGCTGCTCTTCGATTTTTTTGAGATGANNNAGATCGGAA |
| 64338_1 | TATGAGTGTGGAGTCGTTGCTCCGTATAATGCTGCAGAGAACANNNAGATCGGAA |
| 64422_1 | TATGAGTGTGGAGTCGTTGCGTGGATCATTTGAGGTCAGGAGTNNNAGATCGGAA |
| 64452_1 | TATGAGTGTGGAGTCGTTGCCTGACCAAGAGGGACACCTAAATNNNAGATCGGAA |
| 64459_1 | TATGAGTGTGGAGTCGTTGCGAACCTTCATTTATGTCTTCATAGTCANNNAGATCGG |
| 64494_1 | TATGAGTGTGGAGTCGTTGCGTCTTGAACTGTTGGGCTCAAAGNNNAGATCGGAA |
| 64503_1 | TATGAGTGTGGAGTCGTTGCGTAAGCGGCACTAGTAATCATGGNNNAGATCGGAA |

| | |
|---|---|
| 64514_1 | TATGAGTGTGGAGTCGTTGCAGAAAAACAGGAAGGAGTTACCCANNNAGATCGGA |
| 64532_1 | TATGAGTGTGGAGTCGTTGCAGAGAGAGAGAGAGACAGAGAGTNNNAGATCGGA |
| 64538_1 | TATGAGTGTGGAGTCGTTGCTGGAACTGAAAACTATGACACCCTNNNAGATCGGAA |
| 64569_1 | TATGAGTGTGGAGTCGTTGCATTTTTGCCTCTACTCAGTCAGCNNNAGATCGGAAGA |
| 64576_1 | TATGAGTGTGGAGTCGTTGCGTAACCTGGAGAATGAGGGTGTTNNNAGATCGGAA |
| 64585_1 | TATGAGTGTGGAGTCGTTGCTGCAATTACTAGGTGTTTGACTGANNNAGATCGGAA |
| 64605_1 | TATGAGTGTGGAGTCGTTGCTTGCTTCCTATTGGCCTTTCTCTNNNAGATCGGAAGA |
| 64660_1 | TATGAGTGTGGAGTCGTTGCATACAACAGCTGTACTTCCCCTGNNNAGATCGGAAG |
| 64682_1 | TATGAGTGTGGAGTCGTTGCCATGAACGATGGATGCATAATTCTNNNAGATCGGAA |
| 64716_1 | TATGAGTGTGGAGTCGTTGCACTCCTGGACTAAAGTGATCTGCNNNAGATCGGAAG |
| 64730_1 | TATGAGTGTGGAGTCGTTGCTGAAGATTCACTGGGATGCTTCANNNAGATCGGAAG |
| 64735_1 | TATGAGTGTGGAGTCGTTGCCCTAAGGCTCATGAAACAATACATNNNAGATCGGAA |
| 64771_1 | TATGAGTGTGGAGTCGTTGCTGCCTACTATCACATAGACCTCTGNNNAGATCGGAAG |
| 64835_1 | TATGAGTGTGGAGTCGTTGCTGGTGAGGATGTGGAGAAATCAGNNNAGATCGGA |
| 64847_1 | TATGAGTGTGGAGTCGTTGCACCTAAGCTTAGTGACTGGTTGGNNNAGATCGGAAG |
| 64855_1 | TATGAGTGTGGAGTCGTTGCGGTGCAAATGATCTAGCAATCCCNNNAGATCGGAAG |
| 64877_1 | TATGAGTGTGGAGTCGTTGCTGGCCTGTCCTAAATAGCTCATCNNNAGATCGGAAG |
| 64884_1 | TATGAGTGTGGAGTCGTTGCTGGAGCTGAAAAATTCCTATTGCTNNNAGATCGGAA |
| 64889_1 | TATGAGTGTGGAGTCGTTGCGCTCCTGAGATCTCATCTGATACCNNNAGATCGGAAG |
| 64906_1 | TATGAGTGTGGAGTCGTTGCTCCGTTACCACTGATAATGACTGTNNNAGATCGGAA |
| 64941_1 | TATGAGTGTGGAGTCGTTGCCCCATAATCCGTTGATAGACAGGTNNNAGATCGGAA |
| 64942_1 | TATGAGTGTGGAGTCGTTGCTGAAGTAACAGTTTCCACATCCCTNNNAGATCGGAAG |
| 64953_1 | TATGAGTGTGGAGTCGTTGCTGCTCATAACTCTTTCACATTCACTNNNAGATCGGAA |
| 64955_1 | TATGAGTGTGGAGTCGTTGCACACACACTATATACACACACACTNNNAGATCGGAAG |
| 65017_1 | TATGAGTGTGGAGTCGTTGCTCTCCAGCTAACATGTTTCATGGANNNAGATCGGAA |
| 65025_1 | TATGAGTGTGGAGTCGTTGCTGTGTGTGTGTGTATCGAAAAACANNNAGATCGGA |
| 65026_1 | TATGAGTGTGGAGTCGTTGCTAAGGCTGATACACGAGGGAATGNNNAGATCGGAA |
| 65027_1 | TATGAGTGTGGAGTCGTTGCATCTCAGCTCACTGCAACCTCTANNNAGATCGGAAGA |
| 65043_1 | TATGAGTGTGGAGTCGTTGCTTGTAGCTCTGTCTTGGTAGTGGNNNAGATCGGAAG |
| 65061_1 | TATGAGTGTGGAGTCGTTGCGTCCCATTGTTATGAATGCAGACTNNNAGATCGGAA |
| 65089_1 | TATGAGTGTGGAGTCGTTGCCTTCCACATGTTTGGCCAATATANNNAGATCGGAAG |
| 65174_1 | TATGAGTGTGGAGTCGTTGCTTTTTTTGAGACAGAGTCTTGCTCNNNAGATCGGAA |
| 65201_1 | TATGAGTGTGGAGTCGTTGCGGTGTAGTACTGAGTATCATCCCCNNNAGATCGGAA |
| 65226_1 | TATGAGTGTGGAGTCGTTGCTCTCTTCATCTGCTGATTCACACANNNAGATCGGAAG |
| 65228_1 | TATGAGTGTGGAGTCGTTGCTGGCTTCCTCCTTTTCTCAATGTNNNAGATCGGAAGA |
| 65230_1 | TATGAGTGTGGAGTCGTTGCCCCAAATAGTGAGGTTTCATTCCANNNAGATCGGAA |
| 65236_1 | TATGAGTGTGGAGTCGTTGCTCTCTCCTCTCTCTCTCTTTCACANNNAGATCGGAAGA |
| 65319_1 | TATGAGTGTGGAGTCGTTGCTGAATAGTTCTGTCAGTTTGGTGANNNAGATCGGAA |
| 65324_1 | TATGAGTGTGGAGTCGTTGCTGGATGATGATTGTGATATCAGGANNNAGATCGGA |
| 65392_1 | TATGAGTGTGGAGTCGTTGCAGCTCCTTTAGAAGAACTTGACCANNNAGATCGGAA |
| 65456_1 | TATGAGTGTGGAGTCGTTGCGCAGATGTTCAACTCAATATGGGGNNNAGATCGGA |
| 65508_1 | TATGAGTGTGGAGTCGTTGCAGGAGGAAAGAAGAGAGAATGTGTNNNAGATCGG |
| 65541_1 | TATGAGTGTGGAGTCGTTGCACAGTCTGAAGTGACCCTACTATGNNNAGATCGGAA |

| | |
|---|---|
| 65556_1 | TATGAGTGTGGAGTCGTTGCAGTGGGATTGTTGCAATTTATGGTNNNAGATCGGA |
| 65559_1 | TATGAGTGTGGAGTCGTTGCCATTCAGTCCAAGCTCTGTGTTGNNNAGATCGGAAG |
| 65592_1 | TATGAGTGTGGAGTCGTTGCCTAAGAAGTCCCACAGTCTGCTGNNNAGATCGGAAG |
| 65596_1 | TATGAGTGTGGAGTCGTTGCTCAACCAGGAAGCCAGAAGAAAANNNAGATCGGAA |
| 65614_1 | TATGAGTGTGGAGTCGTTGCTAAACACTGCAGGGACTGAATGANNNAGATCGGAA |
| 65626_1 | TATGAGTGTGGAGTCGTTGCTCTATGTTCTGCCATCAGCTTCTNNNAGATCGGAAGA |
| 65647_1 | TATGAGTGTGGAGTCGTTGCGGAAAAGATATGACCATGGTAGGCNNNAGATCGGA |
| 65663_1 | TATGAGTGTGGAGTCGTTGCACTCCATTGGTTCTGTTTCTCTGANNNAGATCGGAAG |
| 65666_1 | TATGAGTGTGGAGTCGTTGCATGACCTAGCCAAGTTGACACATNNNAGATCGGAAG |
| 65686_1 | TATGAGTGTGGAGTCGTTGCGTGTTTTTTTGAGACTGGGCCTTNNNAGATCGGAAG |
| 65687_1 | TATGAGTGTGGAGTCGTTGCACATCACTTTTTGAAAGGACAGAANNNAGATCGGAA |
| 65767_1 | TATGAGTGTGGAGTCGTTGCACCTGTCAGTGGATGCTTAGTTTNNNAGATCGGAAG |
| 65776_1 | TATGAGTGTGGAGTCGTTGCAAATGGAGTCTTGCTATGTTGCCNNNAGATCGGAAG |
| 65780_1 | TATGAGTGTGGAGTCGTTGCAAAAGAACAAAGCTGGAGGCATCNNNAGATCGGAA |
| 65848_1 | TATGAGTGTGGAGTCGTTGCTGTGTGTGTGTGTGTGTATTTTGTNNNAGATCGGA |
| 65857_1 | TATGAGTGTGGAGTCGTTGCAGGGGATATCCATGACTCTGACTNNNAGATCGGAA |
| 65859_1 | TATGAGTGTGGAGTCGTTGCTCGTGTCCTTATGTTATTGTCTCTNNNAGATCGGAAG |
| 65866_1 | TATGAGTGTGGAGTCGTTGCGAGACAGGGTCTTGCTCTGTTGNNNAGATCGGAAG |
| 65874_1 | TATGAGTGTGGAGTCGTTGCACAAGAAAGACTGTAAAGGTACCANNNAGATCGGA |
| 65908_1 | TATGAGTGTGGAGTCGTTGCAGTGGGAATGGAAACTATTTGAGTNNNAGATCGGA |
| 65922_1 | TATGAGTGTGGAGTCGTTGCACCATGGAATACTATGCAGCCATNNNAGATCGGAAG |
| 65986_1 | TATGAGTGTGGAGTCGTTGCGAAGACAGGGAGCCAGAATGATANNNAGATCGGA |
| 65989_1 | TATGAGTGTGGAGTCGTTGCACAATTGCCTAGAAATGAGGGTCTNNNAGATCGGA |
| 66037_1 | TATGAGTGTGGAGTCGTTGCCTGGATTCAAGCAATTCTCCTGCNNNAGATCGGAAG |
| 66066_1 | TATGAGTGTGGAGTCGTTGCACTAAATGACCTCATTCTAGTGCTNNNAGATCGGAA |
| 66067_1 | TATGAGTGTGGAGTCGTTGCTGGCTTCCTCCTTTTCTCAATGTNNNAGATCGGAAGA |
| 66169_1 | TATGAGTGTGGAGTCGTTGCAGTAGAGATTGCCATTCTTCATCTNNNAGATCGGAA |
| 66205_1 | TATGAGTGTGGAGTCGTTGCAGAGAGAGAGAGAGAGAGAGAACANNNAGATCGG |
| 66206_1 | TATGAGTGTGGAGTCGTTGCAGACAAATAGGTTGGGAGTAGATTGTNNNAGATCG |
| 66221_1 | TATGAGTGTGGAGTCGTTGCTGGACTCCTCCTAAACTCTAACTGNNNAGATCGGAA |
| 66224_1 | TATGAGTGTGGAGTCGTTGCACTCAAACCCAGTAAGGAAACTTGNNNAGATCGGAA |
| 66227_1 | TATGAGTGTGGAGTCGTTGCGGGCCACTGAAGAGATAACAGAANNNAGATCGGAA |
| 66241_1 | TATGAGTGTGGAGTCGTTGCACTGACATACCATTAGCACCCAANNNAGATCGGAAG |
| 66245_1 | TATGAGTGTGGAGTCGTTGCTTTACCCAATCCCCAAGTTGTGANNNAGATCGGAAG |
| 66252_1 | TATGAGTGTGGAGTCGTTGCGTCTTGGAAACACACAAGCTACCNNNAGATCGGAAG |
| 66267_1 | TATGAGTGTGGAGTCGTTGCAGAGAGGCAGAGATAGAGTCCTCNNNAGATCGGAA |
| 66319_1 | TATGAGTGTGGAGTCGTTGCATGAGCTCCTTGAAGAAATGGCTNNNAGATCGGAA |
| 66402_1 | TATGAGTGTGGAGTCGTTGCACTCGAAGTTACTTTGGGATCCANNNAGATCGGAAG |
| 66410_1 | TATGAGTGTGGAGTCGTTGCTTTCTCCCTCATCTGGTTATCCTNNNAGATCGGAAGA |
| 66415_1 | TATGAGTGTGGAGTCGTTGCAAAGATGAACTCCCAGCTACTTATNNNAGATCGGAA |
| 66592_1 | TATGAGTGTGGAGTCGTTGCCCTTTCTGTTTACTTTCAGGTCGTNNNAGATCGGAAG |
| 66653_1 | TATGAGTGTGGAGTCGTTGCACCATCACATGACTGGCTATATCCNNNAGATCGGAA |
| 66657_1 | TATGAGTGTGGAGTCGTTGCAGTCTTTCTTTCTAGGCTGAAGGANNNAGATCGGAA |

| | |
|---|---|
| 66658_1 | TATGAGTGTGGAGTCGTTGCCATGGAATGTGTTAGGGAGGGAGNNNAGATCGGA |
| 66691_1 | TATGAGTGTGGAGTCGTTGCCTGGAGCACATTACTACCAACTTNNNAGATCGGAAG |
| 66745_1 | TATGAGTGTGGAGTCGTTGCGCATTAGGCAAAAAGTACGGAACTNNNAGATCGGA |
| 66779_1 | TATGAGTGTGGAGTCGTTGCGCACGCACATAGACACACATATGNNNAGATCGGAAG |
| 66781_1 | TATGAGTGTGGAGTCGTTGCTGAAACAGTGATTACCATGAGTGGNNNAGATCGGA |
| 66789_1 | TATGAGTGTGGAGTCGTTGCTCTTTCCCTGTTTCCCTGTCATCNNNAGATCGGAAGA |
| 66844_1 | TATGAGTGTGGAGTCGTTGCAGGAGAACTACAAAATACTGCTGANNNAGATCGGA |
| 66861_1 | TATGAGTGTGGAGTCGTTGCGCTAACCAAATCCATCAGCACATNNNAGATCGGAAG |
| 66865_1 | TATGAGTGTGGAGTCGTTGCAAGTTGCAGAATACAACATCAACANNNAGATCGGAA |
| 66914_1 | TATGAGTGTGGAGTCGTTGCAGACTTTCTCAATAACTGGTGAGANNNAGATCGGAA |
| 66918_1 | TATGAGTGTGGAGTCGTTGCGCCTATCAATTCTGTCTCTCTGGANNNAGATCGGAAG |
| 66958_1 | TATGAGTGTGGAGTCGTTGCCCAATGGGACAAAATGGAAGTCCNNNAGATCGGAA |
| 67010_1 | TATGAGTGTGGAGTCGTTGCTGGAATAGCAAATCTACGGAGACANNNAGATCGGA |
| 67038_1 | TATGAGTGTGGAGTCGTTGCACTCACCATCATTAGTCTTCAGGGNNNAGATCGGAA |
| 67052_1 | TATGAGTGTGGAGTCGTTGCACCAAAAATAGTTGTTGGCAAACCNNNAGATCGGAA |
| 67114_1 | TATGAGTGTGGAGTCGTTGCTCCTGTTCCAATGCCTTTTTTTCTNNNAGATCGGAAG |
| 67117_1 | TATGAGTGTGGAGTCGTTGCTGTGTATATGTGTGTGTGTGTGTGNNNAGATCGGA |
| 67131_1 | TATGAGTGTGGAGTCGTTGCGGTTGAGACCACAAATCAAAGAGTNNNAGATCGGA |
| 67134_1 | TATGAGTGTGGAGTCGTTGCCCAGAGTCAGCCTGAACATTCTANNNAGATCGGAAG |
| 67149_1 | TATGAGTGTGGAGTCGTTGCTGGCTTCCTCCTTTTCTCAATGTNNNAGATCGGAAGA |
| 67182_1 | TATGAGTGTGGAGTCGTTGCTGCAGCAAAAACGTATATGGGAGNNNAGATCGGAA |
| 67197_1 | TATGAGTGTGGAGTCGTTGCTGATCCCAAGAATCACCATTTGGNNNAGATCGGAA |
| 67235_1 | TATGAGTGTGGAGTCGTTGCCTATAGGTCAGGTAGGCAAGCTGNNNAGATCGGAA |
| 67249_1 | TATGAGTGTGGAGTCGTTGCGGAGAGAGAGACAGAGAGAAGGANNNAGATCGGA |
| 67274_1 | TATGAGTGTGGAGTCGTTGCAAAAGATGGGTACCAGTCTGTGGNNNAGATCGGAA |
| 67284_1 | TATGAGTGTGGAGTCGTTGCTAAATTTGGGGACATTTTGGCCCNNNAGATCGGAAG |
| 67305_1 | TATGAGTGTGGAGTCGTTGCACCTTCACTATTTTGATAGCTGGANNNAGATCGGAA |
| 67315_1 | TATGAGTGTGGAGTCGTTGCACTTGCATTAGGACATTTTGTTGANNNAGATCGGAA |
| 67482_1 | TATGAGTGTGGAGTCGTTGCAGAGAAAAGTTGGGGAGAAATGGTNNNAGATCGG |
| 67505_1 | TATGAGTGTGGAGTCGTTGCCCCAGCAACATCAACATTACCTGNNNAGATCGGAAG |
| 67522_1 | TATGAGTGTGGAGTCGTTGCAAACAGATCTTGCTTAGTACCTGANNNAGATCGGAA |
| 67548_1 | TATGAGTGTGGAGTCGTTGCACTGAAAGTGCTGAGGAAAGAGTNNNAGATCGGAA |
| 67564_1 | TATGAGTGTGGAGTCGTTGCTCTGTTTTGTGAAATGCCTATTCANNNAGATCGGAAG |
| 67585_1 | TATGAGTGTGGAGTCGTTGCTCAGCCAAATCATAAGCACATTGTNNNAGATCGGAA |
| 67740_1 | TATGAGTGTGGAGTCGTTGCTCTCATCTGCTCAGTACTCAGGANNNAGATCGGAAG |
| 67741_1 | TATGAGTGTGGAGTCGTTGCTGTATTCCTCGAGATACACAAGCANNNAGATCGGAA |
| 67743_1 | TATGAGTGTGGAGTCGTTGCTAGACCAATAGTGATGAGAGCGCNNNAGATCGGAA |
| 67797_1 | TATGAGTGTGGAGTCGTTGCTCCAAGAAGAAAAAAAAAAGGCCTNNNAGATCGGA |
| 67798_1 | TATGAGTGTGGAGTCGTTGCCAAAGCACCTCCTATTTGAAACCANNNAGATCGGAAG |
| 67801_1 | TATGAGTGTGGAGTCGTTGCAGCACTCAAACACAAATCTAGACANNNAGATCGGAA |
| 67836_1 | TATGAGTGTGGAGTCGTTGCTCTTTCCCTCTTTCTTGCTCTCTNNNAGATCGGAAGAG |
| 67845_1 | TATGAGTGTGGAGTCGTTGCTTGAAATGTGATCCCCAGTGTCANNNAGATCGGAAG |
| 67846_1 | TATGAGTGTGGAGTCGTTGCTGAAACTCCCAGATTTCTGACACTNNNAGATCGGAA |

| | |
|---|---|
| 67847_1 | TATGAGTGTGGAGTCGTTGCACCCCCAATGTAGAAATATGCTGANNNAGATCGGAA |
| 67865_1 | TATGAGTGTGGAGTCGTTGCTCAGCTGGTATCTAAAAAGGTGGANNNAGATCGGA |
| 67876_1 | TATGAGTGTGGAGTCGTTGCCACCTAGTGATAAACACAGGGACANNNAGATCGGA |
| 67879_1 | TATGAGTGTGGAGTCGTTGCGTCTGCCTAACCGTAGTAATCCCNNNAGATCGGAAG |
| 67883_1 | TATGAGTGTGGAGTCGTTGCACCAGAAGAGATAGAACTCTAGCTNNNAGATCGGA |
| 67895_1 | TATGAGTGTGGAGTCGTTGCTGTAAGGTCAGTGTTCTCCAGAGNNNAGATCGGAA |
| 67944_1 | TATGAGTGTGGAGTCGTTGCAACTTGGATGCTGCTGGTGTATANNNAGATCGGAA |
| 67974_1 | TATGAGTGTGGAGTCGTTGCACAACCAAATAACACATAACACCTNNNAGATCGGAA |
| 67975_1 | TATGAGTGTGGAGTCGTTGCTGAACTCAATAACTTGGAGGGTGTNNNAGATCGGA |
| 67983_1 | TATGAGTGTGGAGTCGTTGCACCTTCTCGTAATATGGACTCAGANNNAGATCGGAA |
| 68010_1 | TATGAGTGTGGAGTCGTTGCTGAGCTCCAGAAAGAGAAAACAGANNNAGATCGGA |
| 68014_1 | TATGAGTGTGGAGTCGTTGCGGAGTGAAATGGCATGATCTTGGNNNAGATCGGAA |
| 68066_1 | TATGAGTGTGGAGTCGTTGCGCGATCTCAAATCACTGCAACTTNNNAGATCGGAAG |
| 68074_1 | TATGAGTGTGGAGTCGTTGCGAGATAGTCTTGCTCTTGTTGCCNNNAGATCGGAAG |
| 68092_1 | TATGAGTGTGGAGTCGTTGCTCTAAAGCCCGTGTAAACATTCTGNNNAGATCGGAA |
| 68126_1 | TATGAGTGTGGAGTCGTTGCAGACTCTGTCACTGTGGTATCATGNNNAGATCGGAA |
| 68155_1 | TATGAGTGTGGAGTCGTTGCCCAATGGCATAGTAGCAAGTATGCNNNAGATCGGA |
| 68173_1 | TATGAGTGTGGAGTCGTTGCTGTAGAAAATTCCAAGTGCTCACCNNNAGATCGGAA |
| 68191_1 | TATGAGTGTGGAGTCGTTGCACCAGTTTTACTTTGTTTGCCCCNNNAGATCGGAAGA |
| 68203_1 | TATGAGTGTGGAGTCGTTGCGTGTGATCATGACTCAACTGCAGNNNAGATCGGAAG |
| 68206_1 | TATGAGTGTGGAGTCGTTGCACTTTATTGAAGAGCTCCTGTCACTNNNAGATCGGA |
| 68241_1 | TATGAGTGTGGAGTCGTTGCAGGATGTAGACAAGTTGGAACCCNNNAGATCGGAA |
| 68285_1 | TATGAGTGTGGAGTCGTTGCACCCTGACTATTTGCAAATACACTNNNAGATCGGAAG |
| 68292_1 | TATGAGTGTGGAGTCGTTGCCCGCATACAATAACTCACAAGAGCNNNAGATCGGAA |
| 68295_1 | TATGAGTGTGGAGTCGTTGCGCTGGGTAAAGGCAGTATACAGANNNAGATCGGAA |
| 68306_1 | TATGAGTGTGGAGTCGTTGCTTCCCCGCAATCTCATTATGACTNNNAGATCGGAAGA |
| 68367_1 | TATGAGTGTGGAGTCGTTGCACACATACACAAACACACATATGTNNNAGATCGGAA |
| 68369_1 | TATGAGTGTGGAGTCGTTGCTGGCATATAATGGGGAAAAATGGCNNNAGATCGGA |
| 68372_1 | TATGAGTGTGGAGTCGTTGCTATACACACACACACACACACACNNNAGATCGGAAGA |
| 68402_1 | TATGAGTGTGGAGTCGTTGCTGTGTGTGTGTGTGTAGGGTTTANNNAGATCGGAA |
| 68421_1 | TATGAGTGTGGAGTCGTTGCCCTTTATATCTTAGCTTGCTGCCCNNNAGATCGGAAG |
| 68430_1 | TATGAGTGTGGAGTCGTTGCTCTTTCTTCTCTTCCTTTCTTCCTNNNAGATCGGAAGA |
| 68479_1 | TATGAGTGTGGAGTCGTTGCGCTTTAGTCAGTTCTTTGTCCCACNNNAGATCGGAAG |
| 68486_1 | TATGAGTGTGGAGTCGTTGCACTGAAGTGCTTATTCCACATCCANNNAGATCGGAA |
| 68491_1 | TATGAGTGTGGAGTCGTTGCTCTTAGGCCAGTTCATTACCAGGNNNAGATCGGAAG |
| 68493_1 | TATGAGTGTGGAGTCGTTGCACATGTAACTTCTTGGTTGAGAAGNNNAGATCGGAA |
| 68518_1 | TATGAGTGTGGAGTCGTTGCTCAGAGCTCAATGGCCATTATTTGNNNAGATCGGAA |
| 68612_1 | TATGAGTGTGGAGTCGTTGCTTTCTTTTTTAGAAATGGGGTCTAACCNNNAGATCGG |
| 68622_1 | TATGAGTGTGGAGTCGTTGCGGAGTGCAGTGGTGTGATCTTAGNNNAGATCGGAA |
| 68636_1 | TATGAGTGTGGAGTCGTTGCTGCATGAAAACTGTCTTTCAGCTTNNNAGATCGGAA |
| 68660_1 | TATGAGTGTGGAGTCGTTGCAGAAAGGAAAGAAAGCAAGCAAGANNNAGATCGG |
| 68800_1 | TATGAGTGTGGAGTCGTTGCGAGAGAGAGAGAGGAAGGTGGACNNNAGATCGGA |
| 68825_1 | TATGAGTGTGGAGTCGTTGCCACACACACACACATACTTACACANNNAGATCGGAAG |

| | |
|---|---|
| 68943_1 | TATGAGTGTGGAGTCGTTGCAGAGACAACAGATAACCCAGGATNNNAGATCGGAA |
| 69006_1 | TATGAGTGTGGAGTCGTTGCTGCAAAAGTGTACTTGTCAGCAGNNNAGATCGGAAG |
| 69038_1 | TATGAGTGTGGAGTCGTTGCTGAATAACAGAGTTCACGTAAGCANNNAGATCGGAA |
| 69104_1 | TATGAGTGTGGAGTCGTTGCACAGCAATTAGTGTTAGAAAGGGTNNNAGATCGGA |
| 69117_1 | TATGAGTGTGGAGTCGTTGCTGCATGTGTGTGTATGTATCTCCTNNNAGATCGGAA |
| 69120_1 | TATGAGTGTGGAGTCGTTGCGTGGTGTGATCTCGCTTACTGTANNNAGATCGGAAG |
| 69179_1 | TATGAGTGTGGAGTCGTTGCACACACACAGAGAGAGAGAGAGANNNAGATCGGAA |
| 69188_1 | TATGAGTGTGGAGTCGTTGCACTGAGGTCATACGGTATTTGACTNNNAGATCGGAA |
| 69302_1 | TATGAGTGTGGAGTCGTTGCGCTACACACGAGGCATTTGATTANNNAGATCGGAAG |
| 69312_1 | TATGAGTGTGGAGTCGTTGCTCTTTCTCTCTGTCTCTCTCTCTCNNNAGATCGGAAGA |
| 69353_1 | TATGAGTGTGGAGTCGTTGCCAAAAAGCTTGAGAACCCAGAGGNNNAGATCGGAA |
| 69371_1 | TATGAGTGTGGAGTCGTTGCATATGGATAAGATTGGCCAGGCANNNAGATCGGAA |
| 69414_1 | TATGAGTGTGGAGTCGTTGCTGGAGCATGCTGACTGAGTTTATNNNAGATCGGAAG |
| 69420_1 | TATGAGTGTGGAGTCGTTGCAGAGAATACCGGTGCATACAGACNNNAGATCGGAA |
| 69437_1 | TATGAGTGTGGAGTCGTTGCTTGACTTCCTGGGCTCAAATGATNNNAGATCGGAAG |
| 69473_1 | TATGAGTGTGGAGTCGTTGCTCAGCTTGAAATTTCAGCAACTGTNNNAGATCGGAA |
| 69591_1 | TATGAGTGTGGAGTCGTTGCTACAAGCCCAGTTGTCTCTCATCNNNAGATCGGAAGA |
| 69710_1 | TATGAGTGTGGAGTCGTTGCTGCACACACACGAATAAAAAATTTNNNAGATCGGAA |
| 69721_1 | TATGAGTGTGGAGTCGTTGCCCTCCCAAGTAGCTGACAAAAAANNNAGATCGGAAG |
| 69737_1 | TATGAGTGTGGAGTCGTTGCTTGAATACACATGCCAGGACATCNNNAGATCGGAAG |
| 69738_1 | TATGAGTGTGGAGTCGTTGCACCTCCTCAATTTTCTGGAATAGTNNNAGATCGGAAG |
| 69756_1 | TATGAGTGTGGAGTCGTTGCTGAGAGAGAGACAGGGACTTATGANNNAGATCGGA |
| 69769_1 | TATGAGTGTGGAGTCGTTGCGGAAAGAACCAAAAGAAAGGGACANNNAGATCGG |
| 69809_1 | TATGAGTGTGGAGTCGTTGCCCTGAGTAGCTGGGATTATAGGCNNNAGATCGGAA |
| 69842_1 | TATGAGTGTGGAGTCGTTGCCTGTTAGCTACATCCCTCTCTCCNNNAGATCGGAAGA |
| 69849_1 | TATGAGTGTGGAGTCGTTGCAGAAAGAATGGAAGGAAGGAAGGANNNAGATCGG |
| 69856_1 | TATGAGTGTGGAGTCGTTGCCCAAGTGGACAATAAAGTAGCACANNNAGATCGGA |
| 69889_1 | TATGAGTGTGGAGTCGTTGCGCCTCCCAGGTTCATTGTAGTTTNNNAGATCGGAAG |
| 69948_1 | TATGAGTGTGGAGTCGTTGCCATAATTCACTTCACTGGCAGGCNNNAGATCGGAAG |
| 69953_1 | TATGAGTGTGGAGTCGTTGCTCAAGCAATTCCTCACCTCAAGANNNAGATCGGAAG |
| 69973_1 | TATGAGTGTGGAGTCGTTGCTCCTGTGTCTCTGAAGAATCATGANNNAGATCGGAA |
| 69987_1 | TATGAGTGTGGAGTCGTTGCTCTTGGTAGCGCATTTCTACTGTNNNAGATCGGAAG |
| 69996_1 | TATGAGTGTGGAGTCGTTGCGAAAAATCGGGTTTACTGGTGCANNNAGATCGGAA |
| 70019_1 | TATGAGTGTGGAGTCGTTGCTGGAGACACCAGTGAATAGATTCTNNNAGATCGGA |
| 70121_1 | TATGAGTGTGGAGTCGTTGCCCACGCCCGGCTAATTATTTTTTNNNAGATCGGAAGA |
| 70242_1 | TATGAGTGTGGAGTCGTTGCGGGAAGAGCTGTTTGTGATGAGANNNAGATCGGA |
| 70257_1 | TATGAGTGTGGAGTCGTTGCAGAAGAAGGAGAAGGAGAAGGAGANNNAGATCG |
| 70286_1 | TATGAGTGTGGAGTCGTTGCGGAGGAAGGTGGAGTAACACAAGNNNAGATCGGA |
| 70313_1 | TATGAGTGTGGAGTCGTTGCTCTTTCTTCTTGACGGAGTTTCACNNNAGATCGGAAG |
| 70314_1 | TATGAGTGTGGAGTCGTTGCACCTGAAATTGCTGTTCAAGAACANNNAGATCGGAA |
| 70324_1 | TATGAGTGTGGAGTCGTTGCACGGAATATCTTGAACAACAGTGCNNNAGATCGGAA |
| 70372_1 | TATGAGTGTGGAGTCGTTGCAGAAATCTTCCCATCTCAGCCTCNNNAGATCGGAAGA |
| 70441_1 | TATGAGTGTGGAGTCGTTGCAGCAATCTTCCAATTAGCAAGCTNNNAGATCGGAAG |

| | |
|---|---|
| 70474_1 | TATGAGTGTGGAGTCGTTGCCATCTCAACTCACTGCAACCTTTNNNAGATCGGAAGA |
| 70479_1 | TATGAGTGTGGAGTCGTTGCTCTCTCTCTCTCCTCTCTCTCTNNNAGATCGGAAGA |
| 70482_1 | TATGAGTGTGGAGTCGTTGCCCTTCCCTTCCCTTTTCTTTCCTNNNAGATCGGAAGAG |
| 70498_1 | TATGAGTGTGGAGTCGTTGCCCTAATTGCTCTGGCTAGGACTTNNNAGATCGGAAG |
| 70525_1 | TATGAGTGTGGAGTCGTTGCCTCCTGGGTTCAAGTGATTCTCANNNAGATCGGAAG |
| 1_1 | TATGAGTGTGGAGTCGTTGCACGATTCAGGTTCAAGTCTTGGTNNNAGATCGGAAG |
| 6_1 | TATGAGTGTGGAGTCGTTGCTACGATTTCCTCTGGAGGTTGGTNNNAGATCGGAAG |
| 9_1 | TATGAGTGTGGAGTCGTTGCGTGTGATGTGCATGCAATTTGGTNNNAGATCGGAA |
| 13_1 | TATGAGTGTGGAGTCGTTGCAACACACCTGGCAGAAGACAGACNNNAGATCGGAA |
| 17_1 | TATGAGTGTGGAGTCGTTGCTACAGCAAGAAGGCATGACAACANNNAGATCGGAA |
| 20_1 | TATGAGTGTGGAGTCGTTGCCAATGCAGCGATGTGATCAGCTCNNNAGATCGGAA |
| 21_1 | TATGAGTGTGGAGTCGTTGCATGACGCGGGAAAAGATACAACANNNAGATCGGAA |
| 22_1 | TATGAGTGTGGAGTCGTTGCTCAGATTTCGTGGCACTTTTGGTNNNAGATCGGAAG |
| 23_1 | TATGAGTGTGGAGTCGTTGCTCCCACGGACAACATTTACAACANNNAGATCGGAAG |
| 29_1 | TATGAGTGTGGAGTCGTTGCCATACACACTGAAGACAGGCCATNNNAGATCGGAAG |
| 30_1 | TATGAGTGTGGAGTCGTTGCTCCAGTCAGTTTCTAAGCCAACANNNAGATCGGAAG |
| 33_1 | TATGAGTGTGGAGTCGTTGCGGAGGCCAGCAAGTCTAAAATGTNNNAGATCGGAA |
| 34_1 | TATGAGTGTGGAGTCGTTGCAATAGAAATGGGCTTGGCCAACANNNAGATCGGAA |
| 36_1 | TATGAGTGTGGAGTCGTTGCAGAGACAGAGAGGGAGCAAATGTNNNAGATCGGA |
| 37_1 | TATGAGTGTGGAGTCGTTGCAACCTGGTATGAAGCAAGCAACANNNAGATCGGAA |
| 38_1 | TATGAGTGTGGAGTCGTTGCGACAGAACAGGTGAGGATGTGANNNAGATCGGAA |
| 41_1 | TATGAGTGTGGAGTCGTTGCGTTGAGGGACAATGGGGAAATGTNNNAGATCGGA |
| 42_1 | TATGAGTGTGGAGTCGTTGCTCTCTCTCCCCACTTGGAAATGTNNNAGATCGGAAGA |
| 45_1 | TATGAGTGTGGAGTCGTTGCTGGTAAAATGGGCAGGACAATGTNNNAGATCGGAA |
| 49_1 | TATGAGTGTGGAGTCGTTGCAACTGGAACAATCTGGGCAATGTNNNAGATCGGAA |
| 50_1 | TATGAGTGTGGAGTCGTTGCTCCACTTTTGAGAGCTGCAATGTNNNAGATCGGAAG |
| 51_1 | TATGAGTGTGGAGTCGTTGCCACTCTCTGTTTGTCCCAGAACANNNAGATCGGAAG |
| 53_1 | TATGAGTGTGGAGTCGTTGCTGGCTCCTGAAGTTTTGGAATGTNNNAGATCGGAA |
| 54_1 | TATGAGTGTGGAGTCGTTGCACTCCCCCTAACTCCATGAATGTNNNAGATCGGAAGA |
| 58_1 | TATGAGTGTGGAGTCGTTGCCATCACCCATTTCCCAAGGAACANNNAGATCGGAAG |
| 59_1 | TATGAGTGTGGAGTCGTTGCACCCAATACATGGCTGAACATGTNNNAGATCGGAAG |
| 61_1 | TATGAGTGTGGAGTCGTTGCCACAGGTTAGGATGCACACATGTNNNAGATCGGAA |
| 64_1 | TATGAGTGTGGAGTCGTTGCGTGTATGTGGGTGTGTCACATGTNNNAGATCGGAA |
| 66_1 | TATGAGTGTGGAGTCGTTGCGGCACTTCAGATCAAGGACATGTNNNAGATCGGAA |
| 72_1 | TATGAGTGTGGAGTCGTTGCATGCAGACACACACAGATGAACANNNAGATCGGAA |
| 74_1 | TATGAGTGTGGAGTCGTTGCATTCTGAGTGCACAAAGGCAGCANNNAGATCGGAA |
| 76_1 | TATGAGTGTGGAGTCGTTGCTCCCCAAGGTATGTGTCTGAACANNNAGATCGGAAG |
| 77_1 | TATGAGTGTGGAGTCGTTGCATCCAAAATCTGCAGGGTGAACANNNAGATCGGAA |
| 79_1 | TATGAGTGTGGAGTCGTTGCGCACTCCATACTTTCCTCCATGTNNNAGATCGGAAGA |
| 84_1 | TATGAGTGTGGAGTCGTTGCACAAGCCCCTAGGTTGAATAACANNNAGATCGGAAG |
| 85_1 | TATGAGTGTGGAGTCGTTGCTCCCCTCTGCCTTGTACATAACANNNAGATCGGAAGA |
| 88_1 | TATGAGTGTGGAGTCGTTGCGTGTGAGGAGTGTGTATGCATGTNNNAGATCGGAA |
| 91_1 | TATGAGTGTGGAGTCGTTGCATTTGTGGCCAGGTTCATCATGTNNNAGATCGGAAG |

| | |
|---|---|
| 92_1 | TATGAGTGTGGAGTCGTTGCCTTTCCTCTGTGTGGCTGTAACANNNAGATCGGAAGA |
| 94_1 | TATGAGTGTGGAGTCGTTGCTGGTCAGCCCCAGTAATTCATGTNNNAGATCGGAAG |
| 97_1 | TATGAGTGTGGAGTCGTTGCACACAGTACACATGCTTTCATGTNNNAGATCGGAAGA |
| 100_1 | TATGAGTGTGGAGTCGTTGCGGGCTTTTCCAAGACACAGATGTNNNAGATCGGAAG |
| 103_1 | TATGAGTGTGGAGTCGTTGCGCTAGAGGCAAAAGCAGAGATGTNNNAGATCGGAA |
| 105_1 | TATGAGTGTGGAGTCGTTGCACTGAAACCACCAAAGGAACACANNNAGATCGGAAG |
| 106_1 | TATGAGTGTGGAGTCGTTGCGAGCTTTGCTAAAGGGGAGATGTNNNAGATCGGAA |
| 108_1 | TATGAGTGTGGAGTCGTTGCGCCTGAGCAGTACCAATAACACANNNAGATCGGAAG |
| 110_1 | TATGAGTGTGGAGTCGTTGCTTCCCCACCTCAGTTCTAACACANNNAGATCGGAAGA |
| 111_1 | TATGAGTGTGGAGTCGTTGCTTTCTGGTTGCATCGAGTTGTCTNNNAGATCGGAAGA |
| 113_1 | TATGAGTGTGGAGTCGTTGCAAACATGCCCCATTTCTGGATGTNNNAGATCGGAAGA |
| 114_1 | TATGAGTGTGGAGTCGTTGCAAATGCCCACTGAGAAACACACANNNAGATCGGAAG |
| 116_1 | TATGAGTGTGGAGTCGTTGCTTTCCAGCTTTGACACACACACANNNAGATCGGAAGA |
| 117_1 | TATGAGTGTGGAGTCGTTGCGGACACCCCCAAATTCATGATGTNNNAGATCGGAAG |
| 118_1 | TATGAGTGTGGAGTCGTTGCCCCAAATATACCGGACACACACANNNAGATCGGAAGA |
| 120_1 | TATGAGTGTGGAGTCGTTGCGATTGCCCTGTTTCCTCTGATGTNNNAGATCGGAAGA |
| 122_1 | TATGAGTGTGGAGTCGTTGCCCCAACAGACATATGGACACACANNNAGATCGGAAG |
| 123_1 | TATGAGTGTGGAGTCGTTGCCCCACTTCCAATTCTGACACACANNNAGATCGGAAGA |
| 125_1 | TATGAGTGTGGAGTCGTTGCCAGCAGTCCCTCGAACAATATGTNNNAGATCGGAAG |
| 130_1 | TATGAGTGTGGAGTCGTTGCCACCACCACCACGAAAAATACAANNNAGATCGGAAGA |
| 131_1 | TATGAGTGTGGAGTCGTTGCTAGTGACTGAGAAGTCCCACACANNNAGATCGGAAG |
| 132_1 | TATGAGTGTGGAGTCGTTGCCGTATCAGACAGTGATGCACACANNNAGATCGGAAG |
| 133_1 | TATGAGTGTGGAGTCGTTGCAGATGGAGGCTCACTATCACACANNNAGATCGGAAG |
| 135_1 | TATGAGTGTGGAGTCGTTGCTGTGTGTGTATGGTATGGTATGTNNNAGATCGGAAG |
| 137_1 | TATGAGTGTGGAGTCGTTGCGGGGGAAGGGGGATAATGTATGTNNNAGATCGGA |
| 139_1 | TATGAGTGTGGAGTCGTTGCCACTTACCCACACACAAGACACANNNAGATCGGAAGA |
| 140_1 | TATGAGTGTGGAGTCGTTGCCCACGAGAGAAGTTCAAGACACANNNAGATCGGAAG |
| 147_1 | TATGAGTGTGGAGTCGTTGCTTGCTATCGCTGCAGGTTTATGTNNNAGATCGGAAG |
| 148_1 | TATGAGTGTGGAGTCGTTGCCCCTGCTTTTTGACTTGTAGGCTNNNAGATCGGAAGA |
| 153_1 | TATGAGTGTGGAGTCGTTGCTTCCCCAATTGCCAACATACACANNNAGATCGGAAGA |
| 154_1 | TATGAGTGTGGAGTCGTTGCTCTGTCGAAACTCATGGAACTGTNNNAGATCGGAAG |
| 155_1 | TATGAGTGTGGAGTCGTTGCGGCTAGAAGTGCACACATACACANNNAGATCGGAAG |
| 156_1 | TATGAGTGTGGAGTCGTTGCAACACAGAGTGTGGACATACACANNNAGATCGGAAG |
| 158_1 | TATGAGTGTGGAGTCGTTGCATGCCTGCTAGGAAGACAAATGTNNNAGATCGGAAG |
| 160_1 | TATGAGTGTGGAGTCGTTGCCAGCCCAAACAGTTAGCTACACANNNAGATCGGAAG |
| 162_1 | TATGAGTGTGGAGTCGTTGCGACCCATTAGGGCTTTCTACACANNNAGATCGGAAGA |
| 163_1 | TATGAGTGTGGAGTCGTTGCTAGCACTTTCTGCTTTCCACTGTNNNAGATCGGAAGA |
| 164_1 | TATGAGTGTGGAGTCGTTGCGGCACAGGAAGATCATGTACACANNNAGATCGGAAG |
| 167_1 | TATGAGTGTGGAGTCGTTGCCTCTGGGCCTCTGTTTTTACACANNNAGATCGGAAGA |
| 168_1 | TATGAGTGTGGAGTCGTTGCTGTGAAATTGCTTTGTGGTGCANNNAGATCGGAAGA |
| 169_1 | TATGAGTGTGGAGTCGTTGCTGTACGCTGATACGTTTCACTGTNNNAGATCGGAAGA |
| 170_1 | TATGAGTGTGGAGTCGTTGCAAACAGGGCTTCCAAGGGACAGTNNNAGATCGGAAG |
| 171_1 | TATGAGTGTGGAGTCGTTGCTCCCTTACAGCCTATTCACCACANNNAGATCGGAAGA |

| | |
|---|---|
| 172_1 | TATGAGTGTGGAGTCGTTGCTTCCTTTCTCTGACCCAGACTGTNNNAGATCGGAAGA |
| 173_1 | TATGAGTGTGGAGTCGTTGCACCACTTGATTTTGAAGACCACANNNAGATCGGAAG |
| 175_1 | TATGAGTGTGGAGTCGTTGCTGTAGAGGATGGCAGTGATGAGANNNAGATCGGA |
| 176_1 | TATGAGTGTGGAGTCGTTGCTGACTTTGTTCTCTCTGACCACANNNAGATCGGAAGA |
| 177_1 | TATGAGTGTGGAGTCGTTGCGCCCACTGCACACTGTATTCAAANNNAGATCGGAAG |
| 179_1 | TATGAGTGTGGAGTCGTTGCATGATGCAAGACGTCACTACTGTNNNAGATCGGAAG |
| 187_1 | TATGAGTGTGGAGTCGTTGCACCTACTGCCTAATAAGCCCACANNNAGATCGGAAG |
| 189_1 | TATGAGTGTGGAGTCGTTGCCCCACTTTCACACTTAGAGGCTCNNNAGATCGGAAGA |
| 190_1 | TATGAGTGTGGAGTCGTTGCAGGACTTCCAGTACATTCCCACANNNAGATCGGAAG |
| 191_1 | TATGAGTGTGGAGTCGTTGCCACACATTTCCCGTCCAGCAAACNNNAGATCGGAAGA |
| 193_1 | TATGAGTGTGGAGTCGTTGCTTATGAATCCCCTCTGAGCCACANNNAGATCGGAAG |
| 194_1 | TATGAGTGTGGAGTCGTTGCCCTCTGTCTCTCTGTGTACCTGTNNNAGATCGGAAGA |
| 196_1 | TATGAGTGTGGAGTCGTTGCCATAAAAGACACTGCCACCCTGTNNNAGATCGGAAG |
| 197_1 | TATGAGTGTGGAGTCGTTGCCTTCCTGATGATTCTTGGCCACANNNAGATCGGAAG |
| 199_1 | TATGAGTGTGGAGTCGTTGCGGCACTAAGCACTCTCTGTACCCNNNAGATCGGAAG |
| 203_1 | TATGAGTGTGGAGTCGTTGCTCCTTTTCATTTTGGATGCCTGTNNNAGATCGGAAGA |
| 207_1 | TATGAGTGTGGAGTCGTTGCCCTACTCTTACCATTGTGCCTGTNNNAGATCGGAAGA |
| 209_1 | TATGAGTGTGGAGTCGTTGCAGGGGTGAATTTTTGGTTCCACANNNAGATCGGAA |
| 210_1 | TATGAGTGTGGAGTCGTTGCGGTTGCTCTGTCGTAAATCCTGTNNNAGATCGGAAG |
| 213_1 | TATGAGTGTGGAGTCGTTGCCGAAAAGGGAGAGAGAAAGCACANNNAGATCGGA |
| 220_1 | TATGAGTGTGGAGTCGTTGCAATGATGGTGGTGAAGACCAGTGNNNAGATCGGAA |
| 221_1 | TATGAGTGTGGAGTCGTTGCCATCCCTTTATAGTGCCAGCACANNNAGATCGGAAG |
| 223_1 | TATGAGTGTGGAGTCGTTGCATGAGAAAAATGGGACCCCAACCNNNAGATCGGAA |
| 224_1 | TATGAGTGTGGAGTCGTTGCGTTAGCATTTGCTTTGCCTGAGANNNAGATCGGAAG |
| 226_1 | TATGAGTGTGGAGTCGTTGCAAACCACCCAGTTGTAAAGCTGTNNNAGATCGGAAG |
| 227_1 | TATGAGTGTGGAGTCGTTGCAACTGATGGGGAAAACAAGCTGTNNNAGATCGGAA |
| 229_1 | TATGAGTGTGGAGTCGTTGCCGCCTAAAAAGTCATCAAGCTGTNNNAGATCGGAAG |
| 231_1 | TATGAGTGTGGAGTCGTTGCCAGACATCATACACACAGGCACANNNAGATCGGAAG |
| 232_1 | TATGAGTGTGGAGTCGTTGCTTCAACTCAAACCCTTTGTCTGGNNNAGATCGGAAGA |
| 235_1 | TATGAGTGTGGAGTCGTTGCAGAGAGAGAGAGAATTCAGGGCACANNNAGATCGGA |
| 238_1 | TATGAGTGTGGAGTCGTTGCATGTTCCCACCATTCTAGGCTGTNNNAGATCGGAAG |
| 239_1 | TATGAGTGTGGAGTCGTTGCTTTGAAGCTGGTAAATGGGCTGTNNNAGATCGGAA |
| 241_1 | TATGAGTGTGGAGTCGTTGCTCTAACGCTCTTTGTGATGCACANNNAGATCGGAAG |
| 242_1 | TATGAGTGTGGAGTCGTTGCATAAGCAGATTCACAGCTGCACANNNAGATCGGAAG |
| 243_1 | TATGAGTGTGGAGTCGTTGCCCTTTCCAGTCATCATCTGCACANNNAGATCGGAAGA |
| 246_1 | TATGAGTGTGGAGTCGTTGCAAGCTGAGGAAATTGTCTGCACANNNAGATCGGAA |
| 247_1 | TATGAGTGTGGAGTCGTTGCGAAGTCCTAAGCTCCCGTACCTCNNNAGATCGGAAG |
| 248_1 | TATGAGTGTGGAGTCGTTGCATGTGGAAACACAGACATGCTGTNNNAGATCGGAA |
| 250_1 | TATGAGTGTGGAGTCGTTGCTCTTCATTTTTGCAGGATGCTGTNNNAGATCGGAAG |
| 252_1 | TATGAGTGTGGAGTCGTTGCTCCTTTCACTTTCAGTGTGCACANNNAGATCGGAAGA |
| 253_1 | TATGAGTGTGGAGTCGTTGCGAGCAGAACAGGATGAGCAATGTNNNAGATCGGAA |
| 255_1 | TATGAGTGTGGAGTCGTTGCCCTTAGCCTTCATTTCCTGCTGTNNNAGATCGGAAGA |
| 256_1 | TATGAGTGTGGAGTCGTTGCTGATTTGTATGCGTAGCTGCTGTNNNAGATCGGAAG |

| | |
|---|---|
| 257_1 | TATGAGTGTGGAGTCGTTGCGCTCTAAGAATGCTGTCTGCTGTNNNAGATCGGAAG |
| 258_1 | TATGAGTGTGGAGTCGTTGCATACCTTACAGACCGGTAGAGAGNNNAGATCGGAA |
| 262_1 | TATGAGTGTGGAGTCGTTGCCCTTGACAGGTTCTGGTATCACANNNAGATCGGAAG |
| 264_1 | TATGAGTGTGGAGTCGTTGCTCTAAAGCATGTAGCCTTCAACCNNNAGATCGGAAG |
| 265_1 | TATGAGTGTGGAGTCGTTGCAGTAGTAACCTGTCCAGCTCACANNNAGATCGGAAG |
| 266_1 | TATGAGTGTGGAGTCGTTGCAGCAGGTATTTCCTAGGCTCACANNNAGATCGGAAG |
| 267_1 | TATGAGTGTGGAGTCGTTGCTTCTAACCTCAAGCCCAGTGAGANNNAGATCGGAAG |
| 268_1 | TATGAGTGTGGAGTCGTTGCAGCGAGAAAGAGAGGATCTCACANNNAGATCGGAA |
| 269_1 | TATGAGTGTGGAGTCGTTGCCCCCAAAGGTGTACATTCTCACANNNAGATCGGAAG |
| 270_1 | TATGAGTGTGGAGTCGTTGCTCCTGGCAGACCTTGATATCTGTNNNAGATCGGAAG |
| 271_1 | TATGAGTGTGGAGTCGTTGCCCTGCTTCATTCTCCAACTCTGTNNNAGATCGGAAGA |
| 274_1 | TATGAGTGTGGAGTCGTTGCGTTCTTCCTTTTGCCTCCTCTGTNNNAGATCGGAAGA |
| 275_1 | TATGAGTGTGGAGTCGTTGCACAGGTATGCACATGGCAGAAAANNNAGATCGGAA |
| 276_1 | TATGAGTGTGGAGTCGTTGCATTCCATGTTTGGTGTGGTCACANNNAGATCGGAAG |
| 277_1 | TATGAGTGTGGAGTCGTTGCAGTTCTGGAGAGATTTGGTCACANNNAGATCGGAA |
| 281_1 | TATGAGTGTGGAGTCGTTGCCTGAAAATGCTATGGGCAGTCACNNNAGATCGGAA |
| 282_1 | TATGAGTGTGGAGTCGTTGCATCAGAACTTGGTCTCTCCCACTNNNAGATCGGAAG |
| 284_1 | TATGAGTGTGGAGTCGTTGCATGTCACTGCCACTATACTGCAGNNNAGATCGGAAG |
| 287_1 | TATGAGTGTGGAGTCGTTGCCCCCAGAAATCTGACAGTTCACANNNAGATCGGAAG |
| 288_1 | TATGAGTGTGGAGTCGTTGCACAAGCAAGCAAGAGAGTTCACANNNAGATCGGAA |
| 290_1 | TATGAGTGTGGAGTCGTTGCTGGTCTCCCCTGTATTAGTCTGTNNNAGATCGGAAG |
| 292_1 | TATGAGTGTGGAGTCGTTGCGGCTCATTCATCCTCAGGAATGTNNNAGATCGGAAG |
| 294_1 | TATGAGTGTGGAGTCGTTGCACATGCATACCTCTTCTGTCTGTNNNAGATCGGAAGA |
| 295_1 | TATGAGTGTGGAGTCGTTGCGTCTATCCCCAGGTTCTGTCTGTNNNAGATCGGAAG |
| 296_1 | TATGAGTGTGGAGTCGTTGCGGAAGTATGTGCCCTGTTTCACANNNAGATCGGAAG |
| 301_1 | TATGAGTGTGGAGTCGTTGCTGAACCACCCACAACAAAAGACANNNAGATCGGAAG |
| 306_1 | TATGAGTGTGGAGTCGTTGCATGGGAGGTCATCTGGAAAGACANNNAGATCGGAA |
| 307_1 | TATGAGTGTGGAGTCGTTGCGGAGGTGTTCTACCCTCTTCTGTNNNAGATCGGAAG |
| 309_1 | TATGAGTGTGGAGTCGTTGCGGTAGAATCAGAGGAGGTAGTGGNNNAGATCGGA |
| 310_1 | TATGAGTGTGGAGTCGTTGCCCTCTTCTCTTTTTGGGAACGTGNNNAGATCGGAAG |
| 311_1 | TATGAGTGTGGAGTCGTTGCTGAAGTGGGTCATTTGATGGGGTNNNAGATCGGAA |
| 313_1 | TATGAGTGTGGAGTCGTTGCTGGGACACTTAGGTTGGTTCTGTNNNAGATCGGAA |
| 314_1 | TATGAGTGTGGAGTCGTTGCTGGTGATAGGCAGAAGGAAGACANNNAGATCGGA |
| 317_1 | TATGAGTGTGGAGTCGTTGCCCAGCATGCCTTACTCTTTCTGTNNNAGATCGGAAGA |
| 319_1 | TATGAGTGTGGAGTCGTTGCCATCAAACCCCCACTTGAAGACANNNAGATCGGAAG |
| 324_1 | TATGAGTGTGGAGTCGTTGCTGTGCATGTATGTGAGCAAGTGTNNNAGATCGGAA |
| 325_1 | TATGAGTGTGGAGTCGTTGCGACAGTTCACTTTATCCCCCAGGNNNAGATCGGAAG |
| 326_1 | TATGAGTGTGGAGTCGTTGCTCCAATTCCGTAACCTCAAGTGTNNNAGATCGGAAG |
| 328_1 | TATGAGTGTGGAGTCGTTGCTGAATTTTGGGATAACCCAGACANNNAGATCGGAA |
| 333_1 | TATGAGTGTGGAGTCGTTGCTGAAAAGCAGGATGGTGAAGTGTNNNAGATCGGAA |
| 334_1 | TATGAGTGTGGAGTCGTTGCACAGAATGTGCGCCAAAGATTCTNNNAGATCGGAA |
| 335_1 | TATGAGTGTGGAGTCGTTGCACCGCTTCTAGGCTACTAAGTGTNNNAGATCGGAAG |
| 336_1 | TATGAGTGTGGAGTCGTTGCTCTTTCCAGGACTTTGTCCACACNNNAGATCGGAAGA |

| | |
|---|---|
| 338_1 | TATGAGTGTGGAGTCGTTGCGAGTGAGGAGACAGACAGAGACANNNAGATCGGA |
| 339_1 | TATGAGTGTGGAGTCGTTGCGAACTTGCCTCCTCATGGACCTGNNNAGATCGGAAG |
| 340_1 | TATGAGTGTGGAGTCGTTGCGGGAATGAAAGTTGTCCCAGTGTNNNAGATCGGAA |
| 346_1 | TATGAGTGTGGAGTCGTTGCGGAGCAGATTTGGAAGTCAGTGTNNNAGATCGGAA |
| 347_1 | TATGAGTGTGGAGTCGTTGCGGGGGTTGAAACTGAATGAGACANNNAGATCGGA |
| 348_1 | TATGAGTGTGGAGTCGTTGCCAGATTGAGCCAAATATCCTGCGNNNAGATCGGAAG |
| 349_1 | TATGAGTGTGGAGTCGTTGCGAAAGATGGAAGGCCTTGAGACANNNAGATCGGA |
| 350_1 | TATGAGTGTGGAGTCGTTGCGGGCGTGTATGTATGTTGAGACANNNAGATCGGAA |
| 354_1 | TATGAGTGTGGAGTCGTTGCGTTGTTGGGGGACACTTTAGACANNNAGATCGGAA |
| 355_1 | TATGAGTGTGGAGTCGTTGCCTATGAACAACGACATTCTCCGGNNNAGATCGGAAG |
| 358_1 | TATGAGTGTGGAGTCGTTGCTGCTGATAGCTTCCTAAAGGACANNNAGATCGGAAG |
| 361_1 | TATGAGTGTGGAGTCGTTGCACAGCCAAGTTATGCATACGTGTNNNAGATCGGAAG |
| 371_1 | TATGAGTGTGGAGTCGTTGCAGATATGAGACAGTGCCAGGTGTNNNAGATCGGAA |
| 377_1 | TATGAGTGTGGAGTCGTTGCTTTCTTTATCAAAAGCCCGGTGTNNNAGATCGGAAG |
| 378_1 | TATGAGTGTGGAGTCGTTGCTGCTCACTTTATAGGCCCTGCGANNNAGATCGGAAG |
| 381_1 | TATGAGTGTGGAGTCGTTGCTATCAAGTGGGAGATGTGGGTGTNNNAGATCGGA |
| 383_1 | TATGAGTGTGGAGTCGTTGCACCCTTCAACATCACAATGGTGTNNNAGATCGGAAG |
| 384_1 | TATGAGTGTGGAGTCGTTGCTATGATGCCAGGAGCTATGGACANNNAGATCGGAA |
| 386_1 | TATGAGTGTGGAGTCGTTGCCCCCAAGAACATATGCAGACCCTNNNAGATCGGAAG |
| 393_1 | TATGAGTGTGGAGTCGTTGCTGCTGTTGATGATGGTGATGTGTNNNAGATCGGAA |
| 394_1 | TATGAGTGTGGAGTCGTTGCAGGGAGGGAGTGTTTACATGACANNNAGATCGGA |
| 395_1 | TATGAGTGTGGAGTCGTTGCTGGACCTGTGAAGACCTATGTGTNNNAGATCGGAA |
| 396_1 | TATGAGTGTGGAGTCGTTGCACCTCTGCCTTTTGTTCATGACANNNAGATCGGAAG |
| 398_1 | TATGAGTGTGGAGTCGTTGCTTAGTGTGCCTGTGTGTATGTGTNNNAGATCGGAA |
| 399_1 | TATGAGTGTGGAGTCGTTGCGTCCAGGTCTTTGGATTATGTGTNNNAGATCGGAAG |
| 400_1 | TATGAGTGTGGAGTCGTTGCCCATAAACACCAGCTGGATGACANNNAGATCGGAAG |
| 401_1 | TATGAGTGTGGAGTCGTTGCTCTTGTGAAGGTTCTCACTGTGTNNNAGATCGGAAG |
| 402_1 | TATGAGTGTGGAGTCGTTGCTAAGCTGCTCCTATGACCTGTGTNNNAGATCGGAAG |
| 406_1 | TATGAGTGTGGAGTCGTTGCGGGAAATGGAATGGTTCCTGTGTNNNAGATCGGAA |
| 407_1 | TATGAGTGTGGAGTCGTTGCGAACAGTGGCAGAATAGCTGACANNNAGATCGGAA |
| 408_1 | TATGAGTGTGGAGTCGTTGCTTTTCTTGAGGGTCTTGCTGTGTNNNAGATCGGAAG |
| 412_1 | TATGAGTGTGGAGTCGTTGCATAGCAGGAGTGTGTCTCTGTGTNNNAGATCGGAA |
| 415_1 | TATGAGTGTGGAGTCGTTGCCTGAGACTAGGAAGCAGGTGACANNNAGATCGGAA |
| 417_1 | TATGAGTGTGGAGTCGTTGCTCCAGTCTCTCTCAGTTTGTGGTNNNAGATCGGAAG |
| 418_1 | TATGAGTGTGGAGTCGTTGCCTGGGTAACTGCAATCTGTGACANNNAGATCGGAA |
| 419_1 | TATGAGTGTGGAGTCGTTGCGAGATATGGCTGAATGGGTGTGTNNNAGATCGGA |
| 421_1 | TATGAGTGTGGAGTCGTTGCTGACAGAATCACCAGCATTGACANNNAGATCGGAAG |
| 422_1 | TATGAGTGTGGAGTCGTTGCTTGTGCAGTCCATAGCATTGACANNNAGATCGGAAG |
| 423_1 | TATGAGTGTGGAGTCGTTGCTGAGGGAGAGTGAGTATGTGTGTNNNAGATCGGA |
| 425_1 | TATGAGTGTGGAGTCGTTGCTGTTTGGCATTTCCCTCTTGACANNNAGATCGGAAG |
| 426_1 | TATGAGTGTGGAGTCGTTGCGCTCTTTCTCCACATCTGTGTGTNNNAGATCGGAAG |
| 428_1 | TATGAGTGTGGAGTCGTTGCAAAGACTTGGGGCATCTTTGACANNNAGATCGGAA |
| 429_1 | TATGAGTGTGGAGTCGTTGCTCGCCAGTCTCCAAGCATATAGANNNAGATCGGAAG |

| | |
|---|---|
| 432_1 | TATGAGTGTGGAGTCGTTGCGGAGCAGTGGTGTGGAAAATACANNNAGATCGGA |
| 434_1 | TATGAGTGTGGAGTCGTTGCATTGAGCCCTCAGAAAGCCTTAGNNNAGATCGGAAG |
| 436_1 | TATGAGTGTGGAGTCGTTGCCGTGGAGAATAGTGTGTGTGTGTNNNAGATCGGAA |
| 437_1 | TATGAGTGTGGAGTCGTTGCTGAGCCAGTTCCATATTGTGTGTNNNAGATCGGAAG |
| 439_1 | TATGAGTGTGGAGTCGTTGCTCTACTGATTGGCCCATTTTACTGANNNAGATCGGA |
| 440_1 | TATGAGTGTGGAGTCGTTGCTCAGCAAACTTTAGTGCTTGTGTNNNAGATCGGAAG |
| 441_1 | TATGAGTGTGGAGTCGTTGCTGGTGACATAAGCACTCCATACANNNAGATCGGAAG |
| 443_1 | TATGAGTGTGGAGTCGTTGCCCAGGACAGGGATAGAGTTGTGTNNNAGATCGGAA |
| 445_1 | TATGAGTGTGGAGTCGTTGCTGTGTTTGGATGTGATGTTGTGTNNNAGATCGGAA |
| 446_1 | TATGAGTGTGGAGTCGTTGCGGAGGGAAAAGAACCTGTTGTGTNNNAGATCGGA |
| 447_1 | TATGAGTGTGGAGTCGTTGCGGGGGTTGTATCTGCATCATACANNNAGATCGGAA |
| 450_1 | TATGAGTGTGGAGTCGTTGCGCCTCATTCTCCCTGTAGATACANNNAGATCGGAAG |
| 455_1 | TATGAGTGTGGAGTCGTTGCCAAGGCACACTGAAGGTGATACANNNAGATCGGAA |
| 465_1 | TATGAGTGTGGAGTCGTTGCCCAGCCTAATCCCTGTACCTACANNNAGATCGGAAG |
| 467_1 | TATGAGTGTGGAGTCGTTGCAACCTAGAATGCTACACCCTACANNNAGATCGGAAG |
| 468_1 | TATGAGTGTGGAGTCGTTGCCTCTTACGGATGCCAGTCATTGTNNNAGATCGGAAG |
| 469_1 | TATGAGTGTGGAGTCGTTGCTACCTAACACACACAGGCCTACANNNAGATCGGAAG |
| 472_1 | TATGAGTGTGGAGTCGTTGCTCTGAGTAAGGTCACTGGATTGTNNNAGATCGGAA |
| 473_1 | TATGAGTGTGGAGTCGTTGCACTGCTTTTGGTTCTTGGCTACANNNAGATCGGAAG |
| 474_1 | TATGAGTGTGGAGTCGTTGCCGGTGCCCTTCAAGGAATATTGTNNNAGATCGGAAG |
| 475_1 | TATGAGTGTGGAGTCGTTGCTTCCTTCAAGCAGAGGTGCTACANNNAGATCGGAAG |
| 480_1 | TATGAGTGTGGAGTCGTTGCGCTGGCAAGTCCCATTATCTACANNNAGATCGGAAG |
| 481_1 | TATGAGTGTGGAGTCGTTGCGCACCAAATGCATTCCAACTTGTNNNAGATCGGAAG |
| 484_1 | TATGAGTGTGGAGTCGTTGCACCTCTGGTATAACCGGTCTACANNNAGATCGGAAG |
| 485_1 | TATGAGTGTGGAGTCGTTGCCCAGGACCCGTAGGATTTCTACANNNAGATCGGAAG |
| 486_1 | TATGAGTGTGGAGTCGTTGCTGAGCAGCAGCATTTTACCTTGTNNNAGATCGGAAG |
| 487_1 | TATGAGTGTGGAGTCGTTGCGCATGTGTCAAGCAATTCTGAAANNNAGATCGGAA |
| 488_1 | TATGAGTGTGGAGTCGTTGCTGCTCAATCCTTGTGTCAGTACANNNAGATCGGAAG |
| 490_1 | TATGAGTGTGGAGTCGTTGCACTTGTCTGGTTCAGCATTTCACNNNAGATCGGAAG |
| 491_1 | TATGAGTGTGGAGTCGTTGCTCTCCTCCAGTGATTAGCCTTGTNNNAGATCGGAAG |
| 492_1 | TATGAGTGTGGAGTCGTTGCATCCCAGGAGGGAACTGAGTACANNNAGATCGGAA |
| 493_1 | TATGAGTGTGGAGTCGTTGCGTGATGCCCAGTAACATCCTTGTNNNAGATCGGAAG |
| 495_1 | TATGAGTGTGGAGTCGTTGCCCGTGAGTTTGCCCATTAGTACANNNAGATCGGAAG |
| 497_1 | TATGAGTGTGGAGTCGTTGCCGCTTTATGAAGTTTTGGCTTGTNNNAGATCGGAAG |
| 498_1 | TATGAGTGTGGAGTCGTTGCGCCTTTCTCTGCTTCACTCACTCNNNAGATCGGAAGA |
| 499_1 | TATGAGTGTGGAGTCGTTGCTCAGGCATTCATGCACATGTACANNNAGATCGGAAG |
| 503_1 | TATGAGTGTGGAGTCGTTGCCTGTTCCACTGATTGCCCTTACANNNAGATCGGAAG |
| 505_1 | TATGAGTGTGGAGTCGTTGCCAGATGACGTGTGTCCTTCTTGTNNNAGATCGGAAG |
| 509_1 | TATGAGTGTGGAGTCGTTGCCCTCCTGGGAAAGCAAAAGTTGTNNNAGATCGGAA |
| 510_1 | TATGAGTGTGGAGTCGTTGCCAAAGGAGACCCGAGAAAGTTGTNNNAGATCGGAA |
| 514_1 | TATGAGTGTGGAGTCGTTGCGGACCAAGCAGCTGGTATTTACANNNAGATCGGAA |
| 515_1 | TATGAGTGTGGAGTCGTTGCCGGCTTGCTGAATCGTTGAGATGNNNAGATCGGAA |
| 516_1 | TATGAGTGTGGAGTCGTTGCACATGGCTGCTGTTTCCTTTACANNNAGATCGGAAG |

| | |
|---|---|
| 517_1 | TATGAGTGTGGAGTCGTTGCCTCTTCCTCTGTCATGCAGTTGTNNNAGATCGGAAGA |
| 518_1 | TATGAGTGTGGAGTCGTTGCAGCCCATCAGAAAATCGAGTTGTNNNAGATCGGAAG |
| 519_1 | TATGAGTGTGGAGTCGTTGCACCAGCTTCTTCTCTGGTTTACANNNAGATCGGAAGA |
| 521_1 | TATGAGTGTGGAGTCGTTGCATTTTGGAGTAGGACCACGTTGTNNNAGATCGGAAG |
| 523_1 | TATGAGTGTGGAGTCGTTGCACAAGCACCTAAACTTCCGTTGTNNNAGATCGGAAGA |
| 524_1 | TATGAGTGTGGAGTCGTTGCAGGGACTAACTAGAAGCGGTTGTNNNAGATCGGAA |
| 525_1 | TATGAGTGTGGAGTCGTTGCTATGAGCCACTACACCTGGTTGTNNNAGATCGGAAG |
| 527_1 | TATGAGTGTGGAGTCGTTGCACAATGATTAGCCTGCCAAACCANNNAGATCGGAAG |
| 528_1 | TATGAGTGTGGAGTCGTTGCTTGTCTGAGTGCATTCCCATTCTNNNAGATCGGAAGA |
| 529_1 | TATGAGTGTGGAGTCGTTGCCACAGTCCCTTTGACAGAAACCANNNAGATCGGAAGA |
| 536_1 | TATGAGTGTGGAGTCGTTGCGCCTTCAGCCTGTGTTTTCTAGANNNAGATCGGAAGA |
| 537_1 | TATGAGTGTGGAGTCGTTGCGCAGGATGAATGTGGCAATTTGTNNNAGATCGGAAG |
| 538_1 | TATGAGTGTGGAGTCGTTGCCTGCAATATTCCCAACAGAACCANNNAGATCGGAAGA |
| 540_1 | TATGAGTGTGGAGTCGTTGCCCCTCATCTATTGGCTGAAGTGGNNNAGATCGGAAG |
| 541_1 | TATGAGTGTGGAGTCGTTGCAGTCTGGTTCTGTCTGGGTAACANNNAGATCGGAAG |
| 542_1 | TATGAGTGTGGAGTCGTTGCAGGAGGCACACAACTTCATTTGTNNNAGATCGGAAG |
| 543_1 | TATGAGTGTGGAGTCGTTGCCATCTGTGCCTGTGTACTCATGTNNNAGATCGGAAGA |
| 544_1 | TATGAGTGTGGAGTCGTTGCGGAGTTTCGCTCTTATAAACGGGNNNAGATCGGAAG |
| 545_1 | TATGAGTGTGGAGTCGTTGCCTCCTCCCTATGGCTTTCTGAGCNNNAGATCGGAAGA |
| 548_1 | TATGAGTGTGGAGTCGTTGCAGCATTGCCTGAAGAAAACACCANNNAGATCGGAAG |
| 549_1 | TATGAGTGTGGAGTCGTTGCGATTTTCTCCAAGGCCATGTGTCNNNAGATCGGAAGA |
| 550_1 | TATGAGTGTGGAGTCGTTGCTCTAGGAAAAAGGAGCACTTTGTNNNAGATCGGAAG |
| 551_1 | TATGAGTGTGGAGTCGTTGCACACACACATCTATCCAGTTCACNNNAGATCGGAAGA |
| 553_1 | TATGAGTGTGGAGTCGTTGCGTGTGTGTGTCTACCACCTTTGTNNNAGATCGGAAGA |
| 554_1 | TATGAGTGTGGAGTCGTTGCTTGCTGGCCAATTATCCCTTTGTNNNAGATCGGAAGA |
| 556_1 | TATGAGTGTGGAGTCGTTGCAGTTCCCCACTCATGCAATGTTGNNNAGATCGGAAGA |
| 559_1 | TATGAGTGTGGAGTCGTTGCCTCGACTACCTGTTAGGACACCANNNAGATCGGAAGA |
| 560_1 | TATGAGTGTGGAGTCGTTGCCTGGGAAGCAAAAACTGCTTTGTNNNAGATCGGAAG |
| 561_1 | TATGAGTGTGGAGTCGTTGCCCCAAACAACCTTACTGCTTTGTNNNAGATCGGAAGA |
| 562_1 | TATGAGTGTGGAGTCGTTGCCTATGTGCCTCTGTCTGCTTTGTNNNAGATCGGAAGA |
| 563_1 | TATGAGTGTGGAGTCGTTGCTGGCACAAGGAAGTTTTACACCANNNAGATCGGAAG |
| 565_1 | TATGAGTGTGGAGTCGTTGCACATCATGAGACTTACCCCACCANNNAGATCGGAAGA |
| 567_1 | TATGAGTGTGGAGTCGTTGCAAAACAACCAACAACCCAGTGTTNNNAGATCGGAAG |
| 568_1 | TATGAGTGTGGAGTCGTTGCGCTCGGAAATAACTGATCCACCANNNAGATCGGAAG |
| 569_1 | TATGAGTGTGGAGTCGTTGCACAGCCACACATAGCAAGTTTGTNNNAGATCGGAAG |
| 570_1 | TATGAGTGTGGAGTCGTTGCAGGATCAGAAATGAGAAGCACCANNNAGATCGGAA |
| 572_1 | TATGAGTGTGGAGTCGTTGCTTTCTCAGCACCCAGTAGTTTGTNNNAGATCGGAAGA |
| 573_1 | TATGAGTGTGGAGTCGTTGCGAGTGTAGAGCATTGATGCACCANNNAGATCGGAAG |
| 575_1 | TATGAGTGTGGAGTCGTTGCGTGATTTGTACAGGACCTCACCANNNAGATCGGAAG |
| 576_1 | TATGAGTGTGGAGTCGTTGCAGCAAAATCTGCATAGCTCACCANNNAGATCGGAAG |
| 577_1 | TATGAGTGTGGAGTCGTTGCAGTGTCTTTGGACTGCTGTTTGTNNNAGATCGGAAG |
| 578_1 | TATGAGTGTGGAGTCGTTGCTTTCTAGTCTCGCTCTGTCACCANNNAGATCGGAAGA |
| 579_1 | TATGAGTGTGGAGTCGTTGCTCCAAAGGGCAATCCTTGTTTGTNNNAGATCGGAAG |

| | |
|---|---|
| 580_1 | TATGAGTGTGGAGTCGTTGCTCTGACACTGAGCACACTATGGANNNAGATCGGAAG |
| 583_1 | TATGAGTGTGGAGTCGTTGCTGACAAGCACGGAATAAAGACCANNNAGATCGGAA |
| 585_1 | TATGAGTGTGGAGTCGTTGCGCAACATGACATACATAAGACCANNNAGATCGGAAG |
| 586_1 | TATGAGTGTGGAGTCGTTGCATGAAAAACCAAGTCCCAGACCANNNAGATCGGAAG |
| 590_1 | TATGAGTGTGGAGTCGTTGCTCTGTCTCAGTGCTTGGTTTTGTNNNAGATCGGAAG |
| 593_1 | TATGAGTGTGGAGTCGTTGCTTTGCACCTGGTCTGATTTTTGTNNNAGATCGGAAG |
| 598_1 | TATGAGTGTGGAGTCGTTGCTAACTGCCTTCTGAAAGGGACCANNNAGATCGGAAG |
| 600_1 | TATGAGTGTGGAGTCGTTGCCAGGTGTTTCTTCCATTGGACCANNNAGATCGGAAG |
| 601_1 | TATGAGTGTGGAGTCGTTGCGGGGTGGTGGTGCTATGAAAATTNNNAGATCGGAA |
| 602_1 | TATGAGTGTGGAGTCGTTGCCATAAGCAAGCAAGCAATGACCANNNAGATCGGAA |
| 603_1 | TATGAGTGTGGAGTCGTTGCGGGGGAATGGAATATCATGACCANNNAGATCGGAA |
| 605_1 | TATGAGTGTGGAGTCGTTGCTAGTGCTTGCAACAACGCAAATTNNNAGATCGGAAG |
| 606_1 | TATGAGTGTGGAGTCGTTGCCCCCACCCTTGTCTCTACAAAATNNNAGATCGGAAGA |
| 608_1 | TATGAGTGTGGAGTCGTTGCATAAGGAAGTCTGGCCCTGTAAGNNNAGATCGGAA |
| 609_1 | TATGAGTGTGGAGTCGTTGCGCACGCAAGAGCTCATTCAAATTNNNAGATCGGAAG |
| 610_1 | TATGAGTGTGGAGTCGTTGCCCCATACATTCCTGCATTGACCANNNAGATCGGAAG |
| 612_1 | TATGAGTGTGGAGTCGTTGCCTGCTTTAGCCCCATGGGAAATTNNNAGATCGGAAG |
| 613_1 | TATGAGTGTGGAGTCGTTGCAAAGCCAGAGAGCAGTGGAAATTNNNAGATCGGAA |
| 615_1 | TATGAGTGTGGAGTCGTTGCCACAGTACAGCAGCCCTACAATTNNNAGATCGGAAG |
| 616_1 | TATGAGTGTGGAGTCGTTGCATGACGCCAATCTGCCTACAATTNNNAGATCGGAAG |
| 620_1 | TATGAGTGTGGAGTCGTTGCGCATCAGAAGACATGCACTACCANNNAGATCGGAAG |
| 623_1 | TATGAGTGTGGAGTCGTTGCAGGTTCAGAGTGAGATGCTACCANNNAGATCGGAA |
| 624_1 | TATGAGTGTGGAGTCGTTGCGAATAACAGTGCCAGTGCTACCANNNAGATCGGAA |
| 626_1 | TATGAGTGTGGAGTCGTTGCTTCACCAGACAGCAGATCTACCANNNAGATCGGAAG |
| 627_1 | TATGAGTGTGGAGTCGTTGCGGAGGAACTGAGACCTTCTACCANNNAGATCGGAA |
| 628_1 | TATGAGTGTGGAGTCGTTGCGAGTGAGAAGGCACTGTCCAATTNNNAGATCGGAA |
| 630_1 | TATGAGTGTGGAGTCGTTGCGGCTTGCAAAAGTGTGAGCAATTNNNAGATCGGAA |
| 633_1 | TATGAGTGTGGAGTCGTTGCTAAGCAACATGGGAACTGCAATTNNNAGATCGGAA |
| 635_1 | TATGAGTGTGGAGTCGTTGCGGGCTTCTGAGGAATCTGCAATTNNNAGATCGGAA |
| 636_1 | TATGAGTGTGGAGTCGTTGCCTGAGGCGTGTGAATTTGCAATTNNNAGATCGGAA |
| 638_1 | TATGAGTGTGGAGTCGTTGCGGCAGTGGGTAGGCTAATCAATTNNNAGATCGGAA |
| 641_1 | TATGAGTGTGGAGTCGTTGCCGTTTCCAGATTCACAAAACCCANNNAGATCGGAAG |
| 644_1 | TATGAGTGTGGAGTCGTTGCTGGGATGAGGAGGGAAAGGAATTNNNAGATCGGA |
| 645_1 | TATGAGTGTGGAGTCGTTGCGTTACAGAGCCGGGAAAGGAATTNNNAGATCGGAA |
| 646_1 | TATGAGTGTGGAGTCGTTGCGTATGGAGGGCCCTGAAGGAATTNNNAGATCGGAA |
| 647_1 | TATGAGTGTGGAGTCGTTGCCATAACCAACCAGTGTCAACCCANNNAGATCGGAAG |
| 648_1 | TATGAGTGTGGAGTCGTTGCCCCTGATAACTCAGCATGCATCTNNNAGATCGGAAG |
| 656_1 | TATGAGTGTGGAGTCGTTGCCTTGCCGAAGGTCAGAGCTAATTNNNAGATCGGAA |
| 659_1 | TATGAGTGTGGAGTCGTTGCTTTGTGGTTTCTTTGTGGACCCANNNAGATCGGAAG |
| 660_1 | TATGAGTGTGGAGTCGTTGCTTTTGGCTGATAGTACTGACCCANNNAGATCGGAAG |
| 662_1 | TATGAGTGTGGAGTCGTTGCATTATTGCCTGGGTCACTACCCANNNAGATCGGAAG |
| 663_1 | TATGAGTGTGGAGTCGTTGCTAAGGCCACAGTTCTGCAACATTNNNAGATCGGAAG |
| 667_1 | TATGAGTGTGGAGTCGTTGCTTGTTCCTGCCTATCTCCCTTGGNNNAGATCGGAAGA |

| | |
|---|---|
| 669_1 | TATGAGTGTGGAGTCGTTGCGTAGGCTGAGCTCCTGTCTTAGANNNAGATCGGAAG |
| 670_1 | TATGAGTGTGGAGTCGTTGCTCATCATTCCCCTCTCCCACATTNNNAGATCGGAAGAG |
| 671_1 | TATGAGTGTGGAGTCGTTGCCCCCTACAGAAAAGGAGCACATTNNNAGATCGGAAG |
| 672_1 | TATGAGTGTGGAGTCGTTGCTGGGAGACTATTGGAGGCACATTNNNAGATCGGAA |
| 674_1 | TATGAGTGTGGAGTCGTTGCCAGCAGTGTGATTTTGGCACATTNNNAGATCGGAAG |
| 675_1 | TATGAGTGTGGAGTCGTTGCGGCTCAATGTGCTTATAACCCCANNNAGATCGGAAG |
| 676_1 | TATGAGTGTGGAGTCGTTGCGGGAACACAGGGAAACTCACATTNNNAGATCGGAA |
| 677_1 | TATGAGTGTGGAGTCGTTGCGGGGGATTTGGTGCTAAGTTAGANNNAGATCGGAA |
| 679_1 | TATGAGTGTGGAGTCGTTGCTCCTCTCTCTCTGAAGTACCCCANNNAGATCGGAAGA |
| 683_1 | TATGAGTGTGGAGTCGTTGCGCAGTCAGCATACCACTGACATTNNNAGATCGGAAG |
| 685_1 | TATGAGTGTGGAGTCGTTGCCTGTCTTTGCTTATGTGTGCAGGNNNAGATCGGAAG |
| 686_1 | TATGAGTGTGGAGTCGTTGCTCATTTCCCATGGATATCCCCCANNNAGATCGGAAGA |
| 687_1 | TATGAGTGTGGAGTCGTTGCTCAAATCCACACTGGTGTACATTNNNAGATCGGAAG |
| 688_1 | TATGAGTGTGGAGTCGTTGCTTTTCTTTCTCCTTCCTCCCCCANNNAGATCGGAAGAG |
| 693_1 | TATGAGTGTGGAGTCGTTGCGACTTTCCACTCATCCACCCATTNNNAGATCGGAAGA |
| 694_1 | TATGAGTGTGGAGTCGTTGCCCCTCCTACCTTATTATGCCCCANNNAGATCGGAAGA |
| 695_1 | TATGAGTGTGGAGTCGTTGCCTACAACACCTCCATCACTTTGCNNNAGATCGGAAGA |
| 696_1 | TATGAGTGTGGAGTCGTTGCCCTGGAGTTTTAGGTTGCCCATTNNNAGATCGGAAG |
| 697_1 | TATGAGTGTGGAGTCGTTGCAGTCATCTTTGTCATCCTCCCCANNNAGATCGGAAGA |
| 698_1 | TATGAGTGTGGAGTCGTTGCACACAGTCATTCACAATGCCATTNNNAGATCGGAAG |
| 700_1 | TATGAGTGTGGAGTCGTTGCGCTAGTCTACATCAGGTTCCCCANNNAGATCGGAAG |
| 701_1 | TATGAGTGTGGAGTCGTTGCTAAGTCCTCACACTTGTTCCCCANNNAGATCGGAAGA |
| 703_1 | TATGAGTGTGGAGTCGTTGCATGGTTTTAGCACTCAGGCTATGNNNAGATCGGAAG |
| 704_1 | TATGAGTGTGGAGTCGTTGCTCTTCACTGACTGTATCAGCCCANNNAGATCGGAAG |
| 707_1 | TATGAGTGTGGAGTCGTTGCTTTTACCAGAAGGTGGCAGCATTNNNAGATCGGAAG |
| 708_1 | TATGAGTGTGGAGTCGTTGCAAAAGACCCCTCACCAGATGAGANNNAGATCGGAAG |
| 710_1 | TATGAGTGTGGAGTCGTTGCCGTGGAAACAAGCATTCAGCATTNNNAGATCGGAAG |
| 711_1 | TATGAGTGTGGAGTCGTTGCAAGACCGACAATAATTCTCTCCANNNAGATCGGAAG |
| 712_1 | TATGAGTGTGGAGTCGTTGCTGTGTGTCTAGGACTTTAGCCCANNNAGATCGGAAG |
| 715_1 | TATGAGTGTGGAGTCGTTGCGGAGAGGGGGTAATCTAGGCATTNNNAGATCGGAA |
| 717_1 | TATGAGTGTGGAGTCGTTGCAAAAGCTACTTCAGCCTGGCATTNNNAGATCGGAAG |
| 718_1 | TATGAGTGTGGAGTCGTTGCCGTGGTAGGTTATGAAATGCCCANNNAGATCGGAA |
| 720_1 | TATGAGTGTGGAGTCGTTGCGTGGGTCATAGGGTGAATGCATTNNNAGATCGGAA |
| 721_1 | TATGAGTGTGGAGTCGTTGCGTGGGAAAACAGAAGTATGCCCANNNAGATCGGAA |
| 722_1 | TATGAGTGTGGAGTCGTTGCAATAAGTGGCCAAAATCTGCCCANNNAGATCGGAAG |
| 723_1 | TATGAGTGTGGAGTCGTTGCCCTTCCTGTGTAAAATCTGCCCANNNAGATCGGAAG |
| 724_1 | TATGAGTGTGGAGTCGTTGCGGGAGTTTCTATTGTTCTGCCCANNNAGATCGGAAG |
| 725_1 | TATGAGTGTGGAGTCGTTGCGCAATGAGTATGTGTAGTGCCCANNNAGATCGGAA |
| 727_1 | TATGAGTGTGGAGTCGTTGCCCCTCCAAGCAGAATACTGCATTNNNAGATCGGAAG |
| 728_1 | TATGAGTGTGGAGTCGTTGCTAACAGGAGAGGGTAATTGCCCANNNAGATCGGAA |
| 729_1 | TATGAGTGTGGAGTCGTTGCGCTTGTGTGTGTTTGTCTGCATTNNNAGATCGGAAG |
| 730_1 | TATGAGTGTGGAGTCGTTGCCTGGAGAAAGGCCTACTTGCATTNNNAGATCGGAAG |
| 731_1 | TATGAGTGTGGAGTCGTTGCAGGTTTCATCTCTGGACATCCCANNNAGATCGGAAG |

| | |
|---|---|
| 732_1 | TATGAGTGTGGAGTCGTTGCTGCCTCCTACCTTAGCCATCATTNNNAGATCGGAAGA |
| 734_1 | TATGAGTGTGGAGTCGTTGCTTGAGCCTTTATCCAACTTCAGCNNNAGATCGGAAGA |
| 735_1 | TATGAGTGTGGAGTCGTTGCTTCAGTCACCATGATGTTTGTGANNNAGATCGGAAG |
| 737_1 | TATGAGTGTGGAGTCGTTGCAGCCTGTCACCCAAAAACTCATTNNNAGATCGGAAGA |
| 743_1 | TATGAGTGTGGAGTCGTTGCAAGCTCATCCTACACGTATCCCANNNAGATCGGAAGA |
| 746_1 | TATGAGTGTGGAGTCGTTGCTTATCAGGTCATGGGAACTCCCANNNAGATCGGAAG |
| 747_1 | TATGAGTGTGGAGTCGTTGCCACTCAATCAGCATGAACTCCCANNNAGATCGGAAGA |
| 749_1 | TATGAGTGTGGAGTCGTTGCTTCTCCATGCATGCTGTCTCATTNNNAGATCGGAAGA |
| 755_1 | TATGAGTGTGGAGTCGTTGCGGGCAACTAAGCAGCTAGTCATTNNNAGATCGGAAG |
| 756_1 | TATGAGTGTGGAGTCGTTGCCAAGAAAGTTCATCCTGCTCCCANNNAGATCGGAAGA |
| 759_1 | TATGAGTGTGGAGTCGTTGCCCATACCAAAGGGCCTCTTCATTNNNAGATCGGAAGA |
| 760_1 | TATGAGTGTGGAGTCGTTGCCTTCTGCCTGTGGCTAGTTCATTNNNAGATCGGAAGA |
| 762_1 | TATGAGTGTGGAGTCGTTGCTAGGTCAAGAAGCATCAGTCCCANNNAGATCGGAAG |
| 765_1 | TATGAGTGTGGAGTCGTTGCGTGGAAGCATTCCTGTTTCATCTNNNAGATCGGAAGA |
| 767_1 | TATGAGTGTGGAGTCGTTGCCTGCTGTGGAAAGGGAGAAGATTNNNAGATCGGAA |
| 769_1 | TATGAGTGTGGAGTCGTTGCTGTGATTGCCTGGGGGTAAGATTNNNAGATCGGAAG |
| 771_1 | TATGAGTGTGGAGTCGTTGCCCCGCTTCGTGAGATTACAGATTNNNAGATCGGAAG |
| 772_1 | TATGAGTGTGGAGTCGTTGCAAGTAGTGAAGGGTGAAGGGTGGNNNAGATCGGA |
| 773_1 | TATGAGTGTGGAGTCGTTGCTTTACACCAGGGATGACTTCCCANNNAGATCGGAAGA |
| 774_1 | TATGAGTGTGGAGTCGTTGCAAAATAGAAGCTTTGGCTTCCCANNNAGATCGGAAG |
| 775_1 | TATGAGTGTGGAGTCGTTGCCGATCAGGGAGAGTAGGCAGATTNNNAGATCGGAA |
| 777_1 | TATGAGTGTGGAGTCGTTGCGTGCTGATTCCCGGACATCTATGNNNAGATCGGAAG |
| 779_1 | TATGAGTGTGGAGTCGTTGCATGTACCAGGAGAATGGTTCCCANNNAGATCGGAAG |
| 780_1 | TATGAGTGTGGAGTCGTTGCAGGGAGGAGAAGCACTTCAGATTNNNAGATCGGAA |
| 781_1 | TATGAGTGTGGAGTCGTTGCCACACATGACCCTGCAAGAGATTNNNAGATCGGAAG |
| 782_1 | TATGAGTGTGGAGTCGTTGCATCCAGACACATCCCCAGAGATTNNNAGATCGGAAGA |
| 785_1 | TATGAGTGTGGAGTCGTTGCTGCTCTTACCTCTGCCTGAGATTNNNAGATCGGAAGA |
| 787_1 | TATGAGTGTGGAGTCGTTGCACCTACACCAGATCAGTTTCCCANNNAGATCGGAAGA |
| 788_1 | TATGAGTGTGGAGTCGTTGCTCCTCCACTCTCAACTTTTCCCANNNAGATCGGAAGAG |
| 789_1 | TATGAGTGTGGAGTCGTTGCGGAGGTGCTAGTCTGGGTAGATTNNNAGATCGGAA |
| 794_1 | TATGAGTGTGGAGTCGTTGCACTCTAACAATGGGCAAAAGCCANNNAGATCGGAAG |
| 795_1 | TATGAGTGTGGAGTCGTTGCAAAGACCCATGAGCAGGGATTCTNNNAGATCGGAAG |
| 798_1 | TATGAGTGTGGAGTCGTTGCTCAGCCTCAGAAAATGAAAGCCANNNAGATCGGAAG |
| 801_1 | TATGAGTGTGGAGTCGTTGCTGAGAGAGAGAAAGAGCAAGCCANNNAGATCGGAA |
| 803_1 | TATGAGTGTGGAGTCGTTGCAAGTAGCAAGTACTGGCAAGGATNNNAGATCGGAA |
| 806_1 | TATGAGTGTGGAGTCGTTGCTCCCACAAATAACCCACTGGATTNNNAGATCGGAAGA |
| 808_1 | TATGAGTGTGGAGTCGTTGCACACCAGACACCAATATAAGCCANNNAGATCGGAAG |
| 809_1 | TATGAGTGTGGAGTCGTTGCAAACAGAGCAAGTGAGTAAGCCANNNAGATCGGAA |
| 811_1 | TATGAGTGTGGAGTCGTTGCAAGCAGTGAACACTTGTAAGCCANNNAGATCGGAAG |
| 817_1 | TATGAGTGTGGAGTCGTTGCAGTTCATCCTGAGGCACCTGATTNNNAGATCGGAAG |
| 818_1 | TATGAGTGTGGAGTCGTTGCTAGCTGGGGTGGTATACCTGATTNNNAGATCGGAAG |
| 819_1 | TATGAGTGTGGAGTCGTTGCTTTTCAGGGCATCCTTCCTGATTNNNAGATCGGAAGA |
| 821_1 | TATGAGTGTGGAGTCGTTGCCGCCATCATCTATCAATCAGCCANNNAGATCGGAAGA |

822_1 TATGAGTGTGGAGTCGTTGCGGACACTTTCTGGTCTGCTGATTNNNAGATCGGAAG
823_1 TATGAGTGTGGAGTCGTTGCGCCTTTGGGTACTGACAGTGATTNNNAGATCGGAAG
825_1 TATGAGTGTGGAGTCGTTGCGGAGCCACCTAATGAAGGTGATTNNNAGATCGGAA
828_1 TATGAGTGTGGAGTCGTTGCGGGTTGCAATGAAACGTGTGATTNNNAGATCGGAA
829_1 TATGAGTGTGGAGTCGTTGCGTTCCATTTTGTCAGGAGAGCCANNNAGATCGGAAG
830_1 TATGAGTGTGGAGTCGTTGCATGCAAACTGCCTAGGTGTGATTNNNAGATCGGAAG
837_1 TATGAGTGTGGAGTCGTTGCTCCTGGTCCTATACACATAGCCANNNAGATCGGAAG
838_1 TATGAGTGTGGAGTCGTTGCCAACTGTTTGGTCCAGATAGCCANNNAGATCGGAAG
840_1 TATGAGTGTGGAGTCGTTGCGCGTGTGTGTGAGTTTGGATATTNNNAGATCGGAA
842_1 TATGAGTGTGGAGTCGTTGCAGGACTAATTCCCTTGCTAGCCANNNAGATCGGAAG
844_1 TATGAGTGTGGAGTCGTTGCGTGTTGGGAAGCCTGTGACTATTNNNAGATCGGAA
846_1 TATGAGTGTGGAGTCGTTGCAGGGTAGCAGTTCTGATTAGCCANNNAGATCGGAA
847_1 TATGAGTGTGGAGTCGTTGCCCCAACCATGGCCTAAACCTATTNNNAGATCGGAAG
848_1 TATGAGTGTGGAGTCGTTGCGTGCTGATGAAGGCAGACCTATTNNNAGATCGGAA
851_1 TATGAGTGTGGAGTCGTTGCGGACAGGGTCTTTGTTTTAGCCANNNAGATCGGAAG
852_1 TATGAGTGTGGAGTCGTTGCCAGCTTCCCACCATCTTGCTATTNNNAGATCGGAAGA
853_1 TATGAGTGTGGAGTCGTTGCTGCTATTCTACAGTCTACCGCCANNNAGATCGGAAG
854_1 TATGAGTGTGGAGTCGTTGCGGAAAGGACTAACTTTCTCGCCANNNAGATCGGAAG
856_1 TATGAGTGTGGAGTCGTTGCGCCCATACACACAGCTGTCTATTNNNAGATCGGAAG
857_1 TATGAGTGTGGAGTCGTTGCGGAAGCATCTAATCTGGCTTGGTNNNAGATCGGAA
858_1 TATGAGTGTGGAGTCGTTGCGATGGGGTGCCATGGTTTCTATTNNNAGATCGGAA
860_1 TATGAGTGTGGAGTCGTTGCCCCACTCCATAGGTCCCAGTATTNNNAGATCGGAAG
862_1 TATGAGTGTGGAGTCGTTGCGGGGCTTTTTACCCTGCAGTATTNNNAGATCGGAAG
863_1 TATGAGTGTGGAGTCGTTGCGCACTTGGAAGAAAGCGCTATTCNNNAGATCGGAA
864_1 TATGAGTGTGGAGTCGTTGCAGCGTTCCTTGGTCATTCGTATTNNNAGATCGGAAG
865_1 TATGAGTGTGGAGTCGTTGCAGGAGAAGCCCAAAGGAGGTATTNNNAGATCGGAA
867_1 TATGAGTGTGGAGTCGTTGCTCCATTCATCTGTAGATGGGCCANNNAGATCGGAAG
868_1 TATGAGTGTGGAGTCGTTGCTATTTGGTGTTTTCACTGGGCCANNNAGATCGGAAG
871_1 TATGAGTGTGGAGTCGTTGCATAAGAGTGATGGAGTTGGGCCANNNAGATCGGAA
872_1 TATGAGTGTGGAGTCGTTGCTCAGGAAAAGTTGGAAATGGCCANNNAGATCGGAA
873_1 TATGAGTGTGGAGTCGTTGCACAGTGCTAATTGCTAATGGCCANNNAGATCGGAAG
874_1 TATGAGTGTGGAGTCGTTGCCTGAAGAAAGGATTTCCTGGCCANNNAGATCGGAA
875_1 TATGAGTGTGGAGTCGTTGCATGCAGGTGGTAGGAGTGTATGTNNNAGATCGGAA
877_1 TATGAGTGTGGAGTCGTTGCATGTCAGTTTGTCTTGTTGGCCANNNAGATCGGAAG
878_1 TATGAGTGTGGAGTCGTTGCGCACGCAATGAGAATGCTTTATTNNNAGATCGGAAG
879_1 TATGAGTGTGGAGTCGTTGCAGGGCCAGTGGAGGATCTTTATTNNNAGATCGGAA
880_1 TATGAGTGTGGAGTCGTTGCGGCAGAAGGCAGACAGGTTTATTNNNAGATCGGAA
881_1 TATGAGTGTGGAGTCGTTGCGGCTGGGTTTCACTGGGTTTATTNNNAGATCGGAAG
882_1 TATGAGTGTGGAGTCGTTGCCACACTCCTCCAGGCAAAAACTTNNNAGATCGGAAG
883_1 TATGAGTGTGGAGTCGTTGCACGAGATGACTTTCTGAATGCCANNNAGATCGGAAG
884_1 TATGAGTGTGGAGTCGTTGCTGGGAAGTCCATAGGTAATGCCANNNAGATCGGAA
885_1 TATGAGTGTGGAGTCGTTGCACACTAAGGATAAGAACATGCCANNNAGATCGGAA
886_1 TATGAGTGTGGAGTCGTTGCCTTCGTTACACAAATGTGAAGGTNNNAGATCGGAAG

| | |
|---|---|
| 888_1 | TATGAGTGTGGAGTCGTTGCGTGAGATAGCCAAATGGATGCCANNNAGATCGGAA |
| 891_1 | TATGAGTGTGGAGTCGTTGCAGACAAGAAGATCCCTGATGCCANNNAGATCGGAAG |
| 892_1 | TATGAGTGTGGAGTCGTTGCCCCACTTGGCTTCTTCTGTTTACNNNAGATCGGAAGA |
| 895_1 | TATGAGTGTGGAGTCGTTGCCAGGTTCTCTAGTGTGTATGCCANNNAGATCGGAAG |
| 896_1 | TATGAGTGTGGAGTCGTTGCGCCAACAGGTCTTCTCAGAACTTNNNAGATCGGAAG |
| 899_1 | TATGAGTGTGGAGTCGTTGCCAGAAATAGAAGCTCTGCTGCCANNNAGATCGGAAG |
| 900_1 | TATGAGTGTGGAGTCGTTGCTGCGAAAGGATTTCAAGGACACANNNAGATCGGAA |
| 901_1 | TATGAGTGTGGAGTCGTTGCCCATCCCCATATTGTTTCTGCCANNNAGATCGGAAGA |
| 902_1 | TATGAGTGTGGAGTCGTTGCGTTTTTCCTCCTCCTGCACACTTNNNAGATCGGAAGA |
| 903_1 | TATGAGTGTGGAGTCGTTGCATTTCCCACACTTTGGACCACTTNNNAGATCGGAAGA |
| 906_1 | TATGAGTGTGGAGTCGTTGCGAAACTACTTGAACCTGGTGCCANNNAGATCGGAAG |
| 908_1 | TATGAGTGTGGAGTCGTTGCTCAACGTACTTACCTTTCCACTTNNNAGATCGGAAGA |
| 910_1 | TATGAGTGTGGAGTCGTTGCTTGGCTATAAGGAAACAGCACTTNNNAGATCGGAAG |
| 911_1 | TATGAGTGTGGAGTCGTTGCAGTTGAATCACACAGAATTGCCANNNAGATCGGAAG |
| 912_1 | TATGAGTGTGGAGTCGTTGCAGCATTAGTACATGGCAGCACTTNNNAGATCGGAAG |
| 914_1 | TATGAGTGTGGAGTCGTTGCTCCTTTTGGAAACTTCAGCACTTNNNAGATCGGAAG |
| 915_1 | TATGAGTGTGGAGTCGTTGCAAGCTCTATTACAGTGGGCACTTNNNAGATCGGAAG |
| 916_1 | TATGAGTGTGGAGTCGTTGCGCCCCTGCCAATAACAATCACTTNNNAGATCGGAAG |
| 917_1 | TATGAGTGTGGAGTCGTTGCCTATGACCACCATGTGATTGCCANNNAGATCGGAAG |
| 918_1 | TATGAGTGTGGAGTCGTTGCCTGTGAAACACCCTCCATCACTTNNNAGATCGGAAGA |
| 924_1 | TATGAGTGTGGAGTCGTTGCAAGCCAACAGCAAGACTTCACTTNNNAGATCGGAAG |
| 930_1 | TATGAGTGTGGAGTCGTTGCTTGAATGGGATTGCAGCAGACTTNNNAGATCGGAA |
| 931_1 | TATGAGTGTGGAGTCGTTGCCTCTGCATCCCATTTCCAATCCANNNAGATCGGAAGA |
| 933_1 | TATGAGTGTGGAGTCGTTGCCTGGCACTTCACGTATCAATCCANNNAGATCGGAAG |
| 935_1 | TATGAGTGTGGAGTCGTTGCTATCCCAAAGAGCTCCAGGACTTNNNAGATCGGAAG |
| 939_1 | TATGAGTGTGGAGTCGTTGCGAGCAAGCAAACTAGGACATCCANNNAGATCGGAA |
| 940_1 | TATGAGTGTGGAGTCGTTGCTTGAGTGAATCAGACATGGACTTNNNAGATCGGAAG |
| 941_1 | TATGAGTGTGGAGTCGTTGCTTCCTTTCATGCTGAAGCATCCANNNAGATCGGAAG |
| 942_1 | TATGAGTGTGGAGTCGTTGCTGAACACACATTCACAGCATCCANNNAGATCGGAAG |
| 943_1 | TATGAGTGTGGAGTCGTTGCCGAAGAACCCCATGATTGGACTTNNNAGATCGGAAG |
| 944_1 | TATGAGTGTGGAGTCGTTGCGTCCTCACCACAAACTTGGACTTNNNAGATCGGAAG |
| 945_1 | TATGAGTGTGGAGTCGTTGCGGAGACAGCTCGATTTTGGACTTNNNAGATCGGAAG |
| 946_1 | TATGAGTGTGGAGTCGTTGCCCAAGCCAGGTTCTGAATGACTTNNNAGATCGGAAG |
| 947_1 | TATGAGTGTGGAGTCGTTGCGACACAGGTTGGTTTCCTGACTTNNNAGATCGGAAG |
| 948_1 | TATGAGTGTGGAGTCGTTGCACACCACCCTATGATGTCATCCANNNAGATCGGAAG |
| 949_1 | TATGAGTGTGGAGTCGTTGCTTCTCATCTCCAAAGGCTGACTTNNNAGATCGGAAG |
| 950_1 | TATGAGTGTGGAGTCGTTGCGAACATCAGAGGCATGGTGACTTNNNAGATCGGAA |
| 951_1 | TATGAGTGTGGAGTCGTTGCCCTTCCTTGCTTTCTGGTGACTTNNNAGATCGGAAGA |
| 953_1 | TATGAGTGTGGAGTCGTTGCTTGTCTCCTCAGCTTCAGATCCANNNAGATCGGAAGA |
| 954_1 | TATGAGTGTGGAGTCGTTGCAGTCCCCAGATGTGAGCATACTTNNNAGATCGGAAG |
| 958_1 | TATGAGTGTGGAGTCGTTGCCACACACACACTTTGCAAAAGGTNNNAGATCGGAAG |
| 959_1 | TATGAGTGTGGAGTCGTTGCTTTGCTGCATGGAATGTGATCCANNNAGATCGGAAG |
| 963_1 | TATGAGTGTGGAGTCGTTGCATAACGTCTTCTGCTCCTATCCANNNAGATCGGAAGA |

| | |
|---|---|
| 967_1 | TATGAGTGTGGAGTCGTTGCTGTTGTCCTTCCCTTGGTATCCANNNAGATCGGAAGA |
| 974_1 | TATGAGTGTGGAGTCGTTGCTTTTTCTGGGGCAATCAACTCCANNNAGATCGGAAG |
| 975_1 | TATGAGTGTGGAGTCGTTGCTCAACTGGGACATGGTTTGTTGGNNNAGATCGGAA |
| 978_1 | TATGAGTGTGGAGTCGTTGCTGTGCATGTCTAAGTGCAACCTTNNNAGATCGGAAG |
| 979_1 | TATGAGTGTGGAGTCGTTGCTCCTAAAGAGGTGCTTCACTCCANNNAGATCGGAAG |
| 982_1 | TATGAGTGTGGAGTCGTTGCCCTTTGACTAGGTAGCTTTGAGCNNNAGATCGGAAG |
| 983_1 | TATGAGTGTGGAGTCGTTGCAGTGCAACCTGCTTTCAAAAACCNNNAGATCGGAAG |
| 984_1 | TATGAGTGTGGAGTCGTTGCTGGTTTGGGCCTATTCTACTCCANNNAGATCGGAAG |
| 985_1 | TATGAGTGTGGAGTCGTTGCTTTCCCCAATTCTGGGTACTCCANNNAGATCGGAAGA |
| 986_1 | TATGAGTGTGGAGTCGTTGCAATCACTTCTTACACCACCTCCANNNAGATCGGAAGA |
| 988_1 | TATGAGTGTGGAGTCGTTGCACAGAAAACGAAGCCTTCACCTTNNNAGATCGGAAG |
| 992_1 | TATGAGTGTGGAGTCGTTGCTCTTACATTCCCATTCCCCTCCANNNAGATCGGAAGA |
| 993_1 | TATGAGTGTGGAGTCGTTGCTCCATAAGTCTAAATGCCCTCCANNNAGATCGGAAGA |
| 994_1 | TATGAGTGTGGAGTCGTTGCTAGCAAAGAAGACTGTCCCTCCANNNAGATCGGAAG |
| 996_1 | TATGAGTGTGGAGTCGTTGCTTGACACATGAATATGGAGCACANNNAGATCGGAA |
| 998_1 | TATGAGTGTGGAGTCGTTGCCTGTGGCTCCTACTGAGTACCTTNNNAGATCGGAAG |
| 999_1 | TATGAGTGTGGAGTCGTTGCACACACAAACATTCCTTCCTCCANNNAGATCGGAAGA |
| 1000_1 | TATGAGTGTGGAGTCGTTGCCACCCCCACAAATTCTGTACCTTNNNAGATCGGAAGA |
| 1001_1 | TATGAGTGTGGAGTCGTTGCGAAGGTCATTTAGGCAAGCTCCANNNAGATCGGAA |
| 1004_1 | TATGAGTGTGGAGTCGTTGCTATTTGTCTCTCTGCCTCCCCTTNNNAGATCGGAAGA |
| 1005_1 | TATGAGTGTGGAGTCGTTGCATACAGCATGCACTTGATCTCCANNNAGATCGGAAG |
| 1006_1 | TATGAGTGTGGAGTCGTTGCACTGACGCATCGAGAAAAATACTNNNAGATCGGAA |
| 1007_1 | TATGAGTGTGGAGTCGTTGCTGAAACAGACCTATGGCTCTCCANNNAGATCGGAAG |
| 1008_1 | TATGAGTGTGGAGTCGTTGCTGATTCCTGTTCCTCAGTCTCCANNNAGATCGGAAGA |
| 1010_1 | TATGAGTGTGGAGTCGTTGCTTCAGTGGTTTGTGTTGTCCCTTNNNAGATCGGAAG |
| 1011_1 | TATGAGTGTGGAGTCGTTGCGTCATGCTCATTCCACTTCCCTTNNNAGATCGGAAGA |
| 1012_1 | TATGAGTGTGGAGTCGTTGCTGGCACATTTCTTCACTTCTCCANNNAGATCGGAAGA |
| 1013_1 | TATGAGTGTGGAGTCGTTGCGCTGTTTGCTCTCTCTTTCCCTTNNNAGATCGGAAGA |
| 1015_1 | TATGAGTGTGGAGTCGTTGCTCCAATTGCCTCCAATTTCTCCANNNAGATCGGAAGA |
| 1017_1 | TATGAGTGTGGAGTCGTTGCAGGGGGAGATACCATCTAGTCCANNNAGATCGGAA |
| 1019_1 | TATGAGTGTGGAGTCGTTGCTTTCAGGATTTGAATGCCGTCCANNNAGATCGGAAG |
| 1020_1 | TATGAGTGTGGAGTCGTTGCGTTTTCAATGAAGGGAGGGCCTTNNNAGATCGGAA |
| 1024_1 | TATGAGTGTGGAGTCGTTGCTTCACAAAGACAAGTACTGCCTTNNNAGATCGGAAG |
| 1025_1 | TATGAGTGTGGAGTCGTTGCCAAGCACATACATACCCTGCCTTNNNAGATCGGAAGA |
| 1027_1 | TATGAGTGTGGAGTCGTTGCGCCTCAGTTTGTGTCTGGGAATANNNAGATCGGAA |
| 1029_1 | TATGAGTGTGGAGTCGTTGCACAACCTCCTTTCAAAGTGCCTTNNNAGATCGGAAGA |
| 1030_1 | TATGAGTGTGGAGTCGTTGCCTGTGAAACACCAGTTCTGTCCANNNAGATCGGAAG |
| 1031_1 | TATGAGTGTGGAGTCGTTGCCGAACTGCTCTCATTTCTGTCCANNNAGATCGGAAGA |
| 1034_1 | TATGAGTGTGGAGTCGTTGCTATGTGTTGCAAAGGTGTGTCCANNNAGATCGGAA |
| 1035_1 | TATGAGTGTGGAGTCGTTGCCACATCCCAACAGACACATCCTTNNNAGATCGGAAGA |
| 1036_1 | TATGAGTGTGGAGTCGTTGCTTTTTCCATGAGCCAGCATCCTTNNNAGATCGGAAGA |
| 1037_1 | TATGAGTGTGGAGTCGTTGCGCATTTGACAGCTCCTCATCCTTNNNAGATCGGAAGA |
| 1039_1 | TATGAGTGTGGAGTCGTTGCGGGTGTTGAAAGTGGGTATCCTTNNNAGATCGGAA |

| | |
|---|---|
| 1042_1 | TATGAGTGTGGAGTCGTTGCTCTTGGCCATATCCATAATTCCANNNAGATCGGAAGA |
| 1046_1 | TATGAGTGTGGAGTCGTTGCGTTTCTGCCCTAACACCATTCCANNNAGATCGGAAGA |
| 1047_1 | TATGAGTGTGGAGTCGTTGCACAAATCAAAAGGGCCCATTCCANNNAGATCGGAAG |
| 1050_1 | TATGAGTGTGGAGTCGTTGCACTTGCCAGCTGTTTGTATTCCANNNAGATCGGAAGA |
| 1051_1 | TATGAGTGTGGAGTCGTTGCGAATGAAATGCTGCATGCACTGTNNNAGATCGGAAG |
| 1052_1 | TATGAGTGTGGAGTCGTTGCCTAACTCCCTGCACAAACTTCCANNNAGATCGGAAGA |
| 1053_1 | TATGAGTGTGGAGTCGTTGCTGAGACCTGATTCCAGTGTCCTTNNNAGATCGGAAGA |
| 1056_1 | TATGAGTGTGGAGTCGTTGCATGTTTGGACCCTGCCGATTTGANNNAGATCGGAAG |
| 1059_1 | TATGAGTGTGGAGTCGTTGCGAGGCTAGTGTTGAACCCCTGGANNNAGATCGGAAG |
| 1060_1 | TATGAGTGTGGAGTCGTTGCAGCAGAAAGATCCAATCCTTCCANNNAGATCGGAAG |
| 1061_1 | TATGAGTGTGGAGTCGTTGCTTCCTCCCATACTCCTCTTCCTTNNNAGATCGGAAGAG |
| 1062_1 | TATGAGTGTGGAGTCGTTGCACTAGGGGTCTTTGTGGTTCCTTNNNAGATCGGAAG |
| 1063_1 | TATGAGTGTGGAGTCGTTGCTTGTTTCTTGGAAAACACCGGCTNNNAGATCGGAAG |
| 1064_1 | TATGAGTGTGGAGTCGTTGCATAGGTCTGGGACGTGTTTCCTTNNNAGATCGGAAG |
| 1065_1 | TATGAGTGTGGAGTCGTTGCAATGTGGGAAGCACCTTTTCCTTNNNAGATCGGAAG |
| 1066_1 | TATGAGTGTGGAGTCGTTGCTTCGTGGTAAGCAGCTTTTCCTTNNNAGATCGGAAGA |
| 1067_1 | TATGAGTGTGGAGTCGTTGCCCCGTAAGGATTCTTTGCTTCCANNNAGATCGGAAGA |
| 1068_1 | TATGAGTGTGGAGTCGTTGCCCTGAGAAGGAGGGAATCTTCCANNNAGATCGGAAG |
| 1070_1 | TATGAGTGTGGAGTCGTTGCATTTCAGATGGTGAGGCAAGCTTNNNAGATCGGAAG |
| 1072_1 | TATGAGTGTGGAGTCGTTGCACACCAAATGGATGCTCAAGCTTNNNAGATCGGAAG |
| 1073_1 | TATGAGTGTGGAGTCGTTGCGTGAGGGTGGAGAAAAGAAGCTTNNNAGATCGGAA |
| 1074_1 | TATGAGTGTGGAGTCGTTGCCAGGGGAAATTTGGAAGGTTCCANNNAGATCGGAA |
| 1077_1 | TATGAGTGTGGAGTCGTTGCCTATGGACGCATGGAATGTTCCANNNAGATCGGAAG |
| 1078_1 | TATGAGTGTGGAGTCGTTGCCATTTGGGACCTGAGCTAAGCTTNNNAGATCGGAAG |
| 1080_1 | TATGAGTGTGGAGTCGTTGCCACCCACAGATTCCTAACAGCTTNNNAGATCGGAAGA |
| 1081_1 | TATGAGTGTGGAGTCGTTGCTCTGTAAGGCAGAGGTACAGCTTNNNAGATCGGAAG |
| 1082_1 | TATGAGTGTGGAGTCGTTGCAATGACTACAAACCTGGCAGCTTNNNAGATCGGAAG |
| 1084_1 | TATGAGTGTGGAGTCGTTGCTTTTCAGCCTGGTTGTTCAGCTTNNNAGATCGGAAGA |
| 1086_1 | TATGAGTGTGGAGTCGTTGCTGTTCATTTGGCATCAGGAGCTTNNNAGATCGGAAG |
| 1087_1 | TATGAGTGTGGAGTCGTTGCTGGAACCCATTTATCTGGAGCTTNNNAGATCGGAAG |
| 1090_1 | TATGAGTGTGGAGTCGTTGCCTGAGGTTTCCACAAGGTTTCCANNNAGATCGGAAG |
| 1091_1 | TATGAGTGTGGAGTCGTTGCGCCCTTCTGGCTCTTTTTCTGATNNNAGATCGGAAGA |
| 1094_1 | TATGAGTGTGGAGTCGTTGCATTACCCCCTTCCTTCCTGCCTGNNNAGATCGGAAGA |
| 1096_1 | TATGAGTGTGGAGTCGTTGCACAGACAGACACTCACAAAAGCANNNAGATCGGAAG |
| 1098_1 | TATGAGTGTGGAGTCGTTGCCCTGGTGAAAAATCAGAGGGCTTNNNAGATCGGAAG |
| 1100_1 | TATGAGTGTGGAGTCGTTGCCAGTCGTACAGTTATTCACCACANNNAGATCGGAAGA |
| 1101_1 | TATGAGTGTGGAGTCGTTGCAGAAAAGTGATCCTGCCAAAGCANNNAGATCGGAAG |
| 1102_1 | TATGAGTGTGGAGTCGTTGCCCCCCAGGATAAAATCCAAAGCANNNAGATCGGAAG |
| 1103_1 | TATGAGTGTGGAGTCGTTGCACTTCTGAAACCACACCTTGTGGNNNAGATCGGAAGA |
| 1104_1 | TATGAGTGTGGAGTCGTTGCTACATTGTCATACCAGCTCAGGCNNNAGATCGGAAGA |
| 1106_1 | TATGAGTGTGGAGTCGTTGCTACAGGCATGCTCAAAGAAAGCANNNAGATCGGAAG |
| 1107_1 | TATGAGTGTGGAGTCGTTGCAGTTCCTGGTTCTGTAAGTCTGGNNNAGATCGGAAG |
| 1109_1 | TATGAGTGTGGAGTCGTTGCGGATCTGGGTACAAAAGTGGCTTNNNAGATCGGAAG |

| | |
|---|---|
| 1110_1 | TATGAGTGTGGAGTCGTTGCATGCAACACACACTCCTAAAGCANNNAGATCGGAAG |
| 1112_1 | TATGAGTGTGGAGTCGTTGCCCAGGACCACATCTGAAGAAGACNNNAGATCGGAA |
| 1113_1 | TATGAGTGTGGAGTCGTTGCTGAGGGGACGATTATCTTGGCTTNNNAGATCGGAA |
| 1116_1 | TATGAGTGTGGAGTCGTTGCCAGTGAAGGGCTGCTTTTCTTTTNNNAGATCGGAAG |
| 1117_1 | TATGAGTGTGGAGTCGTTGCAAAATCTACAGGGTGCACAAGCANNNAGATCGGAA |
| 1118_1 | TATGAGTGTGGAGTCGTTGCGATTTGCGAGTGAGGACAGAGTTNNNAGATCGGAA |
| 1120_1 | TATGAGTGTGGAGTCGTTGCTGGGAGAGCACTGTATACAAGCANNNAGATCGGAA |
| 1121_1 | TATGAGTGTGGAGTCGTTGCTCCCTCTGGTTCATTGTTTGTGGNNNAGATCGGAAG |
| 1123_1 | TATGAGTGTGGAGTCGTTGCCCTGTGGGATGAAAGAACTGCTTNNNAGATCGGAA |
| 1124_1 | TATGAGTGTGGAGTCGTTGCTCATTTCTGAAGGTTCCCAAGCANNNAGATCGGAAG |
| 1127_1 | TATGAGTGTGGAGTCGTTGCGCCAGGGTTCATTTCTACTGCTTNNNAGATCGGAAG |
| 1128_1 | TATGAGTGTGGAGTCGTTGCCCTATGTGTGTGGTCAAGAAGCANNNAGATCGGAA |
| 1131_1 | TATGAGTGTGGAGTCGTTGCGAGGGCACTGTTTATAGCTGCTTNNNAGATCGGAAG |
| 1132_1 | TATGAGTGTGGAGTCGTTGCGTGTCTCCACAGCATTAGAAGCANNNAGATCGGAAG |
| 1133_1 | TATGAGTGTGGAGTCGTTGCCAGAGAGACGGAGATAGGAAGCANNNAGATCGGA |
| 1134_1 | TATGAGTGTGGAGTCGTTGCTCTCACAGAAACATACAGTGCTTNNNAGATCGGAAG |
| 1135_1 | TATGAGTGTGGAGTCGTTGCTTTTTCTGCTGATGTGGGAAGCANNNAGATCGGAAG |
| 1136_1 | TATGAGTGTGGAGTCGTTGCCCAATCAAGTGACTTTCGTGCTTNNNAGATCGGAAG |
| 1139_1 | TATGAGTGTGGAGTCGTTGCATTCCACCTTGACTTTGGTGCTTNNNAGATCGGAAGA |
| 1140_1 | TATGAGTGTGGAGTCGTTGCCCTCACAAGGTGATACTGAAGCANNNAGATCGGAAG |
| 1141_1 | TATGAGTGTGGAGTCGTTGCCAAAGGGAACAAATCACTCGACGNNNAGATCGGAA |
| 1145_1 | TATGAGTGTGGAGTCGTTGCAAACCTGTGAGGTGAAATTGCTTNNNAGATCGGAAG |
| 1147_1 | TATGAGTGTGGAGTCGTTGCACAATACACAGGACAGCCTTTCANNNAGATCGGAAG |
| 1148_1 | TATGAGTGTGGAGTCGTTGCATGGTAAGCAGAAGCACTTGCTTNNNAGATCGGAA |
| 1149_1 | TATGAGTGTGGAGTCGTTGCAATGGGGTCTTGCTTAGTTGCTTNNNAGATCGGAAG |
| 1151_1 | TATGAGTGTGGAGTCGTTGCACACACTCTCACACACTTTGCTTNNNAGATCGGAAGA |
| 1152_1 | TATGAGTGTGGAGTCGTTGCAGATGAGGGGAGTACAAACAGCANNNAGATCGGA |
| 1154_1 | TATGAGTGTGGAGTCGTTGCAGGTAGATGGGGCCTCAAATCTTNNNAGATCGGAA |
| 1155_1 | TATGAGTGTGGAGTCGTTGCGCTGGCGTAGTAAGGGAAATCTTNNNAGATCGGAA |
| 1156_1 | TATGAGTGTGGAGTCGTTGCACAGAACACACACATACACAGCANNNAGATCGGAAG |
| 1158_1 | TATGAGTGTGGAGTCGTTGCCTAATTTTTCACCGTTTGGCCAGNNNAGATCGGAAG |
| 1161_1 | TATGAGTGTGGAGTCGTTGCAACCCATTTCAACTGCCCATCTTNNNAGATCGGAAGA |
| 1164_1 | TATGAGTGTGGAGTCGTTGCGAGAGAGAGAAAACACGTGCTGTNNNAGATCGGAA |
| 1165_1 | TATGAGTGTGGAGTCGTTGCAGGCTCTCAGGTTCCAAGATCTTNNNAGATCGGAAG |
| 1167_1 | TATGAGTGTGGAGTCGTTGCATTTATGTTGCCAAGACCCAGCANNNAGATCGGAAG |
| 1169_1 | TATGAGTGTGGAGTCGTTGCACATTCCTGCATAGCTCTGTGGNNNAGATCGGAAGA |
| 1170_1 | TATGAGTGTGGAGTCGTTGCGTTGTGAGCCACAGGGTTATCTTNNNAGATCGGAA |
| 1171_1 | TATGAGTGTGGAGTCGTTGCTTGGAGGGCTATATGATCCAGCANNNAGATCGGAA |
| 1172_1 | TATGAGTGTGGAGTCGTTGCCGAGATCACAGCTCAATGCAGACNNNAGATCGGAA |
| 1173_1 | TATGAGTGTGGAGTCGTTGCTTTACTGGGCTCTGAGACAAGGANNNAGATCGGAA |
| 1174_1 | TATGAGTGTGGAGTCGTTGCCCACAACTTACTGAAACCCCACANNNAGATCGGAAG |
| 1176_1 | TATGAGTGTGGAGTCGTTGCACACAGGCTCTCCATCTACTCTTNNNAGATCGGAAGA |
| 1179_1 | TATGAGTGTGGAGTCGTTGCGCCTATCCTAACAGCCTCCTCTTNNNAGATCGGAAGA |

| | |
|---|---|
| 1180_1 | TATGAGTGTGGAGTCGTTGCAAGAGCATTTAGATGTGGCAGCANNNAGATCGGAA |
| 1181_1 | TATGAGTGTGGAGTCGTTGCTATTACTGCCAGATGCTGTGTGTNNNAGATCGGAAG |
| 1183_1 | TATGAGTGTGGAGTCGTTGCAGAGAGAGAGTGTGTATGCAGCANNNAGATCGGA |
| 1185_1 | TATGAGTGTGGAGTCGTTGCGTAGGTGGAGTTAGGGAGCTCTTNNNAGATCGGAA |
| 1186_1 | TATGAGTGTGGAGTCGTTGCGGAAGCTTAGACACACATCAGCANNNAGATCGGAA |
| 1187_1 | TATGAGTGTGGAGTCGTTGCTCTTTCTTTGCTTCCCATCAGCANNNAGATCGGAAGA |
| 1188_1 | TATGAGTGTGGAGTCGTTGCGGGTTTGGGGTGTTGAATCTCTTNNNAGATCGGAA |
| 1189_1 | TATGAGTGTGGAGTCGTTGCTGGCTTTCAGGTACATATCAGCANNNAGATCGGAAG |
| 1190_1 | TATGAGTGTGGAGTCGTTGCCACTCCCTTCCCTATGCTCTCTTNNNAGATCGGAAGA |
| 1195_1 | TATGAGTGTGGAGTCGTTGCGGCACCACTACTTTTCCAGTCTTNNNAGATCGGAAG |
| 1197_1 | TATGAGTGTGGAGTCGTTGCAGGGGGTGGTCTTACTAGGTCTTNNNAGATCGGAA |
| 1202_1 | TATGAGTGTGGAGTCGTTGCGCCGACCTTGACTGTAATGTCTTNNNAGATCGGAAG |
| 1203_1 | TATGAGTGTGGAGTCGTTGCAAAGGAATGGACAACCAAGAGCANNNAGATCGGAA |
| 1204_1 | TATGAGTGTGGAGTCGTTGCAACATTTCCAGCACTGGTGTCTTNNNAGATCGGAAG |
| 1206_1 | TATGAGTGTGGAGTCGTTGCTGACTTGGCCTTTTATTCTCCACCNNNAGATCGGAAG |
| 1209_1 | TATGAGTGTGGAGTCGTTGCCAGGGGACAAGACGGAAATTCTTNNNAGATCGGAA |
| 1210_1 | TATGAGTGTGGAGTCGTTGCGAGGTGCCCATTCTTCCATTCTTNNNAGATCGGAAG |
| 1211_1 | TATGAGTGTGGAGTCGTTGCCAGTCCCTCTACTCCTACCCTGGNNNAGATCGGAAGA |
| 1217_1 | TATGAGTGTGGAGTCGTTGCCAGGACACTCACAAGCACTTCTTNNNAGATCGGAAG |
| 1218_1 | TATGAGTGTGGAGTCGTTGCCCGTAACTTTGAGGGCACTTCTTNNNAGATCGGAAG |
| 1220_1 | TATGAGTGTGGAGTCGTTGCAAGCCACTCAGATCTCCCTTCTTNNNAGATCGGAAGA |
| 1221_1 | TATGAGTGTGGAGTCGTTGCCCTAGCGGAATGCTTTCCTTCTTNNNAGATCGGAAG |
| 1222_1 | TATGAGTGTGGAGTCGTTGCATTTCCTCATCTAGGCAGGAGCANNNAGATCGGAAG |
| 1225_1 | TATGAGTGTGGAGTCGTTGCGAAATGGAGAAAGGAAGGGAGCANNNAGATCGGA |
| 1226_1 | TATGAGTGTGGAGTCGTTGCCAGTTGAGAGAGAGTAGGGAGCANNNAGATCGGA |
| 1229_1 | TATGAGTGTGGAGTCGTTGCACAATAGAAGGAAATTGGGAGCANNNAGATCGGAA |
| 1231_1 | TATGAGTGTGGAGTCGTTGCGGTTTTCATCTGTCTTCACATCANNNAGATCGGAAGA |
| 1234_1 | TATGAGTGTGGAGTCGTTGCCTGTGGTGAATCAAGCAATTCTCNNNAGATCGGAAG |
| 1236_1 | TATGAGTGTGGAGTCGTTGCGAGATCCTTGTTGCTAATGAGCANNNAGATCGGAAG |
| 1238_1 | TATGAGTGTGGAGTCGTTGCTGAAACACCTGCTAACATGAGCANNNAGATCGGAAG |
| 1239_1 | TATGAGTGTGGAGTCGTTGCCTGTGGGCTCTGTTAGGACAGGANNNAGATCGGAA |
| 1240_1 | TATGAGTGTGGAGTCGTTGCTGAAAGAAGGCTCAAGATGAGCANNNAGATCGGAA |
| 1243_1 | TATGAGTGTGGAGTCGTTGCAAGTTTCCCTGTGCCTCTTTCTTNNNAGATCGGAAGA |
| 1244_1 | TATGAGTGTGGAGTCGTTGCGAAATGCAACCATGTACTCCCTCNNNAGATCGGAAG |
| 1247_1 | TATGAGTGTGGAGTCGTTGCGTGCATGCGTTTGTGTGTTTCTTNNNAGATCGGAAG |
| 1248_1 | TATGAGTGTGGAGTCGTTGCTCACTCCTGCCTACCTTCATACCNNNAGATCGGAAGA |
| 1250_1 | TATGAGTGTGGAGTCGTTGCTTGTATGTCTGGGGTAGTGAGCANNNAGATCGGAA |
| 1253_1 | TATGAGTGTGGAGTCGTTGCGGCGGAGCAAAATGTATATGAGANNNAGATCGGAA |
| 1254_1 | TATGAGTGTGGAGTCGTTGCTGGAACAAAACCTCTGCCAAGTTNNNAGATCGGAAG |
| 1255_1 | TATGAGTGTGGAGTCGTTGCTTGCAGCATGTGTCATAATAGCANNNAGATCGGAAG |
| 1256_1 | TATGAGTGTGGAGTCGTTGCGAGTTGGGAGAGAGGGGAAAGAANNNAGATCGGA |
| 1263_1 | TATGAGTGTGGAGTCGTTGCTCAGCATCCTCCTTAGAGAAGTANNNAGATCGGAAG |
| 1265_1 | TATGAGTGTGGAGTCGTTGCAGATAAGCGAAAACCCATTGTGGNNNAGATCGGAA |

| | |
|---|---|
| 1266_1 | TATGAGTGTGGAGTCGTTGCATCCCCAAACCAGATCTATAGCANNNAGATCGGAAG |
| 1267_1 | TATGAGTGTGGAGTCGTTGCAGGTTGGGAGGCATTCTATAGCANNNAGATCGGAA |
| 1269_1 | TATGAGTGTGGAGTCGTTGCAGGCAGAGACAGTCGAGTAAGTTNNNAGATCGGAA |
| 1270_1 | TATGAGTGTGGAGTCGTTGCACTCTGGCTACATCCTGTAAGTTNNNAGATCGGAAG |
| 1271_1 | TATGAGTGTGGAGTCGTTGCTAGAGTTTCCACAAGGCCTAGCANNNAGATCGGAAG |
| 1272_1 | TATGAGTGTGGAGTCGTTGCAGTGAACATCATTTGGCCTAGCANNNAGATCGGAAG |
| 1274_1 | TATGAGTGTGGAGTCGTTGCCCAAGATAGCAGCACAGACAGTTNNNAGATCGGAA |
| 1278_1 | TATGAGTGTGGAGTCGTTGCGTGACCATGGGTTGAAAGTAGCANNNAGATCGGAA |
| 1279_1 | TATGAGTGTGGAGTCGTTGCTCTCCCTTTCTTCCACTCCAGTTNNNAGATCGGAAGA |
| 1280_1 | TATGAGTGTGGAGTCGTTGCCTCCCTCTCAAATCCTGGTAGCANNNAGATCGGAAG |
| 1282_1 | TATGAGTGTGGAGTCGTTGCCGTACACATACCAGCATGTAGCANNNAGATCGGAAG |
| 1284_1 | TATGAGTGTGGAGTCGTTGCGTTCACACCTAGCCATAGCAGTTNNNAGATCGGAAG |
| 1285_1 | TATGAGTGTGGAGTCGTTGCGTCTGGAAGCATCCATTGTAGCANNNAGATCGGAA |
| 1286_1 | TATGAGTGTGGAGTCGTTGCCAGGGACTGAACCAGATGATCCTNNNAGATCGGAA |
| 1287_1 | TATGAGTGTGGAGTCGTTGCCAGATGTTTGTCTCAGGGCAGTTNNNAGATCGGAA |
| 1290_1 | TATGAGTGTGGAGTCGTTGCCCGTGCTCAGCCAATACATTATANNNAGATCGGAAG |
| 1291_1 | TATGAGTGTGGAGTCGTTGCTGTCTGAACCTATCTCAGCCTGTNNNAGATCGGAAG |
| 1292_1 | TATGAGTGTGGAGTCGTTGCGCACTCCTTAGGACTTGTTAGCANNNAGATCGGAAG |
| 1295_1 | TATGAGTGTGGAGTCGTTGCCTTTATGCGATTGTGGGATCTGGNNNAGATCGGAA |
| 1297_1 | TATGAGTGTGGAGTCGTTGCGAGAGAGAAACAAGGCTAACGCANNNAGATCGGA |
| 1298_1 | TATGAGTGTGGAGTCGTTGCAGTGATGACAAAGAGAACACGCANNNAGATCGGAA |
| 1299_1 | TATGAGTGTGGAGTCGTTGCTCAAGCATATCAAATCCCACGCANNNAGATCGGAAG |
| 1302_1 | TATGAGTGTGGAGTCGTTGCTGACATGAGTGGAAGGGAGAGTTNNNAGATCGGA |
| 1304_1 | TATGAGTGTGGAGTCGTTGCGCTCAATGTCTTCTTGGTCTGTGNNNAGATCGGAAG |
| 1307_1 | TATGAGTGTGGAGTCGTTGCGAGAGAGACAGAGATAGAGCGCANNNAGATCGGA |
| 1308_1 | TATGAGTGTGGAGTCGTTGCGGTGCTTTGTACAGACTGGAGTTNNNAGATCGGAA |
| 1309_1 | TATGAGTGTGGAGTCGTTGCAGATGCAACAGGTGCAATGAGTTNNNAGATCGGAA |
| 1310_1 | TATGAGTGTGGAGTCGTTGCAGCCAAAATGCCACAGATGAGTTNNNAGATCGGAA |
| 1311_1 | TATGAGTGTGGAGTCGTTGCGTTCATTGTTGTGCATTCTCGCANNNAGATCGGAAG |
| 1312_1 | TATGAGTGTGGAGTCGTTGCTTCCAGAGGACTGACAGTGAGTTNNNAGATCGGAA |
| 1313_1 | TATGAGTGTGGAGTCGTTGCTCCCTTTGTTTTTCCAAAAGGCANNNAGATCGGAAG |
| 1315_1 | TATGAGTGTGGAGTCGTTGCTGAGCAGGAAGAAAGAAAAGGCANNNAGATCGGA |
| 1316_1 | TATGAGTGTGGAGTCGTTGCAGAATGACACATGGGAAAAGGCANNNAGATCGGA |
| 1317_1 | TATGAGTGTGGAGTCGTTGCAACGGGCGTAGATTCAATCAGATNNNAGATCGGAA |
| 1323_1 | TATGAGTGTGGAGTCGTTGCCTGAGGACAAGCCAAAGCTAGTTNNNAGATCGGAA |
| 1326_1 | TATGAGTGTGGAGTCGTTGCGGGGGCATAGAGAGAGCTTAGTTNNNAGATCGGA |
| 1327_1 | TATGAGTGTGGAGTCGTTGCATGTGCTGACAAATGAGAAGGCANNNAGATCGGAA |
| 1329_1 | TATGAGTGTGGAGTCGTTGCTTCAGACTGGGAAGCCCCACTCTNNNAGATCGGAAG |
| 1331_1 | TATGAGTGTGGAGTCGTTGCGTTAGAAGAACTGGCGTGTCTGCNNNAGATCGGAA |
| 1333_1 | TATGAGTGTGGAGTCGTTGCCTGAGAACACACTAAACACCGTTNNNAGATCGGAAG |
| 1335_1 | TATGAGTGTGGAGTCGTTGCGGGAGAAACAGTAAGGAAGCGTTNNNAGATCGGA |
| 1336_1 | TATGAGTGTGGAGTCGTTGCTTTTCAAGATTCACAGCCAGGCANNNAGATCGGAAG |
| 1338_1 | TATGAGTGTGGAGTCGTTGCGTCCTACCTCGTAAACAAATGCTNNNAGATCGGAAG |

| | |
|---|---|
| 1339_1 | TATGAGTGTGGAGTCGTTGCACCCAGCCATCTTCAAATTCGTTNNNAGATCGGAAGA |
| 1340_1 | TATGAGTGTGGAGTCGTTGCCCCAGCATTTCTCAGAGGAGAACNNNAGATCGGAAG |
| 1349_1 | TATGAGTGTGGAGTCGTTGCATAGAGCAAGTAGCTGAGAGGCANNNAGATCGGAA |
| 1351_1 | TATGAGTGTGGAGTCGTTGCAGTGCTCAGTGTAGTCTCAGGTTNNNAGATCGGAAG |
| 1352_1 | TATGAGTGTGGAGTCGTTGCGGTTTCCCTGCTTGCCTCATTCANNNAGATCGGAAGA |
| 1355_1 | TATGAGTGTGGAGTCGTTGCAGAAACAGGATGGTCAGGAGGTTNNNAGATCGGAA |
| 1356_1 | TATGAGTGTGGAGTCGTTGCAGGTGTGAGTTCAATATGAGGCANNNAGATCGGAA |
| 1358_1 | TATGAGTGTGGAGTCGTTGCACCAAGTATGACCAGCAGGCAATNNNAGATCGGAAG |
| 1359_1 | TATGAGTGTGGAGTCGTTGCCAGCTCCATCTTCTGAATAGGCANNNAGATCGGAAGA |
| 1360_1 | TATGAGTGTGGAGTCGTTGCGGTGCATGGAGACACATTAGGTTNNNAGATCGGAAG |
| 1361_1 | TATGAGTGTGGAGTCGTTGCAAGGTTGAACAGATTCTGCGGTTNNNAGATCGGAAG |
| 1363_1 | TATGAGTGTGGAGTCGTTGCCCTGATCGTAAGTTGGTTCGGTTNNNAGATCGGAAG |
| 1364_1 | TATGAGTGTGGAGTCGTTGCGGAAACACACCAGGTAAAGGGTTNNNAGATCGGAA |
| 1365_1 | TATGAGTGTGGAGTCGTTGCGCTTCCAACTTACTGCATCCTGNNNAGATCGGAAGA |
| 1366_1 | TATGAGTGTGGAGTCGTTGCTTGGGGAGATATCCAGTAATGCGNNNAGATCGGAAG |
| 1368_1 | TATGAGTGTGGAGTCGTTGCATGTTCCAAATTGGTTCCGGGTTNNNAGATCGGAAG |
| 1369_1 | TATGAGTGTGGAGTCGTTGCTCTTTTTCCATGGACTGTAGGCANNNAGATCGGAAGA |
| 1371_1 | TATGAGTGTGGAGTCGTTGCTTGAATACCATCCTGGAGGGGTTNNNAGATCGGAAG |
| 1372_1 | TATGAGTGTGGAGTCGTTGCTCTCCCTAGGCATTTCTTAGGCANNNAGATCGGAAGA |
| 1373_1 | TATGAGTGTGGAGTCGTTGCGCAAGTCTATTTTCCAGGGGGTTNNNAGATCGGAAG |
| 1374_1 | TATGAGTGTGGAGTCGTTGCTGATGGGATGTTTTTCTGGGGTTNNNAGATCGGAAG |
| 1376_1 | TATGAGTGTGGAGTCGTTGCCAACAAGAGCCAGAGAATGGGTTNNNAGATCGGAA |
| 1377_1 | TATGAGTGTGGAGTCGTTGCGAGTAAGTGAATTTTGTCCGGCANNNAGATCGGAAG |
| 1379_1 | TATGAGTGTGGAGTCGTTGCGCAACCTCTACAAACTAAGGGCANNNAGATCGGAAG |
| 1380_1 | TATGAGTGTGGAGTCGTTGCATTGGCTAACATTCTACAGGGCANNNAGATCGGAAG |
| 1381_1 | TATGAGTGTGGAGTCGTTGCTTGCTTCCCTAGATTACAGGGCANNNAGATCGGAAG |
| 1382_1 | TATGAGTGTGGAGTCGTTGCATGATAACTTCCTCGAGAGGGCANNNAGATCGGAAG |
| 1384_1 | TATGAGTGTGGAGTCGTTGCGATGGCTATGAATCTGTGTGGTCNNNAGATCGGAAG |
| 1385_1 | TATGAGTGTGGAGTCGTTGCGACAAGTAGCGGTCTTCATGGTTNNNAGATCGGAAG |
| 1387_1 | TATGAGTGTGGAGTCGTTGCCATACTGATTCTGGCTCAAGGTGNNNAGATCGGAAG |
| 1389_1 | TATGAGTGTGGAGTCGTTGCAGAGAGAGAATAATGTGGGGGCANNNAGATCGGAA |
| 1390_1 | TATGAGTGTGGAGTCGTTGCGCAGTTTCTCCAGTTTCCTGGTTNNNAGATCGGAAGA |
| 1391_1 | TATGAGTGTGGAGTCGTTGCTACCATGCCAGAAAAAGCTGGTTNNNAGATCGGAAG |
| 1394_1 | TATGAGTGTGGAGTCGTTGCCCAAGTGAGTAGAGGGTCTGGTTNNNAGATCGGAA |
| 1395_1 | TATGAGTGTGGAGTCGTTGCAACTTTGTGTTGTATCCTGGGCANNNAGATCGGAAG |
| 1397_1 | TATGAGTGTGGAGTCGTTGCAGAGTAGGAGAAAACAGGTGGTTNNNAGATCGGAA |
| 1398_1 | TATGAGTGTGGAGTCGTTGCTGGTCTATGAAGACAGGGTGGTTNNNAGATCGGAA |
| 1403_1 | TATGAGTGTGGAGTCGTTGCGTTTGCAGATGGCATTTGTGGTTNNNAGATCGGAAG |
| 1404_1 | TATGAGTGTGGAGTCGTTGCAAATGCTGTGGGTAAGAATGGCANNNAGATCGGAA |
| 1405_1 | TATGAGTGTGGAGTCGTTGCGTCTACTCACTTCAGCCTCCCAGNNNAGATCGGAAGA |
| 1407_1 | TATGAGTGTGGAGTCGTTGCAGGAGAGAGAAGATGGAATGGCANNNAGATCGGA |
| 1408_1 | TATGAGTGTGGAGTCGTTGCTTGCACGTTACTGAGGTTTGGTTNNNAGATCGGAAG |
| 1409_1 | TATGAGTGTGGAGTCGTTGCGGGCTTGACTAGAAAACATGGCANNNAGATCGGAAG |

| | |
|---|---|
| 1413_1 | TATGAGTGTGGAGTCGTTGCGGGTAGGGGATCGAGGAAATGTTNNNAGATCGGA |
| 1414_1 | TATGAGTGTGGAGTCGTTGCTAAAGAAAGAAGCCACCATGGCANNNAGATCGGAA |
| 1416_1 | TATGAGTGTGGAGTCGTTGCGCACTTGAATCTTACCCATGGCANNNAGATCGGAAG |
| 1417_1 | TATGAGTGTGGAGTCGTTGCGTGTAAGGATGATGGGGAGAGAANNNAGATCGGA |
| 1418_1 | TATGAGTGTGGAGTCGTTGCAGGGTGTGGATTCATGCAATGTTNNNAGATCGGAA |
| 1420_1 | TATGAGTGTGGAGTCGTTGCCCCAGTGGCTCTAGTTCAATGTTNNNAGATCGGAAG |
| 1421_1 | TATGAGTGTGGAGTCGTTGCGAGGGAGGAGCAGAAAGAATGTTNNNAGATCGGA |
| 1422_1 | TATGAGTGTGGAGTCGTTGCTGGGGGAGGGTTATGAGAATGTTNNNAGATCGGA |
| 1423_1 | TATGAGTGTGGAGTCGTTGCAGCGTTTTCCACATATGAATGTTNNNAGATCGGAAG |
| 1425_1 | TATGAGTGTGGAGTCGTTGCGGTGGTGTCTTCTGAACCATGTTNNNAGATCGGAAG |
| 1426_1 | TATGAGTGTGGAGTCGTTGCGAAGCCACTATGGACACCATGTTNNNAGATCGGAAG |
| 1427_1 | TATGAGTGTGGAGTCGTTGCAAGGTGACATGTTCTGCCATGTTNNNAGATCGGAAG |
| 1428_1 | TATGAGTGTGGAGTCGTTGCCTATCGCCAATATGTGACTGGCANNNAGATCGGAAG |
| 1430_1 | TATGAGTGTGGAGTCGTTGCTGACACTTCTAAACAACCTGGCANNNAGATCGGAAG |
| 1431_1 | TATGAGTGTGGAGTCGTTGCTCCAGAGAAAAAGAGATAGCACANNNAGATCGGAA |
| 1434_1 | TATGAGTGTGGAGTCGTTGCTGGACTGAACATTTTTCCTGGCANNNAGATCGGAAG |
| 1435_1 | TATGAGTGTGGAGTCGTTGCTGAGGAAATAGCAAAAGCTGGCANNNAGATCGGAA |
| 1436_1 | TATGAGTGTGGAGTCGTTGCAAGAGGTTGGGATCCCGTATGTTNNNAGATCGGAA |
| 1437_1 | TATGAGTGTGGAGTCGTTGCCAGTTGGCCCAAAAAGTGTGTGTNNNAGATCGGAA |
| 1439_1 | TATGAGTGTGGAGTCGTTGCGCATGTGTTCTCATTTTCTGGCANNNAGATCGGAAG |
| 1442_1 | TATGAGTGTGGAGTCGTTGCTCAAATCACATATGCAGGTGGCANNNAGATCGGAAG |
| 1445_1 | TATGAGTGTGGAGTCGTTGCGGAGCTTACTTTGTGAGCCTGTTNNNAGATCGGAAG |
| 1446_1 | TATGAGTGTGGAGTCGTTGCAGTTTCTTACGGGGCTATTGACTNNNAGATCGGAAG |
| 1448_1 | TATGAGTGTGGAGTCGTTGCTTTGTGCTGTGGTATGTTGCAGTNNNAGATCGGAAG |
| 1449_1 | TATGAGTGTGGAGTCGTTGCAAGGTGGCTCTGAGTATCCTGTTNNNAGATCGGAAG |
| 1450_1 | TATGAGTGTGGAGTCGTTGCGTCACAAAGGACTGTACTTGGCANNNAGATCGGAAG |
| 1452_1 | TATGAGTGTGGAGTCGTTGCATGAACCAGTGGATGCTGTTACCNNNAGATCGGAAG |
| 1454_1 | TATGAGTGTGGAGTCGTTGCGCAGGTATAGAGACTAGTTGGCANNNAGATCGGAA |
| 1456_1 | TATGAGTGTGGAGTCGTTGCACTCCTCTACTTGTACTTTGGCANNNAGATCGGAAGA |
| 1457_1 | TATGAGTGTGGAGTCGTTGCAGAAGTGAGGTTAGACTGCTGTTNNNAGATCGGAA |
| 1459_1 | TATGAGTGTGGAGTCGTTGCATGAAACTGGGGAAAGTTTGGCANNNAGATCGGAA |
| 1461_1 | TATGAGTGTGGAGTCGTTGCCCCAGGAGCATGAACTATCTGTTNNNAGATCGGAAG |
| 1462_1 | TATGAGTGTGGAGTCGTTGCGAGAGAGACAGGCAGACTACCTTNNNAGATCGGAA |
| 1464_1 | TATGAGTGTGGAGTCGTTGCCTTTCTTCCTGCTGCAGTCTGTTNNNAGATCGGAAGA |
| 1466_1 | TATGAGTGTGGAGTCGTTGCTTTGTGTCTCCTTCCCAAATGCANNNAGATCGGAAGA |
| 1469_1 | TATGAGTGTGGAGTCGTTGCTGTGTGTGTCCTGAAACAGTGTTNNNAGATCGGAAG |
| 1470_1 | TATGAGTGTGGAGTCGTTGCCCCAATTACACACACCAAGTCTANNNAGATCGGAAGA |
| 1471_1 | TATGAGTGTGGAGTCGTTGCTAGAGGCAAAAGGCAACAGTGTTNNNAGATCGGAA |
| 1472_1 | TATGAGTGTGGAGTCGTTGCCATCCACTTTGGCTCACAGTGTTNNNAGATCGGAAG |
| 1474_1 | TATGAGTGTGGAGTCGTTGCGGGAAATGACCTTTGGGAGTGTTNNNAGATCGGAA |
| 1475_1 | TATGAGTGTGGAGTCGTTGCATTTCTCTGCTAGTGTGCGTGTTNNNAGATCGGAAG |
| 1476_1 | TATGAGTGTGGAGTCGTTGCAGAAAGCCTGATGTGTCAATGCANNNAGATCGGAA |
| 1479_1 | TATGAGTGTGGAGTCGTTGCTGTCTCTTTCTGTGCAGAATGCANNNAGATCGGAAG |

| | |
|---|---|
| 1480_1 | TATGAGTGTGGAGTCGTTGCAGAGGCTAAGATGAGTGGGTGTTNNNAGATCGGA |
| 1482_1 | TATGAGTGTGGAGTCGTTGCCTTTTGAATCCCATGGCAGCTGTNNNAGATCGGAAG |
| 1484_1 | TATGAGTGTGGAGTCGTTGCCTCAAAGGGGTGGGGAATGTGTTNNNAGATCGGA |
| 1486_1 | TATGAGTGTGGAGTCGTTGCATTCCCCCATCTGTTCATGTGTTNNNAGATCGGAAGA |
| 1487_1 | TATGAGTGTGGAGTCGTTGCGATGAAATCTTACTCTTGCCCAGNNNAGATCGGAAG |
| 1489_1 | TATGAGTGTGGAGTCGTTGCCGTGCCTGGTGCATATATGTGTTNNNAGATCGGAAG |
| 1490_1 | TATGAGTGTGGAGTCGTTGCTGTGTTATGCCTTCCACTGTGTTNNNAGATCGGAAG |
| 1496_1 | TATGAGTGTGGAGTCGTTGCACCCATCCAGGCAGTTATGAAAGNNNAGATCGGAA |
| 1499_1 | TATGAGTGTGGAGTCGTTGCAAGGACCCAGTGAATGGATGAGANNNAGATCGGA |
| 1500_1 | TATGAGTGTGGAGTCGTTGCCAGATCCAAAGGAGGCTTGTGTTNNNAGATCGGAA |
| 1503_1 | TATGAGTGTGGAGTCGTTGCTATTCGCTAAGCCTTTCCATGCANNNAGATCGGAAG |
| 1504_1 | TATGAGTGTGGAGTCGTTGCTTTCCATGTGATTCCAGCATGCANNNAGATCGGAAG |
| 1507_1 | TATGAGTGTGGAGTCGTTGCCACTGCCTGGGTTGTAGATTGTTNNNAGATCGGAAG |
| 1508_1 | TATGAGTGTGGAGTCGTTGCAAAAGTGTGTCTACCTGCATGCANNNAGATCGGAAG |
| 1509_1 | TATGAGTGTGGAGTCGTTGCTGCTGCTTTCTCAACTGATTGTTNNNAGATCGGAAG |
| 1514_1 | TATGAGTGTGGAGTCGTTGCTAGTTCTCTCCGTCTCTCATGCANNNAGATCGGAAGA |
| 1515_1 | TATGAGTGTGGAGTCGTTGCATGGCACACCTAAACAAGATGCANNNAGATCGGAA |
| 1516_1 | TATGAGTGTGGAGTCGTTGCCAGGAAGACCAAGTGAAGATGCANNNAGATCGGAA |
| 1517_1 | TATGAGTGTGGAGTCGTTGCCTGCCAAGGAACCTCATCTCCGNNNAGATCGGAAGA |
| 1518_1 | TATGAGTGTGGAGTCGTTGCGCAATCCCCATCTTACAGATGCANNNAGATCGGAAG |
| 1521_1 | TATGAGTGTGGAGTCGTTGCCGTGGGAGGATGATTCTGTTGTTNNNAGATCGGAA |
| 1522_1 | TATGAGTGTGGAGTCGTTGCGAAACACTGTAGCAGGAGATGCANNNAGATCGGAA |
| 1525_1 | TATGAGTGTGGAGTCGTTGCGCTGCCTTCCTGTTTTGTTTGTTNNNAGATCGGAAG |
| 1526_1 | TATGAGTGTGGAGTCGTTGCCATCAGGAAAGGTGATGGATGCANNNAGATCGGAA |
| 1527_1 | TATGAGTGTGGAGTCGTTGCTTGGAACAAAAGTCCTGGATGCANNNAGATCGGAA |
| 1529_1 | TATGAGTGTGGAGTCGTTGCAGCATACAGAGCTGCCTTTTGTTNNNAGATCGGAAG |
| 1530_1 | TATGAGTGTGGAGTCGTTGCTTCATTTGTTTGCTTGTGATGCANNNAGATCGGAAG |
| 1531_1 | TATGAGTGTGGAGTCGTTGCTGGGGTACATATGAGGTTTTGTTNNNAGATCGGAA |
| 1533_1 | TATGAGTGTGGAGTCGTTGCCGTACTCCCACAACTCATATGCANNNAGATCGGAAG |
| 1534_1 | TATGAGTGTGGAGTCGTTGCTTGATGGAGCACAGACCCAATTTNNNAGATCGGAAG |
| 1536_1 | TATGAGTGTGGAGTCGTTGCATGACCTCAGACAACGCCAATTTNNNAGATCGGAAG |
| 1539_1 | TATGAGTGTGGAGTCGTTGCAAACAAACCCAATGCTCTATGCANNNAGATCGGAAG |
| 1540_1 | TATGAGTGTGGAGTCGTTGCACACGCACACTTAGTTCTATGCANNNAGATCGGAAG |
| 1541_1 | TATGAGTGTGGAGTCGTTGCTGCTTCCACACAATTTCTCAGGANNNAGATCGGAAG |
| 1542_1 | TATGAGTGTGGAGTCGTTGCCAGCAGCTAGGGAACAGGAATTTNNNAGATCGGAA |
| 1543_1 | TATGAGTGTGGAGTCGTTGCTGATTCCACGAGGACTGGAATTTNNNAGATCGGAA |
| 1544_1 | TATGAGTGTGGAGTCGTTGCTGGGTAGGGGGAGAACTGAATTTNNNAGATCGGA |
| 1545_1 | TATGAGTGTGGAGTCGTTGCCCTGGTGGAGGTTTCCTGAATTTNNNAGATCGGAAG |
| 1546_1 | TATGAGTGTGGAGTCGTTGCGCATGGGTGTGTGTATGTATGCANNNAGATCGGAA |
| 1551_1 | TATGAGTGTGGAGTCGTTGCGTGCGGCCATGTATTTTTATGCANNNAGATCGGAAG |
| 1552_1 | TATGAGTGTGGAGTCGTTGCATGATGTGCACAGAGGCACATTTNNNAGATCGGAA |
| 1554_1 | TATGAGTGTGGAGTCGTTGCGTGGAGGTGCTTGTGTTTATGGANNNAGATCGGAA |
| 1557_1 | TATGAGTGTGGAGTCGTTGCGGGAGGGAAGGAGAATAACTGCANNNAGATCGGA |

| | |
|---|---|
| 1558_1 | TATGAGTGTGGAGTCGTTGCTCCATTCTCTGCAGTGGACATTTNNNAGATCGGAAG |
| 1560_1 | TATGAGTGTGGAGTCGTTGCACGACCTAAATTCTAGCACTGCANNNAGATCGGAAG |
| 1561_1 | TATGAGTGTGGAGTCGTTGCAGGGAGGGAGAATGTCACCATTTNNNAGATCGGAA |
| 1562_1 | TATGAGTGTGGAGTCGTTGCGGGTCAGAGGAGAGAGACCATTTNNNAGATCGGA |
| 1564_1 | TATGAGTGTGGAGTCGTTGCAGAACCAATCAGTTGGCCCATTTNNNAGATCGGAAG |
| 1566_1 | TATGAGTGTGGAGTCGTTGCTCATCTCTGCACCAGTCCCATTTNNNAGATCGGAAGA |
| 1567_1 | TATGAGTGTGGAGTCGTTGCAAAGGCAGGTAATACTGACTGCANNNAGATCGGAA |
| 1569_1 | TATGAGTGTGGAGTCGTTGCGTGTACTAGAGTGCCTGCCATTTNNNAGATCGGAAG |
| 1571_1 | TATGAGTGTGGAGTCGTTGCCCAACAGCCTCACATTTGTTCTTNNNAGATCGGAAGA |
| 1574_1 | TATGAGTGTGGAGTCGTTGCTCTCTTTCTCTCCTTTTGGCTGTNNNAGATCGGAAGA |
| 1576_1 | TATGAGTGTGGAGTCGTTGCCTCAGCTTTGTCAGAGAGCATTTNNNAGATCGGAAG |
| 1578_1 | TATGAGTGTGGAGTCGTTGCCCCAAGGCTAGAATCAGGCATTTNNNAGATCGGAAG |
| 1579_1 | TATGAGTGTGGAGTCGTTGCCAGCTAGAGCCCTCATGAAGGAGNNNAGATCGGAA |
| 1580_1 | TATGAGTGTGGAGTCGTTGCTCCAGAAAGCAAGACAATGCTGTNNNAGATCGGAA |
| 1581_1 | TATGAGTGTGGAGTCGTTGCACATTTTCAGTTGACCAGCTGCANNNAGATCGGAAG |
| 1583_1 | TATGAGTGTGGAGTCGTTGCTTTCTTTAGATGAGCAGGCTGCANNNAGATCGGAAG |
| 1584_1 | TATGAGTGTGGAGTCGTTGCCAGCCAAGATGGACCATCATTTNNNAGATCGGAAG |
| 1586_1 | TATGAGTGTGGAGTCGTTGCTCCCTCTATTATTCCTGGCTGCANNNAGATCGGAAGA |
| 1587_1 | TATGAGTGTGGAGTCGTTGCATCTCACATCTCACTCTGTGGCCNNNAGATCGGAAG |
| 1589_1 | TATGAGTGTGGAGTCGTTGCTTCTCTGGTCTTTTGCATCTGCANNNAGATCGGAAG |
| 1590_1 | TATGAGTGTGGAGTCGTTGCTTTGTTGGAGTGGGTGTGGATGTNNNAGATCGGAA |
| 1593_1 | TATGAGTGTGGAGTCGTTGCTCTCCATCCCACTCTCCAGATTTNNNAGATCGGAAGA |
| 1594_1 | TATGAGTGTGGAGTCGTTGCTGGCCTTTGGACTTTGCAGATTTNNNAGATCGGAAG |
| 1595_1 | TATGAGTGTGGAGTCGTTGCCCAAATGCCTGCTCCAGAGATTTNNNAGATCGGAAG |
| 1596_1 | TATGAGTGTGGAGTCGTTGCTTCATGGGCAGACACTGAGATTTNNNAGATCGGAA |
| 1597_1 | TATGAGTGTGGAGTCGTTGCTGGTCAACTTACTAGGCCGATTTNNNAGATCGGAAG |
| 1598_1 | TATGAGTGTGGAGTCGTTGCGAGGCTGCAGGACTTTAGGATTTNNNAGATCGGAA |
| 1599_1 | TATGAGTGTGGAGTCGTTGCCCTGGCAGTCATCTCTCCAACTCNNNAGATCGGAAGA |
| 1600_1 | TATGAGTGTGGAGTCGTTGCAGAGAAGCCAGCAGTTTGGATTTNNNAGATCGGAA |
| 1601_1 | TATGAGTGTGGAGTCGTTGCAGAGAAAAGACACACCAAGTGCANNNAGATCGGAA |
| 1602_1 | TATGAGTGTGGAGTCGTTGCCTAAGCTCTTCCCTGAAGGTCCTNNNAGATCGGAAG |
| 1603_1 | TATGAGTGTGGAGTCGTTGCCCTTGCCTGGACTTGTCTGATTTNNNAGATCGGAAG |
| 1604_1 | TATGAGTGTGGAGTCGTTGCCCCCCATCTTCCCATTCTGATTTNNNAGATCGGAAGA |
| 1605_1 | TATGAGTGTGGAGTCGTTGCAGACAGACGGGAACAGACAAAGGNNNAGATCGGA |
| 1606_1 | TATGAGTGTGGAGTCGTTGCGAGAGAAAGACAGCATGAGTGCANNNAGATCGGA |
| 1607_1 | TATGAGTGTGGAGTCGTTGCACAGTAAGGAACTGATGAGTGCANNNAGATCGGAA |
| 1608_1 | TATGAGTGTGGAGTCGTTGCGTGCTCCTGCTTCAGAGGAACTANNNAGATCGGAAG |
| 1609_1 | TATGAGTGTGGAGTCGTTGCTGGCTTTACCACTTACTAGTGCANNNAGATCGGAAG |
| 1610_1 | TATGAGTGTGGAGTCGTTGCCCGTGACCTGAGATGTGTGATTTNNNAGATCGGAA |
| 1612_1 | TATGAGTGTGGAGTCGTTGCATGGAATGGAACCCCAAAACTGCNNNAGATCGGAA |
| 1614_1 | TATGAGTGTGGAGTCGTTGCTGACATAGAGAGAGAGACGTGCANNNAGATCGGA |
| 1615_1 | TATGAGTGTGGAGTCGTTGCAGCCTGGGAGAGACCACATATTTNNNAGATCGGAA |
| 1616_1 | TATGAGTGTGGAGTCGTTGCGGTATTGGAAGCTGTATGGTGCANNNAGATCGGAA |

| | |
|---|---|
| 1617_1 | TATGAGTGTGGAGTCGTTGCCCCACCGCCTTTCTCTCATATTTNNNAGATCGGAAGA |
| 1618_1 | TATGAGTGTGGAGTCGTTGCAAAGTACAAATCGGTCTGGTGCANNNAGATCGGAA |
| 1619_1 | TATGAGTGTGGAGTCGTTGCTCTTTGCAATATGGTGTGGTGCANNNAGATCGGAA |
| 1620_1 | TATGAGTGTGGAGTCGTTGCAACTGCCCTCTTATTTTGGTGCANNNAGATCGGAAG |
| 1621_1 | TATGAGTGTGGAGTCGTTGCCCACTGTAAGCCACAGGCTATTTNNNAGATCGGAAG |
| 1622_1 | TATGAGTGTGGAGTCGTTGCTGCCCCCTCTAATCAGGCTATTTNNNAGATCGGAAG |
| 1623_1 | TATGAGTGTGGAGTCGTTGCCAGACATTATGGGGGAATGTGCANNNAGATCGGAA |
| 1624_1 | TATGAGTGTGGAGTCGTTGCCAGGTGCCAGGGTAGTTCTATTTNNNAGATCGGAA |
| 1625_1 | TATGAGTGTGGAGTCGTTGCGGGTGTGTGTGTCCTGAGTATTTNNNAGATCGGAA |
| 1627_1 | TATGAGTGTGGAGTCGTTGCGGGAGGTGTACAATTCATGTGCANNNAGATCGGAA |
| 1628_1 | TATGAGTGTGGAGTCGTTGCGAAGGTGGTTGGTGGTGGTATTTNNNAGATCGGA |
| 1630_1 | TATGAGTGTGGAGTCGTTGCGTCACGCCCAGTCATCTGTATTTNNNAGATCGGAAG |
| 1631_1 | TATGAGTGTGGAGTCGTTGCTGGGAATGTTACAAATCTGTGCANNNAGATCGGAA |
| 1632_1 | TATGAGTGTGGAGTCGTTGCCCCCCAGAAGGGTGGATTTATTTNNNAGATCGGAAG |
| 1635_1 | TATGAGTGTGGAGTCGTTGCTCAATCTTGTGAGGTTGTGTGCANNNAGATCGGAA |
| 1636_1 | TATGAGTGTGGAGTCGTTGCACCTCTCCTGAGCTCTGAACTTTNNNAGATCGGAAG |
| 1643_1 | TATGAGTGTGGAGTCGTTGCTCTCCCCATAGCTCTCTCACTTTNNNAGATCGGAAGA |
| 1648_1 | TATGAGTGTGGAGTCGTTGCTCACACATTTCCTGGGAATTGCANNNAGATCGGAAG |
| 1649_1 | TATGAGTGTGGAGTCGTTGCTGAAGGTCAGAGAGTGGGACTTTNNNAGATCGGAA |
| 1653_1 | TATGAGTGTGGAGTCGTTGCAGGGGGTGGTTATGAGTGACTTTNNNAGATCGGAA |
| 1655_1 | TATGAGTGTGGAGTCGTTGCAGTGTTTCCAGCCCTTTGACTTTNNNAGATCGGAAG |
| 1656_1 | TATGAGTGTGGAGTCGTTGCGGCTGGTGGGTACCAAATACTTTNNNAGATCGGAA |
| 1662_1 | TATGAGTGTGGAGTCGTTGCACTTATTTGGTCGGTGCACCTTTNNNAGATCGGAAG |
| 1664_1 | TATGAGTGTGGAGTCGTTGCAACAGGATACTGGAGGACTTGCANNNAGATCGGAA |
| 1665_1 | TATGAGTGTGGAGTCGTTGCTTCACTGAGCAGCAATACTTGCANNNAGATCGGAAG |
| 1668_1 | TATGAGTGTGGAGTCGTTGCAAACCCTTCAATGCCTCTGTGTTNNNAGATCGGAAG |
| 1671_1 | TATGAGTGTGGAGTCGTTGCTTTCCCTGTGGAATCATCTTGCANNNAGATCGGAAG |
| 1672_1 | TATGAGTGTGGAGTCGTTGCAGTGTCTCTAGCAGCTGAAAGTANNNAGATCGGAA |
| 1673_1 | TATGAGTGTGGAGTCGTTGCACATTTGGTGAGGGGTGAGAGATNNNAGATCGGA |
| 1674_1 | TATGAGTGTGGAGTCGTTGCCTCAGCCTTCCTTTTGTCCCTTTNNNAGATCGGAAGA |
| 1677_1 | TATGAGTGTGGAGTCGTTGCGCACACCCTATTTTTGAGTTGCANNNAGATCGGAAG |
| 1678_1 | TATGAGTGTGGAGTCGTTGCTCAGACAGAAAGACTAGGTTGCANNNAGATCGGAA |
| 1680_1 | TATGAGTGTGGAGTCGTTGCTCTCTTTGGGTTCCTCTCTCTNNNAGATCGGAAGAGC |
| 1681_1 | TATGAGTGTGGAGTCGTTGCGTGGGAGTTGTATGCCATCCTTTNNNAGATCGGAAG |
| 1683_1 | TATGAGTGTGGAGTCGTTGCACATCCACCATCTTCCCTCCTTTNNNAGATCGGAAGA |
| 1684_1 | TATGAGTGTGGAGTCGTTGCTGCTGGGAGATGTTTCATTTGCANNNAGATCGGAA |
| 1686_1 | TATGAGTGTGGAGTCGTTGCTATTCGCAGCCTAACACTTTGCANNNAGATCGGAAG |
| 1687_1 | TATGAGTGTGGAGTCGTTGCTCTGGCCTTTCCTCCTTTCCTTTNNNAGATCGGAAGA |
| 1688_1 | TATGAGTGTGGAGTCGTTGCTTCTAGGTGTCAGAGGGCTATCTNNNAGATCGGAA |
| 1689_1 | TATGAGTGTGGAGTCGTTGCAAAGGGCAAGTGAATCAGAGGAANNNAGATCGGA |
| 1691_1 | TATGAGTGTGGAGTCGTTGCTCTTCTGATGGAGTTGCTTTGCANNNAGATCGGAAG |
| 1693_1 | TATGAGTGTGGAGTCGTTGCGCATACCTTGGGAGTTCAGCTTTNNNAGATCGGAAG |
| 1694_1 | TATGAGTGTGGAGTCGTTGCTCCACCCACTGAAATTTTGCTGTNNNAGATCGGAAG |

| | |
|---|---|
| 1695_1 | TATGAGTGTGGAGTCGTTGCAGAGAGCACTGCATTAGTTTGCANNNAGATCGGAA |
| 1696_1 | TATGAGTGTGGAGTCGTTGCGAGTATTTGTTCCAGGACCCCTANNNAGATCGGAAG |
| 1698_1 | TATGAGTGTGGAGTCGTTGCTGTAAATCCCGCATGCTAGCTTTNNNAGATCGGAAG |
| 1699_1 | TATGAGTGTGGAGTCGTTGCAAGTGAGAAGTCTGCTGTTTGCANNNAGATCGGAA |
| 1702_1 | TATGAGTGTGGAGTCGTTGCCAGTTTGTAATCACATTCACGCANNNAGATCGGAAG |
| 1703_1 | TATGAGTGTGGAGTCGTTGCGGCAGGGTGTCTTGATTAGCTTTNNNAGATCGGAA |
| 1704_1 | TATGAGTGTGGAGTCGTTGCGCAGGAAATAAGCAGCTTTTGCANNNAGATCGGAA |
| 1705_1 | TATGAGTGTGGAGTCGTTGCGGGCCTCTAAAGTAGCTTTTGCANNNAGATCGGAAG |
| 1706_1 | TATGAGTGTGGAGTCGTTGCCATAAGGCCACTGGAAAGGCTTTNNNAGATCGGAA |
| 1707_1 | TATGAGTGTGGAGTCGTTGCGAGAGGTGAGGATGGAAGGCTTTNNNAGATCGGA |
| 1709_1 | TATGAGTGTGGAGTCGTTGCATTTTCCCTGTTGCTGAGGCTTTNNNAGATCGGAAG |
| 1710_1 | TATGAGTGTGGAGTCGTTGCGCCTAAGAAAGACAGTCGGCTTTNNNAGATCGGAA |
| 1711_1 | TATGAGTGTGGAGTCGTTGCTGTTGAAGCGAAATGGAAAATCANNNAGATCGGAA |
| 1712_1 | TATGAGTGTGGAGTCGTTGCTAGATGAGAAGGAGCAGGGCTTTNNNAGATCGGA |
| 1713_1 | TATGAGTGTGGAGTCGTTGCTAGGAGCATAATCCCTGGGCTTTNNNAGATCGGAAG |
| 1714_1 | TATGAGTGTGGAGTCGTTGCAGCCACAGATTCTGCTGAAATCANNNAGATCGGAAG |
| 1715_1 | TATGAGTGTGGAGTCGTTGCGCCAACCCTGGTACCATAAATCANNNAGATCGGAAG |
| 1722_1 | TATGAGTGTGGAGTCGTTGCTGCCATCCCATCCAAATACATTTNNNAGATCGGAAGA |
| 1724_1 | TATGAGTGTGGAGTCGTTGCTGTAATTCGGGACAGCTTGCTTTNNNAGATCGGAAG |
| 1725_1 | TATGAGTGTGGAGTCGTTGCTCACTCAGCATTTGCTTTGCTTTNNNAGATCGGAAGA |
| 1726_1 | TATGAGTGTGGAGTCGTTGCCTGTCCCTTTGGCCACATAATCANNNAGATCGGAAG |
| 1728_1 | TATGAGTGTGGAGTCGTTGCAGGGGTTCATGCCTGTAATCTTTNNNAGATCGGAAG |
| 1729_1 | TATGAGTGTGGAGTCGTTGCCATACAGCCTCAGCCACATCTTTNNNAGATCGGAAG |
| 1730_1 | TATGAGTGTGGAGTCGTTGCCCCTCACATACACAGACATCTTTNNNAGATCGGAAGA |
| 1734_1 | TATGAGTGTGGAGTCGTTGCGCGGCCAAAAATAGTCAACATCANNNAGATCGGAA |
| 1735_1 | TATGAGTGTGGAGTCGTTGCGGGGGTGGGGTTGTTTTATCTTTNNNAGATCGGAA |
| 1736_1 | TATGAGTGTGGAGTCGTTGCCTCCATTCCCGTTTCTTTTCCTCNNNAGATCGGAAGA |
| 1737_1 | TATGAGTGTGGAGTCGTTGCCAGTCCCAATAGCCAAAGCTGAANNNAGATCGGAAG |
| 1738_1 | TATGAGTGTGGAGTCGTTGCTCCCACCTCATCTTTCCCTCTTTNNNAGATCGGAAGAG |
| 1739_1 | TATGAGTGTGGAGTCGTTGCATTTTCTCTTGCCTGGCCTCTTTNNNAGATCGGAAGA |
| 1740_1 | TATGAGTGTGGAGTCGTTGCCAGTGTCACCTTGAAGGCTCTTTNNNAGATCGGAAG |
| 1741_1 | TATGAGTGTGGAGTCGTTGCATGCCTGATGCCTAGCTACATCANNNAGATCGGAAG |
| 1743_1 | TATGAGTGTGGAGTCGTTGCACTGTCCTCTTACTGGACCATCANNNAGATCGGAAG |
| 1744_1 | TATGAGTGTGGAGTCGTTGCGCTGTATTGTACCAAACCCATCANNNAGATCGGAAG |
| 1745_1 | TATGAGTGTGGAGTCGTTGCGAATGGTCACCTGCTGTCTCTTTNNNAGATCGGAAG |
| 1746_1 | TATGAGTGTGGAGTCGTTGCGCACACATCCATACATCCCATCANNNAGATCGGAAGA |
| 1747_1 | TATGAGTGTGGAGTCGTTGCGGATGACAGAAGCCACAGTCTTTNNNAGATCGGAA |
| 1748_1 | TATGAGTGTGGAGTCGTTGCTTAGTCTGATACTGGGGCCATCANNNAGATCGGAAG |
| 1751_1 | TATGAGTGTGGAGTCGTTGCGAACCTCTACGCAACCTGTCTTTNNNAGATCGGAAG |
| 1752_1 | TATGAGTGTGGAGTCGTTGCCACCTCCCTCCAAGATCCTAGCNNNAGATCGGAAGA |
| 1754_1 | TATGAGTGTGGAGTCGTTGCGCTACTTTAGAGAGGGGGCATCANNNAGATCGGAA |
| 1755_1 | TATGAGTGTGGAGTCGTTGCTGACTGGCTGAAAGGCATTCTTTNNNAGATCGGAAG |
| 1756_1 | TATGAGTGTGGAGTCGTTGCTGATTCTGGTGCTCTTTGCATCANNNAGATCGGAAG |

| | |
|---|---|
| 1757_1 | TATGAGTGTGGAGTCGTTGCCTCTCCTTCCCTCCACCTTCTTTNNNAGATCGGAAGAG |
| 1758_1 | TATGAGTGTGGAGTCGTTGCTCCCAAAGGGCTTCTACTCATCANNNAGATCGGAAG |
| 1763_1 | TATGAGTGTGGAGTCGTTGCTGACCTCTCCTTCACAGTCATCANNNAGATCGGAAG |
| 1765_1 | TATGAGTGTGGAGTCGTTGCGCCTCAGCCACAGAAATTCATCANNNAGATCGGAAG |
| 1766_1 | TATGAGTGTGGAGTCGTTGCACCTTTTGGTGCTCTGTTCATCANNNAGATCGGAAG |
| 1768_1 | TATGAGTGTGGAGTCGTTGCCAGACCAAAGACAGGCAAGATCANNNAGATCGGAA |
| 1769_1 | TATGAGTGTGGAGTCGTTGCTCTGCAACTGTGGAAACAAGTTTNNNAGATCGGAAG |
| 1770_1 | TATGAGTGTGGAGTCGTTGCGGACTAGGAGGTCAGGGAAGTTTNNNAGATCGGA |
| 1773_1 | TATGAGTGTGGAGTCGTTGCCAAAAGCAACGAGCCCTGTTAGTNNNAGATCGGAA |
| 1776_1 | TATGAGTGTGGAGTCGTTGCTCACATTTGGAACCCCACAGTTTNNNAGATCGGAAG |
| 1777_1 | TATGAGTGTGGAGTCGTTGCAAACGTCAAAATCCACAGGATCANNNAGATCGGAAG |
| 1779_1 | TATGAGTGTGGAGTCGTTGCTTTTTCCCTTGGTTTCTGGATCANNNAGATCGGAAG |
| 1780_1 | TATGAGTGTGGAGTCGTTGCAGTTTGAAGCACAGCCTCAGTTTNNNAGATCGGAAG |
| 1782_1 | TATGAGTGTGGAGTCGTTGCGGGGAACATGTCTGCTTCAGTTTNNNAGATCGGAA |
| 1783_1 | TATGAGTGTGGAGTCGTTGCAGGGGAAGACAGGGAAAGAGTTTNNNAGATCGGA |
| 1784_1 | TATGAGTGTGGAGTCGTTGCTTCTGACAGTCCTCATCTGACCCNNNAGATCGGAAG |
| 1787_1 | TATGAGTGTGGAGTCGTTGCACCCCTTACCATGAGTTGAGTTTNNNAGATCGGAAG |
| 1788_1 | TATGAGTGTGGAGTCGTTGCATCTGGCTGGCTGAAGGATATCANNNAGATCGGAA |
| 1790_1 | TATGAGTGTGGAGTCGTTGCCGCTCCCTTTCCAGTGTTAGTTTNNNAGATCGGAAG |
| 1791_1 | TATGAGTGTGGAGTCGTTGCGTGGCCCGTAGTCTGTTTAGTTTNNNAGATCGGAAG |
| 1793_1 | TATGAGTGTGGAGTCGTTGCCAATTCTCCCCTTGTGTGGAGACNNNAGATCGGAAG |
| 1794_1 | TATGAGTGTGGAGTCGTTGCTTGCCTTACCCTGTCCTGTATCANNNAGATCGGAAG |
| 1799_1 | TATGAGTGTGGAGTCGTTGCTGTTTTTCCCAGCACGTAGGTTTNNNAGATCGGAAG |
| 1800_1 | TATGAGTGTGGAGTCGTTGCGAGCCCCGTGATTTTGTAGGTTTNNNAGATCGGAAG |
| 1801_1 | TATGAGTGTGGAGTCGTTGCCCCAGGTATTATGTGTGCGGTTTNNNAGATCGGAAG |
| 1802_1 | TATGAGTGTGGAGTCGTTGCTGAAGGGCTAGGTTTTTCGGTTTNNNAGATCGGAA |
| 1806_1 | TATGAGTGTGGAGTCGTTGCGATTACAGTCACTGGCAGGGTTTNNNAGATCGGAA |
| 1807_1 | TATGAGTGTGGAGTCGTTGCTTCACTTTTTGGTCTCGGGGTTTNNNAGATCGGAAG |
| 1809_1 | TATGAGTGTGGAGTCGTTGCGGGAGAGGAATTGTTTGGGGTTTNNNAGATCGGA |
| 1810_1 | TATGAGTGTGGAGTCGTTGCCCAATACCTCCTCCCCTAACTCANNNAGATCGGAAGA |
| 1812_1 | TATGAGTGTGGAGTCGTTGCTGCTGGTAACTCATTGTGGGTTTNNNAGATCGGAAG |
| 1814_1 | TATGAGTGTGGAGTCGTTGCGCAGGCTTGTCACCATTATCAATNNNAGATCGGAAG |
| 1816_1 | TATGAGTGTGGAGTCGTTGCTGAGGATATTTAGGGCCCACTCANNNAGATCGGAA |
| 1818_1 | TATGAGTGTGGAGTCGTTGCAATGTAGCTTTCACGGGTGGTTTNNNAGATCGGAA |
| 1821_1 | TATGAGTGTGGAGTCGTTGCTTACTGGGTCGGGGGATTGCGATNNNAGATCGGAA |
| 1823_1 | TATGAGTGTGGAGTCGTTGCTCCCAAGAATGTCAGTTTGGTTTNNNAGATCGGAAG |
| 1824_1 | TATGAGTGTGGAGTCGTTGCCCAGATGTAGGAAAAGTGGCAAANNNAGATCGGAA |
| 1826_1 | TATGAGTGTGGAGTCGTTGCAATAGCCACTTCAGCTTTCATCTNNNAGATCGGAAG |
| 1827_1 | TATGAGTGTGGAGTCGTTGCCCACACTCACCCCTAGAATGTTTNNNAGATCGGAAG |
| 1828_1 | TATGAGTGTGGAGTCGTTGCGCTGCTGGCTAATCAGTTTACCCNNNAGATCGGAAG |
| 1830_1 | TATGAGTGTGGAGTCGTTGCGTGCCTGGCCTATTGATCTAAATNNNAGATCGGAAG |
| 1832_1 | TATGAGTGTGGAGTCGTTGCTGACAGTAGTCTCCATGCTGTTTNNNAGATCGGAAG |
| 1834_1 | TATGAGTGTGGAGTCGTTGCCTCCAAAACAGGCCTCTCTGTTTNNNAGATCGGAAG |

| | |
|---|---|
| 1836_1 | TATGAGTGTGGAGTCGTTGCCTGGGGGAGAAGGAACTTACTCANNNAGATCGGAA |
| 1837_1 | TATGAGTGTGGAGTCGTTGCGAGAATCTTGCTTGTTACCCAGGNNNAGATCGGAAG |
| 1839_1 | TATGAGTGTGGAGTCGTTGCAACAGACACACACACTACCAACCNNNAGATCGGAAG |
| 1840_1 | TATGAGTGTGGAGTCGTTGCGCAGAATACAGTGGCCTGTGTTTNNNAGATCGGAA |
| 1842_1 | TATGAGTGTGGAGTCGTTGCGACTTCCCAACTCCAGTGTGTTTNNNAGATCGGAAG |
| 1843_1 | TATGAGTGTGGAGTCGTTGCCTGGGTCAAGCAATCCTACCTCANNNAGATCGGAAG |
| 1844_1 | TATGAGTGTGGAGTCGTTGCCAAAGCACTAACCATTCCCCTCANNNAGATCGGAAG |
| 1845_1 | TATGAGTGTGGAGTCGTTGCTCTCCATTTCCTTTTCTCCCTCANNNAGATCGGAAGAG |
| 1847_1 | TATGAGTGTGGAGTCGTTGCCCCCAGGAACTTCCCACAATTTTNNNAGATCGGAAG |
| 1852_1 | TATGAGTGTGGAGTCGTTGCCATCTCTCACCACTTTTGCCTCANNNAGATCGGAAGA |
| 1854_1 | TATGAGTGTGGAGTCGTTGCGAAGAAACCCCCAGGGACATTTTNNNAGATCGGAA |
| 1856_1 | TATGAGTGTGGAGTCGTTGCTTATGCTGTCAGGAGTGAACTTGNNNAGATCGGAA |
| 1858_1 | TATGAGTGTGGAGTCGTTGCCAGTTACCAAACCTGCGCATTTTNNNAGATCGGAAG |
| 1862_1 | TATGAGTGTGGAGTCGTTGCTTGCCTGTTATCAGTGTAGTCTTNNNAGATCGGAAG |
| 1864_1 | TATGAGTGTGGAGTCGTTGCGAAAGGGAGGCAGTCAGGATTTTNNNAGATCGGA |
| 1866_1 | TATGAGTGTGGAGTCGTTGCTGCTTGTTGAGGTTGGGGATTTTNNNAGATCGGAA |
| 1868_1 | TATGAGTGTGGAGTCGTTGCCAGATAGTTGCTGCAGTTCCTCANNNAGATCGGAAG |
| 1872_1 | TATGAGTGTGGAGTCGTTGCCACAAGGTAGCTTTCCAAGCTCANNNAGATCGGAAG |
| 1875_1 | TATGAGTGTGGAGTCGTTGCTCATGGTGATCGGCAAAGTACAGNNNAGATCGGAA |
| 1876_1 | TATGAGTGTGGAGTCGTTGCATGCACGCGAAAGGGATTATTTTNNNAGATCGGAA |
| 1878_1 | TATGAGTGTGGAGTCGTTGCCTTTTTCCCTCTGTGTGAGCTCANNNAGATCGGAAG |
| 1881_1 | TATGAGTGTGGAGTCGTTGCACAGCTTGACATTCCCGACTTTTNNNAGATCGGAAG |
| 1883_1 | TATGAGTGTGGAGTCGTTGCGAATGGCTAGAAATGTGACCAAANNNAGATCGGAA |
| 1884_1 | TATGAGTGTGGAGTCGTTGCCACCAACCTGGCCTGCTTTTTNNNAGATCGGAAGAG |
| 1885_1 | TATGAGTGTGGAGTCGTTGCCCTAGTGGGAGGACTAGCCTTTTNNNAGATCGGAA |
| 1887_1 | TATGAGTGTGGAGTCGTTGCGTTGTATTTGTGCCGTGCCTTTTNNNAGATCGGAAG |
| 1889_1 | TATGAGTGTGGAGTCGTTGCTCCCACTCTGCTTCTTTCTCTGTNNNAGATCGGAAGA |
| 1890_1 | TATGAGTGTGGAGTCGTTGCGCTAGTGGCAGTCATGTCCTTTTNNNAGATCGGAAG |
| 1894_1 | TATGAGTGTGGAGTCGTTGCAGCAGACCTTGTTTGTATGCTCANNNAGATCGGAAG |
| 1898_1 | TATGAGTGTGGAGTCGTTGCCATACTGATCTCAGAGCTGCTCANNNAGATCGGAAG |
| 1900_1 | TATGAGTGTGGAGTCGTTGCTGTAGGGAAGCAAGTTGGCTTTTNNNAGATCGGAA |
| 1901_1 | TATGAGTGTGGAGTCGTTGCTGGTGTCTTTCCTTCACAGGGCTNNNAGATCGGAAG |
| 1902_1 | TATGAGTGTGGAGTCGTTGCAACTGCTTCAATGCCATGCTTTTNNNAGATCGGAAG |
| 1903_1 | TATGAGTGTGGAGTCGTTGCATCTAGCACAAAGGAGGTGCTCANNNAGATCGGAA |
| 1904_1 | TATGAGTGTGGAGTCGTTGCCACAATATGGGCAACCTGCTTTTNNNAGATCGGAAG |
| 1905_1 | TATGAGTGTGGAGTCGTTGCAGCAGGAAAGAGAAACTTGCTCANNNAGATCGGAA |
| 1907_1 | TATGAGTGTGGAGTCGTTGCCAGGTACGAGAGATGGTGCTTTTNNNAGATCGGAA |
| 1911_1 | TATGAGTGTGGAGTCGTTGCATTCTCTAGGCAGGCCAATCTCANNNAGATCGGAAG |
| 1913_1 | TATGAGTGTGGAGTCGTTGCGGGGGCGGTTAGAATTCTCTTTTNNNAGATCGGAA |
| 1914_1 | TATGAGTGTGGAGTCGTTGCTTCCCCAAAGAGCAAAGTCTTTTNNNAGATCGGAAG |
| 1915_1 | TATGAGTGTGGAGTCGTTGCCCACCTATGCCTCTGTGTCTTTTNNNAGATCGGAAGA |
| 1916_1 | TATGAGTGTGGAGTCGTTGCGCTCTCCTTCCACAGGTTCTTTTNNNAGATCGGAAGA |
| 1917_1 | TATGAGTGTGGAGTCGTTGCTCGTGTGATCTCCCTTCATCTCANNNAGATCGGAAGA |

| | |
|---|---|
| 1918_1 | TATGAGTGTGGAGTCGTTGCGGGGCAGGGTATGTCTTTCTTTTNNNAGATCGGAAG |
| 1919_1 | TATGAGTGTGGAGTCGTTGCGAGCTATGGCAGAGAGGATCTCANNNAGATCGGAA |
| 1920_1 | TATGAGTGTGGAGTCGTTGCAGGGAGTGGCATAATGGATCTCANNNAGATCGGAA |
| 1921_1 | TATGAGTGTGGAGTCGTTGCGCAGGGAACCAGAGGAAAATGAGNNNAGATCGGA |
| 1922_1 | TATGAGTGTGGAGTCGTTGCGGACAGACCCAACTCCAAGTTTTNNNAGATCGGAAG |
| 1923_1 | TATGAGTGTGGAGTCGTTGCGGTTGTAAGATGCAGCACTCTCANNNAGATCGGAAG |
| 1926_1 | TATGAGTGTGGAGTCGTTGCTAATTGCCCAGACATCCCTCTCANNNAGATCGGAAGA |
| 1927_1 | TATGAGTGTGGAGTCGTTGCGGCATCATCTCTCAGCCAGTTTTNNNAGATCGGAAGA |
| 1928_1 | TATGAGTGTGGAGTCGTTGCGCAATTCAGGAAGACAGCTCTCANNNAGATCGGAAG |
| 1929_1 | TATGAGTGTGGAGTCGTTGCAGACACACAGATACCAGCTCTCANNNAGATCGGAAG |
| 1930_1 | TATGAGTGTGGAGTCGTTGCCATCCAAATCACACTGGCTCTCANNNAGATCGGAAGA |
| 1932_1 | TATGAGTGTGGAGTCGTTGCAAGATGGACGGTGTTAGCTAGGANNNAGATCGGAA |
| 1933_1 | TATGAGTGTGGAGTCGTTGCTGGGCTAGGATGCAAGTAGTTTTNNNAGATCGGAAG |
| 1934_1 | TATGAGTGTGGAGTCGTTGCAGTTCCAGAGGGGTGGTAGTTTTNNNAGATCGGAAG |
| 1935_1 | TATGAGTGTGGAGTCGTTGCCTCCTCTCCTTCACTCTCTCTCANNNAGATCGGAAGAG |
| 1937_1 | TATGAGTGTGGAGTCGTTGCCCGATCCCACAAGATGAGGTTTTNNNAGATCGGAAG |
| 1938_1 | TATGAGTGTGGAGTCGTTGCACCCATATTCCTGGGAAGTCTCANNNAGATCGGAAGA |
| 1941_1 | TATGAGTGTGGAGTCGTTGCGTGTTTGAACTGCAGTGGGTTTTNNNAGATCGGAAG |
| 1943_1 | TATGAGTGTGGAGTCGTTGCTGTGTGTGTGCTTGTATGGTTTTNNNAGATCGGAAG |
| 1944_1 | TATGAGTGTGGAGTCGTTGCACCAAGGAGAATATCTGGTCTCANNNAGATCGGAAG |
| 1945_1 | TATGAGTGTGGAGTCGTTGCGGTAGGGCAACATCTCTGGTTTTNNNAGATCGGAAG |
| 1946_1 | TATGAGTGTGGAGTCGTTGCTGAATAGCCAGCACCTTGGTTTTNNNAGATCGGAAG |
| 1950_1 | TATGAGTGTGGAGTCGTTGCGTGGGGTGAGTGGATCATGTTTTNNNAGATCGGAA |
| 1951_1 | TATGAGTGTGGAGTCGTTGCGTTTCTTGGGCTCACCATTCTCANNNAGATCGGAAGA |
| 1953_1 | TATGAGTGTGGAGTCGTTGCAGGGAAGATGGTGGTCATTCTCANNNAGATCGGAAG |
| 1957_1 | TATGAGTGTGGAGTCGTTGCTTGAAATTGGCAGAGGCTTCTCANNNAGATCGGAAG |
| 1958_1 | TATGAGTGTGGAGTCGTTGCTTAGAATGGGGGCTGTCTTCTCANNNAGATCGGAAG |
| 1959_1 | TATGAGTGTGGAGTCGTTGCGGTGGTTCTTTGCAGTGTGTTTTNNNAGATCGGAAG |
| 1961_1 | TATGAGTGTGGAGTCGTTGCTTTTCCCAAGTGGAGCTTGTTTTNNNAGATCGGAAGA |
| 1963_1 | TATGAGTGTGGAGTCGTTGCTCAGGACAGAGAAGGATTTCTCANNNAGATCGGAAG |
| 1965_1 | TATGAGTGTGGAGTCGTTGCCGCACACAGGCTTAGCAATTTTTNNNAGATCGGAAGA |
| 1968_1 | TATGAGTGTGGAGTCGTTGCTCACCTCTAGCCTGGTTTTCTCANNNAGATCGGAAGA |
| 1969_1 | TATGAGTGTGGAGTCGTTGCCCAGTTTTCCAGCTTTTCTGGCTNNNAGATCGGAAGA |
| 1970_1 | TATGAGTGTGGAGTCGTTGCTAGCAGGGGTGGATTGCATTTTTNNNAGATCGGAAG |
| 1973_1 | TATGAGTGTGGAGTCGTTGCTGGTGGTTGTGGTCAAAAAGACANNNAGATCGGAA |
| 1974_1 | TATGAGTGTGGAGTCGTTGCAGAGAGAAGTAGCGAAGTCCTAGNNNAGATCGGAA |
| 1975_1 | TATGAGTGTGGAGTCGTTGCGCATCCAGCCTGTAATCAAGTCANNNAGATCGGAAG |
| 1976_1 | TATGAGTGTGGAGTCGTTGCCTGGGCTCGGAGGTTTGATTTTTNNNAGATCGGAAG |
| 1977_1 | TATGAGTGTGGAGTCGTTGCCCCCTGGTTTCCACCCTATTTTTNNNAGATCGGAAGAG |
| 1978_1 | TATGAGTGTGGAGTCGTTGCGGGGGTGGGGAGTTTGTATTTTTNNNAGATCGGAA |
| 1980_1 | TATGAGTGTGGAGTCGTTGCTTGCCATTTTTCCCCTGAAGTCANNNAGATCGGAAGA |
| 1982_1 | TATGAGTGTGGAGTCGTTGCTGCCCTCAAGCAGTGTACTTTTTNNNAGATCGGAAGA |
| 1983_1 | TATGAGTGTGGAGTCGTTGCCACTAAGCCCAGCTCAGCATGATNNNAGATCGGAAG |

| | |
|---|---|
| 1986_1 | TATGAGTGTGGAGTCGTTGCTGGAGGTTGAGAAATCCCAGTCANNNAGATCGGAA |
| 1987_1 | TATGAGTGTGGAGTCGTTGCTTTCTCCCAGGGATATGCAGTCANNNAGATCGGAAG |
| 1989_1 | TATGAGTGTGGAGTCGTTGCGTTCTTCCTTTGGTGGCGTTTTTNNNAGATCGGAAG |
| 1991_1 | TATGAGTGTGGAGTCGTTGCCCTGTGTGCTTCCTGTGGTTTTTNNNAGATCGGAAG |
| 1992_1 | TATGAGTGTGGAGTCGTTGCGCCATCAGCAGCAAGATGTTTTTNNNAGATCGGAAG |
| 2007_1 | TATGAGTGTGGAGTCGTTGCAAGAAGAGCAAGTGCAAAGGTCANNNAGATCGGA |
| 2012_1 | TATGAGTGTGGAGTCGTTGCTTTTGTGTCTGTGTGTCCAGCATNNNAGATCGGAAG |
| 2015_1 | TATGAGTGTGGAGTCGTTGCAACAGATTCACACTTCGAGGTCANNNAGATCGGAAG |
| 2018_1 | TATGAGTGTGGAGTCGTTGCTATCTGTTGGCACTCTGAGGTCANNNAGATCGGAAG |
| 2021_1 | TATGAGTGTGGAGTCGTTGCTGTATGCTTCATTGAAAGGGTCANNNAGATCGGAA |
| 2022_1 | TATGAGTGTGGAGTCGTTGCTTCCAGTTCTGAGTTGAGGGCAANNNAGATCGGAA |
| 2025_1 | TATGAGTGTGGAGTCGTTGCAGTAGATTCCACATCGTGGGTCANNNAGATCGGAA |
| 2027_1 | TATGAGTGTGGAGTCGTTGCAAAGTGCGTCCTAATTTGGGTCANNNAGATCGGAA |
| 2028_1 | TATGAGTGTGGAGTCGTTGCGCACAGAACAAGGTACATGGTCANNNAGATCGGAA |
| 2031_1 | TATGAGTGTGGAGTCGTTGCTGCCTGACACCATAAAGTTCTGTNNNAGATCGGAAG |
| 2032_1 | TATGAGTGTGGAGTCGTTGCGAGCTAACCAGTAAACCTGGTCANNNAGATCGGAA |
| 2034_1 | TATGAGTGTGGAGTCGTTGCCAGGGGATAAGAGGAAAGCTGCANNNAGATCGGA |
| 2040_1 | TATGAGTGTGGAGTCGTTGCTCCCTCACACAAAACATTCTTCTNNNAGATCGGAAGA |
| 2044_1 | TATGAGTGTGGAGTCGTTGCACACCTATCCCAAAGGGATGTCANNNAGATCGGAAG |
| 2049_1 | TATGAGTGTGGAGTCGTTGCTCTATTCCGCAGGGTAAGCACTTNNNAGATCGGAAG |
| 2050_1 | TATGAGTGTGGAGTCGTTGCTCTGTTCCTCTGCAGGTATGTCANNNAGATCGGAAG |
| 2051_1 | TATGAGTGTGGAGTCGTTGCATACCAAGTCCACTCTGTTCAGTNNNAGATCGGAAG |
| 2053_1 | TATGAGTGTGGAGTCGTTGCACCTTTTCACCCTCTTACTGTCANNNAGATCGGAAGA |
| 2057_1 | TATGAGTGTGGAGTCGTTGCGCACAGCTTCATACTTCCTGTCANNNAGATCGGAAG |
| 2058_1 | TATGAGTGTGGAGTCGTTGCCACTGGCTGTCTCTTACTAATCCNNNAGATCGGAAGA |
| 2063_1 | TATGAGTGTGGAGTCGTTGCCCTGGATTGCCCTTCATTTCTGTNNNAGATCGGAAG |
| 2065_1 | TATGAGTGTGGAGTCGTTGCCCAGCCCTCCATTGATGAATCCTNNNAGATCGGAAG |
| 2068_1 | TATGAGTGTGGAGTCGTTGCCTGTGTGTCTAAGCCTCACTCTANNNAGATCGGAAG |
| 2071_1 | TATGAGTGTGGAGTCGTTGCCTGAGTGGGGAATGAGTGGGTAANNNAGATCGGA |
| 2080_1 | TATGAGTGTGGAGTCGTTGCCTTTGAGGGCAGGAAACTTGTCANNNAGATCGGAA |
| 2084_1 | TATGAGTGTGGAGTCGTTGCGAATCATGGCCGATCTCTTGTCANNNAGATCGGAAG |
| 2085_1 | TATGAGTGTGGAGTCGTTGCACTGCCCTGATGTGCATAACTTTNNNAGATCGGAAG |
| 2089_1 | TATGAGTGTGGAGTCGTTGCATTACTCCAAGTCCCAAGTCCACNNNAGATCGGAAG |
| 2097_1 | TATGAGTGTGGAGTCGTTGCTATGTGCGTATCCATCCAGAGGGNNNAGATCGGAA |
| 2098_1 | TATGAGTGTGGAGTCGTTGCCCCAAGGTCACCTGAAAATCTCCNNNAGATCGGAAG |
| 2100_1 | TATGAGTGTGGAGTCGTTGCACAGGAGGCCAGATTACCATTCANNNAGATCGGAA |
| 2102_1 | TATGAGTGTGGAGTCGTTGCCCACGGAATAGCCAAACCATAAGNNNAGATCGGAA |
| 2105_1 | TATGAGTGTGGAGTCGTTGCACAAGCATCAAAGAGGGCATTCANNNAGATCGGAA |
| 2108_1 | TATGAGTGTGGAGTCGTTGCACCCTCACACCCATAGTCATTCANNNAGATCGGAAGA |
| 2112_1 | TATGAGTGTGGAGTCGTTGCTCTTGGCACCAACCTTTCATTCANNNAGATCGGAAGA |
| 2114_1 | TATGAGTGTGGAGTCGTTGCTCTTTCCATGCTGGCTAGTACCTNNNAGATCGGAAG |
| 2119_1 | TATGAGTGTGGAGTCGTTGCTTGACATTGTGCTTGGGGATTCANNNAGATCGGAA |
| 2122_1 | TATGAGTGTGGAGTCGTTGCTTAGAGCCCATCCTGGTGATTCANNNAGATCGGAAG |

| | |
|---|---|
| 2123_1 | TATGAGTGTGGAGTCGTTGCTTCTGTGGAGGTCTGAGCCATCANNNAGATCGGAAG |
| 2124_1 | TATGAGTGTGGAGTCGTTGCAAGCAAATGTGGGACAATATTCANNNAGATCGGAAC |
| 2129_1 | TATGAGTGTGGAGTCGTTGCGGACCGTGTGCTGTAGTTATTCANNNAGATCGGAAG |
| 2138_1 | TATGAGTGTGGAGTCGTTGCTGAGGCTAACAGGAGAGGGTAATNNNAGATCGGAA |
| 2142_1 | TATGAGTGTGGAGTCGTTGCGCACAGTTCATCCTGCTACTTCANNNAGATCGGAAGA |
| 2143_1 | TATGAGTGTGGAGTCGTTGCCAGACCTTTGATCCTTTCCAGATNNNAGATCGGAAGA |
| 2147_1 | TATGAGTGTGGAGTCGTTGCCCTTGAAGATGCCTCTCCCTTCANNNAGATCGGAAGA |
| 2151_1 | TATGAGTGTGGAGTCGTTGCTTGTCTGGGGGTAAATGAGAGGGNNNAGATCGGAA |
| 2159_1 | TATGAGTGTGGAGTCGTTGCTACTTTGAGCCTACCAGGCTTCANNNAGATCGGAAGA |
| 2163_1 | TATGAGTGTGGAGTCGTTGCGGAGGGGGTGGTCTTACTAGGTCNNNAGATCGGAA |
| 2170_1 | TATGAGTGTGGAGTCGTTGCCATCTGGTTGACTTCGCTCTTCANNNAGATCGGAAGA |
| 2175_1 | TATGAGTGTGGAGTCGTTGCCCAGTTTCACACAGTGGTCTTCANNNAGATCGGAAGA |
| 2176_1 | TATGAGTGTGGAGTCGTTGCCCTGTCCACAAAACCTGTCTTCANNNAGATCGGAAGA |
| 2182_1 | TATGAGTGTGGAGTCGTTGCCACACCCCACATCTTCAAGTTCANNNAGATCGGAAGA |
| 2183_1 | TATGAGTGTGGAGTCGTTGCCTTCCTCCCCAGATCTCAGTTCANNNAGATCGGAAGA |
| 2191_1 | TATGAGTGTGGAGTCGTTGCTTTAGATGGAGGCTTACGCTGTTNNNAGATCGGAAG |
| 2192_1 | TATGAGTGTGGAGTCGTTGCAGTGTATCTTGAAGACCCTGGGANNNAGATCGGAAC |
| 2198_1 | TATGAGTGTGGAGTCGTTGCAAACAGATACTGGGGGAGGTTCANNNAGATCGGAA |
| 2202_1 | TATGAGTGTGGAGTCGTTGCCTCTAAGGGAAGAGTGGGGTTCANNNAGATCGGAA |
| 2204_1 | TATGAGTGTGGAGTCGTTGCACACAGTTTCACCAATGCACATGNNNAGATCGGAAGA |
| 2214_1 | TATGAGTGTGGAGTCGTTGCGGCAGTGGTACAAACACTGTTCANNNAGATCGGAAC |
| 2226_1 | TATGAGTGTGGAGTCGTTGCAAAACGGGTCCAGTTCCATTTCANNNAGATCGGAAG |
| 2228_1 | TATGAGTGTGGAGTCGTTGCTGGACAAAATTTGGGGTAAGGGANNNAGATCGGAA |
| 2229_1 | TATGAGTGTGGAGTCGTTGCCCTGGGCTGATTGTCAGATTTCANNNAGATCGGAAG |
| 2231_1 | TATGAGTGTGGAGTCGTTGCCGTGTAGACCAGCAGTGATTTCANNNAGATCGGAAG |
| 2243_1 | TATGAGTGTGGAGTCGTTGCTGTCCACCTGTTCTGTGCCTATANNNAGATCGGAAGA |
| 2246_1 | TATGAGTGTGGAGTCGTTGCTTCTCTGCATCTGGCTTCTTTCANNNAGATCGGAAGA |
| 2248_1 | TATGAGTGTGGAGTCGTTGCTGGAAAGTTGACTGGCAGTTTCANNNAGATCGGAAC |
| 2274_1 | TATGAGTGTGGAGTCGTTGCAGAATGCAGCACAGGGAAAAAGANNNAGATCGGAA |
| 2277_1 | TATGAGTGTGGAGTCGTTGCGGGACACAGGAACCAACAAAAGANNNAGATCGGAA |
| 2279_1 | TATGAGTGTGGAGTCGTTGCCAGACCTTGCAGACACCAAAAGANNNAGATCGGAAG |
| 2280_1 | TATGAGTGTGGAGTCGTTGCGTGTGTTTCTCCACTAAGCCAACNNNAGATCGGAAGA |
| 2282_1 | TATGAGTGTGGAGTCGTTGCTCTGGCATGGGGAGTAGAAAAGANNNAGATCGGAA |
| 2287_1 | TATGAGTGTGGAGTCGTTGCCACACCTGGAGTTGGAACAAAGANNNAGATCGGAAC |
| 2292_1 | TATGAGTGTGGAGTCGTTGCTTGAAATCCACCAGCACCAAAGANNNAGATCGGAAG |
| 2294_1 | TATGAGTGTGGAGTCGTTGCATTCAGCACATACCTGCCAAAGANNNAGATCGGAAG |
| 2295_1 | TATGAGTGTGGAGTCGTTGCGACATACACTCCAGGAGAAGCACNNNAGATCGGAAG |
| 2299_1 | TATGAGTGTGGAGTCGTTGCTGTGGTCAACTAATAGGCAGGGANNNAGATCGGAA |
| 2300_1 | TATGAGTGTGGAGTCGTTGCAGCCTGTGTACATTTGGCAAAGANNNAGATCGGAAC |
| 2304_1 | TATGAGTGTGGAGTCGTTGCGGTAGTGGGGAGGGAAAGAAAGANNNAGATCGGA |
| 2310_1 | TATGAGTGTGGAGTCGTTGCGCAACCACAACTGAAACGAAAGANNNAGATCGGAAC |
| 2312_1 | TATGAGTGTGGAGTCGTTGCTGCCTTTGTACATCCAGGAAAGANNNAGATCGGAAG |
| 2314_1 | TATGAGTGTGGAGTCGTTGCTGTAGATCTGTGCAGGGGAAAGANNNAGATCGGAA |

| | |
|---|---|
| 2318_1 | TATGAGTGTGGAGTCGTTGCTTCTTCCTCTGAGCCCTGAAAGANNNAGATCGGAAG |
| 2322_1 | TATGAGTGTGGAGTCGTTGCGTGATGGGCATGGCTTCTAAAGANNNAGATCGGAA |
| 2324_1 | TATGAGTGTGGAGTCGTTGCTTTTTCTGGGTGAGAGGATCCTANNNAGATCGGAAG |
| 2331_1 | TATGAGTGTGGAGTCGTTGCAAAGCAGAAGGAAAGGACCCATGNNNAGATCGGAA |
| 2332_1 | TATGAGTGTGGAGTCGTTGCCAGAGCCACTATGTTGACCAAGANNNAGATCGGAAG |
| 2334_1 | TATGAGTGTGGAGTCGTTGCAAAATCTGGAAGTCACCCCAAGANNNAGATCGGAAG |
| 2336_1 | TATGAGTGTGGAGTCGTTGCTCCAAAGACTAAAGAAGCCAAGANNNAGATCGGAA |
| 2340_1 | TATGAGTGTGGAGTCGTTGCCCGCAGCTTTACATTTGTGTTTGNNNAGATCGGAAG |
| 2341_1 | TATGAGTGTGGAGTCGTTGCGCACACCTGACTATGAAGCAAGANNNAGATCGGAAG |
| 2342_1 | TATGAGTGTGGAGTCGTTGCGTCTGAGTTTTGCCAGAGCAAGANNNAGATCGGAA |
| 2345_1 | TATGAGTGTGGAGTCGTTGCGGCTCAGCATCCTGAAATCAAGANNNAGATCGGAAG |
| 2349_1 | TATGAGTGTGGAGTCGTTGCGTTTCCCATCTGTGATTCCCTCCNNNAGATCGGAAGA |
| 2352_1 | TATGAGTGTGGAGTCGTTGCTCCATCAGAGAGTGGCTTCAAGANNNAGATCGGAA |
| 2355_1 | TATGAGTGTGGAGTCGTTGCGGGCAACAAAAAGCGAAAGAAGANNNAGATCGGA |
| 2358_1 | TATGAGTGTGGAGTCGTTGCGGGCATGCAAGATGAAACATTCCNNNAGATCGGAA |
| 2362_1 | TATGAGTGTGGAGTCGTTGCGCTCCAAACTGTCTTCAAAGGGTNNNAGATCGGAAG |
| 2364_1 | TATGAGTGTGGAGTCGTTGCCTCAACCAGATGCACAAGGAAGANNNAGATCGGAA |
| 2366_1 | TATGAGTGTGGAGTCGTTGCAACACCCTTACCATCCAGGAAGANNNAGATCGGAAG |
| 2371_1 | TATGAGTGTGGAGTCGTTGCGGCTTTTGTGTCAAGAGGGAAGANNNAGATCGGAA |
| 2372_1 | TATGAGTGTGGAGTCGTTGCCTTTTAGTTGAGGCCTGGGAAGANNNAGATCGGAA |
| 2377_1 | TATGAGTGTGGAGTCGTTGCCTGATGTGTCCTTTGCTGGAAGANNNAGATCGGAAG |
| 2378_1 | TATGAGTGTGGAGTCGTTGCCCTAAGAGGCAGGAAGTGGAAGANNNAGATCGGA |
| 2383_1 | TATGAGTGTGGAGTCGTTGCTAAAATACAAGCCAGGACCCCCTNNNAGATCGGAAG |
| 2385_1 | TATGAGTGTGGAGTCGTTGCCATTGCCCTCCGAATTCTGAAGANNNAGATCGGAAG |
| 2386_1 | TATGAGTGTGGAGTCGTTGCTTGCCAGAGAATTGTGTTAGGGCNNNAGATCGGAA |
| 2396_1 | TATGAGTGTGGAGTCGTTGCTTGGGAGAGAAGTGGCACTAAGANNNAGATCGGA |
| 2398_1 | TATGAGTGTGGAGTCGTTGCGTGCATAGCCCACAAACCTAAGANNNAGATCGGAAG |
| 2402_1 | TATGAGTGTGGAGTCGTTGCTCCCATGCCCTTCCAATCTAAGANNNAGATCGGAAGA |
| 2403_1 | TATGAGTGTGGAGTCGTTGCCTGTGGAGCCAGCATTTCTAAGANNNAGATCGGAAG |
| 2407_1 | TATGAGTGTGGAGTCGTTGCTTCATGCTGGGTGTAAGTAGGGANNNAGATCGGAA |
| 2408_1 | TATGAGTGTGGAGTCGTTGCGGCTGAGGGCTTAGAAGGTAAGANNNAGATCGGA |
| 2419_1 | TATGAGTGTGGAGTCGTTGCCCCAACACACACAGAGAAACAGANNNAGATCGGAAG |
| 2420_1 | TATGAGTGTGGAGTCGTTGCAGAGAGCGAGAGCTAGAAACAGANNNAGATCGGA |
| 2426_1 | TATGAGTGTGGAGTCGTTGCAGATGGAGCAGATATCAACGGGANNNAGATCGGAA |
| 2431_1 | TATGAGTGTGGAGTCGTTGCTGAACTAGAAAGACAAGGCCCCTNNNAGATCGGAAG |
| 2436_1 | TATGAGTGTGGAGTCGTTGCTCTTTTCACTGTAATCCCAGCCCNNNAGATCGGAAGA |
| 2439_1 | TATGAGTGTGGAGTCGTTGCATGGACAACTCTGATGGGCTGGGNNNAGATCGGAA |
| 2444_1 | TATGAGTGTGGAGTCGTTGCATCTGGTCAGGGGAAATCACAGANNNAGATCGGAA |
| 2445_1 | TATGAGTGTGGAGTCGTTGCGCATGAAGGGATACCAAGACAGANNNAGATCGGAA |
| 2446_1 | TATGAGTGTGGAGTCGTTGCAAAGAGAGCACTCCCAAGACAGANNNAGATCGGAA |
| 2447_1 | TATGAGTGTGGAGTCGTTGCTGTTGGTAGCATAGGATTCGGGANNNAGATCGGAA |
| 2451_1 | TATGAGTGTGGAGTCGTTGCGGGAGGACTGAGAGAGAGACAGANNNAGATCGGA |
| 2452_1 | TATGAGTGTGGAGTCGTTGCCTCTCTCTACCTCCGGGAAGTTCNNNAGATCGGAAGA |

| | |
|---|---|
| 2456_1 | TATGAGTGTGGAGTCGTTGCTGGTCTATGCTATTGGGGACAGANNNAGATCGGAA |
| 2457_1 | TATGAGTGTGGAGTCGTTGCATCAATGACTGGAAATGGACAGANNNAGATCGGAA |
| 2464_1 | TATGAGTGTGGAGTCGTTGCCCCTAGCCAACCCTGATCTTTTCNNNAGATCGGAAGA |
| 2465_1 | TATGAGTGTGGAGTCGTTGCAAAGCTAATGGGGAGAATACAGANNNAGATCGGAA |
| 2469_1 | TATGAGTGTGGAGTCGTTGCACCAGCTAAATTCAGAAACCAGANNNAGATCGGAAG |
| 2473_1 | TATGAGTGTGGAGTCGTTGCAACAGAAATGGAAGGACACCAGANNNAGATCGGAA |
| 2476_1 | TATGAGTGTGGAGTCGTTGCAGCCAGTTGTAAAGTACAGTGTGNNNAGATCGGAA |
| 2477_1 | TATGAGTGTGGAGTCGTTGCTGCCATGAGATAACCTGACCAGANNNAGATCGGAAG |
| 2482_1 | TATGAGTGTGGAGTCGTTGCTAGTCACCTTTTACGTCCCCAGANNNAGATCGGAAG |
| 2487_1 | TATGAGTGTGGAGTCGTTGCTTTCTTCTGGGTAAAGTCCCAGANNNAGATCGGAAG |
| 2492_1 | TATGAGTGTGGAGTCGTTGCTAGCACTGTCTGGTAACCACCATNNNAGATCGGAAG |
| 2493_1 | TATGAGTGTGGAGTCGTTGCCATCTCCATTTGCACTAGCCAGANNNAGATCGGAAG |
| 2500_1 | TATGAGTGTGGAGTCGTTGCCTACCCATGCGAATTCAGAGGAGNNNAGATCGGAA |
| 2508_1 | TATGAGTGTGGAGTCGTTGCGAGAGGTGGAGGTATTGTCCAGANNNAGATCGGA |
| 2510_1 | TATGAGTGTGGAGTCGTTGCACTGTCTTGTCTTCCCTTCCAGANNNAGATCGGAAGA |
| 2512_1 | TATGAGTGTGGAGTCGTTGCGTCTTGCTGGTCTAGAACTGGCCNNNAGATCGGAAG |
| 2514_1 | TATGAGTGTGGAGTCGTTGCAGTTTGGACTGGTTGCTGTGTAGNNNAGATCGGAA |
| 2516_1 | TATGAGTGTGGAGTCGTTGCAATGCCTGGACACTATACAGCAGNNNAGATCGGAAG |
| 2520_1 | TATGAGTGTGGAGTCGTTGCGAATCACTAACAGCCCAGACTTCNNNAGATCGGAAG |
| 2522_1 | TATGAGTGTGGAGTCGTTGCTTGCTTTCTAACCATGCAGCAGANNNAGATCGGAAG |
| 2523_1 | TATGAGTGTGGAGTCGTTGCATGCAGGAAATTTGCAGAGCAGANNNAGATCGGAA |
| 2524_1 | TATGAGTGTGGAGTCGTTGCTAAAAGCCCATCTGAGGAGCAGANNNAGATCGGAA |
| 2531_1 | TATGAGTGTGGAGTCGTTGCGGAGAGAGAGCTATTCAGGCAGANNNAGATCGGA |
| 2534_1 | TATGAGTGTGGAGTCGTTGCGCTGTGTAAAGATGAAGGGCAGANNNAGATCGGA |
| 2537_1 | TATGAGTGTGGAGTCGTTGCATGTTGTCAGCTTGAATGGCAGANNNAGATCGGAA |
| 2543_1 | TATGAGTGTGGAGTCGTTGCAGGAAAACGTCACATGATGCAGANNNAGATCGGAA |
| 2544_1 | TATGAGTGTGGAGTCGTTGCTGAAAAGAGATGGGGAATGGGGANNNAGATCGGA |
| 2546_1 | TATGAGTGTGGAGTCGTTGCCATCGTGTTTCTGCTACTGCAGANNNAGATCGGAAG |
| 2549_1 | TATGAGTGTGGAGTCGTTGCTTTGTAATTCAGCCAGCTGCAGANNNAGATCGGAAG |
| 2550_1 | TATGAGTGTGGAGTCGTTGCGGTATTCAGCACCTTCCCTTAGGNNNAGATCGGAAG |
| 2553_1 | TATGAGTGTGGAGTCGTTGCTTTCCTTTTCCCTTTCTGGACGGNNNAGATCGGAAGA |
| 2554_1 | TATGAGTGTGGAGTCGTTGCCAGAGGACCAAAGAGGAAAAGGGNNNAGATCGGA |
| 2564_1 | TATGAGTGTGGAGTCGTTGCGACCAGGGAGGGTTCATATCAGANNNAGATCGGAA |
| 2565_1 | TATGAGTGTGGAGTCGTTGCCACGTTGTTTGGGGGATATCAGANNNAGATCGGAA |
| 2566_1 | TATGAGTGTGGAGTCGTTGCAAGAAAAGGCCATCTAGTGGACANNNAGATCGGAA |
| 2570_1 | TATGAGTGTGGAGTCGTTGCTACTTTGGGAGATCATAGCGTCTNNNAGATCGGAAG |
| 2578_1 | TATGAGTGTGGAGTCGTTGCTACAGCAGTCTCACCAAGTCAGANNNAGATCGGAAG |
| 2581_1 | TATGAGTGTGGAGTCGTTGCTCCCTTTGGCAGTCAGTATAGCCNNNAGATCGGAAG |
| 2582_1 | TATGAGTGTGGAGTCGTTGCTGGGAACCAAATTGCTTGTATTCNNNAGATCGGAAG |
| 2585_1 | TATGAGTGTGGAGTCGTTGCGCCAAGTGTCAGAAACCTTCAGANNNAGATCGGAAG |
| 2590_1 | TATGAGTGTGGAGTCGTTGCGTGTATGGGCGTATGTGTTCAGANNNAGATCGGAA |
| 2595_1 | TATGAGTGTGGAGTCGTTGCCTGGGATTCTCTGAGCTTTCAGANNNAGATCGGAAG |
| 2597_1 | TATGAGTGTGGAGTCGTTGCTGGTGCTATTGTGAGGTTTCAGANNNAGATCGGAA |

| | |
|---|---|
| 2599_1 | TATGAGTGTGGAGTCGTTGCACTTTTCACTCCCGTGTAGAATGNNNAGATCGGAAG |
| 2603_1 | TATGAGTGTGGAGTCGTTGCAGAGACCAGAGGCAAGAAAGAGANNNAGATCGGA |
| 2604_1 | TATGAGTGTGGAGTCGTTGCGGGAGAGCTGAGGGTAGATGATGNNNAGATCGGA |
| 2610_1 | TATGAGTGTGGAGTCGTTGCCAAGATAGCAGCACAGACAGTTCNNNAGATCGGAA |
| 2614_1 | TATGAGTGTGGAGTCGTTGCTTCAGTGCAGTGGCATCATCCTTNNNAGATCGGAAG |
| 2616_1 | TATGAGTGTGGAGTCGTTGCAGAGGGAAAAGCACAGACAGAGANNNAGATCGGA |
| 2622_1 | TATGAGTGTGGAGTCGTTGCGTAATGTGCCCTTCAAGCAGAGANNNAGATCGGAA |
| 2632_1 | TATGAGTGTGGAGTCGTTGCATGAGAGAGAGCATGCAGAGAGANNNAGATCGGA |
| 2634_1 | TATGAGTGTGGAGTCGTTGCTCACGTAGTGGGAGAGAGAGAGANNNAGATCGGA |
| 2656_1 | TATGAGTGTGGAGTCGTTGCGGGTGTTACAGTGAGTTCATGCTNNNAGATCGGAA |
| 2657_1 | TATGAGTGTGGAGTCGTTGCTGTTTTCTCGTTTGAGACGGAATNNNAGATCGGAAG |
| 2658_1 | TATGAGTGTGGAGTCGTTGCGGAGAGGAACTGGAAGAAGGAGANNNAGATCGGA |
| 2659_1 | TATGAGTGTGGAGTCGTTGCAGAGAGGGATGAGGAGAAGGAGANNNAGATCGG |
| 2660_1 | TATGAGTGTGGAGTCGTTGCCCCAGGTGAAGAGAGGGAAGAATNNNAGATCGGA |
| 2664_1 | TATGAGTGTGGAGTCGTTGCTATGTTAGGCGACCATCAGGAGANNNAGATCGGAA |
| 2668_1 | TATGAGTGTGGAGTCGTTGCGAAGTGGGGGCAGATGTTTGAATNNNAGATCGGA |
| 2669_1 | TATGAGTGTGGAGTCGTTGCCAGATGGAGAGATGGAGAGGAGANNNAGATCGGA |
| 2671_1 | TATGAGTGTGGAGTCGTTGCAAGAAACTGGACCACTACGGAGANNNAGATCGGAA |
| 2672_1 | TATGAGTGTGGAGTCGTTGCACCCAGTGAAAAGGTGAAACCATNNNAGATCGGAA |
| 2676_1 | TATGAGTGTGGAGTCGTTGCCAGTATCCTCTCATGGAAGCAGCNNNAGATCGGAAG |
| 2678_1 | TATGAGTGTGGAGTCGTTGCTGGCAACAAGGAGAATAGGGAGANNNAGATCGGA |
| 2680_1 | TATGAGTGTGGAGTCGTTGCAGATATTGATGAGTGCCGGGAGANNNAGATCGGA |
| 2682_1 | TATGAGTGTGGAGTCGTTGCTTTTCTCCGAATGGAATGGGAGANNNAGATCGGAA |
| 2683_1 | TATGAGTGTGGAGTCGTTGCCACCATGCTAAGTGACAACGCATNNNAGATCGGAAG |
| 2686_1 | TATGAGTGTGGAGTCGTTGCACCTGGCTCTGTCACCCATACTGNNNAGATCGGAAG |
| 2688_1 | TATGAGTGTGGAGTCGTTGCTCAGACAAACTCTCCCATGGAGANNNAGATCGGAAG |
| 2691_1 | TATGAGTGTGGAGTCGTTGCCCCAACGCATGTATGTATGGAGANNNAGATCGGAA |
| 2692_1 | TATGAGTGTGGAGTCGTTGCATTGGCTCTGTCTTTCCTGGAGANNNAGATCGGAAG |
| 2696_1 | TATGAGTGTGGAGTCGTTGCGCTCCTTTGCTCTAATCCTGACANNNAGATCGGAAG |
| 2697_1 | TATGAGTGTGGAGTCGTTGCAAAGAACCTCCTTCCAGTGGAGANNNAGATCGGAA |
| 2701_1 | TATGAGTGTGGAGTCGTTGCCAGTTGCTCATCCTTTGTGGAGANNNAGATCGGAAG |
| 2708_1 | TATGAGTGTGGAGTCGTTGCTGTGAGAGCTCAGTGTCATGAGANNNAGATCGGAA |
| 2718_1 | TATGAGTGTGGAGTCGTTGCTTTCTAACCAGTACGGCCTGAGANNNAGATCGGAAG |
| 2720_1 | TATGAGTGTGGAGTCGTTGCTGCCTGAACCCTAATTCCTGAGANNNAGATCGGAAG |
| 2722_1 | TATGAGTGTGGAGTCGTTGCACAGGACAGGCTGGATAAGAACTNNNAGATCGGAA |
| 2724_1 | TATGAGTGTGGAGTCGTTGCTGGCTTTGTTTATCATGCTGAGANNNAGATCGGAAG |
| 2728_1 | TATGAGTGTGGAGTCGTTGCTTGGGGCTTAGCTCAGAATAGTTNNNAGATCGGAA |
| 2729_1 | TATGAGTGTGGAGTCGTTGCTGGACTTTGAGGGACTTTCAGGNNNAGATCGGAAG |
| 2735_1 | TATGAGTGTGGAGTCGTTGCTACAGAGGAGTGGGATTACTGGANNNAGATCGGA |
| 2736_1 | TATGAGTGTGGAGTCGTTGCGCCTGTCTGTCTCTTTCTGGACTNNNAGATCGGAAG |
| 2742_1 | TATGAGTGTGGAGTCGTTGCGGATGTTGCATGAGCTCTTGAGANNNAGATCGGAA |
| 2744_1 | TATGAGTGTGGAGTCGTTGCGGTTGGCTGAGTTCTTGTTGAGANNNAGATCGGAA |
| 2745_1 | TATGAGTGTGGAGTCGTTGCTCAGGGTACATTCCCCACATTTTNNNAGATCGGAAG |

| | |
|---|---|
| 2746_1 | TATGAGTGTGGAGTCGTTGCTGTTTTTGTGCACCCATTTGAGANNNAGATCGGAAG |
| 2749_1 | TATGAGTGTGGAGTCGTTGCATTCATCCGGCTCCTCTTTGAGANNNAGATCGGAAG |
| 2755_1 | TATGAGTGTGGAGTCGTTGCTCACGCAGAATGTTGCACATAGANNNAGATCGGAAG |
| 2757_1 | TATGAGTGTGGAGTCGTTGCTGTGTACCTCTCACTGGCATAGANNNAGATCGGAAG |
| 2767_1 | TATGAGTGTGGAGTCGTTGCCCATTCCCCTTCATTCCCCTAGANNNAGATCGGAAGA |
| 2768_1 | TATGAGTGTGGAGTCGTTGCTTCCCATGAAAGCTTCCCCTAGANNNAGATCGGAAG |
| 2770_1 | TATGAGTGTGGAGTCGTTGCACACTTTGGGGAGTTTCCCTAGANNNAGATCGGAAG |
| 2778_1 | TATGAGTGTGGAGTCGTTGCGGAGGCTCAGAATCTCTGCTAGANNNAGATCGGAA |
| 2779_1 | TATGAGTGTGGAGTCGTTGCCACTCAGGAGAAGAGGTGCTAGANNNAGATCGGAA |
| 2780_1 | TATGAGTGTGGAGTCGTTGCTGAGTTACTTCGATGTTTCAGCTNNNAGATCGGAAG |
| 2782_1 | TATGAGTGTGGAGTCGTTGCTACCTGCTACCTCCCCTTCTAGANNNAGATCGGAAGA |
| 2783_1 | TATGAGTGTGGAGTCGTTGCTGTGTGTAACCAGTGCTTCTAGANNNAGATCGGAAG |
| 2792_1 | TATGAGTGTGGAGTCGTTGCGCTCAAATGCTCATGCCTGTAGANNNAGATCGGAAG |
| 2794_1 | TATGAGTGTGGAGTCGTTGCGAGGCCATTGCTGTAGAAGCAGGNNNAGATCGGAA |
| 2798_1 | TATGAGTGTGGAGTCGTTGCGCAGGCACACACAGAAGATTAGANNNAGATCGGAA |
| 2801_1 | TATGAGTGTGGAGTCGTTGCGTGTGTGGTACATTTACTCCTTGNNNAGATCGGAAG |
| 2803_1 | TATGAGTGTGGAGTCGTTGCGGAGTTGTTCCCCAAAGGAAAACNNNAGATCGGAA |
| 2804_1 | TATGAGTGTGGAGTCGTTGCAATGGCTCCACGCTACTCTTAGANNNAGATCGGAAG |
| 2806_1 | TATGAGTGTGGAGTCGTTGCAACTGGACCAGCAGCTTCTTAGANNNAGATCGGAAG |
| 2807_1 | TATGAGTGTGGAGTCGTTGCGCTCTTCTCCCCTCCATTTCTCTNNNAGATCGGAAGAG |
| 2808_1 | TATGAGTGTGGAGTCGTTGCTCCCCTCTCCTTACAAGGACACTNNNAGATCGGAAGA |
| 2809_1 | TATGAGTGTGGAGTCGTTGCTGGCTTTTGGTGGAACAGTTAGANNNAGATCGGAA |
| 2810_1 | TATGAGTGTGGAGTCGTTGCCTGGGACCATAATGGGTGCATGNNNAGATCGGAAG |
| 2812_1 | TATGAGTGTGGAGTCGTTGCAAAAAGGAATGATTACAGCCAGGNNNAGATCGGAA |
| 2813_1 | TATGAGTGTGGAGTCGTTGCCCTGCCTTTCCTCTCCCTTTAGANNNAGATCGGAAGA |
| 2818_1 | TATGAGTGTGGAGTCGTTGCGGAGCCATGCACATCTGTTTAGANNNAGATCGGAAG |
| 2822_1 | TATGAGTGTGGAGTCGTTGCACCTGGCAAAACAATACTAACGANNNAGATCGGAAG |
| 2823_1 | TATGAGTGTGGAGTCGTTGCTGCTCTCTCTTCTGTAGCAAAGTNNNAGATCGGAAG |
| 2825_1 | TATGAGTGTGGAGTCGTTGCCCATTATTCTTTCCCCAGCACGANNNAGATCGGAAGA |
| 2826_1 | TATGAGTGTGGAGTCGTTGCATAGACAACTACAGAGGGCACGANNNAGATCGGAA |
| 2830_1 | TATGAGTGTGGAGTCGTTGCTGAGGGGTTGGTAGAGGTGAGTGNNNAGATCGGA |
| 2831_1 | TATGAGTGTGGAGTCGTTGCTGACCTAGAGCTTGATTGTGGGANNNAGATCGGAA |
| 2833_1 | TATGAGTGTGGAGTCGTTGCGAGTGAAGAGTGGAGAGAGACGANNNAGATCGGA |
| 2836_1 | TATGAGTGTGGAGTCGTTGCACACACACACAAAGCCTAACAGTNNNAGATCGGAAG |
| 2839_1 | TATGAGTGTGGAGTCGTTGCATCACCTTCCATCACCTCTACGANNNAGATCGGAAGA |
| 2842_1 | TATGAGTGTGGAGTCGTTGCAGAGAGAAGGGGTGTTGAAGAGTNNNAGATCGGA |
| 2849_1 | TATGAGTGTGGAGTCGTTGCGTGAGGGGCTATATCGTTTGTTANNNAGATCGGAA |
| 2851_1 | TATGAGTGTGGAGTCGTTGCAGAGAGATGAGGAGTGAAAGCGANNNAGATCGGA |
| 2854_1 | TATGAGTGTGGAGTCGTTGCAAAGCTGCTAAAACTCAGTCCGTNNNAGATCGGAAG |
| 2860_1 | TATGAGTGTGGAGTCGTTGCCGGTCTGAAGACATCCTCATCGANNNAGATCGGAAG |
| 2861_1 | TATGAGTGTGGAGTCGTTGCGACAACACAGGTCAAGTCATCGANNNAGATCGGAA |
| 2862_1 | TATGAGTGTGGAGTCGTTGCGGCTCATCATTGGAACTGCTTCCNNNAGATCGGAAG |
| 2863_1 | TATGAGTGTGGAGTCGTTGCTCTGAAAGGTCACTGGAACTCGANNNAGATCGGAA |

| | |
|---|---|
| 2868_1 | TATGAGTGTGGAGTCGTTGCCCTGAATGGATAAGGAGGGACANNNAGATCGGAAG |
| 2870_1 | TATGAGTGTGGAGTCGTTGCTTGGCTCACACGATTACAAAGGANNNAGATCGGAAG |
| 2872_1 | TATGAGTGTGGAGTCGTTGCGCCTCAAAGTCACTTCCAAAGGANNNAGATCGGAAG |
| 2873_1 | TATGAGTGTGGAGTCGTTGCCTAAAAAGGGGTCTTCCTGAGCTNNNAGATCGGAAG |
| 2874_1 | TATGAGTGTGGAGTCGTTGCGGAGCAGTGGGAAGTCTCATGATNNNAGATCGGAA |
| 2875_1 | TATGAGTGTGGAGTCGTTGCAAGCTGAGGAAGGGATCAAAGGANNNAGATCGGA |
| 2877_1 | TATGAGTGTGGAGTCGTTGCAATAAGGCTGCATGCTCAAAGGANNNAGATCGGAA |
| 2883_1 | TATGAGTGTGGAGTCGTTGCGGAAACACGACACGTCTAAAGGANNNAGATCGGAA |
| 2887_1 | TATGAGTGTGGAGTCGTTGCCCAAGGTGGAGGAAAAACAAGGANNNAGATCGGA |
| 2888_1 | TATGAGTGTGGAGTCGTTGCTAATAAGGGTGAGGCTGCTCTGTNNNAGATCGGAA |
| 2889_1 | TATGAGTGTGGAGTCGTTGCTAGGAAAAGGTGGTGGACAAGGANNNAGATCGGA |
| 2891_1 | TATGAGTGTGGAGTCGTTGCCAGCACTTGTCTAACAGCAAGGANNNAGATCGGAAG |
| 2892_1 | TATGAGTGTGGAGTCGTTGCGAAAACTGAGTACAAGCAGGACAGGNNNAGATCGG |
| 2896_1 | TATGAGTGTGGAGTCGTTGCTAGAGAAGGACTGGGGAGAAGGANNNAGATCGGA |
| 2897_1 | TATGAGTGTGGAGTCGTTGCTAGGGAAGGCAACATGAGAAGGANNNAGATCGGA |
| 2900_1 | TATGAGTGTGGAGTCGTTGCTTGGGGACAGGATAGGGTAGCATNNNAGATCGGA |
| 2903_1 | TATGAGTGTGGAGTCGTTGCGAACAAGCACAGCTAAGGAAGGANNNAGATCGGAA |
| 2910_1 | TATGAGTGTGGAGTCGTTGCTCTTAGCTCTGTGTGGGAAGAACNNNAGATCGGAA |
| 2914_1 | TATGAGTGTGGAGTCGTTGCTCACTCCTAGCAAGAGCTAAGGANNNAGATCGGAAG |
| 2915_1 | TATGAGTGTGGAGTCGTTGCATGAAAGAATGGCACTTTTGGGANNNAGATCGGAA |
| 2918_1 | TATGAGTGTGGAGTCGTTGCAGAGAAGAGAGACGGGAAAGGCANNNAGATCGGA |
| 2920_1 | TATGAGTGTGGAGTCGTTGCGCAATAGTTGGAGGAAGACAGGANNNAGATCGGA |
| 2924_1 | TATGAGTGTGGAGTCGTTGCCAATGGTTTGGTCTATTTGGGGANNNAGATCGGAA |
| 2925_1 | TATGAGTGTGGAGTCGTTGCAGGGCAAGCTTTACTATACAGGANNNAGATCGGAA |
| 2926_1 | TATGAGTGTGGAGTCGTTGCTACTCCCTACAGAGCCCAGAGTTNNNAGATCGGAAG |
| 2928_1 | TATGAGTGTGGAGTCGTTGCAAACCCAGTCTCTAACACCAGGANNNAGATCGGAAG |
| 2929_1 | TATGAGTGTGGAGTCGTTGCATTCCAGTGACAGTACACCAGGANNNAGATCGGAAG |
| 2932_1 | TATGAGTGTGGAGTCGTTGCGTTTTCATCATCCTCAGCCAGGANNNAGATCGGAAG |
| 2933_1 | TATGAGTGTGGAGTCGTTGCGGGATGTTTGCAAGATGGTGCTNNNAGATCGGAAG |
| 2935_1 | TATGAGTGTGGAGTCGTTGCATAGATCCTGTCCTCCTCCAGGANNNAGATCGGAAG |
| 2936_1 | TATGAGTGTGGAGTCGTTGCTTACAGTCTACCACAACCACACCNNNAGATCGGAAGA |
| 2937_1 | TATGAGTGTGGAGTCGTTGCTGCCAAACTTTCAATCAGCAGGANNNAGATCGGAAG |
| 2938_1 | TATGAGTGTGGAGTCGTTGCCACAGTAATAGGAACGCTGCCTTNNNAGATCGGAAG |
| 2940_1 | TATGAGTGTGGAGTCGTTGCATGCAATAAAGTGAAGTGCAGGANNNAGATCGGAA |
| 2942_1 | TATGAGTGTGGAGTCGTTGCCTCCTGTTTTCCACGTGGTTCTCNNNAGATCGGAAGA |
| 2944_1 | TATGAGTGTGGAGTCGTTGCGTGAGAAGGGGAATGTCCTTGAGNNNAGATCGGA |
| 2947_1 | TATGAGTGTGGAGTCGTTGCGGGTGAAGAGAAAGGTGAAAGACNNNAGATCGGA |
| 2952_1 | TATGAGTGTGGAGTCGTTGCAGTGCTTGAACATACCAAGAGGANNNAGATCGGAA |
| 2954_1 | TATGAGTGTGGAGTCGTTGCGTCATGCTTTGGAGTGAAGAGGANNNAGATCGGAA |
| 2959_1 | TATGAGTGTGGAGTCGTTGCAGGAGAGAGACTGACAGAGAGGANNNAGATCGGA |
| 2960_1 | TATGAGTGTGGAGTCGTTGCAAGACATTCATCCTCAGAGAGGANNNAGATCGGAA |
| 2961_1 | TATGAGTGTGGAGTCGTTGCTTTCTGCGTGTTTTGTGAGAGGANNNAGATCGGAA |
| 2963_1 | TATGAGTGTGGAGTCGTTGCATGGGGGATAGATGTCTTGCCAANNNAGATCGGAA |

| | |
|---|---|
| 2964_1 | TATGAGTGTGGAGTCGTTGCGAAAAAGTTCCTGCTCTCGAGGANNNAGATCGGAA |
| 2965_1 | TATGAGTGTGGAGTCGTTGCGTCCTGTGTCTGTAGAAGGAGGANNNAGATCGGAA |
| 2966_1 | TATGAGTGTGGAGTCGTTGCTAACTACAGCAACGGAAGGAGGANNNAGATCGGAA |
| 2967_1 | TATGAGTGTGGAGTCGTTGCAAGGGGGAGATAGAAGAGGAGGANNNAGATCGG |
| 2968_1 | TATGAGTGTGGAGTCGTTGCTGGAAAAATCCCATGAGGGAGGANNNAGATCGGA |
| 2974_1 | TATGAGTGTGGAGTCGTTGCATAAAACCTGCAGAGGTGGAGGANNNAGATCGGA |
| 2975_1 | TATGAGTGTGGAGTCGTTGCTGGTGGGACAAGCATTTAGTTTTNNNAGATCGGAA |
| 2976_1 | TATGAGTGTGGAGTCGTTGCGGACCAAGATTGATTTTGGAGGANNNAGATCGGAA |
| 2982_1 | TATGAGTGTGGAGTCGTTGCTTCCCCTGTTTCATACCTGAGGANNNAGATCGGAAG |
| 2986_1 | TATGAGTGTGGAGTCGTTGCTTTGTCCTGTCCTTTTCTGAGGANNNAGATCGGAAG |
| 2988_1 | TATGAGTGTGGAGTCGTTGCTATGCCTGTGTCTATGGTGAGGANNNAGATCGGAA |
| 2992_1 | TATGAGTGTGGAGTCGTTGCTGTGGTATGGCTCTCAAATAGGANNNAGATCGGAA |
| 2997_1 | TATGAGTGTGGAGTCGTTGCGCCTCCAGAAGGACTACATAGGANNNAGATCGGAA |
| 2998_1 | TATGAGTGTGGAGTCGTTGCAGCCAAATGTGCAAACCATAGGANNNAGATCGGAA |
| 3009_1 | TATGAGTGTGGAGTCGTTGCTTCCTTTCCTACAGTCCCTAGGANNNAGATCGGAAGA |
| 3016_1 | TATGAGTGTGGAGTCGTTGCATGGGATTTCTCTCTTGTCCAGGNNNAGATCGGAAG |
| 3040_1 | TATGAGTGTGGAGTCGTTGCCAGTCAATTGAGAAGGACACGGANNNAGATCGGAA |
| 3060_1 | TATGAGTGTGGAGTCGTTGCGAAAGAGCATTAGAGCTATGCCTNNNAGATCGGAA |
| 3070_1 | TATGAGTGTGGAGTCGTTGCTGAACAGGATGGAAAAGAAGGGANNNAGATCGGA |
| 3073_1 | TATGAGTGTGGAGTCGTTGCCTGGTTAGGCACTAGAGAAGGGANNNAGATCGGA |
| 3076_1 | TATGAGTGTGGAGTCGTTGCTGGCAGAATTCACTAGGAAGGGANNNAGATCGGAA |
| 3082_1 | TATGAGTGTGGAGTCGTTGCGCTGCAAGATTCTTTTGAAGGGANNNAGATCGGAA |
| 3095_1 | TATGAGTGTGGAGTCGTTGCCCAAGGTTTGATTTGGACAGGGANNNAGATCGGAA |
| 3101_1 | TATGAGTGTGGAGTCGTTGCGTCCAGCCAAGTTTTGTTATTGTNNNAGATCGGAAG |
| 3106_1 | TATGAGTGTGGAGTCGTTGCGTCCCTGAAGCATATCCCAGCAGNNNAGATCGGAAG |
| 3109_1 | TATGAGTGTGGAGTCGTTGCAATGGCAAGGGTAATGGCATTGTNNNAGATCGGAA |
| 3111_1 | TATGAGTGTGGAGTCGTTGCAGAGAGAGAGATTTGTGCAGGGANNNAGATCGGA |
| 3124_1 | TATGAGTGTGGAGTCGTTGCGCTGCTTGGAAAATAAGGAGGGANNNAGATCGGA |
| 3129_1 | TATGAGTGTGGAGTCGTTGCTAAAACCCTCTTCCAGTGAGGGANNNAGATCGGAAG |
| 3133_1 | TATGAGTGTGGAGTCGTTGCTGGAGGTCAGTCCCATATAGGGANNNAGATCGGAA |
| 3134_1 | TATGAGTGTGGAGTCGTTGCTCCTAAAATCAGGCATAAGGCAGNNNAGATCGGAA |
| 3136_1 | TATGAGTGTGGAGTCGTTGCCTACCTTTCTGCTTCCTTCAGGANNNAGATCGGAAGA |
| 3142_1 | TATGAGTGTGGAGTCGTTGCAATCTCTTTCCTTCCTAGCGGGANNNAGATCGGAAG |
| 3150_1 | TATGAGTGTGGAGTCGTTGCAGAGGATTTGAGCTAACAGGGGANNNAGATCGGA |
| 3159_1 | TATGAGTGTGGAGTCGTTGCAACTGAGGACTTTTCGTAGGGGANNNAGATCGGAA |
| 3160_1 | TATGAGTGTGGAGTCGTTGCTAGCAATCACCATGTGTAGGGGANNNAGATCGGAA |
| 3175_1 | TATGAGTGTGGAGTCGTTGCTCTAACAGACTGAAAGCTGGGGANNNAGATCGGAA |
| 3184_1 | TATGAGTGTGGAGTCGTTGCACAAACTGGGAAAAGGAATGGGANNNAGATCGGA |
| 3185_1 | TATGAGTGTGGAGTCGTTGCACCAGATGACAGAGTGAATGGGANNNAGATCGGA |
| 3187_1 | TATGAGTGTGGAGTCGTTGCAACACTGGTCCAGACTAATGGGANNNAGATCGGAA |
| 3197_1 | TATGAGTGTGGAGTCGTTGCTGAAGGTCAGGGAGTATATGGGANNNAGATCGGA |
| 3198_1 | TATGAGTGTGGAGTCGTTGCAAATACCCACACGTCCTATGGGANNNAGATCGGAAG |
| 3202_1 | TATGAGTGTGGAGTCGTTGCGGAAGGGAAGGGAGATTATGGGANNNAGATCGGA |

| | |
|---|---|
| 3204_1 | TATGAGTGTGGAGTCGTTGCTCTAACAGATGAGGCAACTGGGANNNAGATCGGAA |
| 3207_1 | TATGAGTGTGGAGTCGTTGCCCTCACTGTCACTTAGACTGGGANNNAGATCGGAAG |
| 3209_1 | TATGAGTGTGGAGTCGTTGCGGACCTATGTGAAAGACCTGGGANNNAGATCGGAA |
| 3211_1 | TATGAGTGTGGAGTCGTTGCTAGGTGTGGAAATAAGCCTGGGANNNAGATCGGAA |
| 3212_1 | TATGAGTGTGGAGTCGTTGCACTGATTAGCTGTAAGCCTGGGANNNAGATCGGAA |
| 3213_1 | TATGAGTGTGGAGTCGTTGCTATCTACTGAGAGAGGCCTGGGANNNAGATCGGAA |
| 3215_1 | TATGAGTGTGGAGTCGTTGCCAATTCTTACGTGGAAGCTGGGANNNAGATCGGAA |
| 3217_1 | TATGAGTGTGGAGTCGTTGCGCACCTATAATCATGCCAGACCANNNAGATCGGAAG |
| 3218_1 | TATGAGTGTGGAGTCGTTGCTATGGCATGGAGAATAGCTGGGANNNAGATCGGAA |
| 3222_1 | TATGAGTGTGGAGTCGTTGCAAGAGCTCAGACATGTTCTGGGANNNAGATCGGAA |
| 3225_1 | TATGAGTGTGGAGTCGTTGCTTTGACCATACTCTGTCGTGGGANNNAGATCGGAAG |
| 3226_1 | TATGAGTGTGGAGTCGTTGCGGTGTAGGTATGTGTAGGTGGGANNNAGATCGGA |
| 3227_1 | TATGAGTGTGGAGTCGTTGCGAACAACGTGAATATGGGTGGGANNNAGATCGGA |
| 3228_1 | TATGAGTGTGGAGTCGTTGCCCAGGAGGAGATTTATGGTGGGANNNAGATCGGA |
| 3229_1 | TATGAGTGTGGAGTCGTTGCTAGATGAAGGTGCTGATGTGGGANNNAGATCGGA |
| 3232_1 | TATGAGTGTGGAGTCGTTGCTGTTAGCCTTCAGCAAATTGGGANNNAGATCGGAA |
| 3237_1 | TATGAGTGTGGAGTCGTTGCTTTGTCATCTCCAGATGTTGGGANNNAGATCGGAAG |
| 3238_1 | TATGAGTGTGGAGTCGTTGCTGTGTGATGAATGCCATTTGGGANNNAGATCGGAA |
| 3246_1 | TATGAGTGTGGAGTCGTTGCTACAAAGTCTCGTGTGTCTGTCTNNNAGATCGGAAG |
| 3247_1 | TATGAGTGTGGAGTCGTTGCCACAGCCTGAATTGGTACATGGANNNAGATCGGAA |
| 3248_1 | TATGAGTGTGGAGTCGTTGCTAGAAGATCATGGTGGCCATGGANNNAGATCGGAA |
| 3249_1 | TATGAGTGTGGAGTCGTTGCCATTGACTAAAGGGGAGGCCAAANNNAGATCGGAA |
| 3254_1 | TATGAGTGTGGAGTCGTTGCGTCAGGGTCTGGAAAGTTTTGGGNNNAGATCGGAA |
| 3259_1 | TATGAGTGTGGAGTCGTTGCTCTTTGGATGGTGGGATGATGGANNNAGATCGGAA |
| 3260_1 | TATGAGTGTGGAGTCGTTGCTTGTCCAGCTATCCATTGATGGANNNAGATCGGAAG |
| 3262_1 | TATGAGTGTGGAGTCGTTGCTGAGGGACACACAGAACTATGGANNNAGATCGGAA |
| 3266_1 | TATGAGTGTGGAGTCGTTGCTCTATGGCTTCCAACTTACTGCANNNAGATCGGAAG |
| 3269_1 | TATGAGTGTGGAGTCGTTGCTATCAGCTATCACCCAGACTGGANNNAGATCGGAAG |
| 3270_1 | TATGAGTGTGGAGTCGTTGCACTCACCATTTTCCCAGACTGGANNNAGATCGGAAG |
| 3271_1 | TATGAGTGTGGAGTCGTTGCTCGTGGACAAAAAGAATACTGGANNNAGATCGGAA |
| 3272_1 | TATGAGTGTGGAGTCGTTGCGCTCTCAGCTTTCTAGATACTGGANNNAGATCGGAA |
| 3275_1 | TATGAGTGTGGAGTCGTTGCTATTACATTCGCAGCAACCTGGANNNAGATCGGAAG |
| 3276_1 | TATGAGTGTGGAGTCGTTGCACACATACTGTAGCAACCCTGGANNNAGATCGGAAG |
| 3278_1 | TATGAGTGTGGAGTCGTTGCAATGACACATGAGAGGTCCTGGANNNAGATCGGAA |
| 3281_1 | TATGAGTGTGGAGTCGTTGCGAAAGATGCAGGAAAGAGCTGGANNNAGATCGGA |
| 3283_1 | TATGAGTGTGGAGTCGTTGCTGCTCCCAAAGTTCACCACTATTNNNAGATCGGAAGA |
| 3287_1 | TATGAGTGTGGAGTCGTTGCTCTGTTGTCTCTGTCTCTCTGGANNNAGATCGGAAG |
| 3288_1 | TATGAGTGTGGAGTCGTTGCCTGAAGGAGTGTCATTGTCTGGANNNAGATCGGAA |
| 3290_1 | TATGAGTGTGGAGTCGTTGCCCAGCACTTTCTTACTTTCTGGANNNAGATCGGAAGA |
| 3291_1 | TATGAGTGTGGAGTCGTTGCGCTGAAGGCACACAACAATTGGTNNNAGATCGGAA |
| 3294_1 | TATGAGTGTGGAGTCGTTGCCTGAAGTGTTTTTCCCCAGTGGANNNAGATCGGAAG |
| 3297_1 | TATGAGTGTGGAGTCGTTGCAACATAGAGTAGGCAGCAGTGGANNNAGATCGGAA |
| 3298_1 | TATGAGTGTGGAGTCGTTGCGTCAGTGTGCTGGCATATGTGTGNNNAGATCGGAA |

| | |
|---|---|
| 3299_1 | TATGAGTGTGGAGTCGTTGCGTGAAGACAGGCTTTTCAGTGGANNNAGATCGGAA |
| 3300_1 | TATGAGTGTGGAGTCGTTGCGTGATTTGTGCACCTAGAGTGGANNNAGATCGGAA |
| 3302_1 | TATGAGTGTGGAGTCGTTGCGGCTTCCTCATCTCTAAGGTGGANNNAGATCGGAAG |
| 3303_1 | TATGAGTGTGGAGTCGTTGCCCCGAGACAGTATCTCACCATATNNNAGATCGGAAG |
| 3304_1 | TATGAGTGTGGAGTCGTTGCAAGTGGGGACATGATTAGGTGGANNNAGATCGGA |
| 3305_1 | TATGAGTGTGGAGTCGTTGCAGAGAATCAAAGGAGTGGGTGGANNNAGATCGGA |
| 3307_1 | TATGAGTGTGGAGTCGTTGCATGGGAAATGAGAAGCATGTGGANNNAGATCGGA |
| 3308_1 | TATGAGTGTGGAGTCGTTGCTAGAATGCGGTGGCTGAATAACGNNNAGATCGGAA |
| 3309_1 | TATGAGTGTGGAGTCGTTGCTTCCTCCTCAACACTCTCCCACCNNNAGATCGGAAGA |
| 3311_1 | TATGAGTGTGGAGTCGTTGCATGGGGTAGGAGAAAAGTGTGGANNNAGATCGGA |
| 3312_1 | TATGAGTGTGGAGTCGTTGCGCATGTATGACTGTGAGTGTGGANNNAGATCGGAA |
| 3317_1 | TATGAGTGTGGAGTCGTTGCAGGTAAAAGAGAAAGGCCAATGTNNNAGATCGGA |
| 3322_1 | TATGAGTGTGGAGTCGTTGCGAGAGGGGTATGGTTACCTTGGANNNAGATCGGA |
| 3324_1 | TATGAGTGTGGAGTCGTTGCACTGGGGAAGATCAAAAACTGGANNNAGATCGGAA |
| 3327_1 | TATGAGTGTGGAGTCGTTGCATTGGGGTCAGAGTGAAAAGAGCNNNAGATCGGA |
| 3328_1 | TATGAGTGTGGAGTCGTTGCCTCTGTGTGTTGGTTATTGGCCTNNNAGATCGGAAG |
| 3332_1 | TATGAGTGTGGAGTCGTTGCCAGAGTTCTAAAGAGCAGGGCTCNNNAGATCGGAA |
| 3333_1 | TATGAGTGTGGAGTCGTTGCTTCTTCCAGGCAAACACTTTGGANNNAGATCGGAAG |
| 3334_1 | TATGAGTGTGGAGTCGTTGCTGAGCTATGGTGAGACCTTTGGANNNAGATCGGAA |
| 3338_1 | TATGAGTGTGGAGTCGTTGCAGAGATAGCGACCAGAAAAAGACNNNAGATCGGAA |
| 3340_1 | TATGAGTGTGGAGTCGTTGCTGTAGTGGCAGATGCAAAAATGANNNAGATCGGAA |
| 3342_1 | TATGAGTGTGGAGTCGTTGCTTGCATTGCTGTCATCACAATGANNNAGATCGGAAG |
| 3344_1 | TATGAGTGTGGAGTCGTTGCCCCACCCAGAAAAGTGACAATGANNNAGATCGGAAG |
| 3345_1 | TATGAGTGTGGAGTCGTTGCTCTACTCAGTCCACAGCCAATGANNNAGATCGGAAG |
| 3350_1 | TATGAGTGTGGAGTCGTTGCATCTTGTCTGACACCTGCTACCANNNAGATCGGAAG |
| 3353_1 | TATGAGTGTGGAGTCGTTGCTGACAGAGTGAAAAGCATCTGGANNNAGATCGGAA |
| 3354_1 | TATGAGTGTGGAGTCGTTGCCACCCATTGACCTGTTCACATGANNNAGATCGGAAG |
| 3357_1 | TATGAGTGTGGAGTCGTTGCTGGGATGAGCTGAGTGTACATGANNNAGATCGGAA |
| 3358_1 | TATGAGTGTGGAGTCGTTGCGGATCTGAGGATGGAGACCATGANNNAGATCGGA |
| 3359_1 | TATGAGTGTGGAGTCGTTGCGGAAGGACATGTTTGGACCATGANNNAGATCGGAA |
| 3360_1 | TATGAGTGTGGAGTCGTTGCCCAGGACTGCATGGATTGGTACANNNAGATCGGAA |
| 3362_1 | TATGAGTGTGGAGTCGTTGCGGAGCATAATATGGTGGCCATGANNNAGATCGGAA |
| 3364_1 | TATGAGTGTGGAGTCGTTGCGGGGGTCCATATGAGTTCCATGANNNAGATCGGAA |
| 3365_1 | TATGAGTGTGGAGTCGTTGCAATCCACTCACTACCCACCAGATNNNAGATCGGAAG |
| 3367_1 | TATGAGTGTGGAGTCGTTGCAGCAAAGAATGGAACTGGCATGANNNAGATCGGAA |
| 3373_1 | TATGAGTGTGGAGTCGTTGCAGAAAATCAAAGGGCTGCATGGANNNAGATCGGAA |
| 3375_1 | TATGAGTGTGGAGTCGTTGCCCCGTAAGACGAAAGTTTCATGANNNAGATCGGAA |
| 3376_1 | TATGAGTGTGGAGTCGTTGCAAAAAGCTGCCACTGGAAGATGANNNAGATCGGAA |
| 3379_1 | TATGAGTGTGGAGTCGTTGCTCCAGAAACCTTAGCTTCTCCAANNNAGATCGGAAG |
| 3380_1 | TATGAGTGTGGAGTCGTTGCGGTGGGGTAAGGGAAAGAGATGANNNAGATCGGA |
| 3381_1 | TATGAGTGTGGAGTCGTTGCAATATGACAGGGCAGGGAAGGAGNNNAGATCGGA |
| 3383_1 | TATGAGTGTGGAGTCGTTGCTGAACCTGGTGGATCCCATTAGANNNAGATCGGAA |
| 3386_1 | TATGAGTGTGGAGTCGTTGCGATGAGAAGCCACTGTTTCACAANNNAGATCGGAA |

| | |
|---|---|
| 3387_1 | TATGAGTGTGGAGTCGTTGCTTACACCTTCTACCGTGGGATGANNNAGATCGGAAG |
| 3389_1 | TATGAGTGTGGAGTCGTTGCAACTCCCATTCATTCATCTGCCCNNNAGATCGGAAGA |
| 3395_1 | TATGAGTGTGGAGTCGTTGCACAAAAACAAAACCCTGTGATGANNNAGATCGGAAG |
| 3396_1 | TATGAGTGTGGAGTCGTTGCGGCTGCACTTCAGAAATTTGGAGNNNAGATCGGAA |
| 3397_1 | TATGAGTGTGGAGTCGTTGCAACAGGAGCTACAGGCCATATGANNNAGATCGGAA |
| 3401_1 | TATGAGTGTGGAGTCGTTGCTCTTTTCCAACAGTCTCCTTCTGNNNAGATCGGAAGA |
| 3402_1 | TATGAGTGTGGAGTCGTTGCGCTTTGCCTTTAGCCATGTATGANNNAGATCGGAAG |
| 3403_1 | TATGAGTGTGGAGTCGTTGCACACGGCCTCTTTTCATGTATGANNNAGATCGGAAG |
| 3404_1 | TATGAGTGTGGAGTCGTTGCAGCAGCAGACACAAACTGTATGANNNAGATCGGAA |
| 3406_1 | TATGAGTGTGGAGTCGTTGCACCTTGGGAGCCTCTACTTATGANNNAGATCGGAAG |
| 3407_1 | TATGAGTGTGGAGTCGTTGCCCCCACCAACAATCACCTTATGANNNAGATCGGAAGA |
| 3409_1 | TATGAGTGTGGAGTCGTTGCAGTTGACAGAAGCACCAAACTGANNNAGATCGGAA |
| 3410_1 | TATGAGTGTGGAGTCGTTGCTAGCTCTGTGTGGGAAGAACTGANNNAGATCGGAA |
| 3411_1 | TATGAGTGTGGAGTCGTTGCACTCAAATCTCTTCCCCAGTGGANNNAGATCGGAAG |
| 3412_1 | TATGAGTGTGGAGTCGTTGCGGTGCATGTAGGTAGCCATTTCANNNAGATCGGAAG |
| 3417_1 | TATGAGTGTGGAGTCGTTGCCACAGGTCAGATTCAAGCACTGANNNAGATCGGAAG |
| 3418_1 | TATGAGTGTGGAGTCGTTGCAGATGAGAGAGAGAGAGCACTGANNNAGATCGGA |
| 3422_1 | TATGAGTGTGGAGTCGTTGCACTTTCCATCCTCAGGACAAGCANNNAGATCGGAAG |
| 3424_1 | TATGAGTGTGGAGTCGTTGCGCCTGTTTTCCATTTGGGACTGANNNAGATCGGAAG |
| 3426_1 | TATGAGTGTGGAGTCGTTGCTGTTTTGTTCAACAGTGGACTGANNNAGATCGGAAG |
| 3429_1 | TATGAGTGTGGAGTCGTTGCGTGTGTGTGTGTGGCTATACTGANNNAGATCGGAA |
| 3430_1 | TATGAGTGTGGAGTCGTTGCCACAGGTACTTCACAAAAGAGGGNNNAGATCGGAA |
| 3431_1 | TATGAGTGTGGAGTCGTTGCAAGCTGTGTCAGGGCAGGAATACNNNAGATCGGAA |
| 3434_1 | TATGAGTGTGGAGTCGTTGCTTTTCTGGGTCTCAGTCACCTGANNNAGATCGGAAG |
| 3435_1 | TATGAGTGTGGAGTCGTTGCACATTCCAACCTATGGGACCTGANNNAGATCGGAAG |
| 3437_1 | TATGAGTGTGGAGTCGTTGCCCTAGAACAATGAAACCCCCTGANNNAGATCGGAAG |
| 3438_1 | TATGAGTGTGGAGTCGTTGCGTGGTGTGGAGCAATATACCGTTNNNAGATCGGAA |
| 3445_1 | TATGAGTGTGGAGTCGTTGCACTTTCTCTGAAAACATGCCTGANNNAGATCGGAAG |
| 3448_1 | TATGAGTGTGGAGTCGTTGCTGGTAAGGAAATGAGGCTCCTGANNNAGATCGGAA |
| 3450_1 | TATGAGTGTGGAGTCGTTGCCCGTGAATTGTATGGCTTCCTGANNNAGATCGGAAG |
| 3453_1 | TATGAGTGTGGAGTCGTTGCTTTGGTAGTGCTGTTTTCCAGGCNNNAGATCGGAAG |
| 3456_1 | TATGAGTGTGGAGTCGTTGCGGAAACCAGGAAACCTATACTGTNNNAGATCGGAAG |
| 3458_1 | TATGAGTGTGGAGTCGTTGCTTTCTCCAACCACTTTCAGCTGANNNAGATCGGAAGA |
| 3460_1 | TATGAGTGTGGAGTCGTTGCTCTGATGAGTGACATGGAGCTGANNNAGATCGGAA |
| 3461_1 | TATGAGTGTGGAGTCGTTGCCCCTATACACACTCCTGAGCTGANNNAGATCGGAAG |
| 3462_1 | TATGAGTGTGGAGTCGTTGCGTTCTCATAGTTCTGGACGCTGANNNAGATCGGAAG |
| 3464_1 | TATGAGTGTGGAGTCGTTGCTCAAAGTAGTTTCTCAAGGCTGANNNAGATCGGAAG |
| 3467_1 | TATGAGTGTGGAGTCGTTGCACCTTTCATACCTGTGAGGCTGANNNAGATCGGAAG |
| 3471_1 | TATGAGTGTGGAGTCGTTGCGTGATCTTTGGGATCTTGGCTGANNNAGATCGGAA |
| 3473_1 | TATGAGTGTGGAGTCGTTGCACGGACAACCATATCCCTTACANNNAGATCGGAAGA |
| 3475_1 | TATGAGTGTGGAGTCGTTGCGAGCATCCCTTGGTTTATGCTGANNNAGATCGGAAG |
| 3476_1 | TATGAGTGTGGAGTCGTTGCCTGTAACACACATACACTGCTGANNNAGATCGGAAG |
| 3479_1 | TATGAGTGTGGAGTCGTTGCACACACAAGCAGGAGAAATCTGANNNAGATCGGAA |

| | |
|---|---|
| 3481_1 | TATGAGTGTGGAGTCGTTGCTCTCACTCACTGAAGCCATCTGANNNAGATCGGAAG |
| 3484_1 | TATGAGTGTGGAGTCGTTGCATAACTTACCCTGTCTCCTCAGTNNNAGATCGGAAGA |
| 3487_1 | TATGAGTGTGGAGTCGTTGCTAGAACATGCCAGGACACTCTGANNNAGATCGGAA |
| 3493_1 | TATGAGTGTGGAGTCGTTGCGAGTGTCTCAGCCTTAGCTCTGANNNAGATCGGAAG |
| 3497_1 | TATGAGTGTGGAGTCGTTGCTCTGCCCATTTTTGATCGTCTGANNNAGATCGGAAG |
| 3499_1 | TATGAGTGTGGAGTCGTTGCTTCTGGTGAGCTCATCCTTCTGANNNAGATCGGAAG |
| 3502_1 | TATGAGTGTGGAGTCGTTGCTAGTCCTGTGAAGTGGGTTCTGANNNAGATCGGAA |
| 3503_1 | TATGAGTGTGGAGTCGTTGCCAGCCTCAAACTCCTAGACAATCNNNAGATCGGAAG |
| 3505_1 | TATGAGTGTGGAGTCGTTGCTGTCTGAAAGTTCCCGTTTCTGANNNAGATCGGAAG |
| 3508_1 | TATGAGTGTGGAGTCGTTGCATTGAGCCAACTGTGGAAAGTGANNNAGATCGGAA |
| 3510_1 | TATGAGTGTGGAGTCGTTGCTGGGAAAGTGTCCACTAAAGTGANNNAGATCGGAA |
| 3512_1 | TATGAGTGTGGAGTCGTTGCGCCTTATGGACCAACAGAAGTGANNNAGATCGGAA |
| 3513_1 | TATGAGTGTGGAGTCGTTGCTTGAAGGAAGAAACGGGAAGTGANNNAGATCGGA |
| 3514_1 | TATGAGTGTGGAGTCGTTGCCAAACTCCTGGCTTGAGCAACCCNNNAGATCGGAAG |
| 3515_1 | TATGAGTGTGGAGTCGTTGCACAGTGAGAGGGAAAGACAGTGANNNAGATCGGA |
| 3517_1 | TATGAGTGTGGAGTCGTTGCAGAGAGACCTTTTGTCCCAGTGANNNAGATCGGAA |
| 3518_1 | TATGAGTGTGGAGTCGTTGCTTGGTGACACAGCTCTTCTTGTTNNNAGATCGGAAG |
| 3521_1 | TATGAGTGTGGAGTCGTTGCTGGAGACTTTATCCTGGCAGTGANNNAGATCGGAA |
| 3522_1 | TATGAGTGTGGAGTCGTTGCTGCCATGTAGTCTCCAATTAGTGNNNAGATCGGAAG |
| 3531_1 | TATGAGTGTGGAGTCGTTGCTGATCTACCCTGTGCTTACGTGANNNAGATCGGAAG |
| 3532_1 | TATGAGTGTGGAGTCGTTGCCTTGGTTGATGTTTGGCATCATCNNNAGATCGGAAG |
| 3533_1 | TATGAGTGTGGAGTCGTTGCTCCACTTGCCATTTCAAAGGTGANNNAGATCGGAAG |
| 3534_1 | TATGAGTGTGGAGTCGTTGCTTTATGGAGAGGCCTCAAGGTGANNNAGATCGGAA |
| 3536_1 | TATGAGTGTGGAGTCGTTGCAACGTAAACACCGCATAAGGTGANNNAGATCGGAA |
| 3537_1 | TATGAGTGTGGAGTCGTTGCCCAGAGTGAGAGTGATCAGGTGANNNAGATCGGA |
| 3538_1 | TATGAGTGTGGAGTCGTTGCCTCCTCACTGTGTTGGCCAGGATNNNAGATCGGAAG |
| 3539_1 | TATGAGTGTGGAGTCGTTGCTCCATTGGTTGACACTCAGGTGANNNAGATCGGAA |
| 3540_1 | TATGAGTGTGGAGTCGTTGCGATGAAGAGAGTGGGAGAGGTGANNNAGATCGG |
| 3542_1 | TATGAGTGTGGAGTCGTTGCAAAAACACTGGGTCTCAGCATGTNNNAGATCGGAA |
| 3543_1 | TATGAGTGTGGAGTCGTTGCGTAACCCTGACACTTGGAGGTGANNNAGATCGGAA |
| 3544_1 | TATGAGTGTGGAGTCGTTGCAATGTATGACAGGGGTGAGGTGANNNAGATCGGA |
| 3546_1 | TATGAGTGTGGAGTCGTTGCGAGAGAAGAGACAGACAGGGTGANNNAGATCGGA |
| 3548_1 | TATGAGTGTGGAGTCGTTGCACAGACCGAGAACTAATGGGTGANNNAGATCGGAA |
| 3550_1 | TATGAGTGTGGAGTCGTTGCAGAGAGAGAAATGGGAATGGTGANNNAGATCGGA |
| 3551_1 | TATGAGTGTGGAGTCGTTGCGGGAGAGACTGAAGAGGAAATGANNNAGATCGGA |
| 3553_1 | TATGAGTGTGGAGTCGTTGCTGTGCATGGTGAGTGACTTTGGANNNAGATCGGAA |
| 3554_1 | TATGAGTGTGGAGTCGTTGCTACTTGCAGGGGAGTTATGGTGANNNAGATCGGAA |
| 3555_1 | TATGAGTGTGGAGTCGTTGCGGAAATGCTTGCTAGACTGGTGANNNAGATCGGAA |
| 3557_1 | TATGAGTGTGGAGTCGTTGCTTGTCCCTCCCTCCCTTATTTGTNNNAGATCGGAAGA |
| 3558_1 | TATGAGTGTGGAGTCGTTGCTGACAGGGTGTGATAAGTGGTGANNNAGATCGGA |
| 3560_1 | TATGAGTGTGGAGTCGTTGCAGGCTTGTTGCTAACTTTGGTGANNNAGATCGGAA |
| 3564_1 | TATGAGTGTGGAGTCGTTGCCACACCAGCATTTGCCTTTCTTTNNNAGATCGGAAGA |
| 3565_1 | TATGAGTGTGGAGTCGTTGCAGTTCAAAATCTCCTGGTCAGGTNNNAGATCGGAAG |

| | |
|---|---|
| 3566_1 | TATGAGTGTGGAGTCGTTGCCTGCTTGGTCTTCCCAGAGTTCANNNAGATCGGAAG |
| 3569_1 | TATGAGTGTGGAGTCGTTGCTGTTTCCAATAGGGTCAAGAGCCNNNAGATCGGAAG |
| 3575_1 | TATGAGTGTGGAGTCGTTGCAATATCCTTCCCATGGTGTCCATNNNAGATCGGAAG |
| 3576_1 | TATGAGTGTGGAGTCGTTGCTCACTGCTGTTTGTTCAGTGTGANNNAGATCGGAAG |
| 3582_1 | TATGAGTGTGGAGTCGTTGCGTGTTCAAGGGTCAACTGTGTGANNNAGATCGGAA |
| 3587_1 | TATGAGTGTGGAGTCGTTGCTGTCCAAAGTACAAGCGTTGTGANNNAGATCGGAAG |
| 3591_1 | TATGAGTGTGGAGTCGTTGCAACAACTCTGTGTAACTGGCTCANNNAGATCGGAAG |
| 3595_1 | TATGAGTGTGGAGTCGTTGCCTGGTCACTTGGATCACCTACACANNNAGATCGGAA |
| 3597_1 | TATGAGTGTGGAGTCGTTGCCAGTGTTGAGTGGAGCACATACNNNAGATCGGAAG |
| 3598_1 | TATGAGTGTGGAGTCGTTGCAGATAAAATGAGGGGGAGGAGCCNNNAGATCGGA |
| 3599_1 | TATGAGTGTGGAGTCGTTGCGATGCTTGAGACCACGACTACTGNNNAGATCGGAAG |
| 3602_1 | TATGAGTGTGGAGTCGTTGCTTACTCTCTCCTGCTGGCTTTGTNNNAGATCGGAAGA |
| 3607_1 | TATGAGTGTGGAGTCGTTGCGCTTGTTTTGGAGGTCTGCTGAANNNAGATCGGAA |
| 3610_1 | TATGAGTGTGGAGTCGTTGCCCACACTGATTTCGGTGACTTGANNNAGATCGGAAG |
| 3611_1 | TATGAGTGTGGAGTCGTTGCACATCTCACACATGCACCTTCTGNNNAGATCGGAAGA |
| 3612_1 | TATGAGTGTGGAGTCGTTGCTTTGGGAGACGCATCTCGTGTATNNNAGATCGGAA |
| 3615_1 | TATGAGTGTGGAGTCGTTGCACCCTCCTTTCGGTCTGTTTGTGNNNAGATCGGAAG |
| 3616_1 | TATGAGTGTGGAGTCGTTGCCCTCTCTCTCACACACTCCTTGANNNAGATCGGAAGA |
| 3618_1 | TATGAGTGTGGAGTCGTTGCGAAGAACCCTCACTAATAGCGGANNNAGATCGGAA |
| 3620_1 | TATGAGTGTGGAGTCGTTGCGGTATGTGTTGTATGTGCATGTGNNNAGATCGGAA |
| 3621_1 | TATGAGTGTGGAGTCGTTGCAGTGTGGTATCAAGGAGGCTTGANNNAGATCGGAA |
| 3623_1 | TATGAGTGTGGAGTCGTTGCTGTACCAGGCATGCAAATCTTGANNNAGATCGGAAG |
| 3624_1 | TATGAGTGTGGAGTCGTTGCGAAACCCGGTCATCACATCTTGANNNAGATCGGAAG |
| 3625_1 | TATGAGTGTGGAGTCGTTGCCTGTGGCTTGGATACCATCTTGANNNAGATCGGAAG |
| 3628_1 | TATGAGTGTGGAGTCGTTGCAGACAGCTAGAGGAGAGACAGAGNNNAGATCGGA |
| 3632_1 | TATGAGTGTGGAGTCGTTGCCCAAAGGTTTCTAAGAGCCACCANNNAGATCGGAAG |
| 3633_1 | TATGAGTGTGGAGTCGTTGCCCAATGTAGCACCAAGAGGTTGANNNAGATCGGAA |
| 3635_1 | TATGAGTGTGGAGTCGTTGCTGGGAAGAGACAACAAAAGCCTACNNNAGATCGGA |
| 3638_1 | TATGAGTGTGGAGTCGTTGCAAGCCAATCATCCAGCTTGTTGANNNAGATCGGAAG |
| 3643_1 | TATGAGTGTGGAGTCGTTGCCCTGATGTTCTGCTGGGATTTGANNNAGATCGGAAG |
| 3644_1 | TATGAGTGTGGAGTCGTTGCTTGCTCTATTGTGCTGCCTTTGANNNAGATCGGAAG |
| 3645_1 | TATGAGTGTGGAGTCGTTGCCTTCTGTGGAATAACGCTTGTGTNNNAGATCGGAAG |
| 3646_1 | TATGAGTGTGGAGTCGTTGCGGGTGCTGTATTACTGGCTTTGANNNAGATCGGAA |
| 3647_1 | TATGAGTGTGGAGTCGTTGCAAACTGTTCCCTTTGCCTCCAATNNNAGATCGGAAGA |
| 3651_1 | TATGAGTGTGGAGTCGTTGCTGGGACTAGAAGGGATGGTTTGANNNAGATCGGA |
| 3654_1 | TATGAGTGTGGAGTCGTTGCACAGGTTCTCTGGTTGAAACCTGNNNAGATCGGAAG |
| 3655_1 | TATGAGTGTGGAGTCGTTGCAGATCCTCAGTTGGTGTGTTTGANNNAGATCGGAA |
| 3658_1 | TATGAGTGTGGAGTCGTTGCTAGCAAGCCTCTTTGCCTTTTGANNNAGATCGGAAG |
| 3660_1 | TATGAGTGTGGAGTCGTTGCATGGACAAAACAACTGCTTTTGANNNAGATCGGAAG |
| 3661_1 | TATGAGTGTGGAGTCGTTGCCTCCTCCTCCACTCCTCTTTTGANNNAGATCGGAAGA |
| 3667_1 | TATGAGTGTGGAGTCGTTGCGGGGACAGGGAACCTGAAAAATANNNAGATCGGA |
| 3668_1 | TATGAGTGTGGAGTCGTTGCAGGAATGCCAAGCCCACAAAATANNNAGATCGGAA |
| 3670_1 | TATGAGTGTGGAGTCGTTGCCGCATACCCCGAAAAAGCAAATANNNAGATCGGAAG |

| | |
|---|---|
| 3671_1 | TATGAGTGTGGAGTCGTTGCGGCAGCATACCACACAGCAAATANNNAGATCGGAAG |
| 3673_1 | TATGAGTGTGGAGTCGTTGCATAGCGGCAGCAGCAAAGAAATANNNAGATCGGAA |
| 3674_1 | TATGAGTGTGGAGTCGTTGCTACCAGCCAAGCACAAGGAAATANNNAGATCGGAAG |
| 3675_1 | TATGAGTGTGGAGTCGTTGCGCACACACGTGAGGAAGGAAATANNNAGATCGGAA |
| 3676_1 | TATGAGTGTGGAGTCGTTGCCCTAATGTGGCCAGCTGGAAATANNNAGATCGGAAG |
| 3677_1 | TATGAGTGTGGAGTCGTTGCTTCTTCGCTCCCTCCCTGAAATANNNAGATCGGAAGA |
| 3678_1 | TATGAGTGTGGAGTCGTTGCAGTTGGAGGGAGCAAAAGTTTGTNNNAGATCGGAA |
| 3683_1 | TATGAGTGTGGAGTCGTTGCGAGTTCTTGGTCGTGCAGAATGTNNNAGATCGGAAG |
| 3686_1 | TATGAGTGTGGAGTCGTTGCAGTGTCTACCCACGGTGTCAATANNNAGATCGGAAG |
| 3687_1 | TATGAGTGTGGAGTCGTTGCTGCGTTGGGGCAGAATTTCAATANNNAGATCGGAAG |
| 3690_1 | TATGAGTGTGGAGTCGTTGCTTGCATGTTTGGTGGCAGGAATANNNAGATCGGAAG |
| 3691_1 | TATGAGTGTGGAGTCGTTGCCATCTGGGAGGAACTTGAAGTGGNNNAGATCGGAA |
| 3692_1 | TATGAGTGTGGAGTCGTTGCTTTGGCCACTCACTGGGTGAATANNNAGATCGGAAG |
| 3693_1 | TATGAGTGTGGAGTCGTTGCGACATTACAGGTGCATGCCATGGNNNAGATCGGAAG |
| 3695_1 | TATGAGTGTGGAGTCGTTGCTCCCCCATGCACCTTCAGTAATANNNAGATCGGAAGA |
| 3696_1 | TATGAGTGTGGAGTCGTTGCCCCCCTGACTTCCCAAAGAACATANNNAGATCGGAAGA |
| 3698_1 | TATGAGTGTGGAGTCGTTGCGCTCCCCTTCTCTCTCACACATANNNAGATCGGAAGAG |
| 3699_1 | TATGAGTGTGGAGTCGTTGCTATCCCAAGTGGTGGCCCACATANNNAGATCGGAAG |
| 3701_1 | TATGAGTGTGGAGTCGTTGCCAGGGAGAGGAAGAGCAGACATANNNAGATCGGAA |
| 3703_1 | TATGAGTGTGGAGTCGTTGCGAAAAGGCTGTTGTGCCAGAAGTNNNAGATCGGAA |
| 3705_1 | TATGAGTGTGGAGTCGTTGCACCATTCTCGACCAGAGACCATANNNAGATCGGAAGA |
| 3706_1 | TATGAGTGTGGAGTCGTTGCATTTCTGCCTCAAAGGGTGGAAGNNNAGATCGGAAG |
| 3707_1 | TATGAGTGTGGAGTCGTTGCTATGGCAGTTCCCTCATGGTCCTNNNAGATCGGAAGA |
| 3709_1 | TATGAGTGTGGAGTCGTTGCGTTTTAGGCCAGTGTGAGCCATANNNAGATCGGAAG |
| 3710_1 | TATGAGTGTGGAGTCGTTGCGAGTTTCTGGGAGGGTAGCCATANNNAGATCGGAA |
| 3711_1 | TATGAGTGTGGAGTCGTTGCGCAAGGAATTCTCACTGGCCATANNNAGATCGGAAG |
| 3713_1 | TATGAGTGTGGAGTCGTTGCCCCCCTAATGCATCCAGCTCCATANNNAGATCGGAAGA |
| 3716_1 | TATGAGTGTGGAGTCGTTGCCCGTCAGCCATCTTTGTTCCATANNNAGATCGGAAGA |
| 3717_1 | TATGAGTGTGGAGTCGTTGCTACATTGTCTTCCCAGCAGTGTTNNNAGATCGGAAGA |
| 3718_1 | TATGAGTGTGGAGTCGTTGCCTGCATGGACTTGAGTGAGCATANNNAGATCGGAAG |
| 3719_1 | TATGAGTGTGGAGTCGTTGCCAGTTCAAGTCGAAGTCCAAAGGNNNAGATCGGAAG |
| 3721_1 | TATGAGTGTGGAGTCGTTGCTCTAAGGATGGCTGAAGGGCATANNNAGATCGGAA |
| 3722_1 | TATGAGTGTGGAGTCGTTGCGTTACACAGCTGAATGGGGCATANNNAGATCGGAAG |
| 3726_1 | TATGAGTGTGGAGTCGTTGCGTGATATGGGATAACCTCCCACCNNNAGATCGGAAG |
| 3727_1 | TATGAGTGTGGAGTCGTTGCCACAGTTGGTGGCATACTGCATANNNAGATCGGAAG |
| 3728_1 | TATGAGTGTGGAGTCGTTGCCAAAGAGAACAGGCAGGTGCATANNNAGATCGGAA |
| 3729_1 | TATGAGTGTGGAGTCGTTGCAAGTGCCAACAGATAGAAGAGGGNNNAGATCGGAA |
| 3731_1 | TATGAGTGTGGAGTCGTTGCCTCCTGCAAATAGCTGGCTCATANNNAGATCGGAAGA |
| 3733_1 | TATGAGTGTGGAGTCGTTGCCCGCAAGCTAAGAGGCAAAGATANNNAGATCGGAAG |
| 3736_1 | TATGAGTGTGGAGTCGTTGCCCTGCGGACCTGATCCTAAGATANNNAGATCGGAAG |
| 3737_1 | TATGAGTGTGGAGTCGTTGCGCCCTACTATGTACCCACAGATANNNAGATCGGAAGA |
| 3738_1 | TATGAGTGTGGAGTCGTTGCCAATGTCAGCCACACACCAGATANNNAGATCGGAAG |
| 3740_1 | TATGAGTGTGGAGTCGTTGCTCCGCGTTGGAAAAACTCAGATANNNAGATCGGAAG |

| | |
|---|---|
| 3742_1 | TATGAGTGTGGAGTCGTTGCTGCGTTTGGATTTGCCAGAGATANNNAGATCGGAA |
| 3743_1 | TATGAGTGTGGAGTCGTTGCAACAACCACTGACCCAGGAGATANNNAGATCGGAA |
| 3744_1 | TATGAGTGTGGAGTCGTTGCGCACAAACCAGAGCAGTGAGATANNNAGATCGGAA |
| 3747_1 | TATGAGTGTGGAGTCGTTGCTCCTTCCACCTTATTACCCCCTGNNNAGATCGGAAGA |
| 3748_1 | TATGAGTGTGGAGTCGTTGCGTTCTTGAGGCTCTTGATGAGCTNNNAGATCGGAAG |
| 3749_1 | TATGAGTGTGGAGTCGTTGCCTGGGTAAGGGAGCAGAAGGATANNNAGATCGGA |
| 3750_1 | TATGAGTGTGGAGTCGTTGCTTGAGGAAGTGGGTCTGGTGTTGNNNAGATCGGA |
| 3752_1 | TATGAGTGTGGAGTCGTTGCGATGAGCCCTGCTTAGGAGGATANNNAGATCGGAA |
| 3753_1 | TATGAGTGTGGAGTCGTTGCGATGGATTTTGACAGGCCGGATANNNAGATCGGAA |
| 3754_1 | TATGAGTGTGGAGTCGTTGCGATGTGGTAGGGGTGAAGGGATANNNAGATCGGA |
| 3755_1 | TATGAGTGTGGAGTCGTTGCGGACACCCCTTAGTTGAGGGATANNNAGATCGGAA |
| 3757_1 | TATGAGTGTGGAGTCGTTGCGGAGTGGATATTGGGGGTGGATANNNAGATCGGA |
| 3758_1 | TATGAGTGTGGAGTCGTTGCAAAAGGTGTTTGATCCCTTCCCTNNNAGATCGGAAG |
| 3761_1 | TATGAGTGTGGAGTCGTTGCGACCACCTTCAGGGATTCTGATANNNAGATCGGAAG |
| 3762_1 | TATGAGTGTGGAGTCGTTGCACCTGACTGCTCCTCTTGTGATANNNAGATCGGAAG |
| 3764_1 | TATGAGTGTGGAGTCGTTGCACCTTGCACCTGCGTGAAATATANNNAGATCGGAAG |
| 3765_1 | TATGAGTGTGGAGTCGTTGCAGGTTCAAGTGGATTGGCTCAACNNNAGATCGGAA |
| 3766_1 | TATGAGTGTGGAGTCGTTGCGGACAGCCACACAGTTGCATATANNNAGATCGGAA |
| 3768_1 | TATGAGTGTGGAGTCGTTGCCAACCCTCCCACCCCCATATATANNNAGATCGGAAGA |
| 3769_1 | TATGAGTGTGGAGTCGTTGCACTGCCACCCAGGTCGATATATANNNAGATCGGAAG |
| 3771_1 | TATGAGTGTGGAGTCGTTGCCCCCTTCCTCAGCCGTTACTATANNNAGATCGGAAGA |
| 3772_1 | TATGAGTGTGGAGTCGTTGCGTTCCTTTGAATTTTGGGGTTTGNNNAGATCGGAAG |
| 3774_1 | TATGAGTGTGGAGTCGTTGCGCAAGGGCAATCCAGAAATCAGANNNAGATCGGAA |
| 3775_1 | TATGAGTGTGGAGTCGTTGCTTCCCACCCTCTTTGCCAGTATANNNAGATCGGAAGA |
| 3776_1 | TATGAGTGTGGAGTCGTTGCATGTCTTTCACTCATGGAGGGAGNNNAGATCGGAA |
| 3777_1 | TATGAGTGTGGAGTCGTTGCTGGGGCAGGAGTGATTACGTATANNNAGATCGGAA |
| 3778_1 | TATGAGTGTGGAGTCGTTGCTAGATTGTCCAGGTTCCAGGAAGNNNAGATCGGAA |
| 3780_1 | TATGAGTGTGGAGTCGTTGCGCCTGCATTTCTGTGCCCTTATANNNAGATCGGAAG |
| 3781_1 | TATGAGTGTGGAGTCGTTGCTAGTTATTGCAGACGGGGCAAATNNNAGATCGGAA |
| 3782_1 | TATGAGTGTGGAGTCGTTGCGGCACAGGCAGTAAGAACAACTANNNAGATCGGAA |
| 3784_1 | TATGAGTGTGGAGTCGTTGCGCCACGTATCAAGCACAGAACTANNNAGATCGGAAG |
| 3785_1 | TATGAGTGTGGAGTCGTTGCGAGTTTATGCTAGGGCAGAACTANNNAGATCGGAA |
| 3786_1 | TATGAGTGTGGAGTCGTTGCGAGGATGGGCTCTAGTCGAACTANNNAGATCGGAA |
| 3787_1 | TATGAGTGTGGAGTCGTTGCGGTCCCAGAGGAGAAAGGAACTANNNAGATCGGA |
| 3788_1 | TATGAGTGTGGAGTCGTTGCACGGGACAGACAACCTGTAACTANNNAGATCGGAA |
| 3790_1 | TATGAGTGTGGAGTCGTTGCCTTGCTCCCAAAGTTCACCACTANNNAGATCGGAAG |
| 3791_1 | TATGAGTGTGGAGTCGTTGCAAGTAACCACTCTCCAGCCACTANNNAGATCGGAAG |
| 3792_1 | TATGAGTGTGGAGTCGTTGCCGTCCACCAAAGATTGAGCACTANNNAGATCGGAAG |
| 3793_1 | TATGAGTGTGGAGTCGTTGCGGGCTGATGAGAAATTGGCACTANNNAGATCGGAA |
| 3794_1 | TATGAGTGTGGAGTCGTTGCCATCACCCAAATGCCCATCACTANNNAGATCGGAAG |
| 3796_1 | TATGAGTGTGGAGTCGTTGCTGCTAAGCCTCTTTCTCAGTCCTNNNAGATCGGAAGA |
| 3797_1 | TATGAGTGTGGAGTCGTTGCCAACCCAAAGTGGTGCAAGACTANNNAGATCGGAA |
| 3800_1 | TATGAGTGTGGAGTCGTTGCAGGGTGGAAAAACAGGCTAATGANNNAGATCGGA |

| | |
|---|---|
| 3801_1 | TATGAGTGTGGAGTCGTTGCACTCCCTTCCATGATGATGACCANNNAGATCGGAAG |
| 3804_1 | TATGAGTGTGGAGTCGTTGCTCCTTCTCTTCCTCTGGTGACTANNNAGATCGGAAGA |
| 3809_1 | TATGAGTGTGGAGTCGTTGCCCTGGCTGATTCCCACTCAAATTNNNAGATCGGAAG |
| 3810_1 | TATGAGTGTGGAGTCGTTGCCTATCTCCTTGTGACTTGGCTGCNNNAGATCGGAAG |
| 3811_1 | TATGAGTGTGGAGTCGTTGCCTTCTCTCACTCATTTGTGTAGCNNNAGATCGGAAGA |
| 3812_1 | TATGAGTGTGGAGTCGTTGCGGGAAGGGTGAGGTGATGTACTANNNAGATCGGA |
| 3813_1 | TATGAGTGTGGAGTCGTTGCATGCAGGAGGTATTGTTATCCCANNNAGATCGGAA |
| 3815_1 | TATGAGTGTGGAGTCGTTGCGATACTTGCTGAGCACCAACCTANNNAGATCGGAAG |
| 3816_1 | TATGAGTGTGGAGTCGTTGCTTTCTGCTGCTACTCTCCACCTANNNAGATCGGAAGA |
| 3819_1 | TATGAGTGTGGAGTCGTTGCTTCCACTTTCAAAGGCCACATGANNNAGATCGGAAG |
| 3821_1 | TATGAGTGTGGAGTCGTTGCGGGGAACAACTGAGCCTTACCTANNNAGATCGGAA |
| 3822_1 | TATGAGTGTGGAGTCGTTGCAGTTGTTTTTCATGAACTGCTCANNNAGATCGGAAG |
| 3823_1 | TATGAGTGTGGAGTCGTTGCCATTTCACTGCATTCCCACCCTANNNAGATCGGAAGA |
| 3826_1 | TATGAGTGTGGAGTCGTTGCTTGTTCGTAGTGTTTTGGTGGGANNNAGATCGGAA |
| 3827_1 | TATGAGTGTGGAGTCGTTGCACCTAGTCATGGAACAAGCCCTANNNAGATCGGAAG |
| 3828_1 | TATGAGTGTGGAGTCGTTGCGATACAAAATGGCCACAGCCCTANNNAGATCGGAAG |
| 3829_1 | TATGAGTGTGGAGTCGTTGCCCACTGACATAGATGAGGCCCTANNNAGATCGGAAG |
| 3830_1 | TATGAGTGTGGAGTCGTTGCACTCATGCCCAGTGTTATCCCTANNNAGATCGGAAG |
| 3832_1 | TATGAGTGTGGAGTCGTTGCCTTTCCTGCCTTGCTTTTCCCTANNNAGATCGGAAGA |
| 3834_1 | TATGAGTGTGGAGTCGTTGCTAGGGTCATGTGCTTTCAGCCTANNNAGATCGGAAG |
| 3836_1 | TATGAGTGTGGAGTCGTTGCAAAAGCACTGGCAGGAAATACCANNNAGATCGGAA |
| 3837_1 | TATGAGTGTGGAGTCGTTGCAAAGACGGAGGACCATATGCCTANNNAGATCGGAA |
| 3838_1 | TATGAGTGTGGAGTCGTTGCAGATTGCTTTTCTTGGCTGCCTANNNAGATCGGAAG |
| 3839_1 | TATGAGTGTGGAGTCGTTGCTGGTCTTTTCCTGTGTTTAGGCCNNNAGATCGGAAG |
| 3841_1 | TATGAGTGTGGAGTCGTTGCCCTTGGTCTGCAAGATTTGCCTANNNAGATCGGAAG |
| 3843_1 | TATGAGTGTGGAGTCGTTGCTCGCCCATGCAGATAGAATCCTANNNAGATCGGAAG |
| 3844_1 | TATGAGTGTGGAGTCGTTGCGCTGCCTTGATTTGTCCATCCTANNNAGATCGGAAG |
| 3847_1 | TATGAGTGTGGAGTCGTTGCCAAGGCTCTGCTGGAAATTCCTANNNAGATCGGAAG |
| 3848_1 | TATGAGTGTGGAGTCGTTGCCCGAGAGATCAGAAGCCTTCCTANNNAGATCGGAAG |
| 3850_1 | TATGAGTGTGGAGTCGTTGCCAGAACTCTAAGATGCTTGGGCANNNAGATCGGAA |
| 3851_1 | TATGAGTGTGGAGTCGTTGCAGAGCCCAGCAAGGTATTTCCTANNNAGATCGGAAG |
| 3852_1 | TATGAGTGTGGAGTCGTTGCCTCCACCTTCCCCTTCTTTCCTANNNAGATCGGAAGAG |
| 3853_1 | TATGAGTGTGGAGTCGTTGCCCAGCATGAGGCAGTTTTTCCTANNNAGATCGGAAG |
| 3854_1 | TATGAGTGTGGAGTCGTTGCGAAAAGCGAATGTAGGGTGGGGANNNAGATCGGA |
| 3857_1 | TATGAGTGTGGAGTCGTTGCATTGTGTGTGTTGCTTCAAGCTANNNAGATCGGAAG |
| 3859_1 | TATGAGTGTGGAGTCGTTGCCTGGGCCTATGCAGCTACTGTTTNNNAGATCGGAAG |
| 3861_1 | TATGAGTGTGGAGTCGTTGCTTCTTTTGCCATTCAGGCAGCTANNNAGATCGGAAG |
| 3864_1 | TATGAGTGTGGAGTCGTTGCTTCCTAGTTTACTTGTGGGGTGTNNNAGATCGGAAG |
| 3866_1 | TATGAGTGTGGAGTCGTTGCTGAGAATTCCTGCTTCCCCAATTNNNAGATCGGAAG |
| 3867_1 | TATGAGTGTGGAGTCGTTGCCCCACCTCTCTGAACAGTAGCTANNNAGATCGGAAG |
| 3868_1 | TATGAGTGTGGAGTCGTTGCCTGGTCCTTGTCTGGAGTAGCTANNNAGATCGGAAG |
| 3870_1 | TATGAGTGTGGAGTCGTTGCCCCTTTCTGTCTCTCTCTCGCTANNNAGATCGGAAGA |
| 3871_1 | TATGAGTGTGGAGTCGTTGCCACATTGCTCCAGGATAAGGCTANNNAGATCGGAAG |

| | |
|---|---|
| 3872_1 | TATGAGTGTGGAGTCGTTGCGAAGTGAATTGGCCTTGAGGCTANNNAGATCGGAA |
| 3873_1 | TATGAGTGTGGAGTCGTTGCGAGACGAGCTTTTGCTTAGGCTANNNAGATCGGAA |
| 3874_1 | TATGAGTGTGGAGTCGTTGCAATGACAATGACCTCCAGGGCTANNNAGATCGGAAG |
| 3878_1 | TATGAGTGTGGAGTCGTTGCATTTACCCCTCCACTGATGGCTANNNAGATCGGAAG |
| 3879_1 | TATGAGTGTGGAGTCGTTGCGAAGTACCAGGTTCCAGCCAATTNNNAGATCGGAAG |
| 3881_1 | TATGAGTGTGGAGTCGTTGCTAAGCAGTATGACAGCCCTACCANNNAGATCGGAAG |
| 3882_1 | TATGAGTGTGGAGTCGTTGCGAAGCCAACTGAAGGAACTGCTANNNAGATCGGAA |
| 3883_1 | TATGAGTGTGGAGTCGTTGCCTCCTTTGTCTTCCAAGCTGCTANNNAGATCGGAAGA |
| 3884_1 | TATGAGTGTGGAGTCGTTGCAAGAGTTAGGGCAACAGCTGCTANNNAGATCGGAA |
| 3885_1 | TATGAGTGTGGAGTCGTTGCCTCTGCATGGTAAAAGGCTGCTANNNAGATCGGAA |
| 3886_1 | TATGAGTGTGGAGTCGTTGCTATGGCACAGACATTTGCTGCTANNNAGATCGGAAG |
| 3888_1 | TATGAGTGTGGAGTCGTTGCGAAGCGAAAAGTAGTGGGTGCTANNNAGATCGGA |
| 3889_1 | TATGAGTGTGGAGTCGTTGCCCCTGATGGGTAGAAATGTGCTANNNAGATCGGAA |
| 3890_1 | TATGAGTGTGGAGTCGTTGCATGATCGCACTGGAAATGTGCTANNNAGATCGGAA |
| 3892_1 | TATGAGTGTGGAGTCGTTGCACCCTCCAGCAAGAAAATTGCTANNNAGATCGGAAG |
| 3893_1 | TATGAGTGTGGAGTCGTTGCATGCTTCTGCTTGGGGTTCTAAGNNNAGATCGGAAG |
| 3894_1 | TATGAGTGTGGAGTCGTTGCAGGAGTACATGAGGCAGTTGCTANNNAGATCGGAA |
| 3895_1 | TATGAGTGTGGAGTCGTTGCGCTGGATGTAGTTGGTGTTGCTANNNAGATCGGAA |
| 3897_1 | TATGAGTGTGGAGTCGTTGCTCCCTGCACCACCAGTAAATCTANNNAGATCGGAAG |
| 3898_1 | TATGAGTGTGGAGTCGTTGCGGGGCACTGCTACCTTACTTTAGNNNAGATCGGAAG |
| 3899_1 | TATGAGTGTGGAGTCGTTGCGGAAGTTTGGCAGGCAGAATCTANNNAGATCGGAA |
| 3900_1 | TATGAGTGTGGAGTCGTTGCGGAGGCGGTAAGTCTGGAATCTANNNAGATCGGAA |
| 3904_1 | TATGAGTGTGGAGTCGTTGCCAGCAGTGGAAGCAGCATATCTANNNAGATCGGAA |
| 3907_1 | TATGAGTGTGGAGTCGTTGCGCTCCCACTGGTGTTGATCTCTANNNAGATCGGAAG |
| 3908_1 | TATGAGTGTGGAGTCGTTGCATAAAACATATGCCTTGGACGGGNNNAGATCGGAA |
| 3911_1 | TATGAGTGTGGAGTCGTTGCGTGTTCAAGGCTAGGAGGGTCTANNNAGATCGGAA |
| 3912_1 | TATGAGTGTGGAGTCGTTGCCACCAATTGCCTTTCTGGGTCTANNNAGATCGGAAG |
| 3915_1 | TATGAGTGTGGAGTCGTTGCGTCTGAACCCTTACCCCTGTCTANNNAGATCGGAAGA |
| 3916_1 | TATGAGTGTGGAGTCGTTGCACCCCAAAGTTACCAGATCACTTNNNAGATCGGAAG |
| 3917_1 | TATGAGTGTGGAGTCGTTGCTCCAATCAGCCTTCTGGTGTCTANNNAGATCGGAAG |
| 3918_1 | TATGAGTGTGGAGTCGTTGCATGTGAGAGCCATTGTGTGTCTANNNAGATCGGAA |
| 3919_1 | TATGAGTGTGGAGTCGTTGCACTGCTTTGTTCCAGCTTGTCTANNNAGATCGGAAG |
| 3920_1 | TATGAGTGTGGAGTCGTTGCTGGGAGCAGAGTGGATTTGTCTANNNAGATCGGAA |
| 3922_1 | TATGAGTGTGGAGTCGTTGCTTGCCCACACATCCATTCCTATGNNNAGATCGGAAGA |
| 3923_1 | TATGAGTGTGGAGTCGTTGCTGTGTGGTCCCAGTGGTATTCTANNNAGATCGGAAG |
| 3924_1 | TATGAGTGTGGAGTCGTTGCGAGCATGTGAGTGCACACTTCTANNNAGATCGGAA |
| 3925_1 | TATGAGTGTGGAGTCGTTGCTCTCCTCGTCCTCCTCACTTCTANNNAGATCGGAAGA |
| 3926_1 | TATGAGTGTGGAGTCGTTGCAAAAATATTAGCTGGGCGCCACCNNNAGATCGGAA |
| 3928_1 | TATGAGTGTGGAGTCGTTGCGTGAAACCCAAGCCACAGTTCTANNNAGATCGGAAG |
| 3929_1 | TATGAGTGTGGAGTCGTTGCTGGGTTATGGAGCCACAGTTCTANNNAGATCGGAA |
| 3930_1 | TATGAGTGTGGAGTCGTTGCGTGCAGATGCCAGTGATGTTCTANNNAGATCGGAA |
| 3931_1 | TATGAGTGTGGAGTCGTTGCATGAGCCCCCAGTAATTGTTATCNNNAGATCGGAAG |
| 3935_1 | TATGAGTGTGGAGTCGTTGCCGTGTGTGTGTACGTTTGCAATTNNNAGATCGGAAG |

| | |
|---|---|
| 3936_1 | TATGAGTGTGGAGTCGTTGCCACCCTGGTCTTCTCGTCAAGTANNNAGATCGGAAGA |
| 3937_1 | TATGAGTGTGGAGTCGTTGCGTGTGTGCGCGCATGTATAAGTANNNAGATCGGAA |
| 3938_1 | TATGAGTGTGGAGTCGTTGCACCCTGGGAAAATTTGATCATGANNNAGATCGGAAG |
| 3939_1 | TATGAGTGTGGAGTCGTTGCAGTGGGGATGTGTGAGAACAGTANNNAGATCGGA |
| 3943_1 | TATGAGTGTGGAGTCGTTGCGAACCAGCATAAGGTTGCCAGTANNNAGATCGGAA |
| 3950_1 | TATGAGTGTGGAGTCGTTGCCTCGGCATTACCTTCAAGGAGTANNNAGATCGGAAG |
| 3952_1 | TATGAGTGTGGAGTCGTTGCCGTCTGTGTAGACTGTGGGAGTANNNAGATCGGAA |
| 3953_1 | TATGAGTGTGGAGTCGTTGCTTGGCTCTCAAGGATGTGGAGTANNNAGATCGGAA |
| 3956_1 | TATGAGTGTGGAGTCGTTGCCAGACGTAAGCATATTCGTGGGGNNNAGATCGGAA |
| 3958_1 | TATGAGTGTGGAGTCGTTGCCAGGTACACACACGTGCTTAGTANNNAGATCGGAAG |
| 3959_1 | TATGAGTGTGGAGTCGTTGCGTGTGTGTGGAGATGGCTTATCTNNNAGATCGGAA |
| 3960_1 | TATGAGTGTGGAGTCGTTGCTCTGGCTGGCATATTTGAACGTANNNAGATCGGAAG |
| 3961_1 | TATGAGTGTGGAGTCGTTGCCTCTACCTCCCAGGCTCAAAGGCNNNAGATCGGAAGA |
| 3962_1 | TATGAGTGTGGAGTCGTTGCTAGTGAACAAAGCAAGCCTTCCTNNNAGATCGGAAG |
| 3966_1 | TATGAGTGTGGAGTCGTTGCGCCTGTGTATCCTTCTTTGCGTANNNAGATCGGAAGA |
| 3970_1 | TATGAGTGTGGAGTCGTTGCGCCAGTAAGCCCAAGAAAAGGTANNNAGATCGGAA |
| 3973_1 | TATGAGTGTGGAGTCGTTGCGGAGTTATTGGGTGTGCAAGGTANNNAGATCGGAA |
| 3974_1 | TATGAGTGTGGAGTCGTTGCAAACCTCTCGATTGGGGAAGGTANNNAGATCGGAA |
| 3977_1 | TATGAGTGTGGAGTCGTTGCACCATCCACAAAGACCTCAGGTANNNAGATCGGAAG |
| 3981_1 | TATGAGTGTGGAGTCGTTGCACCCCCTTTTTCTCCTTCCCTTTNNNAGATCGGAAGAG |
| 3982_1 | TATGAGTGTGGAGTCGTTGCGCATTGTTGAGTTCTGGGAGGTANNNAGATCGGAA |
| 3983_1 | TATGAGTGTGGAGTCGTTGCCAAACTCACACTGTTCCCCTAGANNNAGATCGGAAGA |
| 3984_1 | TATGAGTGTGGAGTCGTTGCGGGCCATGGATTTGTGAATGAAGNNNAGATCGGAA |
| 3985_1 | TATGAGTGTGGAGTCGTTGCGGGGTGATGTTTCTGCCTAGGTANNNAGATCGGAA |
| 3988_1 | TATGAGTGTGGAGTCGTTGCTGTTCCAAGACCCCTGATATCGTNNNAGATCGGAAGA |
| 3989_1 | TATGAGTGTGGAGTCGTTGCCCACAAATGGTAAGCACAGGGTANNNAGATCGGAA |
| 3991_1 | TATGAGTGTGGAGTCGTTGCTTTATGTATGTCTTCCCACGTGTNNNAGATCGGAAGA |
| 3992_1 | TATGAGTGTGGAGTCGTTGCAGGAGGGTGACTACTAAGGGGTANNNAGATCGGA |
| 3994_1 | TATGAGTGTGGAGTCGTTGCCATAGTTGAGTAGGGCAGGGGTANNNAGATCGGA |
| 3995_1 | TATGAGTGTGGAGTCGTTGCTTGAATGGTGACAAAATTGGTGANNNAGATCGGAA |
| 4001_1 | TATGAGTGTGGAGTCGTTGCGGGGTCAACCTTTTTGATGGGTANNNAGATCGGAA |
| 4002_1 | TATGAGTGTGGAGTCGTTGCTATCTTGCCCAGACCTTTCCTCTNNNAGATCGGAAGA |
| 4003_1 | TATGAGTGTGGAGTCGTTGCGGACTGAACATTTTTCCTGGCAGNNNAGATCGGAAG |
| 4005_1 | TATGAGTGTGGAGTCGTTGCCTCCCTCTCTCTCGGAAATGGTANNNAGATCGGAAGA |
| 4006_1 | TATGAGTGTGGAGTCGTTGCATGGTGAGATGGTGGGAATGGTANNNAGATCGGA |
| 4008_1 | TATGAGTGTGGAGTCGTTGCACCAACCTGACAGTGGCTATCTCNNNAGATCGGAAG |
| 4010_1 | TATGAGTGTGGAGTCGTTGCATTGTGAGTTGGGGAGACTGGTANNNAGATCGGAA |
| 4012_1 | TATGAGTGTGGAGTCGTTGCATCCATTTGCACAGATGCTGGTANNNAGATCGGAAG |
| 4014_1 | TATGAGTGTGGAGTCGTTGCGACAACCAAGAATGCCAGTGGTANNNAGATCGGAA |
| 4015_1 | TATGAGTGTGGAGTCGTTGCAAGTCCCAGTCAATGAGGTGGTANNNAGATCGGAA |
| 4017_1 | TATGAGTGTGGAGTCGTTGCGTTCCTGTAGTGGGAGTGTGGTANNNAGATCGGAA |
| 4019_1 | TATGAGTGTGGAGTCGTTGCGCCTGGTTGTGTCTTGATTGGTANNNAGATCGGAAG |
| 4020_1 | TATGAGTGTGGAGTCGTTGCAAGCAGGGAGGGTCATATTGGTANNNAGATCGGAA |

| | |
|---|---|
| 4021_1 | TATGAGTGTGGAGTCGTTGCACCACCATCCTGAATCCTTGGTANNNAGATCGGAAG |
| 4023_1 | TATGAGTGTGGAGTCGTTGCCTTTCCATGGCCTCTACTCCTCANNNAGATCGGAAGA |
| 4024_1 | TATGAGTGTGGAGTCGTTGCGGTCTTTCGTCTGTGCACATGTANNNAGATCGGAAG |
| 4025_1 | TATGAGTGTGGAGTCGTTGCATCTGTCTTTCTGGGGGCATGTANNNAGATCGGAAG |
| 4030_1 | TATGAGTGTGGAGTCGTTGCTTGAGTGACATAGCCCCACTGTANNNAGATCGGAAG |
| 4031_1 | TATGAGTGTGGAGTCGTTGCCCCTAGAAAGTAGGCCCACTGTANNNAGATCGGAAG |
| 4032_1 | TATGAGTGTGGAGTCGTTGCTGGTGTGATAGTCGCTCACTGTANNNAGATCGGAAG |
| 4033_1 | TATGAGTGTGGAGTCGTTGCGGACGGTGGGCATGAGAATAATTNNNAGATCGGAA |
| 4034_1 | TATGAGTGTGGAGTCGTTGCATGGGCCAAAGATTGAGACTGTANNNAGATCGGAA |
| 4035_1 | TATGAGTGTGGAGTCGTTGCGAAGGAAGGCAAGCAAACCTGTANNNAGATCGGAA |
| 4036_1 | TATGAGTGTGGAGTCGTTGCCAAGGCATTTTACCCAGCCTGTANNNAGATCGGAAG |
| 4040_1 | TATGAGTGTGGAGTCGTTGCGCTGTCCCAAAAGGCTTAGTGTANNNAGATCGGAAG |
| 4044_1 | TATGAGTGTGGAGTCGTTGCCAGGTGCATGGTGTATTGGTGTANNNAGATCGGAA |
| 4045_1 | TATGAGTGTGGAGTCGTTGCGTGTGCTGGAGCCAATATGTGTANNNAGATCGGAA |
| 4046_1 | TATGAGTGTGGAGTCGTTGCCTTCACCAGACATTGGCTGTGTANNNAGATCGGAAG |
| 4047_1 | TATGAGTGTGGAGTCGTTGCATGCCTCTATTTCAGCGTGTGTANNNAGATCGGAAG |
| 4049_1 | TATGAGTGTGGAGTCGTTGCCCAATCACCCCAAAGTGTGTGTANNNAGATCGGAAG |
| 4052_1 | TATGAGTGTGGAGTCGTTGCCCAGGATGATGTCATCGCCTCATNNNAGATCGGAAG |
| 4054_1 | TATGAGTGTGGAGTCGTTGCATGGGTGGTCTCAGGAACTTGTANNNAGATCGGAA |
| 4055_1 | TATGAGTGTGGAGTCGTTGCGTTTCACCCCACAGTGACTTGTANNNAGATCGGAAG |
| 4056_1 | TATGAGTGTGGAGTCGTTGCACACACACACAACCTTCCTTGTANNNAGATCGGAAGA |
| 4057_1 | TATGAGTGTGGAGTCGTTGCTACCTCCATGCCCAGTTCTTGTANNNAGATCGGAAGA |
| 4058_1 | TATGAGTGTGGAGTCGTTGCGGGGGAGGAGTGGTGTATTTGTANNNAGATCGGA |
| 4059_1 | TATGAGTGTGGAGTCGTTGCGCCCATGAATGCTGCTGAAATTANNNAGATCGGAAG |
| 4060_1 | TATGAGTGTGGAGTCGTTGCACGCTGTCACGCTCCATAAATTANNNAGATCGGAAG |
| 4061_1 | TATGAGTGTGGAGTCGTTGCGACACAAGAGTGCCACCCAATTANNNAGATCGGAAG |
| 4062_1 | TATGAGTGTGGAGTCGTTGCGGAGTGCAGCATCAAAGCAATTANNNAGATCGGAA |
| 4064_1 | TATGAGTGTGGAGTCGTTGCGGTCCCAAAGTGTAGGCACATTANNNAGATCGGAAG |
| 4065_1 | TATGAGTGTGGAGTCGTTGCCAGCCACTAAAATGGGGCCATTANNNAGATCGGAAG |
| 4066_1 | TATGAGTGTGGAGTCGTTGCCAGTCACTCCCTCCTGTCCATTANNNAGATCGGAAGA |
| 4070_1 | TATGAGTGTGGAGTCGTTGCGTTCTCTCTGAACCTTGGTTCCTNNNAGATCGGAAGA |
| 4072_1 | TATGAGTGTGGAGTCGTTGCATGCAGGCACAGCTTTCAGATTANNNAGATCGGAAG |
| 4073_1 | TATGAGTGTGGAGTCGTTGCTGACCCCATGCCCACATAGATTANNNAGATCGGAAG |
| 4074_1 | TATGAGTGTGGAGTCGTTGCACAAGGGGCTTTCTCTGGGATTANNNAGATCGGAAG |
| 4075_1 | TATGAGTGTGGAGTCGTTGCGGGGGAAAGAGGCAGATGGATTANNNAGATCGGA |
| 4076_1 | TATGAGTGTGGAGTCGTTGCGGCAGGGGTAGGGGAAATGATTANNNAGATCGGA |
| 4077_1 | TATGAGTGTGGAGTCGTTGCGCTTATGGGAACCCCACTGATTANNNAGATCGGAAG |
| 4078_1 | TATGAGTGTGGAGTCGTTGCAGATACATTTTGAGGCACAGAGCNNNAGATCGGAA |
| 4082_1 | TATGAGTGTGGAGTCGTTGCATGGCCTCGTCATCATATACCCANNNAGATCGGAAG |
| 4083_1 | TATGAGTGTGGAGTCGTTGCGACAACCAGCAGCATACCACTTANNNAGATCGGAAG |
| 4085_1 | TATGAGTGTGGAGTCGTTGCCCTGGCACAGTGTTGAGTACTTANNNAGATCGGAAG |
| 4087_1 | TATGAGTGTGGAGTCGTTGCCCATTTCTCCAAAGCCACCCTTANNNAGATCGGAAGA |
| 4088_1 | TATGAGTGTGGAGTCGTTGCCCCCTCCATCTGGTTTACCCTTANNNAGATCGGAAGA |

| | |
|---|---|
| 4089_1 | TATGAGTGTGGAGTCGTTGCTGTCACTCCTTTGTGACCCCTTANNNAGATCGGAAGA |
| 4091_1 | TATGAGTGTGGAGTCGTTGCCCAGGGTTCAACCAATTCCCTTANNNAGATCGGAAG |
| 4095_1 | TATGAGTGTGGAGTCGTTGCAACTTCTCCCCAAGCACTCCTTANNNAGATCGGAAGA |
| 4096_1 | TATGAGTGTGGAGTCGTTGCCTGTGTGTCTGTCGATGTCCTTANNNAGATCGGAAG |
| 4097_1 | TATGAGTGTGGAGTCGTTGCCTCCTTGGCTTCAGGTGTCCTTANNNAGATCGGAAG |
| 4099_1 | TATGAGTGTGGAGTCGTTGCCCTGCACATTGGATCGTTCCTTANNNAGATCGGAAG |
| 4102_1 | TATGAGTGTGGAGTCGTTGCGGAGCATCTCATTCCCAAGCTTANNNAGATCGGAAG |
| 4103_1 | TATGAGTGTGGAGTCGTTGCACGGGGTCATTACAGGAAGCTTANNNAGATCGGAA |
| 4104_1 | TATGAGTGTGGAGTCGTTGCGCCAGGAGGGAGTAATCAGCTTANNNAGATCGGAA |
| 4105_1 | TATGAGTGTGGAGTCGTTGCCACTATGCATGCCCATCAGCTTANNNAGATCGGAAG |
| 4106_1 | TATGAGTGTGGAGTCGTTGCAGGGTGCACAGATCTATGGCTTANNNAGATCGGAA |
| 4108_1 | TATGAGTGTGGAGTCGTTGCGGGGAAGAAACAGGTTTGGCTTANNNAGATCGGA |
| 4109_1 | TATGAGTGTGGAGTCGTTGCTGGAGAAGGAGCAAAAGTCTACTNNNAGATCGGAA |
| 4111_1 | TATGAGTGTGGAGTCGTTGCCCCTTTCTGGGTGAATGTGCTTANNNAGATCGGAAG |
| 4112_1 | TATGAGTGTGGAGTCGTTGCTCCCAACACACCACATTTGCTTANNNAGATCGGAAGA |
| 4113_1 | TATGAGTGTGGAGTCGTTGCGACCTGGGTGCTAGTTTTGCTTANNNAGATCGGAAG |
| 4114_1 | TATGAGTGTGGAGTCGTTGCCCTGCACAAGTGTGTCGATCTTANNNAGATCGGAAG |
| 4115_1 | TATGAGTGTGGAGTCGTTGCGAAAACCAGCCACACACCTCTTANNNAGATCGGAAG |
| 4116_1 | TATGAGTGTGGAGTCGTTGCTGAGGTCTTGCGCAAAATAAAGANNNAGATCGGAA |
| 4117_1 | TATGAGTGTGGAGTCGTTGCGGCACCATGAGTGATATCTTCCTNNNAGATCGGAAG |
| 4118_1 | TATGAGTGTGGAGTCGTTGCTGGTTCTGGAAGCTCCAGTCTTANNNAGATCGGAAG |
| 4120_1 | TATGAGTGTGGAGTCGTTGCTGGGGAGCTGGAGATTGTTCTTANNNAGATCGGAA |
| 4122_1 | TATGAGTGTGGAGTCGTTGCCAGGGTAGCAATGTGCTCAGTTANNNAGATCGGAA |
| 4123_1 | TATGAGTGTGGAGTCGTTGCGTGTGTGAGGGACAGGAGAGTTANNNAGATCGGA |
| 4125_1 | TATGAGTGTGGAGTCGTTGCGTTGCATGGGAAACCCTGAGTTANNNAGATCGGAA |
| 4126_1 | TATGAGTGTGGAGTCGTTGCATTCGCTGCTCATGAGTGAGTTANNNAGATCGGAAG |
| 4127_1 | TATGAGTGTGGAGTCGTTGCTTGAGTGAGGCTTGGGGTAGTTANNNAGATCGGAA |
| 4128_1 | TATGAGTGTGGAGTCGTTGCAGCGTTTAGTCACCTCCAGGTTANNNAGATCGGAAG |
| 4131_1 | TATGAGTGTGGAGTCGTTGCTCAGGGAATGAAGGACAGGGTTANNNAGATCGGA |
| 4132_1 | TATGAGTGTGGAGTCGTTGCTCACCTCTGCCTTATCCTGGTTANNNAGATCGGAAGA |
| 4133_1 | TATGAGTGTGGAGTCGTTGCTACCAGGATCACAAGGCTGGTTANNNAGATCGGAA |
| 4134_1 | TATGAGTGTGGAGTCGTTGCCTTCTAGCCCCTGCTTTTGGTTANNNAGATCGGAAG |
| 4135_1 | TATGAGTGTGGAGTCGTTGCGTGCCTGCCATACAGTCATGTTANNNAGATCGGAAG |
| 4136_1 | TATGAGTGTGGAGTCGTTGCCGCATGGACCCAGAAAACTGTTANNNAGATCGGAA |
| 4137_1 | TATGAGTGTGGAGTCGTTGCTTGTGATTGGGTTGCCACTGTTANNNAGATCGGAAG |
| 4139_1 | TATGAGTGTGGAGTCGTTGCTCCACACCTACCTTCAGCTGTTANNNAGATCGGAAGA |
| 4140_1 | TATGAGTGTGGAGTCGTTGCGGTTGGTGTTGGTCTCAGTGTTANNNAGATCGGAA |
| 4141_1 | TATGAGTGTGGAGTCGTTGCGCAGGCTGTGATCTTGAGTGTTANNNAGATCGGAA |
| 4142_1 | TATGAGTGTGGAGTCGTTGCTTTGAAGGGAGAGGAGGGTGTTANNNAGATCGGA |
| 4143_1 | TATGAGTGTGGAGTCGTTGCTACTGGGGGACATGGATGTGTTANNNAGATCGGAA |
| 4145_1 | TATGAGTGTGGAGTCGTTGCAGTGAGGCCCAGAGAAGTTGTTANNNAGATCGGAA |
| 4146_1 | TATGAGTGTGGAGTCGTTGCGGTGGTGGTGAGAAGTGTTGTTANNNAGATCGGA |
| 4148_1 | TATGAGTGTGGAGTCGTTGCGCCTCAGAGTTGCCCAGAATTTANNNAGATCGGAAG |

| | |
|---|---|
| 4150_1 | TATGAGTGTGGAGTCGTTGCCACACACACACGGCAAACATTTANNNAGATCGGAAG |
| 4151_1 | TATGAGTGTGGAGTCGTTGCGGAGCCCATCACCTGTACATTTANNNAGATCGGAAG |
| 4152_1 | TATGAGTGTGGAGTCGTTGCTGCCATCTGTTTCCTGGCATTTANNNAGATCGGAAG |
| 4154_1 | TATGAGTGTGGAGTCGTTGCAAGGAAGTGCTGCTGTAGGTAAGNNNAGATCGGA |
| 4155_1 | TATGAGTGTGGAGTCGTTGCGGTGGACTGGATGCTTGCATTTANNNAGATCGGAA |
| 4157_1 | TATGAGTGTGGAGTCGTTGCCTGGTCCCATTTTCCTCTCACAGNNNAGATCGGAAGA |
| 4159_1 | TATGAGTGTGGAGTCGTTGCGGACCCGTAGACCAATACCTTTANNNAGATCGGAAG |
| 4161_1 | TATGAGTGTGGAGTCGTTGCTGTCACCAATGTGATGCCCTTTANNNAGATCGGAAG |
| 4164_1 | TATGAGTGTGGAGTCGTTGCGAAATCAGGGTGAGCTGGCTTTANNNAGATCGGAA |
| 4167_1 | TATGAGTGTGGAGTCGTTGCTTCTTTGCGGTGGAGCCTCTTTANNNAGATCGGAAG |
| 4170_1 | TATGAGTGTGGAGTCGTTGCCACTACCACCACCACTGTCTTTANNNAGATCGGAAGA |
| 4172_1 | TATGAGTGTGGAGTCGTTGCGCATGTGTGTTTCCCTAAGTTTANNNAGATCGGAAG |
| 4175_1 | TATGAGTGTGGAGTCGTTGCGTGTGTGTTGTGAGGAGGGTTTANNNAGATCGGA |
| 4177_1 | TATGAGTGTGGAGTCGTTGCGCATTTGTGTGGGTGCATGTTTANNNAGATCGGAA |
| 4179_1 | TATGAGTGTGGAGTCGTTGCTAAGGGATCTGGGAGCGTACATTNNNAGATCGGAA |
| 4180_1 | TATGAGTGTGGAGTCGTTGCCGAGGATCACGTGTGTGTGTTTANNNAGATCGGAA |
| 4182_1 | TATGAGTGTGGAGTCGTTGCTCCCTAGGCCTTACCTAAACTACNNNAGATCGGAAG |
| 4184_1 | TATGAGTGTGGAGTCGTTGCTCTTACTCCAGGGGGCACTTTTANNNAGATCGGAAG |
| 4185_1 | TATGAGTGTGGAGTCGTTGCGGAGGAAGTGGTGATGCCTTTTANNNAGATCGGAA |
| 4187_1 | TATGAGTGTGGAGTCGTTGCGGGATGGGTGATGTTGCCTTTTANNNAGATCGGAA |
| 4188_1 | TATGAGTGTGGAGTCGTTGCTCTCCTCCATGCCCTTCCTTTTANNNAGATCGGAAGA |
| 4190_1 | TATGAGTGTGGAGTCGTTGCGTCTGTCCTCACTTGGGCTTTTANNNAGATCGGAAG |
| 4192_1 | TATGAGTGTGGAGTCGTTGCTCAACTAGGGCCCCTGTCTTTTANNNAGATCGGAAG |
| 4194_1 | TATGAGTGTGGAGTCGTTGCCAGTGCTCACCCTCTTAGTTTTANNNAGATCGGAAG |
| 4195_1 | TATGAGTGTGGAGTCGTTGCGCCTAGTGGGGAGAGAGGTTTTANNNAGATCGGA |
| 4196_1 | TATGAGTGTGGAGTCGTTGCCACCATCATCCACTTAGCAGCTCNNNAGATCGGAAG |
| 4197_1 | TATGAGTGTGGAGTCGTTGCAGAGGCCTGCCATCAGCTTTTTANNNAGATCGGAAG |
| 4199_1 | TATGAGTGTGGAGTCGTTGCACACACACCCTGCACAGTTTTTANNNAGATCGGAAG |
| 4208_1 | TATGAGTGTGGAGTCGTTGCGAGAGAGAGAGACCCTGCAAAACNNNAGATCGGAA |
| 4209_1 | TATGAGTGTGGAGTCGTTGCGGGCAAGGATACATGCAGAAAACNNNAGATCGGAA |
| 4212_1 | TATGAGTGTGGAGTCGTTGCGGTAGCAGGTAAGGGTGGAAAACNNNAGATCGGA |
| 4213_1 | TATGAGTGTGGAGTCGTTGCCCCTTCCCCAGGTGTTCTAAAACNNNAGATCGGAAG |
| 4214_1 | TATGAGTGTGGAGTCGTTGCATGCCTGGCTGACAACTCAATTNNNAGATCGGAAGA |
| 4215_1 | TATGAGTGTGGAGTCGTTGCCAAGGAATGGCAGACACACAAACNNNAGATCGGAA |
| 4216_1 | TATGAGTGTGGAGTCGTTGCTATGTATGCCTCTGGCCACAAACNNNAGATCGGAAG |
| 4217_1 | TATGAGTGTGGAGTCGTTGCGCGTCCCTTCAAAAAGATATGTTNNNAGATCGGAAG |
| 4218_1 | TATGAGTGTGGAGTCGTTGCGCTCAGTCTGGAAAAGCACAAACNNNAGATCGGAA |
| 4219_1 | TATGAGTGTGGAGTCGTTGCACCAGCTGCCTCTAAAGACAAACNNNAGATCGGAAG |
| 4223_1 | TATGAGTGTGGAGTCGTTGCTCTTGCATCTGTGGGTGTTTTCANNNAGATCGGAAG |
| 4226_1 | TATGAGTGTGGAGTCGTTGCATGTGTGTAAACAAGGCCTGTCTNNNAGATCGGAA |
| 4227_1 | TATGAGTGTGGAGTCGTTGCAACTGGCCTTCTGGAGATCAAACNNNAGATCGGAAG |
| 4229_1 | TATGAGTGTGGAGTCGTTGCCAGCAGCAATGGTTCTCTCAAACNNNAGATCGGAAG |
| 4233_1 | TATGAGTGTGGAGTCGTTGCAAGGAAGTGGTGAGACACGAAACNNNAGATCGGA |

| | |
|---|---|
| 4239_1 | TATGAGTGTGGAGTCGTTGCGACAACAAGCAGGGTGTTGAAACNNNAGATCGGAA |
| 4240_1 | TATGAGTGTGGAGTCGTTGCCCAGACCATCAGACAGCCTAAACNNNAGATCGGAAG |
| 4241_1 | TATGAGTGTGGAGTCGTTGCTAGCATCCTTGCCTTCGCTAAACNNNAGATCGGAAG |
| 4243_1 | TATGAGTGTGGAGTCGTTGCAGGCAGGAGTTGCTGTTCTAAACNNNAGATCGGAA |
| 4244_1 | TATGAGTGTGGAGTCGTTGCTACCTCAAAGCAGGGCAGTAAACNNNAGATCGGAA |
| 4246_1 | TATGAGTGTGGAGTCGTTGCGCCAGAGACAAGAGCAAAACAACNNNAGATCGGAA |
| 4248_1 | TATGAGTGTGGAGTCGTTGCGATAGGCTCTTGCTACCCACAACNNNAGATCGGAAG |
| 4249_1 | TATGAGTGTGGAGTCGTTGCAGCCTGTCCAGAATCAAGACAACNNNAGATCGGAAG |
| 4250_1 | TATGAGTGTGGAGTCGTTGCGTCCCACTCTCTGAACAGACAACNNNAGATCGGAAG |
| 4252_1 | TATGAGTGTGGAGTCGTTGCGGGAAGACTGTGCTCTAGACAACNNNAGATCGGAA |
| 4262_1 | TATGAGTGTGGAGTCGTTGCTTTGCTGTAGACAGGTGTCCAACNNNAGATCGGAAG |
| 4263_1 | TATGAGTGTGGAGTCGTTGCCCCAGGAAACTGACAAAAGCAACNNNAGATCGGAA |
| 4264_1 | TATGAGTGTGGAGTCGTTGCGTAAGTGCTGCTTCGTAAGCAACNNNAGATCGGAA |
| 4266_1 | TATGAGTGTGGAGTCGTTGCAAGGCTGTTAGAACACTGGCAACNNNAGATCGGAA |
| 4267_1 | TATGAGTGTGGAGTCGTTGCGTGCAATCCTGGTTTAGTGCAACNNNAGATCGGAAG |
| 4269_1 | TATGAGTGTGGAGTCGTTGCATTGTCACTGGCAGCTCATCAACNNNAGATCGGAAG |
| 4274_1 | TATGAGTGTGGAGTCGTTGCTCCTCTTCCATGTAGCTGTCAACNNNAGATCGGAAGA |
| 4275_1 | TATGAGTGTGGAGTCGTTGCTGGTGGGATGTTTCTCTGTCAACNNNAGATCGGAAG |
| 4277_1 | TATGAGTGTGGAGTCGTTGCCCTTTGCCCCAAAACTCTTCAACNNNAGATCGGAAGA |
| 4279_1 | TATGAGTGTGGAGTCGTTGCAGGGAGGATAAGGAACCCAGAACNNNAGATCGGAA |
| 4281_1 | TATGAGTGTGGAGTCGTTGCTTGGCTCTGTTTGTTTGCAGAACNNNAGATCGGAAG |
| 4282_1 | TATGAGTGTGGAGTCGTTGCGTGGTGGTGGGTAAGAAGAGAACNNNAGATCGGA |
| 4289_1 | TATGAGTGTGGAGTCGTTGCCCATCTCACGATCTCAGCTCACTNNNAGATCGGAAGA |
| 4290_1 | TATGAGTGTGGAGTCGTTGCTTTTGCAGCCACCTGGATAGAACNNNAGATCGGAAG |
| 4291_1 | TATGAGTGTGGAGTCGTTGCGCTGCTTTGGCTAGTGATATGGTNNNAGATCGGAA |
| 4292_1 | TATGAGTGTGGAGTCGTTGCTCATGCAAAGGAAAATGCCGAACNNNAGATCGGAA |
| 4294_1 | TATGAGTGTGGAGTCGTTGCGGGCACGTCCTATTATCAGGAACNNNAGATCGGAA |
| 4297_1 | TATGAGTGTGGAGTCGTTGCAACGGGGAGCAATGCTAAGATTTNNNAGATCGGAA |
| 4299_1 | TATGAGTGTGGAGTCGTTGCTTGATTTTGCCTCTAGGGGGAACNNNAGATCGGAA |
| 4307_1 | TATGAGTGTGGAGTCGTTGCTGGCTGGAACAGACTTGATGAACNNNAGATCGGAA |
| 4308_1 | TATGAGTGTGGAGTCGTTGCCTCTGGGTATCATGGCACTGAACNNNAGATCGGAAG |
| 4309_1 | TATGAGTGTGGAGTCGTTGCCCACAGACGTACTCTCACTGAACNNNAGATCGGAAG |
| 4310_1 | TATGAGTGTGGAGTCGTTGCCTCTACCTGGATGACTTAGCCCANNNAGATCGGAAG |
| 4315_1 | TATGAGTGTGGAGTCGTTGCCACACAGGCACACAGAATTGAACNNNAGATCGGAA |
| 4316_1 | TATGAGTGTGGAGTCGTTGCAAACACCCCATCAGAGGTTGAACNNNAGATCGGAAG |
| 4317_1 | TATGAGTGTGGAGTCGTTGCGCTCCCACAGCACCTAGAATAACNNNAGATCGGAAG |
| 4318_1 | TATGAGTGTGGAGTCGTTGCCATACCTCCCTGCACACGATAACNNNAGATCGGAAG |
| 4319_1 | TATGAGTGTGGAGTCGTTGCCCTACCTTCCATGGGCAACTAACNNNAGATCGGAAG |
| 4321_1 | TATGAGTGTGGAGTCGTTGCCTCATGCCACACTGTCAGCTAACNNNAGATCGGAAG |
| 4325_1 | TATGAGTGTGGAGTCGTTGCATGGGGGTAGTGGGAGAAGTAACNNNAGATCGGA |
| 4327_1 | TATGAGTGTGGAGTCGTTGCCTCCTCTCCCCTCCCTAAGTAACNNNAGATCGGAAGA |
| 4329_1 | TATGAGTGTGGAGTCGTTGCAGTTGCCCAGAATAGCCTGTAACNNNAGATCGGAAG |
| 4330_1 | TATGAGTGTGGAGTCGTTGCGCTTCTTTGGTGCCCTTTGTAACNNNAGATCGGAAG |

| | |
|---|---|
| 4332_1 | TATGAGTGTGGAGTCGTTGCGAGTAGATGCCCAGTGGTGGAATNNNAGATCGGAA |
| 4334_1 | TATGAGTGTGGAGTCGTTGCAACTTGTCCACCAGGCTAAACACNNNAGATCGGAAG |
| 4336_1 | TATGAGTGTGGAGTCGTTGCCACCCATGCAAATACACCAACACNNNAGATCGGAAG |
| 4337_1 | TATGAGTGTGGAGTCGTTGCAAGCAACAATGGTGATATGCCCANNNAGATCGGAA |
| 4338_1 | TATGAGTGTGGAGTCGTTGCTCCATCTCCTGACCTTGTACTGANNNAGATCGGAAG |
| 4339_1 | TATGAGTGTGGAGTCGTTGCAAAAACAGGGGACAGGAGAACACNNNAGATCGGA |
| 4342_1 | TATGAGTGTGGAGTCGTTGCACAAGTGGGTTGGCTAGTAACACNNNAGATCGGAA |
| 4344_1 | TATGAGTGTGGAGTCGTTGCTGAGAGCTGGTTTACCAACACACNNNAGATCGGAAG |
| 4346_1 | TATGAGTGTGGAGTCGTTGCTTAGCTTCCTGTGACACACACACNNNAGATCGGAAG |
| 4347_1 | TATGAGTGTGGAGTCGTTGCGGGGAGAAAGAGTACACACACACNNNAGATCGGAA |
| 4349_1 | TATGAGTGTGGAGTCGTTGCCACTACTCAGCCATGACGAAAATNNNAGATCGGAAG |
| 4350_1 | TATGAGTGTGGAGTCGTTGCTTCTTTCAGCTCTGAGCACACACNNNAGATCGGAAG |
| 4351_1 | TATGAGTGTGGAGTCGTTGCGAGAGAAACATGCATGCACACACNNNAGATCGGAA |
| 4352_1 | TATGAGTGTGGAGTCGTTGCTCACTTCAGCCTCCCAAGATTTTNNNAGATCGGAAGA |
| 4353_1 | TATGAGTGTGGAGTCGTTGCCGGCAAGACTGAAATTCACACACNNNAGATCGGAAG |
| 4354_1 | TATGAGTGTGGAGTCGTTGCTGCTATTGGTGCCTGATACACACNNNAGATCGGAAG |
| 4355_1 | TATGAGTGTGGAGTCGTTGCATTCTGGTAGCCATTAGCCACACNNNAGATCGGAAG |
| 4357_1 | TATGAGTGTGGAGTCGTTGCAAATGAACTTCCTAGCAGCAGCTNNNAGATCGGAAG |
| 4358_1 | TATGAGTGTGGAGTCGTTGCTGTGTTGGGATGAACAGGTCTGANNNAGATCGGAA |
| 4360_1 | TATGAGTGTGGAGTCGTTGCGAGAAAGACACTCAGAGGAAGGANNNAGATCGGA |
| 4366_1 | TATGAGTGTGGAGTCGTTGCCACACACGATCCCAGATAGACACNNNAGATCGGAAG |
| 4367_1 | TATGAGTGTGGAGTCGTTGCAGGGAGAGAGAGAAACAGGACACNNNAGATCGGA |
| 4369_1 | TATGAGTGTGGAGTCGTTGCTATGAAGCGTTTCTCCAGGACACNNNAGATCGGAAG |
| 4371_1 | TATGAGTGTGGAGTCGTTGCTTGCATAGTGTTTGGAGGGACACNNNAGATCGGAA |
| 4375_1 | TATGAGTGTGGAGTCGTTGCGGCTTGAGAGAAGCAGAGTACACNNNAGATCGGAA |
| 4376_1 | TATGAGTGTGGAGTCGTTGCAGCACCTTAGCTCTCTGGTACACNNNAGATCGGAAG |
| 4377_1 | TATGAGTGTGGAGTCGTTGCAACTGCTGGAACTTTGCTTACACNNNAGATCGGAAG |
| 4379_1 | TATGAGTGTGGAGTCGTTGCAGGGAATGACCACTGAAAACCACNNNAGATCGGAA |
| 4380_1 | TATGAGTGTGGAGTCGTTGCGCTATGGGGAAATGTCAAACCACNNNAGATCGGAA |
| 4383_1 | TATGAGTGTGGAGTCGTTGCTGGAAATAAAGGGCCAGAACCACNNNAGATCGGAA |
| 4385_1 | TATGAGTGTGGAGTCGTTGCAGGAGAGGTGGCAAAGTGATAAGNNNAGATCGGA |
| 4387_1 | TATGAGTGTGGAGTCGTTGCTGTGTGTGTGTCTTGAGGGAAAGNNNAGATCGGAA |
| 4388_1 | TATGAGTGTGGAGTCGTTGCTGCCTTTCACCTTCTGTAACCACNNNAGATCGGAAGA |
| 4390_1 | TATGAGTGTGGAGTCGTTGCGGGAGGAGAAATAACACCACCACNNNAGATCGGAA |
| 4391_1 | TATGAGTGTGGAGTCGTTGCGATTTCTGGTGGGAGAAGACCACNNNAGATCGGAA |
| 4392_1 | TATGAGTGTGGAGTCGTTGCGCTATCAGACACAGACCTGGGAGNNNAGATCGGAA |
| 4393_1 | TATGAGTGTGGAGTCGTTGCTGACTTACTGAAGCTGTGACCACNNNAGATCGGAAG |
| 4397_1 | TATGAGTGTGGAGTCGTTGCAAGCAGTGTTCCATCTGTACCACNNNAGATCGGAAG |
| 4398_1 | TATGAGTGTGGAGTCGTTGCAACCCACACATACAAGAACCCACNNNAGATCGGAAG |
| 4399_1 | TATGAGTGTGGAGTCGTTGCGCTCCCTCTTTTCACTAACCCACNNNAGATCGGAAGA |
| 4401_1 | TATGAGTGTGGAGTCGTTGCCTTATTTCTGCTCCTCCACCCACNNNAGATCGGAAGA |
| 4402_1 | TATGAGTGTGGAGTCGTTGCGGTGGGGAGAGAAAATTACCCACNNNAGATCGGAA |
| 4403_1 | TATGAGTGTGGAGTCGTTGCCAATAGCTTTCCTCATCCCCCACNNNAGATCGGAAGA |

| | |
|---|---|
| 4407_1 | TATGAGTGTGGAGTCGTTGCCCAGCACCATGTCAGTGATCATTNNNAGATCGGAAG |
| 4409_1 | TATGAGTGTGGAGTCGTTGCTCCCTGTCTCTATTTCTCCCCACNNNAGATCGGAAGA |
| 4411_1 | TATGAGTGTGGAGTCGTTGCAATAAGGCAAAACACATGCCCACNNNAGATCGGAAG |
| 4413_1 | TATGAGTGTGGAGTCGTTGCAGGGGGAACCACTATAATCCCACNNNAGATCGGAA |
| 4416_1 | TATGAGTGTGGAGTCGTTGCCACTAAGGCATCTTTTGCCCTGANNNAGATCGGAAG |
| 4417_1 | TATGAGTGTGGAGTCGTTGCGCTAAATTACAGTCAAGCGCCACNNNAGATCGGAAG |
| 4418_1 | TATGAGTGTGGAGTCGTTGCGAGAAATTGGGGTATTGAGGAGGNNNAGATCGGA |
| 4419_1 | TATGAGTGTGGAGTCGTTGCACATGAGTTGAGTAACTGGCCACNNNAGATCGGAA |
| 4420_1 | TATGAGTGTGGAGTCGTTGCAGAAGGGCAAGATGAAAACCCAGNNNAGATCGGA |
| 4421_1 | TATGAGTGTGGAGTCGTTGCCTTCCATTTATGGGCATTGCCACNNNAGATCGGAAG |
| 4425_1 | TATGAGTGTGGAGTCGTTGCAACAGGAGGTAAATGCCATCCACNNNAGATCGGAA |
| 4428_1 | TATGAGTGTGGAGTCGTTGCATGATGGAAGGCAACTTCTCCACNNNAGATCGGAAG |
| 4430_1 | TATGAGTGTGGAGTCGTTGCGTTACACACAGACCTCTGTCCACNNNAGATCGGAAG |
| 4432_1 | TATGAGTGTGGAGTCGTTGCACCCAAGGAAACAGGAATTCCACNNNAGATCGGAA |
| 4434_1 | TATGAGTGTGGAGTCGTTGCGCACACATACATGCCATTTCCACNNNAGATCGGAAG |
| 4435_1 | TATGAGTGTGGAGTCGTTGCGCAATCCAATGGAGGAAAAGCACNNNAGATCGGAA |
| 4437_1 | TATGAGTGTGGAGTCGTTGCAAGTTAGCTACCAACCCAAGCACNNNAGATCGGAAG |
| 4438_1 | TATGAGTGTGGAGTCGTTGCAAATCAGTCTGGCTGAGAAGCACNNNAGATCGGAA |
| 4439_1 | TATGAGTGTGGAGTCGTTGCATACACCTCCTTCACAACAGCACNNNAGATCGGAAG |
| 4444_1 | TATGAGTGTGGAGTCGTTGCGTTTGTAGTGTGTGCGTGTTCNNNAGATCGGAAGA |
| 4446_1 | TATGAGTGTGGAGTCGTTGCGCACAGAACCAACTGAAAGAGCTNNNAGATCGGAA |
| 4447_1 | TATGAGTGTGGAGTCGTTGCCTAAAATACACACACCCACGCACNNNAGATCGGAAG |
| 4448_1 | TATGAGTGTGGAGTCGTTGCGATACACTGCAAGAATGACGCACNNNAGATCGGAA |
| 4451_1 | TATGAGTGTGGAGTCGTTGCAGAACAGATCCGAAAAGAGGCACNNNAGATCGGAA |
| 4452_1 | TATGAGTGTGGAGTCGTTGCCCCACTGTAGTACTGGAAGAGCTNNNAGATCGGAA |
| 4454_1 | TATGAGTGTGGAGTCGTTGCGCATTAGCCTGAAAATACGGCACNNNAGATCGGAA |
| 4457_1 | TATGAGTGTGGAGTCGTTGCTGAAAGACCCCATCCTCATACCANNNAGATCGGAAG |
| 4459_1 | TATGAGTGTGGAGTCGTTGCGCTCGCTTGCTCACACCTATAATNNNAGATCGGAAG |
| 4460_1 | TATGAGTGTGGAGTCGTTGCGGTCTAATAAGAGGGCATGGCACNNNAGATCGGAA |
| 4461_1 | TATGAGTGTGGAGTCGTTGCCACTCACATTCACAAACTGGCACNNNAGATCGGAAG |
| 4465_1 | TATGAGTGTGGAGTCGTTGCGAGACAAGGTCACACAAATGCACNNNAGATCGGAA |
| 4467_1 | TATGAGTGTGGAGTCGTTGCACGAATAAGACCTGGTTCTGCACNNNAGATCGGAAG |
| 4468_1 | TATGAGTGTGGAGTCGTTGCATGAAGTGACAAGGAAGGTGCACNNNAGATCGGA |
| 4470_1 | TATGAGTGTGGAGTCGTTGCGAGGGGAAGTGAGGAGAGAGAGANNNAGATCGG |
| 4471_1 | TATGAGTGTGGAGTCGTTGCGCAGGTTGAGAGACAAATTGCACNNNAGATCGGAA |
| 4475_1 | TATGAGTGTGGAGTCGTTGCTGGTGAACAAAAGATGCTTGCACNNNAGATCGGAA |
| 4476_1 | TATGAGTGTGGAGTCGTTGCGCCGCATTTATCTAAGTAGCAGTNNNAGATCGGAAG |
| 4478_1 | TATGAGTGTGGAGTCGTTGCTGGTATAAAAACCTTTTTGCGGCNNNAGATCGGAAG |
| 4479_1 | TATGAGTGTGGAGTCGTTGCAGAGAGGGAGAGAGAGCAATCACNNNAGATCGGA |
| 4480_1 | TATGAGTGTGGAGTCGTTGCCCATCGCCCTTTTGCATAATCACNNNAGATCGGAAGA |
| 4483_1 | TATGAGTGTGGAGTCGTTGCTATGTACTGGCAAAGGCCATCACNNNAGATCGGAAG |
| 4485_1 | TATGAGTGTGGAGTCGTTGCATGTATGTGAGACAGAGCCTCACNNNAGATCGGAA |
| 4487_1 | TATGAGTGTGGAGTCGTTGCTCAGCGCATGTCTATTGGTTTAGNNNAGATCGGAAG |

| | |
|---|---|
| 4488_1 | TATGAGTGTGGAGTCGTTGCATAATGGCGCTACAGCAACTTCANNNAGATCGGAAG |
| 4489_1 | TATGAGTGTGGAGTCGTTGCCCTTCAGTCTTCCGGGTTATCACNNNAGATCGGAAGA |
| 4491_1 | TATGAGTGTGGAGTCGTTGCGCCACACCCTCTGATCTCTTTACNNNAGATCGGAAGA |
| 4492_1 | TATGAGTGTGGAGTCGTTGCAGGCTTTGGAAACAAGCAGATGANNNAGATCGGAA |
| 4495_1 | TATGAGTGTGGAGTCGTTGCGGTTTCAGAAAAATGCCGTGGAANNNAGATCGGAA |
| 4499_1 | TATGAGTGTGGAGTCGTTGCAAATAAGCAAGCCCCTAGCTCACNNNAGATCGGAAG |
| 4500_1 | TATGAGTGTGGAGTCGTTGCGCTATTACGAGGAACTGGCTCACNNNAGATCGGAAG |
| 4502_1 | TATGAGTGTGGAGTCGTTGCCAGGCACTTACCCAGTGTTCATTNNNAGATCGGAAG |
| 4503_1 | TATGAGTGTGGAGTCGTTGCCAGCCAGAGATAAGCACTCTCACNNNAGATCGGAAG |
| 4504_1 | TATGAGTGTGGAGTCGTTGCTTCTCTGTGTGTGAAGCTCTCACNNNAGATCGGAAG |
| 4505_1 | TATGAGTGTGGAGTCGTTGCCGCTCTCACTTTCACTCTCTCACNNNAGATCGGAAGA |
| 4507_1 | TATGAGTGTGGAGTCGTTGCGCCCTGAGCTTATCAGTTCTCACNNNAGATCGGAAG |
| 4508_1 | TATGAGTGTGGAGTCGTTGCACATAATATTCCTGGGGGTCCCANNNAGATCGGAAG |
| 4511_1 | TATGAGTGTGGAGTCGTTGCCAACCCTACAGACACACAGTCACNNNAGATCGGAAG |
| 4514_1 | TATGAGTGTGGAGTCGTTGCGGAATTTTGATGCAGCTGGTCACNNNAGATCGGAA |
| 4517_1 | TATGAGTGTGGAGTCGTTGCTATTGGGCAGTGTTGGAATCATTNNNAGATCGGAA |
| 4519_1 | TATGAGTGTGGAGTCGTTGCTCCATAAATGCCTTGGGTGTCACNNNAGATCGGAAG |
| 4521_1 | TATGAGTGTGGAGTCGTTGCGAGGAGGGGTGTGTACAATTCACNNNAGATCGGAA |
| 4523_1 | TATGAGTGTGGAGTCGTTGCAGAGGAATGGATTGGCGATTCACNNNAGATCGGAA |
| 4524_1 | TATGAGTGTGGAGTCGTTGCAGTGGTTTCACGGGCATATTCACNNNAGATCGGAAG |
| 4525_1 | TATGAGTGTGGAGTCGTTGCTTGCATGTGCGTGTTTACTTCACNNNAGATCGGAAG |
| 4528_1 | TATGAGTGTGGAGTCGTTGCACCCTCTATGGCTCTCTCTTCACNNNAGATCGGAAGA |
| 4529_1 | TATGAGTGTGGAGTCGTTGCACAGGAACTCAGCAGAAGTTCACNNNAGATCGGAA |
| 4530_1 | TATGAGTGTGGAGTCGTTGCCAGACACGAGAAAGCATACCCAANNNAGATCGGAA |
| 4531_1 | TATGAGTGTGGAGTCGTTGCTGTGTGTGTGTCAGAATGAGTAGNNNAGATCGGAA |
| 4532_1 | TATGAGTGTGGAGTCGTTGCGAGGGGTGGTTAGTCCATTTCACNNNAGATCGGAA |
| 4533_1 | TATGAGTGTGGAGTCGTTGCTTCCCTTCCAAGAGTGCTTTCACNNNAGATCGGAAGA |
| 4535_1 | TATGAGTGTGGAGTCGTTGCTATGCAGTGAGACAGGGTTTCACNNNAGATCGGAA |
| 4538_1 | TATGAGTGTGGAGTCGTTGCACCTGCACTGTGGAAGAAAAGACNNNAGATCGGAA |
| 4539_1 | TATGAGTGTGGAGTCGTTGCGGCTCCGAATAGATCCCAAAGACNNNAGATCGGAAG |
| 4540_1 | TATGAGTGTGGAGTCGTTGCTCAGGAAGAGCAAGAGCAAAGACNNNAGATCGGAA |
| 4546_1 | TATGAGTGTGGAGTCGTTGCATAAGACACTGGAACCGCAAGACNNNAGATCGGAA |
| 4547_1 | TATGAGTGTGGAGTCGTTGCAGGGATGTCCTCATTTCCAAACTNNNAGATCGGAAG |
| 4548_1 | TATGAGTGTGGAGTCGTTGCATAGGGTGGTAAAGGGGGAAGACNNNAGATCGGA |
| 4553_1 | TATGAGTGTGGAGTCGTTGCCCAAAGTGGGATCTCTGACAGACNNNAGATCGGAA |
| 4554_1 | TATGAGTGTGGAGTCGTTGCGCCTGGTCATGATGGATACAGACNNNAGATCGGAA |
| 4555_1 | TATGAGTGTGGAGTCGTTGCACTGTCTCCCTCACCTTACAGACNNNAGATCGGAAGA |
| 4556_1 | TATGAGTGTGGAGTCGTTGCTTATGTGCTGACACCAACCAGACNNNAGATCGGAAG |
| 4557_1 | TATGAGTGTGGAGTCGTTGCGGATAATGAGGCACAGACCAGACNNNAGATCGGAA |
| 4558_1 | TATGAGTGTGGAGTCGTTGCCCTACACCTTCTCCCTACCAGACNNNAGATCGGAAGA |
| 4560_1 | TATGAGTGTGGAGTCGTTGCGGCTCACTTGTAGTTTCCCAGACNNNAGATCGGAAG |
| 4561_1 | TATGAGTGTGGAGTCGTTGCGCAGAGAGATCAAGAAGCCAGACNNNAGATCGGAA |
| 4564_1 | TATGAGTGTGGAGTCGTTGCAGTGTCACAGAATCACTGCAGACNNNAGATCGGAA |

| | |
|---|---|
| 4566_1 | TATGAGTGTGGAGTCGTTGCGTTTATTGGGTCAGGCCTCAGACNNNAGATCGGAAG |
| 4575_1 | TATGAGTGTGGAGTCGTTGCCAGACAGACAGACAAGCAGAGACNNNAGATCGGAA |
| 4577_1 | TATGAGTGTGGAGTCGTTGCTCTCACCCTCCCAATGTAGAGACNNNAGATCGGAAGA |
| 4578_1 | TATGAGTGTGGAGTCGTTGCAGGGGTAAGGAGACAAAGGAGACNNNAGATCGGA |
| 4580_1 | TATGAGTGTGGAGTCGTTGCAAAGAGAGAAAGGGGAGGGAGACNNNAGATCGGA |
| 4582_1 | TATGAGTGTGGAGTCGTTGCAACAGGCAGTAAAGCAAACAATTNNNAGATCGGAAG |
| 4583_1 | TATGAGTGTGGAGTCGTTGCAATCGGCTTGATGGAACTGAGACNNNAGATCGGAAG |
| 4586_1 | TATGAGTGTGGAGTCGTTGCCCCAGACTTCAAAGGCCATAGACNNNAGATCGGAAG |
| 4588_1 | TATGAGTGTGGAGTCGTTGCACCTCTTTTCTCCCCCACTAGACNNNAGATCGGAAGAG |
| 4589_1 | TATGAGTGTGGAGTCGTTGCAGGGGTTGGGATGACTTCTAGACNNNAGATCGGAA |
| 4590_1 | TATGAGTGTGGAGTCGTTGCCAGTATGCCCCATGGTTGTAGACNNNAGATCGGAAG |
| 4596_1 | TATGAGTGTGGAGTCGTTGCTATGAAGACATGTGACCAGCGACNNNAGATCGGAAG |
| 4597_1 | TATGAGTGTGGAGTCGTTGCTAGAGAGCTTACCCTTAGGCGACNNNAGATCGGAAG |
| 4599_1 | TATGAGTGTGGAGTCGTTGCAGGAACTGTTTGGAGACAAGGACNNNAGATCGGAA |
| 4600_1 | TATGAGTGTGGAGTCGTTGCCACCTATACTTTGGCAACATGATNNNAGATCGGAAGA |
| 4601_1 | TATGAGTGTGGAGTCGTTGCTTATTTCCTGCCACGTGAAGGACNNNAGATCGGAAG |
| 4602_1 | TATGAGTGTGGAGTCGTTGCGGAAACCAGGAGAGAAACAGGACNNNAGATCGGAA |
| 4603_1 | TATGAGTGTGGAGTCGTTGCCGCACACATGCACAGAGTTATGTNNNAGATCGGAAG |
| 4604_1 | TATGAGTGTGGAGTCGTTGCGGCATCAGACTGGCTTTGAGATTNNNAGATCGGAAG |
| 4605_1 | TATGAGTGTGGAGTCGTTGCACTTTGTGCCAATGAGCATAGCTNNNAGATCGGAAG |
| 4607_1 | TATGAGTGTGGAGTCGTTGCACCTTGGGGGTTATCATAGGGACNNNAGATCGGAAG |
| 4608_1 | TATGAGTGTGGAGTCGTTGCTTGGAGTTTCTCATGGTTAGGGCNNNAGATCGGAAG |
| 4609_1 | TATGAGTGTGGAGTCGTTGCTTAGGAAAGAGAGGACTGGGGACNNNAGATCGGAA |
| 4610_1 | TATGAGTGTGGAGTCGTTGCCCTTTCTTTCTACCCTTGGGGACNNNAGATCGGAAGA |
| 4612_1 | TATGAGTGTGGAGTCGTTGCGTGTTCCTTCCCACCCTCAAATTNNNAGATCGGAAGA |
| 4613_1 | TATGAGTGTGGAGTCGTTGCACTACTTTTCCCTCTCAGCGATTNNNAGATCGGAAGA |
| 4616_1 | TATGAGTGTGGAGTCGTTGCACGGTGGAAGCATGAAGATCTATNNNAGATCGGAAG |
| 4619_1 | TATGAGTGTGGAGTCGTTGCGTCTCCATCCTCTTTCTCTTTGCNNNAGATCGGAAGAG |
| 4621_1 | TATGAGTGTGGAGTCGTTGCAGTGTGTGAAATGTCTGGTGGACNNNAGATCGGAA |
| 4623_1 | TATGAGTGTGGAGTCGTTGCGAACCCATAAGCAGTGGGGTAAGNNNAGATCGGAA |
| 4624_1 | TATGAGTGTGGAGTCGTTGCCACACCAGCTTTGGGAAAATGACNNNAGATCGGAAG |
| 4628_1 | TATGAGTGTGGAGTCGTTGCTTGTTTCCAGGGGACATGATGACNNNAGATCGGAAG |
| 4629_1 | TATGAGTGTGGAGTCGTTGCTATTTGCAGTCACGCCCTATGACNNNAGATCGGAAGA |
| 4630_1 | TATGAGTGTGGAGTCGTTGCACTGGCTCTGCATGATTACTGACNNNAGATCGGAAG |
| 4631_1 | TATGAGTGTGGAGTCGTTGCCTGCTTGTCTAGAGGACCCTGACNNNAGATCGGAAG |
| 4632_1 | TATGAGTGTGGAGTCGTTGCGGATTGACAGAGGGTTTCCTGACNNNAGATCGGAAG |
| 4634_1 | TATGAGTGTGGAGTCGTTGCGTTTGGGTACAGGGACATCTGACNNNAGATCGGAAG |
| 4636_1 | TATGAGTGTGGAGTCGTTGCCTCTTCTGGGAAGCCTTTCTGACNNNAGATCGGAAGA |
| 4637_1 | TATGAGTGTGGAGTCGTTGCGTGTTTCAGAGAGGGGAAGTGACNNNAGATCGGAA |
| 4640_1 | TATGAGTGTGGAGTCGTTGCTAACTCCATACGCCCTCACTTTACNNNAGATCGGAAGA |
| 4641_1 | TATGAGTGTGGAGTCGTTGCGTAACATCCAGGACACCTGTGACNNNAGATCGGAAG |
| 4644_1 | TATGAGTGTGGAGTCGTTGCGAGGCGCAGAGAAATTTTGTGACNNNAGATCGGAA |
| 4645_1 | TATGAGTGTGGAGTCGTTGCACATGAACCAGAACCCACTTGACNNNAGATCGGAAG |

| | |
|---|---|
| 4648_1 | TATGAGTGTGGAGTCGTTGCTGGCAGACAGATTTCAGCTTGACNNNAGATCGGAA |
| 4649_1 | TATGAGTGTGGAGTCGTTGCATAAGCAGCCAAGTGCTCTTGACNNNAGATCGGAAG |
| 4651_1 | TATGAGTGTGGAGTCGTTGCGGGTGGGTCAATTCTTTGTTGACNNNAGATCGGAA |
| 4655_1 | TATGAGTGTGGAGTCGTTGCGGGGAAGGTGGGGAGAAAAATACNNNAGATCGGA |
| 4656_1 | TATGAGTGTGGAGTCGTTGCGTTACAGCATCGTGGCCAAATACNNNAGATCGGAAG |
| 4657_1 | TATGAGTGTGGAGTCGTTGCCTCCTCCTTCACCTGGCAAATACNNNAGATCGGAAGA |
| 4658_1 | TATGAGTGTGGAGTCGTTGCAAGGCTGTTGCTTTTCCATCTGANNNAGATCGGAAG |
| 4659_1 | TATGAGTGTGGAGTCGTTGCTGGCAGCACACTGATGAGAATACNNNAGATCGGAA |
| 4660_1 | TATGAGTGTGGAGTCGTTGCGCGACACCAAACAGGATGAATACNNNAGATCGGAA |
| 4662_1 | TATGAGTGTGGAGTCGTTGCCAGAGCATAGGCAAGACCCATACNNNAGATCGGAA |
| 4664_1 | TATGAGTGTGGAGTCGTTGCCATTAGCCAGTCCTTGGCCATACNNNAGATCGGAAG |
| 4665_1 | TATGAGTGTGGAGTCGTTGCCTGCTCCTATTCCTGTGGCATACNNNAGATCGGAAG |
| 4667_1 | TATGAGTGTGGAGTCGTTGCTACCTACCCATGCGAATTCAGAGNNNAGATCGGAAG |
| 4669_1 | TATGAGTGTGGAGTCGTTGCGCTGCCTCTCTGATTCTGGATACNNNAGATCGGAAG |
| 4671_1 | TATGAGTGTGGAGTCGTTGCCCACCACCACAGAGCACATATACNNNAGATCGGAAG |
| 4672_1 | TATGAGTGTGGAGTCGTTGCCCTTTTCAGGTGGCCGGATATACNNNAGATCGGAAG |
| 4674_1 | TATGAGTGTGGAGTCGTTGCGCCTCAGGTAGCCGTCATTATACNNNAGATCGGAAG |
| 4675_1 | TATGAGTGTGGAGTCGTTGCTGCCAAATCCCTGGCTGTTATACNNNAGATCGGAAG |
| 4676_1 | TATGAGTGTGGAGTCGTTGCCAAAAATGATCACCGAGGGGGTANNNAGATCGGAA |
| 4677_1 | TATGAGTGTGGAGTCGTTGCAGGACATTTGCCAGGGAAACTACNNNAGATCGGAA |
| 4679_1 | TATGAGTGTGGAGTCGTTGCCCACTGCTTCTGTAACCCACTACNNNAGATCGGAAGA |
| 4681_1 | TATGAGTGTGGAGTCGTTGCCTGGAAGACTTGCCCATGACTACNNNAGATCGGAAG |
| 4682_1 | TATGAGTGTGGAGTCGTTGCACCTCTGTTCCCTTCTGGAAGAGNNNAGATCGGAAG |
| 4683_1 | TATGAGTGTGGAGTCGTTGCAATCTGGATCATGTTGCATACACNNNAGATCGGAAG |
| 4684_1 | TATGAGTGTGGAGTCGTTGCCTTGGGGCAGGTCCTTTTACTACNNNAGATCGGAAG |
| 4690_1 | TATGAGTGTGGAGTCGTTGCCTGCAATGGGCGGTTATAGTCTGNNNAGATCGGAA |
| 4691_1 | TATGAGTGTGGAGTCGTTGCCTCTGGGTCTACCAGTTGCCTACNNNAGATCGGAAG |
| 4692_1 | TATGAGTGTGGAGTCGTTGCCTTCCAGGCAAATCACCTCCTACNNNAGATCGGAAGA |
| 4694_1 | TATGAGTGTGGAGTCGTTGCCCATCCCCTGAATTGACAGCTACNNNAGATCGGAAG |
| 4695_1 | TATGAGTGTGGAGTCGTTGCAAACTCCCATTGTGACCAGCTACNNNAGATCGGAAG |
| 4701_1 | TATGAGTGTGGAGTCGTTGCAGACCCCCAGCATACTGATCTACNNNAGATCGGAAG |
| 4702_1 | TATGAGTGTGGAGTCGTTGCCGTAGCAGGAGCATTCACTCTACNNNAGATCGGAAG |
| 4704_1 | TATGAGTGTGGAGTCGTTGCTTCTAGCTGGGTGTCTCGTCTACNNNAGATCGGAAG |
| 4707_1 | TATGAGTGTGGAGTCGTTGCCCCTGCCTCTGGAGTTTTCTTACNNNAGATCGGAAGA |
| 4709_1 | TATGAGTGTGGAGTCGTTGCAGGTGCTAATTGGGCATCAGTACNNNAGATCGGAA |
| 4710_1 | TATGAGTGTGGAGTCGTTGCATCTTGTCTCAGGGCTGGAGTACNNNAGATCGGAA |
| 4712_1 | TATGAGTGTGGAGTCGTTGCAAGATTGCTAGGCTGCAAGGTACNNNAGATCGGAA |
| 4713_1 | TATGAGTGTGGAGTCGTTGCCGTTCTGGTCACAGCTTAGGTACNNNAGATCGGAAG |
| 4717_1 | TATGAGTGTGGAGTCGTTGCGACAGACAATCAGGAGGCTGTACNNNAGATCGGAA |
| 4718_1 | TATGAGTGTGGAGTCGTTGCACATGCTTGGTGTTCCTTCATTTNNNAGATCGGAAG |
| 4719_1 | TATGAGTGTGGAGTCGTTGCGGGGAGAGAGGGAGAGATTGTACNNNAGATCGGA |
| 4720_1 | TATGAGTGTGGAGTCGTTGCGAAGTCCCTCTGCTGAGCATTACNNNAGATCGGAAG |
| 4721_1 | TATGAGTGTGGAGTCGTTGCAGAGACGGAGACGGAGAAGGGAANNNAGATCGGA |

| | |
|---|---|
| 4723_1 | TATGAGTGTGGAGTCGTTGCAGAGAGGGACAGAGAGCACTTACNNNAGATCGGAA |
| 4724_1 | TATGAGTGTGGAGTCGTTGCATGTACGCAGACACACACCTTACNNNAGATCGGAAG |
| 4727_1 | TATGAGTGTGGAGTCGTTGCCTAACTGCCTCTCGAAGGCTTACNNNAGATCGGAAG |
| 4728_1 | TATGAGTGTGGAGTCGTTGCTCCCTATGGCTAGAAGGGCTTACNNNAGATCGGAAG |
| 4730_1 | TATGAGTGTGGAGTCGTTGCTACAACAAATGCCCCCAGGTTACNNNAGATCGGAAG |
| 4732_1 | TATGAGTGTGGAGTCGTTGCTGACTGCCCAAAATACGACACATNNNAGATCGGAAG |
| 4733_1 | TATGAGTGTGGAGTCGTTGCGCAGGGAGTTTAGGAAAAGAGGCNNNAGATCGGA |
| 4735_1 | TATGAGTGTGGAGTCGTTGCGCAGAGACAGCAACACCATTTACNNNAGATCGGAAG |
| 4736_1 | TATGAGTGTGGAGTCGTTGCATAATCCTCCCACCTCAGTGAAGNNNAGATCGGAAG |
| 4737_1 | TATGAGTGTGGAGTCGTTGCTATGTCCAGGCACTGTGCTTTACNNNAGATCGGAAG |
| 4738_1 | TATGAGTGTGGAGTCGTTGCCTTTTGTTCCTCTGGAGAACGCTNNNAGATCGGAAG |
| 4742_1 | TATGAGTGTGGAGTCGTTGCCTCTCTCTCCTTCCAGCAAAACCNNNAGATCGGAAGA |
| 4747_1 | TATGAGTGTGGAGTCGTTGCTTTCACCCAGGATTGAGTGTACTNNNAGATCGGAAG |
| 4749_1 | TATGAGTGTGGAGTCGTTGCACACTCATATCTGTCGGCAAACCNNNAGATCGGAAG |
| 4751_1 | TATGAGTGTGGAGTCGTTGCAGACGTACAATGCTCCTCAAACCNNNAGATCGGAAG |
| 4754_1 | TATGAGTGTGGAGTCGTTGCGAGAAGGTGGTCTGTCTGAAACCNNNAGATCGGAA |
| 4755_1 | TATGAGTGTGGAGTCGTTGCGCCTCTTGAGCAGCTCCAATATTNNNAGATCGGAAG |
| 4756_1 | TATGAGTGTGGAGTCGTTGCAGACGGGGCAAATTCTCTAAACCNNNAGATCGGAA |
| 4757_1 | TATGAGTGTGGAGTCGTTGCGTGCATTTGTTGCAAGGTAAACCNNNAGATCGGAA |
| 4758_1 | TATGAGTGTGGAGTCGTTGCGTTTTGTGAATGGCTCTAGCCTTNNNAGATCGGAAG |
| 4760_1 | TATGAGTGTGGAGTCGTTGCTCTGGCTCACAGCTATGGACTAANNNAGATCGGAAG |
| 4761_1 | TATGAGTGTGGAGTCGTTGCTCCTTCAACTTTCTGTCCCAACCNNNAGATCGGAAGA |
| 4764_1 | TATGAGTGTGGAGTCGTTGCAAAAGAAAGAACCCAACGCAACCNNNAGATCGGAA |
| 4766_1 | TATGAGTGTGGAGTCGTTGCGTCTGTCTGTCTGTGTCTCAACCNNNAGATCGGAAG |
| 4767_1 | TATGAGTGTGGAGTCGTTGCGCCCCAAATGTAGCTGGAACTACNNNAGATCGGAAG |
| 4768_1 | TATGAGTGTGGAGTCGTTGCAGGGGTATGAGGAGACTTCAACCNNNAGATCGGAA |
| 4769_1 | TATGAGTGTGGAGTCGTTGCAGACCCCTTCAATCCACAAACTTNNNAGATCGGAAGA |
| 4770_1 | TATGAGTGTGGAGTCGTTGCACACACACCAGAGAAACAGAACCNNNAGATCGGAA |
| 4772_1 | TATGAGTGTGGAGTCGTTGCCCTCCCTCTCATTGTGTTTCCAGNNNAGATCGGAAGA |
| 4773_1 | TATGAGTGTGGAGTCGTTGCTGGCTCTCTTTCTTAGGAGAACCNNNAGATCGGAAG |
| 4774_1 | TATGAGTGTGGAGTCGTTGCTTTTGTTCACCACCCTGAGAACCNNNAGATCGGAAGA |
| 4775_1 | TATGAGTGTGGAGTCGTTGCGACAGAGAGAAAGAGGGGGATATNNNAGATCGGA |
| 4776_1 | TATGAGTGTGGAGTCGTTGCGGTACTGCATCTGGTTAGGAACCNNNAGATCGGAA |
| 4779_1 | TATGAGTGTGGAGTCGTTGCCTTCCTGATCTGGGTTGTGAACCNNNAGATCGGAAG |
| 4781_1 | TATGAGTGTGGAGTCGTTGCCTCACAGAGCTTGGGTTTGAACCNNNAGATCGGAAG |
| 4782_1 | TATGAGTGTGGAGTCGTTGCAACAGGCCTTGGCTTTCATAACCNNNAGATCGGAAG |
| 4785_1 | TATGAGTGTGGAGTCGTTGCCAAGACTGGGTAATGTATGGAGANNNAGATCGGAA |
| 4786_1 | TATGAGTGTGGAGTCGTTGCACCCCAGAATGCAGAAAGTAACCNNNAGATCGGAA |
| 4787_1 | TATGAGTGTGGAGTCGTTGCATGGGTAGACCAGGGAAGTAACCNNNAGATCGGAA |
| 4788_1 | TATGAGTGTGGAGTCGTTGCATTCTTCACCTGTGAGCGTAACCNNNAGATCGGAAG |
| 4789_1 | TATGAGTGTGGAGTCGTTGCGGATCAGAGTGTTGCAAAACACCNNNAGATCGGAA |
| 4794_1 | TATGAGTGTGGAGTCGTTGCAGACAGACACATGAACACACACCNNNAGATCGGAAG |
| 4796_1 | TATGAGTGTGGAGTCGTTGCACACAGTGAAGGTTTCAGACACCNNNAGATCGGAA |

| | |
|---|---|
| 4797_1 | TATGAGTGTGGAGTCGTTGCGCCCTTTTGAGTTTCTTGACACCNNNAGATCGGAAG |
| 4799_1 | TATGAGTGTGGAGTCGTTGCCGTTCTTGGAGCCTGATTTCTGANNNAGATCGGAAG |
| 4802_1 | TATGAGTGTGGAGTCGTTGCAAGAAATGAGCACACCTACCACCNNNAGATCGGAAG |
| 4805_1 | TATGAGTGTGGAGTCGTTGCAGATTGGGTTACAACAACCCACCNNNAGATCGGAAG |
| 4806_1 | TATGAGTGTGGAGTCGTTGCTTTAGAGAAGTCTTTGCCCCACCNNNAGATCGGAAG |
| 4807_1 | TATGAGTGTGGAGTCGTTGCAGGGTGACTGAAAAATGCTTGCTNNNAGATCGGAA |
| 4808_1 | TATGAGTGTGGAGTCGTTGCCCTTACATGCTACTTTCCCCACCNNNAGATCGGAAGA |
| 4809_1 | TATGAGTGTGGAGTCGTTGCGTTCCAGGCATAAGCTACCATGTNNNAGATCGGAAG |
| 4813_1 | TATGAGTGTGGAGTCGTTGCATCCTAGTTGATCCTCTGCCACCNNNAGATCGGAAGA |
| 4816_1 | TATGAGTGTGGAGTCGTTGCTTACTCCAAGTCCCAAGTCCACCNNNAGATCGGAAGA |
| 4817_1 | TATGAGTGTGGAGTCGTTGCCTGGTCTACCTACCTCTTCCACCNNNAGATCGGAAGA |
| 4818_1 | TATGAGTGTGGAGTCGTTGCACACAGTACTTAGCTCTTCCACCNNNAGATCGGAAGA |
| 4819_1 | TATGAGTGTGGAGTCGTTGCGCTACACTGCATTACAAAGCACCNNNAGATCGGAAG |
| 4820_1 | TATGAGTGTGGAGTCGTTGCTCTCTGGATCATACGGTCATTCTNNNAGATCGGAAG |
| 4823_1 | TATGAGTGTGGAGTCGTTGCTAGCTTAGTGTTTGCAGAGCACCNNNAGATCGGAAG |
| 4825_1 | TATGAGTGTGGAGTCGTTGCTAGCAGGAAACATGCTTAGCACCNNNAGATCGGAA |
| 4826_1 | TATGAGTGTGGAGTCGTTGCTATGTCTCATGGTGTGATGCACCNNNAGATCGGAAG |
| 4827_1 | TATGAGTGTGGAGTCGTTGCTTACTAGCCATACCATCTGGCCTNNNAGATCGGAAG |
| 4829_1 | TATGAGTGTGGAGTCGTTGCTCTCCCACTCCCATATCATCACCNNNAGATCGGAAGA |
| 4830_1 | TATGAGTGTGGAGTCGTTGCCTGCCTTCTGACCCTTTGCTATTNNNAGATCGGAAGA |
| 4831_1 | TATGAGTGTGGAGTCGTTGCTCACTCTCCTGAACTCACTCACCNNNAGATCGGAAGA |
| 4833_1 | TATGAGTGTGGAGTCGTTGCGAACATGGGAAAGAGAGGTCACCNNNAGATCGGAA |
| 4837_1 | TATGAGTGTGGAGTCGTTGCCTGAGAATTACTGCCCCTTCACCNNNAGATCGGAAG |
| 4838_1 | TATGAGTGTGGAGTCGTTGCAGACATCCTCATCCTCCTTCACCNNNAGATCGGAAGA |
| 4839_1 | TATGAGTGTGGAGTCGTTGCTCTTCACTCTCATCCTGCAGTGGNNNAGATCGGAAG |
| 4841_1 | TATGAGTGTGGAGTCGTTGCTACAAGCCTATCTGGGTTTCACCNNNAGATCGGAAG |
| 4844_1 | TATGAGTGTGGAGTCGTTGCATGGGGATTTGGTTCTAGGATCCNNNAGATCGGAA |
| 4845_1 | TATGAGTGTGGAGTCGTTGCAGGTTCATGTACACACCAAGACCNNNAGATCGGAAG |
| 4846_1 | TATGAGTGTGGAGTCGTTGCCTGAAACTGCTTTTGCCAAGACCNNNAGATCGGAAG |
| 4848_1 | TATGAGTGTGGAGTCGTTGCTGAAAAAGGAGGCAGAACAGACCNNNAGATCGGAA |
| 4850_1 | TATGAGTGTGGAGTCGTTGCAGTGAAGACTTAGACCCCAGACCNNNAGATCGGAA |
| 4851_1 | TATGAGTGTGGAGTCGTTGCCTGGTCCTTTTACTCTGCAGACCNNNAGATCGGAAG |
| 4852_1 | TATGAGTGTGGAGTCGTTGCTTCTCCCTGTGCTTACAGAGACCNNNAGATCGGAAG |
| 4853_1 | TATGAGTGTGGAGTCGTTGCAGGGGCATATACATGGAGAGACCNNNAGATCGGAA |
| 4855_1 | TATGAGTGTGGAGTCGTTGCTTTTTCCTGGGTCAATGGAGACCNNNAGATCGGAAG |
| 4857_1 | TATGAGTGTGGAGTCGTTGCAAGGCAGATACCTCAGAAGCAAGNNNAGATCGGAA |
| 4858_1 | TATGAGTGTGGAGTCGTTGCACCCGCACTGGAGTATATAGACCNNNAGATCGGAAG |
| 4859_1 | TATGAGTGTGGAGTCGTTGCACAACCATCACCACTTGTAGACCNNNAGATCGGAAG |
| 4860_1 | TATGAGTGTGGAGTCGTTGCAGTGTGTGGAACATTGAAGGACCNNNAGATCGGAA |
| 4861_1 | TATGAGTGTGGAGTCGTTGCTTTTGACTAGGGTGACCAGGACCNNNAGATCGGAA |
| 4864_1 | TATGAGTGTGGAGTCGTTGCGGTAACACTGAAATGAGGGGACCNNNAGATCGGAA |
| 4865_1 | TATGAGTGTGGAGTCGTTGCAAAACCTAAAAGTGCCCTGGACCNNNAGATCGGAAG |
| 4866_1 | TATGAGTGTGGAGTCGTTGCTATTTCATTTGGGCAGCTGGACCNNNAGATCGGAAG |

| | |
|---|---|
| 4869_1 | TATGAGTGTGGAGTCGTTGCGTGGAGGGTAGACATGGAGAAAANNNAGATCGGA |
| 4871_1 | TATGAGTGTGGAGTCGTTGCGAGGACACTTGGGAATGATGACCNNNAGATCGGAA |
| 4875_1 | TATGAGTGTGGAGTCGTTGCGTGTGTGAGAGAGAGACTTGACCNNNAGATCGGAA |
| 4876_1 | TATGAGTGTGGAGTCGTTGCTCTCTCCAGTGGTGTGTGGTATTNNNAGATCGGAAG |
| 4879_1 | TATGAGTGTGGAGTCGTTGCGCTGCCATTATACCCAGCATACCNNNAGATCGGAAG |
| 4881_1 | TATGAGTGTGGAGTCGTTGCCGCCCCACTACACTAGGCTAATTNNNAGATCGGAAG |
| 4883_1 | TATGAGTGTGGAGTCGTTGCTCTATACCCAGACAAAGCAGGCTNNNAGATCGGAAG |
| 4885_1 | TATGAGTGTGGAGTCGTTGCATGAACCAGCAGTTTCCAATTTNNNAGATCGGAAGA |
| 4887_1 | TATGAGTGTGGAGTCGTTGCTGCTACCTTCCCCTTCTTCTACCNNNAGATCGGAAGA |
| 4889_1 | TATGAGTGTGGAGTCGTTGCTCCCAAGAGCCTAGACTAGTACCNNNAGATCGGAAG |
| 4890_1 | TATGAGTGTGGAGTCGTTGCATGCGTTGAAATGAGGAGGTACCNNNAGATCGGAA |
| 4892_1 | TATGAGTGTGGAGTCGTTGCCAAACACATGCACACTGTGTACCNNNAGATCGGAAG |
| 4893_1 | TATGAGTGTGGAGTCGTTGCCACACACGACTGAATTGTGTACCNNNAGATCGGAAG |
| 4894_1 | TATGAGTGTGGAGTCGTTGCGCTTGCCTGTCCTTTATTGTACCNNNAGATCGGAAGA |
| 4895_1 | TATGAGTGTGGAGTCGTTGCCAAGCCCAAGGCATCAAATTACCNNNAGATCGGAAG |
| 4896_1 | TATGAGTGTGGAGTCGTTGCATACCTACCCTGAGGGCATTACCNNNAGATCGGAAG |
| 4899_1 | TATGAGTGTGGAGTCGTTGCCTCTCTTGGTTGCAGGACTTACCNNNAGATCGGAAG |
| 4901_1 | TATGAGTGTGGAGTCGTTGCATTTGCAGCAACCCAGTTGGAGTNNNAGATCGGAAG |
| 4902_1 | TATGAGTGTGGAGTCGTTGCTGCTGGCTCATTCTTCCTTTACCNNNAGATCGGAAGA |
| 4903_1 | TATGAGTGTGGAGTCGTTGCTACACAGCAACTCCAGGTTAGGANNNAGATCGGAAG |
| 4904_1 | TATGAGTGTGGAGTCGTTGCTACTCCAAAGCCTAGCAAAACCCNNNAGATCGGAAG |
| 4905_1 | TATGAGTGTGGAGTCGTTGCTAGCCAAACATTGCATCAAACCCNNNAGATCGGAAG |
| 4908_1 | TATGAGTGTGGAGTCGTTGCAGTATTGGCAGGATGCTAAACCCNNNAGATCGGAA |
| 4909_1 | TATGAGTGTGGAGTCGTTGCGATGGGTGCGAGGTGACATTTCANNNAGATCGGAA |
| 4910_1 | TATGAGTGTGGAGTCGTTGCACCTGGCTTATTCTCTTCAACCCNNNAGATCGGAAGA |
| 4911_1 | TATGAGTGTGGAGTCGTTGCTCTCTTGCCTTCTCCTCTAACCCNNNAGATCGGAAGA |
| 4917_1 | TATGAGTGTGGAGTCGTTGCCAAGGTTTTGATTGCTGACACCCNNNAGATCGGAAG |
| 4920_1 | TATGAGTGTGGAGTCGTTGCCTGTCTTGAGCATCTTCTCACCCNNNAGATCGGAAGA |
| 4921_1 | TATGAGTGTGGAGTCGTTGCCTGTCTCTGCTCTTTTCTCACCCNNNAGATCGGAAGA |
| 4923_1 | TATGAGTGTGGAGTCGTTGCTAGAATGCAAGTCTGTGAGGGCANNNAGATCGGAA |
| 4925_1 | TATGAGTGTGGAGTCGTTGCAACAAGACATCAAGGCAAGACCCNNNAGATCGGAA |
| 4926_1 | TATGAGTGTGGAGTCGTTGCGAATTGGAGGTAGCCAGTGTCAANNNAGATCGGAA |
| 4927_1 | TATGAGTGTGGAGTCGTTGCATTGCAGAAACATCCTCAGACCCNNNAGATCGGAAG |
| 4928_1 | TATGAGTGTGGAGTCGTTGCGCATATTGTGGGTCAAGAGACCCNNNAGATCGGAA |
| 4929_1 | TATGAGTGTGGAGTCGTTGCTAAGCTCTTCAGCAAAAGGACCCNNNAGATCGGAAG |
| 4930_1 | TATGAGTGTGGAGTCGTTGCGTAAATGGGCATGGGAGCCAATTNNNAGATCGGAA |
| 4931_1 | TATGAGTGTGGAGTCGTTGCAATCCACTTCTCTACCCTGACCCNNNAGATCGGAAGA |
| 4932_1 | TATGAGTGTGGAGTCGTTGCACATGGAGAAAGGACTGGAGTGANNNAGATCGGA |
| 4933_1 | TATGAGTGTGGAGTCGTTGCGGATTGTGTAGAATTGGCGTTGANNNAGATCGGAA |
| 4935_1 | TATGAGTGTGGAGTCGTTGCCCACCGGAGCTATAGCAAAACTTNNNAGATCGGAAG |
| 4936_1 | TATGAGTGTGGAGTCGTTGCGTGGGATGACAGGTATGTACCACNNNAGATCGGAA |
| 4941_1 | TATGAGTGTGGAGTCGTTGCTTGCACACTCCAAAAATGTACCCNNNAGATCGGAAG |
| 4942_1 | TATGAGTGTGGAGTCGTTGCCCTCTGGGAAAGAGACCTGGAAGNNNAGATCGGAA |

| | |
|---|---|
| 4947_1 | TATGAGTGTGGAGTCGTTGCTAGGGGATTGATTTCTCGACCCCNNNAGATCGGAAG |
| 4949_1 | TATGAGTGTGGAGTCGTTGCCATACTGGCTTCATCTTGACCCCNNNAGATCGGAAGA |
| 4952_1 | TATGAGTGTGGAGTCGTTGCTTTTCTTGGCAGTTCCTTACCCCNNNAGATCGGAAGA |
| 4955_1 | TATGAGTGTGGAGTCGTTGCAAATGAGATGCATCAGAACCCCCNNNAGATCGGAAG |
| 4956_1 | TATGAGTGTGGAGTCGTTGCAAGACTACCTAAAAACAGCCCCCNNNAGATCGGAAGA |
| 4961_1 | TATGAGTGTGGAGTCGTTGCTTTGTCTTCAATATGCCTCCCCCNNNAGATCGGAAGAG |
| 4963_1 | TATGAGTGTGGAGTCGTTGCTCAGCAAATACAGCCAGCCCTTTNNNAGATCGGAAGA |
| 4965_1 | TATGAGTGTGGAGTCGTTGCTAGAATAGCAAAAAGCGAGCCCCNNNAGATCGGAAG |
| 4966_1 | TATGAGTGTGGAGTCGTTGCCTTCTTTTCAGGACACTAGCCCCNNNAGATCGGAAGA |
| 4968_1 | TATGAGTGTGGAGTCGTTGCTTATACACAATATGGCTGGCCCCNNNAGATCGGAAGA |
| 4970_1 | TATGAGTGTGGAGTCGTTGCGAATCTTTGAAACCTCATGCCCCNNNAGATCGGAAGA |
| 4972_1 | TATGAGTGTGGAGTCGTTGCAACTCACAGGTTTTCTATGCCCCNNNAGATCGGAAGA |
| 4973_1 | TATGAGTGTGGAGTCGTTGCACTGTCCTCCCAAGTCTCAACTTNNNAGATCGGAAGA |
| 4974_1 | TATGAGTGTGGAGTCGTTGCTTTCCTCTTTCAGCCTTACAGGGNNNAGATCGGAAGA |
| 4975_1 | TATGAGTGTGGAGTCGTTGCTGCTTCTAATTATGGTGTGCCCCNNNAGATCGGAAGA |
| 4976_1 | TATGAGTGTGGAGTCGTTGCTTTTTGTGTATGTGCTTTGCCCCNNNAGATCGGAAGA |
| 4978_1 | TATGAGTGTGGAGTCGTTGCGAAGGCAGGACATTTTACTCCCCNNNAGATCGGAAG |
| 4979_1 | TATGAGTGTGGAGTCGTTGCCCTGGCAGTAAAAGAACGTATGANNNAGATCGGAAG |
| 4981_1 | TATGAGTGTGGAGTCGTTGCCAGAGACCTGGGAACCAGATATGNNNAGATCGGAAG |
| 4983_1 | TATGAGTGTGGAGTCGTTGCACATGGCAAAATCATCTCTCCCCNNNAGATCGGAAGA |
| 4989_1 | TATGAGTGTGGAGTCGTTGCTCTCTTTGGGTTTCATCTTCCCCNNNAGATCGGAAGAG |
| 4991_1 | TATGAGTGTGGAGTCGTTGCTCTCCTTTCGATCATACAGGGCTNNNAGATCGGAAGA |
| 4992_1 | TATGAGTGTGGAGTCGTTGCACACGGACACAAATACAAAGCCCNNNAGATCGGAAG |
| 4995_1 | TATGAGTGTGGAGTCGTTGCTGGAGTTACATTCAGATAAGCCCNNNAGATCGGAAG |
| 4996_1 | TATGAGTGTGGAGTCGTTGCCCACACTAGAACACAGTAAGCCCNNNAGATCGGAAG |
| 4998_1 | TATGAGTGTGGAGTCGTTGCATGTCTTGAGGTTATGCTGGCAGNNNAGATCGGAAG |
| 5002_1 | TATGAGTGTGGAGTCGTTGCTACCTTTGTAGCAATGTGAGCCCNNNAGATCGGAAG |
| 5003_1 | TATGAGTGTGGAGTCGTTGCACAGGGCTCAAGGAGGATAACTTNNNAGATCGGAAG |
| 5004_1 | TATGAGTGTGGAGTCGTTGCAATCTGAAGCTCAATGCTAGCCCNNNAGATCGGAAG |
| 5007_1 | TATGAGTGTGGAGTCGTTGCAGATTTTGAGACTGACAACGCCCNNNAGATCGGAAG |
| 5010_1 | TATGAGTGTGGAGTCGTTGCAAAGTGGTGCCTAATAATGGCCCNNNAGATCGGAAG |
| 5013_1 | TATGAGTGTGGAGTCGTTGCGTTCTGTGTTGACAATTTGGCCCNNNAGATCGGAAG |
| 5014_1 | TATGAGTGTGGAGTCGTTGCGGAGAATGAGGTAGTTTTGGCCCNNNAGATCGGAA |
| 5015_1 | TATGAGTGTGGAGTCGTTGCCTTTGCAGTGATGCTAAATGCCCNNNAGATCGGAAG |
| 5018_1 | TATGAGTGTGGAGTCGTTGCACGATACTTGCAGTTCTATGCCCNNNAGATCGGAAGA |
| 5022_1 | TATGAGTGTGGAGTCGTTGCCTGTTGGGTGAGACAAAACATCTNNNAGATCGGAAG |
| 5023_1 | TATGAGTGTGGAGTCGTTGCCTTGAACAGAGTACATCCTGCCCNNNAGATCGGAAGA |
| 5024_1 | TATGAGTGTGGAGTCGTTGCTTTTTGTATTTGTCGTGCTGCCCNNNAGATCGGAAGA |
| 5025_1 | TATGAGTGTGGAGTCGTTGCTTCCAGATCCATGGTATTCAGTCNNNAGATCGGAAGA |
| 5027_1 | TATGAGTGTGGAGTCGTTGCTGTTACTTGGGGATTTTCTGCCCNNNAGATCGGAAGA |
| 5028_1 | TATGAGTGTGGAGTCGTTGCCAGCCTTGATGTGTTCTAATGGCNNNAGATCGGAAG |
| 5029_1 | TATGAGTGTGGAGTCGTTGCATTTCTGTTCCCCTAGAGTGCCCNNNAGATCGGAAGA |
| 5030_1 | TATGAGTGTGGAGTCGTTGCAGGGGTAAAGCACATTATTGCCCNNNAGATCGGAAG |

| | |
|---|---|
| 5032_1 | TATGAGTGTGGAGTCGTTGCTTGTGGAAGGTGTAGAAGGGTGANNNAGATCGGA |
| 5033_1 | TATGAGTGTGGAGTCGTTGCTGTGTTTTGTCACTTTGTTGCCCNNNAGATCGGAAGA |
| 5034_1 | TATGAGTGTGGAGTCGTTGCATTTCTGGAGACAAGCTTTGCCCNNNAGATCGGAAG |
| 5035_1 | TATGAGTGTGGAGTCGTTGCAGCGGAAGTGATTGAGAAATCCCNNNAGATCGGAA |
| 5036_1 | TATGAGTGTGGAGTCGTTGCAGTCAGATATGGGTGGGAATCCCNNNAGATCGGAA |
| 5039_1 | TATGAGTGTGGAGTCGTTGCTCACTAACCTCTCTTCCCATCCCNNNAGATCGGAAGA |
| 5041_1 | TATGAGTGTGGAGTCGTTGCATCATCCCCAAAGCAATCATCCCNNNAGATCGGAAGA |
| 5042_1 | TATGAGTGTGGAGTCGTTGCTATTTGCAAGCTCCCATCATCCCNNNAGATCGGAAGA |
| 5046_1 | TATGAGTGTGGAGTCGTTGCCTTGTGGGAGCCTTTCTTATCCCNNNAGATCGGAAG |
| 5049_1 | TATGAGTGTGGAGTCGTTGCATTATAGCTCCTCCCACACTCCCNNNAGATCGGAAGA |
| 5053_1 | TATGAGTGTGGAGTCGTTGCTGTATGTATGTGCTTTGCCTCCCNNNAGATCGGAAG |
| 5054_1 | TATGAGTGTGGAGTCGTTGCAGTTTTGAAGCAGACACCTGGAANNNAGATCGGAA |
| 5055_1 | TATGAGTGTGGAGTCGTTGCTGGCCTCTTCATGATATCCTCCCNNNAGATCGGAAGA |
| 5057_1 | TATGAGTGTGGAGTCGTTGCATTTCTTTGCTTCTCCAGCTCCCNNNAGATCGGAAGA |
| 5059_1 | TATGAGTGTGGAGTCGTTGCCCTCCCTATCCAGATATGCTCCCNNNAGATCGGAAGA |
| 5060_1 | TATGAGTGTGGAGTCGTTGCCTTGTACACACATACCTGCTCCCNNNAGATCGGAAGA |
| 5061_1 | TATGAGTGTGGAGTCGTTGCACTCATCAACACATCTTGCTCCCNNNAGATCGGAAGA |
| 5063_1 | TATGAGTGTGGAGTCGTTGCAGGAGGTTCTTCTTTCCTCTCCCNNNAGATCGGAAGA |
| 5066_1 | TATGAGTGTGGAGTCGTTGCCTTACATGCTCACACAGTCTCCCNNNAGATCGGAAGA |
| 5069_1 | TATGAGTGTGGAGTCGTTGCTTGAACCAAACCAGTTGTCTCCCNNNAGATCGGAAG |
| 5070_1 | TATGAGTGTGGAGTCGTTGCATTGATGGTGCTGCTATTCTCCCNNNAGATCGGAAG |
| 5071_1 | TATGAGTGTGGAGTCGTTGCATGCTGGAATGAATGGAAGTCCCNNNAGATCGGAA |
| 5072_1 | TATGAGTGTGGAGTCGTTGCACTCACTCCTTCCTTGAAGTCCCNNNAGATCGGAAGA |
| 5075_1 | TATGAGTGTGGAGTCGTTGCAAACACTTAGATGCCTTGGTCCCNNNAGATCGGAAG |
| 5076_1 | TATGAGTGTGGAGTCGTTGCGAGTTACGAAGCACAGATGTCCCNNNAGATCGGAAG |
| 5077_1 | TATGAGTGTGGAGTCGTTGCCCAGAGTTAGGAAAAGCTGTCCCNNNAGATCGGAAG |
| 5079_1 | TATGAGTGTGGAGTCGTTGCTACAACTTTCAGCCTGCATTCCCNNNAGATCGGAAGA |
| 5084_1 | TATGAGTGTGGAGTCGTTGCAATGTCCTGTTTTCCCACTTCCCNNNAGATCGGAAGA |
| 5085_1 | TATGAGTGTGGAGTCGTTGCAAAGGAACTGAAAATGCTGGTGANNNAGATCGGAA |
| 5086_1 | TATGAGTGTGGAGTCGTTGCCCTGAGAGAATGGAGTTTTGCCANNNAGATCGGAA |
| 5088_1 | TATGAGTGTGGAGTCGTTGCTACAAGCAACACTGTGAATGGCTNNNAGATCGGAAG |
| 5090_1 | TATGAGTGTGGAGTCGTTGCTGTGGCAGGAATTTCTTCTTCCCNNNAGATCGGAAG |
| 5091_1 | TATGAGTGTGGAGTCGTTGCCACCTCAGCATCACTTAGTTCCCNNNAGATCGGAAGA |
| 5092_1 | TATGAGTGTGGAGTCGTTGCCTGCTAACTCAAACCAGGTTCCCNNNAGATCGGAAG |
| 5093_1 | TATGAGTGTGGAGTCGTTGCTCGCTTCTTTTCTCACTGTTCCCNNNAGATCGGAAGA |
| 5097_1 | TATGAGTGTGGAGTCGTTGCTTTCCAGTGCAATTCCAAAAGCCNNNAGATCGGAAG |
| 5098_1 | TATGAGTGTGGAGTCGTTGCATCTCCTGTCTCATGGAAAAGCCNNNAGATCGGAAG |
| 5102_1 | TATGAGTGTGGAGTCGTTGCCGATTTTGCTCTGGTGTAAAGCCNNNAGATCGGAAG |
| 5106_1 | TATGAGTGTGGAGTCGTTGCCTGGAAATAAACCATGCCAAGCCNNNAGATCGGAAG |
| 5107_1 | TATGAGTGTGGAGTCGTTGCACTAGAGCTAAGGACAGCAAGCCNNNAGATCGGAA |
| 5108_1 | TATGAGTGTGGAGTCGTTGCTTTTCTACTCCTTCTGGCAAGCCNNNAGATCGGAAGA |
| 5109_1 | TATGAGTGTGGAGTCGTTGCAACTTTGTGCTCAGAATGAAGCCNNNAGATCGGAAG |
| 5110_1 | TATGAGTGTGGAGTCGTTGCTGTCTATCTGAGGCTCCTAAGCCNNNAGATCGGAAG |

| | |
|---|---|
| 5111_1 | TATGAGTGTGGAGTCGTTGCCCTGACCCCTGAAGCATGAAAGTNNNAGATCGGAAG |
| 5115_1 | TATGAGTGTGGAGTCGTTGCATTATGAAGTTGAGGCCACAGCCNNNAGATCGGAA |
| 5116_1 | TATGAGTGTGGAGTCGTTGCAGCAGAGAAAAACAGAGACAGCCNNNAGATCGGAA |
| 5117_1 | TATGAGTGTGGAGTCGTTGCATCTCTCTTTTCTGCTTCCTGGTNNNAGATCGGAAGA |
| 5118_1 | TATGAGTGTGGAGTCGTTGCTCATCTCTACCTGTAGTCCAGCCNNNAGATCGGAAGA |
| 5120_1 | TATGAGTGTGGAGTCGTTGCGTAGGTAGTTCAGGAAAGCAGCCNNNAGATCGGAA |
| 5123_1 | TATGAGTGTGGAGTCGTTGCCTACACAGAAACAAACTGCAGCCNNNAGATCGGAAG |
| 5125_1 | TATGAGTGTGGAGTCGTTGCACAATGACAGGAGACAATCAGCCNNNAGATCGGAA |
| 5127_1 | TATGAGTGTGGAGTCGTTGCGAGGTTCTCACTGTTGATCAGCCNNNAGATCGGAAG |
| 5128_1 | TATGAGTGTGGAGTCGTTGCATCACCACCATTTTCTCTCAGCCNNNAGATCGGAAGA |
| 5130_1 | TATGAGTGTGGAGTCGTTGCAGCCCACATTTCCTGAAAACTGTNNNAGATCGGAAG |
| 5132_1 | TATGAGTGTGGAGTCGTTGCCCCAGTCATACACATGAAGAGCCNNNAGATCGGAAG |
| 5134_1 | TATGAGTGTGGAGTCGTTGCCTACACAAGTAACACCAGGAGCCNNNAGATCGGAAG |
| 5135_1 | TATGAGTGTGGAGTCGTTGCTGGCCTTACCTTGGGAGAATAGTNNNAGATCGGAA |
| 5136_1 | TATGAGTGTGGAGTCGTTGCGCACGAAATCAGAAATCTGAGCCNNNAGATCGGAA |
| 5139_1 | TATGAGTGTGGAGTCGTTGCGGATTTCTCGACTTCTCTGAGCCNNNAGATCGGAAG |
| 5140_1 | TATGAGTGTGGAGTCGTTGCAGGGATTTACTTTTCGGTGAGCCNNNAGATCGGAA |
| 5143_1 | TATGAGTGTGGAGTCGTTGCAGGTGGAAACGGATATTTGAGCCNNNAGATCGGAA |
| 5144_1 | TATGAGTGTGGAGTCGTTGCAGAGACAGTACCACCTCATAGCCNNNAGATCGGAAG |
| 5146_1 | TATGAGTGTGGAGTCGTTGCGTTACTGAATTGGCTCCCTAGCCNNNAGATCGGAAG |
| 5147_1 | TATGAGTGTGGAGTCGTTGCCAGGCTAGTCATAACTCCTAGCCNNNAGATCGGAAG |
| 5149_1 | TATGAGTGTGGAGTCGTTGCTTTTCCTCTTGCCATGAGTAGCCNNNAGATCGGAAG |
| 5150_1 | TATGAGTGTGGAGTCGTTGCTAGAGGAGGCAGATAACGTAGCCNNNAGATCGGAA |
| 5153_1 | TATGAGTGTGGAGTCGTTGCGGATAGTTCAGGGAAGGTGAGGTNNNAGATCGGA |
| 5154_1 | TATGAGTGTGGAGTCGTTGCCTTGGCTCCTATGGACATTAGCCNNNAGATCGGAAG |
| 5155_1 | TATGAGTGTGGAGTCGTTGCGAGAGAAGCTGTAGCCATTAGCCNNNAGATCGGAA |
| 5157_1 | TATGAGTGTGGAGTCGTTGCAAAAGAACACTCCCCACTTAGCCNNNAGATCGGAAG |
| 5159_1 | TATGAGTGTGGAGTCGTTGCTCACTCCAGAGCTAGTCTTAGCCNNNAGATCGGAAG |
| 5160_1 | TATGAGTGTGGAGTCGTTGCTACGCCAAAGAAAAAGACTGGCTNNNAGATCGGAA |
| 5161_1 | TATGAGTGTGGAGTCGTTGCCAGCATGTATTCCATTTCACGCCNNNAGATCGGAAGA |
| 5162_1 | TATGAGTGTGGAGTCGTTGCAAAGACATGAAAAACCAACCGCCNNNAGATCGGAA |
| 5164_1 | TATGAGTGTGGAGTCGTTGCCAAGTAAATACGGTCCTCTCGCCNNNAGATCGGAAG |
| 5165_1 | TATGAGTGTGGAGTCGTTGCAGGGAATTTGTACTCCATTCGCCNNNAGATCGGAAG |
| 5166_1 | TATGAGTGTGGAGTCGTTGCTCCCTCACATCAAAGAAAAGGCCNNNAGATCGGAAG |
| 5167_1 | TATGAGTGTGGAGTCGTTGCCCTCAGTCTTCTTCAGAAAGGCCNNNAGATCGGAAG |
| 5169_1 | TATGAGTGTGGAGTCGTTGCTTGTGGAACTGGTGAATAAGGCCNNNAGATCGGAA |
| 5170_1 | TATGAGTGTGGAGTCGTTGCGGGCTACTACCTCTGGTCTGTGGNNNAGATCGGAA |
| 5172_1 | TATGAGTGTGGAGTCGTTGCAGTAACATGTATGTCCCCAGGCCNNNAGATCGGAAG |
| 5173_1 | TATGAGTGTGGAGTCGTTGCAGTTTCCCTATTTTTCCCAGGCCNNNAGATCGGAAGA |
| 5174_1 | TATGAGTGTGGAGTCGTTGCATTTACTGTGCATTCTCCAGGCCNNNAGATCGGAAG |
| 5175_1 | TATGAGTGTGGAGTCGTTGCTGATTATCCAAGAACTGCAGGCCNNNAGATCGGAAG |
| 5178_1 | TATGAGTGTGGAGTCGTTGCCATCAGTTCAGATTCTGGAGGCCNNNAGATCGGAAG |
| 5180_1 | TATGAGTGTGGAGTCGTTGCGAAATAACCCCACATCCTAGGCCNNNAGATCGGAAG |

| | |
|---|---|
| 5181_1 | TATGAGTGTGGAGTCGTTGCGGTCAAGAAGCAGAAGCTGACTTNNNAGATCGGAA |
| 5184_1 | TATGAGTGTGGAGTCGTTGCGCCAGTCATAGAATTCTGCACACNNNAGATCGGAAG |
| 5185_1 | TATGAGTGTGGAGTCGTTGCCTAAGAACCTAAAACCTCCGGCCNNNAGATCGGAAG |
| 5186_1 | TATGAGTGTGGAGTCGTTGCCTTAGAGAAGACAACCAAGGGCCNNNAGATCGGAA |
| 5187_1 | TATGAGTGTGGAGTCGTTGCATAAAAACTAACAGGCCAGGGCCNNNAGATCGGAA |
| 5188_1 | TATGAGTGTGGAGTCGTTGCATCACTCTTATTAGGGGAGGGCCNNNAGATCGGAA |
| 5189_1 | TATGAGTGTGGAGTCGTTGCCTTTGGAAACAATTCAAGGGGCCNNNAGATCGGAA |
| 5192_1 | TATGAGTGTGGAGTCGTTGCTTAGTCTTAGAGACAGGTGGGCCNNNAGATCGGAA |
| 5193_1 | TATGAGTGTGGAGTCGTTGCTTTATTGTGCTTTCCTGCTGGCTNNNAGATCGGAAG |
| 5194_1 | TATGAGTGTGGAGTCGTTGCTCTTGATAACTGCCCAAATGGCCNNNAGATCGGAAG |
| 5196_1 | TATGAGTGTGGAGTCGTTGCTTTTCTCTCTCTCATGCATGGCCNNNAGATCGGAAGA |
| 5201_1 | TATGAGTGTGGAGTCGTTGCTGGTCTCTGGTTTCTGCTTGTGTNNNAGATCGGAAG |
| 5202_1 | TATGAGTGTGGAGTCGTTGCCCTGACCTGGAGTGTTTTGACTTNNNAGATCGGAAG |
| 5206_1 | TATGAGTGTGGAGTCGTTGCTTCACAACTACTTTGGTCTGGCCNNNAGATCGGAAG |
| 5208_1 | TATGAGTGTGGAGTCGTTGCCTTGGGTTCTATGAGAAGTGGCCNNNAGATCGGAA |
| 5210_1 | TATGAGTGTGGAGTCGTTGCCCCAGATGATTCTAATGGTGGCCNNNAGATCGGAAG |
| 5211_1 | TATGAGTGTGGAGTCGTTGCTCTATTTAGCATCTGCTGTGGCCNNNAGATCGGAAG |
| 5217_1 | TATGAGTGTGGAGTCGTTGCCTGAATCGTGTCCCTGAAATGCCNNNAGATCGGAAG |
| 5218_1 | TATGAGTGTGGAGTCGTTGCAAGAACGCCTGCCAAATACATCTNNNAGATCGGAAG |
| 5219_1 | TATGAGTGTGGAGTCGTTGCAGGAGGTAGAGATACCCAATGCCNNNAGATCGGAA |
| 5220_1 | TATGAGTGTGGAGTCGTTGCCGGATTTCTAAGGCTTTCTGGCTNNNAGATCGGAAG |
| 5221_1 | TATGAGTGTGGAGTCGTTGCCGGGTCACAGGGAGTTACTACTTNNNAGATCGGAA |
| 5222_1 | TATGAGTGTGGAGTCGTTGCTACGATAGCCACAAGGTACACGTNNNAGATCGGAA |
| 5224_1 | TATGAGTGTGGAGTCGTTGCCCATTGTACTCTCCTAGCATGCCNNNAGATCGGAAGA |
| 5225_1 | TATGAGTGTGGAGTCGTTGCTTTCACATTGTCCCACCTTTCTGNNNAGATCGGAAGA |
| 5226_1 | TATGAGTGTGGAGTCGTTGCTAGGGAAAGGGGATAGAGATGCCNNNAGATCGGA |
| 5229_1 | TATGAGTGTGGAGTCGTTGCAAGGATCATGTGAATGGGATGCCNNNAGATCGGAA |
| 5230_1 | TATGAGTGTGGAGTCGTTGCAGAACTGGTTGATCATGGATGCCNNNAGATCGGAA |
| 5231_1 | TATGAGTGTGGAGTCGTTGCATGTAACCATGTCAGGTGATGCCNNNAGATCGGAA |
| 5232_1 | TATGAGTGTGGAGTCGTTGCTGGTGGAGGTTATAGCATATGCCNNNAGATCGGAA |
| 5233_1 | TATGAGTGTGGAGTCGTTGCGGAACAAGTGGAAGGTGCTACTTNNNAGATCGGAA |
| 5235_1 | TATGAGTGTGGAGTCGTTGCTGTCAAGGCTCTCAGCTCTACTTNNNAGATCGGAAG |
| 5236_1 | TATGAGTGTGGAGTCGTTGCACCAAGATGTGACTGTAACTGCCNNNAGATCGGAAG |
| 5237_1 | TATGAGTGTGGAGTCGTTGCTACACACACACTCACATACTGCCNNNAGATCGGAAGA |
| 5239_1 | TATGAGTGTGGAGTCGTTGCTTTTGATGGGTACAGAACCTGCCNNNAGATCGGAAG |
| 5240_1 | TATGAGTGTGGAGTCGTTGCGGGCCGCTAGAAATGAAGTACTTNNNAGATCGGAA |
| 5241_1 | TATGAGTGTGGAGTCGTTGCTTTGCTCATACTCTTCACCTGCCNNNAGATCGGAAGA |
| 5243_1 | TATGAGTGTGGAGTCGTTGCTATTGTGTTCATATGCCCCTGCCNNNAGATCGGAAG |
| 5244_1 | TATGAGTGTGGAGTCGTTGCTACATTCTGATGCTTTCCCTGCCNNNAGATCGGAAGA |
| 5246_1 | TATGAGTGTGGAGTCGTTGCAAGTTCCTTAGTCTTTGCCTGCCNNNAGATCGGAAG |
| 5249_1 | TATGAGTGTGGAGTCGTTGCGGAAGGTGAGGGCATTTGTACTTNNNAGATCGGAA |
| 5253_1 | TATGAGTGTGGAGTCGTTGCGATGACCCACTGATACTTCTGCCNNNAGATCGGAAG |
| 5255_1 | TATGAGTGTGGAGTCGTTGCTGAAGCACCAACCAAGCTTACTTNNNAGATCGGAAG |

| | |
|---|---|
| 5257_1 | TATGAGTGTGGAGTCGTTGCAACAGCAAGTCAAATTCAGTGCCNNNAGATCGGAAG |
| 5259_1 | TATGAGTGTGGAGTCGTTGCACCTCTCTTTTAGTTGTGGTGCCNNNAGATCGGAAG |
| 5260_1 | TATGAGTGTGGAGTCGTTGCTCAGGTTCTGCTAAAGATGTGCCNNNAGATCGGAAG |
| 5261_1 | TATGAGTGTGGAGTCGTTGCTCAATGTCTTCTTGGTCTGTGCCNNNAGATCGGAAG |
| 5262_1 | TATGAGTGTGGAGTCGTTGCCATTTATTGGATGCCTGTGTGCCNNNAGATCGGAAG |
| 5263_1 | TATGAGTGTGGAGTCGTTGCTTTTAGGATGATCGGTGTGTGCCNNNAGATCGGAA |
| 5264_1 | TATGAGTGTGGAGTCGTTGCAGCATCATCCCTTCCTAATTGCCNNNAGATCGGAAGA |
| 5266_1 | TATGAGTGTGGAGTCGTTGCCCCAGGTACATACACTTCTTGCCNNNAGATCGGAAG |
| 5267_1 | TATGAGTGTGGAGTCGTTGCTTCAAGTGGATTGGCTCAACCTTNNNAGATCGGAAG |
| 5268_1 | TATGAGTGTGGAGTCGTTGCAGTTGCCTTACATCTCTGTTGCCNNNAGATCGGAAG |
| 5269_1 | TATGAGTGTGGAGTCGTTGCATTACTTCAGTCCCAGTGTTGCCNNNAGATCGGAAG |
| 5270_1 | TATGAGTGTGGAGTCGTTGCCTGCAACAGCTTCTAAGTTTGCCNNNAGATCGGAAG |
| 5271_1 | TATGAGTGTGGAGTCGTTGCTCTCTCTCTCTCTCCAGTTTGCCNNNAGATCGGAAGA |
| 5272_1 | TATGAGTGTGGAGTCGTTGCGCCTCTGTTGTTACCATTTTGCCNNNAGATCGGAAGA |
| 5276_1 | TATGAGTGTGGAGTCGTTGCTATCAGGAATGGTTGGCCAATCCNNNAGATCGGAAG |
| 5277_1 | TATGAGTGTGGAGTCGTTGCAGCCCTTGGAACTTCATAAAGGTNNNAGATCGGAAG |
| 5278_1 | TATGAGTGTGGAGTCGTTGCTGGTGGCAGGTCACATGGTTATTNNNAGATCGGAA |
| 5279_1 | TATGAGTGTGGAGTCGTTGCCCCCAGAAAAGTGGCTAGAATCCNNNAGATCGGAAG |
| 5280_1 | TATGAGTGTGGAGTCGTTGCGATGCCCTTCACACCTGTAATCCNNNAGATCGGAAG |
| 5285_1 | TATGAGTGTGGAGTCGTTGCGGGAACTAATGTGACCACCATCCNNNAGATCGGAAG |
| 5286_1 | TATGAGTGTGGAGTCGTTGCCTTATTGGATCAGGGCACCATCCNNNAGATCGGAAG |
| 5287_1 | TATGAGTGTGGAGTCGTTGCGAAGAAACTGGTGGTCACCATCCNNNAGATCGGAA |
| 5288_1 | TATGAGTGTGGAGTCGTTGCTCTCTGTCTGTACCCATCCATCCNNNAGATCGGAAGA |
| 5289_1 | TATGAGTGTGGAGTCGTTGCGGGTGTACCATCTGTTTCCATCCNNNAGATCGGAAG |
| 5291_1 | TATGAGTGTGGAGTCGTTGCTAAGGGAATGACACCATGCATCCNNNAGATCGGAA |
| 5292_1 | TATGAGTGTGGAGTCGTTGCAAACAAGGCTTAGCCAGTCATCCNNNAGATCGGAAG |
| 5296_1 | TATGAGTGTGGAGTCGTTGCTGCTCTAAGGAACTCCCTGATCCNNNAGATCGGAAG |
| 5298_1 | TATGAGTGTGGAGTCGTTGCCTCCATGGTTTCCTCTGTGATCCNNNAGATCGGAAGA |
| 5299_1 | TATGAGTGTGGAGTCGTTGCCCTGCTCTTGAAGGAGTTTGTGANNNAGATCGGAA |
| 5300_1 | TATGAGTGTGGAGTCGTTGCACCCCAAAAGCCACTCAATATCCNNNAGATCGGAAG |
| 5301_1 | TATGAGTGTGGAGTCGTTGCCATCTTGCTGTGTGGGGATATCCNNNAGATCGGAAG |
| 5303_1 | TATGAGTGTGGAGTCGTTGCTGGAGATGATGCCCTAGCTATCCNNNAGATCGGAAG |
| 5305_1 | TATGAGTGTGGAGTCGTTGCTATGGTGATCATGCGCTGTATCCNNNAGATCGGAAG |
| 5306_1 | TATGAGTGTGGAGTCGTTGCCAGGGCCTCCAATTCTTTTGTGANNNAGATCGGAAG |
| 5309_1 | TATGAGTGTGGAGTCGTTGCTCTGTTGCCACATCTTCAACTCCNNNAGATCGGAAGA |
| 5310_1 | TATGAGTGTGGAGTCGTTGCATCATTCTCTGCGGTTCAACTCCNNNAGATCGGAAGA |
| 5311_1 | TATGAGTGTGGAGTCGTTGCCTGCATTCACTCTGGAGAACTCCNNNAGATCGGAAG |
| 5313_1 | TATGAGTGTGGAGTCGTTGCAAAGAAGCAGACAGAGACACTCCNNNAGATCGGAA |
| 5320_1 | TATGAGTGTGGAGTCGTTGCCTTTGCTTCTGCCATTTCCTCCANNNAGATCGGAAGA |
| 5322_1 | TATGAGTGTGGAGTCGTTGCATCTCCTATCAACCTCACCCTCCNNNAGATCGGAAGA |
| 5324_1 | TATGAGTGTGGAGTCGTTGCGGATCAATTTTTGGTTGCCCTCCNNNAGATCGGAAG |
| 5326_1 | TATGAGTGTGGAGTCGTTGCTATGTAGGTTTTCCCTGGCCTCCNNNAGATCGGAAG |
| 5328_1 | TATGAGTGTGGAGTCGTTGCTTTCTTGTCTGTTGCCATCCTCCNNNAGATCGGAAGA |

5329_1 TATGAGTGTGGAGTCGTTGCTGCTTCTCTCTGGGCATTTCTCANNNAGATCGGAAGA
5330_1 TATGAGTGTGGAGTCGTTGCATACTAGGGAAGTGTGCTCCTCCNNNAGATCGGAAG
5331_1 TATGAGTGTGGAGTCGTTGCTGTGAATAAGGGGGTTCTCCTCCNNNAGATCGGAA
5332_1 TATGAGTGTGGAGTCGTTGCCTCCTGCCATTCTACTTTCCTCCNNNAGATCGGAAGA
5334_1 TATGAGTGTGGAGTCGTTGCGTCACCCGAAATACAGTAGCTCCNNNAGATCGGAAG
5335_1 TATGAGTGTGGAGTCGTTGCAACCCAGTACTTCCAACCAACAGNNNAGATCGGAAG
5338_1 TATGAGTGTGGAGTCGTTGCAGGTCAATTTCCACTCTTGCTCCNNNAGATCGGAAGA
5340_1 TATGAGTGTGGAGTCGTTGCCTTGAGCTGTGGGATCAATCTCCNNNAGATCGGAAG
5341_1 TATGAGTGTGGAGTCGTTGCAGGCTCAGAGTTAGGTCATCTCCNNNAGATCGGAAG
5342_1 TATGAGTGTGGAGTCGTTGCAACGGAAAGCACTGAAGATCTCCNNNAGATCGGAA
5343_1 TATGAGTGTGGAGTCGTTGCCTTGTTGCCAGCTATGATCTCCANNNAGATCGGAAG
5345_1 TATGAGTGTGGAGTCGTTGCTTGAGACCCTTGATCTCCTCTCCNNNAGATCGGAAGA
5346_1 TATGAGTGTGGAGTCGTTGCAAAACAACTGGAAGGAGCTCTCCNNNAGATCGGAA
5347_1 TATGAGTGTGGAGTCGTTGCAAGGGAAAAGGGCAAAAGTCTCCNNNAGATCGGAA
5348_1 TATGAGTGTGGAGTCGTTGCTAGTGGAGGGGAGCTTTCATTGANNNAGATCGGAA
5350_1 TATGAGTGTGGAGTCGTTGCAGACTACTTCAGCCTCTGTCTCCNNNAGATCGGAAGA
5351_1 TATGAGTGTGGAGTCGTTGCCACCCAGCACTCCTAAATTCTCCNNNAGATCGGAAGA
5353_1 TATGAGTGTGGAGTCGTTGCAAGACATTCTCTGCTCCTTCTCCNNNAGATCGGAAGA
5354_1 TATGAGTGTGGAGTCGTTGCCTCCTTCTCCACTCTCCTTCTCCNNNAGATCGGAAGAG
5357_1 TATGAGTGTGGAGTCGTTGCTCTGTCTTTTGGTGCCTTTCTCCNNNAGATCGGAAGA
5358_1 TATGAGTGTGGAGTCGTTGCTCCTTTTCCGGTTCTGTTTCTCCNNNAGATCGGAAGA
5359_1 TATGAGTGTGGAGTCGTTGCGTGGGTACAGATCGTGAAAGTCCNNNAGATCGGAA
5360_1 TATGAGTGTGGAGTCGTTGCATGGTTGGCTCATTCTCAAGTCCNNNAGATCGGAAG
5361_1 TATGAGTGTGGAGTCGTTGCACACAGAGCAGAAAGCTAAGTCCNNNAGATCGGAA
5365_1 TATGAGTGTGGAGTCGTTGCTTGACTTGAAGTTGCCTCAGTCCNNNAGATCGGAAG
5366_1 TATGAGTGTGGAGTCGTTGCCAGAGGAAACTTCAGCTCAGTCCNNNAGATCGGAAG
5368_1 TATGAGTGTGGAGTCGTTGCACACACCCCACAAACAATAGTCCNNNAGATCGGAAG
5372_1 TATGAGTGTGGAGTCGTTGCCTTCTTTCTCACCCCAAAGGTCCNNNAGATCGGAAGA
5374_1 TATGAGTGTGGAGTCGTTGCCTTGTAAATGAGCCCAGAGGTCCNNNAGATCGGAA
5377_1 TATGAGTGTGGAGTCGTTGCCTCCTCTCTAGGAAACCTGGTCCNNNAGATCGGAAG
5378_1 TATGAGTGTGGAGTCGTTGCTTTTCCCAGTTTCTCAGTGGTCCNNNAGATCGGAAGA
5381_1 TATGAGTGTGGAGTCGTTGCCCTGGGCATCTCTTTGTATGTCCNNNAGATCGGAAG
5383_1 TATGAGTGTGGAGTCGTTGCATACACACAATCCCACACTGTCCNNNAGATCGGAAGA
5384_1 TATGAGTGTGGAGTCGTTGCCTCGATCATCTTACAGCCTGTCCNNNAGATCGGAAGA
5385_1 TATGAGTGTGGAGTCGTTGCTTGAACTGCTTCTCATCCTGTCCNNNAGATCGGAAGA
5386_1 TATGAGTGTGGAGTCGTTGCGAGATCCAGACAACTCAGTGTCCNNNAGATCGGAAG
5389_1 TATGAGTGTGGAGTCGTTGCCAGGCTCTCCTCTTGGAAATTCCNNNAGATCGGAAG
5390_1 TATGAGTGTGGAGTCGTTGCTGGCTGGTGTTTGTTGAAATTCCNNNAGATCGGAAG
5391_1 TATGAGTGTGGAGTCGTTGCAATCCCATTGCCTGGAGAATTCCNNNAGATCGGAAG
5392_1 TATGAGTGTGGAGTCGTTGCACCAGGGGTATGGGAGTAATTCCNNNAGATCGGAA
5394_1 TATGAGTGTGGAGTCGTTGCGCAAGGACTACCAGACAACTTCCNNNAGATCGGAAG
5395_1 TATGAGTGTGGAGTCGTTGCACTGATCCTCCATCCAGACTTCCNNNAGATCGGAAGA
5398_1 TATGAGTGTGGAGTCGTTGCCTTTGTTGCCTTCATTGCCTTCCNNNAGATCGGAAGA

| | |
|---|---|
| 5401_1 | TATGAGTGTGGAGTCGTTGCGGTGTTTGGGGAATTCACCTTGANNNAGATCGGAA |
| 5402_1 | TATGAGTGTGGAGTCGTTGCATAGGACACTCCCACTTCCTTCCNNNAGATCGGAAGA |
| 5404_1 | TATGAGTGTGGAGTCGTTGCCTTGATGTTGGAAGGTGGCTTCCNNNAGATCGGAAG |
| 5405_1 | TATGAGTGTGGAGTCGTTGCTTCTAGTTGCTCACTTGGCTTCCNNNAGATCGGAAGA |
| 5406_1 | TATGAGTGTGGAGTCGTTGCTCCATTCAGCCATCAATGCTTCCNNNAGATCGGAAGA |
| 5409_1 | TATGAGTGTGGAGTCGTTGCCAAAGACTGCAACTCCTAGCCTTNNNAGATCGGAAG |
| 5415_1 | TATGAGTGTGGAGTCGTTGCCACCAACCACAGTTCATCCTTGANNNAGATCGGAAGA |
| 5417_1 | TATGAGTGTGGAGTCGTTGCTAGTCCTGTTGGATCAGGGTTCCNNNAGATCGGAAG |
| 5418_1 | TATGAGTGTGGAGTCGTTGCCATTGCTGTGAACAACTGGTTCCNNNAGATCGGAAG |
| 5419_1 | TATGAGTGTGGAGTCGTTGCACAGATTCCTAACAACCAGGGGTNNNAGATCGGAAG |
| 5421_1 | TATGAGTGTGGAGTCGTTGCGTTGCTTCTGTTGCTTCTGTTCCNNNAGATCGGAAGA |
| 5422_1 | TATGAGTGTGGAGTCGTTGCTTGCTTGTAGCCATAGGTGTTCCNNNAGATCGGAAG |
| 5424_1 | TATGAGTGTGGAGTCGTTGCTGTTGTCTGATTGCTCAACTGCTNNNAGATCGGAAG |
| 5425_1 | TATGAGTGTGGAGTCGTTGCATCCCCCAAGCTGTACAATTTCCNNNAGATCGGAAGA |
| 5427_1 | TATGAGTGTGGAGTCGTTGCGTCTAGTGGCTTCTCCCATTTCCNNNAGATCGGAAGA |
| 5428_1 | TATGAGTGTGGAGTCGTTGCAGAAAGAGACCTGAAGGGCATCTNNNAGATCGGAA |
| 5431_1 | TATGAGTGTGGAGTCGTTGCATGCCTTTTGTCTGGGGATTTCCNNNAGATCGGAAG |
| 5433_1 | TATGAGTGTGGAGTCGTTGCGGGGAACTAGCAAGTCACTTTCCNNNAGATCGGAAG |
| 5435_1 | TATGAGTGTGGAGTCGTTGCAGGCAAAGTCCATCACTCTTTCCNNNAGATCGGAAG |
| 5436_1 | TATGAGTGTGGAGTCGTTGCCCTCTTGGGAGATAAGGGCATCTNNNAGATCGGAA |
| 5438_1 | TATGAGTGTGGAGTCGTTGCACATGCACCTCTGGTTTCTTTCCNNNAGATCGGAAGA |
| 5439_1 | TATGAGTGTGGAGTCGTTGCGTTTCCACTCCCCTGTAGTTTCCNNNAGATCGGAAGA |
| 5440_1 | TATGAGTGTGGAGTCGTTGCAGCCTTTGGAGTATGTCGTTTCCNNNAGATCGGAAG |
| 5441_1 | TATGAGTGTGGAGTCGTTGCTTTGGCTTTTGCGCACATAATGGNNNAGATCGGAAG |
| 5443_1 | TATGAGTGTGGAGTCGTTGCGAGATTCACCAGGGGGAAAAAGGNNNAGATCGGA |
| 5445_1 | TATGAGTGTGGAGTCGTTGCTGCTGTGTCTGGGTATGTTTTCCNNNAGATCGGAAG |
| 5446_1 | TATGAGTGTGGAGTCGTTGCCTGCTGGGGTTTGGAATTTTTCCNNNAGATCGGAAG |
| 5449_1 | TATGAGTGTGGAGTCGTTGCCCCAAAAGACAGTGTGACAAAGCNNNAGATCGGAA |
| 5450_1 | TATGAGTGTGGAGTCGTTGCATCATCCCCCTTTCATCCAAAGCNNNAGATCGGAAGA |
| 5451_1 | TATGAGTGTGGAGTCGTTGCATCCTCTACTGGGAAAGCAAAGCNNNAGATCGGAAG |
| 5452_1 | TATGAGTGTGGAGTCGTTGCAGTTTCCTGAGGGTAGGCTTTTTNNNAGATCGGAAG |
| 5453_1 | TATGAGTGTGGAGTCGTTGCGTGCTGAACTGCAGGTGTCTCTTNNNAGATCGGAAG |
| 5454_1 | TATGAGTGTGGAGTCGTTGCACCTTGTCCATCACTGTCAAAGCNNNAGATCGGAAG |
| 5455_1 | TATGAGTGTGGAGTCGTTGCAGAGGTTCAGATTTGACCCAGGTNNNAGATCGGAA |
| 5457_1 | TATGAGTGTGGAGTCGTTGCAGGCAAGGAAGATGAAGGAAAGCNNNAGATCGGA |
| 5459_1 | TATGAGTGTGGAGTCGTTGCATGATGACAAGAGGGTGACAAGCNNNAGATCGGAA |
| 5460_1 | TATGAGTGTGGAGTCGTTGCGACATCATGGGTGACATGCATCTNNNAGATCGGAAG |
| 5461_1 | TATGAGTGTGGAGTCGTTGCGCAGAACAATCAGTTATCCCAAGCNNNAGATCGGAA |
| 5462_1 | TATGAGTGTGGAGTCGTTGCTAAATGGAAAGACCAGGCCAAGCNNNAGATCGGAA |
| 5463_1 | TATGAGTGTGGAGTCGTTGCATGAAGTGCTACTATGGCCAAGCNNNAGATCGGAAG |
| 5464_1 | TATGAGTGTGGAGTCGTTGCCAAGAAATCCCACAGAAGCAAGCNNNAGATCGGAAG |
| 5466_1 | TATGAGTGTGGAGTCGTTGCTCGTTTTCTGGGGGAAAAGTTGANNNAGATCGGAA |
| 5467_1 | TATGAGTGTGGAGTCGTTGCCAGTATGTATGCCCTTTGCAAGCNNNAGATCGGAAG |

| | |
|---|---|
| 5468_1 | TATGAGTGTGGAGTCGTTGCGCAACGCAGGCTGAAGTATAATGNNNAGATCGGAA |
| 5469_1 | TATGAGTGTGGAGTCGTTGCGCAGCAACACATAATCCTCAAGCNNNAGATCGGAAG |
| 5470_1 | TATGAGTGTGGAGTCGTTGCTGCTTACCTTCTCACAGTCAAGCNNNAGATCGGAAG |
| 5472_1 | TATGAGTGTGGAGTCGTTGCCAGAGCTTCTTTCCACTTGTCCANNNAGATCGGAAG |
| 5473_1 | TATGAGTGTGGAGTCGTTGCCTGGAAACAATGCAGCTTCAAGCNNNAGATCGGAA |
| 5474_1 | TATGAGTGTGGAGTCGTTGCTCTCTCTCCCAAAGAGTTCAAGCNNNAGATCGGAAG |
| 5475_1 | TATGAGTGTGGAGTCGTTGCAAAGTAGTGATCCTGGTCCTGCTNNNAGATCGGAAG |
| 5476_1 | TATGAGTGTGGAGTCGTTGCACATATAGCAGTGGCACAGAAGCNNNAGATCGGAA |
| 5477_1 | TATGAGTGTGGAGTCGTTGCTTGTTGACCTGTTCTTTCCTGCTNNNAGATCGGAAGA |
| 5479_1 | TATGAGTGTGGAGTCGTTGCCAGAAAGAGAGAGAGCGAGAAGCNNNAGATCGGA |
| 5481_1 | TATGAGTGTGGAGTCGTTGCCGTCCCTAACAGCATCTAGAAGCNNNAGATCGGAAG |
| 5482_1 | TATGAGTGTGGAGTCGTTGCTACCTCTCCACTGGCAAATTCCANNNAGATCGGAAG |
| 5484_1 | TATGAGTGTGGAGTCGTTGCCAGAAAGGAGGTGAGAAGGAAGCNNNAGATCGGA |
| 5485_1 | TATGAGTGTGGAGTCGTTGCATCACAGCAGAAGTGAAGGAAGCNNNAGATCGGAA |
| 5486_1 | TATGAGTGTGGAGTCGTTGCCTCTTCCTTTCAGCCTAGGAAGCNNNAGATCGGAAG |
| 5487_1 | TATGAGTGTGGAGTCGTTGCTGTAAGCTCCAAAAGAGGGAAGCNNNAGATCGGAA |
| 5489_1 | TATGAGTGTGGAGTCGTTGCGTAAGAGACCTGGGAATGGAAGCNNNAGATCGGA |
| 5490_1 | TATGAGTGTGGAGTCGTTGCAACTCCCTCTTCATGTTGGAAGCNNNAGATCGGAAG |
| 5492_1 | TATGAGTGTGGAGTCGTTGCGCTGGGGGTTTCAACATAAAACCNNNAGATCGGAA |
| 5493_1 | TATGAGTGTGGAGTCGTTGCAAGATGCAGAAGGTAGCTGAAGCNNNAGATCGGA |
| 5497_1 | TATGAGTGTGGAGTCGTTGCGAGGTTTTGGAACATGGGTTCCTNNNAGATCGGAA |
| 5499_1 | TATGAGTGTGGAGTCGTTGCCAACACTTGGGAGACCTCTAAGCNNNAGATCGGAAG |
| 5500_1 | TATGAGTGTGGAGTCGTTGCTAATGTTGGAGGGAAGGACTGGTNNNAGATCGGA |
| 5501_1 | TATGAGTGTGGAGTCGTTGCAGAGAGAGAGGGGTTCTGTAAGCNNNAGATCGGA |
| 5502_1 | TATGAGTGTGGAGTCGTTGCGAGAGAGAACTGGCCTTGTAAGCNNNAGATCGGAA |
| 5503_1 | TATGAGTGTGGAGTCGTTGCAGACGACCCATCATAACAACAGCNNNAGATCGGAAG |
| 5505_1 | TATGAGTGTGGAGTCGTTGCATGGTATCGGGTTTCTCAACAGCNNNAGATCGGAAG |
| 5506_1 | TATGAGTGTGGAGTCGTTGCGAGAGAGCATTCCATTCAACAGCNNNAGATCGGAA |
| 5507_1 | TATGAGTGTGGAGTCGTTGCCTGGGATTTGCCTTCAATCTGCTNNNAGATCGGAAG |
| 5509_1 | TATGAGTGTGGAGTCGTTGCGCCCAGTCAGAGAAATCTACAGCNNNAGATCGGAA |
| 5511_1 | TATGAGTGTGGAGTCGTTGCTTCCCAAGAGCTTTCCTTACAGCNNNAGATCGGAAG |
| 5514_1 | TATGAGTGTGGAGTCGTTGCTCTCTTTTCCTCCCTTCCCAGAGNNNAGATCGGAAGA |
| 5517_1 | TATGAGTGTGGAGTCGTTGCCTTGGTGAGTGGCTTGAGATTCCNNNAGATCGGAA |
| 5521_1 | TATGAGTGTGGAGTCGTTGCATTCCCTTGATGAGATTCCCAGCNNNAGATCGGAAG |
| 5522_1 | TATGAGTGTGGAGTCGTTGCTAAAAGACCTCTCCCTTCCCAGCNNNAGATCGGAAG |
| 5523_1 | TATGAGTGTGGAGTCGTTGCCCTGTTTTCCTGTTTCCTCTGCTNNNAGATCGGAAGA |
| 5524_1 | TATGAGTGTGGAGTCGTTGCGAAGTTTTGAAAAGTGGGCCAGCNNNAGATCGGAA |
| 5527_1 | TATGAGTGTGGAGTCGTTGCATTCATGTCAGCAGAGATCCAGCNNNAGATCGGAAG |
| 5528_1 | TATGAGTGTGGAGTCGTTGCTTGCTGGTTTCTGTAGATCCAGCNNNAGATCGGAAG |
| 5533_1 | TATGAGTGTGGAGTCGTTGCCTCCTTCCTTGAGTTCTTCCAGCNNNAGATCGGAAGA |
| 5536_1 | TATGAGTGTGGAGTCGTTGCGAATGGAGTGGCAGGAGTGCNNNAGATCGGAAGA |
| 5537_1 | TATGAGTGTGGAGTCGTTGCTTGGCTGTTTCTCAGGTTGATCCNNNAGATCGGAAG |
| 5539_1 | TATGAGTGTGGAGTCGTTGCAAGAATGAGTTGAGTGGAGCAGCNNNAGATCGGA |

| | |
|---|---|
| 5541_1 | TATGAGTGTGGAGTCGTTGCGAAACTGAGCAAAGAAAGGCAGCNNNAGATCGGA |
| 5542_1 | TATGAGTGTGGAGTCGTTGCATGTTTTCAGGGTAGAAGGCAGCNNNAGATCGGAA |
| 5544_1 | TATGAGTGTGGAGTCGTTGCAGAATGCATGTGGTATAGGCAGCNNNAGATCGGAA |
| 5547_1 | TATGAGTGTGGAGTCGTTGCATTCTTTTCCTCTCCCTGCAGCNNNAGATCGGAAGA |
| 5548_1 | TATGAGTGTGGAGTCGTTGCTTGTCCTGCCCAAACTGTAAGTCNNNAGATCGGAAG |
| 5551_1 | TATGAGTGTGGAGTCGTTGCTCTTTGGTACTACCCATTGCAGCNNNAGATCGGAAG |
| 5554_1 | TATGAGTGTGGAGTCGTTGCTGAACTTCTGTGTGTCCATCAGCNNNAGATCGGAAG |
| 5556_1 | TATGAGTGTGGAGTCGTTGCCCATACCCAGCTCATGTATCAGCNNNAGATCGGAAG |
| 5557_1 | TATGAGTGTGGAGTCGTTGCGGCTGGGCTTTTAGGAAAGTCTTNNNAGATCGGAA |
| 5558_1 | TATGAGTGTGGAGTCGTTGCTGCCATGTAAGATGAAGGCAGAGNNNAGATCGGAA |
| 5562_1 | TATGAGTGTGGAGTCGTTGCTACGCACAGACTTTTGATTCCTTNNNAGATCGGAAG |
| 5567_1 | TATGAGTGTGGAGTCGTTGCCACACAGTGGAAAGCTATTCAGCNNNAGATCGGAA |
| 5568_1 | TATGAGTGTGGAGTCGTTGCAATCAATGAGAGCGCTATTCAGCNNNAGATCGGAA |
| 5571_1 | TATGAGTGTGGAGTCGTTGCGTTCCTCTCCTCCCAATTTCAGCNNNAGATCGGAAGA |
| 5572_1 | TATGAGTGTGGAGTCGTTGCTGTTCCTCACCTTCTCTTTCAGCNNNAGATCGGAAGA |
| 5574_1 | TATGAGTGTGGAGTCGTTGCAGTCGATCCCAAGTGAAAAGAGCNNNAGATCGGAA |
| 5575_1 | TATGAGTGTGGAGTCGTTGCCCCACTCTTTCTCTTCCAAGAGCNNNAGATCGGAAGA |
| 5578_1 | TATGAGTGTGGAGTCGTTGCTGGGGAAATGAAAGACACAGAGCNNNAGATCGGA |
| 5579_1 | TATGAGTGTGGAGTCGTTGCCCGAGGCAGATAGAACATCAGAGNNNAGATCGGAA |
| 5580_1 | TATGAGTGTGGAGTCGTTGCATGGGTAACACACAGACACACAGNNNAGATCGGAA |
| 5581_1 | TATGAGTGTGGAGTCGTTGCTCACTCCCATGAAATCAAGTGCTNNNAGATCGGAAG |
| 5582_1 | TATGAGTGTGGAGTCGTTGCAGGAATCTGTTTGTGAGCAGAGCNNNAGATCGGAA |
| 5585_1 | TATGAGTGTGGAGTCGTTGCCCCTGAATTTCTCTGGTCAGAGCNNNAGATCGGAAG |
| 5590_1 | TATGAGTGTGGAGTCGTTGCGAGAGAGACAGCAACCAAAGCTTNNNAGATCGGAA |
| 5591_1 | TATGAGTGTGGAGTCGTTGCCTGAACTCCAACCTCTTGAGAGCNNNAGATCGGAAG |
| 5593_1 | TATGAGTGTGGAGTCGTTGCTAACCTTGGCTTTTGGGTAGAGCNNNAGATCGGAA |
| 5598_1 | TATGAGTGTGGAGTCGTTGCACACATACACACTTTCCAGGAGCNNNAGATCGGAAG |
| 5599_1 | TATGAGTGTGGAGTCGTTGCACTAGAACCAGAACCAGAGGAGCNNNAGATCGGAA |
| 5601_1 | TATGAGTGTGGAGTCGTTGCGAGAGAGAGGAGAGCATAGGAGCNNNAGATCGGA |
| 5602_1 | TATGAGTGTGGAGTCGTTGCGCCCACAAACAAAACAACTTTCCNNNAGATCGGAAG |
| 5603_1 | TATGAGTGTGGAGTCGTTGCGTAATGGGGTTTTAGGAGGGAGCNNNAGATCGGA |
| 5607_1 | TATGAGTGTGGAGTCGTTGCATAAGGTTGTCAGCAACTGGAGCNNNAGATCGGAA |
| 5608_1 | TATGAGTGTGGAGTCGTTGCTATTCTCTCTGGGTGAGTGGAGCNNNAGATCGGAA |
| 5609_1 | TATGAGTGTGGAGTCGTTGCAAGTGAGAGCACCATATTGGAGCNNNAGATCGGAA |
| 5610_1 | TATGAGTGTGGAGTCGTTGCGAGGGAGAAAATCTGGTTGGAGCNNNAGATCGGA |
| 5611_1 | TATGAGTGTGGAGTCGTTGCTACCCAGTGTGTGTTTTTGGAGCNNNAGATCGGAA |
| 5612_1 | TATGAGTGTGGAGTCGTTGCGGCAACTCCAGGTAAAAATGAGCNNNAGATCGGAA |
| 5613_1 | TATGAGTGTGGAGTCGTTGCAGTGGTAATGGAACCGAATGAGCNNNAGATCGGAA |
| 5614_1 | TATGAGTGTGGAGTCGTTGCCACAAGGAGGGATAACACTGAGCNNNAGATCGGAA |
| 5618_1 | TATGAGTGTGGAGTCGTTGCTGCTAAAAGTGGTGTCTCTGAGCNNNAGATCGGAA |
| 5619_1 | TATGAGTGTGGAGTCGTTGCTACGTTCATCAGACCAAGTGAGCNNNAGATCGGAA |
| 5621_1 | TATGAGTGTGGAGTCGTTGCAGAAAAAGAGAGGAACGGTGAGCNNNAGATCGGA |
| 5623_1 | TATGAGTGTGGAGTCGTTGCCACCACTTCACTTCATTGTGAGCNNNAGATCGGAAG |

| | |
|---|---|
| 5624_1 | TATGAGTGTGGAGTCGTTGCAGCAACTGTCTGCAAACCATGAANNNAGATCGGAAG |
| 5625_1 | TATGAGTGTGGAGTCGTTGCCCACTCCCAAGCTATGTGCTTATNNNAGATCGGAAGA |
| 5628_1 | TATGAGTGTGGAGTCGTTGCAGGCTTTCAGGCCGAATAATAGCNNNAGATCGGAA |
| 5630_1 | TATGAGTGTGGAGTCGTTGCAGGCAGTCTGTACTTTGCATAGCNNNAGATCGGAAG |
| 5631_1 | TATGAGTGTGGAGTCGTTGCATGGAATGCAGTGGTGAGATAGCNNNAGATCGGAA |
| 5632_1 | TATGAGTGTGGAGTCGTTGCAAAAACCTGCCAGCCTAACTAGCNNNAGATCGGAAG |
| 5635_1 | TATGAGTGTGGAGTCGTTGCATGAAAGAGTGAGTGCTGCTAGCNNNAGATCGGAA |
| 5637_1 | TATGAGTGTGGAGTCGTTGCCTTTCCCCCTCTGTGTTTCTAGCNNNAGATCGGAAGA |
| 5638_1 | TATGAGTGTGGAGTCGTTGCCACGCTGGAGTACATACAGTAGCNNNAGATCGGAA |
| 5640_1 | TATGAGTGTGGAGTCGTTGCAAGATCAGTGCCAAGAAGGTAGCNNNAGATCGGAA |
| 5644_1 | TATGAGTGTGGAGTCGTTGCCCCAAATCAGACCTTCCTGTAGCNNNAGATCGGAAG |
| 5647_1 | TATGAGTGTGGAGTCGTTGCCACCTGCCTATTCTCAGAAGCAGNNNAGATCGGAAG |
| 5648_1 | TATGAGTGTGGAGTCGTTGCCCAGCAACTGCAAGTCAATTAGCNNNAGATCGGAAG |
| 5649_1 | TATGAGTGTGGAGTCGTTGCCAGACATCAGGTCCCACATTAGCNNNAGATCGGAAG |
| 5650_1 | TATGAGTGTGGAGTCGTTGCCCCAGCTCAACCAGACTATTAGCNNNAGATCGGAAG |
| 5651_1 | TATGAGTGTGGAGTCGTTGCTGGAGGTGCACAAATAGGTTAGCNNNAGATCGGAA |
| 5652_1 | TATGAGTGTGGAGTCGTTGCTCTGCTGTGAATCTTGGGTTAGCNNNAGATCGGAAG |
| 5654_1 | TATGAGTGTGGAGTCGTTGCGGTATCTCCCAACTGCCTTTAGCNNNAGATCGGAAG |
| 5655_1 | TATGAGTGTGGAGTCGTTGCAGATGGCCTGGTAATGCTTTAGCNNNAGATCGGAA |
| 5657_1 | TATGAGTGTGGAGTCGTTGCAATAGTGTGGATGATTCCCACGCNNNAGATCGGAA |
| 5659_1 | TATGAGTGTGGAGTCGTTGCATCTTCTCAGTGGTTCTTCACGCNNNAGATCGGAAG |
| 5660_1 | TATGAGTGTGGAGTCGTTGCGTTTTGACTTGAGCTTGAGACGCNNNAGATCGGAA |
| 5661_1 | TATGAGTGTGGAGTCGTTGCGAGGATGATAAGCTGAAGGACGCNNNAGATCGGA |
| 5663_1 | TATGAGTGTGGAGTCGTTGCTTTGGTGAACTGCCTATGTACGCNNNAGATCGGAAG |
| 5664_1 | TATGAGTGTGGAGTCGTTGCCTCCAATTCCTCCGTGTTCAAGTNNNAGATCGGAAGA |
| 5665_1 | TATGAGTGTGGAGTCGTTGCAATAGTTGAAGGCAGTCTACCGCNNNAGATCGGAA |
| 5666_1 | TATGAGTGTGGAGTCGTTGCGCTCTAATCTTCCTTAGCTCCGCNNNAGATCGGAAGA |
| 5667_1 | TATGAGTGTGGAGTCGTTGCTGCTTCCTTATGCAATCTGTGCTNNNAGATCGGAAG |
| 5668_1 | TATGAGTGTGGAGTCGTTGCCAAATAGATGACCCCTATGGCGCNNNAGATCGGAAG |
| 5670_1 | TATGAGTGTGGAGTCGTTGCTTCCATTTGAGCTTGATTTGCGCNNNAGATCGGAAG |
| 5671_1 | TATGAGTGTGGAGTCGTTGCGAAAGACAATCCCTCTCTGTCGCNNNAGATCGGAAG |
| 5674_1 | TATGAGTGTGGAGTCGTTGCCACTTCAGTATGCAAAGCTTCGCNNNAGATCGGAAG |
| 5675_1 | TATGAGTGTGGAGTCGTTGCTGTGAGAGGACTGTATGTGTGCTNNNAGATCGGAA |
| 5676_1 | TATGAGTGTGGAGTCGTTGCAAAATCCCCCTTCCAGAAAAGGCNNNAGATCGGAAG |
| 5677_1 | TATGAGTGTGGAGTCGTTGCAAAAGGTAGTGGTACCCAAAGGCNNNAGATCGGAA |
| 5678_1 | TATGAGTGTGGAGTCGTTGCGCAACAGAATAATGGGCAAAGGCNNNAGATCGGAA |
| 5679_1 | TATGAGTGTGGAGTCGTTGCAAGAAAGAAAGGTGGGGAAAGGCNNNAGATCGGA |
| 5680_1 | TATGAGTGTGGAGTCGTTGCGGAGAGCCTTGACTGATAAAGGCNNNAGATCGGAA |
| 5684_1 | TATGAGTGTGGAGTCGTTGCGGACATAAGCAAATCTGCAAGGCNNNAGATCGGAA |
| 5685_1 | TATGAGTGTGGAGTCGTTGCCCATCCTCACATCATCTCAAGGCNNNAGATCGGAAGA |
| 5686_1 | TATGAGTGTGGAGTCGTTGCAGGTGTTAGTGATGCTTCAAGGCNNNAGATCGGAA |
| 5688_1 | TATGAGTGTGGAGTCGTTGCCTAACAGGATGGTTCAGGAAGGCNNNAGATCGGAA |
| 5689_1 | TATGAGTGTGGAGTCGTTGCTAGGAGCAGTATCAGAGGAAGGCNNNAGATCGGA |

| | |
|---|---|
| 5693_1 | TATGAGTGTGGAGTCGTTGCAGAAACCATGTCTACGAACAGGCNNNAGATCGGAA |
| 5694_1 | TATGAGTGTGGAGTCGTTGCCACACACATATCCAAACACAGGCNNNAGATCGGAAG |
| 5696_1 | TATGAGTGTGGAGTCGTTGCGTGTGCATATGCTTGTTTGTGCTNNNAGATCGGAAG |
| 5698_1 | TATGAGTGTGGAGTCGTTGCCTTTTCTGGTTAGTACCCCAGGCNNNAGATCGGAAG |
| 5699_1 | TATGAGTGTGGAGTCGTTGCATTTTCTGGCATGTATCCCAGGCNNNAGATCGGAAG |
| 5702_1 | TATGAGTGTGGAGTCGTTGCCCATAATTGACTGAGGTCCAGGCNNNAGATCGGAA |
| 5705_1 | TATGAGTGTGGAGTCGTTGCCATCCAGGGTCATTCAGTAATGCNNNAGATCGGAAG |
| 5706_1 | TATGAGTGTGGAGTCGTTGCGCAGTGGAAGATGGAATGACTGTNNNAGATCGGA |
| 5707_1 | TATGAGTGTGGAGTCGTTGCTTTCCACAAAGGAGAGATCAGGCNNNAGATCGGAA |
| 5708_1 | TATGAGTGTGGAGTCGTTGCCAGCTAAGGAAGTGGTATCAGGCNNNAGATCGGAA |
| 5709_1 | TATGAGTGTGGAGTCGTTGCAAGAAGGAAATCAGCACTCAGGCNNNAGATCGGAA |
| 5710_1 | TATGAGTGTGGAGTCGTTGCGAGACAGAAAGTAGAGCTCAGGCNNNAGATCGGA |
| 5712_1 | TATGAGTGTGGAGTCGTTGCTATGTAATGCACCAAGGTCAGGCNNNAGATCGGAA |
| 5713_1 | TATGAGTGTGGAGTCGTTGCATTGTGCTTATTGTGTGTCAGGCNNNAGATCGGAAG |
| 5715_1 | TATGAGTGTGGAGTCGTTGCTTGGGAAAATTCAGGGAAGAGGCNNNAGATCGGA |
| 5716_1 | TATGAGTGTGGAGTCGTTGCCCAAATCCCAGATTCTCAGAGGCNNNAGATCGGAAG |
| 5718_1 | TATGAGTGTGGAGTCGTTGCGTGAGTGTGTGTATGAGAGAGGCNNNAGATCGGA |
| 5719_1 | TATGAGTGTGGAGTCGTTGCGTGAAAAAGAGAGAGGGAGAGGCNNNAGATCGG |
| 5720_1 | TATGAGTGTGGAGTCGTTGCTCCCAATCTACGTTTTGAGAGGCNNNAGATCGGAAG |
| 5721_1 | TATGAGTGTGGAGTCGTTGCTGAAACCCTCAGAGAGTAGAGGCNNNAGATCGGAA |
| 5723_1 | TATGAGTGTGGAGTCGTTGCCTTTGCAACAAGATATGGGAGGCNNNAGATCGGAA |
| 5724_1 | TATGAGTGTGGAGTCGTTGCTTGGATTTCAGATGTTGGGAGGCNNNAGATCGGAA |
| 5725_1 | TATGAGTGTGGAGTCGTTGCATCCCAAGAGACTGTGAATTCTCNNNAGATCGGAAG |
| 5726_1 | TATGAGTGTGGAGTCGTTGCATTGTTTCCTCAGTTCTGGAGGCNNNAGATCGGAAG |
| 5727_1 | TATGAGTGTGGAGTCGTTGCAGGTACAGATATCAGGTGGAGGCNNNAGATCGGA |
| 5728_1 | TATGAGTGTGGAGTCGTTGCGAGAAACAGAGACAGAGTGAGGCNNNAGATCGGA |
| 5729_1 | TATGAGTGTGGAGTCGTTGCGGGGTAGGACAATTTCTTGAGGCNNNAGATCGGAA |
| 5730_1 | TATGAGTGTGGAGTCGTTGCTAGGCCGGAGTGCATGATCATAGNNNAGATCGGAA |
| 5731_1 | TATGAGTGTGGAGTCGTTGCGTGATTGTGAGGGCTAACTAGGCNNNAGATCGGAA |
| 5732_1 | TATGAGTGTGGAGTCGTTGCGTGTGTGTTGAGGGGATACACAGNNNAGATCGGA |
| 5735_1 | TATGAGTGTGGAGTCGTTGCTGAAGCATTTGACCCTTATTGCTNNNAGATCGGAAG |
| 5736_1 | TATGAGTGTGGAGTCGTTGCGGGAGGATAGCAGAAAATGGCTTNNNAGATCGGA |
| 5737_1 | TATGAGTGTGGAGTCGTTGCTTCCAAAGGGCACAATTGTAGGCNNNAGATCGGAA |
| 5738_1 | TATGAGTGTGGAGTCGTTGCGCAACCCTTATGTCTCCTTAGGCNNNAGATCGGAAG |
| 5739_1 | TATGAGTGTGGAGTCGTTGCGAGCACATAGCACTCTCTTAGGCNNNAGATCGGAAG |
| 5741_1 | TATGAGTGTGGAGTCGTTGCCAGGAGGGAAGGATGTTTTAGGCNNNAGATCGGA |
| 5743_1 | TATGAGTGTGGAGTCGTTGCGTGCTTTTTGAGTACTTTCCGGCNNNAGATCGGAAG |
| 5744_1 | TATGAGTGTGGAGTCGTTGCCAGTAATAAAACACAAGGGCGGCNNNAGATCGGAA |
| 5745_1 | TATGAGTGTGGAGTCGTTGCCCAAAACTCGAGGATAGAAGGGCNNNAGATCGGAA |
| 5746_1 | TATGAGTGTGGAGTCGTTGCTATGCAGTTTCTGATGACAGGGCNNNAGATCGGAA |
| 5747_1 | TATGAGTGTGGAGTCGTTGCTCAAGATAGTTGCAACAGAGGGCNNNAGATCGGAA |
| 5749_1 | TATGAGTGTGGAGTCGTTGCGGTATGTTTTCTCTCCGTGGCTTNNNAGATCGGAAG |
| 5750_1 | TATGAGTGTGGAGTCGTTGCAAAATGCTTCCCCAATAAGGGGCNNNAGATCGGAA |

| | |
|---|---|
| 5752_1 | TATGAGTGTGGAGTCGTTGCGTAGAGCTGACTTTATGAGGGGCNNNAGATCGGAA |
| 5753_1 | TATGAGTGTGGAGTCGTTGCCCCAAAGGATAAATGTAGGGGGCNNNAGATCGGAA |
| 5754_1 | TATGAGTGTGGAGTCGTTGCCACCCACACTAAGCCATTTTCCTNNNAGATCGGAAGA |
| 5758_1 | TATGAGTGTGGAGTCGTTGCCCAAATTCAAGCACTGTTGGCTTNNNAGATCGGAAG |
| 5759_1 | TATGAGTGTGGAGTCGTTGCGCCACAGTTGACTAAGCAGAAGTNNNAGATCGGAA |
| 5763_1 | TATGAGTGTGGAGTCGTTGCTGCCAGAAAAATGGGACAAGGAANNNAGATCGGA |
| 5764_1 | TATGAGTGTGGAGTCGTTGCTTTTCTTAGAGAGGTTGCTGGGCNNNAGATCGGAA |
| 5766_1 | TATGAGTGTGGAGTCGTTGCTGCTAAGATAGCTCTGTCTGGGCNNNAGATCGGAA |
| 5768_1 | TATGAGTGTGGAGTCGTTGCGGAGTTTTTATCCTCAGGTGGGCNNNAGATCGGAA |
| 5770_1 | TATGAGTGTGGAGTCGTTGCCTCCTCTACATTAGCATGTGGGCNNNAGATCGGAAG |
| 5771_1 | TATGAGTGTGGAGTCGTTGCAAGCCTGAACATCGATATTGGGCNNNAGATCGGAA |
| 5773_1 | TATGAGTGTGGAGTCGTTGCACAAAAGTTTTCCTGAGTTGGGCNNNAGATCGGAA |
| 5774_1 | TATGAGTGTGGAGTCGTTGCCTCCATTTCACCCATTTTTGGGCNNNAGATCGGAAGA |
| 5775_1 | TATGAGTGTGGAGTCGTTGCAAGAAGATGGGTGGTGAAATGGCNNNAGATCGGA |
| 5776_1 | TATGAGTGTGGAGTCGTTGCTACCCAGTGCCTAAAACAATGGCNNNAGATCGGAAG |
| 5777_1 | TATGAGTGTGGAGTCGTTGCTCACTGTCTATTCAGGTCCCCAANNNAGATCGGAAG |
| 5778_1 | TATGAGTGTGGAGTCGTTGCCCTTCGGCAACCATTTAGAAAGCNNNAGATCGGAAG |
| 5779_1 | TATGAGTGTGGAGTCGTTGCAGGTGTGTATCACAGTGAATGGCNNNAGATCGGAA |
| 5780_1 | TATGAGTGTGGAGTCGTTGCGGGGAGAAACTGGAACTAATGGCNNNAGATCGGA |
| 5781_1 | TATGAGTGTGGAGTCGTTGCAAGCCCTCAATGCTAGTAATGGCNNNAGATCGGAAG |
| 5782_1 | TATGAGTGTGGAGTCGTTGCTCTAGCAGGGAAAAGTTGCAAGANNNAGATCGGAA |
| 5785_1 | TATGAGTGTGGAGTCGTTGCCCATTCACTCACAGAATCATGGCNNNAGATCGGAAG |
| 5786_1 | TATGAGTGTGGAGTCGTTGCAAACTCCAAGTCCTGATCATGGCNNNAGATCGGAAG |
| 5787_1 | TATGAGTGTGGAGTCGTTGCGAAGTGTGTGTACTGCCAGCGTTNNNAGATCGGAA |
| 5789_1 | TATGAGTGTGGAGTCGTTGCCTATGCAGCTACTGTTTTGGGCTNNNAGATCGGAAG |
| 5791_1 | TATGAGTGTGGAGTCGTTGCCCATCAGGAGAATAGCAACTGGCNNNAGATCGGAA |
| 5792_1 | TATGAGTGTGGAGTCGTTGCTACGCTGAATTGTCCTAACTGGCNNNAGATCGGAAG |
| 5793_1 | TATGAGTGTGGAGTCGTTGCCTTCTTAGTGATCACCCACTGGCNNNAGATCGGAAG |
| 5795_1 | TATGAGTGTGGAGTCGTTGCTAGAACCCTCCTTCCTTACTGGCNNNAGATCGGAAGA |
| 5799_1 | TATGAGTGTGGAGTCGTTGCAAATCACTTAGGACAAGCCTGGCNNNAGATCGGAA |
| 5801_1 | TATGAGTGTGGAGTCGTTGCCTACGTGAAGATTCTGCCTGCTTNNNAGATCGGAAG |
| 5802_1 | TATGAGTGTGGAGTCGTTGCAAAATCTGTAGGGTATGGCTGGCNNNAGATCGGAA |
| 5804_1 | TATGAGTGTGGAGTCGTTGCCTGGTGATCTTTGTCTCTCTGGCNNNAGATCGGAAG |
| 5805_1 | TATGAGTGTGGAGTCGTTGCAGGGTCAAACACAACTGAGATCTNNNAGATCGGAA |
| 5806_1 | TATGAGTGTGGAGTCGTTGCCAAATGGCTAAAAGCTGTCTGGCNNNAGATCGGAA |
| 5807_1 | TATGAGTGTGGAGTCGTTGCATGGTGTTCACTTGACTTCTGGCNNNAGATCGGAAG |
| 5808_1 | TATGAGTGTGGAGTCGTTGCTTTGGGGTTTTCTCTCTTCTGGCNNNAGATCGGAAG |
| 5810_1 | TATGAGTGTGGAGTCGTTGCCTCTGTCTCACCATACAAGTGGCNNNAGATCGGAAG |
| 5811_1 | TATGAGTGTGGAGTCGTTGCTCCTCTCAAAGCAAGTAAGTGGCNNNAGATCGGAAG |
| 5812_1 | TATGAGTGTGGAGTCGTTGCTGTCTAGGATATAGGGCAGTGGCNNNAGATCGGAA |
| 5814_1 | TATGAGTGTGGAGTCGTTGCAAATGTCCAGAGCTAAGAGTGGCNNNAGATCGGAA |
| 5815_1 | TATGAGTGTGGAGTCGTTGCCCCACAATGAGGACAATCTGCTTNNNAGATCGGAAG |
| 5817_1 | TATGAGTGTGGAGTCGTTGCAAATCTGTTCCGTTCTAGGTGGCNNNAGATCGGAAG |

| | |
|---|---|
| 5818_1 | TATGAGTGTGGAGTCGTTGCAGGTCCAGATAGAAGTAGGTGGCNNNAGATCGGAA |
| 5819_1 | TATGAGTGTGGAGTCGTTGCTGGAAAATAGGCTTGTAGGTGGCNNNAGATCGGAA |
| 5820_1 | TATGAGTGTGGAGTCGTTGCTAATAGAGAGTAGCGAGGGTGGCNNNAGATCGGAA |
| 5822_1 | TATGAGTGTGGAGTCGTTGCTGCTCTGAATCATTACCTGTGGCNNNAGATCGGAAG |
| 5823_1 | TATGAGTGTGGAGTCGTTGCCCAGTAATAACACCCTCTGTGGCNNNAGATCGGAAG |
| 5824_1 | TATGAGTGTGGAGTCGTTGCGGAATGCTATGATAGGGTGTGGCNNNAGATCGGAA |
| 5825_1 | TATGAGTGTGGAGTCGTTGCTATGAAGAGGCAGTGTTTGTGGCNNNAGATCGGAA |
| 5826_1 | TATGAGTGTGGAGTCGTTGCGAGACAGAACGACGAAAATTGGCNNNAGATCGGAA |
| 5829_1 | TATGAGTGTGGAGTCGTTGCCTTCTGTATCCTTGGGGATTGGCNNNAGATCGGAAG |
| 5832_1 | TATGAGTGTGGAGTCGTTGCACATATCCTTGGAACAGCTTGGCNNNAGATCGGAAG |
| 5833_1 | TATGAGTGTGGAGTCGTTGCTGAGTGAGTACATCTTGCTTGGCNNNAGATCGGAAG |
| 5834_1 | TATGAGTGTGGAGTCGTTGCGTGCAGGGTATTCTCAACAGGAGNNNAGATCGGAA |
| 5835_1 | TATGAGTGTGGAGTCGTTGCGCAGGGCATTATGAAATGTTGGCNNNAGATCGGAA |
| 5837_1 | TATGAGTGTGGAGTCGTTGCTTGATTGATTGCAAGGTGTTGGCNNNAGATCGGAA |
| 5838_1 | TATGAGTGTGGAGTCGTTGCTGGGGTTGAATGGAAAATTTGGCNNNAGATCGGAA |
| 5840_1 | TATGAGTGTGGAGTCGTTGCTGGCTACAGAGCATGGTTTCTTTNNNAGATCGGAAG |
| 5841_1 | TATGAGTGTGGAGTCGTTGCCCCACTTTCTTTTCGTCTTTGGCNNNAGATCGGAAGA |
| 5842_1 | TATGAGTGTGGAGTCGTTGCGTGAGATGCTGCTCCTGTTTACCNNNAGATCGGAAG |
| 5845_1 | TATGAGTGTGGAGTCGTTGCCTGTAGTGAAGTGCCCAAAATGCNNNAGATCGGAA |
| 5846_1 | TATGAGTGTGGAGTCGTTGCCCAGGCTGCTTATACGAAAATGCNNNAGATCGGAAG |
| 5847_1 | TATGAGTGTGGAGTCGTTGCTAGAAATGGCCAAGAGCAAATGCNNNAGATCGGAA |
| 5848_1 | TATGAGTGTGGAGTCGTTGCAGTGCTAAGTTTCCTACGCATGGNNNAGATCGGAAG |
| 5849_1 | TATGAGTGTGGAGTCGTTGCCAAGGGGCAAAGACAGATCAAGANNNAGATCGGAA |
| 5850_1 | TATGAGTGTGGAGTCGTTGCCTCTGTAGCCTAGCCTGAAATGCNNNAGATCGGAAG |
| 5852_1 | TATGAGTGTGGAGTCGTTGCTCTGCCTTGAGAAGAGACAATGCNNNAGATCGGAA |
| 5854_1 | TATGAGTGTGGAGTCGTTGCCCTGGCTCATAACTCCTATGGCTNNNAGATCGGAAG |
| 5855_1 | TATGAGTGTGGAGTCGTTGCGCAATCTTCCAGAGCTTCAATGCNNNAGATCGGAAG |
| 5856_1 | TATGAGTGTGGAGTCGTTGCACACACATACTGTGGAACATTTTCNNNAGATCGGAA |
| 5857_1 | TATGAGTGTGGAGTCGTTGCTCGGAAGGCTAACAGAAGAATGCNNNAGATCGGAA |
| 5858_1 | TATGAGTGTGGAGTCGTTGCGAGATGACCCCAGATCAGAATGCNNNAGATCGGAA |
| 5859_1 | TATGAGTGTGGAGTCGTTGCACCAAAGCTGTGACCTAGAATGCNNNAGATCGGAA |
| 5862_1 | TATGAGTGTGGAGTCGTTGCCAATTCAGTCCGTAGCAACATGCNNNAGATCGGAAG |
| 5863_1 | TATGAGTGTGGAGTCGTTGCAGGACGTAAATTGTGACACATGCNNNAGATCGGAA |
| 5864_1 | TATGAGTGTGGAGTCGTTGCAGAGAGAGACTGCATTGACATGCNNNAGATCGGAA |
| 5865_1 | TATGAGTGTGGAGTCGTTGCGCTCACTACCAGCTAGTACATGCNNNAGATCGGAAG |
| 5866_1 | TATGAGTGTGGAGTCGTTGCTAGAGAAGTCCTCAGAGGCAAGCNNNAGATCGGAA |
| 5867_1 | TATGAGTGTGGAGTCGTTGCGCCCACAATTATTTCTGCCATGCNNNAGATCGGAAGA |
| 5869_1 | TATGAGTGTGGAGTCGTTGCAGAGAGAGAGCATACAAGCATGCNNNAGATCGGAA |
| 5870_1 | TATGAGTGTGGAGTCGTTGCGAACCTTGCCCTTTTCCTGTTGANNNAGATCGGAAGA |
| 5871_1 | TATGAGTGTGGAGTCGTTGCATGCCACACATTTACATGCATGCNNNAGATCGGAAG |
| 5872_1 | TATGAGTGTGGAGTCGTTGCAGAGAGAGAAAGGCTATGCATGCNNNAGATCGGAA |
| 5873_1 | TATGAGTGTGGAGTCGTTGCCCTCTGTGTATTTGTGTGCATGCNNNAGATCGGAAG |
| 5874_1 | TATGAGTGTGGAGTCGTTGCTGTGTGTGTGTCTCAGTTTTGCTNNNAGATCGGAAG |

| | |
|---|---|
| 5876_1 | TATGAGTGTGGAGTCGTTGCGGTGGCATTCATTCTGTTCATGCNNNAGATCGGAAG |
| 5880_1 | TATGAGTGTGGAGTCGTTGCACTGGCCGCTTATATACAGATGCNNNAGATCGGAAG |
| 5882_1 | TATGAGTGTGGAGTCGTTGCAGATTGGTGTGGTAACGAGATGCNNNAGATCGGAA |
| 5883_1 | TATGAGTGTGGAGTCGTTGCTGTGTGTTTCAGATGCAGTCTTGNNNAGATCGGAAG |
| 5884_1 | TATGAGTGTGGAGTCGTTGCGAGAGAGAGATGCACATGGATGCNNNAGATCGGA |
| 5885_1 | TATGAGTGTGGAGTCGTTGCATCCCTAACAACACTGGTGATGCNNNAGATCGGAAG |
| 5886_1 | TATGAGTGTGGAGTCGTTGCACACACACTGCTTTTGCATATGCNNNAGATCGGAAG |
| 5887_1 | TATGAGTGTGGAGTCGTTGCGCCAGCCCTTGATTTAGATATGCNNNAGATCGGAAG |
| 5888_1 | TATGAGTGTGGAGTCGTTGCGCATGAGAAGTCCCAGGATATGCNNNAGATCGGAA |
| 5889_1 | TATGAGTGTGGAGTCGTTGCTTGTCAGAAAGGCCTGAGTATGCNNNAGATCGGAA |
| 5891_1 | TATGAGTGTGGAGTCGTTGCCACCTGAGCCTGTAGAGTTATGCNNNAGATCGGAAG |
| 5892_1 | TATGAGTGTGGAGTCGTTGCTTGGGCACGAAGATTTGTTATGCNNNAGATCGGAA |
| 5893_1 | TATGAGTGTGGAGTCGTTGCAGAAGAAGGCAGGGCATTATTGGNNNAGATCGGA |
| 5895_1 | TATGAGTGTGGAGTCGTTGCCCAGAACTGCCCAAACTTTATGCNNNAGATCGGAAG |
| 5896_1 | TATGAGTGTGGAGTCGTTGCCCCAATGGTTGTGAGCTTTATGCNNNAGATCGGAAG |
| 5897_1 | TATGAGTGTGGAGTCGTTGCGCTAGTTCAGGACTGGATTCACANNNAGATCGGAA |
| 5899_1 | TATGAGTGTGGAGTCGTTGCCCCCTCAGTGAATGAGTAACTGCNNNAGATCGGAAG |
| 5900_1 | TATGAGTGTGGAGTCGTTGCCTGGTTTATGAAGGGGACACTGCNNNAGATCGGAA |
| 5901_1 | TATGAGTGTGGAGTCGTTGCACAGAGGGCTAAATTCTCACTGCNNNAGATCGGAAG |
| 5902_1 | TATGAGTGTGGAGTCGTTGCGACCTTTCCATGCATTTCACTGCNNNAGATCGGAAGA |
| 5903_1 | TATGAGTGTGGAGTCGTTGCTCCCTCCCTTGATTTTTCACTGCNNNAGATCGGAAGA |
| 5904_1 | TATGAGTGTGGAGTCGTTGCATCCCAACTCACAATCAGACTGCNNNAGATCGGAAG |
| 5905_1 | TATGAGTGTGGAGTCGTTGCTCAGAGCCTTGTTCCATTACTGCNNNAGATCGGAAG |
| 5906_1 | TATGAGTGTGGAGTCGTTGCGGAAGGGTTGGCTCTATTACTGCNNNAGATCGGAA |
| 5909_1 | TATGAGTGTGGAGTCGTTGCCTCTCTCTCTTTCACCAACCTGCNNNAGATCGGAAGA |
| 5910_1 | TATGAGTGTGGAGTCGTTGCGAAAAATATAACCGCTGCCCTGCNNNAGATCGGAAG |
| 5911_1 | TATGAGTGTGGAGTCGTTGCATTTACTGTGGAACTCAGCCTGCNNNAGATCGGAAG |
| 5913_1 | TATGAGTGTGGAGTCGTTGCCATAAAAAGCAGGCAACTCCTGCNNNAGATCGGAAG |
| 5914_1 | TATGAGTGTGGAGTCGTTGCTCTCTATCACCAACCTCTCCTGCNNNAGATCGGAAGA |
| 5915_1 | TATGAGTGTGGAGTCGTTGCCTGGACCTGTTGACTATTCCTGCNNNAGATCGGAAG |
| 5917_1 | TATGAGTGTGGAGTCGTTGCACAGTCTGGGTTTCATCAGAGGTNNNAGATCGGAA |
| 5918_1 | TATGAGTGTGGAGTCGTTGCCTTGGGAAGCCTCTTTGCTTACTNNNAGATCGGAAG |
| 5920_1 | TATGAGTGTGGAGTCGTTGCAGAACCAGTTGACAGTAAGCTGCNNNAGATCGGAA |
| 5921_1 | TATGAGTGTGGAGTCGTTGCGTGTACTTGTGTCAAACAGCTGCNNNAGATCGGAAG |
| 5922_1 | TATGAGTGTGGAGTCGTTGCTCCCAAGCTTTATTCTCAGCTGCNNNAGATCGGAAGA |
| 5923_1 | TATGAGTGTGGAGTCGTTGCCAGTGGCTAACTAGAAGAGCTGCNNNAGATCGGAA |
| 5924_1 | TATGAGTGTGGAGTCGTTGCTTTGTAGAGCTTCAGTGAGCTGCNNNAGATCGGAA |
| 5925_1 | TATGAGTGTGGAGTCGTTGCAATGTGGAGAACTTACAGGCTGCNNNAGATCGGAA |
| 5926_1 | TATGAGTGTGGAGTCGTTGCGAGTAAATTTTGGAGCAGGCTGCNNNAGATCGGAA |
| 5927_1 | TATGAGTGTGGAGTCGTTGCGTCTTCACACAAAATCAGGCTGCNNNAGATCGGAAG |
| 5929_1 | TATGAGTGTGGAGTCGTTGCGAGGGTGGTGACAAGTTAGAAGCNNNAGATCGGA |
| 5930_1 | TATGAGTGTGGAGTCGTTGCTTCAACAAGTAGCCGTATGCTGCNNNAGATCGGAAG |
| 5932_1 | TATGAGTGTGGAGTCGTTGCGGGATTGTTCCTGCACTTCCATANNNAGATCGGAAG |

| | |
|---|---|
| 5933_1 | TATGAGTGTGGAGTCGTTGCTTGGTGCCAGATTATTCTGCTGCNNNAGATCGGAAG |
| 5934_1 | TATGAGTGTGGAGTCGTTGCCTCTGCTTATTGCTGATTGCTGCNNNAGATCGGAAG |
| 5936_1 | TATGAGTGTGGAGTCGTTGCAATGGGAAAGAAGCCATGAGCAANNNAGATCGGAA |
| 5937_1 | TATGAGTGTGGAGTCGTTGCCTTGTTGCGGACTTTGTATCTGCNNNAGATCGGAAG |
| 5940_1 | TATGAGTGTGGAGTCGTTGCACTGGAACCATCAGCCTGAATCTNNNAGATCGGAAG |
| 5941_1 | TATGAGTGTGGAGTCGTTGCTCAGAAATGAGGTCTCGCTCTGCNNNAGATCGGAAG |
| 5942_1 | TATGAGTGTGGAGTCGTTGCAAAATGCTACCCCTCATCTCTGCNNNAGATCGGAAG |
| 5947_1 | TATGAGTGTGGAGTCGTTGCTTAGAAAATCCCAACGGGTCTGCNNNAGATCGGAAG |
| 5948_1 | TATGAGTGTGGAGTCGTTGCGAAATCTGTGCTTTGGTGTCTGCNNNAGATCGGAAG |
| 5949_1 | TATGAGTGTGGAGTCGTTGCGGGTGGTTGCTAAAATGTTCTGCNNNAGATCGGAA |
| 5953_1 | TATGAGTGTGGAGTCGTTGCGACTGGCTGTGTCATTTTTCTGCNNNAGATCGGAAG |
| 5954_1 | TATGAGTGTGGAGTCGTTGCCTGTCTACCTGTGGGTAAAGTGCNNNAGATCGGAA |
| 5955_1 | TATGAGTGTGGAGTCGTTGCTGGTCGGGGAGCAGTATATTCATNNNAGATCGGAA |
| 5956_1 | TATGAGTGTGGAGTCGTTGCCTGTTACCTCTGTGGAGAAGTGCNNNAGATCGGAA |
| 5957_1 | TATGAGTGTGGAGTCGTTGCCTCCAGGGCTCTTTACTAAGTGCNNNAGATCGGAAG |
| 5958_1 | TATGAGTGTGGAGTCGTTGCAGGCCACCTTTCTATAACAGTGCNNNAGATCGGAAG |
| 5959_1 | TATGAGTGTGGAGTCGTTGCCCAACCTTACACATAACCAGTGCNNNAGATCGGAAG |
| 5960_1 | TATGAGTGTGGAGTCGTTGCACACAGACATACTGAAGCAGTGCNNNAGATCGGAA |
| 5962_1 | TATGAGTGTGGAGTCGTTGCTGTGTCTGTCTGTCTATCAGTGCNNNAGATCGGAAG |
| 5963_1 | TATGAGTGTGGAGTCGTTGCACTCACACAAATCTGTAACAGCTNNNAGATCGGAAG |
| 5964_1 | TATGAGTGTGGAGTCGTTGCGTGTCTATCACAGGCTAGAGTGCNNNAGATCGGAA |
| 5965_1 | TATGAGTGTGGAGTCGTTGCTCCTGTCTTTCCTTAGTGAGTGCNNNAGATCGGAAG |
| 5966_1 | TATGAGTGTGGAGTCGTTGCTAGTAGGTGAGCCCAGATAGTGCNNNAGATCGGAA |
| 5968_1 | TATGAGTGTGGAGTCGTTGCCCATTTCCTTGTCATTAGCGTGCNNNAGATCGGAAG |
| 5969_1 | TATGAGTGTGGAGTCGTTGCAGAGAGAGAAGGGGTAAAGGTGCNNNAGATCGGA |
| 5971_1 | TATGAGTGTGGAGTCGTTGCAACTGATTCACTCCCACCTCTTTNNNAGATCGGAAGA |
| 5972_1 | TATGAGTGTGGAGTCGTTGCAGAAGCCTGGAGCTATTCAGGAANNNAGATCGGAA |
| 5974_1 | TATGAGTGTGGAGTCGTTGCGAATGAAAACAAGAGCTGGGTGCNNNAGATCGGAA |
| 5975_1 | TATGAGTGTGGAGTCGTTGCTTATAGATCCTGTGCCATGGTGCNNNAGATCGGAAG |
| 5980_1 | TATGAGTGTGGAGTCGTTGCCCTCCCAGAATAGGAACATGTGCNNNAGATCGGAAG |
| 5981_1 | TATGAGTGTGGAGTCGTTGCGGGAGCTATCTCTGTACATGTGCNNNAGATCGGAA |
| 5982_1 | TATGAGTGTGGAGTCGTTGCGACTGTACGGAGAAATGGAAGGCNNNAGATCGGAA |
| 5983_1 | TATGAGTGTGGAGTCGTTGCTGTGTGTATGCTCATGGATGTGCNNNAGATCGGAA |
| 5984_1 | TATGAGTGTGGAGTCGTTGCAACGTCAGGCTTAGAGTATGTGCNNNAGATCGGAA |
| 5986_1 | TATGAGTGTGGAGTCGTTGCTCCCTTCAGATGTTGGACCATCTNNNAGATCGGAAG |
| 5987_1 | TATGAGTGTGGAGTCGTTGCAGATAGGACAAGCTTAGCTGTGCNNNAGATCGGAA |
| 5988_1 | TATGAGTGTGGAGTCGTTGCCTAAGCAAAGTATGCTGCTGTGCNNNAGATCGGAA |
| 5989_1 | TATGAGTGTGGAGTCGTTGCTTCAAGTGTATCAGGCTCTGTGCNNNAGATCGGAAG |
| 5990_1 | TATGAGTGTGGAGTCGTTGCCTTTCACACCAAGTGTTCTGTGCNNNAGATCGGAAG |
| 5991_1 | TATGAGTGTGGAGTCGTTGCATAAAGCAGCTGGGTTCAGTAGGNNNAGATCGGAA |
| 5992_1 | TATGAGTGTGGAGTCGTTGCAAAAACCCGTGTAATAAACCTGCNNNAGATCGGAAG |
| 5993_1 | TATGAGTGTGGAGTCGTTGCCCGCTGCCTCTCTTCTATAAGGANNNAGATCGGAAG |
| 5997_1 | TATGAGTGTGGAGTCGTTGCTATTTGGGACCCTGAACTTGTGCNNNAGATCGGAAG |

| | |
|---|---|
| 5998_1 | TATGAGTGTGGAGTCGTTGCCCAAGAGCAAGAGAGCACAGTTTNNNAGATCGGAA |
| 6000_1 | TATGAGTGTGGAGTCGTTGCTTTGTGTGTCTGTCTGTTTGTGCNNNAGATCGGAAG |
| 6002_1 | TATGAGTGTGGAGTCGTTGCACTACAGAACCCCAGGATTCCTCNNNAGATCGGAAG |
| 6003_1 | TATGAGTGTGGAGTCGTTGCCTGTCACTTGCTAGGTCAATTGCNNNAGATCGGAAG |
| 6004_1 | TATGAGTGTGGAGTCGTTGCCACCCTGTTCACTTCAGAATTGCNNNAGATCGGAAG |
| 6005_1 | TATGAGTGTGGAGTCGTTGCTTCTCAGGGTGTGGGCTTCTAATNNNAGATCGGAAG |
| 6006_1 | TATGAGTGTGGAGTCGTTGCTGTTGGGAACTGGTTCAATGGAGNNNAGATCGGAA |
| 6007_1 | TATGAGTGTGGAGTCGTTGCAGGTGCAGGCTAATATACCTGATNNNAGATCGGAA |
| 6008_1 | TATGAGTGTGGAGTCGTTGCATGCTTGAAAAAGGACCCATTGCNNNAGATCGGAA |
| 6009_1 | TATGAGTGTGGAGTCGTTGCAGAGGCAGAGAATACAGCATTGCNNNAGATCGGAA |
| 6010_1 | TATGAGTGTGGAGTCGTTGCCTCCAGCTGTTCCTTTCCTTCAANNNAGATCGGAAGA |
| 6011_1 | TATGAGTGTGGAGTCGTTGCTAGTGGGTCTGCATGGAAAGAGNNNAGATCGGAA |
| 6013_1 | TATGAGTGTGGAGTCGTTGCACCTTCCTGTAGGATACACCCAGNNNAGATCGGAAG |
| 6016_1 | TATGAGTGTGGAGTCGTTGCGCTCTCACTCATTCTCAGCTCAGNNNAGATCGGAAG |
| 6017_1 | TATGAGTGTGGAGTCGTTGCACGGCAAATATTATGGCAGCCTTNNNAGATCGGAAG |
| 6019_1 | TATGAGTGTGGAGTCGTTGCGGAGGAGGAAACCTTAGCCATTGNNNAGATCGGAA |
| 6021_1 | TATGAGTGTGGAGTCGTTGCGTGGGTGTGGGAGAAGTAGATCANNNAGATCGGA |
| 6024_1 | TATGAGTGTGGAGTCGTTGCGGGGAGCTTTGATCTGAGTTTGANNNAGATCGGAA |
| 6025_1 | TATGAGTGTGGAGTCGTTGCAATAGTTTTCTGCAGGTACTTGCNNNAGATCGGAAG |
| 6029_1 | TATGAGTGTGGAGTCGTTGCACCTTTCTGTAAGTGGTCCTTGCNNNAGATCGGAAG |
| 6033_1 | TATGAGTGTGGAGTCGTTGCTGTAGGTCTGTGTATGTGCTTGCNNNAGATCGGAA |
| 6035_1 | TATGAGTGTGGAGTCGTTGCTGAAAAAGCCTTCTTGGAGGAGGNNNAGATCGGAA |
| 6038_1 | TATGAGTGTGGAGTCGTTGCCCTGGTCCTCACTGCTCTAGTAANNNAGATCGGAAG |
| 6039_1 | TATGAGTGTGGAGTCGTTGCTTTCAGTCACTCCATGGTCTTGCNNNAGATCGGAAG |
| 6041_1 | TATGAGTGTGGAGTCGTTGCCAAGCCATTCCAATCCTTCTTGCNNNAGATCGGAAGA |
| 6042_1 | TATGAGTGTGGAGTCGTTGCAGACTGTGTTTCCCTCTTCTTGCNNNAGATCGGAAG |
| 6043_1 | TATGAGTGTGGAGTCGTTGCGGAAAAGCCTAAGGCACTGGTTANNNAGATCGGAA |
| 6044_1 | TATGAGTGTGGAGTCGTTGCACAAGTAAAGACTTCCACCTGCCNNNAGATCGGAAG |
| 6046_1 | TATGAGTGTGGAGTCGTTGCAAAACCCTTCCTTGGGAGTAGCANNNAGATCGGAAG |
| 6047_1 | TATGAGTGTGGAGTCGTTGCGGTTGTATTGTTGTTGCTGTTGCNNNAGATCGGAAG |
| 6048_1 | TATGAGTGTGGAGTCGTTGCGTGTGTGTGTAGTATCCTAGGCANNNAGATCGGAA |
| 6049_1 | TATGAGTGTGGAGTCGTTGCCTCTCTCTATGCCACAGTGTTGCNNNAGATCGGAAG |
| 6050_1 | TATGAGTGTGGAGTCGTTGCCCTAAACATTCAGCAGGTGTTGCNNNAGATCGGAAG |
| 6052_1 | TATGAGTGTGGAGTCGTTGCTGAGCCTATTCTCCTTCTCAGCANNNAGATCGGAAG |
| 6053_1 | TATGAGTGTGGAGTCGTTGCTCCCAAGAATGTGCAGAATTTGCNNNAGATCGGAAG |
| 6055_1 | TATGAGTGTGGAGTCGTTGCCTCTCTTGTGGTGACTGATTTGCNNNAGATCGGAAG |
| 6056_1 | TATGAGTGTGGAGTCGTTGCCTGGTAACAACGGTCTGATTTGCNNNAGATCGGAAG |
| 6057_1 | TATGAGTGTGGAGTCGTTGCCTCTTACCACTCTTCTCCACCCANNNAGATCGGAAGA |
| 6058_1 | TATGAGTGTGGAGTCGTTGCAATATCATGCCCTCTCCCTTTGCNNNAGATCGGAAGA |
| 6059_1 | TATGAGTGTGGAGTCGTTGCACAATGGAGCTGTAAGGTCCTCANNNAGATCGGAA |
| 6060_1 | TATGAGTGTGGAGTCGTTGCGGTCATCAGAATGCATGGTTTGCNNNAGATCGGAA |
| 6061_1 | TATGAGTGTGGAGTCGTTGCCTCCTTCAAACCTTGGTGTTTGCNNNAGATCGGAAG |
| 6063_1 | TATGAGTGTGGAGTCGTTGCGAGCCGTGGTATCTTTTGTTTGCNNNAGATCGGAAG |

| | |
|---|---|
| 6065_1 | TATGAGTGTGGAGTCGTTGCTGTCTTCCCACGTGTTTTGATCTNNNAGATCGGAAGA |
| 6066_1 | TATGAGTGTGGAGTCGTTGCCCCTCCCTAACCTCTTCTTTTGCNNNAGATCGGAAGA |
| 6067_1 | TATGAGTGTGGAGTCGTTGCCCTGCTCCAGTACGATCTCATCTNNNAGATCGGAAGA |
| 6070_1 | TATGAGTGTGGAGTCGTTGCCAGAAGGATAAACTGGCTTGCCTNNNAGATCGGAA |
| 6071_1 | TATGAGTGTGGAGTCGTTGCGCAAGAGCACACTTCTGCAAATCNNNAGATCGGAAG |
| 6075_1 | TATGAGTGTGGAGTCGTTGCGGAAGTGGGGATGTCTCACAATCNNNAGATCGGAA |
| 6076_1 | TATGAGTGTGGAGTCGTTGCCCTGGGCTGGAATTCTACCAATCNNNAGATCGGAAG |
| 6077_1 | TATGAGTGTGGAGTCGTTGCTGAAGGCACAGATCAGAGCAATCNNNAGATCGGAA |
| 6079_1 | TATGAGTGTGGAGTCGTTGCGCCCATGCTGAGGTAGAAGAATCNNNAGATCGGAA |
| 6081_1 | TATGAGTGTGGAGTCGTTGCTTCCTGGATCTCCCCTGAGAATCNNNAGATCGGAAG |
| 6082_1 | TATGAGTGTGGAGTCGTTGCGGACAAAGGTCAGGACAGGAATCNNNAGATCGGAA |
| 6083_1 | TATGAGTGTGGAGTCGTTGCCTTTTTGCTGGCCTTGAGGAATCNNNAGATCGGAAG |
| 6085_1 | TATGAGTGTGGAGTCGTTGCCAATTTCAGACAGGGTGGCTAGGNNNAGATCGGAA |
| 6087_1 | TATGAGTGTGGAGTCGTTGCAGTTGCCTCTGATCAAAGCTGTTNNNAGATCGGAAG |
| 6088_1 | TATGAGTGTGGAGTCGTTGCACCCCATCCCACTCCATCTAATCNNNAGATCGGAAGA |
| 6089_1 | TATGAGTGTGGAGTCGTTGCTCAGGGCAAGGCAAAGTCTAATCNNNAGATCGGAA |
| 6091_1 | TATGAGTGTGGAGTCGTTGCCTTAGGGCTCTGCCACAAACATCNNNAGATCGGAAG |
| 6093_1 | TATGAGTGTGGAGTCGTTGCACCCTTCCTGTAGAGGACACATCNNNAGATCGGAAG |
| 6096_1 | TATGAGTGTGGAGTCGTTGCGCCTCCAGGACATATAGGACATCNNNAGATCGGAAG |
| 6097_1 | TATGAGTGTGGAGTCGTTGCCCCCTGACAGCTGGAAATACATCNNNAGATCGGAAG |
| 6098_1 | TATGAGTGTGGAGTCGTTGCAGAGGCCTGATCCAGGATACATCNNNAGATCGGAA |
| 6099_1 | TATGAGTGTGGAGTCGTTGCTGGTTTGCCCTATTGGCTACATCNNNAGATCGGAAG |
| 6100_1 | TATGAGTGTGGAGTCGTTGCCAGTGCTAAACCCAGGGTACATCNNNAGATCGGAAG |
| 6101_1 | TATGAGTGTGGAGTCGTTGCATTACCGCTTTGAGAGCACCATCNNNAGATCGGAAG |
| 6102_1 | TATGAGTGTGGAGTCGTTGCCTGTGAGGAAGAAGCTCACCATCNNNAGATCGGAA |
| 6104_1 | TATGAGTGTGGAGTCGTTGCCCCTCCTGGATTATCTCCCCATCNNNAGATCGGAAGA |
| 6106_1 | TATGAGTGTGGAGTCGTTGCCGCAGAAAGTAGGTGATGCCATCNNNAGATCGGAA |
| 6107_1 | TATGAGTGTGGAGTCGTTGCCTGCTCACAGCTCTTGATCCATCNNNAGATCGGAAGA |
| 6108_1 | TATGAGTGTGGAGTCGTTGCTTATACCTAGCCCGAGCTCCATCNNNAGATCGGAAGA |
| 6110_1 | TATGAGTGTGGAGTCGTTGCATCTTCCACCAATGCTGTCCATCNNNAGATCGGAAGA |
| 6111_1 | TATGAGTGTGGAGTCGTTGCTTGGGCTTGTCTCTCTTTGTCTTNNNAGATCGGAAGA |
| 6112_1 | TATGAGTGTGGAGTCGTTGCTTTCCTCCTTCCCCTTTTCCATCNNNAGATCGGAAGAG |
| 6113_1 | TATGAGTGTGGAGTCGTTGCTGAATCTAGGCTGTGTTTCCAGCNNNAGATCGGAAG |
| 6116_1 | TATGAGTGTGGAGTCGTTGCCAAGCCAAGGAGAGAGGAGATCTNNNAGATCGGAA |
| 6117_1 | TATGAGTGTGGAGTCGTTGCCTTTAGGTTGGTCCCTGAGCATCNNNAGATCGGAAG |
| 6118_1 | TATGAGTGTGGAGTCGTTGCTCAATGGCACAGTCACTAGCATCNNNAGATCGGAAG |
| 6119_1 | TATGAGTGTGGAGTCGTTGCCCAAGTGATCAAGAATGCGCATCNNNAGATCGGAA |
| 6121_1 | TATGAGTGTGGAGTCGTTGCAGAGAGAGAGAGGAACTGGCATCNNNAGATCGGA |
| 6122_1 | TATGAGTGTGGAGTCGTTGCTCAAGGTTCACTCATGTGGCATCNNNAGATCGGAAG |
| 6123_1 | TATGAGTGTGGAGTCGTTGCAAGATTGAACCCAGGGATGCATCNNNAGATCGGAA |
| 6126_1 | TATGAGTGTGGAGTCGTTGCGGAAAAGCATCAACCTTTGTGCANNNAGATCGGAA |
| 6128_1 | TATGAGTGTGGAGTCGTTGCTCTGCCAGGGTAAAAGCATCATCNNNAGATCGGAAG |
| 6131_1 | TATGAGTGTGGAGTCGTTGCGTTCACCTCCTTCAGAGCTCATCNNNAGATCGGAAGA |

| | |
|---|---|
| 6134_1 | TATGAGTGTGGAGTCGTTGCCATCCACACATGCCCTTCCTATCNNNAGATCGGAAGA |
| 6135_1 | TATGAGTGTGGAGTCGTTGCCATCTGTCTGCCTCTCTGTCATCNNNAGATCGGAAGA |
| 6136_1 | TATGAGTGTGGAGTCGTTGCGTAAGACAGGCCTCGGATTCATCNNNAGATCGGAA |
| 6140_1 | TATGAGTGTGGAGTCGTTGCAGTCTCAGCAACCGAACAAGATCNNNAGATCGGAAG |
| 6141_1 | TATGAGTGTGGAGTCGTTGCAGTGGAACCATACCAGTGGATCTNNNAGATCGGAA |
| 6142_1 | TATGAGTGTGGAGTCGTTGCTTCACACTGGGCTTCAACAGATCNNNAGATCGGAAG |
| 6143_1 | TATGAGTGTGGAGTCGTTGCCAGGGGAGAGAGAGAGACAGATCNNNAGATCGGA |
| 6147_1 | TATGAGTGTGGAGTCGTTGCGCTGGGCCATATTAGTGCAGATCNNNAGATCGGAA |
| 6149_1 | TATGAGTGTGGAGTCGTTGCGGGGACTGAAAAGTGCTGAGATCNNNAGATCGGA |
| 6150_1 | TATGAGTGTGGAGTCGTTGCTGAATTTTCTGGAAGAGGGGCAGNNNAGATCGGAA |
| 6151_1 | TATGAGTGTGGAGTCGTTGCGAAAGCAGAGAGAGACAGGCAAGNNNAGATCGGA |
| 6153_1 | TATGAGTGTGGAGTCGTTGCACAACCACAAGTTCACCAGGATCNNNAGATCGGAAG |
| 6154_1 | TATGAGTGTGGAGTCGTTGCGTCGAGTAAGCACACAGAGGATCNNNAGATCGGAA |
| 6156_1 | TATGAGTGTGGAGTCGTTGCGCCTAACCTGATAGCAGGGTTTCNNNAGATCGGAAG |
| 6157_1 | TATGAGTGTGGAGTCGTTGCTATGACACCAGCCGTATTGGATCNNNAGATCGGAAG |
| 6158_1 | TATGAGTGTGGAGTCGTTGCGGCTTACATCATGGCCAGAGTATNNNAGATCGGAA |
| 6160_1 | TATGAGTGTGGAGTCGTTGCTTTTGGCAGTTAGGCTTGTTCCTNNNAGATCGGAAG |
| 6161_1 | TATGAGTGTGGAGTCGTTGCCGAATGGCTTTTGGTGCCATATCNNNAGATCGGAAG |
| 6163_1 | TATGAGTGTGGAGTCGTTGCCCTGACCCTTTGTCACCACTATCNNNAGATCGGAAGA |
| 6164_1 | TATGAGTGTGGAGTCGTTGCTCCTACTGACTTCTCCCCTTTGGNNNAGATCGGAAGA |
| 6165_1 | TATGAGTGTGGAGTCGTTGCCCCTCTCTGTAACTCTGCTTCTTNNNAGATCGGAAGA |
| 6166_1 | TATGAGTGTGGAGTCGTTGCGCCACAGATTTCCAACATCCATGNNNAGATCGGAAG |
| 6168_1 | TATGAGTGTGGAGTCGTTGCCACCACTTCACCCAGCTTCTATCNNNAGATCGGAAGA |
| 6169_1 | TATGAGTGTGGAGTCGTTGCCTGCTGTGTCCAGCTCAAGTATCNNNAGATCGGAAG |
| 6170_1 | TATGAGTGTGGAGTCGTTGCGGTGAGAGACGATGCACAGTATCNNNAGATCGGAA |
| 6171_1 | TATGAGTGTGGAGTCGTTGCGCATGTTGCTGGTAAAGCGTATCNNNAGATCGGAA |
| 6173_1 | TATGAGTGTGGAGTCGTTGCGACCACTAGGCAACAGGCTTATCNNNAGATCGGAAG |
| 6174_1 | TATGAGTGTGGAGTCGTTGCCACCTGCACATCCACCTCTTATCNNNAGATCGGAAGA |
| 6175_1 | TATGAGTGTGGAGTCGTTGCCCTGACAGTTGGCCCTTCTTATCNNNAGATCGGAAG |
| 6176_1 | TATGAGTGTGGAGTCGTTGCTCCCCATCTAAGTTTCTGGAGGTNNNAGATCGGAAG |
| 6177_1 | TATGAGTGTGGAGTCGTTGCAACATGCCAGTGGAGGAAAACTCNNNAGATCGGAA |
| 6179_1 | TATGAGTGTGGAGTCGTTGCTATGGAGCAGAAGAGGCAAACTCNNNAGATCGGAA |
| 6181_1 | TATGAGTGTGGAGTCGTTGCGCAGTCCCACAGACTAGAAACTCNNNAGATCGGAAG |
| 6182_1 | TATGAGTGTGGAGTCGTTGCAAGGGAGACTGGAAAACCAACTCNNNAGATCGGAA |
| 6185_1 | TATGAGTGTGGAGTCGTTGCCGCACCAGAAAAACCTTCAACTCNNNAGATCGGAAG |
| 6186_1 | TATGAGTGTGGAGTCGTTGCTCAGCTAGCAACTCACAGAACTCNNNAGATCGGAAG |
| 6188_1 | TATGAGTGTGGAGTCGTTGCCATAAAGGGGCCAGTCTGAACTCNNNAGATCGGAA |
| 6189_1 | TATGAGTGTGGAGTCGTTGCTGATGGAACAGGCTCTTGAACTCNNNAGATCGGAA |
| 6190_1 | TATGAGTGTGGAGTCGTTGCCCACCATAACCTACTGCACACTCNNNAGATCGGAAGA |
| 6192_1 | TATGAGTGTGGAGTCGTTGCCAGTAACACAGCACTCAACCAAGNNNAGATCGGAAG |
| 6193_1 | TATGAGTGTGGAGTCGTTGCTGTGTGTGTGAATGTGCTCTACANNNAGATCGGAA |
| 6194_1 | TATGAGTGTGGAGTCGTTGCTGGTATAAATGTGCAGCCCACTCNNNAGATCGGAAG |
| 6195_1 | TATGAGTGTGGAGTCGTTGCTCTAGGATAAGCTCTGGCCACTCNNNAGATCGGAAG |

| | |
|---|---|
| 6196_1 | TATGAGTGTGGAGTCGTTGCATGCTTGAAAGTGGTCTCAGAGCNNNAGATCGGAAC |
| 6197_1 | TATGAGTGTGGAGTCGTTGCTGGCTTCAGATGATTCTGCACTCNNNAGATCGGAAGA |
| 6198_1 | TATGAGTGTGGAGTCGTTGCACACAGCCCTGCCATCTATATCTNNNAGATCGGAAGA |
| 6199_1 | TATGAGTGTGGAGTCGTTGCCTTGTGTGCATGTGTCTCTTTCANNNAGATCGGAAGA |
| 6200_1 | TATGAGTGTGGAGTCGTTGCTGGTAGGTGATTGTGCTTCACTCNNNAGATCGGAAG |
| 6201_1 | TATGAGTGTGGAGTCGTTGCTGGACACAACAGGCATCTCTAGANNNAGATCGGAAG |
| 6204_1 | TATGAGTGTGGAGTCGTTGCGTGCAGGGAAAATGCACCTATCTNNNAGATCGGAAG |
| 6205_1 | TATGAGTGTGGAGTCGTTGCAGGATTTAGAGGCCACCCTACTCNNNAGATCGGAAG |
| 6207_1 | TATGAGTGTGGAGTCGTTGCTGGGTTGGCCTTGAAGTTTACTCNNNAGATCGGAAG |
| 6208_1 | TATGAGTGTGGAGTCGTTGCCAGTTTTGCCCCCTACCTTTCTTNNNAGATCGGAAGAC |
| 6209_1 | TATGAGTGTGGAGTCGTTGCGTCAAAAGCGTCCTTGAAACCTCNNNAGATCGGAAG |
| 6210_1 | TATGAGTGTGGAGTCGTTGCCTCCCTTCCTCCATACCAACCTCNNNAGATCGGAAGAG |
| 6211_1 | TATGAGTGTGGAGTCGTTGCCCTCTGGAACTCCTCAGAACCTCNNNAGATCGGAAGA |
| 6212_1 | TATGAGTGTGGAGTCGTTGCAGTAAATCAGGCACTACCTGTCTNNNAGATCGGAAG |
| 6213_1 | TATGAGTGTGGAGTCGTTGCGTTGGGATGAGTTTCCAGACCTCNNNAGATCGGAAG |
| 6214_1 | TATGAGTGTGGAGTCGTTGCGATGTAGCACTGACCATGACCTCNNNAGATCGGAAG |
| 6215_1 | TATGAGTGTGGAGTCGTTGCTAGAACCCTGCCCTTATGACCTCNNNAGATCGGAAGA |
| 6216_1 | TATGAGTGTGGAGTCGTTGCAAAAAGCACAGACCTCTGACCTCNNNAGATCGGAAG |
| 6217_1 | TATGAGTGTGGAGTCGTTGCATTCTTGGTGAAGGCCATACCTCNNNAGATCGGAAG |
| 6218_1 | TATGAGTGTGGAGTCGTTGCACAGAAGGAGTGTGCTTTACCTCNNNAGATCGGAAG |
| 6222_1 | TATGAGTGTGGAGTCGTTGCAAATGAAAGCAGAGAAGCCCCTCNNNAGATCGGAAC |
| 6223_1 | TATGAGTGTGGAGTCGTTGCCTTTAGGCACCAAGAATCCCCTCNNNAGATCGGAAGA |
| 6224_1 | TATGAGTGTGGAGTCGTTGCCTGAGCAAATCTGGAATCCCCTCNNNAGATCGGAAGA |
| 6227_1 | TATGAGTGTGGAGTCGTTGCGGGAATTTGGGTTACAGTCCCTCNNNAGATCGGAAG |
| 6229_1 | TATGAGTGTGGAGTCGTTGCATCCAGCACTCTTTGTTTCCCTCNNNAGATCGGAAGAC |
| 6230_1 | TATGAGTGTGGAGTCGTTGCGAAACTCAAAAGTCAGCAGCAGCNNNAGATCGGAAC |
| 6233_1 | TATGAGTGTGGAGTCGTTGCCCAATGACACTATGTGAGGCCTCNNNAGATCGGAAG |
| 6234_1 | TATGAGTGTGGAGTCGTTGCCAGAGAAGGCAGGGGGAGAGAAANNNAGATCGGA |
| 6236_1 | TATGAGTGTGGAGTCGTTGCGGCAAGATCTCACTTACTGCCTCNNNAGATCGGAAGA |
| 6238_1 | TATGAGTGTGGAGTCGTTGCTGCTTATGGACACATTCTGCCTCNNNAGATCGGAAGA |
| 6240_1 | TATGAGTGTGGAGTCGTTGCAACTTCCTTCTCCGAATTGCCTCNNNAGATCGGAAGA |
| 6241_1 | TATGAGTGTGGAGTCGTTGCGGTCTGGTTGTCCATGAATCCTCNNNAGATCGGAAG |
| 6243_1 | TATGAGTGTGGAGTCGTTGCAGAAAGTCCCTGAAGACCTCCTCNNNAGATCGGAAG |
| 6244_1 | TATGAGTGTGGAGTCGTTGCCTCATTTTTCCTCTGCCCTCCTCNNNAGATCGGAAGAG |
| 6245_1 | TATGAGTGTGGAGTCGTTGCGCCAGTCTTCTTATTGGCTCCTCNNNAGATCGGAAGA |
| 6246_1 | TATGAGTGTGGAGTCGTTGCTATTGCCCCTGAAAATGCTCCTCNNNAGATCGGAAGA |
| 6248_1 | TATGAGTGTGGAGTCGTTGCATTGTCTGAAAAACGGGGTCCTCNNNAGATCGGAAG |
| 6250_1 | TATGAGTGTGGAGTCGTTGCACCTCTCAGGAACAGGATTCCTCNNNAGATCGGAAGA |
| 6251_1 | TATGAGTGTGGAGTCGTTGCGATGACTGCCTCCTCACTTCCTCNNNAGATCGGAAGA |
| 6252_1 | TATGAGTGTGGAGTCGTTGCACAAAGTGACAAACCCCTTCCTCNNNAGATCGGAAGA |
| 6253_1 | TATGAGTGTGGAGTCGTTGCTTACCTCTGAGTTCCCCTTCCTCNNNAGATCGGAAGAC |
| 6254_1 | TATGAGTGTGGAGTCGTTGCTTTTCCCTGTGTTGCTCTTCCTCNNNAGATCGGAAGAC |
| 6255_1 | TATGAGTGTGGAGTCGTTGCAGAGCAATCACATCTAAGGGACANNNAGATCGGAAC |

6256_1 TATGAGTGTGGAGTCGTTGCTCAAAAATGGCAAGAAGGCAAGCNNNAGATCGGAA
6257_1 TATGAGTGTGGAGTCGTTGCCCTACTGCTCACACACTTTCCTCNNNAGATCGGAAGA
6260_1 TATGAGTGTGGAGTCGTTGCCAGAAACGTTCCCCTAGAAGCTCNNNAGATCGGAAG
6261_1 TATGAGTGTGGAGTCGTTGCTTACGATGCAGAACTGACAGCTCNNNAGATCGGAAG
6262_1 TATGAGTGTGGAGTCGTTGCGGCTGGAGTGAAGATTACAGCTCNNNAGATCGGAA
6264_1 TATGAGTGTGGAGTCGTTGCCCACAGGGTGTATTGAAGAGCTCNNNAGATCGGAA
6266_1 TATGAGTGTGGAGTCGTTGCCACTGTCTACCTTTGCAGAGCTCNNNAGATCGGAAG
6267_1 TATGAGTGTGGAGTCGTTGCACAAAGAGAGAGGGAGAGAGCTCNNNAGATCGGA
6268_1 TATGAGTGTGGAGTCGTTGCCTCTTTGTTCTGGGACTGAGCTCNNNAGATCGGAAG
6270_1 TATGAGTGTGGAGTCGTTGCGCTCATCTTTTTGCCTTGAGCTCNNNAGATCGGAAGA
6271_1 TATGAGTGTGGAGTCGTTGCACATTTGGCAGTGAGCAGAAGTTNNNAGATCGGAA
6272_1 TATGAGTGTGGAGTCGTTGCTGCTGTACGCCTAAGTTTAGCTCNNNAGATCGGAAG
6275_1 TATGAGTGTGGAGTCGTTGCGTTGGTTTTACAGGAGTGGAGGTNNNAGATCGGAA
6277_1 TATGAGTGTGGAGTCGTTGCTTCATGGTCTTCCAGGAGGAAGANNNAGATCGGAA
6279_1 TATGAGTGTGGAGTCGTTGCGCATTGTGTTCTTGATACGGCTCNNNAGATCGGAAG
6280_1 TATGAGTGTGGAGTCGTTGCCCTATGGTTTGTGATGAGGGCTCNNNAGATCGGAAG
6281_1 TATGAGTGTGGAGTCGTTGCGGCTTGGGTTTCAAATATGGCTCNNNAGATCGGAAG
6282_1 TATGAGTGTGGAGTCGTTGCATTCTCCTCAGGACTACTGGCTCNNNAGATCGGAAG
6284_1 TATGAGTGTGGAGTCGTTGCAAAGGTGTGATATGAGCCTGCTCNNNAGATCGGAA
6285_1 TATGAGTGTGGAGTCGTTGCTTCTTAGACAGGGTCTCCTGCTCNNNAGATCGGAAG
6286_1 TATGAGTGTGGAGTCGTTGCCTCCCATCTCTCAGAGTTCAGCTNNNAGATCGGAAGA
6287_1 TATGAGTGTGGAGTCGTTGCACCTTTCTTCTTTCCTCTGCAGCNNNAGATCGGAAGA
6288_1 TATGAGTGTGGAGTCGTTGCGGAATGACATTGTTGAGCTGCTCNNNAGATCGGAAG
6289_1 TATGAGTGTGGAGTCGTTGCCTCCAAAGACCTGCTATCTGCTCNNNAGATCGGAAGA
6290_1 TATGAGTGTGGAGTCGTTGCCAATTCTCCCACCTTCTCTGCTCNNNAGATCGGAAGA
6292_1 TATGAGTGTGGAGTCGTTGCTTGAAGTGTCTGGCATAGTGCTCNNNAGATCGGAAG
6293_1 TATGAGTGTGGAGTCGTTGCATAGTCAGGGGTTCCTAGTGCTCNNNAGATCGGAAG
6294_1 TATGAGTGTGGAGTCGTTGCGCCTGTTGTCTCTGGATGAAGTTNNNAGATCGGAAG
6296_1 TATGAGTGTGGAGTCGTTGCATACACAAGGCAAGGACTTGCTCNNNAGATCGGAAG
6298_1 TATGAGTGTGGAGTCGTTGCATGTATTGAGACGGAGCTTGCTCNNNAGATCGGAAG
6300_1 TATGAGTGTGGAGTCGTTGCGGCTTTGGTTTGAACTCTTGCTCNNNAGATCGGAAG
6301_1 TATGAGTGTGGAGTCGTTGCGTCTGTTGCTTGTCAAGTTGCTCNNNAGATCGGAAG
6302_1 TATGAGTGTGGAGTCGTTGCGCGTGTATGTGTCTTGTTTGCTCNNNAGATCGGAAG
6303_1 TATGAGTGTGGAGTCGTTGCTGGAATGAGCAGCAGGAAATCTCNNNAGATCGGAA
6305_1 TATGAGTGTGGAGTCGTTGCCGGGGTCTTAGCATTCTCATCTCNNNAGATCGGAAG
6306_1 TATGAGTGTGGAGTCGTTGCCCCAGCGCATCTCTATTCATCTCNNNAGATCGGAAGA
6308_1 TATGAGTGTGGAGTCGTTGCTGGTGTGCTTTCTGCTGTATCTCNNNAGATCGGAAG
6309_1 TATGAGTGTGGAGTCGTTGCCGTGCAAAATCCTGGTAGGAAGANNNAGATCGGAA
6310_1 TATGAGTGTGGAGTCGTTGCGCTTGACATACCAGCATGAAGCTNNNAGATCGGAAG
6311_1 TATGAGTGTGGAGTCGTTGCATCATTTGCAAGCGTACCCTCTCNNNAGATCGGAAGA
6312_1 TATGAGTGTGGAGTCGTTGCGAAGGGGCATTATTCAGCCTCTCNNNAGATCGGAAG
6313_1 TATGAGTGTGGAGTCGTTGCCTACTCGCAGCTGATAAGCTCTCNNNAGATCGGAAG
6317_1 TATGAGTGTGGAGTCGTTGCTGCCTGACTGTGTGAATGCATTTNNNAGATCGGAAG

| | |
|---|---|
| 6318_1 | TATGAGTGTGGAGTCGTTGCCCACTCCCATACCTCTCTCTCTCNNNAGATCGGAAGAG |
| 6319_1 | TATGAGTGTGGAGTCGTTGCTAGTGAGGGGTGGATAACTCAGCNNNAGATCGGAA |
| 6321_1 | TATGAGTGTGGAGTCGTTGCTGGCTAAATCACAGCTGTCTGAGNNNAGATCGGAAG |
| 6323_1 | TATGAGTGTGGAGTCGTTGCGTCCTGGGAAATCCTTTGGTCTCNNNAGATCGGAAG |
| 6324_1 | TATGAGTGTGGAGTCGTTGCCGGTCATCTTCTGTCTGTGTCTCNNNAGATCGGAAGA |
| 6325_1 | TATGAGTGTGGAGTCGTTGCACCACATGCAGATGTCTTGTCTCNNNAGATCGGAAGA |
| 6326_1 | TATGAGTGTGGAGTCGTTGCCCTGTCTCACTTGGTGTTGTCTCNNNAGATCGGAAGA |
| 6327_1 | TATGAGTGTGGAGTCGTTGCTGCCTCTCTTGCACTTCATTCTCNNNAGATCGGAAGA |
| 6328_1 | TATGAGTGTGGAGTCGTTGCGTCCCCTTCTGACAACACTTCTCNNNAGATCGGAAGA |
| 6329_1 | TATGAGTGTGGAGTCGTTGCGGCACTTTCCTTCTTCCCTTCTCNNNAGATCGGAAGA |
| 6330_1 | TATGAGTGTGGAGTCGTTGCCCTTTCTCCTCATTGGCCTTCTCNNNAGATCGGAAGA |
| 6331_1 | TATGAGTGTGGAGTCGTTGCCTCCCTTCACAGTCAAGCTTCTCNNNAGATCGGAAGA |
| 6333_1 | TATGAGTGTGGAGTCGTTGCCCTCCCATGACTTGATGCTTCTCNNNAGATCGGAAGA |
| 6334_1 | TATGAGTGTGGAGTCGTTGCGCATTTGTCGGCTTCATGTTCTCNNNAGATCGGAAGA |
| 6335_1 | TATGAGTGTGGAGTCGTTGCAGACTGCTCTGTGGTTTGTTCTCNNNAGATCGGAAG |
| 6336_1 | TATGAGTGTGGAGTCGTTGCCTGCTCTCTGGACCCAATTTCTCNNNAGATCGGAAGA |
| 6337_1 | TATGAGTGTGGAGTCGTTGCTTGGAGCCTCTGCAGAATTTCTCNNNAGATCGGAAG |
| 6338_1 | TATGAGTGTGGAGTCGTTGCCGTAAGGCAAACAAGCCTTTCTCNNNAGATCGGAAG |
| 6339_1 | TATGAGTGTGGAGTCGTTGCTGTCCCCGAGTTTCTGTATTTGCNNNAGATCGGAAGA |
| 6340_1 | TATGAGTGTGGAGTCGTTGCGCCATTCCATTTGCAGGTTTCTCNNNAGATCGGAAGA |
| 6342_1 | TATGAGTGTGGAGTCGTTGCTTACTGTCCTCTTGGCACCACTANNNAGATCGGAAGA |
| 6343_1 | TATGAGTGTGGAGTCGTTGCCCTTTTCAGTTGCCTCTCCAGTTNNNAGATCGGAAGA |
| 6344_1 | TATGAGTGTGGAGTCGTTGCACCCCCTCCTCTCTCTTTTTCTCNNNAGATCGGAAGAG |
| 6345_1 | TATGAGTGTGGAGTCGTTGCTGCCACCAAAAACCCAAAAAGTCNNNAGATCGGAAG |
| 6346_1 | TATGAGTGTGGAGTCGTTGCGGTGGCAAAAGGTGCTAAAAGTCNNNAGATCGGAA |
| 6347_1 | TATGAGTGTGGAGTCGTTGCAGGCAGCCAGAGAAAAGAAAGTCNNNAGATCGGAA |
| 6348_1 | TATGAGTGTGGAGTCGTTGCACCCAGAGGTTTGCTAACTCTTGNNNAGATCGGAAG |
| 6349_1 | TATGAGTGTGGAGTCGTTGCCACTCATCCCTTCTGCTCAAGTCNNNAGATCGGAAGA |
| 6350_1 | TATGAGTGTGGAGTCGTTGCCCTAGAGAACACGTGCAGAAGTCNNNAGATCGGAA |
| 6351_1 | TATGAGTGTGGAGTCGTTGCAGGTCTTCCAAGCTATAGAAGTCNNNAGATCGGAAG |
| 6352_1 | TATGAGTGTGGAGTCGTTGCCCACAAGGATAGCTCAGGAAGTCNNNAGATCGGAA |
| 6353_1 | TATGAGTGTGGAGTCGTTGCGCAGCTAGACGATCAGTGAAGTCNNNAGATCGGAA |
| 6354_1 | TATGAGTGTGGAGTCGTTGCCAAGCACTGACAGGTCCTAAGTCNNNAGATCGGAAG |
| 6356_1 | TATGAGTGTGGAGTCGTTGCTGCTCTGCTTGTTTTGAACAGTCNNNAGATCGGAAG |
| 6357_1 | TATGAGTGTGGAGTCGTTGCAGCAAAAGTAGCCACTGACAGTCNNNAGATCGGAA |
| 6358_1 | TATGAGTGTGGAGTCGTTGCGGGGTAGGGTAGGGAATACAGTCNNNAGATCGGA |
| 6360_1 | TATGAGTGTGGAGTCGTTGCCACAGGCAAAAACTTCACCAGTCNNNAGATCGGAAG |
| 6361_1 | TATGAGTGTGGAGTCGTTGCTGAGGACCAAACAGTTACCAGTCNNNAGATCGGAA |
| 6364_1 | TATGAGTGTGGAGTCGTTGCCTTCTCTTCTGCTCTCTGCAGTCNNNAGATCGGAAGA |
| 6365_1 | TATGAGTGTGGAGTCGTTGCTTCTTGTAACCACCACTGCAGTTNNNAGATCGGAAGA |
| 6366_1 | TATGAGTGTGGAGTCGTTGCATTTTTGGAGGAGAGCTGCAGTTNNNAGATCGGAA |
| 6367_1 | TATGAGTGTGGAGTCGTTGCCAGGGAAGCACATGCCTATAGTCNNNAGATCGGAA |
| 6368_1 | TATGAGTGTGGAGTCGTTGCAGTTCTCTGGTGGCTGTTACTCTNNNAGATCGGAAG |

| | |
|---|---|
| 6370_1 | TATGAGTGTGGAGTCGTTGCGGGGCTGGGGAAGATATCTAGTCNNNAGATCGGAA |
| 6371_1 | TATGAGTGTGGAGTCGTTGCCCCTGGTTCTTCTGACAGTAGTCNNNAGATCGGAAG |
| 6373_1 | TATGAGTGTGGAGTCGTTGCCTGGGAAAATGCAGGAAAACGTCNNNAGATCGGAA |
| 6374_1 | TATGAGTGTGGAGTCGTTGCATGCCTGATTTCTCAACTGCAAANNNAGATCGGAAG |
| 6375_1 | TATGAGTGTGGAGTCGTTGCAAAGACCATATTGGCAGACCGTCNNNAGATCGGAAG |
| 6376_1 | TATGAGTGTGGAGTCGTTGCTAAAACAGCACTCGGGATGTGAGNNNAGATCGGAA |
| 6377_1 | TATGAGTGTGGAGTCGTTGCCAAACAGGAAAAGGAAAGCCGTCNNNAGATCGGAA |
| 6378_1 | TATGAGTGTGGAGTCGTTGCGAAACGTCTTACTGGGGAAGGTCNNNAGATCGGAA |
| 6380_1 | TATGAGTGTGGAGTCGTTGCTGGCAGTGAAGAATTACCAGGTCNNNAGATCGGAA |
| 6381_1 | TATGAGTGTGGAGTCGTTGCGAGCCCTAGAAGCAGTGAGACTANNNAGATCGGAA |
| 6383_1 | TATGAGTGTGGAGTCGTTGCCTTGCTGGTTCTCTTCAGAGGTCNNNAGATCGGAAG |
| 6384_1 | TATGAGTGTGGAGTCGTTGCTGTCTTTCACTCATGGAGGGAGTNNNAGATCGGAAG |
| 6385_1 | TATGAGTGTGGAGTCGTTGCTAGCAGATTCCTCTGGCTCAGTTNNNAGATCGGAAG |
| 6386_1 | TATGAGTGTGGAGTCGTTGCTGTCATTCTTCAGGTGGACCTCTNNNAGATCGGAAG |
| 6387_1 | TATGAGTGTGGAGTCGTTGCCACAGCTTCTGGTGTCTTAGGTCNNNAGATCGGAAG |
| 6388_1 | TATGAGTGTGGAGTCGTTGCTCAGATTTGGTCTTGCAAGGGTCNNNAGATCGGAAG |
| 6389_1 | TATGAGTGTGGAGTCGTTGCGTTGGTCATTCCTCTGATGGGTCNNNAGATCGGAAG |
| 6390_1 | TATGAGTGTGGAGTCGTTGCAAACCTTCAAACCTTCCTGGGTCNNNAGATCGGAAG |
| 6391_1 | TATGAGTGTGGAGTCGTTGCTGAATTGCTGGAGGCTTATGGTCNNNAGATCGGAA |
| 6393_1 | TATGAGTGTGGAGTCGTTGCATTGTGGCTACCTCAGCTGACTANNNAGATCGGAAG |
| 6395_1 | TATGAGTGTGGAGTCGTTGCGGAGAATCAGAGAGGCACATGTCNNNAGATCGGAA |
| 6396_1 | TATGAGTGTGGAGTCGTTGCATCTGCAACTTGGAGGACATGTCNNNAGATCGGAAG |
| 6397_1 | TATGAGTGTGGAGTCGTTGCTGTTCAACTCAGGACACCATGTCNNNAGATCGGAAG |
| 6398_1 | TATGAGTGTGGAGTCGTTGCAAAAGCCTTACCCTGCAGATGTCNNNAGATCGGAAG |
| 6399_1 | TATGAGTGTGGAGTCGTTGCCGCTCAGTGGTTTCATGTGACTANNNAGATCGGAAG |
| 6400_1 | TATGAGTGTGGAGTCGTTGCCTTTTTGCCTGTCGTGTGATGTCNNNAGATCGGAAG |
| 6401_1 | TATGAGTGTGGAGTCGTTGCCTGCCTTTTGCCTTTGCTATGTCNNNAGATCGGAAGA |
| 6402_1 | TATGAGTGTGGAGTCGTTGCGTGTGTAGGAACCTGTGACTGTCNNNAGATCGGAA |
| 6403_1 | TATGAGTGTGGAGTCGTTGCTAAACTGCAGCCAAGTTCCTGTCNNNAGATCGGAAG |
| 6405_1 | TATGAGTGTGGAGTCGTTGCAGAGTGAAAGAAAGCCAGCTGTCNNNAGATCGGAA |
| 6406_1 | TATGAGTGTGGAGTCGTTGCAGCGGTAGGTATAGAGAGCTGTCNNNAGATCGGAA |
| 6407_1 | TATGAGTGTGGAGTCGTTGCCTTTTCAGCCTCTCCCATCTGTCNNNAGATCGGAAGA |
| 6408_1 | TATGAGTGTGGAGTCGTTGCGGCAGTGGCATATGATGAGAGTTNNNAGATCGGAA |
| 6409_1 | TATGAGTGTGGAGTCGTTGCACACACACGAGTTTCACTCTGTCNNNAGATCGGAAG |
| 6410_1 | TATGAGTGTGGAGTCGTTGCTAGAAGCCCTGAGTAGACAGAGCNNNAGATCGGAA |
| 6411_1 | TATGAGTGTGGAGTCGTTGCAGTTGGAGACAAGCAGGCTACTANNNAGATCGGAA |
| 6412_1 | TATGAGTGTGGAGTCGTTGCTTTGTTCTGTGGTCCCTTCTGTCNNNAGATCGGAAGA |
| 6413_1 | TATGAGTGTGGAGTCGTTGCGCAAAAGTAAAGCTGGACGTGTCNNNAGATCGGAA |
| 6416_1 | TATGAGTGTGGAGTCGTTGCCCGAATACAATGTCCCTGGTGTCNNNAGATCGGAAG |
| 6417_1 | TATGAGTGTGGAGTCGTTGCTCAGAGGCAATGCTGTATGTGTCNNNAGATCGGAA |
| 6418_1 | TATGAGTGTGGAGTCGTTGCTAGGGGTCTCACTAAGCTGTGTCNNNAGATCGGAAG |
| 6419_1 | TATGAGTGTGGAGTCGTTGCCAGTGACAGAATCTTGCTGTGTCNNNAGATCGGAAG |
| 6420_1 | TATGAGTGTGGAGTCGTTGCGTGTCCCTGTGTATGTCTGTGTCNNNAGATCGGAAG |

| | |
|---|---|
| 6421_1 | TATGAGTGTGGAGTCGTTGCTATTTGGGGGATGTGTGTGTGTCNNNAGATCGGAA |
| 6422_1 | TATGAGTGTGGAGTCGTTGCATTGTCTTAGCCAGCCACTTGTCNNNAGATCGGAAG |
| 6423_1 | TATGAGTGTGGAGTCGTTGCATGCAGCCAGATTTCATGACAGTNNNAGATCGGAAG |
| 6424_1 | TATGAGTGTGGAGTCGTTGCACAGAGATCCACGATGAGTTGTCNNNAGATCGGAA |
| 6426_1 | TATGAGTGTGGAGTCGTTGCGGGGAGGGGTGTACTCAAAATTCNNNAGATCGGAA |
| 6427_1 | TATGAGTGTGGAGTCGTTGCCGGCAAAGGATGGTTCCAAATTCNNNAGATCGGAA |
| 6428_1 | TATGAGTGTGGAGTCGTTGCGAAACAGCAGCAGGAGGAAATTCNNNAGATCGGAA |
| 6429_1 | TATGAGTGTGGAGTCGTTGCTTCTAGGCAGCACAGACCAATTCNNNAGATCGGAAG |
| 6430_1 | TATGAGTGTGGAGTCGTTGCAAGACAGAGTTGGGCCTGAATTCNNNAGATCGGAA |
| 6431_1 | TATGAGTGTGGAGTCGTTGCTGCCCTCTGGATTGTGAACATTCNNNAGATCGGAAG |
| 6432_1 | TATGAGTGTGGAGTCGTTGCCTGACACACACAACAGCACATTCNNNAGATCGGAAG |
| 6433_1 | TATGAGTGTGGAGTCGTTGCCATAACACCCCTCTTCAGCCTCTNNNAGATCGGAAGA |
| 6434_1 | TATGAGTGTGGAGTCGTTGCTGTCCTGCTAGTGTTGAGCATTCNNNAGATCGGAAG |
| 6437_1 | TATGAGTGTGGAGTCGTTGCAACTCCTCCCTAAGGGGTCATTCNNNAGATCGGAAG |
| 6438_1 | TATGAGTGTGGAGTCGTTGCTTTGGAACCGTGTCCTCAGATTCNNNAGATCGGAAG |
| 6440_1 | TATGAGTGTGGAGTCGTTGCACCTGCTCAAACGTGGTAGATTCNNNAGATCGGAAG |
| 6441_1 | TATGAGTGTGGAGTCGTTGCGGCAAACACAGCTCATTGGATTCNNNAGATCGGAAG |
| 6442_1 | TATGAGTGTGGAGTCGTTGCTGGCAAAGTTTGAAGGCTGATTCNNNAGATCGGAA |
| 6443_1 | TATGAGTGTGGAGTCGTTGCTGATGAGTTTGCAGCCCCTATTCNNNAGATCGGAAG |
| 6444_1 | TATGAGTGTGGAGTCGTTGCATAGCCCTAGTGTGCCCCTATTCNNNAGATCGGAAG |
| 6446_1 | TATGAGTGTGGAGTCGTTGCATTCGTGGGGTCACTTGCTATTCNNNAGATCGGAAG |
| 6447_1 | TATGAGTGTGGAGTCGTTGCGGAATGTGCTTGGGAGGGTATTCNNNAGATCGGAA |
| 6448_1 | TATGAGTGTGGAGTCGTTGCGAAGGCTCAATGTGGCAAACTTCNNNAGATCGGAA |
| 6449_1 | TATGAGTGTGGAGTCGTTGCCCAGAAGGACATTTCCCCACTTCNNNAGATCGGAAG |
| 6450_1 | TATGAGTGTGGAGTCGTTGCGTATAGTGCAGGCCACTCACTTCNNNAGATCGGAAG |
| 6451_1 | TATGAGTGTGGAGTCGTTGCTAGGCTGGTCAGGATGAGACTTCNNNAGATCGGAA |
| 6452_1 | TATGAGTGTGGAGTCGTTGCACATAAAGACCCCTCTTGCCTCTNNNAGATCGGAAGA |
| 6454_1 | TATGAGTGTGGAGTCGTTGCGGGTACTTCCGTGTGTGTACTTCNNNAGATCGGAAG |
| 6456_1 | TATGAGTGTGGAGTCGTTGCTGAAGAGCTTTGGGCATACCTTCNNNAGATCGGAAG |
| 6457_1 | TATGAGTGTGGAGTCGTTGCGCCTTCTCAGTTCAGCTACCTTCNNNAGATCGGAAGA |
| 6458_1 | TATGAGTGTGGAGTCGTTGCATCTTGGTAAGACCCTCCCCTTCNNNAGATCGGAAGA |
| 6459_1 | TATGAGTGTGGAGTCGTTGCGAAATGGATGCTTTGTGCCCTTCNNNAGATCGGAAG |
| 6461_1 | TATGAGTGTGGAGTCGTTGCTTGTTCTAGCACCTTCAGCCTTCNNNAGATCGGAAGA |
| 6462_1 | TATGAGTGTGGAGTCGTTGCGGAAAGACCTCACTTCTGCCTTCNNNAGATCGGAAG |
| 6463_1 | TATGAGTGTGGAGTCGTTGCCCTCATGCTCTGACACATCCTTCNNNAGATCGGAAGA |
| 6464_1 | TATGAGTGTGGAGTCGTTGCTTTGAAGGTGACCAAGCTCCTTCNNNAGATCGGAAG |
| 6466_1 | TATGAGTGTGGAGTCGTTGCAGGCTGCACGAATCTTTTCCTTCNNNAGATCGGAAG |
| 6467_1 | TATGAGTGTGGAGTCGTTGCGTGTGCTTTGTTTCCTCAGCTTCNNNAGATCGGAAG |
| 6470_1 | TATGAGTGTGGAGTCGTTGCTCAAGCTCTCTCATGTGTCCTCTNNNAGATCGGAAGA |
| 6471_1 | TATGAGTGTGGAGTCGTTGCATAATCCCAGGCAATCTGGCTTCNNNAGATCGGAAG |
| 6473_1 | TATGAGTGTGGAGTCGTTGCACGAGGGGTTGTGGATTACAGGANNNAGATCGGA |
| 6474_1 | TATGAGTGTGGAGTCGTTGCCTGAAGCCCCTGTATACTGCTTCNNNAGATCGGAAG |
| 6475_1 | TATGAGTGTGGAGTCGTTGCTGGTCCTTTCTCCTCATTGCTTCNNNAGATCGGAAGA |

| | |
|---|---|
| 6476_1 | TATGAGTGTGGAGTCGTTGCCCCCCTGTTCACTTGCTATCTTCNNNAGATCGGAAGA |
| 6480_1 | TATGAGTGTGGAGTCGTTGCCTTCGCTGCCTAGTATGGTCTTCNNNAGATCGGAAG |
| 6481_1 | TATGAGTGTGGAGTCGTTGCTGGACTGGGTTGAATGGTTCTTCNNNAGATCGGAA |
| 6483_1 | TATGAGTGTGGAGTCGTTGCGAAGAGGGGAGTGCAGTTTCTTCNNNAGATCGGAA |
| 6485_1 | TATGAGTGTGGAGTCGTTGCTGTTTGTGTCTCTCCCGAAGTTCNNNAGATCGGAAG |
| 6486_1 | TATGAGTGTGGAGTCGTTGCTGAAGAAGAGGAAGAATGCGAGCNNNAGATCGGA |
| 6488_1 | TATGAGTGTGGAGTCGTTGCTGCTTTGTTTCAAGCCTCAGTTCNNNAGATCGGAAG |
| 6489_1 | TATGAGTGTGGAGTCGTTGCGAGGAGGAGGTGTGTTTCAGTTCNNNAGATCGGAA |
| 6491_1 | TATGAGTGTGGAGTCGTTGCACCTGTGAGACGAACAGTAGTTCNNNAGATCGGAA |
| 6492_1 | TATGAGTGTGGAGTCGTTGCATACGCCTGCATGAAAAACGTTCNNNAGATCGGAAG |
| 6493_1 | TATGAGTGTGGAGTCGTTGCAGGTAGGAGAGGAACCAAGGTTCNNNAGATCGGA |
| 6495_1 | TATGAGTGTGGAGTCGTTGCAAATCTCGCCTCCATGAAGGTTCNNNAGATCGGAAG |
| 6496_1 | TATGAGTGTGGAGTCGTTGCAAAACAATTGCCTCTGCAGGTTCNNNAGATCGGAAG |
| 6497_1 | TATGAGTGTGGAGTCGTTGCCGATCTCCAAGCTGTAAGGGTTCNNNAGATCGGAAG |
| 6498_1 | TATGAGTGTGGAGTCGTTGCCCAATCTCTGCAGCTTAGGGTTCNNNAGATCGGAAG |
| 6499_1 | TATGAGTGTGGAGTCGTTGCTTCTCACTCCAACATGTGGGTTCNNNAGATCGGAAG |
| 6500_1 | TATGAGTGTGGAGTCGTTGCTTACATATTCCGTGGCCTGGTTCNNNAGATCGGAAG |
| 6501_1 | TATGAGTGTGGAGTCGTTGCTCCTTGTACCTTGCAAGTGGTTCNNNAGATCGGAAG |
| 6503_1 | TATGAGTGTGGAGTCGTTGCCGAGACAGGATCTCACCATGTTCNNNAGATCGGAAG |
| 6504_1 | TATGAGTGTGGAGTCGTTGCAGTCAGCCACAACTACCATGTTCNNNAGATCGGAAG |
| 6506_1 | TATGAGTGTGGAGTCGTTGCAAACATCACTCACTCCAGGGTTGNNNAGATCGGAAG |
| 6507_1 | TATGAGTGTGGAGTCGTTGCAGGACATTGGATTGGAGCTGTTCNNNAGATCGGAA |
| 6508_1 | TATGAGTGTGGAGTCGTTGCCCCTCTCACTCTTTCCAGTGTTCNNNAGATCGGAAGA |
| 6509_1 | TATGAGTGTGGAGTCGTTGCGGAACTGACTGGAGGAGTTGTTCNNNAGATCGGAA |
| 6510_1 | TATGAGTGTGGAGTCGTTGCTGGTTCTGGTGATGAGATGGTTGNNNAGATCGGAA |
| 6511_1 | TATGAGTGTGGAGTCGTTGCGGCAGGGCAGATGACAAAATTTCNNNAGATCGGAA |
| 6512_1 | TATGAGTGTGGAGTCGTTGCGCCACATCACAATCCCCAATTTCNNNAGATCGGAAGA |
| 6513_1 | TATGAGTGTGGAGTCGTTGCCAGCATGACAGCACTGACATTTCNNNAGATCGGAAG |
| 6514_1 | TATGAGTGTGGAGTCGTTGCCATCTCCTCCAGGAAGCCATTTCNNNAGATCGGAAG |
| 6515_1 | TATGAGTGTGGAGTCGTTGCCCCATCTCTGATTGCAGCATTTCNNNAGATCGGAAGA |
| 6517_1 | TATGAGTGTGGAGTCGTTGCCTGTGCCTTCCGGATTGCATTTCNNNAGATCGGAAG |
| 6518_1 | TATGAGTGTGGAGTCGTTGCTTGTGGTCGGTCTAACGGATTTCNNNAGATCGGAAG |
| 6519_1 | TATGAGTGTGGAGTCGTTGCCACTTGCCATGTTCCTGGATTTCNNNAGATCGGAAG |
| 6520_1 | TATGAGTGTGGAGTCGTTGCAACGCTCTCCACTTCCTGATTTCNNNAGATCGGAAGA |
| 6521_1 | TATGAGTGTGGAGTCGTTGCGTCTCGCTCTCCCCTGTTATTTCNNNAGATCGGAAGA |
| 6522_1 | TATGAGTGTGGAGTCGTTGCAGGCTGACTAGAATGTGAGCTCTNNNAGATCGGAA |
| 6523_1 | TATGAGTGTGGAGTCGTTGCCTCTACCTTCCATGCCCACTTTCNNNAGATCGGAAGA |
| 6524_1 | TATGAGTGTGGAGTCGTTGCCAGAACAGGGAGCTGAGACTTTCNNNAGATCGGAA |
| 6525_1 | TATGAGTGTGGAGTCGTTGCCCCACCCTCCCTTGTTTACTTTCNNNAGATCGGAAGA |
| 6527_1 | TATGAGTGTGGAGTCGTTGCCAAATACGGCAAAACCCCCTTTCNNNAGATCGGAAG |
| 6529_1 | TATGAGTGTGGAGTCGTTGCACTGTCATGGGTATGTGCCTTTCNNNAGATCGGAAG |
| 6530_1 | TATGAGTGTGGAGTCGTTGCTCTGAGGGAAAAAGCCTCCTTTCNNNAGATCGGAAG |
| 6531_1 | TATGAGTGTGGAGTCGTTGCTCTGGGGCAAGTTCATAGCTTTCNNNAGATCGGAAG |

| | |
|---|---|
| 6533_1 | TATGAGTGTGGAGTCGTTGCCTGCTTTTCTGCAGATGGCTTTCNNNAGATCGGAAG |
| 6534_1 | TATGAGTGTGGAGTCGTTGCCCTCATGCCCTCATGTCTCTTTCNNNAGATCGGAAGA |
| 6535_1 | TATGAGTGTGGAGTCGTTGCGTCCCCTTTTCAGCCTCAGTTTCNNNAGATCGGAAGA |
| 6536_1 | TATGAGTGTGGAGTCGTTGCAGGGTGTAGCTGGAGAGAGTTTCNNNAGATCGGAA |
| 6538_1 | TATGAGTGTGGAGTCGTTGCCAGCACACACTCATGGAGAAGTTNNNAGATCGGAAG |
| 6539_1 | TATGAGTGTGGAGTCGTTGCGAAAACGTGAATGCCTGGGTTTCNNNAGATCGGAA |
| 6540_1 | TATGAGTGTGGAGTCGTTGCTTTGGGTTTGCAGTTTGGGTTTCNNNAGATCGGAAG |
| 6541_1 | TATGAGTGTGGAGTCGTTGCCCCTCTACTCCTACCCTGGTTTCNNNAGATCGGAAGA |
| 6542_1 | TATGAGTGTGGAGTCGTTGCTCTTGCTTGCTCCTTCTGGTTTCNNNAGATCGGAAGA |
| 6543_1 | TATGAGTGTGGAGTCGTTGCCCTGACCTAACCCACTTGGTTTCNNNAGATCGGAAGA |
| 6544_1 | TATGAGTGTGGAGTCGTTGCCCTTCTCTGGCTCTCCATGTTTCNNNAGATCGGAAGA |
| 6545_1 | TATGAGTGTGGAGTCGTTGCCGTAGAAGCTGGAGGGATGTTTCNNNAGATCGGAA |
| 6547_1 | TATGAGTGTGGAGTCGTTGCTCAAAATACTTCTGGAGTGCCTANNNAGATCGGAAG |
| 6549_1 | TATGAGTGTGGAGTCGTTGCAGCCAAAACAGAGCAGCATTTTCNNNAGATCGGAAG |
| 6550_1 | TATGAGTGTGGAGTCGTTGCTCATGTGTGCCTGTTGCATTTTCNNNAGATCGGAAG |
| 6551_1 | TATGAGTGTGGAGTCGTTGCATCCTGGGTGCACTCTCATTTTCNNNAGATCGGAAG |
| 6553_1 | TATGAGTGTGGAGTCGTTGCGGGGGTGGATTTGGAGGATTTTCNNNAGATCGGAA |
| 6554_1 | TATGAGTGTGGAGTCGTTGCCTTGTCAGCACAGCAGACTTTTCNNNAGATCGGAAG |
| 6555_1 | TATGAGTGTGGAGTCGTTGCCCCATCCAACTCACCTCCTTTTCNNNAGATCGGAAGA |
| 6556_1 | TATGAGTGTGGAGTCGTTGCCACTCTTGGCTCCTCTCCTTTTCNNNAGATCGGAAGA |
| 6558_1 | TATGAGTGTGGAGTCGTTGCATTTTGCATGGGATGGCGTTTTCNNNAGATCGGAAG |
| 6559_1 | TATGAGTGTGGAGTCGTTGCCGCACGCACATATACCCTTTTTCNNNAGATCGGAAGA |
| 6560_1 | TATGAGTGTGGAGTCGTTGCCAGAAGGAACAGGACAAAGCTGTNNNAGATCGGAA |
| 6561_1 | TATGAGTGTGGAGTCGTTGCTGAGTGAGAGGGGTGAGAAAAAGNNNAGATCGGA |
| 6562_1 | TATGAGTGTGGAGTCGTTGCGGGGTGCGATGATGCATAAAAAGNNNAGATCGGA |
| 6563_1 | TATGAGTGTGGAGTCGTTGCGGGAGGAAGAGAGAAGCCAAAAGNNNAGATCGGA |
| 6564_1 | TATGAGTGTGGAGTCGTTGCATGTCACAGCATTGGTGCAAAAGNNNAGATCGGAA |
| 6565_1 | TATGAGTGTGGAGTCGTTGCGGCAAGAGCTGACTTTGCAAAAGNNNAGATCGGAA |
| 6567_1 | TATGAGTGTGGAGTCGTTGCTCATTGTGGCTTCCTCACTCCTANNNAGATCGGAAGA |
| 6568_1 | TATGAGTGTGGAGTCGTTGCGCATTGTTTGCCTCTGGGAAAAGNNNAGATCGGAA |
| 6569_1 | TATGAGTGTGGAGTCGTTGCGCAAGGAAACGTCAGTGGAAAAGNNNAGATCGGA |
| 6570_1 | TATGAGTGTGGAGTCGTTGCTGACAGATTGCAGCACTGAAAAGNNNAGATCGGAA |
| 6571_1 | TATGAGTGTGGAGTCGTTGCGGGGAGCTGGGAGGTAATAAAAGNNNAGATCGGA |
| 6572_1 | TATGAGTGTGGAGTCGTTGCCCCAGCCACCATCTGAGTAAAAGNNNAGATCGGAAG |
| 6573_1 | TATGAGTGTGGAGTCGTTGCGATGGGAATGCCAGGAGGAGAAGNNNAGATCGGA |
| 6574_1 | TATGAGTGTGGAGTCGTTGCCAGTTGGCCTGAAACACACAAAGNNNAGATCGGAA |
| 6575_1 | TATGAGTGTGGAGTCGTTGCCTGAAAGCTGAACAGCACCAAAGNNNAGATCGGAA |
| 6576_1 | TATGAGTGTGGAGTCGTTGCACTGTAAACCAGCTGGACCAAAGNNNAGATCGGAA |
| 6577_1 | TATGAGTGTGGAGTCGTTGCTCTAAAGGCAGCTTTCCCCAAAGNNNAGATCGGAAG |
| 6578_1 | TATGAGTGTGGAGTCGTTGCGAATGAAGTGAAACGGCCCAAAGNNNAGATCGGAA |
| 6579_1 | TATGAGTGTGGAGTCGTTGCGCTTCTCCAGTTTTCTGGAGGTTNNNAGATCGGAAG |
| 6580_1 | TATGAGTGTGGAGTCGTTGCTTGTTCTATTGGGGGCCTCAAAGNNNAGATCGGAA |
| 6581_1 | TATGAGTGTGGAGTCGTTGCTGACACCTCCCCTACTCTCAAAGNNNAGATCGGAAGA |

6582_1 TATGAGTGTGGAGTCGTTGCGCAGCTCTTCTGTGACTTCAAAGNNNAGATCGGAAG
6583_1 TATGAGTGTGGAGTCGTTGCGGGGAGGGGAGAATCAAAGAAAGNNNAGATCGGA
6584_1 TATGAGTGTGGAGTCGTTGCAAACCAGCAAGAGGGGAAGAAAGNNNAGATCGGA
6585_1 TATGAGTGTGGAGTCGTTGCGGAGCGGAAGACCACTAAGAAAGNNNAGATCGGA
6587_1 TATGAGTGTGGAGTCGTTGCACACACGCTTGCTTTTCAGAAAGNNNAGATCGGAAG
6588_1 TATGAGTGTGGAGTCGTTGCAGGACCAGGAGAAGACAGGAAAGNNNAGATCGGA
6589_1 TATGAGTGTGGAGTCGTTGCACCCACAGCAGAATACAGGAAAGNNNAGATCGGAA
6590_1 TATGAGTGTGGAGTCGTTGCAGACAGAGAGATGGGCAGGAAAGNNNAGATCGGA
6593_1 TATGAGTGTGGAGTCGTTGCGCGATCCAAAAATGCAGGGAAAGNNNAGATCGGAA
6594_1 TATGAGTGTGGAGTCGTTGCTGTATGAACCTTCTTGTCTCTCGNNNAGATCGGAAG
6595_1 TATGAGTGTGGAGTCGTTGCACACTCAGGAGACTTTCTGCTCTNNNAGATCGGAAG
6596_1 TATGAGTGTGGAGTCGTTGCTGTCTGGTACTTTGGCTGGAAAGNNNAGATCGGAA
6597_1 TATGAGTGTGGAGTCGTTGCGGGATAGGTGCTTCAGTGGAAAGNNNAGATCGGA
6598_1 TATGAGTGTGGAGTCGTTGCGTTAGACCACTGTGGGTGGAAAGNNNAGATCGGAA
6600_1 TATGAGTGTGGAGTCGTTGCCACCTTCCAGCTGTACCTGAAAGNNNAGATCGGAAG
6601_1 TATGAGTGTGGAGTCGTTGCTTATACCTGCCTGCTCCTGAAAGNNNAGATCGGAAG
6602_1 TATGAGTGTGGAGTCGTTGCTTTGCTGAAGGCTTTGGTGAAAGNNNAGATCGGAA
6603_1 TATGAGTGTGGAGTCGTTGCATGGTCACACCTGCCAAGTAAAGNNNAGATCGGAA
6604_1 TATGAGTGTGGAGTCGTTGCTCCCCTGGACCTTGGTAGTAAAGNNNAGATCGGAAG
6605_1 TATGAGTGTGGAGTCGTTGCGAGTTGGAGGCTGAGAGAACAAGNNNAGATCGGA
6606_1 TATGAGTGTGGAGTCGTTGCAGTGGAAGTAGAGCCACAAGCTANNNAGATCGGAA
6607_1 TATGAGTGTGGAGTCGTTGCCTGGTCACAGGGAGATGAACAAGNNNAGATCGGAA
6608_1 TATGAGTGTGGAGTCGTTGCGTCACACCAAAGCATCACACAAGNNNAGATCGGAAG
6610_1 TATGAGTGTGGAGTCGTTGCGACATAAACTCGCCAAGCACAAGNNNAGATCGGAA
6611_1 TATGAGTGTGGAGTCGTTGCAGTTTTTCTGGCTCCAGCTGTCCNNNAGATCGGAAG
6613_1 TATGAGTGTGGAGTCGTTGCCCCTGGCTGAGAATATCGACAAGNNNAGATCGGAA
6616_1 TATGAGTGTGGAGTCGTTGCAAGACTGTAGGTCTCTGAGGGTTNNNAGATCGGAA
6617_1 TATGAGTGTGGAGTCGTTGCAGTAGTTCCATTCTGGCACCAAGNNNAGATCGGAAG
6618_1 TATGAGTGTGGAGTCGTTGCAACAGCACCTTTCTGAGACCAAGNNNAGATCGGAAG
6619_1 TATGAGTGTGGAGTCGTTGCCCCCCTCTATCCCTAATCCCAAGNNNAGATCGGAAGA
6620_1 TATGAGTGTGGAGTCGTTGCGTTTTGGGGACTGAACTCCCAAGNNNAGATCGGAA
6622_1 TATGAGTGTGGAGTCGTTGCTATTTGGCTCAAGGCTTCCCAAGNNNAGATCGGAAG
6627_1 TATGAGTGTGGAGTCGTTGCATTGAAGGTGACTTCTGGCCAACNNNAGATCGGAA
6631_1 TATGAGTGTGGAGTCGTTGCCGCTATGAAGTTTTTGGGGCAAGNNNAGATCGGAA
6632_1 TATGAGTGTGGAGTCGTTGCAGCATTTGCCTTATGTGGGCAAGNNNAGATCGGAA
6633_1 TATGAGTGTGGAGTCGTTGCCAGAAAGCTGACAAAGTGGCAAGNNNAGATCGGAA
6634_1 TATGAGTGTGGAGTCGTTGCCTGCACTTTCACTTCTTGGCAAGNNNAGATCGGAAG
6635_1 TATGAGTGTGGAGTCGTTGCAAGGTCCCGTTGTCTTTATCTCTNNNAGATCGGAAG
6636_1 TATGAGTGTGGAGTCGTTGCCCTCTTCTGCCATCTGATGCAAGNNNAGATCGGAAG
6637_1 TATGAGTGTGGAGTCGTTGCGTGAGAAGGTATCCACCTGCAAGNNNAGATCGGAA
6638_1 TATGAGTGTGGAGTCGTTGCCCATTTGTCAAGCTTCCTGCAAGNNNAGATCGGAAG
6639_1 TATGAGTGTGGAGTCGTTGCGTGATACGGAGGCTATCTGCAAGNNNAGATCGGAA
6640_1 TATGAGTGTGGAGTCGTTGCGCAAAACTCAAATGTGGTGCAAGNNNAGATCGGAA

| | |
|---|---|
| 6641_1 | TATGAGTGTGGAGTCGTTGCCAGAAACAGTGTGGATGTGCAAGNNNAGATCGGAA |
| 6642_1 | TATGAGTGTGGAGTCGTTGCACACATCCCCCTCACAAATCAAGNNNAGATCGGAAG |
| 6643_1 | TATGAGTGTGGAGTCGTTGCAGGGAATGGCCAGTTGAATCAAGNNNAGATCGGAA |
| 6644_1 | TATGAGTGTGGAGTCGTTGCCAGTGAATCACCTCCCCATCAAGNNNAGATCGGAAG |
| 6645_1 | TATGAGTGTGGAGTCGTTGCATGCCCGCACTCTATTCATCAAGNNNAGATCGGAAG |
| 6648_1 | TATGAGTGTGGAGTCGTTGCGCCCATTCTCATCCACACTCAAGNNNAGATCGGAAGA |
| 6649_1 | TATGAGTGTGGAGTCGTTGCTGCTGCTTCAGGAACTACTCAAGNNNAGATCGGAAG |
| 6650_1 | TATGAGTGTGGAGTCGTTGCACTCCTTTTCCCTTCCACTCTCTNNNAGATCGGAAGAG |
| 6656_1 | TATGAGTGTGGAGTCGTTGCAAGGAGCCTCCAAACACTAGCTANNNAGATCGGAAG |
| 6657_1 | TATGAGTGTGGAGTCGTTGCACCATTCCCGAGTTTCTGTCAAGNNNAGATCGGAAG |
| 6658_1 | TATGAGTGTGGAGTCGTTGCTGGACCTCTACCCAGTCTTCAAGNNNAGATCGGAAG |
| 6659_1 | TATGAGTGTGGAGTCGTTGCCAGTGGTGCATCCTAGGTTCAAGNNNAGATCGGAA |
| 6660_1 | TATGAGTGTGGAGTCGTTGCTGTAAGCTTTGGGAGTGATGGACNNNAGATCGGAA |
| 6662_1 | TATGAGTGTGGAGTCGTTGCATGCTCCCGAAGTCAGAAAGAAGNNNAGATCGGAA |
| 6665_1 | TATGAGTGTGGAGTCGTTGCGTGGGCAGTAAAGAGGGAAGAAGNNNAGATCGGA |
| 6666_1 | TATGAGTGTGGAGTCGTTGCGTGGCTGAGGAAGAGTGAAGAAGNNNAGATCGGA |
| 6670_1 | TATGAGTGTGGAGTCGTTGCAAATCCCTCCATGGGATCAGAAGNNNAGATCGGAA |
| 6672_1 | TATGAGTGTGGAGTCGTTGCGCCCCTACTGTCCAAAAGAGAAGNNNAGATCGGAA |
| 6673_1 | TATGAGTGTGGAGTCGTTGCCGGGAGGCTTTTCAGAAGAGAAGNNNAGATCGGAA |
| 6674_1 | TATGAGTGTGGAGTCGTTGCTTGGAGAAGCACGAACAATGGTTNNNAGATCGGAA |
| 6675_1 | TATGAGTGTGGAGTCGTTGCTAGGAGTACCCAGAACCCTCTCTNNNAGATCGGAAG |
| 6676_1 | TATGAGTGTGGAGTCGTTGCCATTTGGGGGATTGGGAGAGAAGNNNAGATCGGA |
| 6678_1 | TATGAGTGTGGAGTCGTTGCCCATGCAGAGAGAACAGGAGAAGNNNAGATCGGA |
| 6679_1 | TATGAGTGTGGAGTCGTTGCTTGGCACCTGTTTGTTTGTAACTNNNAGATCGGAAG |
| 6681_1 | TATGAGTGTGGAGTCGTTGCCTCTTGTCTCCTTTTTGGAGGGGNNNAGATCGGAAG |
| 6682_1 | TATGAGTGTGGAGTCGTTGCTCCTCCCTTTCTTGGATCTTGGTNNNAGATCGGAAGA |
| 6683_1 | TATGAGTGTGGAGTCGTTGCCAGAATTGAGTAGAGGGCCGAAGNNNAGATCGGA |
| 6686_1 | TATGAGTGTGGAGTCGTTGCGAATGCCAGACTTTACACCAGCTNNNAGATCGGAAG |
| 6688_1 | TATGAGTGTGGAGTCGTTGCACAGCAACTCACCAAGAAGGAAGNNNAGATCGGAA |
| 6689_1 | TATGAGTGTGGAGTCGTTGCGGTTCTTGCCCTTGAGAAGGAAGNNNAGATCGGAA |
| 6690_1 | TATGAGTGTGGAGTCGTTGCAGATTCACAGCTGCACAGGAGAGNNNAGATCGGAA |
| 6692_1 | TATGAGTGTGGAGTCGTTGCTCCCTTTGTCTCTCTCCAGGAAGNNNAGATCGGAAG |
| 6694_1 | TATGAGTGTGGAGTCGTTGCTGGGGAGAAAGAAGGAGAGGAAGNNNAGATCGG |
| 6695_1 | TATGAGTGTGGAGTCGTTGCGAAATGGAGTGCTCAGGAGGAAGNNNAGATCGGA |
| 6697_1 | TATGAGTGTGGAGTCGTTGCCTGGTTGGTATGGGAAAGGGAAGNNNAGATCGGA |
| 6699_1 | TATGAGTGTGGAGTCGTTGCACAGATTTTCTGCCACAGGGAAGNNNAGATCGGAA |
| 6701_1 | TATGAGTGTGGAGTCGTTGCTTAGGGGAACATGAGAGGGGAAGNNNAGATCGGA |
| 6703_1 | TATGAGTGTGGAGTCGTTGCGTCTCCTTTGCTCTCCATGGAAGNNNAGATCGGAAG |
| 6705_1 | TATGAGTGTGGAGTCGTTGCTGCTTATGCCAGTTGACTGGAAGNNNAGATCGGAA |
| 6706_1 | TATGAGTGTGGAGTCGTTGCGGGATCTTATGCAGGACCTAGGTNNNAGATCGGAA |
| 6709_1 | TATGAGTGTGGAGTCGTTGCACAAGAGGGCATGAGTTTGGAAGNNNAGATCGGA |
| 6710_1 | TATGAGTGTGGAGTCGTTGCACCAAGGAAACCCTACTTTCAGANNNAGATCGGAAG |
| 6712_1 | TATGAGTGTGGAGTCGTTGCCAGCCAAAACAGCTTCCATGAAGNNNAGATCGGAAG |

| | |
|---|---|
| 6713_1 | TATGAGTGTGGAGTCGTTGCTCTCTCACACACACAGCATGAAGNNNAGATCGGAAG |
| 6714_1 | TATGAGTGTGGAGTCGTTGCGACCCAAATGTTAGCGCATGAAGNNNAGATCGGAA |
| 6715_1 | TATGAGTGTGGAGTCGTTGCAGCACCCCTTGTCTTTGATGAAGNNNAGATCGGAAG |
| 6717_1 | TATGAGTGTGGAGTCGTTGCACATGATAGGACCTGGCTACCTGNNNAGATCGGAA |
| 6718_1 | TATGAGTGTGGAGTCGTTGCGAAAGGACACCTGCTTCCTGAAGNNNAGATCGGAA |
| 6722_1 | TATGAGTGTGGAGTCGTTGCCCACCAAAGGGAAGAGGTAAGACNNNAGATCGGAA |
| 6727_1 | TATGAGTGTGGAGTCGTTGCTTAGGGACAGGGGGATCTTGAAGNNNAGATCGGA |
| 6729_1 | TATGAGTGTGGAGTCGTTGCATGTGAAGGTGAGCAGGCATAAGNNNAGATCGGA |
| 6732_1 | TATGAGTGTGGAGTCGTTGCGGTGGGGGCGGTGTTATTATAAGNNNAGATCGGA |
| 6733_1 | TATGAGTGTGGAGTCGTTGCAAATTCTAGCAAATCCTGGGGCANNNAGATCGGAA |
| 6734_1 | TATGAGTGTGGAGTCGTTGCGGAGACATGCCAAGGAACCTAAGNNNAGATCGGAA |
| 6735_1 | TATGAGTGTGGAGTCGTTGCGGTCATGGACAAGGTGAGGGGTANNNAGATCGGA |
| 6736_1 | TATGAGTGTGGAGTCGTTGCTGTCCCCTGATGGGGTAAGTAAGNNNAGATCGGAA |
| 6737_1 | TATGAGTGTGGAGTCGTTGCGGGAAGGTGTTACTCAGCGTAAGNNNAGATCGGAA |
| 6738_1 | TATGAGTGTGGAGTCGTTGCGGGGTGATGGTTTTCCAGGTAAGNNNAGATCGGAA |
| 6739_1 | TATGAGTGTGGAGTCGTTGCTAGGCATGGGTGGACACTGTAAGNNNAGATCGGAA |
| 6740_1 | TATGAGTGTGGAGTCGTTGCATGTATGCAAGTGGGCCTGTAAGNNNAGATCGGAA |
| 6743_1 | TATGAGTGTGGAGTCGTTGCACATTTGGGAGACGGTGTGTAAGNNNAGATCGGAA |
| 6744_1 | TATGAGTGTGGAGTCGTTGCCATCAGAAACCATCCCTGCTGGCNNNAGATCGGAAG |
| 6745_1 | TATGAGTGTGGAGTCGTTGCTGTGTCTAGTTGTGCCTTCTCTGNNNAGATCGGAAG |
| 6746_1 | TATGAGTGTGGAGTCGTTGCTAGTTGTTTGCCATCTGAACAGGNNNAGATCGGAAG |
| 6753_1 | TATGAGTGTGGAGTCGTTGCTAGCCATCTGCAACCCAGGAATANNNAGATCGGAAG |
| 6754_1 | TATGAGTGTGGAGTCGTTGCAGGAGAGACAGCATATCAATGGANNNAGATCGGAA |
| 6755_1 | TATGAGTGTGGAGTCGTTGCGCATCCCACCTTTACAGAAACAGNNNAGATCGGAAG |
| 6756_1 | TATGAGTGTGGAGTCGTTGCATGCAGCTATCAGTGGGAAACAGNNNAGATCGGAA |
| 6757_1 | TATGAGTGTGGAGTCGTTGCCAGCCTAAGCCCTTCCTAAACAGNNNAGATCGGAAG |
| 6760_1 | TATGAGTGTGGAGTCGTTGCTTTGTCACCTGGATTCCCAACAGNNNAGATCGGAAG |
| 6761_1 | TATGAGTGTGGAGTCGTTGCCGCACCTGTCTTAGCCTCTGAAANNNAGATCGGAAG |
| 6762_1 | TATGAGTGTGGAGTCGTTGCTCATACACAGAGTGGGGACGTAGNNNAGATCGGAA |
| 6765_1 | TATGAGTGTGGAGTCGTTGCACACATCAGTCTGCAAGGAACAGNNNAGATCGGAA |
| 6766_1 | TATGAGTGTGGAGTCGTTGCGGGGTGTTACTGCTACTGAACAGNNNAGATCGGAA |
| 6767_1 | TATGAGTGTGGAGTCGTTGCATAACCAGAGAGCAGCTGAACAGNNNAGATCGGAA |
| 6768_1 | TATGAGTGTGGAGTCGTTGCGCCTGTTCCTTCTCTCTGAACAGNNNAGATCGGAAG |
| 6769_1 | TATGAGTGTGGAGTCGTTGCTTAGAAGCGGATCGCCATAACAGNNNAGATCGGAA |
| 6770_1 | TATGAGTGTGGAGTCGTTGCTCTCCATTGCCCTTCACTAACAGNNNAGATCGGAAGA |
| 6771_1 | TATGAGTGTGGAGTCGTTGCAGAGCTGAAAATTCTTGTGGGCANNNAGATCGGAA |
| 6772_1 | TATGAGTGTGGAGTCGTTGCACAGCTCACTACAGCCTGGATTCNNNAGATCGGAAG |
| 6773_1 | TATGAGTGTGGAGTCGTTGCCTTGAGCTTCCCATTACCCACTCNNNAGATCGGAAGA |
| 6774_1 | TATGAGTGTGGAGTCGTTGCGATGCACAGAGGTACAGACACAGNNNAGATCGGAA |
| 6776_1 | TATGAGTGTGGAGTCGTTGCACCATCAACATCAGTCACCACAGNNNAGATCGGAAG |
| 6777_1 | TATGAGTGTGGAGTCGTTGCATGATAAGCTGGTGAAGCCACAGNNNAGATCGGAA |
| 6779_1 | TATGAGTGTGGAGTCGTTGCGTCCTGGCACAGAATATCCACAGNNNAGATCGGAAG |
| 6781_1 | TATGAGTGTGGAGTCGTTGCCACAGAGAGACAGCTTTCCACAGNNNAGATCGGAA |

| | |
|---|---|
| 6782_1 | TATGAGTGTGGAGTCGTTGCGCGGTGCCTCAATCTTGGACTTCNNNAGATCGGAAG |
| 6783_1 | TATGAGTGTGGAGTCGTTGCAAGTAACCTGAAGGACTGCACAGNNNAGATCGGAA |
| 6785_1 | TATGAGTGTGGAGTCGTTGCCAGGCACATTCTCTCTTGTGGTTNNNAGATCGGAAG |
| 6786_1 | TATGAGTGTGGAGTCGTTGCAAGTGAGGCACAAATTCCACAGTNNNAGATCGGAA |
| 6787_1 | TATGAGTGTGGAGTCGTTGCCCCCTTCCCCCATTTTATCACAGNNNAGATCGGAAGA |
| 6788_1 | TATGAGTGTGGAGTCGTTGCCCTGCTTCCCCATCATATCCATCNNNAGATCGGAAGA |
| 6789_1 | TATGAGTGTGGAGTCGTTGCACAGATGCTCAAGTTGCTCATACNNNAGATCGGAAG |
| 6790_1 | TATGAGTGTGGAGTCGTTGCGCTCACTTGAAGGTGTCTCACAGNNNAGATCGGAAG |
| 6792_1 | TATGAGTGTGGAGTCGTTGCAGGGGTAGTTTTGCTTGTCACAGNNNAGATCGGAA |
| 6793_1 | TATGAGTGTGGAGTCGTTGCGCATTGTCTCTCCCCATTCACAGNNNAGATCGGAAG |
| 6794_1 | TATGAGTGTGGAGTCGTTGCTCTGTGCCTCTGTTTCTTCACAGNNNAGATCGGAAG |
| 6795_1 | TATGAGTGTGGAGTCGTTGCGGAATATGCTCAGCAGAGGTAGCNNNAGATCGGAA |
| 6798_1 | TATGAGTGTGGAGTCGTTGCTCAACGGCTCACTATCTTGCCCANNNAGATCGGAAG |
| 6800_1 | TATGAGTGTGGAGTCGTTGCCCATGTTCTTCACCCTCAGACAGNNNAGATCGGAAG |
| 6802_1 | TATGAGTGTGGAGTCGTTGCGGAAGACCATGTGAGGGAGACAGNNNAGATCGGA |
| 6803_1 | TATGAGTGTGGAGTCGTTGCCGCTGGCATTTTCTTTGAGACAGNNNAGATCGGAAG |
| 6804_1 | TATGAGTGTGGAGTCGTTGCAGTTTTTGGGGGCAGTTAGACAGNNNAGATCGGAA |
| 6805_1 | TATGAGTGTGGAGTCGTTGCGTGTAGGCAGTCTTGAAGGACAGNNNAGATCGGAA |
| 6808_1 | TATGAGTGTGGAGTCGTTGCAGTGCCATGAAGGCAAATGACAGNNNAGATCGGAA |
| 6809_1 | TATGAGTGTGGAGTCGTTGCCGTTATTTGTCTCCTCTGCAAGGNNNAGATCGGAAG |
| 6812_1 | TATGAGTGTGGAGTCGTTGCTGGGAGAGTGCAAGATAATTCACNNNAGATCGGAA |
| 6814_1 | TATGAGTGTGGAGTCGTTGCTGGCAGGGAAGAGGATACTACAGNNNAGATCGGA |
| 6815_1 | TATGAGTGTGGAGTCGTTGCCACAGCTAGGAGACAAGCTACAGNNNAGATCGGAA |
| 6818_1 | TATGAGTGTGGAGTCGTTGCCACAAGAGCAAGAGCCTGTACAGNNNAGATCGGAA |
| 6819_1 | TATGAGTGTGGAGTCGTTGCTGGCCGTTCTAGTAAGTGTACAGNNNAGATCGGAA |
| 6821_1 | TATGAGTGTGGAGTCGTTGCGGGGATCACGAGGTGAATTACAGNNNAGATCGGA |
| 6823_1 | TATGAGTGTGGAGTCGTTGCAAGCCCCATAGCCATTGTTACAGNNNAGATCGGAAG |
| 6827_1 | TATGAGTGTGGAGTCGTTGCGAGGTTATGGTGAGATTACGCGCNNNAGATCGGAA |
| 6828_1 | TATGAGTGTGGAGTCGTTGCCGATGAGGGATCTTTGGAACCAGNNNAGATCGGAA |
| 6830_1 | TATGAGTGTGGAGTCGTTGCGCGATAGTTTTCTCAACCACCAGNNNAGATCGGAAG |
| 6831_1 | TATGAGTGTGGAGTCGTTGCTTAGCATGGAAGACACTCACCAGNNNAGATCGGAA |
| 6832_1 | TATGAGTGTGGAGTCGTTGCACCATGACCCTGACCTGGAATAANNNAGATCGGAAG |
| 6833_1 | TATGAGTGTGGAGTCGTTGCCCTCCCTCTACCTACTTCACCAGNNNAGATCGGAAGA |
| 6835_1 | TATGAGTGTGGAGTCGTTGCCATAACAAATGGAGGGGGACCAGNNNAGATCGGA |
| 6843_1 | TATGAGTGTGGAGTCGTTGCGTTCTGCTTTAGGTCTGACCCAGNNNAGATCGGAAG |
| 6847_1 | TATGAGTGTGGAGTCGTTGCTTATAGACATGGCAGGTCCCCAGNNNAGATCGGAA |
| 6848_1 | TATGAGTGTGGAGTCGTTGCCTTCCAAAAGAAACATGGCCCAGNNNAGATCGGAAG |
| 6849_1 | TATGAGTGTGGAGTCGTTGCAAAGCACACACTACTATGCCCAGNNNAGATCGGAAG |
| 6850_1 | TATGAGTGTGGAGTCGTTGCAAGGAAAGCCAAAGTAGATCCCANNNAGATCGGAA |
| 6854_1 | TATGAGTGTGGAGTCGTTGCTTCTTTGTGGAGCATTGTCCCAGNNNAGATCGGAAG |
| 6855_1 | TATGAGTGTGGAGTCGTTGCAGGTGAGGTCTAAGAGTTCCCAGNNNAGATCGGAA |
| 6857_1 | TATGAGTGTGGAGTCGTTGCACAGTTTCAGCTTTGTTTCCCAGNNNAGATCGGAAG |
| 6858_1 | TATGAGTGTGGAGTCGTTGCTAAGGAACTGAAGCAGAGATCCTNNNAGATCGGAA |

| | |
|---|---|
| 6860_1 | TATGAGTGTGGAGTCGTTGCGGGTTCTTACTCTGACAAGCCAGNNNAGATCGGAAG |
| 6861_1 | TATGAGTGTGGAGTCGTTGCCTCTCCAGTATGGACCCATTTTTNNNAGATCGGAAGA |
| 6863_1 | TATGAGTGTGGAGTCGTTGCTCCAAATCCACAGATTCAGCCAGNNNAGATCGGAAG |
| 6865_1 | TATGAGTGTGGAGTCGTTGCTATCCAATTCCCAAACAGGCCAGNNNAGATCGGAAG |
| 6866_1 | TATGAGTGTGGAGTCGTTGCGCCTCTCTCCTCCTTCCTTTCCANNNAGATCGGAAGAG |
| 6868_1 | TATGAGTGTGGAGTCGTTGCGAAATAGACCCAATCTGGGCCAGNNNAGATCGGAA |
| 6873_1 | TATGAGTGTGGAGTCGTTGCTTCAAGTCCCAAATTCCCTCCAGNNNAGATCGGAAGA |
| 6875_1 | TATGAGTGTGGAGTCGTTGCTTCTCCTCCCCTTCTTTCTCCAGNNNAGATCGGAAGAG |
| 6876_1 | TATGAGTGTGGAGTCGTTGCTTCCCTTGGACTAAGAGGTCCAGNNNAGATCGGAAG |
| 6880_1 | TATGAGTGTGGAGTCGTTGCAGAAAGGGGATTGTGTGCATGTTNNNAGATCGGAA |
| 6885_1 | TATGAGTGTGGAGTCGTTGCCTCGGGTCATACTGGTTTTCCAGNNNAGATCGGAAG |
| 6886_1 | TATGAGTGTGGAGTCGTTGCTCTGGACCATCCCTCTAGTAGGTNNNAGATCGGAAG |
| 6887_1 | TATGAGTGTGGAGTCGTTGCGCAAAGCAAAGAAAACACAGCAGNNNAGATCGGAA |
| 6888_1 | TATGAGTGTGGAGTCGTTGCCAGCAAAATCCTTTTCCCAGCAGNNNAGATCGGAAG |
| 6889_1 | TATGAGTGTGGAGTCGTTGCCCACAGAATTTCAAAGCCAGCAGNNNAGATCGGAAG |
| 6890_1 | TATGAGTGTGGAGTCGTTGCAAATATGAAGTCCCAGCCAGCAGNNNAGATCGGAA |
| 6891_1 | TATGAGTGTGGAGTCGTTGCATGTACCAGCATTTCTCCAGCAGNNNAGATCGGAAG |
| 6892_1 | TATGAGTGTGGAGTCGTTGCAGGTTCAAGTACAAGAGCAGCAGNNNAGATCGGAA |
| 6893_1 | TATGAGTGTGGAGTCGTTGCACCAATGCCTGGCTCATTTCGTANNNAGATCGGAAG |
| 6895_1 | TATGAGTGTGGAGTCGTTGCGTTGTGAAGTAGGACCTCAGCAGNNNAGATCGGAA |
| 6896_1 | TATGAGTGTGGAGTCGTTGCAAACAGATGAAGAGCCTCAGCAGNNNAGATCGGAA |
| 6898_1 | TATGAGTGTGGAGTCGTTGCAACTTTGATTTGCGTGTCAGCAGNNNAGATCGGAAG |
| 6900_1 | TATGAGTGTGGAGTCGTTGCAAAGAAACCAGACTGCAGAGCAGNNNAGATCGGAA |
| 6902_1 | TATGAGTGTGGAGTCGTTGCCGTACTGCTAGAGAGTTGAGCAGNNNAGATCGGAA |
| 6904_1 | TATGAGTGTGGAGTCGTTGCTCGGTCTTGACATCCAAAGCAAANNNAGATCGGAAG |
| 6905_1 | TATGAGTGTGGAGTCGTTGCGGAAGAAAGTTGGCCTTTAGCAGNNNAGATCGGAA |
| 6906_1 | TATGAGTGTGGAGTCGTTGCATTATAGCCAGAGATGGACGCAGNNNAGATCGGAA |
| 6907_1 | TATGAGTGTGGAGTCGTTGCTGTAGTGTCCATCATTTCCGCAGNNNAGATCGGAAG |
| 6909_1 | TATGAGTGTGGAGTCGTTGCAAGGTCTGCAAAGGAAAAGGCAGNNNAGATCGGAA |
| 6910_1 | TATGAGTGTGGAGTCGTTGCGAGAAGAGACAGTGTGAAGGCAGNNNAGATCGGA |
| 6911_1 | TATGAGTGTGGAGTCGTTGCAGAGAAGGGATAAGTCCAGGCAGNNNAGATCGGA |
| 6912_1 | TATGAGTGTGGAGTCGTTGCCGAGGAAAAAGAGATGGAGGCAGNNNAGATCGGA |
| 6913_1 | TATGAGTGTGGAGTCGTTGCCAGCTTCAAAATTCACAGGGCAGNNNAGATCGGAAG |
| 6914_1 | TATGAGTGTGGAGTCGTTGCGGAGTTTCTCATGGTTAGGGCAGNNNAGATCGGAA |
| 6915_1 | TATGAGTGTGGAGTCGTTGCTAGCAGGGTGTGATAAATGGCAGNNNAGATCGGAA |
| 6916_1 | TATGAGTGTGGAGTCGTTGCACTGATAGGCAAAAGACTGGCAGNNNAGATCGGAA |
| 6918_1 | TATGAGTGTGGAGTCGTTGCGCACATCACCACGACTGGTAATTNNNAGATCGGAAG |
| 6920_1 | TATGAGTGTGGAGTCGTTGCACCTTTCTTCTCCCATTTGGCAGNNNAGATCGGAAGA |
| 6922_1 | TATGAGTGTGGAGTCGTTGCAGTGTCCTATGAACAGCATGCAGNNNAGATCGGAAG |
| 6923_1 | TATGAGTGTGGAGTCGTTGCGTTGGGAGAACAGACCTATGCAGNNNAGATCGGAA |
| 6925_1 | TATGAGTGTGGAGTCGTTGCTCTAGGGAAGATGCAAACTGCAGNNNAGATCGGAA |
| 6926_1 | TATGAGTGTGGAGTCGTTGCCACTGTTTTCCAGGGTGGAGCTTNNNAGATCGGAAG |
| 6929_1 | TATGAGTGTGGAGTCGTTGCCCTCTCTTCCCTCCTTTTGAAGTNNNAGATCGGAAGA |

| | |
|---|---|
| 6930_1 | TATGAGTGTGGAGTCGTTGCTTGAAGCAATCTCTGTAGTGGCANNNAGATCGGAAG |
| 6931_1 | TATGAGTGTGGAGTCGTTGCTTAGAATAGCAAAGGGGCTGCAGNNNAGATCGGAA |
| 6933_1 | TATGAGTGTGGAGTCGTTGCGTGGTTGTGGTGAAATTCTGCAGNNNAGATCGGAA |
| 6935_1 | TATGAGTGTGGAGTCGTTGCTAGGCGGAATTCGATAAGTGCAGNNNAGATCGGAA |
| 6936_1 | TATGAGTGTGGAGTCGTTGCGTTGCCAGGCTGATCTCAAACCTNNNAGATCGGAAG |
| 6938_1 | TATGAGTGTGGAGTCGTTGCGAGTTCTGACTGAGAAGGTGCAGNNNAGATCGGAA |
| 6941_1 | TATGAGTGTGGAGTCGTTGCGCAGGCAGAAGGATAATTTGCAGNNNAGATCGGAA |
| 6942_1 | TATGAGTGTGGAGTCGTTGCTTAGTGGCATTTGGCATTTGCAGNNNAGATCGGAAG |
| 6943_1 | TATGAGTGTGGAGTCGTTGCGATGGGGTTTGGCTCTCTGTTAGNNNAGATCGGAA |
| 6944_1 | TATGAGTGTGGAGTCGTTGCTTCGGCCATCCTTTCTGAATCAGNNNAGATCGGAAG |
| 6945_1 | TATGAGTGTGGAGTCGTTGCTACAGCAAGAAGGCAGTGGTCATNNNAGATCGGAA |
| 6946_1 | TATGAGTGTGGAGTCGTTGCTGGTTTCATTGATCGCTGTCCCTNNNAGATCGGAAG |
| 6948_1 | TATGAGTGTGGAGTCGTTGCTGAATGAGGCACACTTGGATCAGNNNAGATCGGAA |
| 6951_1 | TATGAGTGTGGAGTCGTTGCGAGCAGTAGGCACAGGTTATCAGNNNAGATCGGAA |
| 6952_1 | TATGAGTGTGGAGTCGTTGCGCCTGGTACACACTAACACTCAGNNNAGATCGGAAG |
| 6953_1 | TATGAGTGTGGAGTCGTTGCCTGGCTTTCCAAAAAGCACTCAGNNNAGATCGGAAG |
| 6958_1 | TATGAGTGTGGAGTCGTTGCGCTCTGGAACTTTCATCCCTCAGNNNAGATCGGAAG |
| 6959_1 | TATGAGTGTGGAGTCGTTGCAGAAAACATCAGAGCAGCCTCAGNNNAGATCGGAA |
| 6960_1 | TATGAGTGTGGAGTCGTTGCGGGTAACTGCTAAAGAGCCTCAGNNNAGATCGGAA |
| 6961_1 | TATGAGTGTGGAGTCGTTGCAGTTTGAGACCCAAGGTACTGGGNNNAGATCGGAA |
| 6964_1 | TATGAGTGTGGAGTCGTTGCTTGGTTGTGTCTACAGAGCTCAGNNNAGATCGGAAG |
| 6966_1 | TATGAGTGTGGAGTCGTTGCGATACAAATGTCCCAGGGCTCAGNNNAGATCGGAAG |
| 6967_1 | TATGAGTGTGGAGTCGTTGCCCTGCTTTCTTCTATGGGCTCAGNNNAGATCGGAAG |
| 6970_1 | TATGAGTGTGGAGTCGTTGCACCCAGTGATTCTCCCATCTCAGNNNAGATCGGAAGA |
| 6973_1 | TATGAGTGTGGAGTCGTTGCTTTTGGGGGTTCAGACAAGTCAGNNNAGATCGGAA |
| 6974_1 | TATGAGTGTGGAGTCGTTGCTGTGATGTTATGCACCCAGTCAGNNNAGATCGGAAG |
| 6978_1 | TATGAGTGTGGAGTCGTTGCCCAGGTTCCAGTACAATGGTCAGNNNAGATCGGAAG |
| 6979_1 | TATGAGTGTGGAGTCGTTGCGAGAGCACCTCAGATTTGGTCAGNNNAGATCGGAA |
| 6980_1 | TATGAGTGTGGAGTCGTTGCATCTTCTGAGCACACAGTGAGACNNNAGATCGGAAG |
| 6981_1 | TATGAGTGTGGAGTCGTTGCTTTTGCTAGGGTCTTGCTGTCAGNNNAGATCGGAAG |
| 6982_1 | TATGAGTGTGGAGTCGTTGCCTCTCTCTCTCTCGGAGTGTCAGNNNAGATCGGAAG |
| 6983_1 | TATGAGTGTGGAGTCGTTGCTCCCCTTTCTATGAGGGTGTCAGNNNAGATCGGAAG |
| 6985_1 | TATGAGTGTGGAGTCGTTGCCGGCCTGAACAAAAGACATTCAGNNNAGATCGGAAG |
| 6986_1 | TATGAGTGTGGAGTCGTTGCGCTTCTTACCCTCACCCATTCAGNNNAGATCGGAAGA |
| 6988_1 | TATGAGTGTGGAGTCGTTGCAGCACCTTGTGATGTGGATTCAGNNNAGATCGGAAG |
| 6989_1 | TATGAGTGTGGAGTCGTTGCGTCTCCTCTACCTCCCACTTCAGNNNAGATCGGAAGA |
| 6990_1 | TATGAGTGTGGAGTCGTTGCGTTTCACTGTTGTTGCCCTTCAGNNNAGATCGGAAG |
| 6991_1 | TATGAGTGTGGAGTCGTTGCTGGGTTCATGAGTTCTGCTTCAGNNNAGATCGGAAG |
| 6996_1 | TATGAGTGTGGAGTCGTTGCCCTGTGTCTGCAGTACCTTTCAGNNNAGATCGGAAG |
| 6997_1 | TATGAGTGTGGAGTCGTTGCTGTGAGAAATTCTGTGAGGCTGCNNNAGATCGGAA |
| 6999_1 | TATGAGTGTGGAGTCGTTGCCCCACCCCCAAATCATTTTTCAGNNNAGATCGGAAGA |
| 7000_1 | TATGAGTGTGGAGTCGTTGCGGGGGAGAAACCAAGGAAAAGAGNNNAGATCGGA |
| 7001_1 | TATGAGTGTGGAGTCGTTGCACAGCAGATAGTGGGGAAAAGAGNNNAGATCGGA |

7002_1 TATGAGTGTGGAGTCGTTGCAGAGGGTAGGGCAGATCAAAGAGNNNAGATCGGA
7003_1 TATGAGTGTGGAGTCGTTGCTGGGGGTACAGAGACAGAAAGAGNNNAGATCGGA
7004_1 TATGAGTGTGGAGTCGTTGCGGCGCATGTCTAGAGAGAAAGAGNNNAGATCGGA
7005_1 TATGAGTGTGGAGTCGTTGCTATGCAGGGAGAGGAGGAAAGAGNNNAGATCGGA
7006_1 TATGAGTGTGGAGTCGTTGCGAAGGGAGGGTAGAGGGAAAGAGNNNAGATCGG
7007_1 TATGAGTGTGGAGTCGTTGCGCGGGAAGGGTGATTCTAAAGAGNNNAGATCGGA
7008_1 TATGAGTGTGGAGTCGTTGCGTTTTTGCTAGGGTCTTGCTGTCNNNAGATCGGAAG
7010_1 TATGAGTGTGGAGTCGTTGCGCAAGGCACCCTATACACAAGAGNNNAGATCGGAA
7012_1 TATGAGTGTGGAGTCGTTGCCCCCCTTCCAAAGATGGAGAGTANNNAGATCGGAAG
7013_1 TATGAGTGTGGAGTCGTTGCACAGCTCCAACATACTGCAAGAGNNNAGATCGGAAG
7014_1 TATGAGTGTGGAGTCGTTGCAAGGGGTGGTTGAGGAAGAAGAGNNNAGATCGGA
7017_1 TATGAGTGTGGAGTCGTTGCTGGAAGGGAAGAGGTGAGAAGAGNNNAGATCGG
7023_1 TATGAGTGTGGAGTCGTTGCACACATGAACACAGGCTGAAGAGNNNAGATCGGAA
7025_1 TATGAGTGTGGAGTCGTTGCTCTGTCTTTCTTTGTCTGGCTCTNNNAGATCGGAAGA
7026_1 TATGAGTGTGGAGTCGTTGCGCCTGTCATCCTGTGTAACAGTTNNNAGATCGGAAG
7027_1 TATGAGTGTGGAGTCGTTGCGAACACAGCCCAGTCTAACAGAGNNNAGATCGGAA
7028_1 TATGAGTGTGGAGTCGTTGCATTAGGTGCTGACACACACAGAGNNNAGATCGGAA
7029_1 TATGAGTGTGGAGTCGTTGCCCTACATCCAGCTTGTCACAGAGNNNAGATCGGAAG
7030_1 TATGAGTGTGGAGTCGTTGCCCTGAATGGGGTGACCTCTTTGGNNNAGATCGGAA
7031_1 TATGAGTGTGGAGTCGTTGCATCAAGGTGGCTGCTTTACAGAGNNNAGATCGGAA
7032_1 TATGAGTGTGGAGTCGTTGCTAATGCTGCCATGAACACCAGAGNNNAGATCGGAAG
7035_1 TATGAGTGTGGAGTCGTTGCAACTCCTCTTACCAGAGCCAGAGNNNAGATCGGAAG
7036_1 TATGAGTGTGGAGTCGTTGCCTTTATGGGACTGGATGCCAGAGNNNAGATCGGAA
7037_1 TATGAGTGTGGAGTCGTTGCAAATTGCTGTGGAGGATCCAGAGNNNAGATCGGAA
7038_1 TATGAGTGTGGAGTCGTTGCATGGGCTAGGCTCTTATCCAGAGNNNAGATCGGAA
7039_1 TATGAGTGTGGAGTCGTTGCTACTTCTGGGTGGTGTAATTCGCNNNAGATCGGAAG
7044_1 TATGAGTGTGGAGTCGTTGCCAGTGAGCCATGTGAGTGTANNNAGATCGGAAGAG
7046_1 TATGAGTGTGGAGTCGTTGCAGGGCATAAATCACAGTGCAGAGNNNAGATCGGAA
7048_1 TATGAGTGTGGAGTCGTTGCTGAGGGGATACTGCAGATCAGAGNNNAGATCGGAA
7049_1 TATGAGTGTGGAGTCGTTGCTCCTCCTCTCTTTCACGTCAGAGNNNAGATCGGAAGA
7051_1 TATGAGTGTGGAGTCGTTGCAAGGCAAACGCCTCAACTTTCTANNNAGATCGGAAG
7054_1 TATGAGTGTGGAGTCGTTGCGGGCCTGGAAACAGTAAAGAGAGNNNAGATCGGA
7055_1 TATGAGTGTGGAGTCGTTGCCCAGGAGGTAGGACTGAAGAGAGNNNAGATCGGA
7057_1 TATGAGTGTGGAGTCGTTGCTTGCTTCTCTGTGCACTTCTGTTNNNAGATCGGAAGA
7058_1 TATGAGTGTGGAGTCGTTGCCTGGATCTCTTCCTCTAAAACCCNNNAGATCGGAAGA
7059_1 TATGAGTGTGGAGTCGTTGCGGCAAGGGGAGAAAAAGAGAGAGNNNAGATCGG
7060_1 TATGAGTGTGGAGTCGTTGCTGAGCTGGGAGAGAAAGAGAGAGNNNAGATCGGA
7062_1 TATGAGTGTGGAGTCGTTGCAGACATGCAAGGGAGAGAGAGAGNNNAGATCGGA
7063_1 TATGAGTGTGGAGTCGTTGCACCCATCAATCTAAAGGCGAGAGNNNAGATCGGAA
7065_1 TATGAGTGTGGAGTCGTTGCAGGTGCTGTGTTTTCAAGGAGAGNNNAGATCGGAA
7068_1 TATGAGTGTGGAGTCGTTGCAGCAGAAATACCCAGCTGGAGAGNNNAGATCGGAA
7070_1 TATGAGTGTGGAGTCGTTGCGGAAGAAGTACAGCCACTGAGAGNNNAGATCGGAA
7071_1 TATGAGTGTGGAGTCGTTGCAGCATTTGATGGATGGCTGAGAGNNNAGATCGGAA

| | |
|---|---|
| 7072_1 | TATGAGTGTGGAGTCGTTGCGCTGCTCAGACTTACAGTGAGAGNNNAGATCGGAA |
| 7073_1 | TATGAGTGTGGAGTCGTTGCTTCCCGCTTTGTAACTGTGAGAGNNNAGATCGGAAG |
| 7076_1 | TATGAGTGTGGAGTCGTTGCCGTGTGTGTGAGGGTAAGTAGAGNNNAGATCGGA |
| 7077_1 | TATGAGTGTGGAGTCGTTGCCGTTGAGGTTCCCACTGTTAGAGNNNAGATCGGAA |
| 7081_1 | TATGAGTGTGGAGTCGTTGCCTGATTCCAACTCAACCTCCGAGNNNAGATCGGAAG |
| 7082_1 | TATGAGTGTGGAGTCGTTGCATAAGTTTTAGCCCACAGGCGAGNNNAGATCGGAA |
| 7083_1 | TATGAGTGTGGAGTCGTTGCGGGAAGATAAAGAAGGGTGCGAGNNNAGATCGGA |
| 7084_1 | TATGAGTGTGGAGTCGTTGCCTTGGAGTGCGGTGGCATGATTANNNAGATCGGAA |
| 7085_1 | TATGAGTGTGGAGTCGTTGCTCCAATCAGTTGCCTTCTTCGAGNNNAGATCGGAAG |
| 7086_1 | TATGAGTGTGGAGTCGTTGCAGGCAAAGGAAGACCAAAAGGAGNNNAGATCGGA |
| 7088_1 | TATGAGTGTGGAGTCGTTGCGGATTGGAAGGTTCGAGAAGGAGNNNAGATCGGA |
| 7089_1 | TATGAGTGTGGAGTCGTTGCTGTGTGTGAGTAGAGGGAAGGAGNNNAGATCGGA |
| 7092_1 | TATGAGTGTGGAGTCGTTGCCACCATCATTATCACCACCTGGGNNNAGATCGGAAG |
| 7093_1 | TATGAGTGTGGAGTCGTTGCTCTCTCTCTCTCTTGATCGACAGNNNAGATCGGAAGA |
| 7094_1 | TATGAGTGTGGAGTCGTTGCCAGCTATGGCACTTTCTCAGGAGNNNAGATCGGAAG |
| 7096_1 | TATGAGTGTGGAGTCGTTGCGAGAATTTGGAGGCAAGGAGGAGNNNAGATCGGA |
| 7097_1 | TATGAGTGTGGAGTCGTTGCCTTTCCTGTGCCGATCAACAGTANNNAGATCGGAAG |
| 7099_1 | TATGAGTGTGGAGTCGTTGCTTACCTGTTCCTCACTGGAGGAGNNNAGATCGGAAG |
| 7100_1 | TATGAGTGTGGAGTCGTTGCGTGAAAGAGTTCAGCTGGAGGAGNNNAGATCGGA |
| 7103_1 | TATGAGTGTGGAGTCGTTGCTTACTTGGGGCCTGAGATAGGAGNNNAGATCGGAA |
| 7106_1 | TATGAGTGTGGAGTCGTTGCGGTGCAGGTTTGCTTTTTAGGAGNNNAGATCGGAA |
| 7108_1 | TATGAGTGTGGAGTCGTTGCAAAAAGAAGGGCAGAAGACGGAGNNNAGATCGGA |
| 7109_1 | TATGAGTGTGGAGTCGTTGCAGGTAGAACTCTGATCACCGGAGNNNAGATCGGAA |
| 7110_1 | TATGAGTGTGGAGTCGTTGCTTTGAGTGACTTTCTCTGCGGAGNNNAGATCGGAAG |
| 7112_1 | TATGAGTGTGGAGTCGTTGCAGTGTTGAGTAAGGGACAGGGAGNNNAGATCGGA |
| 7113_1 | TATGAGTGTGGAGTCGTTGCCAGAAACAGAGCTAACCAGGGAGNNNAGATCGGAA |
| 7114_1 | TATGAGTGTGGAGTCGTTGCAGAATAAAATGGCTGGCAGGGAGNNNAGATCGGA |
| 7115_1 | TATGAGTGTGGAGTCGTTGCATGTGGGAAGTGATCCTAGGGAGNNNAGATCGGA |
| 7117_1 | TATGAGTGTGGAGTCGTTGCTGTGGTCAAAGTGGTTTAGGGAGNNNAGATCGGA |
| 7118_1 | TATGAGTGTGGAGTCGTTGCGTTACAGGGTCTTGCTCTTTCCANNNAGATCGGAAG |
| 7119_1 | TATGAGTGTGGAGTCGTTGCCATTTGCCTAAAACAGTGGCTGTNNNAGATCGGAAG |
| 7122_1 | TATGAGTGTGGAGTCGTTGCGGCTTTGATGGTAAGGATGGGAGNNNAGATCGGA |
| 7125_1 | TATGAGTGTGGAGTCGTTGCTGAATGAAGCCCTGAATTGGGAGNNNAGATCGGAA |
| 7127_1 | TATGAGTGTGGAGTCGTTGCGTCTCTCCCCCTTTCACATTTGTNNNAGATCGGAAGA |
| 7128_1 | TATGAGTGTGGAGTCGTTGCAGTTGTGGTAGGCCAAAATGGAGNNNAGATCGGA |
| 7129_1 | TATGAGTGTGGAGTCGTTGCGGAAGAAAATCCCAGCATTCACGNNNAGATCGGAA |
| 7130_1 | TATGAGTGTGGAGTCGTTGCCCACACACTAACTCACCATGGAGNNNAGATCGGAAG |
| 7131_1 | TATGAGTGTGGAGTCGTTGCTGCGAAGCATACGAGAGTTCAAANNNAGATCGGAA |
| 7132_1 | TATGAGTGTGGAGTCGTTGCCACCCTCTTTACACACACTGGAGNNNAGATCGGAAG |
| 7138_1 | TATGAGTGTGGAGTCGTTGCCTTGTTGAGAATGGAGGCTGGAGNNNAGATCGGA |
| 7139_1 | TATGAGTGTGGAGTCGTTGCTACTTTTTGAGACAGGGCTGGAGNNNAGATCGGAA |
| 7142_1 | TATGAGTGTGGAGTCGTTGCCAACATGATAAACTGCAGTGCAANNNAGATCGGAA |
| 7144_1 | TATGAGTGTGGAGTCGTTGCGACCATTTTTGTGGGCAGGGAAGNNNAGATCGGAA |

| | |
|---|---|
| 7145_1 | TATGAGTGTGGAGTCGTTGCAGAGATTGGAAGGGAAGACGTCTNNNAGATCGGAA |
| 7146_1 | TATGAGTGTGGAGTCGTTGCAAAATAGCCTAGCATAGTGCTGCNNNAGATCGGAAG |
| 7147_1 | TATGAGTGTGGAGTCGTTGCAAGAAAATGGACTTGGGGTGGAGNNNAGATCGGAA |
| 7148_1 | TATGAGTGTGGAGTCGTTGCTTCTTGGCCTCGCTCTGTCATTCNNNAGATCGGAAGA |
| 7149_1 | TATGAGTGTGGAGTCGTTGCCTTTCCAGTTGAGGTCTCACGTGNNNAGATCGGAAG |
| 7150_1 | TATGAGTGTGGAGTCGTTGCCCCCTTTTAGCCTCTTTGTGGAGNNNAGATCGGAAGA |
| 7151_1 | TATGAGTGTGGAGTCGTTGCCTGTGAATGCATGTACCCCTCTGNNNAGATCGGAAGA |
| 7152_1 | TATGAGTGTGGAGTCGTTGCCTCTGTCTCTCAGGACAACTAGANNNAGATCGGAAGA |
| 7153_1 | TATGAGTGTGGAGTCGTTGCCGCAGTGAGTGGAAGTAGCCTGANNNAGATCGGAA |
| 7154_1 | TATGAGTGTGGAGTCGTTGCTTTCCAGGGTCTTGCTCAATTGCNNNAGATCGGAAGA |
| 7155_1 | TATGAGTGTGGAGTCGTTGCGATTGCCAGCTTTCGGAAATGAGNNNAGATCGGAAG |
| 7160_1 | TATGAGTGTGGAGTCGTTGCCTGACATAGGTGGATCGCATGAGNNNAGATCGGAAG |
| 7161_1 | TATGAGTGTGGAGTCGTTGCGGGAAAGGCTTCACAGAGATGAGNNNAGATCGGAA |
| 7162_1 | TATGAGTGTGGAGTCGTTGCCTGTGTTGGGAAAGGGAGATGAGNNNAGATCGGAA |
| 7164_1 | TATGAGTGTGGAGTCGTTGCGAGCTACAAATGGTGGGGATGAGNNNAGATCGGAA |
| 7167_1 | TATGAGTGTGGAGTCGTTGCTTATTGTGACCTGGCCCAGAGTANNNAGATCGGAAG |
| 7168_1 | TATGAGTGTGGAGTCGTTGCGTCAAGTGGCTACCAGAACTGAGNNNAGATCGGAAG |
| 7169_1 | TATGAGTGTGGAGTCGTTGCCCAACCCACCCATAACGATCATTNNNAGATCGGAAGA |
| 7172_1 | TATGAGTGTGGAGTCGTTGCCACACCTTCTCCAAGTCACTGAGNNNAGATCGGAAGA |
| 7174_1 | TATGAGTGTGGAGTCGTTGCGACACAGTGAGAAGGCAAGCTAANNNAGATCGGAA |
| 7175_1 | TATGAGTGTGGAGTCGTTGCATGGCAGTTGTAAGCTCCCTGAGNNNAGATCGGAAG |
| 7176_1 | TATGAGTGTGGAGTCGTTGCAAGGGGCATAGATAAGGATGGTCNNNAGATCGGAA |
| 7177_1 | TATGAGTGTGGAGTCGTTGCCTGTTCCAGTGGGATCTCCTGAGNNNAGATCGGAAG |
| 7178_1 | TATGAGTGTGGAGTCGTTGCTTTGGTCCATTGGTGTTCCTGAGNNNAGATCGGAAG |
| 7181_1 | TATGAGTGTGGAGTCGTTGCGTAAACAGAGGTTCTGGGCTGAGNNNAGATCGGAA |
| 7182_1 | TATGAGTGTGGAGTCGTTGCACTGCTCATGACTTCATGCTGAGNNNAGATCGGAAG |
| 7185_1 | TATGAGTGTGGAGTCGTTGCATTCTTACGCTGTGGCATCTGAGNNNAGATCGGAAG |
| 7186_1 | TATGAGTGTGGAGTCGTTGCTTTCTTGGGGCAACTCATCTGAGNNNAGATCGGAAG |
| 7187_1 | TATGAGTGTGGAGTCGTTGCTTGTCCGTGCATGTGTATCTGAGNNNAGATCGGAAG |
| 7190_1 | TATGAGTGTGGAGTCGTTGCATCACACTTCCGTGGTTTCTGAGNNNAGATCGGAAGA |
| 7191_1 | TATGAGTGTGGAGTCGTTGCCCCGAACTGTGGAAGTAAGTGAGNNNAGATCGGAA |
| 7192_1 | TATGAGTGTGGAGTCGTTGCGCAGGAAGACATACCACAGTGAGNNNAGATCGGAA |
| 7193_1 | TATGAGTGTGGAGTCGTTGCTTTTGAATGAGGCCTCGAGTGAGNNNAGATCGGAAG |
| 7194_1 | TATGAGTGTGGAGTCGTTGCCCCTCTGTCCAAGACACAAACTCNNNAGATCGGAAGA |
| 7195_1 | TATGAGTGTGGAGTCGTTGCAAAGCAAGCTGAGATGAGGTGAGNNNAGATCGGAA |
| 7196_1 | TATGAGTGTGGAGTCGTTGCGAAAATCCTGGGCACTAGGTGAGNNNAGATCGGAA |
| 7197_1 | TATGAGTGTGGAGTCGTTGCGAGCCGATCCTTCCACCATTTACNNNAGATCGGAAGA |
| 7198_1 | TATGAGTGTGGAGTCGTTGCCCTGGATCCTCTTGCTTAGATCCNNNAGATCGGAAGA |
| 7199_1 | TATGAGTGTGGAGTCGTTGCTACCTGAACTCTCCAAGGGTGAGNNNAGATCGGAAG |
| 7201_1 | TATGAGTGTGGAGTCGTTGCCAGTAAAGTCAGGGCACTGTGAGNNNAGATCGGAA |
| 7202_1 | TATGAGTGTGGAGTCGTTGCCTCTTGCACAACTTTGCTGTGAGNNNAGATCGGAAGA |
| 7204_1 | TATGAGTGTGGAGTCGTTGCTGGTGGATAGAGGTACGTGTGAGNNNAGATCGGAA |
| 7206_1 | TATGAGTGTGGAGTCGTTGCGTGTGTGTGTGTTGGACAGTTATNNNAGATCGGAAG |

| | |
|---|---|
| 7207_1 | TATGAGTGTGGAGTCGTTGCGAGCTTCCTACCACTGTTGTGAGNNNAGATCGGAAG |
| 7208_1 | TATGAGTGTGGAGTCGTTGCGACTGGAGCAGAGGGAAATTGAGNNNAGATCGGA |
| 7209_1 | TATGAGTGTGGAGTCGTTGCGCCCTCTACTTGGACACATTGAGNNNAGATCGGAAG |
| 7210_1 | TATGAGTGTGGAGTCGTTGCGGTTGGAACTGGAGGTCATTGAGNNNAGATCGGA |
| 7213_1 | TATGAGTGTGGAGTCGTTGCTTCCTACCTGGGAGATCCTTGAGNNNAGATCGGAAG |
| 7217_1 | TATGAGTGTGGAGTCGTTGCGCCCCATCCCCAGATTTAGATAANNNAGATCGGAAG |
| 7218_1 | TATGAGTGTGGAGTCGTTGCGTTTGGTTTTGTTGCTTGCAATNNNAGATCGGAAGA |
| 7220_1 | TATGAGTGTGGAGTCGTTGCGGTTACAGGCTGCTCTCTTTGAGNNNAGATCGGAAG |
| 7221_1 | TATGAGTGTGGAGTCGTTGCCCTCCCTTGACCATCCTTTTGAGNNNAGATCGGAAGA |
| 7223_1 | TATGAGTGTGGAGTCGTTGCCCTCCCACATCGGCCTATCAAATNNNAGATCGGAAG |
| 7224_1 | TATGAGTGTGGAGTCGTTGCTGCTGAACCAGCCTCTCAAATAGNNNAGATCGGAAG |
| 7225_1 | TATGAGTGTGGAGTCGTTGCAGGGGGTCTGGGGAAAGAAATAGNNNAGATCGGA |
| 7230_1 | TATGAGTGTGGAGTCGTTGCCCAACCCATTTACCCTTCTCCTTNNNAGATCGGAAGA |
| 7231_1 | TATGAGTGTGGAGTCGTTGCATTGCTCCTGCATCTCTGCATAGNNNAGATCGGAAG |
| 7234_1 | TATGAGTGTGGAGTCGTTGCGCTCTCCATATTCTGGGGTTCCANNNAGATCGGAAG |
| 7235_1 | TATGAGTGTGGAGTCGTTGCGAGTAGTGCCAAGAAGCGGATAGNNNAGATCGGA |
| 7238_1 | TATGAGTGTGGAGTCGTTGCGCCTGTGGAACTGTTAGTCCTTANNNAGATCGGAAG |
| 7239_1 | TATGAGTGTGGAGTCGTTGCCACAACCACACACCTGGCTATAGNNNAGATCGGAAG |
| 7241_1 | TATGAGTGTGGAGTCGTTGCACTGTGCTGCGAGAATGCTATAGNNNAGATCGGAA |
| 7242_1 | TATGAGTGTGGAGTCGTTGCGGCTGGAGCAAATCCGGTATAGNNNAGATCGGAAG |
| 7243_1 | TATGAGTGTGGAGTCGTTGCAATCCCCAAGAGCCCCTGTATAGNNNAGATCGGAAG |
| 7244_1 | TATGAGTGTGGAGTCGTTGCGGATGGGCTTGGAGACAAACTAGNNNAGATCGGA |
| 7245_1 | TATGAGTGTGGAGTCGTTGCATCGGCAGCTCAAACTCAACTAGNNNAGATCGGAAG |
| 7246_1 | TATGAGTGTGGAGTCGTTGCGGGTGATCAGAGCAGAAACCTAGNNNAGATCGGAA |
| 7247_1 | TATGAGTGTGGAGTCGTTGCTCTTTGGGTAACAGTCCCCCTAGNNNAGATCGGAAG |
| 7248_1 | TATGAGTGTGGAGTCGTTGCGCAAACTCACACTGTTCCCCTAGNNNAGATCGGAAG |
| 7249_1 | TATGAGTGTGGAGTCGTTGCGTGGAAGCTGAAAGCCAGGATAANNNAGATCGGA |
| 7250_1 | TATGAGTGTGGAGTCGTTGCCAGGCTCTCAGATCCCATCCTAGNNNAGATCGGAAG |
| 7253_1 | TATGAGTGTGGAGTCGTTGCTGGTGCTTTTTCCCAAGTCCTAGNNNAGATCGGAAG |
| 7256_1 | TATGAGTGTGGAGTCGTTGCACAGATGAACAGGCACAAGCTAGNNNAGATCGGAA |
| 7257_1 | TATGAGTGTGGAGTCGTTGCCGCTCACATAAATGACTGGTTTGNNNAGATCGGAAG |
| 7258_1 | TATGAGTGTGGAGTCGTTGCCAACTGCCTCTTGGTACAGAGACNNNAGATCGGAAG |
| 7259_1 | TATGAGTGTGGAGTCGTTGCTCATGGCATTCACCAGAATCCTGNNNAGATCGGAAG |
| 7261_1 | TATGAGTGTGGAGTCGTTGCAGTTTTAGGAGCCTGAGGGTTTTNNNAGATCGGAA |
| 7263_1 | TATGAGTGTGGAGTCGTTGCATGAAATGGGATTGGCTGGCTAGNNNAGATCGGAA |
| 7264_1 | TATGAGTGTGGAGTCGTTGCCAGCTTTCTCTGGTTCTGGCTAGNNNAGATCGGAAG |
| 7265_1 | TATGAGTGTGGAGTCGTTGCGCAGGATGAAGAACTGTGGCTAGNNNAGATCGGA |
| 7267_1 | TATGAGTGTGGAGTCGTTGCAACTAGAGCCCTGAGGATGCTAGNNNAGATCGGAA |
| 7270_1 | TATGAGTGTGGAGTCGTTGCTTTAGGAGTATAGCTGCAGTGGGNNNAGATCGGAA |
| 7271_1 | TATGAGTGTGGAGTCGTTGCCACCACCAGCATGTTTTTGCTAGNNNAGATCGGAAG |
| 7273_1 | TATGAGTGTGGAGTCGTTGCAAACAAAAACGGAGGCCCTCTAGNNNAGATCGGAA |
| 7275_1 | TATGAGTGTGGAGTCGTTGCCTGTGCTCCTCTTGCAGTTAGTTNNNAGATCGGAAG |
| 7278_1 | TATGAGTGTGGAGTCGTTGCGGAGGGGTGGTCGATTTTTCTAGNNNAGATCGGAA |

| | |
|---|---|
| 7280_1 | TATGAGTGTGGAGTCGTTGCGCTTCTTTGGGAAGGGGAAGTAGNNNAGATCGGAA |
| 7282_1 | TATGAGTGTGGAGTCGTTGCGCCAACTGTGCTTCTACCAGTAGNNNAGATCGGAAG |
| 7283_1 | TATGAGTGTGGAGTCGTTGCAGTGTGGTAGTATGCGCCAGTAGNNNAGATCGGAA |
| 7286_1 | TATGAGTGTGGAGTCGTTGCGATGGGTGAGGCAATGTCAGTAGNNNAGATCGGA |
| 7289_1 | TATGAGTGTGGAGTCGTTGCACTCCAAATGCTGTAAGGCGTAGNNNAGATCGGAA |
| 7290_1 | TATGAGTGTGGAGTCGTTGCTTTTTGCTGGGGAACAAGATCTTNNNAGATCGGAAG |
| 7291_1 | TATGAGTGTGGAGTCGTTGCAGAAAGCACAAAGGAGCAGGTAGNNNAGATCGGA |
| 7296_1 | TATGAGTGTGGAGTCGTTGCGGCTTTTCAGTGACCACAGCTATNNNAGATCGGAAG |
| 7300_1 | TATGAGTGTGGAGTCGTTGCCCCCCACCACCTTTTATTGGTAGNNNAGATCGGAAG |
| 7301_1 | TATGAGTGTGGAGTCGTTGCGGGCCTGTACTGGTAGCATGTAGNNNAGATCGGAA |
| 7303_1 | TATGAGTGTGGAGTCGTTGCACCGGAAGGAAGTAGGACTGTAGNNNAGATCGGA |
| 7304_1 | TATGAGTGTGGAGTCGTTGCTACACTCTCCCACCCTCCTGTAGNNNAGATCGGAAGA |
| 7306_1 | TATGAGTGTGGAGTCGTTGCCCCCTGCTCTCTACTCTCTGTAGNNNAGATCGGAAGA |
| 7308_1 | TATGAGTGTGGAGTCGTTGCAGGCCCACAGTGAAATACAGTTANNNAGATCGGAA |
| 7310_1 | TATGAGTGTGGAGTCGTTGCATCCTTTCTGCCTTGCCTTGTAGNNNAGATCGGAAG |
| 7314_1 | TATGAGTGTGGAGTCGTTGCCCCCTCCCTTCGAAATCCAATTTNNNAGATCGGAAGA |
| 7315_1 | TATGAGTGTGGAGTCGTTGCATTAGGGCTGGACAGGACCTTAGNNNAGATCGGAA |
| 7316_1 | TATGAGTGTGGAGTCGTTGCCCCTTTGGAGAGGACAAAGTTGCNNNAGATCGGAA |
| 7318_1 | TATGAGTGTGGAGTCGTTGCGGGTCAGAAAAGAGGTGGCTTAGNNNAGATCGGA |
| 7320_1 | TATGAGTGTGGAGTCGTTGCCACCACTGTGGCTTCCTTCTTAGNNNAGATCGGAAG |
| 7321_1 | TATGAGTGTGGAGTCGTTGCGAGAATGGGCAGGTGGAAGTTAGNNNAGATCGGA |
| 7322_1 | TATGAGTGTGGAGTCGTTGCGACCACCAATACAGGCAGGTTAGNNNAGATCGGAA |
| 7323_1 | TATGAGTGTGGAGTCGTTGCCCCCAAAAGCCTTGTATAGACAGNNNAGATCGGAAG |
| 7324_1 | TATGAGTGTGGAGTCGTTGCAAAAACCTCCAGTAGGGCAGTGTNNNAGATCGGAA |
| 7325_1 | TATGAGTGTGGAGTCGTTGCTTGTAATGCCAGGTCTTGCTCTGNNNAGATCGGAAG |
| 7328_1 | TATGAGTGTGGAGTCGTTGCGAGTGAACCAGGAGGTGCTTTAGNNNAGATCGGAA |
| 7329_1 | TATGAGTGTGGAGTCGTTGCTGGGGGAGGTGGAGAATCTTTAGNNNAGATCGGA |
| 7330_1 | TATGAGTGTGGAGTCGTTGCATGCAGAGGCACACAGTCTTTAGNNNAGATCGGAA |
| 7332_1 | TATGAGTGTGGAGTCGTTGCCCAGGGCACAACAAGCATTTTAGNNNAGATCGGAA |
| 7333_1 | TATGAGTGTGGAGTCGTTGCTATTGGGGCTGGAATGCAATGGCNNNAGATCGGAA |
| 7334_1 | TATGAGTGTGGAGTCGTTGCGACATTGCCCTTCCTCAAAAACGNNNAGATCGGAAG |
| 7335_1 | TATGAGTGTGGAGTCGTTGCCCTGTCGTTTGCAGTAAGTCGTANNNAGATCGGAAG |
| 7336_1 | TATGAGTGTGGAGTCGTTGCTCTCTTTGGCTGGAAGACAAACGNNNAGATCGGAA |
| 7337_1 | TATGAGTGTGGAGTCGTTGCAACGAATTGAGGCATGTCAAACGNNNAGATCGGAA |
| 7338_1 | TATGAGTGTGGAGTCGTTGCAATACAAGCATGTGAAAGACCTCNNNAGATCGGAA |
| 7339_1 | TATGAGTGTGGAGTCGTTGCTAAGCTCTTCAAGGGTGGAAACGNNNAGATCGGAA |
| 7340_1 | TATGAGTGTGGAGTCGTTGCCCAGATCTTACCCAAGACCAACGNNNAGATCGGAAG |
| 7341_1 | TATGAGTGTGGAGTCGTTGCGTTCTCAGGGAAGATCTCCAACGNNNAGATCGGAA |
| 7343_1 | TATGAGTGTGGAGTCGTTGCCTGTGCTCCACTAAAATGCAACGNNNAGATCGGAAG |
| 7345_1 | TATGAGTGTGGAGTCGTTGCCCTATTCCAACCATGACAGACCCNNNAGATCGGAAG |
| 7346_1 | TATGAGTGTGGAGTCGTTGCTGTCTTTGAACCACCTGAGAACGNNNAGATCGGAAG |
| 7350_1 | TATGAGTGTGGAGTCGTTGCGCCGAAGTAACTAGGAGAACACGNNNAGATCGGAA |
| 7351_1 | TATGAGTGTGGAGTCGTTGCCACGTTTATGCTGTACACACACGNNNAGATCGGAAG |

| | |
|---|---|
| 7354_1 | TATGAGTGTGGAGTCGTTGCCACACACACAGAGTTATGGTGATNNNAGATCGGAAG |
| 7355_1 | TATGAGTGTGGAGTCGTTGCTTGAGAGTTCAGCCATTACCACGNNNAGATCGGAAG |
| 7356_1 | TATGAGTGTGGAGTCGTTGCCAAAGTAGACTGACCCTTCCACGNNNAGATCGGAAG |
| 7358_1 | TATGAGTGTGGAGTCGTTGCCATACTCAACACTTCTTGGCACGNNNAGATCGGAAGA |
| 7361_1 | TATGAGTGTGGAGTCGTTGCCCTCCCATCACATAACTCTGCATNNNAGATCGGAAGA |
| 7362_1 | TATGAGTGTGGAGTCGTTGCGTGTTTCCAAGTAATGGCTCACGNNNAGATCGGAAG |
| 7363_1 | TATGAGTGTGGAGTCGTTGCCCTCATTCTGATAGGCAGTCACGNNNAGATCGGAAG |
| 7364_1 | TATGAGTGTGGAGTCGTTGCCGATCTCACCTTGTGGCAATCTCNNNAGATCGGAAGA |
| 7367_1 | TATGAGTGTGGAGTCGTTGCGGACATCTCATTGTGGTTTGTGTNNNAGATCGGAAG |
| 7371_1 | TATGAGTGTGGAGTCGTTGCCGAGAACCAAGTTGGCAACATTTNNNAGATCGGAAG |
| 7374_1 | TATGAGTGTGGAGTCGTTGCATTCTGTCCATTCGGGACATACGNNNAGATCGGAAG |
| 7375_1 | TATGAGTGTGGAGTCGTTGCCTCTCTCTCTTTGTGGCCATACGNNNAGATCGGAAGA |
| 7376_1 | TATGAGTGTGGAGTCGTTGCCCTTTGTGAGTGGCTAGACTACGNNNAGATCGGAAG |
| 7377_1 | TATGAGTGTGGAGTCGTTGCAGAGCCAGTGTTCGGTAACATTTNNNAGATCGGAAG |
| 7380_1 | TATGAGTGTGGAGTCGTTGCAGTAATTGGTCACAGCCAGTACGNNNAGATCGGAAG |
| 7381_1 | TATGAGTGTGGAGTCGTTGCAGGGAAAGGAGGGGAGTTCATTTNNNAGATCGGAA |
| 7382_1 | TATGAGTGTGGAGTCGTTGCACAGTCTCCAGAGCTTCTGATCTNNNAGATCGGAAGA |
| 7383_1 | TATGAGTGTGGAGTCGTTGCGTGGGTAGGTGGGAAAAGTTACGNNNAGATCGGAA |
| 7385_1 | TATGAGTGTGGAGTCGTTGCCCCTCTTCCCATTTCAGTAACCGNNNAGATCGGAAGA |
| 7387_1 | TATGAGTGTGGAGTCGTTGCAGCCAACTTACAGAGAGGTCCACNNNAGATCGGAAG |
| 7388_1 | TATGAGTGTGGAGTCGTTGCCTGTTTCTGAGGCATCTAGACCGNNNAGATCGGAAG |
| 7390_1 | TATGAGTGTGGAGTCGTTGCCTGATTGAAACCCCCAGATACCGNNNAGATCGGAAG |
| 7391_1 | TATGAGTGTGGAGTCGTTGCAATGACTTCAGCAGACTGTACCGNNNAGATCGGAAG |
| 7392_1 | TATGAGTGTGGAGTCGTTGCTGAAATGCCCTTCCTCTTTACCGNNNAGATCGGAAGA |
| 7393_1 | TATGAGTGTGGAGTCGTTGCCAAAAGTTTCACCTGGAAACCCGNNNAGATCGGAAG |
| 7394_1 | TATGAGTGTGGAGTCGTTGCTTGTGTATCTGAATCTGCACCCGNNNAGATCGGAAGA |
| 7396_1 | TATGAGTGTGGAGTCGTTGCAAATAAACCTGTCACCATCCCCGNNNAGATCGGAAGA |
| 7397_1 | TATGAGTGTGGAGTCGTTGCTTTTTGAGACAACTCTGTCCCCGNNNAGATCGGAAGA |
| 7398_1 | TATGAGTGTGGAGTCGTTGCACACACAACTTTCAAAAAGCCCGNNNAGATCGGAAGA |
| 7400_1 | TATGAGTGTGGAGTCGTTGCAAATCTGCATAGATGCCATCCCGNNNAGATCGGAAG |
| 7401_1 | TATGAGTGTGGAGTCGTTGCCCAGCATAAGGTGTATGATCCCGNNNAGATCGGAAG |
| 7402_1 | TATGAGTGTGGAGTCGTTGCACTGTAAGATTTATGCCTGGCCGNNNAGATCGGAAG |
| 7403_1 | TATGAGTGTGGAGTCGTTGCACATAAGAAACCACTCTTGGCCGNNNAGATCGGAAG |
| 7404_1 | TATGAGTGTGGAGTCGTTGCACAAAAGCATCATCAGAATGCCGNNNAGATCGGAAG |
| 7405_1 | TATGAGTGTGGAGTCGTTGCTGTGTGTTCAAATTGGAATGCCGNNNAGATCGGAAG |
| 7406_1 | TATGAGTGTGGAGTCGTTGCAGTTGGGGTGGAAAGTAAATCCGNNNAGATCGGAA |
| 7407_1 | TATGAGTGTGGAGTCGTTGCCTCAACCTCTGTGTTCACATCCGNNNAGATCGGAAGA |
| 7408_1 | TATGAGTGTGGAGTCGTTGCACCACTTTCTAAAAGGCCATCCGNNNAGATCGGAAGA |
| 7409_1 | TATGAGTGTGGAGTCGTTGCAAATGGCAGTGAAGGTATCTCCGNNNAGATCGGAAG |
| 7410_1 | TATGAGTGTGGAGTCGTTGCAGGATGAAGAAGGATGGTCTCCGNNNAGATCGGAA |
| 7411_1 | TATGAGTGTGGAGTCGTTGCACAAAACACAAGAGACCAGTCCGNNNAGATCGGAAG |
| 7413_1 | TATGAGTGTGGAGTCGTTGCTGGAGACAGAAGATGGAAAAGCGNNNAGATCGGAA |
| 7414_1 | TATGAGTGTGGAGTCGTTGCACCTGACTACAATTGGAGAAGCGNNNAGATCGGAAG |

| | |
|---|---|
| 7415_1 | TATGAGTGTGGAGTCGTTGCCTGCACCATACAGCATCACTTCTNNNAGATCGGAAGA |
| 7416_1 | TATGAGTGTGGAGTCGTTGCCCTGGCAATAGAAATTCACAGCGNNNAGATCGGAAG |
| 7417_1 | TATGAGTGTGGAGTCGTTGCGCTATTCAACCCCTCTTAGAGCGNNNAGATCGGAAG |
| 7418_1 | TATGAGTGTGGAGTCGTTGCAGCATTCCAAAAGGAGAGTAGCGNNNAGATCGGAA |
| 7419_1 | TATGAGTGTGGAGTCGTTGCGTAAATGATTTGGGTTGGACGCGNNNAGATCGGAA |
| 7420_1 | TATGAGTGTGGAGTCGTTGCTGATGTGATGCTTTTAGGAGGCGNNNAGATCGGAA |
| 7421_1 | TATGAGTGTGGAGTCGTTGCAGAGAGAGAGAGATGAAAGGGCGNNNAGATCGGA |
| 7422_1 | TATGAGTGTGGAGTCGTTGCGTCTAAATCAAAATGCTGGGGCGNNNAGATCGGAA |
| 7423_1 | TATGAGTGTGGAGTCGTTGCGTTGCTGGAGTTGATAAATGGCGNNNAGATCGGAA |
| 7425_1 | TATGAGTGTGGAGTCGTTGCTTGTTCCTTGGAATTGCAATGCGNNNAGATCGGAAG |
| 7428_1 | TATGAGTGTGGAGTCGTTGCGAAACTTGTCACACGGTATTGCGNNNAGATCGGAAG |
| 7429_1 | TATGAGTGTGGAGTCGTTGCTATGCCCATGAGAAGGAGGGAAANNNAGATCGGAA |
| 7430_1 | TATGAGTGTGGAGTCGTTGCTGTATGGGTTCAGTGCTCTACCTNNNAGATCGGAAG |
| 7431_1 | TATGAGTGTGGAGTCGTTGCTCATCAAGGCAGAACTGGAATCGNNNAGATCGGAA |
| 7432_1 | TATGAGTGTGGAGTCGTTGCTGAGCAGAGAAAGGACAAGATCGNNNAGATCGGAA |
| 7434_1 | TATGAGTGTGGAGTCGTTGCATTCTGATACACAGGCTGGATCGNNNAGATCGGAAG |
| 7435_1 | TATGAGTGTGGAGTCGTTGCAAATCCCATGAGAAGGCTGATCGNNNAGATCGGAA |
| 7436_1 | TATGAGTGTGGAGTCGTTGCACCTTCCAATGAGTGAGAACTCGNNNAGATCGGAAG |
| 7437_1 | TATGAGTGTGGAGTCGTTGCACTGCCCTATGCACTTTTACTCGNNNAGATCGGAAGA |
| 7438_1 | TATGAGTGTGGAGTCGTTGCCCAAAGACCAAATACAACCCTCGNNNAGATCGGAAGA |
| 7439_1 | TATGAGTGTGGAGTCGTTGCTTCTGGGTTCTCTCTACTCCTCGNNNAGATCGGAAGA |
| 7440_1 | TATGAGTGTGGAGTCGTTGCTCCTTCTGGCATTTCTTTCCTCGNNNAGATCGGAAGA |
| 7441_1 | TATGAGTGTGGAGTCGTTGCGCAGAAGGTAAGAAGACCTCTCGNNNAGATCGGAA |
| 7443_1 | TATGAGTGTGGAGTCGTTGCTGGACTGAGGTTACAAAGTCTCGNNNAGATCGGAA |
| 7444_1 | TATGAGTGTGGAGTCGTTGCGGTCTCGAACTCTCTAGGTCTCGNNNAGATCGGAAG |
| 7446_1 | TATGAGTGTGGAGTCGTTGCTCTAGCTCTTTTGGGAGCCTTCTNNNAGATCGGAAGA |
| 7447_1 | TATGAGTGTGGAGTCGTTGCCATTGAGCATCCAAGTTGTGTCGNNNAGATCGGAAG |
| 7448_1 | TATGAGTGTGGAGTCGTTGCTCTGTTTTCTGAGGACTGCTTCGNNNAGATCGGAAGA |
| 7449_1 | TATGAGTGTGGAGTCGTTGCCCGCTTGTAGTTCTCTCTCTTCGNNNAGATCGGAAGA |
| 7450_1 | TATGAGTGTGGAGTCGTTGCGTTTGTATGCATGGCTGTCTTCGNNNAGATCGGAAG |
| 7452_1 | TATGAGTGTGGAGTCGTTGCTGTGTGTCTCATGTGTAGGTTCGNNNAGATCGGAAG |
| 7453_1 | TATGAGTGTGGAGTCGTTGCCCCCTCCTCCACTTATTTGTTCGNNNAGATCGGAAGA |
| 7455_1 | TATGAGTGTGGAGTCGTTGCAAGGGGAGCATGAAGGAAAAAGGNNNAGATCGGA |
| 7456_1 | TATGAGTGTGGAGTCGTTGCCCACCTGAAGAAGGAGGAAAAGGNNNAGATCGGAA |
| 7460_1 | TATGAGTGTGGAGTCGTTGCCGCCTGTTCTCTCTATCCAAAGGNNNAGATCGGAAGA |
| 7461_1 | TATGAGTGTGGAGTCGTTGCAGCAGCAGGAGAATAAGCAAAGGNNNAGATCGGAA |
| 7462_1 | TATGAGTGTGGAGTCGTTGCCCAACCTGGTCTATAGGCAAAGGNNNAGATCGGAAG |
| 7463_1 | TATGAGTGTGGAGTCGTTGCTCCAGTACAAAAGTGGGCAAAGGNNNAGATCGGAA |
| 7464_1 | TATGAGTGTGGAGTCGTTGCAAGCCTCCCAAAACTCTCAAAGGNNNAGATCGGAAG |
| 7466_1 | TATGAGTGTGGAGTCGTTGCTCAGCCTCCCAAAGGTCTACAATNNNAGATCGGAAGA |
| 7467_1 | TATGAGTGTGGAGTCGTTGCTCCACGAGATCCCTCCCATGATANNNAGATCGGAAGA |
| 7468_1 | TATGAGTGTGGAGTCGTTGCATGAGCACTCCCTCCTAGAAAGGNNNAGATCGGAAG |
| 7469_1 | TATGAGTGTGGAGTCGTTGCAAGTTAGGGAGGCTAGGGAAAGGNNNAGATCGGA |

| | |
|---|---|
| 7470_1 | TATGAGTGTGGAGTCGTTGCACACTGGACACTGAGTTGAAAGGNNNAGATCGGAA |
| 7471_1 | TATGAGTGTGGAGTCGTTGCTTCATCCATCTTCCAGCACAAGGNNNAGATCGGAAG |
| 7472_1 | TATGAGTGTGGAGTCGTTGCTAGGATTAGGCAACATGCCAAGGNNNAGATCGGAA |
| 7473_1 | TATGAGTGTGGAGTCGTTGCCTGGATCATAAGCTCTGCCAAGGNNNAGATCGGAAG |
| 7474_1 | TATGAGTGTGGAGTCGTTGCAGCAGGAATGACTTCATCCAAGGNNNAGATCGGAA |
| 7475_1 | TATGAGTGTGGAGTCGTTGCTACCCAGTCTCACATCTCCAAGGNNNAGATCGGAAG |
| 7476_1 | TATGAGTGTGGAGTCGTTGCTCAGGGAGTTCAAATCAGCAAGGNNNAGATCGGAA |
| 7477_1 | TATGAGTGTGGAGTCGTTGCAGAGGACAAAACTCAAGGCAAGGNNNAGATCGGA |
| 7478_1 | TATGAGTGTGGAGTCGTTGCTCTTGTGGATTCTTTGGGCTTCTNNNAGATCGGAAG |
| 7479_1 | TATGAGTGTGGAGTCGTTGCGCAGCGAGAGCTATAACTCAAGGNNNAGATCGGAA |
| 7480_1 | TATGAGTGTGGAGTCGTTGCTCTGCCCACCCAGTATAGTCATTNNNAGATCGGAAG |
| 7482_1 | TATGAGTGTGGAGTCGTTGCGTATAAGCCTGAGCCCAAGAAGGNNNAGATCGGAA |
| 7484_1 | TATGAGTGTGGAGTCGTTGCTTCACCCTGGTTTTGGAAGAAGGNNNAGATCGGAA |
| 7485_1 | TATGAGTGTGGAGTCGTTGCGATGTGCTTGAAACTGCAGAAGGNNNAGATCGGAA |
| 7486_1 | TATGAGTGTGGAGTCGTTGCTACGCTGTGATTTGTGGAGAAGGNNNAGATCGGAA |
| 7488_1 | TATGAGTGTGGAGTCGTTGCTGGGTGTTTGGATTACAGGAAGGNNNAGATCGGA |
| 7489_1 | TATGAGTGTGGAGTCGTTGCAGAGTAGCTGGGAATCAGGAAGGNNNAGATCGGA |
| 7490_1 | TATGAGTGTGGAGTCGTTGCAAAGCACATCACCATCAGGAAGGNNNAGATCGGAA |
| 7491_1 | TATGAGTGTGGAGTCGTTGCATTGGCTCAAGCAGTTAGGAAGGNNNAGATCGGAA |
| 7492_1 | TATGAGTGTGGAGTCGTTGCAGTAGAAGATGAAGGGCAAGGGTNNNAGATCGGA |
| 7493_1 | TATGAGTGTGGAGTCGTTGCTTCTTGCATTTCCCTTGGGAAGGNNNAGATCGGAAG |
| 7494_1 | TATGAGTGTGGAGTCGTTGCTGGGATAGAGACAGAGCTGAAGGNNNAGATCGGA |
| 7495_1 | TATGAGTGTGGAGTCGTTGCAGCAACATGAAGTGAGCTGAAGGNNNAGATCGGA |
| 7496_1 | TATGAGTGTGGAGTCGTTGCAGGAGGAGGATTTTTGCTCTTCTNNNAGATCGGAAG |
| 7497_1 | TATGAGTGTGGAGTCGTTGCCACAACTTAGACTGGGCTGAAGGNNNAGATCGGAA |
| 7500_1 | TATGAGTGTGGAGTCGTTGCTAAGGAGGCAAGCATGTTGAAGGNNNAGATCGGA |
| 7502_1 | TATGAGTGTGGAGTCGTTGCCCATCTGCAACCCAGGAATAAGGNNNAGATCGGAAG |
| 7503_1 | TATGAGTGTGGAGTCGTTGCCAGAGTTGGACCTGCTCATAAGGNNNAGATCGGAA |
| 7506_1 | TATGAGTGTGGAGTCGTTGCCACAGAAAAGCCCTGAGGTAAGGNNNAGATCGGAA |
| 7507_1 | TATGAGTGTGGAGTCGTTGCAATAAGGAGTGGGTGGGGTAAGGNNNAGATCGGA |
| 7508_1 | TATGAGTGTGGAGTCGTTGCAAACTTCCAGCCCTACAAACAGGNNNAGATCGGAAG |
| 7512_1 | TATGAGTGTGGAGTCGTTGCAGAAGTAGAGAGGCTCTGACAGGNNNAGATCGGA |
| 7513_1 | TATGAGTGTGGAGTCGTTGCAGAAAGGGTCTGTTGTTGACAGGNNNAGATCGGAA |
| 7514_1 | TATGAGTGTGGAGTCGTTGCGTCAGCAGTGGAAAGACTACAGGNNNAGATCGGAA |
| 7515_1 | TATGAGTGTGGAGTCGTTGCAAAAGAGAGGGGAGGTCTACAGGNNNAGATCGGA |
| 7516_1 | TATGAGTGTGGAGTCGTTGCACCCTGCCTTTGGAAATTACAGGNNNAGATCGGAAG |
| 7517_1 | TATGAGTGTGGAGTCGTTGCTGTAGGGCTTGCAAATAACCAGGNNNAGATCGGAA |
| 7518_1 | TATGAGTGTGGAGTCGTTGCCTCTCACACACTAACACACCAGGNNNAGATCGGAAG |
| 7519_1 | TATGAGTGTGGAGTCGTTGCAGACTATGAAGCCTCTCACCAGGNNNAGATCGGAAG |
| 7520_1 | TATGAGTGTGGAGTCGTTGCTTCAACTACCTATGGGGACCAGGNNNAGATCGGAAG |
| 7521_1 | TATGAGTGTGGAGTCGTTGCAAATGTAACCTGGGTTGACCAGGNNNAGATCGGAA |
| 7522_1 | TATGAGTGTGGAGTCGTTGCAGAGAGTTGTGAGAGGTACCAGGNNNAGATCGGA |
| 7523_1 | TATGAGTGTGGAGTCGTTGCTATTTGCACTCTGGAAACCCAGGNNNAGATCGGAAG |

| | |
|---|---|
| 7524_1 | TATGAGTGTGGAGTCGTTGCTTTAGAATCAGCCTTCACCCAGGNNNAGATCGGAAG |
| 7525_1 | TATGAGTGTGGAGTCGTTGCATTTTCTGTGCTGTGTACCCAGGNNNAGATCGGAAG |
| 7526_1 | TATGAGTGTGGAGTCGTTGCCAGGCAAGTTTTCGGCAAATGTANNNAGATCGGAAG |
| 7527_1 | TATGAGTGTGGAGTCGTTGCATAGAAAAGCAGATCAGCCCAGGNNNAGATCGGAA |
| 7530_1 | TATGAGTGTGGAGTCGTTGCACAGATTATCAGAAAGCGCCAGGNNNAGATCGGAA |
| 7531_1 | TATGAGTGTGGAGTCGTTGCAAAGAATAGAAGCAAGGGCCAGGNNNAGATCGGA |
| 7532_1 | TATGAGTGTGGAGTCGTTGCTAGAATATCTCACAGGGGCCAGGNNNAGATCGGAA |
| 7534_1 | TATGAGTGTGGAGTCGTTGCTTGAAAACATATCTGGGGCCAGGNNNAGATCGGAA |
| 7535_1 | TATGAGTGTGGAGTCGTTGCGTTTGTTTCCAGAAATGGCCAGGNNNAGATCGGAAG |
| 7536_1 | TATGAGTGTGGAGTCGTTGCTATGCAGGAATTCATGTGCCAGGNNNAGATCGGAA |
| 7538_1 | TATGAGTGTGGAGTCGTTGCAAATGTGGTCCGAATCAAGCAGGNNNAGATCGGAA |
| 7539_1 | TATGAGTGTGGAGTCGTTGCGAGAGAGAGGGAAACTAAGCAGGNNNAGATCGGA |
| 7541_1 | TATGAGTGTGGAGTCGTTGCCTGCCTTTGGTTTTGATAGCAGGNNNAGATCGGAAG |
| 7542_1 | TATGAGTGTGGAGTCGTTGCCTCAATGTAGATAAGGGCGCAGGNNNAGATCGGAA |
| 7543_1 | TATGAGTGTGGAGTCGTTGCTGGAGAACTCTGATCAATGCAGGNNNAGATCGGAA |
| 7544_1 | TATGAGTGTGGAGTCGTTGCTATTCCCTGATGCTTCATGCAGGNNNAGATCGGAAG |
| 7545_1 | TATGAGTGTGGAGTCGTTGCCATCTCCTCAAACCCTATGCAGGNNNAGATCGGAAG |
| 7546_1 | TATGAGTGTGGAGTCGTTGCGGCATCCTTAGAGATACTGCAGGNNNAGATCGGAA |
| 7547_1 | TATGAGTGTGGAGTCGTTGCGCGTATATTTTTCTGCCTGCAGGNNNAGATCGGAAG |
| 7548_1 | TATGAGTGTGGAGTCGTTGCTATTCTGTCTAGGGAGCTGCAGGNNNAGATCGGAA |
| 7550_1 | TATGAGTGTGGAGTCGTTGCATTTGGGAACACAGTCACTCAGGNNNAGATCGGAAG |
| 7552_1 | TATGAGTGTGGAGTCGTTGCTAGAGACCACCCTATTGCTCAGGNNNAGATCGGAAG |
| 7553_1 | TATGAGTGTGGAGTCGTTGCAGCCTCTTTTCAGTCAGTTGAGCNNNAGATCGGAAG |
| 7554_1 | TATGAGTGTGGAGTCGTTGCGGAGACCACTGCACTAATTCAGGNNNAGATCGGAAG |
| 7555_1 | TATGAGTGTGGAGTCGTTGCCCCAGATGACCAGTTGGTTGAAANNNAGATCGGAAG |
| 7557_1 | TATGAGTGTGGAGTCGTTGCGAGGGAGCACAAGAGATTTCAGGNNNAGATCGGAA |
| 7559_1 | TATGAGTGTGGAGTCGTTGCGGAGAGAACACAGACCAAAGAGGNNNAGATCGGA |
| 7561_1 | TATGAGTGTGGAGTCGTTGCGCAGGAGGTCAAAGGTAAAGAGGNNNAGATCGGA |
| 7562_1 | TATGAGTGTGGAGTCGTTGCCAGGCTCACTGCAGGTGATATTTNNNAGATCGGAAG |
| 7563_1 | TATGAGTGTGGAGTCGTTGCCCACCTCAACGAGAATGAAGAGGNNNAGATCGGAA |
| 7564_1 | TATGAGTGTGGAGTCGTTGCGTAGGAGGTGGGAGGATAAGAGGNNNAGATCGG |
| 7565_1 | TATGAGTGTGGAGTCGTTGCTACACTGGATGGGTACACAGAGGNNNAGATCGGAA |
| 7566_1 | TATGAGTGTGGAGTCGTTGCGATGGATGCCAAGGTTACAGAGGNNNAGATCGGA |
| 7567_1 | TATGAGTGTGGAGTCGTTGCAGCATCTGCAACTAAACCAGAGGNNNAGATCGGAAG |
| 7568_1 | TATGAGTGTGGAGTCGTTGCAGAATGGTGGATTCATGCAGAGGNNNAGATCGGA |
| 7569_1 | TATGAGTGTGGAGTCGTTGCTACCACCTCTAAGACTGCAGAGGNNNAGATCGGAAG |
| 7570_1 | TATGAGTGTGGAGTCGTTGCACTACAGGAAGCTAGCTCAGAGGNNNAGATCGGAA |
| 7571_1 | TATGAGTGTGGAGTCGTTGCGCTTCTCTGGGGAAATTCAGAGGNNNAGATCGGAA |
| 7572_1 | TATGAGTGTGGAGTCGTTGCACCAGAGCTATCACTTTCCCAGTNNNAGATCGGAAG |
| 7574_1 | TATGAGTGTGGAGTCGTTGCGAGCACTGAACTTGGAAGAGAGGNNNAGATCGGA |
| 7576_1 | TATGAGTGTGGAGTCGTTGCGCACAGCGTCGTTTTGTCTATTTNNNAGATCGGAAG |
| 7577_1 | TATGAGTGTGGAGTCGTTGCTGGGATGACAGAGTAGTGAGAGGNNNAGATCGGA |
| 7578_1 | TATGAGTGTGGAGTCGTTGCGACAGACTTGCTTGGTGTAGAGGNNNAGATCGGAA |

| | |
|---|---|
| 7582_1 | TATGAGTGTGGAGTCGTTGCAGAAAAAGGCATGGACAAGGAGGNNNAGATCGGA |
| 7583_1 | TATGAGTGTGGAGTCGTTGCAAACTCCCCTTAGAGCAAGGAGGNNNAGATCGGAA |
| 7584_1 | TATGAGTGTGGAGTCGTTGCGGTTGGAACTGAAGAGAAGGAGGNNNAGATCGGA |
| 7585_1 | TATGAGTGTGGAGTCGTTGCTGGGGTTAGGGAACATAAGGAGGNNNAGATCGGA |
| 7586_1 | TATGAGTGTGGAGTCGTTGCGTATTGAGTGCTTTCCCAGGAGGNNNAGATCGGAA |
| 7587_1 | TATGAGTGTGGAGTCGTTGCCCATGAATGTGGTTCAGAGGAGGNNNAGATCGGAA |
| 7588_1 | TATGAGTGTGGAGTCGTTGCGAACACAGACTGGGCAAACTAGANNNAGATCGGAA |
| 7589_1 | TATGAGTGTGGAGTCGTTGCTGTCTGTGTTGGTGGATTCAGGGNNNAGATCGGAA |
| 7590_1 | TATGAGTGTGGAGTCGTTGCCCTTAGCTTGCCCAGAGCATAAANNNAGATCGGAAG |
| 7591_1 | TATGAGTGTGGAGTCGTTGCTACCTTTTTCAGTGCAAGGGAGGNNNAGATCGGAA |
| 7593_1 | TATGAGTGTGGAGTCGTTGCTATTTATTTCGGGGGAGGGGAGGNNNAGATCGGA |
| 7594_1 | TATGAGTGTGGAGTCGTTGCGTGACATACGAAAGAGCCATTCGNNNAGATCGGAA |
| 7595_1 | TATGAGTGTGGAGTCGTTGCTAATGGAGAAGAAGGTGGGGAGGNNNAGATCGGA |
| 7597_1 | TATGAGTGTGGAGTCGTTGCTCTTACCTTCTGTGAGTGGGAGGNNNAGATCGGAA |
| 7598_1 | TATGAGTGTGGAGTCGTTGCCTCTCTCTCTCTCAGTTGGGAGGNNNAGATCGGAAG |
| 7599_1 | TATGAGTGTGGAGTCGTTGCCACCCACCTACAAATCATGGAGGNNNAGATCGGAAG |
| 7600_1 | TATGAGTGTGGAGTCGTTGCAACCCCAAAACTGCTTATGGAGGNNNAGATCGGAAG |
| 7601_1 | TATGAGTGTGGAGTCGTTGCTGTCTTTTGGATGGAACTGGAGGNNNAGATCGGAA |
| 7603_1 | TATGAGTGTGGAGTCGTTGCCACTCATACACACTTCCTGGAGGNNNAGATCGGAAG |
| 7604_1 | TATGAGTGTGGAGTCGTTGCAGAGAGAAATGGGTAGCTGGAGGNNNAGATCGGA |
| 7606_1 | TATGAGTGTGGAGTCGTTGCGCTGCTGTTTTGTTATGTGGAGGNNNAGATCGGAA |
| 7609_1 | TATGAGTGTGGAGTCGTTGCAAGTATAGGTGGCGTTTTGGAGGNNNAGATCGGA |
| 7610_1 | TATGAGTGTGGAGTCGTTGCCTGGTGCTTAGATGGGAATGAGGNNNAGATCGGA |
| 7611_1 | TATGAGTGTGGAGTCGTTGCTCAGGGAAGAGGGATAACTGAGGNNNAGATCGGA |
| 7612_1 | TATGAGTGTGGAGTCGTTGCATTCGTAAGCTGTCACACTGAGGNNNAGATCGGAAG |
| 7613_1 | TATGAGTGTGGAGTCGTTGCAAAATCTCTGCTCAGCTCTGAGGNNNAGATCGGAAG |
| 7614_1 | TATGAGTGTGGAGTCGTTGCCAAAGGGTTGGGATTCTCTGAGGNNNAGATCGGAA |
| 7615_1 | TATGAGTGTGGAGTCGTTGCCAGTGTTTTCCCCTCAAGTGAGGNNNAGATCGGAAG |
| 7616_1 | TATGAGTGTGGAGTCGTTGCAGACGGGAGTAACTACAGTGAGGNNNAGATCGGA |
| 7617_1 | TATGAGTGTGGAGTCGTTGCAATAGTCCACAGGTGAGGTGAGGNNNAGATCGGA |
| 7618_1 | TATGAGTGTGGAGTCGTTGCTTCTCATGTTTGTGTGGGTGAGGNNNAGATCGGAA |
| 7619_1 | TATGAGTGTGGAGTCGTTGCACAAAACAACCCTGTCTGTGAGGNNNAGATCGGAAG |
| 7621_1 | TATGAGTGTGGAGTCGTTGCTGTAGGTGTGATAGGGGTTGAGGNNNAGATCGGA |
| 7622_1 | TATGAGTGTGGAGTCGTTGCGCAGTTTCTGGAGGCTTTCTGTANNNAGATCGGAAG |
| 7623_1 | TATGAGTGTGGAGTCGTTGCGCTGCAATGGGCCAAATAATAGGNNNAGATCGGAA |
| 7624_1 | TATGAGTGTGGAGTCGTTGCAGAGAGAGAGCTGGTCACATAGGNNNAGATCGGA |
| 7626_1 | TATGAGTGTGGAGTCGTTGCGCATCTGTAGTGTCTCTTGTGCANNNAGATCGGAAG |
| 7627_1 | TATGAGTGTGGAGTCGTTGCTATTTCACTTCAGCCAGCTGTGCNNNAGATCGGAAG |
| 7630_1 | TATGAGTGTGGAGTCGTTGCTCTGCAATGGGAAGGGTACTAGGNNNAGATCGGAA |
| 7631_1 | TATGAGTGTGGAGTCGTTGCAACTACTGACAGTGGGACCTAGGNNNAGATCGGAA |
| 7633_1 | TATGAGTGTGGAGTCGTTGCAGTGATACAAACTCTTCCAGGGTNNNAGATCGGAAG |
| 7635_1 | TATGAGTGTGGAGTCGTTGCCCTGCCCAAGATTTCTCTCTAGGNNNAGATCGGAAG |
| 7636_1 | TATGAGTGTGGAGTCGTTGCCCCTTTAGCCTGGGTTTCACTTTNNNAGATCGGAAGA |

| | |
|---|---|
| 7638_1 | TATGAGTGTGGAGTCGTTGCCTTGGGTAAAGGTGTGCAGTAGGNNNAGATCGGAA |
| 7639_1 | TATGAGTGTGGAGTCGTTGCTCATGCTTTGTTCACCGAGTAGGNNNAGATCGGAAG |
| 7640_1 | TATGAGTGTGGAGTCGTTGCTGCTAGGTCATGCTAAGGGTGTANNNAGATCGGAA |
| 7641_1 | TATGAGTGTGGAGTCGTTGCGGTAAGCTCTGCATGTGAGTAGGNNNAGATCGGAA |
| 7644_1 | TATGAGTGTGGAGTCGTTGCAAGTAGAAGCTCAGTTGCTCTAANNNAGATCGGAAG |
| 7645_1 | TATGAGTGTGGAGTCGTTGCAGAGACAGCCACTCAGGATTAGGNNNAGATCGGAA |
| 7646_1 | TATGAGTGTGGAGTCGTTGCGAGGCACTTGTTCTGTCCTTAGGNNNAGATCGGAAG |
| 7647_1 | TATGAGTGTGGAGTCGTTGCGCCAGAAAGCAGGGAGATTTAGGNNNAGATCGGAA |
| 7648_1 | TATGAGTGTGGAGTCGTTGCATCCTTTTCACGGTGGCTTTAGGNNNAGATCGGAAG |
| 7649_1 | TATGAGTGTGGAGTCGTTGCTTAGGGGAGGATACAGGAAACGGNNNAGATCGGA |
| 7650_1 | TATGAGTGTGGAGTCGTTGCAACTGTGTTCCCATGCTGACTTTNNNAGATCGGAAG |
| 7651_1 | TATGAGTGTGGAGTCGTTGCGATAATACAGGCAGGTGACACGGNNNAGATCGGAA |
| 7652_1 | TATGAGTGTGGAGTCGTTGCAGCCAGAGCTTCATTTATCACGGNNNAGATCGGAAG |
| 7653_1 | TATGAGTGTGGAGTCGTTGCTGGTATTAGAAACCTGCAGACGGNNNAGATCGGAA |
| 7654_1 | TATGAGTGTGGAGTCGTTGCAGGACAGAAGAGAAGAGAGACGGNNNAGATCGGA |
| 7657_1 | TATGAGTGTGGAGTCGTTGCATTTTAGGCACACAACAGACCGGNNNAGATCGGAA |
| 7659_1 | TATGAGTGTGGAGTCGTTGCTACACAGCAAACATTCTAGCCGGNNNAGATCGGAAG |
| 7660_1 | TATGAGTGTGGAGTCGTTGCTAAAAATGTGCAACATTGGCCGGNNNAGATCGGAA |
| 7661_1 | TATGAGTGTGGAGTCGTTGCTATGTGTTTGTCTTACCTGCCGGNNNAGATCGGAAG |
| 7662_1 | TATGAGTGTGGAGTCGTTGCATCTCTCTCTATGTGGTCTCCGGNNNAGATCGGAAG |
| 7663_1 | TATGAGTGTGGAGTCGTTGCACATCATTCAATACATTGCGCGGNNNAGATCGGAAG |
| 7664_1 | TATGAGTGTGGAGTCGTTGCTTTAGTCAGGTCCTAGTAGGCGGNNNAGATCGGAA |
| 7665_1 | TATGAGTGTGGAGTCGTTGCACATAACCCTCATTTTCTGGCGGNNNAGATCGGAAG |
| 7667_1 | TATGAGTGTGGAGTCGTTGCTCATCAAGCCAAAATGTATGCGGNNNAGATCGGAA |
| 7669_1 | TATGAGTGTGGAGTCGTTGCACCATATGCATGGAAGAACTCGGNNNAGATCGGAA |
| 7670_1 | TATGAGTGTGGAGTCGTTGCATCCATGTTGTAGCCAATCTCGGNNNAGATCGGAAG |
| 7671_1 | TATGAGTGTGGAGTCGTTGCTTGAGGATTGAATGAGCAGTCGGNNNAGATCGGAA |
| 7672_1 | TATGAGTGTGGAGTCGTTGCAACAATTGGCGAATCTAGGTCGGNNNAGATCGGAA |
| 7674_1 | TATGAGTGTGGAGTCGTTGCTCTTCCAAATCCCAGCAATTCGGNNNAGATCGGAAG |
| 7675_1 | TATGAGTGTGGAGTCGTTGCAGGCTTTTGTTGTTGTTGTTCGGNNNAGATCGGAAG |
| 7678_1 | TATGAGTGTGGAGTCGTTGCTCTGCTTTTGAGATGCTCAAGGGNNNAGATCGGAA |
| 7679_1 | TATGAGTGTGGAGTCGTTGCCTATAACAGGGCATGGTCAAGGGNNNAGATCGGAA |
| 7680_1 | TATGAGTGTGGAGTCGTTGCCACACACACATATGGGAGAAGGGNNNAGATCGGAA |
| 7681_1 | TATGAGTGTGGAGTCGTTGCAGGACATCTGAGAAAGGGAAGGGNNNAGATCGGA |
| 7684_1 | TATGAGTGTGGAGTCGTTGCCAATGGAAATCTGAGGGGAAGGGNNNAGATCGGA |
| 7685_1 | TATGAGTGTGGAGTCGTTGCTGGTGTGCGTTAGTATTGAAGGGNNNAGATCGGAA |
| 7686_1 | TATGAGTGTGGAGTCGTTGCCAAAGAGAAGAGCAGGCTAAGGGNNNAGATCGGA |
| 7688_1 | TATGAGTGTGGAGTCGTTGCAACAACCAGAATCTCTCACAGGGNNNAGATCGGAAG |
| 7689_1 | TATGAGTGTGGAGTCGTTGCGGATCACTCCTTTCTTCACAGGGNNNAGATCGGAAG |
| 7690_1 | TATGAGTGTGGAGTCGTTGCGCAGGAAGATGAAAGGTACAGGGNNNAGATCGGA |
| 7691_1 | TATGAGTGTGGAGTCGTTGCTTTTGTTTTAGGCCTCACCAGGGNNNAGATCGGAAG |
| 7693_1 | TATGAGTGTGGAGTCGTTGCCTCACCCATACTAGGATCCAGGGNNNAGATCGGAAG |
| 7696_1 | TATGAGTGTGGAGTCGTTGCAAGAGGTTCAAAGAACAGCAGGGNNNAGATCGGA |

| | |
|---|---|
| 7697_1 | TATGAGTGTGGAGTCGTTGCGAAAAGGGTACTACACAGCAGGGNNNAGATCGGAA |
| 7698_1 | TATGAGTGTGGAGTCGTTGCGTTCATCCTAGGTTACAGCAGGGNNNAGATCGGAAG |
| 7700_1 | TATGAGTGTGGAGTCGTTGCACGCCAGTCCTATTAGATCAGGGNNNAGATCGGAAG |
| 7702_1 | TATGAGTGTGGAGTCGTTGCGACAAGCCTTCAGATGATCAGGGNNNAGATCGGAAG |
| 7704_1 | TATGAGTGTGGAGTCGTTGCTAGAATACCACGCCTACTCAGGGNNNAGATCGGAAG |
| 7705_1 | TATGAGTGTGGAGTCGTTGCTTGCTGGGTCTTTTTAGTCAGGGNNNAGATCGGAAG |
| 7706_1 | TATGAGTGTGGAGTCGTTGCAGACTGGAGGAGTGTTACAGTGANNNAGATCGGAA |
| 7707_1 | TATGAGTGTGGAGTCGTTGCCACTTCTTCCTTCTGTGTCAGGGNNNAGATCGGAAGA |
| 7708_1 | TATGAGTGTGGAGTCGTTGCATTGGATGAAAGCTCGTTCAGGGNNNAGATCGGAAG |
| 7709_1 | TATGAGTGTGGAGTCGTTGCACAGCCTCTTTTCACTTTCAGGGNNNAGATCGGAAGA |
| 7710_1 | TATGAGTGTGGAGTCGTTGCTACCTCGGTCTTCTCTTTCAGGGNNNAGATCGGAAGA |
| 7711_1 | TATGAGTGTGGAGTCGTTGCCCTCCACTTCAGTGATTCTTGCANNNAGATCGGAAGA |
| 7712_1 | TATGAGTGTGGAGTCGTTGCAATAGCCAGGAGACTGAAGAGGGNNNAGATCGGAA |
| 7713_1 | TATGAGTGTGGAGTCGTTGCGATTTCAGCCTAACACCAGAGGGNNNAGATCGGAAG |
| 7715_1 | TATGAGTGTGGAGTCGTTGCTAGGTAAGGCTTCACTGAGAGGGNNNAGATCGGAA |
| 7716_1 | TATGAGTGTGGAGTCGTTGCCTTGAGATAAGTCCCCAGGAGGGNNNAGATCGGAAG |
| 7718_1 | TATGAGTGTGGAGTCGTTGCCTGAGGGTAGGGATTACTGAGGGNNNAGATCGGAA |
| 7720_1 | TATGAGTGTGGAGTCGTTGCAGAGAACTTTGTGTTGCTGAGGGNNNAGATCGGAA |
| 7722_1 | TATGAGTGTGGAGTCGTTGCAATAAGAAGCCCTGGGTTGAGGGNNNAGATCGGAA |
| 7723_1 | TATGAGTGTGGAGTCGTTGCTTACAGAGGCCAGATGGATAGGGNNNAGATCGGAA |
| 7724_1 | TATGAGTGTGGAGTCGTTGCCCAGACCACTCTGTCCTATAGGGNNNAGATCGGAAG |
| 7725_1 | TATGAGTGTGGAGTCGTTGCTGAACTTTGCCTTTGTCCTAGGGNNNAGATCGGAAG |
| 7727_1 | TATGAGTGTGGAGTCGTTGCATAGGACTGAGCAAGGAGTAGGGNNNAGATCGGAA |
| 7728_1 | TATGAGTGTGGAGTCGTTGCCCTTGTAACAGAGCAACGTAGGGNNNAGATCGGAAG |
| 7729_1 | TATGAGTGTGGAGTCGTTGCCAGTCAAAGGGTGAAGTGTAGGGNNNAGATCGGAA |
| 7733_1 | TATGAGTGTGGAGTCGTTGCCCTGGCACTGTTGATTTTTAGGGNNNAGATCGGAAG |
| 7736_1 | TATGAGTGTGGAGTCGTTGCTAAGGATACCAGTCTGCTACGGGNNNAGATCGGAAG |
| 7737_1 | TATGAGTGTGGAGTCGTTGCGAAAGTTATTTTGCTCTCCCGGGNNNAGATCGGAAG |
| 7738_1 | TATGAGTGTGGAGTCGTTGCTAAAGTTGTATACGAAGGCCGGGNNNAGATCGGAA |
| 7739_1 | TATGAGTGTGGAGTCGTTGCTAGAAAATTGCTTTGGGTCCGGGNNNAGATCGGAAG |
| 7740_1 | TATGAGTGTGGAGTCGTTGCAGTGCTATGAATTCTTCAGCGGGNNNAGATCGGAAG |
| 7741_1 | TATGAGTGTGGAGTCGTTGCTAGCTCAGATCTCTCCAAAGGGGNNNAGATCGGAAG |
| 7742_1 | TATGAGTGTGGAGTCGTTGCAAATTCAGATTGAACCCCAGGGGNNNAGATCGGAAG |
| 7743_1 | TATGAGTGTGGAGTCGTTGCTACTGGTGTATAGAGCTCAGGGGNNNAGATCGGAA |
| 7745_1 | TATGAGTGTGGAGTCGTTGCGACATTTGGGTAGGTTTGAGGGGNNNAGATCGGAA |
| 7746_1 | TATGAGTGTGGAGTCGTTGCGGATGTGGGGTATGGTATAGGGGNNNAGATCGGA |
| 7747_1 | TATGAGTGTGGAGTCGTTGCCTGCCTAAAATCCAAGCTAGGGGNNNAGATCGGAAG |
| 7748_1 | TATGAGTGTGGAGTCGTTGCATTGTACCGGCCTTGGTTTTTCTNNNAGATCGGAAGA |
| 7751_1 | TATGAGTGTGGAGTCGTTGCTTTTGGTTTGGCATTAGAGGGGGNNNAGATCGGAA |
| 7752_1 | TATGAGTGTGGAGTCGTTGCTGTTCAGGGCTTTGAGAAAAGCANNNAGATCGGAAG |
| 7753_1 | TATGAGTGTGGAGTCGTTGCGAATAAAGAAGATGCCCTGGGGGNNNAGATCGGAA |
| 7755_1 | TATGAGTGTGGAGTCGTTGCGCACACAAGTAATTCAACTGGGGNNNAGATCGGAAG |
| 7756_1 | TATGAGTGTGGAGTCGTTGCTGTGAGCACTGACTTGTGAACAANNNAGATCGGAAG |

| | |
|---|---|
| 7757_1 | TATGAGTGTGGAGTCGTTGCAGGCATGCAAAGATATTCTGGGGNNNAGATCGGAA |
| 7758_1 | TATGAGTGTGGAGTCGTTGCTCCCAAGTAAGAGAGAAGTGGGGNNNAGATCGGA |
| 7760_1 | TATGAGTGTGGAGTCGTTGCAGGAAGATCTATTGGAGGTGGGGNNNAGATCGGA |
| 7761_1 | TATGAGTGTGGAGTCGTTGCAATACTACCCACTGTAGGTGGGGNNNAGATCGGAA |
| 7762_1 | TATGAGTGTGGAGTCGTTGCGCATCGGGTGTCCTATCAAAAGTNNNAGATCGGAAG |
| 7763_1 | TATGAGTGTGGAGTCGTTGCATGAAAAATCCACAGTGGTGGGGNNNAGATCGGAA |
| 7765_1 | TATGAGTGTGGAGTCGTTGCTCTCTTTCTCCTTCTCTGTGGGGNNNAGATCGGAAG |
| 7766_1 | TATGAGTGTGGAGTCGTTGCAATGGGCTGATGTAATTGTGGGGNNNAGATCGGA |
| 7767_1 | TATGAGTGTGGAGTCGTTGCACGCCACTTTCAGCCTCAATAATNNNAGATCGGAAG |
| 7768_1 | TATGAGTGTGGAGTCGTTGCTTGACTGGATCTCTCTCTTGGGGNNNAGATCGGAAG |
| 7771_1 | TATGAGTGTGGAGTCGTTGCCAGCGGCACAATAGAAAAATGGGNNNAGATCGGAA |
| 7773_1 | TATGAGTGTGGAGTCGTTGCTCAATCATCTGGGAGCAAATGGGNNNAGATCGGAA |
| 7775_1 | TATGAGTGTGGAGTCGTTGCGAGAGAGAAAGCCAGACAATGGGNNNAGATCGGA |
| 7776_1 | TATGAGTGTGGAGTCGTTGCAACAATGAGGCTGAAGGAATGGGNNNAGATCGGA |
| 7779_1 | TATGAGTGTGGAGTCGTTGCAGATCAACTCACCAATCCATGGGNNNAGATCGGAAG |
| 7780_1 | TATGAGTGTGGAGTCGTTGCGTGCTGGGTTATGCTTATATGGCNNNAGATCGGAA |
| 7781_1 | TATGAGTGTGGAGTCGTTGCAGATTGTGATGGCTAAGCATGGGNNNAGATCGGAA |
| 7782_1 | TATGAGTGTGGAGTCGTTGCGCTTCCTTTGCTTGTTCAGCTTTNNNAGATCGGAAGA |
| 7784_1 | TATGAGTGTGGAGTCGTTGCGGGCGTAGATTCAATCAGATGGGNNNAGATCGGAA |
| 7786_1 | TATGAGTGTGGAGTCGTTGCGGATGAAGTTAGCCTTGGATGGGNNNAGATCGGA |
| 7788_1 | TATGAGTGTGGAGTCGTTGCAACCCAGCCTTCTAGACTATGGGNNNAGATCGGAAG |
| 7790_1 | TATGAGTGTGGAGTCGTTGCTAAAGCGTTCCCAGTTCTATGGGNNNAGATCGGAAG |
| 7791_1 | TATGAGTGTGGAGTCGTTGCGGACTGGGAAGAGTTTGAGCTTTNNNAGATCGGAA |
| 7792_1 | TATGAGTGTGGAGTCGTTGCGTCCTCTTCCTGTGCTTTATGGGNNNAGATCGGAAG |
| 7793_1 | TATGAGTGTGGAGTCGTTGCTGATGTGCATTCTCAAGACTGGGNNNAGATCGGAA |
| 7794_1 | TATGAGTGTGGAGTCGTTGCCTTTTCTCTGTGGCTGTACTGGGNNNAGATCGGAAG |
| 7795_1 | TATGAGTGTGGAGTCGTTGCGGGGTAGAGGAAGAGTTACTGGGNNNAGATCGGA |
| 7799_1 | TATGAGTGTGGAGTCGTTGCAAAAGAGATTTCAGCTCCCTGGGNNNAGATCGGAA |
| 7804_1 | TATGAGTGTGGAGTCGTTGCAACATTACCACTGTGAAGCTGGGNNNAGATCGGAA |
| 7805_1 | TATGAGTGTGGAGTCGTTGCCATTTGAATCTGCAACAGCTGGGNNNAGATCGGAAG |
| 7806_1 | TATGAGTGTGGAGTCGTTGCGCGTGAATGAAAATACAGCTGGGNNNAGATCGGAA |
| 7808_1 | TATGAGTGTGGAGTCGTTGCAGAGAAAGAGAGAGTGAGCTGGGNNNAGATCGGA |
| 7810_1 | TATGAGTGTGGAGTCGTTGCAAGACTAAAACACACAGGCTGGGNNNAGATCGGAA |
| 7811_1 | TATGAGTGTGGAGTCGTTGCTAGAAAGAGAAGGACAGGCTGGGNNNAGATCGGA |
| 7812_1 | TATGAGTGTGGAGTCGTTGCGGTTTACAGTTCTTGAGGCTGGGNNNAGATCGGAA |
| 7813_1 | TATGAGTGTGGAGTCGTTGCGGGTATCTATCACCTAGGCTGGGNNNAGATCGGAA |
| 7820_1 | TATGAGTGTGGAGTCGTTGCCTCAAAAATGGGACTAGAAGGGANNNAGATCGGAA |
| 7822_1 | TATGAGTGTGGAGTCGTTGCCTTTGACATCAGATGGGTCTGGGNNNAGATCGGAA |
| 7823_1 | TATGAGTGTGGAGTCGTTGCTAGTGGAGGCTCTAATGTCTGGGNNNAGATCGGAA |
| 7825_1 | TATGAGTGTGGAGTCGTTGCACTGACTTCCTTTGCATTCTGGGNNNAGATCGGAAG |
| 7826_1 | TATGAGTGTGGAGTCGTTGCTTAGCAGGAATGCAACTTCTGGGNNNAGATCGGAA |
| 7827_1 | TATGAGTGTGGAGTCGTTGCAGAAAGATGTGGCAGTCTAGGGTNNNAGATCGGAA |
| 7830_1 | TATGAGTGTGGAGTCGTTGCGAATAGAAGGCACAGACAGTGGGNNNAGATCGGA |

| | |
|---|---|
| 7834_1 | TATGAGTGTGGAGTCGTTGCAGAGAAAGAGAAGTCGGAGTGGGNNNAGATCGGA |
| 7835_1 | TATGAGTGTGGAGTCGTTGCTAAATTCCCTTCTGGGGAGTGGGNNNAGATCGGAA |
| 7836_1 | TATGAGTGTGGAGTCGTTGCTAGGTTTTATGGCTGTTCGTGGGNNNAGATCGGAA |
| 7840_1 | TATGAGTGTGGAGTCGTTGCGAAGTAGAAATGGGGCATGTGGGNNNAGATCGGA |
| 7841_1 | TATGAGTGTGGAGTCGTTGCCGTCACTTATGGGTGTGAAATTCNNNAGATCGGAAG |
| 7842_1 | TATGAGTGTGGAGTCGTTGCGGAGGAGAAAAAGTCACTGTGGGNNNAGATCGGA |
| 7843_1 | TATGAGTGTGGAGTCGTTGCGAGGTGAAATTCCTCAGTGTGGGNNNAGATCGGAA |
| 7844_1 | TATGAGTGTGGAGTCGTTGCTTGGTGGCAAAGGAACAGAAAGTNNNAGATCGGAA |
| 7845_1 | TATGAGTGTGGAGTCGTTGCCTTCAGGATTCCACATGTGTGGGNNNAGATCGGAA |
| 7846_1 | TATGAGTGTGGAGTCGTTGCGTTTTTCACCAGTGATGTGTGGGNNNAGATCGGAA |
| 7847_1 | TATGAGTGTGGAGTCGTTGCAACATCGGTTTCTACTGTGTGGGNNNAGATCGGAA |
| 7848_1 | TATGAGTGTGGAGTCGTTGCTGCTAAGCTTCTGGATTTGTGGGNNNAGATCGGAA |
| 7850_1 | TATGAGTGTGGAGTCGTTGCAGGCTCCCATCACCATAGAAAGTNNNAGATCGGAAG |
| 7851_1 | TATGAGTGTGGAGTCGTTGCTGTGTATACGAGTGACCCTTGGGNNNAGATCGGAA |
| 7853_1 | TATGAGTGTGGAGTCGTTGCCTAACCAGGGCAGAAACCTAAAANNNAGATCGGAA |
| 7854_1 | TATGAGTGTGGAGTCGTTGCAGATGAGGCTAACACTCCTTGGGNNNAGATCGGAA |
| 7856_1 | TATGAGTGTGGAGTCGTTGCCTTTCTCTTCCTTGCTGCCCTTANNNAGATCGGAAGA |
| 7858_1 | TATGAGTGTGGAGTCGTTGCCCTCTTAGACCCCAACATTTGGGNNNAGATCGGAAG |
| 7860_1 | TATGAGTGTGGAGTCGTTGCGCCCTGCTTTCAAATACTTTGGGNNNAGATCGGAAG |
| 7861_1 | TATGAGTGTGGAGTCGTTGCTTTGATGAGGCTTACCCTTTGGGNNNAGATCGGAAG |
| 7862_1 | TATGAGTGTGGAGTCGTTGCTTCCAAAGACTTGTCCCTTTGGGNNNAGATCGGAAG |
| 7863_1 | TATGAGTGTGGAGTCGTTGCATGAGAGCCCACAGATAGCCTTANNNAGATCGGAA |
| 7864_1 | TATGAGTGTGGAGTCGTTGCGAAAAGTCCTGAATGGCTTTGGGNNNAGATCGGAA |
| 7865_1 | TATGAGTGTGGAGTCGTTGCCCTATGCTCGGTGTATCTTTGGGNNNAGATCGGAAG |
| 7867_1 | TATGAGTGTGGAGTCGTTGCAGATCCTGCCTTGGGAAAAATGGNNNAGATCGGAA |
| 7869_1 | TATGAGTGTGGAGTCGTTGCTACATAAGCCCACACCCAAATGGNNNAGATCGGAAG |
| 7870_1 | TATGAGTGTGGAGTCGTTGCGGGCTCAGTAACTCCTCAAATGGNNNAGATCGGAA |
| 7873_1 | TATGAGTGTGGAGTCGTTGCGCTTTCTGTTTCTTGGCCAATGGNNNAGATCGGAAG |
| 7875_1 | TATGAGTGTGGAGTCGTTGCCCAAAAAGGAGTGCCTGGGTAATNNNAGATCGGAA |
| 7876_1 | TATGAGTGTGGAGTCGTTGCTCTAGGGGAAGGGAACAGAATGGNNNAGATCGGA |
| 7877_1 | TATGAGTGTGGAGTCGTTGCAGTGAACAAGTTGAGATGCTGGCNNNAGATCGGAA |
| 7880_1 | TATGAGTGTGGAGTCGTTGCAGCAGCAGCAACAGTAGTAATGGNNNAGATCGGAA |
| 7882_1 | TATGAGTGTGGAGTCGTTGCAACACACAGAACGGGCATAAAGTNNNAGATCGGAA |
| 7883_1 | TATGAGTGTGGAGTCGTTGCATCAGCGTCTAAGCATCACATGGNNNAGATCGGAAG |
| 7884_1 | TATGAGTGTGGAGTCGTTGCGCTAGTCTGCTACAGTCACATGGNNNAGATCGGAAG |
| 7885_1 | TATGAGTGTGGAGTCGTTGCTTTTCCATGATTGCACCAGTGCANNNAGATCGGAAG |
| 7887_1 | TATGAGTGTGGAGTCGTTGCCAGAGTGTCAGGGACCTTCCTTANNNAGATCGGAAG |
| 7893_1 | TATGAGTGTGGAGTCGTTGCAACAAGACTATAGCTGGGCATGGNNNAGATCGGAA |
| 7895_1 | TATGAGTGTGGAGTCGTTGCCAATCTGCAAAAGGACTGCATGGNNNAGATCGGAA |
| 7896_1 | TATGAGTGTGGAGTCGTTGCGAGGGTCTAGCCATTTCCCAAACNNNAGATCGGAAG |
| 7898_1 | TATGAGTGTGGAGTCGTTGCTCCTGATGAAAGCCATCTCATGGNNNAGATCGGAAG |
| 7899_1 | TATGAGTGTGGAGTCGTTGCACATGACAACCCTTCCTTCATGGNNNAGATCGGAAG |
| 7900_1 | TATGAGTGTGGAGTCGTTGCTCAACTCTGGTGGTTTGTGCTTTNNNAGATCGGAAG |

| | |
|---|---|
| 7902_1 | TATGAGTGTGGAGTCGTTGCAAAAGCAATGAGGGTGGAGATGGNNNAGATCGGA |
| 7904_1 | TATGAGTGTGGAGTCGTTGCAATACATCTGGCTGGTACGATGGNNNAGATCGGAA |
| 7905_1 | TATGAGTGTGGAGTCGTTGCTTATCCAGTCCACCATTCGATGGNNNAGATCGGAAG |
| 7907_1 | TATGAGTGTGGAGTCGTTGCAATGTCCCCTTCTCTGGGAATCANNNAGATCGGAAG |
| 7908_1 | TATGAGTGTGGAGTCGTTGCACTTGGGGACAGGTATAGGATGGNNNAGATCGGA |
| 7914_1 | TATGAGTGTGGAGTCGTTGCTTGACTATGCTGGCTTGTGATGGNNNAGATCGGAA |
| 7916_1 | TATGAGTGTGGAGTCGTTGCGCAATCTACCACGACCCATATGGNNNAGATCGGAAG |
| 7917_1 | TATGAGTGTGGAGTCGTTGCAATCCTGCCATCTGTGACTATGGNNNAGATCGGAAG |
| 7920_1 | TATGAGTGTGGAGTCGTTGCGCCCTCTGGTTTTACTGCTATGGNNNAGATCGGAAG |
| 7922_1 | TATGAGTGTGGAGTCGTTGCTTTGCTCTCAGCTCTCCTTATGGNNNAGATCGGAAGA |
| 7924_1 | TATGAGTGTGGAGTCGTTGCAAACCTCCAAAGGGGAAAACTGGNNNAGATCGGAA |
| 7925_1 | TATGAGTGTGGAGTCGTTGCACAAGTGCAAGCATTCAAACTGGNNNAGATCGGAA |
| 7926_1 | TATGAGTGTGGAGTCGTTGCCTGGGAGGCATTCAAAACCAAGTNNNAGATCGGAA |
| 7927_1 | TATGAGTGTGGAGTCGTTGCCACACACACACTCTTTTTCTGGCNNNAGATCGGAAGA |
| 7929_1 | TATGAGTGTGGAGTCGTTGCCTTGAACCTCCAGTCACCACTCTNNNAGATCGGAAGA |
| 7932_1 | TATGAGTGTGGAGTCGTTGCTCTCGTTTGTCAGAACACACTGGNNNAGATCGGAAG |
| 7936_1 | TATGAGTGTGGAGTCGTTGCAGAAGGTCCCCTCACCTATGTTCNNNAGATCGGAAG |
| 7937_1 | TATGAGTGTGGAGTCGTTGCACTGGTCACTGTTTTAGCACTGGNNNAGATCGGAAG |
| 7939_1 | TATGAGTGTGGAGTCGTTGCTTCATTCTGTGAGGCATCACTGGNNNAGATCGGAAG |
| 7940_1 | TATGAGTGTGGAGTCGTTGCTGGCCCTGCAGATGTCTATCTTTNNNAGATCGGAAG |
| 7941_1 | TATGAGTGTGGAGTCGTTGCTCCAGCCCAATAATTGTCACTGGNNNAGATCGGAAG |
| 7945_1 | TATGAGTGTGGAGTCGTTGCCAGGGACTGTCATTTCAGACTGGNNNAGATCGGAA |
| 7947_1 | TATGAGTGTGGAGTCGTTGCTGGGGGAGTGATACCAACTCTTTNNNAGATCGGAA |
| 7948_1 | TATGAGTGTGGAGTCGTTGCCGGTTTCCTTGGATAGTGACTGGNNNAGATCGGAA |
| 7949_1 | TATGAGTGTGGAGTCGTTGCTGCCAAAAGTTACCATTGACTGGNNNAGATCGGAAG |
| 7950_1 | TATGAGTGTGGAGTCGTTGCAGTTCTAGGCTGTCTGCTACTGGNNNAGATCGGAAG |
| 7951_1 | TATGAGTGTGGAGTCGTTGCCTTCCCTGAGGAGGTAGTACTGGNNNAGATCGGAA |
| 7952_1 | TATGAGTGTGGAGTCGTTGCATTCAAACCTGAGGCAAACCTGGNNNAGATCGGAAG |
| 7954_1 | TATGAGTGTGGAGTCGTTGCGAGGGACAAGAATAGCAACCTGGNNNAGATCGGA |
| 7955_1 | TATGAGTGTGGAGTCGTTGCAACCTGGCTTTAGAAACACCTGGNNNAGATCGGAAG |
| 7957_1 | TATGAGTGTGGAGTCGTTGCTGATAGGCAGTTTACCACTGGGTNNNAGATCGGAA |
| 7960_1 | TATGAGTGTGGAGTCGTTGCATAGGAGAGGAGAAACACCCTGGNNNAGATCGGA |
| 7961_1 | TATGAGTGTGGAGTCGTTGCAGGAAGGATGTTTTTGACCCTGGNNNAGATCGGAA |
| 7962_1 | TATGAGTGTGGAGTCGTTGCAACTAGAAAAAGGCAACCCCTGGNNNAGATCGGAA |
| 7964_1 | TATGAGTGTGGAGTCGTTGCTTATGTGTGTGCTTGATCCCTGGNNNAGATCGGAAG |
| 7965_1 | TATGAGTGTGGAGTCGTTGCCGGCAAGGGGAAACAATAAACATNNNAGATCGGAA |
| 7967_1 | TATGAGTGTGGAGTCGTTGCTTCTTGACAAACCACTTCCCTGGNNNAGATCGGAAG |
| 7969_1 | TATGAGTGTGGAGTCGTTGCTCATCAGCAAGATGCTGCTCTTTNNNAGATCGGAAG |
| 7974_1 | TATGAGTGTGGAGTCGTTGCTGATTGGAAAGTGGTGATCCTGGNNNAGATCGGAA |
| 7975_1 | TATGAGTGTGGAGTCGTTGCCAGCAGTGATCTTGAACTCCTGGNNNAGATCGGAAG |
| 7976_1 | TATGAGTGTGGAGTCGTTGCCCAGGCACAAGATTCTGCTCTTANNNAGATCGGAAG |
| 7977_1 | TATGAGTGTGGAGTCGTTGCAGCCGTAAAGCATCATGGGCATANNNAGATCGGAA |
| 7979_1 | TATGAGTGTGGAGTCGTTGCATTTCCTGTGGCTGTTCATCTGTNNNAGATCGGAAG |

| | |
|---|---|
| 7984_1 | TATGAGTGTGGAGTCGTTGCTACATTCACAAGAATGGGGCTGGNNNAGATCGGAA |
| 7985_1 | TATGAGTGTGGAGTCGTTGCTCTCTGTGTTGTAACATGGCTGGNNNAGATCGGAAG |
| 7986_1 | TATGAGTGTGGAGTCGTTGCTTAGTGTACTGCAAACTGGCTGGNNNAGATCGGAAG |
| 7987_1 | TATGAGTGTGGAGTCGTTGCTGGCACATGGATCTTGGACATCANNNAGATCGGAAG |
| 7990_1 | TATGAGTGTGGAGTCGTTGCTGAGGAAAAGGACAGAATGCTGGNNNAGATCGGA |
| 7991_1 | TATGAGTGTGGAGTCGTTGCTTTCTCCAAGGCTAGAATGCTGGNNNAGATCGGAAG |
| 7992_1 | TATGAGTGTGGAGTCGTTGCTTCAAGCACCTGCTATATGCTGGNNNAGATCGGAAG |
| 7996_1 | TATGAGTGTGGAGTCGTTGCACGAGATTTCACTACCTTGCTGGNNNAGATCGGAAG |
| 7998_1 | TATGAGTGTGGAGTCGTTGCTTTCTCTCCCTCCAGTCATCTGGNNNAGATCGGAAGA |
| 8001_1 | TATGAGTGTGGAGTCGTTGCTCCTTACCAGAACTCTGCTCTGGNNNAGATCGGAAG |
| 8002_1 | TATGAGTGTGGAGTCGTTGCTCATGCAAACTCATTTGCTCTGGNNNAGATCGGAAG |
| 8004_1 | TATGAGTGTGGAGTCGTTGCCCGTATTACACAAGCCTCTCTGGNNNAGATCGGAAG |
| 8005_1 | TATGAGTGTGGAGTCGTTGCAACCAGCAAACCTTCTTCTCTGGNNNAGATCGGAAG |
| 8007_1 | TATGAGTGTGGAGTCGTTGCATCTGGTTTTTGCAGAGGTCTGGNNNAGATCGGAA |
| 8008_1 | TATGAGTGTGGAGTCGTTGCTTCCTAACAGTGTGCTTGTCTGGNNNAGATCGGAAG |
| 8009_1 | TATGAGTGTGGAGTCGTTGCAAAGACACCCTGGGAAATTCTGGNNNAGATCGGAA |
| 8010_1 | TATGAGTGTGGAGTCGTTGCGTAGAGATTGGGGCACATTCTGGNNNAGATCGGAA |
| 8011_1 | TATGAGTGTGGAGTCGTTGCTCTGCATGTCATTCACCTTCTGGNNNAGATCGGAAG |
| 8014_1 | TATGAGTGTGGAGTCGTTGCTGTGACTATGGCAGTGTTTCTGGNNNAGATCGGAA |
| 8020_1 | TATGAGTGTGGAGTCGTTGCAGAACACATATCTTGCCCAGTGGNNNAGATCGGAAG |
| 8021_1 | TATGAGTGTGGAGTCGTTGCACAGCAATAAGTTAGGCCAGTGGNNNAGATCGGAA |
| 8022_1 | TATGAGTGTGGAGTCGTTGCCTCTTCTCAATCCACTCCAGTGGNNNAGATCGGAAGA |
| 8023_1 | TATGAGTGTGGAGTCGTTGCTTTTTCTTGCCTTGTTCCAGTGGNNNAGATCGGAAG |
| 8024_1 | TATGAGTGTGGAGTCGTTGCGTGATTTTGGAACCAAGCAGTGGNNNAGATCGGAA |
| 8025_1 | TATGAGTGTGGAGTCGTTGCGGTGATGAATCTCTGAGCAGTGGNNNAGATCGGAA |
| 8029_1 | TATGAGTGTGGAGTCGTTGCCAGAAACTGGGTACACAGAGTGGNNNAGATCGGAA |
| 8031_1 | TATGAGTGTGGAGTCGTTGCAGCACAGTTATGAAGCAGAGTGGNNNAGATCGGAA |
| 8034_1 | TATGAGTGTGGAGTCGTTGCAAGAAGCTGTGGTTCATGAGTGGNNNAGATCGGAA |
| 8035_1 | TATGAGTGTGGAGTCGTTGCTAGAGCCAGGGAAATCTGAGTGGNNNAGATCGGAA |
| 8038_1 | TATGAGTGTGGAGTCGTTGCTGCTGCCTTTGGAATCATAGTGGNNNAGATCGGAAG |
| 8042_1 | TATGAGTGTGGAGTCGTTGCGAGGTTACAGATCTTTCCCGTGGNNNAGATCGGAAG |
| 8043_1 | TATGAGTGTGGAGTCGTTGCTTTCGCCACAATCCTTATCGTGGNNNAGATCGGAAG |
| 8044_1 | TATGAGTGTGGAGTCGTTGCGGTTCTGGTTTAGGGAAAGGTGGNNNAGATCGGA |
| 8047_1 | TATGAGTGTGGAGTCGTTGCGCAAGAATACAGGGGTAAGGTGGNNNAGATCGGA |
| 8048_1 | TATGAGTGTGGAGTCGTTGCGATACCAAGTGCTTCTGAGGTGGNNNAGATCGGAA |
| 8049_1 | TATGAGTGTGGAGTCGTTGCTACCACCCCAATGCAACCTATGNNNAGATCGGAAGA |
| 8051_1 | TATGAGTGTGGAGTCGTTGCTCCAAAAATGTTTCAGCTGGGATNNNAGATCGGAAG |
| 8052_1 | TATGAGTGTGGAGTCGTTGCAGAGAGAGAAGGTAAAGGGGTGGNNNAGATCGGA |
| 8053_1 | TATGAGTGTGGAGTCGTTGCAGGAGAAGAAAGATTGGGGGTGGNNNAGATCGGA |
| 8054_1 | TATGAGTGTGGAGTCGTTGCTCAACATCATGGAAACTGGGTGGNNNAGATCGGAA |
| 8059_1 | TATGAGTGTGGAGTCGTTGCGAACTCCTCATCTTCCATGGTGGNNNAGATCGGAAG |
| 8062_1 | TATGAGTGTGGAGTCGTTGCGATTGGAGGGGATGTTATGGTGGNNNAGATCGGA |
| 8063_1 | TATGAGTGTGGAGTCGTTGCGTTTGCGAATCTTCAACTGGTGGNNNAGATCGGAAG |

| | |
|---|---|
| 8065_1 | TATGAGTGTGGAGTCGTTGCACAGTGAAGGGCTGCTTTTCTTTNNNAGATCGGAAG |
| 8066_1 | TATGAGTGTGGAGTCGTTGCGTGAACAGACAGTTGCAATGTGGNNNAGATCGGAA |
| 8068_1 | TATGAGTGTGGAGTCGTTGCAGCACAATCATAGCTCCATGTGGNNNAGATCGGAAG |
| 8070_1 | TATGAGTGTGGAGTCGTTGCGAGCTTCTAGGTAATGTAGCAGTNNNAGATCGGAA |
| 8073_1 | TATGAGTGTGGAGTCGTTGCCCCTGATTTTGATTTCCCTGTGGNNNAGATCGGAAG |
| 8076_1 | TATGAGTGTGGAGTCGTTGCCTGTAGTGGAGGTTTCTGTGTGGNNNAGATCGGAA |
| 8077_1 | TATGAGTGTGGAGTCGTTGCAGAGTGACATCAGCCAATTGTGGNNNAGATCGGAA |
| 8078_1 | TATGAGTGTGGAGTCGTTGCAACGACCTAAATTCTAGCACTGCNNNAGATCGGAAG |
| 8079_1 | TATGAGTGTGGAGTCGTTGCTGGAAAAGCTTAGGGGATTGTGGNNNAGATCGGA |
| 8085_1 | TATGAGTGTGGAGTCGTTGCCACATGAACACACACCCAGTATANNNAGATCGGAAG |
| 8087_1 | TATGAGTGTGGAGTCGTTGCTCTGTCATCATCACCGTCATTGGNNNAGATCGGAAG |
| 8089_1 | TATGAGTGTGGAGTCGTTGCAGAAAAAGCCCCGTAAGGATTGGNNNAGATCGGAA |
| 8091_1 | TATGAGTGTGGAGTCGTTGCTTGCACACCCTACACTTGATTGGNNNAGATCGGAAG |
| 8095_1 | TATGAGTGTGGAGTCGTTGCTCACCATCACCACTCAAACTTGGNNNAGATCGGAAG |
| 8097_1 | TATGAGTGTGGAGTCGTTGCGCTGTTTTCACCCAAGAACTTGGNNNAGATCGGAAG |
| 8098_1 | TATGAGTGTGGAGTCGTTGCTATGTCTGTGCAATCCCACTTGGNNNAGATCGGAAG |
| 8099_1 | TATGAGTGTGGAGTCGTTGCCGTTGGTTGTCACAAAGACTTGGNNNAGATCGGAA |
| 8100_1 | TATGAGTGTGGAGTCGTTGCGTGTGTGTGTCTGTATCCCTTGGNNNAGATCGGAAG |
| 8101_1 | TATGAGTGTGGAGTCGTTGCGGGGTAGCTGAAATAAGCCTTGGNNNAGATCGGAA |
| 8102_1 | TATGAGTGTGGAGTCGTTGCCCTCACATAATCTCCTGCCTTGGNNNAGATCGGAAGA |
| 8104_1 | TATGAGTGTGGAGTCGTTGCTTTGTTTTCTTCCCTCATCCCCCNNNAGATCGGAAGAG |
| 8105_1 | TATGAGTGTGGAGTCGTTGCAAGGAGAAAAACAGGAGGCTTGGNNNAGATCGGA |
| 8106_1 | TATGAGTGTGGAGTCGTTGCCCTGCAAAAACAGACCATCTTGGNNNAGATCGGAAG |
| 8108_1 | TATGAGTGTGGAGTCGTTGCAAATGGCTGGGGATCATTCTTGGNNNAGATCGGAA |
| 8109_1 | TATGAGTGTGGAGTCGTTGCTAGTAGGGGTTGAGGGTTCTTGGNNNAGATCGGAA |
| 8110_1 | TATGAGTGTGGAGTCGTTGCATGGCAGCTCTAAATGGAGTTGGNNNAGATCGGAA |
| 8111_1 | TATGAGTGTGGAGTCGTTGCAAAAGCTGTGGGATTGGAGTTGGNNNAGATCGGA |
| 8112_1 | TATGAGTGTGGAGTCGTTGCTTATAGCTTCCCCAAGACGTTGGNNNAGATCGGAAG |
| 8114_1 | TATGAGTGTGGAGTCGTTGCAATGTCAACTGTACCGAGGTTGGNNNAGATCGGAA |
| 8115_1 | TATGAGTGTGGAGTCGTTGCACACACATTAGTACAGGGGTTGGNNNAGATCGGAA |
| 8118_1 | TATGAGTGTGGAGTCGTTGCTCTAGGCACCTTGCATATGTTGGNNNAGATCGGAAG |
| 8120_1 | TATGAGTGTGGAGTCGTTGCGATACTACCACTGGGCTTGTTGGNNNAGATCGGAAG |
| 8122_1 | TATGAGTGTGGAGTCGTTGCTTCAGGGTGCATGCTCTATGTTGNNNAGATCGGAAG |
| 8123_1 | TATGAGTGTGGAGTCGTTGCTGATGAGATTCCTGGGCATTTGGNNNAGATCGGAA |
| 8127_1 | TATGAGTGTGGAGTCGTTGCAGCCAAACACTCTGGTACTTTGGNNNAGATCGGAAG |
| 8128_1 | TATGAGTGTGGAGTCGTTGCTAGCTGAACAGAGTGAGCTTTGGNNNAGATCGGAA |
| 8129_1 | TATGAGTGTGGAGTCGTTGCCACACTGTCTTCTTACGGTTTGGNNNAGATCGGAAG |
| 8132_1 | TATGAGTGTGGAGTCGTTGCTGGGAAAACATGTTCCTGTTTGGNNNAGATCGGAA |
| 8134_1 | TATGAGTGTGGAGTCGTTGCAGCCTATGTCTGCCTCATTTTGGNNNAGATCGGAAG |
| 8135_1 | TATGAGTGTGGAGTCGTTGCAGGCTGGGGAAGAAAGATTTTGGNNNAGATCGGA |
| 8136_1 | TATGAGTGTGGAGTCGTTGCCCTGGGACCTGACGATATTTTGGNNNAGATCGGAA |
| 8138_1 | TATGAGTGTGGAGTCGTTGCATAAAACCGCTGACACAACTGGTNNNAGATCGGAAG |
| 8140_1 | TATGAGTGTGGAGTCGTTGCCGTGTGCATGTATCTGGTTTTGGNNNAGATCGGAA |

| | |
|---|---|
| 8141_1 | TATGAGTGTGGAGTCGTTGCCTTGGGGTCATTGAGCTTTTTGGNNNAGATCGGAAG |
| 8142_1 | TATGAGTGTGGAGTCGTTGCTCCCATCATCTGCCCTCAAAATGNNNAGATCGGAAGA |
| 8143_1 | TATGAGTGTGGAGTCGTTGCGCTCTCCTCTGGGTCTCAAAATGNNNAGATCGGAAGA |
| 8145_1 | TATGAGTGTGGAGTCGTTGCTGTCTGTGGTGATAGATAGGGGTNNNAGATCGGAA |
| 8147_1 | TATGAGTGTGGAGTCGTTGCCCCCATCTTCTCTCCCACAAATGNNNAGATCGGAAGA |
| 8148_1 | TATGAGTGTGGAGTCGTTGCACAAAAGCCAGAGTGCACAAATGNNNAGATCGGAA |
| 8149_1 | TATGAGTGTGGAGTCGTTGCAGAATCAGTTTGCTGGCCAAATGNNNAGATCGGAAG |
| 8150_1 | TATGAGTGTGGAGTCGTTGCGGCTGGTCTGCAATGTTCTATCANNNAGATCGGAAG |
| 8152_1 | TATGAGTGTGGAGTCGTTGCGCAGGTCCGCACTAAATCAAATGNNNAGATCGGAAG |
| 8154_1 | TATGAGTGTGGAGTCGTTGCTGTCCCTCTAAGGGGAGGAAATGNNNAGATCGGAA |
| 8155_1 | TATGAGTGTGGAGTCGTTGCCGGGTAGGGAGGTCTTTGAAATGNNNAGATCGGAA |
| 8157_1 | TATGAGTGTGGAGTCGTTGCCTTTAGCAGGACACTGCACAATGNNNAGATCGGAAG |
| 8161_1 | TATGAGTGTGGAGTCGTTGCTGCGTTGATACCAGGGAAGAATGNNNAGATCGGAA |
| 8164_1 | TATGAGTGTGGAGTCGTTGCGTGTTGTGTTGTGCAGGAGAATGNNNAGATCGGAA |
| 8165_1 | TATGAGTGTGGAGTCGTTGCGCTGGGTGCCAAGAAGTAGAATGNNNAGATCGGAA |
| 8166_1 | TATGAGTGTGGAGTCGTTGCGGGAGGAGAGCATACTGAAATGCNNNAGATCGGAA |
| 8167_1 | TATGAGTGTGGAGTCGTTGCCACCAACGTTGCCTTGTAGAATGNNNAGATCGGAAG |
| 8171_1 | TATGAGTGTGGAGTCGTTGCGCTGTACCACTTGAGAGGGAATGNNNAGATCGGAA |
| 8172_1 | TATGAGTGTGGAGTCGTTGCATGCCAGTTTTGCAATGGGAATGNNNAGATCGGAA |
| 8173_1 | TATGAGTGTGGAGTCGTTGCTGTCTCTTCACCAACTGGGAATGNNNAGATCGGAAG |
| 8176_1 | TATGAGTGTGGAGTCGTTGCCCTGGGTATGTGGATGGTGAATGNNNAGATCGGAA |
| 8178_1 | TATGAGTGTGGAGTCGTTGCCTCTGGTGGAAGCCATGCATTTANNNAGATCGGAAG |
| 8181_1 | TATGAGTGTGGAGTCGTTGCCCTTCTGGGGAAATGGGGTAATGNNNAGATCGGAA |
| 8182_1 | TATGAGTGTGGAGTCGTTGCTCATGCAAGCCAACAGAAACATGNNNAGATCGGAAG |
| 8185_1 | TATGAGTGTGGAGTCGTTGCGAGTTTCCCAACATGGCAACATGNNNAGATCGGAAG |
| 8186_1 | TATGAGTGTGGAGTCGTTGCTCTGTGGTGTACGTCGTAACATGNNNAGATCGGAAG |
| 8187_1 | TATGAGTGTGGAGTCGTTGCCAGGGAAAAGACCACACCACATGNNNAGATCGGAA |
| 8190_1 | TATGAGTGTGGAGTCGTTGCATCAAGCACCAGAACCAGACATGNNNAGATCGGAAG |
| 8193_1 | TATGAGTGTGGAGTCGTTGCAAACATGCCTGGACCACTACATGNNNAGATCGGAAG |
| 8202_1 | TATGAGTGTGGAGTCGTTGCTGCTTACTTTTAGAGGCCCCATGNNNAGATCGGAAGA |
| 8203_1 | TATGAGTGTGGAGTCGTTGCACTTATCTTGCAGCATCCCCATGNNNAGATCGGAAGA |
| 8204_1 | TATGAGTGTGGAGTCGTTGCGAGGGCAGTATGATTTCCCCATGNNNAGATCGGAAG |
| 8208_1 | TATGAGTGTGGAGTCGTTGCGCTGCTACTTTTTGGAAGCCATGNNNAGATCGGAAG |
| 8209_1 | TATGAGTGTGGAGTCGTTGCTATTTGGTTCCCAGATGGCCATGNNNAGATCGGAAG |
| 8211_1 | TATGAGTGTGGAGTCGTTGCAGCCAGACTGTGATATTGCCATGNNNAGATCGGAAG |
| 8214_1 | TATGAGTGTGGAGTCGTTGCTTATTCCGTCCTGTTGCTCCATGNNNAGATCGGAAGA |
| 8216_1 | TATGAGTGTGGAGTCGTTGCGGGGACTAAGAGAGAGAAGCATGNNNAGATCGGAA |
| 8218_1 | TATGAGTGTGGAGTCGTTGCGGTAAGGTATCTTGGGTGGCATGNNNAGATCGGAA |
| 8223_1 | TATGAGTGTGGAGTCGTTGCCCAGTTATTGCCCCATTTGCATGNNNAGATCGGAAGA |
| 8224_1 | TATGAGTGTGGAGTCGTTGCCACCAGGCTCACATATTTGCATGNNNAGATCGGAAGA |
| 8225_1 | TATGAGTGTGGAGTCGTTGCAGCAGCAAAACCAAAGCATCATGNNNAGATCGGAAG |
| 8227_1 | TATGAGTGTGGAGTCGTTGCGGGAATGACAGGGTCTCATCATGNNNAGATCGGAA |
| 8229_1 | TATGAGTGTGGAGTCGTTGCGAGCTAAGCCAATGCTGATCATGNNNAGATCGGAAG |

| | |
|---|---|
| 8230_1 | TATGAGTGTGGAGTCGTTGCTCTCTCTCTCTCCAAGGCTCATGNNNAGATCGGAAGA |
| 8232_1 | TATGAGTGTGGAGTCGTTGCGGAGAACCCTGATTGCTCTCATGNNNAGATCGGAAG |
| 8233_1 | TATGAGTGTGGAGTCGTTGCGGCGAAAAGACCAACAAGTCATGNNNAGATCGGAA |
| 8239_1 | TATGAGTGTGGAGTCGTTGCCCCAGCTGACTTCTAGGTTCATGNNNAGATCGGAAG |
| 8240_1 | TATGAGTGTGGAGTCGTTGCGGCTGTCATCATGGCATTTCATGNNNAGATCGGAAG |
| 8243_1 | TATGAGTGTGGAGTCGTTGCCAGGCATAGAGACTGCCAAGATGNNNAGATCGGAA |
| 8244_1 | TATGAGTGTGGAGTCGTTGCTTAGCGATGCAGATGGGAAGATGNNNAGATCGGAA |
| 8245_1 | TATGAGTGTGGAGTCGTTGCTAGAGGGAGCAGGATGGAAGATGNNNAGATCGGA |
| 8246_1 | TATGAGTGTGGAGTCGTTGCGCCAGTGTACTGAGTGGAAGATGNNNAGATCGGAA |
| 8247_1 | TATGAGTGTGGAGTCGTTGCCGGGAGAAGGCTATGTGAAGATGNNNAGATCGGA |
| 8253_1 | TATGAGTGTGGAGTCGTTGCTTCATTTCCCCCACCAAGAGATGNNNAGATCGGAAG |
| 8254_1 | TATGAGTGTGGAGTCGTTGCACTCTCACTCACTCACACACAGTNNNAGATCGGAAGA |
| 8255_1 | TATGAGTGTGGAGTCGTTGCCACTGAGGCTGCTGTTAGAGATGNNNAGATCGGAA |
| 8257_1 | TATGAGTGTGGAGTCGTTGCCCTGACTTTGCCCTTAGGAGATGNNNAGATCGGAAG |
| 8258_1 | TATGAGTGTGGAGTCGTTGCCAAAAGGACCCAGACTGGAGATGNNNAGATCGGAA |
| 8259_1 | TATGAGTGTGGAGTCGTTGCTCATGGGGCTTTTTCCTGAGATGNNNAGATCGGAAG |
| 8260_1 | TATGAGTGTGGAGTCGTTGCTGTAGTAGGTGGAGGCTGAGATGNNNAGATCGGA |
| 8261_1 | TATGAGTGTGGAGTCGTTGCTGGGTGTCTTCAGCAACTAGATGNNNAGATCGGAA |
| 8263_1 | TATGAGTGTGGAGTCGTTGCAAATGGTTGTAGGGCCAAGGATGNNNAGATCGGAA |
| 8264_1 | TATGAGTGTGGAGTCGTTGCTGCTGGCTCTGGTATTTTGGTTTNNNAGATCGGAAG |
| 8266_1 | TATGAGTGTGGAGTCGTTGCTAGTGATTAGGGCCAGGAGGATGNNNAGATCGGA |
| 8269_1 | TATGAGTGTGGAGTCGTTGCCGTCATGATTAGAGAGCGGGATGNNNAGATCGGAA |
| 8270_1 | TATGAGTGTGGAGTCGTTGCGGACAGAATGAATCCTGGGGATGNNNAGATCGGAA |
| 8271_1 | TATGAGTGTGGAGTCGTTGCGGCAATCCACCAGGTATGCTTTANNNAGATCGGAAG |
| 8273_1 | TATGAGTGTGGAGTCGTTGCGAGAGAGAAACAGTGCTGGGATGNNNAGATCGGA |
| 8275_1 | TATGAGTGTGGAGTCGTTGCAAGAAGAGCAGCAAGGATGGATGNNNAGATCGGA |
| 8278_1 | TATGAGTGTGGAGTCGTTGCGCTTGAGAGCTATGCAGTGGATGNNNAGATCGGAA |
| 8280_1 | TATGAGTGTGGAGTCGTTGCTGTGTAGGATTGGCCTTTGGATGNNNAGATCGGAA |
| 8281_1 | TATGAGTGTGGAGTCGTTGCAAACACAGCATCCGCAAATGATGNNNAGATCGGAA |
| 8282_1 | TATGAGTGTGGAGTCGTTGCGGAGTGGGTAGTAAGCCATGATGNNNAGATCGGA |
| 8283_1 | TATGAGTGTGGAGTCGTTGCAGAAAGGGCACCTCATGATGATGNNNAGATCGGAA |
| 8285_1 | TATGAGTGTGGAGTCGTTGCCCCTGGCCATCATCTCCTCTTTANNNAGATCGGAAGA |
| 8287_1 | TATGAGTGTGGAGTCGTTGCTGCCCTTCACATTAGCTCTGATGNNNAGATCGGAAG |
| 8289_1 | TATGAGTGTGGAGTCGTTGCGGTGTTTGTGACAAGCTGTGATGNNNAGATCGGAA |
| 8290_1 | TATGAGTGTGGAGTCGTTGCCAGGAGAACAAGAGGGTGTGATGNNNAGATCGGA |
| 8291_1 | TATGAGTGTGGAGTCGTTGCGGATGGGAGACTGTGACTTGATGNNNAGATCGGAA |
| 8292_1 | TATGAGTGTGGAGTCGTTGCAGCCTGATTGCTGCATTTTGATGNNNAGATCGGAAG |
| 8294_1 | TATGAGTGTGGAGTCGTTGCAGAGAGATTGGGGTGGGGATATGNNNAGATCGGA |
| 8296_1 | TATGAGTGTGGAGTCGTTGCGCTCCCCTGTGTGTGTGTATATGNNNAGATCGGAAG |
| 8297_1 | TATGAGTGTGGAGTCGTTGCCCACCCTACCCTACTCCACTATGNNNAGATCGGAAGA |
| 8299_1 | TATGAGTGTGGAGTCGTTGCGACCTCCAAACTGGGAAGCTATGNNNAGATCGGAA |
| 8303_1 | TATGAGTGTGGAGTCGTTGCAGCATGTGGCAACTAGCAGTATGNNNAGATCGGAA |
| 8305_1 | TATGAGTGTGGAGTCGTTGCGAGCAAGCATATTCTCCGACAGTNNNAGATCGGAAG |

| | |
|---|---|
| 8306_1 | TATGAGTGTGGAGTCGTTGCTGCTCTGGACACTGTTTGGTATGNNNAGATCGGAAG |
| 8307_1 | TATGAGTGTGGAGTCGTTGCGAGTGTGGGCCAGTGTATGTATGNNNAGATCGGAA |
| 8310_1 | TATGAGTGTGGAGTCGTTGCGGCGGGGTCCGATAGTAATTATGNNNAGATCGGAA |
| 8311_1 | TATGAGTGTGGAGTCGTTGCGTGGGTGAGACTGGGAGTTTATGNNNAGATCGGA |
| 8313_1 | TATGAGTGTGGAGTCGTTGCAACCCAAACACCACAGAAAACTGNNNAGATCGGAAG |
| 8314_1 | TATGAGTGTGGAGTCGTTGCATGACACGCTTGTTGACAAACTGNNNAGATCGGAAG |
| 8318_1 | TATGAGTGTGGAGTCGTTGCTGAGACCCACTTCAGGAGAACTGNNNAGATCGGAA |
| 8319_1 | TATGAGTGTGGAGTCGTTGCCATCATTGCTTTGCCAGGAACTGNNNAGATCGGAAG |
| 8320_1 | TATGAGTGTGGAGTCGTTGCAAATGGTCTGCAGCTAGGAACTGNNNAGATCGGAA |
| 8321_1 | TATGAGTGTGGAGTCGTTGCTCCTTACATTCCCACCAACACTGNNNAGATCGGAAGA |
| 8325_1 | TATGAGTGTGGAGTCGTTGCCTAACCAGGAACCAATCCCACTGNNNAGATCGGAAG |
| 8326_1 | TATGAGTGTGGAGTCGTTGCGGGAAATTAGGAGACAGCCACTGNNNAGATCGGAA |
| 8328_1 | TATGAGTGTGGAGTCGTTGCCAGACCCTTGTGTGTATCCACTGNNNAGATCGGAAG |
| 8330_1 | TATGAGTGTGGAGTCGTTGCGACAAGCTTCCTTACCTCCACTGNNNAGATCGGAAG |
| 8332_1 | TATGAGTGTGGAGTCGTTGCGAGGACATTTTGAGTTGGCACTGNNNAGATCGGAA |
| 8333_1 | TATGAGTGTGGAGTCGTTGCAAACCAGGAGAACTTCTGCACTGNNNAGATCGGAA |
| 8339_1 | TATGAGTGTGGAGTCGTTGCATTTGGTCACGATGAAGTCACTGNNNAGATCGGAA |
| 8343_1 | TATGAGTGTGGAGTCGTTGCACGCAGATGATACAGAAATGGGTNNNAGATCGGAA |
| 8352_1 | TATGAGTGTGGAGTCGTTGCGGCATAGTCCTCCATTTGGACTGNNNAGATCGGAAG |
| 8353_1 | TATGAGTGTGGAGTCGTTGCTCACAGTGCAGATTTCCTGACTGNNNAGATCGGAAG |
| 8354_1 | TATGAGTGTGGAGTCGTTGCTCTGCAACATGCTCTAGTGACTGNNNAGATCGGAAG |
| 8356_1 | TATGAGTGTGGAGTCGTTGCGAAGAGACACTGAGCTGTGACTGNNNAGATCGGAA |
| 8357_1 | TATGAGTGTGGAGTCGTTGCAATCTGGGTGGGAGAGGATACTGNNNAGATCGGA |
| 8362_1 | TATGAGTGTGGAGTCGTTGCGGCAGAGTGAAGGCTTTCTACTGNNNAGATCGGAA |
| 8368_1 | TATGAGTGTGGAGTCGTTGCCTCAATGCAATGTCAACCACCTGNNNAGATCGGAAG |
| 8372_1 | TATGAGTGTGGAGTCGTTGCGAGAAAGACTTCAGACCACCCTGNNNAGATCGGAA |
| 8373_1 | TATGAGTGTGGAGTCGTTGCAGAGATAGAGACAGAGCACCCTGNNNAGATCGGAA |
| 8374_1 | TATGAGTGTGGAGTCGTTGCCTGGGTTTTCTTCAGGTACCCTGNNNAGATCGGAAG |
| 8376_1 | TATGAGTGTGGAGTCGTTGCAGGCCATCAACATTTTCACCAGTNNNAGATCGGAAG |
| 8377_1 | TATGAGTGTGGAGTCGTTGCGCAGACCACATCCTGTAGAGGTTNNNAGATCGGAA |
| 8378_1 | TATGAGTGTGGAGTCGTTGCCCTATTCCTGTTTGTTCCCCCTGNNNAGATCGGAAGA |
| 8379_1 | TATGAGTGTGGAGTCGTTGCATTGTATTTCTGCTCCAGCCCTGNNNAGATCGGAAG |
| 8380_1 | TATGAGTGTGGAGTCGTTGCAACATGAAAGGAGCATAGCCCTGNNNAGATCGGAA |
| 8381_1 | TATGAGTGTGGAGTCGTTGCTCTTAGTCACGGATGGATCCCTGNNNAGATCGGAAG |
| 8383_1 | TATGAGTGTGGAGTCGTTGCTGGGAATTGCATAAGTGTCCCTGNNNAGATCGGAA |
| 8384_1 | TATGAGTGTGGAGTCGTTGCATGATGCTGTAAGTGCTTCCCTGNNNAGATCGGAAG |
| 8385_1 | TATGAGTGTGGAGTCGTTGCGGGGATTCACAAAACAAAGCCTGNNNAGATCGGAA |
| 8386_1 | TATGAGTGTGGAGTCGTTGCAACACTGCCACTTAGAAAGCCTGNNNAGATCGGAAG |
| 8387_1 | TATGAGTGTGGAGTCGTTGCGTCTTCACTTTTAGGGGAGCCTGNNNAGATCGGAAG |
| 8388_1 | TATGAGTGTGGAGTCGTTGCAGCATAACTTTGAGTGGAGCCTGNNNAGATCGGAA |
| 8389_1 | TATGAGTGTGGAGTCGTTGCACAGTTTAGTTCATGGAGGCCTGNNNAGATCGGAA |
| 8390_1 | TATGAGTGTGGAGTCGTTGCTCCATCTATCTCAGGATGGCCTGNNNAGATCGGAAG |
| 8391_1 | TATGAGTGTGGAGTCGTTGCAAGAGAATATGTGTGCTGGCCTGNNNAGATCGGAA |

| | |
|---|---|
| 8392_1 | TATGAGTGTGGAGTCGTTGCTATTCTAGGCCCTCCACATCCTGNNNAGATCGGAAG |
| 8393_1 | TATGAGTGTGGAGTCGTTGCGTGGGGAAACACTTGAGATCCTGNNNAGATCGGAA |
| 8394_1 | TATGAGTGTGGAGTCGTTGCAAGTGTTGGCTCACATGATCCTGNNNAGATCGGAAG |
| 8395_1 | TATGAGTGTGGAGTCGTTGCTATTTGCCAGCTCAGAACTCCTGNNNAGATCGGAAG |
| 8397_1 | TATGAGTGTGGAGTCGTTGCATCACAATGGTCTCTCCCAGACANNNAGATCGGAAG |
| 8399_1 | TATGAGTGTGGAGTCGTTGCATGAAAGGCAAAGGTGATGCTCTNNNAGATCGGAA |
| 8401_1 | TATGAGTGTGGAGTCGTTGCCAGCCATAGAATGCCTTGTCCTGNNNAGATCGGAAG |
| 8402_1 | TATGAGTGTGGAGTCGTTGCTGGTAGGGAGTGAAGTCTTCCTGNNNAGATCGGAA |
| 8406_1 | TATGAGTGTGGAGTCGTTGCAGAACTCACAGTGACCAAAGCTGNNNAGATCGGAA |
| 8407_1 | TATGAGTGTGGAGTCGTTGCTAGCTGAGTGCCTTGTAAAGCTGNNNAGATCGGAA |
| 8409_1 | TATGAGTGTGGAGTCGTTGCACGAAAGTTGCATACACAAGCTGNNNAGATCGGAA |
| 8411_1 | TATGAGTGTGGAGTCGTTGCGGTCCAGATGGTTTTACCAGCAANNNAGATCGGAA |
| 8413_1 | TATGAGTGTGGAGTCGTTGCCACAATGGCATGTTACACAGCTGNNNAGATCGGAAG |
| 8415_1 | TATGAGTGTGGAGTCGTTGCACCTATCTACATGAAGGGCCAGTNNNAGATCGGAAG |
| 8416_1 | TATGAGTGTGGAGTCGTTGCCCTGGCCATTACTTATCCAGCTGNNNAGATCGGAAG |
| 8417_1 | TATGAGTGTGGAGTCGTTGCGATACTCCTCCACTTACGCTGGGNNNAGATCGGAAG |
| 8421_1 | TATGAGTGTGGAGTCGTTGCAATGCACACTGATGGTAGAGCTGNNNAGATCGGAA |
| 8422_1 | TATGAGTGTGGAGTCGTTGCAGACTTTGCAATACTGGGAGCTGNNNAGATCGGAA |
| 8423_1 | TATGAGTGTGGAGTCGTTGCTAGGATTGCACAGAAGTGAGCTGNNNAGATCGGAA |
| 8424_1 | TATGAGTGTGGAGTCGTTGCCAATAGAGCCTAGAGCTGCCAGTNNNAGATCGGAA |
| 8426_1 | TATGAGTGTGGAGTCGTTGCTGACAAGGGAGAGAGACTAGCTGNNNAGATCGGA |
| 8427_1 | TATGAGTGTGGAGTCGTTGCGAGAATGCTGCTGGTAGTAGCTGNNNAGATCGGAA |
| 8428_1 | TATGAGTGTGGAGTCGTTGCAGAGAAGTCAACAGCTGTAGCTGNNNAGATCGGAA |
| 8430_1 | TATGAGTGTGGAGTCGTTGCGAGACAGATGGTTCTGAAGGCTGNNNAGATCGGAA |
| 8431_1 | TATGAGTGTGGAGTCGTTGCCGATTAGCCTAAGGTGAAGGCTGNNNAGATCGGAA |
| 8432_1 | TATGAGTGTGGAGTCGTTGCCTTTTCTGCTGTTGTGAAGGCTGNNNAGATCGGAAG |
| 8433_1 | TATGAGTGTGGAGTCGTTGCTACATAGCTACGGGTACAGGCTGNNNAGATCGGAA |
| 8434_1 | TATGAGTGTGGAGTCGTTGCCACCACAGCATGGATGGAAAAACNNNAGATCGGAA |
| 8435_1 | TATGAGTGTGGAGTCGTTGCGGTCCTAGACACTAGAGAGGCTGNNNAGATCGGAA |
| 8438_1 | TATGAGTGTGGAGTCGTTGCGGTCCATGTGATTGTAGAGGCTGNNNAGATCGGAA |
| 8439_1 | TATGAGTGTGGAGTCGTTGCACTTTACGTGGCTCATTAGGCTGNNNAGATCGGAAG |
| 8441_1 | TATGAGTGTGGAGTCGTTGCATCTGTCGAAATCACAAGGGCTGNNNAGATCGGAA |
| 8442_1 | TATGAGTGTGGAGTCGTTGCACAAAATAGGGTTGACTGGGCTGNNNAGATCGGAA |
| 8444_1 | TATGAGTGTGGAGTCGTTGCAAGTTTTGTCTGGAGGATGGCTGNNNAGATCGGAA |
| 8445_1 | TATGAGTGTGGAGTCGTTGCCTCTCTCTCATTCGTGATGGCTGNNNAGATCGGAAG |
| 8446_1 | TATGAGTGTGGAGTCGTTGCACATCAGGTTTTCTAGCTGGCTGNNNAGATCGGAAG |
| 8447_1 | TATGAGTGTGGAGTCGTTGCTTCTTTCTGACAGTGTCTGGCTGNNNAGATCGGAAG |
| 8449_1 | TATGAGTGTGGAGTCGTTGCGCTATCTCCTTGTGACTTGGCTGNNNAGATCGGAAG |
| 8450_1 | TATGAGTGTGGAGTCGTTGCAGGTTCCCTCTTCACAAATGCTGNNNAGATCGGAAG |
| 8451_1 | TATGAGTGTGGAGTCGTTGCTTTCCTCAATGAACCTCCTGCTGNNNAGATCGGAAG |
| 8453_1 | TATGAGTGTGGAGTCGTTGCTGGAGGTGGAAAGTTCATAGCAANNNAGATCGGAA |
| 8455_1 | TATGAGTGTGGAGTCGTTGCTTCTTAGCATGCACTTTGTGCTGNNNAGATCGGAAG |
| 8456_1 | TATGAGTGTGGAGTCGTTGCTTATTCCCACTGCCTGATTGCTGNNNAGATCGGAAG |

| | |
|---|---|
| 8457_1 | TATGAGTGTGGAGTCGTTGCGCGTAAATGAATGCAACTTGCTGNNNAGATCGGAA |
| 8458_1 | TATGAGTGTGGAGTCGTTGCATGTTTATTGCAGCATCTTGCTGNNNAGATCGGAAG |
| 8459_1 | TATGAGTGTGGAGTCGTTGCCTACATTGCCTTGCATCTTGCTGNNNAGATCGGAAG |
| 8460_1 | TATGAGTGTGGAGTCGTTGCCTCAACACAGGCAATTCTTGCTGNNNAGATCGGAAG |
| 8463_1 | TATGAGTGTGGAGTCGTTGCAGGCTGAGGAATCCAACAATCTGNNNAGATCGGAA |
| 8464_1 | TATGAGTGTGGAGTCGTTGCATGGACAAACCTGGGACAATCTGNNNAGATCGGAA |
| 8465_1 | TATGAGTGTGGAGTCGTTGCGTTTGTGTGTGTGCCAGAATCTGNNNAGATCGGAA |
| 8468_1 | TATGAGTGTGGAGTCGTTGCACTTGGAGTGCCTCTTGAATCTGNNNAGATCGGAAG |
| 8469_1 | TATGAGTGTGGAGTCGTTGCATACAGCCAGCCCTTTGAATCTGNNNAGATCGGAAG |
| 8470_1 | TATGAGTGTGGAGTCGTTGCTTGCCTCCACTTCTGAACATCTGNNNAGATCGGAAGA |
| 8471_1 | TATGAGTGTGGAGTCGTTGCAAGGGTGCTGAGTTTCACATCTGNNNAGATCGGAA |
| 8472_1 | TATGAGTGTGGAGTCGTTGCATGCCTGATCCCAAACTCATCTGNNNAGATCGGAAG |
| 8476_1 | TATGAGTGTGGAGTCGTTGCATTTGATGCTCTCGACCACTCTGNNNAGATCGGAAG |
| 8477_1 | TATGAGTGTGGAGTCGTTGCTGCTCAAGGCTTGTCTTACTCTGNNNAGATCGGAAG |
| 8479_1 | TATGAGTGTGGAGTCGTTGCGCTCCACACTCAATAACCCTCTGNNNAGATCGGAAGA |
| 8481_1 | TATGAGTGTGGAGTCGTTGCCAATAAATGCTGGCACAGCTCTGNNNAGATCGGAAG |
| 8482_1 | TATGAGTGTGGAGTCGTTGCTTGCTGAATAGGATGCAGCTCTGNNNAGATCGGAA |
| 8483_1 | TATGAGTGTGGAGTCGTTGCCTGGGTAAGGTCTTTTGGCTCTGNNNAGATCGGAAG |
| 8485_1 | TATGAGTGTGGAGTCGTTGCGGGGTAACATGCCAAACTCTCTGNNNAGATCGGAAG |
| 8489_1 | TATGAGTGTGGAGTCGTTGCTCTCTCTCCAGCCATCTTCTCTGNNNAGATCGGAAGA |
| 8490_1 | TATGAGTGTGGAGTCGTTGCCTTGGTTTGCAATATGGAGGCAGNNNAGATCGGAA |
| 8496_1 | TATGAGTGTGGAGTCGTTGCCAGCAAGGTGTCTTTCCACAAACNNNAGATCGGAAG |
| 8497_1 | TATGAGTGTGGAGTCGTTGCCTCACTACCATTGCTTCCGTCTGNNNAGATCGGAAGA |
| 8502_1 | TATGAGTGTGGAGTCGTTGCTCCTCTCTGTAGCATCTGGTCTGNNNAGATCGGAAG |
| 8503_1 | TATGAGTGTGGAGTCGTTGCTATGGCAATCACTGTGTGGTCTGNNNAGATCGGAAG |
| 8504_1 | TATGAGTGTGGAGTCGTTGCTAGAGGGAGCCTTGAGATGTCTGNNNAGATCGGAA |
| 8505_1 | TATGAGTGTGGAGTCGTTGCCCCATTCCAATACCCACTGTCTGNNNAGATCGGAAGA |
| 8506_1 | TATGAGTGTGGAGTCGTTGCCTTCTCCACATTCCCACTGTCTGNNNAGATCGGAAGA |
| 8507_1 | TATGAGTGTGGAGTCGTTGCCAAACCCCAGGTAAGACTGTCTGNNNAGATCGGAAG |
| 8508_1 | TATGAGTGTGGAGTCGTTGCTAGTGAGGTTTTGTGCCTGTCTGNNNAGATCGGAAG |
| 8509_1 | TATGAGTGTGGAGTCGTTGCAAAGTCCCATCAAGTGCTGTCTGNNNAGATCGGAAG |
| 8511_1 | TATGAGTGTGGAGTCGTTGCGCTAACGGAGTAGAAAAGGGCAANNNAGATCGGA |
| 8518_1 | TATGAGTGTGGAGTCGTTGCGGGAGAGGGAGAGAGGAATTCTGNNNAGATCGGA |
| 8524_1 | TATGAGTGTGGAGTCGTTGCAACAGAGGGTCAAAGCACTTCTGNNNAGATCGGAA |
| 8533_1 | TATGAGTGTGGAGTCGTTGCTCTGGACCTACTACAGGCTTCTGNNNAGATCGGAAG |
| 8535_1 | TATGAGTGTGGAGTCGTTGCCACAGACCCTTGGTGATCTTCTGNNNAGATCGGAAG |
| 8538_1 | TATGAGTGTGGAGTCGTTGCATGCAGGTCTGTTTCCAGTTCTGNNNAGATCGGAAG |
| 8548_1 | TATGAGTGTGGAGTCGTTGCCAGCTTTTCCACAGCACTTTCTGNNNAGATCGGAAGA |
| 8551_1 | TATGAGTGTGGAGTCGTTGCGAGGGGAGAGAGAGAGACAAGTGNNNAGATCGGA |
| 8552_1 | TATGAGTGTGGAGTCGTTGCACCCAAGTACATGATGGAAGGGANNNAGATCGGAA |
| 8556_1 | TATGAGTGTGGAGTCGTTGCCCGTGAAAGTAGGGTACACAGTGNNNAGATCGGAA |
| 8557_1 | TATGAGTGTGGAGTCGTTGCTGCCCTTTGAGAATCTGGCTTTTNNNAGATCGGAAG |
| 8559_1 | TATGAGTGTGGAGTCGTTGCTCCCATGTTTTTACCCACCAGTGNNNAGATCGGAAGA |

| | |
|---|---|
| 8561_1 | TATGAGTGTGGAGTCGTTGCTCCAGCTCCAAATGTAAGCAGTGNNNAGATCGGAAG |
| 8563_1 | TATGAGTGTGGAGTCGTTGCACACAGTGAGAGTAGAGGCAGTGNNNAGATCGGA |
| 8564_1 | TATGAGTGTGGAGTCGTTGCATTACCTGGTTTTGGAGGCAGTGNNNAGATCGGAA |
| 8565_1 | TATGAGTGTGGAGTCGTTGCAAGCATGTAAAGAGAGGGCAGTGNNNAGATCGGA |
| 8566_1 | TATGAGTGTGGAGTCGTTGCTTTGAGATCTGTCCTTGGCAGTGNNNAGATCGGAAG |
| 8567_1 | TATGAGTGTGGAGTCGTTGCACCCAGAAGCTACAACATCAGTGNNNAGATCGGAAG |
| 8569_1 | TATGAGTGTGGAGTCGTTGCTGAGCCAATCATGGAAGTCAGTGNNNAGATCGGAA |
| 8570_1 | TATGAGTGTGGAGTCGTTGCGTCCTTGTACCTGTGTGTCAGTGNNNAGATCGGAAG |
| 8572_1 | TATGAGTGTGGAGTCGTTGCAGGAACTGCAGCATGTAAGAGTGNNNAGATCGGAA |
| 8574_1 | TATGAGTGTGGAGTCGTTGCACACACAAAATTAGGCGGGAGTGNNNAGATCGGAA |
| 8575_1 | TATGAGTGTGGAGTCGTTGCTGATGGACTAGGCCTATGGAGTGNNNAGATCGGAA |
| 8577_1 | TATGAGTGTGGAGTCGTTGCGTGAAGATCTTGGGCTTGGAGTGNNNAGATCGGAA |
| 8583_1 | TATGAGTGTGGAGTCGTTGCTCAAATGGCTACATCCGTGAGTGNNNAGATCGGAA |
| 8587_1 | TATGAGTGTGGAGTCGTTGCCCTGACCACATTTGGGGATAGTGNNNAGATCGGAA |
| 8588_1 | TATGAGTGTGGAGTCGTTGCAAGTTCCAATAGGCTGTCACAAANNNAGATCGGAAG |
| 8589_1 | TATGAGTGTGGAGTCGTTGCACAGGTTTTTCTGCATGTACGTGNNNAGATCGGAAG |
| 8590_1 | TATGAGTGTGGAGTCGTTGCATAACATGACACTGGGAACCGTGNNNAGATCGGAA |
| 8592_1 | TATGAGTGTGGAGTCGTTGCGCAGAGGTTACTAGAGAAGCGTGNNNAGATCGGA |
| 8594_1 | TATGAGTGTGGAGTCGTTGCGATGATAGAGAAGCAGAGGCGTGNNNAGATCGGA |
| 8595_1 | TATGAGTGTGGAGTCGTTGCCTGAGCTTATTCTGTCTTGCGTGNNNAGATCGGAAG |
| 8596_1 | TATGAGTGTGGAGTCGTTGCCCTCGTTGTGGTTTCAATTCGTGNNNAGATCGGAAG |
| 8597_1 | TATGAGTGTGGAGTCGTTGCATTAGCCCATGCTTCACTTCGTGNNNAGATCGGAAG |
| 8600_1 | TATGAGTGTGGAGTCGTTGCCCCCAGTGTTTCAGAGTAAGGTGNNNAGATCGGAA |
| 8604_1 | TATGAGTGTGGAGTCGTTGCATTTTTGCCACATCATGCAGGTGNNNAGATCGGAAG |
| 8605_1 | TATGAGTGTGGAGTCGTTGCTACTCCCCATTAGCCAAACCCAANNNAGATCGGAAGA |
| 8606_1 | TATGAGTGTGGAGTCGTTGCTATCTGTAGGGCAGGCTATGCAANNNAGATCGGAA |
| 8607_1 | TATGAGTGTGGAGTCGTTGCAGAACGTGCATATCCCTGGAAACNNNAGATCGGAA |
| 8608_1 | TATGAGTGTGGAGTCGTTGCTTGAACCAAACAGTGGAGAGGTGNNNAGATCGGAA |
| 8609_1 | TATGAGTGTGGAGTCGTTGCCAAGAGACTGAAGCTGAGAGGTGNNNAGATCGGA |
| 8610_1 | TATGAGTGTGGAGTCGTTGCGCACTTTGTGTCTTTGTGAGGTGNNNAGATCGGAA |
| 8611_1 | TATGAGTGTGGAGTCGTTGCGGGGCTTCTCATGTGTATAGGTGNNNAGATCGGAA |
| 8613_1 | TATGAGTGTGGAGTCGTTGCGCAAGTGTGAAGTTTCAAGGGTGNNNAGATCGGAA |
| 8614_1 | TATGAGTGTGGAGTCGTTGCTAGGATTGGGATTTTGCAGGGTGNNNAGATCGGAA |
| 8615_1 | TATGAGTGTGGAGTCGTTGCTGTTGCCCATACCAGACTGAAACNNNAGATCGGAAG |
| 8616_1 | TATGAGTGTGGAGTCGTTGCTCTAAGTGGTCCAAGGTAGGGTGNNNAGATCGGAA |
| 8618_1 | TATGAGTGTGGAGTCGTTGCAAAGAGAGGTGAGAAACTGGGTGNNNAGATCGGA |
| 8619_1 | TATGAGTGTGGAGTCGTTGCATGTATGTGTGTGTCAGTGGGTGNNNAGATCGGAA |
| 8620_1 | TATGAGTGTGGAGTCGTTGCGGCATATACATACAGGGTGGGTGNNNAGATCGGAA |
| 8621_1 | TATGAGTGTGGAGTCGTTGCAAAAAGAAACGTGTGTGTGGGTGNNNAGATCGGA |
| 8622_1 | TATGAGTGTGGAGTCGTTGCTTCTTCCCATTTCCACTTGGGTGNNNAGATCGGAAGA |
| 8623_1 | TATGAGTGTGGAGTCGTTGCGAAAGACCAGAGAGACTTGGGTGNNNAGATCGGA |
| 8624_1 | TATGAGTGTGGAGTCGTTGCCAGGGGTGACAAAAACCTGGATANNNAGATCGGAA |
| 8625_1 | TATGAGTGTGGAGTCGTTGCGGAAGGAATAGCTCTGTTGGGTGNNNAGATCGGA |

| | |
|---|---|
| 8627_1 | TATGAGTGTGGAGTCGTTGCTTCATCTCTGCCCATGAATGGTGNNNAGATCGGAAG |
| 8628_1 | TATGAGTGTGGAGTCGTTGCTTGCTGCTTTTTATGCCATGGTGNNNAGATCGGAAG |
| 8631_1 | TATGAGTGTGGAGTCGTTGCAGCACCTGCACCTTTTATGTTCCNNNAGATCGGAAGA |
| 8634_1 | TATGAGTGTGGAGTCGTTGCCCTAATATGGAAGCCTGCTGGTGNNNAGATCGGAA |
| 8635_1 | TATGAGTGTGGAGTCGTTGCGAGAGAGAAGGTTCATGGTGGTGNNNAGATCGGA |
| 8636_1 | TATGAGTGTGGAGTCGTTGCACTGCAAATAAGTCCCTGTGGTGNNNAGATCGGAA |
| 8637_1 | TATGAGTGTGGAGTCGTTGCGGAGACAAATGTGTGTTGTGGTGNNNAGATCGGA |
| 8638_1 | TATGAGTGTGGAGTCGTTGCGGAAAACAAGATGCCCATTGGTGNNNAGATCGGAA |
| 8641_1 | TATGAGTGTGGAGTCGTTGCAAGGACACTGCGGTACTAATGTGNNNAGATCGGAA |
| 8644_1 | TATGAGTGTGGAGTCGTTGCGAGATGAAAGGCTCTGGGATGTGNNNAGATCGGA |
| 8648_1 | TATGAGTGTGGAGTCGTTGCGGAGGGAAGTGTGTGTAGGTTTTNNNAGATCGGA |
| 8650_1 | TATGAGTGTGGAGTCGTTGCGCTGTGAGGTTTCTCCTACTGTGNNNAGATCGGAAG |
| 8651_1 | TATGAGTGTGGAGTCGTTGCAGTCCCTTGAATCTTCAGCTGTGNNNAGATCGGAAG |
| 8653_1 | TATGAGTGTGGAGTCGTTGCCTGCACACCATTGTGAATCTGTGNNNAGATCGGAAG |
| 8654_1 | TATGAGTGTGGAGTCGTTGCCAGCTTCATGGGTGTGATCTGTGNNNAGATCGGAA |
| 8656_1 | TATGAGTGTGGAGTCGTTGCCAAGCTACAAACCTGCCTCTGTGNNNAGATCGGAAG |
| 8657_1 | TATGAGTGTGGAGTCGTTGCTTTGTTTTCAGGCTTGCTCTGTGNNNAGATCGGAAG |
| 8658_1 | TATGAGTGTGGAGTCGTTGCAACCTCTGTGTAGTTGCTCTGTGNNNAGATCGGAAG |
| 8659_1 | TATGAGTGTGGAGTCGTTGCTTAGAATGGTGGGCAAGTCTGTGNNNAGATCGGAA |
| 8660_1 | TATGAGTGTGGAGTCGTTGCGGGAGGGTCTAAAGATGTCTGTGNNNAGATCGGA |
| 8661_1 | TATGAGTGTGGAGTCGTTGCGGGGGAGCTCTAAATTGTCTGTGNNNAGATCGGAA |
| 8662_1 | TATGAGTGTGGAGTCGTTGCTGATCCATCCAGCACTTTCTGTGNNNAGATCGGAAG |
| 8663_1 | TATGAGTGTGGAGTCGTTGCGAGAGAGAAGCTGGGTTGGTTTTNNNAGATCGGA |
| 8665_1 | TATGAGTGTGGAGTCGTTGCTTGCTGGAGAACACTCTAGTGTGNNNAGATCGGAA |
| 8666_1 | TATGAGTGTGGAGTCGTTGCAAGCAAAAACCAAGCCAGACAACNNNAGATCGGAA |
| 8667_1 | TATGAGTGTGGAGTCGTTGCTTGTAGTTGTACGTGTGGGTGTGNNNAGATCGGAA |
| 8668_1 | TATGAGTGTGGAGTCGTTGCCCCAGATGAGAGTTCATGGTGTGNNNAGATCGGAA |
| 8669_1 | TATGAGTGTGGAGTCGTTGCTGATTGATGAGGTCGATGGTGTGNNNAGATCGGAA |
| 8670_1 | TATGAGTGTGGAGTCGTTGCACAATAGGAAAGGCACTGGTGTGNNNAGATCGGAA |
| 8671_1 | TATGAGTGTGGAGTCGTTGCATTAGCTGAGACATGGTGGTGTGNNNAGATCGGAA |
| 8672_1 | TATGAGTGTGGAGTCGTTGCAGGGTTTCCACCACAGATGTTTTNNNAGATCGGAAG |
| 8674_1 | TATGAGTGTGGAGTCGTTGCGGACACTTCTCTGTGAGTGTGTGNNNAGATCGGAA |
| 8676_1 | TATGAGTGTGGAGTCGTTGCGTGTATGCTTGCAGATGTGTGTGNNNAGATCGGAA |
| 8677_1 | TATGAGTGTGGAGTCGTTGCTGTATTGTAGCCCAGTGTGTGTGNNNAGATCGGAA |
| 8678_1 | TATGAGTGTGGAGTCGTTGCTAGGAGCCACATGAACTTGTGTGNNNAGATCGGAA |
| 8679_1 | TATGAGTGTGGAGTCGTTGCTGTGTCATTGTGTGTGTTGTGTGNNNAGATCGGAA |
| 8681_1 | TATGAGTGTGGAGTCGTTGCGATGAAAGGTCCGTCTCCTTGTGNNNAGATCGGAA |
| 8684_1 | TATGAGTGTGGAGTCGTTGCCGTGTATGAGAGGCAGGTGTTTTNNNAGATCGGAA |
| 8686_1 | TATGAGTGTGGAGTCGTTGCTACTGATGGGCTTGGATGTTGTGNNNAGATCGGAA |
| 8688_1 | TATGAGTGTGGAGTCGTTGCTGCCCTTCTCTCACTCATTTGTGNNNAGATCGGAAGA |
| 8689_1 | TATGAGTGTGGAGTCGTTGCCCTCTCCATCCTGACGATTTGTGNNNAGATCGGAAG |
| 8690_1 | TATGAGTGTGGAGTCGTTGCAGCCAGTGGAGAAGGAGAATCAANNNAGATCGGA |
| 8691_1 | TATGAGTGTGGAGTCGTTGCTGTACTGGGGAGTTGTGTTTGTGNNNAGATCGGAA |

| | |
|---|---|
| 8693_1 | TATGAGTGTGGAGTCGTTGCGAAGTCAGAAAAATGCCCCCAACNNNAGATCGGAA |
| 8694_1 | TATGAGTGTGGAGTCGTTGCGTGCTAAGGTCTCAGCGAAATTGNNNAGATCGGAA |
| 8695_1 | TATGAGTGTGGAGTCGTTGCGGCAGGGGGTATACAGGAAATTGNNNAGATCGGA |
| 8696_1 | TATGAGTGTGGAGTCGTTGCGCTCTAATGGAGGTTGGCAATTGNNNAGATCGGAA |
| 8697_1 | TATGAGTGTGGAGTCGTTGCTTGGAGGCTGATGATGGGAATTGNNNAGATCGGA |
| 8698_1 | TATGAGTGTGGAGTCGTTGCAGGTGGGTTGATGACAGACATTGNNNAGATCGGAA |
| 8699_1 | TATGAGTGTGGAGTCGTTGCGCCTCTGAATTATCCTGACCTGCNNNAGATCGGAAG |
| 8701_1 | TATGAGTGTGGAGTCGTTGCCACCCCTACTTCTGTCCAGTAGTNNNAGATCGGAAG |
| 8702_1 | TATGAGTGTGGAGTCGTTGCGCTGCATGGTATGCCTTACATTGNNNAGATCGGAAG |
| 8703_1 | TATGAGTGTGGAGTCGTTGCCCACAACCTCCACAATGTTCTCANNNAGATCGGAAGA |
| 8705_1 | TATGAGTGTGGAGTCGTTGCTGGAGTTGAGGAATTTGCCATTGNNNAGATCGGAA |
| 8706_1 | TATGAGTGTGGAGTCGTTGCTCCTTACCAAGCGTGATCCATTGNNNAGATCGGAAG |
| 8707_1 | TATGAGTGTGGAGTCGTTGCCTTGGCTGGCTTAGAAAGCATTGNNNAGATCGGAA |
| 8709_1 | TATGAGTGTGGAGTCGTTGCAATCCAGTTTGTTGGCAGCATTGNNNAGATCGGAAG |
| 8710_1 | TATGAGTGTGGAGTCGTTGCAGAAGCGCTATGAAGGGTCATTGNNNAGATCGGAA |
| 8712_1 | TATGAGTGTGGAGTCGTTGCTCATCTCCTGGGTCCTGAGATTGNNNAGATCGGAAG |
| 8713_1 | TATGAGTGTGGAGTCGTTGCACCATCTTCTGCCCATAGGATTGNNNAGATCGGAAG |
| 8714_1 | TATGAGTGTGGAGTCGTTGCCAGAAGAGCTGAGTGAGGGATTGNNNAGATCGGA |
| 8715_1 | TATGAGTGTGGAGTCGTTGCCAGCATGGGAGGAAAAGTTCAGTNNNAGATCGGAA |
| 8716_1 | TATGAGTGTGGAGTCGTTGCGTGCCAGCCAGTAAAATGGATTGNNNAGATCGGAA |
| 8719_1 | TATGAGTGTGGAGTCGTTGCCCTGTGGCTTTTTGAGCTGATTGNNNAGATCGGAAG |
| 8720_1 | TATGAGTGTGGAGTCGTTGCCAGCCAGAGAGACAGAGTGATTGNNNAGATCGGAA |
| 8722_1 | TATGAGTGTGGAGTCGTTGCCGCTCAGGGAGGGAAGAATATTGNNNAGATCGGA |
| 8723_1 | TATGAGTGTGGAGTCGTTGCTGCTTTGGGCTGGGTCTATATTGNNNAGATCGGAA |
| 8724_1 | TATGAGTGTGGAGTCGTTGCTGAAGGAGAAGAGGCCAAACTTGNNNAGATCGGA |
| 8725_1 | TATGAGTGTGGAGTCGTTGCCTTCCAGTCATCCTCCCAACTTGNNNAGATCGGAAGA |
| 8726_1 | TATGAGTGTGGAGTCGTTGCAGAAGGGGAAATTGGCTCACTTGNNNAGATCGGAA |
| 8727_1 | TATGAGTGTGGAGTCGTTGCCTCAGTTCAGGGTGTCAGACTTGNNNAGATCGGAA |
| 8728_1 | TATGAGTGTGGAGTCGTTGCGACGCAGGCCTATAAGAGACTTGNNNAGATCGGAA |
| 8729_1 | TATGAGTGTGGAGTCGTTGCGCTGCATATGGTCTCCAAAGTCANNNAGATCGGAAG |
| 8730_1 | TATGAGTGTGGAGTCGTTGCAAATGCGTGTGCTTGTGTACTTGNNNAGATCGGAA |
| 8733_1 | TATGAGTGTGGAGTCGTTGCCATTTCTCCCCAGGTTACCCTTGNNNAGATCGGAAGA |
| 8734_1 | TATGAGTGTGGAGTCGTTGCGAAAGAAAAAGAGAGCGCCCTTGNNNAGATCGGAA |
| 8736_1 | TATGAGTGTGGAGTCGTTGCATTTGCTTCACCTCTGAGCCTTGNNNAGATCGGAAG |
| 8737_1 | TATGAGTGTGGAGTCGTTGCCATACAAACACACTGTGGCCTTGNNNAGATCGGAAG |
| 8738_1 | TATGAGTGTGGAGTCGTTGCTGTCCAAACTATCCTTGGCCTTGNNNAGATCGGAAG |
| 8739_1 | TATGAGTGTGGAGTCGTTGCTCTCTAGGTGCCATCAGAACTGGNNNAGATCGGAAG |
| 8740_1 | TATGAGTGTGGAGTCGTTGCACCATTTCCACTTCTGCTCCTTGNNNAGATCGGAAGA |
| 8742_1 | TATGAGTGTGGAGTCGTTGCGACAGAGACATCCCAGAAGCTTGNNNAGATCGGAA |
| 8743_1 | TATGAGTGTGGAGTCGTTGCCATATTGCTGCTTGGGAAGCTTGNNNAGATCGGAA |
| 8744_1 | TATGAGTGTGGAGTCGTTGCGTGCATGAGAGTGAGTGATTTTTNNNAGATCGGAA |
| 8746_1 | TATGAGTGTGGAGTCGTTGCTTTGTAATGGTACCCCAGGCTTGNNNAGATCGGAAG |
| 8747_1 | TATGAGTGTGGAGTCGTTGCTTTTCCTGGATTCCTCAGGCTTGNNNAGATCGGAAG |

| | |
|---|---|
| 8748_1 | TATGAGTGTGGAGTCGTTGCATGTATACCAAGTGCTGGGCTTGNNNAGATCGGAA |
| 8750_1 | TATGAGTGTGGAGTCGTTGCAAGATGACTTCAGAGGCTGCTTGNNNAGATCGGAA |
| 8751_1 | TATGAGTGTGGAGTCGTTGCAAGGTTGATGGGAAGTGTGCTTGNNNAGATCGGA |
| 8752_1 | TATGAGTGTGGAGTCGTTGCGTTTTGAGGCACAGTCTTGCTTGNNNAGATCGGAAG |
| 8755_1 | TATGAGTGTGGAGTCGTTGCAACTCTGGCGCTTGATACTCTTGNNNAGATCGGAAG |
| 8756_1 | TATGAGTGTGGAGTCGTTGCTCTTCATTTCACCCCTGCTCTTGNNNAGATCGGAAGA |
| 8758_1 | TATGAGTGTGGAGTCGTTGCAGGTAGGGGCTTTGTTTGTCTTGNNNAGATCGGAA |
| 8761_1 | TATGAGTGTGGAGTCGTTGCAAGTTTGGTACTGGGGCTTCTTGNNNAGATCGGAA |
| 8763_1 | TATGAGTGTGGAGTCGTTGCCCCTGCAGATTTGCTTGTTCTTGNNNAGATCGGAAG |
| 8764_1 | TATGAGTGTGGAGTCGTTGCCAATTTCCCTGGCTGCTTTCTTGNNNAGATCGGAAG |
| 8766_1 | TATGAGTGTGGAGTCGTTGCTGTGCCAGACAGACACTAAGTTGNNNAGATCGGAA |
| 8769_1 | TATGAGTGTGGAGTCGTTGCGGTTTCACTTTGTCACCCAGTTGNNNAGATCGGAAG |
| 8770_1 | TATGAGTGTGGAGTCGTTGCGCATTTTATCCCTGGTGCAGTTGNNNAGATCGGAAG |
| 8771_1 | TATGAGTGTGGAGTCGTTGCCCCCCTCCAGTGTTGTATCAGTTGNNNAGATCGGAAG |
| 8773_1 | TATGAGTGTGGAGTCGTTGCGAAGACCTGCAACAACAGAGTTGNNNAGATCGGAA |
| 8774_1 | TATGAGTGTGGAGTCGTTGCAGGCACTTCTCATGTGAGAGTTGNNNAGATCGGAA |
| 8775_1 | TATGAGTGTGGAGTCGTTGCACAAAAGAAGGTGAGGGGAGTTGNNNAGATCGGA |
| 8777_1 | TATGAGTGTGGAGTCGTTGCCTTCCAGTTTCTCCTACCCGTTGNNNAGATCGGAAGA |
| 8780_1 | TATGAGTGTGGAGTCGTTGCGCGTGCACATTTATCAAAGGTTGNNNAGATCGGAA |
| 8781_1 | TATGAGTGTGGAGTCGTTGCGGGGAGGGTTTTTCCTTAGGTTGNNNAGATCGGAA |
| 8787_1 | TATGAGTGTGGAGTCGTTGCCTAAGTATCCGCACCTCTGGTTGNNNAGATCGGAAG |
| 8788_1 | TATGAGTGTGGAGTCGTTGCGTTTTCCGGACATGATGTGGTTGNNNAGATCGGAA |
| 8789_1 | TATGAGTGTGGAGTCGTTGCGGGCTCAGAGAAAGGATTGGTTGNNNAGATCGGA |
| 8793_1 | TATGAGTGTGGAGTCGTTGCTTATGCGCCCGTTTGTTATGTTGNNNAGATCGGAAG |
| 8794_1 | TATGAGTGTGGAGTCGTTGCTTAGATGGAGGCTTACGCTGTTGNNNAGATCGGAA |
| 8798_1 | TATGAGTGTGGAGTCGTTGCCCCTCCAGATTGTAGCAGTGTTGNNNAGATCGGAAG |
| 8799_1 | TATGAGTGTGGAGTCGTTGCTTGGGGGAGTAACAATGGTGTTGNNNAGATCGGA |
| 8800_1 | TATGAGTGTGGAGTCGTTGCTATTGCAAGGGCTTTGCTTGTTGNNNAGATCGGAAG |
| 8802_1 | TATGAGTGTGGAGTCGTTGCTACCACCAAATGCAGTCTTGTTGNNNAGATCGGAAG |
| 8803_1 | TATGAGTGTGGAGTCGTTGCTCACCAGTAGGTTCCCTTTGTTGNNNAGATCGGAAG |
| 8805_1 | TATGAGTGTGGAGTCGTTGCAGAACTGTGGGATGGGGAATTTGNNNAGATCGGA |
| 8807_1 | TATGAGTGTGGAGTCGTTGCTTGGGGCCAGAGAAAAACATTTGNNNAGATCGGAA |
| 8808_1 | TATGAGTGTGGAGTCGTTGCTTTCTACCCTGGCTCCACATTTGNNNAGATCGGAAGA |
| 8809_1 | TATGAGTGTGGAGTCGTTGCGCCGGAGGGATTTTTCACATTTGNNNAGATCGGAA |
| 8810_1 | TATGAGTGTGGAGTCGTTGCGTGTGTGTAGGGAGGGTCATTTGNNNAGATCGGA |
| 8811_1 | TATGAGTGTGGAGTCGTTGCTGTTCCTTTGTAGCAGCACTTTGNNNAGATCGGAAG |
| 8814_1 | TATGAGTGTGGAGTCGTTGCTCCTCCATATACTGGCAGCTTTGNNNAGATCGGAAG |
| 8816_1 | TATGAGTGTGGAGTCGTTGCACTGGGTCAGAATCTCTGCTTTGNNNAGATCGGAAG |
| 8817_1 | TATGAGTGTGGAGTCGTTGCCTCAGACAACTCCTGGCTCTTTGNNNAGATCGGAAG |
| 8818_1 | TATGAGTGTGGAGTCGTTGCATTAGTGATGGCTGTCTCTGGGTNNNAGATCGGAA |
| 8819_1 | TATGAGTGTGGAGTCGTTGCGGAGAAGTAGGCAGGTGTCTTTGNNNAGATCGGA |
| 8821_1 | TATGAGTGTGGAGTCGTTGCTGGAAGAGGTGGTGTTGAGTTTGNNNAGATCGGA |
| 8825_1 | TATGAGTGTGGAGTCGTTGCAAGAGGTCACCTTGGATGGTTTGNNNAGATCGGAA |

| | |
|---|---|
| 8826_1 | TATGAGTGTGGAGTCGTTGCTGAGGCACAGTGATAGTGGTTTGNNNAGATCGGAA |
| 8828_1 | TATGAGTGTGGAGTCGTTGCCAGTGGTGGTCCTAGTCTGTTTGNNNAGATCGGAAG |
| 8829_1 | TATGAGTGTGGAGTCGTTGCCTTGGTTTTCCTCGGTGTGTTTGNNNAGATCGGAAG |
| 8830_1 | TATGAGTGTGGAGTCGTTGCCAGCCACAGCACATCAGATTTTGNNNAGATCGGAAG |
| 8831_1 | TATGAGTGTGGAGTCGTTGCAGTTTCCCGTGGTCTGGATTTTGNNNAGATCGGAAG |
| 8832_1 | TATGAGTGTGGAGTCGTTGCGTGAAATCAAGCCCCACCTTTTGNNNAGATCGGAAG |
| 8833_1 | TATGAGTGTGGAGTCGTTGCGGTGCAAGGAAATCACCCTTTTGNNNAGATCGGAAG |
| 8834_1 | TATGAGTGTGGAGTCGTTGCTGGGTCTTTGCTTTTGCCTTTTGNNNAGATCGGAAG |
| 8835_1 | TATGAGTGTGGAGTCGTTGCCCCCTTCACAGACCTTCCTTTTGNNNAGATCGGAAGA |
| 8836_1 | TATGAGTGTGGAGTCGTTGCGCAACCACCGTAAGTAGCTTTTGNNNAGATCGGAAG |
| 8839_1 | TATGAGTGTGGAGTCGTTGCGTGCTTTGCCATTGTTGCTTTTGNNNAGATCGGAAG |
| 8840_1 | TATGAGTGTGGAGTCGTTGCTGTACTCAGCGAGAGCTCTTTTGNNNAGATCGGAAG |
| 8841_1 | TATGAGTGTGGAGTCGTTGCATGCCTCTCCTTTCCCAGTTTTGNNNAGATCGGAAGA |
| 8842_1 | TATGAGTGTGGAGTCGTTGCTGTAAGGAGGGCTCTTCGTTTTGNNNAGATCGGAA |
| 8843_1 | TATGAGTGTGGAGTCGTTGCAACTGCATGTGGTCATGGTTTTGNNNAGATCGGAAG |
| 8847_1 | TATGAGTGTGGAGTCGTTGCTGGAAAATAATTGCACCGTGGGTNNNAGATCGGAA |
| 8848_1 | TATGAGTGTGGAGTCGTTGCGGGAGGGGAAGAGGTTTCAAAATNNNAGATCGGA |
| 8849_1 | TATGAGTGTGGAGTCGTTGCTAGGAGTGCCGAAGCAAGAAAATNNNAGATCGGAA |
| 8850_1 | TATGAGTGTGGAGTCGTTGCCCCACACCATCTCCAGTGAAAATNNNAGATCGGAAG |
| 8853_1 | TATGAGTGTGGAGTCGTTGCCTGCACCACATCAGTAATCGTCANNNAGATCGGAAG |
| 8855_1 | TATGAGTGTGGAGTCGTTGCGCCTGCAAATCCCAATCACAAATNNNAGATCGGAAG |
| 8856_1 | TATGAGTGTGGAGTCGTTGCAGATTGAGTTTCTTCCCAGGTCGNNNAGATCGGAAG |
| 8857_1 | TATGAGTGTGGAGTCGTTGCCAGGTGGTGCATGTAGACCAAATNNNAGATCGGAA |
| 8860_1 | TATGAGTGTGGAGTCGTTGCCCTTGGGATAAGATGGCCCAAATNNNAGATCGGAAG |
| 8861_1 | TATGAGTGTGGAGTCGTTGCCCGACCCAAGTGTGTTTTAGAGTNNNAGATCGGAAG |
| 8862_1 | TATGAGTGTGGAGTCGTTGCGAAAAAGCCCACATCAGCCAAATNNNAGATCGGAAG |
| 8863_1 | TATGAGTGTGGAGTCGTTGCGCTAGAATCAGAGCAGGCCAAATNNNAGATCGGAA |
| 8864_1 | TATGAGTGTGGAGTCGTTGCAGGTCAAACATCCCATGCCAAATNNNAGATCGGAAG |
| 8866_1 | TATGAGTGTGGAGTCGTTGCACACATGCCCTTTCCATCCAAATNNNAGATCGGAAGA |
| 8867_1 | TATGAGTGTGGAGTCGTTGCCGCTCTTCAGTGACCTTCCAAATNNNAGATCGGAAG |
| 8869_1 | TATGAGTGTGGAGTCGTTGCATCCCAACAACACTGGAGCAAATNNNAGATCGGAAG |
| 8871_1 | TATGAGTGTGGAGTCGTTGCGCTCCACTGTGTGTTTTGCAAATNNNAGATCGGAAG |
| 8872_1 | TATGAGTGTGGAGTCGTTGCGGAACCAGCCATTCCCATCAAATNNNAGATCGGAAG |
| 8875_1 | TATGAGTGTGGAGTCGTTGCGCAGAAGGGAGCAGAAGTCAAATNNNAGATCGGA |
| 8876_1 | TATGAGTGTGGAGTCGTTGCCCCCCACAAACCCTAAGTCAAATNNNAGATCGGAAG |
| 8877_1 | TATGAGTGTGGAGTCGTTGCCTTGTGTGTGCAGTGGTTCAAATNNNAGATCGGAA |
| 8879_1 | TATGAGTGTGGAGTCGTTGCGGGGACTGGGAGGATGAAGAAATNNNAGATCGGA |
| 8880_1 | TATGAGTGTGGAGTCGTTGCTGCTGGAACAAGTTTGAAGAAATNNNAGATCGGAA |
| 8882_1 | TATGAGTGTGGAGTCGTTGCGGCACGTCTTTGTCCTGAGAAATNNNAGATCGGAAG |
| 8883_1 | TATGAGTGTGGAGTCGTTGCTCTTTCCCCCACCCACTAGAAATNNNAGATCGGAAGA |
| 8884_1 | TATGAGTGTGGAGTCGTTGCGGGAAACAGCCTGCACTAGAAATNNNAGATCGGAA |
| 8885_1 | TATGAGTGTGGAGTCGTTGCACACACACACGATAGCAGGAAATNNNAGATCGGAA |
| 8886_1 | TATGAGTGTGGAGTCGTTGCAGCAATAGAGAGGTGGGGGAAATNNNAGATCGGA |

| | |
|---|---|
| 8888_1 | TATGAGTGTGGAGTCGTTGCCTCCAAAGCCATCACCTGGAAATNNNAGATCGGAAG |
| 8889_1 | TATGAGTGTGGAGTCGTTGCTAGGTCTGGAGCATGTGGATCATNNNAGATCGGAA |
| 8890_1 | TATGAGTGTGGAGTCGTTGCGTTGCCCCTAGGAACTTGGAAATNNNAGATCGGAAG |
| 8892_1 | TATGAGTGTGGAGTCGTTGCTGGGCAGCTGAATTTCCTGAAATNNNAGATCGGAAG |
| 8893_1 | TATGAGTGTGGAGTCGTTGCGGCTGGAAGGAAGAAGCTGAAATNNNAGATCGGA |
| 8896_1 | TATGAGTGTGGAGTCGTTGCGCTGACTGTACAGGGTGTGAAATNNNAGATCGGAA |
| 8898_1 | TATGAGTGTGGAGTCGTTGCGGAGTTCGACCTGGGTTTGAAATNNNAGATCGGAA |
| 8899_1 | TATGAGTGTGGAGTCGTTGCCTGGAGCACCCCGACTAATAAATNNNAGATCGGAAG |
| 8901_1 | TATGAGTGTGGAGTCGTTGCAAGAGGGCACCGTTTCCCTAAATNNNAGATCGGAAG |
| 8903_1 | TATGAGTGTGGAGTCGTTGCAGTCGGGTCTCACAGAGCTAAATNNNAGATCGGAAG |
| 8904_1 | TATGAGTGTGGAGTCGTTGCACGGCCTGCAATTTTCAAACAATNNNAGATCGGAAG |
| 8905_1 | TATGAGTGTGGAGTCGTTGCGGGTCAACGAGGATTGGAACAATNNNAGATCGGAA |
| 8908_1 | TATGAGTGTGGAGTCGTTGCACTCACAGGGTATTTTGAGGAGTNNNAGATCGGAA |
| 8910_1 | TATGAGTGTGGAGTCGTTGCCCAGACATACCCGTGTTCACAATNNNAGATCGGAAG |
| 8911_1 | TATGAGTGTGGAGTCGTTGCATACGCGCAGACTTTCTGACAATNNNAGATCGGAAG |
| 8913_1 | TATGAGTGTGGAGTCGTTGCAGTGATATGGGGCCAGGTACAATNNNAGATCGGAA |
| 8916_1 | TATGAGTGTGGAGTCGTTGCTGCCCATTGCTCCAATAACCAATNNNAGATCGGAAG |
| 8919_1 | TATGAGTGTGGAGTCGTTGCAGGTGTTGCAAAGTTCTCCCAATNNNAGATCGGAAG |
| 8920_1 | TATGAGTGTGGAGTCGTTGCGGCAAAGAGAACTTGGTCCCAATNNNAGATCGGAA |
| 8922_1 | TATGAGTGTGGAGTCGTTGCCCTTCAGCTGATTACTGGCCAATNNNAGATCGGAAG |
| 8924_1 | TATGAGTGTGGAGTCGTTGCGAAGTCCATCGCAGTGATCCAATNNNAGATCGGAAG |
| 8927_1 | TATGAGTGTGGAGTCGTTGCGGTGAGTGGCTTGAGATTCCAATNNNAGATCGGAA |
| 8929_1 | TATGAGTGTGGAGTCGTTGCCCCATCTTCTGGGAGTTTCCAATNNNAGATCGGAAG |
| 8930_1 | TATGAGTGTGGAGTCGTTGCTCTGCTGGGAAAAGGAAAGCAATNNNAGATCGGAA |
| 8931_1 | TATGAGTGTGGAGTCGTTGCATGTCACAGATGCTCCAAGCAATNNNAGATCGGAAG |
| 8932_1 | TATGAGTGTGGAGTCGTTGCCCACTTCCAGTCACAGAAGCAATNNNAGATCGGAAG |
| 8935_1 | TATGAGTGTGGAGTCGTTGCCAGCTTGGATGCTAAGGAGCAATNNNAGATCGGAA |
| 8937_1 | TATGAGTGTGGAGTCGTTGCCCCACAAAGATCCAAGAGGCAATNNNAGATCGGAAG |
| 8938_1 | TATGAGTGTGGAGTCGTTGCAAGAATGAGGGAGAGGAGGCAATNNNAGATCGGA |
| 8942_1 | TATGAGTGTGGAGTCGTTGCCGCCTCCATACTTACTTGGCAATNNNAGATCGGAAG |
| 8944_1 | TATGAGTGTGGAGTCGTTGCGAGATGTGGGAGGAAACTGCAATNNNAGATCGGA |
| 8948_1 | TATGAGTGTGGAGTCGTTGCGGGGGACTGGTAACACAATCAATNNNAGATCGGAA |
| 8949_1 | TATGAGTGTGGAGTCGTTGCTTGGCACCTGAAGCTGAATCAATNNNAGATCGGAAG |
| 8951_1 | TATGAGTGTGGAGTCGTTGCCAGGTATGCAACAGGTGCTCAATNNNAGATCGGAAG |
| 8952_1 | TATGAGTGTGGAGTCGTTGCGGTACATAAGCCCTGCAGTCAATNNNAGATCGGAAG |
| 8954_1 | TATGAGTGTGGAGTCGTTGCTGCTTCCCCTTCACCCTTTCAATNNNAGATCGGAAGA |
| 8956_1 | TATGAGTGTGGAGTCGTTGCGTCACAGCCTCAGAGTAAAGAATNNNAGATCGGAAG |
| 8957_1 | TATGAGTGTGGAGTCGTTGCACACTCTTTTGGGGGTCAAGAATNNNAGATCGGAAG |
| 8958_1 | TATGAGTGTGGAGTCGTTGCCAGGACCCACAACTTGGAAGAATNNNAGATCGGAAG |
| 8959_1 | TATGAGTGTGGAGTCGTTGCTCCCTATCCTTATGGGTTTTGGGNNNAGATCGGAAG |
| 8962_1 | TATGAGTGTGGAGTCGTTGCGCTGTCACTGATTGGACCAGAATNNNAGATCGGAAG |
| 8964_1 | TATGAGTGTGGAGTCGTTGCTGATTTTCACCTTGGGTTGGTCANNNAGATCGGAAG |
| 8966_1 | TATGAGTGTGGAGTCGTTGCTCCCCTCTGTTTTCCTGGAGAATNNNAGATCGGAAG |

8968_1 TATGAGTGTGGAGTCGTTGCAGGCAGGGGAGGTGAATTAGAATNNNAGATCGGA
8971_1 TATGAGTGTGGAGTCGTTGCGTGGAGGTCACATCCTTCTCTCTNNNAGATCGGAAG
8972_1 TATGAGTGTGGAGTCGTTGCGAGACATACCCCTAGCCAGGAATNNNAGATCGGAAG
8976_1 TATGAGTGTGGAGTCGTTGCTCTGACTAGGTTCTCACCTGGTTNNNAGATCGGAAG
8980_1 TATGAGTGTGGAGTCGTTGCTTGAAGGTGACAGAGGAGGGAATNNNAGATCGGA
8981_1 TATGAGTGTGGAGTCGTTGCTGTGTACTGATGGGAAGGGGAATNNNAGATCGGA
8982_1 TATGAGTGTGGAGTCGTTGCGAAGGCAGGGACAATGATGGAATNNNAGATCGGA
8983_1 TATGAGTGTGGAGTCGTTGCCCACACACATGGGCTTATGGAATNNNAGATCGGAAG
8984_1 TATGAGTGTGGAGTCGTTGCCTTGTGATCTGAGCTGCTGGAATNNNAGATCGGAA
8985_1 TATGAGTGTGGAGTCGTTGCAAGGAGCCTCACTCATCTGGAATNNNAGATCGGAAG
8987_1 TATGAGTGTGGAGTCGTTGCGACAATGGGCAGAGATGTGGAATNNNAGATCGGA
8990_1 TATGAGTGTGGAGTCGTTGCGGGTAGGCAGTGCATTACTGAATNNNAGATCGGAA
8991_1 TATGAGTGTGGAGTCGTTGCGAGAGCGAGTGCACTAAGTGAATNNNAGATCGGAA
8992_1 TATGAGTGTGGAGTCGTTGCGAGTTCAGGCAGAGAACGTGAATNNNAGATCGGAA
8993_1 TATGAGTGTGGAGTCGTTGCTGTGTGCCCATGTGTATGTGAATNNNAGATCGGAA
8994_1 TATGAGTGTGGAGTCGTTGCAATCTCACCCCTCCCACTTGAATNNNAGATCGGAAGA
8995_1 TATGAGTGTGGAGTCGTTGCGGAGAAGAGGGAGGGACTTGAATNNNAGATCGGA
9000_1 TATGAGTGTGGAGTCGTTGCACCTCAGGGGGCAACTGAATAATNNNAGATCGGAA
9001_1 TATGAGTGTGGAGTCGTTGCAGGGAATGTGGGGGAGACATAATNNNAGATCGGA
9002_1 TATGAGTGTGGAGTCGTTGCCACTCCAAGGCAGATGCCATAATNNNAGATCGGAAG
9006_1 TATGAGTGTGGAGTCGTTGCGAGCAACAGCCACCAGCTATAATNNNAGATCGGAAG
9007_1 TATGAGTGTGGAGTCGTTGCAAAGGCCAGGGAGGTGGTATAATNNNAGATCGGA
9010_1 TATGAGTGTGGAGTCGTTGCCCAAAAACCAGGTGAGGCCTAATNNNAGATCGGAA
9011_1 TATGAGTGTGGAGTCGTTGCGGGGAGGGGTCTAAACTCCTAATNNNAGATCGGAA
9013_1 TATGAGTGTGGAGTCGTTGCCATACCATGGCCCACAAGCTAATNNNAGATCGGAAG
9014_1 TATGAGTGTGGAGTCGTTGCTCTCTTCTGCTCCCCAAGCTAATNNNAGATCGGAAGA
9021_1 TATGAGTGTGGAGTCGTTGCCACGTATGCACAGAGACAAACATNNNAGATCGGAA
9023_1 TATGAGTGTGGAGTCGTTGCCTAAGCTAACCGGGCAGAAACATNNNAGATCGGAA
9024_1 TATGAGTGTGGAGTCGTTGCCACTCCGTCACACAGAGAAACATNNNAGATCGGAAG
9027_1 TATGAGTGTGGAGTCGTTGCAACTTTTGGATCCAGGGCAACATNNNAGATCGGAAG
9028_1 TATGAGTGTGGAGTCGTTGCGCTGGAGAGCATGGTATGAACATNNNAGATCGGAA
9029_1 TATGAGTGTGGAGTCGTTGCGAGCCCCCAAATCAGCATAACATNNNAGATCGGAAG
9030_1 TATGAGTGTGGAGTCGTTGCCCTTTTGAGGCTGCCACTAACATNNNAGATCGGAAG
9032_1 TATGAGTGTGGAGTCGTTGCGTTGGGGGAGAAGGAAAACACATNNNAGATCGGA
9033_1 TATGAGTGTGGAGTCGTTGCGGGGTGTGGTGATACAAACACATNNNAGATCGGAA
9034_1 TATGAGTGTGGAGTCGTTGCCAACTGACAAGCAAAGCACACATNNNAGATCGGAAG
9035_1 TATGAGTGTGGAGTCGTTGCGACAAGCAGTACCTGCAGAAGAANNNAGATCGGAA
9036_1 TATGAGTGTGGAGTCGTTGCATGAACCTCTCCCTTCACCACATNNNAGATCGGAAGA
9040_1 TATGAGTGTGGAGTCGTTGCTCAACCACCACACTTTTCCACATNNNAGATCGGAAGA
9041_1 TATGAGTGTGGAGTCGTTGCAAGTTCAACCAATTCTTCCGCCTNNNAGATCGGAAGA
9042_1 TATGAGTGTGGAGTCGTTGCGCAAACAGAATTGACCAGCACATNNNAGATCGGAAG
9045_1 TATGAGTGTGGAGTCGTTGCCAAGGGCACTAATCCCATCACATNNNAGATCGGAAG
9048_1 TATGAGTGTGGAGTCGTTGCAACACTTTGCAACTGCAAGACATNNNAGATCGGAAG

| | |
|---|---|
| 9049_1 | TATGAGTGTGGAGTCGTTGCGGACCTGAGATGGGAGAAGACATNNNAGATCGGA |
| 9051_1 | TATGAGTGTGGAGTCGTTGCTGAAACTACCTACCATCTGGCCANNNAGATCGGAAG |
| 9052_1 | TATGAGTGTGGAGTCGTTGCATGCACAAGTGAGATGCAGACATNNNAGATCGGAA |
| 9053_1 | TATGAGTGTGGAGTCGTTGCGAGAGGTGAGATGGGAGAGACATNNNAGATCGGA |
| 9054_1 | TATGAGTGTGGAGTCGTTGCCTCATATGCCTCCCCTGAGACATNNNAGATCGGAAG |
| 9055_1 | TATGAGTGTGGAGTCGTTGCACCTGTAGATGCAGTACCGACATNNNAGATCGGAAG |
| 9057_1 | TATGAGTGTGGAGTCGTTGCGGAGCAGGAGACAAAGAGGACATNNNAGATCGGA |
| 9058_1 | TATGAGTGTGGAGTCGTTGCAGAGAAAGGAAGCAGGAGGACATNNNAGATCGGA |
| 9059_1 | TATGAGTGTGGAGTCGTTGCGCTTCATGGGACAATACGGACATNNNAGATCGGAA |
| 9060_1 | TATGAGTGTGGAGTCGTTGCGGTGAACTCATTCTTGGGGACATNNNAGATCGGAA |
| 9061_1 | TATGAGTGTGGAGTCGTTGCTGCTACTTTTGCACTTGGGACATNNNAGATCGGAAG |
| 9065_1 | TATGAGTGTGGAGTCGTTGCTCCCGTGGTCATAGCTGATACATNNNAGATCGGAAG |
| 9066_1 | TATGAGTGTGGAGTCGTTGCACTTACGGCACCATCATATACATNNNAGATCGGAAG |
| 9067_1 | TATGAGTGTGGAGTCGTTGCGACACATGCAGACCCATGTACATNNNAGATCGGAAG |
| 9069_1 | TATGAGTGTGGAGTCGTTGCTTGGAGAGGCATGACCTAGGTAGNNNAGATCGGAA |
| 9072_1 | TATGAGTGTGGAGTCGTTGCATGCACAGAAAAGAAACAACCATNNNAGATCGGAA |
| 9073_1 | TATGAGTGTGGAGTCGTTGCGCCCCTTACTGGAATTCAACCATNNNAGATCGGAAG |
| 9075_1 | TATGAGTGTGGAGTCGTTGCGGAGAGAAAGAAAGCAGCACCATNNNAGATCGGAA |
| 9076_1 | TATGAGTGTGGAGTCGTTGCAACAATTTATGACCGGGCACCATNNNAGATCGGAAG |
| 9078_1 | TATGAGTGTGGAGTCGTTGCGCCCCAAAAGAGAATGTGACCATNNNAGATCGGAA |
| 9080_1 | TATGAGTGTGGAGTCGTTGCCCTTTCAAACAGCTACCACCCATNNNAGATCGGAAGA |
| 9081_1 | TATGAGTGTGGAGTCGTTGCCCATCTGTGTCACTTTCACCCATNNNAGATCGGAAGA |
| 9082_1 | TATGAGTGTGGAGTCGTTGCCCAGCAAACACATACAGACCCATNNNAGATCGGAAG |
| 9083_1 | TATGAGTGTGGAGTCGTTGCCATATTACCACCACCTCCCCCATNNNAGATCGGAAGA |
| 9088_1 | TATGAGTGTGGAGTCGTTGCCACACAGAATCAGCACATCCCATNNNAGATCGGAAG |
| 9089_1 | TATGAGTGTGGAGTCGTTGCCTTCCTCTCCTTCTGGATCCCATNNNAGATCGGAAGA |
| 9090_1 | TATGAGTGTGGAGTCGTTGCTGGTGGGGAGTTCAATATCCCATNNNAGATCGGAA |
| 9092_1 | TATGAGTGTGGAGTCGTTGCGGCAGTATCTCCTTAGGTCCCATNNNAGATCGGAAG |
| 9093_1 | TATGAGTGTGGAGTCGTTGCTCCTTCCTTCTTTACGGTCCCATNNNAGATCGGAAGA |
| 9095_1 | TATGAGTGTGGAGTCGTTGCCTGGAGGAGTATGCTGACAACACNNNAGATCGGAA |
| 9097_1 | TATGAGTGTGGAGTCGTTGCGGTTGTTTGGAGTACTCAGCCATNNNAGATCGGAAG |
| 9099_1 | TATGAGTGTGGAGTCGTTGCCTGAATGAGCTGGAAAGAGCCATNNNAGATCGGAA |
| 9101_1 | TATGAGTGTGGAGTCGTTGCCGTGTATGGGTGAAGTTAGCCATNNNAGATCGGAA |
| 9103_1 | TATGAGTGTGGAGTCGTTGCCCGCTCTAAAAATCTGTGGCCATNNNAGATCGGAAG |
| 9104_1 | TATGAGTGTGGAGTCGTTGCCACAGGGAAAGCGAACTACAGAANNNAGATCGGAA |
| 9105_1 | TATGAGTGTGGAGTCGTTGCTCTGACAAGAAATGTGCTGCCATNNNAGATCGGAAG |
| 9106_1 | TATGAGTGTGGAGTCGTTGCCACTACATGATGGCTTCTGCCATNNNAGATCGGAAG |
| 9107_1 | TATGAGTGTGGAGTCGTTGCCATCTCCACACGTCCAAATCCATNNNAGATCGGAAGA |
| 9108_1 | TATGAGTGTGGAGTCGTTGCTCCCAGTGTATGTCACCATCCATNNNAGATCGGAAGA |
| 9112_1 | TATGAGTGTGGAGTCGTTGCCCGCCTTCTCCATCTCTATCCATNNNAGATCGGAAGA |
| 9113_1 | TATGAGTGTGGAGTCGTTGCAGACCTTGCCTTCCCTTATCCATNNNAGATCGGAAGA |
| 9115_1 | TATGAGTGTGGAGTCGTTGCTTCGGTGTTTTGTTCACCTCCATNNNAGATCGGAAGA |
| 9116_1 | TATGAGTGTGGAGTCGTTGCAGCATTGGTTCTCTCTCCTCCATNNNAGATCGGAAGA |

| | |
|---|---|
| 9119_1 | TATGAGTGTGGAGTCGTTGCTACACACATCCTGTGGTCTCCATNNNAGATCGGAAGA |
| 9120_1 | TATGAGTGTGGAGTCGTTGCGCTGGGCATTCCTTATTCTCCATNNNAGATCGGAAGA |
| 9123_1 | TATGAGTGTGGAGTCGTTGCTTTGGCGAGAACAGTGATTCCATNNNAGATCGGAAG |
| 9124_1 | TATGAGTGTGGAGTCGTTGCCAGGGCTTTGGCCTTTATTCCATNNNAGATCGGAAG |
| 9125_1 | TATGAGTGTGGAGTCGTTGCGAAAAGCACCCCATCTCTTGTCANNNAGATCGGAAG |
| 9126_1 | TATGAGTGTGGAGTCGTTGCACTGTGAATTGCCACCTTTCCATNNNAGATCGGAAGA |
| 9128_1 | TATGAGTGTGGAGTCGTTGCCTTGCCTCATGTGAAAACAGCATNNNAGATCGGAAG |
| 9129_1 | TATGAGTGTGGAGTCGTTGCAGTTCTGGCATTTTTCCCAGCATNNNAGATCGGAAG |
| 9131_1 | TATGAGTGTGGAGTCGTTGCAGCATTTGGTCTCCTAAGAGCATNNNAGATCGGAAG |
| 9132_1 | TATGAGTGTGGAGTCGTTGCGTCAGAGATTTGCTCCTGAGCATNNNAGATCGGAAG |
| 9134_1 | TATGAGTGTGGAGTCGTTGCGCACAGAGCTAACAAGCTAGCATNNNAGATCGGAAG |
| 9135_1 | TATGAGTGTGGAGTCGTTGCACATTTTCTGCAAATCCACGCATNNNAGATCGGAAGA |
| 9136_1 | TATGAGTGTGGAGTCGTTGCTCACATTTTGCCTTTCCTCCTCCNNNAGATCGGAAGAG |
| 9137_1 | TATGAGTGTGGAGTCGTTGCAAAGTAAGGTGTTGGCAAGGCATNNNAGATCGGAA |
| 9139_1 | TATGAGTGTGGAGTCGTTGCCACAGGAGAAAGAAGTGAGGCATNNNAGATCGGA |
| 9140_1 | TATGAGTGTGGAGTCGTTGCTCCAGTTTGAAGCTTTAGGGCATNNNAGATCGGAAG |
| 9141_1 | TATGAGTGTGGAGTCGTTGCTTTAGGGATCTTCCAAGGGGCATNNNAGATCGGAAG |
| 9142_1 | TATGAGTGTGGAGTCGTTGCAGAGAGAAAGAGAGTTGGGGCATNNNAGATCGGA |
| 9143_1 | TATGAGTGTGGAGTCGTTGCTGATTTATGCATGGCATGGGCATNNNAGATCGGAAG |
| 9144_1 | TATGAGTGTGGAGTCGTTGCAACATGAAAAGAAGCCTGGGCATNNNAGATCGGAA |
| 9145_1 | TATGAGTGTGGAGTCGTTGCTCACTCACTAACTAGCTGGGCATNNNAGATCGGAAG |
| 9146_1 | TATGAGTGTGGAGTCGTTGCCCTACTTGAGAGCGTTATGGCATNNNAGATCGGAAG |
| 9147_1 | TATGAGTGTGGAGTCGTTGCTTCTTGGAGACTGAGACTGGCATNNNAGATCGGAAG |
| 9148_1 | TATGAGTGTGGAGTCGTTGCTCTCTCTCTCTCTTCGTTGGCATNNNAGATCGGAAGA |
| 9150_1 | TATGAGTGTGGAGTCGTTGCCACACACGCACAGATAAATGCATNNNAGATCGGAAG |
| 9153_1 | TATGAGTGTGGAGTCGTTGCCTCCTACTCAAGTCCCCATGCATNNNAGATCGGAAGA |
| 9155_1 | TATGAGTGTGGAGTCGTTGCAAATCCGAATAAGGCGGATGCATNNNAGATCGGAA |
| 9156_1 | TATGAGTGTGGAGTCGTTGCAACACACACTCACATGGATGCATNNNAGATCGGAAG |
| 9157_1 | TATGAGTGTGGAGTCGTTGCACCTCAATACCCCCAGGAATTCANNNAGATCGGAAGA |
| 9158_1 | TATGAGTGTGGAGTCGTTGCGCAGAGAGATTGGAGTGATGCATNNNAGATCGGAA |
| 9159_1 | TATGAGTGTGGAGTCGTTGCCACGCACTTACGTACATATGCATNNNAGATCGGAAG |
| 9160_1 | TATGAGTGTGGAGTCGTTGCGCCTATAGAGAGGGGAACTGCATNNNAGATCGGAA |
| 9162_1 | TATGAGTGTGGAGTCGTTGCATTGTCTGTGTGATGAGCTGCATNNNAGATCGGAAG |
| 9163_1 | TATGAGTGTGGAGTCGTTGCTATTACCCCCTCTTCTGCTGCATNNNAGATCGGAAGA |
| 9164_1 | TATGAGTGTGGAGTCGTTGCATCGTTTAGGTAGCTTGCTGCATNNNAGATCGGAAG |
| 9165_1 | TATGAGTGTGGAGTCGTTGCGGTCTGAATGGTAAGTGGTGCATNNNAGATCGGAA |
| 9167_1 | TATGAGTGTGGAGTCGTTGCCTCATATGCTTTACGCTGTGCATNNNAGATCGGAAG |
| 9170_1 | TATGAGTGTGGAGTCGTTGCATCACTCTCTGCTTTGGTTGCATNNNAGATCGGAAGA |
| 9171_1 | TATGAGTGTGGAGTCGTTGCTACAGTGTCTGTGCATGTTGCATNNNAGATCGGAAG |
| 9173_1 | TATGAGTGTGGAGTCGTTGCTTCCTGGCTTCTTGACTTTGCATNNNAGATCGGAAGA |
| 9175_1 | TATGAGTGTGGAGTCGTTGCACCCAGCTTCTACCCCAAATCATNNNAGATCGGAAGA |
| 9176_1 | TATGAGTGTGGAGTCGTTGCACCACAAGGCTTACCTAAATCATNNNAGATCGGAAG |
| 9178_1 | TATGAGTGTGGAGTCGTTGCGGGGAGCCTTTCCTTTCAATCATNNNAGATCGGAAG |

| | |
|---|---|
| 9179_1 | TATGAGTGTGGAGTCGTTGCACTGGCTTAGGAGTGTCCATCATNNNAGATCGGAAG |
| 9180_1 | TATGAGTGTGGAGTCGTTGCCGCACTTACTCACCCATCATCATNNNAGATCGGAAGA |
| 9184_1 | TATGAGTGTGGAGTCGTTGCAAAGAACGTGCCTGCCATATCATNNNAGATCGGAAG |
| 9185_1 | TATGAGTGTGGAGTCGTTGCCCTGGGCAGCCTTGATTTATCATNNNAGATCGGAAG |
| 9187_1 | TATGAGTGTGGAGTCGTTGCCATCCTGACCCCTCCTAACTCATNNNAGATCGGAAGA |
| 9188_1 | TATGAGTGTGGAGTCGTTGCCCTCTCTGAACTTTGCCACTCATNNNAGATCGGAAGA |
| 9189_1 | TATGAGTGTGGAGTCGTTGCCCCCACACCTGTTCTCTACTCATNNNAGATCGGAAGA |
| 9191_1 | TATGAGTGTGGAGTCGTTGCGAACACCATTGTCCCAACCTCATNNNAGATCGGAAGA |
| 9193_1 | TATGAGTGTGGAGTCGTTGCACCATGAGATACTCCAGCCTCATNNNAGATCGGAAG |
| 9194_1 | TATGAGTGTGGAGTCGTTGCTTCTGTCTGGTACTCTGCCTCATNNNAGATCGGAAGA |
| 9195_1 | TATGAGTGTGGAGTCGTTGCACTGAAAGAGAGAGGTGCCTCATNNNAGATCGGAA |
| 9196_1 | TATGAGTGTGGAGTCGTTGCCTCTTAGCCATCATGCAGCTCATNNNAGATCGGAAGA |
| 9197_1 | TATGAGTGTGGAGTCGTTGCAGGGAAAATACAGATGTGCTCATNNNAGATCGGAA |
| 9198_1 | TATGAGTGTGGAGTCGTTGCGACCTCTTCTCCAAACATCTCATNNNAGATCGGAAGA |
| 9199_1 | TATGAGTGTGGAGTCGTTGCAGCCTTGACTTCCTCACTCTCATNNNAGATCGGAAGA |
| 9201_1 | TATGAGTGTGGAGTCGTTGCAGCAACAGTGTCTTCCTAGTCATNNNAGATCGGAAG |
| 9202_1 | TATGAGTGTGGAGTCGTTGCGGGGTAGTGGTCATGGTAGTCATNNNAGATCGGAA |
| 9203_1 | TATGAGTGTGGAGTCGTTGCCTGCTATTCCACTGTGACGTCATNNNAGATCGGAAG |
| 9205_1 | TATGAGTGTGGAGTCGTTGCGTTCTGCACAGGTACTTGGTCATNNNAGATCGGAAG |
| 9207_1 | TATGAGTGTGGAGTCGTTGCGCCCACTACAGTTCCTTTGTCATNNNAGATCGGAAGA |
| 9211_1 | TATGAGTGTGGAGTCGTTGCATCAAGCCTGGAAGTTTGTCTCTNNNAGATCGGAAG |
| 9214_1 | TATGAGTGTGGAGTCGTTGCTTGAAAAAGAGGGGCTGCTTCATNNNAGATCGGAA |
| 9217_1 | TATGAGTGTGGAGTCGTTGCCCCACTCTGCAGTTGTTCTTCATNNNAGATCGGAAGA |
| 9218_1 | TATGAGTGTGGAGTCGTTGCCTCTCTCTCTCCACCGAGTTCATNNNAGATCGGAAGA |
| 9220_1 | TATGAGTGTGGAGTCGTTGCGGATAACAACCCCTTCCGTTCATNNNAGATCGGAAG |
| 9221_1 | TATGAGTGTGGAGTCGTTGCCACAACTAACCCGTGAGGTTCATNNNAGATCGGAAG |
| 9222_1 | TATGAGTGTGGAGTCGTTGCGCTAGTTCCACTCTCATGTTCATNNNAGATCGGAAGA |
| 9224_1 | TATGAGTGTGGAGTCGTTGCTGCCAAAGACCTCTTCGTTTCATNNNAGATCGGAAG |
| 9227_1 | TATGAGTGTGGAGTCGTTGCCGTGGTAAGAGCACCTCAAAGATNNNAGATCGGAA |
| 9229_1 | TATGAGTGTGGAGTCGTTGCACAGTCTTGGGAGAGTGCAAGATNNNAGATCGGAA |
| 9232_1 | TATGAGTGTGGAGTCGTTGCTGTAGCAGGGAGACATGGAAGATNNNAGATCGGA |
| 9233_1 | TATGAGTGTGGAGTCGTTGCCGAAAGACTGAGAATGCGCTAGTNNNAGATCGGAA |
| 9234_1 | TATGAGTGTGGAGTCGTTGCGCACCCTGTTGACAGCATAAGATNNNAGATCGGAAG |
| 9238_1 | TATGAGTGTGGAGTCGTTGCACTTTTTAGAGCAGGGAACAGATNNNAGATCGGAA |
| 9240_1 | TATGAGTGTGGAGTCGTTGCGCAGAACCCTGACTATCACAGATNNNAGATCGGAAG |
| 9242_1 | TATGAGTGTGGAGTCGTTGCCCAATTCCATTTGCCTCCCAGATNNNAGATCGGAAGA |
| 9243_1 | TATGAGTGTGGAGTCGTTGCGGAAGGATCCGAGAGATCCAGATNNNAGATCGGAA |
| 9245_1 | TATGAGTGTGGAGTCGTTGCCCTTTATGCCAAGTGCTCCAGATNNNAGATCGGAAG |
| 9246_1 | TATGAGTGTGGAGTCGTTGCGTTGATAGAGCAAGGCAGCAGATNNNAGATCGGAA |
| 9247_1 | TATGAGTGTGGAGTCGTTGCCTGGCATGAGGATGTAGAAAGCTNNNAGATCGGAA |
| 9248_1 | TATGAGTGTGGAGTCGTTGCAGGGAGAGATTCACTTCGCAGATNNNAGATCGGAA |
| 9250_1 | TATGAGTGTGGAGTCGTTGCCTCCCCAACTTGGAATTGCAGATNNNAGATCGGAAG |
| 9251_1 | TATGAGTGTGGAGTCGTTGCTTGTGCACTGTGTACTTGCAGATNNNAGATCGGAAG |

| | |
|---|---|
| 9254_1 | TATGAGTGTGGAGTCGTTGCAAAGCTGGAGGAGTGTCTCAGATNNNAGATCGGAA |
| 9255_1 | TATGAGTGTGGAGTCGTTGCAAAACAATTGCCCTGGGTCAGATNNNAGATCGGAAG |
| 9256_1 | TATGAGTGTGGAGTCGTTGCAAGGGTAGAGTGCCACTTCAGATNNNAGATCGGAA |
| 9257_1 | TATGAGTGTGGAGTCGTTGCTGGCTACTCCTGGGATTTCAGATNNNAGATCGGAAG |
| 9258_1 | TATGAGTGTGGAGTCGTTGCCCCCTTCGAAGTCTGTTTCAGATNNNAGATCGGAAG |
| 9259_1 | TATGAGTGTGGAGTCGTTGCACAGGTATCAGAGCCACAGAGATNNNAGATCGGAA |
| 9260_1 | TATGAGTGTGGAGTCGTTGCGGGCAAGGGAGAAAGAGAGAGATNNNAGATCGGA |
| 9264_1 | TATGAGTGTGGAGTCGTTGCGCATTTCTCTCTCCCTACGAGATNNNAGATCGGAAG |
| 9267_1 | TATGAGTGTGGAGTCGTTGCTGTATCCCAGTTGCTCTGGAGATNNNAGATCGGAAG |
| 9268_1 | TATGAGTGTGGAGTCGTTGCAACCATGACAATTCCGTGGAGATNNNAGATCGGAAG |
| 9270_1 | TATGAGTGTGGAGTCGTTGCTGGAGACCCAGGAAAGATGAGATNNNAGATCGGAA |
| 9272_1 | TATGAGTGTGGAGTCGTTGCTGTAGGGGGAAACCAAGTGAGATNNNAGATCGGA |
| 9274_1 | TATGAGTGTGGAGTCGTTGCGCTTTCCAGATCCAGTGTGAGATNNNAGATCGGAAG |
| 9276_1 | TATGAGTGTGGAGTCGTTGCGAGAGAGAGAGGCTGGTTGAGATNNNAGATCGGA |
| 9277_1 | TATGAGTGTGGAGTCGTTGCGGTTTCCAAGCTGTTGAATAGATNNNAGATCGGAAG |
| 9279_1 | TATGAGTGTGGAGTCGTTGCTGTATTTGAAAACGCCCAGTAGTNNNAGATCGGAAG |
| 9282_1 | TATGAGTGTGGAGTCGTTGCGGACCTACACTGCCCTGTTAGATNNNAGATCGGAAG |
| 9283_1 | TATGAGTGTGGAGTCGTTGCATGTGGTCTTGGACTGGAGTAGTNNNAGATCGGAA |
| 9284_1 | TATGAGTGTGGAGTCGTTGCGCATGTATGTTACTCCTCACGATNNNAGATCGGAAG |
| 9285_1 | TATGAGTGTGGAGTCGTTGCTTCCTTACATTCCCACCAACACTNNNAGATCGGAAGA |
| 9288_1 | TATGAGTGTGGAGTCGTTGCAGAGAGAGGACAGCAACAAGGATNNNAGATCGGA |
| 9289_1 | TATGAGTGTGGAGTCGTTGCTCTGTCCACATTTTCCCAAGGATNNNAGATCGGAAG |
| 9293_1 | TATGAGTGTGGAGTCGTTGCGAGGGAGGTGGTTGTAAGAGGATNNNAGATCGGA |
| 9294_1 | TATGAGTGTGGAGTCGTTGCGAAACCTGGTCATGGGAGAGGATNNNAGATCGGAA |
| 9297_1 | TATGAGTGTGGAGTCGTTGCACCTTTCATAGTGGGCTGAGGATNNNAGATCGGAAG |
| 9298_1 | TATGAGTGTGGAGTCGTTGCGGGACAGTGGGAATAGTGAGGATNNNAGATCGGA |
| 9302_1 | TATGAGTGTGGAGTCGTTGCGGCTATTATGAAGACTGTGGCAANNNAGATCGGAA |
| 9304_1 | TATGAGTGTGGAGTCGTTGCTGGAGTTTCTTGGATCCAGGGATNNNAGATCGGAA |
| 9305_1 | TATGAGTGTGGAGTCGTTGCTACATCTCCCTCCCATCAGGGATNNNAGATCGGAAG |
| 9306_1 | TATGAGTGTGGAGTCGTTGCTGATTTTGGTGGAGGAGAGGGATNNNAGATCGGA |
| 9307_1 | TATGAGTGTGGAGTCGTTGCGTGTGTATGGGTAGGTGAGGGATNNNAGATCGGA |
| 9308_1 | TATGAGTGTGGAGTCGTTGCGTATGAGGAAAAGGGGAGGGGATNNNAGATCGGA |
| 9309_1 | TATGAGTGTGGAGTCGTTGCCTATGGCATCCTAAGAGGGGGATNNNAGATCGGAA |
| 9310_1 | TATGAGTGTGGAGTCGTTGCCAGAAAAGAAACTGGCTGGGGATNNNAGATCGGAA |
| 9311_1 | TATGAGTGTGGAGTCGTTGCCTGAAGGATGGAAATTTGGGGATNNNAGATCGGAA |
| 9312_1 | TATGAGTGTGGAGTCGTTGCCCTGTACTCGGGGATTATGGGATNNNAGATCGGAA |
| 9313_1 | TATGAGTGTGGAGTCGTTGCCCTACAAAGAAGCATGCTGGGATNNNAGATCGGAA |
| 9315_1 | TATGAGTGTGGAGTCGTTGCGTAAATCTGCTGCCTCCACTTCANNNAGATCGGAAG |
| 9316_1 | TATGAGTGTGGAGTCGTTGCTGCCATATTTGTGATTTCATGGATNNNAGATCGGAA |
| 9318_1 | TATGAGTGTGGAGTCGTTGCCCTTCAACCTAGGACACCTGGATNNNAGATCGGAAG |
| 9319_1 | TATGAGTGTGGAGTCGTTGCTGATGGTCTTTCCCATCCTGGATNNNAGATCGGAAG |
| 9322_1 | TATGAGTGTGGAGTCGTTGCCCACTTGAGGGTTTTCTGTGGATNNNAGATCGGAAG |
| 9324_1 | TATGAGTGTGGAGTCGTTGCAGAGAACACAAAGATCCTTGGATNNNAGATCGGAA |

| | |
|---|---|
| 9326_1 | TATGAGTGTGGAGTCGTTGCGGGGGAGCGATTTCTATTTGGATNNNAGATCGGAA |
| 9331_1 | TATGAGTGTGGAGTCGTTGCGGGGTGATGCAAGTGGTAATGATNNNAGATCGGA |
| 9332_1 | TATGAGTGTGGAGTCGTTGCTTGGAAGGTGAGAGGTCCATGATNNNAGATCGGAA |
| 9333_1 | TATGAGTGTGGAGTCGTTGCAGTTTCCCTACCCTATGCATGATNNNAGATCGGAAG |
| 9334_1 | TATGAGTGTGGAGTCGTTGCCCCACCCATCCTTCTCTCATGATNNNAGATCGGAAGA |
| 9335_1 | TATGAGTGTGGAGTCGTTGCAACTGGAGAAATGTGGCAGAGGCNNNAGATCGGA |
| 9336_1 | TATGAGTGTGGAGTCGTTGCGCCTGAATGTGTCCTTGGATGATNNNAGATCGGAAG |
| 9337_1 | TATGAGTGTGGAGTCGTTGCCGTGCGCCACAGAGTAATATGATNNNAGATCGGAAG |
| 9338_1 | TATGAGTGTGGAGTCGTTGCCAAAGCCATTTTCTGGCACTGATNNNAGATCGGAAG |
| 9339_1 | TATGAGTGTGGAGTCGTTGCAAGTGAACCGTACTGGGACTGATNNNAGATCGGAA |
| 9340_1 | TATGAGTGTGGAGTCGTTGCCCCAGACCTTTCTCTCTCCTGATNNNAGATCGGAAGA |
| 9341_1 | TATGAGTGTGGAGTCGTTGCTGTTGCATTTGAGGCTTCCTGATNNNAGATCGGAAG |
| 9342_1 | TATGAGTGTGGAGTCGTTGCACATTGCAAAAGAGGAGGCTGATNNNAGATCGGAA |
| 9343_1 | TATGAGTGTGGAGTCGTTGCATTTCCTAGCCCTGTCTGCTGATNNNAGATCGGAAG |
| 9345_1 | TATGAGTGTGGAGTCGTTGCCCCCGCATTGTTGTTTCTCTGATNNNAGATCGGAAGA |
| 9347_1 | TATGAGTGTGGAGTCGTTGCACTTGGGTGGGACTTACAGTGATNNNAGATCGGAA |
| 9348_1 | TATGAGTGTGGAGTCGTTGCCCTGGTTGTTTCTGTTCCCTTCANNNAGATCGGAAGA |
| 9349_1 | TATGAGTGTGGAGTCGTTGCTGTGCATGTGTGCTTGTAGTGATNNNAGATCGGAAG |
| 9350_1 | TATGAGTGTGGAGTCGTTGCAGGGATGCTATACAGAACGTGATNNNAGATCGGAA |
| 9351_1 | TATGAGTGTGGAGTCGTTGCGCATCTTGCAAACTAGGGGTGATNNNAGATCGGAA |
| 9354_1 | TATGAGTGTGGAGTCGTTGCTACACCCCTTTGGGAGGATTGATNNNAGATCGGAAG |
| 9355_1 | TATGAGTGTGGAGTCGTTGCAAGAAAGGGCAATCAGCCTTGATNNNAGATCGGAA |
| 9356_1 | TATGAGTGTGGAGTCGTTGCGGCATATGGGAAGAGTGCTTGATNNNAGATCGGAA |
| 9357_1 | TATGAGTGTGGAGTCGTTGCGGTGGGTTCTGTGCTTTCTTGATNNNAGATCGGAAG |
| 9358_1 | TATGAGTGTGGAGTCGTTGCCCTCAGCTCACAACAGAGTTGATNNNAGATCGGAAG |
| 9361_1 | TATGAGTGTGGAGTCGTTGCGGTTGCTGAAGAGCACTGTTGATNNNAGATCGGAA |
| 9365_1 | TATGAGTGTGGAGTCGTTGCGGTCTTCTCACATCGCCTTTGATNNNAGATCGGAAG |
| 9366_1 | TATGAGTGTGGAGTCGTTGCCACCACCATGCTGAGATTTTGATNNNAGATCGGAAG |
| 9368_1 | TATGAGTGTGGAGTCGTTGCGAGACAGGGAAATTCAGGCAACANNNAGATCGGAA |
| 9369_1 | TATGAGTGTGGAGTCGTTGCCACGCACGCACATACACAAATATNNNAGATCGGAAG |
| 9370_1 | TATGAGTGTGGAGTCGTTGCGGTGGAGTGGCTGTTCCAAATATNNNAGATCGGAA |
| 9371_1 | TATGAGTGTGGAGTCGTTGCCACCCACGGAAACCCTGAAATATNNNAGATCGGAAG |
| 9372_1 | TATGAGTGTGGAGTCGTTGCTGAGAGGTTGTTCGGGGGAATATNNNAGATCGGA |
| 9374_1 | TATGAGTGTGGAGTCGTTGCCTTGACCTCCTGGGCCTACATATNNNAGATCGGAAG |
| 9377_1 | TATGAGTGTGGAGTCGTTGCGCTGACGCTGTACACATCCATATNNNAGATCGGAAG |
| 9379_1 | TATGAGTGTGGAGTCGTTGCCCAGTCCTGAGAACCTTGCATATNNNAGATCGGAAG |
| 9382_1 | TATGAGTGTGGAGTCGTTGCGGCAAAATCAGGCCCTCAGATATNNNAGATCGGAAG |
| 9383_1 | TATGAGTGTGGAGTCGTTGCGGGACAGAGGGACAGAGAGATATNNNAGATCGGA |
| 9384_1 | TATGAGTGTGGAGTCGTTGCTTTGACCCCTTTCGGGGAGATATNNNAGATCGGAAG |
| 9385_1 | TATGAGTGTGGAGTCGTTGCGGTCCCTCCACATGGCTAGATATNNNAGATCGGAAG |
| 9387_1 | TATGAGTGTGGAGTCGTTGCTCACTCACCCGTAGCTTGGATATNNNAGATCGGAAG |
| 9388_1 | TATGAGTGTGGAGTCGTTGCGCCAGTGACAAACCCCATGATATNNNAGATCGGAAG |
| 9389_1 | TATGAGTGTGGAGTCGTTGCGCTCTGGCATTTGCCACCTATATNNNAGATCGGAAG |

| | |
|---|---|
| 9390_1 | TATGAGTGTGGAGTCGTTGCATGAGTGCCCAGGAAGGCTATATNNNAGATCGGAA |
| 9391_1 | TATGAGTGTGGAGTCGTTGCCACACACACCCCAACACGTATATNNNAGATCGGAAG |
| 9392_1 | TATGAGTGTGGAGTCGTTGCAGCCCCCACCCATGGAATTATATNNNAGATCGGAAG |
| 9394_1 | TATGAGTGTGGAGTCGTTGCCCACCACGACCAACAACAACTATNNNAGATCGGAAG |
| 9395_1 | TATGAGTGTGGAGTCGTTGCGCAAGCACTCTGACTTCCACTATNNNAGATCGGAAG |
| 9396_1 | TATGAGTGTGGAGTCGTTGCGCAGGATACATGGGGGTGACTATNNNAGATCGGAA |
| 9397_1 | TATGAGTGTGGAGTCGTTGCTGGAGGGGGCATATGTTGACTATNNNAGATCGGAA |
| 9398_1 | TATGAGTGTGGAGTCGTTGCCTGGCTCCTCTGCCACTTACTATNNNAGATCGGAAGA |
| 9399_1 | TATGAGTGTGGAGTCGTTGCGCTGGGAAGGAGTAGGAACCTATNNNAGATCGGAA |
| 9400_1 | TATGAGTGTGGAGTCGTTGCTACCCACATTCACTGCCACCTATNNNAGATCGGAAGA |
| 9402_1 | TATGAGTGTGGAGTCGTTGCATACCCTCACTGTCTCTCCCTATNNNAGATCGGAAGA |
| 9404_1 | TATGAGTGTGGAGTCGTTGCCACACTATGTTTCCCTCCCATAANNNAGATCGGAAGA |
| 9405_1 | TATGAGTGTGGAGTCGTTGCGCACACGCAAACCCATATCCTATNNNAGATCGGAAG |
| 9406_1 | TATGAGTGTGGAGTCGTTGCACATTGGAGAACAGCCCTCCTATNNNAGATCGGAAG |
| 9407_1 | TATGAGTGTGGAGTCGTTGCTGCTTCCTTGAGTGCTCTCCTATNNNAGATCGGAAG |
| 9408_1 | TATGAGTGTGGAGTCGTTGCCTTCTCCAGCTTGGGTCTCCTATNNNAGATCGGAAG |
| 9409_1 | TATGAGTGTGGAGTCGTTGCAAGGTCAGCAGTGTGTCTCCTATNNNAGATCGGAAG |
| 9410_1 | TATGAGTGTGGAGTCGTTGCTGTGGGGTCCATCATGTTCCTATNNNAGATCGGAAG |
| 9411_1 | TATGAGTGTGGAGTCGTTGCGGAGGTGACAGAGGCTAAGCTATNNNAGATCGGA |
| 9412_1 | TATGAGTGTGGAGTCGTTGCGGAGGGGAGATGAAGAGAGCTATNNNAGATCGGA |
| 9414_1 | TATGAGTGTGGAGTCGTTGCGCCAGGTGTCTCTAGTGAGCTATNNNAGATCGGAA |
| 9415_1 | TATGAGTGTGGAGTCGTTGCCTATGCAGTTAGCCACACGCTATNNNAGATCGGAAG |
| 9419_1 | TATGAGTGTGGAGTCGTTGCAGATGAGGGGAAGGTTGTGCTATNNNAGATCGGA |
| 9420_1 | TATGAGTGTGGAGTCGTTGCAATGTCATAATGGTGTGGTGGCTNNNAGATCGGAA |
| 9423_1 | TATGAGTGTGGAGTCGTTGCTCCCTCCCTGTCACTCTCTCTATNNNAGATCGGAAGA |
| 9424_1 | TATGAGTGTGGAGTCGTTGCAGCGTATGTCTGTTTCCGTCTATNNNAGATCGGAAG |
| 9426_1 | TATGAGTGTGGAGTCGTTGCTTTGGAGAAAGAAAGGCCATGGTNNNAGATCGGAA |
| 9427_1 | TATGAGTGTGGAGTCGTTGCATGGGTTTTTCAGCGCCTTCTATNNNAGATCGGAAG |
| 9428_1 | TATGAGTGTGGAGTCGTTGCTCAGGACCTCCATTTCTGGACCCNNNAGATCGGAAG |
| 9429_1 | TATGAGTGTGGAGTCGTTGCTTCGCTGTCACGTTGGTTTCTATNNNAGATCGGAAG |
| 9430_1 | TATGAGTGTGGAGTCGTTGCGTGCGTGCATCTTTCTAAAGTATNNNAGATCGGAAG |
| 9433_1 | TATGAGTGTGGAGTCGTTGCCTGCCTTCCCTTTTCTGCAGTATNNNAGATCGGAAGA |
| 9434_1 | TATGAGTGTGGAGTCGTTGCCTGCTGGAGGCAAAACAGAGTATNNNAGATCGGAA |
| 9435_1 | TATGAGTGTGGAGTCGTTGCAGGAAGTGAGTGCTTGGGAGTATNNNAGATCGGA |
| 9437_1 | TATGAGTGTGGAGTCGTTGCTACTCCCCCAACAGAACACGTATNNNAGATCGGAAG |
| 9439_1 | TATGAGTGTGGAGTCGTTGCGGGGAGAAAAGCTTGTCACGTATNNNAGATCGGAA |
| 9440_1 | TATGAGTGTGGAGTCGTTGCTGTATGCGTGTGCATGTACGTATNNNAGATCGGAA |
| 9442_1 | TATGAGTGTGGAGTCGTTGCACTTGCCCTGTATACTGTCGTATNNNAGATCGGAAG |
| 9443_1 | TATGAGTGTGGAGTCGTTGCGGGATATAGTCGGCCCTTCGTATNNNAGATCGGAA |
| 9447_1 | TATGAGTGTGGAGTCGTTGCTGCCACATGGATGGATGGTAGATNNNAGATCGGAA |
| 9448_1 | TATGAGTGTGGAGTCGTTGCCATTTGGCTGGGGTCAATGGTATNNNAGATCGGAA |
| 9450_1 | TATGAGTGTGGAGTCGTTGCTGGGACATAATTGAAGTGCACGTNNNAGATCGGAA |
| 9451_1 | TATGAGTGTGGAGTCGTTGCGTGGTCGCATGTGTGTGATGTATNNNAGATCGGAA |

9452_1 TATGAGTGTGGAGTCGTTGCTGGGACTGGTGTGTGTACTGTATNNNAGATCGGAA
9453_1 TATGAGTGTGGAGTCGTTGCCATTCACAGTGGGCAACCTGTATNNNAGATCGGAAG
9454_1 TATGAGTGTGGAGTCGTTGCGGAATGGCTGCATCTGTCTGTATNNNAGATCGGAAG
9456_1 TATGAGTGTGGAGTCGTTGCCCCCACACACCATACAGGTGTATNNNAGATCGGAAG
9457_1 TATGAGTGTGGAGTCGTTGCTCCACTTTCCCACCTCTGTGTATNNNAGATCGGAAGA
9458_1 TATGAGTGTGGAGTCGTTGCGGTGGGTCCCCAGAAAATTGTATNNNAGATCGGAA
9459_1 TATGAGTGTGGAGTCGTTGCCTGGATCCCTGTCCACTTTGTATNNNAGATCGGAAGA
9461_1 TATGAGTGTGGAGTCGTTGCCCTGGAAGGGTCCCTAGCATTATNNNAGATCGGAAG
9463_1 TATGAGTGTGGAGTCGTTGCGCAGGTGGACAGTGGTTCATTATNNNAGATCGGAA
9465_1 TATGAGTGTGGAGTCGTTGCGGTAGAGGGGCAGGGTAACTTATNNNAGATCGGA
9468_1 TATGAGTGTGGAGTCGTTGCCTGTCCTTTTCGTACTTGGACGTNNNAGATCGGAAG
9469_1 TATGAGTGTGGAGTCGTTGCCCAAGCACTCTATGCAGGCTTATNNNAGATCGGAAG
9470_1 TATGAGTGTGGAGTCGTTGCGCCAAGACCTCTTCCATGCTTATNNNAGATCGGAAG
9471_1 TATGAGTGTGGAGTCGTTGCGAAAGGAAGGCCAAAAGTCTTATNNNAGATCGGAA
9472_1 TATGAGTGTGGAGTCGTTGCGAGGGTGAGGGGAGGATTCTTATNNNAGATCGGA
9473_1 TATGAGTGTGGAGTCGTTGCTGGGAAGGGGCTGAACTTCTTATNNNAGATCGGAA
9474_1 TATGAGTGTGGAGTCGTTGCGGGGAGCAAAGCTGTACAGTTATNNNAGATCGGAA
9476_1 TATGAGTGTGGAGTCGTTGCAGGCATTTGAAGCTGGCTGTTATNNNAGATCGGAA
9477_1 TATGAGTGTGGAGTCGTTGCCACGGACCTGGGTGAGAATTTATNNNAGATCGGAA
9478_1 TATGAGTGTGGAGTCGTTGCGGAGGGCAACGGATACGATTTATNNNAGATCGGAA
9480_1 TATGAGTGTGGAGTCGTTGCTCTGGTGACCCTTGGACCTTTATNNNAGATCGGAAG
9482_1 TATGAGTGTGGAGTCGTTGCGGGGTGGTGACAGAGCATTTTATNNNAGATCGGAA
9484_1 TATGAGTGTGGAGTCGTTGCCCCTCAGCAAAGAGTGGAAAACTNNNAGATCGGAA
9485_1 TATGAGTGTGGAGTCGTTGCTGGTTCTCTTGCTTTGGAAAACTNNNAGATCGGAAG
9487_1 TATGAGTGTGGAGTCGTTGCAGTTTCAGGCACCAGAACAAACTNNNAGATCGGAAG
9489_1 TATGAGTGTGGAGTCGTTGCGAACAACAACCTCTCCCCAAACTNNNAGATCGGAAG
9490_1 TATGAGTGTGGAGTCGTTGCCCACATTCATGCACATGCAAACTNNNAGATCGGAAG
9493_1 TATGAGTGTGGAGTCGTTGCTCTATGTGGGTGGTCAGGAAACTNNNAGATCGGAA
9494_1 TATGAGTGTGGAGTCGTTGCATCCCTCCAAAAGCAATTGGAAGNNNAGATCGGAAG
9495_1 TATGAGTGTGGAGTCGTTGCACGGCAAAGAAAGAAGGGAAACTNNNAGATCGGA
9499_1 TATGAGTGTGGAGTCGTTGCTTGAGTGGTGAAATGCCACAACTNNNAGATCGGAA
9501_1 TATGAGTGTGGAGTCGTTGCTCTCCCTTCTCTGAACACCAACTNNNAGATCGGAAGA
9503_1 TATGAGTGTGGAGTCGTTGCAGAGAGAGATGCTGACACCAACTNNNAGATCGGAA
9504_1 TATGAGTGTGGAGTCGTTGCTCCCAGGCTATCAACTACCAACTNNNAGATCGGAAG
9505_1 TATGAGTGTGGAGTCGTTGCTTATCCTCAGAACCCACCCAACTNNNAGATCGGAAGA
9508_1 TATGAGTGTGGAGTCGTTGCGCACCCATACATACATGGCAACTNNNAGATCGGAAG
9510_1 TATGAGTGTGGAGTCGTTGCACTTCATGTGGGCAAAGTCAACTNNNAGATCGGAAG
9512_1 TATGAGTGTGGAGTCGTTGCCCAGCTACTAATGGCTTTCAACTNNNAGATCGGAAG
9514_1 TATGAGTGTGGAGTCGTTGCCTCTGGTGGCAAGAAGAAGAACTNNNAGATCGGAA
9515_1 TATGAGTGTGGAGTCGTTGCGGGAGTTCCAGGTAGACAGAACTNNNAGATCGGAA
9516_1 TATGAGTGTGGAGTCGTTGCTTTGGCTTGGTTTCTAGGGTTCANNNAGATCGGAAG
9517_1 TATGAGTGTGGAGTCGTTGCAGGACTTAGCCCCATGTAGAACTNNNAGATCGGAAG
9518_1 TATGAGTGTGGAGTCGTTGCGGGGAGAGATTGAAGAGGGAACTNNNAGATCGGA

| | |
|---|---|
| 9519_1 | TATGAGTGTGGAGTCGTTGCTTCATTGTGCAAATTGGGGAACTNNNAGATCGGAAG |
| 9520_1 | TATGAGTGTGGAGTCGTTGCCACAGGAAGGAAACATGGGAACTNNNAGATCGGAA |
| 9521_1 | TATGAGTGTGGAGTCGTTGCGACAGGGACTTAGAGGTGGAACTNNNAGATCGGAA |
| 9522_1 | TATGAGTGTGGAGTCGTTGCGTGGATGAAGCCATCTTGGAACTNNNAGATCGGAAG |
| 9524_1 | TATGAGTGTGGAGTCGTTGCTCTACAGCTTCTGGCCCATAACTNNNAGATCGGAAGA |
| 9526_1 | TATGAGTGTGGAGTCGTTGCGGTTGAATCTGAAGTTTGGGGGANNNAGATCGGAA |
| 9527_1 | TATGAGTGTGGAGTCGTTGCATGCCACAGAGGAAGAGGTAACTNNNAGATCGGAA |
| 9528_1 | TATGAGTGTGGAGTCGTTGCGCGTGATTACATGCAGGGTAACTNNNAGATCGGAAG |
| 9529_1 | TATGAGTGTGGAGTCGTTGCAGGATGGCAAGTGTCAAAACACTNNNAGATCGGAAG |
| 9530_1 | TATGAGTGTGGAGTCGTTGCGGGAATACTATGGGCTGGAGGATNNNAGATCGGAA |
| 9534_1 | TATGAGTGTGGAGTCGTTGCGAAACTGAGGTGCTGCTAACACTNNNAGATCGGAAG |
| 9535_1 | TATGAGTGTGGAGTCGTTGCGATTTTCGCAAAGTGCACACACTNNNAGATCGGAAG |
| 9537_1 | TATGAGTGTGGAGTCGTTGCCAGACAAAGATGACAGCCACACTNNNAGATCGGAAG |
| 9540_1 | TATGAGTGTGGAGTCGTTGCCCAGGCTCAAGAACAAAGACACTNNNAGATCGGAAG |
| 9541_1 | TATGAGTGTGGAGTCGTTGCTATCTGCCAGGTTTTGCTGTTCANNNAGATCGGAAGA |
| 9542_1 | TATGAGTGTGGAGTCGTTGCACCTGCTTATCCACTCAGACACTNNNAGATCGGAAGA |
| 9543_1 | TATGAGTGTGGAGTCGTTGCAAGTCGTGCATTTGTGAGACACTNNNAGATCGGAAG |
| 9545_1 | TATGAGTGTGGAGTCGTTGCTCAGGACCTTCAGTAATGACACTNNNAGATCGGAAG |
| 9548_1 | TATGAGTGTGGAGTCGTTGCTCTGAATGGCTTGGGAGTACACTNNNAGATCGGAAG |
| 9549_1 | TATGAGTGTGGAGTCGTTGCTAACACCATCAGCTGGGTACACTNNNAGATCGGAAG |
| 9550_1 | TATGAGTGTGGAGTCGTTGCCCCACACACATGCTCATTACACTNNNAGATCGGAAGA |
| 9553_1 | TATGAGTGTGGAGTCGTTGCATCTGGGGAAGCTGATAACCACTNNNAGATCGGAAG |
| 9557_1 | TATGAGTGTGGAGTCGTTGCAGGAAAGAGAAGGAAGACCCACTNNNAGATCGGAA |
| 9559_1 | TATGAGTGTGGAGTCGTTGCCTCTCTCCCTTCTTACAGCCACTNNNAGATCGGAAGAG |
| 9560_1 | TATGAGTGTGGAGTCGTTGCTCTCTCTCCCAGTCCTATCCACTNNNAGATCGGAAGAG |
| 9563_1 | TATGAGTGTGGAGTCGTTGCTGTCTCCAACTGACTTCTCCACTNNNAGATCGGAAGA |
| 9564_1 | TATGAGTGTGGAGTCGTTGCAGATCCCTAGCCTTTGTTCCACTNNNAGATCGGAAGA |
| 9565_1 | TATGAGTGTGGAGTCGTTGCGGCACAGTATTTTGAGATAGCGTNNNAGATCGGAAG |
| 9567_1 | TATGAGTGTGGAGTCGTTGCTCTGCCAACTCTATCTCAGCACTNNNAGATCGGAAGA |
| 9570_1 | TATGAGTGTGGAGTCGTTGCAAGAACTCATAGGGTCAGGCACTNNNAGATCGGAAG |
| 9578_1 | TATGAGTGTGGAGTCGTTGCAACCATGGAGGTGAAGGATCACTNNNAGATCGGAAG |
| 9579_1 | TATGAGTGTGGAGTCGTTGCTTCACGGTATCCAGGCTATCACTNNNAGATCGGAAGA |
| 9584_1 | TATGAGTGTGGAGTCGTTGCTGATGACTAGACAGTGGCTCACTNNNAGATCGGAAG |
| 9585_1 | TATGAGTGTGGAGTCGTTGCTGTAGTCTGTCCAGATGCTCACTNNNAGATCGGAAGA |
| 9586_1 | TATGAGTGTGGAGTCGTTGCTACCAACACTAGGAGATCTCACTNNNAGATCGGAAGA |
| 9589_1 | TATGAGTGTGGAGTCGTTGCTTCACACACACACACAAGTCACTNNNAGATCGGAAGA |
| 9590_1 | TATGAGTGTGGAGTCGTTGCTCGTCTACTCTCACACAGTCACTNNNAGATCGGAAGA |
| 9592_1 | TATGAGTGTGGAGTCGTTGCAGCCAGGGACTTATTCTGTCACTNNNAGATCGGAAG |
| 9595_1 | TATGAGTGTGGAGTCGTTGCTACGGTTGTGGGATAGAGGTCCTNNNAGATCGGAAG |
| 9605_1 | TATGAGTGTGGAGTCGTTGCTAACTGATGGTATACCCCAGACTNNNAGATCGGAAG |
| 9606_1 | TATGAGTGTGGAGTCGTTGCCGCAGACTAGCAAAATCCAGACTNNNAGATCGGAAG |
| 9608_1 | TATGAGTGTGGAGTCGTTGCTTCCCCTACAGTCCAGTCAGACTNNNAGATCGGAAGA |
| 9610_1 | TATGAGTGTGGAGTCGTTGCTTCCCAGCAATGGAGTAGAGACTNNNAGATCGGAAG |

| | |
|---|---|
| 9611_1 | TATGAGTGTGGAGTCGTTGCCCCATCCTGATGAAACTGAGACTNNNAGATCGGAAG |
| 9612_1 | TATGAGTGTGGAGTCGTTGCGTGCAGACCAAGATTCTGAGACTNNNAGATCGGAA |
| 9613_1 | TATGAGTGTGGAGTCGTTGCGTGCTTTCCATGATCCAGATCGTNNNAGATCGGAAG |
| 9614_1 | TATGAGTGTGGAGTCGTTGCGACCTGAACCTCCAGCTTAGACTNNNAGATCGGAAG |
| 9618_1 | TATGAGTGTGGAGTCGTTGCACTGGAATGATGAAAGCAGGACTNNNAGATCGGAA |
| 9620_1 | TATGAGTGTGGAGTCGTTGCTGCCGAACTCATAACCTAGGACTNNNAGATCGGAAG |
| 9622_1 | TATGAGTGTGGAGTCGTTGCCTTACCCACCTACATACCGGACTNNNAGATCGGAAG |
| 9626_1 | TATGAGTGTGGAGTCGTTGCGGACAATCTACTCCATGGGGACTNNNAGATCGGAA |
| 9627_1 | TATGAGTGTGGAGTCGTTGCTGGAAGAGATCACCTGTGTGTCANNNAGATCGGAA |
| 9629_1 | TATGAGTGTGGAGTCGTTGCGTGCAGTGGTGCTATAATGGACTNNNAGATCGGAA |
| 9633_1 | TATGAGTGTGGAGTCGTTGCTGACTGGGTGGGGAAATATGACTNNNAGATCGGAA |
| 9637_1 | TATGAGTGTGGAGTCGTTGCAAGGTGGTGCTATGAGAGTGACTNNNAGATCGGAA |
| 9639_1 | TATGAGTGTGGAGTCGTTGCCGCAAAAGCTGAAGTTTGTGACTNNNAGATCGGAA |
| 9641_1 | TATGAGTGTGGAGTCGTTGCCACTCCAAACTGTGCTCTTGACTNNNAGATCGGAAG |
| 9642_1 | TATGAGTGTGGAGTCGTTGCTGGGAGAGTGATACGGTTTGACTNNNAGATCGGAA |
| 9644_1 | TATGAGTGTGGAGTCGTTGCCGAAGGCCCCACTTACAAATACTNNNAGATCGGAAG |
| 9647_1 | TATGAGTGTGGAGTCGTTGCAGTCAAGACAGCCTTACCATACTNNNAGATCGGAAG |
| 9648_1 | TATGAGTGTGGAGTCGTTGCTTGGTGATACCCTGCTCCATACTNNNAGATCGGAAG |
| 9649_1 | TATGAGTGTGGAGTCGTTGCACAAAATGGGGTCATCTCATACTNNNAGATCGGAAG |
| 9652_1 | TATGAGTGTGGAGTCGTTGCATGCCTTCACTTCTCCCACTACTNNNAGATCGGAAGA |
| 9653_1 | TATGAGTGTGGAGTCGTTGCAGGAAGAAGGGATTGCCACTACTNNNAGATCGGAA |
| 9654_1 | TATGAGTGTGGAGTCGTTGCATACACACCTCACCAGCACTACTNNNAGATCGGAAG |
| 9655_1 | TATGAGTGTGGAGTCGTTGCGGGCTCAGGAGAAGATGACTACTNNNAGATCGGAA |
| 9656_1 | TATGAGTGTGGAGTCGTTGCAGATCATCCTGGTTTTGCCTACTNNNAGATCGGAAG |
| 9657_1 | TATGAGTGTGGAGTCGTTGCAGGTTTCTGAAGCAATGGCTACTNNNAGATCGGAA |
| 9658_1 | TATGAGTGTGGAGTCGTTGCGTTGTGTCCATAGCTTGGCTACTNNNAGATCGGAAG |
| 9661_1 | TATGAGTGTGGAGTCGTTGCCCTGGAAGTCTTTCATTGCTACTNNNAGATCGGAAG |
| 9662_1 | TATGAGTGTGGAGTCGTTGCGAGGGAGAAGAGGACTTGCTACTNNNAGATCGGAA |
| 9663_1 | TATGAGTGTGGAGTCGTTGCTTGGTTCTTGTCTGTCTCATGTTNNNAGATCGGAAG |
| 9667_1 | TATGAGTGTGGAGTCGTTGCTACAAAACAACCTGGCCAGTACTNNNAGATCGGAAG |
| 9669_1 | TATGAGTGTGGAGTCGTTGCGACGGACACAGGAGACTAGTACTNNNAGATCGGAA |
| 9671_1 | TATGAGTGTGGAGTCGTTGCTCCAGTTTCCTGAGCAGGTACTTNNNAGATCGGAAG |
| 9672_1 | TATGAGTGTGGAGTCGTTGCTGATGCAAAGCAAACCTGGTACTNNNAGATCGGAA |
| 9673_1 | TATGAGTGTGGAGTCGTTGCTCAGCCTCATTCACAAGTTTCGTNNNAGATCGGAAG |
| 9674_1 | TATGAGTGTGGAGTCGTTGCGTATGCCCGAATGTGCATGTACTNNNAGATCGGAAG |
| 9675_1 | TATGAGTGTGGAGTCGTTGCGCATGATAGCACGTGACTGTACTNNNAGATCGGAA |
| 9676_1 | TATGAGTGTGGAGTCGTTGCTGGAGTTGGGGTTTCTCTGTACTNNNAGATCGGAA |
| 9678_1 | TATGAGTGTGGAGTCGTTGCGAGGCAGCACAATTCCTTGTACTNNNAGATCGGAAG |
| 9679_1 | TATGAGTGTGGAGTCGTTGCAGATGAGAGAGAGGTCTGGGTCTNNNAGATCGGA |
| 9683_1 | TATGAGTGTGGAGTCGTTGCAGAGAGAGAGAGGAGGGCTTACTNNNAGATCGGA |
| 9684_1 | TATGAGTGTGGAGTCGTTGCCTTGGGAAATGGGTGTGCTTACTNNNAGATCGGAA |
| 9685_1 | TATGAGTGTGGAGTCGTTGCTGGGCACTGATGTATTGGTTACTNNNAGATCGGAA |
| 9686_1 | TATGAGTGTGGAGTCGTTGCCTGTGCCTGAATCCTGCTTTACTNNNAGATCGGAAG |

| | |
|---|---|
| 9687_1 | TATGAGTGTGGAGTCGTTGCGTCACATTGGGAGAAACTGAGGGNNNAGATCGGAA |
| 9688_1 | TATGAGTGTGGAGTCGTTGCTACCCACGTTGACAACAAAACCTNNNAGATCGGAAG |
| 9689_1 | TATGAGTGTGGAGTCGTTGCCGAGGCACTGTGAATGAAAACCTNNNAGATCGGAA |
| 9690_1 | TATGAGTGTGGAGTCGTTGCTTCAGGTTTCTGCTTCCAAACCTNNNAGATCGGAAG |
| 9691_1 | TATGAGTGTGGAGTCGTTGCGTGGGAAAGGTCATGGAAAAGGTNNNAGATCGGA |
| 9692_1 | TATGAGTGTGGAGTCGTTGCAGGGGCATCAGAGTTCTAAACCTNNNAGATCGGAA |
| 9696_1 | TATGAGTGTGGAGTCGTTGCAGATGTAGCAATCCCCAGAACCTNNNAGATCGGAAG |
| 9699_1 | TATGAGTGTGGAGTCGTTGCCCCAAAGCAAGAGAAAGGAACCTNNNAGATCGGAA |
| 9700_1 | TATGAGTGTGGAGTCGTTGCCTCCCAGGTTCAAGCGTGATCTCNNNAGATCGGAAG |
| 9701_1 | TATGAGTGTGGAGTCGTTGCAAATGCGAACATTCAGGGAACCTNNNAGATCGGAA |
| 9702_1 | TATGAGTGTGGAGTCGTTGCGGCATTTGCACTGTCTGTTTTCANNNAGATCGGAAG |
| 9706_1 | TATGAGTGTGGAGTCGTTGCAGCACTGCAAAGAGACATAACCTNNNAGATCGGAA |
| 9707_1 | TATGAGTGTGGAGTCGTTGCATCTCACCATTTTGCCCAAAGGTNNNAGATCGGAAG |
| 9709_1 | TATGAGTGTGGAGTCGTTGCACAATCTCAGCTGGCTCTAACCTNNNAGATCGGAAG |
| 9710_1 | TATGAGTGTGGAGTCGTTGCTTCAATCCACTCACAGAACACCTNNNAGATCGGAAG |
| 9711_1 | TATGAGTGTGGAGTCGTTGCCACTATGAACACATGCCACACCTNNNAGATCGGAAG |
| 9712_1 | TATGAGTGTGGAGTCGTTGCCCAAATGCATAACCCTCACACCTNNNAGATCGGAAG |
| 9714_1 | TATGAGTGTGGAGTCGTTGCTAGCTGCGTAGACTTGTACACCTNNNAGATCGGAAG |
| 9715_1 | TATGAGTGTGGAGTCGTTGCAAAAGGTAGAGGTAGCACCACCTNNNAGATCGGAA |
| 9719_1 | TATGAGTGTGGAGTCGTTGCACCATGACTGAAACATTGCACCTNNNAGATCGGAAG |
| 9721_1 | TATGAGTGTGGAGTCGTTGCTTTCCCACTCCATTCTCTCACCTNNNAGATCGGAAGAG |
| 9722_1 | TATGAGTGTGGAGTCGTTGCCTCAGCAACAGTGGAATTCACCTNNNAGATCGGAAG |
| 9723_1 | TATGAGTGTGGAGTCGTTGCAGAGAGAGCCCAATATTCACCTNNNAGATCGGAAG |
| 9724_1 | TATGAGTGTGGAGTCGTTGCCAGGGAAGATTCCAGCAGTTACCNNNAGATCGGAA |
| 9727_1 | TATGAGTGTGGAGTCGTTGCCCCATCATTTGCCTGAAAGACCTNNNAGATCGGAAG |
| 9728_1 | TATGAGTGTGGAGTCGTTGCTTGCTTTTCCATTCAGCAGACCTNNNAGATCGGAAG |
| 9730_1 | TATGAGTGTGGAGTCGTTGCTTGGCTGTAAATGGAAAGGACCTNNNAGATCGGAA |
| 9731_1 | TATGAGTGTGGAGTCGTTGCGTGTGTTTTCCACTCAAGGACCTNNNAGATCGGAAG |
| 9732_1 | TATGAGTGTGGAGTCGTTGCTCAGAGGGACAGAATTTAGCAGANNNAGATCGGAA |
| 9733_1 | TATGAGTGTGGAGTCGTTGCATCTTTAGACCCTACGTGGACCTNNNAGATCGGAAG |
| 9735_1 | TATGAGTGTGGAGTCGTTGCGTGCTTGTTAGAAGTGCTGACCTNNNAGATCGGAA |
| 9738_1 | TATGAGTGTGGAGTCGTTGCTTCTGACTGACTGGCTACAAGGTNNNAGATCGGAAG |
| 9740_1 | TATGAGTGTGGAGTCGTTGCAGTGCCTGTTCAGGTAGATACCTNNNAGATCGGAAG |
| 9741_1 | TATGAGTGTGGAGTCGTTGCGGGTCCTAGCTGATGAACTACCTNNNAGATCGGAAG |
| 9742_1 | TATGAGTGTGGAGTCGTTGCCGCACGTGTCTATCTACCTACCTNNNAGATCGGAAG |
| 9743_1 | TATGAGTGTGGAGTCGTTGCTTGGTGTCAGAATGATGCTACCTNNNAGATCGGAAG |
| 9745_1 | TATGAGTGTGGAGTCGTTGCCGGATCATGAGGTCGAGATAGCNNNAGATCGGAAG |
| 9749_1 | TATGAGTGTGGAGTCGTTGCTTGGGCTGATGGCAAAATTACCTNNNAGATCGGAA |
| 9750_1 | TATGAGTGTGGAGTCGTTGCCACCAAGGCTTCCCAAATTACCTNNNAGATCGGAAG |
| 9751_1 | TATGAGTGTGGAGTCGTTGCACCACACCCTGTGTACATTACCTNNNAGATCGGAAG |
| 9756_1 | TATGAGTGTGGAGTCGTTGCGTAGGGTAGAAAGGTAGCACCCTNNNAGATCGGAA |
| 9757_1 | TATGAGTGTGGAGTCGTTGCTAAGTTTCAGCACATTGCACCCTNNNAGATCGGAAG |
| 9758_1 | TATGAGTGTGGAGTCGTTGCCCTCACGGAGCATTTCGTATGTTNNNAGATCGGAAG |

9760_1 TATGAGTGTGGAGTCGTTGCAGAGCCCAATTACTTTAGACCCTNNNAGATCGGAAG
9773_1 TATGAGTGTGGAGTCGTTGCTTATGAACCGAATGGGTACCCCTNNNAGATCGGAAG
9774_1 TATGAGTGTGGAGTCGTTGCTGGGTGTGGGTTATATGGTGGAANNNAGATCGGAA
9775_1 TATGAGTGTGGAGTCGTTGCAAGAGGAGACAATTCAACCCCCTNNNAGATCGGAAG
9777_1 TATGAGTGTGGAGTCGTTGCAGATCACCTAAAACCTACCCCCTNNNAGATCGGAAG
9779_1 TATGAGTGTGGAGTCGTTGCATTCTCTTCACTTCAGAGCCCCTNNNAGATCGGAAGA
9782_1 TATGAGTGTGGAGTCGTTGCAAAACAAGAGTAGCTCCTCCCCTNNNAGATCGGAAG
9783_1 TATGAGTGTGGAGTCGTTGCATCCACTCTATATCCAGTCCCCTNNNAGATCGGAAGA
9786_1 TATGAGTGTGGAGTCGTTGCGCATCTGTGTGTGTATTTCCCCTNNNAGATCGGAAG
9787_1 TATGAGTGTGGAGTCGTTGCTTGTTTTCCTCCTCTTTTCCCCTNNNAGATCGGAAGAG
9789_1 TATGAGTGTGGAGTCGTTGCGAACCACCCCTCTTTTATTGCCANNNAGATCGGAAGA
9790_1 TATGAGTGTGGAGTCGTTGCTGCCAAACCAAAATCTGGAAGGTNNNAGATCGGAA
9791_1 TATGAGTGTGGAGTCGTTGCTGAGCCTCTAAACTATTGGCCCTNNNAGATCGGAAG
9793_1 TATGAGTGTGGAGTCGTTGCACAAGAACGTTTACCCCCAAAGANNNAGATCGGAAG
9794_1 TATGAGTGTGGAGTCGTTGCGTCTCTCACAACAAATGTGCCCTNNNAGATCGGAAG
9795_1 TATGAGTGTGGAGTCGTTGCTCTACATATCAAGCCTGTGCCCTNNNAGATCGGAAG
9798_1 TATGAGTGTGGAGTCGTTGCAAGATAACCCACTACAGATCCCTNNNAGATCGGAAG
9799_1 TATGAGTGTGGAGTCGTTGCGGAACCTCTGTACCTTGATCCCTNNNAGATCGGAAG
9800_1 TATGAGTGTGGAGTCGTTGCAACGTATTTTTCCTGACCTCCCTNNNAGATCGGAAGA
9801_1 TATGAGTGTGGAGTCGTTGCTTGTCTCTTTTTCCTCCCTCCCTNNNAGATCGGAAGAG
9802_1 TATGAGTGTGGAGTCGTTGCAGTTTCCTCTACTCTCCCTCCCTNNNAGATCGGAAGA
9804_1 TATGAGTGTGGAGTCGTTGCCTTCATGCAACCATTTCCTCCCTNNNAGATCGGAAGA
9807_1 TATGAGTGTGGAGTCGTTGCGTGCATAGTCTACAACTCCATGTNNNAGATCGGAAG
9808_1 TATGAGTGTGGAGTCGTTGCCAATCTGCCCAACTTTTCTCCCTNNNAGATCGGAAGA
9809_1 TATGAGTGTGGAGTCGTTGCAGGCTGTACTCTTTCAAGTCCCTNNNAGATCGGAAG
9810_1 TATGAGTGTGGAGTCGTTGCTCGGAGAAATCAAGTCAGTCCCTNNNAGATCGGAAG
9811_1 TATGAGTGTGGAGTCGTTGCTGGTGTTCTGTATCAGTGTCCCTNNNAGATCGGAAG
9815_1 TATGAGTGTGGAGTCGTTGCGGTGGGCACATGATATTTTCCCTNNNAGATCGGAAG
9818_1 TATGAGTGTGGAGTCGTTGCGAAACTGCAAAAACCCAAAGCCTNNNAGATCGGAAG
9819_1 TATGAGTGTGGAGTCGTTGCGAGAGAGAGAAGAGGGAAAGCCTNNNAGATCGGA
9820_1 TATGAGTGTGGAGTCGTTGCAAGTAGTGCATTCAAGCAAGCCTNNNAGATCGGAA
9821_1 TATGAGTGTGGAGTCGTTGCTGACAAACCAAGAGAAGAAGCCTNNNAGATCGGAA
9823_1 TATGAGTGTGGAGTCGTTGCACTGGTGAAACCTGAATAAGCCTNNNAGATCGGAA
9825_1 TATGAGTGTGGAGTCGTTGCCCTTGATATTGTGCCCAGTCTCCNNNAGATCGGAAG
9826_1 TATGAGTGTGGAGTCGTTGCAGTAGCAGCAGCAGCATATCATCNNNAGATCGGAA
9830_1 TATGAGTGTGGAGTCGTTGCAAGTCAACAATAAGCAGCAGCCTNNNAGATCGGAA
9833_1 TATGAGTGTGGAGTCGTTGCCCACACTGTCTTCAAAAGAGCCTNNNAGATCGGAAG
9834_1 TATGAGTGTGGAGTCGTTGCACTATGAGACTGAAGCCACAGGTNNNAGATCGGAA
9835_1 TATGAGTGTGGAGTCGTTGCTGTTCACTGCTGTACTAGAGCCTNNNAGATCGGAAG
9837_1 TATGAGTGTGGAGTCGTTGCAGACATTTCATCTCCCTGAGCCTNNNAGATCGGAAG
9839_1 TATGAGTGTGGAGTCGTTGCCTTGTTCAGGAAACCTGTAGCCTNNNAGATCGGAAG
9840_1 TATGAGTGTGGAGTCGTTGCAACATATACATACACCCCCGCCTNNNAGATCGGAAGA
9844_1 TATGAGTGTGGAGTCGTTGCTTTCATTCACCTCTTTTTCGCCTNNNAGATCGGAAGA

| | |
|---|---|
| 9845_1 | TATGAGTGTGGAGTCGTTGCTTCCTTGGTGTTTTGAAAGGCCTNNNAGATCGGAAG |
| 9846_1 | TATGAGTGTGGAGTCGTTGCTGCTTTCAACAAGTACAAGGCCTNNNAGATCGGAAG |
| 9847_1 | TATGAGTGTGGAGTCGTTGCAGACAGAAAGGTAGGTAAGGCCTNNNAGATCGGAA |
| 9848_1 | TATGAGTGTGGAGTCGTTGCGGGGCAGAAAAGGAAAGGAAAGANNNAGATCGGA |
| 9852_1 | TATGAGTGTGGAGTCGTTGCGTAAACATTGTCACTGAGGGCCTNNNAGATCGGAAG |
| 9853_1 | TATGAGTGTGGAGTCGTTGCAACAGGAATCACAGCATACAGGTNNNAGATCGGAAG |
| 9855_1 | TATGAGTGTGGAGTCGTTGCAGTAACAAAGATGGTTTGGGCCTNNNAGATCGGAAG |
| 9856_1 | TATGAGTGTGGAGTCGTTGCTATTCGTTGAGGAGAACTGGCCTNNNAGATCGGAAG |
| 9857_1 | TATGAGTGTGGAGTCGTTGCTGTATGACTTCACTCTCTGGCCTNNNAGATCGGAAGA |
| 9858_1 | TATGAGTGTGGAGTCGTTGCTCCTGATATTCTGTGTCTGGCCTNNNAGATCGGAAGA |
| 9859_1 | TATGAGTGTGGAGTCGTTGCGGATAGCTAGTGGTATGTGGCCTNNNAGATCGGAAG |
| 9861_1 | TATGAGTGTGGAGTCGTTGCTGGTTGTTGGAATTTCTTGGCCTNNNAGATCGGAAG |
| 9862_1 | TATGAGTGTGGAGTCGTTGCATCAGAACAAAGTGCTTTGGCCTNNNAGATCGGAAG |
| 9863_1 | TATGAGTGTGGAGTCGTTGCCCTTTTGTTGTTTGCTTTGGCCTNNNAGATCGGAAGA |
| 9866_1 | TATGAGTGTGGAGTCGTTGCCTTGCCTCTCTTGTTGAATGCCTNNNAGATCGGAAGA |
| 9867_1 | TATGAGTGTGGAGTCGTTGCTTCCCCTAAAGCAAAACATGCCTNNNAGATCGGAAGA |
| 9869_1 | TATGAGTGTGGAGTCGTTGCGGGTTTCATCCATCAAGATGCCTNNNAGATCGGAAG |
| 9871_1 | TATGAGTGTGGAGTCGTTGCCTGGAACGTGCTTACATATGCCTNNNAGATCGGAAG |
| 9872_1 | TATGAGTGTGGAGTCGTTGCAGCAGCAACTAATTACTATGCCTNNNAGATCGGAAGA |
| 9874_1 | TATGAGTGTGGAGTCGTTGCTCTGGCATTTTACAAACCTGCCTNNNAGATCGGAAGA |
| 9875_1 | TATGAGTGTGGAGTCGTTGCCATATAAAGCAGAGTGCCTGCCTNNNAGATCGGAAG |
| 9876_1 | TATGAGTGTGGAGTCGTTGCTTGCATCTAAAACCATGCTGCCTNNNAGATCGGAAGA |
| 9877_1 | TATGAGTGTGGAGTCGTTGCATTATCTGTGGACCTGTCTGCCTNNNAGATCGGAAGA |
| 9878_1 | TATGAGTGTGGAGTCGTTGCTGGCTCTTTCAAAACTTCTGCCTNNNAGATCGGAAGA |
| 9881_1 | TATGAGTGTGGAGTCGTTGCTTCAGCATTTCTAGGATGTGCCTNNNAGATCGGAAGA |
| 9882_1 | TATGAGTGTGGAGTCGTTGCTCTTACAATGTAGCTTTGGCCCTNNNAGATCGGAAGA |
| 9885_1 | TATGAGTGTGGAGTCGTTGCCCTTGGGAAACAAGCTATGGTGANNNAGATCGGAAG |
| 9887_1 | TATGAGTGTGGAGTCGTTGCAGAAGGGAGAGAGAGTTTTGCCTNNNAGATCGGAA |
| 9889_1 | TATGAGTGTGGAGTCGTTGCTTCTCCTCTTCCTGGGAAATCCTNNNAGATCGGAAGA |
| 9893_1 | TATGAGTGTGGAGTCGTTGCCTACAGGGAGCTGCTCTAATCCTNNNAGATCGGAAG |
| 9894_1 | TATGAGTGTGGAGTCGTTGCAGCGATATGTGTTCTCTCCAGGTNNNAGATCGGAAG |
| 9896_1 | TATGAGTGTGGAGTCGTTGCGTTGCTTTTCTCTCCACCATCCTNNNAGATCGGAAGAG |
| 9898_1 | TATGAGTGTGGAGTCGTTGCAAGATAGGTGCTCGTGGTATCCTNNNAGATCGGAAG |
| 9901_1 | TATGAGTGTGGAGTCGTTGCAAGCAGCTAGAAGCCCTTATCCTNNNAGATCGGAAG |
| 9902_1 | TATGAGTGTGGAGTCGTTGCTCTTGAGTGTCAGTGGAACTCCTNNNAGATCGGAAG |
| 9905_1 | TATGAGTGTGGAGTCGTTGCTCTTAGGTCATACATCCACTCCTNNNAGATCGGAAGA |
| 9906_1 | TATGAGTGTGGAGTCGTTGCCCCTTATCCTCCTCTCCACTCCTNNNAGATCGGAAGAG |
| 9907_1 | TATGAGTGTGGAGTCGTTGCAAATAGTGGCTGTACACCCTCCTNNNAGATCGGAAGA |
| 9909_1 | TATGAGTGTGGAGTCGTTGCAGTAATGTGGGCTTCATCCTCCTNNNAGATCGGAAGA |
| 9912_1 | TATGAGTGTGGAGTCGTTGCATGACTGGTTTAGGCTTCCTCCTNNNAGATCGGAAGA |
| 9915_1 | TATGAGTGTGGAGTCGTTGCCATTGTGTATTCGCTTTGCTCCTNNNAGATCGGAAGA |
| 9917_1 | TATGAGTGTGGAGTCGTTGCCTCATCACACATTCCCCTCGATCNNNAGATCGGAAGAG |
| 9920_1 | TATGAGTGTGGAGTCGTTGCTCTATGTCCTGGACCTCTCTCCTNNNAGATCGGAAGA |

| | |
|---|---|
| 9921_1 | TATGAGTGTGGAGTCGTTGCGTCTTTCTATCCCCTGTCTCCTNNNAGATCGGAAGA |
| 9924_1 | TATGAGTGTGGAGTCGTTGCGTCTACCTCTGCCAGATTCTCCTNNNAGATCGGAAGA |
| 9925_1 | TATGAGTGTGGAGTCGTTGCGAGAGGCTGGAAGAGATTCTCCTNNNAGATCGGAA |
| 9928_1 | TATGAGTGTGGAGTCGTTGCTACTTGCTGCCTACCTTTCTCCTNNNAGATCGGAAGA |
| 9931_1 | TATGAGTGTGGAGTCGTTGCTGTCTTCCTGAGGCTTTTCTCCTNNNAGATCGGAAGA |
| 9936_1 | TATGAGTGTGGAGTCGTTGCAAATCTTCCACCAGGAAGGTCCTNNNAGATCGGAAG |
| 9937_1 | TATGAGTGTGGAGTCGTTGCCTTGTGGCTTAGAGGTAGGTCCTNNNAGATCGGAA |
| 9938_1 | TATGAGTGTGGAGTCGTTGCATCCTGACAAGCTATTGGGTCCTNNNAGATCGGAAG |
| 9941_1 | TATGAGTGTGGAGTCGTTGCAGAAAAGGAGCTGGAGATGTCCTNNNAGATCGGAA |
| 9943_1 | TATGAGTGTGGAGTCGTTGCTACTAAATGCCAGGCTCTGTCCTNNNAGATCGGAAG |
| 9944_1 | TATGAGTGTGGAGTCGTTGCATTGGCATGTGTGATTATGGGGGNNNAGATCGGAA |
| 9945_1 | TATGAGTGTGGAGTCGTTGCGTCCGCAAAACAATTCTTGTCCTNNNAGATCGGAAG |
| 9946_1 | TATGAGTGTGGAGTCGTTGCCCAGTCTCACAATGCTTTGTCCTNNNAGATCGGAAGA |
| 9949_1 | TATGAGTGTGGAGTCGTTGCGCACCCATTATTCACCCATTCCTNNNAGATCGGAAGA |
| 9955_1 | TATGAGTGTGGAGTCGTTGCAGCCAGGTATGGTGCATATTCCTNNNAGATCGGAAG |
| 9966_1 | TATGAGTGTGGAGTCGTTGCTTCACTTTCTGCTTGCACTTCCTNNNAGATCGGAAGA |
| 9979_1 | TATGAGTGTGGAGTCGTTGCATCTTTCCTTCCACTGCCTTCCTNNNAGATCGGAAGA |
| 9982_1 | TATGAGTGTGGAGTCGTTGCGGTGGGGATAGTAGAAGCTTCCTNNNAGATCGGAA |
| 9988_1 | TATGAGTGTGGAGTCGTTGCGGGCCTTGGTCTAACATAAGCTTNNNAGATCGGAAG |
| 9994_1 | TATGAGTGTGGAGTCGTTGCTCACCTTCCTGGTTTCTCTTCCTNNNAGATCGGAAGA |
| 9998_1 | TATGAGTGTGGAGTCGTTGCTCTGTGGATTCCTGGAAGTTCCTNNNAGATCGGAAG |
| 9999_1 | TATGAGTGTGGAGTCGTTGCTGGTTTGTCAGTTCACAGTTCCTNNNAGATCGGAAG |
| 10011_1 | TATGAGTGTGGAGTCGTTGCAAATTCTTGCAGAGCCTGTTCCTNNNAGATCGGAAG |
| 10012_1 | TATGAGTGTGGAGTCGTTGCCCTGAAAATGCCTCTGTGTTCCTNNNAGATCGGAAG |
| 10052_1 | TATGAGTGTGGAGTCGTTGCACAGACATTCACGTGTCAAAGCTNNNAGATCGGAAG |
| 10059_1 | TATGAGTGTGGAGTCGTTGCGGGGAAACAGACGGAATAAAGCTNNNAGATCGGAA |
| 10073_1 | TATGAGTGTGGAGTCGTTGCTTTTTCCATGAGGATGCCAAGCTNNNAGATCGGAAG |
| 10077_1 | TATGAGTGTGGAGTCGTTGCTGGAAATAGGTCTCCCTCAAGCTNNNAGATCGGAAG |
| 10084_1 | TATGAGTGTGGAGTCGTTGCCCTCATACGTAACCACAGCACTTNNNAGATCGGAAG |
| 10089_1 | TATGAGTGTGGAGTCGTTGCACACCTACAGTGAGATGGAAGCTNNNAGATCGGAA |
| 10096_1 | TATGAGTGTGGAGTCGTTGCAATCTCCCCTTCTCTGTGAAGCTNNNAGATCGGAAG |
| 10105_1 | TATGAGTGTGGAGTCGTTGCAACTTCCCAACAGCTACTAAGCTNNNAGATCGGAAG |
| 10114_1 | TATGAGTGTGGAGTCGTTGCATTTCTCAAGGGCAGAAACAGCTNNNAGATCGGAA |
| 10124_1 | TATGAGTGTGGAGTCGTTGCATTTGGAAACAGGAAGGACAGCTNNNAGATCGGAA |
| 10135_1 | TATGAGTGTGGAGTCGTTGCTGCTCTACTTCTGTAACCCAGCTNNNAGATCGGAAG |
| 10152_1 | TATGAGTGTGGAGTCGTTGCCTCTTGAAACCTTCTGAGCAGCTNNNAGATCGGAAG |
| 10154_1 | TATGAGTGTGGAGTCGTTGCAATATGGAAAGTTTGGGGCAGCTNNNAGATCGGAA |
| 10171_1 | TATGAGTGTGGAGTCGTTGCCACTATTCCTAGGGAGCTCAGCTNNNAGATCGGAAG |
| 10181_1 | TATGAGTGTGGAGTCGTTGCGCCCTAAAATAGCCACTTCAGCTNNNAGATCGGAAG |
| 10199_1 | TATGAGTGTGGAGTCGTTGCTTCAGTGAGTGGGATACAGAGCTNNNAGATCGGAA |
| 10206_1 | TATGAGTGTGGAGTCGTTGCCAGGAGGAGGAGAAAAGAGAGCTNNNAGATCGGA |
| 10218_1 | TATGAGTGTGGAGTCGTTGCCTCCATGGTCCTTGATAGGAGCTNNNAGATCGGAAG |
| 10219_1 | TATGAGTGTGGAGTCGTTGCATCAAAACCTGGGTCTAGGAGCTNNNAGATCGGAA |

| | |
|---|---|
| 10245_1 | TATGAGTGTGGAGTCGTTGCGGGGGAAAATGGTAGTTTGAGCTNNNAGATCGGAA |
| 10248_1 | TATGAGTGTGGAGTCGTTGCGCTCTCAGCAAGTTCCAATAGCTNNNAGATCGGAAG |
| 10277_1 | TATGAGTGTGGAGTCGTTGCGAGCTGTAGTCCTAGCAGTAGCTNNNAGATCGGAA |
| 10288_1 | TATGAGTGTGGAGTCGTTGCGCACTGATGTGATCATGGTAGCTNNNAGATCGGAA |
| 10289_1 | TATGAGTGTGGAGTCGTTGCACATTAGGTCGGGTATGGTAGCTNNNAGATCGGAA |
| 10294_1 | TATGAGTGTGGAGTCGTTGCAAAACAGAAGTGCCAGTGTAGCTNNNAGATCGGAA |
| 10312_1 | TATGAGTGTGGAGTCGTTGCACATCCTGCACATGATAAACGCTNNNAGATCGGAAG |
| 10328_1 | TATGAGTGTGGAGTCGTTGCTGACATCCAAAGGTAATTCCGCTNNNAGATCGGAAG |
| 10345_1 | TATGAGTGTGGAGTCGTTGCACTTTGGAAATGACCTGTTCGCTNNNAGATCGGAAG |
| 10347_1 | TATGAGTGTGGAGTCGTTGCTGGCATCAGTTCTTGAAAAGGCTNNNAGATCGGAA |
| 10360_1 | TATGAGTGTGGAGTCGTTGCAAACAGGAGACAGTAGGAAGGCTNNNAGATCGGAA |
| 10365_1 | TATGAGTGTGGAGTCGTTGCCAATTTCCAGTGAGGAACAGGCTNNNAGATCGGAA |
| 10390_1 | TATGAGTGTGGAGTCGTTGCACCTCATCCTCTTTCTTCAGGCTNNNAGATCGGAAGA |
| 10407_1 | TATGAGTGTGGAGTCGTTGCCCCTTCCCATATTGTCCTAGGCTNNNAGATCGGAAGA |
| 10430_1 | TATGAGTGTGGAGTCGTTGCTGGTATATGTTTCACAACGGGCTNNNAGATCGGAAG |
| 10435_1 | TATGAGTGTGGAGTCGTTGCCATTTATTGAGCATCCTGGGGCTNNNAGATCGGAAG |
| 10438_1 | TATGAGTGTGGAGTCGTTGCATTTATTGATGGGCACATGGGCTNNNAGATCGGAA |
| 10439_1 | TATGAGTGTGGAGTCGTTGCACACAGGAAAAATGACATGGGCTNNNAGATCGGAA |
| 10440_1 | TATGAGTGTGGAGTCGTTGCTACCTTACAAAGATCCCTGGGCTNNNAGATCGGAAG |
| 10448_1 | TATGAGTGTGGAGTCGTTGCGCTGTAGAATACTTGTGTGGGCTNNNAGATCGGAA |
| 10451_1 | TATGAGTGTGGAGTCGTTGCTTCTTGTTCTAGACCATTGGGCTNNNAGATCGGAAG |
| 10468_1 | TATGAGTGTGGAGTCGTTGCGATTTCCGGGTTCACTAATGGCTNNNAGATCGGAAG |
| 10472_1 | TATGAGTGTGGAGTCGTTGCAGAGAGACACAGAGAACATGGCTNNNAGATCGGAA |
| 10509_1 | TATGAGTGTGGAGTCGTTGCTTGCTTTCTTTTCTGTCCTGGCTNNNAGATCGGAAGA |
| 10510_1 | TATGAGTGTGGAGTCGTTGCTTCAAATGACTGAATGGCTGGCTNNNAGATCGGAA |
| 10515_1 | TATGAGTGTGGAGTCGTTGCTGATCTCCTGACCTAATCTGGCTNNNAGATCGGAAG |
| 10521_1 | TATGAGTGTGGAGTCGTTGCAAGTCTGATTACCCTGTCTGGCTNNNAGATCGGAAG |
| 10534_1 | TATGAGTGTGGAGTCGTTGCTGTGTATAGCCTGTATGGTGGCTNNNAGATCGGAA |
| 10563_1 | TATGAGTGTGGAGTCGTTGCAACCTGGTATTCCCTTTTTGGCTNNNAGATCGGAAG |
| 10576_1 | TATGAGTGTGGAGTCGTTGCTTGGTGCTTAGGAAAACAATGCTNNNAGATCGGAA |
| 10577_1 | TATGAGTGTGGAGTCGTTGCAGAAGGTCGAGAGAGACAATGCTNNNAGATCGGAA |
| 10581_1 | TATGAGTGTGGAGTCGTTGCTTGTTCAGAGGCCATAGAATGCTNNNAGATCGGAA |
| 10586_1 | TATGAGTGTGGAGTCGTTGCGGGAAGGGAGAAAGGAGTGAAGANNNAGATCGG |
| 10591_1 | TATGAGTGTGGAGTCGTTGCACAGAAGAGGAGAAGTCCATGCTNNNAGATCGGAA |
| 10597_1 | TATGAGTGTGGAGTCGTTGCTCCTTTACATGCTCATGCATGCTNNNAGATCGGAAGA |
| 10607_1 | TATGAGTGTGGAGTCGTTGCCTGAATTGGGACTGACAGATGCTNNNAGATCGGAA |
| 10609_1 | TATGAGTGTGGAGTCGTTGCTCAAATCCAGCTGTAGAGATGCTNNNAGATCGGAAG |
| 10612_1 | TATGAGTGTGGAGTCGTTGCAGTAGGAACATTACAACGATGCTNNNAGATCGGAA |
| 10642_1 | TATGAGTGTGGAGTCGTTGCTCCAGCCTCAAAATATCACTGCTNNNAGATCGGAAGA |
| 10676_1 | TATGAGTGTGGAGTCGTTGCTTGAAACAATTCCTCTGGCTGCTNNNAGATCGGAAG |
| 10692_1 | TATGAGTGTGGAGTCGTTGCGCTGAGAACAAATAGGCTCTGCTNNNAGATCGGAA |
| 10696_1 | TATGAGTGTGGAGTCGTTGCATGCGTTCAGAAAACTCTCTGCTNNNAGATCGGAAG |
| 10698_1 | TATGAGTGTGGAGTCGTTGCTAGGCAAAATCTGCTTCTCTGCTNNNAGATCGGAAG |

| | |
|---|---|
| 10705_1 | TATGAGTGTGGAGTCGTTGCTTTCTGAGGTTTCAGCTTCTGCTNNNAGATCGGAAG |
| 10731_1 | TATGAGTGTGGAGTCGTTGCGAGAGAGAAAGGGAGAGAGTGCTNNNAGATCGGA |
| 10735_1 | TATGAGTGTGGAGTCGTTGCATGGAAATGTTGGGTCTAGTGCTNNNAGATCGGAA |
| 10739_1 | TATGAGTGTGGAGTCGTTGCTGTAGAAACAACACCTACGTGCTNNNAGATCGGAAG |
| 10740_1 | TATGAGTGTGGAGTCGTTGCGTTTCTACCTCTTGTGTCGTGCTNNNAGATCGGAAG |
| 10742_1 | TATGAGTGTGGAGTCGTTGCCCAACGGGAAAGATAAAGGTGCTNNNAGATCGGAA |
| 10746_1 | TATGAGTGTGGAGTCGTTGCATTGAGTACTGGTAGTGGGTGCTNNNAGATCGGAA |
| 10749_1 | TATGAGTGTGGAGTCGTTGCAGTCTGCCTCTAAATTTGGTGCTNNNAGATCGGAAG |
| 10761_1 | TATGAGTGTGGAGTCGTTGCAATAAGGAATTGGCTCGTGTGCTNNNAGATCGGAA |
| 10762_1 | TATGAGTGTGGAGTCGTTGCCACACTCATATCTGTCGGCAAACNNNAGATCGGAAG |
| 10766_1 | TATGAGTGTGGAGTCGTTGCTTGGCAGGAGTAATTTGTGTGCTNNNAGATCGGAA |
| 10769_1 | TATGAGTGTGGAGTCGTTGCTCTCTCTCACCAAACTACCTGGGNNNAGATCGGAAG |
| 10771_1 | TATGAGTGTGGAGTCGTTGCGAGGAAGAAGACAGTGTTGTGCTNNNAGATCGGAA |
| 10773_1 | TATGAGTGTGGAGTCGTTGCTCTGTGCCATACTTTGTTGTGCTNNNAGATCGGAAG |
| 10807_1 | TATGAGTGTGGAGTCGTTGCGGCTATGTTTGCTTTTGCTTGCTNNNAGATCGGAAG |
| 10824_1 | TATGAGTGTGGAGTCGTTGCGCTATCCCCTTGAACATGTTGCTNNNAGATCGGAAG |
| 10825_1 | TATGAGTGTGGAGTCGTTGCTTTCTGCCTTAGTCCATGTTGCTNNNAGATCGGAAGA |
| 10837_1 | TATGAGTGTGGAGTCGTTGCGTCCCCGAGTTTCTGTATTTGCTNNNAGATCGGAAG |
| 10838_1 | TATGAGTGTGGAGTCGTTGCTACAAAGCCCATCACACTTTGCTNNNAGATCGGAAG |
| 10862_1 | TATGAGTGTGGAGTCGTTGCTAGGCGCTTCTCTTTGTTTTGCTNNNAGATCGGAAG |
| 10869_1 | TATGAGTGTGGAGTCGTTGCGCTGTCAATCCAGCATCAAATCTNNNAGATCGGAAG |
| 10871_1 | TATGAGTGTGGAGTCGTTGCCAAGAACCAGTGGAGCGAAATCTNNNAGATCGGAA |
| 10872_1 | TATGAGTGTGGAGTCGTTGCAAGGCAAAGGACCCAAACAATCTNNNAGATCGGAA |
| 10875_1 | TATGAGTGTGGAGTCGTTGCCCTGCTTGTGGTGTTTACAATCTNNNAGATCGGAAG |
| 10877_1 | TATGAGTGTGGAGTCGTTGCTAGGAGAGAACCAAGGCCAATCTNNNAGATCGGAA |
| 10878_1 | TATGAGTGTGGAGTCGTTGCCAGGGTATCTATCACCTAGGCTGNNNAGATCGGAAG |
| 10879_1 | TATGAGTGTGGAGTCGTTGCGAAAGGAAAGTTACCCTGGGACCNNNAGATCGGAA |
| 10881_1 | TATGAGTGTGGAGTCGTTGCCCCACACCAGAGAATTCCACAAANNNAGATCGGAAG |
| 10882_1 | TATGAGTGTGGAGTCGTTGCACTAAGGATAAACCACTGGCCCANNNAGATCGGAAG |
| 10884_1 | TATGAGTGTGGAGTCGTTGCAGAAAGGGGCAACTGACCAAATTNNNAGATCGGAA |
| 10885_1 | TATGAGTGTGGAGTCGTTGCTCTACACAGAGGGGAAACTAGGGNNNAGATCGGAA |
| 10886_1 | TATGAGTGTGGAGTCGTTGCGGGGAGACAGAGACAAAGACAACNNNAGATCGGA |
| 10888_1 | TATGAGTGTGGAGTCGTTGCAGTGTGTGAAGACAGAAGGCCTANNNAGATCGGAA |
| 10889_1 | TATGAGTGTGGAGTCGTTGCGTCACCTAGGCTGTAGGGAATCTNNNAGATCGGAA |
| 10890_1 | TATGAGTGTGGAGTCGTTGCACCATGTGCTACAAGTGCATCATNNNAGATCGGAAG |
| 10891_1 | TATGAGTGTGGAGTCGTTGCCATTCTGCTCTGCACTGGAATCTNNNAGATCGGAAG |
| 10892_1 | TATGAGTGTGGAGTCGTTGCCACTCCCTCCTCCCCATTATCTTNNNAGATCGGAAGA |
| 10895_1 | TATGAGTGTGGAGTCGTTGCTCCCATTCTGTTTTGATGCTCCANNNAGATCGGAAGA |
| 10896_1 | TATGAGTGTGGAGTCGTTGCTTCTCCCATCCTCCTAAAGGTGCNNNAGATCGGAAGA |
| 10897_1 | TATGAGTGTGGAGTCGTTGCAGGCACCCTAAGATGGTGAATCTNNNAGATCGGAA |
| 10898_1 | TATGAGTGTGGAGTCGTTGCTGATTGACAATGCTGTCAAGGCTNNNAGATCGGAA |
| 10899_1 | TATGAGTGTGGAGTCGTTGCTGCCCTCTACTGGCAACAATATCNNNAGATCGGAAG |
| 10900_1 | TATGAGTGTGGAGTCGTTGCCACTCTACCTCCATGAAGAGCACNNNAGATCGGAAG |

| | |
|---|---|
| 10901_1 | TATGAGTGTGGAGTCGTTGCAGGCTGGACCTGAAGATTAGTCANNNAGATCGGAA |
| 10903_1 | TATGAGTGTGGAGTCGTTGCGCCCTGTATAGACGTACACCAACNNNAGATCGGAAG |
| 10905_1 | TATGAGTGTGGAGTCGTTGCATACACCTGGCTACTAAGCACCANNNAGATCGGAAG |
| 10908_1 | TATGAGTGTGGAGTCGTTGCCGTGTGTGTGTGTATGACACTNNNAGATCGGAA |
| 10910_1 | TATGAGTGTGGAGTCGTTGCAGCCAAACAAAAATGTGTACTGTNNNAGATCGGAA |
| 10912_1 | TATGAGTGTGGAGTCGTTGCGCAATGACAAAAGCAACAGCAACNNNAGATCGGAA |
| 10913_1 | TATGAGTGTGGAGTCGTTGCCAGGGTTTGCATTAGCTTCAGGANNNAGATCGGAA |
| 10914_1 | TATGAGTGTGGAGTCGTTGCAGATTCAGATATCACGCTCCCCANNNAGATCGGAAG |
| 10915_1 | TATGAGTGTGGAGTCGTTGCGTGGTGTGAAGGCCAAAAATCTGNNNAGATCGGAA |
| 10918_1 | TATGAGTGTGGAGTCGTTGCCTCCGGGACACAGTTGAACAATTNNNAGATCGGAAG |
| 10919_1 | TATGAGTGTGGAGTCGTTGCACCAGTTTTCATGTGGATGACCCNNNAGATCGGAAG |
| 10920_1 | TATGAGTGTGGAGTCGTTGCGTAGGAAGCAATGACACCTTCCANNNAGATCGGAA |
| 10923_1 | TATGAGTGTGGAGTCGTTGCGATACCACTCAGTACAGTGGGGANNNAGATCGGAA |
| 10924_1 | TATGAGTGTGGAGTCGTTGCAACTTTGCGCATCTCATTTCCTTNNNAGATCGGAAGA |
| 10927_1 | TATGAGTGTGGAGTCGTTGCAGTCCACAGTATGCTCTGATCCCNNNAGATCGGAAG |
| 10928_1 | TATGAGTGTGGAGTCGTTGCTGTCTTGACTAGCTTTCCCATCTNNNAGATCGGAAGA |
| 10929_1 | TATGAGTGTGGAGTCGTTGCCAGTGTCAGGAGAGATTTGGTTTNNNAGATCGGAA |
| 10930_1 | TATGAGTGTGGAGTCGTTGCTGGAAAAGCAAAGGAAGGAAGCTNNNAGATCGGA |
| 10931_1 | TATGAGTGTGGAGTCGTTGCTGGCAACACAAACAGATCCATCTNNNAGATCGGAAG |
| 10932_1 | TATGAGTGTGGAGTCGTTGCTACCCCCAGACCTATCTCCATCTNNNAGATCGGAAGA |
| 10945_1 | TATGAGTGTGGAGTCGTTGCGGGCCATGCATATGGTAAAAACCNNNAGATCGGAA |
| 10946_1 | TATGAGTGTGGAGTCGTTGCACTTTATCTGCTCTTCCTGCTGCNNNAGATCGGAAGA |
| 10947_1 | TATGAGTGTGGAGTCGTTGCAAAACCCAAGAAACACAGACCCANNNAGATCGGAAG |
| 10951_1 | TATGAGTGTGGAGTCGTTGCGCTTTGCCATCCTCTGACTAGACNNNAGATCGGAAG |
| 10960_1 | TATGAGTGTGGAGTCGTTGCCACAGCTGCTTCAGCATCATCTTNNNAGATCGGAAG |
| 10965_1 | TATGAGTGTGGAGTCGTTGCGGCTCTGGATGTTTAGGTTCTCGNNNAGATCGGAA |
| 10971_1 | TATGAGTGTGGAGTCGTTGCATTGTGTGCCTGTATTCCCAGTGNNNAGATCGGAAG |
| 10974_1 | TATGAGTGTGGAGTCGTTGCTGAGAGGCATTGTTTGAGGAGACNNNAGATCGGAA |
| 10975_1 | TATGAGTGTGGAGTCGTTGCGGTTTAGCCAAGTGATTTGCTGGNNNAGATCGGAA |
| 10976_1 | TATGAGTGTGGAGTCGTTGCTTCAGCAAATGAGTATCCCGGTGNNNAGATCGGAA |
| 10977_1 | TATGAGTGTGGAGTCGTTGCGGGGAATCACAACTTCAGGTTGANNNAGATCGGAA |
| 10979_1 | TATGAGTGTGGAGTCGTTGCATGTTAGTTGAGGAGGAAAGGGTNNNAGATCGGA |
| 10981_1 | TATGAGTGTGGAGTCGTTGCAGGCCGCATAAGATAATTAGGGTNNNAGATCGGAA |
| 10990_1 | TATGAGTGTGGAGTCGTTGCATTTTGTTCCGTTTGCAATGCCANNNAGATCGGAAG |
| 10993_1 | TATGAGTGTGGAGTCGTTGCGAGGGATTTACTTTTCGGTGAGCNNNAGATCGGAA |
| 10994_1 | TATGAGTGTGGAGTCGTTGCAATGTGATTTTGCCACCCACAAANNNAGATCGGAAG |
| 10999_1 | TATGAGTGTGGAGTCGTTGCAGATTTCAGTGCAGCTGTGATGANNNAGATCGGAA |
| 11004_1 | TATGAGTGTGGAGTCGTTGCGAGCTATTTCTTTGCTGACGTGCNNNAGATCGGAAG |
| 11005_1 | TATGAGTGTGGAGTCGTTGCGTTGCCTCACTGAAAAGAAACCGNNNAGATCGGAA |
| 11006_1 | TATGAGTGTGGAGTCGTTGCGCCATAACACGTTTACTGTTCCCNNNAGATCGGAAG |
| 11011_1 | TATGAGTGTGGAGTCGTTGCAGTTCATCTGAGCATTCCAAGCCNNNAGATCGGAAG |
| 11013_1 | TATGAGTGTGGAGTCGTTGCATGATAGTCTTCAAGCCTGAGCCNNNAGATCGGAAG |
| 11014_1 | TATGAGTGTGGAGTCGTTGCAGGCATTTCTCTGTTGCACTGAANNNAGATCGGAAG |

| | |
|---|---|
| 11016_1 | TATGAGTGTGGAGTCGTTGCCTATTTGCCCCTAAAGCCTCTGGNNNAGATCGGAAG |
| 11017_1 | TATGAGTGTGGAGTCGTTGCTCCCAGAAGCAGATGTAGAGGAGNNNAGATCGGAA |
| 11019_1 | TATGAGTGTGGAGTCGTTGCCCATAGTAGCCTGGAACCACAGANNNAGATCGGAAG |
| 11021_1 | TATGAGTGTGGAGTCGTTGCTTCCCCAGCTTTCCTCTCCTATCNNNAGATCGGAAGA |
| 11022_1 | TATGAGTGTGGAGTCGTTGCATTCCACAGCATACTTCTCCCCANNNAGATCGGAAGA |
| 11023_1 | TATGAGTGTGGAGTCGTTGCATCCTGGGTTCACGCTGATATCTNNNAGATCGGAAG |
| 11025_1 | TATGAGTGTGGAGTCGTTGCGAGAGCAGGGAATTCAGTTCAGGNNNAGATCGGAA |
| 11027_1 | TATGAGTGTGGAGTCGTTGCTTATTTCTCCAGCCAGTCACACCNNNAGATCGGAAGA |
| 11028_1 | TATGAGTGTGGAGTCGTTGCAAGCTCAAAAGGCAACAGGAAGANNNAGATCGGAA |
| 11029_1 | TATGAGTGTGGAGTCGTTGCGGAAGAGATCACCAGGCCTATCTNNNAGATCGGAA |
| 11032_1 | TATGAGTGTGGAGTCGTTGCGTCATAGTGTGTGTCAAGTGCCTNNNAGATCGGAAG |
| 11034_1 | TATGAGTGTGGAGTCGTTGCTGTTGGTTTCTGGCCATATGAGGNNNAGATCGGAA |
| 11035_1 | TATGAGTGTGGAGTCGTTGCTGTGTTGGTTCAAGTTGTTTGGTNNNAGATCGGAAG |
| 11036_1 | TATGAGTGTGGAGTCGTTGCAGAGCTGAATCAGGTAGGCATGANNNAGATCGGAA |
| 11037_1 | TATGAGTGTGGAGTCGTTGCCTACTATTCCAGTTGCTTGCCCANNNAGATCGGAAGA |
| 11038_1 | TATGAGTGTGGAGTCGTTGCGCTCTTTGCCCTGGACTGTAATTNNNAGATCGGAAG |
| 11039_1 | TATGAGTGTGGAGTCGTTGCAGTTATTGCATCTGTCTGCCAGGNNNAGATCGGAAG |
| 11040_1 | TATGAGTGTGGAGTCGTTGCACTGAGCCTGGCAGTAAGTATCTNNNAGATCGGAAG |
| 11042_1 | TATGAGTGTGGAGTCGTTGCCGCATTTTCCTGTACCTGTATCTNNNAGATCGGAAGA |
| 11043_1 | TATGAGTGTGGAGTCGTTGCATTCCCTTCTTGGGCCTGTATCTNNNAGATCGGAAGA |
| 11045_1 | TATGAGTGTGGAGTCGTTGCCTGGTCATCTGCCTGTAATTGCTNNNAGATCGGAAG |
| 11046_1 | TATGAGTGTGGAGTCGTTGCCACAAGAACCCTAGCTCAGTGTCNNNAGATCGGAAG |
| 11047_1 | TATGAGTGTGGAGTCGTTGCAGGGAAAACTCAGCCAAAACAGTNNNAGATCGGAA |
| 11048_1 | TATGAGTGTGGAGTCGTTGCATTCTCCCACACATGGAGTTCTGNNNAGATCGGAAG |
| 11049_1 | TATGAGTGTGGAGTCGTTGCAGACTTTTCCTGCACCTTGTGAGNNNAGATCGGAAG |
| 11050_1 | TATGAGTGTGGAGTCGTTGCGATGTGCTGTGGATGGAGATCTGNNNAGATCGGAA |
| 11052_1 | TATGAGTGTGGAGTCGTTGCGTGATCCTCTAGTTCTGTGGGCTNNNAGATCGGAAG |
| 11053_1 | TATGAGTGTGGAGTCGTTGCCCTCCTATGCCTCTAGCTCAGTTNNNAGATCGGAAGA |
| 11055_1 | TATGAGTGTGGAGTCGTTGCCACAAAAACAACTCCCTCTGGGANNNAGATCGGAAG |
| 11057_1 | TATGAGTGTGGAGTCGTTGCAAGCTGGAAATCTCTCTCAGGGTNNNAGATCGGAAG |
| 11058_1 | TATGAGTGTGGAGTCGTTGCACACCATGCATGCCAAACAAAAANNNAGATCGGAAG |
| 11061_1 | TATGAGTGTGGAGTCGTTGCCCTGCCTAATTCCATGCAACTCTNNNAGATCGGAAGA |
| 11062_1 | TATGAGTGTGGAGTCGTTGCGCCGGTCCAAGGGAATACAAAAANNNAGATCGGAA |
| 11065_1 | TATGAGTGTGGAGTCGTTGCCAGCTTGTGTCACTCCCCAAAAANNNAGATCGGAAG |
| 11068_1 | TATGAGTGTGGAGTCGTTGCTTCAGCGGTTTCATTGACACTCTNNNAGATCGGAAG |
| 11069_1 | TATGAGTGTGGAGTCGTTGCATCTCCTCAGTGCCTTGCAAAAANNNAGATCGGAAG |
| 11071_1 | TATGAGTGTGGAGTCGTTGCTGACAGAAGTCTGGTTCCACAGANNNAGATCGGAA |
| 11074_1 | TATGAGTGTGGAGTCGTTGCCCAGAGAGGGTGCAACAGAAAAANNNAGATCGGAA |
| 11075_1 | TATGAGTGTGGAGTCGTTGCAACCGTGTCCAATCCCAGAAAAANNNAGATCGGAAG |
| 11076_1 | TATGAGTGTGGAGTCGTTGCCACCATGAGGTAGGCAGGAAAAANNNAGATCGGAA |
| 11079_1 | TATGAGTGTGGAGTCGTTGCAGAAAGAGATCCCTGTGCACTCTNNNAGATCGGAAG |
| 11080_1 | TATGAGTGTGGAGTCGTTGCTTCTCTGAGCTGTTGGTCACTCTNNNAGATCGGAAG |
| 11081_1 | TATGAGTGTGGAGTCGTTGCGGAGGAGTGTCATGGTGGAAAAANNNAGATCGGA |

11087_1 TATGAGTGTGGAGTCGTTGCCTACCTGCCAGCATGGGTAAAAANNNAGATCGGAAG
11088_1 TATGAGTGTGGAGTCGTTGCCCCTCCCTACCAAAGACACAAAANNNAGATCGGAAGA
11091_1 TATGAGTGTGGAGTCGTTGCGGGTGGGTAAGGGATTCTACTCTNNNAGATCGGAA
11093_1 TATGAGTGTGGAGTCGTTGCTACTGGCTTTGGCTGCTACAAAANNNAGATCGGAAG
11094_1 TATGAGTGTGGAGTCGTTGCCACACACGAACCCAAAACCAAAANNNAGATCGGAAG
11100_1 TATGAGTGTGGAGTCGTTGCCTCTTGCCTGTAGTCCTACCTCTNNNAGATCGGAAGA
11103_1 TATGAGTGTGGAGTCGTTGCACCAGTGTCATAACCTGCCAAAANNNAGATCGGAAG
11105_1 TATGAGTGTGGAGTCGTTGCCATTTTCCTTTGCCTGACCCTCTNNNAGATCGGAAGA
11106_1 TATGAGTGTGGAGTCGTTGCCCAAGCAGGCATGATAAGCAAAANNNAGATCGGAA
11109_1 TATGAGTGTGGAGTCGTTGCACTAACATCAAGCACCTCCCTCTNNNAGATCGGAAGA
11110_1 TATGAGTGTGGAGTCGTTGCACCCTCAGTTCTCACATGCAAAANNNAGATCGGAAGA
11112_1 TATGAGTGTGGAGTCGTTGCCCTGACACAAAGGCACATCAAAANNNAGATCGGAAG
11115_1 TATGAGTGTGGAGTCGTTGCCAGCTTCAATCATCACAGCCTCTNNNAGATCGGAAGA
11116_1 TATGAGTGTGGAGTCGTTGCGCAAACAGCACCTGATGTCAAAANNNAGATCGGAAG
11118_1 TATGAGTGTGGAGTCGTTGCAAGCACTCTTCCTCCAAAGAAAANNNAGATCGGAAGA
11119_1 TATGAGTGTGGAGTCGTTGCGCCAGACTTTGGGGAGAAGAAAANNNAGATCGGAA
11120_1 TATGAGTGTGGAGTCGTTGCCCATTCCACCTGGTCACAGAAAANNNAGATCGGAAGA
11121_1 TATGAGTGTGGAGTCGTTGCGGGGTTCACAGGATTCCAGAAAANNNAGATCGGAA
11122_1 TATGAGTGTGGAGTCGTTGCGGTTGCATCTGTCCAAATCCTCTNNNAGATCGGAAGA
11123_1 TATGAGTGTGGAGTCGTTGCTCTCCAGTTTCTTTGTGACCGACNNNAGATCGGAAGA
11124_1 TATGAGTGTGGAGTCGTTGCCTATTGCTCTGGCTGGCAGAAAANNNAGATCGGAAG
11125_1 TATGAGTGTGGAGTCGTTGCGCTTGTTTCTACTACCCAGTGGGNNNAGATCGGAAG
11126_1 TATGAGTGTGGAGTCGTTGCTTTGCCCTAGATTCTCCTCCTCTNNNAGATCGGAAGA
11127_1 TATGAGTGTGGAGTCGTTGCCTCACCTGGTCTATTGCTCCTCTNNNAGATCGGAAGA
11128_1 TATGAGTGTGGAGTCGTTGCGCAGGTGGTTACCTTTGAGAAAANNNAGATCGGAA
11132_1 TATGAGTGTGGAGTCGTTGCTCTGTCAAGAAACTGCCCGAAAANNNAGATCGGAAG
11133_1 TATGAGTGTGGAGTCGTTGCAGAGACGAGGATGGGAAGGAAAANNNAGATCGGA
11135_1 TATGAGTGTGGAGTCGTTGCTCATTCTCTGACAGCCTTCCTCTNNNAGATCGGAAGA
11136_1 TATGAGTGTGGAGTCGTTGCGGCAATCTCTTCCTCCAGGAAAANNNAGATCGGAAG
11138_1 TATGAGTGTGGAGTCGTTGCATGCAGAAGGCAATCAGGGAAAANNNAGATCGGAA
11139_1 TATGAGTGTGGAGTCGTTGCTGTGAAGTACCTGGGAGGGAAAANNNAGATCGGAA
11141_1 TATGAGTGTGGAGTCGTTGCGCTTCTGTTTCTCTGCAGACAGANNNAGATCGGAAGA
11143_1 TATGAGTGTGGAGTCGTTGCAAAGCAAGAGAGGGCATGGAAAANNNAGATCGGAA
11144_1 TATGAGTGTGGAGTCGTTGCGAGAAGCTGAATGCAAGAGCTCTNNNAGATCGGAA
11145_1 TATGAGTGTGGAGTCGTTGCGGAGAGTGAGGGCTACTGGAAAANNNAGATCGGAA
11146_1 TATGAGTGTGGAGTCGTTGCATGATGCCAGGTCTTGTGGAAAANNNAGATCGGAA
11148_1 TATGAGTGTGGAGTCGTTGCGGGCCTCTTTGAGGAGATGAAAANNNAGATCGGAA
11149_1 TATGAGTGTGGAGTCGTTGCGCCAAACCACATGATGCTGAAAANNNAGATCGGAAG
11150_1 TATGAGTGTGGAGTCGTTGCGCACATGAGATTCTTCAGGCTCTNNNAGATCGGAAG
11151_1 TATGAGTGTGGAGTCGTTGCTTCCTGCCATACCAAAGTGAAAANNNAGATCGGAAG
11155_1 TATGAGTGTGGAGTCGTTGCCCCCATGTGCTCCCATCATAAAANNNAGATCGGAAGA
11156_1 TATGAGTGTGGAGTCGTTGCGCCTCAAAGATAGCAATGGCTCTNNNAGATCGGAAG
11159_1 TATGAGTGTGGAGTCGTTGCAGCCAGCCATCTTGAAGCTAAAANNNAGATCGGAAG

11160_1 TATGAGTGTGGAGTCGTTGCCTTTTATCAGGGCAAGTGGCTCTNNNAGATCGGAAG
11161_1 TATGAGTGTGGAGTCGTTGCGGGTGAGAGAGAGAAGAGTGTGTNNNAGATCGGA
11162_1 TATGAGTGTGGAGTCGTTGCCCAACCCATCTTGGCCTCTAAAANNNAGATCGGAAGA
11164_1 TATGAGTGTGGAGTCGTTGCTGACTATCCCCTTGACATGCTCTNNNAGATCGGAAGA
11167_1 TATGAGTGTGGAGTCGTTGCCGTATTGCTGGGGATTGGTAAAANNNAGATCGGAAG
11168_1 TATGAGTGTGGAGTCGTTGCTGTTTTGCAGATGGTCTTGCTCTNNNAGATCGGAAGA
11172_1 TATGAGTGTGGAGTCGTTGCATGAGCACTAGCAGACCATCTCTNNNAGATCGGAAG
11173_1 TATGAGTGTGGAGTCGTTGCACTCACCGGGGTAGCCAGAAAAGNNNAGATCGGAAG
11175_1 TATGAGTGTGGAGTCGTTGCCTGCCTCCCGGTATTCTATCTCTNNNAGATCGGAAGA
11177_1 TATGAGTGTGGAGTCGTTGCCCTGTCCTTGGCTCTACAACAAANNNAGATCGGAAGA
11178_1 TATGAGTGTGGAGTCGTTGCGTTCTCTGTGGAACCCCAACAAANNNAGATCGGAAG
11179_1 TATGAGTGTGGAGTCGTTGCGGGGGAATTCAGAACATAACAAANNNAGATCGGAA
11180_1 TATGAGTGTGGAGTCGTTGCTGAGAACACAAGACCCACTCTCTNNNAGATCGGAAGA
11181_1 TATGAGTGTGGAGTCGTTGCGAGGCAGATACCACACACACAAANNNAGATCGGAAG
11183_1 TATGAGTGTGGAGTCGTTGCGCACTATCACAAGGTCCCACAAANNNAGATCGGAAG
11184_1 TATGAGTGTGGAGTCGTTGCATCCTGCTTGACTTGACCTCTCTNNNAGATCGGAAGA
11186_1 TATGAGTGTGGAGTCGTTGCTGTTTTTCTCTCCCACCCTCTCTNNNAGATCGGAAGAG
11187_1 TATGAGTGTGGAGTCGTTGCCAAACTGAATGGAGCAGCACAAANNNAGATCGGAAG
11189_1 TATGAGTGTGGAGTCGTTGCACGCTTACCCTCTTTTCCTCTCTNNNAGATCGGAAGAG
11190_1 TATGAGTGTGGAGTCGTTGCGCACATACACGAAGATGCACAAANNNAGATCGGAAG
11191_1 TATGAGTGTGGAGTCGTTGCACTGGCACTGAATCCATCTCTCTNNNAGATCGGAAGA
11192_1 TATGAGTGTGGAGTCGTTGCCCCACCCACTTTTCTCAGACAAANNNAGATCGGAAGA
11193_1 TATGAGTGTGGAGTCGTTGCCTGATTGACGTTGGGGAGACAAANNNAGATCGGAA
11194_1 TATGAGTGTGGAGTCGTTGCTACGTGTGTGTGCGTTAGACAAANNNAGATCGGAAG
11196_1 TATGAGTGTGGAGTCGTTGCGGGCCAAACCTCATTAGGACAAANNNAGATCGGAAG
11204_1 TATGAGTGTGGAGTCGTTGCACGCACACACAGAGCTATACAAANNNAGATCGGAAG
11205_1 TATGAGTGTGGAGTCGTTGCACGCAGCTTTCTGGTTCTACAAANNNAGATCGGAAGA
11206_1 TATGAGTGTGGAGTCGTTGCCTTCAGAGTGGAAGCGGTACAAANNNAGATCGGAAG
11209_1 TATGAGTGTGGAGTCGTTGCTTGTGGTGGCTGGATAAACCAAANNNAGATCGGAAG
11211_1 TATGAGTGTGGAGTCGTTGCACCAGAACAAAGGACCAACCAAANNNAGATCGGAAG
11212_1 TATGAGTGTGGAGTCGTTGCTATCTGGGTGCATCTGTGTCTCTNNNAGATCGGAAGA
11214_1 TATGAGTGTGGAGTCGTTGCTCCCTCTTTTCTTCCAGACCAAANNNAGATCGGAAGAG
11215_1 TATGAGTGTGGAGTCGTTGCATCTGGCAATGTGATGGACCAAANNNAGATCGGAAG
11216_1 TATGAGTGTGGAGTCGTTGCCATAGATTCTGGGGTGGACCAAANNNAGATCGGAAG
11217_1 TATGAGTGTGGAGTCGTTGCGCTTCTGTTGCCTGGAATTCTCTNNNAGATCGGAAGA
11219_1 TATGAGTGTGGAGTCGTTGCTATTCAGGACATGCCAACCCAAANNNAGATCGGAAG
11220_1 TATGAGTGTGGAGTCGTTGCAGTGAAAGTCCTTGGAACCCAAANNNAGATCGGAAG
11222_1 TATGAGTGTGGAGTCGTTGCCACCAACATCTCTGTTCCCCAAANNNAGATCGGAAGA
11223_1 TATGAGTGTGGAGTCGTTGCGCATTCATGTGGGCAATTTCTCTNNNAGATCGGAAGA
11224_1 TATGAGTGTGGAGTCGTTGCATGTTCCCTCACTGTTGCCCAAANNNAGATCGGAAGA
11226_1 TATGAGTGTGGAGTCGTTGCAAACAAGCCCTCTTCTCCGTAGTNNNAGATCGGAAGA
11227_1 TATGAGTGTGGAGTCGTTGCTGCCCTACTGGTTCTCTTTCTCTNNNAGATCGGAAGAG
11228_1 TATGAGTGTGGAGTCGTTGCAGCCTGGAGAACATTATGCCAAANNNAGATCGGAAG

| | |
|---|---|
| 11230_1 | TATGAGTGTGGAGTCGTTGCCCTCTGCTGCCATCTTTTTCTCTNNNAGATCGGAAGA |
| 11231_1 | TATGAGTGTGGAGTCGTTGCCTTTCCCCTCCCCATCAAAGTCTNNNAGATCGGAAGA |
| 11232_1 | TATGAGTGTGGAGTCGTTGCCGCCTTCTGAGACAATCTCCAAANNNAGATCGGAAG |
| 11233_1 | TATGAGTGTGGAGTCGTTGCGTGCTCAAGTTGTCTCTACAACCNNNAGATCGGAAG |
| 11234_1 | TATGAGTGTGGAGTCGTTGCCTTGGCATTTCCCTCTCTCCAAANNNAGATCGGAAGA |
| 11237_1 | TATGAGTGTGGAGTCGTTGCAGAGGCTGTGTATGGTCAAGTCTNNNAGATCGGAA |
| 11240_1 | TATGAGTGTGGAGTCGTTGCGGCTGACTGGAGAGTTTTCCAAANNNAGATCGGAA |
| 11241_1 | TATGAGTGTGGAGTCGTTGCCTGATACGCCAGCAAAAAGCAAANNNAGATCGGAA |
| 11242_1 | TATGAGTGTGGAGTCGTTGCGGCAAAAGCTGGGTAAAAGCAAANNNAGATCGGA |
| 11246_1 | TATGAGTGTGGAGTCGTTGCTTTACACTCCCACTTACAGCAAANNNAGATCGGAAG |
| 11247_1 | TATGAGTGTGGAGTCGTTGCTGTAAGAGATGGGGAGGAGCAAANNNAGATCGGA |
| 11248_1 | TATGAGTGTGGAGTCGTTGCGTTCCATCCCCAACAGTAGCAAANNNAGATCGGAAG |
| 11252_1 | TATGAGTGTGGAGTCGTTGCATTTAGGTGAGGTTGCAGGCAAANNNAGATCGGAA |
| 11253_1 | TATGAGTGTGGAGTCGTTGCTTGGTTCTGTTACTGTCGGCAAANNNAGATCGGAAG |
| 11255_1 | TATGAGTGTGGAGTCGTTGCACACTCACATATTTCCGGGCAAANNNAGATCGGAAG |
| 11259_1 | TATGAGTGTGGAGTCGTTGCCACAGGAGTTTGGCAAATGCAAANNNAGATCGGAA |
| 11260_1 | TATGAGTGTGGAGTCGTTGCAGTTGTCCCTTGAACAATGCAAANNNAGATCGGAAG |
| 11261_1 | TATGAGTGTGGAGTCGTTGCCCTGTCATTCCTCTTCCTGCAAANNNAGATCGGAAGA |
| 11262_1 | TATGAGTGTGGAGTCGTTGCGCTTTCCTTTCTGAAGCTGCAAANNNAGATCGGAAG |
| 11263_1 | TATGAGTGTGGAGTCGTTGCGGGTGGGGGAATAGAATTGCAAANNNAGATCGGA |
| 11264_1 | TATGAGTGTGGAGTCGTTGCTGTAGGACCAAGCAACTTGCAAANNNAGATCGGAA |
| 11266_1 | TATGAGTGTGGAGTCGTTGCGTTTCAGGCCTAGAGTGAGGTCTNNNAGATCGGAA |
| 11267_1 | TATGAGTGTGGAGTCGTTGCTGGTTACAGCTCCAGTTAGGTCTNNNAGATCGGAAG |
| 11269_1 | TATGAGTGTGGAGTCGTTGCTTCTGTGTCAGTAACTGGGGTCTNNNAGATCGGAAG |
| 11270_1 | TATGAGTGTGGAGTCGTTGCTACAGGGAGGATCCTATGGGTCTNNNAGATCGGAA |
| 11271_1 | TATGAGTGTGGAGTCGTTGCGAGGAGGTATCAGCAGCATCAAANNNAGATCGGAA |
| 11273_1 | TATGAGTGTGGAGTCGTTGCTGTGCATGTAGTGCCAGATCAAANNNAGATCGGAA |
| 11274_1 | TATGAGTGTGGAGTCGTTGCTATCTCTGCTTCCTTGTGGGTCTNNNAGATCGGAAG |
| 11275_1 | TATGAGTGTGGAGTCGTTGCCCTCCCAAACCCAAGAACTCAAANNNAGATCGGAAG |
| 11278_1 | TATGAGTGTGGAGTCGTTGCAACATAGAAACCGACCCTGGTCTNNNAGATCGGAAG |
| 11279_1 | TATGAGTGTGGAGTCGTTGCGTCCTTCCTAACTGTCCTGGTCTNNNAGATCGGAAG |
| 11280_1 | TATGAGTGTGGAGTCGTTGCGAGCTTCCTGTACACACGTCAAANNNAGATCGGAAG |
| 11282_1 | TATGAGTGTGGAGTCGTTGCGAGCCCACCTTTTACTCTTCAAANNNAGATCGGAAG |
| 11283_1 | TATGAGTGTGGAGTCGTTGCTTTCAACATGAGGGCAGATGTCTNNNAGATCGGAA |
| 11285_1 | TATGAGTGTGGAGTCGTTGCTGTCCAGCAAGGAAGTCTTTGGGNNNAGATCGGAA |
| 11289_1 | TATGAGTGTGGAGTCGTTGCACGGGTGGGATAGTGGAAAGAAANNNAGATCGGA |
| 11290_1 | TATGAGTGTGGAGTCGTTGCACCCATTGAGCTGCTGAAAGAAANNNAGATCGGAA |
| 11294_1 | TATGAGTGTGGAGTCGTTGCCAGCCTTCTCATGTGGGAAGAAANNNAGATCGGAA |
| 11295_1 | TATGAGTGTGGAGTCGTTGCAGCCAATGGTGATTTTGCTGTCTNNNAGATCGGAAG |
| 11297_1 | TATGAGTGTGGAGTCGTTGCCTTACCCTTGTCGAGCACAGAAANNNAGATCGGAAG |
| 11302_1 | TATGAGTGTGGAGTCGTTGCGTGATGGGTGGTGCTTACAGAAANNNAGATCGGAA |
| 11303_1 | TATGAGTGTGGAGTCGTTGCGTGGGAGGCTTCTTTTCCAGAAANNNAGATCGGAA |
| 11305_1 | TATGAGTGTGGAGTCGTTGCGTGGTGAGAAATCTGGGCAGAAANNNAGATCGGA |

| | |
|---|---|
| 11306_1 | TATGAGTGTGGAGTCGTTGCCGTACTCGTGTGTGTATGTGTCTNNNAGATCGGAAG |
| 11308_1 | TATGAGTGTGGAGTCGTTGCTCAAGCTCTCCTGTCCTCAGAAANNNAGATCGGAAG |
| 11309_1 | TATGAGTGTGGAGTCGTTGCGGCTTTGCTGTCTGAGATTGTCTNNNAGATCGGAAG |
| 11310_1 | TATGAGTGTGGAGTCGTTGCAGAGCCTGCATGTAGAAGAGAAANNNAGATCGGAA |
| 11311_1 | TATGAGTGTGGAGTCGTTGCGGATTGGAAAGGGCACAGAGAAANNNAGATCGGA |
| 11312_1 | TATGAGTGTGGAGTCGTTGCGAAGAGGAGGAGGAGACATTTGGNNNAGATCGGA |
| 11313_1 | TATGAGTGTGGAGTCGTTGCGTGGAGCTCAGAGGAAGGAGAAANNNAGATCGGA |
| 11314_1 | TATGAGTGTGGAGTCGTTGCGGGGCAAAGAACAAGAGGAGAAANNNAGATCGGA |
| 11318_1 | TATGAGTGTGGAGTCGTTGCCTTTCTGGCAACTCCCTTTGTCTNNNAGATCGGAAGA |
| 11319_1 | TATGAGTGTGGAGTCGTTGCGGAAGGGAACCAGAAGTTTGTCTNNNAGATCGGAA |
| 11320_1 | TATGAGTGTGGAGTCGTTGCTGTGTATGGCTCGCAATGAGAAANNNAGATCGGAA |
| 11321_1 | TATGAGTGTGGAGTCGTTGCTGGCACCGGAAAAGAATGAGAAANNNAGATCGGAA |
| 11323_1 | TATGAGTGTGGAGTCGTTGCCTTACCTGCTCTTGCCTGAGAAANNNAGATCGGAAG |
| 11324_1 | TATGAGTGTGGAGTCGTTGCACCACTACCTGGGTCACTAGAAANNNAGATCGGAAG |
| 11328_1 | TATGAGTGTGGAGTCGTTGCGGCGTAGAAGGCACAATTAGAAANNNAGATCGGAA |
| 11329_1 | TATGAGTGTGGAGTCGTTGCGCCTGCTGGGAACTTCTTAGAAANNNAGATCGGAA |
| 11331_1 | TATGAGTGTGGAGTCGTTGCGCATGAACAGGTTCCAGATCCTCNNNAGATCGGAAG |
| 11332_1 | TATGAGTGTGGAGTCGTTGCCCCAGGACCTGTAAGTAACGAAANNNAGATCGGAA |
| 11333_1 | TATGAGTGTGGAGTCGTTGCGGGTATAGTGCACGCTTACGAAANNNAGATCGGAA |
| 11334_1 | TATGAGTGTGGAGTCGTTGCTCCAGCAATGCTCTGTTACGAAANNNAGATCGGAAG |
| 11335_1 | TATGAGTGTGGAGTCGTTGCCCCTAACACTACCAGTAGCGAAANNNAGATCGGAAG |
| 11336_1 | TATGAGTGTGGAGTCGTTGCTATTCATTTTGGGGTGGGCGAAANNNAGATCGGAA |
| 11339_1 | TATGAGTGTGGAGTCGTTGCATCCGCCTCTTTTCATGCATTCTNNNAGATCGGAAGA |
| 11340_1 | TATGAGTGTGGAGTCGTTGCGCCTGAATGGTCCAACTCATTCTNNNAGATCGGAAG |
| 11343_1 | TATGAGTGTGGAGTCGTTGCTCCTGAGGGGTTGATGTCATTCTNNNAGATCGGAAG |
| 11347_1 | TATGAGTGTGGAGTCGTTGCGCCAGGACAGCTAAATCAGGAAANNNAGATCGGAA |
| 11349_1 | TATGAGTGTGGAGTCGTTGCCTCCTTCCGGGTTCAGCTATTCTNNNAGATCGGAAGA |
| 11350_1 | TATGAGTGTGGAGTCGTTGCTGCTATCCAGAGATCTTATGGGTNNNAGATCGGAAG |
| 11351_1 | TATGAGTGTGGAGTCGTTGCGCTGTGACCCATCTACCAATTTGNNNAGATCGGAAG |
| 11352_1 | TATGAGTGTGGAGTCGTTGCCACTAAGGCAGCATCAAGGGAAANNNAGATCGGAA |
| 11353_1 | TATGAGTGTGGAGTCGTTGCTATGTAGCTGGGGTGAAGGGAAANNNAGATCGGA |
| 11355_1 | TATGAGTGTGGAGTCGTTGCTCCTTGTTAGCCCAGTGACTTCTNNNAGATCGGAAG |
| 11356_1 | TATGAGTGTGGAGTCGTTGCAGGATCTGGGCCAATTAGGGAAANNNAGATCGGAA |
| 11358_1 | TATGAGTGTGGAGTCGTTGCCGGAGGACACATTTTCACCTTCTNNNAGATCGGAAG |
| 11359_1 | TATGAGTGTGGAGTCGTTGCTTGAATCCCTTGTCTCCCCTTCTNNNAGATCGGAAGA |
| 11360_1 | TATGAGTGTGGAGTCGTTGCCAGGCTTCTCTTTCTTGGGGAAANNNAGATCGGAAG |
| 11362_1 | TATGAGTGTGGAGTCGTTGCGTTCTGGCGTTTCTTCTCCTTCTNNNAGATCGGAAGA |
| 11365_1 | TATGAGTGTGGAGTCGTTGCATACCAGCAGCAAACACTGGAAANNNAGATCGGAA |
| 11366_1 | TATGAGTGTGGAGTCGTTGCAATGACAAACAGACGCAGCTTCTNNNAGATCGGAAG |
| 11367_1 | TATGAGTGTGGAGTCGTTGCGCCATAGGGGAAAAACCTGGAAANNNAGATCGGAA |
| 11369_1 | TATGAGTGTGGAGTCGTTGCAGCACCACTGTCAAATCTGGAAANNNAGATCGGAAG |
| 11372_1 | TATGAGTGTGGAGTCGTTGCTGCTCACAGAGAAGTAGGCTTCTNNNAGATCGGAA |
| 11373_1 | TATGAGTGTGGAGTCGTTGCTGGCATCATAGAATGCGTGGAAANNNAGATCGGAA |

11374_1 TATGAGTGTGGAGTCGTTGCGATTGATGGGGATTTGGCTTCTNNNAGATCGGAA
11375_1 TATGAGTGTGGAGTCGTTGCATCATCAGGTCAGGCTGTGGAAANNNAGATCGGAA
11376_1 TATGAGTGTGGAGTCGTTGCGGACAATGACAACAGTGGCTTCTNNNAGATCGGAAG
11379_1 TATGAGTGTGGAGTCGTTGCGCCTCCTCATCCCTTCATCTTCTNNNAGATCGGAAGAG
11380_1 TATGAGTGTGGAGTCGTTGCCCCCACAGAGTGTAGGATCTTCTNNNAGATCGGAAG
11381_1 TATGAGTGTGGAGTCGTTGCAGGTCATTGCTGGGATCATGAAANNNAGATCGGAA
11382_1 TATGAGTGTGGAGTCGTTGCACACCCACATGCATCTCATGAAANNNAGATCGGAAG
11383_1 TATGAGTGTGGAGTCGTTGCACTTCCACATCATCCTCTCTTCTNNNAGATCGGAAGAG
11384_1 TATGAGTGTGGAGTCGTTGCTACCTGCTTGCTACCTGTCTTCTNNNAGATCGGAAGA
11386_1 TATGAGTGTGGAGTCGTTGCCTAATGTGTGTATGTGCCTGGCTNNNAGATCGGAAG
11387_1 TATGAGTGTGGAGTCGTTGCCCTCCCAAATGACCCAACTGAAANNNAGATCGGAAG
11389_1 TATGAGTGTGGAGTCGTTGCGCAAAGTGGAGACACACCTGAAANNNAGATCGGAA
11390_1 TATGAGTGTGGAGTCGTTGCTGAGGGTTTTCATTGCCCTGAAANNNAGATCGGAAG
11391_1 TATGAGTGTGGAGTCGTTGCCAATGGGCCTAAACCCAAGTTCTNNNAGATCGGAAG
11392_1 TATGAGTGTGGAGTCGTTGCGGGAGAGGGAAGAAGGAAGTTCTNNNAGATCGGA
11393_1 TATGAGTGTGGAGTCGTTGCCCGTTTCAGCATTGTGTGTGTTCNNNAGATCGGAAG
11394_1 TATGAGTGTGGAGTCGTTGCGTTGTGTCCAAGGCCTAGTGAAANNNAGATCGGAA
11396_1 TATGAGTGTGGAGTCGTTGCCCCAAGCATGAAGAAACGTGAAANNNAGATCGGAA
11399_1 TATGAGTGTGGAGTCGTTGCACACGGCTTGAGAAGATGTGAAANNNAGATCGGAA
11400_1 TATGAGTGTGGAGTCGTTGCTGTCTGCTTTCTTGCATCGTTCTNNNAGATCGGAAGA
11403_1 TATGAGTGTGGAGTCGTTGCGAGCAACTGGGATCTGTTTGAAANNNAGATCGGAA
11404_1 TATGAGTGTGGAGTCGTTGCCACTTACGCTGGGGTTATGTTCTNNNAGATCGGAAG
11405_1 TATGAGTGTGGAGTCGTTGCCCATGCTAGGCCCTGCAAATAAANNNAGATCGGAAG
11407_1 TATGAGTGTGGAGTCGTTGCCATTCTAATCACCCACCTGTGCANNNAGATCGGAAGA
11409_1 TATGAGTGTGGAGTCGTTGCGATACGGTAGTGGGAGGAATAAANNNAGATCGGA
11412_1 TATGAGTGTGGAGTCGTTGCGATGGTGAGAGGGGGAGGATAAANNNAGATCGGA
11414_1 TATGAGTGTGGAGTCGTTGCCCGTGTCTGGTACTGCTGATAAANNNAGATCGGAAG
11416_1 TATGAGTGTGGAGTCGTTGCCCAAGGGCAGCAAGAGCTATAAANNNAGATCGGAA
11418_1 TATGAGTGTGGAGTCGTTGCGGGAAGGTCACAGCACAACTAAANNNAGATCGGAA
11419_1 TATGAGTGTGGAGTCGTTGCGTTTCAAAGGGCAACGCACTAAANNNAGATCGGAAG
11420_1 TATGAGTGTGGAGTCGTTGCGGGGAATCCTGCCAAAGATTTCTNNNAGATCGGAAG
11423_1 TATGAGTGTGGAGTCGTTGCGGGTGGGTGTGTTTGTGATTTCTNNNAGATCGGAA
11425_1 TATGAGTGTGGAGTCGTTGCCCATTCTGCCTGGAACACTTTCTNNNAGATCGGAAGA
11426_1 TATGAGTGTGGAGTCGTTGCCGTGACCCCAGAAGAAAGCTAAANNNAGATCGGAA
11427_1 TATGAGTGTGGAGTCGTTGCAAGAGCGGGGAGACTTAGCTAAANNNAGATCGGAA
11429_1 TATGAGTGTGGAGTCGTTGCCCCCCACACTCTAAATGCTTTCTNNNAGATCGGAAGA
11434_1 TATGAGTGTGGAGTCGTTGCGCTGTGGAAGTCTTGAGGGTAAANNNAGATCGGAA
11435_1 TATGAGTGTGGAGTCGTTGCTGCCTACAGTGAATCTCTCCAGANNNAGATCGGAAG
11439_1 TATGAGTGTGGAGTCGTTGCGTGCTGAGAGAGGGCTGGAAATGNNNAGATCGGA
11441_1 TATGAGTGTGGAGTCGTTGCGCACCTACCCTTTCTCTGTTTCTNNNAGATCGGAAGA
11442_1 TATGAGTGTGGAGTCGTTGCTTGGCTATGACTGCTGTGTTTCTNNNAGATCGGAAG
11443_1 TATGAGTGTGGAGTCGTTGCGGAGGGAGGGAGGAAATTATGGTNNNAGATCGGA
11445_1 TATGAGTGTGGAGTCGTTGCGGCACAGCTGTTGTGAAAAACAANNNAGATCGGAA

| | |
|---|---|
| 11446_1 | TATGAGTGTGGAGTCGTTGCGCAGCATGGGTTTCAGAAAACAANNNAGATCGGAA |
| 11448_1 | TATGAGTGTGGAGTCGTTGCGCCGTTGTCACCACTGATTTTCTNNNAGATCGGAAG |
| 11451_1 | TATGAGTGTGGAGTCGTTGCGAACCTGGCAACACTTCAAACAANNNAGATCGGAAG |
| 11452_1 | TATGAGTGTGGAGTCGTTGCCTGGGAAACATCCAGGGAAACAANNNAGATCGGAA |
| 11455_1 | TATGAGTGTGGAGTCGTTGCGCTCTGGCATCAGGGATAAACAANNNAGATCGGAA |
| 11456_1 | TATGAGTGTGGAGTCGTTGCCCAGGCTCATGTAGTGCTTTTCTNNNAGATCGGAAG |
| 11457_1 | TATGAGTGTGGAGTCGTTGCCCCAAGTGGCATAGCTCTTTTCTNNNAGATCGGAAG |
| 11458_1 | TATGAGTGTGGAGTCGTTGCTTTGGCTCCCTGTTGAGTTTTCTNNNAGATCGGAAG |
| 11460_1 | TATGAGTGTGGAGTCGTTGCGCGTGTCTTATTTGTGGCAACAANNNAGATCGGAAG |
| 11462_1 | TATGAGTGTGGAGTCGTTGCGCATATCACTTGGGGGTCAACAANNNAGATCGGAA |
| 11463_1 | TATGAGTGTGGAGTCGTTGCTGCCTTCCTCAGCAAGTTTTTCTNNNAGATCGGAAGA |
| 11465_1 | TATGAGTGTGGAGTCGTTGCCAAGCCCTTTCAGGAGAGAACAANNNAGATCGGAA |
| 11466_1 | TATGAGTGTGGAGTCGTTGCTGAGCAATGATGGAGAGGAACAANNNAGATCGGA |
| 11467_1 | TATGAGTGTGGAGTCGTTGCTGGCATTTGTTATGGGGGAACAANNNAGATCGGAA |
| 11468_1 | TATGAGTGTGGAGTCGTTGCAACGAGAATTGCTCCCCAAAAGTNNNAGATCGGAAG |
| 11469_1 | TATGAGTGTGGAGTCGTTGCCTAACTTGGGTGGAGCCAAAAGTNNNAGATCGGAA |
| 11472_1 | TATGAGTGTGGAGTCGTTGCCTTTTCAACCCAACACATAGGTCNNNAGATCGGAAG |
| 11474_1 | TATGAGTGTGGAGTCGTTGCCCATCACCTTTCTGGGGAAAAGTNNNAGATCGGAAG |
| 11476_1 | TATGAGTGTGGAGTCGTTGCACTGGCATGTGCATCTGAAAAGTNNNAGATCGGAA |
| 11477_1 | TATGAGTGTGGAGTCGTTGCCTGAGGAGAACACACACACACAANNNAGATCGGAA |
| 11478_1 | TATGAGTGTGGAGTCGTTGCGGAAAACAAATAGCAAGAGGTCCNNNAGATCGGAA |
| 11479_1 | TATGAGTGTGGAGTCGTTGCACCAGATACTGTGCTAGACACAANNNAGATCGGAAG |
| 11480_1 | TATGAGTGTGGAGTCGTTGCCCAAAGCAGTTAGGGGACAAAGTNNNAGATCGGAA |
| 11483_1 | TATGAGTGTGGAGTCGTTGCGTAAGCCCATGCTGATCCAAAGTNNNAGATCGGAAG |
| 11486_1 | TATGAGTGTGGAGTCGTTGCGAGTGCACATAAACCAGCAAAGTNNNAGATCGGAA |
| 11487_1 | TATGAGTGTGGAGTCGTTGCTGGTTTTCCAACTTAGCCCACAANNNAGATCGGAAG |
| 11488_1 | TATGAGTGTGGAGTCGTTGCTGCTTGTATTCACACTCCCACAANNNAGATCGGAAGA |
| 11489_1 | TATGAGTGTGGAGTCGTTGCTTTCCCTAGAATGGTAACGGGCANNNAGATCGGAA |
| 11490_1 | TATGAGTGTGGAGTCGTTGCCTCCAAGCCCCATTTCTCAAAGTNNNAGATCGGAAGA |
| 11491_1 | TATGAGTGTGGAGTCGTTGCGACAGACACACAGACATCCACAANNNAGATCGGAAG |
| 11492_1 | TATGAGTGTGGAGTCGTTGCTCTTGGATTCCGGAAGGGGAAGGNNNAGATCGGA |
| 11493_1 | TATGAGTGTGGAGTCGTTGCTCCCCTTTCTCCATTCTCCACAANNNAGATCGGAAGA |
| 11496_1 | TATGAGTGTGGAGTCGTTGCTCTCCAGTGAGGAAAGGGAAAGTNNNAGATCGGAA |
| 11497_1 | TATGAGTGTGGAGTCGTTGCACGGAAGTTGTAAAGTCTCACAANNNAGATCGGAA |
| 11499_1 | TATGAGTGTGGAGTCGTTGCCCAAGTGAGATGCTGCAAGACAANNNAGATCGGAA |
| 11500_1 | TATGAGTGTGGAGTCGTTGCCTGCAAGGGCCAGATCATAAAGTNNNAGATCGGAA |
| 11502_1 | TATGAGTGTGGAGTCGTTGCCCCAGAGCTCAAAAAGGTAAAGTNNNAGATCGGAA |
| 11503_1 | TATGAGTGTGGAGTCGTTGCCCCTTTCAAATGTGTTCAGACAANNNAGATCGGAAG |
| 11504_1 | TATGAGTGTGGAGTCGTTGCACCCAACTCTTCTCTGGAGACAANNNAGATCGGAAG |
| 11507_1 | TATGAGTGTGGAGTCGTTGCTGTTTTTGACCCTGAGGACAAGTNNNAGATCGGAAG |
| 11510_1 | TATGAGTGTGGAGTCGTTGCCCGCCTTGAAAGAGAGATGACAANNNAGATCGGAA |
| 11511_1 | TATGAGTGTGGAGTCGTTGCGGCTTTTGATCTTCCCCTGACAANNNAGATCGGAAG |
| 11512_1 | TATGAGTGTGGAGTCGTTGCGGACTGACTTTGGAAACCCAAGTNNNAGATCGGAA |

11513_1 TATGAGTGTGGAGTCGTTGCATGTGAATGCATACAGCCCAAGTNNNAGATCGGAAG
11514_1 TATGAGTGTGGAGTCGTTGCTATGCTTTGGTTGACGGTGACAANNNAGATCGGAA
11515_1 TATGAGTGTGGAGTCGTTGCTGCTCTTCCCTACTCTGTGACAANNNAGATCGGAAGA
11517_1 TATGAGTGTGGAGTCGTTGCACTGAACATCACAATGGCCAAGTNNNAGATCGGAAG
11518_1 TATGAGTGTGGAGTCGTTGCAGTAGGAGGGTGTCAGTTGACAANNNAGATCGGAA
11519_1 TATGAGTGTGGAGTCGTTGCTCTTCCCCATCCATACTGTGTGTNNNAGATCGGAAGA
11521_1 TATGAGTGTGGAGTCGTTGCTCATGTGGGAGGTGGGAATACAANNNAGATCGGAA
11522_1 TATGAGTGTGGAGTCGTTGCTTCTGAGACCCCTGATTCCAAGTNNNAGATCGGAAG
11524_1 TATGAGTGTGGAGTCGTTGCACAACTGCAATCAAGGGCTACAANNNAGATCGGAAG
11525_1 TATGAGTGTGGAGTCGTTGCCTCTCTCTCTCTTGATCGACAGGNNNAGATCGGAAG
11526_1 TATGAGTGTGGAGTCGTTGCCCGAATGTGCATCCCTTCTACAANNNAGATCGGAAG
11529_1 TATGAGTGTGGAGTCGTTGCTGCCCATGTTGTGAACTGTACAANNNAGATCGGAAG
11530_1 TATGAGTGTGGAGTCGTTGCGGCGATCTTGAGACTGTGTACAANNNAGATCGGAA
11531_1 TATGAGTGTGGAGTCGTTGCTGTTCCTCCTGGTCACCTTACAANNNAGATCGGAAGA
11534_1 TATGAGTGTGGAGTCGTTGCACTTAGCCTTGTGGAACTCAAGTNNNAGATCGGAAG
11535_1 TATGAGTGTGGAGTCGTTGCGCAACCGAGGAATACACAACCAANNNAGATCGGAA
11537_1 TATGAGTGTGGAGTCGTTGCATCTTTGATGGCTGTCCAACCAANNNAGATCGGAAG
11538_1 TATGAGTGTGGAGTCGTTGCCCAGCTACAACTTTTGCAACCAANNNAGATCGGAAG
11540_1 TATGAGTGTGGAGTCGTTGCGGAAGGGGGAGTAAGGAAGAAGTNNNAGATCGGA
11546_1 TATGAGTGTGGAGTCGTTGCGTAAACATGACCTTGGGAGAAGTNNNAGATCGGAA
11547_1 TATGAGTGTGGAGTCGTTGCGGCTCAGGGCACTTTTCAAAAATNNNAGATCGGAAG
11550_1 TATGAGTGTGGAGTCGTTGCCAGACAAAGAAGCAGGCACCTATNNNAGATCGGAA
11551_1 TATGAGTGTGGAGTCGTTGCTCTCCTGTGCTTCCTGATACCAANNNAGATCGGAAGA
11552_1 TATGAGTGTGGAGTCGTTGCAGGGCAAAATATGCAAAGGAAGTNNNAGATCGGAA
11553_1 TATGAGTGTGGAGTCGTTGCCGAGGGTACTGTGAACTTTACGTNNNAGATCGGAA
11554_1 TATGAGTGTGGAGTCGTTGCTTGTTGGAAACTGAGTGACCCAANNNAGATCGGAA
11555_1 TATGAGTGTGGAGTCGTTGCTTGGGGAATGCTTTTTGACCCAANNNAGATCGGAAG
11557_1 TATGAGTGTGGAGTCGTTGCGTGAGCTTGAAAGAAGACCCCAANNNAGATCGGAA
11558_1 TATGAGTGTGGAGTCGTTGCCCAGGATTTTAGACTTGCCCCAANNNAGATCGGAAG
11559_1 TATGAGTGTGGAGTCGTTGCATTAGCTGTGGGAAAGGGAGACTNNNAGATCGGAA
11561_1 TATGAGTGTGGAGTCGTTGCTTTCAGGCAAATTACAAGCCCAANNNAGATCGGAAG
11562_1 TATGAGTGTGGAGTCGTTGCACTTTCAATCATGCAGAGCCCAANNNAGATCGGAAG
11564_1 TATGAGTGTGGAGTCGTTGCACATTTATTGGCTTCAGGCCCAANNNAGATCGGAAG
11566_1 TATGAGTGTGGAGTCGTTGCGTCACATTCCCAGAACGGAAACTNNNAGATCGGAAG
11569_1 TATGAGTGTGGAGTCGTTGCGACCAGCGATTTGTCTATCCCAANNNAGATCGGAAG
11570_1 TATGAGTGTGGAGTCGTTGCCCACTTGATGTTGGAACTCCCAANNNAGATCGGAAG
11571_1 TATGAGTGTGGAGTCGTTGCAGTCCACCACTAGGGTTTGAAGTNNNAGATCGGAA
11573_1 TATGAGTGTGGAGTCGTTGCTCTTCTTGAGGCGGAATCCATCTNNNAGATCGGAAG
11576_1 TATGAGTGTGGAGTCGTTGCTCAGTAGAGGCAAGTCGCTAAGTNNNAGATCGGAA
11577_1 TATGAGTGTGGAGTCGTTGCCGTGTCTATTTGGAATAAGCCAANNNAGATCGGAAG
11578_1 TATGAGTGTGGAGTCGTTGCAAGGCAGAAGGAGCTTGAGATCCNNNAGATCGGAA
11579_1 TATGAGTGTGGAGTCGTTGCACAGTTCTGGCAAATACAGCCAANNNAGATCGGAAG
11580_1 TATGAGTGTGGAGTCGTTGCGTAGTGCTTGAGCATCAGTAAGTNNNAGATCGGAA

| | |
|---|---|
| 11583_1 | TATGAGTGTGGAGTCGTTGCATTATTGTGCCTCAGTCTCCCCGNNNAGATCGGAAG |
| 11584_1 | TATGAGTGTGGAGTCGTTGCCACTCACTCCCCACCCCATACTANNNAGATCGGAAGA |
| 11585_1 | TATGAGTGTGGAGTCGTTGCGGTGAGAAAGTTTATGGGGCCAANNNAGATCGGAA |
| 11586_1 | TATGAGTGTGGAGTCGTTGCGCAGAGTATTTCGATTGGGCCAANNNAGATCGGAA |
| 11592_1 | TATGAGTGTGGAGTCGTTGCTTGGGACTGTTTTTCTCTGCCAANNNAGATCGGAAG |
| 11596_1 | TATGAGTGTGGAGTCGTTGCATTTTTGTGAGTGTCCCCACAGTNNNAGATCGGAAG |
| 11597_1 | TATGAGTGTGGAGTCGTTGCATTTGTTCTCTTTGGCCCACAGTNNNAGATCGGAAG |
| 11599_1 | TATGAGTGTGGAGTCGTTGCGTGAACGAGTGAGAACCATCCAANNNAGATCGGAA |
| 11600_1 | TATGAGTGTGGAGTCGTTGCCCCCCCTGGTTCAAATTCATCCAANNNAGATCGGAAG |
| 11602_1 | TATGAGTGTGGAGTCGTTGCTGGAGTAAAGGATGAAACATGGTNNNAGATCGGA |
| 11603_1 | TATGAGTGTGGAGTCGTTGCGGGTCAGATGTCACAAAGACAGTNNNAGATCGGAA |
| 11607_1 | TATGAGTGTGGAGTCGTTGCAGAGCACAAATTCACAGCTCCAANNNAGATCGGAAG |
| 11608_1 | TATGAGTGTGGAGTCGTTGCGTGAAAAAGGAAAGGGGGACAGTNNNAGATCGGA |
| 11609_1 | TATGAGTGTGGAGTCGTTGCACATAGGCTTCCCTCATCTCCAANNNAGATCGGAAG |
| 11610_1 | TATGAGTGTGGAGTCGTTGCATGCAGCCAGATTTCATGACAGTNNNAGATCGGAAG |
| 11611_1 | TATGAGTGTGGAGTCGTTGCCCACTTCCACCAGAAAAGTCCAANNNAGATCGGAAG |
| 11614_1 | TATGAGTGTGGAGTCGTTGCCACACTGTTATTCCACTGACAGTNNNAGATCGGAAG |
| 11615_1 | TATGAGTGTGGAGTCGTTGCCACCTTCTCCTACCTCTGTCCAANNNAGATCGGAAGA |
| 11616_1 | TATGAGTGTGGAGTCGTTGCTTACAAGTTCCCTGTGTGACAGTNNNAGATCGGAAG |
| 11619_1 | TATGAGTGTGGAGTCGTTGCCACACCATGAGGTTGTCTACAGTNNNAGATCGGAAG |
| 11620_1 | TATGAGTGTGGAGTCGTTGCGAAACTGCACCCATCAGTACAGTNNNAGATCGGAAG |
| 11621_1 | TATGAGTGTGGAGTCGTTGCGGTCTTTTTCTCTGCCTTTCCAANNNAGATCGGAAGA |
| 11622_1 | TATGAGTGTGGAGTCGTTGCCACTTCCTGGGGTTTCTTTCCAANNNAGATCGGAAG |
| 11623_1 | TATGAGTGTGGAGTCGTTGCGTGGTTTGAACGTGTGTTTCCAANNNAGATCGGAAG |
| 11625_1 | TATGAGTGTGGAGTCGTTGCACACGCCACTGATTCTTTTCCAANNNAGATCGGAAG |
| 11629_1 | TATGAGTGTGGAGTCGTTGCGAGAGTGAGCAGAGACAAAGCAANNNAGATCGGA |
| 11630_1 | TATGAGTGTGGAGTCGTTGCTGGAAGAAACACACAGAAAGCAANNNAGATCGGAA |
| 11633_1 | TATGAGTGTGGAGTCGTTGCCACAAGGCAAACATCACAAGCAANNNAGATCGGAA |
| 11635_1 | TATGAGTGTGGAGTCGTTGCTACCTATTGGGTTTCAGCCCAGTNNNAGATCGGAAG |
| 11639_1 | TATGAGTGTGGAGTCGTTGCCCACTCAGAAGATTTCCCAGCAANNNAGATCGGAAG |
| 11641_1 | TATGAGTGTGGAGTCGTTGCTCACATTGGAAACCCATCAGCAANNNAGATCGGAAG |
| 11642_1 | TATGAGTGTGGAGTCGTTGCATGCAGTAGATTATGCTCAGCAANNNAGATCGGAA |
| 11643_1 | TATGAGTGTGGAGTCGTTGCCGAGGCTTTGAGAGAAAGAGCAANNNAGATCGGA |
| 11646_1 | TATGAGTGTGGAGTCGTTGCCAGGAGAGAGGGTAAGAGAGCAANNNAGATCGGA |
| 11647_1 | TATGAGTGTGGAGTCGTTGCTGTCTTGGTTCCCTTACTCCAGTNNNAGATCGGAAG |
| 11648_1 | TATGAGTGTGGAGTCGTTGCTACTAAACCCACAGAGGGAGCAANNNAGATCGGAA |
| 11649_1 | TATGAGTGTGGAGTCGTTGCGACCTGTTCCCCTTATCTCCAGTNNNAGATCGGAAGA |
| 11651_1 | TATGAGTGTGGAGTCGTTGCGAGCATTTTTGCCGTTCTTACGTNNNAGATCGGAAG |
| 11653_1 | TATGAGTGTGGAGTCGTTGCAGTGAGCATTTGAAGGCTAGCAANNNAGATCGGAA |
| 11654_1 | TATGAGTGTGGAGTCGTTGCGCCACAGAAACCTTAGTTCCAGTNNNAGATCGGAAG |
| 11657_1 | TATGAGTGTGGAGTCGTTGCAACTCCCTAACAGTTCTCGCTTCNNNAGATCGGAAGA |
| 11659_1 | TATGAGTGTGGAGTCGTTGCTGGCTAGTGTGAAAAACAGCAGTNNNAGATCGGAA |
| 11663_1 | TATGAGTGTGGAGTCGTTGCTTTTGCGCAGTTTTATGAGGCAANNNAGATCGGAAG |

| | |
|---|---|
| 11664_1 | TATGAGTGTGGAGTCGTTGCCCTGCTGCTTCCAATATAGGCAANNNAGATCGGAAG |
| 11667_1 | TATGAGTGTGGAGTCGTTGCAGATTTGAAAGAAGACAGGGCAANNNAGATCGGAA |
| 11668_1 | TATGAGTGTGGAGTCGTTGCACCTTACTCTAGTGCCTAGCAGTNNNAGATCGGAAG |
| 11670_1 | TATGAGTGTGGAGTCGTTGCAAGAGGATCAGTATGGAGGGCAANNNAGATCGGA |
| 11672_1 | TATGAGTGTGGAGTCGTTGCAAGGATGTCTTGCATTACGCAGTNNNAGATCGGAA |
| 11674_1 | TATGAGTGTGGAGTCGTTGCATGCGTGTGTTTCATTATGGCAANNNAGATCGGAAG |
| 11675_1 | TATGAGTGTGGAGTCGTTGCTGGACTGAAGTAAAACAGGCAGTNNNAGATCGGAA |
| 11676_1 | TATGAGTGTGGAGTCGTTGCTCAGACCCTATTCTACCTGGCAANNNAGATCGGAAG |
| 11677_1 | TATGAGTGTGGAGTCGTTGCCCAAATAAGAGGCAAGAGGCAGTNNNAGATCGGAA |
| 11678_1 | TATGAGTGTGGAGTCGTTGCTGCCATCCTAGAAATAGGGCAGTNNNAGATCGGAA |
| 11681_1 | TATGAGTGTGGAGTCGTTGCTGGCTTTCTCTATCACTCCGTCANNNAGATCGGAAGA |
| 11682_1 | TATGAGTGTGGAGTCGTTGCTGTGTACTTATGCTGCTTGGCAANNNAGATCGGAAG |
| 11686_1 | TATGAGTGTGGAGTCGTTGCTAATGATGGGAGGGAGGATGCAANNNAGATCGGA |
| 11691_1 | TATGAGTGTGGAGTCGTTGCTGGCTACTCTGATAACACTGCAANNNAGATCGGAAG |
| 11693_1 | TATGAGTGTGGAGTCGTTGCATGGTGTTTGGCACTCAATCAGTNNNAGATCGGAAG |
| 11694_1 | TATGAGTGTGGAGTCGTTGCGGAGATGCACCACATGAGATGTGNNNAGATCGGAA |
| 11695_1 | TATGAGTGTGGAGTCGTTGCGGATCTGAATCTTGGCTCTGCAANNNAGATCGGAAG |
| 11696_1 | TATGAGTGTGGAGTCGTTGCACAGCCACAATACAAGTCTGCAANNNAGATCGGAAG |
| 11697_1 | TATGAGTGTGGAGTCGTTGCATTCTTTGGGAAGCAGTCTGCAANNNAGATCGGAA |
| 11698_1 | TATGAGTGTGGAGTCGTTGCCCAGTAGCAGTGTCTTTCTGCAANNNAGATCGGAAG |
| 11704_1 | TATGAGTGTGGAGTCGTTGCACTTTTCCAAGGACAGAGCTTCTNNNAGATCGGAAG |
| 11705_1 | TATGAGTGTGGAGTCGTTGCCATCTGATTGTGGAGGTGTGCAANNNAGATCGGAA |
| 11709_1 | TATGAGTGTGGAGTCGTTGCGGTCTCCAAAGACATCCTTGCAANNNAGATCGGAAG |
| 11710_1 | TATGAGTGTGGAGTCGTTGCAACTTTCACCCTCATGGTTGCAANNNAGATCGGAAG |
| 11712_1 | TATGAGTGTGGAGTCGTTGCGCCTGCCAAGTTTTTCTTTGCAANNNAGATCGGAAG |
| 11713_1 | TATGAGTGTGGAGTCGTTGCCCCTCAAAAGCTGAATGGTCAGTNNNAGATCGGAAG |
| 11717_1 | TATGAGTGTGGAGTCGTTGCTCTCCATCCTATGCTTCAATCAANNNAGATCGGAAGA |
| 11720_1 | TATGAGTGTGGAGTCGTTGCGGCAGGAAACTGGTCTGAATCAANNNAGATCGGAA |
| 11722_1 | TATGAGTGTGGAGTCGTTGCTTCCAAATTCCTGGCCACATCAANNNAGATCGGAAG |
| 11723_1 | TATGAGTGTGGAGTCGTTGCTTCCTCCTCATCTGTCCTTCAGTNNNAGATCGGAAGA |
| 11724_1 | TATGAGTGTGGAGTCGTTGCTAAAGGAAATGGGGAGAGTGGCTNNNAGATCGGA |
| 11726_1 | TATGAGTGTGGAGTCGTTGCGCCTCATTGGCCTTGGATATCAANNNAGATCGGAAG |
| 11729_1 | TATGAGTGTGGAGTCGTTGCCTGAAGCCCATCCATGAACTCAANNNAGATCGGAAG |
| 11732_1 | TATGAGTGTGGAGTCGTTGCTACCTTCAGCAATGCAACCTCAANNNAGATCGGAAG |
| 11734_1 | TATGAGTGTGGAGTCGTTGCCCTAGAAGACAGGAGGACCTCAANNNAGATCGGAA |
| 11735_1 | TATGAGTGTGGAGTCGTTGCAACTCTCTGGAAGAACCCCTCAANNNAGATCGGAAG |
| 11737_1 | TATGAGTGTGGAGTCGTTGCCTTTTGCAGGACAGCTACTGTCTNNNAGATCGGAAG |
| 11738_1 | TATGAGTGTGGAGTCGTTGCATTGCTCCAGGAGAGAGAAGAGTNNNAGATCGGAA |
| 11739_1 | TATGAGTGTGGAGTCGTTGCATGAGGAACAGCTGGAGAAGAGTNNNAGATCGGA |
| 11743_1 | TATGAGTGTGGAGTCGTTGCGGTCACTCAGGGGATCACAGGTANNNAGATCGGAA |
| 11745_1 | TATGAGTGTGGAGTCGTTGCGTTCAAAGGTGACTGGATCACAANNNAGATCGGAA |
| 11746_1 | TATGAGTGTGGAGTCGTTGCGGCAGAACACTAGAACTGCTCAANNNAGATCGGAA |
| 11747_1 | TATGAGTGTGGAGTCGTTGCTGAGGTAGAATCTTGCTGTCACCNNNAGATCGGAAG |

| | |
|---|---|
| 11748_1 | TATGAGTGTGGAGTCGTTGCCCCCCGCAATAGAAATCTACACTNNNAGATCGGAAG |
| 11749_1 | TATGAGTGTGGAGTCGTTGCGGGCCAGTCTAACTTGATCTCAANNNAGATCGGAAG |
| 11751_1 | TATGAGTGTGGAGTCGTTGCTGGCTATTTCTCCCCACTCTCAANNNAGATCGGAAGA |
| 11752_1 | TATGAGTGTGGAGTCGTTGCGTGCTGCAGATTCTTCCTCTCAANNNAGATCGGAAG |
| 11753_1 | TATGAGTGTGGAGTCGTTGCACAGGAGACTGCCAGAAGTTCTTNNNAGATCGGAA |
| 11754_1 | TATGAGTGTGGAGTCGTTGCGGAGAAGGGATCACCAAGAGAGTNNNAGATCGGA |
| 11755_1 | TATGAGTGTGGAGTCGTTGCATGGGAAGAAGGCTGAAGAGAGTNNNAGATCGGA |
| 11756_1 | TATGAGTGTGGAGTCGTTGCAAAATCAACCAAGTGTCGTGGCTNNNAGATCGGAA |
| 11757_1 | TATGAGTGTGGAGTCGTTGCAATGTATGGTCAGCCAGGAGAGTNNNAGATCGGAA |
| 11759_1 | TATGAGTGTGGAGTCGTTGCGAAACGAAACACATGGGGAGAGTNNNAGATCGGA |
| 11760_1 | TATGAGTGTGGAGTCGTTGCTTTTTGGCCACGTACATGAGAGTNNNAGATCGGAAG |
| 11761_1 | TATGAGTGTGGAGTCGTTGCCAAGCAGGGCTACAATCCTACTTNNNAGATCGGAAG |
| 11762_1 | TATGAGTGTGGAGTCGTTGCTTTTCTCTTTCAGGCCTCAACCTNNNAGATCGGAAGA |
| 11765_1 | TATGAGTGTGGAGTCGTTGCTTGACAGGAGGAAAGAGGGTCAANNNAGATCGGA |
| 11766_1 | TATGAGTGTGGAGTCGTTGCTCATGTAGTCCCACCATGGTCAANNNAGATCGGAAG |
| 11769_1 | TATGAGTGTGGAGTCGTTGCTTCCAAGGCCATGTCTCTAGAGTNNNAGATCGGAAG |
| 11770_1 | TATGAGTGTGGAGTCGTTGCCAGGTTGTCCTTTACCCTGTCAANNNAGATCGGAAG |
| 11771_1 | TATGAGTGTGGAGTCGTTGCCCCAACACCTACATCCTTGTCAANNNAGATCGGAAGA |
| 11772_1 | TATGAGTGTGGAGTCGTTGCGCCAGCTTCTAGGAAAAAGGAGTNNNAGATCGGAA |
| 11774_1 | TATGAGTGTGGAGTCGTTGCACCCACATACATATAGGCATTCAANNNAGATCGGAA |
| 11775_1 | TATGAGTGTGGAGTCGTTGCATGATGCCACAACAAACAGGAGTNNNAGATCGGAA |
| 11776_1 | TATGAGTGTGGAGTCGTTGCCACAAGAGCTCCCGGTTATTCAANNNAGATCGGAAG |
| 11777_1 | TATGAGTGTGGAGTCGTTGCGTTATTATGCAAGCGGCATTGTGNNNAGATCGGAA |
| 11779_1 | TATGAGTGTGGAGTCGTTGCGGAAATTGTTGAGGCTGAGGAGTNNNAGATCGGA |
| 11781_1 | TATGAGTGTGGAGTCGTTGCCCTTGGTTGAGCTCCTTCTTCAANNNAGATCGGAAG |
| 11782_1 | TATGAGTGTGGAGTCGTTGCGAAGGCTAGCTGGGTATAGGAGTNNNAGATCGGA |
| 11783_1 | TATGAGTGTGGAGTCGTTGCCAGGGCCATAATTCCGAGTTCAANNNAGATCGGAAG |
| 11784_1 | TATGAGTGTGGAGTCGTTGCATCAAAACCCTGGGTGTAGGAGTNNNAGATCGGAA |
| 11785_1 | TATGAGTGTGGAGTCGTTGCCATTTGACGAAACGCTTAGGAGTNNNAGATCGGAA |
| 11787_1 | TATGAGTGTGGAGTCGTTGCACTGCTTCAGCTTCTCTGTTCAANNNAGATCGGAAG |
| 11791_1 | TATGAGTGTGGAGTCGTTGCAAACTGCTGACAAGTGAAGTCTANNNAGATCGGAA |
| 11792_1 | TATGAGTGTGGAGTCGTTGCGGGGTTGAGTACTAGCCTTTCAANNNAGATCGGAA |
| 11794_1 | TATGAGTGTGGAGTCGTTGCGAGGCTATGAAAGGACTGGGAGTNNNAGATCGGA |
| 11797_1 | TATGAGTGTGGAGTCGTTGCATCTGGTTCACAGTAGCTGGAGTNNNAGATCGGAA |
| 11798_1 | TATGAGTGTGGAGTCGTTGCCGCCCAAAGACTAAGGCAAAGAANNNAGATCGGAA |
| 11799_1 | TATGAGTGTGGAGTCGTTGCGGATGGGATCAAGAGGCAAAGAANNNAGATCGGA |
| 11800_1 | TATGAGTGTGGAGTCGTTGCCAGCCAAAACAACCTGCAAAGAANNNAGATCGGAA |
| 11802_1 | TATGAGTGTGGAGTCGTTGCAAAATGAGCCACCAGAGAAAGAANNNAGATCGGAA |
| 11803_1 | TATGAGTGTGGAGTCGTTGCGCTGGGTTTTGAGGGAGAAAGAANNNAGATCGGA |
| 11804_1 | TATGAGTGTGGAGTCGTTGCGCTCTCATGACTTCCTTTGGAGTNNNAGATCGGAAG |
| 11808_1 | TATGAGTGTGGAGTCGTTGCCCCACCATGCAAGGATACAAGAANNNAGATCGGAA |
| 11809_1 | TATGAGTGTGGAGTCGTTGCTCTGGGGACACGACTTACAAGAANNNAGATCGGAA |
| 11810_1 | TATGAGTGTGGAGTCGTTGCCTTGGTAAGGTGGTGGCATTTCTNNNAGATCGGAA |

| | |
|---|---|
| 11814_1 | TATGAGTGTGGAGTCGTTGCATCCAGAACAAAGGCAGCAAGAANNNAGATCGGAA |
| 11816_1 | TATGAGTGTGGAGTCGTTGCGACAGCCTGCAGTAACTCAAGAANNNAGATCGGAA |
| 11817_1 | TATGAGTGTGGAGTCGTTGCACCGATACTCTGGCCAAGAAGAANNNAGATCGGAA |
| 11818_1 | TATGAGTGTGGAGTCGTTGCGAAGGGTCCTGGACAGTATGAGTNNNAGATCGGAA |
| 11819_1 | TATGAGTGTGGAGTCGTTGCTGGGACATGAACACTCATTGCTCNNNAGATCGGAAG |
| 11820_1 | TATGAGTGTGGAGTCGTTGCGCAGTGGGTCAGAGAGAGAAGAANNNAGATCGGA |
| 11822_1 | TATGAGTGTGGAGTCGTTGCGGGGATGTTTGGGGTTAGAAGAANNNAGATCGGA |
| 11823_1 | TATGAGTGTGGAGTCGTTGCCGCTGAGAGAGAGAGTACTGAGTNNNAGATCGGAA |
| 11825_1 | TATGAGTGTGGAGTCGTTGCATCCCCTTAGGCTTTTCCTGAGTNNNAGATCGGAAG |
| 11826_1 | TATGAGTGTGGAGTCGTTGCCATTCTGGGACTGCTTTCCAAAANNNAGATCGGAAG |
| 11827_1 | TATGAGTGTGGAGTCGTTGCGCAGCCTAGTAATCCATCTGAGTNNNAGATCGGAAG |
| 11828_1 | TATGAGTGTGGAGTCGTTGCGTTCCTGACACTGTGGCTAAGAANNNAGATCGGAA |
| 11829_1 | TATGAGTGTGGAGTCGTTGCACATCCTGAGTTGAGCTCTGAGTNNNAGATCGGAAG |
| 11830_1 | TATGAGTGTGGAGTCGTTGCGGTAGAGGGCTGTCAGGTAAGAANNNAGATCGGA |
| 11832_1 | TATGAGTGTGGAGTCGTTGCCGTGCGTCTAATGGTTTCTGAGTNNNAGATCGGAAG |
| 11833_1 | TATGAGTGTGGAGTCGTTGCCCCTGAACTGTCTAAAAGTGAGTNNNAGATCGGAAG |
| 11837_1 | TATGAGTGTGGAGTCGTTGCAAATGCTCTAGGCAACGAGATGGNNNAGATCGGAA |
| 11839_1 | TATGAGTGTGGAGTCGTTGCATGGTCAAACCAGATGCACAGAANNNAGATCGGAA |
| 11840_1 | TATGAGTGTGGAGTCGTTGCTCCTGCCACAAATTCTTGTGAGTNNNAGATCGGAAG |
| 11841_1 | TATGAGTGTGGAGTCGTTGCACCAGAGTGTTGATGTTGTGAGTNNNAGATCGGAA |
| 11845_1 | TATGAGTGTGGAGTCGTTGCAAAACCTGCCAGATTGCTTGAGTNNNAGATCGGAAG |
| 11847_1 | TATGAGTGTGGAGTCGTTGCGACAGCATGTGCAATTGTTGAGTNNNAGATCGGAA |
| 11848_1 | TATGAGTGTGGAGTCGTTGCGTCAGCAACGTCCTACTTTGAGTNNNAGATCGGAAG |
| 11849_1 | TATGAGTGTGGAGTCGTTGCATTCCTCTGGTCTTCAGCCAGAANNNAGATCGGAAG |
| 11850_1 | TATGAGTGTGGAGTCGTTGCTTATGCATGTAAGTTCGCCAGAANNNAGATCGGAAG |
| 11851_1 | TATGAGTGTGGAGTCGTTGCGGATCATCAGCTGGGCAAATAGTNNNAGATCGGAA |
| 11857_1 | TATGAGTGTGGAGTCGTTGCTAAGCAGTGTTGGGACCCATAGTNNNAGATCGGAA |
| 11858_1 | TATGAGTGTGGAGTCGTTGCTCCCTTCCTTTCACCTGCATAGTNNNAGATCGGAAGA |
| 11859_1 | TATGAGTGTGGAGTCGTTGCATTCGCAGTTGATGGTGCATAGTNNNAGATCGGAA |
| 11861_1 | TATGAGTGTGGAGTCGTTGCTACCAGGGCAGGAATAGGCAGAANNNAGATCGGAA |
| 11862_1 | TATGAGTGTGGAGTCGTTGCTGAAGAAAGGTGTGGATGCAGAANNNAGATCGGA |
| 11863_1 | TATGAGTGTGGAGTCGTTGCCCTGTGTGCATGAATCTGCAGAANNNAGATCGGAA |
| 11864_1 | TATGAGTGTGGAGTCGTTGCCAAGAGAGCATGATTTCCCTCCCNNNAGATCGGAAG |
| 11865_1 | TATGAGTGTGGAGTCGTTGCGTCTCCCCAAGTCCCTATCAGAANNNAGATCGGAAG |
| 11867_1 | TATGAGTGTGGAGTCGTTGCATGCCTGATTCCCACAGACTAGTNNNAGATCGGAAG |
| 11868_1 | TATGAGTGTGGAGTCGTTGCACTTGCCAGCTTACAGGACTAGTNNNAGATCGGAAG |
| 11869_1 | TATGAGTGTGGAGTCGTTGCCATTCACACAGCCACTCTCAGAANNNAGATCGGAAG |
| 11870_1 | TATGAGTGTGGAGTCGTTGCGAGGTGACCATGAAGGACCTAGTNNNAGATCGGAA |
| 11871_1 | TATGAGTGTGGAGTCGTTGCCTGGCACTGCTTTACTGTCAGAANNNAGATCGGAAG |
| 11873_1 | TATGAGTGTGGAGTCGTTGCGGTTGTTCCTCCCTTCTCCTAGTNNNAGATCGGAAGA |
| 11874_1 | TATGAGTGTGGAGTCGTTGCAAAAAGGCCCAGCTTCTTCAGAANNNAGATCGGAAG |
| 11875_1 | TATGAGTGTGGAGTCGTTGCCTTGACCAGCAGAGCTTTCAGAANNNAGATCGGAAG |
| 11876_1 | TATGAGTGTGGAGTCGTTGCAGATCCTTACGGGCCAAAGAGAANNNAGATCGGAA |

| | |
|---|---|
| 11877_1 | TATGAGTGTGGAGTCGTTGCTTGGCCTTGATTGCTAAAGAGAANNNAGATCGGAA |
| 11878_1 | TATGAGTGTGGAGTCGTTGCAAGACCACTGCTACACTGCTAGTNNNAGATCGGAAG |
| 11880_1 | TATGAGTGTGGAGTCGTTGCGCTACCAAACACAGCACTCTAGTNNNAGATCGGAAG |
| 11881_1 | TATGAGTGTGGAGTCGTTGCCAAGCACAGACACGATCAGAGAANNNAGATCGGAA |
| 11882_1 | TATGAGTGTGGAGTCGTTGCCTGCTCTTGCACATTGGTCTAGTNNNAGATCGGAAG |
| 11884_1 | TATGAGTGTGGAGTCGTTGCAGACAAGAAGCTGGGAAAGTAGTNNNAGATCGGA |
| 11885_1 | TATGAGTGTGGAGTCGTTGCACCCACATAAGCCAGGTAGAGAANNNAGATCGGAA |
| 11886_1 | TATGAGTGTGGAGTCGTTGCCTGCTAAAAGAGACAGCCGAGAANNNAGATCGGAA |
| 11887_1 | TATGAGTGTGGAGTCGTTGCGGCACATGAAGACACAAGGAGAANNNAGATCGGA |
| 11888_1 | TATGAGTGTGGAGTCGTTGCTGGAAAGTGGGGAAGAAGGAGAANNNAGATCGG |
| 11889_1 | TATGAGTGTGGAGTCGTTGCGGATCAAAGCGAGATTCAGTGGTNNNAGATCGGAA |
| 11894_1 | TATGAGTGTGGAGTCGTTGCGACAGAGAGAGAGAGACTGTAGTNNNAGATCGGA |
| 11895_1 | TATGAGTGTGGAGTCGTTGCCTCAAACCCCAGAAAGAGATGCANNNAGATCGGAAG |
| 11897_1 | TATGAGTGTGGAGTCGTTGCCATCCCTTCATTTCGGTGCTCAGNNNAGATCGGAAG |
| 11899_1 | TATGAGTGTGGAGTCGTTGCAGGAGCTCAGAAAAGCCTGAGAANNNAGATCGGAA |
| 11900_1 | TATGAGTGTGGAGTCGTTGCTAGGGGTTGCAGTTAGCTGAGAANNNAGATCGGAA |
| 11909_1 | TATGAGTGTGGAGTCGTTGCCTCCTGGACTAGGGCCTCTTAGTNNNAGATCGGAAG |
| 11910_1 | TATGAGTGTGGAGTCGTTGCCACGCACGAAGGAGATCATAGAANNNAGATCGGAA |
| 11911_1 | TATGAGTGTGGAGTCGTTGCTGGGTGAAAGTCATTGTGTTAGTNNNAGATCGGAA |
| 11912_1 | TATGAGTGTGGAGTCGTTGCTGAGAATTGTGCGGCATATAGAANNNAGATCGGAA |
| 11915_1 | TATGAGTGTGGAGTCGTTGCGTCTGGTCTAGCTCCTGCTAGAANNNAGATCGGAAG |
| 11917_1 | TATGAGTGTGGAGTCGTTGCTGGGGCTACACACAGTTCTAGAANNNAGATCGGAAG |
| 11920_1 | TATGAGTGTGGAGTCGTTGCTTTCATTGTGGTGTGGAGAACGTNNNAGATCGGAA |
| 11921_1 | TATGAGTGTGGAGTCGTTGCGGGAGGGCAATCAAATAGAACGTNNNAGATCGGA |
| 11922_1 | TATGAGTGTGGAGTCGTTGCGCCCAAGCATCATCCCATTAGAANNNAGATCGGAAG |
| 11926_1 | TATGAGTGTGGAGTCGTTGCACAAAGGAGGAGGAGAGAACGAANNNAGATCGGA |
| 11930_1 | TATGAGTGTGGAGTCGTTGCAGGTAGTGCAGCATCAAAAGGAANNNAGATCGGAA |
| 11931_1 | TATGAGTGTGGAGTCGTTGCTAAAAACACCTTCTTCAGCCGGGNNNAGATCGGAAG |
| 11934_1 | TATGAGTGTGGAGTCGTTGCTCAGAAAGAACTGAGCACCCTCTNNNAGATCGGAAG |
| 11935_1 | TATGAGTGTGGAGTCGTTGCGGAAACAGAGCACACAGAAGGAANNNAGATCGGA |
| 11936_1 | TATGAGTGTGGAGTCGTTGCTTGGGATCCAGCAGTAGAAGGAANNNAGATCGGAA |
| 11938_1 | TATGAGTGTGGAGTCGTTGCAGCAAGCCTTAGGAATCTTACGTNNNAGATCGGAAG |
| 11940_1 | TATGAGTGTGGAGTCGTTGCATAAAAACCCAAGTGCCCAGGAANNNAGATCGGAA |
| 11941_1 | TATGAGTGTGGAGTCGTTGCAGTTTAGATAGGCTGGCCAGGAANNNAGATCGGAA |
| 11947_1 | TATGAGTGTGGAGTCGTTGCGCTTCAAATGTGGTCCAGAGGAANNNAGATCGGAA |
| 11948_1 | TATGAGTGTGGAGTCGTTGCTGAGCTGGGATGTTGTAGAGGAANNNAGATCGGA |
| 11952_1 | TATGAGTGTGGAGTCGTTGCCCCTCCACCTTCTTTCACGTTTTNNNAGATCGGAAGA |
| 11955_1 | TATGAGTGTGGAGTCGTTGCTTCCTTACCTTCCTCCATTCCGTNNNAGATCGGAAGA |
| 11956_1 | TATGAGTGTGGAGTCGTTGCGAAGACAAGAGTATTGGCAGCGTNNNAGATCGGAA |
| 11958_1 | TATGAGTGTGGAGTCGTTGCGGACCCCACAATTCCTCTAGGAANNNAGATCGGAAG |
| 11961_1 | TATGAGTGTGGAGTCGTTGCGGGGAAAGATGGGAGAGTAGGAANNNAGATCGG |
| 11963_1 | TATGAGTGTGGAGTCGTTGCTATGCCTTGAAATGTTGAGGCGTNNNAGATCGGAA |
| 11966_1 | TATGAGTGTGGAGTCGTTGCGGACATCACAGATGAAACCGGAANNNAGATCGGAA |

| | |
|---|---|
| 11968_1 | TATGAGTGTGGAGTCGTTGCAGAAATGAGGCATATTGTGGCGTNNNAGATCGGAA |
| 11969_1 | TATGAGTGTGGAGTCGTTGCACTGTGTGTACATGTGAATGCGTNNNAGATCGGAA |
| 11972_1 | TATGAGTGTGGAGTCGTTGCGTTACCACAAAAACTAGGCAACGNNNAGATCGGAAG |
| 11973_1 | TATGAGTGTGGAGTCGTTGCCGCTAAAGACAACACTCAGGGAANNNAGATCGGAA |
| 11974_1 | TATGAGTGTGGAGTCGTTGCGTGAGAGAGAGACACAGAGGGAANNNAGATCGGA |
| 11975_1 | TATGAGTGTGGAGTCGTTGCTGACAAGAGTAGCCATGAGGGAANNNAGATCGGA |
| 11976_1 | TATGAGTGTGGAGTCGTTGCTTTCTCAAGGCCTCACTAGGGAANNNAGATCGGAAG |
| 11978_1 | TATGAGTGTGGAGTCGTTGCTCCCTGACAAACATAGAGGGGAANNNAGATCGGAA |
| 11979_1 | TATGAGTGTGGAGTCGTTGCCATTCTCTGGGGGAATAGGGGAANNNAGATCGGAA |
| 11980_1 | TATGAGTGTGGAGTCGTTGCTAGGGTGAGCCTTGTGAGAGAAANNNAGATCGGA |
| 11981_1 | TATGAGTGTGGAGTCGTTGCTTTTCTCACATCTTGGCTGTCGTNNNAGATCGGAAG |
| 11982_1 | TATGAGTGTGGAGTCGTTGCCCCCACACATATTTCAGGCCTACNNNAGATCGGAAG |
| 11983_1 | TATGAGTGTGGAGTCGTTGCGCAACCTTAGAGGCAAATGGGAANNNAGATCGGAA |
| 11984_1 | TATGAGTGTGGAGTCGTTGCATGGAGCAGAGAGCATATGGGAANNNAGATCGGA |
| 11987_1 | TATGAGTGTGGAGTCGTTGCAAGCATTCAAGACACAGTGGGAANNNAGATCGGAA |
| 11988_1 | TATGAGTGTGGAGTCGTTGCACAAGGATGGGAACTAGTGGGAANNNAGATCGGA |
| 11989_1 | TATGAGTGTGGAGTCGTTGCGGGGGCAGTGGATATATTTTCGTNNNAGATCGGAA |
| 11990_1 | TATGAGTGTGGAGTCGTTGCACACACTGCTTTGTACTTTTCGTNNNAGATCGGAAG |
| 11991_1 | TATGAGTGTGGAGTCGTTGCCCAGGTTTATCAGCCCAAAAGGTNNNAGATCGGAAG |
| 11992_1 | TATGAGTGTGGAGTCGTTGCGGACTAACCCAGAAGTTGCAGTCNNNAGATCGGAA |
| 11993_1 | TATGAGTGTGGAGTCGTTGCCAGGGTCTGGAAAGTTTTGGGAANNNAGATCGGAA |
| 11994_1 | TATGAGTGTGGAGTCGTTGCTCTCTCCCCTCACATGAAAAGGTNNNAGATCGGAAG |
| 11995_1 | TATGAGTGTGGAGTCGTTGCGAGAGAGAGAGGGGTGAATGGAANNNAGATCGGA |
| 11997_1 | TATGAGTGTGGAGTCGTTGCGAGAGAGAAGTGTGCACAAAGGTNNNAGATCGGA |
| 11999_1 | TATGAGTGTGGAGTCGTTGCTCCTAACCCCCAGTTTCATGGAANNNAGATCGGAAG |
| 12001_1 | TATGAGTGTGGAGTCGTTGCTCAGTACAGGTAGGGTGATGGAANNNAGATCGGA |
| 12002_1 | TATGAGTGTGGAGTCGTTGCGCTGGCTTCCTTCCAATATGGAANNNAGATCGGAAG |
| 12003_1 | TATGAGTGTGGAGTCGTTGCGGGGATTTGAAGCTTGACTGGAANNNAGATCGGAA |
| 12004_1 | TATGAGTGTGGAGTCGTTGCAGGTCTTGTCTTGGCTACTGGAANNNAGATCGGAAG |
| 12005_1 | TATGAGTGTGGAGTCGTTGCACATGGAGTTCAAGCAACAAGGTNNNAGATCGGAA |
| 12007_1 | TATGAGTGTGGAGTCGTTGCTTTATTGGGAGAAGCTGGGTGCTNNNAGATCGGAA |
| 12008_1 | TATGAGTGTGGAGTCGTTGCAAGCCAACATCATTAGCCTGGAANNNAGATCGGAAG |
| 12009_1 | TATGAGTGTGGAGTCGTTGCTTCATTGAGACAACTGCCTGGAANNNAGATCGGAAG |
| 12011_1 | TATGAGTGTGGAGTCGTTGCTGCTATGTAATGTGGTGCTGGAANNNAGATCGGAA |
| 12016_1 | TATGAGTGTGGAGTCGTTGCGAGACAGAGAAGCCTTTCAAGGTNNNAGATCGGAA |
| 12017_1 | TATGAGTGTGGAGTCGTTGCTCTGAAGGCAAACTCAAGAAGGTNNNAGATCGGAA |
| 12019_1 | TATGAGTGTGGAGTCGTTGCTCATAGATGCCGCAAAATTGGAANNNAGATCGGAA |
| 12020_1 | TATGAGTGTGGAGTCGTTGCCCCCCTACACATTCCACTTGGAANNNAGATCGGAAG |
| 12021_1 | TATGAGTGTGGAGTCGTTGCGGAAGAGGCAACTTGACTTGGAANNNAGATCGGAA |
| 12022_1 | TATGAGTGTGGAGTCGTTGCAACCCACACACCAATACTTGGAANNNAGATCGGAAG |
| 12023_1 | TATGAGTGTGGAGTCGTTGCACACATCAACCTCCACGAAAGGTNNNAGATCGGAAG |
| 12025_1 | TATGAGTGTGGAGTCGTTGCCACCTACCTACCTCCTTGAAGGTNNNAGATCGGAAG |
| 12028_1 | TATGAGTGTGGAGTCGTTGCGGGAGACAATGGACAGTTTGGAANNNAGATCGGA |

| | |
|---|---|
| 12029_1 | TATGAGTGTGGAGTCGTTGCCCAGAGGACCACGATCAATTGTTNNNAGATCGGAAG |
| 12030_1 | TATGAGTGTGGAGTCGTTGCCGTGTTCTGAATGCCCAAATGAANNNAGATCGGAAG |
| 12033_1 | TATGAGTGTGGAGTCGTTGCGAGCAAGAGCAGACAGGAATGAANNNAGATCGGA |
| 12034_1 | TATGAGTGTGGAGTCGTTGCCTCACTCTGCGGAGAAACATGAANNNAGATCGGAAG |
| 12037_1 | TATGAGTGTGGAGTCGTTGCGCACATCTCTTCCAGAGCATGAANNNAGATCGGAAG |
| 12039_1 | TATGAGTGTGGAGTCGTTGCTCCCCCTCACAAGATCAAAAACANNNAGATCGGAAG |
| 12040_1 | TATGAGTGTGGAGTCGTTGCCTGTGACCCACACACATCATGAANNNAGATCGGAAG |
| 12041_1 | TATGAGTGTGGAGTCGTTGCTGCTTTTTGATGGCATACCAGGTNNNAGATCGGAAG |
| 12047_1 | TATGAGTGTGGAGTCGTTGCTGGAAAACGACACAGTGGATGAANNNAGATCGGAA |
| 12052_1 | TATGAGTGTGGAGTCGTTGCTGGTGGCCAGAGATGAGTATGAANNNAGATCGGAA |
| 12053_1 | TATGAGTGTGGAGTCGTTGCGGAAGCAGGTTTGGTGGTATGAANNNAGATCGGA |
| 12054_1 | TATGAGTGTGGAGTCGTTGCTCAATTTCTCCAGGAAAGCAGGTNNNAGATCGGAAG |
| 12055_1 | TATGAGTGTGGAGTCGTTGCACAATCTCCAAATCGCAGAGCTTNNNAGATCGGAAG |
| 12056_1 | TATGAGTGTGGAGTCGTTGCCAGGAAAAGAATGAGAGGCAGGTNNNAGATCGGA |
| 12057_1 | TATGAGTGTGGAGTCGTTGCTGGGGAGGAGAAAGAAGAACTGGNNNAGATCGGA |
| 12060_1 | TATGAGTGTGGAGTCGTTGCGATTTTTGAGCTTGCATGCAGGTNNNAGATCGGAA |
| 12062_1 | TATGAGTGTGGAGTCGTTGCCAGGCCAACCAAATGAATCAGGTNNNAGATCGGAA |
| 12064_1 | TATGAGTGTGGAGTCGTTGCTATTTCAGGAAACGGCTACTGAANNNAGATCGGAA |
| 12065_1 | TATGAGTGTGGAGTCGTTGCCCAAGGCTAGAATCAGGCATTTCNNNAGATCGGAAG |
| 12066_1 | TATGAGTGTGGAGTCGTTGCAATCACCCAGAATCCTCTCAGGTNNNAGATCGGAAG |
| 12067_1 | TATGAGTGTGGAGTCGTTGCTGCCCAGATAGACTTGACCTGAANNNAGATCGGAAG |
| 12069_1 | TATGAGTGTGGAGTCGTTGCATGTCTGAAGGCAAATGCCTGAANNNAGATCGGAA |
| 12070_1 | TATGAGTGTGGAGTCGTTGCGGTTGCCAAGAAGGATTCTGTCTNNNAGATCGGAA |
| 12075_1 | TATGAGTGTGGAGTCGTTGCACTTGTGGCTAGTGAAAGCTGAANNNAGATCGGAA |
| 12076_1 | TATGAGTGTGGAGTCGTTGCAAATCTAGAGACCCTGCAGAGGTNNNAGATCGGAA |
| 12077_1 | TATGAGTGTGGAGTCGTTGCACATTGAGTAAGCTCTGGAAAGTNNNAGATCGGAA |
| 12078_1 | TATGAGTGTGGAGTCGTTGCGAGGAGCATGTAAACCAGCTGAANNNAGATCGGAA |
| 12082_1 | TATGAGTGTGGAGTCGTTGCCAGCTACGTTTGCATGATAACGCNNNAGATCGGAAG |
| 12085_1 | TATGAGTGTGGAGTCGTTGCGGTCTTGCTGTGTAGATGCTGAANNNAGATCGGAA |
| 12086_1 | TATGAGTGTGGAGTCGTTGCGCTCAGGAGTGTCTTCTAGAGGTNNNAGATCGGAA |
| 12087_1 | TATGAGTGTGGAGTCGTTGCACTGAAGGAAGGGCATTAGAGGTNNNAGATCGGA |
| 12088_1 | TATGAGTGTGGAGTCGTTGCGAGAGGTTCAGCTGATTGCTGAANNNAGATCGGAA |
| 12089_1 | TATGAGTGTGGAGTCGTTGCCAACAAACCAATCGTAACGAGGTNNNAGATCGGAAG |
| 12091_1 | TATGAGTGTGGAGTCGTTGCCTAAAAGGAGCGCCAACTCTGAANNNAGATCGGAA |
| 12092_1 | TATGAGTGTGGAGTCGTTGCTAGCCAAGAGCTTAGCCTCTGAANNNAGATCGGAAG |
| 12093_1 | TATGAGTGTGGAGTCGTTGCACATGCTCACACAAAGCTCTGAANNNAGATCGGAAG |
| 12094_1 | TATGAGTGTGGAGTCGTTGCCGCACAACCTCAGATTCTCTGAANNNAGATCGGAAG |
| 12095_1 | TATGAGTGTGGAGTCGTTGCACATAGAGCAAAGGGAAGGAGGTNNNAGATCGGA |
| 12097_1 | TATGAGTGTGGAGTCGTTGCAAGGGGGAAAGGATGAGAGGAAGNNNAGATCGG |
| 12098_1 | TATGAGTGTGGAGTCGTTGCGCTTTGTAGGGAGTTAGGGAGGTNNNAGATCGGA |
| 12099_1 | TATGAGTGTGGAGTCGTTGCAAGGCATGAATAAGAGGGGAGGTNNNAGATCGGA |
| 12100_1 | TATGAGTGTGGAGTCGTTGCTACACAGGATACGTGGGAACACTNNNAGATCGGAA |
| 12101_1 | TATGAGTGTGGAGTCGTTGCCTGCATCCTCCAAAGTCAGTGAANNNAGATCGGAAG |

| | |
|---|---|
| 12102_1 | TATGAGTGTGGAGTCGTTGCAAAAGGCTGCAGTCGAGAGTGAANNNAGATCGGA |
| 12104_1 | TATGAGTGTGGAGTCGTTGCGACAGAGAACTGGACAGGAGGAGNNNAGATCGGA |
| 12108_1 | TATGAGTGTGGAGTCGTTGCTGGAGGGTGAAGAGAAAGGTGAANNNAGATCGGA |
| 12110_1 | TATGAGTGTGGAGTCGTTGCCTCCCCATCACCCTAAGACCTGTNNNAGATCGGAAGA |
| 12112_1 | TATGAGTGTGGAGTCGTTGCAAGGTCACTCTGCTTACTGAGGTNNNAGATCGGAAG |
| 12113_1 | TATGAGTGTGGAGTCGTTGCCCCATGAAACAAGGCTCTCTCTCNNNAGATCGGAAG |
| 12116_1 | TATGAGTGTGGAGTCGTTGCATAGCCTCTTCCCACATGGTGAANNNAGATCGGAAG |
| 12117_1 | TATGAGTGTGGAGTCGTTGCTTTACTAACCAGCAGGGTGAGGTNNNAGATCGGAA |
| 12120_1 | TATGAGTGTGGAGTCGTTGCAGAACGTACACCCCTTCTGTGATNNNAGATCGGAAG |
| 12122_1 | TATGAGTGTGGAGTCGTTGCATACTACCCAGCTCGGATGTGAANNNAGATCGGAAG |
| 12125_1 | TATGAGTGTGGAGTCGTTGCAGATAGATCATGGGGGCTGTGAANNNAGATCGGA |
| 12127_1 | TATGAGTGTGGAGTCGTTGCTTCTGGTTGGGGTTTGTTGTGAANNNAGATCGGAA |
| 12129_1 | TATGAGTGTGGAGTCGTTGCAGATTGTTGGGTGCAAGATAGGTNNNAGATCGGAA |
| 12130_1 | TATGAGTGTGGAGTCGTTGCCACCATTGGCAATCAGCATTGAANNNAGATCGGAAG |
| 12132_1 | TATGAGTGTGGAGTCGTTGCACTCCTGCTGAGAACAACTTGAANNNAGATCGGAAG |
| 12134_1 | TATGAGTGTGGAGTCGTTGCGGGAAAACCCACTTATTCCTCCANNNAGATCGGAAG |
| 12136_1 | TATGAGTGTGGAGTCGTTGCAGATGTGACATGGTGCTCTTGAANNNAGATCGGAA |
| 12141_1 | TATGAGTGTGGAGTCGTTGCGCACAGAGCCATCCTGATTTGAANNNAGATCGGAAG |
| 12142_1 | TATGAGTGTGGAGTCGTTGCACCACAACCACCACAACTTTGAANNNAGATCGGAAG |
| 12143_1 | TATGAGTGTGGAGTCGTTGCCCACTCACCGTTCTCACTTTGAANNNAGATCGGAAGA |
| 12144_1 | TATGAGTGTGGAGTCGTTGCTAACTGTTGTTCCCCGCTTTGAANNNAGATCGGAAG |
| 12145_1 | TATGAGTGTGGAGTCGTTGCCCCAAAGATCCTCCTGCTTTGAANNNAGATCGGAAG |
| 12148_1 | TATGAGTGTGGAGTCGTTGCATCTGCTTTTGGCATTGGTAGGTNNNAGATCGGAAG |
| 12151_1 | TATGAGTGTGGAGTCGTTGCTGAAAAGGGGCAATTTTGTAGGTNNNAGATCGGAA |
| 12155_1 | TATGAGTGTGGAGTCGTTGCTGCTACCTTTACCTGCCTTAGGTNNNAGATCGGAAG |
| 12156_1 | TATGAGTGTGGAGTCGTTGCGCAAGCACACCTGAGAGCAATAANNNAGATCGGAA |
| 12157_1 | TATGAGTGTGGAGTCGTTGCGCCTCTTGGGTAGGTTGCAATAANNNAGATCGGAA |
| 12158_1 | TATGAGTGTGGAGTCGTTGCGGATGAGGAGACAGACTGTGAGTNNNAGATCGGA |
| 12159_1 | TATGAGTGTGGAGTCGTTGCGCTAGAATGCGGTGGCTGAATAANNNAGATCGGAA |
| 12162_1 | TATGAGTGTGGAGTCGTTGCTGCTCTATGATGTCTCTTCCGGTNNNAGATCGGAAG |
| 12163_1 | TATGAGTGTGGAGTCGTTGCAGGCATCACCCTGGACTATTGTTNNNAGATCGGAAG |
| 12165_1 | TATGAGTGTGGAGTCGTTGCAAACAAGGTTACAAAATTGCGGTNNNAGATCGGAA |
| 12166_1 | TATGAGTGTGGAGTCGTTGCCAGTGGTAGCTTTCGAGGCATAANNNAGATCGGAA |
| 12167_1 | TATGAGTGTGGAGTCGTTGCCTCATCAATTGTGTCCCTTCGGTNNNAGATCGGAAG |
| 12169_1 | TATGAGTGTGGAGTCGTTGCTGATGGATCCTGGGGCAAGATAANNNAGATCGGAA |
| 12171_1 | TATGAGTGTGGAGTCGTTGCAGGCACTGGGGGTATGAAGATAANNNAGATCGGA |
| 12173_1 | TATGAGTGTGGAGTCGTTGCTGGAGGAAGGCTAAGGCAGATAANNNAGATCGGA |
| 12175_1 | TATGAGTGTGGAGTCGTTGCGCCCTTGCCATTGTCAGAGATAANNNAGATCGGAAG |
| 12176_1 | TATGAGTGTGGAGTCGTTGCATTGTGGGAAGGGCAGGAGATAANNNAGATCGGA |
| 12177_1 | TATGAGTGTGGAGTCGTTGCGGCATGGGGAGAGATGGAGATAANNNAGATCGGA |
| 12178_1 | TATGAGTGTGGAGTCGTTGCTGGAAAAGAGACAAGCTTTGGCTNNNAGATCGGAA |
| 12179_1 | TATGAGTGTGGAGTCGTTGCAGGTTTTGACTGTGAGGAAGGGTNNNAGATCGGA |
| 12185_1 | TATGAGTGTGGAGTCGTTGCACACAGCCTAAGCAGAGGGATAANNNAGATCGGAA |

| | |
|---|---|
| 12186_1 | TATGAGTGTGGAGTCGTTGCGAACTCAGGGGATCCTGGGATAANNNAGATCGGAA |
| 12189_1 | TATGAGTGTGGAGTCGTTGCTAGAACCAGTGACTTTCCAGGGTNNNAGATCGGAAG |
| 12190_1 | TATGAGTGTGGAGTCGTTGCGGTCCAGAAAAGGCCACTGATAANNNAGATCGGAA |
| 12191_1 | TATGAGTGTGGAGTCGTTGCAACAGTGGTCAGGGTCCTGATAANNNAGATCGGAA |
| 12193_1 | TATGAGTGTGGAGTCGTTGCGGATCTGCTGGGGTGGACTATAANNNAGATCGGAA |
| 12196_1 | TATGAGTGTGGAGTCGTTGCAGGATGGATGTGAGATCAAGCAGNNNAGATCGGA |
| 12197_1 | TATGAGTGTGGAGTCGTTGCAGGGTCTGCTGCTTCTCCTATAANNNAGATCGGAAG |
| 12199_1 | TATGAGTGTGGAGTCGTTGCTAAATGCAGGTCACTCCTAGGGTNNNAGATCGGAAG |
| 12200_1 | TATGAGTGTGGAGTCGTTGCCCCCTTCAGCTCCTTGAAACTAANNNAGATCGGAAGA |
| 12201_1 | TATGAGTGTGGAGTCGTTGCAGCGTGTGTGTGACATAGAGAGANNNAGATCGGA |
| 12202_1 | TATGAGTGTGGAGTCGTTGCGAGATGCTGGTGGCATCAACTAANNNAGATCGGAA |
| 12204_1 | TATGAGTGTGGAGTCGTTGCAGCAAACAGGACTGAATCACTAANNNAGATCGGAA |
| 12206_1 | TATGAGTGTGGAGTCGTTGCTGAAAATTCTGTGAAGTCCGGGTNNNAGATCGGAA |
| 12207_1 | TATGAGTGTGGAGTCGTTGCTAGAATTTTGTGGGTGGACACGGNNNAGATCGGAA |
| 12209_1 | TATGAGTGTGGAGTCGTTGCGCTTTCCTCCAAGGCCTTACTAANNNAGATCGGAAG |
| 12211_1 | TATGAGTGTGGAGTCGTTGCAAACCTTGAGGCAATGCACCTAANNNAGATCGGAAG |
| 12212_1 | TATGAGTGTGGAGTCGTTGCTGAAACCCCTTCTTCCAAAAAGANNNAGATCGGAAG |
| 12213_1 | TATGAGTGTGGAGTCGTTGCCCCGCGTTCCTTTGAATACCTAANNNAGATCGGAAG |
| 12214_1 | TATGAGTGTGGAGTCGTTGCTTAGACGTTCAGTTACCAGGGGTNNNAGATCGGAA |
| 12218_1 | TATGAGTGTGGAGTCGTTGCGGCAAAGGGTGGTAATTCCAGAANNNAGATCGGAA |
| 12221_1 | TATGAGTGTGGAGTCGTTGCGGTTATTTATCACTTCCGGCTGCNNNAGATCGGAAG |
| 12222_1 | TATGAGTGTGGAGTCGTTGCGAATGGGGAGCTTTTCTGCCTAANNNAGATCGGAA |
| 12223_1 | TATGAGTGTGGAGTCGTTGCTCTCTACAGTCAGCCTTGCCTAANNNAGATCGGAAG |
| 12224_1 | TATGAGTGTGGAGTCGTTGCGACACATGCACACACACTCCTAANNNAGATCGGAAG |
| 12225_1 | TATGAGTGTGGAGTCGTTGCGCTTGTGGCTAACACAGTCCTAANNNAGATCGGAAG |
| 12228_1 | TATGAGTGTGGAGTCGTTGCCACGGATTGCTTGCAAAAGCTAANNNAGATCGGAAG |
| 12229_1 | TATGAGTGTGGAGTCGTTGCTCAGCTTGGGCAAAGAAAGCTAANNNAGATCGGAA |
| 12230_1 | TATGAGTGTGGAGTCGTTGCCCCTGTGGAGTAGGAGAAGCTAANNNAGATCGGAA |
| 12233_1 | TATGAGTGTGGAGTCGTTGCCAAAATCTGACACGGGAGGCTAANNNAGATCGGAA |
| 12234_1 | TATGAGTGTGGAGTCGTTGCGGTAATGAGGGGAGAAGGGCTAANNNAGATCGGA |
| 12235_1 | TATGAGTGTGGAGTCGTTGCCAGGAACCAAGAAGCATGGCTAANNNAGATCGGAA |
| 12237_1 | TATGAGTGTGGAGTCGTTGCTCCTCACCAGTCCTCAATGCTAANNNAGATCGGAAGA |
| 12239_1 | TATGAGTGTGGAGTCGTTGCATCGAGTGGGACAGATATGCTAANNNAGATCGGAA |
| 12240_1 | TATGAGTGTGGAGTCGTTGCGTGGAAATATTAGGTCACTGGGTNNNAGATCGGAA |
| 12241_1 | TATGAGTGTGGAGTCGTTGCACAGAAAAGAAGGCCTACTGGGTNNNAGATCGGAA |
| 12242_1 | TATGAGTGTGGAGTCGTTGCCACAGCCAGTCAGAACTTGCTAANNNAGATCGGAAG |
| 12243_1 | TATGAGTGTGGAGTCGTTGCGCTGCGTACACTTGGAACTCTAANNNAGATCGGAAG |
| 12244_1 | TATGAGTGTGGAGTCGTTGCAGGCATCTTGACTTCACAACAGNNNAGATCGGAAGA |
| 12245_1 | TATGAGTGTGGAGTCGTTGCGTATAAAGTGAACTGGGCTGGGTNNNAGATCGGAA |
| 12246_1 | TATGAGTGTGGAGTCGTTGCGAGATAGCCATTTTTGGCTGGGTNNNAGATCGGAA |
| 12247_1 | TATGAGTGTGGAGTCGTTGCGGAAGGGAGAGAGAAATCTGGGTNNNAGATCGGA |
| 12248_1 | TATGAGTGTGGAGTCGTTGCAGAGAGAGAGGGAGACGCTCTAANNNAGATCGGA |
| 12249_1 | TATGAGTGTGGAGTCGTTGCACTGACAGAAGCTGCCTCTCTAANNNAGATCGGAAG |

| | |
|---|---|
| 12250_1 | TATGAGTGTGGAGTCGTTGCCTGTCTGTCTATTTGGGTGGGTGNNNAGATCGGAA |
| 12253_1 | TATGAGTGTGGAGTCGTTGCAGCCACCTTTGCTTTGGTTCTAANNNAGATCGGAAG |
| 12254_1 | TATGAGTGTGGAGTCGTTGCAGAGGGGTCTTTGAGTTTTCTAANNNAGATCGGAAG |
| 12255_1 | TATGAGTGTGGAGTCGTTGCTCACTCCTTTACCCACTGCATTCNNNAGATCGGAAGA |
| 12257_1 | TATGAGTGTGGAGTCGTTGCTATTTCATGCATGTGTGGTGGGTNNNAGATCGGAAG |
| 12258_1 | TATGAGTGTGGAGTCGTTGCAAAGAGAGAGACAGTCTGTGGGTNNNAGATCGGA |
| 12259_1 | TATGAGTGTGGAGTCGTTGCGTCCTGTCATGGCACTGAAGTAANNNAGATCGGAAG |
| 12260_1 | TATGAGTGTGGAGTCGTTGCGGCATCATCCTTGCTCACAGTAANNNAGATCGGAAG |
| 12261_1 | TATGAGTGTGGAGTCGTTGCTGCTGGGCAACTAAAAGCAGTAANNNAGATCGGAA |
| 12263_1 | TATGAGTGTGGAGTCGTTGCGGAGGGAAGAAGCTGGAGAGTAANNNAGATCGGA |
| 12264_1 | TATGAGTGTGGAGTCGTTGCGCGTCTCTGTCAGTGGAGAGTAANNNAGATCGGAA |
| 12265_1 | TATGAGTGTGGAGTCGTTGCTTCTCTTGCCCAGACTGGAGTAANNNAGATCGGAAG |
| 12266_1 | TATGAGTGTGGAGTCGTTGCTCTGCCCATTACATCTTTTGGGTNNNAGATCGGAAG |
| 12271_1 | TATGAGTGTGGAGTCGTTGCGAAAAGCGAGTAACAGCAACACCNNNAGATCGGAA |
| 12272_1 | TATGAGTGTGGAGTCGTTGCGGGGCACAGTTCCTAAATCGTAANNNAGATCGGAAG |
| 12273_1 | TATGAGTGTGGAGTCGTTGCGGCTTTTGTAGGAGGGAAATGGTNNNAGATCGGA |
| 12277_1 | TATGAGTGTGGAGTCGTTGCCCATCCTGGTTTGAGAAGCAGAANNNAGATCGGAAG |
| 12278_1 | TATGAGTGTGGAGTCGTTGCTGGCTGTGAGCTGTACTAGGTAANNNAGATCGGAA |
| 12280_1 | TATGAGTGTGGAGTCGTTGCACAGACAGACAGACAAGGGGTAANNNAGATCGGA |
| 12281_1 | TATGAGTGTGGAGTCGTTGCTTTTGGTCCCCGAGTTTGTTTTTNNNAGATCGGAAG |
| 12283_1 | TATGAGTGTGGAGTCGTTGCAGAGAGTTGGTTTGTGTGGGTAANNNAGATCGGA |
| 12284_1 | TATGAGTGTGGAGTCGTTGCGGAGGGGTGGAGAATTTGGGTAANNNAGATCGGA |
| 12285_1 | TATGAGTGTGGAGTCGTTGCAGACTTTTCTGCCTCCCTGGTAANNNAGATCGGAAG |
| 12287_1 | TATGAGTGTGGAGTCGTTGCAGATGGAGCCTGTTGGTGTTTAGNNNAGATCGGAA |
| 12289_1 | TATGAGTGTGGAGTCGTTGCGAGGGATGTCCCAGGTTTGGTAANNNAGATCGGAA |
| 12290_1 | TATGAGTGTGGAGTCGTTGCTCCAAAGCCCAGATTTCATGTAANNNAGATCGGAAG |
| 12294_1 | TATGAGTGTGGAGTCGTTGCGTTTGGGATGCTGAGAAGATGGTNNNAGATCGGA |
| 12295_1 | TATGAGTGTGGAGTCGTTGCTCACCCCAATACCCCAACTGTAANNNAGATCGGAAGA |
| 12297_1 | TATGAGTGTGGAGTCGTTGCTCTCAGGTAGGGCAACACTGTAANNNAGATCGGAAG |
| 12299_1 | TATGAGTGTGGAGTCGTTGCATTCTTGTTGGCCAAGCCTGTAANNNAGATCGGAAG |
| 12303_1 | TATGAGTGTGGAGTCGTTGCCCTGCACAATACCACAGGTGTAANNNAGATCGGAAG |
| 12304_1 | TATGAGTGTGGAGTCGTTGCGTGCTCTCCGTATGTGTGTGTAANNNAGATCGGAAG |
| 12305_1 | TATGAGTGTGGAGTCGTTGCCAGGAATGGTGGTGCACTTGTAANNNAGATCGGAA |
| 12306_1 | TATGAGTGTGGAGTCGTTGCAGAGAGAGACAAAAGCTGATGGTNNNAGATCGGA |
| 12307_1 | TATGAGTGTGGAGTCGTTGCCTGCTGATGGTTCCCAGTTGTAANNNAGATCGGAAG |
| 12309_1 | TATGAGTGTGGAGTCGTTGCGTGTAGGTGTTGATGGTTGTGGTNNNAGATCGGAA |
| 12312_1 | TATGAGTGTGGAGTCGTTGCCTGCAAAGACAGACCTTGATGGTNNNAGATCGGAAG |
| 12313_1 | TATGAGTGTGGAGTCGTTGCTTTGAGGATGCCACTTTGATGGTNNNAGATCGGAAG |
| 12315_1 | TATGAGTGTGGAGTCGTTGCCTCTACCTCCCCAGGCTCACATCNNNAGATCGGAAGA |
| 12316_1 | TATGAGTGTGGAGTCGTTGCAGTAGATGGGGCCTTGATATGGTNNNAGATCGGAA |
| 12319_1 | TATGAGTGTGGAGTCGTTGCAGAATGGGGCTTGGTACTATGGTNNNAGATCGGAA |
| 12323_1 | TATGAGTGTGGAGTCGTTGCCTGAGGCCAAAAGGTGTTATGGTNNNAGATCGGAA |
| 12326_1 | TATGAGTGTGGAGTCGTTGCGCTGAGGATCACTTTCAACTGGTNNNAGATCGGAAG |

| | |
|---|---|
| 12327_1 | TATGAGTGTGGAGTCGTTGCACAGCCTTCCTCGAGACACACATNNNAGATCGGAAG |
| 12328_1 | TATGAGTGTGGAGTCGTTGCCCGTTTAGGTGTCCAGACATTGANNNAGATCGGAAG |
| 12330_1 | TATGAGTGTGGAGTCGTTGCGGGACTAAAGATGTATCACTGGTNNNAGATCGGAA |
| 12331_1 | TATGAGTGTGGAGTCGTTGCCCAGAAGCAAAACTCTCACTGGTNNNAGATCGGAAG |
| 12332_1 | TATGAGTGTGGAGTCGTTGCAGTACCCAACTCACTTCACTGGTNNNAGATCGGAAG |
| 12333_1 | TATGAGTGTGGAGTCGTTGCCTACCCCCTCAAGAGTATCACACNNNAGATCGGAAG |
| 12334_1 | TATGAGTGTGGAGTCGTTGCACTGGTACAGAGAAGAACCTGGTNNNAGATCGGAA |
| 12335_1 | TATGAGTGTGGAGTCGTTGCCCCTTCCTACCTCATAGCACCTTNNNAGATCGGAAGA |
| 12336_1 | TATGAGTGTGGAGTCGTTGCGGAAGGACGGGTTACAGAGGAGANNNAGATCGGA |
| 12337_1 | TATGAGTGTGGAGTCGTTGCATCCTTCTGTTCTTTCCCCTGGTNNNAGATCGGAAGA |
| 12338_1 | TATGAGTGTGGAGTCGTTGCTGTCATAGACTGATGAGCCTGGTNNNAGATCGGAA |
| 12339_1 | TATGAGTGTGGAGTCGTTGCAATGACATCCCCTGAACATTGGTNNNAGATCGGAAG |
| 12345_1 | TATGAGTGTGGAGTCGTTGCGAGCCCATCAAAAGACATCTGGTNNNAGATCGGAA |
| 12353_1 | TATGAGTGTGGAGTCGTTGCGCCCTGGTCCAAACATAAAAACANNNAGATCGGAAG |
| 12357_1 | TATGAGTGTGGAGTCGTTGCGAATGCACTTGAGGAACAAAACANNNAGATCGGAA |
| 12361_1 | TATGAGTGTGGAGTCGTTGCCCCTACCTCAAAAGGGCAAAACANNNAGATCGGAAG |
| 12363_1 | TATGAGTGTGGAGTCGTTGCGAAAATGCCACAGAGTGAGTGGTNNNAGATCGGAA |
| 12365_1 | TATGAGTGTGGAGTCGTTGCAGCAGCCACTCAGAAAGAAAACANNNAGATCGGAA |
| 12367_1 | TATGAGTGTGGAGTCGTTGCGAGTATGTATGTGTGCACGTGGTNNNAGATCGGAA |
| 12369_1 | TATGAGTGTGGAGTCGTTGCCCAGTCTAAAACAGATGCAGGCTNNNAGATCGGAA |
| 12373_1 | TATGAGTGTGGAGTCGTTGCAGGTAGAAGCTAAGGGTGCTGCANNNAGATCGGA |
| 12374_1 | TATGAGTGTGGAGTCGTTGCTTTGTGATGAGATTGAGGGTGGTNNNAGATCGGAA |
| 12375_1 | TATGAGTGTGGAGTCGTTGCAGTGAGGATGGGAACATGATGGANNNAGATCGGA |
| 12379_1 | TATGAGTGTGGAGTCGTTGCACCTCCATTTGCTTTCCCAAACANNNAGATCGGAAGA |
| 12380_1 | TATGAGTGTGGAGTCGTTGCGCAGTCTTTGAGATCACTGTGGTNNNAGATCGGAA |
| 12381_1 | TATGAGTGTGGAGTCGTTGCAGTGTTTGTCGGTAGAGCAAACANNNAGATCGGAA |
| 12383_1 | TATGAGTGTGGAGTCGTTGCGAAAGAGGATGTCCAGGCAAACANNNAGATCGGAA |
| 12385_1 | TATGAGTGTGGAGTCGTTGCTGAATGCCTAATGTGTGCAAACANNNAGATCGGAA |
| 12387_1 | TATGAGTGTGGAGTCGTTGCAGAGAGAGAGACTGGCTCAAACANNNAGATCGGAA |
| 12391_1 | TATGAGTGTGGAGTCGTTGCTGCTGTAACTTCTGGTTCAAACANNNAGATCGGAAG |
| 12394_1 | TATGAGTGTGGAGTCGTTGCATGCCTGTGTATGTGTTTGTGGTNNNAGATCGGAA |
| 12395_1 | TATGAGTGTGGAGTCGTTGCTTCTTCCCATCTTCTGGTCTCCCNNNAGATCGGAAGA |
| 12397_1 | TATGAGTGTGGAGTCGTTGCCTTGCTTTCCCTTGGGAATTGGTNNNAGATCGGAAG |
| 12399_1 | TATGAGTGTGGAGTCGTTGCTAGCCCAAATGGAAATGCTTGGTNNNAGATCGGAA |
| 12401_1 | TATGAGTGTGGAGTCGTTGCAGATGAGCATGCCAAAGATTGGTNNNAGATCGGAA |
| 12405_1 | TATGAGTGTGGAGTCGTTGCAACAGAGCAAAGGCAATATTGGTNNNAGATCGGAA |
| 12407_1 | TATGAGTGTGGAGTCGTTGCGGACTAGGAGAGGGAATCTTGGTNNNAGATCGGA |
| 12408_1 | TATGAGTGTGGAGTCGTTGCTCTGCCTGTCTCTAGTTATTGGTNNNAGATCGGAAG |
| 12409_1 | TATGAGTGTGGAGTCGTTGCGCATGTTCTCTCTGAACCTTGGTNNNAGATCGGAAG |
| 12412_1 | TATGAGTGTGGAGTCGTTGCGTGAAGTCTTGGAGAAGCTGGAGNNNAGATCGGA |
| 53437_1 | TATGAGTGTGGAGTCGTTGCAGTACACGATTGGTTACAGTCCCNNNAGATCGGAAG |
| 53439_1 | TATGAGTGTGGAGTCGTTGCCTCTCTCTCTCTCTCTCTCTCGGNNNAGATCGGAAGA |
| 53440_1 | TATGAGTGTGGAGTCGTTGCGAGATGGAGTCTTACTCTTGCCCNNNAGATCGGAAG |

| | |
|---|---|
| 53441_1 | TATGAGTGTGGAGTCGTTGCCTTTCTCCAAGTATTGAGAGCGCNNNAGATCGGAAG |
| 53442_1 | TATGAGTGTGGAGTCGTTGCTACAAGACTGTGTGTGTGTGTGTNNNAGATCGGAAG |
| 53443_1 | TATGAGTGTGGAGTCGTTGCTCTTGTTGCTTATCAGGGATGCTNNNAGATCGGAAG |
| 53445_1 | TATGAGTGTGGAGTCGTTGCAGACAGAGAGACAGAGACAGACANNNAGATCGGAA |
| 53446_1 | TATGAGTGTGGAGTCGTTGCCTCCCTCTCTCCCTCTCTCTCTCNNNAGATCGGAAGAG |
| 53447_1 | TATGAGTGTGGAGTCGTTGCCACACGTGTAATTCCAGCACTTTNNNAGATCGGAAGA |
| 53449_1 | TATGAGTGTGGAGTCGTTGCTGTGTGTGTGTGTGTAGAGAGAGNNNAGATCGGAA |
| 53450_1 | TATGAGTGTGGAGTCGTTGCTGCAGTTTCTGTCTGTCTCTCTCNNNAGATCGGAAGA |
| 53451_1 | TATGAGTGTGGAGTCGTTGCCCTCTCTCTCTCTCTCTCTCTCGNNNAGATCGGAAGAG |
| 53452_1 | TATGAGTGTGGAGTCGTTGCATCTCTCCTTGAATATGCACCCCNNNAGATCGGAAGA |
| 53453_1 | TATGAGTGTGGAGTCGTTGCCTTGGAGTTCTTTCAGGCAACTGNNNAGATCGGAAG |
| 53454_1 | TATGAGTGTGGAGTCGTTGCGCTTGGAAACACTGGTGTGATTCNNNAGATCGGAAG |
| 53455_1 | TATGAGTGTGGAGTCGTTGCGAAATGGAAATGAGCCGAGATCGNNNAGATCGGAA |
| 53456_1 | TATGAGTGTGGAGTCGTTGCATGATCCATACTCAGCCCTCAAGNNNAGATCGGAAGA |
| 53457_1 | TATGAGTGTGGAGTCGTTGCTCCTCTCTCTCTCTCTCTCTTCCNNNAGATCGGAAGAG |
| 53458_1 | TATGAGTGTGGAGTCGTTGCCTGGTTTTGAAACACCCAGGATGNNNAGATCGGAAG |
| 53459_1 | TATGAGTGTGGAGTCGTTGCCCAGTCACTCCAATTCTGAGTCTNNNAGATCGGAAGA |
| 53460_1 | TATGAGTGTGGAGTCGTTGCAGGCAGTACTAGGTTTGCAAGTTNNNAGATCGGAAG |
| 53463_1 | TATGAGTGTGGAGTCGTTGCGACTGCGGTCTAGGAAAAAGTCTNNNAGATCGGAAG |
| 53465_1 | TATGAGTGTGGAGTCGTTGCAGGGAGATAGCTTCAAGACCTGANNNAGATCGGAAG |
| 53467_1 | TATGAGTGTGGAGTCGTTGCGGAGAGAAAATGCCTAGACAGCTNNNAGATCGGAA |
| 53468_1 | TATGAGTGTGGAGTCGTTGCTGTAGTCCCAGGTACTCAAGAGGNNNAGATCGGAAG |
| 53472_1 | TATGAGTGTGGAGTCGTTGCCCCAACATTCACAGGGTCTTTTCNNNAGATCGGAAGA |
| 53473_1 | TATGAGTGTGGAGTCGTTGCTTCCCCAAAGAACTGCTTCCATANNNAGATCGGAAGA |
| 53474_1 | TATGAGTGTGGAGTCGTTGCAAGAAAACAGCAATTGGTCAGGCNNNAGATCGGAAG |
| 53475_1 | TATGAGTGTGGAGTCGTTGCGAAAGCATCACTTGAGGTCAGGANNNAGATCGGAAG |
| 53478_1 | TATGAGTGTGGAGTCGTTGCGTGCAGAGAAATAATCACGGCTCNNNAGATCGGAAG |
| 53479_1 | TATGAGTGTGGAGTCGTTGCGCTGAGAAGCTGAGGATGAATCCNNNAGATCGGAA |
| 53483_1 | TATGAGTGTGGAGTCGTTGCAAATTAGTCAGGCGTAGTGGTGGNNNAGATCGGAA |
| 53484_1 | TATGAGTGTGGAGTCGTTGCGCCACGTGCAACTGTTATACTAANNNAGATCGGAAG |
| 53486_1 | TATGAGTGTGGAGTCGTTGCTGCTGGAAGAGTAGGTGTTTCTCNNNAGATCGGAAG |
| 53487_1 | TATGAGTGTGGAGTCGTTGCGATTGCAGCTCTTTTGCCATCATNNNAGATCGGAAGA |
| 53489_1 | TATGAGTGTGGAGTCGTTGCTAGCTACTTGGGAGGTTGAGGTANNNAGATCGGAA |
| 53490_1 | TATGAGTGTGGAGTCGTTGCAAACAGGCCGAGCATTTAGAAACNNNAGATCGGAAG |
| 53491_1 | TATGAGTGTGGAGTCGTTGCGCTCCTGTAATCCTTGCTACTCANNNAGATCGGAAGA |
| 53492_1 | TATGAGTGTGGAGTCGTTGCCTAGAGTGCAGTGGTGTGATCTCNNNAGATCGGAAG |
| 53493_1 | TATGAGTGTGGAGTCGTTGCCCTCCCTCTCTCTCTCTCTTTCTNNNAGATCGGAAGAG |
| 53494_1 | TATGAGTGTGGAGTCGTTGCTCCTGAATCCTGAAGGTGATTGGNNNAGATCGGAAG |
| 53495_1 | TATGAGTGTGGAGTCGTTGCTCTTTCTCTCTCCCTTCCTTCCTNNNAGATCGGAAGAG |
| 53498_1 | TATGAGTGTGGAGTCGTTGCTCCCAGAGAACAATGAAGGTTCCNNNAGATCGGAAG |
| 53500_1 | TATGAGTGTGGAGTCGTTGCATATTCCACAACAGTACTGGGGGNNNAGATCGGAAG |
| 53501_1 | TATGAGTGTGGAGTCGTTGCCATGTGTGTGACATTCTCAGGACNNNAGATCGGAAG |
| 53502_1 | TATGAGTGTGGAGTCGTTGCAGGATTTCTTGAGCCCAGAAAGTNNNAGATCGGAAG |

| | |
|---|---|
| 53503_1 | TATGAGTGTGGAGTCGTTGCATGTTGGTTAGGCTGGTCTCAAANNNAGATCGGAA |
| 53504_1 | TATGAGTGTGGAGTCGTTGCGAATGAATGGCATTTGCAGCAACNNNAGATCGGAA |
| 53507_1 | TATGAGTGTGGAGTCGTTGCATATAGATAAAGTGGGCCGCCTGNNNAGATCGGAA |
| 53508_1 | TATGAGTGTGGAGTCGTTGCTCTCAAACTCCTGACCTCAAGTGNNNAGATCGGAAG |
| 53509_1 | TATGAGTGTGGAGTCGTTGCGAACAGAGAGAACAAACCTGGGANNNAGATCGGA |
| 53511_1 | TATGAGTGTGGAGTCGTTGCTCTCCCTCTCTCTCTCTCTCTNNNAGATCGGAAGAG |
| 53512_1 | TATGAGTGTGGAGTCGTTGCTAACTGGTCTGGCTGTTCTGTACNNNAGATCGGAAG |
| 53513_1 | TATGAGTGTGGAGTCGTTGCCAGGGAGAAATCCACTGAAGGAANNNAGATCGGAA |
| 53514_1 | TATGAGTGTGGAGTCGTTGCACACACACACACCATACTCTGTTNNNAGATCGGAAG |
| 53516_1 | TATGAGTGTGGAGTCGTTGCAGGCAGGAGAATAATGTGAACCCNNNAGATCGGAA |
| 53517_1 | TATGAGTGTGGAGTCGTTGCGAAAGGGTCTCACTCTGTCACTTNNNAGATCGGAAG |
| 53518_1 | TATGAGTGTGGAGTCGTTGCGGGTCACCATTTGCAAGTAAGTTNNNAGATCGGAA |
| 53519_1 | TATGAGTGTGGAGTCGTTGCTTCATGGTGAATGTCGATGCAAANNNAGATCGGAA |
| 53522_1 | TATGAGTGTGGAGTCGTTGCACCAAGAAGCCTTTAGTTACCCANNNAGATCGGAAG |
| 53523_1 | TATGAGTGTGGAGTCGTTGCAGGATCTTGATGTGTTGCCTAGGNNNAGATCGGAA |
| 53524_1 | TATGAGTGTGGAGTCGTTGCGTCAATGCCTTTATTGGGTCCAGNNNAGATCGGAAG |
| 53527_1 | TATGAGTGTGGAGTCGTTGCATATTGGGGATGGTGTGACACAANNNAGATCGGAA |
| 53528_1 | TATGAGTGTGGAGTCGTTGCAGAAGATTGCCTTGAGAGAAGCANNNAGATCGGAA |
| 53529_1 | TATGAGTGTGGAGTCGTTGCGCTCTGGGGATAGAAAGACTAGCNNNAGATCGGAA |
| 53530_1 | TATGAGTGTGGAGTCGTTGCACCACAAGGAACTATGGGATGTCNNNAGATCGGAA |
| 53531_1 | TATGAGTGTGGAGTCGTTGCGACTAGGTAGTTCTCCAGGGGATNNNAGATCGGAA |
| 53532_1 | TATGAGTGTGGAGTCGTTGCGCATAGCGGCACATACCTATAGANNNAGATCGGAA |
| 53533_1 | TATGAGTGTGGAGTCGTTGCTGCTACATGATCTCTCTCCTGGANNNAGATCGGAAG |
| 53534_1 | TATGAGTGTGGAGTCGTTGCTAAAGTTGGGGGAAACTAGCCAANNNAGATCGGAA |
| 53535_1 | TATGAGTGTGGAGTCGTTGCAAAGTCCAGAAGACACCAAGGTTNNNAGATCGGAA |
| 53536_1 | TATGAGTGTGGAGTCGTTGCCCTCTACTGCTATGTGGCTTCAANNNAGATCGGAAG |
| 53537_1 | TATGAGTGTGGAGTCGTTGCGGTGAGGTTGAGTGAATGTCTCTNNNAGATCGGAA |
| 53538_1 | TATGAGTGTGGAGTCGTTGCTGCTGGGATTACAGGTGTGATTANNNAGATCGGAA |
| 53539_1 | TATGAGTGTGGAGTCGTTGCTGTTTCTCAATCAAGAGGCCAGANNNAGATCGGAAG |
| 53542_1 | TATGAGTGTGGAGTCGTTGCGACTTTAGATCTTTGAGCCCCCANNNAGATCGGAAG |
| 53543_1 | TATGAGTGTGGAGTCGTTGCTCTCTCAACAGGAATAGCCTTGCNNNAGATCGGAAG |
| 53544_1 | TATGAGTGTGGAGTCGTTGCGGGGGCTCTCAATACAAACAAACNNNAGATCGGAA |
| 53545_1 | TATGAGTGTGGAGTCGTTGCCTCTCCACAGCCATAAACAACATNNNAGATCGGAAG |
| 53546_1 | TATGAGTGTGGAGTCGTTGCAAAAAGGCCTGCAGATAACCCTANNNAGATCGGAA |
| 53547_1 | TATGAGTGTGGAGTCGTTGCGAGGCACCTGGTGTTAGTTTTTCNNNAGATCGGAAG |
| 53548_1 | TATGAGTGTGGAGTCGTTGCCGCCTATAATCCCAGCTACTCAGNNNAGATCGGAAG |
| 53549_1 | TATGAGTGTGGAGTCGTTGCCTGTGATCTCAGCTACTTGGGAGNNNAGATCGGAA |
| 53550_1 | TATGAGTGTGGAGTCGTTGCTAGCATGTTCCTGTAGTCTCAGCNNNAGATCGGAAG |
| 53551_1 | TATGAGTGTGGAGTCGTTGCGGGGATACCAAGCAAAAACACTTNNNAGATCGGAA |
| 53552_1 | TATGAGTGTGGAGTCGTTGCCACCCCACATTTATTGAACAGCANNNAGATCGGAAG |
| 53554_1 | TATGAGTGTGGAGTCGTTGCGGAAACCTTTGGAAACCGAGAACNNNAGATCGGAA |
| 53555_1 | TATGAGTGTGGAGTCGTTGCCAGGAAGGTCTTGATCTCCTGACNNNAGATCGGAAG |
| 53556_1 | TATGAGTGTGGAGTCGTTGCCATGGCTGTTTATTTCACCTGGGNNNAGATCGGAAG |

53557_1 TATGAGTGTGGAGTCGTTGCGTAAGGTACTTGACTCACCTGCANNNAGATCGGAAG
53558_1 TATGAGTGTGGAGTCGTTGCAGATTTTGTGTCTGGTGAGGGTTNNNAGATCGGAA
53559_1 TATGAGTGTGGAGTCGTTGCCAGGGTCCACTCCATTCTAGTTGNNNAGATCGGAAG
53561_1 TATGAGTGTGGAGTCGTTGCGAGAGAGAGAGGACGCAGAAAAANNNAGATCGGA
53563_1 TATGAGTGTGGAGTCGTTGCTGTGTGTGTGTAATGCTCCTCATNNNAGATCGGAAG
53564_1 TATGAGTGTGGAGTCGTTGCCACACACACACACACACACATATNNNAGATCGGAAG
53566_1 TATGAGTGTGGAGTCGTTGCTCTCTCTCTCTCTCTCTCTCCCTNNNAGATCGGAAGAG
53568_1 TATGAGTGTGGAGTCGTTGCCAGCTCAGGATTGGGACAATTTCNNNAGATCGGAA
53569_1 TATGAGTGTGGAGTCGTTGCTTTCTTTAGTCCTGCTCTGTGGCNNNAGATCGGAAG
53570_1 TATGAGTGTGGAGTCGTTGCAAAGGAGAGTCAAAGAGGCAGAGNNNAGATCGGA
53571_1 TATGAGTGTGGAGTCGTTGCAACAAGGATAAGACAAGCAAGGANNNAGATCGGA
53572_1 TATGAGTGTGGAGTCGTTGCCAAAAAGTCAGCACCTTGGACAGNNNAGATCGGAA
53573_1 TATGAGTGTGGAGTCGTTGCGCCTGTAGTCCCAACACTAGTTTNNNAGATCGGAAG
53574_1 TATGAGTGTGGAGTCGTTGCGCCTTGCTTTCCCTTCACTTTTTNNNAGATCGGAAGA
53576_1 TATGAGTGTGGAGTCGTTGCGTCTCGATCTCCTGACCTTGTAANNNAGATCGGAAG
53578_1 TATGAGTGTGGAGTCGTTGCTCTTTCCCTGAGACTGGAGTACANNNAGATCGGAAG
53581_1 TATGAGTGTGGAGTCGTTGCGTTCAAGCGATTTCCGGCTAATTNNNAGATCGGAAG
53582_1 TATGAGTGTGGAGTCGTTGCCCCGAGTAGCTAGGATTACAAGCNNNAGATCGGAA
53583_1 TATGAGTGTGGAGTCGTTGCCTGGGTTCAAGTGATTCTTCTGCNNNAGATCGGAAG
53584_1 TATGAGTGTGGAGTCGTTGCTAGGCAGTTTTGCAAGCAGAAATNNNAGATCGGAA
53586_1 TATGAGTGTGGAGTCGTTGCTACAATCTCAGCTCATGGCAACTNNNAGATCGGAAG
53588_1 TATGAGTGTGGAGTCGTTGCTGAGCCAGTGTGAGTTACTATGGNNNAGATCGGAA
53589_1 TATGAGTGTGGAGTCGTTGCAGGTTCAAGTGATTCTTCTGCCTNNNAGATCGGAAG
53591_1 TATGAGTGTGGAGTCGTTGCGCTGCTTGGTTATCTGCTGAAAANNNAGATCGGAAG
53592_1 TATGAGTGTGGAGTCGTTGCGTCCTCTCTGAGACACAAGTCAGNNNAGATCGGAAG
53594_1 TATGAGTGTGGAGTCGTTGCTCTGGTCCAAAACTCCTAAGCTCNNNAGATCGGAAG
53596_1 TATGAGTGTGGAGTCGTTGCCTGAGTAGCTGGGATTACAGGTGNNNAGATCGGAA
53601_1 TATGAGTGTGGAGTCGTTGCATGCAAAAGAAAGGAGGCTCTTGNNNAGATCGGAA
53603_1 TATGAGTGTGGAGTCGTTGCCAATCTCAGCTCACTGCAAACTCNNNAGATCGGAAG
53604_1 TATGAGTGTGGAGTCGTTGCCAGCCACGTGTAAAAGAACCATTNNNAGATCGGAAG
53607_1 TATGAGTGTGGAGTCGTTGCTGAAGAGATTCCCCCTGGAGTATNNNAGATCGGAA
53608_1 TATGAGTGTGGAGTCGTTGCAAGGAAGGAACGAACGAACAAACNNNAGATCGGA
53609_1 TATGAGTGTGGAGTCGTTGCTGCTGGCAACCTGAAATTCTATGNNNAGATCGGAAG
53611_1 TATGAGTGTGGAGTCGTTGCAGGTAAGGCAATCACTTGAACCTNNNAGATCGGAA
53612_1 TATGAGTGTGGAGTCGTTGCGTTCCAAGGGCAATCAGAATTCCNNNAGATCGGAAG
53615_1 TATGAGTGTGGAGTCGTTGCGATCAAGACCAGACTGGGCAATANNNAGATCGGAA
53617_1 TATGAGTGTGGAGTCGTTGCAAAAAACTAGAAAGGCGTGGTGCNNNAGATCGGAA
53619_1 TATGAGTGTGGAGTCGTTGCAAATGTGCAAATGCCAAGTGTCANNNAGATCGGAA
53620_1 TATGAGTGTGGAGTCGTTGCTAGAGGCCAGACTGTGGTAGTAANNNAGATCGGAA
53621_1 TATGAGTGTGGAGTCGTTGCATATGAGCAAGGTGAGCTTAGGGNNNAGATCGGA
53623_1 TATGAGTGTGGAGTCGTTGCTAATCCTAACACTTTGGGAGGCCNNNAGATCGGAAG
53624_1 TATGAGTGTGGAGTCGTTGCTAAGGCATTAGGAAACCCTGGACNNNAGATCGGAA
53626_1 TATGAGTGTGGAGTCGTTGCTGAAGAACATTGCTTCTCTGGGANNNAGATCGGAA

| | |
|---|---|
| 53627_1 | TATGAGTGTGGAGTCGTTGCACTAGCCATGAGAAAAGGGTGTTNNNAGATCGGAA |
| 53629_1 | TATGAGTGTGGAGTCGTTGCCCACTACGCCTGGCTAATTTTTTNNNAGATCGGAAG |
| 53630_1 | TATGAGTGTGGAGTCGTTGCCATTCTCCTGCCTCAGCCTTCNNNAGATCGGAAGAGC |
| 53631_1 | TATGAGTGTGGAGTCGTTGCAGGCCTACATTGCTGCTATTCTTNNNAGATCGGAAG |
| 53632_1 | TATGAGTGTGGAGTCGTTGCCATGATCATGGCTTACTGCAACCNNNAGATCGGAAG |
| 53633_1 | TATGAGTGTGGAGTCGTTGCCCAGGATGGTCTTGATCACTTGANNNAGATCGGAAG |
| 53634_1 | TATGAGTGTGGAGTCGTTGCTTGAAGTGATTCTCCTGCCTCAGNNNAGATCGGAAG |
| 53635_1 | TATGAGTGTGGAGTCGTTGCCTGAAGTGATCCACCTACCTCAGNNNAGATCGGAAG |
| 53637_1 | TATGAGTGTGGAGTCGTTGCGAGAACCCTGTAAGAAGGCAGAANNNAGATCGGAA |
| 53639_1 | TATGAGTGTGGAGTCGTTGCGTTCAAGCAATTCTCTGCCTCAGNNNAGATCGGAAG |
| 53640_1 | TATGAGTGTGGAGTCGTTGCATCACTGCTCCATACAACCTCTGNNNAGATCGGAAGA |
| 53641_1 | TATGAGTGTGGAGTCGTTGCCAGCCTCCCAAGTAGGTGTAATTNNNAGATCGGAAG |
| 53644_1 | TATGAGTGTGGAGTCGTTGCAGACAGAATCTCACTGTGTCACCNNNAGATCGGAAG |
| 53647_1 | TATGAGTGTGGAGTCGTTGCCCTCCTTGGTTCAAGCAATTCTGNNNAGATCGGAAG |
| 53648_1 | TATGAGTGTGGAGTCGTTGCCAATGGCAAAATCTCAGCTCACTNNNAGATCGGAAG |
| 53649_1 | TATGAGTGTGGAGTCGTTGCGCTAGAGTATAGTGGTGGCTCAGNNNAGATCGGAA |
| 53651_1 | TATGAGTGTGGAGTCGTTGCTTCCAGGTTCACACCATTCTCCNNNAGATCGGAAGAG |
| 53653_1 | TATGAGTGTGGAGTCGTTGCAATCTTGGCTCACTACAACCTCCNNNAGATCGGAAGA |
| 53656_1 | TATGAGTGTGGAGTCGTTGCATACACCAGTCTGATGGGAACACNNNAGATCGGAAG |
| 53660_1 | TATGAGTGTGGAGTCGTTGCGGACCACAGACTACACTGAAAGANNNAGATCGGAA |
| 53661_1 | TATGAGTGTGGAGTCGTTGCTGCCTGTAATTCCAGCTACTCAGNNNAGATCGGAAG |
| 53663_1 | TATGAGTGTGGAGTCGTTGCATGCTACTTGGTTCAGAGCGTAANNNAGATCGGAAG |
| 53664_1 | TATGAGTGTGGAGTCGTTGCATAAGGAGTATGGCCAGACATGGNNNAGATCGGAA |
| 53667_1 | TATGAGTGTGGAGTCGTTGCCAGGGTTTCTGGATACCACTGTTNNNAGATCGGAAG |
| 53668_1 | TATGAGTGTGGAGTCGTTGCCCATCCTAAGAACTACGGAGCAANNNAGATCGGAAG |
| 53670_1 | TATGAGTGTGGAGTCGTTGCTCTGTGACCTCAAACTGACTGAGNNNAGATCGGAAG |
| 53672_1 | TATGAGTGTGGAGTCGTTGCGTGGAAGGATTGTTTGAACCCAGNNNAGATCGGAA |
| 53673_1 | TATGAGTGTGGAGTCGTTGCGTGAGACACCAGACAGGATTCATNNNAGATCGGAA |
| 53675_1 | TATGAGTGTGGAGTCGTTGCTGGGCAGAAAAGGATGTAACTGANNNAGATCGGAA |
| 53677_1 | TATGAGTGTGGAGTCGTTGCAACAGGTGTTGACAAAAGTGTGGNNNAGATCGGAA |
| 53678_1 | TATGAGTGTGGAGTCGTTGCAAGAGGTGTTGAGGTCTTTTGGTNNNAGATCGGAA |
| 53681_1 | TATGAGTGTGGAGTCGTTGCTACAACCACTGTGGGAAACTAGCNNNAGATCGGAAG |
| 53683_1 | TATGAGTGTGGAGTCGTTGCGCAAACCAAATCCAACAACACATNNNAGATCGGAAG |
| 53685_1 | TATGAGTGTGGAGTCGTTGCTTCCCCTCTCCTTATCCATGACTNNNAGATCGGAAGA |
| 53688_1 | TATGAGTGTGGAGTCGTTGCGATATGGACTCCCAGGCTATGTGNNNAGATCGGAA |
| 53689_1 | TATGAGTGTGGAGTCGTTGCGAAGATGCTGTGATGATGGAAGCNNNAGATCGGAA |
| 53690_1 | TATGAGTGTGGAGTCGTTGCCTGCCAGTAAAGCTTTGTGTTGTNNNAGATCGGAAG |
| 53691_1 | TATGAGTGTGGAGTCGTTGCGCCCCAGTAAATTCACTTTCCACNNNAGATCGGAAGA |
| 53692_1 | TATGAGTGTGGAGTCGTTGCATAGTATAACGCAGGGCTAACGGNNNAGATCGGAA |
| 53693_1 | TATGAGTGTGGAGTCGTTGCATTTGCCAGTCGACCTTGAGTATNNNAGATCGGAAG |
| 53694_1 | TATGAGTGTGGAGTCGTTGCGTCTGCATTTTTGGATGGTGACANNNAGATCGGAAG |
| 53696_1 | TATGAGTGTGGAGTCGTTGCCATGGAAAGTAGCTATGCCAAGCNNNAGATCGGAA |
| 53698_1 | TATGAGTGTGGAGTCGTTGCTGGCTCACATCTGTAATCCTAGCNNNAGATCGGAAG |

| | |
|---|---|
| 53700_1 | TATGAGTGTGGAGTCGTTGCAGACAAAGTCCATTGGAACCACCNNNAGATCGGAAG |
| 53704_1 | TATGAGTGTGGAGTCGTTGCCAAACAATAGGGGCATTCTTGGGNNNAGATCGGAA |
| 53705_1 | TATGAGTGTGGAGTCGTTGCTGCCAAATATCTCCAGCACTGATNNNAGATCGGAAG |
| 53706_1 | TATGAGTGTGGAGTCGTTGCCTCAAGTGATTCTCCTGCCTCAGNNNAGATCGGAAG |
| 53707_1 | TATGAGTGTGGAGTCGTTGCCCTCCTGGGTTCAAGTGATTCTCNNNAGATCGGAAG |
| 53708_1 | TATGAGTGTGGAGTCGTTGCATCTCCCAGGTTCAAGTGATTCCNNNAGATCGGAAG |
| 53710_1 | TATGAGTGTGGAGTCGTTGCGAGTGCAATGACATGATCTCAGCNNNAGATCGGAA |
| 53711_1 | TATGAGTGTGGAGTCGTTGCATGATCTTGGCTCACTGAAACCTNNNAGATCGGAAG |
| 53712_1 | TATGAGTGTGGAGTCGTTGCGAGTAGTTTCACCCTTGTCACCCNNNAGATCGGAAG |
| 53713_1 | TATGAGTGTGGAGTCGTTGCTTCCTCCTGGGTTCAAGCAATTCNNNAGATCGGAAG |
| 53716_1 | TATGAGTGTGGAGTCGTTGCCTACCGAGTAGCTGGGACTACAANNNAGATCGGAA |
| 53720_1 | TATGAGTGTGGAGTCGTTGCAAGCAATCTGGGATTACAGGCATNNNAGATCGGAA |
| 53722_1 | TATGAGTGTGGAGTCGTTGCTCCTGGTTCAAGCAATTCTCCTTNNNAGATCGGAAGA |
| 53723_1 | TATGAGTGTGGAGTCGTTGCGAGAAGAGAAGTTCTAGGCTGGGNNNAGATCGGA |
| 53724_1 | TATGAGTGTGGAGTCGTTGCGAAAACAAAAAGCACCACACTGTNNNAGATCGGAA |
| 53725_1 | TATGAGTGTGGAGTCGTTGCGTGTAATCACAGCACTTTGGGAGNNNAGATCGGAA |
| 53726_1 | TATGAGTGTGGAGTCGTTGCTGAAAGTGACCTTAGAGATGGCCNNNAGATCGGAA |
| 53728_1 | TATGAGTGTGGAGTCGTTGCATTCTGACTCAGGAACCTTCTGGNNNAGATCGGAAG |
| 53729_1 | TATGAGTGTGGAGTCGTTGCTTCAAGCAACTCTCCTGCCTTAGNNNAGATCGGAAG |
| 53731_1 | TATGAGTGTGGAGTCGTTGCGGTAGACAGGACTATGTGTGGTCNNNAGATCGGAA |
| 53732_1 | TATGAGTGTGGAGTCGTTGCACTGGGTTCTTTGGTCGTAGTTTNNNAGATCGGAAG |
| 53733_1 | TATGAGTGTGGAGTCGTTGCTGATTTCATAGGTCTGGATGGGCNNNAGATCGGAA |
| 53734_1 | TATGAGTGTGGAGTCGTTGCGAGGTCAGGAGATCAAGACCATCNNNAGATCGGAA |
| 53736_1 | TATGAGTGTGGAGTCGTTGCTAAAAACAGCACTAGCCTGAGGTNNNAGATCGGAA |
| 53738_1 | TATGAGTGTGGAGTCGTTGCAAAATTAGCTGGGTGTGATGGTGNNNAGATCGGAA |
| 53739_1 | TATGAGTGTGGAGTCGTTGCTTTGAATCCAGCTGTGTGACCTTNNNAGATCGGAAG |
| 53740_1 | TATGAGTGTGGAGTCGTTGCATATATATGGCTGGGCATGGTGGNNNAGATCGGAA |
| 53741_1 | TATGAGTGTGGAGTCGTTGCTCACTGATCGTTAGGGAAATGCANNNAGATCGGAA |
| 53743_1 | TATGAGTGTGGAGTCGTTGCTGAATCCACAGATTGCTTTGAGCNNNAGATCGGAAG |
| 53744_1 | TATGAGTGTGGAGTCGTTGCGAGATGGTGTTTCGCTCTTGTTGNNNAGATCGGAA |
| 53745_1 | TATGAGTGTGGAGTCGTTGCGGCTTCAGTGAAATGGCATGATTNNNAGATCGGAA |
| 53746_1 | TATGAGTGTGGAGTCGTTGCGGTTCACGCCATTTTCCTACTTCNNNAGATCGGAAGA |
| 53747_1 | TATGAGTGTGGAGTCGTTGCTTCAAGTGATTCTCCTGCCTCAGNNNAGATCGGAAG |
| 53748_1 | TATGAGTGTGGAGTCGTTGCCGTGCTGGAATTACAGGCATAAGNNNAGATCGGAA |
| 53749_1 | TATGAGTGTGGAGTCGTTGCAGAGTCTTGTTATGTCACCCAGGNNNAGATCGGAAG |
| 53750_1 | TATGAGTGTGGAGTCGTTGCCTTCCCAGATTCAAGCGATTCTCNNNAGATCGGAAG |
| 53751_1 | TATGAGTGTGGAGTCGTTGCCTCCTGAGTAGCTGGGACTACAGNNNAGATCGGAA |
| 53753_1 | TATGAGTGTGGAGTCGTTGCGATTACAGGCACACACCAACATGNNNAGATCGGAAG |
| 53754_1 | TATGAGTGTGGAGTCGTTGCCTGAGTAGCTGGGACTACATGTGNNNAGATCGGAA |
| 53755_1 | TATGAGTGTGGAGTCGTTGCAGGTTCAAGCAATTCTCCTGTCTNNNAGATCGGAAG |
| 53757_1 | TATGAGTGTGGAGTCGTTGCACTAGAGACGGGGTTTTACCATGNNNAGATCGGAA |
| 53759_1 | TATGAGTGTGGAGTCGTTGCGTTCAAGCGATTCTCCTACCTCANNNAGATCGGAAG |
| 53760_1 | TATGAGTGTGGAGTCGTTGCGTAGCTGGGACTACAGGCATATGNNNAGATCGGAA |

| | |
|---|---|
| 53761_1 | TATGAGTGTGGAGTCGTTGCGCAGTAGGGCAATCTCAGAATCTNNNAGATCGGAA |
| 53762_1 | TATGAGTGTGGAGTCGTTGCAAGTTCAAGGGATTCTCGTGCTCNNNAGATCGGAAG |
| 53763_1 | TATGAGTGTGGAGTCGTTGCTTCAAGTGATTCTCCTGCCTCAGNNNAGATCGGAAG |
| 53764_1 | TATGAGTGTGGAGTCGTTGCCCTCAGCCTTCCTAGTAGTAGGANNNAGATCGGAAG |
| 53765_1 | TATGAGTGTGGAGTCGTTGCCTTCCTGGGTTCAAGCAATTCTCNNNAGATCGGAAG |
| 53766_1 | TATGAGTGTGGAGTCGTTGCGACTACAGGCCTGGGACTACAGNNNAGATCGGAAG |
| 53767_1 | TATGAGTGTGGAGTCGTTGCTTCCCAGGTTCAAGTGATTCTCCNNNAGATCGGAAG |
| 53768_1 | TATGAGTGTGGAGTCGTTGCTCAATGGCATGATCTTTGCTCACNNNAGATCGGAAG |
| 53772_1 | TATGAGTGTGGAGTCGTTGCCCAAGCACAAAGACCACATATGGNNNAGATCGGAA |
| 53775_1 | TATGAGTGTGGAGTCGTTGCCATGAGAATTGCTTGAACCTGGGNNNAGATCGGAA |
| 53778_1 | TATGAGTGTGGAGTCGTTGCGCTGTTCCAAATCACTGTGAGACNNNAGATCGGAAG |
| 53783_1 | TATGAGTGTGGAGTCGTTGCCTGTAGTCCTAGCACTTTGGGAGNNNAGATCGGAA |
| 53784_1 | TATGAGTGTGGAGTCGTTGCTCAACTTCAACCACCTTCTCCAANNNAGATCGGAAGA |
| 53785_1 | TATGAGTGTGGAGTCGTTGCAAAAGGCTAAAGGAATGGCAACANNNAGATCGGAA |
| 53786_1 | TATGAGTGTGGAGTCGTTGCCAACCTGACCAACATGGAGAAACNNNAGATCGGAA |
| 53787_1 | TATGAGTGTGGAGTCGTTGCCACACCTGTAATCCCAGCACTTTNNNAGATCGGAAGA |
| 53789_1 | TATGAGTGTGGAGTCGTTGCGATCCAGGGAGCCAGTTAGTTTCNNNAGATCGGAA |
| 53790_1 | TATGAGTGTGGAGTCGTTGCGATGTCTGGGTCAAATGCTTGTCNNNAGATCGGAA |
| 53792_1 | TATGAGTGTGGAGTCGTTGCAGACCTTTTTCAAGGCTTTGTCCNNNAGATCGGAAG |
| 53794_1 | TATGAGTGTGGAGTCGTTGCTGATGTGAGGAGTAAGCCCATTGNNNAGATCGGAA |
| 53795_1 | TATGAGTGTGGAGTCGTTGCCAGGTTCAATGAGGGTTTGCTTTNNNAGATCGGAAG |
| 53796_1 | TATGAGTGTGGAGTCGTTGCCACACCTGTAATCCCAGCACTTTNNNAGATCGGAAGA |
| 53797_1 | TATGAGTGTGGAGTCGTTGCAGCAGTGGATCCAAACAAGATGANNNAGATCGGAA |
| 53800_1 | TATGAGTGTGGAGTCGTTGCGGTTCAAGCTATCCTCCCATCTCNNNAGATCGGAAGA |
| 53801_1 | TATGAGTGTGGAGTCGTTGCGATGGGGTTTTACCATGTTGTCCNNNAGATCGGAAG |
| 53802_1 | TATGAGTGTGGAGTCGTTGCGTGTCTCTTCATGTGAGATGGGTNNNAGATCGGAA |
| 53803_1 | TATGAGTGTGGAGTCGTTGCTCTCCTGGGTTCAAGTGATTCTCNNNAGATCGGAAG |
| 53804_1 | TATGAGTGTGGAGTCGTTGCCTCCCAAGTAGCTGGAACTACAGNNNAGATCGGAAG |
| 53805_1 | TATGAGTGTGGAGTCGTTGCTGCCTACTTTGGGAAAAACTGAGNNNAGATCGGAA |
| 53807_1 | TATGAGTGTGGAGTCGTTGCGCCTCCTGAATAGCTGAGACTACNNNAGATCGGAAG |
| 53809_1 | TATGAGTGTGGAGTCGTTGCAATCTCGGCTCACTGCAACATCNNNAGATCGGAAGA |
| 53810_1 | TATGAGTGTGGAGTCGTTGCCCTCCTGGGTTCAAGTGATTCTTNNNAGATCGGAAG |
| 53813_1 | TATGAGTGTGGAGTCGTTGCACTTCCCGGATTCAAGTGATTCTNNNAGATCGGAAG |
| 53814_1 | TATGAGTGTGGAGTCGTTGCGAGTAGCTGAGACTACAGGTGTGNNNAGATCGGAA |
| 53815_1 | TATGAGTGTGGAGTCGTTGCAGTAGTTGGGATTAGAGGCATGCNNNAGATCGGAA |
| 53816_1 | TATGAGTGTGGAGTCGTTGCGTTTCACCATGTTAGCCAGGATGNNNAGATCGGAAG |
| 53817_1 | TATGAGTGTGGAGTCGTTGCAGAGTCTTGCTCAGTCATCCAAGNNNAGATCGGAAG |
| 53818_1 | TATGAGTGTGGAGTCGTTGCCCTCCTGGGTTCAAGTGATTCTCNNNAGATCGGAAG |
| 53819_1 | TATGAGTGTGGAGTCGTTGCGAGATGGAGTCTTGTTCTGTTGCNNNAGATCGGAA |
| 53822_1 | TATGAGTGTGGAGTCGTTGCAGTCTGTAGAGGAAATGCATTGTNNNAGATCGGAA |
| 53824_1 | TATGAGTGTGGAGTCGTTGCTCAATCTCTTGACCTCGTGATCCNNNAGATCGGAAGA |
| 53826_1 | TATGAGTGTGGAGTCGTTGCGGCTAACATCTGGTGAGTGTGTANNNAGATCGGAA |
| 53827_1 | TATGAGTGTGGAGTCGTTGCGAGAAGGGGTTTTGCCATATTGCNNNAGATCGGAA |

| | |
|---|---|
| 53829_1 | TATGAGTGTGGAGTCGTTGCGCTGTTGGACAGAAAAGCAAGAANNNAGATCGGAA |
| 53830_1 | TATGAGTGTGGAGTCGTTGCTCCGCGGCACTTTTCTTTAGATANNNAGATCGGAAG |
| 53831_1 | TATGAGTGTGGAGTCGTTGCGTTCTTCCCATGGATCAGGCTATNNNAGATCGGAAG |
| 53832_1 | TATGAGTGTGGAGTCGTTGCTTCAAGTGATTCTCCTGCCTCAGNNNAGATCGGAAG |
| 53833_1 | TATGAGTGTGGAGTCGTTGCCAATCTCAGCTCACTGCAAACTCNNNAGATCGGAAG |
| 53834_1 | TATGAGTGTGGAGTCGTTGCAGAATCAGCTGAATCCCATTCCANNNAGATCGGAAG |
| 53835_1 | TATGAGTGTGGAGTCGTTGCTCCTAACCTGGGGTAGTCTAAGGNNNAGATCGGAA |
| 53836_1 | TATGAGTGTGGAGTCGTTGCGAGCTGAGATTGTGGTACATTGCNNNAGATCGGAA |
| 53837_1 | TATGAGTGTGGAGTCGTTGCAGTCGACTCATTTCAGTGACCATNNNAGATCGGAAG |
| 53839_1 | TATGAGTGTGGAGTCGTTGCCCGAAGTAAAATACTGACGGGGANNNAGATCGGAA |
| 53840_1 | TATGAGTGTGGAGTCGTTGCTAACTGGTGGCCATTGATAAGCTNNNAGATCGGAA |
| 53841_1 | TATGAGTGTGGAGTCGTTGCCTGACAGAAACCTGAAATCCACANNNAGATCGGAAG |
| 53842_1 | TATGAGTGTGGAGTCGTTGCACTTCTCTGGGTCTTGGTTTTCTNNNAGATCGGAAG |
| 53843_1 | TATGAGTGTGGAGTCGTTGCAGTGAGTGAGAGTCCGTTTCAAANNNAGATCGGAA |
| 53844_1 | TATGAGTGTGGAGTCGTTGCTGTTTGTAGCAGAGCGGTCTTATNNNAGATCGGAA |
| 53845_1 | TATGAGTGTGGAGTCGTTGCATCCAAAAGGCAGGTATGGTTCTNNNAGATCGGAA |
| 53848_1 | TATGAGTGTGGAGTCGTTGCATGAAGCCAAGCATTGTCACAATNNNAGATCGGAAG |
| 53849_1 | TATGAGTGTGGAGTCGTTGCATAGAAAAGCTAGTGGGGTCGTGNNNAGATCGGAA |
| 53850_1 | TATGAGTGTGGAGTCGTTGCCCAGAGTAAGAGGACAAGCAACANNNAGATCGGAA |
| 53851_1 | TATGAGTGTGGAGTCGTTGCGTCAGTAGAACCCCAAGAGTCTTNNNAGATCGGAAG |
| 53852_1 | TATGAGTGTGGAGTCGTTGCTTGTGCAAAGATGAAGGCTGATGNNNAGATCGGAA |
| 53853_1 | TATGAGTGTGGAGTCGTTGCAGACATTCTCAGTCCTCAGCAAGNNNAGATCGGAAG |
| 53855_1 | TATGAGTGTGGAGTCGTTGCACTATAACCTCAAACTCCTGGGCNNNAGATCGGAAG |
| 53856_1 | TATGAGTGTGGAGTCGTTGCCACTTGTAATCCCAGCACTTTGGNNNAGATCGGAAG |
| 53857_1 | TATGAGTGTGGAGTCGTTGCTCTGACACATGCTACAACATGGANNNAGATCGGAAG |
| 53858_1 | TATGAGTGTGGAGTCGTTGCTTAGGACAATTTGAGCACAGCACNNNAGATCGGAA |
| 53859_1 | TATGAGTGTGGAGTCGTTGCCACACCTGTAATCCCAGCACTTTNNNAGATCGGAAGA |
| 53860_1 | TATGAGTGTGGAGTCGTTGCCAAGTGAGCATGGCTAATTCCTGNNNAGATCGGAA |
| 53861_1 | TATGAGTGTGGAGTCGTTGCTGAGTGGCAGTTATTTGGAACAGNNNAGATCGGAA |
| 53862_1 | TATGAGTGTGGAGTCGTTGCCTGAGGCAGAGAATTGCTTGAACNNNAGATCGGAA |
| 53864_1 | TATGAGTGTGGAGTCGTTGCGTGGCTCACACCTGTAATCCTAGNNNAGATCGGAAG |
| 53866_1 | TATGAGTGTGGAGTCGTTGCTAACAGGTGTTGGCAAAGATGTGNNNAGATCGGAA |
| 53869_1 | TATGAGTGTGGAGTCGTTGCTGCCAAGCATTGTAGTAGACACTNNNAGATCGGAAG |
| 53870_1 | TATGAGTGTGGAGTCGTTGCGGCATACCACCTGTATTCATCCANNNAGATCGGAAG |
| 53871_1 | TATGAGTGTGGAGTCGTTGCCCTGTAGTCCCAGCTACTTAGGANNNAGATCGGAAG |
| 53872_1 | TATGAGTGTGGAGTCGTTGCTGGTGGGGGGAAAGAATTTTGTANNNAGATCGGAA |
| 53873_1 | TATGAGTGTGGAGTCGTTGCTCTGACTTGTGTGCGAATGAAACNNNAGATCGGAA |
| 53874_1 | TATGAGTGTGGAGTCGTTGCACCAAACCAAATCTCAGGGATCANNNAGATCGGAAG |
| 53875_1 | TATGAGTGTGGAGTCGTTGCTTGACTGTTGAACATCTGAGTGTNNNAGATCGGAAG |
| 53876_1 | TATGAGTGTGGAGTCGTTGCCAGGGTTTGCAATCCTAGTCTCTNNNAGATCGGAAG |
| 53878_1 | TATGAGTGTGGAGTCGTTGCTACTTGTTTCTTCTCAGCTGCCTNNNAGATCGGAAGA |
| 53880_1 | TATGAGTGTGGAGTCGTTGCCAATTTGGGCTGGCTGTATTTCANNNAGATCGGAAG |
| 53882_1 | TATGAGTGTGGAGTCGTTGCTTTCAGATCCTTGTTCTCCAGGGNNNAGATCGGAAG |

| | |
|---|---|
| 53885_1 | TATGAGTGTGGAGTCGTTGCCACACCTGTAATCCCAGCACTTTNNNAGATCGGAAGA |
| 53886_1 | TATGAGTGTGGAGTCGTTGCTGGTACTCCACACCTTCCAAATTNNNAGATCGGAAGA |
| 53887_1 | TATGAGTGTGGAGTCGTTGCTTACTGAAAGCTTGCTTTGAGCCNNNAGATCGGAAG |
| 53888_1 | TATGAGTGTGGAGTCGTTGCGATCAGAGCCCTACTCAGGAAACNNNAGATCGGAAG |
| 53891_1 | TATGAGTGTGGAGTCGTTGCAGTCTCACTCTGTTACCTAGGCTNNNAGATCGGAAGA |
| 53892_1 | TATGAGTGTGGAGTCGTTGCCAAGTAGCTGGGATTACAGGTGTNNNAGATCGGAAG |
| 53893_1 | TATGAGTGTGGAGTCGTTGCCTCTGGATCTTGACTCACTGCAGNNNAGATCGGAAGA |
| 53894_1 | TATGAGTGTGGAGTCGTTGCGTTCAAGAGATTCTCCTGCCTCANNNAGATCGGAAGA |
| 53895_1 | TATGAGTGTGGAGTCGTTGCGAAGTTTCACCATGTTACCCAGGNNNAGATCGGAAG |
| 53897_1 | TATGAGTGTGGAGTCGTTGCTAGGACAATAGTGGAGGGAAGGTNNNAGATCGGAA |
| 53898_1 | TATGAGTGTGGAGTCGTTGCTCTATTTTCCCTAAGTGTCGGCCNNNAGATCGGAAGA |
| 53899_1 | TATGAGTGTGGAGTCGTTGCTCAAGGGATTCTCTTGCCTTAGCNNNAGATCGGAAGA |
| 53901_1 | TATGAGTGTGGAGTCGTTGCACTCCACTTTCTGAGTTCAAGCANNNAGATCGGAAGA |
| 53902_1 | TATGAGTGTGGAGTCGTTGCTCTTGAACTCCTGACCTCAAGTGNNNAGATCGGAAGA |
| 53904_1 | TATGAGTGTGGAGTCGTTGCCTCCTGAGTAGCTGGGATTACAGNNNAGATCGGAAG |
| 53907_1 | TATGAGTGTGGAGTCGTTGCGGAGTACAGTGGCACAATCACAGNNNAGATCGGAAG |
| 53908_1 | TATGAGTGTGGAGTCGTTGCCCGAGTAGCTATGACTACAGGTGNNNAGATCGGAAG |
| 53909_1 | TATGAGTGTGGAGTCGTTGCTTCAAGCGATTCTCCTGCCTTAGNNNAGATCGGAAGA |
| 53910_1 | TATGAGTGTGGAGTCGTTGCTAGAGATCGGGTTTCGCCATGTTNNNAGATCGGAAG |
| 53911_1 | TATGAGTGTGGAGTCGTTGCGTCTCCTGGGTTCAATGATTCCTNNNAGATCGGAAGA |
| 53914_1 | TATGAGTGTGGAGTCGTTGCGAGACAAGGTCTTGCTTTGTCACNNNAGATCGGAAG |
| 53916_1 | TATGAGTGTGGAGTCGTTGCTTCAAGTGATTCTCCTGCCTCAGNNNAGATCGGAAGA |
| 53918_1 | TATGAGTGTGGAGTCGTTGCCCTCCTGGGTTCAAGTGATTCTCNNNAGATCGGAAGA |
| 53919_1 | TATGAGTGTGGAGTCGTTGCCAAAGTGCTGGGATTACAGGTGTNNNAGATCGGAAG |
| 53920_1 | TATGAGTGTGGAGTCGTTGCTTCTCCTGGACATTCAAGACTCTNNNAGATCGGAAGA |
| 53921_1 | TATGAGTGTGGAGTCGTTGCTTCAAGTGATTCTCCTGCCTCAGNNNAGATCGGAAGA |
| 53922_1 | TATGAGTGTGGAGTCGTTGCCCTCCCAAATAGCTATAGGCACCNNNAGATCGGAAGA |
| 53923_1 | TATGAGTGTGGAGTCGTTGCGCACAATCTCAGCTAACAGCAATNNNAGATCGGAAG |
| 53924_1 | TATGAGTGTGGAGTCGTTGCATGGAGTTTTCGCTGTTATTGCCNNNAGATCGGAAG |
| 53925_1 | TATGAGTGTGGAGTCGTTGCAGTGCTAGTATTACAGGCGTGAGNNNAGATCGGAAG |
| 53926_1 | TATGAGTGTGGAGTCGTTGCAGAAGGAGTCCCACTCTATCACCNNNAGATCGGAAG |
| 53927_1 | TATGAGTGTGGAGTCGTTGCGAGTAGCTGGGATTACAGACGTGNNNAGATCGGAA |
| 53928_1 | TATGAGTGTGGAGTCGTTGCGAGGAGAAATCTCGCTCTGTCTCNNNAGATCGGAAG |
| 53929_1 | TATGAGTGTGGAGTCGTTGCTTCAAGTGATTCTCCTGCCTCAGNNNAGATCGGAAGA |
| 53930_1 | TATGAGTGTGGAGTCGTTGCTTCAAGTGATTCTCCTGCCTCAGNNNAGATCGGAAGA |
| 53931_1 | TATGAGTGTGGAGTCGTTGCGCCACATTCTAATCTCCCCTGAANNNAGATCGGAAGA |
| 53933_1 | TATGAGTGTGGAGTCGTTGCTGGCATCAAAGACCCTTCAAGATNNNAGATCGGAAG |
| 53934_1 | TATGAGTGTGGAGTCGTTGCGCATGCCTATAATCCCAGCTACTNNNAGATCGGAAGA |
| 53937_1 | TATGAGTGTGGAGTCGTTGCGCACTGTGCCTCTTCCTTTTTAGNNNAGATCGGAAGA |
| 53938_1 | TATGAGTGTGGAGTCGTTGCATGAAGGGATAATGGGAGGGAGANNNAGATCGGA |
| 53939_1 | TATGAGTGTGGAGTCGTTGCCCTGTAATTCCAGCTACTTGGGANNNAGATCGGAAG |
| 53940_1 | TATGAGTGTGGAGTCGTTGCTCCAAGTTAGGTGAGTCTCTTGANNNAGATCGGAAG |
| 53941_1 | TATGAGTGTGGAGTCGTTGCCACCTGTAATCCCACCTAGTTGGNNNAGATCGGAAGA |

| | |
|---|---|
| 53943_1 | TATGAGTGTGGAGTCGTTGCATGAAATCTCAAGGGACCCCAAANNNAGATCGGAAG |
| 53944_1 | TATGAGTGTGGAGTCGTTGCATCTATGTGCTTACTCAGGGCTGNNNAGATCGGAAG |
| 53945_1 | TATGAGTGTGGAGTCGTTGCCAAAATCGTCCCCCATTGAGAACNNNAGATCGGAAGA |
| 53946_1 | TATGAGTGTGGAGTCGTTGCAATACTGAACCATTGCCCCTAGGNNNAGATCGGAAG |
| 53947_1 | TATGAGTGTGGAGTCGTTGCCAGAGGCAGGAGAATCACTTGAANNNAGATCGGAA |
| 53948_1 | TATGAGTGTGGAGTCGTTGCGACAAGCCTTTTAGAACTGCTCTNNNAGATCGGAAGA |
| 53950_1 | TATGAGTGTGGAGTCGTTGCTTAGCCAAATGTTGTGGTGCATCNNNAGATCGGAAG |
| 53952_1 | TATGAGTGTGGAGTCGTTGCGCCATGCTTCTTTTTACACTGGANNNAGATCGGAAGA |
| 53953_1 | TATGAGTGTGGAGTCGTTGCTCTGAGAAGTAGTTTGTCCACGGNNNAGATCGGAAG |
| 53954_1 | TATGAGTGTGGAGTCGTTGCGCCACCATTGCAGTTATGAAGAGNNNAGATCGGAAG |
| 53955_1 | TATGAGTGTGGAGTCGTTGCGTAATCTCAGCTACTCAGGAGGCNNNAGATCGGAAG |
| 53956_1 | TATGAGTGTGGAGTCGTTGCAATGTTAGCACTTTCGGAGGCTANNNAGATCGGAAG |
| 53957_1 | TATGAGTGTGGAGTCGTTGCGAAGAGAGACCTGAGCTAGCATGNNNAGATCGGAA |
| 53959_1 | TATGAGTGTGGAGTCGTTGCACACAAACACCAGTTATTCCCCTNNNAGATCGGAAGA |
| 53960_1 | TATGAGTGTGGAGTCGTTGCGGTGACCTTTCATGTTCAAGTGGNNNAGATCGGAAG |
| 53961_1 | TATGAGTGTGGAGTCGTTGCAAGGGGAGAAGAAAAGGAGCAAANNNAGATCGGA |
| 53962_1 | TATGAGTGTGGAGTCGTTGCACTGGCCTGTTCAATATTCACAGNNNAGATCGGAAGA |
| 53964_1 | TATGAGTGTGGAGTCGTTGCGGAAATCCTCTGGAACTTCAGAANNNAGATCGGAAG |
| 53965_1 | TATGAGTGTGGAGTCGTTGCCACACACACACACACACATACATNNNAGATCGGAAGA |
| 53968_1 | TATGAGTGTGGAGTCGTTGCTTGCAGAGATGAGAAAACACACCNNNAGATCGGAAG |
| 53971_1 | TATGAGTGTGGAGTCGTTGCAGAAGAAGGTGTCCTGATTGGTCNNNAGATCGGAAG |
| 53972_1 | TATGAGTGTGGAGTCGTTGCCGGCCATGAAAAACTGCTAGATTNNNAGATCGGAAG |
| 53973_1 | TATGAGTGTGGAGTCGTTGCTTTAGGCAACGAACAGTGAACACNNNAGATCGGAAG |
| 53974_1 | TATGAGTGTGGAGTCGTTGCAAACAAACACACACACACACACANNNAGATCGGAAGA |
| 53975_1 | TATGAGTGTGGAGTCGTTGCTCCCTCATAACCTAGGAGGAAGGNNNAGATCGGAAG |
| 53976_1 | TATGAGTGTGGAGTCGTTGCGAAATGAAAGGGAGTGGGGAAGANNNAGATCGGA |
| 53977_1 | TATGAGTGTGGAGTCGTTGCGTAGTGTTCTTGCTTGGGGAGATNNNAGATCGGAAG |
| 53979_1 | TATGAGTGTGGAGTCGTTGCCAGATCACGAGATCAGGAGATCGNNNAGATCGGAAG |
| 53980_1 | TATGAGTGTGGAGTCGTTGCCAGAACTTTGGGAGTCTGAGACANNNAGATCGGAAG |
| 53981_1 | TATGAGTGTGGAGTCGTTGCGCGCTCGTGGAAATACAATTTTCNNNAGATCGGAAG |
| 53982_1 | TATGAGTGTGGAGTCGTTGCCCTCCTGAGTAGCTGGGACTATANNNAGATCGGAAG |
| 53983_1 | TATGAGTGTGGAGTCGTTGCATGGAGTCTTGTTCTGTTGTCCANNNAGATCGGAAG |
| 53984_1 | TATGAGTGTGGAGTCGTTGCATGGAGTATTACTCTGTTGCCCANNNAGATCGGAAG |
| 53986_1 | TATGAGTGTGGAGTCGTTGCTACCTCTACAGCAAGTGTCTGTGNNNAGATCGGAAG |
| 53988_1 | TATGAGTGTGGAGTCGTTGCGAGGCAGGAGAATTGTTTGAACCNNNAGATCGGAA |
| 53989_1 | TATGAGTGTGGAGTCGTTGCACACACCTGTAATCTCAGCACTTNNNAGATCGGAAGA |
| 53992_1 | TATGAGTGTGGAGTCGTTGCACTGGTGTTTTTTGTCCTGCTTCNNNAGATCGGAAGA |
| 53993_1 | TATGAGTGTGGAGTCGTTGCTAGTAACATGGTCACGTCTTGGGNNNAGATCGGAAG |
| 53994_1 | TATGAGTGTGGAGTCGTTGCCTCAAAACCAAACAGAAAGCCCANNNAGATCGGAAG |
| 53995_1 | TATGAGTGTGGAGTCGTTGCAATAGCCTGGGTTTGTTTCAGGANNNAGATCGGAAG |
| 53998_1 | TATGAGTGTGGAGTCGTTGCATGAGGAAATCTGACTGCTGTGTNNNAGATCGGAAG |
| 53999_1 | TATGAGTGTGGAGTCGTTGCCTGAGACAGGAGGATCACTTGAGNNNAGATCGGAA |
| 54000_1 | TATGAGTGTGGAGTCGTTGCTATTTCCCGAGAGCACATGACAANNNAGATCGGAAG |

| | |
|---|---|
| 54003_1 | TATGAGTGTGGAGTCGTTGCCAAAGTGTTGGGATTACAAGCGTNNNAGATCGGAA |
| 54004_1 | TATGAGTGTGGAGTCGTTGCCTGGGTTCAAATGATTCTCCTGCNNNAGATCGGAAG |
| 54006_1 | TATGAGTGTGGAGTCGTTGCTGGAGTCTCACTTTGTCATCCAGNNNAGATCGGAAG |
| 54007_1 | TATGAGTGTGGAGTCGTTGCTAGCTGGAACTACAGGCCTTTTCNNNAGATCGGAAG |
| 54008_1 | TATGAGTGTGGAGTCGTTGCATCTGCCTCTTGGTTCAAGTGATNNNAGATCGGAAG |
| 54009_1 | TATGAGTGTGGAGTCGTTGCGGAGTTTTGCTCTTGTTGCCTAGNNNAGATCGGAAG |
| 54010_1 | TATGAGTGTGGAGTCGTTGCCCAAGTAGCTGAGATTACAGGCANNNAGATCGGAA |
| 54011_1 | TATGAGTGTGGAGTCGTTGCGAGTGCAGTGTCATGATCTCAGANNNAGATCGGAA |
| 54012_1 | TATGAGTGTGGAGTCGTTGCGTCTCTACAGTTCAGGCGATTCTNNNAGATCGGAAG |
| 54013_1 | TATGAGTGTGGAGTCGTTGCAGTACATCTGTAGCTTTGGGCATNNNAGATCGGAAG |
| 54014_1 | TATGAGTGTGGAGTCGTTGCTGTATCATTCTCTGCAGTCCCAANNNAGATCGGAAG |
| 54015_1 | TATGAGTGTGGAGTCGTTGCAATCTTGTCTCACTGCAACCTCTNNNAGATCGGAAGA |
| 54016_1 | TATGAGTGTGGAGTCGTTGCCTCTTACCTCAGCCTCCCAAGTANNNAGATCGGAAGA |
| 54019_1 | TATGAGTGTGGAGTCGTTGCTTGATCTCAACTCACTGCAACCTNNNAGATCGGAAG |
| 54020_1 | TATGAGTGTGGAGTCGTTGCGAGACGGAGTCTAGTTCTGTCACNNNAGATCGGAA |
| 54021_1 | TATGAGTGTGGAGTCGTTGCTCCTGGTCTCAAGCAATTCTCTCNNNAGATCGGAAG |
| 54022_1 | TATGAGTGTGGAGTCGTTGCGGGTTCAAGCAATTCTGTCTCAGNNNAGATCGGAAG |
| 54023_1 | TATGAGTGTGGAGTCGTTGCAAACTCCCAGATTCAGGTGATCCNNNAGATCGGAAG |
| 54024_1 | TATGAGTGTGGAGTCGTTGCCTTGATCTCAGCTCATGGCAAACNNNAGATCGGAAG |
| 54025_1 | TATGAGTGTGGAGTCGTTGCACTGTTGGTGTACAGAAGAGGTGNNNAGATCGGAA |
| 54026_1 | TATGAGTGTGGAGTCGTTGCTGAATGCTGCTTCCCTACCTTAGNNNAGATCGGAAG |
| 54029_1 | TATGAGTGTGGAGTCGTTGCGGATTTCACCGTATTAGCCAGGANNNAGATCGGAA |
| 54032_1 | TATGAGTGTGGAGTCGTTGCATAGGGTCTAACTCTGTCACCCANNNAGATCGGAAG |
| 54034_1 | TATGAGTGTGGAGTCGTTGCCTCCTGAGTAGCTGGGATTACAGNNNAGATCGGAA |
| 54036_1 | TATGAGTGTGGAGTCGTTGCCTGGGATTACAGGTGTGAGCTACNNNAGATCGGAA |
| 54037_1 | TATGAGTGTGGAGTCGTTGCGTTTCACAGTGTTAGCCAGGATGNNNAGATCGGAA |
| 54038_1 | TATGAGTGTGGAGTCGTTGCGATTCAAGCAATTCTCCTGCCTCNNNAGATCGGAAG |
| 54039_1 | TATGAGTGTGGAGTCGTTGCTATACTCAGTGGCATGATCACGGNNNAGATCGGAA |
| 54040_1 | TATGAGTGTGGAGTCGTTGCTCCTGGGTTCTAGCTATTCTCCTNNNAGATCGGAAG |
| 54041_1 | TATGAGTGTGGAGTCGTTGCATCCCGGGTTCAAGTGATTCTACNNNAGATCGGAAG |
| 54042_1 | TATGAGTGTGGAGTCGTTGCAGCGATTCTCTTACCTCAACCTCNNNAGATCGGAAGA |
| 54043_1 | TATGAGTGTGGAGTCGTTGCCTCCTGAGTAGCTGGGATTACAGNNNAGATCGGAA |
| 54045_1 | TATGAGTGTGGAGTCGTTGCTGATCATAGCTCACTGCAGTCTCNNNAGATCGGAAG |
| 54046_1 | TATGAGTGTGGAGTCGTTGCTCCCAAGTAGTTGGAAGTACAGGNNNAGATCGGAA |
| 54051_1 | TATGAGTGTGGAGTCGTTGCAGCAGGTGTTCGGTACTTACAATNNNAGATCGGAA |
| 54052_1 | TATGAGTGTGGAGTCGTTGCGAGCCTGGATTGGGAGAGAATTANNNAGATCGGAA |
| 54053_1 | TATGAGTGTGGAGTCGTTGCAACATACATACATACCAGGCCGGNNNAGATCGGAAG |
| 54054_1 | TATGAGTGTGGAGTCGTTGCTCATGGAATGATTGGTTGAAGGANNNAGATCGGAA |
| 54055_1 | TATGAGTGTGGAGTCGTTGCGAGTTAGTCTGCTGGAGGTTCATNNNAGATCGGAA |
| 54056_1 | TATGAGTGTGGAGTCGTTGCGTAAACCAAAGCGGGGGAATTAGNNNAGATCGGAA |
| 54057_1 | TATGAGTGTGGAGTCGTTGCGAGACCTCTCTTTCCCTGTCTCCNNNAGATCGGAAGA |
| 54058_1 | TATGAGTGTGGAGTCGTTGCCTCAAGCCTATAATCCCAGCACTNNNAGATCGGAAG |
| 54059_1 | TATGAGTGTGGAGTCGTTGCGAGAGAACAGAGAACAGTGGGAANNNAGATCGGA |

| | |
|---|---|
| 54060_1 | TATGAGTGTGGAGTCGTTGCCAGGAAAGAAGGAGGTGCAGATANNNAGATCGGA |
| 54061_1 | TATGAGTGTGGAGTCGTTGCGAAATAGGCTGTGATGGGATTGCNNNAGATCGGAA |
| 54062_1 | TATGAGTGTGGAGTCGTTGCGGGTACAGAAGTTGTGAGACCATNNNAGATCGGAA |
| 54063_1 | TATGAGTGTGGAGTCGTTGCACTTGATGAGTTTGGAGCGTTTGNNNAGATCGGAA |
| 54064_1 | TATGAGTGTGGAGTCGTTGCCTCACCCCCATATCTGCCAATAANNNAGATCGGAAGA |
| 54065_1 | TATGAGTGTGGAGTCGTTGCAAATAAGCAAAGGCACAGAGTGGNNNAGATCGGAA |
| 54066_1 | TATGAGTGTGGAGTCGTTGCACACATACACACATACAGGCACTNNNAGATCGGAAG |
| 54067_1 | TATGAGTGTGGAGTCGTTGCCTCCCAACAGCCTCTCAATAGAGNNNAGATCGGAAG |
| 54071_1 | TATGAGTGTGGAGTCGTTGCTCGGTGACTTCCAAGAAAACTGANNNAGATCGGAA |
| 54072_1 | TATGAGTGTGGAGTCGTTGCACTTCTGTCCACACAAAAACCTGNNNAGATCGGAAG |
| 54073_1 | TATGAGTGTGGAGTCGTTGCAGTAAATGTGAAGGCCTAGCACANNNAGATCGGAA |
| 54076_1 | TATGAGTGTGGAGTCGTTGCGCTAGACGATTGCTGCCTTATTGNNNAGATCGGAAG |
| 54077_1 | TATGAGTGTGGAGTCGTTGCAACTCAGCAAGCATAGGTAAGGANNNAGATCGGAA |
| 54080_1 | TATGAGTGTGGAGTCGTTGCGACACTCCCAAATAACCCAGGATNNNAGATCGGAAG |
| 54081_1 | TATGAGTGTGGAGTCGTTGCTAAAGCAGCCACACAAACAAGTCNNNAGATCGGAAG |
| 54082_1 | TATGAGTGTGGAGTCGTTGCTCATTTGCTACTAGGAAAACAGCANNNAGATCGGAA |
| 54083_1 | TATGAGTGTGGAGTCGTTGCGAGAATCAACTGACTCCTGGGAGNNNAGATCGGAA |
| 54084_1 | TATGAGTGTGGAGTCGTTGCGAAAGACCCGTTCCTACAGAAGANNNAGATCGGAA |
| 54085_1 | TATGAGTGTGGAGTCGTTGCCCAGGGCAACAGCTTAGTCTATANNNAGATCGGAAG |
| 54086_1 | TATGAGTGTGGAGTCGTTGCGACAAGAAAATTCACCCTTGGCANNNAGATCGGAAG |
| 54087_1 | TATGAGTGTGGAGTCGTTGCCCCTTTGGAATTCAAATTGGGGTNNNAGATCGGAAG |
| 54088_1 | TATGAGTGTGGAGTCGTTGCAGCAACTGGGAGAATAAACAGAANNNAGATCGGAA |
| 54089_1 | TATGAGTGTGGAGTCGTTGCCTCGGTGGCTCTCAATTTGTTTTNNNAGATCGGAAG |
| 54090_1 | TATGAGTGTGGAGTCGTTGCAAGAGTTTTCCTCCAAGGGTCTCNNNAGATCGGAAG |
| 54093_1 | TATGAGTGTGGAGTCGTTGCAAGACTACAGCCAGAATTGAGGGNNNAGATCGGAA |
| 54094_1 | TATGAGTGTGGAGTCGTTGCATGAAACTGGAAGCCATCATCCTNNNAGATCGGAAG |
| 54096_1 | TATGAGTGTGGAGTCGTTGCACCATCCAGTCTTGAGCTTTTCTNNNAGATCGGAAGA |
| 54097_1 | TATGAGTGTGGAGTCGTTGCCATGCCTGTAATCCCAGCTTTTTNNNAGATCGGAAGA |
| 54100_1 | TATGAGTGTGGAGTCGTTGCAAATGTTTATCGTGCAGGAAGCCNNNAGATCGGAA |
| 54101_1 | TATGAGTGTGGAGTCGTTGCCTCCAGCTTGGGTGACATAATGANNNAGATCGGAA |
| 54104_1 | TATGAGTGTGGAGTCGTTGCGGGTTGCAATCCTACTTTCTGACNNNAGATCGGAAG |
| 54106_1 | TATGAGTGTGGAGTCGTTGCGCATTTGAAAAGCTAAGAACAGCCNNNAGATCGGA |
| 54109_1 | TATGAGTGTGGAGTCGTTGCGGACAAATTTGCAGTGGTGATGANNNAGATCGGAA |
| 54110_1 | TATGAGTGTGGAGTCGTTGCAGAGGAATAACTCGTGGAGGAGANNNAGATCGGA |
| 54111_1 | TATGAGTGTGGAGTCGTTGCCATCGTGCCTCAATGAAACTGAANNNAGATCGGAAG |
| 54113_1 | TATGAGTGTGGAGTCGTTGCTGGGATTCGAATGGGATTCAATCTNNNAGATCGGA |
| 54114_1 | TATGAGTGTGGAGTCGTTGCAAACTTGCATAAAAAGGCCTGCTNNNAGATCGGAAG |
| 54115_1 | TATGAGTGTGGAGTCGTTGCTTGCCCCTCATATATTGGACATTNNNAGATCGGAAG |
| 54117_1 | TATGAGTGTGGAGTCGTTGCGAATCACTTGAACCCAGGAGGAANNNAGATCGGAA |
| 54120_1 | TATGAGTGTGGAGTCGTTGCAAAAGAGGGATTGATCAGACCCCNNNAGATCGGAA |
| 54121_1 | TATGAGTGTGGAGTCGTTGCGTGGTCCTCTCCATGATGTACTCNNNAGATCGGAAG |
| 54122_1 | TATGAGTGTGGAGTCGTTGCGGCTACAGTGAGTCAAGATCACANNNAGATCGGAA |
| 54124_1 | TATGAGTGTGGAGTCGTTGCAAAAACTGTGTGTCACGAGGAAGNNNAGATCGGAA |

| | |
|---|---|
| 54125_1 | TATGAGTGTGGAGTCGTTGCCACCTGTAATTCCAGCACTTTGGNNNAGATCGGAAG |
| 54126_1 | TATGAGTGTGGAGTCGTTGCAGTGTCAAATTGGATGCATTGGGNNNAGATCGGAA |
| 54127_1 | TATGAGTGTGGAGTCGTTGCCTAAGGTTTGATCCCCCCAAGAANNNAGATCGGAAG |
| 54130_1 | TATGAGTGTGGAGTCGTTGCTGGAGTGCAATGGTGTTATCTCANNNAGATCGGAA |
| 54131_1 | TATGAGTGTGGAGTCGTTGCGTTCATGCCATTCTCTTGCTTCANNNAGATCGGAAGA |
| 54132_1 | TATGAGTGTGGAGTCGTTGCTCCCACTTCAGCCTCCTAAGTATNNNAGATCGGAAGA |
| 54136_1 | TATGAGTGTGGAGTCGTTGCCTCCTGAGTACCTGGGACTACAGNNNAGATCGGAAG |
| 54137_1 | TATGAGTGTGGAGTCGTTGCAGATGGAGTCTTGATCTGTTGCCNNNAGATCGGAA |
| 54138_1 | TATGAGTGTGGAGTCGTTGCAAGGCTGGACTTGAACTCTTGAGNNNAGATCGGAA |
| 54139_1 | TATGAGTGTGGAGTCGTTGCCTCCTGTCTGGGATGTGTGTATTNNNAGATCGGAAG |
| 54140_1 | TATGAGTGTGGAGTCGTTGCGCTCACTACAACCTTCACCTACTNNNAGATCGGAAGA |
| 54141_1 | TATGAGTGTGGAGTCGTTGCACCAAAGTAGCTGGGATTACAGGNNNAGATCGGAA |
| 54142_1 | TATGAGTGTGGAGTCGTTGCCCTCCTGAGTAGCTGAGACTACANNNAGATCGGAAG |
| 54143_1 | TATGAGTGTGGAGTCGTTGCGAGTCTCATTCTGTTGCCCAAACNNNAGATCGGAAG |
| 54145_1 | TATGAGTGTGGAGTCGTTGCTGAGATGGAGTCTTGTTCTGTCGNNNAGATCGGAA |
| 54148_1 | TATGAGTGTGGAGTCGTTGCTCAAGTGATTCTCCTGCCTTAGCNNNAGATCGGAAG |
| 54149_1 | TATGAGTGTGGAGTCGTTGCCAGCCTCCAGAGTAATTGGGATTNNNAGATCGGAA |
| 54151_1 | TATGAGTGTGGAGTCGTTGCCTGAGTAGCTGCGAGGGACTATANNNAGATCGGAA |
| 54152_1 | TATGAGTGTGGAGTCGTTGCCAGCATCCTGAGTATCTGGGATCNNNAGATCGGAAG |
| 54153_1 | TATGAGTGTGGAGTCGTTGCCCCACCTTAGCCTTCTGAGTAACNNNAGATCGGAAG |
| 54156_1 | TATGAGTGTGGAGTCGTTGCCAGATGGAGTGTCACTCTGTCACNNNAGATCGGAAG |
| 54157_1 | TATGAGTGTGGAGTCGTTGCTAAAAATTAGTCAGGCGTGGTGGNNNAGATCGGAA |
| 54158_1 | TATGAGTGTGGAGTCGTTGCGCCAACAATCACAGGTTTCTACCNNNAGATCGGAAG |
| 54160_1 | TATGAGTGTGGAGTCGTTGCGATCACAAGGTCAGGAGATCGAGNNNAGATCGGAA |
| 54162_1 | TATGAGTGTGGAGTCGTTGCGAGGTCAGGAGATCAAGACCATCNNNAGATCGGAA |
| 54164_1 | TATGAGTGTGGAGTCGTTGCTCTGTAGTCCCAGCTACTTCAGANNNAGATCGGAAG |
| 54166_1 | TATGAGTGTGGAGTCGTTGCAGACACGAGAATCACTTGAACCCNNNAGATCGGAAG |
| 54169_1 | TATGAGTGTGGAGTCGTTGCTCAAGCCTGTACTCCTAGCTACTNNNAGATCGGAAG |
| 54170_1 | TATGAGTGTGGAGTCGTTGCCTACTTAGCTGGAGGTGAAGCTCNNNAGATCGGAA |
| 54172_1 | TATGAGTGTGGAGTCGTTGCTCTCATGCCTGTAATCCTAGCACNNNAGATCGGAAG |
| 54173_1 | TATGAGTGTGGAGTCGTTGCCAGGCGCATCGTTTTGAAAATTANNNAGATCGGAAG |
| 54177_1 | TATGAGTGTGGAGTCGTTGCGATTCCATCTCAAGCTCCCTGAGNNNAGATCGGAAG |
| 54178_1 | TATGAGTGTGGAGTCGTTGCGTGTACTCACATTTGCAAGATGTNNNAGATCGGAAG |
| 54181_1 | TATGAGTGTGGAGTCGTTGCCTGGGTTCGAATCCTACTTCTCCNNNAGATCGGAAG |
| 54182_1 | TATGAGTGTGGAGTCGTTGCTCACATGAGGTCAGGAGTTTGAGNNNAGATCGGAA |
| 54184_1 | TATGAGTGTGGAGTCGTTGCTGAGACAGTCTCTATCACCCAGGNNNAGATCGGAAG |
| 54185_1 | TATGAGTGTGGAGTCGTTGCGGGCTTTTCTTTTCACTCAGTGGNNNAGATCGGAAG |
| 54188_1 | TATGAGTGTGGAGTCGTTGCTCATGAAACTTTTAGAACCGCGCNNNAGATCGGAAG |
| 54189_1 | TATGAGTGTGGAGTCGTTGCAGGACCAAAACCTCTCGAACTTTNNNAGATCGGAAG |
| 54191_1 | TATGAGTGTGGAGTCGTTGCTCCCGTTTCCATTTCTCAAGTCTNNNAGATCGGAAGA |
| 54192_1 | TATGAGTGTGGAGTCGTTGCTTGTGGTTCTAGAATCAGGGTGGNNNAGATCGGAA |
| 54193_1 | TATGAGTGTGGAGTCGTTGCGGATCACGAGGTCAAGAGATCAANNNAGATCGGAA |
| 54196_1 | TATGAGTGTGGAGTCGTTGCTCACACCTGTAATCCCAACACTTNNNAGATCGGAAGA |

54198_1 TATGAGTGTGGAGTCGTTGCTTACCAAATGTCCCTGGAAGAGGNNNAGATCGGAA
54199_1 TATGAGTGTGGAGTCGTTGCCTGAGGTCAGGAGTTCGATACCANNNAGATCGGAA
54200_1 TATGAGTGTGGAGTCGTTGCACTCACACCTATAATCCCAGCACNNNAGATCGGAAG
54201_1 TATGAGTGTGGAGTCGTTGCATACACACACACAGACACACACANNNAGATCGGAAG
54202_1 TATGAGTGTGGAGTCGTTGCACACGGTAAAACCCCATCTCTACNNNAGATCGGAAG
54203_1 TATGAGTGTGGAGTCGTTGCTTGGTCCATGCCTATAATCCCAGNNNAGATCGGAAG
54204_1 TATGAGTGTGGAGTCGTTGCTCCGGTTTCAAGAGATTCTCCTGNNNAGATCGGAAG
54206_1 TATGAGTGTGGAGTCGTTGCTGCAGCCTCGATTGTTCAAATTNNNAGATCGGAAG
54207_1 TATGAGTGTGGAGTCGTTGCCCTCCCAGATTCACGCTATTCTCNNNAGATCGGAAGA
54210_1 TATGAGTGTGGAGTCGTTGCCTATCTCAGCTCACTGCAACCTCNNNAGATCGGAAGA
54211_1 TATGAGTGTGGAGTCGTTGCGCCATAAAAACTCCCAGGCAAAANNNAGATCGGAAG
54212_1 TATGAGTGTGGAGTCGTTGCTAGAGATGGGTTTTCACCGTGTTNNNAGATCGGAA
54213_1 TATGAGTGTGGAGTCGTTGCCCTCCTAGGTTCAGGCAATTCTCNNNAGATCGGAAG
54216_1 TATGAGTGTGGAGTCGTTGCTTCAAGTGATTCTCCTGCCTCAGNNNAGATCGGAAG
54217_1 TATGAGTGTGGAGTCGTTGCCTCAAGTGATCCTCCCACCTTAGNNNAGATCGGAAG
54218_1 TATGAGTGTGGAGTCGTTGCCTGAATAGCTGGGACTACAGGTGNNNAGATCGGAA
54219_1 TATGAGTGTGGAGTCGTTGCGAGTAGCTGGGATTACAGGAGTGNNNAGATCGGA
54220_1 TATGAGTGTGGAGTCGTTGCCGCCTCCCAGATTCAAGAAATTCNNNAGATCGGAAG
54221_1 TATGAGTGTGGAGTCGTTGCTAATCATGACTCACTGCAGCCTTNNNAGATCGGAAG
54223_1 TATGAGTGTGGAGTCGTTGCCATTCTCCTGCCTAATCCTCCTGNNNAGATCGGAAGA
54225_1 TATGAGTGTGGAGTCGTTGCGAGACGGGATTTCGATCTTGTTGNNNAGATCGGAA
54226_1 TATGAGTGTGGAGTCGTTGCCCATTCTCCTGCCTCATCCTCNNNAGATCGGAAGAGC
54228_1 TATGAGTGTGGAGTCGTTGCCTGCATAGCTGGGATTATAGGCANNNAGATCGGAA
54230_1 TATGAGTGTGGAGTCGTTGCAGTAGTTGGGATTAGAGGCATGCNNNAGATCGGAA
54232_1 TATGAGTGTGGAGTCGTTGCCACCTTGCCATCTGTGTCTATCTNNNAGATCGGAAGA
54235_1 TATGAGTGTGGAGTCGTTGCTCCTGAGTAGCTTGGATTACAGGNNNAGATCGGAA
54237_1 TATGAGTGTGGAGTCGTTGCATAGAGTCTCTCCCTGTCATCCCNNNAGATCGGAAGA
54238_1 TATGAGTGTGGAGTCGTTGCCTCACTGCAACCTCCAGCTATATNNNAGATCGGAAGA
54239_1 TATGAGTGTGGAGTCGTTGCACTCACTACAATTTTCGCCTCCTNNNAGATCGGAAGA
54241_1 TATGAGTGTGGAGTCGTTGCCTCAAGCAATTCTCCTGCCTCAGNNNAGATCGGAAGA
54242_1 TATGAGTGTGGAGTCGTTGCTCTCAAACTCCTGACCTCAAGTGNNNAGATCGGAAG
54244_1 TATGAGTGTGGAGTCGTTGCATGCTACCACGACCAGCTAATTTNNNAGATCGGAAG
54245_1 TATGAGTGTGGAGTCGTTGCTTCAAACGATTCTCCTGCCTCAGNNNAGATCGGAAGA
54246_1 TATGAGTGTGGAGTCGTTGCCAGTAGTACCATCTCGGCTCATTNNNAGATCGGAAG
54248_1 TATGAGTGTGGAGTCGTTGCTTTCTTCCTTGAGGTGTGAACCANNNAGATCGGAAG
54249_1 TATGAGTGTGGAGTCGTTGCAGAGGCATGATCATAGTTCAGGGNNNAGATCGGAA
54250_1 TATGAGTGTGGAGTCGTTGCTTCAAGTGATTCTCCTGCCTCAGNNNAGATCGGAAG
54251_1 TATGAGTGTGGAGTCGTTGCGAGACGGAGTTTGACTCTTGTTGNNNAGATCGGAA
54253_1 TATGAGTGTGGAGTCGTTGCTTCAAGTGATTCTCCTGCCTCAGNNNAGATCGGAAG
54254_1 TATGAGTGTGGAGTCGTTGCTGGAGGATACAGCAACAAGGTACNNNAGATCGGAA
54255_1 TATGAGTGTGGAGTCGTTGCAAGTCCTGACAACATGTGCTCAANNNAGATCGGAAG
54256_1 TATGAGTGTGGAGTCGTTGCGGATTGCTGTCTGGACTCCATAANNNAGATCGGAAG
54259_1 TATGAGTGTGGAGTCGTTGCGTACATGCCTGTAATCCCAGCTANNNAGATCGGAAG

| | |
|---|---|
| 54260_1 | TATGAGTGTGGAGTCGTTGCGGCATTGGTCGAGGGTATATCTTNNNAGATCGGAA |
| 54261_1 | TATGAGTGTGGAGTCGTTGCAGGAAGGAAGGAAGAACACGTTTNNNAGATCGGA |
| 54263_1 | TATGAGTGTGGAGTCGTTGCGGGTGAATGAAACCACACCATTGNNNAGATCGGAA |
| 54265_1 | TATGAGTGTGGAGTCGTTGCCCTCACTCGGTATAAGCCTTCAGNNNAGATCGGAAG |
| 54266_1 | TATGAGTGTGGAGTCGTTGCTCTCTCCTCCCAAGTCTCCTAAGNNNAGATCGGAAGA |
| 54269_1 | TATGAGTGTGGAGTCGTTGCTGCTAGATTGGCCTTTACAATGGNNNAGATCGGAAG |
| 54271_1 | TATGAGTGTGGAGTCGTTGCTAGAAATCTTGTCCTCCGAAGCANNNAGATCGGAAG |
| 54274_1 | TATGAGTGTGGAGTCGTTGCTTCCATCACTTACCAGAAGGTGGNNNAGATCGGAAG |
| 54275_1 | TATGAGTGTGGAGTCGTTGCAATAGCCAACTGATCCCTCACAGNNNAGATCGGAAG |
| 54276_1 | TATGAGTGTGGAGTCGTTGCCCTAAACCAGAACTTGCACACTGNNNAGATCGGAAG |
| 54278_1 | TATGAGTGTGGAGTCGTTGCCCAAAAAACCAAGAACCCAGGTTNNNAGATCGGAAG |
| 54279_1 | TATGAGTGTGGAGTCGTTGCGGGAAGAAGGCTGTGTGAAGATANNNAGATCGGA |
| 54280_1 | TATGAGTGTGGAGTCGTTGCGGTTATGCTTGTAATCCCAGCACNNNAGATCGGAAG |
| 54281_1 | TATGAGTGTGGAGTCGTTGCACTGAGAGGCATAAAGAACAGCANNNAGATCGGAA |
| 54282_1 | TATGAGTGTGGAGTCGTTGCACAGAACCCCCAAGAAAATACGTNNNAGATCGGAAG |
| 54283_1 | TATGAGTGTGGAGTCGTTGCCTGTAGTCCCAGCTACTCAAGAGNNNAGATCGGAAG |
| 54284_1 | TATGAGTGTGGAGTCGTTGCGGGGTTGGGAGACAGTAAGAATTNNNAGATCGGA |
| 54285_1 | TATGAGTGTGGAGTCGTTGCAGGCAGGCCTAATTTCTTTCAGANNNAGATCGGAAG |
| 54286_1 | TATGAGTGTGGAGTCGTTGCATGCAAACGATTTGTGAGTTGGTNNNAGATCGGAA |
| 54288_1 | TATGAGTGTGGAGTCGTTGCTCACAAACAAAGGGCTAGACTCCNNNAGATCGGAAG |
| 54289_1 | TATGAGTGTGGAGTCGTTGCTTTACTTGGTGAGGACTAAGGCCNNNAGATCGGAA |
| 54290_1 | TATGAGTGTGGAGTCGTTGCATAAAAATTTGGGGGGAGGGACANNNAGATCGGA |
| 54291_1 | TATGAGTGTGGAGTCGTTGCCCCTAAGTTACCTCTGCATTTGCNNNAGATCGGAAG |
| 54292_1 | TATGAGTGTGGAGTCGTTGCGTAGTCCTAGCTACTCAGGAGGCNNNAGATCGGAA |
| 54294_1 | TATGAGTGTGGAGTCGTTGCCCCTGCATCAGTCAAGACTTTTGNNNAGATCGGAAG |
| 54295_1 | TATGAGTGTGGAGTCGTTGCCACTACAGGAATGTTGAGGTGGANNNAGATCGGAA |
| 54296_1 | TATGAGTGTGGAGTCGTTGCGAATAGGAACAGCTCCCGTCTACNNNAGATCGGAAG |
| 54300_1 | TATGAGTGTGGAGTCGTTGCGTGAATAGGGCCACCTAGTTCAGNNNAGATCGGAA |
| 54301_1 | TATGAGTGTGGAGTCGTTGCTAATCCCAGCACTTTGAGAGACCNNNAGATCGGAAG |
| 54302_1 | TATGAGTGTGGAGTCGTTGCTGAGAAGACTGAGACAGGAGGATNNNAGATCGGA |
| 54305_1 | TATGAGTGTGGAGTCGTTGCTAAAACACTGAGGGAGAAAGCGANNNAGATCGGA |
| 54306_1 | TATGAGTGTGGAGTCGTTGCGATCACAAGGTCAGGAGTTCGAGNNNAGATCGGAA |
| 54307_1 | TATGAGTGTGGAGTCGTTGCCTGAACTCCAGCCTAGGTAACAGNNNAGATCGGAAG |
| 54310_1 | TATGAGTGTGGAGTCGTTGCGGATTCAAGTGATTCTCCTGCCTNNNAGATCGGAAG |
| 54312_1 | TATGAGTGTGGAGTCGTTGCTCTCAGGTTCAAGCAATTCTCCTNNNAGATCGGAAG |
| 54314_1 | TATGAGTGTGGAGTCGTTGCCTTGCCACAGCTATACTTCCAGANNNAGATCGGAAG |
| 54315_1 | TATGAGTGTGGAGTCGTTGCCTTCCCAGGTTCATGCCATTCTCNNNAGATCGGAAGA |
| 54316_1 | TATGAGTGTGGAGTCGTTGCTGGCTTGAACACTTGATGAACACNNNAGATCGGAAG |
| 54318_1 | TATGAGTGTGGAGTCGTTGCGTTCAAGAGATTCTCCTGCCTCANNNAGATCGGAAG |
| 54320_1 | TATGAGTGTGGAGTCGTTGCTCTTGAACTCCTGACCTCAAGTGNNNAGATCGGAAG |
| 54323_1 | TATGAGTGTGGAGTCGTTGCTTCAAGCAATTCTCCTGCCTCAGNNNAGATCGGAAG |
| 54328_1 | TATGAGTGTGGAGTCGTTGCCTAGAGTGCAGTGGTGAGATCTCNNNAGATCGGAA |
| 54329_1 | TATGAGTGTGGAGTCGTTGCACAAGTGAACAAAGGTCTCTGGTNNNAGATCGGAA |

| | |
|---|---|
| 54330_1 | TATGAGTGTGGAGTCGTTGCCTCAAGCAATCCTCCTACCTCAGNNNAGATCGGAAGA |
| 54331_1 | TATGAGTGTGGAGTCGTTGCGTACTGGGATTACAGGTGTGAGCNNNAGATCGGAA |
| 54332_1 | TATGAGTGTGGAGTCGTTGCGATCTTGGCTCACTACAACCTCCNNNAGATCGGAAGA |
| 54335_1 | TATGAGTGTGGAGTCGTTGCGATCTCAGCTCACTGCAATGTCTNNNAGATCGGAAGA |
| 54336_1 | TATGAGTGTGGAGTCGTTGCCTTTAGTTCAGTGGTGCCATCTCNNNAGATCGGAAGA |
| 54337_1 | TATGAGTGTGGAGTCGTTGCCGATCATGGCTTATTGCAACCTTNNNAGATCGGAAGA |
| 54339_1 | TATGAGTGTGGAGTCGTTGCGAGTACAGTGTGATCATGGCTCANNNAGATCGGAAG |
| 54340_1 | TATGAGTGTGGAGTCGTTGCAATCTCAGTTCACTGCAACCTCTNNNAGATCGGAAGA |
| 54342_1 | TATGAGTGTGGAGTCGTTGCCTGGGTTCAAGTGATCTTCCTGANNNAGATCGGAAG |
| 54343_1 | TATGAGTGTGGAGTCGTTGCACTTCCTGGGCTCAAATGATCTTNNNAGATCGGAAGA |
| 54345_1 | TATGAGTGTGGAGTCGTTGCTCCCATTTTCACGACATTCTCCTNNNAGATCGGAAGAG |
| 54347_1 | TATGAGTGTGGAGTCGTTGCGAACTCCTGACCTCAGATGATCCNNNAGATCGGAAGA |
| 54350_1 | TATGAGTGTGGAGTCGTTGCCCACTCTAACAGGTGTGAAGTGANNNAGATCGGAAG |
| 54351_1 | TATGAGTGTGGAGTCGTTGCGATGTAGTCTCACTCTGTCACCCNNNAGATCGGAAGA |
| 54355_1 | TATGAGTGTGGAGTCGTTGCTGGCATGATCTCAGTTACAGCAANNNAGATCGGAAG |
| 54357_1 | TATGAGTGTGGAGTCGTTGCGTGTTTTTGTTCTTGACCCCACANNNAGATCGGAAGA |
| 54359_1 | TATGAGTGTGGAGTCGTTGCGCGATCTTGGCTCATTTCAACTTNNNAGATCGGAAGA |
| 54360_1 | TATGAGTGTGGAGTCGTTGCTCCAGGTTCAAGTGATTCTCCTGNNNAGATCGGAAGA |
| 54363_1 | TATGAGTGTGGAGTCGTTGCGTTTCAAGCGATTCTCCTGCTTCNNNAGATCGGAAGA |
| 54364_1 | TATGAGTGTGGAGTCGTTGCGAACTCCTGACCTCGTGATCCTCNNNAGATCGGAAGA |
| 54365_1 | TATGAGTGTGGAGTCGTTGCATTTGGCAGGGTCATAGGACAATNNNAGATCGGAAG |
| 54368_1 | TATGAGTGTGGAGTCGTTGCCCTCTGAGACATCTTTCCTGGAGNNNAGATCGGAAGA |
| 54369_1 | TATGAGTGTGGAGTCGTTGCCTTGAGCGATTGACCTACCTCAGNNNAGATCGGAAG |
| 54373_1 | TATGAGTGTGGAGTCGTTGCCGGCATGACTCAGTGAATTTGAANNNAGATCGGAAG |
| 54374_1 | TATGAGTGTGGAGTCGTTGCCAGTAGTGCAATCACAGCTCACTNNNAGATCGGAAG |
| 54375_1 | TATGAGTGTGGAGTCGTTGCCAATCTCGGCTCACTACAACCTCNNNAGATCGGAAGA |
| 54377_1 | TATGAGTGTGGAGTCGTTGCCAGTGGCAGGATCTCAGTTTACTNNNAGATCGGAAG |
| 54378_1 | TATGAGTGTGGAGTCGTTGCTTTTCTTAGGGAGGGTACATGGCNNNAGATCGGAAG |
| 54382_1 | TATGAGTGTGGAGTCGTTGCGCCTCCTAAGTAGCTGGGATTACNNNAGATCGGAAG |
| 54385_1 | TATGAGTGTGGAGTCGTTGCCCCAAGTTCAAGCAATTCTCCTGNNNAGATCGGAAGA |
| 54386_1 | TATGAGTGTGGAGTCGTTGCCTCCTGAGTAGCTGCGATTACAGNNNAGATCGGAAG |
| 54387_1 | TATGAGTGTGGAGTCGTTGCGAACATCTGGCCTCAAACAATCCNNNAGATCGGAAGA |
| 54388_1 | TATGAGTGTGGAGTCGTTGCGATCATTTGAGCCTGGGACATTGNNNAGATCGGAAG |
| 54391_1 | TATGAGTGTGGAGTCGTTGCAAGATAGCTTCCCTCATGCCTTCNNNAGATCGGAAGA |
| 54393_1 | TATGAGTGTGGAGTCGTTGCGAACTAGGGGATTATGCTCCAGTNNNAGATCGGAAG |
| 54395_1 | TATGAGTGTGGAGTCGTTGCAAGAACGAGAGCTTCTACTCCACNNNAGATCGGAAG |
| 54396_1 | TATGAGTGTGGAGTCGTTGCCCAGTGCCTTCTTAGAAACTCCANNNAGATCGGAAGA |
| 54397_1 | TATGAGTGTGGAGTCGTTGCATGGTTAGGGCTTATCTCTGCTGNNNAGATCGGAAG |
| 54399_1 | TATGAGTGTGGAGTCGTTGCGTAATCCCAGCTACTTAGGAGGCNNNAGATCGGAAG |
| 54403_1 | TATGAGTGTGGAGTCGTTGCTAATAATTCTGTGGAGGGCCAGGNNNAGATCGGAAG |
| 54404_1 | TATGAGTGTGGAGTCGTTGCACTGGAAAACAATAGCACCTCCTNNNAGATCGGAAG |
| 54405_1 | TATGAGTGTGGAGTCGTTGCATAGACTTGATCCCAAAGGCGAGNNNAGATCGGAAG |
| 54406_1 | TATGAGTGTGGAGTCGTTGCTTTTGAATCTTTTAGTCCTGCTCTAANNNAGATCGGA |

| | |
|---|---|
| 54408_1 | TATGAGTGTGGAGTCGTTGCAGATGGATTCACAGCCAAGTTCTNNNAGATCGGAAG |
| 54411_1 | TATGAGTGTGGAGTCGTTGCAGGAAATGTTTCAGAAGCACTGCNNNAGATCGGAA |
| 54412_1 | TATGAGTGTGGAGTCGTTGCAAACAAAATGCTTTGGTTCAGCCNNNAGATCGGAAG |
| 54413_1 | TATGAGTGTGGAGTCGTTGCAGGAAATTGTGATGCATGCAACANNNAGATCGGAA |
| 54415_1 | TATGAGTGTGGAGTCGTTGCTGATCCTTCTCAGGAAAGAGCACNNNAGATCGGAAG |
| 54417_1 | TATGAGTGTGGAGTCGTTGCGCCCATGCCTGTAATCCTAACTANNNAGATCGGAAG |
| 54418_1 | TATGAGTGTGGAGTCGTTGCCTATGGTTTGGATCTGTGGCCTTNNNAGATCGGAAG |
| 54419_1 | TATGAGTGTGGAGTCGTTGCCCCAGCTTAGTCTGTGATATCCANNNAGATCGGAAG |
| 54421_1 | TATGAGTGTGGAGTCGTTGCGAAGAGGAGAGAAATGGTGAGGTNNNAGATCGGA |
| 54423_1 | TATGAGTGTGGAGTCGTTGCAAGGAAAGGCAAGGCTACATACANNNAGATCGGAA |
| 54425_1 | TATGAGTGTGGAGTCGTTGCTCACTCAAAACCAGAAAGCAGTTNNNAGATCGGAAG |
| 54428_1 | TATGAGTGTGGAGTCGTTGCGGATGAGACTATCACTAGATCCCCNNNAGATCGGAA |
| 54429_1 | TATGAGTGTGGAGTCGTTGCTAGCCCAGCACTGTATTGATGTTNNNAGATCGGAAG |
| 54430_1 | TATGAGTGTGGAGTCGTTGCAGTGGGGTTGGATGAAACTTCTANNNAGATCGGAA |
| 54431_1 | TATGAGTGTGGAGTCGTTGCTCTCCCAACCACTCTGAATCATGNNNAGATCGGAAGA |
| 54432_1 | TATGAGTGTGGAGTCGTTGCCCATCTGTAGGTCACCAACATCANNNAGATCGGAAG |
| 54436_1 | TATGAGTGTGGAGTCGTTGCTGTTAGTCATCCGTGGTCTCTTCNNNAGATCGGAAG |
| 54438_1 | TATGAGTGTGGAGTCGTTGCGCCCATCCTGTCTGTTTACATCTNNNAGATCGGAAGA |
| 54439_1 | TATGAGTGTGGAGTCGTTGCACTTCTGCATTTCCATCTGAGGTNNNAGATCGGAAG |
| 54442_1 | TATGAGTGTGGAGTCGTTGCCAGAAAACAAAGGCAAAAGTGGCNNNAGATCGGAA |
| 54443_1 | TATGAGTGTGGAGTCGTTGCTAGCATGAGAGGCTTGGAAAGAGNNNAGATCGGA |
| 54444_1 | TATGAGTGTGGAGTCGTTGCTATACGTGTGTGTGTGTGTGTGTNNNAGATCGGAA |
| 54447_1 | TATGAGTGTGGAGTCGTTGCTAGCTGTGATTTCTGGATGGTGTNNNAGATCGGAA |
| 54448_1 | TATGAGTGTGGAGTCGTTGCATGCCTGTAGTCCTAGCTACTTGNNNAGATCGGAAG |
| 54449_1 | TATGAGTGTGGAGTCGTTGCATGATGCCCTGTTACTATGTTGCNNNAGATCGGAAG |
| 54450_1 | TATGAGTGTGGAGTCGTTGCCACACACACACACACACATACATNNNAGATCGGAAG |
| 54451_1 | TATGAGTGTGGAGTCGTTGCTCTCTCCTTCCTTTTAGCGCATTNNNAGATCGGAAGA |
| 54453_1 | TATGAGTGTGGAGTCGTTGCTTGAGACCAGTCTGGGTAAATGGNNNAGATCGGAA |
| 54454_1 | TATGAGTGTGGAGTCGTTGCTTCTAATGAACAGCCAGGGTTGTNNNAGATCGGAA |
| 54455_1 | TATGAGTGTGGAGTCGTTGCGTGCAATCTTGGCTCACTGTAAGNNNAGATCGGAAG |
| 54457_1 | TATGAGTGTGGAGTCGTTGCTTTGACACACATTCTCTGAAGCTNNNAGATCGGAAG |
| 54458_1 | TATGAGTGTGGAGTCGTTGCACCCTGTAGTCCTAGCTACTCAGNNNAGATCGGAAG |
| 54459_1 | TATGAGTGTGGAGTCGTTGCAAGGAAAATGTGTTTGCGCTCTTNNNAGATCGGAA |
| 54460_1 | TATGAGTGTGGAGTCGTTGCAAAAACCCCCAAACAAACAAACANNNAGATCGGAAG |
| 54461_1 | TATGAGTGTGGAGTCGTTGCGGTACAGAGAGAAGAGCACACAANNNAGATCGGA |
| 54462_1 | TATGAGTGTGGAGTCGTTGCAGAAAGAGAGAAAGGAAGGACGANNNAGATCGGA |
| 54465_1 | TATGAGTGTGGAGTCGTTGCAAGCATTACTCACAATGGCCATGNNNAGATCGGAAG |
| 54466_1 | TATGAGTGTGGAGTCGTTGCTGCTTCCATGACAAACCAACAAANNNAGATCGGAAG |
| 54468_1 | TATGAGTGTGGAGTCGTTGCCAAGGTCAGGAGTTCGAGATCAGNNNAGATCGGAA |
| 54469_1 | TATGAGTGTGGAGTCGTTGCAGCTGTGTGATTTGTCATCCCTANNNAGATCGGAAG |
| 54470_1 | TATGAGTGTGGAGTCGTTGCGCAGAATGCCCAGAAAAGAAAGANNNAGATCGGAA |
| 54471_1 | TATGAGTGTGGAGTCGTTGCCAGCCAATCCTTCAAGGAATGACNNNAGATCGGAAG |
| 54472_1 | TATGAGTGTGGAGTCGTTGCAAGCCCCCAAACTTCTTATGACTNNNAGATCGGAAG |

| | |
|---|---|
| 54473_1 | TATGAGTGTGGAGTCGTTGCGAAGTCAGGGTGTCAACAGAGATNNNAGATCGGAA |
| 54474_1 | TATGAGTGTGGAGTCGTTGCCCTCATGTTCTCAGAGCTACACTNNNAGATCGGAAG |
| 54475_1 | TATGAGTGTGGAGTCGTTGCTCCTAATTGCCATCAAGTTCCCANNNAGATCGGAAGA |
| 54479_1 | TATGAGTGTGGAGTCGTTGCGCTGATCATGTGTATTTAGCGCANNNAGATCGGAAG |
| 54480_1 | TATGAGTGTGGAGTCGTTGCACAAAAGCATGAAGGAGGAATGCNNNAGATCGGAA |
| 54481_1 | TATGAGTGTGGAGTCGTTGCAAAGACACCAAAAGCAATCGCAANNNAGATCGGAA |
| 54482_1 | TATGAGTGTGGAGTCGTTGCCTAGCGTTACAGGAAGCTCTAGGNNNAGATCGGAA |
| 54483_1 | TATGAGTGTGGAGTCGTTGCCTGTAGTCCCAGCTACTCGGTAGNNNAGATCGGAAG |
| 54484_1 | TATGAGTGTGGAGTCGTTGCTGTGTGGGAGACAATAAAACGGANNNAGATCGGAA |
| 54487_1 | TATGAGTGTGGAGTCGTTGCAAGGAAGGGATCACCATACACACNNNAGATCGGAA |
| 54488_1 | TATGAGTGTGGAGTCGTTGCAGAGAGATAACCCTCACACTTGANNNAGATCGGAAG |
| 54489_1 | TATGAGTGTGGAGTCGTTGCTCGGTTATCTCCTTCAGAAAGCCNNNAGATCGGAAG |
| 54490_1 | TATGAGTGTGGAGTCGTTGCACAAAGAAGTCAGGTGGGTTCTCNNNAGATCGGAA |
| 54491_1 | TATGAGTGTGGAGTCGTTGCAGAAGGTCAGCAGATAAACACGTNNNAGATCGGAA |
| 54493_1 | TATGAGTGTGGAGTCGTTGCACTCCGCCCAGCTAATTTTTGTANNNAGATCGGAAG |
| 54494_1 | TATGAGTGTGGAGTCGTTGCCAAGCAGCTGAGACTACAGACTTNNNAGATCGGAAG |
| 54496_1 | TATGAGTGTGGAGTCGTTGCTCAAGTGATTCTCCTGCCTTAGCNNNAGATCGGAAG |
| 54498_1 | TATGAGTGTGGAGTCGTTGCCTGGCAGGCTCTCATTTTTACCNNNAGATCGGAAGA |
| 54499_1 | TATGAGTGTGGAGTCGTTGCCTCCCGAGTAGCTAGGACTACAGNNNAGATCGGAAG |
| 54500_1 | TATGAGTGTGGAGTCGTTGCCCATCTCAGCTCACTTCAACCTCNNNAGATCGGAAGA |
| 54501_1 | TATGAGTGTGGAGTCGTTGCCTGAGTAGCTGGGATTACAGGTGNNNAGATCGGAA |
| 54503_1 | TATGAGTGTGGAGTCGTTGCTGGAGTGTAGTGGCATGATTTCANNNAGATCGGAA |
| 54505_1 | TATGAGTGTGGAGTCGTTGCGCCTCTCAAGTAGCTGGGATTATNNNAGATCGGAAG |
| 54506_1 | TATGAGTGTGGAGTCGTTGCGCCTCCTAAGTAGCTGGGATTACNNNAGATCGGAAG |
| 54508_1 | TATGAGTGTGGAGTCGTTGCAGAGGCAAAGTTGATACGAAGGANNNAGATCGGAA |
| 54509_1 | TATGAGTGTGGAGTCGTTGCACCAATTCCCCACAACAAAGAACNNNAGATCGGAAG |
| 54510_1 | TATGAGTGTGGAGTCGTTGCGTTCAAGCAATTCTTCTGCCTCANNNAGATCGGAAG |
| 54511_1 | TATGAGTGTGGAGTCGTTGCCTAGGAGGTTCAGTTCATTGGCTNNNAGATCGGAAG |
| 54512_1 | TATGAGTGTGGAGTCGTTGCGGAAGGAATCCAGAGAGGTGTTTNNNAGATCGGAA |
| 54513_1 | TATGAGTGTGGAGTCGTTGCAGCCTGCCATGTAACTGTGATTANNNAGATCGGAAG |
| 54514_1 | TATGAGTGTGGAGTCGTTGCCTCCTGAGTAGCTGGAACTACAGNNNAGATCGGAAG |
| 54515_1 | TATGAGTGTGGAGTCGTTGCGCTCAAGTGATCCTCCTTCTTCANNNAGATCGGAAGA |
| 54518_1 | TATGAGTGTGGAGTCGTTGCTGTGTTCTTATCGTTCCTTGAGANNNAGATCGGAAG |
| 54519_1 | TATGAGTGTGGAGTCGTTGCTTCAAGTGATTCTCCTGCCTCAGNNNAGATCGGAAG |
| 54520_1 | TATGAGTGTGGAGTCGTTGCTGACAATGGCACAATCTTGGTTCNNNAGATCGGAAG |
| 54521_1 | TATGAGTGTGGAGTCGTTGCGAGTACCCAAGTGAAGAGAACGTNNNAGATCGGAA |
| 54522_1 | TATGAGTGTGGAGTCGTTGCGTTCACACTATTCTCCTGCCTCANNNAGATCGGAAGA |
| 54523_1 | TATGAGTGTGGAGTCGTTGCATGATCTCAGCTCACTGCAACTTNNNAGATCGGAAG |
| 54524_1 | TATGAGTGTGGAGTCGTTGCTCCTAGCTGGGACTACAGGTATGNNNAGATCGGAAG |
| 54526_1 | TATGAGTGTGGAGTCGTTGCCTCCCGAGCAGCTGTAATTACAGNNNAGATCGGAAG |
| 54527_1 | TATGAGTGTGGAGTCGTTGCTAGAGTTCAGTGGCACAATCTCGNNNAGATCGGAAG |
| 54528_1 | TATGAGTGTGGAGTCGTTGCTCTGCATCCCAGGTTCAAATGATNNNAGATCGGAAG |
| 54531_1 | TATGAGTGTGGAGTCGTTGCTTCATGCTATTCTCCTGCCTCAGNNNAGATCGGAAGA |

54533_1 TATGAGTGTGGAGTCGTTGCGGTTCCGTGTCATTTGGTTCTTGNNNAGATCGGAAG
54534_1 TATGAGTGTGGAGTCGTTGCTCTTCTCTGGGTTTATGCTGCTTNNNAGATCGGAAG
54535_1 TATGAGTGTGGAGTCGTTGCTGAATATGCTAGATCACAGCCCCNNNAGATCGGAAG
54537_1 TATGAGTGTGGAGTCGTTGCAGAGGCCAGAAGATAACTTGAGCNNNAGATCGGAA
54539_1 TATGAGTGTGGAGTCGTTGCAGAGATGCCTGCTTTATGTAGGGNNNAGATCGGAA
54540_1 TATGAGTGTGGAGTCGTTGCACAATTGGTGGGATGAATCAGGANNNAGATCGGAA
54541_1 TATGAGTGTGGAGTCGTTGCCTGAGGCAGGAGTATCACTTGAGNNNAGATCGGAA
54543_1 TATGAGTGTGGAGTCGTTGCAGGACACTGAGCCAGTTTATCAGNNNAGATCGGAAG
54546_1 TATGAGTGTGGAGTCGTTGCCAACTAGACAGCCTACAGTGCTANNNAGATCGGAAG
54548_1 TATGAGTGTGGAGTCGTTGCGCCAACATGATGGACACGATACTNNNAGATCGGAAG
54549_1 TATGAGTGTGGAGTCGTTGCTGAGACCAAGAGTTTGAGACCAGNNNAGATCGGAA
54551_1 TATGAGTGTGGAGTCGTTGCATCCCTGTACCACCAAACAACTTNNNAGATCGGAAGA
54552_1 TATGAGTGTGGAGTCGTTGCACACCTCAGACCTTACACACAAANNNAGATCGGAAG
54553_1 TATGAGTGTGGAGTCGTTGCTTCTTTGCTAGCTCCATGGTGAANNNAGATCGGAAG
54554_1 TATGAGTGTGGAGTCGTTGCGGATGGACATCACTGAGGAGAATNNNAGATCGGAA
54556_1 TATGAGTGTGGAGTCGTTGCATAGATGAATGCCCTCCTATGGGNNNAGATCGGAAG
54557_1 TATGAGTGTGGAGTCGTTGCCAGATCACAAGGTCAGGAGTTCANNNAGATCGGAA
54559_1 TATGAGTGTGGAGTCGTTGCGTACTTTGTTCTTCAGCAGCACCNNNAGATCGGAAG
54560_1 TATGAGTGTGGAGTCGTTGCAACATATATGGCCGGGTATGGTGNNNAGATCGGAA
54561_1 TATGAGTGTGGAGTCGTTGCTTATGCTAGGTTCACTCTGCCTGNNNAGATCGGAAG
54562_1 TATGAGTGTGGAGTCGTTGCTGCGGTGGCTCATACCTATAATCNNNAGATCGGAAG
54564_1 TATGAGTGTGGAGTCGTTGCACTCTTTATCCATTTGTCCCCCCNNNAGATCGGAAGA
54565_1 TATGAGTGTGGAGTCGTTGCTGAAGGAAGTTAGTGGATGGTGGNNNAGATCGGA
54566_1 TATGAGTGTGGAGTCGTTGCTGAGGTAGGATCTCAGGCTTTTGNNNAGATCGGAA
54567_1 TATGAGTGTGGAGTCGTTGCCAGACAGAAAGCAGCATGTTCAANNNAGATCGGAA
54570_1 TATGAGTGTGGAGTCGTTGCACAAGGTCAAGAGATCGAAACCANNNAGATCGGAA
54571_1 TATGAGTGTGGAGTCGTTGCGAGGTCAGGAGTTTGAGACCAACNNNAGATCGGAA
54572_1 TATGAGTGTGGAGTCGTTGCGACAAGATGGTGACCAAAGGAGANNNAGATCGGA
54573_1 TATGAGTGTGGAGTCGTTGCACAAATGGTACAGAGGGATCCTGNNNAGATCGGAA
54575_1 TATGAGTGTGGAGTCGTTGCGGGGTTTCTTTTTGGAGTGATGGNNNAGATCGGAA
54577_1 TATGAGTGTGGAGTCGTTGCGAGAGAGAGAGAGTCCTACACCANNNAGATCGGAA
54578_1 TATGAGTGTGGAGTCGTTGCGCACCTGTAGACTTCAATGTCCTNNNAGATCGGAAG
54580_1 TATGAGTGTGGAGTCGTTGCTAGAGGAGTGATTCCAGGAAGGGNNNAGATCGGA
54581_1 TATGAGTGTGGAGTCGTTGCTGGCCACTCCTAGTTTTCTTGAANNNAGATCGGAAG
54584_1 TATGAGTGTGGAGTCGTTGCTCCTTCTGTCTACTACCAACTGANNNAGATCGGAAGA
54585_1 TATGAGTGTGGAGTCGTTGCAGAAGAGCTACTCTGAAGTCCCTNNNAGATCGGAAG
54586_1 TATGAGTGTGGAGTCGTTGCTCACTTGAGTCCAGGAGTTCAAGNNNAGATCGGAAG
54587_1 TATGAGTGTGGAGTCGTTGCAAATGGTTCTAATGCTGCCAACCNNNAGATCGGAAG
54588_1 TATGAGTGTGGAGTCGTTGCACATGGTGAAACCTCGTCTCTACNNNAGATCGGAAG
54589_1 TATGAGTGTGGAGTCGTTGCGGAAGAGAAGCACAGAAGTCACTNNNAGATCGGAA
54591_1 TATGAGTGTGGAGTCGTTGCGTAGCTGGGATTACAGGCATGTANNNAGATCGGAA
54592_1 TATGAGTGTGGAGTCGTTGCCTCCTGAGTAGCTGGGACTACAGNNNAGATCGGAA
54593_1 TATGAGTGTGGAGTCGTTGCTAGACAGGGTGTTACTCTGAAGCNNNAGATCGGAA

| | |
|---|---|
| 54595_1 | TATGAGTGTGGAGTCGTTGCAGTAGCTGGGATTACAAGTGCATNNNAGATCGGAA |
| 54598_1 | TATGAGTGTGGAGTCGTTGCTTCACGACATTCTCCTGCCTTAGNNNAGATCGGAAG |
| 54599_1 | TATGAGTGTGGAGTCGTTGCTCCGGGATTCAAGTCATTTTCCTNNNAGATCGGAAG |
| 54600_1 | TATGAGTGTGGAGTCGTTGCCTGGAGTACAATGGCGTGATTTCNNNAGATCGGAA |
| 54602_1 | TATGAGTGTGGAGTCGTTGCTGAATCCCAGCTCTATTTCCTGGNNNAGATCGGAAG |
| 54603_1 | TATGAGTGTGGAGTCGTTGCCTGGGTGAGGTTGATTCTCCTTGNNNAGATCGGAA |
| 54604_1 | TATGAGTGTGGAGTCGTTGCATTATCCTGTCTCAGTCTCCCGANNNAGATCGGAAG |
| 54605_1 | TATGAGTGTGGAGTCGTTGCTCCCGAGTAGCTAGGCCTAATATNNNAGATCGGAAG |
| 54608_1 | TATGAGTGTGGAGTCGTTGCCTGGAGTGCTATGGTGCAATCTTNNNAGATCGGAA |
| 54609_1 | TATGAGTGTGGAGTCGTTGCCTGAGTAGCTGGGATTACAGGTGNNNAGATCGGAA |
| 54610_1 | TATGAGTGTGGAGTCGTTGCGTAGATTCTCACTCTGTTGCCCANNNAGATCGGAAG |
| 54611_1 | TATGAGTGTGGAGTCGTTGCGCAACAGGACGAATGGATTTCTTNNNAGATCGGAA |
| 54612_1 | TATGAGTGTGGAGTCGTTGCTTCAAGTGATTCTCCTGCCTCAGNNNAGATCGGAAG |
| 54616_1 | TATGAGTGTGGAGTCGTTGCCCTTGAAGGAGGGTTTGTACAGANNNAGATCGGAA |
| 54617_1 | TATGAGTGTGGAGTCGTTGCCTGCCCTAGAGATTTGTGGAACTNNNAGATCGGAAG |
| 54618_1 | TATGAGTGTGGAGTCGTTGCCTGTAGTCCCAGCTACTCAGGAGNNNAGATCGGAAG |
| 54619_1 | TATGAGTGTGGAGTCGTTGCAAAAGAAAAAGGCAAGGAGAGGCNNNAGATCGGA |
| 54621_1 | TATGAGTGTGGAGTCGTTGCTAGTGAACTGTGATTGCACCACTNNNAGATCGGAAG |
| 54623_1 | TATGAGTGTGGAGTCGTTGCGAAGGAGAGAAGGAAGGAAGGGANNNAGATCGG |
| 54624_1 | TATGAGTGTGGAGTCGTTGCGGTGGATCATGACGTCAAGAGATNNNAGATCGGAA |
| 54625_1 | TATGAGTGTGGAGTCGTTGCAAAATAAATTCCTGGCTGGGCACNNNAGATCGGAA |
| 54626_1 | TATGAGTGTGGAGTCGTTGCTGCCAAACACTTTTCCAAAATGGNNNAGATCGGAAG |
| 54628_1 | TATGAGTGTGGAGTCGTTGCCTTGGTGGTCCCTGGAGATTATCNNNAGATCGGAAG |
| 54629_1 | TATGAGTGTGGAGTCGTTGCTCACTTGAGTCCAGGAAATCCAGNNNAGATCGGAAG |
| 54630_1 | TATGAGTGTGGAGTCGTTGCTCAATAGTGGTGAGTGAGTGCTCNNNAGATCGGAA |
| 54631_1 | TATGAGTGTGGAGTCGTTGCATATGAAGCCATCACGGAACTGANNNAGATCGGAA |
| 54633_1 | TATGAGTGTGGAGTCGTTGCCCTGCTCCAGCTATGTAAGATGTNNNAGATCGGAAG |
| 54634_1 | TATGAGTGTGGAGTCGTTGCCCCCTGAGCATAGGAAAGTCAAANNNAGATCGGAA |
| 54635_1 | TATGAGTGTGGAGTCGTTGCGATCACTTGAGCCCAGGAATTTGNNNAGATCGGAA |
| 54636_1 | TATGAGTGTGGAGTCGTTGCATCTGTTTACTAAGGGCTGCCTGNNNAGATCGGAAG |
| 54637_1 | TATGAGTGTGGAGTCGTTGCCCAGTCTTCAAAGCAGAGGAGATNNNAGATCGGAA |
| 54638_1 | TATGAGTGTGGAGTCGTTGCATCACCACTTTGTAACCCCCAATNNNAGATCGGAAGA |
| 54639_1 | TATGAGTGTGGAGTCGTTGCTGCCAGTTTATTACGAAGGACATGNNNAGATCGGA |
| 54640_1 | TATGAGTGTGGAGTCGTTGCGAGGTCAGGAGATCAAGACCATCNNNAGATCGGAA |
| 54641_1 | TATGAGTGTGGAGTCGTTGCCCAATCTGACCTGTGTCCTTGTANNNAGATCGGAAG |
| 54642_1 | TATGAGTGTGGAGTCGTTGCATACAAAAATTAGCCGGGTGTGGNNNAGATCGGAA |
| 54643_1 | TATGAGTGTGGAGTCGTTGCATTGGACCCCCTGTATACTGTTGNNNAGATCGGAAG |
| 54644_1 | TATGAGTGTGGAGTCGTTGCAGGCAAAAAGAATTGTTTCTGCANNNAGATCGGAA |
| 54645_1 | TATGAGTGTGGAGTCGTTGCACTTTAGGGACCTGTGAGACTCTNNNAGATCGGAAG |
| 54646_1 | TATGAGTGTGGAGTCGTTGCCCTCCGAAATTCACGTGTTGAAANNNAGATCGGAAG |
| 54648_1 | TATGAGTGTGGAGTCGTTGCATAACAGGGCCAAGAGTTTCCATNNNAGATCGGAA |
| 54650_1 | TATGAGTGTGGAGTCGTTGCTATTCCTGACAACTGGCTACAGGNNNAGATCGGAAG |
| 54651_1 | TATGAGTGTGGAGTCGTTGCTGACACTGTTGCATTGGGAATTCNNNAGATCGGAAG |

| | |
|---|---|
| 54653_1 | TATGAGTGTGGAGTCGTTGCTCAGGGAAATGCAAATCGAAACCNNNAGATCGGAA |
| 54654_1 | TATGAGTGTGGAGTCGTTGCCACAAGCCTACAGTCCTAGCTACNNNAGATCGGAAG |
| 54657_1 | TATGAGTGTGGAGTCGTTGCCTCCTGGGTTTGAGCAATTCTTGNNNAGATCGGAAG |
| 54659_1 | TATGAGTGTGGAGTCGTTGCGGATCACTGCAATCTCTACCTCCNNNAGATCGGAAG |
| 54661_1 | TATGAGTGTGGAGTCGTTGCCCTGAGTAGCTGGGATTATAGGCNNNAGATCGGAA |
| 54663_1 | TATGAGTGTGGAGTCGTTGCTCATGCCTGTAATCCCAACACTTNNNAGATCGGAAG |
| 54664_1 | TATGAGTGTGGAGTCGTTGCGTCTTGAACTCCTGACCTTGTGANNNAGATCGGAAG |
| 54666_1 | TATGAGTGTGGAGTCGTTGCCTCCTGAGTAGCTGGGATTACAGNNNAGATCGGAA |
| 54667_1 | TATGAGTGTGGAGTCGTTGCCACAATCTTGGCTCACTGCAAGNNNAGATCGGAAGA |
| 54668_1 | TATGAGTGTGGAGTCGTTGCTCTTGACTCACTACAACCTCTGCNNNAGATCGGAAGA |
| 54669_1 | TATGAGTGTGGAGTCGTTGCCCCACAGAGCCACAATTTTTCATNNNAGATCGGAAG |
| 54671_1 | TATGAGTGTGGAGTCGTTGCCCACCCATGAGAAAATATGCTGCNNNAGATCGGAAG |
| 54672_1 | TATGAGTGTGGAGTCGTTGCCAAGTAGCTGGGATTACAGGCATNNNAGATCGGAA |
| 54674_1 | TATGAGTGTGGAGTCGTTGCCCGCAGGTGGGTCATTTCTAATTNNNAGATCGGAAG |
| 54676_1 | TATGAGTGTGGAGTCGTTGCAGACAGGGTCTCACTTTGTTACCNNNAGATCGGAAG |
| 54677_1 | TATGAGTGTGGAGTCGTTGCCCTCCTGGGTTCAAGTGATTCTCNNNAGATCGGAAG |
| 54678_1 | TATGAGTGTGGAGTCGTTGCCCTCCTAAAGTGCTGGGATTACANNNAGATCGGAAG |
| 54680_1 | TATGAGTGTGGAGTCGTTGCCCTGAGTAGCTGGGATTATAGGCNNNAGATCGGAA |
| 54681_1 | TATGAGTGTGGAGTCGTTGCCCTCCTGAGTAGCTGAGACTACANNNAGATCGGAAG |
| 54682_1 | TATGAGTGTGGAGTCGTTGCTTTCAAGTGATTCTCCTGCCTCANNNAGATCGGAAG |
| 54683_1 | TATGAGTGTGGAGTCGTTGCGTGGTATGTGCCTGTAGTCCTAGNNNAGATCGGAA |
| 54684_1 | TATGAGTGTGGAGTCGTTGCCCACTAAAAGGGAGAGAACAGCTNNNAGATCGGAA |
| 54686_1 | TATGAGTGTGGAGTCGTTGCACTCTCTCTCTCTTTCTCCCTCCNNNAGATCGGAAGAG |
| 54688_1 | TATGAGTGTGGAGTCGTTGCTGAGCTAATCTTAGGGCCTACCTNNNAGATCGGAAG |
| 54689_1 | TATGAGTGTGGAGTCGTTGCAAGGTGTTATCTGGCACTGTAGANNNAGATCGGAA |
| 54691_1 | TATGAGTGTGGAGTCGTTGCCAGAGGTTACAGTGAGCCAAGATNNNAGATCGGAA |
| 54696_1 | TATGAGTGTGGAGTCGTTGCGAAGGAAGGAAGGAAGGAAGGAANNNAGATCGG |
| 54697_1 | TATGAGTGTGGAGTCGTTGCAGCATCTTGTTTAGGACTAACTACTNNNAGATCGGA |
| 54701_1 | TATGAGTGTGGAGTCGTTGCGTGAGTCAGCAGAACAATTTCCCNNNAGATCGGAAG |
| 54705_1 | TATGAGTGTGGAGTCGTTGCCGGTGATCGTGAAATAGACATGCNNNAGATCGGAA |
| 54706_1 | TATGAGTGTGGAGTCGTTGCTGGGATAATTTGGCCACTTCTGANNNAGATCGGAAG |
| 54707_1 | TATGAGTGTGGAGTCGTTGCGGCGGAGCTTGCAAAAAAAAAAANNNAGATCGGAA |
| 54709_1 | TATGAGTGTGGAGTCGTTGCCATACGTGCAGCCAACAAACATANNNAGATCGGAAG |
| 54710_1 | TATGAGTGTGGAGTCGTTGCTTGCATTTGGGGATTTCTGGAACNNNAGATCGGAA |
| 54711_1 | TATGAGTGTGGAGTCGTTGCCTCACTGTAACCTCCACTTTCCANNNAGATCGGAAGA |
| 54713_1 | TATGAGTGTGGAGTCGTTGCATAAGAGTCTCATTGTGGCCCAGNNNAGATCGGAA |
| 54714_1 | TATGAGTGTGGAGTCGTTGCAATCTCATTCTATTGCCCAGGCTNNNAGATCGGAAG |
| 54715_1 | TATGAGTGTGGAGTCGTTGCATGATATCAGCTCACTGCAACCTNNNAGATCGGAAG |
| 54716_1 | TATGAGTGTGGAGTCGTTGCTAGCAATCCTCCCATTTCCTGACNNNAGATCGGAAGA |
| 54717_1 | TATGAGTGTGGAGTCGTTGCAGCCTTGATCTTCTGGACTCAAGNNNAGATCGGAAG |
| 54721_1 | TATGAGTGTGGAGTCGTTGCTTACTATATTGCCCAGACCGGACNNNAGATCGGAAG |
| 54722_1 | TATGAGTGTGGAGTCGTTGCCACCTCAGCCTCTTGAGTATCTGNNNAGATCGGAAG |
| 54723_1 | TATGAGTGTGGAGTCGTTGCTTGAACTCCTGACCTCGTGATTCNNNAGATCGGAAG |

| | |
|---|---|
| 54725_1 | TATGAGTGTGGAGTCGTTGCTGGAGTACAGTGGTGTGATCATGNNNAGATCGGAA |
| 54726_1 | TATGAGTGTGGAGTCGTTGCATCCAAGCAGATCCAAACCATCTNNNAGATCGGAAG |
| 54727_1 | TATGAGTGTGGAGTCGTTGCAGGTTCAAGCGATTCTCTTGTCTNNNAGATCGGAAG |
| 54728_1 | TATGAGTGTGGAGTCGTTGCGGTTCAAGTGATTCTCCTGCTTCNNNAGATCGGAAG |
| 54729_1 | TATGAGTGTGGAGTCGTTGCCCTGGGTTCAAGCAATTCTTCTGNNNAGATCGGAAG |
| 54730_1 | TATGAGTGTGGAGTCGTTGCAAGCACAGTATTACACCCTTGCTNNNAGATCGGAAG |
| 54732_1 | TATGAGTGTGGAGTCGTTGCAAGTAATTCTCTGCCTCAGCCTCNNNAGATCGGAAG |
| 54734_1 | TATGAGTGTGGAGTCGTTGCCTCAAGCAATTCTCCTGCCTCAGNNNAGATCGGAAG |
| 54736_1 | TATGAGTGTGGAGTCGTTGCCACCATGCCCAGCTAATTTTTGTNNNAGATCGGAAG |
| 54738_1 | TATGAGTGTGGAGTCGTTGCGATTCCAGACGTGAGTCAGAGTTNNNAGATCGGAA |
| 54741_1 | TATGAGTGTGGAGTCGTTGCACCTCATTGCACTCCATGTACATNNNAGATCGGAAGA |
| 54742_1 | TATGAGTGTGGAGTCGTTGCTGGTGCAAATAGGGATCATGAGANNNAGATCGGAA |
| 54743_1 | TATGAGTGTGGAGTCGTTGCATGATCTCAGTTCACAGCAACCTNNNAGATCGGAAG |
| 54748_1 | TATGAGTGTGGAGTCGTTGCACCACTGGAAGCTCGAAGTATTTNNNAGATCGGAAG |
| 54749_1 | TATGAGTGTGGAGTCGTTGCCTCACTGCAAGCTCTGAATTGAGNNNAGATCGGAAG |
| 54750_1 | TATGAGTGTGGAGTCGTTGCTCTGTTGAATTTGTTGCCTGCTTNNNAGATCGGAAG |
| 54751_1 | TATGAGTGTGGAGTCGTTGCACAACATCTGTGGCAACACTTTANNNAGATCGGAAG |
| 54753_1 | TATGAGTGTGGAGTCGTTGCCAGTGTCATAATCTCGGCTCACTNNNAGATCGGAAG |
| 54755_1 | TATGAGTGTGGAGTCGTTGCGAATAGCTGGGAATACAGGCACCNNNAGATCGGAA |
| 54757_1 | TATGAGTGTGGAGTCGTTGCGATCATTCTTGGGTGTTTCTCGCNNNAGATCGGAAG |
| 54758_1 | TATGAGTGTGGAGTCGTTGCCCTGCCTCAGTCAGACCTTTTTANNNAGATCGGAAGA |
| 54760_1 | TATGAGTGTGGAGTCGTTGCCAAGTAGCTGGGATTACAGGCATNNNAGATCGGAA |
| 54762_1 | TATGAGTGTGGAGTCGTTGCACAGAGGACCTAAAATAACCCAANNNAGATCGGAAG |
| 54763_1 | TATGAGTGTGGAGTCGTTGCGTGAGCCATTTTGTTGGGTGTAANNNAGATCGGAA |
| 54764_1 | TATGAGTGTGGAGTCGTTGCCTTCCTTTCCAGAATGCAACCACNNNAGATCGGAAGA |
| 54765_1 | TATGAGTGTGGAGTCGTTGCAAAGGCTGAAACAGATGCAAACANNNAGATCGGAA |
| 54766_1 | TATGAGTGTGGAGTCGTTGCTCTATCCCTCCTCCACTCATCTGNNNAGATCGGAAGA |
| 54767_1 | TATGAGTGTGGAGTCGTTGCAAGGAAAAGGAGTGTTCAATGGCNNNAGATCGGAA |
| 54768_1 | TATGAGTGTGGAGTCGTTGCGAGAGAGATGACAAAGGAGTGCANNNAGATCGGA |
| 54769_1 | TATGAGTGTGGAGTCGTTGCTTGCGTGTAATTGTGTATCCTGCNNNAGATCGGAAG |
| 54770_1 | TATGAGTGTGGAGTCGTTGCACCTTTTGCTCATGCCCTAAGTANNNAGATCGGAAGA |
| 54771_1 | TATGAGTGTGGAGTCGTTGCGAAGGGACTCTTCCTCAACTCATNNNAGATCGGAAG |
| 54773_1 | TATGAGTGTGGAGTCGTTGCGCCAGAGAGTGAGCTTTTCAAAGNNNAGATCGGAA |
| 54774_1 | TATGAGTGTGGAGTCGTTGCAAAGTAATGGCATGCTACTGGGANNNAGATCGGAA |
| 54775_1 | TATGAGTGTGGAGTCGTTGCCTCCAAGGTCTCTTTCAACTCCANNNAGATCGGAAGA |
| 54777_1 | TATGAGTGTGGAGTCGTTGCAGGATGGAATAGGGAGGATCCTTNNNAGATCGGAA |
| 54779_1 | TATGAGTGTGGAGTCGTTGCACCAAAACGTGAGGACCTGATTANNNAGATCGGAA |
| 54782_1 | TATGAGTGTGGAGTCGTTGCGGGAAGAGAAGTGGGTATGACTGNNNAGATCGGA |
| 54784_1 | TATGAGTGTGGAGTCGTTGCCTGAGGCAGGACAATCACTTGAANNNAGATCGGAA |
| 54785_1 | TATGAGTGTGGAGTCGTTGCGGCTTGGGTTTGGTTTTGTGATANNNAGATCGGAA |
| 54786_1 | TATGAGTGTGGAGTCGTTGCAAAATCCTCTGTCCCCTCTGAACNNNAGATCGGAAGA |
| 54787_1 | TATGAGTGTGGAGTCGTTGCAGGCTCACTTTTCCTGAATTGGANNNAGATCGGAAG |
| 54788_1 | TATGAGTGTGGAGTCGTTGCTTTTACAGGTTAGTGCTGCATGGNNNAGATCGGAA |

| | |
|---|---|
| 54792_1 | TATGAGTGTGGAGTCGTTGCGTGGCAATCCTAGTCTCTGACTANNNAGATCGGAAG |
| 54794_1 | TATGAGTGTGGAGTCGTTGCACAGATTCTGGGGAGGATATGGANNNAGATCGGA |
| 54796_1 | TATGAGTGTGGAGTCGTTGCTCAGTGCATAACCAAGGCCATATNNNAGATCGGAAG |
| 54798_1 | TATGAGTGTGGAGTCGTTGCCAGAATCTGGCAGTCCTAGCTAGNNNAGATCGGAA |
| 54801_1 | TATGAGTGTGGAGTCGTTGCGCTACTTGGGAGGCTAAGGTTAGNNNAGATCGGAA |
| 54802_1 | TATGAGTGTGGAGTCGTTGCCAGTCATTGATGCCAGGACAAAGNNNAGATCGGAA |
| 54803_1 | TATGAGTGTGGAGTCGTTGCTCTCTTCATACATTCGCTTCTGCNNNAGATCGGAAGA |
| 54804_1 | TATGAGTGTGGAGTCGTTGCCAAAAATTAGCTGGGCATGGTGGNNNAGATCGGAA |
| 54805_1 | TATGAGTGTGGAGTCGTTGCACTATAAGCCCCCAGTCTCTTCTNNNAGATCGGAAGA |
| 54807_1 | TATGAGTGTGGAGTCGTTGCTGGACCTGATGCAGATCATGAAANNNAGATCGGAA |
| 54808_1 | TATGAGTGTGGAGTCGTTGCTGAATGGTGATGCTAGGCTATCCNNNAGATCGGAA |
| 54809_1 | TATGAGTGTGGAGTCGTTGCAGAGAGACTGACGACAATTTGCTNNNAGATCGGAA |
| 54811_1 | TATGAGTGTGGAGTCGTTGCCGACTGAATTGCTGCAATCTCATNNNAGATCGGAAG |
| 54812_1 | TATGAGTGTGGAGTCGTTGCTTGGCATGAGCAACTAAAAAGGGNNNAGATCGGAA |
| 54813_1 | TATGAGTGTGGAGTCGTTGCACCTTGAGAAAAGTTTCCCGAGANNNAGATCGGAA |
| 54814_1 | TATGAGTGTGGAGTCGTTGCGATGAGTTCATGTCCTTTGCAGGNNNAGATCGGAA |
| 54816_1 | TATGAGTGTGGAGTCGTTGCGACTGGCCAACATAGTGAAAACCNNNAGATCGGAA |
| 54818_1 | TATGAGTGTGGAGTCGTTGCGTAATCCTAGCTACTTGGGAGGCNNNAGATCGGAA |
| 54820_1 | TATGAGTGTGGAGTCGTTGCCCAGGAATTAGATCCCTACTCGCNNNAGATCGGAAG |
| 54821_1 | TATGAGTGTGGAGTCGTTGCGAATTGCTTGAATCTGGGAGGTGNNNAGATCGGAA |
| 54822_1 | TATGAGTGTGGAGTCGTTGCGTAGGAGAATCACTTGAACCCGTNNNAGATCGGAA |
| 54823_1 | TATGAGTGTGGAGTCGTTGCGCAAAGGCAGGATAATTGCTTGANNNAGATCGGAA |
| 54824_1 | TATGAGTGTGGAGTCGTTGCAACTGGGAGAATCAGCTGTGATCNNNAGATCGGAA |
| 54827_1 | TATGAGTGTGGAGTCGTTGCACTGGCTTGAAGATAATGGGGAANNNAGATCGGAA |
| 54829_1 | TATGAGTGTGGAGTCGTTGCAACCTGGTGAAACCCCATCTTTANNNAGATCGGAAG |
| 54830_1 | TATGAGTGTGGAGTCGTTGCTCGGAGATGGCATCTAGTTTGAGNNNAGATCGGAA |
| 54832_1 | TATGAGTGTGGAGTCGTTGCGGAGGAGGAGGAGGAAGAAAAGANNNAGATCGG |
| 54833_1 | TATGAGTGTGGAGTCGTTGCTCAAAGACTCCTGTCAACTGGACNNNAGATCGGAAG |
| 54834_1 | TATGAGTGTGGAGTCGTTGCAGCCAGATAGAAAGTTGCACAGANNNAGATCGGAA |
| 54839_1 | TATGAGTGTGGAGTCGTTGCGTGATCTCAGCTCACTACAACCTNNNAGATCGGAAG |
| 54840_1 | TATGAGTGTGGAGTCGTTGCCACGCCTGGCTAATTTGTTGTATNNNAGATCGGAAG |
| 54842_1 | TATGAGTGTGGAGTCGTTGCTCAGCCTCCCAAAGTTTTCAGATNNNAGATCGGAAG |
| 54843_1 | TATGAGTGTGGAGTCGTTGCCAATGATCTTGGCTCACTGCAACNNNAGATCGGAAG |
| 54845_1 | TATGAGTGTGGAGTCGTTGCCGATCTCCTGACCTCATGATCTGNNNAGATCGGAAG |
| 54846_1 | TATGAGTGTGGAGTCGTTGCCTTCCCTGAGTCAAACCATCCTCNNNAGATCGGAAGA |
| 54847_1 | TATGAGTGTGGAGTCGTTGCCCTCACTTCAACCTCTGTCCTTTNNNAGATCGGAAGA |
| 54848_1 | TATGAGTGTGGAGTCGTTGCGGGTTCAAGTGTTTCTCCTGTCTNNNAGATCGGAAG |
| 54850_1 | TATGAGTGTGGAGTCGTTGCCCTCCCAAGTAGCTGGAATGATGNNNAGATCGGAA |
| 54851_1 | TATGAGTGTGGAGTCGTTGCTCCAGGTTCAAGTGATTCTCCTGNNNAGATCGGAAG |
| 54852_1 | TATGAGTGTGGAGTCGTTGCGTTCAAGAGATTCTCCTGCCTCANNNAGATCGGAAG |
| 54853_1 | TATGAGTGTGGAGTCGTTGCTTGAACTCCTGGGCTCTAGTGATNNNAGATCGGAAG |
| 54856_1 | TATGAGTGTGGAGTCGTTGCCTTGCTTGCTTTGGATGGAGAAANNNAGATCGGAA |
| 54857_1 | TATGAGTGTGGAGTCGTTGCGGTTCAAACAATTCTCCTGCCTCNNNAGATCGGAAG |

| | |
|---|---|
| 54858_1 | TATGAGTGTGGAGTCGTTGCTCTTGTATTGAACTCCTGGGCTCNNNAGATCGGAAG |
| 54861_1 | TATGAGTGTGGAGTCGTTGCTCAGTATCCAAAGGTGAGTCAGCNNNAGATCGGAA |
| 54862_1 | TATGAGTGTGGAGTCGTTGCATCCTGTGTGTTGGGAACCTTTANNNAGATCGGAAG |
| 54863_1 | TATGAGTGTGGAGTCGTTGCGTAGATGGGACTACAGGTGTGTGNNNAGATCGGA |
| 54865_1 | TATGAGTGTGGAGTCGTTGCTGCAATCTCAGCTTACTGGCTTANNNAGATCGGAAG |
| 54868_1 | TATGAGTGTGGAGTCGTTGCGTTCAAGAGATTCTCCTGCCTCANNNAGATCGGAAG |
| 54869_1 | TATGAGTGTGGAGTCGTTGCTCCCACATTCATGCCATTCTCTCNNNAGATCGGAAGA |
| 54870_1 | TATGAGTGTGGAGTCGTTGCAGGTTCAACTGATTCTTCTGCCTNNNAGATCGGAAG |
| 54872_1 | TATGAGTGTGGAGTCGTTGCGTTCAAGCAATTCTCTTGCCTCANNNAGATCGGAAG |
| 54873_1 | TATGAGTGTGGAGTCGTTGCCACTGTCTCTGTCCCTTGCTAATNNNAGATCGGAAGA |
| 54874_1 | TATGAGTGTGGAGTCGTTGCCTTGAACTCCTGGCTGAAGGTATNNNAGATCGGAAG |
| 54875_1 | TATGAGTGTGGAGTCGTTGCTCCTGGATTCAACAGATTCTCCCNNNAGATCGGAAG |
| 54878_1 | TATGAGTGTGGAGTCGTTGCAACCTTTATCTCCCAGGTTCAGGNNNAGATCGGAAG |
| 54880_1 | TATGAGTGTGGAGTCGTTGCAGATGGAGTCTTCCTTTGTCACCNNNAGATCGGAAG |
| 54881_1 | TATGAGTGTGGAGTCGTTGCAAAGGTTCAAGCGATTCTTCTGCNNNAGATCGGAAG |
| 54882_1 | TATGAGTGTGGAGTCGTTGCTCAGGTTCAAACGATTTTCCTGCNNNAGATCGGAAG |
| 54884_1 | TATGAGTGTGGAGTCGTTGCCACAGGCTCTTTTTTGGTTCCATNNNAGATCGGAAG |
| 54885_1 | TATGAGTGTGGAGTCGTTGCGTTCAAGCAATTCTCTGCCTCAGNNNAGATCGGAAG |
| 54886_1 | TATGAGTGTGGAGTCGTTGCTTCAAGTGATTCTCCTGCCTCAGNNNAGATCGGAAG |
| 54887_1 | TATGAGTGTGGAGTCGTTGCCCTCTGGGGTTCAAGAGATTCTCNNNAGATCGGAAG |
| 54890_1 | TATGAGTGTGGAGTCGTTGCAGCCTCCTGAATAGCTGGGATTANNNAGATCGGAA |
| 54891_1 | TATGAGTGTGGAGTCGTTGCACAGTGACATGATCTCAGCTCTCNNNAGATCGGAAG |
| 54894_1 | TATGAGTGTGGAGTCGTTGCTTCAAGTGGTTCTCCTGTCTCAGNNNAGATCGGAAG |
| 54895_1 | TATGAGTGTGGAGTCGTTGCGAATAGCTGGGATTACACGTGTGNNNAGATCGGAA |
| 54896_1 | TATGAGTGTGGAGTCGTTGCTTATGGCTTACTGTGTCCTCGACNNNAGATCGGAAG |
| 54898_1 | TATGAGTGTGGAGTCGTTGCACTGCAGTCTCAACCTTCTAAGGNNNAGATCGGAAG |
| 54899_1 | TATGAGTGTGGAGTCGTTGCACCGCACTCAGCCTAAAAATTTTNNNAGATCGGAAG |
| 54901_1 | TATGAGTGTGGAGTCGTTGCGATTCAAGCGATTCTCCTGCCTCNNNAGATCGGAAG |
| 54902_1 | TATGAGTGTGGAGTCGTTGCGTTGGTGAGGATGCAGAGAAAAGNNNAGATCGGA |
| 54903_1 | TATGAGTGTGGAGTCGTTGCGCCGTGTGCTGAACATTTATCTTNNNAGATCGGAAG |
| 54905_1 | TATGAGTGTGGAGTCGTTGCAAGTGAAGAGACATCAGCAAGGANNNAGATCGGA |
| 54907_1 | TATGAGTGTGGAGTCGTTGCCTGGGGTAACTACACACTAGCTCNNNAGATCGGAAG |
| 54908_1 | TATGAGTGTGGAGTCGTTGCCTGTAGTCCCAGCTACTCAGGAGNNNAGATCGGAAG |
| 54911_1 | TATGAGTGTGGAGTCGTTGCCTAACACCTGTAATCCCAGCACTNNNAGATCGGAAG |
| 54912_1 | TATGAGTGTGGAGTCGTTGCGGAAACGCAAAAGAAGTAGCTGTNNNAGATCGGAA |
| 54915_1 | TATGAGTGTGGAGTCGTTGCCCATCAGGATTTTGAGAGGGACTNNNAGATCGGAA |
| 54916_1 | TATGAGTGTGGAGTCGTTGCAAGAAAGAAAAAAAGGCCAGGCGNNNAGATCGGA |
| 54919_1 | TATGAGTGTGGAGTCGTTGCCAGTGAGCCAAAATCATGCCATTNNNAGATCGGAAG |
| 54920_1 | TATGAGTGTGGAGTCGTTGCCTGTAGTCCCAGCTACTTGAGAGNNNAGATCGGAAG |
| 54922_1 | TATGAGTGTGGAGTCGTTGCACTTTTACACTGTTGGTGGGACTNNNAGATCGGAAG |
| 54923_1 | TATGAGTGTGGAGTCGTTGCAAGCACCTTTTTTCCCTGTCAACNNNAGATCGGAAGA |
| 54927_1 | TATGAGTGTGGAGTCGTTGCCCTTCCCTTCCTTCCACTGTAATNNNAGATCGGAAGA |
| 54929_1 | TATGAGTGTGGAGTCGTTGCGGACTTGAGAACTCTCTAGCCTGNNNAGATCGGAAG |

| | |
|---|---|
| 54932_1 | TATGAGTGTGGAGTCGTTGCGGGCCATTGGAATTTCCTCAAAANNNAGATCGGAAG |
| 54933_1 | TATGAGTGTGGAGTCGTTGCAGTCCCAAACTGGATGGAAACAANNNAGATCGGAA |
| 54934_1 | TATGAGTGTGGAGTCGTTGCTGAGTGCAAATTCAAAACATGCANNNAGATCGGAA |
| 54935_1 | TATGAGTGTGGAGTCGTTGCACTGTAGGCATGTTAGCATCAACNNNAGATCGGAAG |
| 54937_1 | TATGAGTGTGGAGTCGTTGCACTACTTCCATAATGCCTGAGGTNNNAGATCGGAAG |
| 54938_1 | TATGAGTGTGGAGTCGTTGCTCCACAGCTCTCTTTCAATTCCANNNAGATCGGAAGA |
| 54943_1 | TATGAGTGTGGAGTCGTTGCAAAAATTAGGCTGGGTTTGCTGGNNNAGATCGGAA |
| 54944_1 | TATGAGTGTGGAGTCGTTGCCCTGGGAGGTCGAGGTTATAATGNNNAGATCGGAA |
| 54946_1 | TATGAGTGTGGAGTCGTTGCAGCACAATTGTAAAGCATGCACTNNNAGATCGGAAG |
| 54947_1 | TATGAGTGTGGAGTCGTTGCTCTCCAAAGTGTTCTCATGTGGCNNNAGATCGGAAG |
| 54948_1 | TATGAGTGTGGAGTCGTTGCTCACTTGAGAGGTCAGGAATTCGNNNAGATCGGAA |
| 54949_1 | TATGAGTGTGGAGTCGTTGCTTCTGTCAGAGCAGCTCTTGTTANNNAGATCGGAAG |
| 54951_1 | TATGAGTGTGGAGTCGTTGCCCATGAACTTACAACCCAGCTTGNNNAGATCGGAAG |
| 54953_1 | TATGAGTGTGGAGTCGTTGCGGAAATTGTCGAGTTGCTGTGAGNNNAGATCGGAA |
| 54954_1 | TATGAGTGTGGAGTCGTTGCGTAGGAGATAGTAGTTGCCCTGCNNNAGATCGGAA |
| 54955_1 | TATGAGTGTGGAGTCGTTGCTGAACTCTGGATTTTTTTCCCGANNNAGATCGGAA |
| 54956_1 | TATGAGTGTGGAGTCGTTGCACAAGGTGAAACCCAGTCTCTACNNNAGATCGGAAG |
| 54958_1 | TATGAGTGTGGAGTCGTTGCAATCTGATGAAAGCTGTGGACCTNNNAGATCGGAA |
| 54960_1 | TATGAGTGTGGAGTCGTTGCGGTAAGTCACAGAGATCATGGCANNNAGATCGGAA |
| 54961_1 | TATGAGTGTGGAGTCGTTGCCTCCTGTAATCCCAGCACTTAGGNNNAGATCGGAAG |
| 54962_1 | TATGAGTGTGGAGTCGTTGCACTAAAGGTGAGCAGAGGAATGGNNNAGATCGGA |
| 54965_1 | TATGAGTGTGGAGTCGTTGCTTACATTCTGCTTGAAGATGGGCNNNAGATCGGAAG |
| 54966_1 | TATGAGTGTGGAGTCGTTGCAATTGGCTGTGGCAGATACTACANNNAGATCGGAA |
| 54969_1 | TATGAGTGTGGAGTCGTTGCGTGAGCTGAGATCATGCTATTGCNNNAGATCGGAA |
| 54973_1 | TATGAGTGTGGAGTCGTTGCCAACCTCTGTCTCCTCAAGTTCANNNAGATCGGAAGA |
| 54974_1 | TATGAGTGTGGAGTCGTTGCCCTGAGTAGCTGGGATTATAGGCNNNAGATCGGAA |
| 54975_1 | TATGAGTGTGGAGTCGTTGCAGATGGGTTTTCTCCATGTTGGTNNNAGATCGGAAG |
| 54976_1 | TATGAGTGTGGAGTCGTTGCATCAGTTTCTCATCGGAGCTTGGNNNAGATCGGAAG |
| 54977_1 | TATGAGTGTGGAGTCGTTGCTAGTTGGGACTACAGGTGCAAACNNNAGATCGGAA |
| 54978_1 | TATGAGTGTGGAGTCGTTGCCATGATCTCAGCTCATTGCAACCNNNAGATCGGAAG |
| 54979_1 | TATGAGTGTGGAGTCGTTGCTTCAAGCGATTCTCCTGTCTCAGNNNAGATCGGAAG |
| 54981_1 | TATGAGTGTGGAGTCGTTGCACGTACTGGATACATGGGCTTTTNNNAGATCGGAAG |
| 54983_1 | TATGAGTGTGGAGTCGTTGCGGTTTAGGCAATTCTCGTGTCTCNNNAGATCGGAAG |
| 54985_1 | TATGAGTGTGGAGTCGTTGCGATCTGAGCTCACTGTAACCTCCNNNAGATCGGAAG |
| 54986_1 | TATGAGTGTGGAGTCGTTGCACCAAGACAAAGAAAATGCAGAANNNAGATCGGAA |
| 54987_1 | TATGAGTGTGGAGTCGTTGCGGCAGAATCTCACTCTGTCACTTNNNAGATCGGAAG |
| 54988_1 | TATGAGTGTGGAGTCGTTGCCGCCCGGCCAATAGTTATTATAGNNNAGATCGGAAG |
| 54989_1 | TATGAGTGTGGAGTCGTTGCAGCCTCCTGAGTAGCTGGAATTANNNAGATCGGAA |
| 54990_1 | TATGAGTGTGGAGTCGTTGCGATGGAGTCTCACTCTGTCACTCNNNAGATCGGAAG |
| 54991_1 | TATGAGTGTGGAGTCGTTGCAAGTGTTGGGATTACAGGCATGANNNAGATCGGAA |
| 54992_1 | TATGAGTGTGGAGTCGTTGCTTCAAGTGATTCTCCTGCCTCAGNNNAGATCGGAAG |
| 54993_1 | TATGAGTGTGGAGTCGTTGCCTCCTGAGTAGCTGGGATTACAGNNNAGATCGGAA |
| 54994_1 | TATGAGTGTGGAGTCGTTGCTGGTCTCAAATTCCTGAGCTCAANNNAGATCGGAAG |

| | |
|---|---|
| 54996_1 | TATGAGTGTGGAGTCGTTGCAAGGCATTAGGAATACTTGGGCANNNAGATCGGAA |
| 54998_1 | TATGAGTGTGGAGTCGTTGCCTCCTGAGTAGCTGAGATTGCAGNNNAGATCGGAA |
| 54999_1 | TATGAGTGTGGAGTCGTTGCACAGATAGGGTCTCACTGTGTTGNNNAGATCGGAA |
| 55000_1 | TATGAGTGTGGAGTCGTTGCCTTCCAAGTAGCTGGGACAGTAGNNNAGATCGGAA |
| 55002_1 | TATGAGTGTGGAGTCGTTGCAAAGTAACAACTGTTGAAGCGGGNNNAGATCGGAA |
| 55003_1 | TATGAGTGTGGAGTCGTTGCCCTGAGTAGCTGGAATTACAGGTNNNAGATCGGAA |
| 55010_1 | TATGAGTGTGGAGTCGTTGCAGTTTCCAGCCTTGACCACATAANNNAGATCGGAAG |
| 55011_1 | TATGAGTGTGGAGTCGTTGCCCACTGATAGATGCTGGTCTGTANNNAGATCGGAAG |
| 55012_1 | TATGAGTGTGGAGTCGTTGCTTGTGTGGATTCACTGAAAAGCANNNAGATCGGAA |
| 55013_1 | TATGAGTGTGGAGTCGTTGCAAACCTCTGGGACTGGTAGAAACNNNAGATCGGAA |
| 55014_1 | TATGAGTGTGGAGTCGTTGCGGGGAAAAGACACAAAAAGCTGANNNAGATCGGA |
| 55015_1 | TATGAGTGTGGAGTCGTTGCTCCTCCCTCTTCACGTTTTTTCANNNAGATCGGAAGA |
| 55018_1 | TATGAGTGTGGAGTCGTTGCGCAGCAAGCTGGTAAAGAAGTTTNNNAGATCGGAA |
| 55019_1 | TATGAGTGTGGAGTCGTTGCAGACACACACACATACACACACANNNAGATCGGAAG |
| 55020_1 | TATGAGTGTGGAGTCGTTGCTCTAGACACAGACCTTACACCCTNNNAGATCGGAAG |
| 55024_1 | TATGAGTGTGGAGTCGTTGCTGTATGTTCAGCCTTCACCACTGNNNAGATCGGAAG |
| 55027_1 | TATGAGTGTGGAGTCGTTGCGAGGGTGGCAACTAAAACATCACNNNAGATCGGAA |
| 55028_1 | TATGAGTGTGGAGTCGTTGCTGGAAAAAGTCTATCACAGGCCANNNAGATCGGAA |
| 55029_1 | TATGAGTGTGGAGTCGTTGCTGGAGAGAGAAGGAGAAAGAGGTNNNAGATCGGA |
| 55033_1 | TATGAGTGTGGAGTCGTTGCGCTACTTGCTATGTCCCCTAAGGNNNAGATCGGAAG |
| 55036_1 | TATGAGTGTGGAGTCGTTGCGAGCAGAACAGAGATCCCAGAAANNNAGATCGGAA |
| 55038_1 | TATGAGTGTGGAGTCGTTGCTGCAATTCAGAAATACAACAAGCTCTNNNAGATCGG |
| 55043_1 | TATGAGTGTGGAGTCGTTGCGAATGAGGATCTCACTGTGTTGCNNNAGATCGGAA |
| 55044_1 | TATGAGTGTGGAGTCGTTGCTTCACGAGATTCTTCTGCCTCAGNNNAGATCGGAAG |
| 55047_1 | TATGAGTGTGGAGTCGTTGCCATCCCCACAAAATGCTGGAATTNNNAGATCGGAAG |
| 55048_1 | TATGAGTGTGGAGTCGTTGCATGATCTTGGCTCACTGTAAGCTNNNAGATCGGAAG |
| 55049_1 | TATGAGTGTGGAGTCGTTGCTGTTAGCCAGGATGGTCTTGATCNNNAGATCGGAA |
| 55050_1 | TATGAGTGTGGAGTCGTTGCCTGGGTTCAAAAGATCCTTCTGCNNNAGATCGGAAG |
| 55051_1 | TATGAGTGTGGAGTCGTTGCAGACTGGAGTGCAATGGTACAATNNNAGATCGGAA |
| 55053_1 | TATGAGTGTGGAGTCGTTGCAGACTCCTGGGCTATAGAGATCCNNNAGATCGGAAG |
| 55054_1 | TATGAGTGTGGAGTCGTTGCCTCCCGAGTAGCTGTGACTACAGNNNAGATCGGAAG |
| 55057_1 | TATGAGTGTGGAGTCGTTGCCCTCCTGAGTAGCTGGGATTACANNNAGATCGGAAG |
| 55058_1 | TATGAGTGTGGAGTCGTTGCACAAGATCTTGCTCTGTCACTCANNNAGATCGGAAG |
| 55059_1 | TATGAGTGTGGAGTCGTTGCCTCTCAAGTAGCTGGGACTACAGNNNAGATCGGAA |
| 55060_1 | TATGAGTGTGGAGTCGTTGCTTCAAGTGATGCTCCTGCCTTAGNNNAGATCGGAAG |
| 55061_1 | TATGAGTGTGGAGTCGTTGCAGACCCCCAAGTAGCTAGGATTANNNAGATCGGAA |
| 55062_1 | TATGAGTGTGGAGTCGTTGCTTCAAGTGATTCTCCTGCCTCAGNNNAGATCGGAAG |
| 55063_1 | TATGAGTGTGGAGTCGTTGCAGAGGTGTGCAATCATAGCTCAANNNAGATCGGAA |
| 55064_1 | TATGAGTGTGGAGTCGTTGCGTTCCAGTGATTCTCCTGCCTAANNNAGATCGGAAG |
| 55065_1 | TATGAGTGTGGAGTCGTTGCCTGGAATGTAGTGGTGCGATCTCNNNAGATCGGAA |
| 55071_1 | TATGAGTGTGGAGTCGTTGCGTTTCACCATGTTAGCCAGGATGNNNAGATCGGAAG |
| 55072_1 | TATGAGTGTGGAGTCGTTGCTCCAGGTTCAAGTGATTCTCCTGNNNAGATCGGAAG |
| 55073_1 | TATGAGTGTGGAGTCGTTGCGAGTAGCTGGGATTACAGATGCANNNAGATCGGAA |

| | |
|---|---|
| 55075_1 | TATGAGTGTGGAGTCGTTGCCAATGGCGTGATCTCAGCTTACTNNNAGATCGGAAG |
| 55081_1 | TATGAGTGTGGAGTCGTTGCTTCAAGTGAGCCTCTCAACTCAGNNNAGATCGGAAG |
| 55082_1 | TATGAGTGTGGAGTCGTTGCCTCCTGAGTAGCTGGGACTACAGNNNAGATCGGAAG |
| 55084_1 | TATGAGTGTGGAGTCGTTGCAAGTAGCTGGGATTACAGGCATCNNNAGATCGGAA |
| 55086_1 | TATGAGTGTGGAGTCGTTGCTCAGGCTGATCTAGAACTCCTGANNNAGATCGGAAG |
| 55087_1 | TATGAGTGTGGAGTCGTTGCAGTAGAGACGGGGTTTACCAAGNNNAGATCGGAA |
| 55088_1 | TATGAGTGTGGAGTCGTTGCCTGACTCCCAGGTTCAAGCTATTNNNAGATCGGAAG |
| 55089_1 | TATGAGTGTGGAGTCGTTGCTCAAACAATTCTCCTGCTTCAGCNNNAGATCGGAAGA |
| 55091_1 | TATGAGTGTGGAGTCGTTGCCAAGTAGCTGGGATTACCGACATNNNAGATCGGAAG |
| 55093_1 | TATGAGTGTGGAGTCGTTGCCTCACTACAGCTTTGACCTCCTGNNNAGATCGGAAGA |
| 55094_1 | TATGAGTGTGGAGTCGTTGCTGTGTGAATGGGGCATGATAGAANNNAGATCGGAA |
| 55097_1 | TATGAGTGTGGAGTCGTTGCACACAGTGAAACCCCATCTCTACNNNAGATCGGAAG |
| 55100_1 | TATGAGTGTGGAGTCGTTGCTGCAGAAGAAAGAACTCACTCCANNNAGATCGGAAG |
| 55104_1 | TATGAGTGTGGAGTCGTTGCATGTGATGGCCAATCCTGATCATNNNAGATCGGAAG |
| 55105_1 | TATGAGTGTGGAGTCGTTGCAATGTCTTCCTAGAAGTGGGAGCNNNAGATCGGAA |
| 55106_1 | TATGAGTGTGGAGTCGTTGCAGGCTGTACTTTTAGGAGAGAAGCNNNAGATCGGA |
| 55107_1 | TATGAGTGTGGAGTCGTTGCTCCTGCCTGTAATCCCAACATTTNNNAGATCGGAAGA |
| 55108_1 | TATGAGTGTGGAGTCGTTGCGAGACCTCCACTTTCTGTCTGAGNNNAGATCGGAAG |
| 55109_1 | TATGAGTGTGGAGTCGTTGCTCTCAGAGCTCAAAGACTGGTTCNNNAGATCGGAAG |
| 55110_1 | TATGAGTGTGGAGTCGTTGCAGGAAGGACAAGGAAGGAAAGACNNNAGATCGGA |
| 55112_1 | TATGAGTGTGGAGTCGTTGCCACTGGGAATTTTGGTCACCAAGNNNAGATCGGAAG |
| 55113_1 | TATGAGTGTGGAGTCGTTGCTATTGCAGTGAGCTGAAATCACGNNNAGATCGGAAG |
| 55115_1 | TATGAGTGTGGAGTCGTTGCGCTGTACTCTGTTGGGAGACAATNNNAGATCGGAA |
| 55116_1 | TATGAGTGTGGAGTCGTTGCGAAAATATTCGGGCCTAGGCAACNNNAGATCGGAA |
| 55117_1 | TATGAGTGTGGAGTCGTTGCCCCCTATGCCCAAAGTGAACATANNNAGATCGGAAG |
| 55118_1 | TATGAGTGTGGAGTCGTTGCGCTGTACCTGAAACACAAGCTTTNNNAGATCGGAAG |
| 55119_1 | TATGAGTGTGGAGTCGTTGCACGTGTGTATATACGCGCGTATANNNAGATCGGAAG |
| 55120_1 | TATGAGTGTGGAGTCGTTGCTAACTCACACCTGTAATCCCAGCNNNAGATCGGAAGA |
| 55121_1 | TATGAGTGTGGAGTCGTTGCCTGAGACAGGAGGATCACATGAGNNNAGATCGGAA |
| 55124_1 | TATGAGTGTGGAGTCGTTGCTATGCTCCCATGTCATATGCTCCNNNAGATCGGAAGA |
| 55125_1 | TATGAGTGTGGAGTCGTTGCTGCTAATTTGGTGAAAACCCGTCNNNAGATCGGAAG |
| 55126_1 | TATGAGTGTGGAGTCGTTGCATCATCTACATCTTTGCCTCCCTNNNAGATCGGAAGA |
| 55128_1 | TATGAGTGTGGAGTCGTTGCGGAAAATTCTAGCCCCAGGGATTNNNAGATCGGAA |
| 55129_1 | TATGAGTGTGGAGTCGTTGCAAAAGAGACACACACACACTCCANNNAGATCGGAAG |
| 55131_1 | TATGAGTGTGGAGTCGTTGCCGTATGTAACAAACCTGCACGTTNNNAGATCGGAAG |
| 55132_1 | TATGAGTGTGGAGTCGTTGCAGCATGTCAGTAACAATGGCTTTNNNAGATCGGAAG |
| 55134_1 | TATGAGTGTGGAGTCGTTGCGCTAAGAACTGTGGCTAGCAAATNNNAGATCGGAA |
| 55136_1 | TATGAGTGTGGAGTCGTTGCAGGCAGAAGTTGTAGTAAGCCAANNNAGATCGGAA |
| 55138_1 | TATGAGTGTGGAGTCGTTGCGCATCATCATTTCTTCAAGGTGCNNNAGATCGGAAG |
| 55139_1 | TATGAGTGTGGAGTCGTTGCAGAGTGCATGTAAAAACTGGTGANNNAGATCGGAA |
| 55140_1 | TATGAGTGTGGAGTCGTTGCACAAAAAGCAAAAAAAGGGGGGANNNAGATCGGA |
| 55141_1 | TATGAGTGTGGAGTCGTTGCGATCTGCTTCATGGTAAAGACGANNNAGATCGGAA |
| 55143_1 | TATGAGTGTGGAGTCGTTGCACCAAAGTTGCTGAAAGTCTGAANNNAGATCGGAA |

| | |
|---|---|
| 55144_1 | TATGAGTGTGGAGTCGTTGCTTTTTATTTTGGGAGGGTGCTGCNNNAGATCGGAAG |
| 55146_1 | TATGAGTGTGGAGTCGTTGCCACAAAAACCTTCAAGGGGCTTANNNAGATCGGAAG |
| 55147_1 | TATGAGTGTGGAGTCGTTGCAGCACAGGAAAGAATTGATTGCCNNNAGATCGGAAG |
| 55154_1 | TATGAGTGTGGAGTCGTTGCGTCAGGCTTTACATGAACAGCAANNNAGATCGGAAG |
| 55160_1 | TATGAGTGTGGAGTCGTTGCCCAGTGGTCTCTAACTCTTAGGCNNNAGATCGGAAGA |
| 55161_1 | TATGAGTGTGGAGTCGTTGCGCCTCCTGAGTAGCTGAGATTACNNNAGATCGGAAG |
| 55164_1 | TATGAGTGTGGAGTCGTTGCTAAGCACATAGGATGACCACACCNNNAGATCGGAAG |
| 55166_1 | TATGAGTGTGGAGTCGTTGCCAAGATGGTGTCAATGTCCTGACNNNAGATCGGAAG |
| 55167_1 | TATGAGTGTGGAGTCGTTGCAACAGAAGTCAAGGATCAGGCATNNNAGATCGGAAG |
| 55168_1 | TATGAGTGTGGAGTCGTTGCCAAGTAGCTGGGATTACAGGCATNNNAGATCGGAAG |
| 55171_1 | TATGAGTGTGGAGTCGTTGCCTTGGAAGGAGCCTTGAGTTTTGNNNAGATCGGAAG |
| 55172_1 | TATGAGTGTGGAGTCGTTGCCTCACAAGTAGCTGGGACTACAGNNNAGATCGGAAG |
| 55173_1 | TATGAGTGTGGAGTCGTTGCGGAATGGGTAGGAGAAGCATTGANNNAGATCGGAA |
| 55176_1 | TATGAGTGTGGAGTCGTTGCCTGGGTTCAAGTGATTCTCTTGCNNNAGATCGGAAG |
| 55178_1 | TATGAGTGTGGAGTCGTTGCAATTCTCTCATCTCAGCCTTCCGNNNAGATCGGAAGA |
| 55179_1 | TATGAGTGTGGAGTCGTTGCCCAGTGTGAACATCCAAAGCAAANNNAGATCGGAAG |
| 55180_1 | TATGAGTGTGGAGTCGTTGCCTCCTGAGTAGCTGGGACTACAGNNNAGATCGGAAG |
| 55181_1 | TATGAGTGTGGAGTCGTTGCATGGCTGCATAGTATTCCATGGTNNNAGATCGGAAG |
| 55182_1 | TATGAGTGTGGAGTCGTTGCCTGGAGTGAAATGGCACAATCTCNNNAGATCGGAAG |
| 55183_1 | TATGAGTGTGGAGTCGTTGCTCCCAACTAGCTGAGTCTACAGANNNAGATCGGAAG |
| 55187_1 | TATGAGTGTGGAGTCGTTGCTCAAAGTATTCTCCTGCCTCAGCNNNAGATCGGAAGA |
| 55191_1 | TATGAGTGTGGAGTCGTTGCAATAGCTGGGATTACAGGCATGTNNNAGATCGGAAG |
| 55192_1 | TATGAGTGTGGAGTCGTTGCCAATGGCGTGATCTTGGCTTATTNNNAGATCGGAAG |
| 55193_1 | TATGAGTGTGGAGTCGTTGCAACCTCCTGAGTAGCTGAGACTANNNAGATCGGAAG |
| 55194_1 | TATGAGTGTGGAGTCGTTGCCCACCTCACCCAGCTAATTTTTGNNNAGATCGGAAGA |
| 55195_1 | TATGAGTGTGGAGTCGTTGCCACAATATCAGCTCACTGCAACCNNNAGATCGGAAGA |
| 55196_1 | TATGAGTGTGGAGTCGTTGCCTGCCTCCTGGATCCAAGTTATTNNNAGATCGGAAGA |
| 55198_1 | TATGAGTGTGGAGTCGTTGCCTCCCAAGTAGCTGGAACTACAGNNNAGATCGGAAG |
| 55199_1 | TATGAGTGTGGAGTCGTTGCCTCAGTTGATCCTCCTACCTCTGNNNAGATCGGAAGA |
| 55200_1 | TATGAGTGTGGAGTCGTTGCTTCAAGTGATTCTCCTGCCTCAGNNNAGATCGGAAGA |
| 55205_1 | TATGAGTGTGGAGTCGTTGCAGGTTCAAGTGATTCTTCTGCCTNNNAGATCGGAAGA |
| 55206_1 | TATGAGTGTGGAGTCGTTGCCTCAAGCAATCCTCCCACCTTAGNNNAGATCGGAAGA |
| 55208_1 | TATGAGTGTGGAGTCGTTGCCCAAATAGCTGGGACTATGGGTGNNNAGATCGGAAG |
| 55210_1 | TATGAGTGTGGAGTCGTTGCCACTCATAGGTGGGAACTGAACANNNAGATCGGAAG |
| 55211_1 | TATGAGTGTGGAGTCGTTGCGATTCACGTCATTCTCCTGCCTCNNNAGATCGGAAGA |
| 55212_1 | TATGAGTGTGGAGTCGTTGCCCTCCTGGGTTCAAGTGATTCTCNNNAGATCGGAAGA |
| 55213_1 | TATGAGTGTGGAGTCGTTGCGTTCAAGCAATTCTCTGCCTCAGNNNAGATCGGAAGA |
| 55215_1 | TATGAGTGTGGAGTCGTTGCCTGAGTAACTGGGACTACAGGTGNNNAGATCGGAAG |
| 55216_1 | TATGAGTGTGGAGTCGTTGCTCCCAAGTAGCTGGAATTACAGGNNNAGATCGGAAG |
| 55217_1 | TATGAGTGTGGAGTCGTTGCGAGTAGCTGGGACTACAGGTATGNNNAGATCGGAA |
| 55223_1 | TATGAGTGTGGAGTCGTTGCATGCCCTAATCAGCACTCTGTAGNNNAGATCGGAAGA |
| 55224_1 | TATGAGTGTGGAGTCGTTGCTTCCCAATTCTGTGAGGCCAATANNNAGATCGGAAGA |
| 55228_1 | TATGAGTGTGGAGTCGTTGCCCTCTTGCACGGAAAAGCTTTTANNNAGATCGGAAGA |

| | |
|---|---|
| 55229_1 | TATGAGTGTGGAGTCGTTGCAGAATCACTGGCGTAGGTATGACNNNAGATCGGAA |
| 55230_1 | TATGAGTGTGGAGTCGTTGCCAAGGTCATGCAGTAAGTTGGTGNNNAGATCGGAA |
| 55231_1 | TATGAGTGTGGAGTCGTTGCATGGCAATAAAATGGTTCAGGGCNNNAGATCGGAA |
| 55233_1 | TATGAGTGTGGAGTCGTTGCACAAAACAAAGGATCAGCAAAGTNNNAGATCGGAA |
| 55234_1 | TATGAGTGTGGAGTCGTTGCCTTTGAGGTCATCTCCAGACACANNNAGATCGGAAG |
| 55236_1 | TATGAGTGTGGAGTCGTTGCTCACATGAGGTCAGGAGTTCAAGNNNAGATCGGAA |
| 55237_1 | TATGAGTGTGGAGTCGTTGCGTGAGCTATGATCATGCCTCTGANNNAGATCGGAAG |
| 55239_1 | TATGAGTGTGGAGTCGTTGCCACACTCCTACCATACGATCCAGNNNAGATCGGAAG |
| 55240_1 | TATGAGTGTGGAGTCGTTGCCTGTGATTGTGTTACTGCACTCCNNNAGATCGGAAG |
| 55241_1 | TATGAGTGTGGAGTCGTTGCCAACAACAAAAACAAGATGGCCGNNNAGATCGGAA |
| 55243_1 | TATGAGTGTGGAGTCGTTGCAGGACTATTTCCGGCTTGAAAGTNNNAGATCGGAA |
| 55244_1 | TATGAGTGTGGAGTCGTTGCTGGGAGGAAGGAAGGAAATTCTGNNNAGATCGGA |
| 55245_1 | TATGAGTGTGGAGTCGTTGCGGGCTGAGAAAAATGAATAGCCANNNAGATCGGAA |
| 55246_1 | TATGAGTGTGGAGTCGTTGCAGCCAAACGTTTTGGAAAAGTCANNNAGATCGGAA |
| 55248_1 | TATGAGTGTGGAGTCGTTGCTGTAATCCCAGCTACTCAGGAGANNNAGATCGGAAG |
| 55252_1 | TATGAGTGTGGAGTCGTTGCCAAAAATTGATGTTGGCACGGACNNNAGATCGGAA |
| 55253_1 | TATGAGTGTGGAGTCGTTGCGGCAATTTGAGACAACACTTCGANNNAGATCGGAA |
| 55256_1 | TATGAGTGTGGAGTCGTTGCTGAACTCCACAAAATTACTCCCANNNAGATCGGAAG |
| 55258_1 | TATGAGTGTGGAGTCGTTGCTGAGGTGACTGATACACTGTTGTNNNAGATCGGAA |
| 55259_1 | TATGAGTGTGGAGTCGTTGCATAATACCGTGGGTTGACTGGTCNNNAGATCGGAA |
| 55260_1 | TATGAGTGTGGAGTCGTTGCGTCCAGCTACTCAAGAGGCTAAGNNNAGATCGGAA |
| 55263_1 | TATGAGTGTGGAGTCGTTGCTTTCCCAGAAACAGAGCATTTGCNNNAGATCGGAAG |
| 55264_1 | TATGAGTGTGGAGTCGTTGCCCAGAGGTAGTGTTACTGTGCATNNNAGATCGGAA |
| 55265_1 | TATGAGTGTGGAGTCGTTGCCTATGTTGTACTGAGCACTTGCCNNNAGATCGGAAG |
| 55266_1 | TATGAGTGTGGAGTCGTTGCTGTTTTCAAAATGGAGCAGCTGTNNNAGATCGGAA |
| 55268_1 | TATGAGTGTGGAGTCGTTGCCCTCATCAGGCAGTCAAATCAACNNNAGATCGGAAG |
| 55269_1 | TATGAGTGTGGAGTCGTTGCCACACACACACACACACACAATANNNAGATCGGAAG |
| 55270_1 | TATGAGTGTGGAGTCGTTGCAACTCAAGGGGGTGTCAATCATANNNAGATCGGAA |
| 55271_1 | TATGAGTGTGGAGTCGTTGCAGGGCAATCCATAGAGCTGAATTNNNAGATCGGAA |
| 55272_1 | TATGAGTGTGGAGTCGTTGCGACAGAAAGGATACCATGGACCCNNNAGATCGGAA |
| 55273_1 | TATGAGTGTGGAGTCGTTGCAAAAAATTAGCCTGGCATGGTGGNNNAGATCGGAA |
| 55274_1 | TATGAGTGTGGAGTCGTTGCACACAATCCAGAGGAAATGGTCANNNAGATCGGAA |
| 55275_1 | TATGAGTGTGGAGTCGTTGCACTCTTGATTACCTTGAAGCTGGANNNAGATCGGAA |
| 55279_1 | TATGAGTGTGGAGTCGTTGCTTGAGATTTTGGACTGCCAGTCTNNNAGATCGGAAG |
| 55280_1 | TATGAGTGTGGAGTCGTTGCACACAGGTGACTCTAAGGAAAGGNNNAGATCGGAA |
| 55281_1 | TATGAGTGTGGAGTCGTTGCGTGATCCACCCCTATTATCCCAGNNNAGATCGGAAG |
| 55285_1 | TATGAGTGTGGAGTCGTTGCCATAGTGGAAAGAGCTTAGCCCANNNAGATCGGAA |
| 55287_1 | TATGAGTGTGGAGTCGTTGCGCCCCTTCGTTACACACTGAATANNNAGATCGGAAG |
| 55288_1 | TATGAGTGTGGAGTCGTTGCCTCTGTGGCACCTATCACAACTANNNAGATCGGAAG |
| 55289_1 | TATGAGTGTGGAGTCGTTGCTAGATTTGAATGCTGGCCACTCTNNNAGATCGGAAG |
| 55290_1 | TATGAGTGTGGAGTCGTTGCTGCCCTACCTACCTTGTATTGAANNNAGATCGGAAG |
| 55291_1 | TATGAGTGTGGAGTCGTTGCCGAAGAAGAAGAAGGCCTGAATANNNAGATCGGAA |
| 55292_1 | TATGAGTGTGGAGTCGTTGCATAGGGAGTAGGGAGACTAGGGANNNAGATCGGA |

55294_1 TATGAGTGTGGAGTCGTTGCACTTTTTTTTGCAGGGATAGGGCNNNAGATCGGAAG
55296_1 TATGAGTGTGGAGTCGTTGCTCAAGTGATTCTCCTGCCTTAGCNNNAGATCGGAAGA
55297_1 TATGAGTGTGGAGTCGTTGCGGTTCAAGTGATTTTCCTGCCTCNNNAGATCGGAAGA
55299_1 TATGAGTGTGGAGTCGTTGCCTCCAGAGTAGCAGGGATTACAGNNNAGATCGGAAG
55301_1 TATGAGTGTGGAGTCGTTGCGTTCAAGTGATTCTCCTCCCTCANNNAGATCGGAAGA
55302_1 TATGAGTGTGGAGTCGTTGCTAGAAACAGGGTCTCACTCCTTGNNNAGATCGGAAG
55304_1 TATGAGTGTGGAGTCGTTGCCTTCCTGTGTAGCTGGGATTACANNNAGATCGGAAG
55305_1 TATGAGTGTGGAGTCGTTGCCTCACGAGTAGCTGGGATAACAGNNNAGATCGGAAG
55306_1 TATGAGTGTGGAGTCGTTGCCTCCTGAGTAGCTGGGATTACAGNNNAGATCGGAAG
55307_1 TATGAGTGTGGAGTCGTTGCTGCTCTGCTGTCACCTATCATAGNNNAGATCGGAAGA
55308_1 TATGAGTGTGGAGTCGTTGCTACAAATATGAGCCTCCACAGCCNNNAGATCGGAAGA
55309_1 TATGAGTGTGGAGTCGTTGCAGTGATCTTTCCACCTCAACCTCNNNAGATCGGAAGA
55310_1 TATGAGTGTGGAGTCGTTGCGCTAGAGTGTAGTGGTGCAATCTNNNAGATCGGAAG
55311_1 TATGAGTGTGGAGTCGTTGCCCTCCCAGGTTCAAGCAATTCTCNNNAGATCGGAAGA
55312_1 TATGAGTGTGGAGTCGTTGCTTTGGGTTCAAGCAATTCTCCTGNNNAGATCGGAAGA
55313_1 TATGAGTGTGGAGTCGTTGCCCAAGTAGCTGGGATTACAGGTGNNNAGATCGGAAG
55314_1 TATGAGTGTGGAGTCGTTGCGAGATGGAGTTTCGTGCTTGTTGNNNAGATCGGAAG
55316_1 TATGAGTGTGGAGTCGTTGCTTCATGCCATTCTCCTGTCTCAGNNNAGATCGGAAGA
55317_1 TATGAGTGTGGAGTCGTTGCTTCATGCCATTCTCCTGTCTCAGNNNAGATCGGAAGA
55318_1 TATGAGTGTGGAGTCGTTGCCCTAGTAGCTGGGATTACAGGTGNNNAGATCGGAAG
55320_1 TATGAGTGTGGAGTCGTTGCCTGGGTTCACATCATTCTTCTGCNNNAGATCGGAAGA
55321_1 TATGAGTGTGGAGTCGTTGCAGTAGCTGGGATTACAGGTACCTNNNAGATCGGAAG
55322_1 TATGAGTGTGGAGTCGTTGCGAATGCATGGTGTGATCTTTGCTNNNAGATCGGAAG
55323_1 TATGAGTGTGGAGTCGTTGCTCCTGGGTTCTAGTGATTCTCCTNNNAGATCGGAAGA
55327_1 TATGAGTGTGGAGTCGTTGCCTGGAGTGCAATGATGTGATCTCNNNAGATCGGAAG
55328_1 TATGAGTGTGGAGTCGTTGCCAATCTCAGTTCACTGCAACCTCNNNAGATCGGAAGA
55329_1 TATGAGTGTGGAGTCGTTGCGAGACGGAATCTGGCTCTGTTACNNNAGATCGGAAG
55332_1 TATGAGTGTGGAGTCGTTGCAGCAATCTCAGCTCATTTCAAGCNNNAGATCGGAAGA
55333_1 TATGAGTGTGGAGTCGTTGCCAATCTTGACTCACTGCAAGCTCNNNAGATCGGAAGA
55335_1 TATGAGTGTGGAGTCGTTGCGCACAATCTCGGCTCACTATAACNNNAGATCGGAAGA
55338_1 TATGAGTGTGGAGTCGTTGCTAGGTTCAAGCGATTCTCTTGCTNNNAGATCGGAAGA
55339_1 TATGAGTGTGGAGTCGTTGCCAGAGGTTACAGTAAGCTGAGGTNNNAGATCGGAA
55341_1 TATGAGTGTGGAGTCGTTGCAGGTACGTTCCTACAAACTCACCNNNAGATCGGAAGA
55343_1 TATGAGTGTGGAGTCGTTGCGGGTCAGATTCCCTGAGTTCAAANNNAGATCGGAAG
55344_1 TATGAGTGTGGAGTCGTTGCGAAGGAGGTAGTGTTCTCCCAGANNNAGATCGGAAG
55346_1 TATGAGTGTGGAGTCGTTGCGTGGTGAAACCCCATCTCTACTANNNAGATCGGAAGA
55347_1 TATGAGTGTGGAGTCGTTGCAGGGCTAAATGTCATACGCTGATNNNAGATCGGAAG
55348_1 TATGAGTGTGGAGTCGTTGCACCTGTAATCCTAGCACTTTGGGNNNAGATCGGAAG
55349_1 TATGAGTGTGGAGTCGTTGCCACACCTGTAATCCCAGCACTTTNNNAGATCGGAAGA
55351_1 TATGAGTGTGGAGTCGTTGCCACTTGTAATCCCAGCACTTTGGNNNAGATCGGAAGA
55352_1 TATGAGTGTGGAGTCGTTGCCAGAGGCTTCAGTGAGTCAAGATNNNAGATCGGAAG
55353_1 TATGAGTGTGGAGTCGTTGCCACAAATTAGCTAGGCATGGTGGNNNAGATCGGAAG
55354_1 TATGAGTGTGGAGTCGTTGCCATGAGGAAAGTCTCAAGCTACANNNAGATCGGAAG

55355_1 TATGAGTGTGGAGTCGTTGCGGGCACAAAAATCACTTGAACCTNNNAGATCGGAAG
55357_1 TATGAGTGTGGAGTCGTTGCGGAAAATGAAGTCAAGCCAAGCANNNAGATCGGAA
55358_1 TATGAGTGTGGAGTCGTTGCAGTGTTGGGGAAAATGGAGTCATNNNAGATCGGAA
55359_1 TATGAGTGTGGAGTCGTTGCCCAAAGACCACACTCCAGTTTTCNNNAGATCGGAAG
55360_1 TATGAGTGTGGAGTCGTTGCGCATAGCTAGGCAAATCTGTGTGNNNAGATCGGAA
55361_1 TATGAGTGTGGAGTCGTTGCGTTTTCAGTTCCCTACCAAAGGCNNNAGATCGGAAG
55362_1 TATGAGTGTGGAGTCGTTGCTGGCCCATGGTAAATAACATGGANNNAGATCGGAA
55363_1 TATGAGTGTGGAGTCGTTGCCATTTTCCTTGGGGTTTCTGGACNNNAGATCGGAAG
55364_1 TATGAGTGTGGAGTCGTTGCGTCTCCCTGTGTGAGATCTCATCNNNAGATCGGAAG
55365_1 TATGAGTGTGGAGTCGTTGCGAGGAAGGAGGAGGAGAAGGATANNNAGATCGG
55367_1 TATGAGTGTGGAGTCGTTGCAGTTTGGCTGGATATGGTGTCTCNNNAGATCGGAA
55368_1 TATGAGTGTGGAGTCGTTGCAAAGAGACCTAGAACTCAAGGCCNNNAGATCGGAA
55369_1 TATGAGTGTGGAGTCGTTGCCCTGTAATCCTAGCACTTTGGGANNNAGATCGGAAG
55370_1 TATGAGTGTGGAGTCGTTGCGGAAAATGGGCAAAGGACTTGAANNNAGATCGGA
55371_1 TATGAGTGTGGAGTCGTTGCGTTCCCATCTCATTGAAGCCAACNNNAGATCGGAAG
55372_1 TATGAGTGTGGAGTCGTTGCTTGACACTGGAACAACACAGGTTNNNAGATCGGAA
55373_1 TATGAGTGTGGAGTCGTTGCCTGAGGCATGAGAATCACTTGAGNNNAGATCGGAA
55375_1 TATGAGTGTGGAGTCGTTGCGAATGGTGACTCATGCCTGCAATNNNAGATCGGAA
55376_1 TATGAGTGTGGAGTCGTTGCAACTTGATCTTGGAAGCAAAGGCNNNAGATCGGAA
55379_1 TATGAGTGTGGAGTCGTTGCCCCAAAAAAGAAATGACATGGCANNNAGATCGGAA
55380_1 TATGAGTGTGGAGTCGTTGCTCAAGGCTGCAGTGAACTATGATNNNAGATCGGAA
55381_1 TATGAGTGTGGAGTCGTTGCTGGGTTTCTAAGGATTCCCAGTGNNNAGATCGGAA
55382_1 TATGAGTGTGGAGTCGTTGCGGCTTCATTTGGAAAGCCAGATTNNNAGATCGGAA
55383_1 TATGAGTGTGGAGTCGTTGCGATAGGGCTTCGACAGACAGAATNNNAGATCGGAA
55384_1 TATGAGTGTGGAGTCGTTGCCAAAAAAACAGGGCATTTCAGGCNNNAGATCGGAA
55385_1 TATGAGTGTGGAGTCGTTGCTCACAGACAAGATTTACCTGTTGANNNAGATCGGAA
55388_1 TATGAGTGTGGAGTCGTTGCAGCTAAAATCCAGGACAAGGGAGNNNAGATCGGAA
55389_1 TATGAGTGTGGAGTCGTTGCAGAGCAGTGGGAAGAATCTCATGNNNAGATCGGAA
55391_1 TATGAGTGTGGAGTCGTTGCGTAATGTTCTCAACTTCCACGCCNNNAGATCGGAAG
55392_1 TATGAGTGTGGAGTCGTTGCCACGAGGTCAGGATATTGAGACCNNNAGATCGGAA
55394_1 TATGAGTGTGGAGTCGTTGCACATAAAACCCTTTGCCTTCTGCNNNAGATCGGAAG
55395_1 TATGAGTGTGGAGTCGTTGCCTGAGATGGGAGGATCACTTGAGNNNAGATCGGAA
55397_1 TATGAGTGTGGAGTCGTTGCTCAACTTACGCACCTTATCCACANNNAGATCGGAAG
55402_1 TATGAGTGTGGAGTCGTTGCGGTTCAAGTGATTCTTCTGCCTCNNNAGATCGGAAG
55403_1 TATGAGTGTGGAGTCGTTGCCCTTGGAGTGGGATGCTGAATATNNNAGATCGGAA
55405_1 TATGAGTGTGGAGTCGTTGCGGTTCAAGTGATTCTCCTGCTTCNNNAGATCGGAAG
55406_1 TATGAGTGTGGAGTCGTTGCGGAATTCAGGAAGGCTTCATTGGNNNAGATCGGAA
55407_1 TATGAGTGTGGAGTCGTTGCTTTTGATCTTGTTGCTCAGGCTGNNNAGATCGGAAG
55408_1 TATGAGTGTGGAGTCGTTGCAAGAGATTCTCCTACCTCAGCCTNNNAGATCGGAAG
55409_1 TATGAGTGTGGAGTCGTTGCGGGTTCTTGGTCTCACTTCAAGANNNAGATCGGAAG
55413_1 TATGAGTGTGGAGTCGTTGCCTGAGTAGCTGGGATTACAGGTGNNNAGATCGGAA
55414_1 TATGAGTGTGGAGTCGTTGCCCTCCCAAATTCAAGCGATTCTCNNNAGATCGGAAGA
55418_1 TATGAGTGTGGAGTCGTTGCCAACCTCCACCTCTTGAGTTCAANNNAGATCGGAAGA

| | |
|---|---|
| 55419_1 | TATGAGTGTGGAGTCGTTGCATCTCAGATCACTGCAACCTTTGNNNAGATCGGAAG |
| 55420_1 | TATGAGTGTGGAGTCGTTGCTGCCCAGACACAAGATAGTGTAGNNNAGATCGGAA |
| 55422_1 | TATGAGTGTGGAGTCGTTGCGGTTCAAGCAATTATCCTGCCTCNNNAGATCGGAAG |
| 55424_1 | TATGAGTGTGGAGTCGTTGCCCTGGGTTCAAGCAATTCTCCTANNNAGATCGGAAG |
| 55426_1 | TATGAGTGTGGAGTCGTTGCCATTCTCCTGCCTCAGCCTCNNNAGATCGGAAGAGCA |
| 55427_1 | TATGAGTGTGGAGTCGTTGCCTCCTGAGTAGCTGGGATTACAGNNNAGATCGGAA |
| 55428_1 | TATGAGTGTGGAGTCGTTGCGTTCAAGAGATTCTCCTGCCTCANNNAGATCGGAAG |
| 55429_1 | TATGAGTGTGGAGTCGTTGCTCTTGGACTCCTGACCTCAAATGNNNAGATCGGAAG |
| 55431_1 | TATGAGTGTGGAGTCGTTGCCTGGGTTCAAATGATTCTCCTGCNNNAGATCGGAAG |
| 55432_1 | TATGAGTGTGGAGTCGTTGCCAGGCTGGAGTGCTTCTTTTTTTNNNAGATCGGAAG |
| 55434_1 | TATGAGTGTGGAGTCGTTGCAGAGTAGCAGGGATCACAGGTATNNNAGATCGGAA |
| 55435_1 | TATGAGTGTGGAGTCGTTGCCATGGTCTCAGCCTCAAAGTACTNNNAGATCGGAAG |
| 55436_1 | TATGAGTGTGGAGTCGTTGCATTCTACTGCCTCAGCCTTTCAANNNAGATCGGAAGA |
| 55438_1 | TATGAGTGTGGAGTCGTTGCCTCCTGAGTAGCTGGGACTACAGNNNAGATCGGAA |
| 55439_1 | TATGAGTGTGGAGTCGTTGCACTGCAAGCTTCACAATTTCCTCNNNAGATCGGAAGA |
| 55440_1 | TATGAGTGTGGAGTCGTTGCTAGGACAATAGTGGAGGGAAGGTNNNAGATCGGA |
| 55441_1 | TATGAGTGTGGAGTCGTTGCCTTCCAAGTAGCTGGGACTACAGNNNAGATCGGAA |
| 55442_1 | TATGAGTGTGGAGTCGTTGCCTCCTGAGTAGCTGGGACTACAGNNNAGATCGGAA |
| 55443_1 | TATGAGTGTGGAGTCGTTGCGAGACAGGGTTTCACTATGTTGCNNNAGATCGGAA |
| 55445_1 | TATGAGTGTGGAGTCGTTGCTAAAGTGCTGGGATTACAGGCATNNNAGATCGGAA |
| 55446_1 | TATGAGTGTGGAGTCGTTGCGTCTAGCTCTATTGCTCAGGCTGNNNAGATCGGAAG |
| 55448_1 | TATGAGTGTGGAGTCGTTGCAGGTTCAAGCAATTCTCCTTCCTNNNAGATCGGAAG |
| 55450_1 | TATGAGTGTGGAGTCGTTGCCAGCCTCCTAAAGTGCTGAGATTNNNAGATCGGAAG |
| 55452_1 | TATGAGTGTGGAGTCGTTGCCAGAGACCTGATCATGGCTCATTNNNAGATCGGAAG |
| 55453_1 | TATGAGTGTGGAGTCGTTGCTGATCTAGGCTCACTGCAATCTCNNNAGATCGGAAG |
| 55455_1 | TATGAGTGTGGAGTCGTTGCCAGTGGTGTGATCTCAGCTTACTNNNAGATCGGAAG |
| 55456_1 | TATGAGTGTGGAGTCGTTGCACTTCCTGGGTTCAAGTGATTCANNNAGATCGGAAG |
| 55457_1 | TATGAGTGTGGAGTCGTTGCTCCCGGATACAAGTGATTTTCCTNNNAGATCGGAAG |
| 55459_1 | TATGAGTGTGGAGTCGTTGCTCAAGTTATCCTCCCACTTCAGCNNNAGATCGGAAGA |
| 55460_1 | TATGAGTGTGGAGTCGTTGCCAGACTGGGCAACATACTGAGATNNNAGATCGGAA |
| 55466_1 | TATGAGTGTGGAGTCGTTGCAGTGTAATGGCGTGATCTTGTCTNNNAGATCGGAA |
| 55467_1 | TATGAGTGTGGAGTCGTTGCGGTTCAAGCTACTCTCCCATCTCNNNAGATCGGAAGA |
| 55468_1 | TATGAGTGTGGAGTCGTTGCTAGTACAGTGACGCAATCAATGCNNNAGATCGGAA |
| 55469_1 | TATGAGTGTGGAGTCGTTGCCAATGGTATGATCTCGGCTCACTNNNAGATCGGAAG |
| 55470_1 | TATGAGTGTGGAGTCGTTGCCCCAGGTTCAAGCAATTCTCCTANNNAGATCGGAAG |
| 55471_1 | TATGAGTGTGGAGTCGTTGCAGTGCTGAGATTACAGACGTGAGNNNAGATCGGAA |
| 55472_1 | TATGAGTGTGGAGTCGTTGCGGAGTAACTGGGATTACAGGCATNNNAGATCGGAA |
| 55473_1 | TATGAGTGTGGAGTCGTTGCGTTACAAGCAATTCTCCTGCCTCNNNAGATCGGAAG |
| 55474_1 | TATGAGTGTGGAGTCGTTGCTGCCTCTCATTGCCTTTGAAAAGNNNAGATCGGAAG |
| 55475_1 | TATGAGTGTGGAGTCGTTGCGGGCAGGAAAGAGTGTTTTTGAANNNAGATCGGAA |
| 55477_1 | TATGAGTGTGGAGTCGTTGCGGTGAGGCTGTGGAGAAATAAGANNNAGATCGGA |
| 55478_1 | TATGAGTGTGGAGTCGTTGCGGAGAGGCGTTTGTTGTGAATTTNNNAGATCGGAA |
| 55479_1 | TATGAGTGTGGAGTCGTTGCACACACACATATATGGCTCTTGCNNNAGATCGGAAG |

| | |
|---|---|
| 55480_1 | TATGAGTGTGGAGTCGTTGCGACCACAACTATAATTGCGCAGGNNNAGATCGGAAG |
| 55481_1 | TATGAGTGTGGAGTCGTTGCGAGGCTCACATGACCATATCCTTNNNAGATCGGAAGA |
| 55482_1 | TATGAGTGTGGAGTCGTTGCTCTCCTGAGTCCCTTTCACACTANNNAGATCGGAAGA |
| 55483_1 | TATGAGTGTGGAGTCGTTGCTCACAAGGTCAGGAATTCGAGACNNNAGATCGGAAG |
| 55485_1 | TATGAGTGTGGAGTCGTTGCTTACAAAGTCAGAGGAGGATGGCNNNAGATCGGAA |
| 55487_1 | TATGAGTGTGGAGTCGTTGCACTTGAGGTCAGGAGTTCAAGACNNNAGATCGGAAG |
| 55488_1 | TATGAGTGTGGAGTCGTTGCTGATGAGGGGAGAATCTGACTCANNNAGATCGGAA |
| 55490_1 | TATGAGTGTGGAGTCGTTGCGTAATCCCAGCTACTTAGGAGGCNNNAGATCGGAAG |
| 55491_1 | TATGAGTGTGGAGTCGTTGCGAGACTTTTGATTGGCTGGTCAGNNNAGATCGGAAG |
| 55493_1 | TATGAGTGTGGAGTCGTTGCTTTACCAAAGCCTACCTGACCAANNNAGATCGGAAGA |
| 55494_1 | TATGAGTGTGGAGTCGTTGCTCATGCTTGTAATCCCAGCACTTNNNAGATCGGAAGA |
| 55495_1 | TATGAGTGTGGAGTCGTTGCTCTCAACAACACCACTCAGAGAGNNNAGATCGGAAG |
| 55496_1 | TATGAGTGTGGAGTCGTTGCTCTGTAACTCCAGCTACTCAGGANNNAGATCGGAAGA |
| 55499_1 | TATGAGTGTGGAGTCGTTGCCACTCCATTCCCTCACTGAGAAANNNAGATCGGAAGA |
| 55500_1 | TATGAGTGTGGAGTCGTTGCGTAGTCATAGCTACTTGGGAGGCNNNAGATCGGAAG |
| 55503_1 | TATGAGTGTGGAGTCGTTGCCAATAGACTGGAGTGGATGGAGGNNNAGATCGGAA |
| 55509_1 | TATGAGTGTGGAGTCGTTGCCAGAGGTTGTAGTGAGCCAAGATNNNAGATCGGAA |
| 55510_1 | TATGAGTGTGGAGTCGTTGCCCTAAGAGCCAGTGAAGTGTCTTNNNAGATCGGAAG |
| 55511_1 | TATGAGTGTGGAGTCGTTGCGTTTGTGCCTAATCTGTGGAACCNNNAGATCGGAAG |
| 55515_1 | TATGAGTGTGGAGTCGTTGCGATGCATGTTCAGTGAGGTCAAANNNAGATCGGAAG |
| 55517_1 | TATGAGTGTGGAGTCGTTGCGTGAGCGACTGTAATCTCAGCTANNNAGATCGGAAG |
| 55518_1 | TATGAGTGTGGAGTCGTTGCTTTGTTGCTCCTACTTTTGGTGTNNNAGATCGGAAGA |
| 55519_1 | TATGAGTGTGGAGTCGTTGCGTAGAACCACTGCTCTCATCCAANNNAGATCGGAAGA |
| 55520_1 | TATGAGTGTGGAGTCGTTGCACAGGACACAATAGACACGGTTTNNNAGATCGGAAG |
| 55524_1 | TATGAGTGTGGAGTCGTTGCACTGACCACTTTTATTCAGGGCTNNNAGATCGGAAGA |
| 55527_1 | TATGAGTGTGGAGTCGTTGCGTAATCCCAGGTGCATCTACTCGNNNAGATCGGAAG |
| 55528_1 | TATGAGTGTGGAGTCGTTGCAAGATCTGAAGGGATCCAAAGCANNNAGATCGGAAG |
| 55529_1 | TATGAGTGTGGAGTCGTTGCAAAAGATGCCAAAGAGAACCTGCNNNAGATCGGAAG |
| 55530_1 | TATGAGTGTGGAGTCGTTGCCCAACAATGGAAGGCTGATTCAGNNNAGATCGGAAG |
| 55531_1 | TATGAGTGTGGAGTCGTTGCTATGATGGCATGCACCTGGAATTNNNAGATCGGAAG |
| 55532_1 | TATGAGTGTGGAGTCGTTGCCAGTCCCACTCCAATATCTCACANNNAGATCGGAAGA |
| 55534_1 | TATGAGTGTGGAGTCGTTGCTTCTACAGCAGCGGTTAGGTTTANNNAGATCGGAAG |
| 55535_1 | TATGAGTGTGGAGTCGTTGCCAATTCTCACCCATGCCTCATTCNNNAGATCGGAAGAG |
| 55536_1 | TATGAGTGTGGAGTCGTTGCGGAAAGCCCATGAACTTATCACGNNNAGATCGGAAG |
| 55537_1 | TATGAGTGTGGAGTCGTTGCCCTATAGTCCCAGCTTGGAAGACNNNAGATCGGAAG |
| 55538_1 | TATGAGTGTGGAGTCGTTGCAGGAAGTGGATTTTTCTCTGGCTNNNAGATCGGAAG |
| 55540_1 | TATGAGTGTGGAGTCGTTGCGTAATTCCAGCTACTCGGGAGACNNNAGATCGGAAG |
| 55542_1 | TATGAGTGTGGAGTCGTTGCTGGCCCAGAACAGTCAAATTCTANNNAGATCGGAAG |
| 55544_1 | TATGAGTGTGGAGTCGTTGCGAGGTCAGGAGATTGAGACCATCNNNAGATCGGAA |
| 55545_1 | TATGAGTGTGGAGTCGTTGCCTCACGTCTGTAATCTCAGCACTNNNAGATCGGAAGA |
| 55546_1 | TATGAGTGTGGAGTCGTTGCCTTAGAAGGCTGAGGCAGAAGAANNNAGATCGGAA |
| 55550_1 | TATGAGTGTGGAGTCGTTGCGAGAGTGGTGAAGGGAAAGAGAGNNNAGATCGGA |
| 55551_1 | TATGAGTGTGGAGTCGTTGCTGCTATTCTGCTTCTTCCAGAGANNNAGATCGGAAGA |

55552_1 TATGAGTGTGGAGTCGTTGCCGAGACAGGAAGGAAAACACAACNNNAGATCGGAA
55553_1 TATGAGTGTGGAGTCGTTGCTCCAAGAAATCAAAGCTGCAGTGNNNAGATCGGAA
55555_1 TATGAGTGTGGAGTCGTTGCAATCTCCAAGGAAGAGGTGTCACNNNAGATCGGAA
55557_1 TATGAGTGTGGAGTCGTTGCGCTTGGTTGATTTCATGCAGGTANNNAGATCGGAA
55558_1 TATGAGTGTGGAGTCGTTGCATCAGGCTCACATTTGCATGTTCNNNAGATCGGAAG
55559_1 TATGAGTGTGGAGTCGTTGCTGGAGTACAGTGGTGTGATCTTGNNNAGATCGGAA
55560_1 TATGAGTGTGGAGTCGTTGCGAGTAGTTGGGATCACAGGTCTGNNNAGATCGGAA
55561_1 TATGAGTGTGGAGTCGTTGCCCTCCTGGGTTCAAGCAATTCTCNNNAGATCGGAAG
55562_1 TATGAGTGTGGAGTCGTTGCTGTCAGTTTCTGCTTTTGTTGCANNNAGATCGGAAG
55563_1 TATGAGTGTGGAGTCGTTGCGTTGGTTTCCAGGCAAAAGATGANNNAGATCGGAA
55565_1 TATGAGTGTGGAGTCGTTGCTTCAAGCAATTCTCCTGCCTCAGNNNAGATCGGAAG
55566_1 TATGAGTGTGGAGTCGTTGCGGATTATAGGAACCTGCCACCATNNNAGATCGGAAG
55567_1 TATGAGTGTGGAGTCGTTGCGCTCACACCAAAAGACCAAACAANNNAGATCGGAAG
55568_1 TATGAGTGTGGAGTCGTTGCATGCGTGCTCATTGTGTAATCACNNNAGATCGGAAG
55569_1 TATGAGTGTGGAGTCGTTGCGCCTCTCAAAATGCTGGGATTACNNNAGATCGGAAG
55571_1 TATGAGTGTGGAGTCGTTGCTTCAAGTGATTCTCCTGCCTCAGNNNAGATCGGAAG
55572_1 TATGAGTGTGGAGTCGTTGCATCAAGTGATCTGTCTGCCTCAGNNNAGATCGGAAG
55573_1 TATGAGTGTGGAGTCGTTGCCCCTTGGGAGTAAGTGACAGAAANNNAGATCGGAA
55575_1 TATGAGTGTGGAGTCGTTGCATTACAGGCATGAGTCACTGTGTNNNAGATCGGAA
55576_1 TATGAGTGTGGAGTCGTTGCCCGACTCTTTGGTTCAAGCAATTNNNAGATCGGAAG
55577_1 TATGAGTGTGGAGTCGTTGCAAGTGCTGGGATTATAGGTGTGGNNNAGATCGGAA
55578_1 TATGAGTGTGGAGTCGTTGCCTCCTGAGTAGCTAGCTGGTACTNNNAGATCGGAAG
55579_1 TATGAGTGTGGAGTCGTTGCTACACTCACCAGACTATGGAGGTNNNAGATCGGAAG
55581_1 TATGAGTGTGGAGTCGTTGCTGGAGTACAGTGGCATGATCTTGNNNAGATCGGAA
55582_1 TATGAGTGTGGAGTCGTTGCCCTCCCTGGTTTGAGCAATTCTCNNNAGATCGGAAG
55583_1 TATGAGTGTGGAGTCGTTGCATCTCAGCTAACTGCAACCTTGANNNAGATCGGAAG
55584_1 TATGAGTGTGGAGTCGTTGCCATGTACAAGTCAGGAGAGCTCANNNAGATCGGAA
55585_1 TATGAGTGTGGAGTCGTTGCCTTTGAAAGCAGGCATGAGGAAANNNAGATCGGAA
55586_1 TATGAGTGTGGAGTCGTTGCCATGATCTCAGCTCACTGCAAACNNNAGATCGGAAG
55587_1 TATGAGTGTGGAGTCGTTGCAATCTCAGTTCACTGCAAGCTCCNNNAGATCGGAAG
55591_1 TATGAGTGTGGAGTCGTTGCTTCATCAGCAGCATGAAAACTGGNNNAGATCGGAA
55592_1 TATGAGTGTGGAGTCGTTGCCAGGCACATGCTACATACCCTATNNNAGATCGGAAG
55593_1 TATGAGTGTGGAGTCGTTGCAGCTGGGATTACAGGTGTGTAAANNNAGATCGGAA
55595_1 TATGAGTGTGGAGTCGTTGCTTCACTCTTATTGTCCAGGCTGGNNNAGATCGGAAG
55596_1 TATGAGTGTGGAGTCGTTGCACCAAGGTACATATCACTACGCCNNNAGATCGGAAG
55597_1 TATGAGTGTGGAGTCGTTGCCTTCCTGAGTAGCTGGGACTACANNNAGATCGGAAG
55598_1 TATGAGTGTGGAGTCGTTGCGATGAGGACTCTACCGTCATGACNNNAGATCGGAA
55599_1 TATGAGTGTGGAGTCGTTGCCTGGAATTATGGGCACACATCACNNNAGATCGGAAG
55603_1 TATGAGTGTGGAGTCGTTGCCCTCCTGAGTTCATGCCATTCTCNNNAGATCGGAAGA
55604_1 TATGAGTGTGGAGTCGTTGCAGCCTCTGGAAACCACTAATCTGNNNAGATCGGAAG
55610_1 TATGAGTGTGGAGTCGTTGCCCCTTGTCTGTTTCTTACCATCCNNNAGATCGGAAGA
55613_1 TATGAGTGTGGAGTCGTTGCTACCTTCCTGAAAAAGCTCCCTCNNNAGATCGGAAGA
55615_1 TATGAGTGTGGAGTCGTTGCCTATGGTCAATCAGCAAAGCCAGNNNAGATCGGAA

| | |
|---|---|
| 55616_1 | TATGAGTGTGGAGTCGTTGCGAGATCACACCACTGAACTCCAGNNNAGATCGGAAG |
| 55617_1 | TATGAGTGTGGAGTCGTTGCACAGGAAGTCAAGAGGAAATGCTNNNAGATCGGAA |
| 55618_1 | TATGAGTGTGGAGTCGTTGCAGCTGAAAGAGAAAGCTAGGACANNNAGATCGGA |
| 55619_1 | TATGAGTGTGGAGTCGTTGCACAGCCAAATAATGCACTCGTTCNNNAGATCGGAAG |
| 55620_1 | TATGAGTGTGGAGTCGTTGCGCTACTTCACCCAATCAAAAGCGNNNAGATCGGAAG |
| 55622_1 | TATGAGTGTGGAGTCGTTGCACAACGTGAATGTACTTGATGCCNNNAGATCGGAAG |
| 55626_1 | TATGAGTGTGGAGTCGTTGCCAGTTGACAGAAACAGTGGAACANNNAGATCGGAA |
| 55627_1 | TATGAGTGTGGAGTCGTTGCTCCTACCCTCACAGAACTTACCTNNNAGATCGGAAGA |
| 55630_1 | TATGAGTGTGGAGTCGTTGCAAGGTATTTCAGGCAGCACAAACNNNAGATCGGAA |
| 55631_1 | TATGAGTGTGGAGTCGTTGCAGGATGATGGCTGCCTGAAATAANNNAGATCGGAA |
| 55633_1 | TATGAGTGTGGAGTCGTTGCATGGTGGGAATTCATGCCTAGTTNNNAGATCGGAA |
| 55634_1 | TATGAGTGTGGAGTCGTTGCGTGTGCTTACCTATCAGTCGCTANNNAGATCGGAAG |
| 55636_1 | TATGAGTGTGGAGTCGTTGCGGTGATCAACAGGTCATTAGGGTNNNAGATCGGAA |
| 55638_1 | TATGAGTGTGGAGTCGTTGCAGAGTGAGCAGTACTGTCCATTGNNNAGATCGGAA |
| 55639_1 | TATGAGTGTGGAGTCGTTGCGAGGAGCACTGAAGGGTTTTAGANNNAGATCGGA |
| 55641_1 | TATGAGTGTGGAGTCGTTGCGAACAGATCAAGCAGGCAGAAAANNNAGATCGGAA |
| 55642_1 | TATGAGTGTGGAGTCGTTGCGGAAAAGCTTCACCAGAAAGTCCNNNAGATCGGAA |
| 55643_1 | TATGAGTGTGGAGTCGTTGCATAGTGAGCAGAGATCATGCCACNNNAGATCGGAA |
| 55644_1 | TATGAGTGTGGAGTCGTTGCTGAATCAGAACATCGCATAAGCCNNNAGATCGGAAG |
| 55645_1 | TATGAGTGTGGAGTCGTTGCGATCACGAGGTCAGGAGTTCAAGNNNAGATCGGAA |
| 55646_1 | TATGAGTGTGGAGTCGTTGCTGGACAAAATTCCACCAGCAAAANNNAGATCGGAAG |
| 55647_1 | TATGAGTGTGGAGTCGTTGCTGGTAAAGCAGAGACTCTGATGGNNNAGATCGGAA |
| 55648_1 | TATGAGTGTGGAGTCGTTGCTAGAGTTGTGCAACCTCCACTACNNNAGATCGGAAG |
| 55649_1 | TATGAGTGTGGAGTCGTTGCTTGATGGTACATCAGGCTTCTGGNNNAGATCGGAA |
| 55651_1 | TATGAGTGTGGAGTCGTTGCCAGAATCCCAGGTTGGAGTCATCNNNAGATCGGAA |
| 55652_1 | TATGAGTGTGGAGTCGTTGCCGACAGGGCAAGACTCTATCAAANNNAGATCGGAA |
| 55653_1 | TATGAGTGTGGAGTCGTTGCTCAGTGACGACAGTGCTAGTAAGNNNAGATCGGAA |
| 55654_1 | TATGAGTGTGGAGTCGTTGCCATAGGATCACTGCCACTGAACTNNNAGATCGGAAG |
| 55655_1 | TATGAGTGTGGAGTCGTTGCTATGAAAAATGGAGCCCTGGGATNNNAGATCGGAA |
| 55656_1 | TATGAGTGTGGAGTCGTTGCCACCAATAGAAGAGGGGACTACGNNNAGATCGGAA |
| 55657_1 | TATGAGTGTGGAGTCGTTGCGGGAGGGGATTGCTTTATCATCANNNAGATCGGAA |
| 55658_1 | TATGAGTGTGGAGTCGTTGCTGTGTGTTGGTGGAGAGTCTTTTNNNAGATCGGAA |
| 55659_1 | TATGAGTGTGGAGTCGTTGCTCTCCTCCCTTAGTCTTCCAAGTNNNAGATCGGAAGA |
| 55660_1 | TATGAGTGTGGAGTCGTTGCTTCCTCTTACTTGGTGCACTTCCNNNAGATCGGAAGA |
| 55662_1 | TATGAGTGTGGAGTCGTTGCTGTGTAGGAAGAGGGAGAGAGAGNNNAGATCGG |
| 55663_1 | TATGAGTGTGGAGTCGTTGCAGGAGTATCTCTTGGTCCAGGAGNNNAGATCGGAA |
| 55664_1 | TATGAGTGTGGAGTCGTTGCTACACACAATTAGCTGGGAGTGGNNNAGATCGGAA |
| 55665_1 | TATGAGTGTGGAGTCGTTGCTAATCCCAGCTACTCAGGAGTCTNNNAGATCGGAAG |
| 55666_1 | TATGAGTGTGGAGTCGTTGCGGCTGAGGTGGGAGGATCTATTANNNAGATCGGA |
| 55667_1 | TATGAGTGTGGAGTCGTTGCCCTTCCTTGTGTGTGTGTGTTTTNNNAGATCGGAAG |
| 55668_1 | TATGAGTGTGGAGTCGTTGCCACCTGAATAGGCAAGGTCTCTTNNNAGATCGGAAG |
| 55669_1 | TATGAGTGTGGAGTCGTTGCAGAGTTGCTTGATTCCACAAGGANNNAGATCGGAA |
| 55671_1 | TATGAGTGTGGAGTCGTTGCCCAAGGAACTACCAGACACTAGGNNNAGATCGGAA |

55673_1 TATGAGTGTGGAGTCGTTGCCCAGGCATATCTCCAGACATTCANNNAGATCGGAAG
55674_1 TATGAGTGTGGAGTCGTTGCAATGAGGTTGCAGTGAGGAGAAANNNAGATCGGA
55675_1 TATGAGTGTGGAGTCGTTGCCCCAGATTCAAGCGATCCTCTAGNNNAGATCGGAAG
55676_1 TATGAGTGTGGAGTCGTTGCCTTCTAAAGGGAGAGAGCAGCTTNNNAGATCGGAA
55679_1 TATGAGTGTGGAGTCGTTGCGTGTGTGTGTGTGTTTCTTCTNNNAGATCGGAAG
55680_1 TATGAGTGTGGAGTCGTTGCACTAGTAAAGGTAGAGGGCTGGTNNNAGATCGGA
55682_1 TATGAGTGTGGAGTCGTTGCCCTGTAGTCCCAGCTACTTAGGANNNAGATCGGAAG
55683_1 TATGAGTGTGGAGTCGTTGCTCTTGAACTCCTGACCTCAAGTGNNNAGATCGGAAG
55684_1 TATGAGTGTGGAGTCGTTGCAAACAAAGGGAATGTTGACAGCCNNNAGATCGGAA
55685_1 TATGAGTGTGGAGTCGTTGCGAGAGAGAGAGAGAGACACTCCANNNAGATCGGA
55687_1 TATGAGTGTGGAGTCGTTGCATGATCTCAGTTCACTGCAACCTNNNAGATCGGAAG
55689_1 TATGAGTGTGGAGTCGTTGCATTTTTCGAGATGGAGTCTGGGTNNNAGATCGGAA
55690_1 TATGAGTGTGGAGTCGTTGCATCCCTTATCTCCACCTCTGTCCNNNAGATCGGAAGA
55691_1 TATGAGTGTGGAGTCGTTGCATGTGGTGGTACATGCCTGTAATNNNAGATCGGAA
55692_1 TATGAGTGTGGAGTCGTTGCCTCACACCAGTTTTCCTTGGTTCNNNAGATCGGAAGA
55693_1 TATGAGTGTGGAGTCGTTGCACCCTATCAGCCAATGCCTTTATNNNAGATCGGAAGA
55695_1 TATGAGTGTGGAGTCGTTGCCTCAAACTCCTGACCTCACGTAANNNAGATCGGAAG
55696_1 TATGAGTGTGGAGTCGTTGCTAAAAGACACAAAGTGAGCCCAGNNNAGATCGGAA
55697_1 TATGAGTGTGGAGTCGTTGCACAGAGAAACAGAGAGAGCACAGNNNAGATCGGA
55698_1 TATGAGTGTGGAGTCGTTGCCTATCAATGGGCCTTGTTGCTAGNNNAGATCGGAAG
55699_1 TATGAGTGTGGAGTCGTTGCTGACTCCTTCAGACATCAAGAGCNNNAGATCGGAAG
55700_1 TATGAGTGTGGAGTCGTTGCCTGCCCTTACAACAAAACTCCTGNNNAGATCGGAAG
55701_1 TATGAGTGTGGAGTCGTTGCTTTACTGGATCAGACCTGAAGCCNNNAGATCGGAAG
55702_1 TATGAGTGTGGAGTCGTTGCATTGCTCCACCACAAGTGAGTATNNNAGATCGGAAG
55705_1 TATGAGTGTGGAGTCGTTGCCACACAACGTAACTCCAGGTTTCNNNAGATCGGAAG
55707_1 TATGAGTGTGGAGTCGTTGCACAATCAGATCTCGTGGGAACTCNNNAGATCGGAAG
55708_1 TATGAGTGTGGAGTCGTTGCACAGAAATGAAAACTGGGTGCTGNNNAGATCGGAA
55710_1 TATGAGTGTGGAGTCGTTGCATCTGAGGATGGAGGATTGGACTNNNAGATCGGAA
55711_1 TATGAGTGTGGAGTCGTTGCTAAAACAGCTACCCACAGTGGAGNNNAGATCGGAA
55712_1 TATGAGTGTGGAGTCGTTGCGACTATAGCCACGTACCACCATGNNNAGATCGGAAG
55713_1 TATGAGTGTGGAGTCGTTGCCTACAGAAAACATTCGCTGACCCNNNAGATCGGAAG
55714_1 TATGAGTGTGGAGTCGTTGCCAATCCTTCTACCTTAGCCTCCCNNNAGATCGGAAGA
55716_1 TATGAGTGTGGAGTCGTTGCCTCTCTGAGTAGCTGGGACTACANNNAGATCGGAAG
55717_1 TATGAGTGTGGAGTCGTTGCCTCCCAGGTTCAAGAGATTCTCCNNNAGATCGGAAG
55718_1 TATGAGTGTGGAGTCGTTGCCTGAGTAGCTGGGATTACAGGTGNNNAGATCGGAA
55719_1 TATGAGTGTGGAGTCGTTGCTTCCAGGTTCAAGTGATTCTCGTNNNAGATCGGAAG
55720_1 TATGAGTGTGGAGTCGTTGCTTCAAGTGATTCTCCTGCCTCAGNNNAGATCGGAAG
55724_1 TATGAGTGTGGAGTCGTTGCGTAGAAACAGGGTCTAGCTGTGTNNNAGATCGGAA
55727_1 TATGAGTGTGGAGTCGTTGCCCCAGGTTCAAGCAATTCTCTTGNNNAGATCGGAAG
55729_1 TATGAGTGTGGAGTCGTTGCTCAAGTAATTCTCCTGCCTCAGCNNNAGATCGGAAGA
55730_1 TATGAGTGTGGAGTCGTTGCGGTTCAAACGATTCTTCTGCCTCNNNAGATCGGAAG
55731_1 TATGAGTGTGGAGTCGTTGCCAATCTCGGCTCACTACAACCTCNNNAGATCGGAAGA
55732_1 TATGAGTGTGGAGTCGTTGCGATGGAGTTTTGCTTTTGTTGCCNNNAGATCGGAAG

| | |
|---|---|
| 55734_1 | TATGAGTGTGGAGTCGTTGCCAATGTTGGCCTAAGCAGTTTCANNNAGATCGGAAG |
| 55735_1 | TATGAGTGTGGAGTCGTTGCCCCTTCAAATGTCTGACAAGCTGNNNAGATCGGAAG |
| 55736_1 | TATGAGTGTGGAGTCGTTGCTTCAAGTGATTCTCCTGCCTCAGNNNAGATCGGAAG |
| 55737_1 | TATGAGTGTGGAGTCGTTGCCAATCTTTGCTCACCACAACCTCNNNAGATCGGAAGA |
| 55738_1 | TATGAGTGTGGAGTCGTTGCTACTGGTATAAGCCATTGCACCCNNNAGATCGGAAG |
| 55739_1 | TATGAGTGTGGAGTCGTTGCGGGGAGAGACAAATTGAATGCATNNNAGATCGGAA |
| 55741_1 | TATGAGTGTGGAGTCGTTGCTGGGAGCTCATAGTTGTAGAGGANNNAGATCGGAA |
| 55742_1 | TATGAGTGTGGAGTCGTTGCACAGCCAGCTAACATATCACCATNNNAGATCGGAAG |
| 55743_1 | TATGAGTGTGGAGTCGTTGCCTTTGGGAGGCTGAAGTAGGATTNNNAGATCGGAA |
| 55744_1 | TATGAGTGTGGAGTCGTTGCAAAACTCGCCACCCATAAACAAGNNNAGATCGGAAG |
| 55745_1 | TATGAGTGTGGAGTCGTTGCTGGAGGCTTGAAATGAATTCAGCNNNAGATCGGAA |
| 55747_1 | TATGAGTGTGGAGTCGTTGCTTGTTCATGTCCATCACTCCCATNNNAGATCGGAAGA |
| 55752_1 | TATGAGTGTGGAGTCGTTGCCATCTGCTGGTGAATTATGTGCCNNNAGATCGGAAG |
| 55756_1 | TATGAGTGTGGAGTCGTTGCCAACTACTAGGCCCAGAGGTATGNNNAGATCGGAA |
| 55757_1 | TATGAGTGTGGAGTCGTTGCTACCTCTAGCACCTTTCTAGCTGNNNAGATCGGAAGA |
| 55758_1 | TATGAGTGTGGAGTCGTTGCAGCTTCCATTTAGGGCCAGATTANNNAGATCGGAAG |
| 55759_1 | TATGAGTGTGGAGTCGTTGCAGAGGAAAGTGGTGTTATTCCCCNNNAGATCGGAA |
| 55760_1 | TATGAGTGTGGAGTCGTTGCGCCAAACAACATATGGGCTAGAANNNAGATCGGAA |
| 55763_1 | TATGAGTGTGGAGTCGTTGCGTTCAGCCTTGTTGGAAGTTGTTNNNAGATCGGAAG |
| 55764_1 | TATGAGTGTGGAGTCGTTGCAACTGCTCGAGACTGGGTAATTTNNNAGATCGGAAG |
| 55765_1 | TATGAGTGTGGAGTCGTTGCCTTGGGAAGTGGAAGTGGAAGTANNNAGATCGGA |
| 55767_1 | TATGAGTGTGGAGTCGTTGCGTGTGAGGTCATTTGTGAAGGTGNNNAGATCGGAA |
| 55768_1 | TATGAGTGTGGAGTCGTTGCCTGGAAAAGACAGGGCTCTTCTANNNAGATCGGAA |
| 55771_1 | TATGAGTGTGGAGTCGTTGCATGAGGTCCTACTATGTAGCCCANNNAGATCGGAAG |
| 55774_1 | TATGAGTGTGGAGTCGTTGCTGTACAGTCATTGGTCCAGAAGGNNNAGATCGGAA |
| 55775_1 | TATGAGTGTGGAGTCGTTGCTGAGAAAAGGACAAGGGACAGAGNNNAGATCGGA |
| 55778_1 | TATGAGTGTGGAGTCGTTGCAATGTTGGCAGAAACTCTTGCAANNNAGATCGGAAG |
| 55780_1 | TATGAGTGTGGAGTCGTTGCTTCTTCCAGGTTCAGTTTTGGGANNNAGATCGGAAG |
| 55781_1 | TATGAGTGTGGAGTCGTTGCTGAATTGTGTCCCTCATTCTCTTNNNAGATCGGAAGA |
| 55783_1 | TATGAGTGTGGAGTCGTTGCGACACCAGGTATGGAGCTAACTTNNNAGATCGGAAG |
| 55786_1 | TATGAGTGTGGAGTCGTTGCCAAAGTTTTGGGATGACAGGCATNNNAGATCGGAA |
| 55787_1 | TATGAGTGTGGAGTCGTTGCCTGGGTTCAAGTGATTCTTCTGCNNNAGATCGGAAG |
| 55788_1 | TATGAGTGTGGAGTCGTTGCGCTTCATGGGTTCAAGCAATTCTNNNAGATCGGAAG |
| 55791_1 | TATGAGTGTGGAGTCGTTGCGCTGCAAAGTTTGGAGGTTCTTTNNNAGATCGGAAG |
| 55793_1 | TATGAGTGTGGAGTCGTTGCATGTTCAAGTGATTCTCCTGCCTNNNAGATCGGAAG |
| 55794_1 | TATGAGTGTGGAGTCGTTGCCTAAAACCTCTGCCTCCTGGATTNNNAGATCGGAAG |
| 55795_1 | TATGAGTGTGGAGTCGTTGCGGAGTACAGTGGCAGAATCTTGANNNAGATCGGAA |
| 55797_1 | TATGAGTGTGGAGTCGTTGCGATGGAGTCTCATTCTGTCTCCCNNNAGATCGGAAG |
| 55798_1 | TATGAGTGTGGAGTCGTTGCAGACTGGAGTGCAATGGTACAATNNNAGATCGGAA |
| 55800_1 | TATGAGTGTGGAGTCGTTGCCCCATGTTCAAGCAATTCCTCTGNNNAGATCGGAAG |
| 55802_1 | TATGAGTGTGGAGTCGTTGCGATCTCTCCTCACTGCAACCTTTNNNAGATCGGAAGA |
| 55804_1 | TATGAGTGTGGAGTCGTTGCTTCAAGCTATTCTCCTGCCTCAGNNNAGATCGGAAG |
| 55807_1 | TATGAGTGTGGAGTCGTTGCTTCTGTCCTGAGCAAAATAGGCTNNNAGATCGGAAG |

| | |
|---|---|
| 55808_1 | TATGAGTGTGGAGTCGTTGCTTAGACGGAGTCTCAATCTGTCGNNNAGATCGGAAG |
| 55809_1 | TATGAGTGTGGAGTCGTTGCCTATGTTACCCAGGATGGTCTCGNNNAGATCGGAAG |
| 55810_1 | TATGAGTGTGGAGTCGTTGCTCTCTCAGGCTGAAGTACAATGGNNNAGATCGGAAG |
| 55811_1 | TATGAGTGTGGAGTCGTTGCGTTCAAGAGATTCTCCTGCCTCANNNAGATCGGAAGA |
| 55812_1 | TATGAGTGTGGAGTCGTTGCAGACAAAAACAGACACTCTGGGANNNAGATCGGAAG |
| 55813_1 | TATGAGTGTGGAGTCGTTGCCTCCTGAGTAGCTGGGACTACAGNNNAGATCGGAAG |
| 55814_1 | TATGAGTGTGGAGTCGTTGCGATTCTCCTGCCTCAATCTCCTGNNNAGATCGGAAGA |
| 55815_1 | TATGAGTGTGGAGTCGTTGCTGGATTGCAGTGGTGTGATCATANNNAGATCGGAAG |
| 55818_1 | TATGAGTGTGGAGTCGTTGCTTCAAGTGATTCTCCTGCCTCAGNNNAGATCGGAAGA |
| 55820_1 | TATGAGTGTGGAGTCGTTGCGACTGGGTCTTCCTATGTTTCCCNNNAGATCGGAAGA |
| 55823_1 | TATGAGTGTGGAGTCGTTGCCCTGGGTTCAAACAATTCACCTGNNNAGATCGGAAGA |
| 55824_1 | TATGAGTGTGGAGTCGTTGCCCAAGTAGCTGGGATTACAGGTGNNNAGATCGGAAG |
| 55825_1 | TATGAGTGTGGAGTCGTTGCTTCATATCTGTGACACCCTTCCCNNNAGATCGGAAGA |
| 55828_1 | TATGAGTGTGGAGTCGTTGCTTTTTTCATATGTTACGTGCCCANNNAGATCGGAAGA |
| 55829_1 | TATGAGTGTGGAGTCGTTGCCCCAGTTGGTGGTATTCTGTTATNNNAGATCGGAAG |
| 55831_1 | TATGAGTGTGGAGTCGTTGCGCCAGACTTTGAGAAGCTTTTGANNNAGATCGGAAG |
| 55832_1 | TATGAGTGTGGAGTCGTTGCATTCTGTTGATTTCGGGTGGAGANNNAGATCGGAAG |
| 55833_1 | TATGAGTGTGGAGTCGTTGCTATGGTCTGGAACTTCGGCAAATNNNAGATCGGAAG |
| 55834_1 | TATGAGTGTGGAGTCGTTGCATCTTTGTCACCTCAGTAGTGCCNNNAGATCGGAAGA |
| 55839_1 | TATGAGTGTGGAGTCGTTGCATGAGAGAAAACCAGAAGTGTCTNNNAGATCGGAAG |
| 55840_1 | TATGAGTGTGGAGTCGTTGCGCCCAAGGAAACTCTATGCAAATNNNAGATCGGAAG |
| 55841_1 | TATGAGTGTGGAGTCGTTGCTAGTCATGAGCTGGGCATTTTCTNNNAGATCGGAAG |
| 55842_1 | TATGAGTGTGGAGTCGTTGCACACGTATTACTCAGACCTGGTGNNNAGATCGGAAG |
| 55843_1 | TATGAGTGTGGAGTCGTTGCGATGGAGGAGAAACCTACAGACTNNNAGATCGGAA |
| 55844_1 | TATGAGTGTGGAGTCGTTGCTTTATTGCCGCATACCAATCACCNNNAGATCGGAAGA |
| 55845_1 | TATGAGTGTGGAGTCGTTGCGTGCATCTGTAGTTCCAGCTACTNNNAGATCGGAAGA |
| 55846_1 | TATGAGTGTGGAGTCGTTGCCGCCTATAGTCTCAGCTACTTGGNNNAGATCGGAAGA |
| 55847_1 | TATGAGTGTGGAGTCGTTGCGTTGTCCTACATACTCAGGAGGCNNNAGATCGGAAG |
| 55848_1 | TATGAGTGTGGAGTCGTTGCAAAGTTTGAGGATAGAGGCTGCTNNNAGATCGGAA |
| 55849_1 | TATGAGTGTGGAGTCGTTGCACAAAATAACGAGGAACTGCAGGNNNAGATCGGAA |
| 55850_1 | TATGAGTGTGGAGTCGTTGCGACCCCACATGAGCTAATCTTGTNNNAGATCGGAAGA |
| 55852_1 | TATGAGTGTGGAGTCGTTGCTGTGCTTGCAAATTTGGAAGGTTNNNAGATCGGAAG |
| 55853_1 | TATGAGTGTGGAGTCGTTGCCACTGAAAGGGCACTGTCTTTTTNNNAGATCGGAAG |
| 55856_1 | TATGAGTGTGGAGTCGTTGCACATCTCATTGCAGCCACTCTATNNNAGATCGGAAGA |
| 55858_1 | TATGAGTGTGGAGTCGTTGCGACTTGGAGGATGCACATTTGACNNNAGATCGGAAG |
| 55859_1 | TATGAGTGTGGAGTCGTTGCTCAGACTCCAAATAAAACAACTGCANNNAGATCGGAA |
| 55860_1 | TATGAGTGTGGAGTCGTTGCAGAAGGCAATGTTCCTGAGGATTNNNAGATCGGAAG |
| 55862_1 | TATGAGTGTGGAGTCGTTGCGCGGTTCTTTTCAGATCGAATGTNNNAGATCGGAAG |
| 55863_1 | TATGAGTGTGGAGTCGTTGCAAAAAGCAAGGCATCCTCACAAANNNAGATCGGAAG |
| 55865_1 | TATGAGTGTGGAGTCGTTGCTTTTTTGATGGGAACTGGCTGTGNNNAGATCGGAAG |
| 55866_1 | TATGAGTGTGGAGTCGTTGCAGAGTTGTCTGGTGTGTCTTCTTNNNAGATCGGAAG |
| 55867_1 | TATGAGTGTGGAGTCGTTGCCTGTGGCTCTGAGAAATTGGTTTNNNAGATCGGAAG |
| 55869_1 | TATGAGTGTGGAGTCGTTGCCAAAGAAACAGGGCAGGTACAATNNNAGATCGGAA |

55870_1 TATGAGTGTGGAGTCGTTGCGTTTCTTTTGACAGCCACACTGANNNAGATCGGAAG
55871_1 TATGAGTGTGGAGTCGTTGCGCAAAACTCAGGAGCTGACAAATNNNAGATCGGAA
55872_1 TATGAGTGTGGAGTCGTTGCCCAGTAATCCTAGCACTTTGGGANNNAGATCGGAAG
55873_1 TATGAGTGTGGAGTCGTTGCCTGATTTACTGGTTTGGGGATGCNNNAGATCGGAA
55874_1 TATGAGTGTGGAGTCGTTGCTCATTGACTCACAGTTCCACAGGNNNAGATCGGAAG
55875_1 TATGAGTGTGGAGTCGTTGCATGATTTTGAAAAGCACCAGGGTNNNAGATCGGAA
55877_1 TATGAGTGTGGAGTCGTTGCTTCAAGCAATTCTCCTGCCTCAANNNAGATCGGAAGA
55878_1 TATGAGTGTGGAGTCGTTGCTTGGTCTCATTACCGGTGATGTTNNNAGATCGGAAG
55880_1 TATGAGTGTGGAGTCGTTGCGCAGTGATGTGATCTTGGCTTACNNNAGATCGGAAG
55882_1 TATGAGTGTGGAGTCGTTGCGACAGAGTCTCACTGTTACCCAGNNNAGATCGGAAG
55883_1 TATGAGTGTGGAGTCGTTGCGTAGCTGGGATTACAGGCATGTANNNAGATCGGAA
55885_1 TATGAGTGTGGAGTCGTTGCTTCAAGTGATTCCCTTGCCTGAGNNNAGATCGGAAG
55886_1 TATGAGTGTGGAGTCGTTGCCCCAAGTTCAAGCGATTCTCTTGNNNAGATCGGAAG
55887_1 TATGAGTGTGGAGTCGTTGCCTCAATTGACCCTCCCATCTCAGNNNAGATCGGAAGA
55888_1 TATGAGTGTGGAGTCGTTGCACAATCTTGGCTCACTACAACCTNNNAGATCGGAAG
55889_1 TATGAGTGTGGAGTCGTTGCGCCACCATGTCCGACTATTTTTTNNNAGATCGGAAGA
55890_1 TATGAGTGTGGAGTCGTTGCAATCTCGGCTCATTGAAACCTCTNNNAGATCGGAAG
55891_1 TATGAGTGTGGAGTCGTTGCCAGTGGTGTAATCTCAGCTCACTNNNAGATCGGAAG
55892_1 TATGAGTGTGGAGTCGTTGCCACCTTAGCCTCTCAAGTAGCTGNNNAGATCGGAAG
55894_1 TATGAGTGTGGAGTCGTTGCCCTCCAAGTAGCTGGGATTACAGNNNAGATCGGAAG
55896_1 TATGAGTGTGGAGTCGTTGCGGCATGAATATGGCTTACTGCAGNNNAGATCGGAA
55897_1 TATGAGTGTGGAGTCGTTGCACAGCATCTTACTATGTTGCCCANNNAGATCGGAAG
55898_1 TATGAGTGTGGAGTCGTTGCTTCAAGCGATTCTACTGCCTCAGNNNAGATCGGAAG
55899_1 TATGAGTGTGGAGTCGTTGCCTGAGTTCAAGCAATTCTCCTGCNNNAGATCGGAAG
55900_1 TATGAGTGTGGAGTCGTTGCACAATCATAGCTCGCAGTAACCTNNNAGATCGGAAG
55901_1 TATGAGTGTGGAGTCGTTGCTCAAGTGAGCATGTCCAGCTAATNNNAGATCGGAAG
55902_1 TATGAGTGTGGAGTCGTTGCGAGACGGTATCTTGCTCTGTCTCNNNAGATCGGAAG
55903_1 TATGAGTGTGGAGTCGTTGCTTCATGTGATTCTCCTGCCTCAGNNNAGATCGGAAG
55904_1 TATGAGTGTGGAGTCGTTGCTTCGTCATATTGCTCAGGTTGGTNNNAGATCGGAAG
55906_1 TATGAGTGTGGAGTCGTTGCAACAGTACATCTTGCACCCAGATNNNAGATCGGAAG
55907_1 TATGAGTGTGGAGTCGTTGCGCCTCATAGAAAGATGTGCAAAGTNNNAGATCGGA
55908_1 TATGAGTGTGGAGTCGTTGCCTTTGTAAGTTTCCTGCCACCTGNNNAGATCGGAAG
55909_1 TATGAGTGTGGAGTCGTTGCGAACATTGGAAAGGGAATTGCCANNNAGATCGGAA
55910_1 TATGAGTGTGGAGTCGTTGCTGCCCAAACCACTGTTTATAGGANNNAGATCGGAAG
55912_1 TATGAGTGTGGAGTCGTTGCATATCTCAACACTGCCCACTTGTNNNAGATCGGAAG
55913_1 TATGAGTGTGGAGTCGTTGCAACAGGCAAAACCAATTGATGCTNNNAGATCGGAA
55914_1 TATGAGTGTGGAGTCGTTGCGAAACCAGGACTTGAGTCTCCTTNNNAGATCGGAAG
55916_1 TATGAGTGTGGAGTCGTTGCATCCAGCAACAATGTTTCACCAGNNNAGATCGGAAG
55919_1 TATGAGTGTGGAGTCGTTGCACAGTTTGAAAGAGTTCTGTGGTNNNAGATCGGAA
55923_1 TATGAGTGTGGAGTCGTTGCAAGGCGTAGGAAGGGAAATCTACNNNAGATCGGAA
55924_1 TATGAGTGTGGAGTCGTTGCTACGTTTGATCTCCAGGAACAGGNNNAGATCGGAA
55925_1 TATGAGTGTGGAGTCGTTGCCAGCGTCGAAGCATGTGAAATATNNNAGATCGGAA
55926_1 TATGAGTGTGGAGTCGTTGCACCTGTTCCTATTCCTTCACCACNNNAGATCGGAAGA

| | |
|---|---|
| 55927_1 | TATGAGTGTGGAGTCGTTGCGTCTCAAGTGATCATCCTCCCTCNNNAGATCGGAAGA |
| 55928_1 | TATGAGTGTGGAGTCGTTGCGCAAGGGTTGATGAACAGAACAANNNAGATCGGAA |
| 55929_1 | TATGAGTGTGGAGTCGTTGCTCTGCCTTCCAGATTCAAGTGATNNNAGATCGGAAGA |
| 55930_1 | TATGAGTGTGGAGTCGTTGCTCCCAAGTAGATTACAGGTGTGCNNNAGATCGGAAG |
| 55931_1 | TATGAGTGTGGAGTCGTTGCTGATCTCGGCTCATTGTAAGCTCNNNAGATCGGAAGA |
| 55932_1 | TATGAGTGTGGAGTCGTTGCTTTTTCGAGGCTGATCTGTGGTANNNAGATCGGAAG |
| 55933_1 | TATGAGTGTGGAGTCGTTGCCCTGGGTTCAAGCAATTCTCATGNNNAGATCGGAAG |
| 55934_1 | TATGAGTGTGGAGTCGTTGCAAAGAGTCTTGCTGTGTTACCCANNNAGATCGGAAG |
| 55935_1 | TATGAGTGTGGAGTCGTTGCGGAGATAGGTCCCTTTTGGGTTTNNNAGATCGGAAG |
| 55936_1 | TATGAGTGTGGAGTCGTTGCGCAATGGCACTATCATCACTCACNNNAGATCGGAAGA |
| 55937_1 | TATGAGTGTGGAGTCGTTGCTATGGAGTTTTGCTGTTTTGCCCNNNAGATCGGAAGA |
| 55938_1 | TATGAGTGTGGAGTCGTTGCACAGAAATGAATGCATTGGCTGGNNNAGATCGGAAG |
| 55939_1 | TATGAGTGTGGAGTCGTTGCATGGCAGTGGTGTAATCATAGCTNNNAGATCGGAAG |
| 55940_1 | TATGAGTGTGGAGTCGTTGCTTCAAGCAATTTTCCTGCCTCAGNNNAGATCGGAAGA |
| 55942_1 | TATGAGTGTGGAGTCGTTGCCTCCACAGTAGCAGGGATTACAGNNNAGATCGGAAG |
| 55943_1 | TATGAGTGTGGAGTCGTTGCCTCAAGCTATCCTCCCACCTCAGNNNAGATCGGAAGA |
| 55944_1 | TATGAGTGTGGAGTCGTTGCAGTATAGACGGGATTTCACCACGNNNAGATCGGAAG |
| 55945_1 | TATGAGTGTGGAGTCGTTGCCCTCTGGAGTTGAAGCGATTCTCNNNAGATCGGAAG |
| 55947_1 | TATGAGTGTGGAGTCGTTGCGATCACAAGGTCAGGAGTTCGAGNNNAGATCGGAA |
| 55948_1 | TATGAGTGTGGAGTCGTTGCCTGTCAAAACCTTCCCACCTACTNNNAGATCGGAAGA |
| 55952_1 | TATGAGTGTGGAGTCGTTGCCCAAGCAGTTGGGACTATAGGTGNNNAGATCGGAAG |
| 55953_1 | TATGAGTGTGGAGTCGTTGCACCTGAAATGGCTGTCTTCTTGANNNAGATCGGAAG |
| 55954_1 | TATGAGTGTGGAGTCGTTGCGAGTAGCTGGGATTACAGGCATGNNNAGATCGGAA |
| 55956_1 | TATGAGTGTGGAGTCGTTGCCTGAAAGAAAGGAAGGGGATTGCNNNAGATCGGAA |
| 55957_1 | TATGAGTGTGGAGTCGTTGCCACACCTGTAATCCCAGCACTTTNNNAGATCGGAAGA |
| 55959_1 | TATGAGTGTGGAGTCGTTGCCCCAACAAGGAGACATCTCTTGTNNNAGATCGGAAG |
| 55960_1 | TATGAGTGTGGAGTCGTTGCGCAAGCTTTTGATTTGTGTCTGCNNNAGATCGGAAG |
| 55964_1 | TATGAGTGTGGAGTCGTTGCCCAATCCAATTCTGCCCAGTTTTNNNAGATCGGAAGA |
| 55965_1 | TATGAGTGTGGAGTCGTTGCAAACATCACTGGTATGTGCACAANNNAGATCGGAAG |
| 55968_1 | TATGAGTGTGGAGTCGTTGCATGTGATTCAATTCTGGCCAACGNNNAGATCGGAAG |
| 55969_1 | TATGAGTGTGGAGTCGTTGCGGATCATGAGGTCAGGAGTTTGANNNAGATCGGAA |
| 55970_1 | TATGAGTGTGGAGTCGTTGCGGATCATGAGGTCAGGAGTTTGANNNAGATCGGAA |
| 55971_1 | TATGAGTGTGGAGTCGTTGCTGCTAAAGATGGAGGAAGAAACANNNAGATCGGAA |
| 55972_1 | TATGAGTGTGGAGTCGTTGCGATGTGGTGATGCATGCTTGTAGNNNAGATCGGAAG |
| 55975_1 | TATGAGTGTGGAGTCGTTGCCCTAACCTAATCACAGGCATCCTNNNAGATCGGAAGA |
| 55976_1 | TATGAGTGTGGAGTCGTTGCGCTGCTGCTGGGAGTATAAATTGNNNAGATCGGAAG |
| 55977_1 | TATGAGTGTGGAGTCGTTGCATCTCATTTCCTTCCCACGACAANNNAGATCGGAAGA |
| 55978_1 | TATGAGTGTGGAGTCGTTGCGAAATGAACATTCTTGGGCCACANNNAGATCGGAAG |
| 55981_1 | TATGAGTGTGGAGTCGTTGCAGGTGCATGCCTATAATCCTAGCNNNAGATCGGAAG |
| 55982_1 | TATGAGTGTGGAGTCGTTGCGGGGACAGAGTCAGAAACTAGTGNNNAGATCGGAA |
| 55983_1 | TATGAGTGTGGAGTCGTTGCAGTAAGCCAAGATCGCAACATTGNNNAGATCGGAAG |
| 55985_1 | TATGAGTGTGGAGTCGTTGCACTGAGGTAGGAGGATCACTTGANNNAGATCGGAA |
| 55986_1 | TATGAGTGTGGAGTCGTTGCAAAGGTGGAGACTGTAGCAAACANNNAGATCGGAA |

| | |
|---|---|
| 55987_1 | TATGAGTGTGGAGTCGTTGCAGTGACACCATTTTGATTCTGACANNNAGATCGGAA |
| 55988_1 | TATGAGTGTGGAGTCGTTGCGAACTAGGGAAGAGCTAGTGCAANNNAGATCGGAA |
| 55989_1 | TATGAGTGTGGAGTCGTTGCTTCAGGACCTTGCACTATGGTTTNNNAGATCGGAAG |
| 55990_1 | TATGAGTGTGGAGTCGTTGCAATACGGGTTCTTAGGCCTCATCNNNAGATCGGAAG |
| 55998_1 | TATGAGTGTGGAGTCGTTGCCCATCCTTCTACCCTCAGCATACNNNAGATCGGAAGA |
| 55999_1 | TATGAGTGTGGAGTCGTTGCAGCAGGAGAAAAAAAAGTGTCGGNNNAGATCGGA |
| 56000_1 | TATGAGTGTGGAGTCGTTGCTTCAAGCAATTCTCCTGCCTCAGNNNAGATCGGAAG |
| 56001_1 | TATGAGTGTGGAGTCGTTGCTTAGTAGAGGCAGGACTTCTCCANNNAGATCGGAA |
| 56002_1 | TATGAGTGTGGAGTCGTTGCTCAAGCTATTCTCCTGCCTCATCNNNAGATCGGAAGA |
| 56004_1 | TATGAGTGTGGAGTCGTTGCCCCGAGTACCTGGGATTATAGGTNNNAGATCGGAA |
| 56005_1 | TATGAGTGTGGAGTCGTTGCCTGAAATTACAGGTACACGCCACNNNAGATCGGAAG |
| 56007_1 | TATGAGTGTGGAGTCGTTGCTTCCTGGGTTCAAGCAATTTTCCNNNAGATCGGAAG |
| 56009_1 | TATGAGTGTGGAGTCGTTGCTTATTCTTCCTCAGCCTCCCAAGNNNAGATCGGAAGA |
| 56010_1 | TATGAGTGTGGAGTCGTTGCTTCAAGCTATTCTCCTGCCTCAGNNNAGATCGGAAG |
| 56011_1 | TATGAGTGTGGAGTCGTTGCCAGGATGGTCTTGAACTCTTGGANNNAGATCGGAA |
| 56014_1 | TATGAGTGTGGAGTCGTTGCTTTAGTAGAGACGGGGTTTCTGCNNNAGATCGGAA |
| 56018_1 | TATGAGTGTGGAGTCGTTGCTAAAGTGCTGGGATTACAGGCATNNNAGATCGGAA |
| 56019_1 | TATGAGTGTGGAGTCGTTGCCCCTCCCAAGTTCAAGCAATTTTNNNAGATCGGAAG |
| 56021_1 | TATGAGTGTGGAGTCGTTGCGCTCAAGCAATCCTCCTATCTCANNNAGATCGGAAG |
| 56022_1 | TATGAGTGTGGAGTCGTTGCTTCAAGTGATTCTCCTGCCTCAGNNNAGATCGGAAG |
| 56024_1 | TATGAGTGTGGAGTCGTTGCGGAGTATAGTGTCACGATCTCGGNNNAGATCGGAA |
| 56025_1 | TATGAGTGTGGAGTCGTTGCATCACACCGTTGAAGTAGCTCTTNNNAGATCGGAAG |
| 56026_1 | TATGAGTGTGGAGTCGTTGCTTCTGTGTGGTTCAGGAACTCAANNNAGATCGGAAG |
| 56027_1 | TATGAGTGTGGAGTCGTTGCAAAGAAGCAGAACAGTAAGGGCTNNNAGATCGGAA |
| 56028_1 | TATGAGTGTGGAGTCGTTGCCTGGAGTGAAGTGGCACAATCTTNNNAGATCGGAA |
| 56029_1 | TATGAGTGTGGAGTCGTTGCGATTCAAGCGATTCTTCTGCCTCNNNAGATCGGAAG |
| 56030_1 | TATGAGTGTGGAGTCGTTGCTCCAGTGAAATCAATGCTTTGGCNNNAGATCGGAAG |
| 56031_1 | TATGAGTGTGGAGTCGTTGCGCACCTGTTGTTTCCTGACTTTTNNNAGATCGGAAG |
| 56032_1 | TATGAGTGTGGAGTCGTTGCGTTCATGCCATTCTCCTACCTCANNNAGATCGGAAGA |
| 56033_1 | TATGAGTGTGGAGTCGTTGCCCTGGAACACCAATGAGACAGAANNNAGATCGGAA |
| 56034_1 | TATGAGTGTGGAGTCGTTGCTGCTCACTGTAACCTGTAACCTCNNNAGATCGGAAG |
| 56035_1 | TATGAGTGTGGAGTCGTTGCGCTTTAGCCTCTCAAAGTGTTGGNNNAGATCGGAAG |
| 56036_1 | TATGAGTGTGGAGTCGTTGCCACAATCACAGCTCACAATCACANNNAGATCGGAAG |
| 56037_1 | TATGAGTGTGGAGTCGTTGCAGATGGAGTCTTGTTCTGTCACCNNNAGATCGGAAG |
| 56038_1 | TATGAGTGTGGAGTCGTTGCTTCAAGTGATTCTCCTGCCTCAGNNNAGATCGGAAG |
| 56039_1 | TATGAGTGTGGAGTCGTTGCCTTGACTGCAACCTTCGATCTTGNNNAGATCGGAAG |
| 56041_1 | TATGAGTGTGGAGTCGTTGCGAGAGATCTCAGCTCACTGCAACNNNAGATCGGAAG |
| 56043_1 | TATGAGTGTGGAGTCGTTGCGGATGACAGATCCCTGGATACACNNNAGATCGGAA |
| 56044_1 | TATGAGTGTGGAGTCGTTGCGGGTTCAAGTGATTCTCCTGTCTNNNAGATCGGAAG |
| 56045_1 | TATGAGTGTGGAGTCGTTGCAGATAGGGTCTCACTCTGTCACCNNNAGATCGGAAG |
| 56046_1 | TATGAGTGTGGAGTCGTTGCATAAGGGCTCTCTTGGCAAGAAANNNAGATCGGAA |
| 56047_1 | TATGAGTGTGGAGTCGTTGCCAAGACCCTCACCCATCATCTAGNNNAGATCGGAAG |
| 56048_1 | TATGAGTGTGGAGTCGTTGCAGCAATCTTCCCATCTTAGCCTCNNNAGATCGGAAGA |

56049_1 TATGAGTGTGGAGTCGTTGCCAGGATAGATGAAATCCAGGCCANNNAGATCGGAA
56051_1 TATGAGTGTGGAGTCGTTGCGATGGAGTCTTACTCTGTCACCCNNNAGATCGGAAG
56052_1 TATGAGTGTGGAGTCGTTGCCCAACCAAGAATACTGTGTCCAGNNNAGATCGGAAG
56053_1 TATGAGTGTGGAGTCGTTGCGGACCAAAGGAGATAACCGATGANNNAGATCGGAA
56054_1 TATGAGTGTGGAGTCGTTGCTTCTGAGGATCAAGGCACCACTANNNAGATCGGAAG
56055_1 TATGAGTGTGGAGTCGTTGCGGGGGGATATCTTTACCTTCAGGNNNAGATCGGAA
56056_1 TATGAGTGTGGAGTCGTTGCGGGCAACATAGCAAGATCTTGTCNNNAGATCGGAA
56057_1 TATGAGTGTGGAGTCGTTGCGGTGAAACCCCGTCTCTACTAAANNNAGATCGGAAG
56058_1 TATGAGTGTGGAGTCGTTGCAAACAGTGTCTTGCTACATTGCCNNNAGATCGGAAG
56060_1 TATGAGTGTGGAGTCGTTGCGAGGTCTTCCAAGAGAGAACCTGNNNAGATCGGAA
56061_1 TATGAGTGTGGAGTCGTTGCGTAGTACCTCCCCCTTCTCTCTCNNNAGATCGGAAGA
56066_1 TATGAGTGTGGAGTCGTTGCATTACAGGGGAGTAATGAGCAGCNNNAGATCGGAA
56067_1 TATGAGTGTGGAGTCGTTGCGTGGTGTGCTTTTTCAGTAACGTNNNAGATCGGAAG
56070_1 TATGAGTGTGGAGTCGTTGCTAAGTGCTATGGGAAAACAGGGTNNNAGATCGGAA
56071_1 TATGAGTGTGGAGTCGTTGCTGGTGCTGAGACCTGTAAATACCNNNAGATCGGAA
56072_1 TATGAGTGTGGAGTCGTTGCAAAAAGGAAAGAGACCTTCCCCCNNNAGATCGGAA
56075_1 TATGAGTGTGGAGTCGTTGCAGGAAGGAAGGAAACCACCAAATNNNAGATCGGAA
56077_1 TATGAGTGTGGAGTCGTTGCAGGGATATGATCTTGCTGGCAATNNNAGATCGGAA
56078_1 TATGAGTGTGGAGTCGTTGCATTGGCAGTGGTATGAAATCCCTNNNAGATCGGAA
56080_1 TATGAGTGTGGAGTCGTTGCCAGAACCTGAATACTGGAAGCCTNNNAGATCGGAA
56081_1 TATGAGTGTGGAGTCGTTGCAGATAGTAAGGAAAATGGGGCCGNNNAGATCGGA
56082_1 TATGAGTGTGGAGTCGTTGCAAGAAATAACGAGTGGAAACCGCNNNAGATCGGAA
56084_1 TATGAGTGTGGAGTCGTTGCCCTGGTGAGTATCATTGGGGAAANNNAGATCGGAA
56086_1 TATGAGTGTGGAGTCGTTGCTCAAAGGTGAGTAGGATATGGGCNNNAGATCGGAA
56087_1 TATGAGTGTGGAGTCGTTGCACGGTGAAACCCTGTCTCTACTANNNAGATCGGAAG
56088_1 TATGAGTGTGGAGTCGTTGCGCTGCAAGGGAATAACTTGAATGTNNNAGATCGGA
56089_1 TATGAGTGTGGAGTCGTTGCCCAAAACCCACCAAAACCAAGATNNNAGATCGGAAG
56090_1 TATGAGTGTGGAGTCGTTGCGAAGGAGAGAAGGAGGGATCTCTNNNAGATCGGA
56091_1 TATGAGTGTGGAGTCGTTGCGAGGAAGGACAGAACCACATGATNNNAGATCGGAA
56092_1 TATGAGTGTGGAGTCGTTGCCACCTGTAGGCCTAGCTATTCAGNNNAGATCGGAAG
56093_1 TATGAGTGTGGAGTCGTTGCGGCCAGAGAATCACTTGAACTTGNNNAGATCGGAA
56094_1 TATGAGTGTGGAGTCGTTGCAAAAGAGAAAAGGAAAAGCCGCCNNNAGATCGGAA
56095_1 TATGAGTGTGGAGTCGTTGCTTTGGTCACGTTCTCACTTAGGTNNNAGATCGGAAG
56097_1 TATGAGTGTGGAGTCGTTGCCATCCTGGCTAACATGGTGAAACNNNAGATCGGAAG
56099_1 TATGAGTGTGGAGTCGTTGCACTAAGGCATGAGAATCGCTTGANNNAGATCGGAA
56101_1 TATGAGTGTGGAGTCGTTGCACTCCAGGTGTTTTCTCATCAGANNNAGATCGGAAG
56102_1 TATGAGTGTGGAGTCGTTGCTCCTAGCTACTCAGGAGGCTTAGNNNAGATCGGAAG
56104_1 TATGAGTGTGGAGTCGTTGCCACACACACACACACACACATATNNNAGATCGGAAGA
56105_1 TATGAGTGTGGAGTCGTTGCCAGGAGGCTGAAGAAGGAGTATCNNNAGATCGGA
56107_1 TATGAGTGTGGAGTCGTTGCGTGAATTGTTGTTTGCAGGGGAANNNAGATCGGAA
56108_1 TATGAGTGTGGAGTCGTTGCAAGGGTCATACTATGTTGCCCAGNNNAGATCGGAA
56109_1 TATGAGTGTGGAGTCGTTGCTGTCACATGACCCAGTAGTTACANNNAGATCGGAAG
56113_1 TATGAGTGTGGAGTCGTTGCTGGTGCTCTTTTCTTGTCTACCCNNNAGATCGGAAGA

| | |
|---|---|
| 56114_1 | TATGAGTGTGGAGTCGTTGCTTATCCAGGAGTGGTAGTGCATGNNNAGATCGGAA |
| 56115_1 | TATGAGTGTGGAGTCGTTGCCCATTGCAGGTCACATGAAAACTNNNAGATCGGAAG |
| 56116_1 | TATGAGTGTGGAGTCGTTGCCAGAGGTGGGGAAAGACATAATNNNAGATCGGA |
| 56117_1 | TATGAGTGTGGAGTCGTTGCCTATTTGCTCAGGCTGGTCTTGNNNAGATCGGAAG |
| 56121_1 | TATGAGTGTGGAGTCGTTGCAACTCTTGACCTCAGGTAATCCGNNNAGATCGGAAG |
| 56122_1 | TATGAGTGTGGAGTCGTTGCCAGCCTCTCAAGTATCTGGGATCNNNAGATCGGAAG |
| 56125_1 | TATGAGTGTGGAGTCGTTGCTTCACACAATTCTCCTGCCTCAGNNNAGATCGGAAGA |
| 56126_1 | TATGAGTGTGGAGTCGTTGCACCTCCTGAGTTGAAGCAATTCTNNNAGATCGGAAG |
| 56127_1 | TATGAGTGTGGAGTCGTTGCGTTCATTCTCCTGCCTCATCCTCNNNAGATCGGAAGA |
| 56129_1 | TATGAGTGTGGAGTCGTTGCCCCAGGTTCAAGCAATTCTCCTANNNAGATCGGAAG |
| 56130_1 | TATGAGTGTGGAGTCGTTGCTTCAAACCATTCTCCTGCCTCAGNNNAGATCGGAAGA |
| 56131_1 | TATGAGTGTGGAGTCGTTGCCCAAAATGCTGGGATTACAGGTGNNNAGATCGGAA |
| 56132_1 | TATGAGTGTGGAGTCGTTGCCCGAGTAACTGAGATTACAGGCANNNAGATCGGAA |
| 56134_1 | TATGAGTGTGGAGTCGTTGCAGATGGAGTCTCACTATGTTGCCNNNAGATCGGAAG |
| 56135_1 | TATGAGTGTGGAGTCGTTGCCAAATCTCAGCTCACTGCAACTTNNNAGATCGGAAG |
| 56137_1 | TATGAGTGTGGAGTCGTTGCCTGAGTAGCTGGGATTACAGGTGNNNAGATCGGAA |
| 56138_1 | TATGAGTGTGGAGTCGTTGCAAATAGCTGGGATTACAGGCACANNNAGATCGGAA |
| 56139_1 | TATGAGTGTGGAGTCGTTGCCATGGGTTCAAGCGATTCTCTTGNNNAGATCGGAAG |
| 56142_1 | TATGAGTGTGGAGTCGTTGCAGTCTTGCTCTCACCCAGACTATNNNAGATCGGAAG |
| 56144_1 | TATGAGTGTGGAGTCGTTGCTTCAAGCAATTCTCCTGCCTCAGNNNAGATCGGAAG |
| 56146_1 | TATGAGTGTGGAGTCGTTGCTTCATGTCATTCTCCTGCCTCAGNNNAGATCGGAAGA |
| 56148_1 | TATGAGTGTGGAGTCGTTGCCCTTGGTTCAAGCAATTCTCCTGNNNAGATCGGAAG |
| 56149_1 | TATGAGTGTGGAGTCGTTGCGACAGAGTCTCGTTCTTTCACCCNNNAGATCGGAAG |
| 56150_1 | TATGAGTGTGGAGTCGTTGCAAACGGAGTCTCACTCTATTGCCNNNAGATCGGAAG |
| 56151_1 | TATGAGTGTGGAGTCGTTGCCTGAGTAGCTGGGATTATAGGCGNNNAGATCGGAA |
| 56152_1 | TATGAGTGTGGAGTCGTTGCTTCGTGCCATTCTCTTACCTCAGNNNAGATCGGAAGA |
| 56153_1 | TATGAGTGTGGAGTCGTTGCAGATGGAGTCTCATTCTGTTGCCNNNAGATCGGAAG |
| 56154_1 | TATGAGTGTGGAGTCGTTGCGCTAAAGTGTAGTGGTGCTCTCANNNAGATCGGAA |
| 56155_1 | TATGAGTGTGGAGTCGTTGCTGGAGTCTCTCTCTCTCTGTCTCNNNAGATCGGAAGA |
| 56158_1 | TATGAGTGTGGAGTCGTTGCGAGTAAAATGGCTCGATCTTGGCNNNAGATCGGAA |
| 56159_1 | TATGAGTGTGGAGTCGTTGCCCTCCTGGAGTCAAGTGATTCTCNNNAGATCGGAAG |
| 56160_1 | TATGAGTGTGGAGTCGTTGCACATCTCTTGTGCTGCATCAGTANNNAGATCGGAAG |
| 56161_1 | TATGAGTGTGGAGTCGTTGCACTCCTGACCTCAAATGATCCACNNNAGATCGGAAG |
| 56163_1 | TATGAGTGTGGAGTCGTTGCCAATGGCTCAATCTTTGCTCACTNNNAGATCGGAAG |
| 56164_1 | TATGAGTGTGGAGTCGTTGCGAGTGCAATGATGTGATCTCAGCNNNAGATCGGAA |
| 56166_1 | TATGAGTGTGGAGTCGTTGCATGAACAAATGACAACCTCTGCCNNNAGATCGGAAG |
| 56167_1 | TATGAGTGTGGAGTCGTTGCGCTGGGATTACAGACATGATCCTNNNAGATCGGAA |
| 56168_1 | TATGAGTGTGGAGTCGTTGCCCGTGGAAGGTTTGTTTCACTTTNNNAGATCGGAAG |
| 56169_1 | TATGAGTGTGGAGTCGTTGCTTCAAGTGATTCTCCTGCCTCAANNNAGATCGGAAG |
| 56170_1 | TATGAGTGTGGAGTCGTTGCTTTCGTAGAGACGGTGTTTCACCNNNAGATCGGAAG |
| 56171_1 | TATGAGTGTGGAGTCGTTGCATGATCTTGACTCACTGCAACCTNNNAGATCGGAAG |
| 56173_1 | TATGAGTGTGGAGTCGTTGCCCCAGGTTCAAGCAATTCTCCTANNNAGATCGGAAG |
| 56174_1 | TATGAGTGTGGAGTCGTTGCGCAATTCTCCTGCATCAATCTCCNNNAGATCGGAAGA |

| | |
|---|---|
| 56175_1 | TATGAGTGTGGAGTCGTTGCCGAGGAGCTGGGAGTGAATTTTTNNNAGATCGGAA |
| 56177_1 | TATGAGTGTGGAGTCGTTGCGGAATGGGGAGACTTGAAGGAAANNNAGATCGGA |
| 56178_1 | TATGAGTGTGGAGTCGTTGCAGTGCTGAGATTACAGGTGTGAGNNNAGATCGGAA |
| 56179_1 | TATGAGTGTGGAGTCGTTGCGACGAAGTCTTGCTTTGTCTCAGNNNAGATCGGAAG |
| 56180_1 | TATGAGTGTGGAGTCGTTGCGCTGGAGTACAAATGGCATGATCNNNAGATCGGAA |
| 56181_1 | TATGAGTGTGGAGTCGTTGCGAGCTGGGATTTTGCTATGTTGGNNNAGATCGGAA |
| 56182_1 | TATGAGTGTGGAGTCGTTGCCTGGGTTCAGTGTTCTGTTGTTGNNNAGATCGGAAG |
| 56183_1 | TATGAGTGTGGAGTCGTTGCCCTAAGGAAACAGGCCTAATGGTNNNAGATCGGAA |
| 56185_1 | TATGAGTGTGGAGTCGTTGCCTAGCAGAACAAGGCTGAGAAGANNNAGATCGGAA |
| 56186_1 | TATGAGTGTGGAGTCGTTGCGGTGACAAGAGCAAAGTTCCATCNNNAGATCGGAA |
| 56187_1 | TATGAGTGTGGAGTCGTTGCGATTGGAAGAGTCAGGGGAGTTTNNNAGATCGGA |
| 56189_1 | TATGAGTGTGGAGTCGTTGCAGTCTTGCTCTATCTTCAGGCTGNNNAGATCGGAAG |
| 56190_1 | TATGAGTGTGGAGTCGTTGCGGTGGATAACATGAGTCCAGGAGNNNAGATCGGA |
| 56192_1 | TATGAGTGTGGAGTCGTTGCCCGAAGTGAGCTACCAAAAAAGANNNAGATCGGAA |
| 56193_1 | TATGAGTGTGGAGTCGTTGCTGTCTGGAATTGGTTTGGACTGANNNAGATCGGAA |
| 56194_1 | TATGAGTGTGGAGTCGTTGCGTCAAACAGATTGTGAGGGAACGNNNAGATCGGAA |
| 56197_1 | TATGAGTGTGGAGTCGTTGCTCTTATGGCACATTTCTGTTCCANNNAGATCGGAAGA |
| 56200_1 | TATGAGTGTGGAGTCGTTGCGAAGAAAAGACAAACTCCCAGGCNNNAGATCGGAA |
| 56201_1 | TATGAGTGTGGAGTCGTTGCCTCATGCCTGTAATCCCAGAACTNNNAGATCGGAAG |
| 56204_1 | TATGAGTGTGGAGTCGTTGCAGGAAATTCTCTTGATGCAGGGANNNAGATCGGAA |
| 56206_1 | TATGAGTGTGGAGTCGTTGCCTGAGCTCAAACTATCTTGCTGCNNNAGATCGGAAG |
| 56208_1 | TATGAGTGTGGAGTCGTTGCATGTGCTAGGAGGGAAGAGTTTGNNNAGATCGGA |
| 56209_1 | TATGAGTGTGGAGTCGTTGCATTGAAGATGAAGGACTGGGGACNNNAGATCGGA |
| 56210_1 | TATGAGTGTGGAGTCGTTGCTACAAAATTAGCTGGGTGTGGTGNNNAGATCGGAA |
| 56211_1 | TATGAGTGTGGAGTCGTTGCCCTAGATAATTGGGAGGCTGAGGNNNAGATCGGAA |
| 56212_1 | TATGAGTGTGGAGTCGTTGCAACTGTGACTCTCATACAACGCTNNNAGATCGGAAG |
| 56213_1 | TATGAGTGTGGAGTCGTTGCTAGTACCAACTACTCAGGAGGCTNNNAGATCGGAAG |
| 56214_1 | TATGAGTGTGGAGTCGTTGCATTCAGGCTCTGATCTCTACCCTNNNAGATCGGAAGA |
| 56215_1 | TATGAGTGTGGAGTCGTTGCTCATGCCTGTAATCTCTGCACTTNNNAGATCGGAAGA |
| 56216_1 | TATGAGTGTGGAGTCGTTGCATTCAGGCCTAAGGAACACCAATNNNAGATCGGAAG |
| 56219_1 | TATGAGTGTGGAGTCGTTGCGAACCAGATCGCTAAACTTCCAGNNNAGATCGGAAG |
| 56220_1 | TATGAGTGTGGAGTCGTTGCTTGAACTGCATTCCTCCCAAATGNNNAGATCGGAAG |
| 56221_1 | TATGAGTGTGGAGTCGTTGCATATAGTGGGTGTTTGGAGGGGTNNNAGATCGGA |
| 56222_1 | TATGAGTGTGGAGTCGTTGCGGGGAGACATAGAGGAAGGTAGTNNNAGATCGGA |
| 56225_1 | TATGAGTGTGGAGTCGTTGCCTGTAGTTCCAGCTACTCAGGAGNNNAGATCGGAAG |
| 56228_1 | TATGAGTGTGGAGTCGTTGCAATCTCAGTACTTTGGGAGGCTGNNNAGATCGGAA |
| 56229_1 | TATGAGTGTGGAGTCGTTGCTAACCTCACATCTGCCTGATTCCNNNAGATCGGAAGA |
| 56230_1 | TATGAGTGTGGAGTCGTTGCTGTCCAAGGTCACATAGCTAGAANNNAGATCGGAA |
| 56231_1 | TATGAGTGTGGAGTCGTTGCAGGAAACAGCATTCCCCAAACTANNNAGATCGGAAG |
| 56232_1 | TATGAGTGTGGAGTCGTTGCCATTTTCAGCGGGAATACTGCATNNNAGATCGGAAG |
| 56233_1 | TATGAGTGTGGAGTCGTTGCTAGTCCCAGCATTCAGCACTTTANNNAGATCGGAAG |
| 56234_1 | TATGAGTGTGGAGTCGTTGCCCCTGACCCAGATGATGTCTAAGNNNAGATCGGAAG |
| 56235_1 | TATGAGTGTGGAGTCGTTGCGGGGGAGGGAAAAAAGGAGTAATNNNAGATCGGA |

56236_1 TATGAGTGTGGAGTCGTTGCCTGGGGAATTTGGGTAGACAAGANNNAGATCGGAA
56237_1 TATGAGTGTGGAGTCGTTGCAGTGTGTATGTGTGTGTGTGTGTNNNAGATCGGAA
56238_1 TATGAGTGTGGAGTCGTTGCGAGGTAGGAGAACCACTTGAACCNNNAGATCGGAA
56240_1 TATGAGTGTGGAGTCGTTGCAATTAGCCGTGTGTAGTCGTACANNNAGATCGGAAG
56241_1 TATGAGTGTGGAGTCGTTGCACTAAGCTACTAGGGAGGCTGAANNNAGATCGGAA
56242_1 TATGAGTGTGGAGTCGTTGCTGGTGACCATGAACACTACTCACNNNAGATCGGAAG
56243_1 TATGAGTGTGGAGTCGTTGCTGAGCCTGAGTGAGTATATGGGANNNAGATCGGAA
56245_1 TATGAGTGTGGAGTCGTTGCTCTTATCTCCACACTCCATCCTGNNNAGATCGGAAGA
56246_1 TATGAGTGTGGAGTCGTTGCATGTTGAGCAACTGGAACTCTCTNNNAGATCGGAAG
56247_1 TATGAGTGTGGAGTCGTTGCTCTCTAGCCCAAACTTCCTTGTCNNNAGATCGGAAGA
56248_1 TATGAGTGTGGAGTCGTTGCGTACACACCTGTAGCTGCTATCANNNAGATCGGAAGA
56249_1 TATGAGTGTGGAGTCGTTGCGGATAGGTCAACATAACTGCGTGNNNAGATCGGAA
56250_1 TATGAGTGTGGAGTCGTTGCGACTGAAGCAGAAGAGGACTTGANNNAGATCGGAA
56251_1 TATGAGTGTGGAGTCGTTGCATGCGCAGTTTACAATACCGTTCNNNAGATCGGAAGA
56252_1 TATGAGTGTGGAGTCGTTGCAGATGTTTGCTGAGAGGAGGAAGNNNAGATCGGAA
56255_1 TATGAGTGTGGAGTCGTTGCTTTCCGGGTTCAGGTGATTTCTCNNNAGATCGGAAGA
56257_1 TATGAGTGTGGAGTCGTTGCAGAAATACCATTTGACCCAGCCANNNAGATCGGAAG
56258_1 TATGAGTGTGGAGTCGTTGCCTCAAGCGATCCTTCCACCTTAGNNNAGATCGGAAGA
56259_1 TATGAGTGTGGAGTCGTTGCACATGAAATCTCACTCTTGCCCANNNAGATCGGAAGA
56261_1 TATGAGTGTGGAGTCGTTGCGAAGAGACCACCAAACAGGCTTTNNNAGATCGGAAG
56262_1 TATGAGTGTGGAGTCGTTGCCAGCAGTAAATAGCAGACACAGCNNNAGATCGGAA
56263_1 TATGAGTGTGGAGTCGTTGCATTCAAGGGATTTTCCTGCCTCANNNAGATCGGAAGA
56264_1 TATGAGTGTGGAGTCGTTGCAGATCTCAGCTCACTGCAATCTCNNNAGATCGGAAGA
56265_1 TATGAGTGTGGAGTCGTTGCACGCCCGGCTAATTTTTTGTATTNNNAGATCGGAAGA
56266_1 TATGAGTGTGGAGTCGTTGCGGTTCAAGTGATTCTCCTGCTTCNNNAGATCGGAAGA
56268_1 TATGAGTGTGGAGTCGTTGCAGGAAACCCCATAGTCTTGTCACNNNAGATCGGAAG
56271_1 TATGAGTGTGGAGTCGTTGCGTGCCCTATCTCAAGACTGTGTTNNNAGATCGGAAGA
56272_1 TATGAGTGTGGAGTCGTTGCCTAGGTGTGCAGACTGACTTTCTNNNAGATCGGAAG
56273_1 TATGAGTGTGGAGTCGTTGCGGGTAGTGTGTCTGGGTAGTTTTNNNAGATCGGAA
56274_1 TATGAGTGTGGAGTCGTTGCGGTTCAATGCTGAGGGTTTGAAANNNAGATCGGAA
56277_1 TATGAGTGTGGAGTCGTTGCAATCAGGGTCTCACTCTGTTCACNNNAGATCGGAAGA
56278_1 TATGAGTGTGGAGTCGTTGCAGATGGACTCTCACTCTGTCACCNNNAGATCGGAAGA
56279_1 TATGAGTGTGGAGTCGTTGCTTCAAGTGATTCTCCTGCCTCTGNNNAGATCGGAAGA
56280_1 TATGAGTGTGGAGTCGTTGCCTCTGGAGTAGCTGGGATTACAGNNNAGATCGGAA
56283_1 TATGAGTGTGGAGTCGTTGCAATACAGCACACTGATGGGTCTTNNNAGATCGGAAG
56285_1 TATGAGTGTGGAGTCGTTGCTTCTGCTTTAGCCTCCCAAGTAGNNNAGATCGGAAGA
56286_1 TATGAGTGTGGAGTCGTTGCGAAATGGAGTCTTTTTGCTGGCTNNNAGATCGGAAG
56287_1 TATGAGTGTGGAGTCGTTGCTTCAAGCGATTGTCTGAGTAGCTNNNAGATCGGAAG
56288_1 TATGAGTGTGGAGTCGTTGCTGCAGTACAATGGTGTGATCTCANNNAGATCGGAAG
56289_1 TATGAGTGTGGAGTCGTTGCGAGTAGCTGGGATTACAGGTGAGNNNAGATCGGAA
56290_1 TATGAGTGTGGAGTCGTTGCCTCCTGAGTAGCTGGGACTACAGNNNAGATCGGAAG
56294_1 TATGAGTGTGGAGTCGTTGCTTCTCCTACCTAAGCCTCCTGAANNNAGATCGGAAGA
56295_1 TATGAGTGTGGAGTCGTTGCTCTTGGGTTCAGGTGATTCTCTGNNNAGATCGGAAG

| | |
|---|---|
| 56296_1 | TATGAGTGTGGAGTCGTTGCATGGAGTTTCGTTCTATTGCCCANNNAGATCGGAAGA |
| 56298_1 | TATGAGTGTGGAGTCGTTGCCTAGGTTCAAGCGATTCTCTTGCNNNAGATCGGAAGA |
| 56300_1 | TATGAGTGTGGAGTCGTTGCGCTGGTCTCTATCTCCTGACTTCNNNAGATCGGAAGA |
| 56302_1 | TATGAGTGTGGAGTCGTTGCCTGGGTTCAAGCAATTCTACTGCNNNAGATCGGAAG |
| 56303_1 | TATGAGTGTGGAGTCGTTGCTTCAAGCAATTCTCCTGCCTCAGNNNAGATCGGAAGA |
| 56305_1 | TATGAGTGTGGAGTCGTTGCGAGCTTCTCACTCTGTCACCTAGNNNAGATCGGAAGA |
| 56306_1 | TATGAGTGTGGAGTCGTTGCAGTTTTGTTCTTGTTTCCCAGGCNNNAGATCGGAAGA |
| 56307_1 | TATGAGTGTGGAGTCGTTGCATCTCCCAGGCTCAAACAATTCTNNNAGATCGGAAGA |
| 56308_1 | TATGAGTGTGGAGTCGTTGCTAAGCATTCTGATCCCTTGGTGCNNNAGATCGGAAGA |
| 56309_1 | TATGAGTGTGGAGTCGTTGCTACAGTGGTGTAATCATGGCTCANNNAGATCGGAA |
| 56310_1 | TATGAGTGTGGAGTCGTTGCGTGCTGGGATTACAGGGATAAGTNNNAGATCGGAA |
| 56312_1 | TATGAGTGTGGAGTCGTTGCATTCAAGTGATTCTCCTGCCTCANNNAGATCGGAAGA |
| 56313_1 | TATGAGTGTGGAGTCGTTGCCAGGCTGGTCTTGAACTTTTGACNNNAGATCGGAAG |
| 56314_1 | TATGAGTGTGGAGTCGTTGCTGGAGTCTCACTCTGTCATTCAGNNNAGATCGGAAGA |
| 56317_1 | TATGAGTGTGGAGTCGTTGCAGAGGCTGGGATCATAGGTTAGANNNAGATCGGAA |
| 56318_1 | TATGAGTGTGGAGTCGTTGCCTCAAGCAATCCTCCCATCTCAANNNAGATCGGAAGA |
| 56319_1 | TATGAGTGTGGAGTCGTTGCTAGAGTTGGGATTTTGCCATGTGNNNAGATCGGAA |
| 56320_1 | TATGAGTGTGGAGTCGTTGCTTCAAGTGATTCTCCTGCCTCAANNNAGATCGGAAGA |
| 56324_1 | TATGAGTGTGGAGTCGTTGCCAGAATAGCTGGTACTACAGGGGNNNAGATCGGAA |
| 56325_1 | TATGAGTGTGGAGTCGTTGCGAGTAGCTGAGACTACAAGTGCCNNNAGATCGGAA |
| 56326_1 | TATGAGTGTGGAGTCGTTGCCAATTATGGCTCACTGCAACCTCNNNAGATCGGAAGA |
| 56327_1 | TATGAGTGTGGAGTCGTTGCCCTCCCTGATTCATGCCATTCTCNNNAGATCGGAAGA |
| 56328_1 | TATGAGTGTGGAGTCGTTGCCACCACACCTGGCTAATTTTTGTNNNAGATCGGAAGA |
| 56330_1 | TATGAGTGTGGAGTCGTTGCCAGTCTTGGCTCACTGGAATCTCNNNAGATCGGAAGA |
| 56332_1 | TATGAGTGTGGAGTCGTTGCCACCTCAGCCTCCCTAGTAACTANNNAGATCGGAAGA |
| 56333_1 | TATGAGTGTGGAGTCGTTGCTCTCCTGAGTAGCTGGGATTACANNNAGATCGGAAG |
| 56334_1 | TATGAGTGTGGAGTCGTTGCCTAGGTTCAAGCAATTCTCCTGCNNNAGATCGGAAGA |
| 56335_1 | TATGAGTGTGGAGTCGTTGCCCTCCTAGTAGCTGCGATTACAGNNNAGATCGGAAG |
| 56336_1 | TATGAGTGTGGAGTCGTTGCGTGTAATCTCGGCTCATTGCAACNNNAGATCGGAAG |
| 56341_1 | TATGAGTGTGGAGTCGTTGCCTCAAGTGATTCTCCTGCCTCAGNNNAGATCGGAAGA |
| 56343_1 | TATGAGTGTGGAGTCGTTGCCTGGGTTCAAGCTATTCTCCTGTNNNAGATCGGAAGA |
| 56344_1 | TATGAGTGTGGAGTCGTTGCAGGCTGGATTCAAGCTATTCTCCNNNAGATCGGAAG |
| 56345_1 | TATGAGTGTGGAGTCGTTGCCTCCCAGGTTCAGGCCATTATCNNNAGATCGGAAGA |
| 56346_1 | TATGAGTGTGGAGTCGTTGCATGATCTCAACTCACTGCAACCTNNNAGATCGGAAGA |
| 56347_1 | TATGAGTGTGGAGTCGTTGCCCTACCAGGTTCAAGCGATTCTCNNNAGATCGGAAGA |
| 56350_1 | TATGAGTGTGGAGTCGTTGCAGAGATGGGGTCTTGCTATGTTGNNNAGATCGGAA |
| 56351_1 | TATGAGTGTGGAGTCGTTGCGAGACGAAGTTTCACCATGTTGGNNNAGATCGGAA |
| 56352_1 | TATGAGTGTGGAGTCGTTGCATCTCAGTTCCTCACAACCTCTGNNNAGATCGGAAGA |
| 56353_1 | TATGAGTGTGGAGTCGTTGCTTCAAGCAATTTTCCTGCCTCAGNNNAGATCGGAAGA |
| 56354_1 | TATGAGTGTGGAGTCGTTGCGGTTCCAGTGATTCTCCTACCTCNNNAGATCGGAAGA |
| 56356_1 | TATGAGTGTGGAGTCGTTGCAGAGAAGATATAGGGAGAGGGGCNNNAGATCGGA |
| 56357_1 | TATGAGTGTGGAGTCGTTGCCAGGCCAATGGAATGAATGCTAANNNAGATCGGAA |
| 56358_1 | TATGAGTGTGGAGTCGTTGCTGACCATGATAATCACCTCTCCTNNNAGATCGGAAGA |

| | |
|---|---|
| 56361_1 | TATGAGTGTGGAGTCGTTGCTTTTCCTTAGCTCCAGCATTTGCNNNAGATCGGAAGA |
| 56363_1 | TATGAGTGTGGAGTCGTTGCCTTGGAGGCAGAATTGTTGTTCCNNNAGATCGGAAG |
| 56364_1 | TATGAGTGTGGAGTCGTTGCAGACAAGAAACAGCCCTGAGTAGNNNAGATCGGAA |
| 56369_1 | TATGAGTGTGGAGTCGTTGCTAAAACCGTATTTGGCAGCTTGGNNNAGATCGGAAG |
| 56370_1 | TATGAGTGTGGAGTCGTTGCGAAAAGTGGGGGACAATTGTCAGNNNAGATCGGA |
| 56371_1 | TATGAGTGTGGAGTCGTTGCGGCCCAGCATTTGTGAAGTATTTNNNAGATCGGAAG |
| 56373_1 | TATGAGTGTGGAGTCGTTGCCTGAGGTGGGAGGATTACCTGAGNNNAGATCGGA |
| 56374_1 | TATGAGTGTGGAGTCGTTGCCTCCTGTGTTTTTAGCTGGCAAANNNAGATCGGAAG |
| 56375_1 | TATGAGTGTGGAGTCGTTGCTCCATTAGAAAGTTGCTCTGGGCNNNAGATCGGAAG |
| 56376_1 | TATGAGTGTGGAGTCGTTGCTTCCATGTGTCATTCATGTCCCTNNNAGATCGGAAGA |
| 56378_1 | TATGAGTGTGGAGTCGTTGCTGGTGATCTTGAGATAAAGGGGANNNAGATCGGA |
| 56379_1 | TATGAGTGTGGAGTCGTTGCAACACCTTCCATGAAATTGCCAGNNNAGATCGGAAG |
| 56381_1 | TATGAGTGTGGAGTCGTTGCACATCCCTAAAATAACCCCTGCANNNAGATCGGAAG |
| 56383_1 | TATGAGTGTGGAGTCGTTGCGTGAGTAAAACTGGCAAAGACCCNNNAGATCGGAA |
| 56384_1 | TATGAGTGTGGAGTCGTTGCTTCTACCCAAATCTCTTCACCAANNNAGATCGGAAGA |
| 56385_1 | TATGAGTGTGGAGTCGTTGCGTCATGGTGGCACAAACTTGTAGNNNAGATCGGAA |
| 56386_1 | TATGAGTGTGGAGTCGTTGCAACGGATGCTGAAGAGGATGTAGNNNAGATCGGA |
| 56388_1 | TATGAGTGTGGAGTCGTTGCACTCAAAGCTCATGACAGTCCAANNNAGATCGGAAG |
| 56389_1 | TATGAGTGTGGAGTCGTTGCAAGTGTCCAAATACTTCAGCTGTNNNAGATCGGAAG |
| 56390_1 | TATGAGTGTGGAGTCGTTGCCTTAGAGCATCAGCAGATGGAGANNNAGATCGGAA |
| 56391_1 | TATGAGTGTGGAGTCGTTGCGAGGCAGGAGAATTGCTTGAATCNNNAGATCGGAA |
| 56394_1 | TATGAGTGTGGAGTCGTTGCGCATGCCTATAATCCCAGCTACTNNNAGATCGGAAG |
| 56395_1 | TATGAGTGTGGAGTCGTTGCCTTTCCTCTCACAACGCCTAGAGNNNAGATCGGAAG |
| 56397_1 | TATGAGTGTGGAGTCGTTGCAGAGGCTGCAGTAAGCTATGATCNNNAGATCGGAA |
| 56399_1 | TATGAGTGTGGAGTCGTTGCCACACCTGTAATCTCAGCACCTTNNNAGATCGGAAG |
| 56400_1 | TATGAGTGTGGAGTCGTTGCCTGCACACCTATTAGGATGGCTANNNAGATCGGAAG |
| 56401_1 | TATGAGTGTGGAGTCGTTGCCGAATAGGAACAGCTCCAGTCTGNNNAGATCGGAA |
| 56402_1 | TATGAGTGTGGAGTCGTTGCCAAAGGTCACCAACATCGAAGACNNNAGATCGGAAG |
| 56403_1 | TATGAGTGTGGAGTCGTTGCTGAGAAGTTAGCTACCTTGTTGCTNNNAGATCGGAA |
| 56406_1 | TATGAGTGTGGAGTCGTTGCCCTACTGCTTTACACCAAACTGCNNNAGATCGGAAG |
| 56407_1 | TATGAGTGTGGAGTCGTTGCGAGGTGAGAGGATCATTCAGTCCNNNAGATCGGAA |
| 56408_1 | TATGAGTGTGGAGTCGTTGCCATGAGGGATAAGGGAGCATCTCNNNAGATCGGAA |
| 56410_1 | TATGAGTGTGGAGTCGTTGCGGAAGAAGGGGTACAGAGCATAGNNNAGATCGGA |
| 56411_1 | TATGAGTGTGGAGTCGTTGCGGGGAATGAATGGGGAAACATTTNNNAGATCGGA |
| 56413_1 | TATGAGTGTGGAGTCGTTGCTATGAAATCTCTCCACAGACCCCNNNAGATCGGAAG |
| 56414_1 | TATGAGTGTGGAGTCGTTGCGAGTCATCATGTCTTGGAAGCCTNNNAGATCGGAAG |
| 56416_1 | TATGAGTGTGGAGTCGTTGCTTCCGTTATTCCGACTCTCACTGNNNAGATCGGAAGA |
| 56417_1 | TATGAGTGTGGAGTCGTTGCGACTTGAACAACACAAGAAGCCANNNAGATCGGAA |
| 56418_1 | TATGAGTGTGGAGTCGTTGCGGCAACCTGTATGCAATGACTAGNNNAGATCGGAA |
| 56419_1 | TATGAGTGTGGAGTCGTTGCAGGAGAGAGAAAGAGACCTTGGTNNNAGATCGGA |
| 56420_1 | TATGAGTGTGGAGTCGTTGCTGATTCCCCAGAGCAAGCTTATTNNNAGATCGGAAG |
| 56421_1 | TATGAGTGTGGAGTCGTTGCGAGGCTGAGCATCTTTTTTCTCCNNNAGATCGGAAG |
| 56424_1 | TATGAGTGTGGAGTCGTTGCTGAGCAGAATGTGAAGCAAACTGNNNAGATCGGAA |

56425_1 TATGAGTGTGGAGTCGTTGCACTGTCTACTAGCTGAGTGACCTNNNAGATCGGAAG
56426_1 TATGAGTGTGGAGTCGTTGCTGTCTTTTAGGGCCTTGATCCAANNNAGATCGGAAG
56427_1 TATGAGTGTGGAGTCGTTGCGCATCCCTGACCTAGAGAATCAGNNNAGATCGGAAG
56428_1 TATGAGTGTGGAGTCGTTGCGCATTGTCTTCTAAACTGGGCTCNNNAGATCGGAAG
56431_1 TATGAGTGTGGAGTCGTTGCATATGTGGGGAACCAGTGAAGACNNNAGATCGGAA
56432_1 TATGAGTGTGGAGTCGTTGCCGCAGGCATATCTTCGCTTTTATNNNAGATCGGAAG
56433_1 TATGAGTGTGGAGTCGTTGCACTTTTCCAAAACACAGACGCAANNNAGATCGGAAG
56434_1 TATGAGTGTGGAGTCGTTGCTAGTTCAATGTCCCTCTTGCACANNNAGATCGGAAG
56435_1 TATGAGTGTGGAGTCGTTGCCTGAAGCAGGAGAACGGTATGAANNNAGATCGGAA
56436_1 TATGAGTGTGGAGTCGTTGCCTGAGGCAGGAGAATCACTTGAANNNAGATCGGAA
56439_1 TATGAGTGTGGAGTCGTTGCCTTTCTTCACTCGTTCACGGAACNNNAGATCGGAAGA
56440_1 TATGAGTGTGGAGTCGTTGCTGGCTCATACCTGTAATCTCAGCNNNAGATCGGAAG
56441_1 TATGAGTGTGGAGTCGTTGCAAAGGGGAATAACACGTCCAAGGNNNAGATCGGAA
56442_1 TATGAGTGTGGAGTCGTTGCGGCACCTGTAATCCCAACTACTTNNNAGATCGGAAG
56444_1 TATGAGTGTGGAGTCGTTGCCTCAGGAATTCTGGTAGAGGGTGNNNAGATCGGAA
56445_1 TATGAGTGTGGAGTCGTTGCTTGAATTCTCCTTAGCTCTGCCCNNNAGATCGGAAGA
56446_1 TATGAGTGTGGAGTCGTTGCCTGTAATCCCAGCTACTCCACAGNNNAGATCGGAAG
56448_1 TATGAGTGTGGAGTCGTTGCTGCTGGGTTTATCTCAGAACTCCNNNAGATCGGAAG
56449_1 TATGAGTGTGGAGTCGTTGCATTAGGGCAGACCACATATCCACNNNAGATCGGAAG
56450_1 TATGAGTGTGGAGTCGTTGCGCTCTGCTTTTACAAGACCACTGNNNAGATCGGAAG
56451_1 TATGAGTGTGGAGTCGTTGCAAATTATTCCCCTGCCTCAGCTTNNNAGATCGGAAGA
56452_1 TATGAGTGTGGAGTCGTTGCTTCAAGTGATTCTCCTGCCTCAGNNNAGATCGGAAG
56454_1 TATGAGTGTGGAGTCGTTGCCCTCCCAGGTTCAAGTGATTCTGNNNAGATCGGAAG
56455_1 TATGAGTGTGGAGTCGTTGCGCCTCCTTAGTAGCTGGGATTACNNNAGATCGGAAG
56457_1 TATGAGTGTGGAGTCGTTGCCAGAGTCTTGCTATGTCACCCAGNNNAGATCGGAAG
56458_1 TATGAGTGTGGAGTCGTTGCAGAGACAGGGTTTCCTCATGTTGNNNAGATCGGAA
56462_1 TATGAGTGTGGAGTCGTTGCCCTCCCAAGTTCACACCATTCTCNNNAGATCGGAAGA
56464_1 TATGAGTGTGGAGTCGTTGCAGCATTCCTGAGTAGCTGGAATTNNNAGATCGGAAG
56466_1 TATGAGTGTGGAGTCGTTGCCAGTAGCTGGGATTACAGGCATGNNNAGATCGGAA
56467_1 TATGAGTGTGGAGTCGTTGCGTAGCTGGGACTATAGGCATGTGNNNAGATCGGAA
56468_1 TATGAGTGTGGAGTCGTTGCTTTAGTAGAGACGAGGTTTCGCCNNNAGATCGGAA
56469_1 TATGAGTGTGGAGTCGTTGCGTCAAGTGATTCTCCTGCCTCAGNNNAGATCGGAAG
56470_1 TATGAGTGTGGAGTCGTTGCTGCCTCCCAAGTTCAAGAGATTTNNNAGATCGGAAG
56471_1 TATGAGTGTGGAGTCGTTGCGTCAAGAGATTCTCCTTGCCTCANNNAGATCGGAAG
56472_1 TATGAGTGTGGAGTCGTTGCGAGTTTTGCTCTTATTGCCCAGGNNNAGATCGGAAG
56473_1 TATGAGTGTGGAGTCGTTGCCCTCCCAGGTTCAAGCTATTCTGNNNAGATCGGAAG
56474_1 TATGAGTGTGGAGTCGTTGCTGGTGCGAACATAGTTCACTACANNNAGATCGGAAG
56475_1 TATGAGTGTGGAGTCGTTGCCCTCCCAGGATCAAGAGATTCTCNNNAGATCGGAAG
56477_1 TATGAGTGTGGAGTCGTTGCTTCAAGTGATTCTCCTGCCTCAGNNNAGATCGGAAG
56478_1 TATGAGTGTGGAGTCGTTGCCTGAGTAGCTGGGATTACAGGTGNNNAGATCGGAA
56479_1 TATGAGTGTGGAGTCGTTGCCCAAATTGCTGGGATTACAGGTGNNNAGATCGGAA
56480_1 TATGAGTGTGGAGTCGTTGCGAGACAGAATCTTGCTCTTTCGCNNNAGATCGGAAG
56486_1 TATGAGTGTGGAGTCGTTGCGCCTCCTGGGTTCAAGTAATTCTNNNAGATCGGAAG

| | |
|---|---|
| 56487_1 | TATGAGTGTGGAGTCGTTGCTGGGTTCAAACAATTCTCATGCCNNNAGATCGGAAG |
| 56490_1 | TATGAGTGTGGAGTCGTTGCCCATCTCAGCTTGCAGAAACATTNNNAGATCGGAAG |
| 56491_1 | TATGAGTGTGGAGTCGTTGCGAGAGCAAATCCGTACAGAAGGTNNNAGATCGGAA |
| 56492_1 | TATGAGTGTGGAGTCGTTGCTCCGTCCCAAACTGAAAGGATTANNNAGATCGGAAG |
| 56494_1 | TATGAGTGTGGAGTCGTTGCCCAGGCAGTGTAGAGTTCTGAATNNNAGATCGGAA |
| 56496_1 | TATGAGTGTGGAGTCGTTGCTGGCCTTTTATAGGGAAGTTTGCNNNAGATCGGAA |
| 56497_1 | TATGAGTGTGGAGTCGTTGCGGGATCACACAGGTCTACATGTANNNAGATCGGAA |
| 56498_1 | TATGAGTGTGGAGTCGTTGCGTTGTGGTGCATGCCTATAATCCNNNAGATCGGAAG |
| 56499_1 | TATGAGTGTGGAGTCGTTGCGAAATGGGCCGTGATTCATCTTTNNNAGATCGGAAG |
| 56501_1 | TATGAGTGTGGAGTCGTTGCGAAAGAGAGAGGAATGGAAGGGTNNNAGATCGG |
| 56502_1 | TATGAGTGTGGAGTCGTTGCCTACTTTGCAGCTGGAGGGAATANNNAGATCGGAA |
| 56503_1 | TATGAGTGTGGAGTCGTTGCTGAAACCCCGTTGCTACTAAAGANNNAGATCGGAAG |
| 56505_1 | TATGAGTGTGGAGTCGTTGCCCCTTTTCCAATGCCTTATGGTGNNNAGATCGGAAG |
| 56506_1 | TATGAGTGTGGAGTCGTTGCGGAGTTCCCTGGTATCAAGATCANNNAGATCGGAA |
| 56507_1 | TATGAGTGTGGAGTCGTTGCATCATGAGGTCAGGAGATCGAGANNNAGATCGGAA |
| 56508_1 | TATGAGTGTGGAGTCGTTGCTTTGCAGCAACATGAATGAACCTNNNAGATCGGAAG |
| 56509_1 | TATGAGTGTGGAGTCGTTGCATCAGAGACTTGAGCATCCATGGNNNAGATCGGAA |
| 56511_1 | TATGAGTGTGGAGTCGTTGCGTGGGAGAGGCAGATTATGACATNNNAGATCGGA |
| 56512_1 | TATGAGTGTGGAGTCGTTGCGAGGTCAGGAGATCAAGACCATCNNNAGATCGGAA |
| 56513_1 | TATGAGTGTGGAGTCGTTGCGTGCTGATGAGGATCTGGAGAAANNNAGATCGGAA |
| 56514_1 | TATGAGTGTGGAGTCGTTGCTGGATTCAAACTCTCAGCCTCAANNNAGATCGGAAG |
| 56515_1 | TATGAGTGTGGAGTCGTTGCAAGACAGACCATGAAGCACATCTNNNAGATCGGAA |
| 56516_1 | TATGAGTGTGGAGTCGTTGCGTGCAGTGGTGTAATTATGGCTCNNNAGATCGGAA |
| 56517_1 | TATGAGTGTGGAGTCGTTGCGAAAATGAGCAGCTGACATCTCANNNAGATCGGAA |
| 56518_1 | TATGAGTGTGGAGTCGTTGCTAATGAAGTCATTCTGGCCTGGTNNNAGATCGGAAG |
| 56519_1 | TATGAGTGTGGAGTCGTTGCAAGGGATGGATGACAGAGACATGNNNAGATCGGA |
| 56521_1 | TATGAGTGTGGAGTCGTTGCGGAAGTATCATACCATTGGGGCTNNNAGATCGGAA |
| 56524_1 | TATGAGTGTGGAGTCGTTGCAGTACAACTACCCAAGCCTCTTGNNNAGATCGGAAG |
| 56526_1 | TATGAGTGTGGAGTCGTTGCAGGAAGCTGAAGCATGAGAATCANNNAGATCGGAA |
| 56527_1 | TATGAGTGTGGAGTCGTTGCCAACCATTGAATAACCCGTGTCCNNNAGATCGGAAG |
| 56528_1 | TATGAGTGTGGAGTCGTTGCGCTGAAACCCCGTCTCTACTAAANNNAGATCGGAAG |
| 56529_1 | TATGAGTGTGGAGTCGTTGCCCGATTTAGACAGTGTGGTTTGGNNNAGATCGGAA |
| 56530_1 | TATGAGTGTGGAGTCGTTGCAGGAGGAAGGAAGGAAAGAAAGTNNNAGATCGG |
| 56532_1 | TATGAGTGTGGAGTCGTTGCCTCACACCTGTAATCTCAGCACTNNNAGATCGGAAG |
| 56533_1 | TATGAGTGTGGAGTCGTTGCCACTTGACTCCTGTCCACAGAATNNNAGATCGGAAG |
| 56535_1 | TATGAGTGTGGAGTCGTTGCAAGGGCTGGACATTCTTCATAGGNNNAGATCGGAA |
| 56536_1 | TATGAGTGTGGAGTCGTTGCGAGAGAACGAGAGAGATGGAGGANNNAGATCGG |
| 56537_1 | TATGAGTGTGGAGTCGTTGCGGATCACTTGAGGTCAGGAGTTCNNNAGATCGGAA |
| 56538_1 | TATGAGTGTGGAGTCGTTGCGAGGCTGTGTGTATTGTGCATTANNNAGATCGGAA |
| 56539_1 | TATGAGTGTGGAGTCGTTGCCATGGTTGTCCTTCTAGCCTTCTNNNAGATCGGAAG |
| 56541_1 | TATGAGTGTGGAGTCGTTGCAGTTTGAGATCAGTCTGGCCAATNNNAGATCGGAA |
| 56543_1 | TATGAGTGTGGAGTCGTTGCATGGAGAAACCCCGTCTCTACTANNNAGATCGGAAG |
| 56544_1 | TATGAGTGTGGAGTCGTTGCAGAGAAGCCAAAGGATCTTCCAGNNNAGATCGGAA |

| | |
|---|---|
| 56546_1 | TATGAGTGTGGAGTCGTTGCGGAAGGAAACCCAGAGTTCTTCTNNNAGATCGGAA |
| 56547_1 | TATGAGTGTGGAGTCGTTGCCAATACTCCCCTCACAAACTGGANNNAGATCGGAAG |
| 56548_1 | TATGAGTGTGGAGTCGTTGCTGTACTTGACTCTGCTTCTCACCNNNAGATCGGAAGA |
| 56550_1 | TATGAGTGTGGAGTCGTTGCGGGTGATTCAATGGGATAGGGTTNNNAGATCGGA |
| 56551_1 | TATGAGTGTGGAGTCGTTGCGAAAAGAAACGTGGAGGGATTCGNNNAGATCGGA |
| 56552_1 | TATGAGTGTGGAGTCGTTGCGTGGCCCACAAAGTCTAAAACATNNNAGATCGGAAG |
| 56554_1 | TATGAGTGTGGAGTCGTTGCGCCTCCCAAGTAGCTAGGATTACNNNAGATCGGAAG |
| 56557_1 | TATGAGTGTGGAGTCGTTGCAAGTGCTGAGATTACAGGCATGANNNAGATCGGAA |
| 56558_1 | TATGAGTGTGGAGTCGTTGCGACTCCCTGGTTCAAGTGATTCTNNNAGATCGGAAG |
| 56560_1 | TATGAGTGTGGAGTCGTTGCCAGTGTTACAATCTCGGCTCACTNNNAGATCGGAAG |
| 56561_1 | TATGAGTGTGGAGTCGTTGCCCATGCCCGGCTAATTTTTTGTANNNAGATCGGAAG |
| 56562_1 | TATGAGTGTGGAGTCGTTGCGTTAGGTCAAGCAATTCCTCTGCNNNAGATCGGAAG |
| 56564_1 | TATGAGTGTGGAGTCGTTGCCAATCTTGGCTCACTGCAACTTCNNNAGATCGGAAG |
| 56565_1 | TATGAGTGTGGAGTCGTTGCTTCAAGCTATCTTCCTGCCTCAGNNNAGATCGGAAGA |
| 56566_1 | TATGAGTGTGGAGTCGTTGCCTGTATCTTCAAACTCCTGGGCTNNNAGATCGGAAG |
| 56567_1 | TATGAGTGTGGAGTCGTTGCCTGTTGCTCTGTTGCTCTGTTTCNNNAGATCGGAAGA |
| 56568_1 | TATGAGTGTGGAGTCGTTGCGCCTCCCAAGTAGCTAGGATTACNNNAGATCGGAAG |
| 56571_1 | TATGAGTGTGGAGTCGTTGCGTAGAGATGGGGTTTAGCCATGTNNNAGATCGGAA |
| 56573_1 | TATGAGTGTGGAGTCGTTGCCTAGGTTGAAGCAATTCTCCTGCNNNAGATCGGAAG |
| 56574_1 | TATGAGTGTGGAGTCGTTGCTTCAAGTGATTCTCCTGCCTCAGNNNAGATCGGAAG |
| 56575_1 | TATGAGTGTGGAGTCGTTGCTGTTTGGGTAGGAAAGTTGGACTNNNAGATCGGAA |
| 56576_1 | TATGAGTGTGGAGTCGTTGCTTCAAGTGATTCTCCTGCCTCAGNNNAGATCGGAAG |
| 56577_1 | TATGAGTGTGGAGTCGTTGCCAAGTAGCTGGGATTACAGGTGTNNNAGATCGGAA |
| 56580_1 | TATGAGTGTGGAGTCGTTGCGAGAAGGAGTTTTCACTCTTGGTNNNAGATCGGAA |
| 56581_1 | TATGAGTGTGGAGTCGTTGCTTCATGCCATTCTCCTGTCTCAGNNNAGATCGGAAGA |
| 56583_1 | TATGAGTGTGGAGTCGTTGCCTCACCACAACCTCTGCCTCNNNAGATCGGAAGAGCA |
| 56585_1 | TATGAGTGTGGAGTCGTTGCGTTTGACCATGTTAGCCAGGATGNNNAGATCGGAA |
| 56587_1 | TATGAGTGTGGAGTCGTTGCCTCCTGAGTAGCTGGGATTACAGNNNAGATCGGAA |
| 56588_1 | TATGAGTGTGGAGTCGTTGCTGGAGTTTTACTCTTGTCACCCANNNAGATCGGAAG |
| 56590_1 | TATGAGTGTGGAGTCGTTGCAAGTTCATGTGATTCTCCTGCCTNNNAGATCGGAAG |
| 56591_1 | TATGAGTGTGGAGTCGTTGCGCACAATCTCAGCTCACTGTAAGNNNAGATCGGAAG |
| 56592_1 | TATGAGTGTGGAGTCGTTGCAAACTCCTGACCTCAAGTGATCCNNNAGATCGGAAG |
| 56597_1 | TATGAGTGTGGAGTCGTTGCTTCAAGTGATTCTCCTGCCTCAGNNNAGATCGGAAG |
| 56598_1 | TATGAGTGTGGAGTCGTTGCCAATCTCAGCTCACTGCAAACTCNNNAGATCGGAAG |
| 56600_1 | TATGAGTGTGGAGTCGTTGCAAGGGATTGAGTTCCTGATTTGANNNAGATCGGAA |
| 56601_1 | TATGAGTGTGGAGTCGTTGCTGATCCTGGCTTACTGCAACTTCNNNAGATCGGAAG |
| 56603_1 | TATGAGTGTGGAGTCGTTGCTTTCTCCATGTTGATCAGGCTGANNNAGATCGGAAG |
| 56604_1 | TATGAGTGTGGAGTCGTTGCAGAGATTGGGTTTTGCCATGTTGNNNAGATCGGAA |
| 56605_1 | TATGAGTGTGGAGTCGTTGCGCATGATCTCAGGTCATTGGAACNNNAGATCGGAA |
| 56606_1 | TATGAGTGTGGAGTCGTTGCAGCCTCCAAATAGCTGGGATTACNNNAGATCGGAAG |
| 56607_1 | TATGAGTGTGGAGTCGTTGCCCATCTCTGCTCACTGTAGTTCCNNNAGATCGGAAGA |
| 56608_1 | TATGAGTGTGGAGTCGTTGCCCTCAGGCTAGAGTACAGTGATGNNNAGATCGGAA |
| 56609_1 | TATGAGTGTGGAGTCGTTGCCTCCCAGGTTCAAGTGATTCTCANNNAGATCGGAAG |

| | |
|---|---|
| 56610_1 | TATGAGTGTGGAGTCGTTGCTTACTTTGTCCTCTGGAGAAGCCNNNAGATCGGAAGA |
| 56614_1 | TATGAGTGTGGAGTCGTTGCCAATGGCAAGACCTCAGTTCACTNNNAGATCGGAAG |
| 56615_1 | TATGAGTGTGGAGTCGTTGCGCATCTTGCTCTGTCACTACTCANNNAGATCGGAAGA |
| 56617_1 | TATGAGTGTGGAGTCGTTGCGGGAGACGGAGTTTTACTCTTGTNNNAGATCGGAAG |
| 56618_1 | TATGAGTGTGGAGTCGTTGCAACCTCTGTCTCATGGGTTCAAGNNNAGATCGGAAG |
| 56619_1 | TATGAGTGTGGAGTCGTTGCCACCATGCCCAGCTAATTTTTGTNNNAGATCGGAAGA |
| 56622_1 | TATGAGTGTGGAGTCGTTGCTGGAGTACAGTGGCATGATCTTGNNNAGATCGGAAG |
| 56623_1 | TATGAGTGTGGAGTCGTTGCCTGAGTAGCTGGAATTACAGGCANNNAGATCGGAAG |
| 56624_1 | TATGAGTGTGGAGTCGTTGCGTATCTGGGACCAAAGGCACATANNNAGATCGGAAG |
| 56625_1 | TATGAGTGTGGAGTCGTTGCGCTAAGGCAGGAGAATCACTTGANNNAGATCGGAAG |
| 56628_1 | TATGAGTGTGGAGTCGTTGCTCATGCCTGTAATCTCAGCACTTNNNAGATCGGAAGA |
| 56629_1 | TATGAGTGTGGAGTCGTTGCTGGATCCACTTCTCCTGTTTCAGNNNAGATCGGAAGA |
| 56631_1 | TATGAGTGTGGAGTCGTTGCAAAAGACCCTAAATCCTCTTCCANNNAGATCGGAAGA |
| 56632_1 | TATGAGTGTGGAGTCGTTGCTGAAACCCAGCCTCTACTGAAAANNNAGATCGGAAG |
| 56633_1 | TATGAGTGTGGAGTCGTTGCTTGACAATAACGAGGAATGGGGANNNAGATCGGAA |
| 56635_1 | TATGAGTGTGGAGTCGTTGCTAGTTTTGGGTGGGAAACATCGANNNAGATCGGAAG |
| 56637_1 | TATGAGTGTGGAGTCGTTGCATGGACTAGGAAATGCTAAGGCCNNNAGATCGGAAG |
| 56639_1 | TATGAGTGTGGAGTCGTTGCTTCTGCAGACAAGAAACATGCTGNNNAGATCGGAAG |
| 56640_1 | TATGAGTGTGGAGTCGTTGCCTGTGTGTTTGTGGTGAGTCATCNNNAGATCGGAAG |
| 56641_1 | TATGAGTGTGGAGTCGTTGCGGGTTGCAATCCTAGTCTCTGATNNNAGATCGGAAG |
| 56642_1 | TATGAGTGTGGAGTCGTTGCTAAAAACCACTCCCTGTCTCTGGNNNAGATCGGAAGA |
| 56643_1 | TATGAGTGTGGAGTCGTTGCTCATCTTCCTCTCAACCCTTTCCNNNAGATCGGAAGAG |
| 56644_1 | TATGAGTGTGGAGTCGTTGCTCCACTATGCTCCTCATTGTCAANNNAGATCGGAAGA |
| 56645_1 | TATGAGTGTGGAGTCGTTGCCTTCACTTGCCACCATTGGAAAGNNNAGATCGGAAGA |
| 56646_1 | TATGAGTGTGGAGTCGTTGCAAACGCTGGACAAACCTAAACTGNNNAGATCGGAAG |
| 56647_1 | TATGAGTGTGGAGTCGTTGCCTGCAGCAAGGAGAAATCCAAAANNNAGATCGGAAG |
| 56648_1 | TATGAGTGTGGAGTCGTTGCAGCATGTAAAGAGCAGAACTGGANNNAGATCGGAA |
| 56649_1 | TATGAGTGTGGAGTCGTTGCATGGTTTGGATCTGTGTCCCTACNNNAGATCGGAAG |
| 56654_1 | TATGAGTGTGGAGTCGTTGCTGCTCAAAGCTGAACTCTGACTTNNNAGATCGGAAGA |
| 56655_1 | TATGAGTGTGGAGTCGTTGCGGAACAGAATACAGAGCCCAGAANNNAGATCGGAA |
| 56656_1 | TATGAGTGTGGAGTCGTTGCTTGGGGTTCTGGAATGAGAAGAGNNNAGATCGGAA |
| 56658_1 | TATGAGTGTGGAGTCGTTGCGGACCAGCCCATTCACAAATATGNNNAGATCGGAAG |
| 56659_1 | TATGAGTGTGGAGTCGTTGCTGGCTGTTCTGAAAGGGGAATAANNNAGATCGGAA |
| 56660_1 | TATGAGTGTGGAGTCGTTGCGTGTATGGCAACTGAGACTCAGANNNAGATCGGAAG |
| 56663_1 | TATGAGTGTGGAGTCGTTGCCATGATTATGCCTCTGGACTCCANNNAGATCGGAAGA |
| 56666_1 | TATGAGTGTGGAGTCGTTGCGCAGTGATCCCTTTCAACATGTTNNNAGATCGGAAGA |
| 56667_1 | TATGAGTGTGGAGTCGTTGCCTACTCAGCCACTCAGCCATAAANNNAGATCGGAAGA |
| 56669_1 | TATGAGTGTGGAGTCGTTGCAACTCCTGGAATCAAGCAATCCTNNNAGATCGGAAGA |
| 56670_1 | TATGAGTGTGGAGTCGTTGCGTGGACATTGCCAACCGTTATACNNNAGATCGGAAG |
| 56671_1 | TATGAGTGTGGAGTCGTTGCTTTCAGGAGGCTATCTCAAGACANNNAGATCGGAAG |
| 56672_1 | TATGAGTGTGGAGTCGTTGCCTTTGGACGTGGCACGATTTATTNNNAGATCGGAAG |
| 56675_1 | TATGAGTGTGGAGTCGTTGCAGTAAGTCTGTGGGGAAGAAGACNNNAGATCGGAA |
| 56676_1 | TATGAGTGTGGAGTCGTTGCATGTTGCCTAGGCTGATCTGAAANNNAGATCGGAAG |

| | |
|---|---|
| 56678_1 | TATGAGTGTGGAGTCGTTGCGAAAATGACTCAAGAGGCATGGCNNNAGATCGGAA |
| 56680_1 | TATGAGTGTGGAGTCGTTGCTGAGAACCAAGGATGTAGTTGGGNNNAGATCGGA |
| 56681_1 | TATGAGTGTGGAGTCGTTGCACACATAAGCATACAGTCAGAAGANNNAGATCGGA |
| 56682_1 | TATGAGTGTGGAGTCGTTGCAATTAGCTGGACTTGGTGGTGTANNNAGATCGGAA |
| 56684_1 | TATGAGTGTGGAGTCGTTGCGAAGATGAATCTGCTGGGAGTGANNNAGATCGGA |
| 56686_1 | TATGAGTGTGGAGTCGTTGCTCTTGTACGTGTATTCCCAGCAANNNAGATCGGAAG |
| 56687_1 | TATGAGTGTGGAGTCGTTGCTGGATACTCAGTAAGTGGCAACANNNAGATCGGAA |
| 56688_1 | TATGAGTGTGGAGTCGTTGCGGCAACCTACAGAATGGGAGTAANNNAGATCGGAA |
| 56689_1 | TATGAGTGTGGAGTCGTTGCAAGGTAGGAGGATTGTTTGAGCCNNNAGATCGGAA |
| 56690_1 | TATGAGTGTGGAGTCGTTGCGAGGCTGAGTCATGAGAATCACTNNNAGATCGGAA |
| 56691_1 | TATGAGTGTGGAGTCGTTGCCAATTCAGATTTCCCACCAGCAGNNNAGATCGGAAG |
| 56693_1 | TATGAGTGTGGAGTCGTTGCTGTGTACTGAGATGCAGGGTATGNNNAGATCGGAA |
| 56694_1 | TATGAGTGTGGAGTCGTTGCGGATCACTTGAGTCCAGGACTTTNNNAGATCGGAAG |
| 56695_1 | TATGAGTGTGGAGTCGTTGCATTTGGTTAGGGATTGGGAGCATNNNAGATCGGAA |
| 56697_1 | TATGAGTGTGGAGTCGTTGCTTCAGGAGTTTGGATGAGACTGGNNNAGATCGGAA |
| 56698_1 | TATGAGTGTGGAGTCGTTGCCCAGGTCTAAGCTGTACATGAGTNNNAGATCGGAAG |
| 56699_1 | TATGAGTGTGGAGTCGTTGCAGAGGATGGTCATGTGAACTTGGNNNAGATCGGAA |
| 56701_1 | TATGAGTGTGGAGTCGTTGCGGATACACAGATAGGCAGTGGAGNNNAGATCGGA |
| 56702_1 | TATGAGTGTGGAGTCGTTGCAGGAAGATTGCTTGAGACTGCAGNNNAGATCGGAA |
| 56704_1 | TATGAGTGTGGAGTCGTTGCCTCTCAAAGTGCTGGGACTACAANNNAGATCGGAAG |
| 56705_1 | TATGAGTGTGGAGTCGTTGCATAAGATTTTGCCATGTTGCCCANNNAGATCGGAAG |
| 56706_1 | TATGAGTGTGGAGTCGTTGCGGTCATTTTCATCCTTGGCACTCNNNAGATCGGAAG |
| 56707_1 | TATGAGTGTGGAGTCGTTGCCCTTCCAAGTAGCTGGGACTTTANNNAGATCGGAAG |
| 56708_1 | TATGAGTGTGGAGTCGTTGCCCTGAACTCCTAGACTCAAGCAGNNNAGATCGGAAG |
| 56709_1 | TATGAGTGTGGAGTCGTTGCCCAGGATTATAGGTACCTGCCACNNNAGATCGGAAG |
| 56710_1 | TATGAGTGTGGAGTCGTTGCGAGTGCAGTGGTATGATCTCAGTNNNAGATCGGAA |
| 56711_1 | TATGAGTGTGGAGTCGTTGCGAGACTGAGTTTTCCCATGTTGGNNNAGATCGGAAG |
| 56712_1 | TATGAGTGTGGAGTCGTTGCCCTTCTGGGTTCAAGCGATTCTCNNNAGATCGGAAG |
| 56713_1 | TATGAGTGTGGAGTCGTTGCGTTCAAGCAATTTTCCCACCTCANNNAGATCGGAAGA |
| 56714_1 | TATGAGTGTGGAGTCGTTGCGGTGATCTTGGTTCTCTGCAATCNNNAGATCGGAAG |
| 56715_1 | TATGAGTGTGGAGTCGTTGCGAGACAGGTTCTCACTCTATCGCNNNAGATCGGAAG |
| 56716_1 | TATGAGTGTGGAGTCGTTGCGATCTTGGCTCATTGCAATCTCCNNNAGATCGGAAG |
| 56717_1 | TATGAGTGTGGAGTCGTTGCTCCTCCTAAGTAGCTGGGACTACNNNAGATCGGAAG |
| 56718_1 | TATGAGTGTGGAGTCGTTGCCGCTTACCAGGTTCAAGTGATTCNNNAGATCGGAAG |
| 56719_1 | TATGAGTGTGGAGTCGTTGCGATCTCAGCTCATTGCAAGGTCTNNNAGATCGGAAG |
| 56723_1 | TATGAGTGTGGAGTCGTTGCGTTCAAGCAATTCTCTGCCTCAGNNNAGATCGGAAG |
| 56724_1 | TATGAGTGTGGAGTCGTTGCCCAAGTAGCTGGGATTACAGGCNNNAGATCGGAAG |
| 56725_1 | TATGAGTGTGGAGTCGTTGCGGCTAGAGTGCAATGACACAATCNNNAGATCGGAA |
| 56727_1 | TATGAGTGTGGAGTCGTTGCTTTCAAGTGATTCTCCTGCCTCANNNAGATCGGAAGA |
| 56729_1 | TATGAGTGTGGAGTCGTTGCATCTTCTGGGTTCAAGCGATTCTNNNAGATCGGAAG |
| 56730_1 | TATGAGTGTGGAGTCGTTGCGATGGAGTCTTACTCTGTCACCCNNNAGATCGGAAG |
| 56731_1 | TATGAGTGTGGAGTCGTTGCGGGTCTCATTCTGTTACCTAGGCNNNAGATCGGAAG |
| 56733_1 | TATGAGTGTGGAGTCGTTGCCTCACTGCAAACTCTACCTGCTANNNAGATCGGAAGA |

| | |
|---|---|
| 56735_1 | TATGAGTGTGGAGTCGTTGCATTCAAGTGATTCTCCTGCCTCANNNAGATCGGAAG |
| 56736_1 | TATGAGTGTGGAGTCGTTGCCTTCCAGGTTCAAGCAATTCTCGNNNAGATCGGAAG |
| 56737_1 | TATGAGTGTGGAGTCGTTGCGGTTCAAGCAATTCTCCTGACTCNNNAGATCGGAAG |
| 56739_1 | TATGAGTGTGGAGTCGTTGCCAATAGCACAATCTTGGCTCACTNNNAGATCGGAAG |
| 56740_1 | TATGAGTGTGGAGTCGTTGCTTCAAGCAATTTTCCTGCCTCAGNNNAGATCGGAAGA |
| 56743_1 | TATGAGTGTGGAGTCGTTGCCTATCTTGGCTCACTGCAAGCTCNNNAGATCGGAAG |
| 56744_1 | TATGAGTGTGGAGTCGTTGCGGATTACAAGCATGAACCACCACNNNAGATCGGAAG |
| 56745_1 | TATGAGTGTGGAGTCGTTGCTAATCTCAGCTCACTGCAACCTGNNNAGATCGGAAG |
| 56746_1 | TATGAGTGTGGAGTCGTTGCCAAGTAGCTAGGACTACAGGTGCNNNAGATCGGAA |
| 56747_1 | TATGAGTGTGGAGTCGTTGCAAGACAGAGCTCCCATACAAAGGNNNAGATCGGAA |
| 56748_1 | TATGAGTGTGGAGTCGTTGCTACAGACTCTTAGACCCTGTCCANNNAGATCGGAAG |
| 56749_1 | TATGAGTGTGGAGTCGTTGCCCTCCCAGGATCAAGTGATTCTCNNNAGATCGGAAG |
| 56751_1 | TATGAGTGTGGAGTCGTTGCCGATTCCCTGGTTCAAGCAATTCNNNAGATCGGAAG |
| 56753_1 | TATGAGTGTGGAGTCGTTGCATCCTGAGTAGCTGGGACTACAGNNNAGATCGGAA |
| 56757_1 | TATGAGTGTGGAGTCGTTGCGCAATCTTGGCTCATCACAACTTNNNAGATCGGAAG |
| 56758_1 | TATGAGTGTGGAGTCGTTGCCTCCCAAGTAGCTGTGACTACAGNNNAGATCGGAAG |
| 56759_1 | TATGAGTGTGGAGTCGTTGCACTAGATCTCATTCTGTTGCCCTNNNAGATCGGAAGA |
| 56760_1 | TATGAGTGTGGAGTCGTTGCACAAGGTCTCCTTATTTTGCCCANNNAGATCGGAAG |
| 56761_1 | TATGAGTGTGGAGTCGTTGCGACTCCCGAGTAGCTAGGACTATNNNAGATCGGAA |
| 56765_1 | TATGAGTGTGGAGTCGTTGCTTCAAGTGATTCTCCTGCCTCAGNNNAGATCGGAAG |
| 56766_1 | TATGAGTGTGGAGTCGTTGCGCCTCCCAAGTAGCTAGGAATACNNNAGATCGGAAG |
| 56767_1 | TATGAGTGTGGAGTCGTTGCGGAATACAGTGGCACAATCACAGNNNAGATCGGAA |
| 56768_1 | TATGAGTGTGGAGTCGTTGCAAGAGATTCTCCTACCTCAGCCTNNNAGATCGGAAG |
| 56772_1 | TATGAGTGTGGAGTCGTTGCCCTACGCTGGTTTCAAACTTCTGNNNAGATCGGAAG |
| 56773_1 | TATGAGTGTGGAGTCGTTGCGTTCAAGAGATTCTCCTGCCTCANNNAGATCGGAAG |
| 56775_1 | TATGAGTGTGGAGTCGTTGCTACAGTGGTGAGATTATGGCTCANNNAGATCGGAA |
| 56776_1 | TATGAGTGTGGAGTCGTTGCGGAGTGCAGTAGTGTGATCTCAGNNNAGATCGGAA |
| 56777_1 | TATGAGTGTGGAGTCGTTGCTCCTGGGAAGAGGAAGGAACTATNNNAGATCGGAA |
| 56778_1 | TATGAGTGTGGAGTCGTTGCCTTGATCTCACCTCAACTTCCCANNNAGATCGGAAGA |
| 56779_1 | TATGAGTGTGGAGTCGTTGCCTCCTGAGTAGCTGGGACTACAGNNNAGATCGGAAG |
| 56780_1 | TATGAGTGTGGAGTCGTTGCTCTGGGTTCAAGTGATTCTCCTGNNNAGATCGGAAG |
| 56782_1 | TATGAGTGTGGAGTCGTTGCGTTCAAGTGATTCTCCTGCCTTGNNNAGATCGGAAG |
| 56783_1 | TATGAGTGTGGAGTCGTTGCCAGGAGCCTGGAGGACAAATATTNNNAGATCGGAA |
| 56786_1 | TATGAGTGTGGAGTCGTTGCAGACAAGGTCTCACTCTGTCATGNNNAGATCGGAAG |
| 56787_1 | TATGAGTGTGGAGTCGTTGCCATTGGTTATGTGTGTGGCAAGTNNNAGATCGGAA |
| 56788_1 | TATGAGTGTGGAGTCGTTGCAGTGCTGGAATTACAGACCTGAGNNNAGATCGGAA |
| 56791_1 | TATGAGTGTGGAGTCGTTGCTGAGTTCAAGTAATTCCCTGCCTNNNAGATCGGAAG |
| 56792_1 | TATGAGTGTGGAGTCGTTGCAGCAGGAATTGATTGTTTCAGGCNNNAGATCGGAA |
| 56795_1 | TATGAGTGTGGAGTCGTTGCACAATCTTGGCTCACTGTAACCTNNNAGATCGGAAG |
| 56796_1 | TATGAGTGTGGAGTCGTTGCTGGTGTGATCACAGCTCAATGTANNNAGATCGGAA |
| 56798_1 | TATGAGTGTGGAGTCGTTGCATCTTGGCTCACTGTAACCTCTGNNNAGATCGGAAG |
| 56800_1 | TATGAGTGTGGAGTCGTTGCCTCTCAAGTAGCTGGGACTACAGNNNAGATCGGAA |
| 56801_1 | TATGAGTGTGGAGTCGTTGCGCAAGTAGTGAAACCCCATCTCTNNNAGATCGGAAG |

| | |
|---|---|
| 56802_1 | TATGAGTGTGGAGTCGTTGCGGCACAATCTTGTCTCAATGCAANNNAGATCGGAAG |
| 56803_1 | TATGAGTGTGGAGTCGTTGCAGCTGGGATTACAGGAATATGCCNNNAGATCGGAA |
| 56804_1 | TATGAGTGTGGAGTCGTTGCTAGTGGCATGATCAAAGCTCACTNNNAGATCGGAAG |
| 56809_1 | TATGAGTGTGGAGTCGTTGCTGCTTTTCTGTTTTGAGCCAGAGNNNAGATCGGAAG |
| 56812_1 | TATGAGTGTGGAGTCGTTGCCATTCTCCTACCTCATTCTCCCGNNNAGATCGGAAGA |
| 56813_1 | TATGAGTGTGGAGTCGTTGCCTCCCAAGTAGCTGGAACTACAGNNNAGATCGGAAG |
| 56817_1 | TATGAGTGTGGAGTCGTTGCTTCAGGTGATTCTTCTGCCTCAGNNNAGATCGGAAG |
| 56818_1 | TATGAGTGTGGAGTCGTTGCCCTGAGTAGCTGAGACTAAAGGCNNNAGATCGGAA |
| 56819_1 | TATGAGTGTGGAGTCGTTGCAGTGCGTTGTATTTCACTTTGCANNNAGATCGGAAG |
| 56820_1 | TATGAGTGTGGAGTCGTTGCTCCGTCCCAAACTGAAAGGATTANNNAGATCGGAAG |
| 56821_1 | TATGAGTGTGGAGTCGTTGCTCCGTCCCAAACTGAAAGGATTANNNAGATCGGAAG |
| 56822_1 | TATGAGTGTGGAGTCGTTGCAGGAAAGAAGGGAAAGAAAGGGCNNNAGATCGG |
| 56823_1 | TATGAGTGTGGAGTCGTTGCCCCTGATTCAAGAAGCTCTACCANNNAGATCGGAAG |
| 56824_1 | TATGAGTGTGGAGTCGTTGCTACCTGTAATCCCAGCTACTTGGNNNAGATCGGAAG |
| 56825_1 | TATGAGTGTGGAGTCGTTGCGCAGGCAACTGGTTAGAAACAATNNNAGATCGGAA |
| 56826_1 | TATGAGTGTGGAGTCGTTGCCTCCTGAGTAGCTGGGATTACAGNNNAGATCGGAA |
| 56827_1 | TATGAGTGTGGAGTCGTTGCCCAACATGGTGAAACTCCATCTCNNNAGATCGGAAG |
| 56828_1 | TATGAGTGTGGAGTCGTTGCACAGGGCTTTCCACAAATGAGTANNNAGATCGGAAG |
| 56829_1 | TATGAGTGTGGAGTCGTTGCGGGGCTTCTCCTAGGCTTTATTTNNNAGATCGGAAG |
| 56832_1 | TATGAGTGTGGAGTCGTTGCACTAATCATGGGTTTCTGGGCTTNNNAGATCGGAAG |
| 56839_1 | TATGAGTGTGGAGTCGTTGCAGAAATCAAATTGCAGCCTCACGNNNAGATCGGAAG |
| 56840_1 | TATGAGTGTGGAGTCGTTGCCCATTGTTCTTTTTTCCACCCCANNNAGATCGGAAGA |
| 56846_1 | TATGAGTGTGGAGTCGTTGCGAGAGAAACAAAACGGACACACGNNNAGATCGGAA |
| 56847_1 | TATGAGTGTGGAGTCGTTGCCCAGCAGTTTTCCAGAGAATAGGNNNAGATCGGAA |
| 56848_1 | TATGAGTGTGGAGTCGTTGCCATAGGACTAGCGGCTACATGTTNNNAGATCGGAA |
| 56849_1 | TATGAGTGTGGAGTCGTTGCAAGCCAAAGAACAAGTGCAAAGTNNNAGATCGGAA |
| 56850_1 | TATGAGTGTGGAGTCGTTGCTGGGTGGATTGAAGGTTACTGAGNNNAGATCGGAA |
| 56852_1 | TATGAGTGTGGAGTCGTTGCGGGCCCCCTTGATACATCATTATNNNAGATCGGAAG |
| 56853_1 | TATGAGTGTGGAGTCGTTGCTACCAGATCGTAAAGTTTGGGCTNNNAGATCGGAA |
| 56854_1 | TATGAGTGTGGAGTCGTTGCGATCACAAGGTCAGGAGTTCGAGNNNAGATCGGAA |
| 56855_1 | TATGAGTGTGGAGTCGTTGCAACACCAAACAAGACTTTGCACANNNAGATCGGAAG |
| 56856_1 | TATGAGTGTGGAGTCGTTGCCTGTGGAATGAGAGAAGGACACANNNAGATCGGAA |
| 56858_1 | TATGAGTGTGGAGTCGTTGCGTGACCCAAAAAAGACAGACAGGNNNAGATCGGAA |
| 56859_1 | TATGAGTGTGGAGTCGTTGCGTAACCTGTGTCAGTGCTGAAACNNNAGATCGGAA |
| 56860_1 | TATGAGTGTGGAGTCGTTGCTGCATACACAAAGATTTGCCAGGNNNAGATCGGAA |
| 56862_1 | TATGAGTGTGGAGTCGTTGCTGGCTTTGTTTGTTTCTTCTCCCNNNAGATCGGAAGA |
| 56863_1 | TATGAGTGTGGAGTCGTTGCCAGCCAGCATTTTTCAGCGTATANNNAGATCGGAAG |
| 56866_1 | TATGAGTGTGGAGTCGTTGCAGTGTTTGGTAGCAGTATAGGGCNNNAGATCGGAA |
| 56869_1 | TATGAGTGTGGAGTCGTTGCAACACACACACACACACAAAAGTNNNAGATCGGAAG |
| 56870_1 | TATGAGTGTGGAGTCGTTGCCTGAGCCAGGAGAATCACTTGAANNNAGATCGGAA |
| 56873_1 | TATGAGTGTGGAGTCGTTGCCACACCTGTAATCCCAGCACTTTNNNAGATCGGAAGA |
| 56874_1 | TATGAGTGTGGAGTCGTTGCGGCAAGACCACATTCCTACAGAANNNAGATCGGAAG |
| 56876_1 | TATGAGTGTGGAGTCGTTGCTAGTGGCAGGCATTGAAAGAACTNNNAGATCGGAA |

| | |
|---|---|
| 56883_1 | TATGAGTGTGGAGTCGTTGCTGAAATTATGAGGCCTCTGTGCANNNAGATCGGAA |
| 56884_1 | TATGAGTGTGGAGTCGTTGCATAGTGGGCTTTCAATGGTAGCTNNNAGATCGGAA |
| 56885_1 | TATGAGTGTGGAGTCGTTGCCTGAGGTGGGAGGATTACTTGAGNNNAGATCGGA |
| 56887_1 | TATGAGTGTGGAGTCGTTGCGCCTGTAATCTAGCACTTTGGGANNNAGATCGGAAG |
| 56889_1 | TATGAGTGTGGAGTCGTTGCGTGTGATAGCTCAGGTGGATCAANNNAGATCGGA |
| 56890_1 | TATGAGTGTGGAGTCGTTGCTACAATATAAAGCCCCTCCCCTGNNNAGATCGGAAG |
| 56891_1 | TATGAGTGTGGAGTCGTTGCACTCTGTTGGTGGGAGTGAAAATNNNAGATCGGA |
| 56892_1 | TATGAGTGTGGAGTCGTTGCCTTTTCCACTTTCCTGCTGTCAGNNNAGATCGGAAGA |
| 56893_1 | TATGAGTGTGGAGTCGTTGCTTTCTAAGAGGACAACCAAGGCANNNAGATCGGAA |
| 56894_1 | TATGAGTGTGGAGTCGTTGCGCAGGTTTTGAAGATGTGTTGGANNNAGATCGGAA |
| 56895_1 | TATGAGTGTGGAGTCGTTGCACACAAAACCAAGGCATTGCATANNNAGATCGGAAG |
| 56896_1 | TATGAGTGTGGAGTCGTTGCTCACTTGAGGTCAGGAGTTCAAGNNNAGATCGGAA |
| 56897_1 | TATGAGTGTGGAGTCGTTGCGTCAGTTTCCTCAGCCTCTACTCNNNAGATCGGAAGA |
| 56899_1 | TATGAGTGTGGAGTCGTTGCTTAGCTGGTTGTGGTAGTACAGGNNNAGATCGGA |
| 56901_1 | TATGAGTGTGGAGTCGTTGCAAATCATGGGCTCTGGGTTTTTGNNNAGATCGGAA |
| 56903_1 | TATGAGTGTGGAGTCGTTGCCCACTTTGGATGGAAAAGGCATANNNAGATCGGAA |
| 56904_1 | TATGAGTGTGGAGTCGTTGCTCAACCAGGTGTCAAGGTAGAAGNNNAGATCGGA |
| 56905_1 | TATGAGTGTGGAGTCGTTGCAGAGAACCACAAAAGACCCCAAANNNAGATCGGAA |
| 56906_1 | TATGAGTGTGGAGTCGTTGCTTCTGATGTTCCAGTGTGTGAGGNNNAGATCGGAA |
| 56907_1 | TATGAGTGTGGAGTCGTTGCTGGCTAATCTCCAAAATCCTGCTNNNAGATCGGAAG |
| 56909_1 | TATGAGTGTGGAGTCGTTGCAGCAAACAAATTCAAAAGCTCGTNNNAGATCGGAAG |
| 56910_1 | TATGAGTGTGGAGTCGTTGCGAGGGGGAAGTACTTGAAGGAATNNNAGATCGGA |
| 56911_1 | TATGAGTGTGGAGTCGTTGCAGGTTCTTATTGCCTACTCTGCCNNNAGATCGGAAG |
| 56912_1 | TATGAGTGTGGAGTCGTTGCTCGAATTGTGACGCATGACAAAANNNAGATCGGAA |
| 56914_1 | TATGAGTGTGGAGTCGTTGCGTGTGTGCCTCAGAACAATTTGANNNAGATCGGAA |
| 56915_1 | TATGAGTGTGGAGTCGTTGCACAGTGGGCAAAAAACATGAACANNNAGATCGGA |
| 56917_1 | TATGAGTGTGGAGTCGTTGCAATACAAATTGAAGGGACAATGACANNNAGATCGG |
| 56918_1 | TATGAGTGTGGAGTCGTTGCGTGAATGTTGGCCATGTGTGTATNNNAGATCGGAA |
| 56919_1 | TATGAGTGTGGAGTCGTTGCACATGTGGATGGGTGTGGAATTANNNAGATCGGAA |
| 56920_1 | TATGAGTGTGGAGTCGTTGCTTTTTTCTCAAGCCGGAGTCTTGNNNAGATCGGAAG |
| 56921_1 | TATGAGTGTGGAGTCGTTGCAAGAACCAATGAGTTCGGCAATGNNNAGATCGGAA |
| 56922_1 | TATGAGTGTGGAGTCGTTGCGCTCCAGAGGCACTAAACAAAAGNNNAGATCGGAA |
| 56923_1 | TATGAGTGTGGAGTCGTTGCAATCTCAATCTCAGAGTCGGAGCNNNAGATCGGAAG |
| 56924_1 | TATGAGTGTGGAGTCGTTGCAGCCAGGAATACAGGTTCAATGTNNNAGATCGGAA |
| 56926_1 | TATGAGTGTGGAGTCGTTGCGGAACTGCACAGGTCCACTTATANNNAGATCGGAAG |
| 56928_1 | TATGAGTGTGGAGTCGTTGCATTTCATCCACACAGTAGCCCTTNNNAGATCGGAAGA |
| 56929_1 | TATGAGTGTGGAGTCGTTGCGGATAAACACACACACATGCACANNNAGATCGGAAG |
| 56931_1 | TATGAGTGTGGAGTCGTTGCCTGGAGAAGCAAAGTTCTCATGCNNNAGATCGGAA |
| 56933_1 | TATGAGTGTGGAGTCGTTGCGACACCTGTAATCCCAGCACTTTNNNAGATCGGAAG |
| 56935_1 | TATGAGTGTGGAGTCGTTGCGCAGCCTGGAAAAACTAATGCTTNNNAGATCGGAAG |
| 56937_1 | TATGAGTGTGGAGTCGTTGCGAGGTCAAGAGATCGAGACCATCNNNAGATCGGAA |
| 56938_1 | TATGAGTGTGGAGTCGTTGCCAGTACAAACTCCAAAGAGGGCTNNNAGATCGGAA |
| 56940_1 | TATGAGTGTGGAGTCGTTGCTCTACCCCTGTGTTTCTCAAAGGNNNAGATCGGAAG |

| | |
|---|---|
| 56941_1 | TATGAGTGTGGAGTCGTTGCTTCTTCCGCAACCAGATAAAGGANNNAGATCGGAAG |
| 56943_1 | TATGAGTGTGGAGTCGTTGCAGCATACAGGGTCCTTCAAAGTTNNNAGATCGGAAG |
| 56950_1 | TATGAGTGTGGAGTCGTTGCAAAAAACACAAAAAACCCCCAGANNNAGATCGGAAG |
| 56951_1 | TATGAGTGTGGAGTCGTTGCGAGCCCAGACCAATCTAAGTGTCNNNAGATCGGAAG |
| 56952_1 | TATGAGTGTGGAGTCGTTGCATCTCAACTCACTACAACCTCCGNNNAGATCGGAAGA |
| 56953_1 | TATGAGTGTGGAGTCGTTGCTCTGGGCAACATAGGAAGACTTCNNNAGATCGGAAG |
| 56954_1 | TATGAGTGTGGAGTCGTTGCCAATCTTGGCTCATTGCATCCTCNNNAGATCGGAAGA |
| 56955_1 | TATGAGTGTGGAGTCGTTGCAAATGTAGTGGCACAATCTTGGCNNNAGATCGGAAG |
| 56957_1 | TATGAGTGTGGAGTCGTTGCCAATCTTGGCTCATTGCAAGCTCNNNAGATCGGAAGA |
| 56961_1 | TATGAGTGTGGAGTCGTTGCCCTCCTCCCAGATTCAAGGAATTNNNAGATCGGAAGA |
| 56962_1 | TATGAGTGTGGAGTCGTTGCGTAGAGACGGAGTTTCACCATGTNNNAGATCGGAAG |
| 56964_1 | TATGAGTGTGGAGTCGTTGCTCCTAGGCTCAAGCGATTTTCTTNNNAGATCGGAAGA |
| 56966_1 | TATGAGTGTGGAGTCGTTGCGCCTCCTGATGAGCTAGGATTACNNNAGATCGGAAG |
| 56967_1 | TATGAGTGTGGAGTCGTTGCGTTCACACTATTCTCCTGCCTCANNNAGATCGGAAGA |
| 56968_1 | TATGAGTGTGGAGTCGTTGCCCTCCCAAAGTGCTAGGATTACANNNAGATCGGAAG |
| 56969_1 | TATGAGTGTGGAGTCGTTGCTTCAAGCAATTTTCCTGCCTCAGNNNAGATCGGAAGA |
| 56970_1 | TATGAGTGTGGAGTCGTTGCCCTACTGGGTTCAAGCGATTCTCNNNAGATCGGAAGA |
| 56971_1 | TATGAGTGTGGAGTCGTTGCCCTCCCGAGTAGCTGAGATTACANNNAGATCGGAAG |
| 56975_1 | TATGAGTGTGGAGTCGTTGCAGCTAGGCACAGATTCCAGAAAANNNAGATCGGAAG |
| 56978_1 | TATGAGTGTGGAGTCGTTGCAGACCGCAGCAGAGCTATTATAGNNNAGATCGGAAG |
| 56983_1 | TATGAGTGTGGAGTCGTTGCGCTGGAGTGTAATGGTGTGATCTNNNAGATCGGAAG |
| 56985_1 | TATGAGTGTGGAGTCGTTGCTCTCCCTGGTTCAAGTGATTCTCNNNAGATCGGAAGA |
| 56986_1 | TATGAGTGTGGAGTCGTTGCCAAAGTGCTGGGATTAGAGGCATNNNAGATCGGAA |
| 56988_1 | TATGAGTGTGGAGTCGTTGCAGTAGTTGGAATTACAGGCGTGTNNNAGATCGGAAG |
| 56989_1 | TATGAGTGTGGAGTCGTTGCATACAGTGGCATAATCTCGGCTTNNNAGATCGGAAG |
| 56990_1 | TATGAGTGTGGAGTCGTTGCCAGCTCACTGCAATCTCCATTTCNNNAGATCGGAAGA |
| 56991_1 | TATGAGTGTGGAGTCGTTGCAAATTCCTGGGCTCAAGTGATCTNNNAGATCGGAAG |
| 56994_1 | TATGAGTGTGGAGTCGTTGCCTCCTGAGTAGCTGGGATTACACNNNAGATCGGAAG |
| 56995_1 | TATGAGTGTGGAGTCGTTGCATTCATGTGATTCTCCTGCCTCANNNAGATCGGAAGA |
| 56996_1 | TATGAGTGTGGAGTCGTTGCCCTCTTGGGTTCATGCCATTCTCNNNAGATCGGAAGA |
| 56997_1 | TATGAGTGTGGAGTCGTTGCCAGCCTCCTGAGTAATTGGGATTNNNAGATCGGAAG |
| 56999_1 | TATGAGTGTGGAGTCGTTGCTCCTGAGTAGCTGGAACTACAGANNNAGATCGGAAG |
| 57000_1 | TATGAGTGTGGAGTCGTTGCGTTTCACTGTGTTAGCCAGGATGNNNAGATCGGAAG |
| 57001_1 | TATGAGTGTGGAGTCGTTGCCTCTGGAGTAGCTGGGATTACAGNNNAGATCGGAAG |
| 57004_1 | TATGAGTGTGGAGTCGTTGCCCTGAGTAGCTAGGACTACAGGTNNNAGATCGGAAG |
| 57005_1 | TATGAGTGTGGAGTCGTTGCACGCCCGGCTAATTTTTTGTATTNNNAGATCGGAAGA |
| 57010_1 | TATGAGTGTGGAGTCGTTGCCTCTTGAGTAGCTGGGACTACAGNNNAGATCGGAAG |
| 57011_1 | TATGAGTGTGGAGTCGTTGCACCTCCTGAGTTCAAGCAATTCTNNNAGATCGGAAGA |
| 57013_1 | TATGAGTGTGGAGTCGTTGCCCTCCCAAAGTGCTAGGATTACANNNAGATCGGAAG |
| 57014_1 | TATGAGTGTGGAGTCGTTGCCCTCCTGGGTTGAAGCAATTCTTNNNAGATCGGAAGA |
| 57015_1 | TATGAGTGTGGAGTCGTTGCCTCCCATGTAGCTGGAACTACAGNNNAGATCGGAAG |
| 57016_1 | TATGAGTGTGGAGTCGTTGCGAGGGTTGTGAAGTCAGAAAAGCNNNAGATCGGAA |
| 57017_1 | TATGAGTGTGGAGTCGTTGCCTTGGGAAGTTGAGGTGAGAAGANNNAGATCGGAA |

| | |
|---|---|
| 57018_1 | TATGAGTGTGGAGTCGTTGCATTCTCAAGTGGAGTGACTTTGCNNNAGATCGGAAG |
| 57019_1 | TATGAGTGTGGAGTCGTTGCCATTGGTCCTGGCCACTAAAATGNNNAGATCGGAAG |
| 57020_1 | TATGAGTGTGGAGTCGTTGCCTGTAATCCCAGCTACTCAGGAGNNNAGATCGGAAG |
| 57022_1 | TATGAGTGTGGAGTCGTTGCCCTGTAATCTCAGCACTTTGGGANNNAGATCGGAAG |
| 57023_1 | TATGAGTGTGGAGTCGTTGCTGAGGCAGGAGAATTGCTTACATNNNAGATCGGAA |
| 57026_1 | TATGAGTGTGGAGTCGTTGCCAGGAGATTCCCACCTCTTTCAANNNAGATCGGAAG |
| 57028_1 | TATGAGTGTGGAGTCGTTGCACAACTCAGCCTAGAAATGAGGANNNAGATCGGAA |
| 57029_1 | TATGAGTGTGGAGTCGTTGCTCTGTGAATGCTTATTTGTGGCTNNNAGATCGGAAG |
| 57030_1 | TATGAGTGTGGAGTCGTTGCGAGGTAGACATGCAGAGATGGAGNNNAGATCGGA |
| 57034_1 | TATGAGTGTGGAGTCGTTGCTCACTCTCCCCTAGTGGAAATCTNNNAGATCGGAAG |
| 57039_1 | TATGAGTGTGGAGTCGTTGCAAAAGACTGATTGTTCTGTGGCCNNNAGATCGGAA |
| 57041_1 | TATGAGTGTGGAGTCGTTGCAGTGTGAGCGATACACAAAAAGCNNNAGATCGGAA |
| 57043_1 | TATGAGTGTGGAGTCGTTGCAGGGGAGGAAACAAATAGTGACTNNNAGATCGGA |
| 57044_1 | TATGAGTGTGGAGTCGTTGCCATCCTGGCTAACATGGTGAAACNNNAGATCGGAAG |
| 57046_1 | TATGAGTGTGGAGTCGTTGCACACAGTAAAACCCCGTCTCTACNNNAGATCGGAAG |
| 57047_1 | TATGAGTGTGGAGTCGTTGCCACACCTGTAATCCCAGCACTTTNNNAGATCGGAAG |
| 57048_1 | TATGAGTGTGGAGTCGTTGCGCATTTTGAAGTCTGCTCTTGCTNNNAGATCGGAAG |
| 57049_1 | TATGAGTGTGGAGTCGTTGCTTGACAAAGCAAAGTACCAGAGCNNNAGATCGGAA |
| 57050_1 | TATGAGTGTGGAGTCGTTGCACAGTGACTCACACCTGTAATCCNNNAGATCGGAAG |
| 57052_1 | TATGAGTGTGGAGTCGTTGCGAGAACAGTTTGGAGGTTCCTGANNNAGATCGGAA |
| 57053_1 | TATGAGTGTGGAGTCGTTGCCTGAGATCAGGAGTTCGAAACCANNNAGATCGGAA |
| 57054_1 | TATGAGTGTGGAGTCGTTGCTTTTTCACACACACACACACACANNNAGATCGGAAGA |
| 57055_1 | TATGAGTGTGGAGTCGTTGCTGAGTCGGAACCCAATCTAGAACNNNAGATCGGAA |
| 57056_1 | TATGAGTGTGGAGTCGTTGCTGGAGGCTACAGTGAGGTTTTTTNNNAGATCGGAA |
| 57057_1 | TATGAGTGTGGAGTCGTTGCGAGAATCACTTGAACCCAGGAGGNNNAGATCGGAA |
| 57058_1 | TATGAGTGTGGAGTCGTTGCCCTAAGCTAAGTCTGGAACAGCANNNAGATCGGAA |
| 57059_1 | TATGAGTGTGGAGTCGTTGCAAGAAAGAAATCTTGCAGCTGGGNNNAGATCGGAA |
| 57060_1 | TATGAGTGTGGAGTCGTTGCACTGAAAGAAAATTGTGGGCTGGNNNAGATCGGAA |
| 57061_1 | TATGAGTGTGGAGTCGTTGCTGAGAGCAATTTGAGATTTGGCCNNNAGATCGGAA |
| 57062_1 | TATGAGTGTGGAGTCGTTGCAAACATTATGTCCACTCCTCCCCNNNAGATCGGAAGA |
| 57063_1 | TATGAGTGTGGAGTCGTTGCAAGAGGGCAGAGATTCAAGACAGNNNAGATCGGA |
| 57064_1 | TATGAGTGTGGAGTCGTTGCATCTCAGTTCTCACTCCAGACCTNNNAGATCGGAAGA |
| 57068_1 | TATGAGTGTGGAGTCGTTGCTCAGGAAGCTAAGGCACAAGAATNNNAGATCGGAA |
| 57072_1 | TATGAGTGTGGAGTCGTTGCTGTAAAGAGCTTAGACCGGTACCNNNAGATCGGAA |
| 57073_1 | TATGAGTGTGGAGTCGTTGCTCCTTTTTTGACTCTCCAGAGCCNNNAGATCGGAAGA |
| 57074_1 | TATGAGTGTGGAGTCGTTGCCCTTACAAGATACCAACCCTGCTNNNAGATCGGAAG |
| 57077_1 | TATGAGTGTGGAGTCGTTGCAGGTCTCCTCTCACTGTAACTGANNNAGATCGGAAG |
| 57079_1 | TATGAGTGTGGAGTCGTTGCAATGCTTTTACACTGTTGGTGGGNNNAGATCGGAA |
| 57081_1 | TATGAGTGTGGAGTCGTTGCAGAGAAGAGAAGAGAAGGGGAGGNNNAGATCGG |
| 57082_1 | TATGAGTGTGGAGTCGTTGCGAAACAAATAGGCCTCTTCCAGCNNNAGATCGGAAG |
| 57083_1 | TATGAGTGTGGAGTCGTTGCATGGCAACTCAAACAAAACCCAANNNAGATCGGAAG |
| 57084_1 | TATGAGTGTGGAGTCGTTGCAACATGCTCAACATCACCAATCGNNNAGATCGGAAG |
| 57085_1 | TATGAGTGTGGAGTCGTTGCCAGCGCTTGATGAGACTCTGTAANNNAGATCGGAA |

| | |
|---|---|
| 57089_1 | TATGAGTGTGGAGTCGTTGCTTCACACCATTCTCCTGCCTTAGNNNAGATCGGAAGA |
| 57090_1 | TATGAGTGTGGAGTCGTTGCACAGTGTAGTGGTGCAATCATGGNNNAGATCGGAA |
| 57093_1 | TATGAGTGTGGAGTCGTTGCAAGCTTCCGGAGTATCTAGGACTNNNAGATCGGAAG |
| 57094_1 | TATGAGTGTGGAGTCGTTGCCAAGTAGCTGGGATTACAGGCATNNNAGATCGGAA |
| 57095_1 | TATGAGTGTGGAGTCGTTGCCCTCCTGGGTTCAAGTGATTCTGNNNAGATCGGAAG |
| 57098_1 | TATGAGTGTGGAGTCGTTGCGGAAACTCCAAGTCTCCTGTGAANNNAGATCGGAAG |
| 57100_1 | TATGAGTGTGGAGTCGTTGCCCTGCCAAGTAGTTGGGATTACANNNAGATCGGAAG |
| 57102_1 | TATGAGTGTGGAGTCGTTGCATGATTATCCTGCTTCAGCCTCCNNNAGATCGGAAGA |
| 57103_1 | TATGAGTGTGGAGTCGTTGCGATCTTGGCTCACTGCAACCTTTNNNAGATCGGAAG |
| 57105_1 | TATGAGTGTGGAGTCGTTGCGATTCTTCTGCCTGAGACTCCTGNNNAGATCGGAAG |
| 57109_1 | TATGAGTGTGGAGTCGTTGCGAGTAGCTAGGATTACAGGCACCNNNAGATCGGAA |
| 57110_1 | TATGAGTGTGGAGTCGTTGCTGTTGTCCATGCTTGATTCCAACNNNAGATCGGAAG |
| 57112_1 | TATGAGTGTGGAGTCGTTGCGGACTATAGGCATGCACCATCATNNNAGATCGGAAG |
| 57113_1 | TATGAGTGTGGAGTCGTTGCCGGAATGGGTACCTCTGTCATACNNNAGATCGGAAG |
| 57114_1 | TATGAGTGTGGAGTCGTTGCCCTCCAAAGTAGCTGGGATTACANNNAGATCGGAAG |
| 57115_1 | TATGAGTGTGGAGTCGTTGCCTCCCAAGTAGCTGGAACTACAGNNNAGATCGGAAG |
| 57116_1 | TATGAGTGTGGAGTCGTTGCTTCAAGTGATTCTCCTGCCTCAGNNNAGATCGGAAG |
| 57118_1 | TATGAGTGTGGAGTCGTTGCGATTCAAGCAATTCTCCTGCCTCNNNAGATCGGAAG |
| 57119_1 | TATGAGTGTGGAGTCGTTGCGTTCATACCATTCTCCTGCCTCANNNAGATCGGAAGA |
| 57122_1 | TATGAGTGTGGAGTCGTTGCTCATCTGCATGCATCAGAAAGTCNNNAGATCGGAAG |
| 57123_1 | TATGAGTGTGGAGTCGTTGCCAGGATGGTCTCGATCTCTTGACNNNAGATCGGAAG |
| 57124_1 | TATGAGTGTGGAGTCGTTGCGCCTCCCAAGTAGCTAGGATTACNNNAGATCGGAAG |
| 57127_1 | TATGAGTGTGGAGTCGTTGCATCGAGTAGCTGAGACTACAGGTNNNAGATCGGAA |
| 57128_1 | TATGAGTGTGGAGTCGTTGCCTGGGATTATAGGCGTGAGCTACNNNAGATCGGAA |
| 57129_1 | TATGAGTGTGGAGTCGTTGCAGACAGGGTTTCACTTGTTACCCNNNAGATCGGAAG |
| 57130_1 | TATGAGTGTGGAGTCGTTGCCCTCTTGAGTAGCTGGGACTAGANNNAGATCGGAA |
| 57132_1 | TATGAGTGTGGAGTCGTTGCAACAGCTTTCTGGCGTACTTAGANNNAGATCGGAAG |
| 57134_1 | TATGAGTGTGGAGTCGTTGCTGTTTGGAATGGAGACCAGGAAANNNAGATCGGAA |
| 57135_1 | TATGAGTGTGGAGTCGTTGCAGGAGGAGGCTACTTTCCATTTGNNNAGATCGGAA |
| 57137_1 | TATGAGTGTGGAGTCGTTGCATGAAGTTGAACCCCTACCTCACNNNAGATCGGAAG |
| 57138_1 | TATGAGTGTGGAGTCGTTGCAGGCAGGAGAATAGTCTGAAACGNNNAGATCGGAA |
| 57140_1 | TATGAGTGTGGAGTCGTTGCGCATTGTCCTGCTCAAAAACCTTNNNAGATCGGAAG |
| 57141_1 | TATGAGTGTGGAGTCGTTGCGAAAAAAATAGGCTGAGCACGGTNNNAGATCGGAA |
| 57144_1 | TATGAGTGTGGAGTCGTTGCAGGCTGCATTGAACTATGATTGCNNNAGATCGGAA |
| 57146_1 | TATGAGTGTGGAGTCGTTGCCTGTAGTCCCAGCTTCTAGGGAGNNNAGATCGGAA |
| 57147_1 | TATGAGTGTGGAGTCGTTGCTCTGGTCTCAGTTACTTGGAAGCNNNAGATCGGAAG |
| 57148_1 | TATGAGTGTGGAGTCGTTGCCCAAGTTGCTGCAAAAGAGGTTANNNAGATCGGAA |
| 57149_1 | TATGAGTGTGGAGTCGTTGCGCAAGGTTGTGGAGAAAATGGAANNNAGATCGGA |
| 57150_1 | TATGAGTGTGGAGTCGTTGCCACCATGTGAAGTCTCAGTGAGANNNAGATCGGAA |
| 57152_1 | TATGAGTGTGGAGTCGTTGCTGTTGAGATGGGAGGATCACTTGNNNAGATCGGAA |
| 57153_1 | TATGAGTGTGGAGTCGTTGCCTCCTGACCACAATGCCATATCTNNNAGATCGGAAGA |
| 57155_1 | TATGAGTGTGGAGTCGTTGCATAGCCTAGGCCAGTATGTCTGANNNAGATCGGAA |
| 57156_1 | TATGAGTGTGGAGTCGTTGCATACAAAAAAATTAGCCGGGCGTNNNAGATCGGAA |

| | |
|---|---|
| 57157_1 | TATGAGTGTGGAGTCGTTGCGCAAACCTAGGGTGGTAGGTAAANNNAGATCGGAA |
| 57158_1 | TATGAGTGTGGAGTCGTTGCATACGGAAAAAGATGTCTGCCCTNNNAGATCGGAAG |
| 57159_1 | TATGAGTGTGGAGTCGTTGCTGTCATTGTGGATGAACAGCAACNNNAGATCGGAAG |
| 57160_1 | TATGAGTGTGGAGTCGTTGCCTCACAGACAGGGCCATTATGAGNNNAGATCGGAAG |
| 57161_1 | TATGAGTGTGGAGTCGTTGCGCTTTGCTAGAATCCCTCTGAGTNNNAGATCGGAAGA |
| 57162_1 | TATGAGTGTGGAGTCGTTGCATGGAATTGCTGGGTCATATGGTNNNAGATCGGAAG |
| 57163_1 | TATGAGTGTGGAGTCGTTGCCAGTGAGCTGAGATCAAACCTCTNNNAGATCGGAAG |
| 57164_1 | TATGAGTGTGGAGTCGTTGCACATGAGGCTACTAACCTGGAAGNNNAGATCGGAAG |
| 57166_1 | TATGAGTGTGGAGTCGTTGCAAGAAAGAAAGCCAAACCAACGANNNAGATCGGAAG |
| 57167_1 | TATGAGTGTGGAGTCGTTGCACTTTTACACTGTTGGTGGGACTNNNAGATCGGAAG |
| 57168_1 | TATGAGTGTGGAGTCGTTGCCACACCTGTAATCCCAGCACTTTNNNAGATCGGAAGA |
| 57169_1 | TATGAGTGTGGAGTCGTTGCGATTGCTTGATCCAAGGAGGTTGNNNAGATCGGAAG |
| 57171_1 | TATGAGTGTGGAGTCGTTGCATTACCCTGAGGAGTCAGAGGATNNNAGATCGGAAG |
| 57172_1 | TATGAGTGTGGAGTCGTTGCACTGATGGTGTTTACCAATCCCANNNAGATCGGAAGA |
| 57174_1 | TATGAGTGTGGAGTCGTTGCCGCTTGTAATCCCAGCTAACTTGNNNAGATCGGAAGA |
| 57176_1 | TATGAGTGTGGAGTCGTTGCGGATCATCAGCTGAATTGGATGCNNNAGATCGGAAG |
| 57180_1 | TATGAGTGTGGAGTCGTTGCAATTCCTGGCCTCAAGTGATTCTNNNAGATCGGAAGA |
| 57181_1 | TATGAGTGTGGAGTCGTTGCCTCACTATGTTACCCAAGCTGGTNNNAGATCGGAAGA |
| 57182_1 | TATGAGTGTGGAGTCGTTGCCTACCAAGGAGCTGGGACTAGAGNNNAGATCGGAA |
| 57183_1 | TATGAGTGTGGAGTCGTTGCTGTTTCATTTTGCACTGGGATCCNNNAGATCGGAAGA |
| 57184_1 | TATGAGTGTGGAGTCGTTGCCTGAGTAGCTGGGATTACAGGTGNNNAGATCGGAA |
| 57185_1 | TATGAGTGTGGAGTCGTTGCCTGGGACTATAGATGAGCACCACNNNAGATCGGAAG |
| 57186_1 | TATGAGTGTGGAGTCGTTGCTGTAATGGTGTGATCTTGGCTCANNNAGATCGGAAG |
| 57187_1 | TATGAGTGTGGAGTCGTTGCAGATGGTGTTTTGCTCTTGTTGCNNNAGATCGGAAG |
| 57188_1 | TATGAGTGTGGAGTCGTTGCAATCTCAGTTCACTGCAACCTCTNNNAGATCGGAAGA |
| 57190_1 | TATGAGTGTGGAGTCGTTGCTCCCAGACTCAAACCATTCTTCCNNNAGATCGGAAGA |
| 57191_1 | TATGAGTGTGGAGTCGTTGCGTACAGGCTCAAAAGGGAAACACNNNAGATCGGAAG |
| 57194_1 | TATGAGTGTGGAGTCGTTGCGTTCAAGAGATTCTCCTGCCTCANNNAGATCGGAAGA |
| 57196_1 | TATGAGTGTGGAGTCGTTGCACAAACGCCTCTAGAACCTTTCTNNNAGATCGGAAGA |
| 57197_1 | TATGAGTGTGGAGTCGTTGCCTGAAAGCTCTGCCTCCTGTATTNNNAGATCGGAAGA |
| 57198_1 | TATGAGTGTGGAGTCGTTGCTTCTTTGAAGAGCCTGTTGGAGTNNNAGATCGGAAG |
| 57200_1 | TATGAGTGTGGAGTCGTTGCCTGTCCTTACGAAATCCTCCCTCNNNAGATCGGAAGA |
| 57203_1 | TATGAGTGTGGAGTCGTTGCGTCCCGAGTAGCTAGGACTACAGNNNAGATCGGAAG |
| 57204_1 | TATGAGTGTGGAGTCGTTGCGCAAGATCTTGGCTCACTGAAAANNNAGATCGGAAG |
| 57207_1 | TATGAGTGTGGAGTCGTTGCACCTTCATTTCTGTGGGTCACTTNNNAGATCGGAAGA |
| 57208_1 | TATGAGTGTGGAGTCGTTGCCTCCTGAGTAGCTGGGACTACAGNNNAGATCGGAAG |
| 57209_1 | TATGAGTGTGGAGTCGTTGCGGGTTCACACCATTCTGTCTCAGNNNAGATCGGAAGA |
| 57210_1 | TATGAGTGTGGAGTCGTTGCAGTAGCTAGGATTACAGGCATGCNNNAGATCGGAAG |
| 57212_1 | TATGAGTGTGGAGTCGTTGCTCTCCTGGGTTCATGCAATTCTCNNNAGATCGGAAGA |
| 57213_1 | TATGAGTGTGGAGTCGTTGCATGGAAAGCAAGCTGCCATAAAGNNNAGATCGGAAG |
| 57215_1 | TATGAGTGTGGAGTCGTTGCAGATGGAGTCTCTCTCTCTGTCGNNNAGATCGGAAG |
| 57217_1 | TATGAGTGTGGAGTCGTTGCTTCTTGAAGGAGGCAGCTTACTTNNNAGATCGGAAG |
| 57218_1 | TATGAGTGTGGAGTCGTTGCGGAATGGAACTGGGAATCAGAGTNNNAGATCGGAA |

| | |
|---|---|
| 57219_1 | TATGAGTGTGGAGTCGTTGCGCATTACTGTATGATGCAGCAGTNNNAGATCGGAAG |
| 57220_1 | TATGAGTGTGGAGTCGTTGCTCACGCGGGATGAATAAGTCTAGNNNAGATCGGAA |
| 57222_1 | TATGAGTGTGGAGTCGTTGCAAAGGGAACCTGGATCTCAGAACNNNAGATCGGAA |
| 57223_1 | TATGAGTGTGGAGTCGTTGCCCAGCCCCTCTCATGAGAATATTNNNAGATCGGAAG |
| 57224_1 | TATGAGTGTGGAGTCGTTGCGTCAGAAAGAGAGAACACCGTCTNNNAGATCGGAA |
| 57225_1 | TATGAGTGTGGAGTCGTTGCACACTCCTGAACTTTAGGCCTTTNNNAGATCGGAAG |
| 57228_1 | TATGAGTGTGGAGTCGTTGCGATCACACCACTATACTCCAGCCNNNAGATCGGAAG |
| 57229_1 | TATGAGTGTGGAGTCGTTGCTTCACCACGGTATGATTCCACTTNNNAGATCGGAAG |
| 57230_1 | TATGAGTGTGGAGTCGTTGCTAAGGCTGAAGTGAGCTGTGATCNNNAGATCGGAA |
| 57231_1 | TATGAGTGTGGAGTCGTTGCTGTCTGATCTTCGACAAAGCTGANNNAGATCGGAAG |
| 57234_1 | TATGAGTGTGGAGTCGTTGCGCTCAGATTCTATCCTGGGCTTTNNNAGATCGGAAG |
| 57235_1 | TATGAGTGTGGAGTCGTTGCGCTGTCTGACCCTACCTTTTTCTNNNAGATCGGAAGA |
| 57237_1 | TATGAGTGTGGAGTCGTTGCGGTCCATGTGTTTGTTTGTTTGTGNNNAGATCGGAA |
| 57239_1 | TATGAGTGTGGAGTCGTTGCTTACTGAAGTAAGGGAGGGTCCTNNNAGATCGGAA |
| 57240_1 | TATGAGTGTGGAGTCGTTGCTAGTGGAAGAGACCATTTGGCTCNNNAGATCGGAA |
| 57244_1 | TATGAGTGTGGAGTCGTTGCAAATACAAAAATTAGCCGGGCGTNNNAGATCGGAA |
| 57245_1 | TATGAGTGTGGAGTCGTTGCTCCTATGACTCCTTTACTCACTCANNNAGATCGGAAG |
| 57246_1 | TATGAGTGTGGAGTCGTTGCAGAGTGAGGTGTTGCAGAAAAGANNNAGATCGGA |
| 57247_1 | TATGAGTGTGGAGTCGTTGCAGGCGAAGTCAATGTACCAGTTANNNAGATCGGAA |
| 57250_1 | TATGAGTGTGGAGTCGTTGCGGAGGATGGGATTTGAGGTGTTANNNAGATCGGA |
| 57251_1 | TATGAGTGTGGAGTCGTTGCTTCTCATGCATGGGGATTCTTGANNNAGATCGGAAG |
| 57253_1 | TATGAGTGTGGAGTCGTTGCTATGTGAAGTGGAAGAAGCCAGANNNAGATCGGAA |
| 57254_1 | TATGAGTGTGGAGTCGTTGCGCATCTCAATCTAGGAAAACCGCNNNAGATCGGAAG |
| 57258_1 | TATGAGTGTGGAGTCGTTGCCCAGAGATGAAGGATGAAGGCTTNNNAGATCGGAA |
| 57259_1 | TATGAGTGTGGAGTCGTTGCCCCAGCTGTTAGTTTTTCTTGCANNNAGATCGGAAG |
| 57260_1 | TATGAGTGTGGAGTCGTTGCGCATGGCAATGGCAGTATGTTTANNNAGATCGGAA |
| 57261_1 | TATGAGTGTGGAGTCGTTGCATGACTGTTGACTTGGGGAAGTTNNNAGATCGGAA |
| 57262_1 | TATGAGTGTGGAGTCGTTGCCAGACTGAGAGCTGAAGTTTCCANNNAGATCGGAA |
| 57264_1 | TATGAGTGTGGAGTCGTTGCACTGCATTGAGGGTTATGGTGATNNNAGATCGGAA |
| 57266_1 | TATGAGTGTGGAGTCGTTGCCCTGAAAATGTGATTCTGCTGCTNNNAGATCGGAAG |
| 57269_1 | TATGAGTGTGGAGTCGTTGCTGGGATTACAGACATGCACTACCNNNAGATCGGAAG |
| 57270_1 | TATGAGTGTGGAGTCGTTGCCAATGGTGCAATCTCGGCTTATTNNNAGATCGGAAG |
| 57271_1 | TATGAGTGTGGAGTCGTTGCGAAGGAGCTTCATTGAGCAACAGNNNAGATCGGAA |
| 57272_1 | TATGAGTGTGGAGTCGTTGCGGCTGGCTTGAGTTTTTCTTCTTNNNAGATCGGAAG |
| 57274_1 | TATGAGTGTGGAGTCGTTGCGGTTCAGATGATCCTACCTCAGCNNNAGATCGGAAG |
| 57275_1 | TATGAGTGTGGAGTCGTTGCCTTCCAGTCCAAGCAATTCTCCTNNNAGATCGGAAGA |
| 57276_1 | TATGAGTGTGGAGTCGTTGCTTCAAGCTATTCTCCTGCCTCAGNNNAGATCGGAAGA |
| 57280_1 | TATGAGTGTGGAGTCGTTGCTTCAAGTGATTCTCCTGCCTCAGNNNAGATCGGAAG |
| 57281_1 | TATGAGTGTGGAGTCGTTGCAATCTTGGCTCACTACAACCTCCNNNAGATCGGAAGA |
| 57282_1 | TATGAGTGTGGAGTCGTTGCCAATCTTGACTCACTGCAAGCTCNNNAGATCGGAAG |
| 57283_1 | TATGAGTGTGGAGTCGTTGCCTGTAAATTGTGTGGCAGGTTCANNNAGATCGGAA |
| 57285_1 | TATGAGTGTGGAGTCGTTGCCCACCTCCTGAGTTCTAGCAATTNNNAGATCGGAAG |
| 57288_1 | TATGAGTGTGGAGTCGTTGCCAGGATGCTCTCAAAGTGATCCTNNNAGATCGGAAG |

| | |
|---|---|
| 57290_1 | TATGAGTGTGGAGTCGTTGCCTGTGCCTGGTCTGTTCTTTTTTNNNAGATCGGAAGA |
| 57292_1 | TATGAGTGTGGAGTCGTTGCACAGTGGTACGATCTTAGCTCACNNNAGATCGGAAG |
| 57293_1 | TATGAGTGTGGAGTCGTTGCGAATAACTGAGACTACAGGCGCTNNNAGATCGGAAG |
| 57294_1 | TATGAGTGTGGAGTCGTTGCCCTCCCAAGTAGCTGTGATTACANNNAGATCGGAAGA |
| 57297_1 | TATGAGTGTGGAGTCGTTGCGACTACAGGCTTCCACCACTTAGNNNAGATCGGAAGA |
| 57298_1 | TATGAGTGTGGAGTCGTTGCAGTAGCTGGGATTACAGGAATGCNNNAGATCGGAA |
| 57299_1 | TATGAGTGTGGAGTCGTTGCCAAGTTCCACCTCCCAAGTTCATNNNAGATCGGAAGA |
| 57300_1 | TATGAGTGTGGAGTCGTTGCGACCTTCCCACATCAGCTTTCTANNNAGATCGGAAGA |
| 57301_1 | TATGAGTGTGGAGTCGTTGCGGATGTTCTTGCCTCTGACAGTANNNAGATCGGAAG |
| 57303_1 | TATGAGTGTGGAGTCGTTGCTTCCAGTTTGGGGCTATTCTGAANNNAGATCGGAAG |
| 57304_1 | TATGAGTGTGGAGTCGTTGCTTTTTAGTGGAGAGGGCTTCACCNNNAGATCGGAAG |
| 57305_1 | TATGAGTGTGGAGTCGTTGCCTTCAACTGATTCTCCTGCCTCANNNAGATCGGAAGA |
| 57306_1 | TATGAGTGTGGAGTCGTTGCCTCCGATTTCAAGCGATTCTCCTNNNAGATCGGAAGA |
| 57307_1 | TATGAGTGTGGAGTCGTTGCCTCAAGTGATCCTCCTGCCTTAGNNNAGATCGGAAGA |
| 57310_1 | TATGAGTGTGGAGTCGTTGCTGGCATGATCTTGACTCACTGTANNNAGATCGGAAG |
| 57315_1 | TATGAGTGTGGAGTCGTTGCAGTGCTGGGATTATAGGTGTGAGNNNAGATCGGAA |
| 57316_1 | TATGAGTGTGGAGTCGTTGCTTGATTCTACCCATCCATGAGCANNNAGATCGGAAGA |
| 57317_1 | TATGAGTGTGGAGTCGTTGCGATTTCGGTTCACTGCAAGCTCNNNAGATCGGAAGA |
| 57318_1 | TATGAGTGTGGAGTCGTTGCCCTTCTCCTGGGTTCAAGTGATTNNNAGATCGGAAGA |
| 57319_1 | TATGAGTGTGGAGTCGTTGCAAGGAGTAGTTGAAGCTTCCAGGNNNAGATCGGAA |
| 57320_1 | TATGAGTGTGGAGTCGTTGCTCAGAGCTTGTGTGATTATGGCTNNNAGATCGGAAG |
| 57322_1 | TATGAGTGTGGAGTCGTTGCCAAGTAGCTGGGACTATAGGTGCNNNAGATCGGAA |
| 57323_1 | TATGAGTGTGGAGTCGTTGCAGTAGCTAGGATTACAGGCATGCNNNAGATCGGAA |
| 57324_1 | TATGAGTGTGGAGTCGTTGCGTCTTACTGAGCTTTTTGGGGGANNNAGATCGGAAG |
| 57325_1 | TATGAGTGTGGAGTCGTTGCTACAGGCGGGAAACATTTCTCAANNNAGATCGGAAG |
| 57327_1 | TATGAGTGTGGAGTCGTTGCTGAGTAACTGGGATTACAGGCACNNNAGATCGGAA |
| 57330_1 | TATGAGTGTGGAGTCGTTGCCTCCCAGATTCAAGCCATTCTCTNNNAGATCGGAAGA |
| 57332_1 | TATGAGTGTGGAGTCGTTGCATCCAGGAATCCTACTTCTGGGANNNAGATCGGAAG |
| 57333_1 | TATGAGTGTGGAGTCGTTGCGTTGGAAAGGCTTCAGGATTGTCNNNAGATCGGAA |
| 57335_1 | TATGAGTGTGGAGTCGTTGCTAATCTCAGAACTTTGGGAGGCCNNNAGATCGGAAG |
| 57337_1 | TATGAGTGTGGAGTCGTTGCTCCTTGCTCTCCACTTCACATTTNNNAGATCGGAAGA |
| 57338_1 | TATGAGTGTGGAGTCGTTGCCAGAGCTTAGAGTGAGCCAAGATNNNAGATCGGAA |
| 57339_1 | TATGAGTGTGGAGTCGTTGCGGAGTCTTTTGGAGAGGTAGGACNNNAGATCGGAA |
| 57340_1 | TATGAGTGTGGAGTCGTTGCCTGCTTATTCTGCCTAACCATGCNNNAGATCGGAAGA |
| 57341_1 | TATGAGTGTGGAGTCGTTGCGACAAAGCAGTGGAATGTTACCCNNNAGATCGGAA |
| 57342_1 | TATGAGTGTGGAGTCGTTGCGTCAGAGGAATGTGCATGTGTTTNNNAGATCGGAA |
| 57343_1 | TATGAGTGTGGAGTCGTTGCTGCTGATGACCTTTGAAACACTGNNNAGATCGGAAG |
| 57344_1 | TATGAGTGTGGAGTCGTTGCTGTAGCAGAATGTCTTCCAAGGGNNNAGATCGGAA |
| 57346_1 | TATGAGTGTGGAGTCGTTGCAAGCTTTACCCCTCTTTTCAGCTNNNAGATCGGAAGA |
| 57348_1 | TATGAGTGTGGAGTCGTTGCTAGTTTGCACTGGTTGAACACACNNNAGATCGGAAG |
| 57349_1 | TATGAGTGTGGAGTCGTTGCACATGTTCGCTGAAAGATTTGCTNNNAGATCGGAAG |
| 57352_1 | TATGAGTGTGGAGTCGTTGCATACCCCAGGGGAGTGAAAAAAANNNAGATCGGAA |
| 57354_1 | TATGAGTGTGGAGTCGTTGCCCTTGGAGCATAAGACATGAAGCNNNAGATCGGAA |

| | |
|---|---|
| 57357_1 | TATGAGTGTGGAGTCGTTGCGCAGTGGTTGTTGAGTGATTGAANNNAGATCGGAA |
| 57358_1 | TATGAGTGTGGAGTCGTTGCATAAGAATGGAGGCAAGGACACANNNAGATCGGA |
| 57359_1 | TATGAGTGTGGAGTCGTTGCAAAAGAGGGAGGAATGAAAGGGGNNNAGATCGG |
| 57362_1 | TATGAGTGTGGAGTCGTTGCCACATAAGGCACGAAAAACAGGTNNNAGATCGGAA |
| 57364_1 | TATGAGTGTGGAGTCGTTGCGAATTTTCAATTGGAGGCCTGGTNNNAGATCGGAA |
| 57366_1 | TATGAGTGTGGAGTCGTTGCATTGCAGACTTTACTCTCCTGGGNNNAGATCGGAAG |
| 57367_1 | TATGAGTGTGGAGTCGTTGCGTAGTGGTTCACGCATGTAATCCNNNAGATCGGAAG |
| 57369_1 | TATGAGTGTGGAGTCGTTGCAAAGCAATAGAGACTTGGTTCCANNNAGATCGGAA |
| 57371_1 | TATGAGTGTGGAGTCGTTGCTTTTTTCTTTGAACAGCGCTTGCNNNAGATCGGAAG |
| 57372_1 | TATGAGTGTGGAGTCGTTGCAGAATTTGGGTGAGCACAAAGTGNNNAGATCGGAA |
| 57373_1 | TATGAGTGTGGAGTCGTTGCCACAGTGGCTCATGTTTGTAGTCNNNAGATCGGAAG |
| 57374_1 | TATGAGTGTGGAGTCGTTGCCAATGTGGCAAACATCATGGCTANNNAGATCGGAA |
| 57376_1 | TATGAGTGTGGAGTCGTTGCGACTTTCAGACTCTGTTCCTGGTNNNAGATCGGAAG |
| 57377_1 | TATGAGTGTGGAGTCGTTGCTCACACATACTCCAAACCCTCTGNNNAGATCGGAAGA |
| 57385_1 | TATGAGTGTGGAGTCGTTGCATCCCAGTACTTCAGGAGACTGANNNAGATCGGAAG |
| 57388_1 | TATGAGTGTGGAGTCGTTGCGCACCGCTTCCCAGATAATTTTTNNNAGATCGGAAG |
| 57390_1 | TATGAGTGTGGAGTCGTTGCAAAGCCAAAGACGAGGAGAGAATNNNAGATCGGA |
| 57391_1 | TATGAGTGTGGAGTCGTTGCCTGAGGCAGGAGAATCGTTTGAANNNAGATCGGAA |
| 57394_1 | TATGAGTGTGGAGTCGTTGCAGAAATAAGGCCAAGGTGGTGATNNNAGATCGGA |
| 57396_1 | TATGAGTGTGGAGTCGTTGCCCGTTTTCTCACTGCTTTCCTTTNNNAGATCGGAAGA |
| 57397_1 | TATGAGTGTGGAGTCGTTGCCCTCAGTGCAGAAGGAACATCTANNNAGATCGGAA |
| 57399_1 | TATGAGTGTGGAGTCGTTGCTGCAAAGGTCTTAGGACAGGAAANNNAGATCGGAA |
| 57400_1 | TATGAGTGTGGAGTCGTTGCTGATAGTGGAATCAGGAGCCTGANNNAGATCGGAA |
| 57401_1 | TATGAGTGTGGAGTCGTTGCTCTGCTTTCTTCCTTTCATGGGANNNAGATCGGAAGA |
| 57403_1 | TATGAGTGTGGAGTCGTTGCGTAATCCTAGCTACTTGGGAGGCNNNAGATCGGAA |
| 57404_1 | TATGAGTGTGGAGTCGTTGCATCATCAGGGTAAGAGGGAAAGCNNNAGATCGGA |
| 57406_1 | TATGAGTGTGGAGTCGTTGCAAATGTCGGGGACTGCTGATTTANNNAGATCGGAA |
| 57407_1 | TATGAGTGTGGAGTCGTTGCAAGGGAGAGAAAAATGAAAGGCANNNAGATCGGA |
| 57408_1 | TATGAGTGTGGAGTCGTTGCTTAGCCAGTGTCCACTTTCCATTNNNAGATCGGAAGA |
| 57409_1 | TATGAGTGTGGAGTCGTTGCTAGGGATGAGGGAGATGAGAACTNNNAGATCGGA |
| 57411_1 | TATGAGTGTGGAGTCGTTGCTTGCATTTGGCTGACTAGTTGTGNNNAGATCGGAAG |
| 57413_1 | TATGAGTGTGGAGTCGTTGCCCAACGTGCTATGATGAAAAGTANNNAGATCGGAA |
| 57415_1 | TATGAGTGTGGAGTCGTTGCAGGTTCAAGCAATTCTCCTGTCTNNNAGATCGGAAG |
| 57416_1 | TATGAGTGTGGAGTCGTTGCTGTGGCTGTTATTTTTTTCCTGTNNNAGATCGGAAG |
| 57417_1 | TATGAGTGTGGAGTCGTTGCCAAACTCTGCATCCCAGTTTCAANNNAGATCGGAAG |
| 57418_1 | TATGAGTGTGGAGTCGTTGCCTGTACTTTCACTGCAACTTCCGNNNAGATCGGAAGA |
| 57420_1 | TATGAGTGTGGAGTCGTTGCGCCTTCTGGGTTCAAGAGATTCTNNNAGATCGGAAG |
| 57422_1 | TATGAGTGTGGAGTCGTTGCCTAGAGTGCAGTGGTGTGATCTTNNNAGATCGGAA |
| 57423_1 | TATGAGTGTGGAGTCGTTGCCTAGTTCAGTCTTGGGATGCTGTNNNAGATCGGAAG |
| 57425_1 | TATGAGTGTGGAGTCGTTGCTCCCAGTCTGTATTACTCAGGGTNNNAGATCGGAAG |
| 57426_1 | TATGAGTGTGGAGTCGTTGCTTGAGTAGCTGAGATTACAGGCGNNNAGATCGGAA |
| 57428_1 | TATGAGTGTGGAGTCGTTGCTCCTAAACTGTGGGATTACAGGCNNNAGATCGGAAG |
| 57429_1 | TATGAGTGTGGAGTCGTTGCCAATAGCTGGGATTACAGGCATGNNNAGATCGGAA |

| | |
|---|---|
| 57430_1 | TATGAGTGTGGAGTCGTTGCAGTTTTACCGGTTACAACTGGGANNNAGATCGGAAG |
| 57432_1 | TATGAGTGTGGAGTCGTTGCTACAGCACATTGATGGGTCTTGANNNAGATCGGAAG |
| 57433_1 | TATGAGTGTGGAGTCGTTGCGACTACAAACACACACCACCATGNNNAGATCGGAAGA |
| 57434_1 | TATGAGTGTGGAGTCGTTGCGATAGGATCTTGCTCTGTCACCTNNNAGATCGGAAGA |
| 57436_1 | TATGAGTGTGGAGTCGTTGCCACCTGAGTAACTGGGACTACAGNNNAGATCGGAAG |
| 57437_1 | TATGAGTGTGGAGTCGTTGCCTCACTGAAAACTTTGCCTCCTGNNNAGATCGGAAGA |
| 57438_1 | TATGAGTGTGGAGTCGTTGCGTTCAAGCAATTCTCTGCCTCAGNNNAGATCGGAAGA |
| 57441_1 | TATGAGTGTGGAGTCGTTGCGTTTAGCTCTTGGCATGATCTCGNNNAGATCGGAAG |
| 57443_1 | TATGAGTGTGGAGTCGTTGCAGATGGAGTCTCTCTATGTCGCCNNNAGATCGGAAG |
| 57444_1 | TATGAGTGTGGAGTCGTTGCGCCTCTCAAATAGCTGGGACTATNNNAGATCGGAAG |
| 57446_1 | TATGAGTGTGGAGTCGTTGCAGCAATTCTCCTGACTTCTGAGTNNNAGATCGGAAGA |
| 57449_1 | TATGAGTGTGGAGTCGTTGCCTGAAGTGTGAGGTGCAATCATGNNNAGATCGGAAG |
| 57453_1 | TATGAGTGTGGAGTCGTTGCCCAGCTTCAAGTGGTTCTCCTATNNNAGATCGGAAGA |
| 57454_1 | TATGAGTGTGGAGTCGTTGCTGTATTTCCTGGGTTCACACCATNNNAGATCGGAAGA |
| 57455_1 | TATGAGTGTGGAGTCGTTGCCACTACCATGCCCAGCTAATTTTNNNAGATCGGAAGA |
| 57456_1 | TATGAGTGTGGAGTCGTTGCAATCCTGGCTCACTACAACTTCCNNNAGATCGGAAGA |
| 57457_1 | TATGAGTGTGGAGTCGTTGCCTGGGCCTAAGTGATCCTTCTTTNNNAGATCGGAAGA |
| 57458_1 | TATGAGTGTGGAGTCGTTGCTTCAAGTGATTCTCCCACCTCAGNNNAGATCGGAAGA |
| 57459_1 | TATGAGTGTGGAGTCGTTGCACCACCATACCCAGCTGATTTTTNNNAGATCGGAAGA |
| 57460_1 | TATGAGTGTGGAGTCGTTGCCATTTCCCAGTTCAATGGCATGANNNAGATCGGAAGA |
| 57461_1 | TATGAGTGTGGAGTCGTTGCGGTTCAAGCCATTCTCCTGTCTCNNNAGATCGGAAGA |
| 57462_1 | TATGAGTGTGGAGTCGTTGCCGTGATCTAAGCTCTCTGCAAACNNNAGATCGGAAGA |
| 57466_1 | TATGAGTGTGGAGTCGTTGCATCTTGGTTCACTGCAACCTTTGNNNAGATCGGAAGA |
| 57467_1 | TATGAGTGTGGAGTCGTTGCTACACAGGTGATATGGTTTGGCTNNNAGATCGGAAG |
| 57468_1 | TATGAGTGTGGAGTCGTTGCGAGCAACGCATCACACATTTACCNNNAGATCGGAAGA |
| 57469_1 | TATGAGTGTGGAGTCGTTGCGCGATTACCGTCTTAGAAAACTTNNNAGATCGGAAG |
| 57470_1 | TATGAGTGTGGAGTCGTTGCGGGAGCACCTTGTTCTGGATTATNNNAGATCGGAAG |
| 57471_1 | TATGAGTGTGGAGTCGTTGCCAAAGGTGGAAGTGTGTAGCAAANNNAGATCGGAA |
| 57472_1 | TATGAGTGTGGAGTCGTTGCGGGGATGGTCAGTCAGTACAAAANNNAGATCGGAA |
| 57473_1 | TATGAGTGTGGAGTCGTTGCATAAACTGTGGATGCAAGCCCTANNNAGATCGGAAG |
| 57474_1 | TATGAGTGTGGAGTCGTTGCGCAATAAAGCAGGGGAAGTAAAANNNAGATCGGAA |
| 57475_1 | TATGAGTGTGGAGTCGTTGCAGGTCATGAGAATAGAGCCCCTANNNAGATCGGAAG |
| 57477_1 | TATGAGTGTGGAGTCGTTGCAGGCCAGAAGATGAATGTACCAANNNAGATCGGAAG |
| 57478_1 | TATGAGTGTGGAGTCGTTGCAAATGAGGATACTGGTGGGACAGNNNAGATCGGAA |
| 57479_1 | TATGAGTGTGGAGTCGTTGCTGTGTGACAGAATGACAGACACANNNAGATCGGAAG |
| 57480_1 | TATGAGTGTGGAGTCGTTGCGGGGGAGTCAGTATCTTTCACTGNNNAGATCGGAAG |
| 57481_1 | TATGAGTGTGGAGTCGTTGCATTTTTCTTGGGCTCCAGATCCTNNNAGATCGGAAGA |
| 57482_1 | TATGAGTGTGGAGTCGTTGCATCAAAACATCATGTTGCCTGCANNNAGATCGGAAGA |
| 57483_1 | TATGAGTGTGGAGTCGTTGCACATCGACTTCCTACGACCTAGANNNAGATCGGAAGA |
| 57484_1 | TATGAGTGTGGAGTCGTTGCCTGAGGCAGAGAATTGCTTGAACNNNAGATCGGAAG |
| 57489_1 | TATGAGTGTGGAGTCGTTGCCTGAGGCAGGAAAATTGCTTGAANNNAGATCGGAAG |
| 57490_1 | TATGAGTGTGGAGTCGTTGCCTTCTCCAGCATCATCCCATACTNNNAGATCGGAAGAG |
| 57492_1 | TATGAGTGTGGAGTCGTTGCGACTTCAAATTGCACTACAGGGCNNNAGATCGGAAG |

| | |
|---|---|
| 57494_1 | TATGAGTGTGGAGTCGTTGCATCCAACTCACTTCTGCCTAAGGNNNAGATCGGAAG |
| 57496_1 | TATGAGTGTGGAGTCGTTGCTTCTGAGCCCTGTAAACTCTTCCNNNAGATCGGAAG |
| 57500_1 | TATGAGTGTGGAGTCGTTGCTAAACAGACGAATAGTTGGGCCANNNAGATCGGAA |
| 57501_1 | TATGAGTGTGGAGTCGTTGCAGTCCTGAATGGCTGAACTGTAGNNNAGATCGGAA |
| 57507_1 | TATGAGTGTGGAGTCGTTGCGGCAGATTATGTGGTCAGGAGATNNNAGATCGGAA |
| 57509_1 | TATGAGTGTGGAGTCGTTGCACCATCATCATCCACTTTTTCCANNNAGATCGGAAGA |
| 57510_1 | TATGAGTGTGGAGTCGTTGCTCCCAAAACTGTTTGCTTAGCTTNNNAGATCGGAAG |
| 57511_1 | TATGAGTGTGGAGTCGTTGCAAAAACGTCTTTTTGTAAGGTTTGTNNNAGATCGGA |
| 57512_1 | TATGAGTGTGGAGTCGTTGCGATTACACCACTGGACTCTACCCNNNAGATCGGAAG |
| 57515_1 | TATGAGTGTGGAGTCGTTGCGATCAACTTTCACCCCCTTCTCTNNNAGATCGGAAGA |
| 57517_1 | TATGAGTGTGGAGTCGTTGCTGGGGAAGATTCTCCTTTTCACANNNAGATCGGAAG |
| 57518_1 | TATGAGTGTGGAGTCGTTGCAGAGAGCGCGCTAGTGTTTTATANNNAGATCGGAA |
| 57519_1 | TATGAGTGTGGAGTCGTTGCGGCTAAGAAACTTCCAGGACTGANNNAGATCGGAA |
| 57520_1 | TATGAGTGTGGAGTCGTTGCGACTGTAGTCCCAGCTATTGGAGNNNAGATCGGAA |
| 57522_1 | TATGAGTGTGGAGTCGTTGCAAGAGAAAGCTGAGGCACATAGTNNNAGATCGGAA |
| 57523_1 | TATGAGTGTGGAGTCGTTGCACCCGAGGTTAGGAATTTGAGACNNNAGATCGGAA |
| 57524_1 | TATGAGTGTGGAGTCGTTGCATCAGAGCAGAGTAACTGGGTTGNNNAGATCGGAA |
| 57526_1 | TATGAGTGTGGAGTCGTTGCAGGAGGATTGCTTGAGTCACAATNNNAGATCGGAA |
| 57528_1 | TATGAGTGTGGAGTCGTTGCCTGTAGTCCCAGCTACTTGGTACNNNAGATCGGAAG |
| 57531_1 | TATGAGTGTGGAGTCGTTGCAACAAAATACGTGGCTCACATGGNNNAGATCGGAA |
| 57534_1 | TATGAGTGTGGAGTCGTTGCGGAAAAGCTGACTCTCAGGTACTNNNAGATCGGAA |
| 57538_1 | TATGAGTGTGGAGTCGTTGCCTACTAAGGAGGCAGAGGTGGAANNNAGATCGGAA |
| 57540_1 | TATGAGTGTGGAGTCGTTGCAGATTTGGGGTAGGAGCATTTGTNNNAGATCGGAA |
| 57542_1 | TATGAGTGTGGAGTCGTTGCTTCAGCATGCATTAGAATCAGCGNNNAGATCGGAAG |
| 57543_1 | TATGAGTGTGGAGTCGTTGCCTAGGAGAGCTGGGACTCAATTCNNNAGATCGGAA |
| 57544_1 | TATGAGTGTGGAGTCGTTGCTGGCTCATGACTGTAAGCACTTTNNNAGATCGGAAG |
| 57546_1 | TATGAGTGTGGAGTCGTTGCGCAATGCCTTTGATTCTTGGAGTNNNAGATCGGAAG |
| 57547_1 | TATGAGTGTGGAGTCGTTGCGGAGCCCCAAATGTACATCGATANNNAGATCGGAAG |
| 57548_1 | TATGAGTGTGGAGTCGTTGCACTGATCATAGCCAGATTCCTTGTNNNAGATCGGAA |
| 57549_1 | TATGAGTGTGGAGTCGTTGCGTGACCCATGATCAGAGAGACTCNNNAGATCGGAAG |
| 57550_1 | TATGAGTGTGGAGTCGTTGCAACCTGTTTTTGCACCACTGTACNNNAGATCGGAAGA |
| 57551_1 | TATGAGTGTGGAGTCGTTGCTTGATGAAGCAATATTGGCTGCCNNNAGATCGGAAG |
| 57552_1 | TATGAGTGTGGAGTCGTTGCCTCACATCTGTAATCCCAGCACTNNNAGATCGGAAGA |
| 57553_1 | TATGAGTGTGGAGTCGTTGCGGCTTCATTCCTAACAACATTCANNNAGATCGGAAGA |
| 57554_1 | TATGAGTGTGGAGTCGTTGCAGAAACACAAGTCTGGACAGTGANNNAGATCGGAA |
| 57555_1 | TATGAGTGTGGAGTCGTTGCGTGATGGCAAGTTTCTGTGTCTCNNNAGATCGGAAG |
| 57556_1 | TATGAGTGTGGAGTCGTTGCAAGTGTGTGAAAGTGAAAGTGCTNNNAGATCGGAA |
| 57559_1 | TATGAGTGTGGAGTCGTTGCAAGTGGGCAAAGAACATGAAACANNNAGATCGGAA |
| 57562_1 | TATGAGTGTGGAGTCGTTGCCCTCCTGGATTCAAGCGATTCTCNNNAGATCGGAAG |
| 57564_1 | TATGAGTGTGGAGTCGTTGCGAAAAAGTCTTGCTCTGTCACCCNNNAGATCGGAAG |
| 57565_1 | TATGAGTGTGGAGTCGTTGCCTATGTTGCCCACTTTGGTTTGTNNNAGATCGGAAG |
| 57566_1 | TATGAGTGTGGAGTCGTTGCTTCAAGTGATTCTCCTGCCTCAGNNNAGATCGGAAG |
| 57567_1 | TATGAGTGTGGAGTCGTTGCAGATGGAATCTCACTGTGTCACCNNNAGATCGGAAG |

57569_1 TATGAGTGTGGAGTCGTTGCGGAGTACAGTGGTGCAATTTTGGNNNAGATCGGAA
57570_1 TATGAGTGTGGAGTCGTTGCCATCTTAGCTCACTGCAACCTCTNNNAGATCGGAAGA
57571_1 TATGAGTGTGGAGTCGTTGCGGGTTCAAGTGATTCTACTGCCTNNNAGATCGGAAG
57572_1 TATGAGTGTGGAGTCGTTGCAGCAGTCTTTGGTCTTAGGATCTNNNAGATCGGAAG
57574_1 TATGAGTGTGGAGTCGTTGCCCTCCTTGAGTAGCTGGGATTACNNNAGATCGGAAG
57575_1 TATGAGTGTGGAGTCGTTGCAGCAATCCTCTTACTTCAGCCTCNNNAGATCGGAAGA
57576_1 TATGAGTGTGGAGTCGTTGCTCAAGTGATTCTCCTGCCTTAGCNNNAGATCGGAAG
57577_1 TATGAGTGTGGAGTCGTTGCTGAGTAGCTGAGACTACAAGCACNNNAGATCGGAA
57578_1 TATGAGTGTGGAGTCGTTGCCCTCCCTGGTTCAAAAAATTCCCNNNAGATCGGAAG
57582_1 TATGAGTGTGGAGTCGTTGCCTCAAGCAATCCTTTCACCTCAGNNNAGATCGGAAG
57583_1 TATGAGTGTGGAGTCGTTGCCCCAGGTTCAAGCAATTCTCATGNNNAGATCGGAAG
57585_1 TATGAGTGTGGAGTCGTTGCAGACGGTTGAAAAACATTGCCTTNNNAGATCGGAA
57586_1 TATGAGTGTGGAGTCGTTGCCATGATCTTGGCTTACTGCAACCNNNAGATCGGAAG
57587_1 TATGAGTGTGGAGTCGTTGCCCCAGGTTCAAGCAATTCTCTTGNNNAGATCGGAAG
57588_1 TATGAGTGTGGAGTCGTTGCCTTCTGCCATGTGGACTCAAATGNNNAGATCGGAAG
57590_1 TATGAGTGTGGAGTCGTTGCGCATCCAGGCATTTCTATACAGCNNNAGATCGGAAG
57591_1 TATGAGTGTGGAGTCGTTGCGATCTCAGCTCACTGCAACTTCTNNNAGATCGGAAG
57592_1 TATGAGTGTGGAGTCGTTGCAGTGGCACAATCTTGGCTAACTANNNAGATCGGAAG
57596_1 TATGAGTGTGGAGTCGTTGCAGGTTCAAGTGATTCTTCTGCCTNNNAGATCGGAAG
57597_1 TATGAGTGTGGAGTCGTTGCCATCCCGAGTAGCTGGAATTACANNNAGATCGGAAG
57598_1 TATGAGTGTGGAGTCGTTGCATCTCTGTTTCCTGGGCTCAAATNNNAGATCGGAAG
57599_1 TATGAGTGTGGAGTCGTTGCAGCTAGAATTACAGGTGTGCACANNNAGATCGGAA
57600_1 TATGAGTGTGGAGTCGTTGCTTCAAGTGATTCTCCTGCCTCAGNNNAGATCGGAAG
57601_1 TATGAGTGTGGAGTCGTTGCTCTTTCTATGTTGCCCAGGTTGANNNAGATCGGAAG
57604_1 TATGAGTGTGGAGTCGTTGCGAAACGGAGTCTCACTCTGTCACNNNAGATCGGAAG
57606_1 TATGAGTGTGGAGTCGTTGCGATTACAGGGGTTAGCCAGTGTGNNNAGATCGGAA
57609_1 TATGAGTGTGGAGTCGTTGCGGAGATTACAAAGGTGGTCCTGANNNAGATCGGAA
57612_1 TATGAGTGTGGAGTCGTTGCGGAGATTACAAAGGTGGTCCTGANNNAGATCGGAA
57616_1 TATGAGTGTGGAGTCGTTGCAACAGATGGCTTCACAGGAGAATNNNAGATCGGAA
57617_1 TATGAGTGTGGAGTCGTTGCTCAGAAGCATGGGATGTCTTTTCNNNAGATCGGAAG
57618_1 TATGAGTGTGGAGTCGTTGCTATCGTTCTGCCTTCAAGATGCTNNNAGATCGGAAG
57619_1 TATGAGTGTGGAGTCGTTGCCTTTGTAATCCCCACTGCTCACTNNNAGATCGGAAGA
57621_1 TATGAGTGTGGAGTCGTTGCAAAAGCTTAGGACCCTCAACACTNNNAGATCGGAAG
57622_1 TATGAGTGTGGAGTCGTTGCCAAATGTTCTCTGCAGCCTCTTTNNNAGATCGGAAG
57623_1 TATGAGTGTGGAGTCGTTGCGCACAGCTAACAGTGTCTTGTTTNNNAGATCGGAAG
57624_1 TATGAGTGTGGAGTCGTTGCCACACTGACGATGGCTTTTTCTTNNNAGATCGGAAG
57625_1 TATGAGTGTGGAGTCGTTGCCCAGAAAGTTCTCTCATGCCTCTNNNAGATCGGAAG
57627_1 TATGAGTGTGGAGTCGTTGCAGACAGGAGAATCACATGAACCCNNNAGATCGGAA
57628_1 TATGAGTGTGGAGTCGTTGCAGCCTGAAGACGTGACTGAATTANNNAGATCGGAA
57629_1 TATGAGTGTGGAGTCGTTGCTTTCCTTTGAAATGTGTGTCACANNNAGATCGGAAG
57630_1 TATGAGTGTGGAGTCGTTGCGTAATCCCAGCTACTCAGATGGCNNNAGATCGGAAG
57632_1 TATGAGTGTGGAGTCGTTGCTTATGCCTGTAATCCCAGCACTTNNNAGATCGGAAG
57633_1 TATGAGTGTGGAGTCGTTGCCTGTGGTCCTAGCTACTCAGGAANNNAGATCGGAAG

| | |
|---|---|
| 57634_1 | TATGAGTGTGGAGTCGTTGCGTCACCTTTGAATTCAGCAAGGANNNAGATCGGAAG |
| 57635_1 | TATGAGTGTGGAGTCGTTGCCGAGATTGTGTCACTGTACTCCANNNAGATCGGAAG |
| 57636_1 | TATGAGTGTGGAGTCGTTGCATTTCTCAAAAGAAGCCGTGCAANNNAGATCGGAAG |
| 57637_1 | TATGAGTGTGGAGTCGTTGCTAGCCTGCTGGAGAGATGAGATANNNAGATCGGAA |
| 57638_1 | TATGAGTGTGGAGTCGTTGCAAGAAAAGGAGAGGGTGTGGTACNNNAGATCGGA |
| 57639_1 | TATGAGTGTGGAGTCGTTGCGCACCAGTTTCCAGTTACTCAGANNNAGATCGGAAG |
| 57640_1 | TATGAGTGTGGAGTCGTTGCGACTCAAGAGGCCTAAACATGGANNNAGATCGGAA |
| 57642_1 | TATGAGTGTGGAGTCGTTGCCAGGGTCTTGTATATAGCTGCCCNNNAGATCGGAAG |
| 57644_1 | TATGAGTGTGGAGTCGTTGCTGCAAAACTTTTGAGGGAAAGGGNNNAGATCGGAA |
| 57646_1 | TATGAGTGTGGAGTCGTTGCTAAATAAAGCAGGAGTGGGCTCANNNAGATCGGAA |
| 57647_1 | TATGAGTGTGGAGTCGTTGCTCCATTCTCCCTTCACTTTCCAGNNNAGATCGGAAGA |
| 57648_1 | TATGAGTGTGGAGTCGTTGCCTGAGGCAGGAGAATCACTTGAANNNAGATCGGAA |
| 57649_1 | TATGAGTGTGGAGTCGTTGCGATGGCTGTCACTCTCCACATATNNNAGATCGGAAG |
| 57650_1 | TATGAGTGTGGAGTCGTTGCCTGCTCTGTACGTTATTCCCAGANNNAGATCGGAAG |
| 57651_1 | TATGAGTGTGGAGTCGTTGCGAGGTAGAGGCTGCAGTAAACTGNNNAGATCGGAA |
| 57652_1 | TATGAGTGTGGAGTCGTTGCCAACACGCAAGAATTGAGGGAATNNNAGATCGGAA |
| 57653_1 | TATGAGTGTGGAGTCGTTGCGCAGGAGAATTGCTTGAACTCAGNNNAGATCGGAA |
| 57654_1 | TATGAGTGTGGAGTCGTTGCCCATATGGTGCTCTGTGACTTCTNNNAGATCGGAAG |
| 57655_1 | TATGAGTGTGGAGTCGTTGCTGAGTAACCTGAGAGATGGAGGTNNNAGATCGGAA |
| 57656_1 | TATGAGTGTGGAGTCGTTGCATGGAGCTGCCTTACCTATTACGNNNAGATCGGAAG |
| 57658_1 | TATGAGTGTGGAGTCGTTGCGGTATCTGAGAAGCATTGCAAGGNNNAGATCGGAA |
| 57659_1 | TATGAGTGTGGAGTCGTTGCTGGAGTGACAGTTTCTGCAAATGNNNAGATCGGAA |
| 57660_1 | TATGAGTGTGGAGTCGTTGCACAGGACATGTAGCTTGAGAACANNNAGATCGGAA |
| 57661_1 | TATGAGTGTGGAGTCGTTGCCTGTAGTCCCAGCTACTCAGGAGNNNAGATCGGAAG |
| 57662_1 | TATGAGTGTGGAGTCGTTGCACCCAGAACACTCATCCTTCAAANNNAGATCGGAAG |
| 57664_1 | TATGAGTGTGGAGTCGTTGCAGAAACGATGGAGTTGCTGAGATNNNAGATCGGAA |
| 57665_1 | TATGAGTGTGGAGTCGTTGCCTGAAGCTATCCTCCCACCTAAGNNNAGATCGGAAG |
| 57666_1 | TATGAGTGTGGAGTCGTTGCAGTGAGGTCTCATTATGTTGCCANNNAGATCGGAAG |
| 57667_1 | TATGAGTGTGGAGTCGTTGCTAACTGGTACTACAGGTGTGTGCNNNAGATCGGAA |
| 57670_1 | TATGAGTGTGGAGTCGTTGCTTCAAGTGATTCTCCTGCCTCAGNNNAGATCGGAAG |
| 57671_1 | TATGAGTGTGGAGTCGTTGCGCAATCTTGACTCACTGCAACATNNNAGATCGGAAG |
| 57672_1 | TATGAGTGTGGAGTCGTTGCCCTCTTGGGTTCATGCCATTCTCNNNAGATCGGAAGA |
| 57675_1 | TATGAGTGTGGAGTCGTTGCGATCTTTGTTCACTGCAACCTCCNNNAGATCGGAAGA |
| 57676_1 | TATGAGTGTGGAGTCGTTGCGGCCTCTTCAGCCTCTTTTCTAANNNAGATCGGAAGA |
| 57677_1 | TATGAGTGTGGAGTCGTTGCAGTGCTGGGATTACAGGTGTAACNNNAGATCGGAA |
| 57678_1 | TATGAGTGTGGAGTCGTTGCGCATGATCATGGTTACTTCAGCCNNNAGATCGGAAG |
| 57681_1 | TATGAGTGTGGAGTCGTTGCAAATGATTCTCCTGCCTCAGCTTNNNAGATCGGAAG |
| 57682_1 | TATGAGTGTGGAGTCGTTGCTTCAAGTGATTCTCCCGTCTCAGNNNAGATCGGAAG |
| 57683_1 | TATGAGTGTGGAGTCGTTGCCTCCTGGACTCAAGTGATTCTCCNNNAGATCGGAAG |
| 57684_1 | TATGAGTGTGGAGTCGTTGCCTGGAGTATAGTGGTGCGATCTCNNNAGATCGGAA |
| 57685_1 | TATGAGTGTGGAGTCGTTGCTGGTGCGATCTGAGTTCAAGTAANNNAGATCGGAA |
| 57686_1 | TATGAGTGTGGAGTCGTTGCGCAGAGGCTAGGGAAAAGTAACTNNNAGATCGGAA |
| 57687_1 | TATGAGTGTGGAGTCGTTGCAAGACACCAGATTAGGGTTTGGGNNNAGATCGGAA |

| | |
|---|---|
| 57688_1 | TATGAGTGTGGAGTCGTTGCAGGTTCAAATGATTCTCCTGCCTNNNAGATCGGAAG |
| 57692_1 | TATGAGTGTGGAGTCGTTGCGGTTCAAGTGATTTTCCTGCCTCNNNAGATCGGAAG |
| 57693_1 | TATGAGTGTGGAGTCGTTGCCTCCTGAGTAGCTGGGATTACAGNNNAGATCGGAA |
| 57694_1 | TATGAGTGTGGAGTCGTTGCCACACCACCAAGTCTTGCTAATTNNNAGATCGGAAG |
| 57696_1 | TATGAGTGTGGAGTCGTTGCGGCACAATCTTGGTTCACTGAAANNNAGATCGGAAG |
| 57700_1 | TATGAGTGTGGAGTCGTTGCGATAGCATCTCACTCTGTCACCCNNNAGATCGGAAG |
| 57701_1 | TATGAGTGTGGAGTCGTTGCTCAAACTCCCGAAGTGAAGTGATNNNAGATCGGAAG |
| 57703_1 | TATGAGTGTGGAGTCGTTGCATTCAAGTGATTCTCCTGCCTCANNNAGATCGGAAG |
| 57704_1 | TATGAGTGTGGAGTCGTTGCGATTCTCCTTACTCAGCCTCCTGNNNAGATCGGAAGA |
| 57705_1 | TATGAGTGTGGAGTCGTTGCCAATTAGCTGGGATTACAGGTGCNNNAGATCGGAA |
| 57709_1 | TATGAGTGTGGAGTCGTTGCGGAGTACAGTGGTATGATCACGGNNNAGATCGGAA |
| 57710_1 | TATGAGTGTGGAGTCGTTGCCTCCTGAGTAGCTGGGATTACAGNNNAGATCGGAA |
| 57712_1 | TATGAGTGTGGAGTCGTTGCGTGCTGGAATTATAGGCATGAGCNNNAGATCGGAA |
| 57713_1 | TATGAGTGTGGAGTCGTTGCCTGGGTTCAAGTGATTCTTCTGCNNNAGATCGGAAG |
| 57717_1 | TATGAGTGTGGAGTCGTTGCGCCTCCCAAGTAGCTAGGATTACNNNAGATCGGAAG |
| 57719_1 | TATGAGTGTGGAGTCGTTGCTCATGAGCTGTTTCTTCTTCCGANNNAGATCGGAAG |
| 57722_1 | TATGAGTGTGGAGTCGTTGCAGTGCTGAGATTACAGGTGTGAGNNNAGATCGGAA |
| 57723_1 | TATGAGTGTGGAGTCGTTGCCCTCCCAAGTAGCTAGGACTACANNNAGATCGGAAG |
| 57725_1 | TATGAGTGTGGAGTCGTTGCTTAGCCAGGATGGTCTTGAACTCNNNAGATCGGAAG |
| 57727_1 | TATGAGTGTGGAGTCGTTGCGGAATACAGTGGCACCATCTCAGNNNAGATCGGAA |
| 57728_1 | TATGAGTGTGGAGTCGTTGCTGCAGGAAAATGGTAGAAGCATTNNNAGATCGGAA |
| 57729_1 | TATGAGTGTGGAGTCGTTGCGGGTTCAAGTGATTCTCCTGTGTNNNAGATCGGAAG |
| 57730_1 | TATGAGTGTGGAGTCGTTGCTTAGTAGATACAGGGTTTCGCCGNNNAGATCGGAA |
| 57731_1 | TATGAGTGTGGAGTCGTTGCCAGTGGTGTAAACATGGCTCACTNNNAGATCGGAAG |
| 57732_1 | TATGAGTGTGGAGTCGTTGCGCCTCCTGGATTCAAGTGATTCTNNNAGATCGGAAG |
| 57733_1 | TATGAGTGTGGAGTCGTTGCCTCAAGGAATTGTTGTGCCTCAGNNNAGATCGGAAG |
| 57734_1 | TATGAGTGTGGAGTCGTTGCTTTTCCCATGCCTATGTCCTGAANNNAGATCGGAAG |
| 57735_1 | TATGAGTGTGGAGTCGTTGCAATCTCAGCTCACTACAACCTCCNNNAGATCGGAAGA |
| 57736_1 | TATGAGTGTGGAGTCGTTGCACCTAAGGCCACCTCAAGAAATTNNNAGATCGGAAG |
| 57737_1 | TATGAGTGTGGAGTCGTTGCTGTTAGCATCCACTCTCTTCCACNNNAGATCGGAAGA |
| 57738_1 | TATGAGTGTGGAGTCGTTGCGGAGAAAGCAGAGGGTGAAATTGNNNAGATCGGA |
| 57739_1 | TATGAGTGTGGAGTCGTTGCGACAGAGTGAAAAGACAACCTGCNNNAGATCGGAA |
| 57742_1 | TATGAGTGTGGAGTCGTTGCTAGGAGCTTCATTCTTAGCAGGTNNNAGATCGGAAG |
| 57743_1 | TATGAGTGTGGAGTCGTTGCGACCCATTATAGCCAAGGCCTAANNNAGATCGGAAG |
| 57744_1 | TATGAGTGTGGAGTCGTTGCACAAAACCAAAACCACTGGAGTGNNNAGATCGGAA |
| 57747_1 | TATGAGTGTGGAGTCGTTGCCAATGGAGCACAACTTCCACATTNNNAGATCGGAAG |
| 57748_1 | TATGAGTGTGGAGTCGTTGCGATTGCCCACTGATTCCACATTCNNNAGATCGGAAG |
| 57749_1 | TATGAGTGTGGAGTCGTTGCGAGAAACTTCATGTCAACACGGGNNNAGATCGGAA |
| 57750_1 | TATGAGTGTGGAGTCGTTGCTTGAGGTTGCCATGAGCTATGATNNNAGATCGGAA |
| 57751_1 | TATGAGTGTGGAGTCGTTGCCTGAGCTGCATCATTGATAAGGCNNNAGATCGGAA |
| 57753_1 | TATGAGTGTGGAGTCGTTGCACAGAGTCTTGTTATGTTGCCCANNNAGATCGGAAG |
| 57754_1 | TATGAGTGTGGAGTCGTTGCAGTCATCTAGATTGGAAAGGAGCANNNAGATCGGA |
| 57755_1 | TATGAGTGTGGAGTCGTTGCCATGCCTGTAATCCCAGCTACTCNNNAGATCGGAAG |

| | |
|---|---|
| 57761_1 | TATGAGTGTGGAGTCGTTGCTGTTTTGCCCTTATTAGCAGCACNNNAGATCGGAAG |
| 57764_1 | TATGAGTGTGGAGTCGTTGCAGGGGAGAAGGAAGGACTAAAACNNNAGATCGGA |
| 57765_1 | TATGAGTGTGGAGTCGTTGCGCTTATGCCTGTAATCTCAGCACNNNAGATCGGAAG |
| 57767_1 | TATGAGTGTGGAGTCGTTGCATTTGGCTTGGTTTGACAGACTGNNNAGATCGGAAG |
| 57769_1 | TATGAGTGTGGAGTCGTTGCGTGGCTCACTCCTGTAATCTCAGNNNAGATCGGAAG |
| 57770_1 | TATGAGTGTGGAGTCGTTGCGGAAGCTAAGGTGGGAAGATCATNNNAGATCGGA |
| 57771_1 | TATGAGTGTGGAGTCGTTGCTGCTGGAAAATGGGCTCAAAAAANNNAGATCGGAA |
| 57772_1 | TATGAGTGTGGAGTCGTTGCCAAGTCTCACCAGGAGTTCTCTCNNNAGATCGGAAG |
| 57775_1 | TATGAGTGTGGAGTCGTTGCTGAGGCAGGAGAACTACTTGAACNNNAGATCGGAA |
| 57776_1 | TATGAGTGTGGAGTCGTTGCTGAAGCAGGTGGATTACTTGAGGNNNAGATCGGAA |
| 57778_1 | TATGAGTGTGGAGTCGTTGCTAACCTAGCACTGCCCATCTTTTNNNAGATCGGAAGA |
| 57780_1 | TATGAGTGTGGAGTCGTTGCCAGGATGGTCAACATAGCGAAACNNNAGATCGGAA |
| 57781_1 | TATGAGTGTGGAGTCGTTGCTTTTCCTGAGACCAAGTCTTGCTNNNAGATCGGAAG |
| 57782_1 | TATGAGTGTGGAGTCGTTGCGGCGTCTGTAGTCTCAGTTACTCNNNAGATCGGAAG |
| 57787_1 | TATGAGTGTGGAGTCGTTGCTTGGTATCAGGATTATGCTGGCCNNNAGATCGGAAG |
| 57789_1 | TATGAGTGTGGAGTCGTTGCAGAAAAGTCCAGGGTTTAGCGATNNNAGATCGGAA |
| 57791_1 | TATGAGTGTGGAGTCGTTGCATTCACTGGTTCTTTCTCCCCTCNNNAGATCGGAAGA |
| 57792_1 | TATGAGTGTGGAGTCGTTGCTACAAAAAATTAGCCAGGCGTGGNNNAGATCGGAA |
| 57793_1 | TATGAGTGTGGAGTCGTTGCCTGGACACTTTACCTTCCCTCTCNNNAGATCGGAAGA |
| 57797_1 | TATGAGTGTGGAGTCGTTGCTAGTCTCATCTACCTGGAAGGCTNNNAGATCGGAAG |
| 57798_1 | TATGAGTGTGGAGTCGTTGCACCCAAGAGTTCATTCCAGTTGTNNNAGATCGGAAG |
| 57799_1 | TATGAGTGTGGAGTCGTTGCCTGAGGGTGTAATAGTCGCGATTNNNAGATCGGAA |
| 57801_1 | TATGAGTGTGGAGTCGTTGCTTCCACAGTCAATGGTATGGACCNNNAGATCGGAAG |
| 57803_1 | TATGAGTGTGGAGTCGTTGCTAGACCTGCCTTCCAAGAAATGTNNNAGATCGGAAG |
| 57804_1 | TATGAGTGTGGAGTCGTTGCCATGCCTGTAGTCTCAGCTACTTNNNAGATCGGAAG |
| 57806_1 | TATGAGTGTGGAGTCGTTGCAAGGCATTTGGTATCTGATTCCTNNNAGATCGGAAG |
| 57808_1 | TATGAGTGTGGAGTCGTTGCGAGATAACGCCATTGCACTTCAGNNNAGATCGGAAG |
| 57811_1 | TATGAGTGTGGAGTCGTTGCGCTTCTCTGAAGTCTTCACCAAANNNAGATCGGAAG |
| 57813_1 | TATGAGTGTGGAGTCGTTGCGGCCGACCCTTACAAAAATGAAANNNAGATCGGAAG |
| 57814_1 | TATGAGTGTGGAGTCGTTGCTGGTGAGACTGTAAGATTGGTGCNNNAGATCGGAA |
| 57815_1 | TATGAGTGTGGAGTCGTTGCGTGTGTCCTGGGTCATAAGTGAANNNAGATCGGAA |
| 57816_1 | TATGAGTGTGGAGTCGTTGCGCTACTAGGGAGCCTAAGGTAGANNNAGATCGGAA |
| 57817_1 | TATGAGTGTGGAGTCGTTGCCAACCTCTCTACACACCCTGAAANNNAGATCGGAAGA |
| 57819_1 | TATGAGTGTGGAGTCGTTGCTTCTCCTTTTTCCATGCGATGTGNNNAGATCGGAAGA |
| 57820_1 | TATGAGTGTGGAGTCGTTGCACTCTACCCTTTGCCTTTACACTNNNAGATCGGAAGA |
| 57823_1 | TATGAGTGTGGAGTCGTTGCTGCAAAGCATAGTGTGGAATACANNNAGATCGGAA |
| 57826_1 | TATGAGTGTGGAGTCGTTGCAGTCAGCAGCCATCAATACTGAGNNNAGATCGGAAG |
| 57827_1 | TATGAGTGTGGAGTCGTTGCCGAGACTGGGTGATTTCTGAAGANNNAGATCGGAA |
| 57828_1 | TATGAGTGTGGAGTCGTTGCCAAGTGCTGGTGAAAATGGAGAGNNNAGATCGGA |
| 57830_1 | TATGAGTGTGGAGTCGTTGCGTGGAGTGACCTTGGTTCACTATNNNAGATCGGAAG |
| 57831_1 | TATGAGTGTGGAGTCGTTGCGGTTCAAGTGATTTTCCTGCCTCNNNAGATCGGAAG |
| 57832_1 | TATGAGTGTGGAGTCGTTGCGAATATCTGGGAGTACAGGCACCNNNAGATCGGAA |
| 57833_1 | TATGAGTGTGGAGTCGTTGCGAGACGGAGTTTCATCACACTGANNNAGATCGGAAG |

| | |
|---|---|
| 57836_1 | TATGAGTGTGGAGTCGTTGCTTATCTCCACCTCTCGAGTAGCTNNNAGATCGGAAGA |
| 57839_1 | TATGAGTGTGGAGTCGTTGCCACAATCTTAGCTCACTGCAACCNNNAGATCGGAAGA |
| 57840_1 | TATGAGTGTGGAGTCGTTGCTGCAGTGTCACAATCTCGACTTANNNAGATCGGAAGA |
| 57841_1 | TATGAGTGTGGAGTCGTTGCTCTTGGCTCACTACAATCTCCACNNNAGATCGGAAGA |
| 57843_1 | TATGAGTGTGGAGTCGTTGCCTGTTTCAGTCTCCCAAGTAGCTNNNAGATCGGAAGA |
| 57845_1 | TATGAGTGTGGAGTCGTTGCGATCTCAGCTTACTGCAACCTCTNNNAGATCGGAAGA |
| 57847_1 | TATGAGTGTGGAGTCGTTGCATCTCCTGGGTTCAAGCAATTCTNNNAGATCGGAAGA |
| 57848_1 | TATGAGTGTGGAGTCGTTGCTTGTACCTCCCTAGATACAGGCANNNAGATCGGAAGA |
| 57850_1 | TATGAGTGTGGAGTCGTTGCCCACATTCAAGCAATTTTCCTGCNNNAGATCGGAAGA |
| 57851_1 | TATGAGTGTGGAGTCGTTGCTTCTGAGTAGCTGCAACTACAGGNNNAGATCGGAAG |
| 57852_1 | TATGAGTGTGGAGTCGTTGCGAGATGGAGTCTTGCTTTGTCACNNNAGATCGGAAG |
| 57853_1 | TATGAGTGTGGAGTCGTTGCCTCATGTAATCCTCCCACCTCAGNNNAGATCGGAAGA |
| 57854_1 | TATGAGTGTGGAGTCGTTGCTTCAAGTGATTCTCCTGCCTCAGNNNAGATCGGAAGA |
| 57857_1 | TATGAGTGTGGAGTCGTTGCTTTCCCATCTCATTCTTTGCCCTNNNAGATCGGAAGAG |
| 57858_1 | TATGAGTGTGGAGTCGTTGCAAGTGCTGGGATTACAGGTTTGANNNAGATCGGAAG |
| 57859_1 | TATGAGTGTGGAGTCGTTGCAGAGCTTAGAATGCCCATTTGGANNNAGATCGGAAG |
| 57860_1 | TATGAGTGTGGAGTCGTTGCTTGAACTCCTGGACTCAAGCAATNNNAGATCGGAAG |
| 57861_1 | TATGAGTGTGGAGTCGTTGCATCTTGGCTCACTACAACCTCTGNNNAGATCGGAAGA |
| 57863_1 | TATGAGTGTGGAGTCGTTGCAACAGGGTTTCACTTTGTTGCTCNNNAGATCGGAAGA |
| 57864_1 | TATGAGTGTGGAGTCGTTGCCCTGAGTAGCTGGGATTATAGGCNNNAGATCGGAAG |
| 57865_1 | TATGAGTGTGGAGTCGTTGCGACGGAGTTTCATTCCTATTGCCNNNAGATCGGAAGA |
| 57866_1 | TATGAGTGTGGAGTCGTTGCCCGTTTCCAGGTTCAACTGATTCNNNAGATCGGAAGA |
| 57867_1 | TATGAGTGTGGAGTCGTTGCTGTACTGGGATTACAGGCATGAGNNNAGATCGGAAG |
| 57868_1 | TATGAGTGTGGAGTCGTTGCTAGCAAAATCTAGGCTCACTGCANNNAGATCGGAAG |
| 57869_1 | TATGAGTGTGGAGTCGTTGCAAACAGAAGCACTTCAAAGGACGNNNAGATCGGAAG |
| 57873_1 | TATGAGTGTGGAGTCGTTGCATCTCAGCTCGCTGTAACCTAAGNNNAGATCGGAAGA |
| 57874_1 | TATGAGTGTGGAGTCGTTGCGCCAAGCAAACCACACATATGATNNNAGATCGGAAG |
| 57875_1 | TATGAGTGTGGAGTCGTTGCTGGAATGTAGTGTGAGCTTGACTNNNAGATCGGAAG |
| 57877_1 | TATGAGTGTGGAGTCGTTGCTTCAAGTGATTCTCCTGTCTCGGNNNAGATCGGAAGA |
| 57879_1 | TATGAGTGTGGAGTCGTTGCACAGTAGGAGGAGACTACAGGAGNNNAGATCGGAA |
| 57883_1 | TATGAGTGTGGAGTCGTTGCTCCAAGTAAAAGGAGAGCTTCCCNNNAGATCGGAAG |
| 57884_1 | TATGAGTGTGGAGTCGTTGCCTGGTCTCAAACTCCTGAACTCANNNAGATCGGAAGA |
| 57885_1 | TATGAGTGTGGAGTCGTTGCCTGAGTTCAAGTGATTCTCGTGCNNNAGATCGGAAG |
| 57889_1 | TATGAGTGTGGAGTCGTTGCCTCAAGCAATCCTCCTTCCTCAGNNNAGATCGGAAGA |
| 57890_1 | TATGAGTGTGGAGTCGTTGCCACACAGACCCTTCATCTTGAGANNNAGATCGGAAGA |
| 57891_1 | TATGAGTGTGGAGTCGTTGCCTCGAACTCCTGACCTCATGATTNNNAGATCGGAAGA |
| 57892_1 | TATGAGTGTGGAGTCGTTGCTTCTAGCAATTCTCCTGCCTCAGNNNAGATCGGAAGA |
| 57894_1 | TATGAGTGTGGAGTCGTTGCACACAATCTTTCTCTGTCACCCANNNAGATCGGAAGA |
| 57896_1 | TATGAGTGTGGAGTCGTTGCGATTACAAGCACACACAACCACANNNAGATCGGAAG |
| 57897_1 | TATGAGTGTGGAGTCGTTGCCGTGATCTCGGTTCTTTTAGGGTNNNAGATCGGAAG |
| 57899_1 | TATGAGTGTGGAGTCGTTGCGATCTAGGCTCATTGCAACCTCTNNNAGATCGGAAGA |
| 57901_1 | TATGAGTGTGGAGTCGTTGCGGGTTCAAACAATTTTCCTGCCTNNNAGATCGGAAGA |
| 57903_1 | TATGAGTGTGGAGTCGTTGCCAATCTCAGCTCACTGCAAACTCNNNAGATCGGAAGA |

| | |
|---|---|
| 57904_1 | TATGAGTGTGGAGTCGTTGCCTCCCGAGTAGCTGGAACTACAGNNNAGATCGGAAG |
| 57907_1 | TATGAGTGTGGAGTCGTTGCTTCACACCATTCTCTTGCCTCAGNNNAGATCGGAAGA |
| 57910_1 | TATGAGTGTGGAGTCGTTGCAACCTCTTGAGTAGCTGGGACTANNNAGATCGGAAG |
| 57911_1 | TATGAGTGTGGAGTCGTTGCAAGCGATTATCTTGTCTCAGCCTNNNAGATCGGAAG |
| 57912_1 | TATGAGTGTGGAGTCGTTGCATTTTTGCTCTTTCTGTCCAGGCNNNAGATCGGAAGA |
| 57913_1 | TATGAGTGTGGAGTCGTTGCTCTTGGCTTACTGCAACCTCTACNNNAGATCGGAAGA |
| 57914_1 | TATGAGTGTGGAGTCGTTGCGTAGTTGGGATTATAGGCACCCGNNNAGATCGGAA |
| 57916_1 | TATGAGTGTGGAGTCGTTGCTAGAGTGCAATGGTACGATCTCGNNNAGATCGGAA |
| 57917_1 | TATGAGTGTGGAGTCGTTGCACAACTTCTCCAGAAAGACTGCANNNAGATCGGAAG |
| 57918_1 | TATGAGTGTGGAGTCGTTGCACAAATCCTTAGTGCATGGTGGANNNAGATCGGAAG |
| 57919_1 | TATGAGTGTGGAGTCGTTGCAGGGTACACACAGAGAAGGTAGANNNAGATCGGAA |
| 57920_1 | TATGAGTGTGGAGTCGTTGCCGGAGTAGAAGGAAAACTGGAGTNNNAGATCGGA |
| 57924_1 | TATGAGTGTGGAGTCGTTGCACAATATGATCCAGCAACCCACTNNNAGATCGGAAG |
| 57925_1 | TATGAGTGTGGAGTCGTTGCAACTCTGAATCATTTCTTGGGCANNNAGATCGGAAG |
| 57927_1 | TATGAGTGTGGAGTCGTTGCCCAGCCGTTTTTACTCTGAGAACNNNAGATCGGAAG |
| 57928_1 | TATGAGTGTGGAGTCGTTGCTCACCTCCTATCCCAAAACTCTTNNNAGATCGGAAGA |
| 57930_1 | TATGAGTGTGGAGTCGTTGCGGGAGTCACTCATATCTTCCAGGNNNAGATCGGAAG |
| 57931_1 | TATGAGTGTGGAGTCGTTGCCTGACATGTGGGGATTATGGGATNNNAGATCGGAA |
| 57932_1 | TATGAGTGTGGAGTCGTTGCAGGGGGGGAATACATTAGAGGAANNNAGATCGGA |
| 57933_1 | TATGAGTGTGGAGTCGTTGCCATGCCTGTAGTCCTAGCTACTCNNNAGATCGGAAG |
| 57938_1 | TATGAGTGTGGAGTCGTTGCCTGTTGTCTGAACACTTCACCACNNNAGATCGGAAGA |
| 57939_1 | TATGAGTGTGGAGTCGTTGCTCTGACAGCTCATTTCTCTTGCTNNNAGATCGGAAGA |
| 57943_1 | TATGAGTGTGGAGTCGTTGCACCAAAACCAAATTGCACAGCTANNNAGATCGGAAG |
| 57944_1 | TATGAGTGTGGAGTCGTTGCATGGGAAAATATGACCATGCCCANNNAGATCGGAAG |
| 57945_1 | TATGAGTGTGGAGTCGTTGCTGACTGATTAGGCCCAGATGAAGNNNAGATCGGAA |
| 57947_1 | TATGAGTGTGGAGTCGTTGCTGAGAAAGGATATCTGCCTGAGCNNNAGATCGGAA |
| 57948_1 | TATGAGTGTGGAGTCGTTGCGAGAATCTACCCGGGAAGAGTTCNNNAGATCGGAA |
| 57949_1 | TATGAGTGTGGAGTCGTTGCAGTGTTGTTTGCATAAAGCTGGGNNNAGATCGGAA |
| 57950_1 | TATGAGTGTGGAGTCGTTGCCTTGAGAAGCTGAGGTTAGTGGANNNAGATCGGAA |
| 57951_1 | TATGAGTGTGGAGTCGTTGCAATGGAACAACAAAGCCTGGATGNNNAGATCGGAA |
| 57952_1 | TATGAGTGTGGAGTCGTTGCCCCAAAAAACAAACACACACACANNNAGATCGGAAG |
| 57953_1 | TATGAGTGTGGAGTCGTTGCTTTATGTCCTGCCAGAAGAAGCTNNNAGATCGGAAG |
| 57955_1 | TATGAGTGTGGAGTCGTTGCCCAGCTTTAGCACTCAACATGACNNNAGATCGGAAG |
| 57956_1 | TATGAGTGTGGAGTCGTTGCCACCTATAGTCCCAGCTACTTGGNNNAGATCGGAAG |
| 57957_1 | TATGAGTGTGGAGTCGTTGCCCTGGAGAACGGGTGGTATATACNNNAGATCGGAA |
| 57958_1 | TATGAGTGTGGAGTCGTTGCAATTAGCTGAGCTTCAGACCACANNNAGATCGGAAG |
| 57959_1 | TATGAGTGTGGAGTCGTTGCACAATCATAGGCAAGTCAGCAGANNNAGATCGGAA |
| 57960_1 | TATGAGTGTGGAGTCGTTGCCTCACGACTGTAATCTCAGCACTNNNAGATCGGAAGA |
| 57964_1 | TATGAGTGTGGAGTCGTTGCCTTGTGAGGTTATGAAGGCTGATNNNAGATCGGAA |
| 57966_1 | TATGAGTGTGGAGTCGTTGCTTCGCTATGCTTCTTCCTTACCCNNNAGATCGGAAGA |
| 57971_1 | TATGAGTGTGGAGTCGTTGCTATTGTATTGGACAGTGCAGGCTNNNAGATCGGAAG |
| 57973_1 | TATGAGTGTGGAGTCGTTGCTTTAGAGGGCTCAATTGCACAGANNNAGATCGGAAG |
| 57976_1 | TATGAGTGTGGAGTCGTTGCTTTGGTCTGCAGCCTCAATTTTCNNNAGATCGGAAGA |

| | |
|---|---|
| 57977_1 | TATGAGTGTGGAGTCGTTGCAATTAGCCAGAGTAGGGGTCAGANNNAGATCGGAA |
| 57978_1 | TATGAGTGTGGAGTCGTTGCCCTGTAATCCTAGCATGTTGGGANNNAGATCGGAAG |
| 57979_1 | TATGAGTGTGGAGTCGTTGCTAATCCCAGTACTTTGGAAGGCCNNNAGATCGGAAG |
| 57981_1 | TATGAGTGTGGAGTCGTTGCAGGAGGAATGGGAGAAAAACAGTNNNAGATCGGA |
| 57983_1 | TATGAGTGTGGAGTCGTTGCAGGTTGAAGTCTACCGAAAGTGANNNAGATCGGAA |
| 57984_1 | TATGAGTGTGGAGTCGTTGCTCATGCCTGTAATCTCAGCACTTNNNAGATCGGAAG |
| 57988_1 | TATGAGTGTGGAGTCGTTGCAAAAAAAATCAGCATGGACGCAGNNNAGATCGGAA |
| 57990_1 | TATGAGTGTGGAGTCGTTGCAGGTTGAGATTCAAGTTCTGCCTNNNAGATCGGAAG |
| 57991_1 | TATGAGTGTGGAGTCGTTGCGAGGAAGGAAGGAAGGAAGGAAGNNNAGATCGG |
| 57993_1 | TATGAGTGTGGAGTCGTTGCCCTCAGCTATCTCATGCTCTCAGNNNAGATCGGAAGA |
| 57994_1 | TATGAGTGTGGAGTCGTTGCCGCCTGTAATCCTAGCATTTTGGNNNAGATCGGAAG |
| 57996_1 | TATGAGTGTGGAGTCGTTGCACCTTATCCTTTTTGCCCTTCCTNNNAGATCGGAAGA |
| 57997_1 | TATGAGTGTGGAGTCGTTGCGGTGTTAGAGAAGGTGGACACATNNNAGATCGGAA |
| 57998_1 | TATGAGTGTGGAGTCGTTGCCTCCCACTATCCTTAGCACTACANNNAGATCGGAAGA |
| 57999_1 | TATGAGTGTGGAGTCGTTGCCTCAGTGAGGATTCTGGAACACANNNAGATCGGAA |
| 58000_1 | TATGAGTGTGGAGTCGTTGCTGGTAGTACACACCTGTAGTCCTNNNAGATCGGAAG |
| 58002_1 | TATGAGTGTGGAGTCGTTGCAGCTTTCATGGCACATCAAGTTCNNNAGATCGGAAG |
| 58003_1 | TATGAGTGTGGAGTCGTTGCAGCTCCACCAACTTTGAGTTCTANNNAGATCGGAAG |
| 58004_1 | TATGAGTGTGGAGTCGTTGCATTGAGCATTTCCCCAGCATTTGNNNAGATCGGAAG |
| 58006_1 | TATGAGTGTGGAGTCGTTGCTACTGGTTCTTCCTATCCACGAGNNNAGATCGGAAG |
| 58007_1 | TATGAGTGTGGAGTCGTTGCGGGTTCACACCATTCTTCTACCTNNNAGATCGGAAGA |
| 58008_1 | TATGAGTGTGGAGTCGTTGCTGAACCTAGGCTCAAGCAATCTTNNNAGATCGGAAG |
| 58010_1 | TATGAGTGTGGAGTCGTTGCAAGTACTGGCATTATGGGCATGANNNAGATCGGAA |
| 58012_1 | TATGAGTGTGGAGTCGTTGCGTAGAGACGGAGTTTCACCATGTNNNAGATCGGAA |
| 58013_1 | TATGAGTGTGGAGTCGTTGCATACCACTGGGATTACAGCCATCNNNAGATCGGAAG |
| 58014_1 | TATGAGTGTGGAGTCGTTGCTCTCCTGGATTCAAGCAGTTCTCNNNAGATCGGAAG |
| 58015_1 | TATGAGTGTGGAGTCGTTGCGGTTCAAACAATTCTCCTGCCTCNNNAGATCGGAAG |
| 58016_1 | TATGAGTGTGGAGTCGTTGCTCTGATGCATCCCAAAGGTCATANNNAGATCGGAAG |
| 58019_1 | TATGAGTGTGGAGTCGTTGCGCATTACAGGCATTAGGTGGTTCNNNAGATCGGAA |
| 58020_1 | TATGAGTGTGGAGTCGTTGCATCATTCTTGGGTGTTTCTCGGANNNAGATCGGAAG |
| 58021_1 | TATGAGTGTGGAGTCGTTGCCAATCTCCATCTCCCAGGTTCAANNNAGATCGGAAGA |
| 58022_1 | TATGAGTGTGGAGTCGTTGCTTACCAAGACACCAGAGGTTCAGNNNAGATCGGAA |
| 58023_1 | TATGAGTGTGGAGTCGTTGCCTGGGCCCAAGTGATTCTCTTATNNNAGATCGGAAG |
| 58025_1 | TATGAGTGTGGAGTCGTTGCCTTCCGGATTCAAGCAATTCTCCNNNAGATCGGAAG |
| 58026_1 | TATGAGTGTGGAGTCGTTGCAAACTCTTGACCTCAGGTGATCCNNNAGATCGGAAG |
| 58027_1 | TATGAGTGTGGAGTCGTTGCTTCAAGTGATTCTCCTGCCTCAGNNNAGATCGGAAG |
| 58028_1 | TATGAGTGTGGAGTCGTTGCTTTGGGTTCAAGCAATTCTCCTGNNNAGATCGGAAG |
| 58029_1 | TATGAGTGTGGAGTCGTTGCAGACGTAGTTTTGCTCATGTTGCNNNAGATCGGAAG |
| 58030_1 | TATGAGTGTGGAGTCGTTGCATTCAAGTGATTCTCCTGCCTCANNNAGATCGGAAG |
| 58031_1 | TATGAGTGTGGAGTCGTTGCCTCCAAAAGTGCTGGGTTTACAGNNNAGATCGGAA |
| 58032_1 | TATGAGTGTGGAGTCGTTGCCTGAGTAGCTGGGATTACAGGTGNNNAGATCGGAA |
| 58033_1 | TATGAGTGTGGAGTCGTTGCCTGAGTAGCTGGAACTACAGGTGNNNAGATCGGAA |
| 58034_1 | TATGAGTGTGGAGTCGTTGCCCTCCTGGGTTCAAGCAATTCTCNNNAGATCGGAAG |

| | |
|---|---|
| 58035_1 | TATGAGTGTGGAGTCGTTGCAGACAGAGTCTCACTTTGTCACCNNNAGATCGGAAG |
| 58036_1 | TATGAGTGTGGAGTCGTTGCCCAAGTAGCTGGGACTACAGAAGNNNAGATCGGAA |
| 58039_1 | TATGAGTGTGGAGTCGTTGCCAAGTAGCTGAGACTACAGGCATNNNAGATCGGAA |
| 58041_1 | TATGAGTGTGGAGTCGTTGCAAGTGCTGAGATTACAGGCATGANNNAGATCGGAA |
| 58044_1 | TATGAGTGTGGAGTCGTTGCGTGCAATGGTGTGATCTCAGTTCNNNAGATCGGAAG |
| 58046_1 | TATGAGTGTGGAGTCGTTGCAAAGCCACCCTTGTTCAATCATGNNNAGATCGGAAG |
| 58048_1 | TATGAGTGTGGAGTCGTTGCTCCCGAGTTCAAGTGATTCTCTGNNNAGATCGGAAG |
| 58049_1 | TATGAGTGTGGAGTCGTTGCGAATCTCAGCTCTCCACAACCTGNNNAGATCGGAAG |
| 58051_1 | TATGAGTGTGGAGTCGTTGCCCAAGTAGCTGAGATTACAGGCANNNAGATCGGAA |
| 58053_1 | TATGAGTGTGGAGTCGTTGCCCAGTTACTTGGGAGGTTGTCTTNNNAGATCGGAAG |
| 58054_1 | TATGAGTGTGGAGTCGTTGCGTGGATCATGAGGTCAGGAGATCNNNAGATCGGAA |
| 58055_1 | TATGAGTGTGGAGTCGTTGCCTGGAAATGAAACACTTAGGGCGNNNAGATCGGAA |
| 58056_1 | TATGAGTGTGGAGTCGTTGCTAATCTCAGCACTTTGGAAGGCTNNNAGATCGGAAG |
| 58057_1 | TATGAGTGTGGAGTCGTTGCAATTCTAGCTGTGCCACTTACCANNNAGATCGGAAG |
| 58059_1 | TATGAGTGTGGAGTCGTTGCGCAAGACCCTGTCTCAAAAAAAANNNAGATCGGAAG |
| 58060_1 | TATGAGTGTGGAGTCGTTGCTGTTTTCCCTCATTTCTGCCAACNNNAGATCGGAAGA |
| 58062_1 | TATGAGTGTGGAGTCGTTGCCGTGGCGAAACCTGTCTCTATTANNNAGATCGGAAG |
| 58066_1 | TATGAGTGTGGAGTCGTTGCCAAGAGAACTTAGAGGCTGGTGTNNNAGATCGGAA |
| 58067_1 | TATGAGTGTGGAGTCGTTGCCAGATCAGCCTGGACAACATAGTNNNAGATCGGAAG |
| 58068_1 | TATGAGTGTGGAGTCGTTGCTCTAACCCACACTCTCTCTCTCGNNNAGATCGGAAGA |
| 58070_1 | TATGAGTGTGGAGTCGTTGCGGAACTGGCTTGGTGAATTTGAANNNAGATCGGAA |
| 58071_1 | TATGAGTGTGGAGTCGTTGCCAAGGTCAAGAGATGGAGACCATNNNAGATCGGAA |
| 58072_1 | TATGAGTGTGGAGTCGTTGCCAGAATTGCTTCAAACTGGGAGGNNNAGATCGGAA |
| 58073_1 | TATGAGTGTGGAGTCGTTGCAGAGTATGGGTCATGGATGAGGANNNAGATCGGA |
| 58074_1 | TATGAGTGTGGAGTCGTTGCTGCACAGAACACCACTAAGTACTNNNAGATCGGAAG |
| 58075_1 | TATGAGTGTGGAGTCGTTGCATGTGAGGCTCAAGAACAGGAAANNNAGATCGGAA |
| 58076_1 | TATGAGTGTGGAGTCGTTGCCACACCTGTAATCCCAGCACTTTNNNAGATCGGAAGA |
| 58077_1 | TATGAGTGTGGAGTCGTTGCGTCTCTGAAGGATTCCAAGCAGANNNAGATCGGAA |
| 58078_1 | TATGAGTGTGGAGTCGTTGCGTGCTTCAGTGTCCAGAGTTTTTNNNAGATCGGAAG |
| 58079_1 | TATGAGTGTGGAGTCGTTGCAGGTGGAGATCATGAAAATGGCANNNAGATCGGA |
| 58080_1 | TATGAGTGTGGAGTCGTTGCGTGTCGAGAATATGAAGCCCAGANNNAGATCGGAA |
| 58082_1 | TATGAGTGTGGAGTCGTTGCTCTACAACAAAAACAGGGTGGGANNNAGATCGGAA |
| 58083_1 | TATGAGTGTGGAGTCGTTGCTCAGCTCTGAATGTACTGCATCCNNNAGATCGGAAG |
| 58084_1 | TATGAGTGTGGAGTCGTTGCGCATAGACAGAGAGGCAAGTGATNNNAGATCGGA |
| 58085_1 | TATGAGTGTGGAGTCGTTGCCATCCTGGCTAACATGGTGAAACNNNAGATCGGAAG |
| 58086_1 | TATGAGTGTGGAGTCGTTGCAGCTTTCTACTCCCTGGTTCTTGNNNAGATCGGAAGA |
| 58087_1 | TATGAGTGTGGAGTCGTTGCACAGATGTCAGTACAGATGTCGGNNNAGATCGGAA |
| 58089_1 | TATGAGTGTGGAGTCGTTGCCAGGAGGTGGAGCTTGAAGTTAGNNNAGATCGGA |
| 58090_1 | TATGAGTGTGGAGTCGTTGCCAAGGGAAAAAAGAGGGAGAGGTNNNAGATCGGA |
| 58091_1 | TATGAGTGTGGAGTCGTTGCGCAAGCCTGTAGTCCTAGCTATTNNNAGATCGGAAG |
| 58093_1 | TATGAGTGTGGAGTCGTTGCAGATCCAGTAACTGATACCCCCANNNAGATCGGAAG |
| 58094_1 | TATGAGTGTGGAGTCGTTGCGACTGGCAAATCGGATAAAGAGCNNNAGATCGGAA |
| 58095_1 | TATGAGTGTGGAGTCGTTGCTGATGTCCATGATGCATCTCTGTNNNAGATCGGAAG |

58096_1 TATGAGTGTGGAGTCGTTGCATGCTACCACAATGCGGTTTATGNNNAGATCGGAAG
58099_1 TATGAGTGTGGAGTCGTTGCAAGATGTGACTTTGCTCCTCCTTNNNAGATCGGAAG
58100_1 TATGAGTGTGGAGTCGTTGCATGCTGGAAAGAGAGCTGAAAGANNNAGATCGGAA
58101_1 TATGAGTGTGGAGTCGTTGCTACAGTGGCTCAGGAAATATCCCNNNAGATCGGAAG
58102_1 TATGAGTGTGGAGTCGTTGCGAAAAAAAATTGGCCAGGCACAGNNNAGATCGGAA
58103_1 TATGAGTGTGGAGTCGTTGCGTGTAATCCCAGCTACTCAGGAGNNNAGATCGGAA
58104_1 TATGAGTGTGGAGTCGTTGCACGTTTTCAGATGATTGGGATTCANNNAGATCGGAA
58111_1 TATGAGTGTGGAGTCGTTGCTGCTTGACCAGTGATATGGATGTNNNAGATCGGAA
58112_1 TATGAGTGTGGAGTCGTTGCGAGGAAAGTTGGCGAAAGAAGAGNNNAGATCGGA
58113_1 TATGAGTGTGGAGTCGTTGCCAAGGTCAAGAGATGGAGACCATNNNAGATCGGAA
58115_1 TATGAGTGTGGAGTCGTTGCTATGAAATGAGGGCTGTCATCCANNNAGATCGGAA
58116_1 TATGAGTGTGGAGTCGTTGCCCTCCTCCTTTCCCATACTGAGTNNNAGATCGGAAGA
58117_1 TATGAGTGTGGAGTCGTTGCACCAGAGACCAACTTTGAGATCCNNNAGATCGGAAG
58118_1 TATGAGTGTGGAGTCGTTGCCAGGACAGTCTCAGGAGAAAGTCNNNAGATCGGAA
58119_1 TATGAGTGTGGAGTCGTTGCGCAAATTTGGAAGGCAAAACTGCNNNAGATCGGAA
58120_1 TATGAGTGTGGAGTCGTTGCTCCACCTGAGAAAATGTTGGCTANNNAGATCGGAAG
58122_1 TATGAGTGTGGAGTCGTTGCTGGGAGATATTTGGCTTATGGGGNNNAGATCGGAA
58124_1 TATGAGTGTGGAGTCGTTGCAAGTTCAAGCAGAGTGGAAAACGNNNAGATCGGAA
58125_1 TATGAGTGTGGAGTCGTTGCAGAATTGCCTGAACTTGGGAGATNNNAGATCGGAA
58127_1 TATGAGTGTGGAGTCGTTGCTGAGGTAGGAGAATCGCTTGAACNNNAGATCGGAA
58131_1 TATGAGTGTGGAGTCGTTGCCAATCTCTGACTCCCTCGTTCAANNNAGATCGGAAGA
58133_1 TATGAGTGTGGAGTCGTTGCCTCCTGAGTAGCTGGGATTACAGNNNAGATCGGAA
58135_1 TATGAGTGTGGAGTCGTTGCGATCTTGGCTCACTGTAAGCTCCNNNAGATCGGAAG
58136_1 TATGAGTGTGGAGTCGTTGCGCCTCCTGGGTTCATGTAATTCTNNNAGATCGGAAG
58137_1 TATGAGTGTGGAGTCGTTGCGTAGAGATGGGGTTTGGTCATGTNNNAGATCGGAA
58140_1 TATGAGTGTGGAGTCGTTGCTTCAAGGGATTCTTCTGCCTCAGNNNAGATCGGAAG
58143_1 TATGAGTGTGGAGTCGTTGCGAGACAGAGTTTCACCCCTGTTGNNNAGATCGGAAG
58144_1 TATGAGTGTGGAGTCGTTGCATGAAGTCTCACTCTTGTTCCCCNNNAGATCGGAAGA
58145_1 TATGAGTGTGGAGTCGTTGCACTCCTGGGTTCAAATGATCCTCNNNAGATCGGAAG
58146_1 TATGAGTGTGGAGTCGTTGCAGACGGAGTCTTACTCTTTTGCCNNNAGATCGGAAG
58147_1 TATGAGTGTGGAGTCGTTGCAGTAGCTGGGACTACAGGTACATNNNAGATCGGAA
58149_1 TATGAGTGTGGAGTCGTTGCTCAAGTGATTCTCCTGCCTTAGCNNNAGATCGGAAG
58150_1 TATGAGTGTGGAGTCGTTGCCAAGTAGCTGGGATTACAGGTGTNNNAGATCGGAA
58152_1 TATGAGTGTGGAGTCGTTGCCTCCTGAGTAGCTGGGATTACAGNNNAGATCGGAA
58157_1 TATGAGTGTGGAGTCGTTGCATCACTCTATTCTGTCCTGGCCTNNNAGATCGGAAGA
58160_1 TATGAGTGTGGAGTCGTTGCGTGTCACTCAGTCTGGAGTACAGNNNAGATCGGAA
58161_1 TATGAGTGTGGAGTCGTTGCTCTCCTGAGTAGCTGGGATTACANNNAGATCGGAAG
58163_1 TATGAGTGTGGAGTCGTTGCCAGTAGCACAATCTTGGCTCACTNNNAGATCGGAAG
58164_1 TATGAGTGTGGAGTCGTTGCGCTTACTGCAACCTCCATCTTTCNNNAGATCGGAAGA
58165_1 TATGAGTGTGGAGTCGTTGCGAGTTTCGTTCTTATTGCCCAGGNNNAGATCGGAAG
58166_1 TATGAGTGTGGAGTCGTTGCTCTCCTGAGTAGCTGGGATTACANNNAGATCGGAAG
58167_1 TATGAGTGTGGAGTCGTTGCGACGACTTGGAAAAGGTATGCAGNNNAGATCGGAA
58172_1 TATGAGTGTGGAGTCGTTGCTGCCTGGCCCTATGTGATATTTANNNAGATCGGAAG

| | |
|---|---|
| 58173_1 | TATGAGTGTGGAGTCGTTGCCTCAGTGCAGCTTCAAAATCCTGNNNAGATCGGAAG |
| 58174_1 | TATGAGTGTGGAGTCGTTGCCCCAAGACCTAAACATCTCCCAANNNAGATCGGAAGA |
| 58175_1 | TATGAGTGTGGAGTCGTTGCATGGCTAACTGCAGACTTGAACTNNNAGATCGGAAG |
| 58179_1 | TATGAGTGTGGAGTCGTTGCTTTGAGTAGAGATGGGGTTTCACNNNAGATCGGAA |
| 58180_1 | TATGAGTGTGGAGTCGTTGCACATTCAAGTGATTCTCCTGCCTNNNAGATCGGAAGA |
| 58181_1 | TATGAGTGTGGAGTCGTTGCCTAGAGTACAGTGGGGCAATCTCNNNAGATCGGAA |
| 58182_1 | TATGAGTGTGGAGTCGTTGCGCCTCCTGAGTAGCTGAGATTACNNNAGATCGGAAG |
| 58183_1 | TATGAGTGTGGAGTCGTTGCTAGCTTGGATTACAGGCATGTGTNNNAGATCGGAAG |
| 58185_1 | TATGAGTGTGGAGTCGTTGCGATGCTATTCGGGGAAATCTTGCNNNAGATCGGAAG |
| 58186_1 | TATGAGTGTGGAGTCGTTGCCCTGGGTTCAAGCAATTCTCTTGNNNAGATCGGAAG |
| 58188_1 | TATGAGTGTGGAGTCGTTGCACAATCTGGGCTCACTTCAAGATNNNAGATCGGAAG |
| 58189_1 | TATGAGTGTGGAGTCGTTGCTTTCAGCAATTCTCCTGCCTCAGNNNAGATCGGAAGA |
| 58190_1 | TATGAGTGTGGAGTCGTTGCTTGAACTCCTGGACTCAAACGATNNNAGATCGGAAG |
| 58191_1 | TATGAGTGTGGAGTCGTTGCCTGCAACCTCCATCTACCAGATTNNNAGATCGGAAGA |
| 58192_1 | TATGAGTGTGGAGTCGTTGCGTTCAAGAGATTCTCCTGCCTCANNNAGATCGGAAGA |
| 58193_1 | TATGAGTGTGGAGTCGTTGCACCTATGTGGGGGTGGATTTTTTNNNAGATCGGAAG |
| 58194_1 | TATGAGTGTGGAGTCGTTGCTCAAGTGTGATTCTCAGTGGAGGNNNAGATCGGAA |
| 58195_1 | TATGAGTGTGGAGTCGTTGCAGTAGCTAGGACCACAGGTATGTNNNAGATCGGAA |
| 58197_1 | TATGAGTGTGGAGTCGTTGCCCTCCTGAGTAACTGGAAACACANNNAGATCGGAAG |
| 58199_1 | TATGAGTGTGGAGTCGTTGCGTCAGGGAACAAACATTATGGCANNNAGATCGGAA |
| 58200_1 | TATGAGTGTGGAGTCGTTGCGTAGCTGGGACTATAGGCATTCANNNAGATCGGAA |
| 58201_1 | TATGAGTGTGGAGTCGTTGCCCTGAGTAGATGGGATTACAGGCNNNAGATCGGAA |
| 58202_1 | TATGAGTGTGGAGTCGTTGCGAATGCAGTGGTATGATCTTGGCNNNAGATCGGAA |
| 58203_1 | TATGAGTGTGGAGTCGTTGCCTCCTGAGTAGCTGGGACTACAGNNNAGATCGGAAG |
| 58204_1 | TATGAGTGTGGAGTCGTTGCCCTTGGTCTCCGCTTTCAGATTCNNNAGATCGGAAGA |
| 58205_1 | TATGAGTGTGGAGTCGTTGCTAGCTGGGATTACAGATGTGTGCNNNAGATCGGAA |
| 58206_1 | TATGAGTGTGGAGTCGTTGCTTCAAGACATTCTCCTGCCTCAGNNNAGATCGGAAGA |
| 58207_1 | TATGAGTGTGGAGTCGTTGCAGGTTCAAGCAATTCTCCTTCCTNNNAGATCGGAAGA |
| 58210_1 | TATGAGTGTGGAGTCGTTGCTTCAAGTGATTCTCCTGCCTCAGNNNAGATCGGAAGA |
| 58213_1 | TATGAGTGTGGAGTCGTTGCGCTCAAGTGATCTTCCTGCCTTANNNAGATCGGAAGA |
| 58215_1 | TATGAGTGTGGAGTCGTTGCCTGAGTTCACACCATTCTTCTGCNNNAGATCGGAAGA |
| 58216_1 | TATGAGTGTGGAGTCGTTGCGAGTAGCTAGGATTACAGGCACCNNNAGATCGGAA |
| 58217_1 | TATGAGTGTGGAGTCGTTGCGATCTTGTCTCACTGCAACCTCTNNNAGATCGGAAGA |
| 58218_1 | TATGAGTGTGGAGTCGTTGCAGTAGCTGGGATTATAGGCATGCNNNAGATCGGAA |
| 58222_1 | TATGAGTGTGGAGTCGTTGCGTTCAAGCGATTCTCATGCCTCNNNAGATCGGAAGA |
| 58223_1 | TATGAGTGTGGAGTCGTTGCCTCATGAGATTCTCCTGCCTCAGNNNAGATCGGAAGA |
| 58224_1 | TATGAGTGTGGAGTCGTTGCTCTTGAACTCCTAGCCTCCAGTANNNAGATCGGAAGA |
| 58225_1 | TATGAGTGTGGAGTCGTTGCCTGGAGTGTAGTGGTGCAATCTCNNNAGATCGGAA |
| 58226_1 | TATGAGTGTGGAGTCGTTGCCAGGTGTCTGTGTCCAAAAAAAANNNAGATCGGAA |
| 58229_1 | TATGAGTGTGGAGTCGTTGCTTTTCATGTGGACCCTTCAGCTANNNAGATCGGAAGA |
| 58230_1 | TATGAGTGTGGAGTCGTTGCTCTCCTCATGTTTTCAAGAGGGANNNAGATCGGAAG |
| 58231_1 | TATGAGTGTGGAGTCGTTGCGAAGACAGCTATCCAGGAGTCAGNNNAGATCGGAA |
| 58232_1 | TATGAGTGTGGAGTCGTTGCAACTAAGAGGAAGTTGGCAGAGTNNNAGATCGGAA |

| | |
|---|---|
| 58233_1 | TATGAGTGTGGAGTCGTTGCAAAAAGAAGAGCATGCTGTCAGCNNNAGATCGGAA |
| 58235_1 | TATGAGTGTGGAGTCGTTGCCAACTATTCGAGGTGGACATTGCNNNAGATCGGAAG |
| 58241_1 | TATGAGTGTGGAGTCGTTGCGGGTGAGATGATTTTGACACAGCNNNAGATCGGAA |
| 58243_1 | TATGAGTGTGGAGTCGTTGCCTGTATTTCTGGCAGACAGCAACNNNAGATCGGAAG |
| 58244_1 | TATGAGTGTGGAGTCGTTGCTATGTTTTGTGCAAGTTGGTGGANNNAGATCGGAA |
| 58245_1 | TATGAGTGTGGAGTCGTTGCCTGAGCCAGACCATACAAGAACTNNNAGATCGGAAG |
| 58247_1 | TATGAGTGTGGAGTCGTTGCTGGGAGACTAAGAAAGGCAGATGNNNAGATCGGA |
| 58248_1 | TATGAGTGTGGAGTCGTTGCGAGCTTGCAGTGAACCGAGATAGNNNAGATCGGAA |
| 58249_1 | TATGAGTGTGGAGTCGTTGCATAGGAAAGCAGACACAACCAGTNNNAGATCGGAA |
| 58250_1 | TATGAGTGTGGAGTCGTTGCAACTCAGTGGCTGAAGGATAAGGNNNAGATCGGAA |
| 58251_1 | TATGAGTGTGGAGTCGTTGCTTTTCAGAGAGAAAGAGGGAGCTNNNAGATCGGAA |
| 58253_1 | TATGAGTGTGGAGTCGTTGCAGCGTATATTCCTGCATGACCATNNNAGATCGGAAG |
| 58254_1 | TATGAGTGTGGAGTCGTTGCGAGGAAGGAAGGAAGAAAGGGAGNNNAGATCGG |
| 58257_1 | TATGAGTGTGGAGTCGTTGCTTTTTTGCCCCATCCCTTCTTTGNNNAGATCGGAAGA |
| 58258_1 | TATGAGTGTGGAGTCGTTGCGACAAGCAAGAGTCTCTCAGGAANNNAGATCGGAA |
| 58261_1 | TATGAGTGTGGAGTCGTTGCATGCTGGGGATCAGTTGAATACANNNAGATCGGAA |
| 58262_1 | TATGAGTGTGGAGTCGTTGCTTTCCTCCTCCCTACACTCTTCTNNNAGATCGGAAGAG |
| 58263_1 | TATGAGTGTGGAGTCGTTGCTTTCAACCCAGAGTTTGAGGACTNNNAGATCGGAAG |
| 58264_1 | TATGAGTGTGGAGTCGTTGCTCCTGATGGGGTACACAAATCTGNNNAGATCGGAAG |
| 58269_1 | TATGAGTGTGGAGTCGTTGCGATGGCTCCCAATAATCCCTGTANNNAGATCGGAAG |
| 58270_1 | TATGAGTGTGGAGTCGTTGCGCTTTACATCCTCTTCTGGGGAANNNAGATCGGAAG |
| 58273_1 | TATGAGTGTGGAGTCGTTGCCTGAGGCAGGAGAATAGCTTGAANNNAGATCGGAA |
| 58274_1 | TATGAGTGTGGAGTCGTTGCGCCTCTGGATCATTTGCTCATTCNNNAGATCGGAAGA |
| 58276_1 | TATGAGTGTGGAGTCGTTGCAACAACACTACTCCTGCTCTTGGNNNAGATCGGAAGA |
| 58277_1 | TATGAGTGTGGAGTCGTTGCAGGTTGTCACTTTTGTTACCCCTNNNAGATCGGAAGA |
| 58278_1 | TATGAGTGTGGAGTCGTTGCAATAAAGTTTCCCCGCATGCATCNNNAGATCGGAAG |
| 58280_1 | TATGAGTGTGGAGTCGTTGCGATTTATCGCACAGACTTGCCAGNNNAGATCGGAAG |
| 58284_1 | TATGAGTGTGGAGTCGTTGCCCCAGAGAGATCCCTCAACTTTTNNNAGATCGGAAG |
| 58285_1 | TATGAGTGTGGAGTCGTTGCGCAGACGCAAAAGACATACTACANNNAGATCGGAA |
| 58286_1 | TATGAGTGTGGAGTCGTTGCGTTTGCTACAAAAAGTGGAGCCTNNNAGATCGGAA |
| 58289_1 | TATGAGTGTGGAGTCGTTGCCTGAGGCAGGAAAATTGCTTGAANNNAGATCGGAA |
| 58290_1 | TATGAGTGTGGAGTCGTTGCAAGTCCAACTTCTTACCATGACANNNAGATCGGAAG |
| 58292_1 | TATGAGTGTGGAGTCGTTGCACTTCACAGGTGATGATACGTGANNNAGATCGGAA |
| 58294_1 | TATGAGTGTGGAGTCGTTGCGTCTTGCTTTCTTGGCCACATTCNNNAGATCGGAAGA |
| 58295_1 | TATGAGTGTGGAGTCGTTGCAACTTTTGCTCTTCCAAGACACCNNNAGATCGGAAGA |
| 58299_1 | TATGAGTGTGGAGTCGTTGCAACCAGAAAAGGGAAAGGCAAACNNNAGATCGGAA |
| 58300_1 | TATGAGTGTGGAGTCGTTGCAGCTTCTACATTACTAGGCCAGCNNNAGATCGGAAG |
| 58303_1 | TATGAGTGTGGAGTCGTTGCCTTCCTTGTGGAGTTGTTTGGAGNNNAGATCGGAA |
| 58304_1 | TATGAGTGTGGAGTCGTTGCCTGAGGCAGAAGAATTGCTTGAANNNAGATCGGAA |
| 58305_1 | TATGAGTGTGGAGTCGTTGCCTCACACCTATAATCCCAGCACTNNNAGATCGGAAGA |
| 58306_1 | TATGAGTGTGGAGTCGTTGCTGAGGCCCATAAAATTGAATGGCNNNAGATCGGAA |
| 58307_1 | TATGAGTGTGGAGTCGTTGCTGGATCTACTGTCCTTTTGGCTCNNNAGATCGGAAG |
| 58309_1 | TATGAGTGTGGAGTCGTTGCCCCAGCTTATTCAGTTCAGTTGCNNNAGATCGGAAG |

| | |
|---|---|
| 58310_1 | TATGAGTGTGGAGTCGTTGCCAATCTTGGCTCACTGCAACATCNNNAGATCGGAAG |
| 58311_1 | TATGAGTGTGGAGTCGTTGCCCGAGAAAGGAGTACAGTTTCTCNNNAGATCGGAA |
| 58312_1 | TATGAGTGTGGAGTCGTTGCCAACAGAGCAAGACTCCATCTCANNNAGATCGGAAG |
| 58313_1 | TATGAGTGTGGAGTCGTTGCGAAAGAAGGAAGGAAGGAGGGAANNNAGATCGG |
| 58314_1 | TATGAGTGTGGAGTCGTTGCCCAAATCCAGAGCATGAGCATTTNNNAGATCGGAAG |
| 58315_1 | TATGAGTGTGGAGTCGTTGCCCAGATTGCTTCCTTTCTCTCCTNNNAGATCGGAAGA |
| 58317_1 | TATGAGTGTGGAGTCGTTGCATGCATGCTTGTAGTCCCAGTTANNNAGATCGGAAG |
| 58318_1 | TATGAGTGTGGAGTCGTTGCGCCAAGAAGCAAAGGAAAAATGCNNNAGATCGGAA |
| 58319_1 | TATGAGTGTGGAGTCGTTGCTGCTTGGATTGCTGCTAAAGATGNNNAGATCGGAA |
| 58320_1 | TATGAGTGTGGAGTCGTTGCACGTGGGAAGAATCAAAAAGTGANNNAGATCGGA |
| 58321_1 | TATGAGTGTGGAGTCGTTGCGCAAGGAGGAAGATGCATTTCAANNNAGATCGGAA |
| 58322_1 | TATGAGTGTGGAGTCGTTGCTCCAGACAGGACGTATAGGGTTANNNAGATCGGAA |
| 58325_1 | TATGAGTGTGGAGTCGTTGCTGCTGTCTGTATTTTCTGCTGTGNNNAGATCGGAAG |
| 58326_1 | TATGAGTGTGGAGTCGTTGCCTGTAGACCCCACTACTCAGGAGNNNAGATCGGAAG |
| 58328_1 | TATGAGTGTGGAGTCGTTGCGGCGCCTACAGGATCTAGAAAATNNNAGATCGGAA |
| 58330_1 | TATGAGTGTGGAGTCGTTGCGTCAACTGACAAAAATGTGACCANNNAGATCGGAA |
| 58331_1 | TATGAGTGTGGAGTCGTTGCTAACCTTTTGGAATGGGCCTTTGNNNAGATCGGAAG |
| 58332_1 | TATGAGTGTGGAGTCGTTGCCCCAAAGAAACACATGCAGGATTNNNAGATCGGAA |
| 58333_1 | TATGAGTGTGGAGTCGTTGCTTGGGGAGGAGAGATCATATGGTNNNAGATCGGA |
| 58334_1 | TATGAGTGTGGAGTCGTTGCCTTCTGGGCAGAGAGACTTCTTTNNNAGATCGGAAG |
| 58335_1 | TATGAGTGTGGAGTCGTTGCCCTTGTGAAGTCTTGAGCAAAGGNNNAGATCGGAA |
| 58338_1 | TATGAGTGTGGAGTCGTTGCTCACAGATGACTGAAGGGGTTTTNNNAGATCGGAA |
| 58340_1 | TATGAGTGTGGAGTCGTTGCCAAAGAGGTGTGAGTTCTGGAGANNNAGATCGGA |
| 58341_1 | TATGAGTGTGGAGTCGTTGCCTGGTGCTGGTCTTTAGAAGCAGNNNAGATCGGAA |
| 58345_1 | TATGAGTGTGGAGTCGTTGCGCCTCTCAAAGTGCTGTGATTACNNNAGATCGGAAG |
| 58348_1 | TATGAGTGTGGAGTCGTTGCCAAGGTTCAAGTGATTCTCCTGCNNNAGATCGGAAG |
| 58349_1 | TATGAGTGTGGAGTCGTTGCCAAGAGATTCTCCTGACTCACCCNNNAGATCGGAAG |
| 58355_1 | TATGAGTGTGGAGTCGTTGCCCTCCCAGGTTCCAGTGATTATCNNNAGATCGGAAG |
| 58356_1 | TATGAGTGTGGAGTCGTTGCGATCTTGGCTCACTGCAACTTTTNNNAGATCGGAAG |
| 58357_1 | TATGAGTGTGGAGTCGTTGCAGTGCTATTTCGGCTCACTACAANNNAGATCGGAAG |
| 58358_1 | TATGAGTGTGGAGTCGTTGCAATCATAGCTCACTGTAGCCTCGNNNAGATCGGAAG |
| 58360_1 | TATGAGTGTGGAGTCGTTGCACTCCTGAACTCAAGTGATCCACNNNAGATCGGAAG |
| 58361_1 | TATGAGTGTGGAGTCGTTGCTTCAAACAATTCTTCAGCCTCCGNNNAGATCGGAAGA |
| 58362_1 | TATGAGTGTGGAGTCGTTGCCTCTCAGGCTCAAGCAATCCTTANNNAGATCGGAAG |
| 58363_1 | TATGAGTGTGGAGTCGTTGCCTGAGTAGCTGGGATTACAGGTGNNNAGATCGGAA |
| 58365_1 | TATGAGTGTGGAGTCGTTGCCTGGAGTGCAATGGTGTGATCTCNNNAGATCGGAA |
| 58367_1 | TATGAGTGTGGAGTCGTTGCCCATGCCCGGCTAATTTTTTGTANNNAGATCGGAAG |
| 58368_1 | TATGAGTGTGGAGTCGTTGCCTTATTGAAACCTCCACCTCCCANNNAGATCGGAAGA |
| 58370_1 | TATGAGTGTGGAGTCGTTGCGCTTCAAGCAATTCTCCTGTCTCNNNAGATCGGAAGA |
| 58371_1 | TATGAGTGTGGAGTCGTTGCATTCAAGTGATTCTCCTGCCTCANNNAGATCGGAAG |
| 58372_1 | TATGAGTGTGGAGTCGTTGCCCCAAGTAGCTGAGATTACAGGTNNNAGATCGGAA |
| 58373_1 | TATGAGTGTGGAGTCGTTGCATCGGCTCTGTTGTAAATCCTGTNNNAGATCGGAAG |
| 58374_1 | TATGAGTGTGGAGTCGTTGCCTGTCTCCTGGGTTCAAGTGATTNNNAGATCGGAAG |

| | |
|---|---|
| 58375_1 | TATGAGTGTGGAGTCGTTGCACAGAGTCTCACTTTGTTGCTCANNNAGATCGGAAG |
| 58376_1 | TATGAGTGTGGAGTCGTTGCTACTGGTGATGCTGCTTCTAGTCNNNAGATCGGAAG |
| 58378_1 | TATGAGTGTGGAGTCGTTGCGAGACGAAGTCTTGCTCTTGTCCNNNAGATCGGAAG |
| 58379_1 | TATGAGTGTGGAGTCGTTGCATGAGGGATCCACTTCTATTGCCNNNAGATCGGAAG |
| 58380_1 | TATGAGTGTGGAGTCGTTGCCTTCCAGGTTCAGATGATCCTCCNNNAGATCGGAAG |
| 58381_1 | TATGAGTGTGGAGTCGTTGCCAAGTAGCTGGGATTACAGGCATNNNAGATCGGAA |
| 58382_1 | TATGAGTGTGGAGTCGTTGCCACTACAACCTCTACCTCTTGGGNNNAGATCGGAAG |
| 58383_1 | TATGAGTGTGGAGTCGTTGCTTCAAGCAATTCTCCTGCCTCAGNNNAGATCGGAAG |
| 58385_1 | TATGAGTGTGGAGTCGTTGCCTGAGTAGCTGGGATTACAGGTGNNNAGATCGGAA |
| 58386_1 | TATGAGTGTGGAGTCGTTGCCTCCTGAGTAGCTGGGATTACAGNNNAGATCGGAA |
| 58387_1 | TATGAGTGTGGAGTCGTTGCTTATCCAGGCTGGTCTCAAACTCNNNAGATCGGAAG |
| 58389_1 | TATGAGTGTGGAGTCGTTGCCAGGTAGCTGGGATTATAGGCGNNNAGATCGGAAG |
| 58390_1 | TATGAGTGTGGAGTCGTTGCGTTCAAGTGATCCCAAAGTGCTGNNNAGATCGGAA |
| 58391_1 | TATGAGTGTGGAGTCGTTGCGTCACCCAGACTGGAATGATGATNNNAGATCGGAA |
| 58394_1 | TATGAGTGTGGAGTCGTTGCAAACTCTTGACCTCAGGTGATCCNNNAGATCGGAAG |
| 58396_1 | TATGAGTGTGGAGTCGTTGCGAGACAAGGACAAGGTCTTGCTANNNAGATCGGAA |
| 58398_1 | TATGAGTGTGGAGTCGTTGCAATCTCTGCCTCCTGAGTTCAAGNNNAGATCGGAAG |
| 58399_1 | TATGAGTGTGGAGTCGTTGCTTTCATGCCATTCTCCTGACTCANNNAGATCGGAAGA |
| 58401_1 | TATGAGTGTGGAGTCGTTGCCAAGTAGCTGGGATTACAGGCATNNNAGATCGGAA |
| 58404_1 | TATGAGTGTGGAGTCGTTGCCTGGAGTACAAGTGGCATGATCTNNNAGATCGGAA |
| 58405_1 | TATGAGTGTGGAGTCGTTGCAGGCCAAGGGAATTCTAACTCAGNNNAGATCGGAA |
| 58406_1 | TATGAGTGTGGAGTCGTTGCGATCTCCTGACCTCATGATCCACNNNAGATCGGAAG |
| 58408_1 | TATGAGTGTGGAGTCGTTGCTGATCTTCTGGGCTCAAGTGATCNNNAGATCGGAAG |
| 58409_1 | TATGAGTGTGGAGTCGTTGCATTCTCCTGAGCGTCCCAAGTAGNNNAGATCGGAAG |
| 58410_1 | TATGAGTGTGGAGTCGTTGCCTCTCTCTCTTTCGGGAACACAANNNAGATCGGAAG |
| 58413_1 | TATGAGTGTGGAGTCGTTGCCATTCTTCTACCTCAGCCTCCTGNNNAGATCGGAAGA |
| 58414_1 | TATGAGTGTGGAGTCGTTGCCTCCTGAGTACCTGGGACTACAGNNNAGATCGGAAG |
| 58417_1 | TATGAGTGTGGAGTCGTTGCCATTCGTATGGTGTTGATGGACANNNAGATCGGAA |
| 58419_1 | TATGAGTGTGGAGTCGTTGCCTCTAAACACACAAGGCACCAACNNNAGATCGGAAG |
| 58421_1 | TATGAGTGTGGAGTCGTTGCGCTGTCACCTCACACTGAAAATCNNNAGATCGGAAG |
| 58423_1 | TATGAGTGTGGAGTCGTTGCAACCTCACTTAGAGCACTCTTGGNNNAGATCGGAAG |
| 58424_1 | TATGAGTGTGGAGTCGTTGCTTGGACAACATAGCGATTCTCCTNNNAGATCGGAAG |
| 58425_1 | TATGAGTGTGGAGTCGTTGCGAGCTAACACTTTGTCATGGACTNNNAGATCGGAAG |
| 58426_1 | TATGAGTGTGGAGTCGTTGCTGACTTGAGTAGCTCTGTTGATGANNNAGATCGGAA |
| 58430_1 | TATGAGTGTGGAGTCGTTGCTTTAGGGGGGACACAAACATTCANNNAGATCGGAA |
| 58431_1 | TATGAGTGTGGAGTCGTTGCAAACTAAGCCCAAACCAGCAAAANNNAGATCGGAAG |
| 58432_1 | TATGAGTGTGGAGTCGTTGCAAGCTCCAAGCAACTAAAAGGAANNNAGATCGGAA |
| 58434_1 | TATGAGTGTGGAGTCGTTGCTATGTGAGGAAAGGCAATGCAACNNNAGATCGGAA |
| 58435_1 | TATGAGTGTGGAGTCGTTGCTAGATAGCTTCAGGATGAGGGCTNNNAGATCGGAA |
| 58436_1 | TATGAGTGTGGAGTCGTTGCACAAAAAAAACCCCACAGGATCGNNNAGATCGGAAG |
| 58437_1 | TATGAGTGTGGAGTCGTTGCTGGGCTAAAAAGCTAGATCGTGTNNNAGATCGGAA |
| 58438_1 | TATGAGTGTGGAGTCGTTGCTCTGTAATTCCAGCACTTTGGGANNNAGATCGGAAG |
| 58440_1 | TATGAGTGTGGAGTCGTTGCAGACCACCAAAGACAAGAGGATCNNNAGATCGGAA |

| | |
|---|---|
| 58441_1 | TATGAGTGTGGAGTCGTTGCTTCCCGAAGAGAAAGCATGAGAANNNAGATCGGAA |
| 58443_1 | TATGAGTGTGGAGTCGTTGCAGCAGTTAGTGTTAGTGGCAGTTNNNAGATCGGAA |
| 58445_1 | TATGAGTGTGGAGTCGTTGCCTCCTCCCCAAGAAGTACAACTGNNNAGATCGGAAG |
| 58446_1 | TATGAGTGTGGAGTCGTTGCTGTGAAGGCCATAAGAGAGAACCNNNAGATCGGAA |
| 58448_1 | TATGAGTGTGGAGTCGTTGCTACCATCTCATGCCATCTCAGTTNNNAGATCGGAAGA |
| 58449_1 | TATGAGTGTGGAGTCGTTGCCAGAAATGTGAGCACTGAAGCAANNNAGATCGGAA |
| 58450_1 | TATGAGTGTGGAGTCGTTGCGCATACACCATGGGAATGAGACTNNNAGATCGGAA |
| 58451_1 | TATGAGTGTGGAGTCGTTGCTGAAAGAAGCCAATCCAAGAGGANNNAGATCGGAA |
| 58452_1 | TATGAGTGTGGAGTCGTTGCGAGGAAAAAGGAAGCTCTGGAGANNNAGATCGGA |
| 58454_1 | TATGAGTGTGGAGTCGTTGCTGAGGGGGATGGTATATTGCAAGNNNAGATCGGA |
| 58455_1 | TATGAGTGTGGAGTCGTTGCTCTATCTTGTTCAACCACATGGCNNNAGATCGGAAG |
| 58456_1 | TATGAGTGTGGAGTCGTTGCCCCAGGTCAGACTAGCAAATGTANNNAGATCGGAA |
| 58457_1 | TATGAGTGTGGAGTCGTTGCTTCATTAGGTTATAACAAAAGCTGGTNNNAGATCGG |
| 58458_1 | TATGAGTGTGGAGTCGTTGCATCACAAGGTCAGAGTTCGAGACNNNAGATCGGAA |
| 58459_1 | TATGAGTGTGGAGTCGTTGCTCTCATGGCTCCATTTTAGGCTTNNNAGATCGGAAG |
| 58460_1 | TATGAGTGTGGAGTCGTTGCAAGGAAAAGGAAAAGGTACCTGANNNAGATCGGA |
| 58461_1 | TATGAGTGTGGAGTCGTTGCCAGCATGTTTCCGTTGAGATTCANNNAGATCGGAAG |
| 58462_1 | TATGAGTGTGGAGTCGTTGCAAGCAGGAGAATCACTTGAACCTNNNAGATCGGAA |
| 58463_1 | TATGAGTGTGGAGTCGTTGCGCCCGGCCAAGTATATCTTTTTTNNNAGATCGGAAG |
| 58465_1 | TATGAGTGTGGAGTCGTTGCTGACCAGATAACTGATTAGCGCCNNNAGATCGGAAG |
| 58468_1 | TATGAGTGTGGAGTCGTTGCACTTGCTGGCTATGTAACTTCCANNNAGATCGGAAG |
| 58469_1 | TATGAGTGTGGAGTCGTTGCGGCATAGAGGGCAAAAATACAGGNNNAGATCGGA |
| 58470_1 | TATGAGTGTGGAGTCGTTGCACAGAAGTCTCCAGCATATGACANNNAGATCGGAAG |
| 58471_1 | TATGAGTGTGGAGTCGTTGCCCTGTAATTCCAGCTACTTGGGANNNAGATCGGAAG |
| 58472_1 | TATGAGTGTGGAGTCGTTGCTGAGGTCCATATGTTGAACTGGANNNAGATCGGAA |
| 58473_1 | TATGAGTGTGGAGTCGTTGCTGTGTGTGTGTGTGTGATTGTTGNNNAGATCGGAA |
| 58474_1 | TATGAGTGTGGAGTCGTTGCTCTTCAACCTGTGAGATTTGGGTNNNAGATCGGAAG |
| 58475_1 | TATGAGTGTGGAGTCGTTGCCGGTACACTACTTCCTCTTCAGGNNNAGATCGGAAG |
| 58477_1 | TATGAGTGTGGAGTCGTTGCAGATTCCAAATTGTCTTGCAGCTNNNAGATCGGAAG |
| 58478_1 | TATGAGTGTGGAGTCGTTGCTCTTGATGACACAGAGCACAGACNNNAGATCGGAA |
| 58479_1 | TATGAGTGTGGAGTCGTTGCCCACATATCCCATTTGACTCCCTNNNAGATCGGAAGA |
| 58480_1 | TATGAGTGTGGAGTCGTTGCAGCATTGAGGTGGGATAACCAAANNNAGATCGGAA |
| 58481_1 | TATGAGTGTGGAGTCGTTGCCCCAGGCTAGACAGATTCAAACTNNNAGATCGGAAG |
| 58482_1 | TATGAGTGTGGAGTCGTTGCAGAATTTCACCAGATTGGGGTCANNNAGATCGGAA |
| 58483_1 | TATGAGTGTGGAGTCGTTGCGCTGACTCAAAAAGCCACGTAAANNNAGATCGGAA |
| 58484_1 | TATGAGTGTGGAGTCGTTGCGGATCACTTGAGGTCAGGAGTTTNNNAGATCGGAA |
| 58485_1 | TATGAGTGTGGAGTCGTTGCGTAAAAAAAGTCAGGCCTCACCCNNNAGATCGGAAG |
| 58488_1 | TATGAGTGTGGAGTCGTTGCGTGCCTGTATGGTCATAGCAGTANNNAGATCGGAA |
| 58490_1 | TATGAGTGTGGAGTCGTTGCATGTAGTCAGAGCACAGTTTGGTNNNAGATCGGAA |
| 58492_1 | TATGAGTGTGGAGTCGTTGCACCACCCAAGGTAATCTAAGGTGNNNAGATCGGAAG |
| 58496_1 | TATGAGTGTGGAGTCGTTGCACAGTTGAAGTCTCTGGGCTTAGNNNAGATCGGAA |
| 58497_1 | TATGAGTGTGGAGTCGTTGCAGAGGTTGAAGTGAGCCAAGATTNNNAGATCGGAA |
| 58499_1 | TATGAGTGTGGAGTCGTTGCCCCAAGTAGCTGGAATTACAGGTNNNAGATCGGAA |

| | |
|---|---|
| 58500_1 | TATGAGTGTGGAGTCGTTGCATCTCCTGTGTTCAAGCGATTCTNNNAGATCGGAAG |
| 58501_1 | TATGAGTGTGGAGTCGTTGCCTCTGGAGTAACTGGGACCACAGNNNAGATCGGAA |
| 58503_1 | TATGAGTGTGGAGTCGTTGCGAGTAGCCTGGACTACAGATGTGNNNAGATCGGAA |
| 58504_1 | TATGAGTGTGGAGTCGTTGCGGATGTTCGCTATAGTGCTGCTANNNAGATCGGAA |
| 58505_1 | TATGAGTGTGGAGTCGTTGCAAACAATGATTTCAAAGTCCAGGANNNAGATCGGA |
| 58506_1 | TATGAGTGTGGAGTCGTTGCGGAATGACCAAGCTACCTTCAGTNNNAGATCGGAAG |
| 58507_1 | TATGAGTGTGGAGTCGTTGCTGATCTTAGCTCACCACAACCTCNNNAGATCGGAAG |
| 58508_1 | TATGAGTGTGGAGTCGTTGCCGTCTGTCATTGTTACACCCTCANNNAGATCGGAAG |
| 58509_1 | TATGAGTGTGGAGTCGTTGCTGATCTCAACTCACTGCAACCTCNNNAGATCGGAAG |
| 58512_1 | TATGAGTGTGGAGTCGTTGCGATTCAAGCAATTCTCCTGCCTCNNNAGATCGGAAG |
| 58513_1 | TATGAGTGTGGAGTCGTTGCCATGATCTCGGTTCACTGCAAACNNNAGATCGGAAG |
| 58515_1 | TATGAGTGTGGAGTCGTTGCACAAGTGATCCTTCAGTCTCAGCNNNAGATCGGAAG |
| 58516_1 | TATGAGTGTGGAGTCGTTGCCCTCCCCATAAATGCTTTCAGTNNNAGATCGGAAGA |
| 58517_1 | TATGAGTGTGGAGTCGTTGCAGCCTGAGCAACATAATGAGACANNNAGATCGGAA |
| 58520_1 | TATGAGTGTGGAGTCGTTGCGTTCAAGCAATTCTCTGCCTCAGNNNAGATCGGAAG |
| 58522_1 | TATGAGTGTGGAGTCGTTGCGGAAAGGCAGCATATTGTGAAGGNNNAGATCGGA |
| 58524_1 | TATGAGTGTGGAGTCGTTGCAAAGACTTGAAGCATAGACAAAAGANNNAGATCGG |
| 58525_1 | TATGAGTGTGGAGTCGTTGCGACATGACACTGCTGCAGTAAATNNNAGATCGGAA |
| 58527_1 | TATGAGTGTGGAGTCGTTGCCTGGTCTCAAACTTGTGGGTTCTNNNAGATCGGAAG |
| 58529_1 | TATGAGTGTGGAGTCGTTGCCACCTCCATTTCCTCACCTGTAANNNAGATCGGAAGA |
| 58530_1 | TATGAGTGTGGAGTCGTTGCCATCTCAGCTAACTGCAACCTCTNNNAGATCGGAAGA |
| 58531_1 | TATGAGTGTGGAGTCGTTGCGCAAATCACATGCTATCTGGGTANNNAGATCGGAAG |
| 58532_1 | TATGAGTGTGGAGTCGTTGCTGGAATCAACCAAAATGCCCATCNNNAGATCGGAAG |
| 58533_1 | TATGAGTGTGGAGTCGTTGCTCCTTGATCAACCCCTAAATCCCNNNAGATCGGAAGA |
| 58538_1 | TATGAGTGTGGAGTCGTTGCGAAAGGAAGGAAGGAAGGAAGGANNNAGATCGG |
| 58539_1 | TATGAGTGTGGAGTCGTTGCTAAGTCAGTCCACTTTCCTGAGCNNNAGATCGGAAG |
| 58541_1 | TATGAGTGTGGAGTCGTTGCACTAGGTTTCTGCTATTGCCACANNNAGATCGGAAG |
| 58544_1 | TATGAGTGTGGAGTCGTTGCTTGTGGTGGTTCATGCCTGTAATNNNAGATCGGAAG |
| 58545_1 | TATGAGTGTGGAGTCGTTGCCTCCTGAGTAGCTGGGATTACAGNNNAGATCGGAA |
| 58546_1 | TATGAGTGTGGAGTCGTTGCATAAAGTCTTGCTCTGTCACCCANNNAGATCGGAAG |
| 58547_1 | TATGAGTGTGGAGTCGTTGCGTCTCACTCTGTTACTAGGCTGCNNNAGATCGGAAG |
| 58548_1 | TATGAGTGTGGAGTCGTTGCCTCCTGAGTAGCTGGGATTACTGNNNAGATCGGAA |
| 58549_1 | TATGAGTGTGGAGTCGTTGCCACAGCCCTTCTTTGGAGTATCTNNNAGATCGGAAG |
| 58555_1 | TATGAGTGTGGAGTCGTTGCTCTCCCAAAGTGCTGGCATTATANNNAGATCGGAAG |
| 58556_1 | TATGAGTGTGGAGTCGTTGCAGTAGCTAGGATTACAGGCATGCNNNAGATCGGAA |
| 58557_1 | TATGAGTGTGGAGTCGTTGCGTTAGGTGGTTCTCCTGCCTAAGNNNAGATCGGAAG |
| 58560_1 | TATGAGTGTGGAGTCGTTGCGGAGTACAGTGGTGCGATCTCNNNAGATCGGAAGA |
| 58562_1 | TATGAGTGTGGAGTCGTTGCTTTCAAGCGATTCTCCTGTCTCANNNAGATCGGAAGA |
| 58563_1 | TATGAGTGTGGAGTCGTTGCCAAGCTATTCTCCTACCTCAGCCNNNAGATCGGAAGA |
| 58565_1 | TATGAGTGTGGAGTCGTTGCTGGAGTGCAATGGTGTAATCTCANNNAGATCGGAA |
| 58566_1 | TATGAGTGTGGAGTCGTTGCAGTGCTGGGATTATAGGCATGAGNNNAGATCGGAA |
| 58569_1 | TATGAGTGTGGAGTCGTTGCTCTCAGTTCATTGTAGCCTCTGCNNNAGATCGGAAG |
| 58570_1 | TATGAGTGTGGAGTCGTTGCTGAGTAGCTGGGACTATAGGCATNNNAGATCGGAA |

| | |
|---|---|
| 58572_1 | TATGAGTGTGGAGTCGTTGCGACAAGATCTCACTGTCACCCAGNNNAGATCGGAAG |
| 58573_1 | TATGAGTGTGGAGTCGTTGCGTGGTGCAATCATGACTCATAGCNNNAGATCGGAAG |
| 58574_1 | TATGAGTGTGGAGTCGTTGCGAGATGGAATCTTGCTCTGTTGCNNNAGATCGGAAG |
| 58575_1 | TATGAGTGTGGAGTCGTTGCCCACGCCCAGCTAATTTTTGTATNNNAGATCGGAAG |
| 58576_1 | TATGAGTGTGGAGTCGTTGCAGATGGAGTCTCGCTTTGTCATCNNNAGATCGGAAG |
| 58577_1 | TATGAGTGTGGAGTCGTTGCCAATGGTGCAATTTCAGCTCACTNNNAGATCGGAAG |
| 58578_1 | TATGAGTGTGGAGTCGTTGCTCTGGGCATTTTTGTCAGTTGTGNNNAGATCGGAAG |
| 58579_1 | TATGAGTGTGGAGTCGTTGCAAACTCCTGACCTCAAGTGATCCNNNAGATCGGAAG |
| 58580_1 | TATGAGTGTGGAGTCGTTGCCTCTGCTTCTCTGGTTCAAGACANNNAGATCGGAAG |
| 58583_1 | TATGAGTGTGGAGTCGTTGCCAGGCCGATCTATTCCTTTCTGANNNAGATCGGAAG |
| 58587_1 | TATGAGTGTGGAGTCGTTGCGGGTGACAAGTCATACGCTATCTNNNAGATCGGAAG |
| 58588_1 | TATGAGTGTGGAGTCGTTGCCAGTAGTGGCTCACGTCTGTAATNNNAGATCGGAAG |
| 58589_1 | TATGAGTGTGGAGTCGTTGCCCAGTTTGGTGTGCGAGTTTATCNNNAGATCGGAAG |
| 58591_1 | TATGAGTGTGGAGTCGTTGCCCGTGACCTTGAATTAGGCAATGNNNAGATCGGAAG |
| 58593_1 | TATGAGTGTGGAGTCGTTGCGCTATTTGAGAATGGGTGTCAGCNNNAGATCGGAA |
| 58594_1 | TATGAGTGTGGAGTCGTTGCTGACTCAGCAGTAAAACGACAAANNNAGATCGGAA |
| 58596_1 | TATGAGTGTGGAGTCGTTGCCTTCTCCCAGGTTCAAGCAATTCNNNAGATCGGAAG |
| 58597_1 | TATGAGTGTGGAGTCGTTGCAGGCTGATCCAAGGTCTTTCTTTNNNAGATCGGAAG |
| 58598_1 | TATGAGTGTGGAGTCGTTGCTTTAGTAGAGACAGGTTTCGCCANNNAGATCGGAAG |
| 58599_1 | TATGAGTGTGGAGTCGTTGCAAGACAGGATTTCACCATGTTGCNNNAGATCGGAAG |
| 58600_1 | TATGAGTGTGGAGTCGTTGCGTGTGTGCCTATAATCCCAGCTANNNAGATCGGAAG |
| 58604_1 | TATGAGTGTGGAGTCGTTGCAGGGTAATGCTGGCCTCATAAAANNNAGATCGGAA |
| 58605_1 | TATGAGTGTGGAGTCGTTGCCTATAGTCCCAGCTACTCCGGAGNNNAGATCGGAAG |
| 58606_1 | TATGAGTGTGGAGTCGTTGCGCTACTTGGGAGGATCTCTTGAGNNNAGATCGGAA |
| 58609_1 | TATGAGTGTGGAGTCGTTGCTTGAGACAGGTCTTGCTTTGTTGNNNAGATCGGAAG |
| 58610_1 | TATGAGTGTGGAGTCGTTGCGCATAATCATAGCTCACTGCAGCNNNAGATCGGAAG |
| 58613_1 | TATGAGTGTGGAGTCGTTGCTCAACTGTTCTAGCCCCAATTGTNNNAGATCGGAAG |
| 58614_1 | TATGAGTGTGGAGTCGTTGCGGTGAAAGACCTCTACAAGGAAANNNAGATCGGAA |
| 58615_1 | TATGAGTGTGGAGTCGTTGCATGGACATTTAGGTTGCCTCTGTNNNAGATCGGAAG |
| 58617_1 | TATGAGTGTGGAGTCGTTGCTTGTTCGTAATGTCTTTGCCTGGNNNAGATCGGAAG |
| 58618_1 | TATGAGTGTGGAGTCGTTGCAACTATTCATGGTTGGGCACAGTNNNAGATCGGAAG |
| 58619_1 | TATGAGTGTGGAGTCGTTGCGCAGGTGAGTTGTTTCTAGAACCNNNAGATCGGAA |
| 58621_1 | TATGAGTGTGGAGTCGTTGCCACACCGGAATACTATTCAGCCTNNNAGATCGGAAG |
| 58623_1 | TATGAGTGTGGAGTCGTTGCATCTTCTGGGCTCAAACAATCCTNNNAGATCGGAAG |
| 58624_1 | TATGAGTGTGGAGTCGTTGCGCTTCAACACAAGAAATACGCCTNNNAGATCGGAAG |
| 58626_1 | TATGAGTGTGGAGTCGTTGCGAGATAGGTTTTCGCCATGTTGCNNNAGATCGGAAG |
| 58628_1 | TATGAGTGTGGAGTCGTTGCACACACACACACAGAGCTTCTTANNNAGATCGGAAG |
| 58629_1 | TATGAGTGTGGAGTCGTTGCGGTTATTTTTGGCTGAGGCTGTTNNNAGATCGGAAG |
| 58630_1 | TATGAGTGTGGAGTCGTTGCACTACTCAGGAAGTTGAGGCATGNNNAGATCGGAA |
| 58631_1 | TATGAGTGTGGAGTCGTTGCTCAGTCTCCCAAGTTCAAGCAATNNNAGATCGGAAG |
| 58632_1 | TATGAGTGTGGAGTCGTTGCATGGGGACAGAAAAACATCTGGANNNAGATCGGAA |
| 58633_1 | TATGAGTGTGGAGTCGTTGCCCTGTAATCCCAGCACTTTGAGANNNAGATCGGAAG |
| 58634_1 | TATGAGTGTGGAGTCGTTGCTGTGGGTTATGTCCTTGCTTTCANNNAGATCGGAAG |

| | |
|---|---|
| 58635_1 | TATGAGTGTGGAGTCGTTGCCAAGAAAGTGGAGAGAGACCCTCNNNAGATCGGAA |
| 58638_1 | TATGAGTGTGGAGTCGTTGCTCTTCAATCTCCTTCCCTCTCCTNNNAGATCGGAAGAG |
| 58639_1 | TATGAGTGTGGAGTCGTTGCTTCCAGTCGAAAGATCACTCCTCNNNAGATCGGAAG |
| 58640_1 | TATGAGTGTGGAGTCGTTGCATAGGGTTTCACTCTGTTAGCCANNNAGATCGGAAG |
| 58643_1 | TATGAGTGTGGAGTCGTTGCGTTTCACTATGTTTCCCAGGCTGNNNAGATCGGAAG |
| 58645_1 | TATGAGTGTGGAGTCGTTGCTCTCCATGTCACCAAATCCCAATNNNAGATCGGAAGA |
| 58646_1 | TATGAGTGTGGAGTCGTTGCGGGACCTGAAACACAGACAAAAGNNNAGATCGGAA |
| 58647_1 | TATGAGTGTGGAGTCGTTGCAAACAATTTGTCTTCCAGGCTGGNNNAGATCGGAAG |
| 58648_1 | TATGAGTGTGGAGTCGTTGCTGGTTCTGACTTACAGGAGTGACNNNAGATCGGAA |
| 58649_1 | TATGAGTGTGGAGTCGTTGCGTTTCTTTCCCTCTGTCGGTTTGNNNAGATCGGAAG |
| 58651_1 | TATGAGTGTGGAGTCGTTGCGGATTGAAGAAACATCAGCCCACNNNAGATCGGAA |
| 58653_1 | TATGAGTGTGGAGTCGTTGCTCAGGATAAAGCAGACAAAGGGTNNNAGATCGGAA |
| 58654_1 | TATGAGTGTGGAGTCGTTGCTAATAATAACTGCCTCCCCGCTCNNNAGATCGGAAG |
| 58656_1 | TATGAGTGTGGAGTCGTTGCACTTTCAAACAACCAGACCTCCTNNNAGATCGGAAG |
| 58657_1 | TATGAGTGTGGAGTCGTTGCACTTGCTTCCTCTCTTTCTCTGGNNNAGATCGGAAGA |
| 58658_1 | TATGAGTGTGGAGTCGTTGCTGGGTTTTGAAGGTCTTATAGGGANNNAGATCGGA |
| 58659_1 | TATGAGTGTGGAGTCGTTGCGAGAGGGAGAGCAAAAGAAGGAANNNAGATCGG |
| 58660_1 | TATGAGTGTGGAGTCGTTGCCAATCCTATCAGATTAGGGCCCCNNNAGATCGGAAG |
| 58661_1 | TATGAGTGTGGAGTCGTTGCCCCATTATCTGTCCACACTGACANNNAGATCGGAAG |
| 58662_1 | TATGAGTGTGGAGTCGTTGCAGAGAGAAGCCTGAAGCAAATGANNNAGATCGGA |
| 58663_1 | TATGAGTGTGGAGTCGTTGCGAGAGAGAGAGAGAGATCGACCANNNAGATCGGA |
| 58664_1 | TATGAGTGTGGAGTCGTTGCCTTGCCCTCTCTCTCTCTCTTTCNNNAGATCGGAAGAG |
| 58665_1 | TATGAGTGTGGAGTCGTTGCGCTTAGCCGTGGGTTTCTTTTTTNNNAGATCGGAAG |
| 58668_1 | TATGAGTGTGGAGTCGTTGCATATGTGTGTGTGTGTGTGTGTGNNNAGATCGGAA |
| 58669_1 | TATGAGTGTGGAGTCGTTGCGAAGGAGCGAGAGAGAAGAGAAGNNNAGATCGG |
| 58670_1 | TATGAGTGTGGAGTCGTTGCTCACTTGAGGTCAGGAGTTTGAGNNNAGATCGGAA |
| 58673_1 | TATGAGTGTGGAGTCGTTGCGATGGAGTCTCACTCTGTTACCCNNNAGATCGGAAG |
| 58674_1 | TATGAGTGTGGAGTCGTTGCAGCCTCCTGAGTAGCTGAGATTANNNAGATCGGAA |
| 58678_1 | TATGAGTGTGGAGTCGTTGCCAGCCTCCAGAGTAGTTGGAATTNNNAGATCGGAA |
| 58681_1 | TATGAGTGTGGAGTCGTTGCAGATGGAGTCTCACTCTGTCACCNNNAGATCGGAAG |
| 58682_1 | TATGAGTGTGGAGTCGTTGCGAGACAGGGTTTTGTTCTGTTGCNNNAGATCGGAA |
| 58683_1 | TATGAGTGTGGAGTCGTTGCCCCTAGTAGCTGGGATTACAAGCNNNAGATCGGAA |
| 58684_1 | TATGAGTGTGGAGTCGTTGCTCAGGGATTCAACTTCTTCCTGGNNNAGATCGGAAG |
| 58685_1 | TATGAGTGTGGAGTCGTTGCGGTAGAGACAGGGTTTTGTGACCNNNAGATCGGAA |
| 58687_1 | TATGAGTGTGGAGTCGTTGCCTGAGTAGCTGGGATTACAGGTGNNNAGATCGGAA |
| 58692_1 | TATGAGTGTGGAGTCGTTGCCAGCATGATCTTGGTTCATTGCANNNAGATCGGAAG |
| 58693_1 | TATGAGTGTGGAGTCGTTGCGGTTCAAGTCATTTTCCTGCCTCNNNAGATCGGAAG |
| 58696_1 | TATGAGTGTGGAGTCGTTGCGCAGTTGTGTGATATCTTGGCTCNNNAGATCGGAAG |
| 58697_1 | TATGAGTGTGGAGTCGTTGCCCTCCCAGTTTCAAGCAATTCTCNNNAGATCGGAAGA |
| 58698_1 | TATGAGTGTGGAGTCGTTGCGGTTCAAGTGATTCTCTTGCCTCNNNAGATCGGAAG |
| 58701_1 | TATGAGTGTGGAGTCGTTGCGGTTCAAGTGATTCTCCTGTTGCNNNAGATCGGAAG |
| 58702_1 | TATGAGTGTGGAGTCGTTGCCTGGAGTGTAGTGGTGTGATCACNNNAGATCGGAA |
| 58703_1 | TATGAGTGTGGAGTCGTTGCCAAGTAACTGGGACTACAGGCATNNNAGATCGGAA |

58704_1 TATGAGTGTGGAGTCGTTGCCTCCGGAATACCTGGGATTACAGNNNAGATCGGAA
58705_1 TATGAGTGTGGAGTCGTTGCGATTAGCCAGCATGGTCTCCATCNNNAGATCGGAAG
58706_1 TATGAGTGTGGAGTCGTTGCCCATTCTCCTGCCTCAACCTCNNNAGATCGGAAGAGC
58710_1 TATGAGTGTGGAGTCGTTGCGTAATCTCCGCCTCTTGGGTTTANNNAGATCGGAAG
58711_1 TATGAGTGTGGAGTCGTTGCACAATGGTGCAATCATGACTCACNNNAGATCGGAAG
58713_1 TATGAGTGTGGAGTCGTTGCTGTACTAGGCAGCCATTGAAACANNNAGATCGGAA
58715_1 TATGAGTGTGGAGTCGTTGCAAATGTCCAAACCTGTCTGACCTNNNAGATCGGAAG
58718_1 TATGAGTGTGGAGTCGTTGCTGACAAAAGGAGGGAGAATCAGGNNNAGATCGGA
58721_1 TATGAGTGTGGAGTCGTTGCGACAATCTCCATCTACGCCTACCNNNAGATCGGAAGA
58723_1 TATGAGTGTGGAGTCGTTGCGAAAACAAGACCCTCACAACAGCNNNAGATCGGAAG
58724_1 TATGAGTGTGGAGTCGTTGCCACATGCCTGTAGTCCTAGCTACNNNAGATCGGAAG
58725_1 TATGAGTGTGGAGTCGTTGCATGGACAAATCTCTTGGAAAGGTNNNAGATCGGAA
58726_1 TATGAGTGTGGAGTCGTTGCGAAGGATTGGAGCTGAGCCTTATNNNAGATCGGAA
58727_1 TATGAGTGTGGAGTCGTTGCACTTCTTTTGTCCTGTCACCTGANNNAGATCGGAAGA
58728_1 TATGAGTGTGGAGTCGTTGCCCCCAGGGTTCTTGTGTTTAGATNNNAGATCGGAAG
58729_1 TATGAGTGTGGAGTCGTTGCTGGACAACAGTAGCAAAATGCAGNNNAGATCGGAA
58731_1 TATGAGTGTGGAGTCGTTGCACATTTGCCAGTGAGTGAATTGANNNAGATCGGAA
58732_1 TATGAGTGTGGAGTCGTTGCTCACGTATCAACCCAAAAAGACANNNAGATCGGAAG
58733_1 TATGAGTGTGGAGTCGTTGCGTAATACCAGCTACTCAGGAGGCNNNAGATCGGAA
58734_1 TATGAGTGTGGAGTCGTTGCCAGTGAGCTATGATCACACCACTNNNAGATCGGAAG
58739_1 TATGAGTGTGGAGTCGTTGCCTGAGGCAGAGAATTGCTTGAACNNNAGATCGGAA
58740_1 TATGAGTGTGGAGTCGTTGCGGTGACATACAACAAGCAGCTTTNNNAGATCGGAAG
58741_1 TATGAGTGTGGAGTCGTTGCACCACAGTCACCATGATGTAACTNNNAGATCGGAAG
58743_1 TATGAGTGTGGAGTCGTTGCTGCCTGGAAAGAACAAAGTTGTCNNNAGATCGGAA
58744_1 TATGAGTGTGGAGTCGTTGCCAAATGGAAACACATCCCATGCTNNNAGATCGGAAG
58746_1 TATGAGTGTGGAGTCGTTGCCTGGCTAAAACAGTGAAACCCTGNNNAGATCGGAAG
58747_1 TATGAGTGTGGAGTCGTTGCTATATCCATCCCGCATTCAGTGGNNNAGATCGGAAG
58748_1 TATGAGTGTGGAGTCGTTGCCGGAGCTTCATTTTTTTCAAGCANNNAGATCGGAAG
58749_1 TATGAGTGTGGAGTCGTTGCAAGGATTGCACATTGAGTCTGTTNNNAGATCGGAAG
58750_1 TATGAGTGTGGAGTCGTTGCCCTCCTAAGTAGCCAGGACTACTNNNAGATCGGAAG
58752_1 TATGAGTGTGGAGTCGTTGCCCTCCTGGGTTCAAGTGATTCTCNNNAGATCGGAAG
58754_1 TATGAGTGTGGAGTCGTTGCCCCTGTCACACACACACAAATACNNNAGATCGGAAG
58755_1 TATGAGTGTGGAGTCGTTGCATGATCTTGACTCATGTCCCGAGNNNAGATCGGAAG
58756_1 TATGAGTGTGGAGTCGTTGCTAGTGCAGTAGCTATTCACAGGCNNNAGATCGGAAG
58757_1 TATGAGTGTGGAGTCGTTGCAGACAGATTCACACTCCATCACCNNNAGATCGGAAG
58758_1 TATGAGTGTGGAGTCGTTGCGGATTACAGACATGCACCAACACNNNAGATCGGAAG
58761_1 TATGAGTGTGGAGTCGTTGCGCAATGGTATGATCACAGCTCACNNNAGATCGGAAG
58763_1 TATGAGTGTGGAGTCGTTGCGATCTAGACTCACTGCAAGCTCCNNNAGATCGGAAG
58764_1 TATGAGTGTGGAGTCGTTGCGTTCAAGCAATTCTCTGCCTCAGNNNAGATCGGAAG
58765_1 TATGAGTGTGGAGTCGTTGCCTCCGAAGTAGCTGGGATTACAGNNNAGATCGGAA
58766_1 TATGAGTGTGGAGTCGTTGCGCAGTGGTACAATCATGGCTTACNNNAGATCGGAAG
58767_1 TATGAGTGTGGAGTCGTTGCCAGTCTGTTATCTAGGCTGGAGGNNNAGATCGGAA
58768_1 TATGAGTGTGGAGTCGTTGCCCTCCCAAGTAGCTAGGACTACANNNAGATCGGAAG

| | |
|---|---|
| 58771_1 | TATGAGTGTGGAGTCGTTGCCCTCCTGAGTAGCTAGGACTACANNNAGATCGGAAG |
| 58772_1 | TATGAGTGTGGAGTCGTTGCCCTGGGTTCAAGCAATTCTTCTGNNNAGATCGGAAG |
| 58775_1 | TATGAGTGTGGAGTCGTTGCCAAAGTGCTGGGATTAGAGGTGTNNNAGATCGGAA |
| 58776_1 | TATGAGTGTGGAGTCGTTGCCCTCCTGGGGCTTGAATGTTTTANNNAGATCGGAAG |
| 58777_1 | TATGAGTGTGGAGTCGTTGCGTACAATGGTGGTGCAATCTCAGNNNAGATCGGAA |
| 58778_1 | TATGAGTGTGGAGTCGTTGCCTGGTCCTTTTGTGAAGCAAGAANNNAGATCGGAA |
| 58780_1 | TATGAGTGTGGAGTCGTTGCATAGCTGGGAATACAGGTATGCGNNNAGATCGGAA |
| 58781_1 | TATGAGTGTGGAGTCGTTGCCAGTAGCAAGATCTCAGCTCACTNNNAGATCGGAAG |
| 58782_1 | TATGAGTGTGGAGTCGTTGCCTCCTGAGTAGCTGGGACTACAGNNNAGATCGGAA |
| 58783_1 | TATGAGTGTGGAGTCGTTGCCACTCCCAGGTTCATGCTATTCTNNNAGATCGGAAGA |
| 58784_1 | TATGAGTGTGGAGTCGTTGCAGAGACAGTGTCTGGCTATGTTGNNNAGATCGGAA |
| 58786_1 | TATGAGTGTGGAGTCGTTGCCTAACTCATTGGATTCCACCCCANNNAGATCGGAAGA |
| 58789_1 | TATGAGTGTGGAGTCGTTGCTTCAAGCAATTCTCCTGCCTCAGNNNAGATCGGAAG |
| 58791_1 | TATGAGTGTGGAGTCGTTGCATTTTCCTGCCTCCTCAATAGCTNNNAGATCGGAAGA |
| 58792_1 | TATGAGTGTGGAGTCGTTGCTCTGTGCTCATTCTGGTGATTCANNNAGATCGGAAG |
| 58793_1 | TATGAGTGTGGAGTCGTTGCGAGTAGCTGGGATTACAGGTTCCNNNAGATCGGAA |
| 58794_1 | TATGAGTGTGGAGTCGTTGCCAGTGGTACAATCTCAGCTCACTNNNAGATCGGAAG |
| 58795_1 | TATGAGTGTGGAGTCGTTGCATGGAGTCTCACTCTCTCGATCANNNAGATCGGAAG |
| 58796_1 | TATGAGTGTGGAGTCGTTGCCAGTGGTACAATCTCAGCTCACTNNNAGATCGGAAG |
| 58797_1 | TATGAGTGTGGAGTCGTTGCTGAGATTACAGGTATGTGCCACCNNNAGATCGGAA |
| 58798_1 | TATGAGTGTGGAGTCGTTGCCCCAGTAGTTCTCAGCCTCATTTNNNAGATCGGAAGA |
| 58799_1 | TATGAGTGTGGAGTCGTTGCCTGGTCTTGAACTCCTTAGCTCANNNAGATCGGAAG |
| 58800_1 | TATGAGTGTGGAGTCGTTGCATGTTGCCCAAGTTGGTTTTGAANNNAGATCGGAAG |
| 58802_1 | TATGAGTGTGGAGTCGTTGCGCCTCCCAGGTTCAAGTGATTATNNNAGATCGGAAG |
| 58803_1 | TATGAGTGTGGAGTCGTTGCGATTACAGGCATCCACCGCTATANNNAGATCGGAAG |
| 58804_1 | TATGAGTGTGGAGTCGTTGCGTACAATTTCGGCTCACTGCAAGNNNAGATCGGAAG |
| 58805_1 | TATGAGTGTGGAGTCGTTGCTGATCTTGGCTCACTGTAACCTCNNNAGATCGGAAG |
| 58806_1 | TATGAGTGTGGAGTCGTTGCTGGTAGAAATGGTTGCATCCTCANNNAGATCGGAA |
| 58807_1 | TATGAGTGTGGAGTCGTTGCAGCATGGATGACTGAATGAAGGTNNNAGATCGGAA |
| 58811_1 | TATGAGTGTGGAGTCGTTGCGCCTTCTGAGTAGCTGGGATTACNNNAGATCGGAA |
| 58812_1 | TATGAGTGTGGAGTCGTTGCCCTCCCAGATTCATGCCATTCTCNNNAGATCGGAAGA |
| 58813_1 | TATGAGTGTGGAGTCGTTGCCTTCCTGAGTAGCTGGGACTACANNNAGATCGGAAG |
| 58814_1 | TATGAGTGTGGAGTCGTTGCGAATCTCAGCTCACTGCAACCTCNNNAGATCGGAAG |
| 58815_1 | TATGAGTGTGGAGTCGTTGCCCTCCCAGGTTCAAGTGATTCTCNNNAGATCGGAAG |
| 58816_1 | TATGAGTGTGGAGTCGTTGCCTCTTGAGTAGCTGGGACTACAGNNNAGATCGGAA |
| 58817_1 | TATGAGTGTGGAGTCGTTGCATGGCATTCATGGTAACCTGGATNNNAGATCGGAA |
| 58818_1 | TATGAGTGTGGAGTCGTTGCAGGAGGGAGGATCCTGTTCTTATNNNAGATCGGAA |
| 58819_1 | TATGAGTGTGGAGTCGTTGCCTCTTATCGATGTTGTTCCTGCANNNAGATCGGAAG |
| 58820_1 | TATGAGTGTGGAGTCGTTGCGAGGTAAAATTAGCGTGGCAAGGNNNAGATCGGA |
| 58822_1 | TATGAGTGTGGAGTCGTTGCAAAGGAAGGTCTTGAATCTGGCTNNNAGATCGGAA |
| 58824_1 | TATGAGTGTGGAGTCGTTGCCAGTCATGGTGGTAAGAGTCTGTNNNAGATCGGAA |
| 58828_1 | TATGAGTGTGGAGTCGTTGCTCCTAAAAGTCTCCAAGTGCCAGNNNAGATCGGAAG |
| 58829_1 | TATGAGTGTGGAGTCGTTGCTATTGAGCTACTGCACTCCAGTCNNNAGATCGGAAG |

| | |
|---|---|
| 58831_1 | TATGAGTGTGGAGTCGTTGCGAGGGACCTCTTCACCAAAGTACNNNAGATCGGAAG |
| 58832_1 | TATGAGTGTGGAGTCGTTGCGCAACCCAAAAGGATCACTGTAGNNNAGATCGGAA |
| 58833_1 | TATGAGTGTGGAGTCGTTGCACGGAATCAACCTAAGTGTCCATNNNAGATCGGAAG |
| 58835_1 | TATGAGTGTGGAGTCGTTGCACTGTATACTCCCCTTCCCTACCNNNAGATCGGAAGA |
| 58836_1 | TATGAGTGTGGAGTCGTTGCAAAGCAGTTCAATGGAGAAAGGANNNAGATCGGA |
| 58837_1 | TATGAGTGTGGAGTCGTTGCCCACACATATCTCCAGAGAGCTCNNNAGATCGGAAG |
| 58839_1 | TATGAGTGTGGAGTCGTTGCGCAAGCAAGCAAACAAAATGACANNNAGATCGGAA |
| 58841_1 | TATGAGTGTGGAGTCGTTGCAGACACTCTTAGAGGAGCAAAGCNNNAGATCGGAA |
| 58842_1 | TATGAGTGTGGAGTCGTTGCGCAAGCAAGCAAACAAAATGACANNNAGATCGGAA |
| 58848_1 | TATGAGTGTGGAGTCGTTGCAGAAAGAAGTCAATCACCCCCTCNNNAGATCGGAAG |
| 58849_1 | TATGAGTGTGGAGTCGTTGCACTTTGGGATACTGAGGCAGAAGNNNAGATCGGAA |
| 58850_1 | TATGAGTGTGGAGTCGTTGCTAACTCAGTTCTCAACCTTGCCTNNNAGATCGGAAGA |
| 58851_1 | TATGAGTGTGGAGTCGTTGCTGGTGAATCCCCATCTCTACTAANNNAGATCGGAAG |
| 58852_1 | TATGAGTGTGGAGTCGTTGCACACAGTCAGAGGAGACAAAAATNNNAGATCGGAA |
| 58853_1 | TATGAGTGTGGAGTCGTTGCTCACACCTGTAATCCCAACACTTNNNAGATCGGAAGA |
| 58855_1 | TATGAGTGTGGAGTCGTTGCATCTTCTTGGAAAGAGGCCCAAANNNAGATCGGAA |
| 58856_1 | TATGAGTGTGGAGTCGTTGCGAGTGGTGGCACATTCCTATAGTNNNAGATCGGAA |
| 58858_1 | TATGAGTGTGGAGTCGTTGCTTATCAATTGCATGTCCCAGAGCNNNAGATCGGAAG |
| 58860_1 | TATGAGTGTGGAGTCGTTGCGGCTAGTGCAACTAGGGAACTAANNNAGATCGGAA |
| 58861_1 | TATGAGTGTGGAGTCGTTGCTCCAATTACCGAACTTAGGTGGANNNAGATCGGAAG |
| 58862_1 | TATGAGTGTGGAGTCGTTGCACTTGGCAACTGAGACTCTATGTNNNAGATCGGAAG |
| 58863_1 | TATGAGTGTGGAGTCGTTGCCCCAAAAGCAAGAAAAAGGACATNNNAGATCGGAA |
| 58865_1 | TATGAGTGTGGAGTCGTTGCATCTGAAGCTGGTGTCAAAGAGANNNAGATCGGAA |
| 58868_1 | TATGAGTGTGGAGTCGTTGCCTTAGAATTGGCAGTGGAGTCCTNNNAGATCGGAA |
| 58870_1 | TATGAGTGTGGAGTCGTTGCTGGGATGATTCACTAAGCCCAAANNNAGATCGGAA |
| 58872_1 | TATGAGTGTGGAGTCGTTGCTCCCTCAAAAACACCATGAGTTGNNNAGATCGGAAG |
| 58874_1 | TATGAGTGTGGAGTCGTTGCAGACCAAAAAAATGGAGAAGGAAANNNAGATCGG |
| 58875_1 | TATGAGTGTGGAGTCGTTGCCAGTTACCCACAGGATTATGCCTNNNAGATCGGAAG |
| 58876_1 | TATGAGTGTGGAGTCGTTGCAAGATCATACCACTGCACTCCAGNNNAGATCGGAAG |
| 58880_1 | TATGAGTGTGGAGTCGTTGCAAAAAAAATACTGCTCCCTGGGCNNNAGATCGGAAG |
| 58881_1 | TATGAGTGTGGAGTCGTTGCCACAAGGTGGCAAAGAGACAAAANNNAGATCGGAA |
| 58882_1 | TATGAGTGTGGAGTCGTTGCATCTCAGGAAGAGCAGGGAAATGNNNAGATCGGA |
| 58883_1 | TATGAGTGTGGAGTCGTTGCTCTGGGATAGATGAACACACACANNNAGATCGGAA |
| 58884_1 | TATGAGTGTGGAGTCGTTGCAGTGAGCAGAAATAGCTATGCCTNNNAGATCGGAA |
| 58886_1 | TATGAGTGTGGAGTCGTTGCGAGACAGGAGAATAGGGTGAACCNNNAGATCGGA |
| 58887_1 | TATGAGTGTGGAGTCGTTGCAGGAATGCAGTACTGATTCATGCNNNAGATCGGAA |
| 58888_1 | TATGAGTGTGGAGTCGTTGCAAGGCCCCGTTTGTACAAAAAAANNNAGATCGGAA |
| 58889_1 | TATGAGTGTGGAGTCGTTGCGGCAGGCACTTATAATCCAGCTANNNAGATCGGAAG |
| 58890_1 | TATGAGTGTGGAGTCGTTGCAAACAAAACACAGAGGGGAATGGNNNAGATCGGA |
| 58891_1 | TATGAGTGTGGAGTCGTTGCTTCCTGTGGAGATTTTCACTGAANNNAGATCGGAAG |
| 58892_1 | TATGAGTGTGGAGTCGTTGCCTGAAACTCCACGTCCTATTCCANNNAGATCGGAAGA |
| 58895_1 | TATGAGTGTGGAGTCGTTGCGGTGCCTATAATCCCAGCTACTCNNNAGATCGGAAG |
| 58897_1 | TATGAGTGTGGAGTCGTTGCCTCACATCTGTAATCCCAGCACTNNNAGATCGGAAGA |

| | |
|---|---|
| 58901_1 | TATGAGTGTGGAGTCGTTGCTACTTAGGGAGCTGAGGAAGGAANNNAGATCGGAA |
| 58902_1 | TATGAGTGTGGAGTCGTTGCGAGATTCTTGAGGCAGGAGTGAGNNNAGATCGGAA |
| 58903_1 | TATGAGTGTGGAGTCGTTGCAGGATCAGTAACAAGGGTAGCACNNNAGATCGGAA |
| 58904_1 | TATGAGTGTGGAGTCGTTGCGTGTCCCAGGAACTTTAGTCAAGNNNAGATCGGAAG |
| 58905_1 | TATGAGTGTGGAGTCGTTGCTATCCTAGGTTCCAGGCATCCAANNNAGATCGGAAGA |
| 58907_1 | TATGAGTGTGGAGTCGTTGCACTGAATTTTACTGCTGGGTTGGNNNAGATCGGAAG |
| 58908_1 | TATGAGTGTGGAGTCGTTGCGGAGTGGAAATGAGACCCTCTTTNNNAGATCGGAAG |
| 58911_1 | TATGAGTGTGGAGTCGTTGCTTTAGTAGAGACGAGGATTCGCCNNNAGATCGGAAG |
| 58913_1 | TATGAGTGTGGAGTCGTTGCCCACCTCCTGAGTTCAAGTGATTNNNAGATCGGAAGA |
| 58914_1 | TATGAGTGTGGAGTCGTTGCTAGCACTTTAGGAGGCTGAAGTGNNNAGATCGGAAG |
| 58915_1 | TATGAGTGTGGAGTCGTTGCCATGATCTGGGCTCACTGTAACCNNNAGATCGGAAG |
| 58916_1 | TATGAGTGTGGAGTCGTTGCTTCTGAGTTCACGCCATTCTACTNNNAGATCGGAAGA |
| 58917_1 | TATGAGTGTGGAGTCGTTGCCAAGTATCTGGGACTACAGGCTCNNNAGATCGGAAG |
| 58918_1 | TATGAGTGTGGAGTCGTTGCTTCAAGCAATTCTCCTGCCTCAGNNNAGATCGGAAGA |
| 58920_1 | TATGAGTGTGGAGTCGTTGCCTCCTGAGTAGCTGGGATTACAGNNNAGATCGGAAG |
| 58921_1 | TATGAGTGTGGAGTCGTTGCCCTCTCCATTGCACCTGTTGATANNNAGATCGGAAGA |
| 58922_1 | TATGAGTGTGGAGTCGTTGCATAGAATCTCGCTCTGTCTCCCTNNNAGATCGGAAGA |
| 58924_1 | TATGAGTGTGGAGTCGTTGCCACTGTGGGGATAGACCAGAATTNNNAGATCGGAAG |
| 58925_1 | TATGAGTGTGGAGTCGTTGCAGACAGAGTCTCACTTTGTTGCCNNNAGATCGGAAG |
| 58926_1 | TATGAGTGTGGAGTCGTTGCTGGAGTTCAGTGACATGATCTCGNNNAGATCGGAAG |
| 58928_1 | TATGAGTGTGGAGTCGTTGCGTGGCATGATCTCTGTTTACTGCNNNAGATCGGAAG |
| 58929_1 | TATGAGTGTGGAGTCGTTGCCTTCCCGAGTTCACACCATTCTCNNNAGATCGGAAGA |
| 58931_1 | TATGAGTGTGGAGTCGTTGCAGCAATTCTCTGTCTCAGCCTACNNNAGATCGGAAGA |
| 58932_1 | TATGAGTGTGGAGTCGTTGCCTTGGTTTTTGCCTAGGCTGAANNNAGATCGGAAG |
| 58933_1 | TATGAGTGTGGAGTCGTTGCGATCTCGGCTCAATGTAACCTCTNNNAGATCGGAAGA |
| 58934_1 | TATGAGTGTGGAGTCGTTGCTTCACATCATTCTCCTGCCTCAGNNNAGATCGGAAGA |
| 58935_1 | TATGAGTGTGGAGTCGTTGCAGCCTCCTGAGTAGCTAGGATTANNNAGATCGGAAG |
| 58936_1 | TATGAGTGTGGAGTCGTTGCCTTAGCTCACTGTAGCCTCTGTCNNNAGATCGGAAGA |
| 58937_1 | TATGAGTGTGGAGTCGTTGCTTCAAGTGATTCTCCTGCCTCAGNNNAGATCGGAAGA |
| 58938_1 | TATGAGTGTGGAGTCGTTGCCCTCTGAGTTCAGGTGATTCTCCNNNAGATCGGAAGA |
| 58941_1 | TATGAGTGTGGAGTCGTTGCTTCCTGGGTGATTTTCTTGGAGTNNNAGATCGGAAG |
| 58943_1 | TATGAGTGTGGAGTCGTTGCAAAAGCAAAAGTTCCACAGGGTCNNNAGATCGGAAG |
| 58945_1 | TATGAGTGTGGAGTCGTTGCACCTGTAATCCCAGCACTTTAGGNNNAGATCGGAAGA |
| 58946_1 | TATGAGTGTGGAGTCGTTGCAAGGGGAAAACATCTTTGAGGGTNNNAGATCGGAA |
| 58947_1 | TATGAGTGTGGAGTCGTTGCCAGGACTTCTGAGCATAGACAGGNNNAGATCGGAAG |
| 58948_1 | TATGAGTGTGGAGTCGTTGCGGAGGCTAAGACACAAGAATTGCNNNAGATCGGAA |
| 58950_1 | TATGAGTGTGGAGTCGTTGCAGAAAAGAAAAAATCGGCTGGGTNNNAGATCGGAA |
| 58951_1 | TATGAGTGTGGAGTCGTTGCGGCTCTGTCTGTCTCTTCCTAACNNNAGATCGGAAGA |
| 58952_1 | TATGAGTGTGGAGTCGTTGCTGTTTCCTAAGAGCTGACAGGACNNNAGATCGGAAG |
| 58954_1 | TATGAGTGTGGAGTCGTTGCCATGGGAGAAACCTCGTCTCTACNNNAGATCGGAAG |
| 58956_1 | TATGAGTGTGGAGTCGTTGCGGGGCTGTAGTATGCTGTGATTANNNAGATCGGAAG |
| 58961_1 | TATGAGTGTGGAGTCGTTGCTTCACAAAATTAGCCAGGCATGGNNNAGATCGGAAG |
| 58962_1 | TATGAGTGTGGAGTCGTTGCAGGCACATAGATCAAGGGAACAGNNNAGATCGGAA |

| | |
|---|---|
| 58963_1 | TATGAGTGTGGAGTCGTTGCTGTAATAGCCTCATGTTGCCTCGNNNAGATCGGAAG |
| 58964_1 | TATGAGTGTGGAGTCGTTGCAAAAAAGAAATAGGGCCAGTGGCNNNAGATCGGAA |
| 58965_1 | TATGAGTGTGGAGTCGTTGCGCAGATGTCTTGGACAATGACACNNNAGATCGGAA |
| 58968_1 | TATGAGTGTGGAGTCGTTGCCACACCTGTAATCCCAGCATCTTNNNAGATCGGAAG |
| 58969_1 | TATGAGTGTGGAGTCGTTGCACAGAGCGATGTCTTGTCTCAAANNNAGATCGGAAG |
| 58970_1 | TATGAGTGTGGAGTCGTTGCTTGACAGTAATAACACAGCCCCANNNAGATCGGAAG |
| 58974_1 | TATGAGTGTGGAGTCGTTGCCTGAGTCAGGAGAATGGTGTGAANNNAGATCGGAA |
| 58975_1 | TATGAGTGTGGAGTCGTTGCGCACGTTGAAATCACTAGGTTCCNNNAGATCGGAAG |
| 58977_1 | TATGAGTGTGGAGTCGTTGCTTCATGCAAAATCTAACGTGGGTNNNAGATCGGAAG |
| 58978_1 | TATGAGTGTGGAGTCGTTGCGAGTCTCACTTTGTTTTCCAGGCNNNAGATCGGAAG |
| 58980_1 | TATGAGTGTGGAGTCGTTGCCTCAAGTGATACTCCCACCTCAGNNNAGATCGGAAG |
| 58981_1 | TATGAGTGTGGAGTCGTTGCCAGAGCTTGGTGTGCATTTTACANNNAGATCGGAAG |
| 58985_1 | TATGAGTGTGGAGTCGTTGCCAAGTAGCTGGGACTACAGGTTCNNNAGATCGGAA |
| 58986_1 | TATGAGTGTGGAGTCGTTGCCAAGTAGCTGGGATTACAGGCATNNNAGATCGGAA |
| 58987_1 | TATGAGTGTGGAGTCGTTGCAGGATTGGGTAGATAGCTCCACTNNNAGATCGGAA |
| 58988_1 | TATGAGTGTGGAGTCGTTGCCCAGTCTCAGAACATCCTCCATTNNNAGATCGGAAG |
| 58989_1 | TATGAGTGTGGAGTCGTTGCCTCCTGAGTAGCTGGGATTACAGNNNAGATCGGAA |
| 58990_1 | TATGAGTGTGGAGTCGTTGCCAAAGTGCTGGGATTACAGGTGTNNNAGATCGGAA |
| 58991_1 | TATGAGTGTGGAGTCGTTGCCTCCTGAGTAGCTGGGACTACAGNNNAGATCGGAA |
| 58993_1 | TATGAGTGTGGAGTCGTTGCATTCCCTTCTACTAGCACCTTGCNNNAGATCGGAAGA |
| 58994_1 | TATGAGTGTGGAGTCGTTGCCTGATCTCAAACTCCTGGGTTCANNNAGATCGGAAG |
| 58996_1 | TATGAGTGTGGAGTCGTTGCTCAGTAGTGTGATCTTGGCTCACNNNAGATCGGAAG |
| 58997_1 | TATGAGTGTGGAGTCGTTGCAGATGGAGTTCACTCTTGTTCCCNNNAGATCGGAAG |
| 58998_1 | TATGAGTGTGGAGTCGTTGCACTCCTGACCTCAAATGATCCACNNNAGATCGGAAG |
| 58999_1 | TATGAGTGTGGAGTCGTTGCGCCTCCTGATTACCTGGGATTACNNNAGATCGGAAG |
| 59000_1 | TATGAGTGTGGAGTCGTTGCTCCCAGATTCAAGCAATTCTCCTNNNAGATCGGAAG |
| 59001_1 | TATGAGTGTGGAGTCGTTGCGCCTCCCGAAACCCCTATATTTTNNNAGATCGGAAGA |
| 59002_1 | TATGAGTGTGGAGTCGTTGCGTCAAGCACAAAATCAGGCATCTNNNAGATCGGAAG |
| 59003_1 | TATGAGTGTGGAGTCGTTGCAGTGCTAGGATTACAGGCATGAGNNNAGATCGGAA |
| 59005_1 | TATGAGTGTGGAGTCGTTGCCAACCTTTGCCTCCTCAATTCAGNNNAGATCGGAAGA |
| 59006_1 | TATGAGTGTGGAGTCGTTGCCCGGGTTCAAGTGATTATCATGCNNNAGATCGGAAG |
| 59007_1 | TATGAGTGTGGAGTCGTTGCCCGGGTTCAAGTGATTATCATGCNNNAGATCGGAAG |
| 59008_1 | TATGAGTGTGGAGTCGTTGCCTGGGTTCAAATGATTCTCCTGCNNNAGATCGGAAG |
| 59009_1 | TATGAGTGTGGAGTCGTTGCGCCTCCCTAGTAGCTGTGATTACNNNAGATCGGAAG |
| 59010_1 | TATGAGTGTGGAGTCGTTGCAGGTTCAAGCAATTATCCCACCTNNNAGATCGGAAG |
| 59011_1 | TATGAGTGTGGAGTCGTTGCCAGTCTCCCAAGTAGCTGGAATTNNNAGATCGGAAG |
| 59015_1 | TATGAGTGTGGAGTCGTTGCTCAGGAGACTGAGGAAGAAGGATNNNAGATCGGA |
| 59016_1 | TATGAGTGTGGAGTCGTTGCTGAAGAAGAGGTTGACAGTAGGCNNNAGATCGGA |
| 59017_1 | TATGAGTGTGGAGTCGTTGCGCACAATGAACCAAGGGTTTACANNNAGATCGGAA |
| 59018_1 | TATGAGTGTGGAGTCGTTGCACTTAGTGATTCCTGTTGAGGGGNNNAGATCGGAA |
| 59019_1 | TATGAGTGTGGAGTCGTTGCTGCTCCCAGGAAAATGTTTTCACNNNAGATCGGAAG |
| 59022_1 | TATGAGTGTGGAGTCGTTGCATGGTGGTGAATGCCTGTAATCTNNNAGATCGGAA |
| 59025_1 | TATGAGTGTGGAGTCGTTGCCATGTTGCTGGTGGGATTGTAAANNNAGATCGGAA |

| | |
|---|---|
| 59027_1 | TATGAGTGTGGAGTCGTTGCGGAGAATGGAAGTTAGGAGGCAANNNAGATCGGA |
| 59028_1 | TATGAGTGTGGAGTCGTTGCGGGCACAATGGACAATTTCTTCANNNAGATCGGAAG |
| 59029_1 | TATGAGTGTGGAGTCGTTGCAGAAGGGAAATGGACAGTTCTCTNNNAGATCGGAA |
| 59030_1 | TATGAGTGTGGAGTCGTTGCTAGCCAGACCAACTTATCAAGGGNNNAGATCGGAAG |
| 59031_1 | TATGAGTGTGGAGTCGTTGCACACCTTGGGCAAGAAAGTAAGANNNAGATCGGAA |
| 59033_1 | TATGAGTGTGGAGTCGTTGCGCACACACATGGATAATCATGGGNNNAGATCGGAA |
| 59034_1 | TATGAGTGTGGAGTCGTTGCGGGGAATTGGCATAAATGGCATGNNNAGATCGGAA |
| 59035_1 | TATGAGTGTGGAGTCGTTGCGGGATGAAGGAAGGGAACAAACTNNNAGATCGGA |
| 59037_1 | TATGAGTGTGGAGTCGTTGCTTTTTTGTAAAGACGAGGGCCTCNNNAGATCGGAAG |
| 59038_1 | TATGAGTGTGGAGTCGTTGCCTCCCTAGGCCTTTTCATATCCCNNNAGATCGGAAGA |
| 59039_1 | TATGAGTGTGGAGTCGTTGCATCCCTGTGGTGTCATCTAACACNNNAGATCGGAAG |
| 59040_1 | TATGAGTGTGGAGTCGTTGCATCACAGAGTTCAGCTGAAACCANNNAGATCGGAAG |
| 59041_1 | TATGAGTGTGGAGTCGTTGCAGGGTATAGCTGACATTGTTGGCNNNAGATCGGAA |
| 59042_1 | TATGAGTGTGGAGTCGTTGCCACACACACACAAAACTTGACCTNNNAGATCGGAAG |
| 59043_1 | TATGAGTGTGGAGTCGTTGCAGCAGTCTTGGAGAAAAGGAGACNNNAGATCGGAA |
| 59044_1 | TATGAGTGTGGAGTCGTTGCCCTCAGTGAGCAAAGGAAGATCTNNNAGATCGGAA |
| 59046_1 | TATGAGTGTGGAGTCGTTGCAAAGAAGAGGTTGGTTAGGGGAGNNNAGATCGGA |
| 59047_1 | TATGAGTGTGGAGTCGTTGCTAGCTAAGCTAAGCAGACCCCTANNNAGATCGGAAG |
| 59048_1 | TATGAGTGTGGAGTCGTTGCCACGAAGTCAGGAGTTTGAAACCNNNAGATCGGAA |
| 59049_1 | TATGAGTGTGGAGTCGTTGCCAGGGAAAAGCAAGGGCAAAAAANNNAGATCGGA |
| 59051_1 | TATGAGTGTGGAGTCGTTGCGCAGATGAAGTTCAACCTAGGGANNNAGATCGGAA |
| 59052_1 | TATGAGTGTGGAGTCGTTGCATGCCTCTAATCCCAGCTACTTGNNNAGATCGGAAG |
| 59056_1 | TATGAGTGTGGAGTCGTTGCTGCCTTAGCCTCTCATAGTGTTGNNNAGATCGGAAG |
| 59059_1 | TATGAGTGTGGAGTCGTTGCCAAAGTGCTGGCATTATAGGCATNNNAGATCGGAA |
| 59060_1 | TATGAGTGTGGAGTCGTTGCCCTGAGTAGCTGGGATTATAGGCNNNAGATCGGAA |
| 59061_1 | TATGAGTGTGGAGTCGTTGCGATTCAAGTGATTCTGCTGCCTCNNNAGATCGGAAG |
| 59063_1 | TATGAGTGTGGAGTCGTTGCATCTCTTGACCTTGTGATCCACCNNNAGATCGGAAG |
| 59064_1 | TATGAGTGTGGAGTCGTTGCCCCAGGTTCAAGCAATTCTTCTGNNNAGATCGGAAG |
| 59065_1 | TATGAGTGTGGAGTCGTTGCCCATGGCACAATCTTGACTCATCNNNAGATCGGAAG |
| 59068_1 | TATGAGTGTGGAGTCGTTGCGAAGGATGGAGTCTCACTCTGTCNNNAGATCGGAA |
| 59069_1 | TATGAGTGTGGAGTCGTTGCCAGGGTCTCACTATGTTGCCTAGNNNAGATCGGAAG |
| 59070_1 | TATGAGTGTGGAGTCGTTGCTGGGATTACAGGTGAATGCTACCNNNAGATCGGAA |
| 59071_1 | TATGAGTGTGGAGTCGTTGCCCTCCTGAGTAGCTGGGACTATANNNAGATCGGAAG |
| 59072_1 | TATGAGTGTGGAGTCGTTGCCCTGGGCTTTTGTTAGGAGACTTNNNAGATCGGAAG |
| 59073_1 | TATGAGTGTGGAGTCGTTGCAGAATAGCTGGGATTACAGACGCNNNAGATCGGAA |
| 59076_1 | TATGAGTGTGGAGTCGTTGCAGACGGTGTTTCATTCTTGTTGCNNNAGATCGGAAG |
| 59077_1 | TATGAGTGTGGAGTCGTTGCAAAAGGCCTTAGACAGTGGATCCNNNAGATCGGAA |
| 59079_1 | TATGAGTGTGGAGTCGTTGCAACCTCCTGAGTAGCTGGGATTANNNAGATCGGAAG |
| 59080_1 | TATGAGTGTGGAGTCGTTGCGTTCAAGCAATTCTCTGCCTCAGNNNAGATCGGAAG |
| 59081_1 | TATGAGTGTGGAGTCGTTGCTCCTCTCCTTTTCTTTGTCCCTGNNNAGATCGGAAGA |
| 59084_1 | TATGAGTGTGGAGTCGTTGCCTCCTGGTATAACGCCTAAGGTCNNNAGATCGGAAG |
| 59085_1 | TATGAGTGTGGAGTCGTTGCTTGCTTCTATCTCTGTGCTTGGANNNAGATCGGAAG |
| 59087_1 | TATGAGTGTGGAGTCGTTGCGATGTTGGCTTACTGCAACCTTTNNNAGATCGGAAG |

| | |
|---|---|
| 59088_1 | TATGAGTGTGGAGTCGTTGCCCTCCTGAGTAGCTGAGACTACANNNAGATCGGAAG |
| 59089_1 | TATGAGTGTGGAGTCGTTGCGCAATCTCAGCTCACCATATCCTNNNAGATCGGAAGA |
| 59090_1 | TATGAGTGTGGAGTCGTTGCCTCCTGAGTAGCTGGGATTACAGNNNAGATCGGAA |
| 59092_1 | TATGAGTGTGGAGTCGTTGCGAGTAGCTGGGATTACAGGCATGNNNAGATCGGAA |
| 59095_1 | TATGAGTGTGGAGTCGTTGCCGATCTTGGCTCACTGTAACTTCNNNAGATCGGAAG |
| 59096_1 | TATGAGTGTGGAGTCGTTGCTTCAAGCGATTCTTCCGTCTCAGNNNAGATCGGAAG |
| 59097_1 | TATGAGTGTGGAGTCGTTGCGATCACAGCTCATTTGCAGTCTCNNNAGATCGGAAG |
| 59098_1 | TATGAGTGTGGAGTCGTTGCCATTCTTCTACCTCAGCCTCCTGNNNAGATCGGAAGA |
| 59100_1 | TATGAGTGTGGAGTCGTTGCCAATGGCATGATCTGAGCTCACTNNNAGATCGGAAG |
| 59102_1 | TATGAGTGTGGAGTCGTTGCCCTCAAACTCTTTGGGGTCAAACNNNAGATCGGAAG |
| 59103_1 | TATGAGTGTGGAGTCGTTGCCAGTGGTGTAAGTGCAATCAAGGNNNAGATCGGAA |
| 59105_1 | TATGAGTGTGGAGTCGTTGCGCCTCCTAAGTAGCTGGGATTACNNNAGATCGGAAG |
| 59107_1 | TATGAGTGTGGAGTCGTTGCCTCAAGTAATCCTCCCACCTCAGNNNAGATCGGAAGA |
| 59108_1 | TATGAGTGTGGAGTCGTTGCTAGAGTGCAATGGCGTGATCTTANNNAGATCGGAA |
| 59112_1 | TATGAGTGTGGAGTCGTTGCATGCTGGAGTTCAGTGGTACAATNNNAGATCGGAA |
| 59114_1 | TATGAGTGTGGAGTCGTTGCTCCTGAGTAGCTGGGATTACAGANNNAGATCGGAA |
| 59115_1 | TATGAGTGTGGAGTCGTTGCAGTGCTGGGATTATAGGTGTGAGNNNAGATCGGAA |
| 59117_1 | TATGAGTGTGGAGTCGTTGCGCCTTCTGGGTTCAAGAGATTCTNNNAGATCGGAAG |
| 59119_1 | TATGAGTGTGGAGTCGTTGCCTGCCTCAATTTCAGAGCTTGTTNNNAGATCGGAAG |
| 59120_1 | TATGAGTGTGGAGTCGTTGCGATTGTCCTGCCTCAGTCTCCNNNAGATCGGAAGAG |
| 59122_1 | TATGAGTGTGGAGTCGTTGCTCTCAAACTCCTGACCTCAAGTGNNNAGATCGGAAG |
| 59126_1 | TATGAGTGTGGAGTCGTTGCTGGTGCAATCTTGGATCACTGTANNNAGATCGGAAG |
| 59128_1 | TATGAGTGTGGAGTCGTTGCTGATCTCAACTCACTGCAAGCTCNNNAGATCGGAAG |
| 59131_1 | TATGAGTGTGGAGTCGTTGCCCAACTAGCTGGGATTACAGGAGNNNAGATCGGAA |
| 59132_1 | TATGAGTGTGGAGTCGTTGCCATGATCTGGCTCACTGTAACCTNNNAGATCGGAAG |
| 59133_1 | TATGAGTGTGGAGTCGTTGCCATGGCTGTATGACAAGGACTCTNNNAGATCGGAA |
| 59134_1 | TATGAGTGTGGAGTCGTTGCGCCTCTCAGGTATCTGGGACTAANNNAGATCGGAAG |
| 59135_1 | TATGAGTGTGGAGTCGTTGCTTGAGTAGCTGGGATTACAGGTGNNNAGATCGGAA |
| 59136_1 | TATGAGTGTGGAGTCGTTGCTGAGTAGCTGGAACTACAAGTGCNNNAGATCGGAA |
| 59137_1 | TATGAGTGTGGAGTCGTTGCTACTGCACCTGGCCTAGAATTTTNNNAGATCGGAAG |
| 59138_1 | TATGAGTGTGGAGTCGTTGCTCTTCCCCTCTAGAACTGCACTANNNAGATCGGAAGA |
| 59140_1 | TATGAGTGTGGAGTCGTTGCGGGAAATGTAGGGCAAGAGAGAANNNAGATCGGA |
| 59141_1 | TATGAGTGTGGAGTCGTTGCTTCAAGTGATTCTCCTGCCTCAGNNNAGATCGGAAG |
| 59142_1 | TATGAGTGTGGAGTCGTTGCTATCAGCTCATTGCAACCTCCTTNNNAGATCGGAAGA |
| 59143_1 | TATGAGTGTGGAGTCGTTGCTGAACTCCTGGCCTCAAGTAATCNNNAGATCGGAAG |
| 59144_1 | TATGAGTGTGGAGTCGTTGCATGATCTCAACTCACTGCAACCTNNNAGATCGGAAG |
| 59145_1 | TATGAGTGTGGAGTCGTTGCCCTCATGGGTTCAAGTGATCCTCNNNAGATCGGAAG |
| 59146_1 | TATGAGTGTGGAGTCGTTGCATGAGGTCTCATTATGTTGCCCANNNAGATCGGAAG |
| 59149_1 | TATGAGTGTGGAGTCGTTGCTGCCCAAAAAAACCACAGATTCANNNAGATCGGAAG |
| 59152_1 | TATGAGTGTGGAGTCGTTGCGGTTCCAGTTTCTCCGTATCCTTNNNAGATCGGAAGA |
| 59153_1 | TATGAGTGTGGAGTCGTTGCAGCCTCAGCAACATAGTGAGATTNNNAGATCGGAA |
| 59155_1 | TATGAGTGTGGAGTCGTTGCCTAGGTGGGTGGATAACAAGGTCNNNAGATCGGAA |
| 59156_1 | TATGAGTGTGGAGTCGTTGCATGTGGAAGCAACTTTGGAACTGNNNAGATCGGAA |

| | |
|---|---|
| 59157_1 | TATGAGTGTGGAGTCGTTGCAATGAACCCTGAAAACATGCTCGNNNAGATCGGAAG |
| 59158_1 | TATGAGTGTGGAGTCGTTGCTTGAGGTTGCAGTGAGCTATGATNNNAGATCGGAA |
| 59162_1 | TATGAGTGTGGAGTCGTTGCTCCTGGGAACAAAAGCTTCATGANNNAGATCGGAAG |
| 59163_1 | TATGAGTGTGGAGTCGTTGCTGAAAACCTGTCCTCAATGTTGTNNNAGATCGGAAG |
| 59165_1 | TATGAGTGTGGAGTCGTTGCGAAAGCTGCAGTTATTTGCTTGCNNNAGATCGGAAG |
| 59166_1 | TATGAGTGTGGAGTCGTTGCAAGCAAATGACATGAACAGGTGGNNNAGATCGGAA |
| 59167_1 | TATGAGTGTGGAGTCGTTGCTCTTCTTGTCATCTCCCACATCTNNNAGATCGGAAGA |
| 59168_1 | TATGAGTGTGGAGTCGTTGCCGTTATACTGGTAGGAAGCACGANNNAGATCGGAA |
| 59170_1 | TATGAGTGTGGAGTCGTTGCCTAACTAGGGAAACTGAGGCACANNNAGATCGGAA |
| 59172_1 | TATGAGTGTGGAGTCGTTGCAGTTTCAGTCAGAGCATCAGGTTNNNAGATCGGAAG |
| 59173_1 | TATGAGTGTGGAGTCGTTGCCAAAGTGTTGGGATTTCAGGCATNNNAGATCGGAA |
| 59174_1 | TATGAGTGTGGAGTCGTTGCAACAAAAAACGGAAAGAGTGGCANNNAGATCGGA |
| 59177_1 | TATGAGTGTGGAGTCGTTGCTGGAGGAGTTCAGTTGAGTCTTGNNNAGATCGGAA |
| 59178_1 | TATGAGTGTGGAGTCGTTGCGGCAAACCAACTTCCTATTTTGCNNNAGATCGGAAG |
| 59179_1 | TATGAGTGTGGAGTCGTTGCCTGTAATCCCAGCTACTCAGGAGNNNAGATCGGAAG |
| 59183_1 | TATGAGTGTGGAGTCGTTGCCTGTAGTCCTAGCTACTCGGGAGNNNAGATCGGAAG |
| 59184_1 | TATGAGTGTGGAGTCGTTGCTAAACATTAGGTAGCTGGGCACANNNAGATCGGAA |
| 59186_1 | TATGAGTGTGGAGTCGTTGCCCTGGGGAAAGCAGTTTAGAGATNNNAGATCGGAA |
| 59190_1 | TATGAGTGTGGAGTCGTTGCAAGTCTCTCAAGATAGGGGAGAANNNAGATCGGAA |
| 59192_1 | TATGAGTGTGGAGTCGTTGCCATCTCATGCCACTAACCCCAATNNNAGATCGGAAGA |
| 59196_1 | TATGAGTGTGGAGTCGTTGCACTGTAGCCTAATAAGTGCACAANNNAGATCGGAAG |
| 59197_1 | TATGAGTGTGGAGTCGTTGCGTCCTGATCCCTGAGCTAACTTCNNNAGATCGGAAG |
| 59198_1 | TATGAGTGTGGAGTCGTTGCGCAGGTGGACCATAAGTTCAAGANNNAGATCGGAA |
| 59199_1 | TATGAGTGTGGAGTCGTTGCAAAGCAAAGCAAAAGGAAATGGCNNNAGATCGGA |
| 59200_1 | TATGAGTGTGGAGTCGTTGCTTGGTAGTTTGAAACTGGCCATGNNNAGATCGGAAG |
| 59202_1 | TATGAGTGTGGAGTCGTTGCCACACACACACACAGATGACAATNNNAGATCGGAAG |
| 59205_1 | TATGAGTGTGGAGTCGTTGCTCCACTACTGACTCATGTGTTCCNNNAGATCGGAAGA |
| 59207_1 | TATGAGTGTGGAGTCGTTGCCACACACATACACACACACACATNNNAGATCGGAAG |
| 59208_1 | TATGAGTGTGGAGTCGTTGCACTAGAAGATAGCAGAGGCAGGANNNAGATCGGA |
| 59209_1 | TATGAGTGTGGAGTCGTTGCTATGGTCCCTATGATGCAGCAAANNNAGATCGGAAG |
| 59210_1 | TATGAGTGTGGAGTCGTTGCGCAGTGTGCTTTGCTGATTTTAGNNNAGATCGGAAG |
| 59211_1 | TATGAGTGTGGAGTCGTTGCGAAAAACCTGACAAACACTGCCTNNNAGATCGGAAG |
| 59212_1 | TATGAGTGTGGAGTCGTTGCCCTCATCCTGAAGGCTATAGCAGNNNAGATCGGAAG |
| 59214_1 | TATGAGTGTGGAGTCGTTGCCAACTGCAAAGATGGGGGAAAGNNNAGATCGGA |
| 59215_1 | TATGAGTGTGGAGTCGTTGCGGAAAATTTTTGGACCCAAAGCCNNNAGATCGGAAG |
| 59217_1 | TATGAGTGTGGAGTCGTTGCCAGGAGGAGAAGCTCACTCAATTNNNAGATCGGAA |
| 59218_1 | TATGAGTGTGGAGTCGTTGCGAGAGTATCAGGACAGTGTGGTGNNNAGATCGGA |
| 59219_1 | TATGAGTGTGGAGTCGTTGCTTACCCCATTCCCATGGCTTTTANNNAGATCGGAAGA |
| 59221_1 | TATGAGTGTGGAGTCGTTGCCTCATGCCTATAATCCCAGCACTNNNAGATCGGAAGA |
| 59224_1 | TATGAGTGTGGAGTCGTTGCCTCAGGGCCTTTGCATATCCTATNNNAGATCGGAAG |
| 59225_1 | TATGAGTGTGGAGTCGTTGCCCCGATTCAAGTGATTCTCCTGTNNNAGATCGGAAG |
| 59226_1 | TATGAGTGTGGAGTCGTTGCACCTCCTGGATTCAAACGATTCTNNNAGATCGGAAG |
| 59227_1 | TATGAGTGTGGAGTCGTTGCTGTTAGCCAGGATGGTCTTGATCNNNAGATCGGAAG |

| | |
|---|---|
| 59228_1 | TATGAGTGTGGAGTCGTTGCGCTGGTCTTGAATTCCTTGGTTCNNNAGATCGGAAG |
| 59232_1 | TATGAGTGTGGAGTCGTTGCTTTGAAAACGTATGTCCGCACAANNNAGATCGGAAG |
| 59234_1 | TATGAGTGTGGAGTCGTTGCAAGACAGGATTTCACCATGTTGGNNNAGATCGGAA |
| 59235_1 | TATGAGTGTGGAGTCGTTGCGCCTCCTGAGTAGCTGAGATTACNNNAGATCGGAAG |
| 59236_1 | TATGAGTGTGGAGTCGTTGCTAGGACAATAGTGGAGGGAAGGTNNNAGATCGGA |
| 59237_1 | TATGAGTGTGGAGTCGTTGCGGATTTGTACAGGCATGTGACACNNNAGATCGGAA |
| 59238_1 | TATGAGTGTGGAGTCGTTGCAAGTCTCACTCTTGTCATCCAGGNNNAGATCGGAAG |
| 59239_1 | TATGAGTGTGGAGTCGTTGCGGTTCAAGCAATTCTCCTGTCTCNNNAGATCGGAAG |
| 59240_1 | TATGAGTGTGGAGTCGTTGCCCTCCTGAGTAGCTGGGACTATANNNAGATCGGAAG |
| 59242_1 | TATGAGTGTGGAGTCGTTGCGTAGAGACGGAGTTTCACCATGTNNNAGATCGGAA |
| 59243_1 | TATGAGTGTGGAGTCGTTGCCCAGGTTCAAGCAATTCTAGTGCNNNAGATCGGAAG |
| 59247_1 | TATGAGTGTGGAGTCGTTGCCTGTGTTCAAGTGATTCTCCTGCNNNAGATCGGAAG |
| 59248_1 | TATGAGTGTGGAGTCGTTGCCTCAAGTGACCCTACTGTCTCAGNNNAGATCGGAAG |
| 59250_1 | TATGAGTGTGGAGTCGTTGCGATTCAAGCAATTCTCCTGCCTCNNNAGATCGGAAG |
| 59251_1 | TATGAGTGTGGAGTCGTTGCATGTTCAAACAATTCTCCTGCCGNNNAGATCGGAAG |
| 59252_1 | TATGAGTGTGGAGTCGTTGCCTCCTGAGAAGCTGGGATTACAGNNNAGATCGGAA |
| 59253_1 | TATGAGTGTGGAGTCGTTGCATCTTGGCTCACTGTAACCTCTGNNNAGATCGGAAG |
| 59254_1 | TATGAGTGTGGAGTCGTTGCTTCACTTCATTCTCCTGCCTCAGNNNAGATCGGAAGA |
| 59255_1 | TATGAGTGTGGAGTCGTTGCAGATGGATTCTCACTCTGTTGCCNNNAGATCGGAAG |
| 59257_1 | TATGAGTGTGGAGTCGTTGCCCAAGTTCAAGTGATTCTCCTGCNNNAGATCGGAAG |
| 59258_1 | TATGAGTGTGGAGTCGTTGCCCCTAGTAGCTGAGATTACAGGCNNNAGATCGGAAG |
| 59259_1 | TATGAGTGTGGAGTCGTTGCGATCAGGTGATTCTCCCACTTCANNNAGATCGGAAG |
| 59261_1 | TATGAGTGTGGAGTCGTTGCTCTCGCTCTTGTACTATCTTGGCNNNAGATCGGAAG |
| 59263_1 | TATGAGTGTGGAGTCGTTGCCCAAGTAGCTGGGATTACAGGTGNNNAGATCGGAA |
| 59266_1 | TATGAGTGTGGAGTCGTTGCCAATCTTGGCTTACTGCAACCTCNNNAGATCGGAAG |
| 59267_1 | TATGAGTGTGGAGTCGTTGCTTTTGGCTCACTACAACTTCTGCNNNAGATCGGAAG |
| 59271_1 | TATGAGTGTGGAGTCGTTGCCCTCCTGGGTTCACAACATTCTCNNNAGATCGGAAG |
| 59274_1 | TATGAGTGTGGAGTCGTTGCGAATTTGCCTTGGGACACTGAAANNNAGATCGGAA |
| 59275_1 | TATGAGTGTGGAGTCGTTGCTCTCAAACTCCTGACCTCAAGTGNNNAGATCGGAAG |
| 59277_1 | TATGAGTGTGGAGTCGTTGCCAGACCATTTGGGAGATGTACCANNNAGATCGGAA |
| 59278_1 | TATGAGTGTGGAGTCGTTGCAATCTTGACTCACTGCAACCTCTNNNAGATCGGAAG |
| 59279_1 | TATGAGTGTGGAGTCGTTGCTCATGGTTTACTACAGCCTCCACNNNAGATCGGAAG |
| 59280_1 | TATGAGTGTGGAGTCGTTGCCTCCTGAGTAGCTGGGATTACAGNNNAGATCGGAA |
| 59281_1 | TATGAGTGTGGAGTCGTTGCCACTTCCCAGGTTCCAGTGATTANNNAGATCGGAAG |
| 59282_1 | TATGAGTGTGGAGTCGTTGCGTTTCACCATGTTCACCAGGATGNNNAGATCGGAAG |
| 59285_1 | TATGAGTGTGGAGTCGTTGCATGATCTCAGCTCACTGCAACTTNNNAGATCGGAAG |
| 59286_1 | TATGAGTGTGGAGTCGTTGCAGTGATGCAACCTCAGCTCATTANNNAGATCGGAAG |
| 59289_1 | TATGAGTGTGGAGTCGTTGCCTGAGTAGCTGGGATTACAGGTGNNNAGATCGGAA |
| 59290_1 | TATGAGTGTGGAGTCGTTGCCCTCCTGGGTTCAAGCGATTATCNNNAGATCGGAAG |
| 59292_1 | TATGAGTGTGGAGTCGTTGCCCGGGTTCAAGTGATTCTCCTATNNNAGATCGGAAG |
| 59293_1 | TATGAGTGTGGAGTCGTTGCAGTAGCTGGGATTACAGGCATACNNNAGATCGGAA |
| 59294_1 | TATGAGTGTGGAGTCGTTGCATCTTGGCTCACTACAACCTCTGNNNAGATCGGAAG |
| 59296_1 | TATGAGTGTGGAGTCGTTGCCAATTTTGGGTCACTGGAACCTCNNNAGATCGGAAG |

| | |
|---|---|
| 59297_1 | TATGAGTGTGGAGTCGTTGCCTAGAGTAGCTGGGACTACAGGTNNNAGATCGGAA |
| 59298_1 | TATGAGTGTGGAGTCGTTGCCTCTATTGCCCAGGCTGGAATACNNNAGATCGGAAG |
| 59300_1 | TATGAGTGTGGAGTCGTTGCCTGAGGCAGGAGAATCACTTGAANNNAGATCGGAA |
| 59302_1 | TATGAGTGTGGAGTCGTTGCGGAGATGAGCAGCTGACTTATGANNNAGATCGGAA |
| 59303_1 | TATGAGTGTGGAGTCGTTGCGGTGAGTCACTTACATCACCTTGNNNAGATCGGAAG |
| 59304_1 | TATGAGTGTGGAGTCGTTGCTGTAATCCTAGCACTTGTGGGAGNNNAGATCGGAA |
| 59305_1 | TATGAGTGTGGAGTCGTTGCCAATGGCACAGGTTCATCATACCNNNAGATCGGAAG |
| 59309_1 | TATGAGTGTGGAGTCGTTGCCTGCAGAAGTTGAACAAGCCAAANNNAGATCGGAA |
| 59311_1 | TATGAGTGTGGAGTCGTTGCGCTGTGAAGGACCTGGGTTATAANNNAGATCGGAA |
| 59317_1 | TATGAGTGTGGAGTCGTTGCGTAATGGTGCATGCCTGTAATCCNNNAGATCGGAAG |
| 59319_1 | TATGAGTGTGGAGTCGTTGCAAAGAAAAAAAATTGGCCCTGCGNNNAGATCGGAA |
| 59320_1 | TATGAGTGTGGAGTCGTTGCTGCCTTTACCCCCTCTGTATCTANNNAGATCGGAAGA |
| 59322_1 | TATGAGTGTGGAGTCGTTGCTACATACAGCTTTCCATCTGGCCNNNAGATCGGAAG |
| 59323_1 | TATGAGTGTGGAGTCGTTGCGGCTTGCACCTATAATCCCAGAANNNAGATCGGAAG |
| 59324_1 | TATGAGTGTGGAGTCGTTGCAATTTGGTGAAACCCCGTCTCTANNNAGATCGGAAG |
| 59325_1 | TATGAGTGTGGAGTCGTTGCTCTTATGGCACATTTCTGTTCCANNNAGATCGGAAG |
| 59327_1 | TATGAGTGTGGAGTCGTTGCACTCCCTGAGAAATCCTGTTTCCNNNAGATCGGAAG |
| 59328_1 | TATGAGTGTGGAGTCGTTGCTACATTGATTTCTTTCGGGGCCTNNNAGATCGGAAG |
| 59331_1 | TATGAGTGTGGAGTCGTTGCGTAATCCCAGTTACTCAGGAGGCNNNAGATCGGAA |
| 59332_1 | TATGAGTGTGGAGTCGTTGCCAGCTCCAATCAAGGAAGCAAAANNNAGATCGGAA |
| 59333_1 | TATGAGTGTGGAGTCGTTGCGGTCTCTCATTCGGTTCTGAAGTNNNAGATCGGAAG |
| 59334_1 | TATGAGTGTGGAGTCGTTGCGCATGAAACCCAACTCAGGAAAANNNAGATCGGAA |
| 59336_1 | TATGAGTGTGGAGTCGTTGCTACAGTGAAGGGCAGTGTGTTAGNNNAGATCGGAA |
| 59337_1 | TATGAGTGTGGAGTCGTTGCCTATCTCAACCCCTGCCCTAAAANNNAGATCGGAAGA |
| 59338_1 | TATGAGTGTGGAGTCGTTGCTCACACCTGTAATCCCAACACTTNNNAGATCGGAAGA |
| 59340_1 | TATGAGTGTGGAGTCGTTGCACAAGCGCTCTAAGTATGTGAGANNNAGATCGGAA |
| 59341_1 | TATGAGTGTGGAGTCGTTGCTCCAGATGTGCACATTGATACCANNNAGATCGGAAG |
| 59342_1 | TATGAGTGTGGAGTCGTTGCTCTTTCTGTTCTGCCTTCTCAGANNNAGATCGGAAGA |
| 59343_1 | TATGAGTGTGGAGTCGTTGCTGATTCTGTGTTTGGCAACTGTCNNNAGATCGGAAG |
| 59344_1 | TATGAGTGTGGAGTCGTTGCCAGTAAAACTGGCCAGGCATAGTNNNAGATCGGAA |
| 59345_1 | TATGAGTGTGGAGTCGTTGCCCTTCAGGCTTTACTCAGAGTCANNNAGATCGGAAG |
| 59346_1 | TATGAGTGTGGAGTCGTTGCAAAAGGAGTAAACTGAGGCCTGGNNNAGATCGGAA |
| 59347_1 | TATGAGTGTGGAGTCGTTGCTTGGATGCACAGAGGAATCATCTNNNAGATCGGAA |
| 59348_1 | TATGAGTGTGGAGTCGTTGCGTGTCAGGAGGGAGTATGGTAACNNNAGATCGGA |
| 59350_1 | TATGAGTGTGGAGTCGTTGCAAAATTCGCTTTGCACATCCTCANNNAGATCGGAAG |
| 59353_1 | TATGAGTGTGGAGTCGTTGCTCTGCAGAGACCCAAAATCTAGGNNNAGATCGGAA |
| 59354_1 | TATGAGTGTGGAGTCGTTGCCACAGACTCAACATGTGTGTGTGNNNAGATCGGAA |
| 59356_1 | TATGAGTGTGGAGTCGTTGCCATTGGCACCAAAGTGAGACTTTNNNAGATCGGAAG |
| 59359_1 | TATGAGTGTGGAGTCGTTGCTGCCTGTAATTCCAGCTACTCAGNNNAGATCGGAAG |
| 59361_1 | TATGAGTGTGGAGTCGTTGCAAAGTATTCATTGGCTGGGCAAGNNNAGATCGGAA |
| 59362_1 | TATGAGTGTGGAGTCGTTGCACGCCTGTAATCCCAACACTTTANNNAGATCGGAAG |
| 59363_1 | TATGAGTGTGGAGTCGTTGCTGGAAAAGCTGAATGTGCAAGAGNNNAGATCGGA |
| 59364_1 | TATGAGTGTGGAGTCGTTGCTGCCTGTAATCCCACTACTTTGGNNNAGATCGGAAG |

| | |
|---|---|
| 59365_1 | TATGAGTGTGGAGTCGTTGCCATACCTGTAGTTCCAGCTGCTCNNNAGATCGGAAG |
| 59369_1 | TATGAGTGTGGAGTCGTTGCGAGGTCAGGAGATCGAGTCCATCNNNAGATCGGAA |
| 59370_1 | TATGAGTGTGGAGTCGTTGCCAATAAACACAGAAGCATCGCCANNNAGATCGGAAG |
| 59371_1 | TATGAGTGTGGAGTCGTTGCCCTATCCTCCCAACCCCTACTATNNNAGATCGGAAGA |
| 59372_1 | TATGAGTGTGGAGTCGTTGCAGATGTAACCTCAGAGGGTCCTTNNNAGATCGGAAG |
| 59373_1 | TATGAGTGTGGAGTCGTTGCTGAAGTCAAGATCCCCCTGAAACNNNAGATCGGAAG |
| 59374_1 | TATGAGTGTGGAGTCGTTGCAATCAGATTCTGAGGTTCCAGGCNNNAGATCGGAAG |
| 59375_1 | TATGAGTGTGGAGTCGTTGCTGCTCCTAACAATAGATGTGCGTNNNAGATCGGAAG |
| 59377_1 | TATGAGTGTGGAGTCGTTGCCACTGTTTTAGGCATGGTGGTTTNNNAGATCGGAAG |
| 59378_1 | TATGAGTGTGGAGTCGTTGCCAGATCACCTGAGGTCAGAAGTTNNNAGATCGGAA |
| 59379_1 | TATGAGTGTGGAGTCGTTGCCATGAGAATTGCTTGAACCCAGGNNNAGATCGGAA |
| 59380_1 | TATGAGTGTGGAGTCGTTGCTACAAACACACACACACACACACNNNAGATCGGAAG |
| 59382_1 | TATGAGTGTGGAGTCGTTGCAGGGACCCACAATAGCCAAAATANNNAGATCGGAA |
| 59384_1 | TATGAGTGTGGAGTCGTTGCGCCCGTGTCCTATTTTCTACAGANNNAGATCGGAAG |
| 59385_1 | TATGAGTGTGGAGTCGTTGCTAGTTCTGTTTCATCTCCCTCGCNNNAGATCGGAAGA |
| 59391_1 | TATGAGTGTGGAGTCGTTGCCAATCTCGGCTCACTGCAATATCNNNAGATCGGAAG |
| 59393_1 | TATGAGTGTGGAGTCGTTGCAGTGATTCTCTTGCCTTAGCCTCNNNAGATCGGAAG |
| 59394_1 | TATGAGTGTGGAGTCGTTGCATCAAGTGATTCTCCTGCCTCAGNNNAGATCGGAAG |
| 59396_1 | TATGAGTGTGGAGTCGTTGCCTAACCTTGGGGAAAAGTGGTCTNNNAGATCGGAA |
| 59397_1 | TATGAGTGTGGAGTCGTTGCCCTGGGTTCAAGCAATTCTCTTGNNNAGATCGGAAG |
| 59398_1 | TATGAGTGTGGAGTCGTTGCGATTCAAGCAATTCTCCTGCCTCNNNAGATCGGAAG |
| 59399_1 | TATGAGTGTGGAGTCGTTGCGAGAGATGGGGTTTTGCCATACTNNNAGATCGGAA |
| 59400_1 | TATGAGTGTGGAGTCGTTGCCTCCTGAGTAGCTGGGATTACAGNNNAGATCGGAA |
| 59401_1 | TATGAGTGTGGAGTCGTTGCCTCACTACAATCTCTGCCTCCTGNNNAGATCGGAAGA |
| 59406_1 | TATGAGTGTGGAGTCGTTGCCCTCCCAGGTTCAAGTGATTCTCNNNAGATCGGAAG |
| 59407_1 | TATGAGTGTGGAGTCGTTGCTATCACAGTTCACTGCAGCCTTANNNAGATCGGAAG |
| 59408_1 | TATGAGTGTGGAGTCGTTGCCTTGCCAGGTTCAAGCGATTCTTNNNAGATCGGAAG |
| 59411_1 | TATGAGTGTGGAGTCGTTGCGAGCAGTGGATCACACCTGTAATNNNAGATCGGAA |
| 59415_1 | TATGAGTGTGGAGTCGTTGCGACTTGAGGGACCAGGGTAAAATNNNAGATCGGAA |
| 59417_1 | TATGAGTGTGGAGTCGTTGCAGCAGCAAACACAAATGGAAAGANNNAGATCGGAA |
| 59418_1 | TATGAGTGTGGAGTCGTTGCACCATGCCATCTTGCTTCTTCTANNNAGATCGGAAGA |
| 59419_1 | TATGAGTGTGGAGTCGTTGCCAGAGATTGCACTGAGCTGAGATNNNAGATCGGAA |
| 59420_1 | TATGAGTGTGGAGTCGTTGCCACAGTGCTAGTGGGAGAGTAAANNNAGATCGGAA |
| 59421_1 | TATGAGTGTGGAGTCGTTGCAGTTTAGTGTCTGAATGAGGCCCNNNAGATCGGAA |
| 59424_1 | TATGAGTGTGGAGTCGTTGCATCTAGAGGCAGTTGGCAGTTACNNNAGATCGGAA |
| 59427_1 | TATGAGTGTGGAGTCGTTGCGGAAAAGAATGCCAGCCTCTTTTNNNAGATCGGAAG |
| 59428_1 | TATGAGTGTGGAGTCGTTGCCCCAGGTCTGTTTTCCATGTAGANNNAGATCGGAAG |
| 59431_1 | TATGAGTGTGGAGTCGTTGCTGCAAAGCTCAAACTACATGGTGNNNAGATCGGAAG |
| 59432_1 | TATGAGTGTGGAGTCGTTGCAAGGGGCTTGGAGGTAAGAGATANNNAGATCGGA |
| 59433_1 | TATGAGTGTGGAGTCGTTGCGCACATAGCTGTAGTCCCAACTANNNAGATCGGAAG |
| 59436_1 | TATGAGTGTGGAGTCGTTGCTGAAATAGTCCCCCTGTCCCTATNNNAGATCGGAAGA |
| 59438_1 | TATGAGTGTGGAGTCGTTGCACAGAGAGTCCAGAAAGAAACCCNNNAGATCGGAA |
| 59439_1 | TATGAGTGTGGAGTCGTTGCTGGCTCACACTTATAATCCCAGCNNNAGATCGGAAG |

59440_1 TATGAGTGTGGAGTCGTTGCAATAGCTGGGATTACAGGCACATNNNAGATCGGAA
59442_1 TATGAGTGTGGAGTCGTTGCGGTCTTGAACTCCTGGACTCAAGNNNAGATCGGAAG
59443_1 TATGAGTGTGGAGTCGTTGCTCGATCTCTTGACCTTGTGATCCNNNAGATCGGAAG
59444_1 TATGAGTGTGGAGTCGTTGCCTGAGTAGCTGGGATTACAGGTGNNNAGATCGGAA
59445_1 TATGAGTGTGGAGTCGTTGCCTCTCGAGTAGCTGGGACTACAGNNNAGATCGGAA
59446_1 TATGAGTGTGGAGTCGTTGCGTGTGATCTTAGCTCATGGCAACNNNAGATCGGAAG
59447_1 TATGAGTGTGGAGTCGTTGCTTCAAGTGATTCTCCCACCTCAGNNNAGATCGGAAG
59449_1 TATGAGTGTGGAGTCGTTGCATGCTTTTGACAAAGGGCTCATCNNNAGATCGGAAG
59450_1 TATGAGTGTGGAGTCGTTGCTCTCACTCTGTCACCTAGGCTAGNNNAGATCGGAAG
59452_1 TATGAGTGTGGAGTCGTTGCGAACTTTTGACCTCAAGCGATCCNNNAGATCGGAAG
59453_1 TATGAGTGTGGAGTCGTTGCCTCCTGAGTAGCTGGGATTACAGNNNAGATCGGAA
59454_1 TATGAGTGTGGAGTCGTTGCGAGTAGCTAGGATTACAGGCACCNNNAGATCGGAA
59455_1 TATGAGTGTGGAGTCGTTGCAGTGATTCTCCTGCCTTAGCTTCNNNAGATCGGAAG
59457_1 TATGAGTGTGGAGTCGTTGCCCAATCTGGTCTCGAACTCCTAGNNNAGATCGGAAG
59458_1 TATGAGTGTGGAGTCGTTGCACCAAGCTATCCAGTCCTTCTTGNNNAGATCGGAAG
59460_1 TATGAGTGTGGAGTCGTTGCGGATTACAGGTATGAGCCACCATNNNAGATCGGAA
59461_1 TATGAGTGTGGAGTCGTTGCATTCTCAGCTCACTGCAATCTCTNNNAGATCGGAAGA
59463_1 TATGAGTGTGGAGTCGTTGCCAATCTCAGCTCACTGCAACTTCNNNAGATCGGAAGA
59464_1 TATGAGTGTGGAGTCGTTGCGGAGTCTCTAGTCTCGCTCTGTCNNNAGATCGGAAG
59465_1 TATGAGTGTGGAGTCGTTGCTCTGAAACTCCTAGGCTCAAGTGNNNAGATCGGAAG
59466_1 TATGAGTGTGGAGTCGTTGCTTCACACCATTCTCCTGTCTCAGNNNAGATCGGAAGA
59467_1 TATGAGTGTGGAGTCGTTGCTCTCCATCTCCTGACCTCATGATNNNAGATCGGAAGA
59470_1 TATGAGTGTGGAGTCGTTGCCTTGTACCTCAGCATCCCAAGTANNNAGATCGGAAG
59471_1 TATGAGTGTGGAGTCGTTGCCTCCTGGGTAGATGGGATTACAGNNNAGATCGGAA
59473_1 TATGAGTGTGGAGTCGTTGCTCCTGAGTAGCTGGGATTACAGANNNAGATCGGAA
59474_1 TATGAGTGTGGAGTCGTTGCTGTCTTCTGGGCAATTCTCCTACNNNAGATCGGAAG
59477_1 TATGAGTGTGGAGTCGTTGCAGTAGCTCAGATTACAGGCATGCNNNAGATCGGAA
59478_1 TATGAGTGTGGAGTCGTTGCTATGGGAGGCAATAAGGAGATGCNNNAGATCGGAA
59479_1 TATGAGTGTGGAGTCGTTGCGGTTCAAGTGATTCTCCTGCCTCNNNAGATCGGAAG
59480_1 TATGAGTGTGGAGTCGTTGCGTCCCTAGCTAGTCTGCCTTTTTNNNAGATCGGAAGA
59481_1 TATGAGTGTGGAGTCGTTGCGCTGGTCTTGAAATCTTGAGCTCNNNAGATCGGAAG
59482_1 TATGAGTGTGGAGTCGTTGCCAAGTAGCTGGGATTACAGGCATNNNAGATCGGAA
59484_1 TATGAGTGTGGAGTCGTTGCCTCAAGTGATCCTCCCACCTCAGNNNAGATCGGAAG
59485_1 TATGAGTGTGGAGTCGTTGCACAATCTCGGCTCACTGCAANNNAGATCGGAAGAGC
59486_1 TATGAGTGTGGAGTCGTTGCCAAGTAGCTGGGACTACAGACGNNNAGATCGGAAG
59487_1 TATGAGTGTGGAGTCGTTGCTTGAACTCCTGACCTCATGATCCNNNAGATCGGAAG
59490_1 TATGAGTGTGGAGTCGTTGCGTCTCCTGGGTTCAAGTGATTCTNNNAGATCGGAAG
59491_1 TATGAGTGTGGAGTCGTTGCCAATCACATGCAATCTCAGCTCANNNAGATCGGAAG
59493_1 TATGAGTGTGGAGTCGTTGCGTTATTTCCACAAGTCTTGGGCCNNNAGATCGGAAG
59494_1 TATGAGTGTGGAGTCGTTGCGTTTAGCTGGGTGAACACTGTTCNNNAGATCGGAA
59496_1 TATGAGTGTGGAGTCGTTGCTTCATGCTATTCTCCTGCCTCAGNNNAGATCGGAAGA
59497_1 TATGAGTGTGGAGTCGTTGCAGACTTCTGGTGGCAAGATATCCNNNAGATCGGAA
59498_1 TATGAGTGTGGAGTCGTTGCAGTTTCCTTAGCGGTAGAACCAGNNNAGATCGGAA

59499_1 TATGAGTGTGGAGTCGTTGCGAGTAGCTGGGATTATAGGTGCCNNNAGATCGGAA
59500_1 TATGAGTGTGGAGTCGTTGCCTCCTGAGTAGCTGGGACTACAGNNNAGATCGGAA
59501_1 TATGAGTGTGGAGTCGTTGCCCAAGTAGCTGAGATTACAGGCANNNAGATCGGAA
59502_1 TATGAGTGTGGAGTCGTTGCCTCAGCTTCCAGATTACAGGTGTNNNAGATCGGAAG
59505_1 TATGAGTGTGGAGTCGTTGCTATATTGGGGTGAGTGGCATCTGNNNAGATCGGAA
59507_1 TATGAGTGTGGAGTCGTTGCGAGAGGGAGCTTCACTCTTGTTGNNNAGATCGGAA
59508_1 TATGAGTGTGGAGTCGTTGCGGTTCAAGTGATTTTCCTGCCTCNNNAGATCGGAAG
59509_1 TATGAGTGTGGAGTCGTTGCAACAGTTTAGCTCTAGTTGCCCANNNAGATCGGAAG
59510_1 TATGAGTGTGGAGTCGTTGCGTGATTCTCCTACCTCAGCTTCCNNNAGATCGGAAGA
59512_1 TATGAGTGTGGAGTCGTTGCTTCTAGCAATTCTCCTGCCTCAGNNNAGATCGGAAG
59513_1 TATGAGTGTGGAGTCGTTGCCAATCTTGACTCACTGCAAGCTCNNNAGATCGGAAG
59514_1 TATGAGTGTGGAGTCGTTGCTAGCTAGGATTACAGGAACACGCNNNAGATCGGAA
59518_1 TATGAGTGTGGAGTCGTTGCAATTTCAGCTCACTGCAACCTCTNNNAGATCGGAAG
59519_1 TATGAGTGTGGAGTCGTTGCTATCTTGGCTCATTGCAACCTCTNNNAGATCGGAAG
59520_1 TATGAGTGTGGAGTCGTTGCAGACAAGGTCTCATTCTGTCACCNNNAGATCGGAAG
59521_1 TATGAGTGTGGAGTCGTTGCTGATCTCAGCTCACTTCAACCTCNNNAGATCGGAAGA
59524_1 TATGAGTGTGGAGTCGTTGCTCTCCCAGGTTCAACTGATTCTCNNNAGATCGGAAG
59526_1 TATGAGTGTGGAGTCGTTGCCTTCCAAGTAGCTGGGACTACAGNNNAGATCGGAA
59527_1 TATGAGTGTGGAGTCGTTGCTCATTGAAGCCTCAAACTCCTGANNNAGATCGGAAG
59528_1 TATGAGTGTGGAGTCGTTGCCTCCCAAGTAGCTGGAACTACAGNNNAGATCGGAAG
59531_1 TATGAGTGTGGAGTCGTTGCTGGTGACAGAGGAAAGAGACAAGNNNAGATCGGA
59533_1 TATGAGTGTGGAGTCGTTGCTATCTGGGATTACAGGCATGAGCNNNAGATCGGAA
59536_1 TATGAGTGTGGAGTCGTTGCTCTGATCTTTGAGGGTGTGGAAGNNNAGATCGGAA
59537_1 TATGAGTGTGGAGTCGTTGCTTCAAAGAAAGTGGTGTCCCAGANNNAGATCGGAA
59540_1 TATGAGTGTGGAGTCGTTGCACTTGCCCAAGAACACATAGCTANNNAGATCGGAAG
59542_1 TATGAGTGTGGAGTCGTTGCGATGTGGCTAACAAGAAAGGCAANNNAGATCGGAA
59544_1 TATGAGTGTGGAGTCGTTGCGACTGAGGAGTAGAGGCAAAGTCNNNAGATCGGA
59546_1 TATGAGTGTGGAGTCGTTGCCATTCCCCTCTCTTACCTCTGTGNNNAGATCGGAAGA
59547_1 TATGAGTGTGGAGTCGTTGCCTCCCTCACTCTCAAATCACCAANNNAGATCGGAAGA
59548_1 TATGAGTGTGGAGTCGTTGCTTCTCCCATTCCTTCGATTGGAGNNNAGATCGGAAG
59549_1 TATGAGTGTGGAGTCGTTGCTTACCCTTTCTGAGTAAAGCCAANNNAGATCGGAAG
59551_1 TATGAGTGTGGAGTCGTTGCAGGGCCACACAATCTACTAAAGGNNNAGATCGGAA
59556_1 TATGAGTGTGGAGTCGTTGCGCAGCAGCTGGAGACTATAAAAANNNAGATCGGAA
59558_1 TATGAGTGTGGAGTCGTTGCATACAGTCCAGCTTCAGTGATCCNNNAGATCGGAAG
59560_1 TATGAGTGTGGAGTCGTTGCTTCAGGTCATTGCAAATGCCATTNNNAGATCGGAAG
59561_1 TATGAGTGTGGAGTCGTTGCAGAAAAAGAAAACTGCCCATCCCNNNAGATCGGAAG
59562_1 TATGAGTGTGGAGTCGTTGCATGGAGAAACCCCGTCTCTACTANNNAGATCGGAAG
59564_1 TATGAGTGTGGAGTCGTTGCTAACCTCACCGTCGTTTCTCTACNNNAGATCGGAAGA
59565_1 TATGAGTGTGGAGTCGTTGCCTGAGGCAGGAGAATCACTTGAANNNAGATCGGAA
59566_1 TATGAGTGTGGAGTCGTTGCACTCAAGAAAGCAAAATGCAAGCNNNAGATCGGAA
59567_1 TATGAGTGTGGAGTCGTTGCTTTTCTAACTGTCTTGGGAGGGGNNNAGATCGGAA
59569_1 TATGAGTGTGGAGTCGTTGCAGGTCAGCAGATAAGGGGAGTATNNNAGATCGGA
59572_1 TATGAGTGTGGAGTCGTTGCTGAGACCATCTTGTTCTGTCACCNNNAGATCGGAAG

| | |
|---|---|
| 59573_1 | TATGAGTGTGGAGTCGTTGCTGCTAATGCTGCTAAGGAAATGANNNAGATCGGAA |
| 59574_1 | TATGAGTGTGGAGTCGTTGCGGAAGACAAAAAAGGGAAAGGGGNNNAGATCGG |
| 59575_1 | TATGAGTGTGGAGTCGTTGCTCACTTTTGTACCTTAGGGAGCCNNNAGATCGGAAG |
| 59580_1 | TATGAGTGTGGAGTCGTTGCCTGAGGCAGGAGAATCACTTGATNNNAGATCGGAA |
| 59581_1 | TATGAGTGTGGAGTCGTTGCAGTCTAAGTGTCAACGTACCTCTNNNAGATCGGAAG |
| 59582_1 | TATGAGTGTGGAGTCGTTGCTCGCTTCTTGAAGGAAGAGAACANNNAGATCGGAA |
| 59585_1 | TATGAGTGTGGAGTCGTTGCGAGTGACAGAGTGAGGCTCAATCNNNAGATCGGAA |
| 59587_1 | TATGAGTGTGGAGTCGTTGCAATTCTGCAGATTCCAGCAATGGNNNAGATCGGAAG |
| 59592_1 | TATGAGTGTGGAGTCGTTGCAATTTCTGGGCTGCAACTAAAGCNNNAGATCGGAAG |
| 59593_1 | TATGAGTGTGGAGTCGTTGCTGCCACTGTAATTTGTTGTCCTGNNNAGATCGGAAG |
| 59594_1 | TATGAGTGTGGAGTCGTTGCTTACAGTGAGCTGAGATCACGTCNNNAGATCGGAA |
| 59599_1 | TATGAGTGTGGAGTCGTTGCCAGTGAATCATGAGCCAAGGTTGNNNAGATCGGAA |
| 59600_1 | TATGAGTGTGGAGTCGTTGCTATTCATGGGGTTTTCCAAGGGANNNAGATCGGAA |
| 59602_1 | TATGAGTGTGGAGTCGTTGCTAGTGAAGCTTCCTTCGTCCTTCNNNAGATCGGAAG |
| 59603_1 | TATGAGTGTGGAGTCGTTGCTGGCTTACACCTATAATCCCAGCNNNAGATCGGAAG |
| 59604_1 | TATGAGTGTGGAGTCGTTGCAGCATCAGTGAAACAAACTCGTCNNNAGATCGGAAG |
| 59606_1 | TATGAGTGTGGAGTCGTTGCATATAGGGTGGGACTGTTGGCTANNNAGATCGGAA |
| 59607_1 | TATGAGTGTGGAGTCGTTGCCGGAGAAGACCATTTGCAAAGTTNNNAGATCGGAA |
| 59609_1 | TATGAGTGTGGAGTCGTTGCGCTTTGCGCAAAGGACATTACTANNNAGATCGGAAG |
| 59610_1 | TATGAGTGTGGAGTCGTTGCAAAGCCAACTTCAAAACTGCCTCNNNAGATCGGAAG |
| 59611_1 | TATGAGTGTGGAGTCGTTGCAGTGAATGGAAATCTGGCTGTCANNNAGATCGGAA |
| 59614_1 | TATGAGTGTGGAGTCGTTGCCCAGAGTTCAGCAGACAACAAATNNNAGATCGGAA |
| 59617_1 | TATGAGTGTGGAGTCGTTGCCAGGGGGTATAACTGTGTTCTCANNNAGATCGGAA |
| 59619_1 | TATGAGTGTGGAGTCGTTGCTCACTTTAGCCTTGGAGGACAAANNNAGATCGGAAG |
| 59620_1 | TATGAGTGTGGAGTCGTTGCGAGCCTGTAGTCTCAGCTACTTGNNNAGATCGGAAG |
| 59622_1 | TATGAGTGTGGAGTCGTTGCTGTTTCATGCTCAGTAAATCGGCNNNAGATCGGAAG |
| 59623_1 | TATGAGTGTGGAGTCGTTGCGGGTTGCAATCCTAGTCTCTGATNNNAGATCGGAAG |
| 59624_1 | TATGAGTGTGGAGTCGTTGCCAGCTTTTCAAATGTGCAAGCAGNNNAGATCGGAAG |
| 59625_1 | TATGAGTGTGGAGTCGTTGCCCGCTGACATTTTCCACCATAATNNNAGATCGGAAG |
| 59626_1 | TATGAGTGTGGAGTCGTTGCTTTGCAACTAGGGCAGAAAGTTTNNNAGATCGGAA |
| 59627_1 | TATGAGTGTGGAGTCGTTGCGCAAAACCAATGTGTATGACCCANNNAGATCGGAAG |
| 59628_1 | TATGAGTGTGGAGTCGTTGCAAGAAATAGAGGGCTAGGCACAGNNNAGATCGGA |
| 59630_1 | TATGAGTGTGGAGTCGTTGCAGTGATACCAAGTGGCCTAAACTNNNAGATCGGAA |
| 59632_1 | TATGAGTGTGGAGTCGTTGCCCCTGCATGCTCCTCTAGTTTATNNNAGATCGGAAGA |
| 59633_1 | TATGAGTGTGGAGTCGTTGCGGAGAATTAGTGGGGTGGTCTTTNNNAGATCGGAA |
| 59637_1 | TATGAGTGTGGAGTCGTTGCTCCGTGAAGCAAAGGTGTAAGATNNNAGATCGGAA |
| 59638_1 | TATGAGTGTGGAGTCGTTGCGTCTAAGCCCTACCAAGCACATANNNAGATCGGAAG |
| 59639_1 | TATGAGTGTGGAGTCGTTGCAAAAAAGAAAGTGGGGCTAGGTGNNNAGATCGG |
| 59640_1 | TATGAGTGTGGAGTCGTTGCAGAGGACAGGACCCAGAGATTTANNNAGATCGGAA |
| 59641_1 | TATGAGTGTGGAGTCGTTGCGAGTGGGAGTGAGATTCAGGAAGNNNAGATCGG |
| 59642_1 | TATGAGTGTGGAGTCGTTGCGACAGCACCGTCTATGAGATACTNNNAGATCGGAAG |
| 59644_1 | TATGAGTGTGGAGTCGTTGCGAAACAGAAGCAGGTAGTGAGGANNNAGATCGG |
| 59645_1 | TATGAGTGTGGAGTCGTTGCGTTCCTTTTTTTCCCCCAGCAATNNNAGATCGGAAGA |

| | |
|---|---|
| 59647_1 | TATGAGTGTGGAGTCGTTGCGAGATGGAGTCTTGCTGTATTGCNNNAGATCGGAA |
| 59648_1 | TATGAGTGTGGAGTCGTTGCTTCAAGTGATTCTCCTGCCTCAGNNNAGATCGGAAG |
| 59649_1 | TATGAGTGTGGAGTCGTTGCGTTTCAAGCGATTCTGCTACCTCNNNAGATCGGAAG |
| 59651_1 | TATGAGTGTGGAGTCGTTGCGAGTCTTGCTCTGTTGCTTAGGCNNNAGATCGGAAG |
| 59652_1 | TATGAGTGTGGAGTCGTTGCCCTCCTGAGTAGCTGGGATTAGANNNAGATCGGAA |
| 59653_1 | TATGAGTGTGGAGTCGTTGCCCAAGTAGCTGGGATTATAGGCGNNNAGATCGGAA |
| 59654_1 | TATGAGTGTGGAGTCGTTGCCCTACGAGGTTCACACCATTCTCNNNAGATCGGAAG |
| 59655_1 | TATGAGTGTGGAGTCGTTGCAGTGCAGTGATATGATCTTGGCTNNNAGATCGGAA |
| 59657_1 | TATGAGTGTGGAGTCGTTGCGGATCTCACTATGTTACCCAGGCNNNAGATCGGAAG |
| 59658_1 | TATGAGTGTGGAGTCGTTGCTTCAAGTGATTCTCCTGCCTCAGNNNAGATCGGAAG |
| 59659_1 | TATGAGTGTGGAGTCGTTGCCCTCCTGAGCTCAAGAGATTCTCNNNAGATCGGAAG |
| 59660_1 | TATGAGTGTGGAGTCGTTGCGAACTTCCAGGCCTAAGTGATCCNNNAGATCGGAAG |
| 59661_1 | TATGAGTGTGGAGTCGTTGCCGAATTCCCTGGCTCAAGTGATCNNNAGATCGGAAG |
| 59663_1 | TATGAGTGTGGAGTCGTTGCTGGGATCACAAATGTCTACCACCNNNAGATCGGAAG |
| 59664_1 | TATGAGTGTGGAGTCGTTGCCCGAGTAGCTAGGACTACAGGTANNNAGATCGGAA |
| 59666_1 | TATGAGTGTGGAGTCGTTGCCCCAGGTTTGAGCAATTCTCTTGNNNAGATCGGAAG |
| 59667_1 | TATGAGTGTGGAGTCGTTGCGGTCTGTAGCACCTGTCATAACANNNAGATCGGAAG |
| 59668_1 | TATGAGTGTGGAGTCGTTGCCAATCTTGGCTCACTGAAACCTCNNNAGATCGGAAG |
| 59671_1 | TATGAGTGTGGAGTCGTTGCAGCCTTCAAGTAGCTGGGATTATNNNAGATCGGAA |
| 59674_1 | TATGAGTGTGGAGTCGTTGCTTCAAGCAATTCCCCTGCCTTAGNNNAGATCGGAAG |
| 59676_1 | TATGAGTGTGGAGTCGTTGCTCTTGGTTCACTACAACCTCTGCNNNAGATCGGAAGA |
| 59677_1 | TATGAGTGTGGAGTCGTTGCGTTCAAGAGATTCTCCTGCCTCANNNAGATCGGAAG |
| 59680_1 | TATGAGTGTGGAGTCGTTGCAACCTCTGACTTCCTGGTTCAAGNNNAGATCGGAAG |
| 59681_1 | TATGAGTGTGGAGTCGTTGCGGAGAGCAGTGGTACAATCTTGANNNAGATCGGAA |
| 59682_1 | TATGAGTGTGGAGTCGTTGCGCAATCTCAACTCACTGCAACTTNNNAGATCGGAAG |
| 59685_1 | TATGAGTGTGGAGTCGTTGCCGAAATCCTGAGTTCAAGCGATCNNNAGATCGGAAG |
| 59686_1 | TATGAGTGTGGAGTCGTTGCCCTGGGTTCAAGCAATTCTTCTGNNNAGATCGGAAG |
| 59687_1 | TATGAGTGTGGAGTCGTTGCCAAAGTGCTGGGATTACAGGTGTNNNAGATCGGAA |
| 59688_1 | TATGAGTGTGGAGTCGTTGCATCTTGACTCACTGCAATCTCCANNNAGATCGGAAGA |
| 59689_1 | TATGAGTGTGGAGTCGTTGCAAGTGATTCTTGTGTCTCAGCCTNNNAGATCGGAAG |
| 59690_1 | TATGAGTGTGGAGTCGTTGCCTGAATAGCTGGGACTACAGGACNNNAGATCGGAA |
| 59691_1 | TATGAGTGTGGAGTCGTTGCAGCAATTTTCAGCCTCCTGAGTANNNAGATCGGAAG |
| 59692_1 | TATGAGTGTGGAGTCGTTGCCAGGTTCAAGAGATCCTTCTGCTNNNAGATCGGAAG |
| 59696_1 | TATGAGTGTGGAGTCGTTGCTTCAAGTGATTCTCCTGCCTCAGNNNAGATCGGAAG |
| 59697_1 | TATGAGTGTGGAGTCGTTGCCAACCTTGACCTCTCAGGATCAGNNNAGATCGGAAG |
| 59698_1 | TATGAGTGTGGAGTCGTTGCTTCAAGTGATTCTCCTGCCTCAGNNNAGATCGGAAG |
| 59699_1 | TATGAGTGTGGAGTCGTTGCAATCTCAGTTCACTGCAGAGGTTNNNAGATCGGAAG |
| 59700_1 | TATGAGTGTGGAGTCGTTGCCTCCTGAGTAGCTGGGATTACAGNNNAGATCGGAA |
| 59701_1 | TATGAGTGTGGAGTCGTTGCCAGAATCTGGGTAGGAGGTTTGGNNNAGATCGGAA |
| 59703_1 | TATGAGTGTGGAGTCGTTGCGAGAGGGAGTCTCACTTTCTTGCNNNAGATCGGAA |
| 59704_1 | TATGAGTGTGGAGTCGTTGCTTCAAGTGATTCTCCCACCTCTGNNNAGATCGGAAGA |
| 59705_1 | TATGAGTGTGGAGTCGTTGCAGTGCTGGGATTATAGGCATGAGNNNAGATCGGAA |
| 59707_1 | TATGAGTGTGGAGTCGTTGCGACACGATCTCGGTTCACTGTAANNNAGATCGGAAG |

| | |
|---|---|
| 59708_1 | TATGAGTGTGGAGTCGTTGCCTTTCCAGGGTCGAGTGATTCTCNNNAGATCGGAAG |
| 59712_1 | TATGAGTGTGGAGTCGTTGCCCTCCCAGATTCAGGTGATTCTCNNNAGATCGGAAG |
| 59713_1 | TATGAGTGTGGAGTCGTTGCAGCACTTCTTCTTTCTTCTGCCTNNNAGATCGGAAGA |
| 59714_1 | TATGAGTGTGGAGTCGTTGCGAGTAGCTAGGATTACAGGCACCNNNAGATCGGAA |
| 59717_1 | TATGAGTGTGGAGTCGTTGCCAATCTCAGCTCCCTGAAACCTCNNNAGATCGGAAGA |
| 59718_1 | TATGAGTGTGGAGTCGTTGCAGAAGTCTTCCTAGGTTTGGTGGNNNAGATCGGAA |
| 59720_1 | TATGAGTGTGGAGTCGTTGCCAGTTCTGGGAGCTTAGTGAACTNNNAGATCGGAA |
| 59725_1 | TATGAGTGTGGAGTCGTTGCACAACTCCTCCAGCTAAAATCGTNNNAGATCGGAAG |
| 59727_1 | TATGAGTGTGGAGTCGTTGCCAGATGAGGAAACTGAACCCAGANNNAGATCGGAA |
| 59729_1 | TATGAGTGTGGAGTCGTTGCGTCTTCAGAGGATCCAATGGGAGNNNAGATCGGAA |
| 59730_1 | TATGAGTGTGGAGTCGTTGCTTCTACTCTGCCCTCCCATCTTANNNAGATCGGAAGA |
| 59732_1 | TATGAGTGTGGAGTCGTTGCTGCAGCTTGATTTTTCAGGCTACNNNAGATCGGAAG |
| 59733_1 | TATGAGTGTGGAGTCGTTGCGAGGGAAGTTGAGATGGTGTGAANNNAGATCGGA |
| 59736_1 | TATGAGTGTGGAGTCGTTGCGCCTGGGCAAGATAGTAAAAACCNNNAGATCGGAA |
| 59738_1 | TATGAGTGTGGAGTCGTTGCCCATTTCCTTTGTGGCCATGAAANNNAGATCGGAAG |
| 59740_1 | TATGAGTGTGGAGTCGTTGCCATATGCCTGTAGTCCCAGCTACNNNAGATCGGAAG |
| 59741_1 | TATGAGTGTGGAGTCGTTGCTGCTCCACCAACACTTATTGAGTNNNAGATCGGAAG |
| 59742_1 | TATGAGTGTGGAGTCGTTGCCAAGACAAGGATGCTGGAAACAGNNNAGATCGGAA |
| 59743_1 | TATGAGTGTGGAGTCGTTGCCAATAGGAAGAGGGGAGCCATTTNNNAGATCGGAA |
| 59744_1 | TATGAGTGTGGAGTCGTTGCGGGACTCAAAGAAGGACCAGATGNNNAGATCGGAA |
| 59746_1 | TATGAGTGTGGAGTCGTTGCACAATGGCTTACACCTGTAGTCTNNNAGATCGGAAG |
| 59747_1 | TATGAGTGTGGAGTCGTTGCTGGAGAATTGCTTGAACCTGAGANNNAGATCGGAA |
| 59748_1 | TATGAGTGTGGAGTCGTTGCTTTGGGGAATGGAGAGGTTTTCANNNAGATCGGAA |
| 59749_1 | TATGAGTGTGGAGTCGTTGCCCTAGATTACAGAAGGCACCAGGNNNAGATCGGAA |
| 59750_1 | TATGAGTGTGGAGTCGTTGCGTCTGCAGTGTGATTGCTTTTCTNNNAGATCGGAAG |
| 59751_1 | TATGAGTGTGGAGTCGTTGCTTTGCAACAACATGGATGAAGCTNNNAGATCGGAAG |
| 59752_1 | TATGAGTGTGGAGTCGTTGCGGCTGAATGCAACCATACAGAAANNNAGATCGGAA |
| 59753_1 | TATGAGTGTGGAGTCGTTGCTCTTGGGTAGAATTGCACCTTCANNNAGATCGGAAG |
| 59754_1 | TATGAGTGTGGAGTCGTTGCAGCCAATCTGATGCTGGTATTGANNNAGATCGGAA |
| 59756_1 | TATGAGTGTGGAGTCGTTGCAGGAGAATTGTTTGAACCCAGGANNNAGATCGGAA |
| 59758_1 | TATGAGTGTGGAGTCGTTGCGCAAATCCACAGAGGCAGAAAATNNNAGATCGGAA |
| 59759_1 | TATGAGTGTGGAGTCGTTGCAGTAGTCACCAGAAAGTGAACCCNNNAGATCGGAA |
| 59760_1 | TATGAGTGTGGAGTCGTTGCCGGATCACAAGGTAAGGAGTTCANNNAGATCGGAA |
| 59761_1 | TATGAGTGTGGAGTCGTTGCTGTATCTCAGAACTTTGGGAGGCNNNAGATCGGAA |
| 59763_1 | TATGAGTGTGGAGTCGTTGCGTGGGAAGATTGCTTGAGTTCAGNNNAGATCGGAA |
| 59767_1 | TATGAGTGTGGAGTCGTTGCGATTTCTCACCTCTTCCCCAGTCNNNAGATCGGAAGA |
| 59768_1 | TATGAGTGTGGAGTCGTTGCAACTTGTTCACCTCTTCCACCATNNNAGATCGGAAGA |
| 59769_1 | TATGAGTGTGGAGTCGTTGCCCAGAGGAGGCAGAAAAAGAAGTNNNAGATCGGA |
| 59770_1 | TATGAGTGTGGAGTCGTTGCGGCGTGGTAGCTCATGTCTATAANNNAGATCGGAA |
| 59771_1 | TATGAGTGTGGAGTCGTTGCAAAAAGAGGAACCCAGGATGACANNNAGATCGGAA |
| 59772_1 | TATGAGTGTGGAGTCGTTGCTGAGACCTCAGGTGAGTCTAGAGNNNAGATCGGAA |
| 59774_1 | TATGAGTGTGGAGTCGTTGCAAATAATGACCTGGGCCACAAGANNNAGATCGGAA |
| 59775_1 | TATGAGTGTGGAGTCGTTGCTTGCTCTCGAGATGAAAGAACCANNNAGATCGGAA |

| | |
|---|---|
| 59776_1 | TATGAGTGTGGAGTCGTTGCAAAACACTCTTCAATTCCTGCCCNNNAGATCGGAAGA |
| 59778_1 | TATGAGTGTGGAGTCGTTGCATCACGAGGTCAGGAGATGGTAANNNAGATCGGAA |
| 59779_1 | TATGAGTGTGGAGTCGTTGCCTGAGGTCAGGAGTTCAAAACCANNNAGATCGGAA |
| 59780_1 | TATGAGTGTGGAGTCGTTGCGAGAACAAAAAAACCAAACCCCGNNNAGATCGGAA |
| 59781_1 | TATGAGTGTGGAGTCGTTGCCACAATCTCAGCTCACTGCAATCNNNAGATCGGAAG |
| 59783_1 | TATGAGTGTGGAGTCGTTGCTTAGGGGAGGGAGAGCATTATGANNNAGATCGGA |
| 59784_1 | TATGAGTGTGGAGTCGTTGCCGGGAAGGTGCGATATAAGATCANNNAGATCGGAA |
| 59786_1 | TATGAGTGTGGAGTCGTTGCGGCACAGTTACTCACTCCTGTAANNNAGATCGGAAG |
| 59787_1 | TATGAGTGTGGAGTCGTTGCATCACCAAGGAAGCTATCAGTGGNNNAGATCGGAA |
| 59788_1 | TATGAGTGTGGAGTCGTTGCTCAGCCTGTGATACCTACATGTGNNNAGATCGGAAG |
| 59789_1 | TATGAGTGTGGAGTCGTTGCCTGTAATCGCAGGACTTTGGAAGNNNAGATCGGAA |
| 59797_1 | TATGAGTGTGGAGTCGTTGCAGTGGATGTGGCTAAAGGCTTTANNNAGATCGGAA |
| 59798_1 | TATGAGTGTGGAGTCGTTGCTTTTCATTCCTCAGCCTCCTGAANNNAGATCGGAAGA |
| 59799_1 | TATGAGTGTGGAGTCGTTGCCATGGGTTCAAGCAATTCTCCTGNNNAGATCGGAAG |
| 59800_1 | TATGAGTGTGGAGTCGTTGCACAAGTGAACAAAGGTCTCTGGTNNNAGATCGGAA |
| 59803_1 | TATGAGTGTGGAGTCGTTGCGTGGCAGGGTCATAGGATGATAGNNNAGATCGGA |
| 59806_1 | TATGAGTGTGGAGTCGTTGCGTCATCCAGGCTAGAGTGTAGTGNNNAGATCGGAA |
| 59807_1 | TATGAGTGTGGAGTCGTTGCGTCTCCTGGGTTCAAGTGATTCTNNNAGATCGGAAG |
| 59809_1 | TATGAGTGTGGAGTCGTTGCCGAGTTCAAGCAATTCTCCTGTCNNNAGATCGGAAG |
| 59810_1 | TATGAGTGTGGAGTCGTTGCGGTTCTCATAGGTCTGGAGCATTNNNAGATCGGAAG |
| 59812_1 | TATGAGTGTGGAGTCGTTGCACTCTGTCTCAAAAGCTGGAGTGNNNAGATCGGAAG |
| 59813_1 | TATGAGTGTGGAGTCGTTGCCTCTCAAGTAGCTGGGACAACAGNNNAGATCGGAA |
| 59815_1 | TATGAGTGTGGAGTCGTTGCCTGGTCTCAAACTCTTGTCCTCANNNAGATCGGAAGA |
| 59816_1 | TATGAGTGTGGAGTCGTTGCCCTCCTGAGTAGCTGGGATTTTANNNAGATCGGAAG |
| 59817_1 | TATGAGTGTGGAGTCGTTGCCCTGGGTTCAAGTGATTCTCTCANNNAGATCGGAAG |
| 59818_1 | TATGAGTGTGGAGTCGTTGCGGAGTACAGTGGCATGATCTCAGNNNAGATCGGAA |
| 59819_1 | TATGAGTGTGGAGTCGTTGCAGTAGCTGAGACTAAAGGTGCACNNNAGATCGGAA |
| 59821_1 | TATGAGTGTGGAGTCGTTGCACTGACTCCCAACAAAGAATTGTNNNAGATCGGAAG |
| 59822_1 | TATGAGTGTGGAGTCGTTGCTGATCTCAGTTCAGTGCAACCTCNNNAGATCGGAAG |
| 59823_1 | TATGAGTGTGGAGTCGTTGCAAGTAGCTGGGATTATAGGCACGNNNAGATCGGAA |
| 59824_1 | TATGAGTGTGGAGTCGTTGCCTGGGTTCAAGTGATTCTCTTGCNNNAGATCGGAAG |
| 59825_1 | TATGAGTGTGGAGTCGTTGCTTCAAGTGATTCTCCTGGCTCAGNNNAGATCGGAAG |
| 59827_1 | TATGAGTGTGGAGTCGTTGCCTCCCGGGTTCAAGATATTCTCCNNNAGATCGGAAG |
| 59829_1 | TATGAGTGTGGAGTCGTTGCCTCCTGAGCAGCTGGGATTATAGNNNAGATCGGAA |
| 59830_1 | TATGAGTGTGGAGTCGTTGCGGCTGGAGTGCAATTATGTGATCNNNAGATCGGAA |
| 59833_1 | TATGAGTGTGGAGTCGTTGCCTCCTGAGTAGCTGGGACTACAGNNNAGATCGGAA |
| 59835_1 | TATGAGTGTGGAGTCGTTGCCCTCCCAGGCTCAAGTAATTCTCNNNAGATCGGAAG |
| 59836_1 | TATGAGTGTGGAGTCGTTGCGATCTCAGCTCACTGCAAAATCCNNNAGATCGGAAG |
| 59837_1 | TATGAGTGTGGAGTCGTTGCTATAGTGGCACAATCTTGGCTCANNNAGATCGGAAG |
| 59838_1 | TATGAGTGTGGAGTCGTTGCCCTGTGTTCAAGCAATTCTCCTGNNNAGATCGGAAG |
| 59839_1 | TATGAGTGTGGAGTCGTTGCGATCTCAGCTCAGTGCAACCTATNNNAGATCGGAAG |
| 59840_1 | TATGAGTGTGGAGTCGTTGCGCTTCCTGAATAGCTGGGACTATNNNAGATCGGAAG |
| 59842_1 | TATGAGTGTGGAGTCGTTGCCTGCCAAATTCAAGCGATTCTCANNNAGATCGGAAG |

| | |
|---|---|
| 59843_1 | TATGAGTGTGGAGTCGTTGCCCTCCTGAATAGCTGGGACTATGNNNAGATCGGAAG |
| 59845_1 | TATGAGTGTGGAGTCGTTGCCCTCCTAGGTTCATGCCATTCTCNNNAGATCGGAAGA |
| 59846_1 | TATGAGTGTGGAGTCGTTGCTAGCTGGGATTACAGGCATATGCNNNAGATCGGAA |
| 59847_1 | TATGAGTGTGGAGTCGTTGCAGTAGCTGGTACTACAGGGATGTNNNAGATCGGAA |
| 59848_1 | TATGAGTGTGGAGTCGTTGCTGGAGTGCAATGACACAATCTTGNNNAGATCGGAA |
| 59849_1 | TATGAGTGTGGAGTCGTTGCAAACACGATCACAACAGGCAAATNNNAGATCGGAA |
| 59850_1 | TATGAGTGTGGAGTCGTTGCCGATCTTGACTCACTGCAATCTCNNNAGATCGGAAG |
| 59851_1 | TATGAGTGTGGAGTCGTTGCTATCTTGACTCACTGCAACCTCCNNNAGATCGGAAGA |
| 59852_1 | TATGAGTGTGGAGTCGTTGCTTTCCATTCATGTCTTCTGGCCANNNAGATCGGAAGA |
| 59853_1 | TATGAGTGTGGAGTCGTTGCTGGAGTGCAGTGATGTGACAATANNNAGATCGGAA |
| 59854_1 | TATGAGTGTGGAGTCGTTGCCCCTAAACCGGTCTGGCAATATANNNAGATCGGAAG |
| 59855_1 | TATGAGTGTGGAGTCGTTGCAGGCCTGGAGACAGATCAAAAATNNNAGATCGGAA |
| 59856_1 | TATGAGTGTGGAGTCGTTGCGACTGCATACAATGGCACAATCTNNNAGATCGGAAG |
| 59860_1 | TATGAGTGTGGAGTCGTTGCTTCAAGTGATTCTCCTGCCTCAGNNNAGATCGGAAG |
| 59862_1 | TATGAGTGTGGAGTCGTTGCCAGTGGCTCAATCATAGCTCACTNNNAGATCGGAAG |
| 59863_1 | TATGAGTGTGGAGTCGTTGCACCACAGCCGGCTAATTTTTTTTNNNAGATCGGAAG |
| 59864_1 | TATGAGTGTGGAGTCGTTGCCTCCTGAGTAGCTGGGACTACAGNNNAGATCGGAA |
| 59866_1 | TATGAGTGTGGAGTCGTTGCTTCACATCATTCTCCTGCCTCAGNNNAGATCGGAAGA |
| 59869_1 | TATGAGTGTGGAGTCGTTGCATGATCTTAGCTCACTGCAACCTNNNAGATCGGAAG |
| 59870_1 | TATGAGTGTGGAGTCGTTGCGAGTCTTACTCTGTTGCCTAGGCNNNAGATCGGAAG |
| 59873_1 | TATGAGTGTGGAGTCGTTGCAGAGATGGGGTTTCACTCTGTTGNNNAGATCGGAA |
| 59874_1 | TATGAGTGTGGAGTCGTTGCCTCCTGAGTAGCTGGGACTACAGNNNAGATCGGAA |
| 59875_1 | TATGAGTGTGGAGTCGTTGCCAATTCTCCTGCCTCAGTCTTCTNNNAGATCGGAAGA |
| 59876_1 | TATGAGTGTGGAGTCGTTGCTCAAGTAGTTGTGACTCCAGGTGNNNAGATCGGAA |
| 59877_1 | TATGAGTGTGGAGTCGTTGCAGCTTTAGCACCTTCCTTTGGAANNNAGATCGGAAG |
| 59879_1 | TATGAGTGTGGAGTCGTTGCCTCTCAAAGCGCTTGGATTACAGNNNAGATCGGAAG |
| 59881_1 | TATGAGTGTGGAGTCGTTGCTAGTAGAGACAAGGTTTCGCCACNNNAGATCGGAA |
| 59882_1 | TATGAGTGTGGAGTCGTTGCCTAATAGCTGGGACTACAGGTGCNNNAGATCGGAA |
| 59884_1 | TATGAGTGTGGAGTCGTTGCCTGGAGTGCTATGGCTCAATCTCNNNAGATCGGAAG |
| 59885_1 | TATGAGTGTGGAGTCGTTGCCAATCACAGCTTACTACAGCCTGNNNAGATCGGAAG |
| 59886_1 | TATGAGTGTGGAGTCGTTGCTTGGGAAGATACCAAGGATTGCANNNAGATCGGAA |
| 59887_1 | TATGAGTGTGGAGTCGTTGCAACATGGTGAAACCTCTTCTCCANNNAGATCGGAAG |
| 59888_1 | TATGAGTGTGGAGTCGTTGCTGGATGCATAGAGCTGTTTCAGANNNAGATCGGAA |
| 59889_1 | TATGAGTGTGGAGTCGTTGCAGTAATTATCCTGCGTCACCCTCNNNAGATCGGAAG |
| 59890_1 | TATGAGTGTGGAGTCGTTGCTCTTTTCTTCCCCCATCTCTGTGNNNAGATCGGAAGA |
| 59891_1 | TATGAGTGTGGAGTCGTTGCAGGTGCCAAAAGAATTCAATGGANNNAGATCGGAA |
| 59893_1 | TATGAGTGTGGAGTCGTTGCAGGAGACCACAATTGGGAAAGTTNNNAGATCGGAA |
| 59894_1 | TATGAGTGTGGAGTCGTTGCCTCTCTCAACCAATGGAACTCCANNNAGATCGGAAG |
| 59896_1 | TATGAGTGTGGAGTCGTTGCGTGACAGAATGGGACCCTATCTCNNNAGATCGGAA |
| 59897_1 | TATGAGTGTGGAGTCGTTGCATGCCCACACATTAGAATCACCTNNNAGATCGGAAG |
| 59898_1 | TATGAGTGTGGAGTCGTTGCGCTATTGCCCACTTCTCTGTTTCNNNAGATCGGAAGA |
| 59899_1 | TATGAGTGTGGAGTCGTTGCCATTTCAGTTACTCTCCCACCCANNNAGATCGGAAGA |
| 59900_1 | TATGAGTGTGGAGTCGTTGCTGTGTTTTCACCTTGCACCTTTTNNNAGATCGGAAGA |

| | |
|---|---|
| 59901_1 | TATGAGTGTGGAGTCGTTGCAAGCTGGGCATTTGATCCTTCTANNNAGATCGGAAG |
| 59904_1 | TATGAGTGTGGAGTCGTTGCATGGGGAGAGTGGGAAAATAAGGNNNAGATCGGA |
| 59906_1 | TATGAGTGTGGAGTCGTTGCAGATTCTGGAGAGGATGTGGAGANNNAGATCGGA |
| 59907_1 | TATGAGTGTGGAGTCGTTGCATTCCATGCCATCAGTTTGTCACNNNAGATCGGAAG |
| 59908_1 | TATGAGTGTGGAGTCGTTGCATCCTTTGGTTTCCAGGTTCCTTNNNAGATCGGAAG |
| 59913_1 | TATGAGTGTGGAGTCGTTGCTTTGGTTGCTAGGGAAGAAAAGCNNNAGATCGGAA |
| 59916_1 | TATGAGTGTGGAGTCGTTGCTCACGTCTGTAATCCCAACACTTNNNAGATCGGAAG |
| 59917_1 | TATGAGTGTGGAGTCGTTGCGGCCAAGAACAGACTTCTGAGATNNNAGATCGGAA |
| 59918_1 | TATGAGTGTGGAGTCGTTGCTGACTATGAAGGCAAGTGGGATGNNNAGATCGGA |
| 59919_1 | TATGAGTGTGGAGTCGTTGCGGCTGGTGAAACCTCAGGTATTTNNNAGATCGGAA |
| 59922_1 | TATGAGTGTGGAGTCGTTGCATTGAAGCCAGCCAATGAATCTGNNNAGATCGGAA |
| 59924_1 | TATGAGTGTGGAGTCGTTGCTCAGCACATCCCTAATACTTGCANNNAGATCGGAAG |
| 59925_1 | TATGAGTGTGGAGTCGTTGCAAAAGCAGGGAGAGGGGTAAAATNNNAGATCGGA |
| 59927_1 | TATGAGTGTGGAGTCGTTGCACCAGTCTGGGCAACATAGTAAGNNNAGATCGGAA |
| 59928_1 | TATGAGTGTGGAGTCGTTGCTGAGACTAACCTGACCAACATGGNNNAGATCGGAA |
| 59929_1 | TATGAGTGTGGAGTCGTTGCTTACCAAATACCTGGCAATCCCANNNAGATCGGAAG |
| 59932_1 | TATGAGTGTGGAGTCGTTGCTACAAACAATAGTGGGGCACAGANNNAGATCGGAA |
| 59933_1 | TATGAGTGTGGAGTCGTTGCCGATAGCAAATGGAGACGTGAACNNNAGATCGGAA |
| 59935_1 | TATGAGTGTGGAGTCGTTGCTAGAAGCAGAGACCTGAAGCAAGNNNAGATCGGAA |
| 59936_1 | TATGAGTGTGGAGTCGTTGCCATGATGGTATTCTAAGCCCCCANNNAGATCGGAAG |
| 59937_1 | TATGAGTGTGGAGTCGTTGCGCGGTTGCTGTTTGATCAAAAAGNNNAGATCGGAA |
| 59938_1 | TATGAGTGTGGAGTCGTTGCCAAAACACATGACAGTGAGCAGTNNNAGATCGGAA |
| 59939_1 | TATGAGTGTGGAGTCGTTGCGCAAAATTTGGGCCCTATTAGCANNNAGATCGGAAG |
| 59941_1 | TATGAGTGTGGAGTCGTTGCTGAATAAAGCTTGCAGGACTGGANNNAGATCGGAA |
| 59943_1 | TATGAGTGTGGAGTCGTTGCCCTCTTCAAAGACTTTCCTCCCANNNAGATCGGAAGA |
| 59944_1 | TATGAGTGTGGAGTCGTTGCTTTCTGAACTTCCAAAATGGGCTNNNAGATCGGAAG |
| 59945_1 | TATGAGTGTGGAGTCGTTGCTCATGCCTGTAATACCAGCACTTNNNAGATCGGAAG |
| 59946_1 | TATGAGTGTGGAGTCGTTGCTGAGGCAGGAGAATCTCTTGAACNNNAGATCGGAA |
| 59947_1 | TATGAGTGTGGAGTCGTTGCAGCTCTTATACACTGTTGGTGGGNNNAGATCGGAA |
| 59952_1 | TATGAGTGTGGAGTCGTTGCCCTGAAACTCTGCTTTTACAGGCNNNAGATCGGAAG |
| 59954_1 | TATGAGTGTGGAGTCGTTGCGTGATTCCAGAGAAGAGGAGTGGNNNAGATCGGA |
| 59955_1 | TATGAGTGTGGAGTCGTTGCGGCCTCAATTGCTCGATTTTCTCNNNAGATCGGAAG |
| 59958_1 | TATGAGTGTGGAGTCGTTGCTGGTCAATACTAGTGAAGTGGCTNNNAGATCGGAA |
| 59959_1 | TATGAGTGTGGAGTCGTTGCAGGACTATATTGGTTGCTTGGGGNNNAGATCGGAA |
| 59961_1 | TATGAGTGTGGAGTCGTTGCCTGAGGCAGGAGGATTGTGTAAGNNNAGATCGGA |
| 59962_1 | TATGAGTGTGGAGTCGTTGCGGATCACTTGAGTCCAGGAGATCNNNAGATCGGAA |
| 59963_1 | TATGAGTGTGGAGTCGTTGCGGAGCTTTGCTGTAAAGGGAAACNNNAGATCGGAA |
| 59964_1 | TATGAGTGTGGAGTCGTTGCTCACTCACTCACTCACTCATTCGNNNAGATCGGAAGA |
| 59965_1 | TATGAGTGTGGAGTCGTTGCGGATGGTCAGAGTAAGTTTCATGGNNNAGATCGGA |
| 59966_1 | TATGAGTGTGGAGTCGTTGCTGCAACACTGGCTTTTTCCATAGNNNAGATCGGAAG |
| 59967_1 | TATGAGTGTGGAGTCGTTGCTCAGTGAACGTGAAGATAAGTCANNNAGATCGGAA |
| 59968_1 | TATGAGTGTGGAGTCGTTGCCCAAGTAAGAGGACCAAGTACCCNNNAGATCGGAA |
| 59969_1 | TATGAGTGTGGAGTCGTTGCTCGTGTTTATGCCACTAGACTCCNNNAGATCGGAAG |

| | |
|---|---|
| 59971_1 | TATGAGTGTGGAGTCGTTGCTGGGGTCTGTGGAGAAATTTCATNNNAGATCGGAA |
| 59972_1 | TATGAGTGTGGAGTCGTTGCTGGTGGTGAAGGGATTTCAGAATNNNAGATCGGAA |
| 59973_1 | TATGAGTGTGGAGTCGTTGCATGGTGCCTACTTCATTGCCTTANNNAGATCGGAAG |
| 59974_1 | TATGAGTGTGGAGTCGTTGCATGAGTACATCCAGGGAGACAGANNNAGATCGGAA |
| 59975_1 | TATGAGTGTGGAGTCGTTGCGAAGAGAAGTCAGGTGAGTGGTTNNNAGATCGGA |
| 59977_1 | TATGAGTGTGGAGTCGTTGCAATCCTTCTTTCCCCCTCCTTTCNNNAGATCGGAAGAG |
| 59979_1 | TATGAGTGTGGAGTCGTTGCTGAAACCAGTAGCAAGTGACTGANNNAGATCGGAA |
| 59980_1 | TATGAGTGTGGAGTCGTTGCCAACAACTACAAAAACCCCAGCANNNAGATCGGAAG |
| 59982_1 | TATGAGTGTGGAGTCGTTGCCATACACTCAAGCATGACACTGCNNNAGATCGGAAG |
| 59983_1 | TATGAGTGTGGAGTCGTTGCACCTAACTCCTGATGGTGTTTCANNNAGATCGGAAG |
| 59984_1 | TATGAGTGTGGAGTCGTTGCATACTGAAGAAGCCTCCCCTTTGNNNAGATCGGAAG |
| 59985_1 | TATGAGTGTGGAGTCGTTGCTTAGGACCTGAGAAAGCAGCATGNNNAGATCGGAA |
| 59986_1 | TATGAGTGTGGAGTCGTTGCTTGGTCCCACTACATAATTCCCANNNAGATCGGAAGA |
| 59987_1 | TATGAGTGTGGAGTCGTTGCAGTTACAGAGATAGGGCTGGAGTNNNAGATCGGAA |
| 59988_1 | TATGAGTGTGGAGTCGTTGCGGCTTGTGTATATGTGCATCCACNNNAGATCGGAAG |
| 59991_1 | TATGAGTGTGGAGTCGTTGCTGAGGCAGAAGAATTGCTTGAACNNNAGATCGGAA |
| 59992_1 | TATGAGTGTGGAGTCGTTGCCCGAAAGACTCCTGGAACTACATNNNAGATCGGAAG |
| 59993_1 | TATGAGTGTGGAGTCGTTGCTGGGCTGTTTTCGTATTTTTGCANNNAGATCGGAAG |
| 59995_1 | TATGAGTGTGGAGTCGTTGCGATCTCAGCTTACTGCAACCTCCNNNAGATCGGAAG |
| 59996_1 | TATGAGTGTGGAGTCGTTGCAACAGGGTCTTGCTTTGTTTCACNNNAGATCGGAAG |
| 59998_1 | TATGAGTGTGGAGTCGTTGCATCTTGGCTCACTACAACCTCTGNNNAGATCGGAAG |
| 59999_1 | TATGAGTGTGGAGTCGTTGCCCAGGTTCAACCGATTCTCCTATNNNAGATCGGAAG |
| 60000_1 | TATGAGTGTGGAGTCGTTGCGCCCGGCGAATTTTTGTATTTTTNNNAGATCGGAAG |
| 60001_1 | TATGAGTGTGGAGTCGTTGCAGTGGTACCATCTCTGCTTACTGNNNAGATCGGAAG |
| 60004_1 | TATGAGTGTGGAGTCGTTGCCTCAGTGTCCCAAGTAGTTGTGANNNAGATCGGAAG |
| 60005_1 | TATGAGTGTGGAGTCGTTGCGTAGTGGTGCAATCAAAGCTCAANNNAGATCGGAA |
| 60006_1 | TATGAGTGTGGAGTCGTTGCATCTTGGCTCACTACAACCTCTGNNNAGATCGGAAG |
| 60007_1 | TATGAGTGTGGAGTCGTTGCTTCAAGTGATTCTCCTGCCTCAGNNNAGATCGGAAG |
| 60008_1 | TATGAGTGTGGAGTCGTTGCTTCAAACGATTCCCCTACCTCAGNNNAGATCGGAAGA |
| 60010_1 | TATGAGTGTGGAGTCGTTGCCTGGGTTCAAGCAATTATCCTGCNNNAGATCGGAAG |
| 60011_1 | TATGAGTGTGGAGTCGTTGCAGATGGAGTCTCGCTTTTGTCACNNNAGATCGGAAG |
| 60012_1 | TATGAGTGTGGAGTCGTTGCGGGTTGATCTCAGCTCACTACAANNNAGATCGGAAG |
| 60015_1 | TATGAGTGTGGAGTCGTTGCAGGTTCAAGCAATTCTCCTGTCTNNNAGATCGGAAG |
| 60017_1 | TATGAGTGTGGAGTCGTTGCTTCAAGTGATTCTCCTGCCTCAGNNNAGATCGGAAG |
| 60018_1 | TATGAGTGTGGAGTCGTTGCTTCAAGCTATTCTCCTGCCTCAGNNNAGATCGGAAGA |
| 60019_1 | TATGAGTGTGGAGTCGTTGCTGGAGTTCAGTGGTACGATCTTGNNNAGATCGGAA |
| 60020_1 | TATGAGTGTGGAGTCGTTGCTTCACACCATTCTCTTGCCTCAGNNNAGATCGGAAGA |
| 60021_1 | TATGAGTGTGGAGTCGTTGCTTCATGCCATTCTCCTGTCTCAGNNNAGATCGGAAGA |
| 60022_1 | TATGAGTGTGGAGTCGTTGCCAGTCTTGGTTCATTGCAACCTTNNNAGATCGGAAG |
| 60023_1 | TATGAGTGTGGAGTCGTTGCCCTGAGTAGCTGGGAATATAGGCNNNAGATCGGAA |
| 60024_1 | TATGAGTGTGGAGTCGTTGCCCTCCCAACTTCAAGCAATTCTCNNNAGATCGGAAGA |
| 60025_1 | TATGAGTGTGGAGTCGTTGCCTCAATGAAACCTCCACCTACCANNNAGATCGGAAGA |
| 60026_1 | TATGAGTGTGGAGTCGTTGCCACTACAACTTCTGCCTCCTAGGNNNAGATCGGAAG |

| | |
|---|---|
| 60027_1 | TATGAGTGTGGAGTCGTTGCCTGGAGTATGGTGCTGTGATCTCNNNAGATCGGAA |
| 60028_1 | TATGAGTGTGGAGTCGTTGCGAGTCTCACTCTCTCTCCCAGTCNNNAGATCGGAAGA |
| 60029_1 | TATGAGTGTGGAGTCGTTGCGAGATGGAGTCTCTCTGTCACCANNNAGATCGGAAG |
| 60030_1 | TATGAGTGTGGAGTCGTTGCGGTTCAAGCAATTCTCCTGTCTCNNNAGATCGGAAG |
| 60031_1 | TATGAGTGTGGAGTCGTTGCTGAACTCCTGAGCTCAAGTGATTNNNAGATCGGAAG |
| 60032_1 | TATGAGTGTGGAGTCGTTGCGGCACAATTTTGGCTCACTGTAANNNAGATCGGAAG |
| 60033_1 | TATGAGTGTGGAGTCGTTGCTGTTACCTAGGCTGGTCTCAAACNNNAGATCGGAAG |
| 60034_1 | TATGAGTGTGGAGTCGTTGCTGTCATGTTGCCCAGAATGATCTNNNAGATCGGAAG |
| 60035_1 | TATGAGTGTGGAGTCGTTGCCAGCTTACTGCAACTTTGACCTCNNNAGATCGGAAG |
| 60036_1 | TATGAGTGTGGAGTCGTTGCCTATTACCTAGGCTGGAGTGCAGNNNAGATCGGAA |
| 60037_1 | TATGAGTGTGGAGTCGTTGCCAATCTCACCTCACTGCTACCTCNNNAGATCGGAAGA |
| 60038_1 | TATGAGTGTGGAGTCGTTGCAAGCTTTGCTCACAAATTCCTCCNNNAGATCGGAAGA |
| 60039_1 | TATGAGTGTGGAGTCGTTGCGAGATGGGGGTCTCACTATGTTGNNNAGATCGGAA |
| 60040_1 | TATGAGTGTGGAGTCGTTGCCTCAGGCTAGTCTTGAACTCCTGNNNAGATCGGAAG |
| 60041_1 | TATGAGTGTGGAGTCGTTGCTCCAACTCCAGTGTTCAAGTGATNNNAGATCGGAAG |
| 60042_1 | TATGAGTGTGGAGTCGTTGCTGGGTTCAAGCAATTTTCCTGTCNNNAGATCGGAAG |
| 60043_1 | TATGAGTGTGGAGTCGTTGCCACAATCTTAGCTCACTGCAACCNNNAGATCGGAAG |
| 60045_1 | TATGAGTGTGGAGTCGTTGCTTGGAGTACAGTGGCGTTATCTCNNNAGATCGGAA |
| 60046_1 | TATGAGTGTGGAGTCGTTGCCCCAGTGTGCTAGGATTATAGGCNNNAGATCGGAA |
| 60048_1 | TATGAGTGTGGAGTCGTTGCTTCAAGTGATTCTCCTGCCTCAGNNNAGATCGGAAG |
| 60050_1 | TATGAGTGTGGAGTCGTTGCTGATCTGGGCTCATTGCAATTTCNNNAGATCGGAAG |
| 60052_1 | TATGAGTGTGGAGTCGTTGCCTCACTGTAGCCTCAACCTTCTGNNNAGATCGGAAGA |
| 60054_1 | TATGAGTGTGGAGTCGTTGCGAGGCAGAATCTCACTCTGTCTCNNNAGATCGGAAG |
| 60055_1 | TATGAGTGTGGAGTCGTTGCCATGATCTTGGCTCACTGCAATCNNNAGATCGGAAG |
| 60056_1 | TATGAGTGTGGAGTCGTTGCCCTGACCTTGTAATCTACCCACCNNNAGATCGGAAGA |
| 60057_1 | TATGAGTGTGGAGTCGTTGCAGCCTCTTGAGTAGCTAGGACTANNNAGATCGGAAG |
| 60058_1 | TATGAGTGTGGAGTCGTTGCCAAGTAGCTGGGATTACAGGCATNNNAGATCGGAA |
| 60059_1 | TATGAGTGTGGAGTCGTTGCAGACAGGGTCTCACTCACTCTAANNNAGATCGGAAG |
| 60061_1 | TATGAGTGTGGAGTCGTTGCGCACAGGAGCTGGTTTTTTGAAANNNAGATCGGAA |
| 60062_1 | TATGAGTGTGGAGTCGTTGCGGTTCAAGCAATTCTCTTGCCTTNNNAGATCGGAAG |
| 60063_1 | TATGAGTGTGGAGTCGTTGCCAGTGGTGTGAACAGAACTTGTGNNNAGATCGGAA |
| 60064_1 | TATGAGTGTGGAGTCGTTGCTGATCTCAGCTCACTGCAAGCNNNAGATCGGAAGAG |
| 60066_1 | TATGAGTGTGGAGTCGTTGCGCAACCTCCATCTCTCAGATTCANNNAGATCGGAAGA |
| 60069_1 | TATGAGTGTGGAGTCGTTGCAACTGGGATTATAGACGTGAGCCNNNAGATCGGAA |
| 60070_1 | TATGAGTGTGGAGTCGTTGCGCTTCCCTAGTAGCTGTGACTACNNNAGATCGGAAG |
| 60072_1 | TATGAGTGTGGAGTCGTTGCTCTCAGCTCACTACAAACTCTGCNNNAGATCGGAAGA |
| 60075_1 | TATGAGTGTGGAGTCGTTGCGACTCCCGAGTAGCTAGGACTATNNNAGATCGGAAG |
| 60078_1 | TATGAGTGTGGAGTCGTTGCCTAACCTCAGGTGATCCGTGATCNNNAGATCGGAAG |
| 60079_1 | TATGAGTGTGGAGTCGTTGCAAGTGATTCTCATGCCTCAGCTTNNNAGATCGGAAG |
| 60080_1 | TATGAGTGTGGAGTCGTTGCATTCTCTTGCTTCAGCTTCCTGANNNAGATCGGAAGA |
| 60082_1 | TATGAGTGTGGAGTCGTTGCCTCCACCTCCCAGATTCAAGTAGNNNAGATCGGAAG |
| 60085_1 | TATGAGTGTGGAGTCGTTGCGATCTCAGCTCACTCTACCCTCTNNNAGATCGGAAGA |
| 60086_1 | TATGAGTGTGGAGTCGTTGCGGTTCAAGCGATTTTCCTGTCTCNNNAGATCGGAAG |

| | |
|---|---|
| 60087_1 | TATGAGTGTGGAGTCGTTGCACAAGATCTTGCTCGGTTTACCANNNAGATCGGAAG |
| 60088_1 | TATGAGTGTGGAGTCGTTGCTTCCTCCTGGGTTCAAGCAATTCNNNAGATCGGAAG |
| 60089_1 | TATGAGTGTGGAGTCGTTGCGAGTCTCTGTTGTTTAGGCTGGANNNAGATCGGAA |
| 60092_1 | TATGAGTGTGGAGTCGTTGCCTCTGGAGTAGCTGGAACTACAGNNNAGATCGGAA |
| 60093_1 | TATGAGTGTGGAGTCGTTGCGTTTCACTGTATTGGCCAGACTGNNNAGATCGGAAG |
| 60095_1 | TATGAGTGTGGAGTCGTTGCGACAAAGTCAAAGCAGCTGTGATNNNAGATCGGAA |
| 60096_1 | TATGAGTGTGGAGTCGTTGCCCAAGTAGCTGGGATTATAGGCANNNAGATCGGAA |
| 60097_1 | TATGAGTGTGGAGTCGTTGCTCATGACCTCAAATGATCCACCCNNNAGATCGGAAG |
| 60098_1 | TATGAGTGTGGAGTCGTTGCATCCTGACCTCAAATGATCCACCNNNAGATCGGAAG |
| 60099_1 | TATGAGTGTGGAGTCGTTGCGACAGAGTCTCGGTCTGTTGTTCNNNAGATCGGAAG |
| 60100_1 | TATGAGTGTGGAGTCGTTGCCAGTGATTCTCTTGTCTCAGCCTNNNAGATCGGAAG |
| 60104_1 | TATGAGTGTGGAGTCGTTGCTGCAGTGACATGATCTTGACACANNNAGATCGGAAG |
| 60105_1 | TATGAGTGTGGAGTCGTTGCTTCCAGCAATTCTCCTGTCTCAGNNNAGATCGGAAGA |
| 60106_1 | TATGAGTGTGGAGTCGTTGCAAACTCCTGGCCTCAAGTGATTTNNNAGATCGGAAG |
| 60107_1 | TATGAGTGTGGAGTCGTTGCACAAGTGATTCTCCTACCTCAGCNNNAGATCGGAAG |
| 60108_1 | TATGAGTGTGGAGTCGTTGCATGGAGTGCAGTGGTACAATCTTNNNAGATCGGAA |
| 60109_1 | TATGAGTGTGGAGTCGTTGCCAAGTAGCTGGGACTATAGGCACNNNAGATCGGAA |
| 60111_1 | TATGAGTGTGGAGTCGTTGCCACAAAGTTGCTGGGATTACAGGNNNAGATCGGAA |
| 60112_1 | TATGAGTGTGGAGTCGTTGCTTCAAGTGATTCTCCTGCCTCAGNNNAGATCGGAAG |
| 60113_1 | TATGAGTGTGGAGTCGTTGCTTCAAGCTATTCTCCTGCCTCAGNNNAGATCGGAAGA |
| 60114_1 | TATGAGTGTGGAGTCGTTGCCTCCTGAGTAGCTGGGACTACAGNNNAGATCGGAA |
| 60115_1 | TATGAGTGTGGAGTCGTTGCTCAAGAGGTTCTCATGCCTTAGCNNNAGATCGGAAG |
| 60116_1 | TATGAGTGTGGAGTCGTTGCTCCCAAGTTCAAGCAATTCTCCTNNNAGATCGGAAGA |
| 60117_1 | TATGAGTGTGGAGTCGTTGCTCAAGTGATTCTCCTTCCTCAGCNNNAGATCGGAAGA |
| 60118_1 | TATGAGTGTGGAGTCGTTGCCTTTCGGGGTTTTATTGGCAAGTNNNAGATCGGAAG |
| 60119_1 | TATGAGTGTGGAGTCGTTGCGTTCAAGAGATTCTCCTGCCTCANNNAGATCGGAAG |
| 60120_1 | TATGAGTGTGGAGTCGTTGCTTCAAGTGATTCTCCTGCCTCAGNNNAGATCGGAAG |
| 60121_1 | TATGAGTGTGGAGTCGTTGCCCTCCTATGTTCACGCCATTCTCNNNAGATCGGAAGA |
| 60122_1 | TATGAGTGTGGAGTCGTTGCGCCTCCTAAGTAGCTGGGATTACNNNAGATCGGAAG |
| 60123_1 | TATGAGTGTGGAGTCGTTGCTCCTGAGCTCAAACTATCTTCCCNNNAGATCGGAAGA |
| 60124_1 | TATGAGTGTGGAGTCGTTGCGAATGCAGTGGTGGAATCATGTCNNNAGATCGGAA |
| 60126_1 | TATGAGTGTGGAGTCGTTGCTATAAATCCTTCCCCTGTCCCCTNNNAGATCGGAAGA |
| 60127_1 | TATGAGTGTGGAGTCGTTGCAATCTCTGCCTCCCAGGTAAAAGNNNAGATCGGAAG |
| 60128_1 | TATGAGTGTGGAGTCGTTGCTTCAAGTGATTCTCCTGCCTCAGNNNAGATCGGAAG |
| 60129_1 | TATGAGTGTGGAGTCGTTGCAGACAATGTCTCACTCTGTCACCNNNAGATCGGAAG |
| 60131_1 | TATGAGTGTGGAGTCGTTGCTACTGGGTCTCATTCCATTCCACNNNAGATCGGAAGA |
| 60132_1 | TATGAGTGTGGAGTCGTTGCCCCATTCAAGCAATTCTCCTGTCNNNAGATCGGAAGA |
| 60133_1 | TATGAGTGTGGAGTCGTTGCGCTCAAGCAATCCTCCCATATCANNNAGATCGGAAG |
| 60135_1 | TATGAGTGTGGAGTCGTTGCTTTCACCATGTTAGCCAGGATGANNNAGATCGGAAG |
| 60137_1 | TATGAGTGTGGAGTCGTTGCCTGTAGTCCCAGCTACTTGGAAGNNNAGATCGGAAG |
| 60138_1 | TATGAGTGTGGAGTCGTTGCTTTTCTACCTGCCAAGGGACTACNNNAGATCGGAAG |
| 60139_1 | TATGAGTGTGGAGTCGTTGCCCCTTCACAGCAAACATCTGAAGNNNAGATCGGAAG |
| 60142_1 | TATGAGTGTGGAGTCGTTGCATGCCTGTAATCCCAGCACTTTANNNAGATCGGAAG |

| | |
|---|---|
| 60143_1 | TATGAGTGTGGAGTCGTTGCAAGCATATGGACTATTGCAGCACNNNAGATCGGAAG |
| 60144_1 | TATGAGTGTGGAGTCGTTGCAGGCAATGTGAAAAGACTCATCCNNNAGATCGGAAG |
| 60145_1 | TATGAGTGTGGAGTCGTTGCAGGCAATGTGAAAAGACTCATCCNNNAGATCGGAAG |
| 60149_1 | TATGAGTGTGGAGTCGTTGCAGGGCTTTCAAAATCTGGTCTCTNNNAGATCGGAAG |
| 60150_1 | TATGAGTGTGGAGTCGTTGCGAAAACACATGTCCTCAGCACTGNNNAGATCGGAAG |
| 60151_1 | TATGAGTGTGGAGTCGTTGCAACTTTGCAGAGGAGACAGTGATNNNAGATCGGAA |
| 60153_1 | TATGAGTGTGGAGTCGTTGCGAGTGAAGGGAGTGGCATTAGAANNNAGATCGGA |
| 60156_1 | TATGAGTGTGGAGTCGTTGCGGAATTTCACTCTTGTTGCCCAANNNAGATCGGAAG |
| 60158_1 | TATGAGTGTGGAGTCGTTGCGACTAATGGGGACGTGAGTATGGNNNAGATCGGA |
| 60159_1 | TATGAGTGTGGAGTCGTTGCTTTGAGCTGGGAAAATAGAGGCTNNNAGATCGGAA |
| 60160_1 | TATGAGTGTGGAGTCGTTGCAGCATAATACCCAGCTCTTAGGCNNNAGATCGGAAG |
| 60161_1 | TATGAGTGTGGAGTCGTTGCTCTTTCTTCAGGGAGACATGGTGNNNAGATCGGAAG |
| 60163_1 | TATGAGTGTGGAGTCGTTGCTAGTGGCATGATCATAGCTCACCNNNAGATCGGAAG |
| 60165_1 | TATGAGTGTGGAGTCGTTGCCTGTAATCCCAGCACTTTTGGAGNNNAGATCGGAAG |
| 60169_1 | TATGAGTGTGGAGTCGTTGCCGTCTGTAATCCCAGCATTTTGGNNNAGATCGGAAG |
| 60171_1 | TATGAGTGTGGAGTCGTTGCAAAACTGAATCAGAGCCAATGGCNNNAGATCGGAA |
| 60172_1 | TATGAGTGTGGAGTCGTTGCAATGAGATACCACTTCCACACCCNNNAGATCGGAAG |
| 60177_1 | TATGAGTGTGGAGTCGTTGCCCTGGGCCATATTGGAAGAAGAANNNAGATCGGAA |
| 60178_1 | TATGAGTGTGGAGTCGTTGCCAGCGAAAATGTGGAGGAGAAAANNNAGATCGGA |
| 60180_1 | TATGAGTGTGGAGTCGTTGCAAGATGGAGAAGGGATGGAGACTNNNAGATCGGA |
| 60181_1 | TATGAGTGTGGAGTCGTTGCATATGGCACATGGGGGTTTCTAANNNAGATCGGAA |
| 60182_1 | TATGAGTGTGGAGTCGTTGCGTGGCTCACACCTGTAATCCTAGNNNAGATCGGAAG |
| 60183_1 | TATGAGTGTGGAGTCGTTGCGAAGTCAAGGCTGCATTGAGATGNNNAGATCGGAA |
| 60185_1 | TATGAGTGTGGAGTCGTTGCAGATTTAGCCTGAGTCACCCATGNNNAGATCGGAAG |
| 60187_1 | TATGAGTGTGGAGTCGTTGCATGGAACAAGTCATGGACATGGTNNNAGATCGGAA |
| 60189_1 | TATGAGTGTGGAGTCGTTGCGTCCCTCTGAAGAGCCATGATAGNNNAGATCGGAAG |
| 60190_1 | TATGAGTGTGGAGTCGTTGCTGCAAAATAAAGGCTCCCAAAGGNNNAGATCGGAA |
| 60192_1 | TATGAGTGTGGAGTCGTTGCTGATTCACATTGGCCAAGTTGTCNNNAGATCGGAAG |
| 60193_1 | TATGAGTGTGGAGTCGTTGCCAGAGATCCTTGGAGAAATGGCTNNNAGATCGGAA |
| 60194_1 | TATGAGTGTGGAGTCGTTGCGGCAGCAAAAAGGACCAGTTTATNNNAGATCGGAA |
| 60195_1 | TATGAGTGTGGAGTCGTTGCTGTCTTTGCCTCACTACATCTCCNNNAGATCGGAAGA |
| 60198_1 | TATGAGTGTGGAGTCGTTGCCATAAAGGACCGAATGGCCTCTANNNAGATCGGAAG |
| 60203_1 | TATGAGTGTGGAGTCGTTGCCTTCCAGCAAAAAGGTATGTGGGNNNAGATCGGAA |
| 60205_1 | TATGAGTGTGGAGTCGTTGCGTTGCCTGCCTATAGTTCTAGCTNNNAGATCGGAAG |
| 60206_1 | TATGAGTGTGGAGTCGTTGCTAAAGCAGAGAGATGGCTTGTGTNNNAGATCGGAA |
| 60208_1 | TATGAGTGTGGAGTCGTTGCCAGTTAGCTAAGACTGAGCCACANNNAGATCGGAAG |
| 60209_1 | TATGAGTGTGGAGTCGTTGCAAGATGTTCACTGTGGCATTGTCNNNAGATCGGAAG |
| 60210_1 | TATGAGTGTGGAGTCGTTGCGCTTTGGTGAAACTCACATGTGANNNAGATCGGAAG |
| 60211_1 | TATGAGTGTGGAGTCGTTGCAGGCTGTGATGTTTGAGATGGATNNNAGATCGGAA |
| 60212_1 | TATGAGTGTGGAGTCGTTGCGCAGTAAGTGTTCAGTGTTTGCTNNNAGATCGGAAG |
| 60215_1 | TATGAGTGTGGAGTCGTTGCGGAAACATCGGCCTAGCTTACTTNNNAGATCGGAAG |
| 60216_1 | TATGAGTGTGGAGTCGTTGCCTTATGAGTTTGAGGCTGCAGTGNNNAGATCGGAA |
| 60218_1 | TATGAGTGTGGAGTCGTTGCAATGTGTATCTACCCGCAGTTGTNNNAGATCGGAAG |

| | |
|---|---|
| 60219_1 | TATGAGTGTGGAGTCGTTGCGGTGAAACCCCGTCTCTACTAAANNNAGATCGGAAG |
| 60220_1 | TATGAGTGTGGAGTCGTTGCGCTCACATTTGTAATCCCAGCACNNNAGATCGGAAG |
| 60224_1 | TATGAGTGTGGAGTCGTTGCGTGGTTTGAATGTGTTCCCCAAANNNAGATCGGAAG |
| 60225_1 | TATGAGTGTGGAGTCGTTGCTCCTTGACAGCACAAAGAAAGGANNNAGATCGGAA |
| 60226_1 | TATGAGTGTGGAGTCGTTGCCCTGCCTACCCATATCCTTCTTCNNNAGATCGGAAGA |
| 60228_1 | TATGAGTGTGGAGTCGTTGCCAGTGAGCTAGGATAGGACCACTNNNAGATCGGAA |
| 60230_1 | TATGAGTGTGGAGTCGTTGCGAAAGTATTGAGGCCAAGCATGGNNNAGATCGGAA |
| 60231_1 | TATGAGTGTGGAGTCGTTGCCAGGATCCTGAAGCTACAGAGTCNNNAGATCGGAA |
| 60232_1 | TATGAGTGTGGAGTCGTTGCGGCCATTATCTCAAGTCCATTGCNNNAGATCGGAAG |
| 60233_1 | TATGAGTGTGGAGTCGTTGCTTCAGTTTCCCTAGGCCCTAGTANNNAGATCGGAAG |
| 60235_1 | TATGAGTGTGGAGTCGTTGCTTGTCTCTCGTTTCCTCTGAAGGNNNAGATCGGAAG |
| 60236_1 | TATGAGTGTGGAGTCGTTGCCTTGGTTCTTACTGGCCTCCAAGNNNAGATCGGAAG |
| 60237_1 | TATGAGTGTGGAGTCGTTGCCCATTTCTTTGAGGGGAGTGGATNNNAGATCGGAA |
| 60243_1 | TATGAGTGTGGAGTCGTTGCCACACCATCACCCACCTCAATATNNNAGATCGGAAGA |
| 60245_1 | TATGAGTGTGGAGTCGTTGCTGATTTTCAGGTGAAATCCGTGCNNNAGATCGGAAG |
| 60246_1 | TATGAGTGTGGAGTCGTTGCGGTTCCATGACTAGACTACAGCCNNNAGATCGGAAG |
| 60247_1 | TATGAGTGTGGAGTCGTTGCTTGAGGCAGGAGAATTGCTTGAANNNAGATCGGAA |
| 60249_1 | TATGAGTGTGGAGTCGTTGCTTATGCCTGTAATCCCAGCACTTNNNAGATCGGAAG |
| 60250_1 | TATGAGTGTGGAGTCGTTGCGGAATGGGAGGACTTGGATCATTNNNAGATCGGAA |
| 60251_1 | TATGAGTGTGGAGTCGTTGCCACCATCTGCAAAAGGGGAAAATNNNAGATCGGAA |
| 60252_1 | TATGAGTGTGGAGTCGTTGCGAAGACACTGAAAAAGGCTGCTTNNNAGATCGGAA |
| 60253_1 | TATGAGTGTGGAGTCGTTGCCATGCCTGTAATTCCAGCACTTTNNNAGATCGGAAG |
| 60257_1 | TATGAGTGTGGAGTCGTTGCCACAACTAGTTTTTGGCCACAGTNNNAGATCGGAAG |
| 60258_1 | TATGAGTGTGGAGTCGTTGCTGCACAGTTTTTGTATCTGCTGGNNNAGATCGGAAG |
| 60259_1 | TATGAGTGTGGAGTCGTTGCACGGACATGGTTACAGAGAAAGANNNAGATCGGAA |
| 60260_1 | TATGAGTGTGGAGTCGTTGCTTTTTTTAGGTGGAGTCTCGCTCNNNAGATCGGAAG |
| 60261_1 | TATGAGTGTGGAGTCGTTGCTTATGAGCCTGGGTTGAGTCATCNNNAGATCGGAA |
| 60262_1 | TATGAGTGTGGAGTCGTTGCAAGATTGGGACAAGGAGGTATGGNNNAGATCGGA |
| 60263_1 | TATGAGTGTGGAGTCGTTGCTATCCTCCCATAACCTACCCTGGNNNAGATCGGAAGA |
| 60264_1 | TATGAGTGTGGAGTCGTTGCCTGTAATCCCAGCACTTTGTGAGNNNAGATCGGAAG |
| 60265_1 | TATGAGTGTGGAGTCGTTGCAGATAAAGAGCTTCTGCACAGCANNNAGATCGGAA |
| 60266_1 | TATGAGTGTGGAGTCGTTGCATATGGTTTGGCTCTATGTCCCCNNNAGATCGGAAG |
| 60269_1 | TATGAGTGTGGAGTCGTTGCAGGTTCCTGTAGTCTCAGCTACTNNNAGATCGGAAG |
| 60270_1 | TATGAGTGTGGAGTCGTTGCTTTCCTGCTAGAATCCAGCTCTCNNNAGATCGGAAGA |
| 60271_1 | TATGAGTGTGGAGTCGTTGCAACACTGCAAACTCCCCATAGATNNNAGATCGGAAG |
| 60272_1 | TATGAGTGTGGAGTCGTTGCTAGAACCAGATGCAGAGACAACCNNNAGATCGGAA |
| 60273_1 | TATGAGTGTGGAGTCGTTGCTTTTCTGAAACAGAGACTTGCTCNNNAGATCGGAAG |
| 60274_1 | TATGAGTGTGGAGTCGTTGCTGCAAAACCTAAAGGAGGAAAGCNNNAGATCGGAA |
| 60275_1 | TATGAGTGTGGAGTCGTTGCAGGCTTGAAGTGACATTAGCTCTNNNAGATCGGAA |
| 60276_1 | TATGAGTGTGGAGTCGTTGCAATGCAACCGAGAAAGAGAACACNNNAGATCGGAA |
| 60278_1 | TATGAGTGTGGAGTCGTTGCTGGTTCAAGATCTCTCTCTCCCANNNAGATCGGAAGA |
| 60279_1 | TATGAGTGTGGAGTCGTTGCGATTGGGAGCACATTCTCAACAGNNNAGATCGGAA |
| 60280_1 | TATGAGTGTGGAGTCGTTGCACCCAAAGGAGAATACACCTTCTNNNAGATCGGAAG |

| | |
|---|---|
| 60281_1 | TATGAGTGTGGAGTCGTTGCGAGTTTTGCTCTTTTTGCCCAGGNNNAGATCGGAAG |
| 60283_1 | TATGAGTGTGGAGTCGTTGCGGTCTCGAACTCCTAACCTCAGANNNAGATCGGAAG |
| 60285_1 | TATGAGTGTGGAGTCGTTGCTCAAGTGATTCTCCCATCTCAGCNNNAGATCGGAAG |
| 60286_1 | TATGAGTGTGGAGTCGTTGCCCTCCCAAGTAGCTGTGATTACANNNAGATCGGAAG |
| 60287_1 | TATGAGTGTGGAGTCGTTGCATGAAACAGTCTTGCTCTGTCACNNNAGATCGGAAG |
| 60288_1 | TATGAGTGTGGAGTCGTTGCTTGAAACGATTCTCCTGCCTCAGNNNAGATCGGAAG |
| 60289_1 | TATGAGTGTGGAGTCGTTGCTTCAAGCGATTCTCCTGTCTCAGNNNAGATCGGAAG |
| 60290_1 | TATGAGTGTGGAGTCGTTGCAGTAGCTGGGATTATAGGGACCCNNNAGATCGGAA |
| 60291_1 | TATGAGTGTGGAGTCGTTGCGTCATGTAGTTCCCCAGTGGTAGNNNAGATCGGAAG |
| 60292_1 | TATGAGTGTGGAGTCGTTGCTGAGCTCAGGTAATTCTCCCAACNNNAGATCGGAAG |
| 60294_1 | TATGAGTGTGGAGTCGTTGCTCACGGGTTCAAGTGATTCTTCTNNNAGATCGGAAG |
| 60296_1 | TATGAGTGTGGAGTCGTTGCTACAATCTCAGCTCACTGCAACTNNNAGATCGGAAG |
| 60297_1 | TATGAGTGTGGAGTCGTTGCATCTTGGCTCACTACAACCTCTGNNNAGATCGGAAG |
| 60298_1 | TATGAGTGTGGAGTCGTTGCCCTCCCAGGTTCATGTGATTCTCNNNAGATCGGAAG |
| 60299_1 | TATGAGTGTGGAGTCGTTGCAGCTGGGATTACAGGTATGTGTGNNNAGATCGGAA |
| 60300_1 | TATGAGTGTGGAGTCGTTGCGGAGTGCAATGGTGTGATCTCAGNNNAGATCGGAA |
| 60301_1 | TATGAGTGTGGAGTCGTTGCGGTCAAGTGACTCTTCTGCCTTANNNAGATCGGAAG |
| 60302_1 | TATGAGTGTGGAGTCGTTGCCACAATCTCAGCTCACTGCAATCNNNAGATCGGAAG |
| 60304_1 | TATGAGTGTGGAGTCGTTGCGCCTTCCTAGTAGCTGGGATTACNNNAGATCGGAAG |
| 60306_1 | TATGAGTGTGGAGTCGTTGCTCTTTTTGAGACGGAGTCTTGCTNNNAGATCGGAAG |
| 60309_1 | TATGAGTGTGGAGTCGTTGCAGATGGAGTCTCACTCTGTCTCCNNNAGATCGGAAG |
| 60310_1 | TATGAGTGTGGAGTCGTTGCGTTCAAGTGATTCTCCTACCCCANNNAGATCGGAAG |
| 60311_1 | TATGAGTGTGGAGTCGTTGCCTTCCTGAGTATCCAGCTCCATCNNNAGATCGGAAG |
| 60312_1 | TATGAGTGTGGAGTCGTTGCTTCAAGTGATTCTCCTGCCTCAGNNNAGATCGGAAG |
| 60313_1 | TATGAGTGTGGAGTCGTTGCTTCAAGCAATTCTCCTGCCTCAGNNNAGATCGGAAG |
| 60314_1 | TATGAGTGTGGAGTCGTTGCCTCCTGAGGTCAAGGTATATGCCNNNAGATCGGAAG |
| 60315_1 | TATGAGTGTGGAGTCGTTGCCCGAAGTTCAAGCAATTCTCCTGNNNAGATCGGAAG |
| 60316_1 | TATGAGTGTGGAGTCGTTGCGCTTCCTGAGTAGCTGGGATTACNNNAGATCGGAAG |
| 60321_1 | TATGAGTGTGGAGTCGTTGCCTCCTGAGTAGCTGGGATTACAGNNNAGATCGGAA |
| 60323_1 | TATGAGTGTGGAGTCGTTGCTTTTTTTTGAGATGGAGTCCCGCNNNAGATCGGAAG |
| 60325_1 | TATGAGTGTGGAGTCGTTGCCTCAAACTCCTGGACTCAAGTGANNNAGATCGGAAG |
| 60326_1 | TATGAGTGTGGAGTCGTTGCCTGGGATTACAGGACTGTACCACNNNAGATCGGAAG |
| 60327_1 | TATGAGTGTGGAGTCGTTGCAGTACAATGGCACTCACTATCTGTNNNAGATCGGAA |
| 60328_1 | TATGAGTGTGGAGTCGTTGCTTCAAGTGATTCTCCTGCCTCAGNNNAGATCGGAAG |
| 60329_1 | TATGAGTGTGGAGTCGTTGCCGAGTTTTGCTCTTGTTACCCAGNNNAGATCGGAAG |
| 60331_1 | TATGAGTGTGGAGTCGTTGCCTCCTGAGTAGCTGGGATTACAGNNNAGATCGGAA |
| 60332_1 | TATGAGTGTGGAGTCGTTGCTCCGGAGTAGCTGGGATTATAGGNNNAGATCGGAA |
| 60333_1 | TATGAGTGTGGAGTCGTTGCGGAGTACAATGTTGCAATCTCGGNNNAGATCGGAA |
| 61952_1 | TATGAGTGTGGAGTCGTTGCATTCACTGGATAATGCTGTCAGCNNNAGATCGGAAG |
| 61953_1 | TATGAGTGTGGAGTCGTTGCGGTGACTTGAAGAAGCAGGAAACNNNAGATCGGAA |
| 61955_1 | TATGAGTGTGGAGTCGTTGCTGCAGACAGAGAGAGAGAGAGAANNNAGATCGGA |
| 61956_1 | TATGAGTGTGGAGTCGTTGCAACGATACAGGAAAGAAGGCAGTNNNAGATCGGAA |
| 61957_1 | TATGAGTGTGGAGTCGTTGCTGTTACTGGAAAGGGCAGTCAATNNNAGATCGGAA |

| | |
|---|---|
| 61958_1 | TATGAGTGTGGAGTCGTTGCGTAGGGTCACAAGATTGGGAAGANNNAGATCGGA |
| 61959_1 | TATGAGTGTGGAGTCGTTGCGAGGCTGCCTTTTAGAAACAGTGNNNAGATCGGAA |
| 61960_1 | TATGAGTGTGGAGTCGTTGCTTTATTCTGGTCTGTTCCACCCANNNAGATCGGAAGA |
| 61961_1 | TATGAGTGTGGAGTCGTTGCACACATGAGTGAAAGGCTTAGAANNNAGATCGGAA |
| 61962_1 | TATGAGTGTGGAGTCGTTGCCATCAACTGGGAACAAGCACAATNNNAGATCGGAAG |
| 61963_1 | TATGAGTGTGGAGTCGTTGCGCAGGGCTCAGCATTTTATTCTTNNNAGATCGGAAG |
| 61964_1 | TATGAGTGTGGAGTCGTTGCCTCTGAAGTCACTGCTCTGACAGNNNAGATCGGAAG |
| 61965_1 | TATGAGTGTGGAGTCGTTGCCCTGCAAGGCAAGAACCAATAAANNNAGATCGGAA |
| 61966_1 | TATGAGTGTGGAGTCGTTGCAGCACCAAACACAATAACTTGCANNNAGATCGGAAG |
| 61967_1 | TATGAGTGTGGAGTCGTTGCTCTCTGATCCTGCACAAACCATCNNNAGATCGGAAG |
| 61968_1 | TATGAGTGTGGAGTCGTTGCGGGCTAGAGTGGGATCTAGAGAANNNAGATCGGA |
| 61971_1 | TATGAGTGTGGAGTCGTTGCGTGAGTCTACACTATGTTGCCCANNNAGATCGGAAG |
| 61973_1 | TATGAGTGTGGAGTCGTTGCGTGGAAATGTGTCAGCTGTGATTNNNAGATCGGAA |
| 61974_1 | TATGAGTGTGGAGTCGTTGCAGAAAGACAGAGAGACAGAGGGANNNAGATCGGA |
| 61975_1 | TATGAGTGTGGAGTCGTTGCGAAGGAGTTGGAAGTTGGATGGANNNAGATCGGA |
| 61976_1 | TATGAGTGTGGAGTCGTTGCGAGGTTACACATCAGTCACAGGTNNNAGATCGGAA |
| 61977_1 | TATGAGTGTGGAGTCGTTGCACTACTGACCAGAAGCAGTGATGNNNAGATCGGAA |
| 61979_1 | TATGAGTGTGGAGTCGTTGCGCTCTACCTCCTGGACATGTTAGNNNAGATCGGAAG |
| 61980_1 | TATGAGTGTGGAGTCGTTGCAGCTGTAAGAGTAGGAATTGCCANNNAGATCGGAA |
| 61981_1 | TATGAGTGTGGAGTCGTTGCGGGTCATCATTCATGCAGAACACNNNAGATCGGAAG |
| 61982_1 | TATGAGTGTGGAGTCGTTGCGTTGTGGGCTTTACGACTTCATGNNNAGATCGGAAG |
| 61984_1 | TATGAGTGTGGAGTCGTTGCCTCCACTTCAGTGAAAGGAAAGCNNNAGATCGGAAG |
| 61985_1 | TATGAGTGTGGAGTCGTTGCTACATGCCAGACTCCCAAATTCTNNNAGATCGGAAG |
| 61986_1 | TATGAGTGTGGAGTCGTTGCAGTTTCCTGGTGGGGTTCTTTAGNNNAGATCGGAAG |
| 61987_1 | TATGAGTGTGGAGTCGTTGCTCATGCACATCTATGGGAATGTTNNNAGATCGGAAG |
| 61989_1 | TATGAGTGTGGAGTCGTTGCCCAGTCCACATCTAACAGGAGTCNNNAGATCGGAAG |
| 61992_1 | TATGAGTGTGGAGTCGTTGCTTTCCTGGAAATTGCAAGCAGAGNNNAGATCGGAA |
| 61993_1 | TATGAGTGTGGAGTCGTTGCGACTCGACTAACAACAGGACAGTNNNAGATCGGAA |
| 61995_1 | TATGAGTGTGGAGTCGTTGCCTCATGTGGGAGAGAGGTGAAAANNNAGATCGGA |
| 61996_1 | TATGAGTGTGGAGTCGTTGCCAACACAGAAGACGGGTGATTTCNNNAGATCGGAA |
| 61997_1 | TATGAGTGTGGAGTCGTTGCTTCAAGTGATTCTCCTGTCTCGGNNNAGATCGGAAG |
| 61998_1 | TATGAGTGTGGAGTCGTTGCCCTCCAAGTAGCTGGGATTACAGNNNAGATCGGAA |
| 61999_1 | TATGAGTGTGGAGTCGTTGCCAGACCCGTCAAGAAAACAGTTCNNNAGATCGGAAG |
| 62000_1 | TATGAGTGTGGAGTCGTTGCAGTGCTGGAATTACAGGAGTGAGNNNAGATCGGA |
| 62002_1 | TATGAGTGTGGAGTCGTTGCAAATGCTCTATGGGCTGCCTTATNNNAGATCGGAAG |
| 62003_1 | TATGAGTGTGGAGTCGTTGCAAGAGAGAAGAACCAAACGGACANNNAGATCGGA |
| 62004_1 | TATGAGTGTGGAGTCGTTGCAACATGGGCCTTCAAAACAATGANNNAGATCGGAA |
| 62005_1 | TATGAGTGTGGAGTCGTTGCCCTGGGTTGCTTTTCTCCAGATANNNAGATCGGAAG |
| 62006_1 | TATGAGTGTGGAGTCGTTGCAGAGAACGAGCTCAGTGAATTGTNNNAGATCGGAA |
| 62007_1 | TATGAGTGTGGAGTCGTTGCCTCATGTGGTTTTCCTGCCTCAGNNNAGATCGGAAG |
| 62008_1 | TATGAGTGTGGAGTCGTTGCCATGCCCACACACATACACATACNNNAGATCGGAAG |
| 62010_1 | TATGAGTGTGGAGTCGTTGCTAAGAGTGTGTGTGTGTGTGTGTNNNAGATCGGAA |
| 62011_1 | TATGAGTGTGGAGTCGTTGCTTCAAGCAATTCTCCTGCCTCAGNNNAGATCGGAAG |

| | |
|---|---|
| 62012_1 | TATGAGTGTGGAGTCGTTGCCACTCCGTTATTCATTGGAGCACNNNAGATCGGAAG |
| 62013_1 | TATGAGTGTGGAGTCGTTGCTCTCTTATGCTTTGTCACCCAGGNNNAGATCGGAAG |
| 62014_1 | TATGAGTGTGGAGTCGTTGCACACACACACACACACACAAATTNNNAGATCGGAAG |
| 62016_1 | TATGAGTGTGGAGTCGTTGCATGGATTCTCACTCTTTTGCCCANNNAGATCGGAAGA |
| 62017_1 | TATGAGTGTGGAGTCGTTGCTTCAAGCGATTATCCTGCCTCAGNNNAGATCGGAAG |
| 62018_1 | TATGAGTGTGGAGTCGTTGCAGGTGACATGTGAGTGAGTTTGANNNAGATCGGAA |
| 62020_1 | TATGAGTGTGGAGTCGTTGCATCATTCTTGGGTGTTTCTCGGANNNAGATCGGAAG |
| 62021_1 | TATGAGTGTGGAGTCGTTGCTCAGATTTCACCATTTTGCCCAGNNNAGATCGGAAG |
| 62022_1 | TATGAGTGTGGAGTCGTTGCGTACAATGGCACAATCACAGCTCNNNAGATCGGAAG |
| 62023_1 | TATGAGTGTGGAGTCGTTGCGTGAGCCTGTAATTCCTGCTACTNNNAGATCGGAAG |
| 62024_1 | TATGAGTGTGGAGTCGTTGCGTCCTTGAAGGAGAGAGACTGTGNNNAGATCGGAA |
| 62027_1 | TATGAGTGTGGAGTCGTTGCTGCTATTCCCAGCTAAGTCACAGNNNAGATCGGAAG |
| 62028_1 | TATGAGTGTGGAGTCGTTGCCCAGCAGTAACTTCTGTTGTGTGNNNAGATCGGAAG |
| 62029_1 | TATGAGTGTGGAGTCGTTGCTGGCCAAAGCAACTTACAGATTCNNNAGATCGGAAG |
| 62030_1 | TATGAGTGTGGAGTCGTTGCAGGAACTCAGAATACCTCAAGCANNNAGATCGGAAG |
| 62031_1 | TATGAGTGTGGAGTCGTTGCGGCAGGGAAAGGGAATAAGGTAANNNAGATCGGA |
| 62034_1 | TATGAGTGTGGAGTCGTTGCGGCATTTGAAAGGGAAGGGAAAGNNNAGATCGGA |
| 62035_1 | TATGAGTGTGGAGTCGTTGCGAATGTGATCATGTCCCTTGCAGNNNAGATCGGAAG |
| 62036_1 | TATGAGTGTGGAGTCGTTGCGCTCACTACAGCCTCAAACAATCNNNAGATCGGAAG |
| 62039_1 | TATGAGTGTGGAGTCGTTGCGCTATTCTGGAGTCTGAAGTGGGNNNAGATCGGAA |
| 62040_1 | TATGAGTGTGGAGTCGTTGCTCCCAAAGTGCTAGGATTATGGGNNNAGATCGGAA |
| 62042_1 | TATGAGTGTGGAGTCGTTGCCACAAAATTAGCCAGGTGTGGTGNNNAGATCGGAA |
| 62044_1 | TATGAGTGTGGAGTCGTTGCGATCACTTCAGCCCAGGAGTTAGNNNAGATCGGAAG |
| 62045_1 | TATGAGTGTGGAGTCGTTGCAGAAGAGCTCCTATCTGTGAGGTNNNAGATCGGAA |
| 62046_1 | TATGAGTGTGGAGTCGTTGCAGATGACTTCAGAGGTAGGACCANNNAGATCGGAA |
| 62047_1 | TATGAGTGTGGAGTCGTTGCCCTGGGGTTATTGGTCTGATCAANNNAGATCGGAAG |
| 62048_1 | TATGAGTGTGGAGTCGTTGCGGATGCTGAGGAGAAGAGTTTCANNNAGATCGGAA |
| 62050_1 | TATGAGTGTGGAGTCGTTGCACAGACATCATCAGGACACCAAANNNAGATCGGAAG |
| 62051_1 | TATGAGTGTGGAGTCGTTGCGGTTATTTTTGGCTGAGGCTGTTNNNAGATCGGAAG |
| 62053_1 | TATGAGTGTGGAGTCGTTGCTGAGACTGGTCTTGAAATCCTGGNNNAGATCGGAA |
| 62054_1 | TATGAGTGTGGAGTCGTTGCTTCCAGGAAAGGGGAATGAATGTNNNAGATCGGAA |
| 62055_1 | TATGAGTGTGGAGTCGTTGCGAGAGAACCAACACCTTCAATGCNNNAGATCGGAAG |
| 62058_1 | TATGAGTGTGGAGTCGTTGCTTCTAAGTCTCCCCACTGATAGCNNNAGATCGGAAG |
| 62059_1 | TATGAGTGTGGAGTCGTTGCATGACTGGTCTGCAAAGGAGAATNNNAGATCGGAA |
| 62060_1 | TATGAGTGTGGAGTCGTTGCTGGTGGAGAATTTGGCTTCTTTCNNNAGATCGGAAG |
| 62061_1 | TATGAGTGTGGAGTCGTTGCCTACCTCCCAGGCTCAAATGATTNNNAGATCGGAAG |
| 62062_1 | TATGAGTGTGGAGTCGTTGCTTTTTTGAGACGGAGTTTCGCTCNNNAGATCGGAAG |
| 62063_1 | TATGAGTGTGGAGTCGTTGCAAGAGAACAAGAGTCTCTGCCTGNNNAGATCGGAA |
| 62065_1 | TATGAGTGTGGAGTCGTTGCCTGCTGTTGGTTCTGTTTCTCTGNNNAGATCGGAAG |
| 62067_1 | TATGAGTGTGGAGTCGTTGCGTCACAAATTTTGGCTTCGAGGANNNAGATCGGAAG |
| 62068_1 | TATGAGTGTGGAGTCGTTGCCTGGAGTGCAATGGTACGATCTCNNNAGATCGGAA |
| 62070_1 | TATGAGTGTGGAGTCGTTGCCTCTGTCTCAAACACACACACACNNNAGATCGGAAGA |
| 62071_1 | TATGAGTGTGGAGTCGTTGCCACTCCTGTAATCCCAGCACTTTNNNAGATCGGAAGA |

| | |
|---|---|
| 62072_1 | TATGAGTGTGGAGTCGTTGCCTGAGACAGGAGGATCACCTAAGNNNAGATCGGAA |
| 62073_1 | TATGAGTGTGGAGTCGTTGCACAGGTCATCAGGGAATTCTGTCNNNAGATCGGAA |
| 62074_1 | TATGAGTGTGGAGTCGTTGCGATCTTGGCTCACTGAAACCTCTNNNAGATCGGAAG |
| 62075_1 | TATGAGTGTGGAGTCGTTGCAGTCCTAACACTCAAGGAGCATGNNNAGATCGGAA |
| 62077_1 | TATGAGTGTGGAGTCGTTGCAGGGTCAGTCCATCCTATATCCCNNNAGATCGGAAG |
| 62081_1 | TATGAGTGTGGAGTCGTTGCCACACACACACACACATTCACATNNNAGATCGGAAG |
| 62082_1 | TATGAGTGTGGAGTCGTTGCGGGAAATTTTGGCAGTCAAGGAANNNAGATCGGAA |
| 62083_1 | TATGAGTGTGGAGTCGTTGCACTCTGAAAAGCCGAATCTAGCANNNAGATCGGAAG |
| 62084_1 | TATGAGTGTGGAGTCGTTGCGCAGAGAACTCCTGACAGATGTTNNNAGATCGGAA |
| 62085_1 | TATGAGTGTGGAGTCGTTGCATATACACACACACACACACACANNNAGATCGGAAG |
| 62086_1 | TATGAGTGTGGAGTCGTTGCCTTCCTGAGCTCAAGGAATCCTTNNNAGATCGGAAG |
| 62087_1 | TATGAGTGTGGAGTCGTTGCCTCCCAGCTACCATCTTGAACTTNNNAGATCGGAAG |
| 62089_1 | TATGAGTGTGGAGTCGTTGCATCCCATCTTTCCAGCATCATCTNNNAGATCGGAAGA |
| 62090_1 | TATGAGTGTGGAGTCGTTGCTTCCTAATACCTGGGACTGCCTANNNAGATCGGAAG |
| 62091_1 | TATGAGTGTGGAGTCGTTGCTTGCTAGAGGGTAGTGAGGAACTNNNAGATCGGAA |
| 62093_1 | TATGAGTGTGGAGTCGTTGCTCTCTCTCCCTACCTACCTACCTNNNAGATCGGAAGA |
| 62094_1 | TATGAGTGTGGAGTCGTTGCATTCCCCCAAAAAGTTCCCTCATNNNAGATCGGAAG |
| 62095_1 | TATGAGTGTGGAGTCGTTGCCCTCTTGCTTGCTTGCTTGAAAGNNNAGATCGGAAG |
| 62096_1 | TATGAGTGTGGAGTCGTTGCGAAAGGGAGGGGAAGAAAGAGAGNNNAGATCGG |
| 62097_1 | TATGAGTGTGGAGTCGTTGCGGTTCTAGAACAGCAGGTAGCATNNNAGATCGGAA |
| 62098_1 | TATGAGTGTGGAGTCGTTGCTTGTGGTCTACAGCCTAAAGCATNNNAGATCGGAAG |
| 62100_1 | TATGAGTGTGGAGTCGTTGCCTCTCTCTCTCTCACACACACACNNNAGATCGGAAGA |
| 62101_1 | TATGAGTGTGGAGTCGTTGCGACAGGGTCTTACTCTGTTACCCNNNAGATCGGAAG |
| 62102_1 | TATGAGTGTGGAGTCGTTGCGAACCAACCCTCTGGAACTAGAGNNNAGATCGGAA |
| 62103_1 | TATGAGTGTGGAGTCGTTGCTGAGACAATGTCTATTGCCCAGGNNNAGATCGGAA |
| 62104_1 | TATGAGTGTGGAGTCGTTGCGAAAGGAAATTGCAGGCAGTGAANNNAGATCGGA |
| 62105_1 | TATGAGTGTGGAGTCGTTGCCCAAAGCTTTGCATTCCCTGTAANNNAGATCGGAAG |
| 62106_1 | TATGAGTGTGGAGTCGTTGCTTGTGAGACGGTGTCTTACTCTGNNNAGATCGGAAG |
| 62107_1 | TATGAGTGTGGAGTCGTTGCTTCCACACTTTACGGAGGCTAAANNNAGATCGGAAG |
| 62108_1 | TATGAGTGTGGAGTCGTTGCAGAGATGGAGAGAAAGATGGGGANNNAGATCGGA |
| 62109_1 | TATGAGTGTGGAGTCGTTGCAGAGAGAGAGAGAGGGAGAGAGANNNAGATCGG |
| 62111_1 | TATGAGTGTGGAGTCGTTGCGCTTTTCAGCTCCATTCCCAAATNNNAGATCGGAAGA |
| 62113_1 | TATGAGTGTGGAGTCGTTGCCTGAGGCAGGAGAATCACTTGAANNNAGATCGGAA |
| 62115_1 | TATGAGTGTGGAGTCGTTGCGAGGCAAGAAAATCCCTTGAACCNNNAGATCGGAA |
| 62116_1 | TATGAGTGTGGAGTCGTTGCCCATCCCAGATCTGCTTGAAAAGNNNAGATCGGAAG |
| 62117_1 | TATGAGTGTGGAGTCGTTGCTCTCACCTCTTTGTCCTTGCTTTNNNAGATCGGAAGA |
| 62118_1 | TATGAGTGTGGAGTCGTTGCAGCATGGTGCAGACTCTCAATAANNNAGATCGGAA |
| 62119_1 | TATGAGTGTGGAGTCGTTGCACACACATCATGCAAAAGTGGAGNNNAGATCGGAA |
| 62120_1 | TATGAGTGTGGAGTCGTTGCTAGCACCTGGGGATAAATGTGACNNNAGATCGGAA |
| 62121_1 | TATGAGTGTGGAGTCGTTGCACTGGACTCTACAAAAGGCTGAGNNNAGATCGGAA |
| 62122_1 | TATGAGTGTGGAGTCGTTGCTGAGTCAACAAGTGGGAGGAAATNNNAGATCGGA |
| 62123_1 | TATGAGTGTGGAGTCGTTGCCTATCCATCTGTTCAGGGTGAGGNNNAGATCGGAA |
| 62124_1 | TATGAGTGTGGAGTCGTTGCAGCTAAAAATGCGCATGGTGAAANNNAGATCGGAA |

| | |
|---|---|
| 62125_1 | TATGAGTGTGGAGTCGTTGCACTCTGCCACATTTTAGCCTACANNNAGATCGGAAG |
| 62126_1 | TATGAGTGTGGAGTCGTTGCCCTGGGAGGCTCTAGATAGTACANNNAGATCGGAA |
| 62127_1 | TATGAGTGTGGAGTCGTTGCCAGGGTAACAGACAAGTTTGACANNNAGATCGGAA |
| 62128_1 | TATGAGTGTGGAGTCGTTGCATTTAGGCAGGATTTGATGCTGCNNNAGATCGGAA |
| 62129_1 | TATGAGTGTGGAGTCGTTGCTTCCCTGTAGACTCCAAATGCTCNNNAGATCGGAAG |
| 62130_1 | TATGAGTGTGGAGTCGTTGCCACGAGGAAGCTAGAAGAAACCTNNNAGATCGGAA |
| 62132_1 | TATGAGTGTGGAGTCGTTGCCTTGGCATTTGCTCTTCTATGGGNNNAGATCGGAAG |
| 62133_1 | TATGAGTGTGGAGTCGTTGCGAGCTGAGTTGGAGATAAGGGTCNNNAGATCGGA |
| 62134_1 | TATGAGTGTGGAGTCGTTGCCCTTTTCCACGAAGTCGATGAAGNNNAGATCGGAAG |
| 62135_1 | TATGAGTGTGGAGTCGTTGCTTTCAACTAGTGTCTCTGCTGCTNNNAGATCGGAAG |
| 62136_1 | TATGAGTGTGGAGTCGTTGCTTTAGCAACAGATCCAAGCAAGCNNNAGATCGGAAG |
| 62137_1 | TATGAGTGTGGAGTCGTTGCACAGCATGAATGACATCCACTCTNNNAGATCGGAAG |
| 62138_1 | TATGAGTGTGGAGTCGTTGCCATTCAAAGCCTGATCATCCAGCNNNAGATCGGAAG |
| 62139_1 | TATGAGTGTGGAGTCGTTGCGCTTCTGTGGCACTGTCTCTTTANNNAGATCGGAAG |
| 62140_1 | TATGAGTGTGGAGTCGTTGCGGAGGCTTAGCAAGTCCAAAATCNNNAGATCGGAA |
| 62141_1 | TATGAGTGTGGAGTCGTTGCCTCCTGAGTAGCTGGGACTACAGNNNAGATCGGAA |
| 62142_1 | TATGAGTGTGGAGTCGTTGCCCCATCTCAAAAAGTCAAGTGCCNNNAGATCGGAAG |
| 62144_1 | TATGAGTGTGGAGTCGTTGCAAACTCCATTTTATCCCCCTCCCNNNAGATCGGAAGA |
| 62145_1 | TATGAGTGTGGAGTCGTTGCGGAGAGCAGATGAAAGTGACTGANNNAGATCGGA |
| 62146_1 | TATGAGTGTGGAGTCGTTGCGACCCAGACACAGGCAATATCTANNNAGATCGGAAG |
| 62147_1 | TATGAGTGTGGAGTCGTTGCGTGTGTGTGTGTGTTTCTCTCNNNAGATCGGAAG |
| 62148_1 | TATGAGTGTGGAGTCGTTGCGAGACAGAGAGAGATCTGGCATGNNNAGATCGGA |
| 62149_1 | TATGAGTGTGGAGTCGTTGCGAGTAGCTGGGATTATACAGGCGNNNAGATCGGAA |
| 62150_1 | TATGAGTGTGGAGTCGTTGCCTGACTCTGGATTTGCCTCTTCTNNNAGATCGGAAG |
| 62152_1 | TATGAGTGTGGAGTCGTTGCCATCGCCTACTTACCCAGGAAAANNNAGATCGGAAG |
| 62153_1 | TATGAGTGTGGAGTCGTTGCTAGTTTGACTGTCACAGCTTCCANNNAGATCGGAAG |
| 62154_1 | TATGAGTGTGGAGTCGTTGCAGTCATGAAGAGTCAGGATCTGCNNNAGATCGGAA |
| 62155_1 | TATGAGTGTGGAGTCGTTGCAGAGAGAGGAAGAGAAACAGGGANNNAGATCGG |
| 62156_1 | TATGAGTGTGGAGTCGTTGCGGGCTCTGTGAAGAGAATGGTTANNNAGATCGGAA |
| 62157_1 | TATGAGTGTGGAGTCGTTGCCCTCCTGGATTCAAGCGATTCTTNNNAGATCGGAAG |
| 62158_1 | TATGAGTGTGGAGTCGTTGCTCCACAACCATTTCACATCCTCTNNNAGATCGGAAGA |
| 62159_1 | TATGAGTGTGGAGTCGTTGCCAGGTGGTATCATTCCAGTCCAANNNAGATCGGAAG |
| 62160_1 | TATGAGTGTGGAGTCGTTGCGCCTCATTCAAGATGCTAGGAGANNNAGATCGGAA |
| 62161_1 | TATGAGTGTGGAGTCGTTGCCCTGGGACCTAGGCTTGATTTTTNNNAGATCGGAAG |
| 62162_1 | TATGAGTGTGGAGTCGTTGCGGCAGTGTGAGAATCAAGGATCANNNAGATCGGAA |
| 62163_1 | TATGAGTGTGGAGTCGTTGCGTAAGCCACCCCTCTAGTTTCTCNNNAGATCGGAAG |
| 62164_1 | TATGAGTGTGGAGTCGTTGCTTCATGCTATTCTCCTGCCTCAGNNNAGATCGGAAG |
| 62166_1 | TATGAGTGTGGAGTCGTTGCAGGCTGGTCTCAAACACATAACTNNNAGATCGGAAG |
| 62167_1 | TATGAGTGTGGAGTCGTTGCTCTCCAACTCCTTTCCTCTACCTNNNAGATCGGAAGA |
| 62168_1 | TATGAGTGTGGAGTCGTTGCTTTTGGGGTTTTCAGAAGCTGTTNNNAGATCGGAAG |
| 62170_1 | TATGAGTGTGGAGTCGTTGCCGCGCATGTGTGTATGTATGTATNNNAGATCGGAA |
| 62171_1 | TATGAGTGTGGAGTCGTTGCTGAAGTTATGGCTGGCTGATTCTNNNAGATCGGAA |
| 62172_1 | TATGAGTGTGGAGTCGTTGCGCCTGAGCAAGTGAACAATAGAGNNNAGATCGGAA |

| | |
|---|---|
| 62173_1 | TATGAGTGTGGAGTCGTTGCATCACTGAGCTTTCACCCAGAAANNNAGATCGGAAG[illegible] |
| 62175_1 | TATGAGTGTGGAGTCGTTGCTGCTGAGGTTGACACATAAAGGANNNAGATCGGAA[illegible] |
| 62176_1 | TATGAGTGTGGAGTCGTTGCTCAAGAAATAGCCTGCTGTCCTTNNNAGATCGGAAG[illegible] |
| 62177_1 | TATGAGTGTGGAGTCGTTGCAGAAGGATGAGAAGGAGGAGGAANNNAGATCGG[illegible] |
| 62178_1 | TATGAGTGTGGAGTCGTTGCCTCTATCCAGGAGTAAGGCCAGTNNNAGATCGGAAG |
| 62179_1 | TATGAGTGTGGAGTCGTTGCAGTATACCACTGAACCAACCAGCNNNAGATCGGAAG |
| 62180_1 | TATGAGTGTGGAGTCGTTGCTAAATTGACCTGCCAGTGTCAGANNNAGATCGGAAG |
| 62181_1 | TATGAGTGTGGAGTCGTTGCACGAGTTAGCACTAACCTGGAAGNNNAGATCGGAA[illegible] |
| 62182_1 | TATGAGTGTGGAGTCGTTGCTGAACAAGTCCTTGGTTTCTGGTNNNAGATCGGAAG |
| 62183_1 | TATGAGTGTGGAGTCGTTGCAAATGCAGTGAAGAGGAGACTGTNNNAGATCGGA[illegible] |
| 62184_1 | TATGAGTGTGGAGTCGTTGCTTGGCTAGCACAGAGTCATTCTTNNNAGATCGGAAG |
| 62185_1 | TATGAGTGTGGAGTCGTTGCGTTCCCATGCATGAATCTGACTCNNNAGATCGGAAG[illegible] |
| 62186_1 | TATGAGTGTGGAGTCGTTGCCTATTGAACCCCACCTTGTGAGANNNAGATCGGAAG[illegible] |
| 62187_1 | TATGAGTGTGGAGTCGTTGCGCAGTACTTCACTTCTGGTACCANNNAGATCGGAAG[illegible] |
| 62188_1 | TATGAGTGTGGAGTCGTTGCCTCTCGAGTAGCTGGGATTACAGNNNAGATCGGAA[illegible] |
| 62190_1 | TATGAGTGTGGAGTCGTTGCGAGTAGACAGGACTACAGGCATGNNNAGATCGGAA |
| 62191_1 | TATGAGTGTGGAGTCGTTGCAAACCTGTCATCTCTCATTGGCTNNNAGATCGGAAG[illegible] |
| 62193_1 | TATGAGTGTGGAGTCGTTGCTCTGGGTTCAAGTGATTCTCCTGNNNAGATCGGAAG |
| 62196_1 | TATGAGTGTGGAGTCGTTGCGGAAAAGGGAGAGGTGAGAATGTNNNAGATCGGA |
| 62197_1 | TATGAGTGTGGAGTCGTTGCACATGCTTATCCTGCTACAGAGGNNNAGATCGGAAG |
| 62198_1 | TATGAGTGTGGAGTCGTTGCGCAACAAAAAAAGGGCTCCTGTTNNNAGATCGGAA[illegible] |
| 62200_1 | TATGAGTGTGGAGTCGTTGCGGAAGAGGGCAGATTTGGAAAATNNNAGATCGGA[illegible] |
| 62201_1 | TATGAGTGTGGAGTCGTTGCGAAAGAAGAGAGAGACAGGGCAANNNAGATCGGA |
| 62202_1 | TATGAGTGTGGAGTCGTTGCACTGCACTGGAAAATGGAAAATGANNNAGATCGGA[illegible] |
| 62203_1 | TATGAGTGTGGAGTCGTTGCGCTGTTTCTTGTGTTTCAACAGGNNNAGATCGGAAG |
| 62204_1 | TATGAGTGTGGAGTCGTTGCGAAGGTCTTGAATGGCCTAACCANNNAGATCGGAA[illegible] |
| 62205_1 | TATGAGTGTGGAGTCGTTGCTGTTTGAACCCTGGAAGTAGAGGNNNAGATCGGAA[illegible] |
| 62206_1 | TATGAGTGTGGAGTCGTTGCTCACACACTTGGAATCTCACTGTNNNAGATCGGAAG[illegible] |
| 62207_1 | TATGAGTGTGGAGTCGTTGCATCTCCAGGGTTCAAGCAATTCTNNNAGATCGGAAG[illegible] |
| 62208_1 | TATGAGTGTGGAGTCGTTGCTCCCTCTCCTACACTTGCCTATANNNAGATCGGAAGA[illegible] |
| 62209_1 | TATGAGTGTGGAGTCGTTGCATAGTGGAGGGAAGGTCAGAAGANNNAGATCGGA[illegible] |
| 62211_1 | TATGAGTGTGGAGTCGTTGCAGCAACTTGGATAGAACTGGAGGNNNAGATCGGAA |
| 62212_1 | TATGAGTGTGGAGTCGTTGCGTACTTTGGCTTCATGCACTCTGNNNAGATCGGAAG[illegible] |
| 62213_1 | TATGAGTGTGGAGTCGTTGCTCTCTGGACTCACAGCTATCAGANNNAGATCGGAAG[illegible] |
| 62214_1 | TATGAGTGTGGAGTCGTTGCTGGCCCTAGTCTATCCCAACATANNNAGATCGGAAG[illegible] |
| 62216_1 | TATGAGTGTGGAGTCGTTGCATCATCCCAGGCTCTTTTTCTCCNNNAGATCGGAAGA[illegible] |
| 62217_1 | TATGAGTGTGGAGTCGTTGCTGCTTGGGCTGACTAATTTCTTGNNNAGATCGGAAG |
| 62218_1 | TATGAGTGTGGAGTCGTTGCGCGTAGAGAATCTCTGGTGACATNNNAGATCGGAA[illegible] |
| 62219_1 | TATGAGTGTGGAGTCGTTGCCTGAGATCACACCACTGCACTACNNNAGATCGGAAG[illegible] |
| 62220_1 | TATGAGTGTGGAGTCGTTGCAAGGGTAAATCTTCATGGCCTCANNNAGATCGGAAG |
| 62221_1 | TATGAGTGTGGAGTCGTTGCAGAGAACGAGCTCAGTGAATTGTNNNAGATCGGAA[illegible] |
| 62222_1 | TATGAGTGTGGAGTCGTTGCGAGTTTGAGGTCACAGTGAGCTANNNAGATCGGAA[illegible] |
| 62223_1 | TATGAGTGTGGAGTCGTTGCTTCTCCATCCGGTACTGGTAGATNNNAGATCGGAAG[illegible] |

| | |
|---|---|
| 62224_1 | TATGAGTGTGGAGTCGTTGCAATGCTGTGGAGAAAAAGGAAGCNNNAGATCGGA |
| 62226_1 | TATGAGTGTGGAGTCGTTGCTCTCTAACTCTGACCTCAGGTGANNNAGATCGGAAG |
| 62227_1 | TATGAGTGTGGAGTCGTTGCGACACAGTGGCTCCCTTAGAAATNNNAGATCGGAAG |
| 62228_1 | TATGAGTGTGGAGTCGTTGCTTGATTGATGGGCATTTGGACTGNNNAGATCGGAA |
| 62229_1 | TATGAGTGTGGAGTCGTTGCGATCACAAGGTCAGGAGTTCGAGNNNAGATCGGAA |
| 62231_1 | TATGAGTGTGGAGTCGTTGCCTGTAGTCCCAGCTACTCAGGAGNNNAGATCGGAAG |
| 62234_1 | TATGAGTGTGGAGTCGTTGCTGTATTTGTCCCCACCCAAATCTNNNAGATCGGAAGA |
| 62237_1 | TATGAGTGTGGAGTCGTTGCGCTGAAAAGAAAAGGCAGAGGAGNNNAGATCGGA |
| 62239_1 | TATGAGTGTGGAGTCGTTGCCCTCTGTGCTCTACCAGAAACTTNNNAGATCGGAAG |
| 62242_1 | TATGAGTGTGGAGTCGTTGCCCTCAAGTGATCCTCTTGCCTTANNNAGATCGGAAG |
| 62243_1 | TATGAGTGTGGAGTCGTTGCTTTTGGTAGAGGAGTGGTTTCTCNNNAGATCGGAA |
| 62244_1 | TATGAGTGTGGAGTCGTTGCAGTCTCACTCTATCACTCAGGCTNNNAGATCGGAAG |
| 62246_1 | TATGAGTGTGGAGTCGTTGCTAGAGATGGGGTTTATGTTGCCCNNNAGATCGGAA |
| 62248_1 | TATGAGTGTGGAGTCGTTGCGCACACACACACAGACTCAATTTNNNAGATCGGAAG |
| 62249_1 | TATGAGTGTGGAGTCGTTGCTGAGTAGCTGAGATTACATGCGTNNNAGATCGGAA |
| 62250_1 | TATGAGTGTGGAGTCGTTGCTCACAGGCAAAATTACTCGAAGCNNNAGATCGGAAG |
| 62251_1 | TATGAGTGTGGAGTCGTTGCCATAGCCACTTGTCACAAAAGGGNNNAGATCGGAAG |
| 62253_1 | TATGAGTGTGGAGTCGTTGCGGCTGAAGTAGGATTGTTTGAGCNNNAGATCGGAA |
| 62254_1 | TATGAGTGTGGAGTCGTTGCCTGTAGTCCCAGCTACTCAGGAGNNNAGATCGGAAG |
| 62255_1 | TATGAGTGTGGAGTCGTTGCCATTTGGGCTGGTTCCGTATTTTNNNAGATCGGAAG |
| 62256_1 | TATGAGTGTGGAGTCGTTGCTTTTTTGAGACAGAGTCTGGCTCNNNAGATCGGAAG |
| 62257_1 | TATGAGTGTGGAGTCGTTGCGCTCTAGAGTACTGAAACTCCCCNNNAGATCGGAAG |
| 62258_1 | TATGAGTGTGGAGTCGTTGCATGATCCAGAAATCCCTCTGCTGNNNAGATCGGAAG |
| 62259_1 | TATGAGTGTGGAGTCGTTGCATATGTGTGTGTGTGTGTGTGTGNNNAGATCGGAA |
| 62260_1 | TATGAGTGTGGAGTCGTTGCTGGTAACAGAAATAAGGGTCCCANNNAGATCGGAA |
| 62262_1 | TATGAGTGTGGAGTCGTTGCCGGCAGAGGAACATAAATTGTGANNNAGATCGGAA |
| 62263_1 | TATGAGTGTGGAGTCGTTGCTCTCTTCCACATGAGTCAGTGTGNNNAGATCGGAAG |
| 62264_1 | TATGAGTGTGGAGTCGTTGCACAGAAGCATCTCTACACGTGTTNNNAGATCGGAAG |
| 62265_1 | TATGAGTGTGGAGTCGTTGCGATAGAGTTTTGCCATGTTGCCCNNNAGATCGGAAG |
| 62266_1 | TATGAGTGTGGAGTCGTTGCTTTTGGGAGAGAAGTTGTCCTGTNNNAGATCGGAA |
| 62269_1 | TATGAGTGTGGAGTCGTTGCCACTGGGGTGGGTATTATGTTGTNNNAGATCGGAA |
| 62271_1 | TATGAGTGTGGAGTCGTTGCGGAGCTATTCAGTGCCATCCTTANNNAGATCGGAAG |
| 62272_1 | TATGAGTGTGGAGTCGTTGCATCTGTGATGGCTGTTTGAGTCANNNAGATCGGAAG |
| 62273_1 | TATGAGTGTGGAGTCGTTGCCTGGCCATGACTGTGTTCTTTAGNNNAGATCGGAAG |
| 62274_1 | TATGAGTGTGGAGTCGTTGCCAAAAAGTAGGCTGCAACCATGANNNAGATCGGAA |
| 62276_1 | TATGAGTGTGGAGTCGTTGCGTGTGTTGTGTGTGTGTGTTGTANNNAGATCGGAA |
| 62280_1 | TATGAGTGTGGAGTCGTTGCGCCACCAAGCCCAGATAATTTTTNNNAGATCGGAAG |
| 62281_1 | TATGAGTGTGGAGTCGTTGCTACTGGTTCTGTTCCTCTGGAGANNNAGATCGGAAG |
| 62282_1 | TATGAGTGTGGAGTCGTTGCTGAAACAGGATCTCATTCTGCCANNNAGATCGGAAG |
| 62284_1 | TATGAGTGTGGAGTCGTTGCGACAGGTTCTTGCTTTGTTACCCNNNAGATCGGAAG |
| 62285_1 | TATGAGTGTGGAGTCGTTGCGGCACTACAAGATTCTCCAGGATNNNAGATCGGAAG |
| 62286_1 | TATGAGTGTGGAGTCGTTGCGGAGTCCTACTATGTTGCTCAGGNNNAGATCGGAAG |
| 62287_1 | TATGAGTGTGGAGTCGTTGCAGAGATGGGGTTTCTCCATGTTGNNNAGATCGGAA |

| | |
|---|---|
| 62288_1 | TATGAGTGTGGAGTCGTTGCTTCCTCCAAGATACAGCTGTCACNNNAGATCGGAAG |
| 62289_1 | TATGAGTGTGGAGTCGTTGCTCACAGGAAACTAGGACAATGCTNNNAGATCGGAA |
| 62290_1 | TATGAGTGTGGAGTCGTTGCTCTATAACTCTGGCAGGTTGAGCNNNAGATCGGAAG |
| 62291_1 | TATGAGTGTGGAGTCGTTGCCAACTTGGCCTTGGGAGATTTTNNNAGATCGGAAG |
| 62293_1 | TATGAGTGTGGAGTCGTTGCGTTCATGCCATTCTCCTACCTCANNNAGATCGGAAGA |
| 62294_1 | TATGAGTGTGGAGTCGTTGCGGCATCCCCTAATGTCCATCTTTNNNAGATCGGAAG |
| 62295_1 | TATGAGTGTGGAGTCGTTGCTAGAGACTGGGTTTCGCCATTTTNNNAGATCGGAAG |
| 62297_1 | TATGAGTGTGGAGTCGTTGCCCAGTGGGAGGTAATCGAATCATNNNAGATCGGAA |
| 62298_1 | TATGAGTGTGGAGTCGTTGCGAACTCCTGACCTCAAGTGATCCNNNAGATCGGAAG |
| 62299_1 | TATGAGTGTGGAGTCGTTGCTTTCCCCACAATTTATTCGTGCCNNNAGATCGGAAGA |
| 62300_1 | TATGAGTGTGGAGTCGTTGCGGTGTATGTCCCAGCTACTTAGGNNNAGATCGGAAG |
| 62301_1 | TATGAGTGTGGAGTCGTTGCGGATCACTTGAGTCCAGGAGTTTNNNAGATCGGAA |
| 62302_1 | TATGAGTGTGGAGTCGTTGCTACAGGAAGACCATGTACTCAGCNNNAGATCGGAAG |
| 62303_1 | TATGAGTGTGGAGTCGTTGCTGTCTCTGACAGTTTGACGAAGANNNAGATCGGAA |
| 62305_1 | TATGAGTGTGGAGTCGTTGCAGCTCCCTCTTAGTGAAAGACTTNNNAGATCGGAAG |
| 62306_1 | TATGAGTGTGGAGTCGTTGCAAACCAGTGAAGGCTATGTGACTNNNAGATCGGAA |
| 62307_1 | TATGAGTGTGGAGTCGTTGCAAAAAAAAACTGCTGGCTGGACANNNAGATCGGAA |
| 62310_1 | TATGAGTGTGGAGTCGTTGCACTCTTTCTTCATTCCTGCCTCCNNNAGATCGGAAGA |
| 62311_1 | TATGAGTGTGGAGTCGTTGCTGTTATACCTAGCAGGGTGAGGANNNAGATCGGAA |
| 62313_1 | TATGAGTGTGGAGTCGTTGCAATGTGGGGCTGGATCAAATTTGNNNAGATCGGAA |
| 62314_1 | TATGAGTGTGGAGTCGTTGCTGAAACTGCAGTGAGTTGGGATTNNNAGATCGGAA |
| 62315_1 | TATGAGTGTGGAGTCGTTGCTCAGGAAAGAGGTCAACTTTGCTNNNAGATCGGAA |
| 62316_1 | TATGAGTGTGGAGTCGTTGCGAATACCTGTTACCTGGGACTGGNNNAGATCGGAA |
| 62317_1 | TATGAGTGTGGAGTCGTTGCTTACTTCTGGGTGTGTCTGTGAGNNNAGATCGGAAG |
| 62319_1 | TATGAGTGTGGAGTCGTTGCTGACCTTGGAGAAGAGGGAGTATNNNAGATCGGAA |
| 62320_1 | TATGAGTGTGGAGTCGTTGCACTACAAAGACAGTTGTGCATGCNNNAGATCGGAAG |
| 62321_1 | TATGAGTGTGGAGTCGTTGCGAAAAGCTGGTGGTGTCATTTGANNNAGATCGGAA |
| 62322_1 | TATGAGTGTGGAGTCGTTGCATGCCTGTAGTCCTAGCTACTCANNNAGATCGGAAG |
| 62323_1 | TATGAGTGTGGAGTCGTTGCCTTAGAAGGGGCTGGTGTTCTTANNNAGATCGGAA |
| 62324_1 | TATGAGTGTGGAGTCGTTGCATGTTGATCTCATCTGCCAGAGGNNNAGATCGGAAG |
| 62325_1 | TATGAGTGTGGAGTCGTTGCGTGTTTGTCCCTTTCTAATGCCGNNNAGATCGGAAG |
| 62327_1 | TATGAGTGTGGAGTCGTTGCCCCTAGCTACTTGGGAGGCTAAGNNNAGATCGGAA |
| 62328_1 | TATGAGTGTGGAGTCGTTGCTCAAATGAGAGGCTAGGCTCATCNNNAGATCGGAA |
| 62329_1 | TATGAGTGTGGAGTCGTTGCCACACACAATTAGCCTGACATGGNNNAGATCGGAAG |
| 62330_1 | TATGAGTGTGGAGTCGTTGCATAAGGCAATTCCATTCCTGGGTNNNAGATCGGAAG |
| 62331_1 | TATGAGTGTGGAGTCGTTGCCACCGGTAATTACAGCACTTTGGNNNAGATCGGAAG |
| 62332_1 | TATGAGTGTGGAGTCGTTGCTGTTGTTATAGCTTGCTGATTCCCNNNAGATCGGAA |
| 62333_1 | TATGAGTGTGGAGTCGTTGCCCACAAAGTGCTGGGATTACAAGNNNAGATCGGAA |
| 62334_1 | TATGAGTGTGGAGTCGTTGCGAGGAGGACTTCAGGAAACCATTNNNAGATCGGAA |
| 62336_1 | TATGAGTGTGGAGTCGTTGCCCAGATATTCAGGACGCTGAGATNNNAGATCGGAA |
| 62337_1 | TATGAGTGTGGAGTCGTTGCACAAAGCCTGTCTCTGAATTCCANNNAGATCGGAAG |
| 62338_1 | TATGAGTGTGGAGTCGTTGCCCTCTAGAGATTACACAGGGTGGNNNAGATCGGAA |
| 62339_1 | TATGAGTGTGGAGTCGTTGCCCTCTAGAGATTACACAGGGTGGNNNAGATCGGAA |

| | |
|---|---|
| 62340_1 | TATGAGTGTGGAGTCGTTGCCTGTAGTTCCAGCTACTCAGGAGNNNAGATCGGAAG |
| 62342_1 | TATGAGTGTGGAGTCGTTGCACAACCAAGTAGCTAACCCAGAGNNNAGATCGGAAG |
| 62343_1 | TATGAGTGTGGAGTCGTTGCACAAACCTGAGAGAACTGAAAGANNNAGATCGGAA |
| 62344_1 | TATGAGTGTGGAGTCGTTGCCACTGGTGCAATCATAGCTCACTNNNAGATCGGAAG |
| 62345_1 | TATGAGTGTGGAGTCGTTGCCACTCCTGTAATCCCAGCACTTTNNNAGATCGGAAGA |
| 62347_1 | TATGAGTGTGGAGTCGTTGCACACACAGAGACCCACCTACTTANNNAGATCGGAAG |
| 62348_1 | TATGAGTGTGGAGTCGTTGCTGGTCAATCACAGCTCCTCATTTNNNAGATCGGAAG |
| 62349_1 | TATGAGTGTGGAGTCGTTGCGCTTCCCAAAGTGCTGGTATTACNNNAGATCGGAAG |
| 62352_1 | TATGAGTGTGGAGTCGTTGCAACTCCTGACCTCAAGCAATCTTNNNAGATCGGAAG |
| 62353_1 | TATGAGTGTGGAGTCGTTGCAGAGAGAGAGAGAGAGAGAGAGCNNNAGATCGG |
| 62354_1 | TATGAGTGTGGAGTCGTTGCGTGAAGCTAGCACATTGGATTGGNNNAGATCGGAA |
| 62355_1 | TATGAGTGTGGAGTCGTTGCCAGAGTTTTACCATGTTGGCCAGNNNAGATCGGAAG |
| 62356_1 | TATGAGTGTGGAGTCGTTGCAAATGGTCCACAGAGTCAACTCANNNAGATCGGAAG |
| 62357_1 | TATGAGTGTGGAGTCGTTGCGGAGCTGAGGATCAAGTCAAGATNNNAGATCGGAA |
| 62358_1 | TATGAGTGTGGAGTCGTTGCGAGAATTGCTTGAGCGTAGGAGANNNAGATCGGA |
| 62359_1 | TATGAGTGTGGAGTCGTTGCTTTCACAGGCACATCCACAAATCNNNAGATCGGAAG |
| 62360_1 | TATGAGTGTGGAGTCGTTGCACCACAGACAGCCCATCTTTATTNNNAGATCGGAAG |
| 62361_1 | TATGAGTGTGGAGTCGTTGCGTCACCCAGCAAGACTCTTTTTGNNNAGATCGGAAG |
| 62362_1 | TATGAGTGTGGAGTCGTTGCCCCAATACCCCCTAGGCAATTTANNNAGATCGGAAG |
| 62363_1 | TATGAGTGTGGAGTCGTTGCTAATCATAGCTCACTGCAGCCTCNNNAGATCGGAAG |
| 62364_1 | TATGAGTGTGGAGTCGTTGCGGTCTTTCTCTATTGCTCAGGCTNNNAGATCGGAAG |
| 62365_1 | TATGAGTGTGGAGTCGTTGCGAGTTTCGTTCTTGTTACCCAGGNNNAGATCGGAAG |
| 62366_1 | TATGAGTGTGGAGTCGTTGCTCCCAGATATATGAGGGAAGGCTNNNAGATCGGAA |
| 62367_1 | TATGAGTGTGGAGTCGTTGCCATCCCTGTAATCCCAGCACTTTNNNAGATCGGAAGA |
| 62368_1 | TATGAGTGTGGAGTCGTTGCGCCTCCTGGGTTCAAGTGATTATNNNAGATCGGAAG |
| 62371_1 | TATGAGTGTGGAGTCGTTGCTCTCAACCTCTCTCTCTCTCTCANNNAGATCGGAAGA |
| 62372_1 | TATGAGTGTGGAGTCGTTGCTCTCTGCTGGCACTTCATGTAATNNNAGATCGGAAG |
| 62373_1 | TATGAGTGTGGAGTCGTTGCGAGAGGGAGAATACAACCAGTGGNNNAGATCGGA |
| 62375_1 | TATGAGTGTGGAGTCGTTGCTCTGTCTGTCTCTGCACTCATTCNNNAGATCGGAAGA |
| 62376_1 | TATGAGTGTGGAGTCGTTGCATGCAGATTTGGGGGGAAAAATGNNNAGATCGGA |
| 62377_1 | TATGAGTGTGGAGTCGTTGCTCACTCTCACTCTTACCCTCACTNNNAGATCGGAAGA |
| 62381_1 | TATGAGTGTGGAGTCGTTGCTTAGGTGGGTCATTAGGATGCAGNNNAGATCGGAA |
| 62382_1 | TATGAGTGTGGAGTCGTTGCGAGGGAAGAAGGAAGGAAGGAAGNNNAGATCGG |
| 62384_1 | TATGAGTGTGGAGTCGTTGCTTACATCAGAAACCATGGAGGCCNNNAGATCGGAA |
| 62386_1 | TATGAGTGTGGAGTCGTTGCTCTCTGTCTCTGTCTCTCTCTGTNNNAGATCGGAAGA |
| 62387_1 | TATGAGTGTGGAGTCGTTGCCACCACGTCCAGCTAATTTTTGTNNNAGATCGGAAG |
| 62388_1 | TATGAGTGTGGAGTCGTTGCTCTCCTCCTCTTTCTGGAAGTCTNNNAGATCGGAAGA |
| 62389_1 | TATGAGTGTGGAGTCGTTGCCAAACACACACACACACAAACACNNNAGATCGGAAG |
| 62391_1 | TATGAGTGTGGAGTCGTTGCACCAATTGAGTATAGGGAGGGGANNNAGATCGGA |
| 62393_1 | TATGAGTGTGGAGTCGTTGCATTGTAAGTTTCCTGAGGCCTCCNNNAGATCGGAAG |
| 62394_1 | TATGAGTGTGGAGTCGTTGCAGAAGGAGTCTTGTTCTGTCACCNNNAGATCGGAAG |
| 62395_1 | TATGAGTGTGGAGTCGTTGCCCACTTCACAGGGCTGAGATAATNNNAGATCGGAAG |
| 62397_1 | TATGAGTGTGGAGTCGTTGCTCTCTCTCTCCTCTCTCTCCTNNNAGATCGGAAGAG |

| | |
|---|---|
| 62398_1 | TATGAGTGTGGAGTCGTTGCCTCTGTCTGTCTGTCTGTCTGTTNNNAGATCGGAAGA |
| 62399_1 | TATGAGTGTGGAGTCGTTGCCAACCAGGCTAGAGTACAATGGTNNNAGATCGGAAG |
| 62400_1 | TATGAGTGTGGAGTCGTTGCTATATAGGGGAACAGGGTGTGGTNNNAGATCGGAA |
| 62401_1 | TATGAGTGTGGAGTCGTTGCTCCTGAGTAGCTAGGACTACAGGNNNAGATCGGAAG |
| 62402_1 | TATGAGTGTGGAGTCGTTGCCACACACACACACACAGTCATTCNNNAGATCGGAAGA |
| 62403_1 | TATGAGTGTGGAGTCGTTGCCCCCCCTCTCTTTTTCTCTCTTCNNNAGATCGGAAGAG |
| 62404_1 | TATGAGTGTGGAGTCGTTGCCACTGTACTTCACTCTCAGGCTTNNNAGATCGGAAGA |
| 62405_1 | TATGAGTGTGGAGTCGTTGCGCTCTTTGCCTTCTCTGTTTGTTNNNAGATCGGAAGA |
| 62406_1 | TATGAGTGTGGAGTCGTTGCTCCTTTCCTGGAGTGTAGGATCANNNAGATCGGAAG |
| 62408_1 | TATGAGTGTGGAGTCGTTGCATAAAAGTTGGACCAGACCAGGGNNNAGATCGGAA |
| 62409_1 | TATGAGTGTGGAGTCGTTGCTGACCAACATGGAGAAACCTCATNNNAGATCGGAAG |
| 62410_1 | TATGAGTGTGGAGTCGTTGCTCCTTGGTTCAAGTGATTCTCGTNNNAGATCGGAAGA |
| 62411_1 | TATGAGTGTGGAGTCGTTGCCTGAGTAGCTGGGATTACAAGCANNNAGATCGGAAG |
| 62414_1 | TATGAGTGTGGAGTCGTTGCGGTTCTTAGTGTGACAGCGTTTCNNNAGATCGGAAG |
| 62415_1 | TATGAGTGTGGAGTCGTTGCGATTACAAGCATGAGTCACCACGNNNAGATCGGAAG |
| 62416_1 | TATGAGTGTGGAGTCGTTGCAGAACTCATAGGCCATGTTCCAGNNNAGATCGGAAG |
| 62417_1 | TATGAGTGTGGAGTCGTTGCCCTCCTGTGATCCAATTACCTCCNNNAGATCGGAAGA |
| 62418_1 | TATGAGTGTGGAGTCGTTGCCCGCAAGTAGATGCCTTAGTTTGNNNAGATCGGAAG |
| 62422_1 | TATGAGTGTGGAGTCGTTGCTTCAAGCGATTCTCCTGTCTCAANNNAGATCGGAAGA |
| 62423_1 | TATGAGTGTGGAGTCGTTGCCACAGGTTCAAGCAATTCTCCTGNNNAGATCGGAAGA |
| 62425_1 | TATGAGTGTGGAGTCGTTGCGAGTCTTGCTCTATTGCCTGTCTNNNAGATCGGAAGA |
| 62426_1 | TATGAGTGTGGAGTCGTTGCACCACCATGTCCAGCTAAGTTTTNNNAGATCGGAAGA |
| 62429_1 | TATGAGTGTGGAGTCGTTGCTGCCACTCAGCCTACTTCATATCNNNAGATCGGAAGA |
| 62430_1 | TATGAGTGTGGAGTCGTTGCGGCCTCCTAATGTCCAAATGTTGNNNAGATCGGAAG |
| 62431_1 | TATGAGTGTGGAGTCGTTGCTCAGCAAATTTGAGCCACAAAGANNNAGATCGGAAG |
| 62432_1 | TATGAGTGTGGAGTCGTTGCTCCTGTAAGACCAATTCCCTTCCNNNAGATCGGAAGA |
| 62433_1 | TATGAGTGTGGAGTCGTTGCAGAGAGAGAGAGAGAGAGAGAGANNNAGATCGGA |
| 62434_1 | TATGAGTGTGGAGTCGTTGCATTGCATTATTGGTTGGCCTAAANNNAGATCGGAAG |
| 62437_1 | TATGAGTGTGGAGTCGTTGCGTGGCTTATTCTAGCTTTCCCCTNNNAGATCGGAAGA |
| 62440_1 | TATGAGTGTGGAGTCGTTGCGCCTCCTGAATAGCTGGGATTACNNNAGATCGGAAG |
| 62441_1 | TATGAGTGTGGAGTCGTTGCACAATCACAGCTCACTACAACCTNNNAGATCGGAAGA |
| 62443_1 | TATGAGTGTGGAGTCGTTGCGATTGAGGGAAGGAAGGACGTTANNNAGATCGGA |
| 62444_1 | TATGAGTGTGGAGTCGTTGCTATGATGCTGTAAGAAGGCCCTCNNNAGATCGGAAG |
| 62445_1 | TATGAGTGTGGAGTCGTTGCGGGGGGGGGGGATATAAAAATATNNNAGATCGGA |
| 62447_1 | TATGAGTGTGGAGTCGTTGCGTGGATCATGAGGTCAGGAGATCNNNAGATCGGAA |
| 62448_1 | TATGAGTGTGGAGTCGTTGCTGCTTGCTGGAAATACAGATCCTNNNAGATCGGAAG |
| 62449_1 | TATGAGTGTGGAGTCGTTGCCTTCCGAGTAGCTGGGATTACAGNNNAGATCGGAAG |
| 62450_1 | TATGAGTGTGGAGTCGTTGCGAACACATGTGCTGCAACTTACTNNNAGATCGGAAG |
| 62452_1 | TATGAGTGTGGAGTCGTTGCTAGATGATGTGACCTTTGAGGCCNNNAGATCGGAAG |
| 62454_1 | TATGAGTGTGGAGTCGTTGCCCCAGGTTCAAGCAATTCTCTTGNNNAGATCGGAAGA |
| 62456_1 | TATGAGTGTGGAGTCGTTGCTCACTGTCACTTACGCTAGAGTGNNNAGATCGGAAG |
| 62457_1 | TATGAGTGTGGAGTCGTTGCGCCTACTGAGTAGCTGGGATTACNNNAGATCGGAAG |
| 62458_1 | TATGAGTGTGGAGTCGTTGCCTGGTCTTGAACTCCTGGACTTTNNNAGATCGGAAGA |

| | |
|---|---|
| 62459_1 | TATGAGTGTGGAGTCGTTGCGTTCAAGCAATTTCCTGCCTCAGNNNAGATCGGAAG |
| 62460_1 | TATGAGTGTGGAGTCGTTGCTTCAAGTGATTCTCCTGCCTCAGNNNAGATCGGAAG |
| 62461_1 | TATGAGTGTGGAGTCGTTGCGAGTCGGAATCTCACTCTCTCACNNNAGATCGGAAG |
| 62464_1 | TATGAGTGTGGAGTCGTTGCGCCTTGGTCTTCAAATGAGCTTTNNNAGATCGGAAG |
| 62466_1 | TATGAGTGTGGAGTCGTTGCACAGAGTTTCTCCATGTTGGTCANNNAGATCGGAAG |
| 62467_1 | TATGAGTGTGGAGTCGTTGCTGAATAGGTGGAACTACAGGCACNNNAGATCGGAA |
| 62468_1 | TATGAGTGTGGAGTCGTTGCAGCTTATTGCAGCCTCAAACATCNNNAGATCGGAAG |
| 62469_1 | TATGAGTGTGGAGTCGTTGCCCTCCCGAATAGCTGAGATTACANNNAGATCGGAAG |
| 62470_1 | TATGAGTGTGGAGTCGTTGCTAGCTAGGACTACAGGCATATGCNNNAGATCGGAA |
| 62471_1 | TATGAGTGTGGAGTCGTTGCTGATCATAACTCACTGCAGCCTTNNNAGATCGGAAG |
| 62472_1 | TATGAGTGTGGAGTCGTTGCTAATCTCTCTCTGAACTGCAGGCNNNAGATCGGAAG |
| 62473_1 | TATGAGTGTGGAGTCGTTGCCCTGCCTTTACTTCCCATAGTGTNNNAGATCGGAAGA |
| 62475_1 | TATGAGTGTGGAGTCGTTGCTCCTGAGTAGCTGGGATTACAGANNNAGATCGGAA |
| 62478_1 | TATGAGTGTGGAGTCGTTGCGAGGTGGGGTTTCACTATACTGGNNNAGATCGGAA |
| 62479_1 | TATGAGTGTGGAGTCGTTGCTCTGGGTTCAAGTGATTCTCCTGNNNAGATCGGAAG |
| 62482_1 | TATGAGTGTGGAGTCGTTGCGAGTACAGTGGAGAGATCTTGGCNNNAGATCGGAA |
| 62483_1 | TATGAGTGTGGAGTCGTTGCGAGTTTCCCTCTTGTTACCCAGGNNNAGATCGGAAG |
| 62484_1 | TATGAGTGTGGAGTCGTTGCGCTAGTCTCAAACTCCTGGACTCNNNAGATCGGAAG |
| 62486_1 | TATGAGTGTGGAGTCGTTGCTGTTAGCCAGGATGGTCTCAATCNNNAGATCGGAAG |
| 62488_1 | TATGAGTGTGGAGTCGTTGCCTGGTTCTTTGCTTCAGGACAAGNNNAGATCGGAAG |
| 62490_1 | TATGAGTGTGGAGTCGTTGCTGAAGGAGGAAGAAAGGTGATGGNNNAGATCGGA |
| 62491_1 | TATGAGTGTGGAGTCGTTGCTACAGTGGTTTGATCATGGCTCANNNAGATCGGAAG |
| 62492_1 | TATGAGTGTGGAGTCGTTGCCAGGTACCTGGGATCTGTTCATCNNNAGATCGGAAG |
| 62494_1 | TATGAGTGTGGAGTCGTTGCCTGGGTTCAAGTGATTCTTCTGCNNNAGATCGGAAG |
| 62495_1 | TATGAGTGTGGAGTCGTTGCCCTTCTGCAGAACACATTTTGGANNNAGATCGGAAG |
| 62496_1 | TATGAGTGTGGAGTCGTTGCCTCCTGAGTAGCTGGGATTACAGNNNAGATCGGAA |
| 62498_1 | TATGAGTGTGGAGTCGTTGCAGAAACAGGATTTCGCCATGTTGNNNAGATCGGAA |
| 62499_1 | TATGAGTGTGGAGTCGTTGCGGTTGAAGTGATTTTCCTGCCTCNNNAGATCGGAAG |
| 62500_1 | TATGAGTGTGGAGTCGTTGCCCAAGTAGCTGGTATTACAGGCANNNAGATCGGAA |
| 62501_1 | TATGAGTGTGGAGTCGTTGCGTCTCACTCTATTACCCAGGCTGNNNAGATCGGAAG |
| 62502_1 | TATGAGTGTGGAGTCGTTGCGCCTCCTGGATTCAAGTGATTCTNNNAGATCGGAAG |
| 62505_1 | TATGAGTGTGGAGTCGTTGCCTGAGTAGCTGGGATTACAGGTGNNNAGATCGGAA |
| 62506_1 | TATGAGTGTGGAGTCGTTGCTTCAAGCAACTCTCTTCCCTCAGNNNAGATCGGAAGA |
| 62510_1 | TATGAGTGTGGAGTCGTTGCTCAAACTCCTGACCCCAAATGATNNNAGATCGGAAG |
| 62512_1 | TATGAGTGTGGAGTCGTTGCTCTCCTGAGTAGCTGGGATTACANNNAGATCGGAAG |
| 62513_1 | TATGAGTGTGGAGTCGTTGCTCCTGGGTATCTGGGATTACAGTNNNAGATCGGAA |
| 62514_1 | TATGAGTGTGGAGTCGTTGCTAGGGTCTCACTCTCATGACTCANNNAGATCGGAAG |
| 62516_1 | TATGAGTGTGGAGTCGTTGCTTCAAGTGATTCTCCTGCCTCAGNNNAGATCGGAAG |
| 62519_1 | TATGAGTGTGGAGTCGTTGCTTCAAGTGATTCTCCTGCCTCAGNNNAGATCGGAAG |
| 62520_1 | TATGAGTGTGGAGTCGTTGCAAGCAATCTTCTCACCTCAGGTTNNNAGATCGGAAG |
| 62522_1 | TATGAGTGTGGAGTCGTTGCTCATGGCTGTAATCCCAACACTTNNNAGATCGGAAG |
| 62523_1 | TATGAGTGTGGAGTCGTTGCCTCCTGAGTAGCTGGGATTACAGNNNAGATCGGAA |
| 62524_1 | TATGAGTGTGGAGTCGTTGCGAGACAAAGTCTTGCTCTGTTGCNNNAGATCGGAAG |

| | |
|---|---|
| 62525_1 | TATGAGTGTGGAGTCGTTGCTTCAAGCGATTATCCTGCCTCAGNNNAGATCGGAAG |
| 62526_1 | TATGAGTGTGGAGTCGTTGCACTGCAACCTCTGAGATTACAGGNNNAGATCGGAAG |
| 62527_1 | TATGAGTGTGGAGTCGTTGCTTCCAAGTAGCTGGGATTAGTGGNNNAGATCGGAA |
| 62528_1 | TATGAGTGTGGAGTCGTTGCTTCAAGTGATTCTCCTGCCTCAGNNNAGATCGGAAG |
| 62529_1 | TATGAGTGTGGAGTCGTTGCTGATTTTCCTGCCTCAGTCTCTCNNNAGATCGGAAGA |
| 62530_1 | TATGAGTGTGGAGTCGTTGCCAAGTAGCTGGGACTATAGGTGCNNNAGATCGGAA |
| 62531_1 | TATGAGTGTGGAGTCGTTGCCCGGGCTTAGTTTTTGCTTCTAANNNAGATCGGAAG |
| 62532_1 | TATGAGTGTGGAGTCGTTGCCCTCTCAAAGTGCTGGGATTACANNNAGATCGGAAG |
| 62534_1 | TATGAGTGTGGAGTCGTTGCCTCCCGAGTAGCTGAGATTACAGNNNAGATCGGAAG |
| 62535_1 | TATGAGTGTGGAGTCGTTGCTCTCCCAGGTTTGAGTGATTCTCNNNAGATCGGAAG |
| 62540_1 | TATGAGTGTGGAGTCGTTGCGGTTTTTCAGAGTGCCCATAAGGNNNAGATCGGAAG |
| 62542_1 | TATGAGTGTGGAGTCGTTGCGGGCTCTCCTTTGAGGTGATAAANNNAGATCGGAAG |
| 62543_1 | TATGAGTGTGGAGTCGTTGCCTGTAGTCTCAGCTACTTGGGAGNNNAGATCGGAAG |
| 62544_1 | TATGAGTGTGGAGTCGTTGCACAGTAAGAAAGGGCTAGGGAAGNNNAGATCGGA |
| 62545_1 | TATGAGTGTGGAGTCGTTGCGTGATGCTGTCATAAGTCACTGCNNNAGATCGGAAG |
| 62546_1 | TATGAGTGTGGAGTCGTTGCTTTTTTTTGAGACAGGGTCTGGCNNNAGATCGGAAG |
| 62547_1 | TATGAGTGTGGAGTCGTTGCACAACAACAGACAAACAACCTCANNNAGATCGGAAG |
| 62549_1 | TATGAGTGTGGAGTCGTTGCGAATGCCACTAATTGACAAGAGTNNNAGATCGGAAG |
| 62551_1 | TATGAGTGTGGAGTCGTTGCTTATCACCACCTTGGGAAATGGTNNNAGATCGGAAG |
| 62552_1 | TATGAGTGTGGAGTCGTTGCGATTCCAGAAACTCACCCCTCTTNNNAGATCGGAAGA |
| 62553_1 | TATGAGTGTGGAGTCGTTGCAGTGTTAGTTCAGGCTCCAAACTNNNAGATCGGAAG |
| 62554_1 | TATGAGTGTGGAGTCGTTGCCCCATTTTCCAAGCAATAGCCAANNNAGATCGGAAG |
| 62555_1 | TATGAGTGTGGAGTCGTTGCGATGAGATTTCACCATGTTGCCCNNNAGATCGGAAG |
| 62556_1 | TATGAGTGTGGAGTCGTTGCGTAAAGAGAGAACCTGAGGTGCANNNAGATCGGAA |
| 62557_1 | TATGAGTGTGGAGTCGTTGCCGGTCCCTCAGTCATATAGTCCTNNNAGATCGGAAG |
| 62561_1 | TATGAGTGTGGAGTCGTTGCGAGATTGCACTGAGCTGTGATTGNNNAGATCGGAA |
| 62566_1 | TATGAGTGTGGAGTCGTTGCAAAGAGGCACATTTGACACAACCNNNAGATCGGAAG |
| 62567_1 | TATGAGTGTGGAGTCGTTGCGAGAGAAGTCAAGGCTACATGGTNNNAGATCGGAA |
| 62568_1 | TATGAGTGTGGAGTCGTTGCGCATCACAGCTTTGAAAAAAGGCNNNAGATCGGAAG |
| 62570_1 | TATGAGTGTGGAGTCGTTGCGCAGGATCTCAGCTCACTACAAGNNNAGATCGGAAG |
| 62571_1 | TATGAGTGTGGAGTCGTTGCGATGGAGTTTCGCTCATATTGCCNNNAGATCGGAAG |
| 62572_1 | TATGAGTGTGGAGTCGTTGCAGATGGAGTCTCTCTCTGTCACANNNAGATCGGAAG |
| 62573_1 | TATGAGTGTGGAGTCGTTGCCACCACACCTGGCTATTTTCTTTNNNAGATCGGAAGA |
| 62574_1 | TATGAGTGTGGAGTCGTTGCTTCATGCTATTCTCCTGCCTCAGNNNAGATCGGAAGA |
| 62575_1 | TATGAGTGTGGAGTCGTTGCTCCTGGTTCAAGTGATTCTCCTGNNNAGATCGGAAG |
| 62576_1 | TATGAGTGTGGAGTCGTTGCCTCCTGAGTAGCTGGGACTACAGNNNAGATCGGAAG |
| 62579_1 | TATGAGTGTGGAGTCGTTGCACCAGAGGTACAAAGATGAGCTGNNNAGATCGGAA |
| 62581_1 | TATGAGTGTGGAGTCGTTGCTTTCTACACAAAACCCTGCTGGANNNAGATCGGAAG |
| 62582_1 | TATGAGTGTGGAGTCGTTGCTGAGGATGTGAAGCAACTGGAATNNNAGATCGGAA |
| 62584_1 | TATGAGTGTGGAGTCGTTGCCTAAGGCTGGAGAATCGCTTGAANNNAGATCGGAA |
| 62585_1 | TATGAGTGTGGAGTCGTTGCTTCAAGTGATTCTCCTGCCTCAGNNNAGATCGGAAG |
| 62586_1 | TATGAGTGTGGAGTCGTTGCACTACAATCTCCGCCTCTCTAGTNNNAGATCGGAAGA |
| 62587_1 | TATGAGTGTGGAGTCGTTGCGCACCATCATGTCTGGCTAATTTNNNAGATCGGAAG |

| | |
|---|---|
| 62588_1 | TATGAGTGTGGAGTCGTTGCTTCAAGCAATTCTCCTGCCTCAANNNAGATCGGAAGA |
| 62591_1 | TATGAGTGTGGAGTCGTTGCCAAAGTGCTGGGATTACATGCATNNNAGATCGGAAG |
| 62592_1 | TATGAGTGTGGAGTCGTTGCGAGATGGAGTTTCACCCTTGTTGNNNAGATCGGAAG |
| 62593_1 | TATGAGTGTGGAGTCGTTGCCAATGTGTTAGGCTTCTGCTTCCNNNAGATCGGAAGA |
| 62594_1 | TATGAGTGTGGAGTCGTTGCCTGTAATCCCAGCTACTCAGGAGNNNAGATCGGAAG |
| 62595_1 | TATGAGTGTGGAGTCGTTGCAATGTGCAGAAAAGGAAAATCCANNNAGATCGGAAG |
| 62598_1 | TATGAGTGTGGAGTCGTTGCCCCACCAATGACAGACTGGATAANNNAGATCGGAAG |
| 62599_1 | TATGAGTGTGGAGTCGTTGCCAGTAGGGGAAGGATTAGGGAGANNNAGATCGGA |
| 62600_1 | TATGAGTGTGGAGTCGTTGCAAGACTGCAAAAACCACAGCTTTNNNAGATCGGAAG |
| 62602_1 | TATGAGTGTGGAGTCGTTGCCCTCCTGGGTTCAAGTGATTCTCNNNAGATCGGAAGA |
| 62604_1 | TATGAGTGTGGAGTCGTTGCCCTCCTGGGTTCAAGCTATTCTTNNNAGATCGGAAGA |
| 62607_1 | TATGAGTGTGGAGTCGTTGCTCCCAAGTTCAAGCAATTCTCCTNNNAGATCGGAAGA |
| 62609_1 | TATGAGTGTGGAGTCGTTGCACACACACACACACGTACACATANNNAGATCGGAAGA |
| 62610_1 | TATGAGTGTGGAGTCGTTGCCTAAACAGCAGCACTTGAATCCCNNNAGATCGGAAGA |
| 62611_1 | TATGAGTGTGGAGTCGTTGCAAGCTTCCTAGGTGATTCTTGGGNNNAGATCGGAAG |
| 62612_1 | TATGAGTGTGGAGTCGTTGCTTAGACTGCTTGTTACTGTCCCCNNNAGATCGGAAGA |
| 62613_1 | TATGAGTGTGGAGTCGTTGCGGTCATGCCACTTTCACAGTTTTNNNAGATCGGAAGA |
| 62614_1 | TATGAGTGTGGAGTCGTTGCTTCTGGTTTCTTGCCTTTTGGTCNNNAGATCGGAAGA |
| 62615_1 | TATGAGTGTGGAGTCGTTGCTAATCCCAGCTATCCAGGAGACTNNNAGATCGGAAG |
| 62616_1 | TATGAGTGTGGAGTCGTTGCTCTAAAAGCTGGACCAGTATGCTNNNAGATCGGAAG |
| 62619_1 | TATGAGTGTGGAGTCGTTGCGCGATGGAGTAGAGTAGGTTCAGNNNAGATCGGAA |
| 62620_1 | TATGAGTGTGGAGTCGTTGCGATTCTCATGCCTCAGCTTCCTANNNAGATCGGAAGA |
| 62621_1 | TATGAGTGTGGAGTCGTTGCGGATTACAGGAGTGAGCCATCATNNNAGATCGGAAG |
| 62622_1 | TATGAGTGTGGAGTCGTTGCATTCTCAGCTCATTTGCAACGTCNNNAGATCGGAAGA |
| 62624_1 | TATGAGTGTGGAGTCGTTGCCAAAGTGCTAGGACTACAGGTGTNNNAGATCGGAAG |
| 62625_1 | TATGAGTGTGGAGTCGTTGCGCAGTGAGCTATGATTGAACCACNNNAGATCGGAAG |
| 62627_1 | TATGAGTGTGGAGTCGTTGCTGAGCACTTATTGTATGCCAGGTNNNAGATCGGAAG |
| 62628_1 | TATGAGTGTGGAGTCGTTGCAAACTGAGGCAGAAGAATCGCTANNNAGATCGGAAG |
| 62630_1 | TATGAGTGTGGAGTCGTTGCAGGAAGGTACCAGTATGTAGCCTNNNAGATCGGAAG |
| 62631_1 | TATGAGTGTGGAGTCGTTGCGACCCAGATGATTGTGAGCATTTNNNAGATCGGAAG |
| 62634_1 | TATGAGTGTGGAGTCGTTGCAGCCTCTACCTCTCCCAGATATCNNNAGATCGGAAGA |
| 62638_1 | TATGAGTGTGGAGTCGTTGCATTCAAGCAATTCTCCCACCTCANNNAGATCGGAAGA |
| 62644_1 | TATGAGTGTGGAGTCGTTGCTGAGAAACCAACACAGGACTGAANNNAGATCGGAAG |
| 62645_1 | TATGAGTGTGGAGTCGTTGCTTCAGGGCCTATGTGAATGACAGNNNAGATCGGAAG |
| 62647_1 | TATGAGTGTGGAGTCGTTGCCCCAGCCTATGCAGAAAGAGAATNNNAGATCGGAAG |
| 62648_1 | TATGAGTGTGGAGTCGTTGCAACCAAGACTCGATATGTTGCCTNNNAGATCGGAAG |
| 62649_1 | TATGAGTGTGGAGTCGTTGCGGAGTGTAATGGCACAATCTTGGNNNAGATCGGAAG |
| 62651_1 | TATGAGTGTGGAGTCGTTGCGCCCAGCCTCAATTTTCTTTTCTNNNAGATCGGAAGAG |
| 62652_1 | TATGAGTGTGGAGTCGTTGCCTGAAGTGCAGTGACACAGTCATNNNAGATCGGAAG |
| 62654_1 | TATGAGTGTGGAGTCGTTGCAGCTTTTATGCATCGAAGGACACNNNAGATCGGAAG |
| 62655_1 | TATGAGTGTGGAGTCGTTGCAGAAGCTACTGTTTCAACTGCACNNNAGATCGGAAG |
| 62656_1 | TATGAGTGTGGAGTCGTTGCAGTCTTACAAAGGCAGTCTGGTCNNNAGATCGGAAG |
| 62658_1 | TATGAGTGTGGAGTCGTTGCACACAGTGAAACCCCATCTCTACNNNAGATCGGAAGA |

| | |
|---|---|
| 62659_1 | TATGAGTGTGGAGTCGTTGCGAGACACGAGACAGGTAACCATTNNNAGATCGGAA |
| 62663_1 | TATGAGTGTGGAGTCGTTGCCCAATTGTGAATGGGAGTTTGCTNNNAGATCGGAAG |
| 62664_1 | TATGAGTGTGGAGTCGTTGCGAAGACGACAAATAGGCAACAGGNNNAGATCGGAA |
| 62665_1 | TATGAGTGTGGAGTCGTTGCTACCAGACCTACAAAAGAGCTGGNNNAGATCGGAAG |
| 62666_1 | TATGAGTGTGGAGTCGTTGCAGCATACAAATGACAAAGATGAGAGANNNAGATCG |
| 62667_1 | TATGAGTGTGGAGTCGTTGCGTTCTGGGGTTTTGAACTTGGAGNNNAGATCGGAA |
| 62668_1 | TATGAGTGTGGAGTCGTTGCAGTGTAGAGATTGGTTTGCCTGTNNNAGATCGGAA |
| 62671_1 | TATGAGTGTGGAGTCGTTGCCCTCTGGAACATGGAACTGAGAANNNAGATCGGAA |
| 62672_1 | TATGAGTGTGGAGTCGTTGCTCTGAATCTGTCAAGCTGAGGACNNNAGATCGGAAG |
| 62673_1 | TATGAGTGTGGAGTCGTTGCTCATAGCAATCACAGGGGTTCTTNNNAGATCGGAAG |
| 62677_1 | TATGAGTGTGGAGTCGTTGCCTGAGGCAGGAGAATCACTTGACNNNAGATCGGAA |
| 62678_1 | TATGAGTGTGGAGTCGTTGCGCAAAGCTGTCTTTCTGTTGAGGNNNAGATCGGAAG |
| 62680_1 | TATGAGTGTGGAGTCGTTGCCTGAGACGGGAGTATCACTTGAGNNNAGATCGGAA |
| 62681_1 | TATGAGTGTGGAGTCGTTGCTGTAATCACAGCTACTTGGGAGGNNNAGATCGGAA |
| 62682_1 | TATGAGTGTGGAGTCGTTGCACTTTCAGGTGTATGGAACAGCANNNAGATCGGAAG |
| 62683_1 | TATGAGTGTGGAGTCGTTGCTTTGCTAGCACAATAGGGTGACTNNNAGATCGGAAG |
| 62684_1 | TATGAGTGTGGAGTCGTTGCTGTAATCCAAGCTACATGGTGGGNNNAGATCGGAA |
| 62687_1 | TATGAGTGTGGAGTCGTTGCGAGTGTGCCTCAGTAATAACCCANNNAGATCGGAAG |
| 62688_1 | TATGAGTGTGGAGTCGTTGCCTTGTAATCCTCCAACTTCCCCANNNAGATCGGAAGA |
| 62689_1 | TATGAGTGTGGAGTCGTTGCCTCATGCCTATAATCCCAGCACTNNNAGATCGGAAGA |
| 62690_1 | TATGAGTGTGGAGTCGTTGCGGCCTTGCTTCCTAGTTTTTCTGNNNAGATCGGAAG |
| 62691_1 | TATGAGTGTGGAGTCGTTGCAAGGTGAGGAAGACCACAGTTACNNNAGATCGGAA |
| 62692_1 | TATGAGTGTGGAGTCGTTGCTGTTCTTATGCACGATCTCAGCTNNNAGATCGGAAG |
| 62693_1 | TATGAGTGTGGAGTCGTTGCAAGGATCTGTAAACTCCCCCAACNNNAGATCGGAAG |
| 62694_1 | TATGAGTGTGGAGTCGTTGCAGAAAACACGTAGCCTGAGAGAGNNNAGATCGGAA |
| 62695_1 | TATGAGTGTGGAGTCGTTGCTCACTTTGTGGCAGAAGTATCCANNNAGATCGGAAG |
| 62696_1 | TATGAGTGTGGAGTCGTTGCTTGACAAACATGTGCCTTTGTGTNNNAGATCGGAAG |
| 62697_1 | TATGAGTGTGGAGTCGTTGCCCTTTGCTATTTTCTGGGCACAANNNAGATCGGAAG |
| 62698_1 | TATGAGTGTGGAGTCGTTGCATGTTTGCATGCTCTGTAAAGTGNNNAGATCGGAAG |
| 62701_1 | TATGAGTGTGGAGTCGTTGCGTCTCCCAGGAGGTTGTAGAAAGNNNAGATCGGAA |
| 62702_1 | TATGAGTGTGGAGTCGTTGCTTGACCACAGGAAGAACAAGGAANNNAGATCGGAA |
| 62703_1 | TATGAGTGTGGAGTCGTTGCGGAGAATTGCGTGATCCTAGGAGNNNAGATCGGAA |
| 62704_1 | TATGAGTGTGGAGTCGTTGCTAGAGATGCAGCCTCAAATTCCTNNNAGATCGGAAG |
| 62706_1 | TATGAGTGTGGAGTCGTTGCACACAGCAGTGATTCTAGAGCAANNNAGATCGGAAG |
| 62708_1 | TATGAGTGTGGAGTCGTTGCGCAGCAGACCATCACATCATTTCNNNAGATCGGAAG |
| 62709_1 | TATGAGTGTGGAGTCGTTGCGAAAATTGGCAGGAATGGTGACANNNAGATCGGAA |
| 62710_1 | TATGAGTGTGGAGTCGTTGCGATAGCTATTGCCAGCCCCTTATNNNAGATCGGAAG |
| 62711_1 | TATGAGTGTGGAGTCGTTGCCAAGGCTGCAGTGAGTTATGATGNNNAGATCGGAA |
| 62712_1 | TATGAGTGTGGAGTCGTTGCCCAGAAGCAATATTTCCAGCCAGNNNAGATCGGAAG |
| 62713_1 | TATGAGTGTGGAGTCGTTGCTTTTACTTGCATGCCCCAAACTTNNNAGATCGGAAGA |
| 62714_1 | TATGAGTGTGGAGTCGTTGCCCCTTGCAAAAGAAAAGAGAAGCNNNAGATCGGAA |
| 62715_1 | TATGAGTGTGGAGTCGTTGCAGCTGATACTGCTCCTACTGAAANNNAGATCGGAAG |
| 62717_1 | TATGAGTGTGGAGTCGTTGCGTGTGGTGACTCACTCCTGTAATNNNAGATCGGAAG |

| | |
|---|---|
| 62718_1 | TATGAGTGTGGAGTCGTTGCCACACTCGACTGGAAAAGTACCTNNNAGATCGGAAG |
| 62722_1 | TATGAGTGTGGAGTCGTTGCTGGGTGAAAGGAACAATGACAGANNNAGATCGGA |
| 62723_1 | TATGAGTGTGGAGTCGTTGCAGAGAGAAAGAAAGAGGGAGGGANNNAGATCGG |
| 62724_1 | TATGAGTGTGGAGTCGTTGCGGACAGAGAAGTGGAGAAATGGTNNNAGATCGGA |
| 62725_1 | TATGAGTGTGGAGTCGTTGCAAAAAAGAGGCTCAATTGGCTCANNNAGATCGGAA |
| 62726_1 | TATGAGTGTGGAGTCGTTGCCACACACACACACACACACATATNNNAGATCGGAAG |
| 62727_1 | TATGAGTGTGGAGTCGTTGCGAGAGAGATCTGGTTTCCTTGGGNNNAGATCGGAA |
| 62728_1 | TATGAGTGTGGAGTCGTTGCTGTTAGCCTTTCTCACACTCTCCNNNAGATCGGAAGA |
| 62729_1 | TATGAGTGTGGAGTCGTTGCAACAACCTCAGTAGTAACTTGCANNNAGATCGGAAG |
| 62730_1 | TATGAGTGTGGAGTCGTTGCAATTTGCAATCCACCCACAATGTNNNAGATCGGAAG |
| 62731_1 | TATGAGTGTGGAGTCGTTGCGCCGACAGCACAGTAAAAAGATCNNNAGATCGGAA |
| 62733_1 | TATGAGTGTGGAGTCGTTGCGGCATAGAAGCTAAGAGGTTCGANNNAGATCGGAA |
| 62736_1 | TATGAGTGTGGAGTCGTTGCGCCACAGACAGAATTTAGGTGGTNNNAGATCGGAA |
| 62737_1 | TATGAGTGTGGAGTCGTTGCGGTGGTGGCAGTAGAAGAGTAAANNNAGATCGGA |
| 62738_1 | TATGAGTGTGGAGTCGTTGCGTAGACAAGGCTCCCTTTGAGAANNNAGATCGGAA |
| 62740_1 | TATGAGTGTGGAGTCGTTGCTGGCTCTTCAACCATTACAGTCANNNAGATCGGAAG |
| 62742_1 | TATGAGTGTGGAGTCGTTGCTGTTGCATACCTGTAGTCCTAGCNNNAGATCGGAAG |
| 62744_1 | TATGAGTGTGGAGTCGTTGCAAACAGCACCAAATTAGTGGCAANNNAGATCGGAA |
| 62746_1 | TATGAGTGTGGAGTCGTTGCAGTAGGATGTTCTCTTTGCAGGTNNNAGATCGGAAG |
| 62748_1 | TATGAGTGTGGAGTCGTTGCAGGACTTAGGTATACACACGTGANNNAGATCGGAA |
| 62752_1 | TATGAGTGTGGAGTCGTTGCCACAAAAAGCACTACAGCTGTCANNNAGATCGGAAG |
| 62753_1 | TATGAGTGTGGAGTCGTTGCAAAAAAGGGAAGCAACAAGGAGCNNNAGATCGGA |
| 62755_1 | TATGAGTGTGGAGTCGTTGCTAATCCCAGCTACTCAGGAGGTTNNNAGATCGGAAG |
| 62756_1 | TATGAGTGTGGAGTCGTTGCTCTGGCTGGATGACAAGAAGTTCNNNAGATCGGAA |
| 62759_1 | TATGAGTGTGGAGTCGTTGCTCTCCTGACTAGCTGGGATTACANNNAGATCGGAAG |
| 62760_1 | TATGAGTGTGGAGTCGTTGCGGAGGAAATATTCATGCCACACGNNNAGATCGGAA |
| 62761_1 | TATGAGTGTGGAGTCGTTGCCTGAGTTCAAGCGTTTCTCCTGANNNAGATCGGAAG |
| 62762_1 | TATGAGTGTGGAGTCGTTGCGCACCCATCACCTAAGCAGTATANNNAGATCGGAAG |
| 62763_1 | TATGAGTGTGGAGTCGTTGCCTCAAGTGATCCTCCCACCATAGNNNAGATCGGAAG |
| 62765_1 | TATGAGTGTGGAGTCGTTGCCTGAGTAGCTGGGATTACAGGTGNNNAGATCGGAA |
| 62767_1 | TATGAGTGTGGAGTCGTTGCAGACAGGGTCTCATTCTGTTTCCNNNAGATCGGAAG |
| 62769_1 | TATGAGTGTGGAGTCGTTGCTTAGGGTACATGTGCACATTGTGNNNAGATCGGAA |
| 62770_1 | TATGAGTGTGGAGTCGTTGCTTGAACTCCTAAGCTCAAGCGATNNNAGATCGGAAG |
| 62771_1 | TATGAGTGTGGAGTCGTTGCTCTCACTCTATCACCCAGGCTAGNNNAGATCGGAAG |
| 62772_1 | TATGAGTGTGGAGTCGTTGCCGCTTCCTGGATTCAAGTGATTCNNNAGATCGGAAG |
| 62773_1 | TATGAGTGTGGAGTCGTTGCCAAATGCTGGGATTACAAGGCATNNNAGATCGGAA |
| 62774_1 | TATGAGTGTGGAGTCGTTGCTTCAAGTGATTCTCCTGCCTCAGNNNAGATCGGAAG |
| 62775_1 | TATGAGTGTGGAGTCGTTGCAGACTTCCAAGTAGGTGGGACTANNNAGATCGGAA |
| 62776_1 | TATGAGTGTGGAGTCGTTGCTTCACATCATTCTCCTGCCTCAGNNNAGATCGGAAGA |
| 62778_1 | TATGAGTGTGGAGTCGTTGCTCCCAGGTTCAAGTGATTTTCCTNNNAGATCGGAAG |
| 62779_1 | TATGAGTGTGGAGTCGTTGCAGTTTTATTCTTGTTGCCCAGGCNNNAGATCGGAAG |
| 62780_1 | TATGAGTGTGGAGTCGTTGCTCTCCCAGGTTCAATCGATTCTCNNNAGATCGGAAGA |
| 62781_1 | TATGAGTGTGGAGTCGTTGCGCATTTTTAGTAGAGACGGGTTTNNNAGATCGGAA |

| | |
|---|---|
| 62782_1 | TATGAGTGTGGAGTCGTTGCGATTCTCCTGCCTCAGCTTCTAGNNNAGATCGGAAGA |
| 62783_1 | TATGAGTGTGGAGTCGTTGCGAGTGAAGTGGTACAAACTTGGCNNNAGATCGGAA |
| 62784_1 | TATGAGTGTGGAGTCGTTGCCATTTAGGGATCCCCAACTCCAGNNNAGATCGGAAGA |
| 62785_1 | TATGAGTGTGGAGTCGTTGCCCATGTTTGCCAAACTAGTCTCGNNNAGATCGGAAGA |
| 62786_1 | TATGAGTGTGGAGTCGTTGCAGACAGGATTTCTCTCTGTTGGCNNNAGATCGGAAG |
| 62787_1 | TATGAGTGTGGAGTCGTTGCCACTGTAACCTCTGCCTCTTAGGNNNAGATCGGAAGA |
| 62791_1 | TATGAGTGTGGAGTCGTTGCCCTGCTGTCTAAACATAAGCAGCNNNAGATCGGAAG |
| 62793_1 | TATGAGTGTGGAGTCGTTGCATCTCCTGACCTTGTGATCCATGNNNAGATCGGAAGA |
| 62794_1 | TATGAGTGTGGAGTCGTTGCAGATGTTGTCTCACTCTGTCACCNNNAGATCGGAAGA |
| 62795_1 | TATGAGTGTGGAGTCGTTGCCCTCTTGGATTCCAGCGATTCTCNNNAGATCGGAAGA |
| 62797_1 | TATGAGTGTGGAGTCGTTGCGCCTCCTGGATTCAAGTGATTCTNNNAGATCGGAAGA |
| 62798_1 | TATGAGTGTGGAGTCGTTGCGTTCAAGCGATTCTCCTACCTCANNNAGATCGGAAGA |
| 62799_1 | TATGAGTGTGGAGTCGTTGCCTGGGACTATAGGCATGTACCACNNNAGATCGGAAG |
| 62800_1 | TATGAGTGTGGAGTCGTTGCAGAGTAGCTGGGAGTATAGGTGCNNNAGATCGGAA |
| 62801_1 | TATGAGTGTGGAGTCGTTGCAGTGCTGGGATTACAGACATGAGNNNAGATCGGAA |
| 62802_1 | TATGAGTGTGGAGTCGTTGCCGGATTCAAGCAATTCTCCTGTCNNNAGATCGGAAGA |
| 62803_1 | TATGAGTGTGGAGTCGTTGCTTCAAGCGATTCTCCTTCCTCAANNNAGATCGGAAGA |
| 62804_1 | TATGAGTGTGGAGTCGTTGCCTACAGTCTCTATCTCCTGGGCTNNNAGATCGGAAGA |
| 62805_1 | TATGAGTGTGGAGTCGTTGCTTCAAGCAATTTTCCTGCCTCAGNNNAGATCGGAAGA |
| 62806_1 | TATGAGTGTGGAGTCGTTGCTGGAGTGCAGTTGTAAGATCTCGNNNAGATCGGAAG |
| 62807_1 | TATGAGTGTGGAGTCGTTGCGAGTGCAGTGGTATGAACACAACNNNAGATCGGAAG |
| 62808_1 | TATGAGTGTGGAGTCGTTGCCCAGTTTATAGGCGTGAGTCACTNNNAGATCGGAAG |
| 62809_1 | TATGAGTGTGGAGTCGTTGCCTCCTGAATAGCTGGGACTACAGNNNAGATCGGAAG |
| 62810_1 | TATGAGTGTGGAGTCGTTGCGACCTCATGATCCACCTGTTTCTNNNAGATCGGAAGA |
| 62811_1 | TATGAGTGTGGAGTCGTTGCCCACCATGCCCAGCTAATTTTTANNNAGATCGGAAGA |
| 62812_1 | TATGAGTGTGGAGTCGTTGCCAGTGGCATGATCATGGCTTACTNNNAGATCGGAAG |
| 62814_1 | TATGAGTGTGGAGTCGTTGCGATCTTACTGCCTCAGTCTCCAGNNNAGATCGGAAGA |
| 62816_1 | TATGAGTGTGGAGTCGTTGCCCATGTTCAAGCAATTCTCACGTNNNAGATCGGAAGA |
| 62817_1 | TATGAGTGTGGAGTCGTTGCTCCTGACTAGCTGGGATTACAGANNNAGATCGGAAG |
| 62818_1 | TATGAGTGTGGAGTCGTTGCCCAAGTAGCTGGGATTACAGACANNNAGATCGGAAG |
| 62819_1 | TATGAGTGTGGAGTCGTTGCCTCCTGAGTAGCTGGGACTACAGNNNAGATCGGAAG |
| 62820_1 | TATGAGTGTGGAGTCGTTGCAGACAGAGTCTCACTTCTTTGCCNNNAGATCGGAAGA |
| 62821_1 | TATGAGTGTGGAGTCGTTGCCTTTGCCTCCAGCTGAGATTTTTNNNAGATCGGAAGA |
| 62822_1 | TATGAGTGTGGAGTCGTTGCGTTCAAGTGATCCTCCTGCCTTANNNAGATCGGAAGA |
| 62823_1 | TATGAGTGTGGAGTCGTTGCTCTTGCTCTTGTCACTTAGGCTGNNNAGATCGGAAGA |
| 62824_1 | TATGAGTGTGGAGTCGTTGCTGGGATTATCAACATGAGCCACTNNNAGATCGGAAG |
| 62825_1 | TATGAGTGTGGAGTCGTTGCCCCTGTGCCTGGCCTAATTATATNNNAGATCGGAAGA |
| 62826_1 | TATGAGTGTGGAGTCGTTGCCCAGGCTGTGTGTATACCTAACANNNAGATCGGAAG |
| 62827_1 | TATGAGTGTGGAGTCGTTGCCGAGTAGCTGGGATTACAAGTGTNNNAGATCGGAAG |
| 62829_1 | TATGAGTGTGGAGTCGTTGCGATAGCTGAGACTACAGGCATCCNNNAGATCGGAAG |
| 62830_1 | TATGAGTGTGGAGTCGTTGCGTGGCACAATCATCATAGCTCACNNNAGATCGGAAG |
| 62831_1 | TATGAGTGTGGAGTCGTTGCTCAAGCTATTATCCTGCCTCAGCNNNAGATCGGAAGA |
| 62833_1 | TATGAGTGTGGAGTCGTTGCCAATCTTGGCTTACTGCAACCTCNNNAGATCGGAAGA |

62834_1 TATGAGTGTGGAGTCGTTGCTGTAATGTGGAGTTCCTGCAGATNNNAGATCGGAA
62835_1 TATGAGTGTGGAGTCGTTGCATCCTCTGAGATGACAGAAGGGANNNAGATCGGAA
62838_1 TATGAGTGTGGAGTCGTTGCAGGTTCGAGTGATTCTTTCGTCTNNNAGATCGGAAG
62839_1 TATGAGTGTGGAGTCGTTGCCTTCAGCTTCAAGCACTTCTCCTNNNAGATCGGAAGA
62840_1 TATGAGTGTGGAGTCGTTGCGCACCCACTAACTCGTCATCTAGNNNAGATCGGAAG
62842_1 TATGAGTGTGGAGTCGTTGCCTCTCAGGTTCATGCGATTTTCCNNNAGATCGGAAG
62843_1 TATGAGTGTGGAGTCGTTGCAAAGGGTCTCACTCTGTCACCTANNNAGATCGGAAG
62845_1 TATGAGTGTGGAGTCGTTGCCAATCCTCCCACATTATCCCTCCNNNAGATCGGAAGA
62846_1 TATGAGTGTGGAGTCGTTGCCCCAGGTTCAAGCAATTCTTCTGNNNAGATCGGAAG
62847_1 TATGAGTGTGGAGTCGTTGCCTTCCAAAGGGCTGGGATTACAGNNNAGATCGGAA
62848_1 TATGAGTGTGGAGTCGTTGCTGGCATGATCATAGCTCATCGAANNNAGATCGGAAG
62849_1 TATGAGTGTGGAGTCGTTGCTCTCCTGGCTTCAAGTGATTCTCNNNAGATCGGAAG
62851_1 TATGAGTGTGGAGTCGTTGCCTCCTGAGTAGCTGGGACTACAGNNNAGATCGGAA
62853_1 TATGAGTGTGGAGTCGTTGCCCTCCTGAGTAGCTGGGACTATANNNAGATCGGAAG
62854_1 TATGAGTGTGGAGTCGTTGCCCCAAGTAGCTGAAACTACAGGTNNNAGATCGGAA
62855_1 TATGAGTGTGGAGTCGTTGCTCCCAGATTCAAGCAATTCTCCTNNNAGATCGGAAG
62856_1 TATGAGTGTGGAGTCGTTGCCCTGGGTTCAAGCAATTCTCCTANNNAGATCGGAAG
62858_1 TATGAGTGTGGAGTCGTTGCTCCAGGTTCAAGTGATTCTCCTGNNNAGATCGGAAG
62860_1 TATGAGTGTGGAGTCGTTGCTCTTGGGTTCAAACGATTCTCCTNNNAGATCGGAAG
62861_1 TATGAGTGTGGAGTCGTTGCAGATGGGGTTTCTCCATATTGGGNNNAGATCGGAA
62862_1 TATGAGTGTGGAGTCGTTGCCCACCTCCTGAGTTCAAGTGATTNNNAGATCGGAAG
62863_1 TATGAGTGTGGAGTCGTTGCTGCCTCCCAGGTTCAAAAAATTCNNNAGATCGGAAG
62864_1 TATGAGTGTGGAGTCGTTGCCTCAATCCCCGAGTAGTTGAGACNNNAGATCGGAAG
62865_1 TATGAGTGTGGAGTCGTTGCCCTCCTGAGTGACTGGAATTACANNNAGATCGGAAG
62866_1 TATGAGTGTGGAGTCGTTGCCTGTAGTCTCAAATTCCCAGGCTNNNAGATCGGAAG
62867_1 TATGAGTGTGGAGTCGTTGCATCTCCTGGGTTCAAGCAATTCTNNNAGATCGGAAG
62868_1 TATGAGTGTGGAGTCGTTGCCTGGAGTGTAGTGGTGTGATCTCNNNAGATCGGAA
62870_1 TATGAGTGTGGAGTCGTTGCATCCAAGGTGGGTCAATTCAACTNNNAGATCGGAAG
62871_1 TATGAGTGTGGAGTCGTTGCTCAGCTATTGTGAAAGGGGTTGANNNAGATCGGAA
62872_1 TATGAGTGTGGAGTCGTTGCGATGGAGTCTCACTCTAACACCCNNNAGATCGGAAG
62874_1 TATGAGTGTGGAGTCGTTGCGTGTAATCTTGGCTCACTGCAACNNNAGATCGGAAG
62875_1 TATGAGTGTGGAGTCGTTGCCTCCTGAGAAGCTGGGACTACAGNNNAGATCGGAA
62876_1 TATGAGTGTGGAGTCGTTGCCTCCTGAGTAACTGGGATCACAGNNNAGATCGGAAG
62877_1 TATGAGTGTGGAGTCGTTGCGATCTCCTGACCTAGTGATCTGCNNNAGATCGGAAG
62878_1 TATGAGTGTGGAGTCGTTGCATCCTGAGTAGCTGGGACTACAGNNNAGATCGGAA
62879_1 TATGAGTGTGGAGTCGTTGCTTCAAGTGATTCTCCTGCCTCAGNNNAGATCGGAAG
62880_1 TATGAGTGTGGAGTCGTTGCGTGATCTCAGCTCACTACAACCTNNNAGATCGGAAG
62883_1 TATGAGTGTGGAGTCGTTGCGCTCAAGAAATCCTCCCACCTTANNNAGATCGGAAG
62885_1 TATGAGTGTGGAGTCGTTGCCTCTGGAGTAGCTGGGACTACAGNNNAGATCGGAA
62886_1 TATGAGTGTGGAGTCGTTGCCCCAGAAGAGACACAGATCACAANNNAGATCGGAAG
62887_1 TATGAGTGTGGAGTCGTTGCCTGAGTAGCTGGGATTACAGGTGNNNAGATCGGAA
62888_1 TATGAGTGTGGAGTCGTTGCGCGGTACAATCATAACTCACTGCNNNAGATCGGAAG
62891_1 TATGAGTGTGGAGTCGTTGCTCGCTTTGCATGTACTTTGAGACNNNAGATCGGAAG

62892_1 TATGAGTGTGGAGTCGTTGCCTGGGACTATAGGCATGTACCACNNNAGATCGGAAG
62894_1 TATGAGTGTGGAGTCGTTGCAGGATGGTTTCAATCTCCTGACCNNNAGATCGGAAG
62895_1 TATGAGTGTGGAGTCGTTGCCCAATTTGCCAGTCTGTGTCTTTNNNAGATCGGAAGA
62896_1 TATGAGTGTGGAGTCGTTGCACAATTTCACCATTCCCTAGGCTNNNAGATCGGAAGA
62898_1 TATGAGTGTGGAGTCGTTGCCGAACTTCTGGACTCAAGCAATCNNNAGATCGGAAG
62900_1 TATGAGTGTGGAGTCGTTGCATCTCTGCTCACTACAACTTCCGNNNAGATCGGAAGA
62902_1 TATGAGTGTGGAGTCGTTGCAGAGATGACAGGACAGTGAGAGANNNAGATCGGAA
62903_1 TATGAGTGTGGAGTCGTTGCAGACAAACTCTCGCTCTGTCATCNNNAGATCGGAAGA
62904_1 TATGAGTGTGGAGTCGTTGCCCCAGATTCAAGCAATTCTCTGCNNNAGATCGGAAGA
62905_1 TATGAGTGTGGAGTCGTTGCGGTTCAAACATCAAGCAGTCCTCNNNAGATCGGAAG
62906_1 TATGAGTGTGGAGTCGTTGCTTCAAGCTATTGTCCTGCCTCAGNNNAGATCGGAAGA
62907_1 TATGAGTGTGGAGTCGTTGCGCCTCAGCCTCCTTGTAATTTTTNNNAGATCGGAAGA
62910_1 TATGAGTGTGGAGTCGTTGCGATTCTCGCTCTTTTCCCAGACTNNNAGATCGGAAGA
62911_1 TATGAGTGTGGAGTCGTTGCAGTGGCAGGTAGACATCTCAAAGNNNAGATCGGAAG
62914_1 TATGAGTGTGGAGTCGTTGCGAAGTGCAGTGGTGAAATCTCAGNNNAGATCGGAA
62915_1 TATGAGTGTGGAGTCGTTGCTTTTGAATTCCCAGGCTCAAACCNNNAGATCGGAAGA
62916_1 TATGAGTGTGGAGTCGTTGCTCATGGTTCACTGCAGTTTCAACNNNAGATCGGAAGA
62917_1 TATGAGTGTGGAGTCGTTGCCAAGTAGCTGGGACTACAGTCACNNNAGATCGGAAG
62919_1 TATGAGTGTGGAGTCGTTGCTCATGTGATTCTCCTGCCTTAGCNNNAGATCGGAAGA
62920_1 TATGAGTGTGGAGTCGTTGCTTCAAGCAATTTTCCTGCCTCAGNNNAGATCGGAAGA
62921_1 TATGAGTGTGGAGTCGTTGCCAATTCTCCTGTCTCAGACTCCCNNNAGATCGGAAGA
62923_1 TATGAGTGTGGAGTCGTTGCCAAGTAGCTAGGACTACAGGCACNNNAGATCGGAAG
62924_1 TATGAGTGTGGAGTCGTTGCTCTGGTCATTCATCCCTTGTCAGNNNAGATCGGAAGA
62925_1 TATGAGTGTGGAGTCGTTGCTGCCTACTATGCCTACTACAGGANNNAGATCGGAAGA
62928_1 TATGAGTGTGGAGTCGTTGCAGGAAATGAACCATCTGACCACANNNAGATCGGAAG
62930_1 TATGAGTGTGGAGTCGTTGCGCCAACAACACTATTGCTTCCATNNNAGATCGGAAGA
62931_1 TATGAGTGTGGAGTCGTTGCCAGCAGAAAAAACAGACAGCAGANNNAGATCGGAA
62933_1 TATGAGTGTGGAGTCGTTGCCCAATGCCAGGTGTGAATTCAAANNNAGATCGGAAG
62934_1 TATGAGTGTGGAGTCGTTGCCAGAGACTGCAGTGAACTGAGATNNNAGATCGGAAG
62935_1 TATGAGTGTGGAGTCGTTGCAAGTTTTGCCTGGTGGAAAAAGGNNNAGATCGGAA
62936_1 TATGAGTGTGGAGTCGTTGCCCTGTCCTTCTCCCTTTTGTCTTNNNAGATCGGAAGAG
62938_1 TATGAGTGTGGAGTCGTTGCAAGTTGGGAAATAGGCATTTGGCNNNAGATCGGAA
62939_1 TATGAGTGTGGAGTCGTTGCCGTTGGGGGAAGAATAAGCTTACNNNAGATCGGAA
62940_1 TATGAGTGTGGAGTCGTTGCAGGCTGAAAAGAGGAGAAATGGTNNNAGATCGGAA
62941_1 TATGAGTGTGGAGTCGTTGCTGTGAGGAAACTGGAATTGAGCTNNNAGATCGGAA
62943_1 TATGAGTGTGGAGTCGTTGCGGGAAAGAAGTACTGGCAAGGTANNNAGATCGGAA
62946_1 TATGAGTGTGGAGTCGTTGCTGCCCAATCTTATTCCCTGGAAANNNAGATCGGAAGA
62947_1 TATGAGTGTGGAGTCGTTGCACTCCAGGGCTAAGGTGAAAAAANNNAGATCGGAAG
62948_1 TATGAGTGTGGAGTCGTTGCGCTGAGGCAAGAGAATCACTTGANNNAGATCGGAAG
62949_1 TATGAGTGTGGAGTCGTTGCCCATCATAGTAGCACAGGAGCTCNNNAGATCGGAAG
62950_1 TATGAGTGTGGAGTCGTTGCCTCCTGTAGTCCCAGCTACTGAGNNNAGATCGGAAGA
62951_1 TATGAGTGTGGAGTCGTTGCCTCCAAAGATGTGTTGCAATGGTNNNAGATCGGAAG
62952_1 TATGAGTGTGGAGTCGTTGCAACTACAGCAGGCATCTATGAGGNNNAGATCGGAAG

| | |
|---|---|
| 62953_1 | TATGAGTGTGGAGTCGTTGCTGATGCCAGTTCTTCCAGTTTCTNNNAGATCGGAAGA |
| 62954_1 | TATGAGTGTGGAGTCGTTGCTTCCCTCATCTACCTCAGAGAGTNNNAGATCGGAAGA |
| 62955_1 | TATGAGTGTGGAGTCGTTGCAACAAAACCCAAGTGTTAGCTGCNNNAGATCGGAAG |
| 62956_1 | TATGAGTGTGGAGTCGTTGCCCATTTGGCCTAGAACCCTTACTNNNAGATCGGAAGA |
| 62958_1 | TATGAGTGTGGAGTCGTTGCTAGACAGGCTGAACATGAGAACCNNNAGATCGGAAG |
| 62959_1 | TATGAGTGTGGAGTCGTTGCGAGAATTGCTTGAACCCAGGAGANNNAGATCGGAAG |
| 62960_1 | TATGAGTGTGGAGTCGTTGCGCCACGAAAAGACACAGAGAAACNNNAGATCGGAAG |
| 62962_1 | TATGAGTGTGGAGTCGTTGCATGACTAAATCAAGAGGGGGCTGNNNAGATCGGAA |
| 62963_1 | TATGAGTGTGGAGTCGTTGCGATTGTGAGGTCAGGAGATCGAGNNNAGATCGGAA |
| 62964_1 | TATGAGTGTGGAGTCGTTGCTTTACCATGTGGAGATGAGGTGGNNNAGATCGGAA |
| 62965_1 | TATGAGTGTGGAGTCGTTGCAAAACAGCAAACACACAGACACANNNAGATCGGAAG |
| 62967_1 | TATGAGTGTGGAGTCGTTGCTCCCACTCACCACCATGTAAAATNNNAGATCGGAAGA |
| 62969_1 | TATGAGTGTGGAGTCGTTGCATTGTTGCTGCTCCACAATCAAANNNAGATCGGAAGA |
| 62971_1 | TATGAGTGTGGAGTCGTTGCCTGAACACTCATTGAGCCCATTTNNNAGATCGGAAGA |
| 62972_1 | TATGAGTGTGGAGTCGTTGCGAAATGCCCACACTCTTCAGAACNNNAGATCGGAAGA |
| 62973_1 | TATGAGTGTGGAGTCGTTGCTCACAGCAAGATTACAGGTGTGANNNAGATCGGAAG |
| 62974_1 | TATGAGTGTGGAGTCGTTGCCTGTGGTTGTATTTTCCAGCCAANNNAGATCGGAAGA |
| 62977_1 | TATGAGTGTGGAGTCGTTGCATGGTCAATCCTTTAGCCTCTCCNNNAGATCGGAAGA |
| 62978_1 | TATGAGTGTGGAGTCGTTGCATATAATCCAGCAATCCCGCCTCNNNAGATCGGAAGA |
| 62980_1 | TATGAGTGTGGAGTCGTTGCCACACACACACACCCAAGAAATANNNAGATCGGAAGA |
| 62981_1 | TATGAGTGTGGAGTCGTTGCAATAGGTGTGGTTGGTACTGTCCNNNAGATCGGAAG |
| 62982_1 | TATGAGTGTGGAGTCGTTGCACATGCAAATGGGCTGATTGAAANNNAGATCGGAAG |
| 62984_1 | TATGAGTGTGGAGTCGTTGCGCCAGTGAGGAGCCATACTTTATNNNAGATCGGAAG |
| 62986_1 | TATGAGTGTGGAGTCGTTGCCCCAAAAATTCCAGACCCTAAGGNNNAGATCGGAAG |
| 62987_1 | TATGAGTGTGGAGTCGTTGCATAGAGCTAGCCACGAAAGGAAGNNNAGATCGGAA |
| 62988_1 | TATGAGTGTGGAGTCGTTGCTTGCAGAACGTATACACTGAGGGNNNAGATCGGAAG |
| 62992_1 | TATGAGTGTGGAGTCGTTGCACTTCCCTGGGCCTCAATTTTATNNNAGATCGGAAGA |
| 62994_1 | TATGAGTGTGGAGTCGTTGCTCTGGCACATATCATGATCCCTGNNNAGATCGGAAGA |
| 62997_1 | TATGAGTGTGGAGTCGTTGCACATCCAAATGGCAAACACGAATNNNAGATCGGAAG |
| 62998_1 | TATGAGTGTGGAGTCGTTGCACTGGATAATGTGCCCTGAATCANNNAGATCGGAAG |
| 62999_1 | TATGAGTGTGGAGTCGTTGCAGCAGGTTGCAAACACTCTTTTTNNNAGATCGGAAGA |
| 63001_1 | TATGAGTGTGGAGTCGTTGCTGACCTCTTCACTTACTGAAAGCTNNNAGATCGGAAG |
| 63002_1 | TATGAGTGTGGAGTCGTTGCGCCTGTATTCTAACACCTCCCAANNNAGATCGGAAGA |
| 63003_1 | TATGAGTGTGGAGTCGTTGCTTCTGGCACTCACCTTGTAAGAGNNNAGATCGGAAG |
| 63004_1 | TATGAGTGTGGAGTCGTTGCACTTGTGTATGCTGCATAACTGTNNNAGATCGGAAG |
| 63006_1 | TATGAGTGTGGAGTCGTTGCTGTAGTCCCAGCTATTGAGGAGANNNAGATCGGAAG |
| 63007_1 | TATGAGTGTGGAGTCGTTGCACATTTAGCTTCCCACTGTCTGGNNNAGATCGGAAGA |
| 63008_1 | TATGAGTGTGGAGTCGTTGCCAGGGAAAGCTTTACCAGAGAGTNNNAGATCGGAAG |
| 63009_1 | TATGAGTGTGGAGTCGTTGCTCACAAGCTAACCTACACTGGTGNNNAGATCGGAAG |
| 63010_1 | TATGAGTGTGGAGTCGTTGCTGTGGTGGCATGTGTCTATAGTCNNNAGATCGGAAG |
| 63011_1 | TATGAGTGTGGAGTCGTTGCCCTGGGCAACAAGAATGAAACTCNNNAGATCGGAAG |
| 63012_1 | TATGAGTGTGGAGTCGTTGCTCGAGATTATCCTGGCTAACACGNNNAGATCGGAAG |
| 63013_1 | TATGAGTGTGGAGTCGTTGCAATAGGAGAGATGCTCGTTACCCNNNAGATCGGAAG |

| | |
|---|---|
| 63015_1 | TATGAGTGTGGAGTCGTTGCGGAATTTGGTACTGAACAATCCGTNNNAGATCGGAA |
| 63016_1 | TATGAGTGTGGAGTCGTTGCGAGGTCAGGAGATCAAGACCATCNNNAGATCGGAA |
| 63017_1 | TATGAGTGTGGAGTCGTTGCTTTGTAGTGGTAAGGAGTTGGGGNNNAGATCGGAA |
| 63019_1 | TATGAGTGTGGAGTCGTTGCACAAAGACTGAAAGTCCTGTGGTNNNAGATCGGAA |
| 63021_1 | TATGAGTGTGGAGTCGTTGCATCAAAACAATGTGGTACTGCCANNNAGATCGGAAG |
| 63023_1 | TATGAGTGTGGAGTCGTTGCTTTGTTGTCTTCTGGCATGTACGNNNAGATCGGAAG |
| 63025_1 | TATGAGTGTGGAGTCGTTGCTTAGCCCAGTAAGACCTAGGTCANNNAGATCGGAAG |
| 63027_1 | TATGAGTGTGGAGTCGTTGCTTTCTGTAGTCCCAGCTACTTGGNNNAGATCGGAAG |
| 63028_1 | TATGAGTGTGGAGTCGTTGCCTTAGGGGTGTAGTTTCAGGCATNNNAGATCGGAA |
| 63029_1 | TATGAGTGTGGAGTCGTTGCCTGAGATGGGAGGATCACTTGAGNNNAGATCGGAA |
| 63031_1 | TATGAGTGTGGAGTCGTTGCTGTGAGCTCAAGTGTTGGGAATANNNAGATCGGAA |
| 63035_1 | TATGAGTGTGGAGTCGTTGCTTTGGAAGGTAGATAGTTCCCCCNNNAGATCGGAAG |
| 63036_1 | TATGAGTGTGGAGTCGTTGCCTGCACCAAAACAATGGGAAAGTNNNAGATCGGAA |
| 63037_1 | TATGAGTGTGGAGTCGTTGCAAAGCCACCTGAATCAATGAACGNNNAGATCGGAAG |
| 63038_1 | TATGAGTGTGGAGTCGTTGCGGGGAATATCGATGCTTTCAGGANNNAGATCGGAA |
| 63039_1 | TATGAGTGTGGAGTCGTTGCCACATGCCTATAGTCCCAGCTAANNNAGATCGGAAG |
| 63040_1 | TATGAGTGTGGAGTCGTTGCATATTGCACAGGCTATGAGGAGGNNNAGATCGGAA |
| 63041_1 | TATGAGTGTGGAGTCGTTGCGCTAGAGAGAACAGCCTTTCCTTNNNAGATCGGAAG |
| 63043_1 | TATGAGTGTGGAGTCGTTGCTGCAACATCAGTCTTTTTTGTCTNNNAGATCGGAAGA |
| 63044_1 | TATGAGTGTGGAGTCGTTGCACTACATTCACATAGGCAGGTGGNNNAGATCGGAA |
| 63047_1 | TATGAGTGTGGAGTCGTTGCTGTGGCAAGGGGATACAACAATTNNNAGATCGGAA |
| 63051_1 | TATGAGTGTGGAGTCGTTGCATGGTGGGGCAGGAATTCTAATTNNNAGATCGGAA |
| 63053_1 | TATGAGTGTGGAGTCGTTGCACTCTACTTTTTTTGAGCCCCANNNAGATCGGAAGA |
| 63054_1 | TATGAGTGTGGAGTCGTTGCACAACATGGATGAAACAGGAGGANNNAGATCGGAA |
| 63057_1 | TATGAGTGTGGAGTCGTTGCAGTGTAAAATAAAGCCCCTTGTNNNAGATCGGAAG |
| 63058_1 | TATGAGTGTGGAGTCGTTGCGCAGAGAGCACATTAGCTACAGTNNNAGATCGGAA |
| 63059_1 | TATGAGTGTGGAGTCGTTGCTCACACCTGTAATCCCAACACTTNNNAGATCGGAAGA |
| 63064_1 | TATGAGTGTGGAGTCGTTGCAGGTAAAGCAAGATGGCCACATANNNAGATCGGAA |
| 63067_1 | TATGAGTGTGGAGTCGTTGCGTACAGTGTCTAGTCCCATGTCTNNNAGATCGGAAG |
| 63068_1 | TATGAGTGTGGAGTCGTTGCAATACTTCACACAACCTGCAAGCNNNAGATCGGAAG |
| 63071_1 | TATGAGTGTGGAGTCGTTGCGTTACGAATGAGACAGGGGACATNNNAGATCGGAA |
| 63072_1 | TATGAGTGTGGAGTCGTTGCATACCACACTCTCCTCTCACTCANNNAGATCGGAAGA |
| 63073_1 | TATGAGTGTGGAGTCGTTGCTGGAGAATGAAATTTTGGCACCANNNAGATCGGAA |
| 63075_1 | TATGAGTGTGGAGTCGTTGCCCTGTCTTACCACCACACCAATANNNAGATCGGAAGA |
| 63076_1 | TATGAGTGTGGAGTCGTTGCTGCAAGTTTTAGGTTCAGTAGAGANNNAGATCGGA |
| 63077_1 | TATGAGTGTGGAGTCGTTGCAACATCTTCATGTATGTGGGGGGNNNAGATCGGAA |
| 63078_1 | TATGAGTGTGGAGTCGTTGCGAGAAGGTCCACTTACTAGGTGCNNNAGATCGGAA |
| 63079_1 | TATGAGTGTGGAGTCGTTGCAGCAGGAGAATTGTTTGAGCCTANNNAGATCGGAA |
| 63080_1 | TATGAGTGTGGAGTCGTTGCAGTAACTGAAGTGCTTCCTGAGGNNNAGATCGGAA |
| 63082_1 | TATGAGTGTGGAGTCGTTGCAAAAGTGAATGGCAGGTGTGTACNNNAGATCGGAA |
| 63083_1 | TATGAGTGTGGAGTCGTTGCACATGCTTGCTGGATGGGTATTANNNAGATCGGAA |
| 63084_1 | TATGAGTGTGGAGTCGTTGCGGAATGAAGGGAATGGTGGGTAANNNAGATCGGA |
| 63085_1 | TATGAGTGTGGAGTCGTTGCGGTCACCTGAAGTTGGGAATTTGNNNAGATCGGAA |

| | |
|---|---|
| 63086_1 | TATGAGTGTGGAGTCGTTGCGCACAGTGGCACATGCTAATTTTNNNAGATCGGAAG |
| 63088_1 | TATGAGTGTGGAGTCGTTGCGTCAGCACCTCAATTTTACACCCNNNAGATCGGAAGA |
| 63090_1 | TATGAGTGTGGAGTCGTTGCCCCCAGATCACTACAACTGAGAGNNNAGATCGGAAG |
| 63091_1 | TATGAGTGTGGAGTCGTTGCTCCACCTCAAACCTAGAATCAGCNNNAGATCGGAAGA |
| 63092_1 | TATGAGTGTGGAGTCGTTGCAAGAAGTTTCCTGACCTCTGCTTNNNAGATCGGAAGA |
| 63093_1 | TATGAGTGTGGAGTCGTTGCCACACAAAGGAGCCAACAGAAATNNNAGATCGGAAG |
| 63095_1 | TATGAGTGTGGAGTCGTTGCGCTGAGAACCTCTGGGAAAAAACNNNAGATCGGAAG |
| 63096_1 | TATGAGTGTGGAGTCGTTGCACGTTACTGGTTATGAGGGGTTCNNNAGATCGGAAG |
| 63097_1 | TATGAGTGTGGAGTCGTTGCGAAAAGCTTGGCAAACAGAGCTANNNAGATCGGAAG |
| 63099_1 | TATGAGTGTGGAGTCGTTGCCCACCATCACTCTAATGCCAAAANNNAGATCGGAAGA |
| 63100_1 | TATGAGTGTGGAGTCGTTGCAAAACAGTTGGATGTCCAGATGCNNNAGATCGGAAG |
| 63102_1 | TATGAGTGTGGAGTCGTTGCGAATGAGGCTGGCCCTATCAATANNNAGATCGGAAG |
| 63103_1 | TATGAGTGTGGAGTCGTTGCTGCCTAAGACCTAGAATGTTCCCNNNAGATCGGAAGA |
| 63104_1 | TATGAGTGTGGAGTCGTTGCACAGCTTTGTCATCTGATGAGATNNNAGATCGGAAG |
| 63105_1 | TATGAGTGTGGAGTCGTTGCGATAGTGAGAGGTGACAGTGTGCNNNAGATCGGAA |
| 63106_1 | TATGAGTGTGGAGTCGTTGCATTCTCAAGCAATTCTCCTGCCTNNNAGATCGGAAGA |
| 63108_1 | TATGAGTGTGGAGTCGTTGCATCTTGGGTTTTCAAGGTATCCANNNAGATCGGAAG |
| 63109_1 | TATGAGTGTGGAGTCGTTGCTTCTTCTTTTGCCAACACAACGTNNNAGATCGGAAGA |
| 63110_1 | TATGAGTGTGGAGTCGTTGCTTAGCTAGTCCAAGCCATCCATGNNNAGATCGGAAGA |
| 63111_1 | TATGAGTGTGGAGTCGTTGCCAGGATCTTGCGCTGTTACCTAGNNNAGATCGGAAG |
| 63112_1 | TATGAGTGTGGAGTCGTTGCCTGGGATTATAGGTGTCTGCCACNNNAGATCGGAAG |
| 63113_1 | TATGAGTGTGGAGTCGTTGCCAATTCTCCTGCCTCAGTCTCTGNNNAGATCGGAAGA |
| 63114_1 | TATGAGTGTGGAGTCGTTGCCTCACATCTGTAATCCCAGCACTNNNAGATCGGAAGA |
| 63115_1 | TATGAGTGTGGAGTCGTTGCTCCTGAGTAGCTGGGATTACAGANNNAGATCGGAAG |
| 63117_1 | TATGAGTGTGGAGTCGTTGCGTGAAGTGACGCAATCATAGCTCNNNAGATCGGAAG |
| 63119_1 | TATGAGTGTGGAGTCGTTGCCTCCTGAGTAGCTGGGATTACAGNNNAGATCGGAAG |
| 63121_1 | TATGAGTGTGGAGTCGTTGCTGCCTCCCTGGTTCAAATGATTANNNAGATCGGAAGA |
| 63123_1 | TATGAGTGTGGAGTCGTTGCAATTCTCCTGTCTCAGCATCCTGNNNAGATCGGAAGA |
| 63125_1 | TATGAGTGTGGAGTCGTTGCACCCCCATGATTCAATTACCTCCNNNAGATCGGAAGA |
| 63126_1 | TATGAGTGTGGAGTCGTTGCGTGGGTGGGGAAGTTTTGAAAAANNNAGATCGGAA |
| 63127_1 | TATGAGTGTGGAGTCGTTGCTTCTCTCTTTGTAAGTGCCCCTCNNNAGATCGGAAGA |
| 63128_1 | TATGAGTGTGGAGTCGTTGCCATCTCCAGCTTTGACAAATCGGNNNAGATCGGAAGA |
| 63129_1 | TATGAGTGTGGAGTCGTTGCGACAGGAGCATCATTTGAGCTTGNNNAGATCGGAAG |
| 63131_1 | TATGAGTGTGGAGTCGTTGCGACAACTTACAGCATGGGAGAAANNNAGATCGGAAG |
| 63132_1 | TATGAGTGTGGAGTCGTTGCTCCCAGGTTCAAGTGATTTTCCTNNNAGATCGGAAGA |
| 63135_1 | TATGAGTGTGGAGTCGTTGCTGTAGCTAGAGGAGTCAGGAGAGNNNAGATCGGAA |
| 63136_1 | TATGAGTGTGGAGTCGTTGCGGGGGTGCTGCATGTATACAATANNNAGATCGGAAG |
| 63137_1 | TATGAGTGTGGAGTCGTTGCGCTCAAGATATCTGCCTGCATTGNNNAGATCGGAAG |
| 63138_1 | TATGAGTGTGGAGTCGTTGCAATTGGCCATGCAGGAGATATGANNNAGATCGGAAG |
| 63140_1 | TATGAGTGTGGAGTCGTTGCGAGATGGAGCTTGGTTCTTGTTGNNNAGATCGGAAG |
| 63144_1 | TATGAGTGTGGAGTCGTTGCTAGCGGTGCTACTGATTTGTGTANNNAGATCGGAAG |
| 63145_1 | TATGAGTGTGGAGTCGTTGCTTACATTGTGGGTATGGAAGGCANNNAGATCGGAAG |
| 63147_1 | TATGAGTGTGGAGTCGTTGCCCTGCTTCTGCTTAGTATTTGGCNNNAGATCGGAAGA |

| | |
|---|---|
| 63149_1 | TATGAGTGTGGAGTCGTTGCAATGCTGTTCTAGAGGTAGCTGGNNNAGATCGGAA |
| 63150_1 | TATGAGTGTGGAGTCGTTGCGTTGCCAATAGTCTGCTTGCAAGNNNAGATCGGAAG |
| 63152_1 | TATGAGTGTGGAGTCGTTGCAGGCAGGGTATTTATAGCAGTGGNNNAGATCGGAA |
| 63153_1 | TATGAGTGTGGAGTCGTTGCAATTCTCTCCCACTCTGGAAAGGNNNAGATCGGAAG |
| 63154_1 | TATGAGTGTGGAGTCGTTGCTCTCTTCAGCAATTCTAGCAGCANNNAGATCGGAAG |
| 63155_1 | TATGAGTGTGGAGTCGTTGCATAGGAAGAGCTGCAAAGACACANNNAGATCGGAA |
| 63156_1 | TATGAGTGTGGAGTCGTTGCCTTTGGTATGGTCCAGAAAGGCANNNAGATCGGAA |
| 63157_1 | TATGAGTGTGGAGTCGTTGCGCTGGAATACAGTGGCATGAACNNNAGATCGGAAG |
| 63158_1 | TATGAGTGTGGAGTCGTTGCCCAACATGATGAAACCCCATCTCNNNAGATCGGAAG |
| 63159_1 | TATGAGTGTGGAGTCGTTGCATAATTTAGTGCTTGCCTGCAGGNNNAGATCGGAAG |
| 63161_1 | TATGAGTGTGGAGTCGTTGCTCACTTGAGGTCAGGAGTTTGAGNNNAGATCGGAA |
| 63162_1 | TATGAGTGTGGAGTCGTTGCAATGGGCAAGACACATCTGGATANNNAGATCGGAA |
| 63164_1 | TATGAGTGTGGAGTCGTTGCTTGAAGTCTCGTTCTGTCACCTGNNNAGATCGGAAG |
| 63165_1 | TATGAGTGTGGAGTCGTTGCGCCTTGAAAACACCAAGTTTGGANNNAGATCGGAAG |
| 63166_1 | TATGAGTGTGGAGTCGTTGCTCCTGAGTAGCTGGGATTAGACANNNAGATCGGAA |
| 63167_1 | TATGAGTGTGGAGTCGTTGCATGACTATAGTCCAGAGGCGAGANNNAGATCGGAA |
| 63168_1 | TATGAGTGTGGAGTCGTTGCAGTGATTCTCTTGCCTTAGCCTCNNNAGATCGGAAG |
| 63169_1 | TATGAGTGTGGAGTCGTTGCGAGGCAGGAAAATCACTTGAACCNNNAGATCGGAA |
| 63171_1 | TATGAGTGTGGAGTCGTTGCGTGTTGGGATTATAGGCATGAGCNNNAGATCGGAA |
| 63172_1 | TATGAGTGTGGAGTCGTTGCAGGTTCAAATGATTCTCCTGCCTNNNAGATCGGAAG |
| 63173_1 | TATGAGTGTGGAGTCGTTGCGAAACTCCTGCAGTCAAGCAATCNNNAGATCGGAAG |
| 63174_1 | TATGAGTGTGGAGTCGTTGCAGTGTAAAGGGGTCTTGGAAACCNNNAGATCGGAA |
| 63177_1 | TATGAGTGTGGAGTCGTTGCCTGCAGTCTCAGCACTTTGAGAGNNNAGATCGGAAG |
| 63178_1 | TATGAGTGTGGAGTCGTTGCCCACTCTCTACCTCTTCTGCAAANNNAGATCGGAAGA |
| 63179_1 | TATGAGTGTGGAGTCGTTGCATCTCCTGGGTTCAAGCATTTCTNNNAGATCGGAAG |
| 63180_1 | TATGAGTGTGGAGTCGTTGCTCTCAGGAAGAAATGTGGCAGAANNNAGATCGGAA |
| 63181_1 | TATGAGTGTGGAGTCGTTGCGACTGGGAAGTTCAGTGGTCTTANNNAGATCGGAA |
| 63182_1 | TATGAGTGTGGAGTCGTTGCATGTTGAACGTTTGATATGGGCCNNNAGATCGGAAG |
| 63185_1 | TATGAGTGTGGAGTCGTTGCGGTGGGAGCAGAGTTCTAAATCANNNAGATCGGAA |
| 63187_1 | TATGAGTGTGGAGTCGTTGCATCTGCTTCTGTGCTCTTCTTGANNNAGATCGGAAGA |
| 63189_1 | TATGAGTGTGGAGTCGTTGCTCAAACTGCTTCCTGTAACCTCANNNAGATCGGAAGA |
| 63191_1 | TATGAGTGTGGAGTCGTTGCTCTCAAACTCCTGGGTTCACAAGNNNAGATCGGAAG |
| 63193_1 | TATGAGTGTGGAGTCGTTGCATCTGAATCTCGGAATCCACCTGNNNAGATCGGAAG |
| 63194_1 | TATGAGTGTGGAGTCGTTGCTATGTGTGTGTGTGTTTGTGGTGNNNAGATCGGAA |
| 63195_1 | TATGAGTGTGGAGTCGTTGCATACGGTGTGGAAAACAGCAATGNNNAGATCGGAA |
| 63196_1 | TATGAGTGTGGAGTCGTTGCCTGAGACAGGAGAATCGCTTGAANNNAGATCGGAA |
| 63199_1 | TATGAGTGTGGAGTCGTTGCATGAATGAACCAGACCTCTCTCCNNNAGATCGGAAG |
| 63201_1 | TATGAGTGTGGAGTCGTTGCGGTGAAACCCCTTCGCTACTAAANNNAGATCGGAAG |
| 63203_1 | TATGAGTGTGGAGTCGTTGCCTCCTGAGTAGCTGGGATTACAGNNNAGATCGGAA |
| 63204_1 | TATGAGTGTGGAGTCGTTGCCAATGGTGCTATCTAGGCTCACTNNNAGATCGGAAG |
| 63207_1 | TATGAGTGTGGAGTCGTTGCTTCAAGCAATTCTCCCGTCTCAGNNNAGATCGGAAGA |
| 63208_1 | TATGAGTGTGGAGTCGTTGCTCAAACAATTCTTCTGCCTCAGCNNNAGATCGGAAGA |
| 63209_1 | TATGAGTGTGGAGTCGTTGCTCAGCCTCCCGAATAGATAGGATNNNAGATCGGAAG |

| | |
|---|---|
| 63210_1 | TATGAGTGTGGAGTCGTTGCCAGAGTCTTGCTAAGTCACCCATNNNAGATCGGAAG |
| 63211_1 | TATGAGTGTGGAGTCGTTGCGCAGTGGCATAATCTTGGCTTACNNNAGATCGGAAG |
| 63214_1 | TATGAGTGTGGAGTCGTTGCATCCTACTTACACCCAAACACCCNNNAGATCGGAAGA |
| 63215_1 | TATGAGTGTGGAGTCGTTGCGGGTGTGGTCATGAAAATGAAGGNNNAGATCGGA |
| 63216_1 | TATGAGTGTGGAGTCGTTGCAACAAAAATAGCCAGGTGTGGTGNNNAGATCGGAA |
| 63217_1 | TATGAGTGTGGAGTCGTTGCCACGCCTATAACCCCAGTACTTTNNNAGATCGGAAG |
| 63220_1 | TATGAGTGTGGAGTCGTTGCAATGAGTTCCTGATGCTCCACATNNNAGATCGGAAG |
| 63221_1 | TATGAGTGTGGAGTCGTTGCTTCAAGTGATTCTCCTGCCTCAGNNNAGATCGGAAG |
| 63223_1 | TATGAGTGTGGAGTCGTTGCCATGATCTCGACTCACTGCTACCNNNAGATCGGAAG |
| 63226_1 | TATGAGTGTGGAGTCGTTGCCTCCTGAGTAGCTGGGATTACAGNNNAGATCGGAA |
| 63227_1 | TATGAGTGTGGAGTCGTTGCGCCTCCTGAGTAACTGGGATTACNNNAGATCGGAAG |
| 63228_1 | TATGAGTGTGGAGTCGTTGCGTAGAGACAGGGTTTCACTGTGTNNNAGATCGGAA |
| 63229_1 | TATGAGTGTGGAGTCGTTGCGCGATCTCTACTTACCACAAGCTNNNAGATCGGAAG |
| 63231_1 | TATGAGTGTGGAGTCGTTGCGCTCAAGTGATTCTCCTGTCTCANNNAGATCGGAAG |
| 63232_1 | TATGAGTGTGGAGTCGTTGCTTCAAGCAACACAAACAGAGGTGNNNAGATCGGAA |
| 63234_1 | TATGAGTGTGGAGTCGTTGCGAGCATTCAGCATAAAGGGGGTANNNAGATCGGAA |
| 63236_1 | TATGAGTGTGGAGTCGTTGCATCCTGGAGAAAATAGACCCAGCNNNAGATCGGAA |
| 63237_1 | TATGAGTGTGGAGTCGTTGCTCAAGCAATTATTCTGCCTCAGCNNNAGATCGGAAG |
| 63238_1 | TATGAGTGTGGAGTCGTTGCATGGGATAAACCAGCCAACATCANNNAGATCGGAAG |
| 63241_1 | TATGAGTGTGGAGTCGTTGCGTTCAAGCGATTCTCCTACCTCANNNAGATCGGAAG |
| 63242_1 | TATGAGTGTGGAGTCGTTGCGATACAGGGTCTTTCTGTGTTGCNNNAGATCGGAAG |
| 63244_1 | TATGAGTGTGGAGTCGTTGCCCTCTATCCCCTTTTTTGAGGCTNNNAGATCGGAAGA |
| 63245_1 | TATGAGTGTGGAGTCGTTGCCTCCTGAGTAGCTGGGACTACAGNNNAGATCGGAA |
| 63247_1 | TATGAGTGTGGAGTCGTTGCGAGTAGCTGGAACTACAAGCGAGNNNAGATCGGAA |
| 63249_1 | TATGAGTGTGGAGTCGTTGCTCCAGGTTCAAGTGATTCTCCTGNNNAGATCGGAAG |
| 63250_1 | TATGAGTGTGGAGTCGTTGCAGTGGTGGGATTATAGGCATGAGNNNAGATCGGA |
| 63252_1 | TATGAGTGTGGAGTCGTTGCCAATCTTGACTCACTGCAACCTCNNNAGATCGGAAG |
| 63253_1 | TATGAGTGTGGAGTCGTTGCAACAACACACCACAAGTTTGTCANNNAGATCGGAAG |
| 63254_1 | TATGAGTGTGGAGTCGTTGCAAGTCTTCCCAGGCTCAAGTGATNNNAGATCGGAAG |
| 63259_1 | TATGAGTGTGGAGTCGTTGCTGATGAGTCATGAAGTGCAGAACNNNAGATCGGAA |
| 63260_1 | TATGAGTGTGGAGTCGTTGCGGTGCTTCATCTTTTGGTGACATNNNAGATCGGAAG |
| 63261_1 | TATGAGTGTGGAGTCGTTGCGCTGCTCCTTCCAGTCAATAGAANNNAGATCGGAAG |
| 63262_1 | TATGAGTGTGGAGTCGTTGCACAAATGAGTACATCCACAGGCTNNNAGATCGGAAG |
| 63263_1 | TATGAGTGTGGAGTCGTTGCAGGAAATTTCTGAAGCAAGGCACNNNAGATCGGAA |
| 63264_1 | TATGAGTGTGGAGTCGTTGCAGAGAGGGTAAGAGTTGGGAAGANNNAGATCGGA |
| 63265_1 | TATGAGTGTGGAGTCGTTGCGAGAGAGAGAAGGAGGGAGACTTNNNAGATCGGA |
| 63266_1 | TATGAGTGTGGAGTCGTTGCGCAGTGACTCACACCTGTAATCTNNNAGATCGGAAG |
| 63267_1 | TATGAGTGTGGAGTCGTTGCGAAGAGGTCCAAGTCTGAGGTTTNNNAGATCGGAA |
| 63268_1 | TATGAGTGTGGAGTCGTTGCACTTCCTGCAACAAAGATGGAGANNNAGATCGGAA |
| 63269_1 | TATGAGTGTGGAGTCGTTGCCGACAGAGAGAGCCCATTTCAAANNNAGATCGGAA |
| 63270_1 | TATGAGTGTGGAGTCGTTGCAAAGAGACACCAGGTATGCACATNNNAGATCGGAA |
| 63272_1 | TATGAGTGTGGAGTCGTTGCCTGTAGTCTCAGCTACTTGGGAGNNNAGATCGGAA |
| 63273_1 | TATGAGTGTGGAGTCGTTGCTGTACTGGGTAGCTCCATTGAACNNNAGATCGGAAG |

| | |
|---|---|
| 63274_1 | TATGAGTGTGGAGTCGTTGCGCTCTCTAGATTGTGCTACTGACANNNAGATCGGAA |
| 63275_1 | TATGAGTGTGGAGTCGTTGCGCTCCTCTAAGTCCAAGCTACTCNNNAGATCGGAAG |
| 63276_1 | TATGAGTGTGGAGTCGTTGCACCAAGTGCACTAACTCATCAGTNNNAGATCGGAAG |
| 63278_1 | TATGAGTGTGGAGTCGTTGCAACCCAGCATAGCCTATTGTAGGNNNAGATCGGAAG |
| 63279_1 | TATGAGTGTGGAGTCGTTGCTTCATGGATGCAAGCAAAGGTTTNNNAGATCGGAA |
| 63280_1 | TATGAGTGTGGAGTCGTTGCAGTTTCAGTGGCTTACACAGTGANNNAGATCGGAA |
| 63281_1 | TATGAGTGTGGAGTCGTTGCGCAAATCTTGTCATCTCCAGCTCNNNAGATCGGAAG |
| 63283_1 | TATGAGTGTGGAGTCGTTGCGATACAGGGCATGATGAGGTCATNNNAGATCGGAA |
| 63284_1 | TATGAGTGTGGAGTCGTTGCTTCTGGCCTTGTTTTCAAAAGCANNNAGATCGGAAG |
| 63286_1 | TATGAGTGTGGAGTCGTTGCAACACTGACTCTGAGACCATGACNNNAGATCGGAAG |
| 63287_1 | TATGAGTGTGGAGTCGTTGCACACAGAGGAATCATTGTCTGCTNNNAGATCGGAAG |
| 63289_1 | TATGAGTGTGGAGTCGTTGCTTCCCGTTTCATTTCTTCCAAGTNNNAGATCGGAAGA |
| 63290_1 | TATGAGTGTGGAGTCGTTGCAAACCTTTGTTCCTCTCTTGGGTNNNAGATCGGAAG |
| 63291_1 | TATGAGTGTGGAGTCGTTGCATAGCTGGGATTACAGATGCCTGNNNAGATCGGAA |
| 63292_1 | TATGAGTGTGGAGTCGTTGCCCTCTCTCTTCACATCTCCCAAGNNNAGATCGGAAGA |
| 63293_1 | TATGAGTGTGGAGTCGTTGCCAGCGCACAACTATTTCAGAGACNNNAGATCGGAAG |
| 63294_1 | TATGAGTGTGGAGTCGTTGCCTAGCACAGCACATGAAGCATACNNNAGATCGGAAG |
| 63295_1 | TATGAGTGTGGAGTCGTTGCCTCCTGGACTCTATGCCATCCTANNNAGATCGGAAG |
| 63296_1 | TATGAGTGTGGAGTCGTTGCTCCCTCTTGGAAGTTTTTCAAGTNNNAGATCGGAAG |
| 63297_1 | TATGAGTGTGGAGTCGTTGCAGGGTGGGTGTGAAAAAGTAGAGNNNAGATCGGA |
| 63298_1 | TATGAGTGTGGAGTCGTTGCGGAAACCACAGTAAGATCCCACTNNNAGATCGGAAG |
| 63299_1 | TATGAGTGTGGAGTCGTTGCAGAAACAGCAATTTGGAGGAAGCNNNAGATCGGAA |
| 63300_1 | TATGAGTGTGGAGTCGTTGCATAATAGCTCACTGCAGCCTCAANNNAGATCGGAAG |
| 63302_1 | TATGAGTGTGGAGTCGTTGCATTTCTAGGCCCTCTTCCTTCACNNNAGATCGGAAGA |
| 63304_1 | TATGAGTGTGGAGTCGTTGCAAGATCCTGTTCTTTTCCTGGCANNNAGATCGGAAG |
| 63305_1 | TATGAGTGTGGAGTCGTTGCTTCAAGCAATTCTCCCACCTCAGNNNAGATCGGAAG |
| 63306_1 | TATGAGTGTGGAGTCGTTGCGACATGGTTTCATTCTGTTGCCANNNAGATCGGAAG |
| 63307_1 | TATGAGTGTGGAGTCGTTGCGTTAGCCAGGATGGTCTCGTATCNNNAGATCGGAA |
| 63308_1 | TATGAGTGTGGAGTCGTTGCAGGGTTGAGATCTTCCCAGTAGANNNAGATCGGAA |
| 63309_1 | TATGAGTGTGGAGTCGTTGCTAGAGAGTCTCAGCTCTGTCACCNNNAGATCGGAAG |
| 63310_1 | TATGAGTGTGGAGTCGTTGCAGTCCTGACCTCTCCTTCAGTTANNNAGATCGGAAG |
| 63311_1 | TATGAGTGTGGAGTCGTTGCTGTCAACACACAGCAGTCTCTAANNNAGATCGGAAG |
| 63313_1 | TATGAGTGTGGAGTCGTTGCATAGCTAGAGTGATCTCCCTGCANNNAGATCGGAAG |
| 63314_1 | TATGAGTGTGGAGTCGTTGCCTGGAGTGTAGTGGCATGATCTCNNNAGATCGGAA |
| 63315_1 | TATGAGTGTGGAGTCGTTGCCCAAGTAGGTGGGATTACAGACANNNAGATCGGAA |
| 63316_1 | TATGAGTGTGGAGTCGTTGCCATTGCTGACTTTACTGGTGCTCNNNAGATCGGAAG |
| 63317_1 | TATGAGTGTGGAGTCGTTGCTGCATGCATGTGTTATTGACAGGNNNAGATCGGAA |
| 63318_1 | TATGAGTGTGGAGTCGTTGCGAACCTTCCTTCTTACCTGTGGANNNAGATCGGAAG |
| 63319_1 | TATGAGTGTGGAGTCGTTGCCTCTGTTTCCCCTCTCTCTCTTCNNNAGATCGGAAGAG |
| 63320_1 | TATGAGTGTGGAGTCGTTGCCCGAGTAGCTGGGATTGATTACANNNAGATCGGAA |
| 63321_1 | TATGAGTGTGGAGTCGTTGCTCTCAGTTCACTGTAACCTCTGCNNNAGATCGGAAG |
| 63322_1 | TATGAGTGTGGAGTCGTTGCGCAGATCACGAGATCAGGAGATTNNNAGATCGGAA |
| 63323_1 | TATGAGTGTGGAGTCGTTGCTAACCTGGGCCATTGTTGACTTANNNAGATCGGAAG |

63324_1 TATGAGTGTGGAGTCGTTGCACGAGATGACAGACTCTTTGAGGNNNAGATCGGAA
63326_1 TATGAGTGTGGAGTCGTTGCCTCTGGTGGAACTCTGATTGACANNNAGATCGGAAG
63327_1 TATGAGTGTGGAGTCGTTGCAAAGCCCACTGTGGAAGAAAGTANNNAGATCGGAA
63329_1 TATGAGTGTGGAGTCGTTGCACCAGACGCTATTCATCAAGACANNNAGATCGGAAG
63330_1 TATGAGTGTGGAGTCGTTGCGCAGCTAGAATTTGGGAGACCTANNNAGATCGGAA
63331_1 TATGAGTGTGGAGTCGTTGCCTCCCAACCAGCTAGAAGCAATANNNAGATCGGAAG
63332_1 TATGAGTGTGGAGTCGTTGCCAGCTAGTATCTGGGGTGAGATGNNNAGATCGGAA
63333_1 TATGAGTGTGGAGTCGTTGCCTCATTTTGGGTCTGTTTGGGTCNNNAGATCGGAAG
63335_1 TATGAGTGTGGAGTCGTTGCGAGTGTGAGGAAGGAACCAGATGNNNAGATCGGA
63336_1 TATGAGTGTGGAGTCGTTGCACAGAAGGAGGAGTGGTTAGCTANNNAGATCGGA
63337_1 TATGAGTGTGGAGTCGTTGCATGTGGTTACTCTGTGCATGACANNNAGATCGGAAG
63338_1 TATGAGTGTGGAGTCGTTGCGCGAGAAAGAGAGACAGTCATCANNNAGATCGGAA
63339_1 TATGAGTGTGGAGTCGTTGCCTTGAGTGCAGTGGTGTAGTCATNNNAGATCGGAA
63340_1 TATGAGTGTGGAGTCGTTGCTCTTGCTCAATATAGCCTCGACCNNNAGATCGGAAG
63342_1 TATGAGTGTGGAGTCGTTGCTCTTCAACTCCTGGACTCAAGTGNNNAGATCGGAAG
63343_1 TATGAGTGTGGAGTCGTTGCACATGTAAACAAGGAAAGGGAGANNNAGATCGGA
63344_1 TATGAGTGTGGAGTCGTTGCCCTGACTAATGCACTCCCTTTCTNNNAGATCGGAAGA
63345_1 TATGAGTGTGGAGTCGTTGCTCCTGGTTTTTGACAAATGTGCTNNNAGATCGGAAG
63347_1 TATGAGTGTGGAGTCGTTGCTGGCAGAAAATGTCATTTCACCANNNAGATCGGAAG
63348_1 TATGAGTGTGGAGTCGTTGCGAGACCCGAGAAGTGGTATGTTTNNNAGATCGGAA
63349_1 TATGAGTGTGGAGTCGTTGCGTGTGGGTCTCCAAGTGCTTATANNNAGATCGGAAG
63350_1 TATGAGTGTGGAGTCGTTGCAGGCATCCTGAAAACTTGAGACTNNNAGATCGGAAG
63351_1 TATGAGTGTGGAGTCGTTGCTCCTCTGTTTCAGGGCAAAATCTNNNAGATCGGAAG
63354_1 TATGAGTGTGGAGTCGTTGCCACGCCTATAATCCTAGCCCTTTNNNAGATCGGAAGA
63355_1 TATGAGTGTGGAGTCGTTGCGAGTGACACAGACAGAGAGACAGNNNAGATCGGA
63356_1 TATGAGTGTGGAGTCGTTGCCAACTCCACCAGAAACTCTGAGANNNAGATCGGAAG
63357_1 TATGAGTGTGGAGTCGTTGCAGTGATATGGTTTGGCTGTGTTCNNNAGATCGGAA
63358_1 TATGAGTGTGGAGTCGTTGCGAAGAAGAAAGGGAAAGCAGGTGNNNAGATCGGA
63359_1 TATGAGTGTGGAGTCGTTGCGAGTAGCTGGGATTACAGGCTACNNNAGATCGGAA
63360_1 TATGAGTGTGGAGTCGTTGCAGTAAAGTCACTGAGGTAGGCAANNNAGATCGGAA
63361_1 TATGAGTGTGGAGTCGTTGCGGAAAGCCATCCCTCTCCAAATANNNAGATCGGAAG
63362_1 TATGAGTGTGGAGTCGTTGCAAATGTTAGCTCTGTGTGTGGGGNNNAGATCGGAA
63363_1 TATGAGTGTGGAGTCGTTGCATAGCAGGAAAGAGTGGAATGCANNNAGATCGGA
63365_1 TATGAGTGTGGAGTCGTTGCGAGACTGAGGTAGGAGGATCAGTNNNAGATCGGA
63366_1 TATGAGTGTGGAGTCGTTGCAACACTCCTGTGAGCCAAATTTGNNNAGATCGGAAG
63367_1 TATGAGTGTGGAGTCGTTGCCATCTGCTTTCTGGCTTCCAAAANNNAGATCGGAAG
63368_1 TATGAGTGTGGAGTCGTTGCCATTCCACCTCCCAGACCAATAANNNAGATCGGAAG
63369_1 TATGAGTGTGGAGTCGTTGCGATGGAATGTTAGCAACCACACCNNNAGATCGGAAG
63370_1 TATGAGTGTGGAGTCGTTGCTGGCATTTTACCCATGTCACAACNNNAGATCGGAAG
63371_1 TATGAGTGTGGAGTCGTTGCGAGAGACAGAGAGAGAGAGAGGGNNNAGATCGG
63373_1 TATGAGTGTGGAGTCGTTGCGTTCTGGCTACCCTTCTTGCTATNNNAGATCGGAAG
63374_1 TATGAGTGTGGAGTCGTTGCACCTCTCCCAGTCAGAATTTTGGNNNAGATCGGAAG
63375_1 TATGAGTGTGGAGTCGTTGCGTGAGTAGCTGGGACTACATGTGNNNAGATCGGAA

| | |
|---|---|
| 63376_1 | TATGAGTGTGGAGTCGTTGCGGAAGGGAGCACAGAGATATCAGNNNAGATCGGA |
| 63377_1 | TATGAGTGTGGAGTCGTTGCGTTCAAGAGATTCTCCTGCCTCANNNAGATCGGAAG |
| 63378_1 | TATGAGTGTGGAGTCGTTGCGTATGTGGTGTGCTAGCTGGATANNNAGATCGGAA |
| 63379_1 | TATGAGTGTGGAGTCGTTGCAGGAATAACTGACTTGGCAGGAGNNNAGATCGGAA |
| 63380_1 | TATGAGTGTGGAGTCGTTGCCAGTTTGCAGTTTAGCCTTTGGTNNNAGATCGGAAG |
| 63382_1 | TATGAGTGTGGAGTCGTTGCCATCCTTCCAAGTAGCTGGGATTNNNAGATCGGAAG |
| 63383_1 | TATGAGTGTGGAGTCGTTGCCAAAGCAACAACCAAAAAAGGGGNNNAGATCGGAA |
| 63384_1 | TATGAGTGTGGAGTCGTTGCCCGAGTATCTGGAATTACAGGCANNNAGATCGGAA |
| 63385_1 | TATGAGTGTGGAGTCGTTGCCCACCAGTCTCTACATGTAGCATNNNAGATCGGAAG |
| 63386_1 | TATGAGTGTGGAGTCGTTGCTAGGACAATAGTGGAGGGAAGGTNNNAGATCGGA |
| 63387_1 | TATGAGTGTGGAGTCGTTGCTTGTGACAAAGTGCCATCAGATGNNNAGATCGGAA |
| 63388_1 | TATGAGTGTGGAGTCGTTGCTCAGGATTTATGCAAGCTGTGTGNNNAGATCGGAA |
| 63389_1 | TATGAGTGTGGAGTCGTTGCGCACAGACAGAAAAAGCTACAGGNNNAGATCGGAA |
| 63390_1 | TATGAGTGTGGAGTCGTTGCCAACCTCCATCTCCTGAGTTCAANNNAGATCGGAAGA |
| 63392_1 | TATGAGTGTGGAGTCGTTGCCTATCTCGGCTCACTGTAACCTCNNNAGATCGGAAGA |
| 63394_1 | TATGAGTGTGGAGTCGTTGCCTCAAGCAATCCTCCCATCTCATNNNAGATCGGAAGA |
| 63395_1 | TATGAGTGTGGAGTCGTTGCCCTCCTGGGTTCAAGAGATTCTCNNNAGATCGGAAG |
| 63398_1 | TATGAGTGTGGAGTCGTTGCCTAGAGTGCAGTGGAGTGATCTCNNNAGATCGGAA |
| 63399_1 | TATGAGTGTGGAGTCGTTGCATAGAGCTTCCCTGTATTGCTGCNNNAGATCGGAAG |
| 63402_1 | TATGAGTGTGGAGTCGTTGCTTCCCTTCTTTCTTCTCTCTGCCNNNAGATCGGAAGAG |
| 63403_1 | TATGAGTGTGGAGTCGTTGCAAGGGCTTTCTGTCCTCTAAAGGNNNAGATCGGAAG |
| 63404_1 | TATGAGTGTGGAGTCGTTGCCCAGGAGTCTATTGATACTGCCGNNNAGATCGGAAG |
| 63405_1 | TATGAGTGTGGAGTCGTTGCACCATTTCAGCTCACCATAACCTNNNAGATCGGAAGA |
| 63406_1 | TATGAGTGTGGAGTCGTTGCGAGAGAAAGCTACCTTGCTGTAANNNAGATCGGAA |
| 63407_1 | TATGAGTGTGGAGTCGTTGCTTCCTCCATGTAAAATCCCATGTNNNAGATCGGAAGA |
| 63408_1 | TATGAGTGTGGAGTCGTTGCGCAGAGAAGAAGGAATGCTTACGNNNAGATCGGAA |
| 63409_1 | TATGAGTGTGGAGTCGTTGCGTGCCTACTTTCTTAGGAGGGTCNNNAGATCGGAAG |
| 63410_1 | TATGAGTGTGGAGTCGTTGCCAGCAACAAGGCACCATTTTAGANNNAGATCGGAAG |
| 63411_1 | TATGAGTGTGGAGTCGTTGCCTGCATCCATTTCCATCACAGTGNNNAGATCGGAAGA |
| 63412_1 | TATGAGTGTGGAGTCGTTGCTGAGATGAAGTCCTTACCTCCCTNNNAGATCGGAAG |
| 63413_1 | TATGAGTGTGGAGTCGTTGCCACCCTTAGAGGCAATAGATGGTNNNAGATCGGAA |
| 63414_1 | TATGAGTGTGGAGTCGTTGCTCACAAGTGGGACCTGTATTTGTNNNAGATCGGAAG |
| 63415_1 | TATGAGTGTGGAGTCGTTGCCCCTTTCACAGCTAAGAACTCCTNNNAGATCGGAAGA |
| 63416_1 | TATGAGTGTGGAGTCGTTGCTTAGGCAAATACAGGGTCGATGTNNNAGATCGGAA |
| 63417_1 | TATGAGTGTGGAGTCGTTGCCTACTGAGACATCGTCCCATCAGNNNAGATCGGAAG |
| 63418_1 | TATGAGTGTGGAGTCGTTGCTGGGAGAATGGAAGGAAAATGGTNNNAGATCGGA |
| 63419_1 | TATGAGTGTGGAGTCGTTGCAAAAGTGGGGAGAATGTGAGGAGNNNAGATCGGA |
| 63421_1 | TATGAGTGTGGAGTCGTTGCGGAAGGAAGGAACGAAGGAGAAANNNAGATCGGA |
| 63423_1 | TATGAGTGTGGAGTCGTTGCCACAACTCCCTCTTTTGCACTTTNNNAGATCGGAAGA |
| 63424_1 | TATGAGTGTGGAGTCGTTGCTGTATCATGATTGAGGCCATGGGNNNAGATCGGAA |
| 63425_1 | TATGAGTGTGGAGTCGTTGCACAAAAATTAGGTGTGGTGGCAGNNNAGATCGGAA |
| 63426_1 | TATGAGTGTGGAGTCGTTGCGTTGCTACCGTGCTAGATACTGANNNAGATCGGAAG |
| 63427_1 | TATGAGTGTGGAGTCGTTGCTCCTTCAGAACTAGACAACTGGCNNNAGATCGGAAG |

| | |
|---|---|
| 63428_1 | TATGAGTGTGGAGTCGTTGCAGGAGAACTTGGAAAGCAAATGCNNNAGATCGGAA |
| 63429_1 | TATGAGTGTGGAGTCGTTGCCCTGCTTTTTCTCTTGTTCCCAGNNNAGATCGGAAGA |
| 63430_1 | TATGAGTGTGGAGTCGTTGCCCAAACCAAACCAAAATTCTGGCNNNAGATCGGAAG |
| 63431_1 | TATGAGTGTGGAGTCGTTGCAGGACTTCACCATACCCTTTTGGNNNAGATCGGAAG |
| 63435_1 | TATGAGTGTGGAGTCGTTGCCAAGTAGCTCAGACCACAAGTGTNNNAGATCGGAAG |
| 63437_1 | TATGAGTGTGGAGTCGTTGCCTCCTGAGTAGCTGGGACTACAGNNNAGATCGGAA |
| 63438_1 | TATGAGTGTGGAGTCGTTGCGATCTCAGCTCACTGCAAATTCCNNNAGATCGGAAG |
| 63439_1 | TATGAGTGTGGAGTCGTTGCTATATATGGCTGGGTCCAACACCNNNAGATCGGAAG |
| 63440_1 | TATGAGTGTGGAGTCGTTGCATCTCCCAGGTTCAAGCAATTCTNNNAGATCGGAAG |
| 63441_1 | TATGAGTGTGGAGTCGTTGCTCCAATCTTTCTGGCCTGAAGAANNNAGATCGGAAG |
| 63442_1 | TATGAGTGTGGAGTCGTTGCCCTTTGGAGTAACTGGGACTACANNNAGATCGGAAG |
| 63443_1 | TATGAGTGTGGAGTCGTTGCGAGAACCCCAAGTCAGACTACAGNNNAGATCGGAA |
| 63444_1 | TATGAGTGTGGAGTCGTTGCAGGGGGAGAGACAAAGGACTAATNNNAGATCGGA |
| 63445_1 | TATGAGTGTGGAGTCGTTGCTGCTTGTAATCCCAGCTACTCAGNNNAGATCGGAAG |
| 63446_1 | TATGAGTGTGGAGTCGTTGCTTACTGCAGCCTTGACCTTCTAGNNNAGATCGGAAG |
| 63448_1 | TATGAGTGTGGAGTCGTTGCGTACGGTAGCTCACACCTGTAATNNNAGATCGGAAG |
| 63449_1 | TATGAGTGTGGAGTCGTTGCCAAAGTGCTGGGATTACAGGTCTNNNAGATCGGAA |
| 63450_1 | TATGAGTGTGGAGTCGTTGCGCACACCTGTAATCCCAGCTATGNNNAGATCGGAAG |
| 63452_1 | TATGAGTGTGGAGTCGTTGCCCTCACTGATTCCTCTTCTGTGTNNNAGATCGGAAGA |
| 63453_1 | TATGAGTGTGGAGTCGTTGCGGAAGAAAGAAAAGATCTGGGGANNNAGATCGGA |
| 63454_1 | TATGAGTGTGGAGTCGTTGCGGACAACATTGACCTTGAGTGTGNNNAGATCGGAA |
| 63456_1 | TATGAGTGTGGAGTCGTTGCCAGGCTCCCATCCAATTTTTGTTNNNAGATCGGAAGA |
| 63459_1 | TATGAGTGTGGAGTCGTTGCGGGTTCAAGTGATTATCCTGCCTNNNAGATCGGAAG |
| 63460_1 | TATGAGTGTGGAGTCGTTGCCAATCTCAGCTCACTGCAACTTCNNNAGATCGGAAGA |
| 63461_1 | TATGAGTGTGGAGTCGTTGCGATTCTCATACCTCAGCCTCCTGNNNAGATCGGAAGA |
| 63463_1 | TATGAGTGTGGAGTCGTTGCCTGAGTAACTGGGACTACAGGTGNNNAGATCGGAA |
| 63464_1 | TATGAGTGTGGAGTCGTTGCAGATAAGGTCTCACTCTGTTGCCNNNAGATCGGAAG |
| 63465_1 | TATGAGTGTGGAGTCGTTGCTTAGATCCTGACCCCACAAATCCNNNAGATCGGAAGA |
| 63466_1 | TATGAGTGTGGAGTCGTTGCCACACACACACACACATAAGTGGNNNAGATCGGAAG |
| 63467_1 | TATGAGTGTGGAGTCGTTGCCTCAAGCGATCCTCTTACCTCAGNNNAGATCGGAAGA |
| 63469_1 | TATGAGTGTGGAGTCGTTGCCTCCCAAGTAGCTCGGATTACAGNNNAGATCGGAAG |
| 63472_1 | TATGAGTGTGGAGTCGTTGCCTCCTGAGTAGCTGGGATTACAGNNNAGATCGGAA |
| 63473_1 | TATGAGTGTGGAGTCGTTGCGGAGTGCAGTGGAGTGATATCAGNNNAGATCGGA |
| 63474_1 | TATGAGTGTGGAGTCGTTGCTAGTTTCATTCTCGTAGCCCAGGNNNAGATCGGAAG |
| 63476_1 | TATGAGTGTGGAGTCGTTGCGTTCAAGCAATTCTCTTGCCTCANNNAGATCGGAAGA |
| 63478_1 | TATGAGTGTGGAGTCGTTGCAATCTCTCCAGTATGTCAGCTGCNNNAGATCGGAAG |
| 63479_1 | TATGAGTGTGGAGTCGTTGCCTCCCAGTCCAACTTTCTTGACTNNNAGATCGGAAGA |
| 63480_1 | TATGAGTGTGGAGTCGTTGCTCAGATTTTCTTCCCTCCAGGTGNNNAGATCGGAAGA |
| 63481_1 | TATGAGTGTGGAGTCGTTGCCCTAGTATCTGGGATTACAGGCGNNNAGATCGGAA |
| 63482_1 | TATGAGTGTGGAGTCGTTGCATGCACTAATTCCCATCCCTGTTNNNAGATCGGAAGA |
| 63483_1 | TATGAGTGTGGAGTCGTTGCTGGGATTACAAATGTGTGCCAACNNNAGATCGGAA |
| 63484_1 | TATGAGTGTGGAGTCGTTGCAGGCAGGAGAATCACTTGAACTTNNNAGATCGGAA |
| 63487_1 | TATGAGTGTGGAGTCGTTGCGTAGCAGTGAGTCAAGATTGCACNNNAGATCGGAA |

| | |
|---|---|
| 63489_1 | TATGAGTGTGGAGTCGTTGCTCTCAGCTCATCACAACCTACACNNNAGATCGGAAGA |
| 63490_1 | TATGAGTGTGGAGTCGTTGCGAGTAGCTGGGATTACAGGAGTGNNNAGATCGGAA |
| 63491_1 | TATGAGTGTGGAGTCGTTGCAGTAGCTGGGATTACAGGCATACNNNAGATCGGAAG |
| 63492_1 | TATGAGTGTGGAGTCGTTGCGAGTTCAAGACCAGCCTGGATAANNNAGATCGGAAG |
| 63494_1 | TATGAGTGTGGAGTCGTTGCCCCAAGGTCAGTGGAAAAAACAGNNNAGATCGGAAG |
| 63495_1 | TATGAGTGTGGAGTCGTTGCGACCAACCAGAGAAAGTTTGAGCNNNAGATCGGAAG |
| 63499_1 | TATGAGTGTGGAGTCGTTGCGCATGTTCCTGTAGTTCCAGCTANNNAGATCGGAAGA |
| 63500_1 | TATGAGTGTGGAGTCGTTGCAAAAAGGTAACAGGGACTCTCCGNNNAGATCGGAAG |
| 63501_1 | TATGAGTGTGGAGTCGTTGCTTCAAGTGATTCTCCTGCCTCAGNNNAGATCGGAAGA |
| 63504_1 | TATGAGTGTGGAGTCGTTGCATTCTCCTACCTTGGACTCCTGANNNAGATCGGAAGA |
| 63506_1 | TATGAGTGTGGAGTCGTTGCTCCACTGTTGCTCTGACTTTTCTNNNAGATCGGAAGA |
| 63508_1 | TATGAGTGTGGAGTCGTTGCACTGTAGTCAGGGTTTGGGTTTTNNNAGATCGGAAG |
| 63509_1 | TATGAGTGTGGAGTCGTTGCCTCCCAGGTTCAAGTGATTCTCANNNAGATCGGAAGA |
| 63511_1 | TATGAGTGTGGAGTCGTTGCCAATGCAGCGTTTTTGTCATGTTNNNAGATCGGAAGA |
| 63512_1 | TATGAGTGTGGAGTCGTTGCTTTTGCTTACAATGTGCGTGACTNNNAGATCGGAAGA |
| 63513_1 | TATGAGTGTGGAGTCGTTGCTTCAAGCAATTCTCCTGCCTCAGNNNAGATCGGAAGA |
| 63514_1 | TATGAGTGTGGAGTCGTTGCGAATGCAGTGGTGTGATCATAGCNNNAGATCGGAAG |
| 63515_1 | TATGAGTGTGGAGTCGTTGCTCAAACTCATGGCCTCAAGTGATNNNAGATCGGAAG |
| 63517_1 | TATGAGTGTGGAGTCGTTGCCTCCCAGGTTCAAGTGATTCTCANNNAGATCGGAAGA |
| 63518_1 | TATGAGTGTGGAGTCGTTGCATTCTTGTGCTTCAGCCTCCTAANNNAGATCGGAAGA |
| 63519_1 | TATGAGTGTGGAGTCGTTGCTTCAAGTGATTCTCCTGCCTCAGNNNAGATCGGAAGA |
| 63520_1 | TATGAGTGTGGAGTCGTTGCTTGGGATAGCAGCCATTTTCTCTNNNAGATCGGAAGA |
| 63521_1 | TATGAGTGTGGAGTCGTTGCCCTCTCAAAGTGCTGGGATACAGNNNAGATCGGAAG |
| 63522_1 | TATGAGTGTGGAGTCGTTGCGGAATGGGAAGTGAGGTGTTTTGNNNAGATCGGAA |
| 63523_1 | TATGAGTGTGGAGTCGTTGCTGGCCCTAGTCTATCCCAACATANNNAGATCGGAAGA |
| 63524_1 | TATGAGTGTGGAGTCGTTGCAGACAGAGTCTCACTTTGTTGCCNNNAGATCGGAAG |
| 63525_1 | TATGAGTGTGGAGTCGTTGCGCCTCTTGGTTTCAAGCAATTCTNNNAGATCGGAAGA |
| 63526_1 | TATGAGTGTGGAGTCGTTGCGATGGAGTCTCACTCTGTTGTCCNNNAGATCGGAAG |
| 63527_1 | TATGAGTGTGGAGTCGTTGCCTGAGGATATTCACGGCCTACTCNNNAGATCGGAAG |
| 63529_1 | TATGAGTGTGGAGTCGTTGCCCTTCCAAGTGACTCCCTTTGTANNNAGATCGGAAGA |
| 63530_1 | TATGAGTGTGGAGTCGTTGCTCTCTCTCTCTCTCTCTGTGTGTNNNAGATCGGAAGAG |
| 63531_1 | TATGAGTGTGGAGTCGTTGCAAGTCTCTGTTACCTAGGCTGGANNNAGATCGGAAG |
| 63532_1 | TATGAGTGTGGAGTCGTTGCCAGGGTTTCATTATGTTGGCCAGNNNAGATCGGAAG |
| 63533_1 | TATGAGTGTGGAGTCGTTGCTGTATTTGTGCTCTCAGGACACANNNAGATCGGAAG |
| 63535_1 | TATGAGTGTGGAGTCGTTGCAGAGAACGAGCTCAGTGAATTGTNNNAGATCGGAAG |
| 63536_1 | TATGAGTGTGGAGTCGTTGCAAGCTGAAGCAACTTCCAGAAAGNNNAGATCGGAAG |
| 63537_1 | TATGAGTGTGGAGTCGTTGCACACCCACACACATACAAGAAGANNNAGATCGGAAG |
| 63539_1 | TATGAGTGTGGAGTCGTTGCCCTCCTGAGTAGCTGGGACTATANNNAGATCGGAAG |
| 63540_1 | TATGAGTGTGGAGTCGTTGCCGCTAAGTGAAATAAGCCACAGGNNNAGATCGGAAG |
| 63541_1 | TATGAGTGTGGAGTCGTTGCCCGTCTTCCCCTGTAAGGATAGANNNAGATCGGAAG |
| 63542_1 | TATGAGTGTGGAGTCGTTGCAGAATGGTCTTGATCCGATCTCGNNNAGATCGGAAG |
| 63543_1 | TATGAGTGTGGAGTCGTTGCGCTATTGATAGGAGCTGGAACGANNNAGATCGGAA |
| 63544_1 | TATGAGTGTGGAGTCGTTGCAGATGCGGTGGGTTATTTGTACCNNNAGATCGGAAG |

| | |
|---|---|
| 63545_1 | TATGAGTGTGGAGTCGTTGCTGTGTTCACTTTTTCTACCTGCCNNNAGATCGGAAGA |
| 63546_1 | TATGAGTGTGGAGTCGTTGCGGTTCAAGCTATTGTCTTGCCTCNNNAGATCGGAAG |
| 63548_1 | TATGAGTGTGGAGTCGTTGCAAGGAGAAAGAGAAGGAGAGGGANNNAGATCGG |
| 63549_1 | TATGAGTGTGGAGTCGTTGCGCCTCAACCTCCTAGACTCAATCNNNAGATCGGAAGA |
| 63550_1 | TATGAGTGTGGAGTCGTTGCGCCCAGGTTCAAGAGATTCTTCTNNNAGATCGGAAG |
| 63551_1 | TATGAGTGTGGAGTCGTTGCCAATCATAGCTTGTTGCAGCCTTNNNAGATCGGAAG |
| 63552_1 | TATGAGTGTGGAGTCGTTGCTTTGAGACAGGATCTTGCTCTGTNNNAGATCGGAAG |
| 63553_1 | TATGAGTGTGGAGTCGTTGCCCCAGGCTCAAGTGATTCTCTAANNNAGATCGGAAG |
| 63555_1 | TATGAGTGTGGAGTCGTTGCAACTCCTGTGTCAAATGATCCCANNNAGATCGGAAG |
| 63556_1 | TATGAGTGTGGAGTCGTTGCGTGAATCTGGAGATTGCTTTGGGNNNAGATCGGAA |
| 63557_1 | TATGAGTGTGGAGTCGTTGCTCCTGCTGTATGTGTATGTGTGTNNNAGATCGGAAG |
| 63558_1 | TATGAGTGTGGAGTCGTTGCTTTTCAACTGCAGAATGGAGCTGNNNAGATCGGAA |
| 63559_1 | TATGAGTGTGGAGTCGTTGCAAATGTTAGAGCTGGAAGGGTCCNNNAGATCGGAA |
| 63561_1 | TATGAGTGTGGAGTCGTTGCACCAGGTTCAAGTGATTCTCCTGNNNAGATCGGAAG |
| 63563_1 | TATGAGTGTGGAGTCGTTGCTGTTATCCAACACTGAGACTGGGNNNAGATCGGAA |
| 63564_1 | TATGAGTGTGGAGTCGTTGCCTGAGTAGCTAGGATTACAGGCGNNNAGATCGGAA |
| 63565_1 | TATGAGTGTGGAGTCGTTGCTGTGCCTATAGTCCCACCTACTTNNNAGATCGGAAGA |
| 63566_1 | TATGAGTGTGGAGTCGTTGCACACAGAGAGCATACATGAAGCANNNAGATCGGAA |
| 63570_1 | TATGAGTGTGGAGTCGTTGCTGGAGTCATCTTGTTCTGTCACCNNNAGATCGGAAG |
| 63571_1 | TATGAGTGTGGAGTCGTTGCTCTCGCTCTCTCTCTCTATTGGTNNNAGATCGGAAGA |
| 63572_1 | TATGAGTGTGGAGTCGTTGCCCCTGTCTCAAAAAAAAACCCCANNNAGATCGGAAG |
| 63573_1 | TATGAGTGTGGAGTCGTTGCACAAAATGCGGAATCACTGAAGGNNNAGATCGGAA |
| 63574_1 | TATGAGTGTGGAGTCGTTGCGCTTCCCTTGTCAAGTTTTCCTGNNNAGATCGGAAGA |
| 63575_1 | TATGAGTGTGGAGTCGTTGCTCCGCCTCCCGAAAATTCTAATTNNNAGATCGGAAGA |
| 63577_1 | TATGAGTGTGGAGTCGTTGCTGTGATGAAAACTTTCCAGTGGCNNNAGATCGGAA |
| 63578_1 | TATGAGTGTGGAGTCGTTGCATTACAAGATGCTGAGGTGTGGGNNNAGATCGGAA |
| 63579_1 | TATGAGTGTGGAGTCGTTGCCCAGGCCTCTCTGTCATTCTTATNNNAGATCGGAAGA |
| 63580_1 | TATGAGTGTGGAGTCGTTGCTAACAAAGAGAGAATTCCGGCCANNNAGATCGGAA |
| 63581_1 | TATGAGTGTGGAGTCGTTGCATGTGTGTGGAGTCTCCAGTTTTNNNAGATCGGAAG |
| 63582_1 | TATGAGTGTGGAGTCGTTGCTTGTGCCAGGCCTAAGATATCAANNNAGATCGGAAG |
| 63583_1 | TATGAGTGTGGAGTCGTTGCGGATGTGTAATGCTTCTCACAGCNNNAGATCGGAAG |
| 63584_1 | TATGAGTGTGGAGTCGTTGCCAGAAGATAGAAAGGCGGAGAGGNNNAGATCGGA |
| 63585_1 | TATGAGTGTGGAGTCGTTGCTGCTCAAACTTCATTTTCTCCCANNNAGATCGGAAGA |
| 63586_1 | TATGAGTGTGGAGTCGTTGCCTCATCAGGGTGAGGATAAGGTGNNNAGATCGGAA |
| 63588_1 | TATGAGTGTGGAGTCGTTGCAAGCCAACTGAGGATCTTGCTAANNNAGATCGGAA |
| 63589_1 | TATGAGTGTGGAGTCGTTGCCTGGAGTGTAGTGTTGCGATCTCNNNAGATCGGAA |
| 63592_1 | TATGAGTGTGGAGTCGTTGCACAGGAGCACAAGGTGATCTAAGNNNAGATCGGAA |
| 63593_1 | TATGAGTGTGGAGTCGTTGCGGGGAAGATCCTACATGATGGACNNNAGATCGGAA |
| 63594_1 | TATGAGTGTGGAGTCGTTGCGGCCAAGCTGACAGTAAAATTGANNNAGATCGGAA |
| 63595_1 | TATGAGTGTGGAGTCGTTGCCGCTCTTTGCCCTATGAAAACATNNNAGATCGGAAG |
| 63596_1 | TATGAGTGTGGAGTCGTTGCGACTACAGGAGTACACCACCATGNNNAGATCGGAA |
| 63597_1 | TATGAGTGTGGAGTCGTTGCGTTGATACCAGTAGAGTGGGGTGNNNAGATCGGAA |
| 63598_1 | TATGAGTGTGGAGTCGTTGCTGTTTCTCTTACTGGACTCTGGCNNNAGATCGGAAG |

| | |
|---|---|
| 63599_1 | TATGAGTGTGGAGTCGTTGCCACATTCAAGTTTAGCCAAGCCANNNAGATCGGAAG |
| 63600_1 | TATGAGTGTGGAGTCGTTGCATAACACCATTCACAGCAACCTGNNNAGATCGGAAG |
| 63601_1 | TATGAGTGTGGAGTCGTTGCCAACATAGCAAGACCCTGTCTCTNNNAGATCGGAAG |
| 63602_1 | TATGAGTGTGGAGTCGTTGCCTCAAGCAATCCTCCCATCTCAGNNNAGATCGGAAGA |
| 63603_1 | TATGAGTGTGGAGTCGTTGCATCTTGGCTCACTGCAATCTTTGNNNAGATCGGAAG |
| 63605_1 | TATGAGTGTGGAGTCGTTGCCTGGAGTGTAGTGGAGTGATCTCNNNAGATCGGAA |
| 63606_1 | TATGAGTGTGGAGTCGTTGCGCAAGAAAAAGAAAGCAAGCACCNNNAGATCGGAA |
| 63607_1 | TATGAGTGTGGAGTCGTTGCTCTACCACTACACACACACACACNNNAGATCGGAAGA |
| 63608_1 | TATGAGTGTGGAGTCGTTGCACACACACACAAATGACATTGCANNNAGATCGGAAG |
| 63609_1 | TATGAGTGTGGAGTCGTTGCCCCCCCACAAAGAAAGAAAAACANNNAGATCGGAAG |
| 63610_1 | TATGAGTGTGGAGTCGTTGCATTCTTACCCTTTGCTGTCATGCNNNAGATCGGAAGA |
| 63611_1 | TATGAGTGTGGAGTCGTTGCGTTCCATTCGTCTATGTGTTGGTNNNAGATCGGAAG |
| 63612_1 | TATGAGTGTGGAGTCGTTGCATCTATGATGTTGGTTGGGCAGANNNAGATCGGAA |
| 63617_1 | TATGAGTGTGGAGTCGTTGCGTGTGTATGTGTGTGTGTGTGTGNNNAGATCGGAA |
| 63618_1 | TATGAGTGTGGAGTCGTTGCTTGAACTCGCGACCTCAAGTAATNNNAGATCGGAAG |
| 63619_1 | TATGAGTGTGGAGTCGTTGCGTCCTGGGAATACAGCAGTGAATNNNAGATCGGAA |
| 63621_1 | TATGAGTGTGGAGTCGTTGCAGTGCCTGTAATCCCAGCTATTTNNNAGATCGGAAG |
| 63623_1 | TATGAGTGTGGAGTCGTTGCGGCATAGACCACAGCATACCAATNNNAGATCGGAAG |
| 63624_1 | TATGAGTGTGGAGTCGTTGCTTTTTCCAAGAAGGCCTATAATACCTNNNAGATCGGA |
| 63625_1 | TATGAGTGTGGAGTCGTTGCCTATCTGTCTTGGCTTTCCCACTNNNAGATCGGAAGA |
| 63626_1 | TATGAGTGTGGAGTCGTTGCCCTCCCATTGGTTCTGTTTCTCTNNNAGATCGGAAGA |
| 63627_1 | TATGAGTGTGGAGTCGTTGCTAGTCTTGTCCTTGCATTGCTCTNNNAGATCGGAAGA |
| 63629_1 | TATGAGTGTGGAGTCGTTGCCCCACTCCTGACTGACTTCAAAANNNAGATCGGAAGA |
| 63631_1 | TATGAGTGTGGAGTCGTTGCTCTCACTATAGCCTGGAACTCCTNNNAGATCGGAAGA |
| 63632_1 | TATGAGTGTGGAGTCGTTGCATATTGACACCTCGACTGTGGACNNNAGATCGGAAG |
| 63633_1 | TATGAGTGTGGAGTCGTTGCGGCTTACCATAGCATCTACCTCCNNNAGATCGGAAGA |
| 63635_1 | TATGAGTGTGGAGTCGTTGCTTACCCAGAAAGCATCATCCTGGNNNAGATCGGAAG |
| 63637_1 | TATGAGTGTGGAGTCGTTGCGAGAGAGACTCTGTTTGGGTGAANNNAGATCGGAA |
| 63638_1 | TATGAGTGTGGAGTCGTTGCACCAGCACAGGAGAAAGATGTAGNNNAGATCGGAA |
| 63640_1 | TATGAGTGTGGAGTCGTTGCTGTCCATATCCTACTGGTCCTGTNNNAGATCGGAAGA |
| 63642_1 | TATGAGTGTGGAGTCGTTGCGCTTAGGCAGGAAAATTGCTTGANNNAGATCGGAA |
| 63644_1 | TATGAGTGTGGAGTCGTTGCCCTCCACTTCAAGGTATAGCCAANNNAGATCGGAAG |
| 63646_1 | TATGAGTGTGGAGTCGTTGCGTGGCACAATTTCAGCTCACTAANNNAGATCGGAAG |
| 63647_1 | TATGAGTGTGGAGTCGTTGCTCTCATCATGCTTGGTATCTGCANNNAGATCGGAAG |
| 63649_1 | TATGAGTGTGGAGTCGTTGCAGAGCTGTCCTTTCCTTATCTCTNNNAGATCGGAAGA |
| 63651_1 | TATGAGTGTGGAGTCGTTGCAACAGAGTGTGGACTACAGAACCNNNAGATCGGAA |
| 63652_1 | TATGAGTGTGGAGTCGTTGCACAGCATCTGTAAGGAGGATTCCNNNAGATCGGAA |
| 63653_1 | TATGAGTGTGGAGTCGTTGCGTGCTTTGATTACGGGTATGAGCNNNAGATCGGAA |
| 63654_1 | TATGAGTGTGGAGTCGTTGCGGTCAGGGTAGTGAGGTGTTTACNNNAGATCGGAA |
| 63655_1 | TATGAGTGTGGAGTCGTTGCCTCCTGAGTAGCTGGGATTACAGNNNAGATCGGAA |
| 63657_1 | TATGAGTGTGGAGTCGTTGCGTGCCTGTAATCCCAGCATTTTGNNNAGATCGGAAG |
| 63659_1 | TATGAGTGTGGAGTCGTTGCAGAGCGAAAGAGAGAGAGAGAGANNNAGATCGGA |
| 63660_1 | TATGAGTGTGGAGTCGTTGCAACAATGGAATACCCTCCTCAGTNNNAGATCGGAAG |

| | |
|---|---|
| 63661_1 | TATGAGTGTGGAGTCGTTGCAAAAGGCAATCACAACATCTGGANNNAGATCGGAA |
| 63662_1 | TATGAGTGTGGAGTCGTTGCAACACAGGGAGAAGATAACCACCNNNAGATCGGAA |
| 63663_1 | TATGAGTGTGGAGTCGTTGCTGTTGGCATCAGAACAGGACATANNNAGATCGGAA |
| 63664_1 | TATGAGTGTGGAGTCGTTGCTTCCTTTGATTCTGACCACACCANNNAGATCGGAAGA |
| 63665_1 | TATGAGTGTGGAGTCGTTGCAGTAACTGGGATTACAGGCACACNNNAGATCGGAA |
| 63667_1 | TATGAGTGTGGAGTCGTTGCCAAGCCGTATGGTGTTTGAACTTNNNAGATCGGAAG |
| 63668_1 | TATGAGTGTGGAGTCGTTGCAAGTAGACTGCTCTGACATTCCANNNAGATCGGAAG |
| 63669_1 | TATGAGTGTGGAGTCGTTGCAGTAGCTAGGATTACAGGCATGCNNNAGATCGGAA |
| 63670_1 | TATGAGTGTGGAGTCGTTGCTATGGTGTTTGTGTGTGTGTGTGNNNAGATCGGAA |
| 63672_1 | TATGAGTGTGGAGTCGTTGCAAGACTAGAAAAAGCTGGCCAGANNNAGATCGGAA |
| 63674_1 | TATGAGTGTGGAGTCGTTGCGAAGTCAACTGAAAGAGCTTGCANNNAGATCGGAA |
| 63675_1 | TATGAGTGTGGAGTCGTTGCCCCTCACCAATAACGTCTCAAGANNNAGATCGGAAG |
| 63676_1 | TATGAGTGTGGAGTCGTTGCAGTGTTCTCTCTATGTTGCCCAGNNNAGATCGGAAG |
| 63678_1 | TATGAGTGTGGAGTCGTTGCCCAAGTGTTCGAGACTACAGTGANNNAGATCGGAA |
| 63679_1 | TATGAGTGTGGAGTCGTTGCAGATGGGAGAATCACTTCAACCCNNNAGATCGGAAG |
| 63680_1 | TATGAGTGTGGAGTCGTTGCCTCTGTTGGTGAGGTAGGATGTGNNNAGATCGGAA |
| 63681_1 | TATGAGTGTGGAGTCGTTGCCATCCCTGGTAACTGCCATGATANNNAGATCGGAAG |
| 63682_1 | TATGAGTGTGGAGTCGTTGCGTCTCCCTATATTGACCAGGCTGNNNAGATCGGAAG |
| 63684_1 | TATGAGTGTGGAGTCGTTGCCAGAAAGTCACTCTGCTACCACTNNNAGATCGGAAG |
| 63686_1 | TATGAGTGTGGAGTCGTTGCATCTCTTACCTCACCCATCACTGNNNAGATCGGAAGA |
| 63687_1 | TATGAGTGTGGAGTCGTTGCACAGAGTCTTGTTATGTTGCCCANNNAGATCGGAAG |
| 63688_1 | TATGAGTGTGGAGTCGTTGCCCAATGCAAGGCTACTCAATGAANNNAGATCGGAAG |
| 63690_1 | TATGAGTGTGGAGTCGTTGCTCTGGCGGATGAAAATAACCAGANNNAGATCGGAA |
| 63691_1 | TATGAGTGTGGAGTCGTTGCGAGACAAGGTCTTGCTATGTTGCNNNAGATCGGAAG |
| 63692_1 | TATGAGTGTGGAGTCGTTGCCAATCTTGGCTTACTGCAACCTCNNNAGATCGGAAGA |
| 63693_1 | TATGAGTGTGGAGTCGTTGCGGAAGTTCCTGAGACGTGTCATANNNAGATCGGAA |
| 63696_1 | TATGAGTGTGGAGTCGTTGCGCTTGCCTGTAATCCTAGCTACTNNNAGATCGGAAGA |
| 63698_1 | TATGAGTGTGGAGTCGTTGCCCCACAATGGAATACTGTTCAGCNNNAGATCGGAAG |
| 63699_1 | TATGAGTGTGGAGTCGTTGCCCCCCTTTGACCTTGTCTTCTACANNNAGATCGGAAGA |
| 63700_1 | TATGAGTGTGGAGTCGTTGCTACACCAAAACTTGACAAGTGGCNNNAGATCGGAAG |
| 63701_1 | TATGAGTGTGGAGTCGTTGCATGGGAAAGTGAGAGAAGCAGAGNNNAGATCGGA |
| 63703_1 | TATGAGTGTGGAGTCGTTGCATGAGATCTCATTCTGTTGCCCANNNAGATCGGAAG |
| 63704_1 | TATGAGTGTGGAGTCGTTGCGGGGTTCTGCTACTACAAACCAANNNAGATCGGAAG |
| 63705_1 | TATGAGTGTGGAGTCGTTGCTAGGTGTAAGGATCACAGCAGTGNNNAGATCGGAA |
| 63706_1 | TATGAGTGTGGAGTCGTTGCCTTTCTCCCTTTCTCCCTTCCTCNNNAGATCGGAAGAG |
| 63707_1 | TATGAGTGTGGAGTCGTTGCTTGCCTAGACTGGTCTCAAACTCNNNAGATCGGAAG |
| 63708_1 | TATGAGTGTGGAGTCGTTGCGTTCAAGAGATTCTCCTGCCTCANNNAGATCGGAAG |
| 63709_1 | TATGAGTGTGGAGTCGTTGCGAAGAAGGGAAGGAGGGAGAAAGNNNAGATCGG |
| 63710_1 | TATGAGTGTGGAGTCGTTGCAAGAGATTCTCCTACCTCAGCCTNNNAGATCGGAAG |
| 63711_1 | TATGAGTGTGGAGTCGTTGCAGGATCTTGTTATGTTGCCCAGGNNNAGATCGGAAG |
| 63713_1 | TATGAGTGTGGAGTCGTTGCTTCTGTGGCTAAGAGTCCTTTCANNNAGATCGGAAG |
| 63714_1 | TATGAGTGTGGAGTCGTTGCTCAAATGTAAAGCCCAGGAGTGANNNAGATCGGAA |
| 63715_1 | TATGAGTGTGGAGTCGTTGCTGTCATTGTGCTATCAAAGAGCANNNAGATCGGAAG |

| | |
|---|---|
| 63716_1 | TATGAGTGTGGAGTCGTTGCTATTCTTGCAAGGTTAGGAGCCANNNAGATCGGAAG |
| 63717_1 | TATGAGTGTGGAGTCGTTGCAGATGGAGTCTCACTCTGTCACTNNNAGATCGGAAG |
| 63718_1 | TATGAGTGTGGAGTCGTTGCGGCCAGCATTCCAACTTTAGAAANNNAGATCGGAAG |
| 63719_1 | TATGAGTGTGGAGTCGTTGCTGTTGGATCTGTCTCTCTGGAGANNNAGATCGGAAG |
| 63720_1 | TATGAGTGTGGAGTCGTTGCAATCACAGAACGGGCAAATCTTGNNNAGATCGGAAG |
| 63721_1 | TATGAGTGTGGAGTCGTTGCGCTGGGGTTTTGCTACATTTCTTNNNAGATCGGAAGA |
| 63722_1 | TATGAGTGTGGAGTCGTTGCTCAGGGACTGACTCTAACCTGAGNNNAGATCGGAAG |
| 63723_1 | TATGAGTGTGGAGTCGTTGCCTATCTCCTGACCTTGTGATCCGNNNAGATCGGAAGA |
| 63724_1 | TATGAGTGTGGAGTCGTTGCTTGTCAGGGTTTTAGGTTTTGCCNNNAGATCGGAAG |
| 63725_1 | TATGAGTGTGGAGTCGTTGCGTTCCACAGTAGGCAGAACTCTTNNNAGATCGGAAG |
| 63726_1 | TATGAGTGTGGAGTCGTTGCTGTTCGTTCCTCCCCATAATCAGNNNAGATCGGAAGA |
| 63727_1 | TATGAGTGTGGAGTCGTTGCGAGTAGCTGGGACTATAGCCATGNNNAGATCGGAAG |
| 63728_1 | TATGAGTGTGGAGTCGTTGCTGAAAACTCTCTCCACCTAAGGANNNAGATCGGAAGA |
| 63729_1 | TATGAGTGTGGAGTCGTTGCAAACAGAGCCTCTCCTCTCAAAGNNNAGATCGGAAGA |
| 63730_1 | TATGAGTGTGGAGTCGTTGCTCTGATAGGTTGCTCTCAAACCTNNNAGATCGGAAGA |
| 63731_1 | TATGAGTGTGGAGTCGTTGCCTGGACAACAGAGCAAGAATCTGNNNAGATCGGAAG |
| 63733_1 | TATGAGTGTGGAGTCGTTGCGCCAGGGAAGCTAGGAGATAAAGNNNAGATCGGAA |
| 63734_1 | TATGAGTGTGGAGTCGTTGCGACCAAGTCTTGCTTTGTTACCCNNNAGATCGGAAGA |
| 63735_1 | TATGAGTGTGGAGTCGTTGCAGAGTGCTTGACGTGGTTTATCTNNNAGATCGGAAG |
| 63736_1 | TATGAGTGTGGAGTCGTTGCACTCCTGACTTCAAGTGATCCACNNNAGATCGGAAGA |
| 63737_1 | TATGAGTGTGGAGTCGTTGCAGATGGAGTCTCACTTTGTCACCNNNAGATCGGAAG |
| 63738_1 | TATGAGTGTGGAGTCGTTGCGTAGTATGCACCTGTAGTCCCAGNNNAGATCGGAAG |
| 63739_1 | TATGAGTGTGGAGTCGTTGCATGATGGCATACACCTGTAGTCCNNNAGATCGGAAG |
| 63740_1 | TATGAGTGTGGAGTCGTTGCTTTCACAGTAACCTGGATGGAGGNNNAGATCGGAAG |
| 63742_1 | TATGAGTGTGGAGTCGTTGCAACCACAACCCTAATGCAAAACCNNNAGATCGGAAGA |
| 63743_1 | TATGAGTGTGGAGTCGTTGCCACACACACACACATGAGCTTTANNNAGATCGGAAGA |
| 63744_1 | TATGAGTGTGGAGTCGTTGCTTTCTCTCTTCCCTCACCACCTANNNAGATCGGAAGAG |
| 63745_1 | TATGAGTGTGGAGTCGTTGCGGGAAAGAATGCAAGAGGTGAACNNNAGATCGGAA |
| 63747_1 | TATGAGTGTGGAGTCGTTGCTCAGGGTAATGCTGGCTTCATAGNNNAGATCGGAAG |
| 63749_1 | TATGAGTGTGGAGTCGTTGCCCTCCCAGGTTCAAGCAATTCTCNNNAGATCGGAAGA |
| 63750_1 | TATGAGTGTGGAGTCGTTGCCTGGGTTGTAATGTTGGCAAGAANNNAGATCGGAAG |
| 63751_1 | TATGAGTGTGGAGTCGTTGCACTCCTGACCTCAAATGATCCACNNNAGATCGGAAGA |
| 63754_1 | TATGAGTGTGGAGTCGTTGCATCCTCCTGAGTAGCTGGGATTANNNAGATCGGAAG |
| 63756_1 | TATGAGTGTGGAGTCGTTGCGATCACCAAGAGATCCAGACCATNNNAGATCGGAAG |
| 63758_1 | TATGAGTGTGGAGTCGTTGCTCTCTCTCTCTCTCTCTCTCCCTNNNAGATCGGAAGAG |
| 63759_1 | TATGAGTGTGGAGTCGTTGCGCTTGGAATAACAGCACACACTTNNNAGATCGGAAG |
| 63760_1 | TATGAGTGTGGAGTCGTTGCACTTTCATTTGGAAGTGACAGCGNNNAGATCGGAAG |
| 63761_1 | TATGAGTGTGGAGTCGTTGCATCTTTCCTGAAGCAGAAGTCCCNNNAGATCGGAAGA |
| 63763_1 | TATGAGTGTGGAGTCGTTGCATGTTTAGAGGGGGGTAGGGTTANNNAGATCGGAA |
| 63764_1 | TATGAGTGTGGAGTCGTTGCTTTTTCCTGAGACGGAGTCTTGCNNNAGATCGGAAG |
| 63765_1 | TATGAGTGTGGAGTCGTTGCCTTTTGGGATTTGGGCATAACGTNNNAGATCGGAAG |
| 63767_1 | TATGAGTGTGGAGTCGTTGCGTGACGGAGTCTCACTTTGTCTCNNNAGATCGGAAG |
| 63768_1 | TATGAGTGTGGAGTCGTTGCGGTCAGACCTGAAGACAGCATAGNNNAGATCGGAAG |

| | |
|---|---|
| 63770_1 | TATGAGTGTGGAGTCGTTGCTCACAGACAGGATAGGTCAGTCTNNNAGATCGGAAG |
| 63772_1 | TATGAGTGTGGAGTCGTTGCATTGTTTTGTAGCTGGAGGCATGNNNAGATCGGAAG |
| 63773_1 | TATGAGTGTGGAGTCGTTGCCGCCTGTAATATCAGCACTTTGGNNNAGATCGGAAG |
| 63774_1 | TATGAGTGTGGAGTCGTTGCCCTGTAGTCCCAGTTACTAGGGANNNAGATCGGAAG |
| 63776_1 | TATGAGTGTGGAGTCGTTGCTGGATGGAATTGGAGACCAACATNNNAGATCGGAAG |
| 63778_1 | TATGAGTGTGGAGTCGTTGCTTTTGCTCTTGTTTTCCAGGCTGNNNAGATCGGAAGA |
| 63779_1 | TATGAGTGTGGAGTCGTTGCGAGTCCTCCTTTTCAACCGTTTGNNNAGATCGGAAGA |
| 63781_1 | TATGAGTGTGGAGTCGTTGCAAATTCAATCATGAGCAGCAGCANNNAGATCGGAAG |
| 63783_1 | TATGAGTGTGGAGTCGTTGCCCTGTAATCCCAGTGACACAGAANNNAGATCGGAAG |
| 63785_1 | TATGAGTGTGGAGTCGTTGCCTTGCCTGAAGTCTGAGATCCTTNNNAGATCGGAAGA |
| 63787_1 | TATGAGTGTGGAGTCGTTGCTCTGTCTCAGGTTCAAGCGATTCNNNAGATCGGAAGA |
| 63789_1 | TATGAGTGTGGAGTCGTTGCAACTCCTGAGCTCAAGCGATCTANNNAGATCGGAAG |
| 63790_1 | TATGAGTGTGGAGTCGTTGCAGTGTGTGTTTCTGTTTGTGGTGNNNAGATCGGAAG |
| 63791_1 | TATGAGTGTGGAGTCGTTGCTTCCAGTTATTCAGGAGGCTGAGNNNAGATCGGAAG |
| 63792_1 | TATGAGTGTGGAGTCGTTGCATTTGGGGGCATGATTTCATTCCNNNAGATCGGAAG |
| 63793_1 | TATGAGTGTGGAGTCGTTGCCACTCCTGTAATCCCAGCACTTTNNNAGATCGGAAGA |
| 63795_1 | TATGAGTGTGGAGTCGTTGCCAAATTCTCAAGAGGCTGAGCTGNNNAGATCGGAAG |
| 63797_1 | TATGAGTGTGGAGTCGTTGCTCAGAAAGACAGAGCGAGGAAAANNNAGATCGGAA |
| 63798_1 | TATGAGTGTGGAGTCGTTGCACAAATAAAGGTAGGGTCCCGAGNNNAGATCGGAA |
| 63799_1 | TATGAGTGTGGAGTCGTTGCGGTCTAGATTGCAACGTAGGTCANNNAGATCGGAAG |
| 63800_1 | TATGAGTGTGGAGTCGTTGCTTATCACGGCATTATCAGCAGGTNNNAGATCGGAAG |
| 63801_1 | TATGAGTGTGGAGTCGTTGCATCTCGGCTTACTATAACCGCTGNNNAGATCGGAAGA |
| 63802_1 | TATGAGTGTGGAGTCGTTGCGCCTAGGAGCAATAGCCTATACCNNNAGATCGGAAG |
| 63803_1 | TATGAGTGTGGAGTCGTTGCGCCCAAACTCAGAAACTCAGAACNNNAGATCGGAAG |
| 63804_1 | TATGAGTGTGGAGTCGTTGCCACACACACACACACGGTATAAGNNNAGATCGGAAG |
| 63805_1 | TATGAGTGTGGAGTCGTTGCAGATGCAGTGAAGGTGTCATCATNNNAGATCGGAAG |
| 63806_1 | TATGAGTGTGGAGTCGTTGCCAGGAGAATCACTTGAACCTGGANNNAGATCGGAAG |
| 63807_1 | TATGAGTGTGGAGTCGTTGCTAAGCTGGAACAAACTGCCTAGTNNNAGATCGGAAG |
| 63808_1 | TATGAGTGTGGAGTCGTTGCTTTGAGACAAGGTCACACTCTGTNNNAGATCGGAAG |
| 63811_1 | TATGAGTGTGGAGTCGTTGCACAGAGTCTCATTTTGTCGCTCANNNAGATCGGAAGA |
| 63812_1 | TATGAGTGTGGAGTCGTTGCCCATTCCTAAGTCCCCCATCTTCNNNAGATCGGAAGAG |
| 63813_1 | TATGAGTGTGGAGTCGTTGCCCACCTGGGTTTATTAGGCTCTTNNNAGATCGGAAGA |
| 63814_1 | TATGAGTGTGGAGTCGTTGCCTTCTCCTGGGTTCAAGCGATTCNNNAGATCGGAAGA |
| 63815_1 | TATGAGTGTGGAGTCGTTGCAATGAGGTCAGGAGATCGAAACCNNNAGATCGGAA |
| 63816_1 | TATGAGTGTGGAGTCGTTGCCAAATTCCCGAGTCCTTGGAGAGNNNAGATCGGAAG |
| 63818_1 | TATGAGTGTGGAGTCGTTGCTTCAAGTGATTCTCCCACCTCAGNNNAGATCGGAAGA |
| 63819_1 | TATGAGTGTGGAGTCGTTGCCTGTAATCCCAGCTACTCAGGAGNNNAGATCGGAAG |
| 63821_1 | TATGAGTGTGGAGTCGTTGCCCTGTACCAACTCAGCCTTCTTANNNAGATCGGAAGA |
| 63822_1 | TATGAGTGTGGAGTCGTTGCGAGTTATGCAGCTACAAGCGAAGNNNAGATCGGAAG |
| 63823_1 | TATGAGTGTGGAGTCGTTGCCCAGTAAATTCCATTCCCCCTCANNNAGATCGGAAGA |
| 63825_1 | TATGAGTGTGGAGTCGTTGCGGCACCTATAATCCCAGCTACTTNNNAGATCGGAAGA |
| 63826_1 | TATGAGTGTGGAGTCGTTGCTTTGTGGTGTCTGTTGAGTCTCANNNAGATCGGAAG |
| 63827_1 | TATGAGTGTGGAGTCGTTGCTCTCTCTCTCTCTCCCTCTCTNNNAGATCGGAAGAG |

| | |
|---|---|
| 63829_1 | TATGAGTGTGGAGTCGTTGCGGAAAGAACTTCAAGGGATTGGCNNNAGATCGGAA |
| 63830_1 | TATGAGTGTGGAGTCGTTGCAGGAGGAGGATGAGTAGCAGAAANNNAGATCGGA |
| 63831_1 | TATGAGTGTGGAGTCGTTGCCACGAGTAATCCCAGCTACTCAGNNNAGATCGGAAG |
| 63832_1 | TATGAGTGTGGAGTCGTTGCACATGACCTACAGAGGAACAAGGNNNAGATCGGAA |
| 63833_1 | TATGAGTGTGGAGTCGTTGCGCCAAGCAGCCATATTCTCATTANNNAGATCGGAAG |
| 63834_1 | TATGAGTGTGGAGTCGTTGCTCCCAAGTTCAAGCAATTCTCCTNNNAGATCGGAAG |
| 63835_1 | TATGAGTGTGGAGTCGTTGCCAGAAGTCTTAGCAGTGGGATGTNNNAGATCGGAA |
| 63836_1 | TATGAGTGTGGAGTCGTTGCCGTATACACACACACACACACACNNNAGATCGGAAG |
| 63837_1 | TATGAGTGTGGAGTCGTTGCTCTTTCCATTCAGTGTCTGTCCANNNAGATCGGAAGA |
| 63838_1 | TATGAGTGTGGAGTCGTTGCTTCCACGCAGTACTTTTTTTCCCNNNAGATCGGAAGA |
| 63839_1 | TATGAGTGTGGAGTCGTTGCAATTGAAAACCTTCTGGGAGGGANNNAGATCGGAA |
| 63840_1 | TATGAGTGTGGAGTCGTTGCGAGAGAAGATTTCACTGCATGGCNNNAGATCGGAA |
| 63841_1 | TATGAGTGTGGAGTCGTTGCTACTCACTCTGGAAAAGTCAGCCNNNAGATCGGAAG |
| 63843_1 | TATGAGTGTGGAGTCGTTGCCATACACACACACACACACACACNNNAGATCGGAAG |
| 63845_1 | TATGAGTGTGGAGTCGTTGCCACAGTTGTTTTTGGAGCTCTGTNNNAGATCGGAAG |
| 63846_1 | TATGAGTGTGGAGTCGTTGCTTGGGATTTGACATGGATGGCTANNNAGATCGGAA |
| 63847_1 | TATGAGTGTGGAGTCGTTGCAATGGTGCAGGAACTATCTTGGTNNNAGATCGGAA |
| 63849_1 | TATGAGTGTGGAGTCGTTGCGGTGAGGAATTGGAAGGGAAGTTNNNAGATCGGA |
| 63850_1 | TATGAGTGTGGAGTCGTTGCTTGTAATTCTTCTCTCCAGGGCCNNNAGATCGGAAG |
| 63852_1 | TATGAGTGTGGAGTCGTTGCTGTGTGTGTGTGTGTGTGTATAANNNAGATCGGAA |
| 63853_1 | TATGAGTGTGGAGTCGTTGCTGTTACTATATTGTCCCGGCTGGNNNAGATCGGAAG |
| 63854_1 | TATGAGTGTGGAGTCGTTGCAAATGTCAGCAGTGATTGTCTGGNNNAGATCGGAA |
| 63858_1 | TATGAGTGTGGAGTCGTTGCCTTCCTTCCTTCCTCCCTTTCTCNNNAGATCGGAAGAG |
| 63860_1 | TATGAGTGTGGAGTCGTTGCAAAAGTCAATGGATCCAGGAGCTNNNAGATCGGAA |
| 63861_1 | TATGAGTGTGGAGTCGTTGCGAAAGAGAAAGAGAAAGGCAGGCNNNAGATCGGA |
| 63862_1 | TATGAGTGTGGAGTCGTTGCGTTCAAGTGATTGTCCTGCCTCANNNAGATCGGAAG |
| 63863_1 | TATGAGTGTGGAGTCGTTGCACCTCCCATACTCAAGTGATCCTNNNAGATCGGAAG |
| 63864_1 | TATGAGTGTGGAGTCGTTGCGTGTTTGTGTGTGTGTGTGTGTANNNAGATCGGAA |
| 63865_1 | TATGAGTGTGGAGTCGTTGCAAATTGGCCAGCATGTTGATCTTNNNAGATCGGAAG |
| 63866_1 | TATGAGTGTGGAGTCGTTGCCGGGAGAGAGAGAGAGAGAGAAANNNAGATCGG |
| 63867_1 | TATGAGTGTGGAGTCGTTGCCTGTACCTGGGAAGAACAACACTNNNAGATCGGAA |
| 63868_1 | TATGAGTGTGGAGTCGTTGCTTCTTATCTCACTCTGTTGCCCANNNAGATCGGAAGA |
| 63870_1 | TATGAGTGTGGAGTCGTTGCGGGTAATATTTCCAGGGGTGTGTNNNAGATCGGAA |
| 63871_1 | TATGAGTGTGGAGTCGTTGCTTGACCTGGATTCTTCAACCTGTNNNAGATCGGAAG |
| 63872_1 | TATGAGTGTGGAGTCGTTGCTGAAGCAAAGAAAGAACTGTGCANNNAGATCGGAA |
| 63873_1 | TATGAGTGTGGAGTCGTTGCCATTCACCCTAGTATCCACAGCANNNAGATCGGAAG |
| 63874_1 | TATGAGTGTGGAGTCGTTGCAGCTTTGCCCACATGTTTACATCNNNAGATCGGAAG |
| 63876_1 | TATGAGTGTGGAGTCGTTGCCTCCCCCATTTCTCTCTCTGTTCNNNAGATCGGAAGA |
| 63878_1 | TATGAGTGTGGAGTCGTTGCAGAGAGAGAGAGAGAGAGAAAGAAANNNAGATCG |
| 63879_1 | TATGAGTGTGGAGTCGTTGCGAGCTCTCTGGTGTCTCTTCTTANNNAGATCGGAAG |
| 63880_1 | TATGAGTGTGGAGTCGTTGCCCATTGCAGTGACACTGAATCAANNNAGATCGGAAG |
| 63881_1 | TATGAGTGTGGAGTCGTTGCAAAAGCTGGAATGGATGACTTGGNNNAGATCGGAA |
| 63882_1 | TATGAGTGTGGAGTCGTTGCTTTTGTGCTAGGAGTCTCGAGTCNNNAGATCGGAAG |

| | |
|---|---|
| 63883_1 | TATGAGTGTGGAGTCGTTGCTGAAGGAGGAAGGAAAGAAGGTGNNNAGATCGGA |
| 63884_1 | TATGAGTGTGGAGTCGTTGCGTGGGTAGGTACCAGACATTTGTNNNAGATCGGAA |
| 63887_1 | TATGAGTGTGGAGTCGTTGCTATAAACTGCTCAGCCCTCACTCNNNAGATCGGAAGA |
| 63888_1 | TATGAGTGTGGAGTCGTTGCCAGCCACCATACCCAGCTAATTNNNAGATCGGAAGA |
| 63890_1 | TATGAGTGTGGAGTCGTTGCAGGAGAAGCTTTGCCTAATCACANNNAGATCGGAAG |
| 63891_1 | TATGAGTGTGGAGTCGTTGCTCTCATCACTCACTCTGGAGGAANNNAGATCGGAAG |
| 63892_1 | TATGAGTGTGGAGTCGTTGCTCTGTTTTCCTAGGGCTAAGTGCNNNAGATCGGAAG |
| 63893_1 | TATGAGTGTGGAGTCGTTGCGGGTTGCCACAAACTTTCAATTTNNNAGATCGGAAG |
| 63894_1 | TATGAGTGTGGAGTCGTTGCCCTTTGTCTATGTGGGCTCCATANNNAGATCGGAAG |
| 63895_1 | TATGAGTGTGGAGTCGTTGCTAAATTATCTGGGTGTGGTGGCANNNAGATCGGAA |
| 63896_1 | TATGAGTGTGGAGTCGTTGCTTCAGGCTTTCAACTGAGACACTNNNAGATCGGAAG |
| 63897_1 | TATGAGTGTGGAGTCGTTGCCTCTCTGTCTGTCTCTGTCCGTCNNNAGATCGGAAGA |
| 63898_1 | TATGAGTGTGGAGTCGTTGCCTCTCTGTCTGTCTCTGTCCGTCNNNAGATCGGAAGA |
| 63899_1 | TATGAGTGTGGAGTCGTTGCAAAGAGGGAAAGAGAGAAGTGGCNNNAGATCGGA |
| 63900_1 | TATGAGTGTGGAGTCGTTGCGGATCACTTGAGGTCAGGAGTTTNNNAGATCGGAA |
| 63903_1 | TATGAGTGTGGAGTCGTTGCACTGATGCAAAAACCCTCAACAANNNAGATCGGAAG |
| 63904_1 | TATGAGTGTGGAGTCGTTGCCCAGTAGCCCAGGATACTTTCAANNNAGATCGGAAG |
| 63905_1 | TATGAGTGTGGAGTCGTTGCAAAATGTAGACCTGTGCAGTTGCNNNAGATCGGAA |
| 63906_1 | TATGAGTGTGGAGTCGTTGCAGCTAGTTCTTTTGCCAGTTCCTNNNAGATCGGAAGA |
| 63907_1 | TATGAGTGTGGAGTCGTTGCGCCTAGGAAAGATCTAGCAGTCGNNNAGATCGGAA |
| 63908_1 | TATGAGTGTGGAGTCGTTGCACCAGAATCTGACCATGCAAGATNNNAGATCGGAAG |
| 63909_1 | TATGAGTGTGGAGTCGTTGCGCAGAGACAGATGGCAGAGTTATNNNAGATCGGAA |
| 63910_1 | TATGAGTGTGGAGTCGTTGCTTGTCTCTCCGTGTCATGATTGTNNNAGATCGGAAG |
| 63911_1 | TATGAGTGTGGAGTCGTTGCTTTGTTAGGTCTCAGCTCTCCACNNNAGATCGGAAGA |
| 63913_1 | TATGAGTGTGGAGTCGTTGCCTGTAGTCCCAGCTACTCAGGAGNNNAGATCGGAAG |
| 63914_1 | TATGAGTGTGGAGTCGTTGCTCTGTTACAGCACGGTTTACAGANNNAGATCGGAAG |
| 63915_1 | TATGAGTGTGGAGTCGTTGCGATTCAAGCAATTCTCCTGCCTCNNNAGATCGGAAGA |
| 63916_1 | TATGAGTGTGGAGTCGTTGCACTGCCAAACTATGTTCCAGAGTNNNAGATCGGAAG |
| 63917_1 | TATGAGTGTGGAGTCGTTGCGATTACAGGCATGCACTACCATGNNNAGATCGGAAG |
| 63918_1 | TATGAGTGTGGAGTCGTTGCTTATAAGGGTGCAGTGATTGGCTNNNAGATCGGAA |
| 63919_1 | TATGAGTGTGGAGTCGTTGCTCTAGATGTGTTGCTTAGGCTGGNNNAGATCGGAA |
| 63920_1 | TATGAGTGTGGAGTCGTTGCGGTTTCTTAGGTTCCTCTGGGAGNNNAGATCGGAA |
| 63921_1 | TATGAGTGTGGAGTCGTTGCCAAACTCCTGAGCTCAAGCAATCNNNAGATCGGAAG |
| 63922_1 | TATGAGTGTGGAGTCGTTGCCCACCACACCCAGCTAATTTTTTNNNAGATCGGAAGA |
| 63923_1 | TATGAGTGTGGAGTCGTTGCGCTCCTGGGTTCAAACAATTGTTNNNAGATCGGAAG |
| 63924_1 | TATGAGTGTGGAGTCGTTGCAAGGCAGGAGAATCACTTGAACTNNNAGATCGGAA |
| 63925_1 | TATGAGTGTGGAGTCGTTGCAACTGGCCAAGGTGCTAAGTTTCNNNAGATCGGAA |
| 63926_1 | TATGAGTGTGGAGTCGTTGCCTAGAGTGTAGTGGCACGATCTCNNNAGATCGGAA |
| 63927_1 | TATGAGTGTGGAGTCGTTGCAGTACTGGGGACTACAGGTGTATNNNAGATCGGAA |
| 63928_1 | TATGAGTGTGGAGTCGTTGCTTCCAACAGTCACAGTTTTCAGCNNNAGATCGGAAG |
| 63929_1 | TATGAGTGTGGAGTCGTTGCCTCACTTGCTCTATTCTTCCCCANNNAGATCGGAAGA |
| 63930_1 | TATGAGTGTGGAGTCGTTGCCCCAGGTTCAAGAAATTCTCTGCNNNAGATCGGAAG |
| 63933_1 | TATGAGTGTGGAGTCGTTGCGAGATTTGTTGGAATGCCTAGGGNNNAGATCGGAA |

| | |
|---|---|
| 63936_1 | TATGAGTGTGGAGTCGTTGCCCAAGTAGTTGGGATTACAGGCANNNAGATCGGAA |
| 63937_1 | TATGAGTGTGGAGTCGTTGCGCCAGATTTTGTCCTGATACAACCNNNAGATCGGAA |
| 63939_1 | TATGAGTGTGGAGTCGTTGCGAAGGCCAATTCTGCTTTTGGATNNNAGATCGGAAG |
| 63940_1 | TATGAGTGTGGAGTCGTTGCATGCTGAAATCTGGCAGTTAGGTNNNAGATCGGAA |
| 63942_1 | TATGAGTGTGGAGTCGTTGCTCAGCCTCCCAAGTAACTAGGATNNNAGATCGGAAG |
| 63943_1 | TATGAGTGTGGAGTCGTTGCTCTTTTTCCTGTCTACCTCCCAGNNNAGATCGGAAGA |
| 63944_1 | TATGAGTGTGGAGTCGTTGCGCGATAGTCGATGAATGGCAGAANNNAGATCGGAA |
| 63946_1 | TATGAGTGTGGAGTCGTTGCTACACACTCAAAATCGTAGGCCANNNAGATCGGAAG |
| 63948_1 | TATGAGTGTGGAGTCGTTGCATGATGGACAGTTCACAGGGTACNNNAGATCGGAA |
| 63949_1 | TATGAGTGTGGAGTCGTTGCCGACACATACGCACAAAAGAAATNNNAGATCGGAAG |
| 63950_1 | TATGAGTGTGGAGTCGTTGCCCAGGTGTGATCTCAGCTACTTGNNNAGATCGGAAG |
| 63951_1 | TATGAGTGTGGAGTCGTTGCTATAGGAAAGCCTGATGCAGCAANNNAGATCGGAA |
| 63952_1 | TATGAGTGTGGAGTCGTTGCACCAGGAGAGGGACTTGAAAATCNNNAGATCGGAA |
| 63953_1 | TATGAGTGTGGAGTCGTTGCGTATCTTGCAGCAACTTGGATGGNNNAGATCGGAA |
| 63954_1 | TATGAGTGTGGAGTCGTTGCTAAAAATACCAGGCCCACAAGGANNNAGATCGGAA |
| 63955_1 | TATGAGTGTGGAGTCGTTGCTGCTCCCCAAGACTCATTTTCTTNNNAGATCGGAAGA |
| 63957_1 | TATGAGTGTGGAGTCGTTGCCTGGGTTCAAATGATTCTCCTGCNNNAGATCGGAAG |
| 63960_1 | TATGAGTGTGGAGTCGTTGCAAAAATGGAAAGCAACCTCAGCANNNAGATCGGAA |
| 63961_1 | TATGAGTGTGGAGTCGTTGCCTGTAGTCCCAGCTACTCAGGAGNNNAGATCGGAAG |
| 63963_1 | TATGAGTGTGGAGTCGTTGCCGGAAAAACAGTGAGACCAGATGNNNAGATCGGAA |
| 63964_1 | TATGAGTGTGGAGTCGTTGCAGTTGGGAGGATAACTTGAACCCNNNAGATCGGAA |
| 63966_1 | TATGAGTGTGGAGTCGTTGCGAATAGGGTTGAAGGCTCTTGGGNNNAGATCGGAA |
| 63968_1 | TATGAGTGTGGAGTCGTTGCCAGGCCATACTCCAGACCAATAANNNAGATCGGAAG |
| 63969_1 | TATGAGTGTGGAGTCGTTGCCTGTAGTCCCAGCTACTTGAGAGNNNAGATCGGAAG |
| 63970_1 | TATGAGTGTGGAGTCGTTGCCTCCCTTTGCCTCTTTTGACATGNNNAGATCGGAAGA |
| 63971_1 | TATGAGTGTGGAGTCGTTGCGGAGGCATAGGGGAATATGTTGTNNNAGATCGGAA |
| 63972_1 | TATGAGTGTGGAGTCGTTGCGGGTTGCAATCCTAGTCTCTGATNNNAGATCGGAAG |
| 63974_1 | TATGAGTGTGGAGTCGTTGCTGTTACGTGACTTGCTATTACTGANNNAGATCGGAA |
| 63975_1 | TATGAGTGTGGAGTCGTTGCAACTCCAAGACACAAGGCAGTAANNNAGATCGGAA |
| 63978_1 | TATGAGTGTGGAGTCGTTGCTTTCCTCAAACCAATTGCAACCCNNNAGATCGGAAGA |
| 63979_1 | TATGAGTGTGGAGTCGTTGCGGATCATCTGAGGTCAGGAGTTCNNNAGATCGGAA |
| 63982_1 | TATGAGTGTGGAGTCGTTGCACTTCAAAGCCCTGTGCATTTTTNNNAGATCGGAAG |
| 63985_1 | TATGAGTGTGGAGTCGTTGCGGCTGACATATTGGACATTGAGCNNNAGATCGGAA |
| 63987_1 | TATGAGTGTGGAGTCGTTGCCTGGACTTCAAAATGCATCTGGGNNNAGATCGGAA |
| 63991_1 | TATGAGTGTGGAGTCGTTGCGCCTCCTGAGTAGCTGAGATTACNNNAGATCGGAAG |
| 63992_1 | TATGAGTGTGGAGTCGTTGCGACTCTCCCAGGCTTAGATGATCNNNAGATCGGAAG |
| 63993_1 | TATGAGTGTGGAGTCGTTGCAGATGAAGTCTTGCTTTGTTGCCNNNAGATCGGAAG |
| 63994_1 | TATGAGTGTGGAGTCGTTGCCTCCCAGGTTCAAGTGATTCTCANNNAGATCGGAAG |
| 63995_1 | TATGAGTGTGGAGTCGTTGCCCAAGTAGCTGGGATTACAGGTGNNNAGATCGGAA |
| 63996_1 | TATGAGTGTGGAGTCGTTGCTTCAAGCGATTATCCTGCCTCAGNNNAGATCGGAAG |
| 63998_1 | TATGAGTGTGGAGTCGTTGCTAGATAGCCCTTTCCCAGCTTTGNNNAGATCGGAAG |
| 63999_1 | TATGAGTGTGGAGTCGTTGCCCCAAGTTGAAGCAATTCTCCTGNNNAGATCGGAAG |
| 64001_1 | TATGAGTGTGGAGTCGTTGCCAAACACATTGCAAACCCCTAAANNNAGATCGGAAG |

| | |
|---|---|
| 64002_1 | TATGAGTGTGGAGTCGTTGCGCCTCCTGAGTAGTTGGGATTACNNNAGATCGGAAG |
| 64003_1 | TATGAGTGTGGAGTCGTTGCCAATCTCGGCTCTCTACAAGCTCNNNAGATCGGAAGA |
| 64004_1 | TATGAGTGTGGAGTCGTTGCCTCCTGAGTAGCTGTGACTACAGNNNAGATCGGAAG |
| 64005_1 | TATGAGTGTGGAGTCGTTGCAATTTATTCTTGGCCAGACACGGNNNAGATCGGAAG |
| 64007_1 | TATGAGTGTGGAGTCGTTGCCTCCTGAGTAGCTGGGACTACAGNNNAGATCGGAAG |
| 64009_1 | TATGAGTGTGGAGTCGTTGCTTGAGATGGGGTTTCATCCTGTTNNNAGATCGGAAG |
| 64011_1 | TATGAGTGTGGAGTCGTTGCCTAGGACTACAGGCACACACTACNNNAGATCGGAAG |
| 64012_1 | TATGAGTGTGGAGTCGTTGCCTGGAGTGTAGTGGCATGATCTTNNNAGATCGGAA |
| 64015_1 | TATGAGTGTGGAGTCGTTGCCTCTCAAGTAGCTGGGACTACAGNNNAGATCGGAAG |
| 64016_1 | TATGAGTGTGGAGTCGTTGCCTCCTGAGTAGCTGGGACTACAGNNNAGATCGGAA |
| 64018_1 | TATGAGTGTGGAGTCGTTGCGTTCAAGCAATTCTCTGCCTCAGNNNAGATCGGAAG |
| 64019_1 | TATGAGTGTGGAGTCGTTGCCAGTGGTACAATCTCGGATCACTNNNAGATCGGAAG |
| 64020_1 | TATGAGTGTGGAGTCGTTGCGTTCAAGAGATTCTCCTGCCTCANNNAGATCGGAAG |
| 64021_1 | TATGAGTGTGGAGTCGTTGCCTCAGTTGATCCTTCCACCTCAGNNNAGATCGGAAGA |
| 64022_1 | TATGAGTGTGGAGTCGTTGCCCTCCCGGATTCAAGTGATTCTTNNNAGATCGGAAG |
| 64023_1 | TATGAGTGTGGAGTCGTTGCGGGGACAAAATCTCCAAGGCATANNNAGATCGGAA |
| 64024_1 | TATGAGTGTGGAGTCGTTGCTAGATGGGTGCATGATTTGTCCTNNNAGATCGGAAG |
| 64025_1 | TATGAGTGTGGAGTCGTTGCCCAGCACATAGTAGGGACTCAAANNNAGATCGGAAG |
| 64026_1 | TATGAGTGTGGAGTCGTTGCAGGAGAAATGTCTTAGGGGCAAGNNNAGATCGGAA |
| 64027_1 | TATGAGTGTGGAGTCGTTGCGTCCTTCCAAACTGCTGGGATTANNNAGATCGGAAG |
| 64028_1 | TATGAGTGTGGAGTCGTTGCGAGCTCATTGCAATCCTCGATTCNNNAGATCGGAAG |
| 64030_1 | TATGAGTGTGGAGTCGTTGCTTTTATTCAGCAGCTCAGGTCCTNNNAGATCGGAAG |
| 64031_1 | TATGAGTGTGGAGTCGTTGCTAAAACAGCAAAAGCCCCAAACANNNAGATCGGAAG |
| 64032_1 | TATGAGTGTGGAGTCGTTGCCGGCCAAGAAATCTTTGTAGCTTNNNAGATCGGAAG |
| 64033_1 | TATGAGTGTGGAGTCGTTGCTGATGACAAACGTGAGTCACTCTNNNAGATCGGAAG |
| 64034_1 | TATGAGTGTGGAGTCGTTGCCGGTGAAAGATGAGCCTTGTTTTNNNAGATCGGAAG |
| 64035_1 | TATGAGTGTGGAGTCGTTGCCTGAAATTGGGCCAGGTTGAAATNNNAGATCGGAA |
| 64036_1 | TATGAGTGTGGAGTCGTTGCTCCACTCAAGCCCAAAAATTCACNNNAGATCGGAAG |
| 64037_1 | TATGAGTGTGGAGTCGTTGCGGATTGTATATGGTGTGCTTGGGNNNAGATCGGAA |
| 64039_1 | TATGAGTGTGGAGTCGTTGCGACATTCCAGGTAGCAGAGACAANNNAGATCGGAA |
| 64040_1 | TATGAGTGTGGAGTCGTTGCGCACCTGTCCTTGTTTATGCTTANNNAGATCGGAAGA |
| 64041_1 | TATGAGTGTGGAGTCGTTGCGAAAAGAGCACTCACCAGAAACCNNNAGATCGGAAG |
| 64042_1 | TATGAGTGTGGAGTCGTTGCGGATTAGCTCTCGGATTAGCCAANNNAGATCGGAAG |
| 64043_1 | TATGAGTGTGGAGTCGTTGCTTGAGGGAGATATTTTGTGGCCANNNAGATCGGAA |
| 64044_1 | TATGAGTGTGGAGTCGTTGCCAGATGGTTTTGCTTCCATTGGTNNNAGATCGGAAG |
| 64045_1 | TATGAGTGTGGAGTCGTTGCCATGATCTGCCATCTGAAAGCTGNNNAGATCGGAAG |
| 64046_1 | TATGAGTGTGGAGTCGTTGCCACACACACACACACACATACATNNNAGATCGGAAGA |
| 64047_1 | TATGAGTGTGGAGTCGTTGCTGCTTCTTGATCTGTGATGGTTTNNNAGATCGGAAG |
| 64048_1 | TATGAGTGTGGAGTCGTTGCTTTTGGTATGCTCACACCTCAGTNNNAGATCGGAAG |
| 64049_1 | TATGAGTGTGGAGTCGTTGCGGAGATTCTCAGGGTGGGAATTTNNNAGATCGGAA |
| 64050_1 | TATGAGTGTGGAGTCGTTGCGACACACATATGCATACACAGCCNNNAGATCGGAAG |
| 64051_1 | TATGAGTGTGGAGTCGTTGCAGTCGTGTCTGTCTCTCACTAGANNNAGATCGGAAG |
| 64052_1 | TATGAGTGTGGAGTCGTTGCCCTTGTCTCAGGGAACATAACCANNNAGATCGGAAG |

64053_1 TATGAGTGTGGAGTCGTTGCACAGATATTGTGGTTGTTTGGGTNNNAGATCGGAA
64054_1 TATGAGTGTGGAGTCGTTGCATGTGGGACTCTTCTCAATTGCTNNNAGATCGGAAG
64055_1 TATGAGTGTGGAGTCGTTGCATGGGAGACTGGAACATCATGTCNNNAGATCGGAA
64056_1 TATGAGTGTGGAGTCGTTGCTAGCTTGCATCTTCTGTGTCCTTNNNAGATCGGAAGA
64057_1 TATGAGTGTGGAGTCGTTGCCACAGTTGGTTAGCAAATCCCTGNNNAGATCGGAAG
64058_1 TATGAGTGTGGAGTCGTTGCATACAGACTCTCCCGACAGCTATNNNAGATCGGAAG
64059_1 TATGAGTGTGGAGTCGTTGCGGTGGGAGACAGTAAAAGGAGAGNNNAGATCGGA
64060_1 TATGAGTGTGGAGTCGTTGCCTAGCAGAAATGTGACTGGCAAGNNNAGATCGGAA
64061_1 TATGAGTGTGGAGTCGTTGCAAAGTTCAGTAGCTGTTGGGTGANNNAGATCGGAA
64062_1 TATGAGTGTGGAGTCGTTGCAGAAATCTCAGCTTCTCACCTGGNNNAGATCGGAAG
64064_1 TATGAGTGTGGAGTCGTTGCTCCTGTGTTTCCTGAAAATTGGTNNNAGATCGGAAG
64065_1 TATGAGTGTGGAGTCGTTGCCTGCCAATGGTCAAAGCTAGAACNNNAGATCGGAAG
64066_1 TATGAGTGTGGAGTCGTTGCCCCCAGAGTCACACTTTCACTAANNNAGATCGGAAGA
64067_1 TATGAGTGTGGAGTCGTTGCAAGGGCTAACACCTAGATGCAAANNNAGATCGGAA
64069_1 TATGAGTGTGGAGTCGTTGCTCAGTCAACAGACCCTCCTATCTNNNAGATCGGAAGA
64070_1 TATGAGTGTGGAGTCGTTGCTCATGGTCTTATGGGACAGGTTCNNNAGATCGGAAG
64071_1 TATGAGTGTGGAGTCGTTGCTCCATGACCCTGATTCACATGTTNNNAGATCGGAAGA
64072_1 TATGAGTGTGGAGTCGTTGCATCCTGCCTATACAAACACAGCANNNAGATCGGAAG
64073_1 TATGAGTGTGGAGTCGTTGCTAAAAGGGTCCGAGTTCTTGAGGNNNAGATCGGAA
64074_1 TATGAGTGTGGAGTCGTTGCTTACCAACATTTCATCGGTGGGANNNAGATCGGAAG
64075_1 TATGAGTGTGGAGTCGTTGCAAAGTCAGCCCTAGTCCTCTTTGNNNAGATCGGAAG
64076_1 TATGAGTGTGGAGTCGTTGCAATGAGTTCCTGGCAGATGATGTNNNAGATCGGAA
64077_1 TATGAGTGTGGAGTCGTTGCCAATTGTTGGGATACTCAGCTGGNNNAGATCGGAAG
64078_1 TATGAGTGTGGAGTCGTTGCATTATTGGACCCCTTTCACTGGGNNNAGATCGGAAG
64079_1 TATGAGTGTGGAGTCGTTGCTTTCTCTACAGAGGAATGGGCAGNNNAGATCGGAA
64081_1 TATGAGTGTGGAGTCGTTGCAGAGAGAGAGAGAGACCTTCACCNNNAGATCGGAA
64083_1 TATGAGTGTGGAGTCGTTGCTAGACTCATATAACCCCCAGCCANNNAGATCGGAAG
64084_1 TATGAGTGTGGAGTCGTTGCACCAGAACAGCACCTTATGGAATNNNAGATCGGAAG
64085_1 TATGAGTGTGGAGTCGTTGCGACGGAAGAGCTGTCAGTCATAANNNAGATCGGAA
64086_1 TATGAGTGTGGAGTCGTTGCAGCTCCAACTGCTTCTAAGTTTTNNNAGATCGGAAGA
64087_1 TATGAGTGTGGAGTCGTTGCTTGACCAGCCACTAGTGACTTTTNNNAGATCGGAAG
64088_1 TATGAGTGTGGAGTCGTTGCAACGGAGGCATGTCTATAAAGGGNNNAGATCGGAA
64089_1 TATGAGTGTGGAGTCGTTGCCCTCAGACCAAAAGTGGGTATGANNNAGATCGGAA
64090_1 TATGAGTGTGGAGTCGTTGCCACCTCTGGTTTCTAGACCTCACNNNAGATCGGAAGA
64091_1 TATGAGTGTGGAGTCGTTGCTGGACTGACTCATACACTTTGCANNNAGATCGGAAG
64092_1 TATGAGTGTGGAGTCGTTGCAGGACTATTTGTGTTCCAGCAGTNNNAGATCGGAAG
64093_1 TATGAGTGTGGAGTCGTTGCTCTAAGTGGTGAGACACTGATGCNNNAGATCGGAA
64094_1 TATGAGTGTGGAGTCGTTGCTTGTGGTAGGCAGCTTCTAAGAGNNNAGATCGGAA
64096_1 TATGAGTGTGGAGTCGTTGCGAACAACAGAAACTGACTCTGGCNNNAGATCGGAAG
64097_1 TATGAGTGTGGAGTCGTTGCTGTGTTTTCCAGCCAGAAAGTTGNNNAGATCGGAAG
64098_1 TATGAGTGTGGAGTCGTTGCACAGTGACTCACGCCTATAATCCNNNAGATCGGAAG
64099_1 TATGAGTGTGGAGTCGTTGCAGAAAAAGGCCTCCAGTTAGAGGNNNAGATCGGAA
64100_1 TATGAGTGTGGAGTCGTTGCTGGCTTGATTGTTTTGTAGGTCANNNAGATCGGAAG

| | |
|---|---|
| 64102_1 | TATGAGTGTGGAGTCGTTGCGGAGACATATCAAAGCTGGGTCANNNAGATCGGAA |
| 64103_1 | TATGAGTGTGGAGTCGTTGCCCCAGTAACTCAGATCCAATGCTNNNAGATCGGAAG |
| 64104_1 | TATGAGTGTGGAGTCGTTGCTGCCATGTATGCACATAAGGTTTNNNAGATCGGAAG |
| 64105_1 | TATGAGTGTGGAGTCGTTGCTTCAAGCTATTCTCCTGCCTCAGNNNAGATCGGAAG |
| 64106_1 | TATGAGTGTGGAGTCGTTGCTCTCTTGTGTCTCTCTCATCCCTNNNAGATCGGAAGA |
| 64107_1 | TATGAGTGTGGAGTCGTTGCTGGATGTTGCAAATGAGATGTGGNNNAGATCGGAA |
| 64108_1 | TATGAGTGTGGAGTCGTTGCTGGAACTGTACTTTTTGGCCACNNNAGATCGGAAG |
| 64109_1 | TATGAGTGTGGAGTCGTTGCAAATCTCTGGGGCTCTCCTTTTTNNNAGATCGGAAG |
| 64110_1 | TATGAGTGTGGAGTCGTTGCCTTTCACAACACTCCCTACTCCANNNAGATCGGAAGA |
| 64111_1 | TATGAGTGTGGAGTCGTTGCTGTCTGGTGGAGAAACTAGCTTCNNNAGATCGGAA |
| 64112_1 | TATGAGTGTGGAGTCGTTGCGCCATGTGGATATCTTACTGGGTNNNAGATCGGAA |
| 64114_1 | TATGAGTGTGGAGTCGTTGCACACAGAGAGAGAGAGAGAGAGANNNAGATCGGA |
| 64115_1 | TATGAGTGTGGAGTCGTTGCACTGATTGGATGAGGTTCACACANNNAGATCGGAA |
| 64117_1 | TATGAGTGTGGAGTCGTTGCCACATTTTGTTCCCTCTTGTCCCNNNAGATCGGAAGA |
| 64119_1 | TATGAGTGTGGAGTCGTTGCCCAGGACTAGGGAGAACAGAAACNNNAGATCGGAA |
| 64120_1 | TATGAGTGTGGAGTCGTTGCAACCTGGCAGTACAATGTCAGATNNNAGATCGGAA |
| 64121_1 | TATGAGTGTGGAGTCGTTGCATGCCAGAAAGTGTCTTACTGGTNNNAGATCGGAAG |
| 64122_1 | TATGAGTGTGGAGTCGTTGCTTTCCAGGGGTCAGGAAGTAAAGNNNAGATCGGAA |
| 64125_1 | TATGAGTGTGGAGTCGTTGCCATCCCGTGAGTTAGAAGTCCATNNNAGATCGGAAG |
| 64126_1 | TATGAGTGTGGAGTCGTTGCAGCCATTTTATACTCACAGGCCANNNAGATCGGAAG |
| 64127_1 | TATGAGTGTGGAGTCGTTGCCCTAGAAAAGCCCTTGTATCCGTNNNAGATCGGAAG |
| 64128_1 | TATGAGTGTGGAGTCGTTGCCAGAATCTTTCAGGCATGACCACNNNAGATCGGAAG |
| 64129_1 | TATGAGTGTGGAGTCGTTGCTAGAGAAGCTTCCACATAGCCACNNNAGATCGGAAG |
| 64130_1 | TATGAGTGTGGAGTCGTTGCGACTCTGGTCTGGGAAAGGATTTNNNAGATCGGAA |
| 64131_1 | TATGAGTGTGGAGTCGTTGCACTCAAAGTGTGTGTTTTTGGGGNNNAGATCGGAA |
| 64132_1 | TATGAGTGTGGAGTCGTTGCCTCATGGCCTCTTCATGTGGTTANNNAGATCGGAAG |
| 64134_1 | TATGAGTGTGGAGTCGTTGCGGTTTCAGTAACTTGCGGTCATCNNNAGATCGGAAG |
| 64135_1 | TATGAGTGTGGAGTCGTTGCTCTCTCCATCCCCTTAGTCTTGTNNNAGATCGGAAGA |
| 64136_1 | TATGAGTGTGGAGTCGTTGCATTTCCTAGCTCTGTCTGCTGAGNNNAGATCGGAAG |
| 64137_1 | TATGAGTGTGGAGTCGTTGCAGTGGCATGATCTCAGCTAACAANNNAGATCGGAA |
| 64138_1 | TATGAGTGTGGAGTCGTTGCACAAGAAACATCCTGCAATGGTGNNNAGATCGGAA |
| 64139_1 | TATGAGTGTGGAGTCGTTGCTAGGAACACAGGGGGAAATCTTGNNNAGATCGGAA |
| 64140_1 | TATGAGTGTGGAGTCGTTGCCTTACAACACTGGCTGCCTTTTTNNNAGATCGGAAGA |
| 64142_1 | TATGAGTGTGGAGTCGTTGCCTCTACATGGGTATATGGCCACCNNNAGATCGGAAG |
| 64144_1 | TATGAGTGTGGAGTCGTTGCTTTTAGCCATGTGATTCGCGTTTNNNAGATCGGAAG |
| 64145_1 | TATGAGTGTGGAGTCGTTGCGTGTGTGTGAACCTGGAAATCAGNNNAGATCGGAA |
| 64146_1 | TATGAGTGTGGAGTCGTTGCGGGAGGGAAGGAAGAGAAGAATCNNNAGATCGGA |
| 64147_1 | TATGAGTGTGGAGTCGTTGCAACTCTGTGGCATGGACTGAATANNNAGATCGGAA |
| 64148_1 | TATGAGTGTGGAGTCGTTGCGGGAATGTGGGGGAGACATAATANNNAGATCGGA |
| 64149_1 | TATGAGTGTGGAGTCGTTGCTACAAGTTTTGCGATGTTTGCCANNNAGATCGGAAG |
| 64150_1 | TATGAGTGTGGAGTCGTTGCTTCTCTCCAGATTCACACACCAGNNNAGATCGGAAGA |
| 64151_1 | TATGAGTGTGGAGTCGTTGCATACTCCCCTGCTAGACTTGCTANNNAGATCGGAAGA |
| 64152_1 | TATGAGTGTGGAGTCGTTGCATGCAGGAAGACTCAACCTTTCTNNNAGATCGGAAG |

| | |
|---|---|
| 64153_1 | TATGAGTGTGGAGTCGTTGCGACAAAGACAGAGACAGACTCCCNNNAGATCGGAA |
| 64154_1 | TATGAGTGTGGAGTCGTTGCTCCGCTACTATAGACACATGCACNNNAGATCGGAAG |
| 64155_1 | TATGAGTGTGGAGTCGTTGCATGCCTCCACACATTTTCAAAGGNNNAGATCGGAAG |
| 64156_1 | TATGAGTGTGGAGTCGTTGCTTCCATTCCCTCTTGAAACCACTNNNAGATCGGAAGA |
| 64157_1 | TATGAGTGTGGAGTCGTTGCAGAGAGAGAGAGAGAGAGATGCANNNAGATCGGA |
| 64158_1 | TATGAGTGTGGAGTCGTTGCAAGTTCTTGGGCAATAGGGTTCTNNNAGATCGGAA |
| 64161_1 | TATGAGTGTGGAGTCGTTGCTCACCCCTCAATTTCTGTGACTCNNNAGATCGGAAGA |
| 64162_1 | TATGAGTGTGGAGTCGTTGCACATCAAGCACATTTTTCCTCGTNNNAGATCGGAAGA |
| 64165_1 | TATGAGTGTGGAGTCGTTGCGACTTCGGCTGTGTGTTATATGCNNNAGATCGGAAG |
| 64166_1 | TATGAGTGTGGAGTCGTTGCAAGGGGTTTTGGTTTTGTGTGTTNNNAGATCGGAA |
| 64167_1 | TATGAGTGTGGAGTCGTTGCCAAGTGCCAGCTGAAATGACATTNNNAGATCGGAA |
| 64168_1 | TATGAGTGTGGAGTCGTTGCCATTTCCCTGTTGCTTGACTCTGNNNAGATCGGAAGA |
| 64169_1 | TATGAGTGTGGAGTCGTTGCAAGCTCCTAAATGATTCTGCAGCNNNAGATCGGAAG |
| 64170_1 | TATGAGTGTGGAGTCGTTGCTTAGTCATCTCAGGAAGGGGAGTNNNAGATCGGAA |
| 64171_1 | TATGAGTGTGGAGTCGTTGCGGTCCTGCCTCTTTCCTAACTTTNNNAGATCGGAAGA |
| 64172_1 | TATGAGTGTGGAGTCGTTGCTCAGAGTCTGGGGCTATCTTTTGNNNAGATCGGAAG |
| 64173_1 | TATGAGTGTGGAGTCGTTGCCACTTCAGCAAAAAGTCCCTGAGNNNAGATCGGAAG |
| 64174_1 | TATGAGTGTGGAGTCGTTGCGGGGTCTTGCTATGTAACCTAGGNNNAGATCGGAA |
| 64175_1 | TATGAGTGTGGAGTCGTTGCCTCTGTTCCTCTGACTGCTCAAANNNAGATCGGAAGA |
| 64176_1 | TATGAGTGTGGAGTCGTTGCTTAGAAATTCCCCGACTCCCTTCNNNAGATCGGAAGA |
| 64178_1 | TATGAGTGTGGAGTCGTTGCAGATTGAGGCTTCCTAACCATGGNNNAGATCGGAA |
| 64179_1 | TATGAGTGTGGAGTCGTTGCATCAGGTATTTCTCCAAGGAGCCNNNAGATCGGAAG |
| 64181_1 | TATGAGTGTGGAGTCGTTGCATTATGACCACACTCCCCTTCAGNNNAGATCGGAAGA |
| 64182_1 | TATGAGTGTGGAGTCGTTGCTGTGTACATGGGAGACAGAATCCNNNAGATCGGAA |
| 64183_1 | TATGAGTGTGGAGTCGTTGCCCCCATAGAATACAGGAGGAAGCNNNAGATCGGAA |
| 64184_1 | TATGAGTGTGGAGTCGTTGCACTACTGTTCTACCATGCAGCTTNNNAGATCGGAAGA |
| 64185_1 | TATGAGTGTGGAGTCGTTGCAGAGACAGAGAGAAGAGAGGGACNNNAGATCGGA |
| 64186_1 | TATGAGTGTGGAGTCGTTGCTTTTGTTTGCTGCCCAGATACAGNNNAGATCGGAAG |
| 64188_1 | TATGAGTGTGGAGTCGTTGCAAGCAAATCCTGTCTTTCTCCCTNNNAGATCGGAAGA |
| 64189_1 | TATGAGTGTGGAGTCGTTGCTACGGCTAAGGGACTTAGAGTCANNNAGATCGGAA |
| 64192_1 | TATGAGTGTGGAGTCGTTGCCACTGGCCTCTTCCAAAACATTTNNNAGATCGGAAGA |
| 64193_1 | TATGAGTGTGGAGTCGTTGCATGTGTGTGATGGAGTTTTGCTCNNNAGATCGGAA |
| 64194_1 | TATGAGTGTGGAGTCGTTGCCTCAAACTCCTGACCTCGTGATCNNNAGATCGGAAGA |
| 64195_1 | TATGAGTGTGGAGTCGTTGCGCCAGCAAATTCCTGATTAGAGGNNNAGATCGGAA |
| 64196_1 | TATGAGTGTGGAGTCGTTGCAAGGACAGAAAGACAAAGAGGCANNNAGATCGGA |
| 64197_1 | TATGAGTGTGGAGTCGTTGCGATCCTCCTATTTCAGCCTCCAGNNNAGATCGGAAGA |
| 64198_1 | TATGAGTGTGGAGTCGTTGCGATCTCAGCTCACTGCAACATCTNNNAGATCGGAAG |
| 64199_1 | TATGAGTGTGGAGTCGTTGCACACTTGGGTTGATTCCATAGCTNNNAGATCGGAAG |
| 64200_1 | TATGAGTGTGGAGTCGTTGCGGTTGAAGACATCAGGTGTTTGGNNNAGATCGGAA |
| 64201_1 | TATGAGTGTGGAGTCGTTGCTCCGTGCTGTGTTCTGAAATAGTNNNAGATCGGAAG |
| 64202_1 | TATGAGTGTGGAGTCGTTGCACACGCACTCCAAAAAAAGACAANNNAGATCGGAAG |
| 64203_1 | TATGAGTGTGGAGTCGTTGCCACTGATGAGGGAGTAACAGGACNNNAGATCGGAA |
| 64204_1 | TATGAGTGTGGAGTCGTTGCCACTGATGAGGGAGTAACAGGACNNNAGATCGGAA |

| | |
|---|---|
| 64205_1 | TATGAGTGTGGAGTCGTTGCGTAATCCCAGCTACTCAGTAGGCNNNAGATCGGAAG |
| 64206_1 | TATGAGTGTGGAGTCGTTGCCACAGATACCCAAAACATCCCCTNNNAGATCGGAAG |
| 64207_1 | TATGAGTGTGGAGTCGTTGCGATTCAGGCAGAGGGAACAAAAGNNNAGATCGGA |
| 64208_1 | TATGAGTGTGGAGTCGTTGCCCCTACCAAAATGGAAGTCACAGNNNAGATCGGAAG |
| 64209_1 | TATGAGTGTGGAGTCGTTGCCCCTTCTCTTCTCCACCATGAATNNNAGATCGGAAGA |
| 64211_1 | TATGAGTGTGGAGTCGTTGCGGACAGTAGCAATTTGAAGGCTCNNNAGATCGGAA |
| 64213_1 | TATGAGTGTGGAGTCGTTGCCATGCCTGTAATCCCAACACTTGNNNAGATCGGAAG |
| 64214_1 | TATGAGTGTGGAGTCGTTGCCCAATGTCTGTCTTTGGCAACATNNNAGATCGGAAG |
| 64215_1 | TATGAGTGTGGAGTCGTTGCAACATGGTTGGTTCTGGAGATCANNNAGATCGGAA |
| 64216_1 | TATGAGTGTGGAGTCGTTGCGCCAACACTTACTGAACATGAGTNNNAGATCGGAAG |
| 64217_1 | TATGAGTGTGGAGTCGTTGCACAACCCAATTGAAAAATAGGCANNNAGATCGGAA |
| 64218_1 | TATGAGTGTGGAGTCGTTGCCACACACACACACACAATGAGAGNNNAGATCGGAAG |
| 64219_1 | TATGAGTGTGGAGTCGTTGCTATGACACCTATGACAGTTGGGCNNNAGATCGGAA |
| 64220_1 | TATGAGTGTGGAGTCGTTGCGTGTGTGTGTGTGTGTGTGTTTANNNAGATCGGAA |
| 64221_1 | TATGAGTGTGGAGTCGTTGCCCCCTGTCATGAAATCCTTCTCTNNNAGATCGGAAGA |
| 64223_1 | TATGAGTGTGGAGTCGTTGCTACTTGCAATGACTCTTCAGCCTNNNAGATCGGAAG |
| 64225_1 | TATGAGTGTGGAGTCGTTGCGAGGGAAACCTGCATACCAACTANNNAGATCGGAA |
| 64226_1 | TATGAGTGTGGAGTCGTTGCGGCACATCATAAATACCCACAGCNNNAGATCGGAAG |
| 64227_1 | TATGAGTGTGGAGTCGTTGCGAGAGAGAGAGAGAGAGACAGGGNNNAGATCGG |
| 64230_1 | TATGAGTGTGGAGTCGTTGCAAGTGGCCTTGGGATGATTTAGTNNNAGATCGGAA |
| 64231_1 | TATGAGTGTGGAGTCGTTGCGAAATGAGTGTGGTCTTGTGTGGNNNAGATCGGAA |
| 64232_1 | TATGAGTGTGGAGTCGTTGCACCCCTAGCTGTCCTCATTTTTTNNNAGATCGGAAGA |
| 64237_1 | TATGAGTGTGGAGTCGTTGCCCAGGCTTTGGTATCAGGATGATNNNAGATCGGAA |
| 64238_1 | TATGAGTGTGGAGTCGTTGCGTAGAGATGGGGTTTTACCACGTNNNAGATCGGAA |
| 64239_1 | TATGAGTGTGGAGTCGTTGCCTCCCCGATTCAAGTGATTCTCTNNNAGATCGGAAGA |
| 64240_1 | TATGAGTGTGGAGTCGTTGCTTGTCTGCTCAGGACATTTAGGTNNNAGATCGGAAG |
| 64242_1 | TATGAGTGTGGAGTCGTTGCTTTGATCTGGTCTGGAACTCCTGNNNAGATCGGAAG |
| 64243_1 | TATGAGTGTGGAGTCGTTGCCTTATCCACGTGGAGTTGAAGGANNNAGATCGGAA |
| 64244_1 | TATGAGTGTGGAGTCGTTGCCAGACATACACACACACACACACNNNAGATCGGAAG |
| 64245_1 | TATGAGTGTGGAGTCGTTGCCCAGCTAAGAAGGAAGACCAGAGNNNAGATCGGAA |
| 64246_1 | TATGAGTGTGGAGTCGTTGCTGCAGGCCATCTAGAAAGGTATTNNNAGATCGGAA |
| 64250_1 | TATGAGTGTGGAGTCGTTGCTGGAGTGCAGTGTCACAATCATANNNAGATCGGAA |
| 64252_1 | TATGAGTGTGGAGTCGTTGCACTACAACTATGACTGTGCTGGGNNNAGATCGGAA |
| 64254_1 | TATGAGTGTGGAGTCGTTGCCACCCAGATTCACAAAGCAAGTTNNNAGATCGGAAG |
| 64255_1 | TATGAGTGTGGAGTCGTTGCTTCACTAAGTTGCTCAGGTTGGTNNNAGATCGGAAG |
| 64257_1 | TATGAGTGTGGAGTCGTTGCTTCTCCAGTCCTACAGACACAGANNNAGATCGGAAG |
| 64259_1 | TATGAGTGTGGAGTCGTTGCAATTGCCCTCAGGTTCCTAGAAGNNNAGATCGGAAG |
| 64261_1 | TATGAGTGTGGAGTCGTTGCAACCCCCACATTTTCCTAAGTGANNNAGATCGGAAG |
| 64265_1 | TATGAGTGTGGAGTCGTTGCTGAGAACATGCAGCATTTGGTTTNNNAGATCGGAA |
| 64266_1 | TATGAGTGTGGAGTCGTTGCTGCATCTGTCTGTCTATCTTGGGNNNAGATCGGAAG |
| 64267_1 | TATGAGTGTGGAGTCGTTGCGGTCAATCCCACCAGAATAGTGANNNAGATCGGAA |
| 64268_1 | TATGAGTGTGGAGTCGTTGCCTGATAAAACAGACAGCACGGTGNNNAGATCGGAA |
| 64269_1 | TATGAGTGTGGAGTCGTTGCGTGTGTGTCTGTGTGTGTATGTGNNNAGATCGGAA |

| | |
|---|---|
| 64270_1 | TATGAGTGTGGAGTCGTTGCAATTTGCAGCCAGTATCAACCAANNNAGATCGGAAG |
| 64271_1 | TATGAGTGTGGAGTCGTTGCGATATAAGCAGCCGGGTACTCTGNNNAGATCGGAA |
| 64272_1 | TATGAGTGTGGAGTCGTTGCAAAATCCATGAATCCAGGAGCTGNNNAGATCGGAA |
| 64273_1 | TATGAGTGTGGAGTCGTTGCAGCACCAGCAAACTACTAGGATTNNNAGATCGGAAG |
| 64274_1 | TATGAGTGTGGAGTCGTTGCGAGAGAGTGCCGGATGTGATATCNNNAGATCGGAA |
| 64276_1 | TATGAGTGTGGAGTCGTTGCGGAAATGAGGGGGGAAAAAACTGNNNAGATCGGA |
| 64277_1 | TATGAGTGTGGAGTCGTTGCCTGAGGCAGGAGAATCACTTGAANNNAGATCGGAA |
| 64278_1 | TATGAGTGTGGAGTCGTTGCACCTTTCAATCAGATTGGAGGCTNNNAGATCGGAAG |
| 64279_1 | TATGAGTGTGGAGTCGTTGCCCTACTGCTTCTGTTTCTCTGGANNNAGATCGGAAGA |
| 64280_1 | TATGAGTGTGGAGTCGTTGCCAAGTCATAAAAGTGCAGCCCAANNNAGATCGGAAG |
| 64281_1 | TATGAGTGTGGAGTCGTTGCATGGTGCCGAATGACAGAAAAAGNNNAGATCGGAA |
| 64284_1 | TATGAGTGTGGAGTCGTTGCCTCACCAAGTGCTGGGATTACAGNNNAGATCGGAAG |
| 64285_1 | TATGAGTGTGGAGTCGTTGCGCCTCCCAAAGTGCTAGGATTATNNNAGATCGGAAG |
| 64286_1 | TATGAGTGTGGAGTCGTTGCTGTCGTCCAAGCATACAAAAACGNNNAGATCGGAAG |
| 64287_1 | TATGAGTGTGGAGTCGTTGCCTTGAGGGGTTGTTGGAGGATAANNNAGATCGGAA |
| 64288_1 | TATGAGTGTGGAGTCGTTGCCCTCCCAATGTGCTAGGATTACANNNAGATCGGAAG |
| 64290_1 | TATGAGTGTGGAGTCGTTGCACAGGTGCTGAATGAGTTTGATTNNNAGATCGGAA |
| 64292_1 | TATGAGTGTGGAGTCGTTGCGGAGAGAGAGAAGAGGATGAGGTNNNAGATCGG |
| 64293_1 | TATGAGTGTGGAGTCGTTGCATGGTAACCATGTGAGGTGTTGANNNAGATCGGAA |
| 64294_1 | TATGAGTGTGGAGTCGTTGCACTCAAGAAGCCTGAGAGACAAGNNNAGATCGGAA |
| 64295_1 | TATGAGTGTGGAGTCGTTGCACCCCCTCTCCACATATACATGANNNAGATCGGAAGA |
| 64297_1 | TATGAGTGTGGAGTCGTTGCCACACACACACACACACACATATNNNAGATCGGAAG |
| 64298_1 | TATGAGTGTGGAGTCGTTGCCTTCCCAGGTTCAAGCAATTCTCNNNAGATCGGAAG |
| 64299_1 | TATGAGTGTGGAGTCGTTGCTCTCAAACTCCTGAACTCAAGGGNNNAGATCGGAAG |
| 64300_1 | TATGAGTGTGGAGTCGTTGCGCATTTGGGCTGGTTCCATATTTNNNAGATCGGAAG |
| 64302_1 | TATGAGTGTGGAGTCGTTGCAGACTGGAACCTGAAATCCCATCNNNAGATCGGAAG |
| 64303_1 | TATGAGTGTGGAGTCGTTGCGATGGAGTCTCACTCTGTCACTCNNNAGATCGGAAG |
| 64304_1 | TATGAGTGTGGAGTCGTTGCGAGAGAGAGAGAGAGAGACAGGGNNNAGATCGG |
| 64305_1 | TATGAGTGTGGAGTCGTTGCGGATGTGTGTGAATTTGGGAGTGNNNAGATCGGA |
| 64306_1 | TATGAGTGTGGAGTCGTTGCTAGAATGCTGGAGAGGCACTTTTNNNAGATCGGAA |
| 64308_1 | TATGAGTGTGGAGTCGTTGCTGGTTATGAGCAAATTGGCTTGGNNNAGATCGGAA |
| 64311_1 | TATGAGTGTGGAGTCGTTGCACAGTTTCGACAGATGATCACCANNNAGATCGGAAG |
| 64312_1 | TATGAGTGTGGAGTCGTTGCAAGCAAGATGGAGCTGTTTAGGTNNNAGATCGGAA |
| 64313_1 | TATGAGTGTGGAGTCGTTGCAGAACCAGCCCTGTAGATTTTGTNNNAGATCGGAAG |
| 64314_1 | TATGAGTGTGGAGTCGTTGCTCTCAGCCTCAAGGACTTTGTACNNNAGATCGGAAG |
| 64315_1 | TATGAGTGTGGAGTCGTTGCCCTCTCCCTCTCCTCTCTTTTTCNNNAGATCGGAAGAG |
| 64316_1 | TATGAGTGTGGAGTCGTTGCTTTTTGAGACAGTCTTGCTCTGTNNNAGATCGGAAG |
| 64317_1 | TATGAGTGTGGAGTCGTTGCCCAAGATCTGTAGTCGACAAGCTNNNAGATCGGAAG |
| 64318_1 | TATGAGTGTGGAGTCGTTGCTCTTGAACTCCTGACCTCAAGTGNNNAGATCGGAAG |
| 64319_1 | TATGAGTGTGGAGTCGTTGCGGGGAAGAAAGTAAGAAGGGAGGNNNAGATCGG |
| 64320_1 | TATGAGTGTGGAGTCGTTGCCTGGGGAATGTGGTCGTTATCATNNNAGATCGGAA |
| 64321_1 | TATGAGTGTGGAGTCGTTGCGTAATCCCAGCTACTCAGGAAGCNNNAGATCGGAAG |
| 64322_1 | TATGAGTGTGGAGTCGTTGCGTGCCATCTCAAACTCCTCCTTANNNAGATCGGAAGA |

64323_1 TATGAGTGTGGAGTCGTTGCAGGAGGTTGAGACTACAGAAAGCNNNAGATCGGAA
64324_1 TATGAGTGTGGAGTCGTTGCAACAAATTCTCTGCAATCCACCCNNNAGATCGGAAGA
64327_1 TATGAGTGTGGAGTCGTTGCGTTGCCTAGGCTAGTCTCAAACTNNNAGATCGGAAG
64329_1 TATGAGTGTGGAGTCGTTGCTCTCAAGTGTTTCGGATAAGGGANNNAGATCGGAA
64330_1 TATGAGTGTGGAGTCGTTGCTCCTACTTGGGTTTTTATCTGCTNNNAGATCGGAAG
64331_1 TATGAGTGTGGAGTCGTTGCTGCAGCCAGTAGTCTTGTATCTTNNNAGATCGGAAG
64333_1 TATGAGTGTGGAGTCGTTGCAGGGAGAGAGAGAGAGAGAGAGANNNAGATCGGA
64334_1 TATGAGTGTGGAGTCGTTGCCCGCAGAATTTTATGCCCCATTANNNAGATCGGAAG
64335_1 TATGAGTGTGGAGTCGTTGCAATACACACACACACACACACACNNNAGATCGGAAG
64336_1 TATGAGTGTGGAGTCGTTGCAAGTGGCCTTGGGATGATTTAGTNNNAGATCGGAA
64337_1 TATGAGTGTGGAGTCGTTGCCTGAGGCAGGAAAATTGCTTGAANNNAGATCGGAA
64339_1 TATGAGTGTGGAGTCGTTGCCTGGGGTGTTTCTCAACTCCTTANNNAGATCGGAAG
64340_1 TATGAGTGTGGAGTCGTTGCAATAAAAAACACTTGGCCAGGCANNNAGATCGGAAG
64341_1 TATGAGTGTGGAGTCGTTGCTAAAGGAAAAATTCTGGCCAGGCNNNAGATCGGAA
64342_1 TATGAGTGTGGAGTCGTTGCAAGGCAAGATAGGATGAGGTGTGNNNAGATCGGA
64343_1 TATGAGTGTGGAGTCGTTGCTGTTATACCTAGCAGGGTGAGGANNNAGATCGGAA
64344_1 TATGAGTGTGGAGTCGTTGCGCCAATAAGACCCTGAAAACCAGNNNAGATCGGAAG
64345_1 TATGAGTGTGGAGTCGTTGCGGGTTCAAATCCTGACTCTGCTANNNAGATCGGAAG
64346_1 TATGAGTGTGGAGTCGTTGCTCCCAGGTTCAAGCAATTCTTCTNNNAGATCGGAAGA
64347_1 TATGAGTGTGGAGTCGTTGCGGACCATAAGTCAGACAACTGGTNNNAGATCGGAA
64348_1 TATGAGTGTGGAGTCGTTGCCAGAGTCTCAATCTACTGCCCAGNNNAGATCGGAAG
64349_1 TATGAGTGTGGAGTCGTTGCCCTCCCATGTTCGAGCTATTCTCNNNAGATCGGAAGA
64350_1 TATGAGTGTGGAGTCGTTGCTTTTTCCAGACAGAGTCTCACTCNNNAGATCGGAAGA
64351_1 TATGAGTGTGGAGTCGTTGCGGTCCTAATGAAAGAGTGTGGGANNNAGATCGGAA
64352_1 TATGAGTGTGGAGTCGTTGCAGGCTGTAGTGAGTTGTGATCAGNNNAGATCGGAA
64354_1 TATGAGTGTGGAGTCGTTGCGAGAGTGCACAAGAGACAGAGAGNNNAGATCGGA
64355_1 TATGAGTGTGGAGTCGTTGCCTTGCCCAGAAAAGAAGTCATGGNNNAGATCGGAA
64356_1 TATGAGTGTGGAGTCGTTGCTCTCCATCCTGCAAACTTTAGCANNNAGATCGGAAGA
64357_1 TATGAGTGTGGAGTCGTTGCAAGTTGGAGAAAACATTTGGGGCNNNAGATCGGAA
64358_1 TATGAGTGTGGAGTCGTTGCGGCTACCTAGATGCTAACACACANNNAGATCGGAAG
64359_1 TATGAGTGTGGAGTCGTTGCAACTGTTTTTTTCTCCCTCCCCANNNAGATCGGAAGA
64360_1 TATGAGTGTGGAGTCGTTGCTTTTAGTAGAGTCGGGGTTTCGCNNNAGATCGGAAG
64361_1 TATGAGTGTGGAGTCGTTGCCGTGTTAGCATCCTGAAGGTCTANNNAGATCGGAAG
64362_1 TATGAGTGTGGAGTCGTTGCACAAGTTCTTTCTAGGTTGGCCANNNAGATCGGAAG
64363_1 TATGAGTGTGGAGTCGTTGCAGGAGTTCAAGAGGGAAAGTGACNNNAGATCGGA
64364_1 TATGAGTGTGGAGTCGTTGCTTTCTCCATCCAGCACTGTTTCTNNNAGATCGGAAGA
64365_1 TATGAGTGTGGAGTCGTTGCTTCTCACCCAAGAATCTTCTGGGNNNAGATCGGAAG
64366_1 TATGAGTGTGGAGTCGTTGCAATGGGTACAAGGAAGTGGGATCNNNAGATCGGA
64367_1 TATGAGTGTGGAGTCGTTGCAATATCAATGACTCCAGGAGCTGNNNAGATCGGAAG
64368_1 TATGAGTGTGGAGTCGTTGCGTGTGTACGTATGTGTGTGTGTGNNNAGATCGGAA
64369_1 TATGAGTGTGGAGTCGTTGCAGAGAAAATGAACACCGATGGGANNNAGATCGGAA
64370_1 TATGAGTGTGGAGTCGTTGCAACAGCCACCTATAAACAGGACCNNNAGATCGGAAG
64371_1 TATGAGTGTGGAGTCGTTGCGGACATTTAGGCTAGTCCATCCCNNNAGATCGGAAG

64372_1 TATGAGTGTGGAGTCGTTGCAAATAGATCGGACACAGTGGCTCNNNAGATCGGAA
64374_1 TATGAGTGTGGAGTCGTTGCTTCCCACATGACTTCTCCTGAAGNNNAGATCGGAAGA
64375_1 TATGAGTGTGGAGTCGTTGCATTGTGGCTGAGCTTTCCAAAAANNNAGATCGGAAG
64376_1 TATGAGTGTGGAGTCGTTGCCACACTTGCATCTTGGAATATGANNNAGATCGGAAG
64378_1 TATGAGTGTGGAGTCGTTGCTCAGTGGTGTATTATTGGGTGGGNNNAGATCGGAA
64381_1 TATGAGTGTGGAGTCGTTGCGAGAGAGAGAGAGAGAGAGAGGCNNNAGATCGGA
64382_1 TATGAGTGTGGAGTCGTTGCGTGTGTGTGAAATGTAGTCTCGCNNNAGATCGGAA
64383_1 TATGAGTGTGGAGTCGTTGCGCGGCACATACCTATAGTCCTAGNNNAGATCGGAAG
64384_1 TATGAGTGTGGAGTCGTTGCGACCCCCTCACAGTTGAAAATCTNNNAGATCGGAAGA
64385_1 TATGAGTGTGGAGTCGTTGCCACACACACAGAAATTAGCCAGGNNNAGATCGGAAG
64386_1 TATGAGTGTGGAGTCGTTGCCTTCACACCCACTAGGATGACAANNNAGATCGGAAGA
64387_1 TATGAGTGTGGAGTCGTTGCCCCCATGTGGCAGAAGAAAAAAANNNAGATCGGAA
64388_1 TATGAGTGTGGAGTCGTTGCGCCCTCAATGGATTGAATAAGGCNNNAGATCGGAA
64389_1 TATGAGTGTGGAGTCGTTGCACACACATTCCAGAAGTAGCACANNNAGATCGGAAG
64390_1 TATGAGTGTGGAGTCGTTGCTTTCTTCAGACAGGGTCTCACTCNNNAGATCGGAAGA
64391_1 TATGAGTGTGGAGTCGTTGCGGCTCAGTTCAAATCAACAGGTTNNNAGATCGGAAG
64393_1 TATGAGTGTGGAGTCGTTGCATGAGTGACAGACTTCAAGTGCANNNAGATCGGAA
64394_1 TATGAGTGTGGAGTCGTTGCAGGATGATGCTGGCCTCATAAAANNNAGATCGGAA
64395_1 TATGAGTGTGGAGTCGTTGCTCTGTTCAACTGCTGCTTTTGAGNNNAGATCGGAAGA
64396_1 TATGAGTGTGGAGTCGTTGCCCCTAGGACTTGTAACAGGACACNNNAGATCGGAAG
64398_1 TATGAGTGTGGAGTCGTTGCCCCCATCCACAAATCCTGAATCTNNNAGATCGGAAGA
64399_1 TATGAGTGTGGAGTCGTTGCTATTAGGAGTTTTGCAGTGGCCTNNNAGATCGGAAG
64400_1 TATGAGTGTGGAGTCGTTGCCAAAAGAGAATGCAGAGGGTGTGNNNAGATCGGAA
64401_1 TATGAGTGTGGAGTCGTTGCGACTTCCTTCTCTTCCACCCAAANNNAGATCGGAAGA
64402_1 TATGAGTGTGGAGTCGTTGCCTTTCCTCAGGGTCCCTAAGAACNNNAGATCGGAAGA
64403_1 TATGAGTGTGGAGTCGTTGCTTGATGACCAGGAAGGAAGTCAGNNNAGATCGGAA
64405_1 TATGAGTGTGGAGTCGTTGCTGCATACACACATCCTATTGGCTNNNAGATCGGAAGA
64406_1 TATGAGTGTGGAGTCGTTGCTTGACCCTTGAAGTTATGGGCTTNNNAGATCGGAAG
64407_1 TATGAGTGTGGAGTCGTTGCACACTGACAATCAGATCTGGCTTNNNAGATCGGAAG
64408_1 TATGAGTGTGGAGTCGTTGCCTTGAATCCTGGCCTCATCTTCANNNAGATCGGAAGA
64409_1 TATGAGTGTGGAGTCGTTGCACCAAAATCTGTGGATGCTCAAGNNNAGATCGGAAG
64411_1 TATGAGTGTGGAGTCGTTGCACAACAAAGAATGCTAACGTGGGNNNAGATCGGAA
64412_1 TATGAGTGTGGAGTCGTTGCTTGAGTTGGAGTCTTGGTCTGTCNNNAGATCGGAAG
64414_1 TATGAGTGTGGAGTCGTTGCTGTAGTCACAGCCCCATTTACAANNNAGATCGGAAGA
64416_1 TATGAGTGTGGAGTCGTTGCCTTCTCCTAGTCATCTGGGATGCNNNAGATCGGAAGA
64417_1 TATGAGTGTGGAGTCGTTGCTCTGTCTCTCTCTCACCTTGTNNNAGATCGGAAGA
64419_1 TATGAGTGTGGAGTCGTTGCCAGCAACCTGGATGGAATTGAAGNNNAGATCGGAA
64420_1 TATGAGTGTGGAGTCGTTGCTCTCTCTCTCTCTCACCCTTTGTNNNAGATCGGAAGAG
64424_1 TATGAGTGTGGAGTCGTTGCGGGGGTGTTGTTTGTTTGTTTTGNNNAGATCGGAA
64425_1 TATGAGTGTGGAGTCGTTGCAGGTTCAATCAATTCTCCTGCCTNNNAGATCGGAAGA
64426_1 TATGAGTGTGGAGTCGTTGCGTTGAAACAGATCACAGCAACGANNNAGATCGGAA
64427_1 TATGAGTGTGGAGTCGTTGCCTTCAAATGGTTCTCTGCCTCAGNNNAGATCGGAAGA
64429_1 TATGAGTGTGGAGTCGTTGCCTGAACCTGGAACACAAAGCAAANNNAGATCGGAA

| | |
|---|---|
| 64431_1 | TATGAGTGTGGAGTCGTTGCAATGCACACACACACACACAAATNNNAGATCGGAAG |
| 64432_1 | TATGAGTGTGGAGTCGTTGCAACAACACATCAGGTCTTTTGGGNNNAGATCGGAAG |
| 64433_1 | TATGAGTGTGGAGTCGTTGCCCTGTAATCCCAGCACTTTAGGANNNAGATCGGAAG |
| 64434_1 | TATGAGTGTGGAGTCGTTGCTCGCTTTTCCATGAAACAGTGTTNNNAGATCGGAAG |
| 64436_1 | TATGAGTGTGGAGTCGTTGCTTGTAGAGACAGGGTTTTGCCATNNNAGATCGGAA |
| 64437_1 | TATGAGTGTGGAGTCGTTGCGCAAATTCCTGCTTTTTCTTGGCNNNAGATCGGAAG |
| 64438_1 | TATGAGTGTGGAGTCGTTGCCAACAGAACACGAGGAAAAGGTCNNNAGATCGGAA |
| 64439_1 | TATGAGTGTGGAGTCGTTGCGAAGGGCATTCCAAAACATCTGGNNNAGATCGGAA |
| 64440_1 | TATGAGTGTGGAGTCGTTGCACTTATTTGCAGGGGTTCAGGATNNNAGATCGGAA |
| 64441_1 | TATGAGTGTGGAGTCGTTGCCAGATGGGCAAATGTCAACAACTNNNAGATCGGAA |
| 64442_1 | TATGAGTGTGGAGTCGTTGCATAGTCCCACCTACTCAGGAGACNNNAGATCGGAAG |
| 64443_1 | TATGAGTGTGGAGTCGTTGCAGAGAACAATTGCCTCTGAGGAGNNNAGATCGGAA |
| 64444_1 | TATGAGTGTGGAGTCGTTGCGGGACAGGAAGAGAGAAAAGAGGNNNAGATCGG |
| 64445_1 | TATGAGTGTGGAGTCGTTGCAATTGTGTGGGAGTAGCCATAGGNNNAGATCGGAA |
| 64446_1 | TATGAGTGTGGAGTCGTTGCTATCTACTCATTGGCTGATGGGCNNNAGATCGGAAG |
| 64448_1 | TATGAGTGTGGAGTCGTTGCCCCCCCTAGTAATCTTATTCGCCNNNAGATCGGAAGA |
| 64449_1 | TATGAGTGTGGAGTCGTTGCTCCATGTAGTTCCCATTACTGGTNNNAGATCGGAAG |
| 64451_1 | TATGAGTGTGGAGTCGTTGCATTCTCATTCTGTCACCCAAGCTNNNAGATCGGAAGA |
| 64453_1 | TATGAGTGTGGAGTCGTTGCAAAGAGCTTCCAATGACCAAAGCNNNAGATCGGAA |
| 64454_1 | TATGAGTGTGGAGTCGTTGCAATCTCTCTCTATCACCCAGGCTNNNAGATCGGAAGA |
| 64455_1 | TATGAGTGTGGAGTCGTTGCGTCTGTGTTTGTGTGCAGAGATTNNNAGATCGGAA |
| 64457_1 | TATGAGTGTGGAGTCGTTGCCTCCACCTCCTAGCATCACTTTTNNNAGATCGGAAGA |
| 64461_1 | TATGAGTGTGGAGTCGTTGCCTGTTGTCCAGGCTGGAGTATAGNNNAGATCGGAA |
| 64463_1 | TATGAGTGTGGAGTCGTTGCGAGAGAATGGTCCTTTCTGGTGTNNNAGATCGGAA |
| 64464_1 | TATGAGTGTGGAGTCGTTGCTGGTAAACAGGAACTGCTGTCATNNNAGATCGGAA |
| 64465_1 | TATGAGTGTGGAGTCGTTGCGAATTTTCTCCGTGAGGCACATCNNNAGATCGGAAG |
| 64466_1 | TATGAGTGTGGAGTCGTTGCTAGAGAACCACTGAGTTACAGCCNNNAGATCGGAA |
| 64467_1 | TATGAGTGTGGAGTCGTTGCAAATGTTCAGAGAGGCAAATCCANNNAGATCGGAA |
| 64468_1 | TATGAGTGTGGAGTCGTTGCTGTCCAAAACCAAGGTGTTGAAGNNNAGATCGGAA |
| 64469_1 | TATGAGTGTGGAGTCGTTGCTTCAAGTGATTCTCCTGCCTCAGNNNAGATCGGAAG |
| 64470_1 | TATGAGTGTGGAGTCGTTGCTAAGTCTGTGCGTGTGTGTGTATNNNAGATCGGAA |
| 64472_1 | TATGAGTGTGGAGTCGTTGCAATCTCCTTCTGCCTCTCTCCTANNNAGATCGGAAGA |
| 64473_1 | TATGAGTGTGGAGTCGTTGCAGTGTTTGTTCATGTCCTTTGCCNNNAGATCGGAAG |
| 64474_1 | TATGAGTGTGGAGTCGTTGCGATCAGCCTAGGCAATATGGTGANNNAGATCGGAA |
| 64475_1 | TATGAGTGTGGAGTCGTTGCGAGTATGGCCTTGGCATAGGTAANNNAGATCGGAA |
| 64476_1 | TATGAGTGTGGAGTCGTTGCCTCTCTCTCTCTCTCTCAGGGNNNAGATCGGAAGA |
| 64477_1 | TATGAGTGTGGAGTCGTTGCAAATCAGGTCCCAGACTAGATGCNNNAGATCGGAA |
| 64478_1 | TATGAGTGTGGAGTCGTTGCTCACAAGGACTGAAATCACCCTCNNNAGATCGGAAG |
| 64479_1 | TATGAGTGTGGAGTCGTTGCACCATGCTAGTCCACTAAGTTGTNNNAGATCGGAAG |
| 64482_1 | TATGAGTGTGGAGTCGTTGCTCTGTCATGTTGAGAGGCTTTGTNNNAGATCGGAAG |
| 64483_1 | TATGAGTGTGGAGTCGTTGCGAGCAGGAATCTTCAGTGATCCANNNAGATCGGAA |
| 64486_1 | TATGAGTGTGGAGTCGTTGCAAAGAAAAATTAGTGGCCAGGCGNNNAGATCGGAA |
| 64487_1 | TATGAGTGTGGAGTCGTTGCTATATGAAGTAGGCCAGACACGGNNNAGATCGGAA |

64488_1 TATGAGTGTGGAGTCGTTGCTCTTTCTCTCTCTTTCTCCCCCANNNAGATCGGAAGAG
64489_1 TATGAGTGTGGAGTCGTTGCTGCATATGTGTCACATTCACACANNNAGATCGGAAGA
64490_1 TATGAGTGTGGAGTCGTTGCCACACCGCCTTATCTGTATCTCANNNAGATCGGAAGA
64491_1 TATGAGTGTGGAGTCGTTGCGTGTGTGTGTGTGTGTGTGTATGNNNAGATCGGAA
64492_1 TATGAGTGTGGAGTCGTTGCCAAGCTCTTGAAAACAACCAGCTNNNAGATCGGAAG
64493_1 TATGAGTGTGGAGTCGTTGCTGCCTTTTACAACCTAGTCTCCCNNNAGATCGGAAGA
64495_1 TATGAGTGTGGAGTCGTTGCATTTACTCAAAACAAGAGGCCGCNNNAGATCGGAAG
64496_1 TATGAGTGTGGAGTCGTTGCCAGTGAAGCACTCATTCAAGAGCNNNAGATCGGAAG
64497_1 TATGAGTGTGGAGTCGTTGCTCCTATGGCTCTCTTTCTCTGGANNNAGATCGGAAGA
64498_1 TATGAGTGTGGAGTCGTTGCACCAATTGAGTATAGGGAGGGGANNNAGATCGGAA
64499_1 TATGAGTGTGGAGTCGTTGCTCCAGAAGACTGCAAAGAGACTCNNNAGATCGGAAG
64500_1 TATGAGTGTGGAGTCGTTGCAAGAGAAAGAAAGAGGCTCACCANNNAGATCGGAA
64501_1 TATGAGTGTGGAGTCGTTGCCTCCTGCCACATCCCATTATGTANNNAGATCGGAAGA
64502_1 TATGAGTGTGGAGTCGTTGCCACAAGCTCTCTGGTGTCCTATTNNNAGATCGGAAGA
64504_1 TATGAGTGTGGAGTCGTTGCTTCTCCACAATAGGCATGGCTAGNNNAGATCGGAAG
64505_1 TATGAGTGTGGAGTCGTTGCGAGAGGGAGAGAGAGTCATCTCANNNAGATCGGAA
64506_1 TATGAGTGTGGAGTCGTTGCAGAAAATATCCACACCCTTGGGCNNNAGATCGGAAG
64507_1 TATGAGTGTGGAGTCGTTGCCACAACCACGTCCAGCTAATTTTNNNAGATCGGAAGA
64508_1 TATGAGTGTGGAGTCGTTGCACACTTGGATTTTCCCTATGCCANNNAGATCGGAAGA
64509_1 TATGAGTGTGGAGTCGTTGCAAAGATGGTGAAGGGAGATTCCANNNAGATCGGAA
64510_1 TATGAGTGTGGAGTCGTTGCAGAAATGACATGTCCTGGGCATANNNAGATCGGAAG
64511_1 TATGAGTGTGGAGTCGTTGCGATTCAGGAGAGCCAATGGTGTANNNAGATCGGAA
64512_1 TATGAGTGTGGAGTCGTTGCGTAAGTTTCCTGAGGCATCCCTANNNAGATCGGAAG
64513_1 TATGAGTGTGGAGTCGTTGCTCTCTGTCTCTCTCTCTCTCTGCNNNAGATCGGAAGAG
64515_1 TATGAGTGTGGAGTCGTTGCCCCAGGAGGATTATAGACCAACCNNNAGATCGGAAG
64516_1 TATGAGTGTGGAGTCGTTGCCCTCCCTCTGGTAAGTAGTCTCANNNAGATCGGAAGA
64517_1 TATGAGTGTGGAGTCGTTGCCTGGCCCATAAGTCTTGAACTCTNNNAGATCGGAAGA
64518_1 TATGAGTGTGGAGTCGTTGCTGTGTGGTGTGTGCTTTTTTTCTNNNAGATCGGAAGA
64519_1 TATGAGTGTGGAGTCGTTGCATAAGTTTTCTGAGGCTCCCCAGNNNAGATCGGAAG
64520_1 TATGAGTGTGGAGTCGTTGCGAGAGACAGAAAGCAGGAGAGAGNNNAGATCGGA
64521_1 TATGAGTGTGGAGTCGTTGCTGTGTGTGTGTGTGTGTGTTTTANNNAGATCGGAAG
64523_1 TATGAGTGTGGAGTCGTTGCGAAGGGAGGAAGGGAAAGAAGAGNNNAGATCGGA
64525_1 TATGAGTGTGGAGTCGTTGCACATGATTGTAGAGGCTGTCAGGNNNAGATCGGAAG
64526_1 TATGAGTGTGGAGTCGTTGCGCACCTGGCCTTCACTTTTTTTANNNAGATCGGAAGA
64528_1 TATGAGTGTGGAGTCGTTGCAGCTGGTATGGTCTATCAGTCCCNNNAGATCGGAAG
64529_1 TATGAGTGTGGAGTCGTTGCTACCTTTACCATGGCTCCTTGTCNNNAGATCGGAAGA
64531_1 TATGAGTGTGGAGTCGTTGCTCATTCCAACCATCCAAATCCCTNNNAGATCGGAAGAG
64533_1 TATGAGTGTGGAGTCGTTGCTCCCAGTTTCCAGTCCTTTTCTTNNNAGATCGGAAGAG
64534_1 TATGAGTGTGGAGTCGTTGCGGTGACATCATAACCAATCGCACNNNAGATCGGAAG
64535_1 TATGAGTGTGGAGTCGTTGCTGTCCCAACCCCTTTGATAGAAGNNNAGATCGGAAGA
64536_1 TATGAGTGTGGAGTCGTTGCACATCAAAGAATATCAGGCCCCANNNAGATCGGAAG
64537_1 TATGAGTGTGGAGTCGTTGCGGCCGTAAGGAAAAGGTACCATANNNAGATCGGAA
64539_1 TATGAGTGTGGAGTCGTTGCAGAGAGAAGAGCAGGTGAAACTGNNNAGATCGGA

64540_1 TATGAGTGTGGAGTCGTTGCCAGTGCTAGAGACTCAACCCAATNNNAGATCGGAAG
64541_1 TATGAGTGTGGAGTCGTTGCCGAAGTGGCAATGTCTGAAGAAANNNAGATCGGAA
64542_1 TATGAGTGTGGAGTCGTTGCGCTGCTCCTGAATGAAAAGACTCNNNAGATCGGAAG
64544_1 TATGAGTGTGGAGTCGTTGCCAGAACATGTGACAATGTGGGTTNNNAGATCGGAA
64545_1 TATGAGTGTGGAGTCGTTGCTTCATAGCTGAGGACCAGAATGGNNNAGATCGGAA
64546_1 TATGAGTGTGGAGTCGTTGCTACACATTTCTATGCAACCGGGANNNAGATCGGAAG
64547_1 TATGAGTGTGGAGTCGTTGCAGAGAGAGAGAGAGAGAGAGGGANNNAGATCGG
64548_1 TATGAGTGTGGAGTCGTTGCATCCAAGAGGGCACAAACAAATGNNNAGATCGGAA
64549_1 TATGAGTGTGGAGTCGTTGCCTCACAACAAAACTGCTGCAAACNNNAGATCGGAAG
64550_1 TATGAGTGTGGAGTCGTTGCGAGAAGCCAGCTCCATGATTCTANNNAGATCGGAAG
64551_1 TATGAGTGTGGAGTCGTTGCCATCTTATGCACACACACACACANNNAGATCGGAAG
64552_1 TATGAGTGTGGAGTCGTTGCCTGGGTTGGAGCACCTAAGTTATNNNAGATCGGAA
64553_1 TATGAGTGTGGAGTCGTTGCCTGGTGGCTGACTTTAGAGACATNNNAGATCGGAA
64554_1 TATGAGTGTGGAGTCGTTGCTCCTTCACACTTTCTGAACACCANNNAGATCGGAAGA
64555_1 TATGAGTGTGGAGTCGTTGCAGGAGGGTAGATCAGTACCAACTNNNAGATCGGAA
64556_1 TATGAGTGTGGAGTCGTTGCACTCCAAAAACAACAACAACCCANNNAGATCGGAAG
64557_1 TATGAGTGTGGAGTCGTTGCTAGAGTGCAGTGGTAGGATCAGTNNNAGATCGGAA
64558_1 TATGAGTGTGGAGTCGTTGCCGTGACTCATTCTGCTTGACATCNNNAGATCGGAAG
64559_1 TATGAGTGTGGAGTCGTTGCTTTTGCCAGCATCACTGAATCTGNNNAGATCGGAAG
64560_1 TATGAGTGTGGAGTCGTTGCTCTGGAAGCACTGAGGAAGAAAANNNAGATCGGAA
64561_1 TATGAGTGTGGAGTCGTTGCCCCTTTACTCACTGTGTGACCTTNNNAGATCGGAAGA
64562_1 TATGAGTGTGGAGTCGTTGCGTAAGGAATTGGCCTGTGCAATTNNNAGATCGGAA
64563_1 TATGAGTGTGGAGTCGTTGCTGGCAAAGAGCATCAAGAAAGTGNNNAGATCGGAA
64564_1 TATGAGTGTGGAGTCGTTGCTGCTTAGCACAGTTTACATAGCANNNAGATCGGAAG
64565_1 TATGAGTGTGGAGTCGTTGCGGACAATACTCAGACTCCCAGTGNNNAGATCGGAAG
64567_1 TATGAGTGTGGAGTCGTTGCGCTATAAGCTTCATAGGGGCAGTNNNAGATCGGAA
64568_1 TATGAGTGTGGAGTCGTTGCGTTGCCCAGTTTTCAGTTCATGANNNAGATCGGAAG
64570_1 TATGAGTGTGGAGTCGTTGCACCCTTGGCAGCTATAAACAGAANNNAGATCGGAAG
64571_1 TATGAGTGTGGAGTCGTTGCAAGGCCGTCCAGAATTTAGTTCTNNNAGATCGGAAG
64572_1 TATGAGTGTGGAGTCGTTGCGAAGGGCAGAGACGAATGTACATNNNAGATCGGAA
64573_1 TATGAGTGTGGAGTCGTTGCAGGAAAGAGGCAGAAGGAACATTNNNAGATCGGA
64574_1 TATGAGTGTGGAGTCGTTGCAAAGACCGTGAGAGAGGTAGAGANNNAGATCGGA
64575_1 TATGAGTGTGGAGTCGTTGCGAGAAAATGTGTGTGCTTTGGGANNNAGATCGGAA
64577_1 TATGAGTGTGGAGTCGTTGCCCGTATCCTATTGGCATTCACCANNNAGATCGGAAG
64578_1 TATGAGTGTGGAGTCGTTGCGGGTTGAGAGGAGCTTAGTTCAANNNAGATCGGAA
64580_1 TATGAGTGTGGAGTCGTTGCTTGGAGATGAATTGACTGGAGCANNNAGATCGGAA
64581_1 TATGAGTGTGGAGTCGTTGCTGTGTGAGAATATGTGTGCAGGANNNAGATCGGAA
64582_1 TATGAGTGTGGAGTCGTTGCACATTGGTGACTATGTTGCAAATNNNAGATCGGAAG
64583_1 TATGAGTGTGGAGTCGTTGCATAACCCAACTGCCTATCAGGTGNNNAGATCGGAAG
64584_1 TATGAGTGTGGAGTCGTTGCCCACTTATGGGAATGGGATGTGANNNAGATCGGAA
64586_1 TATGAGTGTGGAGTCGTTGCTTCCATAGCTATGAAATGGGCGANNNAGATCGGAA
64587_1 TATGAGTGTGGAGTCGTTGCTCATGCTTGCACATTCACTCATGNNNAGATCGGAAG
64588_1 TATGAGTGTGGAGTCGTTGCAAGGCTGGGCAATAAAGACCTAANNNAGATCGGAA

64589_1 TATGAGTGTGGAGTCGTTGCGTTAGACACCTCCTCATAGCCAGNNNAGATCGGAAG
64590_1 TATGAGTGTGGAGTCGTTGCTTCGTTCTGGGATAGACCAATCGNNNAGATCGGAAG
64591_1 TATGAGTGTGGAGTCGTTGCGGTGCCTGTAGTCATAGCTACTCNNNAGATCGGAAG
64592_1 TATGAGTGTGGAGTCGTTGCTGGAAAGAATGAAAGCTTGCCAGNNNAGATCGGAA
64593_1 TATGAGTGTGGAGTCGTTGCTGTACTGGCATTCAGGTACATCCNNNAGATCGGAAG
64594_1 TATGAGTGTGGAGTCGTTGCTTATTGTGTTCTGGCCAGTGAGANNNAGATCGGAA
64595_1 TATGAGTGTGGAGTCGTTGCAATTAGCATGTCTCTTCCAGGCTNNNAGATCGGAAG
64596_1 TATGAGTGTGGAGTCGTTGCGGACTCACAGAAAACAAGCTGTCNNNAGATCGGAA
64597_1 TATGAGTGTGGAGTCGTTGCTTCCATGTGATTGGCAAGACCTANNNAGATCGGAAG
64598_1 TATGAGTGTGGAGTCGTTGCCTTTTGCCGGGGAAACAATAACANNNAGATCGGAA
64599_1 TATGAGTGTGGAGTCGTTGCATGAACCTTGCAAAAAGAGAGGCNNNAGATCGGAA
64600_1 TATGAGTGTGGAGTCGTTGCTAATTAGCAGGAAGAATGGGCCANNNAGATCGGAA
64601_1 TATGAGTGTGGAGTCGTTGCTCACAAGAAAAAACACTGGCAGCNNNAGATCGGAA
64602_1 TATGAGTGTGGAGTCGTTGCAGAGAACAATGGATGGCACTTCTNNNAGATCGGAA
64603_1 TATGAGTGTGGAGTCGTTGCTTTATTGGGGAGTGACATCTGGGNNNAGATCGGAA
64604_1 TATGAGTGTGGAGTCGTTGCAAGGCATGAGATGGACAATCAGANNNAGATCGGAA
64606_1 TATGAGTGTGGAGTCGTTGCAGAATAAGGAACCCTACCTCCCANNNAGATCGGAAG
64607_1 TATGAGTGTGGAGTCGTTGCTTTTCTGTCCCTGGGCATATCAANNNAGATCGGAAG
64608_1 TATGAGTGTGGAGTCGTTGCGTATATATCCAAACCCACCCGCTNNNAGATCGGAAG
64609_1 TATGAGTGTGGAGTCGTTGCTTTGAGATGGTCACTCTGTCACCNNNAGATCGGAAG
64610_1 TATGAGTGTGGAGTCGTTGCAGTCATCAGGACCTATTTGCCAANNNAGATCGGAAG
64611_1 TATGAGTGTGGAGTCGTTGCAAGAATCATCACACGCACACAAGNNNAGATCGGAAG
64612_1 TATGAGTGTGGAGTCGTTGCTAATGGAGGCTGAGAAATCCCACNNNAGATCGGAA
64613_1 TATGAGTGTGGAGTCGTTGCTCCTAATCCTAACAAGGCTGAGCNNNAGATCGGAAG
64614_1 TATGAGTGTGGAGTCGTTGCGGAACAAGGAAAAGAGCCACTTCNNNAGATCGGAA
64615_1 TATGAGTGTGGAGTCGTTGCTCAGAAGGTCCTGGTAACATGTGNNNAGATCGGAA
64616_1 TATGAGTGTGGAGTCGTTGCGGCCAGATGCAACAACTTATCATNNNAGATCGGAAG
64617_1 TATGAGTGTGGAGTCGTTGCTTGCTGTTGTTCTCTTTGCCATTNNNAGATCGGAAGA
64618_1 TATGAGTGTGGAGTCGTTGCACTTATCACCCATTCCAGCACTTNNNAGATCGGAAGA
64619_1 TATGAGTGTGGAGTCGTTGCCAAGTTTTTGAGCAGGAAGGGAGNNNAGATCGGAA
64620_1 TATGAGTGTGGAGTCGTTGCACCATCTTGGAGTCTCAGCAAATNNNAGATCGGAAG
64621_1 TATGAGTGTGGAGTCGTTGCATTAGTGACTGGTTGCTTCTGGANNNAGATCGGAA
64622_1 TATGAGTGTGGAGTCGTTGCTGCACGCATTTTTTTCTATCCCTNNNAGATCGGAAGA
64623_1 TATGAGTGTGGAGTCGTTGCGAGCCTCACATCACTTGCAAATTNNNAGATCGGAAG
64624_1 TATGAGTGTGGAGTCGTTGCCAACAACCCTCACTCCCATTTTCNNNAGATCGGAAGA
64625_1 TATGAGTGTGGAGTCGTTGCTTCCCTTAGCTGTAACCAAGCATNNNAGATCGGAAG
64626_1 TATGAGTGTGGAGTCGTTGCCATCCATCCATCCAGGGATTCATNNNAGATCGGAAG
64627_1 TATGAGTGTGGAGTCGTTGCTGATCAGTTGGAAATCTGGGACANNNAGATCGGAA
64628_1 TATGAGTGTGGAGTCGTTGCAGGTGTGTGGAGCTGAATGTTTANNNAGATCGGAA
64629_1 TATGAGTGTGGAGTCGTTGCCCAAGATGCAGTAGGAAAAGCAGNNNAGATCGGAA
64630_1 TATGAGTGTGGAGTCGTTGCTAGGCACCAGGCAATTTAGTCTTNNNAGATCGGAAG
64631_1 TATGAGTGTGGAGTCGTTGCTATGCTATCCTCCCTTGGCTTTCNNNAGATCGGAAGA
64632_1 TATGAGTGTGGAGTCGTTGCGGCCCCATTTACCTTTCTTCATGNNNAGATCGGAAGA

| | |
|---|---|
| 64633_1 | TATGAGTGTGGAGTCGTTGCGAGGGATAAACATGTCGGAGTTTNNNAGATCGGAA |
| 64634_1 | TATGAGTGTGGAGTCGTTGCCAATCTGACATTCTGCACACAGGNNNAGATCGGAAG |
| 64635_1 | TATGAGTGTGGAGTCGTTGCTGTCTGCATTCACTTGATGAGAGNNNAGATCGGAAG |
| 64636_1 | TATGAGTGTGGAGTCGTTGCAAATTTAGCCTAGATGCCCCCAANNNAGATCGGAAG |
| 64637_1 | TATGAGTGTGGAGTCGTTGCCACACACGGAGAGAGAGATTGATNNNAGATCGGAA |
| 64638_1 | TATGAGTGTGGAGTCGTTGCTGCGAAGCATTTCTCATCATCATNNNAGATCGGAAG |
| 64639_1 | TATGAGTGTGGAGTCGTTGCCTCAGGGTAGAGGGTTCGTTTTTNNNAGATCGGAA |
| 64640_1 | TATGAGTGTGGAGTCGTTGCTCTGAGAAAAACCCCACAGTGAANNNAGATCGGAA |
| 64641_1 | TATGAGTGTGGAGTCGTTGCGGGCAGCCTCTTGATTCTTTTACNNNAGATCGGAAG |
| 64642_1 | TATGAGTGTGGAGTCGTTGCCACAGAGGAAGGAATGAGCTCATNNNAGATCGGAA |
| 64643_1 | TATGAGTGTGGAGTCGTTGCACAAAGTTGTCTACTCCAGCTGANNNAGATCGGAAG |
| 64644_1 | TATGAGTGTGGAGTCGTTGCTGCTCTTGGACATCAGAATTCCANNNAGATCGGAAG |
| 64645_1 | TATGAGTGTGGAGTCGTTGCTAATGTGGAGTCATCCCAGCAAANNNAGATCGGAA |
| 64646_1 | TATGAGTGTGGAGTCGTTGCGCAATTATGAGCCCTGGAGTCTANNNAGATCGGAA |
| 64647_1 | TATGAGTGTGGAGTCGTTGCCTGACTGTACTGCTAAGTCCCTGNNNAGATCGGAAG |
| 64648_1 | TATGAGTGTGGAGTCGTTGCCATGCTTCTCCACCAACATGATGNNNAGATCGGAAG |
| 64649_1 | TATGAGTGTGGAGTCGTTGCTGGCATAAGGAATAGAGGGCTTCNNNAGATCGGAA |
| 64650_1 | TATGAGTGTGGAGTCGTTGCTCCTGACCCACTAAGTCTCATCTNNNAGATCGGAAGA |
| 64651_1 | TATGAGTGTGGAGTCGTTGCGTCTCTGGACTCTCTGGCTTTTCNNNAGATCGGAAG |
| 64652_1 | TATGAGTGTGGAGTCGTTGCCAGTGGGCTAGAGACCTAAAGAANNNAGATCGGAA |
| 64653_1 | TATGAGTGTGGAGTCGTTGCAAAGAAAAACAGAAGCCCTCAGCNNNAGATCGGAA |
| 64654_1 | TATGAGTGTGGAGTCGTTGCTATACACAAAGACGGTTCTTGCCNNNAGATCGGAAG |
| 64655_1 | TATGAGTGTGGAGTCGTTGCCTCTCCACACAGGTTCATGAAGANNNAGATCGGAAG |
| 64656_1 | TATGAGTGTGGAGTCGTTGCTTCAGGACACTAGGGTTCATGTGNNNAGATCGGAA |
| 64657_1 | TATGAGTGTGGAGTCGTTGCTTATTTGCTCCACGACATTCAGCNNNAGATCGGAAG |
| 64659_1 | TATGAGTGTGGAGTCGTTGCAAGAGCTGCCTGTCTGATCTAACNNNAGATCGGAAG |
| 64661_1 | TATGAGTGTGGAGTCGTTGCTCGGAGCTGTAATGTTGTGAAGANNNAGATCGGAA |
| 64662_1 | TATGAGTGTGGAGTCGTTGCAAAATTACTTCTCTGGCCCCCATNNNAGATCGGAAGA |
| 64663_1 | TATGAGTGTGGAGTCGTTGCCACTTCTGTAGTCCCTGCTACTCNNNAGATCGGAAGA |
| 64664_1 | TATGAGTGTGGAGTCGTTGCATAAGCTCAGTCACCCATTCCAGNNNAGATCGGAAG |
| 64665_1 | TATGAGTGTGGAGTCGTTGCCTTCTTGTACTTCGGGGAAGCAGNNNAGATCGGAA |
| 64666_1 | TATGAGTGTGGAGTCGTTGCTCTCCATGGGAATCCTACTGACTNNNAGATCGGAAG |
| 64667_1 | TATGAGTGTGGAGTCGTTGCGTGTGTGTGTGTGTGTGTGTTATNNNAGATCGGAA |
| 64668_1 | TATGAGTGTGGAGTCGTTGCTCTGACTTTCTCCCTCTTTCCTGNNNAGATCGGAAGA |
| 64669_1 | TATGAGTGTGGAGTCGTTGCAGAAAAGCAAAAGAAGCTGACGANNNAGATCGGAA |
| 64670_1 | TATGAGTGTGGAGTCGTTGCACATGCTTGTCTAACCTGTCAGTNNNAGATCGGAAG |
| 64671_1 | TATGAGTGTGGAGTCGTTGCATCCTAATTATGCACTGCCCCATNNNAGATCGGAAGA |
| 64673_1 | TATGAGTGTGGAGTCGTTGCCCAAATCACCCTGTTGTGAGTTCNNNAGATCGGAAG |
| 64674_1 | TATGAGTGTGGAGTCGTTGCCTTGGAGTCACAGAGTTTTCCCTNNNAGATCGGAAG |
| 64675_1 | TATGAGTGTGGAGTCGTTGCCACACACACACTCTCTCTCTCTCNNNAGATCGGAAGA |
| 64676_1 | TATGAGTGTGGAGTCGTTGCTTCAAATGGTCAGATCCCTCCAGNNNAGATCGGAAG |
| 64678_1 | TATGAGTGTGGAGTCGTTGCACTGGGTACCGAATTCTTAGGTGNNNAGATCGGAA |
| 64679_1 | TATGAGTGTGGAGTCGTTGCGAAATCATTTCCCCGAGAGAGCANNNAGATCGGAA |

| | |
|---|---|
| 64680_1 | TATGAGTGTGGAGTCGTTGCCACACTTTCTTTCTCCTTGGGGANNNAGATCGGAAG |
| 64681_1 | TATGAGTGTGGAGTCGTTGCCATTGAAAACCCCCATTCCTTCNNNAGATCGGAAGA |
| 64683_1 | TATGAGTGTGGAGTCGTTGCCACAGAGCTTCTAAGACCCTTGTNNNAGATCGGAAG |
| 64684_1 | TATGAGTGTGGAGTCGTTGCCATTTCCATGGGTTCACAGTCTGNNNAGATCGGAAG |
| 64685_1 | TATGAGTGTGGAGTCGTTGCACCTGTGGTTCTTGTTCTCTGAANNNAGATCGGAAG |
| 64686_1 | TATGAGTGTGGAGTCGTTGCTTTTGCAGTCAGACAGAGCTTTTNNNAGATCGGAAG |
| 64687_1 | TATGAGTGTGGAGTCGTTGCTATCAACCTGTCATCCCAGCAATNNNAGATCGGAAG |
| 64688_1 | TATGAGTGTGGAGTCGTTGCTTGTTCTCATTCCACAAGACCCTNNNAGATCGGAAGA |
| 64689_1 | TATGAGTGTGGAGTCGTTGCAGAAAGAAAGGCTGGGACTACTGNNNAGATCGGAA |
| 64690_1 | TATGAGTGTGGAGTCGTTGCACTGTCCAGGATGAGAATGAACCNNNAGATCGGAA |
| 64692_1 | TATGAGTGTGGAGTCGTTGCGAACACACTCCTCCTTCCCATTANNNAGATCGGAAGA |
| 64693_1 | TATGAGTGTGGAGTCGTTGCTATTTAGCCCTTGGATCCAGCTGNNNAGATCGGAAG |
| 64694_1 | TATGAGTGTGGAGTCGTTGCCTTCCAGTTTGCCTCTTGAGTCTNNNAGATCGGAAG |
| 64695_1 | TATGAGTGTGGAGTCGTTGCAACTGCTACTTTAGGGCTTTGGTNNNAGATCGGAAG |
| 64696_1 | TATGAGTGTGGAGTCGTTGCAAATCTCAGAACCTCACAGCTGGNNNAGATCGGAAG |
| 64697_1 | TATGAGTGTGGAGTCGTTGCTAGACATGTTCACATAAGGCCGGNNNAGATCGGAA |
| 64698_1 | TATGAGTGTGGAGTCGTTGCCTCCTGTGAATGCTCAGTATGGANNNAGATCGGAAG |
| 64699_1 | TATGAGTGTGGAGTCGTTGCTTATTCAACCATGTGCTAGGTGCNNNAGATCGGAAG |
| 64701_1 | TATGAGTGTGGAGTCGTTGCTATTTCCTTCTGCTCTGACTCCGNNNAGATCGGAAGA |
| 64702_1 | TATGAGTGTGGAGTCGTTGCGAGGGGCTCTGCAAATTGTAATCNNNAGATCGGAA |
| 64703_1 | TATGAGTGTGGAGTCGTTGCGGGGGGATGCTATTTCTCTTTCANNNAGATCGGAAG |
| 64704_1 | TATGAGTGTGGAGTCGTTGCCCAGTACCACAGGGATCATTCAANNNAGATCGGAAG |
| 64705_1 | TATGAGTGTGGAGTCGTTGCTTCTCAACTGCTCCATGGATTGANNNAGATCGGAAG |
| 64706_1 | TATGAGTGTGGAGTCGTTGCTTTCTGCTTGCTTACAGTGAAGCNNNAGATCGGAAG |
| 64707_1 | TATGAGTGTGGAGTCGTTGCAGCATATTAGACAGCACCGAGAGNNNAGATCGGAA |
| 64708_1 | TATGAGTGTGGAGTCGTTGCCTATCGATGTGCAAAGGTATGCGNNNAGATCGGAA |
| 64709_1 | TATGAGTGTGGAGTCGTTGCGGACAGGATGTGCAGCAGATTATNNNAGATCGGAA |
| 64712_1 | TATGAGTGTGGAGTCGTTGCACAACCAGCTATTAGGAAAAGCCNNNAGATCGGAA |
| 64713_1 | TATGAGTGTGGAGTCGTTGCTTTTTTTGCTGGGGGAGGATACTNNNAGATCGGAA |
| 64714_1 | TATGAGTGTGGAGTCGTTGCCTAGGCTTGATTCTGGTGTGACTNNNAGATCGGAAG |
| 64715_1 | TATGAGTGTGGAGTCGTTGCTGACACAACTTCCATGCTTCCTANNNAGATCGGAAGA |
| 64717_1 | TATGAGTGTGGAGTCGTTGCATTCTGGGGCCATAGTTGTCTTTNNNAGATCGGAAG |
| 64718_1 | TATGAGTGTGGAGTCGTTGCAGTTGTCCTTCAGTATCCATGGGNNNAGATCGGAAG |
| 64719_1 | TATGAGTGTGGAGTCGTTGCGAGAGAGAGAGAGAGAGAGACCCNNNAGATCGGA |
| 64720_1 | TATGAGTGTGGAGTCGTTGCCATCCTGAAACCATCTCTGTCCTNNNAGATCGGAAGA |
| 64721_1 | TATGAGTGTGGAGTCGTTGCCAAACCAAACCTGAGTCCTGTCTNNNAGATCGGAAG |
| 64722_1 | TATGAGTGTGGAGTCGTTGCAAAGCTTCTCCTTGATAGTGCCANNNAGATCGGAAG |
| 64723_1 | TATGAGTGTGGAGTCGTTGCTGTGTGTGCTACAGAACATGTCTNNNAGATCGGAAG |
| 64724_1 | TATGAGTGTGGAGTCGTTGCGCAATGCCAAATTTCCTTTGAGGNNNAGATCGGAAG |
| 64725_1 | TATGAGTGTGGAGTCGTTGCCAGGGGTCTTATCTTGGATGGTTNNNAGATCGGAA |
| 64726_1 | TATGAGTGTGGAGTCGTTGCAGTTTGGTAACGAATCTGCTTGCNNNAGATCGGAAG |
| 64727_1 | TATGAGTGTGGAGTCGTTGCGGGAAACCCAGTTCACACAATTCNNNAGATCGGAAG |
| 64728_1 | TATGAGTGTGGAGTCGTTGCGCAGGTTTCTCTCAGGTCTAACANNNAGATCGGAAG |

| | |
|---|---|
| 64729_1 | TATGAGTGTGGAGTCGTTGCGATTGTGTCACAAAGCCCAAGAANNNAGATCGGAAG |
| 64731_1 | TATGAGTGTGGAGTCGTTGCTCCCATCGTTTTGTATCCCTGAGNNNAGATCGGAAGA |
| 64732_1 | TATGAGTGTGGAGTCGTTGCGGATATTTGTTCCCCCCAAGTCTNNNAGATCGGAAGA |
| 64733_1 | TATGAGTGTGGAGTCGTTGCCTGGAAGTACCGTTTCCTAGGTTNNNAGATCGGAAG |
| 64736_1 | TATGAGTGTGGAGTCGTTGCCAGGGTTCTTTCTAGCCTTTCCTNNNAGATCGGAAGA |
| 64737_1 | TATGAGTGTGGAGTCGTTGCATTTAGTCAGCAGTTTGCCAACCNNNAGATCGGAAGA |
| 64738_1 | TATGAGTGTGGAGTCGTTGCAAGGTATGGAGGGTGCTAGGTATNNNAGATCGGAA |
| 64739_1 | TATGAGTGTGGAGTCGTTGCCTGGAATAAGGAGCCAGGAATGANNNAGATCGGAA |
| 64740_1 | TATGAGTGTGGAGTCGTTGCTGCTCAGGTCTGAAGGTTTTTCTNNNAGATCGGAAG |
| 64741_1 | TATGAGTGTGGAGTCGTTGCAGCTTAGCCAGGCTCAATCAATANNNAGATCGGAAG |
| 64742_1 | TATGAGTGTGGAGTCGTTGCCCTCTAACTAGGGTGAGTTTGGCNNNAGATCGGAAG |
| 64743_1 | TATGAGTGTGGAGTCGTTGCGTCAGTGGTTAGAGGTCTGTTCANNNAGATCGGAAG |
| 64745_1 | TATGAGTGTGGAGTCGTTGCGTGTTTGTGTGTGTGGATGTCTTNNNAGATCGGAAG |
| 64746_1 | TATGAGTGTGGAGTCGTTGCGCCACAAGAGCATTAGCAAATTGNNNAGATCGGAAG |
| 64747_1 | TATGAGTGTGGAGTCGTTGCGCACCAACTCTGATCTCTGAAGANNNAGATCGGAAG |
| 64749_1 | TATGAGTGTGGAGTCGTTGCTGCAATGGAGAGTTGGAGATTCTNNNAGATCGGAAG |
| 64750_1 | TATGAGTGTGGAGTCGTTGCCTTTCTTCGCCTATCATGTGTGCNNNAGATCGGAAGA |
| 64751_1 | TATGAGTGTGGAGTCGTTGCTTTCCCTAAAAGTGACCCTTCCCNNNAGATCGGAAGA |
| 64752_1 | TATGAGTGTGGAGTCGTTGCGCTTAGGAGAAACAGGAACCAGANNNAGATCGGAA |
| 64754_1 | TATGAGTGTGGAGTCGTTGCTTCCCCCTCCCTCTCCTTATCTANNNAGATCGGAAGAG |
| 64755_1 | TATGAGTGTGGAGTCGTTGCTGTGACCTATCTTGTTCCACCAGNNNAGATCGGAAGA |
| 64756_1 | TATGAGTGTGGAGTCGTTGCTTCTTATAAGTTTGGGGCTGCCTNNNAGATCGGAAG |
| 64757_1 | TATGAGTGTGGAGTCGTTGCAGAGAGAGAGAGTGTAGTGGGTTNNNAGATCGGA |
| 64758_1 | TATGAGTGTGGAGTCGTTGCAGTGTGGGACACAATAAGTTCCTNNNAGATCGGAAG |
| 64759_1 | TATGAGTGTGGAGTCGTTGCTTCTCTCTGTCTAGTCTCCCCTCNNNAGATCGGAAGAG |
| 64760_1 | TATGAGTGTGGAGTCGTTGCAGGACAGACATCAGCTGTTTCATNNNAGATCGGAAG |
| 64761_1 | TATGAGTGTGGAGTCGTTGCAGTTGATTACCCCAGGAAAGCATNNNAGATCGGAAG |
| 64762_1 | TATGAGTGTGGAGTCGTTGCACAGATGACTATTTCAGCCAGCANNNAGATCGGAAG |
| 64763_1 | TATGAGTGTGGAGTCGTTGCGAGTCTGCAAGGCTGTTCATTTTNNNAGATCGGAAG |
| 64764_1 | TATGAGTGTGGAGTCGTTGCGTTTGGAGGACAATTTCTCGCAGNNNAGATCGGAAG |
| 64765_1 | TATGAGTGTGGAGTCGTTGCGTCAGAAGTCTTCACCGAGCTATNNNAGATCGGAAG |
| 64766_1 | TATGAGTGTGGAGTCGTTGCGGTAAGAGGTTGGGATGGAGAAGNNNAGATCGGA |
| 64767_1 | TATGAGTGTGGAGTCGTTGCTGGCTAATACCTGTAATCCCAGCNNNAGATCGGAAGA |
| 64768_1 | TATGAGTGTGGAGTCGTTGCGATCTGGTCCATCCTTGCTTTCTNNNAGATCGGAAGA |
| 64769_1 | TATGAGTGTGGAGTCGTTGCCAGGATCACCATTCTCCCTTCTCNNNAGATCGGAAGA |
| 64770_1 | TATGAGTGTGGAGTCGTTGCCAATACAGCTTCCTTGGCCATTCNNNAGATCGGAAGA |
| 64773_1 | TATGAGTGTGGAGTCGTTGCGGCTGTTCCTTGCTGAGAAAAATNNNAGATCGGAAG |
| 64774_1 | TATGAGTGTGGAGTCGTTGCAGGATCTCAGTTCACTGCAACTTNNNAGATCGGAAGA |
| 64776_1 | TATGAGTGTGGAGTCGTTGCTGTGTGTTGTGTGTACATGTGTGNNNAGATCGGAAG |
| 64778_1 | TATGAGTGTGGAGTCGTTGCTTCTACCTGGCCCTAGTCTATCCNNNAGATCGGAAGA |
| 64779_1 | TATGAGTGTGGAGTCGTTGCAATTCCTGGGCTCAAGCAATCTANNNAGATCGGAAG |
| 64780_1 | TATGAGTGTGGAGTCGTTGCGTCTCATCTCAGGCCATGTAACANNNAGATCGGAAGA |
| 64781_1 | TATGAGTGTGGAGTCGTTGCTTAGAGAGGTCTTGCTCTGTTGCNNNAGATCGGAAG |

| | |
|---|---|
| 64783_1 | TATGAGTGTGGAGTCGTTGCTCTTGAACTCCTGACCTCAAGTGNNNAGATCGGAAG |
| 64784_1 | TATGAGTGTGGAGTCGTTGCTCATTATGGAGGCTCACAAGACCNNNAGATCGGAAG |
| 64785_1 | TATGAGTGTGGAGTCGTTGCTTGTCCCCCTTTATCCACAGTTTNNNAGATCGGAAGA |
| 64786_1 | TATGAGTGTGGAGTCGTTGCTTCAAGCAATTCTCCCACCTCAGNNNAGATCGGAAGA |
| 64787_1 | TATGAGTGTGGAGTCGTTGCGAGATAGGGTTTTGCCATGTTGGNNNAGATCGGAA |
| 64788_1 | TATGAGTGTGGAGTCGTTGCTCTGAACCTCCTCCTTTTCATGGNNNAGATCGGAAGA |
| 64789_1 | TATGAGTGTGGAGTCGTTGCAGTTGAGATTGCAGTTGTGAAGCNNNAGATCGGAA |
| 64790_1 | TATGAGTGTGGAGTCGTTGCCATTCTTTTCCATTTGGCCCTGTNNNAGATCGGAAGA |
| 64791_1 | TATGAGTGTGGAGTCGTTGCAGCTTCAGTTTTCATCCCTTGACNNNAGATCGGAAGA |
| 64792_1 | TATGAGTGTGGAGTCGTTGCTTGGTGTTTTGATAGGGATTGCANNNAGATCGGAA |
| 64793_1 | TATGAGTGTGGAGTCGTTGCATCTACAATTGGCAAGCTGGAGANNNAGATCGGAA |
| 64794_1 | TATGAGTGTGGAGTCGTTGCAACTCACATGATCACAAGGTCCCNNNAGATCGGAAG |
| 64796_1 | TATGAGTGTGGAGTCGTTGCCTGAAGGACATCTTGGTTGCTTCNNNAGATCGGAAG |
| 64797_1 | TATGAGTGTGGAGTCGTTGCCTTAGCTCTTACTCTTCCCAGCCNNNAGATCGGAAGA |
| 64798_1 | TATGAGTGTGGAGTCGTTGCAAGGTCAGGAGATCGAGACCATCNNNAGATCGGAA |
| 64801_1 | TATGAGTGTGGAGTCGTTGCAAGTACAGCAGTCTCCTTGTTGTNNNAGATCGGAAG |
| 64802_1 | TATGAGTGTGGAGTCGTTGCTGAGGAGTTGGATGGTGATAGTGNNNAGATCGGA |
| 64803_1 | TATGAGTGTGGAGTCGTTGCATAACCCGTTTGTTGAGCATCTGNNNAGATCGGAAG |
| 64805_1 | TATGAGTGTGGAGTCGTTGCGGCTTGTGCTATGGTTTGAATGANNNAGATCGGAA |
| 64806_1 | TATGAGTGTGGAGTCGTTGCCAATGAGCTAGGATCACACCACTNNNAGATCGGAAG |
| 64807_1 | TATGAGTGTGGAGTCGTTGCTTCTACTTCTCTGCAGCTTTGGCNNNAGATCGGAAGA |
| 64808_1 | TATGAGTGTGGAGTCGTTGCAACTCCCCTTTCTGTACCGTAAANNNAGATCGGAAGA |
| 64809_1 | TATGAGTGTGGAGTCGTTGCAAAACCTGGAGAAGACACAACCTNNNAGATCGGAA |
| 64810_1 | TATGAGTGTGGAGTCGTTGCGAGATCAAAATGCAGCTGCCTAGNNNAGATCGGAA |
| 64811_1 | TATGAGTGTGGAGTCGTTGCAGAGAACGAGCTCAGTGAATTGTNNNAGATCGGAA |
| 64813_1 | TATGAGTGTGGAGTCGTTGCAATTCGCTAGACATGGTGGATGGNNNAGATCGGAA |
| 64814_1 | TATGAGTGTGGAGTCGTTGCGAAAATGGAACCCTCACACACTGNNNAGATCGGAAG |
| 64816_1 | TATGAGTGTGGAGTCGTTGCAGGCTCTAGACTTTTCTTGTGCANNNAGATCGGAAG |
| 64817_1 | TATGAGTGTGGAGTCGTTGCCACTCATTGATTGATGGGCGTTTNNNAGATCGGAAG |
| 64818_1 | TATGAGTGTGGAGTCGTTGCTTGATTCTTTTGACTGATGCGGGNNNAGATCGGAAG |
| 64819_1 | TATGAGTGTGGAGTCGTTGCGGCTGGTTCCATATTTTTGCAGTNNNAGATCGGAAG |
| 64820_1 | TATGAGTGTGGAGTCGTTGCTGATGAAACACACACACACACACNNNAGATCGGAAG |
| 64822_1 | TATGAGTGTGGAGTCGTTGCTCTCGATCTCCTGACCTCATGATNNNAGATCGGAAGA |
| 64823_1 | TATGAGTGTGGAGTCGTTGCAGAGGTAGGTATCGTGTCCCAATNNNAGATCGGAA |
| 64824_1 | TATGAGTGTGGAGTCGTTGCAGAGATTCCTCCAGGTAAGAGCTNNNAGATCGGAA |
| 64825_1 | TATGAGTGTGGAGTCGTTGCGTCACTGAGGGACTGAAACTCTCNNNAGATCGGAA |
| 64826_1 | TATGAGTGTGGAGTCGTTGCATATGTGTGTGTGTGTGTGTGTGNNNAGATCGGAA |
| 64827_1 | TATGAGTGTGGAGTCGTTGCTTCATGTGATTCTCCTGCCTCAGNNNAGATCGGAAGA |
| 64828_1 | TATGAGTGTGGAGTCGTTGCTGTATTTCTGCCAGGTTTGGGTANNNAGATCGGAAG |
| 64829_1 | TATGAGTGTGGAGTCGTTGCGAGAGAGAGAGAGAGAGACAGGGNNNAGATCGG |
| 64830_1 | TATGAGTGTGGAGTCGTTGCTGGTGGTGTTGCTGTTGTTATTCNNNAGATCGGAAG |
| 64831_1 | TATGAGTGTGGAGTCGTTGCGGGGAGGGAGGGTATAACTTTTCNNNAGATCGGAA |
| 64832_1 | TATGAGTGTGGAGTCGTTGCTTTCTCTTGCCTGATTGCTTTGGNNNAGATCGGAAGA |

| | |
|---|---|
| 64833_1 | TATGAGTGTGGAGTCGTTGCTGGCTCTGAGAACCAGTTACTTCNNNAGATCGGAAG |
| 64834_1 | TATGAGTGTGGAGTCGTTGCAGGCATGGGAACTACTTCACATTNNNAGATCGGAAG |
| 64837_1 | TATGAGTGTGGAGTCGTTGCTGGTGGTGTAATTCAGTCCAAGTNNNAGATCGGAA |
| 64838_1 | TATGAGTGTGGAGTCGTTGCCCTAGAGGAAGACAGGGAGAAAANNNAGATCGGA |
| 64839_1 | TATGAGTGTGGAGTCGTTGCAAAAAACAAGCTGATGGGAAGCCNNNAGATCGGAA |
| 64840_1 | TATGAGTGTGGAGTCGTTGCTCCCACATTCAAAAATCCCTCCTNNNAGATCGGAAGA |
| 64842_1 | TATGAGTGTGGAGTCGTTGCACACCCAACAGAAATGTGTTCACNNNAGATCGGAAG |
| 64843_1 | TATGAGTGTGGAGTCGTTGCGTAACAGAGTCTCACTTTGTCGCNNNAGATCGGAAG |
| 64844_1 | TATGAGTGTGGAGTCGTTGCCCATATGTGTCTTTCTGTGCCTGNNNAGATCGGAAG |
| 64845_1 | TATGAGTGTGGAGTCGTTGCAAAAGGGGCAAAAGTGTGAAACANNNAGATCGGA |
| 64846_1 | TATGAGTGTGGAGTCGTTGCACAGAAGTCTGCTCATGTGTAGGNNNAGATCGGAA |
| 64848_1 | TATGAGTGTGGAGTCGTTGCATCTCTGTGTGTGTGTGTGTGTANNNAGATCGGAA |
| 64849_1 | TATGAGTGTGGAGTCGTTGCTGAGATGGGAGGATCACTTGAACNNNAGATCGGAA |
| 64850_1 | TATGAGTGTGGAGTCGTTGCTCCCAGTACTTTCAGAAGCCAAGNNNAGATCGGAAG |
| 64851_1 | TATGAGTGTGGAGTCGTTGCCTACTTAGGCTGAGGTGAGAGGANNNAGATCGGAA |
| 64852_1 | TATGAGTGTGGAGTCGTTGCATTTCTTGAGAAGCGGGGAGAAGNNNAGATCGGA |
| 64853_1 | TATGAGTGTGGAGTCGTTGCACACACACACACACACACAAATTNNNAGATCGGAAG |
| 64856_1 | TATGAGTGTGGAGTCGTTGCGCAAAGGGGATCAGAAACAACAANNNAGATCGGAA |
| 64857_1 | TATGAGTGTGGAGTCGTTGCACAAAATAATGGCATTTGCAGCGNNNAGATCGGAA |
| 64858_1 | TATGAGTGTGGAGTCGTTGCTCAGGTTTATGCCATTCTCCTGTNNNAGATCGGAAG |
| 64859_1 | TATGAGTGTGGAGTCGTTGCCCCTGGTTCCTTTTACTGGAGAANNNAGATCGGAAG |
| 64860_1 | TATGAGTGTGGAGTCGTTGCCAAAGCTACATCAGGTGTTTGCANNNAGATCGGAAG |
| 64861_1 | TATGAGTGTGGAGTCGTTGCATCTCAAACTCCTGGCCTCAAATNNNAGATCGGAAG |
| 64862_1 | TATGAGTGTGGAGTCGTTGCGATTCTCTCTCTGTCACTGAGGCNNNAGATCGGAAG |
| 64863_1 | TATGAGTGTGGAGTCGTTGCAGTGTAATCATAGCTCCCTGCAGNNNAGATCGGAAG |
| 64864_1 | TATGAGTGTGGAGTCGTTGCAGACAGGGTTTCACTATGTTGGCNNNAGATCGGAA |
| 64865_1 | TATGAGTGTGGAGTCGTTGCCAGTGGTGTGATCTCAGCTTACTNNNAGATCGGAAG |
| 64866_1 | TATGAGTGTGGAGTCGTTGCTTGTCATGGGCTCTACTCATTGGNNNAGATCGGAAG |
| 64867_1 | TATGAGTGTGGAGTCGTTGCGTGTGTATGTGTGTGTGTCTGTGNNNAGATCGGAA |
| 64868_1 | TATGAGTGTGGAGTCGTTGCTGTGTGTGTGTGTGTGTGTATCTNNNAGATCGGAA |
| 64869_1 | TATGAGTGTGGAGTCGTTGCGGGGTCCCTTCTCAATTTTTTGTNNNAGATCGGAAG |
| 64870_1 | TATGAGTGTGGAGTCGTTGCTGCCTCAGTCCTAAAATCAACCANNNAGATCGGAAG |
| 64871_1 | TATGAGTGTGGAGTCGTTGCGGTGTGTTTGATCCTGTTCATGGNNNAGATCGGAA |
| 64872_1 | TATGAGTGTGGAGTCGTTGCAGTGAGCCTAGATTGTGTCACAGNNNAGATCGGAA |
| 64873_1 | TATGAGTGTGGAGTCGTTGCGCTTTGCTGGGGGAGTAATTTTTNNNAGATCGGAA |
| 64874_1 | TATGAGTGTGGAGTCGTTGCTTGCTTGAATCTGAGAGGTGGAANNNAGATCGGAA |
| 64875_1 | TATGAGTGTGGAGTCGTTGCTGCAGGATACGAAATCAACACACNNNAGATCGGAA |
| 64876_1 | TATGAGTGTGGAGTCGTTGCTCCGACTCAAAACACACAAACACNNNAGATCGGAAG |
| 64878_1 | TATGAGTGTGGAGTCGTTGCCTTCAATCCCTGGTCTCACTTGANNNAGATCGGAAGA |
| 64879_1 | TATGAGTGTGGAGTCGTTGCACATTCATGGGTGATCGTACCTCNNNAGATCGGAAG |
| 64880_1 | TATGAGTGTGGAGTCGTTGCGAGGGTAGGAAAATCACGACTCANNNAGATCGGAA |
| 64881_1 | TATGAGTGTGGAGTCGTTGCAATCCAGGAGCACATCAAAAAGCNNNAGATCGGAA |
| 64882_1 | TATGAGTGTGGAGTCGTTGCTACTCCTGGAGGGTTGCTACTTCNNNAGATCGGAAG |

64883_1 TATGAGTGTGGAGTCGTTGCGGAGACCGCTATTCTAAGTGAGGNNNAGATCGGAA
64885_1 TATGAGTGTGGAGTCGTTGCATTCCATGTCTTTGCTATGCTGCNNNAGATCGGAAG
64886_1 TATGAGTGTGGAGTCGTTGCGTGTAGCCAAGGGACTATCAGTTNNNAGATCGGAA
64887_1 TATGAGTGTGGAGTCGTTGCATTTGGGCCACATAGATCTGGTTNNNAGATCGGAAG
64888_1 TATGAGTGTGGAGTCGTTGCCACCTGTCATCTCAGCTACTCAGNNNAGATCGGAAG
64890_1 TATGAGTGTGGAGTCGTTGCTTCCAGTGATTCTTCTGCCTCAGNNNAGATCGGAAG
64891_1 TATGAGTGTGGAGTCGTTGCGAGGTTGCAGTTACTGTACTCCANNNAGATCGGAAG
64892_1 TATGAGTGTGGAGTCGTTGCACTGTTATGGAGACTGGGAAGTCNNNAGATCGGAA
64893_1 TATGAGTGTGGAGTCGTTGCACAGACATACAGCAGGTAACACCNNNAGATCGGAA
64894_1 TATGAGTGTGGAGTCGTTGCTCAAGGAATGAGATGAATGGCCANNNAGATCGGAA
64896_1 TATGAGTGTGGAGTCGTTGCACTTGAGATCGGGAGTTTGAGACNNNAGATCGGAA
64897_1 TATGAGTGTGGAGTCGTTGCCCCCCCCAAACAAAAACAAAAACNNNAGATCGGAAG
64898_1 TATGAGTGTGGAGTCGTTGCGGAAGATGGAAGAAGTCTGGGTTNNNAGATCGGA
64899_1 TATGAGTGTGGAGTCGTTGCATGATAGTGACCAAAGGAGACGGNNNAGATCGGA
64900_1 TATGAGTGTGGAGTCGTTGCAAGAACAAGCTCATGTCCTTTGCNNNAGATCGGAAG
64901_1 TATGAGTGTGGAGTCGTTGCCCCACAGGAGAAAGAAAGAACCANNNAGATCGGAA
64902_1 TATGAGTGTGGAGTCGTTGCGGGTGACAGAAGGAAACACAAAANNNAGATCGGA
64903_1 TATGAGTGTGGAGTCGTTGCAAAAAAAAAAAAGCCAGGCACGGNNNAGATCGGAA
64904_1 TATGAGTGTGGAGTCGTTGCTTGGGAAATGTCTGTTCAGATCANNNAGATCGGAA
64905_1 TATGAGTGTGGAGTCGTTGCACACACTTCTGACTTTCCAAGGTNNNAGATCGGAAG
64908_1 TATGAGTGTGGAGTCGTTGCCTCAAGCAATGCTCTCACATCAGNNNAGATCGGAAG
64909_1 TATGAGTGTGGAGTCGTTGCATAAGAGTTGGCTGGTACTCAGGNNNAGATCGGAA
64910_1 TATGAGTGTGGAGTCGTTGCATCAGTCAGTAGGGATGCTGAAGNNNAGATCGGAA
64912_1 TATGAGTGTGGAGTCGTTGCTAACATTCCCCAGCATCATTCCANNNAGATCGGAAGA
64913_1 TATGAGTGTGGAGTCGTTGCGAGAAAGATAGTGGTCCAGAGGCNNNAGATCGGA
64914_1 TATGAGTGTGGAGTCGTTGCGGTTCAAGTGATTCTCCCACTCANNNAGATCGGAAG
64916_1 TATGAGTGTGGAGTCGTTGCTCTTTGATGAGAGCCTCAGTTGCNNNAGATCGGAAG
64917_1 TATGAGTGTGGAGTCGTTGCAATTTGCAGCCAGTATCAACCAANNNAGATCGGAAG
64918_1 TATGAGTGTGGAGTCGTTGCCAAGTAGCTAGCATTACGGGTGTNNNAGATCGGAA
64919_1 TATGAGTGTGGAGTCGTTGCAGGATTCATAGGATCATGCTGGGNNNAGATCGGAA
64920_1 TATGAGTGTGGAGTCGTTGCCTGAGAGCTGAGAAACAAGCAAGNNNAGATCGGAA
64922_1 TATGAGTGTGGAGTCGTTGCGCCCAGGTGGTATTTGTTCAAATNNNAGATCGGAAG
64923_1 TATGAGTGTGGAGTCGTTGCTATCCTTACTCCTCCCCAGAGACNNNAGATCGGAAGA
64924_1 TATGAGTGTGGAGTCGTTGCAAAGACGGGAAGCTACTTTGGAANNNAGATCGGAA
64925_1 TATGAGTGTGGAGTCGTTGCAAGTGCAGTAACTGAGGAATGGANNNAGATCGGA
64926_1 TATGAGTGTGGAGTCGTTGCCTGAGGCTGGAGAATCACTTGAGNNNAGATCGGAA
64927_1 TATGAGTGTGGAGTCGTTGCCAAAAATAAGGCATGGTGGCTCANNNAGATCGGAA
64928_1 TATGAGTGTGGAGTCGTTGCGCACACACTTGCTTTATCCAGTTNNNAGATCGGAAG
64929_1 TATGAGTGTGGAGTCGTTGCAGTAGGAGCCTCTAGTTTACCCANNNAGATCGGAAG
64930_1 TATGAGTGTGGAGTCGTTGCGCAGATGAAACCACCTTCTGTTCNNNAGATCGGAAG
64931_1 TATGAGTGTGGAGTCGTTGCCACTCCTACCCACATGTCTGATTNNNAGATCGGAAGA
64932_1 TATGAGTGTGGAGTCGTTGCAGACTTTGCAAGAACCACCATTGNNNAGATCGGAAG
64933_1 TATGAGTGTGGAGTCGTTGCCAATCCCAGAGAGCAGAGGAATTNNNAGATCGGAA

| | |
|---|---|
| 64934_1 | TATGAGTGTGGAGTCGTTGCAACAGCTGCATCCAGAAAATCAGNNNAGATCGGAAG |
| 64936_1 | TATGAGTGTGGAGTCGTTGCAAGTGCTGAGATTACAGGCATGANNNAGATCGGAA |
| 64938_1 | TATGAGTGTGGAGTCGTTGCGTTCCGGGATAAATGTGCAGAACNNNAGATCGGAA |
| 64939_1 | TATGAGTGTGGAGTCGTTGCGCACAGAAACAGAAGAGAAGCAANNNAGATCGGAA |
| 64940_1 | TATGAGTGTGGAGTCGTTGCCTGAGTAGCTGGGATTATAGGCGNNNAGATCGGAA |
| 64943_1 | TATGAGTGTGGAGTCGTTGCCCTCCCTGCCTTCTAATTTGACTNNNAGATCGGAAGA |
| 64944_1 | TATGAGTGTGGAGTCGTTGCCCAAAGTTCAAGCGATTCTCCTGNNNAGATCGGAAG |
| 64945_1 | TATGAGTGTGGAGTCGTTGCGGAATGCAATTCGGGGTAGAATGNNNAGATCGGAA |
| 64947_1 | TATGAGTGTGGAGTCGTTGCGATCCACGGGGAAATTTCTGTTGNNNAGATCGGAAG |
| 64948_1 | TATGAGTGTGGAGTCGTTGCTGTCATTTACAACAACGTGGGTGNNNAGATCGGAAG |
| 64949_1 | TATGAGTGTGGAGTCGTTGCCTTTACCCAACTGATCCCAGGAANNNAGATCGGAAG |
| 64950_1 | TATGAGTGTGGAGTCGTTGCGTTGGCTTTGACTAAAACAGGCANNNAGATCGGAAG |
| 64951_1 | TATGAGTGTGGAGTCGTTGCGAGTAGTACTTCCATGCATGTGCNNNAGATCGGAAG |
| 64952_1 | TATGAGTGTGGAGTCGTTGCGCTTGACTTTTGCCTGGATCAAANNNAGATCGGAAG |
| 64954_1 | TATGAGTGTGGAGTCGTTGCACCTACACACATACACACACACANNNAGATCGGAAGA |
| 64956_1 | TATGAGTGTGGAGTCGTTGCGGTCTTGAACTCCTGACCTCAAGNNNAGATCGGAAG |
| 64957_1 | TATGAGTGTGGAGTCGTTGCGATGGAGTTCAGTGACACGATCTNNNAGATCGGAA |
| 64958_1 | TATGAGTGTGGAGTCGTTGCGAACAACTCCAGCACTGTCTAGANNNAGATCGGAAG |
| 64959_1 | TATGAGTGTGGAGTCGTTGCTGCTGTGGATGAAGGAAAGCATANNNAGATCGGAA |
| 64961_1 | TATGAGTGTGGAGTCGTTGCAAGGAGCCACCATTTTTTCTGTGNNNAGATCGGAAG |
| 64962_1 | TATGAGTGTGGAGTCGTTGCGACAAGATCTCACTGTCACCCAGNNNAGATCGGAAG |
| 64963_1 | TATGAGTGTGGAGTCGTTGCTCCATCAAGGTAGCCAAGAAACANNNAGATCGGAAG |
| 64964_1 | TATGAGTGTGGAGTCGTTGCTTGAGGACAGTGGTGCAATCATANNNAGATCGGAA |
| 64966_1 | TATGAGTGTGGAGTCGTTGCTGGTCAGTTTACATAGCAGAGGANNNAGATCGGAA |
| 64967_1 | TATGAGTGTGGAGTCGTTGCAATTTGCATCTCTCCTGAGCAAANNNAGATCGGAAG |
| 64968_1 | TATGAGTGTGGAGTCGTTGCAGAGAGAGAGAGAAGCTGAGGTTNNNAGATCGGA |
| 64969_1 | TATGAGTGTGGAGTCGTTGCAAGGGAAAAGGAGCTTTGAATGGNNNAGATCGGA |
| 64972_1 | TATGAGTGTGGAGTCGTTGCGAGGTCAGGAGATCAAGACCATCNNNAGATCGGAA |
| 64973_1 | TATGAGTGTGGAGTCGTTGCTGCACACACACACATCTCATTTGNNNAGATCGGAAGA |
| 64974_1 | TATGAGTGTGGAGTCGTTGCTTTGTGTGTGTGTGACAGTTTCANNNAGATCGGAAG |
| 64975_1 | TATGAGTGTGGAGTCGTTGCAGTAACCATCAAAACCCGGAAGANNNAGATCGGAAG |
| 64976_1 | TATGAGTGTGGAGTCGTTGCACATGGCTGGCTTTGAAAATGAANNNAGATCGGAA |
| 64977_1 | TATGAGTGTGGAGTCGTTGCCTGTGTGTATGTGTGTGTGTGTGNNNAGATCGGAA |
| 64978_1 | TATGAGTGTGGAGTCGTTGCTCATTTCTCACAGTTCTGGAGGCNNNAGATCGGAAG |
| 64980_1 | TATGAGTGTGGAGTCGTTGCACTCCTAAGCTCAAGTGATCTGCNNNAGATCGGAAG |
| 64982_1 | TATGAGTGTGGAGTCGTTGCTTTTTTGAGACGGAGTCTTGCTCNNNAGATCGGAAG |
| 64983_1 | TATGAGTGTGGAGTCGTTGCTAAATTCTCAGCTTTGCCACACGNNNAGATCGGAAGA |
| 64984_1 | TATGAGTGTGGAGTCGTTGCAGAGAGAAAGTGTGTGTGTGTGTNNNAGATCGGAA |
| 64985_1 | TATGAGTGTGGAGTCGTTGCAATCCTCATAAGATTCCCTGCCCNNNAGATCGGAAGA |
| 64986_1 | TATGAGTGTGGAGTCGTTGCGTCGAGTTTGTGGACAACCAAAANNNAGATCGGAA |
| 64987_1 | TATGAGTGTGGAGTCGTTGCACTGAGACAGAAGACTTGCTTGANNNAGATCGGAA |
| 64988_1 | TATGAGTGTGGAGTCGTTGCCACACACACATACACACACATGTNNNAGATCGGAAG |
| 64989_1 | TATGAGTGTGGAGTCGTTGCCTGAGATCAGGAGTTCAAGACCANNNAGATCGGAA |

64990_1 TATGAGTGTGGAGTCGTTGCATCCACCTGTACATACACATGCCNNNAGATCGGAGA
64991_1 TATGAGTGTGGAGTCGTTGCAATGGTTTCAGACACTGGACAGANNNAGATCGGAAC
64992_1 TATGAGTGTGGAGTCGTTGCAGCAGTCATTTTCAATTCCCTCCNNNAGATCGGAAGA
64995_1 TATGAGTGTGGAGTCGTTGCGATGTGTGTGTGTTTGTGTGTGTNNNAGATCGGAAC
64996_1 TATGAGTGTGGAGTCGTTGCGCATATGTTCACGATGGTCTGTGNNNAGATCGGAAG
64997_1 TATGAGTGTGGAGTCGTTGCAGGGAGAGAAAGAGACAGAGACANNNAGATCGGA
64999_1 TATGAGTGTGGAGTCGTTGCAGGAGGGAGAGAGAGAGAGAAACNNNAGATCGGA
65000_1 TATGAGTGTGGAGTCGTTGCTGAACAAAACACCAGCAAACCAANNNAGATCGGAAG
65001_1 TATGAGTGTGGAGTCGTTGCTATGGTGGTGCATGCCTATAGTCNNNAGATCGGAAG
65003_1 TATGAGTGTGGAGTCGTTGCCATACACAATGCAAGAGCGGTAGNNNAGATCGGAAC
65004_1 TATGAGTGTGGAGTCGTTGCCTGTAGTCCCAGCTACCTGAGAGNNNAGATCGGAAG
65005_1 TATGAGTGTGGAGTCGTTGCTTGAAGCAACATGGATGGAACTGNNNAGATCGGAAC
65006_1 TATGAGTGTGGAGTCGTTGCATGGTGTAGAAATGCAGTGCTTGNNNAGATCGGAAC
65007_1 TATGAGTGTGGAGTCGTTGCAGAGATCCTAGCCATTCTAGCCANNNAGATCGGAAG
65008_1 TATGAGTGTGGAGTCGTTGCTTGATTCTTTTGACTGATGCGGGNNNAGATCGGAAG
65009_1 TATGAGTGTGGAGTCGTTGCTCACCATCTGTACCTTGGTTCAGNNNAGATCGGAAGA
65011_1 TATGAGTGTGGAGTCGTTGCCTCCCATCCTAGTCCAATCATGGNNNAGATCGGAAGA
65012_1 TATGAGTGTGGAGTCGTTGCGGTTCAAGTGATTCTTCTGCCTCNNNAGATCGGAAGA
65013_1 TATGAGTGTGGAGTCGTTGCCCAAGATTCCCAGCACATCATTCNNNAGATCGGAAGA
65015_1 TATGAGTGTGGAGTCGTTGCTCCATCATCAGTGATCTTGGCAANNNAGATCGGAAGA
65016_1 TATGAGTGTGGAGTCGTTGCATAAGTTTGTCATGGCCACTTGCNNNAGATCGGAAG
65018_1 TATGAGTGTGGAGTCGTTGCGCCTGTTGTGATTTTGATTGGGANNNAGATCGGAAC
65019_1 TATGAGTGTGGAGTCGTTGCTCCCGTAGTGACTAACAGGATCTNNNAGATCGGAAG
65020_1 TATGAGTGTGGAGTCGTTGCCTTCTGGGTTCAAGCGATTCTTCNNNAGATCGGAAGA
65021_1 TATGAGTGTGGAGTCGTTGCCCTTGGGGGAAATGTCAGTTTTCNNNAGATCGGAAG
65022_1 TATGAGTGTGGAGTCGTTGCGCAGTCTTGGTGATAACACACACNNNAGATCGGAAG
65023_1 TATGAGTGTGGAGTCGTTGCAGAAGCAGTCTAACCTTCTGAGCNNNAGATCGGAAG
65024_1 TATGAGTGTGGAGTCGTTGCCTCCACCCCATTAGCAGTAAACTNNNAGATCGGAAGA
65028_1 TATGAGTGTGGAGTCGTTGCGTGATGAAGAGGGGTGTATGTGTNNNAGATCGGAA
65029_1 TATGAGTGTGGAGTCGTTGCTCTCTTTCTTTCTCTCCCTCCCTNNNAGATCGGAAGAG
65030_1 TATGAGTGTGGAGTCGTTGCTGAACTCCTGAGCTCAAGCAATCNNNAGATCGGAAG
65031_1 TATGAGTGTGGAGTCGTTGCCCGAAAAGAGTCAGCAAAAGGAGNNNAGATCGGAA
65034_1 TATGAGTGTGGAGTCGTTGCAAGAAATAGGTCGGGCATAGTGGNNNAGATCGGAA
65035_1 TATGAGTGTGGAGTCGTTGCCCTTGTTGATTTGGGACTTGACCNNNAGATCGGAAG
65036_1 TATGAGTGTGGAGTCGTTGCAAAGCATTCAGAACAATCCTGGCNNNAGATCGGAAG
65037_1 TATGAGTGTGGAGTCGTTGCATGTCTAACTAAGTGGAGGGACANNNAGATCGGAAC
65038_1 TATGAGTGTGGAGTCGTTGCCACATCCACCTTCCCAGATACATNNNAGATCGGAAGA
65039_1 TATGAGTGTGGAGTCGTTGCAATAAATAGCCAGGTGTGGTGGTNNNAGATCGGAA
65040_1 TATGAGTGTGGAGTCGTTGCAGGTAGAAAAGAACTGGCTGAGGNNNAGATCGGAA
65041_1 TATGAGTGTGGAGTCGTTGCCAAAATCCGCAGTCAGAACATGANNNAGATCGGAAG
65042_1 TATGAGTGTGGAGTCGTTGCCACACATGCAGAGAGAGAGAGAGNNNAGATCGGAA
65045_1 TATGAGTGTGGAGTCGTTGCCTCACTGAAAGCTCTGTCTCCTGNNNAGATCGGAAGA
65047_1 TATGAGTGTGGAGTCGTTGCTGGTTTTTTCAGTTGCAGCTTCANNNAGATCGGAAGA

65048_1 TATGAGTGTGGAGTCGTTGCCAGAGTTTTCTTCTTGTTGCCCANNNAGATCGGAAG
65049_1 TATGAGTGTGGAGTCGTTGCACAAAAACCAGCACATTGTTCCANNNAGATCGGAAG
65050_1 TATGAGTGTGGAGTCGTTGCGCCAACCAAGGTGCATGTAAAAANNNAGATCGGAA
65052_1 TATGAGTGTGGAGTCGTTGCGATAGAACCAGCACGGATCAGATNNNAGATCGGAA
65053_1 TATGAGTGTGGAGTCGTTGCAACTTTCCACCCATACAGTCCTCNNNAGATCGGAAGA
65054_1 TATGAGTGTGGAGTCGTTGCCTGTAGTCTGAGCTACTCGGGAGNNNAGATCGGAA
65055_1 TATGAGTGTGGAGTCGTTGCCTGTCAAGGAATCCACTCACAGANNNAGATCGGAAG
65056_1 TATGAGTGTGGAGTCGTTGCAAGCCTCAATTGTTCCACCCTATNNNAGATCGGAAG
65057_1 TATGAGTGTGGAGTCGTTGCTGTTTCTCCACTCCACCCTTTTTNNNAGATCGGAAGA
65059_1 TATGAGTGTGGAGTCGTTGCGTGGTTTTTGATGAGGCTTCTCCNNNAGATCGGAAG
65060_1 TATGAGTGTGGAGTCGTTGCCTCTGAAAATGGCAAAGGCTCAANNNAGATCGGAA
65062_1 TATGAGTGTGGAGTCGTTGCTGAGTTCTTGATTTGACTCAGCTNNNAGATCGGAAG
65063_1 TATGAGTGTGGAGTCGTTGCTTACTTCCCATGAATGCCTGTGTNNNAGATCGGAAG
65064_1 TATGAGTGTGGAGTCGTTGCTATTAGACAAGAAGGTCTGGCCGNNNAGATCGGAA
65065_1 TATGAGTGTGGAGTCGTTGCTGTGTGTGTGTGTGTAGAGATNNNAGATCGGAA
65066_1 TATGAGTGTGGAGTCGTTGCCACAGAAAGGGACCTAACTGCTANNNAGATCGGAA
65067_1 TATGAGTGTGGAGTCGTTGCCTGGTCTGAAACTCCTGAACTCANNNAGATCGGAAG
65068_1 TATGAGTGTGGAGTCGTTGCTGCTTGTAATCCCAGCATTTTGGNNNAGATCGGAAG
65069_1 TATGAGTGTGGAGTCGTTGCACACAATTTTGTATGGCAGGCTCNNNAGATCGGAAG
65070_1 TATGAGTGTGGAGTCGTTGCTGCTTCCTGAATATAAGGGAGCCNNNAGATCGGAA
65071_1 TATGAGTGTGGAGTCGTTGCTAGGGGCAAATAACCCAGTATGGNNNAGATCGGAA
65072_1 TATGAGTGTGGAGTCGTTGCGCAGGAGAGCAAAGAATACGTTCNNNAGATCGGAA
65073_1 TATGAGTGTGGAGTCGTTGCTGTGGTGTATGATGGAGTTTCACNNNAGATCGGAA
65075_1 TATGAGTGTGGAGTCGTTGCGAGACAGACAGACAGACAGACAGNNNAGATCGGAA
65076_1 TATGAGTGTGGAGTCGTTGCCCGTGTCTCTGTGTGATGCTATANNNAGATCGGAAG
65077_1 TATGAGTGTGGAGTCGTTGCGAGAATCGCTTGAACCTGGGAAGNNNAGATCGGAA
65079_1 TATGAGTGTGGAGTCGTTGCTTTTCCCTAGAAGATTGGCAGCTNNNAGATCGGAAG
65080_1 TATGAGTGTGGAGTCGTTGCCTTGTTCCAGGCAACATGCTAAGNNNAGATCGGAAG
65081_1 TATGAGTGTGGAGTCGTTGCTGGAGTGCAGTGTCACAATCATANNNAGATCGGAA
65084_1 TATGAGTGTGGAGTCGTTGCGTTTGTGATGCATGCTTCTACGTNNNAGATCGGAAG
65085_1 TATGAGTGTGGAGTCGTTGCCAGTCTCTAGGCCTTTCTGGTTTNNNAGATCGGAAG
65086_1 TATGAGTGTGGAGTCGTTGCATCACAAGCAGGAAAGAACAGGANNNAGATCGGAA
65087_1 TATGAGTGTGGAGTCGTTGCCACCCCACATTTATTGAACAGCANNNAGATCGGAAG
65088_1 TATGAGTGTGGAGTCGTTGCCACCCCACATTTATTGAACAGCANNNAGATCGGAAG
65090_1 TATGAGTGTGGAGTCGTTGCAATAGAAGAAAGCTGCCCAGACANNNAGATCGGAA
65091_1 TATGAGTGTGGAGTCGTTGCGTGTGACATTGATATCCGCAAGGNNNAGATCGGAA
65092_1 TATGAGTGTGGAGTCGTTGCGAGAAGTGGATGTCGTGGGAATCNNNAGATCGGA
65093_1 TATGAGTGTGGAGTCGTTGCTGTACTTTCTCGGTTCTGGTGAGNNNAGATCGGAAG
65094_1 TATGAGTGTGGAGTCGTTGCGAGCTTAGGAGTCAAGAGACCAGNNNAGATCGGAA
65095_1 TATGAGTGTGGAGTCGTTGCTGTGCCTGTAATCCCAGCTATTTNNNAGATCGGAAG
65096_1 TATGAGTGTGGAGTCGTTGCTGAAAAATACCTACCTTCCCCCGNNNAGATCGGAAG
65097_1 TATGAGTGTGGAGTCGTTGCAGGCACTTACATATTCAGCCACANNNAGATCGGAAG
65098_1 TATGAGTGTGGAGTCGTTGCCTGAAGTCAGGAGTTCAAGACCANNNAGATCGGAA

| | |
|---|---|
| 65099_1 | TATGAGTGTGGAGTCGTTGCACCTCAAACTCATGGGCTAAAGTNNNAGATCGGAAG |
| 65101_1 | TATGAGTGTGGAGTCGTTGCAATGAGGTCTCACTATGTTGCCCNNNAGATCGGAAG |
| 65104_1 | TATGAGTGTGGAGTCGTTGCCTGAGGCAGGAGAATCACTTGAANNNAGATCGGAA |
| 65105_1 | TATGAGTGTGGAGTCGTTGCAATTGCTCTGGCTAGAACTTCCANNNAGATCGGAAG |
| 65106_1 | TATGAGTGTGGAGTCGTTGCTATTGAGACAGGGTCTTGCTCTGNNNAGATCGGAA |
| 65107_1 | TATGAGTGTGGAGTCGTTGCGCCTCCCAAAGCACTAGGATTATNNNAGATCGGAAG |
| 65108_1 | TATGAGTGTGGAGTCGTTGCAGAGACGTGTTTTCACCATGTTGNNNAGATCGGAAG |
| 65110_1 | TATGAGTGTGGAGTCGTTGCGAGCAGCACAGACACACATATGTNNNAGATCGGAA |
| 65111_1 | TATGAGTGTGGAGTCGTTGCAATGCACACACACACACACAAATNNNAGATCGGAAG |
| 65112_1 | TATGAGTGTGGAGTCGTTGCTCCTCATCCCCATTCATGCATAGNNNAGATCGGAAGA |
| 65113_1 | TATGAGTGTGGAGTCGTTGCTTCGATATCCCCAGTCCACAATCNNNAGATCGGAAG |
| 65114_1 | TATGAGTGTGGAGTCGTTGCACTGCAACTTCAACACGATTCAGNNNAGATCGGAAG |
| 65115_1 | TATGAGTGTGGAGTCGTTGCAGACAGACAGAGAGACAGAGACANNNAGATCGGA |
| 65116_1 | TATGAGTGTGGAGTCGTTGCAGAGACAGAGACAGAGACAGACANNNAGATCGGA |
| 65117_1 | TATGAGTGTGGAGTCGTTGCAGCTTGGCAGGAAGAACAAAGATNNNAGATCGGAA |
| 65119_1 | TATGAGTGTGGAGTCGTTGCCAATCTCAGCTCACTGCAACTTCNNNAGATCGGAAG |
| 65121_1 | TATGAGTGTGGAGTCGTTGCTTGTGAGACAGAGTCTTGTTCTGNNNAGATCGGAA |
| 65123_1 | TATGAGTGTGGAGTCGTTGCGTGTGTGTGAGAGAGAGAGAGNNNAGATCGGA |
| 65125_1 | TATGAGTGTGGAGTCGTTGCTACCATTGCCTGATCCAGAATCANNNAGATCGGAAG |
| 65126_1 | TATGAGTGTGGAGTCGTTGCATGTGTTTTGTAGAGACGGGGTTNNNAGATCGGAA |
| 65127_1 | TATGAGTGTGGAGTCGTTGCCCCATCTTATCTGTGAGCTCCTCNNNAGATCGGAAGA |
| 65128_1 | TATGAGTGTGGAGTCGTTGCACACACACACACACACACAAAATNNNAGATCGGAAG |
| 65129_1 | TATGAGTGTGGAGTCGTTGCTTCCCTCAGAGATTGTGCAGAATNNNAGATCGGAAG |
| 65130_1 | TATGAGTGTGGAGTCGTTGCTGAGAGACAGAGAGAGAAGGGAGNNNAGATCGGA |
| 65132_1 | TATGAGTGTGGAGTCGTTGCTTTTACTTCTCAGAGCTGGGCTTNNNAGATCGGAAG |
| 65133_1 | TATGAGTGTGGAGTCGTTGCATCTATAGAACCTGGGAGGCAGANNNAGATCGGAA |
| 65134_1 | TATGAGTGTGGAGTCGTTGCCTACTGTCCTCTAATTCCAGCCCNNNAGATCGGAAGA |
| 65135_1 | TATGAGTGTGGAGTCGTTGCGTTCTTCAAATGCACCAAGCTCTNNNAGATCGGAAG |
| 65136_1 | TATGAGTGTGGAGTCGTTGCTCTACTCCATGCCAGACATTCTGNNNAGATCGGAAG |
| 65137_1 | TATGAGTGTGGAGTCGTTGCTGCAGGTGATTACGATCTCAAGTNNNAGATCGGAA |
| 65138_1 | TATGAGTGTGGAGTCGTTGCGTGTGTGTGTGTGTGATCAATNNNAGATCGGAA |
| 65139_1 | TATGAGTGTGGAGTCGTTGCACGCCAGCCTTCTATTTTGACTANNNAGATCGGAAG |
| 65140_1 | TATGAGTGTGGAGTCGTTGCCCCAAGTAGCTGAGAATACAGGTNNNAGATCGGAA |
| 65141_1 | TATGAGTGTGGAGTCGTTGCAGAGAGAGAGAAAGAAAGCTGCCNNNAGATCGGA |
| 65144_1 | TATGAGTGTGGAGTCGTTGCTCCAGAGAGCCCTGACTAATACANNNAGATCGGAAG |
| 65145_1 | TATGAGTGTGGAGTCGTTGCCCCTACAATCCACAGTTCTGCTANNNAGATCGGAAG |
| 65146_1 | TATGAGTGTGGAGTCGTTGCCACACACATACACACACACACATNNNAGATCGGAAG |
| 65147_1 | TATGAGTGTGGAGTCGTTGCGAGTTCTGGGCTGTAGTGTGTTANNNAGATCGGAA |
| 65148_1 | TATGAGTGTGGAGTCGTTGCACACACACTCGTACTTTTAGCCTNNNAGATCGGAAG |
| 65149_1 | TATGAGTGTGGAGTCGTTGCTTTGGAAATAGTTGCAGCACAGGNNNAGATCGGAA |
| 65150_1 | TATGAGTGTGGAGTCGTTGCCTATTTGTGAGGCTGAGGTGGAANNNAGATCGGAA |
| 65151_1 | TATGAGTGTGGAGTCGTTGCAGCAATTCTCCTACCTCAACCTCNNNAGATCGGAAGA |
| 65152_1 | TATGAGTGTGGAGTCGTTGCTTCAAACGATTCTCCTGCCTCAGNNNAGATCGGAAG |

| | |
|---|---|
| 65153_1 | TATGAGTGTGGAGTCGTTGCGAAGTTTTGCTCTTGTCACCCAGNNNAGATCGGAAG |
| 65156_1 | TATGAGTGTGGAGTCGTTGCGCCTGAACACCCTTTTGAGACTANNNAGATCGGAAG |
| 65157_1 | TATGAGTGTGGAGTCGTTGCGTCTGAAACACTGGTTTGGGAAANNNAGATCGGAA |
| 65158_1 | TATGAGTGTGGAGTCGTTGCCATCCCCACTGCTGTAGGAAAATNNNAGATCGGAAG |
| 65159_1 | TATGAGTGTGGAGTCGTTGCAAAATATATCATGGCCGGGCATGNNNAGATCGGAA |
| 65160_1 | TATGAGTGTGGAGTCGTTGCCTATTCTGACCTCAGCTGATCCCNNNAGATCGGAAG |
| 65161_1 | TATGAGTGTGGAGTCGTTGCTCACAAGGTCAGGAGATCAAGACNNNAGATCGGAA |
| 65162_1 | TATGAGTGTGGAGTCGTTGCTTCTGCGGGTTAGGAAATTGGTANNNAGATCGGAA |
| 65163_1 | TATGAGTGTGGAGTCGTTGCCTCTCTCTCTCTCTCTCTCCANNNAGATCGGAAGAG |
| 65164_1 | TATGAGTGTGGAGTCGTTGCGTGGGAGGCGGATTCTTTTTTTTNNNAGATCGGAA |
| 65165_1 | TATGAGTGTGGAGTCGTTGCCCTCCCAAAGTGCTAGGATTACANNNAGATCGGAAG |
| 65167_1 | TATGAGTGTGGAGTCGTTGCCTCCTGAGTAGCTGGGATTACAGNNNAGATCGGAA |
| 65168_1 | TATGAGTGTGGAGTCGTTGCCACTACAGCCTTGACCTCTCAGNNNAGATCGGAAGA |
| 65169_1 | TATGAGTGTGGAGTCGTTGCTGAGAAGCCAGAACAGAAAAAGANNNAGATCGGA |
| 65170_1 | TATGAGTGTGGAGTCGTTGCTAGAGAAAGAGAGAGAGCGAGCANNNAGATCGGA |
| 65171_1 | TATGAGTGTGGAGTCGTTGCGAGAGAGAGAGAGAGAGAGCGAGNNNAGATCGG |
| 65172_1 | TATGAGTGTGGAGTCGTTGCGACATTTTCAGACTGCTTTCGGTNNNAGATCGGAAG |
| 65173_1 | TATGAGTGTGGAGTCGTTGCCACAACTACAGTCCCAGCTACTCNNNAGATCGGAAG |
| 65175_1 | TATGAGTGTGGAGTCGTTGCTCTGGTCTTTGTGGAATCACCTTNNNAGATCGGAAG |
| 65177_1 | TATGAGTGTGGAGTCGTTGCCATGATCAGCACACTGCAATCTCNNNAGATCGGAAG |
| 65178_1 | TATGAGTGTGGAGTCGTTGCTGTCTGTGTGTGTCAGAGTCTTGNNNAGATCGGAA |
| 65179_1 | TATGAGTGTGGAGTCGTTGCAGACAGAGACAGAGACAGAGACANNNAGATCGGA |
| 65181_1 | TATGAGTGTGGAGTCGTTGCATTTCCACTGCAGCATCATTTGTNNNAGATCGGAAG |
| 65182_1 | TATGAGTGTGGAGTCGTTGCCTTGGGAGACTTAGGCATGAGAANNNAGATCGGAA |
| 65184_1 | TATGAGTGTGGAGTCGTTGCGACAAGGTCTCAGTCTGTAACCCNNNAGATCGGAAG |
| 65185_1 | TATGAGTGTGGAGTCGTTGCAGAGAAAGAGACTCTCCCTCTCCNNNAGATCGGAAG |
| 65186_1 | TATGAGTGTGGAGTCGTTGCCTCCCACCTGATTCTAGGACATGNNNAGATCGGAAG |
| 65187_1 | TATGAGTGTGGAGTCGTTGCACACATACACACACACGCATATGNNNAGATCGGAAG |
| 65188_1 | TATGAGTGTGGAGTCGTTGCTTGTGCTTCAGAATTCTCAGGGANNNAGATCGGAAG |
| 65189_1 | TATGAGTGTGGAGTCGTTGCTCTCCCTCTCTCTCTCTCTCTCTNNNAGATCGGAAGAG |
| 65190_1 | TATGAGTGTGGAGTCGTTGCATCTGGACTCCAAGGTGTGATTCNNNAGATCGGAAG |
| 65191_1 | TATGAGTGTGGAGTCGTTGCAGAGAGACAGAGGAGACAGAACTNNNAGATCGGA |
| 65192_1 | TATGAGTGTGGAGTCGTTGCAGGCAGAAGAAACCATTCCACATNNNAGATCGGAA |
| 65193_1 | TATGAGTGTGGAGTCGTTGCCTAATTATCTCCCAAAGGCCCCANNNAGATCGGAAG |
| 65194_1 | TATGAGTGTGGAGTCGTTGCTGAGAGAGAGTTCAGGTGTCTCTNNNAGATCGGAA |
| 65195_1 | TATGAGTGTGGAGTCGTTGCAAAGACAAAGAGCCTCTGCAAACNNNAGATCGGAA |
| 65196_1 | TATGAGTGTGGAGTCGTTGCTCTCAGGGATGCAGGAATGATTTNNNAGATCGGAA |
| 65197_1 | TATGAGTGTGGAGTCGTTGCAAGACATTGGTTTTGCAGTCCAGNNNAGATCGGAA |
| 65199_1 | TATGAGTGTGGAGTCGTTGCAGAGAGAGAGAGACAAGGTCTGGNNNAGATCGGA |
| 65200_1 | TATGAGTGTGGAGTCGTTGCTGGGGAGAGAGAGAGAGAATTGTNNNAGATCGGA |
| 65202_1 | TATGAGTGTGGAGTCGTTGCTATTTCATTGCAGCCTGGAACTCNNNAGATCGGAAG |
| 65203_1 | TATGAGTGTGGAGTCGTTGCGAGAGAGACAGAGTATGCCCCTANNNAGATCGGAA |
| 65204_1 | TATGAGTGTGGAGTCGTTGCCATGTGAAGAAACTGCCCAACATNNNAGATCGGAAG |

| | |
|---|---|
| 65205_1 | TATGAGTGTGGAGTCGTTGCCAAGGTCCCATCTCCAAATACCANNNAGATCGGAAGA |
| 65206_1 | TATGAGTGTGGAGTCGTTGCCCTGTAGTCCTAGCTACTTGGGANNNAGATCGGAAG |
| 65207_1 | TATGAGTGTGGAGTCGTTGCCTTTGGAATTGAGTGACACCACCNNNAGATCGGAAG |
| 65208_1 | TATGAGTGTGGAGTCGTTGCATAAGCTCTGGTCTCTCTTCCCTNNNAGATCGGAAGA |
| 65209_1 | TATGAGTGTGGAGTCGTTGCCTTCCCCATCACAGACACACATANNNAGATCGGAAGA |
| 65210_1 | TATGAGTGTGGAGTCGTTGCAGATGGATTTTCGCTCTTGTTGCNNNAGATCGGAAG |
| 65211_1 | TATGAGTGTGGAGTCGTTGCGTCCCCTATTTGAAATGGTGCAGNNNAGATCGGAAG |
| 65212_1 | TATGAGTGTGGAGTCGTTGCTCAGTGGAAATGAACAGAGGAGGNNNAGATCGGAA |
| 65213_1 | TATGAGTGTGGAGTCGTTGCGAACAAGAGTCTCTGCCTGGTAANNNAGATCGGAAG |
| 65214_1 | TATGAGTGTGGAGTCGTTGCGAAAAGAAACAAGGGCCACTGTTNNNAGATCGGAAG |
| 65215_1 | TATGAGTGTGGAGTCGTTGCCCACTGGGACTATTGCATCAGATNNNAGATCGGAAG |
| 65217_1 | TATGAGTGTGGAGTCGTTGCTTGGGGGAAAATGAGTCAAGACANNNAGATCGGAA |
| 65218_1 | TATGAGTGTGGAGTCGTTGCTCTCCCTCTCTCTCTCTCTCTCTNNNAGATCGGAAGAG |
| 65219_1 | TATGAGTGTGGAGTCGTTGCCAGTGGCATGAACATAGCTCACTNNNAGATCGGAAG |
| 65220_1 | TATGAGTGTGGAGTCGTTGCTTTAGCCAAATCTTCCCTGTGGTNNNAGATCGGAAGA |
| 65221_1 | TATGAGTGTGGAGTCGTTGCCCTGGGGTGCTTGTCTATATTGTNNNAGATCGGAAG |
| 65222_1 | TATGAGTGTGGAGTCGTTGCGGAGAGAGAGAGAGAGAGAGGAGNNNAGATCGGA |
| 65223_1 | TATGAGTGTGGAGTCGTTGCAACTTCTGTCTCCTGGGTTCAAGNNNAGATCGGAAGA |
| 65225_1 | TATGAGTGTGGAGTCGTTGCGAATTCCAGAACGTTGTCCTGTGNNNAGATCGGAAG |
| 65227_1 | TATGAGTGTGGAGTCGTTGCACAAGCACATACACACACCTACTNNNAGATCGGAAGA |
| 65229_1 | TATGAGTGTGGAGTCGTTGCAACAGAGTTAGGAAGAAGCTGGGNNNAGATCGGAA |
| 65231_1 | TATGAGTGTGGAGTCGTTGCTTTTTTTTGACGGAGTCTCGCTCNNNAGATCGGAAGA |
| 65232_1 | TATGAGTGTGGAGTCGTTGCTCATGGTTCACTGTAGCCTCAAANNNAGATCGGAAGA |
| 65233_1 | TATGAGTGTGGAGTCGTTGCTTCCTTCTAGACCAGGTCTCCAANNNAGATCGGAAGA |
| 65234_1 | TATGAGTGTGGAGTCGTTGCAAGAGCAAGGCAGAAGCTGTATANNNAGATCGGAA |
| 65235_1 | TATGAGTGTGGAGTCGTTGCGGCAATGTTGGGTGAATAACCAANNNAGATCGGAAG |
| 65237_1 | TATGAGTGTGGAGTCGTTGCTATCTCTCTCTGCCAGCTCATTGNNNAGATCGGAAGA |
| 65238_1 | TATGAGTGTGGAGTCGTTGCTTTTGGACTTCTCAGCATCCAGANNNAGATCGGAAGA |
| 65239_1 | TATGAGTGTGGAGTCGTTGCCAGTGGTGCAATCTCTTCTCACTNNNAGATCGGAAGA |
| 65241_1 | TATGAGTGTGGAGTCGTTGCAAGATCACCTGGAAGACAATCCCNNNAGATCGGAAG |
| 65242_1 | TATGAGTGTGGAGTCGTTGCTCTAAGTGAAGTAACTCAGGGGTNNNAGATCGGAAG |
| 65243_1 | TATGAGTGTGGAGTCGTTGCTACACAGCTGAGTCTTGCCTAAGNNNAGATCGGAAG |
| 65244_1 | TATGAGTGTGGAGTCGTTGCAGGAAGGAAGGAAGAAAGAGAGANNNAGATCGGA |
| 65245_1 | TATGAGTGTGGAGTCGTTGCCCAGCCCTCTCATCTCTTTGATANNNAGATCGGAAGA |
| 65247_1 | TATGAGTGTGGAGTCGTTGCCAGTGGGACAAGAAGAGATGGAANNNAGATCGGA |
| 65248_1 | TATGAGTGTGGAGTCGTTGCAATGAAGTGGTGCAATCATAGCGNNNAGATCGGAAG |
| 65249_1 | TATGAGTGTGGAGTCGTTGCAGACCTAGCATAGCAAGAAAGCANNNAGATCGGAAG |
| 65250_1 | TATGAGTGTGGAGTCGTTGCCCTTGTCTGGGAAAACTCTCTGANNNAGATCGGAAG |
| 65251_1 | TATGAGTGTGGAGTCGTTGCTTCTTTCTTTCGTCTCGCTCTGTNNNAGATCGGAAGAG |
| 65252_1 | TATGAGTGTGGAGTCGTTGCCAGTGGTGTGATCTTTCCTCTCTNNNAGATCGGAAGA |
| 65254_1 | TATGAGTGTGGAGTCGTTGCAGATCAAACCACAAAGCCCTCTANNNAGATCGGAAGA |
| 65255_1 | TATGAGTGTGGAGTCGTTGCGGCTCAAGTGATTCTCCTGAGTANNNAGATCGGAAG |
| 65256_1 | TATGAGTGTGGAGTCGTTGCCATCCATCCATCCATCTGTCCATNNNAGATCGGAAGAG |

| | |
|---|---|
| 65257_1 | TATGAGTGTGGAGTCGTTGCCCCTCTCTCCCTTTTTTTGACCTNNNAGATCGGAAGAG |
| 65259_1 | TATGAGTGTGGAGTCGTTGCGTGTGTGTGTGTGTGTCTATGNNNAGATCGGAA |
| 65260_1 | TATGAGTGTGGAGTCGTTGCCAAGCAAGGGAAATGCCAGATAGNNNAGATCGGAA |
| 65261_1 | TATGAGTGTGGAGTCGTTGCAGAGAGAGAGAGAAAGAGAGCCANNNAGATCGGA |
| 65262_1 | TATGAGTGTGGAGTCGTTGCCCCAACCTCCTCTCCTTGATTTTNNNAGATCGGAAGA |
| 65263_1 | TATGAGTGTGGAGTCGTTGCGGAGAGATGGAGACAGGGAGATANNNAGATCGGA |
| 65264_1 | TATGAGTGTGGAGTCGTTGCCTCCAAGCAGGCCAGTTTATTTCNNNAGATCGGAAG |
| 65265_1 | TATGAGTGTGGAGTCGTTGCACGCCCGGCTAATTTTTGTATTTNNNAGATCGGAAG |
| 65266_1 | TATGAGTGTGGAGTCGTTGCCTAGGGGATATTGTTTGAGCCCANNNAGATCGGAA |
| 65268_1 | TATGAGTGTGGAGTCGTTGCCTTCTGCCAAAAAGATCCTTCCANNNAGATCGGAAG |
| 65270_1 | TATGAGTGTGGAGTCGTTGCAAAACTTTTTTCATGGCCACTGTNNNAGATCGGAAG |
| 65271_1 | TATGAGTGTGGAGTCGTTGCCGCAACACTTTCTGACAGGAAATNNNAGATCGGAAG |
| 65272_1 | TATGAGTGTGGAGTCGTTGCGGCTTGCCCATGTTAGTTGAAATNNNAGATCGGAAG |
| 65273_1 | TATGAGTGTGGAGTCGTTGCAACTATGATCCCTCCCTAGGCTTNNNAGATCGGAAG |
| 65274_1 | TATGAGTGTGGAGTCGTTGCATTATCTCGTTTGACCTTGGGGTNNNAGATCGGAAG |
| 65275_1 | TATGAGTGTGGAGTCGTTGCCAACACATATGAAGGATGGCGTCNNNAGATCGGAA |
| 65276_1 | TATGAGTGTGGAGTCGTTGCCTGGAGCCAAGCCAAGAAGTATANNNAGATCGGAA |
| 65277_1 | TATGAGTGTGGAGTCGTTGCGGGGTTCTATGTGGGAGTAGAACNNNAGATCGGAA |
| 65278_1 | TATGAGTGTGGAGTCGTTGCTTTCCCCTCATAGACATCAAGCCNNNAGATCGGAAGA |
| 65279_1 | TATGAGTGTGGAGTCGTTGCTCTAGACACAGTCATGCATTGCTNNNAGATCGGAAG |
| 65280_1 | TATGAGTGTGGAGTCGTTGCTGGAGATAACAAGTGCAAGTGGANNNAGATCGGAA |
| 65281_1 | TATGAGTGTGGAGTCGTTGCCCCTCAGAAGACTCGTTTGCTAANNNAGATCGGAAG |
| 65282_1 | TATGAGTGTGGAGTCGTTGCTACCTCCCATAAGGACAAACAGCNNNAGATCGGAAG |
| 65283_1 | TATGAGTGTGGAGTCGTTGCCTAGAAAAGGGCTAATGCAACGGNNNAGATCGGAA |
| 65284_1 | TATGAGTGTGGAGTCGTTGCTCTCCCCTCATAGAGCTGTAACANNNAGATCGGAAG |
| 65285_1 | TATGAGTGTGGAGTCGTTGCGTTCTAAGCAGATCTGGAGAGGANNNAGATCGGAA |
| 65286_1 | TATGAGTGTGGAGTCGTTGCTTTCCGATCAGTTCCGTCTTCTTNNNAGATCGGAAGA |
| 65287_1 | TATGAGTGTGGAGTCGTTGCTCCTTATAGGACTGTTTAGGGGANNNAGATCGGAA |
| 65288_1 | TATGAGTGTGGAGTCGTTGCTGTTAGTGCTACCACAACTCTGGNNNAGATCGGAAG |
| 65289_1 | TATGAGTGTGGAGTCGTTGCCACATACAGATCAGGGTGGTCTCNNNAGATCGGAA |
| 65290_1 | TATGAGTGTGGAGTCGTTGCGAGGACATCATCTTTCCGAGAGANNNAGATCGGAA |
| 65291_1 | TATGAGTGTGGAGTCGTTGCGTTCTCATGACATGGCAGTTGTCNNNAGATCGGAAG |
| 65292_1 | TATGAGTGTGGAGTCGTTGCAAGAGAACAAGAGTCTCTGCCTGNNNAGATCGGAA |
| 65293_1 | TATGAGTGTGGAGTCGTTGCTCCTCTCCATCTAGTAAGCCAGTNNNAGATCGGAAG |
| 65294_1 | TATGAGTGTGGAGTCGTTGCAGAAAAGGTTGGCAGCATTACACNNNAGATCGGAA |
| 65295_1 | TATGAGTGTGGAGTCGTTGCTCTTACCAAAAAGGATTCAGCCANNNAGATCGGAAG |
| 65296_1 | TATGAGTGTGGAGTCGTTGCACACTGTCATCTTACCGTTTCATNNNAGATCGGAAGA |
| 65297_1 | TATGAGTGTGGAGTCGTTGCGTCTCCAGAGCTAGCACTGTATCNNNAGATCGGAAG |
| 65298_1 | TATGAGTGTGGAGTCGTTGCGTGTGTGTGTGTGTGAATTGANNNAGATCGGAA |
| 65299_1 | TATGAGTGTGGAGTCGTTGCTGTCTTTTTATAGCTCCTCGGCANNNAGATCGGAAG |
| 65300_1 | TATGAGTGTGGAGTCGTTGCTGACACTAAGACTGCAGCATTGTNNNAGATCGGAAG |
| 65301_1 | TATGAGTGTGGAGTCGTTGCATTCAAAATCTGCAAGCTTGGCTNNNAGATCGGAAG |
| 65302_1 | TATGAGTGTGGAGTCGTTGCTGGGATCTAGCATCCAAGCTATTNNNAGATCGGAAG |

| | |
|---|---|
| 65303_1 | TATGAGTGTGGAGTCGTTGCATATTGTGACAGAGGACGCAAGANNNAGATCGGAA |
| 65304_1 | TATGAGTGTGGAGTCGTTGCGCTCATACACGCTCATACACATGNNNAGATCGGAAG |
| 65305_1 | TATGAGTGTGGAGTCGTTGCAGTACTAAAGGGAGCACACACTGNNNAGATCGGAA |
| 65307_1 | TATGAGTGTGGAGTCGTTGCAGAGAGAGAGAGAGGAGAACTCCNNNAGATCGGA |
| 65308_1 | TATGAGTGTGGAGTCGTTGCCACGTTCCATTGTCCTTTCTTCCNNNAGATCGGAAGA |
| 65311_1 | TATGAGTGTGGAGTCGTTGCTCTGAATGAGATTTGTGAGCTGTNNNAGATCGGAA |
| 65312_1 | TATGAGTGTGGAGTCGTTGCCCACTGCTACTACCACTACCTTCNNNAGATCGGAAGA |
| 65313_1 | TATGAGTGTGGAGTCGTTGCACCCATTTATCCTCCAAGACCTGNNNAGATCGGAAGA |
| 65314_1 | TATGAGTGTGGAGTCGTTGCCCTCCTCTAGCTTTGGGAATGAGNNNAGATCGGAAG |
| 65315_1 | TATGAGTGTGGAGTCGTTGCCTTAGTTGGACCACTTCACCAGANNNAGATCGGAAG |
| 65316_1 | TATGAGTGTGGAGTCGTTGCAGGTGACTAGCAAAAGTGTCCTGNNNAGATCGGAA |
| 65317_1 | TATGAGTGTGGAGTCGTTGCACCTGGGTTCAAATGACCTACAANNNAGATCGGAAG |
| 65318_1 | TATGAGTGTGGAGTCGTTGCAGGGCACAAAGAAGGTAGACATTNNNAGATCGGAA |
| 65320_1 | TATGAGTGTGGAGTCGTTGCCTACAAAGGGCCTGTGATAGTGTNNNAGATCGGAA |
| 65321_1 | TATGAGTGTGGAGTCGTTGCGAGAGAGAGAGAGGAGAGGAGTCNNNAGATCGGA |
| 65322_1 | TATGAGTGTGGAGTCGTTGCAGAGAAGGAAAGGGACATAGGGANNNAGATCGGA |
| 65323_1 | TATGAGTGTGGAGTCGTTGCGCACTGCAGGATAATCCCTAAGANNNAGATCGGAA |
| 65325_1 | TATGAGTGTGGAGTCGTTGCGAATGCCTGGGAGCTGTAAGATANNNAGATCGGAA |
| 65326_1 | TATGAGTGTGGAGTCGTTGCTGAATGGAACGAATGAATGGGGANNNAGATCGGA |
| 65327_1 | TATGAGTGTGGAGTCGTTGCAAGGATGACTACAGGAGAGGGAANNNAGATCGGA |
| 65328_1 | TATGAGTGTGGAGTCGTTGCGTCACAGAAAGGGATGGAGTCTTNNNAGATCGGAA |
| 65330_1 | TATGAGTGTGGAGTCGTTGCTCAATTCCAGTGGCCAAAACTAANNNAGATCGGAAG |
| 65331_1 | TATGAGTGTGGAGTCGTTGCGGGTAAAATGCACAGTTAGGCTCNNNAGATCGGAA |
| 65332_1 | TATGAGTGTGGAGTCGTTGCATAAGATGCTAGTGCCAGAAGGGNNNAGATCGGAA |
| 65333_1 | TATGAGTGTGGAGTCGTTGCTTGTGGGGGCTAATAAGTCCAAANNNAGATCGGAA |
| 65334_1 | TATGAGTGTGGAGTCGTTGCATTATCAGGCTAGAGGTGACTGCNNNAGATCGGAA |
| 65335_1 | TATGAGTGTGGAGTCGTTGCTATATGCCCTCTCATCTGTGCACNNNAGATCGGAAGA |
| 65336_1 | TATGAGTGTGGAGTCGTTGCTAGTGGAAGTTCAGCACTGAGTCNNNAGATCGGAA |
| 65338_1 | TATGAGTGTGGAGTCGTTGCACAGTGGAGGGAAAAGAAACAGTNNNAGATCGGA |
| 65339_1 | TATGAGTGTGGAGTCGTTGCACATCTACATGTCTGTGTGTGCTNNNAGATCGGAAG |
| 65340_1 | TATGAGTGTGGAGTCGTTGCCACCTGCAAGAACCATGCTTATGNNNAGATCGGAAG |
| 65341_1 | TATGAGTGTGGAGTCGTTGCTTCTCAGCCATCTCTGGGTACATNNNAGATCGGAAG |
| 65342_1 | TATGAGTGTGGAGTCGTTGCAAGGGAATCACGTCACATGGTAANNNAGATCGGAA |
| 65343_1 | TATGAGTGTGGAGTCGTTGCTTGCTGGACTGCAGTACTTCTAGNNNAGATCGGAAG |
| 65344_1 | TATGAGTGTGGAGTCGTTGCTGTTGGAACCCTGAAAGTGATGANNNAGATCGGAA |
| 65345_1 | TATGAGTGTGGAGTCGTTGCCCCCACCAAGAATTTTTTTGTCGNNNAGATCGGAAGA |
| 65346_1 | TATGAGTGTGGAGTCGTTGCCCTGAGTCTAACCTCTTTGGCTTNNNAGATCGGAAGA |
| 65347_1 | TATGAGTGTGGAGTCGTTGCGTTGTTGTTGTTGTTGAGACGGANNNAGATCGGAA |
| 65348_1 | TATGAGTGTGGAGTCGTTGCACGCTTCTCTAAAGTCCCAGAAGNNNAGATCGGAAG |
| 65349_1 | TATGAGTGTGGAGTCGTTGCATGAATCTCTGGGTAGTAACGCGNNNAGATCGGAA |
| 65350_1 | TATGAGTGTGGAGTCGTTGCTTGTCCAGCTCCATTTCAGATGANNNAGATCGGAAG |
| 65351_1 | TATGAGTGTGGAGTCGTTGCCCTTGTCCAAGTTCTGAGTGACANNNAGATCGGAAG |
| 65352_1 | TATGAGTGTGGAGTCGTTGCATAGAACTTGCCTCTCCCTTGTGNNNAGATCGGAAG |

| | |
|---|---|
| 65353_1 | TATGAGTGTGGAGTCGTTGCTATGCAGATGTTCTTGCCTCTGGNNNAGATCGGAAG |
| 65354_1 | TATGAGTGTGGAGTCGTTGCGCCCCATTTTTCTACCCTAATGCNNNAGATCGGAAGA |
| 65355_1 | TATGAGTGTGGAGTCGTTGCTTGGGTACAAACCACCTTCTTCANNNAGATCGGAAG |
| 65356_1 | TATGAGTGTGGAGTCGTTGCAGATTTATGCTGGAGGTTCACCANNNAGATCGGAA |
| 65357_1 | TATGAGTGTGGAGTCGTTGCCTCCTGAGTAGCTGGGATTACAGNNNAGATCGGAA |
| 65358_1 | TATGAGTGTGGAGTCGTTGCCCGCCATTTTAGAGATGGGAAACNNNAGATCGGAA |
| 65359_1 | TATGAGTGTGGAGTCGTTGCAGGGAGAGAGAGAGAGAGAGAGANNNAGATCGG |
| 65361_1 | TATGAGTGTGGAGTCGTTGCAGATGAGGACAGAATGGCTTTGTNNNAGATCGGAA |
| 65362_1 | TATGAGTGTGGAGTCGTTGCGAGACTCAGTGCACACATCCATANNNAGATCGGAAG |
| 65363_1 | TATGAGTGTGGAGTCGTTGCATTTGCTTGGCTATGTTGTCCACNNNAGATCGGAAG |
| 65364_1 | TATGAGTGTGGAGTCGTTGCCTCAAGAGTCACCCCCAAAAGTANNNAGATCGGAAG |
| 65365_1 | TATGAGTGTGGAGTCGTTGCCCAGGTGGGAAAGAAAACCAAATNNNAGATCGGAA |
| 65367_1 | TATGAGTGTGGAGTCGTTGCCAGGACCATGATTTCAGCCCTATNNNAGATCGGAAG |
| 65369_1 | TATGAGTGTGGAGTCGTTGCCTTTCTCAGGAGGCTCACTAGTGNNNAGATCGGAAG |
| 65370_1 | TATGAGTGTGGAGTCGTTGCCATTCTCAAACTCCAATCCCCCTNNNAGATCGGAAGA |
| 65371_1 | TATGAGTGTGGAGTCGTTGCTCTCCCGTGTTCAAGCATTATCTNNNAGATCGGAAGA |
| 65372_1 | TATGAGTGTGGAGTCGTTGCGTTCTTTCAGTTGCAAGGGCTAGNNNAGATCGGAA |
| 65373_1 | TATGAGTGTGGAGTCGTTGCTTTGGGCAGGCTTTGTACATTTTNNNAGATCGGAAG |
| 65374_1 | TATGAGTGTGGAGTCGTTGCACCGTGAGTCTTCATATGGCATTNNNAGATCGGAAG |
| 65375_1 | TATGAGTGTGGAGTCGTTGCGCCAGAGTGGGACACTTATTACANNNAGATCGGAA |
| 65376_1 | TATGAGTGTGGAGTCGTTGCTCAGAATCCAAGGGCATGCATATNNNAGATCGGAA |
| 65377_1 | TATGAGTGTGGAGTCGTTGCCTGAACTGAAGCCCCATCCTATTNNNAGATCGGAAG |
| 65378_1 | TATGAGTGTGGAGTCGTTGCGAGGCAGAGACAGAGAGAGAAAGNNNAGATCGGA |
| 65379_1 | TATGAGTGTGGAGTCGTTGCAGAGCCCGAAACAAAATAGAGGANNNAGATCGGAA |
| 65380_1 | TATGAGTGTGGAGTCGTTGCATCAAGGAGTGAAGGCCAAATCANNNAGATCGGAA |
| 65381_1 | TATGAGTGTGGAGTCGTTGCATCTTATCTTACTGCACCTCCGCNNNAGATCGGAAGA |
| 65382_1 | TATGAGTGTGGAGTCGTTGCATGAGTCCTCCCTATCCTCAACANNNAGATCGGAAGA |
| 65383_1 | TATGAGTGTGGAGTCGTTGCTTTTGCCAGATTTTCAAAGGGGCNNNAGATCGGAAG |
| 65385_1 | TATGAGTGTGGAGTCGTTGCTCTCCTTCCAGTATGTCTCCAGTNNNAGATCGGAAGA |
| 65386_1 | TATGAGTGTGGAGTCGTTGCTTAGTTGTAAGGCAGGTGGCTAGNNNAGATCGGAA |
| 65387_1 | TATGAGTGTGGAGTCGTTGCGCAGTGATGTGATGGTGATCTTGNNNAGATCGGAA |
| 65388_1 | TATGAGTGTGGAGTCGTTGCGGGAGGAACTCTAAGGAAAGCAANNNAGATCGGA |
| 65389_1 | TATGAGTGTGGAGTCGTTGCCACTCACCAAGAGAGGTTCTGAANNNAGATCGGAA |
| 65390_1 | TATGAGTGTGGAGTCGTTGCTTGTTTTCTGGATTACGTGCACANNNAGATCGGAAG |
| 65391_1 | TATGAGTGTGGAGTCGTTGCGAGGCACACAGTTTTCTGAAACANNNAGATCGGAA |
| 65393_1 | TATGAGTGTGGAGTCGTTGCAGAGAGAGAGAGAGAGAGACGTGNNNAGATCGGA |
| 65397_1 | TATGAGTGTGGAGTCGTTGCAGCTAAATCCAGCACAGATCACANNNAGATCGGAAG |
| 65398_1 | TATGAGTGTGGAGTCGTTGCAGAGAGAGAGGGAGAGAGAGAGANNNAGATCGG |
| 65400_1 | TATGAGTGTGGAGTCGTTGCGGTGTCATCTTATCTAGGGGCTGNNNAGATCGGAA |
| 65401_1 | TATGAGTGTGGAGTCGTTGCTTGCATGAAATTTAGCACCCCTCNNNAGATCGGAAG |
| 65402_1 | TATGAGTGTGGAGTCGTTGCAATTGGATAGGCTAAACCCACCANNNAGATCGGAAG |
| 65403_1 | TATGAGTGTGGAGTCGTTGCGCTCAGCACTTTATAAGCGTGATNNNAGATCGGAAG |
| 65404_1 | TATGAGTGTGGAGTCGTTGCGCTGCTGGACACAGTTTTCATAGNNNAGATCGGAAG |

| | |
|---|---|
| 65405_1 | TATGAGTGTGGAGTCGTTGCCACTGGTGTACAGGATGAAGAGTNNNAGATCGGAA |
| 65407_1 | TATGAGTGTGGAGTCGTTGCCATTGAACGACTTTAGGGGCAAGNNNAGATCGGAA |
| 65408_1 | TATGAGTGTGGAGTCGTTGCTTCTTGCTAAGCCCTCTTTTCCTNNNAGATCGGAAGA |
| 65409_1 | TATGAGTGTGGAGTCGTTGCTAAATTCTCCAATCATGGGGGCANNNAGATCGGAAG |
| 65410_1 | TATGAGTGTGGAGTCGTTGCAAAATTCAGCCTCAGTCTGGTGTNNNAGATCGGAAG |
| 65411_1 | TATGAGTGTGGAGTCGTTGCACTGGGCAGCCTGTAATATTCTTNNNAGATCGGAAG |
| 65412_1 | TATGAGTGTGGAGTCGTTGCTATGTGTGTGTGTGTCTGTCTGTNNNAGATCGGAAG |
| 65413_1 | TATGAGTGTGGAGTCGTTGCGAGGTGCATGAGGTGTCTGTAAANNNAGATCGGAA |
| 65414_1 | TATGAGTGTGGAGTCGTTGCCCTGTGTCCTGTCACCTTCTTTCNNNAGATCGGAAGA |
| 65415_1 | TATGAGTGTGGAGTCGTTGCTCAATGGCAAAAAGAATGCACCTNNNAGATCGGAAG |
| 65416_1 | TATGAGTGTGGAGTCGTTGCCATACTTCCCAGACCTTTTCCCANNNAGATCGGAAGA |
| 65417_1 | TATGAGTGTGGAGTCGTTGCCAAAAGCAAAAGGGAAGGAGGAGNNNAGATCGGA |
| 65418_1 | TATGAGTGTGGAGTCGTTGCTTATAATCAGGGCGTCAGTGGACNNNAGATCGGAA |
| 65419_1 | TATGAGTGTGGAGTCGTTGCAAGGATGCTGGGTACAAGCATAANNNAGATCGGAA |
| 65420_1 | TATGAGTGTGGAGTCGTTGCTCTGTCTGTCTGTCTGTCTCTCTNNNAGATCGGAAGA |
| 65421_1 | TATGAGTGTGGAGTCGTTGCCCTTGCGCATTAGGAACTTCTTANNNAGATCGGAAG |
| 65422_1 | TATGAGTGTGGAGTCGTTGCCTCTTAGTTCTCTGGCTGTGGTCNNNAGATCGGAAG |
| 65423_1 | TATGAGTGTGGAGTCGTTGCATTTACAAGTGCAAATCCCCTCCNNNAGATCGGAAG |
| 65424_1 | TATGAGTGTGGAGTCGTTGCCAAGATGGTGGTTTTTGGAGCATNNNAGATCGGAA |
| 65425_1 | TATGAGTGTGGAGTCGTTGCGAGAAACAGGACCAGTGCATAGANNNAGATCGGAA |
| 65426_1 | TATGAGTGTGGAGTCGTTGCATAATGGCATTCACAGCAACCTGNNNAGATCGGAAG |
| 65427_1 | TATGAGTGTGGAGTCGTTGCTAGCACCCTTAGCTTTTGATCCANNNAGATCGGAAG |
| 65428_1 | TATGAGTGTGGAGTCGTTGCTCTCTGGAGAACCCTGACTAACANNNAGATCGGAAG |
| 65429_1 | TATGAGTGTGGAGTCGTTGCAATCTACCTACTATCACCCAGGTNNNAGATCGGAAG |
| 65430_1 | TATGAGTGTGGAGTCGTTGCAACACTGAAGACTGAACTTTGCANNNAGATCGGAAG |
| 65431_1 | TATGAGTGTGGAGTCGTTGCAACATTCCAAACTCAATGCAGCANNNAGATCGGAAG |
| 65432_1 | TATGAGTGTGGAGTCGTTGCCTCAATCACATCTGCAAAGTCCCNNNAGATCGGAAG |
| 65433_1 | TATGAGTGTGGAGTCGTTGCACAAACACAGGCACACATACATGNNNAGATCGGAAG |
| 65434_1 | TATGAGTGTGGAGTCGTTGCAGACTTGGGCTACTTGTGTGAATNNNAGATCGGAA |
| 65435_1 | TATGAGTGTGGAGTCGTTGCGTCAAAGGGTGTCATCTCAGACTNNNAGATCGGAA |
| 65436_1 | TATGAGTGTGGAGTCGTTGCTGCACACATGTTTACAAGCATCTNNNAGATCGGAAG |
| 65437_1 | TATGAGTGTGGAGTCGTTGCAAGTCCTTGGAACCAGATGTACCNNNAGATCGGAAG |
| 65439_1 | TATGAGTGTGGAGTCGTTGCACATGAAGGAGCAGGGATTTGAANNNAGATCGGA |
| 65440_1 | TATGAGTGTGGAGTCGTTGCGTGTGGGAGCTTTATGTGTGAACNNNAGATCGGAA |
| 65441_1 | TATGAGTGTGGAGTCGTTGCCAAATGTGAGAGGGGACTGGTATNNNAGATCGGA |
| 65443_1 | TATGAGTGTGGAGTCGTTGCTCAAGTGATCTGCCCATCTTTGANNNAGATCGGAAG |
| 65444_1 | TATGAGTGTGGAGTCGTTGCCATCCCTTCCTGATGCCTTCTAGNNNAGATCGGAAGA |
| 65445_1 | TATGAGTGTGGAGTCGTTGCTTTCAGAAAAGGGCCAATTTCCCNNNAGATCGGAAG |
| 65446_1 | TATGAGTGTGGAGTCGTTGCATTATACTCACTTGAGGCTGGGCNNNAGATCGGAAG |
| 65447_1 | TATGAGTGTGGAGTCGTTGCCACAGACTTTGCTTAGCATGCTTNNNAGATCGGAAG |
| 65448_1 | TATGAGTGTGGAGTCGTTGCACAAACCACACTCACTCACTCTTNNNAGATCGGAAGA |
| 65449_1 | TATGAGTGTGGAGTCGTTGCTCAAAGGGAGAAGAAAGGAGGTGNNNAGATCGGA |
| 65450_1 | TATGAGTGTGGAGTCGTTGCAGATACACAAACAGAGACGGAGGNNNAGATCGGA |

| | |
|---|---|
| 65451_1 | TATGAGTGTGGAGTCGTTGCGAGAAATTGCTCCAGAAATGGGGNNNAGATCGGAA |
| 65452_1 | TATGAGTGTGGAGTCGTTGCCCCTTCACCTTCTACTCATCCTTNNNAGATCGGAAGA |
| 65453_1 | TATGAGTGTGGAGTCGTTGCTAGTGAGGTCAGAGATGGTAGGGNNNAGATCGGA |
| 65455_1 | TATGAGTGTGGAGTCGTTGCCACACAATCACAAGGTGAGTTCCNNNAGATCGGAAG |
| 65457_1 | TATGAGTGTGGAGTCGTTGCTGTTTCTCTGGGGAATCTTGACTNNNAGATCGGAAG |
| 65458_1 | TATGAGTGTGGAGTCGTTGCAAACACAAAACTGCTGATCCCACNNNAGATCGGAAG |
| 65459_1 | TATGAGTGTGGAGTCGTTGCCAACACACTGGGCATTTTGATCTNNNAGATCGGAAG |
| 65460_1 | TATGAGTGTGGAGTCGTTGCGAGGAACTGTCATACCCCTTCTGNNNAGATCGGAAG |
| 65461_1 | TATGAGTGTGGAGTCGTTGCGCTGAAATTTGCACACTTTGGGANNNAGATCGGAA |
| 65462_1 | TATGAGTGTGGAGTCGTTGCTGGCCTAGAAGCAATGAGACATTNNNAGATCGGAA |
| 65463_1 | TATGAGTGTGGAGTCGTTGCATCAAGCTCTCCCCAATTTGAGTNNNAGATCGGAAG |
| 65464_1 | TATGAGTGTGGAGTCGTTGCGGCAAGTCTCTAGTTCAGGGATTNNNAGATCGGAA |
| 65465_1 | TATGAGTGTGGAGTCGTTGCAATGCAGTCATCAGAGTCCTCAGNNNAGATCGGAAG |
| 65466_1 | TATGAGTGTGGAGTCGTTGCTGGAAGGATGCTGATATGAGCTGNNNAGATCGGAA |
| 65467_1 | TATGAGTGTGGAGTCGTTGCTGTTCTTCTTTAGAGACAGGGCCNNNAGATCGGAAG |
| 65468_1 | TATGAGTGTGGAGTCGTTGCGTGGGTCTATTTTATCCCTGCCANNNAGATCGGAAG |
| 65469_1 | TATGAGTGTGGAGTCGTTGCTGAAAGTGTACAGCTCAGTGACANNNAGATCGGAA |
| 65470_1 | TATGAGTGTGGAGTCGTTGCTATACTGATCCCACCCTGACTCANNNAGATCGGAAGA |
| 65472_1 | TATGAGTGTGGAGTCGTTGCATTTTCAAAGGCATCAGTGACCCNNNAGATCGGAAG |
| 65473_1 | TATGAGTGTGGAGTCGTTGCCTCAGCTCTGATCAATACCTGCTNNNAGATCGGAAG |
| 65474_1 | TATGAGTGTGGAGTCGTTGCCCAAATGGGCCTGAGATACTGATNNNAGATCGGAA |
| 65475_1 | TATGAGTGTGGAGTCGTTGCAGAGTTGCTACCTCCCTTTCAAGNNNAGATCGGAAG |
| 65476_1 | TATGAGTGTGGAGTCGTTGCGGTGGAAAGTAGGGAAGAGACAANNNAGATCGGA |
| 65477_1 | TATGAGTGTGGAGTCGTTGCGTGTGACAATCAAGGCTGGATTGNNNAGATCGGAA |
| 65478_1 | TATGAGTGTGGAGTCGTTGCTATCTTGCATCCATTGTCCCACTNNNAGATCGGAAGA |
| 65481_1 | TATGAGTGTGGAGTCGTTGCGAACATATCCCCAAAAGAGCACGNNNAGATCGGAA |
| 65483_1 | TATGAGTGTGGAGTCGTTGCTCTGTGCTCAATTGTCCTGATGANNNAGATCGGAAG |
| 65484_1 | TATGAGTGTGGAGTCGTTGCTTGTGGTTCTGTGAGTGTGTTTGNNNAGATCGGAA |
| 65485_1 | TATGAGTGTGGAGTCGTTGCGAGAGAGGAACCCACACCAATAGNNNAGATCGGAA |
| 65486_1 | TATGAGTGTGGAGTCGTTGCAATTATGGAGGCTGACAAGTCCCNNNAGATCGGAA |
| 65487_1 | TATGAGTGTGGAGTCGTTGCGGGGAAGGGTTGGAGGATATTTTNNNAGATCGGA |
| 65488_1 | TATGAGTGTGGAGTCGTTGCACAAGTTTGTACACAGTAGGCCANNNAGATCGGAA |
| 65489_1 | TATGAGTGTGGAGTCGTTGCTACCTGTGCACTCAAAAATGCTGNNNAGATCGGAAG |
| 65490_1 | TATGAGTGTGGAGTCGTTGCGCCATGCCAAGAGATCACAAATTNNNAGATCGGAAG |
| 65491_1 | TATGAGTGTGGAGTCGTTGCTGGCACAAAGGATGGACTCAATANNNAGATCGGAA |
| 65492_1 | TATGAGTGTGGAGTCGTTGCAGAAGAGAACCCTGTTCTTTCCCNNNAGATCGGAAG |
| 65493_1 | TATGAGTGTGGAGTCGTTGCTCCCCATTTTCTGCCAACATTTCNNNAGATCGGAAGA |
| 65495_1 | TATGAGTGTGGAGTCGTTGCTTTGTGGTGATGAGTCTGAGAGGNNNAGATCGGAA |
| 65496_1 | TATGAGTGTGGAGTCGTTGCAAAGACAGGGAGTAGCCATAAGCNNNAGATCGGAA |
| 65497_1 | TATGAGTGTGGAGTCGTTGCTGGACCTAAAACTTGCTGCTTTGNNNAGATCGGAAG |
| 65498_1 | TATGAGTGTGGAGTCGTTGCATTTCCCACTATGTTCCAGGCATNNNAGATCGGAAG |
| 65499_1 | TATGAGTGTGGAGTCGTTGCTGGTGTTTACTGTCTGCATGGTANNNAGATCGGAAG |
| 65500_1 | TATGAGTGTGGAGTCGTTGCCTTTTCCATTATGGGGTTGCCTGNNNAGATCGGAAG |

| | |
|---|---|
| 65501_1 | TATGAGTGTGGAGTCGTTGCTACAAGTCAGCCTGGGTTTCTTTNNNAGATCGGAAG |
| 65503_1 | TATGAGTGTGGAGTCGTTGCCCACCTAGCCCTTTCCTAGTTTTNNNAGATCGGAAGA |
| 65504_1 | TATGAGTGTGGAGTCGTTGCTCACGTACCTGGAGATGCAGATANNNAGATCGGAAG |
| 65505_1 | TATGAGTGTGGAGTCGTTGCGGCAGAGAGTTCCTGACTAGATGNNNAGATCGGAA |
| 65506_1 | TATGAGTGTGGAGTCGTTGCGTTAGGAGGAGATGGAATGGTGGNNNAGATCGGA |
| 65507_1 | TATGAGTGTGGAGTCGTTGCTATGCACAGAGTGGCCAAAAATCNNNAGATCGGAA |
| 65509_1 | TATGAGTGTGGAGTCGTTGCTCATGCTTCTACCAGTTAGGGACNNNAGATCGGAAG |
| 65510_1 | TATGAGTGTGGAGTCGTTGCCCCCTGTGATAATAAGACCCAGGNNNAGATCGGAAG |
| 65511_1 | TATGAGTGTGGAGTCGTTGCCCAATCCCCATAAAAGCACTTGGNNNAGATCGGAAG |
| 65513_1 | TATGAGTGTGGAGTCGTTGCTAACAGAAGCATAAACCAAGCCCNNNAGATCGGAAG |
| 65514_1 | TATGAGTGTGGAGTCGTTGCGATGGCCTCTCAGTAAGCTTCTTNNNAGATCGGAAG |
| 65515_1 | TATGAGTGTGGAGTCGTTGCCCATAGTTTGTCAATGCCTGGTCNNNAGATCGGAAG |
| 65516_1 | TATGAGTGTGGAGTCGTTGCATTCTTTGCCTCTGGAGAATCCTNNNAGATCGGAAG |
| 65517_1 | TATGAGTGTGGAGTCGTTGCGTCTTGGTCCACTTCTCATCCTTNNNAGATCGGAAGA |
| 65518_1 | TATGAGTGTGGAGTCGTTGCTAAGTGTTCTCTGTCCCTTCTCCNNNAGATCGGAAGA |
| 65519_1 | TATGAGTGTGGAGTCGTTGCTTTCCAGCTCATCACTTCCTAGCNNNAGATCGGAAGA |
| 65520_1 | TATGAGTGTGGAGTCGTTGCGACTCCTAAACAAAGGTGTGGGANNNAGATCGGAA |
| 65521_1 | TATGAGTGTGGAGTCGTTGCTCTAGGGACCACAGGCTATTACANNNAGATCGGAAG |
| 65522_1 | TATGAGTGTGGAGTCGTTGCGGGCAAAAGATTGTGTTGTGGTANNNAGATCGGAA |
| 65523_1 | TATGAGTGTGGAGTCGTTGCTCTGTAGGAATCCTGAAGGCCTANNNAGATCGGAAG |
| 65524_1 | TATGAGTGTGGAGTCGTTGCTGTGTTTTTGAAATTGGGTTGCTNNNAGATCGGAAG |
| 65525_1 | TATGAGTGTGGAGTCGTTGCATTGAAAATGGGTCCTGTGCATGNNNAGATCGGAA |
| 65527_1 | TATGAGTGTGGAGTCGTTGCAATGAAGACACCAATGCTCTTGCNNNAGATCGGAAG |
| 65528_1 | TATGAGTGTGGAGTCGTTGCAGACCAGAAGCACATATCACAGGNNNAGATCGGAA |
| 65529_1 | TATGAGTGTGGAGTCGTTGCAAATCATCTGCCAAATGGTCTCTNNNAGATCGGAAG |
| 65530_1 | TATGAGTGTGGAGTCGTTGCTGCATATGCTAATTGGTCTTGGCNNNAGATCGGAAG |
| 65531_1 | TATGAGTGTGGAGTCGTTGCATGTTTGAAATACCAAGCCTGCCNNNAGATCGGAAG |
| 65532_1 | TATGAGTGTGGAGTCGTTGCACCTAGACATCTTTGGGCAATGTNNNAGATCGGAAG |
| 65534_1 | TATGAGTGTGGAGTCGTTGCTCACATGCTGATGCTGATACCATNNNAGATCGGAAG |
| 65535_1 | TATGAGTGTGGAGTCGTTGCAGGGAGGAATAGCTCTTCTGGTTNNNAGATCGGAA |
| 65537_1 | TATGAGTGTGGAGTCGTTGCAAGCACTTTTGTCCAGGCAAGTANNNAGATCGGAAG |
| 65538_1 | TATGAGTGTGGAGTCGTTGCTTGGAGATTCTGCCACATTGGTANNNAGATCGGAAG |
| 65539_1 | TATGAGTGTGGAGTCGTTGCCGTGGCTTTGATGTTAGTTGGTANNNAGATCGGAA |
| 65540_1 | TATGAGTGTGGAGTCGTTGCGAGTCACTCGAACAATGTGTGTGNNNAGATCGGAA |
| 65542_1 | TATGAGTGTGGAGTCGTTGCTGGCTAGTTGTAGCAATGCATTCNNNAGATCGGAAG |
| 65543_1 | TATGAGTGTGGAGTCGTTGCTTCCTGCCTTCTGATTGGATCAANNNAGATCGGAAG |
| 65544_1 | TATGAGTGTGGAGTCGTTGCAACCTGTACAGCTGCACATATCANNNAGATCGGAAG |
| 65545_1 | TATGAGTGTGGAGTCGTTGCAAGCAGTGCTGGATCTTGAGTTANNNAGATCGGAA |
| 65546_1 | TATGAGTGTGGAGTCGTTGCCAGACATATTTGTGGCCAGGATGNNNAGATCGGAA |
| 65548_1 | TATGAGTGTGGAGTCGTTGCCTTCCTCCCCAAATCACATTCCANNNAGATCGGAAGA |
| 65549_1 | TATGAGTGTGGAGTCGTTGCGAAAGAAGTCTGTGGGCTCCATTNNNAGATCGGAA |
| 65550_1 | TATGAGTGTGGAGTCGTTGCTATATCTTGCCACCACACCCATCNNNAGATCGGAAGA |
| 65551_1 | TATGAGTGTGGAGTCGTTGCACCAGAGGCCCTTGAAAATGTAANNNAGATCGGAA |

| | |
|---|---|
| 65552_1 | TATGAGTGTGGAGTCGTTGCTTATAAGGCCTCCAGTCCTACCANNNAGATCGGAAG |
| 65553_1 | TATGAGTGTGGAGTCGTTGCCTTTTGCTGTCTCCCATCAGTTGNNNAGATCGGAAGA |
| 65554_1 | TATGAGTGTGGAGTCGTTGCCTCTACCTCCTCTTATGAAGGCCNNNAGATCGGAAGA |
| 65555_1 | TATGAGTGTGGAGTCGTTGCAGAGAGAGAGTTTTGCAAGGCTTNNNAGATCGGAA |
| 65557_1 | TATGAGTGTGGAGTCGTTGCGTGTGTATCGTTGAGGTAGGAGGNNNAGATCGGAA |
| 65558_1 | TATGAGTGTGGAGTCGTTGCCCATGAATCATGTGGCTTCTCACNNNAGATCGGAAG |
| 65560_1 | TATGAGTGTGGAGTCGTTGCATGACTCTAGTCCCAACACTTGCNNNAGATCGGAAG |
| 65562_1 | TATGAGTGTGGAGTCGTTGCGCCTGAGTCACTTTTGTCAACAGNNNAGATCGGAAG |
| 65564_1 | TATGAGTGTGGAGTCGTTGCACAAAGCTTTGATCTGCCACTTCNNNAGATCGGAAG |
| 65565_1 | TATGAGTGTGGAGTCGTTGCTCCCTACTTCCCTCCCTCTTTAGNNNAGATCGGAAGA |
| 65566_1 | TATGAGTGTGGAGTCGTTGCTTCTCAAAGTCTCCCTGCTTCTGNNNAGATCGGAAGA |
| 65567_1 | TATGAGTGTGGAGTCGTTGCTAATTGGTGGTTTAGAGCTGGCTNNNAGATCGGAA |
| 65568_1 | TATGAGTGTGGAGTCGTTGCTGAGTGTATGTGTGTGTGTGTGANNNAGATCGGAA |
| 65569_1 | TATGAGTGTGGAGTCGTTGCGCCCACCTTCAATATGCAGAATCNNNAGATCGGAAG |
| 65570_1 | TATGAGTGTGGAGTCGTTGCTCACAGAAAGGGTGAACTGCTTANNNAGATCGGAA |
| 65572_1 | TATGAGTGTGGAGTCGTTGCCACTCTGTGTCTAGCTGAAGGTTNNNAGATCGGAAG |
| 65573_1 | TATGAGTGTGGAGTCGTTGCGCGAAAAGAAGTCTCTAGCTCTGNNNAGATCGGAA |
| 65574_1 | TATGAGTGTGGAGTCGTTGCATGTGATTTGCCTTTGTCCCTTGNNNAGATCGGAAG |
| 65575_1 | TATGAGTGTGGAGTCGTTGCGAGGTCAGGAGATTGAGACCATCNNNAGATCGGAA |
| 65576_1 | TATGAGTGTGGAGTCGTTGCTTTATTCTCCTATCTGGCCCCACNNNAGATCGGAAGA |
| 65577_1 | TATGAGTGTGGAGTCGTTGCAATCCAAAAATCAGCCGGACATGNNNAGATCGGAAG |
| 65578_1 | TATGAGTGTGGAGTCGTTGCGGTCTTGAACTCCTAACCTCAGGNNNAGATCGGAAG |
| 65579_1 | TATGAGTGTGGAGTCGTTGCGAGATAGGAGAAATGGGAGCAGCNNNAGATCGGA |
| 65580_1 | TATGAGTGTGGAGTCGTTGCTCTCAAACTCCTGACCTCAAGTGNNNAGATCGGAAG |
| 65581_1 | TATGAGTGTGGAGTCGTTGCGAGACAAGCTCTGACTCTATCGCNNNAGATCGGAAG |
| 65582_1 | TATGAGTGTGGAGTCGTTGCAGATTACAGGTGTGAGCAACCATNNNAGATCGGAA |
| 65583_1 | TATGAGTGTGGAGTCGTTGCTACATTGGCATGGGAAATGACACNNNAGATCGGAA |
| 65584_1 | TATGAGTGTGGAGTCGTTGCTTGTTTGCCTCTGTAGACTGAGGNNNAGATCGGAAG |
| 65585_1 | TATGAGTGTGGAGTCGTTGCCCCTGAATTCTAACCCAGTACCANNNAGATCGGAAGA |
| 65586_1 | TATGAGTGTGGAGTCGTTGCATCTGAAGCAGCCTGATTTCTGANNNAGATCGGAAG |
| 65587_1 | TATGAGTGTGGAGTCGTTGCGTTCAGAAGATGGATGTCCCAGTNNNAGATCGGAA |
| 65588_1 | TATGAGTGTGGAGTCGTTGCGGTTTGCATCATAGATCCTTCGCNNNAGATCGGAAG |
| 65590_1 | TATGAGTGTGGAGTCGTTGCTCTCTTGTCTGATTGCTCTAGCTNNNAGATCGGAAGA |
| 65595_1 | TATGAGTGTGGAGTCGTTGCGCAACATGCCTATGTACATTCCCNNNAGATCGGAAG |
| 65597_1 | TATGAGTGTGGAGTCGTTGCCTAAGAAGAGCCTCTCACCCTTCNNNAGATCGGAAG |
| 65598_1 | TATGAGTGTGGAGTCGTTGCGAGAAAGAGAGAGCAATGACCCANNNAGATCGGAA |
| 65599_1 | TATGAGTGTGGAGTCGTTGCGCCTCCTGGGTTCAAGTTATTCTNNNAGATCGGAAG |
| 65601_1 | TATGAGTGTGGAGTCGTTGCATCAGACTTTCGATTCCAGGTCCNNNAGATCGGAAG |
| 65602_1 | TATGAGTGTGGAGTCGTTGCTATGGTGGCATGTGTCTGTAGTCNNNAGATCGGAA |
| 65603_1 | TATGAGTGTGGAGTCGTTGCTTCCTAGGAACACAGCTAAGCAGNNNAGATCGGAAG |
| 65604_1 | TATGAGTGTGGAGTCGTTGCGGGAAGAAATTCTGCGGAACAAANNNAGATCGGAA |
| 65605_1 | TATGAGTGTGGAGTCGTTGCGGGTAGAGACTTACTCAGCAAGGNNNAGATCGGAA |
| 65606_1 | TATGAGTGTGGAGTCGTTGCTGGTCCTAAGGGTAAACAGCAAGNNNAGATCGGAA |

| | |
|---|---|
| 65607_1 | TATGAGTGTGGAGTCGTTGCTGGTGTATGTTCTGTAATCCACANNNAGATCGGAAG |
| 65608_1 | TATGAGTGTGGAGTCGTTGCCCTGGGTTCAAGCAATTCTCATGNNNAGATCGGAAG |
| 65610_1 | TATGAGTGTGGAGTCGTTGCACATACTGACAGCATTGTCCTGTNNNAGATCGGAAG |
| 65613_1 | TATGAGTGTGGAGTCGTTGCGCCATTTGCAACTACATGGATGANNNAGATCGGAAG |
| 65615_1 | TATGAGTGTGGAGTCGTTGCCCAAAATGCTTGGGACCAGAAATNNNAGATCGGAAG |
| 65616_1 | TATGAGTGTGGAGTCGTTGCGCCTTGGTCTGTCACAGATAGATNNNAGATCGGAAG |
| 65618_1 | TATGAGTGTGGAGTCGTTGCACAGAGGTAGGACAGATCCAGAANNNAGATCGGAA |
| 65619_1 | TATGAGTGTGGAGTCGTTGCTGCATCACATTCCTAGCTGGTAGNNNAGATCGGAAG |
| 65620_1 | TATGAGTGTGGAGTCGTTGCATAATGGCATATGCAGCAACCTGNNNAGATCGGAAG |
| 65622_1 | TATGAGTGTGGAGTCGTTGCTTTTTGTAGAGACAGGGGTTGCTNNNAGATCGGAA |
| 65623_1 | TATGAGTGTGGAGTCGTTGCGCATGCCAAAGAAGAGTGGTAAGNNNAGATCGGAA |
| 65624_1 | TATGAGTGTGGAGTCGTTGCTCAAGCGATTCTCCTATCTCAGCNNNAGATCGGAAG |
| 65625_1 | TATGAGTGTGGAGTCGTTGCTGTGATGTTAGGGTGTCGATTTGNNNAGATCGGAA |
| 65627_1 | TATGAGTGTGGAGTCGTTGCTGGGTACTGCATTGTTTACTCGANNNAGATCGGAAG |
| 65628_1 | TATGAGTGTGGAGTCGTTGCTCTGTGCTACTCTCATGACATGGNNNAGATCGGAAG |
| 65629_1 | TATGAGTGTGGAGTCGTTGCAGAGACAGGGTTTCTCCATGTTGNNNAGATCGGAA |
| 65630_1 | TATGAGTGTGGAGTCGTTGCTTAGGATGCCATCATGGTGGATTNNNAGATCGGAA |
| 65631_1 | TATGAGTGTGGAGTCGTTGCTCCTGACTGGGGTCTAAGAAGATNNNAGATCGGAA |
| 65632_1 | TATGAGTGTGGAGTCGTTGCGAGACAGAAAGAGACAGAGACGGNNNAGATCGGA |
| 65633_1 | TATGAGTGTGGAGTCGTTGCCAAACTCCTTTAGCTCCCAGGATNNNAGATCGGAAG |
| 65634_1 | TATGAGTGTGGAGTCGTTGCACAGACAGAGAGAGAGACAGACANNNAGATCGGAA |
| 65635_1 | TATGAGTGTGGAGTCGTTGCGGTCGTCCAGAAAGCTTGCTATANNNAGATCGGAAG |
| 65637_1 | TATGAGTGTGGAGTCGTTGCTGTCAGGGTAATGGATGTGACTGNNNAGATCGGAA |
| 65638_1 | TATGAGTGTGGAGTCGTTGCACACACACACAGAAAGAGAGAGANNNAGATCGGAA |
| 65639_1 | TATGAGTGTGGAGTCGTTGCTGCATTGCACTTACCAAGTTGAGNNNAGATCGGAAG |
| 65640_1 | TATGAGTGTGGAGTCGTTGCAGATACTTGGGTTTGCATCTCGGNNNAGATCGGAAG |
| 65641_1 | TATGAGTGTGGAGTCGTTGCCCTACTCATGGCTGTTAGGGAAGNNNAGATCGGAA |
| 65642_1 | TATGAGTGTGGAGTCGTTGCAAATCTCACCCCATCCATTAGCANNNAGATCGGAAGA |
| 65643_1 | TATGAGTGTGGAGTCGTTGCATAGTCCTAGCTACTTGGGAGGCNNNAGATCGGAA |
| 65644_1 | TATGAGTGTGGAGTCGTTGCAGTTCCAGTTAGGGATAAGTGCTNNNAGATCGGAA |
| 65646_1 | TATGAGTGTGGAGTCGTTGCCTGTAATCCCAGCTACTCCTGAGNNNAGATCGGAAG |
| 65648_1 | TATGAGTGTGGAGTCGTTGCTTTTGAGGTGTGTTGAGATCCCANNNAGATCGGAAG |
| 65649_1 | TATGAGTGTGGAGTCGTTGCAAACTATGGGCCAATATCTGTGANNNAGATCGGAAG |
| 65650_1 | TATGAGTGTGGAGTCGTTGCTTCTCGACATAAGTGCCCAAAGANNNAGATCGGAAG |
| 65651_1 | TATGAGTGTGGAGTCGTTGCTCACACTCACTATCTGTTTGGGGNNNAGATCGGAAG |
| 65652_1 | TATGAGTGTGGAGTCGTTGCTACACAACGATCTCTTTGTCCCCNNNAGATCGGAAGA |
| 65653_1 | TATGAGTGTGGAGTCGTTGCGAGCAAAGAACACTGCAATGGATNNNAGATCGGAA |
| 65654_1 | TATGAGTGTGGAGTCGTTGCTCTTTTGGCTGCACAGGAAAATTNNNAGATCGGAAG |
| 65655_1 | TATGAGTGTGGAGTCGTTGCGACAGTTGGGACTCTACACACATNNNAGATCGGAAG |
| 65656_1 | TATGAGTGTGGAGTCGTTGCGTTAGAACTGTGGCTGGTAGACANNNAGATCGGAA |
| 65657_1 | TATGAGTGTGGAGTCGTTGCTAAAGTGCAGTGGTACCATCACANNNAGATCGGAAG |
| 65658_1 | TATGAGTGTGGAGTCGTTGCTTCCACCCTTCCTTCTCTCTACTNNNAGATCGGAAGAG |
| 65659_1 | TATGAGTGTGGAGTCGTTGCGATGGAGTCTCACTCTGTCACTCNNNAGATCGGAAG |

| | |
|---|---|
| 65661_1 | TATGAGTGTGGAGTCGTTGCAACTGCCTGCAGAGATGACAATANNNAGATCGGAAG |
| 65662_1 | TATGAGTGTGGAGTCGTTGCATATCCCATGTGCAGGTGTCTTTNNNAGATCGGAAGA |
| 65667_1 | TATGAGTGTGGAGTCGTTGCGACAGAGAATGTGTGTGTGTGTGNNNAGATCGGAA |
| 65668_1 | TATGAGTGTGGAGTCGTTGCTTTGGTAGAGTTGGTGTTTTGCCNNNAGATCGGAAG |
| 65669_1 | TATGAGTGTGGAGTCGTTGCACAGTACAGTACAGTAGTTCGCCNNNAGATCGGAAG |
| 65670_1 | TATGAGTGTGGAGTCGTTGCCCCACACTTCAAGGAGGACATAANNNAGATCGGAAG |
| 65671_1 | TATGAGTGTGGAGTCGTTGCAATCCAGACTCCTCATGATGGTGNNNAGATCGGAAG |
| 65672_1 | TATGAGTGTGGAGTCGTTGCACGAGAACTGGTTGAACCTGTAGNNNAGATCGGAAG |
| 65673_1 | TATGAGTGTGGAGTCGTTGCCAGGTTCTTTTCCTCTCCCAGAANNNAGATCGGAAGA |
| 65674_1 | TATGAGTGTGGAGTCGTTGCGGTGTGCTTATCATGCATGACTCNNNAGATCGGAAG |
| 65675_1 | TATGAGTGTGGAGTCGTTGCCCACTTGTTGAATGATGGGCATTNNNAGATCGGAAG |
| 65676_1 | TATGAGTGTGGAGTCGTTGCAGAAGATGCAGTGAGCCAAGATCNNNAGATCGGAA |
| 65677_1 | TATGAGTGTGGAGTCGTTGCATGTATTTGTTGACCCAGAAGCANNNAGATCGGAAG |
| 65678_1 | TATGAGTGTGGAGTCGTTGCAGAGCTACAAAATGCCACTCAGANNNAGATCGGAAG |
| 65679_1 | TATGAGTGTGGAGTCGTTGCGACAGAAGGCTAATTGGGAAAGCNNNAGATCGGAA |
| 65681_1 | TATGAGTGTGGAGTCGTTGCGAGATTTCATGTCCTCAGCTGGANNNAGATCGGAAG |
| 65682_1 | TATGAGTGTGGAGTCGTTGCGTGACTTTCATCATTGCTGGGAGNNNAGATCGGAAG |
| 65683_1 | TATGAGTGTGGAGTCGTTGCTCTGCCCATTTTGCTTGAATAGTNNNAGATCGGAAGA |
| 65684_1 | TATGAGTGTGGAGTCGTTGCTGGTGGGGGAGAGTTTCAATTTTNNNAGATCGGAAG |
| 65685_1 | TATGAGTGTGGAGTCGTTGCAATGGCTTGGACTAAGGTGGTAGNNNAGATCGGAA |
| 65689_1 | TATGAGTGTGGAGTCGTTGCAATCACTTGAATCAGGGAGGTGGNNNAGATCGGAA |
| 65690_1 | TATGAGTGTGGAGTCGTTGCGAGAGAGAGGGAGAGGAAGAGAGNNNAGATCGGA |
| 65691_1 | TATGAGTGTGGAGTCGTTGCGACAGACAAAGCAAGAACCTGTCNNNAGATCGGAAG |
| 65693_1 | TATGAGTGTGGAGTCGTTGCTGAGTGCATGTAGAACTGGAGAANNNAGATCGGAA |
| 65694_1 | TATGAGTGTGGAGTCGTTGCGGTGAGGGGCTATACCATTTAGGNNNAGATCGGAA |
| 65695_1 | TATGAGTGTGGAGTCGTTGCATCCTTTCTTCCCAACATCACCANNNAGATCGGAAGAG |
| 65697_1 | TATGAGTGTGGAGTCGTTGCAAATGGTGATTAGTTGCTGGGTGNNNAGATCGGAA |
| 65698_1 | TATGAGTGTGGAGTCGTTGCAAACAAGCTCTTTGCCCTTGAAANNNAGATCGGAAGA |
| 65699_1 | TATGAGTGTGGAGTCGTTGCAAACTCTTAGGCTCGAGTGATCCNNNAGATCGGAAG |
| 65700_1 | TATGAGTGTGGAGTCGTTGCTGAGAAGTCCCATATCTGCTGTCNNNAGATCGGAAG |
| 65701_1 | TATGAGTGTGGAGTCGTTGCTCCACCTCCATGACATCAACTTTNNNAGATCGGAAGA |
| 65702_1 | TATGAGTGTGGAGTCGTTGCGCAGCTGCTGATTACACTTTGAANNNAGATCGGAAG |
| 65703_1 | TATGAGTGTGGAGTCGTTGCGCTCCTTTATAAACTCCATGGCCNNNAGATCGGAAGA |
| 65704_1 | TATGAGTGTGGAGTCGTTGCCACAAACCTTCCCCTACCCTTTANNNAGATCGGAAGAG |
| 65705_1 | TATGAGTGTGGAGTCGTTGCTCTTCAGCGATAAGAGTGGACAGNNNAGATCGGAAG |
| 65706_1 | TATGAGTGTGGAGTCGTTGCAATTCGGGAGTTTCTTGGACACTNNNAGATCGGAAG |
| 65707_1 | TATGAGTGTGGAGTCGTTGCTCAACAGTAGACTCAGTCAAGCANNNAGATCGGAAG |
| 65708_1 | TATGAGTGTGGAGTCGTTGCTAGTTCTCCTTGAAGGGGTCCTTNNNAGATCGGAAG |
| 65709_1 | TATGAGTGTGGAGTCGTTGCAGGGTTGCCTAACACAGTATCAGNNNAGATCGGAAG |
| 65710_1 | TATGAGTGTGGAGTCGTTGCTCGAAGAGATACCTGTACTCCCANNNAGATCGGAAG |
| 65712_1 | TATGAGTGTGGAGTCGTTGCGGAGTGTGGTGGTGTGATCTCNNNAGATCGGAAGA |
| 65713_1 | TATGAGTGTGGAGTCGTTGCAGACAGTCTTGCTATGCTACCTGNNNAGATCGGAAG |
| 65714_1 | TATGAGTGTGGAGTCGTTGCGAACAGCATAGAGAAAGGCGTTGNNNAGATCGGAA |

| | |
|---|---|
| 65715_1 | TATGAGTGTGGAGTCGTTGCTGTGTGTATGTCTGTGTCCATGTNNNAGATCGGAAG |
| 65716_1 | TATGAGTGTGGAGTCGTTGCGCCTCCTGAGTTCAAGTGATTCTNNNAGATCGGAAG |
| 65717_1 | TATGAGTGTGGAGTCGTTGCGGTGTGGTTGTGTGCTTTCATTANNNAGATCGGAAG |
| 65719_1 | TATGAGTGTGGAGTCGTTGCACAACATGGATGATTCTGGAGGANNNAGATCGGAA |
| 65720_1 | TATGAGTGTGGAGTCGTTGCACCTTTGGTTCTGTTTCTCTGGANNNAGATCGGAAG |
| 65721_1 | TATGAGTGTGGAGTCGTTGCGTCACTTGTGAATAGCCGTGTTGNNNAGATCGGAAG |
| 65722_1 | TATGAGTGTGGAGTCGTTGCGGCACAATTATAGCTCACTGCAANNNAGATCGGAAG |
| 65723_1 | TATGAGTGTGGAGTCGTTGCCAAAAAACCCACCAGAATAGCCANNNAGATCGGAAG |
| 65724_1 | TATGAGTGTGGAGTCGTTGCCTAGGATGAGCTAACTCTGCCAGNNNAGATCGGAAG |
| 65726_1 | TATGAGTGTGGAGTCGTTGCCTTCTTTCTGCAAATCCCCTTCCNNNAGATCGGAAGA |
| 65727_1 | TATGAGTGTGGAGTCGTTGCAGCTACTTAGAAGGCTGAGGTTGNNNAGATCGGAA |
| 65728_1 | TATGAGTGTGGAGTCGTTGCTGGACAGAAAAGAGGAACTCAGGNNNAGATCGGA |
| 65729_1 | TATGAGTGTGGAGTCGTTGCAAGAAGAGAAAGGACCAGAAGGGNNNAGATCGGA |
| 65730_1 | TATGAGTGTGGAGTCGTTGCGATGGCCTACATAGCAGATCACANNNAGATCGGAAG |
| 65732_1 | TATGAGTGTGGAGTCGTTGCCACTCTGTCCAGCCTTGACTAATNNNAGATCGGAAG |
| 65733_1 | TATGAGTGTGGAGTCGTTGCCATCTGTTGATGGGCACAATCTCNNNAGATCGGAAG |
| 65734_1 | TATGAGTGTGGAGTCGTTGCTGTTTTTGAGATGGAGTCTGCCTNNNAGATCGGAAG |
| 65735_1 | TATGAGTGTGGAGTCGTTGCTGGATGGAGTTTTGCTTCTGTTGNNNAGATCGGAAG |
| 65736_1 | TATGAGTGTGGAGTCGTTGCCACTCATTGGTTGATGGGCATTTNNNAGATCGGAAG |
| 65737_1 | TATGAGTGTGGAGTCGTTGCTGTGTGTGTGTGTGAGAGAGAAANNNAGATCGGA |
| 65738_1 | TATGAGTGTGGAGTCGTTGCACTTGGATGGGAGCAATTATCAANNNAGATCGGAA |
| 65739_1 | TATGAGTGTGGAGTCGTTGCTTCTACAGTCTTACTCTGTCGCCNNNAGATCGGAAGA |
| 65740_1 | TATGAGTGTGGAGTCGTTGCCCCAGGTTGAAGTGATTCTCTCANNNAGATCGGAAG |
| 65741_1 | TATGAGTGTGGAGTCGTTGCCTGGGGAACACAGGCTATATTCANNNAGATCGGAAG |
| 65742_1 | TATGAGTGTGGAGTCGTTGCAGCCATCATGACCTAATCACCTCNNNAGATCGGAAG |
| 65743_1 | TATGAGTGTGGAGTCGTTGCTTCCAGGTTCAAGCAATTTCCTGNNNAGATCGGAAG |
| 65745_1 | TATGAGTGTGGAGTCGTTGCACATCACTTCAGAGGCATGAACTNNNAGATCGGAAG |
| 65746_1 | TATGAGTGTGGAGTCGTTGCTCTCCTATTGGCTGTTTCTCTGGNNNAGATCGGAAG |
| 65748_1 | TATGAGTGTGGAGTCGTTGCCTCTCAGCGATTCAGCTCTTGTCNNNAGATCGGAAG |
| 65749_1 | TATGAGTGTGGAGTCGTTGCACCCCAGAGGAAACAAACATGATNNNAGATCGGAAG |
| 65751_1 | TATGAGTGTGGAGTCGTTGCAGATTGTTGAGGCTCAGAACACTNNNAGATCGGAAG |
| 65752_1 | TATGAGTGTGGAGTCGTTGCACTGTTCTGTAGAGATTCGTGCANNNAGATCGGAAG |
| 65753_1 | TATGAGTGTGGAGTCGTTGCAAATTATGTGAGCCAGGTGAGGTNNNAGATCGGAA |
| 65754_1 | TATGAGTGTGGAGTCGTTGCGGGACATGGCTGGTGAAAAAAAANNNAGATCGGA |
| 65755_1 | TATGAGTGTGGAGTCGTTGCACAGCTGAGCCACATACTACAATNNNAGATCGGAAG |
| 65756_1 | TATGAGTGTGGAGTCGTTGCGCTGGGATTCAGTACCTCAAAGANNNAGATCGGAAG |
| 65757_1 | TATGAGTGTGGAGTCGTTGCCAGCGTAGGATATTTGCCTTGTGNNNAGATCGGAAG |
| 65758_1 | TATGAGTGTGGAGTCGTTGCTGTAACAGGATACACCAAGCACANNNAGATCGGAAG |
| 65759_1 | TATGAGTGTGGAGTCGTTGCAGCCCTCAATGAAATGCAAAGACNNNAGATCGGAAG |
| 65760_1 | TATGAGTGTGGAGTCGTTGCGACCTGATAGAAGGGCATGGAAANNNAGATCGGA |
| 65761_1 | TATGAGTGTGGAGTCGTTGCTGTGTGTATCCTACTGGTTCTGCNNNAGATCGGAAG |
| 65762_1 | TATGAGTGTGGAGTCGTTGCGCTGTCTTATCCCTACATGCTCANNNAGATCGGAAGA |
| 65763_1 | TATGAGTGTGGAGTCGTTGCTCAAGAAATTTGTCCACCTTGGCNNNAGATCGGAAG |

| | |
|---|---|
| 65764_1 | TATGAGTGTGGAGTCGTTGCAAAGAGAGAGACAGAGACAGGGANNNAGATCGGA |
| 65765_1 | TATGAGTGTGGAGTCGTTGCCACAGTGGCTTGCATCTGTAATCNNNAGATCGGAAG |
| 65769_1 | TATGAGTGTGGAGTCGTTGCCTTCCTGGGTTCAAGCAATTCTCNNNAGATCGGAAG |
| 65771_1 | TATGAGTGTGGAGTCGTTGCCCTCCCGCATTCAAATGATTCTCNNNAGATCGGAAGA |
| 65772_1 | TATGAGTGTGGAGTCGTTGCTTGCTATTGTGAGTGTTGCTGTGNNNAGATCGGAA |
| 65774_1 | TATGAGTGTGGAGTCGTTGCCAGAATCCCACTCTTGTCACTCANNNAGATCGGAAG |
| 65775_1 | TATGAGTGTGGAGTCGTTGCACATAAAAAGCACTGAGCCAAGTNNNAGATCGGAA |
| 65777_1 | TATGAGTGTGGAGTCGTTGCATCTAAAAGAAGCTAGGCTGCCCNNNAGATCGGAA |
| 65778_1 | TATGAGTGTGGAGTCGTTGCCTCAAGTGATCTTCCCACCTCAGNNNAGATCGGAAG |
| 65779_1 | TATGAGTGTGGAGTCGTTGCTTCCCCCTAACCCCCTACTATTTNNNAGATCGGAAGA |
| 65781_1 | TATGAGTGTGGAGTCGTTGCCACCAGTAATCCCAGCTACTCAGNNNAGATCGGAAG |
| 65782_1 | TATGAGTGTGGAGTCGTTGCCTTGTATGCATGCACACACATCANNNAGATCGGAAG |
| 65783_1 | TATGAGTGTGGAGTCGTTGCAGAGAAGAGGCAAGCAAAGATGANNNAGATCGGA |
| 65785_1 | TATGAGTGTGGAGTCGTTGCCATTTGTTCATCGACAGGCACTTNNNAGATCGGAAG |
| 65786_1 | TATGAGTGTGGAGTCGTTGCCTGTAGTCCCAGCTACTCAGGAGNNNAGATCGGAAG |
| 65787_1 | TATGAGTGTGGAGTCGTTGCAAGGGACCAGTTTTTTTCACTGCNNNAGATCGGAAG |
| 65788_1 | TATGAGTGTGGAGTCGTTGCCAGAATCCCTTCTATGTCGAGCANNNAGATCGGAAG |
| 65789_1 | TATGAGTGTGGAGTCGTTGCGCCAAAGGTAGGATACAGAGGACNNNAGATCGGA |
| 65790_1 | TATGAGTGTGGAGTCGTTGCAATTACCAGAAGCAGAAGGAGGGNNNAGATCGGA |
| 65791_1 | TATGAGTGTGGAGTCGTTGCCTCTCAGGGTGTAAACCTAGCAGNNNAGATCGGAA |
| 65792_1 | TATGAGTGTGGAGTCGTTGCGAGAAGGAAGGAAGGAAGGAAGGNNNAGATCGG |
| 65793_1 | TATGAGTGTGGAGTCGTTGCCCACAAAGACCGGGAGAATATGANNNAGATCGGAA |
| 65794_1 | TATGAGTGTGGAGTCGTTGCCACCATGGAATACTACTCAGCCANNNAGATCGGAAG |
| 65795_1 | TATGAGTGTGGAGTCGTTGCCAGTGTCACAATCTAGGCTCACTNNNAGATCGGAAG |
| 65796_1 | TATGAGTGTGGAGTCGTTGCAACCAACTGCAAATAGAAAGGGCNNNAGATCGGAA |
| 65797_1 | TATGAGTGTGGAGTCGTTGCGTGAGTACAAGTACAACTGGGGANNNAGATCGGAA |
| 65798_1 | TATGAGTGTGGAGTCGTTGCCAACAACCCAGGTGCCTATAGTCNNNAGATCGGAAG |
| 65799_1 | TATGAGTGTGGAGTCGTTGCTCAGGAGAGGGAAGATTTCAAGCNNNAGATCGGAA |
| 65800_1 | TATGAGTGTGGAGTCGTTGCGTAGGGGCAGTGCATCTAGAAATNNNAGATCGGAA |
| 65801_1 | TATGAGTGTGGAGTCGTTGCTTGACCCAGGACTAGAGATGGATNNNAGATCGGAA |
| 65802_1 | TATGAGTGTGGAGTCGTTGCCCAAATGATATCCACGTTGCCTCNNNAGATCGGAAG |
| 65803_1 | TATGAGTGTGGAGTCGTTGCCCAAGTGTGCAACAACTAGGTTANNNAGATCGGAA |
| 65804_1 | TATGAGTGTGGAGTCGTTGCAACAATGTACCAATGGAGCCAACNNNAGATCGGAA |
| 65805_1 | TATGAGTGTGGAGTCGTTGCTTTCTGGCACTTCAAAATGCTCCNNNAGATCGGAAG |
| 65806_1 | TATGAGTGTGGAGTCGTTGCAAACTCTTGACCTCAGGTGATCCNNNAGATCGGAAG |
| 65807_1 | TATGAGTGTGGAGTCGTTGCCAAAATCCCTTGGAACTCACAGGNNNAGATCGGAAG |
| 65808_1 | TATGAGTGTGGAGTCGTTGCTCCTTTCTGAAAAAGCCTGTTGCNNNAGATCGGAAG |
| 65809_1 | TATGAGTGTGGAGTCGTTGCCTGGAGTGCAATGACACAATCTCNNNAGATCGGAAG |
| 65810_1 | TATGAGTGTGGAGTCGTTGCGCTCAAGTGATTCTCCCATCTCANNNAGATCGGAAG |
| 65812_1 | TATGAGTGTGGAGTCGTTGCACACACACACGGTTTCATAGGTANNNAGATCGGAAG |
| 65813_1 | TATGAGTGTGGAGTCGTTGCGTCCCAGCTATTTCATAGGCTGANNNAGATCGGAAG |
| 65814_1 | TATGAGTGTGGAGTCGTTGCCATGCCTGTAATCCCAGCTACTTNNNAGATCGGAAG |
| 65815_1 | TATGAGTGTGGAGTCGTTGCTCCAGTGCTTTCAAAACAAGACTNNNAGATCGGAAG |

| | |
|---|---|
| 65817_1 | TATGAGTGTGGAGTCGTTGCGAAGACTCACTTTCTGTCCACCTNNNAGATCGGAAG |
| 65819_1 | TATGAGTGTGGAGTCGTTGCTATAGAGCAGAAACTTCGGGTGGNNNAGATCGGAA |
| 65820_1 | TATGAGTGTGGAGTCGTTGCACTGACCTCTAGCAATGATGGTGNNNAGATCGGAA |
| 65821_1 | TATGAGTGTGGAGTCGTTGCACTGACCTCTAGCAATGATGGTGNNNAGATCGGAA |
| 65822_1 | TATGAGTGTGGAGTCGTTGCGTTTCACCGTGTTAGCCAGAATGNNNAGATCGGAAG |
| 65823_1 | TATGAGTGTGGAGTCGTTGCTTGCTGGGTCAAGAAATAGGTGANNNAGATCGGAA |
| 65824_1 | TATGAGTGTGGAGTCGTTGCTCAAGTTATGTTGTTTGCGGCATNNNAGATCGGAAG |
| 65827_1 | TATGAGTGTGGAGTCGTTGCCTGAGGTGGGAGAATCACTTGAANNNAGATCGGAA |
| 65828_1 | TATGAGTGTGGAGTCGTTGCTCCAAAATCAAGGTACCTGGAGGNNNAGATCGGAA |
| 65829_1 | TATGAGTGTGGAGTCGTTGCAATTAGCTGGTCATAGTGGTGGGNNNAGATCGGAA |
| 65831_1 | TATGAGTGTGGAGTCGTTGCATTTTTTCCATCGCCCTTCAAGGNNNAGATCGGAAGA |
| 65832_1 | TATGAGTGTGGAGTCGTTGCGAGGAAAGAGAGAACGGGAGAAGNNNAGATCGG |
| 65833_1 | TATGAGTGTGGAGTCGTTGCAGCTACTCCACGCTACGTATTTTNNNAGATCGGAAG |
| 65835_1 | TATGAGTGTGGAGTCGTTGCACAAATAGGCTCCAGCTTGAAGANNNAGATCGGAA |
| 65836_1 | TATGAGTGTGGAGTCGTTGCGGTAAACACGAAATACACCCAGCNNNAGATCGGAA |
| 65837_1 | TATGAGTGTGGAGTCGTTGCAGTCAACAGAATGTCCTTCCCTCNNNAGATCGGAAG |
| 65838_1 | TATGAGTGTGGAGTCGTTGCTTTCAGTAGAGACGGGGTTTCTCNNNAGATCGGAA |
| 65839_1 | TATGAGTGTGGAGTCGTTGCTCCTTGAGTACTGTGATCTGCTCNNNAGATCGGAAG |
| 65840_1 | TATGAGTGTGGAGTCGTTGCCTGCAGAGTTGCCAAAAGTACCTNNNAGATCGGAAG |
| 65841_1 | TATGAGTGTGGAGTCGTTGCTGGGTTGGTGACATTTACAAGACNNNAGATCGGAA |
| 65842_1 | TATGAGTGTGGAGTCGTTGCCTCCCATCCTAGTCCAATCATGGNNNAGATCGGAAG |
| 65843_1 | TATGAGTGTGGAGTCGTTGCCCTGCCACAGAAATTCTGAACAANNNAGATCGGAAG |
| 65844_1 | TATGAGTGTGGAGTCGTTGCTCCTGGCTGCTTTTGAGATTTTCNNNAGATCGGAAG |
| 65845_1 | TATGAGTGTGGAGTCGTTGCGTGGCTCACACCTATAATCCCAANNNAGATCGGAAG |
| 65849_1 | TATGAGTGTGGAGTCGTTGCACTAAAAAGGTGGGCAGGAGATTNNNAGATCGGA |
| 65850_1 | TATGAGTGTGGAGTCGTTGCCCAAGTAGCTGAGAGTATAGGCGNNNAGATCGGAA |
| 65851_1 | TATGAGTGTGGAGTCGTTGCGAAACTCAGTACATTCAGCTGGCNNNAGATCGGAAG |
| 65852_1 | TATGAGTGTGGAGTCGTTGCTGTCCCATATATCTTGCTCCAGCNNNAGATCGGAAG |
| 65853_1 | TATGAGTGTGGAGTCGTTGCATTTAGAGAGGGTTTGGCAGGAGNNNAGATCGGA |
| 65854_1 | TATGAGTGTGGAGTCGTTGCTTTTTTGAGACAGAGTCTTGCTCNNNAGATCGGAAG |
| 65855_1 | TATGAGTGTGGAGTCGTTGCTTCATGCGATTCTCCTACCTCAGNNNAGATCGGAAG |
| 65856_1 | TATGAGTGTGGAGTCGTTGCATACCCTGAAGAGGAACAGAGGTNNNAGATCGGAA |
| 65858_1 | TATGAGTGTGGAGTCGTTGCAGAGCATGTGCAGAAATAAGGGANNNAGATCGGA |
| 65860_1 | TATGAGTGTGGAGTCGTTGCGTTTCTCAGCAGGCAGACTTTTTNNNAGATCGGAAG |
| 65861_1 | TATGAGTGTGGAGTCGTTGCGAATGTCCATGTTGCGATGTCTTNNNAGATCGGAAG |
| 65862_1 | TATGAGTGTGGAGTCGTTGCGCCATCTGCAAATAAGGAAGTGANNNAGATCGGAA |
| 65863_1 | TATGAGTGTGGAGTCGTTGCGACCACTGTGTTAGAGGACAGAGNNNAGATCGGAA |
| 65865_1 | TATGAGTGTGGAGTCGTTGCCCTTACTCTTTCCTGTGTAGGGGNNNAGATCGGAAG |
| 65867_1 | TATGAGTGTGGAGTCGTTGCGTCTGTCTGTCTGTCTGTGTGTCNNNAGATCGGAAG |
| 65868_1 | TATGAGTGTGGAGTCGTTGCTTCTGCTCCAATTTCTAGGCTTTNNNAGATCGGAAG |
| 65869_1 | TATGAGTGTGGAGTCGTTGCGGCAGGGAATGTTCAGTGAAAAANNNAGATCGGA |
| 65872_1 | TATGAGTGTGGAGTCGTTGCCACCCAGGTTCTGCTACTTTCTANNNAGATCGGAAG |
| 65873_1 | TATGAGTGTGGAGTCGTTGCTTTCTGACCCCCACAACTCTAACNNNAGATCGGAAGA |

| | |
|---|---|
| 65875_1 | TATGAGTGTGGAGTCGTTGCCTGAGGCAGGAGAATCACTTGAANNNAGATCGGAA |
| 65876_1 | TATGAGTGTGGAGTCGTTGCAGATGCCCTCTGATTGACACTTTNNNAGATCGGAAG |
| 65877_1 | TATGAGTGTGGAGTCGTTGCATCACTTGAGGTCAGTTCGAGACNNNAGATCGGAAG |
| 65878_1 | TATGAGTGTGGAGTCGTTGCAACACACAGCTGATTTGAGATGCNNNAGATCGGAAG |
| 65879_1 | TATGAGTGTGGAGTCGTTGCTTCAGAGAACACAGACACACCTGNNNAGATCGGAAG |
| 65881_1 | TATGAGTGTGGAGTCGTTGCTAGCTCACTACAGCCTTGAACTCNNNAGATCGGAAG |
| 65883_1 | TATGAGTGTGGAGTCGTTGCACGGAGTCTTTCTCTTGTCATCCNNNAGATCGGAAG |
| 65884_1 | TATGAGTGTGGAGTCGTTGCTAGACAGGGTCTCACTTTATCGCNNNAGATCGGAAG |
| 65885_1 | TATGAGTGTGGAGTCGTTGCGGCTTGATCTCGGCTCACTATAANNNAGATCGGAAG |
| 65886_1 | TATGAGTGTGGAGTCGTTGCGAGTCATGGGCATATCATGAGGANNNAGATCGGAA |
| 65887_1 | TATGAGTGTGGAGTCGTTGCCTGAGTAGCTGGGATTACAAGCANNNAGATCGGAA |
| 65888_1 | TATGAGTGTGGAGTCGTTGCAGAGAACCCAGAAATAACGCCAANNNAGATCGGAA |
| 65890_1 | TATGAGTGTGGAGTCGTTGCCTGAGACAGGAGAATTGCTGGAANNNAGATCGGAA |
| 65891_1 | TATGAGTGTGGAGTCGTTGCTGCTACAGGACACTAGTCAGAGANNNAGATCGGAA |
| 65892_1 | TATGAGTGTGGAGTCGTTGCACACACATACCCCATCAGAACTCNNNAGATCGGAAG |
| 65893_1 | TATGAGTGTGGAGTCGTTGCAAATCCTGTCATTTGCAGCAACANNNAGATCGGAAG |
| 65894_1 | TATGAGTGTGGAGTCGTTGCTGCTATGGTGAATAGTGCTGTGANNNAGATCGGAA |
| 65895_1 | TATGAGTGTGGAGTCGTTGCGATGGAGTCTCTCTCTGTTGCTGNNNAGATCGGAAG |
| 65896_1 | TATGAGTGTGGAGTCGTTGCGAGGAACCCTAAACCACCCTAAGNNNAGATCGGAAG |
| 65897_1 | TATGAGTGTGGAGTCGTTGCACAAGAGAGAGAGTCCAGGAATGNNNAGATCGGAA |
| 65898_1 | TATGAGTGTGGAGTCGTTGCTATGTGTGTGTCTGTGTTTGTGANNNAGATCGGAAG |
| 65899_1 | TATGAGTGTGGAGTCGTTGCACCGAAGTCTTGAATCCCTCAAANNNAGATCGGAAG |
| 65900_1 | TATGAGTGTGGAGTCGTTGCAAACTCCTGGACTCAAGGGATTCNNNAGATCGGAAG |
| 65901_1 | TATGAGTGTGGAGTCGTTGCCGGGAGGATTTTGGCTTATTTCCNNNAGATCGGAAG |
| 65902_1 | TATGAGTGTGGAGTCGTTGCTTATAGGTAGGAGCCTCCATGCTNNNAGATCGGAAG |
| 65903_1 | TATGAGTGTGGAGTCGTTGCAGATGGAGTTTCGCTTTTGTTGCNNNAGATCGGAAG |
| 65904_1 | TATGAGTGTGGAGTCGTTGCAATGGTACCAGCACAATTACCCANNNAGATCGGAAG |
| 65905_1 | TATGAGTGTGGAGTCGTTGCCAACGTCCTTGAAGCCAAATGAANNNAGATCGGAAG |
| 65906_1 | TATGAGTGTGGAGTCGTTGCACACACACACTAGGTATCCACAANNNAGATCGGAAG |
| 65907_1 | TATGAGTGTGGAGTCGTTGCGTCTCCTGTGAGGCTACAGAAAANNNAGATCGGAAG |
| 65909_1 | TATGAGTGTGGAGTCGTTGCTTCAGTCTTTCTTAGGGCAGGAGNNNAGATCGGAAG |
| 65910_1 | TATGAGTGTGGAGTCGTTGCGTTTTTGAGACAGGGTCACACACNNNAGATCGGAAG |
| 65911_1 | TATGAGTGTGGAGTCGTTGCACAAGAACTACAAGACACTGCTGNNNAGATCGGAAG |
| 65912_1 | TATGAGTGTGGAGTCGTTGCGATTTTGGGGGTCAATGTGAAGGNNNAGATCGGAA |
| 65913_1 | TATGAGTGTGGAGTCGTTGCCTGTAATCCCAGCTACTCAGGAGNNNAGATCGGAAG |
| 65914_1 | TATGAGTGTGGAGTCGTTGCTATGCACACACAATCACACACACNNNAGATCGGAAG |
| 65915_1 | TATGAGTGTGGAGTCGTTGCCCTTGTAGAGATTCGTCACCTCCNNNAGATCGGAAG |
| 65916_1 | TATGAGTGTGGAGTCGTTGCACAAACCTGTCAACTAGGAGCTTNNNAGATCGGAAG |
| 65917_1 | TATGAGTGTGGAGTCGTTGCAATTTATTTTATGGCCCCAGCGGNNNAGATCGGAAG |
| 65918_1 | TATGAGTGTGGAGTCGTTGCACCTACTTCCCAAGCAAGGATTTNNNAGATCGGAAG |
| 65920_1 | TATGAGTGTGGAGTCGTTGCATATGTTGCAGACCAAGAGCTGANNNAGATCGGAA |
| 65923_1 | TATGAGTGTGGAGTCGTTGCAGGCCCACTCATTCAACATTTTGNNNAGATCGGAAG |
| 65925_1 | TATGAGTGTGGAGTCGTTGCACACACACACACACACACAAAATNNNAGATCGGAAG |

| | |
|---|---|
| 65926_1 | TATGAGTGTGGAGTCGTTGCTCTGGGACAATTTAGACATGCCANNNAGATCGGAAG |
| 65927_1 | TATGAGTGTGGAGTCGTTGCTGCCTTGATTAGCTTCCTGTTTGNNNAGATCGGAAG |
| 65928_1 | TATGAGTGTGGAGTCGTTGCCCCTGCTAAGTCTTCTCCCTTTTNNNAGATCGGAAGA |
| 65929_1 | TATGAGTGTGGAGTCGTTGCACTCCATCTCAAAACACACACACNNNAGATCGGAAGA |
| 65930_1 | TATGAGTGTGGAGTCGTTGCTATAGGAGTCTGCAGTGGGAAACNNNAGATCGGAA |
| 65931_1 | TATGAGTGTGGAGTCGTTGCCTACATCTTCTCTCCCTGCAGTCNNNAGATCGGAAGA |
| 65933_1 | TATGAGTGTGGAGTCGTTGCAAGTTCCTGGAGTCCCAAAGATCNNNAGATCGGAAG |
| 65934_1 | TATGAGTGTGGAGTCGTTGCGTCCCAAGATCTATGGTCAGCAANNNAGATCGGAAG |
| 65935_1 | TATGAGTGTGGAGTCGTTGCGTTTTGTTACCCAGGCTGAAGTGNNNAGATCGGAA |
| 65938_1 | TATGAGTGTGGAGTCGTTGCACCACAATGGAATATCTGCCTCANNNAGATCGGAAG |
| 65941_1 | TATGAGTGTGGAGTCGTTGCTTCACTCTCTTACCTAGGCTGGANNNAGATCGGAAG |
| 65943_1 | TATGAGTGTGGAGTCGTTGCTCTGCTCATGTCTTACCAGCTACNNNAGATCGGAAGA |
| 65944_1 | TATGAGTGTGGAGTCGTTGCGGCCTGGTTTCATGAAATCACATNNNAGATCGGAAG |
| 65945_1 | TATGAGTGTGGAGTCGTTGCACTAGGCACATCTGAAATCTGCANNNAGATCGGAAG |
| 65946_1 | TATGAGTGTGGAGTCGTTGCGCTGTTTTGCTGGAATTCAGGATNNNAGATCGGAA |
| 65947_1 | TATGAGTGTGGAGTCGTTGCTTCCCCAGACATTTGGATTTCCANNNAGATCGGAAG |
| 65948_1 | TATGAGTGTGGAGTCGTTGCCCTCTAGAGATTACACAGGGTGGNNNAGATCGGAA |
| 65949_1 | TATGAGTGTGGAGTCGTTGCCACAACCTGGGCTCTCATCTAAANNNAGATCGGAAG |
| 65950_1 | TATGAGTGTGGAGTCGTTGCCCTGTAATCCCAACACTTCAGGANNNAGATCGGAAG |
| 65951_1 | TATGAGTGTGGAGTCGTTGCATTTGGAGATGGAGTCTTGCAGTNNNAGATCGGAA |
| 65952_1 | TATGAGTGTGGAGTCGTTGCATGATCAAGTTGGCTTCATCCCTNNNAGATCGGAAG |
| 65953_1 | TATGAGTGTGGAGTCGTTGCCAGTCTCCTAAAGTGCTGGGATTNNNAGATCGGAAG |
| 65954_1 | TATGAGTGTGGAGTCGTTGCTACGTCTTTGAGCAGATGAAGCTNNNAGATCGGAA |
| 65955_1 | TATGAGTGTGGAGTCGTTGCTCCTTCAAAATCTCTCTCCTGGGNNNAGATCGGAAGA |
| 65958_1 | TATGAGTGTGGAGTCGTTGCTATAAGGGATTGCTGTACCTGGCNNNAGATCGGAA |
| 65959_1 | TATGAGTGTGGAGTCGTTGCAGGCAGGAGAATCATTTGAACCTNNNAGATCGGAA |
| 65960_1 | TATGAGTGTGGAGTCGTTGCTCCAAGTGATTCTTCTGCCTCAGNNNAGATCGGAAG |
| 65961_1 | TATGAGTGTGGAGTCGTTGCTCTTGAACTCCTGACCTCATGTGNNNAGATCGGAAG |
| 65962_1 | TATGAGTGTGGAGTCGTTGCAGAAAGCCCAGACACAGATCATTNNNAGATCGGAA |
| 65963_1 | TATGAGTGTGGAGTCGTTGCGTGGGAAAGGTGGGAACAGAATANNNAGATCGGA |
| 65964_1 | TATGAGTGTGGAGTCGTTGCCGTGTTTTTGAGATGGGGTCTTGNNNAGATCGGAA |
| 65966_1 | TATGAGTGTGGAGTCGTTGCTTACACCCAAATGAGTGCAAAGCNNNAGATCGGAAG |
| 65968_1 | TATGAGTGTGGAGTCGTTGCGCTGTGTGACTTTGTGAAAGACANNNAGATCGGAA |
| 65971_1 | TATGAGTGTGGAGTCGTTGCTGGTAAGTCTGAAATCCACAGGGNNNAGATCGGAA |
| 65974_1 | TATGAGTGTGGAGTCGTTGCCTTGAGTTTGGGAGGTTGAGGTTNNNAGATCGGAA |
| 65975_1 | TATGAGTGTGGAGTCGTTGCTTGTGTGTGTGTGTATGTGTGTGNNNAGATCGGAA |
| 65978_1 | TATGAGTGTGGAGTCGTTGCTAGGTTTGGAAGGGTCAAGAGTGNNNAGATCGGA |
| 65981_1 | TATGAGTGTGGAGTCGTTGCTCTGGGTTCAAGTGATTCTCCTGNNNAGATCGGAAG |
| 65983_1 | TATGAGTGTGGAGTCGTTGCCGAAAAGCTAGGACTACAGGTGTNNNAGATCGGAA |
| 65984_1 | TATGAGTGTGGAGTCGTTGCAAAGATGGTTGTGAGACAAAGCGNNNAGATCGGA |
| 65985_1 | TATGAGTGTGGAGTCGTTGCGCCTCTTGAGTAGCTGGGATTACNNNAGATCGGAA |
| 65988_1 | TATGAGTGTGGAGTCGTTGCAAAAGATCCTCATGCCTCAGCTTNNNAGATCGGAAG |
| 65990_1 | TATGAGTGTGGAGTCGTTGCCTACAACACCTGTTTACCTCCCANNNAGATCGGAAGA |

| | |
|---|---|
| 65991_1 | TATGAGTGTGGAGTCGTTGCTTTGTGTGTTGCAAGGGTTACAGNNNAGATCGGAA |
| 65993_1 | TATGAGTGTGGAGTCGTTGCCTGGGTTCAAGTGATTCTTCTGCNNNAGATCGGAAG |
| 65995_1 | TATGAGTGTGGAGTCGTTGCTTTTGAGGAGTCTCGCTTTGTTGNNNAGATCGGAAG |
| 65996_1 | TATGAGTGTGGAGTCGTTGCGAGGATCACTTGAGCCTAGGAAGNNNAGATCGGAA |
| 65997_1 | TATGAGTGTGGAGTCGTTGCGGCAAAAAAGCAACAGTTCCTTGNNNAGATCGGAA |
| 65998_1 | TATGAGTGTGGAGTCGTTGCGAAACAAACAGCCAAAGGGATCANNNAGATCGGAA |
| 65999_1 | TATGAGTGTGGAGTCGTTGCCAGGAGAATCACTTGAACCCAGANNNAGATCGGAA |
| 66001_1 | TATGAGTGTGGAGTCGTTGCATTCTGAGACAATGGGGACTTGTNNNAGATCGGAA |
| 66002_1 | TATGAGTGTGGAGTCGTTGCTCAGCTCAAGTCCTCAGAGAATGNNNAGATCGGAAG |
| 66004_1 | TATGAGTGTGGAGTCGTTGCAGATGCAGTGAAGGTGTCATCATNNNAGATCGGAA |
| 66005_1 | TATGAGTGTGGAGTCGTTGCCCCCATGGAATACTATTCAGCCANNNAGATCGGAAG |
| 66006_1 | TATGAGTGTGGAGTCGTTGCTCTTTGTCCCAGTTTCTGACACANNNAGATCGGAAGA |
| 66007_1 | TATGAGTGTGGAGTCGTTGCTCTCCTTCAATGAGCAGAACCTTNNNAGATCGGAAG |
| 66009_1 | TATGAGTGTGGAGTCGTTGCTTCAAGTGATTCTCCTGCCTCAGNNNAGATCGGAAG |
| 66010_1 | TATGAGTGTGGAGTCGTTGCTCCAACTGCAAATGACTTTGAGCNNNAGATCGGAAG |
| 66011_1 | TATGAGTGTGGAGTCGTTGCGTTCAAGCGATTCTTCTGCCTCNNNAGATCGGAAGA |
| 66012_1 | TATGAGTGTGGAGTCGTTGCGGAAGATAAAGGAGTTTGGGGGTNNNAGATCGGA |
| 66013_1 | TATGAGTGTGGAGTCGTTGCTTCAAGTGATTCTCCTGCCTCAGNNNAGATCGGAAG |
| 66014_1 | TATGAGTGTGGAGTCGTTGCGGCATGGCAAATACAACACCATANNNAGATCGGAA |
| 66015_1 | TATGAGTGTGGAGTCGTTGCAGTGCCAAGGCTAATTGTAGACANNNAGATCGGAA |
| 66017_1 | TATGAGTGTGGAGTCGTTGCGAACTCCAGACCTCAGGTGATTCNNNAGATCGGAAG |
| 66019_1 | TATGAGTGTGGAGTCGTTGCGCCTGAACACCCTTTTGAGACTANNNAGATCGGAAG |
| 66020_1 | TATGAGTGTGGAGTCGTTGCTTGGTGGCTATAGTTCCCCTCATNNNAGATCGGAAG |
| 66021_1 | TATGAGTGTGGAGTCGTTGCCTACTTCAAGGGTCCATCCTGTGNNNAGATCGGAAG |
| 66023_1 | TATGAGTGTGGAGTCGTTGCTTTCACAGGCACATCCACAAATCNNNAGATCGGAAG |
| 66024_1 | TATGAGTGTGGAGTCGTTGCCAAGCTACTGGAGAAGCTGATGTNNNAGATCGGAA |
| 66025_1 | TATGAGTGTGGAGTCGTTGCCACTGCAGAATCAGGGAAACAAGNNNAGATCGGAA |
| 66026_1 | TATGAGTGTGGAGTCGTTGCTTTCTAGGTTTACGTCCTGCTCCNNNAGATCGGAAGA |
| 66028_1 | TATGAGTGTGGAGTCGTTGCTTTTTTTGAGTTGGAGTCTCGCCNNNAGATCGGAAG |
| 66029_1 | TATGAGTGTGGAGTCGTTGCTTTCACAGGCACATCCACAAATCNNNAGATCGGAAG |
| 66030_1 | TATGAGTGTGGAGTCGTTGCCATTCAAACATTTCTCCCCAGGCNNNAGATCGGAAGA |
| 66031_1 | TATGAGTGTGGAGTCGTTGCTTCCCAGTCCAGTGTTGATATCCNNNAGATCGGAAG |
| 66032_1 | TATGAGTGTGGAGTCGTTGCCCCACAATGAGCACCTTAGGAAANNNAGATCGGAAG |
| 66033_1 | TATGAGTGTGGAGTCGTTGCTCTGAAGTTGAACCTGTTCTCCCNNNAGATCGGAAG |
| 66034_1 | TATGAGTGTGGAGTCGTTGCTGGGGAGGCTAGTATAAAGACCANNNAGATCGGAA |
| 66035_1 | TATGAGTGTGGAGTCGTTGCTTTCTTCAGCTACAAGGAAGCCTNNNAGATCGGAAG |
| 66038_1 | TATGAGTGTGGAGTCGTTGCCCAACTTTGTCTTCCGCTTTGATNNNAGATCGGAAGA |
| 66039_1 | TATGAGTGTGGAGTCGTTGCCGGCAGGAGTTATTTTTAGAGGCNNNAGATCGGAA |
| 66040_1 | TATGAGTGTGGAGTCGTTGCTTCAGAGTCACCTATCTCAGCCTNNNAGATCGGAAG |
| 66042_1 | TATGAGTGTGGAGTCGTTGCTGCCTCCTCTAAATTACAGCTGTNNNAGATCGGAAG |
| 66043_1 | TATGAGTGTGGAGTCGTTGCTTGTGTGTCTTTGGTCCCAACTANNNAGATCGGAAG |
| 66044_1 | TATGAGTGTGGAGTCGTTGCCAGAGAACCCTGGCTAATACAATNNNAGATCGGAAG |
| 66045_1 | TATGAGTGTGGAGTCGTTGCTCTCGAATGATCCACTTGCTCAANNNAGATCGGAAG |

| | |
|---|---|
| 66046_1 | TATGAGTGTGGAGTCGTTGCAGCTCCTTCCTCAGCAAATATGTNNNAGATCGGAAG |
| 66047_1 | TATGAGTGTGGAGTCGTTGCCTGAGGCAGAGAATTGCTTGAACNNNAGATCGGAA |
| 66050_1 | TATGAGTGTGGAGTCGTTGCGAGCCAGTTTCATCATTGGACACNNNAGATCGGAAG |
| 66051_1 | TATGAGTGTGGAGTCGTTGCTTGAGGAGTTTCAGGCTTTCTCANNNAGATCGGAAG |
| 66052_1 | TATGAGTGTGGAGTCGTTGCATCTCAGCTCACTGCAATCTTCANNNAGATCGGAAGA |
| 66053_1 | TATGAGTGTGGAGTCGTTGCCTTCTCATGAAATGCTGCCTCTGNNNAGATCGGAAG |
| 66054_1 | TATGAGTGTGGAGTCGTTGCTGTTGGCCTGGAAAATTTCTTCCNNNAGATCGGAAG |
| 66056_1 | TATGAGTGTGGAGTCGTTGCGCCTGCAACTCTCTCTCTTTTTTNNNAGATCGGAAGA |
| 66057_1 | TATGAGTGTGGAGTCGTTGCGCAGATACACATACACACACGTGNNNAGATCGGAAG |
| 66058_1 | TATGAGTGTGGAGTCGTTGCCTATCAAGCCATACTCCCACCTCNNNAGATCGGAAGA |
| 66059_1 | TATGAGTGTGGAGTCGTTGCGTGGATCACTTGAATCCAGGAGTNNNAGATCGGAA |
| 66060_1 | TATGAGTGTGGAGTCGTTGCATTGATTGCAACTGACTGAGCTGNNNAGATCGGAA |
| 66061_1 | TATGAGTGTGGAGTCGTTGCCTCACCTATGTCCATGGGAGTTCNNNAGATCGGAAG |
| 66062_1 | TATGAGTGTGGAGTCGTTGCGCTAAAAGTGGCTCTAGGTGTCTNNNAGATCGGAA |
| 66063_1 | TATGAGTGTGGAGTCGTTGCGAAAGAAGGAATGGAGGAAGGGANNNAGATCGG |
| 66064_1 | TATGAGTGTGGAGTCGTTGCGTACTTTGGCTTCATGCACTCTGNNNAGATCGGAAG |
| 66068_1 | TATGAGTGTGGAGTCGTTGCGGAACACACACCACTAATGTTCANNNAGATCGGAAG |
| 66069_1 | TATGAGTGTGGAGTCGTTGCCACACACACACTCTCTCTCTGTTNNNAGATCGGAAGA |
| 66070_1 | TATGAGTGTGGAGTCGTTGCGCAACTACTCTTTCTTCTCCCGANNNAGATCGGAAGA |
| 66071_1 | TATGAGTGTGGAGTCGTTGCTCTGAAGATCTCACTAGGGCTCANNNAGATCGGAAG |
| 66072_1 | TATGAGTGTGGAGTCGTTGCTGTGTTCAAATGCACCAATTTCCNNNAGATCGGAAG |
| 66073_1 | TATGAGTGTGGAGTCGTTGCTCCCTCTCTCTCTCTCTCTCTCTNNNAGATCGGAAGAG |
| 66074_1 | TATGAGTGTGGAGTCGTTGCGTCAGGGAAATACTGGCCTTGTANNNAGATCGGAA |
| 66075_1 | TATGAGTGTGGAGTCGTTGCTTCCCTCTCTCTCTCTGCCAATANNNAGATCGGAAGA |
| 66076_1 | TATGAGTGTGGAGTCGTTGCAAGAGAGAGAGAGAGAGACGGAGNNNAGATCGG |
| 66077_1 | TATGAGTGTGGAGTCGTTGCGGAGGCCTGGTGTAGATGATAAGNNNAGATCGGA |
| 66078_1 | TATGAGTGTGGAGTCGTTGCCTCTCTCTCTCTGGCTCTCTCTCNNNAGATCGGAAGA |
| 66080_1 | TATGAGTGTGGAGTCGTTGCTGGGACTGTACTAGGTGAGACTTNNNAGATCGGAA |
| 66081_1 | TATGAGTGTGGAGTCGTTGCGCACTCAGGATCAGAGATGTGAANNNAGATCGGAA |
| 66082_1 | TATGAGTGTGGAGTCGTTGCTAACAGCGCATGACACAAAACTCNNNAGATCGGAAG |
| 66083_1 | TATGAGTGTGGAGTCGTTGCATTCTGTGGGAAATAGAAAAATGAGANNNAGATCG |
| 66084_1 | TATGAGTGTGGAGTCGTTGCAAAATCTGTTATTGTGCTGGGCANNNAGATCGGAA |
| 66085_1 | TATGAGTGTGGAGTCGTTGCAGAGAGAAGCCTGAAGCAAATGANNNAGATCGGA |
| 66087_1 | TATGAGTGTGGAGTCGTTGCAGACTGAAAGTTCTTTGGAGGCANNNAGATCGGAA |
| 66088_1 | TATGAGTGTGGAGTCGTTGCCACACACACATACACACACAGTCNNNAGATCGGAAG |
| 66089_1 | TATGAGTGTGGAGTCGTTGCCTCTCTCTGTGTGTCTCTGGTTCNNNAGATCGGAAGA |
| 66090_1 | TATGAGTGTGGAGTCGTTGCTTCTTTCGAGATGGAGTTTCGCTNNNAGATCGGAAG |
| 66091_1 | TATGAGTGTGGAGTCGTTGCTCCCTCTCAACTGGAACTCGATANNNAGATCGGAAG |
| 66092_1 | TATGAGTGTGGAGTCGTTGCATCTCAGTATGTTGCCTAGGCTGNNNAGATCGGAAG |
| 66093_1 | TATGAGTGTGGAGTCGTTGCCGAGAGAGAGGGAGAGAGTGTATNNNAGATCGGA |
| 66094_1 | TATGAGTGTGGAGTCGTTGCGCCTTCAACTATTTTTCCCCCTGNNNAGATCGGAAGA |
| 66096_1 | TATGAGTGTGGAGTCGTTGCCACTAATCCCATCATGAGGACCCNNNAGATCGGAAG |
| 66097_1 | TATGAGTGTGGAGTCGTTGCGGGTTTTGCTATGTTGTCCAGTCNNNAGATCGGAAG |

| | |
|---|---|
| 66098_1 | TATGAGTGTGGAGTCGTTGCTTCTTTTTGGCAGAGTCTTGCTCNNNAGATCGGAAG |
| 66102_1 | TATGAGTGTGGAGTCGTTGCATCATAGTTCACTGCTGCCTTGANNNAGATCGGAAG |
| 66103_1 | TATGAGTGTGGAGTCGTTGCACCATCTAACCAGGCTAACACAGNNNAGATCGGAAG |
| 66104_1 | TATGAGTGTGGAGTCGTTGCTGAAGAGTTGTTGTTCAAATGCCNNNAGATCGGAA |
| 66107_1 | TATGAGTGTGGAGTCGTTGCGATGTGGTCTCACTCTATCACCCNNNAGATCGGAAG |
| 66108_1 | TATGAGTGTGGAGTCGTTGCGAAGGGAGGAAGGAAGAAAAGGTNNNAGATCGG |
| 66109_1 | TATGAGTGTGGAGTCGTTGCTGGACTTTGATTTTCCCTTTGCTNNNAGATCGGAAGA |
| 66110_1 | TATGAGTGTGGAGTCGTTGCGATCATAACCTCCTGCAATGGGANNNAGATCGGAAG |
| 66111_1 | TATGAGTGTGGAGTCGTTGCTGTAGAAGATGAGGGAAAAGCATGNNNAGATCGG |
| 66112_1 | TATGAGTGTGGAGTCGTTGCGCATAGGCCCCTTCTGAACTTATNNNAGATCGGAAG |
| 66113_1 | TATGAGTGTGGAGTCGTTGCTCTTTTTGAGGTGGAGTCTTGCTNNNAGATCGGAAG |
| 66114_1 | TATGAGTGTGGAGTCGTTGCTCTGAGTAGACTGAAGCTAGGGGNNNAGATCGGAA |
| 66115_1 | TATGAGTGTGGAGTCGTTGCAATCATGACTCACTGCAGACTCANNNAGATCGGAAG |
| 66116_1 | TATGAGTGTGGAGTCGTTGCGTGTGTGTGTGTGTGTTTCTGNNNAGATCGGAA |
| 66118_1 | TATGAGTGTGGAGTCGTTGCAGACAGACAGACAGACAAAGACCNNNAGATCGGAA |
| 66119_1 | TATGAGTGTGGAGTCGTTGCTTGTCTCTGCACCATCCTTACAANNNAGATCGGAAGA |
| 66121_1 | TATGAGTGTGGAGTCGTTGCTATTCTCAGGTCTGCTGGGAAAGNNNAGATCGGAA |
| 66122_1 | TATGAGTGTGGAGTCGTTGCTGAAACTAGACCTCTACCTCACANNNAGATCGGAAG |
| 66123_1 | TATGAGTGTGGAGTCGTTGCAAACCATCATGTCAGCTAAGTGANNNAGATCGGAAG |
| 66125_1 | TATGAGTGTGGAGTCGTTGCTACTATGTTGTGAGCAGCCCAATNNNAGATCGGAAG |
| 66126_1 | TATGAGTGTGGAGTCGTTGCACCCCCTCTACTGTCTCTTCTTTNNNAGATCGGAAGA |
| 66127_1 | TATGAGTGTGGAGTCGTTGCACTTTAGCTGCCTTTAGTCTCCCNNNAGATCGGAAGA |
| 66128_1 | TATGAGTGTGGAGTCGTTGCCAGAATTGGCACAATGTCACACTNNNAGATCGGAAG |
| 66131_1 | TATGAGTGTGGAGTCGTTGCTCCAAGCAATTCTCCTGTCTCAGNNNAGATCGGAAG |
| 66133_1 | TATGAGTGTGGAGTCGTTGCAGAATGACAGAGAGGGAGAGACANNNAGATCGGA |
| 66134_1 | TATGAGTGTGGAGTCGTTGCCCTATTGGATTAGGGTCCCACATNNNAGATCGGAAG |
| 66136_1 | TATGAGTGTGGAGTCGTTGCGAGAGAGAGGGAGAGAGAGAGAANNNAGATCGG |
| 66137_1 | TATGAGTGTGGAGTCGTTGCATTGTAGCTCACTGCAGACTTGANNNAGATCGGAAG |
| 66138_1 | TATGAGTGTGGAGTCGTTGCCTGTAGCCTTGAACTTCTGGACTNNNAGATCGGAAG |
| 66139_1 | TATGAGTGTGGAGTCGTTGCCACACACACACACACACACATATNNNAGATCGGAAG |
| 66140_1 | TATGAGTGTGGAGTCGTTGCCACACACACACACACTCTCTCTCNNNAGATCGGAAGA |
| 66141_1 | TATGAGTGTGGAGTCGTTGCGGAGAGAGAGAGAGACTGGAGAGNNNAGATCGG |
| 66142_1 | TATGAGTGTGGAGTCGTTGCCTCTCTCTCTCTCTCTCTCTCCCNNNAGATCGGAAGAG |
| 66144_1 | TATGAGTGTGGAGTCGTTGCACCTCTGTGGGCTATTTACCAAANNNAGATCGGAAG |
| 66145_1 | TATGAGTGTGGAGTCGTTGCCCGAGTACTGGACCCATTGATAGNNNAGATCGGAA |
| 66146_1 | TATGAGTGTGGAGTCGTTGCAGAGCAGGATTTTTCCCCTTGAANNNAGATCGGAAG |
| 66147_1 | TATGAGTGTGGAGTCGTTGCAGGGATTGGGTGAATGTTGAGAANNNAGATCGGA |
| 66149_1 | TATGAGTGTGGAGTCGTTGCAAAAGGCTAATCTGTCCCTCTCANNNAGATCGGAAG |
| 66150_1 | TATGAGTGTGGAGTCGTTGCATGGGATGAAAACTGACCTGTCANNNAGATCGGAA |
| 66152_1 | TATGAGTGTGGAGTCGTTGCAAAGCATCTGAACATTGACAGGCNNNAGATCGGAA |
| 66154_1 | TATGAGTGTGGAGTCGTTGCGATTGCATCCTGATTGTGTCACCNNNAGATCGGAAG |
| 66155_1 | TATGAGTGTGGAGTCGTTGCCCTATGGATTTGGGGGACTGAATNNNAGATCGGAA |
| 66156_1 | TATGAGTGTGGAGTCGTTGCCCATTCCCCAGCCACATAACTATNNNAGATCGGAAGA |

| | |
|---|---|
| 66157_1 | TATGAGTGTGGAGTCGTTGCAGGCTGCATGACCTTTCATAAGTNNNAGATCGGAAG |
| 66158_1 | TATGAGTGTGGAGTCGTTGCTCTCTGTCTCTCTCTCTGTGTCCNNNAGATCGGAAGA |
| 66159_1 | TATGAGTGTGGAGTCGTTGCCCCCACCTCTTCTACCATGTAAGNNNAGATCGGAAGA |
| 66160_1 | TATGAGTGTGGAGTCGTTGCGCTCCCATTCAACAAGGAAAACANNNAGATCGGAAG |
| 66161_1 | TATGAGTGTGGAGTCGTTGCACATGACATGATAGTGGCAAGGTNNNAGATCGGAA |
| 66162_1 | TATGAGTGTGGAGTCGTTGCAAGTTTTTGAGGGCCAGCATAACNNNAGATCGGAA |
| 66163_1 | TATGAGTGTGGAGTCGTTGCCACACCCACAGATACCACAGATANNNAGATCGGAAG |
| 66164_1 | TATGAGTGTGGAGTCGTTGCAGATAGGATGAGATTTGGCCACANNNAGATCGGAA |
| 66165_1 | TATGAGTGTGGAGTCGTTGCCAGCACATGATGTCTTTCCACAANNNAGATCGGAAG |
| 66166_1 | TATGAGTGTGGAGTCGTTGCAGGCAGCTCTCTGACTTTTTCTTNNNAGATCGGAAG |
| 66167_1 | TATGAGTGTGGAGTCGTTGCGTGAGCTCACCAGCCTATGAATANNNAGATCGGAAG |
| 66168_1 | TATGAGTGTGGAGTCGTTGCGAGAGAGAGACCAAGACAGTTGGNNNAGATCGGA |
| 66170_1 | TATGAGTGTGGAGTCGTTGCCAAACTGAACACTGTCATCAGCCNNNAGATCGGAAG |
| 66173_1 | TATGAGTGTGGAGTCGTTGCGAATGCCTGTTTGAGTTCTGGTTNNNAGATCGGAAG |
| 66174_1 | TATGAGTGTGGAGTCGTTGCCTTCTTCCCCATGTTTTCACCACNNNAGATCGGAAGA |
| 66175_1 | TATGAGTGTGGAGTCGTTGCTTTCTAAGATGCCCAGGGACTTCNNNAGATCGGAAG |
| 66176_1 | TATGAGTGTGGAGTCGTTGCGTGTGTGTGTGTGTGTGTAAANNNAGATCGGAA |
| 66177_1 | TATGAGTGTGGAGTCGTTGCCCCAAAAAAGGTGCATTTCTGGANNNAGATCGGAA |
| 66178_1 | TATGAGTGTGGAGTCGTTGCATCACTGTTTCAACTCCCCCATTNNNAGATCGGAAGA |
| 66179_1 | TATGAGTGTGGAGTCGTTGCTCTCTCTCTCTCTCTCTCCCTNNNAGATCGGAAGAG |
| 66180_1 | TATGAGTGTGGAGTCGTTGCGCAGAGGACAGCTGATAACCTTANNNAGATCGGAA |
| 66181_1 | TATGAGTGTGGAGTCGTTGCAACCTTCTCAGTCTACCCTCCAANNNAGATCGGAAGA |
| 66183_1 | TATGAGTGTGGAGTCGTTGCAACTCAATCTGCTTAGTTCGGGTNNNAGATCGGAAG |
| 66184_1 | TATGAGTGTGGAGTCGTTGCCACACACACACAAGATGCTTGTTNNNAGATCGGAAG |
| 66186_1 | TATGAGTGTGGAGTCGTTGCTTTCTTGGCTGAAGCAAGGAATGNNNAGATCGGAA |
| 66188_1 | TATGAGTGTGGAGTCGTTGCACAAGGGAATCTCCTGTACGTTCNNNAGATCGGAAG |
| 66189_1 | TATGAGTGTGGAGTCGTTGCTGCACCTAAAGTTATGTCAGAGCNNNAGATCGGAAG |
| 66190_1 | TATGAGTGTGGAGTCGTTGCAGAGCATAATTCAGGCTTCCACANNNAGATCGGAAG |
| 66191_1 | TATGAGTGTGGAGTCGTTGCCCAGCTGAGTGAAAGAGAGACTTNNNAGATCGGAA |
| 66192_1 | TATGAGTGTGGAGTCGTTGCGGAGGATGTCAGAGTCAGACATGNNNAGATCGGA |
| 66194_1 | TATGAGTGTGGAGTCGTTGCGGAGGGATTGTGCTTTGGTATTTNNNAGATCGGAA |
| 66195_1 | TATGAGTGTGGAGTCGTTGCATGGTGGACATAGAATGGATGCANNNAGATCGGAA |
| 66197_1 | TATGAGTGTGGAGTCGTTGCTGTATATCCCTGACTCTCTGGCANNNAGATCGGAAG |
| 66198_1 | TATGAGTGTGGAGTCGTTGCAGAAGTCCAAAGACAGAACCTCANNNAGATCGGAA |
| 66199_1 | TATGAGTGTGGAGTCGTTGCCTCACTGAAGAGTAGGGAGATGCNNNAGATCGGAA |
| 66200_1 | TATGAGTGTGGAGTCGTTGCGCGCTATCAGACTGATTGCTATGNNNAGATCGGAAG |
| 66201_1 | TATGAGTGTGGAGTCGTTGCCTTGGCTTCCTGAAGTACTGAGANNNAGATCGGAAG |
| 66204_1 | TATGAGTGTGGAGTCGTTGCTTGTATAGATGTGGGTGAGGCTGNNNAGATCGGAA |
| 66207_1 | TATGAGTGTGGAGTCGTTGCCATGTACAGGCCAATTCTGCAAANNNAGATCGGAAG |
| 66208_1 | TATGAGTGTGGAGTCGTTGCGTAAAAGACCTGTGACCCCGTATNNNAGATCGGAAG |
| 66209_1 | TATGAGTGTGGAGTCGTTGCAGCTAGGTTGGAGACAGTTTGAGNNNAGATCGGAA |
| 66210_1 | TATGAGTGTGGAGTCGTTGCATGGGGTAATGACTCTTTGAGATNNNAGATCGGAA |
| 66211_1 | TATGAGTGTGGAGTCGTTGCGTCCACAAAGCCTAGAAAACACCNNNAGATCGGAAG |

| | |
|---|---|
| 66212_1 | TATGAGTGTGGAGTCGTTGCGATATCTGTAGACTGGGCCTGTGNNNAGATCGGAA |
| 66213_1 | TATGAGTGTGGAGTCGTTGCAAACAGAAATTCCCACAGCACAGNNNAGATCGGAAG |
| 66215_1 | TATGAGTGTGGAGTCGTTGCGGCACGAAGAGACTGATCAGTAANNNAGATCGGAA |
| 66216_1 | TATGAGTGTGGAGTCGTTGCGGGATGGTGACATATCTGAGGACNNNAGATCGGAA |
| 66217_1 | TATGAGTGTGGAGTCGTTGCAGAATGGGACCAGGTCAATATCGNNNAGATCGGAA |
| 66219_1 | TATGAGTGTGGAGTCGTTGCTGATTTGAATGACCTAGCTGCCANNNAGATCGGAAG |
| 66220_1 | TATGAGTGTGGAGTCGTTGCGACACTAACCACTGGGAGTATGGNNNAGATCGGAA |
| 66222_1 | TATGAGTGTGGAGTCGTTGCAACGCATATGTGTGTCTCCTTCTNNNAGATCGGAAG |
| 66223_1 | TATGAGTGTGGAGTCGTTGCAGTCTGAAACATCTCCATGCCTTNNNAGATCGGAAG |
| 66225_1 | TATGAGTGTGGAGTCGTTGCTACCAGGCCAAAAGACATAGGTCNNNAGATCGGAA |
| 66226_1 | TATGAGTGTGGAGTCGTTGCTGGCTCATACCTGTAATCTCAGCNNNAGATCGGAAG |
| 66228_1 | TATGAGTGTGGAGTCGTTGCCATTTTATCCTCACAGCAACCCCNNNAGATCGGAAGA |
| 66229_1 | TATGAGTGTGGAGTCGTTGCGGATCAAAGGAAAGGGAAACTGGNNNAGATCGGA |
| 66230_1 | TATGAGTGTGGAGTCGTTGCCCTCAATCAATCTTACTGCCTCCNNNAGATCGGAAGA |
| 66231_1 | TATGAGTGTGGAGTCGTTGCTGACAGTTGGAACGTGTTACTCTNNNAGATCGGAAG |
| 66232_1 | TATGAGTGTGGAGTCGTTGCTCCCAGCTAGATATCCCTGGAAANNNAGATCGGAAG |
| 66233_1 | TATGAGTGTGGAGTCGTTGCTCCTGGTAACTTCCTGAGAGTCTNNNAGATCGGAAG |
| 66234_1 | TATGAGTGTGGAGTCGTTGCATTGTTCCACTGCACTGTAAAAANNNAGATCGGAAG |
| 66235_1 | TATGAGTGTGGAGTCGTTGCTTGCAGACTCAGTGATGCTTAGTNNNAGATCGGAAG |
| 66236_1 | TATGAGTGTGGAGTCGTTGCATAAACTTTACTGGGCCAGAGCANNNAGATCGGAA |
| 66237_1 | TATGAGTGTGGAGTCGTTGCATGGCAGACCTAAAACCAGTTCANNNAGATCGGAAG |
| 66238_1 | TATGAGTGTGGAGTCGTTGCTACTCAAGAGGATCATTGCCCTGNNNAGATCGGAAG |
| 66239_1 | TATGAGTGTGGAGTCGTTGCGCTGTACCCACTGAGAACACTTANNNAGATCGGAAG |
| 66240_1 | TATGAGTGTGGAGTCGTTGCCACCCAGCTCTCTTCTAACAACTNNNAGATCGGAAGA |
| 66242_1 | TATGAGTGTGGAGTCGTTGCCCTTAGAAGGGAGCCTCGAATTTNNNAGATCGGAA |
| 66244_1 | TATGAGTGTGGAGTCGTTGCTGGTACAATCACAGATCACTGCANNNAGATCGGAAG |
| 66246_1 | TATGAGTGTGGAGTCGTTGCCCTCACAGGCATGCAGATATACANNNAGATCGGAAG |
| 66247_1 | TATGAGTGTGGAGTCGTTGCTCTTGGGTTACTGTAGCCTGAACNNNAGATCGGAAG |
| 66248_1 | TATGAGTGTGGAGTCGTTGCTTGATGTACAAGACTCCAGAGGCNNNAGATCGGAA |
| 66249_1 | TATGAGTGTGGAGTCGTTGCCTCTCGGTGGAGGAATGTATGAGNNNAGATCGGAA |
| 66250_1 | TATGAGTGTGGAGTCGTTGCGGGAGCTGGGGTTTCAAACTATANNNAGATCGGAA |
| 66253_1 | TATGAGTGTGGAGTCGTTGCGCAATTCTAGGCCTCTGTTACCTNNNAGATCGGAAG |
| 66254_1 | TATGAGTGTGGAGTCGTTGCTCTCAAGACCGCTAATTGTACCCNNNAGATCGGAAG |
| 66255_1 | TATGAGTGTGGAGTCGTTGCCACAAAGCCTCAGTCACTAGGATNNNAGATCGGAAG |
| 66256_1 | TATGAGTGTGGAGTCGTTGCCTGTCTCCATCATGTGTAGTCCCNNNAGATCGGAAG |
| 66257_1 | TATGAGTGTGGAGTCGTTGCCCTCTCTCATCAATGGGCTGATTNNNAGATCGGAAG |
| 66258_1 | TATGAGTGTGGAGTCGTTGCCAGTTGGAGGAGGGATGATGTAGNNNAGATCGGA |
| 66259_1 | TATGAGTGTGGAGTCGTTGCGCAATATGATCAGGGAGGGGTAGNNNAGATCGGA |
| 66260_1 | TATGAGTGTGGAGTCGTTGCATCCATAATCTCTCTCAGCCTGCNNNAGATCGGAAGA |
| 66261_1 | TATGAGTGTGGAGTCGTTGCCCATCTCTACATCAGGCATCCATNNNAGATCGGAAGA |
| 66262_1 | TATGAGTGTGGAGTCGTTGCGTCTCTCCCATGGACCATTACTCNNNAGATCGGAAGA |
| 66263_1 | TATGAGTGTGGAGTCGTTGCTCTTATTGGCCAAACTGAAGGGANNNAGATCGGAA |
| 66264_1 | TATGAGTGTGGAGTCGTTGCGTGTGATCTCTGTGTTGTGTGTGNNNAGATCGGAA |

| | |
|---|---|
| 66265_1 | TATGAGTGTGGAGTCGTTGCACCCCTTGAAATGTTCTGCAATGNNNAGATCGGAAG |
| 66266_1 | TATGAGTGTGGAGTCGTTGCTGTTCTCCAGTCTCCAGAGATGTNNNAGATCGGAAG |
| 66268_1 | TATGAGTGTGGAGTCGTTGCCCAAGGATTCCCCATCACACTAANNNAGATCGGAAG |
| 66270_1 | TATGAGTGTGGAGTCGTTGCGAAGGGTCATCATAGCTACCGTCNNNAGATCGGAA |
| 66271_1 | TATGAGTGTGGAGTCGTTGCGTCTTATTGTCACGTCTCCACCTNNNAGATCGGAAGA |
| 66273_1 | TATGAGTGTGGAGTCGTTGCCACAGGAGTTAGGGTCATCAACANNNAGATCGGAA |
| 66274_1 | TATGAGTGTGGAGTCGTTGCTTGAGACAGGAATGAATGGCTGANNNAGATCGGAA |
| 66275_1 | TATGAGTGTGGAGTCGTTGCTGGAATGTTCACAAAGGAGGTGANNNAGATCGGAA |
| 66276_1 | TATGAGTGTGGAGTCGTTGCCTGGTTGGCTATAGCTCTGCATANNNAGATCGGAAG |
| 66277_1 | TATGAGTGTGGAGTCGTTGCATCTCATTTCCAACTGTCTGCCTNNNAGATCGGAAGA |
| 66278_1 | TATGAGTGTGGAGTCGTTGCAGTTGGAAGAAGTCAAAGGGTGTNNNAGATCGGA |
| 66279_1 | TATGAGTGTGGAGTCGTTGCCTTGTCCCTATGTAATCCAGGCANNNAGATCGGAAG |
| 66280_1 | TATGAGTGTGGAGTCGTTGCAGGCCGCTGGTTAGTTCTTATTANNNAGATCGGAAG |
| 66281_1 | TATGAGTGTGGAGTCGTTGCCAGAAAGACACACACACACACACNNNAGATCGGAAG |
| 66282_1 | TATGAGTGTGGAGTCGTTGCGGAGAGACTTTCATCGGCTTGTANNNAGATCGGAA |
| 66283_1 | TATGAGTGTGGAGTCGTTGCAGAGAGAGAGAGAGAGAGGGAGANNNAGATCGG |
| 66284_1 | TATGAGTGTGGAGTCGTTGCTAAAGTTTGAGCACTTGCAGACGNNNAGATCGGAA |
| 66285_1 | TATGAGTGTGGAGTCGTTGCCACTCTGCACGTCTAGATCTCTCNNNAGATCGGAAG |
| 66286_1 | TATGAGTGTGGAGTCGTTGCTTATGAGACAGGGTCTTGCTCTGNNNAGATCGGAA |
| 66287_1 | TATGAGTGTGGAGTCGTTGCGGCAGCCTCTCTGGATCAATAATNNNAGATCGGAAG |
| 66288_1 | TATGAGTGTGGAGTCGTTGCGATTCTTGGGTTTTCAGGTGGTGNNNAGATCGGAA |
| 66289_1 | TATGAGTGTGGAGTCGTTGCAGAGAGAGAGAGTTAGTGGCTGTNNNAGATCGGA |
| 66290_1 | TATGAGTGTGGAGTCGTTGCTCTCTCTCTCTGTTTCTCTCCCCNNNAGATCGGAAGAG |
| 66291_1 | TATGAGTGTGGAGTCGTTGCCCCATCAGGCTGGCTTCATATATNNNAGATCGGAAG |
| 66292_1 | TATGAGTGTGGAGTCGTTGCGACATGCTTAGTCTGTTCACGTGNNNAGATCGGAAG |
| 66293_1 | TATGAGTGTGGAGTCGTTGCGCACACTTTCTTCCCTTGTCTTCNNNAGATCGGAAGA |
| 66295_1 | TATGAGTGTGGAGTCGTTGCCAAGGGAGTATAAAGACGTGGCTNNNAGATCGGAA |
| 66296_1 | TATGAGTGTGGAGTCGTTGCGAGAGAGACAGACAGACACAAGGNNNAGATCGGA |
| 66297_1 | TATGAGTGTGGAGTCGTTGCAGGGGGCCTGATAATTTGTCTTTNNNAGATCGGAA |
| 66298_1 | TATGAGTGTGGAGTCGTTGCGCTTGCATTCTTCTTTGTCCCTTNNNAGATCGGAAGA |
| 66299_1 | TATGAGTGTGGAGTCGTTGCTGCAGGGTTTACATAGTTGCTCTNNNAGATCGGAAG |
| 66300_1 | TATGAGTGTGGAGTCGTTGCTCCTTTCTGAGCTTTGCTTCTCANNNAGATCGGAAGA |
| 66302_1 | TATGAGTGTGGAGTCGTTGCTTCTTCTCAAGGAAACTCCCAGGNNNAGATCGGAAG |
| 66303_1 | TATGAGTGTGGAGTCGTTGCCCATAGGGGAAATAAAACTGCCCNNNAGATCGGAA |
| 66304_1 | TATGAGTGTGGAGTCGTTGCGACAACGTGGAAAGCTTGAAAGANNNAGATCGGAA |
| 66305_1 | TATGAGTGTGGAGTCGTTGCTTGAGTTCCATTTTGTTGCCCAGNNNAGATCGGAAG |
| 66306_1 | TATGAGTGTGGAGTCGTTGCTCCTCAAGACTATAGCCTCCAGTNNNAGATCGGAAG |
| 66307_1 | TATGAGTGTGGAGTCGTTGCAAGCAGAGGAAGAAGCTGTGATTNNNAGATCGGA |
| 66308_1 | TATGAGTGTGGAGTCGTTGCCAGAAGGGGCCTTAGGAATCAATNNNAGATCGGAA |
| 66309_1 | TATGAGTGTGGAGTCGTTGCCCTAGGCATGTCTCAGAGATGACNNNAGATCGGAA |
| 66310_1 | TATGAGTGTGGAGTCGTTGCCGCATTTTGAGGCTGAAGTAACANNNAGATCGGAA |
| 66311_1 | TATGAGTGTGGAGTCGTTGCCACGAAGGATCTGGAAGCCTAAGNNNAGATCGGAA |
| 66312_1 | TATGAGTGTGGAGTCGTTGCCTGAAGCAGAATCTTGGGGAGATNNNAGATCGGAA |

| | |
|---|---|
| 66313_1 | TATGAGTGTGGAGTCGTTGCTTCTACAGAATGCCTCGAGTAGCNNNAGATCGGAAG |
| 66314_1 | TATGAGTGTGGAGTCGTTGCGCTAGGAAGAGTCAAGGAAGGAGNNNAGATCGGA |
| 66315_1 | TATGAGTGTGGAGTCGTTGCAGAAACAAACAGGACCCAGTCTTNNNAGATCGGAAG |
| 66316_1 | TATGAGTGTGGAGTCGTTGCCCTCCTCTAGGGTTTCAACACTGNNNAGATCGGAAG |
| 66317_1 | TATGAGTGTGGAGTCGTTGCATATGAAGCTCCCTCTCTCTCCANNNAGATCGGAAGA |
| 66318_1 | TATGAGTGTGGAGTCGTTGCGGTGGCTTGTAGGGAAAAGATTCNNNAGATCGGAA |
| 66320_1 | TATGAGTGTGGAGTCGTTGCCAAAGTGAGAAAGCAGCTTCCTTNNNAGATCGGAAG |
| 66321_1 | TATGAGTGTGGAGTCGTTGCTCATATTGAGGAGCAAAGGCAGANNNAGATCGGAA |
| 66322_1 | TATGAGTGTGGAGTCGTTGCCTCACTGCAAACTCTGTCTCTCANNNAGATCGGAAGA |
| 66323_1 | TATGAGTGTGGAGTCGTTGCGTGTGTGTGTGTGTGTAGTCTTGNNNAGATCGGAA |
| 66324_1 | TATGAGTGTGGAGTCGTTGCGAGTCAGGGAAACGTTTTGGAACNNNAGATCGGAA |
| 66325_1 | TATGAGTGTGGAGTCGTTGCCCATCAGAGGACAACACAAGAGANNNAGATCGGAA |
| 66327_1 | TATGAGTGTGGAGTCGTTGCACTGTCCACAAAGAGTCAGAGTCNNNAGATCGGAAG |
| 66328_1 | TATGAGTGTGGAGTCGTTGCTTGTTTACCCCTAAGATCTGGCCNNNAGATCGGAAG |
| 66329_1 | TATGAGTGTGGAGTCGTTGCGACAAAGAAGAGGGATGAGTCCANNNAGATCGGA |
| 66330_1 | TATGAGTGTGGAGTCGTTGCGAAAGTGAGGCAGGAGTGTTTTCNNNAGATCGGAA |
| 66331_1 | TATGAGTGTGGAGTCGTTGCGGGTGAGGAGAGAATATGGCAAANNNAGATCGGA |
| 66332_1 | TATGAGTGTGGAGTCGTTGCCCAAGGTACAGACTGCCTAGATGNNNAGATCGGAA |
| 66333_1 | TATGAGTGTGGAGTCGTTGCTGCAACACAGGTAGGAAAAGTCTNNNAGATCGGAA |
| 66334_1 | TATGAGTGTGGAGTCGTTGCTCAGGCTGTTCATTGAGCATCTANNNAGATCGGAAG |
| 66335_1 | TATGAGTGTGGAGTCGTTGCTGGGTGGTACTAAGTCATTCACGNNNAGATCGGAA |
| 66336_1 | TATGAGTGTGGAGTCGTTGCCTGATGCCACATTTTTACACCGANNNAGATCGGAAG |
| 66337_1 | TATGAGTGTGGAGTCGTTGCGCACAAGCATTTGTCTGATGTGTNNNAGATCGGAAG |
| 66339_1 | TATGAGTGTGGAGTCGTTGCAGGACCAATTCTATAGGCCGAACNNNAGATCGGAAG |
| 66340_1 | TATGAGTGTGGAGTCGTTGCGGTGCGATCATGTATACCCACATNNNAGATCGGAAG |
| 66341_1 | TATGAGTGTGGAGTCGTTGCACAGAAAAGACGTGGCCTAGTAGNNNAGATCGGAA |
| 66342_1 | TATGAGTGTGGAGTCGTTGCCCATCTTCACAACAGTCCTACCANNNAGATCGGAAGA |
| 66343_1 | TATGAGTGTGGAGTCGTTGCCACAATCACAAAATCACAGGGCTNNNAGATCGGAAG |
| 66344_1 | TATGAGTGTGGAGTCGTTGCTGTTTCCACAGGATTACAGCTGANNNAGATCGGAAG |
| 66345_1 | TATGAGTGTGGAGTCGTTGCCAGTGCACGATGTCTTCCTCTATNNNAGATCGGAAG |
| 66346_1 | TATGAGTGTGGAGTCGTTGCAGTGGTTCAATCTCAGCTCAACTNNNAGATCGGAAG |
| 66347_1 | TATGAGTGTGGAGTCGTTGCGCCTTGACTCAGCAATCACAATTNNNAGATCGGAAG |
| 66348_1 | TATGAGTGTGGAGTCGTTGCCCGAAACATCCATGGAAAACACANNNAGATCGGAAG |
| 66349_1 | TATGAGTGTGGAGTCGTTGCAGACTGGGATCGGAGTTTGAAAANNNAGATCGGAA |
| 66350_1 | TATGAGTGTGGAGTCGTTGCAAAGAAAGGACTGAGGATGAGGGNNNAGATCGGA |
| 66351_1 | TATGAGTGTGGAGTCGTTGCTTTTTCCTCTGGATCAGGTAGGGNNNAGATCGGAAG |
| 66352_1 | TATGAGTGTGGAGTCGTTGCGGGGAGTAAAAAAATCCCAGTGCNNNAGATCGGAA |
| 66353_1 | TATGAGTGTGGAGTCGTTGCGAGAGAGAGAGAGAGAGACCTCANNNAGATCGGA |
| 66354_1 | TATGAGTGTGGAGTCGTTGCCGCGTGATCTATTTACGTTTGCTNNNAGATCGGAAG |
| 66355_1 | TATGAGTGTGGAGTCGTTGCAGAGGTCTGCTAGATGGTACACTNNNAGATCGGAA |
| 66356_1 | TATGAGTGTGGAGTCGTTGCTTGCCTAGACTGGTCTCAAACTCNNNAGATCGGAAG |
| 66357_1 | TATGAGTGTGGAGTCGTTGCAGGGAGAGAGAGAGTCATGGAAANNNAGATCGGA |
| 66358_1 | TATGAGTGTGGAGTCGTTGCCAAATGTGCATTCCTCTTGTGGTNNNAGATCGGAAG |

66359_1 TATGAGTGTGGAGTCGTTGCTTAGTGGCCTAGAGAGGACAGAANNNAGATCGGAA
66360_1 TATGAGTGTGGAGTCGTTGCCATCTCCTTCATTATCCACCCCANNNAGATCGGAAGA
66361_1 TATGAGTGTGGAGTCGTTGCATCAGCGATCTGGGGAATTTCATNNNAGATCGGAAG
66362_1 TATGAGTGTGGAGTCGTTGCAGTAATTGCCACTCACTCCACATNNNAGATCGGAAGA
66363_1 TATGAGTGTGGAGTCGTTGCTAGGAATGAGCTGTGGAAATGCTNNNAGATCGGAA
66364_1 TATGAGTGTGGAGTCGTTGCCTTGACTTCCCAGGCTAAGTGATNNNAGATCGGAAG
66365_1 TATGAGTGTGGAGTCGTTGCTTCAAATGGAAGAGCCTTGGTTGNNNAGATCGGAA
66366_1 TATGAGTGTGGAGTCGTTGCCAAGTCCAAAATCTGCAGACCAGNNNAGATCGGAAG
66367_1 TATGAGTGTGGAGTCGTTGCTCCTACTGTTTCTGTTTGGAGCTNNNAGATCGGAAG
66368_1 TATGAGTGTGGAGTCGTTGCACACTAATGGCTAATCGGGTGAANNNAGATCGGAA
66369_1 TATGAGTGTGGAGTCGTTGCTGGGTAGATCAGTGAGTCTCCTGNNNAGATCGGAA
66370_1 TATGAGTGTGGAGTCGTTGCACTTAGTGTGGGTGGCTCATTAGNNNAGATCGGAA
66371_1 TATGAGTGTGGAGTCGTTGCCTGTGTGTTGTGACAAGTGTGAGNNNAGATCGGAA
66373_1 TATGAGTGTGGAGTCGTTGCCTTTCAGGACTTTGGAAACCACCNNNAGATCGGAAG
66374_1 TATGAGTGTGGAGTCGTTGCGCACAGTGTGAGTCTACAGAGAANNNAGATCGGAA
66375_1 TATGAGTGTGGAGTCGTTGCTTGCCCTGTGAGGAATTTTGTTCNNNAGATCGGAAG
66376_1 TATGAGTGTGGAGTCGTTGCTACCATCACCTTACAATGGAGCTNNNAGATCGGAAG
66377_1 TATGAGTGTGGAGTCGTTGCGGTATTTCAAGGTCAACAGCACCNNNAGATCGGAAG
66378_1 TATGAGTGTGGAGTCGTTGCATCAGCTGGTGGAGTGAAGAATTNNNAGATCGGAA
66379_1 TATGAGTGTGGAGTCGTTGCGGGAATGGGAGGTAATATGACCCNNNAGATCGGAA
66380_1 TATGAGTGTGGAGTCGTTGCTGTGGTCCTAATAGCTAAAGCCCNNNAGATCGGAAG
66381_1 TATGAGTGTGGAGTCGTTGCCCTCACGCTCTTCTTCATGTAGTNNNAGATCGGAAGA
66382_1 TATGAGTGTGGAGTCGTTGCATACCAATGGCTTTCCAGGATGTNNNAGATCGGAAG
66383_1 TATGAGTGTGGAGTCGTTGCATCTCAAGCAACCTGTCCAAAACNNNAGATCGGAAG
66384_1 TATGAGTGTGGAGTCGTTGCGCCACAATTCCATAGGGACAATGNNNAGATCGGAAG
66385_1 TATGAGTGTGGAGTCGTTGCATTGCCTCTCTCTGGGAAGACTANNNAGATCGGAAG
66386_1 TATGAGTGTGGAGTCGTTGCTAACACACAAGCGCATGCATTTANNNAGATCGGAAG
66387_1 TATGAGTGTGGAGTCGTTGCGCTTTCTTGTTGTGGCTTCCATANNNAGATCGGAAG
66388_1 TATGAGTGTGGAGTCGTTGCTATGTGGTGGACTCTTCTGTGTGNNNAGATCGGAAG
66389_1 TATGAGTGTGGAGTCGTTGCTGGTTTTACTCCTCCTTCTCAGCNNNAGATCGGAAGA
66390_1 TATGAGTGTGGAGTCGTTGCGGAGGGAGGGATAGAGGAAAGAANNNAGATCGG
66391_1 TATGAGTGTGGAGTCGTTGCGACCAGTGTAGGATGCAAGGATCNNNAGATCGGAA
66393_1 TATGAGTGTGGAGTCGTTGCAGCTGTGCTCCAAGTGTTAGTTANNNAGATCGGAAG
66394_1 TATGAGTGTGGAGTCGTTGCGTGATATCAAGGTAGCAGGACCCNNNAGATCGGAA
66395_1 TATGAGTGTGGAGTCGTTGCGGAGTCTCTACAGCGAGGTAAAGNNNAGATCGGAA
66396_1 TATGAGTGTGGAGTCGTTGCTATGACAGACAGACACACACACANNNAGATCGGAAG
66397_1 TATGAGTGTGGAGTCGTTGCGCGCTGTGCTTCTTCTGTTTATTNNNAGATCGGAAG
66398_1 TATGAGTGTGGAGTCGTTGCTTTGAAACTGGAGGAAGGGTCAANNNAGATCGGAA
66399_1 TATGAGTGTGGAGTCGTTGCATCTTTGGGATGAAGGACTCCACNNNAGATCGGAAG
66400_1 TATGAGTGTGGAGTCGTTGCTGAAGCAATGGTTTCCTGACTCTNNNAGATCGGAAG
66401_1 TATGAGTGTGGAGTCGTTGCATGGGGTCTCACTACACTGACTANNNAGATCGGAAG
66403_1 TATGAGTGTGGAGTCGTTGCTGCTACACAACACAGATGAACCTNNNAGATCGGAAG
66404_1 TATGAGTGTGGAGTCGTTGCTCTGCGGTTTTCTTCACTAACCTNNNAGATCGGAAGA

| | |
|---|---|
| 66405_1 | TATGAGTGTGGAGTCGTTGCTCCTCTGGAGAACCCTGTCTAATNNNAGATCGGAAG |
| 66406_1 | TATGAGTGTGGAGTCGTTGCTGAGAGCAAGTAGTCCCCTCTTANNNAGATCGGAAG |
| 66407_1 | TATGAGTGTGGAGTCGTTGCTTGTATCACGTGTAATGGTGGGTNNNAGATCGGAA |
| 66409_1 | TATGAGTGTGGAGTCGTTGCAAAGCCAAAAGAACAGGACCCTANNNAGATCGGAA |
| 66411_1 | TATGAGTGTGGAGTCGTTGCTTTAGAGACAGGGTCTTGCTCTGNNNAGATCGGAAG |
| 66412_1 | TATGAGTGTGGAGTCGTTGCTGATGTGATACCCCTGCTTTCTCNNNAGATCGGAAG |
| 66413_1 | TATGAGTGTGGAGTCGTTGCTGCTGGTATTGAAAGCTGGAGAANNNAGATCGGAA |
| 66414_1 | TATGAGTGTGGAGTCGTTGCTGGTCTAATTCAACTTGCTGTGANNNAGATCGGAAG |
| 66416_1 | TATGAGTGTGGAGTCGTTGCGGAGAAAACCTTATTGCTGAGGGNNNAGATCGGAA |
| 66417_1 | TATGAGTGTGGAGTCGTTGCAGAGAGATTTTGGTTGGAAGCCANNNAGATCGGAA |
| 66418_1 | TATGAGTGTGGAGTCGTTGCACCTGTAAAGCCCAGACAATGATNNNAGATCGGAAG |
| 66419_1 | TATGAGTGTGGAGTCGTTGCCCCTTCTTGTCATGTCTCTGCTANNNAGATCGGAAGA |
| 66420_1 | TATGAGTGTGGAGTCGTTGCACAAAAAGTGTGTTGTGACATCTNNNAGATCGGAAG |
| 66421_1 | TATGAGTGTGGAGTCGTTGCCTGTCTCTCTCTCTCACACACACNNNAGATCGGAAGA |
| 66422_1 | TATGAGTGTGGAGTCGTTGCTTATCTATGACATGGAGGGTGCGNNNAGATCGGAA |
| 66423_1 | TATGAGTGTGGAGTCGTTGCTCTCATCTGGGCTTGAAAAACTCNNNAGATCGGAAG |
| 66424_1 | TATGAGTGTGGAGTCGTTGCCCTTCAGTCTCTGACTCTTGCATNNNAGATCGGAAGA |
| 66425_1 | TATGAGTGTGGAGTCGTTGCTTTTTGAAGACAGCGTGTGACTCNNNAGATCGGAAG |
| 66426_1 | TATGAGTGTGGAGTCGTTGCTCTCTGGTCTCTGATAACTGGTTNNNAGATCGGAAG |
| 66427_1 | TATGAGTGTGGAGTCGTTGCTCCCTCTGGTGACTCTTATCTCTNNNAGATCGGAAGA |
| 66428_1 | TATGAGTGTGGAGTCGTTGCTATAACTGTCAGAGCAGGTGCAGNNNAGATCGGAA |
| 66430_1 | TATGAGTGTGGAGTCGTTGCTGGAATTGGTTCTTAGGGAGGTCNNNAGATCGGAA |
| 66431_1 | TATGAGTGTGGAGTCGTTGCTCAGCAAGTGAAATTTCCTGTGTNNNAGATCGGAAG |
| 66432_1 | TATGAGTGTGGAGTCGTTGCTCCATCCAAGCTTACCAGAAGTCNNNAGATCGGAAG |
| 66433_1 | TATGAGTGTGGAGTCGTTGCGGACTAAAATGGTCATGGCAGTGNNNAGATCGGAA |
| 66434_1 | TATGAGTGTGGAGTCGTTGCTGTCTGAGAAACAGCAAAACAGGNNNAGATCGGAA |
| 66435_1 | TATGAGTGTGGAGTCGTTGCTCTTTCTCCATGAGCAGGTCAATNNNAGATCGGAAG |
| 66436_1 | TATGAGTGTGGAGTCGTTGCTCAGAGTACCAGTTCCCGTATCTNNNAGATCGGAAG |
| 66437_1 | TATGAGTGTGGAGTCGTTGCTAGAGCTTGTTCATGAGGGGATCNNNAGATCGGAA |
| 66438_1 | TATGAGTGTGGAGTCGTTGCTGGAGAGAGAAAGAGAGATGTCANNNAGATCGGA |
| 66439_1 | TATGAGTGTGGAGTCGTTGCTTGCTCTCCTTACGTGTAAAGGANNNAGATCGGAAG |
| 66440_1 | TATGAGTGTGGAGTCGTTGCTCATCAGTTCCTATGCCACATGTNNNAGATCGGAAG |
| 66441_1 | TATGAGTGTGGAGTCGTTGCTTCAAGTTGTTGGGAATGTGAGCNNNAGATCGGAA |
| 66442_1 | TATGAGTGTGGAGTCGTTGCTTCCAGTCTGTGGTGTTAGGATGNNNAGATCGGAAG |
| 66443_1 | TATGAGTGTGGAGTCGTTGCAGCAGGTAAGGTTTTTGGACTGTNNNAGATCGGAA |
| 66444_1 | TATGAGTGTGGAGTCGTTGCTCTCTGTACCTTCTCTTGGGTCTNNNAGATCGGAAGA |
| 66445_1 | TATGAGTGTGGAGTCGTTGCACTATCCCAACTCTGTTTGTCCANNNAGATCGGAAGA |
| 66446_1 | TATGAGTGTGGAGTCGTTGCCCTGAGTTATAGGTTGCTGGTGTNNNAGATCGGAAG |
| 66447_1 | TATGAGTGTGGAGTCGTTGCTTCATGCATGTGTGTGTGTGTACNNNAGATCGGAAG |
| 66448_1 | TATGAGTGTGGAGTCGTTGCAAGCCAGCTGACTTTTGATTAGGNNNAGATCGGAAG |
| 66449_1 | TATGAGTGTGGAGTCGTTGCTGCTGCCAAGTAAGAGGTATTGTNNNAGATCGGAA |
| 66450_1 | TATGAGTGTGGAGTCGTTGCGGAAAACACATCTCTAACTCGCGNNNAGATCGGAAG |
| 66451_1 | TATGAGTGTGGAGTCGTTGCAAAATTTGGAAAAGGCTGGTCGTNNNAGATCGGAA |

| | |
|---|---|
| 66452_1 | TATGAGTGTGGAGTCGTTGCGTGTTGTTGTAATGGCTGTCACANNNAGATCGGAAG |
| 66453_1 | TATGAGTGTGGAGTCGTTGCTGGGATCCTGTCAAGAAAGTTTGNNNAGATCGGAAG |
| 66455_1 | TATGAGTGTGGAGTCGTTGCTAATGGATACGTAGAAGGTGGCGNNNAGATCGGAA |
| 66456_1 | TATGAGTGTGGAGTCGTTGCTCAGAGACACACTCAGGTTTGACNNNAGATCGGAAG |
| 66457_1 | TATGAGTGTGGAGTCGTTGCGGGAGTAGGTGGTAAGTTCCAAGNNNAGATCGGAA |
| 66458_1 | TATGAGTGTGGAGTCGTTGCTGGATGGATGTCTGACCTCTACTNNNAGATCGGAAG |
| 66459_1 | TATGAGTGTGGAGTCGTTGCAGTTCTAGTCTCAGACTCTGCCTNNNAGATCGGAAGA |
| 66460_1 | TATGAGTGTGGAGTCGTTGCTTTTAGGAGACAGTCTCGCTCTGNNNAGATCGGAAG |
| 66461_1 | TATGAGTGTGGAGTCGTTGCAAACTGCTTTCCTCACAACACTGNNNAGATCGGAAGA |
| 66462_1 | TATGAGTGTGGAGTCGTTGCTGTGTGCCTTGTTTTGTTTGAGTNNNAGATCGGAAG |
| 66463_1 | TATGAGTGTGGAGTCGTTGCTGTTCCCACATTCTACACACAGANNNAGATCGGAAGA |
| 66465_1 | TATGAGTGTGGAGTCGTTGCATTTTTCTTCTTCCTGCCTCCCTNNNAGATCGGAAGAG |
| 66466_1 | TATGAGTGTGGAGTCGTTGCGGGAGAAGTTGGGTTACTCACATNNNAGATCGGAAG |
| 66467_1 | TATGAGTGTGGAGTCGTTGCGTTGCCTCCTGAGAGTAGAAGAGNNNAGATCGGAAG |
| 66468_1 | TATGAGTGTGGAGTCGTTGCTCTCAAGCTGATTCTGAAGACGANNNAGATCGGAAG |
| 66469_1 | TATGAGTGTGGAGTCGTTGCCCAACAGCACAGAGAATGTTTCCNNNAGATCGGAAG |
| 66470_1 | TATGAGTGTGGAGTCGTTGCAATGTAGAGAAGGGCATTCCAGGNNNAGATCGGAA |
| 66471_1 | TATGAGTGTGGAGTCGTTGCGCTGAAGTTGAGGTCTAGTGGAGNNNAGATCGGAA |
| 66472_1 | TATGAGTGTGGAGTCGTTGCAGCTCTAAGTTTCTGAGGGCTTCNNNAGATCGGAAG |
| 66473_1 | TATGAGTGTGGAGTCGTTGCAATTTCTCTGGCCAGAACTTCCANNNAGATCGGAAGA |
| 66474_1 | TATGAGTGTGGAGTCGTTGCAGACCTGGTTTTGAACTCTGGTTNNNAGATCGGAAG |
| 66475_1 | TATGAGTGTGGAGTCGTTGCAGCATTGAGTAGTGAACAGCTGANNNAGATCGGAAG |
| 66476_1 | TATGAGTGTGGAGTCGTTGCGTTGAGAATCAGTGGTCTAGGCANNNAGATCGGAAG |
| 66477_1 | TATGAGTGTGGAGTCGTTGCAAGTGTTGGGGGTCAAGTTTCATNNNAGATCGGAAG |
| 66479_1 | TATGAGTGTGGAGTCGTTGCATTCAGTTTGAATCTGCTTGGGCNNNAGATCGGAAG |
| 66481_1 | TATGAGTGTGGAGTCGTTGCACTGAGCTAATCTGGTCTTGTGGNNNAGATCGGAAG |
| 66482_1 | TATGAGTGTGGAGTCGTTGCCCACCACATCGGGATAATTTTCGNNNAGATCGGAAGA |
| 66483_1 | TATGAGTGTGGAGTCGTTGCAGTGCTGAGATTACAGGTGTGAGNNNAGATCGGAA |
| 66485_1 | TATGAGTGTGGAGTCGTTGCATCTGAGTTTGCTCTCTGTCCAGNNNAGATCGGAAGA |
| 66486_1 | TATGAGTGTGGAGTCGTTGCATCCCTAGGTTAGGCCAGGTAATNNNAGATCGGAAG |
| 66487_1 | TATGAGTGTGGAGTCGTTGCGATCCTCCTATTTCAGCCTCCAGNNNAGATCGGAAGA |
| 66488_1 | TATGAGTGTGGAGTCGTTGCCACATTGTGCGGGTTAGTTACATNNNAGATCGGAAG |
| 66490_1 | TATGAGTGTGGAGTCGTTGCCCTGCTAAATTGAGGAGGACACTNNNAGATCGGAAG |
| 66492_1 | TATGAGTGTGGAGTCGTTGCGTAGAGATGGGGTTTACCGTGTNNNAGATCGGAAG |
| 66493_1 | TATGAGTGTGGAGTCGTTGCACTCCTGACCTCAAATGATCCACNNNAGATCGGAAGA |
| 66494_1 | TATGAGTGTGGAGTCGTTGCGACAAGTTCTGCATCCATTTCCCNNNAGATCGGAAGA |
| 66495_1 | TATGAGTGTGGAGTCGTTGCCTGGAGTGCAATGATGTGATCTCNNNAGATCGGAAG |
| 66496_1 | TATGAGTGTGGAGTCGTTGCTTTTTTTTTTTGGGGGGAGACGGNNNAGATCGGAAG |
| 66497_1 | TATGAGTGTGGAGTCGTTGCGGCCTCCTAAAGTGTTGGGAATANNNAGATCGGAAG |
| 66498_1 | TATGAGTGTGGAGTCGTTGCAATCTCAGTTCACTGCAACCTCTNNNAGATCGGAAGA |
| 66499_1 | TATGAGTGTGGAGTCGTTGCGCTTGAGTTTCATTGGGCTTCTTNNNAGATCGGAAGA |
| 66500_1 | TATGAGTGTGGAGTCGTTGCCTTTGGGTGGTTGGTTTGAATCANNNAGATCGGAAG |
| 66501_1 | TATGAGTGTGGAGTCGTTGCTTTTTGAGACAGGGTCTTGCTCTNNNAGATCGGAAG |

| | |
|---|---|
| 66502_1 | TATGAGTGTGGAGTCGTTGCCAAAGGAAACGCAGAATGAGTGTNNNAGATCGGAA |
| 66503_1 | TATGAGTGTGGAGTCGTTGCGTGTGTGTATTTTGCTCGGAGACNNNAGATCGGAA |
| 66504_1 | TATGAGTGTGGAGTCGTTGCAAATAAGTACCCCTATGGCCACCNNNAGATCGGAAG |
| 66505_1 | TATGAGTGTGGAGTCGTTGCTGCACCTGGCCTGATAATACTTTNNNAGATCGGAAG |
| 66506_1 | TATGAGTGTGGAGTCGTTGCATCTTGGCTCATTGCAACCTCTANNNAGATCGGAAG |
| 66507_1 | TATGAGTGTGGAGTCGTTGCAGGACTCACTGTGGTTTCTTGTTNNNAGATCGGAAG |
| 66508_1 | TATGAGTGTGGAGTCGTTGCATCCACTTATCTGCTGTTGGACANNNAGATCGGAAG |
| 66509_1 | TATGAGTGTGGAGTCGTTGCTAGGTGAGTGCTTCAAGCTGAAANNNAGATCGGAA |
| 66510_1 | TATGAGTGTGGAGTCGTTGCAAGAAGAAATGGCCTCTCATGGGNNNAGATCGGAA |
| 66511_1 | TATGAGTGTGGAGTCGTTGCGATCCTCCTATTTCAGCCTCCAGNNNAGATCGGAAG |
| 66512_1 | TATGAGTGTGGAGTCGTTGCTTTGAGATGGAGACTTGCTGTGTNNNAGATCGGAA |
| 66513_1 | TATGAGTGTGGAGTCGTTGCTAACAGAGTGATGTATGCCTGGCNNNAGATCGGAA |
| 66514_1 | TATGAGTGTGGAGTCGTTGCTTTTTTTTTTTGAGACGGGGTCTNNNAGATCGGAAG |
| 66516_1 | TATGAGTGTGGAGTCGTTGCCAGCTCACTGAAATCTCCACCTCNNNAGATCGGAAG |
| 66517_1 | TATGAGTGTGGAGTCGTTGCTAAAAATGGCCAGTGTCACTTGCNNNAGATCGGAAG |
| 66518_1 | TATGAGTGTGGAGTCGTTGCCACACCAGAATTGGGTTGTTCTCNNNAGATCGGAAG |
| 66521_1 | TATGAGTGTGGAGTCGTTGCGACCTAATCTGGGGTGGAATACCNNNAGATCGGAA |
| 66522_1 | TATGAGTGTGGAGTCGTTGCATGGTGGCCTGATTTATTGAGGCNNNAGATCGGAA |
| 66523_1 | TATGAGTGTGGAGTCGTTGCCATCCATGGTCTCAGTGCTTACTNNNAGATCGGAAG |
| 66524_1 | TATGAGTGTGGAGTCGTTGCTGTATTTGTGCTCTCAGGACACANNNAGATCGGAAG |
| 66526_1 | TATGAGTGTGGAGTCGTTGCGGCCAAAGAGCAGAATACTTTCCNNNAGATCGGAAG |
| 66527_1 | TATGAGTGTGGAGTCGTTGCAGACAAGGTCTCACTTTGTCACCNNNAGATCGGAAG |
| 66529_1 | TATGAGTGTGGAGTCGTTGCGCCACTCACTAGCTACAAGTCTTNNNAGATCGGAAG |
| 66530_1 | TATGAGTGTGGAGTCGTTGCCTGAGGTCAGAAGTTTGAGACCANNNAGATCGGAA |
| 66531_1 | TATGAGTGTGGAGTCGTTGCCAGAATGAGTTCCCTGTCTCACANNNAGATCGGAAG |
| 66532_1 | TATGAGTGTGGAGTCGTTGCTATTTCACAGTACCACAGGACCCNNNAGATCGGAAG |
| 66534_1 | TATGAGTGTGGAGTCGTTGCAAGGAAAACACACACACACACACNNNAGATCGGAAG |
| 66535_1 | TATGAGTGTGGAGTCGTTGCTCACACTCTCTCTAGTGGTGACTNNNAGATCGGAAG |
| 66536_1 | TATGAGTGTGGAGTCGTTGCACAGTATCGAAGACACACACACANNNAGATCGGAAG |
| 66537_1 | TATGAGTGTGGAGTCGTTGCCATTCAGCCTCAGACTGAGAGTTNNNAGATCGGAAG |
| 66538_1 | TATGAGTGTGGAGTCGTTGCGGGATGGAAGGAATAAAGCAACANNNAGATCGGAA |
| 66539_1 | TATGAGTGTGGAGTCGTTGCTAATGGCATTTGCAGAAACCTGGNNNAGATCGGAA |
| 66540_1 | TATGAGTGTGGAGTCGTTGCCTGAGGCTTGGACAAAAAAGGACNNNAGATCGGAA |
| 66541_1 | TATGAGTGTGGAGTCGTTGCAGGCATAAACTGGTAGGAAGCAANNNAGATCGGAA |
| 66542_1 | TATGAGTGTGGAGTCGTTGCTTGTGATCTAGTTTCCTCTGGCCNNNAGATCGGAAG |
| 66543_1 | TATGAGTGTGGAGTCGTTGCTCGGAGATATTGTGGGTTTGGTTNNNAGATCGGAA |
| 66544_1 | TATGAGTGTGGAGTCGTTGCTGATGCTGTTCCTTTTGGAAACCNNNAGATCGGAAG |
| 66545_1 | TATGAGTGTGGAGTCGTTGCAGACAGACAGAGAGACAGAGACANNNAGATCGGAA |
| 66546_1 | TATGAGTGTGGAGTCGTTGCCTGTGTGTTGTGTGTAGTGTGTGNNNAGATCGGAA |
| 66547_1 | TATGAGTGTGGAGTCGTTGCATGAGAGAAGAGGAGACTCCACANNNAGATCGGAA |
| 66548_1 | TATGAGTGTGGAGTCGTTGCTCTAAGCAACTTGAGTGGAGGTCNNNAGATCGGAA |
| 66549_1 | TATGAGTGTGGAGTCGTTGCAAGACATGGTTATGTGTGACGGANNNAGATCGGAA |
| 66550_1 | TATGAGTGTGGAGTCGTTGCAATCTTGGCTCACTACAACCTCCNNNAGATCGGAAG |

66551_1 TATGAGTGTGGAGTCGTTGCGACTCCTTTGTGCTACCCCTTTANNNAGATCGGAAGA
66552_1 TATGAGTGTGGAGTCGTTGCCGCCTGACCAGTCCTTTTTATTGNNNAGATCGGAAG
66553_1 TATGAGTGTGGAGTCGTTGCTGGGATTGCATTCTTGATTTGCTNNNAGATCGGAAG
66554_1 TATGAGTGTGGAGTCGTTGCAGCCATTCTAACAGGTGTAAGGTNNNAGATCGGAA
66555_1 TATGAGTGTGGAGTCGTTGCTAGCTGAAGGCATCACATTAGCTNNNAGATCGGAAG
66556_1 TATGAGTGTGGAGTCGTTGCCCTCTCTCATCCCCATATGCAAANNNAGATCGGAAGA
66557_1 TATGAGTGTGGAGTCGTTGCCCTTCCAAAGCTCTGGGATTACANNNAGATCGGAAG
66558_1 TATGAGTGTGGAGTCGTTGCCTATCTCAGCTCACTGCAACCTCNNNAGATCGGAAGA
66560_1 TATGAGTGTGGAGTCGTTGCTTCGGACTGGACTTCGTGTTATTNNNAGATCGGAAG
66561_1 TATGAGTGTGGAGTCGTTGCGGAACTGCATTCTTGATTTGGCTNNNAGATCGGAAG
66562_1 TATGAGTGTGGAGTCGTTGCTGTAGGGTGGGGGTTTCTTATTGNNNAGATCGGAA
66563_1 TATGAGTGTGGAGTCGTTGCCAGCTCCAGAAAAGAGAAACTGCNNNAGATCGGAA
66564_1 TATGAGTGTGGAGTCGTTGCTTTAGTAGAGATGGGGTTTCGCCNNNAGATCGGAA
66565_1 TATGAGTGTGGAGTCGTTGCGCATTTGGGCTGGTTCCATATTTNNNAGATCGGAAG
66566_1 TATGAGTGTGGAGTCGTTGCTCTCTCTCTCATTTTCTGCCAGCNNNAGATCGGAAGA
66569_1 TATGAGTGTGGAGTCGTTGCCCACTTGTCAATTGATGGGCATTNNNAGATCGGAAG
66570_1 TATGAGTGTGGAGTCGTTGCAGATATGAACAGTGCAGGCAGAANNNAGATCGGAA
66571_1 TATGAGTGTGGAGTCGTTGCAGTGCTAGGATTACAGGCATGAGNNNAGATCGGAA
66572_1 TATGAGTGTGGAGTCGTTGCGAGAGAAAGGGAGGAAGACACTGNNNAGATCGGA
66573_1 TATGAGTGTGGAGTCGTTGCTTCAAGTGATTCTCCTGCCTCAGNNNAGATCGGAAG
66574_1 TATGAGTGTGGAGTCGTTGCAGTGACAGGGTCTTACTCTGTTGNNNAGATCGGAA
66575_1 TATGAGTGTGGAGTCGTTGCAGAGTCCCACAGAATCAAACCAANNNAGATCGGAAG
66577_1 TATGAGTGTGGAGTCGTTGCCTGACCTCACTTACTTCCTGGTCNNNAGATCGGAAGA
66579_1 TATGAGTGTGGAGTCGTTGCGGTGCCTGTAATACCAGCTACTTNNNAGATCGGAAG
66580_1 TATGAGTGTGGAGTCGTTGCTGAATCGAGGAGCTTGTTTCTTGNNNAGATCGGAA
66581_1 TATGAGTGTGGAGTCGTTGCTCCTCAACAAAATACGGGCAAACNNNAGATCGGAAG
66582_1 TATGAGTGTGGAGTCGTTGCGGAGTACAATGGCACAATCTTGGNNNAGATCGGAA
66583_1 TATGAGTGTGGAGTCGTTGCACACAATGAAGCCCTAAAAAGGCNNNAGATCGGAA
66586_1 TATGAGTGTGGAGTCGTTGCGGTTCAAGTGATTCTCTTGCCTCNNNAGATCGGAAG
66587_1 TATGAGTGTGGAGTCGTTGCAAGAAAAGATGATCCAGGCTGGGNNNAGATCGGAA
66588_1 TATGAGTGTGGAGTCGTTGCGAAATACAGCTAGCCAAGGAGGTNNNAGATCGGAA
66589_1 TATGAGTGTGGAGTCGTTGCCATCCACATAGTCTCAGAACCCANNNAGATCGGAAG
66590_1 TATGAGTGTGGAGTCGTTGCCAGCCTGGTCAATATGGTGAAACNNNAGATCGGAA
66591_1 TATGAGTGTGGAGTCGTTGCTGTGACACAGGCATTCAGGAATANNNAGATCGGAA
66593_1 TATGAGTGTGGAGTCGTTGCATGGGATGTTTGAGCTGAATCCTNNNAGATCGGAA
66594_1 TATGAGTGTGGAGTCGTTGCTCTCATTTTTGACAAAGGCACCANNNAGATCGGAAG
66595_1 TATGAGTGTGGAGTCGTTGCGCCTGGGTAAGTCCAAACTAGAANNNAGATCGGAA
66596_1 TATGAGTGTGGAGTCGTTGCGAAGCAAAGATTACAGTGGTGCANNNAGATCGGAA
66598_1 TATGAGTGTGGAGTCGTTGCCTTGATCCTGTGTCCTCCTTCAANNNAGATCGGAAGA
66599_1 TATGAGTGTGGAGTCGTTGCATCTCCTTTTCTGCATGTCGACTNNNAGATCGGAAGA
66601_1 TATGAGTGTGGAGTCGTTGCGCCATGAGTTTTTCTGGGGAAAANNNAGATCGGAA
66602_1 TATGAGTGTGGAGTCGTTGCTAATGGCATTTGCAGAAACCTGGNNNAGATCGGAA
66603_1 TATGAGTGTGGAGTCGTTGCCATACACACAAACACCACACACANNNAGATCGGAAG

66604_1 TATGAGTGTGGAGTCGTTGCCACACAGAGCAAGAGAGAGAGAGNNNAGATCGGA
66605_1 TATGAGTGTGGAGTCGTTGCATTTTTCATCATGGCACCGCTACNNNAGATCGGAAG
66606_1 TATGAGTGTGGAGTCGTTGCATTGGCCCCTAAGAAGAGTTACCNNNAGATCGGAAG
66607_1 TATGAGTGTGGAGTCGTTGCACATCCCAATTCTTCAGGTAGCANNNAGATCGGAAG
66608_1 TATGAGTGTGGAGTCGTTGCACACACACCACAGTTTATCCACTNNNAGATCGGAAG
66609_1 TATGAGTGTGGAGTCGTTGCGTTTCTCTGAAGAACCCTGGCTANNNAGATCGGAAG
66610_1 TATGAGTGTGGAGTCGTTGCCAGTGGAGTGTTCTTAGCTCACTNNNAGATCGGAAG
66612_1 TATGAGTGTGGAGTCGTTGCGTGGCATAATCTTGGCTCATTGANNNAGATCGGAAG
66613_1 TATGAGTGTGGAGTCGTTGCCCAGTCAGATTCAATTAGGGCCTNNNAGATCGGAAG
66614_1 TATGAGTGTGGAGTCGTTGCCTTGAGTGTGAAGACAGTCTGGANNNAGATCGGAA
66615_1 TATGAGTGTGGAGTCGTTGCTTCTTCATGTTGGTCAGACTGGTNNNAGATCGGAAG
66616_1 TATGAGTGTGGAGTCGTTGCTCGATTGCTGGTAATGTTCCTGANNNAGATCGGAAG
66617_1 TATGAGTGTGGAGTCGTTGCTATACACCACACACACACACACANNNAGATCGGAAG
66618_1 TATGAGTGTGGAGTCGTTGCACACATACACACACAGCTTCCTANNNAGATCGGAAG
66619_1 TATGAGTGTGGAGTCGTTGCGGCTGTGGTGTGGATATTTTTCCNNNAGATCGGAAG
66621_1 TATGAGTGTGGAGTCGTTGCACACAATATAGGCCTGCAGTAGGNNNAGATCGGAA
66622_1 TATGAGTGTGGAGTCGTTGCTCTGTTTCTATGAAAAGCGCTGCNNNAGATCGGAAG
66623_1 TATGAGTGTGGAGTCGTTGCGGAATTGGCTCACATAATCGCAANNNAGATCGGAAG
66624_1 TATGAGTGTGGAGTCGTTGCGATGGGAGTTATTCATTGCAGGCNNNAGATCGGAA
66625_1 TATGAGTGTGGAGTCGTTGCTCCTATGTAAGACTGCCCTCTCANNNAGATCGGAAG
66626_1 TATGAGTGTGGAGTCGTTGCATGGAGTCTCACTTTGCTGACAANNNAGATCGGAAG
66628_1 TATGAGTGTGGAGTCGTTGCGCGCACAAAGAATAAAGCATTGGNNNAGATCGGAA
66629_1 TATGAGTGTGGAGTCGTTGCATACAGCTACGAAGCCTCTCATGNNNAGATCGGAAG
66630_1 TATGAGTGTGGAGTCGTTGCCATTTGAACCACAGCAACTCCATNNNAGATCGGAAG
66631_1 TATGAGTGTGGAGTCGTTGCTTTTTTGATTCGGAGTCTCGCTGNNNAGATCGGAAG
66632_1 TATGAGTGTGGAGTCGTTGCGAGGTGGGAGGATCATCTGAAAANNNAGATCGGA
66633_1 TATGAGTGTGGAGTCGTTGCGCGCAGAGGAGATAAAGGATGTANNNAGATCGGA
66634_1 TATGAGTGTGGAGTCGTTGCAGTTGTGGCAACCAAAAATGTCTNNNAGATCGGAAG
66636_1 TATGAGTGTGGAGTCGTTGCCTGTGACTCACACCTGTAATCACNNNAGATCGGAAG
66637_1 TATGAGTGTGGAGTCGTTGCAAGCAGGCAGCCTCAAAATTATGNNNAGATCGGAAG
66638_1 TATGAGTGTGGAGTCGTTGCGCACTTAGAAAATGCCAGGCAAGNNNAGATCGGAA
66639_1 TATGAGTGTGGAGTCGTTGCTTGAGTGGAGTTTTGGAGAGTGTNNNAGATCGGAA
66640_1 TATGAGTGTGGAGTCGTTGCAGGATACTGAATGTCCCCAACACNNNAGATCGGAAG
66641_1 TATGAGTGTGGAGTCGTTGCCTGACAAAAACAAGCAATGGGGANNNAGATCGGAA
66643_1 TATGAGTGTGGAGTCGTTGCCAGAGTCAGACTGTCCTTGAGTCNNNAGATCGGAAG
66644_1 TATGAGTGTGGAGTCGTTGCAAGGAAACCTGCTCTCAGAATGTNNNAGATCGGAAG
66645_1 TATGAGTGTGGAGTCGTTGCACAGCCACAGAGAATTCCAAGATNNNAGATCGGAAG
66647_1 TATGAGTGTGGAGTCGTTGCCCCTACAACCATCCTGATCCTTCNNNAGATCGGAAGA
66648_1 TATGAGTGTGGAGTCGTTGCCCAGTGGGAAATACGGATGGAAANNNAGATCGGAA
66649_1 TATGAGTGTGGAGTCGTTGCTTTGCATGTCTGATGTATGTGCCNNNAGATCGGAAG
66650_1 TATGAGTGTGGAGTCGTTGCTGACTCTAGAATTCTGGCTTGCANNNAGATCGGAAG
66652_1 TATGAGTGTGGAGTCGTTGCGTTCAAGCAATTCTCATGCCTCANNNAGATCGGAAG
66655_1 TATGAGTGTGGAGTCGTTGCGTGTGTGTATCAAGCCAGAAAGCNNNAGATCGGAA

| | |
|---|---|
| 66656_1 | TATGAGTGTGGAGTCGTTGCTGATCTCGGCTAACTGCAACTTCNNNAGATCGGAAG |
| 66659_1 | TATGAGTGTGGAGTCGTTGCTTGACTCTCCTTTTCTCTTGGGGNNNAGATCGGAAG |
| 66660_1 | TATGAGTGTGGAGTCGTTGCGGAAAATGTGCCCACTACTGTTTNNNAGATCGGAAG |
| 66661_1 | TATGAGTGTGGAGTCGTTGCTAGTGTAATCTCAGCTTGCTGCANNNAGATCGGAAG |
| 66662_1 | TATGAGTGTGGAGTCGTTGCCAGTGGTATGATCACAGCTCACTNNNAGATCGGAAG |
| 66663_1 | TATGAGTGTGGAGTCGTTGCATGAAATCCGGAGAGTTCTAGGCNNNAGATCGGAA |
| 66665_1 | TATGAGTGTGGAGTCGTTGCTAGAAGAGAAAGAGAGGAGGGCANNNAGATCGGA |
| 66666_1 | TATGAGTGTGGAGTCGTTGCGCAGCTTGGAGAAGTCTCATCTANNNAGATCGGAA |
| 66667_1 | TATGAGTGTGGAGTCGTTGCAGATCTCAGCTCACTGCATACTCNNNAGATCGGAAG |
| 66668_1 | TATGAGTGTGGAGTCGTTGCGAAGTTTCACTCTTGTTGCCCAGNNNAGATCGGAAG |
| 66669_1 | TATGAGTGTGGAGTCGTTGCAGCACCGCATGAGTTAGAAAAACNNNAGATCGGAA |
| 66670_1 | TATGAGTGTGGAGTCGTTGCATATGTGTGTGTGTGTGTGTGTGNNNAGATCGGAA |
| 66671_1 | TATGAGTGTGGAGTCGTTGCAAAAATGAAATCATGGCTGGGCGNNNAGATCGGAA |
| 66672_1 | TATGAGTGTGGAGTCGTTGCATTCCTTTGCCATTCTCTTCCCTNNNAGATCGGAAGA |
| 66673_1 | TATGAGTGTGGAGTCGTTGCCTCTCTTCCTCCCTTCTGGCTTANNNAGATCGGAAGA |
| 66674_1 | TATGAGTGTGGAGTCGTTGCCACCCAATACTGGAGCTGTGATANNNAGATCGGAAG |
| 66675_1 | TATGAGTGTGGAGTCGTTGCCAATCTTGAAGCCAGACAGCAAANNNAGATCGGAA |
| 66676_1 | TATGAGTGTGGAGTCGTTGCGGGGTTACTTCCTGGAAAGACATNNNAGATCGGAA |
| 66677_1 | TATGAGTGTGGAGTCGTTGCAATACTTCTTCAAGGCCATCTGCNNNAGATCGGAAG |
| 66678_1 | TATGAGTGTGGAGTCGTTGCCTGAGGTGGAAGGATCAGTTGAGNNNAGATCGGA |
| 66680_1 | TATGAGTGTGGAGTCGTTGCACAGACATACAGCAGGTAACACCNNNAGATCGGAA |
| 66681_1 | TATGAGTGTGGAGTCGTTGCACTCAAACACACACACACACATTNNNAGATCGGAAG |
| 66682_1 | TATGAGTGTGGAGTCGTTGCACAGCTAACTGGGAAGGTGAAAANNNAGATCGGAA |
| 66683_1 | TATGAGTGTGGAGTCGTTGCAACTACAAAACACTGCTGAGGGANNNAGATCGGAA |
| 66684_1 | TATGAGTGTGGAGTCGTTGCTCTCTGTAGAACCCAAACACAGTNNNAGATCGGAAG |
| 66685_1 | TATGAGTGTGGAGTCGTTGCCACACACTCTCACACTCCACATANNNAGATCGGAAGA |
| 66686_1 | TATGAGTGTGGAGTCGTTGCTACAGGCCAATGACTTTCAACCTNNNAGATCGGAAG |
| 66689_1 | TATGAGTGTGGAGTCGTTGCTGTCTCCACACTCAAGGTAATGGNNNAGATCGGAAG |
| 66690_1 | TATGAGTGTGGAGTCGTTGCGTGTGTGAGAGACAGTCTTGCTANNNAGATCGGAA |
| 66693_1 | TATGAGTGTGGAGTCGTTGCCAATGGAAGTGTCTAGGGATGCTNNNAGATCGGAA |
| 66694_1 | TATGAGTGTGGAGTCGTTGCGGGAATCTGGGAATATTGGTGCTNNNAGATCGGAA |
| 66695_1 | TATGAGTGTGGAGTCGTTGCCTGGAGTGTAGTGGCATGATCTCNNNAGATCGGAA |
| 66696_1 | TATGAGTGTGGAGTCGTTGCGTGCTGGCAAGTCTAAAATGTGTNNNAGATCGGAA |
| 66697_1 | TATGAGTGTGGAGTCGTTGCGCACACATAAACACACACACGTANNNAGATCGGAAG |
| 66698_1 | TATGAGTGTGGAGTCGTTGCTTTGAAACGGAGTCTCACTCTGTNNNAGATCGGAAG |
| 66700_1 | TATGAGTGTGGAGTCGTTGCGCTCAAGCAATTCTCCCTCTTTGNNNAGATCGGAAGA |
| 66701_1 | TATGAGTGTGGAGTCGTTGCGAAGAGTCGATGCTACAGCTCAANNNAGATCGGAA |
| 66702_1 | TATGAGTGTGGAGTCGTTGCAATTTGCAGCCAGTATCAACCAANNNAGATCGGAAG |
| 66705_1 | TATGAGTGTGGAGTCGTTGCATGCTACATGGATATGAGCAGCCNNNAGATCGGAA |
| 66706_1 | TATGAGTGTGGAGTCGTTGCTATTCCATTGTGTGAATGTGCCANNNAGATCGGAAG |
| 66707_1 | TATGAGTGTGGAGTCGTTGCGGGTAATAATGATGGTGAGGGCTNNNAGATCGGA |
| 66708_1 | TATGAGTGTGGAGTCGTTGCTCTCCTGAGTAGCTGGGATTACANNNAGATCGGAAG |
| 66709_1 | TATGAGTGTGGAGTCGTTGCCAGAATTTGCAGTGAGCCAAGATNNNAGATCGGAA |

| | |
|---|---|
| 66710_1 | TATGAGTGTGGAGTCGTTGCGCACACGCACATCTTTTTTTTTNNNAGATCGGAAGA |
| 66711_1 | TATGAGTGTGGAGTCGTTGCACACCAACACTAGCCATCTTGATNNNAGATCGGAAG |
| 66712_1 | TATGAGTGTGGAGTCGTTGCAATACCTGCAAACCAAATCCAGCNNNAGATCGGAAG |
| 66713_1 | TATGAGTGTGGAGTCGTTGCCTAAGAAGAGCCTCTCACCCTTCNNNAGATCGGAAG |
| 66714_1 | TATGAGTGTGGAGTCGTTGCTGACACCCTCTTGTGAACATAGANNNAGATCGGAAG |
| 66716_1 | TATGAGTGTGGAGTCGTTGCTCCACATAGAGAGAGCAGAGTCANNNAGATCGGAA |
| 66718_1 | TATGAGTGTGGAGTCGTTGCTGAAATAGTGGCATTTGCAGCAANNNAGATCGGAA |
| 66719_1 | TATGAGTGTGGAGTCGTTGCATCTCCCAGAATTCCCATGTGTTNNNAGATCGGAAG |
| 66720_1 | TATGAGTGTGGAGTCGTTGCGCTGATTGGCAATGTTTTTGGGANNNAGATCGGAA |
| 66721_1 | TATGAGTGTGGAGTCGTTGCCCTGTGTCAGAGGTCTTTGGATANNNAGATCGGAA |
| 66722_1 | TATGAGTGTGGAGTCGTTGCAGGTTCAAGCAATTCTCCTGACTNNNAGATCGGAAG |
| 66723_1 | TATGAGTGTGGAGTCGTTGCCTGCCTCCATGGTTCTAGTGATTNNNAGATCGGAAG |
| 66724_1 | TATGAGTGTGGAGTCGTTGCTTTAGTGTCTGTCAGTACCAGGCNNNAGATCGGAAG |
| 66725_1 | TATGAGTGTGGAGTCGTTGCCACCTGTTTTCACTGCTGTCATGNNNAGATCGGAAG |
| 66726_1 | TATGAGTGTGGAGTCGTTGCCTCGAACTCCTGGGCTAAATGATNNNAGATCGGAAG |
| 66727_1 | TATGAGTGTGGAGTCGTTGCAGAGGAAGAGAGAGACAGAGAGGNNNAGATCGG |
| 66728_1 | TATGAGTGTGGAGTCGTTGCTACAGGCAGATGCATGTTTCTCTNNNAGATCGGAAG |
| 66730_1 | TATGAGTGTGGAGTCGTTGCTACTGGAGTGTAATGGTGCGATCNNNAGATCGGAA |
| 66731_1 | TATGAGTGTGGAGTCGTTGCAATTCCTAGCCCATCCCATCTTCNNNAGATCGGAAGA |
| 66732_1 | TATGAGTGTGGAGTCGTTGCTGGCTTGAGGATTTCCATTTGTGNNNAGATCGGAAG |
| 66733_1 | TATGAGTGTGGAGTCGTTGCCCCTTTCTCTGGGAGAGTACTCTNNNAGATCGGAAG |
| 66734_1 | TATGAGTGTGGAGTCGTTGCCCCACTTTCTTTCCACCTCTCATNNNAGATCGGAAGA |
| 66735_1 | TATGAGTGTGGAGTCGTTGCAGACAGCAAACAAGTACAAAAACANNNAGATCGGA |
| 66737_1 | TATGAGTGTGGAGTCGTTGCTTTGAGACAGAGTCTTGCTGTGTNNNAGATCGGAA |
| 66738_1 | TATGAGTGTGGAGTCGTTGCTGTGATCATACCACTGCATTCCANNNAGATCGGAAG |
| 66739_1 | TATGAGTGTGGAGTCGTTGCTATTATGGCCTTCGTGTATTGGCNNNAGATCGGAAG |
| 66741_1 | TATGAGTGTGGAGTCGTTGCCTGTAAGGCTGTCCCACTACAATNNNAGATCGGAAG |
| 66743_1 | TATGAGTGTGGAGTCGTTGCGAGGATTGCTTGAGCTCAGTAGTNNNAGATCGGAA |
| 66744_1 | TATGAGTGTGGAGTCGTTGCTCAAATTTTTCCCCAGTTGGAGCNNNAGATCGGAAG |
| 66746_1 | TATGAGTGTGGAGTCGTTGCCCCTCTCCTGAAATCACACTTCANNNAGATCGGAAGA |
| 66747_1 | TATGAGTGTGGAGTCGTTGCACACACTTCATACTCCTGTGCTGNNNAGATCGGAAG |
| 66748_1 | TATGAGTGTGGAGTCGTTGCTTGTTGTACACTGGATCACACCTNNNAGATCGGAAG |
| 66749_1 | TATGAGTGTGGAGTCGTTGCACTAACAGCATTTGCAATGACCCNNNAGATCGGAAG |
| 66750_1 | TATGAGTGTGGAGTCGTTGCAATCACATCTGAACTTTCCCCCANNNAGATCGGAAGA |
| 66752_1 | TATGAGTGTGGAGTCGTTGCCAAGGTCTGAAGGGAATAGCCTTNNNAGATCGGAA |
| 66753_1 | TATGAGTGTGGAGTCGTTGCAAATCCTTTCTTCATGGGGACGGNNNAGATCGGAAG |
| 66755_1 | TATGAGTGTGGAGTCGTTGCCCTGGTTCAAGTGATTCTCCTGTNNNAGATCGGAAG |
| 66756_1 | TATGAGTGTGGAGTCGTTGCATCTCATCTCACTGCAACCTCTGNNNAGATCGGAAGA |
| 66757_1 | TATGAGTGTGGAGTCGTTGCTAAAGTGTGTGTGTGTGTGTGTGNNNAGATCGGAA |
| 66758_1 | TATGAGTGTGGAGTCGTTGCTTTTCCTCTGTTGCTTGCAAGTTNNNAGATCGGAAGA |
| 66760_1 | TATGAGTGTGGAGTCGTTGCATATGTGTGCACGAAACACTTGGNNNAGATCGGAA |
| 66762_1 | TATGAGTGTGGAGTCGTTGCCACCCACTATAACAGCAGGACTTNNNAGATCGGAAG |
| 66763_1 | TATGAGTGTGGAGTCGTTGCTATCCCACTGTGACCTCATCTCANNNAGATCGGAAGA |

| | |
|---|---|
| 66764_1 | TATGAGTGTGGAGTCGTTGCACATGCTTTTTTTGGGAGACTGANNNAGATCGGAAG |
| 66765_1 | TATGAGTGTGGAGTCGTTGCCCTTCCCAGGGAGAAACTCATTTNNNAGATCGGAAG |
| 66766_1 | TATGAGTGTGGAGTCGTTGCTTGCTGCAAAAACTCCTAACTGGNNNAGATCGGAAG |
| 66767_1 | TATGAGTGTGGAGTCGTTGCACAAAACAATAGCCAGAAGGTGGNNNAGATCGGAA |
| 66769_1 | TATGAGTGTGGAGTCGTTGCGGAGGCTGAAGTGAGCTATGATTNNNAGATCGGAA |
| 66770_1 | TATGAGTGTGGAGTCGTTGCCTAGAAACATGGCCTTCCTCTGTNNNAGATCGGAAG |
| 66771_1 | TATGAGTGTGGAGTCGTTGCGCAGAAAAGCCAGTGTTGTACTTNNNAGATCGGAA |
| 66772_1 | TATGAGTGTGGAGTCGTTGCACTTTTCGAAGGTTGTCCTACCANNNAGATCGGAAG |
| 66775_1 | TATGAGTGTGGAGTCGTTGCCTAGATGAAGGTCACCTTTCCCANNNAGATCGGAAG |
| 66776_1 | TATGAGTGTGGAGTCGTTGCTGCTTTCTTATAAACCCCCCAGTNNNAGATCGGAAGA |
| 66777_1 | TATGAGTGTGGAGTCGTTGCGCACAGCCTTGACTAGTGAAAAGNNNAGATCGGAA |
| 66778_1 | TATGAGTGTGGAGTCGTTGCAGCATCACATTATTCCTGGGGTTNNNAGATCGGAAG |
| 66780_1 | TATGAGTGTGGAGTCGTTGCGGCATGGAACAGATTCTCCCTAANNNAGATCGGAAG |
| 66782_1 | TATGAGTGTGGAGTCGTTGCATAATGCTGCAAAAGGAACCAGGNNNAGATCGGAA |
| 66783_1 | TATGAGTGTGGAGTCGTTGCCTTGAAGGAGGAACCAACACCTANNNAGATCGGAA |
| 66784_1 | TATGAGTGTGGAGTCGTTGCCGGAGAGACAGAGAGAAAGAGAGNNNAGATCGGA |
| 66785_1 | TATGAGTGTGGAGTCGTTGCAAAGCTGGCATAATGTGAGCAAANNNAGATCGGAA |
| 66786_1 | TATGAGTGTGGAGTCGTTGCGAGTTTCGCTCTTTGTTGCCCNNNAGATCGGAAGAG |
| 66787_1 | TATGAGTGTGGAGTCGTTGCGTGTGTGTGTCTGTTTGAGATGGNNNAGATCGGAA |
| 66788_1 | TATGAGTGTGGAGTCGTTGCTGGTCCTCAGCCTGAACATTATTNNNAGATCGGAAG |
| 66790_1 | TATGAGTGTGGAGTCGTTGCTGCAGTAGCTCACACCTGTAATTNNNAGATCGGAAG |
| 66791_1 | TATGAGTGTGGAGTCGTTGCTTCCCAGGTTCAAGAGATTCTGGNNNAGATCGGAAG |
| 66792_1 | TATGAGTGTGGAGTCGTTGCGGAATACCATTCAGCCACAAAAANNNAGATCGGAAG |
| 66793_1 | TATGAGTGTGGAGTCGTTGCCGTCTGTGTGTCTGGTATTGACTNNNAGATCGGAAG |
| 66794_1 | TATGAGTGTGGAGTCGTTGCGTTTGCTCTGGGTTTGCTAGTTTNNNAGATCGGAAG |
| 66795_1 | TATGAGTGTGGAGTCGTTGCAGACCATGGGGAGCAACTTTATTNNNAGATCGGAA |
| 66796_1 | TATGAGTGTGGAGTCGTTGCCAATGGCACAATCTTGGTTCACCNNNAGATCGGAAG |
| 66797_1 | TATGAGTGTGGAGTCGTTGCGAGTCTCATTATGTCGCCAAAGCNNNAGATCGGAAG |
| 66798_1 | TATGAGTGTGGAGTCGTTGCCAGGTGAGTAGCCCATTGGATAANNNAGATCGGAA |
| 66799_1 | TATGAGTGTGGAGTCGTTGCAATCCCAAGTCCAGTCAGAAGACNNNAGATCGGAAG |
| 66800_1 | TATGAGTGTGGAGTCGTTGCGAGTTTAGCCATGACTCCAGGAANNNAGATCGGAA |
| 66801_1 | TATGAGTGTGGAGTCGTTGCTTTTCTGCCAATGTGGGATTCTGNNNAGATCGGAAG |
| 66802_1 | TATGAGTGTGGAGTCGTTGCGGAGTTGCAACTTACAAGCAGAGNNNAGATCGGAA |
| 66803_1 | TATGAGTGTGGAGTCGTTGCCAAACACACACACACAAAAAGCCNNNAGATCGGAAG |
| 66805_1 | TATGAGTGTGGAGTCGTTGCCTGAATACCAACTGCGCACTAAGNNNAGATCGGAAG |
| 66806_1 | TATGAGTGTGGAGTCGTTGCCTAAAGAAGTAGCCAAGGGGTCANNNAGATCGGAA |
| 66807_1 | TATGAGTGTGGAGTCGTTGCTAATGTCATCTGCAGCAACATGGNNNAGATCGGAAG |
| 66808_1 | TATGAGTGTGGAGTCGTTGCATGCCACACATCTACAACGATCTNNNAGATCGGAAG |
| 66810_1 | TATGAGTGTGGAGTCGTTGCACTGCTGTGATAAGTTTTGGGGANNNAGATCGGAA |
| 66811_1 | TATGAGTGTGGAGTCGTTGCGAATCACAGCTTCCAACACAGACNNNAGATCGGAAG |
| 66812_1 | TATGAGTGTGGAGTCGTTGCTTGAGTCTTATCGAAAGGCCCAANNNAGATCGGAAG |
| 66813_1 | TATGAGTGTGGAGTCGTTGCTGTGTCTATCAGCCTGAGGAGTANNNAGATCGGAA |
| 66814_1 | TATGAGTGTGGAGTCGTTGCACTCGACAGAAGAAAAGTAGCGTNNNAGATCGGAA |

66815_1 TATGAGTGTGGAGTCGTTGCTCTCTCTCTCTCTCTCTCTCAGCNNNAGATCGGAAGA
66816_1 TATGAGTGTGGAGTCGTTGCATTCACAGTGACCTGGATGGAATNNNAGATCGGAA
66817_1 TATGAGTGTGGAGTCGTTGCCTGAATTCTTTCTGTGCCTGCTCNNNAGATCGGAAG
66818_1 TATGAGTGTGGAGTCGTTGCCCACACACCCACTCAAAAAGAAANNNAGATCGGAAG
66820_1 TATGAGTGTGGAGTCGTTGCTTATACCTCCTGCAATTCCCACANNNAGATCGGAAGA
66821_1 TATGAGTGTGGAGTCGTTGCAGTGCTAGGATTACAGCCATGAGNNNAGATCGGAA
66822_1 TATGAGTGTGGAGTCGTTGCAAGTAGCTGGAATTACAGGTGCANNNAGATCGGAA
66824_1 TATGAGTGTGGAGTCGTTGCAGGTTGCAGTGAAATGGAGTGATNNNAGATCGGA
66825_1 TATGAGTGTGGAGTCGTTGCCCTGTGAGGCTAATGAGTAACCANNNAGATCGGAA
66826_1 TATGAGTGTGGAGTCGTTGCGGCATCCTGGCTTCTTATGGTAANNNAGATCGGAAG
66827_1 TATGAGTGTGGAGTCGTTGCACTCACCCTTCTTGTGTTGATGTNNNAGATCGGAAG
66828_1 TATGAGTGTGGAGTCGTTGCTGTGTGTGTGTGTTTGTGAATNNNAGATCGGAA
66829_1 TATGAGTGTGGAGTCGTTGCTAGGGGACAATGACAACAGACATNNNAGATCGGAA
66830_1 TATGAGTGTGGAGTCGTTGCTATTTGGGGGGGTTGTTTGTTTGNNNAGATCGGAA
66831_1 TATGAGTGTGGAGTCGTTGCTGTGTGTGTGGAAAGAGAGAGAGNNNAGATCGGA
66832_1 TATGAGTGTGGAGTCGTTGCTGTGTATGTAACTGCCCTTGGTTNNNAGATCGGAAG
66833_1 TATGAGTGTGGAGTCGTTGCCACTCTGGAGGTTAGGAATGGTTNNNAGATCGGAA
66834_1 TATGAGTGTGGAGTCGTTGCGTACCCCAAAGTTCATGTGTTGGNNNAGATCGGAAG
66835_1 TATGAGTGTGGAGTCGTTGCCACCAATAACAGAAGCACCACAGNNNAGATCGGAAG
66836_1 TATGAGTGTGGAGTCGTTGCGGAGGCTGGTTCAGGATAATCATNNNAGATCGGAA
66837_1 TATGAGTGTGGAGTCGTTGCTCTATTTTCGGGCAACTTGCTTCNNNAGATCGGAAG
66839_1 TATGAGTGTGGAGTCGTTGCAATAATGGCATTTGCAGCAACCTNNNAGATCGGAAG
66840_1 TATGAGTGTGGAGTCGTTGCCCAGAGGTCAAGGTAGCATTCTTNNNAGATCGGAA
66841_1 TATGAGTGTGGAGTCGTTGCTTGCAGCAACCTGTATGAAGTTGNNNAGATCGGAA
66843_1 TATGAGTGTGGAGTCGTTGCCGTGACAACCAAAAATGTCTCCANNNAGATCGGAAG
66845_1 TATGAGTGTGGAGTCGTTGCTCTGTAATCCCAGCTACTCAGGANNNAGATCGGAAG
66846_1 TATGAGTGTGGAGTCGTTGCAACTCCTGACCTCAAGTGATTGGNNNAGATCGGAAG
66847_1 TATGAGTGTGGAGTCGTTGCTACCCTGCCATTGGTCATTTACANNNAGATCGGAAG
66848_1 TATGAGTGTGGAGTCGTTGCCAATAGATCTTCTGGCACCCTCANNNAGATCGGAAG
66849_1 TATGAGTGTGGAGTCGTTGCTACTACCCCATCCCAACCTTAGTNNNAGATCGGAAGA
66851_1 TATGAGTGTGGAGTCGTTGCGTTCAAGTGATTCTCCTGCCTCANNNAGATCGGAAG
66853_1 TATGAGTGTGGAGTCGTTGCCTCCCATCCTAGTCCAATCATGGNNNAGATCGGAAG
66854_1 TATGAGTGTGGAGTCGTTGCTGTACTTTTTTGCCCACCTTTTTNNNAGATCGGAAGA
66855_1 TATGAGTGTGGAGTCGTTGCAATGCTACAATGAACATGGGAGCNNNAGATCGGAA
66856_1 TATGAGTGTGGAGTCGTTGCCAGGGTCTCACTCTGTTGCTTAGNNNAGATCGGAAG
66859_1 TATGAGTGTGGAGTCGTTGCATAAGTTTGTCATGGCCACTTGCNNNAGATCGGAAG
66860_1 TATGAGTGTGGAGTCGTTGCCTCCCTCTTTCTTCCATCCCATTNNNAGATCGGAAGAG
66862_1 TATGAGTGTGGAGTCGTTGCCATATGCCTGTAGTCCCAGCTACNNNAGATCGGAAG
66863_1 TATGAGTGTGGAGTCGTTGCGATGCTTGAGCTCAGGAGTTCTANNNAGATCGGAA
66864_1 TATGAGTGTGGAGTCGTTGCACATAGCGTCATCTTCCATCTCCNNNAGATCGGAAGA
66866_1 TATGAGTGTGGAGTCGTTGCCAGCCTGTAATCCCAGCAATTTGNNNAGATCGGAAG
66867_1 TATGAGTGTGGAGTCGTTGCACCAATGCACAAAAATCAGTTGTNNNAGATCGGAAG
66868_1 TATGAGTGTGGAGTCGTTGCATGGAGCACCCTAGTTATGAACANNNAGATCGGAA

| | |
|---|---|
| 66869_1 | TATGAGTGTGGAGTCGTTGCGAATGGCATTAGCAGCAACCTAGNNNAGATCGGAA |
| 66871_1 | TATGAGTGTGGAGTCGTTGCAAGAAATGCAATCGGGAGTACCTNNNAGATCGGAA |
| 66872_1 | TATGAGTGTGGAGTCGTTGCTGTGTGTTTATAGTCCCAGCCTCNNNAGATCGGAAG |
| 66873_1 | TATGAGTGTGGAGTCGTTGCCACACCTGTGATCTCAGCTACTTNNNAGATCGGAAG |
| 66874_1 | TATGAGTGTGGAGTCGTTGCGCAACATGGATGGAACTAGATGCNNNAGATCGGAA |
| 66875_1 | TATGAGTGTGGAGTCGTTGCCCCCCCAACAAAAAACCTACAAANNNAGATCGGAAG |
| 66876_1 | TATGAGTGTGGAGTCGTTGCACACTCTCTCTCTCTCTCTCTCTNNNAGATCGGAAGAG |
| 66879_1 | TATGAGTGTGGAGTCGTTGCACTGCAGTCTTGGTGATAACACANNNAGATCGGAAG |
| 66880_1 | TATGAGTGTGGAGTCGTTGCACTGATGCTGACTATAAGGGCTGNNNAGATCGGAA |
| 66882_1 | TATGAGTGTGGAGTCGTTGCTACGTGTGTATGTGTATGTGTGTNNNAGATCGGAA |
| 66883_1 | TATGAGTGTGGAGTCGTTGCGGACAACTTGAAGCATACCTGTGNNNAGATCGGAA |
| 66884_1 | TATGAGTGTGGAGTCGTTGCCTCAAGCAATCCTCCTTCCTCATNNNAGATCGGAAGA |
| 66885_1 | TATGAGTGTGGAGTCGTTGCTCACAGTTCATAAAAGCGGAAGCNNNAGATCGGAA |
| 66886_1 | TATGAGTGTGGAGTCGTTGCACTCTGGAGAAAGGTGGAGTAGANNNAGATCGGA |
| 66887_1 | TATGAGTGTGGAGTCGTTGCTTTAGTAGAGACGGGGTTTCAGCNNNAGATCGGAA |
| 66888_1 | TATGAGTGTGGAGTCGTTGCACTGCCGAAATGAATTGATGGTGNNNAGATCGGAA |
| 66889_1 | TATGAGTGTGGAGTCGTTGCTAGCTCACTGTAACCTCAAGCTCNNNAGATCGGAAG |
| 66890_1 | TATGAGTGTGGAGTCGTTGCCTGGGGAACCACAAAGAAGTCTANNNAGATCGGAA |
| 66892_1 | TATGAGTGTGGAGTCGTTGCTTACCTAGGCTGGTGTTGAACTCNNNAGATCGGAAG |
| 66893_1 | TATGAGTGTGGAGTCGTTGCGATGGAGTCTCACTCTGTCACTCNNNAGATCGGAAG |
| 66894_1 | TATGAGTGTGGAGTCGTTGCGCACACACACGCACAAATATTTANNNAGATCGGAAG |
| 66895_1 | TATGAGTGTGGAGTCGTTGCATGTGTGATGAGATGCAGCAAACNNNAGATCGGAA |
| 66897_1 | TATGAGTGTGGAGTCGTTGCTTGGGTCCTAAGCATCCCTAAACNNNAGATCGGAAG |
| 66898_1 | TATGAGTGTGGAGTCGTTGCGAGAGAGATAGGGCCTCACTCTGNNNAGATCGGAA |
| 66899_1 | TATGAGTGTGGAGTCGTTGCCGCGCATGCATATAGACAAATGANNNAGATCGGAA |
| 66900_1 | TATGAGTGTGGAGTCGTTGCGTGTACCTGTGTGTTGTCTTGTGNNNAGATCGGAAG |
| 66901_1 | TATGAGTGTGGAGTCGTTGCTTAGAATAGACTTGTCTGGGCCGNNNAGATCGGAA |
| 66902_1 | TATGAGTGTGGAGTCGTTGCCACATGTAATCCCAGCTACTCGANNNAGATCGGAAG |
| 66903_1 | TATGAGTGTGGAGTCGTTGCGGCTAGCCTACAATGATACAGCANNNAGATCGGAA |
| 66904_1 | TATGAGTGTGGAGTCGTTGCTCCATCAGTAAGTGAGAGGGGATNNNAGATCGGAA |
| 66905_1 | TATGAGTGTGGAGTCGTTGCTGCAATTATGGAGGCTGAGAAGTNNNAGATCGGAA |
| 66906_1 | TATGAGTGTGGAGTCGTTGCGGAAGAGAGAGAGACACACACAGNNNAGATCGGA |
| 66907_1 | TATGAGTGTGGAGTCGTTGCAGAAAAGCTGCACAATGACAACANNNAGATCGGAA |
| 66908_1 | TATGAGTGTGGAGTCGTTGCGCTCATGTGATTACAGAGGCTGANNNAGATCGGAA |
| 66909_1 | TATGAGTGTGGAGTCGTTGCTGTATACACACTCATGGTCCACANNNAGATCGGAAG |
| 66910_1 | TATGAGTGTGGAGTCGTTGCACACACACCTCCACAACTGATTANNNAGATCGGAAG |
| 66911_1 | TATGAGTGTGGAGTCGTTGCGCACTCGAGCAGGATTTTGTAAGNNNAGATCGGAA |
| 66912_1 | TATGAGTGTGGAGTCGTTGCTGAGTTCAAGTTCCCCATGTCTTNNNAGATCGGAAG |
| 66915_1 | TATGAGTGTGGAGTCGTTGCATGGCATTCACAGTAACCTGGATNNNAGATCGGAAG |
| 66916_1 | TATGAGTGTGGAGTCGTTGCCAAGGGAACAGAGGATGACTCAANNNAGATCGGAA |
| 66917_1 | TATGAGTGTGGAGTCGTTGCACCTTACCAGCAAGTTCTAACGTNNNAGATCGGAAG |
| 66919_1 | TATGAGTGTGGAGTCGTTGCACAGTTTCAAGGGACTGACTTGTNNNAGATCGGAAG |
| 66920_1 | TATGAGTGTGGAGTCGTTGCAGATGGGATCTCACTCTGTCTCCNNNAGATCGGAAG |

| | |
|---|---|
| 66921_1 | TATGAGTGTGGAGTCGTTGCTGTTTTGGGTGTGTCTTTGACTGNNNAGATCGGAAG |
| 66923_1 | TATGAGTGTGGAGTCGTTGCTTCCCTCCCCTCATGAAAGAGTANNNAGATCGGAAG |
| 66924_1 | TATGAGTGTGGAGTCGTTGCTGGCTACAATATTTTGCGGATCCNNNAGATCGGAAG |
| 66925_1 | TATGAGTGTGGAGTCGTTGCCCAAAGTACTGGGATTACAGGCANNNAGATCGGAA |
| 66926_1 | TATGAGTGTGGAGTCGTTGCTTGAGATGAAGTTGCTAGCCTGTNNNAGATCGGAA |
| 66927_1 | TATGAGTGTGGAGTCGTTGCTGCAGTATCATGTTCTCAGCTCANNNAGATCGGAAG |
| 66928_1 | TATGAGTGTGGAGTCGTTGCTTCAAGTGATTCTCCAGCCTCAGNNNAGATCGGAAG |
| 66929_1 | TATGAGTGTGGAGTCGTTGCCACACCTGTAATCCCAGCACTTTNNNAGATCGGAAGA |
| 66931_1 | TATGAGTGTGGAGTCGTTGCTACACACAGAGTAGAAGTGGCACNNNAGATCGGAA |
| 66932_1 | TATGAGTGTGGAGTCGTTGCTTGGACAGTGTATGGTATGGGTGNNNAGATCGGAA |
| 66933_1 | TATGAGTGTGGAGTCGTTGCCCTTGTGTACTCCCAAACAATGGNNNAGATCGGAAG |
| 66934_1 | TATGAGTGTGGAGTCGTTGCCAGGTGGATCATGAGATCAGGAGNNNAGATCGGAA |
| 66935_1 | TATGAGTGTGGAGTCGTTGCGCAGAGTCAAAGGAATGCAACATNNNAGATCGGAA |
| 66936_1 | TATGAGTGTGGAGTCGTTGCTGTAGTCCCAGCTACTCAGAAGANNNAGATCGGAAG |
| 66938_1 | TATGAGTGTGGAGTCGTTGCCATTCCACATCTCAGAGCATGTCNNNAGATCGGAAG |
| 66939_1 | TATGAGTGTGGAGTCGTTGCTGACACCACCCTTGTTAGTACTGNNNAGATCGGAAG |
| 66940_1 | TATGAGTGTGGAGTCGTTGCCTCTCTCTCTCTTCCTCCCTCTCNNNAGATCGGAAGAG |
| 66941_1 | TATGAGTGTGGAGTCGTTGCACATTGCTACTCACAGCCAGTTANNNAGATCGGAAG |
| 66942_1 | TATGAGTGTGGAGTCGTTGCGGATGGAGCTGGAGACCATTATTNNNAGATCGGAA |
| 66943_1 | TATGAGTGTGGAGTCGTTGCAAATGAATGAAATTGGCTGGGTGNNNAGATCGGAA |
| 66944_1 | TATGAGTGTGGAGTCGTTGCTAGATGGAGTCTCACTCTGTTGCNNNAGATCGGAAG |
| 66945_1 | TATGAGTGTGGAGTCGTTGCAGAGGTAGAAAGGGACCAGATCANNNAGATCGGAA |
| 66946_1 | TATGAGTGTGGAGTCGTTGCTTCTTGTGAGAGCCTCAAAGTCANNNAGATCGGAAG |
| 66947_1 | TATGAGTGTGGAGTCGTTGCTGTGTGGTGTATGTGTGTGCATANNNAGATCGGAA |
| 66949_1 | TATGAGTGTGGAGTCGTTGCCAGGCTGGTCTAGAGAACTCTTGNNNAGATCGGAA |
| 66950_1 | TATGAGTGTGGAGTCGTTGCTTCAGTTTGGATGTTGCTTGTGTNNNAGATCGGAAG |
| 66951_1 | TATGAGTGTGGAGTCGTTGCTAGAAAGTCAGAGGTGCATGGACNNNAGATCGGAA |
| 66952_1 | TATGAGTGTGGAGTCGTTGCACGTTTCACTTCTGGTAGGTTGTNNNAGATCGGAAG |
| 66953_1 | TATGAGTGTGGAGTCGTTGCCTCCTGCTTTCAAGGCTTTACTGNNNAGATCGGAAG |
| 66954_1 | TATGAGTGTGGAGTCGTTGCTTCAAGTGATTCTCCTGCCTCAGNNNAGATCGGAAG |
| 66955_1 | TATGAGTGTGGAGTCGTTGCCCACCAAAATAACCGGCATCATTNNNAGATCGGAAG |
| 66956_1 | TATGAGTGTGGAGTCGTTGCATTGGGAGAGAAGGAAGGCTTTTNNNAGATCGGAA |
| 66957_1 | TATGAGTGTGGAGTCGTTGCTTGAAGGAAGTTGCCCTTAGTGTNNNAGATCGGAA |
| 66959_1 | TATGAGTGTGGAGTCGTTGCTGTAAGTTGAGCCAAACGCAAAANNNAGATCGGAA |
| 66961_1 | TATGAGTGTGGAGTCGTTGCAAATGTGCCCTCACCATAGCTAANNNAGATCGGAAG |
| 66962_1 | TATGAGTGTGGAGTCGTTGCAGGGAGAGAGAGAGAGAGAGAGANNNAGATCGG |
| 66964_1 | TATGAGTGTGGAGTCGTTGCTTTTTTCCTCTGCACTGTGTGTGNNNAGATCGGAAG |
| 66965_1 | TATGAGTGTGGAGTCGTTGCGGTGATTGGGCATCTGAGATCTANNNAGATCGGAA |
| 66966_1 | TATGAGTGTGGAGTCGTTGCTCTGTGTGTGTGAGAAGATGAGGNNNAGATCGGAA |
| 66967_1 | TATGAGTGTGGAGTCGTTGCTGTTAGTTCTTTCTTGAGCCCCANNNAGATCGGAAG |
| 66968_1 | TATGAGTGTGGAGTCGTTGCGGAAAGCCTCACTCAGAAGATGANNNAGATCGGAA |
| 66969_1 | TATGAGTGTGGAGTCGTTGCTGTCAATGGTGTTGCAATGAACANNNAGATCGGAA |
| 66970_1 | TATGAGTGTGGAGTCGTTGCAAACTCCTGGGATCAAGCAATCTNNNAGATCGGAAG |

66971_1 TATGAGTGTGGAGTCGTTGCTGGAGTATAGTGGTGCCATCATGNNNAGATCGGAA
66973_1 TATGAGTGTGGAGTCGTTGCGAGCCAGTTTCATCATTGGACACNNNAGATCGGAAG
66974_1 TATGAGTGTGGAGTCGTTGCTGAGCCATGTGAACATCTTGAGANNNAGATCGGAA
66976_1 TATGAGTGTGGAGTCGTTGCGGAAAGGGAAGAAGGGAAAAAGCNNNAGATCGG
66977_1 TATGAGTGTGGAGTCGTTGCGTTCAAGCAGTTCTCCTACCTCANNNAGATCGGAAG
66978_1 TATGAGTGTGGAGTCGTTGCCTGTAATTGCAGCTACTCGGGAGNNNAGATCGGAA
66979_1 TATGAGTGTGGAGTCGTTGCGAGATCATACCACTGCACTCCAGNNNAGATCGGAAG
66980_1 TATGAGTGTGGAGTCGTTGCTTTTTTTTTTTGAGACGGCGTCTNNNAGATCGGAAG
66981_1 TATGAGTGTGGAGTCGTTGCTCTTTTGAGATGGAGTTTCGCTCNNNAGATCGGAAG
66982_1 TATGAGTGTGGAGTCGTTGCCCAGAAGCAGATGCACAAATCTGNNNAGATCGGAA
66983_1 TATGAGTGTGGAGTCGTTGCTCACTTTAGCCCAGGAGTTCAAGNNNAGATCGGAAG
66984_1 TATGAGTGTGGAGTCGTTGCTTTGTATCCCTGCCCAAATCTCANNNAGATCGGAAGA
66985_1 TATGAGTGTGGAGTCGTTGCCATGGACACACAGAGACAAACACNNNAGATCGGAA
66986_1 TATGAGTGTGGAGTCGTTGCAGAGAGAAAGACAGAGGATCCCANNNAGATCGGA
66987_1 TATGAGTGTGGAGTCGTTGCTTGTGCATCATTGTCTTGCTCAGNNNAGATCGGAAG
66989_1 TATGAGTGTGGAGTCGTTGCCAAGTCTGGCCTCTAGCAAATCTNNNAGATCGGAAG
66990_1 TATGAGTGTGGAGTCGTTGCTGAAGAAAAAGCCAAATGCCCTTNNNAGATCGGAA
66991_1 TATGAGTGTGGAGTCGTTGCCCATTACTGTGGGGACTCTAACCNNNAGATCGGAAG
66992_1 TATGAGTGTGGAGTCGTTGCAGTAACTCAGGAGCAGAAAACCANNNAGATCGGAA
66993_1 TATGAGTGTGGAGTCGTTGCATATTTCTAGAACAGGGCAGGGGNNNAGATCGGAA
66994_1 TATGAGTGTGGAGTCGTTGCTTTTAGTTTGCAGCCAGGTGAAGNNNAGATCGGAA
66995_1 TATGAGTGTGGAGTCGTTGCTCTCTGGAAAACCCTGACTGATGNNNAGATCGGAAG
66996_1 TATGAGTGTGGAGTCGTTGCCCCTCATGGATGTGATATCAGCANNNAGATCGGAAG
66998_1 TATGAGTGTGGAGTCGTTGCCTCACTGCAACCTTCACCTCTAANNNAGATCGGAAGA
66999_1 TATGAGTGTGGAGTCGTTGCGGGGCAAGTGTACTATTTTCTGCNNNAGATCGGAA
67000_1 TATGAGTGTGGAGTCGTTGCGAGACAGAGAGAGAGAGAGAGGANNNAGATCGG
67002_1 TATGAGTGTGGAGTCGTTGCAGTACCTGGAGAGATGGGACTACNNNAGATCGGAA
67003_1 TATGAGTGTGGAGTCGTTGCTCAATCTAAGTGCCCATCAACCANNNAGATCGGAAG
67004_1 TATGAGTGTGGAGTCGTTGCTCACTAGAAGACAAGCAAAGCCTNNNAGATCGGAA
67005_1 TATGAGTGTGGAGTCGTTGCCAGCCATTGTGTTCTGCTAAACANNNAGATCGGAAG
67006_1 TATGAGTGTGGAGTCGTTGCGTGCCTGTAATCTCAGCTACTCANNNAGATCGGAAG
67007_1 TATGAGTGTGGAGTCGTTGCAGACAGGAAGATTGCTTGAACCTNNNAGATCGGAA
67008_1 TATGAGTGTGGAGTCGTTGCAAGAACAAGCACCTAGTTTCCCTNNNAGATCGGAAG
67009_1 TATGAGTGTGGAGTCGTTGCGCTACTTGAGAGGGAGTGTGTAGNNNAGATCGGA
67011_1 TATGAGTGTGGAGTCGTTGCGAGAAGGGCTTATGTGCTACACTNNNAGATCGGAA
67013_1 TATGAGTGTGGAGTCGTTGCGCTCACACCGCAATTTCAATACTNNNAGATCGGAAG
67014_1 TATGAGTGTGGAGTCGTTGCCTCTCTCTCTCTGCCCATCTTTCNNNAGATCGGAAGA
67016_1 TATGAGTGTGGAGTCGTTGCGAGTGTGAGGAACGGACTAGATGNNNAGATCGG
67020_1 TATGAGTGTGGAGTCGTTGCGAATCCCCATTCTTTCACCAACCNNNAGATCGGAAGA
67021_1 TATGAGTGTGGAGTCGTTGCTGCCTGTGTATATAAGGGATCGCNNNAGATCGGAA
67022_1 TATGAGTGTGGAGTCGTTGCATGTGTATGTGCAGTGTCCATCTNNNAGATCGGAAG
67024_1 TATGAGTGTGGAGTCGTTGCTTATGTGTGTTATGGGGATGGGGNNNAGATCGGA
67025_1 TATGAGTGTGGAGTCGTTGCGCAGCAGGAAGAAAATGAAAGGTNNNAGATCGG

| | |
|---|---|
| 67026_1 | TATGAGTGTGGAGTCGTTGCACACACACACACACACATTTTTTNNNAGATCGGAAGA |
| 67027_1 | TATGAGTGTGGAGTCGTTGCCCCAAAGTGCTAGGATTACAGGTNNNAGATCGGAA |
| 67028_1 | TATGAGTGTGGAGTCGTTGCTCATTGATTGATGGGCATTTGGGNNNAGATCGGAA |
| 67029_1 | TATGAGTGTGGAGTCGTTGCTCAAGCCCAGGACTAAATGAGACNNNAGATCGGAA |
| 67030_1 | TATGAGTGTGGAGTCGTTGCGGTTGAAGCTATTCTTGTGCCTCNNNAGATCGGAAG |
| 67031_1 | TATGAGTGTGGAGTCGTTGCCTCCTTGAGTATGAGACCTTGCANNNAGATCGGAAG |
| 67032_1 | TATGAGTGTGGAGTCGTTGCGTGGGAAAGGTGGGAACAGAATANNNAGATCGGA |
| 67033_1 | TATGAGTGTGGAGTCGTTGCTAGTAAAAATGGGGAGGTGGTGGNNNAGATCGGA |
| 67035_1 | TATGAGTGTGGAGTCGTTGCCTGTAATTCCAGCACTTTGGGAGNNNAGATCGGAAG |
| 67036_1 | TATGAGTGTGGAGTCGTTGCCAGACTGGCTTTTGAAGGAAGTGNNNAGATCGGAA |
| 67037_1 | TATGAGTGTGGAGTCGTTGCATTTTTTTGGCTTGGAGTCTCGCNNNAGATCGGAAG |
| 67039_1 | TATGAGTGTGGAGTCGTTGCCACTCCTGTAATCCCAGCACTTTNNNAGATCGGAAGA |
| 67040_1 | TATGAGTGTGGAGTCGTTGCGCAAGTTGTTCCTTGGTGGTTTANNNAGATCGGAAG |
| 67041_1 | TATGAGTGTGGAGTCGTTGCAGCTGAAATTCCTCAGTTGGAGTNNNAGATCGGAA |
| 67043_1 | TATGAGTGTGGAGTCGTTGCTGGCTCACATCTGTAATCCTAGCNNNAGATCGGAAG |
| 67046_1 | TATGAGTGTGGAGTCGTTGCACTCAGCATAGCACTGTTCATGANNNAGATCGGAAG |
| 67047_1 | TATGAGTGTGGAGTCGTTGCTTTTTGAGACACAGTTTCGCTCGNNNAGATCGGAAG |
| 67048_1 | TATGAGTGTGGAGTCGTTGCTCTCTAACACACACACACACACANNNAGATCGGAAGA |
| 67049_1 | TATGAGTGTGGAGTCGTTGCCCCTGATGAACAGAGATGCAAAANNNAGATCGGAA |
| 67053_1 | TATGAGTGTGGAGTCGTTGCGGTTCAAGCAATTCTCACACCTCNNNAGATCGGAAG |
| 67055_1 | TATGAGTGTGGAGTCGTTGCTCTCTATCAGGTTGGTGCAAAAANNNAGATCGGAAG |
| 67056_1 | TATGAGTGTGGAGTCGTTGCTGTAGAGATGGAGCTTCACCATGNNNAGATCGGAA |
| 67058_1 | TATGAGTGTGGAGTCGTTGCAAACTCCTGACCTCAAGTGATCCNNNAGATCGGAAG |
| 67059_1 | TATGAGTGTGGAGTCGTTGCGAGACAGGGTTTTGCTATGTTGCNNNAGATCGGAA |
| 67060_1 | TATGAGTGTGGAGTCGTTGCTAGTGGCACCGAATACATTGACANNNAGATCGGAA |
| 67061_1 | TATGAGTGTGGAGTCGTTGCGCCAAAGCTATCCTGAGCAAAAANNNAGATCGGAA |
| 67062_1 | TATGAGTGTGGAGTCGTTGCCAAGCCTGAGATTCAGAGAAGGTNNNAGATCGGAA |
| 67063_1 | TATGAGTGTGGAGTCGTTGCAATTAGCTGAGTGTAGTGGTGGGNNNAGATCGGAA |
| 67067_1 | TATGAGTGTGGAGTCGTTGCATGCACAGCTTGGAATTGTCAATNNNAGATCGGAAG |
| 67069_1 | TATGAGTGTGGAGTCGTTGCAAGGGATAACAAGTGTAGGCCAGNNNAGATCGGAA |
| 67070_1 | TATGAGTGTGGAGTCGTTGCTTATCCAGTCCACCATTGAAGGGNNNAGATCGGAAG |
| 67073_1 | TATGAGTGTGGAGTCGTTGCAAGAGCTGCTACAGTGATGGTACNNNAGATCGGAA |
| 67074_1 | TATGAGTGTGGAGTCGTTGCAAGGTATTTGCCAACCTCAGTCTNNNAGATCGGAAG |
| 67075_1 | TATGAGTGTGGAGTCGTTGCTGTTTTTTTCTGTTGGGTGCAGTNNNAGATCGGAAG |
| 67076_1 | TATGAGTGTGGAGTCGTTGCTTCTGAAATTTGAGTGGTTGGCCNNNAGATCGGAA |
| 67077_1 | TATGAGTGTGGAGTCGTTGCACAGAGTGGAAGAAAATCTGGGANNNAGATCGGAA |
| 67079_1 | TATGAGTGTGGAGTCGTTGCGGAAAGGGGTGGATACTCAAACTNNNAGATCGGAA |
| 67080_1 | TATGAGTGTGGAGTCGTTGCCCTGCTGTGGTTTTGAATGGAATNNNAGATCGGAA |
| 67081_1 | TATGAGTGTGGAGTCGTTGCATACACACACACCCCTGCATATTNNNAGATCGGAAGA |
| 67082_1 | TATGAGTGTGGAGTCGTTGCCCAAACTGGTCAGGTTTTTGCTTNNNAGATCGGAAG |
| 67083_1 | TATGAGTGTGGAGTCGTTGCTATTACCCAGGCTGGTCTTGAACNNNAGATCGGAAG |
| 67084_1 | TATGAGTGTGGAGTCGTTGCCACAGTGGCTCATGCTTGTAATCNNNAGATCGGAAG |
| 67085_1 | TATGAGTGTGGAGTCGTTGCCAAGTAGCTGGGATTACAGGCATNNNAGATCGGAA |

| | |
|---|---|
| 67086_1 | TATGAGTGTGGAGTCGTTGCATGTGTGTGTGTTTGTGTGTTNNNAGATCGGAAG |
| 67088_1 | TATGAGTGTGGAGTCGTTGCGATGGGAAGATGACTAACTGGGGNNNAGATCGGAA |
| 67089_1 | TATGAGTGTGGAGTCGTTGCGATCTCCTGACCTCATGATCCACNNNAGATCGGAAGA |
| 67090_1 | TATGAGTGTGGAGTCGTTGCGTGTATGTTTCTGGGTATGTGCGNNNAGATCGGAAG |
| 67091_1 | TATGAGTGTGGAGTCGTTGCTCTTTGCAATAAAACCAGGCAGGNNNAGATCGGAAG |
| 67092_1 | TATGAGTGTGGAGTCGTTGCAGTCACAGAGAGATACGGAGGAANNNAGATCGGAA |
| 67094_1 | TATGAGTGTGGAGTCGTTGCCCTGTGTCCATGTGTTCTCATTGNNNAGATCGGAAGA |
| 67095_1 | TATGAGTGTGGAGTCGTTGCCATGTGTGTACAGACAGGTGTTGNNNAGATCGGAAG |
| 67096_1 | TATGAGTGTGGAGTCGTTGCGAGGGTGAATCATCAGAGGCTATNNNAGATCGGAA |
| 67097_1 | TATGAGTGTGGAGTCGTTGCTTAGCTGGTTGTGTTACTCAGGGNNNAGATCGGAAG |
| 67098_1 | TATGAGTGTGGAGTCGTTGCGTGATCTCAGCTCACTGTAACCTNNNAGATCGGAAGA |
| 67100_1 | TATGAGTGTGGAGTCGTTGCATGGTTGTACACACCTGTAGTCCNNNAGATCGGAAG |
| 67101_1 | TATGAGTGTGGAGTCGTTGCGCTGAAGGACACTTGTGAGTTTGNNNAGATCGGAAG |
| 67102_1 | TATGAGTGTGGAGTCGTTGCAATTGAGGTCAGGAGTTCGAGACNNNAGATCGGAA |
| 67103_1 | TATGAGTGTGGAGTCGTTGCTATATACACACAGCCCCTGGACCNNNAGATCGGAAGA |
| 67107_1 | TATGAGTGTGGAGTCGTTGCATCCTCACAAAGTGCTGGGATTANNNAGATCGGAAG |
| 67108_1 | TATGAGTGTGGAGTCGTTGCCTCCCTTCTCTCTCTGTCTCTCANNNAGATCGGAAGAG |
| 67110_1 | TATGAGTGTGGAGTCGTTGCTGCACTTTGTAGAGGTTGTACCANNNAGATCGGAAG |
| 67111_1 | TATGAGTGTGGAGTCGTTGCGAGAATTGCTTGAACTTGGGAGGNNNAGATCGGAA |
| 67112_1 | TATGAGTGTGGAGTCGTTGCTTGACTCACTGGATCAACCTCTGNNNAGATCGGAAGA |
| 67113_1 | TATGAGTGTGGAGTCGTTGCTAACAAGGCTGCCTTTTTGCTAGNNNAGATCGGAAG |
| 67115_1 | TATGAGTGTGGAGTCGTTGCGAGTCCAGGGCAAGAACATCTTANNNAGATCGGAAG |
| 67118_1 | TATGAGTGTGGAGTCGTTGCGCATCTGTCATCTTGGCTTTCTCNNNAGATCGGAAGA |
| 67121_1 | TATGAGTGTGGAGTCGTTGCCTCTTGCTACACCACACTGATCTNNNAGATCGGAAGA |
| 67122_1 | TATGAGTGTGGAGTCGTTGCTGGCTGACACTTCATTATCCACANNNAGATCGGAAGA |
| 67123_1 | TATGAGTGTGGAGTCGTTGCTAGTTTCAGCTGAGTCCCTTTGGNNNAGATCGGAAG |
| 67124_1 | TATGAGTGTGGAGTCGTTGCGATATATAGGGCTGGGTGCAGTGNNNAGATCGGAA |
| 67125_1 | TATGAGTGTGGAGTCGTTGCATGCCTGGACAGAACTAGACAAGNNNAGATCGGAAG |
| 67126_1 | TATGAGTGTGGAGTCGTTGCCTCAACAGGAGAGCAGGAATCTTNNNAGATCGGAAG |
| 67127_1 | TATGAGTGTGGAGTCGTTGCGCTGATGCCTATAATCCCAACACNNNAGATCGGAAGA |
| 67128_1 | TATGAGTGTGGAGTCGTTGCTGGCCAGATTTTGGAACAGGATANNNAGATCGGAAG |
| 67129_1 | TATGAGTGTGGAGTCGTTGCTCAGCATTTTCTCCTCACATGGANNNAGATCGGAAGA |
| 67130_1 | TATGAGTGTGGAGTCGTTGCCTCATCTACTCAACTCAGGAGGCNNNAGATCGGAAGA |
| 67132_1 | TATGAGTGTGGAGTCGTTGCGAAACCAGAAGGATTCCTGGGTCNNNAGATCGGAAG |
| 67133_1 | TATGAGTGTGGAGTCGTTGCCTCTGTCTCTGTCTCTCATGCTGNNNAGATCGGAAGA |
| 67136_1 | TATGAGTGTGGAGTCGTTGCGAAAGAAACCTGGTTCCATGCTCNNNAGATCGGAAG |
| 67137_1 | TATGAGTGTGGAGTCGTTGCGCTTCAACAATGGCCATATGGTTNNNAGATCGGAAG |
| 67139_1 | TATGAGTGTGGAGTCGTTGCCACACACACACACATACACATNNNAGATCGGAAGA |
| 67140_1 | TATGAGTGTGGAGTCGTTGCCTCATCACAAACTGACCATGCTGNNNAGATCGGAAGA |
| 67141_1 | TATGAGTGTGGAGTCGTTGCGGAGAGACAGAGAGAGAGCAAAGNNNAGATCGGA |
| 67142_1 | TATGAGTGTGGAGTCGTTGCGAATGACCATTTCCAGAGCCATGNNNAGATCGGAAG |
| 67144_1 | TATGAGTGTGGAGTCGTTGCTACAATTCCTTGTGCCAGAGACANNNAGATCGGAAG |
| 67146_1 | TATGAGTGTGGAGTCGTTGCCCATCTCAACTCACTGCAACTTCNNNAGATCGGAAGA |

| | |
|---|---|
| 67147_1 | TATGAGTGTGGAGTCGTTGCGAGGCTAGTGTCTTGACAAGGAANNNAGATCGGAA |
| 67148_1 | TATGAGTGTGGAGTCGTTGCTCTCTAAGTGCAGACAGTAAGGCNNNAGATCGGAA |
| 67150_1 | TATGAGTGTGGAGTCGTTGCACTCTCTCTGTTACCCACATTCTNNNAGATCGGAAGA |
| 67151_1 | TATGAGTGTGGAGTCGTTGCGAAGAGAAGTGTCTGTGCTGGTANNNAGATCGGAA |
| 67152_1 | TATGAGTGTGGAGTCGTTGCGACAGACAGACAGACAGACAGAGNNNAGATCGGAA |
| 67153_1 | TATGAGTGTGGAGTCGTTGCTATCACAAGAACAGCAATGGGGANNNAGATCGGAA |
| 67154_1 | TATGAGTGTGGAGTCGTTGCGGAGCTAAGCAAAACAACCAGATNNNAGATCGGAA |
| 67155_1 | TATGAGTGTGGAGTCGTTGCTGGTGAAGTTTGTCCTTGGATGANNNAGATCGGAA |
| 67156_1 | TATGAGTGTGGAGTCGTTGCCTGAGTTCAAACGATTCTCCTGCNNNAGATCGGAAGA |
| 67157_1 | TATGAGTGTGGAGTCGTTGCTTGGATCTTCTGCAGATGACAGTNNNAGATCGGAAG |
| 67158_1 | TATGAGTGTGGAGTCGTTGCTTCTTATCCTTTTGCTGGCATGCNNNAGATCGGAAGA |
| 67159_1 | TATGAGTGTGGAGTCGTTGCCCCGCTCATGTTACAAGGAAAAANNNAGATCGGAAG |
| 67160_1 | TATGAGTGTGGAGTCGTTGCCTTACCAGGTTAGGGTCCCATTCNNNAGATCGGAAGA |
| 67161_1 | TATGAGTGTGGAGTCGTTGCACCCTATTTTTTTCTGCAGTGGCNNNAGATCGGAAGA |
| 67162_1 | TATGAGTGTGGAGTCGTTGCAACAGAACAGAGCCCTCAGAAATNNNAGATCGGAA |
| 67163_1 | TATGAGTGTGGAGTCGTTGCACTATCATGAGAACAGCAAGGGGNNNAGATCGGAA |
| 67164_1 | TATGAGTGTGGAGTCGTTGCTTTAGATGTGGGGGGTACATGTGNNNAGATCGGAA |
| 67166_1 | TATGAGTGTGGAGTCGTTGCAGTAAACACTGCTTCCCAAGACANNNAGATCGGAAG |
| 67167_1 | TATGAGTGTGGAGTCGTTGCTCCCATCACCTGAAAATTTCCCTNNNAGATCGGAAGA |
| 67168_1 | TATGAGTGTGGAGTCGTTGCTCTTCTCTCGGACTTCTCAGTCTNNNAGATCGGAAGA |
| 67169_1 | TATGAGTGTGGAGTCGTTGCTCTCTCTCTCTCTTCTGGTCTCCNNNAGATCGGAAGA |
| 67171_1 | TATGAGTGTGGAGTCGTTGCCCCCAAGGATACTTCTGAAGGTCNNNAGATCGGAAG |
| 67172_1 | TATGAGTGTGGAGTCGTTGCGCAGAAAAAGAGGCCAATTCAGTNNNAGATCGGAA |
| 67173_1 | TATGAGTGTGGAGTCGTTGCCACACACACACACAAATCAGCTGNNNAGATCGGAAGA |
| 67174_1 | TATGAGTGTGGAGTCGTTGCATACACCTGCTATGTACCCACAANNNAGATCGGAAGA |
| 67175_1 | TATGAGTGTGGAGTCGTTGCTTCCTTCAACAGTCTCCAACACANNNAGATCGGAAGA |
| 67177_1 | TATGAGTGTGGAGTCGTTGCATTTCCCAAGTGATGACCTGTCANNNAGATCGGAAGA |
| 67179_1 | TATGAGTGTGGAGTCGTTGCCTCTAGTTATCTGCCCACCTTGGNNNAGATCGGAAGA |
| 67180_1 | TATGAGTGTGGAGTCGTTGCGCACAGAAGCCACAGTCTTTTTANNNAGATCGGAAG |
| 67181_1 | TATGAGTGTGGAGTCGTTGCAGCTGGCTGAATTAGAGAGGATGNNNAGATCGGAA |
| 67183_1 | TATGAGTGTGGAGTCGTTGCATGTATGTGTGTGTGTGTGTGTGNNNAGATCGGAA |
| 67184_1 | TATGAGTGTGGAGTCGTTGCTCTCTCTTTCCTCCTCCCATCATNNNAGATCGGAAGAG |
| 67185_1 | TATGAGTGTGGAGTCGTTGCGCAGTTTTCTGGCAGAAATCCTTNNNAGATCGGAAG |
| 67186_1 | TATGAGTGTGGAGTCGTTGCGGCCACTCTGAAGGACAATTTTTNNNAGATCGGAAG |
| 67187_1 | TATGAGTGTGGAGTCGTTGCTACACACACACACTGAGGTTTGANNNAGATCGGAAG |
| 67188_1 | TATGAGTGTGGAGTCGTTGCTTTGAAGATGGAGTGTCGCTTTGNNNAGATCGGAA |
| 67189_1 | TATGAGTGTGGAGTCGTTGCGAATACAGTGGCATGATCTTGGCNNNAGATCGGAA |
| 67190_1 | TATGAGTGTGGAGTCGTTGCGCTATCCTTGGTCTTAGGGCTTTNNNAGATCGGAAGA |
| 67191_1 | TATGAGTGTGGAGTCGTTGCCCATGGCCTTACCTAAATGCAAGNNNAGATCGGAAG |
| 67193_1 | TATGAGTGTGGAGTCGTTGCAGCCTGATTCTCATCTGATCCTCNNNAGATCGGAAGA |
| 67194_1 | TATGAGTGTGGAGTCGTTGCTTCCTCCCAAGAGAATTTACCCCNNNAGATCGGAAGA |
| 67195_1 | TATGAGTGTGGAGTCGTTGCATCATCCTAGAGCTGTGACTCCTNNNAGATCGGAAGA |
| 67196_1 | TATGAGTGTGGAGTCGTTGCTGGCAGCATAATTCACAAATGCANNNAGATCGGAAG |

| | |
|---|---|
| 67198_1 | TATGAGTGTGGAGTCGTTGCTTGAGACGGTATCTTGCTCTGTCNNNAGATCGGAAG |
| 67199_1 | TATGAGTGTGGAGTCGTTGCATTCAGGTAAGAGGTGAGGCTTGNNNAGATCGGAA |
| 67200_1 | TATGAGTGTGGAGTCGTTGCAGAGAGGGAGAGAGAGAGACCTANNNAGATCGGA |
| 67201_1 | TATGAGTGTGGAGTCGTTGCTCCCTCTGTTTCTCTCTCTTCCTNNNAGATCGGAAGAG |
| 67202_1 | TATGAGTGTGGAGTCGTTGCTGAGAAATTCTTTGGTGGGAGCTNNNAGATCGGAA |
| 67203_1 | TATGAGTGTGGAGTCGTTGCCCACACTGTTGAGAAACTGTCAGNNNAGATCGGAAG |
| 67204_1 | TATGAGTGTGGAGTCGTTGCCTACTGTCCTCTAATTCCAGCCCNNNAGATCGGAAGA |
| 67205_1 | TATGAGTGTGGAGTCGTTGCAGTTAGAGACTGCCTCAAAAGGGNNNAGATCGGAA |
| 67206_1 | TATGAGTGTGGAGTCGTTGCTTCTGAGCTCAAGAGATTCACCCNNNAGATCGGAAG |
| 67207_1 | TATGAGTGTGGAGTCGTTGCCTCTCCCATGGATACTCAAGCTCNNNAGATCGGAAG |
| 67208_1 | TATGAGTGTGGAGTCGTTGCGAGAAAGGAGAGGAAAGCACAGANNNAGATCGGA |
| 67209_1 | TATGAGTGTGGAGTCGTTGCTGAGGTCTGTGTTACCCAGATTGNNNAGATCGGAA |
| 67210_1 | TATGAGTGTGGAGTCGTTGCTTTTTTCAGTCGGAGTCTTGCTCNNNAGATCGGAAG |
| 67211_1 | TATGAGTGTGGAGTCGTTGCCTCTCTTTCTGTCTCTGTCCGTCNNNAGATCGGAAGA |
| 67212_1 | TATGAGTGTGGAGTCGTTGCACAATCTGCTGTTCCTTAGGGTTNNNAGATCGGAAG |
| 67213_1 | TATGAGTGTGGAGTCGTTGCCTCTCGCTGTCTCTGTCTCATTTNNNAGATCGGAAGA |
| 67215_1 | TATGAGTGTGGAGTCGTTGCTCCTTTCCAGTGAGATCGACATGNNNAGATCGGAAG |
| 67217_1 | TATGAGTGTGGAGTCGTTGCCTCTCTCTCCCCTGATCTCTAGGNNNAGATCGGAAGA |
| 67218_1 | TATGAGTGTGGAGTCGTTGCTGTGCTCTATTATCAGGTCGTCGNNNAGATCGGAAG |
| 67219_1 | TATGAGTGTGGAGTCGTTGCCTTCAATCCCTCAATCCCTGGTTNNNAGATCGGAAGA |
| 67220_1 | TATGAGTGTGGAGTCGTTGCAAGAGAGAGAGCACCTACTATGCNNNAGATCGGAA |
| 67221_1 | TATGAGTGTGGAGTCGTTGCATAATTGTCAGGGCACAGACCTTNNNAGATCGGAAG |
| 67222_1 | TATGAGTGTGGAGTCGTTGCAGAAGTCCCACAATATGCCATCTNNNAGATCGGAAG |
| 67223_1 | TATGAGTGTGGAGTCGTTGCGCCCAGCCATACCATGATAAAGANNNAGATCGGAAG |
| 67224_1 | TATGAGTGTGGAGTCGTTGCGCCCCACAGCAGATTCAAAAATANNNAGATCGGAAG |
| 67225_1 | TATGAGTGTGGAGTCGTTGCGGAAGCTTTCTCTAGGTGTAGCANNNAGATCGGAA |
| 67226_1 | TATGAGTGTGGAGTCGTTGCACATGCTGGATTCAAGACTGAGTNNNAGATCGGAA |
| 67227_1 | TATGAGTGTGGAGTCGTTGCCTGTGTCCTTGGGAAAAGCAAAANNNAGATCGGAA |
| 67229_1 | TATGAGTGTGGAGTCGTTGCGCATTCCTTCCACACAATGACTTNNNAGATCGGAAG |
| 67230_1 | TATGAGTGTGGAGTCGTTGCCTGACGAGATTTGGCCTTACAACNNNAGATCGGAAG |
| 67231_1 | TATGAGTGTGGAGTCGTTGCATCCCTCCTGAACTCATAGACCANNNAGATCGGAAG |
| 67232_1 | TATGAGTGTGGAGTCGTTGCCCCCATTTCATCTCTCACTCCTCNNNAGATCGGAAGAG |
| 67233_1 | TATGAGTGTGGAGTCGTTGCGGTAGGAAGATGCACAGAGGAATNNNAGATCGGA |
| 67234_1 | TATGAGTGTGGAGTCGTTGCTTCTACACTGGCTGGCTGTAATTNNNAGATCGGAAG |
| 67236_1 | TATGAGTGTGGAGTCGTTGCCATGTTTTTAGACTCGGCCGTTTNNNAGATCGGAAG |
| 67237_1 | TATGAGTGTGGAGTCGTTGCCTCAGAAGCAGAGAAGTTCCCATNNNAGATCGGAA |
| 67238_1 | TATGAGTGTGGAGTCGTTGCGCTGCTACATGGTGTTCTAGGAANNNAGATCGGAA |
| 67239_1 | TATGAGTGTGGAGTCGTTGCCACCTGAGAAACCTAAGGAACGANNNAGATCGGAA |
| 67240_1 | TATGAGTGTGGAGTCGTTGCTTAGTGCTTGGATGGGAGACTTCNNNAGATCGGAA |
| 67242_1 | TATGAGTGTGGAGTCGTTGCGCCACTGGGAAAGATGATGAAAGNNNAGATCGGAA |
| 67243_1 | TATGAGTGTGGAGTCGTTGCAGGATAGGGAAGACTCACCTGAANNNAGATCGGAA |
| 67244_1 | TATGAGTGTGGAGTCGTTGCCAGCCAGCACCAAGGAAAATAAANNNAGATCGGAA |
| 67245_1 | TATGAGTGTGGAGTCGTTGCAGAGGGCCTGCAATCAGAATAAGNNNAGATCGGAA |

| | |
|---|---|
| 67246_1 | TATGAGTGTGGAGTCGTTGCAGGAGCTCTGCTAATTACACCACNNNAGATCGGAAG |
| 67247_1 | TATGAGTGTGGAGTCGTTGCGTCTTAGCCTCTAACCGGTGATTNNNAGATCGGAAG |
| 67248_1 | TATGAGTGTGGAGTCGTTGCGGGTTTTCTTACTTGCCAGCAATNNNAGATCGGAAG |
| 67250_1 | TATGAGTGTGGAGTCGTTGCGTGCACACACGCCTTAGTTTTANNNAGATCGGAAG |
| 67251_1 | TATGAGTGTGGAGTCGTTGCAGTGATACAGCACACAGAGTGTTNNNAGATCGGAA |
| 67252_1 | TATGAGTGTGGAGTCGTTGCTAAGGAAACTGAGGCATGGAACANNNAGATCGGAA |
| 67253_1 | TATGAGTGTGGAGTCGTTGCCCACACAAACAAAAGACACAGGTNNNAGATCGGAA |
| 67254_1 | TATGAGTGTGGAGTCGTTGCCTCTGTTTCTGGCTAGGAAGACANNNAGATCGGAAG |
| 67255_1 | TATGAGTGTGGAGTCGTTGCACCCGGCCTATACAACCTAAATCNNNAGATCGGAAG |
| 67256_1 | TATGAGTGTGGAGTCGTTGCGGAAGAACAAACAGAGTGACACGNNNAGATCGGAA |
| 67257_1 | TATGAGTGTGGAGTCGTTGCATCAGAGAACACCTTGACCAAGCNNNAGATCGGAAG |
| 67258_1 | TATGAGTGTGGAGTCGTTGCCATCAGGGCAACACATTGGTTTANNNAGATCGGAAG |
| 67259_1 | TATGAGTGTGGAGTCGTTGCCAGACTGGGACAATGTAGACACANNNAGATCGGAA |
| 67260_1 | TATGAGTGTGGAGTCGTTGCTGTGTGTGTGTGGTTTCCTCTTANNNAGATCGGAAG |
| 67261_1 | TATGAGTGTGGAGTCGTTGCGCAAGTAATACATGCTCACGGTGNNNAGATCGGAA |
| 67262_1 | TATGAGTGTGGAGTCGTTGCTGGGTGGAAATGTCCTTGAGAATNNNAGATCGGAA |
| 67263_1 | TATGAGTGTGGAGTCGTTGCGACAACACATAAAGAGCAAGGCCNNNAGATCGGAA |
| 67264_1 | TATGAGTGTGGAGTCGTTGCTATCTCTCCATCCCTCTGTTGGTNNNAGATCGGAAGA |
| 67265_1 | TATGAGTGTGGAGTCGTTGCAGTGGGAATACGAATCTTGTCCANNNAGATCGGAA |
| 67266_1 | TATGAGTGTGGAGTCGTTGCTCCATGGGTTCCAGGAAGAATTTNNNAGATCGGAAG |
| 67267_1 | TATGAGTGTGGAGTCGTTGCTGGAGAATCCGCAAACTGATACANNNAGATCGGAA |
| 67268_1 | TATGAGTGTGGAGTCGTTGCCTATGGTTTTCCCCCACATACCANNNAGATCGGAAGA |
| 67269_1 | TATGAGTGTGGAGTCGTTGCAGTATGAAGACGACGCAGATGTTNNNAGATCGGAA |
| 67270_1 | TATGAGTGTGGAGTCGTTGCAGAGACAGTGGGAGTATAGACGTNNNAGATCGGAA |
| 67271_1 | TATGAGTGTGGAGTCGTTGCAAGTGCACATTTTCGAGAACTGGNNNAGATCGGAA |
| 67272_1 | TATGAGTGTGGAGTCGTTGCGTGAGACTCCTTCACTGACTGAANNNAGATCGGAAG |
| 67273_1 | TATGAGTGTGGAGTCGTTGCAACTAGGAAATGGGGTTGTCACANNNAGATCGGAA |
| 67275_1 | TATGAGTGTGGAGTCGTTGCAAGCTCTGTGTGGCCTGTTTATTNNNAGATCGGAAG |
| 67276_1 | TATGAGTGTGGAGTCGTTGCCCAGTGGTATCCTTGGCTACTTTNNNAGATCGGAAG |
| 67277_1 | TATGAGTGTGGAGTCGTTGCAACAAAATACCACAGGCAAGGTGNNNAGATCGGAA |
| 67278_1 | TATGAGTGTGGAGTCGTTGCGACCCTGCACTTCCAATTCAATCNNNAGATCGGAAGA |
| 67279_1 | TATGAGTGTGGAGTCGTTGCTGGGTTCTTGGCTCATGAATCATNNNAGATCGGAAG |
| 67280_1 | TATGAGTGTGGAGTCGTTGCTGCAGTGAAGGAAAGATTGTTGCNNNAGATCGGAA |
| 67281_1 | TATGAGTGTGGAGTCGTTGCACTTTGGGAGGAAGTCTCATTCANNNAGATCGGAA |
| 67282_1 | TATGAGTGTGGAGTCGTTGCTTGCCGCTGTGATAAAATCTGTCNNNAGATCGGAAG |
| 67283_1 | TATGAGTGTGGAGTCGTTGCTGCAGCTTTCTAAGACCAGTCATNNNAGATCGGAAG |
| 67285_1 | TATGAGTGTGGAGTCGTTGCATAGAAGTTCCTCCCTCTGGTCANNNAGATCGGAAG |
| 67286_1 | TATGAGTGTGGAGTCGTTGCAACACCTTTCAGATCAGCACAGANNNAGATCGGAAG |
| 67287_1 | TATGAGTGTGGAGTCGTTGCAGAGAGAGAGAGAGAGAGAGGGANNNAGATCGG |
| 67288_1 | TATGAGTGTGGAGTCGTTGCTTCATCAGTTACTGCTTGCAGAANNNAGATCGGAAG |
| 67290_1 | TATGAGTGTGGAGTCGTTGCGAAAATTGTGAGAGCTTGGCCTTNNNAGATCGGAA |
| 67291_1 | TATGAGTGTGGAGTCGTTGCCGAGAAGGGTTCCAAGTTTTGTGNNNAGATCGGAA |
| 67292_1 | TATGAGTGTGGAGTCGTTGCTCAAAAACTGACTGGACCCATGANNNAGATCGGAAG |

| | |
|---|---|
| 67293_1 | TATGAGTGTGGAGTCGTTGCGGACAGCTAAAACTCCCAGAGAANNNAGATCGGAA |
| 67294_1 | TATGAGTGTGGAGTCGTTGCAGCTACAGAGAGAGAGAGTGTGTNNNAGATCGGA |
| 67295_1 | TATGAGTGTGGAGTCGTTGCAGTAGAACAAGGTGAGGGGGATANNNAGATCGGA |
| 67297_1 | TATGAGTGTGGAGTCGTTGCGAGAGAGAGAGAGCACCAATGAGNNNAGATCGGA |
| 67298_1 | TATGAGTGTGGAGTCGTTGCGGAGGAGAGAAAGGAAAGGAAGGNNNAGATCGG |
| 67299_1 | TATGAGTGTGGAGTCGTTGCCATCACTTGAGCAAAAGAAGCCANNNAGATCGGAA |
| 67300_1 | TATGAGTGTGGAGTCGTTGCTGAGTCCATTGATGTACAAGGGANNNAGATCGGAA |
| 67301_1 | TATGAGTGTGGAGTCGTTGCTGACTTTTCACAGGGAAGGTGAANNNAGATCGGAA |
| 67302_1 | TATGAGTGTGGAGTCGTTGCATTTGGAGATAATGAGCCCAGGGNNNAGATCGGAA |
| 67303_1 | TATGAGTGTGGAGTCGTTGCTGTGTATGGGCCCATAAACTGAANNNAGATCGGAA |
| 67304_1 | TATGAGTGTGGAGTCGTTGCGCCTAGATGGTGAGATTTGGGAANNNAGATCGGAA |
| 67306_1 | TATGAGTGTGGAGTCGTTGCGAGTCTCTCTGTATTGCTCAGGCNNNAGATCGGAAG |
| 67307_1 | TATGAGTGTGGAGTCGTTGCAAGCCTTCTTTTTCTAGTGCACANNNAGATCGGAAG |
| 67308_1 | TATGAGTGTGGAGTCGTTGCTGCACACATAGAGTTCTCCCAAANNNAGATCGGAAG |
| 67309_1 | TATGAGTGTGGAGTCGTTGCCCAGGGACCATACTACTCCTGTANNNAGATCGGAAG |
| 67310_1 | TATGAGTGTGGAGTCGTTGCGTGAAGAGGCTCACAAATGACTCNNNAGATCGGAA |
| 67311_1 | TATGAGTGTGGAGTCGTTGCAATCTGTGGCTTTCCTCCTTCATNNNAGATCGGAAGA |
| 67314_1 | TATGAGTGTGGAGTCGTTGCTACAGTGAGTGATGGAGGTGTTGNNNAGATCGGA |
| 67316_1 | TATGAGTGTGGAGTCGTTGCACTCACGGTCAATCATATGCTCANNNAGATCGGAAG |
| 67317_1 | TATGAGTGTGGAGTCGTTGCCAACTGGCACCTTGATTTTGAGTNNNAGATCGGAAG |
| 67318_1 | TATGAGTGTGGAGTCGTTGCGACCCCAAGGTATAGATTCCACCNNNAGATCGGAAG |
| 67319_1 | TATGAGTGTGGAGTCGTTGCTCACAGACATTCTCACAGTGGTTNNNAGATCGGAAG |
| 67320_1 | TATGAGTGTGGAGTCGTTGCTCATCTTCCTGAGCAGCTGATACNNNAGATCGGAAG |
| 67321_1 | TATGAGTGTGGAGTCGTTGCTCCCTGGATATGTGACTGAGAGANNNAGATCGGAA |
| 67322_1 | TATGAGTGTGGAGTCGTTGCTGTGTGTGTGTGTGTGTGTATTGNNNAGATCGGAA |
| 67323_1 | TATGAGTGTGGAGTCGTTGCTGTATATGGCAGGAAGTCACCAGNNNAGATCGGAA |
| 67324_1 | TATGAGTGTGGAGTCGTTGCATGGATGCCCTGACTTGGATAAANNNAGATCGGAA |
| 67325_1 | TATGAGTGTGGAGTCGTTGCCCTACCTCCAGAGCTTGTTGATANNNAGATCGGAAG |
| 67326_1 | TATGAGTGTGGAGTCGTTGCGTGTGTGTTTTGGAAGCAGCTATNNNAGATCGGAA |
| 67327_1 | TATGAGTGTGGAGTCGTTGCCCTGAGGGTTTCTTGGTAGCTAANNNAGATCGGAA |
| 67328_1 | TATGAGTGTGGAGTCGTTGCGTTCCCTTAGGTCTTGGGCTAAANNNAGATCGGAAG |
| 67329_1 | TATGAGTGTGGAGTCGTTGCGATCTGAGGGGCACTGATGTTATNNNAGATCGGAA |
| 67330_1 | TATGAGTGTGGAGTCGTTGCATGGTTCTCACTCTATTGCCCAGNNNAGATCGGAAG |
| 67331_1 | TATGAGTGTGGAGTCGTTGCTCTGTTTTTCTGGAGGATGCTGANNNAGATCGGAAG |
| 67332_1 | TATGAGTGTGGAGTCGTTGCCTTATGAGCTCTAGGACCATGGGNNNAGATCGGAA |
| 67333_1 | TATGAGTGTGGAGTCGTTGCGCCCTTCGTTTTTGGTCTAAACANNNAGATCGGAAG |
| 67334_1 | TATGAGTGTGGAGTCGTTGCATGCAAAGTAAGCAAATGGGGAGNNNAGATCGGA |
| 67335_1 | TATGAGTGTGGAGTCGTTGCTGAACTATGATTGTGCCACTGGANNNAGATCGGAA |
| 67336_1 | TATGAGTGTGGAGTCGTTGCTTTCGTGTATTTCAAGTGAGCCCNNNAGATCGGAAG |
| 67337_1 | TATGAGTGTGGAGTCGTTGCGAGCATGTGAAAAGAGAGGGAGANNNAGATCGGA |
| 67338_1 | TATGAGTGTGGAGTCGTTGCATAAACCGCAAGGGAAGGAAAGANNNAGATCGGA |
| 67339_1 | TATGAGTGTGGAGTCGTTGCGCGCGTTGCCCAAAGTTATATAANNNAGATCGGAAG |
| 67341_1 | TATGAGTGTGGAGTCGTTGCCCCTGCGTGTTCTAAACCTAGATNNNAGATCGGAAG |

| | |
|---|---|
| 67342_1 | TATGAGTGTGGAGTCGTTGCACACACACACACACAGAGACATTNNNAGATCGGAAG |
| 67344_1 | TATGAGTGTGGAGTCGTTGCAACCAGATCCATTTTCTCCCTGTNNNAGATCGGAAGA |
| 67345_1 | TATGAGTGTGGAGTCGTTGCACCACATTGGCCAAGAACATTTTNNNAGATCGGAAG |
| 67347_1 | TATGAGTGTGGAGTCGTTGCATCTTTCAGCCCTGAGTGAGTTTNNNAGATCGGAAG |
| 67348_1 | TATGAGTGTGGAGTCGTTGCTGTAAGATTGCCTTCCAGCACTANNNAGATCGGAAG |
| 67349_1 | TATGAGTGTGGAGTCGTTGCGAGAGCGAATGTGAGCATGAAAGNNNAGATCGGA |
| 67350_1 | TATGAGTGTGGAGTCGTTGCTTTGTGCTTTTACCCAAATCGCANNNAGATCGGAAG |
| 67352_1 | TATGAGTGTGGAGTCGTTGCTAGCACATCCAAAACTCACAGGANNNAGATCGGAAG |
| 67353_1 | TATGAGTGTGGAGTCGTTGCCTCTGTTTCTCTGGAGAGTCACCNNNAGATCGGAAG |
| 67354_1 | TATGAGTGTGGAGTCGTTGCGAAACAGAGGGGGATCAGGTAAGNNNAGATCGGA |
| 67355_1 | TATGAGTGTGGAGTCGTTGCATTGCTGCTGTAACAAGTTACCANNNAGATCGGAAG |
| 67356_1 | TATGAGTGTGGAGTCGTTGCGTCTCCGAACCATTTTGTGTCAGNNNAGATCGGAAG |
| 67359_1 | TATGAGTGTGGAGTCGTTGCTAGCTATATCCCACCAGCTCTCANNNAGATCGGAAG |
| 67360_1 | TATGAGTGTGGAGTCGTTGCTAGATGAGGAAACAGGCACTTCCNNNAGATCGGAA |
| 67361_1 | TATGAGTGTGGAGTCGTTGCTTGCTGAACTGAATATGGGACCANNNAGATCGGAA |
| 67362_1 | TATGAGTGTGGAGTCGTTGCCATCAGTCCTTCACCCAGAGAAANNNAGATCGGAAG |
| 67363_1 | TATGAGTGTGGAGTCGTTGCGTAACAGAGCTAGGACCAGAACCNNNAGATCGGAA |
| 67364_1 | TATGAGTGTGGAGTCGTTGCTGGGTGATCAGGAGTGATTCTTGNNNAGATCGGAA |
| 67365_1 | TATGAGTGTGGAGTCGTTGCATGGAGTCTCATCTGTACCTGGANNNAGATCGGAAG |
| 67366_1 | TATGAGTGTGGAGTCGTTGCAGCCAGATTCTTACCCAGGTTTTNNNAGATCGGAAG |
| 67367_1 | TATGAGTGTGGAGTCGTTGCACAGGGGAATCTCTCTCTCTCTCNNNAGATCGGAAG |
| 67368_1 | TATGAGTGTGGAGTCGTTGCGAGGGGTGTGTGTAAGTGTGTTANNNAGATCGGA |
| 67369_1 | TATGAGTGTGGAGTCGTTGCGACAGGCAGCCTAGTAGAAGAATNNNAGATCGGAA |
| 67370_1 | TATGAGTGTGGAGTCGTTGCCAATGTCAGAGTGAGAGAGTCCCNNNAGATCGGAA |
| 67371_1 | TATGAGTGTGGAGTCGTTGCGCCAGTAAACATCCTAAAGTGCCNNNAGATCGGAAG |
| 67372_1 | TATGAGTGTGGAGTCGTTGCGCCCCATTCTCAACACACAATAANNNAGATCGGAAG |
| 67373_1 | TATGAGTGTGGAGTCGTTGCCACCCTTGCACTCCATCTTTTAGNNNAGATCGGAAGA |
| 67374_1 | TATGAGTGTGGAGTCGTTGCCCAGGTTTCAGAGTTTTACAGGCNNNAGATCGGAAG |
| 67375_1 | TATGAGTGTGGAGTCGTTGCACCAGAATCTCAAAAGCCCATCTNNNAGATCGGAAG |
| 67376_1 | TATGAGTGTGGAGTCGTTGCAAATGTACCAATGGCTTCCCTCANNNAGATCGGAAG |
| 67377_1 | TATGAGTGTGGAGTCGTTGCTGTGGTTATTGGTTGGTTTTCCCNNNAGATCGGAAG |
| 67378_1 | TATGAGTGTGGAGTCGTTGCATTTCCTTGGGGTAGGTCATACCNNNAGATCGGAAG |
| 67379_1 | TATGAGTGTGGAGTCGTTGCCATGAAATTTCCCCAGATGCCAANNNAGATCGGAAG |
| 67380_1 | TATGAGTGTGGAGTCGTTGCCAAATCTGGAGTGACAGAGTCCANNNAGATCGGAA |
| 67381_1 | TATGAGTGTGGAGTCGTTGCGTCATTCTTCTCGGTCCCTACTCNNNAGATCGGAAGA |
| 67382_1 | TATGAGTGTGGAGTCGTTGCAGGAGCTCTCTTCCTCTCTTGTCNNNAGATCGGAAG |
| 67383_1 | TATGAGTGTGGAGTCGTTGCGCAGAAGGGTATGAACAGTCAGANNNAGATCGGA |
| 67384_1 | TATGAGTGTGGAGTCGTTGCACAGGAGTGATAAGCTGAGAAGCNNNAGATCGGA |
| 67385_1 | TATGAGTGTGGAGTCGTTGCGCCATACAGCTGTGTACCTCAATNNNAGATCGGAAG |
| 67386_1 | TATGAGTGTGGAGTCGTTGCCAGGAGTATCTCTTACCGTGGTGNNNAGATCGGAA |
| 67387_1 | TATGAGTGTGGAGTCGTTGCCTCAATTACCTACCAAACCCCCANNNAGATCGGAAGA |
| 67388_1 | TATGAGTGTGGAGTCGTTGCAGAGAGGAAGAGAGAGAGAGAGCNNNAGATCGG |
| 67389_1 | TATGAGTGTGGAGTCGTTGCACCAAAGCTGAGCAATTTTGTCANNNAGATCGGAAG |

| | |
|---|---|
| 67390_1 | TATGAGTGTGGAGTCGTTGCTTGTAGTGCAGTAAATGAGGGCTNNNAGATCGGAA |
| 67392_1 | TATGAGTGTGGAGTCGTTGCAGAGAAAACTCAGGCAGTACAGGNNNAGATCGGAA |
| 67393_1 | TATGAGTGTGGAGTCGTTGCAGGCCATCCCTTACATCTTTGAGNNNAGATCGGAAG |
| 67394_1 | TATGAGTGTGGAGTCGTTGCTAACTAATACAGGTGCCTGGCTCNNNAGATCGGAAG |
| 67395_1 | TATGAGTGTGGAGTCGTTGCGCTGTTTCTTAGCACAACTTTGCNNNAGATCGGAAG |
| 67396_1 | TATGAGTGTGGAGTCGTTGCCTGAGGAGTCCCATGATACGATGNNNAGATCGGAA |
| 67397_1 | TATGAGTGTGGAGTCGTTGCGATTGTAGCCTCCTTGAGTTCCANNNAGATCGGAAG |
| 67398_1 | TATGAGTGTGGAGTCGTTGCCTGGATAACGAGTATGGCAGGAANNNAGATCGGAA |
| 67399_1 | TATGAGTGTGGAGTCGTTGCGCCTAGGAATCCACACTGAATCANNNAGATCGGAAG |
| 67400_1 | TATGAGTGTGGAGTCGTTGCAGAGTGCCATATACACTGCTCTGNNNAGATCGGAAG |
| 67401_1 | TATGAGTGTGGAGTCGTTGCATAGATCTGGACAGGGGATCACANNNAGATCGGAA |
| 67402_1 | TATGAGTGTGGAGTCGTTGCATACTCTGAGATCCCTCTAGCCCNNNAGATCGGAAG |
| 67403_1 | TATGAGTGTGGAGTCGTTGCAGAGTCTAGAGTGCCCAAAGAGANNNAGATCGGAA |
| 67404_1 | TATGAGTGTGGAGTCGTTGCGATTCTGAGCTCCATAGGAAGGGNNNAGATCGGAA |
| 67405_1 | TATGAGTGTGGAGTCGTTGCTAGTCTCCTCCCACCTAGGAAACNNNAGATCGGAAG |
| 67407_1 | TATGAGTGTGGAGTCGTTGCAGGGCAGGGACTATGTCTTTTTCNNNAGATCGGAA |
| 67408_1 | TATGAGTGTGGAGTCGTTGCAAGCCAGGTTGAATCTAACTCCTNNNAGATCGGAAG |
| 67409_1 | TATGAGTGTGGAGTCGTTGCCTGAATGCCTGTGCTATTGCTACNNNAGATCGGAAG |
| 67410_1 | TATGAGTGTGGAGTCGTTGCCTATCCCCTGTTCCAAGCTCATTNNNAGATCGGAAGA |
| 67411_1 | TATGAGTGTGGAGTCGTTGCACCACGCTAAGTTCTGTAGGAATNNNAGATCGGAAG |
| 67412_1 | TATGAGTGTGGAGTCGTTGCGGACCTGATGGAGGACATTTTCTNNNAGATCGGAA |
| 67413_1 | TATGAGTGTGGAGTCGTTGCACCTGACCTGAGTCTCCAAGATANNNAGATCGGAAG |
| 67414_1 | TATGAGTGTGGAGTCGTTGCTCAAAGAACGCATCTGAAGGAGANNNAGATCGGAA |
| 67415_1 | TATGAGTGTGGAGTCGTTGCTTCAGACGGAAAGATGGGGAAATNNNAGATCGGAA |
| 67416_1 | TATGAGTGTGGAGTCGTTGCCCAGGCCCATGGTTTATGAAAGANNNAGATCGGAA |
| 67417_1 | TATGAGTGTGGAGTCGTTGCGGGGAAACAATAAATGCTCCCTGNNNAGATCGGAA |
| 67418_1 | TATGAGTGTGGAGTCGTTGCCATGTTCGAGTTGCTTGCCTATTNNNAGATCGGAAG |
| 67419_1 | TATGAGTGTGGAGTCGTTGCGGTTATCCATGACCCTTCCTCTCNNNAGATCGGAAGA |
| 67420_1 | TATGAGTGTGGAGTCGTTGCTCTCAAGAAGTGTTTCCTTGGCTNNNAGATCGGAAG |
| 67421_1 | TATGAGTGTGGAGTCGTTGCGATGTGACAAATCCGAGATGCTGNNNAGATCGGAA |
| 67422_1 | TATGAGTGTGGAGTCGTTGCATCCAAATGTAATGGCAGCATGGNNNAGATCGGAA |
| 67423_1 | TATGAGTGTGGAGTCGTTGCGTGTGTGCTCTCCTCTTCTTGTANNNAGATCGGAAG |
| 67424_1 | TATGAGTGTGGAGTCGTTGCCTGGCAAAGGGATCAACAGATTGNNNAGATCGGAA |
| 67425_1 | TATGAGTGTGGAGTCGTTGCGAGAGATTGGGTCAACTCCGTAANNNAGATCGGAA |
| 67426_1 | TATGAGTGTGGAGTCGTTGCACAATTGGCAAGTAGGCTACTGANNNAGATCGGAA |
| 67427_1 | TATGAGTGTGGAGTCGTTGCTCAAAGATCTGACAGGCAATGGANNNAGATCGGAA |
| 67428_1 | TATGAGTGTGGAGTCGTTGCAAGTGGGTGTAGTGCGATTGTAANNNAGATCGGAA |
| 67429_1 | TATGAGTGTGGAGTCGTTGCTGACCAACGCAGATTGACTATCTNNNAGATCGGAAG |
| 67430_1 | TATGAGTGTGGAGTCGTTGCCATACATCAGCCTGCAGGAAAACNNNAGATCGGAAG |
| 67431_1 | TATGAGTGTGGAGTCGTTGCTCAATGTCTCCAGAGTCATCAGANNNAGATCGGAAG |
| 67432_1 | TATGAGTGTGGAGTCGTTGCTACAAAGATGGCACTGTGACTCANNNAGATCGGAA |
| 67433_1 | TATGAGTGTGGAGTCGTTGCAGAAAAACACACAGAGAGAGGCANNNAGATCGGAA |
| 67434_1 | TATGAGTGTGGAGTCGTTGCTTTCTTTCCTTGCTTTTTGGCCANNNAGATCGGAAGA |

| | |
|---|---|
| 67435_1 | TATGAGTGTGGAGTCGTTGCCGTGCAAGTGAGAGAGAGAGAATNNNAGATCGGA |
| 67436_1 | TATGAGTGTGGAGTCGTTGCAATTGGAGTCTGAGAAGAGCTGGNNNAGATCGGAA |
| 67437_1 | TATGAGTGTGGAGTCGTTGCCAGCTATGTTGTGGGACAAAGTGNNNAGATCGGAA |
| 67438_1 | TATGAGTGTGGAGTCGTTGCTCTGATGTTTGCATTGCACTGAGNNNAGATCGGAAG |
| 67439_1 | TATGAGTGTGGAGTCGTTGCGCTTTCAGGAGTTAGACACCAGANNNAGATCGGAA |
| 67440_1 | TATGAGTGTGGAGTCGTTGCTGATCATGTTTCTGGAGGGTCTCNNNAGATCGGAAG |
| 67441_1 | TATGAGTGTGGAGTCGTTGCTGGAAGGACCTGTTATCTCTCCTNNNAGATCGGAAG |
| 67442_1 | TATGAGTGTGGAGTCGTTGCGAAATGAGAGAGGCCCAATCTCANNNAGATCGGAA |
| 67443_1 | TATGAGTGTGGAGTCGTTGCATTCCCTTCTTATGAGGTCAGGCNNNAGATCGGAAG |
| 67444_1 | TATGAGTGTGGAGTCGTTGCACCACAGTGTTTAGTTTGGGACANNNAGATCGGAA |
| 67445_1 | TATGAGTGTGGAGTCGTTGCTTCTTACATAACCGCAAGGTGGTNNNAGATCGGAAG |
| 67447_1 | TATGAGTGTGGAGTCGTTGCTGCCAAGTTCTGCTTTTTAGAGCNNNAGATCGGAAG |
| 67448_1 | TATGAGTGTGGAGTCGTTGCATCACTACAAGAGCCCCATTCAGNNNAGATCGGAAG |
| 67449_1 | TATGAGTGTGGAGTCGTTGCACCTGGACTTTCTGACTGAATCCNNNAGATCGGAAG |
| 67450_1 | TATGAGTGTGGAGTCGTTGCTAAAGTGAGCTTCCAAGAGGTCCNNNAGATCGGAA |
| 67451_1 | TATGAGTGTGGAGTCGTTGCCTCTCTCTCTGACCATTTCCTGGNNNAGATCGGAAGA |
| 67453_1 | TATGAGTGTGGAGTCGTTGCACCTTTCCAACTCTCCTTCATCCNNNAGATCGGAAGA |
| 67454_1 | TATGAGTGTGGAGTCGTTGCCCATGGGAGCAGAGAATTTTGTCNNNAGATCGGAA |
| 67456_1 | TATGAGTGTGGAGTCGTTGCTGGAGTGCAGTGGTGAAATCATANNNAGATCGGAA |
| 67457_1 | TATGAGTGTGGAGTCGTTGCCAAGAGAGACTGGGAAAGGCTAANNNAGATCGGA |
| 67458_1 | TATGAGTGTGGAGTCGTTGCGGGGACCAGATTCTAGGTGATTCNNNAGATCGGAA |
| 67459_1 | TATGAGTGTGGAGTCGTTGCTGCTCTCTCTCTCTCTCTCTCTCNNNAGATCGGAAGAG |
| 67460_1 | TATGAGTGTGGAGTCGTTGCATCTTTTCCTGAGCTGTGGGATTNNNAGATCGGAAG |
| 67461_1 | TATGAGTGTGGAGTCGTTGCTGGAGGTGAGGTCCAGTAACTATNNNAGATCGGAA |
| 67462_1 | TATGAGTGTGGAGTCGTTGCATTCCTCCCTCACATCTCCCTAANNNAGATCGGAAGA |
| 67463_1 | TATGAGTGTGGAGTCGTTGCTCTGATTCTGTGAGTCTCTCCACNNNAGATCGGAAG |
| 67465_1 | TATGAGTGTGGAGTCGTTGCCTTGAAGAGATCACTGGGGACAANNNAGATCGGAA |
| 67467_1 | TATGAGTGTGGAGTCGTTGCGGCTAATCAGAGTCATTCCCCAANNNAGATCGGAAG |
| 67469_1 | TATGAGTGTGGAGTCGTTGCACCAACAGTAAACTTCTGGGCTANNNAGATCGGAAG |
| 67470_1 | TATGAGTGTGGAGTCGTTGCGGCATAAATCCACCAAGACATGGNNNAGATCGGAA |
| 67471_1 | TATGAGTGTGGAGTCGTTGCTCCTCTTGATGGTTCTGTTTCCCNNNAGATCGGAAGA |
| 67472_1 | TATGAGTGTGGAGTCGTTGCAGGCAGCCTCAATAAACTGTGTANNNAGATCGGAA |
| 67473_1 | TATGAGTGTGGAGTCGTTGCCAACAACAAAACTGTGGATGGGANNNAGATCGGAA |
| 67474_1 | TATGAGTGTGGAGTCGTTGCTAGTCAGCTGAAGCAATTCCACTNNNAGATCGGAAG |
| 67475_1 | TATGAGTGTGGAGTCGTTGCCTTTGCGGCAGTTCTACTTCTTCNNNAGATCGGAAGA |
| 67476_1 | TATGAGTGTGGAGTCGTTGCCACAGGCTTCAAACTTTCTGGAANNNAGATCGGAAG |
| 67477_1 | TATGAGTGTGGAGTCGTTGCTCAGACTCAGAGGCAATGGAAAANNNAGATCGGAA |
| 67478_1 | TATGAGTGTGGAGTCGTTGCGACTAATACACCAATCCCTGGGGNNNAGATCGGAAG |
| 67479_1 | TATGAGTGTGGAGTCGTTGCTAAAAACAGTGAATAGCCCCCCANNNAGATCGGAAG |
| 67480_1 | TATGAGTGTGGAGTCGTTGCGTATCAAGAGCCTGTAGTCCCAGNNNAGATCGGAA |
| 67483_1 | TATGAGTGTGGAGTCGTTGCCCTAGAGCCCATCCAAGTTTTCTNNNAGATCGGAAG |
| 67484_1 | TATGAGTGTGGAGTCGTTGCGAGCACAGAGGAAGGACAAGTAANNNAGATCGGA |
| 67485_1 | TATGAGTGTGGAGTCGTTGCAGAGAGAGAGGAGGCACTAAGTTNNNAGATCGGA |

| | |
|---|---|
| 67487_1 | TATGAGTGTGGAGTCGTTGCAACAGCGAGACTAGTGAAGAGTCNNNAGATCGGAA |
| 67488_1 | TATGAGTGTGGAGTCGTTGCTGCTCAGAGATGGGAAAGACATCNNNAGATCGGAA |
| 67489_1 | TATGAGTGTGGAGTCGTTGCATGTGTGAGTGAGGAAGAAGGTGNNNAGATCGGA |
| 67490_1 | TATGAGTGTGGAGTCGTTGCTCCTATGTTATGCTTCCTGAGGCNNNAGATCGGAAG |
| 67491_1 | TATGAGTGTGGAGTCGTTGCTGTACATGCACACACAGACTTCTNNNAGATCGGAAG |
| 67492_1 | TATGAGTGTGGAGTCGTTGCAACAGCATTGCTTGGCTTCAAATNNNAGATCGGAAG |
| 67493_1 | TATGAGTGTGGAGTCGTTGCGAATACCCACTCTGCCTGTCTAGNNNAGATCGGAAG |
| 67495_1 | TATGAGTGTGGAGTCGTTGCCAAACTCAAGCTCTGCCACTAACNNNAGATCGGAAG |
| 67496_1 | TATGAGTGTGGAGTCGTTGCACAGAGCCTTTACCTGTCTTGTCNNNAGATCGGAAG |
| 67497_1 | TATGAGTGTGGAGTCGTTGCCCAAGCTTAGGCAATAATACGGCNNNAGATCGGAA |
| 67498_1 | TATGAGTGTGGAGTCGTTGCCGCTAGTTGTCAGGTCAAAGAGANNNAGATCGGAA |
| 67499_1 | TATGAGTGTGGAGTCGTTGCCTATGCTCAGTTTTGTTGCCCAANNNAGATCGGAAG |
| 67500_1 | TATGAGTGTGGAGTCGTTGCACTCAAGCTACACTCATTGACGTNNNAGATCGGAAG |
| 67502_1 | TATGAGTGTGGAGTCGTTGCATGGAAGCATCTGGCCTAAATGANNNAGATCGGAA |
| 67504_1 | TATGAGTGTGGAGTCGTTGCAGAGAGAGAGAGAGAGTGTCTGGNNNAGATCGGA |
| 67506_1 | TATGAGTGTGGAGTCGTTGCGGGGTGCTCCTTATCAACATCTTNNNAGATCGGAAG |
| 67507_1 | TATGAGTGTGGAGTCGTTGCACTCAGTTCTCCTTGTGCACATTNNNAGATCGGAAGA |
| 67508_1 | TATGAGTGTGGAGTCGTTGCCACACACACACACACCCTTTATCNNNAGATCGGAAGA |
| 67509_1 | TATGAGTGTGGAGTCGTTGCGAACCCTGACTCATACAGATCCCNNNAGATCGGAAG |
| 67510_1 | TATGAGTGTGGAGTCGTTGCGATTAGGGCCATGATGAGACCATNNNAGATCGGAA |
| 67511_1 | TATGAGTGTGGAGTCGTTGCAGAGAGAAAGAGACTGACCTCCANNNAGATCGGAA |
| 67512_1 | TATGAGTGTGGAGTCGTTGCTTCCGGTAACAATGAACACCTCTNNNAGATCGGAAG |
| 67513_1 | TATGAGTGTGGAGTCGTTGCCTCCAGTTGGTTGAAGGAGGTAANNNAGATCGGAA |
| 67514_1 | TATGAGTGTGGAGTCGTTGCGAAAGACTGGAGGGAAATGCATGNNNAGATCGGA |
| 67515_1 | TATGAGTGTGGAGTCGTTGCACTGCCAGTCTAACCTCATACTGNNNAGATCGGAAG |
| 67517_1 | TATGAGTGTGGAGTCGTTGCACGAGACCCAAAAGCTTTGTTTTNNNAGATCGGAAG |
| 67518_1 | TATGAGTGTGGAGTCGTTGCTTTGTAGCTGGCAATGAAATGCANNNAGATCGGAA |
| 67519_1 | TATGAGTGTGGAGTCGTTGCAATAAAAAACACAGAAGGGCCGGNNNAGATCGGAA |
| 67520_1 | TATGAGTGTGGAGTCGTTGCTCAAGCCATTTTTTTGTCTGCCANNNAGATCGGAAGA |
| 67521_1 | TATGAGTGTGGAGTCGTTGCCTTAGCCTGCATTTGTGGTCTCANNNAGATCGGAAG |
| 67523_1 | TATGAGTGTGGAGTCGTTGCCTCCCAGGAACACCTAGAACTTTNNNAGATCGGAAG |
| 67524_1 | TATGAGTGTGGAGTCGTTGCGGGTGGAGTAACAAGTATCCCTGNNNAGATCGGAA |
| 67525_1 | TATGAGTGTGGAGTCGTTGCAGAGAGAACGAGAAACTCCAAGGNNNAGATCGGAA |
| 67526_1 | TATGAGTGTGGAGTCGTTGCCACTTCAGTGACCTTCTTGTCCTNNNAGATCGGAAGA |
| 67527_1 | TATGAGTGTGGAGTCGTTGCTGGCTATGCTAATTTGCTTGACTNNNAGATCGGAAG |
| 67528_1 | TATGAGTGTGGAGTCGTTGCTACTGGGAAACAGCTTGATGGTTNNNAGATCGGAA |
| 67529_1 | TATGAGTGTGGAGTCGTTGCGTGTGATAACTGCCAAACCCATCNNNAGATCGGAAG |
| 67531_1 | TATGAGTGTGGAGTCGTTGCCTTCACATGAGAGGGTGTCTTGANNNAGATCGGAA |
| 67532_1 | TATGAGTGTGGAGTCGTTGCCCTACTGAGTCAAAGTCAGCAGTNNNAGATCGGAAG |
| 67533_1 | TATGAGTGTGGAGTCGTTGCCCAGGAGAAAGCAGAGATATCCCNNNAGATCGGAA |
| 67534_1 | TATGAGTGTGGAGTCGTTGCGTAAATCCTCAAGTCTCCCCCTCNNNAGATCGGAAGA |
| 67535_1 | TATGAGTGTGGAGTCGTTGCATAAAAGCACAGCCCTCAAGACTNNNAGATCGGAAG |
| 67537_1 | TATGAGTGTGGAGTCGTTGCCCACATCCTCAAGGGTATCCATCNNNAGATCGGAAG |

| | |
|---|---|
| 67538_1 | TATGAGTGTGGAGTCGTTGCGCCTAGAAAGAGAAAAATGGGGCNNNAGATCGGAA |
| 67539_1 | TATGAGTGTGGAGTCGTTGCTTTTCCAGTTCAGGCCAACTAGTNNNAGATCGGAAGA |
| 67540_1 | TATGAGTGTGGAGTCGTTGCTGCGTGTAAGTCTCCTAGGAAACNNNAGATCGGAAG |
| 67542_1 | TATGAGTGTGGAGTCGTTGCTATGGGGCTCTCTTAGACCTTGANNNAGATCGGAAG |
| 67543_1 | TATGAGTGTGGAGTCGTTGCATCAAAGTTCATAAGTCCGCCAGNNNAGATCGGAAG |
| 67544_1 | TATGAGTGTGGAGTCGTTGCAAAGGAAACAGTAGGCTGAGTGTNNNAGATCGGAA |
| 67545_1 | TATGAGTGTGGAGTCGTTGCTTTCACTGAAATTTGTCCTGCCTNNNAGATCGGAAGA |
| 67546_1 | TATGAGTGTGGAGTCGTTGCTTGCAGTGATGAAAATGCTTTCANNNAGATCGGAAG |
| 67547_1 | TATGAGTGTGGAGTCGTTGCGCCCAATCTGCATAAACCATGAANNNAGATCGGAAG |
| 67549_1 | TATGAGTGTGGAGTCGTTGCGCCTCCCAAAGTGCTAGGATTATNNNAGATCGGAAG |
| 67550_1 | TATGAGTGTGGAGTCGTTGCTAGCCTGATCTACCCTCACTTGANNNAGATCGGAAGA |
| 67552_1 | TATGAGTGTGGAGTCGTTGCCTTGAGCACATATACCCCCATGTNNNAGATCGGAAGA |
| 67554_1 | TATGAGTGTGGAGTCGTTGCTCCCCTTCCTCTATCACCACATTNNNAGATCGGAAGAG |
| 67555_1 | TATGAGTGTGGAGTCGTTGCATGCCAACTTTCCTCTTTCCTCTNNNAGATCGGAAGAG |
| 67556_1 | TATGAGTGTGGAGTCGTTGCAAGTTTTGTTCTTCCCCCTCCTTNNNAGATCGGAAGAG |
| 67557_1 | TATGAGTGTGGAGTCGTTGCAGTACAGAACTGAGAACTGAGGCNNNAGATCGGAA |
| 67558_1 | TATGAGTGTGGAGTCGTTGCCGACACAGCACTGAGAAAAACAANNNAGATCGGAAG |
| 67559_1 | TATGAGTGTGGAGTCGTTGCCCAATAGCCAGTAAGGACCTGAGNNNAGATCGGAAG |
| 67560_1 | TATGAGTGTGGAGTCGTTGCATTGTCTCTTCAACCCTGGAGTCNNNAGATCGGAAGA |
| 67561_1 | TATGAGTGTGGAGTCGTTGCTTAGTATGGAACACGGCTCTGTCNNNAGATCGGAAG |
| 67562_1 | TATGAGTGTGGAGTCGTTGCACCACAATTTACTTTGGCTTCCANNNAGATCGGAAGA |
| 67563_1 | TATGAGTGTGGAGTCGTTGCCACAGAGAAGGGAAGTTCGAAGANNNAGATCGGAA |
| 67565_1 | TATGAGTGTGGAGTCGTTGCCCCTTTTGGAAGTAGGTACCAGTNNNAGATCGGAAG |
| 67566_1 | TATGAGTGTGGAGTCGTTGCTCCAGTACGCATTGCATTTCTTGNNNAGATCGGAAGA |
| 67567_1 | TATGAGTGTGGAGTCGTTGCCTTGTGGTCTCAGGAACATCTCANNNAGATCGGAAG |
| 67568_1 | TATGAGTGTGGAGTCGTTGCAAAACTAAGTGGAAACATGCCCGNNNAGATCGGAAG |
| 67569_1 | TATGAGTGTGGAGTCGTTGCTGAGGCAGGATGGAACACAATAANNNAGATCGGAA |
| 67570_1 | TATGAGTGTGGAGTCGTTGCTGACCACCTATGTTGTGACAAACNNNAGATCGGAAG |
| 67571_1 | TATGAGTGTGGAGTCGTTGCGTGGAGAGAGGTCCCAGTTTTAGNNNAGATCGGAA |
| 67572_1 | TATGAGTGTGGAGTCGTTGCGGCTTCAAGGAAAGGCTAGTAGTNNNAGATCGGAA |
| 67573_1 | TATGAGTGTGGAGTCGTTGCTTCATCTTCTGGAAGTGCTGAGTNNNAGATCGGAAG |
| 67574_1 | TATGAGTGTGGAGTCGTTGCCTCTCTCCTTTCTCTCTCTCCCANNNAGATCGGAAGAG |
| 67575_1 | TATGAGTGTGGAGTCGTTGCGTCGAGATTCTGGTGTACCTACCNNNAGATCGGAAG |
| 67576_1 | TATGAGTGTGGAGTCGTTGCAAATGTTGCTGTCTTCATCGTGGNNNAGATCGGAAG |
| 67577_1 | TATGAGTGTGGAGTCGTTGCAGCCTGAATATTTGTGTTCCCCTNNNAGATCGGAAGA |
| 67578_1 | TATGAGTGTGGAGTCGTTGCTCAGCCACCAAAAAAATACAGGTNNNAGATCGGAAG |
| 67579_1 | TATGAGTGTGGAGTCGTTGCGATTTGTCTCTGGAGGACCTGTTNNNAGATCGGAAG |
| 67580_1 | TATGAGTGTGGAGTCGTTGCTCACTTTGTTGCCTATGGTAGCTNNNAGATCGGAAGA |
| 67581_1 | TATGAGTGTGGAGTCGTTGCCTCCCCACATACACACACACATANNNAGATCGGAAGA |
| 67582_1 | TATGAGTGTGGAGTCGTTGCCCTTGAGTTGAGCTTTCGTTCAGNNNAGATCGGAAG |
| 67584_1 | TATGAGTGTGGAGTCGTTGCTGTCACGTTACGAAAACAATCCANNNAGATCGGAAG |
| 67586_1 | TATGAGTGTGGAGTCGTTGCAGTGAGAAAGCTGGCTTCCATTANNNAGATCGGAAG |
| 67587_1 | TATGAGTGTGGAGTCGTTGCTGTTGCCCTTTGAGTCTTGTAGTNNNAGATCGGAAGA |

| | |
|---|---|
| 67588_1 | TATGAGTGTGGAGTCGTTGCATTTGTCTGCTTTCCCCCATAGTNNNAGATCGGAAGA |
| 67589_1 | TATGAGTGTGGAGTCGTTGCCGTCTGTGGGAAGAGTCAAATTGNNNAGATCGGAA |
| 67590_1 | TATGAGTGTGGAGTCGTTGCTGGTCAGGCGAGAATTATTCCAANNNAGATCGGAA |
| 67592_1 | TATGAGTGTGGAGTCGTTGCTTTCGCTCACCATTTGCATTTCTNNNAGATCGGAAGA |
| 67593_1 | TATGAGTGTGGAGTCGTTGCTGTGTGTGGTGTACCTCATTCTTNNNAGATCGGAAG |
| 67594_1 | TATGAGTGTGGAGTCGTTGCTATGTGAAAGGGTGACAGGGAAGNNNAGATCGGA |
| 67595_1 | TATGAGTGTGGAGTCGTTGCGGAAGGGAAGTTGATGTTGATGCNNNAGATCGGA |
| 67596_1 | TATGAGTGTGGAGTCGTTGCGTTCCATTTTGCATGCCTTTTGCNNNAGATCGGAAGA |
| 67597_1 | TATGAGTGTGGAGTCGTTGCTGGATGTACCTCTTTCTTCTGGGNNNAGATCGGAAG |
| 67598_1 | TATGAGTGTGGAGTCGTTGCTTCTCTCCTCCAATCCCAGAAACNNNAGATCGGAAGA |
| 67599_1 | TATGAGTGTGGAGTCGTTGCCGTGTGTGTGTGTGTATGCATATNNNAGATCGGAA |
| 67600_1 | TATGAGTGTGGAGTCGTTGCTGCTCTAAAAAATGGCAATGGCANNNAGATCGGAA |
| 67601_1 | TATGAGTGTGGAGTCGTTGCCACTCTGTCACCTAGGCTGAAATNNNAGATCGGAAG |
| 67602_1 | TATGAGTGTGGAGTCGTTGCGACCAAGCAGTTTTCCCTCTTTCNNNAGATCGGAAGA |
| 67603_1 | TATGAGTGTGGAGTCGTTGCATGATCACAAACAACCCTGCTTGNNNAGATCGGAAG |
| 67604_1 | TATGAGTGTGGAGTCGTTGCATTTCTGCCTCGACATCCTCAATNNNAGATCGGAAGA |
| 67605_1 | TATGAGTGTGGAGTCGTTGCGGAATCAGCTTAGGAGGGTTTCANNNAGATCGGAA |
| 67606_1 | TATGAGTGTGGAGTCGTTGCGGAGGGAGGGAAATATGGTTCAGNNNAGATCGGA |
| 67608_1 | TATGAGTGTGGAGTCGTTGCGTGGTAATAGCTGACACCTCTGTNNNAGATCGGAAG |
| 67609_1 | TATGAGTGTGGAGTCGTTGCATTCAGGGATTCTGCAACTTCCTNNNAGATCGGAAG |
| 67610_1 | TATGAGTGTGGAGTCGTTGCGTTTCTGGCTTTTGGAGGAAGTCNNNAGATCGGAA |
| 67611_1 | TATGAGTGTGGAGTCGTTGCTTTTGAATGACCAGTGCCATCACNNNAGATCGGAAG |
| 67612_1 | TATGAGTGTGGAGTCGTTGCCTGAGGTTTCCGCCATCTTTTTTNNNAGATCGGAAGA |
| 67614_1 | TATGAGTGTGGAGTCGTTGCGCCTCTGCCCAAAGACTAATTCTNNNAGATCGGAAG |
| 67615_1 | TATGAGTGTGGAGTCGTTGCAAATACAAAAATTAGCCGGGCGTNNNAGATCGGAA |
| 67616_1 | TATGAGTGTGGAGTCGTTGCATTTATAGCCTAGTCACTGCGGGNNNAGATCGGAAG |
| 67617_1 | TATGAGTGTGGAGTCGTTGCCTCTTCCTCCTCTTGTCTTCTGGNNNAGATCGGAAGA |
| 67618_1 | TATGAGTGTGGAGTCGTTGCCGCTGGAATCTCTCATTGCAATTNNNAGATCGGAAG |
| 67620_1 | TATGAGTGTGGAGTCGTTGCGGACAACAACACTCAATCCCAAGNNNAGATCGGAAG |
| 67621_1 | TATGAGTGTGGAGTCGTTGCTTCCTATCTCTGTGTCCTAAGCGNNNAGATCGGAAG |
| 67622_1 | TATGAGTGTGGAGTCGTTGCCACACTGTGGAAGCTTTGTTCTTNNNAGATCGGAAG |
| 67623_1 | TATGAGTGTGGAGTCGTTGCTGATTCTGGAGGCTGAAAGTTCANNNAGATCGGAA |
| 67624_1 | TATGAGTGTGGAGTCGTTGCGGCTGAAGGCTTTTGTTCAGATTNNNAGATCGGAA |
| 67625_1 | TATGAGTGTGGAGTCGTTGCTTCAAGTGATTCTCCTGTCTCGGNNNAGATCGGAAG |
| 67627_1 | TATGAGTGTGGAGTCGTTGCCAGAGAAACCAGAGGAACCAGAANNNAGATCGGAA |
| 67629_1 | TATGAGTGTGGAGTCGTTGCTACACCCACAAACATACTCCCTGNNNAGATCGGAAGA |
| 67630_1 | TATGAGTGTGGAGTCGTTGCACAGAGAGCTGATTGGTCTGTTTNNNAGATCGGAA |
| 67631_1 | TATGAGTGTGGAGTCGTTGCTGTGCTAGACTTTTTCATGTGGCNNNAGATCGGAAG |
| 67632_1 | TATGAGTGTGGAGTCGTTGCTTTGTTTTGTTTTGAGGCAGGGTNNNAGATCGGAA |
| 67633_1 | TATGAGTGTGGAGTCGTTGCTAGCTAGGATTACAGGCACATGCNNNAGATCGGAA |
| 67634_1 | TATGAGTGTGGAGTCGTTGCAGAGAAAAAGAGGGAGGGAGAGANNNAGATCGG |
| 67635_1 | TATGAGTGTGGAGTCGTTGCCAATCTTGGCTTACTGCAACCTCNNNAGATCGGAAG |
| 67637_1 | TATGAGTGTGGAGTCGTTGCCAGACTAGAGTGTAGTGGCACAGNNNAGATCGGAA |

| | |
|---|---|
| 67638_1 | TATGAGTGTGGAGTCGTTGCTGTGTGTCAGAGAAGGCTTTCTANNNAGATCGGAA |
| 67639_1 | TATGAGTGTGGAGTCGTTGCCTTTTTGAGACAGGGTTTCGCTGNNNAGATCGGAAG |
| 67640_1 | TATGAGTGTGGAGTCGTTGCACCTGTGATTGTCAGCATGGTAANNNAGATCGGAAG |
| 67641_1 | TATGAGTGTGGAGTCGTTGCATTCAAGTGATTCTCCTGCCTCANNNAGATCGGAAG |
| 67642_1 | TATGAGTGTGGAGTCGTTGCTGATAGGTATTTGGGCTGGTTCCNNNAGATCGGAA |
| 67643_1 | TATGAGTGTGGAGTCGTTGCTGTGTTTCCTATCTAGGCCATGGNNNAGATCGGAAG |
| 67644_1 | TATGAGTGTGGAGTCGTTGCCTAGAGTGCAGTGGCTATTCACANNNAGATCGGAAG |
| 67645_1 | TATGAGTGTGGAGTCGTTGCAGAGAGAGGGAAAGAAGGAAGGANNNAGATCGG |
| 67646_1 | TATGAGTGTGGAGTCGTTGCTCTGTCCCAACATGAAGGCAATANNNAGATCGGAAG |
| 67647_1 | TATGAGTGTGGAGTCGTTGCCATTCTTTTCCATTTGGCCCTGTNNNAGATCGGAAGA |
| 67648_1 | TATGAGTGTGGAGTCGTTGCTCTGTCCCAACATGAAGGCAATANNNAGATCGGAAG |
| 67649_1 | TATGAGTGTGGAGTCGTTGCTTTGAGACAGCGTCTGTGTTAGTNNNAGATCGGAAG |
| 67650_1 | TATGAGTGTGGAGTCGTTGCTTTGTTTAGTGTGTGTGTGTGCANNNAGATCGGAAG |
| 67651_1 | TATGAGTGTGGAGTCGTTGCATGGACATTTGGGCTAGTTCTGTNNNAGATCGGAAG |
| 67652_1 | TATGAGTGTGGAGTCGTTGCAATGTCTCTCTGGAAATAGGCCCNNNAGATCGGAAG |
| 67653_1 | TATGAGTGTGGAGTCGTTGCTGTAATGCCCGTTTTACTGGAGANNNAGATCGGAAG |
| 67654_1 | TATGAGTGTGGAGTCGTTGCAACACACTAGGCATCCACATCAANNNAGATCGGAAG |
| 67657_1 | TATGAGTGTGGAGTCGTTGCCTTGGTTCTGTTCCTCTGGAGAANNNAGATCGGAAG |
| 67659_1 | TATGAGTGTGGAGTCGTTGCATGCCTGGACAGAACTAGACAAGNNNAGATCGGAA |
| 67660_1 | TATGAGTGTGGAGTCGTTGCCACATCTCACTCTATCACCCAGGNNNAGATCGGAAGA |
| 67661_1 | TATGAGTGTGGAGTCGTTGCGCAGAGTCCTTTTGTGTGGTAACNNNAGATCGGAAG |
| 67662_1 | TATGAGTGTGGAGTCGTTGCCATCATCTCCTCCTCTTTTCCCCNNNAGATCGGAAGAG |
| 67663_1 | TATGAGTGTGGAGTCGTTGCCAGAGAGAGAGAGACTGCAACAGNNNAGATCGGAA |
| 67664_1 | TATGAGTGTGGAGTCGTTGCGAATAAAACAGAGATGCTGCCCCNNNAGATCGGAAG |
| 67666_1 | TATGAGTGTGGAGTCGTTGCACAATCTTGGCTCCCTACAACTTNNNAGATCGGAAGA |
| 67668_1 | TATGAGTGTGGAGTCGTTGCAGGCAGGAGAATAATGTGAACCCNNNAGATCGGAA |
| 67669_1 | TATGAGTGTGGAGTCGTTGCTATACACACACACCCTAAGCACCNNNAGATCGGAAGA |
| 67670_1 | TATGAGTGTGGAGTCGTTGCCTACCTTTGCCAACCTGAAAAGGNNNAGATCGGAAG |
| 67671_1 | TATGAGTGTGGAGTCGTTGCCGCCTATAATCCCAGCTACTCAGNNNAGATCGGAAG |
| 67672_1 | TATGAGTGTGGAGTCGTTGCGAGGAACTCCCCACAAAGAAGTANNNAGATCGGAA |
| 67674_1 | TATGAGTGTGGAGTCGTTGCCCTGGGTGACTGAAACATTTGTCNNNAGATCGGAAG |
| 67675_1 | TATGAGTGTGGAGTCGTTGCGTTTCTGTAACTGGTCACATGGCNNNAGATCGGAAG |
| 67676_1 | TATGAGTGTGGAGTCGTTGCCCTCCTGTGAGAGATAGCCCTAANNNAGATCGGAAG |
| 67677_1 | TATGAGTGTGGAGTCGTTGCTGATAGATTGGGTTATGTCTCTGGANNNAGATCGG |
| 67678_1 | TATGAGTGTGGAGTCGTTGCGGTAGAAGAATCATCTGAGCCCANNNAGATCGGAA |
| 67679_1 | TATGAGTGTGGAGTCGTTGCCCACAGCAAGGATATTTGTGTGGNNNAGATCGGAA |
| 67680_1 | TATGAGTGTGGAGTCGTTGCTATTGTCTAGCATCCTTGGCTCCNNNAGATCGGAAG |
| 67681_1 | TATGAGTGTGGAGTCGTTGCATATTGCAGCAACATGGATGCAANNNAGATCGGAAG |
| 67682_1 | TATGAGTGTGGAGTCGTTGCCTGGAGTGAAGTGGCATGATCTCNNNAGATCGGAA |
| 67683_1 | TATGAGTGTGGAGTCGTTGCCACCATGTCACTACTCCACTCTTNNNAGATCGGAAGA |
| 67684_1 | TATGAGTGTGGAGTCGTTGCATATCGAATATGGCTGGGCAACANNNAGATCGGAA |
| 67685_1 | TATGAGTGTGGAGTCGTTGCGCATGTGCCTGTAGTTCTAGCTANNNAGATCGGAAG |
| 67686_1 | TATGAGTGTGGAGTCGTTGCTGCTCTCAAATTCCTGGGATCAANNNAGATCGGAAG |

| | |
|---|---|
| 67687_1 | TATGAGTGTGGAGTCGTTGCAGTTAGCTGGGCATGGTAGTATGNNNAGATCGGAA |
| 67690_1 | TATGAGTGTGGAGTCGTTGCATATACACCCACACACACAGACANNNAGATCGGAAG |
| 67691_1 | TATGAGTGTGGAGTCGTTGCATCTAGTGAGTAGAGGCCAAGGANNNAGATCGGAA |
| 67692_1 | TATGAGTGTGGAGTCGTTGCTCATGGATTTTTTTCACCGCACANNNAGATCGGAAG |
| 67693_1 | TATGAGTGTGGAGTCGTTGCTAATCCTAGCTACTCAGGAGGCTNNNAGATCGGAAG |
| 67694_1 | TATGAGTGTGGAGTCGTTGCGGATTATACCACTCCCCTCCAACNNNAGATCGGAAG |
| 67695_1 | TATGAGTGTGGAGTCGTTGCTCAAAACAACACACACACACACANNNAGATCGGAAG |
| 67697_1 | TATGAGTGTGGAGTCGTTGCGCCAAACTCACTTCCTTCCAAATNNNAGATCGGAAGA |
| 67698_1 | TATGAGTGTGGAGTCGTTGCCAAAGCTCATCAAAACCACTCCCNNNAGATCGGAAG |
| 67699_1 | TATGAGTGTGGAGTCGTTGCGCCAAAAATAAGCGAATGGGGATNNNAGATCGGAA |
| 67700_1 | TATGAGTGTGGAGTCGTTGCCAAACCTTTGTCGTTCAGTCTGGNNNAGATCGGAAG |
| 67701_1 | TATGAGTGTGGAGTCGTTGCGCCAACATGATGAAACCTTCCTTNNNAGATCGGAAG |
| 67702_1 | TATGAGTGTGGAGTCGTTGCACTAAAACACATTTCCCCACCCTNNNAGATCGGAAGA |
| 67703_1 | TATGAGTGTGGAGTCGTTGCCTCAAACTCCTGACCTCCTGATCNNNAGATCGGAAGA |
| 67704_1 | TATGAGTGTGGAGTCGTTGCTGGGGAATGTTTCTAAGTCCAGGNNNAGATCGGAA |
| 67705_1 | TATGAGTGTGGAGTCGTTGCCACACATACACACACACACACATNNNAGATCGGAAG |
| 67709_1 | TATGAGTGTGGAGTCGTTGCTAGACCGCATATACAACAGCAGTNNNAGATCGGAAG |
| 67710_1 | TATGAGTGTGGAGTCGTTGCGGTCATCTGAGTCCTCCAACTTTNNNAGATCGGAAG |
| 67711_1 | TATGAGTGTGGAGTCGTTGCAGAATTGCTTTGGCTATTCAGGCNNNAGATCGGAAG |
| 67714_1 | TATGAGTGTGGAGTCGTTGCGCTCAAGTGATTCTTCTGCCTCANNNAGATCGGAAG |
| 67716_1 | TATGAGTGTGGAGTCGTTGCTACATTCCCCATTCTAGTGGCACNNNAGATCGGAAG |
| 67717_1 | TATGAGTGTGGAGTCGTTGCAAAATCCCAGACCCTTCACAGTTNNNAGATCGGAAG |
| 67718_1 | TATGAGTGTGGAGTCGTTGCCTCTTGGCTGATTATCTCCCTGGNNNAGATCGGAAG |
| 67719_1 | TATGAGTGTGGAGTCGTTGCTGTGTGTGTGTGTGTGTGGTTATNNNAGATCGGAA |
| 67721_1 | TATGAGTGTGGAGTCGTTGCAAGGCCAGAGAGATATACAAGCGNNNAGATCGGAA |
| 67722_1 | TATGAGTGTGGAGTCGTTGCTACAGAGGCATTTTGTAGGCTCANNNAGATCGGAAG |
| 67723_1 | TATGAGTGTGGAGTCGTTGCAAGTGCTGAGATTACAGGCATGANNNAGATCGGAA |
| 67724_1 | TATGAGTGTGGAGTCGTTGCCACTTCAATAAATCTGGGGGGGANNNAGATCGGAA |
| 67725_1 | TATGAGTGTGGAGTCGTTGCCTGTAGTCCCAGCTACTCAGGAGNNNAGATCGGAAG |
| 67726_1 | TATGAGTGTGGAGTCGTTGCGTTAGGGGGAAGAAGCTCAAGAANNNAGATCGGA |
| 67727_1 | TATGAGTGTGGAGTCGTTGCACCAGAAATCCTTCACTTCTCCCNNNAGATCGGAAGA |
| 67728_1 | TATGAGTGTGGAGTCGTTGCAATAATGGCATTTGCAGCAACCTNNNAGATCGGAAG |
| 67729_1 | TATGAGTGTGGAGTCGTTGCAGGAATACAGCTAACCAGGGAGANNNAGATCGGAA |
| 67730_1 | TATGAGTGTGGAGTCGTTGCTTTTTTTTGAGACGGAGTCTGGCNNNAGATCGGAAG |
| 67732_1 | TATGAGTGTGGAGTCGTTGCCTGTAGTCCCAGCTACTCAGGAGNNNAGATCGGAAG |
| 67733_1 | TATGAGTGTGGAGTCGTTGCGAGTTGAGGGAGGAATGGAAAGTNNNAGATCGGA |
| 67734_1 | TATGAGTGTGGAGTCGTTGCGTCACCCAGGATAGAGTGCAATGNNNAGATCGGAA |
| 67736_1 | TATGAGTGTGGAGTCGTTGCTCCGGGGAGCATATATAAGGCTANNNAGATCGGAA |
| 67737_1 | TATGAGTGTGGAGTCGTTGCATTTCTCCAAGGAGCCCTGATTTNNNAGATCGGAAG |
| 67738_1 | TATGAGTGTGGAGTCGTTGCGCCTTCCAGATTCAAGCAATTCTNNNAGATCGGAAG |
| 67742_1 | TATGAGTGTGGAGTCGTTGCCCCATGAGAACTCTTCCTCCAAANNNAGATCGGAAG |
| 67744_1 | TATGAGTGTGGAGTCGTTGCAAAGAGTTTTGTGGTCAGGCATGNNNAGATCGGAA |
| 67745_1 | TATGAGTGTGGAGTCGTTGCCTCTCTCTCTCTCACACACACACNNNAGATCGGAAGA |

| | |
|---|---|
| 67746_1 | TATGAGTGTGGAGTCGTTGCAGAGAGACACAGAGACAGAGACANNNAGATCGGAA |
| 67747_1 | TATGAGTGTGGAGTCGTTGCTGTCTGTAGTCCCAGCTACTCAGNNNAGATCGGAAG |
| 67748_1 | TATGAGTGTGGAGTCGTTGCCTGAGGTAGGAGAGTCACTGGAANNNAGATCGGAA |
| 67749_1 | TATGAGTGTGGAGTCGTTGCTAGGGTTTTCTGACAAGGAAGGGNNNAGATCGGAA |
| 67750_1 | TATGAGTGTGGAGTCGTTGCAGGATCCCACTGCTATAGGAACANNNAGATCGGAAG |
| 67751_1 | TATGAGTGTGGAGTCGTTGCAATCTCTGCGATCTCACTAACCCNNNAGATCGGAAGA |
| 67752_1 | TATGAGTGTGGAGTCGTTGCAAACACTTGAACTCAGGAGGTGTNNNAGATCGGAAG |
| 67753_1 | TATGAGTGTGGAGTCGTTGCACGGAGACAAGGATTTATGGTCCNNNAGATCGGAAG |
| 67754_1 | TATGAGTGTGGAGTCGTTGCGCACACACACACAACTGTATCTGNNNAGATCGGAAGA |
| 67755_1 | TATGAGTGTGGAGTCGTTGCTTCACCAGCTGCTAAGAAAGTCTNNNAGATCGGAAG |
| 67756_1 | TATGAGTGTGGAGTCGTTGCACACATACACATGAGTTCAGGCTNNNAGATCGGAAG |
| 67757_1 | TATGAGTGTGGAGTCGTTGCCATATGCCCATAGTCCCAGCTACNNNAGATCGGAAGA |
| 67758_1 | TATGAGTGTGGAGTCGTTGCTGGACACAGAAAATCTAACCCGANNNAGATCGGAAG |
| 67759_1 | TATGAGTGTGGAGTCGTTGCGCAAAAACAAAAACCAGCCTCAGNNNAGATCGGAAG |
| 67761_1 | TATGAGTGTGGAGTCGTTGCAACAACCTGGATGGGAGAGATTCNNNAGATCGGAAG |
| 67762_1 | TATGAGTGTGGAGTCGTTGCAAGCAGCTTCTTGAGAGGAGAAANNNAGATCGGAA |
| 67763_1 | TATGAGTGTGGAGTCGTTGCTTCATTGGAGGTGGAATTCTGGTNNNAGATCGGAAG |
| 67765_1 | TATGAGTGTGGAGTCGTTGCGCACCACATCCTGCTAATTTCTGNNNAGATCGGAAGA |
| 67767_1 | TATGAGTGTGGAGTCGTTGCTTGAGAGTTCAGTTCCAGACCACNNNAGATCGGAAG |
| 67768_1 | TATGAGTGTGGAGTCGTTGCATGTTGTTATGGTGGTGGAACCTNNNAGATCGGAAG |
| 67769_1 | TATGAGTGTGGAGTCGTTGCCCAAGCAAGATCCCCTGTAGAATNNNAGATCGGAAG |
| 67771_1 | TATGAGTGTGGAGTCGTTGCCCACAAACACACCCCATTCATACNNNAGATCGGAAGA |
| 67772_1 | TATGAGTGTGGAGTCGTTGCGGAGGGAGAGAAAGAGAGAGAGTNNNAGATCGGA |
| 67773_1 | TATGAGTGTGGAGTCGTTGCCTGCTCTGACCAACAAAACTAGCNNNAGATCGGAAGA |
| 67774_1 | TATGAGTGTGGAGTCGTTGCAGGAATCCTGGCTCTGTAACTTGNNNAGATCGGAAG |
| 67775_1 | TATGAGTGTGGAGTCGTTGCAGATGAATTCAGCAGGGGTTAGGNNNAGATCGGAA |
| 67777_1 | TATGAGTGTGGAGTCGTTGCGTGCCTGCAGTATCAACCATAACNNNAGATCGGAAG |
| 67779_1 | TATGAGTGTGGAGTCGTTGCCTAGGCTAGAGTGCAGTGGTATGNNNAGATCGGAA |
| 67780_1 | TATGAGTGTGGAGTCGTTGCAAATAAAGCAAGAGGAAAGGGCGNNNAGATCGGA |
| 67781_1 | TATGAGTGTGGAGTCGTTGCTGTATTTGCCAGGATGGTCTCAANNNAGATCGGAAG |
| 67782_1 | TATGAGTGTGGAGTCGTTGCGTGCTGGAATTACAGATGTGAGCNNNAGATCGGAAG |
| 67783_1 | TATGAGTGTGGAGTCGTTGCATTTGGCGGTATAGCATTGCTTANNNAGATCGGAAG |
| 67784_1 | TATGAGTGTGGAGTCGTTGCTGCATGTATAGTAGCCCTTACCTNNNAGATCGGAAGA |
| 67785_1 | TATGAGTGTGGAGTCGTTGCACGCAGCCATAAAAAGGAATGAANNNAGATCGGAAG |
| 67787_1 | TATGAGTGTGGAGTCGTTGCAACCTGTCATCTGAATCCTCACCNNNAGATCGGAAGA |
| 67788_1 | TATGAGTGTGGAGTCGTTGCCTGCTGCTCCTAAATTGACAACCNNNAGATCGGAAGA |
| 67789_1 | TATGAGTGTGGAGTCGTTGCTCTTCTGTGATTTCGAGCATTGCNNNAGATCGGAAGA |
| 67790_1 | TATGAGTGTGGAGTCGTTGCTTCAAACGATTCTCCTGCCTCAGNNNAGATCGGAAGA |
| 67791_1 | TATGAGTGTGGAGTCGTTGCCGCGCACATGTATGTAGAGAAAANNNAGATCGGAAG |
| 67792_1 | TATGAGTGTGGAGTCGTTGCTGATCTTGAACTCTTGGGCTCAANNNAGATCGGAAG |
| 67793_1 | TATGAGTGTGGAGTCGTTGCAGGAGTCTTGCTTTATCACCCAGNNNAGATCGGAAG |
| 67794_1 | TATGAGTGTGGAGTCGTTGCACACACACACGCACACATATTTTNNNAGATCGGAAGA |
| 67795_1 | TATGAGTGTGGAGTCGTTGCCTCCCATTCCGAAGAGAAAGGAANNNAGATCGGAAG |

| | |
|---|---|
| 67799_1 | TATGAGTGTGGAGTCGTTGCTGAATGGCTTTGACCAAAATGCTNNNAGATCGGAAG |
| 67800_1 | TATGAGTGTGGAGTCGTTGCCGCAGATGAAGCTACCGTCTATANNNAGATCGGAAG |
| 67802_1 | TATGAGTGTGGAGTCGTTGCAATGCTGACAGAGTGATGGGATTNNNAGATCGGAA |
| 67803_1 | TATGAGTGTGGAGTCGTTGCCTGCAGTGAGTCATGATAGAGCANNNAGATCGGAAG |
| 67804_1 | TATGAGTGTGGAGTCGTTGCGAGCGAGACTGTCTCAAAAACACNNNAGATCGGAAG |
| 67806_1 | TATGAGTGTGGAGTCGTTGCCCCTACAGCCTAACCAAGAAAGTNNNAGATCGGAAG |
| 67807_1 | TATGAGTGTGGAGTCGTTGCGTTACCCTTTGATTCCATGCAGGNNNAGATCGGAAGA |
| 67808_1 | TATGAGTGTGGAGTCGTTGCCACACACACACACAGAGGAAAATNNNAGATCGGAAG |
| 67809_1 | TATGAGTGTGGAGTCGTTGCAAGAACCAAAAACAGGTGAGCACNNNAGATCGGAAG |
| 67811_1 | TATGAGTGTGGAGTCGTTGCAGGGAATCAGAAACGTGTGTTCTNNNAGATCGGAAG |
| 67812_1 | TATGAGTGTGGAGTCGTTGCATAGACACACACACACACACACANNNAGATCGGAAGA |
| 67813_1 | TATGAGTGTGGAGTCGTTGCCTGTCTCTACACACACACACACANNNAGATCGGAAGA |
| 67814_1 | TATGAGTGTGGAGTCGTTGCTTTTGACTTTGCATCCAGAGCTGNNNAGATCGGAAGA |
| 67815_1 | TATGAGTGTGGAGTCGTTGCGTCTTATCCAGCTGCAGATCTCANNNAGATCGGAAGA |
| 67816_1 | TATGAGTGTGGAGTCGTTGCTTGCATGCATTTCTTTTGGGGATNNNAGATCGGAAG |
| 67817_1 | TATGAGTGTGGAGTCGTTGCTTTGACCATGGCTCAAGGAAAACNNNAGATCGGAAG |
| 67818_1 | TATGAGTGTGGAGTCGTTGCTTCTTTAGCCAATCCACTGGTGANNNAGATCGGAAGA |
| 67819_1 | TATGAGTGTGGAGTCGTTGCGCTCCAGTCTTAGGAAACCAGAANNNAGATCGGAAG |
| 67820_1 | TATGAGTGTGGAGTCGTTGCCAGTGACCAGGAGGACTTATGTCNNNAGATCGGAAG |
| 67822_1 | TATGAGTGTGGAGTCGTTGCCAAGTCATCCTCCTATCTCAGCCNNNAGATCGGAAGA |
| 67824_1 | TATGAGTGTGGAGTCGTTGCTCTCAGCTCATTGCAACCTCTACNNNAGATCGGAAGA |
| 67825_1 | TATGAGTGTGGAGTCGTTGCATGTTGGTCATGCTGGTCTTGAANNNAGATCGGAAG |
| 67826_1 | TATGAGTGTGGAGTCGTTGCTCCATAGGGGAAAGGTGAAAAGGNNNAGATCGGAA |
| 67828_1 | TATGAGTGTGGAGTCGTTGCGTGTGTGTGTGTGTGTGTGTAAANNNAGATCGGAA |
| 67830_1 | TATGAGTGTGGAGTCGTTGCGGGTCTCTAAAAAGGAGAGTGGTNNNAGATCGGAA |
| 67831_1 | TATGAGTGTGGAGTCGTTGCTCTTTCACTTTCCCTGTCTAGCCNNNAGATCGGAAGAG |
| 67832_1 | TATGAGTGTGGAGTCGTTGCTGATCTCTTGACCTCTTGATCCGNNNAGATCGGAAGA |
| 67833_1 | TATGAGTGTGGAGTCGTTGCTGGAGTGCAGTGTCACAATCATANNNAGATCGGAAG |
| 67834_1 | TATGAGTGTGGAGTCGTTGCGAGAGAGAGAGAGAGAGTTGGGANNNAGATCGGA |
| 67835_1 | TATGAGTGTGGAGTCGTTGCTGCCTGGCATGTAGAAGATACTCNNNAGATCGGAAG |
| 67837_1 | TATGAGTGTGGAGTCGTTGCGCAAGGGCTATAGGAACTTCTTTNNNAGATCGGAAG |
| 67838_1 | TATGAGTGTGGAGTCGTTGCAACAGAATGAGCATAGTAGGGGCNNNAGATCGGAA |
| 67839_1 | TATGAGTGTGGAGTCGTTGCGAGGGATGGAAAGGCAGTAAGAANNNAGATCGGA |
| 67840_1 | TATGAGTGTGGAGTCGTTGCCAGGAGGCTGAGGTACAAGAATTNNNAGATCGGAA |
| 67841_1 | TATGAGTGTGGAGTCGTTGCCTTTGTGCAGCAGCTGTGATTATNNNAGATCGGAAG |
| 67843_1 | TATGAGTGTGGAGTCGTTGCGAGTCACACATTGCCCAAATTCANNNAGATCGGAAGA |
| 67844_1 | TATGAGTGTGGAGTCGTTGCACCTTTATTGGCTGTCTTCCCTTNNNAGATCGGAAGA |
| 67848_1 | TATGAGTGTGGAGTCGTTGCAGCAACATGGATGGAACTAGAGGNNNAGATCGGAA |
| 67849_1 | TATGAGTGTGGAGTCGTTGCAACAAGCTGATCAAAAGATGGGCNNNAGATCGGAAG |
| 67850_1 | TATGAGTGTGGAGTCGTTGCACAAAATGGATGGAATTGGAGGCNNNAGATCGGAA |
| 67851_1 | TATGAGTGTGGAGTCGTTGCTAAGGCCTAGTGCAAGCATTGTANNNAGATCGGAAG |
| 67852_1 | TATGAGTGTGGAGTCGTTGCACAGACATACAGCAGGTAACACCNNNAGATCGGAAG |
| 67853_1 | TATGAGTGTGGAGTCGTTGCACAGACATACAGCAGGTAACACCNNNAGATCGGAAG |

| | |
|---|---|
| 67854_1 | TATGAGTGTGGAGTCGTTGCATGCCTGTAATACCAGCTACTCGNNNAGATCGGAAG |
| 67855_1 | TATGAGTGTGGAGTCGTTGCTTTCACCAATTCATGGAGCCAACNNNAGATCGGAAG |
| 67856_1 | TATGAGTGTGGAGTCGTTGCACCTTCTGTGTACCTACAACAGGNNNAGATCGGAAG |
| 67857_1 | TATGAGTGTGGAGTCGTTGCCGCACGCAAAAAGAAAGAAGACTNNNAGATCGGAA |
| 67858_1 | TATGAGTGTGGAGTCGTTGCAGAATTGAACTGAACGGTTTGGCNNNAGATCGGAA |
| 67859_1 | TATGAGTGTGGAGTCGTTGCCCATGGGATACTATGCAGCCATANNNAGATCGGAAG |
| 67860_1 | TATGAGTGTGGAGTCGTTGCGAGCTAGAGGGGATCCTGAATTGNNNAGATCGGAA |
| 67861_1 | TATGAGTGTGGAGTCGTTGCTACCGTACCCTACATCACCTAGGNNNAGATCGGAAG |
| 67862_1 | TATGAGTGTGGAGTCGTTGCAAATAGGATTGCAACTGCTGTGCNNNAGATCGGAA |
| 67863_1 | TATGAGTGTGGAGTCGTTGCGAGAAATTTGGGGGCTGATGAAGNNNAGATCGGA |
| 67864_1 | TATGAGTGTGGAGTCGTTGCAATAATGGCATTTGCAGCAACCTNNNAGATCGGAAG |
| 67866_1 | TATGAGTGTGGAGTCGTTGCACACACACATATATGTACACATACACNNNAGATCGGA |
| 67867_1 | TATGAGTGTGGAGTCGTTGCTGTGATGGAATACTACTCAGCCANNNAGATCGGAAG |
| 67868_1 | TATGAGTGTGGAGTCGTTGCAGATATTTATTCCCCCTCCCCCTNNNAGATCGGAAGA |
| 67869_1 | TATGAGTGTGGAGTCGTTGCAGGGTTAGTACCATTTGCCTCTGNNNAGATCGGAAG |
| 67870_1 | TATGAGTGTGGAGTCGTTGCGCAGTTGTTGAGTGCTTTTCTCTNNNAGATCGGAAG |
| 67872_1 | TATGAGTGTGGAGTCGTTGCGGACTAATTTCCTTCCATTGGCANNNAGATCGGAAG |
| 67873_1 | TATGAGTGTGGAGTCGTTGCCTATCCCCTAAGAGCAAAGGCATNNNAGATCGGAAG |
| 67875_1 | TATGAGTGTGGAGTCGTTGCTGTGTGTGGTGCATTGTGATTTCNNNAGATCGGAA |
| 67877_1 | TATGAGTGTGGAGTCGTTGCTGAGGAAAAACGTAGAAGCTGGTNNNAGATCGGAA |
| 67880_1 | TATGAGTGTGGAGTCGTTGCTGAGACAGAGTCTAGCTGTTGTCNNNAGATCGGAA |
| 67881_1 | TATGAGTGTGGAGTCGTTGCACCTGCTCCTAAATGACTTTTGGNNNAGATCGGAAG |
| 67882_1 | TATGAGTGTGGAGTCGTTGCTATGTCTGATAACGCCTAGGCACNNNAGATCGGAAG |
| 67884_1 | TATGAGTGTGGAGTCGTTGCAGGTGATGGATATGCCAATTTGCNNNAGATCGGAA |
| 67885_1 | TATGAGTGTGGAGTCGTTGCATCAGGTGATCCAGCAATTCCATNNNAGATCGGAAG |
| 67887_1 | TATGAGTGTGGAGTCGTTGCAGAGAGAGAGAGACCAGTACAGANNNAGATCGGA |
| 67888_1 | TATGAGTGTGGAGTCGTTGCCACACACACATACACTCACACACNNNAGATCGGAAGA |
| 67889_1 | TATGAGTGTGGAGTCGTTGCGGCTAGAGTACAGTGGTGTGATCNNNAGATCGGAA |
| 67890_1 | TATGAGTGTGGAGTCGTTGCCTAGCTATGTAGGAAGCTGAGGCNNNAGATCGGAA |
| 67891_1 | TATGAGTGTGGAGTCGTTGCCTCTCTCTCTCTCTCGCCAATTCNNNAGATCGGAAGA |
| 67893_1 | TATGAGTGTGGAGTCGTTGCAGGCAGGAGAATCATTCAAACCTNNNAGATCGGAA |
| 67894_1 | TATGAGTGTGGAGTCGTTGCAAACCAAAAGATGCTGGTGAGACNNNAGATCGGAA |
| 67896_1 | TATGAGTGTGGAGTCGTTGCTTTCACTCTTGTAGTCCAGGCTGNNNAGATCGGAAG |
| 67897_1 | TATGAGTGTGGAGTCGTTGCCGTGACCTTGACATGTTCTTTCCNNNAGATCGGAAGA |
| 67898_1 | TATGAGTGTGGAGTCGTTGCTCTCTGTCACTTTCGAGCATTCANNNAGATCGGAAGA |
| 67899_1 | TATGAGTGTGGAGTCGTTGCGACAAACACACAGTGGAAGACACNNNAGATCGGAA |
| 67900_1 | TATGAGTGTGGAGTCGTTGCAGAGAGAGAAGTGAGTGAGGAGANNNAGATCGGA |
| 67902_1 | TATGAGTGTGGAGTCGTTGCGACATGATCATATGCCCATCCCTNNNAGATCGGAAG |
| 67903_1 | TATGAGTGTGGAGTCGTTGCCTCTCTCTCTCTCACACACACACNNNAGATCGGAAGA |
| 67904_1 | TATGAGTGTGGAGTCGTTGCGAGATGGATTCTTGCTGTGTTGCNNNAGATCGGAA |
| 67906_1 | TATGAGTGTGGAGTCGTTGCAGGTAGCTGATGGGAACATTTGTNNNAGATCGGAA |
| 67908_1 | TATGAGTGTGGAGTCGTTGCGCTGTCAGCTGATATCCAAATGCNNNAGATCGGAAG |
| 67909_1 | TATGAGTGTGGAGTCGTTGCTTGCCAAAATCCCTTTACTGAGANNNAGATCGGAAG |

| | |
|---|---|
| 67910_1 | TATGAGTGTGGAGTCGTTGCAGGTGCTGCTAAACATCCTACAANNNAGATCGGAAG |
| 67911_1 | TATGAGTGTGGAGTCGTTGCGACAAGGGGTTCTGAGAACTCATNNNAGATCGGAA |
| 67912_1 | TATGAGTGTGGAGTCGTTGCCAGGTCTCAAACTTCTGGACTCANNNAGATCGGAAG |
| 67913_1 | TATGAGTGTGGAGTCGTTGCTTGAGATGGAGTCTGCTCTGTCANNNAGATCGGAAG |
| 67915_1 | TATGAGTGTGGAGTCGTTGCTTTCCCCTTCTTTTCACAGGTGTNNNAGATCGGAAGA |
| 67916_1 | TATGAGTGTGGAGTCGTTGCCTGGTTTCAAGCAATTCTCCCACNNNAGATCGGAAG |
| 67917_1 | TATGAGTGTGGAGTCGTTGCTGTTGTAAGGGAAAGGGATGAGGNNNAGATCGGA |
| 67918_1 | TATGAGTGTGGAGTCGTTGCAAAGGGTGGAAAGGAGAAGTGAGNNNAGATCGGA |
| 67919_1 | TATGAGTGTGGAGTCGTTGCAGTGAGGAAATGAGGTTAGAGCANNNAGATCGGA |
| 67920_1 | TATGAGTGTGGAGTCGTTGCCATGATCATACCACTGCTCTCCANNNAGATCGGAAGA |
| 67921_1 | TATGAGTGTGGAGTCGTTGCGGGTCTTTGCAAACAGGTTCTACNNNAGATCGGAAG |
| 67922_1 | TATGAGTGTGGAGTCGTTGCAATCTCAGCTCACTGGAACCTTCNNNAGATCGGAAG |
| 67923_1 | TATGAGTGTGGAGTCGTTGCACAGAGACAGGAAGAAACACGAANNNAGATCGGAA |
| 67924_1 | TATGAGTGTGGAGTCGTTGCTGCAATTTAGAGACACCTGGCTANNNAGATCGGAAG |
| 67926_1 | TATGAGTGTGGAGTCGTTGCCAAGTGAAATCTTACCAGGCAGTNNNAGATCGGAAG |
| 67928_1 | TATGAGTGTGGAGTCGTTGCATGGAAAAGCATCCCATCCTCATNNNAGATCGGAAG |
| 67929_1 | TATGAGTGTGGAGTCGTTGCGTAATCCCAGCTACTTACTCGGGNNNAGATCGGAAG |
| 67930_1 | TATGAGTGTGGAGTCGTTGCAACCTGTATCTTGCTTCTAGCCANNNAGATCGGAAGA |
| 67931_1 | TATGAGTGTGGAGTCGTTGCTGTTGAAGATCACAGGGCAGTAANNNAGATCGGAA |
| 67933_1 | TATGAGTGTGGAGTCGTTGCACGGAAACCAAACCAAAACAAGTNNNAGATCGGAAG |
| 67934_1 | TATGAGTGTGGAGTCGTTGCCACCATGGAATACTACACAGCCANNNAGATCGGAAG |
| 67935_1 | TATGAGTGTGGAGTCGTTGCAAAAGAAAAACAGCGGCTCACTTNNNAGATCGGAAG |
| 67936_1 | TATGAGTGTGGAGTCGTTGCTATAAAGCTTCCCCCAGTGAAGGNNNAGATCGGAAG |
| 67938_1 | TATGAGTGTGGAGTCGTTGCCAGACAGCATTTGGATCAGCTTTNNNAGATCGGAAG |
| 67939_1 | TATGAGTGTGGAGTCGTTGCGGCTGTTCTATGCACTGTAGGATNNNAGATCGGAAG |
| 67940_1 | TATGAGTGTGGAGTCGTTGCGGCCTACAGCTTCATCCAAATTGNNNAGATCGGAAG |
| 67941_1 | TATGAGTGTGGAGTCGTTGCTGGAGTTCAGTGGTGTGATCTTGNNNAGATCGGAA |
| 67942_1 | TATGAGTGTGGAGTCGTTGCTGCCCAAATAAATTCAGGTCCTCNNNAGATCGGAAG |
| 67943_1 | TATGAGTGTGGAGTCGTTGCAGGGTCATTTGCAGTGACACTAANNNAGATCGGAAG |
| 67945_1 | TATGAGTGTGGAGTCGTTGCACAGTCTGTTTGTGAGAGTGACTNNNAGATCGGAAG |
| 67946_1 | TATGAGTGTGGAGTCGTTGCGGTTGGAGTGTAATGGTGCAATCNNNAGATCGGAA |
| 67947_1 | TATGAGTGTGGAGTCGTTGCTCTTTATTTCAGGGAGAGGGCAGNNNAGATCGGAA |
| 67948_1 | TATGAGTGTGGAGTCGTTGCTGGAACATTATCAGGGAATGGCANNNAGATCGGAA |
| 67949_1 | TATGAGTGTGGAGTCGTTGCCCTTCCTCTCTCCCCCCATATATNNNAGATCGGAAGAG |
| 67950_1 | TATGAGTGTGGAGTCGTTGCTTGGCAAGTATGAAATCTGCAGGNNNAGATCGGAA |
| 67951_1 | TATGAGTGTGGAGTCGTTGCGATCTCGCTGGCTGTTTTTGTAANNNAGATCGGAAG |
| 67952_1 | TATGAGTGTGGAGTCGTTGCCAAGGCTGGTCTCAAACTCCTATNNNAGATCGGAAG |
| 67953_1 | TATGAGTGTGGAGTCGTTGCCACAGGCAGAGCAACCTAATTTTNNNAGATCGGAAG |
| 67954_1 | TATGAGTGTGGAGTCGTTGCTTTAGGCAAATCCACGGTTACTGNNNAGATCGGAAG |
| 67955_1 | TATGAGTGTGGAGTCGTTGCTGTGTGTGTGTGTGTTTTGGTTTNNNAGATCGGAAG |
| 67956_1 | TATGAGTGTGGAGTCGTTGCCACGAGCATGCGATTGGAATATTNNNAGATCGGAAG |
| 67957_1 | TATGAGTGTGGAGTCGTTGCGTAGTTGGAATTACAGGTGCACGNNNAGATCGGAA |
| 67958_1 | TATGAGTGTGGAGTCGTTGCAACACCCCCAAAACTAAATTGCTNNNAGATCGGAAG |

| | |
|---|---|
| 67959_1 | TATGAGTGTGGAGTCGTTGCAGCCAACTGAAAGAGATCCGAATNNNAGATCGGAA |
| 67960_1 | TATGAGTGTGGAGTCGTTGCCCTGAATCCTCCTATGAACTGGGNNNAGATCGGAAG |
| 67961_1 | TATGAGTGTGGAGTCGTTGCATAGCTCGCTGAAATGGCTTTTCNNNAGATCGGAAG |
| 67962_1 | TATGAGTGTGGAGTCGTTGCGATGATGTGGGAGGATCACTTGANNNAGATCGGAA |
| 67963_1 | TATGAGTGTGGAGTCGTTGCACACACACACACACACAACAAAANNNAGATCGGAAG |
| 67964_1 | TATGAGTGTGGAGTCGTTGCAAACATCTGCCTGTCCACTAGAGNNNAGATCGGAAG |
| 67965_1 | TATGAGTGTGGAGTCGTTGCCCAGGACAAGGAAACACAGGATANNNAGATCGGAA |
| 67966_1 | TATGAGTGTGGAGTCGTTGCGACAGAAAGAGCATGCAAGTGTTNNNAGATCGGAA |
| 67967_1 | TATGAGTGTGGAGTCGTTGCGGACCGAGAGTACACTTGGATTTNNNAGATCGGAA |
| 67968_1 | TATGAGTGTGGAGTCGTTGCCACAAACACTCATACAGCAGCTTNNNAGATCGGAAG |
| 67970_1 | TATGAGTGTGGAGTCGTTGCCAAAACACAAGGGAATAGGCCAGNNNAGATCGGAA |
| 67972_1 | TATGAGTGTGGAGTCGTTGCGGGTTTGGGGTGACCTTATCTAANNNAGATCGGAA |
| 67977_1 | TATGAGTGTGGAGTCGTTGCTTGGGAGGCAGAGACATAAGAACNNNAGATCGGAA |
| 67978_1 | TATGAGTGTGGAGTCGTTGCTCTTGAACTCCTGACCTCAAGTGNNNAGATCGGAAG |
| 67979_1 | TATGAGTGTGGAGTCGTTGCTGATACTCTGTGTGTGTGTGTGTNNNAGATCGGAA |
| 67980_1 | TATGAGTGTGGAGTCGTTGCTAGTATATCGCCCAGACTGGAGTNNNAGATCGGAA |
| 67981_1 | TATGAGTGTGGAGTCGTTGCCTCACTACAGCCTCAACATCCTGNNNAGATCGGAAGA |
| 67982_1 | TATGAGTGTGGAGTCGTTGCATTATTATCAGGCTGGGCATGGTNNNAGATCGGAA |
| 67984_1 | TATGAGTGTGGAGTCGTTGCTGTATGTTGAAATCCCAACCCCTNNNAGATCGGAAG |
| 67987_1 | TATGAGTGTGGAGTCGTTGCTTCAGCTTGAATGCTATTGGTGTNNNAGATCGGAAG |
| 67988_1 | TATGAGTGTGGAGTCGTTGCTGAGACAGCATAGCAGGTAATGGNNNAGATCGGAA |
| 67990_1 | TATGAGTGTGGAGTCGTTGCGGCACAATTATAGCTCACTGCAGNNNAGATCGGAAG |
| 67991_1 | TATGAGTGTGGAGTCGTTGCCAACTTTCTGCAAATGTTCGTGCNNNAGATCGGAAG |
| 67993_1 | TATGAGTGTGGAGTCGTTGCGCTTCATTGTTCTCTGGCTGAAANNNAGATCGGAAG |
| 67994_1 | TATGAGTGTGGAGTCGTTGCGAGAGGGATGACAGAAGTCTCAANNNAGATCGGAA |
| 67995_1 | TATGAGTGTGGAGTCGTTGCTGGCTTCAGACCATTTATCTGCANNNAGATCGGAAG |
| 67996_1 | TATGAGTGTGGAGTCGTTGCGACTGAGACAGAGAATGGCTCAANNNAGATCGGAA |
| 67997_1 | TATGAGTGTGGAGTCGTTGCACACACACACGGTTTCATAGGTANNNAGATCGGAAG |
| 67998_1 | TATGAGTGTGGAGTCGTTGCACACACAGGGTTTCATAGGTGAANNNAGATCGGAA |
| 67999_1 | TATGAGTGTGGAGTCGTTGCCATGGATGCTCAAGTCCCTCATANNNAGATCGGAAG |
| 68000_1 | TATGAGTGTGGAGTCGTTGCATTATGTGGGTTCATGAGGAGGGNNNAGATCGGAA |
| 68001_1 | TATGAGTGTGGAGTCGTTGCACATTCCATTTGAGCCTGAAAGCNNNAGATCGGAAG |
| 68003_1 | TATGAGTGTGGAGTCGTTGCAGCTGAACACCAATAACTGATGCNNNAGATCGGAAG |
| 68004_1 | TATGAGTGTGGAGTCGTTGCGCACATGGTACAAAAGAGCCAAANNNAGATCGGAA |
| 68005_1 | TATGAGTGTGGAGTCGTTGCCCTAGGCTGGTTTCAAACTCATGNNNAGATCGGAAG |
| 68008_1 | TATGAGTGTGGAGTCGTTGCAAGCTTTGTCCTGGATGCTTAGANNNAGATCGGAAG |
| 68009_1 | TATGAGTGTGGAGTCGTTGCGCAACCCAAGTGTCCATCAATAGNNNAGATCGGAAG |
| 68011_1 | TATGAGTGTGGAGTCGTTGCGAAGTTTTCCTGAGGCTTCCCTANNNAGATCGGAAG |
| 68012_1 | TATGAGTGTGGAGTCGTTGCCCAGAGCTAAATGACAAGCCATGNNNAGATCGGAA |
| 68013_1 | TATGAGTGTGGAGTCGTTGCCGGTAACAAATACTGAAGGGCTTNNNAGATCGGAA |
| 68015_1 | TATGAGTGTGGAGTCGTTGCGTGACCTCTGAGCTTCCTGTATCNNNAGATCGGAAG |
| 68018_1 | TATGAGTGTGGAGTCGTTGCTGTCACACCTAACTTGTTGCTTTNNNAGATCGGAAGA |
| 68019_1 | TATGAGTGTGGAGTCGTTGCCAAGTAGCTGGGATTACAGGGTCNNNAGATCGGAA |

| | |
|---|---|
| 68020_1 | TATGAGTGTGGAGTCGTTGCCAAAGTGCTGGAATTACAGGCATNNNAGATCGGAA |
| 68021_1 | TATGAGTGTGGAGTCGTTGCACTGAAAGGAAGTGGTAACGTCANNNAGATCGGAA |
| 68022_1 | TATGAGTGTGGAGTCGTTGCCATTGATTGATGGGCATTTGTGCNNNAGATCGGAAG |
| 68023_1 | TATGAGTGTGGAGTCGTTGCACTTTTGCTTCCTGAGTTCAAGCNNNAGATCGGAAG |
| 68024_1 | TATGAGTGTGGAGTCGTTGCTTTTGGATAAGGGGGTGAGGAAGNNNAGATCGGA |
| 68025_1 | TATGAGTGTGGAGTCGTTGCAAGTAGCTGAGATTACAGGTGCCNNNAGATCGGAA |
| 68027_1 | TATGAGTGTGGAGTCGTTGCTCCAGTTGCTCCATATCTTCACCNNNAGATCGGAAGA |
| 68028_1 | TATGAGTGTGGAGTCGTTGCTGATTCTCCTACCTCAGTCTCCCNNNAGATCGGAAGA |
| 68029_1 | TATGAGTGTGGAGTCGTTGCGCAGGTCACATTTGTAAAGCACTNNNAGATCGGAAG |
| 68030_1 | TATGAGTGTGGAGTCGTTGCGGAGAGGAGATGATGCATTCAGANNNAGATCGGA |
| 68031_1 | TATGAGTGTGGAGTCGTTGCTGAGAAATGTGGAGATACGCCTTNNNAGATCGGAA |
| 68033_1 | TATGAGTGTGGAGTCGTTGCGATCTCCGCTCACTATAAGCTCCNNNAGATCGGAAG |
| 68034_1 | TATGAGTGTGGAGTCGTTGCGTGTGTGTGTGTGTGTTGTATNNNAGATCGGAA |
| 68035_1 | TATGAGTGTGGAGTCGTTGCGCCTCTTTGCAGTAATCAATGGGNNNAGATCGGAAG |
| 68036_1 | TATGAGTGTGGAGTCGTTGCTGCTTGACTGCAGTAATCATTCCNNNAGATCGGAAG |
| 68037_1 | TATGAGTGTGGAGTCGTTGCAGGAAATTTACTCGAACCCGGAGNNNAGATCGGAA |
| 68039_1 | TATGAGTGTGGAGTCGTTGCGAGAACAGAGTGAGTGACCACATNNNAGATCGGAA |
| 68040_1 | TATGAGTGTGGAGTCGTTGCCAAGATCGTGTCACTGTACTCCANNNAGATCGGAAG |
| 68041_1 | TATGAGTGTGGAGTCGTTGCCGTTCCCTCTAGTCCCTGAGAAGNNNAGATCGGAAG |
| 68042_1 | TATGAGTGTGGAGTCGTTGCTTTCAGAAAACCTTCACGTCAGCNNNAGATCGGAAG |
| 68043_1 | TATGAGTGTGGAGTCGTTGCGCAAAAGGAATTCACCAGCACATNNNAGATCGGAAG |
| 68044_1 | TATGAGTGTGGAGTCGTTGCCACCACCACCACAACAACAATATNNNAGATCGGAAG |
| 68045_1 | TATGAGTGTGGAGTCGTTGCTCCAGATTGCAGTAAACACCTGANNNAGATCGGAAG |
| 68047_1 | TATGAGTGTGGAGTCGTTGCGGCATGAGTAGTTCAGGCATTTCNNNAGATCGGAAG |
| 68048_1 | TATGAGTGTGGAGTCGTTGCATCCACACACACACATACAGACANNNAGATCGGAAG |
| 68049_1 | TATGAGTGTGGAGTCGTTGCCCCAATGATCCAGTGGTACTGAANNNAGATCGGAAG |
| 68051_1 | TATGAGTGTGGAGTCGTTGCGTGGTGAGAGAAGAAGGTGAGAANNNAGATCGGA |
| 68052_1 | TATGAGTGTGGAGTCGTTGCTCTCGAACTCCTGACCTCTAGTGNNNAGATCGGAAG |
| 68053_1 | TATGAGTGTGGAGTCGTTGCTGCTGTCAAACAAGACTGAGAGANNNAGATCGGAA |
| 68056_1 | TATGAGTGTGGAGTCGTTGCTTTTTTTTTGAGCCGGAGTCTCGNNNAGATCGGAAG |
| 68057_1 | TATGAGTGTGGAGTCGTTGCGAAATGGGGAGAGCATCTGTAGTNNNAGATCGGA |
| 68058_1 | TATGAGTGTGGAGTCGTTGCCTCCCATCCTAGTCCAATCATGGNNNAGATCGGAAG |
| 68059_1 | TATGAGTGTGGAGTCGTTGCGGACCATAAGTCAGACAACTGGTNNNAGATCGGAA |
| 68060_1 | TATGAGTGTGGAGTCGTTGCATATTGGTCAGGCTGGTCTTGAANNNAGATCGGAAG |
| 68062_1 | TATGAGTGTGGAGTCGTTGCTATGTCAGTTGAGTCCAAAGGCANNNAGATCGGAAG |
| 68064_1 | TATGAGTGTGGAGTCGTTGCCTGGGTTCAAGTGATTCTCTTGCNNNAGATCGGAAG |
| 68065_1 | TATGAGTGTGGAGTCGTTGCTGGCATAGTTTCAGCTCACTGTANNNAGATCGGAAG |
| 68067_1 | TATGAGTGTGGAGTCGTTGCTCAGGCTCTCAGACAGACTAGAANNNAGATCGGAAG |
| 68068_1 | TATGAGTGTGGAGTCGTTGCCAAGGCTGTTTATTTCACCTGGGNNNAGATCGGAAG |
| 68069_1 | TATGAGTGTGGAGTCGTTGCGGTTCAAGCGATTCTCTTTCCTCNNNAGATCGGAAG |
| 68070_1 | TATGAGTGTGGAGTCGTTGCTGTGTATGTGTGTGTGTGAATGCNNNAGATCGGAA |
| 68071_1 | TATGAGTGTGGAGTCGTTGCAAAGCATGCTGTGTCACAAATCANNNAGATCGGAAG |
| 68072_1 | TATGAGTGTGGAGTCGTTGCTATGAAGGCAATTGCAAAGAGCCNNNAGATCGGAA |

68073_1 TATGAGTGTGGAGTCGTTGCCCTCTGTATCTGTGGGTTCTGAANNNAGATCGGAAG
68075_1 TATGAGTGTGGAGTCGTTGCGTCTGTCCAGCCTTGACTAATCANNNAGATCGGAAG
68076_1 TATGAGTGTGGAGTCGTTGCCACTCTTCCAGCTTACTGCAGTANNNAGATCGGAAGA
68078_1 TATGAGTGTGGAGTCGTTGCGATGTTTTGGCACACCTCTAGTGNNNAGATCGGAAG
68079_1 TATGAGTGTGGAGTCGTTGCCTGGGTATGAGATGTGTGTGTGTNNNAGATCGGAA
68080_1 TATGAGTGTGGAGTCGTTGCCAGTGAGCTATGATCATGCCACTNNNAGATCGGAAG
68081_1 TATGAGTGTGGAGTCGTTGCCCATCTCTGTGTGTCCGGAATATNNNAGATCGGAAG
68082_1 TATGAGTGTGGAGTCGTTGCCCAACATGGTGAAACCCAATCTCNNNAGATCGGAAG
68083_1 TATGAGTGTGGAGTCGTTGCGTACTGGAACAGTTGGACATCCANNNAGATCGGAA
68084_1 TATGAGTGTGGAGTCGTTGCCAGGAGAATGGCTTGAAAACCTGNNNAGATCGGAA
68085_1 TATGAGTGTGGAGTCGTTGCAACATTCCCAACCCTTGATGGTANNNAGATCGGAAG
68087_1 TATGAGTGTGGAGTCGTTGCAGATCATTAGATAGGGAGGGGCANNNAGATCGGA
68088_1 TATGAGTGTGGAGTCGTTGCATCAATCCTTACAGCATGCATGCNNNAGATCGGAAG
68089_1 TATGAGTGTGGAGTCGTTGCCCCCCCAGCTACAGGAATTATTTNNNAGATCGGAAG
68090_1 TATGAGTGTGGAGTCGTTGCCACAGGGAGGAAGAAGAGACTTCNNNAGATCGGAA
68091_1 TATGAGTGTGGAGTCGTTGCGGTAGTCCCAGCTATTAGGGAGGNNNAGATCGGAA
68093_1 TATGAGTGTGGAGTCGTTGCTCCCAGGTAATTGAATCATGGGGNNNAGATCGGAA
68094_1 TATGAGTGTGGAGTCGTTGCAAGATCATGTCCTTTGCAGCAACNNNAGATCGGAAG
68095_1 TATGAGTGTGGAGTCGTTGCAGCTCCACCCCAATTCCTAAATTNNNAGATCGGAAGA
68096_1 TATGAGTGTGGAGTCGTTGCACACACACACACACACACAAAATNNNAGATCGGAAG
68097_1 TATGAGTGTGGAGTCGTTGCCAGAAACACAGACACACACAGACNNNAGATCGGAAG
68098_1 TATGAGTGTGGAGTCGTTGCACCCTAATACTGAACCTTCCAGCNNNAGATCGGAAGA
68099_1 TATGAGTGTGGAGTCGTTGCGCTCAGGGAAGCAACTAAACTCANNNAGATCGGAA
68100_1 TATGAGTGTGGAGTCGTTGCAGGAGTAAGCCAGAATCCAAGTGNNNAGATCGGAA
68101_1 TATGAGTGTGGAGTCGTTGCGATCCTCCTATTTCAGCCTCCAGNNNAGATCGGAAGA
68103_1 TATGAGTGTGGAGTCGTTGCGGTTTTGCCATTGAAATTGAGGCNNNAGATCGGAA
68104_1 TATGAGTGTGGAGTCGTTGCGCTCTGGTTTGGCCTAAATGTTTNNNAGATCGGAAG
68105_1 TATGAGTGTGGAGTCGTTGCTCTCCTGTCATCATCTGGAAACANNNAGATCGGAAG
68106_1 TATGAGTGTGGAGTCGTTGCTGGGGTGTGGAGAGTAAGATGTANNNAGATCGGA
68107_1 TATGAGTGTGGAGTCGTTGCGGAGAGAGGAAATTTCAGGTCGTNNNAGATCGGAA
68108_1 TATGAGTGTGGAGTCGTTGCACACACACACACGGAACAAATTTNNNAGATCGGAAG
68109_1 TATGAGTGTGGAGTCGTTGCCATTTACTTGTGCTCTTCAGGGCNNNAGATCGGAAG
68110_1 TATGAGTGTGGAGTCGTTGCATAGCTTCAGATCTGAGGCCAAGNNNAGATCGGAA
68111_1 TATGAGTGTGGAGTCGTTGCAAGTGCTGAGATGACAGACGTAGNNNAGATCGGAA
68112_1 TATGAGTGTGGAGTCGTTGCAACATGTCTTCCTAGCTCCACTGNNNAGATCGGAAGA
68113_1 TATGAGTGTGGAGTCGTTGCGTAGTCTAGATCCCACTCCAGGTNNNAGATCGGAAG
68114_1 TATGAGTGTGGAGTCGTTGCGGAGTCTTGCTTTTATTGCCCAGNNNAGATCGGAAG
68115_1 TATGAGTGTGGAGTCGTTGCCTCTAAAAAGCCTCTCCCTCTGGNNNAGATCGGAAGA
68116_1 TATGAGTGTGGAGTCGTTGCACACTCACCTCATTCCCATTGTTNNNAGATCGGAAGA
68117_1 TATGAGTGTGGAGTCGTTGCTACATACATACACACCCTGCCTCNNNAGATCGGAAGA
68118_1 TATGAGTGTGGAGTCGTTGCTGAGTAGCTGGTATGAAGCAGTCNNNAGATCGGAA
68119_1 TATGAGTGTGGAGTCGTTGCGTGGCTGTTCAACGTCAGATAAGNNNAGATCGGAA
68120_1 TATGAGTGTGGAGTCGTTGCTCTCAAGTCAACTTCATCCCTGGNNNAGATCGGAAGA

68121_1 TATGAGTGTGGAGTCGTTGCGTGTGTGTGTGCATGTACATANNNAGATCGGAA
68124_1 TATGAGTGTGGAGTCGTTGCATGCAACAAAGTGATGATGTCTTNNNAGATCGGAA
68125_1 TATGAGTGTGGAGTCGTTGCCAGATGCCTGTAGTCCCAGTTACNNNAGATCGGAAG
68127_1 TATGAGTGTGGAGTCGTTGCGGAGGTAGGGACAATAGCTAAGCNNNAGATCGGA
68128_1 TATGAGTGTGGAGTCGTTGCGAATAGCCCTGCTCTGTCTATGGNNNAGATCGGAAG
68129_1 TATGAGTGTGGAGTCGTTGCATGGAATTTTGCTCTTGTTGCCCNNNAGATCGGAAG
68130_1 TATGAGTGTGGAGTCGTTGCCTGAGGTGGGAAGATCACTTGAGNNNAGATCGGAA
68131_1 TATGAGTGTGGAGTCGTTGCCTCCCAACTGCACCAAAACATAANNNAGATCGGAAG
68133_1 TATGAGTGTGGAGTCGTTGCGCATTTGCTACCAACAAGCTCTANNNAGATCGGAAG
68134_1 TATGAGTGTGGAGTCGTTGCAGAAGAAATGATCGTCACCTTCTNNNAGATCGGAAG
68135_1 TATGAGTGTGGAGTCGTTGCGGCACACACATATCCTCAGACTTNNNAGATCGGAAG
68136_1 TATGAGTGTGGAGTCGTTGCAATACGGAAAGGAGCACAAGAGANNNAGATCGGA
68137_1 TATGAGTGTGGAGTCGTTGCAGAAGGACGTGCTCAAATGTACTNNNAGATCGGAA
68138_1 TATGAGTGTGGAGTCGTTGCCACACAGACACACATACACACACNNNAGATCGGAAG
68139_1 TATGAGTGTGGAGTCGTTGCAGGTCTATTTTCACCACTGGCTTNNNAGATCGGAAG
68140_1 TATGAGTGTGGAGTCGTTGCATGCCTGGACAGAACTAGACAAGNNNAGATCGGAA
68141_1 TATGAGTGTGGAGTCGTTGCGCTCAAGTGATACTCCTGCCTTANNNAGATCGGAAG
68142_1 TATGAGTGTGGAGTCGTTGCGTGAAAAGACACATACGAAGGCCNNNAGATCGGAA
68143_1 TATGAGTGTGGAGTCGTTGCGTCTCTGTCTCCTGGGTTCATTTNNNAGATCGGAAG
68144_1 TATGAGTGTGGAGTCGTTGCTACGTGTGTGTGTGTATACGTGTNNNAGATCGGAA
68146_1 TATGAGTGTGGAGTCGTTGCCATGGATGAAGCTGGAAACCATCNNNAGATCGGAA
68147_1 TATGAGTGTGGAGTCGTTGCGGTAAATGCTGCAGCTACAAACANNNAGATCGGAA
68148_1 TATGAGTGTGGAGTCGTTGCGGAAGCTTTTGCACGTGTGTATTNNNAGATCGGAA
68149_1 TATGAGTGTGGAGTCGTTGCTTCAAGCTATTCTCCTGCCTCAGNNNAGATCGGAAG
68150_1 TATGAGTGTGGAGTCGTTGCGTCACACACATACATGCACTCACNNNAGATCGGAAG
68151_1 TATGAGTGTGGAGTCGTTGCCACACTCACATAACAAACCTGCANNNAGATCGGAAG
68152_1 TATGAGTGTGGAGTCGTTGCAGTCTCGCTTTGTTGTCAGACTANNNAGATCGGAAG
68154_1 TATGAGTGTGGAGTCGTTGCTGGTAGGAAAAGGAGAAAGCACTNNNAGATCGGA
68156_1 TATGAGTGTGGAGTCGTTGCGACTCAGCCAAAGAACAAAGACCNNNAGATCGGAA
68157_1 TATGAGTGTGGAGTCGTTGCTATGGACCAGGAAAGTGGAGAACNNNAGATCGGA
68158_1 TATGAGTGTGGAGTCGTTGCAAGTTCAAGGCTTCAGTGAGCTANNNAGATCGGAA
68159_1 TATGAGTGTGGAGTCGTTGCGTTTGCGCTCCTCCAATCTATTCNNNAGATCGGAAG
68160_1 TATGAGTGTGGAGTCGTTGCTTGTGGGTCAGTGATGCTTATCANNNAGATCGGAA
68161_1 TATGAGTGTGGAGTCGTTGCATTCTTTTGAAGGTGGGCCAATGNNNAGATCGGAA
68162_1 TATGAGTGTGGAGTCGTTGCTGGAGGACCCTGACTAATACACTNNNAGATCGGAAG
68163_1 TATGAGTGTGGAGTCGTTGCTCTCCACAGATGACCACTATCCTNNNAGATCGGAAG
68164_1 TATGAGTGTGGAGTCGTTGCTGGAGGAGAGAGACAGAGATCAGNNNAGATCGGA
68165_1 TATGAGTGTGGAGTCGTTGCCACAACACCCTCTGATACTTCCTNNNAGATCGGAAGA
68166_1 TATGAGTGTGGAGTCGTTGCCCTGTAATACCAGCACTTTGGGANNNAGATCGGAAG
68167_1 TATGAGTGTGGAGTCGTTGCAGATGGAGTCTCTCTCTGTCGTCNNNAGATCGGAAG
68169_1 TATGAGTGTGGAGTCGTTGCGGGTTTTGATTCATTCTGGCCAANNNAGATCGGAAG
68170_1 TATGAGTGTGGAGTCGTTGCGACCACCCTAGGATTTGTGCTATNNNAGATCGGAAG
68171_1 TATGAGTGTGGAGTCGTTGCATAATGGCATTGGCAGAAACCTGNNNAGATCGGAA

| | |
|---|---|
| 68172_1 | TATGAGTGTGGAGTCGTTGCAGGAAAGCATCTAACCAAGGAGGNNNAGATCGGAA |
| 68174_1 | TATGAGTGTGGAGTCGTTGCCAAGAGGGGGTGAACGGATATTTNNNAGATCGGAA |
| 68175_1 | TATGAGTGTGGAGTCGTTGCGTTGGATTTATGCTGCCACAAGTNNNAGATCGGAAG |
| 68176_1 | TATGAGTGTGGAGTCGTTGCACTACTACAACTTCCCCAGGACANNNAGATCGGAAGA |
| 68178_1 | TATGAGTGTGGAGTCGTTGCTCTCTCTCCCCCCCTTACTATANNNAGATCGGAAGA |
| 68179_1 | TATGAGTGTGGAGTCGTTGCTTAGGAGAAATATGGCCAGGAGCNNNAGATCGGAA |
| 68180_1 | TATGAGTGTGGAGTCGTTGCCAAAGTGCTAGGATGACAGGTGTNNNAGATCGGAA |
| 68181_1 | TATGAGTGTGGAGTCGTTGCTTTTTTGAGACTGAGTCTTGCTCNNNAGATCGGAAGA |
| 68182_1 | TATGAGTGTGGAGTCGTTGCGGTGTCCAAAGGATTGAAGACAGNNNAGATCGGAA |
| 68183_1 | TATGAGTGTGGAGTCGTTGCGTGTGTGTTTCAGGGAGAGTCTCNNNAGATCGGAA |
| 68184_1 | TATGAGTGTGGAGTCGTTGCATTTGATGTTCTTGTGGGGAGGANNNAGATCGGAA |
| 68185_1 | TATGAGTGTGGAGTCGTTGCCTGCTTTATGAACCTGGGTGTTCNNNAGATCGGAAG |
| 68186_1 | TATGAGTGTGGAGTCGTTGCAGGGAAAACATTTTTTGAGGCCCNNNAGATCGGAA |
| 68187_1 | TATGAGTGTGGAGTCGTTGCGTTGCAGGATACAAGCCAAACATNNNAGATCGGAA |
| 68188_1 | TATGAGTGTGGAGTCGTTGCTCCAGCCCTCTTTAGCCTTTATGNNNAGATCGGAAGA |
| 68189_1 | TATGAGTGTGGAGTCGTTGCCATGCTCACGGATTGGAAGACTANNNAGATCGGAA |
| 68190_1 | TATGAGTGTGGAGTCGTTGCTCACTTGAGGTCAGAAGTTCGAGNNNAGATCGGAA |
| 68192_1 | TATGAGTGTGGAGTCGTTGCACTCTCTAGCACCTCCATACCTTNNNAGATCGGAAGA |
| 68193_1 | TATGAGTGTGGAGTCGTTGCCAGACGCCTGTAATAGCTACTCTNNNAGATCGGAAGA |
| 68195_1 | TATGAGTGTGGAGTCGTTGCCATTAGACAGAGGGAACCACACANNNAGATCGGAA |
| 68196_1 | TATGAGTGTGGAGTCGTTGCTACCCAACTGAGCATATGCAACTNNNAGATCGGAAGA |
| 68197_1 | TATGAGTGTGGAGTCGTTGCCCCCAACCACCCTATCAAAAGTANNNAGATCGGAAGA |
| 68198_1 | TATGAGTGTGGAGTCGTTGCGCCTTTCCCATCTCTGTAACCTTNNNAGATCGGAAGA |
| 68201_1 | TATGAGTGTGGAGTCGTTGCGATCTCAGCTTACTGCAACCTCTNNNAGATCGGAAGA |
| 68202_1 | TATGAGTGTGGAGTCGTTGCAAAATGCATTCTATGGGCACTGGNNNAGATCGGAA |
| 68204_1 | TATGAGTGTGGAGTCGTTGCGGTTCAAGCGATTCTCCTACTCANNNAGATCGGAAGA |
| 68205_1 | TATGAGTGTGGAGTCGTTGCTATCCTCCCAAAATGCATGTCCANNNAGATCGGAAGA |
| 68208_1 | TATGAGTGTGGAGTCGTTGCGATGTGGCCAGCAAAACCTAAAANNNAGATCGGAA |
| 68209_1 | TATGAGTGTGGAGTCGTTGCCTTTCCAGTGACCACAGAGTCAANNNAGATCGGAAG |
| 68210_1 | TATGAGTGTGGAGTCGTTGCACAAATCCACAAGAGGCATTTGGNNNAGATCGGAA |
| 68211_1 | TATGAGTGTGGAGTCGTTGCACTGGAAACCATTATCCCATGCANNNAGATCGGAAGA |
| 68212_1 | TATGAGTGTGGAGTCGTTGCTCATTTGGGTAGGTTCCAGGTTTNNNAGATCGGAAG |
| 68213_1 | TATGAGTGTGGAGTCGTTGCGCACACCAGAATAACACTGCAAANNNAGATCGGAAG |
| 68214_1 | TATGAGTGTGGAGTCGTTGCTCTCAAAACACACACACACACACNNNAGATCGGAAGA |
| 68215_1 | TATGAGTGTGGAGTCGTTGCCTAGAAGCAGGAGAATCGCTTGANNNAGATCGGAA |
| 68217_1 | TATGAGTGTGGAGTCGTTGCTGGGTAGGAGATGGAGTCACATANNNAGATCGGAA |
| 68218_1 | TATGAGTGTGGAGTCGTTGCTTTCTTTTTCTGGAAAGTGCAGANNNAGATCGGAAG |
| 68219_1 | TATGAGTGTGGAGTCGTTGCGGAGTGAATGTCAGTAGGAAGCANNNAGATCGGAA |
| 68220_1 | TATGAGTGTGGAGTCGTTGCATGTTGCTAGGGTGGTCAAACTCNNNAGATCGGAA |
| 68221_1 | TATGAGTGTGGAGTCGTTGCAGTTTTCAGTTTGGGAGAGGGAGNNNAGATCGGAA |
| 68223_1 | TATGAGTGTGGAGTCGTTGCGGAACAGTGTTGGGGGAAGATATNNNAGATCGGAA |
| 68225_1 | TATGAGTGTGGAGTCGTTGCTGGATCAGGAGGTCAGGAGATAANNNAGATCGGAA |
| 68227_1 | TATGAGTGTGGAGTCGTTGCCCAGAGGGAACCACAATCCATAANNNAGATCGGAA |

| | |
|---|---|
| 68228_1 | TATGAGTGTGGAGTCGTTGCATCCAGCAATTCCAGTTATGGGTNNNAGATCGGAAG |
| 68229_1 | TATGAGTGTGGAGTCGTTGCTGGAGTAAAGTGGTGCAATCTCANNNAGATCGGAA |
| 68230_1 | TATGAGTGTGGAGTCGTTGCAACTCCTGAGCTCAAGTGATCTGNNNAGATCGGAAG |
| 68231_1 | TATGAGTGTGGAGTCGTTGCAGAGATGGTGTTTTGCCATGTTGNNNAGATCGGAA |
| 68232_1 | TATGAGTGTGGAGTCGTTGCTGAGATGGACTCTTGCTCTGTTGNNNAGATCGGAAG |
| 68233_1 | TATGAGTGTGGAGTCGTTGCTGTTAGCCAGGATGGTCTTGATCNNNAGATCGGAA |
| 68234_1 | TATGAGTGTGGAGTCGTTGCTGTGTGTGAGTGTATCAGTGGAGNNNAGATCGGAA |
| 68236_1 | TATGAGTGTGGAGTCGTTGCTCTCCCCAGCATCTAGTTTCATGNNNAGATCGGAAG |
| 68237_1 | TATGAGTGTGGAGTCGTTGCTTTGGGAGGTCAAAGTTGCAATGNNNAGATCGGAA |
| 68238_1 | TATGAGTGTGGAGTCGTTGCTCAATCTAAGTGCCCATCAACCANNNAGATCGGAAG |
| 68240_1 | TATGAGTGTGGAGTCGTTGCGAGTGTGAGGAAGGGACCAGATCNNNAGATCGGA |
| 68242_1 | TATGAGTGTGGAGTCGTTGCCAGGAGAATCTGAGTCCAGGAACNNNAGATCGGAA |
| 68243_1 | TATGAGTGTGGAGTCGTTGCCTGAGGTCAGGATTCGAGACTAGNNNAGATCGGAA |
| 68244_1 | TATGAGTGTGGAGTCGTTGCGCCCAAAAAAATGTGTCCCCTTANNNAGATCGGAAG |
| 68245_1 | TATGAGTGTGGAGTCGTTGCCACACACACACACACGCATATATNNNAGATCGGAAG |
| 68246_1 | TATGAGTGTGGAGTCGTTGCCTGTAATCCCAGCTACTCAGGAGNNNAGATCGGAAG |
| 68247_1 | TATGAGTGTGGAGTCGTTGCATGGTGTTATGTGCCTGTAGTCCNNNAGATCGGAAG |
| 68248_1 | TATGAGTGTGGAGTCGTTGCACAAAAACCAATGGCATTTCTGTNNNAGATCGGAAG |
| 68250_1 | TATGAGTGTGGAGTCGTTGCCAAGAGAAAGCTAACCCCTGACANNNAGATCGGAA |
| 68251_1 | TATGAGTGTGGAGTCGTTGCTGGTTCCTTCCCAATCTATGCTTNNNAGATCGGAAGA |
| 68252_1 | TATGAGTGTGGAGTCGTTGCCGTATACACACACACACACACACNNNAGATCGGAAG |
| 68253_1 | TATGAGTGTGGAGTCGTTGCCACACACACACATACACACACACNNNAGATCGGAAG |
| 68254_1 | TATGAGTGTGGAGTCGTTGCTGAGAGGGCTGGCTACTATTCTANNNAGATCGGAA |
| 68255_1 | TATGAGTGTGGAGTCGTTGCTGGAGTACAGTGGGACAATCATGNNNAGATCGGAA |
| 68258_1 | TATGAGTGTGGAGTCGTTGCCATGATTTTTCCTTGTGGCCCTTNNNAGATCGGAAG |
| 68259_1 | TATGAGTGTGGAGTCGTTGCTGGTAACACACGGATCTTGCTAANNNAGATCGGAAG |
| 68260_1 | TATGAGTGTGGAGTCGTTGCGCCTTTGTAACCAAGGAAAGGACNNNAGATCGGAA |
| 68261_1 | TATGAGTGTGGAGTCGTTGCTCTCTCTCTCTCTCTCTCTCTCANNNAGATCGGAAGAG |
| 68262_1 | TATGAGTGTGGAGTCGTTGCGAGACAGGTCTTACTCTATCGCCNNNAGATCGGAAG |
| 68263_1 | TATGAGTGTGGAGTCGTTGCGGGGTGAGAGAGAGAAAGAGAGANNNAGATCGG |
| 68264_1 | TATGAGTGTGGAGTCGTTGCGCGGAACACTCAGAGAGACATTANNNAGATCGGAA |
| 68265_1 | TATGAGTGTGGAGTCGTTGCATGTATGTGTGTGTGTGTGTGTGNNNAGATCGGAA |
| 68266_1 | TATGAGTGTGGAGTCGTTGCAGTGCTAGGATTACAGGCATGAGNNNAGATCGGAA |
| 68267_1 | TATGAGTGTGGAGTCGTTGCCAAGTAATCCTCCAACTCAGCCTNNNAGATCGGAAG |
| 68269_1 | TATGAGTGTGGAGTCGTTGCTAGAAAATGTGGCTGGCTCATCTNNNAGATCGGAA |
| 68271_1 | TATGAGTGTGGAGTCGTTGCCTTCAAACGGTTCTCTGCCTCAGNNNAGATCGGAAG |
| 68272_1 | TATGAGTGTGGAGTCGTTGCTGCTGGCACCTTGATTTTGAATTNNNAGATCGGAAG |
| 68273_1 | TATGAGTGTGGAGTCGTTGCCCTTTGGAGATGAGAGCTTGGAANNNAGATCGGAA |
| 68275_1 | TATGAGTGTGGAGTCGTTGCGTGTACCTAGGCAGTCAACACAGNNNAGATCGGAA |
| 68276_1 | TATGAGTGTGGAGTCGTTGCACTACCCCTACCATTCCTCTTCANNNAGATCGGAAGA |
| 68277_1 | TATGAGTGTGGAGTCGTTGCTTCAAGTGATTCTCCTGCCTCAGNNNAGATCGGAAG |
| 68278_1 | TATGAGTGTGGAGTCGTTGCGAGCAGCACAGACACACATATGTNNNAGATCGGAA |
| 68279_1 | TATGAGTGTGGAGTCGTTGCTTTTCCATGAAGGCACCAAGAACNNNAGATCGGAAG |

| | |
|---|---|
| 68280_1 | TATGAGTGTGGAGTCGTTGCTGGAGTGCAGTGATGTGATCTTGNNNAGATCGGAA |
| 68281_1 | TATGAGTGTGGAGTCGTTGCACAACATGGATGGAACTAGAGGANNNAGATCGGAA |
| 68283_1 | TATGAGTGTGGAGTCGTTGCAGTAAAACTCGCTCAAGGACAGTNNNAGATCGGAA |
| 68284_1 | TATGAGTGTGGAGTCGTTGCGAAAATGAGGCTGAGACTTGCTGNNNAGATCGGAA |
| 68286_1 | TATGAGTGTGGAGTCGTTGCCCTGTAGTCTCAGCTACTCAGGANNNAGATCGGAAG |
| 68287_1 | TATGAGTGTGGAGTCGTTGCCACACAAGAATGACATGAGGCTGNNNAGATCGGAA |
| 68288_1 | TATGAGTGTGGAGTCGTTGCCCAGGAGACCACAGATACACAAGNNNAGATCGGAA |
| 68289_1 | TATGAGTGTGGAGTCGTTGCGATGGGTCTCTACTGGGTTGAAGNNNAGATCGGAA |
| 68290_1 | TATGAGTGTGGAGTCGTTGCCTCAAACCATTGCACAGGTCATTNNNAGATCGGAAG |
| 68291_1 | TATGAGTGTGGAGTCGTTGCTGCAAGAATAGCCAAGGCAATTCNNNAGATCGGAA |
| 68294_1 | TATGAGTGTGGAGTCGTTGCAGACTTTCCTAACAATAACCCCANNNAGATCGGAAG |
| 68297_1 | TATGAGTGTGGAGTCGTTGCTATAGACAGTGTCTCGCTCTGTGNNNAGATCGGAAG |
| 68298_1 | TATGAGTGTGGAGTCGTTGCCCTCCAACCTACACCACAGTAAANNNAGATCGGAAG |
| 68299_1 | TATGAGTGTGGAGTCGTTGCAGACAGGCCTAAATGCAAAGGTANNNAGATCGGAA |
| 68300_1 | TATGAGTGTGGAGTCGTTGCATGCCTGGACAGAACTAGACAAGNNNAGATCGGAA |
| 68301_1 | TATGAGTGTGGAGTCGTTGCCTGGATCAGATGCCGATAAGTGANNNAGATCGGAA |
| 68302_1 | TATGAGTGTGGAGTCGTTGCTCACTTGAGTCCAGGAGTTCAAGNNNAGATCGGAA |
| 68303_1 | TATGAGTGTGGAGTCGTTGCGTGTCTCTGTGTGCAAAAGTGTTNNNAGATCGGAA |
| 68304_1 | TATGAGTGTGGAGTCGTTGCTTGAGGTACAGGTGGTATTTGGTNNNAGATCGGAA |
| 68305_1 | TATGAGTGTGGAGTCGTTGCGGTGTGGTATCTCATGCCTGTAANNNAGATCGGAA |
| 68307_1 | TATGAGTGTGGAGTCGTTGCCACACACACAACCTTTCAGCTAGNNNAGATCGGAAG |
| 68308_1 | TATGAGTGTGGAGTCGTTGCGAGCCTCAGTTTCCTCATCAGAANNNAGATCGGAAG |
| 68309_1 | TATGAGTGTGGAGTCGTTGCGAGAATTGCTTGAATGTGGGAGGNNNAGATCGGA |
| 68311_1 | TATGAGTGTGGAGTCGTTGCGAGTTCTAGACCCTCCAAAGAGCNNNAGATCGGAA |
| 68312_1 | TATGAGTGTGGAGTCGTTGCAACCCTCTCAAACCATCCAACTTNNNAGATCGGAAGA |
| 68313_1 | TATGAGTGTGGAGTCGTTGCTCATGAGGTCAGGAGTTTGAGACNNNAGATCGGAA |
| 68314_1 | TATGAGTGTGGAGTCGTTGCCTCCTATCTGGCCTCCATTGTTTNNNAGATCGGAAGA |
| 68315_1 | TATGAGTGTGGAGTCGTTGCTCACCATCTGTACCTTGGTTCAGNNNAGATCGGAAG |
| 68316_1 | TATGAGTGTGGAGTCGTTGCAGATGGAGTCTCACTCTGTCACCNNNAGATCGGAAG |
| 68317_1 | TATGAGTGTGGAGTCGTTGCATAATGGGAGAGACAGGAGTCCANNNAGATCGGA |
| 68318_1 | TATGAGTGTGGAGTCGTTGCAAACTCCTGACCTCAAGTGATCCNNNAGATCGGAAG |
| 68320_1 | TATGAGTGTGGAGTCGTTGCTCTTTTTGCAGGGTTGTGGTTTTNNNAGATCGGAAG |
| 68321_1 | TATGAGTGTGGAGTCGTTGCTTGCAAAGACTAGCAGCTTTTCCNNNAGATCGGAAG |
| 68322_1 | TATGAGTGTGGAGTCGTTGCGGAGTACAGTGGTGCCATCATAGNNNAGATCGGAA |
| 68324_1 | TATGAGTGTGGAGTCGTTGCGCAGTGTAGTCTGGAAGAAGTGANNNAGATCGGA |
| 68325_1 | TATGAGTGTGGAGTCGTTGCCACTCGTTGATTGATGGGCATTTNNNAGATCGGAAG |
| 68327_1 | TATGAGTGTGGAGTCGTTGCGCTAAAGGAATCCTCCCACTTCANNNAGATCGGAAG |
| 68328_1 | TATGAGTGTGGAGTCGTTGCATAAAAAGAGAGGCAGTGGAGGGNNNAGATCGGA |
| 68331_1 | TATGAGTGTGGAGTCGTTGCCCTCTCTTGGACTCTGAATGCATNNNAGATCGGAAG |
| 68332_1 | TATGAGTGTGGAGTCGTTGCCCTCCTGGGTTCAAGTGATTCTCNNNAGATCGGAAG |
| 68333_1 | TATGAGTGTGGAGTCGTTGCTAGACATGATCATTGTGCACTGCNNNAGATCGGAAG |
| 68334_1 | TATGAGTGTGGAGTCGTTGCGTGTGTGTGTGTGTGGTCTTTTANNNAGATCGGAA |
| 68335_1 | TATGAGTGTGGAGTCGTTGCATGAAGTATACGGCAGGGATGTCNNNAGATCGGAA |

| | |
|---|---|
| 68338_1 | TATGAGTGTGGAGTCGTTGCCTCAGCTGGGATTACAGGCATAANNNAGATCGGAA |
| 68339_1 | TATGAGTGTGGAGTCGTTGCGTGTCCAAACTCAGTGACTAGGTNNNAGATCGGAA |
| 68340_1 | TATGAGTGTGGAGTCGTTGCGGCACCTGTAATCCCAGTTACTTNNNAGATCGGAAG |
| 68341_1 | TATGAGTGTGGAGTCGTTGCGAGATATCTTAGGGATGGCACCCNNNAGATCGGAA |
| 68342_1 | TATGAGTGTGGAGTCGTTGCTGGAGTAAAATGGCCCAATCTCANNNAGATCGGAA |
| 68343_1 | TATGAGTGTGGAGTCGTTGCAATACCCAAGGTGATATCAGGGTNNNAGATCGGAA |
| 68344_1 | TATGAGTGTGGAGTCGTTGCACACACACATGCACATTTAGTGGNNNAGATCGGAAG |
| 68345_1 | TATGAGTGTGGAGTCGTTGCTAAACACACACACACACACACACNNNAGATCGGAAG |
| 68346_1 | TATGAGTGTGGAGTCGTTGCTTGAAAATCTACCTACTGGGGGCNNNAGATCGGAA |
| 68347_1 | TATGAGTGTGGAGTCGTTGCTTCTTCAGTGAATACCAGGTGGGNNNAGATCGGAA |
| 68349_1 | TATGAGTGTGGAGTCGTTGCGAGGTCTCACTATGTTGTCCAGGNNNAGATCGGAA |
| 68350_1 | TATGAGTGTGGAGTCGTTGCTCCTTTCAGTGTCTACGTCTACGNNNAGATCGGAAG |
| 68351_1 | TATGAGTGTGGAGTCGTTGCGTGACTCATGCCTGTAATCCCAGNNNAGATCGGAAG |
| 68352_1 | TATGAGTGTGGAGTCGTTGCTGTAGTGGCATGATCTTGGTTCANNNAGATCGGAA |
| 68353_1 | TATGAGTGTGGAGTCGTTGCCCAGCATCCACCTCTGTATCAAANNNAGATCGGAAG |
| 68354_1 | TATGAGTGTGGAGTCGTTGCTCTCCATGAACATTGGGACTTCCNNNAGATCGGAAG |
| 68355_1 | TATGAGTGTGGAGTCGTTGCGAGGGCCTGAGTTTTCTTCATCANNNAGATCGGAAG |
| 68356_1 | TATGAGTGTGGAGTCGTTGCTCCAAGAAGAATACTGTGGGGTGNNNAGATCGGAA |
| 68357_1 | TATGAGTGTGGAGTCGTTGCCTCCTTTGAATTCCTATGGCCCTNNNAGATCGGAAGA |
| 68358_1 | TATGAGTGTGGAGTCGTTGCATGTGTTTGGGAGGAGCAGAAAGNNNAGATCGGA |
| 68359_1 | TATGAGTGTGGAGTCGTTGCCTGGGGCTTTTGGTTGTATGTTTNNNAGATCGGAAG |
| 68360_1 | TATGAGTGTGGAGTCGTTGCCTGCATGCCAGTAAAAACTCTGTNNNAGATCGGAAG |
| 68361_1 | TATGAGTGTGGAGTCGTTGCCCCACTCCAAGGCTGTTTATGAANNNAGATCGGAAG |
| 68362_1 | TATGAGTGTGGAGTCGTTGCACCTAAGATGCCAATGAGTAGCCNNNAGATCGGAA |
| 68363_1 | TATGAGTGTGGAGTCGTTGCGAGTACAGTGGTAGGATCACAGCNNNAGATCGGAA |
| 68366_1 | TATGAGTGTGGAGTCGTTGCCTTGAATTTCTGGGCTCAAGCAANNNAGATCGGAAG |
| 68368_1 | TATGAGTGTGGAGTCGTTGCTACAGTATCAACCCAGCAGCAATNNNAGATCGGAAG |
| 68370_1 | TATGAGTGTGGAGTCGTTGCGCCCTTCTGAATTCCCTCTACAANNNAGATCGGAAGA |
| 68371_1 | TATGAGTGTGGAGTCGTTGCTCTTCCTTGTCCCTTCTACCTCTNNNAGATCGGAAGA |
| 68374_1 | TATGAGTGTGGAGTCGTTGCCTGCTTCTCCTACCACCATGATTNNNAGATCGGAAGA |
| 68375_1 | TATGAGTGTGGAGTCGTTGCCCTAGCGGACCTTGATTTACGTCNNNAGATCGGAAG |
| 68376_1 | TATGAGTGTGGAGTCGTTGCATGGCAGGCGTAATAACACTTTGNNNAGATCGGAA |
| 68377_1 | TATGAGTGTGGAGTCGTTGCAAAAATGGTCAACTGTGCGAAGTNNNAGATCGGAA |
| 68378_1 | TATGAGTGTGGAGTCGTTGCATGGAGTGCAGTGGTATGATCTCNNNAGATCGGAA |
| 68380_1 | TATGAGTGTGGAGTCGTTGCGCAAGTCCAAAAATCTGCAGAGTNNNAGATCGGAA |
| 68381_1 | TATGAGTGTGGAGTCGTTGCGAGGGATTGTGTATGCATGTGTGNNNAGATCGGAA |
| 68383_1 | TATGAGTGTGGAGTCGTTGCGAGGAGGTGTCACTGAAAGTCTCNNNAGATCGGAA |
| 68384_1 | TATGAGTGTGGAGTCGTTGCATATATGTGTGTGCGTGTGTGTGNNNAGATCGGAA |
| 68385_1 | TATGAGTGTGGAGTCGTTGCCCTATCCAAGACTACCAAGGTGGNNNAGATCGGAAG |
| 68386_1 | TATGAGTGTGGAGTCGTTGCAGGAGAATGACACCATACGTAGCNNNAGATCGGAA |
| 68387_1 | TATGAGTGTGGAGTCGTTGCTGCCAGGGATCTTGAGGAAAATTNNNAGATCGGAA |
| 68388_1 | TATGAGTGTGGAGTCGTTGCAGACCCCACCTCTCCAAAAAAATNNNAGATCGGAAG |
| 68389_1 | TATGAGTGTGGAGTCGTTGCTTATAAGATACATGAGGCCGGGCNNNAGATCGGAA |

| | |
|---|---|
| 68390_1 | TATGAGTGTGGAGTCGTTGCCTCTCTCTCTGTCTCTCTCCCTCNNNAGATCGGAAGAG |
| 68391_1 | TATGAGTGTGGAGTCGTTGCAGAAAACATGAGCTCTATGGCCTNNNAGATCGGAAG |
| 68394_1 | TATGAGTGTGGAGTCGTTGCATCTCAACTCACTGCAACTTCCANNNAGATCGGAAGA |
| 68395_1 | TATGAGTGTGGAGTCGTTGCTGACGTTGGGTTCTCTAGTTCACNNNAGATCGGAAG |
| 68396_1 | TATGAGTGTGGAGTCGTTGCTCTCTCTATCTCTTGACCTGCCTNNNAGATCGGAAGA |
| 68397_1 | TATGAGTGTGGAGTCGTTGCTTTTTTGGAGACAGAGTCTGGCANNNAGATCGGAAG |
| 68398_1 | TATGAGTGTGGAGTCGTTGCTTGTAAGTTTCCTGAGGCTTCCCNNNAGATCGGAAGA |
| 68399_1 | TATGAGTGTGGAGTCGTTGCATGTGGAAAGTCTCATCCCTCACNNNAGATCGGAAGA |
| 68400_1 | TATGAGTGTGGAGTCGTTGCACTCTGGACTTTCTTCCTCTTCTNNNAGATCGGAAGA |
| 68401_1 | TATGAGTGTGGAGTCGTTGCGAAGGAAGGAAGGAAGGAAGGAANNNAGATCGGA |
| 68403_1 | TATGAGTGTGGAGTCGTTGCTGTCTCTGTCTCTGTCTCTCTGTNNNAGATCGGAAGA |
| 68404_1 | TATGAGTGTGGAGTCGTTGCTACTGGGTTCAAGCAATTCTCCTNNNAGATCGGAAGA |
| 68405_1 | TATGAGTGTGGAGTCGTTGCCACGAGAGTGAGAGAGAGAGAGANNNAGATCGGA |
| 68407_1 | TATGAGTGTGGAGTCGTTGCGAACAATGTCCTAATGTGGGGGANNNAGATCGGAA |
| 68408_1 | TATGAGTGTGGAGTCGTTGCCGCACAAGCAATTCATATTCACANNNAGATCGGAAGA |
| 68410_1 | TATGAGTGTGGAGTCGTTGCTCTTACCCTTTTTGCTGGCATTCNNNAGATCGGAAGA |
| 68412_1 | TATGAGTGTGGAGTCGTTGCGAACTCATCGAGTACCAGGAGAGNNNAGATCGGAAG |
| 68413_1 | TATGAGTGTGGAGTCGTTGCTCAAAAGTTTAGGTGGCTGCTTGNNNAGATCGGAAG |
| 68414_1 | TATGAGTGTGGAGTCGTTGCGAGAGAGAGAGAGTGTGTGTGTGNNNAGATCGGA |
| 68415_1 | TATGAGTGTGGAGTCGTTGCCTTGTTCAACACATCCAGCTAGGNNNAGATCGGAAGA |
| 68416_1 | TATGAGTGTGGAGTCGTTGCTTGCTCATTATTACCTGACTGCANNNAGATCGGAAGA |
| 68417_1 | TATGAGTGTGGAGTCGTTGCATTAGCTTCCAAAGGCCAACCTANNNAGATCGGAAGA |
| 68418_1 | TATGAGTGTGGAGTCGTTGCTAGGGTCTCACTATGTTGCCCAGNNNAGATCGGAAG |
| 68420_1 | TATGAGTGTGGAGTCGTTGCAGAGAGAGAAAGGAAATGGCCAGNNNAGATCGGAA |
| 68423_1 | TATGAGTGTGGAGTCGTTGCCTCCCTCTCTCTCTCTCTCTCTCNNNAGATCGGAAGAG |
| 68424_1 | TATGAGTGTGGAGTCGTTGCGACAGAGAAGAGAGAGAATGGCCNNNAGATCGGAA |
| 68427_1 | TATGAGTGTGGAGTCGTTGCCAGGCTCGTAGAGATGGGAATTANNNAGATCGGAA |
| 68428_1 | TATGAGTGTGGAGTCGTTGCATTTTGCCCATTAGCTCCAATGTNNNAGATCGGAAGA |
| 68429_1 | TATGAGTGTGGAGTCGTTGCCAATCTCGACTCATTGCAACCTCNNNAGATCGGAAGA |
| 68431_1 | TATGAGTGTGGAGTCGTTGCCCCTGGCTTCCTAGTTTTCTCAANNNAGATCGGAAGA |
| 68433_1 | TATGAGTGTGGAGTCGTTGCAAGAAGACCAGAGGCAAGAGATGNNNAGATCGGAA |
| 68434_1 | TATGAGTGTGGAGTCGTTGCCCCCTTCTCTGCTCTGAAGAATANNNAGATCGGAAGA |
| 68436_1 | TATGAGTGTGGAGTCGTTGCACAAAGACAAGATTGAGAGCGGANNNAGATCGGAA |
| 68437_1 | TATGAGTGTGGAGTCGTTGCACAACATGACCATCTATAGCCCCNNNAGATCGGAAGA |
| 68438_1 | TATGAGTGTGGAGTCGTTGCCATATGCACACACACACACACATNNNAGATCGGAAGA |
| 68441_1 | TATGAGTGTGGAGTCGTTGCCAACAAACTGCAAACCAGGAAGANNNAGATCGGAAG |
| 68443_1 | TATGAGTGTGGAGTCGTTGCCTCCACACCTAATCCATCACCAANNNAGATCGGAAGA |
| 68445_1 | TATGAGTGTGGAGTCGTTGCGAGAAGGGAGAGGAGAGAGAAGANNNAGATCGGA |
| 68446_1 | TATGAGTGTGGAGTCGTTGCCAGCAACCTGGATTCAATTGGAGNNNAGATCGGAAG |
| 68447_1 | TATGAGTGTGGAGTCGTTGCACCCCTAACCAGAATCTTGCAATNNNAGATCGGAAGA |
| 68448_1 | TATGAGTGTGGAGTCGTTGCGAGAGAGCGCATACATCTCTGTTNNNAGATCGGAAG |
| 68450_1 | TATGAGTGTGGAGTCGTTGCACCCTGAATGTTCTGAGATCCTGNNNAGATCGGAAG |
| 68451_1 | TATGAGTGTGGAGTCGTTGCGGCAGGAAATCCAGGGTGATTATNNNAGATCGGAA |

| | |
|---|---|
| 68452_1 | TATGAGTGTGGAGTCGTTGCAAGCAATACAAGGCCAGAGGTAANNNAGATCGGAA |
| 68453_1 | TATGAGTGTGGAGTCGTTGCCTCTTCCTGATGGCTGTAGAAGGNNNAGATCGGAAG |
| 68454_1 | TATGAGTGTGGAGTCGTTGCAGAACAGAGAGCTCCAAAACTCTNNNAGATCGGAAG |
| 68455_1 | TATGAGTGTGGAGTCGTTGCAGCTCTGGTCTCTCTCTTCTGATNNNAGATCGGAAG |
| 68456_1 | TATGAGTGTGGAGTCGTTGCACACTCAAGAATGCTTTCTGTGCNNNAGATCGGAAG |
| 68457_1 | TATGAGTGTGGAGTCGTTGCCTATTCTAACCGCTCACCTGTGTNNNAGATCGGAAG |
| 68458_1 | TATGAGTGTGGAGTCGTTGCTTCATTTCCCCGCAAGATTTCACNNNAGATCGGAAGA |
| 68460_1 | TATGAGTGTGGAGTCGTTGCAACGTCTTGTGGAACACTGTAGANNNAGATCGGAA |
| 68461_1 | TATGAGTGTGGAGTCGTTGCGGTGGTGGTGTTCATGAATCATGNNNAGATCGGAA |
| 68462_1 | TATGAGTGTGGAGTCGTTGCTCTCTCCCTCTCTCTCTCTCTCTNNNAGATCGGAAGAG |
| 68464_1 | TATGAGTGTGGAGTCGTTGCGAAAAAACCTACCAGCATGCCTTNNNAGATCGGAAG |
| 68465_1 | TATGAGTGTGGAGTCGTTGCTCAAGGTACAACTGCTAGCAGACNNNAGATCGGAA |
| 68466_1 | TATGAGTGTGGAGTCGTTGCGAGAGAGAGAGAGCACACATCAGNNNAGATCGGA |
| 68467_1 | TATGAGTGTGGAGTCGTTGCCCTCATCAACAGGACTATGGAGGNNNAGATCGGAA |
| 68468_1 | TATGAGTGTGGAGTCGTTGCAAGTGATTATCATGCCTCAGCCTNNNAGATCGGAAG |
| 68469_1 | TATGAGTGTGGAGTCGTTGCCTCTCTTTCTGTCTCTGTCCGTCNNNAGATCGGAAGA |
| 68470_1 | TATGAGTGTGGAGTCGTTGCTTTTTTGAGACGGAGTCTTGCTCNNNAGATCGGAAG |
| 68473_1 | TATGAGTGTGGAGTCGTTGCCACAATCTCAGCTTACTGCAACCNNNAGATCGGAAG |
| 68474_1 | TATGAGTGTGGAGTCGTTGCGAGGGAAGGGAAGGAAAGAAAGANNNAGATCGG |
| 68475_1 | TATGAGTGTGGAGTCGTTGCCGTGGGAAAAATGCTTCTCAGAANNNAGATCGGAA |
| 68476_1 | TATGAGTGTGGAGTCGTTGCTATGCAAGGAGCCAGATAAGCAANNNAGATCGGAA |
| 68478_1 | TATGAGTGTGGAGTCGTTGCTTTGGGAGAGGGGTAAAAAGAGGNNNAGATCGGA |
| 68480_1 | TATGAGTGTGGAGTCGTTGCCTCCTTCTCTCACCAGTCCAATCNNNAGATCGGAAGA |
| 68481_1 | TATGAGTGTGGAGTCGTTGCACTCCAAGGGTTCCAAACTCAATNNNAGATCGGAAG |
| 68482_1 | TATGAGTGTGGAGTCGTTGCACCAGAGGGAAAACAATCCTGAANNNAGATCGGAA |
| 68483_1 | TATGAGTGTGGAGTCGTTGCAATCAGTAGGCACCTTGATCCTGNNNAGATCGGAAG |
| 68484_1 | TATGAGTGTGGAGTCGTTGCTAGGCCCAGTCCTGTAATCCTAANNNAGATCGGAAG |
| 68485_1 | TATGAGTGTGGAGTCGTTGCCTCTGATCTCGTGGAAACACAGANNNAGATCGGAAG |
| 68487_1 | TATGAGTGTGGAGTCGTTGCTCGGTCATTTACATCAAGTCCCCNNNAGATCGGAAG |
| 68488_1 | TATGAGTGTGGAGTCGTTGCGTTCCATGTTTTCTTCTGAGCCCNNNAGATCGGAAG |
| 68489_1 | TATGAGTGTGGAGTCGTTGCTTCTACTTCCATGACATCCCTGCNNNAGATCGGAAGA |
| 68490_1 | TATGAGTGTGGAGTCGTTGCTTGATCTCCACTCCTAGATCCCCNNNAGATCGGAAGA |
| 68492_1 | TATGAGTGTGGAGTCGTTGCACCATTGTCTCTTGCACTACTGTNNNAGATCGGAAG |
| 68494_1 | TATGAGTGTGGAGTCGTTGCTGTGTGTCTCTCTCCCTGTCTATNNNAGATCGGAAGA |
| 68496_1 | TATGAGTGTGGAGTCGTTGCCGCTATGTTGTGAGTAAGCCAAGNNNAGATCGGAA |
| 68497_1 | TATGAGTGTGGAGTCGTTGCAGGGAAGACAAAAGGACAAGAGGNNNAGATCGGA |
| 68498_1 | TATGAGTGTGGAGTCGTTGCACAGAATTGTCCTGCTCCTTGATNNNAGATCGGAAG |
| 68499_1 | TATGAGTGTGGAGTCGTTGCCCACCCGCATATAAAGGTACCATNNNAGATCGGAAG |
| 68500_1 | TATGAGTGTGGAGTCGTTGCCTCTAATTCCCAAAGCTCCCACTNNNAGATCGGAAGA |
| 68501_1 | TATGAGTGTGGAGTCGTTGCCTCCTTGAATGAGGCAGAGTTCTNNNAGATCGGAAG |
| 68502_1 | TATGAGTGTGGAGTCGTTGCCTCTCTCATCTCCACAAGGATGGNNNAGATCGGAAG |
| 68503_1 | TATGAGTGTGGAGTCGTTGCGGCTGCCATGATAATCCTTTTCCNNNAGATCGGAAG |
| 68504_1 | TATGAGTGTGGAGTCGTTGCCTCAGAAAAGAGCTGAAGTTGCCNNNAGATCGGAA |

| | |
|---|---|
| 68505_1 | TATGAGTGTGGAGTCGTTGCGGCTAGCTTGGGTTCATCATAGANNNAGATCGGAAC |
| 68506_1 | TATGAGTGTGGAGTCGTTGCCACACAAACTCACATTCCTCCACNNNAGATCGGAAGA |
| 68507_1 | TATGAGTGTGGAGTCGTTGCCTATTGGCCAAAGCAAGTCACAANNNAGATCGGAAG |
| 68508_1 | TATGAGTGTGGAGTCGTTGCGGGAGAGGAAACTTGATGCTTCTNNNAGATCGGAAC |
| 68509_1 | TATGAGTGTGGAGTCGTTGCAGAAGGAACAGATTGGAGCAGAGNNNAGATCGGAA |
| 68512_1 | TATGAGTGTGGAGTCGTTGCCCTTCTGAGTCTATGGCTGGAAANNNAGATCGGAAG |
| 68513_1 | TATGAGTGTGGAGTCGTTGCCGTTTGAGGGGATAGGTAAGCATNNNAGATCGGAA |
| 68514_1 | TATGAGTGTGGAGTCGTTGCCAGTATCTGTGAGGGACTGGTTCNNNAGATCGGAAC |
| 68515_1 | TATGAGTGTGGAGTCGTTGCGGACCCTCATGTCATACTCCTTCNNNAGATCGGAAGA |
| 68516_1 | TATGAGTGTGGAGTCGTTGCATTTCTACTCTTCAGCTGTGGCANNNAGATCGGAAGA |
| 68517_1 | TATGAGTGTGGAGTCGTTGCTAGACGAATCAATTGCTTCCCCANNNAGATCGGAAGA |
| 68519_1 | TATGAGTGTGGAGTCGTTGCCTGATGTCTCTGACCAATAGCCANNNAGATCGGAAGA |
| 68520_1 | TATGAGTGTGGAGTCGTTGCTCCTACCCCATCCTCAATACTCTNNNAGATCGGAAGAC |
| 68521_1 | TATGAGTGTGGAGTCGTTGCATGCCAGTCATTCTAGGCAAGATNNNAGATCGGAAG |
| 68522_1 | TATGAGTGTGGAGTCGTTGCAGATCTCAGGCAATGAAGATGCTNNNAGATCGGAAC |
| 68523_1 | TATGAGTGTGGAGTCGTTGCCAGACCCCAGTTCCAGTAATTCANNNAGATCGGAAGA |
| 68524_1 | TATGAGTGTGGAGTCGTTGCATCTAGGGTGTTTCTTTTGCCGTNNNAGATCGGAAGA |
| 68525_1 | TATGAGTGTGGAGTCGTTGCTCAGGAGAAGAAGGAAGATTCGCNNNAGATCGGAA |
| 68526_1 | TATGAGTGTGGAGTCGTTGCTGTAATTACTCTGGCAACGTCCANNNAGATCGGAAGA |
| 68527_1 | TATGAGTGTGGAGTCGTTGCCTCCTGAGTAGCTGGGACTACAGNNNAGATCGGAAC |
| 68529_1 | TATGAGTGTGGAGTCGTTGCTTCAACCCATGAACATCAATGGCNNNAGATCGGAAGA |
| 68530_1 | TATGAGTGTGGAGTCGTTGCTCATGTGCATCTAGCTCCTTCAGNNNAGATCGGAAGA |
| 68531_1 | TATGAGTGTGGAGTCGTTGCAGGGGTTTCCACTTTTCCTTCTTNNNAGATCGGAAGA |
| 68532_1 | TATGAGTGTGGAGTCGTTGCGTGGCGTAGCTTTTCTTTCCTTCNNNAGATCGGAAGA |
| 68533_1 | TATGAGTGTGGAGTCGTTGCGGATTGGTGTGCTGTAGATCTGANNNAGATCGGAAC |
| 68534_1 | TATGAGTGTGGAGTCGTTGCTCCCTTTCTATTTCCAAACAGCCNNNAGATCGGAAGAC |
| 68535_1 | TATGAGTGTGGAGTCGTTGCCTCAAAGTGTTGGGATTACAGGCNNNAGATCGGAAC |
| 68536_1 | TATGAGTGTGGAGTCGTTGCAGACCTGACAACCAATTCCATGTNNNAGATCGGAAGA |
| 68537_1 | TATGAGTGTGGAGTCGTTGCGTGTGCGAAGAGGCAGTATTTTTNNNAGATCGGAAC |
| 68539_1 | TATGAGTGTGGAGTCGTTGCCCAAGTACCTGGACATTGGTTTCNNNAGATCGGAAG |
| 68541_1 | TATGAGTGTGGAGTCGTTGCTGAGGAACCTGAGTACTCATCCTNNNAGATCGGAAG |
| 68542_1 | TATGAGTGTGGAGTCGTTGCAGTAATCATTTGGGATAGCGGCTNNNAGATCGGAAC |
| 68543_1 | TATGAGTGTGGAGTCGTTGCTTCCCTCGTATTACGGAAAAGCANNNAGATCGGAAG |
| 68544_1 | TATGAGTGTGGAGTCGTTGCACTCCAGTCTGTCAGATCCTGTANNNAGATCGGAAGA |
| 68545_1 | TATGAGTGTGGAGTCGTTGCCAGGGGAGAGGAGAGAGAAGTAANNNAGATCGGA |
| 68546_1 | TATGAGTGTGGAGTCGTTGCGAGAATCTCTGCTAAGCTCCCAGNNNAGATCGGAAG |
| 68548_1 | TATGAGTGTGGAGTCGTTGCCTGAGTAGCTGGGATTACAGGTGNNNAGATCGGAA |
| 68549_1 | TATGAGTGTGGAGTCGTTGCCAGCTGCCATCCAATACCTACTGNNNAGATCGGAAGA |
| 68550_1 | TATGAGTGTGGAGTCGTTGCCAGAGTTTCACTTTTGTTGCCCANNNAGATCGGAAGA |
| 68551_1 | TATGAGTGTGGAGTCGTTGCCCCCTCCTTCCTTCTGAAATGTTNNNAGATCGGAAGAC |
| 68552_1 | TATGAGTGTGGAGTCGTTGCCAATACCGGGAATGAAACAGTGGNNNAGATCGGAA |
| 68553_1 | TATGAGTGTGGAGTCGTTGCCCCATGGTTTCTCACGTTCTCTANNNAGATCGGAAGA |
| 68554_1 | TATGAGTGTGGAGTCGTTGCTCTTCTTTGGTCTCCATTGCTGANNNAGATCGGAAGA |

| | |
|---|---|
| 68555_1 | TATGAGTGTGGAGTCGTTGCTGATCTTAGCTCACTGCAACGTCNNNAGATCGGAAG |
| 68556_1 | TATGAGTGTGGAGTCGTTGCCCTAGCGGACCTTGATTTACGTCNNNAGATCGGAAG |
| 68558_1 | TATGAGTGTGGAGTCGTTGCTTGTGCAGTTTTCTTGGGGTTTTNNNAGATCGGAAG |
| 68559_1 | TATGAGTGTGGAGTCGTTGCAGAGAGGGAGAGAGAGAGAGAGANNNAGATCGG |
| 68560_1 | TATGAGTGTGGAGTCGTTGCTTGTGCACACATATATGCTCAGCNNNAGATCGGAAG |
| 68561_1 | TATGAGTGTGGAGTCGTTGCACACACACACACACACAGACTAANNNAGATCGGAAG |
| 68562_1 | TATGAGTGTGGAGTCGTTGCACTAATCCCATTCATGAGGGCTCNNNAGATCGGAAG |
| 68563_1 | TATGAGTGTGGAGTCGTTGCGGTTTCCACAGCCTGCTAGTTTANNNAGATCGGAAG |
| 68564_1 | TATGAGTGTGGAGTCGTTGCAGAGAGAGCAGCATGTGATTTCANNNAGATCGGAA |
| 68566_1 | TATGAGTGTGGAGTCGTTGCTGCATGTAGATACCCAGTTGTCCNNNAGATCGGAAG |
| 68568_1 | TATGAGTGTGGAGTCGTTGCACTCAAGTTCTAGAACCAGCAGGNNNAGATCGGAA |
| 68569_1 | TATGAGTGTGGAGTCGTTGCAAGAGAGAGAATGAGGGCAATCCNNNAGATCGGA |
| 68570_1 | TATGAGTGTGGAGTCGTTGCATCTGGTCCATTCTCAAGGGAAGNNNAGATCGGAA |
| 68571_1 | TATGAGTGTGGAGTCGTTGCCAAGTGGATAGACAGAGTGACGTNNNAGATCGGAA |
| 68572_1 | TATGAGTGTGGAGTCGTTGCATAGGCTGTCTACAAGCTTGAGGNNNAGATCGGAA |
| 68573_1 | TATGAGTGTGGAGTCGTTGCGACTGAGACTGTGTGCAGAAAACNNNAGATCGGAA |
| 68575_1 | TATGAGTGTGGAGTCGTTGCCTTTCTCCAGCAGTAGCCTTCTTNNNAGATCGGAAGA |
| 68576_1 | TATGAGTGTGGAGTCGTTGCTTCAAAGCACCTGTTTTGAGGAANNNAGATCGGAAG |
| 68577_1 | TATGAGTGTGGAGTCGTTGCTGAGGCCATATGATTAGCTCAGGNNNAGATCGGAA |
| 68578_1 | TATGAGTGTGGAGTCGTTGCAAGTGCTGAGATTACAGGCATGANNNAGATCGGAA |
| 68579_1 | TATGAGTGTGGAGTCGTTGCTGATCTCCTGGCTTCAAGTGATCNNNAGATCGGAAG |
| 68580_1 | TATGAGTGTGGAGTCGTTGCAGTCTTGCTCTATTGCCCAGAATNNNAGATCGGAAG |
| 68581_1 | TATGAGTGTGGAGTCGTTGCGTGCATTGTATGTCTACACCACANNNAGATCGGAAG |
| 68582_1 | TATGAGTGTGGAGTCGTTGCGAGAGCAGCCAAAAAGAAAGAGGNNNAGATCGGA |
| 68583_1 | TATGAGTGTGGAGTCGTTGCGAGGAAGGAAGGAAGGAAGGAAGNNNAGATCGG |
| 68584_1 | TATGAGTGTGGAGTCGTTGCCAGAGAGTAGAGGGAGGTAGGAGNNNAGATCGGA |
| 68585_1 | TATGAGTGTGGAGTCGTTGCGCTACTGTTGTTTTCTCTGCCTGNNNAGATCGGAAG |
| 68586_1 | TATGAGTGTGGAGTCGTTGCGGTCTCTCTGTCTCTGTCTCTCTNNNAGATCGGAAGA |
| 68587_1 | TATGAGTGTGGAGTCGTTGCCCTCACATGACCTAGTGACCAAANNNAGATCGGAAG |
| 68589_1 | TATGAGTGTGGAGTCGTTGCCCTCTCCTCTCTGTCTCCGAATANNNAGATCGGAAGA |
| 68592_1 | TATGAGTGTGGAGTCGTTGCCATGTGTGTTTGTGTGTGTGTGTNNNAGATCGGAA |
| 68593_1 | TATGAGTGTGGAGTCGTTGCTCCCATTTCTTCTCTCCCTCTCTNNNAGATCGGAAGAG |
| 68594_1 | TATGAGTGTGGAGTCGTTGCAAGAACCACCAACTGACTGACTTNNNAGATCGGAAG |
| 68595_1 | TATGAGTGTGGAGTCGTTGCACACACACATACACATACTCCCCNNNAGATCGGAAGA |
| 68596_1 | TATGAGTGTGGAGTCGTTGCGAAGGAAGGAAGGAAGGAAGGAANNNAGATCGG |
| 68597_1 | TATGAGTGTGGAGTCGTTGCGGAAAGAGGAAGGAAGGAAGGAGNNNAGATCGG |
| 68598_1 | TATGAGTGTGGAGTCGTTGCTGATCCAATAACCTCCCAACAGGNNNAGATCGGAAG |
| 68599_1 | TATGAGTGTGGAGTCGTTGCGGGAAGAGAACAAGAGTGTCCATNNNAGATCGGA |
| 68600_1 | TATGAGTGTGGAGTCGTTGCGAAACCACACCCATGATCCAATCNNNAGATCGGAAG |
| 68601_1 | TATGAGTGTGGAGTCGTTGCTCTAGAAAATGGCTGCTTGTTGANNNAGATCGGAA |
| 68602_1 | TATGAGTGTGGAGTCGTTGCTGCAGTGACATGATCATAGCTCANNNAGATCGGAAG |
| 68603_1 | TATGAGTGTGGAGTCGTTGCGGTTCAAGCAATTCTTCTGCCTTNNNAGATCGGAAG |
| 68604_1 | TATGAGTGTGGAGTCGTTGCAGCCTCCTGAGTAGCTGAGATTANNNAGATCGGAA |

| | |
|---|---|
| 68605_1 | TATGAGTGTGGAGTCGTTGCGCATTGAACAGGGTCTCTAGAGTNNNAGATCGGAA |
| 68606_1 | TATGAGTGTGGAGTCGTTGCTCTCTCTCTCTCTCCATGGTCTCNNNAGATCGGAAGA |
| 68607_1 | TATGAGTGTGGAGTCGTTGCAGAGAGAGAAACCTGGAAACCTGNNNAGATCGGAA |
| 68608_1 | TATGAGTGTGGAGTCGTTGCTAAGTAAATTGGAAACTGCGGGCNNNAGATCGGAA |
| 68609_1 | TATGAGTGTGGAGTCGTTGCCTAACCTCCAGGATTCAAGCAGTNNNAGATCGGAAG |
| 68610_1 | TATGAGTGTGGAGTCGTTGCGAAGGAAGGAAGGAAGGAAGGAANNNAGATCGG |
| 68613_1 | TATGAGTGTGGAGTCGTTGCGAAGGACAAACTGGCTTCATGTCNNNAGATCGGAA |
| 68614_1 | TATGAGTGTGGAGTCGTTGCTACGCATCCATCTACCTGTGATCNNNAGATCGGAAG |
| 68615_1 | TATGAGTGTGGAGTCGTTGCGTTGACCAGACAGATCTCGAACTNNNAGATCGGAAG |
| 68616_1 | TATGAGTGTGGAGTCGTTGCGGGTCCTATTCCCAGAGTTTCTGNNNAGATCGGAAG |
| 68617_1 | TATGAGTGTGGAGTCGTTGCCAATCTTGGCTCACTGCAATCTCNNNAGATCGGAAG |
| 68618_1 | TATGAGTGTGGAGTCGTTGCGGAGGGGGAGAGAAAGAAAGAAANNNAGATCGG |
| 68620_1 | TATGAGTGTGGAGTCGTTGCCTTGATCCTGGACTTCTAGCCTCNNNAGATCGGAAG |
| 68621_1 | TATGAGTGTGGAGTCGTTGCATGCTGAATAGTAGAGCAAGGGGNNNAGATCGGA |
| 68623_1 | TATGAGTGTGGAGTCGTTGCCACACACACACACACCCTATCTANNNAGATCGGAAGA |
| 68626_1 | TATGAGTGTGGAGTCGTTGCCGGATATAAGCTCTCCAGCCTTTNNNAGATCGGAAG |
| 68627_1 | TATGAGTGTGGAGTCGTTGCTCTTACTCTATCACCAGCCTGGANNNAGATCGGAAG |
| 68628_1 | TATGAGTGTGGAGTCGTTGCCACACACACACACACACATACAANNNAGATCGGAAG |
| 68629_1 | TATGAGTGTGGAGTCGTTGCCAAACTCCTGGGTTCAAGCAATCNNNAGATCGGAAG |
| 68630_1 | TATGAGTGTGGAGTCGTTGCTGGCTGGAAAGTTCAAAATTGGGNNNAGATCGGAA |
| 68631_1 | TATGAGTGTGGAGTCGTTGCAGAGGGAGAGAGAGAGAGAGAGANNNAGATCGG |
| 68632_1 | TATGAGTGTGGAGTCGTTGCCTGTCTGTCTCTGTGTCTGTCTCNNNAGATCGGAAGA |
| 68633_1 | TATGAGTGTGGAGTCGTTGCCTATGTATGGGAACTGGTCTGGGNNNAGATCGGAA |
| 68635_1 | TATGAGTGTGGAGTCGTTGCGGGAGAGAGAGAGAGAGAGAGAGNNNAGATCGG |
| 68637_1 | TATGAGTGTGGAGTCGTTGCTGTTCATCACCACTAGGACTGTGNNNAGATCGGAAG |
| 68638_1 | TATGAGTGTGGAGTCGTTGCCCCCAAACAAGAATCTGCTCTTGNNNAGATCGGAAG |
| 68639_1 | TATGAGTGTGGAGTCGTTGCTTTCTCTCTGCTGCTGGGTAATTNNNAGATCGGAAG |
| 68640_1 | TATGAGTGTGGAGTCGTTGCCCCCTGGGGATCTTGATAAGATGNNNAGATCGGAA |
| 68641_1 | TATGAGTGTGGAGTCGTTGCTTTGCCAACAACAGCTATGTGAGNNNAGATCGGAAG |
| 68644_1 | TATGAGTGTGGAGTCGTTGCCTCTCTCTCTCTTTCTCGTTGGGNNNAGATCGGAAGA |
| 68645_1 | TATGAGTGTGGAGTCGTTGCCTGGCCTGAAATGATCAACTCACNNNAGATCGGAAG |
| 68646_1 | TATGAGTGTGGAGTCGTTGCAGTGTCCAAAAGCAATTGCAACANNNAGATCGGAA |
| 68647_1 | TATGAGTGTGGAGTCGTTGCATTGAACTCCTGGACTCAAGCAANNNAGATCGGAAG |
| 68648_1 | TATGAGTGTGGAGTCGTTGCTGTGTGTGTGTGTGATGTACTCTNNNAGATCGGAA |
| 68649_1 | TATGAGTGTGGAGTCGTTGCACACACACACAGTCATTCTACCANNNAGATCGGAAG |
| 68650_1 | TATGAGTGTGGAGTCGTTGCTTGCTTGGAACTGGAGGAGAAATNNNAGATCGGAA |
| 68651_1 | TATGAGTGTGGAGTCGTTGCTCCCTCTCTCTCTCTCTCTCTCTNNNAGATCGGAAGAG |
| 68653_1 | TATGAGTGTGGAGTCGTTGCTTGGAGCAGACAGTGATTCAAGANNNAGATCGGAA |
| 68654_1 | TATGAGTGTGGAGTCGTTGCTGAAGCCCTTCAACCATATGGATNNNAGATCGGAAG |
| 68655_1 | TATGAGTGTGGAGTCGTTGCTTAGTTGTGACACCCTGGTATGGNNNAGATCGGAA |
| 68656_1 | TATGAGTGTGGAGTCGTTGCGGACACCATGTGCTTTGATTCAANNNAGATCGGAAG |
| 68657_1 | TATGAGTGTGGAGTCGTTGCCTAATTCCACTTCAGAGCCCTGTNNNAGATCGGAAG |
| 68658_1 | TATGAGTGTGGAGTCGTTGCCAGTGAGTCCTGTTTCTAGTCCCNNNAGATCGGAAG |

| | |
|---|---|
| 68659_1 | TATGAGTGTGGAGTCGTTGCGGATGAATACCTGGAGGCAAGAANNNAGATCGGAA |
| 68661_1 | TATGAGTGTGGAGTCGTTGCGGGCCTAGAGTCACACATTTAGTNNNAGATCGGAAG |
| 68662_1 | TATGAGTGTGGAGTCGTTGCACAGAGAGAGAGAAGGTCTTCCTNNNAGATCGGAA |
| 68663_1 | TATGAGTGTGGAGTCGTTGCTCCCCTCATTGGTTCTGTTTCTCNNNAGATCGGAAGA |
| 68664_1 | TATGAGTGTGGAGTCGTTGCACAAGAGTTCCAGTTGGTAGCAANNNAGATCGGAA |
| 68665_1 | TATGAGTGTGGAGTCGTTGCACCAGGTAATGGATCATGTGCTTNNNAGATCGGAAG |
| 68666_1 | TATGAGTGTGGAGTCGTTGCCTCGGAACTCAGGAAAGCAAAAGNNNAGATCGGAA |
| 68667_1 | TATGAGTGTGGAGTCGTTGCGTAGCAAGTGACAACTGGAAACCNNNAGATCGGAA |
| 68668_1 | TATGAGTGTGGAGTCGTTGCTCCTTAGTGGTGTTCTTCAGGTGNNNAGATCGGAAG |
| 68669_1 | TATGAGTGTGGAGTCGTTGCGGGAGAGAGAGAGAGAGAGAGAGNNNAGATCGG |
| 68670_1 | TATGAGTGTGGAGTCGTTGCAATCATTGTTGGCTGATTGAGGCNNNAGATCGGAAG |
| 68671_1 | TATGAGTGTGGAGTCGTTGCTGAGACGGAGTCTATCTCTGTCANNNAGATCGGAAG |
| 68672_1 | TATGAGTGTGGAGTCGTTGCTCTCTTTCTGTCTCTCTCCGTCTNNNAGATCGGAAGAG |
| 68673_1 | TATGAGTGTGGAGTCGTTGCAACAATCTTGGCTTACTGCAACCNNNAGATCGGAAG |
| 68674_1 | TATGAGTGTGGAGTCGTTGCAAAGAAAGAGAAAAAAGGGCCGGNNNAGATCGGA |
| 68675_1 | TATGAGTGTGGAGTCGTTGCTTGAAAAAGTTCCCAGGTGACAGNNNAGATCGGAA |
| 68679_1 | TATGAGTGTGGAGTCGTTGCCCAAGGAAGTATGGTTCAGCATCNNNAGATCGGAAG |
| 68680_1 | TATGAGTGTGGAGTCGTTGCCCCTTCATCAGTTTCCAGCAATCNNNAGATCGGAAGA |
| 68681_1 | TATGAGTGTGGAGTCGTTGCGAGGGAGGGGCTATAAAATGGAGNNNAGATCGGA |
| 68682_1 | TATGAGTGTGGAGTCGTTGCCTCGAGCTCTGGTCTCTTTCTTTNNNAGATCGGAAGA |
| 68683_1 | TATGAGTGTGGAGTCGTTGCGTTCAAGCAATTCTCATGCCTCANNNAGATCGGAAG |
| 68684_1 | TATGAGTGTGGAGTCGTTGCGGCAGCTTATTTCCAAGGTGAAGNNNAGATCGGAA |
| 68685_1 | TATGAGTGTGGAGTCGTTGCACCTAGCAATGTGTATGGAACGTNNNAGATCGGAAG |
| 68686_1 | TATGAGTGTGGAGTCGTTGCATACACATACACACACACACACANNNAGATCGGAAG |
| 68687_1 | TATGAGTGTGGAGTCGTTGCTCACTATATGCCAACAACACCCANNNAGATCGGAAG |
| 68688_1 | TATGAGTGTGGAGTCGTTGCCCACAGAAAGCCACATCATTGAGNNNAGATCGGAAG |
| 68689_1 | TATGAGTGTGGAGTCGTTGCTGTGGTAAGGAATGAAAGTCCCTNNNAGATCGGAA |
| 68690_1 | TATGAGTGTGGAGTCGTTGCTCCACCTCCAAATAGACACAGTGNNNAGATCGGAAG |
| 68691_1 | TATGAGTGTGGAGTCGTTGCACCAATCCTGTTAGCACCTTGATNNNAGATCGGAAG |
| 68692_1 | TATGAGTGTGGAGTCGTTGCAGACTGTACGTGCATAACTGTGTNNNAGATCGGAAG |
| 68693_1 | TATGAGTGTGGAGTCGTTGCTGCCCTTATAACCATGAGAGAGCNNNAGATCGGAAG |
| 68694_1 | TATGAGTGTGGAGTCGTTGCTATCTCTCACACACACACACACANNNAGATCGGAAGA |
| 68695_1 | TATGAGTGTGGAGTCGTTGCAGGGCAGGGTAGTCTTGTTTATGNNNAGATCGGAA |
| 68696_1 | TATGAGTGTGGAGTCGTTGCCACAGGCACTTACTGTATGACCANNNAGATCGGAAG |
| 68697_1 | TATGAGTGTGGAGTCGTTGCCTAGGAAAAAATCGGAACCAGGCNNNAGATCGGAA |
| 68699_1 | TATGAGTGTGGAGTCGTTGCCCCTTGCGTCTCTTCTTCTTCTTNNNAGATCGGAAGAG |
| 68700_1 | TATGAGTGTGGAGTCGTTGCAGTGAAAAAGGCACTGAAGCATCNNNAGATCGGAA |
| 68701_1 | TATGAGTGTGGAGTCGTTGCGTCTTGTTGTTTGGTTCCTGCTTNNNAGATCGGAAG |
| 68702_1 | TATGAGTGTGGAGTCGTTGCAGAAAGAAAACGAGGGAGAGCATNNNAGATCGGA |
| 68703_1 | TATGAGTGTGGAGTCGTTGCCCATTGAATTGGCTCTATGGCTGNNNAGATCGGAAG |
| 68704_1 | TATGAGTGTGGAGTCGTTGCGTGTGTGTGTGTGTTTGCCATATNNNAGATCGGAAG |
| 68705_1 | TATGAGTGTGGAGTCGTTGCTACTGTGACTGAGCTAGGTCAGANNNAGATCGGAA |
| 68706_1 | TATGAGTGTGGAGTCGTTGCTGTGTATGGATGTTTGTTCGTCANNNAGATCGGAAG |

68707_1 TATGAGTGTGGAGTCGTTGCATGGATGGTATGACAGAAGCTGTNNNAGATCGGAA
68708_1 TATGAGTGTGGAGTCGTTGCCATTGCTCAGTACTGACAGCTTGNNNAGATCGGAAG
68709_1 TATGAGTGTGGAGTCGTTGCGAGACCCCTCCATCTGTGTATTCNNNAGATCGGAAG
68710_1 TATGAGTGTGGAGTCGTTGCATACACACACACAACACACACACNNNAGATCGGAAG
68711_1 TATGAGTGTGGAGTCGTTGCAGAAGGAAGGGACAGATGCAAATNNNAGATCGGA
68712_1 TATGAGTGTGGAGTCGTTGCTCAAGGGTGACAAGTGCTAATGANNNAGATCGGAA
68713_1 TATGAGTGTGGAGTCGTTGCCTGCTTGGCAATCCAGGAAATTANNNAGATCGGAAG
68714_1 TATGAGTGTGGAGTCGTTGCTTTCTGACTCTGCTTCCTTGTGTNNNAGATCGGAAGA
68715_1 TATGAGTGTGGAGTCGTTGCGTGTGTGTGTGTGTCTGTATGNNNAGATCGGAA
68717_1 TATGAGTGTGGAGTCGTTGCTGCTGTGAGGTATAGAGGGAGAANNNAGATCGGA
68720_1 TATGAGTGTGGAGTCGTTGCCTGAGGCAGGAAAATTGCTTGAANNNAGATCGGAA
68721_1 TATGAGTGTGGAGTCGTTGCGGAGGAAGACAGAATGATGCTCTNNNAGATCGGAA
68722_1 TATGAGTGTGGAGTCGTTGCATTATGTCTCACTGCAGCCTTGANNNAGATCGGAAG
68723_1 TATGAGTGTGGAGTCGTTGCTTCCCTGAGACAGTTTTCCTCTGNNNAGATCGGAAG
68724_1 TATGAGTGTGGAGTCGTTGCAGTCCTTCACTTTCCCTGTTCTCNNNAGATCGGAAGA
68725_1 TATGAGTGTGGAGTCGTTGCGTTTGAAAGACTATGATGGGCCGNNNAGATCGGAA
68726_1 TATGAGTGTGGAGTCGTTGCATTCTAGCTCTGATTCACCAGGCNNNAGATCGGAAG
68727_1 TATGAGTGTGGAGTCGTTGCGGACATGGAAATGGCTGAACATTNNNAGATCGGAA
68729_1 TATGAGTGTGGAGTCGTTGCGGAATAAAAGCAGGCTGCAATCTNNNAGATCGGAA
68732_1 TATGAGTGTGGAGTCGTTGCGTTTTACCATGTTCGTCAGGCTGNNNAGATCGGAAG
68734_1 TATGAGTGTGGAGTCGTTGCGGTCAAAAACTTCCCAAACGTGANNNAGATCGGAAG
68737_1 TATGAGTGTGGAGTCGTTGCTCCCTCAATATCCACTCTTCCCTNNNAGATCGGAAGA
68739_1 TATGAGTGTGGAGTCGTTGCTCTTGTTGTACTAACAGCGTCCANNNAGATCGGAAG
68740_1 TATGAGTGTGGAGTCGTTGCTAGGTCCAAGGAACAACACAACTNNNAGATCGGAA
68741_1 TATGAGTGTGGAGTCGTTGCCAAAGTACTGGGGTTATCGGTGTNNNAGATCGGAA
68742_1 TATGAGTGTGGAGTCGTTGCTTTTTTTAGAGTCGGGGTTTGGCNNNAGATCGGAA
68744_1 TATGAGTGTGGAGTCGTTGCAGCATTTGGGAGAAAAGTGTGTGNNNAGATCGGA
68746_1 TATGAGTGTGGAGTCGTTGCGCTTCCGATACTAGGTCACAGAGNNNAGATCGGAA
68747_1 TATGAGTGTGGAGTCGTTGCCCGGTTCAAGTTTGTCATCAACTNNNAGATCGGAAG
68748_1 TATGAGTGTGGAGTCGTTGCCTTGCAAGTTCTCTGCTAAGCTGNNNAGATCGGAAG
68749_1 TATGAGTGTGGAGTCGTTGCTAGGCTCTCTGGGTTTCTAGTGANNNAGATCGGAAG
68751_1 TATGAGTGTGGAGTCGTTGCTGTAAGGATATCTAAGCCCCCACNNNAGATCGGAAG
68752_1 TATGAGTGTGGAGTCGTTGCATACAAAAATTAGCCGGGCATGGNNNAGATCGGAA
68753_1 TATGAGTGTGGAGTCGTTGCGTGTAAAGAGAACCAAGCCAACCNNNAGATCGGAA
68754_1 TATGAGTGTGGAGTCGTTGCCGAGACTCCCACTCACAAAAAAANNNAGATCGGAAG
68756_1 TATGAGTGTGGAGTCGTTGCAGTGTGGGTTTTTGGTTTTGAGGNNNAGATCGGAA
68758_1 TATGAGTGTGGAGTCGTTGCTCTGCTTTCTGTGTTACCCATGANNNAGATCGGAAG
68759_1 TATGAGTGTGGAGTCGTTGCCTCACTACAGACTCAAACACCCANNNAGATCGGAAG
68760_1 TATGAGTGTGGAGTCGTTGCAGAAAGCAGCATCCTATAAGCCANNNAGATCGGAA
68761_1 TATGAGTGTGGAGTCGTTGCGTGTGTGTATATGTGTGTGGTCGNNNAGATCGGAA
68762_1 TATGAGTGTGGAGTCGTTGCAGGAGCAAAAGCCTATATAGCCCNNNAGATCGGAA
68763_1 TATGAGTGTGGAGTCGTTGCTCTCTGTCTCCCTCCCATTGTATNNNAGATCGGAAGA
68764_1 TATGAGTGTGGAGTCGTTGCCACACACACACACACACACAAATNNNAGATCGGAAG

| | |
|---|---|
| 68765_1 | TATGAGTGTGGAGTCGTTGCGAGAGGCAGGGAACTTAGATTGTNNNAGATCGGAA |
| 68766_1 | TATGAGTGTGGAGTCGTTGCACTCAGTGGGTAAAAGCCAAAGANNNAGATCGGAA |
| 68767_1 | TATGAGTGTGGAGTCGTTGCGCTGTAATGGAGTCGTGTCTAGTNNNAGATCGGAA |
| 68768_1 | TATGAGTGTGGAGTCGTTGCAACACTCTAACTTCCCGTGGTTTNNNAGATCGGAAG |
| 68769_1 | TATGAGTGTGGAGTCGTTGCCTCTGGTATCCTGTTTCTTCCCCNNNAGATCGGAAGA |
| 68770_1 | TATGAGTGTGGAGTCGTTGCCCATGACCTAATGACCTCCCAAANNNAGATCGGAAG |
| 68771_1 | TATGAGTGTGGAGTCGTTGCGACACAAAGAGACAGAGAGAGCANNNAGATCGGAA |
| 68772_1 | TATGAGTGTGGAGTCGTTGCGCATGATCATAGTTCACTGCAGCNNNAGATCGGAAG |
| 68773_1 | TATGAGTGTGGAGTCGTTGCAAAGAAAAAGCAAGCAAGCAAGCNNNAGATCGGAA |
| 68774_1 | TATGAGTGTGGAGTCGTTGCTGAAATGTGAAGAGAAGGGGTGGNNNAGATCGGA |
| 68775_1 | TATGAGTGTGGAGTCGTTGCTGAAATGTGAAGAGAAGGGGTGGNNNAGATCGGA |
| 68776_1 | TATGAGTGTGGAGTCGTTGCGAGAGAGAGAGAGAGAGACGGAANNNAGATCGG |
| 68777_1 | TATGAGTGTGGAGTCGTTGCGGAGCAAGAGAGAGAGAGAGAGANNNAGATCGG |
| 68778_1 | TATGAGTGTGGAGTCGTTGCATTTTGGGGGTTTTTGTGTGTGTNNNAGATCGGAA |
| 68780_1 | TATGAGTGTGGAGTCGTTGCAGAGAGAGAGAGAAAAGCTTGGCNNNAGATCGGA |
| 68781_1 | TATGAGTGTGGAGTCGTTGCGTGTCTTGCTCTCACTCTCTCAGNNNAGATCGGAAG |
| 68782_1 | TATGAGTGTGGAGTCGTTGCCTGCCTCCTAGGTTCAAGTGATTNNNAGATCGGAAG |
| 68783_1 | TATGAGTGTGGAGTCGTTGCTGGGTTCTATAACTTGGACCAGCNNNAGATCGGAAG |
| 68785_1 | TATGAGTGTGGAGTCGTTGCGTATCCAGCCCATTCGGTGATTANNNAGATCGGAAG |
| 68786_1 | TATGAGTGTGGAGTCGTTGCAGCCTGCATGCAAATTCTCTTTTNNNAGATCGGAAG |
| 68787_1 | TATGAGTGTGGAGTCGTTGCGATGACGAAATGCACCATCTGAGNNNAGATCGGAA |
| 68789_1 | TATGAGTGTGGAGTCGTTGCCGTCCTGGGTCTTAGAAGCTAAGNNNAGATCGGAA |
| 68790_1 | TATGAGTGTGGAGTCGTTGCTGGTTCTCCTTATCTGGTTCGACNNNAGATCGGAAG |
| 68791_1 | TATGAGTGTGGAGTCGTTGCCTGCATGATGTCTGCCCATATTGNNNAGATCGGAAG |
| 68792_1 | TATGAGTGTGGAGTCGTTGCACAGATGCTCAAGTCCCTGATTTNNNAGATCGGAAG |
| 68793_1 | TATGAGTGTGGAGTCGTTGCAACCATCCATCTGAATCAAGGCTNNNAGATCGGAAG |
| 68794_1 | TATGAGTGTGGAGTCGTTGCACACACACACACACTACTGACAANNNAGATCGGAAG |
| 68795_1 | TATGAGTGTGGAGTCGTTGCACACACACACACAAAACACCTTTNNNAGATCGGAAG |
| 68796_1 | TATGAGTGTGGAGTCGTTGCTAAATCTGCCTTGCCTGTACCTTNNNAGATCGGAAG |
| 68797_1 | TATGAGTGTGGAGTCGTTGCTTCCAAATTCTGTAAGGCTCCCANNNAGATCGGAAG |
| 68798_1 | TATGAGTGTGGAGTCGTTGCATGTACCCAGTCTTTACTCCCTGNNNAGATCGGAAG |
| 68799_1 | TATGAGTGTGGAGTCGTTGCCAGAGATTTGGTGCTTCATCTGCNNNAGATCGGAAG |
| 68801_1 | TATGAGTGTGGAGTCGTTGCAAGAAGAAGGAAGAGGAGGAGGANNNAGATCGG |
| 68802_1 | TATGAGTGTGGAGTCGTTGCCCTGTCTCTGTGTTTGTGTGTTTNNNAGATCGGAAG |
| 68804_1 | TATGAGTGTGGAGTCGTTGCCAGGCTGAAGTCATAGCTCACTGNNNAGATCGGAA |
| 68805_1 | TATGAGTGTGGAGTCGTTGCTGTAAATTTCCTGAAGCCTCCCANNNAGATCGGAAG |
| 68806_1 | TATGAGTGTGGAGTCGTTGCCCACATGGTGAGGAACTGAAAACNNNAGATCGGAA |
| 68808_1 | TATGAGTGTGGAGTCGTTGCGCATGGGAAAAGAACGTTCAGAANNNAGATCGGAA |
| 68809_1 | TATGAGTGTGGAGTCGTTGCATCTTCACTGCCTTTCTGATCGTNNNAGATCGGAAGA |
| 68811_1 | TATGAGTGTGGAGTCGTTGCGCTCAACTGATTCTCCCACATCTNNNAGATCGGAAGA |
| 68812_1 | TATGAGTGTGGAGTCGTTGCTACTCTCTCCTTCTGTCTCCCTTNNNAGATCGGAAGAG |
| 68813_1 | TATGAGTGTGGAGTCGTTGCTTCCTTCCTTCCTTCCTTCCTTCNNNAGATCGGAAGAG |
| 68814_1 | TATGAGTGTGGAGTCGTTGCCTACGCCAGAGAGAGACAAAGACNNNAGATCGGAA |

| | |
|---|---|
| 68815_1 | TATGAGTGTGGAGTCGTTGCCTCACACACACACATACACACACNNNAGATCGGAAGA |
| 68816_1 | TATGAGTGTGGAGTCGTTGCAGACAGAGAGACAGAGAGAGTCANNNAGATCGGA |
| 68817_1 | TATGAGTGTGGAGTCGTTGCTTTGGCCAAAACGTTTTGAGACTNNNAGATCGGAAG |
| 68818_1 | TATGAGTGTGGAGTCGTTGCCCTACCCTCAGGACCTCATCTAANNNAGATCGGAAGA |
| 68819_1 | TATGAGTGTGGAGTCGTTGCCCCCTGAATGGTTCACTAACAATGNNNAGATCGGAA |
| 68821_1 | TATGAGTGTGGAGTCGTTGCCACTACAGGCACAGATCACCATANNNAGATCGGAAG |
| 68822_1 | TATGAGTGTGGAGTCGTTGCCTCCTGAGTAGCTGGAATGACAGNNNAGATCGGAA |
| 68823_1 | TATGAGTGTGGAGTCGTTGCATACACACACACACACAACTACANNNAGATCGGAAGA |
| 68824_1 | TATGAGTGTGGAGTCGTTGCCCTCTCTCCTCTCTCTCTGTCTCNNNAGATCGGAAGAG |
| 68826_1 | TATGAGTGTGGAGTCGTTGCCATAGACAACAGTGGCTCAAAGCNNNAGATCGGAA |
| 68827_1 | TATGAGTGTGGAGTCGTTGCGGAGCTGGGACACTCTTTTTTTCNNNAGATCGGAAG |
| 68828_1 | TATGAGTGTGGAGTCGTTGCTCAAGTGATTCTCCTGCCTTAGCNNNAGATCGGAAGA |
| 68830_1 | TATGAGTGTGGAGTCGTTGCATACTGAGATGTGGAAGTGAGGCNNNAGATCGGAA |
| 68831_1 | TATGAGTGTGGAGTCGTTGCCACCACATCCGGCTAATTTTTGTNNNAGATCGGAAGA |
| 68833_1 | TATGAGTGTGGAGTCGTTGCTCTGTGAGATTTGTCCATGTGGTNNNAGATCGGAAG |
| 68834_1 | TATGAGTGTGGAGTCGTTGCAGAGAGAGAGAGAAAGCATGCTGNNNAGATCGGA |
| 68835_1 | TATGAGTGTGGAGTCGTTGCACTGTGAGAATGTGAGAAGCTGGNNNAGATCGGAA |
| 68836_1 | TATGAGTGTGGAGTCGTTGCCAGCTACTAGGGAGGGTGAGATANNNAGATCGGAA |
| 68837_1 | TATGAGTGTGGAGTCGTTGCGTCTTGCCCAAAGCTCACTCTAANNNAGATCGGAAGA |
| 68838_1 | TATGAGTGTGGAGTCGTTGCTCCTGGTCTTTGGGAGATCTTTGNNNAGATCGGAAG |
| 68839_1 | TATGAGTGTGGAGTCGTTGCTCTGTCTCTGTCTCTCTCTGTCCNNNAGATCGGAAGA |
| 68840_1 | TATGAGTGTGGAGTCGTTGCGCTTCCACATCAGACTCCAGTAANNNAGATCGGAAGA |
| 68842_1 | TATGAGTGTGGAGTCGTTGCAGAGCCAACAGCAAGTACTAACANNNAGATCGGAA |
| 68843_1 | TATGAGTGTGGAGTCGTTGCTACTTTTGCTGCCTAATTGCTGCNNNAGATCGGAAGA |
| 68844_1 | TATGAGTGTGGAGTCGTTGCATTCTCCAGTCTTCACAGACACCNNNAGATCGGAAGA |
| 68845_1 | TATGAGTGTGGAGTCGTTGCACCAGAAACCAACTTGGACTTCTNNNAGATCGGAAG |
| 68847_1 | TATGAGTGTGGAGTCGTTGCAGAGAGAGAGAGAGAGAGGGAGANNNAGATCGG |
| 68848_1 | TATGAGTGTGGAGTCGTTGCAGCTCACTGCAACCTTTTTTGTTNNNAGATCGGAAGA |
| 68849_1 | TATGAGTGTGGAGTCGTTGCGTCCTTTCAGAAATTCGGCCTTCNNNAGATCGGAAGA |
| 68850_1 | TATGAGTGTGGAGTCGTTGCCCATGTCAACCCTCCAGATGATTNNNAGATCGGAAGA |
| 68851_1 | TATGAGTGTGGAGTCGTTGCTCAATGATCGTGGGATCTTCTGGNNNAGATCGGAA |
| 68852_1 | TATGAGTGTGGAGTCGTTGCAAGACTTGTCCTGGAGAGAAACCNNNAGATCGGAA |
| 68853_1 | TATGAGTGTGGAGTCGTTGCGGGTTCATTGCCACTACTTCATGNNNAGATCGGAAG |
| 68854_1 | TATGAGTGTGGAGTCGTTGCGGGAGAGAGAGAGAGTCAAGGTTNNNAGATCGGA |
| 68855_1 | TATGAGTGTGGAGTCGTTGCTGTTTGGGGAAAGTGAGAAGTGANNNAGATCGGA |
| 68856_1 | TATGAGTGTGGAGTCGTTGCCACACACACACACACAAAATCCTNNNAGATCGGAAGA |
| 68858_1 | TATGAGTGTGGAGTCGTTGCACAGAAAAATCCCTGGTCCCATTNNNAGATCGGAAG |
| 68859_1 | TATGAGTGTGGAGTCGTTGCATACGAAAGCAACTGCAGGTTTCNNNAGATCGGAA |
| 68861_1 | TATGAGTGTGGAGTCGTTGCATGTCCCTTGCAACTAGAATGGTNNNAGATCGGAAG |
| 68862_1 | TATGAGTGTGGAGTCGTTGCCCCTTCACTACCCCTTTTGTCTTNNNAGATCGGAAGA |
| 68863_1 | TATGAGTGTGGAGTCGTTGCTCTGTGGTTACATTCCCAGGAAGNNNAGATCGGAAG |
| 68864_1 | TATGAGTGTGGAGTCGTTGCCCTGTAATTCTAGCACTGTGGGANNNAGATCGGAAG |
| 68865_1 | TATGAGTGTGGAGTCGTTGCGCAGGCTGATTGGTCTTTGATTTNNNAGATCGGAAG |

| | |
|---|---|
| 68866_1 | TATGAGTGTGGAGTCGTTGCTGCTCAGCCTTATTCTCCATCTCNNNAGATCGGAAGA |
| 68867_1 | TATGAGTGTGGAGTCGTTGCTGGAAGGGGACACCATAAACAATNNNAGATCGGAA |
| 68869_1 | TATGAGTGTGGAGTCGTTGCGCGACAATCATGACAACACAGAANNNAGATCGGAAG |
| 68870_1 | TATGAGTGTGGAGTCGTTGCACCTAGACAGAGCCTACCATTCTNNNAGATCGGAAG |
| 68871_1 | TATGAGTGTGGAGTCGTTGCAGAACAGAGGGAAGAGATGGAGANNNAGATCGGA |
| 68872_1 | TATGAGTGTGGAGTCGTTGCGAACCAGGAAGAGAGGGAGAAAGNNNAGATCGGA |
| 68873_1 | TATGAGTGTGGAGTCGTTGCCTCCCTCTCTCTCTCTCCATGNNNAGATCGGAAGAG |
| 68874_1 | TATGAGTGTGGAGTCGTTGCGCCTTCCACCATGGGATAATGTANNNAGATCGGAAG |
| 68875_1 | TATGAGTGTGGAGTCGTTGCAAACAATTCTCCTGACTCAGCCTNNNAGATCGGAAG |
| 68876_1 | TATGAGTGTGGAGTCGTTGCAGCCAGGTTTAGTTGGAATCGATNNNAGATCGGAA |
| 68877_1 | TATGAGTGTGGAGTCGTTGCGATCATAGCTCACTGCAGTCTCANNNAGATCGGAAG |
| 68878_1 | TATGAGTGTGGAGTCGTTGCTGTGCCAGGCTAGATTTTACAGANNNAGATCGGAAG |
| 68879_1 | TATGAGTGTGGAGTCGTTGCGCCTGTATTCCCAACTCCAACTANNNAGATCGGAAG |
| 68880_1 | TATGAGTGTGGAGTCGTTGCATGACCACACAAAAATCCTGCAGNNNAGATCGGAAG |
| 68881_1 | TATGAGTGTGGAGTCGTTGCACGTAATTGTAGAGGAGTGCTCGNNNAGATCGGAA |
| 68884_1 | TATGAGTGTGGAGTCGTTGCAATGGGGGAGGTGTGATTCTTTTNNNAGATCGGAA |
| 68885_1 | TATGAGTGTGGAGTCGTTGCGAGAGAGAGAGAGAGAGAGAGCGNNNAGATCGG |
| 68887_1 | TATGAGTGTGGAGTCGTTGCACTCAGGACAGTCACAGAACATGNNNAGATCGGAA |
| 68888_1 | TATGAGTGTGGAGTCGTTGCGAAAGGAAGGAAGGAAGGAAGGANNNAGATCGG |
| 68889_1 | TATGAGTGTGGAGTCGTTGCGAATAGCACAAGAAAGACCTGCCNNNAGATCGGAA |
| 68891_1 | TATGAGTGTGGAGTCGTTGCCTTTTTCCATGGGGCCAAAATCTNNNAGATCGGAAG |
| 68892_1 | TATGAGTGTGGAGTCGTTGCCTGCATGTCTTCCATCTGTCTCTNNNAGATCGGAAG |
| 68893_1 | TATGAGTGTGGAGTCGTTGCTCTTCCCAAACTTCACAAGGAAANNNAGATCGGAAG |
| 68894_1 | TATGAGTGTGGAGTCGTTGCGTGTGTGTGTGTGTGTGTGTATTNNNAGATCGGAA |
| 68895_1 | TATGAGTGTGGAGTCGTTGCACACACACACACACACACAATTTNNNAGATCGGAAG |
| 68896_1 | TATGAGTGTGGAGTCGTTGCAGAGAGAGAGAGAGAGGCAAGTTNNNAGATCGGA |
| 68897_1 | TATGAGTGTGGAGTCGTTGCATATGGCAACATGACAAAACCCCNNNAGATCGGAAG |
| 68898_1 | TATGAGTGTGGAGTCGTTGCTGTTCTCCATCACCATTGGTTCTNNNAGATCGGAAG |
| 68899_1 | TATGAGTGTGGAGTCGTTGCACACACACAATGGGATCTAGCATNNNAGATCGGAAG |
| 68900_1 | TATGAGTGTGGAGTCGTTGCGTCTGGTTGTCGTTCTCAAACTGNNNAGATCGGAAG |
| 68901_1 | TATGAGTGTGGAGTCGTTGCTATACACACACACACACACACACNNNAGATCGGAAG |
| 68902_1 | TATGAGTGTGGAGTCGTTGCTTGGGAAGTCCTCTAAGCTTGTCNNNAGATCGGAAG |
| 68903_1 | TATGAGTGTGGAGTCGTTGCAATATGCAGGGCAAGTTGTCAACNNNAGATCGGAA |
| 68904_1 | TATGAGTGTGGAGTCGTTGCTTCCAAATCTTCCTAGGGCCTTCNNNAGATCGGAAG |
| 68905_1 | TATGAGTGTGGAGTCGTTGCTCTTGCCTGTCAAACTTCAGAGANNNAGATCGGAAG |
| 68906_1 | TATGAGTGTGGAGTCGTTGCTGAACCACCTAAAACTTGGGGAANNNAGATCGGAAG |
| 68907_1 | TATGAGTGTGGAGTCGTTGCAAAAAAACCAGATAAGCGTGCGANNNAGATCGGAA |
| 68908_1 | TATGAGTGTGGAGTCGTTGCCCAACAGACAGTTTTCTCTCCCTNNNAGATCGGAAG |
| 68909_1 | TATGAGTGTGGAGTCGTTGCAATAAGGTGCACAGAAGCTCTGANNNAGATCGGAA |
| 68910_1 | TATGAGTGTGGAGTCGTTGCAGTCTTACTCTATCGGCTAGGCTNNNAGATCGGAAG |
| 68912_1 | TATGAGTGTGGAGTCGTTGCGCATGACAGAACTGGGGTTACTANNNAGATCGGAA |
| 68913_1 | TATGAGTGTGGAGTCGTTGCTCCAGGCTCAAGAAATAGCAAGTNNNAGATCGGAAG |
| 68914_1 | TATGAGTGTGGAGTCGTTGCAGACCTTACCATGAGTCAAACCCNNNAGATCGGAAG |

| | |
|---|---|
| 68915_1 | TATGAGTGTGGAGTCGTTGCGGTTCAAGTGATTTTCCTGCCTCNNNAGATCGGAAG |
| 68916_1 | TATGAGTGTGGAGTCGTTGCCCCAGAAGACACCAAGATTCAGANNNAGATCGGAAG |
| 68917_1 | TATGAGTGTGGAGTCGTTGCATCTGTCTTTGAGAGAGTGCCAGNNNAGATCGGAAG |
| 68918_1 | TATGAGTGTGGAGTCGTTGCCACACACACACATGAGAGAGAGANNNAGATCGGAA |
| 68919_1 | TATGAGTGTGGAGTCGTTGCGAGCGAGGCCAATTTTGTGTATANNNAGATCGGAA |
| 68920_1 | TATGAGTGTGGAGTCGTTGCTTTCTAGCCTCCAGAACTGTGAGNNNAGATCGGAAG |
| 68921_1 | TATGAGTGTGGAGTCGTTGCTGTTATGGAGTCTCTCTGTGTCGNNNAGATCGGAAG |
| 68922_1 | TATGAGTGTGGAGTCGTTGCCTGCAGATGCCTGAAACTCAAAANNNAGATCGGAAG |
| 68923_1 | TATGAGTGTGGAGTCGTTGCAATGAGAACTTCTGCGGAGACATNNNAGATCGGAA |
| 68924_1 | TATGAGTGTGGAGTCGTTGCGACTGAGTCACAAGACCAGCTTANNNAGATCGGAAG |
| 68925_1 | TATGAGTGTGGAGTCGTTGCTCACAGAACACTGGACTAGAAGCNNNAGATCGGAAG |
| 68926_1 | TATGAGTGTGGAGTCGTTGCAACTGTGTGTGATCAGAGATGCANNNAGATCGGAA |
| 68927_1 | TATGAGTGTGGAGTCGTTGCCCAGGAGTACAATGAGTGAGGACNNNAGATCGGAA |
| 68928_1 | TATGAGTGTGGAGTCGTTGCGGGGTATTGATTTTCTTCCCGGANNNAGATCGGAAG |
| 68929_1 | TATGAGTGTGGAGTCGTTGCCAAAGAAAATGGGAGAACGGGACNNNAGATCGGA |
| 68930_1 | TATGAGTGTGGAGTCGTTGCCACCTCCAAATGCCATTACACTGNNNAGATCGGAAG |
| 68931_1 | TATGAGTGTGGAGTCGTTGCATTTCTGTGGTTGCATCCTGACTNNNAGATCGGAAG |
| 68932_1 | TATGAGTGTGGAGTCGTTGCTCAACCTGCTTTATCTCTAGGCCNNNAGATCGGAAG |
| 68933_1 | TATGAGTGTGGAGTCGTTGCCTTGACTTCCCAGGCTAAGTGATNNNAGATCGGAAG |
| 68934_1 | TATGAGTGTGGAGTCGTTGCCACACACACACACACACAGAAAANNNAGATCGGAAG |
| 68936_1 | TATGAGTGTGGAGTCGTTGCCTCTCTCTCTCTCACACACACACNNNAGATCGGAAGA |
| 68937_1 | TATGAGTGTGGAGTCGTTGCACTACCATTGAGACTGCAGTGAGNNNAGATCGGAAG |
| 68938_1 | TATGAGTGTGGAGTCGTTGCTGCGAATTTGAGAAAGCGTCATTNNNAGATCGGAAG |
| 68939_1 | TATGAGTGTGGAGTCGTTGCTGTGTGTGTGTGTGTGTAGTGATNNNAGATCGGAA |
| 68940_1 | TATGAGTGTGGAGTCGTTGCCCTTCAGATGACCCCATTCTCAANNNAGATCGGAAG |
| 68941_1 | TATGAGTGTGGAGTCGTTGCCCCCTTGGCTAAAACAACAGTTTNNNAGATCGGAAG |
| 68942_1 | TATGAGTGTGGAGTCGTTGCACATTGAGTTGCAAAGACCATCCNNNAGATCGGAAG |
| 68944_1 | TATGAGTGTGGAGTCGTTGCATGCCCTTGAAGCACATCAGATANNNAGATCGGAAG |
| 68945_1 | TATGAGTGTGGAGTCGTTGCATTTAGTCAGCAGTTTGCCAACCNNNAGATCGGAAG |
| 68946_1 | TATGAGTGTGGAGTCGTTGCAAGCATTCAAACATACATGCACANNNAGATCGGAAG |
| 68947_1 | TATGAGTGTGGAGTCGTTGCTGAGTCTAGGTGGGAAAACTTGTNNNAGATCGGAA |
| 68948_1 | TATGAGTGTGGAGTCGTTGCGAGAAACACATACCCCACCCTTANNNAGATCGGAAG |
| 68949_1 | TATGAGTGTGGAGTCGTTGCGACATCAGGGTTAGGGTATGTGGNNNAGATCGGA |
| 68950_1 | TATGAGTGTGGAGTCGTTGCGCCCTCTGGTGGACAGTTTATTTNNNAGATCGGAAG |
| 68951_1 | TATGAGTGTGGAGTCGTTGCAGGGAAGGGTAGGTATCTATGCTNNNAGATCGGAA |
| 68952_1 | TATGAGTGTGGAGTCGTTGCACACACATCCAGTTAGCTCCAATNNNAGATCGGAAG |
| 68953_1 | TATGAGTGTGGAGTCGTTGCGGATAAACCAGTGTGTGTGTGTGNNNAGATCGGAA |
| 68955_1 | TATGAGTGTGGAGTCGTTGCCTGTGTGGGAAAGCTGTTGTAAGNNNAGATCGGAA |
| 68956_1 | TATGAGTGTGGAGTCGTTGCAAAAACAAAGCCAGAAACCCACANNNAGATCGGAAG |
| 68958_1 | TATGAGTGTGGAGTCGTTGCCGCCATCACACCCAACTAATTTTNNNAGATCGGAAGA |
| 68961_1 | TATGAGTGTGGAGTCGTTGCAGTGGTTTTCCCTCCCTTTTACCNNNAGATCGGAAGA |
| 68964_1 | TATGAGTGTGGAGTCGTTGCCCGGGGGGAAAATCTATGCATATNNNAGATCGGAA |
| 68965_1 | TATGAGTGTGGAGTCGTTGCGTTAGTTTGCATGCCTTCCGTAGNNNAGATCGGAAG |

| | |
|---|---|
| 68966_1 | TATGAGTGTGGAGTCGTTGCTAGGAGTAAGCAACTGCATCCAGNNNAGATCGGAA |
| 68967_1 | TATGAGTGTGGAGTCGTTGCGGCGGTGCTATTACTCCTCATATNNNAGATCGGAAG |
| 68969_1 | TATGAGTGTGGAGTCGTTGCTTCTACCCTCTTTGTACCCTGGANNNAGATCGGAAGA |
| 68970_1 | TATGAGTGTGGAGTCGTTGCGGTTCAAGCGATTATCTTGCCTCNNNAGATCGGAAG |
| 68973_1 | TATGAGTGTGGAGTCGTTGCGGACGTAGAGCAGAGAAGAGAACNNNAGATCGGA |
| 68974_1 | TATGAGTGTGGAGTCGTTGCTCTGATTGAGGGAATGAACACGTNNNAGATCGGAA |
| 68975_1 | TATGAGTGTGGAGTCGTTGCTATGCCTTATCTGGGTCTCATGCNNNAGATCGGAAG |
| 68976_1 | TATGAGTGTGGAGTCGTTGCCATGACATACTGAGACACCAGCTNNNAGATCGGAAG |
| 68977_1 | TATGAGTGTGGAGTCGTTGCAGCCTACCAAAATGCTGGGATTANNNAGATCGGAAG |
| 68980_1 | TATGAGTGTGGAGTCGTTGCTTCAGTAAACTCTGGGTGACTCGNNNAGATCGGAAG |
| 68982_1 | TATGAGTGTGGAGTCGTTGCATACTTTGCTTTGCCTCCCAAACNNNAGATCGGAAGA |
| 68983_1 | TATGAGTGTGGAGTCGTTGCCAGGATGGTCTCGATCTCTTGACNNNAGATCGGAAG |
| 68985_1 | TATGAGTGTGGAGTCGTTGCCAATGGCTTGGTCTTGTCTCACTNNNAGATCGGAAG |
| 68987_1 | TATGAGTGTGGAGTCGTTGCCTTCACTCACTCACACTTTCCCTNNNAGATCGGAAGA |
| 68988_1 | TATGAGTGTGGAGTCGTTGCTAGGAGCAGCAATTTATCCGTGTNNNAGATCGGAAG |
| 68989_1 | TATGAGTGTGGAGTCGTTGCACCCTGCAAAAAAAGGAAACCAANNNAGATCGGAA |
| 68991_1 | TATGAGTGTGGAGTCGTTGCCCATCATGCCCGGCTAATTTTTTNNNAGATCGGAAGA |
| 68992_1 | TATGAGTGTGGAGTCGTTGCTATGCCCTTTGTGAGAGTTCCAANNNAGATCGGAAG |
| 68995_1 | TATGAGTGTGGAGTCGTTGCTTCTAAACCAGACGAGAGAAGGCNNNAGATCGGAA |
| 68996_1 | TATGAGTGTGGAGTCGTTGCTCTACTGCTGTCCTTTGTACCTGNNNAGATCGGAAGA |
| 68997_1 | TATGAGTGTGGAGTCGTTGCTCTGGTTTCACAGGCAATAGTCANNNAGATCGGAAG |
| 68998_1 | TATGAGTGTGGAGTCGTTGCAGGGGAAGCTCTTTGATTTCGATNNNAGATCGGAA |
| 69000_1 | TATGAGTGTGGAGTCGTTGCATCTGTTTCTGAAAGGCAGGGATNNNAGATCGGAA |
| 69001_1 | TATGAGTGTGGAGTCGTTGCAGGTAGCAATAAAGGCCCTGAAANNNAGATCGGAA |
| 69003_1 | TATGAGTGTGGAGTCGTTGCCTCTGCTTAGATGAATGTGTGCGNNNAGATCGGAAG |
| 69004_1 | TATGAGTGTGGAGTCGTTGCCCCTGAGACATGAATCACCAGAANNNAGATCGGAAG |
| 69005_1 | TATGAGTGTGGAGTCGTTGCGCCTCCTGAGTAGCTGAGATTACNNNAGATCGGAAG |
| 69007_1 | TATGAGTGTGGAGTCGTTGCGGTAAACATGAAGCACAAAGGCTNNNAGATCGGAA |
| 69008_1 | TATGAGTGTGGAGTCGTTGCCAAGCTGTGAAATGGGGACAATGNNNAGATCGGAA |
| 69009_1 | TATGAGTGTGGAGTCGTTGCCAGAGCTGAATTGTCTGCATGAGNNNAGATCGGAA |
| 69011_1 | TATGAGTGTGGAGTCGTTGCTGTGTGTTTCCCAGTAGACCAAANNNAGATCGGAAG |
| 69013_1 | TATGAGTGTGGAGTCGTTGCCAAGCTGTGCTGAGAAACTCTTCNNNAGATCGGAAG |
| 69014_1 | TATGAGTGTGGAGTCGTTGCTCTCCAAGTTTTCAGTGTGGTGANNNAGATCGGAAG |
| 69015_1 | TATGAGTGTGGAGTCGTTGCCTCCGGAGTAGCTGAGATTACAGNNNAGATCGGAA |
| 69016_1 | TATGAGTGTGGAGTCGTTGCACTCAGTGGGTAAAAGCCAAAGANNNAGATCGGAA |
| 69017_1 | TATGAGTGTGGAGTCGTTGCGGAGAGAGAGACTGACTGACTCTNNNAGATCGGAA |
| 69018_1 | TATGAGTGTGGAGTCGTTGCCTTCTTAGTCTGTTCCACTGGCTNNNAGATCGGAAGA |
| 69019_1 | TATGAGTGTGGAGTCGTTGCATGGCTTCATAGACTGTCCAGTGNNNAGATCGGAAG |
| 69020_1 | TATGAGTGTGGAGTCGTTGCCAGGGACTAGGGCCACTAAAAAANNNAGATCGGAA |
| 69021_1 | TATGAGTGTGGAGTCGTTGCCACCTCCAAACATGCTCTCTTTGNNNAGATCGGAAGA |
| 69022_1 | TATGAGTGTGGAGTCGTTGCGAAACAAGTCATAGCAACACCGGNNNAGATCGGAA |
| 69023_1 | TATGAGTGTGGAGTCGTTGCTTGCCTCTCTCTCTCTCTCTCTCNNNAGATCGGAAGAG |
| 69024_1 | TATGAGTGTGGAGTCGTTGCTCTCTCTCTCTCCTCATCAGCNNNAGATCGGAAGA |

| | |
|---|---|
| 69025_1 | TATGAGTGTGGAGTCGTTGCATCAAGCAACTCTATCAGTGCCANNNAGATCGGAAG |
| 69026_1 | TATGAGTGTGGAGTCGTTGCCACACACACACACACACACATACNNNAGATCGGAAG |
| 69028_1 | TATGAGTGTGGAGTCGTTGCAGAGAGAGAGAGAGAGAGACTGGNNNAGATCGG |
| 69031_1 | TATGAGTGTGGAGTCGTTGCCAGAACCATGAGCCAAATAAGCCNNNAGATCGGAAG |
| 69032_1 | TATGAGTGTGGAGTCGTTGCTCACTGGGGAAATTTTGGCAATCNNNAGATCGGAAG |
| 69033_1 | TATGAGTGTGGAGTCGTTGCAAGAGAGAGAGCACTGAGGTTTGNNNAGATCGGA |
| 69035_1 | TATGAGTGTGGAGTCGTTGCCCCAATACACAGAGTGGTTCTCANNNAGATCGGAAG |
| 69037_1 | TATGAGTGTGGAGTCGTTGCCCACACCCTTACGACCTCTTTTANNNAGATCGGAAGA |
| 69039_1 | TATGAGTGTGGAGTCGTTGCGAGAGACCAAGGCCATACTAACANNNAGATCGGAA |
| 69040_1 | TATGAGTGTGGAGTCGTTGCTCTATCTCTCCATCCCTCTGTCCNNNAGATCGGAAGA |
| 69041_1 | TATGAGTGTGGAGTCGTTGCGCCTTCAAGTCTTGTTCAGTGTCNNNAGATCGGAAG |
| 69042_1 | TATGAGTGTGGAGTCGTTGCTTCAGAATGTCATTTGTGGTGGCNNNAGATCGGAAG |
| 69043_1 | TATGAGTGTGGAGTCGTTGCTCTGTGTCTCTCTCTGTCTCACANNNAGATCGGAAGA |
| 69044_1 | TATGAGTGTGGAGTCGTTGCATGATTCCAAAGAGCTCCTGTGTNNNAGATCGGAAG |
| 69045_1 | TATGAGTGTGGAGTCGTTGCACCTGCTTCTTACTGCCTTAGAANNNAGATCGGAAG |
| 69046_1 | TATGAGTGTGGAGTCGTTGCTCCCTCACCAGATATGGACTCTTNNNAGATCGGAAG |
| 69047_1 | TATGAGTGTGGAGTCGTTGCTCTTTCTTGACGGAGTCTCACTCNNNAGATCGGAAG |
| 69048_1 | TATGAGTGTGGAGTCGTTGCGTAGAGCAGAAGATTGGACCACANNNAGATCGGAA |
| 69049_1 | TATGAGTGTGGAGTCGTTGCCACACACACACTCTCTCTCTCTCNNNAGATCGGAAGA |
| 69051_1 | TATGAGTGTGGAGTCGTTGCCACACACACACACAAAACAACACNNNAGATCGGAAG |
| 69052_1 | TATGAGTGTGGAGTCGTTGCAGCACAAAATATTTGCAGCTGGTNNNAGATCGGAAG |
| 69053_1 | TATGAGTGTGGAGTCGTTGCACCTAATGACCTCTGCCTGTACTNNNAGATCGGAAG |
| 69054_1 | TATGAGTGTGGAGTCGTTGCGAAAGAGATAGGCAGGGAGGAAGNNNAGATCGGA |
| 69055_1 | TATGAGTGTGGAGTCGTTGCAAATGTGTTGACTGAGGATGGGTNNNAGATCGGAA |
| 69056_1 | TATGAGTGTGGAGTCGTTGCTACCACAGAATTGCAGCAATAGCNNNAGATCGGAAG |
| 69057_1 | TATGAGTGTGGAGTCGTTGCGGTAGGCCGACCAGAGATAATTTNNNAGATCGGAA |
| 69058_1 | TATGAGTGTGGAGTCGTTGCTAAGATGACAGCAAGCAAAGTGCNNNAGATCGGAA |
| 69060_1 | TATGAGTGTGGAGTCGTTGCTGATGGTCCAAGAGAAGTCAAGGNNNAGATCGGAA |
| 69061_1 | TATGAGTGTGGAGTCGTTGCCTCCCTCGTAAGACAGTGGTAAGNNNAGATCGGAAG |
| 69062_1 | TATGAGTGTGGAGTCGTTGCCCCTCTCCCTCTCCCATCTATAANNNAGATCGGAAGA |
| 69063_1 | TATGAGTGTGGAGTCGTTGCAGAGACAGAGAGAGAGATAGGGCNNNAGATCGGA |
| 69064_1 | TATGAGTGTGGAGTCGTTGCTTCTTCCAGACCTTGTCCAAGACNNNAGATCGGAAG |
| 69065_1 | TATGAGTGTGGAGTCGTTGCGTGTGTGTGTGTGTGTGTATGTTNNNAGATCGGAA |
| 69066_1 | TATGAGTGTGGAGTCGTTGCAGGGAGAGAGAGAGAGAGAGAGANNNAGATCGG |
| 69067_1 | TATGAGTGTGGAGTCGTTGCTGTGTGTGTGTGTATACAGGCATNNNAGATCGGAA |
| 69068_1 | TATGAGTGTGGAGTCGTTGCCACACACACACACACACACATATNNNAGATCGGAAG |
| 69070_1 | TATGAGTGTGGAGTCGTTGCTGTGCCTACCCTAAAGCTAACTGNNNAGATCGGAAG |
| 69071_1 | TATGAGTGTGGAGTCGTTGCCCCCGAGTACACCTTGATGAATTNNNAGATCGGAAG |
| 69072_1 | TATGAGTGTGGAGTCGTTGCTCTTCAGTATCTTCAGTAACGGCNNNAGATCGGAAG |
| 69073_1 | TATGAGTGTGGAGTCGTTGCCCTAATGGTGACTTGGAATTGCCNNNAGATCGGAAG |
| 69074_1 | TATGAGTGTGGAGTCGTTGCTCAGAGATGGAGAGACAGCAAAGNNNAGATCGGA |
| 69075_1 | TATGAGTGTGGAGTCGTTGCATTCCCACACCTGCAGAACTTATNNNAGATCGGAAG |
| 69076_1 | TATGAGTGTGGAGTCGTTGCGAGAGAGAACCCCAAGTTTCAGANNNAGATCGGAA |

| | |
|---|---|
| 69079_1 | TATGAGTGTGGAGTCGTTGCGCCTGCCTGCCTATCTATCTATCNNNAGATCGGAAGA |
| 69080_1 | TATGAGTGTGGAGTCGTTGCTCTGAGTCACCTAAACCTAGGGANNNAGATCGGAAG |
| 69082_1 | TATGAGTGTGGAGTCGTTGCATCTCTCTCTCTCTTTCTCCCCCNNNAGATCGGAAGAG |
| 69083_1 | TATGAGTGTGGAGTCGTTGCATCCCTTCATTGCTGCTGTCTAANNNAGATCGGAAGA |
| 69084_1 | TATGAGTGTGGAGTCGTTGCAGATGGGCAGATGATGAAGTCAGNNNAGATCGGA |
| 69085_1 | TATGAGTGTGGAGTCGTTGCGTGTGTGTGTGTGTGTGTGTTTANNNAGATCGGAA |
| 69086_1 | TATGAGTGTGGAGTCGTTGCTCTCTCTCTCTCTCTCTCTGTGCNNNAGATCGGAAGA |
| 69087_1 | TATGAGTGTGGAGTCGTTGCCTCCTGAGTAGGTGGGATTACAGNNNAGATCGGAA |
| 69089_1 | TATGAGTGTGGAGTCGTTGCATTCATCTGCTTTTATGGCCCAGNNNAGATCGGAAG |
| 69090_1 | TATGAGTGTGGAGTCGTTGCACACAATGCCCGTCAGTTTTATTNNNAGATCGGAAG |
| 69091_1 | TATGAGTGTGGAGTCGTTGCCTCTCACCTCTCTCTCGGTCATANNNAGATCGGAAGA |
| 69092_1 | TATGAGTGTGGAGTCGTTGCGCTTCCTCACTTTCTCCTGTGTANNNAGATCGGAAGA |
| 69093_1 | TATGAGTGTGGAGTCGTTGCACACTTCCAGCTTCCATAACCATNNNAGATCGGAAGA |
| 69095_1 | TATGAGTGTGGAGTCGTTGCCCATTGGATCTCGTGAGAACTCANNNAGATCGGAAG |
| 69096_1 | TATGAGTGTGGAGTCGTTGCTCTTGCAGGTTTTTGTACAGAGANNNAGATCGGAAG |
| 69097_1 | TATGAGTGTGGAGTCGTTGCAAGAAGGAAGGAAGGAGGAAAGANNNAGATCGG |
| 69098_1 | TATGAGTGTGGAGTCGTTGCTTGAATGGCACCTTAGGTTACGANNNAGATCGGAA |
| 69099_1 | TATGAGTGTGGAGTCGTTGCCCTCACCTAAGCCCAATTACCTTNNNAGATCGGAAGA |
| 69100_1 | TATGAGTGTGGAGTCGTTGCTTCTGTGCTGTAATGGCAGTTTGNNNAGATCGGAAG |
| 69101_1 | TATGAGTGTGGAGTCGTTGCACACACACACACACAAATGCATANNNAGATCGGAAG |
| 69102_1 | TATGAGTGTGGAGTCGTTGCATATTGCATAGCAGGAAGAGGGGNNNAGATCGGA |
| 69103_1 | TATGAGTGTGGAGTCGTTGCAGAGAGGGAGAAAGACAAAGACANNNAGATCGGA |
| 69105_1 | TATGAGTGTGGAGTCGTTGCCACATAGGGTTCTCAGCCTCTTTNNNAGATCGGAAG |
| 69106_1 | TATGAGTGTGGAGTCGTTGCAGAGAGAGAGAGAGATGTGTGCTNNNAGATCGGA |
| 69107_1 | TATGAGTGTGGAGTCGTTGCCCCACTGTCATGACCTCACTAAANNNAGATCGGAAG |
| 69108_1 | TATGAGTGTGGAGTCGTTGCCTTTTGTCATTGCTGCAGGGTAANNNAGATCGGAAG |
| 69109_1 | TATGAGTGTGGAGTCGTTGCGCTCAATCGATCTTCCCACCTTANNNAGATCGGAAGA |
| 69111_1 | TATGAGTGTGGAGTCGTTGCATTTCCCCTTCATCTTCTGCCATNNNAGATCGGAAGA |
| 69113_1 | TATGAGTGTGGAGTCGTTGCTTGGAGAGTTAGTTTCTGGCTCCNNNAGATCGGAAG |
| 69114_1 | TATGAGTGTGGAGTCGTTGCCTGGGGTCTCTTTCTCGTCTTATNNNAGATCGGAAG |
| 69115_1 | TATGAGTGTGGAGTCGTTGCGGAGGAAGTGAGAGAAGAAGAGGNNNAGATCGG |
| 69116_1 | TATGAGTGTGGAGTCGTTGCAGTGCAATGATACAGTCTCTGCTNNNAGATCGGAAG |
| 69119_1 | TATGAGTGTGGAGTCGTTGCCTGTGTTTGCCAGTATTGGACATNNNAGATCGGAAG |
| 69121_1 | TATGAGTGTGGAGTCGTTGCTGAATAGGCTGAGTGGGGTAGTANNNAGATCGGA |
| 69122_1 | TATGAGTGTGGAGTCGTTGCTGAGATGGGAATTTTTTAGGGGANNNAGATCGGAA |
| 69123_1 | TATGAGTGTGGAGTCGTTGCTGCACAGAGAGGCCAAATAAGATNNNAGATCGGAA |
| 69124_1 | TATGAGTGTGGAGTCGTTGCATTCACTGTGGTCTTGAACTCCANNNAGATCGGAAG |
| 69125_1 | TATGAGTGTGGAGTCGTTGCCATCCAGACTAGAGTATGGTGGCNNNAGATCGGAA |
| 69127_1 | TATGAGTGTGGAGTCGTTGCGGTAAGGGAGCAGAGAAGATAGCNNNAGATCGGA |
| 69128_1 | TATGAGTGTGGAGTCGTTGCAGTGTTTTCAGGTGTGAGTCAGANNNAGATCGGAA |
| 69129_1 | TATGAGTGTGGAGTCGTTGCGGCCATCTGCTTTACTCAGTCTANNNAGATCGGAAG |
| 69130_1 | TATGAGTGTGGAGTCGTTGCAAGAAGACAATGGAGGTGACTGGNNNAGATCGGA |
| 69131_1 | TATGAGTGTGGAGTCGTTGCGGAAGGAAGGAAGAAAGGAAGGANNNAGATCGG |

| | |
|---|---|
| 69133_1 | TATGAGTGTGGAGTCGTTGCATCAGTGCGCAAACATTTCCTACNNNAGATCGGAAG |
| 69134_1 | TATGAGTGTGGAGTCGTTGCAAGAAAGGCGTTTGGCTTTAGACNNNAGATCGGAA |
| 69135_1 | TATGAGTGTGGAGTCGTTGCTTCAAGTGATTCTCCTGCCTCAGNNNAGATCGGAAG |
| 69136_1 | TATGAGTGTGGAGTCGTTGCTCCAAAATAACGAACCCACTCCTNNNAGATCGGAAG |
| 69137_1 | TATGAGTGTGGAGTCGTTGCCCTAGGGTGGCAAAGTGAGTTATNNNAGATCGGAA |
| 69138_1 | TATGAGTGTGGAGTCGTTGCGATAGGGAGCCCATCTTCAGATGNNNAGATCGGAA |
| 69139_1 | TATGAGTGTGGAGTCGTTGCACACACACACACACACACAAATTNNNAGATCGGAAG |
| 69140_1 | TATGAGTGTGGAGTCGTTGCTAAGAGGGAACCGACATCAACTGNNNAGATCGGAA |
| 69141_1 | TATGAGTGTGGAGTCGTTGCCTAGACGTTGTCCAAGACCATCANNNAGATCGGAAG |
| 69142_1 | TATGAGTGTGGAGTCGTTGCTACGTTGGGGAAAACCTACTCAGNNNAGATCGGAA |
| 69143_1 | TATGAGTGTGGAGTCGTTGCGAGAACAAGAGTCTGCCTGGTAANNNAGATCGGAA |
| 69145_1 | TATGAGTGTGGAGTCGTTGCTTCTTTGAGATGGAGTCTTGCTCNNNAGATCGGAAG |
| 69147_1 | TATGAGTGTGGAGTCGTTGCAGGAAGAAACTCCAGCATACCAGNNNAGATCGGAA |
| 69148_1 | TATGAGTGTGGAGTCGTTGCAGATGCAAAGTGAGGTATCGAGGNNNAGATCGGA |
| 69149_1 | TATGAGTGTGGAGTCGTTGCAAGAGAAGAGAAAAGGCCTGGTTNNNAGATCGGA |
| 69150_1 | TATGAGTGTGGAGTCGTTGCCAAACTCAGCCTCCTAAGTAGCTNNNAGATCGGAAG |
| 69151_1 | TATGAGTGTGGAGTCGTTGCAAGTGATTCTCCTGCCTCAGTTTNNNAGATCGGAAG |
| 69153_1 | TATGAGTGTGGAGTCGTTGCTGATCTGGTTAGGGTTGTGGATGNNNAGATCGGAA |
| 69155_1 | TATGAGTGTGGAGTCGTTGCACAAACACACACCCCTACTCATTNNNAGATCGGAAGA |
| 69156_1 | TATGAGTGTGGAGTCGTTGCCCTCTCTCCCTCTCTCTCTCTTCNNNAGATCGGAAGAG |
| 69157_1 | TATGAGTGTGGAGTCGTTGCCTCGCCTCTTCCTCATTAGAGTCNNNAGATCGGAAGA |
| 69158_1 | TATGAGTGTGGAGTCGTTGCTGCAGGCATTTATGTCCAGGATANNNAGATCGGAAG |
| 69160_1 | TATGAGTGTGGAGTCGTTGCTCTGTATCTTGTCACCCAGGTTGNNNAGATCGGAAG |
| 69161_1 | TATGAGTGTGGAGTCGTTGCGATACAGGGCTTTTCTCTGTTGCNNNAGATCGGAAG |
| 69162_1 | TATGAGTGTGGAGTCGTTGCAAACTCTGCCTCCTAGGTTCAAGNNNAGATCGGAAG |
| 69163_1 | TATGAGTGTGGAGTCGTTGCAGAGAGAGAGAGATCCTTCACCCNNNAGATCGGAA |
| 69167_1 | TATGAGTGTGGAGTCGTTGCCATGAGAGAGAGTGAGTGAGCAANNNAGATCGGA |
| 69168_1 | TATGAGTGTGGAGTCGTTGCATCACCCCAGCTCTGATTATGACNNNAGATCGGAAG |
| 69169_1 | TATGAGTGTGGAGTCGTTGCGCCCTGAACTTACAAAATGAGGGNNNAGATCGGAA |
| 69170_1 | TATGAGTGTGGAGTCGTTGCCAAAGCACACACACACACAAAACNNNAGATCGGAAG |
| 69171_1 | TATGAGTGTGGAGTCGTTGCGGAGGGAGAGAGAGAGATGAAGTNNNAGATCGGA |
| 69172_1 | TATGAGTGTGGAGTCGTTGCCCCTGCCTCCTTTCTTTCTCTATNNNAGATCGGAAGAG |
| 69173_1 | TATGAGTGTGGAGTCGTTGCCAAGCACGTACACATTCACACATNNNAGATCGGAAG |
| 69174_1 | TATGAGTGTGGAGTCGTTGCAGTTATTGAAAATAGCGCTGGCCNNNAGATCGGAAG |
| 69175_1 | TATGAGTGTGGAGTCGTTGCCCCACGGACAACATTTACAACAANNNAGATCGGAAG |
| 69176_1 | TATGAGTGTGGAGTCGTTGCACCATGGTGAGACAAAGCCTATTNNNAGATCGGAAG |
| 69178_1 | TATGAGTGTGGAGTCGTTGCTCTCCCACCTTCTTTGTTCTAGCNNNAGATCGGAAGA |
| 69180_1 | TATGAGTGTGGAGTCGTTGCCTAAGCCTCAGATGAGACCTCAGNNNAGATCGGAAG |
| 69183_1 | TATGAGTGTGGAGTCGTTGCTCAATTGGGAGACCTATCATGGTNNNAGATCGGAAG |
| 69184_1 | TATGAGTGTGGAGTCGTTGCCACACACACACACAAGATCCAAANNNAGATCGGAAG |
| 69185_1 | TATGAGTGTGGAGTCGTTGCGCATAAATTTTTGGGTGCTGGTGNNNAGATCGGAA |
| 69186_1 | TATGAGTGTGGAGTCGTTGCGTGTGTGTGTGTGTGTGTGTATTNNNAGATCGGAA |
| 69187_1 | TATGAGTGTGGAGTCGTTGCTGAAAATTACTGGACCTGCCCTTNNNAGATCGGAAG |

| | |
|---|---|
| 69189_1 | TATGAGTGTGGAGTCGTTGCATTATGACTCACTGCAGCCTCAANNNAGATCGGAAG |
| 69190_1 | TATGAGTGTGGAGTCGTTGCTCACATGCATGAGTCAAACTTGGNNNAGATCGGAAG |
| 69192_1 | TATGAGTGTGGAGTCGTTGCAGGCAGCCCCATATATTTCCTCNNNAGATCGGAAGA |
| 69193_1 | TATGAGTGTGGAGTCGTTGCAACCAGGTAGACTGAAAGTTTGTNNNAGATCGGAA |
| 69194_1 | TATGAGTGTGGAGTCGTTGCACCCAACTAACCAGCATCTTGATNNNAGATCGGAAG |
| 69195_1 | TATGAGTGTGGAGTCGTTGCCAAGCATCACCTTCTGCAATCATNNNAGATCGGAAG |
| 69197_1 | TATGAGTGTGGAGTCGTTGCTGCTTTGGTCCTGAAGATAGACCNNNAGATCGGAAG |
| 69198_1 | TATGAGTGTGGAGTCGTTGCTTTGGACCTCCAGTTGTAATGCTNNNAGATCGGAAG |
| 69199_1 | TATGAGTGTGGAGTCGTTGCATCTACTATTCGGCCCAGCAATTNNNAGATCGGAAG |
| 69200_1 | TATGAGTGTGGAGTCGTTGCCAGACTCAGCTCAGAATCTTCCANNNAGATCGGAAG |
| 69201_1 | TATGAGTGTGGAGTCGTTGCGGTTGTTTCCCTTCTTTCCTGTGNNNAGATCGGAAGA |
| 69202_1 | TATGAGTGTGGAGTCGTTGCTGCTTTCTCAGGACTCACACTTTNNNAGATCGGAAGA |
| 69203_1 | TATGAGTGTGGAGTCGTTGCTAGCCTTGGTCAATTCTCCACAANNNAGATCGGAAG |
| 69204_1 | TATGAGTGTGGAGTCGTTGCTCCAGGTCAAGTTGTCCTTTGAANNNAGATCGGAAG |
| 69205_1 | TATGAGTGTGGAGTCGTTGCGTGTTGACTGCAGAACCAATACCNNNAGATCGGAAG |
| 69207_1 | TATGAGTGTGGAGTCGTTGCCCCATAACCAACTCCATCCTCAANNNAGATCGGAAGA |
| 69208_1 | TATGAGTGTGGAGTCGTTGCGTGGCTCACACCTGTAATCCCNNNAGATCGGAAGAG |
| 69209_1 | TATGAGTGTGGAGTCGTTGCCTGAGAACATGGCAACCTGAAAGNNNAGATCGGAA |
| 69210_1 | TATGAGTGTGGAGTCGTTGCATTACCCCTTTCAACACTGCCTANNNAGATCGGAAGA |
| 69211_1 | TATGAGTGTGGAGTCGTTGCGTATGAAAGCAGCAGGGCTATTGNNNAGATCGGAA |
| 69212_1 | TATGAGTGTGGAGTCGTTGCCTCTGGCCCTTTCTTTCCTGTATNNNAGATCGGAAGA |
| 69213_1 | TATGAGTGTGGAGTCGTTGCAAGGCAGAATGCAGTAAGGATCANNNAGATCGGAA |
| 69214_1 | TATGAGTGTGGAGTCGTTGCCCCTCTGTCTACGTGGAGAATTCNNNAGATCGGAAG |
| 69215_1 | TATGAGTGTGGAGTCGTTGCTGTTCAACTGTGCAAATGGTGAGNNNAGATCGGAA |
| 69216_1 | TATGAGTGTGGAGTCGTTGCATGCGATGTACCTGTTTCCTCTTNNNAGATCGGAAG |
| 69217_1 | TATGAGTGTGGAGTCGTTGCTGTGTGTGTGTGTGTGTTAGAGANNNAGATCGGAA |
| 69218_1 | TATGAGTGTGGAGTCGTTGCAGAAATGGAAGCTCAGAGAGGTGNNNAGATCGGA |
| 69220_1 | TATGAGTGTGGAGTCGTTGCCCCTATCTCCCTTGCAATCTGATNNNAGATCGGAAGA |
| 69223_1 | TATGAGTGTGGAGTCGTTGCACGAAGGAAACTCGGAAAGTGTANNNAGATCGGAA |
| 69224_1 | TATGAGTGTGGAGTCGTTGCCCCACTCAACTGGACTTTGTTTCNNNAGATCGGAAGA |
| 69228_1 | TATGAGTGTGGAGTCGTTGCAAAGAAAGGAGAGATCCGCAAGTNNNAGATCGGAA |
| 69229_1 | TATGAGTGTGGAGTCGTTGCAGAAGAGAGGTGAGGTTAGGTGANNNAGATCGGA |
| 69231_1 | TATGAGTGTGGAGTCGTTGCGATCCTTCAACTTTCACACAGGCNNNAGATCGGAAG |
| 69232_1 | TATGAGTGTGGAGTCGTTGCGAGACAGGGTTTCATCATGTTGGNNNAGATCGGAA |
| 69233_1 | TATGAGTGTGGAGTCGTTGCGTTCAAGAGATTCTCCTGCCTCANNNAGATCGGAAG |
| 69234_1 | TATGAGTGTGGAGTCGTTGCTACAAAAAATTAGCCAGGCGTGGNNNAGATCGGAA |
| 69235_1 | TATGAGTGTGGAGTCGTTGCTTCCTGGGTTCATGTCATTCTCCNNNAGATCGGAAGA |
| 69236_1 | TATGAGTGTGGAGTCGTTGCCACCGTGCCCAACATTCTTTATTNNNAGATCGGAAGA |
| 69239_1 | TATGAGTGTGGAGTCGTTGCAGGGAGACGTTACAAGCAATACCNNNAGATCGGAA |
| 69240_1 | TATGAGTGTGGAGTCGTTGCTTCCTGTCTTTGATGTCTCCCTGNNNAGATCGGAAGA |
| 69241_1 | TATGAGTGTGGAGTCGTTGCTCTCCCAAAGTGCTGGAATTACANNNAGATCGGAAG |
| 69242_1 | TATGAGTGTGGAGTCGTTGCGACCTTGCTGAATAAAGGCTCCTNNNAGATCGGAAG |
| 69243_1 | TATGAGTGTGGAGTCGTTGCTGGCATCTTCCCTGCATCTTTATNNNAGATCGGAAGA |

| | |
|---|---|
| 69244_1 | TATGAGTGTGGAGTCGTTGCTTTCTGAGTTTCAAATGCCCGTCNNNAGATCGGAAG |
| 69245_1 | TATGAGTGTGGAGTCGTTGCTCTCAGTAGCCTCCTTAGGAACANNNAGATCGGAAG |
| 69246_1 | TATGAGTGTGGAGTCGTTGCTGCCATAGAGATGAGGAGGTACTNNNAGATCGGAA |
| 69247_1 | TATGAGTGTGGAGTCGTTGCGCATTATCCTGCCTACCACAGTANNNAGATCGGAAG |
| 69248_1 | TATGAGTGTGGAGTCGTTGCCCTAAGTTCGTACACTGAGCACTNNNAGATCGGAAG |
| 69250_1 | TATGAGTGTGGAGTCGTTGCGAAACAGATGGCACAAACTCTGGNNNAGATCGGAA |
| 69251_1 | TATGAGTGTGGAGTCGTTGCAAAATGGGTATGTTTGGGTTGGCNNNAGATCGGAA |
| 69252_1 | TATGAGTGTGGAGTCGTTGCCTGGGGGAAAAGAGCAAGACTTCNNNAGATCGGAA |
| 69253_1 | TATGAGTGTGGAGTCGTTGCGGACTATTGGCACATCTTGGAGANNNAGATCGGAA |
| 69254_1 | TATGAGTGTGGAGTCGTTGCAAGGGTCTGCCAGGAGTTTTTATNNNAGATCGGAA |
| 69255_1 | TATGAGTGTGGAGTCGTTGCGCCACACCCAGCTAATTTTTGTANNNAGATCGGAAG |
| 69256_1 | TATGAGTGTGGAGTCGTTGCGCTTTGTTCTTTTGCTGTTGCTCNNNAGATCGGAAGA |
| 69257_1 | TATGAGTGTGGAGTCGTTGCGCCTTTCCTGGTCTGCTAAGTAGNNNAGATCGGAAG |
| 69258_1 | TATGAGTGTGGAGTCGTTGCACCTCTTTCTCCCTAGACTTCCANNNAGATCGGAAGA |
| 69259_1 | TATGAGTGTGGAGTCGTTGCGATGGACTGAGTAAGGACGTCTGNNNAGATCGGAA |
| 69260_1 | TATGAGTGTGGAGTCGTTGCTTCTCATGAAAGGTTGGGCACTANNNAGATCGGAA |
| 69261_1 | TATGAGTGTGGAGTCGTTGCTGCCTATGCTGGTCTTGAATTCTNNNAGATCGGAAG |
| 69262_1 | TATGAGTGTGGAGTCGTTGCACTACTCGGGAAACTGTAGCAAGNNNAGATCGGAA |
| 69263_1 | TATGAGTGTGGAGTCGTTGCTTCAGGGTAAACACATTGGATGANNNAGATCGGAA |
| 69264_1 | TATGAGTGTGGAGTCGTTGCAAGTCTTCATCTAACACCGTCCCNNNAGATCGGAAGA |
| 69268_1 | TATGAGTGTGGAGTCGTTGCAACCTGCTTTCTTCAGAGAAGCANNNAGATCGGAAG |
| 69269_1 | TATGAGTGTGGAGTCGTTGCGGCATTTGTCAGAAGCTACACAGNNNAGATCGGAA |
| 69270_1 | TATGAGTGTGGAGTCGTTGCTCACTTGAAGTCAGGAGTTCGAGNNNAGATCGGAA |
| 69272_1 | TATGAGTGTGGAGTCGTTGCATCTAAACCATGGCACCTCACTTNNNAGATCGGAAGA |
| 69273_1 | TATGAGTGTGGAGTCGTTGCGGTACTCTCTCCTCTTTTGCACANNNAGATCGGAAGA |
| 69274_1 | TATGAGTGTGGAGTCGTTGCTTCAAGTGATTCTCCTGTCTCGGNNNAGATCGGAAG |
| 69275_1 | TATGAGTGTGGAGTCGTTGCTTCAAGTGATTCTCCTGTCTCGGNNNAGATCGGAAG |
| 69276_1 | TATGAGTGTGGAGTCGTTGCAGGTTGTACCCAATGACTGACTCNNNAGATCGGAAG |
| 69277_1 | TATGAGTGTGGAGTCGTTGCCTTTACTGTGCATGGTCCCAGAANNNAGATCGGAAG |
| 69279_1 | TATGAGTGTGGAGTCGTTGCAACTTGGTTACGGCATTAGTCCTNNNAGATCGGAAG |
| 69281_1 | TATGAGTGTGGAGTCGTTGCATTCATCAGGCTGGTCTCAAACTNNNAGATCGGAAG |
| 69282_1 | TATGAGTGTGGAGTCGTTGCCAGGAGAAAACAGAAAGTGCCAGNNNAGATCGGAA |
| 69284_1 | TATGAGTGTGGAGTCGTTGCGGGCTCCCTCTCATTATTCTTGANNNAGATCGGAAGA |
| 69285_1 | TATGAGTGTGGAGTCGTTGCCTCTGGAGCTGCCTGTAGATTACNNNAGATCGGAAG |
| 69286_1 | TATGAGTGTGGAGTCGTTGCACAATGCCTGCAACCATTTACAANNNAGATCGGAAG |
| 69287_1 | TATGAGTGTGGAGTCGTTGCTAAGCAGGGTGATGTAGAACAGGNNNAGATCGGAA |
| 69288_1 | TATGAGTGTGGAGTCGTTGCCAAAGAGAGGTCAGGTTTTGCTGNNNAGATCGGAA |
| 69289_1 | TATGAGTGTGGAGTCGTTGCGAACTCCTGACCTTAGGTGATCCNNNAGATCGGAAG |
| 69290_1 | TATGAGTGTGGAGTCGTTGCGATTTCTTGCACAGTGGTCACTCNNNAGATCGGAAG |
| 69291_1 | TATGAGTGTGGAGTCGTTGCCTACCAATAAAGGAAGGGGAGGGNNNAGATCGGA |
| 69292_1 | TATGAGTGTGGAGTCGTTGCTAAATCAGAGTGCCAAGAGGGACNNNAGATCGGAA |
| 69293_1 | TATGAGTGTGGAGTCGTTGCCATACGTCACACCTGAGAGAGAANNNAGATCGGAA |
| 69295_1 | TATGAGTGTGGAGTCGTTGCGAAAGTGAAGGCCTGAGAGGTTANNNAGATCGGA |

| | |
|---|---|
| 69296_1 | TATGAGTGTGGAGTCGTTGCAACCTGTGTGTCTTATTGCAACGNNNAGATCGGAAG |
| 69300_1 | TATGAGTGTGGAGTCGTTGCCACCAGGCCCCTCTTTTCTTTATNNNAGATCGGAAGA |
| 69301_1 | TATGAGTGTGGAGTCGTTGCTTCTCCTCCAGACTACAAACCAGNNNAGATCGGAAG |
| 69304_1 | TATGAGTGTGGAGTCGTTGCTAAATCTTGCTGCTGCTCACTCTNNNAGATCGGAAG |
| 69307_1 | TATGAGTGTGGAGTCGTTGCGAAACGTCACAAGATGCAGGATTNNNAGATCGGAA |
| 69308_1 | TATGAGTGTGGAGTCGTTGCCAGAGACCACAAGAGTATGGCAGNNNAGATCGGAA |
| 69309_1 | TATGAGTGTGGAGTCGTTGCCTGAGTAGCTGGAATTACAGGCANNNAGATCGGAA |
| 69310_1 | TATGAGTGTGGAGTCGTTGCCCGCTAAGTGGATAGCATGAGATNNNAGATCGGAA |
| 69311_1 | TATGAGTGTGGAGTCGTTGCGAGTGCAGTGATGTTATCATGGCNNNAGATCGGAA |
| 69313_1 | TATGAGTGTGGAGTCGTTGCTTTTTGAGACAGTCTTGCTCTGTNNNAGATCGGAAG |
| 69314_1 | TATGAGTGTGGAGTCGTTGCTCTCTCTCTCTTTCTTCCTCCCTNNNAGATCGGAAGAG |
| 69315_1 | TATGAGTGTGGAGTCGTTGCCCAAAGTGTTGTGTTTAGTGGCANNNAGATCGGAA |
| 69316_1 | TATGAGTGTGGAGTCGTTGCGTGTGTGTGTGTGTGTGTAAANNNAGATCGGAA |
| 69317_1 | TATGAGTGTGGAGTCGTTGCTCTAGGTTAGAAGCCCTGAGAGGNNNAGATCGGAA |
| 69319_1 | TATGAGTGTGGAGTCGTTGCGAAAGGAAGGAAGGAAGGAAGGANNNAGATCGG |
| 69320_1 | TATGAGTGTGGAGTCGTTGCAAGAGAGAGAGGGAGACCTTGTTNNNAGATCGGA |
| 69321_1 | TATGAGTGTGGAGTCGTTGCTGCTGCTATTCTGTTGGTCTGATNNNAGATCGGAAG |
| 69322_1 | TATGAGTGTGGAGTCGTTGCCTATCACATTAGGGCCACACCTTNNNAGATCGGAAG |
| 69324_1 | TATGAGTGTGGAGTCGTTGCGGGAAAAGAACAAGAGTCTCTGCNNNAGATCGGAA |
| 69325_1 | TATGAGTGTGGAGTCGTTGCAGGGAGAGAGAGAGAGAGAGAGANNNAGATCGG |
| 69326_1 | TATGAGTGTGGAGTCGTTGCGTGTGTGTGTGTGTGTGTTTANNNAGATCGGAA |
| 69327_1 | TATGAGTGTGGAGTCGTTGCGGAGAGAGAGAGAGAGACAGAGANNNAGATCGG |
| 69328_1 | TATGAGTGTGGAGTCGTTGCACATAGTCTCACTGCAATGCCTANNNAGATCGGAAG |
| 69329_1 | TATGAGTGTGGAGTCGTTGCGGGATCAAAGTTGCAGGTGATTCNNNAGATCGGAA |
| 69330_1 | TATGAGTGTGGAGTCGTTGCTTCAGGCTCTGTGACTTTCATCANNNAGATCGGAAG |
| 69331_1 | TATGAGTGTGGAGTCGTTGCTTCTCTCCTCTGTCTTTTCCCACNNNAGATCGGAAGA |
| 69332_1 | TATGAGTGTGGAGTCGTTGCGGTAGATACAGATGCAGCATCCANNNAGATCGGAA |
| 69333_1 | TATGAGTGTGGAGTCGTTGCGTGTGTGTGTGATCTCTCTCTNNNAGATCGGAAG |
| 69334_1 | TATGAGTGTGGAGTCGTTGCAGAGAGAGAGAGAGAGAGAAGGCNNNAGATCGG |
| 69335_1 | TATGAGTGTGGAGTCGTTGCGGGACACAGAGAGAAGACAACATNNNAGATCGGA |
| 69336_1 | TATGAGTGTGGAGTCGTTGCGCCTGACTTATAATGGCTCGACTNNNAGATCGGAAG |
| 69337_1 | TATGAGTGTGGAGTCGTTGCAGATTGAGATCATCCCGGCTAACNNNAGATCGGAAG |
| 69338_1 | TATGAGTGTGGAGTCGTTGCAGGATTGTGATGTTTTCCTGTCANNNAGATCGGAAG |
| 69339_1 | TATGAGTGTGGAGTCGTTGCATGTTTGGAATCAGGACTCCGAANNNAGATCGGAA |
| 69340_1 | TATGAGTGTGGAGTCGTTGCTCCACAGATAAGCAGGTGACTACNNNAGATCGGAA |
| 69341_1 | TATGAGTGTGGAGTCGTTGCCCTCCTAAGCTCAAGTGATCCTCNNNAGATCGGAAG |
| 69342_1 | TATGAGTGTGGAGTCGTTGCTGAAGGAAGGAAGGAGGAAAGAGNNNAGATCGG |
| 69344_1 | TATGAGTGTGGAGTCGTTGCTCTCTGTCTCTCACTCTCTGACANNNAGATCGGAAGA |
| 69345_1 | TATGAGTGTGGAGTCGTTGCTGCCCCTGGAGAGGAAAATATTTNNNAGATCGGAA |
| 69346_1 | TATGAGTGTGGAGTCGTTGCTCTGAAAACATATGCCTTGGGGANNNAGATCGGAA |
| 69349_1 | TATGAGTGTGGAGTCGTTGCAGGAGAGAGAGAGAGAAGGCAATNNNAGATCGGA |
| 69350_1 | TATGAGTGTGGAGTCGTTGCCAATCCTCTTACCTCAACCTCCCNNNAGATCGGAAGA |
| 69351_1 | TATGAGTGTGGAGTCGTTGCAGAGAGGGAGAGAGAGAGAGAGANNNAGATCGG |

| | |
|---|---|
| 69352_1 | TATGAGTGTGGAGTCGTTGCGGATATGTCTCAGGCGTCTTCTCNNNAGATCGGAAG |
| 69354_1 | TATGAGTGTGGAGTCGTTGCGCTGGTCTCAGACTCCTAAGTTCNNNAGATCGGAAG |
| 69355_1 | TATGAGTGTGGAGTCGTTGCCCTATTGGGTCCGATACTCACATNNNAGATCGGAAG |
| 69356_1 | TATGAGTGTGGAGTCGTTGCTCTATTGGTTCCGTTTCTCTGGANNNAGATCGGAAG |
| 69359_1 | TATGAGTGTGGAGTCGTTGCGTCTGTCCAGCCTTGACTAATCANNNAGATCGGAAG |
| 69360_1 | TATGAGTGTGGAGTCGTTGCACCCTACTCTAGTTCCCATCCATNNNAGATCGGAAGA |
| 69361_1 | TATGAGTGTGGAGTCGTTGCCAGGGAAGAGAGGAAAGGAGAACNNNAGATCGGA |
| 69362_1 | TATGAGTGTGGAGTCGTTGCGACAGACACTCATGCAGGTTTTCNNNAGATCGGAAG |
| 69363_1 | TATGAGTGTGGAGTCGTTGCTACTTGCTGCTTACCATGTGCTANNNAGATCGGAAG |
| 69364_1 | TATGAGTGTGGAGTCGTTGCCTTGATCCAATCACTTCCCTCCANNNAGATCGGAAGA |
| 69365_1 | TATGAGTGTGGAGTCGTTGCAAATAGGTGTGTGTGTGTGTGTGNNNAGATCGGAA |
| 69366_1 | TATGAGTGTGGAGTCGTTGCAAACCACCTCCATGATCCAATCANNNAGATCGGAAG |
| 69367_1 | TATGAGTGTGGAGTCGTTGCGAGCATGAGAGCCAGATGATACANNNAGATCGGAA |
| 69368_1 | TATGAGTGTGGAGTCGTTGCAGCTGGAGAACAAGGAAAGTTGANNNAGATCGGA |
| 69369_1 | TATGAGTGTGGAGTCGTTGCTGGGCCATAGAAACAACTTGGATNNNAGATCGGAA |
| 69370_1 | TATGAGTGTGGAGTCGTTGCTTCTTTTTCACGGAGTTTTGCTCNNNAGATCGGAAG |
| 69372_1 | TATGAGTGTGGAGTCGTTGCGTGTGTGTGTGTGTGTGTGTATTNNNAGATCGGAA |
| 69374_1 | TATGAGTGTGGAGTCGTTGCTCTGAACACAGCTCTAGTGCATTNNNAGATCGGAAG |
| 69375_1 | TATGAGTGTGGAGTCGTTGCAAGGATGTAGTGCAGTTCAGCATNNNAGATCGGAA |
| 69376_1 | TATGAGTGTGGAGTCGTTGCTGACGGTGAAATTACACAAGCACNNNAGATCGGAA |
| 69378_1 | TATGAGTGTGGAGTCGTTGCGAGGTTCTGGTCTGAGTCTGAAGNNNAGATCGGAA |
| 69379_1 | TATGAGTGTGGAGTCGTTGCGCGTTGTTGCCTTATGGAGTTTTNNNAGATCGGAAG |
| 69380_1 | TATGAGTGTGGAGTCGTTGCACGATCATGAGGACAGTACCAACNNNAGATCGGAA |
| 69383_1 | TATGAGTGTGGAGTCGTTGCCCTAGGGAATTTGTGACCTGTGTNNNAGATCGGAAG |
| 69384_1 | TATGAGTGTGGAGTCGTTGCATGTAAACAAGTGGGCATGGTTGNNNAGATCGGAA |
| 69386_1 | TATGAGTGTGGAGTCGTTGCAGAGAGAAGAGAGAGGGGGAAAANNNAGATCGG |
| 69387_1 | TATGAGTGTGGAGTCGTTGCAGTTAGGAAACACAGGGCTTCATNNNAGATCGGAA |
| 69388_1 | TATGAGTGTGGAGTCGTTGCACCTCTGGTGGCTTGGATTTTATNNNAGATCGGAAG |
| 69389_1 | TATGAGTGTGGAGTCGTTGCTGACTCTCTCTCTGTCTCTCTCGNNNAGATCGGAAGA |
| 69390_1 | TATGAGTGTGGAGTCGTTGCAGAGGAAGAAGAGAAGGAAGGGANNNAGATCGG |
| 69391_1 | TATGAGTGTGGAGTCGTTGCCCCTCATTGTACATCTCAGGGAGNNNAGATCGGAAG |
| 69392_1 | TATGAGTGTGGAGTCGTTGCCGGGAAGTAGATCATCACTGAGCNNNAGATCGGAA |
| 69394_1 | TATGAGTGTGGAGTCGTTGCGTAAGGGTTTGACATTGTGAGCCNNNAGATCGGAA |
| 69395_1 | TATGAGTGTGGAGTCGTTGCGTGTGTGTGTGTGTGTGTTTGTANNNAGATCGGAA |
| 69396_1 | TATGAGTGTGGAGTCGTTGCGGTGTGATCTGGGCTTACTACAANNNAGATCGGAA |
| 69398_1 | TATGAGTGTGGAGTCGTTGCATCAACTCGACCTCCTCTTGTTCNNNAGATCGGAAGA |
| 69400_1 | TATGAGTGTGGAGTCGTTGCCTCCTTCTCCCTGGTTCAAATGANNNAGATCGGAAG |
| 69401_1 | TATGAGTGTGGAGTCGTTGCTCTTTTACCTCTTGGCTTTGAGANNNAGATCGGAAG |
| 69402_1 | TATGAGTGTGGAGTCGTTGCTATTCCTGTGTAGCTCTGCATCCNNNAGATCGGAAG |
| 69404_1 | TATGAGTGTGGAGTCGTTGCATAGCATCACATTGGGAGCTTCANNNAGATCGGAAG |
| 69405_1 | TATGAGTGTGGAGTCGTTGCAAAGCAGCTAACGAGTCACTACTNNNAGATCGGAAG |
| 69406_1 | TATGAGTGTGGAGTCGTTGCAACACAGAAGCTAGGGAGAAAGGNNNAGATCGGA |
| 69407_1 | TATGAGTGTGGAGTCGTTGCGGAGGGTAGTGATGTGCTCATAGNNNAGATCGGAA |

| | |
|---|---|
| 69408_1 | TATGAGTGTGGAGTCGTTGCCTCCTGCCCTTGATATCAGACTCNNNAGATCGGAAG |
| 69409_1 | TATGAGTGTGGAGTCGTTGCTTGTAGGAAGGTTTGGGAGTGAGNNNAGATCGGA |
| 69410_1 | TATGAGTGTGGAGTCGTTGCTGCAGCATTTGACAAACAGTAGANNNAGATCGGAA |
| 69411_1 | TATGAGTGTGGAGTCGTTGCGATCTCAGGCTCACTACAACCTCNNNAGATCGGAAG |
| 69412_1 | TATGAGTGTGGAGTCGTTGCTAAGTTCCTGCCATCAAGAAGCTNNNAGATCGGAAG |
| 69413_1 | TATGAGTGTGGAGTCGTTGCTTCAAGTGATTCTCCTGCCTCAGNNNAGATCGGAAG |
| 69415_1 | TATGAGTGTGGAGTCGTTGCCACACACACACACACTCTTTANNNAGATCGGAAGA |
| 69416_1 | TATGAGTGTGGAGTCGTTGCTGTCTGGCTTCTCTAGTGGAAACNNNAGATCGGAAG |
| 69417_1 | TATGAGTGTGGAGTCGTTGCCACACACACACAGAGTTGTTANNNAGATCGGAAG |
| 69418_1 | TATGAGTGTGGAGTCGTTGCCATTTGGGAAAACAGCTTTTGCANNNAGATCGGAAG |
| 69421_1 | TATGAGTGTGGAGTCGTTGCATGTGCTTCTCCCATGGATCTTTNNNAGATCGGAAG |
| 69422_1 | TATGAGTGTGGAGTCGTTGCCCCTTCTCTCCCTTTCTCTCTCTNNNAGATCGGAAGAG |
| 69423_1 | TATGAGTGTGGAGTCGTTGCGCTCCGGTCTGTTTCTCTTTCTANNNAGATCGGAAGA |
| 69424_1 | TATGAGTGTGGAGTCGTTGCGTGTGAAGATGCCATCATGGAAGNNNAGATCGGAA |
| 69425_1 | TATGAGTGTGGAGTCGTTGCCCCACTCTACACAAGGTCTTGATNNNAGATCGGAAG |
| 69426_1 | TATGAGTGTGGAGTCGTTGCGGAGAGAGAGAAGGAAGGAAGGANNNAGATCGG |
| 69427_1 | TATGAGTGTGGAGTCGTTGCCTACTGGATTTGTCTCTCGGGAGNNNAGATCGGAAG |
| 69428_1 | TATGAGTGTGGAGTCGTTGCAAAGCCTAATCACACAAATGCCANNNAGATCGGAAG |
| 69429_1 | TATGAGTGTGGAGTCGTTGCAGCAGTTGCCAAGACAAAAGATCNNNAGATCGGAA |
| 69431_1 | TATGAGTGTGGAGTCGTTGCCATGGAAAGAGTTGCTTGAAGCTNNNAGATCGGAA |
| 69432_1 | TATGAGTGTGGAGTCGTTGCTCCCTCTCTCTCTCTCTCTCTCTNNNAGATCGGAAGAG |
| 69433_1 | TATGAGTGTGGAGTCGTTGCAAGAAGTGGCATGTAAGCTGAGANNNAGATCGGA |
| 69434_1 | TATGAGTGTGGAGTCGTTGCAGTCTGAATGTTCATGTCCCACANNNAGATCGGAAG |
| 69435_1 | TATGAGTGTGGAGTCGTTGCCCATTAGAGGGATTCACCTGCAANNNAGATCGGAAG |
| 69436_1 | TATGAGTGTGGAGTCGTTGCCATGAAGGGATAAGAGACAGCGANNNAGATCGGA |
| 69438_1 | TATGAGTGTGGAGTCGTTGCTGACGGTTTCCTTATAGGCAAAANNNAGATCGGAAG |
| 69439_1 | TATGAGTGTGGAGTCGTTGCAGGCTCTGCTAAGACTCTGTCTANNNAGATCGGAAG |
| 69440_1 | TATGAGTGTGGAGTCGTTGCTATGAAAAGAGGGTGCCAGAGAGNNNAGATCGGA |
| 69441_1 | TATGAGTGTGGAGTCGTTGCTCTCTTTCTCTCTCTCTCACCCTNNNAGATCGGAAGAG |
| 69442_1 | TATGAGTGTGGAGTCGTTGCTATCATGGCTCAACCCAGTGAAANNNAGATCGGAAG |
| 69445_1 | TATGAGTGTGGAGTCGTTGCTACAAGTCTGCAAGAACCACAGTNNNAGATCGGAAG |
| 69446_1 | TATGAGTGTGGAGTCGTTGCAGAAACATCACCGGGGGAAAATANNNAGATCGGAA |
| 69447_1 | TATGAGTGTGGAGTCGTTGCAGAGAGAGAGAGAGGGAGAGAGANNNAGATCGG |
| 69448_1 | TATGAGTGTGGAGTCGTTGCACAGAACCACTCCTGAGATAACGNNNAGATCGGAAG |
| 69449_1 | TATGAGTGTGGAGTCGTTGCTGTGTGTGTGTGTGTGTAGATNNNAGATCGGAA |
| 69450_1 | TATGAGTGTGGAGTCGTTGCGATCCTCCTACTTCAGCCTTCTGNNNAGATCGGAAGA |
| 69451_1 | TATGAGTGTGGAGTCGTTGCGAAGGAAAGGAAGGAAGGAAGGANNNAGATCGG |
| 69452_1 | TATGAGTGTGGAGTCGTTGCTTTGGGATGTACAGAGACAGTCCNNNAGATCGGAA |
| 69453_1 | TATGAGTGTGGAGTCGTTGCCTCACTGCAGCATTGAACTTCTGNNNAGATCGGAAG |
| 69454_1 | TATGAGTGTGGAGTCGTTGCGAGAGAGAGCGAGCAACTAGTACNNNAGATCGGA |
| 69455_1 | TATGAGTGTGGAGTCGTTGCGTGTGTGTGTGTGTGTGTAAANNNAGATCGGAA |
| 69456_1 | TATGAGTGTGGAGTCGTTGCCCAGGTACTATGGCTGAAAACCTNNNAGATCGGAAG |
| 69457_1 | TATGAGTGTGGAGTCGTTGCGAGAACAGCTGCCTAGTGAAGATNNNAGATCGGAA |

| | |
|---|---|
| 69459_1 | TATGAGTGTGGAGTCGTTGCTCAATCTAAGTGCCCATCAACCANNNAGATCGGAAG |
| 69460_1 | TATGAGTGTGGAGTCGTTGCACACAGTCTGTTTGGGAGAAAGTNNNAGATCGGAA |
| 69462_1 | TATGAGTGTGGAGTCGTTGCCAGTAGCTGGGACTACAGACATGNNNAGATCGGAA |
| 69463_1 | TATGAGTGTGGAGTCGTTGCCTTTCCCCTTCACAGACACACTANNNAGATCGGAAGA |
| 69464_1 | TATGAGTGTGGAGTCGTTGCCAGAGACATTCAGGCCATCAATGNNNAGATCGGAAG |
| 69465_1 | TATGAGTGTGGAGTCGTTGCTTCCTTCCTTCCTTCCTTCCTTCNNNAGATCGGAAGAG |
| 69467_1 | TATGAGTGTGGAGTCGTTGCGGAATGGCTCAGAGAAGGTAAGTNNNAGATCGGA |
| 69468_1 | TATGAGTGTGGAGTCGTTGCTCTACGCTAACACTGAGCAAGTCNNNAGATCGGAAG |
| 69469_1 | TATGAGTGTGGAGTCGTTGCAATGCTTGGCACATAGGAGCTATNNNAGATCGGAA |
| 69470_1 | TATGAGTGTGGAGTCGTTGCCCTGTAATCCCAGCACTTTGAGANNNAGATCGGAAG |
| 69471_1 | TATGAGTGTGGAGTCGTTGCACTAGAGTCGGGAGATCATTGGANNNAGATCGGAA |
| 69472_1 | TATGAGTGTGGAGTCGTTGCCTTGGAGGTGGGAGAAAATCAGANNNAGATCGGA |
| 69474_1 | TATGAGTGTGGAGTCGTTGCGAGGAGATGAGAGGAGAGGAGAANNNAGATCGG |
| 69475_1 | TATGAGTGTGGAGTCGTTGCACGCGCATTTACTATGGACAGATNNNAGATCGGAAG |
| 69476_1 | TATGAGTGTGGAGTCGTTGCCCTCCTAAAGTGCTGGGATTACANNNAGATCGGAAG |
| 69477_1 | TATGAGTGTGGAGTCGTTGCACTAGCTTTCTGGGTAAAGGGTGNNNAGATCGGAA |
| 69478_1 | TATGAGTGTGGAGTCGTTGCGGGAGACTTCAATACCCCACTTTNNNAGATCGGAAG |
| 69479_1 | TATGAGTGTGGAGTCGTTGCGAGGGAAGAGAGCAAAATACCCANNNAGATCGGA |
| 69480_1 | TATGAGTGTGGAGTCGTTGCAGGCTTCTGGGTTTCTTTTCTCTNNNAGATCGGAAG |
| 69481_1 | TATGAGTGTGGAGTCGTTGCACAGAGAGACTGAGAGAGACAGANNNAGATCGGA |
| 69482_1 | TATGAGTGTGGAGTCGTTGCGATTGTTTTTGCCTAGCTGCTGANNNAGATCGGAAG |
| 69483_1 | TATGAGTGTGGAGTCGTTGCCACACACACACACACACACATATNNNAGATCGGAAG |
| 69484_1 | TATGAGTGTGGAGTCGTTGCACCCTAACACATGGAGCATGAATNNNAGATCGGAAG |
| 69485_1 | TATGAGTGTGGAGTCGTTGCGTGTGTGTGTGTGTCTACATGNNNAGATCGGAA |
| 69486_1 | TATGAGTGTGGAGTCGTTGCACTGAGGTTCTCAAGAGCTTCTGNNNAGATCGGAAG |
| 69487_1 | TATGAGTGTGGAGTCGTTGCAACTCTAATCCCATCATGGTGCCNNNAGATCGGAAG |
| 69488_1 | TATGAGTGTGGAGTCGTTGCCTTTGTGGAAAGCAAAACTCCCANNNAGATCGGAAG |
| 69489_1 | TATGAGTGTGGAGTCGTTGCTCCTATTTGTTCTGTCTCTCTGGNNNAGATCGGAAGA |
| 69490_1 | TATGAGTGTGGAGTCGTTGCGAGTGCAGTGGTCCAAATACAACNNNAGATCGGAA |
| 69492_1 | TATGAGTGTGGAGTCGTTGCGAGATGAGAGAGATGGGGAGAGANNNAGATCGG |
| 69493_1 | TATGAGTGTGGAGTCGTTGCCTGAGACAGGAGGAACACTTGAGNNNAGATCGGAA |
| 69495_1 | TATGAGTGTGGAGTCGTTGCCTCATGTCTGTAATCCCAGCACTNNNAGATCGGAAG |
| 69496_1 | TATGAGTGTGGAGTCGTTGCGGAGGGGAAGGGAGAAAAAGAAANNNAGATCGG |
| 69497_1 | TATGAGTGTGGAGTCGTTGCTCTGTTTGAGAGAAAGCAAGGGANNNAGATCGGAA |
| 69498_1 | TATGAGTGTGGAGTCGTTGCTTTTTTCTGAGAGAGTCTCGCCCNNNAGATCGGAAG |
| 69499_1 | TATGAGTGTGGAGTCGTTGCCACTCACACACACTCTCTCTCTCNNNAGATCGGAAGA |
| 69500_1 | TATGAGTGTGGAGTCGTTGCTCTGTCACCCATGATTGTAGTGGNNNAGATCGGAAG |
| 69501_1 | TATGAGTGTGGAGTCGTTGCACCAGAAACAATCCCCTTTCTCANNNAGATCGGAAGA |
| 69502_1 | TATGAGTGTGGAGTCGTTGCAAGTTAGGTTACGTAGAGGCTGCNNNAGATCGGAA |
| 69503_1 | TATGAGTGTGGAGTCGTTGCATTACACACACACACACACACACNNNAGATCGGAAG |
| 69504_1 | TATGAGTGTGGAGTCGTTGCAGAGAGAGAGAGAGAGAGGGAGANNNAGATCGG |
| 69505_1 | TATGAGTGTGGAGTCGTTGCGTGGTGCTAATACACTGGGAAGANNNAGATCGGAA |
| 69506_1 | TATGAGTGTGGAGTCGTTGCTCATCCTTCCTCTGCATTTTTGCNNNAGATCGGAAGA |

| | |
|---|---|
| 69507_1 | TATGAGTGTGGAGTCGTTGCTGGAAGAGAGAAAAACTGGAGCANNNAGATCGGA |
| 69508_1 | TATGAGTGTGGAGTCGTTGCAGAGAGAGAGGAGGAGAGAGAGANNNAGATCGG |
| 69509_1 | TATGAGTGTGGAGTCGTTGCCAAGGGTCAAAACAAGCATGAGTNNNAGATCGGAA |
| 69511_1 | TATGAGTGTGGAGTCGTTGCTGTGAGGTCAGGGCCTATAAGTANNNAGATCGGAA |
| 69512_1 | TATGAGTGTGGAGTCGTTGCCACACACACTCTCCCTTTCTGTANNNAGATCGGAAGA |
| 69513_1 | TATGAGTGTGGAGTCGTTGCCTCTCTCCATACACACACACACANNNAGATCGGAAGA |
| 69514_1 | TATGAGTGTGGAGTCGTTGCAATTGTGGTTTCTCAGTGCCTTGNNNAGATCGGAAG |
| 69515_1 | TATGAGTGTGGAGTCGTTGCCACACACACACACAGAGTGAAATNNNAGATCGGAAG |
| 69516_1 | TATGAGTGTGGAGTCGTTGCCCAAGTTTTTTCTCCAGCTCAGCNNNAGATCGGAAGA |
| 69517_1 | TATGAGTGTGGAGTCGTTGCTGAAGAATGGCAAAACACGTCTCNNNAGATCGGAAG |
| 69518_1 | TATGAGTGTGGAGTCGTTGCCCAGCTGAGTGAAAGAGAGACTTNNNAGATCGGAA |
| 69519_1 | TATGAGTGTGGAGTCGTTGCGTGTGTGTGTGTGCATATGTGTANNNAGATCGGAA |
| 69520_1 | TATGAGTGTGGAGTCGTTGCCTTCCTCAACGGGTTATAGAGCANNNAGATCGGAAG |
| 69522_1 | TATGAGTGTGGAGTCGTTGCTACAAAAGGGGGTTTATCAGGGGNNNAGATCGGA |
| 69524_1 | TATGAGTGTGGAGTCGTTGCGGAGGAGAGAAAGGAAAGGAAGGNNNAGATCGG |
| 69525_1 | TATGAGTGTGGAGTCGTTGCTCGTGGACAAAAAGAATACTGGANNNAGATCGGAA |
| 69526_1 | TATGAGTGTGGAGTCGTTGCCTTGGCTTACCTCTTGGAACTCTNNNAGATCGGAAG |
| 69527_1 | TATGAGTGTGGAGTCGTTGCGGAGCCCTGTCTGTCTCTAACTANNNAGATCGGAAG |
| 69528_1 | TATGAGTGTGGAGTCGTTGCTCTAAGAAGAGGCTGCTAGACCTNNNAGATCGGAA |
| 69529_1 | TATGAGTGTGGAGTCGTTGCAGATTAGTCTCTTTGCCTGCGAGNNNAGATCGGAAG |
| 69530_1 | TATGAGTGTGGAGTCGTTGCGTGTGTCCCTTTCCTCCAAGTTANNNAGATCGGAAG |
| 69532_1 | TATGAGTGTGGAGTCGTTGCGTCCCTGGATTACAATGTCAGGTNNNAGATCGGAAG |
| 69533_1 | TATGAGTGTGGAGTCGTTGCGAGATCATTGCTTGAGTCCTGGANNNAGATCGGAA |
| 69534_1 | TATGAGTGTGGAGTCGTTGCCATCTAAACCTAACTAGCCCCCCNNNAGATCGGAAGA |
| 69535_1 | TATGAGTGTGGAGTCGTTGCATCCTGGGTATCCACAACTGTTCNNNAGATCGGAAG |
| 69536_1 | TATGAGTGTGGAGTCGTTGCGGAAGCAGAATGTAATGGCCAAGNNNAGATCGGA |
| 69537_1 | TATGAGTGTGGAGTCGTTGCGATCAGGGTACCATCCTCATGACNNNAGATCGGAAG |
| 69539_1 | TATGAGTGTGGAGTCGTTGCTCCTTTTCTCTACAGCACTGACCNNNAGATCGGAAGA |
| 69540_1 | TATGAGTGTGGAGTCGTTGCGCTCCCTCTGCAGTTTCTAGATTNNNAGATCGGAAG |
| 69542_1 | TATGAGTGTGGAGTCGTTGCTCTCATACCATACCCTCTCAGCTNNNAGATCGGAAGA |
| 69543_1 | TATGAGTGTGGAGTCGTTGCCACACAGAGCAAGAGAGAGAGAGNNNAGATCGGA |
| 69545_1 | TATGAGTGTGGAGTCGTTGCCCTTGTCCACCAATTACTCTGGANNNAGATCGGAAG |
| 69546_1 | TATGAGTGTGGAGTCGTTGCACTATGGGACTCCAGGACTTACANNNAGATCGGAAG |
| 69547_1 | TATGAGTGTGGAGTCGTTGCCCTCCTCCCAGATAACAGACAAGNNNAGATCGGAAG |
| 69548_1 | TATGAGTGTGGAGTCGTTGCAGCATGGTGTCTAGATTTACCCTNNNAGATCGGAAG |
| 69549_1 | TATGAGTGTGGAGTCGTTGCGCTGGCACCTTGATCTTTGATTTNNNAGATCGGAAG |
| 69550_1 | TATGAGTGTGGAGTCGTTGCACCCAAGAAAGCCAGTGGTATAGNNNAGATCGGAA |
| 69551_1 | TATGAGTGTGGAGTCGTTGCGAATCAATGGAAAGTGGGAAGGCNNNAGATCGGA |
| 69552_1 | TATGAGTGTGGAGTCGTTGCTCTACTATTCTGGGTGCTGGAGANNNAGATCGGAA |
| 69553_1 | TATGAGTGTGGAGTCGTTGCATAACAAACATCCTGGGTGCTCTNNNAGATCGGAAG |
| 69555_1 | TATGAGTGTGGAGTCGTTGCAGTATCATAGGCTGTGAGGCAACNNNAGATCGGAA |
| 69556_1 | TATGAGTGTGGAGTCGTTGCATAGATATGGACACAGGGAGTCANNNAGATCGGAA |
| 69557_1 | TATGAGTGTGGAGTCGTTGCGAAGAGCAGAGGCAATCCAAAACNNNAGATCGGAA |

| | |
|---|---|
| 69559_1 | TATGAGTGTGGAGTCGTTGCACACACACAGAGAGAGAGAGAGANNNAGATCGGAA |
| 69560_1 | TATGAGTGTGGAGTCGTTGCTTTTCCAGTTCAGGCCAACTAGTNNNAGATCGGAAGA |
| 69561_1 | TATGAGTGTGGAGTCGTTGCAAACCTACATTGCCTCAAATGCCNNNAGATCGGAAGA |
| 69562_1 | TATGAGTGTGGAGTCGTTGCTCATGAGAATCACTTGAGGCCAGNNNAGATCGGAAG |
| 69563_1 | TATGAGTGTGGAGTCGTTGCAGGTATCCACATGTCACATGTCANNNAGATCGGAAG |
| 69565_1 | TATGAGTGTGGAGTCGTTGCTTTTTTTGCCAAGGGGGTGAAATNNNAGATCGGAAG |
| 69566_1 | TATGAGTGTGGAGTCGTTGCAACTTCCTGATCATTTCAAGGGTNNNAGATCGGAAGA |
| 69567_1 | TATGAGTGTGGAGTCGTTGCGGAGGGGGTAAGCTAGGTAATGTNNNAGATCGGAA |
| 69568_1 | TATGAGTGTGGAGTCGTTGCGACCTTGGTGTTGAGGGATATGTNNNAGATCGGAAG |
| 69569_1 | TATGAGTGTGGAGTCGTTGCTTATTCCAAGACGTGTGAAGAGTNNNAGATCGGAAG |
| 69570_1 | TATGAGTGTGGAGTCGTTGCTCAGGAACAAGAGAATCAGGTGGNNNAGATCGGAA |
| 69571_1 | TATGAGTGTGGAGTCGTTGCATACCCAAGAAATGTCGGAACCANNNAGATCGGAAG |
| 69572_1 | TATGAGTGTGGAGTCGTTGCTTGGCTGACATGACTATGTAGGCNNNAGATCGGAAG |
| 69573_1 | TATGAGTGTGGAGTCGTTGCAGGGGAACTGCACAATTAGTGAANNNAGATCGGAA |
| 69574_1 | TATGAGTGTGGAGTCGTTGCTTGGCAACAAAATCACAGTTCTTNNNAGATCGGAAGA |
| 69576_1 | TATGAGTGTGGAGTCGTTGCCCATGCAAAGAGAGTGAACCAATNNNAGATCGGAAG |
| 69577_1 | TATGAGTGTGGAGTCGTTGCATTTCCCCTCTTAGGTTCTGAGCNNNAGATCGGAAGA |
| 69578_1 | TATGAGTGTGGAGTCGTTGCACAAAAATTAGCCAGGTGTGGTGNNNAGATCGGAAG |
| 69579_1 | TATGAGTGTGGAGTCGTTGCCCCACTCTAGCTACCTGGAGATANNNAGATCGGAAGA |
| 69580_1 | TATGAGTGTGGAGTCGTTGCATGACCATGGCATTTCTTTTGCANNNAGATCGGAAGA |
| 69583_1 | TATGAGTGTGGAGTCGTTGCTGCTTGGCACTTTCTTATGTGTGNNNAGATCGGAAGA |
| 69584_1 | TATGAGTGTGGAGTCGTTGCTCTGGTTTCAGGAGATTCGAGTCNNNAGATCGGAAG |
| 69585_1 | TATGAGTGTGGAGTCGTTGCTTAGGAACAACAGAAGAGTCGGGNNNAGATCGGAA |
| 69586_1 | TATGAGTGTGGAGTCGTTGCGGAGGAAGGCATTCAGTTTTCACNNNAGATCGGAAG |
| 69587_1 | TATGAGTGTGGAGTCGTTGCCTGCATTCTCTTTGATGACCACCNNNAGATCGGAAGA |
| 69588_1 | TATGAGTGTGGAGTCGTTGCTTGCAAAAGGAGGCATAATGTGGNNNAGATCGGAA |
| 69589_1 | TATGAGTGTGGAGTCGTTGCACAGGAGCCTTTCTCTCACATTTNNNAGATCGGAAGA |
| 69590_1 | TATGAGTGTGGAGTCGTTGCGAAGAGGAATTTGATGTCAGCCGNNNAGATCGGAA |
| 69592_1 | TATGAGTGTGGAGTCGTTGCAAGCACATACCACAACTCACTCTNNNAGATCGGAAGA |
| 69593_1 | TATGAGTGTGGAGTCGTTGCGTCCTTTTGTACACTGTGGAAGCNNNAGATCGGAAG |
| 69594_1 | TATGAGTGTGGAGTCGTTGCAGCTTATGGAATTGCAGCAGTTGNNNAGATCGGAAG |
| 69596_1 | TATGAGTGTGGAGTCGTTGCGAGAGGCTATACTCCTTCCTCAGNNNAGATCGGAAG |
| 69598_1 | TATGAGTGTGGAGTCGTTGCAACCACCACCTTGAAAGATCCATNNNAGATCGGAAGA |
| 69599_1 | TATGAGTGTGGAGTCGTTGCGAGTAAACTGGGTGAGGAGTCAGNNNAGATCGGAA |
| 69602_1 | TATGAGTGTGGAGTCGTTGCCTTTATGGCTATCTGTGGGTGGTNNNAGATCGGAAG |
| 69603_1 | TATGAGTGTGGAGTCGTTGCATATCATCCTTCAGCAAAGCCCANNNAGATCGGAAGA |
| 69604_1 | TATGAGTGTGGAGTCGTTGCACAAAAATTAGCCTGGTGTGGTGNNNAGATCGGAAG |
| 69605_1 | TATGAGTGTGGAGTCGTTGCACTCACACTCATTTGGTCACAGANNNAGATCGGAAGA |
| 69607_1 | TATGAGTGTGGAGTCGTTGCCCTATGAGCTAAGACGGGAAGTCNNNAGATCGGAAG |
| 69608_1 | TATGAGTGTGGAGTCGTTGCAACAATGTCCCCAGGTCAATACANNNAGATCGGAAG |
| 69609_1 | TATGAGTGTGGAGTCGTTGCGGTCAGCTTTCATGCAATGCTTANNNAGATCGGAAG |
| 69610_1 | TATGAGTGTGGAGTCGTTGCATCTTGCCTGTGATTACCTGTCANNNAGATCGGAAGA |
| 69611_1 | TATGAGTGTGGAGTCGTTGCGAGAGGAGAAAAGCTTCCTGGAANNNAGATCGGAA |

69612_1 TATGAGTGTGGAGTCGTTGCCTCCTGAGTAGCTGGGACTACAGNNNAGATCGGAAG
69613_1 TATGAGTGTGGAGTCGTTGCAAGAGAGGCGTGAGAAAAGAGTTNNNAGATCGGAA
69614_1 TATGAGTGTGGAGTCGTTGCGAGGTGACTACAGAGGCTGAAAANNNAGATCGGAA
69617_1 TATGAGTGTGGAGTCGTTGCAGCATCAATGTTGACCCTAAGGTNNNAGATCGGAAG
69618_1 TATGAGTGTGGAGTCGTTGCTGACCTATACCTGCTCCTACTCCNNNAGATCGGAAGA
69621_1 TATGAGTGTGGAGTCGTTGCATTACAGAGACCAAAGCAAGGGGNNNAGATCGGAA
69623_1 TATGAGTGTGGAGTCGTTGCACTCATCTTGTTCACACCCTAGTNNNAGATCGGAAGA
69624_1 TATGAGTGTGGAGTCGTTGCGAGGAATGTCGAGGTTTGGTTTGNNNAGATCGGAA
69625_1 TATGAGTGTGGAGTCGTTGCCCCGCTTTTTGAGATAGGGTCTTNNNAGATCGGAAG
69626_1 TATGAGTGTGGAGTCGTTGCTTCAAGTGATTCTCCTGCCTCAGNNNAGATCGGAAGA
69627_1 TATGAGTGTGGAGTCGTTGCGGTCCAATGAGTACGAGTCCAATNNNAGATCGGAAG
69628_1 TATGAGTGTGGAGTCGTTGCCCTAAGTTCAGGAGTTCGAGACCNNNAGATCGGAAG
69631_1 TATGAGTGTGGAGTCGTTGCGAGAGAGCAAGAACAACCCAAACNNNAGATCGGAAG
69632_1 TATGAGTGTGGAGTCGTTGCATTCTCTTTCCCAGGGATGTTCCNNNAGATCGGAAGA
69634_1 TATGAGTGTGGAGTCGTTGCCATAAAGGTTCTCCACTTCCCCANNNAGATCGGAAGA
69636_1 TATGAGTGTGGAGTCGTTGCAGGGAATAAATCAATGACCTGGTNNNAGATCGGAAG
69638_1 TATGAGTGTGGAGTCGTTGCGCTGCTGACTTGAGGAGAAATGANNNAGATCGGAA
69639_1 TATGAGTGTGGAGTCGTTGCGTGGAATTCACTCTGTCACTCCTNNNAGATCGGAAGA
69640_1 TATGAGTGTGGAGTCGTTGCATCCCATAATACACAGGATGGCCNNNAGATCGGAAG
69641_1 TATGAGTGTGGAGTCGTTGCACAGAAAAATTAGCTGGGCATGGNNNAGATCGGAA
69642_1 TATGAGTGTGGAGTCGTTGCCTTGGGGGGAAATGTTCTTTGACNNNAGATCGGAAG
69643_1 TATGAGTGTGGAGTCGTTGCACTCTGAGTTTTGTTTCCCGAGANNNAGATCGGAAGA
69644_1 TATGAGTGTGGAGTCGTTGCAACCCATCCGAAATGCACTTTTTNNNAGATCGGAAGA
69646_1 TATGAGTGTGGAGTCGTTGCTAAGTAGCTAGGACTACAGGCCCNNNAGATCGGAAG
69647_1 TATGAGTGTGGAGTCGTTGCTCCTAACCTACCACAGAGCTACANNNAGATCGGAAGA
69648_1 TATGAGTGTGGAGTCGTTGCGATGTAGGGGAAACTCCAGCTTCNNNAGATCGGAAG
69650_1 TATGAGTGTGGAGTCGTTGCCAGCATTCAATTCAGCCTTGGAANNNAGATCGGAAG
69651_1 TATGAGTGTGGAGTCGTTGCAATACAAAAGTAAGCCAGGCGTGNNNAGATCGGAAG
69652_1 TATGAGTGTGGAGTCGTTGCTTTTCCCTCTGATTTCTCCAGGGNNNAGATCGGAAGA
69653_1 TATGAGTGTGGAGTCGTTGCACTGAACTGGTTTTCTCCTTGGTNNNAGATCGGAAGA
69655_1 TATGAGTGTGGAGTCGTTGCGAAGCAGAGGAAGGAGAGAGAAGNNNAGATCGGA
69656_1 TATGAGTGTGGAGTCGTTGCGTGGGACTATAGGCATGTACCACNNNAGATCGGAAG
69657_1 TATGAGTGTGGAGTCGTTGCTTCTTCACACATCCATGTCCCTTNNNAGATCGGAAGAG
69659_1 TATGAGTGTGGAGTCGTTGCTCACGCTTGTAATCCTAGCAGTTNNNAGATCGGAAGA
69661_1 TATGAGTGTGGAGTCGTTGCAACACATTTTTTGGTTGGGGGAGNNNAGATCGGAAG
69662_1 TATGAGTGTGGAGTCGTTGCCTGGGATTACAGGTGTGAGTCACNNNAGATCGGAAG
69663_1 TATGAGTGTGGAGTCGTTGCTGTGGAAGCTTTGTTCTTTTGCTNNNAGATCGGAAGA
69667_1 TATGAGTGTGGAGTCGTTGCGCCCGGCCTGAATTTAGATCTTANNNAGATCGGAAG
69668_1 TATGAGTGTGGAGTCGTTGCTTCATGTCATTCTCCTGCCTCAGNNNAGATCGGAAGA
69669_1 TATGAGTGTGGAGTCGTTGCGCAGTGAACACAAATTGTCAGGANNNAGATCGGAAG
69671_1 TATGAGTGTGGAGTCGTTGCCCTTTCTTCTGCACACAGCTTTTNNNAGATCGGAAGAG
69672_1 TATGAGTGTGGAGTCGTTGCGGTGAAAAGTAGCCTCGATGTGANNNAGATCGGAA
69673_1 TATGAGTGTGGAGTCGTTGCGGATTCTGCTGGTTGACTTTCTGNNNAGATCGGAAG

| | |
|---|---|
| 69674_1 | TATGAGTGTGGAGTCGTTGCTTTCTTCAGTCTCCCTCTGTTGCNNNAGATCGGAAGA( |
| 69675_1 | TATGAGTGTGGAGTCGTTGCTGAGCAAGACTCTGTCTCAAACANNNAGATCGGAAG |
| 69676_1 | TATGAGTGTGGAGTCGTTGCAAGCATGAGTATCACAAGTGGGTNNNAGATCGGAA( |
| 69677_1 | TATGAGTGTGGAGTCGTTGCACCTCAGACCATTTGCCCATATTNNNAGATCGGAAGA |
| 69678_1 | TATGAGTGTGGAGTCGTTGCCTACTGTGTAGAGAACAGCCTGGNNNAGATCGGAA( |
| 69679_1 | TATGAGTGTGGAGTCGTTGCTGAGAGCAGAATGTAGAGTTGGGNNNAGATCGGA/ |
| 69680_1 | TATGAGTGTGGAGTCGTTGCGTAACAGGGTCTCACTCTGTCACNNNAGATCGGAAG |
| 69681_1 | TATGAGTGTGGAGTCGTTGCGAGTGTGTGTTCTTGCAACTTGANNNAGATCGGAAG |
| 69683_1 | TATGAGTGTGGAGTCGTTGCAATCTTTCAGCTCTACGACGGTTNNNAGATCGGAAG/ |
| 69685_1 | TATGAGTGTGGAGTCGTTGCATTCCCAACAGCACTCTGAGAAANNNAGATCGGAAG |
| 69686_1 | TATGAGTGTGGAGTCGTTGCGCTTGGACTACAGGCATATACCANNNAGATCGGAAG |
| 69687_1 | TATGAGTGTGGAGTCGTTGCCTGTTGCTCAGGCTAGAGTGTAANNNAGATCGGAA( |
| 69688_1 | TATGAGTGTGGAGTCGTTGCCTTCAGCCTTCTACCATGGGATGNNNAGATCGGAAG/ |
| 69689_1 | TATGAGTGTGGAGTCGTTGCCATTCTTTTCCATTTGGCCCTGTNNNAGATCGGAAGA |
| 69691_1 | TATGAGTGTGGAGTCGTTGCTCTCTCTCTTTTTTTTCCCCGGNNNAGATCGGAAGA( |
| 69692_1 | TATGAGTGTGGAGTCGTTGCGGGCAAAGCAGGAAAATATGGTTNNNAGATCGGAA |
| 69693_1 | TATGAGTGTGGAGTCGTTGCTTGAGACAAGGTCTTACTCTGCCNNNAGATCGGAAG |
| 69694_1 | TATGAGTGTGGAGTCGTTGCGGCCAAGGTTGATCAATGATGAGNNNAGATCGGAA |
| 69695_1 | TATGAGTGTGGAGTCGTTGCTTTCCCAATTCAGAGTCTTGCTCNNNAGATCGGAAGA |
| 69696_1 | TATGAGTGTGGAGTCGTTGCAATGATGGAGTAATTGACACCCTNNNAGATCGGAAG |
| 69697_1 | TATGAGTGTGGAGTCGTTGCTTCTGCAGTTTGATCACATGCTCNNNAGATCGGAAG/ |
| 69698_1 | TATGAGTGTGGAGTCGTTGCGACATGGTAGCTCACACCTGTAANNNAGATCGGAAG |
| 69699_1 | TATGAGTGTGGAGTCGTTGCGAAGTCTCTATCTGCAGAAGGGGNNNAGATCGGAA( |
| 69700_1 | TATGAGTGTGGAGTCGTTGCATTTGATCTTCGAGAGAGCGGTTNNNAGATCGGAAG |
| 69701_1 | TATGAGTGTGGAGTCGTTGCCACTGATGAGGGAGTAACAGGACNNNAGATCGGAA |
| 69702_1 | TATGAGTGTGGAGTCGTTGCCATTTGTACTCCACGCATGATCCNNNAGATCGGAAG/ |
| 69703_1 | TATGAGTGTGGAGTCGTTGCTTAGACCCAGGTCAGTTTGGAAGNNNAGATCGGAA( |
| 69704_1 | TATGAGTGTGGAGTCGTTGCGTGTGTGTGTGTGTGTGTGTATANNNAGATCGGAA( |
| 69705_1 | TATGAGTGTGGAGTCGTTGCCATGCTGACACCCTGATTTCAAANNNAGATCGGAAG/ |
| 69706_1 | TATGAGTGTGGAGTCGTTGCTCCTTGCAAGACAGACAATAGGTNNNAGATCGGAAG |
| 69707_1 | TATGAGTGTGGAGTCGTTGCTACACACACAAACACACACACACNNNAGATCGGAAG/ |
| 69708_1 | TATGAGTGTGGAGTCGTTGCTAGTTCTGTGACTGAGGCAACTCNNNAGATCGGAAG |
| 69709_1 | TATGAGTGTGGAGTCGTTGCTAAAACCCAGAACCCATGTCCATNNNAGATCGGAAG/ |
| 69711_1 | TATGAGTGTGGAGTCGTTGCACACACAAACACACACACACAATNNNAGATCGGAAG/ |
| 69712_1 | TATGAGTGTGGAGTCGTTGCGCCATGTTGTGAGAACAGTTAGGNNNAGATCGGAA( |
| 69715_1 | TATGAGTGTGGAGTCGTTGCAGAGGTACTATGGCACAGTAGTTNNNAGATCGGAA( |
| 69716_1 | TATGAGTGTGGAGTCGTTGCGGTCTCTTGAGTCCATGTAGCATNNNAGATCGGAAG |
| 69717_1 | TATGAGTGTGGAGTCGTTGCTCTGTCCTGTCTGCAAATGAAGTNNNAGATCGGAAG |
| 69718_1 | TATGAGTGTGGAGTCGTTGCTGACAGGCTAGAAACACAGGAAANNNAGATCGGAA |
| 69719_1 | TATGAGTGTGGAGTCGTTGCAGCCTCCCAAATAGCTAGGACTANNNAGATCGGAAG |
| 69720_1 | TATGAGTGTGGAGTCGTTGCAGATGGAGTCTCACTCTGTAGCCNNNAGATCGGAAG |
| 69722_1 | TATGAGTGTGGAGTCGTTGCATAGAAACGGGAAATGTCCAAGCNNNAGATCGGAA( |
| 69723_1 | TATGAGTGTGGAGTCGTTGCACCACCAGAAGACAAAGGGTAAGNNNAGATCGGAA |

| | |
|---|---|
| 69724_1 | TATGAGTGTGGAGTCGTTGCTGGAAAAATATGGCTTCCCCTGANNNAGATCGGAAG |
| 69725_1 | TATGAGTGTGGAGTCGTTGCTCTCTTTCTCTGTGTGTGTGTGTNNNAGATCGGAAG |
| 69726_1 | TATGAGTGTGGAGTCGTTGCCCTCCCAGCACCTTTTTTTCTTTNNNAGATCGGAAGA |
| 69727_1 | TATGAGTGTGGAGTCGTTGCCACACACATTCTCAGCAGGTCTANNNAGATCGGAAG |
| 69729_1 | TATGAGTGTGGAGTCGTTGCCAATCTCGGCTCACTGTAACCTCNNNAGATCGGAAG |
| 69730_1 | TATGAGTGTGGAGTCGTTGCTCCATGTCCTTCAAACTCCATGTNNNAGATCGGAAGA |
| 69731_1 | TATGAGTGTGGAGTCGTTGCTAATGAAACACAGGCATGATGGCNNNAGATCGGAA |
| 69732_1 | TATGAGTGTGGAGTCGTTGCCCTTTGAGATGCTTGCAGATGTTNNNAGATCGGAAG |
| 69733_1 | TATGAGTGTGGAGTCGTTGCTCTCCTGGGTTCAAGTGATTCTCNNNAGATCGGAAG |
| 69734_1 | TATGAGTGTGGAGTCGTTGCACAGAGACAGGAAGCAATACCAGNNNAGATCGGAA |
| 69735_1 | TATGAGTGTGGAGTCGTTGCCCACCATGTCCCAAATCCATAGANNNAGATCGGAAG |
| 69736_1 | TATGAGTGTGGAGTCGTTGCCCAGATCTCATGAGAACTCACCCNNNAGATCGGAAG |
| 69739_1 | TATGAGTGTGGAGTCGTTGCTGACCACCTGGATCTAATCACACNNNAGATCGGAAG |
| 69740_1 | TATGAGTGTGGAGTCGTTGCACTTCCTGCCATTTCCATGTACTNNNAGATCGGAAGA |
| 69742_1 | TATGAGTGTGGAGTCGTTGCGGGCAAGAGAAGATACCAGAGAGNNNAGATCGGA |
| 69744_1 | TATGAGTGTGGAGTCGTTGCTTCTTGTTGGGGAGGAAATCAGTNNNAGATCGGAA |
| 69745_1 | TATGAGTGTGGAGTCGTTGCCCTCCAGAACTGTGAGAATGTGANNNAGATCGGAA |
| 69746_1 | TATGAGTGTGGAGTCGTTGCAAGAGCATAAACAGCAGGCAATGNNNAGATCGGAA |
| 69749_1 | TATGAGTGTGGAGTCGTTGCATTCCGTATCTTGGTGGTTGTGANNNAGATCGGAAG |
| 69750_1 | TATGAGTGTGGAGTCGTTGCCCCACAGAACTGTGAGGTGATTANNNAGATCGGAA |
| 69751_1 | TATGAGTGTGGAGTCGTTGCCAGACAGTCGGATGAGATGCATANNNAGATCGGAA |
| 69752_1 | TATGAGTGTGGAGTCGTTGCCAATCTTCAGAGAGGCTGTCCAANNNAGATCGGAAG |
| 69753_1 | TATGAGTGTGGAGTCGTTGCTCTGGTTGTGTAAGATGTGGCATNNNAGATCGGAA |
| 69755_1 | TATGAGTGTGGAGTCGTTGCGAGAGTCTCCCTTTGTCACTCAGNNNAGATCGGAAG |
| 69757_1 | TATGAGTGTGGAGTCGTTGCTCCAGATCACCACAGTAAAGCAANNNAGATCGGAAG |
| 69758_1 | TATGAGTGTGGAGTCGTTGCTTCCATGTGACTTTTCCAATGCCNNNAGATCGGAAG |
| 69760_1 | TATGAGTGTGGAGTCGTTGCAGAGACAGGTCTTGCTATGTTGCNNNAGATCGGAA |
| 69761_1 | TATGAGTGTGGAGTCGTTGCTGACTGTCCAATTGTTTCAGCACNNNAGATCGGAAG |
| 69763_1 | TATGAGTGTGGAGTCGTTGCGCTCTGATGAAGCAAGTTACCACNNNAGATCGGAAG |
| 69764_1 | TATGAGTGTGGAGTCGTTGCTTCCCCTCTCTCCCTCTCTTATCNNNAGATCGGAAGAG |
| 69766_1 | TATGAGTGTGGAGTCGTTGCCAATCATGGCTCACTGCAATCTCNNNAGATCGGAAG |
| 69767_1 | TATGAGTGTGGAGTCGTTGCTAGGACTCCATTTCCTCTTCTGCNNNAGATCGGAAGA |
| 69768_1 | TATGAGTGTGGAGTCGTTGCCCATTACAGAAAGAGGAAGCAGCNNNAGATCGGAA |
| 69770_1 | TATGAGTGTGGAGTCGTTGCCTCTCAAGGTGTCCAGAGTCTTCNNNAGATCGGAAG |
| 69771_1 | TATGAGTGTGGAGTCGTTGCTAGGAGAAGGAAACAGGAGTGGANNNAGATCGGA |
| 69772_1 | TATGAGTGTGGAGTCGTTGCCAAGACCTCCAAGATGGTACCTCNNNAGATCGGAAG |
| 69774_1 | TATGAGTGTGGAGTCGTTGCTTCAAGTGATTCTCCTGCCTCTCNNNAGATCGGAAGA |
| 69775_1 | TATGAGTGTGGAGTCGTTGCCAAGAGCAAAACATTCCTAGGGCNNNAGATCGGAA |
| 69776_1 | TATGAGTGTGGAGTCGTTGCAGAGAGACAGAAAGGACTCCAGANNNAGATCGGAA |
| 69777_1 | TATGAGTGTGGAGTCGTTGCTCTTCCAGGCTCAAGTGATCTTCNNNAGATCGGAAG |
| 69778_1 | TATGAGTGTGGAGTCGTTGCAAACAGAGGGGCTTTACTCTCTGNNNAGATCGGAA |
| 69779_1 | TATGAGTGTGGAGTCGTTGCTTCTTTCTTTCTTGAGACGGGGTNNNAGATCGGAAG |
| 69780_1 | TATGAGTGTGGAGTCGTTGCAATGAACTATCCTACCCTTGGGCNNNAGATCGGAAG |

| | |
|---|---|
| 69781_1 | TATGAGTGTGGAGTCGTTGCGTGTCAGCATGTTAGATGGAAGCNNNAGATCGGAA |
| 69782_1 | TATGAGTGTGGAGTCGTTGCGCATGATCTCTAGTCACTGCAGANNNAGATCGGAAG |
| 69784_1 | TATGAGTGTGGAGTCGTTGCTTCCCCTTCTGCCATGATTGTAANNNAGATCGGAAG |
| 69785_1 | TATGAGTGTGGAGTCGTTGCGAGTGCAATGATGTGATCTCAGCNNNAGATCGGAA |
| 69786_1 | TATGAGTGTGGAGTCGTTGCTATAACTGCCCCTTTCCCATCTCNNNAGATCGGAAGA |
| 69787_1 | TATGAGTGTGGAGTCGTTGCGAATGGCACACATCCAGTAATGGNNNAGATCGGAA |
| 69788_1 | TATGAGTGTGGAGTCGTTGCGGAGGACATCCATGTTGAGTGAANNNAGATCGGAA |
| 69790_1 | TATGAGTGTGGAGTCGTTGCGTGGCTAAAAAGTCATGAGGCTGNNNAGATCGGAA |
| 69791_1 | TATGAGTGTGGAGTCGTTGCTGTAGGATTTCTGCACTGAGTCCNNNAGATCGGAAG |
| 69792_1 | TATGAGTGTGGAGTCGTTGCGGAGAATCTGCTGATCCCAGAATNNNAGATCGGAA |
| 69793_1 | TATGAGTGTGGAGTCGTTGCGGGAGAGAGAGAGAGAGAGAGAGNNNAGATCGG |
| 69794_1 | TATGAGTGTGGAGTCGTTGCCTAGGCAGAGGTAGAAGCTATGCNNNAGATCGGAA |
| 69795_1 | TATGAGTGTGGAGTCGTTGCCTTTCCTCCTCTCAGCACTGTAGNNNAGATCGGAAGA |
| 69797_1 | TATGAGTGTGGAGTCGTTGCTCTCCTTCTCTGATTCTGACCCTNNNAGATCGGAAGA |
| 69798_1 | TATGAGTGTGGAGTCGTTGCACCTCTTGTCCATCTATCTGGGANNNAGATCGGAAG |
| 69799_1 | TATGAGTGTGGAGTCGTTGCTTCATTCAGGGAGGTCAAGACTGNNNAGATCGGAA |
| 69800_1 | TATGAGTGTGGAGTCGTTGCTGGTTTCTGAGACACAAAGGTCTNNNAGATCGGAA |
| 69801_1 | TATGAGTGTGGAGTCGTTGCCAGAGTACAGTGGCAGAATCACTNNNAGATCGGAA |
| 69803_1 | TATGAGTGTGGAGTCGTTGCAAGAAAGGAAGCAGGAAAGGAGGNNNAGATCGG |
| 69804_1 | TATGAGTGTGGAGTCGTTGCCCAGGAAGTCAGAATCAGGACATNNNAGATCGGAA |
| 69805_1 | TATGAGTGTGGAGTCGTTGCAAAGGGACAGGTAGTCAGGAAACNNNAGATCGGA |
| 69806_1 | TATGAGTGTGGAGTCGTTGCATAGATAACTATTGCCGGGCGTTNNNAGATCGGAAG |
| 69807_1 | TATGAGTGTGGAGTCGTTGCTTGAGGTCAGGAGTTCAAAACCANNNAGATCGGAA |
| 69810_1 | TATGAGTGTGGAGTCGTTGCCTCTCTCTCTCTCTCTCTCTCCCNNNAGATCGGAAGAG |
| 69811_1 | TATGAGTGTGGAGTCGTTGCGGAAAAGCCTTCCACAGAAACTGNNNAGATCGGAA |
| 69812_1 | TATGAGTGTGGAGTCGTTGCCTTCCCCTTTGCTTTCCATCATGNNNAGATCGGAAGA |
| 69813_1 | TATGAGTGTGGAGTCGTTGCTTTTTAGGCTCTATCCCAGGACCNNNAGATCGGAAG |
| 69814_1 | TATGAGTGTGGAGTCGTTGCGATCGTTGGCTTGGGGTTTATTTNNNAGATCGGAAG |
| 69816_1 | TATGAGTGTGGAGTCGTTGCGGAGAAACGAGCATAGAGAGAGGNNNAGATCGGA |
| 69817_1 | TATGAGTGTGGAGTCGTTGCTGGGACAGCAATTGGGATTCTTANNNAGATCGGAA |
| 69818_1 | TATGAGTGTGGAGTCGTTGCCACTCTACTTGGAATCAGCCACTNNNAGATCGGAAG |
| 69819_1 | TATGAGTGTGGAGTCGTTGCATTCAGAAACAGACTGGGGACAGNNNAGATCGGAA |
| 69821_1 | TATGAGTGTGGAGTCGTTGCTCTCACTATGTTGGTTAGGCTGGNNNAGATCGGAAG |
| 69822_1 | TATGAGTGTGGAGTCGTTGCGAGGGAGGAAGGAAGGAAAGAAGNNNAGATCGG |
| 69823_1 | TATGAGTGTGGAGTCGTTGCCTGAGTAAGAAAGACCCACCCTCNNNAGATCGGAAG |
| 69824_1 | TATGAGTGTGGAGTCGTTGCTCCTGTAGGAGTGCTTCATTCTTNNNAGATCGGAAG |
| 69825_1 | TATGAGTGTGGAGTCGTTGCACACACACACACAAATGCAAAGANNNAGATCGGAAG |
| 69826_1 | TATGAGTGTGGAGTCGTTGCTGAAGGTTTGGGAATCACATCCTNNNAGATCGGAA |
| 69828_1 | TATGAGTGTGGAGTCGTTGCATCTCTTTCTCACTCCCTTCTCCNNNAGATCGGAAGAG |
| 69829_1 | TATGAGTGTGGAGTCGTTGCAAAAATTCAGAATGCAGCTGGCTNNNAGATCGGAA |
| 69830_1 | TATGAGTGTGGAGTCGTTGCCTCTCTCTCGCCTCTAAGTGTTCNNNAGATCGGAAGA |
| 69832_1 | TATGAGTGTGGAGTCGTTGCTAGATGGAGTCTCACTCTGTTGCNNNAGATCGGAAG |
| 69833_1 | TATGAGTGTGGAGTCGTTGCAGAAATGCACATTCTCATGCCACNNNAGATCGGAAG |

| | |
|---|---|
| 69834_1 | TATGAGTGTGGAGTCGTTGCTAAGGACACCAGCCTATCAGACTNNNAGATCGGAAG |
| 69835_1 | TATGAGTGTGGAGTCGTTGCTCAAAAGGCAATTCCGGAGACTANNNAGATCGGAAG |
| 69836_1 | TATGAGTGTGGAGTCGTTGCAGAGAGAGAGAGAGAGAAAGCGANNNAGATCGGA |
| 69837_1 | TATGAGTGTGGAGTCGTTGCCTCACAAGTAGCTGGGACTACAGNNNAGATCGGAAG |
| 69838_1 | TATGAGTGTGGAGTCGTTGCTCACACAAACAAAGACTGAGGGANNNAGATCGGAAG |
| 69839_1 | TATGAGTGTGGAGTCGTTGCGAAGGAGTGAGAAAAATGTGCCCNNNAGATCGGAA |
| 69840_1 | TATGAGTGTGGAGTCGTTGCGGGAAAGTTGAAGTTGGGTGAAGNNNAGATCGGA |
| 69841_1 | TATGAGTGTGGAGTCGTTGCGCAGAGACAGAGAGACAGAAACANNNAGATCGGAA |
| 69844_1 | TATGAGTGTGGAGTCGTTGCGGAGAGAGAGAGACTGACTCAATNNNAGATCGGAA |
| 69845_1 | TATGAGTGTGGAGTCGTTGCTAGGGAGGGAGGGATTACCTTTTNNNAGATCGGAA |
| 69846_1 | TATGAGTGTGGAGTCGTTGCAGAGAGGAGAGAGAGAGAGGAGANNNAGATCGGA |
| 69847_1 | TATGAGTGTGGAGTCGTTGCTATCTCTGCCCACTAGTTCCAACNNNAGATCGGAAGA |
| 69848_1 | TATGAGTGTGGAGTCGTTGCAATATACACATCCTTGGGCTGGGNNNAGATCGGAAG |
| 69851_1 | TATGAGTGTGGAGTCGTTGCTCTCTCTAAGGGCAACATGCATTNNNAGATCGGAAGA |
| 69852_1 | TATGAGTGTGGAGTCGTTGCAGAGATGCTAGAGAGGCTTCTGANNNAGATCGGAA |
| 69853_1 | TATGAGTGTGGAGTCGTTGCAGATTCCAAAGTCTCACCCTGTCNNNAGATCGGAAGA |
| 69854_1 | TATGAGTGTGGAGTCGTTGCCAATGGTGCAATCTTGGTTCACTNNNAGATCGGAAG |
| 69857_1 | TATGAGTGTGGAGTCGTTGCTGCAGTGGTATGATCACAGTTCANNNAGATCGGAAG |
| 69858_1 | TATGAGTGTGGAGTCGTTGCCACCCAAGGAATTTCCCTCTACANNNAGATCGGAAGA |
| 69860_1 | TATGAGTGTGGAGTCGTTGCCTCTCTCTGGTTCTCTGTTTCCCNNNAGATCGGAAGAG |
| 69861_1 | TATGAGTGTGGAGTCGTTGCCTCTATCTCTCTCTCACCTCCCCNNNAGATCGGAAGAG |
| 69862_1 | TATGAGTGTGGAGTCGTTGCCTTACCCTCCCAGAAAAGTCTCCNNNAGATCGGAAGA |
| 69863_1 | TATGAGTGTGGAGTCGTTGCTCTCTCTCTCTCTCCCCAGAATGNNNAGATCGGAAGAG |
| 69864_1 | TATGAGTGTGGAGTCGTTGCTTCTCTTCAGGGCCATTCAAAGANNNAGATCGGAAGA |
| 69866_1 | TATGAGTGTGGAGTCGTTGCTTCCATACCACCTCATTCACCAANNNAGATCGGAAGAG |
| 69867_1 | TATGAGTGTGGAGTCGTTGCCAGTGGCACAATCATGACTCACNNNAGATCGGAAGA |
| 69868_1 | TATGAGTGTGGAGTCGTTGCTCCATGAGGAGCCAAAGATTCAANNNAGATCGGAAG |
| 69870_1 | TATGAGTGTGGAGTCGTTGCTTTTTTGAGACGGAGTCTTGCTCNNNAGATCGGAAG |
| 69871_1 | TATGAGTGTGGAGTCGTTGCCACACACACACGCTTTTATAACCTNNNAGATCGGAAG |
| 69872_1 | TATGAGTGTGGAGTCGTTGCGAGTGGTACCATCATGGCTCATTNNNAGATCGGAAG |
| 69873_1 | TATGAGTGTGGAGTCGTTGCACTGCAAGGGGGAAATAAGAGAGNNNAGATCGGA |
| 69874_1 | TATGAGTGTGGAGTCGTTGCTGTCTCTCTCTCTGTCTCTCTGTNNNAGATCGGAAGAG |
| 69875_1 | TATGAGTGTGGAGTCGTTGCCTCCATACACTCCCTCTCTCTCANNNAGATCGGAAGAG |
| 69876_1 | TATGAGTGTGGAGTCGTTGCTGAAGATATGCTTGCTTCCCCTTNNNAGATCGGAAGA |
| 69877_1 | TATGAGTGTGGAGTCGTTGCGTGATTCTCAGTGTTGGATGTGGNNNAGATCGGAAG |
| 69878_1 | TATGAGTGTGGAGTCGTTGCGGAATGTAGTGGCGCAATCATTGNNNAGATCGGAAG |
| 69879_1 | TATGAGTGTGGAGTCGTTGCACACATGACAAAATGCCCAACATNNNAGATCGGAAG |
| 69880_1 | TATGAGTGTGGAGTCGTTGCTTCTCTTTCTCCCTCCTTCCCTCNNNAGATCGGAAGAG |
| 69881_1 | TATGAGTGTGGAGTCGTTGCGGTGTGATCTTGGTTCATTGCAANNNAGATCGGAAG |
| 69882_1 | TATGAGTGTGGAGTCGTTGCTCTCTCTCTCTCTCTCTCTCCCTNNNAGATCGGAAGAG |
| 69883_1 | TATGAGTGTGGAGTCGTTGCGTTCAAGAGATTCTCCTGCCTCANNNAGATCGGAAGA |
| 69884_1 | TATGAGTGTGGAGTCGTTGCAAGTAAGTGATACTGCATGCCCANNNAGATCGGAAG |
| 69885_1 | TATGAGTGTGGAGTCGTTGCAGAAACCTCATGTCATGTCCAGANNNAGATCGGAAG |

69886_1 TATGAGTGTGGAGTCGTTGCACCAAAATCCCAGTGATAGAGGGNNNAGATCGGAA
69887_1 TATGAGTGTGGAGTCGTTGCAAGGAAGGAGCAAGAAAGGAAGANNNAGATCGGA
69888_1 TATGAGTGTGGAGTCGTTGCTGGACACCATGTGCTATGATTCANNNAGATCGGAAG
69890_1 TATGAGTGTGGAGTCGTTGCTTCAAGTGATTCTCCTGCCTCAGNNNAGATCGGAAG
69891_1 TATGAGTGTGGAGTCGTTGCCCCAGGACAGTATTTCCAGTTTGNNNAGATCGGAAG
69892_1 TATGAGTGTGGAGTCGTTGCAGCATAAAAATGGGCCAACAGTCNNNAGATCGGAA
69893_1 TATGAGTGTGGAGTCGTTGCCAGTGAGTCCTGTTTCTAGTCCCNNNAGATCGGAAG
69894_1 TATGAGTGTGGAGTCGTTGCTTGTGGTGTGTAAAAAGCACTCCNNNAGATCGGAA
69895_1 TATGAGTGTGGAGTCGTTGCTTAGTCAAGTGAAGCAGTGGGAGNNNAGATCGGA
69896_1 TATGAGTGTGGAGTCGTTGCACACACACACACACAAACAAACANNNAGATCGGAAG
69897_1 TATGAGTGTGGAGTCGTTGCGACAATTTTTCTACAGAGCGGGGNNNAGATCGGAA
69898_1 TATGAGTGTGGAGTCGTTGCCTCGCCTCTTCCTCATTAGAGTCNNNAGATCGGAAGA
69899_1 TATGAGTGTGGAGTCGTTGCCTCTCAGAGAACATAGCCTCAGTNNNAGATCGGAAG
69900_1 TATGAGTGTGGAGTCGTTGCCTTACACGAATTTAGGAGCTGCGNNNAGATCGGAA
69902_1 TATGAGTGTGGAGTCGTTGCTGGAGTGCAGTGACGTTATCTTGNNNAGATCGGAA
69903_1 TATGAGTGTGGAGTCGTTGCAGAGAGAGAGAGAAAAGGACGGANNNAGATCGG
69905_1 TATGAGTGTGGAGTCGTTGCGCCATTCAAAGCACTCAGAGAAANNNAGATCGGAA
69906_1 TATGAGTGTGGAGTCGTTGCGAGCAACTCAGAGGTTACTGGATNNNAGATCGGAA
69907_1 TATGAGTGTGGAGTCGTTGCACTTCTCCTTCTGCCATGATTGTNNNAGATCGGAAGA
69908_1 TATGAGTGTGGAGTCGTTGCCACACACACACACACACATTCATNNNAGATCGGAAGA
69909_1 TATGAGTGTGGAGTCGTTGCCCATGCCCGGCTAATTTTTGTATNNNAGATCGGAAG
69910_1 TATGAGTGTGGAGTCGTTGCAGACAGAGAGAGAGAGAGACACCNNNAGATCGGA
69911_1 TATGAGTGTGGAGTCGTTGCAAGAGAGGGAAGGCTCAATTCAGNNNAGATCGGA
69912_1 TATGAGTGTGGAGTCGTTGCTTGTGCCCATCAACAGAAGATTGNNNAGATCGGAAG
69913_1 TATGAGTGTGGAGTCGTTGCATGGAGTCTCACTTAGTCACCCANNNAGATCGGAAG
69915_1 TATGAGTGTGGAGTCGTTGCAATTAGTCAGGCATGGTAGTGCANNNAGATCGGAA
69916_1 TATGAGTGTGGAGTCGTTGCGTCAGTCAGTGTACAGGAGGAAGNNNAGATCGGA
69917_1 TATGAGTGTGGAGTCGTTGCAGAGGGAGAGAATTTGGGAGGTANNNAGATCGGA
69918_1 TATGAGTGTGGAGTCGTTGCGTCGCCCTCTCTTTCTATCTCTGNNNAGATCGGAAGA
69919_1 TATGAGTGTGGAGTCGTTGCGTCGCCCTCTCTTTCTATCTCTGNNNAGATCGGAAGA
69920_1 TATGAGTGTGGAGTCGTTGCCTGAGATCAGAGGGAAGACAGAGNNNAGATCGGA
69921_1 TATGAGTGTGGAGTCGTTGCTCTCTCTCACACACACACAAACANNNAGATCGGAAGA
69922_1 TATGAGTGTGGAGTCGTTGCGGTGATTGAGGTCCAGAGAAAGTNNNAGATCGGA
69924_1 TATGAGTGTGGAGTCGTTGCATCTTCCTGAGTAGCTGGGACTANNNAGATCGGAAG
69925_1 TATGAGTGTGGAGTCGTTGCCCTTCACCTTCCGAGATGATCATNNNAGATCGGAAG
69926_1 TATGAGTGTGGAGTCGTTGCTTAGGGGTGGGGAGATAATCTGANNNAGATCGGA
69927_1 TATGAGTGTGGAGTCGTTGCGGAGTCCCAGGAAAGTTGATGATNNNAGATCGGAA
69928_1 TATGAGTGTGGAGTCGTTGCAAGGAAAGCAAAGAGAGAGGCTTNNNAGATCGGA
69929_1 TATGAGTGTGGAGTCGTTGCGAGAACTTGCGGTGTTTGGTAAANNNAGATCGGAA
69930_1 TATGAGTGTGGAGTCGTTGCGAGTGTGTGTGTGTGTGTGTATGNNNAGATCGGAA
69931_1 TATGAGTGTGGAGTCGTTGCAGAGAGACTAGAGAGAGCTCCTGNNNAGATCGGAA
69932_1 TATGAGTGTGGAGTCGTTGCTGTGTGTGTGTTTGTGTGTGTACNNNAGATCGGAA
69933_1 TATGAGTGTGGAGTCGTTGCTTTATGACCTAGCCTTGGCAGTCNNNAGATCGGAAG

| | |
|---|---|
| 69934_1 | TATGAGTGTGGAGTCGTTGCAAGAAGGACATGTTTGCTTCTGCNNNAGATCGGAAG |
| 69935_1 | TATGAGTGTGGAGTCGTTGCTCTGCCATGATTGTAAGCTTCCTNNNAGATCGGAAGA |
| 69936_1 | TATGAGTGTGGAGTCGTTGCCACACACACAAACCCATGAGTACNNNAGATCGGAAGA |
| 69938_1 | TATGAGTGTGGAGTCGTTGCATCCCGGTACCTCTCCATTCTTANNNAGATCGGAAGA |
| 69939_1 | TATGAGTGTGGAGTCGTTGCCAGGCTAGCTAGAGTTACAGGTGNNNAGATCGGAA |
| 69940_1 | TATGAGTGTGGAGTCGTTGCGCCTAAAGTTTTGACGGGCATTANNNAGATCGGAAG |
| 69941_1 | TATGAGTGTGGAGTCGTTGCCAGAAAGTGGTGAGAAAGGGCTANNNAGATCGGAA |
| 69943_1 | TATGAGTGTGGAGTCGTTGCTGTCCCTAGCAAAGAGAGAGAGANNNAGATCGGAA |
| 69944_1 | TATGAGTGTGGAGTCGTTGCTTATCTGTCCTTCCCAATTGCCTNNNAGATCGGAAGA |
| 69945_1 | TATGAGTGTGGAGTCGTTGCCCCCTCTTTTTCCAGTTTGCAAANNNAGATCGGAAGA |
| 69946_1 | TATGAGTGTGGAGTCGTTGCAACACTGAAGCATTTGACAGAGCNNNAGATCGGAAG |
| 69947_1 | TATGAGTGTGGAGTCGTTGCGGAGTGATAGCCATTTATCGAGCNNNAGATCGGAA |
| 69949_1 | TATGAGTGTGGAGTCGTTGCACAGAAGTTTTGCAGGCTCAAAGNNNAGATCGGAA |
| 69950_1 | TATGAGTGTGGAGTCGTTGCGGAAGGAGGGGGAAAAAACCTAANNNAGATCGGA |
| 69951_1 | TATGAGTGTGGAGTCGTTGCGCAATTATTCTGCCTCAACCTCCNNNAGATCGGAAGA |
| 69952_1 | TATGAGTGTGGAGTCGTTGCCTGCCCCAAAATCTACAGCATTTNNNAGATCGGAAGA |
| 69955_1 | TATGAGTGTGGAGTCGTTGCCACACACAGATGCATACATGGACNNNAGATCGGAAG |
| 69956_1 | TATGAGTGTGGAGTCGTTGCCAATTGCCCAGTCTCAAGTGATCNNNAGATCGGAAGA |
| 69957_1 | TATGAGTGTGGAGTCGTTGCACACAGTTTACACAGTCACAGGTNNNAGATCGGAAG |
| 69958_1 | TATGAGTGTGGAGTCGTTGCCACTCGGTAGATAAGAGGTTGGGNNNAGATCGGAA |
| 69959_1 | TATGAGTGTGGAGTCGTTGCAGAGAGAGAGAAAGAGAGACGCTNNNAGATCGGA |
| 69960_1 | TATGAGTGTGGAGTCGTTGCGTTGAGAAACAGTAAAGGCACCCNNNAGATCGGAA |
| 69961_1 | TATGAGTGTGGAGTCGTTGCTTCCCACTCTCACTACATACCCTNNNAGATCGGAAGA |
| 69962_1 | TATGAGTGTGGAGTCGTTGCCATTGCTATCACTGAGGCACTTGNNNAGATCGGAAG |
| 69965_1 | TATGAGTGTGGAGTCGTTGCCAGAGTGTGTCAAGAGAACAGGANNNAGATCGGAA |
| 69967_1 | TATGAGTGTGGAGTCGTTGCCCTGTGACCTAATGACCTCCAAANNNAGATCGGAAGA |
| 69968_1 | TATGAGTGTGGAGTCGTTGCGAGAGATGTAGAAGCAAGCTCCANNNAGATCGGAA |
| 69969_1 | TATGAGTGTGGAGTCGTTGCTGTGTGTGTGTGTTTCCTACAGANNNAGATCGGAAG |
| 69970_1 | TATGAGTGTGGAGTCGTTGCACACACACACACACACACAAAATNNNAGATCGGAAGA |
| 69971_1 | TATGAGTGTGGAGTCGTTGCAGCAAGAATTTTGTCACCAGCTCNNNAGATCGGAAG |
| 69972_1 | TATGAGTGTGGAGTCGTTGCATAAGCAGAGCATTTGGGTCTGANNNAGATCGGAA |
| 69974_1 | TATGAGTGTGGAGTCGTTGCACAACCACAGTCATGGAGCTTATNNNAGATCGGAAG |
| 69975_1 | TATGAGTGTGGAGTCGTTGCCACACACACACACACACACAATANNNAGATCGGAAGA |
| 69976_1 | TATGAGTGTGGAGTCGTTGCTGTGTGTGTGTGTGTGTTTCATTNNNAGATCGGAAG |
| 69977_1 | TATGAGTGTGGAGTCGTTGCTTCTCCAAGGAAAGCTGACTTCCNNNAGATCGGAAGA |
| 69979_1 | TATGAGTGTGGAGTCGTTGCGGTATTCTTTCGGTGTTCAGTGCNNNAGATCGGAAG |
| 69980_1 | TATGAGTGTGGAGTCGTTGCCATTCGTAGCTTATTGCAGCCTCNNNAGATCGGAAGA |
| 69981_1 | TATGAGTGTGGAGTCGTTGCTAGAAGGGAAAACTCGACTTGGGNNNAGATCGGAA |
| 69982_1 | TATGAGTGTGGAGTCGTTGCCCTGTATGTGTGCCTGTATCTGTNNNAGATCGGAAGA |
| 69983_1 | TATGAGTGTGGAGTCGTTGCAAAATACCCTCCATATCTCGGCCNNNAGATCGGAAGA |
| 69984_1 | TATGAGTGTGGAGTCGTTGCGACATAAGCACTCCAAGCTCTCTNNNAGATCGGAAGA |
| 69985_1 | TATGAGTGTGGAGTCGTTGCGATTTCCTCTCTTCCCAACCCTTNNNAGATCGGAAGA |
| 69986_1 | TATGAGTGTGGAGTCGTTGCTTCCCCTGCACAGTCTCTAATTCNNNAGATCGGAAGA |

| | |
|---|---|
| 69988_1 | TATGAGTGTGGAGTCGTTGCGCAAGCTAGCAATGTGCAGAATANNNAGATCGGAA |
| 69989_1 | TATGAGTGTGGAGTCGTTGCATTTTTCTGAGTTAGGCCAGGCTNNNAGATCGGAAG |
| 69990_1 | TATGAGTGTGGAGTCGTTGCGAGCTAGGAGTCTGTGTCTTCACNNNAGATCGGAA |
| 69991_1 | TATGAGTGTGGAGTCGTTGCAATTATGGAGGCTGAGAAGTCCCNNNAGATCGGAA |
| 69992_1 | TATGAGTGTGGAGTCGTTGCTCCACTCTCTCAGCCATAACCTANNNAGATCGGAAGA |
| 69993_1 | TATGAGTGTGGAGTCGTTGCCACCTCATCCATCACTCATCCTTNNNAGATCGGAAGA |
| 69994_1 | TATGAGTGTGGAGTCGTTGCAAAGACTGTAATAAGGGCTGGGGNNNAGATCGGA |
| 69995_1 | TATGAGTGTGGAGTCGTTGCAGTGGGGAATCCTGACAAATTCANNNAGATCGGAA |
| 69997_1 | TATGAGTGTGGAGTCGTTGCCCCAGATTCTAGTCAGAGTGCTCNNNAGATCGGAAG |
| 69998_1 | TATGAGTGTGGAGTCGTTGCCTTGAAAGGCATCAGGGAAATCCNNNAGATCGGAA |
| 69999_1 | TATGAGTGTGGAGTCGTTGCCTGATAGTGTCGTTCCAGTCCAANNNAGATCGGAAG |
| 70001_1 | TATGAGTGTGGAGTCGTTGCTGAATCAGTAACAGAGGTTGGGGNNNAGATCGGA |
| 70002_1 | TATGAGTGTGGAGTCGTTGCAAGCGTGTATGTGTATCTCTCGANNNAGATCGGAAG |
| 70003_1 | TATGAGTGTGGAGTCGTTGCATGCAGCAAGAGAGAGGAATGAANNNAGATCGGA |
| 70004_1 | TATGAGTGTGGAGTCGTTGCTTTCTCACTCTCCATGTCTGCTCNNNAGATCGGAAGA |
| 70005_1 | TATGAGTGTGGAGTCGTTGCTATGTGCCTGGGAAAAAGATGGANNNAGATCGGAA |
| 70006_1 | TATGAGTGTGGAGTCGTTGCACTGGTGGTGTAATTCTGTGTGANNNAGATCGGAA |
| 70007_1 | TATGAGTGTGGAGTCGTTGCACAAGATTTACAAGGGAGGCTGANNNAGATCGGAA |
| 70008_1 | TATGAGTGTGGAGTCGTTGCGTGTGTGTGTTCCAGGTATGGTANNNAGATCGGAA |
| 70009_1 | TATGAGTGTGGAGTCGTTGCTAATCACCCAGCAAGTTCTTCCTNNNAGATCGGAAGA |
| 70010_1 | TATGAGTGTGGAGTCGTTGCGAAGAGAGGAAAAGACTGCTGGANNNAGATCGGA |
| 70011_1 | TATGAGTGTGGAGTCGTTGCTGACCTAGCTTTAGAAGTCCCACNNNAGATCGGAAG |
| 70012_1 | TATGAGTGTGGAGTCGTTGCCACAGAGTAGGGGGACACATTTTNNNAGATCGGAA |
| 70014_1 | TATGAGTGTGGAGTCGTTGCTTGAGGTCTAAAGACAGGAGCACNNNAGATCGGAA |
| 70015_1 | TATGAGTGTGGAGTCGTTGCAAGTGCTAGCCTTAGACTTGCTTNNNAGATCGGAAG |
| 70016_1 | TATGAGTGTGGAGTCGTTGCCACAGAGTGAGAGCAAGAGACTCNNNAGATCGGAA |
| 70017_1 | TATGAGTGTGGAGTCGTTGCTGGTGTCTCTTCTTATAAGGGCGNNNAGATCGGAAG |
| 70018_1 | TATGAGTGTGGAGTCGTTGCCTGTAATGACAAGGAGGACACCANNNAGATCGGAA |
| 70020_1 | TATGAGTGTGGAGTCGTTGCACCTTTCCCAAACAGATGATGGANNNAGATCGGAAG |
| 70021_1 | TATGAGTGTGGAGTCGTTGCTGTGTGTGTGTGTACTTACCANNNAGATCGGAAG |
| 70022_1 | TATGAGTGTGGAGTCGTTGCGGCCAAATGTCAGATACTAAAGCCNNNAGATCGGA |
| 70023_1 | TATGAGTGTGGAGTCGTTGCACACACACACAGAGAGGAAAAGANNNAGATCGGAA |
| 70024_1 | TATGAGTGTGGAGTCGTTGCGGAAGGGAATACTGAAGGTCTGANNNAGATCGGA |
| 70025_1 | TATGAGTGTGGAGTCGTTGCAGGGGAGAGAAATAATCCTGGGANNNAGATCGGA |
| 70026_1 | TATGAGTGTGGAGTCGTTGCTTCTCCACAAGAAACACTTCCCANNNAGATCGGAAGA |
| 70028_1 | TATGAGTGTGGAGTCGTTGCCTCCTATTGGTTCTACTGCCCTGNNNAGATCGGAAGA |
| 70029_1 | TATGAGTGTGGAGTCGTTGCTGTGGTTTCTACATCCTCTGTGTNNNAGATCGGAAG |
| 70030_1 | TATGAGTGTGGAGTCGTTGCATTACGTGATTTGGGTGGAGACANNNAGATCGGAA |
| 70031_1 | TATGAGTGTGGAGTCGTTGCCACACACACACACATAGACACACNNNAGATCGGAAG |
| 70032_1 | TATGAGTGTGGAGTCGTTGCGGAGGTCGTTTGCTTCTTTGTTTNNNAGATCGGAAG |
| 70033_1 | TATGAGTGTGGAGTCGTTGCTCTCACGGCTGTTCAACTGATTTNNNAGATCGGAAG |
| 70034_1 | TATGAGTGTGGAGTCGTTGCGGTTGTGTGTGTGTGTGTATGTTNNNAGATCGGAA |
| 70035_1 | TATGAGTGTGGAGTCGTTGCCTCTGAGGAGTGACTCATCCCTANNNAGATCGGAAG |

| | |
|---|---|
| 70036_1 | TATGAGTGTGGAGTCGTTGCAGAGACCAGTTGTGACAAGTCTCNNNAGATCGGAAG |
| 70037_1 | TATGAGTGTGGAGTCGTTGCGAAACAAAGGCTTGCTGGAGATTNNNAGATCGGAA |
| 70038_1 | TATGAGTGTGGAGTCGTTGCATCTGAGCTAAGAGCCCTCACTANNNAGATCGGAAG |
| 70040_1 | TATGAGTGTGGAGTCGTTGCAGTTAGTAGGATCTCCCTGGCTTNNNAGATCGGAAG |
| 70041_1 | TATGAGTGTGGAGTCGTTGCTGAGACTGTCTTTGTCATGCTGTNNNAGATCGGAAG |
| 70042_1 | TATGAGTGTGGAGTCGTTGCCCCTGCTTTCATTTGTTTGGGATNNNAGATCGGAAGA |
| 70043_1 | TATGAGTGTGGAGTCGTTGCGCTAGGTCAGAATGAGAGACTGGNNNAGATCGGAA |
| 70044_1 | TATGAGTGTGGAGTCGTTGCCAATCCCCAAGCTGCTTATCTTGNNNAGATCGGAAGA |
| 70045_1 | TATGAGTGTGGAGTCGTTGCACTGTCAATATCGCAGGGAGTTTNNNAGATCGGAAG |
| 70046_1 | TATGAGTGTGGAGTCGTTGCCAGGTGGGTTGGGGTAGAAAATANNNAGATCGGAA |
| 70047_1 | TATGAGTGTGGAGTCGTTGCAACACTGTACACCATTTTTCCCANNNAGATCGGAAGA |
| 70048_1 | TATGAGTGTGGAGTCGTTGCGTACAGTGTTTTTCCTGGGAAGCNNNAGATCGGAAG |
| 70049_1 | TATGAGTGTGGAGTCGTTGCGCCTTTTTCTTTTCCCAGACAGGNNNAGATCGGAAGA |
| 70051_1 | TATGAGTGTGGAGTCGTTGCCTCCCAAAGACTCTGACCCTTTTNNNAGATCGGAAGA |
| 70052_1 | TATGAGTGTGGAGTCGTTGCAATATGAAAATTGGGCAGGCAGGNNNAGATCGGAA |
| 70053_1 | TATGAGTGTGGAGTCGTTGCCTGGAAAGGGAAGCGCTTAGATANNNAGATCGGAA |
| 70055_1 | TATGAGTGTGGAGTCGTTGCGCCCAGCATCATGTTTTTGAGATNNNAGATCGGAAG |
| 70056_1 | TATGAGTGTGGAGTCGTTGCACAAAAATTAGCCAGGTGTGGTGNNNAGATCGGAA |
| 70057_1 | TATGAGTGTGGAGTCGTTGCGGCTTTCCAGAATGCAGTTCATTNNNAGATCGGAAG |
| 70059_1 | TATGAGTGTGGAGTCGTTGCGCCCAAGAAGATAGGACTACAGGNNNAGATCGGAA |
| 70060_1 | TATGAGTGTGGAGTCGTTGCAATACCAAAAGAATGGGTGTGCCNNNAGATCGGAA |
| 70061_1 | TATGAGTGTGGAGTCGTTGCGAAGAGATGTGCACAATCAGCTCNNNAGATCGGAA |
| 70062_1 | TATGAGTGTGGAGTCGTTGCTTTTCTAGTGGAGGAAACTGCCANNNAGATCGGAAG |
| 70064_1 | TATGAGTGTGGAGTCGTTGCAGTGAATGGTTCTCTGGCCTTAGNNNAGATCGGAAG |
| 70065_1 | TATGAGTGTGGAGTCGTTGCGTGGATGCTCAAGTCCCTGATATNNNAGATCGGAAG |
| 70066_1 | TATGAGTGTGGAGTCGTTGCCTCTGTACTTAGGGCCCAATGTTNNNAGATCGGAAG |
| 70067_1 | TATGAGTGTGGAGTCGTTGCTCAAGGTTCATCCACGTTGTACTNNNAGATCGGAAG |
| 70068_1 | TATGAGTGTGGAGTCGTTGCCCTGCTGTCTTAGAAGAACTGGTNNNAGATCGGAAG |
| 70069_1 | TATGAGTGTGGAGTCGTTGCTAACCACTGGACAAGCAACCTTANNNAGATCGGAAG |
| 70073_1 | TATGAGTGTGGAGTCGTTGCGTAATCCCACCTACTCATGAGGCNNNAGATCGGAAG |
| 70074_1 | TATGAGTGTGGAGTCGTTGCATGACAGGCCCAGATCTCATTTTNNNAGATCGGAAG |
| 70075_1 | TATGAGTGTGGAGTCGTTGCACGTCCTTCTAGGCAAGTGATTTNNNAGATCGGAAG |
| 70076_1 | TATGAGTGTGGAGTCGTTGCGAGGGGACTGTGCTCCAAAATAANNNAGATCGGAA |
| 70077_1 | TATGAGTGTGGAGTCGTTGCCTAGTCCGCCTTCATGTTTCATCNNNAGATCGGAAGA |
| 70078_1 | TATGAGTGTGGAGTCGTTGCGAATCATCTCGGGGCTGTAGAATNNNAGATCGGAA |
| 70079_1 | TATGAGTGTGGAGTCGTTGCAATACCAGTGACCCAGTGATCACNNNAGATCGGAAG |
| 70080_1 | TATGAGTGTGGAGTCGTTGCATTGATTGGAGGCCTCACTGAATNNNAGATCGGAAG |
| 70081_1 | TATGAGTGTGGAGTCGTTGCTGAAACAGGGTCTCACTTTGTCANNNAGATCGGAAG |
| 70082_1 | TATGAGTGTGGAGTCGTTGCACACCCCTCCTGTGATATTGTTCNNNAGATCGGAAGA |
| 70083_1 | TATGAGTGTGGAGTCGTTGCAAGAGATTCTCCTACCTCAGCCTNNNAGATCGGAAGA |
| 70084_1 | TATGAGTGTGGAGTCGTTGCCTTGAGAAAAAACGGGCTTGGAANNNAGATCGGAA |
| 70085_1 | TATGAGTGTGGAGTCGTTGCGATCACAAGGTCAGGAGTTCGAGNNNAGATCGGAA |
| 70086_1 | TATGAGTGTGGAGTCGTTGCTCCTTTTCTACCACTTCTACGGCNNNAGATCGGAAGA |

| | |
|---|---|
| 70087_1 | TATGAGTGTGGAGTCGTTGCATCTGCAAAGGATTCACCAGTCTNNNAGATCGGAAG |
| 70088_1 | TATGAGTGTGGAGTCGTTGCAGCTGGACCACTGTTCACTATTTNNNAGATCGGAAG |
| 70089_1 | TATGAGTGTGGAGTCGTTGCTGAGCCCCTATTTTTTGTTATGGGNNNAGATCGGAA |
| 70090_1 | TATGAGTGTGGAGTCGTTGCTAAGAGTACCAGCCACCAGAATGNNNAGATCGGAA |
| 70091_1 | TATGAGTGTGGAGTCGTTGCAAAGATTGTGCCACTGTACTCCANNNAGATCGGAAG |
| 70092_1 | TATGAGTGTGGAGTCGTTGCAATCTGTGATTGTTGTCAGCAGCNNNAGATCGGAAG |
| 70093_1 | TATGAGTGTGGAGTCGTTGCAACTGCAGATAGTACTGAACCCTNNNAGATCGGAAG |
| 70094_1 | TATGAGTGTGGAGTCGTTGCAACTTGGTTAGGCTATCGTGGTTNNNAGATCGGAA |
| 70095_1 | TATGAGTGTGGAGTCGTTGCGGACTGGAAGTAAGAGGGAAAGGNNNAGATCGGA |
| 70097_1 | TATGAGTGTGGAGTCGTTGCAAGGAAGGGGAAAGTCATTGTGANNNAGATCGGA |
| 70098_1 | TATGAGTGTGGAGTCGTTGCAGGAGAGAGTGGGGAAGGATTTANNNAGATCGGA |
| 70099_1 | TATGAGTGTGGAGTCGTTGCAGACAAACATATCCAGGGGACACNNNAGATCGGAA |
| 70100_1 | TATGAGTGTGGAGTCGTTGCGACTTAGACAGGTTGGGAGGAAGNNNAGATCGGA |
| 70101_1 | TATGAGTGTGGAGTCGTTGCTATGAGAGGGGCCTTCCTAGAAANNNAGATCGGAA |
| 70102_1 | TATGAGTGTGGAGTCGTTGCTTCCCAATTGCCAAACAGAACAANNNAGATCGGAAG |
| 70103_1 | TATGAGTGTGGAGTCGTTGCTTTGCAATTGTCCCTGATTCTGGNNNAGATCGGAAG |
| 70106_1 | TATGAGTGTGGAGTCGTTGCGATTCAAGCAATTCTCCTGCCTCNNNAGATCGGAAG |
| 70107_1 | TATGAGTGTGGAGTCGTTGCCTGAATTGCATCCCTCCCAATTCNNNAGATCGGAAGA |
| 70108_1 | TATGAGTGTGGAGTCGTTGCCCATGTTCTCACTTATTTGCGGGNNNAGATCGGAAG |
| 70109_1 | TATGAGTGTGGAGTCGTTGCTAGACCAATTCAGCCCACTTCTGNNNAGATCGGAAG |
| 70110_1 | TATGAGTGTGGAGTCGTTGCACTGCAGTCACCCAAGTAACTAGNNNAGATCGGAAG |
| 70112_1 | TATGAGTGTGGAGTCGTTGCCAGAGGGTCAGGAAACCTTGAAGNNNAGATCGGAA |
| 70114_1 | TATGAGTGTGGAGTCGTTGCCTGGCCACATTGATTGACTCTTCNNNAGATCGGAAG |
| 70117_1 | TATGAGTGTGGAGTCGTTGCAAAGTGCTGGGATTACAGGCATANNNAGATCGGAA |
| 70118_1 | TATGAGTGTGGAGTCGTTGCTATGAACTCTCTGAGCTCTGGGANNNAGATCGGAAG |
| 70119_1 | TATGAGTGTGGAGTCGTTGCGTAGCGTTCGAGCTCTTCAAAAGNNNAGATCGGAA |
| 70120_1 | TATGAGTGTGGAGTCGTTGCGTTGGGCCTATAGTTACACACCANNNAGATCGGAAG |
| 70122_1 | TATGAGTGTGGAGTCGTTGCTCTATGAGGCTAGCATCATCCTGNNNAGATCGGAAG |
| 70124_1 | TATGAGTGTGGAGTCGTTGCTTTTTTATGGATGCAGGCACTGGNNNAGATCGGAA |
| 70125_1 | TATGAGTGTGGAGTCGTTGCCCAATCTCTCGCTCTCTGAAACTNNNAGATCGGAAGA |
| 70126_1 | TATGAGTGTGGAGTCGTTGCGTTGGCATTATCTGACCAGCAATNNNAGATCGGAAG |
| 70127_1 | TATGAGTGTGGAGTCGTTGCACAGTGCTAGCCATTTACCTCTCNNNAGATCGGAAG |
| 70128_1 | TATGAGTGTGGAGTCGTTGCCCACAGAATTTCAGCTTCCCAAGNNNAGATCGGAAG |
| 70129_1 | TATGAGTGTGGAGTCGTTGCTTTTTTTTGGAGGCAGGTCTCACNNNAGATCGGAAG |
| 70130_1 | TATGAGTGTGGAGTCGTTGCAGATGCAAAATGGTCTAAAGGCANNNAGATCGGAA |
| 70131_1 | TATGAGTGTGGAGTCGTTGCGTGCACACCTGGCTGATTTATTTNNNAGATCGGAAG |
| 70132_1 | TATGAGTGTGGAGTCGTTGCCTGTGCACCAAAAGCAGTCATTANNNAGATCGGAAG |
| 70133_1 | TATGAGTGTGGAGTCGTTGCACACCCCCTACGATATTGAGAGTNNNAGATCGGAAG |
| 70135_1 | TATGAGTGTGGAGTCGTTGCAAAAAAAGCCCTGGACCAAATGGNNNAGATCGGAA |
| 70136_1 | TATGAGTGTGGAGTCGTTGCACTGAAGAAAGTAAGTGCCCTGTNNNAGATCGGAA |
| 70137_1 | TATGAGTGTGGAGTCGTTGCCATGCTTGAGGAAGGAGAGACTTNNNAGATCGGAA |
| 70138_1 | TATGAGTGTGGAGTCGTTGCTCATTTATCCTCCTTCAGGGCTGNNNAGATCGGAAG |
| 70139_1 | TATGAGTGTGGAGTCGTTGCTTCTAGTTTTCTGCCTGGACACTNNNAGATCGGAAG |

| | |
|---|---|
| 70141_1 | TATGAGTGTGGAGTCGTTGCAAGTGGACTCACACAGGATTTGTNNNAGATCGGAA |
| 70143_1 | TATGAGTGTGGAGTCGTTGCGAACTCCTGACCTCAGATGATCCNNNAGATCGGAAG |
| 70144_1 | TATGAGTGTGGAGTCGTTGCCCTTGGGATCTGGCTCATGATAANNNAGATCGGAAG |
| 70145_1 | TATGAGTGTGGAGTCGTTGCGCAAGTAGAAAGAGGGAAGGACANNNAGATCGGA |
| 70146_1 | TATGAGTGTGGAGTCGTTGCAAAATACAAAATCAGCTGGGCGTNNNAGATCGGAA |
| 70148_1 | TATGAGTGTGGAGTCGTTGCATAAAGGTGACCCAAGAAGGCATNNNAGATCGGAA |
| 70149_1 | TATGAGTGTGGAGTCGTTGCACGCACACATGAGTCCTAATCATNNNAGATCGGAAG |
| 70151_1 | TATGAGTGTGGAGTCGTTGCGGCATTGACTTCCTGCGAATATCNNNAGATCGGAAG |
| 70152_1 | TATGAGTGTGGAGTCGTTGCAAGAGAAGCCTTGGAAAGACCTCNNNAGATCGGAA |
| 70153_1 | TATGAGTGTGGAGTCGTTGCTCATGTCTCTCTTGCCTCCAAAANNNAGATCGGAAGA |
| 70154_1 | TATGAGTGTGGAGTCGTTGCTCAGTTCCCCATTTGACTTGTGANNNAGATCGGAAG |
| 70156_1 | TATGAGTGTGGAGTCGTTGCTAATATCCAGAGGGCGAGAGGATNNNAGATCGGAA |
| 70157_1 | TATGAGTGTGGAGTCGTTGCCCAAGCAGAGACCTTCTAAGGAGNNNAGATCGGAA |
| 70158_1 | TATGAGTGTGGAGTCGTTGCGACGGTGACAATGTTTACAAGCANNNAGATCGGAA |
| 70159_1 | TATGAGTGTGGAGTCGTTGCACTTGAAGAAAGCAAGGGAGTGANNNAGATCGGA |
| 70160_1 | TATGAGTGTGGAGTCGTTGCAGGAAAAACCTGTGCAAAAAGCTNNNAGATCGGAA |
| 70161_1 | TATGAGTGTGGAGTCGTTGCAACATCCTCCCAGGAAACTTACCNNNAGATCGGAAG |
| 70162_1 | TATGAGTGTGGAGTCGTTGCCATTCAGCAACTCAAAAAGGGGTNNNAGATCGGAA |
| 70163_1 | TATGAGTGTGGAGTCGTTGCCATTCTCAGATCTCCAGCTGTGTNNNAGATCGGAAG |
| 70164_1 | TATGAGTGTGGAGTCGTTGCAGAATGGACTTAGAGGCACGAAGNNNAGATCGGAA |
| 70165_1 | TATGAGTGTGGAGTCGTTGCGTAAGTAGGAGGACAGGTATGCCNNNAGATCGGAA |
| 70166_1 | TATGAGTGTGGAGTCGTTGCTAGTCACAGAGCAGGTAAGACCANNNAGATCGGAA |
| 70167_1 | TATGAGTGTGGAGTCGTTGCCCTCCAGCTTCTTCTTAGTGGTTNNNAGATCGGAAGA |
| 70168_1 | TATGAGTGTGGAGTCGTTGCGACTGCATATGATCGCCAAAGAANNNAGATCGGAA |
| 70169_1 | TATGAGTGTGGAGTCGTTGCCCCCTCGTGGATATTAGGAACAGNNNAGATCGGAA |
| 70171_1 | TATGAGTGTGGAGTCGTTGCCTCCCGGCTACTTTTTCTTTCTTNNNAGATCGGAAGA |
| 70172_1 | TATGAGTGTGGAGTCGTTGCGGCAGAAATGCAAAAGTAGCCTTNNNAGATCGGAA |
| 70173_1 | TATGAGTGTGGAGTCGTTGCGCATACCTGTGATGAAAAGGAGCNNNAGATCGGAA |
| 70174_1 | TATGAGTGTGGAGTCGTTGCTGAGCCTGGTAAATCATGTACCCNNNAGATCGGAAG |
| 70176_1 | TATGAGTGTGGAGTCGTTGCAATAAGGCTGTGGTCCCTATCACNNNAGATCGGAAG |
| 70177_1 | TATGAGTGTGGAGTCGTTGCGCCAGCAAAAGGGGAAAAAAAAANNNAGATCGGAA |
| 70180_1 | TATGAGTGTGGAGTCGTTGCCTGGACCTACAATCCCTTTTGGANNNAGATCGGAAG |
| 70181_1 | TATGAGTGTGGAGTCGTTGCAATGTCAGGGGTTAGCTTTCTCTNNNAGATCGGAAG |
| 70182_1 | TATGAGTGTGGAGTCGTTGCCATCTTGGGGAAGGCGTTCTATANNNAGATCGGAA |
| 70183_1 | TATGAGTGTGGAGTCGTTGCCTTGATGAGACCTTCACAGGGAANNNAGATCGGAA |
| 70185_1 | TATGAGTGTGGAGTCGTTGCCTGTCCCTTGGCTGAGAATCAAANNNAGATCGGAAG |
| 70186_1 | TATGAGTGTGGAGTCGTTGCCTAGGAATCTTACGGGGCAAGAGNNNAGATCGGAA |
| 70187_1 | TATGAGTGTGGAGTCGTTGCAGTTTCCATAGCAACTGCAGAGANNNAGATCGGAA |
| 70189_1 | TATGAGTGTGGAGTCGTTGCTAAGAGCTAGCGTTTCCTCGTTTNNNAGATCGGAAG |
| 70190_1 | TATGAGTGTGGAGTCGTTGCAAGAGCCAGAACTTTCCTTAGCANNNAGATCGGAAG |
| 70193_1 | TATGAGTGTGGAGTCGTTGCGGACCCAAAACCAGAAATCAAGGNNNAGATCGGAA |
| 70194_1 | TATGAGTGTGGAGTCGTTGCGCAGGAAAGTAACTTCACACTCGNNNAGATCGGAA |
| 70195_1 | TATGAGTGTGGAGTCGTTGCGGGAGGGGGTCATACGATATTTGNNNAGATCGGAA |

| | |
|---|---|
| 70197_1 | TATGAGTGTGGAGTCGTTGCATAAAGTCTCTGCAGCACTGTGANNNAGATCGGAAG |
| 70198_1 | TATGAGTGTGGAGTCGTTGCGAGGTTTTGACTCCCGGGATTTANNNAGATCGGAAG |
| 70199_1 | TATGAGTGTGGAGTCGTTGCGAAATAGTTTATGGGCCATGGGCNNNAGATCGGAA |
| 70200_1 | TATGAGTGTGGAGTCGTTGCGGTCTCTGTCAACACAATTCAGCNNNAGATCGGAAG |
| 70201_1 | TATGAGTGTGGAGTCGTTGCGTTTGTGTTTTGGAGACAGGGTCNNNAGATCGGAA |
| 70202_1 | TATGAGTGTGGAGTCGTTGCGAGGCTAAGACAGGAGAATTGCTNNNAGATCGGAA |
| 70203_1 | TATGAGTGTGGAGTCGTTGCCCAATTCTCCTTCCTGATCTGCANNNAGATCGGAAGA |
| 70204_1 | TATGAGTGTGGAGTCGTTGCCCGTGGCCCTCAAATGTATTTTTNNNAGATCGGAAG |
| 70205_1 | TATGAGTGTGGAGTCGTTGCGCGTTGGTTGGAAAATAAGGTCANNNAGATCGGAA |
| 70206_1 | TATGAGTGTGGAGTCGTTGCAGATTAGTGTCAACATGTGGGCTNNNAGATCGGAA |
| 70207_1 | TATGAGTGTGGAGTCGTTGCAGATGTAGCCAGGGAAGGAAAAGNNNAGATCGGA |
| 70209_1 | TATGAGTGTGGAGTCGTTGCGTAGGGAGTGGATGCAGACATAGNNNAGATCGGA |
| 70210_1 | TATGAGTGTGGAGTCGTTGCACCCATTCTGTATTCCCTTTGCTNNNAGATCGGAAGA |
| 70211_1 | TATGAGTGTGGAGTCGTTGCCTAAGCGTGCTTTCAGGATGATGNNNAGATCGGAA |
| 70212_1 | TATGAGTGTGGAGTCGTTGCGAGACGTGGCTAAACAATCACACNNNAGATCGGAA |
| 70214_1 | TATGAGTGTGGAGTCGTTGCAGAACCTTAGTGATGTGACGCTTNNNAGATCGGAA |
| 70215_1 | TATGAGTGTGGAGTCGTTGCCACCTGTAGTCCCAGTTACTTGGNNNAGATCGGAAG |
| 70218_1 | TATGAGTGTGGAGTCGTTGCTGGAGAGAGATAAAGGGGAGAGGNNNAGATCGG |
| 70219_1 | TATGAGTGTGGAGTCGTTGCTCTTTACCCGTGATCACTGAGTGNNNAGATCGGAAG |
| 70220_1 | TATGAGTGTGGAGTCGTTGCCAAAATCCGGGTTCTGCCAAAATNNNAGATCGGAAG |
| 70221_1 | TATGAGTGTGGAGTCGTTGCGGTGAAACCCTGTCTCCACTAAANNNAGATCGGAAG |
| 70222_1 | TATGAGTGTGGAGTCGTTGCCACTGATCCCATCATGAGGACTCNNNAGATCGGAAG |
| 70223_1 | TATGAGTGTGGAGTCGTTGCGTGTGTGTGTGTGTGTGTGTATANNNAGATCGGAA |
| 70225_1 | TATGAGTGTGGAGTCGTTGCTACTTGCTCTCCTGTCAATAGGCNNNAGATCGGAAG |
| 70226_1 | TATGAGTGTGGAGTCGTTGCCCAGTCTTGCTTTCCTTTCTCCTNNNAGATCGGAAGA |
| 70227_1 | TATGAGTGTGGAGTCGTTGCGGTGTACTGAATGAGGCTTTTGCNNNAGATCGGAA |
| 70228_1 | TATGAGTGTGGAGTCGTTGCTCAGCGACACTCTCATCTCTCTANNNAGATCGGAAGA |
| 70231_1 | TATGAGTGTGGAGTCGTTGCGGACAGTGTGCCTTTCTTTAGGANNNAGATCGGAAG |
| 70232_1 | TATGAGTGTGGAGTCGTTGCCGACAAAGCAGTGGTAGAAACAANNNAGATCGGAA |
| 70233_1 | TATGAGTGTGGAGTCGTTGCACCTGTAATCCTAGCACTTTGGGNNNAGATCGGAAG |
| 70234_1 | TATGAGTGTGGAGTCGTTGCCTGAGGTGGAAGGATTGTGTGAGNNNAGATCGGA |
| 70235_1 | TATGAGTGTGGAGTCGTTGCGCTGAGTCTCTAACAGATGCAGTNNNAGATCGGAAG |
| 70238_1 | TATGAGTGTGGAGTCGTTGCCATTCTTTTCCATTTGGCCCTGTNNNAGATCGGAAGA |
| 70239_1 | TATGAGTGTGGAGTCGTTGCGCTGAAGTGATCCTCTTGCCTTANNNAGATCGGAAG |
| 70240_1 | TATGAGTGTGGAGTCGTTGCCTGCAAAACGGAGAAGAAGGAAGNNNAGATCGGA |
| 70241_1 | TATGAGTGTGGAGTCGTTGCTTCCTGGAACCAAAAGGAACTCANNNAGATCGGAAG |
| 70243_1 | TATGAGTGTGGAGTCGTTGCGCTCTCTGTCTCACTCTCTCTCTNNNAGATCGGAAGA |
| 70244_1 | TATGAGTGTGGAGTCGTTGCAGTGTGGGTTTTTGGTTTTGAGGNNNAGATCGGAA |
| 70245_1 | TATGAGTGTGGAGTCGTTGCACACAACAACAATGATGGCTTGTNNNAGATCGGAAG |
| 70246_1 | TATGAGTGTGGAGTCGTTGCTCTCTTCATTGTCTGGATCCAGGNNNAGATCGGAAG |
| 70247_1 | TATGAGTGTGGAGTCGTTGCTCTCCCTCTCTCTCTCTCTCTCTNNNAGATCGGAAGAG |
| 70248_1 | TATGAGTGTGGAGTCGTTGCCTCTTACCGTTCCAGGGAGTATGNNNAGATCGGAAG |
| 70249_1 | TATGAGTGTGGAGTCGTTGCATGTGAGTGAAAGTGTGCATGTGNNNAGATCGGAA |

| | |
|---|---|
| 70250_1 | TATGAGTGTGGAGTCGTTGCTGGGAGAGAGATGAGAGAGAGAGNNNAGATCGGA |
| 70251_1 | TATGAGTGTGGAGTCGTTGCTTACTGGGATGACCAAGTCCTCANNNAGATCGGAAG |
| 70252_1 | TATGAGTGTGGAGTCGTTGCCAAGCTCTGGTCTCTCTTCCTTTNNNAGATCGGAAGA |
| 70253_1 | TATGAGTGTGGAGTCGTTGCCCAAGCTGCATCACAAAGGAAAANNNAGATCGGAAG |
| 70254_1 | TATGAGTGTGGAGTCGTTGCTTGACAGTCTTGTTCTTGTCACCNNNAGATCGGAAGA |
| 70255_1 | TATGAGTGTGGAGTCGTTGCTCTCACCAAACCTAGCCTTCATCNNNAGATCGGAAGA |
| 70256_1 | TATGAGTGTGGAGTCGTTGCAACACAGAGTGACACAGATACCCNNNAGATCGGAAG |
| 70258_1 | TATGAGTGTGGAGTCGTTGCTCAAGACACCTGGTCGTACAATTNNNAGATCGGAAG |
| 70259_1 | TATGAGTGTGGAGTCGTTGCATTCATGTTGGTGGAGCCCTTATNNNAGATCGGAAG |
| 70260_1 | TATGAGTGTGGAGTCGTTGCAGGGAGATGGAAAGGGGATATCANNNAGATCGGA |
| 70261_1 | TATGAGTGTGGAGTCGTTGCCCTGTATGGAAATTCACTGGGGANNNAGATCGGAA |
| 70264_1 | TATGAGTGTGGAGTCGTTGCGTTTCTTGGTCCCTTTGAAGCTCNNNAGATCGGAAGA |
| 70265_1 | TATGAGTGTGGAGTCGTTGCGGAGGAGGTACCAGGTTCTTTTTNNNAGATCGGAA |
| 70268_1 | TATGAGTGTGGAGTCGTTGCATATGTGTGTGTGTGTGTGTGTGNNNAGATCGGAA |
| 70269_1 | TATGAGTGTGGAGTCGTTGCCTTAGCCTCCCTAGTAGCTGAGANNNAGATCGGAAG |
| 70270_1 | TATGAGTGTGGAGTCGTTGCCTCACACACACACACACATACACNNNAGATCGGAAGA |
| 70271_1 | TATGAGTGTGGAGTCGTTGCGCTGTGTTGAAAATGAAGGCTCTNNNAGATCGGAA |
| 70272_1 | TATGAGTGTGGAGTCGTTGCTTTTTTGAGACGGAGTCTTGCTCNNNAGATCGGAAG |
| 70273_1 | TATGAGTGTGGAGTCGTTGCCACACACACACACACTTGTTANNNAGATCGGAAGA |
| 70274_1 | TATGAGTGTGGAGTCGTTGCACACACATACACACCCCTTACTCNNNAGATCGGAAGA |
| 70275_1 | TATGAGTGTGGAGTCGTTGCCACACACACATGTTCCTTGACTTNNNAGATCGGAAGA |
| 70277_1 | TATGAGTGTGGAGTCGTTGCCCTGTTCTTTACTGTCCAGAGCTNNNAGATCGGAAGA |
| 70278_1 | TATGAGTGTGGAGTCGTTGCCAGAGAGACACCCAAGACTTCAANNNAGATCGGAAG |
| 70279_1 | TATGAGTGTGGAGTCGTTGCCTAATCCCATCTGATCAGGACCCNNNAGATCGGAAGA |
| 70281_1 | TATGAGTGTGGAGTCGTTGCACCCTATCAGCCAATGCCTTTATNNNAGATCGGAAGA |
| 70282_1 | TATGAGTGTGGAGTCGTTGCTGTGGCCCTCATGTTGCTAAATANNNAGATCGGAAG |
| 70283_1 | TATGAGTGTGGAGTCGTTGCAGAGAGAGAGAGAGAGAGAGGGANNNAGATCGGA |
| 70284_1 | TATGAGTGTGGAGTCGTTGCGGAGAGACAGAGAGTGAGAGACANNNAGATCGGA |
| 70285_1 | TATGAGTGTGGAGTCGTTGCACAATTCAGCATGAGATTTGGGCNNNAGATCGGAAG |
| 70287_1 | TATGAGTGTGGAGTCGTTGCGCTTACATGGTTATGGAGGGTGANNNAGATCGGAA |
| 70288_1 | TATGAGTGTGGAGTCGTTGCTGGCCTAATATACCTGCATGTCCNNNAGATCGGAAGA |
| 70289_1 | TATGAGTGTGGAGTCGTTGCCCTCCCACCTTTCTTTCTCTCTCNNNAGATCGGAAGAG |
| 70290_1 | TATGAGTGTGGAGTCGTTGCGTTCATGTGTCCCAAGTCTCAAGNNNAGATCGGAAG |
| 70291_1 | TATGAGTGTGGAGTCGTTGCCTCTCTCTCTCCTGTTCTGCAATNNNAGATCGGAAGA |
| 70292_1 | TATGAGTGTGGAGTCGTTGCCTCCCTATTCCCTGAGACACAAGNNNAGATCGGAAGA |
| 70293_1 | TATGAGTGTGGAGTCGTTGCATTTCCCTCACATTTCTTGGGCTNNNAGATCGGAAGA |
| 70294_1 | TATGAGTGTGGAGTCGTTGCCCTGCCTTAGCCTCCTAAGTAACNNNAGATCGGAAGA |
| 70295_1 | TATGAGTGTGGAGTCGTTGCCTCTCTCATCCCTGAGCTCTTTCNNNAGATCGGAAGA |
| 70296_1 | TATGAGTGTGGAGTCGTTGCGGCATATGGACACTTGGACTGTANNNAGATCGGAA |
| 70298_1 | TATGAGTGTGGAGTCGTTGCCCACCTCCTGTTTTCAAGCAATTNNNAGATCGGAAGA |
| 70299_1 | TATGAGTGTGGAGTCGTTGCGGGAGGGAAGGAAAGAAGGAAAANNNAGATCGGA |
| 70301_1 | TATGAGTGTGGAGTCGTTGCCAAGTCCCAAATTTGCGGAGTAGNNNAGATCGGAAG |
| 70302_1 | TATGAGTGTGGAGTCGTTGCTGTCCTTCCACGGTTATTTCACANNNAGATCGGAAGA |

| | |
|---|---|
| 70303_1 | TATGAGTGTGGAGTCGTTGCGAGAGAGAGAGCAGGGTACTGTANNNAGATCGGA |
| 70304_1 | TATGAGTGTGGAGTCGTTGCACTGAGGGTAATGGCTTTCTTCANNNAGATCGGAAG |
| 70306_1 | TATGAGTGTGGAGTCGTTGCAGAGAGAGAGACAGAGTTGGAGGNNNAGATCGGA |
| 70307_1 | TATGAGTGTGGAGTCGTTGCATGAGAACAGGAAGGGGGAAATCNNNAGATCGGA |
| 70308_1 | TATGAGTGTGGAGTCGTTGCGTGTGTGTGTGTTTGTGTGTGTANNNAGATCGGAA |
| 70309_1 | TATGAGTGTGGAGTCGTTGCAGGATGTTTGGCTGATTCTCTGTNNNAGATCGGAAG |
| 70310_1 | TATGAGTGTGGAGTCGTTGCGGCAGAAAGAGAGAGAGTCCATCNNNAGATCGGAA |
| 70311_1 | TATGAGTGTGGAGTCGTTGCCCATGTCGAGATCAAAAAGGCAANNNAGATCGGAA |
| 70312_1 | TATGAGTGTGGAGTCGTTGCATCTTCCTCTTCCTCTCTCTCCCNNNAGATCGGAAGAG |
| 70315_1 | TATGAGTGTGGAGTCGTTGCCCCTACCATCCAGGAGAGGATATNNNAGATCGGAAG |
| 70316_1 | TATGAGTGTGGAGTCGTTGCTCTCAAGGAAGAGATTTGGCTCCNNNAGATCGGAAG |
| 70317_1 | TATGAGTGTGGAGTCGTTGCTAGGAAGGATGACTGTGTGTGTGNNNAGATCGGAA |
| 70318_1 | TATGAGTGTGGAGTCGTTGCTCCAAAGAGGAAAGTACGATGGGNNNAGATCGGAA |
| 70319_1 | TATGAGTGTGGAGTCGTTGCTAAAGCATTAGAAACCACCCCCANNNAGATCGGAAG |
| 70320_1 | TATGAGTGTGGAGTCGTTGCGTGTGAGAGAGCCAGTGCATATANNNAGATCGGAA |
| 70321_1 | TATGAGTGTGGAGTCGTTGCTGGTGGTCACGTTGTTTAGAAGANNNAGATCGGAA |
| 70322_1 | TATGAGTGTGGAGTCGTTGCAAAGAGAAAGAGCAAGCTCTGGTNNNAGATCGGAA |
| 70323_1 | TATGAGTGTGGAGTCGTTGCAGCACAAGTGAATGATCTGGACTNNNAGATCGGAA |
| 70325_1 | TATGAGTGTGGAGTCGTTGCATGAGAGATTGGCTCACACCATTNNNAGATCGGAAG |
| 70326_1 | TATGAGTGTGGAGTCGTTGCACAGCATAGGGCAATTACCTTGANNNAGATCGGAAG |
| 70327_1 | TATGAGTGTGGAGTCGTTGCATTACTCAGCTCCAAGACAGAGGNNNAGATCGGAAG |
| 70328_1 | TATGAGTGTGGAGTCGTTGCATAGCCGAATGTCTCTAGGAGGTNNNAGATCGGAA |
| 70329_1 | TATGAGTGTGGAGTCGTTGCGAGTAGCTGGGATTACAGGCATCNNNAGATCGGAA |
| 70331_1 | TATGAGTGTGGAGTCGTTGCGCTGTATACCTAAATCGGGAGGGNNNAGATCGGAA |
| 70332_1 | TATGAGTGTGGAGTCGTTGCGTCTTCCACCATAATTGTGAGGCNNNAGATCGGAAG |
| 70333_1 | TATGAGTGTGGAGTCGTTGCTTCCTTCCTTCCTTCCTCTCTCTNNNAGATCGGAAGAG |
| 70334_1 | TATGAGTGTGGAGTCGTTGCGCTGTATACCTAAATCGGGAGGGNNNAGATCGGAA |
| 70335_1 | TATGAGTGTGGAGTCGTTGCCTTTCTTTCTTTGCGACAGGGTCNNNAGATCGGAAG |
| 70336_1 | TATGAGTGTGGAGTCGTTGCCACACACACAACACACACATACANNNAGATCGGAAG |
| 70337_1 | TATGAGTGTGGAGTCGTTGCGAACAGCAAAGGGAAAATCCTCCNNNAGATCGGAA |
| 70338_1 | TATGAGTGTGGAGTCGTTGCTCCAGGTTCAAGTGATTCTCCTGNNNAGATCGGAAG |
| 70339_1 | TATGAGTGTGGAGTCGTTGCATACCCTGAGTTTGGTAGAAGCCNNNAGATCGGAAG |
| 70340_1 | TATGAGTGTGGAGTCGTTGCATACTCCTTGCCTTCTTCCAAGGNNNAGATCGGAAGA |
| 70341_1 | TATGAGTGTGGAGTCGTTGCACGCTTCTGTTTTTTCCCTAACANNNAGATCGGAAGA |
| 70342_1 | TATGAGTGTGGAGTCGTTGCGGAGAGAGGAGAGAGAGAGACAGNNNAGATCGG |
| 70344_1 | TATGAGTGTGGAGTCGTTGCATGGAACAGTGGTAACAGAGCATNNNAGATCGGAA |
| 70346_1 | TATGAGTGTGGAGTCGTTGCATAGCTGGAATGACAGACGTGAGNNNAGATCGGAA |
| 70347_1 | TATGAGTGTGGAGTCGTTGCTCTCTCTGTCTCTCTCTCTCTCTNNNAGATCGGAAGAG |
| 70348_1 | TATGAGTGTGGAGTCGTTGCATGTATACCTACACACACACACANNNAGATCGGAAG |
| 70350_1 | TATGAGTGTGGAGTCGTTGCTTTTGGAAGTCTCACTGTGTTGCNNNAGATCGGAAG |
| 70351_1 | TATGAGTGTGGAGTCGTTGCGCACAATCTCAGCTCACTACAACNNNAGATCGGAAG |
| 70352_1 | TATGAGTGTGGAGTCGTTGCGTGAACAGCTTAGGGCAAATGAANNNAGATCGGAA |
| 70353_1 | TATGAGTGTGGAGTCGTTGCAGGTCAGTTGCTGAGCTTTTCTANNNAGATCGGAAG |

| | |
|---|---|
| 70354_1 | TATGAGTGTGGAGTCGTTGCACAGTGGCTCACACCTATAATCCNNNAGATCGGAAG |
| 70355_1 | TATGAGTGTGGAGTCGTTGCGGCAGGGTTTATTGGGTGAAAAGNNNAGATCGGAA |
| 70356_1 | TATGAGTGTGGAGTCGTTGCGCTCACACCTGTAATCCCAGTACNNNAGATCGGAAG |
| 70357_1 | TATGAGTGTGGAGTCGTTGCGGGAAGAGAACAAGAGTCTCTGGNNNAGATCGGAA |
| 70358_1 | TATGAGTGTGGAGTCGTTGCTCCATTGGATGCAGCTTCTAAGANNNAGATCGGAAG |
| 70359_1 | TATGAGTGTGGAGTCGTTGCAGATGTGTGAGTGAAGCCATCTTNNNAGATCGGAA |
| 70360_1 | TATGAGTGTGGAGTCGTTGCGTGCAAAGGGGGAAATGTATAGCNNNAGATCGGAA |
| 70361_1 | TATGAGTGTGGAGTCGTTGCATTCCCACACCTGCAGAACTTATNNNAGATCGGAAG |
| 70362_1 | TATGAGTGTGGAGTCGTTGCAACCAGAAGGTAAGACCAACCTGNNNAGATCGGAA |
| 70363_1 | TATGAGTGTGGAGTCGTTGCCCTCAAACCTAATCACCTCCCAANNNAGATCGGAAGA |
| 70364_1 | TATGAGTGTGGAGTCGTTGCAACCAGGTAGAAGTTGCAGTCTTNNNAGATCGGAA |
| 70365_1 | TATGAGTGTGGAGTCGTTGCGTAGCTGGGACTGTAAGTGTTCANNNAGATCGGAA |
| 70366_1 | TATGAGTGTGGAGTCGTTGCTCACTTGGATTTTCCCTATGCCANNNAGATCGGAAG |
| 70367_1 | TATGAGTGTGGAGTCGTTGCTGTCTCTCTCTCCCTTCTTTCCTNNNAGATCGGAAGAG |
| 70368_1 | TATGAGTGTGGAGTCGTTGCGTTTCCCTTTCATTCTGCTTCCCNNNAGATCGGAAGA |
| 70369_1 | TATGAGTGTGGAGTCGTTGCTCTTCCCTTCTAGCAAAAGGAAANNNAGATCGGAAG |
| 70370_1 | TATGAGTGTGGAGTCGTTGCATGGCCTTCAGTAAGTGAATGGTNNNAGATCGGAA |
| 70371_1 | TATGAGTGTGGAGTCGTTGCTCTGGAGAATTCTGACCAATGCANNNAGATCGGAAG |
| 70373_1 | TATGAGTGTGGAGTCGTTGCACACACACACACACACAAAGAAGNNNAGATCGGAAG |
| 70374_1 | TATGAGTGTGGAGTCGTTGCCACAGTGGCTCACACCTGTAATANNNAGATCGGAAG |
| 70375_1 | TATGAGTGTGGAGTCGTTGCTTTGAACCCAAGAATGACAAGCANNNAGATCGGAAG |
| 70376_1 | TATGAGTGTGGAGTCGTTGCAAGAAAGAAAGCCCCACCAAATGNNNAGATCGGAA |
| 70377_1 | TATGAGTGTGGAGTCGTTGCTCATGAAACCACCCAATCTAGCANNNAGATCGGAAG |
| 70378_1 | TATGAGTGTGGAGTCGTTGCGAAAGGAAGGAAGGAAGGAGAGANNNAGATCGG |
| 70379_1 | TATGAGTGTGGAGTCGTTGCATGGTGAGGATACTGTCTTGCAGNNNAGATCGGAA |
| 70380_1 | TATGAGTGTGGAGTCGTTGCGGACAGAGAGAAACCAGTAGCAANNNAGATCGGAA |
| 70381_1 | TATGAGTGTGGAGTCGTTGCAGCCATAAAAAATTAGCTGGGCANNNAGATCGGAA |
| 70382_1 | TATGAGTGTGGAGTCGTTGCTGAGAAAGAGAGAGAGAGAGGCTNNNAGATCGGA |
| 70383_1 | TATGAGTGTGGAGTCGTTGCAGAGAGAGAGAGACGGAGTTTCANNNAGATCGGA |
| 70384_1 | TATGAGTGTGGAGTCGTTGCTTTTTTGGTGTGAGGAGACAGGANNNAGATCGGAA |
| 70385_1 | TATGAGTGTGGAGTCGTTGCCACCTCCCAGACTCAAGTGATTCNNNAGATCGGAAG |
| 70386_1 | TATGAGTGTGGAGTCGTTGCCCCTACCCCCGTCTCTTTTATTTNNNAGATCGGAAGA |
| 70387_1 | TATGAGTGTGGAGTCGTTGCTTGTGAAATGCCCTTCATCTGTGNNNAGATCGGAAG |
| 70388_1 | TATGAGTGTGGAGTCGTTGCGCCACAACATCTCGCTTATTCTCNNNAGATCGGAAGA |
| 70389_1 | TATGAGTGTGGAGTCGTTGCTCATAGCTCCCAGTCTCCCATATNNNAGATCGGAAGA |
| 70390_1 | TATGAGTGTGGAGTCGTTGCCTCTCCCTCTTCATCATGTGAGGNNNAGATCGGAAG |
| 70391_1 | TATGAGTGTGGAGTCGTTGCCACACAGACACACACACACATACNNNAGATCGGAAG |
| 70392_1 | TATGAGTGTGGAGTCGTTGCGAGAGAGAGAGAGATCTTTGCGGNNNAGATCGGA |
| 70393_1 | TATGAGTGTGGAGTCGTTGCGGGGGGGAAGAAAGAAAGAAAGANNNAGATCGG |
| 70395_1 | TATGAGTGTGGAGTCGTTGCAGAGAGAGACAGAGAGGCTTCAGNNNAGATCGGA |
| 70396_1 | TATGAGTGTGGAGTCGTTGCAAAGAGATGGACTCCAGCAACATNNNAGATCGGAA |
| 70397_1 | TATGAGTGTGGAGTCGTTGCTTTATGTAGGGAAAGGTGCCACANNNAGATCGGAA |
| 70398_1 | TATGAGTGTGGAGTCGTTGCACAAAAATTAGCCAGGTGTGGTGNNNAGATCGGAA |

| | |
|---|---|
| 70399_1 | TATGAGTGTGGAGTCGTTGCGTGTGTGTGTGTGTGTGTAATNNNAGATCGGAA |
| 70400_1 | TATGAGTGTGGAGTCGTTGCCAATGGCATGATCTCAGGTCACTNNNAGATCGGAAG |
| 70402_1 | TATGAGTGTGGAGTCGTTGCCATTACCAATCTAACCGGGACCANNNAGATCGGAAG |
| 70403_1 | TATGAGTGTGGAGTCGTTGCCCACCTCTCTCTCTCTCTCTCTCNNNAGATCGGAAGAG |
| 70405_1 | TATGAGTGTGGAGTCGTTGCCATGGGAGCACCTAACTCTGTAANNNAGATCGGAAG |
| 70406_1 | TATGAGTGTGGAGTCGTTGCAAGAAAACTGTGAGGGAAGAGGGNNNAGATCGGA |
| 70407_1 | TATGAGTGTGGAGTCGTTGCAGAGAGAAAGAAACACCTAGGCCNNNAGATCGGAA |
| 70408_1 | TATGAGTGTGGAGTCGTTGCAGTCTTTGCAACTAGCTGAAGGANNNAGATCGGAA |
| 70409_1 | TATGAGTGTGGAGTCGTTGCTATGCCCCAGAGTCTGAAAACATNNNAGATCGGAAG |
| 70410_1 | TATGAGTGTGGAGTCGTTGCGAATCCTGGCACCAAATATGAGCNNNAGATCGGAA |
| 70411_1 | TATGAGTGTGGAGTCGTTGCAAGAGAAGTACAAAAGGCCAGGTNNNAGATCGGA |
| 70412_1 | TATGAGTGTGGAGTCGTTGCCAGATCTATAATGAGGGCCCCACNNNAGATCGGAAG |
| 70413_1 | TATGAGTGTGGAGTCGTTGCGTGTGTGTGTGTTTGTGTGTGTANNNAGATCGGAA |
| 70416_1 | TATGAGTGTGGAGTCGTTGCTGCACATCCTCCTGTTTCCTTTANNNAGATCGGAAGA |
| 70417_1 | TATGAGTGTGGAGTCGTTGCATAGCCTGCAGAATCATGAACCANNNAGATCGGAAG |
| 70418_1 | TATGAGTGTGGAGTCGTTGCTGGAGACACAGAAAAGCTGAAGTNNNAGATCGGAA |
| 70420_1 | TATGAGTGTGGAGTCGTTGCTTCACCATTCCCCTGTCTAACAGNNNAGATCGGAAGA |
| 70421_1 | TATGAGTGTGGAGTCGTTGCGAAAGGAAGGAAGGAAGGAAGGANNNAGATCGG |
| 70422_1 | TATGAGTGTGGAGTCGTTGCTACCATCTGGCCTTTCACAGAAANNNAGATCGGAAG |
| 70423_1 | TATGAGTGTGGAGTCGTTGCCCATTGCGCCCATTTACATGTAANNNAGATCGGAAG |
| 70424_1 | TATGAGTGTGGAGTCGTTGCTACTGAGAACATGTGCCCTACACNNNAGATCGGAAG |
| 70426_1 | TATGAGTGTGGAGTCGTTGCCTCACTACAGCCTCAAACTCCTGNNNAGATCGGAAG |
| 70427_1 | TATGAGTGTGGAGTCGTTGCTGTGATCTTGGTTCATTGCAACCNNNAGATCGGAAG |
| 70428_1 | TATGAGTGTGGAGTCGTTGCATATGTGTGTGTGTGTGTGTGTGNNNAGATCGGAA |
| 70429_1 | TATGAGTGTGGAGTCGTTGCTCCAATGTGCACTTATCATCTGTNNNAGATCGGAAG |
| 70430_1 | TATGAGTGTGGAGTCGTTGCTACCTTCCTCTTCCTACTCTGGGNNNAGATCGGAAGA |
| 70431_1 | TATGAGTGTGGAGTCGTTGCTCTGTTTTCTGCTTAGGTGTCCTNNNAGATCGGAAG |
| 70432_1 | TATGAGTGTGGAGTCGTTGCTCTCTCTTTCTCTCTCCCTGTCCNNNAGATCGGAAGAG |
| 70433_1 | TATGAGTGTGGAGTCGTTGCGGTGGTGTCTTTGGGAGGTAATTNNNAGATCGGAA |
| 70434_1 | TATGAGTGTGGAGTCGTTGCTGTCTCTCTCTCCATCCATCCATNNNAGATCGGAAGA |
| 70435_1 | TATGAGTGTGGAGTCGTTGCCCTGGTTTTGAGGCTTTTGAACTNNNAGATCGGAAG |
| 70436_1 | TATGAGTGTGGAGTCGTTGCAAGACCCTAACAATAGGCCATGGNNNAGATCGGAA |
| 70437_1 | TATGAGTGTGGAGTCGTTGCCTCCTTCTCAGCTCACTCATCTGNNNAGATCGGAAGA |
| 70438_1 | TATGAGTGTGGAGTCGTTGCTTCCATCACATTGGGGGTTAGAGNNNAGATCGGAA |
| 70439_1 | TATGAGTGTGGAGTCGTTGCAATAAATGTGATCAGGCTGCACGNNNAGATCGGAA |
| 70442_1 | TATGAGTGTGGAGTCGTTGCTTCAGCTGATGAGAGAAGTTTGTNNNAGATCGGAA |
| 70443_1 | TATGAGTGTGGAGTCGTTGCTTGTCTTTGAAGCTTTTGCCACANNNAGATCGGAAG |
| 70444_1 | TATGAGTGTGGAGTCGTTGCGTCTCTCTCTCTCTCTTTCCCCTNNNAGATCGGAAGAG |
| 70445_1 | TATGAGTGTGGAGTCGTTGCGTGTGTGTGTGTGTGTATGTGTGNNNAGATCGGAA |
| 70446_1 | TATGAGTGTGGAGTCGTTGCCTCTTCTCTCCTTTCCCCTGAACNNNAGATCGGAAGA |
| 70447_1 | TATGAGTGTGGAGTCGTTGCCCCCCCCCAAGATCTTTGTTAGATNNNAGATCGGAAG |
| 70448_1 | TATGAGTGTGGAGTCGTTGCATCAGAGTACCTGGAAAACACCCNNNAGATCGGAAG |
| 70449_1 | TATGAGTGTGGAGTCGTTGCACTTCCCAGAGTCTCTCCTGTAANNNAGATCGGAAG |

| | |
|---|---|
| 70451_1 | TATGAGTGTGGAGTCGTTGCAGAGAGAGAGACTACCAGCCAATNNNAGATCGGAA |
| 70452_1 | TATGAGTGTGGAGTCGTTGCGACAGACAGACAGACAGACAGAGNNNAGATCGGAA |
| 70453_1 | TATGAGTGTGGAGTCGTTGCAGCACAGTCATAGTTACCACCTGNNNAGATCGGAAG |
| 70454_1 | TATGAGTGTGGAGTCGTTGCATCTTCCCCAAGCAAGAACATCTNNNAGATCGGAAG |
| 70455_1 | TATGAGTGTGGAGTCGTTGCCAGGAAGAGGGAGAAGAGAGAGANNNAGATCGG |
| 70456_1 | TATGAGTGTGGAGTCGTTGCGGACAGAGAGAGCTTCTTGCATANNNAGATCGGAA |
| 70457_1 | TATGAGTGTGGAGTCGTTGCAGAAAGGAAGGAAGGAAGGAAGANNNAGATCGG |
| 70459_1 | TATGAGTGTGGAGTCGTTGCAGAGACAAAGAGAGACACAGAGANNNAGATCGGA |
| 70460_1 | TATGAGTGTGGAGTCGTTGCGAAGGGAATACAACTGAGCATGCNNNAGATCGGAA |
| 70461_1 | TATGAGTGTGGAGTCGTTGCTTCAAGCAATTCTCATGCCTCAGNNNAGATCGGAAG |
| 70463_1 | TATGAGTGTGGAGTCGTTGCCTCTGTCTCTCTCTCTCTCCCNNNAGATCGGAAGAG |
| 70464_1 | TATGAGTGTGGAGTCGTTGCAATGTCACTCTTGTTGTTCAGGCNNNAGATCGGAAG |
| 70465_1 | TATGAGTGTGGAGTCGTTGCAGAAAGGAAGCCAGGAAGATAGCNNNAGATCGGA |
| 70466_1 | TATGAGTGTGGAGTCGTTGCTTGAGACATCTTAGACGCTCCACNNNAGATCGGAAG |
| 70467_1 | TATGAGTGTGGAGTCGTTGCCACACACACACACACAACTTCNNNAGATCGGAAGA |
| 70468_1 | TATGAGTGTGGAGTCGTTGCAAGGGAAAGAAGGGAAGGAAAGGNNNAGATCGG |
| 70469_1 | TATGAGTGTGGAGTCGTTGCCAATCAATGGCCACTGTGGAAATNNNAGATCGGAA |
| 70470_1 | TATGAGTGTGGAGTCGTTGCAAAAAAAAGGCCACAGAGAGCTCNNNAGATCGGAA |
| 70471_1 | TATGAGTGTGGAGTCGTTGCCACTGGCTGGATTTTGTTCATCANNNAGATCGGAAG |
| 70472_1 | TATGAGTGTGGAGTCGTTGCCTTTTGGAGGGAAATTGCAGAGGNNNAGATCGGAA |
| 70473_1 | TATGAGTGTGGAGTCGTTGCTTCTGTCTTAGGAAAACGGAGCANNNAGATCGGAA |
| 70475_1 | TATGAGTGTGGAGTCGTTGCATGATCTCAGTTCACTGCAACCTNNNAGATCGGAAG |
| 70476_1 | TATGAGTGTGGAGTCGTTGCCTCACAACCAGCTGAGAAGTTTCNNNAGATCGGAAG |
| 70478_1 | TATGAGTGTGGAGTCGTTGCTTCCCCTTCTGCCATGATTGTAANNNAGATCGGAAGA |
| 70480_1 | TATGAGTGTGGAGTCGTTGCCTTACACAGTTAGGGAGGCTGAGNNNAGATCGGAA |
| 70481_1 | TATGAGTGTGGAGTCGTTGCAATTTCTCTGGAGGTGGAGTTGGNNNAGATCGGAA |
| 70483_1 | TATGAGTGTGGAGTCGTTGCTACAAAGTGAAAGAGGGCAACCTNNNAGATCGGAA |
| 70484_1 | TATGAGTGTGGAGTCGTTGCATAATAGGGGGTGTACCTGGTCANNNAGATCGGAA |
| 70486_1 | TATGAGTGTGGAGTCGTTGCGTCTCTGTGGCTGTCTCTATCTGNNNAGATCGGAAG |
| 70487_1 | TATGAGTGTGGAGTCGTTGCTAAGGACCACTGTGATTACACCGNNNAGATCGGAAG |
| 70488_1 | TATGAGTGTGGAGTCGTTGCCAATGGCACAATCTCAGCACATTNNNAGATCGGAAG |
| 70489_1 | TATGAGTGTGGAGTCGTTGCCTCCCATAGCTCAAAAATTGCCCNNNAGATCGGAAG |
| 70490_1 | TATGAGTGTGGAGTCGTTGCCTCATATCCCCTTTCTTGCCCTTNNNAGATCGGAAGA |
| 70491_1 | TATGAGTGTGGAGTCGTTGCTTCTCTGTCTCTCCCTCTCTCTGNNNAGATCGGAAGA |
| 70492_1 | TATGAGTGTGGAGTCGTTGCTTCTCTCCTCTGTCTTTTCCCACNNNAGATCGGAAGA |
| 70493_1 | TATGAGTGTGGAGTCGTTGCAGGGTGATTGTCCACTTACAACANNNAGATCGGAAG |
| 70494_1 | TATGAGTGTGGAGTCGTTGCAGGCTAAAGTACATGCACTGTCANNNAGATCGGAA |
| 70495_1 | TATGAGTGTGGAGTCGTTGCGGGAGAGAGAGAGAGAGAGAGAGNNNAGATCGG |
| 70496_1 | TATGAGTGTGGAGTCGTTGCAATGAGAGAGAGAGAGGAGGCTTNNNAGATCGGA |
| 70497_1 | TATGAGTGTGGAGTCGTTGCGAGAGAGGGAGAGAGGGAGAGAGNNNAGATCGG |
| 70499_1 | TATGAGTGTGGAGTCGTTGCCACACACACAGACACACATACACNNNAGATCGGAAG |
| 70501_1 | TATGAGTGTGGAGTCGTTGCGTACCTCTATGAGTCCGCATGAGNNNAGATCGGAAG |
| 70502_1 | TATGAGTGTGGAGTCGTTGCGGTTGTATGACAAAGACGGCATTNNNAGATCGGAA |

| | |
|---|---|
| 70503_1 | TATGAGTGTGGAGTCGTTGCCTGAGTAGCTGGGACTACAAGTGNNNAGATCGGAA |
| 70504_1 | TATGAGTGTGGAGTCGTTGCTATGGCTCTCATCCCTCTACTCCNNNAGATCGGAAGA |
| 70505_1 | TATGAGTGTGGAGTCGTTGCAGAATCATTTGAGCCCAGGAGTTNNNAGATCGGAA |
| 70506_1 | TATGAGTGTGGAGTCGTTGCGAAGTGGCTAATGCAACGATGAANNNAGATCGGAA |
| 70507_1 | TATGAGTGTGGAGTCGTTGCAGAGAAGAAAGGAGGCCAGATTGNNNAGATCGGA |
| 70508_1 | TATGAGTGTGGAGTCGTTGCTTCCTTTTTTGTTGGGACAGAGTNNNAGATCGGAAG |
| 70510_1 | TATGAGTGTGGAGTCGTTGCTGGAGACCGACAGTAAGTTTCAGNNNAGATCGGAA |
| 70511_1 | TATGAGTGTGGAGTCGTTGCTCACTCTCCCCAACCATTGAATCNNNAGATCGGAAGA |
| 70512_1 | TATGAGTGTGGAGTCGTTGCACACATGCATTTCTGGAGTACCTNNNAGATCGGAAG |
| 70514_1 | TATGAGTGTGGAGTCGTTGCTGAAACCCCGTCTCCACTAAAAANNNAGATCGGAAG |
| 70515_1 | TATGAGTGTGGAGTCGTTGCGGCTCACCAAAATATGTTACCGGNNNAGATCGGAAG |
| 70516_1 | TATGAGTGTGGAGTCGTTGCGGTAGCACAGCCAAGAGTCTAAGNNNAGATCGGAA |
| 70517_1 | TATGAGTGTGGAGTCGTTGCCTCCTCTTTCTCTCTCCCTCTCANNNAGATCGGAAGAG |
| 70518_1 | TATGAGTGTGGAGTCGTTGCAGAGTAGCTGGGACTACAGATGTNNNAGATCGGAA |
| 70519_1 | TATGAGTGTGGAGTCGTTGCTTACTCACTATCCCAAGAGCAGCNNNAGATCGGAAG |
| 70520_1 | TATGAGTGTGGAGTCGTTGCGTCTTGCTGTTTAGTGAAGCTGGNNNAGATCGGAA |
| 70521_1 | TATGAGTGTGGAGTCGTTGCGAGGGAGGAAAGGAAGGAAAGAANNNAGATCGG |
| 70522_1 | TATGAGTGTGGAGTCGTTGCGGTAGCATGGCTCAGTTCAAATCNNNAGATCGGAAG |
| 70523_1 | TATGAGTGTGGAGTCGTTGCAACATGCCCATTTTTCCTGTCTGNNNAGATCGGAAGA |
| 70524_1 | TATGAGTGTGGAGTCGTTGCAATGTGCCAAGACTCGTAATGTGNNNAGATCGGAA |
| 70528_1 | TATGAGTGTGGAGTCGTTGCTCGAAGCTGCAGTGAGTTATGATNNNAGATCGGAA |
| 70529_1 | TATGAGTGTGGAGTCGTTGCTATTGACCTGATGCCACTGGTTTNNNAGATCGGAAG |
| 70530_1 | TATGAGTGTGGAGTCGTTGCAGGGAGGTAGGTAGAGAGAGAGANNNAGATCGG |
| 70531_1 | TATGAGTGTGGAGTCGTTGCGTGTGTGACAGAGTCTCAGTCTGNNNAGATCGGAA |
| 70532_1 | TATGAGTGTGGAGTCGTTGCGGAAGGAAGGGGAGAGAGAGAAANNNAGATCGG |
| 35_1 | TATGAGTGTGGAGTCGTTGCCCGTATCCATTCCAAGCAAATGTNNNAGATCGGAAG |
| 73_1 | TATGAGTGTGGAGTCGTTGCCCAACCAGCCCTTCTTCCCATTNNNAGATCGGAAGAG |
| 121_1 | TATGAGTGTGGAGTCGTTGCAAGAGGAAGCCTTGACCCCCTTNNNAGATCGGAAGA |
| 249_1 | TATGAGTGTGGAGTCGTTGCTCACATACACAGTAGCATGCTGTNNNAGATCGGAAG |
| 259_1 | TATGAGTGTGGAGTCGTTGCTGATTTTTGTCCACAGACCCCANNNAGATCGGAAGA |
| 279_1 | TATGAGTGTGGAGTCGTTGCGGGAAGCAGGATGGAGATTAGAATNNNAGATCGG |
| 344_1 | TATGAGTGTGGAGTCGTTGCTTCAGGAGCTGGTGGGACACATNNNAGATCGGAAG |
| 360_1 | TATGAGTGTGGAGTCGTTGCAGTCTGTGTGTGTGTTCACGTGTNNNAGATCGGAA |
| 372_1 | TATGAGTGTGGAGTCGTTGCCCAAGAGGGGTGACAAGATCTCNNNAGATCGGAAG |
| 433_1 | TATGAGTGTGGAGTCGTTGCAGTTCTCAGATCATGCGTCTGTNNNAGATCGGAAGA |
| 464_1 | TATGAGTGTGGAGTCGTTGCAGAAAAGCCACCTCACCTCCCTNNNAGATCGGAAGA |
| 520_1 | TATGAGTGTGGAGTCGTTGCTCCTTCCTTTTCAAGGCCCCGCNNNAGATCGGAAGAG |
| 584_1 | TATGAGTGTGGAGTCGTTGCAATGGAACAAATAAGGCGCGGTNNNAGATCGGAAG |
| 632_1 | TATGAGTGTGGAGTCGTTGCGAATGAGGCACCCTGGAAGTGGNNNAGATCGGAA |
| 634_1 | TATGAGTGTGGAGTCGTTGCTACCAGCCTCCTACTGTCCTGCNNNAGATCGGAAGAG |
| 691_1 | TATGAGTGTGGAGTCGTTGCCTCTTTTGGCAGAATAAGCCCCANNNAGATCGGAAG |
| 705_1 | TATGAGTGTGGAGTCGTTGCCAAAAGAGTAAGTGCGGCCTGANNNAGATCGGAAG |
| 709_1 | TATGAGTGTGGAGTCGTTGCCTACCCTCCGGGCTCACTTCGANNNAGATCGGAAGA |

| | |
|---|---|
| 719_1 | TATGAGTGTGGAGTCGTTGCGCGTGCGTGTTTGAAATGCATTNNNAGATCGGAAGA |
| 738_1 | TATGAGTGTGGAGTCGTTGCGAGACATCTTGGCGAGGCTCAANNNAGATCGGAAG |
| 744_1 | TATGAGTGTGGAGTCGTTGCACACGAAGAGAAAGTTGCTCATTNNNAGATCGGAA |
| 752_1 | TATGAGTGTGGAGTCGTTGCAGAGAGAGAGAGAATCCCTCCCANNNAGATCGGAA |
| 764_1 | TATGAGTGTGGAGTCGTTGCATACCCACACACACATAGTCCCANNNAGATCGGAAG |
| 778_1 | TATGAGTGTGGAGTCGTTGCCCTACTTGGATTCTCCTGCTCGNNNAGATCGGAAGA |
| 869_1 | TATGAGTGTGGAGTCGTTGCGTGCTCGCAGTAGATTCCAAGTNNNAGATCGGAAG |
| 932_1 | TATGAGTGTGGAGTCGTTGCGGTGGGAATGGGTAGAGAGACTTNNNAGATCGGA |
| 938_1 | TATGAGTGTGGAGTCGTTGCTCTTGCTCAGTTTCCTGGCAGGNNNAGATCGGAAGA |
| 956_1 | TATGAGTGTGGAGTCGTTGCCTTGGGTAAGCCTTGGTTTGTCNNNAGATCGGAAG |
| 1002_1 | TATGAGTGTGGAGTCGTTGCAACCTTTGCCTTTCTACCGGACNNNAGATCGGAAGA |
| 1032_1 | TATGAGTGTGGAGTCGTTGCTGTGTCTCCATCTGTGTTCACCNNNAGATCGGAAGA |
| 1071_1 | TATGAGTGTGGAGTCGTTGCTTGGGGAGCAAGCAAAGTTTCANNNAGATCGGAAG |
| 1095_1 | TATGAGTGTGGAGTCGTTGCCTCTCTCTCCCCCTCTTTTTCCANNNAGATCGGAAGAG |
| 1108_1 | TATGAGTGTGGAGTCGTTGCCTGTAACCTCACACTGGGCTTANNNAGATCGGAAGA |
| 1219_1 | TATGAGTGTGGAGTCGTTGCAGTTATTGGGTGAGCTATGCCCNNNAGATCGGAAG |
| 1224_1 | TATGAGTGTGGAGTCGTTGCTCAGCACCATGGAAAAGGTACCNNNAGATCGGAAG |
| 1252_1 | TATGAGTGTGGAGTCGTTGCACAGGAGTTTGGCCAGAAAAGTTNNNAGATCGGAA |
| 1273_1 | TATGAGTGTGGAGTCGTTGCATTGGATCGTCTGAAGCCTTGGNNNAGATCGGAAG |
| 1289_1 | TATGAGTGTGGAGTCGTTGCTGTGGGAGGTGGGAATACAAACNNNAGATCGGAA |
| 1296_1 | TATGAGTGTGGAGTCGTTGCTAAGAGGAGGAGAGGCCCGAACNNNAGATCGGAA |
| 1324_1 | TATGAGTGTGGAGTCGTTGCACCAGGGCATCATTTTCCACCTNNNAGATCGGAAGA |
| 1325_1 | TATGAGTGTGGAGTCGTTGCCCTGCTGCCTTAGTATGAGCAANNNAGATCGGAAGA |
| 1334_1 | TATGAGTGTGGAGTCGTTGCTTTCTCTCTAATGGCTAGGGGGNNNAGATCGGAAG |
| 1345_1 | TATGAGTGTGGAGTCGTTGCGTTCTGGTGAGATTGCAGGACTNNNAGATCGGAAG |
| 1357_1 | TATGAGTGTGGAGTCGTTGCAGGAAGCAAGGAAGCAATCGGNNNAGATCGGAAG |
| 1400_1 | TATGAGTGTGGAGTCGTTGCCCCACTGCAAGCTTTGTCTCTCNNNAGATCGGAAGA |
| 1402_1 | TATGAGTGTGGAGTCGTTGCTGGTGGGTGAGAGTGGTTAGAATNNNAGATCGGA |
| 1410_1 | TATGAGTGTGGAGTCGTTGCTTCGGTCTGTTTGTGGCATATCNNNAGATCGGAAGA |
| 1429_1 | TATGAGTGTGGAGTCGTTGCATGGCAGTGGGATCTGGATCTGNNNAGATCGGAAG |
| 1440_1 | TATGAGTGTGGAGTCGTTGCCATCCTGTCACCTTGTAGCGTTNNNAGATCGGAAGA |
| 1444_1 | TATGAGTGTGGAGTCGTTGCTGAGCATATGTGTGTGTGTGGCNNNAGATCGGAAG |
| 1495_1 | TATGAGTGTGGAGTCGTTGCGTTGTAAAGCCGGGAGTGTGTTNNNAGATCGGAAG |
| 1510_1 | TATGAGTGTGGAGTCGTTGCACGATGTACTGGCTGGAAGGAANNNAGATCGGAAG |
| 1572_1 | TATGAGTGTGGAGTCGTTGCACCTTTTCCTTCTCCATGAAGCNNNAGATCGGAAGAG |
| 1575_1 | TATGAGTGTGGAGTCGTTGCCTCAAGACGGGCTACAGCATTTNNNAGATCGGAAG |
| 1652_1 | TATGAGTGTGGAGTCGTTGCCTGTGGTGTGAGGTATGTGTGGNNNAGATCGGAA |
| 1670_1 | TATGAGTGTGGAGTCGTTGCACAAGGAGGATTCTGGGGGATTNNNAGATCGGAA |
| 1774_1 | TATGAGTGTGGAGTCGTTGCTCCCCTCATCCTGTGGTAAGTTTNNNAGATCGGAAG |
| 1781_1 | TATGAGTGTGGAGTCGTTGCACTCCTCTTACCAGAGCCAGAGNNNAGATCGGAAGA |
| 1815_1 | TATGAGTGTGGAGTCGTTGCTGTAGGCAGTCTTGAAGGACAGNNNAGATCGGAAG |
| 1817_1 | TATGAGTGTGGAGTCGTTGCCACAGAGAAGGGAGAGCTTGCTNNNAGATCGGAAG |
| 1841_1 | TATGAGTGTGGAGTCGTTGCTGCCTGGCCTCCAATTCTTAGTAANNNAGATCGGAA |

| | |
|---|---|
| 1855_1 | TATGAGTGTGGAGTCGTTGCACCGTGCCTAGCTAATCCATTTNNNAGATCGGAAGA |
| 1869_1 | TATGAGTGTGGAGTCGTTGCTGCCCAGCCTTTCTTCTTCTTTTTNNNAGATCGGAAG |
| 1873_1 | TATGAGTGTGGAGTCGTTGCTCCGTGTGTATTATGGGATGGCNNNAGATCGGAAG |
| 1874_1 | TATGAGTGTGGAGTCGTTGCAGCCAAATGTTTCTTCTGGCATNNNAGATCGGAAGA |
| 1880_1 | TATGAGTGTGGAGTCGTTGCCCATTCTCTCCTCAGCCTCACTNNNAGATCGGAAGAG |
| 1891_1 | TATGAGTGTGGAGTCGTTGCGCTCAAACAGTCCACCTACCCANNNAGATCGGAAGA |
| 1910_1 | TATGAGTGTGGAGTCGTTGCTGAAGGATGACCAGGTGGAAAGNNNAGATCGGAA |
| 1940_1 | TATGAGTGTGGAGTCGTTGCCTGTGTCAGGAATGGGTAGGGANNNAGATCGGAA |
| 1948_1 | TATGAGTGTGGAGTCGTTGCCTCACAAGTACCTAAGTGGCCTNNNAGATCGGAAGA |
| 1952_1 | TATGAGTGTGGAGTCGTTGCACCACACTCAGCTACCTTGTTGNNNAGATCGGAAGA |
| 1962_1 | TATGAGTGTGGAGTCGTTGCTGTAGAAGGGGTCTTTCTGTGTNNNAGATCGGAAG |
| 1967_1 | TATGAGTGTGGAGTCGTTGCCTTGCCACACGTGTCCATTTTTNNNAGATCGGAAGA |
| 1988_1 | TATGAGTGTGGAGTCGTTGCAGATGCTGTAGGCTCACCCTGNNNAGATCGGAAGA |
| 2009_1 | TATGAGTGTGGAGTCGTTGCCCACTCCAAGGCAAAACAGCAGNNNAGATCGGAAG |
| 2019_1 | TATGAGTGTGGAGTCGTTGCTGTTAGGTTCCGTGTGTGTGTGNNNAGATCGGAAG |
| 2086_1 | TATGAGTGTGGAGTCGTTGCGAGGTTAGGTGTAGCCACACTTNNNAGATCGGAAG |
| 2091_1 | TATGAGTGTGGAGTCGTTGCTATGTGCTGACACCAACCAGACNNNAGATCGGAAGA |
| 2117_1 | TATGAGTGTGGAGTCGTTGCGAGAAACAGGTGACACGGATTCANNNAGATCGGAA |
| 2136_1 | TATGAGTGTGGAGTCGTTGCGTGTCTCCCAATGTCATGGTGGNNNAGATCGGAAG |
| 2187_1 | TATGAGTGTGGAGTCGTTGCACGGTTGGTTTCCTCTCTACACNNNAGATCGGAAGA |
| 2210_1 | TATGAGTGTGGAGTCGTTGCGCCTGGTGAAGCATCTGCTGAGNNNAGATCGGAAG |
| 2233_1 | TATGAGTGTGGAGTCGTTGCTCACAGAGAAAGCACCACTTTCANNNAGATCGGAAG |
| 2255_1 | TATGAGTGTGGAGTCGTTGCGCTCCTGGGCTTTTGAACTCCTNNNAGATCGGAAGA |
| 2260_1 | TATGAGTGTGGAGTCGTTGCATGTCTGTGGACCTGAATGTGCNNNAGATCGGAAG |
| 2273_1 | TATGAGTGTGGAGTCGTTGCGGAGAAGCAAGCAAGCAAGAAGNNNAGATCGGAA |
| 2293_1 | TATGAGTGTGGAGTCGTTGCGTTTAGGCTCCAAAGGCAGAGGNNNAGATCGGAAG |
| 2382_1 | TATGAGTGTGGAGTCGTTGCTGAAAATGCCTACCACCTCCCTNNNAGATCGGAAGA |
| 2415_1 | TATGAGTGTGGAGTCGTTGCAACCTTTCCCCCAGTGGTTTCNNNAGATCGGAAGAG |
| 2422_1 | TATGAGTGTGGAGTCGTTGCGCCCAGCCCAGTTTATCCTTTTNNNAGATCGGAAGA |
| 2430_1 | TATGAGTGTGGAGTCGTTGCCTGTAGGAGCACAGCTTCAACCNNNAGATCGGAAGA |
| 2435_1 | TATGAGTGTGGAGTCGTTGCAACCTCCTATGCCTCAGCAATTNNNAGATCGGAAGA |
| 2450_1 | TATGAGTGTGGAGTCGTTGCATCTTGGCTCACTGCTCATTTTNNNAGATCGGAAGA |
| 2517_1 | TATGAGTGTGGAGTCGTTGCTCTGACCACAAAAGGCTGAAGGNNNAGATCGGAAG |
| 2601_1 | TATGAGTGTGGAGTCGTTGCAGGCGAAAAGGGAGAGAGAAAGNNNAGATCGGAA |
| 2605_1 | TATGAGTGTGGAGTCGTTGCGTGTACTGCAAACTGGCTGGTTNNNAGATCGGAAG |
| 2608_1 | TATGAGTGTGGAGTCGTTGCATTCTGGAGCTAAGGAGGGCAANNNAGATCGGAAG |
| 2633_1 | TATGAGTGTGGAGTCGTTGCGAAGAGGCCTGGAATGGTCACANNNAGATCGGAAG |
| 2638_1 | TATGAGTGTGGAGTCGTTGCCTCTTGGAGACAGAGTGTCACTNNNAGATCGGAAG |
| 2650_1 | TATGAGTGTGGAGTCGTTGCGAGAATGATGCCATACACAGCTNNNAGATCGGAAG |
| 2715_1 | TATGAGTGTGGAGTCGTTGCATGCAATTGTTGAGGCTTGCAANNNAGATCGGAAG |
| 2732_1 | TATGAGTGTGGAGTCGTTGCTGGTGTGCAAGTGTTGTTGGAGNNNAGATCGGAAG |
| 2753_1 | TATGAGTGTGGAGTCGTTGCATTCCTCCCTCCCCTTTTTGTCNNNAGATCGGAAGAG |
| 2759_1 | TATGAGTGTGGAGTCGTTGCTTAGGCTCTGGACTGAAACCCANNNAGATCGGAAGA |

| | |
|---|---|
| 2763_1 | TATGAGTGTGGAGTCGTTGCTGGGTATGTAGGTGGCTTAGGCNNNAGATCGGAAG |
| 2771_1 | TATGAGTGTGGAGTCGTTGCCAGAAGCAAGTGCTGCCATTTGNNNAGATCGGAAG |
| 2777_1 | TATGAGTGTGGAGTCGTTGCATGTGTGTGTATCCTTTCAGCANNNAGATCGGAAGA |
| 2838_1 | TATGAGTGTGGAGTCGTTGCGTGTGTTCTTTGGCCTTGTTTTNNNAGATCGGAAGA |
| 2850_1 | TATGAGTGTGGAGTCGTTGCCCATGATTTCTCAGAGGGAGTGNNNAGATCGGAAG |
| 2901_1 | TATGAGTGTGGAGTCGTTGCGGGCAGGATCACCCAGAGAAAGNNNAGATCGGAAG |
| 2948_1 | TATGAGTGTGGAGTCGTTGCTAATTGGAATGGAGCCCAATGCNNNAGATCGGAAG |
| 2969_1 | TATGAGTGTGGAGTCGTTGCATAGGGTTGGGGAAAGATGGGANNNAGATCGGAA |
| 3035_1 | TATGAGTGTGGAGTCGTTGCGTGGGAGTAGGGGTATGAAATCNNNAGATCGGAA |
| 3132_1 | TATGAGTGTGGAGTCGTTGCATGTGGTCAAAACTCCCAAAAGNNNAGATCGGAAG |
| 3148_1 | TATGAGTGTGGAGTCGTTGCGTGACTGTGAATTGTGCTGCTTNNNAGATCGGAAG |
| 3206_1 | TATGAGTGTGGAGTCGTTGCATAGAGACCTCAGGGGGCAACTNNNAGATCGGAAG |
| 3239_1 | TATGAGTGTGGAGTCGTTGCACATCTGGAAAGTGACCAGGGANNNAGATCGGAAG |
| 3240_1 | TATGAGTGTGGAGTCGTTGCAGAGAGACTACAGCTGTTTGGGANNNAGATCGGAA |
| 3244_1 | TATGAGTGTGGAGTCGTTGCTCTACATATCAAGCCTGTGCCCNNNAGATCGGAAGA |
| 3253_1 | TATGAGTGTGGAGTCGTTGCAGGCGGAATTCGATAAGTGCAGNNNAGATCGGAAG |
| 3258_1 | TATGAGTGTGGAGTCGTTGCTGCCAACTCTATCTCAGCACTGNNNAGATCGGAAGA |
| 3263_1 | TATGAGTGTGGAGTCGTTGCTGACTTAGAGGTAGCCCCCTCNNNAGATCGGAAGAG |
| 3277_1 | TATGAGTGTGGAGTCGTTGCAAAATCTGGAAGTCACCCCAAGNNNAGATCGGAAG |
| 3280_1 | TATGAGTGTGGAGTCGTTGCGCAGAGGGAAAGGAAGGTTGCANNNAGATCGGAA |
| 3282_1 | TATGAGTGTGGAGTCGTTGCAGGGATGTTGTTGAGGAGGAGGNNNAGATCGGAA |
| 3301_1 | TATGAGTGTGGAGTCGTTGCCCGAGGGCCAACCATACTATGTNNNAGATCGGAAGA |
| 3316_1 | TATGAGTGTGGAGTCGTTGCCTGCTCCTGCACATGTTTTGAANNNAGATCGGAAGA |
| 3323_1 | TATGAGTGTGGAGTCGTTGCTGTGTCTAAGTAAGGGGCTTGGANNNAGATCGGAA |
| 3337_1 | TATGAGTGTGGAGTCGTTGCACCAAGTAAAGGCCATGCCAGTNNNAGATCGGAAG |
| 3356_1 | TATGAGTGTGGAGTCGTTGCCATCCTGCCAAATTCTCCCGAGNNNAGATCGGAAGA |
| 3370_1 | TATGAGTGTGGAGTCGTTGCACATGACAATTCTAAGCCCCTTTNNNAGATCGGAAG |
| 3385_1 | TATGAGTGTGGAGTCGTTGCCTTTTCATCTCTGGTGTGCCACNNNAGATCGGAAGA |
| 3423_1 | TATGAGTGTGGAGTCGTTGCTAGCGGTTGCTTCCTGGAGAAANNNAGATCGGAAG |
| 3432_1 | TATGAGTGTGGAGTCGTTGCACACACAAAAAGCAACCCTGCCNNNAGATCGGAAGA |
| 3446_1 | TATGAGTGTGGAGTCGTTGCGGGGGTAATTTTTGGTATCCTGANNNAGATCGGAA |
| 3454_1 | TATGAGTGTGGAGTCGTTGCTCACCGTAAACGCCCCAAGATCNNNAGATCGGAAGA |
| 3463_1 | TATGAGTGTGGAGTCGTTGCTGGTCCTGTTGGAAGAGGAGTTANNNAGATCGGAA |
| 3468_1 | TATGAGTGTGGAGTCGTTGCAGTCCACAGTTCCCAGGGTTTTNNNAGATCGGAAGA |
| 3474_1 | TATGAGTGTGGAGTCGTTGCGTGCGCCTCCATCGATGCATATNNNAGATCGGAAGA |
| 3482_1 | TATGAGTGTGGAGTCGTTGCTTTGTGTGGATTGGGGCTGTTANNNAGATCGGAAG |
| 3485_1 | TATGAGTGTGGAGTCGTTGCTTGGTTTGCATTTCTCCCCTTCNNNAGATCGGAAGAG |
| 3486_1 | TATGAGTGTGGAGTCGTTGCCTCCCAGAACACGTCAGTAACANNNAGATCGGAAGA |
| 3495_1 | TATGAGTGTGGAGTCGTTGCCTTTGGCCCTGGATCAGTCTGANNNAGATCGGAAGA |
| 3500_1 | TATGAGTGTGGAGTCGTTGCTGACTATGGAGATGTGGGAGGGNNNAGATCGGAA |
| 3525_1 | TATGAGTGTGGAGTCGTTGCGTAGAGACGGGGGAACTCACCANNNAGATCGGAAG |
| 3530_1 | TATGAGTGTGGAGTCGTTGCAGCAGAGCTCCCAGTCACATCCNNNAGATCGGAAGA |
| 3547_1 | TATGAGTGTGGAGTCGTTGCACAGAGGAAGTGACCAGAGCAANNNAGATCGGAAG |

| | |
|---|---|
| 3586_1 | TATGAGTGTGGAGTCGTTGCCCAGCACCCCAAAAAGATCCTGNNNAGATCGGAAGA |
| 3609_1 | TATGAGTGTGGAGTCGTTGCGTGTGTGTGTCTTGAAAGCTCANNNAGATCGGAAG |
| 3614_1 | TATGAGTGTGGAGTCGTTGCAGACGAGGGTTCCATGACAGAGNNNAGATCGGAAG |
| 3627_1 | TATGAGTGTGGAGTCGTTGCAGAGAGAGGAAGATGGTGTGGGNNNAGATCGGAA |
| 3631_1 | TATGAGTGTGGAGTCGTTGCGCTTGTTGTAGACTGGAGGTGCNNNAGATCGGAAG |
| 3641_1 | TATGAGTGTGGAGTCGTTGCTCAAGAGGACACATGGTCAGGGNNNAGATCGGAAG |
| 3659_1 | TATGAGTGTGGAGTCGTTGCAGCGGGGAGACTTAGCTAAAAANNNAGATCGGAAG |
| 3672_1 | TATGAGTGTGGAGTCGTTGCGAATAGGCTGTGGTGGGCAAATANNNAGATCGGAA |
| 3720_1 | TATGAGTGTGGAGTCGTTGCGGTGGAGCAGAAACTGAATTTGNNNAGATCGGAAG |
| 3730_1 | TATGAGTGTGGAGTCGTTGCACTCTGTACTGCCTGCTTGATTNNNAGATCGGAAGA |
| 3741_1 | TATGAGTGTGGAGTCGTTGCGGGGGCAGGGAGCAATTGATTANNNAGATCGGAA |
| 3745_1 | TATGAGTGTGGAGTCGTTGCGGCATTTGTCGGCTTCATGTTCNNNAGATCGGAAGA |
| 3759_1 | TATGAGTGTGGAGTCGTTGCGGGCAGATGCTGGGAAATGGAANNNAGATCGGAA |
| 3807_1 | TATGAGTGTGGAGTCGTTGCGTGGCTGGCAAACAAGTGTTGGNNNAGATCGGAAG |
| 3808_1 | TATGAGTGTGGAGTCGTTGCAGGGGTTGATGTCATTCTCCCCNNNAGATCGGAAGA |
| 3849_1 | TATGAGTGTGGAGTCGTTGCCTTTGCCTTTGTCCTAGGGAGCNNNAGATCGGAAGA |
| 3903_1 | TATGAGTGTGGAGTCGTTGCGCCACTGTTTAGTAGACTGCTTNNNAGATCGGAAGA |
| 3913_1 | TATGAGTGTGGAGTCGTTGCAGTGGTTACTTTCCCAGGGTGANNNAGATCGGAAG |
| 3965_1 | TATGAGTGTGGAGTCGTTGCTTTGGCAGGTCACTGGTTTGAANNNAGATCGGAAG |
| 4011_1 | TATGAGTGTGGAGTCGTTGCCTGGCACTTCAGATCAAGGACANNNAGATCGGAAG |
| 4048_1 | TATGAGTGTGGAGTCGTTGCTTGTATGTCTCCGAGGGTGTGTANNNAGATCGGAA |
| 4071_1 | TATGAGTGTGGAGTCGTTGCAGAGGCTTGATCATTACAGGGTNNNAGATCGGAAG |
| 4079_1 | TATGAGTGTGGAGTCGTTGCGGGTTCATGCCTGTAATCTTTTNNNAGATCGGAAGA |
| 4090_1 | TATGAGTGTGGAGTCGTTGCATCCTACTGAGCCCATCTGCAANNNAGATCGGAAGA |
| 4098_1 | TATGAGTGTGGAGTCGTTGCCTTGCCTACTTCATCTTTGCTGNNNAGATCGGAAGAG |
| 4129_1 | TATGAGTGTGGAGTCGTTGCAAACACAAGGGGGCTTCCTTTTNNNAGATCGGAAGA |
| 4138_1 | TATGAGTGTGGAGTCGTTGCATAAATGTGCAGCCCACTCAGGNNNAGATCGGAAG |
| 4158_1 | TATGAGTGTGGAGTCGTTGCGAGAGACTGAGAGTGTCAGGCANNNAGATCGGAA |
| 4160_1 | TATGAGTGTGGAGTCGTTGCCCCTGCAGAACCTGAGTATACCNNNAGATCGGAAGA |
| 4173_1 | TATGAGTGTGGAGTCGTTGCAAGGCCCCAAAGAGACCACTAANNNAGATCGGAAG |
| 4181_1 | TATGAGTGTGGAGTCGTTGCACAGGTGATCCTCCTCAGCCACNNNAGATCGGAAGA |
| 4191_1 | TATGAGTGTGGAGTCGTTGCTGTTGGTGTGTGTTGGGTATGTNNNAGATCGGAAG |
| 4200_1 | TATGAGTGTGGAGTCGTTGCTAATGAGTTGGGGAGGAGTGTTNNNAGATCGGAA |
| 4251_1 | TATGAGTGTGGAGTCGTTGCCACACGCACCCACACAGTTATGNNNAGATCGGAAGA |
| 4259_1 | TATGAGTGTGGAGTCGTTGCGCATAGAACTGGGCTTCCCACACNNNAGATCGGAAG |
| 4285_1 | TATGAGTGTGGAGTCGTTGCGTGTGTGTGTGTGATTCATAGGANNNAGATCGGAA |
| 4293_1 | TATGAGTGTGGAGTCGTTGCATTTACTCACCGATGCCCAGAANNNAGATCGGAAGA |
| 4296_1 | TATGAGTGTGGAGTCGTTGCTTTATGGCATTTACAGCGACCNNNAGATCGGAAGAG |
| 4298_1 | TATGAGTGTGGAGTCGTTGCTGCCTTCTTTGAGCTTGATGATNNNAGATCGGAAGA |
| 4348_1 | TATGAGTGTGGAGTCGTTGCTACCAAGGGCAGAGACTGGATNNNAGATCGGAAGA |
| 4356_1 | TATGAGTGTGGAGTCGTTGCGAAAGGCATCCTCCCCTTGTGTCANNNAGATCGGAA |
| 4365_1 | TATGAGTGTGGAGTCGTTGCGGGGGAAGCTAGGAATGAACTGNNNAGATCGGAA |
| 4368_1 | TATGAGTGTGGAGTCGTTGCGGGATTGTGGGGACCTATGTGTNNNAGATCGGAAG |

| | |
|---|---|
| 4370_1 | TATGAGTGTGGAGTCGTTGCGTATAAGGGAGCCCAGGGACACNNNAGATCGGAAG |
| 4415_1 | TATGAGTGTGGAGTCGTTGCTCTATCCCTCCCCTGACAAGACNNNAGATCGGAAGAG |
| 4423_1 | TATGAGTGTGGAGTCGTTGCGGGTAGGGGCTGTAAGTAAACTNNNAGATCGGAAG |
| 4424_1 | TATGAGTGTGGAGTCGTTGCGCTTATCCAGGGTCCCTTTTGGNNNAGATCGGAAGA |
| 4431_1 | TATGAGTGTGGAGTCGTTGCATCCGTCTGCCTCAGGTTACCANNNAGATCGGAAGA |
| 4443_1 | TATGAGTGTGGAGTCGTTGCAAAGCTGGGAGAGAGGAGACTGNNNAGATCGGAA |
| 4464_1 | TATGAGTGTGGAGTCGTTGCATGTCACCAGTTTCTCCCCCTTNNNAGATCGGAAGAG |
| 4466_1 | TATGAGTGTGGAGTCGTTGCGGATAAGTTTTACGTGGCTGCACNNNAGATCGGAA |
| 4469_1 | TATGAGTGTGGAGTCGTTGCGATTGCATGCTTGTAGGTGCACNNNAGATCGGAAG |
| 4484_1 | TATGAGTGTGGAGTCGTTGCTCCATCCACTCGTTCATCCAATNNNAGATCGGAAGAG |
| 4501_1 | TATGAGTGTGGAGTCGTTGCCGTTGCCTCAGAGAAGATTCCCNNNAGATCGGAAGA |
| 4510_1 | TATGAGTGTGGAGTCGTTGCCTGGGAGACAAAGCGGAAATCANNNAGATCGGAAG |
| 4515_1 | TATGAGTGTGGAGTCGTTGCCGTGAGATTTCTAGGATGGTTCCCNNNAGATCGGAA |
| 4537_1 | TATGAGTGTGGAGTCGTTGCATAGCAGCCCTTTGCAAACAAANNNAGATCGGAAGA |
| 4541_1 | TATGAGTGTGGAGTCGTTGCATAAAGTCCATACCAGGCCAGGNNNAGATCGGAAG |
| 4542_1 | TATGAGTGTGGAGTCGTTGCTACAGACATGGCCGGCTAATTTNNNAGATCGGAAGA |
| 4550_1 | TATGAGTGTGGAGTCGTTGCTGTGCATGTGTGCTTGTAGTGANNNAGATCGGAAG |
| 4552_1 | TATGAGTGTGGAGTCGTTGCAGAGTGGTGAGGAGAAGACAGACNNNAGATCGGA |
| 4584_1 | TATGAGTGTGGAGTCGTTGCGGTGTGTTACAGTGTCAGTGGGNNNAGATCGGAAG |
| 4620_1 | TATGAGTGTGGAGTCGTTGCACAAGAGGCTTCTGCCTATCCCNNNAGATCGGAAGA |
| 4639_1 | TATGAGTGTGGAGTCGTTGCGGATTCTGAGTGTGTGCTTTGCNNNAGATCGGAAG |
| 4643_1 | TATGAGTGTGGAGTCGTTGCCCCCCGCCAATAGGTACATTTTNNNAGATCGGAAGA |
| 4654_1 | TATGAGTGTGGAGTCGTTGCCTAAGCATCAGGAGAGGCAGTANNNAGATCGGAAG |
| 4685_1 | TATGAGTGTGGAGTCGTTGCACAGTCTGCTGGAGAAGTCCTTNNNAGATCGGAAG |
| 4686_1 | TATGAGTGTGGAGTCGTTGCCACACACCCCCAAACACTTGTGNNNAGATCGGAAGA |
| 4689_1 | TATGAGTGTGGAGTCGTTGCCACCTGCTGTTTTCTCTCCCTACNNNAGATCGGAAGA |
| 4750_1 | TATGAGTGTGGAGTCGTTGCGATTGTCTAGTGGGGAGGACTGNNNAGATCGGAAG |
| 4759_1 | TATGAGTGTGGAGTCGTTGCGTCTAATGCCAAAGGGCAATATTNNNAGATCGGAAG |
| 4783_1 | TATGAGTGTGGAGTCGTTGCCTGGCAATGGAGGTCGATAACCNNNAGATCGGAAG |
| 4792_1 | TATGAGTGTGGAGTCGTTGCTGGGGCAGGTGGTATTTCAAAANNNAGATCGGAAG |
| 4800_1 | TATGAGTGTGGAGTCGTTGCGTTGTTGTTGTTGTCCACCACCNNNAGATCGGAAGA |
| 4804_1 | TATGAGTGTGGAGTCGTTGCCAGATAGCACTGGAACCAGGACNNNAGATCGGAAG |
| 4822_1 | TATGAGTGTGGAGTCGTTGCAAAGGTCTCAAAAGGCAGCACCNNNAGATCGGAAG |
| 4834_1 | TATGAGTGTGGAGTCGTTGCAAAGACAGCACTTTTCCTGCCCNNNAGATCGGAAGA |
| 4873_1 | TATGAGTGTGGAGTCGTTGCATTATGCCTGGGTTCCATCCCTNNNAGATCGGAAGA |
| 4878_1 | TATGAGTGTGGAGTCGTTGCACTGGAAGGCATGGGGTAATGANNNAGATCGGAA |
| 4918_1 | TATGAGTGTGGAGTCGTTGCAGTTCTTCCAAGCATCAACCCCNNNAGATCGGAAGA |
| 4938_1 | TATGAGTGTGGAGTCGTTGCCTGTCACACCCATGGTCTACCCNNNAGATCGGAAGA |
| 4944_1 | TATGAGTGTGGAGTCGTTGCCAAAATCCAGAGGGGCAGTCCCNNNAGATCGGAAG |
| 4988_1 | TATGAGTGTGGAGTCGTTGCATTCCCTCTTCCTCAAAGGACGNNNAGATCGGAAGA |
| 4994_1 | TATGAGTGTGGAGTCGTTGCTGAAAAGGTATTTGTGGCCGGGNNNAGATCGGAAG |
| 5005_1 | TATGAGTGTGGAGTCGTTGCTCTGTCTGTTGTATGTGTGCGANNNAGATCGGAAGA |
| 5012_1 | TATGAGTGTGGAGTCGTTGCTACTACGATTTCCTAGGTGGCCCNNNAGATCGGAAG |

| | |
|---|---|
| 5019_1 | TATGAGTGTGGAGTCGTTGCCCTTCACTAGAACACACTGCCCNNNAGATCGGAAGA |
| 5051_1 | TATGAGTGTGGAGTCGTTGCGAAACAATCGGCTTCACTGCAGNNNAGATCGGAAG |
| 5058_1 | TATGAGTGTGGAGTCGTTGCATTATGGAGGTTGAGGGCTCCCNNNAGATCGGAAG |
| 5064_1 | TATGAGTGTGGAGTCGTTGCCCTTGAATCCTTTGCTCTCTCCCNNNAGATCGGAAGA |
| 5065_1 | TATGAGTGTGGAGTCGTTGCACAGGGTCTACTCAGGTTTCCCNNNAGATCGGAAGA |
| 5067_1 | TATGAGTGTGGAGTCGTTGCACTCTCCCTTTTTCCCCTCTTCNNNAGATCGGAAGAG |
| 5082_1 | TATGAGTGTGGAGTCGTTGCATAACACCCCATGCAACTTCCCNNNAGATCGGAAGA |
| 5099_1 | TATGAGTGTGGAGTCGTTGCCACACACTCACATCCACCTTCCNNNAGATCGGAAGAG |
| 5101_1 | TATGAGTGTGGAGTCGTTGCGGACCATCTCTAGCCCATGTGANNNAGATCGGAAGA |
| 5105_1 | TATGAGTGTGGAGTCGTTGCTTGGTGCCCTTTGTAACCCTTCNNNAGATCGGAAGA |
| 5114_1 | TATGAGTGTGGAGTCGTTGCAGAAGTCACAGTGCTCCCATCTNNNAGATCGGAAGA |
| 5121_1 | TATGAGTGTGGAGTCGTTGCGAGAGAAGGCTGATTTCTGCTGNNNAGATCGGAAG |
| 5145_1 | TATGAGTGTGGAGTCGTTGCAGGAAGCATGGAAGCATAGCAGNNNAGATCGGAA |
| 5148_1 | TATGAGTGTGGAGTCGTTGCCGGAAGGGAAGTAGGTTGTCATNNNAGATCGGAA |
| 5152_1 | TATGAGTGTGGAGTCGTTGCGAAGGCACTGCTGGCACAATATNNNAGATCGGAAG |
| 5168_1 | TATGAGTGTGGAGTCGTTGCAAAGTCATCTGTTGTTGGGCACNNNAGATCGGAAG |
| 5177_1 | TATGAGTGTGGAGTCGTTGCCGCTCAGAGGCTTTCACACAAANNNAGATCGGAAGA |
| 5179_1 | TATGAGTGTGGAGTCGTTGCGAGAACAGAGTAGGCCTCACCCNNNAGATCGGAAG |
| 5190_1 | TATGAGTGTGGAGTCGTTGCGTGGGGTGGAGATGGACAGCAAANNNAGATCGGA |
| 5203_1 | TATGAGTGTGGAGTCGTTGCCTTGCGTCAAATCGAAAGCTGTNNNAGATCGGAAG |
| 5227_1 | TATGAGTGTGGAGTCGTTGCTCCCCTCAGCAAAGATGCCAATNNNAGATCGGAAGA |
| 5256_1 | TATGAGTGTGGAGTCGTTGCACAGACCTATGGGGCCTGGAAGNNNAGATCGGAAG |
| 5295_1 | TATGAGTGTGGAGTCGTTGCAAGGTCAGATGTCTGGGAGATCCNNNAGATCGGAA |
| 5316_1 | TATGAGTGTGGAGTCGTTGCTGTCGGAGAGCCAAGGAATGATNNNAGATCGGAAG |
| 5319_1 | TATGAGTGTGGAGTCGTTGCGCTCAGTCAATAGCACACCTCCNNNAGATCGGAAGA |
| 5363_1 | TATGAGTGTGGAGTCGTTGCAGCACCCCAGAGATATACAGTCCNNNAGATCGGAAG |
| 5411_1 | TATGAGTGTGGAGTCGTTGCTACACAGCAAACATTCTAGCCGNNNAGATCGGAAGA |
| 5430_1 | TATGAGTGTGGAGTCGTTGCAGATGGTTGCTGACGGATTTCCNNNAGATCGGAAG |
| 5434_1 | TATGAGTGTGGAGTCGTTGCTGCCAGATACTTGCTTCCTTTCCNNNAGATCGGAAGA |
| 5498_1 | TATGAGTGTGGAGTCGTTGCCGTGATGTCCAGATTCTCACAATNNNAGATCGGAAG |
| 5560_1 | TATGAGTGTGGAGTCGTTGCCTTTCTCACCAACCTCCTCAGCNNNAGATCGGAAGAG |
| 5653_1 | TATGAGTGTGGAGTCGTTGCAAGAAATCCTGTGACCTGCTGTNNNAGATCGGAAG |
| 5691_1 | TATGAGTGTGGAGTCGTTGCGGTCAGGTCTTTTGAGCCCTTGNNNAGATCGGAAG |
| 5697_1 | TATGAGTGTGGAGTCGTTGCGGACAACAAACATGCACCAACGNNNAGATCGGAAG |
| 5701_1 | TATGAGTGTGGAGTCGTTGCAAAATAAAGGTCTGGGCCAGGCNNNAGATCGGAAG |
| 5816_1 | TATGAGTGTGGAGTCGTTGCTCCTCCTCTCTCAGTATGCTGANNNAGATCGGAAGA |
| 5995_1 | TATGAGTGTGGAGTCGTTGCGCTTCTCATGGACGAAATTGTGCNNNAGATCGGAAG |
| 6078_1 | TATGAGTGTGGAGTCGTTGCCCTTTGCTTTTCCGCTGCAATCNNNAGATCGGAAGAG |
| 6095_1 | TATGAGTGTGGAGTCGTTGCGCCTGTAAGGAGACCCATACTTNNNAGATCGGAAG |
| 6133_1 | TATGAGTGTGGAGTCGTTGCTGATGAGAAGGGTTTGAGGCTCTNNNAGATCGGAA |
| 6145_1 | TATGAGTGTGGAGTCGTTGCCTTTGCAGACATTCCCCCAGATCNNNAGATCGGAAG |
| 6202_1 | TATGAGTGTGGAGTCGTTGCCTTAGGAGGGTGAGGAATGGCCNNNAGATCGGAA |
| 6203_1 | TATGAGTGTGGAGTCGTTGCAAGCAGGCCCAACGTAAACATANNNAGATCGGAAG |

| | |
|---|---|
| 6219_1 | TATGAGTGTGGAGTCGTTGCAGAGCAGCTGAGAAAGGCTAAANNNAGATCGGAAG |
| 6232_1 | TATGAGTGTGGAGTCGTTGCCACGATCTTCTGCTTCAGCCTCNNNAGATCGGAAGAG |
| 6239_1 | TATGAGTGTGGAGTCGTTGCCCCCACAGCCATGTTGGTTGNNNAGATCGGAAGAGC |
| 6258_1 | TATGAGTGTGGAGTCGTTGCGTCAAGAGGGAGACACAAGCTCNNNAGATCGGAAG |
| 6269_1 | TATGAGTGTGGAGTCGTTGCCTATTTCCTGGGCTGTGAGCTCNNNAGATCGGAAGA |
| 6299_1 | TATGAGTGTGGAGTCGTTGCGTGTAGTGGCAGGATCTTGCTCNNNAGATCGGAAG |
| 6307_1 | TATGAGTGTGGAGTCGTTGCCTTCCCGCTTCCCGTCTATCTCNNNAGATCGGAAGAG |
| 6341_1 | TATGAGTGTGGAGTCGTTGCTCTCAGGTCTTGGGGGTTTCTCNNNAGATCGGAAGA |
| 6469_1 | TATGAGTGTGGAGTCGTTGCCTTCTGGGGTCTTCGTAGCTTCNNNAGATCGGAAGA |
| 6478_1 | TATGAGTGTGGAGTCGTTGCATGCTCACAGGAAAGCCTCTTCNNNAGATCGGAAGA |
| 6505_1 | TATGAGTGTGGAGTCGTTGCACATGCGCACACAATCATGTTCNNNAGATCGGAAGA |
| 6546_1 | TATGAGTGTGGAGTCGTTGCGTTGCACATGGATGGACTGTTTCNNNAGATCGGAAG |
| 6557_1 | TATGAGTGTGGAGTCGTTGCTCTTCCCACTCCACATCTCCAANNNAGATCGGAAGAG |
| 6612_1 | TATGAGTGTGGAGTCGTTGCGTATCCGAAAGGGCCAGACAAGNNNAGATCGGAAG |
| 6614_1 | TATGAGTGTGGAGTCGTTGCATCACAGTTTGACCAAGGCGAANNNAGATCGGAAG |
| 6624_1 | TATGAGTGTGGAGTCGTTGCCTTACAGCCTCCTCCCTCCAAGNNNAGATCGGAAGAG |
| 6704_1 | TATGAGTGTGGAGTCGTTGCGGTGACAAGTTAGAAGCGCTTGNNNAGATCGGAAG |
| 6711_1 | TATGAGTGTGGAGTCGTTGCCCTGAAAAGCTGCCCCATGAAGNNNAGATCGGAAG |
| 6719_1 | TATGAGTGTGGAGTCGTTGCTGAGGAGGAGACGGGTACGTACNNNAGATCGGAA |
| 6723_1 | TATGAGTGTGGAGTCGTTGCGAGGATCACTGGGGAGGTGAAGNNNAGATCGGAA |
| 6730_1 | TATGAGTGTGGAGTCGTTGCTCTGTCTCTGGCCAGTTCTCATNNNAGATCGGAAGA |
| 6759_1 | TATGAGTGTGGAGTCGTTGCCCTTGCCTTTCTGTCTTGACCANNNAGATCGGAAGAG |
| 6775_1 | TATGAGTGTGGAGTCGTTGCAGGTCATCCACACTGTCCACACNNNAGATCGGAAGA |
| 6778_1 | TATGAGTGTGGAGTCGTTGCTCTCTCCCCTTTCTCTCCAGTCNNNAGATCGGAAGAG |
| 6810_1 | TATGAGTGTGGAGTCGTTGCAGGCTTCTTCTGTTTCTCAGGCNNNAGATCGGAAGA |
| 6838_1 | TATGAGTGTGGAGTCGTTGCTCCTGCTGTCACTTCTGCATGTNNNAGATCGGAAGA |
| 6839_1 | TATGAGTGTGGAGTCGTTGCAAATGAGGCCTCTGCCTCACAANNNAGATCGGAAGA |
| 6845_1 | TATGAGTGTGGAGTCGTTGCGCTCTGTTCCAAATCCTACCCAGNNNAGATCGGAAG |
| 6852_1 | TATGAGTGTGGAGTCGTTGCTCTCTCTCTTGAAAGGGGTGCCNNNAGATCGGAAGA |
| 6871_1 | TATGAGTGTGGAGTCGTTGCCTCACTCTGTTGTGCAGGCCTGNNNAGATCGGAAGA |
| 6897_1 | TATGAGTGTGGAGTCGTTGCAAACCTCCCTGAGTACTCCCCNNNAGATCGGAAGAG |
| 6908_1 | TATGAGTGTGGAGTCGTTGCATCTGCTGCACTGTTATCGCAGNNNAGATCGGAAGA |
| 6919_1 | TATGAGTGTGGAGTCGTTGCCAGAGCCTAAAGTGAGTGGCAGNNNAGATCGGAAG |
| 6937_1 | TATGAGTGTGGAGTCGTTGCGGAAAACGACACAGTGGATGAANNNAGATCGGAAG |
| 6956_1 | TATGAGTGTGGAGTCGTTGCGTTGCAGGGAAAAGACTGGGGTNNNAGATCGGAA |
| 6968_1 | TATGAGTGTGGAGTCGTTGCAAGTGGATCTTGAGCCTGGTGCNNNAGATCGGAAG |
| 6992_1 | TATGAGTGTGGAGTCGTTGCCTCCCTCCCTCCCATTTGTAGANNNAGATCGGAAGAG |
| 6993_1 | TATGAGTGTGGAGTCGTTGCTTCTCACTAGGCTTTTCCCCCANNNAGATCGGAAGAG |
| 7011_1 | TATGAGTGTGGAGTCGTTGCCTCTCACATCCCTTGCTCTGCTNNNAGATCGGAAGAG |
| 7043_1 | TATGAGTGTGGAGTCGTTGCGAGATGGCTTCTTCTACAGGCCNNNAGATCGGAAGA |
| 7066_1 | TATGAGTGTGGAGTCGTTGCGTCCTTCTCATCTTAGCACCTGNNNAGATCGGAAGAG |
| 7075_1 | TATGAGTGTGGAGTCGTTGCGGATCAAGGGGCCTCTCTAGAGNNNAGATCGGAAG |
| 7078_1 | TATGAGTGTGGAGTCGTTGCCTGCTTCTGGTGTGGAGAGTTANNNAGATCGGAAG |

| | |
|---|---|
| 7087_1 | TATGAGTGTGGAGTCGTTGCTCTCGGTGGGAAAACCAAATGGNNNAGATCGGAAG |
| 7091_1 | TATGAGTGTGGAGTCGTTGCGGCAATGCTCACAGCTAAGGAGNNNAGATCGGAAG |
| 7101_1 | TATGAGTGTGGAGTCGTTGCTGCTTGATTCTACTTCTGCCCANNNAGATCGGAAGA |
| 7104_1 | TATGAGTGTGGAGTCGTTGCCTGCAAGTGGCCAGATCTCTGTNNNAGATCGGAAG |
| 7111_1 | TATGAGTGTGGAGTCGTTGCCAATGAAGAGATGAGCTGCGCCNNNAGATCGGAAG |
| 7116_1 | TATGAGTGTGGAGTCGTTGCTCTAGGATATAGGGCAGTGGCTNNNAGATCGGAAG |
| 7141_1 | TATGAGTGTGGAGTCGTTGCTCACTCAGCTGCCTCTTTAGACNNNAGATCGGAAGA |
| 7157_1 | TATGAGTGTGGAGTCGTTGCGAAGACAGGCCAAGGAACATGAGNNNAGATCGGA |
| 7158_1 | TATGAGTGTGGAGTCGTTGCACCAGCATTTGCCTTTCTTTCANNNAGATCGGAAGAG |
| 7170_1 | TATGAGTGTGGAGTCGTTGCGGCAGTGGTACAAACACTGTTCNNNAGATCGGAAG |
| 7183_1 | TATGAGTGTGGAGTCGTTGCATAGGTAACCCCCAGTGCTGAGNNNAGATCGGAAG |
| 7188_1 | TATGAGTGTGGAGTCGTTGCCAACAGCATCAAGGCTGAACATNNNAGATCGGAAG |
| 7189_1 | TATGAGTGTGGAGTCGTTGCCTGTCTGCCCTTTTCTGTCTGGNNNAGATCGGAAGA |
| 7216_1 | TATGAGTGTGGAGTCGTTGCAGAAAACACACGGTTTCGTTGAGNNNAGATCGGAA |
| 7219_1 | TATGAGTGTGGAGTCGTTGCCCAGCATGAGCACAGAGAAAAGNNNAGATCGGAAG |
| 7222_1 | TATGAGTGTGGAGTCGTTGCTCTGCTGGAAAAGCTTGATTCCNNNAGATCGGAAGA |
| 7232_1 | TATGAGTGTGGAGTCGTTGCATAAAGCCCCACTTTTCTGGGGNNNAGATCGGAAGA |
| 7262_1 | TATGAGTGTGGAGTCGTTGCTTCCGCCATCTGATCTTGGACTNNNAGATCGGAAGA |
| 7272_1 | TATGAGTGTGGAGTCGTTGCATTAGCTTAGGGCTCCACCCTTANNNAGATCGGAAG |
| 7281_1 | TATGAGTGTGGAGTCGTTGCAGTCAGGCCCTATGTAACAATGNNNAGATCGGAAG |
| 7298_1 | TATGAGTGTGGAGTCGTTGCACCCTAGCAATCTTCCCACCCCNNNAGATCGGAAGAG |
| 7312_1 | TATGAGTGTGGAGTCGTTGCAGTTCAGTGGTGCAATCCTCCCNNNAGATCGGAAGA |
| 7319_1 | TATGAGTGTGGAGTCGTTGCTCTTTCCTTGTCTCCCCTCTCTNNNAGATCGGAAGAG |
| 7357_1 | TATGAGTGTGGAGTCGTTGCAGCACCTAAACCCCTTTTGCTCNNNAGATCGGAAGA |
| 7359_1 | TATGAGTGTGGAGTCGTTGCACAATGACACACTGATCTGCACGNNNAGATCGGAAG |
| 7366_1 | TATGAGTGTGGAGTCGTTGCGTCTGGAATGCAGTGTTGTGGCNNNAGATCGGAAG |
| 7369_1 | TATGAGTGTGGAGTCGTTGCGGTGGCTGATGTAGGAAGATTTNNNAGATCGGAAG |
| 7424_1 | TATGAGTGTGGAGTCGTTGCATTACTCATGGAAGCCATGGCGNNNAGATCGGAAG |
| 7433_1 | TATGAGTGTGGAGTCGTTGCACTGTGGCATGACATGAGATCGNNNAGATCGGAAG |
| 7575_1 | TATGAGTGTGGAGTCGTTGCCTGGTGCAGAGTTAGGGAGAGGNNNAGATCGGAA |
| 7607_1 | TATGAGTGTGGAGTCGTTGCACATCTTTCTCCTCCCTGGGACNNNAGATCGGAAGA |
| 7676_1 | TATGAGTGTGGAGTCGTTGCGCCTCAAATTATTCTGGCTTTCGGNNNAGATCGGAA |
| 7683_1 | TATGAGTGTGGAGTCGTTGCTTTCAAGGAGGGTAGGGAAGGGNNNAGATCGGAA |
| 7687_1 | TATGAGTGTGGAGTCGTTGCCATTCACAACTTCTTTGGTGGGCNNNAGATCGGAAG |
| 7694_1 | TATGAGTGTGGAGTCGTTGCCATTCAGCATTCTAGCCCGGGANNNAGATCGGAAGA |
| 7734_1 | TATGAGTGTGGAGTCGTTGCATAATTGCTTACCTGCCACGGGNNNAGATCGGAAGA |
| 7735_1 | TATGAGTGTGGAGTCGTTGCTTTTTCTTTCCGTCGAGACGGGNNNAGATCGGAAGA |
| 7754_1 | TATGAGTGTGGAGTCGTTGCTAGCACTTGGAAATGAGCCTCANNNAGATCGGAAG |
| 7859_1 | TATGAGTGTGGAGTCGTTGCACAGCAAGACAGCTAGATTTGGGNNNAGATCGGAA |
| 7872_1 | TATGAGTGTGGAGTCGTTGCTCCTTCCTACGCAGATACAATGGNNNAGATCGGAAG |
| 7903_1 | TATGAGTGTGGAGTCGTTGCAGAGGCTTGGGGTGTATAGATGGNNNAGATCGGA |
| 7963_1 | TATGAGTGTGGAGTCGTTGCTTTTTCTGATGCTCTCCCCTGGNNNAGATCGGAAGA |
| 8061_1 | TATGAGTGTGGAGTCGTTGCACAGTGAGGGAGTAGTATGGTGGNNNAGATCGGA |

| | |
|---|---|
| 8092_1 | TATGAGTGTGGAGTCGTTGCCCCCCAAAACATCTGCTATTGGNNNAGATCGGAAGA |
| 8169_1 | TATGAGTGTGGAGTCGTTGCCGCTGCATGAATACACTCGAATGNNNAGATCGGAAG |
| 8198_1 | TATGAGTGTGGAGTCGTTGCGGCACCAAACAGAATTCACCATGNNNAGATCGGAAG |
| 8200_1 | TATGAGTGTGGAGTCGTTGCGCAGGAAGAGACAAGACCCATGNNNAGATCGGAAG |
| 8316_1 | TATGAGTGTGGAGTCGTTGCGGATAGAAGCTGCCCAGAAACTGNNNAGATCGGAA |
| 8371_1 | TATGAGTGTGGAGTCGTTGCCATTGTCAGGGGTGTGTACCTGNNNAGATCGGAAG |
| 8382_1 | TATGAGTGTGGAGTCGTTGCTGTACCTGGCAAGCTGTAGACNNNAGATCGGAAGA |
| 8404_1 | TATGAGTGTGGAGTCGTTGCCCACATCAAGGCTGGGGTTGATNNNAGATCGGAAG |
| 8425_1 | TATGAGTGTGGAGTCGTTGCGGGATTACCTGGTCCAGCCTTCNNNAGATCGGAAGA |
| 8429_1 | TATGAGTGTGGAGTCGTTGCCCGTCCCTAGTCAAATTTGCAANNNAGATCGGAAGA |
| 8454_1 | TATGAGTGTGGAGTCGTTGCGTAGAGGGAGGAGGATGTGCTGNNNAGATCGGAA |
| 8467_1 | TATGAGTGTGGAGTCGTTGCTACAGTTGCAGGATGAGAATCTGNNNAGATCGGAA |
| 8474_1 | TATGAGTGTGGAGTCGTTGCACTTTGTGCCAGGTACTGATCTGNNNAGATCGGAAG |
| 8488_1 | TATGAGTGTGGAGTCGTTGCCGGTGACCACTCAACTTCTCTGNNNAGATCGGAAGA |
| 8521_1 | TATGAGTGTGGAGTCGTTGCCTTCTCCCACGTGTGGATTCTGNNNAGATCGGAAGA |
| 8546_1 | TATGAGTGTGGAGTCGTTGCTTTGATGCTGTCCCTTTGTCTCNNNAGATCGGAAGA |
| 8549_1 | TATGAGTGTGGAGTCGTTGCGTCCGTCAGTCCTATGCTTTCTGNNNAGATCGGAAG |
| 8581_1 | TATGAGTGTGGAGTCGTTGCGGATGCACAGTCAGAGTGAGTGNNNAGATCGGAA |
| 8585_1 | TATGAGTGTGGAGTCGTTGCGTGAAAGTGGTGGCTGTGAGTGNNNAGATCGGAA |
| 8633_1 | TATGAGTGTGGAGTCGTTGCCTTCTTTTAGGCCAGCCTGGTGNNNAGATCGGAAGA |
| 8682_1 | TATGAGTGTGGAGTCGTTGCTATTCACATGCTGGCAGTTGTGNNNAGATCGGAAG |
| 8685_1 | TATGAGTGTGGAGTCGTTGCATGTGTGCCCCTTCATGTTGTGNNNAGATCGGAAGA |
| 8717_1 | TATGAGTGTGGAGTCGTTGCAGTGTCTCTCCCACCTGGATTGNNNAGATCGGAAGA |
| 8844_1 | TATGAGTGTGGAGTCGTTGCTCTGGCTGTGCTTTTGTGTTTTGNNNAGATCGGAAG |
| 8912_1 | TATGAGTGTGGAGTCGTTGCGCCAGAAGCATACATGTAGGAGTNNNAGATCGGAA |
| 8918_1 | TATGAGTGTGGAGTCGTTGCTCTGTTCTGTGTCTCCTCCCAATNNNAGATCGGAAGA |
| 8973_1 | TATGAGTGTGGAGTCGTTGCCTATTCCCTCGGCATCCAGGAATNNNAGATCGGAAG |
| 9026_1 | TATGAGTGTGGAGTCGTTGCGCTTTGTGGCTATTGCCAACATNNNAGATCGGAAGA |
| 9043_1 | TATGAGTGTGGAGTCGTTGCAGAGGTATTCCATGTCAGCACATNNNAGATCGGAAG |
| 9152_1 | TATGAGTGTGGAGTCGTTGCCGTCTGCACCATAGACATGCATNNNAGATCGGAAGA |
| 9177_1 | TATGAGTGTGGAGTCGTTGCCAGTGGCACAAAGCACAATCATNNNAGATCGGAAG |
| 9558_1 | TATGAGTGTGGAGTCGTTGCCTCAGACCCCAAACTCTCCCAGNNNAGATCGGAAGA |
| 9591_1 | TATGAGTGTGGAGTCGTTGCGCCCACCCAGAGAATGTTCTCGNNNAGATCGGAAGA |
| 9634_1 | TATGAGTGTGGAGTCGTTGCATGTAGCTTTGTTTTGCCTGACTNNNAGATCGGAAG |
| 9643_1 | TATGAGTGTGGAGTCGTTGCAGTCATCTTGAATTGGGGCCCCNNNAGATCGGAAG |
| 9748_1 | TATGAGTGTGGAGTCGTTGCTACAAGGGACCAGGCTTTTTCCNNNAGATCGGAAGA |
| 9868_1 | TATGAGTGTGGAGTCGTTGCAGAGAGATACAATGTGCATGCCTNNNAGATCGGAA |
| 9884_1 | TATGAGTGTGGAGTCGTTGCACATTTCCAGGTCATAGGGCATNNNAGATCGGAAG |
| 9904_1 | TATGAGTGTGGAGTCGTTGCAACCTGAGGGAACAGAGACAAGANNNAGATCGGA |
| 9951_1 | TATGAGTGTGGAGTCGTTGCTTGAGAGGGACACCTTCATTCCTNNNAGATCGGAAG |
| 9971_1 | TATGAGTGTGGAGTCGTTGCCCAGGATACTTTCCCACTGAGCNNNAGATCGGAAGA |
| 10596_1 | TATGAGTGTGGAGTCGTTGCTACATTGCGGTAAACTCAGGGCNNNAGATCGGAAG |
| 10870_1 | TATGAGTGTGGAGTCGTTGCCGGAAGACACTGCTACTAGCCTNNNAGATCGGAAG |

| | |
|---|---|
| 10902_1 | TATGAGTGTGGAGTCGTTGCGCTATCCCAGCCTCTTCTCTGAANNNAGATCGGAAGA |
| 10939_1 | TATGAGTGTGGAGTCGTTGCTGGGATCTAAACTCTATGCCTGCNNNAGATCGGAAG |
| 10986_1 | TATGAGTGTGGAGTCGTTGCCTCTCACTGGGGAAGGTGACTAGNNNAGATCGGAA |
| 11008_1 | TATGAGTGTGGAGTCGTTGCATCAGGGTCAGTGTGCAGACATNNNAGATCGGAAGA |
| 11063_1 | TATGAGTGTGGAGTCGTTGCCGTGCCTGGCCTGTGATACATANNNAGATCGGAAGA |
| 11064_1 | TATGAGTGTGGAGTCGTTGCTCCAGCCATTTTTGCACCCATTNNNAGATCGGAAGAG |
| 11108_1 | TATGAGTGTGGAGTCGTTGCATACTCACACTCACACCCCCTCNNNAGATCGGAAGAG |
| 11137_1 | TATGAGTGTGGAGTCGTTGCTACCTCGGTCTTCTCTTTCAGGNNNAGATCGGAAGA |
| 11142_1 | TATGAGTGTGGAGTCGTTGCAGAGACAGAAAGAGCAAGAGCTNNNAGATCGGAA |
| 11169_1 | TATGAGTGTGGAGTCGTTGCGACTTTCCAAGGAGCTACTGCTNNNAGATCGGAAGA |
| 11218_1 | TATGAGTGTGGAGTCGTTGCGCCCCGCCCCAATAAGTCTGTCNNNAGATCGGAAGA |
| 11256_1 | TATGAGTGTGGAGTCGTTGCTTACGGCTTACACAGTCCCCTTGNNNAGATCGGAAGA |
| 11315_1 | TATGAGTGTGGAGTCGTTGCTGCCTGTATTTGCTGGAGAACTNNNAGATCGGAAGA |
| 11325_1 | TATGAGTGTGGAGTCGTTGCACCCTGCTCCTCTTTTCTCAGGNNNAGATCGGAAGAG |
| 11395_1 | TATGAGTGTGGAGTCGTTGCCACAAAACTCAGCCCAGCCAGGNNNAGATCGGAAGA |
| 11398_1 | TATGAGTGTGGAGTCGTTGCCACCCATTGTGACCTGGAGCTANNNAGATCGGAAGA |
| 11410_1 | TATGAGTGTGGAGTCGTTGCCTGTCTCTGTGGATCGGAGGACNNNAGATCGGAAGA |
| 11432_1 | TATGAGTGTGGAGTCGTTGCTCAGAGGGACATGGGGTGATTGNNNAGATCGGAA |
| 11440_1 | TATGAGTGTGGAGTCGTTGCTGTCTGCACTCCTGATTCCAATGNNNAGATCGGAAGA |
| 11461_1 | TATGAGTGTGGAGTCGTTGCCTTCCAATTTAGTTGGGCCAGCCNNNAGATCGGAAGA |
| 11482_1 | TATGAGTGTGGAGTCGTTGCCTTCCACATCCCCAGCATCCATNNNAGATCGGAAGAG |
| 11565_1 | TATGAGTGTGGAGTCGTTGCCTTGACACAGCTGCCTGGGAATNNNAGATCGGAAGA |
| 11582_1 | TATGAGTGTGGAGTCGTTGCGGCTCCACTGCTTCTCTTTCTCNNNAGATCGGAAGAG |
| 11598_1 | TATGAGTGTGGAGTCGTTGCTTCTTCTTGGAAACCTGCATGCANNNAGATCGGAAGA |
| 11636_1 | TATGAGTGTGGAGTCGTTGCCCCCATTCATTCTTTCAAGCTGNNNAGATCGGAAGAG |
| 11637_1 | TATGAGTGTGGAGTCGTTGCCCCACCGGTCTTTCATAAAGGANNNAGATCGGAAGA |
| 11701_1 | TATGAGTGTGGAGTCGTTGCCCCCTCGCCCAATCTCTACAAANNNAGATCGGAAGAG |
| 11750_1 | TATGAGTGTGGAGTCGTTGCTGTCTGGGATGGTGGAAGTCTNNNAGATCGGAAGA |
| 11789_1 | TATGAGTGTGGAGTCGTTGCCTGGTTAGTTTCTGCTTGTGCGNNNAGATCGGAAGA |
| 11793_1 | TATGAGTGTGGAGTCGTTGCTGACATTCCGAGTTCCTGGAAGNNNAGATCGGAAGA |
| 11821_1 | TATGAGTGTGGAGTCGTTGCCACACCCTACACTTGATTGGCTNNNAGATCGGAAGA |
| 11854_1 | TATGAGTGTGGAGTCGTTGCCTTAGTTCCTCCCTCCACCCCCNNNAGATCGGAAGAG |
| 11901_1 | TATGAGTGTGGAGTCGTTGCCCTCCTATTGGCTTCTTGTGGTNNNAGATCGGAAGA |
| 11908_1 | TATGAGTGTGGAGTCGTTGCGGGGGAAATCTAGGGGTTGAGAANNNAGATCGGA |
| 11933_1 | TATGAGTGTGGAGTCGTTGCAAGAGACTTCACTGGGCACAGTNNNAGATCGGAAGA |
| 11962_1 | TATGAGTGTGGAGTCGTTGCCTCCCTTGACCTTCCTCCAGCTNNNAGATCGGAAGAG |
| 11985_1 | TATGAGTGTGGAGTCGTTGCAGTTGTTCTAGGTCTGGGTTCGTNNNAGATCGGAA |
| 12036_1 | TATGAGTGTGGAGTCGTTGCGATTGGAGAAGGCTTCACGGGTNNNAGATCGGAAG |
| 12058_1 | TATGAGTGTGGAGTCGTTGCGACCTTATTCCTTGGGGTCTGANNNAGATCGGAAGA |
| 12061_1 | TATGAGTGTGGAGTCGTTGCTTGCTGCCATTACTGTGGCTCANNNAGATCGGAAGA |
| 12081_1 | TATGAGTGTGGAGTCGTTGCGTCTAGTGATGCTTGGCTACCCNNNAGATCGGAAGA |
| 12111_1 | TATGAGTGTGGAGTCGTTGCATTTCCAATTGCTGAAGCTGGAGANNNAGATCGGAA |
| 12118_1 | TATGAGTGTGGAGTCGTTGCCACACACACTCACAGCCATCTCNNNAGATCGGAAGA |

| | |
|---|---|
| 12121_1 | TATGAGTGTGGAGTCGTTGCAGCAGTTCCGAGTAGTTAGTGGTNNNAGATCGGAA |
| 12182_1 | TATGAGTGTGGAGTCGTTGCTTAGCGCCCTCTCTTATGTGGGNNNAGATCGGAAGA |
| 12183_1 | TATGAGTGTGGAGTCGTTGCGCAATAAGTTAGGCCAGTGGTTNNNAGATCGGAAG |
| 12203_1 | TATGAGTGTGGAGTCGTTGCTTTCCAGCATGTGGGTGTATGTNNNAGATCGGAAG |
| 12262_1 | TATGAGTGTGGAGTCGTTGCAAAACCCAAAAACCGTTCGGCANNNAGATCGGAAG |
| 12318_1 | TATGAGTGTGGAGTCGTTGCCCCAGTTCACTGAAAGGATCCTNNNAGATCGGAAGA |
| 12342_1 | TATGAGTGTGGAGTCGTTGCTGTACTCTTGCACAACTTTGCTNNNAGATCGGAAGA |
| 12343_1 | TATGAGTGTGGAGTCGTTGCGCTCATACCTGCATTTGGTGGTNNNAGATCGGAAGA |
| 12344_1 | TATGAGTGTGGAGTCGTTGCGCTAAGGCTCTCAGAACTCCCGNNNAGATCGGAAGA |
| 12389_1 | TATGAGTGTGGAGTCGTTGCCCACCCTTTTCTCAACCTTTGCNNNAGATCGGAAGAG |
| 53438_1 | TATGAGTGTGGAGTCGTTGCCCAAGAGCTCTGCTAGGGAAAGNNNAGATCGGAAG |
| 53448_1 | TATGAGTGTGGAGTCGTTGCCACCACTACACACCTCTCACTCNNNAGATCGGAAGAG |
| 53477_1 | TATGAGTGTGGAGTCGTTGCGAAACAGAGGGATACGGGGATACNNNAGATCGGA |
| 53480_1 | TATGAGTGTGGAGTCGTTGCCTGTTCCTGTCTTTGCTCACACNNNAGATCGGAAGAG |
| 53481_1 | TATGAGTGTGGAGTCGTTGCTTGCACAAGTAGCTCCAAGTTTGNNNAGATCGGAAG |
| 53485_1 | TATGAGTGTGGAGTCGTTGCGCTTCTGTAGTCCCAGCTACTCNNNAGATCGGAAGA |
| 53496_1 | TATGAGTGTGGAGTCGTTGCCTCCCTCCTCCTCTCTCTTCTGNNNAGATCGGAAGAG |
| 53497_1 | TATGAGTGTGGAGTCGTTGCCTGGAGTACAATGGTGCGATCTNNNAGATCGGAAG |
| 53499_1 | TATGAGTGTGGAGTCGTTGCCATTCTGCCTTGAATCTGGTGCNNNAGATCGGAAGA |
| 53521_1 | TATGAGTGTGGAGTCGTTGCCTCTTTCTGAGGATGGGAAGGAANNNAGATCGGAA |
| 53541_1 | TATGAGTGTGGAGTCGTTGCAGGGAACAGAAATTGCTGAGGATNNNAGATCGGAA |
| 53553_1 | TATGAGTGTGGAGTCGTTGCGACCTCAGTGATCACACCTCAGNNNAGATCGGAAGA |
| 53560_1 | TATGAGTGTGGAGTCGTTGCAGACGAGGTCTCATCACTTTGCNNNAGATCGGAAGA |
| 53562_1 | TATGAGTGTGGAGTCGTTGCTCACATTTAGCCCTGGTAAGCATNNNAGATCGGAAG |
| 53565_1 | TATGAGTGTGGAGTCGTTGCCAGCTAGGAGGAGTTGTCCAACNNNAGATCGGAAG |
| 53567_1 | TATGAGTGTGGAGTCGTTGCCAATGTCGTGATCTCGGCTCANNNAGATCGGAAGAG |
| 53577_1 | TATGAGTGTGGAGTCGTTGCTGGAGTACAGTGGCACAATCTCNNNAGATCGGAAG |
| 53587_1 | TATGAGTGTGGAGTCGTTGCCTCCCAGGTTCATGCCATTCTCNNNAGATCGGAAGAG |
| 53595_1 | TATGAGTGTGGAGTCGTTGCCTGATCTCAAACTCCCGACCTCNNNAGATCGGAAGAG |
| 53598_1 | TATGAGTGTGGAGTCGTTGCCAGTAGCTGGGACTACAAGAGCNNNAGATCGGAAG |
| 53599_1 | TATGAGTGTGGAGTCGTTGCGTCTCCTGAGTAGCTGGGACTANNNAGATCGGAAG |
| 53605_1 | TATGAGTGTGGAGTCGTTGCGAAGGGGAGGGTTATATGGATGGNNNAGATCGGA |
| 53614_1 | TATGAGTGTGGAGTCGTTGCAGCAAAGAGTTAGAATCTAGGCANNNAGATCGGAA |
| 53622_1 | TATGAGTGTGGAGTCGTTGCATGGGAATTTTGGTTGAGGGAGANNNAGATCGGA |
| 53625_1 | TATGAGTGTGGAGTCGTTGCTGTGAGGACTTGGCGTTTCTATANNNAGATCGGAAG |
| 53628_1 | TATGAGTGTGGAGTCGTTGCGATGATCTTGGCTCACTGCAACNNNAGATCGGAAGA |
| 53645_1 | TATGAGTGTGGAGTCGTTGCCAAGTAGCTGGGACTACAGACGNNNAGATCGGAAG |
| 53646_1 | TATGAGTGTGGAGTCGTTGCCTCCTGGGTTCATGCCATTCTCNNNAGATCGGAAGA |
| 53650_1 | TATGAGTGTGGAGTCGTTGCGCTGATTCAAGCGATTCTCCTGNNNAGATCGGAAGA |
| 53652_1 | TATGAGTGTGGAGTCGTTGCCTCCCAGGTTCAAGCTGTTCTCNNNAGATCGGAAGA |
| 53654_1 | TATGAGTGTGGAGTCGTTGCAACCTTCACCTCCTGCATTCAGNNNAGATCGGAAGAG |
| 53655_1 | TATGAGTGTGGAGTCGTTGCGTAGTGGTGTGATCACAGCTCAANNNAGATCGGAA |
| 53657_1 | TATGAGTGTGGAGTCGTTGCATACACAACCAGAGGGAATGCATNNNAGATCGGAA |

53658_1 TATGAGTGTGGAGTCGTTGCGCCTTTACTGCCAGTTACCAATCNNNAGATCGGAAGA
53665_1 TATGAGTGTGGAGTCGTTGCCCAGGGAAATGCAAATCAAAACCNNNAGATCGGAAG
53671_1 TATGAGTGTGGAGTCGTTGCACTCACTGTGTCAACCTTGTTTANNNAGATCGGAAGA
53680_1 TATGAGTGTGGAGTCGTTGCAGTCATGAATGCCAAAACCAGTGNNNAGATCGGAAG
53682_1 TATGAGTGTGGAGTCGTTGCTTTCAACTTTGTTTGGACACCCCNNNAGATCGGAAGA
53702_1 TATGAGTGTGGAGTCGTTGCTAGAGAGCAGTGGCGTGATCTCNNNAGATCGGAAG
53703_1 TATGAGTGTGGAGTCGTTGCCTCCCAGGTTCATGCCATTCTCNNNAGATCGGAAGAG
53715_1 TATGAGTGTGGAGTCGTTGCCTCTGGGGTTCATGCCATTCTCNNNAGATCGGAAGA
53727_1 TATGAGTGTGGAGTCGTTGCCCTGCCTCTGAGATGTGGTATTTNNNAGATCGGAAG
53730_1 TATGAGTGTGGAGTCGTTGCAGAGGTTTCAGTGAGCCAAGATCNNNAGATCGGAA
53735_1 TATGAGTGTGGAGTCGTTGCCGTCTGTAGTCCCAGCTACTTGNNNAGATCGGAAGA
53742_1 TATGAGTGTGGAGTCGTTGCGTTCAAGCAATTCTCCTGCCTCNNNAGATCGGAAGAG
53752_1 TATGAGTGTGGAGTCGTTGCGCACGATCTCAGCTCATTGTAACNNNAGATCGGAAG
53769_1 TATGAGTGTGGAGTCGTTGCTTAGCTGGAAGTGGTGCTATGGNNNAGATCGGAAG
53771_1 TATGAGTGTGGAGTCGTTGCCAACAAACTAGGCATTGAAGGCANNNAGATCGGAA
53773_1 TATGAGTGTGGAGTCGTTGCGGCTGAGGAAGGATAAGCACTTNNNAGATCGGAAG
53776_1 TATGAGTGTGGAGTCGTTGCCCCAGGCTGAAAATGAAACTCAANNNAGATCGGAAG
53779_1 TATGAGTGTGGAGTCGTTGCAGAAAAAAAAGAAAGTGGCCCGGNNNAGATCGGA
53780_1 TATGAGTGTGGAGTCGTTGCAGCTGTAGATATTTTGTGTCCTGGNNNAGATCGGAA
53782_1 TATGAGTGTGGAGTCGTTGCGGGGCAAGAGACAGAGAGAATAGNNNAGATCGGA
53788_1 TATGAGTGTGGAGTCGTTGCCACCTTGCTAAGTCCCCATAGATNNNAGATCGGAAG
53791_1 TATGAGTGTGGAGTCGTTGCCATGCCTGTAATCCCAGCAATTTNNNAGATCGGAAG
53798_1 TATGAGTGTGGAGTCGTTGCCTCCTGGGTTCATGCGACTATCNNNAGATCGGAAGA
53812_1 TATGAGTGTGGAGTCGTTGCGATCTCTGCTCACCACAAGCTCNNNAGATCGGAAGA
53828_1 TATGAGTGTGGAGTCGTTGCCTCCCAGGTTCAAGCGATTCTCNNNAGATCGGAAGA
53863_1 TATGAGTGTGGAGTCGTTGCTGAGCAAATCAAAGTGAGGCCTANNNAGATCGGAA
53865_1 TATGAGTGTGGAGTCGTTGCTCTGTAGCACTCTGGAGAATGTGNNNAGATCGGAA
53877_1 TATGAGTGTGGAGTCGTTGCTATACACACACACACACACACACNNNAGATCGGAAGA
53879_1 TATGAGTGTGGAGTCGTTGCCAGCCTGAGCAACATGATGTAACNNNAGATCGGAAG
53881_1 TATGAGTGTGGAGTCGTTGCCTTGAGCCCTCAGGAGTTTGAGNNNAGATCGGAAGA
53890_1 TATGAGTGTGGAGTCGTTGCGATCTTGGCTCATTGCAACCTCNNNAGATCGGAAGA
53903_1 TATGAGTGTGGAGTCGTTGCCATCGAGGCTCACTGTAACCTCNNNAGATCGGAAGA
53906_1 TATGAGTGTGGAGTCGTTGCGAGATGGTGTCTTGCTCTGTTGNNNAGATCGGAAGA
53912_1 TATGAGTGTGGAGTCGTTGCGAATGCAGTGGTATGATCTTGGCNNNAGATCGGAA
53913_1 TATGAGTGTGGAGTCGTTGCCTCCTGGGTTCAAGCGATTCTCNNNAGATCGGAAGA
53917_1 TATGAGTGTGGAGTCGTTGCGATCTCAGCTCACTGCAACCTCNNNAGATCGGAAGA
53932_1 TATGAGTGTGGAGTCGTTGCCCTGTGGAGTAACCTTTTCTCGTNNNAGATCGGAAG
53935_1 TATGAGTGTGGAGTCGTTGCTCTTATGTTGTGGGCATGGCTTANNNAGATCGGAAG
53936_1 TATGAGTGTGGAGTCGTTGCCAGGTAGTGGCAGAGTGAAGTCNNNAGATCGGAAG
53942_1 TATGAGTGTGGAGTCGTTGCGGGTTAGGCGAATGGCATAAAATNNNAGATCGGAA
53951_1 TATGAGTGTGGAGTCGTTGCTGTAGCCTCAAAGCACAAATGTGNNNAGATCGGAAG
53963_1 TATGAGTGTGGAGTCGTTGCAGAATTACCTCTGAATGGCTCCANNNAGATCGGAAG
53996_1 TATGAGTGTGGAGTCGTTGCTACGGAGTCCTTCTCAGACCATANNNAGATCGGAAG

| | |
|---|---|
| 53997_1 | TATGAGTGTGGAGTCGTTGCCTGATTCCAGTTGTAGCCTTCTTNNNAGATCGGAAGA |
| 54005_1 | TATGAGTGTGGAGTCGTTGCATCCCAGGTTCAAGCGATTCTCNNNAGATCGGAAGA |
| 54018_1 | TATGAGTGTGGAGTCGTTGCGAGATAATCTCGCTCTGTCGCCNNNAGATCGGAAGA |
| 54028_1 | TATGAGTGTGGAGTCGTTGCGAATAGCTGGGACTACGGGTGNNNAGATCGGAAGA |
| 54044_1 | TATGAGTGTGGAGTCGTTGCGAGTTTCGCTCTTGTTGTAGGCNNNAGATCGGAAGA |
| 54047_1 | TATGAGTGTGGAGTCGTTGCGTTTCATCGTGTTAGCCAGGATGNNNAGATCGGAAG |
| 54048_1 | TATGAGTGTGGAGTCGTTGCCTGCCTCAGCCTCCTGAATATCNNNAGATCGGAAGA |
| 54049_1 | TATGAGTGTGGAGTCGTTGCCTCCAGGGTTCACACCATTCTCNNNAGATCGGAAGA |
| 54050_1 | TATGAGTGTGGAGTCGTTGCCACCATGCCCAGCTAATTTTGTNNNAGATCGGAAGA |
| 54074_1 | TATGAGTGTGGAGTCGTTGCAGCCAAGCTACGTGTATCAATCANNNAGATCGGAAG |
| 54075_1 | TATGAGTGTGGAGTCGTTGCATTTCAGCCACACAGTCTACCTCNNNAGATCGGAAGA |
| 54095_1 | TATGAGTGTGGAGTCGTTGCAAGAGATCAAAGCAAGCACACTGNNNAGATCGGAA |
| 54098_1 | TATGAGTGTGGAGTCGTTGCCCACCCCCTAATGGTTTTCAAACNNNAGATCGGAAGA |
| 54108_1 | TATGAGTGTGGAGTCGTTGCCATGCCTGTAATCCCAGCTACCNNNAGATCGGAAGA |
| 54116_1 | TATGAGTGTGGAGTCGTTGCATAAAGAATGATGGCTCGTGGGTNNNAGATCGGAA |
| 54119_1 | TATGAGTGTGGAGTCGTTGCTGAGATTTGCCAAACACCTTTCCNNNAGATCGGAAG |
| 54123_1 | TATGAGTGTGGAGTCGTTGCAAGGAATGAAGATGCTGGGTTCTNNNAGATCGGAA |
| 54128_1 | TATGAGTGTGGAGTCGTTGCAGATGGAGTCTCACTCTGTCACCNNNAGATCGGAAG |
| 54133_1 | TATGAGTGTGGAGTCGTTGCCCGGATTCAAATGATTCTCCTGCNNNAGATCGGAAG |
| 54134_1 | TATGAGTGTGGAGTCGTTGCATGATCTTGGCTCACTGGAACCNNNAGATCGGAAGA |
| 54144_1 | TATGAGTGTGGAGTCGTTGCGATCTAGGCTCACTGCAAGCTCNNNAGATCGGAAGA |
| 54147_1 | TATGAGTGTGGAGTCGTTGCGAGTTTTGCTCTTCTTCCGCAGNNNAGATCGGAAGA |
| 54150_1 | TATGAGTGTGGAGTCGTTGCGGTCTTGCTCTGTCTTTCCATTCNNNAGATCGGAAGA |
| 54155_1 | TATGAGTGTGGAGTCGTTGCGATCTCGGCTCACTACAACCTCNNNAGATCGGAAGA |
| 54161_1 | TATGAGTGTGGAGTCGTTGCCATCTCCTGGGCTCAAGTGATCNNNAGATCGGAAGA |
| 54163_1 | TATGAGTGTGGAGTCGTTGCATAAGCAAATTGCAACGGGAAGTNNNAGATCGGAA |
| 54167_1 | TATGAGTGTGGAGTCGTTGCACACGGTGAAACCTCATCTCTACNNNAGATCGGAAG |
| 54168_1 | TATGAGTGTGGAGTCGTTGCTCTCACTGTCCTTTGCTATTTGCNNNAGATCGGAAGA |
| 54174_1 | TATGAGTGTGGAGTCGTTGCAGCACTTTGGGAGGCTGTAATCNNNAGATCGGAAG |
| 54179_1 | TATGAGTGTGGAGTCGTTGCTCACTAAGTCCTTGGGATTTGCANNNAGATCGGAAG |
| 54180_1 | TATGAGTGTGGAGTCGTTGCACTCCTAGATAGGTTGGCTCTGANNNAGATCGGAAG |
| 54183_1 | TATGAGTGTGGAGTCGTTGCTTCTGCACATAAAATGCTGAGCCNNNAGATCGGAAG |
| 54186_1 | TATGAGTGTGGAGTCGTTGCATGTGAGAAGGCCAACATCCAANNNAGATCGGAAG |
| 54187_1 | TATGAGTGTGGAGTCGTTGCAAATAAAGAGTGAGCCCTCCCAGNNNAGATCGGAA |
| 54190_1 | TATGAGTGTGGAGTCGTTGCCAGGAAATAGCAACTGCAGCAAGNNNAGATCGGAA |
| 54195_1 | TATGAGTGTGGAGTCGTTGCGCTTGGGGAAAAGGAAAATGTGANNNAGATCGGA |
| 54197_1 | TATGAGTGTGGAGTCGTTGCAATTCCAAGGGATTTAGGAGCCCNNNAGATCGGAA |
| 54205_1 | TATGAGTGTGGAGTCGTTGCGAGACGGAGTCTTACTCTGTCGNNNAGATCGGAAG |
| 54215_1 | TATGAGTGTGGAGTCGTTGCGATCTCGGCTCACTACAACCTCNNNAGATCGGAAGA |
| 54233_1 | TATGAGTGTGGAGTCGTTGCCACAATCTCAGCTCACTGCAAGNNNAGATCGGAAGA |
| 54234_1 | TATGAGTGTGGAGTCGTTGCGTAGTTTAGGCTGGAGTGCTGTNNNAGATCGGAAG |
| 54236_1 | TATGAGTGTGGAGTCGTTGCGGAGGAGTAGCTGGGACTACAGNNNAGATCGGAA |
| 54243_1 | TATGAGTGTGGAGTCGTTGCGAGTAGCTGGGACTATAGGCGTNNNAGATCGGAAG |

| | |
|---|---|
| 54252_1 | TATGAGTGTGGAGTCGTTGCAATCTCGGCTCACTACAAGCTCNNNAGATCGGAAGA |
| 54257_1 | TATGAGTGTGGAGTCGTTGCGGCTGTAAGGGCTTTGTTGTTANNNAGATCGGAA |
| 54272_1 | TATGAGTGTGGAGTCGTTGCATCACCTGAGGTCACAAGTTCAANNNAGATCGGAAG |
| 54277_1 | TATGAGTGTGGAGTCGTTGCGCAGATAGGTTGAGCTCAGAAGTNNNAGATCGGAA |
| 54287_1 | TATGAGTGTGGAGTCGTTGCGAACCCAGGAGGCAGAACTTACNNNAGATCGGAAG |
| 54293_1 | TATGAGTGTGGAGTCGTTGCCACACACACAGTCACATGTCAATNNNAGATCGGAAG |
| 54297_1 | TATGAGTGTGGAGTCGTTGCTCAGTCAGCAGCTGTCAAAATTGNNNAGATCGGAAG |
| 54298_1 | TATGAGTGTGGAGTCGTTGCTGGAAATAGACAGAGGTGGTTGTNNNAGATCGGAA |
| 54304_1 | TATGAGTGTGGAGTCGTTGCTCTGTTTAGGACCCGTGTTACTGNNNAGATCGGAAG |
| 54311_1 | TATGAGTGTGGAGTCGTTGCCTCCTGGGTTCAAGCAGTTTTCNNNAGATCGGAAGA |
| 54313_1 | TATGAGTGTGGAGTCGTTGCCTTCTCTCGAGTAGCTGGGACTANNNAGATCGGAAG |
| 54319_1 | TATGAGTGTGGAGTCGTTGCCTCCTGGGTTCAAGCGATTCTCNNNAGATCGGAAGA |
| 54321_1 | TATGAGTGTGGAGTCGTTGCAGGTTCAAGCGATTCTCCTGTCNNNAGATCGGAAGA |
| 54324_1 | TATGAGTGTGGAGTCGTTGCAGATAGTCTCCCTCTGTTGTCCANNNAGATCGGAAG |
| 54326_1 | TATGAGTGTGGAGTCGTTGCGATCTTGGCTCACTGCAACCTCNNNAGATCGGAAGA |
| 54327_1 | TATGAGTGTGGAGTCGTTGCCCCTAGTAGCTGGGACTACAGGNNNAGATCGGAAG |
| 54334_1 | TATGAGTGTGGAGTCGTTGCCTCCTGGGTTCATGGGATTCTCNNNAGATCGGAAGA |
| 54344_1 | TATGAGTGTGGAGTCGTTGCCTCCTAGGTTCACGCCATTCTCNNNAGATCGGAAGAG |
| 54352_1 | TATGAGTGTGGAGTCGTTGCCTCAGGTTCAAGTGATTCTCCCANNNAGATCGGAAG |
| 54353_1 | TATGAGTGTGGAGTCGTTGCGAGATGGAGACTCACTCTGTCGNNNAGATCGGAAG |
| 54362_1 | TATGAGTGTGGAGTCGTTGCGATCTCAGCTCACTGCAACCTCNNNAGATCGGAAGA |
| 54370_1 | TATGAGTGTGGAGTCGTTGCCTCCTGGGTTCACACCATTGTCNNNAGATCGGAAGA |
| 54372_1 | TATGAGTGTGGAGTCGTTGCAGAGTCTCTCTCAGTCACCCAGNNNAGATCGGAAGA |
| 54379_1 | TATGAGTGTGGAGTCGTTGCGTCTCCTGAGTAGCTGGGACTANNNAGATCGGAAG |
| 54380_1 | TATGAGTGTGGAGTCGTTGCGTTCACGCTATTCTCCTGCCTCNNNAGATCGGAAGAG |
| 54381_1 | TATGAGTGTGGAGTCGTTGCCTCCCAGGTTCAAGCGATTCTCNNNAGATCGGAAGA |
| 54384_1 | TATGAGTGTGGAGTCGTTGCGCTGGAGTACAATGGTGTGATCTNNNAGATCGGAA |
| 54389_1 | TATGAGTGTGGAGTCGTTGCCAAAGAACCATGACCCAACGATGNNNAGATCGGAAG |
| 54390_1 | TATGAGTGTGGAGTCGTTGCCTGCGTGACAGAGTAAGAACCTANNNAGATCGGAA |
| 54401_1 | TATGAGTGTGGAGTCGTTGCTACTCTTTTCCATAGTGGCTGCANNNAGATCGGAAG |
| 54407_1 | TATGAGTGTGGAGTCGTTGCCCAGCAGCACATCAAAAAGCTTANNNAGATCGGAAG |
| 54410_1 | TATGAGTGTGGAGTCGTTGCGGGTCTTGTAAGGAAAGCTCTGANNNAGATCGGAA |
| 54416_1 | TATGAGTGTGGAGTCGTTGCTTCTGGCTTGTAGAGTTTCTGCTNNNAGATCGGAAG |
| 54426_1 | TATGAGTGTGGAGTCGTTGCTGCTACAAATCCAGAAAAATAAGCCNNNAGATCGGA |
| 54434_1 | TATGAGTGTGGAGTCGTTGCGAGTCAGTGGCCCAAAAGAAAAANNNAGATCGGAA |
| 54440_1 | TATGAGTGTGGAGTCGTTGCGCCTGTAATCCCAGCTACTTGGNNNAGATCGGAAGA |
| 54441_1 | TATGAGTGTGGAGTCGTTGCTGAACTATTTAGGCCAGGACTCCNNNAGATCGGAAG |
| 54445_1 | TATGAGTGTGGAGTCGTTGCACCTTTCTGCTCTTGTCAGACAANNNAGATCGGAAG |
| 54452_1 | TATGAGTGTGGAGTCGTTGCGTGCCTGTAATCCCAGCTACTTNNNAGATCGGAAGA |
| 54467_1 | TATGAGTGTGGAGTCGTTGCTTCTGTGTTGGGTACTGGCAATANNNAGATCGGAAG |
| 54477_1 | TATGAGTGTGGAGTCGTTGCACCACGGCATATTTCTGATCTCANNNAGATCGGAAG |
| 54486_1 | TATGAGTGTGGAGTCGTTGCGCTTGTAATCAGAATGGTGGTCGNNNAGATCGGAA |
| 54497_1 | TATGAGTGTGGAGTCGTTGCCTAGTTGCTGGGACTACAGACGNNNAGATCGGAAG |

| | |
|---|---|
| 54504_1 | TATGAGTGTGGAGTCGTTGCGATTTCGGCTCACTGCAAACTCNNNAGATCGGAAGA |
| 54507_1 | TATGAGTGTGGAGTCGTTGCCTCCTGGGTTCATGCCATTCTCNNNAGATCGGAAGA |
| 54516_1 | TATGAGTGTGGAGTCGTTGCCAGTGGTGCAATCATAGCTCACNNNAGATCGGAAGA |
| 54525_1 | TATGAGTGTGGAGTCGTTGCAGACAGAATCTCGCTCTTTCGCNNNAGATCGGAAGA |
| 54532_1 | TATGAGTGTGGAGTCGTTGCCAATGGTGCAATCTCAGCTCACNNNAGATCGGAAGA |
| 54542_1 | TATGAGTGTGGAGTCGTTGCCCTGGGTGACTAAGCAAGACTCNNNAGATCGGAAG |
| 54547_1 | TATGAGTGTGGAGTCGTTGCATCTTCAGCTCTCCCTTTGGAAGNNNAGATCGGAAG |
| 54563_1 | TATGAGTGTGGAGTCGTTGCACAAAAATTAGCAGGGTGTGGTGNNNAGATCGGAA |
| 54568_1 | TATGAGTGTGGAGTCGTTGCTTGCAACATACATGAAAAGGCCANNNAGATCGGAA |
| 54569_1 | TATGAGTGTGGAGTCGTTGCAGAAGAATCACACTGTTGCTGAANNNAGATCGGAA |
| 54583_1 | TATGAGTGTGGAGTCGTTGCTGTCCTTGAAAAGTCATCAGGGTNNNAGATCGGAA |
| 54594_1 | TATGAGTGTGGAGTCGTTGCCTCCCAGGTTCATGGCATTCTCNNNAGATCGGAAGA |
| 54601_1 | TATGAGTGTGGAGTCGTTGCCAGGTTCAAGCAATTCTCCTGCNNNAGATCGGAAGA |
| 54614_1 | TATGAGTGTGGAGTCGTTGCCAGGTTCAAGCAATTCTCCTGCNNNAGATCGGAAGA |
| 54620_1 | TATGAGTGTGGAGTCGTTGCAGCAAAGGAAAAGGCATTCAACANNNAGATCGGAA |
| 54622_1 | TATGAGTGTGGAGTCGTTGCTGCACTTGACTGAAGAGTTGTCANNNAGATCGGAA |
| 54632_1 | TATGAGTGTGGAGTCGTTGCGTGCTATAGCCATCCATCAGTCANNNAGATCGGAAG |
| 54658_1 | TATGAGTGTGGAGTCGTTGCTTCAAGTGATTCTCCTGCCTCAGNNNAGATCGGAAG |
| 54665_1 | TATGAGTGTGGAGTCGTTGCCAATGGTGCAAACTCAGCTCACNNNAGATCGGAAGA |
| 54673_1 | TATGAGTGTGGAGTCGTTGCCTCCTGGGTTCATGCCATTTTCNNNAGATCGGAAGA |
| 54675_1 | TATGAGTGTGGAGTCGTTGCCAAGTAGCTGGGACTACAGGTGNNNAGATCGGAAG |
| 54679_1 | TATGAGTGTGGAGTCGTTGCCTGGGTTCATGTGATTCTTCTGCNNNAGATCGGAAG |
| 54690_1 | TATGAGTGTGGAGTCGTTGCAATTCTTCCCATTGCTGGTTTGGNNNAGATCGGAAG |
| 54692_1 | TATGAGTGTGGAGTCGTTGCGAAGTTCCTCAGGTACCTTTTTCTNNNAGATCGGAA |
| 54694_1 | TATGAGTGTGGAGTCGTTGCACACACCATCCCATTATGACAAANNNAGATCGGAAG |
| 54699_1 | TATGAGTGTGGAGTCGTTGCTGAACAAGGGTTTTCTGCAAAGGNNNAGATCGGAA |
| 54704_1 | TATGAGTGTGGAGTCGTTGCAGAGTTGCCCTTTTATCTGCTGANNNAGATCGGAAG |
| 54718_1 | TATGAGTGTGGAGTCGTTGCCTCCCAGGTTCACACCATTCTCNNNAGATCGGAAGAG |
| 54735_1 | TATGAGTGTGGAGTCGTTGCCTTCAGCCTCCTGAGTAGTTGGNNNAGATCGGAAGA |
| 54739_1 | TATGAGTGTGGAGTCGTTGCCAAGTAGCTGGGACTACAGGCNNNAGATCGGAAGA |
| 54752_1 | TATGAGTGTGGAGTCGTTGCGAGTAGCTGGGATTACAGGAGCNNNAGATCGGAA |
| 54754_1 | TATGAGTGTGGAGTCGTTGCCTCCCAGGTTCATGCCATTCTCNNNAGATCGGAAGA |
| 54756_1 | TATGAGTGTGGAGTCGTTGCCCTCCTGAGTAGCTGTCAAGTCNNNAGATCGGAAGA |
| 54800_1 | TATGAGTGTGGAGTCGTTGCGTGCCCTGATTTTGATGCATTCTNNNAGATCGGAAG |
| 54810_1 | TATGAGTGTGGAGTCGTTGCCAACGTAAAACGAATCAATGTGTTNNNAGATCGGAA |
| 54817_1 | TATGAGTGTGGAGTCGTTGCACACCATGCAAAGGATGAATTGCNNNAGATCGGAA |
| 54825_1 | TATGAGTGTGGAGTCGTTGCCTGTGAAGCCAGTATCACCCTAANNNAGATCGGAAG |
| 54826_1 | TATGAGTGTGGAGTCGTTGCTGTCTATTATGTGAGCATGCCCANNNAGATCGGAAG |
| 54828_1 | TATGAGTGTGGAGTCGTTGCCTGAGTAGCTGGGATTACAGGTGNNNAGATCGGAA |
| 54831_1 | TATGAGTGTGGAGTCGTTGCGATGAGTAGGCAGTACACAGGAGNNNAGATCGGAA |
| 54836_1 | TATGAGTGTGGAGTCGTTGCTGTTAGCCAGGATGGTCTTGATCNNNAGATCGGAA |
| 54844_1 | TATGAGTGTGGAGTCGTTGCAGATGGAATCTCGCTCTGTCACNNNAGATCGGAAGA |
| 54849_1 | TATGAGTGTGGAGTCGTTGCGATCTCGGCTCACTGCAAACTCNNNAGATCGGAAGA |

| | |
|---|---|
| 54883_1 | TATGAGTGTGGAGTCGTTGCGATCTCGACTCACTGCAAGCTCNNNAGATCGGAAGA |
| 54888_1 | TATGAGTGTGGAGTCGTTGCGAGTTTTGCTCTCATTGCCCAGNNNAGATCGGAAGA |
| 54889_1 | TATGAGTGTGGAGTCGTTGCATTCTCCTGCCTCAACTTCCAGNNNAGATCGGAAGAG |
| 54893_1 | TATGAGTGTGGAGTCGTTGCGAGTAGCCGGGATTACAGGAGNNNAGATCGGAAG |
| 54906_1 | TATGAGTGTGGAGTCGTTGCGGACCTCTGCTTTTCTGCTTTTANNNAGATCGGAAGA |
| 54914_1 | TATGAGTGTGGAGTCGTTGCAGAGAAGAGGCAAGTAAGACACANNNAGATCGGA |
| 54921_1 | TATGAGTGTGGAGTCGTTGCGAGACAGAGGTTGCAGTGAGTTNNNAGATCGGAA |
| 54930_1 | TATGAGTGTGGAGTCGTTGCTGTTGGGTTTGGTTTATTCTTGCTNNNAGATCGGAA |
| 54942_1 | TATGAGTGTGGAGTCGTTGCCTAGCAAGCTTCTGAGTCATTGCNNNAGATCGGAAG |
| 54952_1 | TATGAGTGTGGAGTCGTTGCACTCTCAAAACCTCAGATAGCCTNNNAGATCGGAAG |
| 54963_1 | TATGAGTGTGGAGTCGTTGCAAAGCAGGTAGAACTCAAGGAGGNNNAGATCGGA |
| 54964_1 | TATGAGTGTGGAGTCGTTGCGGGCAAGAGAGAACCATAGATGGNNNAGATCGGA |
| 54967_1 | TATGAGTGTGGAGTCGTTGCCAGTACTTACTCCTTGCCACCATNNNAGATCGGAAGA |
| 54968_1 | TATGAGTGTGGAGTCGTTGCCATAGGACAAATGAGGCTTTGCCNNNAGATCGGAA |
| 54970_1 | TATGAGTGTGGAGTCGTTGCAACAAAACCAGAGGACCCAGTTNNNAGATCGGAAG |
| 54971_1 | TATGAGTGTGGAGTCGTTGCGACTCCCTGGTTCAAGTGATTCTNNNAGATCGGAAG |
| 54984_1 | TATGAGTGTGGAGTCGTTGCGTTCAAGCGATTCTTCTGCCTCNNNAGATCGGAAGA |
| 55001_1 | TATGAGTGTGGAGTCGTTGCGAGAAAGAGTCTCGGCTCTGTCNNNAGATCGGAAG |
| 55004_1 | TATGAGTGTGGAGTCGTTGCCTCCTGGGTTCACACCATTCTCNNNAGATCGGAAGA |
| 55006_1 | TATGAGTGTGGAGTCGTTGCCCTCCTGGACTCATGCATTCTCNNNAGATCGGAAGA |
| 55007_1 | TATGAGTGTGGAGTCGTTGCCTGTCTGGAATTCCTAGGCTCAANNNAGATCGGAAG |
| 55021_1 | TATGAGTGTGGAGTCGTTGCGACAGTCATGCAAGCTTCAGTTTNNNAGATCGGAAG |
| 55026_1 | TATGAGTGTGGAGTCGTTGCACAAGAACAAAAGGACCAAAGGCNNNAGATCGGAA |
| 55031_1 | TATGAGTGTGGAGTCGTTGCGCAACCTGCATTGGAGTATGATGNNNAGATCGGAA |
| 55035_1 | TATGAGTGTGGAGTCGTTGCAGTATTTTCAGCTTTCCTCCCCANNNAGATCGGAAGA |
| 55037_1 | TATGAGTGTGGAGTCGTTGCGGGTTTGATTGGAAGCACAATGANNNAGATCGGAA |
| 55041_1 | TATGAGTGTGGAGTCGTTGCACACTGTGAAGGTAAAGCACATTNNNAGATCGGAA |
| 55042_1 | TATGAGTGTGGAGTCGTTGCCTCACTGCAGCTTTGACTTCCTNNNAGATCGGAAGA |
| 55068_1 | TATGAGTGTGGAGTCGTTGCCCAGGTTCAAGCAATTCTCGTGNNNAGATCGGAAGA |
| 55083_1 | TATGAGTGTGGAGTCGTTGCGTTCATGCTATTCCCCTGCCTCNNNAGATCGGAAGAG |
| 55092_1 | TATGAGTGTGGAGTCGTTGCCTCTATGTTGTCCAAGCTGGAGTNNNAGATCGGAAG |
| 55095_1 | TATGAGTGTGGAGTCGTTGCTCCATTCTACAGTTGGAGAAGCCNNNAGATCGGAAG |
| 55096_1 | TATGAGTGTGGAGTCGTTGCGGGGCATGAAGCACATTTATGTTNNNAGATCGGAA |
| 55098_1 | TATGAGTGTGGAGTCGTTGCACACACAAAAATTCTGAAGCCGANNNAGATCGGAAG |
| 55099_1 | TATGAGTGTGGAGTCGTTGCTAAAAAGGCAACCCACAGAATGGNNNAGATCGGAA |
| 55111_1 | TATGAGTGTGGAGTCGTTGCATCTGCAGTTTGTGGATTGTCCANNNAGATCGGAAG |
| 55127_1 | TATGAGTGTGGAGTCGTTGCTGTGTTGTACTCTTGAAGACTGCNNNAGATCGGAAG |
| 55130_1 | TATGAGTGTGGAGTCGTTGCTGTGTGTGCCTCATATCTCCTTCNNNAGATCGGAAGA |
| 55148_1 | TATGAGTGTGGAGTCGTTGCTGGTGCACACTTATAGTTCCAGGNNNAGATCGGAAG |
| 55150_1 | TATGAGTGTGGAGTCGTTGCGCAAATCCAGCTACAGAACACGNNNAGATCGGAAG |
| 55153_1 | TATGAGTGTGGAGTCGTTGCTCCCATTTGTAGTTGTAGAAGTGGANNNAGATCGGA |
| 55155_1 | TATGAGTGTGGAGTCGTTGCATGGTGGTGTGCATTTGTAATCCNNNAGATCGGAA |
| 55156_1 | TATGAGTGTGGAGTCGTTGCCAGGTAGATCACAAGGTCAGGAGNNNAGATCGGAA |

| | |
|---|---|
| 55157_1 | TATGAGTGTGGAGTCGTTGCTTATTTGAGGTCAGGAGATCGGCNNNAGATCGGAA |
| 55159_1 | TATGAGTGTGGAGTCGTTGCAGCACTTTGTGTTGCAACTTAGGNNNAGATCGGAAG |
| 55163_1 | TATGAGTGTGGAGTCGTTGCAAGTAGCTGGGACTACAGGCATNNNAGATCGGAAG |
| 55169_1 | TATGAGTGTGGAGTCGTTGCGAGTAGCTGGGACTACAGGCATTNNNAGATCGGAA |
| 55175_1 | TATGAGTGTGGAGTCGTTGCGGAGTGCAGTAGTGTGATCTCGNNNAGATCGGAAG |
| 55184_1 | TATGAGTGTGGAGTCGTTGCCCTCCTGGGTTTATGCCATTCTCNNNAGATCGGAAG |
| 55197_1 | TATGAGTGTGGAGTCGTTGCTCTGGAGTGCAGTGACACAATCNNNAGATCGGAAG |
| 55203_1 | TATGAGTGTGGAGTCGTTGCTGGAGTACAGTGGTGCAATCTCNNNAGATCGGAAG |
| 55204_1 | TATGAGTGTGGAGTCGTTGCACCACACCCAGCCAATCATTATNNNAGATCGGAAGAG |
| 55221_1 | TATGAGTGTGGAGTCGTTGCTTCTCATGGATCTGGCATAGGACNNNAGATCGGAAG |
| 55235_1 | TATGAGTGTGGAGTCGTTGCCCTCCCAGAAGAACCCAGTAAATNNNAGATCGGAAG |
| 55238_1 | TATGAGTGTGGAGTCGTTGCTAGAGCTAGTACCACTGACAGCTNNNAGATCGGAAG |
| 55242_1 | TATGAGTGTGGAGTCGTTGCGCTTAGAAAACAGGCAAAAAGCGNNNAGATCGGAA |
| 55247_1 | TATGAGTGTGGAGTCGTTGCCAGTCATGGAGGAGAACAAGGAANNNAGATCGGA |
| 55251_1 | TATGAGTGTGGAGTCGTTGCACAGGGAGTTCACAGAAACACATNNNAGATCGGAA |
| 55257_1 | TATGAGTGTGGAGTCGTTGCTGCAGAAATGTTCGAACAAAGCTNNNAGATCGGAA |
| 55278_1 | TATGAGTGTGGAGTCGTTGCTCTCACAACTTGTAATCCCAGCANNNAGATCGGAAG |
| 55286_1 | TATGAGTGTGGAGTCGTTGCAATAGAGAAGCTTTTTGTGGGCGNNNAGATCGGAA |
| 55293_1 | TATGAGTGTGGAGTCGTTGCCGATCTCGGCTCACTGCAAGNNNAGATCGGAAGAG |
| 55295_1 | TATGAGTGTGGAGTCGTTGCCTCCTGGGTTCAAGCGATTCTCNNNAGATCGGAAGA |
| 55300_1 | TATGAGTGTGGAGTCGTTGCAATCTCGGCTCACTACAAGCTCNNNAGATCGGAAGA |
| 55315_1 | TATGAGTGTGGAGTCGTTGCCACAATCTCAGCTCACTGCAACNNNAGATCGGAAGA |
| 55319_1 | TATGAGTGTGGAGTCGTTGCCACGATCTTGGTTCACTGCAACNNNAGATCGGAAGA |
| 55325_1 | TATGAGTGTGGAGTCGTTGCATTCTCGAACTCCTGACCTCATGNNNAGATCGGAAG |
| 55334_1 | TATGAGTGTGGAGTCGTTGCGAGACAGAGTTTTGCCTTGTCGNNNAGATCGGAAG |
| 55336_1 | TATGAGTGTGGAGTCGTTGCGAGTTTAGCTCTTGTTGCCCAGNNNAGATCGGAAGA |
| 55340_1 | TATGAGTGTGGAGTCGTTGCGACCAGCCTGACCAACATGTATANNNAGATCGGAAG |
| 55356_1 | TATGAGTGTGGAGTCGTTGCCTGAGTAGCTGGAACTACAGGTGNNNAGATCGGAA |
| 55366_1 | TATGAGTGTGGAGTCGTTGCGTGGATCACTTGAGGTCAGGATTNNNAGATCGGAA |
| 55377_1 | TATGAGTGTGGAGTCGTTGCCCCTCTTTTTCACCCACCAAATCNNNAGATCGGAAGA |
| 55378_1 | TATGAGTGTGGAGTCGTTGCTGGGTGGTACAAGGAGAGGATTANNNAGATCGGA |
| 55387_1 | TATGAGTGTGGAGTCGTTGCAGATGGAGTCTCACTCTGTTGCNNNAGATCGGAAGA |
| 55390_1 | TATGAGTGTGGAGTCGTTGCTGCAGTGAGCTATGATTGTGCTANNNAGATCGGAA |
| 55393_1 | TATGAGTGTGGAGTCGTTGCGCCTGAGCAACAAGAGTGAAATTNNNAGATCGGAA |
| 55396_1 | TATGAGTGTGGAGTCGTTGCACCCCTTGTCCTTCTCGTAAATCNNNAGATCGGAAGA |
| 55398_1 | TATGAGTGTGGAGTCGTTGCTATATACACACCGCACAGAGTCCNNNAGATCGGAAG |
| 55400_1 | TATGAGTGTGGAGTCGTTGCCCCATAGAGCATACTTCCCTGTCNNNAGATCGGAAG |
| 55404_1 | TATGAGTGTGGAGTCGTTGCCACGATCTCAGTTCACTGCAAGNNNAGATCGGAAGA |
| 55410_1 | TATGAGTGTGGAGTCGTTGCCTCCCAGGTTCATGCCATTCTCNNNAGATCGGAAGAG |
| 55415_1 | TATGAGTGTGGAGTCGTTGCCTCCCGAGTAGCTGAGACTACANNNAGATCGGAAGA |
| 55416_1 | TATGAGTGTGGAGTCGTTGCCAAGTAGCTGGGACAAAAGGTGNNNAGATCGGAAG |
| 55417_1 | TATGAGTGTGGAGTCGTTGCACCTCCCAGGCTCAAATGATTCNNNAGATCGGAAGA |
| 55449_1 | TATGAGTGTGGAGTCGTTGCCCAGCTTCAAGCGATTCTCTTGNNNAGATCGGAAGA |

| | |
|---|---|
| 55451_1 | TATGAGTGTGGAGTCGTTGCAGTCTCCTGAGTAGCTGGGATTANNNAGATCGGAA |
| 55454_1 | TATGAGTGTGGAGTCGTTGCCATAGTGCGATCTCTGCTCACTNNNAGATCGGAAGA |
| 55462_1 | TATGAGTGTGGAGTCGTTGCCTCACTGTAGCCTCGACTTCTCNNNAGATCGGAAGAG |
| 55476_1 | TATGAGTGTGGAGTCGTTGCGCACAATTATGCGAAGACAGTGTNNNAGATCGGAA |
| 55489_1 | TATGAGTGTGGAGTCGTTGCTTGCTCACAGTTAGTAATTGGCANNNAGATCGGAAG |
| 55504_1 | TATGAGTGTGGAGTCGTTGCTTCTTCTTCAATTGCAGTGAATGTNNNAGATCGGAA |
| 55505_1 | TATGAGTGTGGAGTCGTTGCCAGCTGCCCAGTTAGATACATGANNNAGATCGGAAG |
| 55508_1 | TATGAGTGTGGAGTCGTTGCCCTACAGTCCTAGCTATGCAGGANNNAGATCGGAAG |
| 55512_1 | TATGAGTGTGGAGTCGTTGCTTGTCCCTTACTGAAACTGTCCTNNNAGATCGGAAG |
| 55513_1 | TATGAGTGTGGAGTCGTTGCTTGCACATTCACTCTTCAGTTCANNNAGATCGGAAGA |
| 55525_1 | TATGAGTGTGGAGTCGTTGCGATGTTGGGCACTGAGAAACTTTNNNAGATCGGAA |
| 55539_1 | TATGAGTGTGGAGTCGTTGCATACTCACCAACAGTAGAACGGANNNAGATCGGAA |
| 55547_1 | TATGAGTGTGGAGTCGTTGCAGAAGTGGCCAAATTTGAGAGATNNNAGATCGGAA |
| 55549_1 | TATGAGTGTGGAGTCGTTGCTCCTCAACTCTGGACATTTCTTTNNNAGATCGGAAGA |
| 55556_1 | TATGAGTGTGGAGTCGTTGCCAAGCATAGCATCTCAAGCAACANNNAGATCGGAAG |
| 55564_1 | TATGAGTGTGGAGTCGTTGCGACCACTTGTTCTGAAGCTGGGNNNAGATCGGAAG |
| 55580_1 | TATGAGTGTGGAGTCGTTGCCAGGTTCAAGCAATTCTCCTGCNNNAGATCGGAAGA |
| 55590_1 | TATGAGTGTGGAGTCGTTGCCAGGTTCAAGCAATTCTCCTGCNNNAGATCGGAAGA |
| 55594_1 | TATGAGTGTGGAGTCGTTGCGATCTCAGCTCACTGCAACCTCNNNAGATCGGAAGA |
| 55600_1 | TATGAGTGTGGAGTCGTTGCCCAGGTTCAAGCAATTGTCCTGNNNAGATCGGAAGA |
| 55607_1 | TATGAGTGTGGAGTCGTTGCCAACCTAGAGATGTCCATCCAGGNNNAGATCGGAAG |
| 55608_1 | TATGAGTGTGGAGTCGTTGCGGGTGCTAGGTGTAAGCTCATTNNNAGATCGGAAG |
| 55609_1 | TATGAGTGTGGAGTCGTTGCTAGAATTCTGGAGGTATGCGACCNNNAGATCGGAA |
| 55611_1 | TATGAGTGTGGAGTCGTTGCGAGAGAGAGAGAGAAAAGGCTGCNNNAGATCGGA |
| 55614_1 | TATGAGTGTGGAGTCGTTGCTCACCGGAGAAAAAAAGCAATGTNNNAGATCGGAA |
| 55625_1 | TATGAGTGTGGAGTCGTTGCCTCAATGCAAACTTGACCTCCAGNNNAGATCGGAAG |
| 55650_1 | TATGAGTGTGGAGTCGTTGCATTGCATGGGAGAGAAGAAAGGTNNNAGATCGGA |
| 55661_1 | TATGAGTGTGGAGTCGTTGCTTGTCTTTCTTGGTTCATGTGCCNNNAGATCGGAAG |
| 55672_1 | TATGAGTGTGGAGTCGTTGCTGTTCTCAAACTCCTGTTTTGCCNNNAGATCGGAAGA |
| 55677_1 | TATGAGTGTGGAGTCGTTGCGAAGTCACCCTGCAAATAGGAGANNNAGATCGGAA |
| 55678_1 | TATGAGTGTGGAGTCGTTGCGGAGTACAGTGGTGTGATCTCGNNNAGATCGGAAG |
| 55688_1 | TATGAGTGTGGAGTCGTTGCAGACAGAGTCTCGCTGTGTTACNNNAGATCGGAAG |
| 55694_1 | TATGAGTGTGGAGTCGTTGCAAGATAGGGTCTGGCTGTCTGGNNNAGATCGGAAG |
| 55704_1 | TATGAGTGTGGAGTCGTTGCAAGACTGAATAGACTGCCAGTGGNNNAGATCGGAA |
| 55715_1 | TATGAGTGTGGAGTCGTTGCCTGGAGTATAGTGGCGTGATCTCNNNAGATCGGAA |
| 55748_1 | TATGAGTGTGGAGTCGTTGCCAGCTGAAACTGTAAGCCAAATTNNNAGATCGGAAG |
| 55749_1 | TATGAGTGTGGAGTCGTTGCCTGAGGCAGGAGAATCACTTGAANNNAGATCGGAA |
| 55754_1 | TATGAGTGTGGAGTCGTTGCCAGAAGAACTGCTTGAAGCCAGNNNAGATCGGAAG |
| 55755_1 | TATGAGTGTGGAGTCGTTGCTGCATCTTTGTCATGATAGTGGTNNNAGATCGGAAG |
| 55762_1 | TATGAGTGTGGAGTCGTTGCGAACAAACAGCTACAGAGTGTGCNNNAGATCGGAA |
| 55769_1 | TATGAGTGTGGAGTCGTTGCCAATGGCATGATCAAGGCTCACNNNAGATCGGAAG |
| 55770_1 | TATGAGTGTGGAGTCGTTGCCAAGTAGCTGGGACTACAGGTGNNNAGATCGGAAG |
| 55773_1 | TATGAGTGTGGAGTCGTTGCCATGATCTTGGCTCACTGCAACNNNAGATCGGAAGA |

55796_1 TATGAGTGTGGAGTCGTTGCGTTCAAGCGATTCTCTTGCCTCNNNAGATCGGAAGA
55799_1 TATGAGTGTGGAGTCGTTGCGATGGTGTCTCACTCTGTCACCNNNAGATCGGAAGA
55816_1 TATGAGTGTGGAGTCGTTGCCTAGGCTGGTCTCAAACTCCTGNNNAGATCGGAAGA
55830_1 TATGAGTGTGGAGTCGTTGCCATCCTGGCTAACATGGTGAAACNNNAGATCGGAAG
55836_1 TATGAGTGTGGAGTCGTTGCTCCACTTTCATCCATGTTGTTGCNNNAGATCGGAAGA
55855_1 TATGAGTGTGGAGTCGTTGCGAAGAGAAAGAGCCCGGGTTACNNNAGATCGGAAG
55868_1 TATGAGTGTGGAGTCGTTGCAACACGTAGCCTTTTTTTCCTGGNNNAGATCGGAAG
55876_1 TATGAGTGTGGAGTCGTTGCGACCAAGAGGAGCGAGAGAAACNNNAGATCGGAA
55884_1 TATGAGTGTGGAGTCGTTGCTAGAGATGGGGTTTTGCCATGTTNNNAGATCGGAA
55893_1 TATGAGTGTGGAGTCGTTGCGAAATGTCAGCTCACTGCAACCNNNAGATCGGAAGA
55905_1 TATGAGTGTGGAGTCGTTGCATGTCCACACAAAAACTTGCACANNNAGATCGGAAG
55911_1 TATGAGTGTGGAGTCGTTGCAAGGAGGCTACACTGAGACTTTGNNNAGATCGGAA
55917_1 TATGAGTGTGGAGTCGTTGCAAAGAGAAAAAAGTCCGGGCATGNNNAGATCGGAA
55922_1 TATGAGTGTGGAGTCGTTGCGAGAGGGGTCTCTGACTCCTAATNNNAGATCGGAA
55946_1 TATGAGTGTGGAGTCGTTGCCATGATCTCGGTTCACTGCAAGNNNAGATCGGAAGA
55951_1 TATGAGTGTGGAGTCGTTGCGGGTTCAAGTGATTCTCCTGTCTNNNAGATCGGAAG
55955_1 TATGAGTGTGGAGTCGTTGCTAGAAATTAGCTGAGCGTGGTGANNNAGATCGGAA
55958_1 TATGAGTGTGGAGTCGTTGCGAGCTACCAGATGAGTTTTGGGTNNNAGATCGGAA
55962_1 TATGAGTGTGGAGTCGTTGCTTCATATTATTTGGTCAACGCCCNNNAGATCGGAAG
55974_1 TATGAGTGTGGAGTCGTTGCCCAGAGATGGCTTCCAAACAAAGNNNAGATCGGAA
55979_1 TATGAGTGTGGAGTCGTTGCACTGGTCTTACATGAACTGCTCANNNAGATCGGAAG
55984_1 TATGAGTGTGGAGTCGTTGCCCATTCGAGTCAAGTGCCTAAGANNNAGATCGGAAG
55993_1 TATGAGTGTGGAGTCGTTGCGAGTAGCTCGGATTACAGGAACCNNNAGATCGGAA
55997_1 TATGAGTGTGGAGTCGTTGCGAGTGCAGTGGTACAGTCATGANNNAGATCGGAAG
56003_1 TATGAGTGTGGAGTCGTTGCTCTCCGAGTAGCTGGGACTACNNNAGATCGGAAGA
56006_1 TATGAGTGTGGAGTCGTTGCCTCCTGGGTTCAAGCGATTCTCNNNAGATCGGAAGA
56017_1 TATGAGTGTGGAGTCGTTGCCAAGTAGCTGGGACCATAGGTGNNNAGATCGGAAG
56042_1 TATGAGTGTGGAGTCGTTGCGTTCAAGCGATTCTTCTGCCTCNNNAGATCGGAAGA
56050_1 TATGAGTGTGGAGTCGTTGCGAGTGCAGTGGTGTGATCTCAGNNNAGATCGGAAG
56068_1 TATGAGTGTGGAGTCGTTGCACAACTATAAAGTGCTGGAGCCANNNAGATCGGAA
56083_1 TATGAGTGTGGAGTCGTTGCGCACTGAGATGAAACTGCACAAANNNAGATCGGAA
56085_1 TATGAGTGTGGAGTCGTTGCGACCATCAGGGGTTGTTTGAATCNNNAGATCGGAA
56096_1 TATGAGTGTGGAGTCGTTGCTAATGCCACAATTTCACAGCAGGNNNAGATCGGAAG
56112_1 TATGAGTGTGGAGTCGTTGCAAAGAGGTTTGTTGGTTCATGGCNNNAGATCGGAA
56118_1 TATGAGTGTGGAGTCGTTGCCCAAATTGCTGGGATTACAGGCNNNAGATCGGAAG
56119_1 TATGAGTGTGGAGTCGTTGCCATGATTGTGGCTCACTGCAACNNNAGATCGGAAGA
56120_1 TATGAGTGTGGAGTCGTTGCGATCTTGGCTCACTGCAAGCTCNNNAGATCGGAAGA
56123_1 TATGAGTGTGGAGTCGTTGCAAACTCCTGACCTTGGGTGATCNNNAGATCGGAAGA
56124_1 TATGAGTGTGGAGTCGTTGCGGTTCAAGTGATTCTCCTGCCTCNNNAGATCGGAAG
56128_1 TATGAGTGTGGAGTCGTTGCCAAAGTGTTGGGATTACAGGCGNNNAGATCGGAAG
56133_1 TATGAGTGTGGAGTCGTTGCGTAGAGATGTTTCCCAGGCTGGNNNAGATCGGAAG
56140_1 TATGAGTGTGGAGTCGTTGCCTCCCAGGTTCACACCATTCTCNNNAGATCGGAAGAG
56141_1 TATGAGTGTGGAGTCGTTGCCTCCTGGGTTCATGCCATTCTCNNNAGATCGGAAGA

| | |
|---|---|
| 56143_1 | TATGAGTGTGGAGTCGTTGCCTCCTGAGTAGCTGGGATTACAGNNNAGATCGGAA |
| 56147_1 | TATGAGTGTGGAGTCGTTGCGACAGAGTCTCACTGTGTCACCNNNAGATCGGAAGA |
| 56156_1 | TATGAGTGTGGAGTCGTTGCGTTCACGCCATTCTCCTACCTCNNNAGATCGGAAGAG |
| 56157_1 | TATGAGTGTGGAGTCGTTGCCTCTCGGGTTTGCACCATTCTCNNNAGATCGGAAGA |
| 56162_1 | TATGAGTGTGGAGTCGTTGCCTCACTGCAACTTCTACCTCCCNNNAGATCGGAAGAG |
| 56165_1 | TATGAGTGTGGAGTCGTTGCTCAAGCTTTTCTCCCACCTCAGNNNAGATCGGAAGAG |
| 56195_1 | TATGAGTGTGGAGTCGTTGCATGTCTGATCAAGTGTAGGGTGGNNNAGATCGGAA |
| 56199_1 | TATGAGTGTGGAGTCGTTGCCACACCTGTAATCCCAGCACTTTNNNAGATCGGAAG |
| 56202_1 | TATGAGTGTGGAGTCGTTGCGAACTGGCTTGATGTGCATTCCNNNAGATCGGAAG |
| 56217_1 | TATGAGTGTGGAGTCGTTGCGATTATGGTCTCAGCCCCTCACNNNAGATCGGAAGA |
| 56218_1 | TATGAGTGTGGAGTCGTTGCGATAGGTTGAGAGCTCGACTCACNNNAGATCGGAA |
| 56226_1 | TATGAGTGTGGAGTCGTTGCCCCATATTCATAGTGCCTGTTGCNNNAGATCGGAAG |
| 56239_1 | TATGAGTGTGGAGTCGTTGCCTCACATCTGTAGCACTTTGGGANNNAGATCGGAAG |
| 56256_1 | TATGAGTGTGGAGTCGTTGCAGACGGAGTCTCACTGTCTCTCNNNAGATCGGAAGA |
| 56275_1 | TATGAGTGTGGAGTCGTTGCCCTAACACTTTCCCTCTAGCACANNNAGATCGGAAGA |
| 56276_1 | TATGAGTGTGGAGTCGTTGCGACTTCCGCTCCGTCTGTAATATNNNAGATCGGAAG |
| 56281_1 | TATGAGTGTGGAGTCGTTGCCGAGTAGCTGGGATTACAGGTGNNNAGATCGGAAG |
| 56282_1 | TATGAGTGTGGAGTCGTTGCCAATCTTGGCTCACTGCAACCNNNAGATCGGAAGAG |
| 56284_1 | TATGAGTGTGGAGTCGTTGCGAGTTTTGCTCTTGTTGCCCAGNNNAGATCGGAAGA |
| 56291_1 | TATGAGTGTGGAGTCGTTGCTGCAGTGCAGTGGTATAACCTCNNNAGATCGGAAG |
| 56292_1 | TATGAGTGTGGAGTCGTTGCCAAGTAGCTGGGACTACAGGTGNNNAGATCGGAAG |
| 56297_1 | TATGAGTGTGGAGTCGTTGCTTAGCCTCCTAAGTAGCTGGGATNNNAGATCGGAAG |
| 56301_1 | TATGAGTGTGGAGTCGTTGCGAGTCTTGCTCTGTCTCCTAGGNNNAGATCGGAAGA |
| 56315_1 | TATGAGTGTGGAGTCGTTGCGGGTTCACACCATCCTCCTGNNNAGATCGGAAGAGC |
| 56316_1 | TATGAGTGTGGAGTCGTTGCAATCTCGACTCACTGCAAGCTCNNNAGATCGGAAGA |
| 56321_1 | TATGAGTGTGGAGTCGTTGCAATGTTGGCTCACTGCAACTTCNNNAGATCGGAAGA |
| 56322_1 | TATGAGTGTGGAGTCGTTGCGAGTATCTGGGACTACAGGCACNNNAGATCGGAAG |
| 56338_1 | TATGAGTGTGGAGTCGTTGCTCTTAGTAGAGACGGGGTTTCACNNNAGATCGGAA |
| 56339_1 | TATGAGTGTGGAGTCGTTGCTGGAGTACAGTGGTGCAATCTCNNNAGATCGGAAG |
| 56342_1 | TATGAGTGTGGAGTCGTTGCGAGTTTCGCTCTTATTGCCCAGNNNAGATCGGAAGA |
| 56348_1 | TATGAGTGTGGAGTCGTTGCAGCTAGCTCCATCTCTCTTCTCANNNAGATCGGAAGA |
| 56355_1 | TATGAGTGTGGAGTCGTTGCTGGTAGAGTCTGCTACACACCTANNNAGATCGGAAG |
| 56362_1 | TATGAGTGTGGAGTCGTTGCATTTCAAAGCCAGCAGACATTCCNNNAGATCGGAAG |
| 56366_1 | TATGAGTGTGGAGTCGTTGCCGTTCTGCTACCAGGAGAAGTAGNNNAGATCGGAA |
| 56368_1 | TATGAGTGTGGAGTCGTTGCCCCATTTTCCTCTCAGTCCAGATNNNAGATCGGAAGA |
| 56372_1 | TATGAGTGTGGAGTCGTTGCACGACTGTAATCCCAGCACTTTNNNAGATCGGAAGA |
| 56380_1 | TATGAGTGTGGAGTCGTTGCCAATGCACTCCAGTCTGGGTAANNNAGATCGGAAG |
| 56382_1 | TATGAGTGTGGAGTCGTTGCGGATCCGTCCCATGAGAGATAACNNNAGATCGGAA |
| 56392_1 | TATGAGTGTGGAGTCGTTGCGGTGAAACCCCGTCTCTACTAAANNNAGATCGGAAG |
| 56396_1 | TATGAGTGTGGAGTCGTTGCGTTTGTGAGACTGAGTTTCGCTCNNNAGATCGGAAG |
| 56398_1 | TATGAGTGTGGAGTCGTTGCAAGCAGAGGGATTTCTGGTAGCNNNAGATCGGAAG |
| 56404_1 | TATGAGTGTGGAGTCGTTGCAGAATGTCAGACCTTCAAGGACCNNNAGATCGGAA |
| 56405_1 | TATGAGTGTGGAGTCGTTGCATGCCTGTAATCCCAGCACTTTNNNAGATCGGAAGA |

| | |
|---|---|
| 56423_1 | TATGAGTGTGGAGTCGTTGCTGTAATGGAGATGGAACAGTGCCNNNAGATCGGAA |
| 56429_1 | TATGAGTGTGGAGTCGTTGCCTGCAACACCCAAGGAATTATGGNNNAGATCGGAAG |
| 56437_1 | TATGAGTGTGGAGTCGTTGCTGAGGATACAGCAAGAAGGCATTNNNAGATCGGAA |
| 56443_1 | TATGAGTGTGGAGTCGTTGCCTCCTGGGTTCAAGCGATTCTCNNNAGATCGGAAGA |
| 56459_1 | TATGAGTGTGGAGTCGTTGCGTTCACTCCATTCTCCTGCCTCNNNAGATCGGAAGAG |
| 56460_1 | TATGAGTGTGGAGTCGTTGCCTCCCAGGTTCATGCCATTCTCNNNAGATCGGAAGAG |
| 56465_1 | TATGAGTGTGGAGTCGTTGCAGACAGGGTCTCACTTTGTCACNNNAGATCGGAAGA |
| 56481_1 | TATGAGTGTGGAGTCGTTGCCTAGAGCGATTCTTCTGCCTCAGNNNAGATCGGAAG |
| 56483_1 | TATGAGTGTGGAGTCGTTGCCAGGCTTTGAGCAATTCTCCTGNNNAGATCGGAAGA |
| 56485_1 | TATGAGTGTGGAGTCGTTGCCTTCCGAGTAGCTGGGACTACANNNAGATCGGAAG |
| 56493_1 | TATGAGTGTGGAGTCGTTGCCCGAGTTAGAGGCTACAGTGAGNNNAGATCGGAAG |
| 56495_1 | TATGAGTGTGGAGTCGTTGCGAAAAGTGTTCACCCAGGAGGCNNNAGATCGGAAG |
| 56504_1 | TATGAGTGTGGAGTCGTTGCTTTTCTGAGATGGAGTTTCGCTCNNNAGATCGGAAG |
| 56510_1 | TATGAGTGTGGAGTCGTTGCGAATCCCTTTTGTCCAGGCAAGNNNAGATCGGAAGA |
| 56525_1 | TATGAGTGTGGAGTCGTTGCACCCATAAAGCCATCTGATCCTGNNNAGATCGGAAG |
| 56531_1 | TATGAGTGTGGAGTCGTTGCAATCCAAAACATGCTGACAGTGGNNNAGATCGGAA |
| 56534_1 | TATGAGTGTGGAGTCGTTGCATGTTGCAATAAAGTTGTGGGGGNNNAGATCGGAA |
| 56542_1 | TATGAGTGTGGAGTCGTTGCCATCCTGGCTAACATGGTGAAACNNNAGATCGGAAG |
| 56545_1 | TATGAGTGTGGAGTCGTTGCTAAATGGACACAGACCCAGACAGNNNAGATCGGAA |
| 56553_1 | TATGAGTGTGGAGTCGTTGCCAGGTTCAAGCAATTCTCCTGCNNNAGATCGGAAGA |
| 56555_1 | TATGAGTGTGGAGTCGTTGCCTCAATCTCCTGACCTCGTGATCNNNAGATCGGAAG |
| 56556_1 | TATGAGTGTGGAGTCGTTGCGATCTTGGCTCACTGCAACCTCNNNAGATCGGAAGA |
| 56559_1 | TATGAGTGTGGAGTCGTTGCGATTTGGCTCACTGCAACCTCTNNNAGATCGGAAGA |
| 56563_1 | TATGAGTGTGGAGTCGTTGCTAGTTGGGACTACAGGCACTCANNNAGATCGGAAG |
| 56570_1 | TATGAGTGTGGAGTCGTTGCTTGATCTCCTGACCTTGTGATCCNNNAGATCGGAAG |
| 56578_1 | TATGAGTGTGGAGTCGTTGCCAAGTAGCTGGGACTACAGGTGNNNAGATCGGAAG |
| 56582_1 | TATGAGTGTGGAGTCGTTGCGAGCTGGAGTCTCTCTGTCAACNNNAGATCGGAAG |
| 56586_1 | TATGAGTGTGGAGTCGTTGCCTCCTGGGTTCACACCATTCTCNNNAGATCGGAAGAG |
| 56593_1 | TATGAGTGTGGAGTCGTTGCCAAGTCCAAGCAATTCTCCTGCNNNAGATCGGAAGA |
| 56594_1 | TATGAGTGTGGAGTCGTTGCGGATCTCGGCTCACTGCAAGNNNAGATCGGAAGAG |
| 56595_1 | TATGAGTGTGGAGTCGTTGCCCAAAATGGTCTCGATCTCCTGANNNAGATCGGAAG |
| 56599_1 | TATGAGTGTGGAGTCGTTGCGATCTTGGCTCACTGCAAGCTCNNNAGATCGGAAGA |
| 56611_1 | TATGAGTGTGGAGTCGTTGCCTCCCAGGTTCATGCCATTCTCNNNAGATCGGAAGAG |
| 56612_1 | TATGAGTGTGGAGTCGTTGCGTTCAGTGATTCTCCTGCCTCANNNAGATCGGAAGA |
| 56613_1 | TATGAGTGTGGAGTCGTTGCCAATGGCATGATCTTGGCTCACNNNAGATCGGAAGA |
| 56616_1 | TATGAGTGTGGAGTCGTTGCATCTCCCAGGTTCACACCATTCNNNAGATCGGAAGAG |
| 56626_1 | TATGAGTGTGGAGTCGTTGCAAGACAGTGTGGTGATTCCTCAANNNAGATCGGAA |
| 56630_1 | TATGAGTGTGGAGTCGTTGCTCCCAGGTTCACGCTATTCTCNNNAGATCGGAAGAG |
| 56650_1 | TATGAGTGTGGAGTCGTTGCCAGGAGATCGAGACCATTCTAGCNNNAGATCGGAA |
| 56651_1 | TATGAGTGTGGAGTCGTTGCCACCTGTAGTCCTGGCTACTTGNNNAGATCGGAAGA |
| 56652_1 | TATGAGTGTGGAGTCGTTGCGATCACTTCAGCCCAGGAGTTCNNNAGATCGGAAGA |
| 56653_1 | TATGAGTGTGGAGTCGTTGCGAGGCCAATATCAACACGAACAGNNNAGATCGGAA |
| 56657_1 | TATGAGTGTGGAGTCGTTGCATTTGGCTGATGTGGATATCCCANNNAGATCGGAAG |

| | |
|---|---|
| 56665_1 | TATGAGTGTGGAGTCGTTGCGGGTTGTTTGTATTTCTGTGGGGNNNAGATCGGAA |
| 56674_1 | TATGAGTGTGGAGTCGTTGCAGCACTACCTATTTGCCTGGTACNNNAGATCGGAAG |
| 56679_1 | TATGAGTGTGGAGTCGTTGCAACCATGCACATCAATTTGGGAGNNNAGATCGGAAG |
| 56692_1 | TATGAGTGTGGAGTCGTTGCCCCATGTGCTAGAGACAACTACANNNAGATCGGAAG |
| 56696_1 | TATGAGTGTGGAGTCGTTGCGAATCACTGCCCCAGAATCCATANNNAGATCGGAAG |
| 56700_1 | TATGAGTGTGGAGTCGTTGCGAGGTCAGGAGTTCGAGACATGNNNAGATCGGAAG |
| 56703_1 | TATGAGTGTGGAGTCGTTGCCTCTGGAGTAGCTGGGACTACAGNNNAGATCGGAA |
| 56720_1 | TATGAGTGTGGAGTCGTTGCCCAAGTAGCTGGGATTACAGGTGNNNAGATCGGAA |
| 56732_1 | TATGAGTGTGGAGTCGTTGCGAGTCTTGCTCTGTTGCCTAGGNNNAGATCGGAAGA |
| 56741_1 | TATGAGTGTGGAGTCGTTGCCTCCCAGGTTCATGCGATTCTCNNNAGATCGGAAGA |
| 56755_1 | TATGAGTGTGGAGTCGTTGCCTCAATCTCAGCTCACTGCAACNNNAGATCGGAAGA |
| 56756_1 | TATGAGTGTGGAGTCGTTGCAAACTCCCAGGCTCAAGTGATCNNNAGATCGGAAGA |
| 56770_1 | TATGAGTGTGGAGTCGTTGCGAGTCTCACTCTTTCACCCAGGNNNAGATCGGAAGA |
| 56784_1 | TATGAGTGTGGAGTCGTTGCAGATGGAGTCTCACTGTCACCCNNNAGATCGGAAGA |
| 56785_1 | TATGAGTGTGGAGTCGTTGCCATCTCAGCTCACTGCAACTTCNNNAGATCGGAAGAG |
| 56789_1 | TATGAGTGTGGAGTCGTTGCCGTAGCACGATCTTGGCTTACTNNNAGATCGGAAGA |
| 56797_1 | TATGAGTGTGGAGTCGTTGCGTGATCTTGGTTCACTGCAACCNNNAGATCGGAAGA |
| 56799_1 | TATGAGTGTGGAGTCGTTGCCTCCCAGGTTCACACCATTCTCNNNAGATCGGAAGAG |
| 56810_1 | TATGAGTGTGGAGTCGTTGCCTCAATCTCCTGACCTCGTGATCNNNAGATCGGAAGA |
| 56811_1 | TATGAGTGTGGAGTCGTTGCGTTCAAGCGATTCTCCTGCTTCNNNAGATCGGAAGA |
| 56830_1 | TATGAGTGTGGAGTCGTTGCTACTTCTCCCCTCCTCCAGAATTNNNAGATCGGAAGA |
| 56833_1 | TATGAGTGTGGAGTCGTTGCCCTCCGAGGGCTATTGATTTCTTNNNAGATCGGAAG |
| 56836_1 | TATGAGTGTGGAGTCGTTGCAGAACTGCCAGAAGAAAGGGAAANNNAGATCGGAA |
| 56837_1 | TATGAGTGTGGAGTCGTTGCGGGTGGGATCATTTTTTCTCCACNNNAGATCGGAAG |
| 56843_1 | TATGAGTGTGGAGTCGTTGCGCCAAAAAAATTGCCTCCAGGTANNNAGATCGGAAG |
| 56844_1 | TATGAGTGTGGAGTCGTTGCATCACGTCAAAAGATCGAGACCANNNAGATCGGAAG |
| 56851_1 | TATGAGTGTGGAGTCGTTGCGCAACAAGAAAGAAACTCTGCCTNNNAGATCGGAAG |
| 56861_1 | TATGAGTGTGGAGTCGTTGCATGAAGTCTTGAGTGGAAAGGCTNNNAGATCGGAA |
| 56865_1 | TATGAGTGTGGAGTCGTTGCGATGATGTCTTCTTGCTGCATCCNNNAGATCGGAAG |
| 56872_1 | TATGAGTGTGGAGTCGTTGCCTGGTGGAGTGGATATAAGCCTCNNNAGATCGGAA |
| 56875_1 | TATGAGTGTGGAGTCGTTGCGAAAGATACCATGCAAAGGCCTGNNNAGATCGGAA |
| 56877_1 | TATGAGTGTGGAGTCGTTGCCAAAGTATGTCCAGGGAAGACCANNNAGATCGGAA |
| 56880_1 | TATGAGTGTGGAGTCGTTGCAGAGAGACGTATGGAGGGAAGAANNNAGATCGGA |
| 56888_1 | TATGAGTGTGGAGTCGTTGCAGAGATGTACAAAAGCTGACCGANNNAGATCGGAA |
| 56898_1 | TATGAGTGTGGAGTCGTTGCATGAGCAGCCATGTTATCATCCANNNAGATCGGAAG |
| 56916_1 | TATGAGTGTGGAGTCGTTGCCTGAGGTGGGAGGATTGTTTGAGNNNAGATCGGAA |
| 56927_1 | TATGAGTGTGGAGTCGTTGCTTTGGGAGTTGTCAGTGCATAGANNNAGATCGGAA |
| 56932_1 | TATGAGTGTGGAGTCGTTGCTATCTACGGAAAACAACGTCCCANNNAGATCGGAAG |
| 56934_1 | TATGAGTGTGGAGTCGTTGCGAGTAGAGAGGAAGGAGAGTGGTNNNAGATCGGA |
| 56942_1 | TATGAGTGTGGAGTCGTTGCCTAACCTGAAAGCACACCCAAGNNNAGATCGGAAGA |
| 56946_1 | TATGAGTGTGGAGTCGTTGCCTCACATCTGTAATCCCAGCACTNNNAGATCGGAAGA |
| 56956_1 | TATGAGTGTGGAGTCGTTGCCTGGTCTCAAACTCCTGACATCANNNAGATCGGAAG |
| 56958_1 | TATGAGTGTGGAGTCGTTGCCAGGTTCAAGCAATTCTCCTGCNNNAGATCGGAAGA |

56959_1 TATGAGTGTGGAGTCGTTGCTCAACCTCCTGAGTAGCTGAGANNNAGATCGGAAGA
56960_1 TATGAGTGTGGAGTCGTTGCGTTCAAGCGATTCTCATGCCTCNNNAGATCGGAAGA
56965_1 TATGAGTGTGGAGTCGTTGCAGCATGCCCGGCTATTTTTTTTNNNAGATCGGAAGA
56973_1 TATGAGTGTGGAGTCGTTGCCTTTCTTTGTTGCCCAGGCTACNNNAGATCGGAAGA
56974_1 TATGAGTGTGGAGTCGTTGCTTCAAGTGATTCTCTCCCTCAGCNNNAGATCGGAAGA
56979_1 TATGAGTGTGGAGTCGTTGCATGATCTCAGGTCACTGCAACCNNNAGATCGGAAGA
56980_1 TATGAGTGTGGAGTCGTTGCGTTCACCATTCTCCTGCCTCAGNNNAGATCGGAAGAG
56987_1 TATGAGTGTGGAGTCGTTGCCCAAGTGATCCTCCCACCTTAGNNNAGATCGGAAGA
56993_1 TATGAGTGTGGAGTCGTTGCCTCCTGGGTTCAAGCGATTCTCNNNAGATCGGAAGA
57002_1 TATGAGTGTGGAGTCGTTGCCAGGCTGGTCTCAAACTCTTGANNNAGATCGGAAGA
57003_1 TATGAGTGTGGAGTCGTTGCAACTACAGCTCACTGCAACCTCNNNAGATCGGAAGA
57021_1 TATGAGTGTGGAGTCGTTGCAACTTCAGCCCTGTAGCAGATACNNNAGATCGGAAG
57025_1 TATGAGTGTGGAGTCGTTGCGGGCTGTTTAGCAACACAAAGTTNNNAGATCGGAA
57027_1 TATGAGTGTGGAGTCGTTGCCACCCATAATCCCAGCTACTCGNNNAGATCGGAAGA
57035_1 TATGAGTGTGGAGTCGTTGCAGCTTTTGGAGTGGTGTACTTCTNNNAGATCGGAAG
57042_1 TATGAGTGTGGAGTCGTTGCTTCAGATCCCAGCACATGTCATTNNNAGATCGGAAGA
57069_1 TATGAGTGTGGAGTCGTTGCCATGCTTGTAATCCCAGCACTTTNNNAGATCGGAAGA
57071_1 TATGAGTGTGGAGTCGTTGCTAGTGGGCAAAACTCTAAGGAGGNNNAGATCGGAA
57075_1 TATGAGTGTGGAGTCGTTGCAGTTGTCAAAGGATATGGACCCCNNNAGATCGGAA
57078_1 TATGAGTGTGGAGTCGTTGCGGGATGACACATGCCTGAAATTTNNNAGATCGGAA
57080_1 TATGAGTGTGGAGTCGTTGCGTCACTAACTGGCTTGATCCTCANNNAGATCGGAAGA
57086_1 TATGAGTGTGGAGTCGTTGCCTGTTCTTGTGTCTCAGCCTCCNNNAGATCGGAAGAG
57091_1 TATGAGTGTGGAGTCGTTGCGCAATGGCACAATCTCTACTCACNNNAGATCGGAAGA
57107_1 TATGAGTGTGGAGTCGTTGCGTCTTGAACTCCTGACCTTGTGANNNAGATCGGAAGA
57111_1 TATGAGTGTGGAGTCGTTGCAGCCTCCCAAAGTGCTGATATTANNNAGATCGGAAG
57117_1 TATGAGTGTGGAGTCGTTGCTCCTGAGTAGCTGGGATTACAGANNNAGATCGGAA
57120_1 TATGAGTGTGGAGTCGTTGCCTCTCAAGTAGCTGGGACTACAGNNNAGATCGGAAG
57121_1 TATGAGTGTGGAGTCGTTGCCATGATCTCAGCTCACTGCAACNNNAGATCGGAAGA
57125_1 TATGAGTGTGGAGTCGTTGCGAGTTTCACTGTTGTTGCCCAGNNNAGATCGGAAGA
57126_1 TATGAGTGTGGAGTCGTTGCGTTTCACCATGTTAGCCAGGATGNNNAGATCGGAAG
57133_1 TATGAGTGTGGAGTCGTTGCAAATATCTCACCCCACTCACCAGNNNAGATCGGAAGA
57139_1 TATGAGTGTGGAGTCGTTGCATTTGCTGGGGGAACACTATGATNNNAGATCGGAA
57142_1 TATGAGTGTGGAGTCGTTGCGGACGCAATCATACAAAAACAGCNNNAGATCGGAA
57143_1 TATGAGTGTGGAGTCGTTGCCTGAGATCACGCCACTGAACTCNNNAGATCGGAAGA
57154_1 TATGAGTGTGGAGTCGTTGCCTTGAGCCCAGGAGTTCAAGACNNNAGATCGGAAGA
57165_1 TATGAGTGTGGAGTCGTTGCTTGCAGTGAGCCAAGATCATACTNNNAGATCGGAAG
57170_1 TATGAGTGTGGAGTCGTTGCAGGCTTCCATTAGAGTGTAGTCGNNNAGATCGGAA
57178_1 TATGAGTGTGGAGTCGTTGCTTCAAGCAATTCTCCTGCCTCTNNNAGATCGGAAGAG
57189_1 TATGAGTGTGGAGTCGTTGCGATCTTGGCTCACTGCAAGCTCNNNAGATCGGAAGA
57192_1 TATGAGTGTGGAGTCGTTGCTTCACGTCATTCTCCTGCCTCNNNAGATCGGAAGAGC
57199_1 TATGAGTGTGGAGTCGTTGCGACAAGTGTCTTGCTCTGTTGCNNNAGATCGGAAGA
57205_1 TATGAGTGTGGAGTCGTTGCCATTCTCCTGCCTCAGCCTCNNNAGATCGGAAGAGCA
57206_1 TATGAGTGTGGAGTCGTTGCCCTCCTGAGTTCATGCCATTCTNNNAGATCGGAAGA

| | |
|---|---|
| 57221_1 | TATGAGTGTGGAGTCGTTGCACCTCTACATATGCACACAGAAAANNNAGATCGGAA |
| 57236_1 | TATGAGTGTGGAGTCGTTGCCGACAAAAATGACGTTACCACCANNNAGATCGGAAG |
| 57241_1 | TATGAGTGTGGAGTCGTTGCAGCTGAGTACATGATGATGCTGANNNAGATCGGAA |
| 57242_1 | TATGAGTGTGGAGTCGTTGCTTCAGCCTTAGAGGAAGTTGACANNNAGATCGGAA |
| 57243_1 | TATGAGTGTGGAGTCGTTGCTTCTCACTACATTTCCAGTGTGTNNNAGATCGGAAG |
| 57249_1 | TATGAGTGTGGAGTCGTTGCTGTGCCATGCATGTGAATATCATNNNAGATCGGAAG |
| 57255_1 | TATGAGTGTGGAGTCGTTGCCCCCAGGAGGAATCAGACATTAGNNNAGATCGGAA |
| 57256_1 | TATGAGTGTGGAGTCGTTGCCACCTGTAATCCGAGCTACTCAANNNAGATCGGAAG |
| 57265_1 | TATGAGTGTGGAGTCGTTGCCCAAAATGCTGGGATTACAGGTGNNNAGATCGGAA |
| 57277_1 | TATGAGTGTGGAGTCGTTGCCACAATCTTGGCTCACTGCAACNNNAGATCGGAAGA |
| 57287_1 | TATGAGTGTGGAGTCGTTGCGTTCACACCATACTCCTGCCTCNNNAGATCGGAAGAG |
| 57291_1 | TATGAGTGTGGAGTCGTTGCTGGAGTACAGTGGTGCAATCTCNNNAGATCGGAAG |
| 57295_1 | TATGAGTGTGGAGTCGTTGCGATTGGGTCTCACTCTGTTGCCNNNAGATCGGAAGA |
| 57313_1 | TATGAGTGTGGAGTCGTTGCCGAGATCTTGGCTCACTGCAAGNNNAGATCGGAAG |
| 57314_1 | TATGAGTGTGGAGTCGTTGCCTCCCAGGTTCATGCCATTCTCNNNAGATCGGAAGA |
| 57326_1 | TATGAGTGTGGAGTCGTTGCCACAATCTTGGCTCACTGCAACNNNAGATCGGAAGA |
| 57328_1 | TATGAGTGTGGAGTCGTTGCGATCCTGTCTCACTGCAAGCTCNNNAGATCGGAAGA |
| 57334_1 | TATGAGTGTGGAGTCGTTGCAGGCAGAAATCAGAGTGACTCAGNNNAGATCGGAA |
| 57336_1 | TATGAGTGTGGAGTCGTTGCGTGATGTAGCAAACCCAAGACACNNNAGATCGGAA |
| 57360_1 | TATGAGTGTGGAGTCGTTGCGACTTGTTGAATGGCTTTGACCANNNAGATCGGAAG |
| 57368_1 | TATGAGTGTGGAGTCGTTGCCATCTGTGGTCCCAGCTACTTGNNNAGATCGGAAGA |
| 57384_1 | TATGAGTGTGGAGTCGTTGCGGCTGGAGGTTCACAGAGTATTTNNNAGATCGGAA |
| 57387_1 | TATGAGTGTGGAGTCGTTGCGACGGTTCTTTCTCAGTCCTCATNNNAGATCGGAAG |
| 57393_1 | TATGAGTGTGGAGTCGTTGCCATTGTCTCACAAAGGTCATCCANNNAGATCGGAAG |
| 57398_1 | TATGAGTGTGGAGTCGTTGCTTTCTGCTCAATGGTGCATTCAGNNNAGATCGGAAG |
| 57405_1 | TATGAGTGTGGAGTCGTTGCGACTCACACTTCTGCCCATGATNNNAGATCGGAAGA |
| 57414_1 | TATGAGTGTGGAGTCGTTGCTCCTAGGATAAGCCCACCTTAGCNNNAGATCGGAAG |
| 57421_1 | TATGAGTGTGGAGTCGTTGCGATCTTGGCTCACTGCAAGCTCNNNAGATCGGAAGA |
| 57427_1 | TATGAGTGTGGAGTCGTTGCCAAGTAGCTGGGACTACAGGTGNNNAGATCGGAAG |
| 57435_1 | TATGAGTGTGGAGTCGTTGCCTCCTGGGTTCATGCCATTCTCNNNAGATCGGAAGA |
| 57439_1 | TATGAGTGTGGAGTCGTTGCAAGTAGCTGGGATTACAGGCACNNNAGATCGGAAG |
| 57440_1 | TATGAGTGTGGAGTCGTTGCCTTCTGGGTTCACGCCATTCTCNNNAGATCGGAAGA |
| 57463_1 | TATGAGTGTGGAGTCGTTGCGTACAATCTCGGCTCACTGCAANNNAGATCGGAAGA |
| 57464_1 | TATGAGTGTGGAGTCGTTGCGTTTCACCATGTTAGCCAGGATGNNNAGATCGGAAG |
| 57465_1 | TATGAGTGTGGAGTCGTTGCAACTTCTGGCCTCAAGCAGTTCNNNAGATCGGAAGA |
| 57487_1 | TATGAGTGTGGAGTCGTTGCCAGGAGGATCACTTGAACCCAGNNNAGATCGGAAG |
| 57498_1 | TATGAGTGTGGAGTCGTTGCGCTGCCAAGTTAGCAATGAATGTNNNAGATCGGAA |
| 57499_1 | TATGAGTGTGGAGTCGTTGCGGTCTGGGTGAATGTAGTTCTGTNNNAGATCGGAA |
| 57504_1 | TATGAGTGTGGAGTCGTTGCACAGCGCACCAGGAGATTATATCNNNAGATCGGAA |
| 57508_1 | TATGAGTGTGGAGTCGTTGCGTGATTGCATGCTTGGAGATAGCNNNAGATCGGAA |
| 57514_1 | TATGAGTGTGGAGTCGTTGCACAAGGTTACCTCAGGCATAAGANNNAGATCGGAA |
| 57521_1 | TATGAGTGTGGAGTCGTTGCGCAGCAGGAAAGAGAAAAGAACANNNAGATCGGA |
| 57525_1 | TATGAGTGTGGAGTCGTTGCAGCAGAAGGTTTTGCATTTCTCTNNNAGATCGGAAG |

| | |
|---|---|
| 57529_1 | TATGAGTGTGGAGTCGTTGCGGGCATGAGAATTGCTTGAACATNNNAGATCGGAA |
| 57532_1 | TATGAGTGTGGAGTCGTTGCGGTTGCAGTGAGCTGAAATCAGNNNAGATCGGAAG |
| 57535_1 | TATGAGTGTGGAGTCGTTGCTGATCACACCACTGCAGTTCAGNNNAGATCGGAAGA |
| 57536_1 | TATGAGTGTGGAGTCGTTGCAGAAATTCTGGGAGAGGATTCCANNNAGATCGGAA |
| 57539_1 | TATGAGTGTGGAGTCGTTGCAGAGGTGGTTGTTTCTACACACTNNNAGATCGGAAG |
| 57541_1 | TATGAGTGTGGAGTCGTTGCACCAAAATTGCTTCATCCAGACANNNAGATCGGAAG |
| 57557_1 | TATGAGTGTGGAGTCGTTGCATGCCTGTAATCCCAGCACTTTNNNAGATCGGAAGA |
| 57558_1 | TATGAGTGTGGAGTCGTTGCCGCCTATAATCCCAGCTACTCGNNNAGATCGGAAGA |
| 57589_1 | TATGAGTGTGGAGTCGTTGCTTTGGCTTTGGGAAGTGCTTTGNNNAGATCGGAAG |
| 57594_1 | TATGAGTGTGGAGTCGTTGCCTCCCAGGTTTACGCCATTCTCNNNAGATCGGAAGAG |
| 57595_1 | TATGAGTGTGGAGTCGTTGCCCTGATTTCAGCTCACTGCAAGNNNAGATCGGAAGA |
| 57602_1 | TATGAGTGTGGAGTCGTTGCCTCCCAGGTTCATGCCATTCTCNNNAGATCGGAAGAG |
| 57605_1 | TATGAGTGTGGAGTCGTTGCATGATCTCGGCTCACTGCAATCNNNAGATCGGAAGA |
| 57607_1 | TATGAGTGTGGAGTCGTTGCAGATGGAGTCTCGCTTTGTCTCNNNAGATCGGAAGA |
| 57608_1 | TATGAGTGTGGAGTCGTTGCGTGCCTGTAATCTCAGCTACTCANNNAGATCGGAAG |
| 57610_1 | TATGAGTGTGGAGTCGTTGCACTTGGTGAAACCTCGTCTCTACNNNAGATCGGAAG |
| 57614_1 | TATGAGTGTGGAGTCGTTGCGGGAGAGGAGTGAAAGGAGAGTANNNAGATCGG |
| 57615_1 | TATGAGTGTGGAGTCGTTGCTTTTTGAGACAGAATCTTGCGGCNNNAGATCGGAAG |
| 57626_1 | TATGAGTGTGGAGTCGTTGCCCAGCCTAGGGGTTGTTGTAATNNNAGATCGGAAG |
| 57641_1 | TATGAGTGTGGAGTCGTTGCATTGCATTGTTGTACCCAGTGAGNNNAGATCGGAAG |
| 57657_1 | TATGAGTGTGGAGTCGTTGCCATGCTTGTAGTCCCAGCTACTCNNNAGATCGGAAG |
| 57663_1 | TATGAGTGTGGAGTCGTTGCCATCAGCAGGGAAAACACCATTTNNNAGATCGGAAG |
| 57669_1 | TATGAGTGTGGAGTCGTTGCCACGATCTTGGCTCACTGAAACNNNAGATCGGAAGA |
| 57680_1 | TATGAGTGTGGAGTCGTTGCCTCCCATGTTCGAGCGATTCTCNNNAGATCGGAAGA |
| 57695_1 | TATGAGTGTGGAGTCGTTGCCCTCCCAGGTTCAAGAGATTCTCNNNAGATCGGAAG |
| 57697_1 | TATGAGTGTGGAGTCGTTGCCATCTAGGCTCACTGCAAGCTCNNNAGATCGGAAGA |
| 57698_1 | TATGAGTGTGGAGTCGTTGCCAGTGGGGTGATCTCAGTTTCCNNNAGATCGGAAGA |
| 57702_1 | TATGAGTGTGGAGTCGTTGCCATATCGGCTAACTGCAAGCTCNNNAGATCGGAAGA |
| 57706_1 | TATGAGTGTGGAGTCGTTGCCTCCCAGGTTCACACCATTCTCNNNAGATCGGAAGAG |
| 57708_1 | TATGAGTGTGGAGTCGTTGCCGACATACATATGGCCCAGGACNNNAGATCGGAAGA |
| 57715_1 | TATGAGTGTGGAGTCGTTGCCAAGTAGCTGGGACTACAGGTGNNNAGATCGGAAG |
| 57720_1 | TATGAGTGTGGAGTCGTTGCGATCTTGGCTCACTGCAAGCTCNNNAGATCGGAAGA |
| 57726_1 | TATGAGTGTGGAGTCGTTGCGACGGAGTCTCATTCTGTCACCNNNAGATCGGAAGA |
| 57740_1 | TATGAGTGTGGAGTCGTTGCGATCACAAGGTCAGGAGATCGAGNNNAGATCGGAA |
| 57741_1 | TATGAGTGTGGAGTCGTTGCTTTCTTACAGTTCTGGAGGCTGGNNNAGATCGGAAG |
| 57756_1 | TATGAGTGTGGAGTCGTTGCAGAACCTCACTTTCCATGAGCAANNNAGATCGGAAG |
| 57757_1 | TATGAGTGTGGAGTCGTTGCGCCACATAGCACAGTACCAGATANNNAGATCGGAAG |
| 57760_1 | TATGAGTGTGGAGTCGTTGCAAATGTGGCCTAATTCTGAACGCNNNAGATCGGAAG |
| 57762_1 | TATGAGTGTGGAGTCGTTGCAGAGTGTCAGTCTCAAAACGACTNNNAGATCGGAAG |
| 57786_1 | TATGAGTGTGGAGTCGTTGCAGTGAACTAAGATGAGCCATGGANNNAGATCGGAA |
| 57788_1 | TATGAGTGTGGAGTCGTTGCATTCTACACCTCGTGGAAATGCTNNNAGATCGGAAG |
| 57810_1 | TATGAGTGTGGAGTCGTTGCCAATACTTTCAATGGAGCAGGGCNNNAGATCGGAA |
| 57818_1 | TATGAGTGTGGAGTCGTTGCATTCAAAGGGAGCTCTGGAATGTNNNAGATCGGAA |

| | |
|---|---|
| 57822_1 | TATGAGTGTGGAGTCGTTGCGTCAGTGTCAGACATCTCACCATNNNAGATCGGAAG |
| 57825_1 | TATGAGTGTGGAGTCGTTGCATGCCTGTAATCCCAGCACTTTNNNAGATCGGAAGA |
| 57838_1 | TATGAGTGTGGAGTCGTTGCGATCTCGGCTTACTGCAACCTCNNNAGATCGGAAGA |
| 57844_1 | TATGAGTGTGGAGTCGTTGCCTCTCTGAGTAGCTGGGACTACANNNAGATCGGAAG |
| 57855_1 | TATGAGTGTGGAGTCGTTGCCTCCTGAGTAGCTGGGACTACAGNNNAGATCGGAA |
| 57856_1 | TATGAGTGTGGAGTCGTTGCGATCTTGGCTCACTGCAAGCTCNNNAGATCGGAAGA |
| 57862_1 | TATGAGTGTGGAGTCGTTGCAATCTCAGCTCACTGCAACCTCNNNAGATCGGAAGA |
| 57871_1 | TATGAGTGTGGAGTCGTTGCCTCCCAAGTTCAGGCCATTCTCNNNAGATCGGAAGA |
| 57886_1 | TATGAGTGTGGAGTCGTTGCCTCCTGAGTAGCTGGGACTACANNNAGATCGGAAG |
| 57888_1 | TATGAGTGTGGAGTCGTTGCGGAGTCTCTCTCTGTCGCCTAGNNNAGATCGGAAGA |
| 57902_1 | TATGAGTGTGGAGTCGTTGCCAAGTAGCTGGGACTACAGGCNNNAGATCGGAAGA |
| 57905_1 | TATGAGTGTGGAGTCGTTGCAATCTCGGCTCACTGCAACTTCNNNAGATCGGAAGA |
| 57915_1 | TATGAGTGTGGAGTCGTTGCCATCTTGGCTCACTGCAACTTCNNNAGATCGGAAGA |
| 57921_1 | TATGAGTGTGGAGTCGTTGCCTTCAGTTGTCACCTCCATGTCTNNNAGATCGGAAGA |
| 57922_1 | TATGAGTGTGGAGTCGTTGCCAGGCTGGGAGAGTTTCCATATTNNNAGATCGGAA |
| 57923_1 | TATGAGTGTGGAGTCGTTGCCACCGTAATCCCAGCACTTTTGNNNAGATCGGAAGA |
| 57929_1 | TATGAGTGTGGAGTCGTTGCGAGAATGGCATGAACCTGGGAGNNNAGATCGGAA |
| 57936_1 | TATGAGTGTGGAGTCGTTGCTGGAGTGCAGTAACGTGATCTTGNNNAGATCGGAA |
| 57946_1 | TATGAGTGTGGAGTCGTTGCTACACACTTTGCATAAACTCGCANNNAGATCGGAAG |
| 57954_1 | TATGAGTGTGGAGTCGTTGCAGGTAGGAGAATTGCTTGAACCCNNNAGATCGGAA |
| 57962_1 | TATGAGTGTGGAGTCGTTGCTGAAAAGAGTGTGCTTTTTTTGAGTNNNAGATCGG |
| 57968_1 | TATGAGTGTGGAGTCGTTGCGTGCTTAGATTTCACAACTTCCGANNNAGATCGGAA |
| 57969_1 | TATGAGTGTGGAGTCGTTGCCTGGGTTCTAGACACAGTTGGAANNNAGATCGGAA |
| 57972_1 | TATGAGTGTGGAGTCGTTGCTTGCATGCTTAGCTCACCTCTTANNNAGATCGGAAGA |
| 57974_1 | TATGAGTGTGGAGTCGTTGCTCTTTCTTTCTCCTCTCTCTCTCNNNAGATCGGAAGA |
| 57982_1 | TATGAGTGTGGAGTCGTTGCCTACTTCTGAGGCAGGAGAATCGNNNAGATCGGAA |
| 57985_1 | TATGAGTGTGGAGTCGTTGCGGCATTCAGATCATCTCAACAGCNNNAGATCGGAAG |
| 57986_1 | TATGAGTGTGGAGTCGTTGCGTCAGCACACATCTAGGACATGANNNAGATCGGAA |
| 57992_1 | TATGAGTGTGGAGTCGTTGCGCCATCCCCAAACCTACTGTTATNNNAGATCGGAAGA |
| 57995_1 | TATGAGTGTGGAGTCGTTGCTAGCTGAGTGTGCTGGTACATGNNNAGATCGGAAG |
| 58005_1 | TATGAGTGTGGAGTCGTTGCCACACACACACACACATATGACCNNNAGATCGGAAG |
| 58045_1 | TATGAGTGTGGAGTCGTTGCTGAGACGGAGTCTTGCTTTGTCNNNAGATCGGAAG |
| 58050_1 | TATGAGTGTGGAGTCGTTGCGAGACGGAATCTTGCTCTGTCANNNAGATCGGAAG |
| 58052_1 | TATGAGTGTGGAGTCGTTGCAGTGCTAGGATTACAGGCATGAGNNNAGATCGGAA |
| 58064_1 | TATGAGTGTGGAGTCGTTGCATCAGAGAAACTCACTTGGCAGTNNNAGATCGGAA |
| 58097_1 | TATGAGTGTGGAGTCGTTGCGTAAAAAAGCAAGCGTACGGAGTNNNAGATCGGAA |
| 58098_1 | TATGAGTGTGGAGTCGTTGCCTGAGTTCAGGAGTTCGAGTCCNNNAGATCGGAAG |
| 58105_1 | TATGAGTGTGGAGTCGTTGCTGGATAGCAGAAGGTAGCTCATCNNNAGATCGGAA |
| 58106_1 | TATGAGTGTGGAGTCGTTGCATGAAATGGAAGCAGGTCTCTGTNNNAGATCGGAA |
| 58107_1 | TATGAGTGTGGAGTCGTTGCGAGGCAGAAGAATCGCTTGAATCNNNAGATCGGAA |
| 58123_1 | TATGAGTGTGGAGTCGTTGCTTTGGAAATGTGTAACAGGCAGGNNNAGATCGGAA |
| 58132_1 | TATGAGTGTGGAGTCGTTGCGATCTCCACTCACTGCAACCTCNNNAGATCGGAAGAG |
| 58138_1 | TATGAGTGTGGAGTCGTTGCGAGTAGCTGGGATTACAGGCACNNNAGATCGGAAG |

| | |
|---|---|
| 58142_1 | TATGAGTGTGGAGTCGTTGCCTCCTGGGTTCAAGCAGTTCTCNNNAGATCGGAAGA |
| 58153_1 | TATGAGTGTGGAGTCGTTGCGAGACAGTCTCACTCTTGTCGCNNNAGATCGGAAGA |
| 58154_1 | TATGAGTGTGGAGTCGTTGCTTACCCAGGCTGGAGTTCAATGNNNAGATCGGAAG |
| 58155_1 | TATGAGTGTGGAGTCGTTGCTTCCCAGGTTCAAGCTATTCTCCNNNAGATCGGAAG |
| 58156_1 | TATGAGTGTGGAGTCGTTGCGATTCAAGCGATTCTCTTGCCTCNNNAGATCGGAAG |
| 58158_1 | TATGAGTGTGGAGTCGTTGCGATCTTGGCTCACTGCAACCTCNNNAGATCGGAAGA |
| 58187_1 | TATGAGTGTGGAGTCGTTGCGATCTCGGCTCAATGCAAGCTCNNNAGATCGGAAGA |
| 58198_1 | TATGAGTGTGGAGTCGTTGCCCATTCTCCTGCCTCAGTCTTCNNNAGATCGGAAGAG |
| 58211_1 | TATGAGTGTGGAGTCGTTGCGAGTCTCGCTCTTTCACCTAGGNNNAGATCGGAAGA |
| 58220_1 | TATGAGTGTGGAGTCGTTGCCTCCCAGGTTCAAGTGATCCTCNNNAGATCGGAAGA |
| 58221_1 | TATGAGTGTGGAGTCGTTGCGAGATGGAGTCTGCTCTGTTGCNNNAGATCGGAAG |
| 58228_1 | TATGAGTGTGGAGTCGTTGCTGAACTCTCTAACCACTGCTCACNNNAGATCGGAAG |
| 58237_1 | TATGAGTGTGGAGTCGTTGCGGTAGGAAGGATAGCGTCTTGTTNNNAGATCGGAA |
| 58239_1 | TATGAGTGTGGAGTCGTTGCGTCTCACTATGTTGCTCAGGCTNNNAGATCGGAAGA |
| 58240_1 | TATGAGTGTGGAGTCGTTGCATGCCTGTAATCCCAGCACTTTNNNAGATCGGAAGA |
| 58246_1 | TATGAGTGTGGAGTCGTTGCAATAAAACCAACTCCTCGGTGGTNNNAGATCGGAAG |
| 58252_1 | TATGAGTGTGGAGTCGTTGCAGGGTTGTGGAAAAGGGATTTCANNNAGATCGGA |
| 58255_1 | TATGAGTGTGGAGTCGTTGCACCATGACGAAATCCAAAACCTGNNNAGATCGGAAG |
| 58267_1 | TATGAGTGTGGAGTCGTTGCTTCACAACCTCCTTGAATCTGGGNNNAGATCGGAAG |
| 58275_1 | TATGAGTGTGGAGTCGTTGCAAACAGGAACAGAATTGGTTGGANNNAGATCGGAA |
| 58279_1 | TATGAGTGTGGAGTCGTTGCTGGTGAGAACGTGGAGAAATAGGNNNAGATCGGA |
| 58282_1 | TATGAGTGTGGAGTCGTTGCTATAGAACTTGACCCACCACCTGNNNAGATCGGAAG |
| 58283_1 | TATGAGTGTGGAGTCGTTGCAAGAAATGCATCTCTCGCTCTCTNNNAGATCGGAAG |
| 58301_1 | TATGAGTGTGGAGTCGTTGCGAGAGGGTAACCTATGATGCAGGNNNAGATCGGA |
| 58316_1 | TATGAGTGTGGAGTCGTTGCACCTCATCAGAATGTCCTTTACTGTNNNAGATCGGAA |
| 58323_1 | TATGAGTGTGGAGTCGTTGCGGGCAACCTGTTGTCAGAATTTTNNNAGATCGGAAG |
| 58329_1 | TATGAGTGTGGAGTCGTTGCCCCCTCTATGTTGACTTGGTCATNNNAGATCGGAAG |
| 58336_1 | TATGAGTGTGGAGTCGTTGCGAGAGGGGCGGGATCTAAAAATANNNAGATCGGA |
| 58337_1 | TATGAGTGTGGAGTCGTTGCGATCCAGCATAAAGGGTAGAGGGNNNAGATCGGA |
| 58339_1 | TATGAGTGTGGAGTCGTTGCTGTGGGCCAGAAACTCAGATTTANNNAGATCGGAA |
| 58343_1 | TATGAGTGTGGAGTCGTTGCGAAAACGGGCTAAGGACATGAACNNNAGATCGGAA |
| 58344_1 | TATGAGTGTGGAGTCGTTGCCACTTGAGGTCAGGAGTTTGAGANNNAGATCGGAA |
| 58350_1 | TATGAGTGTGGAGTCGTTGCAAGTGTTGGGATTACAGGTGTGANNNAGATCGGAA |
| 58364_1 | TATGAGTGTGGAGTCGTTGCCTCACTGCAATCTCTGTCTCCCNNNAGATCGGAAGAG |
| 58366_1 | TATGAGTGTGGAGTCGTTGCAAGTAGCTGGGATTACAGGCACNNNAGATCGGAAG |
| 58369_1 | TATGAGTGTGGAGTCGTTGCCAGGTTCAAGCAATTCTCCTGCNNNAGATCGGAAGA |
| 58377_1 | TATGAGTGTGGAGTCGTTGCGATATTGGCTCACTGCAAGTTCCNNNAGATCGGAAG |
| 58393_1 | TATGAGTGTGGAGTCGTTGCCCCTAGTAGCTGGGACTACAGGNNNAGATCGGAAG |
| 58397_1 | TATGAGTGTGGAGTCGTTGCCTCCAGGGTTCAAGTGAGTCTCNNNAGATCGGAAG |
| 58400_1 | TATGAGTGTGGAGTCGTTGCCAAGTAGCTGGGACTACAGGTGNNNAGATCGGAAG |
| 58402_1 | TATGAGTGTGGAGTCGTTGCTGATCTCGGCTCACTGCAAGNNNAGATCGGAAGAG |
| 58416_1 | TATGAGTGTGGAGTCGTTGCTCTTGAGGCCAGGAGTTCTAAACNNNAGATCGGAA |
| 58420_1 | TATGAGTGTGGAGTCGTTGCTGGCATTTGTAGCAACTTGGATGNNNAGATCGGAA |

| | |
|---|---|
| 58429_1 | TATGAGTGTGGAGTCGTTGCGAGATTGCAGTCAGCCGAAATCNNNAGATCGGAAG |
| 58444_1 | TATGAGTGTGGAGTCGTTGCAGCACTTTCTAACGATGCCCATANNNAGATCGGAAG |
| 58486_1 | TATGAGTGTGGAGTCGTTGCCTCCCGGGTTCAAGAGATTCTCNNNAGATCGGAAGA |
| 58494_1 | TATGAGTGTGGAGTCGTTGCAGATGAAGTCTCACTGTGTTGCCNNNAGATCGGAAG |
| 58511_1 | TATGAGTGTGGAGTCGTTGCCTCAGGTCCTCAGCTGGGATTANNNAGATCGGAAGA |
| 58523_1 | TATGAGTGTGGAGTCGTTGCCCCGGGATGACCTGAACTAATTANNNAGATCGGAAG |
| 58536_1 | TATGAGTGTGGAGTCGTTGCGGTAGGCAGCTGTTCTTTTTTGTNNNAGATCGGAAG |
| 58542_1 | TATGAGTGTGGAGTCGTTGCGAGGCAAGAGAATGGCATGAACNNNAGATCGGAA |
| 58543_1 | TATGAGTGTGGAGTCGTTGCCCAGGAGTTCAAGGCTACAATGANNNAGATCGGAA |
| 58550_1 | TATGAGTGTGGAGTCGTTGCCGAATAGCTGGGACTACAGGTGNNNAGATCGGAAG |
| 58553_1 | TATGAGTGTGGAGTCGTTGCAATCTCAGCTCACTGCAACCTCNNNAGATCGGAAGA |
| 58559_1 | TATGAGTGTGGAGTCGTTGCCAGGTTCAAGCGATTCTCTTGCNNNAGATCGGAAGA |
| 58561_1 | TATGAGTGTGGAGTCGTTGCCAATCTCGGCTCACTGCAAGATNNNAGATCGGAAGA |
| 58567_1 | TATGAGTGTGGAGTCGTTGCCACAATCTTGGCTCACTGCAAGNNNAGATCGGAAGA |
| 58585_1 | TATGAGTGTGGAGTCGTTGCAGGAAGGTTTGGGAAAAATGTGANNNAGATCGGA |
| 58586_1 | TATGAGTGTGGAGTCGTTGCTCTCCATTCTATTGCTGATGGGCNNNAGATCGGAAG |
| 58592_1 | TATGAGTGTGGAGTCGTTGCGTGCACCTGTAGTCTCAGCTATTNNNAGATCGGAAG |
| 58607_1 | TATGAGTGTGGAGTCGTTGCCTGAGGCAGGAGAATCGCTATCNNNAGATCGGAAG |
| 58627_1 | TATGAGTGTGGAGTCGTTGCGAGACAGAGTCTTGCTGTGTCANNNAGATCGGAAG |
| 58636_1 | TATGAGTGTGGAGTCGTTGCGAGCCACCTCAGCTTTCTTTTTTNNNAGATCGGAAGA |
| 58637_1 | TATGAGTGTGGAGTCGTTGCCACTGCTGAGTTCAGGCTATCAGNNNAGATCGGAAG |
| 58642_1 | TATGAGTGTGGAGTCGTTGCATTGAGGGAGAAAAGGGGAAGACNNNAGATCGGA |
| 58644_1 | TATGAGTGTGGAGTCGTTGCATCTCGTATAAGCTCAGTGGCTGNNNAGATCGGAAG |
| 58652_1 | TATGAGTGTGGAGTCGTTGCGCCTTTGTTGGGAGAATGAAGAGNNNAGATCGGAA |
| 58667_1 | TATGAGTGTGGAGTCGTTGCCAGTCGGAAAGCAAAGGAGGATNNNAGATCGGAA |
| 58679_1 | TATGAGTGTGGAGTCGTTGCGAAATTTTACCACGTGGCCCAGNNNAGATCGGAAGA |
| 58686_1 | TATGAGTGTGGAGTCGTTGCAACCTTTGCCTCTTGGGTTCAANNNAGATCGGAAGA |
| 58688_1 | TATGAGTGTGGAGTCGTTGCGTTCATGCCATTCTCCTGCTTCNNNAGATCGGAAGAG |
| 58694_1 | TATGAGTGTGGAGTCGTTGCCTCCCAGGTTCACACCATTCTCNNNAGATCGGAAGAG |
| 58699_1 | TATGAGTGTGGAGTCGTTGCCTCCCAGGTTCAAGCGATTCTCNNNAGATCGGAAGA |
| 58700_1 | TATGAGTGTGGAGTCGTTGCATGAGAGATGGGGTTTTGCCATNNNAGATCGGAAG |
| 58712_1 | TATGAGTGTGGAGTCGTTGCATTTTCCTGGGAGGTAAGCATGTNNNAGATCGGAA |
| 58716_1 | TATGAGTGTGGAGTCGTTGCGAATGGACAACATAAGAAGGCCCNNNAGATCGGAA |
| 58717_1 | TATGAGTGTGGAGTCGTTGCAACTCTCCTATTCATGGTCTGCCNNNAGATCGGAAGA |
| 58730_1 | TATGAGTGTGGAGTCGTTGCCAACAAGAACCAAACTGGGAGTCNNNAGATCGGAA |
| 58735_1 | TATGAGTGTGGAGTCGTTGCGAGGATCACTTGAACCTGGGAGNNNAGATCGGAAG |
| 58736_1 | TATGAGTGTGGAGTCGTTGCGAGGATTGCTTGAGACTGGGAGNNNAGATCGGAA |
| 58751_1 | TATGAGTGTGGAGTCGTTGCCTGGGTTCACACCATTCTCCTGNNNAGATCGGAAGA |
| 58760_1 | TATGAGTGTGGAGTCGTTGCCCCGAGTAGCTGGGACTACAGNNNAGATCGGAAGA |
| 58769_1 | TATGAGTGTGGAGTCGTTGCCCAAGAAGCTGGGACTACAGGNNNAGATCGGAAGA |
| 58787_1 | TATGAGTGTGGAGTCGTTGCGAATAGCTCGGACTATAGGCGCNNNAGATCGGAAG |
| 58801_1 | TATGAGTGTGGAGTCGTTGCGATTACAGGCACACACCACAACNNNAGATCGGAAGA |
| 58809_1 | TATGAGTGTGGAGTCGTTGCGATCTCGGCTCATTGCAAGCTCNNNAGATCGGAAGA |

| | |
|---|---|
| 58825_1 | TATGAGTGTGGAGTCGTTGCGAAGGTGAAAGGCATCTCACATGNNNAGATCGGAA |
| 58830_1 | TATGAGTGTGGAGTCGTTGCCAACACTGGTCTACTCTCTGCTTNNNAGATCGGAAGA |
| 58834_1 | TATGAGTGTGGAGTCGTTGCTTCCCCTTGCTGGCATGTATAATNNNAGATCGGAAG |
| 58838_1 | TATGAGTGTGGAGTCGTTGCTCAGCCTCTTTTTTTCCACATGCNNNAGATCGGAAGA |
| 58854_1 | TATGAGTGTGGAGTCGTTGCAAATAAAAAACGGGGCCTTGTGTNNNAGATCGGAA |
| 58857_1 | TATGAGTGTGGAGTCGTTGCATGCACCCCACACTAAACTGTATNNNAGATCGGAAG |
| 58866_1 | TATGAGTGTGGAGTCGTTGCCCGTATACCCACTCCTAGATCCTNNNAGATCGGAAGA |
| 58867_1 | TATGAGTGTGGAGTCGTTGCTCTCAGTATTGCAGCTACAGTGTNNNAGATCGGAAG |
| 58869_1 | TATGAGTGTGGAGTCGTTGCTCAAAAAACTGTGGATAAGCCCANNNAGATCGGAAG |
| 58873_1 | TATGAGTGTGGAGTCGTTGCTTAGTCAGGTGTGATGGTGCACNNNAGATCGGAAG |
| 58877_1 | TATGAGTGTGGAGTCGTTGCACTTATGTTATTGTGCATCAGCCCNNNAGATCGGAA |
| 58893_1 | TATGAGTGTGGAGTCGTTGCCCTGTAGTCCCAGCTACTCTCCNNNAGATCGGAAGAG |
| 58899_1 | TATGAGTGTGGAGTCGTTGCCTCAGTGGCTCACACCTGTAATCNNNAGATCGGAAG |
| 58900_1 | TATGAGTGTGGAGTCGTTGCAAAAGAAAGAAAGGGCCTGAAGCNNNAGATCGGA |
| 58912_1 | TATGAGTGTGGAGTCGTTGCGAGTCTCACTCTGTCGCTTAGGNNNAGATCGGAAGA |
| 58919_1 | TATGAGTGTGGAGTCGTTGCAGACGGAATCTCACTCTTTCGCNNNAGATCGGAAGA |
| 58923_1 | TATGAGTGTGGAGTCGTTGCCTCCCGGGTTCAAGCTATTCTCNNNAGATCGGAAGA |
| 58930_1 | TATGAGTGTGGAGTCGTTGCCGAGTAGCTGGGATTACAGGTGNNNAGATCGGAAG |
| 58939_1 | TATGAGTGTGGAGTCGTTGCCAGGAAGTGCAAAGACACACAGNNNAGATCGGAAG |
| 58942_1 | TATGAGTGTGGAGTCGTTGCCTGAGGTCAGGAGTTCGAGAACNNNAGATCGGAAG |
| 58955_1 | TATGAGTGTGGAGTCGTTGCCTTGGCCAGGGTGGTATATCAGNNNAGATCGGAAG |
| 58971_1 | TATGAGTGTGGAGTCGTTGCGCCAAGAGATCGAGACTATCCTGNNNAGATCGGAA |
| 58972_1 | TATGAGTGTGGAGTCGTTGCTACTCGTGGGTTTGGTGACAAGNNNAGATCGGAAG |
| 58976_1 | TATGAGTGTGGAGTCGTTGCCTCAATCTCCTGACCTGGTGATCNNNAGATCGGAAG |
| 58995_1 | TATGAGTGTGGAGTCGTTGCCTGAATAGCTGGGACTACAGGTGNNNAGATCGGAA |
| 59012_1 | TATGAGTGTGGAGTCGTTGCAAGTGAAAGAAAGTGGGGGGAATNNNAGATCGGA |
| 59013_1 | TATGAGTGTGGAGTCGTTGCGGAAGATTGGGGACCAGTATCTCNNNAGATCGGAA |
| 59014_1 | TATGAGTGTGGAGTCGTTGCATCATCAGGACCTAGGGCTTAGTNNNAGATCGGAAG |
| 59023_1 | TATGAGTGTGGAGTCGTTGCTTATGGAACCTTTACCACCAGCANNNAGATCGGAAG |
| 59026_1 | TATGAGTGTGGAGTCGTTGCAGCCTCAAAATATTCACCTGTCTNNNAGATCGGAAGA |
| 59032_1 | TATGAGTGTGGAGTCGTTGCCACACACACACACACACACAATANNNAGATCGGAAG |
| 59036_1 | TATGAGTGTGGAGTCGTTGCCTCCTGACCTCAAGTAATCCACCNNNAGATCGGAAGA |
| 59045_1 | TATGAGTGTGGAGTCGTTGCCAGAGCCAGATTCCGTCTCAAANNNAGATCGGAAGA |
| 59053_1 | TATGAGTGTGGAGTCGTTGCTGGTCCATGTGCAATAGGAATCANNNAGATCGGAA |
| 59054_1 | TATGAGTGTGGAGTCGTTGCAGAACTACCTGAGACTGGGAAGTNNNAGATCGGAA |
| 59055_1 | TATGAGTGTGGAGTCGTTGCATTGGAGCAACATGAATTCTGCTNNNAGATCGGAAG |
| 59058_1 | TATGAGTGTGGAGTCGTTGCGTTCAAGCGATTCTTCTGCCTCNNNAGATCGGAAGA |
| 59062_1 | TATGAGTGTGGAGTCGTTGCGTTCAAGCGATTCTTCTGCCTCNNNAGATCGGAAGA |
| 59067_1 | TATGAGTGTGGAGTCGTTGCCTACTGGGTTCACGCCATTCTCNNNAGATCGGAAGA |
| 59075_1 | TATGAGTGTGGAGTCGTTGCGATCTCCACTCACTGCAAGCTCNNNAGATCGGAAGA |
| 59099_1 | TATGAGTGTGGAGTCGTTGCCATGATCTTGGCTCACTGCAAGNNNAGATCGGAAGA |
| 59101_1 | TATGAGTGTGGAGTCGTTGCGATCTTGGCTCACTGCAAGCTANNNAGATCGGAAGA |
| 59104_1 | TATGAGTGTGGAGTCGTTGCTTGCCCAGTTGATCTCAGACTCNNNAGATCGGAAGA |

| | |
|---|---|
| 59106_1 | TATGAGTGTGGAGTCGTTGCAAACTCCCAGGTTCAAGCCATCNNNAGATCGGAAGA |
| 59109_1 | TATGAGTGTGGAGTCGTTGCGACTGCAGTGGTGCTATCTCTGNNNAGATCGGAAG |
| 59110_1 | TATGAGTGTGGAGTCGTTGCCATGGAGCTGGGATTACAGGTACNNNAGATCGGAA |
| 59113_1 | TATGAGTGTGGAGTCGTTGCCTACCGGGTTCATGCCATTCTCNNNAGATCGGAAGA |
| 59118_1 | TATGAGTGTGGAGTCGTTGCGAGTAACTGGGACTACAGGCACNNNAGATCGGAAG |
| 59121_1 | TATGAGTGTGGAGTCGTTGCAGAAGGAGTCTCACTCTGTTGCNNNAGATCGGAAG |
| 59123_1 | TATGAGTGTGGAGTCGTTGCCCCTGCGAGGTAGTCCTATTATTNNNAGATCGGAAG |
| 59127_1 | TATGAGTGTGGAGTCGTTGCCGTAATCTCGGCTCACTGCAAGNNNAGATCGGAAGA |
| 59130_1 | TATGAGTGTGGAGTCGTTGCGGAATGCAGTGGTGTGATCTTGNNNAGATCGGAAG |
| 59139_1 | TATGAGTGTGGAGTCGTTGCCTCCTAGGTTCACGCCATTCTCNNNAGATCGGAAGAG |
| 59148_1 | TATGAGTGTGGAGTCGTTGCCTGTAGTCCCAGCTACTAGGGANNNAGATCGGAAG |
| 59161_1 | TATGAGTGTGGAGTCGTTGCTCTTCTCAGTGGGTTGTACAGTTNNNAGATCGGAAG |
| 59164_1 | TATGAGTGTGGAGTCGTTGCTGTAGGCAATTCATCTGGGAACTNNNAGATCGGAAG |
| 59180_1 | TATGAGTGTGGAGTCGTTGCGAGATGGGGGTTTCATCATGTTGNNNAGATCGGAA |
| 59181_1 | TATGAGTGTGGAGTCGTTGCCATTGGTGGGGAATGGGTAAATTNNNAGATCGGAA |
| 59185_1 | TATGAGTGTGGAGTCGTTGCTCACACCTGTAATCCCAACACTTNNNAGATCGGAAGA |
| 59187_1 | TATGAGTGTGGAGTCGTTGCTGAGACAGAGTCTTGCTACCTTGNNNAGATCGGAAG |
| 59188_1 | TATGAGTGTGGAGTCGTTGCGAAGCTATCTCTAATTCCCGCCANNNAGATCGGAAG |
| 59189_1 | TATGAGTGTGGAGTCGTTGCTTCAGAGCTTCTTTCCAGTCACANNNAGATCGGAAG |
| 59191_1 | TATGAGTGTGGAGTCGTTGCTGGTGACAAGCCTGGATAAATGANNNAGATCGGAA |
| 59206_1 | TATGAGTGTGGAGTCGTTGCCATGTGTAATCCCAGTGCATTGGNNNAGATCGGAAG |
| 59222_1 | TATGAGTGTGGAGTCGTTGCTTCAAGCGATTCTCAGCCTCAGNNNAGATCGGAAGA |
| 59229_1 | TATGAGTGTGGAGTCGTTGCTGGAGTGCAGTGGTGAAATCTTNNNAGATCGGAAG |
| 59230_1 | TATGAGTGTGGAGTCGTTGCAGATGGAGTCTCGTTCTGTTGCNNNAGATCGGAAG |
| 59233_1 | TATGAGTGTGGAGTCGTTGCGATCTCCACTCACTGCAAGCTCNNNAGATCGGAAGA |
| 59244_1 | TATGAGTGTGGAGTCGTTGCCTTCTGGGTTCACGCCATTCTCNNNAGATCGGAAGAG |
| 59256_1 | TATGAGTGTGGAGTCGTTGCGAGATGGGGTTTCGCTCTTGTTNNNAGATCGGAAG |
| 59260_1 | TATGAGTGTGGAGTCGTTGCCAAGTAGCTGGGACTACAGGCNNNAGATCGGAAGA |
| 59268_1 | TATGAGTGTGGAGTCGTTGCGTTCAAGCGATTCTCTTGCTTCANNNAGATCGGAAG |
| 59270_1 | TATGAGTGTGGAGTCGTTGCCTCCAGGGTTCAAGCCATTCTCNNNAGATCGGAAGA |
| 59272_1 | TATGAGTGTGGAGTCGTTGCCTGGGTTCACGCCATTCTTTTGNNNAGATCGGAAGA |
| 59288_1 | TATGAGTGTGGAGTCGTTGCGAGTACAGTGGTGCGATCTCAGNNNAGATCGGAAG |
| 59307_1 | TATGAGTGTGGAGTCGTTGCGCAGCTTAGAAAGGTGACTATAGTNNNAGATCGGA |
| 59312_1 | TATGAGTGTGGAGTCGTTGCTCCACTACACAAGAACAGTGTGGNNNAGATCGGAAG |
| 59315_1 | TATGAGTGTGGAGTCGTTGCATCACTTGAACCCAGAAGGCATANNNAGATCGGAAG |
| 59318_1 | TATGAGTGTGGAGTCGTTGCTTTGGAATGCACCCATGAAGTTCNNNAGATCGGAAG |
| 59330_1 | TATGAGTGTGGAGTCGTTGCTAGCTGAGGGATGGAAAAATGCTNNNAGATCGGAA |
| 59335_1 | TATGAGTGTGGAGTCGTTGCAGTGAACTGAGATTGCCACTACANNNAGATCGGAAG |
| 59351_1 | TATGAGTGTGGAGTCGTTGCCGTTTGGAGATGCATGAATTCCANNNAGATCGGAAG |
| 59366_1 | TATGAGTGTGGAGTCGTTGCCAGTCAGTCAGTCTTGGGGAATTNNNAGATCGGAA |
| 59367_1 | TATGAGTGTGGAGTCGTTGCATGCCTGTAATCCCAGCACTTTNNNAGATCGGAAGA |
| 59376_1 | TATGAGTGTGGAGTCGTTGCCTGGGCAACAGAGTGAGATTCTNNNAGATCGGAAG |
| 59381_1 | TATGAGTGTGGAGTCGTTGCGTATACACACACACACACACACGNNNAGATCGGAAG |

| | |
|---|---|
| 59383_1 | TATGAGTGTGGAGTCGTTGCACATGGTCTGGGATGACATAACANNNAGATCGGAA |
| 59386_1 | TATGAGTGTGGAGTCGTTGCGAAGTCTGACCCAGGTCTCATGNNNAGATCGGAAGA |
| 59387_1 | TATGAGTGTGGAGTCGTTGCCATCTCTGAGCATTGACTTGCGNNNAGATCGGAAGA |
| 59389_1 | TATGAGTGTGGAGTCGTTGCCTCCAGGGTACATGCCATTCTCNNNAGATCGGAAGA |
| 59392_1 | TATGAGTGTGGAGTCGTTGCCAATCTCGGCTCACTGCAAGNNNAGATCGGAAGAGC |
| 59395_1 | TATGAGTGTGGAGTCGTTGCTCTCGATCTCCTGACCTCATGATNNNAGATCGGAAGA |
| 59403_1 | TATGAGTGTGGAGTCGTTGCCATTCTCCTGCCTCAGTCTCTCNNNAGATCGGAAGAG |
| 59404_1 | TATGAGTGTGGAGTCGTTGCGAGTAGCTGGAACTACAGGCACNNNAGATCGGAAG |
| 59414_1 | TATGAGTGTGGAGTCGTTGCTCCCATGTTCCTGTGAAGCTTATNNNAGATCGGAAGA |
| 59422_1 | TATGAGTGTGGAGTCGTTGCACCATAAGAACAGCATGAGGGAANNNAGATCGGAA |
| 59430_1 | TATGAGTGTGGAGTCGTTGCAAGGGGTAGGACTGGAAATACCTNNNAGATCGGAA |
| 59434_1 | TATGAGTGTGGAGTCGTTGCACCAGGAAGAGAAGGGGATAAGANNNAGATCGGA |
| 59435_1 | TATGAGTGTGGAGTCGTTGCTAAATGTTCCCTGTCAACCACACNNNAGATCGGAAGA |
| 59441_1 | TATGAGTGTGGAGTCGTTGCGAGACGAAAACTCACTGTTGCCNNNAGATCGGAAGA |
| 59448_1 | TATGAGTGTGGAGTCGTTGCGAGTAGCTGGGACTATTGGCACNNNAGATCGGAAG |
| 59459_1 | TATGAGTGTGGAGTCGTTGCGTTGGTCAGGATGGTCTCAATCTNNNAGATCGGAAG |
| 59472_1 | TATGAGTGTGGAGTCGTTGCCCCAAGTACCTGGGATTACAGGNNNAGATCGGAAGA |
| 59475_1 | TATGAGTGTGGAGTCGTTGCCACAATCTTGGGTCACTGCAAGNNNAGATCGGAAGA |
| 59476_1 | TATGAGTGTGGAGTCGTTGCGTTCAAGCGATTCTCCTGCTTCNNNAGATCGGAAGA |
| 59483_1 | TATGAGTGTGGAGTCGTTGCCGAGTAGCTGGGTTTACAGGTGNNNAGATCGGAAG |
| 59489_1 | TATGAGTGTGGAGTCGTTGCATCTTGGCTCACTGCAAGCTCNNNAGATCGGAAGAG |
| 59492_1 | TATGAGTGTGGAGTCGTTGCGACAGAGTCTCATTCTCTCGCCNNNAGATCGGAAGA |
| 59495_1 | TATGAGTGTGGAGTCGTTGCCTCACTGCAAACTCCACCTCCNNNAGATCGGAAGAGC |
| 59504_1 | TATGAGTGTGGAGTCGTTGCCTGGGTTCTCGCCATTCTACTGNNNAGATCGGAAGA |
| 59515_1 | TATGAGTGTGGAGTCGTTGCGCGAGTAGCTGGTACTACAGACNNNAGATCGGAAG |
| 59517_1 | TATGAGTGTGGAGTCGTTGCGTTCAAGCGATTCTCTTGCCTCNNNAGATCGGAAGA |
| 59525_1 | TATGAGTGTGGAGTCGTTGCCTCCCTGGTTCAAGCGATTCTCNNNAGATCGGAAGA |
| 59532_1 | TATGAGTGTGGAGTCGTTGCCTCCTGGGTTCATGCCATTCTCNNNAGATCGGAAGAG |
| 59534_1 | TATGAGTGTGGAGTCGTTGCCCAGAGATGGACATGGCATATGANNNAGATCGGAA |
| 59535_1 | TATGAGTGTGGAGTCGTTGCTCCAGTTTCCCCAACACCTTTTANNNAGATCGGAAGA |
| 59541_1 | TATGAGTGTGGAGTCGTTGCCTGAATGCAGTGGGGTAGACTTNNNAGATCGGAAG |
| 59550_1 | TATGAGTGTGGAGTCGTTGCAATTTCCCTCTTGAAGATGCCCTNNNAGATCGGAAGA |
| 59554_1 | TATGAGTGTGGAGTCGTTGCAGGAAGAAACAGTGTCTATGCCANNNAGATCGGAA |
| 59559_1 | TATGAGTGTGGAGTCGTTGCCAGCCTAGCTCAGTGAGAAACTNNNAGATCGGAAGA |
| 59563_1 | TATGAGTGTGGAGTCGTTGCAAGGTAAGGGATAAAGGCCTGTGNNNAGATCGGAA |
| 59568_1 | TATGAGTGTGGAGTCGTTGCTGCTTGCATCTTCTCCTTCTCTTNNNAGATCGGAAGA |
| 59571_1 | TATGAGTGTGGAGTCGTTGCCCTGTAATCCCATAATCCCAGCANNNAGATCGGAAGA |
| 59576_1 | TATGAGTGTGGAGTCGTTGCCACACACACACACACACACATATNNNAGATCGGAAGA |
| 59578_1 | TATGAGTGTGGAGTCGTTGCGATATAAGAGGCAGGGCAGGATTNNNAGATCGGAA |
| 59589_1 | TATGAGTGTGGAGTCGTTGCCACAGGAGGAGTCATCACCTTTANNNAGATCGGAAG |
| 59591_1 | TATGAGTGTGGAGTCGTTGCTATGAAGAGACACCCAACCTCACNNNAGATCGGAAG |
| 59608_1 | TATGAGTGTGGAGTCGTTGCGGAATAACCACCTCTGCAAGGATNNNAGATCGGAAG |
| 59616_1 | TATGAGTGTGGAGTCGTTGCACCAATCCACCATCATGATACGTNNNAGATCGGAAGA |

| | |
|---|---|
| 59621_1 | TATGAGTGTGGAGTCGTTGCTGCAAACAGAGGGACAATTTGACNNNAGATCGGAA |
| 59635_1 | TATGAGTGTGGAGTCGTTGCGAGCAAACAAACCCCAAAGCTATNNNAGATCGGAAG |
| 59636_1 | TATGAGTGTGGAGTCGTTGCTGCGGCTTCTTCCATATTCCATANNNAGATCGGAAGA |
| 59646_1 | TATGAGTGTGGAGTCGTTGCTTTGCAGAAGGGGAAGGAAAAACNNNAGATCGGA |
| 59656_1 | TATGAGTGTGGAGTCGTTGCGAACTCTGGGCCTCAAGTGATCNNNAGATCGGAAG |
| 59669_1 | TATGAGTGTGGAGTCGTTGCGTTCAAACCATTTTCCTGCCTCANNNAGATCGGAAGA |
| 59670_1 | TATGAGTGTGGAGTCGTTGCCAGTGTTGCAATCATCGCTCACNNNAGATCGGAAGA |
| 59673_1 | TATGAGTGTGGAGTCGTTGCATCTCCTGGGTTCAAGCACTTCNNNAGATCGGAAGA |
| 59675_1 | TATGAGTGTGGAGTCGTTGCTACTGAGCCCGGCAAATATCAATNNNAGATCGGAAG |
| 59679_1 | TATGAGTGTGGAGTCGTTGCTTTCACCGTGTTAGCCAGGATAGNNNAGATCGGAAG |
| 59693_1 | TATGAGTGTGGAGTCGTTGCCTCCTGGGTTCACACCATTCTCNNNAGATCGGAAGAG |
| 59694_1 | TATGAGTGTGGAGTCGTTGCCTCCCAGGTTCCAGCTATTCTCNNNAGATCGGAAGAG |
| 59709_1 | TATGAGTGTGGAGTCGTTGCGATCTCGACTCACTGCAAGCTCNNNAGATCGGAAGA |
| 59711_1 | TATGAGTGTGGAGTCGTTGCGACAAAGTCTCACCCTGTCTCCNNNAGATCGGAAGA |
| 59715_1 | TATGAGTGTGGAGTCGTTGCCTCCCTGGTTCAAGCGATTCTCNNNAGATCGGAAGA |
| 59722_1 | TATGAGTGTGGAGTCGTTGCCTGTAGTCCTAGCTGCTTGGAAANNNAGATCGGAAG |
| 59723_1 | TATGAGTGTGGAGTCGTTGCGTCACTCATGGTCAGGGATTTCTNNNAGATCGGAAG |
| 59724_1 | TATGAGTGTGGAGTCGTTGCAATCTCAGTACTTTGGGAGGCTGNNNAGATCGGAA |
| 59726_1 | TATGAGTGTGGAGTCGTTGCCAATAGAGCTAGACGCCGTCTCNNNAGATCGGAAGA |
| 59728_1 | TATGAGTGTGGAGTCGTTGCCACACCTGTAATCCCAGCACTTTNNNAGATCGGAAGA |
| 59731_1 | TATGAGTGTGGAGTCGTTGCCTTCCTCCTTTCCTCTGACCACNNNAGATCGGAAGAG |
| 59737_1 | TATGAGTGTGGAGTCGTTGCTTGAAGGATGAGCAAGCGAATTCNNNAGATCGGAA |
| 59739_1 | TATGAGTGTGGAGTCGTTGCAGAGAGATTAGCTGAGCATGGTGNNNAGATCGGA |
| 59745_1 | TATGAGTGTGGAGTCGTTGCCTCTGGGCTACACAGTAAGATCCNNNAGATCGGAAG |
| 59757_1 | TATGAGTGTGGAGTCGTTGCAACAGCTGCCTCCTCTATTCAAANNNAGATCGGAAG |
| 59762_1 | TATGAGTGTGGAGTCGTTGCCCATTGAACGCTCAAAGACACAANNNAGATCGGAAG |
| 59764_1 | TATGAGTGTGGAGTCGTTGCTCCATTGCATAACTACACCACCANNNAGATCGGAAGA |
| 59765_1 | TATGAGTGTGGAGTCGTTGCTAATCTCAGCACTTTGTGAGGCTNNNAGATCGGAAG |
| 59766_1 | TATGAGTGTGGAGTCGTTGCAGACAAGCGGGAAGGTAAAAAGANNNAGATCGGA |
| 59773_1 | TATGAGTGTGGAGTCGTTGCATTTGAGGTCAGGAGTTCGAGACNNNAGATCGGAA |
| 59777_1 | TATGAGTGTGGAGTCGTTGCTGGGTGACAAAAGAAAGCTCCTANNNAGATCGGAA |
| 59782_1 | TATGAGTGTGGAGTCGTTGCCTGGGCATGTAACGAACTAAACANNNAGATCGGAA |
| 59785_1 | TATGAGTGTGGAGTCGTTGCGGATCATCTGAGGTCAGGAGTTCNNNAGATCGGAA |
| 59792_1 | TATGAGTGTGGAGTCGTTGCGCAGTGAGAGGTAGTTGTGAGAANNNAGATCGGA |
| 59794_1 | TATGAGTGTGGAGTCGTTGCGTATCTCTGCTGTGCTGAAGGTNNNAGATCGGAAG |
| 59802_1 | TATGAGTGTGGAGTCGTTGCCTCGTGGATTGAAGCGATTCTCNNNAGATCGGAAGA |
| 59808_1 | TATGAGTGTGGAGTCGTTGCCAATGGTGCAATCTCAGCTCACNNNAGATCGGAAGA |
| 59811_1 | TATGAGTGTGGAGTCGTTGCCTCCCAGGTTCAAGCGATTCTCNNNAGATCGGAAGA |
| 59820_1 | TATGAGTGTGGAGTCGTTGCCGAGTTGCTGGTACTACAGGTGNNNAGATCGGAAG |
| 59826_1 | TATGAGTGTGGAGTCGTTGCGAGAGATGGGGTTTCATCACCTTNNNAGATCGGAA |
| 59828_1 | TATGAGTGTGGAGTCGTTGCGAGTATTGCTCTGTCGCCTAGGNNNAGATCGGAAG |
| 59831_1 | TATGAGTGTGGAGTCGTTGCCTCCCAGGTTCACACCATTCTCNNNAGATCGGAAGAG |
| 59844_1 | TATGAGTGTGGAGTCGTTGCTGCGATCTCAGCTTACTGCAAGNNNAGATCGGAAGA |

| | |
|---|---|
| 59857_1 | TATGAGTGTGGAGTCGTTGCCTTCCGGGTTCACACCATTCTCNNNAGATCGGAAGA |
| 59858_1 | TATGAGTGTGGAGTCGTTGCACCTCCCAGATTCAAGCAATTCTNNNAGATCGGAAG |
| 59859_1 | TATGAGTGTGGAGTCGTTGCAATCTCAGCTCGCTTCAACCTCNNNAGATCGGAAGA |
| 59861_1 | TATGAGTGTGGAGTCGTTGCCTCCTGGGTTCAAGCAATTCTGNNNAGATCGGAAGA |
| 59868_1 | TATGAGTGTGGAGTCGTTGCGACTGTGTCTCACTCTGTCGTCNNNAGATCGGAAGA |
| 59871_1 | TATGAGTGTGGAGTCGTTGCCAGGTTCAAGCAATTCTCCTGCNNNAGATCGGAAGA |
| 59872_1 | TATGAGTGTGGAGTCGTTGCCAAGATCTTGGCTCACTGCAACNNNAGATCGGAAGA |
| 59878_1 | TATGAGTGTGGAGTCGTTGCCCTCAGCTCACTACAACCTCAGNNNAGATCGGAAGA |
| 59880_1 | TATGAGTGTGGAGTCGTTGCCAGGCTGGAGTACAGTGACATAANNNAGATCGGAA |
| 59883_1 | TATGAGTGTGGAGTCGTTGCCTCCTGGGTTCACGACATTCTCNNNAGATCGGAAGA |
| 59909_1 | TATGAGTGTGGAGTCGTTGCCTTTCCTCCTTTCTTTGGCTGTGNNNAGATCGGAAGA |
| 59911_1 | TATGAGTGTGGAGTCGTTGCATTAGCCTTGCCTGCTGATGATANNNAGATCGGAAG |
| 59914_1 | TATGAGTGTGGAGTCGTTGCTGCAGAATTGTCCTTGACTGTTGNNNAGATCGGAAG |
| 59915_1 | TATGAGTGTGGAGTCGTTGCCTGAGGCAGGAGAATCACTTGAANNNAGATCGGAA |
| 59930_1 | TATGAGTGTGGAGTCGTTGCCAGCACCGCTATCCTCTTCTAGNNNAGATCGGAAGA |
| 59953_1 | TATGAGTGTGGAGTCGTTGCACAATTTGCCAGAGGAAAACCAGNNNAGATCGGAA |
| 59970_1 | TATGAGTGTGGAGTCGTTGCGGAGAATCGCTTGAACTCAGGAGNNNAGATCGGAA |
| 59978_1 | TATGAGTGTGGAGTCGTTGCGTCCTTTGCAACAACATGGATGANNNAGATCGGAAG |
| 59989_1 | TATGAGTGTGGAGTCGTTGCAGTGACATTCTGGAAAGGGTCAANNNAGATCGGAA |
| 59990_1 | TATGAGTGTGGAGTCGTTGCGCAACTGGGGAATAATAGAATGCANNNAGATCGGA |
| 60002_1 | TATGAGTGTGGAGTCGTTGCCATGATCTTGGCTCACTGCAACNNNAGATCGGAAGA |
| 60051_1 | TATGAGTGTGGAGTCGTTGCGCCTCCAAAGTAGCTGGGATTATNNNAGATCGGAA |
| 60053_1 | TATGAGTGTGGAGTCGTTGCTTGGGTTCAAGCAATTCTCGTGNNNAGATCGGAAG |
| 60071_1 | TATGAGTGTGGAGTCGTTGCGTTCATGCCATTCTCCTGCTTCNNNAGATCGGAAGAG |
| 60073_1 | TATGAGTGTGGAGTCGTTGCCACAATCTCGGCTCATTGCAAGNNNAGATCGGAAGA |
| 60074_1 | TATGAGTGTGGAGTCGTTGCCTCTCTGGTTCAGGCGATTCTCNNNAGATCGGAAGA |
| 60081_1 | TATGAGTGTGGAGTCGTTGCCACGATCTCAGCTCAATGCAACNNNAGATCGGAAGA |
| 60084_1 | TATGAGTGTGGAGTCGTTGCTAGAGATGGGGGTCTCACTGTGNNNAGATCGGAAG |
| 60091_1 | TATGAGTGTGGAGTCGTTGCCTAGTAGCTGGGATTACAGGCGNNNAGATCGGAAG |
| 60125_1 | TATGAGTGTGGAGTCGTTGCCCCGAGTAGATGGGACTACAGGNNNAGATCGGAAG |
| 60136_1 | TATGAGTGTGGAGTCGTTGCGATCTTGGCTCATTGCAACCTCNNNAGATCGGAAGA |
| 60141_1 | TATGAGTGTGGAGTCGTTGCGAGCAGAGAGACTCCACCTTTCNNNAGATCGGAAG |
| 60148_1 | TATGAGTGTGGAGTCGTTGCGCCTGTAATCCCAGCATTTTGGNNNAGATCGGAAGA |
| 60155_1 | TATGAGTGTGGAGTCGTTGCCAGAAGACAGGCTCTATGAGGACNNNAGATCGGAA |
| 60157_1 | TATGAGTGTGGAGTCGTTGCCTGCAACTCAGATTCTGAAAGCANNNAGATCGGAAG |
| 60170_1 | TATGAGTGTGGAGTCGTTGCTAGCAGAAGAAAGCATGGTGAGTNNNAGATCGGA |
| 60175_1 | TATGAGTGTGGAGTCGTTGCTCCACTAAATGCCAAGAAAGGGANNNAGATCGGAA |
| 60176_1 | TATGAGTGTGGAGTCGTTGCGAAGTAGGCGGATCTTCTGAGGNNNAGATCGGAAG |
| 60179_1 | TATGAGTGTGGAGTCGTTGCACATACGAGCGTAGGGTACTAGANNNAGATCGGAA |
| 60184_1 | TATGAGTGTGGAGTCGTTGCTGTATTCCATTTCCCTCCACTCCNNNAGATCGGAAGA |
| 60188_1 | TATGAGTGTGGAGTCGTTGCCTTGCCCAAATCCTGTGTCAATCNNNAGATCGGAAG |
| 60191_1 | TATGAGTGTGGAGTCGTTGCCCAGGATGTCGCCTATCTGTATANNNAGATCGGAAG |
| 60196_1 | TATGAGTGTGGAGTCGTTGCCTGAGGTGGGATTCTACAGATGGNNNAGATCGGAA |

| | |
|---|---|
| 60199_1 | TATGAGTGTGGAGTCGTTGCCAACTTGCATATTTGAGCCTGGTNNNAGATCGGAAG |
| 60213_1 | TATGAGTGTGGAGTCGTTGCAAAAAGTTCAGGAGTTCCAGACCNNNAGATCGGAA |
| 60217_1 | TATGAGTGTGGAGTCGTTGCAGGTAGACAGTGAAGAGATCCCTNNNAGATCGGAA |
| 60221_1 | TATGAGTGTGGAGTCGTTGCGCTGGTGTTGATGTAAAGCTGTTNNNAGATCGGAA |
| 60222_1 | TATGAGTGTGGAGTCGTTGCCAAATGAGGTCAGAGAGGTTGGANNNAGATCGGA |
| 60223_1 | TATGAGTGTGGAGTCGTTGCTTTGGTTCCGTTTACATGGGAGANNNAGATCGGAAG |
| 60227_1 | TATGAGTGTGGAGTCGTTGCAAAATGTGGAGGTCGTGAGAGATNNNAGATCGGA |
| 60234_1 | TATGAGTGTGGAGTCGTTGCTGTTCTAGAAAGATGAGGGGTGCNNNAGATCGGAA |
| 60238_1 | TATGAGTGTGGAGTCGTTGCTTTCCCTAAACCTCACCAACGTTNNNAGATCGGAAGA |
| 60244_1 | TATGAGTGTGGAGTCGTTGCCCCTTGAACAACCTGGTTTTGAANNNAGATCGGAAG |
| 60254_1 | TATGAGTGTGGAGTCGTTGCGTGTGATGTAGTAGGCAGAGGAGNNNAGATCGGA |
| 60256_1 | TATGAGTGTGGAGTCGTTGCAAGAACATCCTGGCTAACATGGTNNNAGATCGGAAG |
| 60277_1 | TATGAGTGTGGAGTCGTTGCTGGATCACGAGGTCAGGAGATTNNNAGATCGGAAG |
| 60282_1 | TATGAGTGTGGAGTCGTTGCGAGTCTTGCTCTGTCACCTAGGNNNAGATCGGAAGA |
| 60284_1 | TATGAGTGTGGAGTCGTTGCCACTATCTCGGCTCTCTGCAACNNNAGATCGGAAGAG |
| 60305_1 | TATGAGTGTGGAGTCGTTGCCAAGTAGCTGAGACCACAGGAACNNNAGATCGGAA |
| 60307_1 | TATGAGTGTGGAGTCGTTGCGACTCCTGAGTAGCTGGGACTANNNAGATCGGAAG |
| 60317_1 | TATGAGTGTGGAGTCGTTGCCAATTCTTCTGCTTCAGCCTCCTNNNAGATCGGAAGA |
| 60318_1 | TATGAGTGTGGAGTCGTTGCAGTGCTAGGATTACAGGCATGAGNNNAGATCGGAA |
| 60320_1 | TATGAGTGTGGAGTCGTTGCCGATTAGCTGGGACTACAGGTGNNNAGATCGGAAG |
| 60322_1 | TATGAGTGTGGAGTCGTTGCCCAGGTTCACACCATTCTCCTGNNNAGATCGGAAGAG |
| 60324_1 | TATGAGTGTGGAGTCGTTGCCTCCTGGGTTCATGCCATTCTCNNNAGATCGGAAGAG |
| 61954_1 | TATGAGTGTGGAGTCGTTGCTTATCTTCAAGAGCGTCCAGGAGNNNAGATCGGAAG |
| 61969_1 | TATGAGTGTGGAGTCGTTGCCCTATGGTCCGTATGATCCAGGANNNAGATCGGAAG |
| 61970_1 | TATGAGTGTGGAGTCGTTGCGCCAAGGCCAGAAGAAGTAAAGANNNAGATCGGA |
| 61972_1 | TATGAGTGTGGAGTCGTTGCGCTGGCCTCCTTTTCTGTAATTCNNNAGATCGGAAGA |
| 61978_1 | TATGAGTGTGGAGTCGTTGCCAGCACATCCAGTGACTTCTACANNNAGATCGGAAG |
| 61983_1 | TATGAGTGTGGAGTCGTTGCCCTATTCATGTACCCGCGATGTANNNAGATCGGAAG |
| 62001_1 | TATGAGTGTGGAGTCGTTGCGATCTCGGCTCACTGCAAACTCNNNAGATCGGAAGA |
| 62009_1 | TATGAGTGTGGAGTCGTTGCGTTTCACCATATTAGCCAGGCTGNNNAGATCGGAAG |
| 62019_1 | TATGAGTGTGGAGTCGTTGCGTAGCTCCTGATTCAGCCTCTTANNNAGATCGGAAG |
| 62033_1 | TATGAGTGTGGAGTCGTTGCCTCAGGTTCATGCCATTCTCCTNNNAGATCGGAAGAG |
| 62037_1 | TATGAGTGTGGAGTCGTTGCCAAGTAGCTGGGATTACAGGCANNNAGATCGGAAG |
| 62041_1 | TATGAGTGTGGAGTCGTTGCCACAATCTTGGCTCACTGCAACNNNAGATCGGAAGA |
| 62057_1 | TATGAGTGTGGAGTCGTTGCGTAATCCCAGCTACTCCAGAAGGNNNAGATCGGAAG |
| 62069_1 | TATGAGTGTGGAGTCGTTGCTTCAAGCAATTCCCTGCCTCAGNNNAGATCGGAAGA |
| 62078_1 | TATGAGTGTGGAGTCGTTGCCTGTGGTCCTAGCTTCTTGAGAGNNNAGATCGGAAG |
| 62092_1 | TATGAGTGTGGAGTCGTTGCTCTCTCTCTCTCTCCTCCTCTCTNNNAGATCGGAAGAG |
| 62110_1 | TATGAGTGTGGAGTCGTTGCCTGAGGTGGGAGGATTACTTGAGNNNAGATCGGA |
| 62112_1 | TATGAGTGTGGAGTCGTTGCGCCAACTTAGCTGTATTCTTCTGGNNNAGATCGGAA |
| 62143_1 | TATGAGTGTGGAGTCGTTGCGAGTTTTGCTCTTGTTGCCCAGNNNAGATCGGAAGA |
| 62151_1 | TATGAGTGTGGAGTCGTTGCGTACCAGGGTATGAGAACCAGCNNNAGATCGGAAG |
| 62169_1 | TATGAGTGTGGAGTCGTTGCCTAAGCGAACCAGGAAGACTGTNNNAGATCGGAAG |

| | |
|---|---|
| 62174_1 | TATGAGTGTGGAGTCGTTGCCATGCAGACAGAGCTAGACAGAANNNAGATCGGAA |
| 62189_1 | TATGAGTGTGGAGTCGTTGCGATCTCAGCTCACTGCAACCTCNNNAGATCGGAAGA |
| 62192_1 | TATGAGTGTGGAGTCGTTGCGTTCAAGCGATTCTTCTGCCTCNNNAGATCGGAAGA |
| 62195_1 | TATGAGTGTGGAGTCGTTGCCTGGGACTACAGGCACATACTACNNNAGATCGGAAG |
| 62210_1 | TATGAGTGTGGAGTCGTTGCCACAATCTCAGCTCACTGCAACNNNAGATCGGAAGA |
| 62225_1 | TATGAGTGTGGAGTCGTTGCCTGCCTCTGGGTTCAAGTGATTNNNAGATCGGAAGA |
| 62238_1 | TATGAGTGTGGAGTCGTTGCTGAGGATGTGTAGGTCAAATGCANNNAGATCGGAA |
| 62241_1 | TATGAGTGTGGAGTCGTTGCTGCAATACTGTTTTTCCCAGTGGNNNAGATCGGAAG |
| 62245_1 | TATGAGTGTGGAGTCGTTGCCAATGGCACAATCTCAGCTCACNNNAGATCGGAAGA |
| 62247_1 | TATGAGTGTGGAGTCGTTGCGGGATGCTTCCAAATACAATCGGNNNAGATCGGAA |
| 62252_1 | TATGAGTGTGGAGTCGTTGCATTCAATCCTGGGAGACAGTGAGNNNAGATCGGAA |
| 62261_1 | TATGAGTGTGGAGTCGTTGCCTCCTGGGTTCACACCATTCTCNNNAGATCGGAAGA |
| 62275_1 | TATGAGTGTGGAGTCGTTGCCAAGACCACCTCAGACTTCATGANNNAGATCGGAAG |
| 62278_1 | TATGAGTGTGGAGTCGTTGCCTCTCAGGTTCACGCCATTCTCNNNAGATCGGAAGA |
| 62283_1 | TATGAGTGTGGAGTCGTTGCATCATGGCTTACTGCAGTCTCAANNNAGATCGGAAG |
| 62292_1 | TATGAGTGTGGAGTCGTTGCACTAACAGAACGAACTTGGCTGANNNAGATCGGAA |
| 62296_1 | TATGAGTGTGGAGTCGTTGCATGCCTGTAATCCCAGCACTTTNNNAGATCGGAAGA |
| 62304_1 | TATGAGTGTGGAGTCGTTGCGAGATGGGGTTTTGCCATATTGGNNNAGATCGGAA |
| 62309_1 | TATGAGTGTGGAGTCGTTGCTGAGCATGAACACATAGCAGTGANNNAGATCGGAA |
| 62326_1 | TATGAGTGTGGAGTCGTTGCTACCCAGGCTGATCTCAAACTTCNNNAGATCGGAAG |
| 62341_1 | TATGAGTGTGGAGTCGTTGCCTGCAGTGAGCCATGATTGTTCNNNAGATCGGAAG |
| 62351_1 | TATGAGTGTGGAGTCGTTGCGCAATAAGACAGATTCGACCCATNNNAGATCGGAAG |
| 62378_1 | TATGAGTGTGGAGTCGTTGCCTCTCTCTTTCTCTCCCTCCCTCNNNAGATCGGAAGAG |
| 62380_1 | TATGAGTGTGGAGTCGTTGCGTTCCCGATTCCTCTTTTTGCATNNNAGATCGGAAGA |
| 62385_1 | TATGAGTGTGGAGTCGTTGCTATTGGTAGAGGGCTTCAGCTGNNNAGATCGGAAG |
| 62407_1 | TATGAGTGTGGAGTCGTTGCATGCCTGTAATCCCAGCACTTTNNNAGATCGGAAGA |
| 62428_1 | TATGAGTGTGGAGTCGTTGCGATGTAGTCTCGCTCTGTCACCNNNAGATCGGAAGA |
| 62436_1 | TATGAGTGTGGAGTCGTTGCCAGCTACTTGGAGGCTAAGGTGNNNAGATCGGAAG |
| 62446_1 | TATGAGTGTGGAGTCGTTGCCAGGTGGTAGAGCATCAGACTTTNNNAGATCGGAA |
| 62455_1 | TATGAGTGTGGAGTCGTTGCCGATCTCCTGACCTCATGATCTGNNNAGATCGGAAG |
| 62462_1 | TATGAGTGTGGAGTCGTTGCCTCCCAGGTTCAAGCGATTCTCNNNAGATCGGAAGA |
| 62465_1 | TATGAGTGTGGAGTCGTTGCGTGATACGATCTCGGCTCACTGNNNAGATCGGAAGA |
| 62476_1 | TATGAGTGTGGAGTCGTTGCCAAGTAGCTGGGACTACAGGTGNNNAGATCGGAAG |
| 62481_1 | TATGAGTGTGGAGTCGTTGCGAAACGGAGTCTCACTCTGTCGNNNAGATCGGAAG |
| 62489_1 | TATGAGTGTGGAGTCGTTGCGATCTTGGCTCACTGCAAGCTCNNNAGATCGGAAGA |
| 62504_1 | TATGAGTGTGGAGTCGTTGCGAGTGCAGAGGTGTGATCATGANNNAGATCGGAA |
| 62507_1 | TATGAGTGTGGAGTCGTTGCCTCCCTGGCTCTAGCAATTCTCNNNAGATCGGAAGA |
| 62508_1 | TATGAGTGTGGAGTCGTTGCCAAGTAGCTGGGACTACAGGTGNNNAGATCGGAAG |
| 62509_1 | TATGAGTGTGGAGTCGTTGCGTTCAAGCGATTCTTCTGCCTCNNNAGATCGGAAGA |
| 62511_1 | TATGAGTGTGGAGTCGTTGCCTTACTGCAACCTTCACCTCCTNNNAGATCGGAAGAG |
| 62515_1 | TATGAGTGTGGAGTCGTTGCTGAACTCTTGACCTCGTGATCTGNNNAGATCGGAAG |
| 62517_1 | TATGAGTGTGGAGTCGTTGCCTCCCAGGTTCAAGCGATTCTCNNNAGATCGGAAGA |
| 62518_1 | TATGAGTGTGGAGTCGTTGCCAATTCTCCTGCCTCAGTCTCCNNNAGATCGGAAGAG |

| | |
|---|---|
| 62521_1 | TATGAGTGTGGAGTCGTTGCCTCCTGGGTTCATGCCATTCTCNNNAGATCGGAAGA |
| 62537_1 | TATGAGTGTGGAGTCGTTGCGAGATTGCAGTGAGCCGAGATTNNNAGATCGGAAG |
| 62538_1 | TATGAGTGTGGAGTCGTTGCCTGCAAAGCTGAAGGTACATGAGNNNAGATCGGAA |
| 62539_1 | TATGAGTGTGGAGTCGTTGCAACTCCTCACCCCATTACATGACNNNAGATCGGAAGA |
| 62550_1 | TATGAGTGTGGAGTCGTTGCCGATGGTACTACTGCACTCCAGNNNAGATCGGAAGA |
| 62589_1 | TATGAGTGTGGAGTCGTTGCGAGTGCAGTGACGTGATCTCAGNNNAGATCGGAAG |
| 62590_1 | TATGAGTGTGGAGTCGTTGCGAGTCTCCCTTTGTCTCCCAAGNNNAGATCGGAAGA |
| 62596_1 | TATGAGTGTGGAGTCGTTGCGAAAAGACAAATCTCCCACGCAANNNAGATCGGAAG |
| 62603_1 | TATGAGTGTGGAGTCGTTGCCTCCCAGGTTCACGCTATTCTCNNNAGATCGGAAGAG |
| 62606_1 | TATGAGTGTGGAGTCGTTGCGATGGAGTCTCACTATGTCGCCNNNAGATCGGAAG |
| 62618_1 | TATGAGTGTGGAGTCGTTGCGTGCAATAAGTAGAGGAAGGAAACANNNAGATCGG |
| 62629_1 | TATGAGTGTGGAGTCGTTGCACCATCTCACACCATTCAGAATGNNNAGATCGGAAG |
| 62643_1 | TATGAGTGTGGAGTCGTTGCGGACAGAGGTCATCTTATCACCCNNNAGATCGGAAG |
| 62646_1 | TATGAGTGTGGAGTCGTTGCACTGGCGTTGTATTCTGAGCTTANNNAGATCGGAAG |
| 62661_1 | TATGAGTGTGGAGTCGTTGCAAATGTTCCCTCATCTTCAGAGGNNNAGATCGGAAG |
| 62670_1 | TATGAGTGTGGAGTCGTTGCTGTCACAACAGAATGGTGGAAGANNNAGATCGGAA |
| 62679_1 | TATGAGTGTGGAGTCGTTGCCTGAGACACCAGGATCACTTGAANNNAGATCGGAAG |
| 62685_1 | TATGAGTGTGGAGTCGTTGCGTTGCAGTAAGCCGAGATTGTACNNNAGATCGGAA |
| 62699_1 | TATGAGTGTGGAGTCGTTGCTTGTGAGGCTGAAGTGGGAAATTNNNAGATCGGAA |
| 62700_1 | TATGAGTGTGGAGTCGTTGCGGTTGCAATCCTAGTCTCTGACANNNAGATCGGAAG |
| 62716_1 | TATGAGTGTGGAGTCGTTGCCTCTCTTGCCCTTCTGCCTTTTNNNAGATCGGAAGAG |
| 62735_1 | TATGAGTGTGGAGTCGTTGCCTCTTTCTCTGGTCCAACATGGANNNAGATCGGAAG |
| 62743_1 | TATGAGTGTGGAGTCGTTGCGATTGCTTGAACCCAGGAGTTTGNNNAGATCGGAA |
| 62747_1 | TATGAGTGTGGAGTCGTTGCAAACCCACCCAAACATAGTACCTNNNAGATCGGAAG |
| 62749_1 | TATGAGTGTGGAGTCGTTGCTACATTACAATCCCATCCGTGCTNNNAGATCGGAAGA |
| 62750_1 | TATGAGTGTGGAGTCGTTGCCCTCAGCCTCCTAAAGTGTTTGANNNAGATCGGAAG |
| 62751_1 | TATGAGTGTGGAGTCGTTGCCTGAAGCTGGAGGATCTCTTGAGNNNAGATCGGAA |
| 62768_1 | TATGAGTGTGGAGTCGTTGCCAGTGTCTTACTCTGTCACCCANNNAGATCGGAAGA |
| 62792_1 | TATGAGTGTGGAGTCGTTGCCTCCTGAGTAGCTGGGACTACANNNAGATCGGAAG |
| 62796_1 | TATGAGTGTGGAGTCGTTGCAATCTCAGCTCACTGCAACCTCNNNAGATCGGAAGA |
| 62813_1 | TATGAGTGTGGAGTCGTTGCGTAATATGACTGTGGCCCACCANNNAGATCGGAAG |
| 62832_1 | TATGAGTGTGGAGTCGTTGCGTCTCCTGAGTAGCTGGGACTANNNAGATCGGAAG |
| 62836_1 | TATGAGTGTGGAGTCGTTGCGAGTCATGACCTCGCTCTTTCANNNAGATCGGAAGA |
| 62841_1 | TATGAGTGTGGAGTCGTTGCCGAGTAGCTGGGATTACAGGTGNNNAGATCGGAAG |
| 62850_1 | TATGAGTGTGGAGTCGTTGCTGGAGTGCAGTGGTACAATCTCNNNAGATCGGAAG |
| 62857_1 | TATGAGTGTGGAGTCGTTGCCTCCTGGGTTCAAGCCATTCTCNNNAGATCGGAAGA |
| 62859_1 | TATGAGTGTGGAGTCGTTGCCTCTATTGCCTAGGCTGGAGTGNNNAGATCGGAAG |
| 62869_1 | TATGAGTGTGGAGTCGTTGCGATCTCGACTCACTGCAAGCTCNNNAGATCGGAAGA |
| 62873_1 | TATGAGTGTGGAGTCGTTGCAGACAGAGTCTCACTATGTTGCCNNNAGATCGGAAG |
| 62882_1 | TATGAGTGTGGAGTCGTTGCCTCCCAGGTTCATGCCATTCTCNNNAGATCGGAAGA |
| 62890_1 | TATGAGTGTGGAGTCGTTGCAATTTCCCTACACATTTGTCGCCNNNAGATCGGAAGA |
| 62893_1 | TATGAGTGTGGAGTCGTTGCCAGTGGCACTATCTTGGCTCATNNNAGATCGGAAGA |
| 62897_1 | TATGAGTGTGGAGTCGTTGCCTCCTGGGTTCATGCCATTCTCNNNAGATCGGAAGA |

| | |
|---|---|
| 62908_1 | TATGAGTGTGGAGTCGTTGCCTCCCAGGTTCATGCCATTCTCNNNAGATCGGAAGAG |
| 62909_1 | TATGAGTGTGGAGTCGTTGCCTCCTGGGTTCATGCCATTCTCNNNAGATCGGAAGA |
| 62912_1 | TATGAGTGTGGAGTCGTTGCCAAGTTCCGTCTCCCAAGTTCANNNAGATCGGAAGA |
| 62913_1 | TATGAGTGTGGAGTCGTTGCGATCTTGGCTCACTGCAAGCTCNNNAGATCGGAAGA |
| 62929_1 | TATGAGTGTGGAGTCGTTGCGAAACGAATTGCGCAATCTTGTTNNNAGATCGGAAG |
| 62957_1 | TATGAGTGTGGAGTCGTTGCGTGCCTGTAATCCCAGCTACTTNNNAGATCGGAAGA |
| 62961_1 | TATGAGTGTGGAGTCGTTGCGTTACCAGATGCCACGTGTTTGNNNAGATCGGAAGA |
| 62968_1 | TATGAGTGTGGAGTCGTTGCGAGATGACAATAGGCTGGGCATNNNAGATCGGAAG |
| 62990_1 | TATGAGTGTGGAGTCGTTGCTCTAATCCTCTCCAGCATCCTGTNNNAGATCGGAAGA |
| 62995_1 | TATGAGTGTGGAGTCGTTGCCATGGAACTCGAAGCTACAAAGTNNNAGATCGGAA |
| 62996_1 | TATGAGTGTGGAGTCGTTGCTACCGTTTATTCTGAGCCAGCTTNNNAGATCGGAAG |
| 63024_1 | TATGAGTGTGGAGTCGTTGCCCTCTACCCTCAAGGCGTTTACNNNAGATCGGAAGA |
| 63042_1 | TATGAGTGTGGAGTCGTTGCCACATAAAAGCTGTGGTCAATCTTNNNAGATCGGAA |
| 63046_1 | TATGAGTGTGGAGTCGTTGCTGAAACTGGAGGACAAGTTCACANNNAGATCGGAA |
| 63048_1 | TATGAGTGTGGAGTCGTTGCGAGAATCACTTGAACCCAGGAGGNNNAGATCGGAA |
| 63049_1 | TATGAGTGTGGAGTCGTTGCTGCAGGGAGAGGAAGAGTATACANNNAGATCGGA |
| 63050_1 | TATGAGTGTGGAGTCGTTGCCTGAGGTCAGGAGTTCAAGACCNNNAGATCGGAAG |
| 63055_1 | TATGAGTGTGGAGTCGTTGCGAGGTAGGAGAATGGCGTGAACNNNAGATCGGAA |
| 63056_1 | TATGAGTGTGGAGTCGTTGCTTTGCCATAAGACCTCCTTTGGTNNNAGATCGGAAG |
| 63060_1 | TATGAGTGTGGAGTCGTTGCTCAAAGGCGTTATCCTGATTTCTNNNAGATCGGAAG |
| 63061_1 | TATGAGTGTGGAGTCGTTGCTGAAATTCTGAAGGATGCATGGCNNNAGATCGGAA |
| 63062_1 | TATGAGTGTGGAGTCGTTGCTGTGATTTAGAAGCAGGAAGGCTNNNAGATCGGAA |
| 63065_1 | TATGAGTGTGGAGTCGTTGCCCTCTCAAGACATCCCAGTTCAGNNNAGATCGGAAG |
| 63069_1 | TATGAGTGTGGAGTCGTTGCGGCCATTCCAGAAGAATTTGAGGNNNAGATCGGAA |
| 63089_1 | TATGAGTGTGGAGTCGTTGCAGAAACCTTTAGCCAAAACTGCCNNNAGATCGGAAG |
| 63098_1 | TATGAGTGTGGAGTCGTTGCAAGTAAACAGCTCAGCAAGGAGTNNNAGATCGGAA |
| 63107_1 | TATGAGTGTGGAGTCGTTGCTGTGACAGATAAACTGATGAGGTNNNAGATCGGAA |
| 63120_1 | TATGAGTGTGGAGTCGTTGCGAGGAAGCAGAGCACAAAAGTTTNNNAGATCGGAA |
| 63122_1 | TATGAGTGTGGAGTCGTTGCAAGCCTGTAATCCCAGCACTTTNNNAGATCGGAAGA |
| 63124_1 | TATGAGTGTGGAGTCGTTGCCACATCACACACACACACACACNNNAGATCGGAAGAG |
| 63133_1 | TATGAGTGTGGAGTCGTTGCACATGATGAAACCCCGTCTCTACNNNAGATCGGAAG |
| 63141_1 | TATGAGTGTGGAGTCGTTGCCCCACCGTTGAGAATGTACTTTGNNNAGATCGGAAG |
| 63143_1 | TATGAGTGTGGAGTCGTTGCACGATCTCAGCTCACTGTAAGCNNNAGATCGGAAGA |
| 63146_1 | TATGAGTGTGGAGTCGTTGCCATGAACATGGCTCACTGCATCNNNAGATCGGAAGA |
| 63163_1 | TATGAGTGTGGAGTCGTTGCTGCTCAACATCAATTCTCCTTTCCNNNAGATCGGAAG |
| 63170_1 | TATGAGTGTGGAGTCGTTGCCTAGTAGCTGGGACTACAGGCANNNAGATCGGAAG |
| 63175_1 | TATGAGTGTGGAGTCGTTGCGTTCAAGCGATTCTTCTGCCTCNNNAGATCGGAAGA |
| 63176_1 | TATGAGTGTGGAGTCGTTGCTCTACTCTCCCAATAGTGCTCCANNNAGATCGGAAGA |
| 63183_1 | TATGAGTGTGGAGTCGTTGCTTGGGATTCCACGAAGATGACATNNNAGATCGGAA |
| 63197_1 | TATGAGTGTGGAGTCGTTGCGGCACCAAGACACTCCAAATAACNNNAGATCGGAAG |
| 63198_1 | TATGAGTGTGGAGTCGTTGCCTATTTGCCTGTCTCTCTGGTGGNNNAGATCGGAAG |
| 63202_1 | TATGAGTGTGGAGTCGTTGCGGACGACAGGTGTGCATCTATGNNNAGATCGGAAG |
| 63206_1 | TATGAGTGTGGAGTCGTTGCCAAGTAGCTGGGACTACAGGTGNNNAGATCGGAAG |

| | |
|---|---|
| 63213_1 | TATGAGTGTGGAGTCGTTGCGTAATCCCAGCCACTTGAGAGACNNNAGATCGGAAG |
| 63218_1 | TATGAGTGTGGAGTCGTTGCCAATGTGCTGAGATTGTGCCACNNNAGATCGGAAG |
| 63222_1 | TATGAGTGTGGAGTCGTTGCGAGTAGCTGGGATTACAGGCAGNNNAGATCGGAA |
| 63233_1 | TATGAGTGTGGAGTCGTTGCCACATGACCATGCTTTTGTCACTNNNAGATCGGAAG |
| 63235_1 | TATGAGTGTGGAGTCGTTGCGCCATGTTGCTATCAAATCCAGTNNNAGATCGGAAG |
| 63239_1 | TATGAGTGTGGAGTCGTTGCCCACACATTCTCTGTTGCAACTANNNAGATCGGAAG |
| 63240_1 | TATGAGTGTGGAGTCGTTGCACTATTTCCTAGCTGGCCAAAGTNNNAGATCGGAAG |
| 63312_1 | TATGAGTGTGGAGTCGTTGCCTGTTTTCTTGCCCACACACACNNNAGATCGGAAGAG |
| 63334_1 | TATGAGTGTGGAGTCGTTGCGTGAAACCCCCGTCTCTACTAAANNNAGATCGGAAG |
| 63341_1 | TATGAGTGTGGAGTCGTTGCCAAAAAGACAGAGCCTTGTGTGTNNNAGATCGGAA |
| 63346_1 | TATGAGTGTGGAGTCGTTGCGAAGTGGAGTTGGCTTGTTAGTGNNNAGATCGGAA |
| 63353_1 | TATGAGTGTGGAGTCGTTGCTCTTCCTTTGTTGCCCAGACTGNNNAGATCGGAAGA |
| 63381_1 | TATGAGTGTGGAGTCGTTGCATACATAAAGCTCACCCATCGCCNNNAGATCGGAAG |
| 63432_1 | TATGAGTGTGGAGTCGTTGCTCTCCAAGCCATGAATCAACTCANNNAGATCGGAAG |
| 63433_1 | TATGAGTGTGGAGTCGTTGCTCTGCTTGGTTAGACTTCTGAGCNNNAGATCGGAAG |
| 63447_1 | TATGAGTGTGGAGTCGTTGCAAGGCACACCTACAAGATCTAGANNNAGATCGGAA |
| 63451_1 | TATGAGTGTGGAGTCGTTGCTTCCTATGAACTTCCCACCCTTGNNNAGATCGGAAGA |
| 63455_1 | TATGAGTGTGGAGTCGTTGCAGAAGTGAGAGTAGGTGGAGGTTNNNAGATCGGA |
| 63458_1 | TATGAGTGTGGAGTCGTTGCCTCCTGGGTTCATGCCATTCTCNNNAGATCGGAAGA |
| 63468_1 | TATGAGTGTGGAGTCGTTGCAGACGGAGTCTCACTTTGTTCCNNNAGATCGGAAGA |
| 63471_1 | TATGAGTGTGGAGTCGTTGCTGGTGGTGTTTTGTTCTTTTTGCNNNAGATCGGAAG |
| 63475_1 | TATGAGTGTGGAGTCGTTGCCTCCCAGGTTCATGCCATTCTCNNNAGATCGGAAGA |
| 63485_1 | TATGAGTGTGGAGTCGTTGCCTCCCAGGTTCAAGCGATTCTCNNNAGATCGGAAGA |
| 63486_1 | TATGAGTGTGGAGTCGTTGCCTCCCAGGTTCACACCATTCTCNNNAGATCGGAAGAG |
| 63497_1 | TATGAGTGTGGAGTCGTTGCATTGAGCCGAGATGCCAAGATCNNNAGATCGGAAG |
| 63498_1 | TATGAGTGTGGAGTCGTTGCCATCCGGAATAGCTGGGACTACNNNAGATCGGAAG |
| 63502_1 | TATGAGTGTGGAGTCGTTGCCTTCTGGAGTAGCTGGGACTACANNNAGATCGGAA |
| 63505_1 | TATGAGTGTGGAGTCGTTGCCTCCTGGGTTCAAGTGATCCTCNNNAGATCGGAAGA |
| 63507_1 | TATGAGTGTGGAGTCGTTGCCTTCAACCTCCCCAGGATCATGNNNAGATCGGAAGA |
| 63516_1 | TATGAGTGTGGAGTCGTTGCGTGTGATCTCGGCTAACTGCAANNNAGATCGGAAG |
| 63528_1 | TATGAGTGTGGAGTCGTTGCCCTGAGGCATGATGTCTGAGTGNNNAGATCGGAAG |
| 63554_1 | TATGAGTGTGGAGTCGTTGCGATCTCGACTCACTGCAAGCTCNNNAGATCGGAAGA |
| 63562_1 | TATGAGTGTGGAGTCGTTGCTAGGCATGAGAATCGCTTGAACTNNNAGATCGGAA |
| 63567_1 | TATGAGTGTGGAGTCGTTGCCCCCCTCCCCCATTATTTTCATTNNNAGATCGGAAGA |
| 63569_1 | TATGAGTGTGGAGTCGTTGCCAGGGAAGCTGCATGAATCAACNNNAGATCGGAAG |
| 63604_1 | TATGAGTGTGGAGTCGTTGCACCTTCCTGTGGCATTAGCATTANNNAGATCGGAAG |
| 63622_1 | TATGAGTGTGGAGTCGTTGCCTCACTGCAACATCCATCTCCTNNNAGATCGGAAGAG |
| 63628_1 | TATGAGTGTGGAGTCGTTGCGGTGTTGTCAGTGTAAGTGTGTGNNNAGATCGGAA |
| 63630_1 | TATGAGTGTGGAGTCGTTGCTGTTCCAATGGAGTTGTCATTGCNNNAGATCGGAAG |
| 63634_1 | TATGAGTGTGGAGTCGTTGCTTCATGGCTGGTAGGAACGTAAANNNAGATCGGAA |
| 63636_1 | TATGAGTGTGGAGTCGTTGCGAGTCTTGCTCTGTCTGGAGTGNNNAGATCGGAAG |
| 63656_1 | TATGAGTGTGGAGTCGTTGCCATACAGCCCAGATCTGACCTCNNNAGATCGGAAGA |
| 63658_1 | TATGAGTGTGGAGTCGTTGCCATGGTGGTGTATGCCTGTAGTANNNAGATCGGAA |

| | |
|---|---|
| 63689_1 | TATGAGTGTGGAGTCGTTGCGGAGCCTGAGTGACAAGAATGAANNNAGATCGGAA |
| 63695_1 | TATGAGTGTGGAGTCGTTGCAGACAAGGTCTCACTCTGTTGCNNNAGATCGGAAGA |
| 63697_1 | TATGAGTGTGGAGTCGTTGCAAGCAACCCTAACACTACTGAGGNNNAGATCGGAAG |
| 63702_1 | TATGAGTGTGGAGTCGTTGCGCTTGACTGTTGTCCTAGCTACTNNNAGATCGGAAGA |
| 63732_1 | TATGAGTGTGGAGTCGTTGCCTCCTGAGTAGCTGGGACTACANNNAGATCGGAAGA |
| 63746_1 | TATGAGTGTGGAGTCGTTGCGTTCATGCCATTCTCCTCCCTCNNNAGATCGGAAGAG |
| 63748_1 | TATGAGTGTGGAGTCGTTGCGGACCTCAGGCTATCCAAATGATNNNAGATCGGAAG |
| 63752_1 | TATGAGTGTGGAGTCGTTGCCACACACACACACACAAGTTTCTNNNAGATCGGAAGA |
| 63753_1 | TATGAGTGTGGAGTCGTTGCATCTCAGATCACTGCAAGCTCCNNNAGATCGGAAGAG |
| 63755_1 | TATGAGTGTGGAGTCGTTGCCTCCCATGCTCAAGTGATCCTCNNNAGATCGGAAGAG |
| 63769_1 | TATGAGTGTGGAGTCGTTGCGTTCAAGCGATTCTTCTGCCTCNNNAGATCGGAAGAG |
| 63782_1 | TATGAGTGTGGAGTCGTTGCTCATCTCATGGACTGTCAGCAAANNNAGATCGGAAG |
| 63788_1 | TATGAGTGTGGAGTCGTTGCCACACACACACACACACTCTTGNNNAGATCGGAAGAG |
| 63794_1 | TATGAGTGTGGAGTCGTTGCCCTTTCTCTGTCACCTAGGCTGNNNAGATCGGAAGAG |
| 63796_1 | TATGAGTGTGGAGTCGTTGCAATTAGCTGGGTGTGGTAGTGGNNNAGATCGGAAG |
| 63820_1 | TATGAGTGTGGAGTCGTTGCGAGGCAGAAGAATCGCTTGAACNNNAGATCGGAAG |
| 63824_1 | TATGAGTGTGGAGTCGTTGCCTTGCAGTGAGCTGAGATTGTGNNNAGATCGGAAGA |
| 63848_1 | TATGAGTGTGGAGTCGTTGCTTCCTGTGGATCTACGATAGGGANNNAGATCGGAAG |
| 63855_1 | TATGAGTGTGGAGTCGTTGCCCTCCTGAGTAGCTAGGACTACANNNAGATCGGAAG |
| 63877_1 | TATGAGTGTGGAGTCGTTGCCTCCCAGGTTCATGCCATTCTCNNNAGATCGGAAGAG |
| 63889_1 | TATGAGTGTGGAGTCGTTGCAAGCTGCTGAGGGATTTCAAGTANNNAGATCGGAAG |
| 63912_1 | TATGAGTGTGGAGTCGTTGCCATACAGGGCACAGGACAAAAACNNNAGATCGGAAG |
| 63934_1 | TATGAGTGTGGAGTCGTTGCCTCCCAGGTTCATGCCATTCTCNNNAGATCGGAAGAG |
| 63938_1 | TATGAGTGTGGAGTCGTTGCATGCCTGTAATCCCAGCACTTTNNNAGATCGGAAGAG |
| 63956_1 | TATGAGTGTGGAGTCGTTGCTACTGCTTCTCTGTCTCCCACTANNNAGATCGGAAGAG |
| 63962_1 | TATGAGTGTGGAGTCGTTGCGAAGAAAAAACAGGGCCATCAGGNNNAGATCGGAA |
| 63976_1 | TATGAGTGTGGAGTCGTTGCTTGTTTTGAGTCAGAGTCTCGCTNNNAGATCGGAAG |
| 63980_1 | TATGAGTGTGGAGTCGTTGCTGTTAGCCAGGATGGTCTCAATCNNNAGATCGGAAG |
| 63984_1 | TATGAGTGTGGAGTCGTTGCCCCTGTTGGCTAATTTTTCTCAGGNNNAGATCGGAAG |
| 63997_1 | TATGAGTGTGGAGTCGTTGCGTCTCAAACTCCTGGGCTCAAGNNNAGATCGGAAGA |
| 64000_1 | TATGAGTGTGGAGTCGTTGCCAACAGCCTGAAGTAGTCGGAANNNAGATCGGAAGA |
| 64006_1 | TATGAGTGTGGAGTCGTTGCTCCTGAGTAGCTGGGATTACAGANNNAGATCGGAAG |
| 64010_1 | TATGAGTGTGGAGTCGTTGCCTCCTGGGTTCACACCATTCTCNNNAGATCGGAAGAG |
| 64013_1 | TATGAGTGTGGAGTCGTTGCAGTGCTACGATTACAGGTGTGAGNNNAGATCGGAAG |
| 64068_1 | TATGAGTGTGGAGTCGTTGCCTCAGACACAGGAACAGACCTAGNNNAGATCGGAAG |
| 64113_1 | TATGAGTGTGGAGTCGTTGCTGGGGGAAGAGTGAGAAAATGAGNNNAGATCGGA |
| 64116_1 | TATGAGTGTGGAGTCGTTGCCCTCAAGGTAAGATGCTTCCTGTNNNAGATCGGAAG |
| 64118_1 | TATGAGTGTGGAGTCGTTGCCTCCCAGAAGCTAGATGCACAGNNNAGATCGGAAGA |
| 64123_1 | TATGAGTGTGGAGTCGTTGCAAACCCCCAGTGTTCAGATGAGNNNAGATCGGAAGA |
| 64124_1 | TATGAGTGTGGAGTCGTTGCGAGGAGGAGAGGAACAGGAGAGNNNAGATCGGAA |
| 64141_1 | TATGAGTGTGGAGTCGTTGCAGACATCTGTGGGTGAGACAGANNNAGATCGGAAG |
| 64177_1 | TATGAGTGTGGAGTCGTTGCGAAGAAACCTGCCCACTCTCTCNNNAGATCGGAAGAG |
| 64180_1 | TATGAGTGTGGAGTCGTTGCATCACCCTCCCAATTTTGCACATNNNAGATCGGAAGA |

| | |
|---|---|
| 64210_1 | TATGAGTGTGGAGTCGTTGCGTGCAATCTCAGCTCACTGTAACNNNAGATCGGAAG |
| 64224_1 | TATGAGTGTGGAGTCGTTGCGCTTCCCAGAGTGAAATTGCTCNNNAGATCGGAAGA |
| 64234_1 | TATGAGTGTGGAGTCGTTGCGTGCCTCATCTACAAGACAAGGANNNAGATCGGAAG |
| 64241_1 | TATGAGTGTGGAGTCGTTGCGATCTTGGCTCATTGCAACCTCNNNAGATCGGAAGA |
| 64249_1 | TATGAGTGTGGAGTCGTTGCCTCGAGTTTTCTGCCAGAAATGGNNNAGATCGGAAG |
| 64262_1 | TATGAGTGTGGAGTCGTTGCAAAGGGCAGAACATCATGAGGAANNNAGATCGGAA |
| 64263_1 | TATGAGTGTGGAGTCGTTGCTTTGAGACGGAGTCTTGTTCTGTNNNAGATCGGAAG |
| 64289_1 | TATGAGTGTGGAGTCGTTGCACACACACAGAGAGAGAGAGAGANNNAGATCGGAA |
| 64291_1 | TATGAGTGTGGAGTCGTTGCTACTGGGAGTTAGAGCTTCAACGNNNAGATCGGAA |
| 64310_1 | TATGAGTGTGGAGTCGTTGCTATATATATGCGATCGAGCCCGGNNNAGATCGGAAG |
| 64325_1 | TATGAGTGTGGAGTCGTTGCGATCCTCCTCCTCCCTCTCTTGNNNAGATCGGAAGAG |
| 64328_1 | TATGAGTGTGGAGTCGTTGCCACAATCATGGCTCACTGCAACNNNAGATCGGAAGA |
| 64332_1 | TATGAGTGTGGAGTCGTTGCGCAGCCAAAGGATCTGGATTTGNNNAGATCGGAAG |
| 64373_1 | TATGAGTGTGGAGTCGTTGCTGTTTCACTTCCTACCAAGTCCTNNNAGATCGGAAGA |
| 64377_1 | TATGAGTGTGGAGTCGTTGCGGAAAAAGACTGGTTTCGAGACANNNAGATCGGAA |
| 64380_1 | TATGAGTGTGGAGTCGTTGCCTCCCAGGTTCAAGTGACTCTCNNNAGATCGGAAGA |
| 64397_1 | TATGAGTGTGGAGTCGTTGCGGAATGTAGTGGCGCAATCTTGNNNAGATCGGAAG |
| 64404_1 | TATGAGTGTGGAGTCGTTGCAGCCCATCTCAGTTACTGATTTCANNNAGATCGGAAG |
| 64410_1 | TATGAGTGTGGAGTCGTTGCCCTAGGTAGCTGGGACCACTATNNNAGATCGGAAGA |
| 64421_1 | TATGAGTGTGGAGTCGTTGCGAGTGTGAAGAACGGACCAGATNNNAGATCGGAAG |
| 64423_1 | TATGAGTGTGGAGTCGTTGCGAGTTTTGCTCTTGTTGCCCAGNNNAGATCGGAAGA |
| 64430_1 | TATGAGTGTGGAGTCGTTGCGATGCAGATGGAGGTCTCTGAAANNNAGATCGGAA |
| 64447_1 | TATGAGTGTGGAGTCGTTGCCTCAATCTCGACTCCCTGCAACNNNAGATCGGAAGAG |
| 64462_1 | TATGAGTGTGGAGTCGTTGCCCCAGGCAGAAGAGTAAAATCACNNNAGATCGGAA |
| 64480_1 | TATGAGTGTGGAGTCGTTGCGATGAAGGCTGCTCAGAGAGAGNNNAGATCGGAAG |
| 64485_1 | TATGAGTGTGGAGTCGTTGCTAATCCCATCTACTCAGGAGGCTNNNAGATCGGAAG |
| 64524_1 | TATGAGTGTGGAGTCGTTGCCTGTAGTCGCTGCTACTCAGAAGNNNAGATCGGAAG |
| 64527_1 | TATGAGTGTGGAGTCGTTGCGAGAGAGAGAGAGTGTGTGTGTGNNNAGATCGGA |
| 64777_1 | TATGAGTGTGGAGTCGTTGCGTTCAAGCGATTCTTCTGCCTCNNNAGATCGGAAGA |
| 64800_1 | TATGAGTGTGGAGTCGTTGCGAGAAAACACACACACACACACANNNAGATCGGAAG |
| 64804_1 | TATGAGTGTGGAGTCGTTGCAGATGGAGTCTCACTCTGTTGCNNNAGATCGGAAGA |
| 64812_1 | TATGAGTGTGGAGTCGTTGCGACGCTCCCATTTCATTTCCAGNNNAGATCGGAAGAG |
| 64836_1 | TATGAGTGTGGAGTCGTTGCCTCAATCCAAAGGTGTGTTGCATNNNAGATCGGAAG |
| 64854_1 | TATGAGTGTGGAGTCGTTGCCCTTGATGTCTCCCTCTCCATGNNNAGATCGGAAGAG |
| 64907_1 | TATGAGTGTGGAGTCGTTGCATGTTTTGAGACAGGGTCCCACNNNAGATCGGAAGA |
| 64915_1 | TATGAGTGTGGAGTCGTTGCCTCTGACTCCCAAGTTCAAGTGANNNAGATCGGAAG |
| 64921_1 | TATGAGTGTGGAGTCGTTGCGCACATCACAGTCACCATAAACANNNAGATCGGAAG |
| 64937_1 | TATGAGTGTGGAGTCGTTGCCTCACTCTGTCACCAAGGCTAGNNNAGATCGGAAGA |
| 64960_1 | TATGAGTGTGGAGTCGTTGCGAGACAAGGTGTTGCTCTGTTGNNNAGATCGGAAG |
| 64965_1 | TATGAGTGTGGAGTCGTTGCAAGATGCTGTATGGAAAAGGGGTNNNAGATCGGAA |
| 64970_1 | TATGAGTGTGGAGTCGTTGCGTCAAACCAGGGCAAAACAGACNNNAGATCGGAAG |
| 64971_1 | TATGAGTGTGGAGTCGTTGCTGACCTACATCACCCACATAAGCNNNAGATCGGAAGA |
| 65002_1 | TATGAGTGTGGAGTCGTTGCTGAGCATGAACACATAGCAGTGANNNAGATCGGAA |

| | |
|---|---|
| 65014_1 | TATGAGTGTGGAGTCGTTGCCAGAGTCTTGATCTGTTGCCCANNNAGATCGGAAGA |
| 65032_1 | TATGAGTGTGGAGTCGTTGCAGGTGGATTATCTGAGGTCAGGANNNAGATCGGAA |
| 65033_1 | TATGAGTGTGGAGTCGTTGCGAGAATGGTGTGAACCCAGGAGNNNAGATCGGAA |
| 65044_1 | TATGAGTGTGGAGTCGTTGCGGAGTGCAGTGGTGTGATCATANNNAGATCGGAAG |
| 65051_1 | TATGAGTGTGGAGTCGTTGCCTAAGCTGCATGTACACACCCANNNAGATCGGAAGA |
| 65058_1 | TATGAGTGTGGAGTCGTTGCCCCAAACATAATGCCCTGTTCCNNNAGATCGGAAGA |
| 65078_1 | TATGAGTGTGGAGTCGTTGCTCTGTGTTGCTTCTATCCCATGTNNNAGATCGGAAG |
| 65083_1 | TATGAGTGTGGAGTCGTTGCACACACACACGGGATAAGAGACNNNAGATCGGAAG |
| 65100_1 | TATGAGTGTGGAGTCGTTGCGAGTGTGAAGAACGGACCAGATNNNAGATCGGAA |
| 65103_1 | TATGAGTGTGGAGTCGTTGCCAGAATCTCACTCTGTTGCCCANNNAGATCGGAAGA |
| 65109_1 | TATGAGTGTGGAGTCGTTGCGAGTCTTGCTCTTGTCGTCCAANNNAGATCGGAAGA |
| 65120_1 | TATGAGTGTGGAGTCGTTGCAAGGCTCAGGTCTCTACCGAAANNNAGATCGGAAG |
| 65122_1 | TATGAGTGTGGAGTCGTTGCCCCAAGAAATCTTCTGGCCTCANNNAGATCGGAAGA |
| 65142_1 | TATGAGTGTGGAGTCGTTGCGAGATGGGGTTTCACTTTGTTGCNNNAGATCGGAA |
| 65154_1 | TATGAGTGTGGAGTCGTTGCGGAGTCCTGGGCCACTTTATTTANNNAGATCGGAAG |
| 65155_1 | TATGAGTGTGGAGTCGTTGCCTCCCAGGTTCAAGCGATTCTCNNNAGATCGGAAGA |
| 65166_1 | TATGAGTGTGGAGTCGTTGCCACAGGCTGATCTCAAACTCCTNNNAGATCGGAAGA |
| 65180_1 | TATGAGTGTGGAGTCGTTGCTTGTCTGGTGCAGTGTTTTATGGNNNAGATCGGAA |
| 65183_1 | TATGAGTGTGGAGTCGTTGCGGTGCCTATAATCCCAGCTACTCNNNAGATCGGAAG |
| 65198_1 | TATGAGTGTGGAGTCGTTGCAGATGGAATCTCGCTCTGTCACNNNAGATCGGAAGA |
| 65224_1 | TATGAGTGTGGAGTCGTTGCACAGGTAGCTGGTTGATGGATGNNNAGATCGGAAG |
| 65246_1 | TATGAGTGTGGAGTCGTTGCCACACACACACACACAAGAAANNNAGATCGGAAG |
| 65258_1 | TATGAGTGTGGAGTCGTTGCTTTTTTGAGATGGAGTCTTGCTCNNNAGATCGGAAG |
| 65309_1 | TATGAGTGTGGAGTCGTTGCAGGGGGATCAGTTTACAAAAGCANNNAGATCGGAA |
| 65329_1 | TATGAGTGTGGAGTCGTTGCCCAACTCTGAACTCAGCTCTGANNNAGATCGGAAGA |
| 65337_1 | TATGAGTGTGGAGTCGTTGCCAAGTGCTGGGATTGTAGGCATNNNAGATCGGAAG |
| 65384_1 | TATGAGTGTGGAGTCGTTGCGACGTTTGGGATGATAAACTGGCNNNAGATCGGAA |
| 65406_1 | TATGAGTGTGGAGTCGTTGCGTGCATTGGTGTGTGGATCTTGNNNAGATCGGAAG |
| 65442_1 | TATGAGTGTGGAGTCGTTGCCTAAAGCCTGCTGTAGGACCACNNNAGATCGGAAG |
| 65454_1 | TATGAGTGTGGAGTCGTTGCCTGGGCAGTGGACACTAAACTTANNNAGATCGGAA |
| 65480_1 | TATGAGTGTGGAGTCGTTGCTCCACCAACAAAGACACCACAGNNNAGATCGGAAGA |
| 65494_1 | TATGAGTGTGGAGTCGTTGCGAAGCAGCCACTTTTACATCCTCNNNAGATCGGAAG |
| 65502_1 | TATGAGTGTGGAGTCGTTGCTGAACTTTTGACCTGGTCATGGANNNAGATCGGAAG |
| 65512_1 | TATGAGTGTGGAGTCGTTGCGCAAGAACCACAGCATTATTGGTNNNAGATCGGAA |
| 65547_1 | TATGAGTGTGGAGTCGTTGCAGGCAAACTGCACATAGACAGTANNNAGATCGGAA |
| 65571_1 | TATGAGTGTGGAGTCGTTGCGCAAGTGACCGTGTGTGTAAAGNNNAGATCGGAAG |
| 65591_1 | TATGAGTGTGGAGTCGTTGCCTACCGGGTTCAGGCAATTCTCNNNAGATCGGAAGA |
| 65594_1 | TATGAGTGTGGAGTCGTTGCATGGAGTCTCACTCTTGTTGCCNNNAGATCGGAAGA |
| 65600_1 | TATGAGTGTGGAGTCGTTGCATTGATCCTGGGCCTATGCAAGNNNAGATCGGAAG |
| 65609_1 | TATGAGTGTGGAGTCGTTGCAGGGATCAAAGTATGGCCAAACANNNAGATCGGAA |
| 65611_1 | TATGAGTGTGGAGTCGTTGCGGTTTCAGTGAGCCGAGATCATNNNAGATCGGAAG |
| 65617_1 | TATGAGTGTGGAGTCGTTGCATCCTGAATGTGTCAAGAGCAGTNNNAGATCGGAA |
| 65645_1 | TATGAGTGTGGAGTCGTTGCGAGGCAAGAGAATCGCTTGAATCNNNAGATCGGAA |

| | |
|---|---|
| 65660_1 | TATGAGTGTGGAGTCGTTGCCAGAGTGCTGGGATTACAGGAGNNNAGATCGGAA |
| 65688_1 | TATGAGTGTGGAGTCGTTGCCGATCACACACACACAGAACAAGNNNAGATCGGAAG |
| 65692_1 | TATGAGTGTGGAGTCGTTGCGGAAGCTATACATGGGGTACTCGNNNAGATCGGAA |
| 65718_1 | TATGAGTGTGGAGTCGTTGCCTCCTGGGTTCAAGCCATTCTCNNNAGATCGGAAGA |
| 65725_1 | TATGAGTGTGGAGTCGTTGCGGCCAGAAAAAAAGGCAAAACTGNNNAGATCGGAA |
| 65747_1 | TATGAGTGTGGAGTCGTTGCACACACACATGCATACACACAAANNNAGATCGGAAG |
| 65768_1 | TATGAGTGTGGAGTCGTTGCATAAATTGAGGAGAAGGCAGGGGNNNAGATCGGA |
| 65773_1 | TATGAGTGTGGAGTCGTTGCGAGTTTTGCTCTTGTTGCCCAGNNNAGATCGGAAGA |
| 65784_1 | TATGAGTGTGGAGTCGTTGCAGAAGTGGTGAACAGGTTGCATANNNAGATCGGAA |
| 65818_1 | TATGAGTGTGGAGTCGTTGCGTGCGTTCTTGTCTCACTACAACNNNAGATCGGAAG |
| 65864_1 | TATGAGTGTGGAGTCGTTGCATGCCTATTTCCCTGGTTTTCCTNNNAGATCGGAAGA |
| 65871_1 | TATGAGTGTGGAGTCGTTGCTTTCACTCTTGTTGCCCAGGTTNNNAGATCGGAAGA |
| 65882_1 | TATGAGTGTGGAGTCGTTGCCAGAATTTCACCGTGTTAGCCAGNNNAGATCGGAAG |
| 65889_1 | TATGAGTGTGGAGTCGTTGCATGATCACACCACTACACTCCACNNNAGATCGGAAGA |
| 65921_1 | TATGAGTGTGGAGTCGTTGCTCTAGAGAGTCAGGGGGAAACTTNNNAGATCGGAA |
| 65924_1 | TATGAGTGTGGAGTCGTTGCCACACACACACGGAGGATATGANNNAGATCGGAAG |
| 65932_1 | TATGAGTGTGGAGTCGTTGCCACACACACACACTACAGCAACNNNAGATCGGAAGA |
| 65936_1 | TATGAGTGTGGAGTCGTTGCCATGATCTTGGCTCACTGCAACNNNAGATCGGAAGA |
| 65939_1 | TATGAGTGTGGAGTCGTTGCAATTTCGGCTCATCACAAACTCCNNNAGATCGGAAGA |
| 65940_1 | TATGAGTGTGGAGTCGTTGCTCATCTCATGGACTGTCAGCAAANNNAGATCGGAAG |
| 65956_1 | TATGAGTGTGGAGTCGTTGCCTACACACACACACACACACACNNNAGATCGGAAGA |
| 65965_1 | TATGAGTGTGGAGTCGTTGCGGAGTACAGTGGTGTGATCTCGNNNAGATCGGAAG |
| 65970_1 | TATGAGTGTGGAGTCGTTGCGTGACTCACTCCTGTAATCCCAGNNNAGATCGGAAG |
| 65979_1 | TATGAGTGTGGAGTCGTTGCCTCCTGGGTTCACACCATTCTCNNNAGATCGGAAGA |
| 65980_1 | TATGAGTGTGGAGTCGTTGCAAAGTGCTGGGATTACAGGCATNNNAGATCGGAAG |
| 65982_1 | TATGAGTGTGGAGTCGTTGCGGACAGGAAACAGAGGGGTAAATNNNAGATCGGA |
| 65987_1 | TATGAGTGTGGAGTCGTTGCAATCTCCACTCACCACAACCTCNNNAGATCGGAAGAG |
| 65992_1 | TATGAGTGTGGAGTCGTTGCGAGAGAGAGATAGGGTCTCGCTNNNAGATCGGAA |
| 66008_1 | TATGAGTGTGGAGTCGTTGCTGTGGAGGAAATGGAAGACTCAGNNNAGATCGGAA |
| 66018_1 | TATGAGTGTGGAGTCGTTGCCACAATCTTGACCCACTGCAACNNNAGATCGGAAGA |
| 66022_1 | TATGAGTGTGGAGTCGTTGCCCCAGGGTGTTTCTCTCTTGATTNNNAGATCGGAAG |
| 66055_1 | TATGAGTGTGGAGTCGTTGCGACAAACACACACACACACACANNNAGATCGGAAGA |
| 66095_1 | TATGAGTGTGGAGTCGTTGCCTCTCTCTCTCTCTCTCCACGGNNNAGATCGGAAGAG |
| 66105_1 | TATGAGTGTGGAGTCGTTGCTGTTGTTCAGGTTAGAGTGCAGTNNNAGATCGGAA |
| 66130_1 | TATGAGTGTGGAGTCGTTGCCCCCACTCCTACATAGGTTTTGTNNNAGATCGGAAGA |
| 66135_1 | TATGAGTGTGGAGTCGTTGCCGTGAGAGAGAGACAGAAGTCGNNNAGATCGGAA |
| 66171_1 | TATGAGTGTGGAGTCGTTGCGCCAGGCACTCTTGAAAGAATACNNNAGATCGGAA |
| 66196_1 | TATGAGTGTGGAGTCGTTGCTTTGCAACAGTTGTGTTATTGCTNNNAGATCGGAAG |
| 66203_1 | TATGAGTGTGGAGTCGTTGCCTCAGGGTGGATGGTAGGAGAGNNNAGATCGGAA |
| 66294_1 | TATGAGTGTGGAGTCGTTGCGAGATGGCAGAGGTTCTGGTAGNNNAGATCGGAA |
| 66301_1 | TATGAGTGTGGAGTCGTTGCAGTGGCACAATCTCCTGCTTTCNNNAGATCGGAAGA |
| 66326_1 | TATGAGTGTGGAGTCGTTGCCAGATAGGGAACAGAGGTGCAGNNNAGATCGGAA |
| 66464_1 | TATGAGTGTGGAGTCGTTGCGTTCAAGCAATTCTCCTGCCTCNNNAGATCGGAAGA |

| | |
|---|---|
| 66489_1 | TATGAGTGTGGAGTCGTTGCAGTGTGTGTGTTTGTATGTGTGTNNNAGATCGGAA |
| 66491_1 | TATGAGTGTGGAGTCGTTGCGAGTACAGTGGCGTGATCTCAGNNNAGATCGGAAG |
| 66519_1 | TATGAGTGTGGAGTCGTTGCCCCTAGTCATTCTCTGTTGCTCCNNNAGATCGGAAGA |
| 66528_1 | TATGAGTGTGGAGTCGTTGCTCTCTTCCTCCTCTCCTTCACTTNNNAGATCGGAAGAG |
| 66559_1 | TATGAGTGTGGAGTCGTTGCGTTTCACCTTGTTGGCCAGATGNNNAGATCGGAAGA |
| 66584_1 | TATGAGTGTGGAGTCGTTGCATGCCTGTAATCCCAGCACTTTNNNAGATCGGAAGA |
| 66585_1 | TATGAGTGTGGAGTCGTTGCATGTGCCTGTAATCCCAGCTACNNNAGATCGGAAGA |
| 66611_1 | TATGAGTGTGGAGTCGTTGCGACACTCAGAGAGGCTTCAACTGNNNAGATCGGAA |
| 66635_1 | TATGAGTGTGGAGTCGTTGCAAAGTAAGTGGGAACCATGGGAANNNAGATCGGA |
| 66642_1 | TATGAGTGTGGAGTCGTTGCAGATGGAGTCTCACTGTCACCCNNNAGATCGGAAGA |
| 66646_1 | TATGAGTGTGGAGTCGTTGCGATTGAGGTGGGAGGATTGCTTANNNAGATCGGA |
| 66687_1 | TATGAGTGTGGAGTCGTTGCGATGTACAGGACACAAAGCCAACNNNAGATCGGAA |
| 66692_1 | TATGAGTGTGGAGTCGTTGCGCGCTATAAGGAAAAATGTGTGCNNNAGATCGGAA |
| 66699_1 | TATGAGTGTGGAGTCGTTGCTTTTTTGAGACGCAGTCACTCTGNNNAGATCGGAAG |
| 66704_1 | TATGAGTGTGGAGTCGTTGCAGCTCTGATTTTTGCACTGCAGNNNAGATCGGAAGA |
| 66754_1 | TATGAGTGTGGAGTCGTTGCGAGTTTTGCTCTTGTTGCCCAGNNNAGATCGGAAGA |
| 66773_1 | TATGAGTGTGGAGTCGTTGCCAGGAAAACTGCATGAACCCAGNNNAGATCGGAAG |
| 66809_1 | TATGAGTGTGGAGTCGTTGCAGAATTATCTGCAGTGCTTCCCANNNAGATCGGAAG |
| 66823_1 | TATGAGTGTGGAGTCGTTGCCAGGTTGCAGTGAAATGGAGTGNNNAGATCGGAAG |
| 66838_1 | TATGAGTGTGGAGTCGTTGCGCACACAGAGAGAGATATGCCTANNNAGATCGGAA |
| 66842_1 | TATGAGTGTGGAGTCGTTGCCAGAGCTTGCAATGATCCCAGATNNNAGATCGGAAG |
| 66852_1 | TATGAGTGTGGAGTCGTTGCCCAAAATGCTGGGATTATAGGCANNNAGATCGGAA |
| 66857_1 | TATGAGTGTGGAGTCGTTGCCTCCGAGGTTCAAGCGATTCTCNNNAGATCGGAAGA |
| 66858_1 | TATGAGTGTGGAGTCGTTGCGTTCAAGAGATTCTCCTGCCTCANNNAGATCGGAAG |
| 66870_1 | TATGAGTGTGGAGTCGTTGCCTGTAGTCTCAGCTACTTGGGAGNNNAGATCGGAAG |
| 66877_1 | TATGAGTGTGGAGTCGTTGCGAGGTTGCAGTGAGTCGAGATCNNNAGATCGGAAG |
| 66878_1 | TATGAGTGTGGAGTCGTTGCAAAAATTAGCCAGGTGTGGTTCCNNNAGATCGGAAG |
| 66891_1 | TATGAGTGTGGAGTCGTTGCTTCAAGTGATTCTCCTGCCTCAGNNNAGATCGGAAG |
| 66913_1 | TATGAGTGTGGAGTCGTTGCGCTTGACTCAGAGCTACAGGAAGNNNAGATCGGAA |
| 66922_1 | TATGAGTGTGGAGTCGTTGCGTCATTCTCCTTTGACCTGTCGANNNAGATCGGAAG |
| 66930_1 | TATGAGTGTGGAGTCGTTGCGCCTGTAATCCCAGCTACTTGGNNNAGATCGGAAGA |
| 66988_1 | TATGAGTGTGGAGTCGTTGCCAGAGGACATGAGACTCCCACTNNNAGATCGGAAG |
| 67001_1 | TATGAGTGTGGAGTCGTTGCCGCATGTGTGTGTGTGTATTTNNNAGATCGGAAG |
| 67012_1 | TATGAGTGTGGAGTCGTTGCTATACTTTTACCCCTGGACTGGGNNNAGATCGGAAG |
| 67015_1 | TATGAGTGTGGAGTCGTTGCGAGTGTGAAGAACGGACCAGATNNNAGATCGGAAG |
| 67018_1 | TATGAGTGTGGAGTCGTTGCCAAGATCTTGGCTCACTGCAACNNNAGATCGGAAGA |
| 67019_1 | TATGAGTGTGGAGTCGTTGCCTCCTGGGTTCAAGCGATTCTCNNNAGATCGGAAGA |
| 67042_1 | TATGAGTGTGGAGTCGTTGCGAGACAGGAGAATCGCTGGAACNNNAGATCGGAAG |
| 67051_1 | TATGAGTGTGGAGTCGTTGCGATGGAGTTTCCCTCTTGTCGCNNNAGATCGGAAGA |
| 67054_1 | TATGAGTGTGGAGTCGTTGCGTGTGTGTGTGAAGGTCTTGNNNAGATCGGAAG |
| 67057_1 | TATGAGTGTGGAGTCGTTGCGGAGAGGCTCACGGAAATTCTTTNNNAGATCGGAA |
| 67064_1 | TATGAGTGTGGAGTCGTTGCAATCTCCACTCACTGCAACCTCNNNAGATCGGAAGAG |
| 67065_1 | TATGAGTGTGGAGTCGTTGCAACGGAGTATGATTGCACCACTNNNAGATCGGAAG |

| | |
|---|---|
| 67066_1 | TATGAGTGTGGAGTCGTTGCAACGGAGTATGATTGCACCACTNNNAGATCGGAAGA |
| 67087_1 | TATGAGTGTGGAGTCGTTGCAACAGAGACACAGAGAGAGCAACNNNAGATCGGAA |
| 67093_1 | TATGAGTGTGGAGTCGTTGCGTGCAATCATGGCTCACTTCAGNNNAGATCGGAAGA |
| 67099_1 | TATGAGTGTGGAGTCGTTGCAAAATGTCAGCCAATGGTGAGTCNNNAGATCGGAA |
| 67105_1 | TATGAGTGTGGAGTCGTTGCCCTCATGGGAGCAACTGGTAATNNNAGATCGGAAGA |
| 67116_1 | TATGAGTGTGGAGTCGTTGCGGGTGTTTACAAACTCAGGAACCNNNAGATCGGAA |
| 67119_1 | TATGAGTGTGGAGTCGTTGCAGCCAGTTCTCACATTGCTATGANNNAGATCGGAAG |
| 67135_1 | TATGAGTGTGGAGTCGTTGCTCTGTCTGTCTGTCTTTGAGGCNNNAGATCGGAAGA |
| 67138_1 | TATGAGTGTGGAGTCGTTGCGTCCTCCTTCTCCCAGCAATAAANNNAGATCGGAAGA |
| 67143_1 | TATGAGTGTGGAGTCGTTGCTGATCTTAGCTCACTGCAACGTNNNAGATCGGAAGA |
| 67170_1 | TATGAGTGTGGAGTCGTTGCGTGCAATCATAGCTCACTGCAGNNNAGATCGGAAGA |
| 67176_1 | TATGAGTGTGGAGTCGTTGCAACCTCCTGGGCTAAAGCAATCNNNAGATCGGAAGA |
| 67192_1 | TATGAGTGTGGAGTCGTTGCTTTCCAAGACCCCTTTGCAGATANNNAGATCGGAAGA |
| 67214_1 | TATGAGTGTGGAGTCGTTGCCACACACAGACACACACACAGANNNAGATCGGAAGA |
| 67216_1 | TATGAGTGTGGAGTCGTTGCAGTGGCATGATCTCAGCTCACNNNAGATCGGAAGAG |
| 67340_1 | TATGAGTGTGGAGTCGTTGCGTAAAAGGCAAAGGCATCTGGTCNNNAGATCGGAA |
| 67343_1 | TATGAGTGTGGAGTCGTTGCAGAAGGGAAAGACTCAGAGAAGCNNNAGATCGGA |
| 67391_1 | TATGAGTGTGGAGTCGTTGCCTTTTTGAGATGGCTCTGTCGCNNNAGATCGGAAGA |
| 67452_1 | TATGAGTGTGGAGTCGTTGCTATCCCGATAACTGCTTCACTGGNNNAGATCGGAAG |
| 67464_1 | TATGAGTGTGGAGTCGTTGCATGTCAGGTGTCAAGACTGAACANNNAGATCGGAA |
| 67503_1 | TATGAGTGTGGAGTCGTTGCGAAGCGGTAGCAGAGATCTGAATNNNAGATCGGAA |
| 67516_1 | TATGAGTGTGGAGTCGTTGCTAGTCCTTGCTGAAACCATGCCNNNAGATCGGAAGA |
| 67536_1 | TATGAGTGTGGAGTCGTTGCTGGCTAATTACCAGGTGCAGAGNNNAGATCGGAAG |
| 67551_1 | TATGAGTGTGGAGTCGTTGCTACAGTGAAAGGTCTGGTAACCCNNNAGATCGGAA |
| 67607_1 | TATGAGTGTGGAGTCGTTGCCGACCCCATTCTTTCTCTTTCACNNNAGATCGGAAGA |
| 67655_1 | TATGAGTGTGGAGTCGTTGCTGTTAGCCAGGATGGTCTCAATCNNNAGATCGGAAG |
| 67656_1 | TATGAGTGTGGAGTCGTTGCCTCCTGGGTTCACACCATTCTCNNNAGATCGGAAGAG |
| 67688_1 | TATGAGTGTGGAGTCGTTGCAAGTTGGGGGGCAGAAATATTGANNNAGATCGGA |
| 67706_1 | TATGAGTGTGGAGTCGTTGCGACAAGGTCTCACTCTGTCACCNNNAGATCGGAAGA |
| 67707_1 | TATGAGTGTGGAGTCGTTGCCACAATCTCAGCTCACTGCAACNNNAGATCGGAAGA |
| 67708_1 | TATGAGTGTGGAGTCGTTGCAAACTCCTGACCTCAAGTGATCCNNNAGATCGGAAG |
| 67712_1 | TATGAGTGTGGAGTCGTTGCAAGGGGGTTGATTTGGCAAATCNNNAGATCGGAAG |
| 67713_1 | TATGAGTGTGGAGTCGTTGCGTCTCTCCCTGTCTCTGCAAAGNNNAGATCGGAAGA |
| 67715_1 | TATGAGTGTGGAGTCGTTGCCAAAGTCTCACTCTGTCACCCANNNAGATCGGAAGAG |
| 67735_1 | TATGAGTGTGGAGTCGTTGCGTCCTGTGTACGTGCAATTCTTTNNNAGATCGGAAG |
| 67766_1 | TATGAGTGTGGAGTCGTTGCAACAACTGGCACACTACATAAGCNNNAGATCGGAAG |
| 67776_1 | TATGAGTGTGGAGTCGTTGCGATCTCAGCTCACTGCAACCTCNNNAGATCGGAAGA |
| 67786_1 | TATGAGTGTGGAGTCGTTGCGAGTCTCGCTCTTTTTGCCCAGNNNAGATCGGAAGA |
| 67805_1 | TATGAGTGTGGAGTCGTTGCGAGTTTCGCTCTTGTTGCCTAGNNNAGATCGGAAGA |
| 67821_1 | TATGAGTGTGGAGTCGTTGCCCAATTACTTGGGAGGGTAAGGTNNNAGATCGGAA |
| 67823_1 | TATGAGTGTGGAGTCGTTGCCAGGGTTCTAGCAATTCTCTTGCNNNAGATCGGAAG |
| 67842_1 | TATGAGTGTGGAGTCGTTGCTTGAAAACTTCCCACACTCCAGTNNNAGATCGGAAGA |
| 67874_1 | TATGAGTGTGGAGTCGTTGCGATTCTCCTGCCTCAGACTCCTNNNAGATCGGAAGAG |

| | |
|---|---|
| 67878_1 | TATGAGTGTGGAGTCGTTGCGTTGCAGTGAGACGAGTTCTTGNNNAGATCGGAAG |
| 67886_1 | TATGAGTGTGGAGTCGTTGCTTTGATGACCATCCAGTCCACANNNAGATCGGAAG |
| 67892_1 | TATGAGTGTGGAGTCGTTGCGCATAGTGGTGTATGCCTGTAGTNNNAGATCGGAA |
| 67901_1 | TATGAGTGTGGAGTCGTTGCGAGGCTAAGGCATGAGAATCACTNNNAGATCGGAA |
| 67914_1 | TATGAGTGTGGAGTCGTTGCCCAACTGCAGTGGTGCTATCTCNNNAGATCGGAAGA |
| 67932_1 | TATGAGTGTGGAGTCGTTGCTTCCCATTTCAGCCTTCCAAGTANNNAGATCGGAAGA |
| 67937_1 | TATGAGTGTGGAGTCGTTGCCACACACACACACACACACATATNNNAGATCGGAAG |
| 67971_1 | TATGAGTGTGGAGTCGTTGCCCAGTCTCCTTGGGTACAGAATANNNAGATCGGAAG |
| 67989_1 | TATGAGTGTGGAGTCGTTGCCCAGGCTAGTCTTGAACTCCTGNNNAGATCGGAAGA |
| 67992_1 | TATGAGTGTGGAGTCGTTGCTCAAACTCCTGGGTTCAAGTGATNNNAGATCGGAAG |
| 68006_1 | TATGAGTGTGGAGTCGTTGCTGAGTTCTGTGAACCACTCTAGCNNNAGATCGGAAG |
| 68026_1 | TATGAGTGTGGAGTCGTTGCAGTCTCAGCTAACTGCAACCTCNNNAGATCGGAAGA |
| 68050_1 | TATGAGTGTGGAGTCGTTGCATTCCAGCTGCTCTGCCTTTATCNNNAGATCGGAAGA |
| 68055_1 | TATGAGTGTGGAGTCGTTGCGTTCAAGCAATTCTCGTGCCTCNNNAGATCGGAAGA |
| 68061_1 | TATGAGTGTGGAGTCGTTGCGCTTCAGTGGTTTCAGGTCCTAANNNAGATCGGAAG |
| 68063_1 | TATGAGTGTGGAGTCGTTGCCAAAGTGCTGGGATTATAGGCCTNNNAGATCGGAA |
| 68086_1 | TATGAGTGTGGAGTCGTTGCGTGTGTCTGTAGTCCCAGCTACNNNAGATCGGAAGA |
| 68102_1 | TATGAGTGTGGAGTCGTTGCGAGACGGTCTCACTCTGTTTCCNNNAGATCGGAAGA |
| 68122_1 | TATGAGTGTGGAGTCGTTGCTCTTGTTTGCTTTGTGAACGGTTNNNAGATCGGAAG |
| 68145_1 | TATGAGTGTGGAGTCGTTGCTTTGAGATGGAGTCTTGCCGTGNNNAGATCGGAAG |
| 68199_1 | TATGAGTGTGGAGTCGTTGCGTGTGTGTGTGTGTGTATTCCTGNNNAGATCGGAA |
| 68200_1 | TATGAGTGTGGAGTCGTTGCGTTTCACTCTTGTTGTCCAGGCNNNAGATCGGAAGA |
| 68207_1 | TATGAGTGTGGAGTCGTTGCTTCTACATCTAGAGGGGGGTTCTNNNAGATCGGAA |
| 68226_1 | TATGAGTGTGGAGTCGTTGCATTCTCCTTTGTCAGCTTCCCAANNNAGATCGGAAGA |
| 68239_1 | TATGAGTGTGGAGTCGTTGCCACACAATGCCCAACAGCAAATNNNAGATCGGAAGA |
| 68249_1 | TATGAGTGTGGAGTCGTTGCCTGTAGTCCCAGCTACTCAGGAGNNNAGATCGGAAG |
| 68257_1 | TATGAGTGTGGAGTCGTTGCCCAAGCTACTTGAGAGGCTGAGNNNAGATCGGAAG |
| 68270_1 | TATGAGTGTGGAGTCGTTGCGCGTGTGTGTGTGTATAATGGAGNNNAGATCGGAA |
| 68274_1 | TATGAGTGTGGAGTCGTTGCGATCTGGGCTCACTACAAGCTCNNNAGATCGGAAGA |
| 68296_1 | TATGAGTGTGGAGTCGTTGCCCTCCTGGGTTCAAGCAATTCTNNNAGATCGGAAGA |
| 68319_1 | TATGAGTGTGGAGTCGTTGCCTGGAACTCCTGACAGGTGATCNNNAGATCGGAAG |
| 68329_1 | TATGAGTGTGGAGTCGTTGCTCATACTGAATGGGCAAAAGCTGNNNAGATCGGAA |
| 68348_1 | TATGAGTGTGGAGTCGTTGCTCAAGAAAACAGCAGGAAAAGCANNNAGATCGGAA |
| 68365_1 | TATGAGTGTGGAGTCGTTGCAGGCAGAGTGGACAAAAAGAGAANNNAGATCGGA |
| 68382_1 | TATGAGTGTGGAGTCGTTGCAGAGAGAGGAGAGAGAGGAGAGANNNAGATCGG |
| 68409_1 | TATGAGTGTGGAGTCGTTGCCATCTCAGCTCACTGCAACTTCNNNAGATCGGAAGAG |
| 68411_1 | TATGAGTGTGGAGTCGTTGCAGCTGAGAAGTGCCAACATCTGNNNAGATCGGAAG |
| 68419_1 | TATGAGTGTGGAGTCGTTGCCGAACTCTTGACTGACACAGTTGNNNAGATCGGAAG |
| 68432_1 | TATGAGTGTGGAGTCGTTGCCAACCTGTTTGGACTGTGGAAAGNNNAGATCGGAA |
| 68439_1 | TATGAGTGTGGAGTCGTTGCTATAAAAAAGAACTGCCGGACGCNNNAGATCGGAA |
| 68442_1 | TATGAGTGTGGAGTCGTTGCAATGGCGTGATCTCAGCTCATTNNNAGATCGGAAGA |
| 68463_1 | TATGAGTGTGGAGTCGTTGCTTGGAAAAGCAGAGCAGAGAGATNNNAGATCGGAA |
| 68471_1 | TATGAGTGTGGAGTCGTTGCCACACACAGACACACACACAGANNNAGATCGGAAGA |

| | |
|---|---|
| 68472_1 | TATGAGTGTGGAGTCGTTGCCACACACAGACACACACACAGANNNAGATCGGAAGA |
| 68510_1 | TATGAGTGTGGAGTCGTTGCCTTACCTGTGTGGATGCCAATGNNNAGATCGGAAG |
| 68538_1 | TATGAGTGTGGAGTCGTTGCGATTCCACACCTGCTTCAGATGANNNAGATCGGAAG |
| 68565_1 | TATGAGTGTGGAGTCGTTGCTTTTTTTTCATGGCTGGGTCAGGNNNAGATCGGAAG |
| 68574_1 | TATGAGTGTGGAGTCGTTGCCATCCTTCTTTTGTTCAGGGTGCNNNAGATCGGAAG |
| 68590_1 | TATGAGTGTGGAGTCGTTGCGTTCAAGCGATTCTCTTGCCTCNNNAGATCGGAAGA |
| 68619_1 | TATGAGTGTGGAGTCGTTGCAGGGAGGGAAAGGAAAAGAAAGGNNNAGATCGG |
| 68643_1 | TATGAGTGTGGAGTCGTTGCACACACACACACACACACTAAAGNNNAGATCGGAAG |
| 68676_1 | TATGAGTGTGGAGTCGTTGCCACACACAGACACACACACAGANNNAGATCGGAAGA |
| 68677_1 | TATGAGTGTGGAGTCGTTGCCACACACAGACACACACACAGANNNAGATCGGAAGA |
| 68678_1 | TATGAGTGTGGAGTCGTTGCCACACACACACACTGAGTTTCAGNNNAGATCGGAAG |
| 68698_1 | TATGAGTGTGGAGTCGTTGCAAGAACAAAAGTCTTTGCCTGGTNNNAGATCGGAA |
| 68718_1 | TATGAGTGTGGAGTCGTTGCCTTTCGCCTCTCCCCAAATTTTGNNNAGATCGGAAGA |
| 68719_1 | TATGAGTGTGGAGTCGTTGCCCAGTCACCCGCTTGATTTTTCNNNAGATCGGAAGA |
| 68728_1 | TATGAGTGTGGAGTCGTTGCCATGTCGTGACTAGCCCTATGGNNNAGATCGGAAG |
| 68757_1 | TATGAGTGTGGAGTCGTTGCGAGTTTTGCTCTAATTGCCCAGGNNNAGATCGGAAG |
| 68807_1 | TATGAGTGTGGAGTCGTTGCATAAGAACAGCATAGGGGAAGCCNNNAGATCGGAA |
| 68810_1 | TATGAGTGTGGAGTCGTTGCCAATCTCAGCTCACTGCAACTTCNNNAGATCGGAAG |
| 68832_1 | TATGAGTGTGGAGTCGTTGCCCTCTCTCTACCCCTCTCTCTCNNNAGATCGGAAGAG |
| 68857_1 | TATGAGTGTGGAGTCGTTGCACTGGTGCTTACAGACAGATGAGNNNAGATCGGAA |
| 68860_1 | TATGAGTGTGGAGTCGTTGCGCCTAATAATCCCAGCACTTTGGNNNAGATCGGAAG |
| 68868_1 | TATGAGTGTGGAGTCGTTGCATCAGCAACGTCAAGTGTGTTACNNNAGATCGGAAG |
| 68882_1 | TATGAGTGTGGAGTCGTTGCCACACACAGACACACACACAGANNNAGATCGGAAGA |
| 68883_1 | TATGAGTGTGGAGTCGTTGCCACACACAGACACACACACAGANNNAGATCGGAAGA |
| 68886_1 | TATGAGTGTGGAGTCGTTGCACACAAGATGCAAGATCAATGCCNNNAGATCGGAA |
| 68911_1 | TATGAGTGTGGAGTCGTTGCATGAATCTCTGGGTAGTAACGCGNNNAGATCGGAA |
| 68935_1 | TATGAGTGTGGAGTCGTTGCCTTCCTGGCACAAGCAATTCTTNNNAGATCGGAAGA |
| 68957_1 | TATGAGTGTGGAGTCGTTGCTCTTGTCTTCTTCATGTCCAGGGNNNAGATCGGAAG |
| 68963_1 | TATGAGTGTGGAGTCGTTGCATTGCGGGGATAAACAATGTGTCNNNAGATCGGAA |
| 68971_1 | TATGAGTGTGGAGTCGTTGCCCTCCTGGTTCACACCATTCTCNNNAGATCGGAAGAG |
| 68981_1 | TATGAGTGTGGAGTCGTTGCCAAGTGTGTTTCGGCTGTAGTGNNNAGATCGGAAG |
| 69012_1 | TATGAGTGTGGAGTCGTTGCTGGACTGCAGTGACACAATCTCNNNAGATCGGAAGA |
| 69036_1 | TATGAGTGTGGAGTCGTTGCGAGTGCAGTGGTGCGATCTGNNNAGATCGGAAGAG |
| 69069_1 | TATGAGTGTGGAGTCGTTGCTTTTTTTTTTTTCTCGCCCAGGCNNNAGATCGGAAGA |
| 69112_1 | TATGAGTGTGGAGTCGTTGCCAGCAGGTATTGGGAACAATTGGNNNAGATCGGAA |
| 69118_1 | TATGAGTGTGGAGTCGTTGCACAGGAAGGAACTGACAACATGTNNNAGATCGGAA |
| 69126_1 | TATGAGTGTGGAGTCGTTGCCTTGGACATTAGAACTCCAGGCTNNNAGATCGGAAG |
| 69132_1 | TATGAGTGTGGAGTCGTTGCCTCTCTCTCCCCCTCACTGTTANNNAGATCGGAAGAG |
| 69146_1 | TATGAGTGTGGAGTCGTTGCAGAAGAAAAAGACCCCAGAGAGCNNNAGATCGGAA |
| 69164_1 | TATGAGTGTGGAGTCGTTGCCACACACAGACACACACACAGANNNAGATCGGAAGA |
| 69165_1 | TATGAGTGTGGAGTCGTTGCCACACACAGACACACACACAGANNNAGATCGGAAGA |
| 69166_1 | TATGAGTGTGGAGTCGTTGCCACACACAGACACACACACAGANNNAGATCGGAAGA |
| 69181_1 | TATGAGTGTGGAGTCGTTGCAGTCAACGTTATCTAGAGGCACANNNAGATCGGAA |

| | |
|---|---|
| 69196_1 | TATGAGTGTGGAGTCGTTGCTAATGGTCCCTTCATGAGTCTGCNNNAGATCGGAAG |
| 69225_1 | TATGAGTGTGGAGTCGTTGCGCCAAAAACAAAAAACCCACCAGNNNAGATCGGAAG |
| 69226_1 | TATGAGTGTGGAGTCGTTGCACGTCCTTGTCATGTTAGCCATNNNAGATCGGAAGA |
| 69238_1 | TATGAGTGTGGAGTCGTTGCCAGGGTTCACACCATTCTCCTGNNNAGATCGGAAGA |
| 69265_1 | TATGAGTGTGGAGTCGTTGCGATTGCAGTGAGCCGAGATCATNNNAGATCGGAAG |
| 69266_1 | TATGAGTGTGGAGTCGTTGCCAAGCTGTATGATCCTGGGAGAGNNNAGATCGGAA |
| 69271_1 | TATGAGTGTGGAGTCGTTGCACCTGACATTGCAGCTTACCTTCNNNAGATCGGAAGA |
| 69294_1 | TATGAGTGTGGAGTCGTTGCTTGATATCCCACCTACCCCCTTCNNNAGATCGGAAGA |
| 69297_1 | TATGAGTGTGGAGTCGTTGCGCTATAAAATGGGTGCTGGCTTCNNNAGATCGGAAG |
| 69298_1 | TATGAGTGTGGAGTCGTTGCTGAGATCTTGGCTCACTGCAAGNNNAGATCGGAAGA |
| 69299_1 | TATGAGTGTGGAGTCGTTGCGTCTCTGGCCTTCTGGTCTCTANNNAGATCGGAAGA |
| 69303_1 | TATGAGTGTGGAGTCGTTGCGGTGAGATCATGGAACCTCTGTTNNNAGATCGGAAG |
| 69305_1 | TATGAGTGTGGAGTCGTTGCGTTTCACTGTTAGTTGCCCAGGNNNAGATCGGAAGA |
| 69306_1 | TATGAGTGTGGAGTCGTTGCTTTCACTAGTCAGGCGACACAGNNNAGATCGGAAGA |
| 69323_1 | TATGAGTGTGGAGTCGTTGCAAAAGAACAAAAGCCTCTGCCTGNNNAGATCGGAAG |
| 69357_1 | TATGAGTGTGGAGTCGTTGCCTTCGCCCTTTCTCTCTCTCTCNNNAGATCGGAAGAG |
| 69381_1 | TATGAGTGTGGAGTCGTTGCCACAATCTTGGCTCACTGCAACNNNAGATCGGAAGA |
| 69385_1 | TATGAGTGTGGAGTCGTTGCCTCTCTCTCTCACACACACACACNNNAGATCGGAAGA |
| 69403_1 | TATGAGTGTGGAGTCGTTGCTGTCTCATGCCTGTAAACCCAGNNNAGATCGGAAGA |
| 69430_1 | TATGAGTGTGGAGTCGTTGCGGGGAGAGAGAGAGAGAGAGAACNNNAGATCGGA |
| 69461_1 | TATGAGTGTGGAGTCGTTGCGAATTGCAATCCCCAGTTTTGGANNNAGATCGGAAG |
| 69494_1 | TATGAGTGTGGAGTCGTTGCCATGACGAACACACACGTACAGNNNAGATCGGAAGA |
| 69510_1 | TATGAGTGTGGAGTCGTTGCCAGAGGGAGGAAAAGGAAAAGGANNNAGATCGGA |
| 69521_1 | TATGAGTGTGGAGTCGTTGCAAAGGGTGGAAAGTGGGAAGAGNNNAGATCGGAA |
| 69531_1 | TATGAGTGTGGAGTCGTTGCGAAGTCGATGGCTTCAGCATTGNNNAGATCGGAAG |
| 69544_1 | TATGAGTGTGGAGTCGTTGCGTGATGGGTTTGCTGAGTTACAGNNNAGATCGGAA |
| 69554_1 | TATGAGTGTGGAGTCGTTGCAAGAGGCATAGTAGTTGATTGCTNNNAGATCGGAA |
| 69564_1 | TATGAGTGTGGAGTCGTTGCCACATACACATACACAGTTGGCCNNNAGATCGGAAG |
| 69575_1 | TATGAGTGTGGAGTCGTTGCAAGGCCCTTTCTGTTTTGTTTCCNNNAGATCGGAAGA |
| 69597_1 | TATGAGTGTGGAGTCGTTGCCGTGTCCTGTTCTTCCTGTTTCNNNAGATCGGAAGAG |
| 69601_1 | TATGAGTGTGGAGTCGTTGCCCAGGTTCACACCATCCTTCTGNNNAGATCGGAAGAG |
| 69630_1 | TATGAGTGTGGAGTCGTTGCCTCCTGAGTTCACGCCATTCTCNNNAGATCGGAAGAG |
| 69633_1 | TATGAGTGTGGAGTCGTTGCCACAAGGAAAGTGCTCGAGAAGNNNAGATCGGAAG |
| 69649_1 | TATGAGTGTGGAGTCGTTGCGCGTAAGAACAAGTATTGGCCAGNNNAGATCGGAA |
| 69654_1 | TATGAGTGTGGAGTCGTTGCGTTCCATTCCCAAAAAGGAGCAANNNAGATCGGAAG |
| 69660_1 | TATGAGTGTGGAGTCGTTGCCCTGATCCACTAGAGAAATGGGGNNNAGATCGGAA |
| 69670_1 | TATGAGTGTGGAGTCGTTGCTTTGCCCTCTCTTACTGTTCCAANNNAGATCGGAAGA |
| 69684_1 | TATGAGTGTGGAGTCGTTGCGACCCTCTCAGTTCTTGGGAATTNNNAGATCGGAAG |
| 69741_1 | TATGAGTGTGGAGTCGTTGCATCACTTGCTTCCTTGCTTCCTANNNAGATCGGAAGA |
| 69747_1 | TATGAGTGTGGAGTCGTTGCCTTGACCTACTGGGCTCATGTGNNNAGATCGGAAGA |
| 69748_1 | TATGAGTGTGGAGTCGTTGCGCAACACCATGTACCTAATAGGCNNNAGATCGGAAG |
| 69783_1 | TATGAGTGTGGAGTCGTTGCACCCATTCATCTGTCTGTTGACANNNAGATCGGAAGA |
| 69789_1 | TATGAGTGTGGAGTCGTTGCGATCATAGCTCACTGCAGCCTCNNNAGATCGGAAGA |

| | |
|---|---|
| 69843_1 | TATGAGTGTGGAGTCGTTGCAACTTCCCAGACTCAAGCCATCNNNAGATCGGAAGA |
| 69859_1 | TATGAGTGTGGAGTCGTTGCGAACTCCTGGGCTCAAGTGATCNNNAGATCGGAAG |
| 69923_1 | TATGAGTGTGGAGTCGTTGCCCATGACGAACACACACGTACANNNAGATCGGAAGA |
| 69937_1 | TATGAGTGTGGAGTCGTTGCCAGGGAGTTTAGAAGCAACCCTNNNAGATCGGAAG |
| 69966_1 | TATGAGTGTGGAGTCGTTGCCTTGGCATCAGAACTCTGGGTTNNNAGATCGGAAG |
| 69978_1 | TATGAGTGTGGAGTCGTTGCCGTGGATACTGCAGCTTCTCTGNNNAGATCGGAAGA |
| 70000_1 | TATGAGTGTGGAGTCGTTGCATCTCAGTCTCTCCTATCCTCCTNNNAGATCGGAAGA |
| 70039_1 | TATGAGTGTGGAGTCGTTGCCTCTACTGAGCGCCCTTCCTAGNNNAGATCGGAAGA |
| 70071_1 | TATGAGTGTGGAGTCGTTGCCCTTTCCCGGTCTGCTAAGTAGNNNAGATCGGAAGA |
| 70072_1 | TATGAGTGTGGAGTCGTTGCTATCCCATATATCCTGCTTGGCNNNAGATCGGAAG |
| 70096_1 | TATGAGTGTGGAGTCGTTGCCGTCACCTTTTCTCGGTCTCTCNNNAGATCGGAAGAG |
| 70123_1 | TATGAGTGTGGAGTCGTTGCGACCCTGTCTCAAAAACGCAAAANNNAGATCGGAAG |
| 70140_1 | TATGAGTGTGGAGTCGTTGCTCATCGGGCACATCTGTAAAAGANNNAGATCGGAA |
| 70147_1 | TATGAGTGTGGAGTCGTTGCCACACAATCATGGTGGCCATAAGNNNAGATCGGAA |
| 70230_1 | TATGAGTGTGGAGTCGTTGCTGCAGTGGCTAGTGTCTGTAATCNNNAGATCGGAA |
| 70263_1 | TATGAGTGTGGAGTCGTTGCCCATATTGCCTCTCTTGGTCAGTNNNAGATCGGAAG |
| 70276_1 | TATGAGTGTGGAGTCGTTGCCTCCCGGGTTCAAACGATTCTCNNNAGATCGGAAGA |
| 70297_1 | TATGAGTGTGGAGTCGTTGCAGAGAGAGAGAGAGTTCCTGACCNNNAGATCGGAA |
| 70401_1 | TATGAGTGTGGAGTCGTTGCCTCATTGCAGCCTCAACTTCTCNNNAGATCGGAAGAG |
| 70404_1 | TATGAGTGTGGAGTCGTTGCCACAATCTCAGCTCACTGCAACNNNAGATCGGAAGA |
| 70415_1 | TATGAGTGTGGAGTCGTTGCGATCTTGGCTCACTGCAACTTCNNNAGATCGGAAGA |
| 70462_1 | TATGAGTGTGGAGTCGTTGCTTATATGGGAACGTGTGCAGACANNNAGATCGGAA |
| 70485_1 | TATGAGTGTGGAGTCGTTGCGGGAATGCAATGGCTTGATCTCNNNAGATCGGAAG |
| 70526_1 | TATGAGTGTGGAGTCGTTGCTGAAGAGACATTTCACTGAAGCANNNAGATCGGAA |
| 70527_1 | TATGAGTGTGGAGTCGTTGCTCTCTGTCTTTCTCTCTTCCCCTNNNAGATCGGAAGAG |
| 19_1 | TATGAGTGTGGAGTCGTTGCAACTACTGAACCGCCAGGCATNNNAGATCGGAAGAG |
| 62_1 | TATGAGTGTGGAGTCGTTGCGAGTGTGTTGGTCGAGAGCAATCNNNAGATCGGAA |
| 109_1 | TATGAGTGTGGAGTCGTTGCAGCCATCTGTGTGATGGGACCNNNAGATCGGAAGA |
| 178_1 | TATGAGTGTGGAGTCGTTGCGAGGGACTTCAGGCCACTGACNNNAGATCGGAAGA |
| 182_1 | TATGAGTGTGGAGTCGTTGCTTTGGCCATTCTTCGCCTGACNNNAGATCGGAAGAG |
| 222_1 | TATGAGTGTGGAGTCGTTGCTCCACCTTCTAGAGAAGCCGCNNNAGATCGGAAGAG |
| 234_1 | TATGAGTGTGGAGTCGTTGCTTTAGGGGTCCCAGACCTAGGNNNAGATCGGAAGA |
| 331_1 | TATGAGTGTGGAGTCGTTGCCTCTATGCTCCCAGGCTCACCNNNAGATCGGAAGAG |
| 368_1 | TATGAGTGTGGAGTCGTTGCCCAGGGAGGTGAGTTCAGACTNNNAGATCGGAAGA |
| 414_1 | TATGAGTGTGGAGTCGTTGCGAGGTTTGCTCTCACACAGGCNNNAGATCGGAAGA |
| 449_1 | TATGAGTGTGGAGTCGTTGCGGGTCTCATTCGGCGTCTCCCNNNAGATCGGAAGAG |
| 489_1 | TATGAGTGTGGAGTCGTTGCAGGCTTAGAGGGCAAGTGGAGNNNAGATCGGAAG |
| 535_1 | TATGAGTGTGGAGTCGTTGCTATCTCACTCCGGGCCTCCTGNNNAGATCGGAAGAG |
| 555_1 | TATGAGTGTGGAGTCGTTGCAATGTGTCCACTGATGCGCAGNNNAGATCGGAAGA |
| 588_1 | TATGAGTGTGGAGTCGTTGCTCAAGCCACATTCTGGGTTTTNNNAGATCGGAAGAG |
| 621_1 | TATGAGTGTGGAGTCGTTGCAGGGAAGCAGGATGGAGATTAGANNNAGATCGGA |
| 653_1 | TATGAGTGTGGAGTCGTTGCTGGTTGCCACTGTGAGTTCAGNNNAGATCGGAAGA |
| 666_1 | TATGAGTGTGGAGTCGTTGCCGCACCCAATAAAGCGAATGGNNNAGATCGGAAGA |

| | |
|---|---|
| 668_1 | TATGAGTGTGGAGTCGTTGCAACCCTTCTGGCTTTGTCCTTNNNAGATCGGAAGAG |
| 726_1 | TATGAGTGTGGAGTCGTTGCAGAGTTCCTGCTGCCTCAGACNNNAGATCGGAAGAG |
| 757_1 | TATGAGTGTGGAGTCGTTGCGGAGGAGTTCACAGATCCGCTNNNAGATCGGAAGA |
| 799_1 | TATGAGTGTGGAGTCGTTGCAGTCTCCCTCTCTTGTGGAGANNNAGATCGGAAGAG |
| 841_1 | TATGAGTGTGGAGTCGTTGCGTGCCAGGCTGCTCTCTGAAATNNNAGATCGGAAGA |
| 1028_1 | TATGAGTGTGGAGTCGTTGCGAGGTCGTGAGAGTGGTCTTCNNNAGATCGGAAGA |
| 1040_1 | TATGAGTGTGGAGTCGTTGCTATCGGTGATGTAAGGGGGCGNNNAGATCGGAAG |
| 1055_1 | TATGAGTGTGGAGTCGTTGCTTTCCACTTGGCCAACATTCCTTNNNAGATCGGAAGA |
| 1058_1 | TATGAGTGTGGAGTCGTTGCCAGTGGGTCTGCCCTTGTAGANNNAGATCGGAAGA |
| 1126_1 | TATGAGTGTGGAGTCGTTGCTTGGAAACCTCCAGAGTTGCANNNAGATCGGAAGA |
| 1159_1 | TATGAGTGTGGAGTCGTTGCGCACCAGTCCACTGGAAAATCNNNAGATCGGAAGAG |
| 1193_1 | TATGAGTGTGGAGTCGTTGCTCCGAAAGCCTTTGCTGTGTTCNNNAGATCGGAAGA |
| 1268_1 | TATGAGTGTGGAGTCGTTGCCTCTGTTGCCAGTGTGTGTGTNNNAGATCGGAAGAG |
| 1438_1 | TATGAGTGTGGAGTCGTTGCTGGAAAGCTGGAGGTTCAGGANNNAGATCGGAAG |
| 1441_1 | TATGAGTGTGGAGTCGTTGCTACCTGCCCTCCTTGAATGACACNNNAGATCGGAAG |
| 1468_1 | TATGAGTGTGGAGTCGTTGCCATCAGTGGCATCATCTTGGCNNNAGATCGGAAGAG |
| 1719_1 | TATGAGTGTGGAGTCGTTGCGAGGGGGCGGTTAGAATTCTCNNNAGATCGGAAGA |
| 1749_1 | TATGAGTGTGGAGTCGTTGCGCAGGAGGGTCTCTCAGAACTNNNAGATCGGAAGA |
| 1753_1 | TATGAGTGTGGAGTCGTTGCAGGGTTTTGCTCTTGTGGCACNNNAGATCGGAAGA |
| 1771_1 | TATGAGTGTGGAGTCGTTGCCAGGCTCGATCTCCCAAGGTCNNNAGATCGGAAGAG |
| 1797_1 | TATGAGTGTGGAGTCGTTGCCATTGTTGTGTCCAACACCANNNAGATCGGAAGAGC |
| 1897_1 | TATGAGTGTGGAGTCGTTGCCGGACAGCTCAAGAACAAGGTNNNAGATCGGAAGA |
| 1939_1 | TATGAGTGTGGAGTCGTTGCTCCTCTAGCCTCTGCAGTCTCNNNAGATCGGAAGAG |
| 1942_1 | TATGAGTGTGGAGTCGTTGCCACACCCAGTACTCCAGCCTTATNNNAGATCGGAAGA |
| 1990_1 | TATGAGTGTGGAGTCGTTGCTTCGGCAGCGTTAGGGTTTTTNNNAGATCGGAAGA |
| 2004_1 | TATGAGTGTGGAGTCGTTGCCATCTGTGCATTCAACCTTGGNNNAGATCGGAAGAG |
| 2052_1 | TATGAGTGTGGAGTCGTTGCTAGCAAAGACACGGGGTTCACNNNAGATCGGAAGA |
| 2130_1 | TATGAGTGTGGAGTCGTTGCTGGGACACACTGTGGTTGTCANNNAGATCGGAAGA |
| 2223_1 | TATGAGTGTGGAGTCGTTGCGGGTTGGGGATTGGGGTAATGNNNAGATCGGAAG |
| 2235_1 | TATGAGTGTGGAGTCGTTGCAAATCAACTGAGCAGCCCTTTCANNNAGATCGGAAG |
| 2329_1 | TATGAGTGTGGAGTCGTTGCCCACCATCCCCCACCCTTATCNNNAGATCGGAAGAGC |
| 2434_1 | TATGAGTGTGGAGTCGTTGCAGGCCGTGTATCTGTCACTCTNNNAGATCGGAAGAG |
| 2442_1 | TATGAGTGTGGAGTCGTTGCAGGCTCACTATCACACAGGCTNNNAGATCGGAAGAG |
| 2463_1 | TATGAGTGTGGAGTCGTTGCGGGAAACAGACCTCCCAGTTCNNNAGATCGGAAGA |
| 2661_1 | TATGAGTGTGGAGTCGTTGCGTGCTTCTGAGACCCAGGAGANNNAGATCGGAAGA |
| 2709_1 | TATGAGTGTGGAGTCGTTGCTAGTGTCTGTGCGGGTCTCTGNNNAGATCGGAAGA |
| 2738_1 | TATGAGTGTGGAGTCGTTGCGGGTTGGGGTGAGCAAAGTTTNNNAGATCGGAAG |
| 2784_1 | TATGAGTGTGGAGTCGTTGCCCTCTGTCCTCCCCTATGGGANNNAGATCGGAAGAG |
| 2816_1 | TATGAGTGTGGAGTCGTTGCTGGCAGCAAAACAAGTCACCTNNNAGATCGGAAGA |
| 2893_1 | TATGAGTGTGGAGTCGTTGCGCACCCGTGATGGTGTACTTCNNNAGATCGGAAGAG |
| 2930_1 | TATGAGTGTGGAGTCGTTGCCACACCACTGTTGGCCTGTTTNNNAGATCGGAAGAG |
| 2934_1 | TATGAGTGTGGAGTCGTTGCTACAGCTGCTTGACAACAGAANNNAGATCGGAAGA |
| 2939_1 | TATGAGTGTGGAGTCGTTGCCACCAGGCAGTTCTGCTTTCANNNAGATCGGAAGAG |

| | |
|---|---|
| 3027_1 | TATGAGTGTGGAGTCGTTGCGGGGGCATCACTCAACATTCANNNAGATCGGAAGA |
| 3170_1 | TATGAGTGTGGAGTCGTTGCATGTGTGAGGCGTTTACAGGGNNNAGATCGGAAG |
| 3251_1 | TATGAGTGTGGAGTCGTTGCAGGAAAAGGTGGTGGACAAGGNNNAGATCGGAAG |
| 3331_1 | TATGAGTGTGGAGTCGTTGCTTCTGGCCGTGTTCTGGTTAGNNNAGATCGGAAGA |
| 3335_1 | TATGAGTGTGGAGTCGTTGCAAAAGGCAAATGTGGGGACCCNNNAGATCGGAAG |
| 3377_1 | TATGAGTGTGGAGTCGTTGCCTCACCTTTCCATGCAGGCACNNNAGATCGGAAGAG |
| 3420_1 | TATGAGTGTGGAGTCGTTGCGCCAACGAGGGAGATAAGGTGNNNAGATCGGAAG |
| 3592_1 | TATGAGTGTGGAGTCGTTGCGCCTATTGTGCGGGACTACTCNNNAGATCGGAAGA |
| 3608_1 | TATGAGTGTGGAGTCGTTGCTGGGTGCAGGTGATAAGGACANNNAGATCGGAAG |
| 3642_1 | TATGAGTGTGGAGTCGTTGCTGTGGTGCAATTCTGTGCTGTNNNAGATCGGAAGA |
| 3652_1 | TATGAGTGTGGAGTCGTTGCTGACCCACTGATACTTCTGCCNNNAGATCGGAAGAG |
| 3662_1 | TATGAGTGTGGAGTCGTTGCCTGACCCTCTATGTGTCGGGGNNNAGATCGGAAGA |
| 3715_1 | TATGAGTGTGGAGTCGTTGCGGGTAGGAGAGTTGGGCATGANNNAGATCGGAAG |
| 3724_1 | TATGAGTGTGGAGTCGTTGCTAGGACCTGACCCTGTAGCTTNNNAGATCGGAAGAG |
| 3725_1 | TATGAGTGTGGAGTCGTTGCATGCTCGTGTGGATATGCATANNNAGATCGGAAGA |
| 3869_1 | TATGAGTGTGGAGTCGTTGCTTCTGCTGAGGAACCTTAGGCNNNAGATCGGAAGA |
| 3877_1 | TATGAGTGTGGAGTCGTTGCACACTACGTGACTTCTGGGTGNNNAGATCGGAAGA |
| 3964_1 | TATGAGTGTGGAGTCGTTGCGCTCACCATCCTCTGTTTGCANNNAGATCGGAAGAG |
| 4000_1 | TATGAGTGTGGAGTCGTTGCAAGATACATGAAACGGAGGCANNNAGATCGGAAG |
| 4038_1 | TATGAGTGTGGAGTCGTTGCCTATCGGTGATGTAAGGGGGCNNNAGATCGGAAG |
| 4080_1 | TATGAGTGTGGAGTCGTTGCTGTTTCCATGCCCCACTTCTTNNNAGATCGGAAGAGC |
| 4201_1 | TATGAGTGTGGAGTCGTTGCGGCTAATGGTTTCGGCATGGANNNAGATCGGAAGA |
| 4207_1 | TATGAGTGTGGAGTCGTTGCTACACATGGTGGCTGTGTCTGNNNAGATCGGAAGA |
| 4221_1 | TATGAGTGTGGAGTCGTTGCTGTGTATGGTCTGGGTTGGCANNNAGATCGGAAGA |
| 4230_1 | TATGAGTGTGGAGTCGTTGCCACCACTCACACACCTTCCTGNNNAGATCGGAAGAGC |
| 4232_1 | TATGAGTGTGGAGTCGTTGCGTGTGTTGGGGGAGGGTTCAGNNNAGATCGGAAG |
| 4238_1 | TATGAGTGTGGAGTCGTTGCGTGATGAGGAACTGACCTGACNNNAGATCGGAAGA |
| 4302_1 | TATGAGTGTGGAGTCGTTGCGGGTCTCAGTCTGGTGCTTTTNNNAGATCGGAAGA |
| 4498_1 | TATGAGTGTGGAGTCGTTGCACAACCTGCCCTCACCCAAATNNNAGATCGGAAGAG |
| 4559_1 | TATGAGTGTGGAGTCGTTGCTTCACCGAACCTCACCCAGACNNNAGATCGGAAGAG |
| 4571_1 | TATGAGTGTGGAGTCGTTGCCTCCAAGGTCAAGTCAGCCACNNNAGATCGGAAGAG |
| 4633_1 | TATGAGTGTGGAGTCGTTGCAAATACAAGTGGCCTCAGCTGACNNNAGATCGGAA |
| 4708_1 | TATGAGTGTGGAGTCGTTGCCCTGTGTGCTTCCTGTGGTTTNNNAGATCGGAAGAG |
| 4745_1 | TATGAGTGTGGAGTCGTTGCAGGTATGGGAAGAACCCTCCTNNNAGATCGGAAGA |
| 4882_1 | TATGAGTGTGGAGTCGTTGCTGTGTCAGGTGGGTCTCACTCNNNAGATCGGAAGA |
| 4940_1 | TATGAGTGTGGAGTCGTTGCCTGACTAGGGGAGTGGTACCCNNNAGATCGGAAGA |
| 4977_1 | TATGAGTGTGGAGTCGTTGCGTCACTCAGCACCCAGGCTAGNNNAGATCGGAAGA |
| 4980_1 | TATGAGTGTGGAGTCGTTGCGTTGGGAGGAACCAGGGCTATNNNAGATCGGAAG |
| 4984_1 | TATGAGTGTGGAGTCGTTGCTGAGAAGCATACCAGGTCCCCNNNAGATCGGAAGA |
| 4997_1 | TATGAGTGTGGAGTCGTTGCAGTGAAGTGCAGCCAGTCTAGANNNAGATCGGAAG |
| 5043_1 | TATGAGTGTGGAGTCGTTGCGACCCCCACACACAGACATGCNNNAGATCGGAAGAG |
| 5096_1 | TATGAGTGTGGAGTCGTTGCAAACATATCTGGGGCCAGGCCNNNAGATCGGAAGA |
| 5122_1 | TATGAGTGTGGAGTCGTTGCTTGGCAGGGTTTCACAATGTCNNNAGATCGGAAGA |

| | |
|---|---|
| 5215_1 | TATGAGTGTGGAGTCGTTGCGTGTCAGCCTTTGTTTTGGCCNNNAGATCGGAAGAG |
| 5323_1 | TATGAGTGTGGAGTCGTTGCGAACCCCAGGCAGACAGAATCNNNAGATCGGAAGA |
| 5325_1 | TATGAGTGTGGAGTCGTTGCTCACATCAGTCCCATCCCTCCNNNAGATCGGAAGAGC |
| 5352_1 | TATGAGTGTGGAGTCGTTGCCTCACTTCATCCCCGAATTCTCCNNNAGATCGGAAGA |
| 5379_1 | TATGAGTGTGGAGTCGTTGCTGAGTTTCAGAAAGCCTTGGTCCNNNAGATCGGAAG |
| 5545_1 | TATGAGTGTGGAGTCGTTGCCCCCATTTGTCTGAAGTGCTGNNNAGATCGGAAGAG |
| 5605_1 | TATGAGTGTGGAGTCGTTGCGCTAACACAACCATGGCTTGNNNAGATCGGAAGAG |
| 5722_1 | TATGAGTGTGGAGTCGTTGCGTTATTCCAGCAAGGAGGAGGCNNNAGATCGGAAG |
| 5733_1 | TATGAGTGTGGAGTCGTTGCGCCTCCTGGCCTATTCTTTAGTNNNAGATCGGAAGA |
| 6080_1 | TATGAGTGTGGAGTCGTTGCGTTCAGAAAGGCGTGCACTTCNNNAGATCGGAAGA |
| 6274_1 | TATGAGTGTGGAGTCGTTGCCTTGTCTGTCCCCACTCGCTCNNNAGATCGGAAGAG |
| 6487_1 | TATGAGTGTGGAGTCGTTGCCCAGAAAAGCCCCCATCAGTTCNNNAGATCGGAAGA |
| 6528_1 | TATGAGTGTGGAGTCGTTGCCCACAGAGGAAACGGCCTTTCNNNAGATCGGAAGA |
| 6651_1 | TATGAGTGTGGAGTCGTTGCAGATTCCCATGCCATAGCTGCNNNAGATCGGAAGAG |
| 6708_1 | TATGAGTGTGGAGTCGTTGCCTTTAGGGCTGTCCCTCCTGGNNNAGATCGGAAGAG |
| 6742_1 | TATGAGTGTGGAGTCGTTGCCCTGACTAATGTGTGGTTTGCAGNNNAGATCGGAA |
| 6763_1 | TATGAGTGTGGAGTCGTTGCGAAAAGGCCAGAGCTTCTGCCNNNAGATCGGAAGA |
| 6780_1 | TATGAGTGTGGAGTCGTTGCACTTCTGCAGCTTCATGTGCCNNNAGATCGGAAGAG |
| 6881_1 | TATGAGTGTGGAGTCGTTGCGATTGGAGTTCAGTGGCGGAGNNNAGATCGGAAG |
| 6949_1 | TATGAGTGTGGAGTCGTTGCCTCTCACTTCGCTTCCTGATCAGNNNAGATCGGAAGA |
| 6963_1 | TATGAGTGTGGAGTCGTTGCCCACGTCTCTGAACAGCTCAGNNNAGATCGGAAGAG |
| 6971_1 | TATGAGTGTGGAGTCGTTGCGCCACTGTCATCTGGTGGTTANNNAGATCGGAAGAG |
| 7098_1 | TATGAGTGTGGAGTCGTTGCGGTAGGTCCGTAGGTTTCAGCNNNAGATCGGAAGA |
| 7107_1 | TATGAGTGTGGAGTCGTTGCTTTCCGCACCCTATTTGCTTGNNNAGATCGGAAGAG |
| 7121_1 | TATGAGTGTGGAGTCGTTGCCGGCTGCAGAGCTCAAACTACNNNAGATCGGAAGA |
| 7205_1 | TATGAGTGTGGAGTCGTTGCCCCCCAAAGTCAGCTACTCAANNNAGATCGGAAGAG |
| 7211_1 | TATGAGTGTGGAGTCGTTGCATGTGCTGAACTACAGAAGCANNNAGATCGGAAGA |
| 7229_1 | TATGAGTGTGGAGTCGTTGCACCACTGCCTCCTCAGCATAGNNNAGATCGGAAGAG |
| 7255_1 | TATGAGTGTGGAGTCGTTGCCGTCGGCATCTACACCAATGTNNNAGATCGGAAGAG |
| 7309_1 | TATGAGTGTGGAGTCGTTGCTTTCACTCACCCGTAGCTTGGNNNAGATCGGAAGAG |
| 7353_1 | TATGAGTGTGGAGTCGTTGCACACACACATTTTTCTGACACACGNNNAGATCGGAA |
| 7504_1 | TATGAGTGTGGAGTCGTTGCTCAGGGATCAGCATACCATGGNNNAGATCGGAAGA |
| 7682_1 | TATGAGTGTGGAGTCGTTGCGTGTTCTTAGCCCCACTCGACNNNAGATCGGAAGAG |
| 7695_1 | TATGAGTGTGGAGTCGTTGCCCAAACTGCCCATTTCCAGGGNNNAGATCGGAAGAG |
| 7714_1 | TATGAGTGTGGAGTCGTTGCTTGGTCCTCATGATCCCTTTTCTNNNAGATCGGAAGA |
| 7789_1 | TATGAGTGTGGAGTCGTTGCCTCCGGGTTCAAAGCTATGGGNNNAGATCGGAAGA |
| 7942_1 | TATGAGTGTGGAGTCGTTGCGAGAGCACACAGTCATTCACTGGNNNAGATCGGAA |
| 8013_1 | TATGAGTGTGGAGTCGTTGCGTCCATGCTGCAGGTTTCTGGNNNAGATCGGAAGA |
| 8045_1 | TATGAGTGTGGAGTCGTTGCCAACGGAGCAATCTCCCCTCANNNAGATCGGAAGAG |
| 8304_1 | TATGAGTGTGGAGTCGTTGCCCCCCAAGAAAGTGACCCAAGNNNAGATCGGAAGA |
| 8331_1 | TATGAGTGTGGAGTCGTTGCCAAACAATGGGGCAAGCACTGNNNAGATCGGAAGA |
| 8369_1 | TATGAGTGTGGAGTCGTTGCGGCTCCGAAGAAAGAGACCTGNNNAGATCGGAAGA |
| 8396_1 | TATGAGTGTGGAGTCGTTGCCTTCACATCACAGCTGGCGGANNNAGATCGGAAGA |

| | |
|---|---|
| 8452_1 | TATGAGTGTGGAGTCGTTGCATGCTTCATGCTTCCTCTGCTGNNNAGATCGGAAGA |
| 8512_1 | TATGAGTGTGGAGTCGTTGCACTACGTGACTTCTGGGTGTCTGNNNAGATCGGAAG |
| 8553_1 | TATGAGTGTGGAGTCGTTGCTTAGTTGGGTGCTGTGGCAGANNNAGATCGGAAGA |
| 8591_1 | TATGAGTGTGGAGTCGTTGCTCATCTCTGTTCTTTGCTCCGTGNNNAGATCGGAAGA |
| 8643_1 | TATGAGTGTGGAGTCGTTGCGTTCACAGCTCCACCCATGTGNNNAGATCGGAAGAG |
| 8673_1 | TATGAGTGTGGAGTCGTTGCTCCCAGAAATCCAGCCCTGAGNNNAGATCGGAAGAG |
| 8812_1 | TATGAGTGTGGAGTCGTTGCTACCAGATGTGGGTGCATCTTNNNAGATCGGAAGA |
| 8936_1 | TATGAGTGTGGAGTCGTTGCCCTGACCTCATTGCGCAAAGGNNNAGATCGGAAGA |
| 9241_1 | TATGAGTGTGGAGTCGTTGCCTGAAGGACGGCTACCCAGATNNNAGATCGGAAGA |
| 9286_1 | TATGAGTGTGGAGTCGTTGCCCACCAAGGGAATTCAATGGGNNNAGATCGGAAGA |
| 9327_1 | TATGAGTGTGGAGTCGTTGCCAGACCTAGGGGCAGACAATTNNNAGATCGGAAGA |
| 9422_1 | TATGAGTGTGGAGTCGTTGCGGTTGTTTACCCAGCAGGTGGNNNAGATCGGAAGA |
| 9438_1 | TATGAGTGTGGAGTCGTTGCCTACCCAGCCCCCTCACACGTATNNNAGATCGGAAGA |
| 9460_1 | TATGAGTGTGGAGTCGTTGCCTATCGAGCACTCTTGGGGAGNNNAGATCGGAAGA |
| 9753_1 | TATGAGTGTGGAGTCGTTGCAGCACACTGAGTGAGAACCCTNNNAGATCGGAAGA |
| 9828_1 | TATGAGTGTGGAGTCGTTGCGTGGTGGATCTCCCTTGTTCTNNNAGATCGGAAGAG |
| 10348_1 | TATGAGTGTGGAGTCGTTGCACTCCTACAAGAACGGCACACNNNAGATCGGAAGAG |
| 10893_1 | TATGAGTGTGGAGTCGTTGCTAGACCAGTTGGGGGTGTGTGNNNAGATCGGAAG |
| 10917_1 | TATGAGTGTGGAGTCGTTGCAGGAAAATGACCCTCCCCTCCNNNAGATCGGAAGAG |
| 11077_1 | TATGAGTGTGGAGTCGTTGCACTTGCTGGGAACAAGGGAAAAANNNAGATCGGAA |
| 11078_1 | TATGAGTGTGGAGTCGTTGCGTGGCCTGGTAGGGGGAAAAANNNAGATCGGAAG |
| 11238_1 | TATGAGTGTGGAGTCGTTGCTCAAACCCCAACTCTGGGCTCNNNAGATCGGAAGAG |
| 11258_1 | TATGAGTGTGGAGTCGTTGCAAAGCTACACCCACTCCCCACNNNAGATCGGAAGAG |
| 11326_1 | TATGAGTGTGGAGTCGTTGCGATACGGGGTTCCAGGAAAGGNNNAGATCGGAAG |
| 11377_1 | TATGAGTGTGGAGTCGTTGCATAGCAGGCAGGAAATCTGGCNNNAGATCGGAAGA |
| 11378_1 | TATGAGTGTGGAGTCGTTGCGCTGGCCTTCTCTCTAGCTCTNNNAGATCGGAAGAG |
| 11484_1 | TATGAGTGTGGAGTCGTTGCTAGCAGGTGTCTCTGGCTTCTNNNAGATCGGAAGAG |
| 11658_1 | TATGAGTGTGGAGTCGTTGCCATCCACTCTCCATGGTGTGGNNNAGATCGGAAGAG |
| 11790_1 | TATGAGTGTGGAGTCGTTGCGAGTAGAGGCTGAGGCCTGGANNNAGATCGGAAG |
| 11853_1 | TATGAGTGTGGAGTCGTTGCTGTCTCCCTCTCCACTTCTGGNNNAGATCGGAAGAG |
| 11913_1 | TATGAGTGTGGAGTCGTTGCCCACCCTCCACCTGCAGATTTTANNNAGATCGGAAGA |
| 11959_1 | TATGAGTGTGGAGTCGTTGCTAACCAGGGCACAGAGACACANNNAGATCGGAAGA |
| 12010_1 | TATGAGTGTGGAGTCGTTGCTGATTGGCACGCACCTCTTTTNNNAGATCGGAAGAG |
| 12154_1 | TATGAGTGTGGAGTCGTTGCCAGCTTCACACTTCAGCCCATNNNAGATCGGAAGAG |
| 12164_1 | TATGAGTGTGGAGTCGTTGCCTCTGGCACGCTGCTTATCAGNNNAGATCGGAAGAG |
| 12205_1 | TATGAGTGTGGAGTCGTTGCCTCCTAACCTGTGCTCGTGTGNNNAGATCGGAAGAG |
| 12232_1 | TATGAGTGTGGAGTCGTTGCGAGTACCATCTCGGCTCACCGNNNAGATCGGAAGAG |
| 12268_1 | TATGAGTGTGGAGTCGTTGCGATGCAGGGGGTTGGTTTCAGNNNAGATCGGAAG |
| 12279_1 | TATGAGTGTGGAGTCGTTGCCTTATGTAACCGAGCCCGGGTAANNNAGATCGGAA |
| 12340_1 | TATGAGTGTGGAGTCGTTGCGCAAAGCACAACCTCAGCTCANNNAGATCGGAAGAG |
| 53435_1 | TATGAGTGTGGAGTCGTTGCCCCCCGCCAAAAAAAAGAGAAATNNNAGATCGGAA |
| 53444_1 | TATGAGTGTGGAGTCGTTGCGGGTAACACAAGCACCTCTGTNNNAGATCGGAAGA |
| 53462_1 | TATGAGTGTGGAGTCGTTGCCCAGAAGAGCTTCCTCTGCTAAGNNNAGATCGGAAG |

| | |
|---|---|
| 53469_1 | TATGAGTGTGGAGTCGTTGCAATAAAAAGAGAGCTCTGGGCCANNNAGATCGGAA |
| 53471_1 | TATGAGTGTGGAGTCGTTGCCTGGAGAGGGGTGACACAAACNNNAGATCGGAAGA |
| 53505_1 | TATGAGTGTGGAGTCGTTGCGGCTTTTCTGGTGCTGTTCACNNNAGATCGGAAGAG |
| 53506_1 | TATGAGTGTGGAGTCGTTGCATGTATGTATATATGCCGGGCGCNNNAGATCGGAA |
| 53525_1 | TATGAGTGTGGAGTCGTTGCAGAATTCCTGGCTGTCTTGCTTANNNAGATCGGAAG |
| 53526_1 | TATGAGTGTGGAGTCGTTGCGAGCAGATAATGGCAGGGTGTNNNAGATCGGAAG |
| 53580_1 | TATGAGTGTGGAGTCGTTGCTTAGCCAGGATGGTCTCGATTTCNNNAGATCGGAAG |
| 53593_1 | TATGAGTGTGGAGTCGTTGCCAAGTAGCTGGGACTACAGGCNNNAGATCGGAAGA |
| 53597_1 | TATGAGTGTGGAGTCGTTGCCCATTCTCCTGCCTCAGCTTCNNNAGATCGGAAGAGC |
| 53602_1 | TATGAGTGTGGAGTCGTTGCCTCATTGGCTCACACCTGAAATCNNNAGATCGGAAG |
| 53606_1 | TATGAGTGTGGAGTCGTTGCTTTGCCTGTTTGGTTGTTTAGCANNNAGATCGGAAG |
| 53618_1 | TATGAGTGTGGAGTCGTTGCGTCTCTGAGGCTTGGTGTATGAANNNAGATCGGAA |
| 53638_1 | TATGAGTGTGGAGTCGTTGCAGAGTAGCTGGGACTACAGGCNNNAGATCGGAAGA |
| 53643_1 | TATGAGTGTGGAGTCGTTGCCAAGTAGCTGGGACTACAGGCNNNAGATCGGAAGA |
| 53669_1 | TATGAGTGTGGAGTCGTTGCTTTGGGTTTGCAGAAGGTTGTTTNNNAGATCGGAA |
| 53684_1 | TATGAGTGTGGAGTCGTTGCAAAACTGGCACAAGACAAGGATGNNNAGATCGGAA |
| 53709_1 | TATGAGTGTGGAGTCGTTGCGGGGAAAGAGGGAAGAGGAAAGNNNAGATCGGA |
| 53774_1 | TATGAGTGTGGAGTCGTTGCGATGGACATTGGGGTTCTGCTNNNAGATCGGAAGA |
| 53777_1 | TATGAGTGTGGAGTCGTTGCCTGAGAAAGCCCATCAGAGTCTTNNNAGATCGGAAG |
| 53806_1 | TATGAGTGTGGAGTCGTTGCTGCATCTCAGCTCACTGCAAGNNNAGATCGGAAGAG |
| 53811_1 | TATGAGTGTGGAGTCGTTGCTCCCAAAGTGCTGCAATTACATGNNNAGATCGGAAG |
| 53821_1 | TATGAGTGTGGAGTCGTTGCCTCACTGAAAGCTCCACCTCCNNNAGATCGGAAGAG |
| 53823_1 | TATGAGTGTGGAGTCGTTGCCAAGTAGCTGGGACTACAGGCNNNAGATCGGAAGA |
| 53825_1 | TATGAGTGTGGAGTCGTTGCATCTCGGCTCACTGAAAGCTCNNNAGATCGGAAGAG |
| 53854_1 | TATGAGTGTGGAGTCGTTGCGTAGTCCCAGGTAAATCAGGAGGNNNAGATCGGAA |
| 53867_1 | TATGAGTGTGGAGTCGTTGCATACCAGGCTTGTTTCTCCCAATNNNAGATCGGAAG |
| 53889_1 | TATGAGTGTGGAGTCGTTGCGAGACGGAGTCTTGCTCTGTCNNNAGATCGGAAGA |
| 53896_1 | TATGAGTGTGGAGTCGTTGCGGAGTATAGTGGCGCTGTCTCNNNAGATCGGAAGA |
| 53905_1 | TATGAGTGTGGAGTCGTTGCTCCCAGGTTCAAACGATTCTCTTNNNAGATCGGAAG |
| 53966_1 | TATGAGTGTGGAGTCGTTGCTCCACTGTCCCTAACTCTCTTCTNNNAGATCGGAAGA |
| 53970_1 | TATGAGTGTGGAGTCGTTGCGCTTGCAGTGAGCTGAGATTGNNNAGATCGGAAGA |
| 53991_1 | TATGAGTGTGGAGTCGTTGCTCCCTTGTTACCTTGAAAGGAGCNNNAGATCGGAAG |
| 54030_1 | TATGAGTGTGGAGTCGTTGCCCATTCTCCTGCCTCAGACTCNNNAGATCGGAAGAG |
| 54035_1 | TATGAGTGTGGAGTCGTTGCCAAGTAGCTGGGACTACAGGCNNNAGATCGGAAGA |
| 54068_1 | TATGAGTGTGGAGTCGTTGCTTCCTTGAACACCTACTACCTGCNNNAGATCGGAAGA |
| 54069_1 | TATGAGTGTGGAGTCGTTGCTTAGCACCAAAGAACATAGCCCANNNAGATCGGAAG |
| 54091_1 | TATGAGTGTGGAGTCGTTGCGGAGGGAGGAAGAAAAGACCAAANNNAGATCGGA |
| 54135_1 | TATGAGTGTGGAGTCGTTGCCAAGTAGCTGGGACTACAGGCNNNAGATCGGAAGA |
| 54146_1 | TATGAGTGTGGAGTCGTTGCCAGAGTGCTGGGATTACAGGCNNNAGATCGGAAGA |
| 54159_1 | TATGAGTGTGGAGTCGTTGCCAACTTCCTTTGCATATGAGGCCNNNAGATCGGAAG |
| 54165_1 | TATGAGTGTGGAGTCGTTGCACTTGAGATCGGGAGTTCAAGACNNNAGATCGGAA |
| 54208_1 | TATGAGTGTGGAGTCGTTGCCAGTGGTGTGATCTTGGCTCANNNAGATCGGAAGA |
| 54209_1 | TATGAGTGTGGAGTCGTTGCCAAGTAGCTGGGACTACAGGCNNNAGATCGGAAGA |

| | |
|---|---|
| 54229_1 | TATGAGTGTGGAGTCGTTGCGGAGTACAGTGGTGCGATCTCNNNAGATCGGAAGA |
| 54231_1 | TATGAGTGTGGAGTCGTTGCCAAGTAGCTGGGACTACAGGCNNNAGATCGGAAGA |
| 54258_1 | TATGAGTGTGGAGTCGTTGCCATAAGAGAAGGGGAAAGGGGAGNNNAGATCGGA |
| 54262_1 | TATGAGTGTGGAGTCGTTGCCATCGAGGCCAACTTCAAAGACNNNAGATCGGAAGA |
| 54264_1 | TATGAGTGTGGAGTCGTTGCATCAATTACCTGTGACCCCCAAANNNAGATCGGAAG |
| 54270_1 | TATGAGTGTGGAGTCGTTGCAGAGTAGGCCAATAGACTCCAGTNNNAGATCGGAA |
| 54299_1 | TATGAGTGTGGAGTCGTTGCGAGATTCCGAAACTGGGAAAACCNNNAGATCGGAA |
| 54303_1 | TATGAGTGTGGAGTCGTTGCCAATCAATTGCCTGTGATGGTGANNNAGATCGGAAG |
| 54325_1 | TATGAGTGTGGAGTCGTTGCCAAGTAGCTGGGACTACAGGCNNNAGATCGGAAGA |
| 54338_1 | TATGAGTGTGGAGTCGTTGCCAAGTAGCTGGGACTACAGGCNNNAGATCGGAAGA |
| 54361_1 | TATGAGTGTGGAGTCGTTGCCTCCTGAGTAGCTGGAATCACAGNNNAGATCGGAAG |
| 54376_1 | TATGAGTGTGGAGTCGTTGCGTCTCGAACTCCTGACCTCAGNNNAGATCGGAAGAG |
| 54392_1 | TATGAGTGTGGAGTCGTTGCTTCAGAGGGAACATGTGGGGAANNNAGATCGGAAG |
| 54398_1 | TATGAGTGTGGAGTCGTTGCAAGAGCAGAGAGAAAAGGAACGGNNNAGATCGGA |
| 54400_1 | TATGAGTGTGGAGTCGTTGCGTTGCAGTGAACTGAGATCGTGNNNAGATCGGAAG |
| 54402_1 | TATGAGTGTGGAGTCGTTGCTTAGCCAGGCGTAGTGTCATGNNNAGATCGGAAGA |
| 54409_1 | TATGAGTGTGGAGTCGTTGCTTAGCTGTGTGTGGTGGTGTGNNNAGATCGGAAGA |
| 54424_1 | TATGAGTGTGGAGTCGTTGCAATTTTTGCAGTCTGGGCTTGTTNNNAGATCGGAAG |
| 54446_1 | TATGAGTGTGGAGTCGTTGCCCAAAGGTCTATCAACAGGGGAANNNAGATCGGAA |
| 54463_1 | TATGAGTGTGGAGTCGTTGCTGTCTTGAGTGCTACCTTGTACCNNNAGATCGGAAG |
| 54485_1 | TATGAGTGTGGAGTCGTTGCTGTCAATCAAGACTGCTGATCCANNNAGATCGGAAG |
| 54492_1 | TATGAGTGTGGAGTCGTTGCGGGTTCATGCCATTCTTCTGCNNNAGATCGGAAGAG |
| 54495_1 | TATGAGTGTGGAGTCGTTGCCTGACCTCAGGCAATCTACCCNNNAGATCGGAAGAG |
| 54502_1 | TATGAGTGTGGAGTCGTTGCGCAATCTCAGCTCACTGCAAGNNNAGATCGGAAGAG |
| 54530_1 | TATGAGTGTGGAGTCGTTGCAGAGTAGCTGGGACTACAGGCNNNAGATCGGAAGA |
| 54536_1 | TATGAGTGTGGAGTCGTTGCTTACGAGTCAAATTTGCACAGGCNNNAGATCGGAAG |
| 54550_1 | TATGAGTGTGGAGTCGTTGCGCTGTTTCAGATCAGTGGCATACNNNAGATCGGAAG |
| 54555_1 | TATGAGTGTGGAGTCGTTGCTGCCTTGCTGATTATACACAGTGNNNAGATCGGAAG |
| 54574_1 | TATGAGTGTGGAGTCGTTGCGAACTATGTTGGGCTTGCATTGTNNNAGATCGGAAG |
| 54576_1 | TATGAGTGTGGAGTCGTTGCCAGACTCCCTTCACTCCATTCCNNNAGATCGGAAGAG |
| 54590_1 | TATGAGTGTGGAGTCGTTGCCCATTCTCCTGCCTCAGCTTCNNNAGATCGGAAGAGC |
| 54607_1 | TATGAGTGTGGAGTCGTTGCCTCTGTTGCCTACGTTGGAGTNNNAGATCGGAAGAG |
| 54613_1 | TATGAGTGTGGAGTCGTTGCACAATCTCGGCTCACTGCAAGNNNAGATCGGAAGAG |
| 54615_1 | TATGAGTGTGGAGTCGTTGCGGTCCCTCACAGACGAAAGTGNNNAGATCGGAAGA |
| 54652_1 | TATGAGTGTGGAGTCGTTGCGAATGATACGGCCCTCTCCTGNNNAGATCGGAAGAG |
| 54655_1 | TATGAGTGTGGAGTCGTTGCGAATGGAAAACTGTACACACGCANNNAGATCGGAA |
| 54660_1 | TATGAGTGTGGAGTCGTTGCGAGACGGAGTCTTGCTCTGTCNNNAGATCGGAAGA |
| 54670_1 | TATGAGTGTGGAGTCGTTGCCAAGTAGCTGGGACTACAGGCNNNAGATCGGAAGA |
| 54698_1 | TATGAGTGTGGAGTCGTTGCTCCTCAGCATACAACTATTCTGCTNNNAGATCGGAAG |
| 54702_1 | TATGAGTGTGGAGTCGTTGCCTGAATCTATTCCCAGGCCTCTCNNNAGATCGGAAGA |
| 54703_1 | TATGAGTGTGGAGTCGTTGCAAGCACCCATGAGGAATACACATNNNAGATCGGAAG |
| 54712_1 | TATGAGTGTGGAGTCGTTGCAAGTCTCGCTTTGTCACCCAGNNNAGATCGGAAGAG |
| 54731_1 | TATGAGTGTGGAGTCGTTGCCAAGTAGCTGGGACTACAGGCNNNAGATCGGAAGA |

| | |
|---|---|
| 54740_1 | TATGAGTGTGGAGTCGTTGCATGATCTCGGCTCACTGCAAGNNNAGATCGGAAGAG |
| 54745_1 | TATGAGTGTGGAGTCGTTGCCAAGTAGCTGGGACTACAGGCNNNAGATCGGAAGA |
| 54747_1 | TATGAGTGTGGAGTCGTTGCCAAGTAGCTGGGACTACAGGCNNNAGATCGGAAGA |
| 54778_1 | TATGAGTGTGGAGTCGTTGCGAAGACATACAAGCAGCCAACAGNNNAGATCGGAA |
| 54781_1 | TATGAGTGTGGAGTCGTTGCCTTGCAGTGAGCCAAGATCACNNNAGATCGGAAGAG |
| 54791_1 | TATGAGTGTGGAGTCGTTGCTCTGCACAAGCCTTCTCTTTTTGNNNAGATCGGAAGA |
| 54793_1 | TATGAGTGTGGAGTCGTTGCGGCAACACCTAAGTACAGGAACANNNAGATCGGAA |
| 54837_1 | TATGAGTGTGGAGTCGTTGCGTCTCGAACTCCTGACCTCAGNNNAGATCGGAAGAG |
| 54838_1 | TATGAGTGTGGAGTCGTTGCATTACAGGCATGAGACGGAATCTNNNAGATCGGAA |
| 54841_1 | TATGAGTGTGGAGTCGTTGCAGACGGAGTCTGTCTCTGTCANNNAGATCGGAAGA |
| 54859_1 | TATGAGTGTGGAGTCGTTGCGAGACGGAGTCTTGCTCTGTCNNNAGATCGGAAGA |
| 54871_1 | TATGAGTGTGGAGTCGTTGCGCAATCTTGGCTCACTGCAAGNNNAGATCGGAAGAG |
| 54877_1 | TATGAGTGTGGAGTCGTTGCACCTCGCATTCATTGGCTTTTTTNNNAGATCGGAAGA |
| 54897_1 | TATGAGTGTGGAGTCGTTGCCATTCTCCTGCCTCAGCCTCNNNAGATCGGAAGAGCA |
| 54910_1 | TATGAGTGTGGAGTCGTTGCCTTGCAGTGAGCCGAGATCTCNNNAGATCGGAAGAG |
| 54917_1 | TATGAGTGTGGAGTCGTTGCAGCCTTTTCTGAGACCTTGTAGGNNNAGATCGGAAG |
| 54924_1 | TATGAGTGTGGAGTCGTTGCAGGTCTCTCTACACTCAGGCTAGNNNAGATCGGAAG |
| 54926_1 | TATGAGTGTGGAGTCGTTGCGGAGGCTGAAGCAGGAGAATGNNNAGATCGGAAG |
| 54931_1 | TATGAGTGTGGAGTCGTTGCGCTTACCAAGCATCTAACGTTAGTNNNAGATCGGAA |
| 54936_1 | TATGAGTGTGGAGTCGTTGCAAAATCAATAATGAGGGCCCAGCNNNAGATCGGAA |
| 54939_1 | TATGAGTGTGGAGTCGTTGCGGAAAGAAGGAAGGAAGGAAGGANNNAGATCGG |
| 54982_1 | TATGAGTGTGGAGTCGTTGCATCTCGGCTTACTGCAACTCCNNNAGATCGGAAGAG |
| 55016_1 | TATGAGTGTGGAGTCGTTGCCAGGCCTATCTGTACCCCTACTANNNAGATCGGAAG |
| 55040_1 | TATGAGTGTGGAGTCGTTGCACAAATTTGAAAAGCTAGCAGCANNNAGATCGGAA |
| 55056_1 | TATGAGTGTGGAGTCGTTGCCAAGTAGCTGGGACTACAGGCNNNAGATCGGAAGA |
| 55066_1 | TATGAGTGTGGAGTCGTTGCCAAGTAGCTGGGACTACAGGCNNNAGATCGGAAGA |
| 55070_1 | TATGAGTGTGGAGTCGTTGCGCGATCTCAGCTCACTGCAAGNNNAGATCGGAAGAG |
| 55076_1 | TATGAGTGTGGAGTCGTTGCCAAGTAGCTGGGACTACAGGCNNNAGATCGGAAGA |
| 55078_1 | TATGAGTGTGGAGTCGTTGCGACAGAGTCTCATCCCGTTGCNNNAGATCGGAAGAG |
| 55079_1 | TATGAGTGTGGAGTCGTTGCTTCAAGTGGTTCTCCTGCCTCNNNAGATCGGAAGAG |
| 55080_1 | TATGAGTGTGGAGTCGTTGCGCTATCTCGGCTCACTGCAAGNNNAGATCGGAAGAG |
| 55102_1 | TATGAGTGTGGAGTCGTTGCACCAGAGAGAATGTTCAGCAATCTNNNAGATCGGAA |
| 55103_1 | TATGAGTGTGGAGTCGTTGCGAGGCAGGAGAATGGTGTGAANNNAGATCGGAAG |
| 55114_1 | TATGAGTGTGGAGTCGTTGCATGAATTAGCCTAGTGTGGTGGCNNNAGATCGGAA |
| 55123_1 | TATGAGTGTGGAGTCGTTGCATGGGAAATGCTGCAAGAACATTNNNAGATCGGAA |
| 55135_1 | TATGAGTGTGGAGTCGTTGCAAGGTATTTTGCAGTGCTAACCANNNAGATCGGAAG |
| 55145_1 | TATGAGTGTGGAGTCGTTGCAGTGCTCAACTCCAAATGGTAGANNNAGATCGGAAG |
| 55149_1 | TATGAGTGTGGAGTCGTTGCGAGGCAGGAGAATCGCTTGAANNNAGATCGGAAGA |
| 55158_1 | TATGAGTGTGGAGTCGTTGCGAGATTCTGTTTTGACGAAGGCANNNAGATCGGAA |
| 55165_1 | TATGAGTGTGGAGTCGTTGCGAGACGGAGTCTTGCTCTGTCNNNAGATCGGAAGA |
| 55170_1 | TATGAGTGTGGAGTCGTTGCCTCCCAGGTTCAGGCAAATCTNNNAGATCGGAAGAG |
| 55177_1 | TATGAGTGTGGAGTCGTTGCTTCAAGCGATTCTCCTGCCTCNNNAGATCGGAAGAG |
| 55185_1 | TATGAGTGTGGAGTCGTTGCCTGTGGCACAATCTCAGCTCANNNAGATCGGAAGAG |

| | |
|---|---|
| 55188_1 | TATGAGTGTGGAGTCGTTGCAGTGCAGTGGAGTGATCTTGGNNNAGATCGGAAGA |
| 55189_1 | TATGAGTGTGGAGTCGTTGCCAAGTAGCTGGGACTACAGGCNNNAGATCGGAAGA |
| 55190_1 | TATGAGTGTGGAGTCGTTGCGGAGTGCAGTGGTGCTATCTCNNNAGATCGGAAGA |
| 55202_1 | TATGAGTGTGGAGTCGTTGCGCAATCTGGGCTTACTGCAAGNNNAGATCGGAAGA |
| 55207_1 | TATGAGTGTGGAGTCGTTGCCGATCTCGGCTCACTGCAGGNNNAGATCGGAAGAG |
| 55219_1 | TATGAGTGTGGAGTCGTTGCGGCTAAATGGAAAGAGCAAGGACNNNAGATCGGA |
| 55220_1 | TATGAGTGTGGAGTCGTTGCAGTTCTTCAAAGGGCAGACTACANNNAGATCGGAAG |
| 55227_1 | TATGAGTGTGGAGTCGTTGCCAGTTTGAGTGGCAAGACGTGNNNAGATCGGAAGA |
| 55249_1 | TATGAGTGTGGAGTCGTTGCTTGTGCTGTCATAGAACTCAACANNNAGATCGGAAG |
| 55250_1 | TATGAGTGTGGAGTCGTTGCAAGACATAAATGGCAAAGGCAGANNNAGATCGGAA |
| 55254_1 | TATGAGTGTGGAGTCGTTGCGGGGCTAGAGTTTGATGGAGAAANNNAGATCGGA |
| 55262_1 | TATGAGTGTGGAGTCGTTGCATAACTGAATCATGGCGGAGGTTNNNAGATCGGAA |
| 55267_1 | TATGAGTGTGGAGTCGTTGCGAGGGTGCATTCCAAGTGTTCNNNAGATCGGAAGA |
| 55283_1 | TATGAGTGTGGAGTCGTTGCTTTGAGGGTTGTTGTTATTTGCTNNNAGATCGGAAG |
| 55324_1 | TATGAGTGTGGAGTCGTTGCGGAGTCTTGCTGTGTCTCCAGNNNAGATCGGAAGAG |
| 55326_1 | TATGAGTGTGGAGTCGTTGCGAGTCTCACTCTTTCGCCCAGNNNAGATCGGAAGAG |
| 55331_1 | TATGAGTGTGGAGTCGTTGCCAAGTAGCTGGGACTACAGGCNNNAGATCGGAAGA |
| 55337_1 | TATGAGTGTGGAGTCGTTGCCAATCACAGCTCACTGCAACCNNNAGATCGGAAGAG |
| 55401_1 | TATGAGTGTGGAGTCGTTGCAGATCCCTGAAGACAGATCTGTGNNNAGATCGGAA |
| 55411_1 | TATGAGTGTGGAGTCGTTGCGACGGAGTCTCACTCTGTTGCNNNAGATCGGAAGAG |
| 55412_1 | TATGAGTGTGGAGTCGTTGCGGAGTACAGTGGCGTGATCTCNNNAGATCGGAAGA |
| 55421_1 | TATGAGTGTGGAGTCGTTGCCAAGTAGCTGGGACTACAGGCNNNAGATCGGAAGA |
| 55423_1 | TATGAGTGTGGAGTCGTTGCGTGATCTCACTGCAACCTCCANNNAGATCGGAAGAG |
| 55430_1 | TATGAGTGTGGAGTCGTTGCCTCACTGCAAGCTCCATCTCCNNNAGATCGGAAGAG |
| 55444_1 | TATGAGTGTGGAGTCGTTGCCAAGTAGCTGGGACTACAGGCNNNAGATCGGAAGA |
| 55447_1 | TATGAGTGTGGAGTCGTTGCCAAGTAGCTGGGACTACAGGCNNNAGATCGGAAGA |
| 55463_1 | TATGAGTGTGGAGTCGTTGCTAGCACCTATGTCTTCAGGAAGCNNNAGATCGGAAG |
| 55465_1 | TATGAGTGTGGAGTCGTTGCTTCACACCATTCTCCTGCCTCNNNAGATCGGAAGAGC |
| 55486_1 | TATGAGTGTGGAGTCGTTGCTGACTTGATACTACCACCGGTTCNNNAGATCGGAAG |
| 55506_1 | TATGAGTGTGGAGTCGTTGCCAGGTGTAGTGCCCTCTGTTATTNNNAGATCGGAAG |
| 55521_1 | TATGAGTGTGGAGTCGTTGCTCTTCAAACGAAAAAAGCATGCTNNNAGATCGGAAG |
| 55522_1 | TATGAGTGTGGAGTCGTTGCCTTCCTAGAAAAGCTCCTCTGGGNNNAGATCGGAAG |
| 55523_1 | TATGAGTGTGGAGTCGTTGCACAGGACAAGGGAATAGTGGATGNNNAGATCGGA |
| 55533_1 | TATGAGTGTGGAGTCGTTGCAGGCTCTACCAGAAAATTGTGCTNNNAGATCGGAAG |
| 55541_1 | TATGAGTGTGGAGTCGTTGCCTGAAGCAACAGCAGTCAAACANNNAGATCGGAAG |
| 55548_1 | TATGAGTGTGGAGTCGTTGCGAGGCAGAAGAATGGGGTGAANNNAGATCGGAAG |
| 55554_1 | TATGAGTGTGGAGTCGTTGCTTCCAGCGTATTGACATTTCACTNNNAGATCGGAAG |
| 55605_1 | TATGAGTGTGGAGTCGTTGCATTCTGTCCTCACAAGACATGCANNNAGATCGGAAG |
| 55621_1 | TATGAGTGTGGAGTCGTTGCTGTGAAACATTGTGCTAGCAGTGNNNAGATCGGAA |
| 55623_1 | TATGAGTGTGGAGTCGTTGCGAGTTCAGTGAACAGACATTGCCNNNAGATCGGAA |
| 55632_1 | TATGAGTGTGGAGTCGTTGCCTTTCCTCAGGTATATGGCCACANNNAGATCGGAAG |
| 55635_1 | TATGAGTGTGGAGTCGTTGCCCTGTAGTCCCAGCTACTCGGNNNAGATCGGAAGAG |
| 55706_1 | TATGAGTGTGGAGTCGTTGCTCGCTCTCTTTCTCTCTCTCTNNNAGATCGGAAGAG |

| | |
|---|---|
| 55721_1 | TATGAGTGTGGAGTCGTTGCCAAGTAGCTGGGACTACAGGCNNNAGATCGGAAGA |
| 55726_1 | TATGAGTGTGGAGTCGTTGCCTGGGTTCACGCCATTCTTCTNNNAGATCGGAAGAG |
| 55728_1 | TATGAGTGTGGAGTCGTTGCCTCACTGCAAGCTCCACTTCCNNNAGATCGGAAGAG |
| 55746_1 | TATGAGTGTGGAGTCGTTGCGCAAGTAGCCTAATGTGACTGGANNNAGATCGGAA |
| 55750_1 | TATGAGTGTGGAGTCGTTGCTTCTTTTTGGAGGGGATCACACANNNAGATCGGAAG |
| 55753_1 | TATGAGTGTGGAGTCGTTGCATTTTCACAGAAGGAAGTACCCTNNNAGATCGGAAG |
| 55772_1 | TATGAGTGTGGAGTCGTTGCCAAATGCTGGGATTACAGGCGNNNAGATCGGAAGA |
| 55776_1 | TATGAGTGTGGAGTCGTTGCCAAGTAGCTGGGACTACAGGCNNNAGATCGGAAGA |
| 55779_1 | TATGAGTGTGGAGTCGTTGCTAGAGACGGGGTTTCACATTGTTNNNAGATCGGAA |
| 55782_1 | TATGAGTGTGGAGTCGTTGCTTCAAGCGATTCTCCTGCCTTAGNNNAGATCGGAAG |
| 55785_1 | TATGAGTGTGGAGTCGTTGCCTCTGTTGGCTCAACTGCAACNNNAGATCGGAAGAG |
| 55790_1 | TATGAGTGTGGAGTCGTTGCGGAGTCTCACTCTGTCACCCANNNAGATCGGAAGAG |
| 55806_1 | TATGAGTGTGGAGTCGTTGCTTCAAGCGATTCTCCTGCCTCNNNAGATCGGAAGAG |
| 55819_1 | TATGAGTGTGGAGTCGTTGCAGACGGAGTCTTTCTGTTTCTCCNNNAGATCGGAAG |
| 55838_1 | TATGAGTGTGGAGTCGTTGCACTAACACTTTGGGCAAAATTCGTNNNAGATCGGAA |
| 55864_1 | TATGAGTGTGGAGTCGTTGCCAAAGGAAACACCTGCACAGAGNNNAGATCGGAAG |
| 55881_1 | TATGAGTGTGGAGTCGTTGCTTCAAGCGATTCTCCTGCCTCNNNAGATCGGAAGAG |
| 55895_1 | TATGAGTGTGGAGTCGTTGCCAAGTAGCTGGGACTACAGGCNNNAGATCGGAAGA |
| 55915_1 | TATGAGTGTGGAGTCGTTGCTGGAAGGAAAATTCCCAAGTCCTNNNAGATCGGAA |
| 55918_1 | TATGAGTGTGGAGTCGTTGCCCATATCCTAGCCTTGGAACCTGNNNAGATCGGAAG |
| 55920_1 | TATGAGTGTGGAGTCGTTGCTAATAAACACAACCTCAGGGCCTNNNAGATCGGAAG |
| 55921_1 | TATGAGTGTGGAGTCGTTGCGAGGATCTGTTGAACCCTGGAGNNNAGATCGGAAG |
| 55950_1 | TATGAGTGTGGAGTCGTTGCCAAGTAGCTGGGACTACAGGCNNNAGATCGGAAGA |
| 55994_1 | TATGAGTGTGGAGTCGTTGCGAGACGGAGTCTTGCTCTGTCNNNAGATCGGAAGA |
| 55996_1 | TATGAGTGTGGAGTCGTTGCAGACGGAGTCTCACTCTGTCGNNNAGATCGGAAGA |
| 56013_1 | TATGAGTGTGGAGTCGTTGCTTCACACCATTCTCCTGCCTCNNNAGATCGGAAGAGC |
| 56015_1 | TATGAGTGTGGAGTCGTTGCGAGACGGAGTCTTGCTCTGTCNNNAGATCGGAAGA |
| 56023_1 | TATGAGTGTGGAGTCGTTGCCAAGTAGCTGGGACTACAGGCNNNAGATCGGAAGA |
| 56059_1 | TATGAGTGTGGAGTCGTTGCTGAGGCTATCAACCAGTGAAGACNNNAGATCGGAA |
| 56103_1 | TATGAGTGTGGAGTCGTTGCACAGGTGGATCACTTTAGCTCAGNNNAGATCGGAA |
| 56111_1 | TATGAGTGTGGAGTCGTTGCGGTAACCCTAGCACTTTGGGAGNNNAGATCGGAAG |
| 56136_1 | TATGAGTGTGGAGTCGTTGCAGACGGGGTCTCACTCTGTAGNNNAGATCGGAAGA |
| 56145_1 | TATGAGTGTGGAGTCGTTGCGCAATCTTGGCTCACTGCAAGNNNAGATCGGAAGA |
| 56172_1 | TATGAGTGTGGAGTCGTTGCCAAGTAGCTGGGACTACAGGCNNNAGATCGGAAGA |
| 56176_1 | TATGAGTGTGGAGTCGTTGCTTCACGCCATTCTCCTGTCTCNNNAGATCGGAAGAGC |
| 56188_1 | TATGAGTGTGGAGTCGTTGCCGTGGCAAGAATTCAGTGAAAGTNNNAGATCGGAA |
| 56191_1 | TATGAGTGTGGAGTCGTTGCGGAGAAATCACTGGCCTACTTGANNNAGATCGGAA |
| 56196_1 | TATGAGTGTGGAGTCGTTGCGGTGAAACCCCGTCTCTACTAAANNNAGATCGGAAG |
| 56198_1 | TATGAGTGTGGAGTCGTTGCAAGAAAGAAATACTCGGCCAGGTNNNAGATCGGAA |
| 56223_1 | TATGAGTGTGGAGTCGTTGCGAAAATTCAGGACCCCAGAGTGGNNNAGATCGGAA |
| 56244_1 | TATGAGTGTGGAGTCGTTGCAACAGGTAGCTAATGGGAACAGGNNNAGATCGGAA |
| 56253_1 | TATGAGTGTGGAGTCGTTGCCTGAATGTGTCAAGGTCCAGGTANNNAGATCGGAA |
| 56269_1 | TATGAGTGTGGAGTCGTTGCAAACCAAGGCAGGATAGAACACANNNAGATCGGAA |

| | |
|---|---|
| 56270_1 | TATGAGTGTGGAGTCGTTGCCCTTCTAGGTAGGTCATCCTCCANNNAGATCGGAAG |
| 56293_1 | TATGAGTGTGGAGTCGTTGCGAGACGGAGTCTTGCTCTGTCNNNAGATCGGAAGA |
| 56299_1 | TATGAGTGTGGAGTCGTTGCCAAGTAGCTGGGACTACAGGCNNNAGATCGGAAGA |
| 56349_1 | TATGAGTGTGGAGTCGTTGCGAGTAGCTGGGACTACAGGCANNNAGATCGGAAG |
| 56367_1 | TATGAGTGTGGAGTCGTTGCTGCAATGGGAAATGCAAGTTACCNNNAGATCGGAA |
| 56377_1 | TATGAGTGTGGAGTCGTTGCAGAACAGAGACTGACACAAGGACNNNAGATCGGAA |
| 56430_1 | TATGAGTGTGGAGTCGTTGCCTACTCAAGAGGCTGAGGCAGNNNAGATCGGAAGA |
| 56456_1 | TATGAGTGTGGAGTCGTTGCCAAGTAGCTGGGACTACAGGCNNNAGATCGGAAGA |
| 56461_1 | TATGAGTGTGGAGTCGTTGCACAGGGTCTCATTCTGTCACCNNNAGATCGGAAGAG |
| 56463_1 | TATGAGTGTGGAGTCGTTGCACTCCTGGCCTCAAGAGATCTNNNAGATCGGAAGAG |
| 56482_1 | TATGAGTGTGGAGTCGTTGCGAGTCTCGCTCTGTTGCCTAGNNNAGATCGGAAGAG |
| 56484_1 | TATGAGTGTGGAGTCGTTGCATCTTGGCTCACTGCAACCTCNNNAGATCGGAAGAG |
| 56572_1 | TATGAGTGTGGAGTCGTTGCCCTCCCAGGTTCACACCATTCNNNAGATCGGAAGAG |
| 56596_1 | TATGAGTGTGGAGTCGTTGCCATGATCTTGGCTCACTGCAACNNNAGATCGGAAGA |
| 56620_1 | TATGAGTGTGGAGTCGTTGCCAATCTTGGCTCACTGCAAGCNNNAGATCGGAAGAG |
| 56636_1 | TATGAGTGTGGAGTCGTTGCCCACTCTCTTCTGGCTTGTAGAGNNNAGATCGGAAG |
| 56664_1 | TATGAGTGTGGAGTCGTTGCACGTATCCATGTTGTCAGAGAAANNNAGATCGGAAG |
| 56685_1 | TATGAGTGTGGAGTCGTTGCACGATGAAACCCCCTCTCTACTANNNAGATCGGAAG |
| 56722_1 | TATGAGTGTGGAGTCGTTGCGTTCAAGCGATTCTTCTGCCTCNNNAGATCGGAAGA |
| 56726_1 | TATGAGTGTGGAGTCGTTGCACAATCTCGGCTCACTGCAAGNNNAGATCGGAAGAG |
| 56728_1 | TATGAGTGTGGAGTCGTTGCCTCCTGGGTTCACGCTATTCTCNNNAGATCGGAAGA |
| 56738_1 | TATGAGTGTGGAGTCGTTGCGCAATCTCAGCTCACTGCAAGNNNAGATCGGAAGAG |
| 56752_1 | TATGAGTGTGGAGTCGTTGCTTCACGTCATTCTCCTGCCTCNNNAGATCGGAAGAGC |
| 56754_1 | TATGAGTGTGGAGTCGTTGCCTCCTGGGTTCACACCATTCTNNNAGATCGGAAGAG |
| 56762_1 | TATGAGTGTGGAGTCGTTGCCGATTAGCTGGGACTACAGGCNNNAGATCGGAAGA |
| 56763_1 | TATGAGTGTGGAGTCGTTGCTTTCAAGCGATTCTCCTGCCTNNNAGATCGGAAGAG |
| 56781_1 | TATGAGTGTGGAGTCGTTGCATGAACTCGGCTCACTGCAANNNAGATCGGAAGAG |
| 56790_1 | TATGAGTGTGGAGTCGTTGCCCAAGTAGCTGGGACTACAGGNNNAGATCGGAAGA |
| 56794_1 | TATGAGTGTGGAGTCGTTGCTGATCTCAGCTCACTGCAAGCNNNAGATCGGAAGAG |
| 56806_1 | TATGAGTGTGGAGTCGTTGCCAAGTAGCTGGGACTACAGGCNNNAGATCGGAAGA |
| 56808_1 | TATGAGTGTGGAGTCGTTGCTTCAAGCGATTCTCCTGCCTCNNNAGATCGGAAGAG |
| 56831_1 | TATGAGTGTGGAGTCGTTGCGGTAAACGGCTCCTTTTGTTCAANNNAGATCGGAAG |
| 56838_1 | TATGAGTGTGGAGTCGTTGCTGGCCCCAGAATATGTACGAAATNNNAGATCGGAA |
| 56842_1 | TATGAGTGTGGAGTCGTTGCACAGCAGAAATTCAAGAATCCCCNNNAGATCGGAAG |
| 56857_1 | TATGAGTGTGGAGTCGTTGCGAGAAGGGCCTGAGACAGAAATCNNNAGATCGGA |
| 56878_1 | TATGAGTGTGGAGTCGTTGCTATCTCCTAATGCTTCCCTCCCCNNNAGATCGGAAGA |
| 56886_1 | TATGAGTGTGGAGTCGTTGCATTAGAACCACAACACCATGCTGNNNAGATCGGAAG |
| 56913_1 | TATGAGTGTGGAGTCGTTGCACAGTAAGTCATAGACAGCAAGCANNNAGATCGGA |
| 56945_1 | TATGAGTGTGGAGTCGTTGCTAGCCACACTTATGATCCAACCCNNNAGATCGGAAG |
| 56948_1 | TATGAGTGTGGAGTCGTTGCGGAGGATCACTTGAGGCTAAGAGNNNAGATCGGA |
| 56949_1 | TATGAGTGTGGAGTCGTTGCCCTCTCAGACAGACACAGAGAAGNNNAGATCGGAA |
| 56976_1 | TATGAGTGTGGAGTCGTTGCAGACGGAGTCTCACTCTGTCGNNNAGATCGGAAGA |
| 56977_1 | TATGAGTGTGGAGTCGTTGCCAAGTAGCTGGGACTACAGGCNNNAGATCGGAAGA |

| | |
|---|---|
| 56982_1 | TATGAGTGTGGAGTCGTTGCCTCACGCCATTCTCCTACCTCNNNAGATCGGAAGAGC |
| 57007_1 | TATGAGTGTGGAGTCGTTGCCCCGAGTAGCTGGGACTACAGNNNAGATCGGAAGA |
| 57009_1 | TATGAGTGTGGAGTCGTTGCCCATTCTCCTGCCTTAGCCTCNNNAGATCGGAAGAGC |
| 57032_1 | TATGAGTGTGGAGTCGTTGCACATAGACCATTTTTCAGCCCCANNNAGATCGGAAGA |
| 57037_1 | TATGAGTGTGGAGTCGTTGCTGTAGGTCTTTTCAGTGGACAGANNNAGATCGGAA |
| 57045_1 | TATGAGTGTGGAGTCGTTGCCACCAGTCCTCGTTGTGTTTATTNNNAGATCGGAAGA |
| 57065_1 | TATGAGTGTGGAGTCGTTGCCTGAGGCTGGAGAATCACTTGAANNNAGATCGGAA |
| 57067_1 | TATGAGTGTGGAGTCGTTGCATTGCCACTGTCATGCTTGAATTNNNAGATCGGAAG |
| 57087_1 | TATGAGTGTGGAGTCGTTGCCCCGAGTAGCTGGGACTACAGNNNAGATCGGAAGA |
| 57097_1 | TATGAGTGTGGAGTCGTTGCCAAGTAGCTGGGACTACAGGCNNNAGATCGGAAGA |
| 57104_1 | TATGAGTGTGGAGTCGTTGCCCCGAGTAGCTGGGATTACAGNNNAGATCGGAAGA |
| 57106_1 | TATGAGTGTGGAGTCGTTGCCAAGTAGCTGGGACTACAGGCNNNAGATCGGAAGA |
| 57177_1 | TATGAGTGTGGAGTCGTTGCAGCATTGTGAAATGTTCTTGGGGNNNAGATCGGAA |
| 57211_1 | TATGAGTGTGGAGTCGTTGCCAATCTCAGCCAACTGCAACCNNNAGATCGGAAGAG |
| 57226_1 | TATGAGTGTGGAGTCGTTGCATCTGGAGGAGTGGAGGTAAGAANNNAGATCGGA |
| 57248_1 | TATGAGTGTGGAGTCGTTGCTGCAAGTAATCCATGTGTCTCAGANNNAGATCGGAA |
| 57268_1 | TATGAGTGTGGAGTCGTTGCTTCACACCATTCTCCTGCCTCNNNAGATCGGAAGAGC |
| 57284_1 | TATGAGTGTGGAGTCGTTGCCCCGAGTAGATGAGATTACAGGCNNNAGATCGGAA |
| 57308_1 | TATGAGTGTGGAGTCGTTGCCAAGTAGCTGGGACTACAGGCNNNAGATCGGAAGA |
| 57309_1 | TATGAGTGTGGAGTCGTTGCGAGACGGAGTCTTGCTCTGTCNNNAGATCGGAAGA |
| 57312_1 | TATGAGTGTGGAGTCGTTGCCAATCTTGGCTCACTGCAACCNNNAGATCGGAAGAG |
| 57345_1 | TATGAGTGTGGAGTCGTTGCTCCTGAGCTGGAGAAGACTATGANNNAGATCGGAA |
| 57350_1 | TATGAGTGTGGAGTCGTTGCAGCCAAGTCACTGTGTACTCAATNNNAGATCGGAAG |
| 57351_1 | TATGAGTGTGGAGTCGTTGCTGAGACAGTCAGGGAGAAACAAANNNAGATCGGAA |
| 57355_1 | TATGAGTGTGGAGTCGTTGCCAGCCATAGTCCCAGTTACTCAGNNNAGATCGGAAG |
| 57356_1 | TATGAGTGTGGAGTCGTTGCCGTGACAGGGTGAACTACAGTTANNNAGATCGGAA |
| 57370_1 | TATGAGTGTGGAGTCGTTGCATGGATGCTAGCAGAAGACACAANNNAGATCGGAA |
| 57375_1 | TATGAGTGTGGAGTCGTTGCTGTTTTTGGGGTATAAGTTGAGCANNNAGATCGGA |
| 57380_1 | TATGAGTGTGGAGTCGTTGCGAGATAGTGTGAGCCATAGCCATNNNAGATCGGAA |
| 57395_1 | TATGAGTGTGGAGTCGTTGCAGTAGTGTCTTAGCTGGGTGTTTNNNAGATCGGAA |
| 57402_1 | TATGAGTGTGGAGTCGTTGCACCTCAAGAGCATCATCCTAAGCNNNAGATCGGAAG |
| 57410_1 | TATGAGTGTGGAGTCGTTGCGGAGACTGTAGGTATATGCCACCNNNAGATCGGAA |
| 57412_1 | TATGAGTGTGGAGTCGTTGCAAGCCTACAATGGACACACAGATNNNAGATCGGAA |
| 57431_1 | TATGAGTGTGGAGTCGTTGCAGTCTCACTCTTGTTGCCCAGNNNAGATCGGAAGAG |
| 57445_1 | TATGAGTGTGGAGTCGTTGCGGAGTCCAGTGGCGTAATCTCNNNAGATCGGAAGA |
| 57447_1 | TATGAGTGTGGAGTCGTTGCCAAGTAGCTGGGACTACAGGCNNNAGATCGGAAGA |
| 57450_1 | TATGAGTGTGGAGTCGTTGCCAATGGCACGATCTCTGCTCANNNAGATCGGAAGAG |
| 57451_1 | TATGAGTGTGGAGTCGTTGCCCCTAGTAGCTGGGACTACAGGNNNAGATCGGAAG |
| 57476_1 | TATGAGTGTGGAGTCGTTGCTTGCTAAGGAACCCAAAGGAAAANNNAGATCGGAA |
| 57493_1 | TATGAGTGTGGAGTCGTTGCCCATTCCAAATTTGACAACTCAGCNNNAGATCGGAA |
| 57495_1 | TATGAGTGTGGAGTCGTTGCAGTGTTTACAAGAGTACCTGGCANNNAGATCGGAA |
| 57505_1 | TATGAGTGTGGAGTCGTTGCAGAAATCAAACACCTCAAATGTGANNNAGATCGGAA |
| 57516_1 | TATGAGTGTGGAGTCGTTGCGCACCTGTAGTCCCAAATACTTGNNNAGATCGGAAG |

| | |
|---|---|
| 57527_1 | TATGAGTGTGGAGTCGTTGCGGCTTTTCTCTGTCCTTTGCATTNNNAGATCGGAAGA |
| 57533_1 | TATGAGTGTGGAGTCGTTGCGCTGGAGAGGATGTGGAGAAATANNNAGATCGGA |
| 57561_1 | TATGAGTGTGGAGTCGTTGCATTCTCCTGCCTCCCGAGTAGNNNAGATCGGAAGAG |
| 57563_1 | TATGAGTGTGGAGTCGTTGCCTCCGGAGTAGCTGGGACTACNNNAGATCGGAAGA |
| 57568_1 | TATGAGTGTGGAGTCGTTGCCCCGAGTAGCTGGGACTACAGNNNAGATCGGAAGA |
| 57573_1 | TATGAGTGTGGAGTCGTTGCCAAGTAGCTGGGACTACAGGCNNNAGATCGGAAGA |
| 57579_1 | TATGAGTGTGGAGTCGTTGCCATCTCGCCTCACTACAAGCTCNNNAGATCGGAAGAG |
| 57580_1 | TATGAGTGTGGAGTCGTTGCCCTGGGGTTTACGCCATTCTCNNNAGATCGGAAGAG |
| 57581_1 | TATGAGTGTGGAGTCGTTGCCAAGTAGCTGGGACTACAGGCNNNAGATCGGAAGA |
| 57584_1 | TATGAGTGTGGAGTCGTTGCAGTGGCATGATCTCTGCTCACNNNAGATCGGAAGAG |
| 57603_1 | TATGAGTGTGGAGTCGTTGCTTCAAGCTATTCTCCTGCCTCAGNNNAGATCGGAAGA |
| 57613_1 | TATGAGTGTGGAGTCGTTGCCACCCAAAATTTTGACGGATGGANNNAGATCGGAAG |
| 57631_1 | TATGAGTGTGGAGTCGTTGCTGGTATTGGAAAGCAGAAAAGCCNNNAGATCGGAA |
| 57643_1 | TATGAGTGTGGAGTCGTTGCGAGTCAGGAGAATGGCGTGAANNNAGATCGGAAG |
| 57699_1 | TATGAGTGTGGAGTCGTTGCGTTCAAGCGATTCTCCTGCATCNNNAGATCGGAAGA |
| 57707_1 | TATGAGTGTGGAGTCGTTGCCAAGTAGCTGGGACTACAGGCNNNAGATCGGAAGA |
| 57716_1 | TATGAGTGTGGAGTCGTTGCGACAGGATCTCGCTGTTGTCCNNNAGATCGGAAGAG |
| 57718_1 | TATGAGTGTGGAGTCGTTGCGAGTAGCTGGGACTAACAGGCNNNAGATCGGAAGA |
| 57721_1 | TATGAGTGTGGAGTCGTTGCTTCACACCATTCTCCTGCCTCNNNAGATCGGAAGAGC |
| 57724_1 | TATGAGTGTGGAGTCGTTGCCTTCCGGGTTTACGCCATTCTNNNAGATCGGAAGAG |
| 57745_1 | TATGAGTGTGGAGTCGTTGCTCCGGACTATTGGCACTTTCTACNNNAGATCGGAAG |
| 57763_1 | TATGAGTGTGGAGTCGTTGCTGGAAAGTGACAAAGAAGGAAGGANNNAGATCGG |
| 57766_1 | TATGAGTGTGGAGTCGTTGCACTCCAGCCTAGGTGACAGAGNNNAGATCGGAAGA |
| 57773_1 | TATGAGTGTGGAGTCGTTGCCTTCCACTACTTCCGGGAAACCNNNAGATCGGAAGA |
| 57784_1 | TATGAGTGTGGAGTCGTTGCTTCTGAGTGCTGAGGAAAGAGAGNNNAGATCGGAA |
| 57794_1 | TATGAGTGTGGAGTCGTTGCCAGGGAACGGAGACTCTTTCAATNNNAGATCGGAAG |
| 57795_1 | TATGAGTGTGGAGTCGTTGCGTGACAGAGCAAGACTCATCTCANNNAGATCGGAAG |
| 57796_1 | TATGAGTGTGGAGTCGTTGCGATCCCTAGGCAGCCAAATTAGANNNAGATCGGAAG |
| 57802_1 | TATGAGTGTGGAGTCGTTGCTAGATTTGGTTGTGTGGTGTGTGNNNAGATCGGAA |
| 57809_1 | TATGAGTGTGGAGTCGTTGCTTGGGGAATTCCTACTAACACCGNNNAGATCGGAAG |
| 57812_1 | TATGAGTGTGGAGTCGTTGCGGAGCAAAGGCAACCATCTAATGNNNAGATCGGAA |
| 57824_1 | TATGAGTGTGGAGTCGTTGCGGAAGTCCTGGCTCTAAGCATATNNNAGATCGGAAG |
| 57829_1 | TATGAGTGTGGAGTCGTTGCGCAATCTTGGCTCACTGCAAGNNNAGATCGGAAGAG |
| 57834_1 | TATGAGTGTGGAGTCGTTGCCAAGTAGCTGGGACTACAGGCNNNAGATCGGAAGA |
| 57872_1 | TATGAGTGTGGAGTCGTTGCGAGTAGCTGGGACTAGAGGCANNNAGATCGGAAGA |
| 57876_1 | TATGAGTGTGGAGTCGTTGCAGACGGAGTCTCACTCTGTCGNNNAGATCGGAAGAG |
| 57882_1 | TATGAGTGTGGAGTCGTTGCCTTCTGAGCAGCTGGGACTACNNNAGATCGGAAGAG |
| 57887_1 | TATGAGTGTGGAGTCGTTGCGGGTTCATGCCATTCTTCTGCNNNAGATCGGAAGAG |
| 57895_1 | TATGAGTGTGGAGTCGTTGCAGGGAGTCTCACTCTGTCTCCNNNAGATCGGAAGAG |
| 57926_1 | TATGAGTGTGGAGTCGTTGCTTTCATTCAGGGAGCATCTGTCANNNAGATCGGAAG |
| 57941_1 | TATGAGTGTGGAGTCGTTGCGTGGTGAGGTGTGAGGATCAGNNNAGATCGGAAG |
| 57967_1 | TATGAGTGTGGAGTCGTTGCACAAATGGTGCTGGAGAAATTGGNNNAGATCGGAA |
| 57987_1 | TATGAGTGTGGAGTCGTTGCATGTGACTCTGCTTCTACACACANNNAGATCGGAAG |

| | |
|---|---|
| 58001_1 | TATGAGTGTGGAGTCGTTGCCAACTCTGACACTGTACCTTCCANNNAGATCGGAAGA |
| 58009_1 | TATGAGTGTGGAGTCGTTGCGAGACGGAGTCTTGCTCTGTCNNNAGATCGGAAGAG |
| 58011_1 | TATGAGTGTGGAGTCGTTGCCAAGTAGCTGGGACTACAGGCNNNAGATCGGAAGA |
| 58018_1 | TATGAGTGTGGAGTCGTTGCCAAGTAGCTGGGACTACAGGCNNNAGATCGGAAGA |
| 58037_1 | TATGAGTGTGGAGTCGTTGCAGAGTAGCTGGGACTACAGGCNNNAGATCGGAAGA |
| 58043_1 | TATGAGTGTGGAGTCGTTGCCAAGTAGCTGGGACTACAGGCNNNAGATCGGAAGA |
| 58063_1 | TATGAGTGTGGAGTCGTTGCTCAAGGAGCAGCTAGATTTGGCNNNAGATCGGAAGA |
| 58069_1 | TATGAGTGTGGAGTCGTTGCAGATTTGCATCCCCTTCATTCATNNNAGATCGGAAGA |
| 58092_1 | TATGAGTGTGGAGTCGTTGCGCCACTGACTGGGAGAAAAAAAANNNAGATCGGAA |
| 58109_1 | TATGAGTGTGGAGTCGTTGCATCAAGAGGTCAGGAGATCGAGANNNAGATCGGAA |
| 58110_1 | TATGAGTGTGGAGTCGTTGCGGAAGTCCTGAGCTACATTACTGTNNNAGATCGGAA |
| 58126_1 | TATGAGTGTGGAGTCGTTGCGGATCATCTGAGGTCAGGAGTTGNNNAGATCGGAA |
| 58128_1 | TATGAGTGTGGAGTCGTTGCGAAATCATCATGTTTGGGCCTGGNNNAGATCGGAAG |
| 58130_1 | TATGAGTGTGGAGTCGTTGCCAAGTAGCTGGGACTACAGGCNNNAGATCGGAAGA |
| 58159_1 | TATGAGTGTGGAGTCGTTGCGACTAGCTTCGGACTACAGGCNNNAGATCGGAAGAG |
| 58170_1 | TATGAGTGTGGAGTCGTTGCCCCGAGTAGGTGGGACTACAGNNNAGATCGGAAGA |
| 58171_1 | TATGAGTGTGGAGTCGTTGCGATCTCGGCTCACTGCAACTTNNNAGATCGGAAGAG |
| 58177_1 | TATGAGTGTGGAGTCGTTGCCAAGTAGCTGGGACTACAGGCNNNAGATCGGAAGA |
| 58184_1 | TATGAGTGTGGAGTCGTTGCCTCTGGGGTAGTTGGGATTACAGNNNAGATCGGAA |
| 58208_1 | TATGAGTGTGGAGTCGTTGCTTCACACCATTCTCCTGCCTCNNNAGATCGGAAGAGCA |
| 58209_1 | TATGAGTGTGGAGTCGTTGCGTCTCGAACTCCTGACCTCAAGNNNAGATCGGAAGA |
| 58212_1 | TATGAGTGTGGAGTCGTTGCACGATCTCAGCTCACTGCAAGNNNAGATCGGAAGAG |
| 58214_1 | TATGAGTGTGGAGTCGTTGCCAAGTAGCTGGGACTACAGGCNNNAGATCGGAAGA |
| 58219_1 | TATGAGTGTGGAGTCGTTGCACGATCTCAGCTCACTGCAAGNNNAGATCGGAAGAG |
| 58234_1 | TATGAGTGTGGAGTCGTTGCAATCTGGAGGATGGAGAAAACCCNNNAGATCGGAA |
| 58238_1 | TATGAGTGTGGAGTCGTTGCATCTCAGCACTTTTAGAGGCTGANNNAGATCGGAAG |
| 58242_1 | TATGAGTGTGGAGTCGTTGCGGAGCATTTAGTCTAGAAGCGCTNNNAGATCGGAAG |
| 58256_1 | TATGAGTGTGGAGTCGTTGCGTGACTTGTGCCTGTAGTCCTAANNNAGATCGGAAG |
| 58272_1 | TATGAGTGTGGAGTCGTTGCAAAACAAAAAAAACACTGGGCGCNNNAGATCGGAAG |
| 58288_1 | TATGAGTGTGGAGTCGTTGCCCAACCAAAAAGAGTACAGGACCNNNAGATCGGAAG |
| 58302_1 | TATGAGTGTGGAGTCGTTGCTTCCTAGCCCATATGGTTTCCACNNNAGATCGGAAGA |
| 58327_1 | TATGAGTGTGGAGTCGTTGCGCATGCATTTCATTCCAACCCAANNNAGATCGGAAGA |
| 58342_1 | TATGAGTGTGGAGTCGTTGCGGACAGAAGATTTAGGAAGGGGGNNNAGATCGGA |
| 58347_1 | TATGAGTGTGGAGTCGTTGCCCTCCTCAGTACGAGCGATTCNNNAGATCGGAAGAG |
| 58351_1 | TATGAGTGTGGAGTCGTTGCCATTCTCCTGCCTCAGCCTCNNNAGATCGGAAGAGCA |
| 58384_1 | TATGAGTGTGGAGTCGTTGCGGGATCTCAGCTCACTCCAAGNNNAGATCGGAAGAG |
| 58388_1 | TATGAGTGTGGAGTCGTTGCATCTCCTGACCTTGTGATTCACCNNNAGATCGGAAGA |
| 58392_1 | TATGAGTGTGGAGTCGTTGCCAAGTAGCTGGGACTACAGGCNNNAGATCGGAAGA |
| 58411_1 | TATGAGTGTGGAGTCGTTGCCGATTAGCTGGGACTACAGGCNNNAGATCGGAAGA |
| 58412_1 | TATGAGTGTGGAGTCGTTGCGAGACGGAGTCTTGCTCTGTCNNNAGATCGGAAGAG |
| 58442_1 | TATGAGTGTGGAGTCGTTGCTTTGTCAATCAAAAGAGCCCACCNNNAGATCGGAAGA |
| 58447_1 | TATGAGTGTGGAGTCGTTGCTGGACCCTGCCTGTTTTCTAAAANNNAGATCGGAAGA |
| 58466_1 | TATGAGTGTGGAGTCGTTGCGTGCTCAGCAGGTGGTTAGATATNNNAGATCGGAAG |

| | |
|---|---|
| 58476_1 | TATGAGTGTGGAGTCGTTGCCGAGTTGGGAGTGACTCACTGNNNAGATCGGAAGA |
| 58487_1 | TATGAGTGTGGAGTCGTTGCGGAGTACAGTGGTGCGATCTCNNNAGATCGGAAGA |
| 58491_1 | TATGAGTGTGGAGTCGTTGCTTCACACCATTCTCCTGCCTCNNNAGATCGGAAGAGC |
| 58493_1 | TATGAGTGTGGAGTCGTTGCGGAGGAAAACAGGCCAGTGTATANNNAGATCGGA |
| 58495_1 | TATGAGTGTGGAGTCGTTGCTTCAAGCGATTCTCCTGCCTCNNNAGATCGGAAGAG |
| 58498_1 | TATGAGTGTGGAGTCGTTGCATGAACCTGGGAGACGGAGTTAGNNNAGATCGGA |
| 58514_1 | TATGAGTGTGGAGTCGTTGCGATCACAAGGTCAGGAGATCGAGNNNAGATCGGAA |
| 58518_1 | TATGAGTGTGGAGTCGTTGCAGGTCTCTTGATGCCTCTTTCAGNNNAGATCGGAAG |
| 58521_1 | TATGAGTGTGGAGTCGTTGCGAGACGGAGTCTTGCTCTGTCNNNAGATCGGAAGA |
| 58537_1 | TATGAGTGTGGAGTCGTTGCCATCCTGGCTAACATGGTGAAACNNNAGATCGGAAG |
| 58540_1 | TATGAGTGTGGAGTCGTTGCCAGTTCCAGAGGTGTTTCAGGAANNNAGATCGGAA |
| 58551_1 | TATGAGTGTGGAGTCGTTGCCTCCTGAGTAGCTGGGATTACAGNNNAGATCGGAA |
| 58552_1 | TATGAGTGTGGAGTCGTTGCTAGCTGGGACTACAGGCATCTNNNAGATCGGAAGA |
| 58558_1 | TATGAGTGTGGAGTCGTTGCCCATTCTCCTGCCTCAACCTCNNNAGATCGGAAGAGC |
| 58564_1 | TATGAGTGTGGAGTCGTTGCCTGGAGTGCAATGGCCAATCTNNNAGATCGGAAGA |
| 58582_1 | TATGAGTGTGGAGTCGTTGCCTTGTCCTTTCTGCCATGTGAAGNNNAGATCGGAAG |
| 58595_1 | TATGAGTGTGGAGTCGTTGCGAGTGCAGTGACGCAATCTTGNNNAGATCGGAAGA |
| 58602_1 | TATGAGTGTGGAGTCGTTGCACACTTTGGGAAGCTGAGGTAGNNNAGATCGGAAG |
| 58603_1 | TATGAGTGTGGAGTCGTTGCGTAGAGACAGGGTTTCGCCATATNNNAGATCGGAA |
| 58612_1 | TATGAGTGTGGAGTCGTTGCCAACCCCATGTCATACTTACCCANNNAGATCGGAAGA |
| 58620_1 | TATGAGTGTGGAGTCGTTGCGCTCCTGTAATCCCAGCTACTGNNNAGATCGGAAGA |
| 58650_1 | TATGAGTGTGGAGTCGTTGCAATGTCTGGGTGGAGACTGAAAANNNAGATCGGAA |
| 58676_1 | TATGAGTGTGGAGTCGTTGCCAGGTTCATGCCATCCTCCTGNNNAGATCGGAAGAG |
| 58677_1 | TATGAGTGTGGAGTCGTTGCCATTCTCCTGCCTCAGCCTCNNNAGATCGGAAGAGCA |
| 58680_1 | TATGAGTGTGGAGTCGTTGCCATGATCTTGGCTCACTGCAGNNNAGATCGGAAGAG |
| 58689_1 | TATGAGTGTGGAGTCGTTGCGAGATCTCGGCTCACTGCAAGNNNAGATCGGAAGA |
| 58690_1 | TATGAGTGTGGAGTCGTTGCCAGTAGCTGGTCCTACAGGTGNNNAGATCGGAAGA |
| 58709_1 | TATGAGTGTGGAGTCGTTGCCCGGGTTCATGCCATTATCCTNNNAGATCGGAAGAG |
| 58720_1 | TATGAGTGTGGAGTCGTTGCGAGAAAATTTCGAGAGGAGGGGANNNAGATCGGA |
| 58722_1 | TATGAGTGTGGAGTCGTTGCCGAGTGACAAAAGTTGGGTCTTCNNNAGATCGGAA |
| 58742_1 | TATGAGTGTGGAGTCGTTGCAAAAATCCTCCACGGTGATTGTGNNNAGATCGGAAG |
| 58753_1 | TATGAGTGTGGAGTCGTTGCCTCTCGGGTTCACTTCTCCTGNNNAGATCGGAAGAG |
| 58773_1 | TATGAGTGTGGAGTCGTTGCCCACTAGGTGTAGGGTTTCATCCNNNAGATCGGAAG |
| 58774_1 | TATGAGTGTGGAGTCGTTGCCCATTCTCCTGCCTTAGCCTCNNNAGATCGGAAGAGC |
| 58790_1 | TATGAGTGTGGAGTCGTTGCGATCCTCTCCTGACCTGGTGTNNNAGATCGGAAGAG |
| 58808_1 | TATGAGTGTGGAGTCGTTGCTTCAAGCGATTCTCCTGCCTCNNNAGATCGGAAGAG |
| 58810_1 | TATGAGTGTGGAGTCGTTGCGAGACGGAGTCTTGCTCTGTCNNNAGATCGGAAGA |
| 58821_1 | TATGAGTGTGGAGTCGTTGCAGGGACCTGACCTCAAGTAGTTANNNAGATCGGAA |
| 58823_1 | TATGAGTGTGGAGTCGTTGCAGGTGCAAAGATGTCTGAGTGATNNNAGATCGGAA |
| 58826_1 | TATGAGTGTGGAGTCGTTGCGGTCATAAGTTTGTCAGGCAAGTNNNAGATCGGAA |
| 58827_1 | TATGAGTGTGGAGTCGTTGCACTAAGAATCATGTGAGCCAGCANNNAGATCGGAA |
| 58840_1 | TATGAGTGTGGAGTCGTTGCCTGGATGGTGTCTTTGGAGGAAANNNAGATCGGAA |
| 58843_1 | TATGAGTGTGGAGTCGTTGCCTGGATGGTGTCTTTGGAGGAAANNNAGATCGGAA |

| | |
|---|---|
| 58845_1 | TATGAGTGTGGAGTCGTTGCGTAGTCCCAGCTACTCCGGAGNNNAGATCGGAAGA |
| 58847_1 | TATGAGTGTGGAGTCGTTGCTAAAACCTCCTGATTGCCCATCANNNAGATCGGAAG |
| 58859_1 | TATGAGTGTGGAGTCGTTGCTAAATACATCAGGAGTGAGCCGGNNNAGATCGGAA |
| 58894_1 | TATGAGTGTGGAGTCGTTGCGAAAGATGCTGGCTTGGAAGCNNNAGATCGGAAG |
| 58896_1 | TATGAGTGTGGAGTCGTTGCGTACAGCTACTTGGGAGGCTGNNNAGATCGGAAGA |
| 58909_1 | TATGAGTGTGGAGTCGTTGCCCATTCTTCTGCCTCAGCCTCNNNAGATCGGAAGAGC |
| 58927_1 | TATGAGTGTGGAGTCGTTGCTAGCTGGGACTACAGGTGCTTNNNAGATCGGAAGA |
| 58957_1 | TATGAGTGTGGAGTCGTTGCCCTTCCTTCAGCTCATTTTAGGCNNNAGATCGGAAGA |
| 58959_1 | TATGAGTGTGGAGTCGTTGCCTGTAGTCCCAGCTACTCAGGAGNNNAGATCGGAAG |
| 58960_1 | TATGAGTGTGGAGTCGTTGCTTGACTTAGGGCCATAGAAAGGGNNNAGATCGGAA |
| 58966_1 | TATGAGTGTGGAGTCGTTGCTTTGAGTTTTGTGGTCAACCAGGNNNAGATCGGAA |
| 58967_1 | TATGAGTGTGGAGTCGTTGCGATCACAGATCATCTCCTCCAGGNNNAGATCGGAAG |
| 58982_1 | TATGAGTGTGGAGTCGTTGCCCAGGTTCAAGCGATTCTCCTNNNAGATCGGAAGAG |
| 58983_1 | TATGAGTGTGGAGTCGTTGCCCTCAACCTCCCAAGCTCAAGNNNAGATCGGAAGAG |
| 58984_1 | TATGAGTGTGGAGTCGTTGCCAAGTAGCTGGGACTACAGGCNNNAGATCGGAAGA |
| 58992_1 | TATGAGTGTGGAGTCGTTGCCAAGTAGCTGGGACTACAGGCNNNAGATCGGAAGA |
| 59004_1 | TATGAGTGTGGAGTCGTTGCTTCACACCATTCTCCTGCCTCNNNAGATCGGAAGAGC |
| 59024_1 | TATGAGTGTGGAGTCGTTGCTAACCCAGGAGTAAGGTTGTGACNNNAGATCGGAA |
| 59057_1 | TATGAGTGTGGAGTCGTTGCCCATTATCCTGCCTCAGCCTCNNNAGATCGGAAGAGC |
| 59082_1 | TATGAGTGTGGAGTCGTTGCCAAGTAGCTGGGACTACAGGCNNNAGATCGGAAGA |
| 59083_1 | TATGAGTGTGGAGTCGTTGCCAAGTAGCTGGGACTACAGGCNNNAGATCGGAAGA |
| 59094_1 | TATGAGTGTGGAGTCGTTGCTGCTGGGATTCCTTCTGTGTATTNNNAGATCGGAAG |
| 59111_1 | TATGAGTGTGGAGTCGTTGCCTCACTGAAAGCTCCTCCTCCNNNAGATCGGAAGAG |
| 59176_1 | TATGAGTGTGGAGTCGTTGCTGGGCAAAAGAGGTGAATAGACANNNAGATCGGA |
| 59182_1 | TATGAGTGTGGAGTCGTTGCGGTCTACTGTTCTCTGCTCTGTTNNNAGATCGGAAG |
| 59195_1 | TATGAGTGTGGAGTCGTTGCCTGATTTTGGGTGTGTGGGTTTTNNNAGATCGGAA |
| 59213_1 | TATGAGTGTGGAGTCGTTGCTTTGTTTTAGAAGCAGGGCCTTGNNNAGATCGGAA |
| 59220_1 | TATGAGTGTGGAGTCGTTGCGAACGGACTGAAATTTCTAACGTNNNAGATCGGAA |
| 59241_1 | TATGAGTGTGGAGTCGTTGCCATTCTCCTGCCTCAGCCTCNNNAGATCGGAAGAGCA |
| 59246_1 | TATGAGTGTGGAGTCGTTGCCCAAGTAGCTGGGACTACAGGNNNAGATCGGAAGA |
| 59262_1 | TATGAGTGTGGAGTCGTTGCGGAGTGCAGTGACACGATCTCNNNAGATCGGAAGA |
| 59264_1 | TATGAGTGTGGAGTCGTTGCCATGAACATGGCTCACTGCAGNNNAGATCGGAAGA |
| 59265_1 | TATGAGTGTGGAGTCGTTGCCCATTCTTCTGCCTCAGCCTCNNNAGATCGGAAGAGC |
| 59269_1 | TATGAGTGTGGAGTCGTTGCCAAGTAGCTGGGACTACAGGCNNNAGATCGGAAGA |
| 59283_1 | TATGAGTGTGGAGTCGTTGCACAATCTCGGCTCACTGCAAGNNNAGATCGGAAGAG |
| 59295_1 | TATGAGTGTGGAGTCGTTGCCAGTGGTGTGATCTTGGCTCANNNAGATCGGAAGA |
| 59299_1 | TATGAGTGTGGAGTCGTTGCTACTTCCAGAGATGATTCCTGGCNNNAGATCGGAAG |
| 59306_1 | TATGAGTGTGGAGTCGTTGCAAGACAAGATAGTAGGCTGGGTGNNNAGATCGGA |
| 59308_1 | TATGAGTGTGGAGTCGTTGCCAGCTACTGGGGAAACTGAGGNNNAGATCGGAAG |
| 59313_1 | TATGAGTGTGGAGTCGTTGCAAGGCCAATCCCTTTTTTTCTCTNNNAGATCGGAAGA |
| 59314_1 | TATGAGTGTGGAGTCGTTGCGAGGCTGAGACAGGAGAATCGNNNAGATCGGAAG |
| 59321_1 | TATGAGTGTGGAGTCGTTGCACGGAAGAGGAAAATCAGAACCTNNNAGATCGGAA |
| 59339_1 | TATGAGTGTGGAGTCGTTGCGAGGTTGCAATGAGCCATGATTGNNNAGATCGGAA |

| | |
|---|---|
| 59349_1 | TATGAGTGTGGAGTCGTTGCCAATCCCAGCTACTCAGGACGNNNAGATCGGAAGAG |
| 59451_1 | TATGAGTGTGGAGTCGTTGCAGACGGTGTCTCACTCTTTCGNNNAGATCGGAAGAG |
| 59468_1 | TATGAGTGTGGAGTCGTTGCAGATGGAGTCTCACTCTGTTGCNNNAGATCGGAAGA |
| 59488_1 | TATGAGTGTGGAGTCGTTGCAGAGTAGCTGGGACTACAGGCNNNAGATCGGAAGA |
| 59516_1 | TATGAGTGTGGAGTCGTTGCGAGTAGCTGGGACTACAGGCANNNAGATCGGAAGA |
| 59522_1 | TATGAGTGTGGAGTCGTTGCCATTCTCCTGCCTCAGCCTCNNNAGATCGGAAGAGCA |
| 59523_1 | TATGAGTGTGGAGTCGTTGCACAATCTCGGCTCACTGCAAGNNNAGATCGGAAGAG |
| 59538_1 | TATGAGTGTGGAGTCGTTGCCATGGAGATTCCCTAGCTGTGTTNNNAGATCGGAAG |
| 59539_1 | TATGAGTGTGGAGTCGTTGCCATGCTGGATTAGGGCGAGTTATNNNAGATCGGAAG |
| 59543_1 | TATGAGTGTGGAGTCGTTGCGATGCCACAATATTACAAGGCCCNNNAGATCGGAAG |
| 59552_1 | TATGAGTGTGGAGTCGTTGCGTTGCAGTTTTACAGGTAAGCCANNNAGATCGGAAG |
| 59553_1 | TATGAGTGTGGAGTCGTTGCCAAAGCTTCAGATCCAGGTTGTGNNNAGATCGGAAG |
| 59557_1 | TATGAGTGTGGAGTCGTTGCCAGCCTGACCAAAATGGAGAAACNNNAGATCGGAAG |
| 59577_1 | TATGAGTGTGGAGTCGTTGCACAAGAACCGGAACTAAGAGTCANNNAGATCGGAAG |
| 59590_1 | TATGAGTGTGGAGTCGTTGCGTGAAGGCATGGGATAGGTTAGTNNNAGATCGGAA |
| 59597_1 | TATGAGTGTGGAGTCGTTGCTGGAGAACCAAGATACTGTTTGCNNNAGATCGGAAG |
| 59601_1 | TATGAGTGTGGAGTCGTTGCTAATTGCTACTACCAGTTGGGCANNNAGATCGGAAG |
| 59605_1 | TATGAGTGTGGAGTCGTTGCGGAAGACAGGCCTGAGGAAATTANNNAGATCGGAA |
| 59612_1 | TATGAGTGTGGAGTCGTTGCCTCCGTTGCGCCTCTTTTATAAANNNAGATCGGAAGA |
| 59613_1 | TATGAGTGTGGAGTCGTTGCGGCCCAAGTAAAAAGAACAAGGTNNNAGATCGGAA |
| 59650_1 | TATGAGTGTGGAGTCGTTGCGAGTAGCTGGGACTACAGGCANNNAGATCGGAAGA |
| 59662_1 | TATGAGTGTGGAGTCGTTGCAGACAGAGTCTCGCTTTGTCACNNNAGATCGGAAGA |
| 59678_1 | TATGAGTGTGGAGTCGTTGCTTCAAGCGATTCTCCTGCCTCNNNAGATCGGAAGAGC |
| 59683_1 | TATGAGTGTGGAGTCGTTGCCTCCCAGGTTCAAGCGATTCTCNNNAGATCGGAAGAG |
| 59684_1 | TATGAGTGTGGAGTCGTTGCCCAGGTTCAGGCCATTCTCTTNNNAGATCGGAAGAGC |
| 59695_1 | TATGAGTGTGGAGTCGTTGCGAGTCTCGCTCTTTCACCCAGNNNAGATCGGAAGAGC |
| 59706_1 | TATGAGTGTGGAGTCGTTGCCAAGTAGCTGGGACTACAGGCNNNAGATCGGAAGA |
| 59710_1 | TATGAGTGTGGAGTCGTTGCCCATTCTCCTGCCTCAGACTCNNNAGATCGGAAGAGC |
| 59719_1 | TATGAGTGTGGAGTCGTTGCTCCTGGATTCTTAGTCAAGGTGCNNNAGATCGGAAG |
| 59721_1 | TATGAGTGTGGAGTCGTTGCCATAAAAGCCCCAGATTGGAACCNNNAGATCGGAAG |
| 59734_1 | TATGAGTGTGGAGTCGTTGCCCTGTAGTCCCAGCTACTTGGNNNAGATCGGAAGAG |
| 59735_1 | TATGAGTGTGGAGTCGTTGCGGGCAAAAGACCTGAAAGACATTNNNAGATCGGAAG |
| 59790_1 | TATGAGTGTGGAGTCGTTGCACTTACTGGGGCATTTCTTCTGTNNNAGATCGGAAGA |
| 59791_1 | TATGAGTGTGGAGTCGTTGCGCTGTAGTGAGCCGTGATCATNNNAGATCGGAAGAG |
| 59793_1 | TATGAGTGTGGAGTCGTTGCGCCATGTATTTGGTACGCAAGTTNNNAGATCGGAAG |
| 59795_1 | TATGAGTGTGGAGTCGTTGCGCAATCTCAGCTCACTGCAAGNNNAGATCGGAAGAG |
| 59796_1 | TATGAGTGTGGAGTCGTTGCCACAATCTTGGCTCACTGCAAGNNNAGATCGGAAGA |
| 59832_1 | TATGAGTGTGGAGTCGTTGCGTGATCTTGGCTTGCTGCAACNNNAGATCGGAAGAG |
| 59834_1 | TATGAGTGTGGAGTCGTTGCGGAGTTCAGTGGCACGATCTCNNNAGATCGGAAGAG |
| 59841_1 | TATGAGTGTGGAGTCGTTGCTTCACACCATTCTCCTGCCTCNNNAGATCGGAAGAGCA |
| 59865_1 | TATGAGTGTGGAGTCGTTGCATGATCTCGGCTCACTGCAAGNNNAGATCGGAAGAG |
| 59902_1 | TATGAGTGTGGAGTCGTTGCGAGATACCTGCACCCCATGTTTANNNAGATCGGAAGA |
| 59905_1 | TATGAGTGTGGAGTCGTTGCAGCCATATGCAGAAAGCTGAAACNNNAGATCGGAAG |

| | |
|---|---|
| 59910_1 | TATGAGTGTGGAGTCGTTGCTGAGCAGTTTCATGCAAAAGGTTNNNAGATCGGAA |
| 59912_1 | TATGAGTGTGGAGTCGTTGCCAGCTCCCAATTTGAGATGCTTCNNNAGATCGGAAG |
| 59920_1 | TATGAGTGTGGAGTCGTTGCAGATATCCCCCTTCACCTCTGTTNNNAGATCGGAAGA |
| 59921_1 | TATGAGTGTGGAGTCGTTGCTAGCTACCATTTACTGAAGGCCTNNNAGATCGGAAG |
| 59931_1 | TATGAGTGTGGAGTCGTTGCTTTCACTCTTATTGCCCAGGCTNNNAGATCGGAAGAG |
| 59942_1 | TATGAGTGTGGAGTCGTTGCTGCTAGGCAACAGGCATCATTATNNNAGATCGGAAG |
| 59949_1 | TATGAGTGTGGAGTCGTTGCACAAAAAACATTTAGCCAGGCGTNNNAGATCGGAAG |
| 59951_1 | TATGAGTGTGGAGTCGTTGCCTCACTGCAACTTCTGTCTCCCNNNAGATCGGAAGAG |
| 59956_1 | TATGAGTGTGGAGTCGTTGCTGAGGTCAGAAGTTTGAGACCAGNNNAGATCGGAA |
| 59957_1 | TATGAGTGTGGAGTCGTTGCACATGTCTTTGAACCCAGCTACTNNNAGATCGGAAG |
| 59960_1 | TATGAGTGTGGAGTCGTTGCGGGACTAGTGGTCACAAAAATGCNNNAGATCGGAA |
| 59976_1 | TATGAGTGTGGAGTCGTTGCTTATCAGTGTACCCTGGAGACTGNNNAGATCGGAAG |
| 59997_1 | TATGAGTGTGGAGTCGTTGCGAGTCTCACTCCATCACCCAGNNNAGATCGGAAGAG |
| 60003_1 | TATGAGTGTGGAGTCGTTGCCAATCTTGGCTCACTGCAACCNNNAGATCGGAAGAG |
| 60016_1 | TATGAGTGTGGAGTCGTTGCGACGGAGTCTCACTCTGTTGCNNNAGATCGGAAGAG |
| 60044_1 | TATGAGTGTGGAGTCGTTGCCAAGTAGCTGGGACTACAGGCNNNAGATCGGAAGA |
| 60076_1 | TATGAGTGTGGAGTCGTTGCCAAGTAGCTGGGACTACAGGCNNNAGATCGGAAGA |
| 60077_1 | TATGAGTGTGGAGTCGTTGCCAAGTAGCTGGGACTACAGGCNNNAGATCGGAAGA |
| 60083_1 | TATGAGTGTGGAGTCGTTGCCAAGTAGCTGGGACTACAGGCNNNAGATCGGAAGA |
| 60090_1 | TATGAGTGTGGAGTCGTTGCTTCAAGCGATTCTCCTGCCTCNNNAGATCGGAAGAG |
| 60094_1 | TATGAGTGTGGAGTCGTTGCCAAGTAGCTGGGACTACAGGCNNNAGATCGGAAGA |
| 60102_1 | TATGAGTGTGGAGTCGTTGCGAGACGGAGTCTTGCTCTGTCNNNAGATCGGAAGA |
| 60103_1 | TATGAGTGTGGAGTCGTTGCGACTACAGGCGTCCGAAGTTCNNNAGATCGGAAGA |
| 60110_1 | TATGAGTGTGGAGTCGTTGCGAGACGGAGTCTTGCTCTGTCNNNAGATCGGAAGA |
| 60130_1 | TATGAGTGTGGAGTCGTTGCCTGATCTCGGCTCACTGCAAGNNNAGATCGGAAGAG |
| 60146_1 | TATGAGTGTGGAGTCGTTGCATAAATTAGTGAGCAGGGGAGGGNNNAGATCGGA |
| 60154_1 | TATGAGTGTGGAGTCGTTGCGAGGTCAAGAGATCGAGACCATCNNNAGATCGGAA |
| 60168_1 | TATGAGTGTGGAGTCGTTGCGGGGAGGAGAAGGCATTACAAANNNAGATCGGAA |
| 60174_1 | TATGAGTGTGGAGTCGTTGCGCCAGAGTCTGCAAGAGTAAGAANNNAGATCGGAA |
| 60197_1 | TATGAGTGTGGAGTCGTTGCTATTCCCATCGCTTCCAAACATGNNNAGATCGGAAGA |
| 60201_1 | TATGAGTGTGGAGTCGTTGCTACTTGGTGGGAGAATGACCTTGNNNAGATCGGAA |
| 60229_1 | TATGAGTGTGGAGTCGTTGCTGTTTCCCCCTTTCCCACAATAANNNAGATCGGAAGA |
| 60241_1 | TATGAGTGTGGAGTCGTTGCGGTTGGAGAAGGTGACTATCTGGNNNAGATCGGAA |
| 60248_1 | TATGAGTGTGGAGTCGTTGCAGGAAAAACAAATAGGCCTTCCCNNNAGATCGGAAG |
| 60293_1 | TATGAGTGTGGAGTCGTTGCCGGGTTCATGCCATTCTCCTNNNAGATCGGAAGAGC |
| 61994_1 | TATGAGTGTGGAGTCGTTGCAAAGTGGGCAGATCACCTGAGNNNAGATCGGAAGA |
| 62015_1 | TATGAGTGTGGAGTCGTTGCTACTAGTCTATCCTGTGAGGGCCNNNAGATCGGAAG |
| 62088_1 | TATGAGTGTGGAGTCGTTGCCCATTCAAGGGATCCCAGAAGAGNNNAGATCGGAA |
| 62165_1 | TATGAGTGTGGAGTCGTTGCCTTCCCACTCCTGGAGAAACTGNNNAGATCGGAAGA |
| 62194_1 | TATGAGTGTGGAGTCGTTGCCAAGTAGCTGGGACTACAGGCNNNAGATCGGAAGA |
| 62215_1 | TATGAGTGTGGAGTCGTTGCGCCTGATCTTAGCTCAATGCAACNNNAGATCGGAAG |
| 62230_1 | TATGAGTGTGGAGTCGTTGCGCAACCTGGAACAAGTAAGTTGGNNNAGATCGGAA |
| 62270_1 | TATGAGTGTGGAGTCGTTGCAAACACTGGTCCTGCTCAGAGNNNAGATCGGAAGA |

| | |
|---|---|
| 62312_1 | TATGAGTGTGGAGTCGTTGCGTGGGTGTATGTCATGCACTAACNNNAGATCGGAAG |
| 62318_1 | TATGAGTGTGGAGTCGTTGCGAGTCTCGCTCTGTCATCAGGNNNAGATCGGAAGAG |
| 62346_1 | TATGAGTGTGGAGTCGTTGCAAAAAGACTGCACTTTCAGAGGCNNNAGATCGGAAG |
| 62374_1 | TATGAGTGTGGAGTCGTTGCAAGGGTCTTTGCCTTGTAATCCANNNAGATCGGAAG |
| 62420_1 | TATGAGTGTGGAGTCGTTGCGAGTAGCTGGGACTACAGGCANNNAGATCGGAAGA |
| 62421_1 | TATGAGTGTGGAGTCGTTGCCTTCCAAGTAGCTGGGACTACAGNNNAGATCGGAAG |
| 62424_1 | TATGAGTGTGGAGTCGTTGCCAAGTAGCTGGGACTACAGGCNNNAGATCGGAAGA |
| 62427_1 | TATGAGTGTGGAGTCGTTGCGAGTGATCCTCCTAGCACAGCNNNAGATCGGAAGAG |
| 62435_1 | TATGAGTGTGGAGTCGTTGCCGATGGCTTGCACTTGTAATTCTNNNAGATCGGAAGA |
| 62442_1 | TATGAGTGTGGAGTCGTTGCCTCCCGGGTTCAAGCAATTCTNNNAGATCGGAAGAG |
| 62477_1 | TATGAGTGTGGAGTCGTTGCTTCACACCATTCTCCTGCCTCNNNAGATCGGAAGAGCA |
| 62480_1 | TATGAGTGTGGAGTCGTTGCCAAGTAGCTGGGACTACAGGCNNNAGATCGGAAGA |
| 62485_1 | TATGAGTGTGGAGTCGTTGCGATGGAGTCTCACTCTGTGGCNNNAGATCGGAAGAG |
| 62493_1 | TATGAGTGTGGAGTCGTTGCGCGATCTCAGCTCACTGCAAGNNNAGATCGGAAGAG |
| 62503_1 | TATGAGTGTGGAGTCGTTGCGCAATCTCAGCTCACTGCAAGNNNAGATCGGAAGAG |
| 62536_1 | TATGAGTGTGGAGTCGTTGCTAGGGTAATGAGGGTTTGCCAAANNNAGATCGGAA |
| 62541_1 | TATGAGTGTGGAGTCGTTGCTAAACAAAGACACATTGGCCAGGNNNAGATCGGAAG |
| 62564_1 | TATGAGTGTGGAGTCGTTGCTGATGTTCTTCACTTGTGAGAGANNNAGATCGGAAG |
| 62577_1 | TATGAGTGTGGAGTCGTTGCAAGTAAAAGTTCGTCCGGGAATGNNNAGATCGGAAG |
| 62578_1 | TATGAGTGTGGAGTCGTTGCTCTTACTTTTTGCGGTGTCCTCTNNNAGATCGGAAGA |
| 62580_1 | TATGAGTGTGGAGTCGTTGCCAACTGTCTCACCTGTCCTACTCNNNAGATCGGAAGA |
| 62583_1 | TATGAGTGTGGAGTCGTTGCATACAAAAATTAGCCAGGCGTGGNNNAGATCGGAAG |
| 62601_1 | TATGAGTGTGGAGTCGTTGCGGTGTGTGCCTGTAGTCCAAGNNNAGATCGGAAGAG |
| 62623_1 | TATGAGTGTGGAGTCGTTGCCAAGTAGCTGGGACTACAGGCNNNAGATCGGAAGA |
| 62632_1 | TATGAGTGTGGAGTCGTTGCTTCTTTCTTGAAGACACCACTGCNNNAGATCGGAAGA |
| 62640_1 | TATGAGTGTGGAGTCGTTGCTTGGAGTGCAGTGGTGTGATCNNNAGATCGGAAGA |
| 62641_1 | TATGAGTGTGGAGTCGTTGCCAAGTAGCTGGGACTACAGGCNNNAGATCGGAAGA |
| 62650_1 | TATGAGTGTGGAGTCGTTGCCTCCGGAGTAGCTGGGACTACNNNAGATCGGAAGAG |
| 62657_1 | TATGAGTGTGGAGTCGTTGCTCCTACGCTGCACTAATTCTCTCNNNAGATCGGAAGA |
| 62660_1 | TATGAGTGTGGAGTCGTTGCCAAGTAGCTGGGACTACAGGCNNNAGATCGGAAGA |
| 62662_1 | TATGAGTGTGGAGTCGTTGCAGGAAACAATAGGTGCTGGAGAGNNNAGATCGGAA |
| 62686_1 | TATGAGTGTGGAGTCGTTGCATTAGCAATGAACAACTGGAAGTTNNNAGATCGGAA |
| 62705_1 | TATGAGTGTGGAGTCGTTGCACTGGAAGCCCAAAGGTCAATTANNNAGATCGGAAG |
| 62719_1 | TATGAGTGTGGAGTCGTTGCTATTACCCAGGAGAACTTCCCCANNNAGATCGGAAGA |
| 62720_1 | TATGAGTGTGGAGTCGTTGCGGATCACTTGAGGTCAGGAGTTTNNNAGATCGGAAG |
| 62721_1 | TATGAGTGTGGAGTCGTTGCTCAACTCATTACTGTGAGGAGGGNNNAGATCGGAAG |
| 62741_1 | TATGAGTGTGGAGTCGTTGCAATTTAGGCAGTGACCTAGCACTNNNAGATCGGAAG |
| 62754_1 | TATGAGTGTGGAGTCGTTGCGTTGCTTCCGGATTCAGTACATTNNNAGATCGGAAGA |
| 62764_1 | TATGAGTGTGGAGTCGTTGCCACCACTACGCCCAGCTAATTNNNAGATCGGAAGAGC |
| 62789_1 | TATGAGTGTGGAGTCGTTGCCAGTGGTGCAATCTCAGCTCANNNAGATCGGAAGAG |
| 62844_1 | TATGAGTGTGGAGTCGTTGCAGGATAGTCTCGATCTCCTGACCNNNAGATCGGAAG |
| 62901_1 | TATGAGTGTGGAGTCGTTGCGCAAGTAGCTGGGACTACAGGNNNAGATCGGAAGA |
| 62922_1 | TATGAGTGTGGAGTCGTTGCCAAGTAGCTGGGACTACAGGCNNNAGATCGGAAGA |

| | |
|---|---|
| 62945_1 | TATGAGTGTGGAGTCGTTGCCTTGAAAGGCTTCTGTTGCGTAGNNNAGATCGGAAG |
| 62970_1 | TATGAGTGTGGAGTCGTTGCCATTGGTGCAATCTCAGCTCACNNNAGATCGGAAGA |
| 62979_1 | TATGAGTGTGGAGTCGTTGCCTGAGGCAGGAAAATTGCTTGAANNNAGATCGGAA |
| 62985_1 | TATGAGTGTGGAGTCGTTGCAGGTACTTCTGGATCTCAGGACANNNAGATCGGAAG |
| 62989_1 | TATGAGTGTGGAGTCGTTGCCACACCTTCTGGAAATAAAGGACANNNAGATCGGAA |
| 62993_1 | TATGAGTGTGGAGTCGTTGCACAAGCAGCAAGAGAAAATTGACTNNNAGATCGGA |
| 63005_1 | TATGAGTGTGGAGTCGTTGCTCCTTTCCTTACTTCCTTCATTCCNNNAGATCGGAAGA |
| 63014_1 | TATGAGTGTGGAGTCGTTGCATTCAATGGAGAAAGGAGTGCCTNNNAGATCGGAA |
| 63018_1 | TATGAGTGTGGAGTCGTTGCATGGACATGACTCCAGCAAAAGANNNAGATCGGAA |
| 63022_1 | TATGAGTGTGGAGTCGTTGCCCAGTGGCTTTCTATCTCTGATGTNNNAGATCGGAAG |
| 63026_1 | TATGAGTGTGGAGTCGTTGCCCTGTGGAGGCTGATATAGGAACNNNAGATCGGAA |
| 63033_1 | TATGAGTGTGGAGTCGTTGCAGGCCTTTCAACTCTGACTTTGANNNAGATCGGAAG |
| 63052_1 | TATGAGTGTGGAGTCGTTGCTGTAGTCCCAGCTACTCAGGAGNNNAGATCGGAAGA |
| 63081_1 | TATGAGTGTGGAGTCGTTGCGAGGCAGGAGAATGGCATGAANNNAGATCGGAAG |
| 63094_1 | TATGAGTGTGGAGTCGTTGCTACACTTCTGCAGTCTCTAACGGNNNAGATCGGAAG |
| 63101_1 | TATGAGTGTGGAGTCGTTGCTATGGGCATGTGACACAATTCTGNNNAGATCGGAAG |
| 63118_1 | TATGAGTGTGGAGTCGTTGCCAAGTAGCTGGGACTACAGGCNNNAGATCGGAAGA |
| 63148_1 | TATGAGTGTGGAGTCGTTGCGGAGACTGACAGTAGAAGGATGGNNNAGATCGGA |
| 63160_1 | TATGAGTGTGGAGTCGTTGCCCATGTGGTACATTGTCTTTGCANNNAGATCGGAAG |
| 63192_1 | TATGAGTGTGGAGTCGTTGCGTAGTCCCAGCTACTCAGGAGANNNAGATCGGAAGA |
| 63205_1 | TATGAGTGTGGAGTCGTTGCGAGTAGCTGGGACTACAGGCANNNAGATCGGAAGA |
| 63212_1 | TATGAGTGTGGAGTCGTTGCTTCCAGATGTGCCACAGGAAATANNNAGATCGGAAG |
| 63224_1 | TATGAGTGTGGAGTCGTTGCGAGTAGCTGGGACCACAGTTGNNNAGATCGGAAGA |
| 63246_1 | TATGAGTGTGGAGTCGTTGCCAAGTAGCTGGGACTACAGGCNNNAGATCGGAAGA |
| 63248_1 | TATGAGTGTGGAGTCGTTGCCAGAGTCTCGTTCTGTCGTCCNNNAGATCGGAAGAG |
| 63251_1 | TATGAGTGTGGAGTCGTTGCCCTCAGCCTCCCATGTAGATGNNNAGATCGGAAGAG |
| 63257_1 | TATGAGTGTGGAGTCGTTGCAGACGGGATCTCACTCTGTCANNNAGATCGGAAGAG |
| 63282_1 | TATGAGTGTGGAGTCGTTGCCTGTAATCCCAGCTACTCAGGAGNNNAGATCGGAAG |
| 63328_1 | TATGAGTGTGGAGTCGTTGCTCCTTATCTACTTGCAGGTGCTGNNNAGATCGGAAG |
| 63372_1 | TATGAGTGTGGAGTCGTTGCTGGGAATATGCTTGTGATCCTGTNNNAGATCGGAAG |
| 63393_1 | TATGAGTGTGGAGTCGTTGCGATCTCCACTCACTACAAGCTCCNNNAGATCGGAAGA |
| 63397_1 | TATGAGTGTGGAGTCGTTGCGAGTACAGTGGCTCCATCTCGNNNAGATCGGAAGAG |
| 63434_1 | TATGAGTGTGGAGTCGTTGCTGTGGTGATGTGTGCCTATAGTCNNNAGATCGGAAG |
| 63457_1 | TATGAGTGTGGAGTCGTTGCCTCACTGCAAGCTCCTCTTCCNNNAGATCGGAAGAGC |
| 63462_1 | TATGAGTGTGGAGTCGTTGCCTCCTGGGTTCACACTGTTCTCNNNAGATCGGAAGAG |
| 63470_1 | TATGAGTGTGGAGTCGTTGCCCAAAGTGCTGGGACTACAGGNNNAGATCGGAAGA |
| 63503_1 | TATGAGTGTGGAGTCGTTGCAAACGAAGGCTCCCTAAGTCTTCNNNAGATCGGAAG |
| 63534_1 | TATGAGTGTGGAGTCGTTGCAGAGTGCTGGAGTGCTCACTNNNAGATCGGAAGAG |
| 63538_1 | TATGAGTGTGGAGTCGTTGCGCCTGTAGTCCCAGCTAATTGGNNNAGATCGGAAGA |
| 63590_1 | TATGAGTGTGGAGTCGTTGCAAAAAAAATTAGCCGGGTGTGGTNNNAGATCGGAA |
| 63591_1 | TATGAGTGTGGAGTCGTTGCTAACATCTGAGATTCCACTGCCCNNNAGATCGGAAGA |
| 63620_1 | TATGAGTGTGGAGTCGTTGCAGCACAAATCATACTCTCAGCATNNNAGATCGGAAGA |
| 63643_1 | TATGAGTGTGGAGTCGTTGCCTAGAGTGCAGTGGCGAGATCNNNAGATCGGAAGA |

| | |
|---|---|
| 63650_1 | TATGAGTGTGGAGTCGTTGCGTTTCACTGTGTTAGCCAGGATGNNNAGATCGGAAG |
| 63683_1 | TATGAGTGTGGAGTCGTTGCCTGAGGTCGGGAGTTCAAGACNNNAGATCGGAAGA |
| 63685_1 | TATGAGTGTGGAGTCGTTGCAATGTGTATCTACCCGCAGTTGTNNNAGATCGGAAG |
| 63694_1 | TATGAGTGTGGAGTCGTTGCCTTGCAGTGAGCCAAGATCACNNNAGATCGGAAGAG |
| 63762_1 | TATGAGTGTGGAGTCGTTGCGAGACGGAGTCTTGCTCTGTCNNNAGATCGGAAGAG |
| 63777_1 | TATGAGTGTGGAGTCGTTGCCTATGTGTGTGTGTGTGTGTCCNNNAGATCGGAAGA |
| 63784_1 | TATGAGTGTGGAGTCGTTGCCCAAGTAGCTGGGACTACAGGNNNAGATCGGAAGA |
| 63786_1 | TATGAGTGTGGAGTCGTTGCACCCCAAAGTCTCAGTCCAAAATNNNAGATCGGAAGA |
| 63809_1 | TATGAGTGTGGAGTCGTTGCTGTACACTGCTCAGAAATCACCANNNAGATCGGAAGA |
| 63817_1 | TATGAGTGTGGAGTCGTTGCGGAAGGGAAGGCAGAAAGAAAGANNNAGATCGGA |
| 63828_1 | TATGAGTGTGGAGTCGTTGCCTGGGGACCTTCTGATTCCAGNNNAGATCGGAAGAG |
| 63842_1 | TATGAGTGTGGAGTCGTTGCTGTATGGAGCAGTGAAGGCATTTNNNAGATCGGAAG |
| 63886_1 | TATGAGTGTGGAGTCGTTGCAGTGACCACTAAGTACAACCCACNNNAGATCGGAAG |
| 63941_1 | TATGAGTGTGGAGTCGTTGCGACATGTGACCAAATATTGCCAANNNAGATCGGAAG |
| 63945_1 | TATGAGTGTGGAGTCGTTGCGCCTGTAGTCCCAGCTACTTGNNNAGATCGGAAGAG |
| 63958_1 | TATGAGTGTGGAGTCGTTGCGCAACTTCTGGTTCCTAAATGTGTNNNAGATCGGAAG |
| 63977_1 | TATGAGTGTGGAGTCGTTGCTGATCTCAGCTCACTGCAAGCNNNAGATCGGAAGAG |
| 63990_1 | TATGAGTGTGGAGTCGTTGCGATGACTGCGTGAAAATGCTTTCNNNAGATCGGAAG |
| 64017_1 | TATGAGTGTGGAGTCGTTGCGAGTGTAGTGGCGCGATCTTNNNAGATCGGAAGAG |
| 64095_1 | TATGAGTGTGGAGTCGTTGCAGGAGACAGGACCACTCCATCNNNAGATCGGAAGAG |
| 64159_1 | TATGAGTGTGGAGTCGTTGCCAGGGTCTGGAAGGAACTGTCNNNAGATCGGAAGA |
| 64160_1 | TATGAGTGTGGAGTCGTTGCGAGACAGTGATCCAGTGGGTTCNNNAGATCGGAAG |
| 64212_1 | TATGAGTGTGGAGTCGTTGCCCTGGTATTCAGCTGCGAAGGNNNAGATCGGAAGAG |
| 64251_1 | TATGAGTGTGGAGTCGTTGCCTGTGTCTGTCGATGAGTCAGAANNNAGATCGGAAG |
| 64283_1 | TATGAGTGTGGAGTCGTTGCCTTGATCTCCTGACCTCGTGATCNNNAGATCGGAAGA |
| 64353_1 | TATGAGTGTGGAGTCGTTGCCTACACCCCGTCCATAAGAAGTCNNNAGATCGGAAGA |
| 64379_1 | TATGAGTGTGGAGTCGTTGCGTCTTGGCTCACTGCAAACTCNNNAGATCGGAAGAG |
| 64392_1 | TATGAGTGTGGAGTCGTTGCATGGGAACGAGAAATAGCAATGGNNNAGATCGGAA |
| 64413_1 | TATGAGTGTGGAGTCGTTGCTTTCACCGTGTTAGCCAGGATGNNNAGATCGGAAGA |
| 64418_1 | TATGAGTGTGGAGTCGTTGCCTCCAGAGGGCCAGCTGTTGNNNAGATCGGAAGAGC |
| 64458_1 | TATGAGTGTGGAGTCGTTGCTTGTTCACTGCACAAAGGAATCCNNNAGATCGGAAG |
| 64471_1 | TATGAGTGTGGAGTCGTTGCGTGTGTGTGTGTGTGTGTGTGTCNNNAGATCGGAAGA |
| 64481_1 | TATGAGTGTGGAGTCGTTGCTGCAGCTCAGTGTATGTGTGTATNNNAGATCGGAAG |
| 64658_1 | TATGAGTGTGGAGTCGTTGCCCTCTGGGCTCTCTTCTCTCCNNNAGATCGGAAGAGC |
| 64672_1 | TATGAGTGTGGAGTCGTTGCGGTGAAGGTGTTGAGCTCCTTNNNAGATCGGAAGAG |
| 64748_1 | TATGAGTGTGGAGTCGTTGCCCAGACTGGGAACAAGTCAAATGNNNAGATCGGAAG |
| 64753_1 | TATGAGTGTGGAGTCGTTGCTAGAATTTGTAGGGCTTGGGGACNNNAGATCGGAA |
| 64795_1 | TATGAGTGTGGAGTCGTTGCCAGTGGTGTGATCTTGGCTCANNNAGATCGGAAGAG |
| 64821_1 | TATGAGTGTGGAGTCGTTGCGTTAGCCAGGATGGTCTCGATCNNNAGATCGGAAGA |
| 64841_1 | TATGAGTGTGGAGTCGTTGCGAGTCTCACTCTGCCATCCAGNNNAGATCGGAAGAG |
| 64895_1 | TATGAGTGTGGAGTCGTTGCTTGCGACAAAAATATTTGCCCANNNAGATCGGAAGA |
| 64935_1 | TATGAGTGTGGAGTCGTTGCGAGACGGAGTCTTGCTCTGTCNNNAGATCGGAAGAG |
| 64994_1 | TATGAGTGTGGAGTCGTTGCGCGATCTCAGCTCACTGCAAGNNNAGATCGGAAGAG |

| | |
|---|---|
| 65082_1 | TATGAGTGTGGAGTCGTTGCAATAAAAAGAAAAGGGGCCCTCGNNNAGATCGGAA |
| 65118_1 | TATGAGTGTGGAGTCGTTGCTACCCATCTCTTCCTTCTTCCCTNNNAGATCGGAAGAG |
| 65124_1 | TATGAGTGTGGAGTCGTTGCAGACTCTCACTCTGTTGCCCANNNAGATCGGAAGAG |
| 65216_1 | TATGAGTGTGGAGTCGTTGCTCTTTCTTTTTTGAGACGGAGTCTNNNAGATCGGAA |
| 65240_1 | TATGAGTGTGGAGTCGTTGCGAGAGAGAGAGAGAGAGAGGTGGNNNAGATCGG |
| 65253_1 | TATGAGTGTGGAGTCGTTGCTGAAGCCCTTCAACCATATGGATNNNAGATCGGAAG |
| 65399_1 | TATGAGTGTGGAGTCGTTGCCCTCAGTCACTGACCTACGTGNNNAGATCGGAAGAG |
| 65482_1 | TATGAGTGTGGAGTCGTTGCGGATGTGCGTGTATGTGATGTGNNNAGATCGGAAG |
| 65536_1 | TATGAGTGTGGAGTCGTTGCCAGTGTGGGGAACATCTGAGGNNNAGATCGGAAG |
| 65561_1 | TATGAGTGTGGAGTCGTTGCGCAATGTGAGCCTGGCAAAAGNNNAGATCGGAAGA |
| 65593_1 | TATGAGTGTGGAGTCGTTGCGATCTTGGGTCACTGCAACCANNNAGATCGGAAGA |
| 65612_1 | TATGAGTGTGGAGTCGTTGCACACACCCAGCACATATAATTGCNNNAGATCGGAAG |
| 65680_1 | TATGAGTGTGGAGTCGTTGCTCACTTGAGGTCAGGAGTTCATGNNNAGATCGGAA |
| 65731_1 | TATGAGTGTGGAGTCGTTGCAGTCACTTCACCTCTCAGAGACTNNNAGATCGGAAG |
| 65750_1 | TATGAGTGTGGAGTCGTTGCCTTGCAGTGAGCCGAGATTGTNNNAGATCGGAAGA |
| 65811_1 | TATGAGTGTGGAGTCGTTGCAAGCCATGAAAAGACTTGGAGGANNNAGATCGGAA |
| 65816_1 | TATGAGTGTGGAGTCGTTGCCTGTACTCCCAGCTACTCAGGAGNNNAGATCGGAAG |
| 65830_1 | TATGAGTGTGGAGTCGTTGCACCAGACACCCATACTCAGGTNNNAGATCGGAAGAG |
| 65834_1 | TATGAGTGTGGAGTCGTTGCACACACACACACACACTCTCTTANNNAGATCGGAAGA |
| 65847_1 | TATGAGTGTGGAGTCGTTGCGTTCCACCTTTGAAACTGTAGGCNNNAGATCGGAAG |
| 65919_1 | TATGAGTGTGGAGTCGTTGCCACAATCATGGCTCACTGCAGNNNAGATCGGAAGAG |
| 65937_1 | TATGAGTGTGGAGTCGTTGCTTCAGTAGAGATGGGCTTTCACCNNNAGATCGGAAG |
| 65957_1 | TATGAGTGTGGAGTCGTTGCCCAGGTTCAAGCGATTCTCCTNNNAGATCGGAAGAG |
| 65976_1 | TATGAGTGTGGAGTCGTTGCACACTTGGCTTATATCCCCTGTGNNNAGATCGGAAG |
| 66003_1 | TATGAGTGTGGAGTCGTTGCCCATGAGAGCAGGGTTTTTTTGTNNNAGATCGGAA |
| 66036_1 | TATGAGTGTGGAGTCGTTGCATTGGCACTATCTCAGCTCGCNNNAGATCGGAAGAG |
| 66041_1 | TATGAGTGTGGAGTCGTTGCTCTTCTCAGAACAAGTTCCTGCANNNAGATCGGAAG |
| 66086_1 | TATGAGTGTGGAGTCGTTGCATCTTGGCTCACTGCAACCTCNNNAGATCGGAAGAG |
| 66117_1 | TATGAGTGTGGAGTCGTTGCGTGTGTGTGTGAGAGAGAGAGAGNNNAGATCGGA |
| 66120_1 | TATGAGTGTGGAGTCGTTGCTACCCCAAAAATAACAGAGCCGTNNNAGATCGGAAG |
| 66129_1 | TATGAGTGTGGAGTCGTTGCCAGTCTCGTTCTGTCGCCTAGNNNAGATCGGAAGAG |
| 66153_1 | TATGAGTGTGGAGTCGTTGCCCTGAGCTTTGGTCTGGTATCTTNNNAGATCGGAAG |
| 66269_1 | TATGAGTGTGGAGTCGTTGCATCATTTCCTGACCCCAGCTCNNNAGATCGGAAGAG |
| 66372_1 | TATGAGTGTGGAGTCGTTGCCACACACACACACACACACATATNNNAGATCGGAAG |
| 66484_1 | TATGAGTGTGGAGTCGTTGCGGCACTCAGAACAGGAAGAATCTNNNAGATCGGAA |
| 66525_1 | TATGAGTGTGGAGTCGTTGCGCGCCTGTAATCCCAACTACTTNNNAGATCGGAAGA |
| 66533_1 | TATGAGTGTGGAGTCGTTGCTGTAATCCTAGCACTTTGAGGGGNNNAGATCGGAA |
| 66567_1 | TATGAGTGTGGAGTCGTTGCCAATGGTGCTATCTCGGCTCANNNAGATCGGAAGAG |
| 66578_1 | TATGAGTGTGGAGTCGTTGCCCTGCAATCCCAGCTACTCAGNNNAGATCGGAAGAG |
| 66654_1 | TATGAGTGTGGAGTCGTTGCCTGTCACTCAGGCTGCAGTACNNNAGATCGGAAGAG |
| 66664_1 | TATGAGTGTGGAGTCGTTGCTTTTCAATGCATGTTTTCCGGGTNNNAGATCGGAAG |
| 66751_1 | TATGAGTGTGGAGTCGTTGCTACACGGAGTTTCACAACCCCNNNAGATCGGAAGAG |
| 66759_1 | TATGAGTGTGGAGTCGTTGCCAGTGGCACAATCTCTGCTCANNNAGATCGGAAGAG |

| | |
|---|---|
| 66761_1 | TATGAGTGTGGAGTCGTTGCGTGTTTCACCATGTTGGCCAGNNNAGATCGGAAGAG |
| 66819_1 | TATGAGTGTGGAGTCGTTGCCAATCTTGGCTCACTGCAACCNNNAGATCGGAAGAG |
| 66948_1 | TATGAGTGTGGAGTCGTTGCTATATAATAGGGCACCAGTGGCCNNNAGATCGGAA |
| 66960_1 | TATGAGTGTGGAGTCGTTGCGATCAATTGAGCCTGGGAGACAGNNNAGATCGGAA |
| 66963_1 | TATGAGTGTGGAGTCGTTGCGATGGAGTCTCCCTCTGTTGCNNNAGATCGGAAGAG |
| 67023_1 | TATGAGTGTGGAGTCGTTGCATGTATATGTGTGTGTGTGTGTGTNNNAGATCGGA |
| 67034_1 | TATGAGTGTGGAGTCGTTGCTACGAACAGAAAGCAGGCCAGNNNAGATCGGAAGA |
| 67068_1 | TATGAGTGTGGAGTCGTTGCTATGTACACACACAGAGCGAGACNNNAGATCGGAA |
| 67104_1 | TATGAGTGTGGAGTCGTTGCGAGTTTTGCTCTTGTTCCCCAGNNNAGATCGGAAGA |
| 67106_1 | TATGAGTGTGGAGTCGTTGCGTGCACAGTGACTCACACCTATANNNAGATCGGAAG |
| 67109_1 | TATGAGTGTGGAGTCGTTGCATCCTCAAAGCAAGAAAGCAAGGNNNAGATCGGAA |
| 67228_1 | TATGAGTGTGGAGTCGTTGCGAGGTGGGAGGAAGGGATGTCNNNAGATCGGAAG |
| 67241_1 | TATGAGTGTGGAGTCGTTGCGTTAGTTGTGCCGTTCTGCTGNNNAGATCGGAAGAG |
| 67289_1 | TATGAGTGTGGAGTCGTTGCGTTCCAGTCTCAGAAGGGCTGNNNAGATCGGAAGA |
| 67358_1 | TATGAGTGTGGAGTCGTTGCAACTACTGCCATATGTGTGACCANNNAGATCGGAAG |
| 67406_1 | TATGAGTGTGGAGTCGTTGCCCACATACACACATGGCATGCNNNAGATCGGAAGAG |
| 67446_1 | TATGAGTGTGGAGTCGTTGCAGAGGAGAGAGATAGAGATCTGGTNNNAGATCGG |
| 67468_1 | TATGAGTGTGGAGTCGTTGCCTGAGGCAGGAGGATCACTTGNNNAGATCGGAAGA |
| 67486_1 | TATGAGTGTGGAGTCGTTGCCAGGGTGGTAGACAGAGACAGTNNNAGATCGGAA |
| 67530_1 | TATGAGTGTGGAGTCGTTGCCAGTTTTTCTGGAGAACCACGACNNNAGATCGGAAG |
| 67591_1 | TATGAGTGTGGAGTCGTTGCGAGGGGTTTGGGAAGCTTTTCNNNAGATCGGAAGA |
| 67613_1 | TATGAGTGTGGAGTCGTTGCTCCTAGAGGACAGAGCCCATGNNNAGATCGGAAGA |
| 67619_1 | TATGAGTGTGGAGTCGTTGCGCTGGTACTGAAACTGCTCCTATNNNAGATCGGAAG |
| 67626_1 | TATGAGTGTGGAGTCGTTGCGAGAGGGAGAGCTTGAGTTGATTNNNAGATCGGA |
| 67658_1 | TATGAGTGTGGAGTCGTTGCGGAGTCTCACTCTGTCACCCANNNAGATCGGAAGAG |
| 67665_1 | TATGAGTGTGGAGTCGTTGCCATCTTCCTACTCAATCCCCACCNNNAGATCGGAAGA |
| 67667_1 | TATGAGTGTGGAGTCGTTGCGGCCTGGAATTCACTGGTGAANNNAGATCGGAAGA |
| 67689_1 | TATGAGTGTGGAGTCGTTGCGAGTTTTGCTCTTGTTGCCCANNNAGATCGGAAGAG |
| 67739_1 | TATGAGTGTGGAGTCGTTGCCAGGAAGAGTCTCCCTCCCTGNNNAGATCGGAAGAG |
| 67778_1 | TATGAGTGTGGAGTCGTTGCGGACTGCAGTAGTGCGATCTTNNNAGATCGGAAGA |
| 67969_1 | TATGAGTGTGGAGTCGTTGCGACTGAAGAAGGGTGGGGAATGNNNAGATCGGAA |
| 67973_1 | TATGAGTGTGGAGTCGTTGCCTCTCTCTCTCTCTCTGCGGATANNNAGATCGGAAGA |
| 67976_1 | TATGAGTGTGGAGTCGTTGCACACACACACACCCATTCCTTCNNNAGATCGGAAGAG |
| 68002_1 | TATGAGTGTGGAGTCGTTGCAAGAGAACAAGAGACTCTGCCTGNNNAGATCGGAA |
| 68017_1 | TATGAGTGTGGAGTCGTTGCGAGTACAGTGGTGAGATCTCAGCNNNAGATCGGAA |
| 68038_1 | TATGAGTGTGGAGTCGTTGCTGAGAGGAAGAGGGAAGATCCTGNNNAGATCGGA |
| 68046_1 | TATGAGTGTGGAGTCGTTGCACACACACACACACACTCTCTTANNNAGATCGGAAGA |
| 68168_1 | TATGAGTGTGGAGTCGTTGCCACCTCTTCTTTGTGGCTGTTCNNNAGATCGGAAGAG |
| 68224_1 | TATGAGTGTGGAGTCGTTGCCTGATCTCTGGTGATCTGTCCACNNNAGATCGGAAG |
| 68235_1 | TATGAGTGTGGAGTCGTTGCTGGGGTTGACTTTCCTTACTTGTNNNAGATCGGAAG |
| 68293_1 | TATGAGTGTGGAGTCGTTGCCTGGAGTGTAGTGGCGTGATCNNNAGATCGGAAGA |
| 68323_1 | TATGAGTGTGGAGTCGTTGCCATAGTCCCAGCTACTCGGAAGNNNAGATCGGAAGA |
| 68330_1 | TATGAGTGTGGAGTCGTTGCATTTCACTTTATTGCCCAGGCTGNNNAGATCGGAAG |

| | |
|---|---|
| 68337_1 | TATGAGTGTGGAGTCGTTGCGAGCAAGACCCTGTCTCAAAGAANNNAGATCGGAAG |
| 68373_1 | TATGAGTGTGGAGTCGTTGCGGCCACCAGAAATACCTGTACTANNNAGATCGGAAG |
| 68379_1 | TATGAGTGTGGAGTCGTTGCTGCATTATGGCCATTGTCATTGANNNAGATCGGAAG |
| 68422_1 | TATGAGTGTGGAGTCGTTGCAGATGGAGTCTCACTGTTGCCNNNAGATCGGAAGAG |
| 68425_1 | TATGAGTGTGGAGTCGTTGCTCTAGAAATCTCTGAGTTGGGGANNNAGATCGGAAG |
| 68435_1 | TATGAGTGTGGAGTCGTTGCCAATCTTGGCTCACTGCAACCNNNAGATCGGAAGAG |
| 68459_1 | TATGAGTGTGGAGTCGTTGCATATATATATCTCGGCTGGGCGCNNNAGATCGGAAG |
| 68495_1 | TATGAGTGTGGAGTCGTTGCTCAGAAAGAAGTCCTCTGCTTCCNNNAGATCGGAAGA |
| 68557_1 | TATGAGTGTGGAGTCGTTGCCTGGGTGACAGAGTGAGACTCNNNAGATCGGAAGA |
| 68567_1 | TATGAGTGTGGAGTCGTTGCTCCAGGCTGGAGTGAAGTGGNNNAGATCGGAAGAG |
| 68588_1 | TATGAGTGTGGAGTCGTTGCAAACTGAGGAGACGTGACAACAANNNAGATCGGAA |
| 68591_1 | TATGAGTGTGGAGTCGTTGCATTACCTCTCTCCCTATCTCCCTNNNAGATCGGAAGAG |
| 68624_1 | TATGAGTGTGGAGTCGTTGCGCCTGTAATCCCAGCACTTTGNNNAGATCGGAAGAG |
| 68733_1 | TATGAGTGTGGAGTCGTTGCCATCTCCTACGACAAGATGGTCANNNAGATCGGAAG |
| 68735_1 | TATGAGTGTGGAGTCGTTGCCAGAGTGCTGGGATTACAGGCNNNAGATCGGAAGA |
| 68738_1 | TATGAGTGTGGAGTCGTTGCCCACTCCCTTGCATTTCCTTCTCNNNAGATCGGAAGAG |
| 68846_1 | TATGAGTGTGGAGTCGTTGCAGAAAGAAAGGAAGGAAGGAAGGANNNAGATCGG |
| 68954_1 | TATGAGTGTGGAGTCGTTGCCAAGTGACCGTGCGTGTAAAGNNNAGATCGGAAGA |
| 68960_1 | TATGAGTGTGGAGTCGTTGCGTGACCTTCTTCCCTGAGGACNNNAGATCGGAAGAG |
| 68972_1 | TATGAGTGTGGAGTCGTTGCCTGTAAGTGTGGTTCCTCCCATGNNNAGATCGGAAG |
| 68990_1 | TATGAGTGTGGAGTCGTTGCCAAGGCTGAGGATAAGGAGTGGNNNAGATCGGAAG |
| 68993_1 | TATGAGTGTGGAGTCGTTGCCTGTCCCTGGGTCATCTCAACNNNAGATCGGAAGAG |
| 69027_1 | TATGAGTGTGGAGTCGTTGCCACACACACACACACACACAAATNNNAGATCGGAAGA |
| 69110_1 | TATGAGTGTGGAGTCGTTGCCAGTGGCATGATCTCAGGTCANNNAGATCGGAAGAG |
| 69222_1 | TATGAGTGTGGAGTCGTTGCAAAGATGGGTAGTTCCAAAGGGGNNNAGATCGGAA |
| 69227_1 | TATGAGTGTGGAGTCGTTGCGAAGCCGAGGAAGAGCGTTTTNNNAGATCGGAAGA |
| 69230_1 | TATGAGTGTGGAGTCGTTGCTAGACACTCATTCAGGCAGATGGNNNAGATCGGAAG |
| 69249_1 | TATGAGTGTGGAGTCGTTGCCTCAGTCCCACAAGACTCACCNNNAGATCGGAAGAG |
| 69280_1 | TATGAGTGTGGAGTCGTTGCAGCATGAGACACTTGGTAGCCNNNAGATCGGAAGAG |
| 69347_1 | TATGAGTGTGGAGTCGTTGCTGAATGAATGAGGGAGGACAAGGNNNAGATCGGAA |
| 69393_1 | TATGAGTGTGGAGTCGTTGCGAGTCTCACTCTGTTGCCGAGNNNAGATCGGAAGAG |
| 69444_1 | TATGAGTGTGGAGTCGTTGCAAAGGAAGGAAGGAACGAAGGAANNNAGATCGGA |
| 69466_1 | TATGAGTGTGGAGTCGTTGCCACGACAGGGGTTCTTCACTATTNNNAGATCGGAAG |
| 69538_1 | TATGAGTGTGGAGTCGTTGCCGTGTCCAGAAGGGAGTCTTCNNNAGATCGGAAGAG |
| 69581_1 | TATGAGTGTGGAGTCGTTGCCTGGCAGGGGCTCTTCTCATCNNNAGATCGGAAGAG |
| 69582_1 | TATGAGTGTGGAGTCGTTGCCTGGGAGGGCCAAATGTCTAANNNAGATCGGAAGA |
| 69595_1 | TATGAGTGTGGAGTCGTTGCCTGGCAGGGAGTGTCATACTCNNNAGATCGGAAGAG |
| 69600_1 | TATGAGTGTGGAGTCGTTGCGGGTGTTAGGGAGAGAGGGAGNNNAGATCGGAAG |
| 69606_1 | TATGAGTGTGGAGTCGTTGCCCTCAACTTAGCAGGGGTAGAACNNNAGATCGGAAG |
| 69616_1 | TATGAGTGTGGAGTCGTTGCGGCTCAGACCTTCAGGGATTCNNNAGATCGGAAGAG |
| 69620_1 | TATGAGTGTGGAGTCGTTGCACCAAGGGAAAAAAAATAGGAGCANNNAGATCGGA |
| 69622_1 | TATGAGTGTGGAGTCGTTGCTTCCACAAACCCTTCATGAAAGCNNNAGATCGGAAGA |
| 69658_1 | TATGAGTGTGGAGTCGTTGCCAGTCCTCCATCCTCTTCTGGANNNAGATCGGAAGAG |

| | |
|---|---|
| 69666_1 | TATGAGTGTGGAGTCGTTGCTTCCTTGGCTGATGCTGAATCTANNNAGATCGGAAG |
| 69682_1 | TATGAGTGTGGAGTCGTTGCCTACTGTGGTTACACTGCCGGNNNAGATCGGAAGAG |
| 69815_1 | TATGAGTGTGGAGTCGTTGCGACCTCCTGGCTCAAGTGATCNNNAGATCGGAAGAG |
| 69831_1 | TATGAGTGTGGAGTCGTTGCGATGCAGTCTCGCTCTCTCTCNNNAGATCGGAAGAG |
| 69855_1 | TATGAGTGTGGAGTCGTTGCCCTGAAAGGCTGTGTACCAAAATNNNAGATCGGAAG |
| 69904_1 | TATGAGTGTGGAGTCGTTGCAACATTGTAACACAGCAGGCAAANNNAGATCGGAA |
| 70050_1 | TATGAGTGTGGAGTCGTTGCTGTCATCCGCCATTTGTTGTTTTNNNAGATCGGAAGA |
| 70063_1 | TATGAGTGTGGAGTCGTTGCGGGTAGATCAGGAGGAGATCTGANNNAGATCGGA |
| 70104_1 | TATGAGTGTGGAGTCGTTGCTCTGTTCATTCCTTCTGGGTCTGNNNAGATCGGAAG |
| 70105_1 | TATGAGTGTGGAGTCGTTGCTCTCAGCGGAAAACCAATCATTGNNNAGATCGGAAG |
| 70113_1 | TATGAGTGTGGAGTCGTTGCACATCGCTATAGCAACCGGTGNNNAGATCGGAAGA |
| 70115_1 | TATGAGTGTGGAGTCGTTGCGTGTGGGTACCTGCCTTATCCNNNAGATCGGAAGAG |
| 70170_1 | TATGAGTGTGGAGTCGTTGCAGGATAAGTTCTCACCAGAGGGANNNAGATCGGAA |
| 70178_1 | TATGAGTGTGGAGTCGTTGCCACCAACTCTCTGGAACCCACNNNAGATCGGAAGAG |
| 70179_1 | TATGAGTGTGGAGTCGTTGCAGAAACACCAAAGGGCAATCAAGNNNAGATCGGAA |
| 70184_1 | TATGAGTGTGGAGTCGTTGCGTGTGGAGCTGGGATGTTCTCNNNAGATCGGAAGA |
| 70188_1 | TATGAGTGTGGAGTCGTTGCTCTTTGGTACCTGTGTTCGTCAGNNNAGATCGGAAG |
| 70213_1 | TATGAGTGTGGAGTCGTTGCGAACTGTGGATGGGGGGACTCNNNAGATCGGAAG |
| 70216_1 | TATGAGTGTGGAGTCGTTGCGAGCTAGATCCCCACTGCAAGNNNAGATCGGAAGA |
| 70305_1 | TATGAGTGTGGAGTCGTTGCGCTTGCAGTGAGCTGAGATTGNNNAGATCGGAAGA |
| 70330_1 | TATGAGTGTGGAGTCGTTGCTTCACACCATTCTCCTGCCTCNNNAGATCGGAAGAGC |
| 70394_1 | TATGAGTGTGGAGTCGTTGCCAGAGAGAACCTATGATGGACATNNNAGATCGGAA |
| 70419_1 | TATGAGTGTGGAGTCGTTGCGCTCACTGCTATCTCCACCTCNNNAGATCGGAAGAGC |
| 70458_1 | TATGAGTGTGGAGTCGTTGCGGAGGGAGGGAGGGAGGAAGNNNAGATCGGAAG |
| 70477_1 | TATGAGTGTGGAGTCGTTGCGGAGTGCAGTGGTACGATCTCNNNAGATCGGAAGA |
| 70500_1 | TATGAGTGTGGAGTCGTTGCCACACACACACACACACACACNNNAGATCGGAAGAG |
| 70513_1 | TATGAGTGTGGAGTCGTTGCGCCTATAATCCCACCCCTTTGGNNNAGATCGGAAGA |
| 186_1 | TATGAGTGTGGAGTCGTTGCGACCATCCATGCCGTTGTCCNNNAGATCGGAAGAGC |
| 280_1 | TATGAGTGTGGAGTCGTTGCACCCCAGCCACCATGAGTAGNNNAGATCGGAAGAGC |
| 293_1 | TATGAGTGTGGAGTCGTTGCTCTTCCCTCCCACCTTGCACNNNAGATCGGAAGAGCA |
| 389_1 | TATGAGTGTGGAGTCGTTGCAGGGGGCGACAGGCCAATCTNNNAGATCGGAAGAG |
| 889_1 | TATGAGTGTGGAGTCGTTGCTCCTGGCATTCGAAGTTCCCNNNAGATCGGAAGAGC |
| 909_1 | TATGAGTGTGGAGTCGTTGCAAATAGTCCGCAGGCTGGGTNNNAGATCGGAAGAG |
| 1150_1 | TATGAGTGTGGAGTCGTTGCCCCTCTCTGTCCCAATCCCANNNAGATCGGAAGAGCA |
| 1485_1 | TATGAGTGTGGAGTCGTTGCGTCCCTTTGGCCTTTGCCTTNNNAGATCGGAAGAGC |
| 1537_1 | TATGAGTGTGGAGTCGTTGCTGTTGTTGTTGTCCACCACCNNNAGATCGGAAGAGC |
| 1931_1 | TATGAGTGTGGAGTCGTTGCCCCGGCTTTTAGCATCAAAANNNAGATCGGAAGAGC |
| 1993_1 | TATGAGTGTGGAGTCGTTGCGGATCTCTGGCTCTGGTACCNNNAGATCGGAAGAG |
| 2568_1 | TATGAGTGTGGAGTCGTTGCGGAGGCCGAATCGCATGAGANNNAGATCGGAAGA |
| 3341_1 | TATGAGTGTGGAGTCGTTGCATGGCGGTGCCTTAGGTTCCNNNAGATCGGAAGAG |
| 3366_1 | TATGAGTGTGGAGTCGTTGCCTCAGCCCTCCCTAGCCCTCNNNAGATCGGAAGAGCA |
| 3391_1 | TATGAGTGTGGAGTCGTTGCCTCACCCTCCAATTACCGGGTNNNAGATCGGAAGAG |
| 3478_1 | TATGAGTGTGGAGTCGTTGCAGTGGCTCATGGGCTGTTTCNNNAGATCGGAAGAG |

| | |
|---|---|
| 3657_1 | TATGAGTGTGGAGTCGTTGCGAAACGCGAGTGACTGGGTCTGNNNAGATCGGAAG |
| 3855_1 | TATGAGTGTGGAGTCGTTGCACCACCCTATTGCTCAGGCTNNNAGATCGGAAGAGC |
| 3914_1 | TATGAGTGTGGAGTCGTTGCTACACCGCTGGGCATGGTAGNNNAGATCGGAAGAG |
| 4171_1 | TATGAGTGTGGAGTCGTTGCGGCACACTGAAGCTGACAACNNNAGATCGGAAGAG |
| 4278_1 | TATGAGTGTGGAGTCGTTGCAATGCCGGGTCACAGCAACCNNNAGATCGGAAGAG |
| 4569_1 | TATGAGTGTGGAGTCGTTGCATGGGGACCAGGCAGACAATNNNAGATCGGAAGAG |
| 4740_1 | TATGAGTGTGGAGTCGTTGCGTGGAGAATGTTGGGGAGGTNNNAGATCGGAAGA |
| 4790_1 | TATGAGTGTGGAGTCGTTGCCAGGGCAAAAGGAAGAAAACACCNNNAGATCGGAA |
| 4828_1 | TATGAGTGTGGAGTCGTTGCCGTGTCTTTCAGCATGGGGANNNAGATCGGAAGAG |
| 4856_1 | TATGAGTGTGGAGTCGTTGCTGCGAGGAAGACTGGAGACCNNNAGATCGGAAGAG |
| 4958_1 | TATGAGTGTGGAGTCGTTGCTATGTTTGTCTCCTCACTCCCCCNNNAGATCGGAAGA |
| 5199_1 | TATGAGTGTGGAGTCGTTGCTGCCCTAATGGAACTGGCCANNNAGATCGGAAGAG |
| 5483_1 | TATGAGTGTGGAGTCGTTGCCTCCCCTAATCTCTACAGCTGCTNNNAGATCGGAAGA |
| 5711_1 | TATGAGTGTGGAGTCGTTGCCCACACACACAGGAAGAAGATCTNNNAGATCGGAAG |
| 5877_1 | TATGAGTGTGGAGTCGTTGCCCTTCTCATAGCTGCCCCCGNNNAGATCGGAAGAGC |
| 6155_1 | TATGAGTGTGGAGTCGTTGCCTTTGTCCTCACTCCTCGTGTCNNNAGATCGGAAGAG |
| 6221_1 | TATGAGTGTGGAGTCGTTGCCACACGACTTGCCTGCCTCANNNAGATCGGAAGAGC |
| 6362_1 | TATGAGTGTGGAGTCGTTGCGGACACCCATCATGCCAGTCNNNAGATCGGAAGAGC |
| 6369_1 | TATGAGTGTGGAGTCGTTGCGGGCAAGGTAAGAGCTGCAANNNAGATCGGAAGA |
| 6382_1 | TATGAGTGTGGAGTCGTTGCTCCACCAAACTCCCCAGGAAANNNAGATCGGAAGAG |
| 6453_1 | TATGAGTGTGGAGTCGTTGCAAGCCCCGTAAGGATTGGGANNNAGATCGGAAGAG |
| 6822_1 | TATGAGTGTGGAGTCGTTGCTCAGAGCTGGGTGCCTTACAGNNNAGATCGGAAGA |
| 6853_1 | TATGAGTGTGGAGTCGTTGCGAAACCCCAGACCAGGTCCANNNAGATCGGAAGAG |
| 6901_1 | TATGAGTGTGGAGTCGTTGCATGGACCCACCCACATGCTCNNNAGATCGGAAGAGC |
| 7079_1 | TATGAGTGTGGAGTCGTTGCGTTTCGGGACTGCAGCCGAGNNNAGATCGGAAGAG |
| 7137_1 | TATGAGTGTGGAGTCGTTGCGGAAAAGTCCATGACGCTGGAGNNNAGATCGGAAG |
| 7228_1 | TATGAGTGTGGAGTCGTTGCACTCCTCCCCACCACCGTGGNNNAGATCGGAAGAGC |
| 7395_1 | TATGAGTGTGGAGTCGTTGCCAATGAAGTCTCGCCTCGCTNNNAGATCGGAAGAGC |
| 7620_1 | TATGAGTGTGGAGTCGTTGCGAGAAATGGAGGATCGTGTGAGGNNNAGATCGGA |
| 7699_1 | TATGAGTGTGGAGTCGTTGCAAGTGCCTCCTGGTGCACCGNNNAGATCGGAAGAG |
| 8197_1 | TATGAGTGTGGAGTCGTTGCGCAGAGTGGTTGTGCATTACATGNNNAGATCGGAA |
| 8206_1 | TATGAGTGTGGAGTCGTTGCGTTAGTGGAGTAGCAGCCCATGNNNAGATCGGAAG |
| 8398_1 | TATGAGTGTGGAGTCGTTGCTCCTTTAGGTGAGTGAGGTCCTGNNNAGATCGGAAG |
| 8639_1 | TATGAGTGTGGAGTCGTTGCCTGGGCTTGAGGTGTTGGTGNNNAGATCGGAAGAG |
| 9353_1 | TATGAGTGTGGAGTCGTTGCTTTTTGACCCTGGGTACTCCNNNAGATCGGAAGAGC |
| 10791_1 | TATGAGTGTGGAGTCGTTGCACCCAGAGTAGGAGGGAGGANNNAGATCGGAAGA |
| 11111_1 | TATGAGTGTGGAGTCGTTGCCAGACGCCATCTCTCCCTCTNNNAGATCGGAAGAGCA |
| 11236_1 | TATGAGTGTGGAGTCGTTGCCGCCCATTCCCCCATTCCACNNNAGATCGGAAGAGCA |
| 11307_1 | TATGAGTGTGGAGTCGTTGCACCACCACCACTGTCAGAGCNNNAGATCGGAAGAGC |
| 11370_1 | TATGAGTGTGGAGTCGTTGCCCTCCCCGCCGTCTCTAGTANNNAGATCGGAAGAGC |
| 11417_1 | TATGAGTGTGGAGTCGTTGCGCCAGGGGCTCTAAGAGTGANNNAGATCGGAAGAG |
| 11590_1 | TATGAGTGTGGAGTCGTTGCGTCAGAGCTAAGCAAAGCCCCNNNAGATCGGAAGA |
| 11801_1 | TATGAGTGTGGAGTCGTTGCCACCGGCAAAACCATCACGANNNAGATCGGAAGAGC |

| | |
|---|---|
| 11903_1 | TATGAGTGTGGAGTCGTTGCCAAAGCAGCCTCTCCTTCCGNNNAGATCGGAAGAGC |
| 12038_1 | TATGAGTGTGGAGTCGTTGCCCCACCCATCCACATAGCCGNNNAGATCGGAAGAGC |
| 12292_1 | TATGAGTGTGGAGTCGTTGCAAGGCTCCGTCTGTTGATGCNNNAGATCGGAAGAG |
| 53461_1 | TATGAGTGTGGAGTCGTTGCTGACCATGGCTCACTGTAGCNNNAGATCGGAAGAG |
| 53466_1 | TATGAGTGTGGAGTCGTTGCGTTAGCCAGGATGGTCTCGATCNNNAGATCGGAAG |
| 53470_1 | TATGAGTGTGGAGTCGTTGCGTGTGTGTGTGTAGTGTGTGNNNAGATCGGAAG |
| 53510_1 | TATGAGTGTGGAGTCGTTGCCCTCCACCCCACATTTACAGTAANNNAGATCGGAAGA |
| 53520_1 | TATGAGTGTGGAGTCGTTGCTGACCTCAGCTCACTGCAAGNNNAGATCGGAAGAG |
| 53540_1 | TATGAGTGTGGAGTCGTTGCCAGGCTGAGGCAGGAGAATGNNNAGATCGGAAGA |
| 53575_1 | TATGAGTGTGGAGTCGTTGCCATTCTCCTGCCTCAGCCTCNNNAGATCGGAAGAGCA |
| 53579_1 | TATGAGTGTGGAGTCGTTGCCATTCTCCTGCCTCAGCCTCNNNAGATCGGAAGAGCA |
| 53585_1 | TATGAGTGTGGAGTCGTTGCCAATCTCGGCTCACTGCAAGNNNAGATCGGAAGAG |
| 53590_1 | TATGAGTGTGGAGTCGTTGCCATTCTCCTGCCTCAGCCTCNNNAGATCGGAAGAGCA |
| 53610_1 | TATGAGTGTGGAGTCGTTGCTTCTCCGTGGTTTACATCCTGAANNNAGATCGGAAG |
| 53613_1 | TATGAGTGTGGAGTCGTTGCTTTTTCTCTGCTACCACACACCTNNNAGATCGGAAGA |
| 53686_1 | TATGAGTGTGGAGTCGTTGCTGATGGACAGAGACCTTCGTTACNNNAGATCGGAA |
| 53687_1 | TATGAGTGTGGAGTCGTTGCCTGTAATCTCAGCTACCCAGGAGNNNAGATCGGAAG |
| 53695_1 | TATGAGTGTGGAGTCGTTGCCACCTGTAAGCCTAGCTACTCAGNNNAGATCGGAAG |
| 53701_1 | TATGAGTGTGGAGTCGTTGCTGAGGTATGATGTATGCACAGCTNNNAGATCGGAA |
| 53721_1 | TATGAGTGTGGAGTCGTTGCCAAGTAGCTGGGACTACAGGCNNNAGATCGGAAGA |
| 53737_1 | TATGAGTGTGGAGTCGTTGCCCCTTCTCTGCCCTCAAATTACANNNAGATCGGAAGA |
| 53756_1 | TATGAGTGTGGAGTCGTTGCCAATCTCGGCTCACTGCAAGNNNAGATCGGAAGAG |
| 53758_1 | TATGAGTGTGGAGTCGTTGCCATTCTCCTGCCTCAGCCTCNNNAGATCGGAAGAGCA |
| 53793_1 | TATGAGTGTGGAGTCGTTGCTGGGCAAAGGATATGAACAGACANNNAGATCGGAA |
| 53808_1 | TATGAGTGTGGAGTCGTTGCCAATCTCGGCTCACTGCAAGNNNAGATCGGAAGAG |
| 53846_1 | TATGAGTGTGGAGTCGTTGCTGGCTGCCTAGACTGTTATGAAGNNNAGATCGGAA |
| 53847_1 | TATGAGTGTGGAGTCGTTGCCTATGTGCTCGCCAGAGTAGTAGNNNAGATCGGAA |
| 53868_1 | TATGAGTGTGGAGTCGTTGCCACACACACAACAACAACAACAANNNAGATCGGAAG |
| 53883_1 | TATGAGTGTGGAGTCGTTGCCTCAGCTCAGAGATGTACATCCCNNNAGATCGGAAG |
| 53915_1 | TATGAGTGTGGAGTCGTTGCCATTCTCCTGCCTCAGCCTCNNNAGATCGGAAGAGCA |
| 53949_1 | TATGAGTGTGGAGTCGTTGCCATTTTGGATCACCTTTGGCTGANNNAGATCGGAAG |
| 53967_1 | TATGAGTGTGGAGTCGTTGCCTTTTGTGCACGAAGGGACATTANNNAGATCGGAA |
| 53969_1 | TATGAGTGTGGAGTCGTTGCACATCACCTGTTCCCTCAAAACTNNNAGATCGGAAGA |
| 53985_1 | TATGAGTGTGGAGTCGTTGCCATTCTCCTGCCTCAGCCTCNNNAGATCGGAAGAGCA |
| 53987_1 | TATGAGTGTGGAGTCGTTGCGCTTGTAATGTAGGAAGGGCTCTNNNAGATCGGAA |
| 54002_1 | TATGAGTGTGGAGTCGTTGCCATTCTCCTGCCTCAGCCTCNNNAGATCGGAAGAGCA |
| 54017_1 | TATGAGTGTGGAGTCGTTGCCAATCTCGGCTCACTGCAAGNNNAGATCGGAAGAG |
| 54027_1 | TATGAGTGTGGAGTCGTTGCCAGGTTCACGCCATTCTCCTNNNAGATCGGAAGAGC |
| 54031_1 | TATGAGTGTGGAGTCGTTGCACAATCTCGGCTCACTGCAANNNAGATCGGAAGAGC |
| 54033_1 | TATGAGTGTGGAGTCGTTGCCATTCTCCTGCCTCAGCCTCNNNAGATCGGAAGAGCA |
| 54078_1 | TATGAGTGTGGAGTCGTTGCAACAGGTGAGTGATAACAGGCTTNNNAGATCGGAA |
| 54092_1 | TATGAGTGTGGAGTCGTTGCACCCATTTTAGTTTCTGTGTCCTNNNAGATCGGAAGA |
| 54099_1 | TATGAGTGTGGAGTCGTTGCCAAGTGCCAAAGCCATAGATCAGNNNAGATCGGAA |

| | |
|---|---|
| 54102_1 | TATGAGTGTGGAGTCGTTGCTGTCTACAACACACAGGGAACTTNNNAGATCGGAAG |
| 54105_1 | TATGAGTGTGGAGTCGTTGCTGTGGGATCAGTCCTGATATCCTNNNAGATCGGAAG |
| 54107_1 | TATGAGTGTGGAGTCGTTGCGAGTTAGACTCTGCATTGCCAACNNNAGATCGGAAG |
| 54129_1 | TATGAGTGTGGAGTCGTTGCCAATCTCGGCTCACTGCAAGNNNAGATCGGAAGAGC |
| 54154_1 | TATGAGTGTGGAGTCGTTGCCATTCTCCTGCCTCAGCCTCNNNAGATCGGAAGAGCA |
| 54176_1 | TATGAGTGTGGAGTCGTTGCAGCTATAGGGTTTTTGGGTGTGGNNNAGATCGGAA |
| 54214_1 | TATGAGTGTGGAGTCGTTGCCATTCTCCTGCCTCAGCCTCNNNAGATCGGAAGAGCA |
| 54222_1 | TATGAGTGTGGAGTCGTTGCCATTCTCCTGCCTCAGCCTCNNNAGATCGGAAGAGCA |
| 54227_1 | TATGAGTGTGGAGTCGTTGCCGATCTCGGCTCACCACAACNNNAGATCGGAAGAGC |
| 54247_1 | TATGAGTGTGGAGTCGTTGCAATCTCAGCTCACTGCAACCTCNNNAGATCGGAAGA |
| 54273_1 | TATGAGTGTGGAGTCGTTGCAGACGGAGTCTCGCTCTGTCNNNAGATCGGAAGAG |
| 54317_1 | TATGAGTGTGGAGTCGTTGCCAGGTTCACGCCATTCTCCTNNNAGATCGGAAGAGC |
| 54341_1 | TATGAGTGTGGAGTCGTTGCCATTCTCCTGCCTCAGCCTCNNNAGATCGGAAGAGCA |
| 54358_1 | TATGAGTGTGGAGTCGTTGCCAGGTTCACGCTGTTCTCCTNNNAGATCGGAAGAGC |
| 54371_1 | TATGAGTGTGGAGTCGTTGCCGGGTTCATGCCATTCTCCTNNNAGATCGGAAGAGC |
| 54383_1 | TATGAGTGTGGAGTCGTTGCCATTCTCCTGCCTCAGCCTCNNNAGATCGGAAGAGCA |
| 54414_1 | TATGAGTGTGGAGTCGTTGCTAAATTGTCCCTCCTTGCAGACANNNAGATCGGAAG |
| 54422_1 | TATGAGTGTGGAGTCGTTGCAGGAAAAAAAGAATGAGGCCAGCNNNAGATCGGA |
| 54433_1 | TATGAGTGTGGAGTCGTTGCAAAGGGGATATCACCACCAATCCNNNAGATCGGAAG |
| 54437_1 | TATGAGTGTGGAGTCGTTGCATGCTGTGTTCTTCTCAAGCAACNNNAGATCGGAAG |
| 54456_1 | TATGAGTGTGGAGTCGTTGCGCAAACCAACGCCTGAAATACTANNNAGATCGGAAG |
| 54476_1 | TATGAGTGTGGAGTCGTTGCCTGTACAGCTTACTGGAGGGAAGNNNAGATCGGAA |
| 54517_1 | TATGAGTGTGGAGTCGTTGCCATTCTCCTGCCTCAGCCTCNNNAGATCGGAAGAGCA |
| 54538_1 | TATGAGTGTGGAGTCGTTGCAACTAAGCATCACACAATTGGGCNNNAGATCGGAAG |
| 54649_1 | TATGAGTGTGGAGTCGTTGCCTGTGACAGGAGGATTGCTTGAANNNAGATCGGAA |
| 54685_1 | TATGAGTGTGGAGTCGTTGCAGAAGTATGCAGGACCTGTAGGANNNAGATCGGAA |
| 54693_1 | TATGAGTGTGGAGTCGTTGCAGGGACTCATGATTTCCACAACANNNAGATCGGAAG |
| 54700_1 | TATGAGTGTGGAGTCGTTGCCCCTCAAGATTCATTATGCTCCGNNNAGATCGGAAG |
| 54708_1 | TATGAGTGTGGAGTCGTTGCAAGGTCTGCTTATTTTGTCAGCANNNAGATCGGAAG |
| 54733_1 | TATGAGTGTGGAGTCGTTGCCATTCTCCTGCCTCAGCCTCNNNAGATCGGAAGAGCA |
| 54737_1 | TATGAGTGTGGAGTCGTTGCCATTCTCCTGCCTCAGCCTCNNNAGATCGGAAGAGCA |
| 54746_1 | TATGAGTGTGGAGTCGTTGCCAATCTCGGCTCACTGCAAGNNNAGATCGGAAGAGC |
| 54759_1 | TATGAGTGTGGAGTCGTTGCCATTCTCCTGCCTCAGCCTCNNNAGATCGGAAGAGCA |
| 54761_1 | TATGAGTGTGGAGTCGTTGCCGGGTTCATGCCATTCTCCTNNNAGATCGGAAGAGC |
| 54772_1 | TATGAGTGTGGAGTCGTTGCATGCTCAGTACTTGGGTGATGAGNNNAGATCGGAA |
| 54776_1 | TATGAGTGTGGAGTCGTTGCCTGCTCTGTTTCTGCGATAACTCNNNAGATCGGAAG |
| 54780_1 | TATGAGTGTGGAGTCGTTGCTCCCCTTCTCTTAGTTGTCTGGANNNAGATCGGAAG |
| 54783_1 | TATGAGTGTGGAGTCGTTGCTCAAAGTTGGAGTCAGTCCTCTGNNNAGATCGGAAG |
| 54789_1 | TATGAGTGTGGAGTCGTTGCATGGAGAGTGTAATGGGAAGCTGNNNAGATCGGA |
| 54790_1 | TATGAGTGTGGAGTCGTTGCTTTGAGGAAGTGAAGCTGTGACTNNNAGATCGGAA |
| 54797_1 | TATGAGTGTGGAGTCGTTGCAATTCCAGCACTTTGGGAAGTCANNNAGATCGGAAG |
| 54799_1 | TATGAGTGTGGAGTCGTTGCACTCAACTGGGAGAAAAGCTTGANNNAGATCGGAA |
| 54806_1 | TATGAGTGTGGAGTCGTTGCTTCCCTCCTGTTTGAGTTACTGGNNNAGATCGGAAG |

| | |
|---|---|
| 54819_1 | TATGAGTGTGGAGTCGTTGCTGCTCTATTTGTGACTGGCATCTNNNAGATCGGAAG |
| 54854_1 | TATGAGTGTGGAGTCGTTGCCGGGTTCATGCCATTCTCCTNNNAGATCGGAAGAGC |
| 54864_1 | TATGAGTGTGGAGTCGTTGCCAATCTCGGCTCACTGCAAGNNNAGATCGGAAGAG |
| 54879_1 | TATGAGTGTGGAGTCGTTGCGGATCTCGGCTCACTGCAAGNNNAGATCGGAAGAG |
| 54892_1 | TATGAGTGTGGAGTCGTTGCCATTCTCCTGCCTCAGCCTCNNNAGATCGGAAGAGCA |
| 54900_1 | TATGAGTGTGGAGTCGTTGCCATTCTCCTGCCTCAGCCTCNNNAGATCGGAAGAGCA |
| 54904_1 | TATGAGTGTGGAGTCGTTGCGAGGCTTTGGATATTTGGCTGACNNNAGATCGGAA |
| 54918_1 | TATGAGTGTGGAGTCGTTGCTGATGCCTGAAGAAAAAAGTGCANNNAGATCGGAA |
| 54941_1 | TATGAGTGTGGAGTCGTTGCCAGGAGCCTTATGTCATGACACTNNNAGATCGGAAG |
| 54945_1 | TATGAGTGTGGAGTCGTTGCTCAAACATGTCAGGAATTTGGGCNNNAGATCGGAA |
| 54959_1 | TATGAGTGTGGAGTCGTTGCGGGTCACCACTAAAGCAGATTCANNNAGATCGGAAG |
| 54972_1 | TATGAGTGTGGAGTCGTTGCCATTCTCCTGCCTCAGCCTCNNNAGATCGGAAGAGCA |
| 54980_1 | TATGAGTGTGGAGTCGTTGCCATTCTCCTGCCTCAGCCTCNNNAGATCGGAAGAGCA |
| 54995_1 | TATGAGTGTGGAGTCGTTGCGTGTTAGCCAGGATGATCTCGATNNNAGATCGGAA |
| 54997_1 | TATGAGTGTGGAGTCGTTGCCAGTGGCGTGATCTCTGCTCNNNAGATCGGAAGAG |
| 55005_1 | TATGAGTGTGGAGTCGTTGCGAACTCACGACCTCAGGTGATCNNNAGATCGGAAGA |
| 55009_1 | TATGAGTGTGGAGTCGTTGCCCTGCAATCCCAGCCACTAGNNNAGATCGGAAGAGC |
| 55017_1 | TATGAGTGTGGAGTCGTTGCAAAATACTGGCAAACCGAATGCANNNAGATCGGAA |
| 55022_1 | TATGAGTGTGGAGTCGTTGCTGATTTCATGCAAAGTCCCACAGNNNAGATCGGAAG |
| 55023_1 | TATGAGTGTGGAGTCGTTGCTTAGTGTCTACTCTTTGCTGCCANNNAGATCGGAAGA |
| 55030_1 | TATGAGTGTGGAGTCGTTGCTTTTTTGAGACAGAGTGTCGCTCNNNAGATCGGAAG |
| 55039_1 | TATGAGTGTGGAGTCGTTGCCTCACAGACACATCAGGACTAGCNNNAGATCGGAAG |
| 55045_1 | TATGAGTGTGGAGTCGTTGCACTCAGGTTGGAGTGCAGTGNNNAGATCGGAAGAG |
| 55055_1 | TATGAGTGTGGAGTCGTTGCCAGGTTCACGCCATTCTCCTNNNAGATCGGAAGAGC |
| 55069_1 | TATGAGTGTGGAGTCGTTGCACGTAGCTGGGACTACAGGTNNNAGATCGGAAGAG |
| 55074_1 | TATGAGTGTGGAGTCGTTGCGAGTGCAGTGGCGTGATCTTNNNAGATCGGAAGAG |
| 55090_1 | TATGAGTGTGGAGTCGTTGCGAGTCTCACTCTGTCCCCCANNNAGATCGGAAGAGC |
| 55101_1 | TATGAGTGTGGAGTCGTTGCTTCTCTTGGTCTTCAATCCAGGTNNNAGATCGGAAGA |
| 55122_1 | TATGAGTGTGGAGTCGTTGCCTCCCGGGTTCAAGTGATTCTCNNNAGATCGGAAGA |
| 55137_1 | TATGAGTGTGGAGTCGTTGCCCTTAGCATGCCCTACCTTAGAGNNNAGATCGGAAG |
| 55162_1 | TATGAGTGTGGAGTCGTTGCCATTCTCCTGCCTCAGCCTCNNNAGATCGGAAGAGCA |
| 55186_1 | TATGAGTGTGGAGTCGTTGCCATTCTCCTGCCTCAGCCTCNNNAGATCGGAAGAGCA |
| 55209_1 | TATGAGTGTGGAGTCGTTGCCAATCTCGGCTCACTGCAAGNNNAGATCGGAAGAG |
| 55214_1 | TATGAGTGTGGAGTCGTTGCCATTCTCCTGCCTCAGCCTCNNNAGATCGGAAGAGCA |
| 55222_1 | TATGAGTGTGGAGTCGTTGCTGCTTTCATTTTGGTGAACAGCANNNAGATCGGAAG |
| 55225_1 | TATGAGTGTGGAGTCGTTGCAGTGACTCTTTGAGGATGATGCANNNAGATCGGAA |
| 55226_1 | TATGAGTGTGGAGTCGTTGCATAGGCCATCACAATCTACTCCTNNNAGATCGGAAGA |
| 55282_1 | TATGAGTGTGGAGTCGTTGCCTAAGGTAGGGCTCATGTACGTGNNNAGATCGGAA |
| 55284_1 | TATGAGTGTGGAGTCGTTGCACATTTGTGGGACAAAAGGCAAANNNAGATCGGAA |
| 55330_1 | TATGAGTGTGGAGTCGTTGCGGATCTCGGCTCACTGCAAGNNNAGATCGGAAGAG |
| 55345_1 | TATGAGTGTGGAGTCGTTGCTACATTTTTAGTGGGGGGTGGAGNNNAGATCGGAA |
| 55425_1 | TATGAGTGTGGAGTCGTTGCCATTCTCCTGCCTCAGCCTCNNNAGATCGGAAGAGCA |
| 55437_1 | TATGAGTGTGGAGTCGTTGCCATTCTCCTGCCTCAGCCTCNNNAGATCGGAAGAGCA |

| | |
|---|---|
| 55464_1 | TATGAGTGTGGAGTCGTTGCCATTCTCCTGCCTCAGCCTCNNNAGATCGGAAGAGCA |
| 55484_1 | TATGAGTGTGGAGTCGTTGCCGCTTGTAGTCCTAGCTACTCAGNNNAGATCGGAAG |
| 55497_1 | TATGAGTGTGGAGTCGTTGCCCTGTAATCCCAGCACTTTGAGANNNAGATCGGAAG |
| 55501_1 | TATGAGTGTGGAGTCGTTGCGCAAAGTGCAGAGAAAGAGAGAANNNAGATCGGA |
| 55507_1 | TATGAGTGTGGAGTCGTTGCAACAATTTGGGAAGCTGAAGTGGNNNAGATCGGAA |
| 55514_1 | TATGAGTGTGGAGTCGTTGCGTGCCTATAGTCCCAGCTACTTGNNNAGATCGGAAG |
| 55516_1 | TATGAGTGTGGAGTCGTTGCAAACCTGTAGAATGAATGGGGGANNNAGATCGGAA |
| 55526_1 | TATGAGTGTGGAGTCGTTGCTTCAGACAAGACAAATTGCCGTCNNNAGATCGGAAG |
| 55570_1 | TATGAGTGTGGAGTCGTTGCCATTCTCCTGCCTCAGCCTCNNNAGATCGGAAGAGCA |
| 55606_1 | TATGAGTGTGGAGTCGTTGCCGGAGGGAAATTAGACACTAGGGNNNAGATCGGA |
| 55612_1 | TATGAGTGTGGAGTCGTTGCGAGTCTTACTCTCTCGCCCAGGNNNAGATCGGAAGA |
| 55703_1 | TATGAGTGTGGAGTCGTTGCGGGGTGTGTGGTGTGTGTAGNNNAGATCGGAAGA |
| 55722_1 | TATGAGTGTGGAGTCGTTGCCCAGGTAGCTGCGACTACAGNNNAGATCGGAAGAG |
| 55725_1 | TATGAGTGTGGAGTCGTTGCCAATCTCGGCTCACTGCAAGNNNAGATCGGAAGAG |
| 55733_1 | TATGAGTGTGGAGTCGTTGCCATTCTCCTGCCTCAGCCTCNNNAGATCGGAAGAGCA |
| 55740_1 | TATGAGTGTGGAGTCGTTGCAGGAGGACTCTGTCGACATAGAANNNAGATCGGAA |
| 55751_1 | TATGAGTGTGGAGTCGTTGCAGGAAAAAGCATGAAACAAACCANNNAGATCGGAA |
| 55761_1 | TATGAGTGTGGAGTCGTTGCTGGGGTCAACCTAATTGTGTTCANNNAGATCGGAAG |
| 55777_1 | TATGAGTGTGGAGTCGTTGCCATTCTCCTGCCTCAGCCTCNNNAGATCGGAAGAGCA |
| 55784_1 | TATGAGTGTGGAGTCGTTGCCAGGTTCACGCCATTCTCCTNNNAGATCGGAAGAGC |
| 55789_1 | TATGAGTGTGGAGTCGTTGCGTTTCACCATGTTAGCCAGGATGNNNAGATCGGAAG |
| 55792_1 | TATGAGTGTGGAGTCGTTGCCATTCTCCTGCCTCAGCCTCNNNAGATCGGAAGAGCA |
| 55817_1 | TATGAGTGTGGAGTCGTTGCACAATCCCAGCTCACTGCAANNNAGATCGGAAGAGC |
| 55826_1 | TATGAGTGTGGAGTCGTTGCGCTGGTGTTCTGGAAAGATCAACNNNAGATCGGAA |
| 55827_1 | TATGAGTGTGGAGTCGTTGCAAGTTCACCATGGCAATGCAATTNNNAGATCGGAAG |
| 55837_1 | TATGAGTGTGGAGTCGTTGCAATTCCCTTCCAGACATCTCAGGNNNAGATCGGAAG |
| 55854_1 | TATGAGTGTGGAGTCGTTGCAGCAGGTAGGGAAAGAATCTGTGNNNAGATCGGA |
| 55861_1 | TATGAGTGTGGAGTCGTTGCGCCTGTATGTAGTTCCAGCTACTNNNAGATCGGAAG |
| 55879_1 | TATGAGTGTGGAGTCGTTGCCATTCTCCTGCCTCAGCCTCNNNAGATCGGAAGAGCA |
| 55941_1 | TATGAGTGTGGAGTCGTTGCCATTCTCCTGCCTCAGCCTCNNNAGATCGGAAGAGCA |
| 55949_1 | TATGAGTGTGGAGTCGTTGCCAATCTCGGCTCACTGCAAGNNNAGATCGGAAGAG |
| 55963_1 | TATGAGTGTGGAGTCGTTGCAAATACAACCTCTGCCTCCTGAGNNNAGATCGGAAG |
| 55967_1 | TATGAGTGTGGAGTCGTTGCGGTGAAACCCCGTCTGTACTAAANNNAGATCGGAAG |
| 55973_1 | TATGAGTGTGGAGTCGTTGCTACATGGTGGTGTGCATCTGTAANNNAGATCGGAA |
| 55980_1 | TATGAGTGTGGAGTCGTTGCGCATGTCACGGTGAGTAATCAAGNNNAGATCGGAA |
| 55991_1 | TATGAGTGTGGAGTCGTTGCAGAAAACTGGTGTTCATACCGGANNNAGATCGGAA |
| 55992_1 | TATGAGTGTGGAGTCGTTGCGTATGGGGATGGGAAAGACCTTCNNNAGATCGGAA |
| 55995_1 | TATGAGTGTGGAGTCGTTGCTTCAAGCCATTCTCCTGTGTCAGNNNAGATCGGAAG |
| 56008_1 | TATGAGTGTGGAGTCGTTGCTTCACACCATTCTCCCTGCCNNNAGATCGGAAGAGCA |
| 56012_1 | TATGAGTGTGGAGTCGTTGCCAATCTCGGCTCACTGCAAGNNNAGATCGGAAGAG |
| 56020_1 | TATGAGTGTGGAGTCGTTGCCTCATGGCAACCTCTACCTCCNNNAGATCGGAAGAG |
| 56040_1 | TATGAGTGTGGAGTCGTTGCGACGGAGTCTCGTTCTGTCGNNNAGATCGGAAGAG |
| 56062_1 | TATGAGTGTGGAGTCGTTGCGGGGACCTTTATGTCACAGACTTNNNAGATCGGAAG |

| | |
|---|---|
| 56064_1 | TATGAGTGTGGAGTCGTTGCTGTTAGAGCTACTTCCCTCATGCNNNAGATCGGAAGA |
| 56073_1 | TATGAGTGTGGAGTCGTTGCACTCACTGACCAGGGAATTTACCNNNAGATCGGAAG |
| 56074_1 | TATGAGTGTGGAGTCGTTGCGCAACAAAGAGAGAGAACTGTGTNNNAGATCGGAA |
| 56079_1 | TATGAGTGTGGAGTCGTTGCTGAGCATCTGCAGATTTGGGTATNNNAGATCGGAAG |
| 56106_1 | TATGAGTGTGGAGTCGTTGCTCACTAAAGGGCAACTTACACGANNNAGATCGGAAG |
| 56184_1 | TATGAGTGTGGAGTCGTTGCATGGCAGGAAAAAGTTGACCAAGNNNAGATCGGAA |
| 56205_1 | TATGAGTGTGGAGTCGTTGCAATTGAGGCATACCACCCTCAAANNNAGATCGGAAG |
| 56254_1 | TATGAGTGTGGAGTCGTTGCAGTTTGTCTGTCCTGGCCAGNNNAGATCGGAAGAGC |
| 56267_1 | TATGAGTGTGGAGTCGTTGCCTTCCAGCCATGAGGTACTACTGNNNAGATCGGAAG |
| 56304_1 | TATGAGTGTGGAGTCGTTGCCGATCTCGGCTCACTGCAAGNNNAGATCGGAAGAGC |
| 56323_1 | TATGAGTGTGGAGTCGTTGCGGTTCACGCTGTTCTGCCTCNNNAGATCGGAAGAGCA |
| 56329_1 | TATGAGTGTGGAGTCGTTGCCATTCTCCTGCCTCAGCCTCNNNAGATCGGAAGAGCA |
| 56331_1 | TATGAGTGTGGAGTCGTTGCCAATCTCGGCTCACTGCAAGNNNAGATCGGAAGAGC |
| 56340_1 | TATGAGTGTGGAGTCGTTGCCAATCTCGGCTCACTGCAACNNNAGATCGGAAGAGCA |
| 56359_1 | TATGAGTGTGGAGTCGTTGCGGAGAAACCCCGTGTCTACTAAANNNAGATCGGAAG |
| 56360_1 | TATGAGTGTGGAGTCGTTGCCCGAAGCAGTTGGATCACCTNNNAGATCGGAAGAGC |
| 56387_1 | TATGAGTGTGGAGTCGTTGCACCCAGTGATCTTTTGCCTACAANNNAGATCGGAAGA |
| 56393_1 | TATGAGTGTGGAGTCGTTGCCGAAACAGCAGTCGGAGTTATTGNNNAGATCGGAAC |
| 56409_1 | TATGAGTGTGGAGTCGTTGCAAATGGGCATCTTGGCTTATTCTNNNAGATCGGAAG |
| 56476_1 | TATGAGTGTGGAGTCGTTGCCCAAAGTGCTGGGATTACAAGCNNNAGATCGGAAGA |
| 56489_1 | TATGAGTGTGGAGTCGTTGCAGACGGAGTCTCGCTGTCACNNNAGATCGGAAGAGC |
| 56500_1 | TATGAGTGTGGAGTCGTTGCGATTGCTTGAGCCCAGGAGTTCNNNAGATCGGAAGA |
| 56549_1 | TATGAGTGTGGAGTCGTTGCTTTTGCAAGGTTCTGTTGCCTTTNNNAGATCGGAAGA |
| 56579_1 | TATGAGTGTGGAGTCGTTGCCATTCTCCTGCCTCAGCCTCNNNAGATCGGAAGAGCA |
| 56589_1 | TATGAGTGTGGAGTCGTTGCTGTTAGCCAGGATGGTCTCAATCNNNAGATCGGAAG |
| 56602_1 | TATGAGTGTGGAGTCGTTGCCATTCTCCTGCCTCAGCCTCNNNAGATCGGAAGAGCA |
| 56621_1 | TATGAGTGTGGAGTCGTTGCCAATCTCGGCTCACTGCAACNNNAGATCGGAAGAGCA |
| 56627_1 | TATGAGTGTGGAGTCGTTGCATAGGGTCTCATTTTGTTGCCCANNNAGATCGGAAGA |
| 56638_1 | TATGAGTGTGGAGTCGTTGCCGCCTGTAGTCCCAGCTACTNNNAGATCGGAAGAGCA |
| 56677_1 | TATGAGTGTGGAGTCGTTGCACAAAATGGAGGCAGCAGGTATANNNAGATCGGAA |
| 56683_1 | TATGAGTGTGGAGTCGTTGCTTCAAATCACTGACTAGCCACCANNNAGATCGGAAGA |
| 56721_1 | TATGAGTGTGGAGTCGTTGCCATTCTCCTGCCTCAGCCTCNNNAGATCGGAAGAGCA |
| 56734_1 | TATGAGTGTGGAGTCGTTGCCATTCTCCTGCCTCAGCCTCNNNAGATCGGAAGAGCA |
| 56742_1 | TATGAGTGTGGAGTCGTTGCCATTCTCCTGCCTCAGCCTCNNNAGATCGGAAGAGCA |
| 56769_1 | TATGAGTGTGGAGTCGTTGCCATTCTCCTGCCTCAGCCTCNNNAGATCGGAAGAGCA |
| 56771_1 | TATGAGTGTGGAGTCGTTGCTGATCTCGGCTCACTGCAAGNNNAGATCGGAAGAGC |
| 56774_1 | TATGAGTGTGGAGTCGTTGCCATTCTCCTGCCTCAGCCTCNNNAGATCGGAAGAGCA |
| 56793_1 | TATGAGTGTGGAGTCGTTGCGAGTGCAGTGGTGCGATCTCNNNAGATCGGAAGAG |
| 56805_1 | TATGAGTGTGGAGTCGTTGCGAATGCAGTGGTGCGATGTCNNNAGATCGGAAGAG |
| 56807_1 | TATGAGTGTGGAGTCGTTGCAGACGGAGTCTCGCTCTGTCNNNAGATCGGAAGAGC |
| 56815_1 | TATGAGTGTGGAGTCGTTGCCTTGATCTCCTGACCTCGTGATCNNNAGATCGGAAGA |
| 56816_1 | TATGAGTGTGGAGTCGTTGCCGATCTCGGCTCACTGCAAGNNNAGATCGGAAGAGC |
| 56834_1 | TATGAGTGTGGAGTCGTTGCCAGAAGAATCCACCTCTCACCTTNNNAGATCGGAAGA |

| | |
|---|---|
| 56835_1 | TATGAGTGTGGAGTCGTTGCGCTTGGGAATCTGCCTAAGTTTGNNNAGATCGGAAG |
| 56841_1 | TATGAGTGTGGAGTCGTTGCAAACTGCCGTGGGTCCTTATAATNNNAGATCGGAAG |
| 56845_1 | TATGAGTGTGGAGTCGTTGCGATTTTCGCATGGATCTCACTGGNNNAGATCGGAAG |
| 56871_1 | TATGAGTGTGGAGTCGTTGCTGCTGTTATTTGGCCATTACTGANNNAGATCGGAAGA |
| 56879_1 | TATGAGTGTGGAGTCGTTGCACACTACAGGACATTGATCTGGGNNNAGATCGGAAG |
| 56900_1 | TATGAGTGTGGAGTCGTTGCCAACTTGTTTTATGAGGCCAGCANNNAGATCGGAAG |
| 56902_1 | TATGAGTGTGGAGTCGTTGCACTTGAGACTTGATGGCTTCACTNNNAGATCGGAAG |
| 56908_1 | TATGAGTGTGGAGTCGTTGCAGATCTGAGCTGTTTGTGAGACANNNAGATCGGAAG |
| 56925_1 | TATGAGTGTGGAGTCGTTGCTTACCTGATTGAGAGAGGGTGGANNNAGATCGGAA |
| 56936_1 | TATGAGTGTGGAGTCGTTGCATCGAGTGTTTGGAGGTTAGGACNNNAGATCGGAA |
| 56984_1 | TATGAGTGTGGAGTCGTTGCCATTCTCCTGCCTCAGCCTCNNNAGATCGGAAGAGCA |
| 56998_1 | TATGAGTGTGGAGTCGTTGCCATTCTCCTGCCTCAGCCTCNNNAGATCGGAAGAGCA |
| 57006_1 | TATGAGTGTGGAGTCGTTGCCATTCTCCTGCCTCAGCCTCNNNAGATCGGAAGAGCA |
| 57012_1 | TATGAGTGTGGAGTCGTTGCCATTCTCCTGCCTCAGCCTCNNNAGATCGGAAGAGCA |
| 57024_1 | TATGAGTGTGGAGTCGTTGCTGTGTGTGTGTGTCTGTGTATGANNNAGATCGGAAG |
| 57033_1 | TATGAGTGTGGAGTCGTTGCTAAAAGAAGCCTGCCAGGTAACCNNNAGATCGGAAG |
| 57038_1 | TATGAGTGTGGAGTCGTTGCTGATCTTATTGCCCTGGCTATCCNNNAGATCGGAAGA |
| 57040_1 | TATGAGTGTGGAGTCGTTGCGCAAACCACTATGGCACATGTATNNNAGATCGGAAG |
| 57088_1 | TATGAGTGTGGAGTCGTTGCCATTCTCCTGCCTCAGCCTCNNNAGATCGGAAGAGCA |
| 57092_1 | TATGAGTGTGGAGTCGTTGCCAATCTCGGCTCACTGCAAGNNNAGATCGGAAGAGC |
| 57096_1 | TATGAGTGTGGAGTCGTTGCCATTCTCCTGCCTCAGCCTCNNNAGATCGGAAGAGCA |
| 57108_1 | TATGAGTGTGGAGTCGTTGCCAATCTCGGCTCACTGCAACNNNAGATCGGAAGAGCA |
| 57136_1 | TATGAGTGTGGAGTCGTTGCTCTGATATGGTTTGGCTCTGTGTNNNAGATCGGAAG |
| 57173_1 | TATGAGTGTGGAGTCGTTGCATACTTAGAAGAGGGCCAAGCAGNNNAGATCGGAA |
| 57179_1 | TATGAGTGTGGAGTCGTTGCCATTCTCCTGCCTCAGCCTCNNNAGATCGGAAGAGCA |
| 57193_1 | TATGAGTGTGGAGTCGTTGCGAGTCTTTCTCTGTCGCCCANNNAGATCGGAAGAGCA |
| 57195_1 | TATGAGTGTGGAGTCGTTGCCAATCTCGGCTCACTGCAAGNNNAGATCGGAAGAGC |
| 57201_1 | TATGAGTGTGGAGTCGTTGCGGGTTCACGCCATTCATGAANNNAGATCGGAAGAGC |
| 57202_1 | TATGAGTGTGGAGTCGTTGCCATTCTCCTGCCTCAGCCTCNNNAGATCGGAAGAGCA |
| 57216_1 | TATGAGTGTGGAGTCGTTGCTCATGCATACACATACACACACANNNAGATCGGAAGA |
| 57227_1 | TATGAGTGTGGAGTCGTTGCATCTGTGGACACTTCATTCCTCCNNNAGATCGGAAGA |
| 57238_1 | TATGAGTGTGGAGTCGTTGCGTGTCGAATGAAAGTAGTGGCTGNNNAGATCGGAA |
| 57252_1 | TATGAGTGTGGAGTCGTTGCCAGGCAGATCACAAGGTCAAAAGNNNAGATCGGAAG |
| 57257_1 | TATGAGTGTGGAGTCGTTGCACCCAGAGATTTTGTCCAAGTCANNNAGATCGGAAG |
| 57263_1 | TATGAGTGTGGAGTCGTTGCCAGGTTCACGCCATTCTCCTNNNAGATCGGAAGAGCA |
| 57267_1 | TATGAGTGTGGAGTCGTTGCCATTCTCCTGCCTCAGCCTCNNNAGATCGGAAGAGCA |
| 57273_1 | TATGAGTGTGGAGTCGTTGCCAATCTCGGCTCACTGCAAGNNNAGATCGGAAGAGC |
| 57286_1 | TATGAGTGTGGAGTCGTTGCCATTCTCCTGCCTCAGCCTCNNNAGATCGGAAGAGCA |
| 57302_1 | TATGAGTGTGGAGTCGTTGCCAATCTCGGCTCACTGCAAGNNNAGATCGGAAGAGC |
| 57311_1 | TATGAGTGTGGAGTCGTTGCCATTCTCCTGCCTCAGCCTCNNNAGATCGGAAGAGCA |
| 57321_1 | TATGAGTGTGGAGTCGTTGCGCCATCATTCTTTTCCCCAAAANNNAGATCGGAAGA |
| 57329_1 | TATGAGTGTGGAGTCGTTGCCATTCTCCTGCCTCAGCCTCNNNAGATCGGAAGAGCA |
| 57331_1 | TATGAGTGTGGAGTCGTTGCGATTCTCCTGCCTCAGCCTCNNNAGATCGGAAGAGCA |

| | |
|---|---|
| 57347_1 | TATGAGTGTGGAGTCGTTGCCCTGTCCTAGCAGAGTTTGTGTTNNNAGATCGGAAG |
| 57361_1 | TATGAGTGTGGAGTCGTTGCACCCCATACCTCTAAAAGTAGCTNNNAGATCGGAAG |
| 57379_1 | TATGAGTGTGGAGTCGTTGCACCCATTTTAGGCTATGAAACGANNNAGATCGGAAG |
| 57381_1 | TATGAGTGTGGAGTCGTTGCTGATCTTTTCTCGTAGGTGCTCCNNNAGATCGGAAG |
| 57383_1 | TATGAGTGTGGAGTCGTTGCCTCTCTGTGAAGATGGCACTGTANNNAGATCGGAAG |
| 57424_1 | TATGAGTGTGGAGTCGTTGCCAATCTCGGCTCACTGCAAGNNNAGATCGGAAGAGC |
| 57442_1 | TATGAGTGTGGAGTCGTTGCCATTCTCCTGCCTCAGCCTCNNNAGATCGGAAGAGCA |
| 57452_1 | TATGAGTGTGGAGTCGTTGCCATTCTCCTGCCTCAGCCTCNNNAGATCGGAAGAGCA |
| 57503_1 | TATGAGTGTGGAGTCGTTGCAACAGGCAAAATAACCAGCTAGCNNNAGATCGGAA |
| 57537_1 | TATGAGTGTGGAGTCGTTGCATATGGAATAAGCAGGTGGAGCANNNAGATCGGA |
| 57545_1 | TATGAGTGTGGAGTCGTTGCTAACAAAGTAGCAAAATGCCCGTNNNAGATCGGAA |
| 57560_1 | TATGAGTGTGGAGTCGTTGCGTGGAGGAATTGGAACCCACATANNNAGATCGGAA |
| 57593_1 | TATGAGTGTGGAGTCGTTGCGAGTGTAGTGGCGCCATCTCNNNAGATCGGAAGAG |
| 57611_1 | TATGAGTGTGGAGTCGTTGCTTTCATGCAAACAAACAGGAACTNNNAGATCGGAAG |
| 57668_1 | TATGAGTGTGGAGTCGTTGCGGGTTCACGCCATTCTCCTGNNNAGATCGGAAGAGC |
| 57673_1 | TATGAGTGTGGAGTCGTTGCGATCTCGGCTCACTGCAACCNNNAGATCGGAAGAGC |
| 57679_1 | TATGAGTGTGGAGTCGTTGCCATTCTCCTGCCTCAGCCTCNNNAGATCGGAAGAGCA |
| 57711_1 | TATGAGTGTGGAGTCGTTGCAATGGCACAATCTCGGCTCANNNAGATCGGAAGAGC |
| 57746_1 | TATGAGTGTGGAGTCGTTGCTGCTCTGGAAAAGGAGATAGGTTNNNAGATCGGAA |
| 57752_1 | TATGAGTGTGGAGTCGTTGCTATGGATCTCTCCTCTTGGGGAANNNAGATCGGAAG |
| 57777_1 | TATGAGTGTGGAGTCGTTGCGCTTTGTGGTCCTCAGTTGTGNNNAGATCGGAAGA |
| 57785_1 | TATGAGTGTGGAGTCGTTGCTTGGTTTTACGAGAAGGGTTGGANNNAGATCGGAA |
| 57800_1 | TATGAGTGTGGAGTCGTTGCCTGTAGTTCCAGCTACTCAGGAGNNNAGATCGGAAG |
| 57821_1 | TATGAGTGTGGAGTCGTTGCGATCACAAGGTCAGGAGATGGAGNNNAGATCGGA |
| 57837_1 | TATGAGTGTGGAGTCGTTGCCATTCTCCTGCCTCAGCCTCNNNAGATCGGAAGAGCA |
| 57842_1 | TATGAGTGTGGAGTCGTTGCCAATCTCGGCTCACTGCAAGNNNAGATCGGAAGAGC |
| 57846_1 | TATGAGTGTGGAGTCGTTGCCATTCTCCTGCCTCAGCCTCNNNAGATCGGAAGAGCA |
| 57870_1 | TATGAGTGTGGAGTCGTTGCCATTCTCCTGCCTCAGCCTCNNNAGATCGGAAGAGCA |
| 57881_1 | TATGAGTGTGGAGTCGTTGCCATTCTCCTGCCTCAGCCTCNNNAGATCGGAAGAGCA |
| 57893_1 | TATGAGTGTGGAGTCGTTGCCATTCTCCTGCCTCAGCCTCNNNAGATCGGAAGAGCA |
| 57898_1 | TATGAGTGTGGAGTCGTTGCCATTCTCCTGCCTCAGCCTCNNNAGATCGGAAGAGCA |
| 57908_1 | TATGAGTGTGGAGTCGTTGCCATTCTCCTGCCTCAGCCTCNNNAGATCGGAAGAGCA |
| 57937_1 | TATGAGTGTGGAGTCGTTGCCAAGTAAAAAGCAGACAGAGGGTNNNAGATCGGA |
| 57940_1 | TATGAGTGTGGAGTCGTTGCTCCCCACCTTATCTACAAGGGATNNNAGATCGGAAG |
| 57942_1 | TATGAGTGTGGAGTCGTTGCCATAGGAAGGGGATAAAGAGGCCNNNAGATCGGA |
| 57961_1 | TATGAGTGTGGAGTCGTTGCACGATCAAAGAGCCTGATTCAGTNNNAGATCGGAA |
| 57963_1 | TATGAGTGTGGAGTCGTTGCTAATTCTGCTCTGCATTCCACCTNNNAGATCGGAAGA |
| 57965_1 | TATGAGTGTGGAGTCGTTGCGGTGGGGTTCTTGAGGTTCTTATNNNAGATCGGAA |
| 57975_1 | TATGAGTGTGGAGTCGTTGCAGAGGGTAGTTATTAGGCCTGGTNNNAGATCGGAA |
| 57980_1 | TATGAGTGTGGAGTCGTTGCTACCTCCTCACTCTGAGATTCCANNNAGATCGGAAGA |
| 58024_1 | TATGAGTGTGGAGTCGTTGCCATTCTCCTGCCTCAGCCTCNNNAGATCGGAAGAGCA |
| 58038_1 | TATGAGTGTGGAGTCGTTGCCATTCTCCTGCCTCAGCCTCNNNAGATCGGAAGAGCA |
| 58040_1 | TATGAGTGTGGAGTCGTTGCCATTCTCCTGCCTCAGCCTCNNNAGATCGGAAGAGCA |

58061_1 TATGAGTGTGGAGTCGTTGCGTGTAATCTCAGCTACTCGGGAGNNNAGATCGGAA
58081_1 TATGAGTGTGGAGTCGTTGCCAAACGCTTCATCTTTCAAAGCANNNAGATCGGAAG
58088_1 TATGAGTGTGGAGTCGTTGCAAGATCAGGTGTATTCAGGGTGGNNNAGATCGGAA
58114_1 TATGAGTGTGGAGTCGTTGCGACAGATAAAGGGTGACTCAGGGNNNAGATCGGA
58129_1 TATGAGTGTGGAGTCGTTGCCATTCTCCTGCCTCAGCCTCNNNAGATCGGAAGAGCA
58139_1 TATGAGTGTGGAGTCGTTGCAGAGTCTCTCTCAGTCGCCCNNNAGATCGGAAGAGC
58148_1 TATGAGTGTGGAGTCGTTGCCGATCTCGGCTCACTGCAAGNNNAGATCGGAAGAG
58162_1 TATGAGTGTGGAGTCGTTGCTAGAGTGCAGTGGCGCGATCNNNAGATCGGAAGAG
58168_1 TATGAGTGTGGAGTCGTTGCTGATCACAGCTCACTGCAGCNNNAGATCGGAAGAG
58178_1 TATGAGTGTGGAGTCGTTGCCGATCTCGGCTCACTGCAAGNNNAGATCGGAAGAG
58196_1 TATGAGTGTGGAGTCGTTGCGAGTGCAGTAGCGGGATCTCNNNAGATCGGAAGAG
58227_1 TATGAGTGTGGAGTCGTTGCTAAAGAAACTAGGCAGGCTCAGGNNNAGATCGGAA
58236_1 TATGAGTGTGGAGTCGTTGCCTGTTCTAATTGCCTACTCCCCANNNAGATCGGAAGA
58259_1 TATGAGTGTGGAGTCGTTGCGGTGCATACCTGTAGTCCCAGNNNAGATCGGAAGA
58260_1 TATGAGTGTGGAGTCGTTGCGTGTGTGTGTGTGGACATTTCATNNNAGATCGGAA
58271_1 TATGAGTGTGGAGTCGTTGCTCTCCAAAGAAGCAGAACAGACANNNAGATCGGAA
58281_1 TATGAGTGTGGAGTCGTTGCAGGAGAAAATGTTCACCCAATGCNNNAGATCGGAA
58287_1 TATGAGTGTGGAGTCGTTGCAGTTATGGTGGTCCTAGCAAACTNNNAGATCGGAAG
58293_1 TATGAGTGTGGAGTCGTTGCCTGAGGCAGGAGAATCACTTGAANNNAGATCGGAA
58352_1 TATGAGTGTGGAGTCGTTGCCATTCTCCTGCCTCAGCCTCNNNAGATCGGAAGAGCA
58353_1 TATGAGTGTGGAGTCGTTGCGAGTCTCTCTCTGTCGCCCANNNAGATCGGAAGAGC
58359_1 TATGAGTGTGGAGTCGTTGCCAGGTTCACGCCATTCTCCTNNNAGATCGGAAGAGC
58395_1 TATGAGTGTGGAGTCGTTGCCATTCTCCTGCCTCAGCCTCNNNAGATCGGAAGAGCA
58403_1 TATGAGTGTGGAGTCGTTGCCATTCTCCTGCCTCAGCCTCNNNAGATCGGAAGAGCA
58428_1 TATGAGTGTGGAGTCGTTGCACACAGAGAACAAACCATCAGGANNNAGATCGGAA
58453_1 TATGAGTGTGGAGTCGTTGCTTCTTCCACAGGAGTGACTATGCNNNAGATCGGAAG
58467_1 TATGAGTGTGGAGTCGTTGCACCATGGAGGCCAAAAGAAAATGNNNAGATCGGAA
58489_1 TATGAGTGTGGAGTCGTTGCGAGGAGACTGGGGCACATAAAAGNNNAGATCGGA
58502_1 TATGAGTGTGGAGTCGTTGCCGATCTCGGCTCACTGCAAGNNNAGATCGGAAGAG
58519_1 TATGAGTGTGGAGTCGTTGCGGTTGCAGAGAGCCGAGATTNNNAGATCGGAAGAG
58528_1 TATGAGTGTGGAGTCGTTGCCGGGGCCAGATGTGATATATCAANNNAGATCGGAA
58534_1 TATGAGTGTGGAGTCGTTGCATCTGGCATGTATGGGTGCTTATNNNAGATCGGAAG
58535_1 TATGAGTGTGGAGTCGTTGCTAATGTGAAGCAAGGCCCTAACTNNNAGATCGGAAG
58554_1 TATGAGTGTGGAGTCGTTGCCGATCTCGGCTCACTGCAAGNNNAGATCGGAAGAG
58571_1 TATGAGTGTGGAGTCGTTGCCTCACTGCAAGCTCCACCTCNNNAGATCGGAAGAGC
58581_1 TATGAGTGTGGAGTCGTTGCCATTCTCCTGCCTCAGCCTCNNNAGATCGGAAGAGCA
58590_1 TATGAGTGTGGAGTCGTTGCTTTAGACATGAATCGCATCCCCANNNAGATCGGAAG
58601_1 TATGAGTGTGGAGTCGTTGCAAACAAACTGAAGGCTCTAAGGCNNNAGATCGGAA
58611_1 TATGAGTGTGGAGTCGTTGCTGGGACCAAAAAACTCAGAGGAANNNAGATCGGAA
58625_1 TATGAGTGTGGAGTCGTTGCGAGTCTTGCTCTGTTGCCCANNNAGATCGGAAGAGC
58671_1 TATGAGTGTGGAGTCGTTGCCAGGTTCACGCCATTCTCCTNNNAGATCGGAAGAGC
58708_1 TATGAGTGTGGAGTCGTTGCGAGTGCAGTGGCCCAATCTCNNNAGATCGGAAGAG
58714_1 TATGAGTGTGGAGTCGTTGCCCAACATGTAGTCAAAGCGTTGTNNNAGATCGGAAG

58745_1 TATGAGTGTGGAGTCGTTGCCTGAGGCAGGGAATCACTTGAANNNAGATCGGAAG
58759_1 TATGAGTGTGGAGTCGTTGCCATTCTCCTGCCTCAGCCTCNNNAGATCGGAAGAGCA
58762_1 TATGAGTGTGGAGTCGTTGCGACGGAGTCTCGCTATGTCGNNNAGATCGGAAGAG
58785_1 TATGAGTGTGGAGTCGTTGCCATTCTCCTGCCTCAGCCTCNNNAGATCGGAAGAGCA
58844_1 TATGAGTGTGGAGTCGTTGCAATTGCAGGGATACCAGTTTGAANNNAGATCGGAA
58898_1 TATGAGTGTGGAGTCGTTGCATTCCTTCCAGAAAGTGACCTCCNNNAGATCGGAAG
58910_1 TATGAGTGTGGAGTCGTTGCCATTCTCCTGCCTCAGCCTCNNNAGATCGGAAGAGCA
58940_1 TATGAGTGTGGAGTCGTTGCCACGGTTTGAGACTGTCACAAAANNNAGATCGGAA
58958_1 TATGAGTGTGGAGTCGTTGCACCAGCATGGCACATGTATACATNNNAGATCGGAAG
58979_1 TATGAGTGTGGAGTCGTTGCCATTCTCCTGCCTCAGCCTCNNNAGATCGGAAGAGCA
59066_1 TATGAGTGTGGAGTCGTTGCCAATCTCGGCTCACTGCAAGNNNAGATCGGAAGAG
59078_1 TATGAGTGTGGAGTCGTTGCGATTTCGCCTTACTGCAGCCTCNNNAGATCGGAAGA
59086_1 TATGAGTGTGGAGTCGTTGCCATTCTCCTGCCTCAGCCTCNNNAGATCGGAAGAGCA
59091_1 TATGAGTGTGGAGTCGTTGCCATTCTCCTGCCTCAGCCTCNNNAGATCGGAAGAGCA
59093_1 TATGAGTGTGGAGTCGTTGCCCCAGGCTGAAGTGCAATTGNNNAGATCGGAAGAG
59124_1 TATGAGTGTGGAGTCGTTGCCATTCTCCTGCCTCAGCCTCNNNAGATCGGAAGAGCA
59147_1 TATGAGTGTGGAGTCGTTGCAGACGGAGTCTCGCTCTGTCNNNAGATCGGAAGAG
59194_1 TATGAGTGTGGAGTCGTTGCGAGCTGAACTGAAGGAGAGAGACNNNAGATCGGA
59204_1 TATGAGTGTGGAGTCGTTGCGACCACACTACTCAAAGCAATCTNNNAGATCGGAAG
59216_1 TATGAGTGTGGAGTCGTTGCCTGTAACCTCAGCTACTTGGGAGNNNAGATCGGAAG
59249_1 TATGAGTGTGGAGTCGTTGCCATTCTCCTGCCTCAGCCTCNNNAGATCGGAAGAGCA
59273_1 TATGAGTGTGGAGTCGTTGCCGGGTTCAAGCCATTCTCCTNNNAGATCGGAAGAGC
59287_1 TATGAGTGTGGAGTCGTTGCCTCTGCCTCCTGGGTTCAAGNNNAGATCGGAAGAGC
59291_1 TATGAGTGTGGAGTCGTTGCCGAGTAGCTGGGATTACTGGCNNNAGATCGGAAGA
59301_1 TATGAGTGTGGAGTCGTTGCGAAAATGCAGCAGTGGTTAGGAGNNNAGATCGGA
59310_1 TATGAGTGTGGAGTCGTTGCCTGGGAAAAATATTTGCGACCCANNNAGATCGGAA
59316_1 TATGAGTGTGGAGTCGTTGCGGTTTGTCAACATCCCAAGTGAANNNAGATCGGAAG
59326_1 TATGAGTGTGGAGTCGTTGCCAGGAAGCTGAGGCAGAAAAATCNNNAGATCGGAA
59329_1 TATGAGTGTGGAGTCGTTGCCCATACACACCCCAAGGGATCNNNAGATCGGAAGAG
59352_1 TATGAGTGTGGAGTCGTTGCCACAAAATAGATTCTGGCTGGGCNNNAGATCGGAA
59358_1 TATGAGTGTGGAGTCGTTGCTTGGTTTTGGAGACAGGACAAGANNNAGATCGGAA
59360_1 TATGAGTGTGGAGTCGTTGCCTGATCAGGGACTTGTAACCAGANNNAGATCGGAA
59368_1 TATGAGTGTGGAGTCGTTGCAAATTAGCTTGGCATAGTGGTGGNNNAGATCGGAA
59390_1 TATGAGTGTGGAGTCGTTGCCCGAGTAGCTGGGACAACAGNNNAGATCGGAAGAG
59409_1 TATGAGTGTGGAGTCGTTGCGTGGGTCTCAGAGCAGAGATTAGNNNAGATCGGAA
59413_1 TATGAGTGTGGAGTCGTTGCTGGTGTCTGGTGTTCTCATGAATNNNAGATCGGAAG
59429_1 TATGAGTGTGGAGTCGTTGCGGACACATGTTCTTGGGATCTCTNNNAGATCGGAAG
59437_1 TATGAGTGTGGAGTCGTTGCTAGATTGCATAGCCCTCATCAGGNNNAGATCGGAAG
59456_1 TATGAGTGTGGAGTCGTTGCCAGGTTCACGCCATTCTCCTNNNAGATCGGAAGAGC
59511_1 TATGAGTGTGGAGTCGTTGCCTTGGCTCACTGCAACCTCTNNNAGATCGGAAGAGC
59529_1 TATGAGTGTGGAGTCGTTGCCATTCTCCTGCCTCAGCCTCNNNAGATCGGAAGAGCA
59555_1 TATGAGTGTGGAGTCGTTGCACTCACAGACCAATGGAACAGAANNNAGATCGGAA
59570_1 TATGAGTGTGGAGTCGTTGCCCAGCCTCAACATAGAGCAATTCNNNAGATCGGAAG

| | |
|---|---|
| 59595_1 | TATGAGTGTGGAGTCGTTGCTTCAGTCTCAACCTCTAGCTCAGNNNAGATCGGAAG |
| 59598_1 | TATGAGTGTGGAGTCGTTGCCCAAGAAAGGCTAGGAGAACCATNNNAGATCGGAA |
| 59629_1 | TATGAGTGTGGAGTCGTTGCCTCCCAACAAAGAAAAGCTCAGCNNNAGATCGGAAG |
| 59631_1 | TATGAGTGTGGAGTCGTTGCAAGAGTAGGAAGGCCTAGGATGANNNAGATCGGA |
| 59634_1 | TATGAGTGTGGAGTCGTTGCGAGGCTGAGGCAGGAGAATGNNNAGATCGGAAGA |
| 59643_1 | TATGAGTGTGGAGTCGTTGCGTGGTGTAGGTCAGTGGTCTAAGNNNAGATCGGAA |
| 59665_1 | TATGAGTGTGGAGTCGTTGCCATTCTCCTGCCTCAGCCTCNNNAGATCGGAAGAGCA |
| 59672_1 | TATGAGTGTGGAGTCGTTGCGTCTCCTGAGTAGCTGGGACTANNNAGATCGGAAG |
| 59702_1 | TATGAGTGTGGAGTCGTTGCCTGTAGCTGGGACCACAGGTNNNAGATCGGAAGAG |
| 59801_1 | TATGAGTGTGGAGTCGTTGCCATTCTCCTGCCTCAGCCTCNNNAGATCGGAAGAGCA |
| 59804_1 | TATGAGTGTGGAGTCGTTGCCTCTGCTTCCCGGGTTCATGNNNAGATCGGAAGAGC |
| 59805_1 | TATGAGTGTGGAGTCGTTGCCATTCTCCTGCCTCAGCCTCNNNAGATCGGAAGAGCA |
| 59814_1 | TATGAGTGTGGAGTCGTTGCCAATCTCGGCTCACTGCAAGNNNAGATCGGAAGAGC |
| 59867_1 | TATGAGTGTGGAGTCGTTGCCATTCTCCTGCCTCAGCCTCNNNAGATCGGAAGAGCA |
| 59892_1 | TATGAGTGTGGAGTCGTTGCCTACCAGAAAAGGACCTCACTGGNNNAGATCGGAAG |
| 59895_1 | TATGAGTGTGGAGTCGTTGCAAGACTGAGAAAAGACCACTGGANNNAGATCGGAA |
| 59903_1 | TATGAGTGTGGAGTCGTTGCATAGGTTTCAGGTCCACTTGAGGNNNAGATCGGAAG |
| 59923_1 | TATGAGTGTGGAGTCGTTGCGGAGAGAGAGTGGGGTTGAATTTNNNAGATCGGA |
| 59981_1 | TATGAGTGTGGAGTCGTTGCAGTTTCACCATGTTAGCCAGGATNNNAGATCGGAAG |
| 59994_1 | TATGAGTGTGGAGTCGTTGCCATTCTCCTGCCTCAGCCTCNNNAGATCGGAAGAGCA |
| 60009_1 | TATGAGTGTGGAGTCGTTGCCCAAATAGCTGGGACTACAGGCNNNAGATCGGAAG |
| 60013_1 | TATGAGTGTGGAGTCGTTGCCATTCTCCTGCCTCAGCCTCNNNAGATCGGAAGAGCA |
| 60014_1 | TATGAGTGTGGAGTCGTTGCGAGTCTTGCTCTGTTGCCCANNNAGATCGGAAGAGC |
| 60060_1 | TATGAGTGTGGAGTCGTTGCCAGGTTCACGCCATTCTCCTNNNAGATCGGAAGAGC |
| 60067_1 | TATGAGTGTGGAGTCGTTGCGAGTATGGTGGCGGGATCTCNNNAGATCGGAAGAG |
| 60068_1 | TATGAGTGTGGAGTCGTTGCCTCCTGGGTTCACGCCATTCNNNAGATCGGAAGAGC |
| 60140_1 | TATGAGTGTGGAGTCGTTGCAATCATGGCCCTGAACCAAAAACNNNAGATCGGAAG |
| 60147_1 | TATGAGTGTGGAGTCGTTGCAACTAGCTGGAAGACTGGAACAGNNNAGATCGGAA |
| 60152_1 | TATGAGTGTGGAGTCGTTGCTCTGATTTTTTTTTGCTGGGGGGNNNAGATCGGAAG |
| 60162_1 | TATGAGTGTGGAGTCGTTGCATCCTGGCTATCATGGTCAAACCNNNAGATCGGAAG |
| 60164_1 | TATGAGTGTGGAGTCGTTGCAAAACACTCTACCTTTTGCTGGGNNNAGATCGGAAG |
| 60186_1 | TATGAGTGTGGAGTCGTTGCGTATAGTACCTGGACCAGTGTGCNNNAGATCGGAAG |
| 60202_1 | TATGAGTGTGGAGTCGTTGCGTTCTTAGGGAAACTGGAGAGGTNNNAGATCGGAA |
| 60207_1 | TATGAGTGTGGAGTCGTTGCCACCAATTGGAACCAGCAAAGATNNNAGATCGGAAG |
| 60214_1 | TATGAGTGTGGAGTCGTTGCTAACAGTTTGGAAGGTGGCCTTANNNAGATCGGAA |
| 60239_1 | TATGAGTGTGGAGTCGTTGCGAACAAGGGCTAGCTAAGTTCTTNNNAGATCGGAAG |
| 60242_1 | TATGAGTGTGGAGTCGTTGCTTGTGCCCAGATTTCTTTTGTCCNNNAGATCGGAAGA |
| 60255_1 | TATGAGTGTGGAGTCGTTGCCATATGCCTGTAGTCCCAGCTACNNNAGATCGGAAG |
| 60267_1 | TATGAGTGTGGAGTCGTTGCAAGAGGATTGGGAGGGATATTGCNNNAGATCGGA |
| 60303_1 | TATGAGTGTGGAGTCGTTGCCATTCTCCTGCCTCAGCCTCNNNAGATCGGAAGAGCA |
| 60330_1 | TATGAGTGTGGAGTCGTTGCGAGTCTTGCTCTGTTGCCCANNNAGATCGGAAGAGC |
| 62038_1 | TATGAGTGTGGAGTCGTTGCCTTGGGACGAGCTACAGGAAGNNNAGATCGGAAGA |
| 62080_1 | TATGAGTGTGGAGTCGTTGCACCCAGAATTTCATATCCAGCCANNNAGATCGGAAG |

| | |
|---|---|
| 62199_1 | TATGAGTGTGGAGTCGTTGCCATTGCCCTTTACACATCCATCCNNNAGATCGGAAGA |
| 62235_1 | TATGAGTGTGGAGTCGTTGCTCTACCTGGCTCCATGACCANNNAGATCGGAAGAGC |
| 62236_1 | TATGAGTGTGGAGTCGTTGCATTTGGAGACACAGTCTTGCTCTNNNAGATCGGAAG |
| 62277_1 | TATGAGTGTGGAGTCGTTGCCAGGTTCACGCCATTCTCCTNNNAGATCGGAAGAGC |
| 62350_1 | TATGAGTGTGGAGTCGTTGCCGATCTCGGCTCACTGCAAGNNNAGATCGGAAGAG |
| 62396_1 | TATGAGTGTGGAGTCGTTGCAGACCACCAAGGCAGTACATTTANNNAGATCGGAAG |
| 62413_1 | TATGAGTGTGGAGTCGTTGCAAGTGGCATGGTATCAGCTTTTGNNNAGATCGGAA |
| 62419_1 | TATGAGTGTGGAGTCGTTGCCATTCTCCTGCCTCAGCCTCNNNAGATCGGAAGAGCA |
| 62438_1 | TATGAGTGTGGAGTCGTTGCGGCAGCTGGTCACAAAATCATAGNNNAGATCGGAA |
| 62453_1 | TATGAGTGTGGAGTCGTTGCCAAGTAGCTGGGACTACAGGCNNNAGATCGGAAGA |
| 62463_1 | TATGAGTGTGGAGTCGTTGCCATTCTCCTGCCTCAGCCTCNNNAGATCGGAAGAGCA |
| 62487_1 | TATGAGTGTGGAGTCGTTGCCATTCTCCTGCCTCAGCCTCNNNAGATCGGAAGAGCA |
| 62548_1 | TATGAGTGTGGAGTCGTTGCACACCTGTAACCCCAACACTTTNNNAGATCGGAAGAG |
| 62563_1 | TATGAGTGTGGAGTCGTTGCATGATCTGTTTCCTACCCACCTCNNNAGATCGGAAGA |
| 62569_1 | TATGAGTGTGGAGTCGTTGCAAAGAAGGAGAGAGTGGAAGCACNNNAGATCGGA |
| 62605_1 | TATGAGTGTGGAGTCGTTGCCTCACTGCAAGCTCCACCTCNNNAGATCGGAAGAGC |
| 62626_1 | TATGAGTGTGGAGTCGTTGCTGTGTGTGTGTGTTTTGTGTGTTNNNAGATCGGAA |
| 62636_1 | TATGAGTGTGGAGTCGTTGCGTGAGACTCAAGGCACGATTTACNNNAGATCGGAA |
| 62637_1 | TATGAGTGTGGAGTCGTTGCCATTCTCCTGCCTCAGCCTCNNNAGATCGGAAGAGCA |
| 62639_1 | TATGAGTGTGGAGTCGTTGCGAGGGCAGTGGCGTGATCTTNNNAGATCGGAAGAG |
| 62642_1 | TATGAGTGTGGAGTCGTTGCCTGGAGCTGCAGTGATCTCGNNNAGATCGGAAGAG |
| 62653_1 | TATGAGTGTGGAGTCGTTGCCATTCTCCTGCCTCAGCCTCNNNAGATCGGAAGAGCA |
| 62674_1 | TATGAGTGTGGAGTCGTTGCTTATTGATGGGCATTTGGGTTGGNNNAGATCGGAA |
| 62675_1 | TATGAGTGTGGAGTCGTTGCCTACCTGTGCTCCAAAAAAGGTGNNNAGATCGGAAG |
| 62732_1 | TATGAGTGTGGAGTCGTTGCAAGAAAGAGCTATGATGTGGGCANNNAGATCGGA |
| 62739_1 | TATGAGTGTGGAGTCGTTGCTGTAGTTGTCCTGATGTTGCAGTNNNAGATCGGAAG |
| 62745_1 | TATGAGTGTGGAGTCGTTGCAAAACTGTCACAGGCTAGAGGAGNNNAGATCGGAA |
| 62757_1 | TATGAGTGTGGAGTCGTTGCCATTCTCCTGCCTCAGCCTCNNNAGATCGGAAGAGCA |
| 62758_1 | TATGAGTGTGGAGTCGTTGCCATTCTCCTGCCTCAGCCTCNNNAGATCGGAAGAGCA |
| 62766_1 | TATGAGTGTGGAGTCGTTGCGGTTCACGCCATTCTCCTGANNNAGATCGGAAGAGC |
| 62777_1 | TATGAGTGTGGAGTCGTTGCACAATCTGGGCTCACTGCAANNNAGATCGGAAGAG |
| 62788_1 | TATGAGTGTGGAGTCGTTGCCCAAGTAGCTGGGTCCACAGNNNAGATCGGAAGAG |
| 62828_1 | TATGAGTGTGGAGTCGTTGCCATTCTCCTGCCTCAGCCTCNNNAGATCGGAAGAGCA |
| 62852_1 | TATGAGTGTGGAGTCGTTGCCATTCTCCTGCCTCAGCCTCNNNAGATCGGAAGAGCA |
| 62884_1 | TATGAGTGTGGAGTCGTTGCCATTCTCCTGCCTCAGCCTCNNNAGATCGGAAGAGCA |
| 62927_1 | TATGAGTGTGGAGTCGTTGCTCATGTCTTCCTGATTACGTTGCNNNAGATCGGAAGA |
| 62966_1 | TATGAGTGTGGAGTCGTTGCTATTTTGTTGGAGTGCACTGGTGNNNAGATCGGAA |
| 62975_1 | TATGAGTGTGGAGTCGTTGCAAGGATGGATGGATGGATGGATGNNNAGATCGGA |
| 62991_1 | TATGAGTGTGGAGTCGTTGCGGTGAAACCCCGTCTCTACTAAANNNAGATCGGAAG |
| 63032_1 | TATGAGTGTGGAGTCGTTGCGTAAAGAAACAAGGCCGGCCNNNAGATCGGAAGAG |
| 63063_1 | TATGAGTGTGGAGTCGTTGCCAATAAATATGGCCGGGCGCNNNAGATCGGAAGAG |
| 63066_1 | TATGAGTGTGGAGTCGTTGCCCACTCCAGTGAGAAGTGATCTTNNNAGATCGGAAG |
| 63070_1 | TATGAGTGTGGAGTCGTTGCGCTTCCTGAGTAGCTGAGATGACNNNAGATCGGAA |

| | |
|---|---|
| 63087_1 | TATGAGTGTGGAGTCGTTGCCTTCCGTGGAGAGCAGAGTGNNNAGATCGGAAGAG |
| 63116_1 | TATGAGTGTGGAGTCGTTGCCATTCTCCTGCCTCAGCCTCNNNAGATCGGAAGAGCA |
| 63130_1 | TATGAGTGTGGAGTCGTTGCTTTCACTGTGCATCAGAAACACCNNNAGATCGGAAG |
| 63139_1 | TATGAGTGTGGAGTCGTTGCCTTCTTTCCTCCTTGGGGTACTCNNNAGATCGGAAGA |
| 63151_1 | TATGAGTGTGGAGTCGTTGCAGTTCCTGACTGTAACTAGCTGGNNNAGATCGGAAG |
| 63184_1 | TATGAGTGTGGAGTCGTTGCTGTCATGTACACTCCAGTATCCANNNAGATCGGAAG |
| 63190_1 | TATGAGTGTGGAGTCGTTGCTGGTCACAATTCTACTTCGGTGTNNNAGATCGGAAG |
| 63200_1 | TATGAGTGTGGAGTCGTTGCCATAAAAAGTCCAGCTACCTCGCNNNAGATCGGAAG |
| 63255_1 | TATGAGTGTGGAGTCGTTGCCTGGTTCACGCCATTCTCCTNNNAGATCGGAAGAGC |
| 63352_1 | TATGAGTGTGGAGTCGTTGCCATTCTCCTGCCTCAGCCTCNNNAGATCGGAAGAGCA |
| 63496_1 | TATGAGTGTGGAGTCGTTGCTGAGATAAAGCTCAACCTCCACANNNAGATCGGAAG |
| 63547_1 | TATGAGTGTGGAGTCGTTGCAGACGGAGTCTCGCTCTGTCNNNAGATCGGAAGAG |
| 63613_1 | TATGAGTGTGGAGTCGTTGCTGGTACAGGTGAGAAAACTGACANNNAGATCGGAA |
| 63614_1 | TATGAGTGTGGAGTCGTTGCTCTTCTTTCCTTCGTTCCTTCCTNNNAGATCGGAAGAG |
| 63615_1 | TATGAGTGTGGAGTCGTTGCCAAACCTTGACACCTGCTATGTGNNNAGATCGGAAG |
| 63639_1 | TATGAGTGTGGAGTCGTTGCCATTCTCCTGCCTCAGCCTCNNNAGATCGGAAGAGCA |
| 63645_1 | TATGAGTGTGGAGTCGTTGCGGGTGGACAGCTATGTAGAGATGNNNAGATCGGA |
| 63648_1 | TATGAGTGTGGAGTCGTTGCCATTCTCCTGCCTCAGCCTCNNNAGATCGGAAGAGCA |
| 63677_1 | TATGAGTGTGGAGTCGTTGCTCATTCACTGAGGCTGCTCTTAGNNNAGATCGGAAG |
| 63712_1 | TATGAGTGTGGAGTCGTTGCTCAGAAAATCGTTTTTAGGCCGGNNNAGATCGGAAG |
| 63741_1 | TATGAGTGTGGAGTCGTTGCGGATTCTCGCTGTCACCCAANNNAGATCGGAAGAGC |
| 63775_1 | TATGAGTGTGGAGTCGTTGCCTATGGCAATGCTGAACACACATNNNAGATCGGAAG |
| 63810_1 | TATGAGTGTGGAGTCGTTGCGTGTTCACGCCATTCTCCTGNNNAGATCGGAAGAGC |
| 63844_1 | TATGAGTGTGGAGTCGTTGCCCTGTGGTTCCAGCTACCAGNNNAGATCGGAAGAGC |
| 63901_1 | TATGAGTGTGGAGTCGTTGCCTGTGTCATGGGGAGAGAATGAANNNAGATCGGA |
| 63902_1 | TATGAGTGTGGAGTCGTTGCTTGAATGAATCACTGGGTCTGGANNNAGATCGGAA |
| 63931_1 | TATGAGTGTGGAGTCGTTGCTGATCTCGGCTCACTGCAAGNNNAGATCGGAAGAG |
| 63935_1 | TATGAGTGTGGAGTCGTTGCTCATGTGAACTGTTATCCCAGCANNNAGATCGGAAG |
| 63947_1 | TATGAGTGTGGAGTCGTTGCCAGGAGAATGGCGTGAACCTNNNAGATCGGAAGAG |
| 63959_1 | TATGAGTGTGGAGTCGTTGCGAGATCACCCCACTGCACTCNNNAGATCGGAAGAGC |
| 63965_1 | TATGAGTGTGGAGTCGTTGCGCCAAATTTCTCTCCATAGGTGCNNNAGATCGGAAG |
| 63986_1 | TATGAGTGTGGAGTCGTTGCACAAACCATGTAATGCCACTGTGNNNAGATCGGAAG |
| 63988_1 | TATGAGTGTGGAGTCGTTGCCCCTGCCTCTCTTTCATTTCCTTNNNAGATCGGAAGAG |
| 63989_1 | TATGAGTGTGGAGTCGTTGCTCTTTGTGAAAAGTGAGGCACACNNNAGATCGGAA |
| 64008_1 | TATGAGTGTGGAGTCGTTGCCATTCTCCTGCCTCAGCCTCNNNAGATCGGAAGAGCA |
| 64014_1 | TATGAGTGTGGAGTCGTTGCCATTCTCCTGCCTCAGCCTCNNNAGATCGGAAGAGCA |
| 64187_1 | TATGAGTGTGGAGTCGTTGCCTCATCATAGAGCGCTGTTTTGGNNNAGATCGGAAG |
| 64190_1 | TATGAGTGTGGAGTCGTTGCGGGAAAGTGGGAGAGAATCAAGTNNNAGATCGGA |
| 64256_1 | TATGAGTGTGGAGTCGTTGCAGACGGAGTCTCGCTCTGTCNNNAGATCGGAAGAG |
| 64309_1 | TATGAGTGTGGAGTCGTTGCTAGTCCCAGCCACTCGGAAGNNNAGATCGGAAGAG |
| 64428_1 | TATGAGTGTGGAGTCGTTGCCCGATTCTGCACCTCCGTCCNNNAGATCGGAAGAGC |
| 64435_1 | TATGAGTGTGGAGTCGTTGCGAGGCTGAGGCAGGAGAATGNNNAGATCGGAAGA |
| 64450_1 | TATGAGTGTGGAGTCGTTGCCAGGCTGAGGCAGGAGAATGNNNAGATCGGAAGA |

| | |
|---|---|
| 64460_1 | TATGAGTGTGGAGTCGTTGCTTTCACCATGTTAGCCAGGCTNNNAGATCGGAAGAG |
| 64484_1 | TATGAGTGTGGAGTCGTTGCGGGGAGGCCTGGGTTTATATATTNNNAGATCGGAA |
| 64530_1 | TATGAGTGTGGAGTCGTTGCCAACCCAGCAGAAACTTCGGANNNAGATCGGAAGA |
| 64543_1 | TATGAGTGTGGAGTCGTTGCGGGTGTCTGTCTGAATAGAGCTGNNNAGATCGGAA |
| 64691_1 | TATGAGTGTGGAGTCGTTGCCTAGTAGGCAGCATGGGGCANNNAGATCGGAAGA |
| 64772_1 | TATGAGTGTGGAGTCGTTGCGTTTTCCTCTCCGTCGGGAAGNNNAGATCGGAAGAG |
| 64775_1 | TATGAGTGTGGAGTCGTTGCATGCATATGTGTGTGTTGTGTGTNNNAGATCGGAA |
| 64782_1 | TATGAGTGTGGAGTCGTTGCCAGAATGGTCTCGATCTCCTGACNNNAGATCGGAAG |
| 64998_1 | TATGAGTGTGGAGTCGTTGCGTGTGTGTGTGTGTGTGTATTNNNAGATCGGAA |
| 65074_1 | TATGAGTGTGGAGTCGTTGCCCAGAGTGCTAGGATTACAGGCNNNAGATCGGAAG |
| 65396_1 | TATGAGTGTGGAGTCGTTGCGTCAGATGGGGAGGAGCATGNNNAGATCGGAAGA |
| 65438_1 | TATGAGTGTGGAGTCGTTGCTAGTTGGGTCCTCTTGGAAAAGGNNNAGATCGGAA |
| 65589_1 | TATGAGTGTGGAGTCGTTGCGAGTGCAGTGGAGCGATCTTNNNAGATCGGAAGAG |
| 65636_1 | TATGAGTGTGGAGTCGTTGCCTGTGTGTGTGTGTGTGTGNNNAGATCGGAAGA |
| 65744_1 | TATGAGTGTGGAGTCGTTGCGAGGCTGAGGCAGGAGAATGNNNAGATCGGAAGA |
| 65770_1 | TATGAGTGTGGAGTCGTTGCCAATGTCGGCTCACTGCAAGNNNAGATCGGAAGAG |
| 65967_1 | TATGAGTGTGGAGTCGTTGCTCTCCCCAGAAGCATCAATATTTNNNAGATCGGAAG |
| 65972_1 | TATGAGTGTGGAGTCGTTGCCACACACACACACACATACACACNNNAGATCGGAAG |
| 65973_1 | TATGAGTGTGGAGTCGTTGCCTCTCTCTCTCTCACACACACACNNNAGATCGGAAGA |
| 65977_1 | TATGAGTGTGGAGTCGTTGCCTGGGTTCAAGCGATTCTCTTGNNNAGATCGGAAGA |
| 66000_1 | TATGAGTGTGGAGTCGTTGCCTTGAGGCTAGGAGTTGGAGATCNNNAGATCGGAA |
| 66016_1 | TATGAGTGTGGAGTCGTTGCGATGCAGTCTCGCTCTGTCACNNNAGATCGGAAGAG |
| 66027_1 | TATGAGTGTGGAGTCGTTGCAGACAGGGTCTCGCTCTGTGNNNAGATCGGAAGAG |
| 66049_1 | TATGAGTGTGGAGTCGTTGCGTGGGGAGGTTTTGTTCACAAACNNNAGATCGGAA |
| 66099_1 | TATGAGTGTGGAGTCGTTGCGGTGCGATCTCCACTCACTGNNNAGATCGGAAGAG |
| 66100_1 | TATGAGTGTGGAGTCGTTGCCATCCCTCTCTCCCTCCCTCNNNAGATCGGAAGAGCA |
| 66148_1 | TATGAGTGTGGAGTCGTTGCAGCCAGCTTCATACACTCTCAAANNNAGATCGGAAG |
| 66600_1 | TATGAGTGTGGAGTCGTTGCATTTGACTTCAGACCCTGTTCCCNNNAGATCGGAAG |
| 66740_1 | TATGAGTGTGGAGTCGTTGCCGGGTTCATGCAGTTCTTCTGNNNAGATCGGAAGAG |
| 66742_1 | TATGAGTGTGGAGTCGTTGCTAGCATCCCTGTTGTCTTTTGGTNNNAGATCGGAAG |
| 66768_1 | TATGAGTGTGGAGTCGTTGCGGAATGAGGTACAGTAATCCCCCNNNAGATCGGAA |
| 66850_1 | TATGAGTGTGGAGTCGTTGCTGATCTCGGCTCACTGCAAGNNNAGATCGGAAGAG |
| 66972_1 | TATGAGTGTGGAGTCGTTGCGAGCTTCCAGGACGACTTGAAGNNNAGATCGGAAG |
| 66997_1 | TATGAGTGTGGAGTCGTTGCCACAGCCTCAGCAGAAGCATNNNAGATCGGAAGAG |
| 67045_1 | TATGAGTGTGGAGTCGTTGCCTGCGGTCTGGAGAGGAGAGNNNAGATCGGAAGA |
| 67050_1 | TATGAGTGTGGAGTCGTTGCATTTGAACTACACAGGGAACACANNNAGATCGGAA |
| 67120_1 | TATGAGTGTGGAGTCGTTGCAAATGCTTCATCAACAAAGGGCGNNNAGATCGGAA |
| 67628_1 | TATGAGTGTGGAGTCGTTGCGAAGTGGAGGGTCTACACGAAGNNNAGATCGGAA |
| 67810_1 | TATGAGTGTGGAGTCGTTGCAACACACACACTGCCCTAGCNNNAGATCGGAAGAGC |
| 67905_1 | TATGAGTGTGGAGTCGTTGCGATTCTCCTGCCTCAGCCTCNNNAGATCGGAAGAGC |
| 68077_1 | TATGAGTGTGGAGTCGTTGCCTTGCAGTGAGCCAAGATCACNNNAGATCGGAAGA |
| 68132_1 | TATGAGTGTGGAGTCGTTGCCAGGGCTCTTTTCACAGATTCAGNNNAGATCGGAAG |
| 68406_1 | TATGAGTGTGGAGTCGTTGCCACAATCTCGGCTCACTGCANNNAGATCGGAAGAGC |

| | |
|---|---|
| 68426_1 | TATGAGTGTGGAGTCGTTGCACGGAGGGAAAAGAGGAAAGAAANNNAGATCGGA |
| 68440_1 | TATGAGTGTGGAGTCGTTGCGCCCAGCCAACATGGTGAAGNNNAGATCGGAAGAG |
| 68547_1 | TATGAGTGTGGAGTCGTTGCACTGACGACTCTTTCACAAGACANNNAGATCGGAAG |
| 68736_1 | TATGAGTGTGGAGTCGTTGCGATGGGTATGTGGGAGCAGGNNNAGATCGGAAGA |
| 68743_1 | TATGAGTGTGGAGTCGTTGCTCTTTGGTACCTGTGTTCGTCAGNNNAGATCGGAAG |
| 68745_1 | TATGAGTGTGGAGTCGTTGCCCTCTGATCACTCTGCCCAAGNNNAGATCGGAAGAG |
| 68750_1 | TATGAGTGTGGAGTCGTTGCGCCGACTGTTCATTCTCTCTCTCNNNAGATCGGAAGA |
| 68755_1 | TATGAGTGTGGAGTCGTTGCGAAGTTCAGCCAAAGGGGAGAGNNNAGATCGGAA |
| 68962_1 | TATGAGTGTGGAGTCGTTGCGCCTGAATTTTTACGGATCCCTTNNNAGATCGGAAG |
| 68978_1 | TATGAGTGTGGAGTCGTTGCCTTTTTGCCCATCGCGGTTTTTTNNNAGATCGGAAGA |
| 68986_1 | TATGAGTGTGGAGTCGTTGCGTTAGCCAGGATGGTCTTGATCTNNNAGATCGGAA |
| 68994_1 | TATGAGTGTGGAGTCGTTGCCCCCTTGTGGTTCTCTGTGACNNNAGATCGGAAGAG |
| 68999_1 | TATGAGTGTGGAGTCGTTGCAACAGTGACATCGACTTGAGCAANNNAGATCGGAA |
| 69002_1 | TATGAGTGTGGAGTCGTTGCCCTTCATCTTAGCGAACACCAAGNNNAGATCGGAAG |
| 69010_1 | TATGAGTGTGGAGTCGTTGCCAGTGGCACAATCTTGGCTCNNNAGATCGGAAGAG |
| 69219_1 | TATGAGTGTGGAGTCGTTGCCAGGAATCAGCTTGCTCCCAGNNNAGATCGGAAGA |
| 69237_1 | TATGAGTGTGGAGTCGTTGCGAAGTGACGGTTCCGGCTCTTNNNAGATCGGAAGA |
| 69267_1 | TATGAGTGTGGAGTCGTTGCGATGACGTCGCTGCAAGTTCTTCNNNAGATCGGAAG |
| 69283_1 | TATGAGTGTGGAGTCGTTGCTAAAGCGGGGATTTGATTTCGTGNNNAGATCGGAA |
| 69348_1 | TATGAGTGTGGAGTCGTTGCTGACCTCAGCTCACTGCAACNNNAGATCGGAAGAGC |
| 69397_1 | TATGAGTGTGGAGTCGTTGCCAGGCTGGTCTGCAATGGTGNNNAGATCGGAAGAG |
| 69443_1 | TATGAGTGTGGAGTCGTTGCAGAAAGAAGGAAGCAAGGAAGGANNNAGATCGGA |
| 69615_1 | TATGAGTGTGGAGTCGTTGCAGAAAAAACAGAAGCATGCCTCANNNAGATCGGAA |
| 69619_1 | TATGAGTGTGGAGTCGTTGCCATGCGCAGAGACACTGGTCNNNAGATCGGAAGAG |
| 69629_1 | TATGAGTGTGGAGTCGTTGCCTGTGTAGAGGGACTCAGACTCCNNNAGATCGGAA |
| 69635_1 | TATGAGTGTGGAGTCGTTGCGGTGCAAAAAGTCGCGACAGNNNAGATCGGAAGA |
| 69645_1 | TATGAGTGTGGAGTCGTTGCCGAAAACGTCCCAGACCTGGNNNAGATCGGAAGAG |
| 69664_1 | TATGAGTGTGGAGTCGTTGCAGAGCATGCCGAACTCTGACNNNAGATCGGAAGAG |
| 69759_1 | TATGAGTGTGGAGTCGTTGCAGACGGAGTCTCACCCTGTCNNNAGATCGGAAGAG |
| 69914_1 | TATGAGTGTGGAGTCGTTGCCAAGCTGGATTGCAATGGCANNNAGATCGGAAGAG |
| 69954_1 | TATGAGTGTGGAGTCGTTGCCCAAGTGGGTGGAAAGAAATCAGNNNAGATCGGA |
| 70054_1 | TATGAGTGTGGAGTCGTTGCCCACCTCGAGTCCAAGTTCGNNNAGATCGGAAGAGC |
| 70134_1 | TATGAGTGTGGAGTCGTTGCCTTCCAGGAGACCCTCTGCANNNAGATCGGAAGAGC |
| 70142_1 | TATGAGTGTGGAGTCGTTGCAACCCACATCACAGTGTCCCNNNAGATCGGAAGAGC |
| 70155_1 | TATGAGTGTGGAGTCGTTGCCGTTTCCATTTCTCTCTGGCATTNNNAGATCGGAAGA |
| 70191_1 | TATGAGTGTGGAGTCGTTGCCTGACAGTGCCAGGAGATGGNNNAGATCGGAAGA |
| 70196_1 | TATGAGTGTGGAGTCGTTGCCTTGGCTTTTTCCACCCTGCNNNAGATCGGAAGAGC |
| 70217_1 | TATGAGTGTGGAGTCGTTGCCACGCATCCCTGTCGATAGACNNNAGATCGGAAGAG |
| 70229_1 | TATGAGTGTGGAGTCGTTGCAAAGCAGGGCCAGTGAGGAAGNNNAGATCGGAAG |
| 70345_1 | TATGAGTGTGGAGTCGTTGCCTCAAGCTGGAGTGCAGTGGNNNAGATCGGAAGAG |
| 70533_1 | TATGAGTGTGGAGTCGTTGCGAGTCTCACTCTGTCGCCCANNNAGATCGGAAGAGC |
| 424_1 | TATGAGTGTGGAGTCGTTGCCAGAACACACCCCCTACACACNNNAGATCGGAAGAG |
| 3216_1 | TATGAGTGTGGAGTCGTTGCTCTTTGCCTGACTAGCCACCAANNNAGATCGGAAGA |

| | |
|---|---|
| 3405_1 | TATGAGTGTGGAGTCGTTGCGAGCGGATGCTATTGCCTACCNNNAGATCGGAAGA |
| 3979_1 | TATGAGTGTGGAGTCGTTGCGCCAGGTTTGCCTCAGAGGTANNNAGATCGGAAGA |
| 4189_1 | TATGAGTGTGGAGTCGTTGCAGTGATTTTCCTGCTTCTGCGNNNAGATCGGAAGAG |
| 6609_1 | TATGAGTGTGGAGTCGTTGCCATAAACCCGCCCCCACAAGNNNAGATCGGAAGAGC |
| 7140_1 | TATGAGTGTGGAGTCGTTGCAGGTTTCCTTCCGGTCAGTCANNNAGATCGGAAGAG |
| 7159_1 | TATGAGTGTGGAGTCGTTGCGATGTGGGTGGGGTGGTGACANNNAGATCGGAAG |
| 9769_1 | TATGAGTGTGGAGTCGTTGCTCTCTGTAGGGTGAGACCCCTNNNAGATCGGAAGA |
| 11906_1 | TATGAGTGTGGAGTCGTTGCGTTCAGGCCTCTCGTCGCCGNNNAGATCGGAAGAG |
| 12210_1 | TATGAGTGTGGAGTCGTTGCTCAGATGTGGAGTTGCTGCTGNNNAGATCGGAAGA |
| 12377_1 | TATGAGTGTGGAGTCGTTGCCCCACAACGGGAGTGGCATCNNNAGATCGGAAGAG |
| 53464_1 | TATGAGTGTGGAGTCGTTGCCCAAATACCTGAGGAAGAGGGGNNNAGATCGGAA |
| 53642_1 | TATGAGTGTGGAGTCGTTGCCATTCTCCTGCCTCAGCCTCNNNAGATCGGAAGAGCA |
| 53717_1 | TATGAGTGTGGAGTCGTTGCCATTCTCCTGCCTCAGCCTCNNNAGATCGGAAGAGCA |
| 54175_1 | TATGAGTGTGGAGTCGTTGCGAGGTTGTAGTGAGCCGAGATCNNNAGATCGGAAG |
| 54194_1 | TATGAGTGTGGAGTCGTTGCGGGAACAGCATGTGCAAAGACNNNAGATCGGAAGA |
| 54308_1 | TATGAGTGTGGAGTCGTTGCGAGGCTGAGGCAGGAGAATGNNNAGATCGGAAGA |
| 54420_1 | TATGAGTGTGGAGTCGTTGCGAAGCAAGAGAATGGCGTGAACNNNAGATCGGAA |
| 55374_1 | TATGAGTGTGGAGTCGTTGCGAGAAGAGGAGCAGAAGCCACNNNAGATCGGAAG |
| 55399_1 | TATGAGTGTGGAGTCGTTGCCAGAATCTCACTCTGTTGCCCANNNAGATCGGAAGA |
| 55670_1 | TATGAGTGTGGAGTCGTTGCCGCTCTTTGTAAGTCGGAGGTNNNAGATCGGAAGA |
| 56453_1 | TATGAGTGTGGAGTCGTTGCGTTCAGGCCATTCTCCTACCTCNNNAGATCGGAAGA |
| 56930_1 | TATGAGTGTGGAGTCGTTGCCAAAAGGGAGAGCCAGAGCCNNNAGATCGGAAGA |
| 58047_1 | TATGAGTGTGGAGTCGTTGCCATTCTCCTGCCTCAGCCTCNNNAGATCGGAAGAGCA |
| 58265_1 | TATGAGTGTGGAGTCGTTGCCCGTGTTCAAGCAATTCTCCTGNNNAGATCGGAAGA |
| 58666_1 | TATGAGTGTGGAGTCGTTGCGAAGGATGCTTCCAAGTGTGAGNNNAGATCGGAAG |
| 59125_1 | TATGAGTGTGGAGTCGTTGCTTCACGCCATTCTCCTGCTTNNNAGATCGGAAGAGCA |
| 59402_1 | TATGAGTGTGGAGTCGTTGCCATTCTCCTGCCTCAGCCTCNNNAGATCGGAAGAGCA |
| 62064_1 | TATGAGTGTGGAGTCGTTGCAGCAAGAAGGCAGGTCTTGGNNNAGATCGGAAGA |
| 62079_1 | TATGAGTGTGGAGTCGTTGCCTGGAGTGTAGTGGTGTGATCCNNNAGATCGGAAG |
| 63587_1 | TATGAGTGTGGAGTCGTTGCGAGGCTGAGGCAGGAGAATGNNNAGATCGGAAGA |
| 63875_1 | TATGAGTGTGGAGTCGTTGCGTTAGCCAGGATGGTCTCGATCNNNAGATCGGAAG |
| 64326_1 | TATGAGTGTGGAGTCGTTGCGTTTCACCATGTTGTCCAGGCNNNAGATCGGAAGAG |
| 65176_1 | TATGAGTGTGGAGTCGTTGCGTCAACCAGGCCAGAGTACAGNNNAGATCGGAAGA |
| 66048_1 | TATGAGTGTGGAGTCGTTGCTGGAGCTTGTTGTTTCCGGATCNNNAGATCGGAAG |
| 66143_1 | TATGAGTGTGGAGTCGTTGCGTAGGGAGCACGGATTCTGACNNNAGATCGGAAGA |
| 66478_1 | TATGAGTGTGGAGTCGTTGCGGAGAGACGAAGAGGAAGGGGNNNAGATCGGAA |
| 67017_1 | TATGAGTGTGGAGTCGTTGCGGGCTCCATTTCCCTCTCAGNNNAGATCGGAAGAGC |
| 68716_1 | TATGAGTGTGGAGTCGTTGCGGTGCTCACACTCATGAGGAGNNNAGATCGGAAGA |
| 68784_1 | TATGAGTGTGGAGTCGTTGCGATCTTGCTCTGTTGCTCCATGNNNAGATCGGAAGA |
| 68959_1 | TATGAGTGTGGAGTCGTTGCGCTCCTCACTTTCCAGTAGGGNNNAGATCGGAAGAG |
| 68968_1 | TATGAGTGTGGAGTCGTTGCCACCAGGTCCCATCAACCACNNNAGATCGGAAGAGC |
| 68979_1 | TATGAGTGTGGAGTCGTTGCGGACACCCTAGGAGACTCCAAGNNNAGATCGGAAG |
| 68984_1 | TATGAGTGTGGAGTCGTTGCGAAACAGTGAAGGGACACACCTNNNAGATCGGAAG |

| | |
|---|---|
| 69278_1 | TATGAGTGTGGAGTCGTTGCGGTGAGGGGTAGCTCAGGATAGNNNAGATCGGAA |
| 69491_1 | TATGAGTGTGGAGTCGTTGCCCCCTTCTCTCCTACCTCCTCNNNAGATCGGAAGAGC |
| 69637_1 | TATGAGTGTGGAGTCGTTGCCTGCCATTGAGGGGAGAAGAGNNNAGATCGGAAG |
| 69665_1 | TATGAGTGTGGAGTCGTTGCCAGTAGCCAAAGTGCTTCACAGNNNAGATCGGAAG |
| 70111_1 | TATGAGTGTGGAGTCGTTGCCTACTGGGAAGTGAGGACCCCNNNAGATCGGAAGA |
| 70150_1 | TATGAGTGTGGAGTCGTTGCCCTCCCTGTAAGGTCAGCTACCNNNAGATCGGAAGA |
| 70175_1 | TATGAGTGTGGAGTCGTTGCCCACTGAAAAACCCCTGAGCTNNNAGATCGGAAGAG |
| 70224_1 | TATGAGTGTGGAGTCGTTGCCAGAGAAGGTGGTGACCAGCNNNAGATCGGAAGA |
| 649_1 | TATGAGTGTGGAGTCGTTGCAAGAGGGGCTGGCTGTCCCTNNNAGATCGGAAGAG |
| 937_1 | TATGAGTGTGGAGTCGTTGCGGTCCCCCTACCGGTGGAGGNNNAGATCGGAAGAG |
| 9018_1 | TATGAGTGTGGAGTCGTTGCGGACGCTGATGTGCACAGTTNNNAGATCGGAAGAG |
| 9583_1 | TATGAGTGTGGAGTCGTTGCCCAAGGCCCTTCTGTCCTGGTNNNAGATCGGAAGAG |
| 11342_1 | TATGAGTGTGGAGTCGTTGCTTGTAGGTTTCGTGGTCGGGNNNAGATCGGAAGAG |
| 12131_1 | TATGAGTGTGGAGTCGTTGCCCCAAGGAACACCCGCCCACNNNAGATCGGAAGAGC |
| 53978_1 | TATGAGTGTGGAGTCGTTGCGATGGAGTCTCGCTTTGTTGCNNNAGATCGGAAGA |
| 57070_1 | TATGAGTGTGGAGTCGTTGCGAGGCAGGAGAATGGTGTGAANNNAGATCGGAAG |
| 57530_1 | TATGAGTGTGGAGTCGTTGCCCCGTAATCCCAGCACTTTGGNNNAGATCGGAAGAG |
| 58042_1 | TATGAGTGTGGAGTCGTTGCCAAGTAGCTGGGACTACAGGCNNNAGATCGGAAGA |
| 58641_1 | TATGAGTGTGGAGTCGTTGCGTGGTGGCCTCCGAATTCTCNNNAGATCGGAAGAG |
| 58906_1 | TATGAGTGTGGAGTCGTTGCCCTCGACCCACAGAACAACAGNNNAGATCGGAAGAG |
| 62066_1 | TATGAGTGTGGAGTCGTTGCACTGTCTCCTAGGCGACCAGNNNAGATCGGAAGAG |
| 66124_1 | TATGAGTGTGGAGTCGTTGCCACACACACACACACACACAGNNNAGATCGGAAGAG |
| 66338_1 | TATGAGTGTGGAGTCGTTGCGGGGCCCAGGAAGGAATAAATNNNAGATCGGAAG |
| 67357_1 | TATGAGTGTGGAGTCGTTGCGTCAGGCGCTCTCCTCCTTCNNNAGATCGGAAGAGC |
| 68528_1 | TATGAGTGTGGAGTCGTTGCGGGCTCCTCACTTCCCAGTAGNNNAGATCGGAAGAG |
| 68540_1 | TATGAGTGTGGAGTCGTTGCGAGGCCCTGCGAGAATTCAAGNNNAGATCGGAAGA |
| 68730_1 | TATGAGTGTGGAGTCGTTGCCAATTGTGGCCCTGCTGTTGNNNAGATCGGAAGAG |
| 68731_1 | TATGAGTGTGGAGTCGTTGCCTACGCTCTCATAACTGGGCGNNNAGATCGGAAGAG |
| 69221_1 | TATGAGTGTGGAGTCGTTGCTCGTTACCATGACGACCCCCNNNAGATCGGAAGAGC |
| 70058_1 | TATGAGTGTGGAGTCGTTGCGGTAGCGCGGGAGCAAAAAGNNNAGATCGGAAGA |
| 70070_1 | TATGAGTGTGGAGTCGTTGCAGAAGCGTGAGTGGAACAGAGNNNAGATCGGAAG |
| 70116_1 | TATGAGTGTGGAGTCGTTGCCTGTGCTCTGCTGGTCAAAAGNNNAGATCGGAAGA |
| 3293_1 | TATGAGTGTGGAGTCGTTGCCCTGGAGCTGCGCTGCTGATNNNAGATCGGAAGAG |
| 6932_1 | TATGAGTGTGGAGTCGTTGCATCCCCCTGGACACGACCACNNNAGATCGGAAGAGC |
| 9572_1 | TATGAGTGTGGAGTCGTTGCGGTCCCCTCTGGAATGCACTNNNAGATCGGAAGAG |
| 11239_1 | TATGAGTGTGGAGTCGTTGCGTGCCAGCCATGGAGCCTCCNNNAGATCGGAAGAG |
| 11660_1 | TATGAGTGTGGAGTCGTTGCCTGTGCCCGCACCCTCATCCNNNAGATCGGAAGAGC |
| 11742_1 | TATGAGTGTGGAGTCGTTGCGGCACCTCTAGCCCGGGGATNNNAGATCGGAAGAG |
| 12195_1 | TATGAGTGTGGAGTCGTTGCCCCCCGCTACCACGGCTTTTNNNAGATCGGAAGAGC |
| 66480_1 | TATGAGTGTGGAGTCGTTGCCCATGTTGTCAGGCCAGGTGNNNAGATCGGAAGAG |
| 70208_1 | TATGAGTGTGGAGTCGTTGCGAGAGCCGAGACCTCAGTTCNNNAGATCGGAAGAG |
| 46553_1 | TGAGTGTGGAGTCGTTGCTGTCAGTTTGAAATACTGTATGTACGTNNNAGATCGGA |
| 48667_1 | TGAGTGTGGAGTCGTTGCATATTTTCCAGTTATACGTACCTCATCNNNAGATCGGAA |

| | |
|---|---|
| 39579_1 | TGAGTGTGGAGTCGTTGCTGCTTTCAAAAAAAGTTCTCCAGAGANNNAGATCGGAA |
| 28785_1 | ATGAGTGTGGAGTCGTTGCTCACATTTTCTTTATCCACTCATTGANNNAGATCGGAA |
| 28827_1 | ATGAGTGTGGAGTCGTTGCACACACACATATAAGCAGGTTATATCTNNNAGATCGG |
| 30741_1 | ATGAGTGTGGAGTCGTTGCACTGGATTTTTTTTCTGTTGTATCGANNNAGATCGGA |
| 30826_1 | ATGAGTGTGGAGTCGTTGCACACTTTTTTTCAATAGGGACTTCTGANNNAGATCGG |
| 30900_1 | ATGAGTGTGGAGTCGTTGCGAGAGTACAATCTCTAGAACAATCAAANNNAGATCGG |
| 31043_1 | ATGAGTGTGGAGTCGTTGCTCTAGTGAAGTATAGAGACCTTACCANNNAGATCGGA |
| 37088_1 | ATGAGTGTGGAGTCGTTGCAAATTCCCAGATAACCAAGATTGAATNNNAGATCGGA |
| 42892_1 | ATGAGTGTGGAGTCGTTGCTGTATATACACGTATACACATATGTGCNNNAGATCGG |
| 44538_1 | ATGAGTGTGGAGTCGTTGCTTTTGGAAAACATTATAGAGGTTACCTNNNAGATCGG |
| 48789_1 | ATGAGTGTGGAGTCGTTGCAAGAAAGAAGAAAAGAGAGAAGATTCANNNAGATC |
| 28888_1 | ATGAGTGTGGAGTCGTTGCAATTCATATATGTGTATGTCTGTGCANNNAGATCGGA |
| 28897_1 | ATGAGTGTGGAGTCGTTGCGTCCTTACCCTTTAGTCACTGTGTNNNAGATCGGAAG |
| 30782_1 | ATGAGTGTGGAGTCGTTGCTGGGATGTGTATCTCTGACTAATACANNNAGATCGGA |
| 31725_1 | ATGAGTGTGGAGTCGTTGCAGATTATCTCTCTCTCTCTCTACANNNAGATCGGAA |
| 35436_1 | ATGAGTGTGGAGTCGTTGCAACAAACAAAAAAAAAACAGGAGGANNNAGATCGG |
| 35763_1 | ATGAGTGTGGAGTCGTTGCACTGGTAATTTTGCAGAAGTGTGANNNAGATCGGAA |
| 43021_1 | ATGAGTGTGGAGTCGTTGCATTCATAAATATTTCCTTTGGGGGAANNNAGATCGGA |
| 43044_1 | ATGAGTGTGGAGTCGTTGCGAAGTGGAATTGCTGGATATATGACNNNAGATCGGA |
| 43537_1 | ATGAGTGTGGAGTCGTTGCAGGTATTTTTTCAATCCTCGTCTTCCNNNAGATCGGAA |
| 341_1 | TATGAGTGTGGAGTCGTTGCTCTCCCTCATTCATATCAGGTACTNNNAGATCGGAAG |
| 622_1 | TATGAGTGTGGAGTCGTTGCATTTTAGAAACCACTGCTCTAAAGAACNNNAGATCG |
| 766_1 | TATGAGTGTGGAGTCGTTGCCCCATACATAAGGAACAGTGAAGTGANNNAGATCG |
| 1198_1 | TATGAGTGTGGAGTCGTTGCCACACATTTACCCCTAAATATTCAGCANNNAGATCGG |
| 1640_1 | TATGAGTGTGGAGTCGTTGCCAAAACACTGATGAAACAATTGAAGATNNNAGATCG |
| 1805_1 | TATGAGTGTGGAGTCGTTGCAGGGGTAAGGTATTATTTCTCCTCANNNAGATCGGA |
| 1912_1 | TATGAGTGTGGAGTCGTTGCTAGTGATGAGATTGCTGGGTCATATNNNAGATCGG |
| 2010_1 | TATGAGTGTGGAGTCGTTGCGTGGGTAGAATGCCTTTGGGAAANNNAGATCGGAA |
| 2079_1 | TATGAGTGTGGAGTCGTTGCCTCTGCATCTGTATTTATCTCTCTACANNNAGATCGG |
| 2150_1 | TATGAGTGTGGAGTCGTTGCTAGATCATTTCTGTTACATGTTCTTCANNNAGATCGG |
| 2220_1 | TATGAGTGTGGAGTCGTTGCATTACCATTTGCTCTCAGCCTGANNNAGATCGGAAG |
| 2242_1 | TATGAGTGTGGAGTCGTTGCCGTGTCATCAATGGGCACACTATANNNAGATCGGAA |
| 2253_1 | TATGAGTGTGGAGTCGTTGCGATGGTGGGGCTATGGCTAATTTTTANNNAGATCG |
| 2281_1 | TATGAGTGTGGAGTCGTTGCTGTTGAATAAACTGTTGTCAGATCAACNNNAGATCG |
| 2353_1 | TATGAGTGTGGAGTCGTTGCTGGGTTATAAGGGTGCTTTCTCCNNNAGATCGGAAG |
| 2394_1 | TATGAGTGTGGAGTCGTTGCCCTGGCATATAGACACATAGTGGCNNNAGATCGGA |
| 2423_1 | TATGAGTGTGGAGTCGTTGCACCATGTACTCAACACTCGCATCNNNAGATCGGAAG |
| 2547_1 | TATGAGTGTGGAGTCGTTGCTGTCCAACTCAAATTCAGGAATATAGGNNNAGATCG |
| 2642_1 | TATGAGTGTGGAGTCGTTGCTCCCCCAGGCTCAGATAAGATTGNNNAGATCGGAAG |
| 2712_1 | TATGAGTGTGGAGTCGTTGCTGCCCATGCCTGTCTTTTACTATNNNAGATCGGAAGA |
| 2740_1 | TATGAGTGTGGAGTCGTTGCACACATGAACACCTATATACACATACTNNNAGATCGG |
| 2819_1 | TATGAGTGTGGAGTCGTTGCAAATAGAGAGCTACCTGAAATAGTTCTNNNAGATCG |
| 3511_1 | TATGAGTGTGGAGTCGTTGCGTTGTTGTTGTAGAGATGAGGTCTAACNNNAGATC |

| | |
|---|---|
| 3519_1 | TATGAGTGTGGAGTCGTTGCAGACACATCAAGTCTCAGTTTCTTCATNNNAGATCGG |
| 4536_1 | TATGAGTGTGGAGTCGTTGCCAAAAGGTAGCATAGTATACACACAGCNNNAGATCG |
| 4562_1 | TATGAGTGTGGAGTCGTTGCTCTGTTTCCTCTCACTTGCTCAAATNNNAGATCGGAA |
| 4670_1 | TATGAGTGTGGAGTCGTTGCAAGATAGAATAACCTAGAATACCACCANNNAGATCG |
| 4698_1 | TATGAGTGTGGAGTCGTTGCTCGTTCGTGTGTTTGGATAGATANNNAGATCGGAA |
| 4964_1 | TATGAGTGTGGAGTCGTTGCCTGCCTATCTGGCTGCATAAACANNNAGATCGGAAG |
| 5089_1 | TATGAGTGTGGAGTCGTTGCACACTCCTCCACTATCATCCCATNNNAGATCGGAAGA |
| 5656_1 | TATGAGTGTGGAGTCGTTGCGACTTTCCTTTGGAGCATTATTGGGNNNAGATCGGA |
| 5703_1 | TATGAGTGTGGAGTCGTTGCCTCAGCATCCTTGAAAGCTCTCANNNAGATCGGAAG |
| 6747_1 | TATGAGTGTGGAGTCGTTGCTCTCTAAAAAGGATTTGTGAGGCTAGTNNNAGATCG |
| 7126_1 | TATGAGTGTGGAGTCGTTGCTCTCCTCTTTTGATCTACAATTTTCGGNNNAGATCGG |
| 7596_1 | TATGAGTGTGGAGTCGTTGCTCTCCTGTGTTGGTCCGGATGATATCANNNAGATCG |
| 7605_1 | TATGAGTGTGGAGTCGTTGCCTCTGCTTTCAGTTCTGTTCACTATNNNAGATCGGAA |
| 7819_1 | TATGAGTGTGGAGTCGTTGCATGAATGAAAACAAACAGAGCTTCAGANNNAGATC |
| 7824_1 | TATGAGTGTGGAGTCGTTGCTCTACCTCAGCCTGTCAAGTAGGTNNNAGATCGGAA |
| 7906_1 | TATGAGTGTGGAGTCGTTGCTCTCTGATAAAACTGGCTTTACACCANNNAGATCGG |
| 8039_1 | TATGAGTGTGGAGTCGTTGCTAGGCAAGTCATACGGAAAACATTTTCNNNAGATCG |
| 8213_1 | TATGAGTGTGGAGTCGTTGCATTAGTGACAGACATTTTAGATGTGATNNNAGATCG |
| 8347_1 | TATGAGTGTGGAGTCGTTGCTGAACCTTATTTATTGTGCTTCTACTGNNNAGATCGG |
| 8785_1 | TATGAGTGTGGAGTCGTTGCATACTGAAAACCATGATATCGTCTGTTNNNAGATCG |
| 9109_1 | TATGAGTGTGGAGTCGTTGCTCCTTCAAACAAATTTCAAAGCAACTNNNAGATCGGA |
| 9169_1 | TATGAGTGTGGAGTCGTTGCTGTGTGAGAGAGATCAAGCACATNNNAGATCGGAA |
| 9266_1 | TATGAGTGTGGAGTCGTTGCTTCTGTGGCTATTTCTCAGCCTGNNNAGATCGGAAG |
| 9378_1 | TATGAGTGTGGAGTCGTTGCGCTGTAAGGTACTAATGGGCACANNNAGATCGGAA |
| 9436_1 | TATGAGTGTGGAGTCGTTGCTTCCTGGGCATGAGTTTCTTGAGNNNAGATCGGAAG |
| 9778_1 | TATGAGTGTGGAGTCGTTGCTTTCAAGTGTACGTGGAGCGTTANNNAGATCGGAA |
| 9803_1 | TATGAGTGTGGAGTCGTTGCAGGGCAACAAGGTGGCTATACTANNNAGATCGGAA |
| 9959_1 | TATGAGTGTGGAGTCGTTGCTTGCTTTGTTTTCAGTTGGAATAAAAGNNNAGATCG |
| 10008_1 | TATGAGTGTGGAGTCGTTGCGCAACAGCCGCATACTTACTATGNNNAGATCGGAAG |
| 10013_1 | TATGAGTGTGGAGTCGTTGCGTGCCAACTCATACAGCACCTAANNNAGATCGGAAG |
| 10274_1 | TATGAGTGTGGAGTCGTTGCAAGTGTGGCTCCTGGAATTTGTANNNAGATCGGAA |
| 10304_1 | TATGAGTGTGGAGTCGTTGCGTAATGGGCCTCCTTGTCTTCTTNNNAGATCGGAAG |
| 10327_1 | TATGAGTGTGGAGTCGTTGCACCTCTCTGAATTTATGTCTTCATTTGNNNAGATCGG |
| 10341_1 | TATGAGTGTGGAGTCGTTGCACAGGCGTTTATGATAGATTTCTAAAGNNNAGATCG |
| 10460_1 | TATGAGTGTGGAGTCGTTGCACTCTTTTGAGCTTCTTTCATTCAAGANNNAGATCGG |
| 10541_1 | TATGAGTGTGGAGTCGTTGCGCATATTGTATTCTGTACCTTGTTTTTNNNAGATCGG |
| 10564_1 | TATGAGTGTGGAGTCGTTGCTGGTAGGTGTTTTTCTTTTGTATTTTTNNNAGATCGG |
| 10629_1 | TATGAGTGTGGAGTCGTTGCACCCCAATAAAAATGGGTGGAAANNNAGATCGGAA |
| 10679_1 | TATGAGTGTGGAGTCGTTGCTGTGTGTCATTCTAGGAGCATGCNNNAGATCGGAA |
| 10747_1 | TATGAGTGTGGAGTCGTTGCGCATGAGAAAAACTCTAAATTCAATGTNNNAGATCG |
| 10852_1 | TATGAGTGTGGAGTCGTTGCCAGGAAACAGTATTTGTAGTTGACATGNNNAGATCG |
| 10934_1 | TATGAGTGTGGAGTCGTTGCGAAACACCAGCTAGATTTATGCATGACNNNAGATCG |
| 11182_1 | TATGAGTGTGGAGTCGTTGCCCTCCCTAGCTGGAATTACAGATGNNNAGATCGGAA |

| | |
|---|---|
| 11337_1 | TATGAGTGTGGAGTCGTTGCCTGCTACTGTTTTTGCATGTTGATTNNNAGATCGGAA |
| 11361_1 | TATGAGTGTGGAGTCGTTGCAGGACACATAATTACAAACACTGGCTGNNNAGATCG |
| 11627_1 | TATGAGTGTGGAGTCGTTGCGATTCTCTAGGTATTTTGGCCGATGATNNNAGATCG |
| 11946_1 | TATGAGTGTGGAGTCGTTGCTCAACAATGTATAACCACACCAGCATNNNAGATCGG |
| 12105_1 | TATGAGTGTGGAGTCGTTGCTCTGACATTGGAGAGAAGATGGAAATNNNAGATCG |
| 12160_1 | TATGAGTGTGGAGTCGTTGCAGTAATGACCACAGATATTTTTGTTTGNNNAGATCG |
| 28781_1 | TATGAGTGTGGAGTCGTTGCTGGTTTCAAACTCTTGAGCTCAANNNAGATCGGAAG |
| 28799_1 | TATGAGTGTGGAGTCGTTGCACACACACACATATACACATATACACANNNAGATCGG |
| 28806_1 | TATGAGTGTGGAGTCGTTGCACACATACTATGTATACACACACTACANNNAGATCGG |
| 28826_1 | TATGAGTGTGGAGTCGTTGCCGTATATATATATGTGTGTGTGTGTGTNNNAGATCG |
| 28974_1 | TATGAGTGTGGAGTCGTTGCTGGAATACTACTCAGCCATAAATATGANNNAGATCG |
| 28995_1 | TATGAGTGTGGAGTCGTTGCTTTGTCATGTCAGAAATTTTGAAGAAANNNAGATCG |
| 29064_1 | TATGAGTGTGGAGTCGTTGCTCTGTACAAGAATAAGGAAACTCCANNNAGATCGGA |
| 29077_1 | TATGAGTGTGGAGTCGTTGCTGGAATTATGCCTCTCTAGTGAGANNNAGATCGGAA |
| 29146_1 | TATGAGTGTGGAGTCGTTGCATTCTGGAGTAAAATGGGTAGTTATTTNNNAGATCG |
| 29202_1 | TATGAGTGTGGAGTCGTTGCAGCATCCAAATAAACTTGCACTAACTNNNAGATCGG |
| 29234_1 | TATGAGTGTGGAGTCGTTGCAGATAGATAGATGAGATGGATGATGGANNNAGAT |
| 29290_1 | TATGAGTGTGGAGTCGTTGCACTTTCTTCTTTTGTATCTTTGAGACCNNNAGATCGG |
| 30736_1 | TATGAGTGTGGAGTCGTTGCAGGTTTCTTTTTGGGGTGTTGAANNNAGATCGGAA |
| 30771_1 | TATGAGTGTGGAGTCGTTGCTGTCTCTTCTTACAGTTTTTGTCTTGANNNAGATCGG |
| 30785_1 | TATGAGTGTGGAGTCGTTGCTGAACAATTTATACAAAAGGTTGTCCTNNNAGATCG |
| 30911_1 | TATGAGTGTGGAGTCGTTGCTGTTCACTGGTCAGAAGACTCAGNNNAGATCGGAA |
| 31074_1 | TATGAGTGTGGAGTCGTTGCACATGATACATACACATGATACATGCANNNAGATCG |
| 31295_1 | TATGAGTGTGGAGTCGTTGCATGTAGCCATGAAAAGGAATGCGNNNAGATCGGAA |
| 31305_1 | TATGAGTGTGGAGTCGTTGCGAGAAAGAGAGAAGAAAGAAAGAAAGANNNAGAT |
| 31420_1 | TATGAGTGTGGAGTCGTTGCACTGAGTGTACCCTCCTCTACTCNNNAGATCGGAAG |
| 31496_1 | TATGAGTGTGGAGTCGTTGCTCAGTGCCATTGTAAATGTAAATGACTNNNAGATCG |
| 31518_1 | TATGAGTGTGGAGTCGTTGCTGTTATGTTTTGATGCAAGCATACANNNAGATCGGA |
| 31741_1 | TATGAGTGTGGAGTCGTTGCTGACAGTTACAGTCCTCAGTTGGNNNAGATCGGAAG |
| 31819_1 | TATGAGTGTGGAGTCGTTGCTGTGTGTGTGTGTATACATATATCCCTNNNAGATCG |
| 31822_1 | TATGAGTGTGGAGTCGTTGCTCTCTATCATCTTTTCTCCTCATGTCANNNAGATCGGA |
| 31832_1 | TATGAGTGTGGAGTCGTTGCAAGAACTTTGCAGTGTTGTTGGGNNNAGATCGGAA |
| 31854_1 | TATGAGTGTGGAGTCGTTGCTCCTGGAAATGTACAACCTCACANNNAGATCGGAAG |
| 31971_1 | TATGAGTGTGGAGTCGTTGCGCAGATAAATGACTATGTGTACTAGAANNNAGATCG |
| 31973_1 | TATGAGTGTGGAGTCGTTGCGGTGTTTTTTGTGTTGATATTCTGTCCNNNAGATCG |
| 31987_1 | TATGAGTGTGGAGTCGTTGCTGTTTGTCTCCTAAGCCCATTCANNNAGATCGGAAG |
| 32014_1 | TATGAGTGTGGAGTCGTTGCACCACATAAATGTTCTAGAAGTCAACANNNAGATCG |
| 32016_1 | TATGAGTGTGGAGTCGTTGCTCAATCAAGTGAAGAGAAAACCTATGANNNAGATCG |
| 32134_1 | TATGAGTGTGGAGTCGTTGCCCACAGGTATCAAAGCAATACTAGCNNNAGATCGGA |
| 32163_1 | TATGAGTGTGGAGTCGTTGCGATCTCTGCTCACTTCAAGCTCTNNNAGATCGGAAG |
| 32339_1 | TATGAGTGTGGAGTCGTTGCAATTCCAGAACAGGAGCAGAGACNNNAGATCGGAA |
| 32407_1 | TATGAGTGTGGAGTCGTTGCACACACCCACACATATATACATATACANNNAGATCGG |
| 32681_1 | TATGAGTGTGGAGTCGTTGCCCCAGGTTCAAGTGATTCTCTCANNNAGATCGGAAG |

| | |
|---|---|
| 32761_1 | TATGAGTGTGGAGTCGTTGCAGAGGAGACAAATTCAAAAGCTAGCNNNAGATCGG |
| 33461_1 | TATGAGTGTGGAGTCGTTGCCCATACCAAGGAGCTGGATGTAANNNAGATCGGAA |
| 33588_1 | TATGAGTGTGGAGTCGTTGCCTTCCTGGGTTCAAGCAATTCTCNNNAGATCGGAAG |
| 34747_1 | TATGAGTGTGGAGTCGTTGCGCCCGGCCAATTTCTGTATTTTTNNNAGATCGGAAG |
| 34876_1 | TATGAGTGTGGAGTCGTTGCGCCTCCTAAGTAGCTGGGATTACNNNAGATCGGAAG |
| 35037_1 | TATGAGTGTGGAGTCGTTGCCTGTCTCAGCTCACTGTAACCTCNNNAGATCGGAAG |
| 35475_1 | TATGAGTGTGGAGTCGTTGCAGGTCAAATCTCACATATTTCATAAGTNNNAGATCG |
| 35619_1 | TATGAGTGTGGAGTCGTTGCGACAATATGCGTTGGTGAACTCCNNNAGATCGGAA |
| 35734_1 | TATGAGTGTGGAGTCGTTGCTCTTTACGGATTTCAAGGGAGCANNNAGATCGGAA |
| 35790_1 | TATGAGTGTGGAGTCGTTGCTCTTTACGGATTTCAAGGGAGCANNNAGATCGGAA |
| 35832_1 | TATGAGTGTGGAGTCGTTGCCTCCCTGGTTCAAGTGATTCTCANNNAGATCGGAAG |
| 35849_1 | TATGAGTGTGGAGTCGTTGCCTTCAAGTGATTCTCCTGCCTCANNNAGATCGGAAG |
| 36765_1 | TATGAGTGTGGAGTCGTTGCATTCATTGCTAGGATAGTGTATTTTGANNNAGATCG |
| 36814_1 | TATGAGTGTGGAGTCGTTGCCTAACATTATGCCCTACTATCTTTTGTNNNAGATCGG |
| 36887_1 | TATGAGTGTGGAGTCGTTGCAGGTTATCTGTGTGTTCATTGAATACTNNNAGATCG |
| 37011_1 | TATGAGTGTGGAGTCGTTGCGGAGAATAGCATGAGTGGGACATNNNAGATCGGA |
| 37031_1 | TATGAGTGTGGAGTCGTTGCACAAGTTAGTCAGAAAACCAGAAATCTNNNAGATCG |
| 37152_1 | TATGAGTGTGGAGTCGTTGCCCTGCTCAAAAAAAATTCCATTCACANNNAGATCGGA |
| 37166_1 | TATGAGTGTGGAGTCGTTGCTGTCATGGAAAAAGTAAAAGTCTTTGCNNNAGATCG |
| 37195_1 | TATGAGTGTGGAGTCGTTGCCCAGATTGAGGATATTAGAGGAAAACANNNAGATC |
| 37216_1 | TATGAGTGTGGAGTCGTTGCTGTGCTACCACTATGAACATATTTTTNNNAGATCGGA |
| 37349_1 | TATGAGTGTGGAGTCGTTGCGCATTCCCATACACATATATGTATACANNNAGATCGG |
| 37426_1 | TATGAGTGTGGAGTCGTTGCTCCTTCAAAAAAACACATAACCAAAGANNNAGATCG |
| 37607_1 | TATGAGTGTGGAGTCGTTGCTTGTAATAAAATTCAGGAATCAGCTCTNNNAGATCG |
| 37855_1 | TATGAGTGTGGAGTCGTTGCTGAGAAAATATTTGTCATCTCCTTCGANNNAGATCG |
| 38290_1 | TATGAGTGTGGAGTCGTTGCTCTAGGAATCACATTACCTAACTTCANNNAGATCGGA |
| 38400_1 | TATGAGTGTGGAGTCGTTGCACTAGGAAGTGATTGATTAGTTTTGAGNNNAGATC |
| 38703_1 | TATGAGTGTGGAGTCGTTGCAGTATATATGCACCCAACACTGGANNNAGATCGGAA |
| 38830_1 | TATGAGTGTGGAGTCGTTGCAGAAAAGTTGCATAAATGTTCAATGGANNNAGATC |
| 39062_1 | TATGAGTGTGGAGTCGTTGCTCCTGGAAATGTACAACCTCACANNNAGATCGGAAG |
| 39106_1 | TATGAGTGTGGAGTCGTTGCAGGAAGCTAGGAATTGAAAGATACAGANNNAGATC |
| 39188_1 | TATGAGTGTGGAGTCGTTGCAGGTCATAGAAATTTTGAAGTGTTGACNNNAGATCG |
| 39449_1 | TATGAGTGTGGAGTCGTTGCCTGGATAGGAGGAATAAATTCAAAACANNNAGATC |
| 39450_1 | TATGAGTGTGGAGTCGTTGCGTGGATTACGAGGTCAGGAGATCNNNAGATCGGAA |
| 39727_1 | TATGAGTGTGGAGTCGTTGCAATTCCAATGACATTTTTGCCGANNNAGATCGGAAG |
| 39781_1 | TATGAGTGTGGAGTCGTTGCACCTAAAAAGAAAACTAAATACTGGGGNNNAGATC |
| 39788_1 | TATGAGTGTGGAGTCGTTGCGCCAGAGCACACAGACAAGATATNNNAGATCGGAA |
| 40379_1 | TATGAGTGTGGAGTCGTTGCAGGAACTTAGATGCAAAATTCTCAACANNNAGATCG |
| 40420_1 | TATGAGTGTGGAGTCGTTGCTGCAAATCAACTGTAAGAGAAAATAGANNNAGATC |
| 40724_1 | TATGAGTGTGGAGTCGTTGCTGTTCATTGCAGCACTATTCACANNNAGATCGGAAG |
| 40935_1 | TATGAGTGTGGAGTCGTTGCGGAAAGAAATAATTGAAAACCTCCAGANNNAGATC |
| 41581_1 | TATGAGTGTGGAGTCGTTGCACATCTTTTATTTCTTGTAAAGAGGGGNNNAGATCG |
| 41796_1 | TATGAGTGTGGAGTCGTTGCGCATAATCTCGGCTCACTGAAACNNNAGATCGGAAG |

| | |
|---|---|
| 42169_1 | TATGAGTGTGGAGTCGTTGCACTCCTGCCATTTTGTTGTTTGTNNNAGATCGGAAG |
| 42243_1 | TATGAGTGTGGAGTCGTTGCTGGTGTGCGTATATATATATGGTGTGTNNNAGATCG |
| 42254_1 | TATGAGTGTGGAGTCGTTGCACACATATATACATATACACACACGTGNNNAGATCGG |
| 42258_1 | TATGAGTGTGGAGTCGTTGCACACATATATACATATACACACACGTGNNNAGATCGG |
| 42295_1 | TATGAGTGTGGAGTCGTTGCTGGCATGATCTCAACTAACTGCTNNNAGATCGGAAG |
| 42303_1 | TATGAGTGTGGAGTCGTTGCACCCCAAAATAGGCAAAATGCATNNNAGATCGGAAG |
| 42310_1 | TATGAGTGTGGAGTCGTTGCGGAGTAGCTGTTCTTTCACCACTNNNAGATCGGAAG |
| 42359_1 | TATGAGTGTGGAGTCGTTGCTGTTTGTCTCCTAAGCCCATTCANNNAGATCGGAAG |
| 42384_1 | TATGAGTGTGGAGTCGTTGCTTATGCTGCAATAGACATGCGTGNNNAGATCGGAAG |
| 42390_1 | TATGAGTGTGGAGTCGTTGCAGGGAAAAGTCAAGCTGGTAACTNNNAGATCGGAA |
| 42707_1 | TATGAGTGTGGAGTCGTTGCACCCCAAAATAGGCAAAATGCATNNNAGATCGGAAG |
| 42761_1 | TATGAGTGTGGAGTCGTTGCCTGTTGTACAGGGTGATAAAAGAAANNNAGATCGG |
| 42764_1 | TATGAGTGTGGAGTCGTTGCTCAGAAATGTAAAGAACAAAGAGGTCCNNNAGATC |
| 42814_1 | TATGAGTGTGGAGTCGTTGCTGCGTATGTAATCAAATACATTGTTGCNNNAGATCG |
| 42843_1 | TATGAGTGTGGAGTCGTTGCATTGAACACCTAGCATTTGACAGNNNAGATCGGAAG |
| 42868_1 | TATGAGTGTGGAGTCGTTGCCCTCCAACATTTATTATGAAAATGGGCNNNAGATCG |
| 42890_1 | TATGAGTGTGGAGTCGTTGCTGGAAGCAAATTGGAGACATCACNNNAGATCGGAA |
| 42897_1 | TATGAGTGTGGAGTCGTTGCTCTGTGTGTGTATATATATGTGTGTGTNNNAGATCG |
| 42917_1 | TATGAGTGTGGAGTCGTTGCCTCCAGTGTTTCTTCTCAAGGGANNNAGATCGGAAG |
| 42991_1 | TATGAGTGTGGAGTCGTTGCTGTGTATGTGTATGCAAATATGTGTGTNNNAGATCG |
| 43289_1 | TATGAGTGTGGAGTCGTTGCACACACACACATCTATATATACACACGNNNAGATCGG |
| 43317_1 | TATGAGTGTGGAGTCGTTGCCTCTCTTTTCTTCTTTATCAGTCTAGCNNNAGATCGGA |
| 43550_1 | TATGAGTGTGGAGTCGTTGCGCAAATATTTTCTCCCATTATTTCTGGNNNAGATCGG |
| 43786_1 | TATGAGTGTGGAGTCGTTGCAATAGAATAAAAATATTCGGGGCAGANNNAGATCG |
| 43800_1 | TATGAGTGTGGAGTCGTTGCGGAACAAAATACCAAACTGAAATTTGCNNNAGATCG |
| 44455_1 | TATGAGTGTGGAGTCGTTGCGCTCAAGTGATTCTCGTACCTCANNNAGATCGGAAG |
| 44522_1 | TATGAGTGTGGAGTCGTTGCACACCCACACATATATACATATACACANNNAGATCGG |
| 44561_1 | TATGAGTGTGGAGTCGTTGCACCAAGAGAGATAAAATGAAAACATGTNNNAGATC |
| 44856_1 | TATGAGTGTGGAGTCGTTGCTCTGGTATTGTCGGAATATGGGCNNNAGATCGGAA |
| 45087_1 | TATGAGTGTGGAGTCGTTGCTGTAGATCATATAACATGTTTGTGGGTNNNAGATCG |
| 45359_1 | TATGAGTGTGGAGTCGTTGCTCCTGTAGAGTTCTTTGAACTTCTNNNAGATCGGAA |
| 45784_1 | TATGAGTGTGGAGTCGTTGCCCATCCTGGGACTTAGAAGAAAGTNNNAGATCGGAA |
| 46000_1 | TATGAGTGTGGAGTCGTTGCCTCCAAACCTAGCTCCCAAGTAGNNNAGATCGGAAG |
| 46791_1 | TATGAGTGTGGAGTCGTTGCACACTACACAAATGTTTTGTATACAGANNNAGATCG |
| 46935_1 | TATGAGTGTGGAGTCGTTGCGTGCTGGGAAATGTGGGTTTTTANNNAGATCGGAA |
| 47199_1 | TATGAGTGTGGAGTCGTTGCCCTCCTGGGTTCAAGTGATTCTCNNNAGATCGGAAG |
| 47654_1 | TATGAGTGTGGAGTCGTTGCGCCTCCTAAGTAGCTGGGATTACNNNAGATCGGAAG |
| 47788_1 | TATGAGTGTGGAGTCGTTGCCCAAGTAGCTGGGATTACAGGTGNNNAGATCGGAA |
| 48177_1 | TATGAGTGTGGAGTCGTTGCCTGGGTTCAAGCAATTCTACTGCNNNAGATCGGAAG |
| 48749_1 | TATGAGTGTGGAGTCGTTGCGCATTTTTCGAGCCAAACCAAAGNNNAGATCGGAAG |
| 48756_1 | TATGAGTGTGGAGTCGTTGCACTGGTATGTTCATTGCACCACTNNNAGATCGGAAG |
| 48763_1 | TATGAGTGTGGAGTCGTTGCCACCAAGCCCAGCTAATTTTTGTNNNAGATCGGAAG |
| 48788_1 | TATGAGTGTGGAGTCGTTGCTCTTTTCATGACTGTGTCCTTTGNNNAGATCGGAAG |

| | |
|---|---|
| 48856_1 | TATGAGTGTGGAGTCGTTGCTGGAGTGCAATGGTGTGATCTTANNNAGATCGGAA |
| 49019_1 | TATGAGTGTGGAGTCGTTGCGCCCGGCCAATTTCTGTATTTTTNNNAGATCGGAAG |
| 49068_1 | TATGAGTGTGGAGTCGTTGCGTGCAAACTCATGGGGAAGAAATNNNAGATCGGAA |
| 49137_1 | TATGAGTGTGGAGTCGTTGCAAACCTGAGTTTCCATCCAGTGTNNNAGATCGGAAG |
| 49240_1 | TATGAGTGTGGAGTCGTTGCGTGAAATTCATGGAAAGAACCCTNNNAGATCGGAA |
| 49342_1 | TATGAGTGTGGAGTCGTTGCGCAAGTCATAGAAAAAAACACAAAAGCNNNAGATC |
| 49378_1 | TATGAGTGTGGAGTCGTTGCGTAGCTGGAATTACACACATGCGNNNAGATCGGAA |
| 49489_1 | TATGAGTGTGGAGTCGTTGCACTAAAAACATCTCAAATTTCCAGAGANNNAGATCG |
| 49538_1 | TATGAGTGTGGAGTCGTTGCCTCCTGAGTAGCTGGGACTACAGNNNAGATCGGAA |
| 49539_1 | TATGAGTGTGGAGTCGTTGCTTCAAGTGATTCTCCTGCCTCAGNNNAGATCGGAAG |
| 49568_1 | TATGAGTGTGGAGTCGTTGCTGTGTTGAATAACAGTGGTGACANNNAGATCGGAA |
| 49718_1 | TATGAGTGTGGAGTCGTTGCACGCCCGGCTAATTTTTTTGTATNNNAGATCGGAAG |
| 49772_1 | TATGAGTGTGGAGTCGTTGCAATGTTGGGCTAGAAGGAACACANNNAGATCGGAA |
| 50850_1 | TATGAGTGTGGAGTCGTTGCAGTATCATGTGTGAATGCAGAGTNNNAGATCGGAA |
| 51342_1 | TATGAGTGTGGAGTCGTTGCTGGACAGAAAATACCTATGAAGAAGGTNNNAGATC |
| 52930_1 | TATGAGTGTGGAGTCGTTGCAGATGTAGATGAAATACTACTCAGCCANNNAGATCG |
| 53127_1 | TATGAGTGTGGAGTCGTTGCGCAGATAAATGACTATGTGTACTAGAANNNAGATCG |
| 53185_1 | TATGAGTGTGGAGTCGTTGCCCATCCCCTCAAGCATTTATCCTNNNAGATCGGAAGA |
| 53196_1 | TATGAGTGTGGAGTCGTTGCCAGAGAGAGAGAAATAGATTAGGAAGANNNAGAT |
| 53333_1 | TATGAGTGTGGAGTCGTTGCTGTGAATTTATATGTGTGTGTGTGTGTNNNAGATCG |
| 53385_1 | TATGAGTGTGGAGTCGTTGCTTACAGATTCAGTGTCTGGTGGGNNNAGATCGGAA |
| 104_1 | TATGAGTGTGGAGTCGTTGCTGTGATTGAATGTGATTTGAAGGCAGNNNAGATCG |
| 146_1 | TATGAGTGTGGAGTCGTTGCCCCCACTAGTAGATTATAAGTTCCTGNNNAGATCGG |
| 212_1 | TATGAGTGTGGAGTCGTTGCAGTAGTGGTTACTGGAATGCTCACANNNAGATCGG |
| 283_1 | TATGAGTGTGGAGTCGTTGCCTGTCCTCCTTGTACTCAACTTGATTNNNAGATCGGA |
| 366_1 | TATGAGTGTGGAGTCGTTGCGCACACATGCAAAAATACAGGACANNNAGATCGGA |
| 392_1 | TATGAGTGTGGAGTCGTTGCTAGCCTTTCTGATAGCTCTGTGGTAANNNAGATCGG |
| 566_1 | TATGAGTGTGGAGTCGTTGCAAAAAGGAATGTCAGGGTTGTGGTAANNNAGATCG |
| 796_1 | TATGAGTGTGGAGTCGTTGCCATTTCACACTCTACATCCTTGGGTANNNAGATCGGA |
| 839_1 | TATGAGTGTGGAGTCGTTGCTGTAAAGAACTCTGAAAGCTCAGTNNNAGATCGGA |
| 929_1 | TATGAGTGTGGAGTCGTTGCCAAAGTTTCAGCTCAGATTGATCACCNNNAGATCGG |
| 1093_1 | TATGAGTGTGGAGTCGTTGCAGGTTCTTTCCAATGCTTTTCCANNNAGATCGGAAG |
| 1208_1 | TATGAGTGTGGAGTCGTTGCTGTGTTTTCATGTGTCTGCCAGATTTNNNAGATCGG |
| 1660_1 | TATGAGTGTGGAGTCGTTGCGTGTGTGTGTATAATTGCCTTTATTCNNNAGATCGG |
| 1690_1 | TATGAGTGTGGAGTCGTTGCAAGAACATGCATGCTGAGAAGTCANNNAGATCGGA |
| 2055_1 | TATGAGTGTGGAGTCGTTGCAGAGAAAGCAGAAGCTTGGAAACANNNAGATCGG |
| 2059_1 | TATGAGTGTGGAGTCGTTGCGTGTGTGTGTGACTAAAGCTAACANNNAGATCGGA |
| 2069_1 | TATGAGTGTGGAGTCGTTGCAAGCTGGAGTGCAGAGGTATAATTTNNNAGATCGG |
| 2113_1 | TATGAGTGTGGAGTCGTTGCATCATTTCCATCAGCATGAGGCANNNAGATCGGAAG |
| 2139_1 | TATGAGTGTGGAGTCGTTGCTGGGTTCAATAACATCATACTTCTCANNNAGATCGG |
| 2199_1 | TATGAGTGTGGAGTCGTTGCACAGGAAATTGACAGATAGTTGTGGANNNAGATCG |
| 2232_1 | TATGAGTGTGGAGTCGTTGCTGTGACGTTTTGGTGTAAATGTTTGANNNAGATCGG |
| 2406_1 | TATGAGTGTGGAGTCGTTGCTCAGGCAGAATAAATTTTCCATCTGCNNNAGATCGG |

| | |
|---|---|
| 2417_1 | TATGAGTGTGGAGTCGTTGCTGGTGGTTTCCGCTATAATATTTGCNNNAGATCGG |
| 2460_1 | TATGAGTGTGGAGTCGTTGCCACCCCGCTCTTTTTCTTTTTCNNNAGATCGGAAGA |
| 2588_1 | TATGAGTGTGGAGTCGTTGCTGGGAAGTTTTGAATAAGTTTGGTGGNNNAGATCG |
| 2747_1 | TATGAGTGTGGAGTCGTTGCGACAGATGCATAAGCTTTTGAAACCTNNNAGATCGG |
| 2905_1 | TATGAGTGTGGAGTCGTTGCCTCAAGATTTCATGTCATTTCATTGTNNNAGATCGGA |
| 3441_1 | TATGAGTGTGGAGTCGTTGCAACTGCAGAATGACAGAAGCCAANNNAGATCGGAA |
| 3449_1 | TATGAGTGTGGAGTCGTTGCGGAGTTTTGGTGTTTCACCCCGGNNNAGATCGGAAG |
| 3905_1 | TATGAGTGTGGAGTCGTTGCTGCCCTTACAATATTTTTCCTTTTGANNNAGATCGGA |
| 4063_1 | TATGAGTGTGGAGTCGTTGCACCCATATTTCAAAGTGATGTCATGTNNNAGATCGG |
| 4178_1 | TATGAGTGTGGAGTCGTTGCTGACTAAAAACTGCATTATGTTGTGTNNNAGATCGG |
| 4286_1 | TATGAGTGTGGAGTCGTTGCATACCAACACCACAATCAAGATATGGNNNAGATCGG |
| 4482_1 | TATGAGTGTGGAGTCGTTGCTTGAAACCTTATGGCCCCATCACNNNAGATCGGAAG |
| 4886_1 | TATGAGTGTGGAGTCGTTGCGTCAAACTTCCCCGTTATCCCAANNNAGATCGGAAGA |
| 4913_1 | TATGAGTGTGGAGTCGTTGCTCATATCCGTGGTTTCCCCATCTNNNAGATCGGAAGA |
| 5318_1 | TATGAGTGTGGAGTCGTTGCTTGGAAAGGAAAAGCCAGAATCTNNNAGATCGGAA |
| 5636_1 | TATGAGTGTGGAGTCGTTGCGGAACTATGCGAGGTAGTATGTATGTNNNAGATCG |
| 5700_1 | TATGAGTGTGGAGTCGTTGCGGTTGATTTTCTGCTAATATCAGGGGNNNAGATCGG |
| 5813_1 | TATGAGTGTGGAGTCGTTGCCCATAGAAATTTGTACCTGTGGAGGANNNAGATCG |
| 5878_1 | TATGAGTGTGGAGTCGTTGCAACATTATCTGTAGCAGTGAGGAAAANNNAGATCG |
| 6064_1 | TATGAGTGTGGAGTCGTTGCGGGAGAGAGACTTATGCATGTGTNNNAGATCGGAA |
| 6092_1 | TATGAGTGTGGAGTCGTTGCTTCTTCTGGCAATAGCATCATCATTGNNNAGATCGGA |
| 6750_1 | TATGAGTGTGGAGTCGTTGCAGAAATGAAACATATCCCATGCTCGTNNNAGATCGG |
| 6820_1 | TATGAGTGTGGAGTCGTTGCTGTCTTTATTCATCCACTGATGGGANNNAGATCGGA |
| 7133_1 | TATGAGTGTGGAGTCGTTGCTCTCCTCTCCTTGAGCTTTCTGTACNNNAGATCGGAA |
| 7785_1 | TATGAGTGTGGAGTCGTTGCCCTCGCTGAGCATCTTTCTATATCAANNNAGATCGGA |
| 7817_1 | TATGAGTGTGGAGTCGTTGCACAACACATCAAAAAGATAGCATCCCNNNAGATCGG |
| 7909_1 | TATGAGTGTGGAGTCGTTGCCCGATAACCAATGATTCACAAAGCCTNNNAGATCGG |
| 7995_1 | TATGAGTGTGGAGTCGTTGCTGCTTGTGGTGTGTACTTCTAATTANNNAGATCGGA |
| 8050_1 | TATGAGTGTGGAGTCGTTGCGGGGAGATGAAATGATATTCTAAACCNNNAGATCG |
| 8080_1 | TATGAGTGTGGAGTCGTTGCATAGAACTTAGACAAGTGGCTCCTGANNNAGATCGG |
| 8096_1 | TATGAGTGTGGAGTCGTTGCCTCCTGGTGAAACCTTATTCATCCAGNNNAGATCGGA |
| 8207_1 | TATGAGTGTGGAGTCGTTGCGCTCAAGTAATTATCCTGTCTCAGCANNNAGATCGG |
| 8249_1 | TATGAGTGTGGAGTCGTTGCAACTGCAATCTCTTACTTCTGGGTTCNNNAGATCGGA |
| 8767_1 | TATGAGTGTGGAGTCGTTGCTGGAGTGTAGTGATTTGATCCCTTCANNNAGATCGG |
| 8915_1 | TATGAGTGTGGAGTCGTTGCTCAGTCCTAAAACAAACTTATGAGGTNNNAGATCGG |
| 8986_1 | TATGAGTGTGGAGTCGTTGCGCATAGTCCTTTATTTAGCTGGTCTNNNAGATCGGA |
| 9079_1 | TATGAGTGTGGAGTCGTTGCAGTGGATTTCTTACTTGTAGTCACCTNNNAGATCGG |
| 9381_1 | TATGAGTGTGGAGTCGTTGCATACAAATGTTAGCAGGCTATAAACANNNAGATCGG |
| 9525_1 | TATGAGTGTGGAGTCGTTGCCAAAATAAAAGCCAAAGTGCACTACCNNNAGATCGG |
| 9621_1 | TATGAGTGTGGAGTCGTTGCTTGTTTTCAAAGTGCTAGCCAGCNNNAGATCGGAAG |
| 9841_1 | TATGAGTGTGGAGTCGTTGCTGTCTACTAGTTCAATCTATTGCTGANNNAGATCGGA |
| 9849_1 | TATGAGTGTGGAGTCGTTGCACAGAACAAAACAAGCAAAACACTGANNNAGATCG |
| 10079_1 | TATGAGTGTGGAGTCGTTGCATCCAGTATGCAAGGGTCCTTATNNNAGATCGGAAG |

| | |
|---|---|
| 10083_1 | TATGAGTGTGGAGTCGTTGCACACACCCCTACCTGTATATGCANNNAGATCGGAAG |
| 10101_1 | TATGAGTGTGGAGTCGTTGCACCTTTGACCGAGATCTGAAGGANNNAGATCGGAA |
| 10132_1 | TATGAGTGTGGAGTCGTTGCAAGGGGAAATGGTAATAGTAATAGCANNNAGATCG |
| 10151_1 | TATGAGTGTGGAGTCGTTGCTGCTGCCTTTTTCCTTTATTTTCTCCNNNAGATCGGAA |
| 10205_1 | TATGAGTGTGGAGTCGTTGCAGTTTCTCCATCCTCCTTTTCAGCNNNAGATCGGAAG |
| 10220_1 | TATGAGTGTGGAGTCGTTGCGCCATGAAAGAGCAGATGGTACANNNAGATCGGAA |
| 10233_1 | TATGAGTGTGGAGTCGTTGCAGAGAAAGAAACCTTGGATTCATGGTNNNAGATCG |
| 10244_1 | TATGAGTGTGGAGTCGTTGCTCTTCTGAATGCCGGGTTCCTATNNNAGATCGGAAG |
| 10254_1 | TATGAGTGTGGAGTCGTTGCAATGCCTCTTAGAAGCTAGAGGGGNNNAGATCGGA |
| 10264_1 | TATGAGTGTGGAGTCGTTGCTTATAGCAGGCATGGATTCTGGCNNNAGATCGGAA |
| 10280_1 | TATGAGTGTGGAGTCGTTGCATAAATGCAGGACTGGGCTAGGANNNAGATCGGAA |
| 10293_1 | TATGAGTGTGGAGTCGTTGCAGGAATAAGATGTTTTAGGAGGCACANNNAGATCG |
| 10314_1 | TATGAGTGTGGAGTCGTTGCAGGGAAAAATAAACATGCAGAAGGTNNNAGATCGG |
| 10325_1 | TATGAGTGTGGAGTCGTTGCCCCAAATAAATGGTGTATACCTACATNNNAGATCGG |
| 10335_1 | TATGAGTGTGGAGTCGTTGCTAAGAGCATGTCATCTACGAAATGAGGNNNAGATCG |
| 10394_1 | TATGAGTGTGGAGTCGTTGCACAGGCAGATCTAAAGTAAATGAGGTNNNAGATCG |
| 10415_1 | TATGAGTGTGGAGTCGTTGCACAAAATTCAGGTCAGTTGTGGTGNNNAGATCGGA |
| 10454_1 | TATGAGTGTGGAGTCGTTGCATGGCGGGGTGGAGTTATAATGANNNAGATCGGA |
| 10503_1 | TATGAGTGTGGAGTCGTTGCCTGGTCCATGAGAAATGACAGTGTNNNAGATCGGA |
| 10574_1 | TATGAGTGTGGAGTCGTTGCATTTTCAGCATACCCAAGGGTGGNNNAGATCGGAA |
| 10637_1 | TATGAGTGTGGAGTCGTTGCAGGACACTTCATCCATGTTGCTGNNNAGATCGGAAG |
| 10748_1 | TATGAGTGTGGAGTCGTTGCAGGAGGTGAGTTTCATATTTTTGGTNNNAGATCGG |
| 10751_1 | TATGAGTGTGGAGTCGTTGCTGTCATCGTCAAATAGGGAAAGGTNNNAGATCGGA |
| 10786_1 | TATGAGTGTGGAGTCGTTGCCACACACACTCCTATTGTTGCAGNNNAGATCGGAAG |
| 10839_1 | TATGAGTGTGGAGTCGTTGCTTTTGAGGTGAATTGAATGCATGCANNNAGATCGG |
| 10921_1 | TATGAGTGTGGAGTCGTTGCTCCCCCATTTTGATATAGCATAGCCTNNNAGATCGGA |
| 10938_1 | TATGAGTGTGGAGTCGTTGCATTGTTGTATCCTAAGTGTCCAGTAGNNNAGATCGG |
| 10978_1 | TATGAGTGTGGAGTCGTTGCAGTCCATGACAACGTAGAATGCTGANNNAGATCGG |
| 10998_1 | TATGAGTGTGGAGTCGTTGCAGAAATAACACAAAGGCAAAGTACCANNNAGATCG |
| 11202_1 | TATGAGTGTGGAGTCGTTGCGCAGTTACCCAGAGTTTTTGAAGGTANNNAGATCGG |
| 11235_1 | TATGAGTGTGGAGTCGTTGCGCTTCTTTTTGGGGTGATATGTTCTANNNAGATCGG |
| 11338_1 | TATGAGTGTGGAGTCGTTGCTTGGGTCCTTATGACAGTTTGCTTANNNAGATCGGA |
| 11354_1 | TATGAGTGTGGAGTCGTTGCGGCTGTGGTTGTAGACATCTGAANNNAGATCGGAA |
| 11464_1 | TATGAGTGTGGAGTCGTTGCGAGTGACTACCTCAACATTATCACAGNNNAGATCGG |
| 11572_1 | TATGAGTGTGGAGTCGTTGCACACACACACGGAGACAAATGAANNNAGATCGGAA |
| 11587_1 | TATGAGTGTGGAGTCGTTGCCTCAACAATTATCAACTTGTGGCCAANNNAGATCGG |
| 11632_1 | TATGAGTGTGGAGTCGTTGCCCTCCTGAATTTGTGCAATTTTTGAANNNAGATCGGA |
| 11634_1 | TATGAGTGTGGAGTCGTTGCTCACAGAAAATGAAAAGTTACCCTGANNNAGATCGG |
| 11715_1 | TATGAGTGTGGAGTCGTTGCTTCCAGTATATATCCCACTCCCTTTGNNNAGATCGGA |
| 11716_1 | TATGAGTGTGGAGTCGTTGCAGTGGCTCCTTCTCTATCATCTCTTTNNNAGATCGGA |
| 11795_1 | TATGAGTGTGGAGTCGTTGCTCCTGTCATTTTGTTATTTTGTTGTTNNNAGATCGGA |
| 11890_1 | TATGAGTGTGGAGTCGTTGCTGAACATGCAGTCTCTCTTGGAGATGNNNAGATCGG |
| 11964_1 | TATGAGTGTGGAGTCGTTGCAAGCCTGGAACGGGGACTTTTGAAAATNNNAGATCG |

12147_1 TATGAGTGTGGAGTCGTTGCGGTGGGAAGAGTAATTTCAGAGGTAGNNNAGATC
12322_1 TATGAGTGTGGAGTCGTTGCGGATCTCAGGTTTAGAACTCCTAGTCNNNAGATCGG
12350_1 TATGAGTGTGGAGTCGTTGCAGCCTGAAAAAGAGGTAATTTCTGGTNNNAGATCG
28780_1 TATGAGTGTGGAGTCGTTGCTGGAATACTACTCAACCATACAAGGANNNAGATCGG
28788_1 TATGAGTGTGGAGTCGTTGCTCACATTTTCTTTATCCTTTCATCCANNNAGATCGGAA
28793_1 TATGAGTGTGGAGTCGTTGCAAAACCTTCTAGTGGCTTTCTGTNNNAGATCGGAAG
28813_1 TATGAGTGTGGAGTCGTTGCTGAGTTGGTTCCACATTTTTGCANNNAGATCGGAAG
28815_1 TATGAGTGTGGAGTCGTTGCTGTTGGTTGAAGAGTAAACTGTTTGTNNNAGATCG
28896_1 TATGAGTGTGGAGTCGTTGCACCCAAAAATTTCTCTACAAATGCCTNNNAGATCGGA
28956_1 TATGAGTGTGGAGTCGTTGCTGTCTATGTGTATATACTGCATGTGTNNNAGATCGG
28960_1 TATGAGTGTGGAGTCGTTGCTGCCTAACGATGGATTTCTCAGANNNAGATCGGAAG
29069_1 TATGAGTGTGGAGTCGTTGCTGCTTATCGTAGAAAATTGGGAAAGTNNNAGATCG
29124_1 TATGAGTGTGGAGTCGTTGCAGACTGTGGTTAGTTTTACTTTGATTNNNAGATCGG
29182_1 TATGAGTGTGGAGTCGTTGCACCTGGAAACAAATACACTAAAACCCNNNAGATCGG
29198_1 TATGAGTGTGGAGTCGTTGCAGGAACATACATATGTGCACACACNNNAGATCGGAA
29232_1 TATGAGTGTGGAGTCGTTGCTTCTCTGTATATCCTGTACATTGGATNNNAGATCGGA
29300_1 TATGAGTGTGGAGTCGTTGCAGTGGGTAAGGGAGAAAATTCTCNNNAGATCGGAA
29357_1 TATGAGTGTGGAGTCGTTGCGCTTGCTCACAAATCACACATCTNNNAGATCGGAAG
30751_1 TATGAGTGTGGAGTCGTTGCTGCAAAAGGAAATGCAGGAAGAANNNAGATCGGA
30766_1 TATGAGTGTGGAGTCGTTGCACACACACACATTTTGTAGATATGGANNNAGATCGG
30802_1 TATGAGTGTGGAGTCGTTGCACAGAAGTTCTTTGTTTCTTCCAACTNNNAGATCGGA
31062_1 TATGAGTGTGGAGTCGTTGCGATTCTAGCAGTGGGAAAAGTGCNNNAGATCGGAA
31181_1 TATGAGTGTGGAGTCGTTGCTCTTTGGCTTTTAGGATGCAATGANNNAGATCGGAA
31252_1 TATGAGTGTGGAGTCGTTGCGAGTCATTGTGGCAGAAACAGACNNNAGATCGGAA
31372_1 TATGAGTGTGGAGTCGTTGCGATTATAATGGTGTTGGCCAGGCNNNAGATCGGAA
31406_1 TATGAGTGTGGAGTCGTTGCACCATCATTTCTTTTCACATCAGGTNNNAGATCGGAA
31407_1 TATGAGTGTGGAGTCGTTGCACTCTGTATTTGACTTCTTCCAAGGANNNAGATCGGA
31587_1 TATGAGTGTGGAGTCGTTGCTGCAATATAGTTATTTGACACATGGANNNAGATCGG
31656_1 TATGAGTGTGGAGTCGTTGCTCCCTATAGGCTAATATCTCTAGGCANNNAGATCGGA
31696_1 TATGAGTGTGGAGTCGTTGCAGTTCTCTTTCTGGGATGGAAAANNNAGATCGGAAG
31698_1 TATGAGTGTGGAGTCGTTGCTCAGTTTTCTCAGGGGATTGGTTNNNAGATCGGAAG
31729_1 TATGAGTGTGGAGTCGTTGCTGCCATTTTGGTTGGTTCAATAGTNNNAGATCGGAA
31827_1 TATGAGTGTGGAGTCGTTGCAGTGCAGATATGCCTTCAACATATTGNNNAGATCGG
31828_1 TATGAGTGTGGAGTCGTTGCGGGATCAAATGACCCTCTTGTCTNNNAGATCGGAAG
31906_1 TATGAGTGTGGAGTCGTTGCACATGAAAAAAAGCTCCACTGATCANNNAGATCGGA
31941_1 TATGAGTGTGGAGTCGTTGCACATGTATATGTGTGCGTATATACACNNNAGATCGG
31948_1 TATGAGTGTGGAGTCGTTGCACTCTGGTGATTATTTGCTGATTTCCNNNAGATCGGA
32162_1 TATGAGTGTGGAGTCGTTGCCAGGTTCAAGCAATTCTCCTGCNNNAGATCGGAAGA
32321_1 TATGAGTGTGGAGTCGTTGCCCACACACACATATATACACAACACANNNAGATCGGA
32459_1 TATGAGTGTGGAGTCGTTGCAGGAAATCTGACAATATCTGCTCTCANNNAGATCGG
33077_1 TATGAGTGTGGAGTCGTTGCCGGGTTCAAGCAATTCTGTTTCANNNAGATCGGAAG
33138_1 TATGAGTGTGGAGTCGTTGCAGGAAAGAGATGTAACAACTAACTGTNNNAGATCG
33258_1 TATGAGTGTGGAGTCGTTGCTTCAAGCGTTTTTCCTGCTTCAGNNNAGATCGGAAG

| | |
|---|---|
| 33338_1 | TATGAGTGTGGAGTCGTTGCGATCTCAGCTCACTGCAATCTCTNNNAGATCGGAAGA |
| 33365_1 | TATGAGTGTGGAGTCGTTGCTCATAACAGTCAAGCTTTGGAAGNNNAGATCGGAAG |
| 33434_1 | TATGAGTGTGGAGTCGTTGCTCCCCTTCCATTTCTTTATAATACGTNNNAGATCGGAA |
| 33603_1 | TATGAGTGTGGAGTCGTTGCGCTGATCAGCTAATTTTGAATTCCAANNNAGATCGGA |
| 33605_1 | TATGAGTGTGGAGTCGTTGCTGGCCGGGATTTCTAGTATGATGNNNAGATCGGAAG |
| 33805_1 | TATGAGTGTGGAGTCGTTGCAGTAGACAGATTCAATGCAAATTCCANNNAGATCGG |
| 33855_1 | TATGAGTGTGGAGTCGTTGCAAATAAAGGTCCAATATCCCAAACATNNNAGATCGGA |
| 34121_1 | TATGAGTGTGGAGTCGTTGCGGCAGACATAAATCCAACTATATCAANNNAGATCGGA |
| 35095_1 | TATGAGTGTGGAGTCGTTGCAGCAACTTTTGGATTTGTTTGATCTNNNAGATCGGAA |
| 35366_1 | TATGAGTGTGGAGTCGTTGCCTCCTGAGTAGGTGGGACTACANNNAGATCGGAAGA |
| 35381_1 | TATGAGTGTGGAGTCGTTGCTGTCCTCTTTTTTTTTTTGAGACGGANNNAGATCGGA |
| 35424_1 | TATGAGTGTGGAGTCGTTGCCACAAGCCAATGTCATATAGTTCACANNNAGATCGGA |
| 35467_1 | TATGAGTGTGGAGTCGTTGCAGTTTCTCATTTATTTGTCCTCATTGTNNNAGATCGGA |
| 35487_1 | TATGAGTGTGGAGTCGTTGCCTCAAGCAATTCTTCTGCCTCAGNNNAGATCGGAAGA |
| 35506_1 | TATGAGTGTGGAGTCGTTGCTTCTAGTGGCACAATCTCTCCACNNNAGATCGGAAGA |
| 35549_1 | TATGAGTGTGGAGTCGTTGCTCTAACTGCCAAGCTGAACTGATNNNAGATCGGAAG |
| 35608_1 | TATGAGTGTGGAGTCGTTGCGTCTCAGCCTCCCAAGTAGACTANNNAGATCGGAAGA |
| 35637_1 | TATGAGTGTGGAGTCGTTGCCCTTGTAGCGTACTTCTTCAATGANNNAGATCGGAAG |
| 35729_1 | TATGAGTGTGGAGTCGTTGCGCACACTACAAACTTCACTTCCCNNNAGATCGGAAGA |
| 35736_1 | TATGAGTGTGGAGTCGTTGCACCTCCCAGATTCAAGCAATTCTNNNAGATCGGAAGA |
| 35834_1 | TATGAGTGTGGAGTCGTTGCGCCTTCCTAGTAGCTGGGATTACNNNAGATCGGAAG |
| 36054_1 | TATGAGTGTGGAGTCGTTGCAGCTTTGGTTCTTTGATGTTCTATCCNNNAGATCGGA |
| 36164_1 | TATGAGTGTGGAGTCGTTGCAGTCAAACATACTTCTGCTTTATCCANNNAGATCGGA |
| 36194_1 | TATGAGTGTGGAGTCGTTGCTGCATTATTCTTACTGTATGTGGAGANNNAGATCGGA |
| 36222_1 | TATGAGTGTGGAGTCGTTGCAGCTACATACATGGATAGATAGAAAGANNNAGATCG |
| 36292_1 | TATGAGTGTGGAGTCGTTGCGTGGCCTGGCAATAAAAATAAAATCANNNAGATCGG |
| 36395_1 | TATGAGTGTGGAGTCGTTGCACATGTACCACTCCATTACTAAATCANNNAGATCGGA |
| 36472_1 | TATGAGTGTGGAGTCGTTGCAGAACTTTTGAAAACACAGAACATGANNNAGATCGG |
| 36760_1 | TATGAGTGTGGAGTCGTTGCTCAAGAACTTCAGTAACGAATATGGTNNNAGATCGG |
| 36781_1 | TATGAGTGTGGAGTCGTTGCTCTTTTGTAACATAGAGGAGAAAGACANNNAGATCG |
| 36801_1 | TATGAGTGTGGAGTCGTTGCTCCTGATGAAGATGCAGAGAAACTNNNAGATCGGAA |
| 36838_1 | TATGAGTGTGGAGTCGTTGCTGGCATCTACTACTTCTGAATTGTCANNNAGATCGGA |
| 36997_1 | TATGAGTGTGGAGTCGTTGCACGGAGAAATGAAATACAATCTCTGTNNNAGATCGG |
| 37054_1 | TATGAGTGTGGAGTCGTTGCAAACTTGTAAAAAGAGAACTACCACANNNAGATCGG |
| 37058_1 | TATGAGTGTGGAGTCGTTGCGAGGAAAGTAGAGTATCATTTCAGTGTNNNAGATCG |
| 37086_1 | TATGAGTGTGGAGTCGTTGCGAGTCAGAATTGGTAAATGATGAATCTNNNAGATCG |
| 37117_1 | TATGAGTGTGGAGTCGTTGCACCACACTCTAATACAAAACAAAGTTNNNAGATCGGA |
| 37124_1 | TATGAGTGTGGAGTCGTTGCACCACAGAAAAAGATGTACTAAAAGGANNNAGATCG |
| 37183_1 | TATGAGTGTGGAGTCGTTGCTCTGTAGTCCTAGAACTATTTGCTCANNNAGATCGGA |
| 37184_1 | TATGAGTGTGGAGTCGTTGCTGGAAAAAGAAAGAAATGCAACAATGNNNAGATCG |
| 37454_1 | TATGAGTGTGGAGTCGTTGCAGAAAAGGAAAAAAAGCTAGGAAACTNNNAGATCG |
| 37500_1 | TATGAGTGTGGAGTCGTTGCAGAGCTCCTTTTATACTCAGGAAACTNNNAGATCGGA |
| 38231_1 | TATGAGTGTGGAGTCGTTGCACCCACATAAACAAACAAAAAACTCANNNAGATCGGA |

| | |
|---|---|
| 38232_1 | TATGAGTGTGGAGTCGTTGCAGTGGTTATTTCTGATTATGTGTGTTNNNAGATCGG |
| 38265_1 | TATGAGTGTGGAGTCGTTGCAGCAAAGCTAACCTTCAGAAATGANNNAGATCGGAA |
| 38272_1 | TATGAGTGTGGAGTCGTTGCTGTGTATGTCTATAAAGCTTAGCTCANNNAGATCGG |
| 38289_1 | TATGAGTGTGGAGTCGTTGCGTTGCCAAGAACATAAACTGGGGNNNAGATCGGAA |
| 38346_1 | TATGAGTGTGGAGTCGTTGCTGACAGACCATTAGTGAGATTATCCANNNAGATCGG |
| 38354_1 | TATGAGTGTGGAGTCGTTGCGTCTCTTCAAGAAATGGTGCTGGNNNAGATCGGAA |
| 38822_1 | TATGAGTGTGGAGTCGTTGCTGTACCACTTTCAAAAAAATGAGAGTNNNAGATCGG |
| 38837_1 | TATGAGTGTGGAGTCGTTGCACTGAACAGATCAATCATGAATTCCANNNAGATCGG |
| 38969_1 | TATGAGTGTGGAGTCGTTGCAGCTTATCTTGCTTCATATCCTTCCTNNNAGATCGGA |
| 38986_1 | TATGAGTGTGGAGTCGTTGCTCACAAAAATCTGGAAAGATTACAGTNNNAGATCGG |
| 39004_1 | TATGAGTGTGGAGTCGTTGCTCAAATGCTTTTTTTCTCACCTACTGANNNAGATCGG |
| 39034_1 | TATGAGTGTGGAGTCGTTGCTCCATATAGGAAAAGAGGAAGTCACANNNAGATCG |
| 39240_1 | TATGAGTGTGGAGTCGTTGCTTCAGTGACTTTCAATCTATGTTTCTNNNAGATCGGA |
| 39289_1 | TATGAGTGTGGAGTCGTTGCAGTACTCTCAGTGAAGAAATCTAAGTNNNAGATCGG |
| 39480_1 | TATGAGTGTGGAGTCGTTGCTGGATCAACTACAGCTATCTGCANNNAGATCGGAAG |
| 39488_1 | TATGAGTGTGGAGTCGTTGCACAAAGCTGTTTGATAACTGAATGGANNNAGATCG |
| 39514_1 | TATGAGTGTGGAGTCGTTGCAGAAAGAATACAAGTTGAAAGGAGCTNNNAGATCG |
| 39523_1 | TATGAGTGTGGAGTCGTTGCCATGACTGCAAACTGTCACTTCCNNNAGATCGGAAG |
| 39547_1 | TATGAGTGTGGAGTCGTTGCTGAAAGGACAAAAAGATAAATCCAGTNNNAGATCG |
| 39548_1 | TATGAGTGTGGAGTCGTTGCTGAAAAGTCAAAGAACTTAGAATAGCTNNNAGATCG |
| 39713_1 | TATGAGTGTGGAGTCGTTGCGGGAAGAAATGAGGAGATGTAGGTNNNAGATCGG |
| 39730_1 | TATGAGTGTGGAGTCGTTGCTGGTTCACTTACCTCAATTAGTTTCCNNNAGATCGGA |
| 40040_1 | TATGAGTGTGGAGTCGTTGCAGGAAAAACATTAGAGAAATCATCGANNNAGATCG |
| 40048_1 | TATGAGTGTGGAGTCGTTGCGATGAAGGTAAAATAACTGTGGATGANNNAGATCG |
| 40121_1 | TATGAGTGTGGAGTCGTTGCATCACGTCTATTGAACAAACTCATAANNNAGATCGG |
| 40301_1 | TATGAGTGTGGAGTCGTTGCTGACAATACATATCAGCATGAATGAANNNAGATCGG |
| 40623_1 | TATGAGTGTGGAGTCGTTGCTTGGTAGAAAAAAGTAAAGCCTAGTTNNNAGATCG |
| 40708_1 | TATGAGTGTGGAGTCGTTGCTGAACCCTTATTTCATACCATACACANNNAGATCGGA |
| 40864_1 | TATGAGTGTGGAGTCGTTGCACCAGAAAGTTTATATCTTGGTCAAANNNAGATCGG |
| 40872_1 | TATGAGTGTGGAGTCGTTGCGGAACAGTTTGGTACTCGAGTCTNNNAGATCGGAA |
| 40877_1 | TATGAGTGTGGAGTCGTTGCGCTACTGCCAATGATAATACTTGACANNNAGATCGG |
| 41035_1 | TATGAGTGTGGAGTCGTTGCGTACAGTGGCAAAATCATGGCTTNNNAGATCGGAA |
| 41056_1 | TATGAGTGTGGAGTCGTTGCATTCATTGCTGGTGGAAATGCAANNNAGATCGGAA |
| 41169_1 | TATGAGTGTGGAGTCGTTGCACACAACACATGTCTAAATAATGGGTNNNAGATCGG |
| 41182_1 | TATGAGTGTGGAGTCGTTGCAGTCAGTGAACTTGAAGATAGAGCANNNAGATCGG |
| 41328_1 | TATGAGTGTGGAGTCGTTGCACAGTCAAAGGACAAATATCTGTTCTNNNAGATCGG |
| 41360_1 | TATGAGTGTGGAGTCGTTGCGGGTGAGGAACTGACAGAATTCANNNAGATCGGAA |
| 41449_1 | TATGAGTGTGGAGTCGTTGCTGTCCTGTGATAAACCTCAGTATTGTNNNAGATCGG |
| 41520_1 | TATGAGTGTGGAGTCGTTGCAGACAGAAAGACAAGAAAGAAAGACANNNAGATCG |
| 41637_1 | TATGAGTGTGGAGTCGTTGCATGGAGTACAGTGGCACAATCTTNNNAGATCGGAA |
| 41684_1 | TATGAGTGTGGAGTCGTTGCACCACACCCAGCTCATTTTTTTTNNNAGATCGGAAGA |
| 41816_1 | TATGAGTGTGGAGTCGTTGCGATCTCGGCTCACTGCAACTTCNNNAGATCGGAAGA |
| 41881_1 | TATGAGTGTGGAGTCGTTGCTCATAGCTATCTAGTGATTGGACAGANNNAGATCGG |

| | |
|---|---|
| 41918_1 | TATGAGTGTGGAGTCGTTGCCGAGTAGCTGGAATTACAGGTGTNNNAGATCGGAA |
| 42263_1 | TATGAGTGTGGAGTCGTTGCTGTTTTTATTGCCTAATTGCTCTGGTNNNAGATCGGA |
| 42317_1 | TATGAGTGTGGAGTCGTTGCCACATAGAAGTCACTGGTCCCAANNNAGATCGGAAG |
| 42339_1 | TATGAGTGTGGAGTCGTTGCAGAGTTTCATCAATTTGGTCAAACTNNNAGATCGGA |
| 42347_1 | TATGAGTGTGGAGTCGTTGCTATACATGTGTTCCCTCCGCAGNNNAGATCGGAAGA |
| 42418_1 | TATGAGTGTGGAGTCGTTGCATCATGGCTAACTGAAGTCTGCANNNAGATCGGAAG |
| 42433_1 | TATGAGTGTGGAGTCGTTGCAGAAAGGGAGAAAGAGAGAAGAAAGANNNAGATC |
| 42500_1 | TATGAGTGTGGAGTCGTTGCTATCACTCTTTCACCCAGGCTGNNNAGATCGGAAGA |
| 42535_1 | TATGAGTGTGGAGTCGTTGCACAGAATTCAAAAGAAAAGGGCANNNAGATCGGAA |
| 42559_1 | TATGAGTGTGGAGTCGTTGCTTCCACTAACAGGAGAACTTCAANNNAGATCGGAAG |
| 42602_1 | TATGAGTGTGGAGTCGTTGCTGTTTTTATTGCCTAATTGCTCTGGTNNNAGATCGGA |
| 42652_1 | TATGAGTGTGGAGTCGTTGCTTTTAGGTGCAGGGTCTTGTTCTNNNAGATCGGAAG |
| 42876_1 | TATGAGTGTGGAGTCGTTGCTCCAGTCAACCACTAATAGGAATGTNNNAGATCGGA |
| 42878_1 | TATGAGTGTGGAGTCGTTGCACACCATGGAGTACTATTCAGTCANNNAGATCGGAA |
| 42884_1 | TATGAGTGTGGAGTCGTTGCTGATGCAACATGTTATATGTAGTGCTNNNAGATCGG |
| 43153_1 | TATGAGTGTGGAGTCGTTGCATCTTCTCTTTTGGTCATTTGAGTAANNNAGATCGGA |
| 43185_1 | TATGAGTGTGGAGTCGTTGCTCTGTGAAGAGAGATAGTTTTCCCTNNNAGATCGGA |
| 43281_1 | TATGAGTGTGGAGTCGTTGCGTCATCAGGTTTCATTTATTGGAGGTNNNAGATCGG |
| 43296_1 | TATGAGTGTGGAGTCGTTGCACACCATAGCATACTAATCAGCCANNNAGATCGGAA |
| 43300_1 | TATGAGTGTGGAGTCGTTGCTTTTTGTGGTTACCATAAGGATTACANNNAGATCGG |
| 43302_1 | TATGAGTGTGGAGTCGTTGCTTCAGAAAGGAAACACAAACCACNNNAGATCGGAAG |
| 43380_1 | TATGAGTGTGGAGTCGTTGCAGGTTCAAGCAATTCTCCTGTCTNNNAGATCGGAAG |
| 43435_1 | TATGAGTGTGGAGTCGTTGCCACGTACGGCTAATTTTTTGTGCNNNAGATCGGAAG |
| 43916_1 | TATGAGTGTGGAGTCGTTGCTGCACATTCAATAACATTCTCAGAGANNNAGATCGG |
| 44224_1 | TATGAGTGTGGAGTCGTTGCTCAAATGACATAAACCAGGAGGGANNNAGATCGGA |
| 44450_1 | TATGAGTGTGGAGTCGTTGCATCTTTGCAGAAATTATATCAGTGAGANNNAGATCG |
| 44499_1 | TATGAGTGTGGAGTCGTTGCTGATAGCATTCGTGTCTATATAGTGTNNNAGATCGG |
| 44501_1 | TATGAGTGTGGAGTCGTTGCACAAGGCTATAATAACCAAAACAGCANNNAGATCGG |
| 44521_1 | TATGAGTGTGGAGTCGTTGCACCTCTGGGAAGTGGAATTCTATGNNNAGATCGGA |
| 44532_1 | TATGAGTGTGGAGTCGTTGCAGAGAGAGAGAGAGAGAGAGAATATTCANNNAGATC |
| 44535_1 | TATGAGTGTGGAGTCGTTGCTGCTGCTATAAACATGCATGTGCNNNAGATCGGAAG |
| 44539_1 | TATGAGTGTGGAGTCGTTGCACACATATAGTGGAATACTACTCAGTNNNAGATCGG |
| 44549_1 | TATGAGTGTGGAGTCGTTGCCACACACCACATTTTGTTAGTATTCANNNAGATCGGA |
| 44552_1 | TATGAGTGTGGAGTCGTTGCCAATGCCACGTTTCTTCTTGCTANNNAGATCGGAAGA |
| 44984_1 | TATGAGTGTGGAGTCGTTGCACACATGGTTCTTTTACAAAAATGCGNNNAGATCGG |
| 45102_1 | TATGAGTGTGGAGTCGTTGCACTGCTAGTTGTCTGTGTTTGGANNNAGATCGGAAG |
| 45741_1 | TATGAGTGTGGAGTCGTTGCAAAAGGATGCCAAGGGGATTTTGNNNAGATCGGAA |
| 45795_1 | TATGAGTGTGGAGTCGTTGCGAGCGTGAGAGAGAGAGAGAAAGNNNAGATCGGA |
| 45804_1 | TATGAGTGTGGAGTCGTTGCCCTCCTAAGCTCAAGTGATCCTCNNNAGATCGGAAGA |
| 45926_1 | TATGAGTGTGGAGTCGTTGCACCTAAAGAACTAAAATGCAAGTGCTNNNAGATCGG |
| 46485_1 | TATGAGTGTGGAGTCGTTGCGGTTCAAGTGATTCTGTTGCCTCNNNAGATCGGAAG |
| 46487_1 | TATGAGTGTGGAGTCGTTGCAGGCTGGTCTCAAACTTCTGATCNNNAGATCGGAAG |
| 46531_1 | TATGAGTGTGGAGTCGTTGCGGACATTCTAATAGGTGTGTAGTGGTNNNAGATCG |

| | |
|---|---|
| 46670_1 | TATGAGTGTGGAGTCGTTGCTCAGTCTCCCAAGTAGCTAGGATNNNAGATCGGAAG |
| 47128_1 | TATGAGTGTGGAGTCGTTGCGATCTCAGCTCACTGCAACCTTANNNAGATCGGAAGA |
| 47194_1 | TATGAGTGTGGAGTCGTTGCAGGTTCAAGTGATTCTCGTGTCTNNNAGATCGGAAG |
| 47709_1 | TATGAGTGTGGAGTCGTTGCACGTGCAGCCAAACAAAACTAATNNNAGATCGGAAG |
| 47807_1 | TATGAGTGTGGAGTCGTTGCCAGGTTCAAGCAATTCTCCTGCNNNAGATCGGAAGA |
| 47877_1 | TATGAGTGTGGAGTCGTTGCCCAAGTAGCTGAGACTACATGCANNNAGATCGGAAG |
| 48088_1 | TATGAGTGTGGAGTCGTTGCGTTCAAGGGATTATCCTGCCTCANNNAGATCGGAAG |
| 48311_1 | TATGAGTGTGGAGTCGTTGCAAAGGCAAAGCATCTTTGTACCCNNNAGATCGGAAG |
| 48395_1 | TATGAGTGTGGAGTCGTTGCCACCACACCCAGCTAATTTTTGTNNNAGATCGGAAGA |
| 48576_1 | TATGAGTGTGGAGTCGTTGCAGCCACAGTTTTTTTTTTTTCCTTCCNNNAGATCGGAA |
| 48657_1 | TATGAGTGTGGAGTCGTTGCTGATTCTCACCTAGAACAATATTGGTNNNAGATCGGA |
| 48855_1 | TATGAGTGTGGAGTCGTTGCGGTTCAAGTGATTCTTCTGCCTCNNNAGATCGGAAGA |
| 48926_1 | TATGAGTGTGGAGTCGTTGCAGGCAAGTCAGATTAGCTTGTGANNNAGATCGGAA |
| 48977_1 | TATGAGTGTGGAGTCGTTGCCTTCCCTGGATAGTGAGCTTCTCNNNAGATCGGAAGA |
| 49055_1 | TATGAGTGTGGAGTCGTTGCACTGATGCCATCATAGCTCACTGNNNAGATCGGAAGA |
| 49328_1 | TATGAGTGTGGAGTCGTTGCAAAGATAGTCATGGCCAGTGGTCNNNAGATCGGAA |
| 49497_1 | TATGAGTGTGGAGTCGTTGCTTTTGGTTCTTTATCCTGCTGGCNNNAGATCGGAAGA |
| 49499_1 | TATGAGTGTGGAGTCGTTGCAGCCTCAGGAGTAGATGGGATTANNNAGATCGGAA |
| 49939_1 | TATGAGTGTGGAGTCGTTGCCATGGGTGAGAATGGGGAAGTAANNNAGATCGGAA |
| 50039_1 | TATGAGTGTGGAGTCGTTGCACTTCTCAGCCATAAAATACAATGAANNNAGATCGGA |
| 50151_1 | TATGAGTGTGGAGTCGTTGCTGAGGATGATGTGCAAGCTTACTNNNAGATCGGAA |
| 50841_1 | TATGAGTGTGGAGTCGTTGCACCTACATTTAGTACACAAGCAGTCTNNNAGATCGGA |
| 50880_1 | TATGAGTGTGGAGTCGTTGCTTTTCTGCTAGGTTCATGGGTGTNNNAGATCGGAAG |
| 51018_1 | TATGAGTGTGGAGTCGTTGCAGTGAAGAAACACAAGAACTATAGCANNNAGATCG |
| 51120_1 | TATGAGTGTGGAGTCGTTGCTATCTCTCTCAGCGTATGGCCTANNNAGATCGGAAGA |
| 51309_1 | TATGAGTGTGGAGTCGTTGCGTCTGTGTATTGGGGAGGTAAGGNNNAGATCGGAA |
| 51731_1 | TATGAGTGTGGAGTCGTTGCTGCTATTGAGAAACAGATAAGTTGCTNNNAGATCGG |
| 52023_1 | TATGAGTGTGGAGTCGTTGCTGCCTTGTATTATGTAACCTTGATGGNNNAGATCGGA |
| 53133_1 | TATGAGTGTGGAGTCGTTGCAGGTGTTTTGAAAACATATGGAACCTNNNAGATCGG |
| 53195_1 | TATGAGTGTGGAGTCGTTGCCTATCATGGGTCACTACAGCCTCNNNAGATCGGAAGA |
| 53376_1 | TATGAGTGTGGAGTCGTTGCACATCTACATGCATATATGTGTGTGTNNNAGATCGGA |
| 53420_1 | TATGAGTGTGGAGTCGTTGCAGAAAGAGAGAGAAAGAAAGAAAGGANNNAGATC |
| 7_1 | TATGAGTGTGGAGTCGTTGCGACCAGAATGGATCACAGATGATCGNNNAGATCGGA |
| 67_1 | TATGAGTGTGGAGTCGTTGCGCATCTATTCACAGCTAAGCAGCTCNNNAGATCGGAA |
| 261_1 | TATGAGTGTGGAGTCGTTGCCTTGAACTTCTCCCAGAAAATGGTTNNNAGATCGGAA |
| 302_1 | TATGAGTGTGGAGTCGTTGCGATTCCTAATTGCCAGCCAGTANNNAGATCGGAAGA |
| 365_1 | TATGAGTGTGGAGTCGTTGCGAGTTTTGATTTGCTGATTGCCAGANNNAGATCGGA |
| 413_1 | TATGAGTGTGGAGTCGTTGCTATCCATTAGCCAATTCTGTGAGCTNNNAGATCGGAA |
| 452_1 | TATGAGTGTGGAGTCGTTGCCCCAAAAGTAGGGCTCTAAAGAAGCNNNAGATCGGA |
| 532_1 | TATGAGTGTGGAGTCGTTGCGAATACAACAATTCCTGCCATCATGNNNAGATCGGAA |
| 599_1 | TATGAGTGTGGAGTCGTTGCTGTTGTCTTCCAGTATGGGACCANNNAGATCGGAAG |
| 736_1 | TATGAGTGTGGAGTCGTTGCGACAGGTGATTTCTGCATTGTGAACNNNAGATCGGA |
| 797_1 | TATGAGTGTGGAGTCGTTGCAGAACAGGATGGGATAATCCTAAGTNNNAGATCGGA |

| | |
|---|---|
| 920_1 | TATGAGTGTGGAGTCGTTGCCTCCACTAGACCTATTCTCTGTGGTNNNAGATCGGAA |
| 1478_1 | TATGAGTGTGGAGTCGTTGCATCAATGCATAGGCTTAGCCCTTNNNAGATCGGAAG |
| 2056_1 | TATGAGTGTGGAGTCGTTGCTGCCAGGAAATTGAAATCAACAACANNNAGATCGGA |
| 2094_1 | TATGAGTGTGGAGTCGTTGCCCATGAGTTCATACTGTTGTCTCCANNNAGATCGGAA |
| 2101_1 | TATGAGTGTGGAGTCGTTGCTACTTACAGTTCCTTCTTTACAGCANNNAGATCGGAA |
| 2107_1 | TATGAGTGTGGAGTCGTTGCATTGACAGCATCACAGCATAGCANNNAGATCGGAAG |
| 2125_1 | TATGAGTGTGGAGTCGTTGCCTGGGGTTACTATAAGGCTTGCANNNAGATCGGAA |
| 2145_1 | TATGAGTGTGGAGTCGTTGCTCACGTTCCATATTTTCCTGTCANNNAGATCGGAAGA |
| 2197_1 | TATGAGTGTGGAGTCGTTGCTCTTTGGGCATTGAAACTACTGGANNNAGATCGGAA |
| 2222_1 | TATGAGTGTGGAGTCGTTGCGCAACTTGGATTGAGCTATAGGTGANNNAGATCGG |
| 2252_1 | TATGAGTGTGGAGTCGTTGCCGCCCAGCCTTGATTTCTATTTTTATNNNAGATCGGA |
| 2256_1 | TATGAGTGTGGAGTCGTTGCCAAGTCCCAAGATCTGCAGTCTANNNAGATCGGAAG |
| 2264_1 | TATGAGTGTGGAGTCGTTGCTCATAATTTCAAGCATTTCCCCTGCNNNAGATCGGAA |
| 2266_1 | TATGAGTGTGGAGTCGTTGCGTTCAGCACCACAGGTGTAGTTANNNAGATCGGAA |
| 2301_1 | TATGAGTGTGGAGTCGTTGCAACATGCACACATCCCTACTCACNNNAGATCGGAAGA |
| 2321_1 | TATGAGTGTGGAGTCGTTGCTTTCTCATCGATTCATCCATCAGTGNNNAGATCGGAA |
| 2413_1 | TATGAGTGTGGAGTCGTTGCCGCTGAGGGTACATAAGATGTGCNNNAGATCGGAA |
| 2418_1 | TATGAGTGTGGAGTCGTTGCACCAATGTCATTCTCTTGCTTTTGCNNNAGATCGGAA |
| 2511_1 | TATGAGTGTGGAGTCGTTGCGAGCTGGATTGAGTTGAGGCTAGNNNAGATCGGAA |
| 2727_1 | TATGAGTGTGGAGTCGTTGCTTCTCTCTCTCAAACTCACCACTNNNAGATCGGAAGA |
| 2758_1 | TATGAGTGTGGAGTCGTTGCAGCCAAGAAGTTACCAATTTCCCTNNNAGATCGGAA |
| 2772_1 | TATGAGTGTGGAGTCGTTGCTGTTCCCAGTGTTTTGTAGTCCTNNNAGATCGGAAG |
| 2774_1 | TATGAGTGTGGAGTCGTTGCCTATTCTTGCACCTTCGCTTCCTNNNAGATCGGAAGA |
| 2853_1 | TATGAGTGTGGAGTCGTTGCACATTTCAGACAAAACCCTGTTAGTNNNAGATCGGA |
| 2876_1 | TATGAGTGTGGAGTCGTTGCCAATGTGTACCTCTCCTGACATGTNNNAGATCGGAA |
| 2890_1 | TATGAGTGTGGAGTCGTTGCTGTAACTATGAAACCAGCTGTCTGTNNNAGATCGGA |
| 3313_1 | TATGAGTGTGGAGTCGTTGCTCTGTCCTCTATAACACCTGAATGANNNAGATCGGA |
| 3419_1 | TATGAGTGTGGAGTCGTTGCCTCCAGGACAGAAATTAGCAAGCTANNNAGATCGG |
| 3887_1 | TATGAGTGTGGAGTCGTTGCCAATACTCATCCTCCTCCTACCCTCNNNAGATCGGAAG |
| 3891_1 | TATGAGTGTGGAGTCGTTGCGAAAAACTGGGAGGTGGTCATAGGANNNAGATCG |
| 3901_1 | TATGAGTGTGGAGTCGTTGCGGTTCTCTACTTCTTTTCTCTGTGGNNNAGATCGGAA |
| 4276_1 | TATGAGTGTGGAGTCGTTGCGCAAGGTCAACAGAAACTTGTAAAGNNNAGATCGG |
| 4872_1 | TATGAGTGTGGAGTCGTTGCATGGATTGGTTCTGTTTCTCTGAAANNNAGATCGGA |
| 4950_1 | TATGAGTGTGGAGTCGTTGCGACACCATAAAGAGCAAGCAACANNNAGATCGGAA |
| 5282_1 | TATGAGTGTGGAGTCGTTGCCCTCTTTCCATTCTAAGCACTCTCANNNAGATCGGAA |
| 5312_1 | TATGAGTGTGGAGTCGTTGCCACTTAGGATCGCAGGAATCAGGNNNAGATCGGAA |
| 5314_1 | TATGAGTGTGGAGTCGTTGCACACACACACACTAAACAGAGCCNNNAGATCGGAAG |
| 5315_1 | TATGAGTGTGGAGTCGTTGCGCCATACCTCTCTGAGTATTTAGCTNNNAGATCGGA |
| 5513_1 | TATGAGTGTGGAGTCGTTGCTGTCAAATATCTCCTGTGTGTGTGTNNNAGATCGGA |
| 5576_1 | TATGAGTGTGGAGTCGTTGCAGCATACCAAAAGATTGTAACATCANNNAGATCGGA |
| 5645_1 | TATGAGTGTGGAGTCGTTGCTCCATTGTCTTTTTGTTTGCATGGTNNNAGATCGGAA |
| 5704_1 | TATGAGTGTGGAGTCGTTGCAACTATTGTCACCATGCTCTCCGNNNAGATCGGAAG |
| 5827_1 | TATGAGTGTGGAGTCGTTGCGAAATCTTGATGGCCTCATTGCANNNAGATCGGAAG |

| | |
|---|---|
| 5853_1 | TATGAGTGTGGAGTCGTTGCTTACATCATGATCAGCCATGGTCACNNNAGATCGGA |
| 6037_1 | TATGAGTGTGGAGTCGTTGCATGGGATGATCTTGTCTCACTGCTANNNAGATCGGA |
| 6172_1 | TATGAGTGTGGAGTCGTTGCTCATTTGTGTAAAGGGGATGGGGAANNNAGATCGG |
| 6599_1 | TATGAGTGTGGAGTCGTTGCTATTCCTCCTTTTGCATCTGGCTTTNNNAGATCGGAA |
| 6716_1 | TATGAGTGTGGAGTCGTTGCAAAATAATGCGTGAAAAGTGCTTGGNNNAGATCGG |
| 6826_1 | TATGAGTGTGGAGTCGTTGCGCCTAGTTCTCCTCTGTTCTTTGTNNNAGATCGGAAG |
| 6829_1 | TATGAGTGTGGAGTCGTTGCCTTTTTGGAGCTCCCTTGTATGATNNNAGATCGGAA |
| 6957_1 | TATGAGTGTGGAGTCGTTGCCACCACCACTCGTATTACACTGTGNNNAGATCGGAA |
| 7020_1 | TATGAGTGTGGAGTCGTTGCCAGCACATCTGGCAGAAAAATTCTTNNNAGATCGGA |
| 7331_1 | TATGAGTGTGGAGTCGTTGCAGGATTCGTACTTTACCTTTGGAGGNNNAGATCGGA |
| 7549_1 | TATGAGTGTGGAGTCGTTGCAGATGTAAGATGTACATGGTGCAGGNNNAGATCGG |
| 7637_1 | TATGAGTGTGGAGTCGTTGCTCAGTGGTTATGGCCAATAGCTNNNAGATCGGAAG |
| 7666_1 | TATGAGTGTGGAGTCGTTGCGGGATTATAGGTGATTCTCCTGCCTNNNAGATCGGA |
| 7677_1 | TATGAGTGTGGAGTCGTTGCGAATGCCTCCTTCTCTCTGATGTTCNNNAGATCGGAA |
| 7703_1 | TATGAGTGTGGAGTCGTTGCCCCTTACCTGATTTTGGTATCAGGGNNNAGATCGGA |
| 7802_1 | TATGAGTGTGGAGTCGTTGCTTTGGTAGAGATGGAGTTTTGCATGNNNAGATCGG |
| 7807_1 | TATGAGTGTGGAGTCGTTGCTCAATTTCAGTCTTGGGAGGGTGTANNNAGATCGG |
| 7889_1 | TATGAGTGTGGAGTCGTTGCGACATCAAGTGTTTGACTGAAGGACNNNAGATCGG |
| 7911_1 | TATGAGTGTGGAGTCGTTGCCCTCATTATTTTTGGAGTGCTGTGTNNNAGATCGGA |
| 7931_1 | TATGAGTGTGGAGTCGTTGCAAAATGCTGTTGGAGACTGGCAAAGNNNAGATCGG |
| 8015_1 | TATGAGTGTGGAGTCGTTGCCAAGCAATAACTGATAAAGGCCAAANNNAGATCGG |
| 8018_1 | TATGAGTGTGGAGTCGTTGCACGAGACCTGAATTATTCTGCTACCNNNAGATCGGA |
| 8030_1 | TATGAGTGTGGAGTCGTTGCTCAGCTAGGCTCTGGACTACAGTNNNAGATCGGAAG |
| 8037_1 | TATGAGTGTGGAGTCGTTGCACTGCCTTCATGATTTAGGATGGACNNNAGATCGGA |
| 8055_1 | TATGAGTGTGGAGTCGTTGCTACAAGAGGTGCAAAGAAGAACAGGNNNAGATCG |
| 8125_1 | TATGAGTGTGGAGTCGTTGCTCGGATTACAGGTGTGAGGAGTTACNNNAGATCGG |
| 8236_1 | TATGAGTGTGGAGTCGTTGCTGAAGTTTTGTTCTTGTTGCTGAGGNNNAGATCGGA |
| 8241_1 | TATGAGTGTGGAGTCGTTGCCATTGGTTTCCTAGGCCTGTTAGAANNNAGATCGGA |
| 8251_1 | TATGAGTGTGGAGTCGTTGCGGGTCTCCCGTTCAAAGTGTTGNNNAGATCGGAAG |
| 8276_1 | TATGAGTGTGGAGTCGTTGCGGTTTGTTTTTGTATTTGTCTCGCTNNNAGATCGGAA |
| 8286_1 | TATGAGTGTGGAGTCGTTGCCATCATACCAATTGCGTCTTTCTGANNNAGATCGGAA |
| 8293_1 | TATGAGTGTGGAGTCGTTGCTGTGATTGGTAACAGGTCAGGAAAGNNNAGATCGG |
| 8300_1 | TATGAGTGTGGAGTCGTTGCTCCAAATGTTTCCAAAACCGCCTTTNNNAGATCGGAA |
| 8301_1 | TATGAGTGTGGAGTCGTTGCAACACCTGTGGGACAGAACTATTCANNNAGATCGGA |
| 8336_1 | TATGAGTGTGGAGTCGTTGCTTTTATTTTGTTTCCCTCGCTACANNNAGATCGGAAG |
| 8359_1 | TATGAGTGTGGAGTCGTTGCTCAGTGGTCCTCCCACAATCANNNAGATCGGAAGAG |
| 8361_1 | TATGAGTGTGGAGTCGTTGCACAAAATGGATGGACTTCAAGCCAGNNNAGATCGG |
| 8412_1 | TATGAGTGTGGAGTCGTTGCACAGACTTCAAATCAAGAACAGCTGNNNAGATCGGA |
| 8776_1 | TATGAGTGTGGAGTCGTTGCACAGTCCCCTTGAGTGAAGAGTTNNNAGATCGGAA |
| 8782_1 | TATGAGTGTGGAGTCGTTGCCACATCTCCCTTTTCTTCATTAGGGNNNAGATCGGAA |
| 8820_1 | TATGAGTGTGGAGTCGTTGCGGGCCAAATGTGTAGGGCTAATTNNNAGATCGGAA |
| 8945_1 | TATGAGTGTGGAGTCGTTGCACATTTGGTTCCTGTGCAAGAGTNNNAGATCGGAAG |
| 8979_1 | TATGAGTGTGGAGTCGTTGCTGTGCTTGTACTTCTTCCTCATTCTNNNAGATCGGAA |

| | |
|---|---|
| 9174_1 | TATGAGTGTGGAGTCGTTGCGCAATCAAGAATGAACAAGGAACATNNNAGATCGG |
| 9209_1 | TATGAGTGTGGAGTCGTTGCAAACAAATTCTGCAAAGTTGCAAGTNNNAGATCGGA |
| 9226_1 | TATGAGTGTGGAGTCGTTGCTCTTTGTCACTCAGGCAGATGTGNNNAGATCGGAAG |
| 9239_1 | TATGAGTGTGGAGTCGTTGCCTGTGCTTGGCTGGAAATGTTGTAANNNAGATCGG |
| 9417_1 | TATGAGTGTGGAGTCGTTGCTGTGTGTGAATGTTTGTATGCACGNNNAGATCGGA |
| 9551_1 | TATGAGTGTGGAGTCGTTGCCCCCTGGTTTTCTACCATGGATCNNNAGATCGGAAG |
| 9577_1 | TATGAGTGTGGAGTCGTTGCGCAGTGTGAACCTGAAGTAATGCNNNAGATCGGAA |
| 9594_1 | TATGAGTGTGGAGTCGTTGCCAGAGAACTTATAGGACCAGAGGGTNNNAGATCGG |
| 9650_1 | TATGAGTGTGGAGTCGTTGCGTTCATTGGTACAATGCTTAGTGCCNNNAGATCGGA |
| 9737_1 | TATGAGTGTGGAGTCGTTGCGACCTTGAAGGAGTACTCAGCACNNNAGATCGGAA |
| 9814_1 | TATGAGTGTGGAGTCGTTGCCCCTCTCCCTCTCTCTGTTCATANNNAGATCGGAAGA |
| 9829_1 | TATGAGTGTGGAGTCGTTGCACACCTCTTGCAATACAGATCTTGANNNAGATCGGA |
| 9860_1 | TATGAGTGTGGAGTCGTTGCTTATTGGCAACTTCCAAATGTTGGANNNAGATCGGA |
| 9935_1 | TATGAGTGTGGAGTCGTTGCTTTCAGTTGTGTTCTGGGCAAGANNNAGATCGGAA |
| 9990_1 | TATGAGTGTGGAGTCGTTGCTGTTGTATGTTCTGTGGGTTTGACANNNAGATCGGA |
| 10177_1 | TATGAGTGTGGAGTCGTTGCTCTGAAGCATGCAAAGTTCATCCTNNNAGATCGGAA |
| 10185_1 | TATGAGTGTGGAGTCGTTGCTGTCACTCCAAATTATGTTGCACATNNNAGATCGGAA |
| 10211_1 | TATGAGTGTGGAGTCGTTGCAGGAGCCAAATTGAAACTCAAGAACNNNAGATCGG |
| 10217_1 | TATGAGTGTGGAGTCGTTGCAGTATTAGTGGATGGGAGCAGCANNNAGATCGGAA |
| 10227_1 | TATGAGTGTGGAGTCGTTGCATCTACAGAATACTCCACCAACCTTNNNAGATCGGAA |
| 10235_1 | TATGAGTGTGGAGTCGTTGCAAGAAGGCACGTGGAAAGTTTCTNNNAGATCGGAA |
| 10269_1 | TATGAGTGTGGAGTCGTTGCAAATTGGGTTGGAAAGAACTGATACNNNAGATCGG |
| 10281_1 | TATGAGTGTGGAGTCGTTGCAGGAAGACAAGGAATAAAGGAAGGANNNAGATCG |
| 10329_1 | TATGAGTGTGGAGTCGTTGCTCCTAGTGGTATCTGTGCATTTGTGNNNAGATCGGA |
| 10330_1 | TATGAGTGTGGAGTCGTTGCTGAATGTAGCATATTTGCAACTCTGNNNAGATCGGA |
| 10342_1 | TATGAGTGTGGAGTCGTTGCCCTGGCTAGAGGTTTGTCAGTTTCNNNAGATCGGAA |
| 10358_1 | TATGAGTGTGGAGTCGTTGCTCTTATTCAATCATCTCTGCAGTGANNNAGATCGGAA |
| 10393_1 | TATGAGTGTGGAGTCGTTGCCACGTATCTAGCATGATTTTAGGGTNNNAGATCGGA |
| 10396_1 | TATGAGTGTGGAGTCGTTGCGGAGGGAGTAAAGCCAAGAGAGTNNNAGATCGGA |
| 10409_1 | TATGAGTGTGGAGTCGTTGCAGCCTCATGTTGTGGACCTCTATNNNAGATCGGAAG |
| 10443_1 | TATGAGTGTGGAGTCGTTGCTGGTACATCAAAAGAATTGCACACCNNNAGATCGGA |
| 10444_1 | TATGAGTGTGGAGTCGTTGCGCTCACACAGATATTGAAGGTTGACNNNAGATCGG |
| 10445_1 | TATGAGTGTGGAGTCGTTGCGGGGGAAGAGAAAGAGACTTATCACNNNAGATCGG |
| 10467_1 | TATGAGTGTGGAGTCGTTGCTCAGAGTTTCAAGTCACAGAAATCANNNAGATCGGA |
| 10473_1 | TATGAGTGTGGAGTCGTTGCGCTCTACATTTCAGATGTGCCCANNNAGATCGGAAG |
| 10478_1 | TATGAGTGTGGAGTCGTTGCACAACAATCTGCAGCTCAAGACANNNAGATCGGAAG |
| 10482_1 | TATGAGTGTGGAGTCGTTGCTTTGAGGAGAGGGAGCAAGAGAANNNAGATCGGA |
| 10492_1 | TATGAGTGTGGAGTCGTTGCGCTTCAAGGCCAAACTTCAGTTTNNNAGATCGGAAG |
| 10497_1 | TATGAGTGTGGAGTCGTTGCATCTGAGTTTGGGGCCAACATTTNNNAGATCGGAAG |
| 10507_1 | TATGAGTGTGGAGTCGTTGCCATTTGGGGAATCAAGTGAAAGGTNNNAGATCGGA |
| 10511_1 | TATGAGTGTGGAGTCGTTGCTGTTTAGGGAGTTCTCATATGGAGTNNNAGATCGG |
| 10542_1 | TATGAGTGTGGAGTCGTTGCTTGCACAGAACAACGCATTTCAANNNAGATCGGAAG |
| 10549_1 | TATGAGTGTGGAGTCGTTGCACACACAGTATGCTCCCGATTTTNNNAGATCGGAAG |

| | |
|---|---|
| 10588_1 | TATGAGTGTGGAGTCGTTGCATCTGGAAGTGAAAGCCATCGTGNNNAGATCGGAA |
| 10668_1 | TATGAGTGTGGAGTCGTTGCTACAGGTGCCAAGTAACAGCTTCNNNAGATCGGAAG |
| 10675_1 | TATGAGTGTGGAGTCGTTGCGCAGGTTGACTAATCTCTTTGTGCNNNAGATCGGAA |
| 10677_1 | TATGAGTGTGGAGTCGTTGCGGACAATAATTGGGTTACTTGCTGCNNNAGATCGG |
| 10757_1 | TATGAGTGTGGAGTCGTTGCCATCTTTGTTGACATACGCAGCTNNNAGATCGGAAG |
| 10763_1 | TATGAGTGTGGAGTCGTTGCTCAAATTACAGGTCAAACACCAACTNNNAGATCGGA |
| 10782_1 | TATGAGTGTGGAGTCGTTGCACCACACACAATAAGCAACTAACGNNNAGATCGGAA |
| 10792_1 | TATGAGTGTGGAGTCGTTGCTTGTTTCCTTGTCTTCACCCTGGNNNAGATCGGAAGA |
| 10823_1 | TATGAGTGTGGAGTCGTTGCTTTTGATGGTGGATGAACAGAGAGGNNNAGATCGG |
| 10831_1 | TATGAGTGTGGAGTCGTTGCTCTGACATAACCTGGAATTTGTTGANNNAGATCGGA |
| 10835_1 | TATGAGTGTGGAGTCGTTGCGCACACACAAGCATTTTCTATTTCANNNAGATCGGAA |
| 10845_1 | TATGAGTGTGGAGTCGTTGCTCACCAAAAGAGCTCCTGATTTACANNNAGATCGGA |
| 10907_1 | TATGAGTGTGGAGTCGTTGCGGTCAGGATTTGAACCCATAGAGTNNNAGATCGGA |
| 10926_1 | TATGAGTGTGGAGTCGTTGCACGATCTCTCGGCTCATTGTAACNNNAGATCGGAAG |
| 10943_1 | TATGAGTGTGGAGTCGTTGCTATCAGTAAAACCCACTTGGAAGGCNNNAGATCGGA |
| 10952_1 | TATGAGTGTGGAGTCGTTGCGATTGTTTACCTCAGCCTCCTGAGTNNNAGATCGGA |
| 10956_1 | TATGAGTGTGGAGTCGTTGCTGGCTGCTTTTGCACTGTAGTTGNNNAGATCGGAAG |
| 10961_1 | TATGAGTGTGGAGTCGTTGCCCTAACTAACAGACTGTACTCCCACNNNAGATCGGAA |
| 10962_1 | TATGAGTGTGGAGTCGTTGCGGGTCCAATCTAGACTTTTACAAGCNNNAGATCGGA |
| 11000_1 | TATGAGTGTGGAGTCGTTGCGACAGGGTTTCAGTTTGTCACTCATNNNAGATCGGA |
| 11009_1 | TATGAGTGTGGAGTCGTTGCCCAACACTCCTGGCTAAAACTTTTTNNNAGATCGGAA |
| 11082_1 | TATGAGTGTGGAGTCGTTGCCCCTGAGCTTCAGTTTCTGTGATTGNNNAGATCGGA |
| 11084_1 | TATGAGTGTGGAGTCGTTGCAGTGCAGAGACATGATCTAGGTTCANNNAGATCGG |
| 11098_1 | TATGAGTGTGGAGTCGTTGCATTTGTAACAGTGATTGGCCCATACNNNAGATCGGA |
| 11104_1 | TATGAGTGTGGAGTCGTTGCAGGACCCCTGGTATCAAACAACCNNNAGATCGGAAG |
| 11140_1 | TATGAGTGTGGAGTCGTTGCCAGGGCTATTGCAGTAGGTAAAAAGNNNAGATCGG |
| 11197_1 | TATGAGTGTGGAGTCGTTGCGACTGGAGTGGAGTGATGTAGTGATNNNAGATCGG |
| 11225_1 | TATGAGTGTGGAGTCGTTGCTTTTACAGAACCAATACCTGGGCTCNNNAGATCGGA |
| 11402_1 | TATGAGTGTGGAGTCGTTGCGTGGAATTCTGTTTGTAATCAGCCGNNNAGATCGGA |
| 11475_1 | TATGAGTGTGGAGTCGTTGCTTCTTCACTGCCCCATTTCATCTAGNNNAGATCGGAA |
| 11560_1 | TATGAGTGTGGAGTCGTTGCTGGGAAATTATAAAGTGTGTGTGTGTNNNAGATCG |
| 11640_1 | TATGAGTGTGGAGTCGTTGCATTTCACTTTTGTTGCCCAAGAGTGNNNAGATCGGA |
| 11831_1 | TATGAGTGTGGAGTCGTTGCAAATTGAGAGAAACCAGCTGATGATNNNAGATCGG |
| 11835_1 | TATGAGTGTGGAGTCGTTGCAGACAGGATTTTACCGTGTTAGACTNNNAGATCGGA |
| 11971_1 | TATGAGTGTGGAGTCGTTGCGAATACCTGCTTTGCAAATGTCTTTNNNAGATCGGA |
| 11986_1 | TATGAGTGTGGAGTCGTTGCAGAAAGATGGATGAGTTGAAGCTTCNNNAGATCGG |
| 12153_1 | TATGAGTGTGGAGTCGTTGCAGGCCAGTAAACTTCCATCTTAGACNNNAGATCGGA |
| 12293_1 | TATGAGTGTGGAGTCGTTGCTGTAGCAATAATAGAAAGCACCCAANNNAGATCGG |
| 12321_1 | TATGAGTGTGGAGTCGTTGCGGATATATTGAGTACGGACCTCCCCNNNAGATCGGA |
| 12341_1 | TATGAGTGTGGAGTCGTTGCAGACTGAAACAACCACTAAAACAGANNNAGATCGG |
| 12349_1 | TATGAGTGTGGAGTCGTTGCCATCCTTCCTCCTCTTTCCTCCGCANNNAGATCGGAAG |
| 12390_1 | TATGAGTGTGGAGTCGTTGCGAGAAGAGAGAGAGGGTCTCTGTNNNAGATCGGA |
| 28775_1 | TATGAGTGTGGAGTCGTTGCGGGTGTGTATTTACAACAATCTTGCNNNAGATCGGA |

| | |
|---|---|
| 28786_1 | TATGAGTGTGGAGTCGTTGCAACTCAGCCATAATAAGCAATGAATNNNAGATCGGA |
| 28810_1 | TATGAGTGTGGAGTCGTTGCAGATCTAGCTTGTAATTGTGTTGTTNNNAGATCGGA |
| 28831_1 | TATGAGTGTGGAGTCGTTGCGAGGGAGAGAGAGAGAGAGAGAGNNNAGATCGG |
| 28927_1 | TATGAGTGTGGAGTCGTTGCGAAGGCATTGTGTAGGTTGCATTNNNAGATCGGAA |
| 28994_1 | TATGAGTGTGGAGTCGTTGCTGCATAAGATATTGGCCAGGTGTNNNAGATCGGAA |
| 29017_1 | TATGAGTGTGGAGTCGTTGCGAGAGAAAGTAGCAGGTAATTAGGTNNNAGATCG |
| 29042_1 | TATGAGTGTGGAGTCGTTGCCCAAACTGTGTAATCTTCAACTTGCNNNAGATCGGA |
| 29055_1 | TATGAGTGTGGAGTCGTTGCTTCAAGTTGCTTTTCCATGATGANNNAGATCGGAAG |
| 29066_1 | TATGAGTGTGGAGTCGTTGCTGGTTGTTTTCACCTTTGAACTGTNNNAGATCGGAA |
| 29088_1 | TATGAGTGTGGAGTCGTTGCTGTCATGTTATATGCCTCCTTCTCTNNNAGATCGGAA |
| 29108_1 | TATGAGTGTGGAGTCGTTGCTGCAGTTTACTCAAAAGGTGAACANNNAGATCGGA |
| 29110_1 | TATGAGTGTGGAGTCGTTGCTCTGAGGGAAATTAGAAGAGTGCTNNNAGATCGGA |
| 29137_1 | TATGAGTGTGGAGTCGTTGCGGAAGGGAAAAAACAATAGGATGGTNNNAGATCG |
| 29143_1 | TATGAGTGTGGAGTCGTTGCCACCATGGAATACTACTCAGCCANNNAGATCGGAAG |
| 29176_1 | TATGAGTGTGGAGTCGTTGCAGCCCTATTCCTTATATTGTGCCTNNNAGATCGGAAG |
| 29188_1 | TATGAGTGTGGAGTCGTTGCCACACACACATATATACACACACACANNNAGATCGGA |
| 29213_1 | TATGAGTGTGGAGTCGTTGCTGAGTTAGGTCTCCTCCCTCATTNNNAGATCGGAAG |
| 29302_1 | TATGAGTGTGGAGTCGTTGCAAAAAAGGAGCTTTCTGGGCTTCNNNAGATCGGAA |
| 29363_1 | TATGAGTGTGGAGTCGTTGCTGGTTCATGGGATACTGTGTACCNNNAGATCGGAA |
| 30792_1 | TATGAGTGTGGAGTCGTTGCACACACACAAATACTCTTTTCAGGGNNNAGATCGGA |
| 30794_1 | TATGAGTGTGGAGTCGTTGCACTCCTAGCTAAGAAATGCAGGGNNNAGATCGGAA |
| 30797_1 | TATGAGTGTGGAGTCGTTGCTGTTTTGGAGACTCAAGGGGAAANNNAGATCGGAA |
| 30805_1 | TATGAGTGTGGAGTCGTTGCTAAAAATAAGTGGAAGCAGCCGGNNNAGATCGGAA |
| 30835_1 | TATGAGTGTGGAGTCGTTGCTGGAGAATTTGGGTAAAGCAGGANNNAGATCGGA |
| 30932_1 | TATGAGTGTGGAGTCGTTGCTCGAAAATCTTACCATTTCAGCACANNNAGATCGGA |
| 30937_1 | TATGAGTGTGGAGTCGTTGCCCTCTCTCTCCCTCTTATTCTACCANNNAGATCGGAAG |
| 31056_1 | TATGAGTGTGGAGTCGTTGCAGCAACTTAGAGCTCATTGACCANNNAGATCGGAAG |
| 31073_1 | TATGAGTGTGGAGTCGTTGCTTCACTGCCTCTTCTCATGTTGTNNNAGATCGGAAGA |
| 31076_1 | TATGAGTGTGGAGTCGTTGCAGCTTGATACCTGACAGAATATCCTNNNAGATCGGA |
| 31108_1 | TATGAGTGTGGAGTCGTTGCACCTAACTGAGTCACTGAAATCACTNNNAGATCGGA |
| 31182_1 | TATGAGTGTGGAGTCGTTGCTCTCCCACACTAAGCAATTCTCCNNNAGATCGGAAGA |
| 31277_1 | TATGAGTGTGGAGTCGTTGCACACAAGACTATCCAGGAACCACNNNAGATCGGAAG |
| 31280_1 | TATGAGTGTGGAGTCGTTGCACAGTCTAGTATCTATGTCAAGCAANNNAGATCGGA |
| 31301_1 | TATGAGTGTGGAGTCGTTGCCAGAACAGTAGGTGACCACGTAANNNAGATCGGAA |
| 31310_1 | TATGAGTGTGGAGTCGTTGCTGGACATCCCTCCCTCAATCTAANNNAGATCGGAAG |
| 31347_1 | TATGAGTGTGGAGTCGTTGCTGGGTAGCAAAATTTGATAGATGCANNNAGATCGG |
| 31422_1 | TATGAGTGTGGAGTCGTTGCCCTTGGTTAGTACAGATTTTGGTCCNNNAGATCGGA |
| 31425_1 | TATGAGTGTGGAGTCGTTGCTGGTTCTGTTTCTCTGTAGAACTCTNNNAGATCGGA |
| 31450_1 | TATGAGTGTGGAGTCGTTGCGTTTGGGGCATGGATATTCATGGNNNAGATCGGAA |
| 31481_1 | TATGAGTGTGGAGTCGTTGCAGCACCATTTTTCCAATAGCATGTNNNAGATCGGAA |
| 31495_1 | TATGAGTGTGGAGTCGTTGCCTTCTCACCACTGGCAATAATGCNNNAGATCGGAAG |
| 31516_1 | TATGAGTGTGGAGTCGTTGCAGCAACCACTAGGAAAACTATACGANNNAGATCGGA |
| 31558_1 | TATGAGTGTGGAGTCGTTGCTGTAAATGAGTTAGGGTTTGCTACANNNAGATCGG |

31617_1 TATGAGTGTGGAGTCGTTGCAAGAGTTATCAGAGGCTGTGCTTNNNAGATCGGAA
31736_1 TATGAGTGTGGAGTCGTTGCACATGTGCAGGTTTGTTACATGGNNNAGATCGGAA
31739_1 TATGAGTGTGGAGTCGTTGCAGAGAGAGAGAGAAAGAGAGAGAGANNNAGATC
31794_1 TATGAGTGTGGAGTCGTTGCAGAAGGAAGAAAAGGAAAGGAAGGANNNAGATCG
31805_1 TATGAGTGTGGAGTCGTTGCTGTGTGTGTGTATATATGTGTGTGTNNNAGATCGG
31884_1 TATGAGTGTGGAGTCGTTGCTGGCTAAGTAGGAGGTTTATGTTGANNNAGATCGG
31885_1 TATGAGTGTGGAGTCGTTGCACCATCTCTTGAAGTGAACATCAGANNNAGATCGGA
31897_1 TATGAGTGTGGAGTCGTTGCAGAAGTATACGTGGCAAATTGATGTNNNAGATCGG
31903_1 TATGAGTGTGGAGTCGTTGCAAGAAGCAGGATGTCTTGGTCATNNNAGATCGGAA
31905_1 TATGAGTGTGGAGTCGTTGCTGACATTGACTCATTTTTGTGTGGTNNNAGATCGGA
31992_1 TATGAGTGTGGAGTCGTTGCACATGATGCTACATCTTGGAGATCTNNNAGATCGGA
32094_1 TATGAGTGTGGAGTCGTTGCTCAGAGGAGCTAAAAAAACACAGGNNNAGATCGGA
32282_1 TATGAGTGTGGAGTCGTTGCTGCAACAACTGATGGACCTGTATNNNAGATCGGAAG
32492_1 TATGAGTGTGGAGTCGTTGCTTCAGCTTGGCTATTGGCGTATANNNAGATCGGAAG
32504_1 TATGAGTGTGGAGTCGTTGCGGTACAATCTTGGCTCATGCAACNNNAGATCGGAAG
32556_1 TATGAGTGTGGAGTCGTTGCGAATTCCTGAGCTCAAGCAATCCNNNAGATCGGAAG
32595_1 TATGAGTGTGGAGTCGTTGCTCAGACTCCATGCATAAAATAGGGANNNAGATCGGA
32628_1 TATGAGTGTGGAGTCGTTGCGCAAGGTACAACATAAACAAACTGCNNNAGATCGGA
32659_1 TATGAGTGTGGAGTCGTTGCGTTCGAGCGATTATTTTGCCTCANNNAGATCGGAAG
32673_1 TATGAGTGTGGAGTCGTTGCAATACAGCACACTGATGGGTCTTNNNAGATCGGAAG
32742_1 TATGAGTGTGGAGTCGTTGCCCATATAACCCAGCAATTCCATTCCNNNAGATCGGAA
32833_1 TATGAGTGTGGAGTCGTTGCATTCATGTGATTCTCCTGCCTCANNNAGATCGGAAGA
32838_1 TATGAGTGTGGAGTCGTTGCGGGTTTTGTAGTTGGTGTGCATANNNAGATCGGAA
32861_1 TATGAGTGTGGAGTCGTTGCGATCTCAGCTCACTGCAACCTATNNNAGATCGGAAG
32900_1 TATGAGTGTGGAGTCGTTGCAGAGTACAACATAGTTTCAGTTGCTNNNAGATCGGA
32977_1 TATGAGTGTGGAGTCGTTGCACAGAAAACAGAAGTGAGGTACAGANNNAGATCGG
32997_1 TATGAGTGTGGAGTCGTTGCACAAAGAAACAAACAAAACCCCCNNNAGATCGGAAG
33195_1 TATGAGTGTGGAGTCGTTGCACCTATGGCCATTTGATCTCTTACANNNAGATCGGAA
33216_1 TATGAGTGTGGAGTCGTTGCTGACTTTGTTTATGTTCTGGTTGCANNNAGATCGGA
33218_1 TATGAGTGTGGAGTCGTTGCCAAAACGTTGGAATAAGCAGAAAGTNNNAGATCGG
33288_1 TATGAGTGTGGAGTCGTTGCAGGGAATGGATATAACACACTTTGTNNNAGATCGG
33310_1 TATGAGTGTGGAGTCGTTGCGGAAGAAAGGCCAGCAACATTTTNNNAGATCGGAA
33392_1 TATGAGTGTGGAGTCGTTGCAGAGAGAGAGAGAGAGAGAGAAAGANNNAGATC
33470_1 TATGAGTGTGGAGTCGTTGCGGAGTAAATGCAGTGTATAGTGAAANNNAGATCGG
33474_1 TATGAGTGTGGAGTCGTTGCACACATGAAAAGATTAGAGGGAACANNNAGATCGG
33513_1 TATGAGTGTGGAGTCGTTGCTGCTGGTAGAGTAAACATTCAGTGANNNAGATCGG
33628_1 TATGAGTGTGGAGTCGTTGCTACTGGCAGCGTAGAATGAGTTTNNNAGATCGGAA
33812_1 TATGAGTGTGGAGTCGTTGCACCAATTCAAATTTGTTACCCTTGANNNAGATCGGAA
33866_1 TATGAGTGTGGAGTCGTTGCAGTACGTCAGAATTCTCTAGTGAAANNNAGATCGGA
33949_1 TATGAGTGTGGAGTCGTTGCTCAAGTGATTCTCCTGCCTTAGCNNNAGATCGGAAG
33951_1 TATGAGTGTGGAGTCGTTGCACTATGCCCGGCGAATTTTTTCNNNAGATCGGAAGA
34148_1 TATGAGTGTGGAGTCGTTGCAGTGATCTATTGCACTATAGGGTGANNNAGATCGG
34207_1 TATGAGTGTGGAGTCGTTGCAGATGGGACTAACAGCAAATTAGGTNNNAGATCGG

34253_1 TATGAGTGTGGAGTCGTTGCCTATTGGAAAAGATGGGCAGCACNNNAGATCGGAA
34415_1 TATGAGTGTGGAGTCGTTGCTCAGTGGTTATAGGATGAATTACCTNNNAGATCGGA
34479_1 TATGAGTGTGGAGTCGTTGCACGGATACCAAATTCTGAGGATTCTNNNAGATCGGA
34668_1 TATGAGTGTGGAGTCGTTGCAGAAGCCTTGTTTTTTTCACACTNNNAGATCGGAAG
34792_1 TATGAGTGTGGAGTCGTTGCTCTGCTTCCTGAGTTCAAGTGATNNNAGATCGGAAG
34941_1 TATGAGTGTGGAGTCGTTGCCACCACACCTGGCTAATTTTTGTNNNAGATCGGAAG
35044_1 TATGAGTGTGGAGTCGTTGCTTCAAGTGATTCTCCTGCCTCAGNNNAGATCGGAAG
35077_1 TATGAGTGTGGAGTCGTTGCTTCAAGCAATTCTCCTGCCTCAGNNNAGATCGGAAG
35092_1 TATGAGTGTGGAGTCGTTGCCACAATCTCAGCTCACTGCAACNNNAGATCGGAAGA
35104_1 TATGAGTGTGGAGTCGTTGCGATCTCGGCTCACTGCAACTTNNNAGATCGGAAGAG
35154_1 TATGAGTGTGGAGTCGTTGCAGTTGTTCCTAATAGTCTCCGATGTNNNAGATCGGA
35245_1 TATGAGTGTGGAGTCGTTGCACAATCTCGGCTCATTGTAACCTNNNAGATCGGAAG
35272_1 TATGAGTGTGGAGTCGTTGCAGAGTAGCTGGGACTATAGGCANNNAGATCGGAAG
35376_1 TATGAGTGTGGAGTCGTTGCCAGGTTCAAGCAATTCTCCTGCNNNAGATCGGAAGA
35543_1 TATGAGTGTGGAGTCGTTGCTTCAAGCTATTCTCCTGCCTCAGNNNAGATCGGAAGA
35567_1 TATGAGTGTGGAGTCGTTGCGTATCATAGCTGGGACTACAGGCNNNAGATCGGAA
35640_1 TATGAGTGTGGAGTCGTTGCAGGATTTTTATCAGGAAGAGATGCTNNNAGATCGG
36015_1 TATGAGTGTGGAGTCGTTGCTGGCTAAGTAGGAGGTTTATGTTGANNNAGATCGG
36019_1 TATGAGTGTGGAGTCGTTGCACCATCTCTTGAAGTGAACATCAGANNNAGATCGGA
36090_1 TATGAGTGTGGAGTCGTTGCAGAAGTATACGTGGCAAATTGATGTNNNAGATCGG
36109_1 TATGAGTGTGGAGTCGTTGCTGGTTACTCAATTCATTGAAGGTGTNNNAGATCGGA
36182_1 TATGAGTGTGGAGTCGTTGCGAAGACGATGACGAAAAAAAAAGAANNNAGATCG
36188_1 TATGAGTGTGGAGTCGTTGCAGAAAGAAAGTTGTCATCTAGTTACCNNNAGATCGG
36291_1 TATGAGTGTGGAGTCGTTGCGGAAATCTGAGAAAATCTGTTTCCANNNAGATCGGA
36665_1 TATGAGTGTGGAGTCGTTGCTGCAATGCAAAAATAAAGAGTGACANNNAGATCGG
36691_1 TATGAGTGTGGAGTCGTTGCAGAAAGAAAGGAGAAAGAAAGAGGANNNAGATCG
36699_1 TATGAGTGTGGAGTCGTTGCAGACTGGTTAGAGGTCATTTATGCANNNAGATCGG
36775_1 TATGAGTGTGGAGTCGTTGCGAGGAACTCCCTGTAGTGAAAAANNNAGATCGGAA
36855_1 TATGAGTGTGGAGTCGTTGCCCCCTTCTCTTATCTACTAGTTACCCNNNAGATCGGAA
36989_1 TATGAGTGTGGAGTCGTTGCAGAGGAAACAGTAAGGAGTGAAAGTNNNAGATCG
37051_1 TATGAGTGTGGAGTCGTTGCACATGTTGAGTGCTTACAATATGCTNNNAGATCGGA
37066_1 TATGAGTGTGGAGTCGTTGCACAAACTTTTGGACGATATTGGTGANNNAGATCGGA
37095_1 TATGAGTGTGGAGTCGTTGCTCCTATTACCTAGCCCTTGATTTCTNNNAGATCGGAA
37190_1 TATGAGTGTGGAGTCGTTGCTCAAATCAGTGGATAGAGATGGACTNNNAGATCGG
37191_1 TATGAGTGTGGAGTCGTTGCAGTCAGTTGAGATAGTGAAGATCCANNNAGATCGG
37207_1 TATGAGTGTGGAGTCGTTGCAGATGGTAAATGTCTGAGAAACACANNNAGATCGG
37214_1 TATGAGTGTGGAGTCGTTGCACAAAGACTTGAGAAACATTGCTGTNNNAGATCGGA
37252_1 TATGAGTGTGGAGTCGTTGCACAAAAAATTCTGAAGTGTTGAAGAGANNNAGATC
37264_1 TATGAGTGTGGAGTCGTTGCTTCTGTGGAAGAGTTATACAGCTATNNNAGATCGGA
37269_1 TATGAGTGTGGAGTCGTTGCCCCTTCTTCAGTTTTAGGGGTTANNNAGATCGGAAG
37276_1 TATGAGTGTGGAGTCGTTGCTGACAAACCTTTAGACAGACCATCTNNNAGATCGGA
37345_1 TATGAGTGTGGAGTCGTTGCTGTCTGTGTGTATATGTGTGTCTGTNNNAGATCGGA
37528_1 TATGAGTGTGGAGTCGTTGCAGACATGTGTATTCAGTTCCACAGANNNAGATCGGA

| | |
|---|---|
| 37543_1 | TATGAGTGTGGAGTCGTTGCGGGTAATACTTTTCGTGTAGTGTTTNNNAGATCGGA |
| 37602_1 | TATGAGTGTGGAGTCGTTGCTGTAACAAGGTGTTGAGGGCTTANNNAGATCGGAA |
| 37603_1 | TATGAGTGTGGAGTCGTTGCAGAGAGAGAAAGAAGAAAGACAGACANNNAGATC |
| 37757_1 | TATGAGTGTGGAGTCGTTGCATTTCAAGAGTTGTAATTCGTCTGANNNAGATCGGA |
| 37781_1 | TATGAGTGTGGAGTCGTTGCAGTGGATGGGTATATATGGTGTTCANNNAGATCGG |
| 37853_1 | TATGAGTGTGGAGTCGTTGCGCCAAAAAGACAAATTTAGTGTCAGGNNNAGATCG |
| 37962_1 | TATGAGTGTGGAGTCGTTGCAGAGAGAGAGAGAGGAAAGAAAGGANNNAGATC |
| 37993_1 | TATGAGTGTGGAGTCGTTGCACTGTAATGCTATTTTGTTCACATGACNNNAGATCGG |
| 38028_1 | TATGAGTGTGGAGTCGTTGCACTGGGTAAAAAAAAGCAAGAACANNNAGATCGGA |
| 38069_1 | TATGAGTGTGGAGTCGTTGCTCAACATACAAAAATCAGTAGCACTNNNAGATCGGA |
| 38075_1 | TATGAGTGTGGAGTCGTTGCTGGCCCAATCAAAAATTACACTTCTNNNAGATCGGA |
| 38313_1 | TATGAGTGTGGAGTCGTTGCAGATCAAAAAGAACAAAGAAAGGCANNNAGATCGG |
| 38379_1 | TATGAGTGTGGAGTCGTTGCTGTTGACTAGTATTCCATTGTCTGTNNNAGATCGGA |
| 38382_1 | TATGAGTGTGGAGTCGTTGCAGTGTCAACCTAACATAACAGACCANNNAGATCGGA |
| 38402_1 | TATGAGTGTGGAGTCGTTGCTGCATCACTTCCATTTTATCTGTCANNNAGATCGGAA |
| 38453_1 | TATGAGTGTGGAGTCGTTGCGCAGACAAATCTTTTTTTGAGACGGNNNAGATCGGA |
| 38555_1 | TATGAGTGTGGAGTCGTTGCACCAAAATTATGAAAACACCCAACANNNAGATCGGA |
| 38656_1 | TATGAGTGTGGAGTCGTTGCTGTCACTCAACTAAAACTATGAAACTNNNAGATCGG |
| 38758_1 | TATGAGTGTGGAGTCGTTGCGCCAGTAGCATTGATGTAGTATTTTNNNAGATCGGA |
| 38785_1 | TATGAGTGTGGAGTCGTTGCACCTGCGAAAAAAATCAAGGAGGNNNAGATCGGAA |
| 38804_1 | TATGAGTGTGGAGTCGTTGCTCAAAGATGACACAAACAAATGGGTNNNAGATCGG |
| 38812_1 | TATGAGTGTGGAGTCGTTGCACCAAGAATACACTACACCAAGTCANNNAGATCGGA |
| 38826_1 | TATGAGTGTGGAGTCGTTGCCGAAAGAAGGGTGTTACATAATGGTNNNAGATCGG |
| 38835_1 | TATGAGTGTGGAGTCGTTGCACGAAGTTTCAGAAGAGAGTTCTCANNNAGATCGG |
| 38854_1 | TATGAGTGTGGAGTCGTTGCTGAAACAAAAAGCTGGTTCTTGGANNNAGATCGGA |
| 38894_1 | TATGAGTGTGGAGTCGTTGCAGACAACTTACATAGCGGAAGAAAANNNAGATCGG |
| 38897_1 | TATGAGTGTGGAGTCGTTGCTGAACTAAAGGAGATTGAGACATGANNNAGATCGG |
| 38944_1 | TATGAGTGTGGAGTCGTTGCAGCAGAACTGAAGGAGATAGAAACANNNAGATCGG |
| 50853_1 | TATGAGTGTGGAGTCGTTGCTTGGTCAACCTACAGAATGGGAGNNNAGATCGGAA |
| 50853_2 | TATGAGTGTGGAGTCGTTGCTTGGTCAACCTACAGAATGGGAGNNNAGATCGGAA |
| 50853_3 | TATGAGTGTGGAGTCGTTGCTTGGTCAACCTACAGAATGGGAGNNNAGATCGGAA |
| 50853_4 | TATGAGTGTGGAGTCGTTGCTTGGTCAACCTACAGAATGGGAGNNNAGATCGGAA |
| 50853_5 | TATGAGTGTGGAGTCGTTGCTTGGTCAACCTACAGAATGGGAGNNNAGATCGGAA |
| 50853_6 | TATGAGTGTGGAGTCGTTGCTTGGTCAACCTACAGAATGGGAGNNNAGATCGGAA |
| 50853_7 | TATGAGTGTGGAGTCGTTGCTTGGTCAACCTACAGAATGGGAGNNNAGATCGGAA |
| 50853_8 | TATGAGTGTGGAGTCGTTGCTTGGTCAACCTACAGAATGGGAGNNNAGATCGGAA |
| 46567_1 | TATGAGTGTGGAGTCGTTGCGGAGTGTAGTGGCGTGATCTCNNNAGATCGGAAGA |
| 46567_2 | TATGAGTGTGGAGTCGTTGCGGAGTGTAGTGGCGTGATCTCNNNAGATCGGAAGA |
| 46567_3 | TATGAGTGTGGAGTCGTTGCGGAGTGTAGTGGCGTGATCTCNNNAGATCGGAAGA |
| 46567_4 | TATGAGTGTGGAGTCGTTGCGGAGTGTAGTGGCGTGATCTCNNNAGATCGGAAGA |
| 46567_5 | TATGAGTGTGGAGTCGTTGCGGAGTGTAGTGGCGTGATCTCNNNAGATCGGAAGA |
| 46567_6 | TATGAGTGTGGAGTCGTTGCGGAGTGTAGTGGCGTGATCTCNNNAGATCGGAAGA |
| 46567_7 | TATGAGTGTGGAGTCGTTGCGGAGTGTAGTGGCGTGATCTCNNNAGATCGGAAGA |

| | |
|---|---|
| 46567_8 | TATGAGTGTGGAGTCGTTGCGGAGTGTAGTGGCGTGATCTCNNNAGATCGGAAGA |
| 33407_1 | TATGAGTGTGGAGTCGTTGCGATTTGACAACTGTACATCGGCCNNNAGATCGGAAG |
| 33407_2 | TATGAGTGTGGAGTCGTTGCGATTTGACAACTGTACATCGGCCNNNAGATCGGAAG |
| 33407_3 | TATGAGTGTGGAGTCGTTGCGATTTGACAACTGTACATCGGCCNNNAGATCGGAAG |
| 33407_4 | TATGAGTGTGGAGTCGTTGCGATTTGACAACTGTACATCGGCCNNNAGATCGGAAG |
| 33407_5 | TATGAGTGTGGAGTCGTTGCGATTTGACAACTGTACATCGGCCNNNAGATCGGAAG |
| 33407_6 | TATGAGTGTGGAGTCGTTGCGATTTGACAACTGTACATCGGCCNNNAGATCGGAAG |
| 33407_7 | TATGAGTGTGGAGTCGTTGCGATTTGACAACTGTACATCGGCCNNNAGATCGGAAG |
| 33407_8 | TATGAGTGTGGAGTCGTTGCGATTTGACAACTGTACATCGGCCNNNAGATCGGAAG |
| 32413_1 | TATGAGTGTGGAGTCGTTGCTGTTTGCTGTATGACCTTGGACTNNNAGATCGGAAG |
| 32413_2 | TATGAGTGTGGAGTCGTTGCTGTTTGCTGTATGACCTTGGACTNNNAGATCGGAAG |
| 32413_3 | TATGAGTGTGGAGTCGTTGCTGTTTGCTGTATGACCTTGGACTNNNAGATCGGAAG |
| 32413_4 | TATGAGTGTGGAGTCGTTGCTGTTTGCTGTATGACCTTGGACTNNNAGATCGGAAG |
| 32413_5 | TATGAGTGTGGAGTCGTTGCTGTTTGCTGTATGACCTTGGACTNNNAGATCGGAAG |
| 32413_6 | TATGAGTGTGGAGTCGTTGCTGTTTGCTGTATGACCTTGGACTNNNAGATCGGAAG |
| 32413_7 | TATGAGTGTGGAGTCGTTGCTGTTTGCTGTATGACCTTGGACTNNNAGATCGGAAG |
| 32413_8 | TATGAGTGTGGAGTCGTTGCTGTTTGCTGTATGACCTTGGACTNNNAGATCGGAAG |
| 37927_1 | TATGAGTGTGGAGTCGTTGCCAGTCTCAGTGCCTCAGTTGAGNNNAGATCGGAAGA |
| 37927_2 | TATGAGTGTGGAGTCGTTGCCAGTCTCAGTGCCTCAGTTGAGNNNAGATCGGAAGA |
| 37927_3 | TATGAGTGTGGAGTCGTTGCCAGTCTCAGTGCCTCAGTTGAGNNNAGATCGGAAGA |
| 37927_4 | TATGAGTGTGGAGTCGTTGCCAGTCTCAGTGCCTCAGTTGAGNNNAGATCGGAAGA |
| 37927_5 | TATGAGTGTGGAGTCGTTGCCAGTCTCAGTGCCTCAGTTGAGNNNAGATCGGAAGA |
| 37927_6 | TATGAGTGTGGAGTCGTTGCCAGTCTCAGTGCCTCAGTTGAGNNNAGATCGGAAGA |
| 37927_7 | TATGAGTGTGGAGTCGTTGCCAGTCTCAGTGCCTCAGTTGAGNNNAGATCGGAAGA |
| 37927_8 | TATGAGTGTGGAGTCGTTGCCAGTCTCAGTGCCTCAGTTGAGNNNAGATCGGAAGA |
| 41720_1 | TATGAGTGTGGAGTCGTTGCGTAGCTGGGATTACAGGGGTATGNNNAGATCGGAA |
| 41720_2 | TATGAGTGTGGAGTCGTTGCGTAGCTGGGATTACAGGGGTATGNNNAGATCGGAA |
| 41720_3 | TATGAGTGTGGAGTCGTTGCGTAGCTGGGATTACAGGGGTATGNNNAGATCGGAA |
| 41720_4 | TATGAGTGTGGAGTCGTTGCGTAGCTGGGATTACAGGGGTATGNNNAGATCGGAA |
| 41720_5 | TATGAGTGTGGAGTCGTTGCGTAGCTGGGATTACAGGGGTATGNNNAGATCGGAA |
| 41720_6 | TATGAGTGTGGAGTCGTTGCGTAGCTGGGATTACAGGGGTATGNNNAGATCGGAA |
| 41720_7 | TATGAGTGTGGAGTCGTTGCGTAGCTGGGATTACAGGGGTATGNNNAGATCGGAA |
| 41720_8 | TATGAGTGTGGAGTCGTTGCGTAGCTGGGATTACAGGGGTATGNNNAGATCGGAA |
| 48955_1 | TATGAGTGTGGAGTCGTTGCCTGACAGGTGACACACTTTACCTNNNAGATCGGAAG |
| 48955_2 | TATGAGTGTGGAGTCGTTGCCTGACAGGTGACACACTTTACCTNNNAGATCGGAAG |
| 48955_3 | TATGAGTGTGGAGTCGTTGCCTGACAGGTGACACACTTTACCTNNNAGATCGGAAG |
| 48955_4 | TATGAGTGTGGAGTCGTTGCCTGACAGGTGACACACTTTACCTNNNAGATCGGAAG |
| 48955_5 | TATGAGTGTGGAGTCGTTGCCTGACAGGTGACACACTTTACCTNNNAGATCGGAAG |
| 48955_6 | TATGAGTGTGGAGTCGTTGCCTGACAGGTGACACACTTTACCTNNNAGATCGGAAG |
| 48955_7 | TATGAGTGTGGAGTCGTTGCCTGACAGGTGACACACTTTACCTNNNAGATCGGAAG |
| 48955_8 | TATGAGTGTGGAGTCGTTGCCTGACAGGTGACACACTTTACCTNNNAGATCGGAAG |
| 45527_1 | TATGAGTGTGGAGTCGTTGCAGCCTCATCTGAGAAACCCAAATNNNAGATCGGAAG |
| 45527_2 | TATGAGTGTGGAGTCGTTGCAGCCTCATCTGAGAAACCCAAATNNNAGATCGGAAG |

| | |
|---|---|
| 45527_3 | TATGAGTGTGGAGTCGTTGCAGCCTCATCTGAGAAACCCAAATNNNAGATCGGAAG |
| 45527_4 | TATGAGTGTGGAGTCGTTGCAGCCTCATCTGAGAAACCCAAATNNNAGATCGGAAG |
| 45527_5 | TATGAGTGTGGAGTCGTTGCAGCCTCATCTGAGAAACCCAAATNNNAGATCGGAAG |
| 45527_6 | TATGAGTGTGGAGTCGTTGCAGCCTCATCTGAGAAACCCAAATNNNAGATCGGAAG |
| 45527_7 | TATGAGTGTGGAGTCGTTGCAGCCTCATCTGAGAAACCCAAATNNNAGATCGGAAG |
| 45527_8 | TATGAGTGTGGAGTCGTTGCAGCCTCATCTGAGAAACCCAAATNNNAGATCGGAAG |
| 43421_1 | TATGAGTGTGGAGTCGTTGCTCTGATTCCCAGGTTCAAGCAATNNNAGATCGGAAG |
| 43421_2 | TATGAGTGTGGAGTCGTTGCTCTGATTCCCAGGTTCAAGCAATNNNAGATCGGAAG |
| 43421_3 | TATGAGTGTGGAGTCGTTGCTCTGATTCCCAGGTTCAAGCAATNNNAGATCGGAAG |
| 43421_4 | TATGAGTGTGGAGTCGTTGCTCTGATTCCCAGGTTCAAGCAATNNNAGATCGGAAG |
| 43421_5 | TATGAGTGTGGAGTCGTTGCTCTGATTCCCAGGTTCAAGCAATNNNAGATCGGAAG |
| 43421_6 | TATGAGTGTGGAGTCGTTGCTCTGATTCCCAGGTTCAAGCAATNNNAGATCGGAAG |
| 43421_7 | TATGAGTGTGGAGTCGTTGCTCTGATTCCCAGGTTCAAGCAATNNNAGATCGGAAG |
| 43421_8 | TATGAGTGTGGAGTCGTTGCTCTGATTCCCAGGTTCAAGCAATNNNAGATCGGAAG |
| 37885_1 | TATGAGTGTGGAGTCGTTGCATCACTTGAACCTGGGAGATGTCNNNAGATCGGAAG |
| 37885_2 | TATGAGTGTGGAGTCGTTGCATCACTTGAACCTGGGAGATGTCNNNAGATCGGAAG |
| 37885_3 | TATGAGTGTGGAGTCGTTGCATCACTTGAACCTGGGAGATGTCNNNAGATCGGAAG |
| 37885_4 | TATGAGTGTGGAGTCGTTGCATCACTTGAACCTGGGAGATGTCNNNAGATCGGAAG |
| 37885_5 | TATGAGTGTGGAGTCGTTGCATCACTTGAACCTGGGAGATGTCNNNAGATCGGAAG |
| 37885_6 | TATGAGTGTGGAGTCGTTGCATCACTTGAACCTGGGAGATGTCNNNAGATCGGAAG |
| 37885_7 | TATGAGTGTGGAGTCGTTGCATCACTTGAACCTGGGAGATGTCNNNAGATCGGAAG |
| 37885_8 | TATGAGTGTGGAGTCGTTGCATCACTTGAACCTGGGAGATGTCNNNAGATCGGAAG |
| 38987_1 | TATGAGTGTGGAGTCGTTGCACCCAACAGAGCAGACAACATTANNNAGATCGGAAG |
| 38987_2 | TATGAGTGTGGAGTCGTTGCACCCAACAGAGCAGACAACATTANNNAGATCGGAAG |
| 38987_3 | TATGAGTGTGGAGTCGTTGCACCCAACAGAGCAGACAACATTANNNAGATCGGAAG |
| 38987_4 | TATGAGTGTGGAGTCGTTGCACCCAACAGAGCAGACAACATTANNNAGATCGGAAG |
| 38987_5 | TATGAGTGTGGAGTCGTTGCACCCAACAGAGCAGACAACATTANNNAGATCGGAAG |
| 38987_6 | TATGAGTGTGGAGTCGTTGCACCCAACAGAGCAGACAACATTANNNAGATCGGAAG |
| 38987_7 | TATGAGTGTGGAGTCGTTGCACCCAACAGAGCAGACAACATTANNNAGATCGGAAG |
| 38987_8 | TATGAGTGTGGAGTCGTTGCACCCAACAGAGCAGACAACATTANNNAGATCGGAAG |
| 47282_1 | TATGAGTGTGGAGTCGTTGCGGGTTCAAGTGATTCTCCTGTCTNNNAGATCGGAAG |
| 47282_2 | TATGAGTGTGGAGTCGTTGCGGGTTCAAGTGATTCTCCTGTCTNNNAGATCGGAAG |
| 47282_3 | TATGAGTGTGGAGTCGTTGCGGGTTCAAGTGATTCTCCTGTCTNNNAGATCGGAAG |
| 47282_4 | TATGAGTGTGGAGTCGTTGCGGGTTCAAGTGATTCTCCTGTCTNNNAGATCGGAAG |
| 47282_5 | TATGAGTGTGGAGTCGTTGCGGGTTCAAGTGATTCTCCTGTCTNNNAGATCGGAAG |
| 47282_6 | TATGAGTGTGGAGTCGTTGCGGGTTCAAGTGATTCTCCTGTCTNNNAGATCGGAAG |
| 47282_7 | TATGAGTGTGGAGTCGTTGCGGGTTCAAGTGATTCTCCTGTCTNNNAGATCGGAAG |
| 47282_8 | TATGAGTGTGGAGTCGTTGCGGGTTCAAGTGATTCTCCTGTCTNNNAGATCGGAAG |
| 38260_1 | TATGAGTGTGGAGTCGTTGCCCTGGAGTCAGCAAAGTGGATATNNNAGATCGGAA |
| 38260_2 | TATGAGTGTGGAGTCGTTGCCCTGGAGTCAGCAAAGTGGATATNNNAGATCGGAA |
| 38260_3 | TATGAGTGTGGAGTCGTTGCCCTGGAGTCAGCAAAGTGGATATNNNAGATCGGAA |
| 38260_4 | TATGAGTGTGGAGTCGTTGCCCTGGAGTCAGCAAAGTGGATATNNNAGATCGGAA |
| 38260_5 | TATGAGTGTGGAGTCGTTGCCCTGGAGTCAGCAAAGTGGATATNNNAGATCGGAA |

| | |
|---|---|
| 38260_6 | TATGAGTGTGGAGTCGTTGCCCTGGAGTCAGCAAAGTGGATATNNNAGATCGGAA |
| 38260_7 | TATGAGTGTGGAGTCGTTGCCCTGGAGTCAGCAAAGTGGATATNNNAGATCGGAA |
| 38260_8 | TATGAGTGTGGAGTCGTTGCCCTGGAGTCAGCAAAGTGGATATNNNAGATCGGAA |
| 36602_1 | TATGAGTGTGGAGTCGTTGCAAGCATAGAGAAATGGGAGACCCNNNAGATCGGAA |
| 36602_2 | TATGAGTGTGGAGTCGTTGCAAGCATAGAGAAATGGGAGACCCNNNAGATCGGAA |
| 36602_3 | TATGAGTGTGGAGTCGTTGCAAGCATAGAGAAATGGGAGACCCNNNAGATCGGAA |
| 36602_4 | TATGAGTGTGGAGTCGTTGCAAGCATAGAGAAATGGGAGACCCNNNAGATCGGAA |
| 36602_5 | TATGAGTGTGGAGTCGTTGCAAGCATAGAGAAATGGGAGACCCNNNAGATCGGAA |
| 36602_6 | TATGAGTGTGGAGTCGTTGCAAGCATAGAGAAATGGGAGACCCNNNAGATCGGAA |
| 36602_7 | TATGAGTGTGGAGTCGTTGCAAGCATAGAGAAATGGGAGACCCNNNAGATCGGAA |
| 36602_8 | TATGAGTGTGGAGTCGTTGCAAGCATAGAGAAATGGGAGACCCNNNAGATCGGAA |
| 13793_1 | TATGAGTGTGGAGTCGTTGCTGGAAGAGGTATAGTTAAAGATCNNNAGATCGGAA |
| 13793_2 | TATGAGTGTGGAGTCGTTGCTGGAAGAGGTATAGTTAAAGATCNNNAGATCGGAA |
| 13793_3 | TATGAGTGTGGAGTCGTTGCTGGAAGAGGTATAGTTAAAGATCNNNAGATCGGAA |
| 13793_4 | TATGAGTGTGGAGTCGTTGCTGGAAGAGGTATAGTTAAAGATCNNNAGATCGGAA |
| 13793_5 | TATGAGTGTGGAGTCGTTGCTGGAAGAGGTATAGTTAAAGATCNNNAGATCGGAA |
| 13793_6 | TATGAGTGTGGAGTCGTTGCTGGAAGAGGTATAGTTAAAGATCNNNAGATCGGAA |
| 13793_7 | TATGAGTGTGGAGTCGTTGCTGGAAGAGGTATAGTTAAAGATCNNNAGATCGGAA |
| 13793_8 | TATGAGTGTGGAGTCGTTGCTGGAAGAGGTATAGTTAAAGATCNNNAGATCGGAA |
| 2574_1 | TATGAGTGTGGAGTCGTTGCTTCTAATCTGCACACTGGTTGGGNNNAGATCGGAAG |
| 2574_2 | TATGAGTGTGGAGTCGTTGCTTCTAATCTGCACACTGGTTGGGNNNAGATCGGAAG |
| 2574_3 | TATGAGTGTGGAGTCGTTGCTTCTAATCTGCACACTGGTTGGGNNNAGATCGGAAG |
| 2574_4 | TATGAGTGTGGAGTCGTTGCTTCTAATCTGCACACTGGTTGGGNNNAGATCGGAAG |
| 2574_5 | TATGAGTGTGGAGTCGTTGCTTCTAATCTGCACACTGGTTGGGNNNAGATCGGAAG |
| 2574_6 | TATGAGTGTGGAGTCGTTGCTTCTAATCTGCACACTGGTTGGGNNNAGATCGGAAG |
| 2574_7 | TATGAGTGTGGAGTCGTTGCTTCTAATCTGCACACTGGTTGGGNNNAGATCGGAAG |
| 2574_8 | TATGAGTGTGGAGTCGTTGCTTCTAATCTGCACACTGGTTGGGNNNAGATCGGAAG |
| 47334_1 | TATGAGTGTGGAGTCGTTGCCAAGAAGCAGCTCAGATTTAGGCNNNAGATCGGAA |
| 47334_2 | TATGAGTGTGGAGTCGTTGCCAAGAAGCAGCTCAGATTTAGGCNNNAGATCGGAA |
| 47334_3 | TATGAGTGTGGAGTCGTTGCCAAGAAGCAGCTCAGATTTAGGCNNNAGATCGGAA |
| 47334_4 | TATGAGTGTGGAGTCGTTGCCAAGAAGCAGCTCAGATTTAGGCNNNAGATCGGAA |
| 47334_5 | TATGAGTGTGGAGTCGTTGCCAAGAAGCAGCTCAGATTTAGGCNNNAGATCGGAA |
| 47334_6 | TATGAGTGTGGAGTCGTTGCCAAGAAGCAGCTCAGATTTAGGCNNNAGATCGGAA |
| 47334_7 | TATGAGTGTGGAGTCGTTGCCAAGAAGCAGCTCAGATTTAGGCNNNAGATCGGAA |
| 47334_8 | TATGAGTGTGGAGTCGTTGCCAAGAAGCAGCTCAGATTTAGGCNNNAGATCGGAA |
| 8279_1 | TATGAGTGTGGAGTCGTTGCAGACGCACTTTCACTGTGTCNNNAGATCGGAAGAGC |
| 8279_2 | TATGAGTGTGGAGTCGTTGCAGACGCACTTTCACTGTGTCNNNAGATCGGAAGAGC |
| 8279_3 | TATGAGTGTGGAGTCGTTGCAGACGCACTTTCACTGTGTCNNNAGATCGGAAGAGC |
| 8279_4 | TATGAGTGTGGAGTCGTTGCAGACGCACTTTCACTGTGTCNNNAGATCGGAAGAGC |
| 8279_5 | TATGAGTGTGGAGTCGTTGCAGACGCACTTTCACTGTGTCNNNAGATCGGAAGAGC |
| 8279_6 | TATGAGTGTGGAGTCGTTGCAGACGCACTTTCACTGTGTCNNNAGATCGGAAGAGC |
| 8279_7 | TATGAGTGTGGAGTCGTTGCAGACGCACTTTCACTGTGTCNNNAGATCGGAAGAGC |
| 8279_8 | TATGAGTGTGGAGTCGTTGCAGACGCACTTTCACTGTGTCNNNAGATCGGAAGAGC |

| | |
|---|---|
| 48281_1 | TATGAGTGTGGAGTCGTTGCCAGCCTTGATCTCCTGAGTTCAANNNAGATCGGAAG |
| 48281_2 | TATGAGTGTGGAGTCGTTGCCAGCCTTGATCTCCTGAGTTCAANNNAGATCGGAAG |
| 48281_3 | TATGAGTGTGGAGTCGTTGCCAGCCTTGATCTCCTGAGTTCAANNNAGATCGGAAG |
| 48281_4 | TATGAGTGTGGAGTCGTTGCCAGCCTTGATCTCCTGAGTTCAANNNAGATCGGAAG |
| 48281_5 | TATGAGTGTGGAGTCGTTGCCAGCCTTGATCTCCTGAGTTCAANNNAGATCGGAAG |
| 48281_6 | TATGAGTGTGGAGTCGTTGCCAGCCTTGATCTCCTGAGTTCAANNNAGATCGGAAG |
| 48281_7 | TATGAGTGTGGAGTCGTTGCCAGCCTTGATCTCCTGAGTTCAANNNAGATCGGAAG |
| 48281_8 | TATGAGTGTGGAGTCGTTGCCAGCCTTGATCTCCTGAGTTCAANNNAGATCGGAAG |
| 34456_1 | TATGAGTGTGGAGTCGTTGCCCCATCTCCTCCCCTCTCACNNNAGATCGGAAGAGCA |
| 34456_2 | TATGAGTGTGGAGTCGTTGCCCCATCTCCTCCCCTCTCACNNNAGATCGGAAGAGCA |
| 34456_3 | TATGAGTGTGGAGTCGTTGCCCCATCTCCTCCCCTCTCACNNNAGATCGGAAGAGCA |
| 34456_4 | TATGAGTGTGGAGTCGTTGCCCCATCTCCTCCCCTCTCACNNNAGATCGGAAGAGCA |
| 34456_5 | TATGAGTGTGGAGTCGTTGCCCCATCTCCTCCCCTCTCACNNNAGATCGGAAGAGCA |
| 34456_6 | TATGAGTGTGGAGTCGTTGCCCCATCTCCTCCCCTCTCACNNNAGATCGGAAGAGCA |
| 34456_7 | TATGAGTGTGGAGTCGTTGCCCCATCTCCTCCCCTCTCACNNNAGATCGGAAGAGCA |
| 34456_8 | TATGAGTGTGGAGTCGTTGCCCCATCTCCTCCCCTCTCACNNNAGATCGGAAGAGCA |
| 31458_1 | TATGAGTGTGGAGTCGTTGCTGTAAGTTTTGAGCTAGGACTGCTNNNAGATCGGAA |
| 31458_2 | TATGAGTGTGGAGTCGTTGCTGTAAGTTTTGAGCTAGGACTGCTNNNAGATCGGAA |
| 31458_3 | TATGAGTGTGGAGTCGTTGCTGTAAGTTTTGAGCTAGGACTGCTNNNAGATCGGAA |
| 31458_4 | TATGAGTGTGGAGTCGTTGCTGTAAGTTTTGAGCTAGGACTGCTNNNAGATCGGAA |
| 31458_5 | TATGAGTGTGGAGTCGTTGCTGTAAGTTTTGAGCTAGGACTGCTNNNAGATCGGAA |
| 31458_6 | TATGAGTGTGGAGTCGTTGCTGTAAGTTTTGAGCTAGGACTGCTNNNAGATCGGAA |
| 31458_7 | TATGAGTGTGGAGTCGTTGCTGTAAGTTTTGAGCTAGGACTGCTNNNAGATCGGAA |
| 31458_8 | TATGAGTGTGGAGTCGTTGCTGTAAGTTTTGAGCTAGGACTGCTNNNAGATCGGAA |
| 6030_1 | TATGAGTGTGGAGTCGTTGCTAACACAGGACACAGCACCCAAANNNAGATCGGAAG |
| 6030_2 | TATGAGTGTGGAGTCGTTGCTAACACAGGACACAGCACCCAAANNNAGATCGGAAG |
| 6030_3 | TATGAGTGTGGAGTCGTTGCTAACACAGGACACAGCACCCAAANNNAGATCGGAAG |
| 6030_4 | TATGAGTGTGGAGTCGTTGCTAACACAGGACACAGCACCCAAANNNAGATCGGAAG |
| 6030_5 | TATGAGTGTGGAGTCGTTGCTAACACAGGACACAGCACCCAAANNNAGATCGGAAG |
| 6030_6 | TATGAGTGTGGAGTCGTTGCTAACACAGGACACAGCACCCAAANNNAGATCGGAAG |
| 6030_7 | TATGAGTGTGGAGTCGTTGCTAACACAGGACACAGCACCCAAANNNAGATCGGAAG |
| 6030_8 | TATGAGTGTGGAGTCGTTGCTAACACAGGACACAGCACCCAAANNNAGATCGGAAG |
| 42378_1 | TATGAGTGTGGAGTCGTTGCCACCACACCTGGCTAATTTTTGTNNNAGATCGGAAG |
| 42378_2 | TATGAGTGTGGAGTCGTTGCCACCACACCTGGCTAATTTTTGTNNNAGATCGGAAG |
| 42378_3 | TATGAGTGTGGAGTCGTTGCCACCACACCTGGCTAATTTTTGTNNNAGATCGGAAG |
| 42378_4 | TATGAGTGTGGAGTCGTTGCCACCACACCTGGCTAATTTTTGTNNNAGATCGGAAG |
| 42378_5 | TATGAGTGTGGAGTCGTTGCCACCACACCTGGCTAATTTTTGTNNNAGATCGGAAG |
| 42378_6 | TATGAGTGTGGAGTCGTTGCCACCACACCTGGCTAATTTTTGTNNNAGATCGGAAG |
| 42378_7 | TATGAGTGTGGAGTCGTTGCCACCACACCTGGCTAATTTTTGTNNNAGATCGGAAG |
| 42378_8 | TATGAGTGTGGAGTCGTTGCCACCACACCTGGCTAATTTTTGTNNNAGATCGGAAG |
| 43065_1 | TATGAGTGTGGAGTCGTTGCTCTCTCTCTCTCTCTCTCACACCNNNAGATCGGAAGAG |
| 43065_2 | TATGAGTGTGGAGTCGTTGCTCTCTCTCTCTCTCTCTCACACCNNNAGATCGGAAGAG |
| 43065_3 | TATGAGTGTGGAGTCGTTGCTCTCTCTCTCTCTCTCTCACACCNNNAGATCGGAAGAG |

| | |
|---|---|
| 43065_4 | TATGAGTGTGGAGTCGTTGCTCTCTCTCTCTCTCTCACACCNNNAGATCGGAAGAG |
| 43065_5 | TATGAGTGTGGAGTCGTTGCTCTCTCTCTCTCTCTCACACCNNNAGATCGGAAGAG |
| 43065_6 | TATGAGTGTGGAGTCGTTGCTCTCTCTCTCTCTCTCACACCNNNAGATCGGAAGAG |
| 43065_7 | TATGAGTGTGGAGTCGTTGCTCTCTCTCTCTCTCTCACACCNNNAGATCGGAAGAG |
| 43065_8 | TATGAGTGTGGAGTCGTTGCTCTCTCTCTCTCTCTCACACCNNNAGATCGGAAGAG |
| 39554_1 | TATGAGTGTGGAGTCGTTGCAAGGAAGTTTGGCAAGATCATGANNNAGATCGGAA |
| 39554_2 | TATGAGTGTGGAGTCGTTGCAAGGAAGTTTGGCAAGATCATGANNNAGATCGGAA |
| 39554_3 | TATGAGTGTGGAGTCGTTGCAAGGAAGTTTGGCAAGATCATGANNNAGATCGGAA |
| 39554_4 | TATGAGTGTGGAGTCGTTGCAAGGAAGTTTGGCAAGATCATGANNNAGATCGGAA |
| 39554_5 | TATGAGTGTGGAGTCGTTGCAAGGAAGTTTGGCAAGATCATGANNNAGATCGGAA |
| 39554_6 | TATGAGTGTGGAGTCGTTGCAAGGAAGTTTGGCAAGATCATGANNNAGATCGGAA |
| 39554_7 | TATGAGTGTGGAGTCGTTGCAAGGAAGTTTGGCAAGATCATGANNNAGATCGGAA |
| 39554_8 | TATGAGTGTGGAGTCGTTGCAAGGAAGTTTGGCAAGATCATGANNNAGATCGGAA |
| 30178_1 | TATGAGTGTGGAGTCGTTGCTCCAAAATTTCATAGACTTTTGCCCTNNNAGATCGGA |
| 30178_2 | TATGAGTGTGGAGTCGTTGCTCCAAAATTTCATAGACTTTTGCCCTNNNAGATCGGA |
| 30178_3 | TATGAGTGTGGAGTCGTTGCTCCAAAATTTCATAGACTTTTGCCCTNNNAGATCGGA |
| 30178_4 | TATGAGTGTGGAGTCGTTGCTCCAAAATTTCATAGACTTTTGCCCTNNNAGATCGGA |
| 30178_5 | TATGAGTGTGGAGTCGTTGCTCCAAAATTTCATAGACTTTTGCCCTNNNAGATCGGA |
| 30178_6 | TATGAGTGTGGAGTCGTTGCTCCAAAATTTCATAGACTTTTGCCCTNNNAGATCGGA |
| 30178_7 | TATGAGTGTGGAGTCGTTGCTCCAAAATTTCATAGACTTTTGCCCTNNNAGATCGGA |
| 30178_8 | TATGAGTGTGGAGTCGTTGCTCCAAAATTTCATAGACTTTTGCCCTNNNAGATCGGA |
| 44859_1 | TATGAGTGTGGAGTCGTTGCACACTGAAGACCAATGATGGAGGNNNAGATCGGAA |
| 44859_2 | TATGAGTGTGGAGTCGTTGCACACTGAAGACCAATGATGGAGGNNNAGATCGGAA |
| 44859_3 | TATGAGTGTGGAGTCGTTGCACACTGAAGACCAATGATGGAGGNNNAGATCGGAA |
| 44859_4 | TATGAGTGTGGAGTCGTTGCACACTGAAGACCAATGATGGAGGNNNAGATCGGAA |
| 44859_5 | TATGAGTGTGGAGTCGTTGCACACTGAAGACCAATGATGGAGGNNNAGATCGGAA |
| 44859_6 | TATGAGTGTGGAGTCGTTGCACACTGAAGACCAATGATGGAGGNNNAGATCGGAA |
| 44859_7 | TATGAGTGTGGAGTCGTTGCACACTGAAGACCAATGATGGAGGNNNAGATCGGAA |
| 44859_8 | TATGAGTGTGGAGTCGTTGCACACTGAAGACCAATGATGGAGGNNNAGATCGGAA |
| 35537_1 | TATGAGTGTGGAGTCGTTGCGTGGGATCTCTACTCACTGCAAGNNNAGATCGGAAG |
| 35537_2 | TATGAGTGTGGAGTCGTTGCGTGGGATCTCTACTCACTGCAAGNNNAGATCGGAAG |
| 35537_3 | TATGAGTGTGGAGTCGTTGCGTGGGATCTCTACTCACTGCAAGNNNAGATCGGAAG |
| 35537_4 | TATGAGTGTGGAGTCGTTGCGTGGGATCTCTACTCACTGCAAGNNNAGATCGGAAG |
| 35537_5 | TATGAGTGTGGAGTCGTTGCGTGGGATCTCTACTCACTGCAAGNNNAGATCGGAAG |
| 35537_6 | TATGAGTGTGGAGTCGTTGCGTGGGATCTCTACTCACTGCAAGNNNAGATCGGAAG |
| 35537_7 | TATGAGTGTGGAGTCGTTGCGTGGGATCTCTACTCACTGCAAGNNNAGATCGGAAG |
| 35537_8 | TATGAGTGTGGAGTCGTTGCGTGGGATCTCTACTCACTGCAAGNNNAGATCGGAAG |
| 31130_1 | TATGAGTGTGGAGTCGTTGCATGCAGCTTTTATATCGCTAGCTNNNAGATCGGAAG |
| 31130_2 | TATGAGTGTGGAGTCGTTGCATGCAGCTTTTATATCGCTAGCTNNNAGATCGGAAG |
| 31130_3 | TATGAGTGTGGAGTCGTTGCATGCAGCTTTTATATCGCTAGCTNNNAGATCGGAAG |
| 31130_4 | TATGAGTGTGGAGTCGTTGCATGCAGCTTTTATATCGCTAGCTNNNAGATCGGAAG |
| 31130_5 | TATGAGTGTGGAGTCGTTGCATGCAGCTTTTATATCGCTAGCTNNNAGATCGGAAG |
| 31130_6 | TATGAGTGTGGAGTCGTTGCATGCAGCTTTTATATCGCTAGCTNNNAGATCGGAAG |

| | |
|---|---|
| 31130_7 | TATGAGTGTGGAGTCGTTGCATGCAGCTTTTATATCGCTAGCTNNNAGATCGGAAG |
| 31130_8 | TATGAGTGTGGAGTCGTTGCATGCAGCTTTTATATCGCTAGCTNNNAGATCGGAAG |
| 31415_1 | TATGAGTGTGGAGTCGTTGCATTAGGCCACAGTCCTCTTTGAGNNNAGATCGGAAG |
| 31415_2 | TATGAGTGTGGAGTCGTTGCATTAGGCCACAGTCCTCTTTGAGNNNAGATCGGAAG |
| 31415_3 | TATGAGTGTGGAGTCGTTGCATTAGGCCACAGTCCTCTTTGAGNNNAGATCGGAAG |
| 31415_4 | TATGAGTGTGGAGTCGTTGCATTAGGCCACAGTCCTCTTTGAGNNNAGATCGGAAG |
| 31415_5 | TATGAGTGTGGAGTCGTTGCATTAGGCCACAGTCCTCTTTGAGNNNAGATCGGAAG |
| 31415_6 | TATGAGTGTGGAGTCGTTGCATTAGGCCACAGTCCTCTTTGAGNNNAGATCGGAAG |
| 31415_7 | TATGAGTGTGGAGTCGTTGCATTAGGCCACAGTCCTCTTTGAGNNNAGATCGGAAG |
| 31415_8 | TATGAGTGTGGAGTCGTTGCATTAGGCCACAGTCCTCTTTGAGNNNAGATCGGAAG |
| 37384_1 | TATGAGTGTGGAGTCGTTGCCCAGGAACCAAGTGAGCATCTAANNNAGATCGGAAG |
| 37384_2 | TATGAGTGTGGAGTCGTTGCCCAGGAACCAAGTGAGCATCTAANNNAGATCGGAAG |
| 37384_3 | TATGAGTGTGGAGTCGTTGCCCAGGAACCAAGTGAGCATCTAANNNAGATCGGAAG |
| 37384_4 | TATGAGTGTGGAGTCGTTGCCCAGGAACCAAGTGAGCATCTAANNNAGATCGGAAG |
| 37384_5 | TATGAGTGTGGAGTCGTTGCCCAGGAACCAAGTGAGCATCTAANNNAGATCGGAAG |
| 37384_6 | TATGAGTGTGGAGTCGTTGCCCAGGAACCAAGTGAGCATCTAANNNAGATCGGAAG |
| 37384_7 | TATGAGTGTGGAGTCGTTGCCCAGGAACCAAGTGAGCATCTAANNNAGATCGGAAG |
| 37384_8 | TATGAGTGTGGAGTCGTTGCCCAGGAACCAAGTGAGCATCTAANNNAGATCGGAAG |
| 42757_1 | TATGAGTGTGGAGTCGTTGCAGAGAGAGGGAATTGGAGAATGCNNNAGATCGGA |
| 42757_2 | TATGAGTGTGGAGTCGTTGCAGAGAGAGGGAATTGGAGAATGCNNNAGATCGGA |
| 42757_3 | TATGAGTGTGGAGTCGTTGCAGAGAGAGGGAATTGGAGAATGCNNNAGATCGGA |
| 42757_4 | TATGAGTGTGGAGTCGTTGCAGAGAGAGGGAATTGGAGAATGCNNNAGATCGGA |
| 42757_5 | TATGAGTGTGGAGTCGTTGCAGAGAGAGGGAATTGGAGAATGCNNNAGATCGGA |
| 42757_6 | TATGAGTGTGGAGTCGTTGCAGAGAGAGGGAATTGGAGAATGCNNNAGATCGGA |
| 42757_7 | TATGAGTGTGGAGTCGTTGCAGAGAGAGGGAATTGGAGAATGCNNNAGATCGGA |
| 42757_8 | TATGAGTGTGGAGTCGTTGCAGAGAGAGGGAATTGGAGAATGCNNNAGATCGGA |
| 30892_1 | TATGAGTGTGGAGTCGTTGCCTGTCTCAGAATGACGGAAATGCNNNAGATCGGAAG |
| 30892_2 | TATGAGTGTGGAGTCGTTGCCTGTCTCAGAATGACGGAAATGCNNNAGATCGGAAG |
| 30892_3 | TATGAGTGTGGAGTCGTTGCCTGTCTCAGAATGACGGAAATGCNNNAGATCGGAAG |
| 30892_4 | TATGAGTGTGGAGTCGTTGCCTGTCTCAGAATGACGGAAATGCNNNAGATCGGAAG |
| 30892_5 | TATGAGTGTGGAGTCGTTGCCTGTCTCAGAATGACGGAAATGCNNNAGATCGGAAG |
| 30892_6 | TATGAGTGTGGAGTCGTTGCCTGTCTCAGAATGACGGAAATGCNNNAGATCGGAAG |
| 30892_7 | TATGAGTGTGGAGTCGTTGCCTGTCTCAGAATGACGGAAATGCNNNAGATCGGAAG |
| 30892_8 | TATGAGTGTGGAGTCGTTGCCTGTCTCAGAATGACGGAAATGCNNNAGATCGGAAG |
| 33756_1 | TATGAGTGTGGAGTCGTTGCTTTTTAGGGCATCCAGAAGCAAGNNNAGATCGGAAG |
| 33756_2 | TATGAGTGTGGAGTCGTTGCTTTTTAGGGCATCCAGAAGCAAGNNNAGATCGGAAG |
| 33756_3 | TATGAGTGTGGAGTCGTTGCTTTTTAGGGCATCCAGAAGCAAGNNNAGATCGGAAG |
| 33756_4 | TATGAGTGTGGAGTCGTTGCTTTTTAGGGCATCCAGAAGCAAGNNNAGATCGGAAG |
| 33756_5 | TATGAGTGTGGAGTCGTTGCTTTTTAGGGCATCCAGAAGCAAGNNNAGATCGGAAG |
| 33756_6 | TATGAGTGTGGAGTCGTTGCTTTTTAGGGCATCCAGAAGCAAGNNNAGATCGGAAG |
| 33756_7 | TATGAGTGTGGAGTCGTTGCTTTTTAGGGCATCCAGAAGCAAGNNNAGATCGGAAG |
| 33756_8 | TATGAGTGTGGAGTCGTTGCTTTTTAGGGCATCCAGAAGCAAGNNNAGATCGGAAG |
| 39355_1 | TATGAGTGTGGAGTCGTTGCGGGAGCCAGAGGAAGAAAGTAAANNNAGATCGGA |

| | |
|---|---|
| 39355_2 | TATGAGTGTGGAGTCGTTGCGGGAGCCAGAGGAAGAAAGTAAANNNAGATCGGA |
| 39355_3 | TATGAGTGTGGAGTCGTTGCGGGAGCCAGAGGAAGAAAGTAAANNNAGATCGGA |
| 39355_4 | TATGAGTGTGGAGTCGTTGCGGGAGCCAGAGGAAGAAAGTAAANNNAGATCGGA |
| 39355_5 | TATGAGTGTGGAGTCGTTGCGGGAGCCAGAGGAAGAAAGTAAANNNAGATCGGA |
| 39355_6 | TATGAGTGTGGAGTCGTTGCGGGAGCCAGAGGAAGAAAGTAAANNNAGATCGGA |
| 39355_7 | TATGAGTGTGGAGTCGTTGCGGGAGCCAGAGGAAGAAAGTAAANNNAGATCGGA |
| 39355_8 | TATGAGTGTGGAGTCGTTGCGGGAGCCAGAGGAAGAAAGTAAANNNAGATCGGA |
| 35897_1 | TATGAGTGTGGAGTCGTTGCAGTCAAAGGCAACGAGGATTCTANNNAGATCGGAA |
| 35897_2 | TATGAGTGTGGAGTCGTTGCAGTCAAAGGCAACGAGGATTCTANNNAGATCGGAA |
| 35897_3 | TATGAGTGTGGAGTCGTTGCAGTCAAAGGCAACGAGGATTCTANNNAGATCGGAA |
| 35897_4 | TATGAGTGTGGAGTCGTTGCAGTCAAAGGCAACGAGGATTCTANNNAGATCGGAA |
| 35897_5 | TATGAGTGTGGAGTCGTTGCAGTCAAAGGCAACGAGGATTCTANNNAGATCGGAA |
| 35897_6 | TATGAGTGTGGAGTCGTTGCAGTCAAAGGCAACGAGGATTCTANNNAGATCGGAA |
| 35897_7 | TATGAGTGTGGAGTCGTTGCAGTCAAAGGCAACGAGGATTCTANNNAGATCGGAA |
| 35897_8 | TATGAGTGTGGAGTCGTTGCAGTCAAAGGCAACGAGGATTCTANNNAGATCGGAA |
| 43951_1 | TATGAGTGTGGAGTCGTTGCTCAACAGAAGAAAGGAGGAGAGCNNNAGATCGGA |
| 43951_2 | TATGAGTGTGGAGTCGTTGCTCAACAGAAGAAAGGAGGAGAGCNNNAGATCGGA |
| 43951_3 | TATGAGTGTGGAGTCGTTGCTCAACAGAAGAAAGGAGGAGAGCNNNAGATCGGA |
| 43951_4 | TATGAGTGTGGAGTCGTTGCTCAACAGAAGAAAGGAGGAGAGCNNNAGATCGGA |
| 43951_5 | TATGAGTGTGGAGTCGTTGCTCAACAGAAGAAAGGAGGAGAGCNNNAGATCGGA |
| 43951_6 | TATGAGTGTGGAGTCGTTGCTCAACAGAAGAAAGGAGGAGAGCNNNAGATCGGA |
| 43951_7 | TATGAGTGTGGAGTCGTTGCTCAACAGAAGAAAGGAGGAGAGCNNNAGATCGGA |
| 43951_8 | TATGAGTGTGGAGTCGTTGCTCAACAGAAGAAAGGAGGAGAGCNNNAGATCGGA |
| 48338_1 | TATGAGTGTGGAGTCGTTGCCCTCCCAAAGTGCTAGGATTACANNNAGATCGGAAG |
| 48338_2 | TATGAGTGTGGAGTCGTTGCCCTCCCAAAGTGCTAGGATTACANNNAGATCGGAAG |
| 48338_3 | TATGAGTGTGGAGTCGTTGCCCTCCCAAAGTGCTAGGATTACANNNAGATCGGAAG |
| 48338_4 | TATGAGTGTGGAGTCGTTGCCCTCCCAAAGTGCTAGGATTACANNNAGATCGGAAG |
| 48338_5 | TATGAGTGTGGAGTCGTTGCCCTCCCAAAGTGCTAGGATTACANNNAGATCGGAAG |
| 48338_6 | TATGAGTGTGGAGTCGTTGCCCTCCCAAAGTGCTAGGATTACANNNAGATCGGAAG |
| 48338_7 | TATGAGTGTGGAGTCGTTGCCCTCCCAAAGTGCTAGGATTACANNNAGATCGGAAG |
| 48338_8 | TATGAGTGTGGAGTCGTTGCCCTCCCAAAGTGCTAGGATTACANNNAGATCGGAAG |
| 36220_1 | TATGAGTGTGGAGTCGTTGCTATCTTTGAAGACACGGGACACANNNAGATCGGAA |
| 36220_2 | TATGAGTGTGGAGTCGTTGCTATCTTTGAAGACACGGGACACANNNAGATCGGAA |
| 36220_3 | TATGAGTGTGGAGTCGTTGCTATCTTTGAAGACACGGGACACANNNAGATCGGAA |
| 36220_4 | TATGAGTGTGGAGTCGTTGCTATCTTTGAAGACACGGGACACANNNAGATCGGAA |
| 36220_5 | TATGAGTGTGGAGTCGTTGCTATCTTTGAAGACACGGGACACANNNAGATCGGAA |
| 36220_6 | TATGAGTGTGGAGTCGTTGCTATCTTTGAAGACACGGGACACANNNAGATCGGAA |
| 36220_7 | TATGAGTGTGGAGTCGTTGCTATCTTTGAAGACACGGGACACANNNAGATCGGAA |
| 36220_8 | TATGAGTGTGGAGTCGTTGCTATCTTTGAAGACACGGGACACANNNAGATCGGAA |
| 13614_1 | TATGAGTGTGGAGTCGTTGCCCCATATGCTATCCCTTATATTTNNNAGATCGGAAGA |
| 13614_2 | TATGAGTGTGGAGTCGTTGCCCCATATGCTATCCCTTATATTTNNNAGATCGGAAGA |
| 13614_3 | TATGAGTGTGGAGTCGTTGCCCCATATGCTATCCCTTATATTTNNNAGATCGGAAGA |
| 13614_4 | TATGAGTGTGGAGTCGTTGCCCCATATGCTATCCCTTATATTTNNNAGATCGGAAGA |

| | |
|---|---|
| 13614_5 | TATGAGTGTGGAGTCGTTGCCCCATATGCTATCCCTTATATTTNNNAGATCGGAAGA |
| 13614_6 | TATGAGTGTGGAGTCGTTGCCCCATATGCTATCCCTTATATTTNNNAGATCGGAAGA |
| 13614_7 | TATGAGTGTGGAGTCGTTGCCCCATATGCTATCCCTTATATTTNNNAGATCGGAAGA |
| 13614_8 | TATGAGTGTGGAGTCGTTGCCCCATATGCTATCCCTTATATTTNNNAGATCGGAAGA |
| 44271_1 | TATGAGTGTGGAGTCGTTGCAACCAGCTTACCCTTGAATTTGCNNNAGATCGGAAG |
| 44271_2 | TATGAGTGTGGAGTCGTTGCAACCAGCTTACCCTTGAATTTGCNNNAGATCGGAAG |
| 44271_3 | TATGAGTGTGGAGTCGTTGCAACCAGCTTACCCTTGAATTTGCNNNAGATCGGAAG |
| 44271_4 | TATGAGTGTGGAGTCGTTGCAACCAGCTTACCCTTGAATTTGCNNNAGATCGGAAG |
| 44271_5 | TATGAGTGTGGAGTCGTTGCAACCAGCTTACCCTTGAATTTGCNNNAGATCGGAAG |
| 44271_6 | TATGAGTGTGGAGTCGTTGCAACCAGCTTACCCTTGAATTTGCNNNAGATCGGAAG |
| 44271_7 | TATGAGTGTGGAGTCGTTGCAACCAGCTTACCCTTGAATTTGCNNNAGATCGGAAG |
| 44271_8 | TATGAGTGTGGAGTCGTTGCAACCAGCTTACCCTTGAATTTGCNNNAGATCGGAAG |
| 12973_1 | TATGAGTGTGGAGTCGTTGCCCTAAAGAAGATCTTATGGAAACNNNAGATCGGAAG |
| 12973_2 | TATGAGTGTGGAGTCGTTGCCCTAAAGAAGATCTTATGGAAACNNNAGATCGGAAG |
| 12973_3 | TATGAGTGTGGAGTCGTTGCCCTAAAGAAGATCTTATGGAAACNNNAGATCGGAAG |
| 12973_4 | TATGAGTGTGGAGTCGTTGCCCTAAAGAAGATCTTATGGAAACNNNAGATCGGAAG |
| 12973_5 | TATGAGTGTGGAGTCGTTGCCCTAAAGAAGATCTTATGGAAACNNNAGATCGGAAG |
| 12973_6 | TATGAGTGTGGAGTCGTTGCCCTAAAGAAGATCTTATGGAAACNNNAGATCGGAAG |
| 12973_7 | TATGAGTGTGGAGTCGTTGCCCTAAAGAAGATCTTATGGAAACNNNAGATCGGAAG |
| 12973_8 | TATGAGTGTGGAGTCGTTGCCCTAAAGAAGATCTTATGGAAACNNNAGATCGGAAG |
| 42767_1 | TATGAGTGTGGAGTCGTTGCGTTACAGCCTGGGTTACAGAGTGNNNAGATCGGAA |
| 42767_2 | TATGAGTGTGGAGTCGTTGCGTTACAGCCTGGGTTACAGAGTGNNNAGATCGGAA |
| 42767_3 | TATGAGTGTGGAGTCGTTGCGTTACAGCCTGGGTTACAGAGTGNNNAGATCGGAA |
| 42767_4 | TATGAGTGTGGAGTCGTTGCGTTACAGCCTGGGTTACAGAGTGNNNAGATCGGAA |
| 42767_5 | TATGAGTGTGGAGTCGTTGCGTTACAGCCTGGGTTACAGAGTGNNNAGATCGGAA |
| 42767_6 | TATGAGTGTGGAGTCGTTGCGTTACAGCCTGGGTTACAGAGTGNNNAGATCGGAA |
| 42767_7 | TATGAGTGTGGAGTCGTTGCGTTACAGCCTGGGTTACAGAGTGNNNAGATCGGAA |
| 42767_8 | TATGAGTGTGGAGTCGTTGCGTTACAGCCTGGGTTACAGAGTGNNNAGATCGGAA |
| 32717_1 | TATGAGTGTGGAGTCGTTGCGGAACTTCCAGAGCAATCTTACGNNNAGATCGGAAG |
| 32717_2 | TATGAGTGTGGAGTCGTTGCGGAACTTCCAGAGCAATCTTACGNNNAGATCGGAAG |
| 32717_3 | TATGAGTGTGGAGTCGTTGCGGAACTTCCAGAGCAATCTTACGNNNAGATCGGAAG |
| 32717_4 | TATGAGTGTGGAGTCGTTGCGGAACTTCCAGAGCAATCTTACGNNNAGATCGGAAG |
| 32717_5 | TATGAGTGTGGAGTCGTTGCGGAACTTCCAGAGCAATCTTACGNNNAGATCGGAAG |
| 32717_6 | TATGAGTGTGGAGTCGTTGCGGAACTTCCAGAGCAATCTTACGNNNAGATCGGAAG |
| 32717_7 | TATGAGTGTGGAGTCGTTGCGGAACTTCCAGAGCAATCTTACGNNNAGATCGGAAG |
| 32717_8 | TATGAGTGTGGAGTCGTTGCGGAACTTCCAGAGCAATCTTACGNNNAGATCGGAAG |
| 34416_1 | TATGAGTGTGGAGTCGTTGCACCAGTCCTAAACACCAGATTGGNNNAGATCGGAAG |
| 34416_2 | TATGAGTGTGGAGTCGTTGCACCAGTCCTAAACACCAGATTGGNNNAGATCGGAAG |
| 34416_3 | TATGAGTGTGGAGTCGTTGCACCAGTCCTAAACACCAGATTGGNNNAGATCGGAAG |
| 34416_4 | TATGAGTGTGGAGTCGTTGCACCAGTCCTAAACACCAGATTGGNNNAGATCGGAAG |
| 34416_5 | TATGAGTGTGGAGTCGTTGCACCAGTCCTAAACACCAGATTGGNNNAGATCGGAAG |
| 34416_6 | TATGAGTGTGGAGTCGTTGCACCAGTCCTAAACACCAGATTGGNNNAGATCGGAAG |
| 34416_7 | TATGAGTGTGGAGTCGTTGCACCAGTCCTAAACACCAGATTGGNNNAGATCGGAAG |

| | |
|---|---|
| 34416_8 | TATGAGTGTGGAGTCGTTGCACCAGTCCTAAACACCAGATTGGNNNAGATCGGAAG |
| 8440_1 | TATGAGTGTGGAGTCGTTGCGTCTGGTCTCACACTTCTGGATACNNNAGATCGGAA |
| 8440_2 | TATGAGTGTGGAGTCGTTGCGTCTGGTCTCACACTTCTGGATACNNNAGATCGGAA |
| 8440_3 | TATGAGTGTGGAGTCGTTGCGTCTGGTCTCACACTTCTGGATACNNNAGATCGGAA |
| 8440_4 | TATGAGTGTGGAGTCGTTGCGTCTGGTCTCACACTTCTGGATACNNNAGATCGGAA |
| 8440_5 | TATGAGTGTGGAGTCGTTGCGTCTGGTCTCACACTTCTGGATACNNNAGATCGGAA |
| 8440_6 | TATGAGTGTGGAGTCGTTGCGTCTGGTCTCACACTTCTGGATACNNNAGATCGGAA |
| 8440_7 | TATGAGTGTGGAGTCGTTGCGTCTGGTCTCACACTTCTGGATACNNNAGATCGGAA |
| 8440_8 | TATGAGTGTGGAGTCGTTGCGTCTGGTCTCACACTTCTGGATACNNNAGATCGGAA |
| 39191_1 | TATGAGTGTGGAGTCGTTGCCCCTCCCTTCCTCTTCTTTTCAGNNNAGATCGGAAGA |
| 39191_2 | TATGAGTGTGGAGTCGTTGCCCCTCCCTTCCTCTTCTTTTCAGNNNAGATCGGAAGA |
| 39191_3 | TATGAGTGTGGAGTCGTTGCCCCTCCCTTCCTCTTCTTTTCAGNNNAGATCGGAAGA |
| 39191_4 | TATGAGTGTGGAGTCGTTGCCCCTCCCTTCCTCTTCTTTTCAGNNNAGATCGGAAGA |
| 39191_5 | TATGAGTGTGGAGTCGTTGCCCCTCCCTTCCTCTTCTTTTCAGNNNAGATCGGAAGA |
| 39191_6 | TATGAGTGTGGAGTCGTTGCCCCTCCCTTCCTCTTCTTTTCAGNNNAGATCGGAAGA |
| 39191_7 | TATGAGTGTGGAGTCGTTGCCCCTCCCTTCCTCTTCTTTTCAGNNNAGATCGGAAGA |
| 39191_8 | TATGAGTGTGGAGTCGTTGCCCCTCCCTTCCTCTTCTTTTCAGNNNAGATCGGAAGA |
| 33335_1 | TATGAGTGTGGAGTCGTTGCCTCCTGAGTAGCTGGGACTACAGNNNAGATCGGAA |
| 33335_2 | TATGAGTGTGGAGTCGTTGCCTCCTGAGTAGCTGGGACTACAGNNNAGATCGGAA |
| 33335_3 | TATGAGTGTGGAGTCGTTGCCTCCTGAGTAGCTGGGACTACAGNNNAGATCGGAA |
| 33335_4 | TATGAGTGTGGAGTCGTTGCCTCCTGAGTAGCTGGGACTACAGNNNAGATCGGAA |
| 33335_5 | TATGAGTGTGGAGTCGTTGCCTCCTGAGTAGCTGGGACTACAGNNNAGATCGGAA |
| 33335_6 | TATGAGTGTGGAGTCGTTGCCTCCTGAGTAGCTGGGACTACAGNNNAGATCGGAA |
| 33335_7 | TATGAGTGTGGAGTCGTTGCCTCCTGAGTAGCTGGGACTACAGNNNAGATCGGAA |
| 33335_8 | TATGAGTGTGGAGTCGTTGCCTCCTGAGTAGCTGGGACTACAGNNNAGATCGGAA |
| 3494_1 | TATGAGTGTGGAGTCGTTGCCCAACTCATTTCTGCTGCCCANNNAGATCGGAAGAG |
| 3494_2 | TATGAGTGTGGAGTCGTTGCCCAACTCATTTCTGCTGCCCANNNAGATCGGAAGAG |
| 3494_3 | TATGAGTGTGGAGTCGTTGCCCAACTCATTTCTGCTGCCCANNNAGATCGGAAGAG |
| 3494_4 | TATGAGTGTGGAGTCGTTGCCCAACTCATTTCTGCTGCCCANNNAGATCGGAAGAG |
| 3494_5 | TATGAGTGTGGAGTCGTTGCCCAACTCATTTCTGCTGCCCANNNAGATCGGAAGAG |
| 3494_6 | TATGAGTGTGGAGTCGTTGCCCAACTCATTTCTGCTGCCCANNNAGATCGGAAGAG |
| 3494_7 | TATGAGTGTGGAGTCGTTGCCCAACTCATTTCTGCTGCCCANNNAGATCGGAAGAG |
| 3494_8 | TATGAGTGTGGAGTCGTTGCCCAACTCATTTCTGCTGCCCANNNAGATCGGAAGAG |
| 34698_1 | TATGAGTGTGGAGTCGTTGCAGGGTCTTACCATATTGCCCATGNNNAGATCGGAAG |
| 34698_2 | TATGAGTGTGGAGTCGTTGCAGGGTCTTACCATATTGCCCATGNNNAGATCGGAAG |
| 34698_3 | TATGAGTGTGGAGTCGTTGCAGGGTCTTACCATATTGCCCATGNNNAGATCGGAAG |
| 34698_4 | TATGAGTGTGGAGTCGTTGCAGGGTCTTACCATATTGCCCATGNNNAGATCGGAAG |
| 34698_5 | TATGAGTGTGGAGTCGTTGCAGGGTCTTACCATATTGCCCATGNNNAGATCGGAAG |
| 34698_6 | TATGAGTGTGGAGTCGTTGCAGGGTCTTACCATATTGCCCATGNNNAGATCGGAAG |
| 34698_7 | TATGAGTGTGGAGTCGTTGCAGGGTCTTACCATATTGCCCATGNNNAGATCGGAAG |
| 34698_8 | TATGAGTGTGGAGTCGTTGCAGGGTCTTACCATATTGCCCATGNNNAGATCGGAAG |
| 50969_1 | TATGAGTGTGGAGTCGTTGCCACAGTAAATGGCAATTCCCTGTNNNAGATCGGAAG |
| 50969_2 | TATGAGTGTGGAGTCGTTGCCACAGTAAATGGCAATTCCCTGTNNNAGATCGGAAG |

| | |
|---|---|
| 50969_3 | TATGAGTGTGGAGTCGTTGCCACAGTAAATGGCAATTCCCTGTNNNAGATCGGAAG |
| 50969_4 | TATGAGTGTGGAGTCGTTGCCACAGTAAATGGCAATTCCCTGTNNNAGATCGGAAG |
| 50969_5 | TATGAGTGTGGAGTCGTTGCCACAGTAAATGGCAATTCCCTGTNNNAGATCGGAAG |
| 50969_6 | TATGAGTGTGGAGTCGTTGCCACAGTAAATGGCAATTCCCTGTNNNAGATCGGAAG |
| 50969_7 | TATGAGTGTGGAGTCGTTGCCACAGTAAATGGCAATTCCCTGTNNNAGATCGGAAG |
| 50969_8 | TATGAGTGTGGAGTCGTTGCCACAGTAAATGGCAATTCCCTGTNNNAGATCGGAAG |
| 30476_1 | TATGAGTGTGGAGTCGTTGCGAAGGGGAGGGGGTTTAGATTGANNNAGATCGGA |
| 30476_2 | TATGAGTGTGGAGTCGTTGCGAAGGGGAGGGGGTTTAGATTGANNNAGATCGGA |
| 30476_3 | TATGAGTGTGGAGTCGTTGCGAAGGGGAGGGGGTTTAGATTGANNNAGATCGGA |
| 30476_4 | TATGAGTGTGGAGTCGTTGCGAAGGGGAGGGGGTTTAGATTGANNNAGATCGGA |
| 30476_5 | TATGAGTGTGGAGTCGTTGCGAAGGGGAGGGGGTTTAGATTGANNNAGATCGGA |
| 30476_6 | TATGAGTGTGGAGTCGTTGCGAAGGGGAGGGGGTTTAGATTGANNNAGATCGGA |
| 30476_7 | TATGAGTGTGGAGTCGTTGCGAAGGGGAGGGGGTTTAGATTGANNNAGATCGGA |
| 30476_8 | TATGAGTGTGGAGTCGTTGCGAAGGGGAGGGGGTTTAGATTGANNNAGATCGGA |
| 40775_1 | TATGAGTGTGGAGTCGTTGCGTGTGTGTGTGTGTGAACTCAAANNNAGATCGGAA[illegible] |
| 40775_2 | TATGAGTGTGGAGTCGTTGCGTGTGTGTGTGTGTGAACTCAAANNNAGATCGGAA[illegible] |
| 40775_3 | TATGAGTGTGGAGTCGTTGCGTGTGTGTGTGTGTGAACTCAAANNNAGATCGGAA[illegible] |
| 40775_4 | TATGAGTGTGGAGTCGTTGCGTGTGTGTGTGTGTGAACTCAAANNNAGATCGGAA[illegible] |
| 40775_5 | TATGAGTGTGGAGTCGTTGCGTGTGTGTGTGTGTGAACTCAAANNNAGATCGGAA[illegible] |
| 40775_6 | TATGAGTGTGGAGTCGTTGCGTGTGTGTGTGTGTGAACTCAAANNNAGATCGGAA[illegible] |
| 40775_7 | TATGAGTGTGGAGTCGTTGCGTGTGTGTGTGTGTGAACTCAAANNNAGATCGGAA[illegible] |
| 40775_8 | TATGAGTGTGGAGTCGTTGCGTGTGTGTGTGTGTGAACTCAAANNNAGATCGGAA[illegible] |
| 48889_1 | TATGAGTGTGGAGTCGTTGCCCATGCCTATGTCCTGAATGGTANNNAGATCGGAAG |
| 48889_2 | TATGAGTGTGGAGTCGTTGCCCATGCCTATGTCCTGAATGGTANNNAGATCGGAAG |
| 48889_3 | TATGAGTGTGGAGTCGTTGCCCATGCCTATGTCCTGAATGGTANNNAGATCGGAAG |
| 48889_4 | TATGAGTGTGGAGTCGTTGCCCATGCCTATGTCCTGAATGGTANNNAGATCGGAAG |
| 48889_5 | TATGAGTGTGGAGTCGTTGCCCATGCCTATGTCCTGAATGGTANNNAGATCGGAAG |
| 48889_6 | TATGAGTGTGGAGTCGTTGCCCATGCCTATGTCCTGAATGGTANNNAGATCGGAAG |
| 48889_7 | TATGAGTGTGGAGTCGTTGCCCATGCCTATGTCCTGAATGGTANNNAGATCGGAAG |
| 48889_8 | TATGAGTGTGGAGTCGTTGCCCATGCCTATGTCCTGAATGGTANNNAGATCGGAAG |
| 38813_1 | TATGAGTGTGGAGTCGTTGCCCTGTAATCCCATAATCCCAGCANNNAGATCGGAAG[illegible] |
| 38813_2 | TATGAGTGTGGAGTCGTTGCCCTGTAATCCCATAATCCCAGCANNNAGATCGGAAG[illegible] |
| 38813_3 | TATGAGTGTGGAGTCGTTGCCCTGTAATCCCATAATCCCAGCANNNAGATCGGAAG[illegible] |
| 38813_4 | TATGAGTGTGGAGTCGTTGCCCTGTAATCCCATAATCCCAGCANNNAGATCGGAAG[illegible] |
| 38813_5 | TATGAGTGTGGAGTCGTTGCCCTGTAATCCCATAATCCCAGCANNNAGATCGGAAG[illegible] |
| 38813_6 | TATGAGTGTGGAGTCGTTGCCCTGTAATCCCATAATCCCAGCANNNAGATCGGAAG[illegible] |
| 38813_7 | TATGAGTGTGGAGTCGTTGCCCTGTAATCCCATAATCCCAGCANNNAGATCGGAAG[illegible] |
| 38813_8 | TATGAGTGTGGAGTCGTTGCCCTGTAATCCCATAATCCCAGCANNNAGATCGGAAG[illegible] |
| 50448_1 | TATGAGTGTGGAGTCGTTGCCTTTCCCAACAAAAGCCCTTTCCNNNAGATCGGAAGA |
| 50448_2 | TATGAGTGTGGAGTCGTTGCCTTTCCCAACAAAAGCCCTTTCCNNNAGATCGGAAGA |
| 50448_3 | TATGAGTGTGGAGTCGTTGCCTTTCCCAACAAAAGCCCTTTCCNNNAGATCGGAAGA |
| 50448_4 | TATGAGTGTGGAGTCGTTGCCTTTCCCAACAAAAGCCCTTTCCNNNAGATCGGAAGA |
| 50448_5 | TATGAGTGTGGAGTCGTTGCCTTTCCCAACAAAAGCCCTTTCCNNNAGATCGGAAGA |

| | |
|---|---|
| 50448_6 | TATGAGTGTGGAGTCGTTGCCTTTCCCAACAAAAGCCCTTTCCNNNAGATCGGAAGA |
| 50448_7 | TATGAGTGTGGAGTCGTTGCCTTTCCCAACAAAAGCCCTTTCCNNNAGATCGGAAGA |
| 50448_8 | TATGAGTGTGGAGTCGTTGCCTTTCCCAACAAAAGCCCTTTCCNNNAGATCGGAAGA |
| 12700_1 | TATGAGTGTGGAGTCGTTGCTATTCGACAAATATGTATGATGCNNNAGATCGGAAG |
| 12700_2 | TATGAGTGTGGAGTCGTTGCTATTCGACAAATATGTATGATGCNNNAGATCGGAAG |
| 12700_3 | TATGAGTGTGGAGTCGTTGCTATTCGACAAATATGTATGATGCNNNAGATCGGAAG |
| 12700_4 | TATGAGTGTGGAGTCGTTGCTATTCGACAAATATGTATGATGCNNNAGATCGGAAG |
| 12700_5 | TATGAGTGTGGAGTCGTTGCTATTCGACAAATATGTATGATGCNNNAGATCGGAAG |
| 12700_6 | TATGAGTGTGGAGTCGTTGCTATTCGACAAATATGTATGATGCNNNAGATCGGAAG |
| 12700_7 | TATGAGTGTGGAGTCGTTGCTATTCGACAAATATGTATGATGCNNNAGATCGGAAG |
| 12700_8 | TATGAGTGTGGAGTCGTTGCTATTCGACAAATATGTATGATGCNNNAGATCGGAAG |
| 42828_1 | TATGAGTGTGGAGTCGTTGCTTCAAGTGATTCTCCTGCCTCAGNNNAGATCGGAAG |
| 42828_2 | TATGAGTGTGGAGTCGTTGCTTCAAGTGATTCTCCTGCCTCAGNNNAGATCGGAAG |
| 42828_3 | TATGAGTGTGGAGTCGTTGCTTCAAGTGATTCTCCTGCCTCAGNNNAGATCGGAAG |
| 42828_4 | TATGAGTGTGGAGTCGTTGCTTCAAGTGATTCTCCTGCCTCAGNNNAGATCGGAAG |
| 42828_5 | TATGAGTGTGGAGTCGTTGCTTCAAGTGATTCTCCTGCCTCAGNNNAGATCGGAAG |
| 42828_6 | TATGAGTGTGGAGTCGTTGCTTCAAGTGATTCTCCTGCCTCAGNNNAGATCGGAAG |
| 42828_7 | TATGAGTGTGGAGTCGTTGCTTCAAGTGATTCTCCTGCCTCAGNNNAGATCGGAAG |
| 42828_8 | TATGAGTGTGGAGTCGTTGCTTCAAGTGATTCTCCTGCCTCAGNNNAGATCGGAAG |
| 42160_1 | TATGAGTGTGGAGTCGTTGCGCCACACACCCTCTTTTTATGTNNNAGATCGGAAGA |
| 42160_2 | TATGAGTGTGGAGTCGTTGCGCCACACACCCTCTTTTTATGTNNNAGATCGGAAGA |
| 42160_3 | TATGAGTGTGGAGTCGTTGCGCCACACACCCTCTTTTTATGTNNNAGATCGGAAGA |
| 42160_4 | TATGAGTGTGGAGTCGTTGCGCCACACACCCTCTTTTTATGTNNNAGATCGGAAGA |
| 42160_5 | TATGAGTGTGGAGTCGTTGCGCCACACACCCTCTTTTTATGTNNNAGATCGGAAGA |
| 42160_6 | TATGAGTGTGGAGTCGTTGCGCCACACACCCTCTTTTTATGTNNNAGATCGGAAGA |
| 42160_7 | TATGAGTGTGGAGTCGTTGCGCCACACACCCTCTTTTTATGTNNNAGATCGGAAGA |
| 42160_8 | TATGAGTGTGGAGTCGTTGCGCCACACACCCTCTTTTTATGTNNNAGATCGGAAGA |
| 40422_1 | TATGAGTGTGGAGTCGTTGCCTCAACGTCATCAGTCATTTGGGNNNAGATCGGAAG |
| 40422_2 | TATGAGTGTGGAGTCGTTGCCTCAACGTCATCAGTCATTTGGGNNNAGATCGGAAG |
| 40422_3 | TATGAGTGTGGAGTCGTTGCCTCAACGTCATCAGTCATTTGGGNNNAGATCGGAAG |
| 40422_4 | TATGAGTGTGGAGTCGTTGCCTCAACGTCATCAGTCATTTGGGNNNAGATCGGAAG |
| 40422_5 | TATGAGTGTGGAGTCGTTGCCTCAACGTCATCAGTCATTTGGGNNNAGATCGGAAG |
| 40422_6 | TATGAGTGTGGAGTCGTTGCCTCAACGTCATCAGTCATTTGGGNNNAGATCGGAAG |
| 40422_7 | TATGAGTGTGGAGTCGTTGCCTCAACGTCATCAGTCATTTGGGNNNAGATCGGAAG |
| 40422_8 | TATGAGTGTGGAGTCGTTGCCTCAACGTCATCAGTCATTTGGGNNNAGATCGGAAG |
| 35961_1 | TATGAGTGTGGAGTCGTTGCAAAAAAGAGGGAGAGAGCTTCCCNNNAGATCGGAA |
| 35961_2 | TATGAGTGTGGAGTCGTTGCAAAAAAGAGGGAGAGAGCTTCCCNNNAGATCGGAA |
| 35961_3 | TATGAGTGTGGAGTCGTTGCAAAAAAGAGGGAGAGAGCTTCCCNNNAGATCGGAA |
| 35961_4 | TATGAGTGTGGAGTCGTTGCAAAAAAGAGGGAGAGAGCTTCCCNNNAGATCGGAA |
| 35961_5 | TATGAGTGTGGAGTCGTTGCAAAAAAGAGGGAGAGAGCTTCCCNNNAGATCGGAA |
| 35961_6 | TATGAGTGTGGAGTCGTTGCAAAAAAGAGGGAGAGAGCTTCCCNNNAGATCGGAA |
| 35961_7 | TATGAGTGTGGAGTCGTTGCAAAAAAGAGGGAGAGAGCTTCCCNNNAGATCGGAA |
| 35961_8 | TATGAGTGTGGAGTCGTTGCAAAAAAGAGGGAGAGAGCTTCCCNNNAGATCGGAA |

| | |
|---|---|
| 53345_1 | TATGAGTGTGGAGTCGTTGCGTGGAGAAAATGGAACCCTTGTGNNNAGATCGGAA |
| 53345_2 | TATGAGTGTGGAGTCGTTGCGTGGAGAAAATGGAACCCTTGTGNNNAGATCGGAA |
| 53345_3 | TATGAGTGTGGAGTCGTTGCGTGGAGAAAATGGAACCCTTGTGNNNAGATCGGAA |
| 53345_4 | TATGAGTGTGGAGTCGTTGCGTGGAGAAAATGGAACCCTTGTGNNNAGATCGGAA |
| 53345_5 | TATGAGTGTGGAGTCGTTGCGTGGAGAAAATGGAACCCTTGTGNNNAGATCGGAA |
| 53345_6 | TATGAGTGTGGAGTCGTTGCGTGGAGAAAATGGAACCCTTGTGNNNAGATCGGAA |
| 53345_7 | TATGAGTGTGGAGTCGTTGCGTGGAGAAAATGGAACCCTTGTGNNNAGATCGGAA |
| 53345_8 | TATGAGTGTGGAGTCGTTGCGTGGAGAAAATGGAACCCTTGTGNNNAGATCGGAA |
| 41595_1 | TATGAGTGTGGAGTCGTTGCGATTACAGGCATGCCCCACTATANNNAGATCGGAAG |
| 41595_2 | TATGAGTGTGGAGTCGTTGCGATTACAGGCATGCCCCACTATANNNAGATCGGAAG |
| 41595_3 | TATGAGTGTGGAGTCGTTGCGATTACAGGCATGCCCCACTATANNNAGATCGGAAG |
| 41595_4 | TATGAGTGTGGAGTCGTTGCGATTACAGGCATGCCCCACTATANNNAGATCGGAAG |
| 41595_5 | TATGAGTGTGGAGTCGTTGCGATTACAGGCATGCCCCACTATANNNAGATCGGAAG |
| 41595_6 | TATGAGTGTGGAGTCGTTGCGATTACAGGCATGCCCCACTATANNNAGATCGGAAG |
| 41595_7 | TATGAGTGTGGAGTCGTTGCGATTACAGGCATGCCCCACTATANNNAGATCGGAAG |
| 41595_8 | TATGAGTGTGGAGTCGTTGCGATTACAGGCATGCCCCACTATANNNAGATCGGAAG |
| 36346_1 | TATGAGTGTGGAGTCGTTGCTCACGTGGCTTTCTGATCCTTTANNNAGATCGGAAG |
| 36346_2 | TATGAGTGTGGAGTCGTTGCTCACGTGGCTTTCTGATCCTTTANNNAGATCGGAAG |
| 36346_3 | TATGAGTGTGGAGTCGTTGCTCACGTGGCTTTCTGATCCTTTANNNAGATCGGAAG |
| 36346_4 | TATGAGTGTGGAGTCGTTGCTCACGTGGCTTTCTGATCCTTTANNNAGATCGGAAG |
| 36346_5 | TATGAGTGTGGAGTCGTTGCTCACGTGGCTTTCTGATCCTTTANNNAGATCGGAAG |
| 36346_6 | TATGAGTGTGGAGTCGTTGCTCACGTGGCTTTCTGATCCTTTANNNAGATCGGAAG |
| 36346_7 | TATGAGTGTGGAGTCGTTGCTCACGTGGCTTTCTGATCCTTTANNNAGATCGGAAG |
| 36346_8 | TATGAGTGTGGAGTCGTTGCTCACGTGGCTTTCTGATCCTTTANNNAGATCGGAAG |
| 32280_1 | TATGAGTGTGGAGTCGTTGCTGGTGCTCAGAACAGATCTATGGNNNAGATCGGAA |
| 32280_2 | TATGAGTGTGGAGTCGTTGCTGGTGCTCAGAACAGATCTATGGNNNAGATCGGAA |
| 32280_3 | TATGAGTGTGGAGTCGTTGCTGGTGCTCAGAACAGATCTATGGNNNAGATCGGAA |
| 32280_4 | TATGAGTGTGGAGTCGTTGCTGGTGCTCAGAACAGATCTATGGNNNAGATCGGAA |
| 32280_5 | TATGAGTGTGGAGTCGTTGCTGGTGCTCAGAACAGATCTATGGNNNAGATCGGAA |
| 32280_6 | TATGAGTGTGGAGTCGTTGCTGGTGCTCAGAACAGATCTATGGNNNAGATCGGAA |
| 32280_7 | TATGAGTGTGGAGTCGTTGCTGGTGCTCAGAACAGATCTATGGNNNAGATCGGAA |
| 32280_8 | TATGAGTGTGGAGTCGTTGCTGGTGCTCAGAACAGATCTATGGNNNAGATCGGAA |
| 35813_1 | TATGAGTGTGGAGTCGTTGCGGAGTCTTACTCTATTGCCCAGGNNNAGATCGGAAG |
| 35813_2 | TATGAGTGTGGAGTCGTTGCGGAGTCTTACTCTATTGCCCAGGNNNAGATCGGAAG |
| 35813_3 | TATGAGTGTGGAGTCGTTGCGGAGTCTTACTCTATTGCCCAGGNNNAGATCGGAAG |
| 35813_4 | TATGAGTGTGGAGTCGTTGCGGAGTCTTACTCTATTGCCCAGGNNNAGATCGGAAG |
| 35813_5 | TATGAGTGTGGAGTCGTTGCGGAGTCTTACTCTATTGCCCAGGNNNAGATCGGAAG |
| 35813_6 | TATGAGTGTGGAGTCGTTGCGGAGTCTTACTCTATTGCCCAGGNNNAGATCGGAAG |
| 35813_7 | TATGAGTGTGGAGTCGTTGCGGAGTCTTACTCTATTGCCCAGGNNNAGATCGGAAG |
| 35813_8 | TATGAGTGTGGAGTCGTTGCGGAGTCTTACTCTATTGCCCAGGNNNAGATCGGAAG |
| 52267_1 | TATGAGTGTGGAGTCGTTGCACACTGGGGGTCAGAGAGTAGNNNAGATCGGAAG |
| 52267_2 | TATGAGTGTGGAGTCGTTGCACACTGGGGGTCAGAGAGTAGNNNAGATCGGAAG |
| 52267_3 | TATGAGTGTGGAGTCGTTGCACACTGGGGGTCAGAGAGTAGNNNAGATCGGAAG |

| | |
|---|---|
| 52267_4 | TATGAGTGTGGAGTCGTTGCACACTGGGGGTCAGAGAGTAGNNNAGATCGGAAG |
| 52267_5 | TATGAGTGTGGAGTCGTTGCACACTGGGGGTCAGAGAGTAGNNNAGATCGGAAG |
| 52267_6 | TATGAGTGTGGAGTCGTTGCACACTGGGGGTCAGAGAGTAGNNNAGATCGGAAG |
| 52267_7 | TATGAGTGTGGAGTCGTTGCACACTGGGGGTCAGAGAGTAGNNNAGATCGGAAG |
| 52267_8 | TATGAGTGTGGAGTCGTTGCACACTGGGGGTCAGAGAGTAGNNNAGATCGGAAG |
| 39767_1 | TATGAGTGTGGAGTCGTTGCTGGCATTCCTGAGAAAGAAAGAGANNNAGATCGGA |
| 39767_2 | TATGAGTGTGGAGTCGTTGCTGGCATTCCTGAGAAAGAAAGAGANNNAGATCGGA |
| 39767_3 | TATGAGTGTGGAGTCGTTGCTGGCATTCCTGAGAAAGAAAGAGANNNAGATCGGA |
| 39767_4 | TATGAGTGTGGAGTCGTTGCTGGCATTCCTGAGAAAGAAAGAGANNNAGATCGGA |
| 39767_5 | TATGAGTGTGGAGTCGTTGCTGGCATTCCTGAGAAAGAAAGAGANNNAGATCGGA |
| 39767_6 | TATGAGTGTGGAGTCGTTGCTGGCATTCCTGAGAAAGAAAGAGANNNAGATCGGA |
| 39767_7 | TATGAGTGTGGAGTCGTTGCTGGCATTCCTGAGAAAGAAAGAGANNNAGATCGGA |
| 39767_8 | TATGAGTGTGGAGTCGTTGCTGGCATTCCTGAGAAAGAAAGAGANNNAGATCGGA |
| 38740_1 | TATGAGTGTGGAGTCGTTGCTGCAGATTCCTGGCTTCATCTAANNNAGATCGGAAG |
| 38740_2 | TATGAGTGTGGAGTCGTTGCTGCAGATTCCTGGCTTCATCTAANNNAGATCGGAAG |
| 38740_3 | TATGAGTGTGGAGTCGTTGCTGCAGATTCCTGGCTTCATCTAANNNAGATCGGAAG |
| 38740_4 | TATGAGTGTGGAGTCGTTGCTGCAGATTCCTGGCTTCATCTAANNNAGATCGGAAG |
| 38740_5 | TATGAGTGTGGAGTCGTTGCTGCAGATTCCTGGCTTCATCTAANNNAGATCGGAAG |
| 38740_6 | TATGAGTGTGGAGTCGTTGCTGCAGATTCCTGGCTTCATCTAANNNAGATCGGAAG |
| 38740_7 | TATGAGTGTGGAGTCGTTGCTGCAGATTCCTGGCTTCATCTAANNNAGATCGGAAG |
| 38740_8 | TATGAGTGTGGAGTCGTTGCTGCAGATTCCTGGCTTCATCTAANNNAGATCGGAAG |
| 42312_1 | TATGAGTGTGGAGTCGTTGCAACAAAAGTTCCAGAATGGGCAGNNNAGATCGGAA |
| 42312_2 | TATGAGTGTGGAGTCGTTGCAACAAAAGTTCCAGAATGGGCAGNNNAGATCGGAA |
| 42312_3 | TATGAGTGTGGAGTCGTTGCAACAAAAGTTCCAGAATGGGCAGNNNAGATCGGAA |
| 42312_4 | TATGAGTGTGGAGTCGTTGCAACAAAAGTTCCAGAATGGGCAGNNNAGATCGGAA |
| 42312_5 | TATGAGTGTGGAGTCGTTGCAACAAAAGTTCCAGAATGGGCAGNNNAGATCGGAA |
| 42312_6 | TATGAGTGTGGAGTCGTTGCAACAAAAGTTCCAGAATGGGCAGNNNAGATCGGAA |
| 42312_7 | TATGAGTGTGGAGTCGTTGCAACAAAAGTTCCAGAATGGGCAGNNNAGATCGGAA |
| 42312_8 | TATGAGTGTGGAGTCGTTGCAACAAAAGTTCCAGAATGGGCAGNNNAGATCGGAA |
| 39281_1 | TATGAGTGTGGAGTCGTTGCAGCCTAGTTGGACAGACATTAGCNNNAGATCGGAA |
| 39281_2 | TATGAGTGTGGAGTCGTTGCAGCCTAGTTGGACAGACATTAGCNNNAGATCGGAA |
| 39281_3 | TATGAGTGTGGAGTCGTTGCAGCCTAGTTGGACAGACATTAGCNNNAGATCGGAA |
| 39281_4 | TATGAGTGTGGAGTCGTTGCAGCCTAGTTGGACAGACATTAGCNNNAGATCGGAA |
| 39281_5 | TATGAGTGTGGAGTCGTTGCAGCCTAGTTGGACAGACATTAGCNNNAGATCGGAA |
| 39281_6 | TATGAGTGTGGAGTCGTTGCAGCCTAGTTGGACAGACATTAGCNNNAGATCGGAA |
| 39281_7 | TATGAGTGTGGAGTCGTTGCAGCCTAGTTGGACAGACATTAGCNNNAGATCGGAA |
| 39281_8 | TATGAGTGTGGAGTCGTTGCAGCCTAGTTGGACAGACATTAGCNNNAGATCGGAA |
| 47804_1 | TATGAGTGTGGAGTCGTTGCCAAAGGCATGATCTTGGCTCACNNNAGATCGGAAGA |
| 47804_2 | TATGAGTGTGGAGTCGTTGCCAAAGGCATGATCTTGGCTCACNNNAGATCGGAAGA |
| 47804_3 | TATGAGTGTGGAGTCGTTGCCAAAGGCATGATCTTGGCTCACNNNAGATCGGAAGA |
| 47804_4 | TATGAGTGTGGAGTCGTTGCCAAAGGCATGATCTTGGCTCACNNNAGATCGGAAGA |
| 47804_5 | TATGAGTGTGGAGTCGTTGCCAAAGGCATGATCTTGGCTCACNNNAGATCGGAAGA |
| 47804_6 | TATGAGTGTGGAGTCGTTGCCAAAGGCATGATCTTGGCTCACNNNAGATCGGAAGA |

| | |
|---|---|
| 47804_7 | TATGAGTGTGGAGTCGTTGCCAAAGGCATGATCTTGGCTCACNNNAGATCGGAAGA |
| 47804_8 | TATGAGTGTGGAGTCGTTGCCAAAGGCATGATCTTGGCTCACNNNAGATCGGAAGA |
| 42627_1 | TATGAGTGTGGAGTCGTTGCCCTCGAGCAATCCTTCTACATCANNNAGATCGGAAGA |
| 42627_2 | TATGAGTGTGGAGTCGTTGCCCTCGAGCAATCCTTCTACATCANNNAGATCGGAAGA |
| 42627_3 | TATGAGTGTGGAGTCGTTGCCCTCGAGCAATCCTTCTACATCANNNAGATCGGAAGA |
| 42627_4 | TATGAGTGTGGAGTCGTTGCCCTCGAGCAATCCTTCTACATCANNNAGATCGGAAGA |
| 42627_5 | TATGAGTGTGGAGTCGTTGCCCTCGAGCAATCCTTCTACATCANNNAGATCGGAAGA |
| 42627_6 | TATGAGTGTGGAGTCGTTGCCCTCGAGCAATCCTTCTACATCANNNAGATCGGAAGA |
| 42627_7 | TATGAGTGTGGAGTCGTTGCCCTCGAGCAATCCTTCTACATCANNNAGATCGGAAGA |
| 42627_8 | TATGAGTGTGGAGTCGTTGCCCTCGAGCAATCCTTCTACATCANNNAGATCGGAAGA |
| 12702_1 | TATGAGTGTGGAGTCGTTGCTTGGCTTTCTTTCTTTTTCTTTANNNAGATCGGAAGA |
| 12702_2 | TATGAGTGTGGAGTCGTTGCTTGGCTTTCTTTCTTTTTCTTTANNNAGATCGGAAGA |
| 12702_3 | TATGAGTGTGGAGTCGTTGCTTGGCTTTCTTTCTTTTTCTTTANNNAGATCGGAAGA |
| 12702_4 | TATGAGTGTGGAGTCGTTGCTTGGCTTTCTTTCTTTTTCTTTANNNAGATCGGAAGA |
| 12702_5 | TATGAGTGTGGAGTCGTTGCTTGGCTTTCTTTCTTTTTCTTTANNNAGATCGGAAGA |
| 12702_6 | TATGAGTGTGGAGTCGTTGCTTGGCTTTCTTTCTTTTTCTTTANNNAGATCGGAAGA |
| 12702_7 | TATGAGTGTGGAGTCGTTGCTTGGCTTTCTTTCTTTTTCTTTANNNAGATCGGAAGA |
| 12702_8 | TATGAGTGTGGAGTCGTTGCTTGGCTTTCTTTCTTTTTCTTTANNNAGATCGGAAGA |
| 38679_1 | TATGAGTGTGGAGTCGTTGCGCTCATCTAGGCTTTTTTCTAGGCNNNAGATCGGAA |
| 38679_2 | TATGAGTGTGGAGTCGTTGCGCTCATCTAGGCTTTTTTCTAGGCNNNAGATCGGAA |
| 38679_3 | TATGAGTGTGGAGTCGTTGCGCTCATCTAGGCTTTTTTCTAGGCNNNAGATCGGAA |
| 38679_4 | TATGAGTGTGGAGTCGTTGCGCTCATCTAGGCTTTTTTCTAGGCNNNAGATCGGAA |
| 38679_5 | TATGAGTGTGGAGTCGTTGCGCTCATCTAGGCTTTTTTCTAGGCNNNAGATCGGAA |
| 38679_6 | TATGAGTGTGGAGTCGTTGCGCTCATCTAGGCTTTTTTCTAGGCNNNAGATCGGAA |
| 38679_7 | TATGAGTGTGGAGTCGTTGCGCTCATCTAGGCTTTTTTCTAGGCNNNAGATCGGAA |
| 38679_8 | TATGAGTGTGGAGTCGTTGCGCTCATCTAGGCTTTTTTCTAGGCNNNAGATCGGAA |
| 39584_1 | TATGAGTGTGGAGTCGTTGCGTTTTTGAGAAACATCCCCTGGGNNNAGATCGGAAG |
| 39584_2 | TATGAGTGTGGAGTCGTTGCGTTTTTGAGAAACATCCCCTGGGNNNAGATCGGAAG |
| 39584_3 | TATGAGTGTGGAGTCGTTGCGTTTTTGAGAAACATCCCCTGGGNNNAGATCGGAAG |
| 39584_4 | TATGAGTGTGGAGTCGTTGCGTTTTTGAGAAACATCCCCTGGGNNNAGATCGGAAG |
| 39584_5 | TATGAGTGTGGAGTCGTTGCGTTTTTGAGAAACATCCCCTGGGNNNAGATCGGAAG |
| 39584_6 | TATGAGTGTGGAGTCGTTGCGTTTTTGAGAAACATCCCCTGGGNNNAGATCGGAAG |
| 39584_7 | TATGAGTGTGGAGTCGTTGCGTTTTTGAGAAACATCCCCTGGGNNNAGATCGGAAG |
| 39584_8 | TATGAGTGTGGAGTCGTTGCGTTTTTGAGAAACATCCCCTGGGNNNAGATCGGAAG |
| 44723_1 | TATGAGTGTGGAGTCGTTGCAGGGGAGGAAGGAAAGTCATAGANNNAGATCGGA |
| 44723_2 | TATGAGTGTGGAGTCGTTGCAGGGGAGGAAGGAAAGTCATAGANNNAGATCGGA |
| 44723_3 | TATGAGTGTGGAGTCGTTGCAGGGGAGGAAGGAAAGTCATAGANNNAGATCGGA |
| 44723_4 | TATGAGTGTGGAGTCGTTGCAGGGGAGGAAGGAAAGTCATAGANNNAGATCGGA |
| 44723_5 | TATGAGTGTGGAGTCGTTGCAGGGGAGGAAGGAAAGTCATAGANNNAGATCGGA |
| 44723_6 | TATGAGTGTGGAGTCGTTGCAGGGGAGGAAGGAAAGTCATAGANNNAGATCGGA |
| 44723_7 | TATGAGTGTGGAGTCGTTGCAGGGGAGGAAGGAAAGTCATAGANNNAGATCGGA |
| 44723_8 | TATGAGTGTGGAGTCGTTGCAGGGGAGGAAGGAAAGTCATAGANNNAGATCGGA |
| 41988_1 | TATGAGTGTGGAGTCGTTGCGTTCATGCCATTCTCCTGCTTCNNNAGATCGGAAGAG |

| | |
|---|---|
| 41988_2 | TATGAGTGTGGAGTCGTTGCGTTCATGCCATTCTCCTGCTTCNNNAGATCGGAAGAG |
| 41988_3 | TATGAGTGTGGAGTCGTTGCGTTCATGCCATTCTCCTGCTTCNNNAGATCGGAAGAG |
| 41988_4 | TATGAGTGTGGAGTCGTTGCGTTCATGCCATTCTCCTGCTTCNNNAGATCGGAAGAG |
| 41988_5 | TATGAGTGTGGAGTCGTTGCGTTCATGCCATTCTCCTGCTTCNNNAGATCGGAAGAG |
| 41988_6 | TATGAGTGTGGAGTCGTTGCGTTCATGCCATTCTCCTGCTTCNNNAGATCGGAAGAG |
| 41988_7 | TATGAGTGTGGAGTCGTTGCGTTCATGCCATTCTCCTGCTTCNNNAGATCGGAAGAG |
| 41988_8 | TATGAGTGTGGAGTCGTTGCGTTCATGCCATTCTCCTGCTTCNNNAGATCGGAAGAG |
| 33883_1 | TATGAGTGTGGAGTCGTTGCCCTCCTGAGTAACTGGGATTACANNNAGATCGGAAG |
| 33883_2 | TATGAGTGTGGAGTCGTTGCCCTCCTGAGTAACTGGGATTACANNNAGATCGGAAG |
| 33883_3 | TATGAGTGTGGAGTCGTTGCCCTCCTGAGTAACTGGGATTACANNNAGATCGGAAG |
| 33883_4 | TATGAGTGTGGAGTCGTTGCCCTCCTGAGTAACTGGGATTACANNNAGATCGGAAG |
| 33883_5 | TATGAGTGTGGAGTCGTTGCCCTCCTGAGTAACTGGGATTACANNNAGATCGGAAG |
| 33883_6 | TATGAGTGTGGAGTCGTTGCCCTCCTGAGTAACTGGGATTACANNNAGATCGGAAG |
| 33883_7 | TATGAGTGTGGAGTCGTTGCCCTCCTGAGTAACTGGGATTACANNNAGATCGGAAG |
| 33883_8 | TATGAGTGTGGAGTCGTTGCCCTCCTGAGTAACTGGGATTACANNNAGATCGGAAG |
| 47231_1 | TATGAGTGTGGAGTCGTTGCCAATCTCGGCTCACTGTAACCTCNNNAGATCGGAAGA |
| 47231_2 | TATGAGTGTGGAGTCGTTGCCAATCTCGGCTCACTGTAACCTCNNNAGATCGGAAGA |
| 47231_3 | TATGAGTGTGGAGTCGTTGCCAATCTCGGCTCACTGTAACCTCNNNAGATCGGAAGA |
| 47231_4 | TATGAGTGTGGAGTCGTTGCCAATCTCGGCTCACTGTAACCTCNNNAGATCGGAAGA |
| 47231_5 | TATGAGTGTGGAGTCGTTGCCAATCTCGGCTCACTGTAACCTCNNNAGATCGGAAGA |
| 47231_6 | TATGAGTGTGGAGTCGTTGCCAATCTCGGCTCACTGTAACCTCNNNAGATCGGAAGA |
| 47231_7 | TATGAGTGTGGAGTCGTTGCCAATCTCGGCTCACTGTAACCTCNNNAGATCGGAAGA |
| 47231_8 | TATGAGTGTGGAGTCGTTGCCAATCTCGGCTCACTGTAACCTCNNNAGATCGGAAGA |
| 40391_1 | TATGAGTGTGGAGTCGTTGCAGCCATAAATGAGATGCAGGAGTNNNAGATCGGAA |
| 40391_2 | TATGAGTGTGGAGTCGTTGCAGCCATAAATGAGATGCAGGAGTNNNAGATCGGAA |
| 40391_3 | TATGAGTGTGGAGTCGTTGCAGCCATAAATGAGATGCAGGAGTNNNAGATCGGAA |
| 40391_4 | TATGAGTGTGGAGTCGTTGCAGCCATAAATGAGATGCAGGAGTNNNAGATCGGAA |
| 40391_5 | TATGAGTGTGGAGTCGTTGCAGCCATAAATGAGATGCAGGAGTNNNAGATCGGAA |
| 40391_6 | TATGAGTGTGGAGTCGTTGCAGCCATAAATGAGATGCAGGAGTNNNAGATCGGAA |
| 40391_7 | TATGAGTGTGGAGTCGTTGCAGCCATAAATGAGATGCAGGAGTNNNAGATCGGAA |
| 40391_8 | TATGAGTGTGGAGTCGTTGCAGCCATAAATGAGATGCAGGAGTNNNAGATCGGAA |
| 50896_1 | TATGAGTGTGGAGTCGTTGCCTCCAGGCATTTAGTGTCCTCTTNNNAGATCGGAAGA |
| 50896_2 | TATGAGTGTGGAGTCGTTGCCTCCAGGCATTTAGTGTCCTCTTNNNAGATCGGAAGA |
| 50896_3 | TATGAGTGTGGAGTCGTTGCCTCCAGGCATTTAGTGTCCTCTTNNNAGATCGGAAGA |
| 50896_4 | TATGAGTGTGGAGTCGTTGCCTCCAGGCATTTAGTGTCCTCTTNNNAGATCGGAAGA |
| 50896_5 | TATGAGTGTGGAGTCGTTGCCTCCAGGCATTTAGTGTCCTCTTNNNAGATCGGAAGA |
| 50896_6 | TATGAGTGTGGAGTCGTTGCCTCCAGGCATTTAGTGTCCTCTTNNNAGATCGGAAGA |
| 50896_7 | TATGAGTGTGGAGTCGTTGCCTCCAGGCATTTAGTGTCCTCTTNNNAGATCGGAAGA |
| 50896_8 | TATGAGTGTGGAGTCGTTGCCTCCAGGCATTTAGTGTCCTCTTNNNAGATCGGAAGA |

AGCACACGTCTGAACTCTTTCCCTACACGACGCTCTTCCGATCTNNNAAGTGATAGAAATCGATCAGTT
AGCACACGTCTGAACTCTTTCCCTACACGACGCTCTTCCGATCTNNNAAGTGATAGAAATCGATCAGTT
AGCACACGTCTGAACTCTTTCCCTACACGACGCTCTTCCGATCTNNNAAGTGATAGAAATCGATCAGTT
AGCACACGTCTGAACTCTTTCCCTACACGACGCTCTTCCGATCTNNNAAGTGATAGAAATCGATCAGTT
AGCACACGTCTGAACTCTTTCCCTACACGACGCTCTTCCGATCTNNNAAGTGATAGAAATCGATCAGTT
AGCACACGTCTGAACTCTTTCCCTACACGACGCTCTTCCGATCTNNNAAGTGATAGAAATCGATCAGTT
AGCACACGTCTGAACTCTTTCCCTACACGACGCTCTTCCGATCTNNNAAGTGATAGAAATCGATCAGTT
AGCACACGTCTGAACTCTTTCCCTACACGACGCTCTTCCGATCTNNNCATTCCCACCAACAGTGTGTAAC
AGCACACGTCTGAACTCTTTCCCTACACGACGCTCTTCCGATCTNNNCATTCCCACCAACAGTGTGTAAC
AGCACACGTCTGAACTCTTTCCCTACACGACGCTCTTCCGATCTNNNCATTCCCACCAACAGTGTGTAAC
AGCACACGTCTGAACTCTTTCCCTACACGACGCTCTTCCGATCTNNNCATTCCCACCAACAGTGTGTAAC
AGCACACGTCTGAACTCTTTCCCTACACGACGCTCTTCCGATCTNNNCATTCCCACCAACAGTGTGTAAC
AGCACACGTCTGAACTCTTTCCCTACACGACGCTCTTCCGATCTNNNCATTCCCACCAACAGTGTGTAAC
AGCACACGTCTGAACTCTTTCCCTACACGACGCTCTTCCGATCTNNNCATTCCCACCAACAGTGTGTAAC
AGCACACGTCTGAACTCTTTCCCTACACGACGCTCTTCCGATCTNNNCACGGATGGCAGAGTTTCTGTA
AGCACACGTCTGAACTCTTTCCCTACACGACGCTCTTCCGATCTNNNCACGGATGGCAGAGTTTCTGTA
AGCACACGTCTGAACTCTTTCCCTACACGACGCTCTTCCGATCTNNNCACGGATGGCAGAGTTTCTGTA
AGCACACGTCTGAACTCTTTCCCTACACGACGCTCTTCCGATCTNNNCACGGATGGCAGAGTTTCTGTA
AGCACACGTCTGAACTCTTTCCCTACACGACGCTCTTCCGATCTNNNCACGGATGGCAGAGTTTCTGTA
AGCACACGTCTGAACTCTTTCCCTACACGACGCTCTTCCGATCTNNNCACGGATGGCAGAGTTTCTGTA
AGCACACGTCTGAACTCTTTCCCTACACGACGCTCTTCCGATCTNNNCACGGATGGCAGAGTTTCTGTA
GAGCACACGTCTGAACTCTTTCCCTACACGACGCTCTTCCGATCTNNNGCTCCAACCACCAGTTTAGATC
GAGCACACGTCTGAACTCTTTCCCTACACGACGCTCTTCCGATCTNNNGCTCCAACCACCAGTTTAGATC
GAGCACACGTCTGAACTCTTTCCCTACACGACGCTCTTCCGATCTNNNGCTCCAACCACCAGTTTAGATC
GAGCACACGTCTGAACTCTTTCCCTACACGACGCTCTTCCGATCTNNNGCTCCAACCACCAGTTTAGATC
GAGCACACGTCTGAACTCTTTCCCTACACGACGCTCTTCCGATCTNNNGCTCCAACCACCAGTTTAGATC
GAGCACACGTCTGAACTCTTTCCCTACACGACGCTCTTCCGATCTNNNGCTCCAACCACCAGTTTAGATC
GAGCACACGTCTGAACTCTTTCCCTACACGACGCTCTTCCGATCTNNNGCTCCAACCACCAGTTTAGATC
AGCACACGTCTGAACTCTTTCCCTACACGACGCTCTTCCGATCTNNNTGTAGTTCTAGCTACTCGGGAG
AGCACACGTCTGAACTCTTTCCCTACACGACGCTCTTCCGATCTNNNTGTAGTTCTAGCTACTCGGGAG
AGCACACGTCTGAACTCTTTCCCTACACGACGCTCTTCCGATCTNNNTGTAGTTCTAGCTACTCGGGAG
AGCACACGTCTGAACTCTTTCCCTACACGACGCTCTTCCGATCTNNNTGTAGTTCTAGCTACTCGGGAG
AGCACACGTCTGAACTCTTTCCCTACACGACGCTCTTCCGATCTNNNTGTAGTTCTAGCTACTCGGGAG
AGCACACGTCTGAACTCTTTCCCTACACGACGCTCTTCCGATCTNNNTGTAGTTCTAGCTACTCGGGAG
AGCACACGTCTGAACTCTTTCCCTACACGACGCTCTTCCGATCTNNNTGTAGTTCTAGCTACTCGGGAG
AGCACACGTCTGAACTCTTTCCCTACACGACGCTCTTCCGATCTNNNGGTAGCTTGAATCCAGGAGTTC
AGCACACGTCTGAACTCTTTCCCTACACGACGCTCTTCCGATCTNNNGGTAGCTTGAATCCAGGAGTTC
AGCACACGTCTGAACTCTTTCCCTACACGACGCTCTTCCGATCTNNNGGTAGCTTGAATCCAGGAGTTC
AGCACACGTCTGAACTCTTTCCCTACACGACGCTCTTCCGATCTNNNGGTAGCTTGAATCCAGGAGTTC
AGCACACGTCTGAACTCTTTCCCTACACGACGCTCTTCCGATCTNNNGGTAGCTTGAATCCAGGAGTTC
AGCACACGTCTGAACTCTTTCCCTACACGACGCTCTTCCGATCTNNNGGTAGCTTGAATCCAGGAGTTC
AGCACACGTCTGAACTCTTTCCCTACACGACGCTCTTCCGATCTNNNGGTAGCTTGAATCCAGGAGTTC

GAGCACACGTCTGAACTCTTTCCCTACACGACGCTCTTCCGATCTNNNAAAAAAACAATGATGGGGT
GAGCACACGTCTGAACTCTTTCCCTACACGACGCTCTTCCGATCTNNNAAAAAAACAATGATGGGGT
GAGCACACGTCTGAACTCTTTCCCTACACGACGCTCTTCCGATCTNNNAAAAAAACAATGATGGGGT
GAGCACACGTCTGAACTCTTTCCCTACACGACGCTCTTCCGATCTNNNAAAAAAACAATGATGGGGT
GAGCACACGTCTGAACTCTTTCCCTACACGACGCTCTTCCGATCTNNNAAAAAAACAATGATGGGGT
GAGCACACGTCTGAACTCTTTCCCTACACGACGCTCTTCCGATCTNNNAAAAAAACAATGATGGGGT
GAGCACACGTCTGAACTCTTTCCCTACACGACGCTCTTCCGATCTNNNAAAAAAACAATGATGGGGT
GAGCACACGTCTGAACTCTTTCCCTACACGACGCTCTTCCGATCTNNNCATGAGAATTGCTTGAACCC
GAGCACACGTCTGAACTCTTTCCCTACACGACGCTCTTCCGATCTNNNCATGAGAATTGCTTGAACCC
GAGCACACGTCTGAACTCTTTCCCTACACGACGCTCTTCCGATCTNNNCATGAGAATTGCTTGAACCC
GAGCACACGTCTGAACTCTTTCCCTACACGACGCTCTTCCGATCTNNNCATGAGAATTGCTTGAACCC
GAGCACACGTCTGAACTCTTTCCCTACACGACGCTCTTCCGATCTNNNCATGAGAATTGCTTGAACCC
GAGCACACGTCTGAACTCTTTCCCTACACGACGCTCTTCCGATCTNNNCATGAGAATTGCTTGAACCC
GAGCACACGTCTGAACTCTTTCCCTACACGACGCTCTTCCGATCTNNNCATGAGAATTGCTTGAACCC
AGCACACGTCTGAACTCTTTCCCTACACGACGCTCTTCCGATCTNNNTGTAAGAAAATGCAGCAGCTC
AGCACACGTCTGAACTCTTTCCCTACACGACGCTCTTCCGATCTNNNTGTAAGAAAATGCAGCAGCTC
AGCACACGTCTGAACTCTTTCCCTACACGACGCTCTTCCGATCTNNNTGTAAGAAAATGCAGCAGCTC
AGCACACGTCTGAACTCTTTCCCTACACGACGCTCTTCCGATCTNNNTGTAAGAAAATGCAGCAGCTC
AGCACACGTCTGAACTCTTTCCCTACACGACGCTCTTCCGATCTNNNTGTAAGAAAATGCAGCAGCTC
AGCACACGTCTGAACTCTTTCCCTACACGACGCTCTTCCGATCTNNNTGTAAGAAAATGCAGCAGCTC
AGCACACGTCTGAACTCTTTCCCTACACGACGCTCTTCCGATCTNNNTGTAAGAAAATGCAGCAGCTC
GCACACGTCTGAACTCTTTCCCTACACGACGCTCTTCCGATCTNNNGCCATGCTCTCTTCATGCAATTC
GCACACGTCTGAACTCTTTCCCTACACGACGCTCTTCCGATCTNNNGCCATGCTCTCTTCATGCAATTC
GCACACGTCTGAACTCTTTCCCTACACGACGCTCTTCCGATCTNNNGCCATGCTCTCTTCATGCAATTC
GCACACGTCTGAACTCTTTCCCTACACGACGCTCTTCCGATCTNNNGCCATGCTCTCTTCATGCAATTC
GCACACGTCTGAACTCTTTCCCTACACGACGCTCTTCCGATCTNNNGCCATGCTCTCTTCATGCAATTC
GCACACGTCTGAACTCTTTCCCTACACGACGCTCTTCCGATCTNNNGCCATGCTCTCTTCATGCAATTC
GCACACGTCTGAACTCTTTCCCTACACGACGCTCTTCCGATCTNNNGCCATGCTCTCTTCATGCAATTC
AGCACACGTCTGAACTCTTTCCCTACACGACGCTCTTCCGATCTNNNTGATTTCTCCGTAGAAGCATT
AGCACACGTCTGAACTCTTTCCCTACACGACGCTCTTCCGATCTNNNTGATTTCTCCGTAGAAGCATT
AGCACACGTCTGAACTCTTTCCCTACACGACGCTCTTCCGATCTNNNTGATTTCTCCGTAGAAGCATT
AGCACACGTCTGAACTCTTTCCCTACACGACGCTCTTCCGATCTNNNTGATTTCTCCGTAGAAGCATT
AGCACACGTCTGAACTCTTTCCCTACACGACGCTCTTCCGATCTNNNTGATTTCTCCGTAGAAGCATT
AGCACACGTCTGAACTCTTTCCCTACACGACGCTCTTCCGATCTNNNTGATTTCTCCGTAGAAGCATT
AGCACACGTCTGAACTCTTTCCCTACACGACGCTCTTCCGATCTNNNTGATTTCTCCGTAGAAGCATT
GCACACGTCTGAACTCTTTCCCTACACGACGCTCTTCCGATCTNNNGGAAGAAATATCCACCAGCTC
GCACACGTCTGAACTCTTTCCCTACACGACGCTCTTCCGATCTNNNGGAAGAAATATCCACCAGCTC
GCACACGTCTGAACTCTTTCCCTACACGACGCTCTTCCGATCTNNNGGAAGAAATATCCACCAGCTC
GCACACGTCTGAACTCTTTCCCTACACGACGCTCTTCCGATCTNNNGGAAGAAATATCCACCAGCTC
GCACACGTCTGAACTCTTTCCCTACACGACGCTCTTCCGATCTNNNGGAAGAAATATCCACCAGCTC
GCACACGTCTGAACTCTTTCCCTACACGACGCTCTTCCGATCTNNNGGAAGAAATATCCACCAGCTC
GCACACGTCTGAACTCTTTCCCTACACGACGCTCTTCCGATCTNNNGGAAGAAATATCCACCAGCTC
AGCACACGTCTGAACTCTTTCCCTACACGACGCTCTTCCGATCTNNNAGATCACACACCATTTCACTCC

AGCACACGTCTGAACTCTTTCCCTACACGACGCTCTTCCGATCTNNNAGATCACACACCATTTCACTCC
AGCACACGTCTGAACTCTTTCCCTACACGACGCTCTTCCGATCTNNNAGATCACACACCATTTCACTCC
AGCACACGTCTGAACTCTTTCCCTACACGACGCTCTTCCGATCTNNNAGATCACACACCATTTCACTCC
AGCACACGTCTGAACTCTTTCCCTACACGACGCTCTTCCGATCTNNNAGATCACACACCATTTCACTCC
AGCACACGTCTGAACTCTTTCCCTACACGACGCTCTTCCGATCTNNNAGATCACACACCATTTCACTCC
AGCACACGTCTGAACTCTTTCCCTACACGACGCTCTTCCGATCTNNNAGATCACACACCATTTCACTCC
AGCACACGTCTGAACTCTTTCCCTACACGACGCTCTTCCGATCTNNNGACGAATAGGGACAGGAAG
AGCACACGTCTGAACTCTTTCCCTACACGACGCTCTTCCGATCTNNNGACGAATAGGGACAGGAAG
AGCACACGTCTGAACTCTTTCCCTACACGACGCTCTTCCGATCTNNNGACGAATAGGGACAGGAAG
AGCACACGTCTGAACTCTTTCCCTACACGACGCTCTTCCGATCTNNNGACGAATAGGGACAGGAAG
AGCACACGTCTGAACTCTTTCCCTACACGACGCTCTTCCGATCTNNNGACGAATAGGGACAGGAAG
AGCACACGTCTGAACTCTTTCCCTACACGACGCTCTTCCGATCTNNNGACGAATAGGGACAGGAAG
AGCACACGTCTGAACTCTTTCCCTACACGACGCTCTTCCGATCTNNNGACGAATAGGGACAGGAAG
GAGCACACGTCTGAACTCTTTCCCTACACGACGCTCTTCCGATCTNNNCTCTGTGTTGAAGAGGTAGC
GAGCACACGTCTGAACTCTTTCCCTACACGACGCTCTTCCGATCTNNNCTCTGTGTTGAAGAGGTAGC
GAGCACACGTCTGAACTCTTTCCCTACACGACGCTCTTCCGATCTNNNCTCTGTGTTGAAGAGGTAGC
GAGCACACGTCTGAACTCTTTCCCTACACGACGCTCTTCCGATCTNNNCTCTGTGTTGAAGAGGTAGC
GAGCACACGTCTGAACTCTTTCCCTACACGACGCTCTTCCGATCTNNNCTCTGTGTTGAAGAGGTAGC
GAGCACACGTCTGAACTCTTTCCCTACACGACGCTCTTCCGATCTNNNCTCTGTGTTGAAGAGGTAGC
GAGCACACGTCTGAACTCTTTCCCTACACGACGCTCTTCCGATCTNNNCTCTGTGTTGAAGAGGTAGC
AGCACACGTCTGAACTCTTTCCCTACACGACGCTCTTCCGATCTNNNCACCTGTAATTCCAGCACTTTG
AGCACACGTCTGAACTCTTTCCCTACACGACGCTCTTCCGATCTNNNCACCTGTAATTCCAGCACTTTG
AGCACACGTCTGAACTCTTTCCCTACACGACGCTCTTCCGATCTNNNCACCTGTAATTCCAGCACTTTG
AGCACACGTCTGAACTCTTTCCCTACACGACGCTCTTCCGATCTNNNCACCTGTAATTCCAGCACTTTG
AGCACACGTCTGAACTCTTTCCCTACACGACGCTCTTCCGATCTNNNCACCTGTAATTCCAGCACTTTG
AGCACACGTCTGAACTCTTTCCCTACACGACGCTCTTCCGATCTNNNCACCTGTAATTCCAGCACTTTG
AGCACACGTCTGAACTCTTTCCCTACACGACGCTCTTCCGATCTNNNCACCTGTAATTCCAGCACTTTG
GAGCACACGTCTGAACTCTTTCCCTACACGACGCTCTTCCGATCTNNNCTGCCACTCTGACATACATAG
GAGCACACGTCTGAACTCTTTCCCTACACGACGCTCTTCCGATCTNNNCTGCCACTCTGACATACATAG
GAGCACACGTCTGAACTCTTTCCCTACACGACGCTCTTCCGATCTNNNCTGCCACTCTGACATACATAG
GAGCACACGTCTGAACTCTTTCCCTACACGACGCTCTTCCGATCTNNNCTGCCACTCTGACATACATAG
GAGCACACGTCTGAACTCTTTCCCTACACGACGCTCTTCCGATCTNNNCTGCCACTCTGACATACATAG
GAGCACACGTCTGAACTCTTTCCCTACACGACGCTCTTCCGATCTNNNCTGCCACTCTGACATACATAG
GAGCACACGTCTGAACTCTTTCCCTACACGACGCTCTTCCGATCTNNNCTGCCACTCTGACATACATAG
GAGCACACGTCTGAACTCTTTCCCTACACGACGCTCTTCCGATCTNNNGTTAGCTGCAGGAGTAGCTT
GAGCACACGTCTGAACTCTTTCCCTACACGACGCTCTTCCGATCTNNNGTTAGCTGCAGGAGTAGCTT
GAGCACACGTCTGAACTCTTTCCCTACACGACGCTCTTCCGATCTNNNGTTAGCTGCAGGAGTAGCTT
GAGCACACGTCTGAACTCTTTCCCTACACGACGCTCTTCCGATCTNNNGTTAGCTGCAGGAGTAGCTT
GAGCACACGTCTGAACTCTTTCCCTACACGACGCTCTTCCGATCTNNNGTTAGCTGCAGGAGTAGCTT
GAGCACACGTCTGAACTCTTTCCCTACACGACGCTCTTCCGATCTNNNGTTAGCTGCAGGAGTAGCTT
GAGCACACGTCTGAACTCTTTCCCTACACGACGCTCTTCCGATCTNNNGTTAGCTGCAGGAGTAGCTT
GAGCACACGTCTGAACTCTTTCCCTACACGACGCTCTTCCGATCTNNNCTACTGCCCTCTCCCATGATTT
GAGCACACGTCTGAACTCTTTCCCTACACGACGCTCTTCCGATCTNNNCTACTGCCCTCTCCCATGATTT

GAGCACACGTCTGAACTCTTTCCCTACACGACGCTCTTCCGATCTNNNCTACTGCCCTCTCCCATGATTT
GAGCACACGTCTGAACTCTTTCCCTACACGACGCTCTTCCGATCTNNNCTACTGCCCTCTCCCATGATTT
GAGCACACGTCTGAACTCTTTCCCTACACGACGCTCTTCCGATCTNNNCTACTGCCCTCTCCCATGATTT
GAGCACACGTCTGAACTCTTTCCCTACACGACGCTCTTCCGATCTNNNCTACTGCCCTCTCCCATGATTT
GAGCACACGTCTGAACTCTTTCCCTACACGACGCTCTTCCGATCTNNNCTACTGCCCTCTCCCATGATTT
AGCACACGTCTGAACTCTTTCCCTACACGACGCTCTTCCGATCTNNNGGGCAACAAAATGAGACTCTG
AGCACACGTCTGAACTCTTTCCCTACACGACGCTCTTCCGATCTNNNGGGCAACAAAATGAGACTCTG
AGCACACGTCTGAACTCTTTCCCTACACGACGCTCTTCCGATCTNNNGGGCAACAAAATGAGACTCTG
AGCACACGTCTGAACTCTTTCCCTACACGACGCTCTTCCGATCTNNNGGGCAACAAAATGAGACTCTG
AGCACACGTCTGAACTCTTTCCCTACACGACGCTCTTCCGATCTNNNGGGCAACAAAATGAGACTCTG
AGCACACGTCTGAACTCTTTCCCTACACGACGCTCTTCCGATCTNNNGGGCAACAAAATGAGACTCTG
AGCACACGTCTGAACTCTTTCCCTACACGACGCTCTTCCGATCTNNNGGGCAACAAAATGAGACTCTG
AGCACACGTCTGAACTCTTTCCCTACACGACGCTCTTCCGATCTNNNCACAGTGCCTGGGGTATAGTAA
AGCACACGTCTGAACTCTTTCCCTACACGACGCTCTTCCGATCTNNNCACAGTGCCTGGGGTATAGTAA
AGCACACGTCTGAACTCTTTCCCTACACGACGCTCTTCCGATCTNNNCACAGTGCCTGGGGTATAGTAA
AGCACACGTCTGAACTCTTTCCCTACACGACGCTCTTCCGATCTNNNCACAGTGCCTGGGGTATAGTAA
AGCACACGTCTGAACTCTTTCCCTACACGACGCTCTTCCGATCTNNNCACAGTGCCTGGGGTATAGTAA
AGCACACGTCTGAACTCTTTCCCTACACGACGCTCTTCCGATCTNNNCACAGTGCCTGGGGTATAGTAA
AGCACACGTCTGAACTCTTTCCCTACACGACGCTCTTCCGATCTNNNCACAGTGCCTGGGGTATAGTAA
AGCACACGTCTGAACTCTTTCCCTACACGACGCTCTTCCGATCTNNNCACCTGTAATCCCAGCTACTCA
AGCACACGTCTGAACTCTTTCCCTACACGACGCTCTTCCGATCTNNNCACCTGTAATCCCAGCTACTCA
AGCACACGTCTGAACTCTTTCCCTACACGACGCTCTTCCGATCTNNNCACCTGTAATCCCAGCTACTCA
AGCACACGTCTGAACTCTTTCCCTACACGACGCTCTTCCGATCTNNNCACCTGTAATCCCAGCTACTCA
AGCACACGTCTGAACTCTTTCCCTACACGACGCTCTTCCGATCTNNNCACCTGTAATCCCAGCTACTCA
AGCACACGTCTGAACTCTTTCCCTACACGACGCTCTTCCGATCTNNNCACCTGTAATCCCAGCTACTCA
AGCACACGTCTGAACTCTTTCCCTACACGACGCTCTTCCGATCTNNNCACCTGTAATCCCAGCTACTCA
ACACGTCTGAACTCTTTCCCTACACGACGCTCTTCCGATCTNNNACATCCCTGTACCCAATGCATTCCAT
ACACGTCTGAACTCTTTCCCTACACGACGCTCTTCCGATCTNNNACATCCCTGTACCCAATGCATTCCAT
ACACGTCTGAACTCTTTCCCTACACGACGCTCTTCCGATCTNNNACATCCCTGTACCCAATGCATTCCAT
ACACGTCTGAACTCTTTCCCTACACGACGCTCTTCCGATCTNNNACATCCCTGTACCCAATGCATTCCAT
ACACGTCTGAACTCTTTCCCTACACGACGCTCTTCCGATCTNNNACATCCCTGTACCCAATGCATTCCAT
ACACGTCTGAACTCTTTCCCTACACGACGCTCTTCCGATCTNNNACATCCCTGTACCCAATGCATTCCAT
ACACGTCTGAACTCTTTCCCTACACGACGCTCTTCCGATCTNNNACATCCCTGTACCCAATGCATTCCAT
AGCACACGTCTGAACTCTTTCCCTACACGACGCTCTTCCGATCTNNNTGAGAATCACTTGAACCCAGGA
AGCACACGTCTGAACTCTTTCCCTACACGACGCTCTTCCGATCTNNNTGAGAATCACTTGAACCCAGGA
AGCACACGTCTGAACTCTTTCCCTACACGACGCTCTTCCGATCTNNNTGAGAATCACTTGAACCCAGGA
AGCACACGTCTGAACTCTTTCCCTACACGACGCTCTTCCGATCTNNNTGAGAATCACTTGAACCCAGGA
AGCACACGTCTGAACTCTTTCCCTACACGACGCTCTTCCGATCTNNNTGAGAATCACTTGAACCCAGGA
AGCACACGTCTGAACTCTTTCCCTACACGACGCTCTTCCGATCTNNNTGAGAATCACTTGAACCCAGGA
AGCACACGTCTGAACTCTTTCCCTACACGACGCTCTTCCGATCTNNNTGAGAATCACTTGAACCCAGGA
AGCACACGTCTGAACTCTTTCCCTACACGACGCTCTTCCGATCTNNNTGGCCTTCTTTGTCTCTTGTGA
AGCACACGTCTGAACTCTTTCCCTACACGACGCTCTTCCGATCTNNNTGGCCTTCTTTGTCTCTTGTGA
AGCACACGTCTGAACTCTTTCCCTACACGACGCTCTTCCGATCTNNNTGGCCTTCTTTGTCTCTTGTGA

AGCACACGTCTGAACTCTTTCCCTACACGACGCTCTTCCGATCTNNNTGGCCTTCTTTGTCTCTTGTGA
AGCACACGTCTGAACTCTTTCCCTACACGACGCTCTTCCGATCTNNNTGGCCTTCTTTGTCTCTTGTGA
AGCACACGTCTGAACTCTTTCCCTACACGACGCTCTTCCGATCTNNNTGGCCTTCTTTGTCTCTTGTGA
AGCACACGTCTGAACTCTTTCCCTACACGACGCTCTTCCGATCTNNNTGGCCTTCTTTGTCTCTTGTGA
GAGCACACGTCTGAACTCTTTCCCTACACGACGCTCTTCCGATCTNNNTAACAGGCCTGCTGTTAGCTT
GAGCACACGTCTGAACTCTTTCCCTACACGACGCTCTTCCGATCTNNNTAACAGGCCTGCTGTTAGCTT
GAGCACACGTCTGAACTCTTTCCCTACACGACGCTCTTCCGATCTNNNTAACAGGCCTGCTGTTAGCTT
GAGCACACGTCTGAACTCTTTCCCTACACGACGCTCTTCCGATCTNNNTAACAGGCCTGCTGTTAGCTT
GAGCACACGTCTGAACTCTTTCCCTACACGACGCTCTTCCGATCTNNNTAACAGGCCTGCTGTTAGCTT
GAGCACACGTCTGAACTCTTTCCCTACACGACGCTCTTCCGATCTNNNTAACAGGCCTGCTGTTAGCTT
GAGCACACGTCTGAACTCTTTCCCTACACGACGCTCTTCCGATCTNNNTAACAGGCCTGCTGTTAGCTT
GAGCACACGTCTGAACTCTTTCCCTACACGACGCTCTTCCGATCTNNNCTTGAGGCTAGGCACAGTTC
GAGCACACGTCTGAACTCTTTCCCTACACGACGCTCTTCCGATCTNNNCTTGAGGCTAGGCACAGTTC
GAGCACACGTCTGAACTCTTTCCCTACACGACGCTCTTCCGATCTNNNCTTGAGGCTAGGCACAGTTC
GAGCACACGTCTGAACTCTTTCCCTACACGACGCTCTTCCGATCTNNNCTTGAGGCTAGGCACAGTTC
GAGCACACGTCTGAACTCTTTCCCTACACGACGCTCTTCCGATCTNNNCTTGAGGCTAGGCACAGTTC
GAGCACACGTCTGAACTCTTTCCCTACACGACGCTCTTCCGATCTNNNCTTGAGGCTAGGCACAGTTC
GAGCACACGTCTGAACTCTTTCCCTACACGACGCTCTTCCGATCTNNNCTTGAGGCTAGGCACAGTTC
GAGCACACGTCTGAACTCTTTCCCTACACGACGCTCTTCCGATCTNNNCAAGATCACACCACTGCACTA
GAGCACACGTCTGAACTCTTTCCCTACACGACGCTCTTCCGATCTNNNCAAGATCACACCACTGCACTA
GAGCACACGTCTGAACTCTTTCCCTACACGACGCTCTTCCGATCTNNNCAAGATCACACCACTGCACTA
GAGCACACGTCTGAACTCTTTCCCTACACGACGCTCTTCCGATCTNNNCAAGATCACACCACTGCACTA
GAGCACACGTCTGAACTCTTTCCCTACACGACGCTCTTCCGATCTNNNCAAGATCACACCACTGCACTA
GAGCACACGTCTGAACTCTTTCCCTACACGACGCTCTTCCGATCTNNNCAAGATCACACCACTGCACTA
GAGCACACGTCTGAACTCTTTCCCTACACGACGCTCTTCCGATCTNNNCAAGATCACACCACTGCACTA
AGCACACGTCTGAACTCTTTCCCTACACGACGCTCTTCCGATCTNNNCGTGATTGTACCACTGCACTCT
AGCACACGTCTGAACTCTTTCCCTACACGACGCTCTTCCGATCTNNNCGTGATTGTACCACTGCACTCT
AGCACACGTCTGAACTCTTTCCCTACACGACGCTCTTCCGATCTNNNCGTGATTGTACCACTGCACTCT
AGCACACGTCTGAACTCTTTCCCTACACGACGCTCTTCCGATCTNNNCGTGATTGTACCACTGCACTCT
AGCACACGTCTGAACTCTTTCCCTACACGACGCTCTTCCGATCTNNNCGTGATTGTACCACTGCACTCT
AGCACACGTCTGAACTCTTTCCCTACACGACGCTCTTCCGATCTNNNCGTGATTGTACCACTGCACTCT
AGCACACGTCTGAACTCTTTCCCTACACGACGCTCTTCCGATCTNNNCGTGATTGTACCACTGCACTCT
GAGCACACGTCTGAACTCTTTCCCTACACGACGCTCTTCCGATCTNNNCCACTCTGTTGACTAGGATGG
GAGCACACGTCTGAACTCTTTCCCTACACGACGCTCTTCCGATCTNNNCCACTCTGTTGACTAGGATGG
GAGCACACGTCTGAACTCTTTCCCTACACGACGCTCTTCCGATCTNNNCCACTCTGTTGACTAGGATGG
GAGCACACGTCTGAACTCTTTCCCTACACGACGCTCTTCCGATCTNNNCCACTCTGTTGACTAGGATGG
GAGCACACGTCTGAACTCTTTCCCTACACGACGCTCTTCCGATCTNNNCCACTCTGTTGACTAGGATGG
GAGCACACGTCTGAACTCTTTCCCTACACGACGCTCTTCCGATCTNNNCCACTCTGTTGACTAGGATGG
GAGCACACGTCTGAACTCTTTCCCTACACGACGCTCTTCCGATCTNNNCCACTCTGTTGACTAGGATGG
GCACACGTCTGAACTCTTTCCCTACACGACGCTCTTCCGATCTNNNGCAGGAGAATTGCTTGAACTCA
GCACACGTCTGAACTCTTTCCCTACACGACGCTCTTCCGATCTNNNGCAGGAGAATTGCTTGAACTCA
GCACACGTCTGAACTCTTTCCCTACACGACGCTCTTCCGATCTNNNGCAGGAGAATTGCTTGAACTCA
GCACACGTCTGAACTCTTTCCCTACACGACGCTCTTCCGATCTNNNGCAGGAGAATTGCTTGAACTCA

AGCACACGTCTGAACTCTTTCCCTACACGACGCTCTTCCGATCTNNNGCAGGAGAATTGCTTGAACTCA
AGCACACGTCTGAACTCTTTCCCTACACGACGCTCTTCCGATCTNNNGCAGGAGAATTGCTTGAACTCA
AGCACACGTCTGAACTCTTTCCCTACACGACGCTCTTCCGATCTNNNGCAGGAGAATTGCTTGAACTCA
GAGCACACGTCTGAACTCTTTCCCTACACGACGCTCTTCCGATCTNNNATTCACAAATGCAGTGCGGTA
GAGCACACGTCTGAACTCTTTCCCTACACGACGCTCTTCCGATCTNNNATTCACAAATGCAGTGCGGTA
GAGCACACGTCTGAACTCTTTCCCTACACGACGCTCTTCCGATCTNNNATTCACAAATGCAGTGCGGTA
GAGCACACGTCTGAACTCTTTCCCTACACGACGCTCTTCCGATCTNNNATTCACAAATGCAGTGCGGTA
GAGCACACGTCTGAACTCTTTCCCTACACGACGCTCTTCCGATCTNNNATTCACAAATGCAGTGCGGTA
GAGCACACGTCTGAACTCTTTCCCTACACGACGCTCTTCCGATCTNNNATTCACAAATGCAGTGCGGTA
GAGCACACGTCTGAACTCTTTCCCTACACGACGCTCTTCCGATCTNNNATTCACAAATGCAGTGCGGTA
GAGCACACGTCTGAACTCTTTCCCTACACGACGCTCTTCCGATCTNNNGTTCATCTGAGCTTGGGAAGT
GAGCACACGTCTGAACTCTTTCCCTACACGACGCTCTTCCGATCTNNNGTTCATCTGAGCTTGGGAAGT
GAGCACACGTCTGAACTCTTTCCCTACACGACGCTCTTCCGATCTNNNGTTCATCTGAGCTTGGGAAGT
GAGCACACGTCTGAACTCTTTCCCTACACGACGCTCTTCCGATCTNNNGTTCATCTGAGCTTGGGAAGT
GAGCACACGTCTGAACTCTTTCCCTACACGACGCTCTTCCGATCTNNNGTTCATCTGAGCTTGGGAAGT
GAGCACACGTCTGAACTCTTTCCCTACACGACGCTCTTCCGATCTNNNGTTCATCTGAGCTTGGGAAGT
GAGCACACGTCTGAACTCTTTCCCTACACGACGCTCTTCCGATCTNNNGTTCATCTGAGCTTGGGAAGT
GAGCACACGTCTGAACTCTTTCCCTACACGACGCTCTTCCGATCTNNNGATCCAAATACCCATCGACAG
GAGCACACGTCTGAACTCTTTCCCTACACGACGCTCTTCCGATCTNNNGATCCAAATACCCATCGACAG
GAGCACACGTCTGAACTCTTTCCCTACACGACGCTCTTCCGATCTNNNGATCCAAATACCCATCGACAG
GAGCACACGTCTGAACTCTTTCCCTACACGACGCTCTTCCGATCTNNNGATCCAAATACCCATCGACAG
GAGCACACGTCTGAACTCTTTCCCTACACGACGCTCTTCCGATCTNNNGATCCAAATACCCATCGACAG
GAGCACACGTCTGAACTCTTTCCCTACACGACGCTCTTCCGATCTNNNGATCCAAATACCCATCGACAG
GAGCACACGTCTGAACTCTTTCCCTACACGACGCTCTTCCGATCTNNNGATCCAAATACCCATCGACAG
GAGCACACGTCTGAACTCTTTCCCTACACGACGCTCTTCCGATCTNNNTGGGGTACAGGTAGTATTTG
GAGCACACGTCTGAACTCTTTCCCTACACGACGCTCTTCCGATCTNNNTGGGGTACAGGTAGTATTTG
GAGCACACGTCTGAACTCTTTCCCTACACGACGCTCTTCCGATCTNNNTGGGGTACAGGTAGTATTTG
GAGCACACGTCTGAACTCTTTCCCTACACGACGCTCTTCCGATCTNNNTGGGGTACAGGTAGTATTTG
GAGCACACGTCTGAACTCTTTCCCTACACGACGCTCTTCCGATCTNNNTGGGGTACAGGTAGTATTTG
GAGCACACGTCTGAACTCTTTCCCTACACGACGCTCTTCCGATCTNNNTGGGGTACAGGTAGTATTTG
GAGCACACGTCTGAACTCTTTCCCTACACGACGCTCTTCCGATCTNNNTGGGGTACAGGTAGTATTTG
GAGCACACGTCTGAACTCTTTCCCTACACGACGCTCTTCCGATCTNNNATATAAAAAGTTAGCCGGGCC
GAGCACACGTCTGAACTCTTTCCCTACACGACGCTCTTCCGATCTNNNATATAAAAAGTTAGCCGGGCC
GAGCACACGTCTGAACTCTTTCCCTACACGACGCTCTTCCGATCTNNNATATAAAAAGTTAGCCGGGCC
GAGCACACGTCTGAACTCTTTCCCTACACGACGCTCTTCCGATCTNNNATATAAAAAGTTAGCCGGGCC
GAGCACACGTCTGAACTCTTTCCCTACACGACGCTCTTCCGATCTNNNATATAAAAAGTTAGCCGGGCC
GAGCACACGTCTGAACTCTTTCCCTACACGACGCTCTTCCGATCTNNNATATAAAAAGTTAGCCGGGCC
GAGCACACGTCTGAACTCTTTCCCTACACGACGCTCTTCCGATCTNNNATATAAAAAGTTAGCCGGGCC
AGAGCACACGTCTGAACTCTTTCCCTACACGACGCTCTTCCGATCTNNNATCACTTGAACCCAGTAGG
AGAGCACACGTCTGAACTCTTTCCCTACACGACGCTCTTCCGATCTNNNATCACTTGAACCCAGTAGG
AGAGCACACGTCTGAACTCTTTCCCTACACGACGCTCTTCCGATCTNNNATCACTTGAACCCAGTAGG
AGAGCACACGTCTGAACTCTTTCCCTACACGACGCTCTTCCGATCTNNNATCACTTGAACCCAGTAGG
AGAGCACACGTCTGAACTCTTTCCCTACACGACGCTCTTCCGATCTNNNATCACTTGAACCCAGTAGG

AGAGCACACGTCTGAACTCTTTCCCTACACGACGCTCTTCCGATCTNNNATCACTTGAACCCAGTAGG
AGAGCACACGTCTGAACTCTTTCCCTACACGACGCTCTTCCGATCTNNNATCACTTGAACCCAGTAGG
AGCACACGTCTGAACTCTTTCCCTACACGACGCTCTTCCGATCTNNNGAGGCTGAGGCAGGAGAATG
AGCACACGTCTGAACTCTTTCCCTACACGACGCTCTTCCGATCTNNNGAGGCTGAGGCAGGAGAATG
AGCACACGTCTGAACTCTTTCCCTACACGACGCTCTTCCGATCTNNNGAGGCTGAGGCAGGAGAATG
AGCACACGTCTGAACTCTTTCCCTACACGACGCTCTTCCGATCTNNNGAGGCTGAGGCAGGAGAATG
AGCACACGTCTGAACTCTTTCCCTACACGACGCTCTTCCGATCTNNNGAGGCTGAGGCAGGAGAATG
AGCACACGTCTGAACTCTTTCCCTACACGACGCTCTTCCGATCTNNNGAGGCTGAGGCAGGAGAATG
AGCACACGTCTGAACTCTTTCCCTACACGACGCTCTTCCGATCTNNNGAGGCTGAGGCAGGAGAATG
GCACACGTCTGAACTCTTTCCCTACACGACGCTCTTCCGATCTNNNGCTCTCTCTCTTTGCCTCTTTCTC
GCACACGTCTGAACTCTTTCCCTACACGACGCTCTTCCGATCTNNNGCTCTCTCTCTTTGCCTCTTTCTC
GCACACGTCTGAACTCTTTCCCTACACGACGCTCTTCCGATCTNNNGCTCTCTCTCTTTGCCTCTTTCTC
GCACACGTCTGAACTCTTTCCCTACACGACGCTCTTCCGATCTNNNGCTCTCTCTCTTTGCCTCTTTCTC
GCACACGTCTGAACTCTTTCCCTACACGACGCTCTTCCGATCTNNNGCTCTCTCTCTTTGCCTCTTTCTC
GCACACGTCTGAACTCTTTCCCTACACGACGCTCTTCCGATCTNNNGCTCTCTCTCTTTGCCTCTTTCTC
GCACACGTCTGAACTCTTTCCCTACACGACGCTCTTCCGATCTNNNGCTCTCTCTCTTTGCCTCTTTCTC
GAGCACACGTCTGAACTCTTTCCCTACACGACGCTCTTCCGATCTNNNTCAGTCTTTGAGGTAGGTTT
GAGCACACGTCTGAACTCTTTCCCTACACGACGCTCTTCCGATCTNNNTCAGTCTTTGAGGTAGGTTT
GAGCACACGTCTGAACTCTTTCCCTACACGACGCTCTTCCGATCTNNNTCAGTCTTTGAGGTAGGTTT
GAGCACACGTCTGAACTCTTTCCCTACACGACGCTCTTCCGATCTNNNTCAGTCTTTGAGGTAGGTTT
GAGCACACGTCTGAACTCTTTCCCTACACGACGCTCTTCCGATCTNNNTCAGTCTTTGAGGTAGGTTT
GAGCACACGTCTGAACTCTTTCCCTACACGACGCTCTTCCGATCTNNNTCAGTCTTTGAGGTAGGTTT
GAGCACACGTCTGAACTCTTTCCCTACACGACGCTCTTCCGATCTNNNTCAGTCTTTGAGGTAGGTTT
GAGCACACGTCTGAACTCTTTCCCTACACGACGCTCTTCCGATCTNNNCCCTGTAGTCCCAGCTACTCC
GAGCACACGTCTGAACTCTTTCCCTACACGACGCTCTTCCGATCTNNNCCCTGTAGTCCCAGCTACTCC
GAGCACACGTCTGAACTCTTTCCCTACACGACGCTCTTCCGATCTNNNCCCTGTAGTCCCAGCTACTCC
GAGCACACGTCTGAACTCTTTCCCTACACGACGCTCTTCCGATCTNNNCCCTGTAGTCCCAGCTACTCC
GAGCACACGTCTGAACTCTTTCCCTACACGACGCTCTTCCGATCTNNNCCCTGTAGTCCCAGCTACTCC
GAGCACACGTCTGAACTCTTTCCCTACACGACGCTCTTCCGATCTNNNCCCTGTAGTCCCAGCTACTCC
GAGCACACGTCTGAACTCTTTCCCTACACGACGCTCTTCCGATCTNNNCCCTGTAGTCCCAGCTACTCC
AGCACACGTCTGAACTCTTTCCCTACACGACGCTCTTCCGATCTNNNAGTGTGTGTATATGTGTGTG
AGCACACGTCTGAACTCTTTCCCTACACGACGCTCTTCCGATCTNNNAGTGTGTGTATATGTGTGTG
AGCACACGTCTGAACTCTTTCCCTACACGACGCTCTTCCGATCTNNNAGTGTGTGTATATGTGTGTG
AGCACACGTCTGAACTCTTTCCCTACACGACGCTCTTCCGATCTNNNAGTGTGTGTATATGTGTGTG
AGCACACGTCTGAACTCTTTCCCTACACGACGCTCTTCCGATCTNNNAGTGTGTGTATATGTGTGTG
AGCACACGTCTGAACTCTTTCCCTACACGACGCTCTTCCGATCTNNNAGTGTGTGTATATGTGTGTG
AGCACACGTCTGAACTCTTTCCCTACACGACGCTCTTCCGATCTNNNAGTGTGTGTATATGTGTGTG
GAGCACACGTCTGAACTCTTTCCCTACACGACGCTCTTCCGATCTNNNACTAATACAGATGCCAAGCA
GAGCACACGTCTGAACTCTTTCCCTACACGACGCTCTTCCGATCTNNNACTAATACAGATGCCAAGCA
GAGCACACGTCTGAACTCTTTCCCTACACGACGCTCTTCCGATCTNNNACTAATACAGATGCCAAGCA
GAGCACACGTCTGAACTCTTTCCCTACACGACGCTCTTCCGATCTNNNACTAATACAGATGCCAAGCA
GAGCACACGTCTGAACTCTTTCCCTACACGACGCTCTTCCGATCTNNNACTAATACAGATGCCAAGCA
GAGCACACGTCTGAACTCTTTCCCTACACGACGCTCTTCCGATCTNNNACTAATACAGATGCCAAGCA

GAGCACACGTCTGAACTCTTTCCCTACACGACGCTCTTCCGATCTNNNACTAATACAGATGCCAAGCA
AGCACACGTCTGAACTCTTTCCCTACACGACGCTCTTCCGATCTNNNGTAATACCAGCTACTCAGGAG
AGCACACGTCTGAACTCTTTCCCTACACGACGCTCTTCCGATCTNNNGTAATACCAGCTACTCAGGAG
AGCACACGTCTGAACTCTTTCCCTACACGACGCTCTTCCGATCTNNNGTAATACCAGCTACTCAGGAG
AGCACACGTCTGAACTCTTTCCCTACACGACGCTCTTCCGATCTNNNGTAATACCAGCTACTCAGGAG
AGCACACGTCTGAACTCTTTCCCTACACGACGCTCTTCCGATCTNNNGTAATACCAGCTACTCAGGAG
AGCACACGTCTGAACTCTTTCCCTACACGACGCTCTTCCGATCTNNNGTAATACCAGCTACTCAGGAG
AGCACACGTCTGAACTCTTTCCCTACACGACGCTCTTCCGATCTNNNGTAATACCAGCTACTCAGGAG
GCACACGTCTGAACTCTTTCCCTACACGACGCTCTTCCGATCTNNNTTTAGTGCTCTAGGGGATTGC
GCACACGTCTGAACTCTTTCCCTACACGACGCTCTTCCGATCTNNNTTTAGTGCTCTAGGGGATTGC
GCACACGTCTGAACTCTTTCCCTACACGACGCTCTTCCGATCTNNNTTTAGTGCTCTAGGGGATTGC
GCACACGTCTGAACTCTTTCCCTACACGACGCTCTTCCGATCTNNNTTTAGTGCTCTAGGGGATTGC
GCACACGTCTGAACTCTTTCCCTACACGACGCTCTTCCGATCTNNNTTTAGTGCTCTAGGGGATTGC
GCACACGTCTGAACTCTTTCCCTACACGACGCTCTTCCGATCTNNNTTTAGTGCTCTAGGGGATTGC
GCACACGTCTGAACTCTTTCCCTACACGACGCTCTTCCGATCTNNNTTTAGTGCTCTAGGGGATTGC
GAGCACACGTCTGAACTCTTTCCCTACACGACGCTCTTCCGATCTNNNGAGGAGCAGTGATTATGGC
GAGCACACGTCTGAACTCTTTCCCTACACGACGCTCTTCCGATCTNNNGAGGAGCAGTGATTATGGC
GAGCACACGTCTGAACTCTTTCCCTACACGACGCTCTTCCGATCTNNNGAGGAGCAGTGATTATGGC
GAGCACACGTCTGAACTCTTTCCCTACACGACGCTCTTCCGATCTNNNGAGGAGCAGTGATTATGGC
GAGCACACGTCTGAACTCTTTCCCTACACGACGCTCTTCCGATCTNNNGAGGAGCAGTGATTATGGC
GAGCACACGTCTGAACTCTTTCCCTACACGACGCTCTTCCGATCTNNNGAGGAGCAGTGATTATGGC
GAGCACACGTCTGAACTCTTTCCCTACACGACGCTCTTCCGATCTNNNGAGGAGCAGTGATTATGGC
AGCACACGTCTGAACTCTTTCCCTACACGACGCTCTTCCGATCTNNNTTCTCCCAGTTTGTAATGTCG
AGCACACGTCTGAACTCTTTCCCTACACGACGCTCTTCCGATCTNNNTTCTCCCAGTTTGTAATGTCG
AGCACACGTCTGAACTCTTTCCCTACACGACGCTCTTCCGATCTNNNTTCTCCCAGTTTGTAATGTCG
AGCACACGTCTGAACTCTTTCCCTACACGACGCTCTTCCGATCTNNNTTCTCCCAGTTTGTAATGTCG
AGCACACGTCTGAACTCTTTCCCTACACGACGCTCTTCCGATCTNNNTTCTCCCAGTTTGTAATGTCG
AGCACACGTCTGAACTCTTTCCCTACACGACGCTCTTCCGATCTNNNTTCTCCCAGTTTGTAATGTCG
AGCACACGTCTGAACTCTTTCCCTACACGACGCTCTTCCGATCTNNNTTCTCCCAGTTTGTAATGTCG
GCACACGTCTGAACTCTTTCCCTACACGACGCTCTTCCGATCTNNNCAGAAAAGTGCAAGACAGGAC
GCACACGTCTGAACTCTTTCCCTACACGACGCTCTTCCGATCTNNNCAGAAAAGTGCAAGACAGGAC
GCACACGTCTGAACTCTTTCCCTACACGACGCTCTTCCGATCTNNNCAGAAAAGTGCAAGACAGGAC
GCACACGTCTGAACTCTTTCCCTACACGACGCTCTTCCGATCTNNNCAGAAAAGTGCAAGACAGGAC
GCACACGTCTGAACTCTTTCCCTACACGACGCTCTTCCGATCTNNNCAGAAAAGTGCAAGACAGGAC
GCACACGTCTGAACTCTTTCCCTACACGACGCTCTTCCGATCTNNNCAGAAAAGTGCAAGACAGGAC
GCACACGTCTGAACTCTTTCCCTACACGACGCTCTTCCGATCTNNNCAGAAAAGTGCAAGACAGGAC
GCACACGTCTGAACTCTTTCCCTACACGACGCTCTTCCGATCTNNNCCACACAGGGTACAACTCAGAG
GCACACGTCTGAACTCTTTCCCTACACGACGCTCTTCCGATCTNNNCCACACAGGGTACAACTCAGAG
GCACACGTCTGAACTCTTTCCCTACACGACGCTCTTCCGATCTNNNCCACACAGGGTACAACTCAGAG
GCACACGTCTGAACTCTTTCCCTACACGACGCTCTTCCGATCTNNNCCACACAGGGTACAACTCAGAG
GCACACGTCTGAACTCTTTCCCTACACGACGCTCTTCCGATCTNNNCCACACAGGGTACAACTCAGAG
GCACACGTCTGAACTCTTTCCCTACACGACGCTCTTCCGATCTNNNCCACACAGGGTACAACTCAGAG
GCACACGTCTGAACTCTTTCCCTACACGACGCTCTTCCGATCTNNNCCACACAGGGTACAACTCAGAG

AGCACACGTCTGAACTCTTTCCCTACACGACGCTCTTCCGATCTNNNTGGGCACCTTAGAAGTTCCTCTT
AGCACACGTCTGAACTCTTTCCCTACACGACGCTCTTCCGATCTNNNTGGGCACCTTAGAAGTTCCTCTT
AGCACACGTCTGAACTCTTTCCCTACACGACGCTCTTCCGATCTNNNTGGGCACCTTAGAAGTTCCTCTT
AGCACACGTCTGAACTCTTTCCCTACACGACGCTCTTCCGATCTNNNTGGGCACCTTAGAAGTTCCTCTT
AGCACACGTCTGAACTCTTTCCCTACACGACGCTCTTCCGATCTNNNTGGGCACCTTAGAAGTTCCTCTT
AGCACACGTCTGAACTCTTTCCCTACACGACGCTCTTCCGATCTNNNTGGGCACCTTAGAAGTTCCTCTT
AGCACACGTCTGAACTCTTTCCCTACACGACGCTCTTCCGATCTNNNTGGGCACCTTAGAAGTTCCTCTT
AGCACACGTCTGAACTCTTTCCCTACACGACGCTCTTCCGATCTNNNAATCTATTCGTCTGACAGAGGG
AGCACACGTCTGAACTCTTTCCCTACACGACGCTCTTCCGATCTNNNAATCTATTCGTCTGACAGAGGG
AGCACACGTCTGAACTCTTTCCCTACACGACGCTCTTCCGATCTNNNAATCTATTCGTCTGACAGAGGG
AGCACACGTCTGAACTCTTTCCCTACACGACGCTCTTCCGATCTNNNAATCTATTCGTCTGACAGAGGG
AGCACACGTCTGAACTCTTTCCCTACACGACGCTCTTCCGATCTNNNAATCTATTCGTCTGACAGAGGG
AGCACACGTCTGAACTCTTTCCCTACACGACGCTCTTCCGATCTNNNAATCTATTCGTCTGACAGAGGG
AGCACACGTCTGAACTCTTTCCCTACACGACGCTCTTCCGATCTNNNAATCTATTCGTCTGACAGAGGG
GCACACGTCTGAACTCTTTCCCTACACGACGCTCTTCCGATCTNNNGTTTGCTTTCTCAGAACCTCCCCA
GCACACGTCTGAACTCTTTCCCTACACGACGCTCTTCCGATCTNNNGTTTGCTTTCTCAGAACCTCCCCA
GCACACGTCTGAACTCTTTCCCTACACGACGCTCTTCCGATCTNNNGTTTGCTTTCTCAGAACCTCCCCA
GCACACGTCTGAACTCTTTCCCTACACGACGCTCTTCCGATCTNNNGTTTGCTTTCTCAGAACCTCCCCA
GCACACGTCTGAACTCTTTCCCTACACGACGCTCTTCCGATCTNNNGTTTGCTTTCTCAGAACCTCCCCA
GCACACGTCTGAACTCTTTCCCTACACGACGCTCTTCCGATCTNNNGTTTGCTTTCTCAGAACCTCCCCA
GCACACGTCTGAACTCTTTCCCTACACGACGCTCTTCCGATCTNNNGTTTGCTTTCTCAGAACCTCCCCA
GAGCACACGTCTGAACTCTTTCCCTACACGACGCTCTTCCGATCTNNNTTAAGTGCACGGTGTGTGACT
GAGCACACGTCTGAACTCTTTCCCTACACGACGCTCTTCCGATCTNNNTTAAGTGCACGGTGTGTGACT
GAGCACACGTCTGAACTCTTTCCCTACACGACGCTCTTCCGATCTNNNTTAAGTGCACGGTGTGTGACT
GAGCACACGTCTGAACTCTTTCCCTACACGACGCTCTTCCGATCTNNNTTAAGTGCACGGTGTGTGACT
GAGCACACGTCTGAACTCTTTCCCTACACGACGCTCTTCCGATCTNNNTTAAGTGCACGGTGTGTGACT
GAGCACACGTCTGAACTCTTTCCCTACACGACGCTCTTCCGATCTNNNTTAAGTGCACGGTGTGTGACT
GAGCACACGTCTGAACTCTTTCCCTACACGACGCTCTTCCGATCTNNNTTAAGTGCACGGTGTGTGACT
AGCACACGTCTGAACTCTTTCCCTACACGACGCTCTTCCGATCTNNNTAGAGTCCCCCAGAAAACTGAG
AGCACACGTCTGAACTCTTTCCCTACACGACGCTCTTCCGATCTNNNTAGAGTCCCCCAGAAAACTGAG
AGCACACGTCTGAACTCTTTCCCTACACGACGCTCTTCCGATCTNNNTAGAGTCCCCCAGAAAACTGAG
AGCACACGTCTGAACTCTTTCCCTACACGACGCTCTTCCGATCTNNNTAGAGTCCCCCAGAAAACTGAG
AGCACACGTCTGAACTCTTTCCCTACACGACGCTCTTCCGATCTNNNTAGAGTCCCCCAGAAAACTGAG
AGCACACGTCTGAACTCTTTCCCTACACGACGCTCTTCCGATCTNNNTAGAGTCCCCCAGAAAACTGAG
AGCACACGTCTGAACTCTTTCCCTACACGACGCTCTTCCGATCTNNNTAGAGTCCCCCAGAAAACTGAG
AGCACACGTCTGAACTCTTTCCCTACACGACGCTCTTCCGATCTNNNGAGGCTGAGGCAGGAGAATGC
AGCACACGTCTGAACTCTTTCCCTACACGACGCTCTTCCGATCTNNNGAGGCTGAGGCAGGAGAATGC
AGCACACGTCTGAACTCTTTCCCTACACGACGCTCTTCCGATCTNNNGAGGCTGAGGCAGGAGAATGC
AGCACACGTCTGAACTCTTTCCCTACACGACGCTCTTCCGATCTNNNGAGGCTGAGGCAGGAGAATGC
AGCACACGTCTGAACTCTTTCCCTACACGACGCTCTTCCGATCTNNNGAGGCTGAGGCAGGAGAATGC
AGCACACGTCTGAACTCTTTCCCTACACGACGCTCTTCCGATCTNNNGAGGCTGAGGCAGGAGAATGC
AGCACACGTCTGAACTCTTTCCCTACACGACGCTCTTCCGATCTNNNGAGGCTGAGGCAGGAGAATGC
AGCACACGTCTGAACTCTTTCCCTACACGACGCTCTTCCGATCTNNNGGGCAACAAGAGTGAACAAAA

AGCACACGTCTGAACTCTTTCCCTACACGACGCTCTTCCGATCTNNNGGGCAACAAGAGTGAACAAAA
AGCACACGTCTGAACTCTTTCCCTACACGACGCTCTTCCGATCTNNNGGGCAACAAGAGTGAACAAAA
AGCACACGTCTGAACTCTTTCCCTACACGACGCTCTTCCGATCTNNNGGGCAACAAGAGTGAACAAAA
AGCACACGTCTGAACTCTTTCCCTACACGACGCTCTTCCGATCTNNNGGGCAACAAGAGTGAACAAAA
AGCACACGTCTGAACTCTTTCCCTACACGACGCTCTTCCGATCTNNNGGGCAACAAGAGTGAACAAAA
AGCACACGTCTGAACTCTTTCCCTACACGACGCTCTTCCGATCTNNNGGGCAACAAGAGTGAACAAAA
GAGCACACGTCTGAACTCTTTCCCTACACGACGCTCTTCCGATCTNNNGTGTCCAAACATTTGAGGAAG
GAGCACACGTCTGAACTCTTTCCCTACACGACGCTCTTCCGATCTNNNGTGTCCAAACATTTGAGGAAG
GAGCACACGTCTGAACTCTTTCCCTACACGACGCTCTTCCGATCTNNNGTGTCCAAACATTTGAGGAAG
GAGCACACGTCTGAACTCTTTCCCTACACGACGCTCTTCCGATCTNNNGTGTCCAAACATTTGAGGAAG
GAGCACACGTCTGAACTCTTTCCCTACACGACGCTCTTCCGATCTNNNGTGTCCAAACATTTGAGGAAG
GAGCACACGTCTGAACTCTTTCCCTACACGACGCTCTTCCGATCTNNNGTGTCCAAACATTTGAGGAAG
GAGCACACGTCTGAACTCTTTCCCTACACGACGCTCTTCCGATCTNNNGTGTCCAAACATTTGAGGAAG
GAGCACACGTCTGAACTCTTTCCCTACACGACGCTCTTCCGATCTNNNTTGTTCCTTCATTCTGGGTTCC
GAGCACACGTCTGAACTCTTTCCCTACACGACGCTCTTCCGATCTNNNTTGTTCCTTCATTCTGGGTTCC
GAGCACACGTCTGAACTCTTTCCCTACACGACGCTCTTCCGATCTNNNTTGTTCCTTCATTCTGGGTTCC
GAGCACACGTCTGAACTCTTTCCCTACACGACGCTCTTCCGATCTNNNTTGTTCCTTCATTCTGGGTTCC
GAGCACACGTCTGAACTCTTTCCCTACACGACGCTCTTCCGATCTNNNTTGTTCCTTCATTCTGGGTTCC
GAGCACACGTCTGAACTCTTTCCCTACACGACGCTCTTCCGATCTNNNTTGTTCCTTCATTCTGGGTTCC
GAGCACACGTCTGAACTCTTTCCCTACACGACGCTCTTCCGATCTNNNTTGTTCCTTCATTCTGGGTTCC
GCACACGTCTGAACTCTTTCCCTACACGACGCTCTTCCGATCTNNNCACTCAGTTCGAATTCCTCCTCACA
GCACACGTCTGAACTCTTTCCCTACACGACGCTCTTCCGATCTNNNCACTCAGTTCGAATTCCTCCTCACA
GCACACGTCTGAACTCTTTCCCTACACGACGCTCTTCCGATCTNNNCACTCAGTTCGAATTCCTCCTCACA
GCACACGTCTGAACTCTTTCCCTACACGACGCTCTTCCGATCTNNNCACTCAGTTCGAATTCCTCCTCACA
GCACACGTCTGAACTCTTTCCCTACACGACGCTCTTCCGATCTNNNCACTCAGTTCGAATTCCTCCTCACA
GCACACGTCTGAACTCTTTCCCTACACGACGCTCTTCCGATCTNNNCACTCAGTTCGAATTCCTCCTCACA
GCACACGTCTGAACTCTTTCCCTACACGACGCTCTTCCGATCTNNNCACTCAGTTCGAATTCCTCCTCACA
GAGCACACGTCTGAACTCTTTCCCTACACGACGCTCTTCCGATCTNNNAGAGGAATCAAATGGGCATG
GAGCACACGTCTGAACTCTTTCCCTACACGACGCTCTTCCGATCTNNNAGAGGAATCAAATGGGCATG
GAGCACACGTCTGAACTCTTTCCCTACACGACGCTCTTCCGATCTNNNAGAGGAATCAAATGGGCATG
GAGCACACGTCTGAACTCTTTCCCTACACGACGCTCTTCCGATCTNNNAGAGGAATCAAATGGGCATG
GAGCACACGTCTGAACTCTTTCCCTACACGACGCTCTTCCGATCTNNNAGAGGAATCAAATGGGCATG
GAGCACACGTCTGAACTCTTTCCCTACACGACGCTCTTCCGATCTNNNAGAGGAATCAAATGGGCATG
GAGCACACGTCTGAACTCTTTCCCTACACGACGCTCTTCCGATCTNNNAGAGGAATCAAATGGGCATG
GCACACGTCTGAACTCTTTCCCTACACGACGCTCTTCCGATCTNNNCTACACTTTGACCTGCACACATCCA
GCACACGTCTGAACTCTTTCCCTACACGACGCTCTTCCGATCTNNNCTACACTTTGACCTGCACACATCCA
GCACACGTCTGAACTCTTTCCCTACACGACGCTCTTCCGATCTNNNCTACACTTTGACCTGCACACATCCA
GCACACGTCTGAACTCTTTCCCTACACGACGCTCTTCCGATCTNNNCTACACTTTGACCTGCACACATCCA
GCACACGTCTGAACTCTTTCCCTACACGACGCTCTTCCGATCTNNNCTACACTTTGACCTGCACACATCCA
GCACACGTCTGAACTCTTTCCCTACACGACGCTCTTCCGATCTNNNCTACACTTTGACCTGCACACATCCA
GCACACGTCTGAACTCTTTCCCTACACGACGCTCTTCCGATCTNNNCTACACTTTGACCTGCACACATCCA
AGAGCACACGTCTGAACTCTTTCCCTACACGACGCTCTTCCGATCTNNNATGACAATTAGCTGGGTGT
AGAGCACACGTCTGAACTCTTTCCCTACACGACGCTCTTCCGATCTNNNATGACAATTAGCTGGGTGT

AGAGCACACGTCTGAACTCTTTCCCTACACGACGCTCTTCCGATCTNNNATGACAATTAGCTGGGTGT
AGAGCACACGTCTGAACTCTTTCCCTACACGACGCTCTTCCGATCTNNNATGACAATTAGCTGGGTGT
AGAGCACACGTCTGAACTCTTTCCCTACACGACGCTCTTCCGATCTNNNATGACAATTAGCTGGGTGT
AGAGCACACGTCTGAACTCTTTCCCTACACGACGCTCTTCCGATCTNNNATGACAATTAGCTGGGTGT
AGAGCACACGTCTGAACTCTTTCCCTACACGACGCTCTTCCGATCTNNNATGACAATTAGCTGGGTGT
AAGAGCACACGTCTGAACTCTTTCCCTACACGACGCTCTTCCGATCTNNNAGAAAAATCACATGCAAAA
AAGAGCACACGTCTGAACTCTTTCCCTACACGACGCTCTTCCGATCTNNNAGAAAAATCACATGCAAAA
AAGAGCACACGTCTGAACTCTTTCCCTACACGACGCTCTTCCGATCTNNNAGAAAAATCACATGCAAAA
AAGAGCACACGTCTGAACTCTTTCCCTACACGACGCTCTTCCGATCTNNNAGAAAAATCACATGCAAAA
AAGAGCACACGTCTGAACTCTTTCCCTACACGACGCTCTTCCGATCTNNNAGAAAAATCACATGCAAAA
AAGAGCACACGTCTGAACTCTTTCCCTACACGACGCTCTTCCGATCTNNNAGAAAAATCACATGCAAAA
AAGAGCACACGTCTGAACTCTTTCCCTACACGACGCTCTTCCGATCTNNNAGAAAAATCACATGCAAAA
AGAGCACACGTCTGAACTCTTTCCCTACACGACGCTCTTCCGATCTNNNGAGGCAATAGAATTGCTTG
AGAGCACACGTCTGAACTCTTTCCCTACACGACGCTCTTCCGATCTNNNGAGGCAATAGAATTGCTTG
AGAGCACACGTCTGAACTCTTTCCCTACACGACGCTCTTCCGATCTNNNGAGGCAATAGAATTGCTTG
AGAGCACACGTCTGAACTCTTTCCCTACACGACGCTCTTCCGATCTNNNGAGGCAATAGAATTGCTTG
AGAGCACACGTCTGAACTCTTTCCCTACACGACGCTCTTCCGATCTNNNGAGGCAATAGAATTGCTTG
AGAGCACACGTCTGAACTCTTTCCCTACACGACGCTCTTCCGATCTNNNGAGGCAATAGAATTGCTTG
AGAGCACACGTCTGAACTCTTTCCCTACACGACGCTCTTCCGATCTNNNGAGGCAATAGAATTGCTTG
AGCACACGTCTGAACTCTTTCCCTACACGACGCTCTTCCGATCTNNNGAACCCCCTATCTCTACCTTCCCCA
AGCACACGTCTGAACTCTTTCCCTACACGACGCTCTTCCGATCTNNNGAACCCCCTATCTCTACCTTCCCCA
AGCACACGTCTGAACTCTTTCCCTACACGACGCTCTTCCGATCTNNNGAACCCCCTATCTCTACCTTCCCCA
AGCACACGTCTGAACTCTTTCCCTACACGACGCTCTTCCGATCTNNNGAACCCCCTATCTCTACCTTCCCCA
AGCACACGTCTGAACTCTTTCCCTACACGACGCTCTTCCGATCTNNNGAACCCCCTATCTCTACCTTCCCCA
AGCACACGTCTGAACTCTTTCCCTACACGACGCTCTTCCGATCTNNNGAACCCCCTATCTCTACCTTCCCCA
AGCACACGTCTGAACTCTTTCCCTACACGACGCTCTTCCGATCTNNNGAACCCCCTATCTCTACCTTCCCCA
CACACGTCTGAACTCTTTCCCTACACGACGCTCTTCCGATCTNNNCACAACGACCACATCCTCATAGACA
CACACGTCTGAACTCTTTCCCTACACGACGCTCTTCCGATCTNNNCACAACGACCACATCCTCATAGACA
CACACGTCTGAACTCTTTCCCTACACGACGCTCTTCCGATCTNNNCACAACGACCACATCCTCATAGACA
CACACGTCTGAACTCTTTCCCTACACGACGCTCTTCCGATCTNNNCACAACGACCACATCCTCATAGACA
CACACGTCTGAACTCTTTCCCTACACGACGCTCTTCCGATCTNNNCACAACGACCACATCCTCATAGACA
CACACGTCTGAACTCTTTCCCTACACGACGCTCTTCCGATCTNNNCACAACGACCACATCCTCATAGACA
CACACGTCTGAACTCTTTCCCTACACGACGCTCTTCCGATCTNNNCACAACGACCACATCCTCATAGACA
AGCACACGTCTGAACTCTTTCCCTACACGACGCTCTTCCGATCTNNNGCAGAAGAATCACTTGAACCCA
AGCACACGTCTGAACTCTTTCCCTACACGACGCTCTTCCGATCTNNNGCAGAAGAATCACTTGAACCCA
AGCACACGTCTGAACTCTTTCCCTACACGACGCTCTTCCGATCTNNNGCAGAAGAATCACTTGAACCCA
AGCACACGTCTGAACTCTTTCCCTACACGACGCTCTTCCGATCTNNNGCAGAAGAATCACTTGAACCCA
AGCACACGTCTGAACTCTTTCCCTACACGACGCTCTTCCGATCTNNNGCAGAAGAATCACTTGAACCCA
AGCACACGTCTGAACTCTTTCCCTACACGACGCTCTTCCGATCTNNNGCAGAAGAATCACTTGAACCCA
AGCACACGTCTGAACTCTTTCCCTACACGACGCTCTTCCGATCTNNNGCAGAAGAATCACTTGAACCCA
GAGCACACGTCTGAACTCTTTCCCTACACGACGCTCTTCCGATCTNNNCCTTTGGATAAAATGTGGGCC
GAGCACACGTCTGAACTCTTTCCCTACACGACGCTCTTCCGATCTNNNCCTTTGGATAAAATGTGGGCC
GAGCACACGTCTGAACTCTTTCCCTACACGACGCTCTTCCGATCTNNNCCTTTGGATAAAATGTGGGCC

GAGCACACGTCTGAACTCTTTCCCTACACGACGCTCTTCCGATCTNNNCCTTTGGATAAAATGTGGGC
GAGCACACGTCTGAACTCTTTCCCTACACGACGCTCTTCCGATCTNNNCCTTTGGATAAAATGTGGGC
GAGCACACGTCTGAACTCTTTCCCTACACGACGCTCTTCCGATCTNNNCCTTTGGATAAAATGTGGGC
GAGCACACGTCTGAACTCTTTCCCTACACGACGCTCTTCCGATCTNNNCCTTTGGATAAAATGTGGGC
GAGCACACGTCTGAACTCTTTCCCTACACGACGCTCTTCCGATCTNNNCCTCACTAATATTCCCCTGGT
GAGCACACGTCTGAACTCTTTCCCTACACGACGCTCTTCCGATCTNNNCCTCACTAATATTCCCCTGGT
GAGCACACGTCTGAACTCTTTCCCTACACGACGCTCTTCCGATCTNNNCCTCACTAATATTCCCCTGGT
GAGCACACGTCTGAACTCTTTCCCTACACGACGCTCTTCCGATCTNNNCCTCACTAATATTCCCCTGGT
GAGCACACGTCTGAACTCTTTCCCTACACGACGCTCTTCCGATCTNNNCCTCACTAATATTCCCCTGGT
GAGCACACGTCTGAACTCTTTCCCTACACGACGCTCTTCCGATCTNNNCCTCACTAATATTCCCCTGGT
GAGCACACGTCTGAACTCTTTCCCTACACGACGCTCTTCCGATCTNNNCCTCACTAATATTCCCCTGGT
AGCACACGTCTGAACTCTTTCCCTACACGACGCTCTTCCGATCTNNNCCAGCTTCTGTGCATGTTTTGTT
AGCACACGTCTGAACTCTTTCCCTACACGACGCTCTTCCGATCTNNNCCAGCTTCTGTGCATGTTTTGTT
AGCACACGTCTGAACTCTTTCCCTACACGACGCTCTTCCGATCTNNNCCAGCTTCTGTGCATGTTTTGTT
AGCACACGTCTGAACTCTTTCCCTACACGACGCTCTTCCGATCTNNNCCAGCTTCTGTGCATGTTTTGTT
AGCACACGTCTGAACTCTTTCCCTACACGACGCTCTTCCGATCTNNNCCAGCTTCTGTGCATGTTTTGTT
AGCACACGTCTGAACTCTTTCCCTACACGACGCTCTTCCGATCTNNNCCAGCTTCTGTGCATGTTTTGTT
AGCACACGTCTGAACTCTTTCCCTACACGACGCTCTTCCGATCTNNNCCAGCTTCTGTGCATGTTTTGTT
AGCACACGTCTGAACTCTTTCCCTACACGACGCTCTTCCGATCTNNNAAGTAAAAGTTTCAAAGGTACT
AGCACACGTCTGAACTCTTTCCCTACACGACGCTCTTCCGATCTNNNAAGTAAAAGTTTCAAAGGTACT
AGCACACGTCTGAACTCTTTCCCTACACGACGCTCTTCCGATCTNNNAAGTAAAAGTTTCAAAGGTACT
AGCACACGTCTGAACTCTTTCCCTACACGACGCTCTTCCGATCTNNNAAGTAAAAGTTTCAAAGGTACT
AGCACACGTCTGAACTCTTTCCCTACACGACGCTCTTCCGATCTNNNAAGTAAAAGTTTCAAAGGTACT
AGCACACGTCTGAACTCTTTCCCTACACGACGCTCTTCCGATCTNNNAAGTAAAAGTTTCAAAGGTACT
AGCACACGTCTGAACTCTTTCCCTACACGACGCTCTTCCGATCTNNNAAGTAAAAGTTTCAAAGGTACT
AGCACACGTCTGAACTCTTTCCCTACACGACGCTCTTCCGATCTNNNTGACCACACAACTTTGCAACTT
AGCACACGTCTGAACTCTTTCCCTACACGACGCTCTTCCGATCTNNNTGACCACACAACTTTGCAACTT
AGCACACGTCTGAACTCTTTCCCTACACGACGCTCTTCCGATCTNNNTGACCACACAACTTTGCAACTT
AGCACACGTCTGAACTCTTTCCCTACACGACGCTCTTCCGATCTNNNTGACCACACAACTTTGCAACTT
AGCACACGTCTGAACTCTTTCCCTACACGACGCTCTTCCGATCTNNNTGACCACACAACTTTGCAACTT
AGCACACGTCTGAACTCTTTCCCTACACGACGCTCTTCCGATCTNNNTGACCACACAACTTTGCAACTT
AGCACACGTCTGAACTCTTTCCCTACACGACGCTCTTCCGATCTNNNTGACCACACAACTTTGCAACTT
AGCACACGTCTGAACTCTTTCCCTACACGACGCTCTTCCGATCTNNNAGGAAAATCACTTGAACTCGGG
AGCACACGTCTGAACTCTTTCCCTACACGACGCTCTTCCGATCTNNNAGGAAAATCACTTGAACTCGGG
AGCACACGTCTGAACTCTTTCCCTACACGACGCTCTTCCGATCTNNNAGGAAAATCACTTGAACTCGGG
AGCACACGTCTGAACTCTTTCCCTACACGACGCTCTTCCGATCTNNNAGGAAAATCACTTGAACTCGGG
AGCACACGTCTGAACTCTTTCCCTACACGACGCTCTTCCGATCTNNNAGGAAAATCACTTGAACTCGGG
AGCACACGTCTGAACTCTTTCCCTACACGACGCTCTTCCGATCTNNNAGGAAAATCACTTGAACTCGGG
AGCACACGTCTGAACTCTTTCCCTACACGACGCTCTTCCGATCTNNNAGGAAAATCACTTGAACTCGGG
GAGCACACGTCTGAACTCTTTCCCTACACGACGCTCTTCCGATCTNNNGAGAATCGCTTGAACCCTGG
GAGCACACGTCTGAACTCTTTCCCTACACGACGCTCTTCCGATCTNNNGAGAATCGCTTGAACCCTGG
GAGCACACGTCTGAACTCTTTCCCTACACGACGCTCTTCCGATCTNNNGAGAATCGCTTGAACCCTGG
GAGCACACGTCTGAACTCTTTCCCTACACGACGCTCTTCCGATCTNNNGAGAATCGCTTGAACCCTGG

GAGCACACGTCTGAACTCTTTCCCTACACGACGCTCTTCCGATCTNNNGAGAATCGCTTGAACCCTGG
GAGCACACGTCTGAACTCTTTCCCTACACGACGCTCTTCCGATCTNNNGAGAATCGCTTGAACCCTGG
GAGCACACGTCTGAACTCTTTCCCTACACGACGCTCTTCCGATCTNNNGAGAATCGCTTGAACCCTGG
AGAGCACACGTCTGAACTCTTTCCCTACACGACGCTCTTCCGATCTNNNATGCATTCAAGGAGAATCG
AGAGCACACGTCTGAACTCTTTCCCTACACGACGCTCTTCCGATCTNNNATGCATTCAAGGAGAATCG
AGAGCACACGTCTGAACTCTTTCCCTACACGACGCTCTTCCGATCTNNNATGCATTCAAGGAGAATCG
AGAGCACACGTCTGAACTCTTTCCCTACACGACGCTCTTCCGATCTNNNATGCATTCAAGGAGAATCG
AGAGCACACGTCTGAACTCTTTCCCTACACGACGCTCTTCCGATCTNNNATGCATTCAAGGAGAATCG
AGAGCACACGTCTGAACTCTTTCCCTACACGACGCTCTTCCGATCTNNNATGCATTCAAGGAGAATCG
AGAGCACACGTCTGAACTCTTTCCCTACACGACGCTCTTCCGATCTNNNATGCATTCAAGGAGAATCG
AGCACACGTCTGAACTCTTTCCCTACACGACGCTCTTCCGATCTNNNCACCCCCAGCTGTATTTCTTTTG
AGCACACGTCTGAACTCTTTCCCTACACGACGCTCTTCCGATCTNNNCACCCCCAGCTGTATTTCTTTTG
AGCACACGTCTGAACTCTTTCCCTACACGACGCTCTTCCGATCTNNNCACCCCCAGCTGTATTTCTTTTG
AGCACACGTCTGAACTCTTTCCCTACACGACGCTCTTCCGATCTNNNCACCCCCAGCTGTATTTCTTTTG
AGCACACGTCTGAACTCTTTCCCTACACGACGCTCTTCCGATCTNNNCACCCCCAGCTGTATTTCTTTTG
AGCACACGTCTGAACTCTTTCCCTACACGACGCTCTTCCGATCTNNNCACCCCCAGCTGTATTTCTTTTG
AGCACACGTCTGAACTCTTTCCCTACACGACGCTCTTCCGATCTNNNCACCCCCAGCTGTATTTCTTTTG
GAGCACACGTCTGAACTCTTTCCCTACACGACGCTCTTCCGATCTNNNCAGCCTCCAACAAAGTGGTA
GAGCACACGTCTGAACTCTTTCCCTACACGACGCTCTTCCGATCTNNNCAGCCTCCAACAAAGTGGTA
GAGCACACGTCTGAACTCTTTCCCTACACGACGCTCTTCCGATCTNNNCAGCCTCCAACAAAGTGGTA
GAGCACACGTCTGAACTCTTTCCCTACACGACGCTCTTCCGATCTNNNCAGCCTCCAACAAAGTGGTA
GAGCACACGTCTGAACTCTTTCCCTACACGACGCTCTTCCGATCTNNNCAGCCTCCAACAAAGTGGTA
GAGCACACGTCTGAACTCTTTCCCTACACGACGCTCTTCCGATCTNNNCAGCCTCCAACAAAGTGGTA
GAGCACACGTCTGAACTCTTTCCCTACACGACGCTCTTCCGATCTNNNCAGCCTCCAACAAAGTGGTA
AGCACACGTCTGAACTCTTTCCCTACACGACGCTCTTCCGATCTNNNCTGAAGCAGGAGAATGGTGT
AGCACACGTCTGAACTCTTTCCCTACACGACGCTCTTCCGATCTNNNCTGAAGCAGGAGAATGGTGT
AGCACACGTCTGAACTCTTTCCCTACACGACGCTCTTCCGATCTNNNCTGAAGCAGGAGAATGGTGT
AGCACACGTCTGAACTCTTTCCCTACACGACGCTCTTCCGATCTNNNCTGAAGCAGGAGAATGGTGT
AGCACACGTCTGAACTCTTTCCCTACACGACGCTCTTCCGATCTNNNCTGAAGCAGGAGAATGGTGT
AGCACACGTCTGAACTCTTTCCCTACACGACGCTCTTCCGATCTNNNCTGAAGCAGGAGAATGGTGT
AGCACACGTCTGAACTCTTTCCCTACACGACGCTCTTCCGATCTNNNCTGAAGCAGGAGAATGGTGT
AGAGCACACGTCTGAACTCTTTCCCTACACGACGCTCTTCCGATCTNNNGCAAACTGAAGCAAATATC
AGAGCACACGTCTGAACTCTTTCCCTACACGACGCTCTTCCGATCTNNNGCAAACTGAAGCAAATATC
AGAGCACACGTCTGAACTCTTTCCCTACACGACGCTCTTCCGATCTNNNGCAAACTGAAGCAAATATC
AGAGCACACGTCTGAACTCTTTCCCTACACGACGCTCTTCCGATCTNNNGCAAACTGAAGCAAATATC
AGAGCACACGTCTGAACTCTTTCCCTACACGACGCTCTTCCGATCTNNNGCAAACTGAAGCAAATATC
AGAGCACACGTCTGAACTCTTTCCCTACACGACGCTCTTCCGATCTNNNGCAAACTGAAGCAAATATC
AGAGCACACGTCTGAACTCTTTCCCTACACGACGCTCTTCCGATCTNNNGCAAACTGAAGCAAATATC
CACACGTCTGAACTCTTTCCCTACACGACGCTCTTCCGATCTNNNTGTAATCCTAGCACTTTGGGAGGC
CACACGTCTGAACTCTTTCCCTACACGACGCTCTTCCGATCTNNNTGTAATCCTAGCACTTTGGGAGGC
CACACGTCTGAACTCTTTCCCTACACGACGCTCTTCCGATCTNNNTGTAATCCTAGCACTTTGGGAGGC
CACACGTCTGAACTCTTTCCCTACACGACGCTCTTCCGATCTNNNTGTAATCCTAGCACTTTGGGAGGC
CACACGTCTGAACTCTTTCCCTACACGACGCTCTTCCGATCTNNNTGTAATCCTAGCACTTTGGGAGGC

CACACGTCTGAACTCTTTCCCTACACGACGCTCTTCCGATCTNNNTGTAATCCTAGCACTTTGGGAGGC
CACACGTCTGAACTCTTTCCCTACACGACGCTCTTCCGATCTNNNTGTAATCCTAGCACTTTGGGAGGC
GCACACGTCTGAACTCTTTCCCTACACGACGCTCTTCCGATCTNNNGCCTTCCTTCCTTCCTTTCTTCTCA
GCACACGTCTGAACTCTTTCCCTACACGACGCTCTTCCGATCTNNNGCCTTCCTTCCTTCCTTTCTTCTCA
GCACACGTCTGAACTCTTTCCCTACACGACGCTCTTCCGATCTNNNGCCTTCCTTCCTTCCTTTCTTCTCA
GCACACGTCTGAACTCTTTCCCTACACGACGCTCTTCCGATCTNNNGCCTTCCTTCCTTCCTTTCTTCTCA
GCACACGTCTGAACTCTTTCCCTACACGACGCTCTTCCGATCTNNNGCCTTCCTTCCTTCCTTTCTTCTCA
GCACACGTCTGAACTCTTTCCCTACACGACGCTCTTCCGATCTNNNGCCTTCCTTCCTTCCTTTCTTCTCA
GCACACGTCTGAACTCTTTCCCTACACGACGCTCTTCCGATCTNNNGCCTTCCTTCCTTCCTTTCTTCTCA
GAGCACACGTCTGAACTCTTTCCCTACACGACGCTCTTCCGATCTNNNGGATGAGAGAGGATGAGA
GAGCACACGTCTGAACTCTTTCCCTACACGACGCTCTTCCGATCTNNNGGATGAGAGAGGATGAGA
GAGCACACGTCTGAACTCTTTCCCTACACGACGCTCTTCCGATCTNNNGGATGAGAGAGGATGAGA
GAGCACACGTCTGAACTCTTTCCCTACACGACGCTCTTCCGATCTNNNGGATGAGAGAGGATGAGA
GAGCACACGTCTGAACTCTTTCCCTACACGACGCTCTTCCGATCTNNNGGATGAGAGAGGATGAGA
GAGCACACGTCTGAACTCTTTCCCTACACGACGCTCTTCCGATCTNNNGGATGAGAGAGGATGAGA
GAGCACACGTCTGAACTCTTTCCCTACACGACGCTCTTCCGATCTNNNGGATGAGAGAGGATGAGA
GCACACGTCTGAACTCTTTCCCTACACGACGCTCTTCCGATCTNNNTGAGAATTGTCGATTAGCCCAC
GCACACGTCTGAACTCTTTCCCTACACGACGCTCTTCCGATCTNNNTGAGAATTGTCGATTAGCCCAC
GCACACGTCTGAACTCTTTCCCTACACGACGCTCTTCCGATCTNNNTGAGAATTGTCGATTAGCCCAC
GCACACGTCTGAACTCTTTCCCTACACGACGCTCTTCCGATCTNNNTGAGAATTGTCGATTAGCCCAC
GCACACGTCTGAACTCTTTCCCTACACGACGCTCTTCCGATCTNNNTGAGAATTGTCGATTAGCCCAC
GCACACGTCTGAACTCTTTCCCTACACGACGCTCTTCCGATCTNNNTGAGAATTGTCGATTAGCCCAC
GCACACGTCTGAACTCTTTCCCTACACGACGCTCTTCCGATCTNNNTGAGAATTGTCGATTAGCCCAC
AGCACACGTCTGAACTCTTTCCCTACACGACGCTCTTCCGATCTNNNTAGGCAGGAAAAGAGCAGTA
AGCACACGTCTGAACTCTTTCCCTACACGACGCTCTTCCGATCTNNNTAGGCAGGAAAAGAGCAGTA
AGCACACGTCTGAACTCTTTCCCTACACGACGCTCTTCCGATCTNNNTAGGCAGGAAAAGAGCAGTA
AGCACACGTCTGAACTCTTTCCCTACACGACGCTCTTCCGATCTNNNTAGGCAGGAAAAGAGCAGTA
AGCACACGTCTGAACTCTTTCCCTACACGACGCTCTTCCGATCTNNNTAGGCAGGAAAAGAGCAGTA
AGCACACGTCTGAACTCTTTCCCTACACGACGCTCTTCCGATCTNNNTAGGCAGGAAAAGAGCAGTA
AGCACACGTCTGAACTCTTTCCCTACACGACGCTCTTCCGATCTNNNTAGGCAGGAAAAGAGCAGTA
GAGCACACGTCTGAACTCTTTCCCTACACGACGCTCTTCCGATCTNNNGGGGGGTAGATCTAACCAA
GAGCACACGTCTGAACTCTTTCCCTACACGACGCTCTTCCGATCTNNNGGGGGGTAGATCTAACCAA
GAGCACACGTCTGAACTCTTTCCCTACACGACGCTCTTCCGATCTNNNGGGGGGTAGATCTAACCAA
GAGCACACGTCTGAACTCTTTCCCTACACGACGCTCTTCCGATCTNNNGGGGGGTAGATCTAACCAA
GAGCACACGTCTGAACTCTTTCCCTACACGACGCTCTTCCGATCTNNNGGGGGGTAGATCTAACCAA
GAGCACACGTCTGAACTCTTTCCCTACACGACGCTCTTCCGATCTNNNGGGGGGTAGATCTAACCAA
GAGCACACGTCTGAACTCTTTCCCTACACGACGCTCTTCCGATCTNNNGGGGGGTAGATCTAACCAA
GCACACGTCTGAACTCTTTCCCTACACGACGCTCTTCCGATCTNNNAAGATTTAATGACCTCAGGTTT
GCACACGTCTGAACTCTTTCCCTACACGACGCTCTTCCGATCTNNNAAGATTTAATGACCTCAGGTTT
GCACACGTCTGAACTCTTTCCCTACACGACGCTCTTCCGATCTNNNAAGATTTAATGACCTCAGGTTT
GCACACGTCTGAACTCTTTCCCTACACGACGCTCTTCCGATCTNNNAAGATTTAATGACCTCAGGTTT
GCACACGTCTGAACTCTTTCCCTACACGACGCTCTTCCGATCTNNNAAGATTTAATGACCTCAGGTTT
GCACACGTCTGAACTCTTTCCCTACACGACGCTCTTCCGATCTNNNAAGATTTAATGACCTCAGGTTT

AGCACACGTCTGAACTCTTTCCCTACACGACGCTCTTCCGATCTNNNAAGATTTAATGACCTCAGGTTT
GAGCACACGTCTGAACTCTTTCCCTACACGACGCTCTTCCGATCTNNNTCTGAACTTTGGACATACCCA
GAGCACACGTCTGAACTCTTTCCCTACACGACGCTCTTCCGATCTNNNTCTGAACTTTGGACATACCCA
GAGCACACGTCTGAACTCTTTCCCTACACGACGCTCTTCCGATCTNNNTCTGAACTTTGGACATACCCA
GAGCACACGTCTGAACTCTTTCCCTACACGACGCTCTTCCGATCTNNNTCTGAACTTTGGACATACCCA
GAGCACACGTCTGAACTCTTTCCCTACACGACGCTCTTCCGATCTNNNTCTGAACTTTGGACATACCCA
GAGCACACGTCTGAACTCTTTCCCTACACGACGCTCTTCCGATCTNNNTCTGAACTTTGGACATACCCA
GAGCACACGTCTGAACTCTTTCCCTACACGACGCTCTTCCGATCTNNNTCTGAACTTTGGACATACCCA
AGCACACGTCTGAACTCTTTCCCTACACGACGCTCTTCCGATCTNNNTACAAAAAATTAGCTGGGCGT
AGCACACGTCTGAACTCTTTCCCTACACGACGCTCTTCCGATCTNNNTACAAAAAATTAGCTGGGCGT
AGCACACGTCTGAACTCTTTCCCTACACGACGCTCTTCCGATCTNNNTACAAAAAATTAGCTGGGCGT
AGCACACGTCTGAACTCTTTCCCTACACGACGCTCTTCCGATCTNNNTACAAAAAATTAGCTGGGCGT
AGCACACGTCTGAACTCTTTCCCTACACGACGCTCTTCCGATCTNNNTACAAAAAATTAGCTGGGCGT
AGCACACGTCTGAACTCTTTCCCTACACGACGCTCTTCCGATCTNNNTACAAAAAATTAGCTGGGCGT
AGCACACGTCTGAACTCTTTCCCTACACGACGCTCTTCCGATCTNNNTACAAAAAATTAGCTGGGCGT
AGCACACGTCTGAACTCTTTCCCTACACGACGCTCTTCCGATCTNNNCAACAGTTCTCTCTGCAGTAGGT
AGCACACGTCTGAACTCTTTCCCTACACGACGCTCTTCCGATCTNNNCAACAGTTCTCTCTGCAGTAGGT
AGCACACGTCTGAACTCTTTCCCTACACGACGCTCTTCCGATCTNNNCAACAGTTCTCTCTGCAGTAGGT
AGCACACGTCTGAACTCTTTCCCTACACGACGCTCTTCCGATCTNNNCAACAGTTCTCTCTGCAGTAGGT
AGCACACGTCTGAACTCTTTCCCTACACGACGCTCTTCCGATCTNNNCAACAGTTCTCTCTGCAGTAGGT
AGCACACGTCTGAACTCTTTCCCTACACGACGCTCTTCCGATCTNNNCAACAGTTCTCTCTGCAGTAGGT
AGCACACGTCTGAACTCTTTCCCTACACGACGCTCTTCCGATCTNNNCAACAGTTCTCTCTGCAGTAGGT
GAGCACACGTCTGAACTCTTTCCCTACACGACGCTCTTCCGATCTNNNTACTTTGATGCACACACACACA
GAGCACACGTCTGAACTCTTTCCCTACACGACGCTCTTCCGATCTNNNTACTTTGATGCACACACACACA
GAGCACACGTCTGAACTCTTTCCCTACACGACGCTCTTCCGATCTNNNTACTTTGATGCACACACACACA
GAGCACACGTCTGAACTCTTTCCCTACACGACGCTCTTCCGATCTNNNTACTTTGATGCACACACACACA
GAGCACACGTCTGAACTCTTTCCCTACACGACGCTCTTCCGATCTNNNTACTTTGATGCACACACACACA
GAGCACACGTCTGAACTCTTTCCCTACACGACGCTCTTCCGATCTNNNTACTTTGATGCACACACACACA
GAGCACACGTCTGAACTCTTTCCCTACACGACGCTCTTCCGATCTNNNTACTTTGATGCACACACACACA
GAGCACACGTCTGAACTCTTTCCCTACACGACGCTCTTCCGATCTNNNAATACAAAAAATTAGCCGGG
GAGCACACGTCTGAACTCTTTCCCTACACGACGCTCTTCCGATCTNNNAATACAAAAAATTAGCCGGG
GAGCACACGTCTGAACTCTTTCCCTACACGACGCTCTTCCGATCTNNNAATACAAAAAATTAGCCGGG
GAGCACACGTCTGAACTCTTTCCCTACACGACGCTCTTCCGATCTNNNAATACAAAAAATTAGCCGGG
GAGCACACGTCTGAACTCTTTCCCTACACGACGCTCTTCCGATCTNNNAATACAAAAAATTAGCCGGG
GAGCACACGTCTGAACTCTTTCCCTACACGACGCTCTTCCGATCTNNNAATACAAAAAATTAGCCGGG
GAGCACACGTCTGAACTCTTTCCCTACACGACGCTCTTCCGATCTNNNAATACAAAAAATTAGCCGGG
AGAGCACACGTCTGAACTCTTTCCCTACACGACGCTCTTCCGATCTNNNTAAAAAAGCAGGATCACGTT
AGAGCACACGTCTGAACTCTTTCCCTACACGACGCTCTTCCGATCTNNNTAAAAAAGCAGGATCACGTT
AGAGCACACGTCTGAACTCTTTCCCTACACGACGCTCTTCCGATCTNNNTAAAAAAGCAGGATCACGTT
AGAGCACACGTCTGAACTCTTTCCCTACACGACGCTCTTCCGATCTNNNTAAAAAAGCAGGATCACGTT
AGAGCACACGTCTGAACTCTTTCCCTACACGACGCTCTTCCGATCTNNNTAAAAAAGCAGGATCACGTT
AGAGCACACGTCTGAACTCTTTCCCTACACGACGCTCTTCCGATCTNNNTAAAAAAGCAGGATCACGTT
AGAGCACACGTCTGAACTCTTTCCCTACACGACGCTCTTCCGATCTNNNTAAAAAAGCAGGATCACGTT

GAGCACACGTCTGAACTCTTTCCCTACACGACGCTCTTCCGATCTNNNCAAGATAGCACCACTGCACT
GAGCACACGTCTGAACTCTTTCCCTACACGACGCTCTTCCGATCTNNNCAAGATAGCACCACTGCACT
GAGCACACGTCTGAACTCTTTCCCTACACGACGCTCTTCCGATCTNNNCAAGATAGCACCACTGCACT
GAGCACACGTCTGAACTCTTTCCCTACACGACGCTCTTCCGATCTNNNCAAGATAGCACCACTGCACT
GAGCACACGTCTGAACTCTTTCCCTACACGACGCTCTTCCGATCTNNNCAAGATAGCACCACTGCACT
GAGCACACGTCTGAACTCTTTCCCTACACGACGCTCTTCCGATCTNNNCAAGATAGCACCACTGCACT
GAGCACACGTCTGAACTCTTTCCCTACACGACGCTCTTCCGATCTNNNCAAGATAGCACCACTGCACT
GCACACGTCTGAACTCTTTCCCTACACGACGCTCTTCCGATCTNNNAGTTTTTCTGGTGTCTTTGGGG
GCACACGTCTGAACTCTTTCCCTACACGACGCTCTTCCGATCTNNNAGTTTTTCTGGTGTCTTTGGGG
GCACACGTCTGAACTCTTTCCCTACACGACGCTCTTCCGATCTNNNAGTTTTTCTGGTGTCTTTGGGG
GCACACGTCTGAACTCTTTCCCTACACGACGCTCTTCCGATCTNNNAGTTTTTCTGGTGTCTTTGGGG
GCACACGTCTGAACTCTTTCCCTACACGACGCTCTTCCGATCTNNNAGTTTTTCTGGTGTCTTTGGGG
GCACACGTCTGAACTCTTTCCCTACACGACGCTCTTCCGATCTNNNAGTTTTTCTGGTGTCTTTGGGG
GCACACGTCTGAACTCTTTCCCTACACGACGCTCTTCCGATCTNNNAGTTTTTCTGGTGTCTTTGGGG
ACACGTCTGAACTCTTTCCCTACACGACGCTCTTCCGATCTNNNCAGTGCTGCCATTTTCTTCTGATCAT
ACACGTCTGAACTCTTTCCCTACACGACGCTCTTCCGATCTNNNCAGTGCTGCCATTTTCTTCTGATCAT
ACACGTCTGAACTCTTTCCCTACACGACGCTCTTCCGATCTNNNCAGTGCTGCCATTTTCTTCTGATCAT
ACACGTCTGAACTCTTTCCCTACACGACGCTCTTCCGATCTNNNCAGTGCTGCCATTTTCTTCTGATCAT
ACACGTCTGAACTCTTTCCCTACACGACGCTCTTCCGATCTNNNCAGTGCTGCCATTTTCTTCTGATCAT
ACACGTCTGAACTCTTTCCCTACACGACGCTCTTCCGATCTNNNCAGTGCTGCCATTTTCTTCTGATCAT
ACACGTCTGAACTCTTTCCCTACACGACGCTCTTCCGATCTNNNCAGTGCTGCCATTTTCTTCTGATCAT
GAGCACACGTCTGAACTCTTTCCCTACACGACGCTCTTCCGATCTNNNTACGACTTTCTCCCTAGGCAA
GAGCACACGTCTGAACTCTTTCCCTACACGACGCTCTTCCGATCTNNNTACGACTTTCTCCCTAGGCAA
GAGCACACGTCTGAACTCTTTCCCTACACGACGCTCTTCCGATCTNNNTACGACTTTCTCCCTAGGCAA
GAGCACACGTCTGAACTCTTTCCCTACACGACGCTCTTCCGATCTNNNTACGACTTTCTCCCTAGGCAA
GAGCACACGTCTGAACTCTTTCCCTACACGACGCTCTTCCGATCTNNNTACGACTTTCTCCCTAGGCAA
GAGCACACGTCTGAACTCTTTCCCTACACGACGCTCTTCCGATCTNNNTACGACTTTCTCCCTAGGCAA
GAGCACACGTCTGAACTCTTTCCCTACACGACGCTCTTCCGATCTNNNTACGACTTTCTCCCTAGGCAA
AGCACACGTCTGAACTCTTTCCCTACACGACGCTCTTCCGATCTNNNTCACACCTATGTTTGTGTGTGT
AGCACACGTCTGAACTCTTTCCCTACACGACGCTCTTCCGATCTNNNTCACACCTATGTTTGTGTGTGT
AGCACACGTCTGAACTCTTTCCCTACACGACGCTCTTCCGATCTNNNTCACACCTATGTTTGTGTGTGT
AGCACACGTCTGAACTCTTTCCCTACACGACGCTCTTCCGATCTNNNTCACACCTATGTTTGTGTGTGT
AGCACACGTCTGAACTCTTTCCCTACACGACGCTCTTCCGATCTNNNTCACACCTATGTTTGTGTGTGT
AGCACACGTCTGAACTCTTTCCCTACACGACGCTCTTCCGATCTNNNTCACACCTATGTTTGTGTGTGT
AGCACACGTCTGAACTCTTTCCCTACACGACGCTCTTCCGATCTNNNTCACACCTATGTTTGTGTGTGT
GAGCACACGTCTGAACTCTTTCCCTACACGACGCTCTTCCGATCTNNNACCTTGGGCAAAACAAGCAA
GAGCACACGTCTGAACTCTTTCCCTACACGACGCTCTTCCGATCTNNNACCTTGGGCAAAACAAGCAA
GAGCACACGTCTGAACTCTTTCCCTACACGACGCTCTTCCGATCTNNNACCTTGGGCAAAACAAGCAA
GAGCACACGTCTGAACTCTTTCCCTACACGACGCTCTTCCGATCTNNNACCTTGGGCAAAACAAGCAA
GAGCACACGTCTGAACTCTTTCCCTACACGACGCTCTTCCGATCTNNNACCTTGGGCAAAACAAGCAA
GAGCACACGTCTGAACTCTTTCCCTACACGACGCTCTTCCGATCTNNNACCTTGGGCAAAACAAGCAA
GAGCACACGTCTGAACTCTTTCCCTACACGACGCTCTTCCGATCTNNNACCTTGGGCAAAACAAGCAA
GCACACGTCTGAACTCTTTCCCTACACGACGCTCTTCCGATCTNNNGGACTGAGTGGAGATTTCACC

AGCACACGTCTGAACTCTTTCCCTACACGACGCTCTTCCGATCTNNNGGACTGAGTGGAGATTTCACCT

AGCACACGTCTGAACTCTTTCCCTACACGACGCTCTTCCGATCTNNNGGACTGAGTGGAGATTTCACCT

AGCACACGTCTGAACTCTTTCCCTACACGACGCTCTTCCGATCTNNNGGACTGAGTGGAGATTTCACCT

AGCACACGTCTGAACTCTTTCCCTACACGACGCTCTTCCGATCTNNNGGACTGAGTGGAGATTTCACCT

AGCACACGTCTGAACTCTTTCCCTACACGACGCTCTTCCGATCTNNNGGACTGAGTGGAGATTTCACCT

AGCACACGTCTGAACTCTTTCCCTACACGACGCTCTTCCGATCTNNNGGACTGAGTGGAGATTTCACCT

AGCACACGTCTGAACTCTTTCCCTACACGACGCTCTTCCGATCTNNNGAGGTTGTGGTGAGCAGAGA

AGCACACGTCTGAACTCTTTCCCTACACGACGCTCTTCCGATCTNNNGAGGTTGTGGTGAGCAGAGA

AGCACACGTCTGAACTCTTTCCCTACACGACGCTCTTCCGATCTNNNGAGGTTGTGGTGAGCAGAGA

AGCACACGTCTGAACTCTTTCCCTACACGACGCTCTTCCGATCTNNNGAGGTTGTGGTGAGCAGAGA

AGCACACGTCTGAACTCTTTCCCTACACGACGCTCTTCCGATCTNNNGAGGTTGTGGTGAGCAGAGA

AGCACACGTCTGAACTCTTTCCCTACACGACGCTCTTCCGATCTNNNGAGGTTGTGGTGAGCAGAGA

AGCACACGTCTGAACTCTTTCCCTACACGACGCTCTTCCGATCTNNNGAGGTTGTGGTGAGCAGAGA

AGCACACGTCTGAACTCTTTCCCTACACGACGCTCTTCCGATCTNNNGCACGATGGAGCAGTGTTTAT

AGCACACGTCTGAACTCTTTCCCTACACGACGCTCTTCCGATCTNNNGCACGATGGAGCAGTGTTTAT

AGCACACGTCTGAACTCTTTCCCTACACGACGCTCTTCCGATCTNNNGCACGATGGAGCAGTGTTTAT

AGCACACGTCTGAACTCTTTCCCTACACGACGCTCTTCCGATCTNNNGCACGATGGAGCAGTGTTTAT

AGCACACGTCTGAACTCTTTCCCTACACGACGCTCTTCCGATCTNNNGCACGATGGAGCAGTGTTTAT

AGCACACGTCTGAACTCTTTCCCTACACGACGCTCTTCCGATCTNNNGCACGATGGAGCAGTGTTTAT

AGCACACGTCTGAACTCTTTCCCTACACGACGCTCTTCCGATCTNNNGCACGATGGAGCAGTGTTTAT

ACACGTCTGAACTCTTTCCCTACACGACGCTCTTCCGATCTNNNCCTTTGTCTCCAGCCCCTTTCCATCG

ACACGTCTGAACTCTTTCCCTACACGACGCTCTTCCGATCTNNNCCTTTGTCTCCAGCCCCTTTCCATCG

ACACGTCTGAACTCTTTCCCTACACGACGCTCTTCCGATCTNNNCCTTTGTCTCCAGCCCCTTTCCATCG

ACACGTCTGAACTCTTTCCCTACACGACGCTCTTCCGATCTNNNCCTTTGTCTCCAGCCCCTTTCCATCG

ACACGTCTGAACTCTTTCCCTACACGACGCTCTTCCGATCTNNNCCTTTGTCTCCAGCCCCTTTCCATCG

ACACGTCTGAACTCTTTCCCTACACGACGCTCTTCCGATCTNNNCCTTTGTCTCCAGCCCCTTTCCATCG

ACACGTCTGAACTCTTTCCCTACACGACGCTCTTCCGATCTNNNCCTTTGTCTCCAGCCCCTTTCCATCG

GAGCACACGTCTGAACTCTTTCCCTACACGACGCTCTTCCGATCTNNNGAGAACAGCTGGATATGCAC

GAGCACACGTCTGAACTCTTTCCCTACACGACGCTCTTCCGATCTNNNGAGAACAGCTGGATATGCAC

GAGCACACGTCTGAACTCTTTCCCTACACGACGCTCTTCCGATCTNNNGAGAACAGCTGGATATGCAC

GAGCACACGTCTGAACTCTTTCCCTACACGACGCTCTTCCGATCTNNNGAGAACAGCTGGATATGCAC

GAGCACACGTCTGAACTCTTTCCCTACACGACGCTCTTCCGATCTNNNGAGAACAGCTGGATATGCAC

GAGCACACGTCTGAACTCTTTCCCTACACGACGCTCTTCCGATCTNNNGAGAACAGCTGGATATGCAC

GAGCACACGTCTGAACTCTTTCCCTACACGACGCTCTTCCGATCTNNNGAGAACAGCTGGATATGCAC

GCACACGTCTGAACTCTTTCCCTACACGACGCTCTTCCGATCTNNNGCTTCCTTGTGCTTAGAGTTGGT

GCACACGTCTGAACTCTTTCCCTACACGACGCTCTTCCGATCTNNNGCTTCCTTGTGCTTAGAGTTGGT

GCACACGTCTGAACTCTTTCCCTACACGACGCTCTTCCGATCTNNNGCTTCCTTGTGCTTAGAGTTGGT

GCACACGTCTGAACTCTTTCCCTACACGACGCTCTTCCGATCTNNNGCTTCCTTGTGCTTAGAGTTGGT

GCACACGTCTGAACTCTTTCCCTACACGACGCTCTTCCGATCTNNNGCTTCCTTGTGCTTAGAGTTGGT

GCACACGTCTGAACTCTTTCCCTACACGACGCTCTTCCGATCTNNNGCTTCCTTGTGCTTAGAGTTGGT

GCACACGTCTGAACTCTTTCCCTACACGACGCTCTTCCGATCTNNNGCTTCCTTGTGCTTAGAGTTGGT

GAGCACACGTCTGAACTCTTTCCCTACACGACGCTCTTCCGATCTNNNACAAAACTTAGCTGGGCATG

GAGCACACGTCTGAACTCTTTCCCTACACGACGCTCTTCCGATCTNNNACAAAACTTAGCTGGGCATG

GAGCACACGTCTGAACTCTTTCCCTACACGACGCTCTTCCGATCTNNNACAAAACTTAGCTGGGCATG
GAGCACACGTCTGAACTCTTTCCCTACACGACGCTCTTCCGATCTNNNACAAAACTTAGCTGGGCATG
GAGCACACGTCTGAACTCTTTCCCTACACGACGCTCTTCCGATCTNNNACAAAACTTAGCTGGGCATG
GAGCACACGTCTGAACTCTTTCCCTACACGACGCTCTTCCGATCTNNNACAAAACTTAGCTGGGCATG
GAGCACACGTCTGAACTCTTTCCCTACACGACGCTCTTCCGATCTNNNACAAAACTTAGCTGGGCATG
AGCACACGTCTGAACTCTTTCCCTACACGACGCTCTTCCGATCTNNNATTTTCCTCCCTCTTGGACTTC
AGCACACGTCTGAACTCTTTCCCTACACGACGCTCTTCCGATCTNNNATTTTCCTCCCTCTTGGACTTC
AGCACACGTCTGAACTCTTTCCCTACACGACGCTCTTCCGATCTNNNATTTTCCTCCCTCTTGGACTTC
AGCACACGTCTGAACTCTTTCCCTACACGACGCTCTTCCGATCTNNNATTTTCCTCCCTCTTGGACTTC
AGCACACGTCTGAACTCTTTCCCTACACGACGCTCTTCCGATCTNNNATTTTCCTCCCTCTTGGACTTC
AGCACACGTCTGAACTCTTTCCCTACACGACGCTCTTCCGATCTNNNATTTTCCTCCCTCTTGGACTTC
AGCACACGTCTGAACTCTTTCCCTACACGACGCTCTTCCGATCTNNNATTTTCCTCCCTCTTGGACTTC
GCACACGTCTGAACTCTTTCCCTACACGACGCTCTTCCGATCTNNNACTAACCACCTTGTCTTTCTGGT
GCACACGTCTGAACTCTTTCCCTACACGACGCTCTTCCGATCTNNNACTAACCACCTTGTCTTTCTGGT
GCACACGTCTGAACTCTTTCCCTACACGACGCTCTTCCGATCTNNNACTAACCACCTTGTCTTTCTGGT
GCACACGTCTGAACTCTTTCCCTACACGACGCTCTTCCGATCTNNNACTAACCACCTTGTCTTTCTGGT
GCACACGTCTGAACTCTTTCCCTACACGACGCTCTTCCGATCTNNNACTAACCACCTTGTCTTTCTGGT
GCACACGTCTGAACTCTTTCCCTACACGACGCTCTTCCGATCTNNNACTAACCACCTTGTCTTTCTGGT
GCACACGTCTGAACTCTTTCCCTACACGACGCTCTTCCGATCTNNNACTAACCACCTTGTCTTTCTGGT
GCACACGTCTGAACTCTTTCCCTACACGACGCTCTTCCGATCTNNNTGCTGCAGTGAGTCATGTATGTC
GCACACGTCTGAACTCTTTCCCTACACGACGCTCTTCCGATCTNNNTGCTGCAGTGAGTCATGTATGTC
GCACACGTCTGAACTCTTTCCCTACACGACGCTCTTCCGATCTNNNTGCTGCAGTGAGTCATGTATGTC
GCACACGTCTGAACTCTTTCCCTACACGACGCTCTTCCGATCTNNNTGCTGCAGTGAGTCATGTATGTC
GCACACGTCTGAACTCTTTCCCTACACGACGCTCTTCCGATCTNNNTGCTGCAGTGAGTCATGTATGTC
GCACACGTCTGAACTCTTTCCCTACACGACGCTCTTCCGATCTNNNTGCTGCAGTGAGTCATGTATGTC
GCACACGTCTGAACTCTTTCCCTACACGACGCTCTTCCGATCTNNNTGCTGCAGTGAGTCATGTATGTC
GCACACGTCTGAACTCTTTCCCTACACGACGCTCTTCCGATCTNNNTTTCACATCGCTCTTCTCTCACAC
GCACACGTCTGAACTCTTTCCCTACACGACGCTCTTCCGATCTNNNTTTCACATCGCTCTTCTCTCACAC
GCACACGTCTGAACTCTTTCCCTACACGACGCTCTTCCGATCTNNNTTTCACATCGCTCTTCTCTCACAC
GCACACGTCTGAACTCTTTCCCTACACGACGCTCTTCCGATCTNNNTTTCACATCGCTCTTCTCTCACAC
GCACACGTCTGAACTCTTTCCCTACACGACGCTCTTCCGATCTNNNTTTCACATCGCTCTTCTCTCACAC
GCACACGTCTGAACTCTTTCCCTACACGACGCTCTTCCGATCTNNNTTTCACATCGCTCTTCTCTCACAC
GCACACGTCTGAACTCTTTCCCTACACGACGCTCTTCCGATCTNNNTTTCACATCGCTCTTCTCTCACAC
AGCACACGTCTGAACTCTTTCCCTACACGACGCTCTTCCGATCTNNNCTTTGCATTTGGAGACTGGGA
AGCACACGTCTGAACTCTTTCCCTACACGACGCTCTTCCGATCTNNNCTTTGCATTTGGAGACTGGGA
AGCACACGTCTGAACTCTTTCCCTACACGACGCTCTTCCGATCTNNNCTTTGCATTTGGAGACTGGGA
AGCACACGTCTGAACTCTTTCCCTACACGACGCTCTTCCGATCTNNNCTTTGCATTTGGAGACTGGGA
AGCACACGTCTGAACTCTTTCCCTACACGACGCTCTTCCGATCTNNNCTTTGCATTTGGAGACTGGGA
AGCACACGTCTGAACTCTTTCCCTACACGACGCTCTTCCGATCTNNNCTTTGCATTTGGAGACTGGGA
AGCACACGTCTGAACTCTTTCCCTACACGACGCTCTTCCGATCTNNNCTTTGCATTTGGAGACTGGGA
AGCACACGTCTGAACTCTTTCCCTACACGACGCTCTTCCGATCTNNNTATATGTGTGTGGGTGTGTGT
AGCACACGTCTGAACTCTTTCCCTACACGACGCTCTTCCGATCTNNNTATATGTGTGTGGGTGTGTGT
AGCACACGTCTGAACTCTTTCCCTACACGACGCTCTTCCGATCTNNNTATATGTGTGTGGGTGTGTGT

AGCACACGTCTGAACTCTTTCCCTACACGACGCTCTTCCGATCTNNNTATATGTGTGTGGGTGTGTGT

AGCACACGTCTGAACTCTTTCCCTACACGACGCTCTTCCGATCTNNNTATATGTGTGTGGGTGTGTGT

AGCACACGTCTGAACTCTTTCCCTACACGACGCTCTTCCGATCTNNNTATATGTGTGTGGGTGTGTGT

AGCACACGTCTGAACTCTTTCCCTACACGACGCTCTTCCGATCTNNNTATATGTGTGTGGGTGTGTGT

GAGCACACGTCTGAACTCTTTCCCTACACGACGCTCTTCCGATCTNNNGACCAATGTCTGAGACCCATTC

GAGCACACGTCTGAACTCTTTCCCTACACGACGCTCTTCCGATCTNNNGACCAATGTCTGAGACCCATTC

GAGCACACGTCTGAACTCTTTCCCTACACGACGCTCTTCCGATCTNNNGACCAATGTCTGAGACCCATTC

GAGCACACGTCTGAACTCTTTCCCTACACGACGCTCTTCCGATCTNNNGACCAATGTCTGAGACCCATTC

GAGCACACGTCTGAACTCTTTCCCTACACGACGCTCTTCCGATCTNNNGACCAATGTCTGAGACCCATTC

GAGCACACGTCTGAACTCTTTCCCTACACGACGCTCTTCCGATCTNNNGACCAATGTCTGAGACCCATTC

GAGCACACGTCTGAACTCTTTCCCTACACGACGCTCTTCCGATCTNNNGACCAATGTCTGAGACCCATTC

AGAGCACACGTCTGAACTCTTTCCCTACACGACGCTCTTCCGATCTNNNACACAGAGCCAAACTGTATC

AGAGCACACGTCTGAACTCTTTCCCTACACGACGCTCTTCCGATCTNNNACACAGAGCCAAACTGTATC

AGAGCACACGTCTGAACTCTTTCCCTACACGACGCTCTTCCGATCTNNNACACAGAGCCAAACTGTATC

AGAGCACACGTCTGAACTCTTTCCCTACACGACGCTCTTCCGATCTNNNACACAGAGCCAAACTGTATC

AGAGCACACGTCTGAACTCTTTCCCTACACGACGCTCTTCCGATCTNNNACACAGAGCCAAACTGTATC

AGAGCACACGTCTGAACTCTTTCCCTACACGACGCTCTTCCGATCTNNNACACAGAGCCAAACTGTATC

AGAGCACACGTCTGAACTCTTTCCCTACACGACGCTCTTCCGATCTNNNACACAGAGCCAAACTGTATC

AGCACACGTCTGAACTCTTTCCCTACACGACGCTCTTCCGATCTNNNTCTATTCATCGCCATTGGAACCA

AGCACACGTCTGAACTCTTTCCCTACACGACGCTCTTCCGATCTNNNTCTATTCATCGCCATTGGAACCA

AGCACACGTCTGAACTCTTTCCCTACACGACGCTCTTCCGATCTNNNTCTATTCATCGCCATTGGAACCA

AGCACACGTCTGAACTCTTTCCCTACACGACGCTCTTCCGATCTNNNTCTATTCATCGCCATTGGAACCA

AGCACACGTCTGAACTCTTTCCCTACACGACGCTCTTCCGATCTNNNTCTATTCATCGCCATTGGAACCA

AGCACACGTCTGAACTCTTTCCCTACACGACGCTCTTCCGATCTNNNTCTATTCATCGCCATTGGAACCA

AGCACACGTCTGAACTCTTTCCCTACACGACGCTCTTCCGATCTNNNTCTATTCATCGCCATTGGAACCA

AGCACACGTCTGAACTCTTTCCCTACACGACGCTCTTCCGATCTNNNAGCAAAGAAAAAAACAGGGAG

AGCACACGTCTGAACTCTTTCCCTACACGACGCTCTTCCGATCTNNNAGCAAAGAAAAAAACAGGGAG

AGCACACGTCTGAACTCTTTCCCTACACGACGCTCTTCCGATCTNNNAGCAAAGAAAAAAACAGGGAG

AGCACACGTCTGAACTCTTTCCCTACACGACGCTCTTCCGATCTNNNAGCAAAGAAAAAAACAGGGAG

AGCACACGTCTGAACTCTTTCCCTACACGACGCTCTTCCGATCTNNNAGCAAAGAAAAAAACAGGGAG

AGCACACGTCTGAACTCTTTCCCTACACGACGCTCTTCCGATCTNNNAGCAAAGAAAAAAACAGGGAG

AGCACACGTCTGAACTCTTTCCCTACACGACGCTCTTCCGATCTNNNAGCAAAGAAAAAAACAGGGAG

GAGCACACGTCTGAACTCTTTCCCTACACGACGCTCTTCCGATCTNNNTTCTTGGGCCACATTTTCTTTC

GAGCACACGTCTGAACTCTTTCCCTACACGACGCTCTTCCGATCTNNNTTCTTGGGCCACATTTTCTTTC

GAGCACACGTCTGAACTCTTTCCCTACACGACGCTCTTCCGATCTNNNTTCTTGGGCCACATTTTCTTTC

GAGCACACGTCTGAACTCTTTCCCTACACGACGCTCTTCCGATCTNNNTTCTTGGGCCACATTTTCTTTC

GAGCACACGTCTGAACTCTTTCCCTACACGACGCTCTTCCGATCTNNNTTCTTGGGCCACATTTTCTTTC

GAGCACACGTCTGAACTCTTTCCCTACACGACGCTCTTCCGATCTNNNTTCTTGGGCCACATTTTCTTTC

GAGCACACGTCTGAACTCTTTCCCTACACGACGCTCTTCCGATCTNNNTTCTTGGGCCACATTTTCTTTC

AGCACACGTCTGAACTCTTTCCCTACACGACGCTCTTCCGATCTNNNAGGAGAATGGCGTGAACCTGCA

AGCACACGTCTGAACTCTTTCCCTACACGACGCTCTTCCGATCTNNNAGGAGAATGGCGTGAACCTGCA

AGCACACGTCTGAACTCTTTCCCTACACGACGCTCTTCCGATCTNNNAGGAGAATGGCGTGAACCTGCA

AGCACACGTCTGAACTCTTTCCCTACACGACGCTCTTCCGATCTNNNAGGAGAATGGCGTGAACCTGCA

AGCACACGTCTGAACTCTTTCCCTACACGACGCTCTTCCGATCTNNNAGGAGAATGGCGTGAACCTGCA
AGCACACGTCTGAACTCTTTCCCTACACGACGCTCTTCCGATCTNNNAGGAGAATGGCGTGAACCTGCA
AGCACACGTCTGAACTCTTTCCCTACACGACGCTCTTCCGATCTNNNAGGAGAATGGCGTGAACCTGCA
AGCACACGTCTGAACTCTTTCCCTACACGACGCTCTTCCGATCTNNNGATCACACCACTGCACTCCACAT
AGCACACGTCTGAACTCTTTCCCTACACGACGCTCTTCCGATCTNNNGATCACACCACTGCACTCCACAT
AGCACACGTCTGAACTCTTTCCCTACACGACGCTCTTCCGATCTNNNGATCACACCACTGCACTCCACAT
AGCACACGTCTGAACTCTTTCCCTACACGACGCTCTTCCGATCTNNNGATCACACCACTGCACTCCACAT
AGCACACGTCTGAACTCTTTCCCTACACGACGCTCTTCCGATCTNNNGATCACACCACTGCACTCCACAT
AGCACACGTCTGAACTCTTTCCCTACACGACGCTCTTCCGATCTNNNGATCACACCACTGCACTCCACAT
AGCACACGTCTGAACTCTTTCCCTACACGACGCTCTTCCGATCTNNNGATCACACCACTGCACTCCACAT
GCACACGTCTGAACTCTTTCCCTACACGACGCTCTTCCGATCTNNNGCTATACTTTATGACACTAAGGAC
GCACACGTCTGAACTCTTTCCCTACACGACGCTCTTCCGATCTNNNGCTATACTTTATGACACTAAGGAC
GCACACGTCTGAACTCTTTCCCTACACGACGCTCTTCCGATCTNNNGCTATACTTTATGACACTAAGGAC
GCACACGTCTGAACTCTTTCCCTACACGACGCTCTTCCGATCTNNNGCTATACTTTATGACACTAAGGAC
GCACACGTCTGAACTCTTTCCCTACACGACGCTCTTCCGATCTNNNGCTATACTTTATGACACTAAGGAC
GCACACGTCTGAACTCTTTCCCTACACGACGCTCTTCCGATCTNNNGCTATACTTTATGACACTAAGGAC
GCACACGTCTGAACTCTTTCCCTACACGACGCTCTTCCGATCTNNNGCTATACTTTATGACACTAAGGAC
GCACACGTCTGAACTCTTTCCCTACACGACGCTCTTCCGATCTNNNAACTTGTGCTTCAATGCCTGTTTC
GCACACGTCTGAACTCTTTCCCTACACGACGCTCTTCCGATCTNNNAACTTGTGCTTCAATGCCTGTTTC
GCACACGTCTGAACTCTTTCCCTACACGACGCTCTTCCGATCTNNNAACTTGTGCTTCAATGCCTGTTTC
GCACACGTCTGAACTCTTTCCCTACACGACGCTCTTCCGATCTNNNAACTTGTGCTTCAATGCCTGTTTC
GCACACGTCTGAACTCTTTCCCTACACGACGCTCTTCCGATCTNNNAACTTGTGCTTCAATGCCTGTTTC
GCACACGTCTGAACTCTTTCCCTACACGACGCTCTTCCGATCTNNNAACTTGTGCTTCAATGCCTGTTTC
GCACACGTCTGAACTCTTTCCCTACACGACGCTCTTCCGATCTNNNAACTTGTGCTTCAATGCCTGTTTC
GAGCACACGTCTGAACTCTTTCCCTACACGACGCTCTTCCGATCTNNNAAATATGGAACCAGCCCAAAT
GAGCACACGTCTGAACTCTTTCCCTACACGACGCTCTTCCGATCTNNNAAATATGGAACCAGCCCAAAT
GAGCACACGTCTGAACTCTTTCCCTACACGACGCTCTTCCGATCTNNNAAATATGGAACCAGCCCAAAT
GAGCACACGTCTGAACTCTTTCCCTACACGACGCTCTTCCGATCTNNNAAATATGGAACCAGCCCAAAT
GAGCACACGTCTGAACTCTTTCCCTACACGACGCTCTTCCGATCTNNNAAATATGGAACCAGCCCAAAT
GAGCACACGTCTGAACTCTTTCCCTACACGACGCTCTTCCGATCTNNNAAATATGGAACCAGCCCAAAT
GAGCACACGTCTGAACTCTTTCCCTACACGACGCTCTTCCGATCTNNNAAATATGGAACCAGCCCAAAT
GCACACGTCTGAACTCTTTCCCTACACGACGCTCTTCCGATCTNNNTTGCGTGTTTTCTGTTGTTTCCTCA
GCACACGTCTGAACTCTTTCCCTACACGACGCTCTTCCGATCTNNNTTGCGTGTTTTCTGTTGTTTCCTCA
GCACACGTCTGAACTCTTTCCCTACACGACGCTCTTCCGATCTNNNTTGCGTGTTTTCTGTTGTTTCCTCA
GCACACGTCTGAACTCTTTCCCTACACGACGCTCTTCCGATCTNNNTTGCGTGTTTTCTGTTGTTTCCTCA
GCACACGTCTGAACTCTTTCCCTACACGACGCTCTTCCGATCTNNNTTGCGTGTTTTCTGTTGTTTCCTCA
GCACACGTCTGAACTCTTTCCCTACACGACGCTCTTCCGATCTNNNTTGCGTGTTTTCTGTTGTTTCCTCA
GCACACGTCTGAACTCTTTCCCTACACGACGCTCTTCCGATCTNNNTTGCGTGTTTTCTGTTGTTTCCTCA
ACACGTCTGAACTCTTTCCCTACACGACGCTCTTCCGATCTNNNTCGGTTTTGAAGGATAAGGGTGACA
ACACGTCTGAACTCTTTCCCTACACGACGCTCTTCCGATCTNNNTCGGTTTTGAAGGATAAGGGTGACA
ACACGTCTGAACTCTTTCCCTACACGACGCTCTTCCGATCTNNNTCGGTTTTGAAGGATAAGGGTGACA
ACACGTCTGAACTCTTTCCCTACACGACGCTCTTCCGATCTNNNTCGGTTTTGAAGGATAAGGGTGACA
ACACGTCTGAACTCTTTCCCTACACGACGCTCTTCCGATCTNNNTCGGTTTTGAAGGATAAGGGTGACA

ACACGTCTGAACTCTTTCCCTACACGACGCTCTTCCGATCTNNNTCGGTTTTGAAGGATAAGGGTGACA
ACACGTCTGAACTCTTTCCCTACACGACGCTCTTCCGATCTNNNTCGGTTTTGAAGGATAAGGGTGACA
GAGCACACGTCTGAACTCTTTCCCTACACGACGCTCTTCCGATCTNNNGAGACAGGTTTTCGCCATGTT
GAGCACACGTCTGAACTCTTTCCCTACACGACGCTCTTCCGATCTNNNGAGACAGGTTTTCGCCATGTT
GAGCACACGTCTGAACTCTTTCCCTACACGACGCTCTTCCGATCTNNNGAGACAGGTTTTCGCCATGTT
GAGCACACGTCTGAACTCTTTCCCTACACGACGCTCTTCCGATCTNNNGAGACAGGTTTTCGCCATGTT
GAGCACACGTCTGAACTCTTTCCCTACACGACGCTCTTCCGATCTNNNGAGACAGGTTTTCGCCATGTT
GAGCACACGTCTGAACTCTTTCCCTACACGACGCTCTTCCGATCTNNNGAGACAGGTTTTCGCCATGTT
GAGCACACGTCTGAACTCTTTCCCTACACGACGCTCTTCCGATCTNNNGAGACAGGTTTTCGCCATGTT
AGAGCACACGTCTGAACTCTTTCCCTACACGACGCTCTTCCGATCTNNNGCAAGAGGAAGAGAGAGCA
AGAGCACACGTCTGAACTCTTTCCCTACACGACGCTCTTCCGATCTNNNGCAAGAGGAAGAGAGAGCA
AGAGCACACGTCTGAACTCTTTCCCTACACGACGCTCTTCCGATCTNNNGCAAGAGGAAGAGAGAGCA
AGAGCACACGTCTGAACTCTTTCCCTACACGACGCTCTTCCGATCTNNNGCAAGAGGAAGAGAGAGCA
AGAGCACACGTCTGAACTCTTTCCCTACACGACGCTCTTCCGATCTNNNGCAAGAGGAAGAGAGAGCA
AGAGCACACGTCTGAACTCTTTCCCTACACGACGCTCTTCCGATCTNNNGCAAGAGGAAGAGAGAGCA
AGAGCACACGTCTGAACTCTTTCCCTACACGACGCTCTTCCGATCTNNNGCAAGAGGAAGAGAGAGCA
AGCACACGTCTGAACTCTTTCCCTACACGACGCTCTTCCGATCTNNNGAGCTCAGAGATACATGGATG
AGCACACGTCTGAACTCTTTCCCTACACGACGCTCTTCCGATCTNNNGAGCTCAGAGATACATGGATG
AGCACACGTCTGAACTCTTTCCCTACACGACGCTCTTCCGATCTNNNGAGCTCAGAGATACATGGATG
AGCACACGTCTGAACTCTTTCCCTACACGACGCTCTTCCGATCTNNNGAGCTCAGAGATACATGGATG
AGCACACGTCTGAACTCTTTCCCTACACGACGCTCTTCCGATCTNNNGAGCTCAGAGATACATGGATG
AGCACACGTCTGAACTCTTTCCCTACACGACGCTCTTCCGATCTNNNGAGCTCAGAGATACATGGATG
AGCACACGTCTGAACTCTTTCCCTACACGACGCTCTTCCGATCTNNNGAGCTCAGAGATACATGGATG
GCACACGTCTGAACTCTTTCCCTACACGACGCTCTTCCGATCTNNNCTCAACAATAGATGGAAATTTCA
GCACACGTCTGAACTCTTTCCCTACACGACGCTCTTCCGATCTNNNCTCAACAATAGATGGAAATTTCA
GCACACGTCTGAACTCTTTCCCTACACGACGCTCTTCCGATCTNNNCTCAACAATAGATGGAAATTTCA
GCACACGTCTGAACTCTTTCCCTACACGACGCTCTTCCGATCTNNNCTCAACAATAGATGGAAATTTCA
GCACACGTCTGAACTCTTTCCCTACACGACGCTCTTCCGATCTNNNCTCAACAATAGATGGAAATTTCA
GCACACGTCTGAACTCTTTCCCTACACGACGCTCTTCCGATCTNNNCTCAACAATAGATGGAAATTTCA
GCACACGTCTGAACTCTTTCCCTACACGACGCTCTTCCGATCTNNNCTCAACAATAGATGGAAATTTCA
GAGCACACGTCTGAACTCTTTCCCTACACGACGCTCTTCCGATCTNNNGAGATCACACCACTACACTCCA
GAGCACACGTCTGAACTCTTTCCCTACACGACGCTCTTCCGATCTNNNGAGATCACACCACTACACTCCA
GAGCACACGTCTGAACTCTTTCCCTACACGACGCTCTTCCGATCTNNNGAGATCACACCACTACACTCCA
GAGCACACGTCTGAACTCTTTCCCTACACGACGCTCTTCCGATCTNNNGAGATCACACCACTACACTCCA
GAGCACACGTCTGAACTCTTTCCCTACACGACGCTCTTCCGATCTNNNGAGATCACACCACTACACTCCA
GAGCACACGTCTGAACTCTTTCCCTACACGACGCTCTTCCGATCTNNNGAGATCACACCACTACACTCCA
GAGCACACGTCTGAACTCTTTCCCTACACGACGCTCTTCCGATCTNNNGAGATCACACCACTACACTCCA
GAGCACACGTCTGAACTCTTTCCCTACACGACGCTCTTCCGATCTNNNGCAGGAGTTGTACACACTTATC
GAGCACACGTCTGAACTCTTTCCCTACACGACGCTCTTCCGATCTNNNGCAGGAGTTGTACACACTTATC
GAGCACACGTCTGAACTCTTTCCCTACACGACGCTCTTCCGATCTNNNGCAGGAGTTGTACACACTTATC
GAGCACACGTCTGAACTCTTTCCCTACACGACGCTCTTCCGATCTNNNGCAGGAGTTGTACACACTTATC
GAGCACACGTCTGAACTCTTTCCCTACACGACGCTCTTCCGATCTNNNGCAGGAGTTGTACACACTTATC
GAGCACACGTCTGAACTCTTTCCCTACACGACGCTCTTCCGATCTNNNGCAGGAGTTGTACACACTTATC

AGCACACGTCTGAACTCTTTCCCTACACGACGCTCTTCCGATCTNNNGCAGGAGTTGTACACACTTAT
GAGCACACGTCTGAACTCTTTCCCTACACGACGCTCTTCCGATCTNNNTTTCTATCTCCTACTCCACCCCC
GAGCACACGTCTGAACTCTTTCCCTACACGACGCTCTTCCGATCTNNNTTTCTATCTCCTACTCCACCCCC
GAGCACACGTCTGAACTCTTTCCCTACACGACGCTCTTCCGATCTNNNTTTCTATCTCCTACTCCACCCCC
GAGCACACGTCTGAACTCTTTCCCTACACGACGCTCTTCCGATCTNNNTTTCTATCTCCTACTCCACCCCC
GAGCACACGTCTGAACTCTTTCCCTACACGACGCTCTTCCGATCTNNNTTTCTATCTCCTACTCCACCCCC
GAGCACACGTCTGAACTCTTTCCCTACACGACGCTCTTCCGATCTNNNTTTCTATCTCCTACTCCACCCCC
GAGCACACGTCTGAACTCTTTCCCTACACGACGCTCTTCCGATCTNNNTTTCTATCTCCTACTCCACCCCC
AGCACACGTCTGAACTCTTTCCCTACACGACGCTCTTCCGATCTNNNGTTTCTGAGGCTGTGAAATGT
AGCACACGTCTGAACTCTTTCCCTACACGACGCTCTTCCGATCTNNNGTTTCTGAGGCTGTGAAATGT
AGCACACGTCTGAACTCTTTCCCTACACGACGCTCTTCCGATCTNNNGTTTCTGAGGCTGTGAAATGT
AGCACACGTCTGAACTCTTTCCCTACACGACGCTCTTCCGATCTNNNGTTTCTGAGGCTGTGAAATGT
AGCACACGTCTGAACTCTTTCCCTACACGACGCTCTTCCGATCTNNNGTTTCTGAGGCTGTGAAATGT
AGCACACGTCTGAACTCTTTCCCTACACGACGCTCTTCCGATCTNNNGTTTCTGAGGCTGTGAAATGT
AGCACACGTCTGAACTCTTTCCCTACACGACGCTCTTCCGATCTNNNGTTTCTGAGGCTGTGAAATGT
AGCACACGTCTGAACTCTTTCCCTACACGACGCTCTTCCGATCTNNNCAGAGGTTACAGTGAGCCAA
AGCACACGTCTGAACTCTTTCCCTACACGACGCTCTTCCGATCTNNNCAGAGGTTACAGTGAGCCAA
AGCACACGTCTGAACTCTTTCCCTACACGACGCTCTTCCGATCTNNNCAGAGGTTACAGTGAGCCAA
AGCACACGTCTGAACTCTTTCCCTACACGACGCTCTTCCGATCTNNNCAGAGGTTACAGTGAGCCAA
AGCACACGTCTGAACTCTTTCCCTACACGACGCTCTTCCGATCTNNNCAGAGGTTACAGTGAGCCAA
AGCACACGTCTGAACTCTTTCCCTACACGACGCTCTTCCGATCTNNNCAGAGGTTACAGTGAGCCAA
AGCACACGTCTGAACTCTTTCCCTACACGACGCTCTTCCGATCTNNNCAGAGGTTACAGTGAGCCAA
GAGCACACGTCTGAACTCTTTCCCTACACGACGCTCTTCCGATCTNNNAAGGCATGAGAATTGCTTGA
GAGCACACGTCTGAACTCTTTCCCTACACGACGCTCTTCCGATCTNNNAAGGCATGAGAATTGCTTGA
GAGCACACGTCTGAACTCTTTCCCTACACGACGCTCTTCCGATCTNNNAAGGCATGAGAATTGCTTGA
GAGCACACGTCTGAACTCTTTCCCTACACGACGCTCTTCCGATCTNNNAAGGCATGAGAATTGCTTGA
GAGCACACGTCTGAACTCTTTCCCTACACGACGCTCTTCCGATCTNNNAAGGCATGAGAATTGCTTGA
GAGCACACGTCTGAACTCTTTCCCTACACGACGCTCTTCCGATCTNNNAAGGCATGAGAATTGCTTGA
GAGCACACGTCTGAACTCTTTCCCTACACGACGCTCTTCCGATCTNNNAAGGCATGAGAATTGCTTGA
AGAGCACACGTCTGAACTCTTTCCCTACACGACGCTCTTCCGATCTNNNAGCATTCTCCAGAGTTGTCC
AGAGCACACGTCTGAACTCTTTCCCTACACGACGCTCTTCCGATCTNNNAGCATTCTCCAGAGTTGTCC
AGAGCACACGTCTGAACTCTTTCCCTACACGACGCTCTTCCGATCTNNNAGCATTCTCCAGAGTTGTCC
AGAGCACACGTCTGAACTCTTTCCCTACACGACGCTCTTCCGATCTNNNAGCATTCTCCAGAGTTGTCC
AGAGCACACGTCTGAACTCTTTCCCTACACGACGCTCTTCCGATCTNNNAGCATTCTCCAGAGTTGTCC
AGAGCACACGTCTGAACTCTTTCCCTACACGACGCTCTTCCGATCTNNNAGCATTCTCCAGAGTTGTCC
AGAGCACACGTCTGAACTCTTTCCCTACACGACGCTCTTCCGATCTNNNAGCATTCTCCAGAGTTGTCC
GAGCACACGTCTGAACTCTTTCCCTACACGACGCTCTTCCGATCTNNNGCACACACCTGTAATCCCAGTT
GAGCACACGTCTGAACTCTTTCCCTACACGACGCTCTTCCGATCTNNNGCACACACCTGTAATCCCAGTT
GAGCACACGTCTGAACTCTTTCCCTACACGACGCTCTTCCGATCTNNNGCACACACCTGTAATCCCAGTT
GAGCACACGTCTGAACTCTTTCCCTACACGACGCTCTTCCGATCTNNNGCACACACCTGTAATCCCAGTT
GAGCACACGTCTGAACTCTTTCCCTACACGACGCTCTTCCGATCTNNNGCACACACCTGTAATCCCAGTT
GAGCACACGTCTGAACTCTTTCCCTACACGACGCTCTTCCGATCTNNNGCACACACCTGTAATCCCAGTT
GAGCACACGTCTGAACTCTTTCCCTACACGACGCTCTTCCGATCTNNNGCACACACCTGTAATCCCAGTT

AGCACACGTCTGAACTCTTTCCCTACACGACGCTCTTCCGATCTNNNAAAAATTATCCAGGCGTGGTT
AGCACACGTCTGAACTCTTTCCCTACACGACGCTCTTCCGATCTNNNAAAAATTATCCAGGCGTGGTT
AGCACACGTCTGAACTCTTTCCCTACACGACGCTCTTCCGATCTNNNAAAAATTATCCAGGCGTGGTT
AGCACACGTCTGAACTCTTTCCCTACACGACGCTCTTCCGATCTNNNAAAAATTATCCAGGCGTGGTT
AGCACACGTCTGAACTCTTTCCCTACACGACGCTCTTCCGATCTNNNAAAAATTATCCAGGCGTGGTT
AGCACACGTCTGAACTCTTTCCCTACACGACGCTCTTCCGATCTNNNAAAAATTATCCAGGCGTGGTT
AGCACACGTCTGAACTCTTTCCCTACACGACGCTCTTCCGATCTNNNAAAAATTATCCAGGCGTGGTT
AGCACACGTCTGAACTCTTTCCCTACACGACGCTCTTCCGATCTNNNCAGCATTACTCTGTGTGTGTG
AGCACACGTCTGAACTCTTTCCCTACACGACGCTCTTCCGATCTNNNCAGCATTACTCTGTGTGTGTG
AGCACACGTCTGAACTCTTTCCCTACACGACGCTCTTCCGATCTNNNCAGCATTACTCTGTGTGTGTG
AGCACACGTCTGAACTCTTTCCCTACACGACGCTCTTCCGATCTNNNCAGCATTACTCTGTGTGTGTG
AGCACACGTCTGAACTCTTTCCCTACACGACGCTCTTCCGATCTNNNCAGCATTACTCTGTGTGTGTG
AGCACACGTCTGAACTCTTTCCCTACACGACGCTCTTCCGATCTNNNCAGCATTACTCTGTGTGTGTG
AGCACACGTCTGAACTCTTTCCCTACACGACGCTCTTCCGATCTNNNCAGCATTACTCTGTGTGTGTG
GAGCACACGTCTGAACTCTTTCCCTACACGACGCTCTTCCGATCTNNNGCACTCCAATCTGAATGCCAG
GAGCACACGTCTGAACTCTTTCCCTACACGACGCTCTTCCGATCTNNNGCACTCCAATCTGAATGCCAG
GAGCACACGTCTGAACTCTTTCCCTACACGACGCTCTTCCGATCTNNNGCACTCCAATCTGAATGCCAG
GAGCACACGTCTGAACTCTTTCCCTACACGACGCTCTTCCGATCTNNNGCACTCCAATCTGAATGCCAG
GAGCACACGTCTGAACTCTTTCCCTACACGACGCTCTTCCGATCTNNNGCACTCCAATCTGAATGCCAG
GAGCACACGTCTGAACTCTTTCCCTACACGACGCTCTTCCGATCTNNNGCACTCCAATCTGAATGCCAG
GAGCACACGTCTGAACTCTTTCCCTACACGACGCTCTTCCGATCTNNNGCACTCCAATCTGAATGCCAG
GCACACGTCTGAACTCTTTCCCTACACGACGCTCTTCCGATCTNNNACACGGTGAAATCCCATCTCTA
GCACACGTCTGAACTCTTTCCCTACACGACGCTCTTCCGATCTNNNACACGGTGAAATCCCATCTCTA
GCACACGTCTGAACTCTTTCCCTACACGACGCTCTTCCGATCTNNNACACGGTGAAATCCCATCTCTA
GCACACGTCTGAACTCTTTCCCTACACGACGCTCTTCCGATCTNNNACACGGTGAAATCCCATCTCTA
GCACACGTCTGAACTCTTTCCCTACACGACGCTCTTCCGATCTNNNACACGGTGAAATCCCATCTCTA
GCACACGTCTGAACTCTTTCCCTACACGACGCTCTTCCGATCTNNNACACGGTGAAATCCCATCTCTA
GCACACGTCTGAACTCTTTCCCTACACGACGCTCTTCCGATCTNNNACACGGTGAAATCCCATCTCTA
AGCACACGTCTGAACTCTTTCCCTACACGACGCTCTTCCGATCTNNNGGCTCACCCAAAGACGTTATTT
AGCACACGTCTGAACTCTTTCCCTACACGACGCTCTTCCGATCTNNNGGCTCACCCAAAGACGTTATTT
AGCACACGTCTGAACTCTTTCCCTACACGACGCTCTTCCGATCTNNNGGCTCACCCAAAGACGTTATTT
AGCACACGTCTGAACTCTTTCCCTACACGACGCTCTTCCGATCTNNNGGCTCACCCAAAGACGTTATTT
AGCACACGTCTGAACTCTTTCCCTACACGACGCTCTTCCGATCTNNNGGCTCACCCAAAGACGTTATTT
AGCACACGTCTGAACTCTTTCCCTACACGACGCTCTTCCGATCTNNNGGCTCACCCAAAGACGTTATTT
AGCACACGTCTGAACTCTTTCCCTACACGACGCTCTTCCGATCTNNNGGCTCACCCAAAGACGTTATTT
AGCACACGTCTGAACTCTTTCCCTACACGACGCTCTTCCGATCTNNNCTGCAAAGGACATGAACTCAT
AGCACACGTCTGAACTCTTTCCCTACACGACGCTCTTCCGATCTNNNCTGCAAAGGACATGAACTCAT
AGCACACGTCTGAACTCTTTCCCTACACGACGCTCTTCCGATCTNNNCTGCAAAGGACATGAACTCAT
AGCACACGTCTGAACTCTTTCCCTACACGACGCTCTTCCGATCTNNNCTGCAAAGGACATGAACTCAT
AGCACACGTCTGAACTCTTTCCCTACACGACGCTCTTCCGATCTNNNCTGCAAAGGACATGAACTCAT
AGCACACGTCTGAACTCTTTCCCTACACGACGCTCTTCCGATCTNNNCTGCAAAGGACATGAACTCAT
AGCACACGTCTGAACTCTTTCCCTACACGACGCTCTTCCGATCTNNNCTGCAAAGGACATGAACTCAT
AGCACACGTCTGAACTCTTTCCCTACACGACGCTCTTCCGATCTNNNCTGTCTATGGCGTTGTTGTTC

AGCACACGTCTGAACTCTTTCCCTACACGACGCTCTTCCGATCTNNNCTGTCTATGGCGTTGTTGTTCT
AGCACACGTCTGAACTCTTTCCCTACACGACGCTCTTCCGATCTNNNCTGTCTATGGCGTTGTTGTTCT
AGCACACGTCTGAACTCTTTCCCTACACGACGCTCTTCCGATCTNNNGCCACCTGTATTACCAGCTACTT
AGCACACGTCTGAACTCTTTCCCTACACGACGCTCTTCCGATCTNNNGCCACCTGTATTACCAGCTACTT
AGCACACGTCTGAACTCTTTCCCTACACGACGCTCTTCCGATCTNNNGCCACCTGTATTACCAGCTACTT
GCACACGTCTGAACTCTTTCCCTACACGACGCTCTTCCGATCTNNNTGGACAGATATGTGTGTGCCTC
GCACACGTCTGAACTCTTTCCCTACACGACGCTCTTCCGATCTNNNTGGACAGATATGTGTGTGCCTC
GCACACGTCTGAACTCTTTCCCTACACGACGCTCTTCCGATCTNNNTGGACAGATATGTGTGTGCCTC
GAGCACACGTCTGAACTCTTTCCCTACACGACGCTCTTCCGATCTNNNGAGTGACACCTCCACAAGAC
GAGCACACGTCTGAACTCTTTCCCTACACGACGCTCTTCCGATCTNNNGAGTGACACCTCCACAAGAC
GAGCACACGTCTGAACTCTTTCCCTACACGACGCTCTTCCGATCTNNNGAGTGACACCTCCACAAGAC
AGCACACGTCTGAACTCTTTCCCTACACGACGCTCTTCCGATCTNNNATAGGCCCCATGCAAGTCTAA
AGCACACGTCTGAACTCTTTCCCTACACGACGCTCTTCCGATCTNNNATAGGCCCCATGCAAGTCTAA
AGCACACGTCTGAACTCTTTCCCTACACGACGCTCTTCCGATCTNNNATAGGCCCCATGCAAGTCTAA
GAGCACACGTCTGAACTCTTTCCCTACACGACGCTCTTCCGATCTNNNCTAGGACTGGAGGATTGTG
GAGCACACGTCTGAACTCTTTCCCTACACGACGCTCTTCCGATCTNNNCTAGGACTGGAGGATTGTG
GAGCACACGTCTGAACTCTTTCCCTACACGACGCTCTTCCGATCTNNNCTAGGACTGGAGGATTGTG
AGCACACGTCTGAACTCTTTCCCTACACGACGCTCTTCCGATCTNNNACTGCAAGTGGTAAACGTGAG
AGCACACGTCTGAACTCTTTCCCTACACGACGCTCTTCCGATCTNNNACTGCAAGTGGTAAACGTGAG
AGCACACGTCTGAACTCTTTCCCTACACGACGCTCTTCCGATCTNNNACTGCAAGTGGTAAACGTGAG
GAGCACACGTCTGAACTCTTTCCCTACACGACGCTCTTCCGATCTNNNGTAGGAAAAGCAATGACAG
GAGCACACGTCTGAACTCTTTCCCTACACGACGCTCTTCCGATCTNNNGTAGGAAAAGCAATGACAG
GAGCACACGTCTGAACTCTTTCCCTACACGACGCTCTTCCGATCTNNNGTAGGAAAAGCAATGACAG
GCACACGTCTGAACTCTTTCCCTACACGACGCTCTTCCGATCTNNNATTCTCTAAAATAACCAAAGCAG
GCACACGTCTGAACTCTTTCCCTACACGACGCTCTTCCGATCTNNNATTCTCTAAAATAACCAAAGCAG
GCACACGTCTGAACTCTTTCCCTACACGACGCTCTTCCGATCTNNNATTCTCTAAAATAACCAAAGCAG
AGCACACGTCTGAACTCTTTCCCTACACGACGCTCTTCCGATCTNNNGGTGAAGCCCCATCTCTACTA
AGCACACGTCTGAACTCTTTCCCTACACGACGCTCTTCCGATCTNNNGGTGAAGCCCCATCTCTACTA
AGCACACGTCTGAACTCTTTCCCTACACGACGCTCTTCCGATCTNNNGGTGAAGCCCCATCTCTACTA
AGAGCACACGTCTGAACTCTTTCCCTACACGACGCTCTTCCGATCTNNNATACCTGGCATTTCTTAGCT
AGAGCACACGTCTGAACTCTTTCCCTACACGACGCTCTTCCGATCTNNNATACCTGGCATTTCTTAGCT
AGAGCACACGTCTGAACTCTTTCCCTACACGACGCTCTTCCGATCTNNNATACCTGGCATTTCTTAGCT
CACACGTCTGAACTCTTTCCCTACACGACGCTCTTCCGATCTNNNATGGGCATTTAGGGTGATACCAT
CACACGTCTGAACTCTTTCCCTACACGACGCTCTTCCGATCTNNNATGGGCATTTAGGGTGATACCAT
CACACGTCTGAACTCTTTCCCTACACGACGCTCTTCCGATCTNNNATGGGCATTTAGGGTGATACCAT
AGCACACGTCTGAACTCTTTCCCTACACGACGCTCTTCCGATCTNNNTCCTAGAGTACCCAGATGCAG
AGCACACGTCTGAACTCTTTCCCTACACGACGCTCTTCCGATCTNNNTCCTAGAGTACCCAGATGCAG
AGCACACGTCTGAACTCTTTCCCTACACGACGCTCTTCCGATCTNNNTCCTAGAGTACCCAGATGCAG
AGCACACGTCTGAACTCTTTCCCTACACGACGCTCTTCCGATCTNNNGAGGTTGCAATGAGCCAAGA
AGCACACGTCTGAACTCTTTCCCTACACGACGCTCTTCCGATCTNNNGAGGTTGCAATGAGCCAAGA
AGCACACGTCTGAACTCTTTCCCTACACGACGCTCTTCCGATCTNNNGAGGTTGCAATGAGCCAAGA
ACACGTCTGAACTCTTTCCCTACACGACGCTCTTCCGATCTNNNACAGCATGAAAATCTGGCAAAGTC
ACACGTCTGAACTCTTTCCCTACACGACGCTCTTCCGATCTNNNACAGCATGAAAATCTGGCAAAGTC

ACACGTCTGAACTCTTTCCCTACACGACGCTCTTCCGATCTNNNACAGCATGAAAATCTGGCAAAGTC
CACACGTCTGAACTCTTTCCCTACACGACGCTCTTCCGATCTNNNAATGGGAAACTGATTTTGGTGG
CACACGTCTGAACTCTTTCCCTACACGACGCTCTTCCGATCTNNNAATGGGAAACTGATTTTGGTGG
CACACGTCTGAACTCTTTCCCTACACGACGCTCTTCCGATCTNNNAATGGGAAACTGATTTTGGTGG
AGCACACGTCTGAACTCTTTCCCTACACGACGCTCTTCCGATCTNNNGAGATTCTCATGCCTCAGACT
AGCACACGTCTGAACTCTTTCCCTACACGACGCTCTTCCGATCTNNNGAGATTCTCATGCCTCAGACT
AGCACACGTCTGAACTCTTTCCCTACACGACGCTCTTCCGATCTNNNGAGATTCTCATGCCTCAGACT
AGCACACGTCTGAACTCTTTCCCTACACGACGCTCTTCCGATCTNNNACAAAAATTAGCTGGGCATGG
AGCACACGTCTGAACTCTTTCCCTACACGACGCTCTTCCGATCTNNNACAAAAATTAGCTGGGCATGG
AGCACACGTCTGAACTCTTTCCCTACACGACGCTCTTCCGATCTNNNACAAAAATTAGCTGGGCATGG
AGCACACGTCTGAACTCTTTCCCTACACGACGCTCTTCCGATCTNNNCTTAGAGGACGGGTCAATAG
AGCACACGTCTGAACTCTTTCCCTACACGACGCTCTTCCGATCTNNNCTTAGAGGACGGGTCAATAG
AGCACACGTCTGAACTCTTTCCCTACACGACGCTCTTCCGATCTNNNCTTAGAGGACGGGTCAATAG
CACACGTCTGAACTCTTTCCCTACACGACGCTCTTCCGATCTNNNTCAGGACACAATTGCCCCACCATC
CACACGTCTGAACTCTTTCCCTACACGACGCTCTTCCGATCTNNNTCAGGACACAATTGCCCCACCATC
CACACGTCTGAACTCTTTCCCTACACGACGCTCTTCCGATCTNNNTCAGGACACAATTGCCCCACCATC
AGCACACGTCTGAACTCTTTCCCTACACGACGCTCTTCCGATCTNNNTGACCACTATGACACACTCAAG
AGCACACGTCTGAACTCTTTCCCTACACGACGCTCTTCCGATCTNNNTGACCACTATGACACACTCAAG
AGCACACGTCTGAACTCTTTCCCTACACGACGCTCTTCCGATCTNNNTGACCACTATGACACACTCAAG
GCACACGTCTGAACTCTTTCCCTACACGACGCTCTTCCGATCTNNNAGACATGGAAAGGGAGTCATCC
GCACACGTCTGAACTCTTTCCCTACACGACGCTCTTCCGATCTNNNAGACATGGAAAGGGAGTCATCC
GCACACGTCTGAACTCTTTCCCTACACGACGCTCTTCCGATCTNNNAGACATGGAAAGGGAGTCATCC
AGCACACGTCTGAACTCTTTCCCTACACGACGCTCTTCCGATCTNNNTGTGTAGAGGTGTGAGTGTT
AGCACACGTCTGAACTCTTTCCCTACACGACGCTCTTCCGATCTNNNTGTGTAGAGGTGTGAGTGTT
AGCACACGTCTGAACTCTTTCCCTACACGACGCTCTTCCGATCTNNNTGTGTAGAGGTGTGAGTGTT
GAGCACACGTCTGAACTCTTTCCCTACACGACGCTCTTCCGATCTNNNCTCCTGAGTAGCTGGGATTA
GAGCACACGTCTGAACTCTTTCCCTACACGACGCTCTTCCGATCTNNNCTCCTGAGTAGCTGGGATTA
GAGCACACGTCTGAACTCTTTCCCTACACGACGCTCTTCCGATCTNNNCTCCTGAGTAGCTGGGATTA
GAGCACACGTCTGAACTCTTTCCCTACACGACGCTCTTCCGATCTNNNCCACCAGTACAGCTATTCCTC
GAGCACACGTCTGAACTCTTTCCCTACACGACGCTCTTCCGATCTNNNCCACCAGTACAGCTATTCCTC
GAGCACACGTCTGAACTCTTTCCCTACACGACGCTCTTCCGATCTNNNCCACCAGTACAGCTATTCCTC
GAGCACACGTCTGAACTCTTTCCCTACACGACGCTCTTCCGATCTNNNGAATTTTACGACCAGAATTAT
GAGCACACGTCTGAACTCTTTCCCTACACGACGCTCTTCCGATCTNNNGAATTTTACGACCAGAATTAT
GAGCACACGTCTGAACTCTTTCCCTACACGACGCTCTTCCGATCTNNNGAATTTTACGACCAGAATTAT
GAGCACACGTCTGAACTCTTTCCCTACACGACGCTCTTCCGATCTNNNGCAGAGGGAAATTGAGGA
GAGCACACGTCTGAACTCTTTCCCTACACGACGCTCTTCCGATCTNNNGCAGAGGGAAATTGAGGA
GAGCACACGTCTGAACTCTTTCCCTACACGACGCTCTTCCGATCTNNNGCAGAGGGAAATTGAGGA
AGCACACGTCTGAACTCTTTCCCTACACGACGCTCTTCCGATCTNNNTGACACCTGTAGAAGCAGAACT
AGCACACGTCTGAACTCTTTCCCTACACGACGCTCTTCCGATCTNNNTGACACCTGTAGAAGCAGAACT
AGCACACGTCTGAACTCTTTCCCTACACGACGCTCTTCCGATCTNNNTGACACCTGTAGAAGCAGAACT
GAGCACACGTCTGAACTCTTTCCCTACACGACGCTCTTCCGATCTNNNTACCAGTATGTGTTCAACCA
GAGCACACGTCTGAACTCTTTCCCTACACGACGCTCTTCCGATCTNNNTACCAGTATGTGTTCAACCA
GAGCACACGTCTGAACTCTTTCCCTACACGACGCTCTTCCGATCTNNNTACCAGTATGTGTTCAACCA

GAGCACACGTCTGAACTCTTTCCCTACACGACGCTCTTCCGATCTNNNCACCTGTAATCCCAGCTACTT
GAGCACACGTCTGAACTCTTTCCCTACACGACGCTCTTCCGATCTNNNCACCTGTAATCCCAGCTACTT
GAGCACACGTCTGAACTCTTTCCCTACACGACGCTCTTCCGATCTNNNCACCTGTAATCCCAGCTACTT
AGCACACGTCTGAACTCTTTCCCTACACGACGCTCTTCCGATCTNNNAGGAGAATCGCTTGAACCTGG
AGCACACGTCTGAACTCTTTCCCTACACGACGCTCTTCCGATCTNNNAGGAGAATCGCTTGAACCTGG
AGCACACGTCTGAACTCTTTCCCTACACGACGCTCTTCCGATCTNNNAGGAGAATCGCTTGAACCTGG
AGCACACGTCTGAACTCTTTCCCTACACGACGCTCTTCCGATCTNNNTGAGGCAGGAGAATCTCTTGA
AGCACACGTCTGAACTCTTTCCCTACACGACGCTCTTCCGATCTNNNTGAGGCAGGAGAATCTCTTGA
AGCACACGTCTGAACTCTTTCCCTACACGACGCTCTTCCGATCTNNNTGAGGCAGGAGAATCTCTTGA
GCACACGTCTGAACTCTTTCCCTACACGACGCTCTTCCGATCTNNNAAACCACACACACAAAGAGCAA
GCACACGTCTGAACTCTTTCCCTACACGACGCTCTTCCGATCTNNNAAACCACACACACAAAGAGCAA
GCACACGTCTGAACTCTTTCCCTACACGACGCTCTTCCGATCTNNNAAACCACACACACAAAGAGCAA
AGCACACGTCTGAACTCTTTCCCTACACGACGCTCTTCCGATCTNNNTCAGACAAGCTTAGCGTTCCAA
AGCACACGTCTGAACTCTTTCCCTACACGACGCTCTTCCGATCTNNNTCAGACAAGCTTAGCGTTCCAA
AGCACACGTCTGAACTCTTTCCCTACACGACGCTCTTCCGATCTNNNTCAGACAAGCTTAGCGTTCCAA
GCACACGTCTGAACTCTTTCCCTACACGACGCTCTTCCGATCTNNNTCCACTTGTTGTAAAGAATTGA
GCACACGTCTGAACTCTTTCCCTACACGACGCTCTTCCGATCTNNNTCCACTTGTTGTAAAGAATTGA
GCACACGTCTGAACTCTTTCCCTACACGACGCTCTTCCGATCTNNNTCCACTTGTTGTAAAGAATTGA
AGCACACGTCTGAACTCTTTCCCTACACGACGCTCTTCCGATCTNNNAGAGCCTCTTTTTCTGAGCTTA
AGCACACGTCTGAACTCTTTCCCTACACGACGCTCTTCCGATCTNNNAGAGCCTCTTTTTCTGAGCTTA
AGCACACGTCTGAACTCTTTCCCTACACGACGCTCTTCCGATCTNNNAGAGCCTCTTTTTCTGAGCTTA
GCACACGTCTGAACTCTTTCCCTACACGACGCTCTTCCGATCTNNNTCCAGGATTTCAACATTCCAGGA
GCACACGTCTGAACTCTTTCCCTACACGACGCTCTTCCGATCTNNNTCCAGGATTTCAACATTCCAGGA
GCACACGTCTGAACTCTTTCCCTACACGACGCTCTTCCGATCTNNNTCCAGGATTTCAACATTCCAGGA
AGCACACGTCTGAACTCTTTCCCTACACGACGCTCTTCCGATCTNNNCAAACCATAGCCTGCCTTTTCA
AGCACACGTCTGAACTCTTTCCCTACACGACGCTCTTCCGATCTNNNCAAACCATAGCCTGCCTTTTCA
AGCACACGTCTGAACTCTTTCCCTACACGACGCTCTTCCGATCTNNNCAAACCATAGCCTGCCTTTTCA
AGCACACGTCTGAACTCTTTCCCTACACGACGCTCTTCCGATCTNNNAGGGGCAGTAATCCAGATTT
AGCACACGTCTGAACTCTTTCCCTACACGACGCTCTTCCGATCTNNNAGGGGCAGTAATCCAGATTT
AGCACACGTCTGAACTCTTTCCCTACACGACGCTCTTCCGATCTNNNAGGGGCAGTAATCCAGATTT
GCACACGTCTGAACTCTTTCCCTACACGACGCTCTTCCGATCTNNNGAGAGAGAGAGAGAGAGAGG
GCACACGTCTGAACTCTTTCCCTACACGACGCTCTTCCGATCTNNNGAGAGAGAGAGAGAGAGAGG
GCACACGTCTGAACTCTTTCCCTACACGACGCTCTTCCGATCTNNNGAGAGAGAGAGAGAGAGAGG
AGCACACGTCTGAACTCTTTCCCTACACGACGCTCTTCCGATCTNNNAGGAATGAGGAGGATGTTTG
AGCACACGTCTGAACTCTTTCCCTACACGACGCTCTTCCGATCTNNNAGGAATGAGGAGGATGTTTG
AGCACACGTCTGAACTCTTTCCCTACACGACGCTCTTCCGATCTNNNAGGAATGAGGAGGATGTTTG
AGCACACGTCTGAACTCTTTCCCTACACGACGCTCTTCCGATCTNNNGACTCACCTTGGGAGAGAACT
AGCACACGTCTGAACTCTTTCCCTACACGACGCTCTTCCGATCTNNNGACTCACCTTGGGAGAGAACT
AGCACACGTCTGAACTCTTTCCCTACACGACGCTCTTCCGATCTNNNGACTCACCTTGGGAGAGAACT
AGCACACGTCTGAACTCTTTCCCTACACGACGCTCTTCCGATCTNNNCTTGAGCACAAGCAGAGCA
AGCACACGTCTGAACTCTTTCCCTACACGACGCTCTTCCGATCTNNNCTTGAGCACAAGCAGAGCA
AGCACACGTCTGAACTCTTTCCCTACACGACGCTCTTCCGATCTNNNCTTGAGCACAAGCAGAGCA
AGCACACGTCTGAACTCTTTCCCTACACGACGCTCTTCCGATCTNNNAGCTTCACCCAGGAGAGCTAA

AGCACACGTCTGAACTCTTTCCCTACACGACGCTCTTCCGATCTNNNAGCTTCACCCAGGAGAGCTAA
AGCACACGTCTGAACTCTTTCCCTACACGACGCTCTTCCGATCTNNNAGCTTCACCCAGGAGAGCTAA
CACACGTCTGAACTCTTTCCCTACACGACGCTCTTCCGATCTNNNCCTGGGCTAGCAAAACATTGTAG
CACACGTCTGAACTCTTTCCCTACACGACGCTCTTCCGATCTNNNCCTGGGCTAGCAAAACATTGTAG
CACACGTCTGAACTCTTTCCCTACACGACGCTCTTCCGATCTNNNCCTGGGCTAGCAAAACATTGTAG
GAGCACACGTCTGAACTCTTTCCCTACACGACGCTCTTCCGATCTNNNGATGTTACAGTTCATGCTCT
GAGCACACGTCTGAACTCTTTCCCTACACGACGCTCTTCCGATCTNNNGATGTTACAGTTCATGCTCT
GAGCACACGTCTGAACTCTTTCCCTACACGACGCTCTTCCGATCTNNNGATGTTACAGTTCATGCTCT
AGCACACGTCTGAACTCTTTCCCTACACGACGCTCTTCCGATCTNNNCAAGTGACAGAGTGAGAACCT
AGCACACGTCTGAACTCTTTCCCTACACGACGCTCTTCCGATCTNNNCAAGTGACAGAGTGAGAACCT
AGCACACGTCTGAACTCTTTCCCTACACGACGCTCTTCCGATCTNNNCAAGTGACAGAGTGAGAACCT
AGCACACGTCTGAACTCTTTCCCTACACGACGCTCTTCCGATCTNNNAAAGGGGCAGATAACACCCAA
AGCACACGTCTGAACTCTTTCCCTACACGACGCTCTTCCGATCTNNNAAAGGGGCAGATAACACCCAA
AGCACACGTCTGAACTCTTTCCCTACACGACGCTCTTCCGATCTNNNAAAGGGGCAGATAACACCCAA
ACACGTCTGAACTCTTTCCCTACACGACGCTCTTCCGATCTNNNTGTCCATGGGGTTCATGTAGTCTC
ACACGTCTGAACTCTTTCCCTACACGACGCTCTTCCGATCTNNNTGTCCATGGGGTTCATGTAGTCTC
ACACGTCTGAACTCTTTCCCTACACGACGCTCTTCCGATCTNNNTGTCCATGGGGTTCATGTAGTCTC
AGCACACGTCTGAACTCTTTCCCTACACGACGCTCTTCCGATCTNNNTGTTCATAGACAGTGCCTCCTT
AGCACACGTCTGAACTCTTTCCCTACACGACGCTCTTCCGATCTNNNTGTTCATAGACAGTGCCTCCTT
AGCACACGTCTGAACTCTTTCCCTACACGACGCTCTTCCGATCTNNNTGTTCATAGACAGTGCCTCCTT
GCACACGTCTGAACTCTTTCCCTACACGACGCTCTTCCGATCTNNNGGTCTCTTGCTTCTGTCCTACAA
GCACACGTCTGAACTCTTTCCCTACACGACGCTCTTCCGATCTNNNGGTCTCTTGCTTCTGTCCTACAA
GCACACGTCTGAACTCTTTCCCTACACGACGCTCTTCCGATCTNNNGGTCTCTTGCTTCTGTCCTACAA
AGCACACGTCTGAACTCTTTCCCTACACGACGCTCTTCCGATCTNNNTCTCCCTCTACGAACTGCTTTTC
AGCACACGTCTGAACTCTTTCCCTACACGACGCTCTTCCGATCTNNNTCTCCCTCTACGAACTGCTTTTC
AGCACACGTCTGAACTCTTTCCCTACACGACGCTCTTCCGATCTNNNTCTCCCTCTACGAACTGCTTTTC
AGCACACGTCTGAACTCTTTCCCTACACGACGCTCTTCCGATCTNNNTTTGTAGTCAGCCTTCAGGTCT
AGCACACGTCTGAACTCTTTCCCTACACGACGCTCTTCCGATCTNNNTTTGTAGTCAGCCTTCAGGTCT
AGCACACGTCTGAACTCTTTCCCTACACGACGCTCTTCCGATCTNNNTTTGTAGTCAGCCTTCAGGTCT
GCACACGTCTGAACTCTTTCCCTACACGACGCTCTTCCGATCTNNNAAAGCTCTCCAATGGCCTGATTC
GCACACGTCTGAACTCTTTCCCTACACGACGCTCTTCCGATCTNNNAAAGCTCTCCAATGGCCTGATTC
GCACACGTCTGAACTCTTTCCCTACACGACGCTCTTCCGATCTNNNAAAGCTCTCCAATGGCCTGATTC
GAGCACACGTCTGAACTCTTTCCCTACACGACGCTCTTCCGATCTNNNTGTCTCCCAGGAAGAATTGA
GAGCACACGTCTGAACTCTTTCCCTACACGACGCTCTTCCGATCTNNNTGTCTCCCAGGAAGAATTGA
GAGCACACGTCTGAACTCTTTCCCTACACGACGCTCTTCCGATCTNNNTGTCTCCCAGGAAGAATTGA
AGCACACGTCTGAACTCTTTCCCTACACGACGCTCTTCCGATCTNNNAGAAGTTGCAGTGAGTCGAG
AGCACACGTCTGAACTCTTTCCCTACACGACGCTCTTCCGATCTNNNAGAAGTTGCAGTGAGTCGAG
AGCACACGTCTGAACTCTTTCCCTACACGACGCTCTTCCGATCTNNNAGAAGTTGCAGTGAGTCGAG
GCACACGTCTGAACTCTTTCCCTACACGACGCTCTTCCGATCTNNNGTAATCCTAGCTACTTGGGAGG
GCACACGTCTGAACTCTTTCCCTACACGACGCTCTTCCGATCTNNNGTAATCCTAGCTACTTGGGAGG
GCACACGTCTGAACTCTTTCCCTACACGACGCTCTTCCGATCTNNNGTAATCCTAGCTACTTGGGAGG
AGCACACGTCTGAACTCTTTCCCTACACGACGCTCTTCCGATCTNNNATACAAAAAAATTAGCCGGGC
AGCACACGTCTGAACTCTTTCCCTACACGACGCTCTTCCGATCTNNNATACAAAAAAATTAGCCGGGC

AGCACACGTCTGAACTCTTTCCCTACACGACGCTCTTCCGATCTNNNATACAAAAAAATTAGCCGGGC
AGCACACGTCTGAACTCTTTCCCTACACGACGCTCTTCCGATCTNNNTAGAGTCCTAAATCTCGAACAA
AGCACACGTCTGAACTCTTTCCCTACACGACGCTCTTCCGATCTNNNTAGAGTCCTAAATCTCGAACAA
AGCACACGTCTGAACTCTTTCCCTACACGACGCTCTTCCGATCTNNNTAGAGTCCTAAATCTCGAACAA
GCACACGTCTGAACTCTTTCCCTACACGACGCTCTTCCGATCTNNNCTGAGGTGGGAGTATCAGCTAAG
GCACACGTCTGAACTCTTTCCCTACACGACGCTCTTCCGATCTNNNCTGAGGTGGGAGTATCAGCTAAG
GCACACGTCTGAACTCTTTCCCTACACGACGCTCTTCCGATCTNNNCTGAGGTGGGAGTATCAGCTAAG
GCACACGTCTGAACTCTTTCCCTACACGACGCTCTTCCGATCTNNNCTCTCAGCCTCCTGAAAAGAGAG
GCACACGTCTGAACTCTTTCCCTACACGACGCTCTTCCGATCTNNNCTCTCAGCCTCCTGAAAAGAGAG
GCACACGTCTGAACTCTTTCCCTACACGACGCTCTTCCGATCTNNNCTCTCAGCCTCCTGAAAAGAGAG
GAGCACACGTCTGAACTCTTTCCCTACACGACGCTCTTCCGATCTNNNGTCAGTTGTTTGCTGATTTTAT
GAGCACACGTCTGAACTCTTTCCCTACACGACGCTCTTCCGATCTNNNGTCAGTTGTTTGCTGATTTTAT
GAGCACACGTCTGAACTCTTTCCCTACACGACGCTCTTCCGATCTNNNGTCAGTTGTTTGCTGATTTTAT
GAGCACACGTCTGAACTCTTTCCCTACACGACGCTCTTCCGATCTNNNGGTGCCTGTAATCCCAACTACT
GAGCACACGTCTGAACTCTTTCCCTACACGACGCTCTTCCGATCTNNNGGTGCCTGTAATCCCAACTACT
GAGCACACGTCTGAACTCTTTCCCTACACGACGCTCTTCCGATCTNNNGGTGCCTGTAATCCCAACTACT
AGAGCACACGTCTGAACTCTTTCCCTACACGACGCTCTTCCGATCTNNNAAAAATTAGCTGGATGTGG
AGAGCACACGTCTGAACTCTTTCCCTACACGACGCTCTTCCGATCTNNNAAAAATTAGCTGGATGTGG
AGAGCACACGTCTGAACTCTTTCCCTACACGACGCTCTTCCGATCTNNNAAAAATTAGCTGGATGTGG
GAGCACACGTCTGAACTCTTTCCCTACACGACGCTCTTCCGATCTNNNAGGAGAATCGCTTGAACTCAT
GAGCACACGTCTGAACTCTTTCCCTACACGACGCTCTTCCGATCTNNNAGGAGAATCGCTTGAACTCAT
GAGCACACGTCTGAACTCTTTCCCTACACGACGCTCTTCCGATCTNNNAGGAGAATCGCTTGAACTCAT
AGAGCACACGTCTGAACTCTTTCCCTACACGACGCTCTTCCGATCTNNNGAGAAGGCCCTGCTTTCAGA
AGAGCACACGTCTGAACTCTTTCCCTACACGACGCTCTTCCGATCTNNNGAGAAGGCCCTGCTTTCAGA
AGAGCACACGTCTGAACTCTTTCCCTACACGACGCTCTTCCGATCTNNNGAGAAGGCCCTGCTTTCAGA
GAGCACACGTCTGAACTCTTTCCCTACACGACGCTCTTCCGATCTNNNATTCTTTTGTGCGTCCGATTGT
GAGCACACGTCTGAACTCTTTCCCTACACGACGCTCTTCCGATCTNNNATTCTTTTGTGCGTCCGATTGT
GAGCACACGTCTGAACTCTTTCCCTACACGACGCTCTTCCGATCTNNNATTCTTTTGTGCGTCCGATTGT
GAGCACACGTCTGAACTCTTTCCCTACACGACGCTCTTCCGATCTNNNAAAAGGACCCAAGACTCGGAT
GAGCACACGTCTGAACTCTTTCCCTACACGACGCTCTTCCGATCTNNNAAAAGGACCCAAGACTCGGAT
GAGCACACGTCTGAACTCTTTCCCTACACGACGCTCTTCCGATCTNNNAAAAGGACCCAAGACTCGGAT
AAGAGCACACGTCTGAACTCTTTCCCTACACGACGCTCTTCCGATCTNNNTCTATTTCCTCAATCCCAAG
AAGAGCACACGTCTGAACTCTTTCCCTACACGACGCTCTTCCGATCTNNNTCTATTTCCTCAATCCCAAG
AAGAGCACACGTCTGAACTCTTTCCCTACACGACGCTCTTCCGATCTNNNTCTATTTCCTCAATCCCAAG
GAGCACACGTCTGAACTCTTTCCCTACACGACGCTCTTCCGATCTNNNCTCCACTGGAACTTGCTCTGTA
GAGCACACGTCTGAACTCTTTCCCTACACGACGCTCTTCCGATCTNNNCTCCACTGGAACTTGCTCTGTA
GAGCACACGTCTGAACTCTTTCCCTACACGACGCTCTTCCGATCTNNNCTCCACTGGAACTTGCTCTGTA
AGCACACGTCTGAACTCTTTCCCTACACGACGCTCTTCCGATCTNNNGAATGATTTGGTATGCTGGAGC
AGCACACGTCTGAACTCTTTCCCTACACGACGCTCTTCCGATCTNNNGAATGATTTGGTATGCTGGAGC
AGCACACGTCTGAACTCTTTCCCTACACGACGCTCTTCCGATCTNNNGAATGATTTGGTATGCTGGAGC
AGCACACGTCTGAACTCTTTCCCTACACGACGCTCTTCCGATCTNNNAACCTGAGAAGCAGAGGTTGTA
AGCACACGTCTGAACTCTTTCCCTACACGACGCTCTTCCGATCTNNNAACCTGAGAAGCAGAGGTTGTA
AGCACACGTCTGAACTCTTTCCCTACACGACGCTCTTCCGATCTNNNAACCTGAGAAGCAGAGGTTGTA

AGCACACGTCTGAACTCTTTCCCTACACGACGCTCTTCCGATCTNNNACAGCACAATTCACAACTGCAA
AGCACACGTCTGAACTCTTTCCCTACACGACGCTCTTCCGATCTNNNACAGCACAATTCACAACTGCAA
AGCACACGTCTGAACTCTTTCCCTACACGACGCTCTTCCGATCTNNNACAGCACAATTCACAACTGCAA
GAGCACACGTCTGAACTCTTTCCCTACACGACGCTCTTCCGATCTNNNTTCCAGTCCTTGCCAGATCAA
GAGCACACGTCTGAACTCTTTCCCTACACGACGCTCTTCCGATCTNNNTTCCAGTCCTTGCCAGATCAA
GAGCACACGTCTGAACTCTTTCCCTACACGACGCTCTTCCGATCTNNNTTCCAGTCCTTGCCAGATCAA
AGAGCACACGTCTGAACTCTTTCCCTACACGACGCTCTTCCGATCTNNNATGCCTGTAATCCCAGCTAC
AGAGCACACGTCTGAACTCTTTCCCTACACGACGCTCTTCCGATCTNNNATGCCTGTAATCCCAGCTAC
AGAGCACACGTCTGAACTCTTTCCCTACACGACGCTCTTCCGATCTNNNATGCCTGTAATCCCAGCTAC
GAGCACACGTCTGAACTCTTTCCCTACACGACGCTCTTCCGATCTNNNATGAAATCAAGACGCTCCTG
GAGCACACGTCTGAACTCTTTCCCTACACGACGCTCTTCCGATCTNNNATGAAATCAAGACGCTCCTG
GAGCACACGTCTGAACTCTTTCCCTACACGACGCTCTTCCGATCTNNNATGAAATCAAGACGCTCCTG
GCACACGTCTGAACTCTTTCCCTACACGACGCTCTTCCGATCTNNNAGTCTAAAGTTAGATCTGTCTT
GCACACGTCTGAACTCTTTCCCTACACGACGCTCTTCCGATCTNNNAGTCTAAAGTTAGATCTGTCTT
GCACACGTCTGAACTCTTTCCCTACACGACGCTCTTCCGATCTNNNAGTCTAAAGTTAGATCTGTCTT
AGCACACGTCTGAACTCTTTCCCTACACGACGCTCTTCCGATCTNNNGGTTCAAGCAATTCTTCTGCCT
AGCACACGTCTGAACTCTTTCCCTACACGACGCTCTTCCGATCTNNNGGTTCAAGCAATTCTTCTGCCT
AGCACACGTCTGAACTCTTTCCCTACACGACGCTCTTCCGATCTNNNGGTTCAAGCAATTCTTCTGCCT
AGCACACGTCTGAACTCTTTCCCTACACGACGCTCTTCCGATCTNNNAGTTGGGAAGAAGAAAAAGG
AGCACACGTCTGAACTCTTTCCCTACACGACGCTCTTCCGATCTNNNAGTTGGGAAGAAGAAAAAGG
AGCACACGTCTGAACTCTTTCCCTACACGACGCTCTTCCGATCTNNNAGTTGGGAAGAAGAAAAAGG
GCACACGTCTGAACTCTTTCCCTACACGACGCTCTTCCGATCTNNNGAGAAGTGTCTTGGGGTGCTTT
GCACACGTCTGAACTCTTTCCCTACACGACGCTCTTCCGATCTNNNGAGAAGTGTCTTGGGGTGCTTT
GCACACGTCTGAACTCTTTCCCTACACGACGCTCTTCCGATCTNNNGAGAAGTGTCTTGGGGTGCTTT
AGCACACGTCTGAACTCTTTCCCTACACGACGCTCTTCCGATCTNNNTCTCATTCTCCCTTCATCCCT
AGCACACGTCTGAACTCTTTCCCTACACGACGCTCTTCCGATCTNNNTCTCATTCTCCCTTCATCCCT
AGCACACGTCTGAACTCTTTCCCTACACGACGCTCTTCCGATCTNNNTCTCATTCTCCCTTCATCCCT
AGCACACGTCTGAACTCTTTCCCTACACGACGCTCTTCCGATCTNNNTCTCTCTCTCTCTCTCCCTCTC
AGCACACGTCTGAACTCTTTCCCTACACGACGCTCTTCCGATCTNNNTCTCTCTCTCTCTCTCCCTCTC
AGCACACGTCTGAACTCTTTCCCTACACGACGCTCTTCCGATCTNNNTCTCTCTCTCTCTCTCCCTCTC
AGCACACGTCTGAACTCTTTCCCTACACGACGCTCTTCCGATCTNNNTGGCTGTTTCTCTAGATGCCTT
AGCACACGTCTGAACTCTTTCCCTACACGACGCTCTTCCGATCTNNNTGGCTGTTTCTCTAGATGCCTT
AGCACACGTCTGAACTCTTTCCCTACACGACGCTCTTCCGATCTNNNTGGCTGTTTCTCTAGATGCCTT
CACACGTCTGAACTCTTTCCCTACACGACGCTCTTCCGATCTNNNAGTTGACACACACAGTTACAGTG
CACACGTCTGAACTCTTTCCCTACACGACGCTCTTCCGATCTNNNAGTTGACACACACAGTTACAGTG
CACACGTCTGAACTCTTTCCCTACACGACGCTCTTCCGATCTNNNAGTTGACACACACAGTTACAGTG
CACACGTCTGAACTCTTTCCCTACACGACGCTCTTCCGATCTNNNATAGCCCCTAGTTTGCCTGTTTTC
CACACGTCTGAACTCTTTCCCTACACGACGCTCTTCCGATCTNNNATAGCCCCTAGTTTGCCTGTTTTC
CACACGTCTGAACTCTTTCCCTACACGACGCTCTTCCGATCTNNNATAGCCCCTAGTTTGCCTGTTTTC
GAGCACACGTCTGAACTCTTTCCCTACACGACGCTCTTCCGATCTNNNTGGAAGAGATGAGCAATGT
GAGCACACGTCTGAACTCTTTCCCTACACGACGCTCTTCCGATCTNNNTGGAAGAGATGAGCAATGT
GAGCACACGTCTGAACTCTTTCCCTACACGACGCTCTTCCGATCTNNNTGGAAGAGATGAGCAATGT
GAGCACACGTCTGAACTCTTTCCCTACACGACGCTCTTCCGATCTNNNCCATGTGGTCGTTAGAGACA

GAGCACACGTCTGAACTCTTTCCCTACACGACGCTCTTCCGATCTNNNCCATGTGGTCGTTAGAGACA
GAGCACACGTCTGAACTCTTTCCCTACACGACGCTCTTCCGATCTNNNCCATGTGGTCGTTAGAGACA
AGCACACGTCTGAACTCTTTCCCTACACGACGCTCTTCCGATCTNNNAGCTTCTCTCAGTCTCTCTCAG
AGCACACGTCTGAACTCTTTCCCTACACGACGCTCTTCCGATCTNNNAGCTTCTCTCAGTCTCTCTCAG
AGCACACGTCTGAACTCTTTCCCTACACGACGCTCTTCCGATCTNNNAGCTTCTCTCAGTCTCTCTCAG
AGCACACGTCTGAACTCTTTCCCTACACGACGCTCTTCCGATCTNNNGAGTGCCTGTAATTCCAGCTA
AGCACACGTCTGAACTCTTTCCCTACACGACGCTCTTCCGATCTNNNGAGTGCCTGTAATTCCAGCTA
AGCACACGTCTGAACTCTTTCCCTACACGACGCTCTTCCGATCTNNNGAGTGCCTGTAATTCCAGCTA
AGCACACGTCTGAACTCTTTCCCTACACGACGCTCTTCCGATCTNNNAGTGGAAAACAGATCACAGG
AGCACACGTCTGAACTCTTTCCCTACACGACGCTCTTCCGATCTNNNAGTGGAAAACAGATCACAGG
AGCACACGTCTGAACTCTTTCCCTACACGACGCTCTTCCGATCTNNNAGTGGAAAACAGATCACAGG
GAGCACACGTCTGAACTCTTTCCCTACACGACGCTCTTCCGATCTNNNGAGCCCGTATTCTACAAGAG
GAGCACACGTCTGAACTCTTTCCCTACACGACGCTCTTCCGATCTNNNGAGCCCGTATTCTACAAGAG
GAGCACACGTCTGAACTCTTTCCCTACACGACGCTCTTCCGATCTNNNGAGCCCGTATTCTACAAGAG
AGCACACGTCTGAACTCTTTCCCTACACGACGCTCTTCCGATCTNNNGAGGCTGAGGCAGGAGAATG
AGCACACGTCTGAACTCTTTCCCTACACGACGCTCTTCCGATCTNNNGAGGCTGAGGCAGGAGAATG
AGCACACGTCTGAACTCTTTCCCTACACGACGCTCTTCCGATCTNNNGAGGCTGAGGCAGGAGAATG
GAGCACACGTCTGAACTCTTTCCCTACACGACGCTCTTCCGATCTNNNGAAGGATTATGGTTGCCTCT
GAGCACACGTCTGAACTCTTTCCCTACACGACGCTCTTCCGATCTNNNGAAGGATTATGGTTGCCTCT
GAGCACACGTCTGAACTCTTTCCCTACACGACGCTCTTCCGATCTNNNGAAGGATTATGGTTGCCTCT
GAGCACACGTCTGAACTCTTTCCCTACACGACGCTCTTCCGATCTNNNAGATCAAGCCATTGCACTCAA
GAGCACACGTCTGAACTCTTTCCCTACACGACGCTCTTCCGATCTNNNAGATCAAGCCATTGCACTCAA
GAGCACACGTCTGAACTCTTTCCCTACACGACGCTCTTCCGATCTNNNAGATCAAGCCATTGCACTCAA
AGCACACGTCTGAACTCTTTCCCTACACGACGCTCTTCCGATCTNNNAGGCAGAAGAATCACTTGAA
AGCACACGTCTGAACTCTTTCCCTACACGACGCTCTTCCGATCTNNNAGGCAGAAGAATCACTTGAA
AGCACACGTCTGAACTCTTTCCCTACACGACGCTCTTCCGATCTNNNAGGCAGAAGAATCACTTGAA
AGCACACGTCTGAACTCTTTCCCTACACGACGCTCTTCCGATCTNNNTATCACTCATCACTCTCCCCTCCC
AGCACACGTCTGAACTCTTTCCCTACACGACGCTCTTCCGATCTNNNTATCACTCATCACTCTCCCCTCCC
AGCACACGTCTGAACTCTTTCCCTACACGACGCTCTTCCGATCTNNNTATCACTCATCACTCTCCCCTCCC
AGCACACGTCTGAACTCTTTCCCTACACGACGCTCTTCCGATCTNNNCATACACACACACACACACACA
AGCACACGTCTGAACTCTTTCCCTACACGACGCTCTTCCGATCTNNNCATACACACACACACACACACA
AGCACACGTCTGAACTCTTTCCCTACACGACGCTCTTCCGATCTNNNCATACACACACACACACACACA
GAGCACACGTCTGAACTCTTTCCCTACACGACGCTCTTCCGATCTNNNGGCTCCTGTAATTCCAGCTAC
GAGCACACGTCTGAACTCTTTCCCTACACGACGCTCTTCCGATCTNNNGGCTCCTGTAATTCCAGCTAC
GAGCACACGTCTGAACTCTTTCCCTACACGACGCTCTTCCGATCTNNNGGCTCCTGTAATTCCAGCTAC
GAGCACACGTCTGAACTCTTTCCCTACACGACGCTCTTCCGATCTNNNGATAGAGCATCAGTGGGGA
GAGCACACGTCTGAACTCTTTCCCTACACGACGCTCTTCCGATCTNNNGATAGAGCATCAGTGGGGA
GAGCACACGTCTGAACTCTTTCCCTACACGACGCTCTTCCGATCTNNNGATAGAGCATCAGTGGGGA
GAGCACACGTCTGAACTCTTTCCCTACACGACGCTCTTCCGATCTNNNAAAATTAGCTGGGGTATGGT
GAGCACACGTCTGAACTCTTTCCCTACACGACGCTCTTCCGATCTNNNAAAATTAGCTGGGGTATGGT
GAGCACACGTCTGAACTCTTTCCCTACACGACGCTCTTCCGATCTNNNAAAATTAGCTGGGGTATGGT
CACACGTCTGAACTCTTTCCCTACACGACGCTCTTCCGATCTNNNCTGTCCCTGTTTCTGTCTCTACCCA
CACACGTCTGAACTCTTTCCCTACACGACGCTCTTCCGATCTNNNCTGTCCCTGTTTCTGTCTCTACCCA

GCACACGTCTGAACTCTTTCCCTACACGACGCTCTTCCGATCTNNNCTGTCCCTGTTTCTGTCTCTACCCA

AGCACACGTCTGAACTCTTTCCCTACACGACGCTCTTCCGATCTNNNTCCTCTCTCTCCTCTCTCTCTCA

AGCACACGTCTGAACTCTTTCCCTACACGACGCTCTTCCGATCTNNNTCCTCTCTCTCCTCTCTCTCTCA

AGCACACGTCTGAACTCTTTCCCTACACGACGCTCTTCCGATCTNNNTCCTCTCTCTCCTCTCTCTCTCA

CACGTCTGAACTCTTTCCCTACACGACGCTCTTCCGATCTNNNTAAGGTGTGTCCAAACTCCCATCCAT

CACGTCTGAACTCTTTCCCTACACGACGCTCTTCCGATCTNNNTAAGGTGTGTCCAAACTCCCATCCAT

CACGTCTGAACTCTTTCCCTACACGACGCTCTTCCGATCTNNNTAAGGTGTGTCCAAACTCCCATCCAT

AGCACACGTCTGAACTCTTTCCCTACACGACGCTCTTCCGATCTNNNACACAAGGAAACCCTGTTTAGGA

AGCACACGTCTGAACTCTTTCCCTACACGACGCTCTTCCGATCTNNNACACAAGGAAACCCTGTTTAGGA

AGCACACGTCTGAACTCTTTCCCTACACGACGCTCTTCCGATCTNNNACACAAGGAAACCCTGTTTAGGA

AGCACACGTCTGAACTCTTTCCCTACACGACGCTCTTCCGATCTNNNTTTCCTTCTGCTCTCTGGAAAGG

AGCACACGTCTGAACTCTTTCCCTACACGACGCTCTTCCGATCTNNNTTTCCTTCTGCTCTCTGGAAAGG

AGCACACGTCTGAACTCTTTCCCTACACGACGCTCTTCCGATCTNNNTTTCCTTCTGCTCTCTGGAAAGG

AGCACACGTCTGAACTCTTTCCCTACACGACGCTCTTCCGATCTNNNGATAAATCAGACGTGATGCTG

AGCACACGTCTGAACTCTTTCCCTACACGACGCTCTTCCGATCTNNNGATAAATCAGACGTGATGCTG

AGCACACGTCTGAACTCTTTCCCTACACGACGCTCTTCCGATCTNNNGATAAATCAGACGTGATGCTG

GAGCACACGTCTGAACTCTTTCCCTACACGACGCTCTTCCGATCTNNNTTTGCCCTTTTGGAGCCATAGT

GAGCACACGTCTGAACTCTTTCCCTACACGACGCTCTTCCGATCTNNNTTTGCCCTTTTGGAGCCATAGT

GAGCACACGTCTGAACTCTTTCCCTACACGACGCTCTTCCGATCTNNNTTTGCCCTTTTGGAGCCATAGT

AGCACACGTCTGAACTCTTTCCCTACACGACGCTCTTCCGATCTNNNAATGGCAATGAAGTCAAGCACT

AGCACACGTCTGAACTCTTTCCCTACACGACGCTCTTCCGATCTNNNAATGGCAATGAAGTCAAGCACT

AGCACACGTCTGAACTCTTTCCCTACACGACGCTCTTCCGATCTNNNAATGGCAATGAAGTCAAGCACT

GAGCACACGTCTGAACTCTTTCCCTACACGACGCTCTTCCGATCTNNNGGTTGCAGTGAGCCAAGATT

GAGCACACGTCTGAACTCTTTCCCTACACGACGCTCTTCCGATCTNNNGGTTGCAGTGAGCCAAGATT

GAGCACACGTCTGAACTCTTTCCCTACACGACGCTCTTCCGATCTNNNGGTTGCAGTGAGCCAAGATT

GCACACGTCTGAACTCTTTCCCTACACGACGCTCTTCCGATCTNNNTACGTCACCATTGATTGTGTCCTC

GCACACGTCTGAACTCTTTCCCTACACGACGCTCTTCCGATCTNNNTACGTCACCATTGATTGTGTCCTC

GCACACGTCTGAACTCTTTCCCTACACGACGCTCTTCCGATCTNNNTACGTCACCATTGATTGTGTCCTC

AGAGCACACGTCTGAACTCTTTCCCTACACGACGCTCTTCCGATCTNNNGGGGTAGAGAGTTGGGGC

AGAGCACACGTCTGAACTCTTTCCCTACACGACGCTCTTCCGATCTNNNGGGGTAGAGAGTTGGGGC

AGAGCACACGTCTGAACTCTTTCCCTACACGACGCTCTTCCGATCTNNNGGGGTAGAGAGTTGGGGC

AGCACACGTCTGAACTCTTTCCCTACACGACGCTCTTCCGATCTNNNGGTGCACCAATACAGTATGCTT

AGCACACGTCTGAACTCTTTCCCTACACGACGCTCTTCCGATCTNNNGGTGCACCAATACAGTATGCTT

AGCACACGTCTGAACTCTTTCCCTACACGACGCTCTTCCGATCTNNNGGTGCACCAATACAGTATGCTT

AGAGCACACGTCTGAACTCTTTCCCTACACGACGCTCTTCCGATCTNNNGCAAATGTTAGTGGCACGT

AGAGCACACGTCTGAACTCTTTCCCTACACGACGCTCTTCCGATCTNNNGCAAATGTTAGTGGCACGT

AGAGCACACGTCTGAACTCTTTCCCTACACGACGCTCTTCCGATCTNNNGCAAATGTTAGTGGCACGT

CACACGTCTGAACTCTTTCCCTACACGACGCTCTTCCGATCTNNNGCCTCTGTCTCTCAGTGACTAGACA

CACACGTCTGAACTCTTTCCCTACACGACGCTCTTCCGATCTNNNGCCTCTGTCTCTCAGTGACTAGACA

CACACGTCTGAACTCTTTCCCTACACGACGCTCTTCCGATCTNNNGCCTCTGTCTCTCAGTGACTAGACA

GAGCACACGTCTGAACTCTTTCCCTACACGACGCTCTTCCGATCTNNNGAGTGAGTCCCATGCTGACA

GAGCACACGTCTGAACTCTTTCCCTACACGACGCTCTTCCGATCTNNNGAGTGAGTCCCATGCTGACA

GAGCACACGTCTGAACTCTTTCCCTACACGACGCTCTTCCGATCTNNNGAGTGAGTCCCATGCTGACA

AGCACACGTCTGAACTCTTTCCCTACACGACGCTCTTCCGATCTNNNCAAGCTGAGAGCCAAAACAAG
AGCACACGTCTGAACTCTTTCCCTACACGACGCTCTTCCGATCTNNNCAAGCTGAGAGCCAAAACAAG
AGCACACGTCTGAACTCTTTCCCTACACGACGCTCTTCCGATCTNNNCAAGCTGAGAGCCAAAACAAG
AGCACACGTCTGAACTCTTTCCCTACACGACGCTCTTCCGATCTNNNAAATTAGGCAGATGTAGTGGC
AGCACACGTCTGAACTCTTTCCCTACACGACGCTCTTCCGATCTNNNAAATTAGGCAGATGTAGTGGC
AGCACACGTCTGAACTCTTTCCCTACACGACGCTCTTCCGATCTNNNAAATTAGGCAGATGTAGTGGC
CACGTCTGAACTCTTTCCCTACACGACGCTCTTCCGATCTNNNGATGAGCAGATGGCACAAATCTCCA
CACGTCTGAACTCTTTCCCTACACGACGCTCTTCCGATCTNNNGATGAGCAGATGGCACAAATCTCCA
CACGTCTGAACTCTTTCCCTACACGACGCTCTTCCGATCTNNNGATGAGCAGATGGCACAAATCTCCA
GAGCACACGTCTGAACTCTTTCCCTACACGACGCTCTTCCGATCTNNNGGTCCTATATGTTTCACACCT
GAGCACACGTCTGAACTCTTTCCCTACACGACGCTCTTCCGATCTNNNGGTCCTATATGTTTCACACCT
GAGCACACGTCTGAACTCTTTCCCTACACGACGCTCTTCCGATCTNNNGGTCCTATATGTTTCACACCT
GAGCACACGTCTGAACTCTTTCCCTACACGACGCTCTTCCGATCTNNNTTTAGTAGAGACGGGGTTTC
GAGCACACGTCTGAACTCTTTCCCTACACGACGCTCTTCCGATCTNNNTTTAGTAGAGACGGGGTTTC
GAGCACACGTCTGAACTCTTTCCCTACACGACGCTCTTCCGATCTNNNTTTAGTAGAGACGGGGTTTC
GCACACGTCTGAACTCTTTCCCTACACGACGCTCTTCCGATCTNNNGGCTCTCAGGTTCTAGCAATTCT
GCACACGTCTGAACTCTTTCCCTACACGACGCTCTTCCGATCTNNNGGCTCTCAGGTTCTAGCAATTCT
GCACACGTCTGAACTCTTTCCCTACACGACGCTCTTCCGATCTNNNGGCTCTCAGGTTCTAGCAATTCT
AGCACACGTCTGAACTCTTTCCCTACACGACGCTCTTCCGATCTNNNAAGAGACTTAGACTCCCACAC
AGCACACGTCTGAACTCTTTCCCTACACGACGCTCTTCCGATCTNNNAAGAGACTTAGACTCCCACAC
AGCACACGTCTGAACTCTTTCCCTACACGACGCTCTTCCGATCTNNNAAGAGACTTAGACTCCCACAC
GCACACGTCTGAACTCTTTCCCTACACGACGCTCTTCCGATCTNNNATATGAAAAAGTACTGGTAGAC
GCACACGTCTGAACTCTTTCCCTACACGACGCTCTTCCGATCTNNNATATGAAAAAGTACTGGTAGAC
GCACACGTCTGAACTCTTTCCCTACACGACGCTCTTCCGATCTNNNATATGAAAAAGTACTGGTAGAC
AGCACACGTCTGAACTCTTTCCCTACACGACGCTCTTCCGATCTNNNGGTGTGTGGATACTAGCTAGC
AGCACACGTCTGAACTCTTTCCCTACACGACGCTCTTCCGATCTNNNGGTGTGTGGATACTAGCTAGC
AGCACACGTCTGAACTCTTTCCCTACACGACGCTCTTCCGATCTNNNGGTGTGTGGATACTAGCTAGC
GAGCACACGTCTGAACTCTTTCCCTACACGACGCTCTTCCGATCTNNNTTTTGAGACCGAATCTTGCTG
GAGCACACGTCTGAACTCTTTCCCTACACGACGCTCTTCCGATCTNNNTTTTGAGACCGAATCTTGCTG
GAGCACACGTCTGAACTCTTTCCCTACACGACGCTCTTCCGATCTNNNTTTTGAGACCGAATCTTGCTG
GAGCACACGTCTGAACTCTTTCCCTACACGACGCTCTTCCGATCTNNNCTTCCAGACCATTGACATCGA
GAGCACACGTCTGAACTCTTTCCCTACACGACGCTCTTCCGATCTNNNCTTCCAGACCATTGACATCGA
GAGCACACGTCTGAACTCTTTCCCTACACGACGCTCTTCCGATCTNNNCTTCCAGACCATTGACATCGA
AGAGCACACGTCTGAACTCTTTCCCTACACGACGCTCTTCCGATCTNNNGCAGACTTCAATGAGACTTG
AGAGCACACGTCTGAACTCTTTCCCTACACGACGCTCTTCCGATCTNNNGCAGACTTCAATGAGACTTG
AGAGCACACGTCTGAACTCTTTCCCTACACGACGCTCTTCCGATCTNNNGCAGACTTCAATGAGACTTG
GAGCACACGTCTGAACTCTTTCCCTACACGACGCTCTTCCGATCTNNNGTGCTTCCCTCTTCTTTCACTCT
GAGCACACGTCTGAACTCTTTCCCTACACGACGCTCTTCCGATCTNNNGTGCTTCCCTCTTCTTTCACTCT
GAGCACACGTCTGAACTCTTTCCCTACACGACGCTCTTCCGATCTNNNGTGCTTCCCTCTTCTTTCACTCT
GAAGAGCACACGTCTGAACTCTTTCCCTACACGACGCTCTTCCGATCTNNNGAGGCTGAGGCAGGAG
GAAGAGCACACGTCTGAACTCTTTCCCTACACGACGCTCTTCCGATCTNNNGAGGCTGAGGCAGGAG
GAAGAGCACACGTCTGAACTCTTTCCCTACACGACGCTCTTCCGATCTNNNGAGGCTGAGGCAGGAG
AGCACACGTCTGAACTCTTTCCCTACACGACGCTCTTCCGATCTNNNAAAGACACAGACTGGCAAATT

AGCACACGTCTGAACTCTTTCCCTACACGACGCTCTTCCGATCTNNNAAAGACACAGACTGGCAAATT

AGCACACGTCTGAACTCTTTCCCTACACGACGCTCTTCCGATCTNNNAAAGACACAGACTGGCAAATT

AGCACACGTCTGAACTCTTTCCCTACACGACGCTCTTCCGATCTNNNCTGAGGCAGGAGAATTGCTT

AGCACACGTCTGAACTCTTTCCCTACACGACGCTCTTCCGATCTNNNCTGAGGCAGGAGAATTGCTT

AGCACACGTCTGAACTCTTTCCCTACACGACGCTCTTCCGATCTNNNCTGAGGCAGGAGAATTGCTT

GAGCACACGTCTGAACTCTTTCCCTACACGACGCTCTTCCGATCTNNNGTGGCAACAGCTTTCCTGAT

GAGCACACGTCTGAACTCTTTCCCTACACGACGCTCTTCCGATCTNNNGTGGCAACAGCTTTCCTGAT

GAGCACACGTCTGAACTCTTTCCCTACACGACGCTCTTCCGATCTNNNGTGGCAACAGCTTTCCTGAT

AGCACACGTCTGAACTCTTTCCCTACACGACGCTCTTCCGATCTNNNGAATGCAGTGGTGTGATCATA

AGCACACGTCTGAACTCTTTCCCTACACGACGCTCTTCCGATCTNNNGAATGCAGTGGTGTGATCATA

AGCACACGTCTGAACTCTTTCCCTACACGACGCTCTTCCGATCTNNNGAATGCAGTGGTGTGATCATA

AGCACACGTCTGAACTCTTTCCCTACACGACGCTCTTCCGATCTNNNGCCAAGATTCTGCCATTGTACT

AGCACACGTCTGAACTCTTTCCCTACACGACGCTCTTCCGATCTNNNGCCAAGATTCTGCCATTGTACT

AGCACACGTCTGAACTCTTTCCCTACACGACGCTCTTCCGATCTNNNGCCAAGATTCTGCCATTGTACT

AGCACACGTCTGAACTCTTTCCCTACACGACGCTCTTCCGATCTNNNTCTAGTCATGGTATTGCAGAA

AGCACACGTCTGAACTCTTTCCCTACACGACGCTCTTCCGATCTNNNTCTAGTCATGGTATTGCAGAA

AGCACACGTCTGAACTCTTTCCCTACACGACGCTCTTCCGATCTNNNTCTAGTCATGGTATTGCAGAA

AGCACACGTCTGAACTCTTTCCCTACACGACGCTCTTCCGATCTNNNTCATTTTTCCAGGCCCAAATT

AGCACACGTCTGAACTCTTTCCCTACACGACGCTCTTCCGATCTNNNTCATTTTTCCAGGCCCAAATT

AGCACACGTCTGAACTCTTTCCCTACACGACGCTCTTCCGATCTNNNTCATTTTTCCAGGCCCAAATT

GCACACGTCTGAACTCTTTCCCTACACGACGCTCTTCCGATCTNNNTGGCCTTCTTTGTCTCTTGATCT

GCACACGTCTGAACTCTTTCCCTACACGACGCTCTTCCGATCTNNNTGGCCTTCTTTGTCTCTTGATCT

GCACACGTCTGAACTCTTTCCCTACACGACGCTCTTCCGATCTNNNTGGCCTTCTTTGTCTCTTGATCT

AGAGCACACGTCTGAACTCTTTCCCTACACGACGCTCTTCCGATCTNNNGTGTGTGTGAGAGAG

AGAGCACACGTCTGAACTCTTTCCCTACACGACGCTCTTCCGATCTNNNGTGTGTGTGAGAGAG

AGAGCACACGTCTGAACTCTTTCCCTACACGACGCTCTTCCGATCTNNNGTGTGTGTGAGAGAG

AGCACACGTCTGAACTCTTTCCCTACACGACGCTCTTCCGATCTNNNAAGTGGGAGTCGAACAATGA

AGCACACGTCTGAACTCTTTCCCTACACGACGCTCTTCCGATCTNNNAAGTGGGAGTCGAACAATGA

AGCACACGTCTGAACTCTTTCCCTACACGACGCTCTTCCGATCTNNNAAGTGGGAGTCGAACAATGA

AGCACACGTCTGAACTCTTTCCCTACACGACGCTCTTCCGATCTNNNTACTCTTACTGAGACCTCACG

AGCACACGTCTGAACTCTTTCCCTACACGACGCTCTTCCGATCTNNNTACTCTTACTGAGACCTCACG

AGCACACGTCTGAACTCTTTCCCTACACGACGCTCTTCCGATCTNNNTACTCTTACTGAGACCTCACG

GAGCACACGTCTGAACTCTTTCCCTACACGACGCTCTTCCGATCTNNNATTTCTTGAGGCCAGGAGTT

GAGCACACGTCTGAACTCTTTCCCTACACGACGCTCTTCCGATCTNNNATTTCTTGAGGCCAGGAGTT

GAGCACACGTCTGAACTCTTTCCCTACACGACGCTCTTCCGATCTNNNATTTCTTGAGGCCAGGAGTT

GCACACGTCTGAACTCTTTCCCTACACGACGCTCTTCCGATCTNNNAGTTAGCCAGAACCGATTTTTG

GCACACGTCTGAACTCTTTCCCTACACGACGCTCTTCCGATCTNNNAGTTAGCCAGAACCGATTTTTG

GCACACGTCTGAACTCTTTCCCTACACGACGCTCTTCCGATCTNNNAGTTAGCCAGAACCGATTTTTG

GCACACGTCTGAACTCTTTCCCTACACGACGCTCTTCCGATCTNNNCGACAGAGTGAGACTCCGTCT

GCACACGTCTGAACTCTTTCCCTACACGACGCTCTTCCGATCTNNNCGACAGAGTGAGACTCCGTCT

GCACACGTCTGAACTCTTTCCCTACACGACGCTCTTCCGATCTNNNCGACAGAGTGAGACTCCGTCT

AGCACACGTCTGAACTCTTTCCCTACACGACGCTCTTCCGATCTNNNTAAAGAGCAGAAATGGTCAG

AGCACACGTCTGAACTCTTTCCCTACACGACGCTCTTCCGATCTNNNTAAAGAGCAGAAATGGTCAG

GAGCACACGTCTGAACTCTTTCCCTACACGACGCTCTTCCGATCTNNNTAAAGAGCAGAAATGGTCAG

GAGCACACGTCTGAACTCTTTCCCTACACGACGCTCTTCCGATCTNNNGCCAGCCTAGGTTGAGGAAT

GAGCACACGTCTGAACTCTTTCCCTACACGACGCTCTTCCGATCTNNNGCCAGCCTAGGTTGAGGAAT

GAGCACACGTCTGAACTCTTTCCCTACACGACGCTCTTCCGATCTNNNGCCAGCCTAGGTTGAGGAAT

GCACACGTCTGAACTCTTTCCCTACACGACGCTCTTCCGATCTNNNTCCAGCTTCATCTTCCACAATCAC

GCACACGTCTGAACTCTTTCCCTACACGACGCTCTTCCGATCTNNNTCCAGCTTCATCTTCCACAATCAC

GCACACGTCTGAACTCTTTCCCTACACGACGCTCTTCCGATCTNNNTCCAGCTTCATCTTCCACAATCAC

AGCACACGTCTGAACTCTTTCCCTACACGACGCTCTTCCGATCTNNNGAAGGAGCCTGGGGTTGGAG

AGCACACGTCTGAACTCTTTCCCTACACGACGCTCTTCCGATCTNNNGAAGGAGCCTGGGGTTGGAG

AGCACACGTCTGAACTCTTTCCCTACACGACGCTCTTCCGATCTNNNGAAGGAGCCTGGGGTTGGAG

AGCACACGTCTGAACTCTTTCCCTACACGACGCTCTTCCGATCTNNNAACAAGCTCTGAAATTGAGGC

AGCACACGTCTGAACTCTTTCCCTACACGACGCTCTTCCGATCTNNNAACAAGCTCTGAAATTGAGGC

AGCACACGTCTGAACTCTTTCCCTACACGACGCTCTTCCGATCTNNNAACAAGCTCTGAAATTGAGGC

AGCACACGTCTGAACTCTTTCCCTACACGACGCTCTTCCGATCTNNNCCTTCAGGATGGGTATCAGCT

AGCACACGTCTGAACTCTTTCCCTACACGACGCTCTTCCGATCTNNNCCTTCAGGATGGGTATCAGCT

AGCACACGTCTGAACTCTTTCCCTACACGACGCTCTTCCGATCTNNNCCTTCAGGATGGGTATCAGCT

GAGCACACGTCTGAACTCTTTCCCTACACGACGCTCTTCCGATCTNNNCGTGCATGTTGAACCATTATT

GAGCACACGTCTGAACTCTTTCCCTACACGACGCTCTTCCGATCTNNNCGTGCATGTTGAACCATTATT

GAGCACACGTCTGAACTCTTTCCCTACACGACGCTCTTCCGATCTNNNCGTGCATGTTGAACCATTATT

GAGCACACGTCTGAACTCTTTCCCTACACGACGCTCTTCCGATCTNNNCATCCTGGCTAACACAGTGAA

GAGCACACGTCTGAACTCTTTCCCTACACGACGCTCTTCCGATCTNNNCATCCTGGCTAACACAGTGAA

GAGCACACGTCTGAACTCTTTCCCTACACGACGCTCTTCCGATCTNNNCATCCTGGCTAACACAGTGAA

AGCACACGTCTGAACTCTTTCCCTACACGACGCTCTTCCGATCTNNNCTTCTTCCCAGAAAGTGCACAAC

AGCACACGTCTGAACTCTTTCCCTACACGACGCTCTTCCGATCTNNNCTTCTTCCCAGAAAGTGCACAAC

AGCACACGTCTGAACTCTTTCCCTACACGACGCTCTTCCGATCTNNNCTTCTTCCCAGAAAGTGCACAAC

AGCACACGTCTGAACTCTTTCCCTACACGACGCTCTTCCGATCTNNNGGACCCCTTTATACTTGGCTCTC

AGCACACGTCTGAACTCTTTCCCTACACGACGCTCTTCCGATCTNNNGGACCCCTTTATACTTGGCTCTC

AGCACACGTCTGAACTCTTTCCCTACACGACGCTCTTCCGATCTNNNGGACCCCTTTATACTTGGCTCTC

AGCACACGTCTGAACTCTTTCCCTACACGACGCTCTTCCGATCTNNNTAGCTTTCCTGACTCACTTGCAA

AGCACACGTCTGAACTCTTTCCCTACACGACGCTCTTCCGATCTNNNTAGCTTTCCTGACTCACTTGCAA

AGCACACGTCTGAACTCTTTCCCTACACGACGCTCTTCCGATCTNNNTAGCTTTCCTGACTCACTTGCAA

GAGCACACGTCTGAACTCTTTCCCTACACGACGCTCTTCCGATCTNNNTGAAAAGTGTCGGCATCCAG

GAGCACACGTCTGAACTCTTTCCCTACACGACGCTCTTCCGATCTNNNTGAAAAGTGTCGGCATCCAG

GAGCACACGTCTGAACTCTTTCCCTACACGACGCTCTTCCGATCTNNNTGAAAAGTGTCGGCATCCAG

GCACACGTCTGAACTCTTTCCCTACACGACGCTCTTCCGATCTNNNGGCCACCAATAATGTATCTAAAG

GCACACGTCTGAACTCTTTCCCTACACGACGCTCTTCCGATCTNNNGGCCACCAATAATGTATCTAAAG

GCACACGTCTGAACTCTTTCCCTACACGACGCTCTTCCGATCTNNNGGCCACCAATAATGTATCTAAAG

GAGCACACGTCTGAACTCTTTCCCTACACGACGCTCTTCCGATCTNNNAAGGAAGACGAGGAATTCAG

GAGCACACGTCTGAACTCTTTCCCTACACGACGCTCTTCCGATCTNNNAAGGAAGACGAGGAATTCAG

GAGCACACGTCTGAACTCTTTCCCTACACGACGCTCTTCCGATCTNNNAAGGAAGACGAGGAATTCAG

GAGCACACGTCTGAACTCTTTCCCTACACGACGCTCTTCCGATCTNNNTAGGATGGGGAGAGACATTG

GAGCACACGTCTGAACTCTTTCCCTACACGACGCTCTTCCGATCTNNNTAGGATGGGGAGAGACATTG

GAGCACACGTCTGAACTCTTTCCCTACACGACGCTCTTCCGATCTNNNTAGGATGGGGAGAGACATTG

AGCACACGTCTGAACTCTTTCCCTACACGACGCTCTTCCGATCTNNNACAGTAGCTGCTAATGCTGAG
AGCACACGTCTGAACTCTTTCCCTACACGACGCTCTTCCGATCTNNNACAGTAGCTGCTAATGCTGAG
AGCACACGTCTGAACTCTTTCCCTACACGACGCTCTTCCGATCTNNNACAGTAGCTGCTAATGCTGAG
AGCACACGTCTGAACTCTTTCCCTACACGACGCTCTTCCGATCTNNNGCTAAGAGAGATCCTGGCTCA
AGCACACGTCTGAACTCTTTCCCTACACGACGCTCTTCCGATCTNNNGCTAAGAGAGATCCTGGCTCA
AGCACACGTCTGAACTCTTTCCCTACACGACGCTCTTCCGATCTNNNGCTAAGAGAGATCCTGGCTCA
GCACACGTCTGAACTCTTTCCCTACACGACGCTCTTCCGATCTNNNACTGTTGAGTCAACTTAGACGC
GCACACGTCTGAACTCTTTCCCTACACGACGCTCTTCCGATCTNNNACTGTTGAGTCAACTTAGACGC
GCACACGTCTGAACTCTTTCCCTACACGACGCTCTTCCGATCTNNNACTGTTGAGTCAACTTAGACGC
AGCACACGTCTGAACTCTTTCCCTACACGACGCTCTTCCGATCTNNNAGGAAAACTGCTTGAACCTAG
AGCACACGTCTGAACTCTTTCCCTACACGACGCTCTTCCGATCTNNNAGGAAAACTGCTTGAACCTAG
AGCACACGTCTGAACTCTTTCCCTACACGACGCTCTTCCGATCTNNNAGGAAAACTGCTTGAACCTAG
AGCACACGTCTGAACTCTTTCCCTACACGACGCTCTTCCGATCTNNNCTAACCTTTTGTGTCTTGTATG
AGCACACGTCTGAACTCTTTCCCTACACGACGCTCTTCCGATCTNNNCTAACCTTTTGTGTCTTGTATG
AGCACACGTCTGAACTCTTTCCCTACACGACGCTCTTCCGATCTNNNCTAACCTTTTGTGTCTTGTATG
CACACGTCTGAACTCTTTCCCTACACGACGCTCTTCCGATCTNNNTGAAAACAAACTTGAATAAACTG
CACACGTCTGAACTCTTTCCCTACACGACGCTCTTCCGATCTNNNTGAAAACAAACTTGAATAAACTG
CACACGTCTGAACTCTTTCCCTACACGACGCTCTTCCGATCTNNNTGAAAACAAACTTGAATAAACTG
AGCACACGTCTGAACTCTTTCCCTACACGACGCTCTTCCGATCTNNNTTTTGGGCAGTAACTGGGAA
AGCACACGTCTGAACTCTTTCCCTACACGACGCTCTTCCGATCTNNNTTTTGGGCAGTAACTGGGAA
AGCACACGTCTGAACTCTTTCCCTACACGACGCTCTTCCGATCTNNNTTTTGGGCAGTAACTGGGAA
AGCACACGTCTGAACTCTTTCCCTACACGACGCTCTTCCGATCTNNNGGTGACAGAGTGAGACTACG
AGCACACGTCTGAACTCTTTCCCTACACGACGCTCTTCCGATCTNNNGGTGACAGAGTGAGACTACG
AGCACACGTCTGAACTCTTTCCCTACACGACGCTCTTCCGATCTNNNGGTGACAGAGTGAGACTACG
AGCACACGTCTGAACTCTTTCCCTACACGACGCTCTTCCGATCTNNNCTTGTACTAAAGACCTGACAAC
AGCACACGTCTGAACTCTTTCCCTACACGACGCTCTTCCGATCTNNNCTTGTACTAAAGACCTGACAAC
AGCACACGTCTGAACTCTTTCCCTACACGACGCTCTTCCGATCTNNNCTTGTACTAAAGACCTGACAAC
AGCACACGTCTGAACTCTTTCCCTACACGACGCTCTTCCGATCTNNNCCCAAAGCAATGGAGGCATTT
AGCACACGTCTGAACTCTTTCCCTACACGACGCTCTTCCGATCTNNNCCCAAAGCAATGGAGGCATTT
AGCACACGTCTGAACTCTTTCCCTACACGACGCTCTTCCGATCTNNNCCCAAAGCAATGGAGGCATTT
GAGCACACGTCTGAACTCTTTCCCTACACGACGCTCTTCCGATCTNNNAAAAATTAGCTGGGCGTGG
GAGCACACGTCTGAACTCTTTCCCTACACGACGCTCTTCCGATCTNNNAAAAATTAGCTGGGCGTGG
GAGCACACGTCTGAACTCTTTCCCTACACGACGCTCTTCCGATCTNNNAAAAATTAGCTGGGCGTGG
AGCACACGTCTGAACTCTTTCCCTACACGACGCTCTTCCGATCTNNNGGCACTTGTAATCCCAGCTAT
AGCACACGTCTGAACTCTTTCCCTACACGACGCTCTTCCGATCTNNNGGCACTTGTAATCCCAGCTAT
AGCACACGTCTGAACTCTTTCCCTACACGACGCTCTTCCGATCTNNNGGCACTTGTAATCCCAGCTAT
GAGCACACGTCTGAACTCTTTCCCTACACGACGCTCTTCCGATCTNNNCAGCGTTCTGTGTTGTTTCTA
GAGCACACGTCTGAACTCTTTCCCTACACGACGCTCTTCCGATCTNNNCAGCGTTCTGTGTTGTTTCTA
GAGCACACGTCTGAACTCTTTCCCTACACGACGCTCTTCCGATCTNNNCAGCGTTCTGTGTTGTTTCTA
CACACGTCTGAACTCTTTCCCTACACGACGCTCTTCCGATCTNNNGCTATCCCACTCTCTCTAGCTACA
CACACGTCTGAACTCTTTCCCTACACGACGCTCTTCCGATCTNNNGCTATCCCACTCTCTCTAGCTACA
CACACGTCTGAACTCTTTCCCTACACGACGCTCTTCCGATCTNNNGCTATCCCACTCTCTCTAGCTACA
AGCACACGTCTGAACTCTTTCCCTACACGACGCTCTTCCGATCTNNNCAAGTCAGAGGCCTTACATCCG

AGCACACGTCTGAACTCTTTCCCTACACGACGCTCTTCCGATCTNNNCAAGTCAGAGGCCTTACATCCG

AGCACACGTCTGAACTCTTTCCCTACACGACGCTCTTCCGATCTNNNCAAGTCAGAGGCCTTACATCCG

GAGCACACGTCTGAACTCTTTCCCTACACGACGCTCTTCCGATCTNNNCTATACAAGTCAGCCTCCAGA

GAGCACACGTCTGAACTCTTTCCCTACACGACGCTCTTCCGATCTNNNCTATACAAGTCAGCCTCCAGA

GAGCACACGTCTGAACTCTTTCCCTACACGACGCTCTTCCGATCTNNNCTATACAAGTCAGCCTCCAGA

GAGCACACGTCTGAACTCTTTCCCTACACGACGCTCTTCCGATCTNNNGGTTCCTTTAGCCCATTTGCTT

GAGCACACGTCTGAACTCTTTCCCTACACGACGCTCTTCCGATCTNNNGGTTCCTTTAGCCCATTTGCTT

GAGCACACGTCTGAACTCTTTCCCTACACGACGCTCTTCCGATCTNNNGGTTCCTTTAGCCCATTTGCTT

GCACACGTCTGAACTCTTTCCCTACACGACGCTCTTCCGATCTNNNGAAGGCCTTATACGTCACAAAG

GCACACGTCTGAACTCTTTCCCTACACGACGCTCTTCCGATCTNNNGAAGGCCTTATACGTCACAAAG

GCACACGTCTGAACTCTTTCCCTACACGACGCTCTTCCGATCTNNNGAAGGCCTTATACGTCACAAAG

AGCACACGTCTGAACTCTTTCCCTACACGACGCTCTTCCGATCTNNNCTGACTTCACACAGAAACACAC

AGCACACGTCTGAACTCTTTCCCTACACGACGCTCTTCCGATCTNNNCTGACTTCACACAGAAACACAC

AGCACACGTCTGAACTCTTTCCCTACACGACGCTCTTCCGATCTNNNCTGACTTCACACAGAAACACAC

GAGCACACGTCTGAACTCTTTCCCTACACGACGCTCTTCCGATCTNNNACTCCAAATTGAGGTTTCTTC

GAGCACACGTCTGAACTCTTTCCCTACACGACGCTCTTCCGATCTNNNACTCCAAATTGAGGTTTCTTC

GAGCACACGTCTGAACTCTTTCCCTACACGACGCTCTTCCGATCTNNNACTCCAAATTGAGGTTTCTTC

GAGCACACGTCTGAACTCTTTCCCTACACGACGCTCTTCCGATCTNNNAGGGTTTCAGGGAGACACT

GAGCACACGTCTGAACTCTTTCCCTACACGACGCTCTTCCGATCTNNNAGGGTTTCAGGGAGACACT

GAGCACACGTCTGAACTCTTTCCCTACACGACGCTCTTCCGATCTNNNAGGGTTTCAGGGAGACACT

GAGCACACGTCTGAACTCTTTCCCTACACGACGCTCTTCCGATCTNNNCTTCATTAGTTCAACAGCTTG

GAGCACACGTCTGAACTCTTTCCCTACACGACGCTCTTCCGATCTNNNCTTCATTAGTTCAACAGCTTG

GAGCACACGTCTGAACTCTTTCCCTACACGACGCTCTTCCGATCTNNNCTTCATTAGTTCAACAGCTTG

GCACACGTCTGAACTCTTTCCCTACACGACGCTCTTCCGATCTNNNGAGTTCAAGACCAGTCTAGGCA

GCACACGTCTGAACTCTTTCCCTACACGACGCTCTTCCGATCTNNNGAGTTCAAGACCAGTCTAGGCA

GCACACGTCTGAACTCTTTCCCTACACGACGCTCTTCCGATCTNNNGAGTTCAAGACCAGTCTAGGCA

CACACGTCTGAACTCTTTCCCTACACGACGCTCTTCCGATCTNNNCTCACCTGCAATGATGAACCAGTCA

CACACGTCTGAACTCTTTCCCTACACGACGCTCTTCCGATCTNNNCTCACCTGCAATGATGAACCAGTCA

CACACGTCTGAACTCTTTCCCTACACGACGCTCTTCCGATCTNNNCTCACCTGCAATGATGAACCAGTCA

CACACGTCTGAACTCTTTCCCTACACGACGCTCTTCCGATCTNNNCTGTAATCCCAGCACTTTTGGAGC

CACACGTCTGAACTCTTTCCCTACACGACGCTCTTCCGATCTNNNCTGTAATCCCAGCACTTTTGGAGC

CACACGTCTGAACTCTTTCCCTACACGACGCTCTTCCGATCTNNNCTGTAATCCCAGCACTTTTGGAGC

AGCACACGTCTGAACTCTTTCCCTACACGACGCTCTTCCGATCTNNNATTTCAGCCAAGGATTCAGGT

AGCACACGTCTGAACTCTTTCCCTACACGACGCTCTTCCGATCTNNNATTTCAGCCAAGGATTCAGGT

AGCACACGTCTGAACTCTTTCCCTACACGACGCTCTTCCGATCTNNNATTTCAGCCAAGGATTCAGGT

AGCACACGTCTGAACTCTTTCCCTACACGACGCTCTTCCGATCTNNNTTCCAAAGATCCTCAGAGCATC

AGCACACGTCTGAACTCTTTCCCTACACGACGCTCTTCCGATCTNNNTTCCAAAGATCCTCAGAGCATC

AGCACACGTCTGAACTCTTTCCCTACACGACGCTCTTCCGATCTNNNTTCCAAAGATCCTCAGAGCATC

GAGCACACGTCTGAACTCTTTCCCTACACGACGCTCTTCCGATCTNNNGGATTGGTCCAAATAACCTGA

GAGCACACGTCTGAACTCTTTCCCTACACGACGCTCTTCCGATCTNNNGGATTGGTCCAAATAACCTGA

GAGCACACGTCTGAACTCTTTCCCTACACGACGCTCTTCCGATCTNNNGGATTGGTCCAAATAACCTGA

AGCACACGTCTGAACTCTTTCCCTACACGACGCTCTTCCGATCTNNNGGTGGAGCTAGGAAAGTCAGA

AGCACACGTCTGAACTCTTTCCCTACACGACGCTCTTCCGATCTNNNGGTGGAGCTAGGAAAGTCAGA

AGCACACGTCTGAACTCTTTCCCTACACGACGCTCTTCCGATCTNNNGGTGGAGCTAGGAAAGTCAGA
GAGCACACGTCTGAACTCTTTCCCTACACGACGCTCTTCCGATCTNNNTGAGCCGAGATTGCACCATTC
GAGCACACGTCTGAACTCTTTCCCTACACGACGCTCTTCCGATCTNNNTGAGCCGAGATTGCACCATTC
GAGCACACGTCTGAACTCTTTCCCTACACGACGCTCTTCCGATCTNNNTGAGCCGAGATTGCACCATTC
AGCACACGTCTGAACTCTTTCCCTACACGACGCTCTTCCGATCTNNNACAAAGAAACTTCTCCAACTCCT
AGCACACGTCTGAACTCTTTCCCTACACGACGCTCTTCCGATCTNNNACAAAGAAACTTCTCCAACTCCT
AGCACACGTCTGAACTCTTTCCCTACACGACGCTCTTCCGATCTNNNACAAAGAAACTTCTCCAACTCCT
GAGCACACGTCTGAACTCTTTCCCTACACGACGCTCTTCCGATCTNNNTCCTCTTGCTGAATTGACCTCTT
GAGCACACGTCTGAACTCTTTCCCTACACGACGCTCTTCCGATCTNNNTCCTCTTGCTGAATTGACCTCTT
GAGCACACGTCTGAACTCTTTCCCTACACGACGCTCTTCCGATCTNNNTCCTCTTGCTGAATTGACCTCTT
AGCACACGTCTGAACTCTTTCCCTACACGACGCTCTTCCGATCTNNNGAGGCTGAGGCAGGAGAATGC
AGCACACGTCTGAACTCTTTCCCTACACGACGCTCTTCCGATCTNNNGAGGCTGAGGCAGGAGAATGC
AGCACACGTCTGAACTCTTTCCCTACACGACGCTCTTCCGATCTNNNGAGGCTGAGGCAGGAGAATGC
AGCACACGTCTGAACTCTTTCCCTACACGACGCTCTTCCGATCTNNNACAGAGAGGCCAAGAAGAATCT
AGCACACGTCTGAACTCTTTCCCTACACGACGCTCTTCCGATCTNNNACAGAGAGGCCAAGAAGAATCT
AGCACACGTCTGAACTCTTTCCCTACACGACGCTCTTCCGATCTNNNACAGAGAGGCCAAGAAGAATCT
AGAGCACACGTCTGAACTCTTTCCCTACACGACGCTCTTCCGATCTNNNGATGAGCTGAAGGTGACAG
AGAGCACACGTCTGAACTCTTTCCCTACACGACGCTCTTCCGATCTNNNGATGAGCTGAAGGTGACAG
AGAGCACACGTCTGAACTCTTTCCCTACACGACGCTCTTCCGATCTNNNGATGAGCTGAAGGTGACAG
GAGCACACGTCTGAACTCTTTCCCTACACGACGCTCTTCCGATCTNNNCTTGCAGTGAGCTGAGATCTT
GAGCACACGTCTGAACTCTTTCCCTACACGACGCTCTTCCGATCTNNNCTTGCAGTGAGCTGAGATCTT
GAGCACACGTCTGAACTCTTTCCCTACACGACGCTCTTCCGATCTNNNCTTGCAGTGAGCTGAGATCTT
AGCACACGTCTGAACTCTTTCCCTACACGACGCTCTTCCGATCTNNNGAGAATCGCTTGAACCTGGGA
AGCACACGTCTGAACTCTTTCCCTACACGACGCTCTTCCGATCTNNNGAGAATCGCTTGAACCTGGGA
AGCACACGTCTGAACTCTTTCCCTACACGACGCTCTTCCGATCTNNNGAGAATCGCTTGAACCTGGGA
GAGCACACGTCTGAACTCTTTCCCTACACGACGCTCTTCCGATCTNNNGTTGCAGTGAGCTGAGATCAT
GAGCACACGTCTGAACTCTTTCCCTACACGACGCTCTTCCGATCTNNNGTTGCAGTGAGCTGAGATCAT
GAGCACACGTCTGAACTCTTTCCCTACACGACGCTCTTCCGATCTNNNGTTGCAGTGAGCTGAGATCAT
AGAGCACACGTCTGAACTCTTTCCCTACACGACGCTCTTCCGATCTNNNAAAAAATTAGCCAGGTGTGG
AGAGCACACGTCTGAACTCTTTCCCTACACGACGCTCTTCCGATCTNNNAAAAAATTAGCCAGGTGTGG
AGAGCACACGTCTGAACTCTTTCCCTACACGACGCTCTTCCGATCTNNNAAAAAATTAGCCAGGTGTGG
GCACACGTCTGAACTCTTTCCCTACACGACGCTCTTCCGATCTNNNATCGCGCCAAAAACAAAACAAAA
GCACACGTCTGAACTCTTTCCCTACACGACGCTCTTCCGATCTNNNATCGCGCCAAAAACAAAACAAAA
GCACACGTCTGAACTCTTTCCCTACACGACGCTCTTCCGATCTNNNATCGCGCCAAAAACAAAACAAAA
AGAGCACACGTCTGAACTCTTTCCCTACACGACGCTCTTCCGATCTNNNTCATCAGAGAGAACAGACAA
AGAGCACACGTCTGAACTCTTTCCCTACACGACGCTCTTCCGATCTNNNTCATCAGAGAGAACAGACAA
AGAGCACACGTCTGAACTCTTTCCCTACACGACGCTCTTCCGATCTNNNTCATCAGAGAGAACAGACAA
GAGCACACGTCTGAACTCTTTCCCTACACGACGCTCTTCCGATCTNNNCTGAGATCACGCCACTGTACTC
GAGCACACGTCTGAACTCTTTCCCTACACGACGCTCTTCCGATCTNNNCTGAGATCACGCCACTGTACTC
GAGCACACGTCTGAACTCTTTCCCTACACGACGCTCTTCCGATCTNNNCTGAGATCACGCCACTGTACTC
GAGCACACGTCTGAACTCTTTCCCTACACGACGCTCTTCCGATCTNNNTCAATGGGATTTGGGGAGAA
GAGCACACGTCTGAACTCTTTCCCTACACGACGCTCTTCCGATCTNNNTCAATGGGATTTGGGGAGAA
GAGCACACGTCTGAACTCTTTCCCTACACGACGCTCTTCCGATCTNNNTCAATGGGATTTGGGGAGAA

AGCACACGTCTGAACTCTTTCCCTACACGACGCTCTTCCGATCTNNNTCCACTAGGGAGTAGGGCAAT
AGCACACGTCTGAACTCTTTCCCTACACGACGCTCTTCCGATCTNNNTCCACTAGGGAGTAGGGCAAT
AGCACACGTCTGAACTCTTTCCCTACACGACGCTCTTCCGATCTNNNTCCACTAGGGAGTAGGGCAAT
AGCACACGTCTGAACTCTTTCCCTACACGACGCTCTTCCGATCTNNNGAGAATCACTTGAACCCAGGA
AGCACACGTCTGAACTCTTTCCCTACACGACGCTCTTCCGATCTNNNGAGAATCACTTGAACCCAGGA
AGCACACGTCTGAACTCTTTCCCTACACGACGCTCTTCCGATCTNNNGAGAATCACTTGAACCCAGGA
AGCACACGTCTGAACTCTTTCCCTACACGACGCTCTTCCGATCTNNNGCTTCTATCCCTCACCTACAAG
AGCACACGTCTGAACTCTTTCCCTACACGACGCTCTTCCGATCTNNNGCTTCTATCCCTCACCTACAAG
AGCACACGTCTGAACTCTTTCCCTACACGACGCTCTTCCGATCTNNNGCTTCTATCCCTCACCTACAAG
GAGCACACGTCTGAACTCTTTCCCTACACGACGCTCTTCCGATCTNNNTTGAGGCAGGAGAATCACTT
GAGCACACGTCTGAACTCTTTCCCTACACGACGCTCTTCCGATCTNNNTTGAGGCAGGAGAATCACTT
GAGCACACGTCTGAACTCTTTCCCTACACGACGCTCTTCCGATCTNNNTTGAGGCAGGAGAATCACTT
GCACACGTCTGAACTCTTTCCCTACACGACGCTCTTCCGATCTNNNGGGTGTAGAATTTTAGGCTGGG
GCACACGTCTGAACTCTTTCCCTACACGACGCTCTTCCGATCTNNNGGGTGTAGAATTTTAGGCTGGG
GCACACGTCTGAACTCTTTCCCTACACGACGCTCTTCCGATCTNNNGGGTGTAGAATTTTAGGCTGGG
AGCACACGTCTGAACTCTTTCCCTACACGACGCTCTTCCGATCTNNNTAGAGGTTACAGTGACCCGAG
AGCACACGTCTGAACTCTTTCCCTACACGACGCTCTTCCGATCTNNNTAGAGGTTACAGTGACCCGAG
AGCACACGTCTGAACTCTTTCCCTACACGACGCTCTTCCGATCTNNNTAGAGGTTACAGTGACCCGAG
AGCACACGTCTGAACTCTTTCCCTACACGACGCTCTTCCGATCTNNNACAAAAGCTTTCTATCCCCAG
AGCACACGTCTGAACTCTTTCCCTACACGACGCTCTTCCGATCTNNNACAAAAGCTTTCTATCCCCAG
AGCACACGTCTGAACTCTTTCCCTACACGACGCTCTTCCGATCTNNNACAAAAGCTTTCTATCCCCAG
GCACACGTCTGAACTCTTTCCCTACACGACGCTCTTCCGATCTNNNTTTCTGCCCTGTAAACCTCAAGA
GCACACGTCTGAACTCTTTCCCTACACGACGCTCTTCCGATCTNNNTTTCTGCCCTGTAAACCTCAAGA
GCACACGTCTGAACTCTTTCCCTACACGACGCTCTTCCGATCTNNNTTTCTGCCCTGTAAACCTCAAGA
GCACACGTCTGAACTCTTTCCCTACACGACGCTCTTCCGATCTNNNTTGAACTTGTGTTTGCCTCTTGG
GCACACGTCTGAACTCTTTCCCTACACGACGCTCTTCCGATCTNNNTTGAACTTGTGTTTGCCTCTTGG
GCACACGTCTGAACTCTTTCCCTACACGACGCTCTTCCGATCTNNNTTGAACTTGTGTTTGCCTCTTGG
GCACACGTCTGAACTCTTTCCCTACACGACGCTCTTCCGATCTNNNAGCTCCAGTGTTCAGTGCATAT
GCACACGTCTGAACTCTTTCCCTACACGACGCTCTTCCGATCTNNNAGCTCCAGTGTTCAGTGCATAT
GCACACGTCTGAACTCTTTCCCTACACGACGCTCTTCCGATCTNNNAGCTCCAGTGTTCAGTGCATAT
AGCACACGTCTGAACTCTTTCCCTACACGACGCTCTTCCGATCTNNNTAGCTGTTGGGTTTCACTCAG
AGCACACGTCTGAACTCTTTCCCTACACGACGCTCTTCCGATCTNNNTAGCTGTTGGGTTTCACTCAG
AGCACACGTCTGAACTCTTTCCCTACACGACGCTCTTCCGATCTNNNTAGCTGTTGGGTTTCACTCAG
GAGCACACGTCTGAACTCTTTCCCTACACGACGCTCTTCCGATCTNNNGATCATGAGGTCAGGAGTTC
GAGCACACGTCTGAACTCTTTCCCTACACGACGCTCTTCCGATCTNNNGATCATGAGGTCAGGAGTTC
GAGCACACGTCTGAACTCTTTCCCTACACGACGCTCTTCCGATCTNNNGATCATGAGGTCAGGAGTTC
AGCACACGTCTGAACTCTTTCCCTACACGACGCTCTTCCGATCTNNNGAGACCTAGGCTTTCTTCAGT
AGCACACGTCTGAACTCTTTCCCTACACGACGCTCTTCCGATCTNNNGAGACCTAGGCTTTCTTCAGT
AGCACACGTCTGAACTCTTTCCCTACACGACGCTCTTCCGATCTNNNGAGACCTAGGCTTTCTTCAGT
GCACACGTCTGAACTCTTTCCCTACACGACGCTCTTCCGATCTNNNGTTTGGTTTTTCTGTTTGGTTGG
GCACACGTCTGAACTCTTTCCCTACACGACGCTCTTCCGATCTNNNGTTTGGTTTTTCTGTTTGGTTGG
GCACACGTCTGAACTCTTTCCCTACACGACGCTCTTCCGATCTNNNGTTTGGTTTTTCTGTTTGGTTGG
AGCACACGTCTGAACTCTTTCCCTACACGACGCTCTTCCGATCTNNNGAAGGTAGGGGCGCAGTTAT

GAGCACACGTCTGAACTCTTTCCCTACACGACGCTCTTCCGATCTNNNGAAGGTAGGGGCGCAGTTAT

GAGCACACGTCTGAACTCTTTCCCTACACGACGCTCTTCCGATCTNNNGAAGGTAGGGGCGCAGTTAT

AGCACACGTCTGAACTCTTTCCCTACACGACGCTCTTCCGATCTNNNAAGACTGAAGCCAAGAAGGTG

AGCACACGTCTGAACTCTTTCCCTACACGACGCTCTTCCGATCTNNNAAGACTGAAGCCAAGAAGGTG

AGCACACGTCTGAACTCTTTCCCTACACGACGCTCTTCCGATCTNNNAAGACTGAAGCCAAGAAGGTG

AGCACACGTCTGAACTCTTTCCCTACACGACGCTCTTCCGATCTNNNGATCTGTGCTGTACCTCAGTGA

AGCACACGTCTGAACTCTTTCCCTACACGACGCTCTTCCGATCTNNNGATCTGTGCTGTACCTCAGTGA

AGCACACGTCTGAACTCTTTCCCTACACGACGCTCTTCCGATCTNNNGATCTGTGCTGTACCTCAGTGA

GCACACGTCTGAACTCTTTCCCTACACGACGCTCTTCCGATCTNNNGGTCTTCAGTTTCACCCACATTGC

GCACACGTCTGAACTCTTTCCCTACACGACGCTCTTCCGATCTNNNGGTCTTCAGTTTCACCCACATTGC

GCACACGTCTGAACTCTTTCCCTACACGACGCTCTTCCGATCTNNNGGTCTTCAGTTTCACCCACATTGC

GAGCACACGTCTGAACTCTTTCCCTACACGACGCTCTTCCGATCTNNNTGTGTGTGTGTATATATGTGT

GAGCACACGTCTGAACTCTTTCCCTACACGACGCTCTTCCGATCTNNNTGTGTGTGTGTATATATGTGT

GAGCACACGTCTGAACTCTTTCCCTACACGACGCTCTTCCGATCTNNNTGTGTGTGTGTATATATGTGT

AGAGCACACGTCTGAACTCTTTCCCTACACGACGCTCTTCCGATCTNNNGAGATTGCGCCATTGCACT

AGAGCACACGTCTGAACTCTTTCCCTACACGACGCTCTTCCGATCTNNNGAGATTGCGCCATTGCACT

AGAGCACACGTCTGAACTCTTTCCCTACACGACGCTCTTCCGATCTNNNGAGATTGCGCCATTGCACT

AGCACACGTCTGAACTCTTTCCCTACACGACGCTCTTCCGATCTNNNTTGGGACATTGTCATTGAGGT

AGCACACGTCTGAACTCTTTCCCTACACGACGCTCTTCCGATCTNNNTTGGGACATTGTCATTGAGGT

AGCACACGTCTGAACTCTTTCCCTACACGACGCTCTTCCGATCTNNNTTGGGACATTGTCATTGAGGT

AGAGCACACGTCTGAACTCTTTCCCTACACGACGCTCTTCCGATCTNNNAACAACTTTGGCAAAGGTTCA

AGAGCACACGTCTGAACTCTTTCCCTACACGACGCTCTTCCGATCTNNNAACAACTTTGGCAAAGGTTCA

AGAGCACACGTCTGAACTCTTTCCCTACACGACGCTCTTCCGATCTNNNAACAACTTTGGCAAAGGTTCA

AGCACACGTCTGAACTCTTTCCCTACACGACGCTCTTCCGATCTNNNTTCCTTGCCCTTTTCAGCTTTCTCA

AGCACACGTCTGAACTCTTTCCCTACACGACGCTCTTCCGATCTNNNTTCCTTGCCCTTTTCAGCTTTCTCA

AGCACACGTCTGAACTCTTTCCCTACACGACGCTCTTCCGATCTNNNTTCCTTGCCCTTTTCAGCTTTCTCA

GCACACGTCTGAACTCTTTCCCTACACGACGCTCTTCCGATCTNNNACATCACTCTTCTCTTGCGAACACA

GCACACGTCTGAACTCTTTCCCTACACGACGCTCTTCCGATCTNNNACATCACTCTTCTCTTGCGAACACA

GCACACGTCTGAACTCTTTCCCTACACGACGCTCTTCCGATCTNNNACATCACTCTTCTCTTGCGAACACA

GAGCACACGTCTGAACTCTTTCCCTACACGACGCTCTTCCGATCTNNNAATATTGCAGCACTCCACTCCA

GAGCACACGTCTGAACTCTTTCCCTACACGACGCTCTTCCGATCTNNNAATATTGCAGCACTCCACTCCA

GAGCACACGTCTGAACTCTTTCCCTACACGACGCTCTTCCGATCTNNNAATATTGCAGCACTCCACTCCA

AGAGCACACGTCTGAACTCTTTCCCTACACGACGCTCTTCCGATCTNNNGTAATCCCAGCTACTTAGGA

AGAGCACACGTCTGAACTCTTTCCCTACACGACGCTCTTCCGATCTNNNGTAATCCCAGCTACTTAGGA

AGAGCACACGTCTGAACTCTTTCCCTACACGACGCTCTTCCGATCTNNNGTAATCCCAGCTACTTAGGA

CACACGTCTGAACTCTTTCCCTACACGACGCTCTTCCGATCTNNNCAGGGTCAGGCTTGCCTGAGCATC

CACACGTCTGAACTCTTTCCCTACACGACGCTCTTCCGATCTNNNCAGGGTCAGGCTTGCCTGAGCATC

CACACGTCTGAACTCTTTCCCTACACGACGCTCTTCCGATCTNNNCAGGGTCAGGCTTGCCTGAGCATC

AGCACACGTCTGAACTCTTTCCCTACACGACGCTCTTCCGATCTNNNTTGTAGTCAGCCGAGATCACAC

AGCACACGTCTGAACTCTTTCCCTACACGACGCTCTTCCGATCTNNNTTGTAGTCAGCCGAGATCACAC

AGCACACGTCTGAACTCTTTCCCTACACGACGCTCTTCCGATCTNNNTTGTAGTCAGCCGAGATCACAC

AGCACACGTCTGAACTCTTTCCCTACACGACGCTCTTCCGATCTNNNCTCCAGCTACTGTTGTATTGGGT

AGCACACGTCTGAACTCTTTCCCTACACGACGCTCTTCCGATCTNNNCTCCAGCTACTGTTGTATTGGGT

GCACACGTCTGAACTCTTTCCCTACACGACGCTCTTCCGATCTNNNCTCCAGCTACTGTTGTATTGGGT

GCACACGTCTGAACTCTTTCCCTACACGACGCTCTTCCGATCTNNNTAGCTGGGACTATACATGTGTG

GCACACGTCTGAACTCTTTCCCTACACGACGCTCTTCCGATCTNNNTAGCTGGGACTATACATGTGTG

GCACACGTCTGAACTCTTTCCCTACACGACGCTCTTCCGATCTNNNTAGCTGGGACTATACATGTGTG

GAGCACACGTCTGAACTCTTTCCCTACACGACGCTCTTCCGATCTNNNGGATGTGGGCTTTCAAGTTT

GAGCACACGTCTGAACTCTTTCCCTACACGACGCTCTTCCGATCTNNNGGATGTGGGCTTTCAAGTTT

GAGCACACGTCTGAACTCTTTCCCTACACGACGCTCTTCCGATCTNNNGGATGTGGGCTTTCAAGTTT

AGCACACGTCTGAACTCTTTCCCTACACGACGCTCTTCCGATCTNNNCAGGGGGCCGTTTTCTTTATTT

AGCACACGTCTGAACTCTTTCCCTACACGACGCTCTTCCGATCTNNNCAGGGGGCCGTTTTCTTTATTT

AGCACACGTCTGAACTCTTTCCCTACACGACGCTCTTCCGATCTNNNCAGGGGGCCGTTTTCTTTATTT

AGCACACGTCTGAACTCTTTCCCTACACGACGCTCTTCCGATCTNNNAATATTTCTAGAGGGAAGGGG

AGCACACGTCTGAACTCTTTCCCTACACGACGCTCTTCCGATCTNNNAATATTTCTAGAGGGAAGGGG

AGCACACGTCTGAACTCTTTCCCTACACGACGCTCTTCCGATCTNNNAATATTTCTAGAGGGAAGGGG

AGCACACGTCTGAACTCTTTCCCTACACGACGCTCTTCCGATCTNNNCTGAGGCAGGAGAATAGTGT

AGCACACGTCTGAACTCTTTCCCTACACGACGCTCTTCCGATCTNNNCTGAGGCAGGAGAATAGTGT

AGCACACGTCTGAACTCTTTCCCTACACGACGCTCTTCCGATCTNNNCTGAGGCAGGAGAATAGTGT

GAGCACACGTCTGAACTCTTTCCCTACACGACGCTCTTCCGATCTNNNCCTAGGAAGCAGAGGTTACA

GAGCACACGTCTGAACTCTTTCCCTACACGACGCTCTTCCGATCTNNNCCTAGGAAGCAGAGGTTACA

GAGCACACGTCTGAACTCTTTCCCTACACGACGCTCTTCCGATCTNNNCCTAGGAAGCAGAGGTTACA

GCACACGTCTGAACTCTTTCCCTACACGACGCTCTTCCGATCTNNNTCTGGTTTCTTGTGGGTAAATGC

GCACACGTCTGAACTCTTTCCCTACACGACGCTCTTCCGATCTNNNTCTGGTTTCTTGTGGGTAAATGC

GCACACGTCTGAACTCTTTCCCTACACGACGCTCTTCCGATCTNNNTCTGGTTTCTTGTGGGTAAATGC

GAGCACACGTCTGAACTCTTTCCCTACACGACGCTCTTCCGATCTNNNGTTGGTCGCTGGGTCCTTTT

GAGCACACGTCTGAACTCTTTCCCTACACGACGCTCTTCCGATCTNNNGTTGGTCGCTGGGTCCTTTT

GAGCACACGTCTGAACTCTTTCCCTACACGACGCTCTTCCGATCTNNNGTTGGTCGCTGGGTCCTTTT

AGAGCACACGTCTGAACTCTTTCCCTACACGACGCTCTTCCGATCTNNNAGGAGCCTATAATCCCAGCT

AGAGCACACGTCTGAACTCTTTCCCTACACGACGCTCTTCCGATCTNNNAGGAGCCTATAATCCCAGCT

AGAGCACACGTCTGAACTCTTTCCCTACACGACGCTCTTCCGATCTNNNAGGAGCCTATAATCCCAGCT

GCACACGTCTGAACTCTTTCCCTACACGACGCTCTTCCGATCTNNNCTTTCCCTGCAGTTGTTTACCAG

GCACACGTCTGAACTCTTTCCCTACACGACGCTCTTCCGATCTNNNCTTTCCCTGCAGTTGTTTACCAG

GCACACGTCTGAACTCTTTCCCTACACGACGCTCTTCCGATCTNNNCTTTCCCTGCAGTTGTTTACCAG

GCACACGTCTGAACTCTTTCCCTACACGACGCTCTTCCGATCTNNNGCAGAAGTGAACTTTGGTGCAT

GCACACGTCTGAACTCTTTCCCTACACGACGCTCTTCCGATCTNNNGCAGAAGTGAACTTTGGTGCAT

GCACACGTCTGAACTCTTTCCCTACACGACGCTCTTCCGATCTNNNGCAGAAGTGAACTTTGGTGCAT

GAGCACACGTCTGAACTCTTTCCCTACACGACGCTCTTCCGATCTNNNCGTATATCTGGGGGAAGCA

GAGCACACGTCTGAACTCTTTCCCTACACGACGCTCTTCCGATCTNNNCGTATATCTGGGGGAAGCA

GAGCACACGTCTGAACTCTTTCCCTACACGACGCTCTTCCGATCTNNNCGTATATCTGGGGGAAGCA

CACACGTCTGAACTCTTTCCCTACACGACGCTCTTCCGATCTNNNTTGACATTCTCCACCCATCTTGTCA

CACACGTCTGAACTCTTTCCCTACACGACGCTCTTCCGATCTNNNTTGACATTCTCCACCCATCTTGTCA

CACACGTCTGAACTCTTTCCCTACACGACGCTCTTCCGATCTNNNTTGACATTCTCCACCCATCTTGTCA

GAGCACACGTCTGAACTCTTTCCCTACACGACGCTCTTCCGATCTNNNCTGAGGCAGAAGAATTGCTT

GAGCACACGTCTGAACTCTTTCCCTACACGACGCTCTTCCGATCTNNNCTGAGGCAGAAGAATTGCTT

GAGCACACGTCTGAACTCTTTCCCTACACGACGCTCTTCCGATCTNNNCTGAGGCAGAAGAATTGCTT

AGCACACGTCTGAACTCTTTCCCTACACGACGCTCTTCCGATCTNNNGGGAAGCATGGCAAGAAAGT
AGCACACGTCTGAACTCTTTCCCTACACGACGCTCTTCCGATCTNNNGGGAAGCATGGCAAGAAAGT
AGCACACGTCTGAACTCTTTCCCTACACGACGCTCTTCCGATCTNNNGGGAAGCATGGCAAGAAAGT
AGCACACGTCTGAACTCTTTCCCTACACGACGCTCTTCCGATCTNNNAAATCACTTGAACTCAGGAGG
AGCACACGTCTGAACTCTTTCCCTACACGACGCTCTTCCGATCTNNNAAATCACTTGAACTCAGGAGG
AGCACACGTCTGAACTCTTTCCCTACACGACGCTCTTCCGATCTNNNAAATCACTTGAACTCAGGAGG
AGCACACGTCTGAACTCTTTCCCTACACGACGCTCTTCCGATCTNNNGTGTCTTTGGGGAAAAAGAG
AGCACACGTCTGAACTCTTTCCCTACACGACGCTCTTCCGATCTNNNGTGTCTTTGGGGAAAAAGAG
AGCACACGTCTGAACTCTTTCCCTACACGACGCTCTTCCGATCTNNNGTGTCTTTGGGGAAAAAGAG
AGCACACGTCTGAACTCTTTCCCTACACGACGCTCTTCCGATCTNNNAATGTCTGGCTTACTGTCAAC
AGCACACGTCTGAACTCTTTCCCTACACGACGCTCTTCCGATCTNNNAATGTCTGGCTTACTGTCAAC
AGCACACGTCTGAACTCTTTCCCTACACGACGCTCTTCCGATCTNNNAATGTCTGGCTTACTGTCAAC
AGCACACGTCTGAACTCTTTCCCTACACGACGCTCTTCCGATCTNNNTTTTGTTTTAAATACCGTTCAG
AGCACACGTCTGAACTCTTTCCCTACACGACGCTCTTCCGATCTNNNTTTTGTTTTAAATACCGTTCAG
AGCACACGTCTGAACTCTTTCCCTACACGACGCTCTTCCGATCTNNNTTTTGTTTTAAATACCGTTCAG
GCACACGTCTGAACTCTTTCCCTACACGACGCTCTTCCGATCTNNNCTGAGGCAGGAGAATCACTTG
GCACACGTCTGAACTCTTTCCCTACACGACGCTCTTCCGATCTNNNCTGAGGCAGGAGAATCACTTG
GCACACGTCTGAACTCTTTCCCTACACGACGCTCTTCCGATCTNNNCTGAGGCAGGAGAATCACTTG
GCACACGTCTGAACTCTTTCCCTACACGACGCTCTTCCGATCTNNNAATAGGTACCAGGGATGTCCGT
GCACACGTCTGAACTCTTTCCCTACACGACGCTCTTCCGATCTNNNAATAGGTACCAGGGATGTCCGT
GCACACGTCTGAACTCTTTCCCTACACGACGCTCTTCCGATCTNNNAATAGGTACCAGGGATGTCCGT
GCACACGTCTGAACTCTTTCCCTACACGACGCTCTTCCGATCTNNNGAGGTTGCAGTGAGCCAAGAT
GCACACGTCTGAACTCTTTCCCTACACGACGCTCTTCCGATCTNNNGAGGTTGCAGTGAGCCAAGAT
GCACACGTCTGAACTCTTTCCCTACACGACGCTCTTCCGATCTNNNGAGGTTGCAGTGAGCCAAGAT
GAGCACACGTCTGAACTCTTTCCCTACACGACGCTCTTCCGATCTNNNGCATGGACGAACCTTGTAAG
GAGCACACGTCTGAACTCTTTCCCTACACGACGCTCTTCCGATCTNNNGCATGGACGAACCTTGTAAG
GAGCACACGTCTGAACTCTTTCCCTACACGACGCTCTTCCGATCTNNNGCATGGACGAACCTTGTAAG
GAGCACACGTCTGAACTCTTTCCCTACACGACGCTCTTCCGATCTNNNAGCAAAATGGACACCAGAGA
GAGCACACGTCTGAACTCTTTCCCTACACGACGCTCTTCCGATCTNNNAGCAAAATGGACACCAGAGA
GAGCACACGTCTGAACTCTTTCCCTACACGACGCTCTTCCGATCTNNNAGCAAAATGGACACCAGAGA
GCACACGTCTGAACTCTTTCCCTACACGACGCTCTTCCGATCTNNNTGGCAATGAGTTTTGTGTTTG
GCACACGTCTGAACTCTTTCCCTACACGACGCTCTTCCGATCTNNNTGGCAATGAGTTTTGTGTTTG
GCACACGTCTGAACTCTTTCCCTACACGACGCTCTTCCGATCTNNNTGGCAATGAGTTTTGTGTTTG
AGCACACGTCTGAACTCTTTCCCTACACGACGCTCTTCCGATCTNNNCAACAAGGAGAGAGATGCTGT
AGCACACGTCTGAACTCTTTCCCTACACGACGCTCTTCCGATCTNNNCAACAAGGAGAGAGATGCTGT
AGCACACGTCTGAACTCTTTCCCTACACGACGCTCTTCCGATCTNNNCAACAAGGAGAGAGATGCTGT
AGCACACGTCTGAACTCTTTCCCTACACGACGCTCTTCCGATCTNNNACACACACAGAGAGAGAGAGA
AGCACACGTCTGAACTCTTTCCCTACACGACGCTCTTCCGATCTNNNACACACACAGAGAGAGAGAGA
AGCACACGTCTGAACTCTTTCCCTACACGACGCTCTTCCGATCTNNNACACACACAGAGAGAGAGAGA
AGCACACGTCTGAACTCTTTCCCTACACGACGCTCTTCCGATCTNNNGATAGCTTCCACCCGTCCCCCA
AGCACACGTCTGAACTCTTTCCCTACACGACGCTCTTCCGATCTNNNGATAGCTTCCACCCGTCCCCCA
AGCACACGTCTGAACTCTTTCCCTACACGACGCTCTTCCGATCTNNNGATAGCTTCCACCCGTCCCCCA
GCACACGTCTGAACTCTTTCCCTACACGACGCTCTTCCGATCTNNNAAAATCACAAAACACAGTTATTT

GCACACGTCTGAACTCTTTCCCTACACGACGCTCTTCCGATCTNNNAAAATCACAAAACACAGTTATTT
GCACACGTCTGAACTCTTTCCCTACACGACGCTCTTCCGATCTNNNAAAATCACAAAACACAGTTATTT
AGCACACGTCTGAACTCTTTCCCTACACGACGCTCTTCCGATCTNNNTCGGCAAATCACTATGTCCTAGC
AGCACACGTCTGAACTCTTTCCCTACACGACGCTCTTCCGATCTNNNTCGGCAAATCACTATGTCCTAGC
AGCACACGTCTGAACTCTTTCCCTACACGACGCTCTTCCGATCTNNNTCGGCAAATCACTATGTCCTAGC
AGCACACGTCTGAACTCTTTCCCTACACGACGCTCTTCCGATCTNNNTGTTTTTCTGGATGGTTGCAG
AGCACACGTCTGAACTCTTTCCCTACACGACGCTCTTCCGATCTNNNTGTTTTTCTGGATGGTTGCAG
AGCACACGTCTGAACTCTTTCCCTACACGACGCTCTTCCGATCTNNNTGTTTTTCTGGATGGTTGCAG
AGCACACGTCTGAACTCTTTCCCTACACGACGCTCTTCCGATCTNNNGAAGTACAGGTGTGAACCACTC
AGCACACGTCTGAACTCTTTCCCTACACGACGCTCTTCCGATCTNNNGAAGTACAGGTGTGAACCACTC
AGCACACGTCTGAACTCTTTCCCTACACGACGCTCTTCCGATCTNNNGAAGTACAGGTGTGAACCACTC
AGCACACGTCTGAACTCTTTCCCTACACGACGCTCTTCCGATCTNNNCTGAGGCAGGAGAATCACTTG
AGCACACGTCTGAACTCTTTCCCTACACGACGCTCTTCCGATCTNNNCTGAGGCAGGAGAATCACTTG
AGCACACGTCTGAACTCTTTCCCTACACGACGCTCTTCCGATCTNNNCTGAGGCAGGAGAATCACTTG
GAGCACACGTCTGAACTCTTTCCCTACACGACGCTCTTCCGATCTNNNGGCAAAAGAATCACTTGAAC
GAGCACACGTCTGAACTCTTTCCCTACACGACGCTCTTCCGATCTNNNGGCAAAAGAATCACTTGAAC
GAGCACACGTCTGAACTCTTTCCCTACACGACGCTCTTCCGATCTNNNGGCAAAAGAATCACTTGAAC
GAGCACACGTCTGAACTCTTTCCCTACACGACGCTCTTCCGATCTNNNCACCTGTAATCCCAGCTACTCT
GAGCACACGTCTGAACTCTTTCCCTACACGACGCTCTTCCGATCTNNNCACCTGTAATCCCAGCTACTCT
GAGCACACGTCTGAACTCTTTCCCTACACGACGCTCTTCCGATCTNNNCACCTGTAATCCCAGCTACTCT
GAGCACACGTCTGAACTCTTTCCCTACACGACGCTCTTCCGATCTNNNTTCTGAGTAGTGGGGACTACA
GAGCACACGTCTGAACTCTTTCCCTACACGACGCTCTTCCGATCTNNNTTCTGAGTAGTGGGGACTACA
GAGCACACGTCTGAACTCTTTCCCTACACGACGCTCTTCCGATCTNNNTTCTGAGTAGTGGGGACTACA
AGCACACGTCTGAACTCTTTCCCTACACGACGCTCTTCCGATCTNNNATCTGGATGGCTTCAAGCTCTTT
AGCACACGTCTGAACTCTTTCCCTACACGACGCTCTTCCGATCTNNNATCTGGATGGCTTCAAGCTCTTT
AGCACACGTCTGAACTCTTTCCCTACACGACGCTCTTCCGATCTNNNATCTGGATGGCTTCAAGCTCTTT
GCACACGTCTGAACTCTTTCCCTACACGACGCTCTTCCGATCTNNNTCTCAGATGAGCATGGAACATTC
GCACACGTCTGAACTCTTTCCCTACACGACGCTCTTCCGATCTNNNTCTCAGATGAGCATGGAACATTC
GCACACGTCTGAACTCTTTCCCTACACGACGCTCTTCCGATCTNNNTCTCAGATGAGCATGGAACATTC
AGCACACGTCTGAACTCTTTCCCTACACGACGCTCTTCCGATCTNNNCAATCCTTCTACCTCAGCCTACCC
AGCACACGTCTGAACTCTTTCCCTACACGACGCTCTTCCGATCTNNNCAATCCTTCTACCTCAGCCTACCC
AGCACACGTCTGAACTCTTTCCCTACACGACGCTCTTCCGATCTNNNCAATCCTTCTACCTCAGCCTACCC
GAGCACACGTCTGAACTCTTTCCCTACACGACGCTCTTCCGATCTNNNTCTTGGAACATCAAGAAGGAA
GAGCACACGTCTGAACTCTTTCCCTACACGACGCTCTTCCGATCTNNNTCTTGGAACATCAAGAAGGAA
GAGCACACGTCTGAACTCTTTCCCTACACGACGCTCTTCCGATCTNNNTCTTGGAACATCAAGAAGGAA
AGCACACGTCTGAACTCTTTCCCTACACGACGCTCTTCCGATCTNNNAAAAATGCAATGCCTGGATCCA
AGCACACGTCTGAACTCTTTCCCTACACGACGCTCTTCCGATCTNNNAAAAATGCAATGCCTGGATCCA
AGCACACGTCTGAACTCTTTCCCTACACGACGCTCTTCCGATCTNNNAAAAATGCAATGCCTGGATCCA
GAGCACACGTCTGAACTCTTTCCCTACACGACGCTCTTCCGATCTNNNGTTCTTATCTAGAGAAGAGCTA
GAGCACACGTCTGAACTCTTTCCCTACACGACGCTCTTCCGATCTNNNGTTCTTATCTAGAGAAGAGCTA
GAGCACACGTCTGAACTCTTTCCCTACACGACGCTCTTCCGATCTNNNGTTCTTATCTAGAGAAGAGCTA
GAGCACACGTCTGAACTCTTTCCCTACACGACGCTCTTCCGATCTNNNGCACCTGTAGTCTCAGCTACTC
GAGCACACGTCTGAACTCTTTCCCTACACGACGCTCTTCCGATCTNNNGCACCTGTAGTCTCAGCTACTC

GAGCACACGTCTGAACTCTTTCCCTACACGACGCTCTTCCGATCTNNNGCACCTGTAGTCTCAGCTACTC

AGCACACGTCTGAACTCTTTCCCTACACGACGCTCTTCCGATCTNNNCTGAGGCAGGAGAATCACTTG

AGCACACGTCTGAACTCTTTCCCTACACGACGCTCTTCCGATCTNNNCTGAGGCAGGAGAATCACTTG

AGCACACGTCTGAACTCTTTCCCTACACGACGCTCTTCCGATCTNNNCTGAGGCAGGAGAATCACTTG

GAGCACACGTCTGAACTCTTTCCCTACACGACGCTCTTCCGATCTNNNGAGGCAGGAGAATCGCTTGA

GAGCACACGTCTGAACTCTTTCCCTACACGACGCTCTTCCGATCTNNNGAGGCAGGAGAATCGCTTGA

GAGCACACGTCTGAACTCTTTCCCTACACGACGCTCTTCCGATCTNNNGAGGCAGGAGAATCGCTTGA

AGCACACGTCTGAACTCTTTCCCTACACGACGCTCTTCCGATCTNNNCCTCCAGAACTGTACGGCAATA

AGCACACGTCTGAACTCTTTCCCTACACGACGCTCTTCCGATCTNNNCCTCCAGAACTGTACGGCAATA

AGCACACGTCTGAACTCTTTCCCTACACGACGCTCTTCCGATCTNNNCCTCCAGAACTGTACGGCAATA

AGCACACGTCTGAACTCTTTCCCTACACGACGCTCTTCCGATCTNNNTCCTAGAGATTGGAAAACGGCT

AGCACACGTCTGAACTCTTTCCCTACACGACGCTCTTCCGATCTNNNTCCTAGAGATTGGAAAACGGCT

AGCACACGTCTGAACTCTTTCCCTACACGACGCTCTTCCGATCTNNNTCCTAGAGATTGGAAAACGGCT

ACACGTCTGAACTCTTTCCCTACACGACGCTCTTCCGATCTNNNAGCAATATTCAGTTGTTTTGGTTTGA

ACACGTCTGAACTCTTTCCCTACACGACGCTCTTCCGATCTNNNAGCAATATTCAGTTGTTTTGGTTTGA

ACACGTCTGAACTCTTTCCCTACACGACGCTCTTCCGATCTNNNAGCAATATTCAGTTGTTTTGGTTTGA

GAGCACACGTCTGAACTCTTTCCCTACACGACGCTCTTCCGATCTNNNAGCAAGACTCCACCTCAAAGAA

GAGCACACGTCTGAACTCTTTCCCTACACGACGCTCTTCCGATCTNNNAGCAAGACTCCACCTCAAAGAA

GAGCACACGTCTGAACTCTTTCCCTACACGACGCTCTTCCGATCTNNNAGCAAGACTCCACCTCAAAGAA

GCACACGTCTGAACTCTTTCCCTACACGACGCTCTTCCGATCTNNNCCCACCTTCGACCAATACACTAAC

GCACACGTCTGAACTCTTTCCCTACACGACGCTCTTCCGATCTNNNCCCACCTTCGACCAATACACTAAC

GCACACGTCTGAACTCTTTCCCTACACGACGCTCTTCCGATCTNNNCCCACCTTCGACCAATACACTAAC

GAGCACACGTCTGAACTCTTTCCCTACACGACGCTCTTCCGATCTNNNGCCCGTGGATATTATCTCCTTC

GAGCACACGTCTGAACTCTTTCCCTACACGACGCTCTTCCGATCTNNNGCCCGTGGATATTATCTCCTTC

GAGCACACGTCTGAACTCTTTCCCTACACGACGCTCTTCCGATCTNNNGCCCGTGGATATTATCTCCTTC

GAGCACACGTCTGAACTCTTTCCCTACACGACGCTCTTCCGATCTNNNCTGAGGCAGGAGAATCACTT

GAGCACACGTCTGAACTCTTTCCCTACACGACGCTCTTCCGATCTNNNCTGAGGCAGGAGAATCACTT

GAGCACACGTCTGAACTCTTTCCCTACACGACGCTCTTCCGATCTNNNCTGAGGCAGGAGAATCACTT

GAGCACACGTCTGAACTCTTTCCCTACACGACGCTCTTCCGATCTNNNTTTGAGACAAAGTCACGCTCT

GAGCACACGTCTGAACTCTTTCCCTACACGACGCTCTTCCGATCTNNNTTTGAGACAAAGTCACGCTCT

GAGCACACGTCTGAACTCTTTCCCTACACGACGCTCTTCCGATCTNNNTTTGAGACAAAGTCACGCTCT

AGCACACGTCTGAACTCTTTCCCTACACGACGCTCTTCCGATCTNNNATGGAGGCCAGCATTCATACAT

AGCACACGTCTGAACTCTTTCCCTACACGACGCTCTTCCGATCTNNNATGGAGGCCAGCATTCATACAT

AGCACACGTCTGAACTCTTTCCCTACACGACGCTCTTCCGATCTNNNATGGAGGCCAGCATTCATACAT

GCACACGTCTGAACTCTTTCCCTACACGACGCTCTTCCGATCTNNNTAAATTAGAGGTAGCTGGGGCAC

GCACACGTCTGAACTCTTTCCCTACACGACGCTCTTCCGATCTNNNTAAATTAGAGGTAGCTGGGGCAC

GCACACGTCTGAACTCTTTCCCTACACGACGCTCTTCCGATCTNNNTAAATTAGAGGTAGCTGGGGCAC

AGCACACGTCTGAACTCTTTCCCTACACGACGCTCTTCCGATCTNNNGTGCCTGGCTTATTTCTCTCTCT

AGCACACGTCTGAACTCTTTCCCTACACGACGCTCTTCCGATCTNNNGTGCCTGGCTTATTTCTCTCTCT

AGCACACGTCTGAACTCTTTCCCTACACGACGCTCTTCCGATCTNNNGTGCCTGGCTTATTTCTCTCTCT

AGCACACGTCTGAACTCTTTCCCTACACGACGCTCTTCCGATCTNNNTAAAGAAGGTCACATGCTGAGG

AGCACACGTCTGAACTCTTTCCCTACACGACGCTCTTCCGATCTNNNTAAAGAAGGTCACATGCTGAGG

AGCACACGTCTGAACTCTTTCCCTACACGACGCTCTTCCGATCTNNNTAAAGAAGGTCACATGCTGAGG

GCACACGTCTGAACTCTTTCCCTACACGACGCTCTTCCGATCTNNNAGTAACTGGGATTACAGGCATG
GCACACGTCTGAACTCTTTCCCTACACGACGCTCTTCCGATCTNNNAGTAACTGGGATTACAGGCATG
GCACACGTCTGAACTCTTTCCCTACACGACGCTCTTCCGATCTNNNAGTAACTGGGATTACAGGCATG
AGCACACGTCTGAACTCTTTCCCTACACGACGCTCTTCCGATCTNNNCTGTAGTCCCAGCTACTTGGAA
AGCACACGTCTGAACTCTTTCCCTACACGACGCTCTTCCGATCTNNNCTGTAGTCCCAGCTACTTGGAA
AGCACACGTCTGAACTCTTTCCCTACACGACGCTCTTCCGATCTNNNCTGTAGTCCCAGCTACTTGGAA
CACACGTCTGAACTCTTTCCCTACACGACGCTCTTCCGATCTNNNGTGTGTGTGCCCCATATAGAT
CACACGTCTGAACTCTTTCCCTACACGACGCTCTTCCGATCTNNNGTGTGTGTGCCCCATATAGAT
CACACGTCTGAACTCTTTCCCTACACGACGCTCTTCCGATCTNNNGTGTGTGTGCCCCATATAGAT
GAGCACACGTCTGAACTCTTTCCCTACACGACGCTCTTCCGATCTNNNGACAAAGAGTCAAGCCTTGA
GAGCACACGTCTGAACTCTTTCCCTACACGACGCTCTTCCGATCTNNNGACAAAGAGTCAAGCCTTGA
GAGCACACGTCTGAACTCTTTCCCTACACGACGCTCTTCCGATCTNNNGACAAAGAGTCAAGCCTTGA
AGCACACGTCTGAACTCTTTCCCTACACGACGCTCTTCCGATCTNNNCTCTGGCGCCTCATTTCCTCCAT
AGCACACGTCTGAACTCTTTCCCTACACGACGCTCTTCCGATCTNNNCTCTGGCGCCTCATTTCCTCCAT
AGCACACGTCTGAACTCTTTCCCTACACGACGCTCTTCCGATCTNNNCTCTGGCGCCTCATTTCCTCCAT
AGCACACGTCTGAACTCTTTCCCTACACGACGCTCTTCCGATCTNNNTGAGCATCTCAAATCTGAACTGA
AGCACACGTCTGAACTCTTTCCCTACACGACGCTCTTCCGATCTNNNTGAGCATCTCAAATCTGAACTGA
AGCACACGTCTGAACTCTTTCCCTACACGACGCTCTTCCGATCTNNNTGAGCATCTCAAATCTGAACTGA
GAGCACACGTCTGAACTCTTTCCCTACACGACGCTCTTCCGATCTNNNCTTGAACCTGGGAGATGGAG
GAGCACACGTCTGAACTCTTTCCCTACACGACGCTCTTCCGATCTNNNCTTGAACCTGGGAGATGGAG
GAGCACACGTCTGAACTCTTTCCCTACACGACGCTCTTCCGATCTNNNCTTGAACCTGGGAGATGGAG
AGCACACGTCTGAACTCTTTCCCTACACGACGCTCTTCCGATCTNNNAAGTGCAGTGAACTAGGCCGT
AGCACACGTCTGAACTCTTTCCCTACACGACGCTCTTCCGATCTNNNAAGTGCAGTGAACTAGGCCGT
AGCACACGTCTGAACTCTTTCCCTACACGACGCTCTTCCGATCTNNNAAGTGCAGTGAACTAGGCCGT
AGAGCACACGTCTGAACTCTTTCCCTACACGACGCTCTTCCGATCTNNNGGGCAAGACTTCGTCAAAA
AGAGCACACGTCTGAACTCTTTCCCTACACGACGCTCTTCCGATCTNNNGGGCAAGACTTCGTCAAAA
AGAGCACACGTCTGAACTCTTTCCCTACACGACGCTCTTCCGATCTNNNGGGCAAGACTTCGTCAAAA
GCACACGTCTGAACTCTTTCCCTACACGACGCTCTTCCGATCTNNNTGTCAATCAGGCTGCAGTCTTAT
GCACACGTCTGAACTCTTTCCCTACACGACGCTCTTCCGATCTNNNTGTCAATCAGGCTGCAGTCTTAT
GCACACGTCTGAACTCTTTCCCTACACGACGCTCTTCCGATCTNNNTGTCAATCAGGCTGCAGTCTTAT
GAGCACACGTCTGAACTCTTTCCCTACACGACGCTCTTCCGATCTNNNCTACCTCAGCCTCCTAAGTACC
GAGCACACGTCTGAACTCTTTCCCTACACGACGCTCTTCCGATCTNNNCTACCTCAGCCTCCTAAGTACC
GAGCACACGTCTGAACTCTTTCCCTACACGACGCTCTTCCGATCTNNNCTACCTCAGCCTCCTAAGTACC
GAGCACACGTCTGAACTCTTTCCCTACACGACGCTCTTCCGATCTNNNGCAACACAGCGAGACTCTATC
GAGCACACGTCTGAACTCTTTCCCTACACGACGCTCTTCCGATCTNNNGCAACACAGCGAGACTCTATC
GAGCACACGTCTGAACTCTTTCCCTACACGACGCTCTTCCGATCTNNNGCAACACAGCGAGACTCTATC
AAGAGCACACGTCTGAACTCTTTCCCTACACGACGCTCTTCCGATCTNNNTGCAAATTTTGAAACAATT
AAGAGCACACGTCTGAACTCTTTCCCTACACGACGCTCTTCCGATCTNNNTGCAAATTTTGAAACAATT
AAGAGCACACGTCTGAACTCTTTCCCTACACGACGCTCTTCCGATCTNNNTGCAAATTTTGAAACAATT
AGCACACGTCTGAACTCTTTCCCTACACGACGCTCTTCCGATCTNNNCCCAAATTTGAGGACTGTGACA
AGCACACGTCTGAACTCTTTCCCTACACGACGCTCTTCCGATCTNNNCCCAAATTTGAGGACTGTGACA
AGCACACGTCTGAACTCTTTCCCTACACGACGCTCTTCCGATCTNNNCCCAAATTTGAGGACTGTGACA
GAGCACACGTCTGAACTCTTTCCCTACACGACGCTCTTCCGATCTNNNTCTGACCTAGTAACCCATTTGA

GAGCACACGTCTGAACTCTTTCCCTACACGACGCTCTTCCGATCTNNNTCTGACCTAGTAACCCATTTGA
GAGCACACGTCTGAACTCTTTCCCTACACGACGCTCTTCCGATCTNNNTCTGACCTAGTAACCCATTTGA
GCACACGTCTGAACTCTTTCCCTACACGACGCTCTTCCGATCTNNNAGAGATTTTACTGCCTCAGCCTCC
GCACACGTCTGAACTCTTTCCCTACACGACGCTCTTCCGATCTNNNAGAGATTTTACTGCCTCAGCCTCC
GCACACGTCTGAACTCTTTCCCTACACGACGCTCTTCCGATCTNNNAGAGATTTTACTGCCTCAGCCTCC
AGCACACGTCTGAACTCTTTCCCTACACGACGCTCTTCCGATCTNNNCTGAGATGGGAGGATCGATTGA
AGCACACGTCTGAACTCTTTCCCTACACGACGCTCTTCCGATCTNNNCTGAGATGGGAGGATCGATTGA
AGCACACGTCTGAACTCTTTCCCTACACGACGCTCTTCCGATCTNNNCTGAGATGGGAGGATCGATTGA
AGAGCACACGTCTGAACTCTTTCCCTACACGACGCTCTTCCGATCTNNNGAACCAGTAGGATGTGTGTG
AGAGCACACGTCTGAACTCTTTCCCTACACGACGCTCTTCCGATCTNNNGAACCAGTAGGATGTGTGTG
AGAGCACACGTCTGAACTCTTTCCCTACACGACGCTCTTCCGATCTNNNGAACCAGTAGGATGTGTGTG
AAGAGCACACGTCTGAACTCTTTCCCTACACGACGCTCTTCCGATCTNNNGAGGCTAAGGAAGGAGAA
AAGAGCACACGTCTGAACTCTTTCCCTACACGACGCTCTTCCGATCTNNNGAGGCTAAGGAAGGAGAA
AAGAGCACACGTCTGAACTCTTTCCCTACACGACGCTCTTCCGATCTNNNGAGGCTAAGGAAGGAGAA
AGCACACGTCTGAACTCTTTCCCTACACGACGCTCTTCCGATCTNNNGTATCCCTGGACACCACTGAAA
AGCACACGTCTGAACTCTTTCCCTACACGACGCTCTTCCGATCTNNNGTATCCCTGGACACCACTGAAA
AGCACACGTCTGAACTCTTTCCCTACACGACGCTCTTCCGATCTNNNGTATCCCTGGACACCACTGAAA
AGCACACGTCTGAACTCTTTCCCTACACGACGCTCTTCCGATCTNNNTCTCTTCCCAAACACAGTTCTGCC
AGCACACGTCTGAACTCTTTCCCTACACGACGCTCTTCCGATCTNNNTCTCTTCCCAAACACAGTTCTGCC
AGCACACGTCTGAACTCTTTCCCTACACGACGCTCTTCCGATCTNNNTCTCTTCCCAAACACAGTTCTGCC
AGCACACGTCTGAACTCTTTCCCTACACGACGCTCTTCCGATCTNNNGCGACAAAGTGAGAGTCCATCT
AGCACACGTCTGAACTCTTTCCCTACACGACGCTCTTCCGATCTNNNGCGACAAAGTGAGAGTCCATCT
AGCACACGTCTGAACTCTTTCCCTACACGACGCTCTTCCGATCTNNNGCGACAAAGTGAGAGTCCATCT
AGCACACGTCTGAACTCTTTCCCTACACGACGCTCTTCCGATCTNNNGAGTCATGGCACTGGACTAGA
AGCACACGTCTGAACTCTTTCCCTACACGACGCTCTTCCGATCTNNNGAGTCATGGCACTGGACTAGA
AGCACACGTCTGAACTCTTTCCCTACACGACGCTCTTCCGATCTNNNGAGTCATGGCACTGGACTAGA
GAGCACACGTCTGAACTCTTTCCCTACACGACGCTCTTCCGATCTNNNTCAGACACACACTCACCAAGA
GAGCACACGTCTGAACTCTTTCCCTACACGACGCTCTTCCGATCTNNNTCAGACACACACTCACCAAGA
GAGCACACGTCTGAACTCTTTCCCTACACGACGCTCTTCCGATCTNNNTCAGACACACACTCACCAAGA
AGAGCACACGTCTGAACTCTTTCCCTACACGACGCTCTTCCGATCTNNNAGCAGTCTCTCTCTCTCTCT
AGAGCACACGTCTGAACTCTTTCCCTACACGACGCTCTTCCGATCTNNNAGCAGTCTCTCTCTCTCTCT
AGAGCACACGTCTGAACTCTTTCCCTACACGACGCTCTTCCGATCTNNNAGCAGTCTCTCTCTCTCTCT
CACACGTCTGAACTCTTTCCCTACACGACGCTCTTCCGATCTNNNCTTACAAAGTAGTAGAAGGCATCA
CACACGTCTGAACTCTTTCCCTACACGACGCTCTTCCGATCTNNNCTTACAAAGTAGTAGAAGGCATCA
CACACGTCTGAACTCTTTCCCTACACGACGCTCTTCCGATCTNNNCTTACAAAGTAGTAGAAGGCATCA
AGCACACGTCTGAACTCTTTCCCTACACGACGCTCTTCCGATCTNNNGAGGTTACAGTGAGCGGAGAT
AGCACACGTCTGAACTCTTTCCCTACACGACGCTCTTCCGATCTNNNGAGGTTACAGTGAGCGGAGAT
AGCACACGTCTGAACTCTTTCCCTACACGACGCTCTTCCGATCTNNNGAGGTTACAGTGAGCGGAGAT
AGCACACGTCTGAACTCTTTCCCTACACGACGCTCTTCCGATCTNNNGAGGCAGGAGAATGGTGTGAA
AGCACACGTCTGAACTCTTTCCCTACACGACGCTCTTCCGATCTNNNGAGGCAGGAGAATGGTGTGAA
AGCACACGTCTGAACTCTTTCCCTACACGACGCTCTTCCGATCTNNNGAGGCAGGAGAATGGTGTGAA
AGCACACGTCTGAACTCTTTCCCTACACGACGCTCTTCCGATCTNNNGCAGGAGAATTGCTTGAACCTG
AGCACACGTCTGAACTCTTTCCCTACACGACGCTCTTCCGATCTNNNGCAGGAGAATTGCTTGAACCTG

AGCACACGTCTGAACTCTTTCCCTACACGACGCTCTTCCGATCTNNNGCAGGAGAATTGCTTGAACCTG
AGCACACGTCTGAACTCTTTCCCTACACGACGCTCTTCCGATCTNNNGCACATCTGTAATCCCAGCTACT
AGCACACGTCTGAACTCTTTCCCTACACGACGCTCTTCCGATCTNNNGCACATCTGTAATCCCAGCTACT
AGCACACGTCTGAACTCTTTCCCTACACGACGCTCTTCCGATCTNNNGCACATCTGTAATCCCAGCTACT
GAGCACACGTCTGAACTCTTTCCCTACACGACGCTCTTCCGATCTNNNAAGACCTTCTTTTGCCAGGGTT
GAGCACACGTCTGAACTCTTTCCCTACACGACGCTCTTCCGATCTNNNAAGACCTTCTTTTGCCAGGGTT
GAGCACACGTCTGAACTCTTTCCCTACACGACGCTCTTCCGATCTNNNAAGACCTTCTTTTGCCAGGGTT
GAGCACACGTCTGAACTCTTTCCCTACACGACGCTCTTCCGATCTNNNTGACCATTAATGTCTGCATCA
GAGCACACGTCTGAACTCTTTCCCTACACGACGCTCTTCCGATCTNNNTGACCATTAATGTCTGCATCA
GAGCACACGTCTGAACTCTTTCCCTACACGACGCTCTTCCGATCTNNNTGACCATTAATGTCTGCATCA
AGCACACGTCTGAACTCTTTCCCTACACGACGCTCTTCCGATCTNNNCCAATAAGCACATGAACAGGTG
AGCACACGTCTGAACTCTTTCCCTACACGACGCTCTTCCGATCTNNNCCAATAAGCACATGAACAGGTG
AGCACACGTCTGAACTCTTTCCCTACACGACGCTCTTCCGATCTNNNCCAATAAGCACATGAACAGGTG
AGCACACGTCTGAACTCTTTCCCTACACGACGCTCTTCCGATCTNNNCTCAGTCTCTGTCCTTGTTTCCC
AGCACACGTCTGAACTCTTTCCCTACACGACGCTCTTCCGATCTNNNCTCAGTCTCTGTCCTTGTTTCCC
AGCACACGTCTGAACTCTTTCCCTACACGACGCTCTTCCGATCTNNNCTCAGTCTCTGTCCTTGTTTCCC
GAGCACACGTCTGAACTCTTTCCCTACACGACGCTCTTCCGATCTNNNTTACAGTGAGTTGAGATCCTG
GAGCACACGTCTGAACTCTTTCCCTACACGACGCTCTTCCGATCTNNNTTACAGTGAGTTGAGATCCTG
GAGCACACGTCTGAACTCTTTCCCTACACGACGCTCTTCCGATCTNNNTTACAGTGAGTTGAGATCCTG
AGCACACGTCTGAACTCTTTCCCTACACGACGCTCTTCCGATCTNNNCTCCAAGAACCACCAGAGTAAC
AGCACACGTCTGAACTCTTTCCCTACACGACGCTCTTCCGATCTNNNCTCCAAGAACCACCAGAGTAAC
AGCACACGTCTGAACTCTTTCCCTACACGACGCTCTTCCGATCTNNNCTCCAAGAACCACCAGAGTAAC
AGAGCACACGTCTGAACTCTTTCCCTACACGACGCTCTTCCGATCTNNNAAAGAAACTCAGGGTCCTCA
AGAGCACACGTCTGAACTCTTTCCCTACACGACGCTCTTCCGATCTNNNAAAGAAACTCAGGGTCCTCA
AGAGCACACGTCTGAACTCTTTCCCTACACGACGCTCTTCCGATCTNNNAAAGAAACTCAGGGTCCTCA
GAGCACACGTCTGAACTCTTTCCCTACACGACGCTCTTCCGATCTNNNGTGATCTCTCCTCACTGCAATC
GAGCACACGTCTGAACTCTTTCCCTACACGACGCTCTTCCGATCTNNNGTGATCTCTCCTCACTGCAATC
GAGCACACGTCTGAACTCTTTCCCTACACGACGCTCTTCCGATCTNNNGTGATCTCTCCTCACTGCAATC
GAGCACACGTCTGAACTCTTTCCCTACACGACGCTCTTCCGATCTNNNCCCGCTTTTTGCCATTCTTTTTC
GAGCACACGTCTGAACTCTTTCCCTACACGACGCTCTTCCGATCTNNNCCCGCTTTTTGCCATTCTTTTTC
GAGCACACGTCTGAACTCTTTCCCTACACGACGCTCTTCCGATCTNNNCCCGCTTTTTGCCATTCTTTTTC
GCACACGTCTGAACTCTTTCCCTACACGACGCTCTTCCGATCTNNNGAAACCTAGGGATGTCACTTGCT
GCACACGTCTGAACTCTTTCCCTACACGACGCTCTTCCGATCTNNNGAAACCTAGGGATGTCACTTGCT
GCACACGTCTGAACTCTTTCCCTACACGACGCTCTTCCGATCTNNNGAAACCTAGGGATGTCACTTGCT
AGCACACGTCTGAACTCTTTCCCTACACGACGCTCTTCCGATCTNNNAGTGCTGCAATCTAAGCTCACT
AGCACACGTCTGAACTCTTTCCCTACACGACGCTCTTCCGATCTNNNAGTGCTGCAATCTAAGCTCACT
AGCACACGTCTGAACTCTTTCCCTACACGACGCTCTTCCGATCTNNNAGTGCTGCAATCTAAGCTCACT
GAGCACACGTCTGAACTCTTTCCCTACACGACGCTCTTCCGATCTNNNCTCTGGGTTGACTAGAAGAGC
GAGCACACGTCTGAACTCTTTCCCTACACGACGCTCTTCCGATCTNNNCTCTGGGTTGACTAGAAGAGC
GAGCACACGTCTGAACTCTTTCCCTACACGACGCTCTTCCGATCTNNNCTCTGGGTTGACTAGAAGAGC
GCACACGTCTGAACTCTTTCCCTACACGACGCTCTTCCGATCTNNNGGACACTTCAGAGCACACAAAAA
GCACACGTCTGAACTCTTTCCCTACACGACGCTCTTCCGATCTNNNGGACACTTCAGAGCACACAAAAA
GCACACGTCTGAACTCTTTCCCTACACGACGCTCTTCCGATCTNNNGGACACTTCAGAGCACACAAAAA

GAGCACACGTCTGAACTCTTTCCCTACACGACGCTCTTCCGATCTNNNGCACCCTCACCAGCATCTATTA

GAGCACACGTCTGAACTCTTTCCCTACACGACGCTCTTCCGATCTNNNGCACCCTCACCAGCATCTATTA

GAGCACACGTCTGAACTCTTTCCCTACACGACGCTCTTCCGATCTNNNGCACCCTCACCAGCATCTATTA

GAGCACACGTCTGAACTCTTTCCCTACACGACGCTCTTCCGATCTNNNAAAAAATTAGCTGGGCATGGT

GAGCACACGTCTGAACTCTTTCCCTACACGACGCTCTTCCGATCTNNNAAAAAATTAGCTGGGCATGGT

GAGCACACGTCTGAACTCTTTCCCTACACGACGCTCTTCCGATCTNNNAAAAAATTAGCTGGGCATGGT

AGCACACGTCTGAACTCTTTCCCTACACGACGCTCTTCCGATCTNNNGGTACACTGGACTTTGGGGAAT

AGCACACGTCTGAACTCTTTCCCTACACGACGCTCTTCCGATCTNNNGGTACACTGGACTTTGGGGAAT

AGCACACGTCTGAACTCTTTCCCTACACGACGCTCTTCCGATCTNNNGGTACACTGGACTTTGGGGAAT

AGCACACGTCTGAACTCTTTCCCTACACGACGCTCTTCCGATCTNNNAATCACATCTACTTGGGAGGCT

AGCACACGTCTGAACTCTTTCCCTACACGACGCTCTTCCGATCTNNNAATCACATCTACTTGGGAGGCT

AGCACACGTCTGAACTCTTTCCCTACACGACGCTCTTCCGATCTNNNAATCACATCTACTTGGGAGGCT

GCACACGTCTGAACTCTTTCCCTACACGACGCTCTTCCGATCTNNNCAGGTTTAGAAGGGGTGTCTGG

GCACACGTCTGAACTCTTTCCCTACACGACGCTCTTCCGATCTNNNCAGGTTTAGAAGGGGTGTCTGG

GCACACGTCTGAACTCTTTCCCTACACGACGCTCTTCCGATCTNNNCAGGTTTAGAAGGGGTGTCTGG

GAGCACACGTCTGAACTCTTTCCCTACACGACGCTCTTCCGATCTNNNCTAGCCATTGGTCATCACCTTC

GAGCACACGTCTGAACTCTTTCCCTACACGACGCTCTTCCGATCTNNNCTAGCCATTGGTCATCACCTTC

GAGCACACGTCTGAACTCTTTCCCTACACGACGCTCTTCCGATCTNNNCTAGCCATTGGTCATCACCTTC

GAGCACACGTCTGAACTCTTTCCCTACACGACGCTCTTCCGATCTNNNACACAAGGCTCCTACAGAGTA

GAGCACACGTCTGAACTCTTTCCCTACACGACGCTCTTCCGATCTNNNACACAAGGCTCCTACAGAGTA

GAGCACACGTCTGAACTCTTTCCCTACACGACGCTCTTCCGATCTNNNACACAAGGCTCCTACAGAGTA

AGAGCACACGTCTGAACTCTTTCCCTACACGACGCTCTTCCGATCTNNNGAGAATGGCATGAACCCAGC

AGAGCACACGTCTGAACTCTTTCCCTACACGACGCTCTTCCGATCTNNNGAGAATGGCATGAACCCAGC

AGAGCACACGTCTGAACTCTTTCCCTACACGACGCTCTTCCGATCTNNNGAGAATGGCATGAACCCAGC

GAGCACACGTCTGAACTCTTTCCCTACACGACGCTCTTCCGATCTNNNCACCCTTACTTCTTTCTGGGGA

GAGCACACGTCTGAACTCTTTCCCTACACGACGCTCTTCCGATCTNNNCACCCTTACTTCTTTCTGGGGA

GAGCACACGTCTGAACTCTTTCCCTACACGACGCTCTTCCGATCTNNNCACCCTTACTTCTTTCTGGGGA

AGCACACGTCTGAACTCTTTCCCTACACGACGCTCTTCCGATCTNNNTTTCTCCTTCCTTCCTTCCTTCCCA

AGCACACGTCTGAACTCTTTCCCTACACGACGCTCTTCCGATCTNNNTTTCTCCTTCCTTCCTTCCTTCCCA

AGCACACGTCTGAACTCTTTCCCTACACGACGCTCTTCCGATCTNNNTTTCTCCTTCCTTCCTTCCTTCCCA

AGCACACGTCTGAACTCTTTCCCTACACGACGCTCTTCCGATCTNNNCTGAGGCAGGAGAATGACATG

AGCACACGTCTGAACTCTTTCCCTACACGACGCTCTTCCGATCTNNNCTGAGGCAGGAGAATGACATG

AGCACACGTCTGAACTCTTTCCCTACACGACGCTCTTCCGATCTNNNCTGAGGCAGGAGAATGACATG

GAGCACACGTCTGAACTCTTTCCCTACACGACGCTCTTCCGATCTNNNGAGTTTGCAGTGAGCTGAGAT

GAGCACACGTCTGAACTCTTTCCCTACACGACGCTCTTCCGATCTNNNGAGTTTGCAGTGAGCTGAGAT

GAGCACACGTCTGAACTCTTTCCCTACACGACGCTCTTCCGATCTNNNGAGTTTGCAGTGAGCTGAGAT

AGCACACGTCTGAACTCTTTCCCTACACGACGCTCTTCCGATCTNNNCTGTAGCCTCAAACTTCTGGGAT

AGCACACGTCTGAACTCTTTCCCTACACGACGCTCTTCCGATCTNNNCTGTAGCCTCAAACTTCTGGGAT

AGCACACGTCTGAACTCTTTCCCTACACGACGCTCTTCCGATCTNNNCTGTAGCCTCAAACTTCTGGGAT

ACACGTCTGAACTCTTTCCCTACACGACGCTCTTCCGATCTNNNACCCATCCTCCCGTAGGCCCCATCGG

ACACGTCTGAACTCTTTCCCTACACGACGCTCTTCCGATCTNNNACCCATCCTCCCGTAGGCCCCATCGG

ACACGTCTGAACTCTTTCCCTACACGACGCTCTTCCGATCTNNNACCCATCCTCCCGTAGGCCCCATCGG

AGCACACGTCTGAACTCTTTCCCTACACGACGCTCTTCCGATCTNNNTTTTCTCACAGCTCTGGACATCAC

GCACACGTCTGAACTCTTTCCCTACACGACGCTCTTCCGATCTNNNTTTTCTCACAGCTCTGGACATCA
GCACACGTCTGAACTCTTTCCCTACACGACGCTCTTCCGATCTNNNTTTTCTCACAGCTCTGGACATCA
GCACACGTCTGAACTCTTTCCCTACACGACGCTCTTCCGATCTNNNTCCATCTGTCTCTCTCTCTCCCTCA
GCACACGTCTGAACTCTTTCCCTACACGACGCTCTTCCGATCTNNNTCCATCTGTCTCTCTCTCTCCCTCA
GCACACGTCTGAACTCTTTCCCTACACGACGCTCTTCCGATCTNNNTCCATCTGTCTCTCTCTCTCCCTCA
GCACACGTCTGAACTCTTTCCCTACACGACGCTCTTCCGATCTNNNGCAGACATTCTCTCATTTCCTGC
GCACACGTCTGAACTCTTTCCCTACACGACGCTCTTCCGATCTNNNGCAGACATTCTCTCATTTCCTGC
GCACACGTCTGAACTCTTTCCCTACACGACGCTCTTCCGATCTNNNGCAGACATTCTCTCATTTCCTGC
AGAGCACACGTCTGAACTCTTTCCCTACACGACGCTCTTCCGATCTNNNGAGAGAGAGAGAGACCATT
AGAGCACACGTCTGAACTCTTTCCCTACACGACGCTCTTCCGATCTNNNGAGAGAGAGAGAGACCATT
AGAGCACACGTCTGAACTCTTTCCCTACACGACGCTCTTCCGATCTNNNGAGAGAGAGAGAGACCATT
AGAGCACACGTCTGAACTCTTTCCCTACACGACGCTCTTCCGATCTNNNCCCTGGAGGCCATAATTAC
AGAGCACACGTCTGAACTCTTTCCCTACACGACGCTCTTCCGATCTNNNCCCTGGAGGCCATAATTAC
AGAGCACACGTCTGAACTCTTTCCCTACACGACGCTCTTCCGATCTNNNCCCTGGAGGCCATAATTAC
AGCACACGTCTGAACTCTTTCCCTACACGACGCTCTTCCGATCTNNNACAGCTACTGAGACAACCCAA
AGCACACGTCTGAACTCTTTCCCTACACGACGCTCTTCCGATCTNNNACAGCTACTGAGACAACCCAA
AGCACACGTCTGAACTCTTTCCCTACACGACGCTCTTCCGATCTNNNACAGCTACTGAGACAACCCAA
AGCACACGTCTGAACTCTTTCCCTACACGACGCTCTTCCGATCTNNNCATTTGAACACCGGTTTCCATG
AGCACACGTCTGAACTCTTTCCCTACACGACGCTCTTCCGATCTNNNCATTTGAACACCGGTTTCCATG
AGCACACGTCTGAACTCTTTCCCTACACGACGCTCTTCCGATCTNNNCATTTGAACACCGGTTTCCATG
GAGCACACGTCTGAACTCTTTCCCTACACGACGCTCTTCCGATCTNNNTTTTCTGAGGTGTTCTCTGTCG
GAGCACACGTCTGAACTCTTTCCCTACACGACGCTCTTCCGATCTNNNTTTTCTGAGGTGTTCTCTGTCG
GAGCACACGTCTGAACTCTTTCCCTACACGACGCTCTTCCGATCTNNNTTTTCTGAGGTGTTCTCTGTCG
AGCACACGTCTGAACTCTTTCCCTACACGACGCTCTTCCGATCTNNNGTGAAGGAGCGAGAGTCCAT
AGCACACGTCTGAACTCTTTCCCTACACGACGCTCTTCCGATCTNNNGTGAAGGAGCGAGAGTCCAT
AGCACACGTCTGAACTCTTTCCCTACACGACGCTCTTCCGATCTNNNGTGAAGGAGCGAGAGTCCAT
GAGCACACGTCTGAACTCTTTCCCTACACGACGCTCTTCCGATCTNNNGAATCGTTTGAACCCAGGAAG
GAGCACACGTCTGAACTCTTTCCCTACACGACGCTCTTCCGATCTNNNGAATCGTTTGAACCCAGGAAG
GAGCACACGTCTGAACTCTTTCCCTACACGACGCTCTTCCGATCTNNNGAATCGTTTGAACCCAGGAAG
GAGCACACGTCTGAACTCTTTCCCTACACGACGCTCTTCCGATCTNNNGCTTGTATCAGACCCTCATCCA
GAGCACACGTCTGAACTCTTTCCCTACACGACGCTCTTCCGATCTNNNGCTTGTATCAGACCCTCATCCA
GAGCACACGTCTGAACTCTTTCCCTACACGACGCTCTTCCGATCTNNNGCTTGTATCAGACCCTCATCCA
AGCACACGTCTGAACTCTTTCCCTACACGACGCTCTTCCGATCTNNNTTTCTCCGTCTTCCCATCTTCAG
AGCACACGTCTGAACTCTTTCCCTACACGACGCTCTTCCGATCTNNNTTTCTCCGTCTTCCCATCTTCAG
AGCACACGTCTGAACTCTTTCCCTACACGACGCTCTTCCGATCTNNNTTTCTCCGTCTTCCCATCTTCAG
AAGAGCACACGTCTGAACTCTTTCCCTACACGACGCTCTTCCGATCTNNNTGTGATGGTGGGTGCCTA
AAGAGCACACGTCTGAACTCTTTCCCTACACGACGCTCTTCCGATCTNNNTGTGATGGTGGGTGCCTA
AAGAGCACACGTCTGAACTCTTTCCCTACACGACGCTCTTCCGATCTNNNTGTGATGGTGGGTGCCTA
GCACACGTCTGAACTCTTTCCCTACACGACGCTCTTCCGATCTNNNCTGCAGTGTTGTGTTCCTGTAA
GCACACGTCTGAACTCTTTCCCTACACGACGCTCTTCCGATCTNNNCTGCAGTGTTGTGTTCCTGTAA
GCACACGTCTGAACTCTTTCCCTACACGACGCTCTTCCGATCTNNNCTGCAGTGTTGTGTTCCTGTAA
CACACGTCTGAACTCTTTCCCTACACGACGCTCTTCCGATCTNNNGACATTCAGTCCCTCGTAGACAGC
CACACGTCTGAACTCTTTCCCTACACGACGCTCTTCCGATCTNNNGACATTCAGTCCCTCGTAGACAGC

CACACGTCTGAACTCTTTCCCTACACGACGCTCTTCCGATCTNNNGACATTCAGTCCCTCGTAGACAGC
GCACACGTCTGAACTCTTTCCCTACACGACGCTCTTCCGATCTNNNGTATTTTTAGTAGAGACAGGGT
GCACACGTCTGAACTCTTTCCCTACACGACGCTCTTCCGATCTNNNGTATTTTTAGTAGAGACAGGGT
GCACACGTCTGAACTCTTTCCCTACACGACGCTCTTCCGATCTNNNGTATTTTTAGTAGAGACAGGGT
AAGAGCACACGTCTGAACTCTTTCCCTACACGACGCTCTTCCGATCTNNNGAAAAACCCAGCCAAGT
AAGAGCACACGTCTGAACTCTTTCCCTACACGACGCTCTTCCGATCTNNNGAAAAACCCAGCCAAGT
AAGAGCACACGTCTGAACTCTTTCCCTACACGACGCTCTTCCGATCTNNNGAAAAACCCAGCCAAGT
CACACGTCTGAACTCTTTCCCTACACGACGCTCTTCCGATCTNNNCTGTACCCTTGTTGGGTTGTCTGC
CACACGTCTGAACTCTTTCCCTACACGACGCTCTTCCGATCTNNNCTGTACCCTTGTTGGGTTGTCTGC
CACACGTCTGAACTCTTTCCCTACACGACGCTCTTCCGATCTNNNCTGTACCCTTGTTGGGTTGTCTGC
GCACACGTCTGAACTCTTTCCCTACACGACGCTCTTCCGATCTNNNCCTGCTGCTTTCTGTTTTGGTAA
GCACACGTCTGAACTCTTTCCCTACACGACGCTCTTCCGATCTNNNCCTGCTGCTTTCTGTTTTGGTAA
GCACACGTCTGAACTCTTTCCCTACACGACGCTCTTCCGATCTNNNCCTGCTGCTTTCTGTTTTGGTAA
GCACACGTCTGAACTCTTTCCCTACACGACGCTCTTCCGATCTNNNGGGTTCCTCAGAGAAGGAGAA
GCACACGTCTGAACTCTTTCCCTACACGACGCTCTTCCGATCTNNNGGGTTCCTCAGAGAAGGAGAA
GCACACGTCTGAACTCTTTCCCTACACGACGCTCTTCCGATCTNNNGGGTTCCTCAGAGAAGGAGAA
GAGCACACGTCTGAACTCTTTCCCTACACGACGCTCTTCCGATCTNNNCTGAGGCATGAGAATTGC
GAGCACACGTCTGAACTCTTTCCCTACACGACGCTCTTCCGATCTNNNCTGAGGCATGAGAATTGC
GAGCACACGTCTGAACTCTTTCCCTACACGACGCTCTTCCGATCTNNNCTGAGGCATGAGAATTGC
GAGCACACGTCTGAACTCTTTCCCTACACGACGCTCTTCCGATCTNNNCTCTGCTCTGCTAGTCTTTCCA
GAGCACACGTCTGAACTCTTTCCCTACACGACGCTCTTCCGATCTNNNCTCTGCTCTGCTAGTCTTTCCA
GAGCACACGTCTGAACTCTTTCCCTACACGACGCTCTTCCGATCTNNNCTCTGCTCTGCTAGTCTTTCCA
AGCACACGTCTGAACTCTTTCCCTACACGACGCTCTTCCGATCTNNNAGCCTGTAATCCCAGTACTTTG
AGCACACGTCTGAACTCTTTCCCTACACGACGCTCTTCCGATCTNNNAGCCTGTAATCCCAGTACTTTG
AGCACACGTCTGAACTCTTTCCCTACACGACGCTCTTCCGATCTNNNAGCCTGTAATCCCAGTACTTTG
GCACACGTCTGAACTCTTTCCCTACACGACGCTCTTCCGATCTNNNCCTGCTGCTTTCTGTTTTGGTAA
GCACACGTCTGAACTCTTTCCCTACACGACGCTCTTCCGATCTNNNCCTGCTGCTTTCTGTTTTGGTAA
GCACACGTCTGAACTCTTTCCCTACACGACGCTCTTCCGATCTNNNCCTGCTGCTTTCTGTTTTGGTAA
AGCACACGTCTGAACTCTTTCCCTACACGACGCTCTTCCGATCTNNNTGCCAACGTTCCATTTTAGTG
AGCACACGTCTGAACTCTTTCCCTACACGACGCTCTTCCGATCTNNNTGCCAACGTTCCATTTTAGTG
AGCACACGTCTGAACTCTTTCCCTACACGACGCTCTTCCGATCTNNNTGCCAACGTTCCATTTTAGTG
AGCACACGTCTGAACTCTTTCCCTACACGACGCTCTTCCGATCTNNNTTCTGTAGAGGCGGTATCTCT
AGCACACGTCTGAACTCTTTCCCTACACGACGCTCTTCCGATCTNNNTTCTGTAGAGGCGGTATCTCT
AGCACACGTCTGAACTCTTTCCCTACACGACGCTCTTCCGATCTNNNTTCTGTAGAGGCGGTATCTCT
GCACACGTCTGAACTCTTTCCCTACACGACGCTCTTCCGATCTNNNTGACTGTCAACTTGTCCATCTGT
GCACACGTCTGAACTCTTTCCCTACACGACGCTCTTCCGATCTNNNTGACTGTCAACTTGTCCATCTGT
GCACACGTCTGAACTCTTTCCCTACACGACGCTCTTCCGATCTNNNTGACTGTCAACTTGTCCATCTGT
AGCACACGTCTGAACTCTTTCCCTACACGACGCTCTTCCGATCTNNNGAGCTGGTACCATTCCTTCTGA
AGCACACGTCTGAACTCTTTCCCTACACGACGCTCTTCCGATCTNNNGAGCTGGTACCATTCCTTCTGA
AGCACACGTCTGAACTCTTTCCCTACACGACGCTCTTCCGATCTNNNGAGCTGGTACCATTCCTTCTGA
GAGCACACGTCTGAACTCTTTCCCTACACGACGCTCTTCCGATCTNNNCTTCTATCCTCATCACCCCCAA
GAGCACACGTCTGAACTCTTTCCCTACACGACGCTCTTCCGATCTNNNCTTCTATCCTCATCACCCCCAA
GAGCACACGTCTGAACTCTTTCCCTACACGACGCTCTTCCGATCTNNNCTTCTATCCTCATCACCCCCAA

GAGCACACGTCTGAACTCTTTCCCTACACGACGCTCTTCCGATCTNNNCTTTACTGCAAAGTCAGGGTT
GAGCACACGTCTGAACTCTTTCCCTACACGACGCTCTTCCGATCTNNNCTTTACTGCAAAGTCAGGGTT
GAGCACACGTCTGAACTCTTTCCCTACACGACGCTCTTCCGATCTNNNCTTTACTGCAAAGTCAGGGTT
AGCACACGTCTGAACTCTTTCCCTACACGACGCTCTTCCGATCTNNNAAAGTTTCTATTCTCCCGGACCC
AGCACACGTCTGAACTCTTTCCCTACACGACGCTCTTCCGATCTNNNAAAGTTTCTATTCTCCCGGACCC
AGCACACGTCTGAACTCTTTCCCTACACGACGCTCTTCCGATCTNNNAAAGTTTCTATTCTCCCGGACCC
AGCACACGTCTGAACTCTTTCCCTACACGACGCTCTTCCGATCTNNNAAGGGAAACGATCACCTCTCTTC
AGCACACGTCTGAACTCTTTCCCTACACGACGCTCTTCCGATCTNNNAAGGGAAACGATCACCTCTCTTC
AGCACACGTCTGAACTCTTTCCCTACACGACGCTCTTCCGATCTNNNAAGGGAAACGATCACCTCTCTTC
GAGCACACGTCTGAACTCTTTCCCTACACGACGCTCTTCCGATCTNNNCACACCACCAATCACGCAAATA
GAGCACACGTCTGAACTCTTTCCCTACACGACGCTCTTCCGATCTNNNCACACCACCAATCACGCAAATA
GAGCACACGTCTGAACTCTTTCCCTACACGACGCTCTTCCGATCTNNNCACACCACCAATCACGCAAATA
GAGCACACGTCTGAACTCTTTCCCTACACGACGCTCTTCCGATCTNNNCTGAGACAGAGGAACTACAG
GAGCACACGTCTGAACTCTTTCCCTACACGACGCTCTTCCGATCTNNNCTGAGACAGAGGAACTACAG
GAGCACACGTCTGAACTCTTTCCCTACACGACGCTCTTCCGATCTNNNCTGAGACAGAGGAACTACAG
GAGCACACGTCTGAACTCTTTCCCTACACGACGCTCTTCCGATCTNNNAGCAATTACCATTTTTTTTTG
GAGCACACGTCTGAACTCTTTCCCTACACGACGCTCTTCCGATCTNNNAGCAATTACCATTTTTTTTTG
GAGCACACGTCTGAACTCTTTCCCTACACGACGCTCTTCCGATCTNNNAGCAATTACCATTTTTTTTTG
AGCACACGTCTGAACTCTTTCCCTACACGACGCTCTTCCGATCTNNNATCCCTTTACCAGTGACCCTCTA
AGCACACGTCTGAACTCTTTCCCTACACGACGCTCTTCCGATCTNNNATCCCTTTACCAGTGACCCTCTA
AGCACACGTCTGAACTCTTTCCCTACACGACGCTCTTCCGATCTNNNATCCCTTTACCAGTGACCCTCTA
GCACACGTCTGAACTCTTTCCCTACACGACGCTCTTCCGATCTNNNAAAATGCCTTGGACACTACCCAGC
GCACACGTCTGAACTCTTTCCCTACACGACGCTCTTCCGATCTNNNAAAATGCCTTGGACACTACCCAGC
GCACACGTCTGAACTCTTTCCCTACACGACGCTCTTCCGATCTNNNAAAATGCCTTGGACACTACCCAGC
CACACGTCTGAACTCTTTCCCTACACGACGCTCTTCCGATCTNNNCGGGAAGCAAGTTTCTTTATCCTCA
CACACGTCTGAACTCTTTCCCTACACGACGCTCTTCCGATCTNNNCGGGAAGCAAGTTTCTTTATCCTCA
CACACGTCTGAACTCTTTCCCTACACGACGCTCTTCCGATCTNNNCGGGAAGCAAGTTTCTTTATCCTCA
GCACACGTCTGAACTCTTTCCCTACACGACGCTCTTCCGATCTNNNCCTTCCTGGAGTTCCTTGTTGTAC
GCACACGTCTGAACTCTTTCCCTACACGACGCTCTTCCGATCTNNNCCTTCCTGGAGTTCCTTGTTGTAC
GCACACGTCTGAACTCTTTCCCTACACGACGCTCTTCCGATCTNNNCCTTCCTGGAGTTCCTTGTTGTAC
AGCACACGTCTGAACTCTTTCCCTACACGACGCTCTTCCGATCTNNNATGGGGGAGAGATCAGAGATT
AGCACACGTCTGAACTCTTTCCCTACACGACGCTCTTCCGATCTNNNATGGGGGAGAGATCAGAGATT
AGCACACGTCTGAACTCTTTCCCTACACGACGCTCTTCCGATCTNNNATGGGGGAGAGATCAGAGATT
AGCACACGTCTGAACTCTTTCCCTACACGACGCTCTTCCGATCTNNNACAATCAAGCAGAAGAAGGC
AGCACACGTCTGAACTCTTTCCCTACACGACGCTCTTCCGATCTNNNACAATCAAGCAGAAGAAGGC
AGCACACGTCTGAACTCTTTCCCTACACGACGCTCTTCCGATCTNNNACAATCAAGCAGAAGAAGGC
AGCACACGTCTGAACTCTTTCCCTACACGACGCTCTTCCGATCTNNNGGCAACAAGATGGAAACTCAGT
AGCACACGTCTGAACTCTTTCCCTACACGACGCTCTTCCGATCTNNNGGCAACAAGATGGAAACTCAGT
AGCACACGTCTGAACTCTTTCCCTACACGACGCTCTTCCGATCTNNNGGCAACAAGATGGAAACTCAGT
AGCACACGTCTGAACTCTTTCCCTACACGACGCTCTTCCGATCTNNNCTGTCGGAAGCAGAGGTCAGCA
AGCACACGTCTGAACTCTTTCCCTACACGACGCTCTTCCGATCTNNNCTGTCGGAAGCAGAGGTCAGCA
AGCACACGTCTGAACTCTTTCCCTACACGACGCTCTTCCGATCTNNNCTGTCGGAAGCAGAGGTCAGCA
GAGCACACGTCTGAACTCTTTCCCTACACGACGCTCTTCCGATCTNNNACCTAGAGCAAGAGTTGATGG

GAGCACACGTCTGAACTCTTTCCCTACACGACGCTCTTCCGATCTNNNACCTAGAGCAAGAGTTGATG
GAGCACACGTCTGAACTCTTTCCCTACACGACGCTCTTCCGATCTNNNACCTAGAGCAAGAGTTGATG
AGCACACGTCTGAACTCTTTCCCTACACGACGCTCTTCCGATCTNNNCAACATAGCACGATCCTGTCTC
AGCACACGTCTGAACTCTTTCCCTACACGACGCTCTTCCGATCTNNNCAACATAGCACGATCCTGTCTC
AGCACACGTCTGAACTCTTTCCCTACACGACGCTCTTCCGATCTNNNCAACATAGCACGATCCTGTCTC
GCACACGTCTGAACTCTTTCCCTACACGACGCTCTTCCGATCTNNNTTGCTGGAGAAGAGTTGTGAT
GCACACGTCTGAACTCTTTCCCTACACGACGCTCTTCCGATCTNNNTTGCTGGAGAAGAGTTGTGAT
GCACACGTCTGAACTCTTTCCCTACACGACGCTCTTCCGATCTNNNTTGCTGGAGAAGAGTTGTGAT
GAGCACACGTCTGAACTCTTTCCCTACACGACGCTCTTCCGATCTNNNTCCACTTGGCTGTTAGAACAC
GAGCACACGTCTGAACTCTTTCCCTACACGACGCTCTTCCGATCTNNNTCCACTTGGCTGTTAGAACAC
GAGCACACGTCTGAACTCTTTCCCTACACGACGCTCTTCCGATCTNNNTCCACTTGGCTGTTAGAACAC
GAGCACACGTCTGAACTCTTTCCCTACACGACGCTCTTCCGATCTNNNAGGAGAATGGCGTGAACCTG
GAGCACACGTCTGAACTCTTTCCCTACACGACGCTCTTCCGATCTNNNAGGAGAATGGCGTGAACCTG
GAGCACACGTCTGAACTCTTTCCCTACACGACGCTCTTCCGATCTNNNAGGAGAATGGCGTGAACCTG
GAGCACACGTCTGAACTCTTTCCCTACACGACGCTCTTCCGATCTNNNTGTAGTCCCAGCTACTCAGGA
GAGCACACGTCTGAACTCTTTCCCTACACGACGCTCTTCCGATCTNNNTGTAGTCCCAGCTACTCAGGA
GAGCACACGTCTGAACTCTTTCCCTACACGACGCTCTTCCGATCTNNNTGTAGTCCCAGCTACTCAGGA
GAGCACACGTCTGAACTCTTTCCCTACACGACGCTCTTCCGATCTNNNATCATTTGTGGTCAGGAGTT
GAGCACACGTCTGAACTCTTTCCCTACACGACGCTCTTCCGATCTNNNATCATTTGTGGTCAGGAGTT
GAGCACACGTCTGAACTCTTTCCCTACACGACGCTCTTCCGATCTNNNATCATTTGTGGTCAGGAGTT
AGCACACGTCTGAACTCTTTCCCTACACGACGCTCTTCCGATCTNNNTATGATAGCACCACTGCATTCCA
AGCACACGTCTGAACTCTTTCCCTACACGACGCTCTTCCGATCTNNNTATGATAGCACCACTGCATTCCA
AGCACACGTCTGAACTCTTTCCCTACACGACGCTCTTCCGATCTNNNTATGATAGCACCACTGCATTCCA
GCACACGTCTGAACTCTTTCCCTACACGACGCTCTTCCGATCTNNNTCAAATGATGTACTCCCGTGTGT
GCACACGTCTGAACTCTTTCCCTACACGACGCTCTTCCGATCTNNNTCAAATGATGTACTCCCGTGTGT
GCACACGTCTGAACTCTTTCCCTACACGACGCTCTTCCGATCTNNNTCAAATGATGTACTCCCGTGTGT
AGCACACGTCTGAACTCTTTCCCTACACGACGCTCTTCCGATCTNNNCACTCTGATCATGTGCTCCCTT
AGCACACGTCTGAACTCTTTCCCTACACGACGCTCTTCCGATCTNNNCACTCTGATCATGTGCTCCCTT
AGCACACGTCTGAACTCTTTCCCTACACGACGCTCTTCCGATCTNNNCACTCTGATCATGTGCTCCCTT
AGAGCACACGTCTGAACTCTTTCCCTACACGACGCTCTTCCGATCTNNNGCTTGTTCTTTCTGCCTAAG
AGAGCACACGTCTGAACTCTTTCCCTACACGACGCTCTTCCGATCTNNNGCTTGTTCTTTCTGCCTAAG
AGAGCACACGTCTGAACTCTTTCCCTACACGACGCTCTTCCGATCTNNNGCTTGTTCTTTCTGCCTAAG
GAGCACACGTCTGAACTCTTTCCCTACACGACGCTCTTCCGATCTNNNCAGATGCTACATGAGTCTGGG
GAGCACACGTCTGAACTCTTTCCCTACACGACGCTCTTCCGATCTNNNCAGATGCTACATGAGTCTGGG
GAGCACACGTCTGAACTCTTTCCCTACACGACGCTCTTCCGATCTNNNCAGATGCTACATGAGTCTGGG
AGCACACGTCTGAACTCTTTCCCTACACGACGCTCTTCCGATCTNNNCTGTAATCCCAGCTACTCAGGA
AGCACACGTCTGAACTCTTTCCCTACACGACGCTCTTCCGATCTNNNCTGTAATCCCAGCTACTCAGGA
AGCACACGTCTGAACTCTTTCCCTACACGACGCTCTTCCGATCTNNNCTGTAATCCCAGCTACTCAGGA
GCACACGTCTGAACTCTTTCCCTACACGACGCTCTTCCGATCTNNNTTATTCACCCACGTTTCAGAGGTC
GCACACGTCTGAACTCTTTCCCTACACGACGCTCTTCCGATCTNNNTTATTCACCCACGTTTCAGAGGTC
GCACACGTCTGAACTCTTTCCCTACACGACGCTCTTCCGATCTNNNTTATTCACCCACGTTTCAGAGGTC
AGCACACGTCTGAACTCTTTCCCTACACGACGCTCTTCCGATCTNNNAAGTGATTCTCATGCCTCAACC
AGCACACGTCTGAACTCTTTCCCTACACGACGCTCTTCCGATCTNNNAAGTGATTCTCATGCCTCAACC

AGCACACGTCTGAACTCTTTCCCTACACGACGCTCTTCCGATCTNNNAAGTGATTCTCATGCCTCAACCT
GAGCACACGTCTGAACTCTTTCCCTACACGACGCTCTTCCGATCTNNNTACAAAAAATTAGCAGGGCGT
GAGCACACGTCTGAACTCTTTCCCTACACGACGCTCTTCCGATCTNNNTACAAAAAATTAGCAGGGCGT
GAGCACACGTCTGAACTCTTTCCCTACACGACGCTCTTCCGATCTNNNTACAAAAAATTAGCAGGGCGT
AGCACACGTCTGAACTCTTTCCCTACACGACGCTCTTCCGATCTNNNCACCTGTAGTCCCAGTTACTTG
AGCACACGTCTGAACTCTTTCCCTACACGACGCTCTTCCGATCTNNNCACCTGTAGTCCCAGTTACTTG
AGCACACGTCTGAACTCTTTCCCTACACGACGCTCTTCCGATCTNNNCACCTGTAGTCCCAGTTACTTG
AGCACACGTCTGAACTCTTTCCCTACACGACGCTCTTCCGATCTNNNTATAAAACCATCAGATCTCATGA
AGCACACGTCTGAACTCTTTCCCTACACGACGCTCTTCCGATCTNNNTATAAAACCATCAGATCTCATGA
AGCACACGTCTGAACTCTTTCCCTACACGACGCTCTTCCGATCTNNNTATAAAACCATCAGATCTCATGA
GAGCACACGTCTGAACTCTTTCCCTACACGACGCTCTTCCGATCTNNNGTCGCTCATAGTTTGACTCGG
GAGCACACGTCTGAACTCTTTCCCTACACGACGCTCTTCCGATCTNNNGTCGCTCATAGTTTGACTCGG
GAGCACACGTCTGAACTCTTTCCCTACACGACGCTCTTCCGATCTNNNGTCGCTCATAGTTTGACTCGG
AGCACACGTCTGAACTCTTTCCCTACACGACGCTCTTCCGATCTNNNGATCCCAGGTGTATCCTTTTCCA
AGCACACGTCTGAACTCTTTCCCTACACGACGCTCTTCCGATCTNNNGATCCCAGGTGTATCCTTTTCCA
AGCACACGTCTGAACTCTTTCCCTACACGACGCTCTTCCGATCTNNNGATCCCAGGTGTATCCTTTTCCA
GAGCACACGTCTGAACTCTTTCCCTACACGACGCTCTTCCGATCTNNNGTTGCAGTGAGCTGAGATCAT
GAGCACACGTCTGAACTCTTTCCCTACACGACGCTCTTCCGATCTNNNGTTGCAGTGAGCTGAGATCAT
GAGCACACGTCTGAACTCTTTCCCTACACGACGCTCTTCCGATCTNNNGTTGCAGTGAGCTGAGATCAT
AGCACACGTCTGAACTCTTTCCCTACACGACGCTCTTCCGATCTNNNCTGGGCAACAAGAGCAAAACT
AGCACACGTCTGAACTCTTTCCCTACACGACGCTCTTCCGATCTNNNCTGGGCAACAAGAGCAAAACT
AGCACACGTCTGAACTCTTTCCCTACACGACGCTCTTCCGATCTNNNCTGGGCAACAAGAGCAAAACT
GCACACGTCTGAACTCTTTCCCTACACGACGCTCTTCCGATCTNNNCCACGAAAACCCTAGTGAATCACC
GCACACGTCTGAACTCTTTCCCTACACGACGCTCTTCCGATCTNNNCCACGAAAACCCTAGTGAATCACC
GCACACGTCTGAACTCTTTCCCTACACGACGCTCTTCCGATCTNNNCCACGAAAACCCTAGTGAATCACC
AGAGCACACGTCTGAACTCTTTCCCTACACGACGCTCTTCCGATCTNNNTCAAATCTTGTGTTTGCTACC
AGAGCACACGTCTGAACTCTTTCCCTACACGACGCTCTTCCGATCTNNNTCAAATCTTGTGTTTGCTACC
AGAGCACACGTCTGAACTCTTTCCCTACACGACGCTCTTCCGATCTNNNTCAAATCTTGTGTTTGCTACC
AGCACACGTCTGAACTCTTTCCCTACACGACGCTCTTCCGATCTNNNTTCACTCCTTTTATTGCAGTGCC
AGCACACGTCTGAACTCTTTCCCTACACGACGCTCTTCCGATCTNNNTTCACTCCTTTTATTGCAGTGCC
AGCACACGTCTGAACTCTTTCCCTACACGACGCTCTTCCGATCTNNNTTCACTCCTTTTATTGCAGTGCC
AGCACACGTCTGAACTCTTTCCCTACACGACGCTCTTCCGATCTNNNATTTCACGAGTCACCCATCACAG
AGCACACGTCTGAACTCTTTCCCTACACGACGCTCTTCCGATCTNNNATTTCACGAGTCACCCATCACAG
AGCACACGTCTGAACTCTTTCCCTACACGACGCTCTTCCGATCTNNNATTTCACGAGTCACCCATCACAG
AGCACACGTCTGAACTCTTTCCCTACACGACGCTCTTCCGATCTNNNACGTGCACACATAGAAAATCAT
AGCACACGTCTGAACTCTTTCCCTACACGACGCTCTTCCGATCTNNNACGTGCACACATAGAAAATCAT
AGCACACGTCTGAACTCTTTCCCTACACGACGCTCTTCCGATCTNNNACGTGCACACATAGAAAATCAT
GAGCACACGTCTGAACTCTTTCCCTACACGACGCTCTTCCGATCTNNNGCCGGAGTAACAACAGCAAA
GAGCACACGTCTGAACTCTTTCCCTACACGACGCTCTTCCGATCTNNNGCCGGAGTAACAACAGCAAA
GAGCACACGTCTGAACTCTTTCCCTACACGACGCTCTTCCGATCTNNNGCCGGAGTAACAACAGCAAA
AGCACACGTCTGAACTCTTTCCCTACACGACGCTCTTCCGATCTNNNGTGGACCAATACCAGGGATAAC
AGCACACGTCTGAACTCTTTCCCTACACGACGCTCTTCCGATCTNNNGTGGACCAATACCAGGGATAAC
AGCACACGTCTGAACTCTTTCCCTACACGACGCTCTTCCGATCTNNNGTGGACCAATACCAGGGATAAC

GAGCACACGTCTGAACTCTTTCCCTACACGACGCTCTTCCGATCTNNNTTCTTGCTTCTTTGTCATTTGG
GAGCACACGTCTGAACTCTTTCCCTACACGACGCTCTTCCGATCTNNNTTCTTGCTTCTTTGTCATTTGG
GAGCACACGTCTGAACTCTTTCCCTACACGACGCTCTTCCGATCTNNNTTCTTGCTTCTTTGTCATTTGG
AGCACACGTCTGAACTCTTTCCCTACACGACGCTCTTCCGATCTNNNTTTGTGTGTGTGTGTGTGTGTA
AGCACACGTCTGAACTCTTTCCCTACACGACGCTCTTCCGATCTNNNTTTGTGTGTGTGTGTGTGTGTA
AGCACACGTCTGAACTCTTTCCCTACACGACGCTCTTCCGATCTNNNTTTGTGTGTGTGTGTGTGTGTA
AGAGCACACGTCTGAACTCTTTCCCTACACGACGCTCTTCCGATCTNNNGGGTCTGGGGTGGTCCTAA
AGAGCACACGTCTGAACTCTTTCCCTACACGACGCTCTTCCGATCTNNNGGGTCTGGGGTGGTCCTAA
AGAGCACACGTCTGAACTCTTTCCCTACACGACGCTCTTCCGATCTNNNGGGTCTGGGGTGGTCCTAA
AGCACACGTCTGAACTCTTTCCCTACACGACGCTCTTCCGATCTNNNGAGAATTGCTTGAACCTGGAG
AGCACACGTCTGAACTCTTTCCCTACACGACGCTCTTCCGATCTNNNGAGAATTGCTTGAACCTGGAG
AGCACACGTCTGAACTCTTTCCCTACACGACGCTCTTCCGATCTNNNGAGAATTGCTTGAACCTGGAG
AGCACACGTCTGAACTCTTTCCCTACACGACGCTCTTCCGATCTNNNTGGATGGTGCAACTAAAAACCA
AGCACACGTCTGAACTCTTTCCCTACACGACGCTCTTCCGATCTNNNTGGATGGTGCAACTAAAAACCA
AGCACACGTCTGAACTCTTTCCCTACACGACGCTCTTCCGATCTNNNTGGATGGTGCAACTAAAAACCA
AGCACACGTCTGAACTCTTTCCCTACACGACGCTCTTCCGATCTNNNTTATCACACCACTGTACTCCAG
AGCACACGTCTGAACTCTTTCCCTACACGACGCTCTTCCGATCTNNNTTATCACACCACTGTACTCCAG
AGCACACGTCTGAACTCTTTCCCTACACGACGCTCTTCCGATCTNNNTTATCACACCACTGTACTCCAG
GAGCACACGTCTGAACTCTTTCCCTACACGACGCTCTTCCGATCTNNNGGTGTGATGTTCTTTCTTTGT
GAGCACACGTCTGAACTCTTTCCCTACACGACGCTCTTCCGATCTNNNGGTGTGATGTTCTTTCTTTGT
GAGCACACGTCTGAACTCTTTCCCTACACGACGCTCTTCCGATCTNNNGGTGTGATGTTCTTTCTTTGT
GAGCACACGTCTGAACTCTTTCCCTACACGACGCTCTTCCGATCTNNNAGCAAGAGAGTACGACATTG
GAGCACACGTCTGAACTCTTTCCCTACACGACGCTCTTCCGATCTNNNAGCAAGAGAGTACGACATTG
GAGCACACGTCTGAACTCTTTCCCTACACGACGCTCTTCCGATCTNNNAGCAAGAGAGTACGACATTG
GAGCACACGTCTGAACTCTTTCCCTACACGACGCTCTTCCGATCTNNNGTAGGAGATTATAGGTGCCT
GAGCACACGTCTGAACTCTTTCCCTACACGACGCTCTTCCGATCTNNNGTAGGAGATTATAGGTGCCT
GAGCACACGTCTGAACTCTTTCCCTACACGACGCTCTTCCGATCTNNNGTAGGAGATTATAGGTGCCT
GAGCACACGTCTGAACTCTTTCCCTACACGACGCTCTTCCGATCTNNNTGAGGCAGGAAAATCGTTTGA
GAGCACACGTCTGAACTCTTTCCCTACACGACGCTCTTCCGATCTNNNTGAGGCAGGAAAATCGTTTGA
GAGCACACGTCTGAACTCTTTCCCTACACGACGCTCTTCCGATCTNNNTGAGGCAGGAAAATCGTTTGA
GAGCACACGTCTGAACTCTTTCCCTACACGACGCTCTTCCGATCTNNNAGCCAAGAACACAGAATGCAA
GAGCACACGTCTGAACTCTTTCCCTACACGACGCTCTTCCGATCTNNNAGCCAAGAACACAGAATGCAA
GAGCACACGTCTGAACTCTTTCCCTACACGACGCTCTTCCGATCTNNNAGCCAAGAACACAGAATGCAA
GCACACGTCTGAACTCTTTCCCTACACGACGCTCTTCCGATCTNNNTCGCAATAATCGTGTCCATCAAA
GCACACGTCTGAACTCTTTCCCTACACGACGCTCTTCCGATCTNNNTCGCAATAATCGTGTCCATCAAA
GCACACGTCTGAACTCTTTCCCTACACGACGCTCTTCCGATCTNNNTCGCAATAATCGTGTCCATCAAA
GAGCACACGTCTGAACTCTTTCCCTACACGACGCTCTTCCGATCTNNNAGAGTATTGCTTGAACCCAG
GAGCACACGTCTGAACTCTTTCCCTACACGACGCTCTTCCGATCTNNNAGAGTATTGCTTGAACCCAG
GAGCACACGTCTGAACTCTTTCCCTACACGACGCTCTTCCGATCTNNNAGAGTATTGCTTGAACCCAG
GCACACGTCTGAACTCTTTCCCTACACGACGCTCTTCCGATCTNNNTTCTTCTTCACTTCTGTTATAGTC
GCACACGTCTGAACTCTTTCCCTACACGACGCTCTTCCGATCTNNNTTCTTCTTCACTTCTGTTATAGTC
GCACACGTCTGAACTCTTTCCCTACACGACGCTCTTCCGATCTNNNTTCTTCTTCACTTCTGTTATAGTC
GAGCACACGTCTGAACTCTTTCCCTACACGACGCTCTTCCGATCTNNNCCCTGTGTCAGCACTCTCTTTA

GAGCACACGTCTGAACTCTTTCCCTACACGACGCTCTTCCGATCTNNNCCCTGTGTCAGCACTCTCTTTA
GAGCACACGTCTGAACTCTTTCCCTACACGACGCTCTTCCGATCTNNNCCCTGTGTCAGCACTCTCTTTA
GAGCACACGTCTGAACTCTTTCCCTACACGACGCTCTTCCGATCTNNNGTAATCTTGAACTCCTGGGCT
GAGCACACGTCTGAACTCTTTCCCTACACGACGCTCTTCCGATCTNNNGTAATCTTGAACTCCTGGGCT
GAGCACACGTCTGAACTCTTTCCCTACACGACGCTCTTCCGATCTNNNGTAATCTTGAACTCCTGGGCT
GAGCACACGTCTGAACTCTTTCCCTACACGACGCTCTTCCGATCTNNNGAGGTTGCAGTGAGTCGAGA
GAGCACACGTCTGAACTCTTTCCCTACACGACGCTCTTCCGATCTNNNGAGGTTGCAGTGAGTCGAGA
GAGCACACGTCTGAACTCTTTCCCTACACGACGCTCTTCCGATCTNNNGAGGTTGCAGTGAGTCGAGA
AGCACACGTCTGAACTCTTTCCCTACACGACGCTCTTCCGATCTNNNCCAGAAAACCAGCATTGCAGTA
AGCACACGTCTGAACTCTTTCCCTACACGACGCTCTTCCGATCTNNNCCAGAAAACCAGCATTGCAGTA
AGCACACGTCTGAACTCTTTCCCTACACGACGCTCTTCCGATCTNNNCCAGAAAACCAGCATTGCAGTA
AGCACACGTCTGAACTCTTTCCCTACACGACGCTCTTCCGATCTNNNCCAGGACATGAGTTCCCATAGA
AGCACACGTCTGAACTCTTTCCCTACACGACGCTCTTCCGATCTNNNCCAGGACATGAGTTCCCATAGA
AGCACACGTCTGAACTCTTTCCCTACACGACGCTCTTCCGATCTNNNCCAGGACATGAGTTCCCATAGA
GCACACGTCTGAACTCTTTCCCTACACGACGCTCTTCCGATCTNNNGAGGAACGAAAAGTGTCCAGTT
GCACACGTCTGAACTCTTTCCCTACACGACGCTCTTCCGATCTNNNGAGGAACGAAAAGTGTCCAGTT
GCACACGTCTGAACTCTTTCCCTACACGACGCTCTTCCGATCTNNNGAGGAACGAAAAGTGTCCAGTT
GAGCACACGTCTGAACTCTTTCCCTACACGACGCTCTTCCGATCTNNNAGTGAGCCAAGATAGTGCCA
GAGCACACGTCTGAACTCTTTCCCTACACGACGCTCTTCCGATCTNNNAGTGAGCCAAGATAGTGCCA
GAGCACACGTCTGAACTCTTTCCCTACACGACGCTCTTCCGATCTNNNAGTGAGCCAAGATAGTGCCA
GCACACGTCTGAACTCTTTCCCTACACGACGCTCTTCCGATCTNNNTTCACGGAAAATAAAAGTGGCC
GCACACGTCTGAACTCTTTCCCTACACGACGCTCTTCCGATCTNNNTTCACGGAAAATAAAAGTGGCC
GCACACGTCTGAACTCTTTCCCTACACGACGCTCTTCCGATCTNNNTTCACGGAAAATAAAAGTGGCC
AGCACACGTCTGAACTCTTTCCCTACACGACGCTCTTCCGATCTNNNCTCTTGGGCTTTGTAGGGTACTT
AGCACACGTCTGAACTCTTTCCCTACACGACGCTCTTCCGATCTNNNCTCTTGGGCTTTGTAGGGTACTT
AGCACACGTCTGAACTCTTTCCCTACACGACGCTCTTCCGATCTNNNCTCTTGGGCTTTGTAGGGTACTT
AGCACACGTCTGAACTCTTTCCCTACACGACGCTCTTCCGATCTNNNGTGAACAGTTGCTTGGGAAGAA
AGCACACGTCTGAACTCTTTCCCTACACGACGCTCTTCCGATCTNNNGTGAACAGTTGCTTGGGAAGAA
AGCACACGTCTGAACTCTTTCCCTACACGACGCTCTTCCGATCTNNNGTGAACAGTTGCTTGGGAAGAA
AGAGCACACGTCTGAACTCTTTCCCTACACGACGCTCTTCCGATCTNNNAGTAGACATCTGCTATTTTG
AGAGCACACGTCTGAACTCTTTCCCTACACGACGCTCTTCCGATCTNNNAGTAGACATCTGCTATTTTG
AGAGCACACGTCTGAACTCTTTCCCTACACGACGCTCTTCCGATCTNNNAGTAGACATCTGCTATTTTG
GAGCACACGTCTGAACTCTTTCCCTACACGACGCTCTTCCGATCTNNNAAGTGGCAGAACCTAGAGAG
GAGCACACGTCTGAACTCTTTCCCTACACGACGCTCTTCCGATCTNNNAAGTGGCAGAACCTAGAGAG
GAGCACACGTCTGAACTCTTTCCCTACACGACGCTCTTCCGATCTNNNAAGTGGCAGAACCTAGAGAG
AGCACACGTCTGAACTCTTTCCCTACACGACGCTCTTCCGATCTNNNTGCCCAGCTAATGTTTGTGTTTT
AGCACACGTCTGAACTCTTTCCCTACACGACGCTCTTCCGATCTNNNTGCCCAGCTAATGTTTGTGTTTT
AGCACACGTCTGAACTCTTTCCCTACACGACGCTCTTCCGATCTNNNTGCCCAGCTAATGTTTGTGTTTT
AGCACACGTCTGAACTCTTTCCCTACACGACGCTCTTCCGATCTNNNCTCTTTGAAGGAATTTGCAGGC
AGCACACGTCTGAACTCTTTCCCTACACGACGCTCTTCCGATCTNNNCTCTTTGAAGGAATTTGCAGGC
AGCACACGTCTGAACTCTTTCCCTACACGACGCTCTTCCGATCTNNNCTCTTTGAAGGAATTTGCAGGC
AGCACACGTCTGAACTCTTTCCCTACACGACGCTCTTCCGATCTNNNGCCCTAAGTGATGCCATGAGAT
AGCACACGTCTGAACTCTTTCCCTACACGACGCTCTTCCGATCTNNNGCCCTAAGTGATGCCATGAGAT

AGCACACGTCTGAACTCTTTCCCTACACGACGCTCTTCCGATCTNNNGCCCTAAGTGATGCCATGAGA
GAGCACACGTCTGAACTCTTTCCCTACACGACGCTCTTCCGATCTNNNATACCCAAGTTACCCTACCTG
GAGCACACGTCTGAACTCTTTCCCTACACGACGCTCTTCCGATCTNNNATACCCAAGTTACCCTACCTG
GAGCACACGTCTGAACTCTTTCCCTACACGACGCTCTTCCGATCTNNNATACCCAAGTTACCCTACCTG
GAGCACACGTCTGAACTCTTTCCCTACACGACGCTCTTCCGATCTNNNCACATCTGTGGTATTCTTCCC
GAGCACACGTCTGAACTCTTTCCCTACACGACGCTCTTCCGATCTNNNCACATCTGTGGTATTCTTCCC
GAGCACACGTCTGAACTCTTTCCCTACACGACGCTCTTCCGATCTNNNCACATCTGTGGTATTCTTCCC
AGCACACGTCTGAACTCTTTCCCTACACGACGCTCTTCCGATCTNNNGGCATGTACTGACGATTATGG
AGCACACGTCTGAACTCTTTCCCTACACGACGCTCTTCCGATCTNNNGGCATGTACTGACGATTATGG
AGCACACGTCTGAACTCTTTCCCTACACGACGCTCTTCCGATCTNNNGGCATGTACTGACGATTATGG
GAGCACACGTCTGAACTCTTTCCCTACACGACGCTCTTCCGATCTNNNTGAGACTCGTTATTTGTGGTC
GAGCACACGTCTGAACTCTTTCCCTACACGACGCTCTTCCGATCTNNNTGAGACTCGTTATTTGTGGTC
GAGCACACGTCTGAACTCTTTCCCTACACGACGCTCTTCCGATCTNNNTGAGACTCGTTATTTGTGGTC
AGCACACGTCTGAACTCTTTCCCTACACGACGCTCTTCCGATCTNNNTAACACCAGTGACTGTGTGCTTA
AGCACACGTCTGAACTCTTTCCCTACACGACGCTCTTCCGATCTNNNTAACACCAGTGACTGTGTGCTTA
AGCACACGTCTGAACTCTTTCCCTACACGACGCTCTTCCGATCTNNNTAACACCAGTGACTGTGTGCTTA
GCACACGTCTGAACTCTTTCCCTACACGACGCTCTTCCGATCTNNNTATACACACACACACACACACC
GCACACGTCTGAACTCTTTCCCTACACGACGCTCTTCCGATCTNNNTATACACACACACACACACACC
GCACACGTCTGAACTCTTTCCCTACACGACGCTCTTCCGATCTNNNTATACACACACACACACACACC
ACACGTCTGAACTCTTTCCCTACACGACGCTCTTCCGATCTNNNCTTCCTTCCTTCCTCATCACCAGCATC
ACACGTCTGAACTCTTTCCCTACACGACGCTCTTCCGATCTNNNCTTCCTTCCTTCCTCATCACCAGCATC
ACACGTCTGAACTCTTTCCCTACACGACGCTCTTCCGATCTNNNCTTCCTTCCTTCCTCATCACCAGCATC
AGCACACGTCTGAACTCTTTCCCTACACGACGCTCTTCCGATCTNNNATGTCTCATTATGTGCTCAGGC
AGCACACGTCTGAACTCTTTCCCTACACGACGCTCTTCCGATCTNNNATGTCTCATTATGTGCTCAGGC
AGCACACGTCTGAACTCTTTCCCTACACGACGCTCTTCCGATCTNNNATGTCTCATTATGTGCTCAGGC
AGCACACGTCTGAACTCTTTCCCTACACGACGCTCTTCCGATCTNNNCTGTAATCCCAGCTACTCAGGA
AGCACACGTCTGAACTCTTTCCCTACACGACGCTCTTCCGATCTNNNCTGTAATCCCAGCTACTCAGGA
AGCACACGTCTGAACTCTTTCCCTACACGACGCTCTTCCGATCTNNNCTGTAATCCCAGCTACTCAGGA
GCACACGTCTGAACTCTTTCCCTACACGACGCTCTTCCGATCTNNNGGAACTGCAACTATCAACACCAT
GCACACGTCTGAACTCTTTCCCTACACGACGCTCTTCCGATCTNNNGGAACTGCAACTATCAACACCAT
GCACACGTCTGAACTCTTTCCCTACACGACGCTCTTCCGATCTNNNGGAACTGCAACTATCAACACCAT
AGCACACGTCTGAACTCTTTCCCTACACGACGCTCTTCCGATCTNNNCCGGATAACTTCCACTGCTAAC
AGCACACGTCTGAACTCTTTCCCTACACGACGCTCTTCCGATCTNNNCCGGATAACTTCCACTGCTAAC
AGCACACGTCTGAACTCTTTCCCTACACGACGCTCTTCCGATCTNNNCCGGATAACTTCCACTGCTAAC
GAGCACACGTCTGAACTCTTTCCCTACACGACGCTCTTCCGATCTNNNCTGGAGTTCAGTGGTGTGAT
GAGCACACGTCTGAACTCTTTCCCTACACGACGCTCTTCCGATCTNNNCTGGAGTTCAGTGGTGTGAT
GAGCACACGTCTGAACTCTTTCCCTACACGACGCTCTTCCGATCTNNNCTGGAGTTCAGTGGTGTGAT
GAGCACACGTCTGAACTCTTTCCCTACACGACGCTCTTCCGATCTNNNTATAGTCCCAGCTACTCAGGA
GAGCACACGTCTGAACTCTTTCCCTACACGACGCTCTTCCGATCTNNNTATAGTCCCAGCTACTCAGGA
GAGCACACGTCTGAACTCTTTCCCTACACGACGCTCTTCCGATCTNNNTATAGTCCCAGCTACTCAGGA
GCACACGTCTGAACTCTTTCCCTACACGACGCTCTTCCGATCTNNNAACCCCTTGCATAATGAAGAGG
GCACACGTCTGAACTCTTTCCCTACACGACGCTCTTCCGATCTNNNAACCCCTTGCATAATGAAGAGG
GCACACGTCTGAACTCTTTCCCTACACGACGCTCTTCCGATCTNNNAACCCCTTGCATAATGAAGAGG

GAGCACACGTCTGAACTCTTTCCCTACACGACGCTCTTCCGATCTNNNCTGAGGCAGGAGAATCGTTT

GAGCACACGTCTGAACTCTTTCCCTACACGACGCTCTTCCGATCTNNNCTGAGGCAGGAGAATCGTTT

GAGCACACGTCTGAACTCTTTCCCTACACGACGCTCTTCCGATCTNNNCTGAGGCAGGAGAATCGTTT

AGCACACGTCTGAACTCTTTCCCTACACGACGCTCTTCCGATCTNNNTGTGCTGTTTTGGTTACTGTAG

AGCACACGTCTGAACTCTTTCCCTACACGACGCTCTTCCGATCTNNNTGTGCTGTTTTGGTTACTGTAG

AGCACACGTCTGAACTCTTTCCCTACACGACGCTCTTCCGATCTNNNTGTGCTGTTTTGGTTACTGTAG

AGCACACGTCTGAACTCTTTCCCTACACGACGCTCTTCCGATCTNNNGGATGTTTGTGGTCTAACAAG

AGCACACGTCTGAACTCTTTCCCTACACGACGCTCTTCCGATCTNNNGGATGTTTGTGGTCTAACAAG

AGCACACGTCTGAACTCTTTCCCTACACGACGCTCTTCCGATCTNNNGGATGTTTGTGGTCTAACAAG

AGCACACGTCTGAACTCTTTCCCTACACGACGCTCTTCCGATCTNNNAGCATTTATCACCCACTAGCACA

AGCACACGTCTGAACTCTTTCCCTACACGACGCTCTTCCGATCTNNNAGCATTTATCACCCACTAGCACA

AGCACACGTCTGAACTCTTTCCCTACACGACGCTCTTCCGATCTNNNAGCATTTATCACCCACTAGCACA

CACACGTCTGAACTCTTTCCCTACACGACGCTCTTCCGATCTNNNGCCTCCCGATTTCTAAGAAGGCAT

CACACGTCTGAACTCTTTCCCTACACGACGCTCTTCCGATCTNNNGCCTCCCGATTTCTAAGAAGGCAT

CACACGTCTGAACTCTTTCCCTACACGACGCTCTTCCGATCTNNNGCCTCCCGATTTCTAAGAAGGCAT

AGCACACGTCTGAACTCTTTCCCTACACGACGCTCTTCCGATCTNNNTTCTTTTCAGGGAGACTCACCA

AGCACACGTCTGAACTCTTTCCCTACACGACGCTCTTCCGATCTNNNTTCTTTTCAGGGAGACTCACCA

AGCACACGTCTGAACTCTTTCCCTACACGACGCTCTTCCGATCTNNNTTCTTTTCAGGGAGACTCACCA

AGCACACGTCTGAACTCTTTCCCTACACGACGCTCTTCCGATCTNNNGCCGTGAGAAAAGATCATGTG

AGCACACGTCTGAACTCTTTCCCTACACGACGCTCTTCCGATCTNNNGCCGTGAGAAAAGATCATGTG

AGCACACGTCTGAACTCTTTCCCTACACGACGCTCTTCCGATCTNNNGCCGTGAGAAAAGATCATGTG

AGCACACGTCTGAACTCTTTCCCTACACGACGCTCTTCCGATCTNNNCCCATCTCTGCAAAAACACAA

AGCACACGTCTGAACTCTTTCCCTACACGACGCTCTTCCGATCTNNNCCCATCTCTGCAAAAACACAA

AGCACACGTCTGAACTCTTTCCCTACACGACGCTCTTCCGATCTNNNCCCATCTCTGCAAAAACACAA

AGCACACGTCTGAACTCTTTCCCTACACGACGCTCTTCCGATCTNNNCCAGTTGGGTGGTTACAAGTA

AGCACACGTCTGAACTCTTTCCCTACACGACGCTCTTCCGATCTNNNCCAGTTGGGTGGTTACAAGTA

AGCACACGTCTGAACTCTTTCCCTACACGACGCTCTTCCGATCTNNNCCAGTTGGGTGGTTACAAGTA

GAGCACACGTCTGAACTCTTTCCCTACACGACGCTCTTCCGATCTNNNGAGACATGAGAATCGCTTCA

GAGCACACGTCTGAACTCTTTCCCTACACGACGCTCTTCCGATCTNNNGAGACATGAGAATCGCTTCA

GAGCACACGTCTGAACTCTTTCCCTACACGACGCTCTTCCGATCTNNNGAGACATGAGAATCGCTTCA

AGCACACGTCTGAACTCTTTCCCTACACGACGCTCTTCCGATCTNNNGCAAGAGAATGGCATGAACCT

AGCACACGTCTGAACTCTTTCCCTACACGACGCTCTTCCGATCTNNNGCAAGAGAATGGCATGAACCT

AGCACACGTCTGAACTCTTTCCCTACACGACGCTCTTCCGATCTNNNGCAAGAGAATGGCATGAACCT

AGCACACGTCTGAACTCTTTCCCTACACGACGCTCTTCCGATCTNNNGTACTTGAGGTGGTGGAGCTA

AGCACACGTCTGAACTCTTTCCCTACACGACGCTCTTCCGATCTNNNGTACTTGAGGTGGTGGAGCTA

AGCACACGTCTGAACTCTTTCCCTACACGACGCTCTTCCGATCTNNNGTACTTGAGGTGGTGGAGCTA

GAGCACACGTCTGAACTCTTTCCCTACACGACGCTCTTCCGATCTNNNTACAAAAATTAGCCAGCGTG

GAGCACACGTCTGAACTCTTTCCCTACACGACGCTCTTCCGATCTNNNTACAAAAATTAGCCAGCGTG

GAGCACACGTCTGAACTCTTTCCCTACACGACGCTCTTCCGATCTNNNTACAAAAATTAGCCAGCGTG

GCACACGTCTGAACTCTTTCCCTACACGACGCTCTTCCGATCTNNNATATAATCTGCCCCCTTGTGGAC

GCACACGTCTGAACTCTTTCCCTACACGACGCTCTTCCGATCTNNNATATAATCTGCCCCCTTGTGGAC

GCACACGTCTGAACTCTTTCCCTACACGACGCTCTTCCGATCTNNNATATAATCTGCCCCCTTGTGGAC

GAGCACACGTCTGAACTCTTTCCCTACACGACGCTCTTCCGATCTNNNATACCTATCCCCAGCAGCTAG

GAGCACACGTCTGAACTCTTTCCCTACACGACGCTCTTCCGATCTNNNATACCTATCCCCAGCAGCTAG

GAGCACACGTCTGAACTCTTTCCCTACACGACGCTCTTCCGATCTNNNATACCTATCCCCAGCAGCTAG

AGCACACGTCTGAACTCTTTCCCTACACGACGCTCTTCCGATCTNNNTATCTTGGCCTTGCAAGCTAG

AGCACACGTCTGAACTCTTTCCCTACACGACGCTCTTCCGATCTNNNTATCTTGGCCTTGCAAGCTAG

AGCACACGTCTGAACTCTTTCCCTACACGACGCTCTTCCGATCTNNNTATCTTGGCCTTGCAAGCTAG

AGCACACGTCTGAACTCTTTCCCTACACGACGCTCTTCCGATCTNNNGCAGGCTGGAGAGAATGTTT

AGCACACGTCTGAACTCTTTCCCTACACGACGCTCTTCCGATCTNNNGCAGGCTGGAGAGAATGTTT

AGCACACGTCTGAACTCTTTCCCTACACGACGCTCTTCCGATCTNNNGCAGGCTGGAGAGAATGTTT

AGCACACGTCTGAACTCTTTCCCTACACGACGCTCTTCCGATCTNNNTCCCAACTAGCTGGAATTACA

AGCACACGTCTGAACTCTTTCCCTACACGACGCTCTTCCGATCTNNNTCCCAACTAGCTGGAATTACA

AGCACACGTCTGAACTCTTTCCCTACACGACGCTCTTCCGATCTNNNTCCCAACTAGCTGGAATTACA

AGCACACGTCTGAACTCTTTCCCTACACGACGCTCTTCCGATCTNNNATTGTCACCACACTCACCCATC

AGCACACGTCTGAACTCTTTCCCTACACGACGCTCTTCCGATCTNNNATTGTCACCACACTCACCCATC

AGCACACGTCTGAACTCTTTCCCTACACGACGCTCTTCCGATCTNNNATTGTCACCACACTCACCCATC

CACACGTCTGAACTCTTTCCCTACACGACGCTCTTCCGATCTNNNCTGAGAACATGGAGGGTCCGCAT

CACACGTCTGAACTCTTTCCCTACACGACGCTCTTCCGATCTNNNCTGAGAACATGGAGGGTCCGCAT

CACACGTCTGAACTCTTTCCCTACACGACGCTCTTCCGATCTNNNCTGAGAACATGGAGGGTCCGCAT

AGCACACGTCTGAACTCTTTCCCTACACGACGCTCTTCCGATCTNNNCACAAAACAAACAAAAGGCCA

AGCACACGTCTGAACTCTTTCCCTACACGACGCTCTTCCGATCTNNNCACAAAACAAACAAAAGGCCA

AGCACACGTCTGAACTCTTTCCCTACACGACGCTCTTCCGATCTNNNCACAAAACAAACAAAAGGCCA

AGCACACGTCTGAACTCTTTCCCTACACGACGCTCTTCCGATCTNNNACAAAAATTAGCTGGGTGTG

AGCACACGTCTGAACTCTTTCCCTACACGACGCTCTTCCGATCTNNNACAAAAATTAGCTGGGTGTG

AGCACACGTCTGAACTCTTTCCCTACACGACGCTCTTCCGATCTNNNACAAAAATTAGCTGGGTGTG

AGCACACGTCTGAACTCTTTCCCTACACGACGCTCTTCCGATCTNNNCTCATGCCTGTAATCCCAGCTA

AGCACACGTCTGAACTCTTTCCCTACACGACGCTCTTCCGATCTNNNCTCATGCCTGTAATCCCAGCTA

AGCACACGTCTGAACTCTTTCCCTACACGACGCTCTTCCGATCTNNNCTCATGCCTGTAATCCCAGCTA

AGCACACGTCTGAACTCTTTCCCTACACGACGCTCTTCCGATCTNNNGACTGGACCACATCTACCCAAA

AGCACACGTCTGAACTCTTTCCCTACACGACGCTCTTCCGATCTNNNGACTGGACCACATCTACCCAAA

AGCACACGTCTGAACTCTTTCCCTACACGACGCTCTTCCGATCTNNNGACTGGACCACATCTACCCAAA

GCACACGTCTGAACTCTTTCCCTACACGACGCTCTTCCGATCTNNNGGGAAGCGACTCTTAGACCTAA

GCACACGTCTGAACTCTTTCCCTACACGACGCTCTTCCGATCTNNNGGGAAGCGACTCTTAGACCTAA

GCACACGTCTGAACTCTTTCCCTACACGACGCTCTTCCGATCTNNNGGGAAGCGACTCTTAGACCTAA

GCACACGTCTGAACTCTTTCCCTACACGACGCTCTTCCGATCTNNNTTTTTGGCTGCATAAGTGTCTTC

GCACACGTCTGAACTCTTTCCCTACACGACGCTCTTCCGATCTNNNTTTTTGGCTGCATAAGTGTCTTC

GCACACGTCTGAACTCTTTCCCTACACGACGCTCTTCCGATCTNNNTTTTTGGCTGCATAAGTGTCTTC

GAGCACACGTCTGAACTCTTTCCCTACACGACGCTCTTCCGATCTNNNGTGGTTGCTTTGAAGAGTG

GAGCACACGTCTGAACTCTTTCCCTACACGACGCTCTTCCGATCTNNNGTGGTTGCTTTGAAGAGTG

GAGCACACGTCTGAACTCTTTCCCTACACGACGCTCTTCCGATCTNNNGTGGTTGCTTTGAAGAGTG

GAGCACACGTCTGAACTCTTTCCCTACACGACGCTCTTCCGATCTNNNAGACTGCATCTTTCCTCTTGG

GAGCACACGTCTGAACTCTTTCCCTACACGACGCTCTTCCGATCTNNNAGACTGCATCTTTCCTCTTGG

GAGCACACGTCTGAACTCTTTCCCTACACGACGCTCTTCCGATCTNNNAGACTGCATCTTTCCTCTTGG

GCACACGTCTGAACTCTTTCCCTACACGACGCTCTTCCGATCTNNNCACATCTGCATGGTCTCCTCTGC

GCACACGTCTGAACTCTTTCCCTACACGACGCTCTTCCGATCTNNNCACATCTGCATGGTCTCCTCTGC

AGCACACGTCTGAACTCTTTCCCTACACGACGCTCTTCCGATCTNNNCACATCTGCATGGTCTCCTCTGC
GAGCACACGTCTGAACTCTTTCCCTACACGACGCTCTTCCGATCTNNNGCATGCGTACGAGATCTTTTG
GAGCACACGTCTGAACTCTTTCCCTACACGACGCTCTTCCGATCTNNNGCATGCGTACGAGATCTTTTG
GAGCACACGTCTGAACTCTTTCCCTACACGACGCTCTTCCGATCTNNNGCATGCGTACGAGATCTTTTG
AGCACACGTCTGAACTCTTTCCCTACACGACGCTCTTCCGATCTNNNTGCCTTCTGACTTAGGTTCTGAC
AGCACACGTCTGAACTCTTTCCCTACACGACGCTCTTCCGATCTNNNTGCCTTCTGACTTAGGTTCTGAC
AGCACACGTCTGAACTCTTTCCCTACACGACGCTCTTCCGATCTNNNTGCCTTCTGACTTAGGTTCTGAC
GAGCACACGTCTGAACTCTTTCCCTACACGACGCTCTTCCGATCTNNNAGAGGTAGGAGAATTGCTTG
GAGCACACGTCTGAACTCTTTCCCTACACGACGCTCTTCCGATCTNNNAGAGGTAGGAGAATTGCTTG
GAGCACACGTCTGAACTCTTTCCCTACACGACGCTCTTCCGATCTNNNAGAGGTAGGAGAATTGCTTG
AGCACACGTCTGAACTCTTTCCCTACACGACGCTCTTCCGATCTNNNGATAGCTTCCTTGCTTTCTGCTG
AGCACACGTCTGAACTCTTTCCCTACACGACGCTCTTCCGATCTNNNGATAGCTTCCTTGCTTTCTGCTG
AGCACACGTCTGAACTCTTTCCCTACACGACGCTCTTCCGATCTNNNGATAGCTTCCTTGCTTTCTGCTG
GAGCACACGTCTGAACTCTTTCCCTACACGACGCTCTTCCGATCTNNNACCCATCAGCCATTTTGTCAAT
GAGCACACGTCTGAACTCTTTCCCTACACGACGCTCTTCCGATCTNNNACCCATCAGCCATTTTGTCAAT
GAGCACACGTCTGAACTCTTTCCCTACACGACGCTCTTCCGATCTNNNACCCATCAGCCATTTTGTCAAT
AGAGCACACGTCTGAACTCTTTCCCTACACGACGCTCTTCCGATCTNNNGGAAACTGAAATGGAAAGG
AGAGCACACGTCTGAACTCTTTCCCTACACGACGCTCTTCCGATCTNNNGGAAACTGAAATGGAAAGG
AGAGCACACGTCTGAACTCTTTCCCTACACGACGCTCTTCCGATCTNNNGGAAACTGAAATGGAAAGG
GAGCACACGTCTGAACTCTTTCCCTACACGACGCTCTTCCGATCTNNNTAAACTGTGTGCGGGCCTAG
GAGCACACGTCTGAACTCTTTCCCTACACGACGCTCTTCCGATCTNNNTAAACTGTGTGCGGGCCTAG
GAGCACACGTCTGAACTCTTTCCCTACACGACGCTCTTCCGATCTNNNTAAACTGTGTGCGGGCCTAG
GAGCACACGTCTGAACTCTTTCCCTACACGACGCTCTTCCGATCTNNNCAGAAATTCAGGGACTCCACA
GAGCACACGTCTGAACTCTTTCCCTACACGACGCTCTTCCGATCTNNNCAGAAATTCAGGGACTCCACA
GAGCACACGTCTGAACTCTTTCCCTACACGACGCTCTTCCGATCTNNNCAGAAATTCAGGGACTCCACA
GAGCACACGTCTGAACTCTTTCCCTACACGACGCTCTTCCGATCTNNNAGAGGTTGCAGTGAGCCTAAA
GAGCACACGTCTGAACTCTTTCCCTACACGACGCTCTTCCGATCTNNNAGAGGTTGCAGTGAGCCTAAA
GAGCACACGTCTGAACTCTTTCCCTACACGACGCTCTTCCGATCTNNNAGAGGTTGCAGTGAGCCTAAA
AGAGCACACGTCTGAACTCTTTCCCTACACGACGCTCTTCCGATCTNNNTAGTCCCAACTACTCAGGAG
AGAGCACACGTCTGAACTCTTTCCCTACACGACGCTCTTCCGATCTNNNTAGTCCCAACTACTCAGGAG
AGAGCACACGTCTGAACTCTTTCCCTACACGACGCTCTTCCGATCTNNNTAGTCCCAACTACTCAGGAG
AGCACACGTCTGAACTCTTTCCCTACACGACGCTCTTCCGATCTNNNATTACAACTTGGGGTTTGCTGT
AGCACACGTCTGAACTCTTTCCCTACACGACGCTCTTCCGATCTNNNATTACAACTTGGGGTTTGCTGT
AGCACACGTCTGAACTCTTTCCCTACACGACGCTCTTCCGATCTNNNATTACAACTTGGGGTTTGCTGT
AGAGCACACGTCTGAACTCTTTCCCTACACGACGCTCTTCCGATCTNNNGTGTTGGAGTTGAAGAAGG
AGAGCACACGTCTGAACTCTTTCCCTACACGACGCTCTTCCGATCTNNNGTGTTGGAGTTGAAGAAGG
AGAGCACACGTCTGAACTCTTTCCCTACACGACGCTCTTCCGATCTNNNGTGTTGGAGTTGAAGAAGG
AGCACACGTCTGAACTCTTTCCCTACACGACGCTCTTCCGATCTNNNAGGATTTTTCTTCCACGTCTGGA
AGCACACGTCTGAACTCTTTCCCTACACGACGCTCTTCCGATCTNNNAGGATTTTTCTTCCACGTCTGGA
AGCACACGTCTGAACTCTTTCCCTACACGACGCTCTTCCGATCTNNNAGGATTTTTCTTCCACGTCTGGA
GAGCACACGTCTGAACTCTTTCCCTACACGACGCTCTTCCGATCTNNNCTCTTCCCCTTGGTTTGTAACAT
GAGCACACGTCTGAACTCTTTCCCTACACGACGCTCTTCCGATCTNNNCTCTTCCCCTTGGTTTGTAACAT
GAGCACACGTCTGAACTCTTTCCCTACACGACGCTCTTCCGATCTNNNCTCTTCCCCTTGGTTTGTAACAT

AGAGCACACGTCTGAACTCTTTCCCTACACGACGCTCTTCCGATCTNNNCAGTCCGAGATGACAGGTA
AGAGCACACGTCTGAACTCTTTCCCTACACGACGCTCTTCCGATCTNNNCAGTCCGAGATGACAGGTA
AGAGCACACGTCTGAACTCTTTCCCTACACGACGCTCTTCCGATCTNNNCAGTCCGAGATGACAGGTA
GCACACGTCTGAACTCTTTCCCTACACGACGCTCTTCCGATCTNNNTCTTTGGATTACCAGGCATGGA
GCACACGTCTGAACTCTTTCCCTACACGACGCTCTTCCGATCTNNNTCTTTGGATTACCAGGCATGGA
GCACACGTCTGAACTCTTTCCCTACACGACGCTCTTCCGATCTNNNTCTTTGGATTACCAGGCATGGA
AGCACACGTCTGAACTCTTTCCCTACACGACGCTCTTCCGATCTNNNCATGATCTCAGCTCACTGCAAG
AGCACACGTCTGAACTCTTTCCCTACACGACGCTCTTCCGATCTNNNCATGATCTCAGCTCACTGCAAG
AGCACACGTCTGAACTCTTTCCCTACACGACGCTCTTCCGATCTNNNCATGATCTCAGCTCACTGCAAG
AGCACACGTCTGAACTCTTTCCCTACACGACGCTCTTCCGATCTNNNTATATGAAAAAGGCACCTGCA
AGCACACGTCTGAACTCTTTCCCTACACGACGCTCTTCCGATCTNNNTATATGAAAAAGGCACCTGCA
AGCACACGTCTGAACTCTTTCCCTACACGACGCTCTTCCGATCTNNNTATATGAAAAAGGCACCTGCA
AGCACACGTCTGAACTCTTTCCCTACACGACGCTCTTCCGATCTNNNGTCCAATTTCAGTGGAAGCAG
AGCACACGTCTGAACTCTTTCCCTACACGACGCTCTTCCGATCTNNNGTCCAATTTCAGTGGAAGCAG
AGCACACGTCTGAACTCTTTCCCTACACGACGCTCTTCCGATCTNNNGTCCAATTTCAGTGGAAGCAG
AGAGCACACGTCTGAACTCTTTCCCTACACGACGCTCTTCCGATCTNNNAGAGTATGATCCCTGTTTGA
AGAGCACACGTCTGAACTCTTTCCCTACACGACGCTCTTCCGATCTNNNAGAGTATGATCCCTGTTTGA
AGAGCACACGTCTGAACTCTTTCCCTACACGACGCTCTTCCGATCTNNNAGAGTATGATCCCTGTTTGA
GCACACGTCTGAACTCTTTCCCTACACGACGCTCTTCCGATCTNNNGTAGCCAGCAATGCATCAATAC
GCACACGTCTGAACTCTTTCCCTACACGACGCTCTTCCGATCTNNNGTAGCCAGCAATGCATCAATAC
GCACACGTCTGAACTCTTTCCCTACACGACGCTCTTCCGATCTNNNGTAGCCAGCAATGCATCAATAC
AGCACACGTCTGAACTCTTTCCCTACACGACGCTCTTCCGATCTNNNCACTCCGACACGAGGGAATAG
AGCACACGTCTGAACTCTTTCCCTACACGACGCTCTTCCGATCTNNNCACTCCGACACGAGGGAATAG
AGCACACGTCTGAACTCTTTCCCTACACGACGCTCTTCCGATCTNNNCACTCCGACACGAGGGAATAG
AGCACACGTCTGAACTCTTTCCCTACACGACGCTCTTCCGATCTNNNGAAGAACGGAGCCACCTTCCC
AGCACACGTCTGAACTCTTTCCCTACACGACGCTCTTCCGATCTNNNGAAGAACGGAGCCACCTTCCC
AGCACACGTCTGAACTCTTTCCCTACACGACGCTCTTCCGATCTNNNGAAGAACGGAGCCACCTTCCC
AGCACACGTCTGAACTCTTTCCCTACACGACGCTCTTCCGATCTNNNCTGTCCACAAATACCATGATAT
AGCACACGTCTGAACTCTTTCCCTACACGACGCTCTTCCGATCTNNNCTGTCCACAAATACCATGATAT
AGCACACGTCTGAACTCTTTCCCTACACGACGCTCTTCCGATCTNNNCTGTCCACAAATACCATGATAT
AGCACACGTCTGAACTCTTTCCCTACACGACGCTCTTCCGATCTNNNTGGTGATGGACCTGCTCAATAT
AGCACACGTCTGAACTCTTTCCCTACACGACGCTCTTCCGATCTNNNTGGTGATGGACCTGCTCAATAT
AGCACACGTCTGAACTCTTTCCCTACACGACGCTCTTCCGATCTNNNTGGTGATGGACCTGCTCAATAT
AGCACACGTCTGAACTCTTTCCCTACACGACGCTCTTCCGATCTNNNCACACACACACACACACACATAT
AGCACACGTCTGAACTCTTTCCCTACACGACGCTCTTCCGATCTNNNCACACACACACACACACACATAT
AGCACACGTCTGAACTCTTTCCCTACACGACGCTCTTCCGATCTNNNCACACACACACACACACACATAT
AGCACACGTCTGAACTCTTTCCCTACACGACGCTCTTCCGATCTNNNACGTAGTGTGATGCTTCCAGA
AGCACACGTCTGAACTCTTTCCCTACACGACGCTCTTCCGATCTNNNACGTAGTGTGATGCTTCCAGA
AGCACACGTCTGAACTCTTTCCCTACACGACGCTCTTCCGATCTNNNACGTAGTGTGATGCTTCCAGA
AGCACACGTCTGAACTCTTTCCCTACACGACGCTCTTCCGATCTNNNTGTCCTTTTTGTAGAGACACGG
AGCACACGTCTGAACTCTTTCCCTACACGACGCTCTTCCGATCTNNNTGTCCTTTTTGTAGAGACACGG
AGCACACGTCTGAACTCTTTCCCTACACGACGCTCTTCCGATCTNNNTGTCCTTTTTGTAGAGACACGG
AGCACACGTCTGAACTCTTTCCCTACACGACGCTCTTCCGATCTNNNAACCTCCTGAGTAGCTGGAACT

AGCACACGTCTGAACTCTTTCCCTACACGACGCTCTTCCGATCTNNNAACCTCCTGAGTAGCTGGAACT
AGCACACGTCTGAACTCTTTCCCTACACGACGCTCTTCCGATCTNNNAACCTCCTGAGTAGCTGGAACT
GAGCACACGTCTGAACTCTTTCCCTACACGACGCTCTTCCGATCTNNNCTCCTGAGTAGCTGGGATTAC
GAGCACACGTCTGAACTCTTTCCCTACACGACGCTCTTCCGATCTNNNCTCCTGAGTAGCTGGGATTAC
GAGCACACGTCTGAACTCTTTCCCTACACGACGCTCTTCCGATCTNNNCTCCTGAGTAGCTGGGATTAC
GAGCACACGTCTGAACTCTTTCCCTACACGACGCTCTTCCGATCTNNNTGAGGGAGGAGAATCACTTGA
GAGCACACGTCTGAACTCTTTCCCTACACGACGCTCTTCCGATCTNNNTGAGGGAGGAGAATCACTTGA
GAGCACACGTCTGAACTCTTTCCCTACACGACGCTCTTCCGATCTNNNTGAGGGAGGAGAATCACTTGA
GAGCACACGTCTGAACTCTTTCCCTACACGACGCTCTTCCGATCTNNNGAGAATCGCTTGAACCTGGGA
GAGCACACGTCTGAACTCTTTCCCTACACGACGCTCTTCCGATCTNNNGAGAATCGCTTGAACCTGGGA
GAGCACACGTCTGAACTCTTTCCCTACACGACGCTCTTCCGATCTNNNGAGAATCGCTTGAACCTGGGA
AGAGCACACGTCTGAACTCTTTCCCTACACGACGCTCTTCCGATCTNNNACATGACCCCTTTCTTTGTGCA
AGAGCACACGTCTGAACTCTTTCCCTACACGACGCTCTTCCGATCTNNNACATGACCCCTTTCTTTGTGCA
AGAGCACACGTCTGAACTCTTTCCCTACACGACGCTCTTCCGATCTNNNACATGACCCCTTTCTTTGTGCA
AGCACACGTCTGAACTCTTTCCCTACACGACGCTCTTCCGATCTNNNGATTGCTTGAGTCCAGGAGGT
AGCACACGTCTGAACTCTTTCCCTACACGACGCTCTTCCGATCTNNNGATTGCTTGAGTCCAGGAGGT
AGCACACGTCTGAACTCTTTCCCTACACGACGCTCTTCCGATCTNNNGATTGCTTGAGTCCAGGAGGT
AGCACACGTCTGAACTCTTTCCCTACACGACGCTCTTCCGATCTNNNAGTGGATGTTCTTGGCTCATCAT
AGCACACGTCTGAACTCTTTCCCTACACGACGCTCTTCCGATCTNNNAGTGGATGTTCTTGGCTCATCAT
AGCACACGTCTGAACTCTTTCCCTACACGACGCTCTTCCGATCTNNNAGTGGATGTTCTTGGCTCATCAT
AGCACACGTCTGAACTCTTTCCCTACACGACGCTCTTCCGATCTNNNGAGGCAGGAGAATTGCTTGAAC
AGCACACGTCTGAACTCTTTCCCTACACGACGCTCTTCCGATCTNNNGAGGCAGGAGAATTGCTTGAAC
AGCACACGTCTGAACTCTTTCCCTACACGACGCTCTTCCGATCTNNNGAGGCAGGAGAATTGCTTGAAC
GAGCACACGTCTGAACTCTTTCCCTACACGACGCTCTTCCGATCTNNNGAGATCATGCCACTTCACTCCA
GAGCACACGTCTGAACTCTTTCCCTACACGACGCTCTTCCGATCTNNNGAGATCATGCCACTTCACTCCA
GAGCACACGTCTGAACTCTTTCCCTACACGACGCTCTTCCGATCTNNNGAGATCATGCCACTTCACTCCA
AGCACACGTCTGAACTCTTTCCCTACACGACGCTCTTCCGATCTNNNGGGTCAGGAAGAAGAGGGAAA
AGCACACGTCTGAACTCTTTCCCTACACGACGCTCTTCCGATCTNNNGGGTCAGGAAGAAGAGGGAAA
AGCACACGTCTGAACTCTTTCCCTACACGACGCTCTTCCGATCTNNNGGGTCAGGAAGAAGAGGGAAA
GAGCACACGTCTGAACTCTTTCCCTACACGACGCTCTTCCGATCTNNNTCTGCAGATGTGGAATGTGAC
GAGCACACGTCTGAACTCTTTCCCTACACGACGCTCTTCCGATCTNNNTCTGCAGATGTGGAATGTGAC
GAGCACACGTCTGAACTCTTTCCCTACACGACGCTCTTCCGATCTNNNTCTGCAGATGTGGAATGTGAC
GAGCACACGTCTGAACTCTTTCCCTACACGACGCTCTTCCGATCTNNNAAGAAATTGCCCCAGAGGAG
GAGCACACGTCTGAACTCTTTCCCTACACGACGCTCTTCCGATCTNNNAAGAAATTGCCCCAGAGGAG
GAGCACACGTCTGAACTCTTTCCCTACACGACGCTCTTCCGATCTNNNAAGAAATTGCCCCAGAGGAG
GAGCACACGTCTGAACTCTTTCCCTACACGACGCTCTTCCGATCTNNNTGATGTACACCTGTGATACCA
GAGCACACGTCTGAACTCTTTCCCTACACGACGCTCTTCCGATCTNNNTGATGTACACCTGTGATACCA
GAGCACACGTCTGAACTCTTTCCCTACACGACGCTCTTCCGATCTNNNTGATGTACACCTGTGATACCA
GCACACGTCTGAACTCTTTCCCTACACGACGCTCTTCCGATCTNNNGTACAAGGGTTCCCTTTTCTCCAC
GCACACGTCTGAACTCTTTCCCTACACGACGCTCTTCCGATCTNNNGTACAAGGGTTCCCTTTTCTCCAC
GCACACGTCTGAACTCTTTCCCTACACGACGCTCTTCCGATCTNNNGTACAAGGGTTCCCTTTTCTCCAC
AGCACACGTCTGAACTCTTTCCCTACACGACGCTCTTCCGATCTNNNCCGGAAAATGCCTAGTTTGACAA
AGCACACGTCTGAACTCTTTCCCTACACGACGCTCTTCCGATCTNNNCCGGAAAATGCCTAGTTTGACAA

GCACACGTCTGAACTCTTTCCCTACACGACGCTCTTCCGATCTNNNCCGGAAAATGCCTAGTTTGACA
GCACACGTCTGAACTCTTTCCCTACACGACGCTCTTCCGATCTNNNAAGCAGGAGAATCACTTGAAC
GCACACGTCTGAACTCTTTCCCTACACGACGCTCTTCCGATCTNNNAAGCAGGAGAATCACTTGAAC
GCACACGTCTGAACTCTTTCCCTACACGACGCTCTTCCGATCTNNNAAGCAGGAGAATCACTTGAAC
GAGCACACGTCTGAACTCTTTCCCTACACGACGCTCTTCCGATCTNNNGAGAAAAGGAGTGCGCTCTT
GAGCACACGTCTGAACTCTTTCCCTACACGACGCTCTTCCGATCTNNNGAGAAAAGGAGTGCGCTCTT
GAGCACACGTCTGAACTCTTTCCCTACACGACGCTCTTCCGATCTNNNGAGAAAAGGAGTGCGCTCTT
AGCACACGTCTGAACTCTTTCCCTACACGACGCTCTTCCGATCTNNNGTACAAACGTTGTCATACCTGT
AGCACACGTCTGAACTCTTTCCCTACACGACGCTCTTCCGATCTNNNGTACAAACGTTGTCATACCTGT
AGCACACGTCTGAACTCTTTCCCTACACGACGCTCTTCCGATCTNNNGTACAAACGTTGTCATACCTGT
AGCACACGTCTGAACTCTTTCCCTACACGACGCTCTTCCGATCTNNNTGTGACCACTGATGTTGAGCA
AGCACACGTCTGAACTCTTTCCCTACACGACGCTCTTCCGATCTNNNTGTGACCACTGATGTTGAGCA
AGCACACGTCTGAACTCTTTCCCTACACGACGCTCTTCCGATCTNNNTGTGACCACTGATGTTGAGCA
GCACACGTCTGAACTCTTTCCCTACACGACGCTCTTCCGATCTNNNACCAAAACTGAGCAAGAGTAG
GCACACGTCTGAACTCTTTCCCTACACGACGCTCTTCCGATCTNNNACCAAAACTGAGCAAGAGTAG
GCACACGTCTGAACTCTTTCCCTACACGACGCTCTTCCGATCTNNNACCAAAACTGAGCAAGAGTAG
CACACGTCTGAACTCTTTCCCTACACGACGCTCTTCCGATCTNNNGAAAACTGCCATGTTGATCTCCG
CACACGTCTGAACTCTTTCCCTACACGACGCTCTTCCGATCTNNNGAAAACTGCCATGTTGATCTCCG
CACACGTCTGAACTCTTTCCCTACACGACGCTCTTCCGATCTNNNGAAAACTGCCATGTTGATCTCCG
AGCACACGTCTGAACTCTTTCCCTACACGACGCTCTTCCGATCTNNNCTCTCAGCTACTCGGGACTTTA
AGCACACGTCTGAACTCTTTCCCTACACGACGCTCTTCCGATCTNNNCTCTCAGCTACTCGGGACTTTA
AGCACACGTCTGAACTCTTTCCCTACACGACGCTCTTCCGATCTNNNCTCTCAGCTACTCGGGACTTTA
GCACACGTCTGAACTCTTTCCCTACACGACGCTCTTCCGATCTNNNATAAACTAATTCTCAGCCGGGCA
GCACACGTCTGAACTCTTTCCCTACACGACGCTCTTCCGATCTNNNATAAACTAATTCTCAGCCGGGCA
GCACACGTCTGAACTCTTTCCCTACACGACGCTCTTCCGATCTNNNATAAACTAATTCTCAGCCGGGCA
GCACACGTCTGAACTCTTTCCCTACACGACGCTCTTCCGATCTNNNCCTGCATGTCTGTCAAATCCATC
GCACACGTCTGAACTCTTTCCCTACACGACGCTCTTCCGATCTNNNCCTGCATGTCTGTCAAATCCATC
GCACACGTCTGAACTCTTTCCCTACACGACGCTCTTCCGATCTNNNCCTGCATGTCTGTCAAATCCATC
GAGCACACGTCTGAACTCTTTCCCTACACGACGCTCTTCCGATCTNNNGCACATACCTGTAATCCCAG
GAGCACACGTCTGAACTCTTTCCCTACACGACGCTCTTCCGATCTNNNGCACATACCTGTAATCCCAG
GAGCACACGTCTGAACTCTTTCCCTACACGACGCTCTTCCGATCTNNNGCACATACCTGTAATCCCAG
AGCACACGTCTGAACTCTTTCCCTACACGACGCTCTTCCGATCTNNNATAAGCCTGGCTAACATGGTG
AGCACACGTCTGAACTCTTTCCCTACACGACGCTCTTCCGATCTNNNATAAGCCTGGCTAACATGGTG
AGCACACGTCTGAACTCTTTCCCTACACGACGCTCTTCCGATCTNNNATAAGCCTGGCTAACATGGTG
AGCACACGTCTGAACTCTTTCCCTACACGACGCTCTTCCGATCTNNNGGGGTTCAGGAATTTTTTGCA
AGCACACGTCTGAACTCTTTCCCTACACGACGCTCTTCCGATCTNNNGGGGTTCAGGAATTTTTTGCA
AGCACACGTCTGAACTCTTTCCCTACACGACGCTCTTCCGATCTNNNGGGGTTCAGGAATTTTTTGCA
CACACGTCTGAACTCTTTCCCTACACGACGCTCTTCCGATCTNNNCATGGGAAGACATTTCCGACAAG
CACACGTCTGAACTCTTTCCCTACACGACGCTCTTCCGATCTNNNCATGGGAAGACATTTCCGACAAG
CACACGTCTGAACTCTTTCCCTACACGACGCTCTTCCGATCTNNNCATGGGAAGACATTTCCGACAAG
GCACACGTCTGAACTCTTTCCCTACACGACGCTCTTCCGATCTNNNTAATGAGTACCTGGAATGTTAC
GCACACGTCTGAACTCTTTCCCTACACGACGCTCTTCCGATCTNNNTAATGAGTACCTGGAATGTTAC
GCACACGTCTGAACTCTTTCCCTACACGACGCTCTTCCGATCTNNNTAATGAGTACCTGGAATGTTAC

GAGCACACGTCTGAACTCTTTCCCTACACGACGCTCTTCCGATCTNNNATCTTCCAGTCATGAAGCCAG
GAGCACACGTCTGAACTCTTTCCCTACACGACGCTCTTCCGATCTNNNATCTTCCAGTCATGAAGCCAG
GAGCACACGTCTGAACTCTTTCCCTACACGACGCTCTTCCGATCTNNNATCTTCCAGTCATGAAGCCAG
AAGAGCACACGTCTGAACTCTTTCCCTACACGACGCTCTTCCGATCTNNNTCATCTACAAGGAAGGCT
AAGAGCACACGTCTGAACTCTTTCCCTACACGACGCTCTTCCGATCTNNNTCATCTACAAGGAAGGCT
AAGAGCACACGTCTGAACTCTTTCCCTACACGACGCTCTTCCGATCTNNNTCATCTACAAGGAAGGCT
AGCACACGTCTGAACTCTTTCCCTACACGACGCTCTTCCGATCTNNNTCAACATCACACGGATAAGGGT
AGCACACGTCTGAACTCTTTCCCTACACGACGCTCTTCCGATCTNNNTCAACATCACACGGATAAGGGT
AGCACACGTCTGAACTCTTTCCCTACACGACGCTCTTCCGATCTNNNTCAACATCACACGGATAAGGGT
AGCACACGTCTGAACTCTTTCCCTACACGACGCTCTTCCGATCTNNNGCTTTTTACACATCACTGCGGA
AGCACACGTCTGAACTCTTTCCCTACACGACGCTCTTCCGATCTNNNGCTTTTTACACATCACTGCGGA
AGCACACGTCTGAACTCTTTCCCTACACGACGCTCTTCCGATCTNNNGCTTTTTACACATCACTGCGGA
AGCACACGTCTGAACTCTTTCCCTACACGACGCTCTTCCGATCTNNNATGATGGAAGCAGAGAATCCT
AGCACACGTCTGAACTCTTTCCCTACACGACGCTCTTCCGATCTNNNATGATGGAAGCAGAGAATCCT
AGCACACGTCTGAACTCTTTCCCTACACGACGCTCTTCCGATCTNNNATGATGGAAGCAGAGAATCCT
CACACGTCTGAACTCTTTCCCTACACGACGCTCTTCCGATCTNNNGGTCCTTTTCAAGCTTCGCAGCAT
CACACGTCTGAACTCTTTCCCTACACGACGCTCTTCCGATCTNNNGGTCCTTTTCAAGCTTCGCAGCAT
CACACGTCTGAACTCTTTCCCTACACGACGCTCTTCCGATCTNNNGGTCCTTTTCAAGCTTCGCAGCAT
GAGCACACGTCTGAACTCTTTCCCTACACGACGCTCTTCCGATCTNNNCCTCTGTCCCAAAATGCTAGGA
GAGCACACGTCTGAACTCTTTCCCTACACGACGCTCTTCCGATCTNNNCCTCTGTCCCAAAATGCTAGGA
GAGCACACGTCTGAACTCTTTCCCTACACGACGCTCTTCCGATCTNNNCCTCTGTCCCAAAATGCTAGGA
AGCACACGTCTGAACTCTTTCCCTACACGACGCTCTTCCGATCTNNNGCTTAACGGAACTTATAAAACA
AGCACACGTCTGAACTCTTTCCCTACACGACGCTCTTCCGATCTNNNGCTTAACGGAACTTATAAAACA
AGCACACGTCTGAACTCTTTCCCTACACGACGCTCTTCCGATCTNNNGCTTAACGGAACTTATAAAACA
AGCACACGTCTGAACTCTTTCCCTACACGACGCTCTTCCGATCTNNNAGTGTCCACATTCCAGCTTTTTG
AGCACACGTCTGAACTCTTTCCCTACACGACGCTCTTCCGATCTNNNAGTGTCCACATTCCAGCTTTTTG
AGCACACGTCTGAACTCTTTCCCTACACGACGCTCTTCCGATCTNNNAGTGTCCACATTCCAGCTTTTTG
GAGCACACGTCTGAACTCTTTCCCTACACGACGCTCTTCCGATCTNNNAAAATTAGCTGGATGTGGTG
GAGCACACGTCTGAACTCTTTCCCTACACGACGCTCTTCCGATCTNNNAAAATTAGCTGGATGTGGTG
GAGCACACGTCTGAACTCTTTCCCTACACGACGCTCTTCCGATCTNNNAAAATTAGCTGGATGTGGTG
AGCACACGTCTGAACTCTTTCCCTACACGACGCTCTTCCGATCTNNNAGAATCACTTGAACTCGGGAG
AGCACACGTCTGAACTCTTTCCCTACACGACGCTCTTCCGATCTNNNAGAATCACTTGAACTCGGGAG
AGCACACGTCTGAACTCTTTCCCTACACGACGCTCTTCCGATCTNNNAGAATCACTTGAACTCGGGAG
GAGCACACGTCTGAACTCTTTCCCTACACGACGCTCTTCCGATCTNNNTTGAACAAAACGTGAAGCAG
GAGCACACGTCTGAACTCTTTCCCTACACGACGCTCTTCCGATCTNNNTTGAACAAAACGTGAAGCAG
GAGCACACGTCTGAACTCTTTCCCTACACGACGCTCTTCCGATCTNNNTTGAACAAAACGTGAAGCAG
AGCACACGTCTGAACTCTTTCCCTACACGACGCTCTTCCGATCTNNNACAAGTTTTCAAGTCTGGTTCC
AGCACACGTCTGAACTCTTTCCCTACACGACGCTCTTCCGATCTNNNACAAGTTTTCAAGTCTGGTTCC
AGCACACGTCTGAACTCTTTCCCTACACGACGCTCTTCCGATCTNNNACAAGTTTTCAAGTCTGGTTCC
AGCACACGTCTGAACTCTTTCCCTACACGACGCTCTTCCGATCTNNNCAGAAGGCAGAAGTTTCAGTG
AGCACACGTCTGAACTCTTTCCCTACACGACGCTCTTCCGATCTNNNCAGAAGGCAGAAGTTTCAGTG
AGCACACGTCTGAACTCTTTCCCTACACGACGCTCTTCCGATCTNNNCAGAAGGCAGAAGTTTCAGTG
CACACGTCTGAACTCTTTCCCTACACGACGCTCTTCCGATCTNNNTGAGTTACACTAAATGGCCACCAC

CACACGTCTGAACTCTTTCCCTACACGACGCTCTTCCGATCTNNNTGAGTTACACTAAATGGCCACCAC
CACACGTCTGAACTCTTTCCCTACACGACGCTCTTCCGATCTNNNTGAGTTACACTAAATGGCCACCAC
AGAGCACACGTCTGAACTCTTTCCCTACACGACGCTCTTCCGATCTNNNACAAGATGCCACAGGTCTCT
AGAGCACACGTCTGAACTCTTTCCCTACACGACGCTCTTCCGATCTNNNACAAGATGCCACAGGTCTCT
AGAGCACACGTCTGAACTCTTTCCCTACACGACGCTCTTCCGATCTNNNACAAGATGCCACAGGTCTCT
GAGCACACGTCTGAACTCTTTCCCTACACGACGCTCTTCCGATCTNNNTTGGATTGGAGTCATGGAGT
GAGCACACGTCTGAACTCTTTCCCTACACGACGCTCTTCCGATCTNNNTTGGATTGGAGTCATGGAGT
GAGCACACGTCTGAACTCTTTCCCTACACGACGCTCTTCCGATCTNNNTTGGATTGGAGTCATGGAGT
GAGCACACGTCTGAACTCTTTCCCTACACGACGCTCTTCCGATCTNNNGGGGGTAATTCTTGACACAG
GAGCACACGTCTGAACTCTTTCCCTACACGACGCTCTTCCGATCTNNNGGGGGTAATTCTTGACACAG
GAGCACACGTCTGAACTCTTTCCCTACACGACGCTCTTCCGATCTNNNGGGGGTAATTCTTGACACAG
GCACACGTCTGAACTCTTTCCCTACACGACGCTCTTCCGATCTNNNGTTTGGGTCATTGCAGCCATATT
GCACACGTCTGAACTCTTTCCCTACACGACGCTCTTCCGATCTNNNGTTTGGGTCATTGCAGCCATATT
GCACACGTCTGAACTCTTTCCCTACACGACGCTCTTCCGATCTNNNGTTTGGGTCATTGCAGCCATATT
GAGCACACGTCTGAACTCTTTCCCTACACGACGCTCTTCCGATCTNNNAGAGGTTGCAGTGAGTTGAG
GAGCACACGTCTGAACTCTTTCCCTACACGACGCTCTTCCGATCTNNNAGAGGTTGCAGTGAGTTGAG
GAGCACACGTCTGAACTCTTTCCCTACACGACGCTCTTCCGATCTNNNAGAGGTTGCAGTGAGTTGAG
AAGAGCACACGTCTGAACTCTTTCCCTACACGACGCTCTTCCGATCTNNNGCCTAGGGGAAGAGAGTA
AAGAGCACACGTCTGAACTCTTTCCCTACACGACGCTCTTCCGATCTNNNGCCTAGGGGAAGAGAGTA
AAGAGCACACGTCTGAACTCTTTCCCTACACGACGCTCTTCCGATCTNNNGCCTAGGGGAAGAGAGTA
GAGCACACGTCTGAACTCTTTCCCTACACGACGCTCTTCCGATCTNNNTTCCCTTTGGAATGATTCAGCT
GAGCACACGTCTGAACTCTTTCCCTACACGACGCTCTTCCGATCTNNNTTCCCTTTGGAATGATTCAGCT
GAGCACACGTCTGAACTCTTTCCCTACACGACGCTCTTCCGATCTNNNTTCCCTTTGGAATGATTCAGCT
GAGCACACGTCTGAACTCTTTCCCTACACGACGCTCTTCCGATCTNNNTGTAATGTTAGCACTTCGGGA
GAGCACACGTCTGAACTCTTTCCCTACACGACGCTCTTCCGATCTNNNTGTAATGTTAGCACTTCGGGA
GAGCACACGTCTGAACTCTTTCCCTACACGACGCTCTTCCGATCTNNNTGTAATGTTAGCACTTCGGGA
AGCACACGTCTGAACTCTTTCCCTACACGACGCTCTTCCGATCTNNNATACCTCAAGCTGCCTTTGACAC
AGCACACGTCTGAACTCTTTCCCTACACGACGCTCTTCCGATCTNNNATACCTCAAGCTGCCTTTGACAC
AGCACACGTCTGAACTCTTTCCCTACACGACGCTCTTCCGATCTNNNATACCTCAAGCTGCCTTTGACAC
AGCACACGTCTGAACTCTTTCCCTACACGACGCTCTTCCGATCTNNNCAAAAACTTAGCCGAGCATGGT
AGCACACGTCTGAACTCTTTCCCTACACGACGCTCTTCCGATCTNNNCAAAAACTTAGCCGAGCATGGT
AGCACACGTCTGAACTCTTTCCCTACACGACGCTCTTCCGATCTNNNCAAAAACTTAGCCGAGCATGGT
AGAGCACACGTCTGAACTCTTTCCCTACACGACGCTCTTCCGATCTNNNGACAGGACACAGCTCCTCGA
AGAGCACACGTCTGAACTCTTTCCCTACACGACGCTCTTCCGATCTNNNGACAGGACACAGCTCCTCGA
AGAGCACACGTCTGAACTCTTTCCCTACACGACGCTCTTCCGATCTNNNGACAGGACACAGCTCCTCGA
GAGCACACGTCTGAACTCTTTCCCTACACGACGCTCTTCCGATCTNNNAACCTGAAAATCGATTTCCAG
GAGCACACGTCTGAACTCTTTCCCTACACGACGCTCTTCCGATCTNNNAACCTGAAAATCGATTTCCAG
GAGCACACGTCTGAACTCTTTCCCTACACGACGCTCTTCCGATCTNNNAACCTGAAAATCGATTTCCAG
AGCACACGTCTGAACTCTTTCCCTACACGACGCTCTTCCGATCTNNNGGTTTTCATCCCTTTCCTCTCCC
AGCACACGTCTGAACTCTTTCCCTACACGACGCTCTTCCGATCTNNNGGTTTTCATCCCTTTCCTCTCCC
AGCACACGTCTGAACTCTTTCCCTACACGACGCTCTTCCGATCTNNNGGTTTTCATCCCTTTCCTCTCCC
GCACACGTCTGAACTCTTTCCCTACACGACGCTCTTCCGATCTNNNGGTTGCAGTGAGTCTATATTGC
GCACACGTCTGAACTCTTTCCCTACACGACGCTCTTCCGATCTNNNGGTTGCAGTGAGTCTATATTGC

GCACACGTCTGAACTCTTTCCCTACACGACGCTCTTCCGATCTNNNGGTTGCAGTGAGTCTATATTGC
AGCACACGTCTGAACTCTTTCCCTACACGACGCTCTTCCGATCTNNNGGAGGTTACAGTGAGCTGAG
AGCACACGTCTGAACTCTTTCCCTACACGACGCTCTTCCGATCTNNNGGAGGTTACAGTGAGCTGAG
AGCACACGTCTGAACTCTTTCCCTACACGACGCTCTTCCGATCTNNNGGAGGTTACAGTGAGCTGAG
AGCACACGTCTGAACTCTTTCCCTACACGACGCTCTTCCGATCTNNNTGGGATCGATCTCTCTCTCT
AGCACACGTCTGAACTCTTTCCCTACACGACGCTCTTCCGATCTNNNTGGGATCGATCTCTCTCTCT
AGCACACGTCTGAACTCTTTCCCTACACGACGCTCTTCCGATCTNNNTGGGATCGATCTCTCTCTCT
AGCACACGTCTGAACTCTTTCCCTACACGACGCTCTTCCGATCTNNNCTCCAGGCAGTAAGAAAAACG
AGCACACGTCTGAACTCTTTCCCTACACGACGCTCTTCCGATCTNNNCTCCAGGCAGTAAGAAAAACG
AGCACACGTCTGAACTCTTTCCCTACACGACGCTCTTCCGATCTNNNCTCCAGGCAGTAAGAAAAACG
AGCACACGTCTGAACTCTTTCCCTACACGACGCTCTTCCGATCTNNNGCATTATGGTGCATGCCTGTA
AGCACACGTCTGAACTCTTTCCCTACACGACGCTCTTCCGATCTNNNGCATTATGGTGCATGCCTGTA
AGCACACGTCTGAACTCTTTCCCTACACGACGCTCTTCCGATCTNNNGCATTATGGTGCATGCCTGTA
AGCACACGTCTGAACTCTTTCCCTACACGACGCTCTTCCGATCTNNNTAAGTCCATATGAGCACCTAGG
AGCACACGTCTGAACTCTTTCCCTACACGACGCTCTTCCGATCTNNNTAAGTCCATATGAGCACCTAGG
AGCACACGTCTGAACTCTTTCCCTACACGACGCTCTTCCGATCTNNNTAAGTCCATATGAGCACCTAGG
AGCACACGTCTGAACTCTTTCCCTACACGACGCTCTTCCGATCTNNNGGATTACTTGACCCTGGGAGT
AGCACACGTCTGAACTCTTTCCCTACACGACGCTCTTCCGATCTNNNGGATTACTTGACCCTGGGAGT
AGCACACGTCTGAACTCTTTCCCTACACGACGCTCTTCCGATCTNNNGGATTACTTGACCCTGGGAGT
AGCACACGTCTGAACTCTTTCCCTACACGACGCTCTTCCGATCTNNNATGTGCTGTCTCACCTCCTTTA
AGCACACGTCTGAACTCTTTCCCTACACGACGCTCTTCCGATCTNNNATGTGCTGTCTCACCTCCTTTA
AGCACACGTCTGAACTCTTTCCCTACACGACGCTCTTCCGATCTNNNATGTGCTGTCTCACCTCCTTTA
AGCACACGTCTGAACTCTTTCCCTACACGACGCTCTTCCGATCTNNNTGCCTGTAATCCCAGCTAAACA
AGCACACGTCTGAACTCTTTCCCTACACGACGCTCTTCCGATCTNNNTGCCTGTAATCCCAGCTAAACA
AGCACACGTCTGAACTCTTTCCCTACACGACGCTCTTCCGATCTNNNTGCCTGTAATCCCAGCTAAACA
AGCACACGTCTGAACTCTTTCCCTACACGACGCTCTTCCGATCTNNNCAAATCTCAAAAGCTTACACGC
AGCACACGTCTGAACTCTTTCCCTACACGACGCTCTTCCGATCTNNNCAAATCTCAAAAGCTTACACGC
AGCACACGTCTGAACTCTTTCCCTACACGACGCTCTTCCGATCTNNNCAAATCTCAAAAGCTTACACGC
AGCACACGTCTGAACTCTTTCCCTACACGACGCTCTTCCGATCTNNNAGAACAGTGCCTGACCTAAAA
AGCACACGTCTGAACTCTTTCCCTACACGACGCTCTTCCGATCTNNNAGAACAGTGCCTGACCTAAAA
AGCACACGTCTGAACTCTTTCCCTACACGACGCTCTTCCGATCTNNNAGAACAGTGCCTGACCTAAAA
CACACGTCTGAACTCTTTCCCTACACGACGCTCTTCCGATCTNNNCACAGCTTTTCACCTGTCAGTCAC
CACACGTCTGAACTCTTTCCCTACACGACGCTCTTCCGATCTNNNCACAGCTTTTCACCTGTCAGTCAC
CACACGTCTGAACTCTTTCCCTACACGACGCTCTTCCGATCTNNNCACAGCTTTTCACCTGTCAGTCAC
GCACACGTCTGAACTCTTTCCCTACACGACGCTCTTCCGATCTNNNCATACCCAGTGTTAGCAAGGAG
GCACACGTCTGAACTCTTTCCCTACACGACGCTCTTCCGATCTNNNCATACCCAGTGTTAGCAAGGAG
GCACACGTCTGAACTCTTTCCCTACACGACGCTCTTCCGATCTNNNCATACCCAGTGTTAGCAAGGAG
AGCACACGTCTGAACTCTTTCCCTACACGACGCTCTTCCGATCTNNNTCTTTATCTCATAGTTTTTGCA
AGCACACGTCTGAACTCTTTCCCTACACGACGCTCTTCCGATCTNNNTCTTTATCTCATAGTTTTTGCA
AGCACACGTCTGAACTCTTTCCCTACACGACGCTCTTCCGATCTNNNTCTTTATCTCATAGTTTTTGCA
GCACACGTCTGAACTCTTTCCCTACACGACGCTCTTCCGATCTNNNGGCAAAACACCTCTTTCAGTCTT
GCACACGTCTGAACTCTTTCCCTACACGACGCTCTTCCGATCTNNNGGCAAAACACCTCTTTCAGTCTT
GCACACGTCTGAACTCTTTCCCTACACGACGCTCTTCCGATCTNNNGGCAAAACACCTCTTTCAGTCTT

CACACGTCTGAACTCTTTCCCTACACGACGCTCTTCCGATCTNNNACATTTTCTGTTTCTGGAAATGCTCA

CACACGTCTGAACTCTTTCCCTACACGACGCTCTTCCGATCTNNNACATTTTCTGTTTCTGGAAATGCTCA

CACACGTCTGAACTCTTTCCCTACACGACGCTCTTCCGATCTNNNACATTTTCTGTTTCTGGAAATGCTCA

GAGCACACGTCTGAACTCTTTCCCTACACGACGCTCTTCCGATCTNNNGATGCAGAGATGGATGGATG

GAGCACACGTCTGAACTCTTTCCCTACACGACGCTCTTCCGATCTNNNGATGCAGAGATGGATGGATG

GAGCACACGTCTGAACTCTTTCCCTACACGACGCTCTTCCGATCTNNNGATGCAGAGATGGATGGATG

GAGCACACGTCTGAACTCTTTCCCTACACGACGCTCTTCCGATCTNNNCCCTGGAATGTTAGTTGCAGCA

GAGCACACGTCTGAACTCTTTCCCTACACGACGCTCTTCCGATCTNNNCCCTGGAATGTTAGTTGCAGCA

GAGCACACGTCTGAACTCTTTCCCTACACGACGCTCTTCCGATCTNNNCCCTGGAATGTTAGTTGCAGCA

GAGCACACGTCTGAACTCTTTCCCTACACGACGCTCTTCCGATCTNNNTTCGTGTGAGTTATCAGGACT

GAGCACACGTCTGAACTCTTTCCCTACACGACGCTCTTCCGATCTNNNTTCGTGTGAGTTATCAGGACT

GAGCACACGTCTGAACTCTTTCCCTACACGACGCTCTTCCGATCTNNNTTCGTGTGAGTTATCAGGACT

GAGCACACGTCTGAACTCTTTCCCTACACGACGCTCTTCCGATCTNNNGATGCATCCACTAAGTTGAAG

GAGCACACGTCTGAACTCTTTCCCTACACGACGCTCTTCCGATCTNNNGATGCATCCACTAAGTTGAAG

GAGCACACGTCTGAACTCTTTCCCTACACGACGCTCTTCCGATCTNNNGATGCATCCACTAAGTTGAAG

GAGCACACGTCTGAACTCTTTCCCTACACGACGCTCTTCCGATCTNNNTAGTAGAGACAGGGTTTTGC

GAGCACACGTCTGAACTCTTTCCCTACACGACGCTCTTCCGATCTNNNTAGTAGAGACAGGGTTTTGC

GAGCACACGTCTGAACTCTTTCCCTACACGACGCTCTTCCGATCTNNNTAGTAGAGACAGGGTTTTGC

AGCACACGTCTGAACTCTTTCCCTACACGACGCTCTTCCGATCTNNNCCCGGCTTTTTTTGTGTTTTGTT

AGCACACGTCTGAACTCTTTCCCTACACGACGCTCTTCCGATCTNNNCCCGGCTTTTTTTGTGTTTTGTT

AGCACACGTCTGAACTCTTTCCCTACACGACGCTCTTCCGATCTNNNCCCGGCTTTTTTTGTGTTTTGTT

AGCACACGTCTGAACTCTTTCCCTACACGACGCTCTTCCGATCTNNNAATACATGCTCAGGCATACCCA

AGCACACGTCTGAACTCTTTCCCTACACGACGCTCTTCCGATCTNNNAATACATGCTCAGGCATACCCA

AGCACACGTCTGAACTCTTTCCCTACACGACGCTCTTCCGATCTNNNAATACATGCTCAGGCATACCCA

GCACACGTCTGAACTCTTTCCCTACACGACGCTCTTCCGATCTNNNAAGCTGAGGTGGAAGAATCACT

GCACACGTCTGAACTCTTTCCCTACACGACGCTCTTCCGATCTNNNAAGCTGAGGTGGAAGAATCACT

GCACACGTCTGAACTCTTTCCCTACACGACGCTCTTCCGATCTNNNAAGCTGAGGTGGAAGAATCACT

AGCACACGTCTGAACTCTTTCCCTACACGACGCTCTTCCGATCTNNNTGTGAAACACTGACCCTCAAAC

AGCACACGTCTGAACTCTTTCCCTACACGACGCTCTTCCGATCTNNNTGTGAAACACTGACCCTCAAAC

AGCACACGTCTGAACTCTTTCCCTACACGACGCTCTTCCGATCTNNNTGTGAAACACTGACCCTCAAAC

AGCACACGTCTGAACTCTTTCCCTACACGACGCTCTTCCGATCTNNNGAGAATGGCATGAACCTGGGA

AGCACACGTCTGAACTCTTTCCCTACACGACGCTCTTCCGATCTNNNGAGAATGGCATGAACCTGGGA

AGCACACGTCTGAACTCTTTCCCTACACGACGCTCTTCCGATCTNNNGAGAATGGCATGAACCTGGGA

AGCACACGTCTGAACTCTTTCCCTACACGACGCTCTTCCGATCTNNNATTTCCATTTCCCAACTCCCCTTCA

AGCACACGTCTGAACTCTTTCCCTACACGACGCTCTTCCGATCTNNNATTTCCATTTCCCAACTCCCCTTCA

AGCACACGTCTGAACTCTTTCCCTACACGACGCTCTTCCGATCTNNNATTTCCATTTCCCAACTCCCCTTCA

GAGCACACGTCTGAACTCTTTCCCTACACGACGCTCTTCCGATCTNNNGTGAGGGAATGAGTTGCAG

GAGCACACGTCTGAACTCTTTCCCTACACGACGCTCTTCCGATCTNNNGTGAGGGAATGAGTTGCAG

GAGCACACGTCTGAACTCTTTCCCTACACGACGCTCTTCCGATCTNNNGTGAGGGAATGAGTTGCAG

AGAGCACACGTCTGAACTCTTTCCCTACACGACGCTCTTCCGATCTNNNTCTGCCTTCTAGGACATTCA

AGAGCACACGTCTGAACTCTTTCCCTACACGACGCTCTTCCGATCTNNNTCTGCCTTCTAGGACATTCA

AGAGCACACGTCTGAACTCTTTCCCTACACGACGCTCTTCCGATCTNNNTCTGCCTTCTAGGACATTCA

GAGCACACGTCTGAACTCTTTCCCTACACGACGCTCTTCCGATCTNNNACCAAATGGGATGACATGGA

AGCACACGTCTGAACTCTTTCCCTACACGACGCTCTTCCGATCTNNNACCAAATGGGATGACATGGA
AGCACACGTCTGAACTCTTTCCCTACACGACGCTCTTCCGATCTNNNACCAAATGGGATGACATGGA
ACACGTCTGAACTCTTTCCCTACACGACGCTCTTCCGATCTNNNCCATGAGCTGTAGGTCCTTTGGCA
ACACGTCTGAACTCTTTCCCTACACGACGCTCTTCCGATCTNNNCCATGAGCTGTAGGTCCTTTGGCA
ACACGTCTGAACTCTTTCCCTACACGACGCTCTTCCGATCTNNNCCATGAGCTGTAGGTCCTTTGGCA
AGCACACGTCTGAACTCTTTCCCTACACGACGCTCTTCCGATCTNNNGAGGTTGCAGTGAGCTGAGA
AGCACACGTCTGAACTCTTTCCCTACACGACGCTCTTCCGATCTNNNGAGGTTGCAGTGAGCTGAGA
AGCACACGTCTGAACTCTTTCCCTACACGACGCTCTTCCGATCTNNNGAGGTTGCAGTGAGCTGAGA
AGCACACGTCTGAACTCTTTCCCTACACGACGCTCTTCCGATCTNNNTGAACATTTCCAGGGGACTTTC
AGCACACGTCTGAACTCTTTCCCTACACGACGCTCTTCCGATCTNNNTGAACATTTCCAGGGGACTTTC
AGCACACGTCTGAACTCTTTCCCTACACGACGCTCTTCCGATCTNNNTGAACATTTCCAGGGGACTTTC
GCACACGTCTGAACTCTTTCCCTACACGACGCTCTTCCGATCTNNNCTGAGGCAGGAGAATCACTTC
GCACACGTCTGAACTCTTTCCCTACACGACGCTCTTCCGATCTNNNCTGAGGCAGGAGAATCACTTC
GCACACGTCTGAACTCTTTCCCTACACGACGCTCTTCCGATCTNNNCTGAGGCAGGAGAATCACTTC
AGCACACGTCTGAACTCTTTCCCTACACGACGCTCTTCCGATCTNNNACTGGAATCCCTGCCAAAATA
AGCACACGTCTGAACTCTTTCCCTACACGACGCTCTTCCGATCTNNNACTGGAATCCCTGCCAAAATA
AGCACACGTCTGAACTCTTTCCCTACACGACGCTCTTCCGATCTNNNACTGGAATCCCTGCCAAAATA
GAGCACACGTCTGAACTCTTTCCCTACACGACGCTCTTCCGATCTNNNGATACTCTACCTCAGCCTCCT
GAGCACACGTCTGAACTCTTTCCCTACACGACGCTCTTCCGATCTNNNGATACTCTACCTCAGCCTCCT
GAGCACACGTCTGAACTCTTTCCCTACACGACGCTCTTCCGATCTNNNGATACTCTACCTCAGCCTCCT
GCACACGTCTGAACTCTTTCCCTACACGACGCTCTTCCGATCTNNNGACCGTGAGTTTGGTTTGCAAA
GCACACGTCTGAACTCTTTCCCTACACGACGCTCTTCCGATCTNNNGACCGTGAGTTTGGTTTGCAAA
GCACACGTCTGAACTCTTTCCCTACACGACGCTCTTCCGATCTNNNGACCGTGAGTTTGGTTTGCAAA
GAGCACACGTCTGAACTCTTTCCCTACACGACGCTCTTCCGATCTNNNAGTTGCTGGAGAATGTGTAC
GAGCACACGTCTGAACTCTTTCCCTACACGACGCTCTTCCGATCTNNNAGTTGCTGGAGAATGTGTAC
GAGCACACGTCTGAACTCTTTCCCTACACGACGCTCTTCCGATCTNNNAGTTGCTGGAGAATGTGTAC
AGCACACGTCTGAACTCTTTCCCTACACGACGCTCTTCCGATCTNNNCTGGGGGTAACAGAGAAAG
AGCACACGTCTGAACTCTTTCCCTACACGACGCTCTTCCGATCTNNNCTGGGGGTAACAGAGAAAG
AGCACACGTCTGAACTCTTTCCCTACACGACGCTCTTCCGATCTNNNCTGGGGGTAACAGAGAAAG
GAGCACACGTCTGAACTCTTTCCCTACACGACGCTCTTCCGATCTNNNCACTTCTGTTTTCCTGCCCAA
GAGCACACGTCTGAACTCTTTCCCTACACGACGCTCTTCCGATCTNNNCACTTCTGTTTTCCTGCCCAA
GAGCACACGTCTGAACTCTTTCCCTACACGACGCTCTTCCGATCTNNNCACTTCTGTTTTCCTGCCCAA
AGCACACGTCTGAACTCTTTCCCTACACGACGCTCTTCCGATCTNNNAGACAATAACAAGTGCTGCT
AGCACACGTCTGAACTCTTTCCCTACACGACGCTCTTCCGATCTNNNAGACAATAACAAGTGCTGCT
AGCACACGTCTGAACTCTTTCCCTACACGACGCTCTTCCGATCTNNNAGACAATAACAAGTGCTGCT
AGCACACGTCTGAACTCTTTCCCTACACGACGCTCTTCCGATCTNNNACACACACACATTCCCAGCTTA
AGCACACGTCTGAACTCTTTCCCTACACGACGCTCTTCCGATCTNNNACACACACACATTCCCAGCTTA
AGCACACGTCTGAACTCTTTCCCTACACGACGCTCTTCCGATCTNNNACACACACACATTCCCAGCTTA
CACACGTCTGAACTCTTTCCCTACACGACGCTCTTCCGATCTNNNCCCAGGAATGGCGTAAGGGGCAT
CACACGTCTGAACTCTTTCCCTACACGACGCTCTTCCGATCTNNNCCCAGGAATGGCGTAAGGGGCAT
CACACGTCTGAACTCTTTCCCTACACGACGCTCTTCCGATCTNNNCCCAGGAATGGCGTAAGGGGCAT
GCACACGTCTGAACTCTTTCCCTACACGACGCTCTTCCGATCTNNNTTGAGGCAGGAGAATTGCTTG
GCACACGTCTGAACTCTTTCCCTACACGACGCTCTTCCGATCTNNNTTGAGGCAGGAGAATTGCTTG

AGCACACGTCTGAACTCTTTCCCTACACGACGCTCTTCCGATCTNNNTTGAGGCAGGAGAATTGCTTG
GAGCACACGTCTGAACTCTTTCCCTACACGACGCTCTTCCGATCTNNNGAGACTGGAGAATTGCTTG
GAGCACACGTCTGAACTCTTTCCCTACACGACGCTCTTCCGATCTNNNGAGACTGGAGAATTGCTTG
GAGCACACGTCTGAACTCTTTCCCTACACGACGCTCTTCCGATCTNNNGAGACTGGAGAATTGCTTG
GAGCACACGTCTGAACTCTTTCCCTACACGACGCTCTTCCGATCTNNNAATCGCTTGAACCCGAGATT
GAGCACACGTCTGAACTCTTTCCCTACACGACGCTCTTCCGATCTNNNAATCGCTTGAACCCGAGATT
GAGCACACGTCTGAACTCTTTCCCTACACGACGCTCTTCCGATCTNNNAATCGCTTGAACCCGAGATT
AGCACACGTCTGAACTCTTTCCCTACACGACGCTCTTCCGATCTNNNGGAGAAGTCCACTTTCCAACT
AGCACACGTCTGAACTCTTTCCCTACACGACGCTCTTCCGATCTNNNGGAGAAGTCCACTTTCCAACT
AGCACACGTCTGAACTCTTTCCCTACACGACGCTCTTCCGATCTNNNGGAGAAGTCCACTTTCCAACT
AGCACACGTCTGAACTCTTTCCCTACACGACGCTCTTCCGATCTNNNAATACCAGTACTTTGGGAGGC
AGCACACGTCTGAACTCTTTCCCTACACGACGCTCTTCCGATCTNNNAATACCAGTACTTTGGGAGGC
AGCACACGTCTGAACTCTTTCCCTACACGACGCTCTTCCGATCTNNNAATACCAGTACTTTGGGAGGC
GCACACGTCTGAACTCTTTCCCTACACGACGCTCTTCCGATCTNNNACGTTGCAGAGAATAGGATGT
GCACACGTCTGAACTCTTTCCCTACACGACGCTCTTCCGATCTNNNACGTTGCAGAGAATAGGATGT
GCACACGTCTGAACTCTTTCCCTACACGACGCTCTTCCGATCTNNNACGTTGCAGAGAATAGGATGT
AGAGCACACGTCTGAACTCTTTCCCTACACGACGCTCTTCCGATCTNNNCACCTCCATCTCTCTCTCTC
AGAGCACACGTCTGAACTCTTTCCCTACACGACGCTCTTCCGATCTNNNCACCTCCATCTCTCTCTCTC
AGAGCACACGTCTGAACTCTTTCCCTACACGACGCTCTTCCGATCTNNNCACCTCCATCTCTCTCTCTC
AGCACACGTCTGAACTCTTTCCCTACACGACGCTCTTCCGATCTNNNACACACACCTCTATCTTTCCAC
AGCACACGTCTGAACTCTTTCCCTACACGACGCTCTTCCGATCTNNNACACACACCTCTATCTTTCCAC
AGCACACGTCTGAACTCTTTCCCTACACGACGCTCTTCCGATCTNNNACACACACCTCTATCTTTCCAC
GAGCACACGTCTGAACTCTTTCCCTACACGACGCTCTTCCGATCTNNNCTTGAGACCAGTGGTTCGA
GAGCACACGTCTGAACTCTTTCCCTACACGACGCTCTTCCGATCTNNNCTTGAGACCAGTGGTTCGA
GAGCACACGTCTGAACTCTTTCCCTACACGACGCTCTTCCGATCTNNNCTTGAGACCAGTGGTTCGA
GAGCACACGTCTGAACTCTTTCCCTACACGACGCTCTTCCGATCTNNNTATAGTCCCAGATGCTCAGG
GAGCACACGTCTGAACTCTTTCCCTACACGACGCTCTTCCGATCTNNNTATAGTCCCAGATGCTCAGG
GAGCACACGTCTGAACTCTTTCCCTACACGACGCTCTTCCGATCTNNNTATAGTCCCAGATGCTCAGG
AGCACACGTCTGAACTCTTTCCCTACACGACGCTCTTCCGATCTNNNTTGCTATTCTTTTCACTTGTGC
AGCACACGTCTGAACTCTTTCCCTACACGACGCTCTTCCGATCTNNNTTGCTATTCTTTTCACTTGTGC
AGCACACGTCTGAACTCTTTCCCTACACGACGCTCTTCCGATCTNNNTTGCTATTCTTTTCACTTGTGC
AGCACACGTCTGAACTCTTTCCCTACACGACGCTCTTCCGATCTNNNTTCTGGACAGTGAAGGCTTAG
AGCACACGTCTGAACTCTTTCCCTACACGACGCTCTTCCGATCTNNNTTCTGGACAGTGAAGGCTTAG
AGCACACGTCTGAACTCTTTCCCTACACGACGCTCTTCCGATCTNNNTTCTGGACAGTGAAGGCTTAG
GCACACGTCTGAACTCTTTCCCTACACGACGCTCTTCCGATCTNNNTTCTGTCTCTATGAACAGGGCA
GCACACGTCTGAACTCTTTCCCTACACGACGCTCTTCCGATCTNNNTTCTGTCTCTATGAACAGGGCA
GCACACGTCTGAACTCTTTCCCTACACGACGCTCTTCCGATCTNNNTTCTGTCTCTATGAACAGGGCA
CACACGTCTGAACTCTTTCCCTACACGACGCTCTTCCGATCTNNNCCCCAAGGATTATGCTAGAGTCC
CACACGTCTGAACTCTTTCCCTACACGACGCTCTTCCGATCTNNNCCCCAAGGATTATGCTAGAGTCC
CACACGTCTGAACTCTTTCCCTACACGACGCTCTTCCGATCTNNNCCCCAAGGATTATGCTAGAGTCC
AGCACACGTCTGAACTCTTTCCCTACACGACGCTCTTCCGATCTNNNCACCTGTTGTCCCAGCTACTCC
AGCACACGTCTGAACTCTTTCCCTACACGACGCTCTTCCGATCTNNNCACCTGTTGTCCCAGCTACTCC
AGCACACGTCTGAACTCTTTCCCTACACGACGCTCTTCCGATCTNNNCACCTGTTGTCCCAGCTACTCC

GAGCACACGTCTGAACTCTTTCCCTACACGACGCTCTTCCGATCTNNNGAGGCAGGAGAATTGTTTGA
GAGCACACGTCTGAACTCTTTCCCTACACGACGCTCTTCCGATCTNNNGAGGCAGGAGAATTGTTTGA
GAGCACACGTCTGAACTCTTTCCCTACACGACGCTCTTCCGATCTNNNGAGGCAGGAGAATTGTTTGA
AGCACACGTCTGAACTCTTTCCCTACACGACGCTCTTCCGATCTNNNAGGTCTGATTGGGTAGGTTCAT
AGCACACGTCTGAACTCTTTCCCTACACGACGCTCTTCCGATCTNNNAGGTCTGATTGGGTAGGTTCAT
AGCACACGTCTGAACTCTTTCCCTACACGACGCTCTTCCGATCTNNNAGGTCTGATTGGGTAGGTTCAT
GAGCACACGTCTGAACTCTTTCCCTACACGACGCTCTTCCGATCTNNNACACCTGTAATCCCAGCTACTTG
GAGCACACGTCTGAACTCTTTCCCTACACGACGCTCTTCCGATCTNNNACACCTGTAATCCCAGCTACTTG
GAGCACACGTCTGAACTCTTTCCCTACACGACGCTCTTCCGATCTNNNACACCTGTAATCCCAGCTACTTG
GCACACGTCTGAACTCTTTCCCTACACGACGCTCTTCCGATCTNNNACGCCCGGCTAATTTTTGTATTTG
GCACACGTCTGAACTCTTTCCCTACACGACGCTCTTCCGATCTNNNACGCCCGGCTAATTTTTGTATTTG
GCACACGTCTGAACTCTTTCCCTACACGACGCTCTTCCGATCTNNNACGCCCGGCTAATTTTTGTATTTG
GAGCACACGTCTGAACTCTTTCCCTACACGACGCTCTTCCGATCTNNNAGTTTGAGTCATTATCCTATTG
GAGCACACGTCTGAACTCTTTCCCTACACGACGCTCTTCCGATCTNNNAGTTTGAGTCATTATCCTATTG
GAGCACACGTCTGAACTCTTTCCCTACACGACGCTCTTCCGATCTNNNAGTTTGAGTCATTATCCTATTG
GAGCACACGTCTGAACTCTTTCCCTACACGACGCTCTTCCGATCTNNNAGTGGAGGTGAAGGAAAGAG
GAGCACACGTCTGAACTCTTTCCCTACACGACGCTCTTCCGATCTNNNAGTGGAGGTGAAGGAAAGAG
GAGCACACGTCTGAACTCTTTCCCTACACGACGCTCTTCCGATCTNNNAGTGGAGGTGAAGGAAAGAG
AGCACACGTCTGAACTCTTTCCCTACACGACGCTCTTCCGATCTNNNCTGAGATGGGAGGACTGATTG
AGCACACGTCTGAACTCTTTCCCTACACGACGCTCTTCCGATCTNNNCTGAGATGGGAGGACTGATTG
AGCACACGTCTGAACTCTTTCCCTACACGACGCTCTTCCGATCTNNNCTGAGATGGGAGGACTGATTG
GAGCACACGTCTGAACTCTTTCCCTACACGACGCTCTTCCGATCTNNNGGAGTTGAGGTGGGAGAATG
GAGCACACGTCTGAACTCTTTCCCTACACGACGCTCTTCCGATCTNNNGGAGTTGAGGTGGGAGAATG
GAGCACACGTCTGAACTCTTTCCCTACACGACGCTCTTCCGATCTNNNGGAGTTGAGGTGGGAGAATG
GCACACGTCTGAACTCTTTCCCTACACGACGCTCTTCCGATCTNNNTTACAGTTGTCGGATTTGTCCATC
GCACACGTCTGAACTCTTTCCCTACACGACGCTCTTCCGATCTNNNTTACAGTTGTCGGATTTGTCCATC
GCACACGTCTGAACTCTTTCCCTACACGACGCTCTTCCGATCTNNNTTACAGTTGTCGGATTTGTCCATC
AGAGCACACGTCTGAACTCTTTCCCTACACGACGCTCTTCCGATCTNNNTCAGAGAGGATTTTGGTGTG
AGAGCACACGTCTGAACTCTTTCCCTACACGACGCTCTTCCGATCTNNNTCAGAGAGGATTTTGGTGTG
AGAGCACACGTCTGAACTCTTTCCCTACACGACGCTCTTCCGATCTNNNTCAGAGAGGATTTTGGTGTG
AGCACACGTCTGAACTCTTTCCCTACACGACGCTCTTCCGATCTNNNCAGAGGTTACAGTGAGCCAAG
AGCACACGTCTGAACTCTTTCCCTACACGACGCTCTTCCGATCTNNNCAGAGGTTACAGTGAGCCAAG
AGCACACGTCTGAACTCTTTCCCTACACGACGCTCTTCCGATCTNNNCAGAGGTTACAGTGAGCCAAG
AGCACACGTCTGAACTCTTTCCCTACACGACGCTCTTCCGATCTNNNGGTTCAAGTGATTTTCCTGCCTG
AGCACACGTCTGAACTCTTTCCCTACACGACGCTCTTCCGATCTNNNGGTTCAAGTGATTTTCCTGCCTG
AGCACACGTCTGAACTCTTTCCCTACACGACGCTCTTCCGATCTNNNGGTTCAAGTGATTTTCCTGCCTG
AGCACACGTCTGAACTCTTTCCCTACACGACGCTCTTCCGATCTNNNCCCCTTTCAACTTCTTCTCTCTTCA
AGCACACGTCTGAACTCTTTCCCTACACGACGCTCTTCCGATCTNNNCCCCTTTCAACTTCTTCTCTCTTCA
AGCACACGTCTGAACTCTTTCCCTACACGACGCTCTTCCGATCTNNNCCCCTTTCAACTTCTTCTCTCTTCA
AGCACACGTCTGAACTCTTTCCCTACACGACGCTCTTCCGATCTNNNGGTTTCCATATTGAGTTCCTGT
AGCACACGTCTGAACTCTTTCCCTACACGACGCTCTTCCGATCTNNNGGTTTCCATATTGAGTTCCTGT
AGCACACGTCTGAACTCTTTCCCTACACGACGCTCTTCCGATCTNNNGGTTTCCATATTGAGTTCCTGT
GAGCACACGTCTGAACTCTTTCCCTACACGACGCTCTTCCGATCTNNNAGAGAGGAATAAAGACAGAT

GAGCACACGTCTGAACTCTTTCCCTACACGACGCTCTTCCGATCTNNNAGAGAGGAATAAAGACAGAT
GAGCACACGTCTGAACTCTTTCCCTACACGACGCTCTTCCGATCTNNNAGAGAGGAATAAAGACAGAT
AGCACACGTCTGAACTCTTTCCCTACACGACGCTCTTCCGATCTNNNTGGTGACATACTCAGGAATCTGC
AGCACACGTCTGAACTCTTTCCCTACACGACGCTCTTCCGATCTNNNTGGTGACATACTCAGGAATCTGC
AGCACACGTCTGAACTCTTTCCCTACACGACGCTCTTCCGATCTNNNTGGTGACATACTCAGGAATCTGC
AGCACACGTCTGAACTCTTTCCCTACACGACGCTCTTCCGATCTNNNGAGATCACACCACTGCATTCCACA
AGCACACGTCTGAACTCTTTCCCTACACGACGCTCTTCCGATCTNNNGAGATCACACCACTGCATTCCACA
AGCACACGTCTGAACTCTTTCCCTACACGACGCTCTTCCGATCTNNNGAGATCACACCACTGCATTCCACA
GAGCACACGTCTGAACTCTTTCCCTACACGACGCTCTTCCGATCTNNNGGGGTCCATTTCTGAGTTGTTT
GAGCACACGTCTGAACTCTTTCCCTACACGACGCTCTTCCGATCTNNNGGGGTCCATTTCTGAGTTGTTT
GAGCACACGTCTGAACTCTTTCCCTACACGACGCTCTTCCGATCTNNNGGGGTCCATTTCTGAGTTGTTT
AGAGCACACGTCTGAACTCTTTCCCTACACGACGCTCTTCCGATCTNNNCTGGGCTACAAGAGTAAGAC
AGAGCACACGTCTGAACTCTTTCCCTACACGACGCTCTTCCGATCTNNNCTGGGCTACAAGAGTAAGAC
AGAGCACACGTCTGAACTCTTTCCCTACACGACGCTCTTCCGATCTNNNCTGGGCTACAAGAGTAAGAC
AGAGCACACGTCTGAACTCTTTCCCTACACGACGCTCTTCCGATCTNNNGCAGGAGAATTGCTTGATCC
AGAGCACACGTCTGAACTCTTTCCCTACACGACGCTCTTCCGATCTNNNGCAGGAGAATTGCTTGATCC
AGAGCACACGTCTGAACTCTTTCCCTACACGACGCTCTTCCGATCTNNNGCAGGAGAATTGCTTGATCC
AGCACACGTCTGAACTCTTTCCCTACACGACGCTCTTCCGATCTNNNGTCTCTCTGATGGTTCACCTTCT
AGCACACGTCTGAACTCTTTCCCTACACGACGCTCTTCCGATCTNNNGTCTCTCTGATGGTTCACCTTCT
AGCACACGTCTGAACTCTTTCCCTACACGACGCTCTTCCGATCTNNNGTCTCTCTGATGGTTCACCTTCT
AGCACACGTCTGAACTCTTTCCCTACACGACGCTCTTCCGATCTNNNGAGTCTTGCTCTGTTGCCCAGC
AGCACACGTCTGAACTCTTTCCCTACACGACGCTCTTCCGATCTNNNGAGTCTTGCTCTGTTGCCCAGC
AGCACACGTCTGAACTCTTTCCCTACACGACGCTCTTCCGATCTNNNGAGTCTTGCTCTGTTGCCCAGC
AAGAGCACACGTCTGAACTCTTTCCCTACACGACGCTCTTCCGATCTNNNCTCGGGGCCAGTTTCCTTAT
AAGAGCACACGTCTGAACTCTTTCCCTACACGACGCTCTTCCGATCTNNNCTCGGGGCCAGTTTCCTTAT
AAGAGCACACGTCTGAACTCTTTCCCTACACGACGCTCTTCCGATCTNNNCTCGGGGCCAGTTTCCTTAT
AGAGCACACGTCTGAACTCTTTCCCTACACGACGCTCTTCCGATCTNNNGAGGTTGCAGTAAGCCGAGA
AGAGCACACGTCTGAACTCTTTCCCTACACGACGCTCTTCCGATCTNNNGAGGTTGCAGTAAGCCGAGA
AGAGCACACGTCTGAACTCTTTCCCTACACGACGCTCTTCCGATCTNNNGAGGTTGCAGTAAGCCGAGA
AGCACACGTCTGAACTCTTTCCCTACACGACGCTCTTCCGATCTNNNCAGGTCTGCCTTCCTTCATCACCA
AGCACACGTCTGAACTCTTTCCCTACACGACGCTCTTCCGATCTNNNCAGGTCTGCCTTCCTTCATCACCA
AGCACACGTCTGAACTCTTTCCCTACACGACGCTCTTCCGATCTNNNCAGGTCTGCCTTCCTTCATCACCA
AGCACACGTCTGAACTCTTTCCCTACACGACGCTCTTCCGATCTNNNCTCTCTTGGCTTGAAGGGAATG
AGCACACGTCTGAACTCTTTCCCTACACGACGCTCTTCCGATCTNNNCTCTCTTGGCTTGAAGGGAATG
AGCACACGTCTGAACTCTTTCCCTACACGACGCTCTTCCGATCTNNNCTCTCTTGGCTTGAAGGGAATG
GAGCACACGTCTGAACTCTTTCCCTACACGACGCTCTTCCGATCTNNNCATCCATGTGCACGTTTATGCC
GAGCACACGTCTGAACTCTTTCCCTACACGACGCTCTTCCGATCTNNNCATCCATGTGCACGTTTATGCC
GAGCACACGTCTGAACTCTTTCCCTACACGACGCTCTTCCGATCTNNNCATCCATGTGCACGTTTATGCC
AGCACACGTCTGAACTCTTTCCCTACACGACGCTCTTCCGATCTNNNGGTCTCAGCCACTCCAGATTTTC
AGCACACGTCTGAACTCTTTCCCTACACGACGCTCTTCCGATCTNNNGGTCTCAGCCACTCCAGATTTTC
AGCACACGTCTGAACTCTTTCCCTACACGACGCTCTTCCGATCTNNNGGTCTCAGCCACTCCAGATTTTC
AGCACACGTCTGAACTCTTTCCCTACACGACGCTCTTCCGATCTNNNTGGCCTCTATAATTATGTGAGCC
AGCACACGTCTGAACTCTTTCCCTACACGACGCTCTTCCGATCTNNNTGGCCTCTATAATTATGTGAGCC

GCACACGTCTGAACTCTTTCCCTACACGACGCTCTTCCGATCTNNNTGGCCTCTATAATTATGTGAGC
AGCACACGTCTGAACTCTTTCCCTACACGACGCTCTTCCGATCTNNNCAGTCTGTATGTCATTCCTGCA
AGCACACGTCTGAACTCTTTCCCTACACGACGCTCTTCCGATCTNNNCAGTCTGTATGTCATTCCTGCA
AGCACACGTCTGAACTCTTTCCCTACACGACGCTCTTCCGATCTNNNCAGTCTGTATGTCATTCCTGCA
GAGCACACGTCTGAACTCTTTCCCTACACGACGCTCTTCCGATCTNNNGGGAGAAGAGAAATAACTG
GAGCACACGTCTGAACTCTTTCCCTACACGACGCTCTTCCGATCTNNNGGGAGAAGAGAAATAACTG
GAGCACACGTCTGAACTCTTTCCCTACACGACGCTCTTCCGATCTNNNGGGAGAAGAGAAATAACTG
AGCACACGTCTGAACTCTTTCCCTACACGACGCTCTTCCGATCTNNNCACCCAGGTAGTGAGCATAAC
AGCACACGTCTGAACTCTTTCCCTACACGACGCTCTTCCGATCTNNNCACCCAGGTAGTGAGCATAAC
AGCACACGTCTGAACTCTTTCCCTACACGACGCTCTTCCGATCTNNNCACCCAGGTAGTGAGCATAAC
CACACGTCTGAACTCTTTCCCTACACGACGCTCTTCCGATCTNNNCAAGAGCGATGGAAAACCATTCC
CACACGTCTGAACTCTTTCCCTACACGACGCTCTTCCGATCTNNNCAAGAGCGATGGAAAACCATTCC
CACACGTCTGAACTCTTTCCCTACACGACGCTCTTCCGATCTNNNCAAGAGCGATGGAAAACCATTCC
AGCACACGTCTGAACTCTTTCCCTACACGACGCTCTTCCGATCTNNNTGAACTCCTGACCTCAAGTGAT
AGCACACGTCTGAACTCTTTCCCTACACGACGCTCTTCCGATCTNNNTGAACTCCTGACCTCAAGTGAT
AGCACACGTCTGAACTCTTTCCCTACACGACGCTCTTCCGATCTNNNTGAACTCCTGACCTCAAGTGAT
GAGCACACGTCTGAACTCTTTCCCTACACGACGCTCTTCCGATCTNNNGAACAGAGGGTAGAAAATG
GAGCACACGTCTGAACTCTTTCCCTACACGACGCTCTTCCGATCTNNNGAACAGAGGGTAGAAAATG
GAGCACACGTCTGAACTCTTTCCCTACACGACGCTCTTCCGATCTNNNGAACAGAGGGTAGAAAATG
GAGCACACGTCTGAACTCTTTCCCTACACGACGCTCTTCCGATCTNNNAGCCATCTCATCAGCCATTTT
GAGCACACGTCTGAACTCTTTCCCTACACGACGCTCTTCCGATCTNNNAGCCATCTCATCAGCCATTTT
GAGCACACGTCTGAACTCTTTCCCTACACGACGCTCTTCCGATCTNNNAGCCATCTCATCAGCCATTTT
AGCACACGTCTGAACTCTTTCCCTACACGACGCTCTTCCGATCTNNNCTGGAGTGCAGTAGTACCATC
AGCACACGTCTGAACTCTTTCCCTACACGACGCTCTTCCGATCTNNNCTGGAGTGCAGTAGTACCATC
AGCACACGTCTGAACTCTTTCCCTACACGACGCTCTTCCGATCTNNNCTGGAGTGCAGTAGTACCATC
AGCACACGTCTGAACTCTTTCCCTACACGACGCTCTTCCGATCTNNNTTGGGGAGAAAGGATAAACA
AGCACACGTCTGAACTCTTTCCCTACACGACGCTCTTCCGATCTNNNTTGGGGAGAAAGGATAAACA
AGCACACGTCTGAACTCTTTCCCTACACGACGCTCTTCCGATCTNNNTTGGGGAGAAAGGATAAACA
AGAGCACACGTCTGAACTCTTTCCCTACACGACGCTCTTCCGATCTNNNGCTCAAAACCTAGACATCTA
AGAGCACACGTCTGAACTCTTTCCCTACACGACGCTCTTCCGATCTNNNGCTCAAAACCTAGACATCTA
AGAGCACACGTCTGAACTCTTTCCCTACACGACGCTCTTCCGATCTNNNGCTCAAAACCTAGACATCTA
GCACACGTCTGAACTCTTTCCCTACACGACGCTCTTCCGATCTNNNTGTGGAAGGTGACATCAAGTT
GCACACGTCTGAACTCTTTCCCTACACGACGCTCTTCCGATCTNNNTGTGGAAGGTGACATCAAGTT
GCACACGTCTGAACTCTTTCCCTACACGACGCTCTTCCGATCTNNNTGTGGAAGGTGACATCAAGTT
AGCACACGTCTGAACTCTTTCCCTACACGACGCTCTTCCGATCTNNNAGGCAACCTACAAAATGGGA
AGCACACGTCTGAACTCTTTCCCTACACGACGCTCTTCCGATCTNNNAGGCAACCTACAAAATGGGA
AGCACACGTCTGAACTCTTTCCCTACACGACGCTCTTCCGATCTNNNAGGCAACCTACAAAATGGGA
GAGCACACGTCTGAACTCTTTCCCTACACGACGCTCTTCCGATCTNNNCTTCATTCACCAGTTACAGCT
GAGCACACGTCTGAACTCTTTCCCTACACGACGCTCTTCCGATCTNNNCTTCATTCACCAGTTACAGCT
GAGCACACGTCTGAACTCTTTCCCTACACGACGCTCTTCCGATCTNNNCTTCATTCACCAGTTACAGCT
AGCACACGTCTGAACTCTTTCCCTACACGACGCTCTTCCGATCTNNNCTTTCTATGAGCCAAGATCGCA
AGCACACGTCTGAACTCTTTCCCTACACGACGCTCTTCCGATCTNNNCTTTCTATGAGCCAAGATCGCA
AGCACACGTCTGAACTCTTTCCCTACACGACGCTCTTCCGATCTNNNCTTTCTATGAGCCAAGATCGCA

AGCACACGTCTGAACTCTTTCCCTACACGACGCTCTTCCGATCTNNNCTCACACTGTATCTCCTCCCATGC
AGCACACGTCTGAACTCTTTCCCTACACGACGCTCTTCCGATCTNNNCTCACACTGTATCTCCTCCCATGC
AGCACACGTCTGAACTCTTTCCCTACACGACGCTCTTCCGATCTNNNCTCACACTGTATCTCCTCCCATGC
AGCACACGTCTGAACTCTTTCCCTACACGACGCTCTTCCGATCTNNNAACAATAACCGGCACACACAAA
AGCACACGTCTGAACTCTTTCCCTACACGACGCTCTTCCGATCTNNNAACAATAACCGGCACACACAAA
AGCACACGTCTGAACTCTTTCCCTACACGACGCTCTTCCGATCTNNNAACAATAACCGGCACACACAAA
AGAGCACACGTCTGAACTCTTTCCCTACACGACGCTCTTCCGATCTNNNTGGAACTGGGAAAGTCTAA
AGAGCACACGTCTGAACTCTTTCCCTACACGACGCTCTTCCGATCTNNNTGGAACTGGGAAAGTCTAA
AGAGCACACGTCTGAACTCTTTCCCTACACGACGCTCTTCCGATCTNNNTGGAACTGGGAAAGTCTAA
GAGCACACGTCTGAACTCTTTCCCTACACGACGCTCTTCCGATCTNNNGCAAACATGATGAAACCCTGT
GAGCACACGTCTGAACTCTTTCCCTACACGACGCTCTTCCGATCTNNNGCAAACATGATGAAACCCTGT
GAGCACACGTCTGAACTCTTTCCCTACACGACGCTCTTCCGATCTNNNGCAAACATGATGAAACCCTGT
AGCACACGTCTGAACTCTTTCCCTACACGACGCTCTTCCGATCTNNNCCAGGTGTTTCTGTGCCTGTAAA
AGCACACGTCTGAACTCTTTCCCTACACGACGCTCTTCCGATCTNNNCCAGGTGTTTCTGTGCCTGTAAA
AGCACACGTCTGAACTCTTTCCCTACACGACGCTCTTCCGATCTNNNCCAGGTGTTTCTGTGCCTGTAAA
GAGCACACGTCTGAACTCTTTCCCTACACGACGCTCTTCCGATCTNNNAGTACCCCCTGGAATACTGAA
GAGCACACGTCTGAACTCTTTCCCTACACGACGCTCTTCCGATCTNNNAGTACCCCCTGGAATACTGAA
GAGCACACGTCTGAACTCTTTCCCTACACGACGCTCTTCCGATCTNNNAGTACCCCCTGGAATACTGAA
AGCACACGTCTGAACTCTTTCCCTACACGACGCTCTTCCGATCTNNNACACCTGTAGTCCCAGCTACTCC
AGCACACGTCTGAACTCTTTCCCTACACGACGCTCTTCCGATCTNNNACACCTGTAGTCCCAGCTACTCC
AGCACACGTCTGAACTCTTTCCCTACACGACGCTCTTCCGATCTNNNACACCTGTAGTCCCAGCTACTCC
GAGCACACGTCTGAACTCTTTCCCTACACGACGCTCTTCCGATCTNNNCCTGGCAACAAGAGTGAAA
GAGCACACGTCTGAACTCTTTCCCTACACGACGCTCTTCCGATCTNNNCCTGGCAACAAGAGTGAAA
GAGCACACGTCTGAACTCTTTCCCTACACGACGCTCTTCCGATCTNNNCCTGGCAACAAGAGTGAAA
AGCACACGTCTGAACTCTTTCCCTACACGACGCTCTTCCGATCTNNNGTTACAGTGAGCTGAGAATGT
AGCACACGTCTGAACTCTTTCCCTACACGACGCTCTTCCGATCTNNNGTTACAGTGAGCTGAGAATGT
AGCACACGTCTGAACTCTTTCCCTACACGACGCTCTTCCGATCTNNNGTTACAGTGAGCTGAGAATGT
AGAGCACACGTCTGAACTCTTTCCCTACACGACGCTCTTCCGATCTNNNATCACTCCACTGTACTCTAGC
AGAGCACACGTCTGAACTCTTTCCCTACACGACGCTCTTCCGATCTNNNATCACTCCACTGTACTCTAGC
AGAGCACACGTCTGAACTCTTTCCCTACACGACGCTCTTCCGATCTNNNATCACTCCACTGTACTCTAGC
GAGCACACGTCTGAACTCTTTCCCTACACGACGCTCTTCCGATCTNNNTTCACCCGAAAAACCTCCTGTA
GAGCACACGTCTGAACTCTTTCCCTACACGACGCTCTTCCGATCTNNNTTCACCCGAAAAACCTCCTGTA
GAGCACACGTCTGAACTCTTTCCCTACACGACGCTCTTCCGATCTNNNTTCACCCGAAAAACCTCCTGTA
AGCACACGTCTGAACTCTTTCCCTACACGACGCTCTTCCGATCTNNNCAAAGTGTGTTCAGCTATGACG
AGCACACGTCTGAACTCTTTCCCTACACGACGCTCTTCCGATCTNNNCAAAGTGTGTTCAGCTATGACG
AGCACACGTCTGAACTCTTTCCCTACACGACGCTCTTCCGATCTNNNCAAAGTGTGTTCAGCTATGACG
AGAGCACACGTCTGAACTCTTTCCCTACACGACGCTCTTCCGATCTNNNAAATTGAACTTCCCTCCCTCC
AGAGCACACGTCTGAACTCTTTCCCTACACGACGCTCTTCCGATCTNNNAAATTGAACTTCCCTCCCTCC
AGAGCACACGTCTGAACTCTTTCCCTACACGACGCTCTTCCGATCTNNNAAATTGAACTTCCCTCCCTCC
AGCACACGTCTGAACTCTTTCCCTACACGACGCTCTTCCGATCTNNNCCCAAAACAGAGTGAACTTTGCT
AGCACACGTCTGAACTCTTTCCCTACACGACGCTCTTCCGATCTNNNCCCAAAACAGAGTGAACTTTGCT
AGCACACGTCTGAACTCTTTCCCTACACGACGCTCTTCCGATCTNNNCCCAAAACAGAGTGAACTTTGCT
GAGCACACGTCTGAACTCTTTCCCTACACGACGCTCTTCCGATCTNNNCACGTTCCATGAGCACAGATA

GAGCACACGTCTGAACTCTTTCCCTACACGACGCTCTTCCGATCTNNNCACGTTCCATGAGCACAGATA

GAGCACACGTCTGAACTCTTTCCCTACACGACGCTCTTCCGATCTNNNCACGTTCCATGAGCACAGATA

GAGCACACGTCTGAACTCTTTCCCTACACGACGCTCTTCCGATCTNNNTGGGAATCACTTGAACCTGAG

GAGCACACGTCTGAACTCTTTCCCTACACGACGCTCTTCCGATCTNNNTGGGAATCACTTGAACCTGAG

GAGCACACGTCTGAACTCTTTCCCTACACGACGCTCTTCCGATCTNNNTGGGAATCACTTGAACCTGAG

CACGTCTGAACTCTTTCCCTACACGACGCTCTTCCGATCTNNNAGACTCCGTCTCAAACAAAACAACAT

CACGTCTGAACTCTTTCCCTACACGACGCTCTTCCGATCTNNNAGACTCCGTCTCAAACAAAACAACAT

CACGTCTGAACTCTTTCCCTACACGACGCTCTTCCGATCTNNNAGACTCCGTCTCAAACAAAACAACAT

AGCACACGTCTGAACTCTTTCCCTACACGACGCTCTTCCGATCTNNNAGGCAGGAGAATAGCTTGAA

AGCACACGTCTGAACTCTTTCCCTACACGACGCTCTTCCGATCTNNNAGGCAGGAGAATAGCTTGAA

AGCACACGTCTGAACTCTTTCCCTACACGACGCTCTTCCGATCTNNNAGGCAGGAGAATAGCTTGAA

GAGCACACGTCTGAACTCTTTCCCTACACGACGCTCTTCCGATCTNNNAATTGCAGACACAGGAAAAG

GAGCACACGTCTGAACTCTTTCCCTACACGACGCTCTTCCGATCTNNNAATTGCAGACACAGGAAAAG

GAGCACACGTCTGAACTCTTTCCCTACACGACGCTCTTCCGATCTNNNAATTGCAGACACAGGAAAAG

AGCACACGTCTGAACTCTTTCCCTACACGACGCTCTTCCGATCTNNNCAAAACCGGCAGAATCAGGAG

AGCACACGTCTGAACTCTTTCCCTACACGACGCTCTTCCGATCTNNNCAAAACCGGCAGAATCAGGAG

AGCACACGTCTGAACTCTTTCCCTACACGACGCTCTTCCGATCTNNNCAAAACCGGCAGAATCAGGAG

AGAGCACACGTCTGAACTCTTTCCCTACACGACGCTCTTCCGATCTNNNTTACAGTGAGCCAAGATCG

AGAGCACACGTCTGAACTCTTTCCCTACACGACGCTCTTCCGATCTNNNTTACAGTGAGCCAAGATCG

AGAGCACACGTCTGAACTCTTTCCCTACACGACGCTCTTCCGATCTNNNTTACAGTGAGCCAAGATCG

AGAGCACACGTCTGAACTCTTTCCCTACACGACGCTCTTCCGATCTNNNCCGTAGTTTTTCTTGGCAGT

AGAGCACACGTCTGAACTCTTTCCCTACACGACGCTCTTCCGATCTNNNCCGTAGTTTTTCTTGGCAGT

AGAGCACACGTCTGAACTCTTTCCCTACACGACGCTCTTCCGATCTNNNCCGTAGTTTTTCTTGGCAGT

GAGCACACGTCTGAACTCTTTCCCTACACGACGCTCTTCCGATCTNNNAGTATCCATTTGCCCCTCATTG

GAGCACACGTCTGAACTCTTTCCCTACACGACGCTCTTCCGATCTNNNAGTATCCATTTGCCCCTCATTG

GAGCACACGTCTGAACTCTTTCCCTACACGACGCTCTTCCGATCTNNNAGTATCCATTTGCCCCTCATTG

AGCACACGTCTGAACTCTTTCCCTACACGACGCTCTTCCGATCTNNNGAGCGTCTGGGGCACTTTAGC

AGCACACGTCTGAACTCTTTCCCTACACGACGCTCTTCCGATCTNNNGAGCGTCTGGGGCACTTTAGC

AGCACACGTCTGAACTCTTTCCCTACACGACGCTCTTCCGATCTNNNGAGCGTCTGGGGCACTTTAGC

AGAGCACACGTCTGAACTCTTTCCCTACACGACGCTCTTCCGATCTNNNAGCATTCTCCAGAGTTGTCC

AGAGCACACGTCTGAACTCTTTCCCTACACGACGCTCTTCCGATCTNNNAGCATTCTCCAGAGTTGTCC

AGAGCACACGTCTGAACTCTTTCCCTACACGACGCTCTTCCGATCTNNNAGCATTCTCCAGAGTTGTCC

AGAGCACACGTCTGAACTCTTTCCCTACACGACGCTCTTCCGATCTNNNTCTTGGGACTTCTCCAGATTC

AGAGCACACGTCTGAACTCTTTCCCTACACGACGCTCTTCCGATCTNNNTCTTGGGACTTCTCCAGATTC

AGAGCACACGTCTGAACTCTTTCCCTACACGACGCTCTTCCGATCTNNNTCTTGGGACTTCTCCAGATTC

GAGCACACGTCTGAACTCTTTCCCTACACGACGCTCTTCCGATCTNNNGGAGATTGCAGTGAACTGGG

GAGCACACGTCTGAACTCTTTCCCTACACGACGCTCTTCCGATCTNNNGGAGATTGCAGTGAACTGGG

GAGCACACGTCTGAACTCTTTCCCTACACGACGCTCTTCCGATCTNNNGGAGATTGCAGTGAACTGGG

GCACACGTCTGAACTCTTTCCCTACACGACGCTCTTCCGATCTNNNAGTATTTGTGTTGTCAGGCAAG

GCACACGTCTGAACTCTTTCCCTACACGACGCTCTTCCGATCTNNNAGTATTTGTGTTGTCAGGCAAG

GCACACGTCTGAACTCTTTCCCTACACGACGCTCTTCCGATCTNNNAGTATTTGTGTTGTCAGGCAAG

GAGCACACGTCTGAACTCTTTCCCTACACGACGCTCTTCCGATCTNNNAGATGGTTTTAGTATTCATCA

GAGCACACGTCTGAACTCTTTCCCTACACGACGCTCTTCCGATCTNNNAGATGGTTTTAGTATTCATCA

GAGCACACGTCTGAACTCTTTCCCTACACGACGCTCTTCCGATCTNNNAGATGGTTTTAGTATTCATCAT
GAGCACACGTCTGAACTCTTTCCCTACACGACGCTCTTCCGATCTNNNTTTGTTTGCCCAGATTTTCTGC
GAGCACACGTCTGAACTCTTTCCCTACACGACGCTCTTCCGATCTNNNTTTGTTTGCCCAGATTTTCTGC
GAGCACACGTCTGAACTCTTTCCCTACACGACGCTCTTCCGATCTNNNTTTGTTTGCCCAGATTTTCTGC
AGAGCACACGTCTGAACTCTTTCCCTACACGACGCTCTTCCGATCTNNNGAGTTTGAAACTGCAGTGAA
AGAGCACACGTCTGAACTCTTTCCCTACACGACGCTCTTCCGATCTNNNGAGTTTGAAACTGCAGTGAA
AGAGCACACGTCTGAACTCTTTCCCTACACGACGCTCTTCCGATCTNNNGAGTTTGAAACTGCAGTGAA
AGCACACGTCTGAACTCTTTCCCTACACGACGCTCTTCCGATCTNNNTCAATCTTTGCATGTTCCCACCA
AGCACACGTCTGAACTCTTTCCCTACACGACGCTCTTCCGATCTNNNTCAATCTTTGCATGTTCCCACCA
AGCACACGTCTGAACTCTTTCCCTACACGACGCTCTTCCGATCTNNNTCAATCTTTGCATGTTCCCACCA
AGCACACGTCTGAACTCTTTCCCTACACGACGCTCTTCCGATCTNNNCATACGTATACATGTGCCATGCC
AGCACACGTCTGAACTCTTTCCCTACACGACGCTCTTCCGATCTNNNCATACGTATACATGTGCCATGCC
AGCACACGTCTGAACTCTTTCCCTACACGACGCTCTTCCGATCTNNNCATACGTATACATGTGCCATGCC
AGCACACGTCTGAACTCTTTCCCTACACGACGCTCTTCCGATCTNNNAGCAATATCCACACCTTGGCTAA
AGCACACGTCTGAACTCTTTCCCTACACGACGCTCTTCCGATCTNNNAGCAATATCCACACCTTGGCTAA
AGCACACGTCTGAACTCTTTCCCTACACGACGCTCTTCCGATCTNNNAGCAATATCCACACCTTGGCTAA
AGCACACGTCTGAACTCTTTCCCTACACGACGCTCTTCCGATCTNNNGGGTTTCCAATATCCCTGAGT
AGCACACGTCTGAACTCTTTCCCTACACGACGCTCTTCCGATCTNNNGGGTTTCCAATATCCCTGAGT
AGCACACGTCTGAACTCTTTCCCTACACGACGCTCTTCCGATCTNNNGGGTTTCCAATATCCCTGAGT
GAGCACACGTCTGAACTCTTTCCCTACACGACGCTCTTCCGATCTNNNGAGAATGGCATGTAACCAGG
GAGCACACGTCTGAACTCTTTCCCTACACGACGCTCTTCCGATCTNNNGAGAATGGCATGTAACCAGG
GAGCACACGTCTGAACTCTTTCCCTACACGACGCTCTTCCGATCTNNNGAGAATGGCATGTAACCAGG
AGCACACGTCTGAACTCTTTCCCTACACGACGCTCTTCCGATCTNNNGTAATCCCAGCTACTATGGAGG
AGCACACGTCTGAACTCTTTCCCTACACGACGCTCTTCCGATCTNNNGTAATCCCAGCTACTATGGAGG
AGCACACGTCTGAACTCTTTCCCTACACGACGCTCTTCCGATCTNNNGTAATCCCAGCTACTATGGAGG
GAGCACACGTCTGAACTCTTTCCCTACACGACGCTCTTCCGATCTNNNCCAGGCTCTGAAGATCAAGTTC
GAGCACACGTCTGAACTCTTTCCCTACACGACGCTCTTCCGATCTNNNCCAGGCTCTGAAGATCAAGTTC
GAGCACACGTCTGAACTCTTTCCCTACACGACGCTCTTCCGATCTNNNCCAGGCTCTGAAGATCAAGTTC
GAGCACACGTCTGAACTCTTTCCCTACACGACGCTCTTCCGATCTNNNTTGGACCAACTGGAATGAACA
GAGCACACGTCTGAACTCTTTCCCTACACGACGCTCTTCCGATCTNNNTTGGACCAACTGGAATGAACA
GAGCACACGTCTGAACTCTTTCCCTACACGACGCTCTTCCGATCTNNNTTGGACCAACTGGAATGAACA
GAGCACACGTCTGAACTCTTTCCCTACACGACGCTCTTCCGATCTNNNTGCATTTTGGAGTGGTCGTATC
GAGCACACGTCTGAACTCTTTCCCTACACGACGCTCTTCCGATCTNNNTGCATTTTGGAGTGGTCGTATC
GAGCACACGTCTGAACTCTTTCCCTACACGACGCTCTTCCGATCTNNNTGCATTTTGGAGTGGTCGTATC
GAGCACACGTCTGAACTCTTTCCCTACACGACGCTCTTCCGATCTNNNATAAATTCCTGGACACATGCAC
GAGCACACGTCTGAACTCTTTCCCTACACGACGCTCTTCCGATCTNNNATAAATTCCTGGACACATGCAC
GAGCACACGTCTGAACTCTTTCCCTACACGACGCTCTTCCGATCTNNNATAAATTCCTGGACACATGCAC
GAGCACACGTCTGAACTCTTTCCCTACACGACGCTCTTCCGATCTNNNGATTTGGTTTATCCCGAAATAT
GAGCACACGTCTGAACTCTTTCCCTACACGACGCTCTTCCGATCTNNNGATTTGGTTTATCCCGAAATAT
GAGCACACGTCTGAACTCTTTCCCTACACGACGCTCTTCCGATCTNNNGATTTGGTTTATCCCGAAATAT
AGCACACGTCTGAACTCTTTCCCTACACGACGCTCTTCCGATCTNNNTCTTGAAGAGAACACCCCCTTTTC
AGCACACGTCTGAACTCTTTCCCTACACGACGCTCTTCCGATCTNNNTCTTGAAGAGAACACCCCCTTTTC
AGCACACGTCTGAACTCTTTCCCTACACGACGCTCTTCCGATCTNNNTCTTGAAGAGAACACCCCCTTTTC

GAGCACACGTCTGAACTCTTTCCCTACACGACGCTCTTCCGATCTNNNAGAACTACATTCTATAGGAAC
GAGCACACGTCTGAACTCTTTCCCTACACGACGCTCTTCCGATCTNNNAGAACTACATTCTATAGGAAC
GAGCACACGTCTGAACTCTTTCCCTACACGACGCTCTTCCGATCTNNNAGAACTACATTCTATAGGAAC
AGCACACGTCTGAACTCTTTCCCTACACGACGCTCTTCCGATCTNNNGGTGGGTAATTTGAGGTCAC
AGCACACGTCTGAACTCTTTCCCTACACGACGCTCTTCCGATCTNNNGGTGGGTAATTTGAGGTCAC
AGCACACGTCTGAACTCTTTCCCTACACGACGCTCTTCCGATCTNNNGGTGGGTAATTTGAGGTCAC
GAGCACACGTCTGAACTCTTTCCCTACACGACGCTCTTCCGATCTNNNAAGTTTTGTTTAAAAAATGTG
GAGCACACGTCTGAACTCTTTCCCTACACGACGCTCTTCCGATCTNNNAAGTTTTGTTTAAAAAATGTG
GAGCACACGTCTGAACTCTTTCCCTACACGACGCTCTTCCGATCTNNNAAGTTTTGTTTAAAAAATGTG
AAGAGCACACGTCTGAACTCTTTCCCTACACGACGCTCTTCCGATCTNNNTTGTGGAACACTTGGAA
AAGAGCACACGTCTGAACTCTTTCCCTACACGACGCTCTTCCGATCTNNNTTGTGGAACACTTGGAA
AAGAGCACACGTCTGAACTCTTTCCCTACACGACGCTCTTCCGATCTNNNTTGTGGAACACTTGGAA
CACACGTCTGAACTCTTTCCCTACACGACGCTCTTCCGATCTNNNACTGTCAAGTCTGGATGATTCCC
CACACGTCTGAACTCTTTCCCTACACGACGCTCTTCCGATCTNNNACTGTCAAGTCTGGATGATTCCC
CACACGTCTGAACTCTTTCCCTACACGACGCTCTTCCGATCTNNNACTGTCAAGTCTGGATGATTCCC
AGCACACGTCTGAACTCTTTCCCTACACGACGCTCTTCCGATCTNNNGGCCCCGGGAATTATTTCTTT
AGCACACGTCTGAACTCTTTCCCTACACGACGCTCTTCCGATCTNNNGGCCCCGGGAATTATTTCTTT
AGCACACGTCTGAACTCTTTCCCTACACGACGCTCTTCCGATCTNNNGGCCCCGGGAATTATTTCTTT
GAGCACACGTCTGAACTCTTTCCCTACACGACGCTCTTCCGATCTNNNTGTAGAAAGATTTCAGTGC
GAGCACACGTCTGAACTCTTTCCCTACACGACGCTCTTCCGATCTNNNTGTAGAAAGATTTCAGTGC
GAGCACACGTCTGAACTCTTTCCCTACACGACGCTCTTCCGATCTNNNTGTAGAAAGATTTCAGTGC
AGCACACGTCTGAACTCTTTCCCTACACGACGCTCTTCCGATCTNNNGTGGGAGATGGAATTATGCA
AGCACACGTCTGAACTCTTTCCCTACACGACGCTCTTCCGATCTNNNGTGGGAGATGGAATTATGCA
AGCACACGTCTGAACTCTTTCCCTACACGACGCTCTTCCGATCTNNNGTGGGAGATGGAATTATGCA
GCACACGTCTGAACTCTTTCCCTACACGACGCTCTTCCGATCTNNNCCCCCTTAGTTCCTCCCTACTTCC
GCACACGTCTGAACTCTTTCCCTACACGACGCTCTTCCGATCTNNNCCCCCTTAGTTCCTCCCTACTTCC
GCACACGTCTGAACTCTTTCCCTACACGACGCTCTTCCGATCTNNNCCCCCTTAGTTCCTCCCTACTTCC
GCACACGTCTGAACTCTTTCCCTACACGACGCTCTTCCGATCTNNNGAGTCTTGCTCTGTCACCCAGC
GCACACGTCTGAACTCTTTCCCTACACGACGCTCTTCCGATCTNNNGAGTCTTGCTCTGTCACCCAGC
GCACACGTCTGAACTCTTTCCCTACACGACGCTCTTCCGATCTNNNGAGTCTTGCTCTGTCACCCAGC
AGCACACGTCTGAACTCTTTCCCTACACGACGCTCTTCCGATCTNNNGGCCACAGAAAGCCCAGATCC
AGCACACGTCTGAACTCTTTCCCTACACGACGCTCTTCCGATCTNNNGGCCACAGAAAGCCCAGATCC
AGCACACGTCTGAACTCTTTCCCTACACGACGCTCTTCCGATCTNNNGGCCACAGAAAGCCCAGATCC
AGAGCACACGTCTGAACTCTTTCCCTACACGACGCTCTTCCGATCTNNNCGGCGGGGAAATAGGTTT
AGAGCACACGTCTGAACTCTTTCCCTACACGACGCTCTTCCGATCTNNNCGGCGGGGAAATAGGTTT
AGAGCACACGTCTGAACTCTTTCCCTACACGACGCTCTTCCGATCTNNNCGGCGGGGAAATAGGTTT
GAGCACACGTCTGAACTCTTTCCCTACACGACGCTCTTCCGATCTNNNGAACATAGTACAACCGCTCT
GAGCACACGTCTGAACTCTTTCCCTACACGACGCTCTTCCGATCTNNNGAACATAGTACAACCGCTCT
GAGCACACGTCTGAACTCTTTCCCTACACGACGCTCTTCCGATCTNNNGAACATAGTACAACCGCTCT
AGCACACGTCTGAACTCTTTCCCTACACGACGCTCTTCCGATCTNNNCTGTAATCTCAGCTGCTTGGG
AGCACACGTCTGAACTCTTTCCCTACACGACGCTCTTCCGATCTNNNCTGTAATCTCAGCTGCTTGGG
AGCACACGTCTGAACTCTTTCCCTACACGACGCTCTTCCGATCTNNNCTGTAATCTCAGCTGCTTGGG
GAGCACACGTCTGAACTCTTTCCCTACACGACGCTCTTCCGATCTNNNTGAGAAGTTCAACAAGGAA

AGAGCACACGTCTGAACTCTTTCCCTACACGACGCTCTTCCGATCTNNNTGAGAAGTTCAACAAGGAAA

AGAGCACACGTCTGAACTCTTTCCCTACACGACGCTCTTCCGATCTNNNTGAGAAGTTCAACAAGGAAA

AGCACACGTCTGAACTCTTTCCCTACACGACGCTCTTCCGATCTNNNGGGAAAGATTCTAATCCCCAGCT

AGCACACGTCTGAACTCTTTCCCTACACGACGCTCTTCCGATCTNNNGGGAAAGATTCTAATCCCCAGCT

AGCACACGTCTGAACTCTTTCCCTACACGACGCTCTTCCGATCTNNNGGGAAAGATTCTAATCCCCAGCT

GAGCACACGTCTGAACTCTTTCCCTACACGACGCTCTTCCGATCTNNNACTTGAGAGATGAGATGTAA

GAGCACACGTCTGAACTCTTTCCCTACACGACGCTCTTCCGATCTNNNACTTGAGAGATGAGATGTAA

GAGCACACGTCTGAACTCTTTCCCTACACGACGCTCTTCCGATCTNNNACTTGAGAGATGAGATGTAA

AGCACACGTCTGAACTCTTTCCCTACACGACGCTCTTCCGATCTNNNCGTACACTGAAACAGAACTGCA

AGCACACGTCTGAACTCTTTCCCTACACGACGCTCTTCCGATCTNNNCGTACACTGAAACAGAACTGCA

AGCACACGTCTGAACTCTTTCCCTACACGACGCTCTTCCGATCTNNNCGTACACTGAAACAGAACTGCA

AGCACACGTCTGAACTCTTTCCCTACACGACGCTCTTCCGATCTNNNAGTACCATCAGTCAACTAGGTG

AGCACACGTCTGAACTCTTTCCCTACACGACGCTCTTCCGATCTNNNAGTACCATCAGTCAACTAGGTG

AGCACACGTCTGAACTCTTTCCCTACACGACGCTCTTCCGATCTNNNAGTACCATCAGTCAACTAGGTG

GAGCACACGTCTGAACTCTTTCCCTACACGACGCTCTTCCGATCTNNNCCAGTTGTCCGAGGCTTGTTTA

GAGCACACGTCTGAACTCTTTCCCTACACGACGCTCTTCCGATCTNNNCCAGTTGTCCGAGGCTTGTTTA

GAGCACACGTCTGAACTCTTTCCCTACACGACGCTCTTCCGATCTNNNCCAGTTGTCCGAGGCTTGTTTA

AGCACACGTCTGAACTCTTTCCCTACACGACGCTCTTCCGATCTNNNCCTGGTAATTCTCATGAAAAATA

AGCACACGTCTGAACTCTTTCCCTACACGACGCTCTTCCGATCTNNNCCTGGTAATTCTCATGAAAAATA

AGCACACGTCTGAACTCTTTCCCTACACGACGCTCTTCCGATCTNNNCCTGGTAATTCTCATGAAAAATA

GAGCACACGTCTGAACTCTTTCCCTACACGACGCTCTTCCGATCTNNNGGTCAACGTTATCCTAAGGCTG

GAGCACACGTCTGAACTCTTTCCCTACACGACGCTCTTCCGATCTNNNGGTCAACGTTATCCTAAGGCTG

GAGCACACGTCTGAACTCTTTCCCTACACGACGCTCTTCCGATCTNNNGGTCAACGTTATCCTAAGGCTG

GAGCACACGTCTGAACTCTTTCCCTACACGACGCTCTTCCGATCTNNNAGAGGGCTTGAATCCAGTAGA

GAGCACACGTCTGAACTCTTTCCCTACACGACGCTCTTCCGATCTNNNAGAGGGCTTGAATCCAGTAGA

GAGCACACGTCTGAACTCTTTCCCTACACGACGCTCTTCCGATCTNNNAGAGGGCTTGAATCCAGTAGA

AGCACACGTCTGAACTCTTTCCCTACACGACGCTCTTCCGATCTNNNACGCCTGTAATCCCAGTACTTTG

AGCACACGTCTGAACTCTTTCCCTACACGACGCTCTTCCGATCTNNNACGCCTGTAATCCCAGTACTTTG

AGCACACGTCTGAACTCTTTCCCTACACGACGCTCTTCCGATCTNNNACGCCTGTAATCCCAGTACTTTG

GAGCACACGTCTGAACTCTTTCCCTACACGACGCTCTTCCGATCTNNNCTGAGGTGGGAAAATCGTTTG

GAGCACACGTCTGAACTCTTTCCCTACACGACGCTCTTCCGATCTNNNCTGAGGTGGGAAAATCGTTTG

GAGCACACGTCTGAACTCTTTCCCTACACGACGCTCTTCCGATCTNNNCTGAGGTGGGAAAATCGTTTG

AGCACACGTCTGAACTCTTTCCCTACACGACGCTCTTCCGATCTNNNCTCCTGATTAGCTGGGACTACAG

AGCACACGTCTGAACTCTTTCCCTACACGACGCTCTTCCGATCTNNNCTCCTGATTAGCTGGGACTACAG

AGCACACGTCTGAACTCTTTCCCTACACGACGCTCTTCCGATCTNNNCTCCTGATTAGCTGGGACTACAG

AGCACACGTCTGAACTCTTTCCCTACACGACGCTCTTCCGATCTNNNCCCTCTTGATAGGGACATCCTGT

AGCACACGTCTGAACTCTTTCCCTACACGACGCTCTTCCGATCTNNNCCCTCTTGATAGGGACATCCTGT

AGCACACGTCTGAACTCTTTCCCTACACGACGCTCTTCCGATCTNNNCCCTCTTGATAGGGACATCCTGT

GAGCACACGTCTGAACTCTTTCCCTACACGACGCTCTTCCGATCTNNNCTGAGGCAGGAGAATCACTTG

GAGCACACGTCTGAACTCTTTCCCTACACGACGCTCTTCCGATCTNNNCTGAGGCAGGAGAATCACTTG

GAGCACACGTCTGAACTCTTTCCCTACACGACGCTCTTCCGATCTNNNCTGAGGCAGGAGAATCACTTG

GAGCACACGTCTGAACTCTTTCCCTACACGACGCTCTTCCGATCTNNNCTCCTGGGTTCGAGTGATTCA

GAGCACACGTCTGAACTCTTTCCCTACACGACGCTCTTCCGATCTNNNCTCCTGGGTTCGAGTGATTCA

GAGCACACGTCTGAACTCTTTCCCTACACGACGCTCTTCCGATCTNNNCTCCTGGGTTCGAGTGATTCA

AGCACACGTCTGAACTCTTTCCCTACACGACGCTCTTCCGATCTNNNGTGACAGAGCGAGATTCCATCT

AGCACACGTCTGAACTCTTTCCCTACACGACGCTCTTCCGATCTNNNGTGACAGAGCGAGATTCCATCT

AGCACACGTCTGAACTCTTTCCCTACACGACGCTCTTCCGATCTNNNGTGACAGAGCGAGATTCCATCT

GAGCACACGTCTGAACTCTTTCCCTACACGACGCTCTTCCGATCTNNNCAGGAGAATTGCTTGAACCTT

GAGCACACGTCTGAACTCTTTCCCTACACGACGCTCTTCCGATCTNNNCAGGAGAATTGCTTGAACCTT

GAGCACACGTCTGAACTCTTTCCCTACACGACGCTCTTCCGATCTNNNCAGGAGAATTGCTTGAACCTT

ACACGTCTGAACTCTTTCCCTACACGACGCTCTTCCGATCTNNNTTCTCAATGAGCTGTTGGGTACACA

ACACGTCTGAACTCTTTCCCTACACGACGCTCTTCCGATCTNNNTTCTCAATGAGCTGTTGGGTACACA

ACACGTCTGAACTCTTTCCCTACACGACGCTCTTCCGATCTNNNTTCTCAATGAGCTGTTGGGTACACA

AGCACACGTCTGAACTCTTTCCCTACACGACGCTCTTCCGATCTNNNTGTGGCTAGCACAATCAAGAAA

AGCACACGTCTGAACTCTTTCCCTACACGACGCTCTTCCGATCTNNNTGTGGCTAGCACAATCAAGAAA

AGCACACGTCTGAACTCTTTCCCTACACGACGCTCTTCCGATCTNNNTGTGGCTAGCACAATCAAGAAA

AGCACACGTCTGAACTCTTTCCCTACACGACGCTCTTCCGATCTNNNGTAACCTTGGGTAGATCATGG

AGCACACGTCTGAACTCTTTCCCTACACGACGCTCTTCCGATCTNNNGTAACCTTGGGTAGATCATGG

AGCACACGTCTGAACTCTTTCCCTACACGACGCTCTTCCGATCTNNNGTAACCTTGGGTAGATCATGG

AGCACACGTCTGAACTCTTTCCCTACACGACGCTCTTCCGATCTNNNGCACTCACACAATTTGACTGGAA

AGCACACGTCTGAACTCTTTCCCTACACGACGCTCTTCCGATCTNNNGCACTCACACAATTTGACTGGAA

AGCACACGTCTGAACTCTTTCCCTACACGACGCTCTTCCGATCTNNNGCACTCACACAATTTGACTGGAA

AGAGCACACGTCTGAACTCTTTCCCTACACGACGCTCTTCCGATCTNNNTTCAGGTAGCCTTGGTATAG

AGAGCACACGTCTGAACTCTTTCCCTACACGACGCTCTTCCGATCTNNNTTCAGGTAGCCTTGGTATAG

AGAGCACACGTCTGAACTCTTTCCCTACACGACGCTCTTCCGATCTNNNTTCAGGTAGCCTTGGTATAG

AGCACACGTCTGAACTCTTTCCCTACACGACGCTCTTCCGATCTNNNTCCTTCTGTTTTTGTGCTTTTCAC

AGCACACGTCTGAACTCTTTCCCTACACGACGCTCTTCCGATCTNNNTCCTTCTGTTTTTGTGCTTTTCAC

AGCACACGTCTGAACTCTTTCCCTACACGACGCTCTTCCGATCTNNNTCCTTCTGTTTTTGTGCTTTTCAC

GAGCACACGTCTGAACTCTTTCCCTACACGACGCTCTTCCGATCTNNNGCAACAAGAGCGAAATTCCAT

GAGCACACGTCTGAACTCTTTCCCTACACGACGCTCTTCCGATCTNNNGCAACAAGAGCGAAATTCCAT

GAGCACACGTCTGAACTCTTTCCCTACACGACGCTCTTCCGATCTNNNGCAACAAGAGCGAAATTCCAT

GAGCACACGTCTGAACTCTTTCCCTACACGACGCTCTTCCGATCTNNNCTGAGGCAGCAGAATCACTTC

GAGCACACGTCTGAACTCTTTCCCTACACGACGCTCTTCCGATCTNNNCTGAGGCAGCAGAATCACTTC

GAGCACACGTCTGAACTCTTTCCCTACACGACGCTCTTCCGATCTNNNCTGAGGCAGCAGAATCACTTC

GCACACGTCTGAACTCTTTCCCTACACGACGCTCTTCCGATCTNNNTGTCCCCACACAAATCTCATCTTCA

GCACACGTCTGAACTCTTTCCCTACACGACGCTCTTCCGATCTNNNTGTCCCCACACAAATCTCATCTTCA

GCACACGTCTGAACTCTTTCCCTACACGACGCTCTTCCGATCTNNNTGTCCCCACACAAATCTCATCTTCA

AGCACACGTCTGAACTCTTTCCCTACACGACGCTCTTCCGATCTNNNCTGAGGCAGGAGAATCAGTTG

AGCACACGTCTGAACTCTTTCCCTACACGACGCTCTTCCGATCTNNNCTGAGGCAGGAGAATCAGTTG

AGCACACGTCTGAACTCTTTCCCTACACGACGCTCTTCCGATCTNNNCTGAGGCAGGAGAATCAGTTG

AAGAGCACACGTCTGAACTCTTTCCCTACACGACGCTCTTCCGATCTNNNGAGGTTGCAGTGAGCTGA

AAGAGCACACGTCTGAACTCTTTCCCTACACGACGCTCTTCCGATCTNNNGAGGTTGCAGTGAGCTGA

AAGAGCACACGTCTGAACTCTTTCCCTACACGACGCTCTTCCGATCTNNNGAGGTTGCAGTGAGCTGA

AGCACACGTCTGAACTCTTTCCCTACACGACGCTCTTCCGATCTNNNCTGATCTTGGCTCACTGTAACCT

AGCACACGTCTGAACTCTTTCCCTACACGACGCTCTTCCGATCTNNNCTGATCTTGGCTCACTGTAACCT

AGCACACGTCTGAACTCTTTCCCTACACGACGCTCTTCCGATCTNNNCTGATCTTGGCTCACTGTAACCT

GAGCACACGTCTGAACTCTTTCCCTACACGACGCTCTTCCGATCTNNNTAGGTAATCACACCACTGCA

GAGCACACGTCTGAACTCTTTCCCTACACGACGCTCTTCCGATCTNNNTAGGTAATCACACCACTGCA

GAGCACACGTCTGAACTCTTTCCCTACACGACGCTCTTCCGATCTNNNTAGGTAATCACACCACTGCA

GAGCACACGTCTGAACTCTTTCCCTACACGACGCTCTTCCGATCTNNNGAGAAAGGGGATGAATAG

GAGCACACGTCTGAACTCTTTCCCTACACGACGCTCTTCCGATCTNNNGAGAAAGGGGATGAATAG

GAGCACACGTCTGAACTCTTTCCCTACACGACGCTCTTCCGATCTNNNGAGAAAGGGGATGAATAG

GCACACGTCTGAACTCTTTCCCTACACGACGCTCTTCCGATCTNNNATAGCCTGGACAACATGGTGA

GCACACGTCTGAACTCTTTCCCTACACGACGCTCTTCCGATCTNNNATAGCCTGGACAACATGGTGA

GCACACGTCTGAACTCTTTCCCTACACGACGCTCTTCCGATCTNNNATAGCCTGGACAACATGGTGA

GAGCACACGTCTGAACTCTTTCCCTACACGACGCTCTTCCGATCTNNNATGACTCTGATGATCATCCAG

GAGCACACGTCTGAACTCTTTCCCTACACGACGCTCTTCCGATCTNNNATGACTCTGATGATCATCCAG

GAGCACACGTCTGAACTCTTTCCCTACACGACGCTCTTCCGATCTNNNATGACTCTGATGATCATCCAG

AGCACACGTCTGAACTCTTTCCCTACACGACGCTCTTCCGATCTNNNGCCATAGGACTTGTGAGAGA

AGCACACGTCTGAACTCTTTCCCTACACGACGCTCTTCCGATCTNNNGCCATAGGACTTGTGAGAGA

AGCACACGTCTGAACTCTTTCCCTACACGACGCTCTTCCGATCTNNNGCCATAGGACTTGTGAGAGA

GCACACGTCTGAACTCTTTCCCTACACGACGCTCTTCCGATCTNNNTAGTCCAGGTGAGAGTCACTG

GCACACGTCTGAACTCTTTCCCTACACGACGCTCTTCCGATCTNNNTAGTCCAGGTGAGAGTCACTG

GCACACGTCTGAACTCTTTCCCTACACGACGCTCTTCCGATCTNNNTAGTCCAGGTGAGAGTCACTG

AGCACACGTCTGAACTCTTTCCCTACACGACGCTCTTCCGATCTNNNAGTAGTGCTTGCTTTATAAATC

AGCACACGTCTGAACTCTTTCCCTACACGACGCTCTTCCGATCTNNNAGTAGTGCTTGCTTTATAAATC

AGCACACGTCTGAACTCTTTCCCTACACGACGCTCTTCCGATCTNNNAGTAGTGCTTGCTTTATAAATC

AGCACACGTCTGAACTCTTTCCCTACACGACGCTCTTCCGATCTNNNCGATCTTGCACAAAACACATC

AGCACACGTCTGAACTCTTTCCCTACACGACGCTCTTCCGATCTNNNCGATCTTGCACAAAACACATC

AGCACACGTCTGAACTCTTTCCCTACACGACGCTCTTCCGATCTNNNCGATCTTGCACAAAACACATC

GCACACGTCTGAACTCTTTCCCTACACGACGCTCTTCCGATCTNNNAGATTGTACCACTGCACTCTAG

GCACACGTCTGAACTCTTTCCCTACACGACGCTCTTCCGATCTNNNAGATTGTACCACTGCACTCTAG

GCACACGTCTGAACTCTTTCCCTACACGACGCTCTTCCGATCTNNNAGATTGTACCACTGCACTCTAG

AGCACACGTCTGAACTCTTTCCCTACACGACGCTCTTCCGATCTNNNAAAGGTGTGTATGTGTGGGG

AGCACACGTCTGAACTCTTTCCCTACACGACGCTCTTCCGATCTNNNAAAGGTGTGTATGTGTGGGG

AGCACACGTCTGAACTCTTTCCCTACACGACGCTCTTCCGATCTNNNAAAGGTGTGTATGTGTGGGG

GCACACGTCTGAACTCTTTCCCTACACGACGCTCTTCCGATCTNNNACTTTTCTCAGTGTGCCACAATG

GCACACGTCTGAACTCTTTCCCTACACGACGCTCTTCCGATCTNNNACTTTTCTCAGTGTGCCACAATG

GCACACGTCTGAACTCTTTCCCTACACGACGCTCTTCCGATCTNNNACTTTTCTCAGTGTGCCACAATG

GCACACGTCTGAACTCTTTCCCTACACGACGCTCTTCCGATCTNNNTTGTCTTTTTCACCACATCGTGA

GCACACGTCTGAACTCTTTCCCTACACGACGCTCTTCCGATCTNNNTTGTCTTTTTCACCACATCGTGA

GCACACGTCTGAACTCTTTCCCTACACGACGCTCTTCCGATCTNNNTTGTCTTTTTCACCACATCGTGA

AGCACACGTCTGAACTCTTTCCCTACACGACGCTCTTCCGATCTNNNCAGAATTCCTTCCCTTCCCTCAA

AGCACACGTCTGAACTCTTTCCCTACACGACGCTCTTCCGATCTNNNCAGAATTCCTTCCCTTCCCTCAA

AGCACACGTCTGAACTCTTTCCCTACACGACGCTCTTCCGATCTNNNCAGAATTCCTTCCCTTCCCTCAA

AGCACACGTCTGAACTCTTTCCCTACACGACGCTCTTCCGATCTNNNACTATGTGTCTTGGAGATGG

AGCACACGTCTGAACTCTTTCCCTACACGACGCTCTTCCGATCTNNNACTATGTGTCTTGGAGATGG

AGCACACGTCTGAACTCTTTCCCTACACGACGCTCTTCCGATCTNNNACTATGTGTCTTGGAGATGG

AGCACACGTCTGAACTCTTTCCCTACACGACGCTCTTCCGATCTNNNAGATAATTTGAATCATGCGGG

AGCACACGTCTGAACTCTTTCCCTACACGACGCTCTTCCGATCTNNNAGATAATTTGAATCATGCGGG
AGCACACGTCTGAACTCTTTCCCTACACGACGCTCTTCCGATCTNNNAGATAATTTGAATCATGCGGG
GAGCACACGTCTGAACTCTTTCCCTACACGACGCTCTTCCGATCTNNNTTATGCTTGAGGTGTTCTGA
GAGCACACGTCTGAACTCTTTCCCTACACGACGCTCTTCCGATCTNNNTTATGCTTGAGGTGTTCTGA
GAGCACACGTCTGAACTCTTTCCCTACACGACGCTCTTCCGATCTNNNTTATGCTTGAGGTGTTCTGA
AGCACACGTCTGAACTCTTTCCCTACACGACGCTCTTCCGATCTNNNTGAGGCACGAAAATCACTTG
AGCACACGTCTGAACTCTTTCCCTACACGACGCTCTTCCGATCTNNNTGAGGCACGAAAATCACTTG
AGCACACGTCTGAACTCTTTCCCTACACGACGCTCTTCCGATCTNNNTGAGGCACGAAAATCACTTG
AGCACACGTCTGAACTCTTTCCCTACACGACGCTCTTCCGATCTNNNAGCTTCTCTCACTTGGCATGTT
AGCACACGTCTGAACTCTTTCCCTACACGACGCTCTTCCGATCTNNNAGCTTCTCTCACTTGGCATGTT
AGCACACGTCTGAACTCTTTCCCTACACGACGCTCTTCCGATCTNNNAGCTTCTCTCACTTGGCATGTT
GCACACGTCTGAACTCTTTCCCTACACGACGCTCTTCCGATCTNNNAAGGGAGTGCTTAGATCATGGT
GCACACGTCTGAACTCTTTCCCTACACGACGCTCTTCCGATCTNNNAAGGGAGTGCTTAGATCATGGT
GCACACGTCTGAACTCTTTCCCTACACGACGCTCTTCCGATCTNNNAAGGGAGTGCTTAGATCATGGT
GCACACGTCTGAACTCTTTCCCTACACGACGCTCTTCCGATCTNNNAGGGGGCAATTTTCAGTTTCAG
GCACACGTCTGAACTCTTTCCCTACACGACGCTCTTCCGATCTNNNAGGGGGCAATTTTCAGTTTCAG
GCACACGTCTGAACTCTTTCCCTACACGACGCTCTTCCGATCTNNNAGGGGGCAATTTTCAGTTTCAG
CACACGTCTGAACTCTTTCCCTACACGACGCTCTTCCGATCTNNNGCCCCTTTAGTAGCTCCACTTAGC
CACACGTCTGAACTCTTTCCCTACACGACGCTCTTCCGATCTNNNGCCCCTTTAGTAGCTCCACTTAGC
CACACGTCTGAACTCTTTCCCTACACGACGCTCTTCCGATCTNNNGCCCCTTTAGTAGCTCCACTTAGC
GAGCACACGTCTGAACTCTTTCCCTACACGACGCTCTTCCGATCTNNNCTCTTCTCACTCTGTCTGTCAC
GAGCACACGTCTGAACTCTTTCCCTACACGACGCTCTTCCGATCTNNNCTCTTCTCACTCTGTCTGTCAC
GAGCACACGTCTGAACTCTTTCCCTACACGACGCTCTTCCGATCTNNNCTCTTCTCACTCTGTCTGTCAC
AGCACACGTCTGAACTCTTTCCCTACACGACGCTCTTCCGATCTNNNGCCTGGTCAACATAACAAATC
AGCACACGTCTGAACTCTTTCCCTACACGACGCTCTTCCGATCTNNNGCCTGGTCAACATAACAAATC
AGCACACGTCTGAACTCTTTCCCTACACGACGCTCTTCCGATCTNNNGCCTGGTCAACATAACAAATC
GCACACGTCTGAACTCTTTCCCTACACGACGCTCTTCCGATCTNNNGGCTCGCAACTAATTTCCTGTTT
GCACACGTCTGAACTCTTTCCCTACACGACGCTCTTCCGATCTNNNGGCTCGCAACTAATTTCCTGTTT
GCACACGTCTGAACTCTTTCCCTACACGACGCTCTTCCGATCTNNNGGCTCGCAACTAATTTCCTGTTT
AGCACACGTCTGAACTCTTTCCCTACACGACGCTCTTCCGATCTNNNGTGCGATCTTGGTTTACTGCA
AGCACACGTCTGAACTCTTTCCCTACACGACGCTCTTCCGATCTNNNGTGCGATCTTGGTTTACTGCA
AGCACACGTCTGAACTCTTTCCCTACACGACGCTCTTCCGATCTNNNGTGCGATCTTGGTTTACTGCA
AGAGCACACGTCTGAACTCTTTCCCTACACGACGCTCTTCCGATCTNNNCAAGACGTGCTGTGGTGA
AGAGCACACGTCTGAACTCTTTCCCTACACGACGCTCTTCCGATCTNNNCAAGACGTGCTGTGGTGA
AGAGCACACGTCTGAACTCTTTCCCTACACGACGCTCTTCCGATCTNNNCAAGACGTGCTGTGGTGA
AGCACACGTCTGAACTCTTTCCCTACACGACGCTCTTCCGATCTNNNGTTGTCTCCTATCACTGCGAGA
AGCACACGTCTGAACTCTTTCCCTACACGACGCTCTTCCGATCTNNNGTTGTCTCCTATCACTGCGAGA
AGCACACGTCTGAACTCTTTCCCTACACGACGCTCTTCCGATCTNNNGTTGTCTCCTATCACTGCGAGA
AGCACACGTCTGAACTCTTTCCCTACACGACGCTCTTCCGATCTNNNTCCCCTCTCTCTCTTTACATGC
AGCACACGTCTGAACTCTTTCCCTACACGACGCTCTTCCGATCTNNNTCCCCTCTCTCTCTTTACATGC
AGCACACGTCTGAACTCTTTCCCTACACGACGCTCTTCCGATCTNNNTCCCCTCTCTCTCTTTACATGC
GCACACGTCTGAACTCTTTCCCTACACGACGCTCTTCCGATCTNNNTCTCCCTCTCTCTCTCTCTCTCA
GCACACGTCTGAACTCTTTCCCTACACGACGCTCTTCCGATCTNNNTCTCCCTCTCTCTCTCTCTCTCA

AGCACACGTCTGAACTCTTTCCCTACACGACGCTCTTCCGATCTNNNTCTCCCTCTCTCTCTCTCTCTCTCA
AGCACACGTCTGAACTCTTTCCCTACACGACGCTCTTCCGATCTNNNGATTGAGACCATCCTGGCTAAC
AGCACACGTCTGAACTCTTTCCCTACACGACGCTCTTCCGATCTNNNGATTGAGACCATCCTGGCTAAC
AGCACACGTCTGAACTCTTTCCCTACACGACGCTCTTCCGATCTNNNGATTGAGACCATCCTGGCTAAC
GAGCACACGTCTGAACTCTTTCCCTACACGACGCTCTTCCGATCTNNNTAGTTTGAAGGTGTGTGAG
GAGCACACGTCTGAACTCTTTCCCTACACGACGCTCTTCCGATCTNNNTAGTTTGAAGGTGTGTGAG
GAGCACACGTCTGAACTCTTTCCCTACACGACGCTCTTCCGATCTNNNTAGTTTGAAGGTGTGTGAG
CACACGTCTGAACTCTTTCCCTACACGACGCTCTTCCGATCTNNNTCATTGGAAGTGGAGAACTGTCT
CACACGTCTGAACTCTTTCCCTACACGACGCTCTTCCGATCTNNNTCATTGGAAGTGGAGAACTGTCT
CACACGTCTGAACTCTTTCCCTACACGACGCTCTTCCGATCTNNNTCATTGGAAGTGGAGAACTGTCT
AGAGCACACGTCTGAACTCTTTCCCTACACGACGCTCTTCCGATCTNNNGAGGATCACTTGAGACTGT
AGAGCACACGTCTGAACTCTTTCCCTACACGACGCTCTTCCGATCTNNNGAGGATCACTTGAGACTGT
AGAGCACACGTCTGAACTCTTTCCCTACACGACGCTCTTCCGATCTNNNGAGGATCACTTGAGACTGT
GAGCACACGTCTGAACTCTTTCCCTACACGACGCTCTTCCGATCTNNNCTTGTATCCAGGAGTTTGAG
GAGCACACGTCTGAACTCTTTCCCTACACGACGCTCTTCCGATCTNNNCTTGTATCCAGGAGTTTGAG
GAGCACACGTCTGAACTCTTTCCCTACACGACGCTCTTCCGATCTNNNCTTGTATCCAGGAGTTTGAG
GAGCACACGTCTGAACTCTTTCCCTACACGACGCTCTTCCGATCTNNNCCTTTACACACACACACACACT
GAGCACACGTCTGAACTCTTTCCCTACACGACGCTCTTCCGATCTNNNCCTTTACACACACACACACACT
GAGCACACGTCTGAACTCTTTCCCTACACGACGCTCTTCCGATCTNNNCCTTTACACACACACACACACT
AGCACACGTCTGAACTCTTTCCCTACACGACGCTCTTCCGATCTNNNCCTGGCCCATGTGAAGGTTTTA
AGCACACGTCTGAACTCTTTCCCTACACGACGCTCTTCCGATCTNNNCCTGGCCCATGTGAAGGTTTTA
AGCACACGTCTGAACTCTTTCCCTACACGACGCTCTTCCGATCTNNNCCTGGCCCATGTGAAGGTTTTA
GAGCACACGTCTGAACTCTTTCCCTACACGACGCTCTTCCGATCTNNNAGAAGTTGCAGTGAGCCAAG
GAGCACACGTCTGAACTCTTTCCCTACACGACGCTCTTCCGATCTNNNAGAAGTTGCAGTGAGCCAAG
GAGCACACGTCTGAACTCTTTCCCTACACGACGCTCTTCCGATCTNNNAGAAGTTGCAGTGAGCCAAG
AGCACACGTCTGAACTCTTTCCCTACACGACGCTCTTCCGATCTNNNGCAGATCATTTGAGGTCAGGA
AGCACACGTCTGAACTCTTTCCCTACACGACGCTCTTCCGATCTNNNGCAGATCATTTGAGGTCAGGA
AGCACACGTCTGAACTCTTTCCCTACACGACGCTCTTCCGATCTNNNGCAGATCATTTGAGGTCAGGA
GAGCACACGTCTGAACTCTTTCCCTACACGACGCTCTTCCGATCTNNNAAAACAGTGAAATGCTGTCC
GAGCACACGTCTGAACTCTTTCCCTACACGACGCTCTTCCGATCTNNNAAAACAGTGAAATGCTGTCC
GAGCACACGTCTGAACTCTTTCCCTACACGACGCTCTTCCGATCTNNNAAAACAGTGAAATGCTGTCC
GAGCACACGTCTGAACTCTTTCCCTACACGACGCTCTTCCGATCTNNNAGACGAAAATGGTCAGAAGT
GAGCACACGTCTGAACTCTTTCCCTACACGACGCTCTTCCGATCTNNNAGACGAAAATGGTCAGAAGT
GAGCACACGTCTGAACTCTTTCCCTACACGACGCTCTTCCGATCTNNNAGACGAAAATGGTCAGAAGT
AGCACACGTCTGAACTCTTTCCCTACACGACGCTCTTCCGATCTNNNAAGGTGTAAGGGGGTATAAG
AGCACACGTCTGAACTCTTTCCCTACACGACGCTCTTCCGATCTNNNAAGGTGTAAGGGGGTATAAG
AGCACACGTCTGAACTCTTTCCCTACACGACGCTCTTCCGATCTNNNAAGGTGTAAGGGGGTATAAG
AGCACACGTCTGAACTCTTTCCCTACACGACGCTCTTCCGATCTNNNTTTAGTCTTCCTAGTGGAGCTG
AGCACACGTCTGAACTCTTTCCCTACACGACGCTCTTCCGATCTNNNTTTAGTCTTCCTAGTGGAGCTG
AGCACACGTCTGAACTCTTTCCCTACACGACGCTCTTCCGATCTNNNTTTAGTCTTCCTAGTGGAGCTG
AGCACACGTCTGAACTCTTTCCCTACACGACGCTCTTCCGATCTNNNACCTATAGTCCCAGCTACTAGG
AGCACACGTCTGAACTCTTTCCCTACACGACGCTCTTCCGATCTNNNACCTATAGTCCCAGCTACTAGG
AGCACACGTCTGAACTCTTTCCCTACACGACGCTCTTCCGATCTNNNACCTATAGTCCCAGCTACTAGG

GAGCACACGTCTGAACTCTTTCCCTACACGACGCTCTTCCGATCTNNNCGGTCAGGTAGGATCTCATGT
GAGCACACGTCTGAACTCTTTCCCTACACGACGCTCTTCCGATCTNNNCGGTCAGGTAGGATCTCATGT
GAGCACACGTCTGAACTCTTTCCCTACACGACGCTCTTCCGATCTNNNCGGTCAGGTAGGATCTCATGT
GAGCACACGTCTGAACTCTTTCCCTACACGACGCTCTTCCGATCTNNNTGCACTCTATGTGTTTTGTGGT
GAGCACACGTCTGAACTCTTTCCCTACACGACGCTCTTCCGATCTNNNTGCACTCTATGTGTTTTGTGGT
GAGCACACGTCTGAACTCTTTCCCTACACGACGCTCTTCCGATCTNNNTGCACTCTATGTGTTTTGTGGT
GAGCACACGTCTGAACTCTTTCCCTACACGACGCTCTTCCGATCTNNNCTTCTTCCCTTCCTTCCTTCCTC
GAGCACACGTCTGAACTCTTTCCCTACACGACGCTCTTCCGATCTNNNCTTCTTCCCTTCCTTCCTTCCTC
GAGCACACGTCTGAACTCTTTCCCTACACGACGCTCTTCCGATCTNNNCTTCTTCCCTTCCTTCCTTCCTC
AGCACACGTCTGAACTCTTTCCCTACACGACGCTCTTCCGATCTNNNGGACTTTCACTTTGGAAATGGC
AGCACACGTCTGAACTCTTTCCCTACACGACGCTCTTCCGATCTNNNGGACTTTCACTTTGGAAATGGC
AGCACACGTCTGAACTCTTTCCCTACACGACGCTCTTCCGATCTNNNGGACTTTCACTTTGGAAATGGC
AGCACACGTCTGAACTCTTTCCCTACACGACGCTCTTCCGATCTNNNCACACACAGAAATGCAAAGCAA
AGCACACGTCTGAACTCTTTCCCTACACGACGCTCTTCCGATCTNNNCACACACAGAAATGCAAAGCAA
AGCACACGTCTGAACTCTTTCCCTACACGACGCTCTTCCGATCTNNNCACACACAGAAATGCAAAGCAA
AGCACACGTCTGAACTCTTTCCCTACACGACGCTCTTCCGATCTNNNTTTCTTCCTTCCTTCCCTCCTTCCA
AGCACACGTCTGAACTCTTTCCCTACACGACGCTCTTCCGATCTNNNTTTCTTCCTTCCTTCCCTCCTTCCA
AGCACACGTCTGAACTCTTTCCCTACACGACGCTCTTCCGATCTNNNTTTCTTCCTTCCTTCCCTCCTTCCA
AGCACACGTCTGAACTCTTTCCCTACACGACGCTCTTCCGATCTNNNATTCAGTCACATCTCTGGGTTCC
AGCACACGTCTGAACTCTTTCCCTACACGACGCTCTTCCGATCTNNNATTCAGTCACATCTCTGGGTTCC
AGCACACGTCTGAACTCTTTCCCTACACGACGCTCTTCCGATCTNNNATTCAGTCACATCTCTGGGTTCC
AGCACACGTCTGAACTCTTTCCCTACACGACGCTCTTCCGATCTNNNATACCTGTAATCCCAGCACTGTG
AGCACACGTCTGAACTCTTTCCCTACACGACGCTCTTCCGATCTNNNATACCTGTAATCCCAGCACTGTG
AGCACACGTCTGAACTCTTTCCCTACACGACGCTCTTCCGATCTNNNATACCTGTAATCCCAGCACTGTG
AGCACACGTCTGAACTCTTTCCCTACACGACGCTCTTCCGATCTNNNCGTCTACTACGTGTTACCTCTCA
AGCACACGTCTGAACTCTTTCCCTACACGACGCTCTTCCGATCTNNNCGTCTACTACGTGTTACCTCTCA
AGCACACGTCTGAACTCTTTCCCTACACGACGCTCTTCCGATCTNNNCGTCTACTACGTGTTACCTCTCA
GCACACGTCTGAACTCTTTCCCTACACGACGCTCTTCCGATCTNNNAATTCAAGGACGCTCAGAAGGTA
GCACACGTCTGAACTCTTTCCCTACACGACGCTCTTCCGATCTNNNAATTCAAGGACGCTCAGAAGGTA
GCACACGTCTGAACTCTTTCCCTACACGACGCTCTTCCGATCTNNNAATTCAAGGACGCTCAGAAGGTA
AGAGCACACGTCTGAACTCTTTCCCTACACGACGCTCTTCCGATCTNNNGGATGAGAGAGCGAGAGA
AGAGCACACGTCTGAACTCTTTCCCTACACGACGCTCTTCCGATCTNNNGGATGAGAGAGCGAGAGA
AGAGCACACGTCTGAACTCTTTCCCTACACGACGCTCTTCCGATCTNNNGGATGAGAGAGCGAGAGA
GAGCACACGTCTGAACTCTTTCCCTACACGACGCTCTTCCGATCTNNNAAAGCATTCTGTCAGTTCACTG
GAGCACACGTCTGAACTCTTTCCCTACACGACGCTCTTCCGATCTNNNAAAGCATTCTGTCAGTTCACTG
GAGCACACGTCTGAACTCTTTCCCTACACGACGCTCTTCCGATCTNNNAAAGCATTCTGTCAGTTCACTG
AGCACACGTCTGAACTCTTTCCCTACACGACGCTCTTCCGATCTNNNGGTAAGATTTCTTGAGCCTGGG
AGCACACGTCTGAACTCTTTCCCTACACGACGCTCTTCCGATCTNNNGGTAAGATTTCTTGAGCCTGGG
AGCACACGTCTGAACTCTTTCCCTACACGACGCTCTTCCGATCTNNNGGTAAGATTTCTTGAGCCTGGG
GAGCACACGTCTGAACTCTTTCCCTACACGACGCTCTTCCGATCTNNNAGGAGAATGGCGTGAACCTG
GAGCACACGTCTGAACTCTTTCCCTACACGACGCTCTTCCGATCTNNNAGGAGAATGGCGTGAACCTG
GAGCACACGTCTGAACTCTTTCCCTACACGACGCTCTTCCGATCTNNNAGGAGAATGGCGTGAACCTG
GAGCACACGTCTGAACTCTTTCCCTACACGACGCTCTTCCGATCTNNNGAGGCAGGAGAGTTGTTTGA

GAGCACACGTCTGAACTCTTTCCCTACACGACGCTCTTCCGATCTNNNGAGGCAGGAGAGTTGTTTGA

GAGCACACGTCTGAACTCTTTCCCTACACGACGCTCTTCCGATCTNNNGAGGCAGGAGAGTTGTTTGA

AGAGCACACGTCTGAACTCTTTCCCTACACGACGCTCTTCCGATCTNNNGCAATGGCCCCTTTGTCATTA

AGAGCACACGTCTGAACTCTTTCCCTACACGACGCTCTTCCGATCTNNNGCAATGGCCCCTTTGTCATTA

AGAGCACACGTCTGAACTCTTTCCCTACACGACGCTCTTCCGATCTNNNGCAATGGCCCCTTTGTCATTA

AGCACACGTCTGAACTCTTTCCCTACACGACGCTCTTCCGATCTNNNCAACAACGCAGAGCCTCAAATAT

AGCACACGTCTGAACTCTTTCCCTACACGACGCTCTTCCGATCTNNNCAACAACGCAGAGCCTCAAATAT

AGCACACGTCTGAACTCTTTCCCTACACGACGCTCTTCCGATCTNNNCAACAACGCAGAGCCTCAAATAT

GAGCACACGTCTGAACTCTTTCCCTACACGACGCTCTTCCGATCTNNNCTGAAGAGTCAGCAGCTTTTG

GAGCACACGTCTGAACTCTTTCCCTACACGACGCTCTTCCGATCTNNNCTGAAGAGTCAGCAGCTTTTG

GAGCACACGTCTGAACTCTTTCCCTACACGACGCTCTTCCGATCTNNNCTGAAGAGTCAGCAGCTTTTG

GAGCACACGTCTGAACTCTTTCCCTACACGACGCTCTTCCGATCTNNNGAAGGAGGGAGTGAATCACC

GAGCACACGTCTGAACTCTTTCCCTACACGACGCTCTTCCGATCTNNNGAAGGAGGGAGTGAATCACC

GAGCACACGTCTGAACTCTTTCCCTACACGACGCTCTTCCGATCTNNNGAAGGAGGGAGTGAATCACC

AGCACACGTCTGAACTCTTTCCCTACACGACGCTCTTCCGATCTNNNTGAGGCAGGAGAATCTCTTGAA

AGCACACGTCTGAACTCTTTCCCTACACGACGCTCTTCCGATCTNNNTGAGGCAGGAGAATCTCTTGAA

AGCACACGTCTGAACTCTTTCCCTACACGACGCTCTTCCGATCTNNNTGAGGCAGGAGAATCTCTTGAA

AGCACACGTCTGAACTCTTTCCCTACACGACGCTCTTCCGATCTNNNCCCATTATGTGACCAGCTTTTCC

AGCACACGTCTGAACTCTTTCCCTACACGACGCTCTTCCGATCTNNNCCCATTATGTGACCAGCTTTTCC

AGCACACGTCTGAACTCTTTCCCTACACGACGCTCTTCCGATCTNNNCCCATTATGTGACCAGCTTTTCC

GCACACGTCTGAACTCTTTCCCTACACGACGCTCTTCCGATCTNNNATGAATGAATTTGAGACGGAACA

GCACACGTCTGAACTCTTTCCCTACACGACGCTCTTCCGATCTNNNATGAATGAATTTGAGACGGAACA

GCACACGTCTGAACTCTTTCCCTACACGACGCTCTTCCGATCTNNNATGAATGAATTTGAGACGGAACA

AGCACACGTCTGAACTCTTTCCCTACACGACGCTCTTCCGATCTNNNCCTCTCTCTCCTCTCATCTTCCAC

AGCACACGTCTGAACTCTTTCCCTACACGACGCTCTTCCGATCTNNNCCTCTCTCTCCTCTCATCTTCCAC

AGCACACGTCTGAACTCTTTCCCTACACGACGCTCTTCCGATCTNNNCCTCTCTCTCCTCTCATCTTCCAC

CACACGTCTGAACTCTTTCCCTACACGACGCTCTTCCGATCTNNNGCTGACTCAACTGGAAATTTCATCG

CACACGTCTGAACTCTTTCCCTACACGACGCTCTTCCGATCTNNNGCTGACTCAACTGGAAATTTCATCG

CACACGTCTGAACTCTTTCCCTACACGACGCTCTTCCGATCTNNNGCTGACTCAACTGGAAATTTCATCG

GAGCACACGTCTGAACTCTTTCCCTACACGACGCTCTTCCGATCTNNNGATCTCTTGACCTCTTGAGCCA

GAGCACACGTCTGAACTCTTTCCCTACACGACGCTCTTCCGATCTNNNGATCTCTTGACCTCTTGAGCCA

GAGCACACGTCTGAACTCTTTCCCTACACGACGCTCTTCCGATCTNNNGATCTCTTGACCTCTTGAGCCA

AGCACACGTCTGAACTCTTTCCCTACACGACGCTCTTCCGATCTNNNAGTAGAGGGGCTCAACTGACA

AGCACACGTCTGAACTCTTTCCCTACACGACGCTCTTCCGATCTNNNAGTAGAGGGGCTCAACTGACA

AGCACACGTCTGAACTCTTTCCCTACACGACGCTCTTCCGATCTNNNAGTAGAGGGGCTCAACTGACA

AGCACACGTCTGAACTCTTTCCCTACACGACGCTCTTCCGATCTNNNAGTATCTCTCCATCTTGTGTTGTT

AGCACACGTCTGAACTCTTTCCCTACACGACGCTCTTCCGATCTNNNAGTATCTCTCCATCTTGTGTTGTT

AGCACACGTCTGAACTCTTTCCCTACACGACGCTCTTCCGATCTNNNAGTATCTCTCCATCTTGTGTTGTT

AGCACACGTCTGAACTCTTTCCCTACACGACGCTCTTCCGATCTNNNCCCAACACTATACCTTGGGCCCAT

AGCACACGTCTGAACTCTTTCCCTACACGACGCTCTTCCGATCTNNNCCCAACACTATACCTTGGGCCCAT

AGCACACGTCTGAACTCTTTCCCTACACGACGCTCTTCCGATCTNNNCCCAACACTATACCTTGGGCCCAT

GAGCACACGTCTGAACTCTTTCCCTACACGACGCTCTTCCGATCTNNNAGATTAGTTGAGACACTAGTT

GAGCACACGTCTGAACTCTTTCCCTACACGACGCTCTTCCGATCTNNNAGATTAGTTGAGACACTAGTT

GAGCACACGTCTGAACTCTTTCCCTACACGACGCTCTTCCGATCTNNNAGATTAGTTGAGACACTAGT
AGCACACGTCTGAACTCTTTCCCTACACGACGCTCTTCCGATCTNNNGCCAAGATCACACCACTGTACT
AGCACACGTCTGAACTCTTTCCCTACACGACGCTCTTCCGATCTNNNGCCAAGATCACACCACTGTACT
AGCACACGTCTGAACTCTTTCCCTACACGACGCTCTTCCGATCTNNNGCCAAGATCACACCACTGTACT
AAGAGCACACGTCTGAACTCTTTCCCTACACGACGCTCTTCCGATCTNNNTACATTTCACAGTCCTTGCC
AAGAGCACACGTCTGAACTCTTTCCCTACACGACGCTCTTCCGATCTNNNTACATTTCACAGTCCTTGCC
AAGAGCACACGTCTGAACTCTTTCCCTACACGACGCTCTTCCGATCTNNNTACATTTCACAGTCCTTGCC
AGCACACGTCTGAACTCTTTCCCTACACGACGCTCTTCCGATCTNNNAATGTTATAGACAGCTTTTTCT
AGCACACGTCTGAACTCTTTCCCTACACGACGCTCTTCCGATCTNNNAATGTTATAGACAGCTTTTTCT
AGCACACGTCTGAACTCTTTCCCTACACGACGCTCTTCCGATCTNNNAATGTTATAGACAGCTTTTTCT
AGCACACGTCTGAACTCTTTCCCTACACGACGCTCTTCCGATCTNNNGGGTAAAACACTGGTGAAACA
AGCACACGTCTGAACTCTTTCCCTACACGACGCTCTTCCGATCTNNNGGGTAAAACACTGGTGAAACA
AGCACACGTCTGAACTCTTTCCCTACACGACGCTCTTCCGATCTNNNGGGTAAAACACTGGTGAAACA
CACACGTCTGAACTCTTTCCCTACACGACGCTCTTCCGATCTNNNGAATGTGTTCCTGGCAAGTGGCA
CACACGTCTGAACTCTTTCCCTACACGACGCTCTTCCGATCTNNNGAATGTGTTCCTGGCAAGTGGCA
CACACGTCTGAACTCTTTCCCTACACGACGCTCTTCCGATCTNNNGAATGTGTTCCTGGCAAGTGGCA
GAGCACACGTCTGAACTCTTTCCCTACACGACGCTCTTCCGATCTNNNCAGGAGAATCACTTGAACCCA
GAGCACACGTCTGAACTCTTTCCCTACACGACGCTCTTCCGATCTNNNCAGGAGAATCACTTGAACCCA
GAGCACACGTCTGAACTCTTTCCCTACACGACGCTCTTCCGATCTNNNCAGGAGAATCACTTGAACCCA
GAGCACACGTCTGAACTCTTTCCCTACACGACGCTCTTCCGATCTNNNGGGAAATGGGGAGTAACTG
GAGCACACGTCTGAACTCTTTCCCTACACGACGCTCTTCCGATCTNNNGGGAAATGGGGAGTAACTG
GAGCACACGTCTGAACTCTTTCCCTACACGACGCTCTTCCGATCTNNNGGGAAATGGGGAGTAACTG
AGCACACGTCTGAACTCTTTCCCTACACGACGCTCTTCCGATCTNNNGGGGTATATACCTAGGAGTGG
AGCACACGTCTGAACTCTTTCCCTACACGACGCTCTTCCGATCTNNNGGGGTATATACCTAGGAGTGG
AGCACACGTCTGAACTCTTTCCCTACACGACGCTCTTCCGATCTNNNGGGGTATATACCTAGGAGTGG
GAGCACACGTCTGAACTCTTTCCCTACACGACGCTCTTCCGATCTNNNGAGTCTCACTCTGTCACCCAGC
GAGCACACGTCTGAACTCTTTCCCTACACGACGCTCTTCCGATCTNNNGAGTCTCACTCTGTCACCCAGC
GAGCACACGTCTGAACTCTTTCCCTACACGACGCTCTTCCGATCTNNNGAGTCTCACTCTGTCACCCAGC
GAGCACACGTCTGAACTCTTTCCCTACACGACGCTCTTCCGATCTNNNTGAGTAACAGAACGAGAACCT
GAGCACACGTCTGAACTCTTTCCCTACACGACGCTCTTCCGATCTNNNTGAGTAACAGAACGAGAACCT
GAGCACACGTCTGAACTCTTTCCCTACACGACGCTCTTCCGATCTNNNTGAGTAACAGAACGAGAACCT
GAGCACACGTCTGAACTCTTTCCCTACACGACGCTCTTCCGATCTNNNGATCTTCAGAATTTTCTTAGC
GAGCACACGTCTGAACTCTTTCCCTACACGACGCTCTTCCGATCTNNNGATCTTCAGAATTTTCTTAGC
GAGCACACGTCTGAACTCTTTCCCTACACGACGCTCTTCCGATCTNNNGATCTTCAGAATTTTCTTAGC
AGCACACGTCTGAACTCTTTCCCTACACGACGCTCTTCCGATCTNNNGCATGAAACGAGAATGAACCC
AGCACACGTCTGAACTCTTTCCCTACACGACGCTCTTCCGATCTNNNGCATGAAACGAGAATGAACCC
AGCACACGTCTGAACTCTTTCCCTACACGACGCTCTTCCGATCTNNNGCATGAAACGAGAATGAACCC
AGCACACGTCTGAACTCTTTCCCTACACGACGCTCTTCCGATCTNNNTGTGGGTCTATTTACAGTGTCT
AGCACACGTCTGAACTCTTTCCCTACACGACGCTCTTCCGATCTNNNTGTGGGTCTATTTACAGTGTCT
AGCACACGTCTGAACTCTTTCCCTACACGACGCTCTTCCGATCTNNNTGTGGGTCTATTTACAGTGTCT
AGAGCACACGTCTGAACTCTTTCCCTACACGACGCTCTTCCGATCTNNNTCTAGTTCCTGTAGGTGTGA
AGAGCACACGTCTGAACTCTTTCCCTACACGACGCTCTTCCGATCTNNNTCTAGTTCCTGTAGGTGTGA
AGAGCACACGTCTGAACTCTTTCCCTACACGACGCTCTTCCGATCTNNNTCTAGTTCCTGTAGGTGTGA

AGCACACGTCTGAACTCTTTCCCTACACGACGCTCTTCCGATCTNNNTACAAAAAATTAGCCGGGCAT
AGCACACGTCTGAACTCTTTCCCTACACGACGCTCTTCCGATCTNNNTACAAAAAATTAGCCGGGCAT
AGCACACGTCTGAACTCTTTCCCTACACGACGCTCTTCCGATCTNNNTACAAAAAATTAGCCGGGCAT
GAGCACACGTCTGAACTCTTTCCCTACACGACGCTCTTCCGATCTNNNGTAGACCGGAGCTGTTCCTA
GAGCACACGTCTGAACTCTTTCCCTACACGACGCTCTTCCGATCTNNNGTAGACCGGAGCTGTTCCTA
GAGCACACGTCTGAACTCTTTCCCTACACGACGCTCTTCCGATCTNNNGTAGACCGGAGCTGTTCCTA
GAGCACACGTCTGAACTCTTTCCCTACACGACGCTCTTCCGATCTNNNTGCCAATCAAATTCCTCCTCT
GAGCACACGTCTGAACTCTTTCCCTACACGACGCTCTTCCGATCTNNNTGCCAATCAAATTCCTCCTCT
GAGCACACGTCTGAACTCTTTCCCTACACGACGCTCTTCCGATCTNNNTGCCAATCAAATTCCTCCTCT
GCACACGTCTGAACTCTTTCCCTACACGACGCTCTTCCGATCTNNNAGAGGTTCCTCTTCTGATGGTT
GCACACGTCTGAACTCTTTCCCTACACGACGCTCTTCCGATCTNNNAGAGGTTCCTCTTCTGATGGTT
GCACACGTCTGAACTCTTTCCCTACACGACGCTCTTCCGATCTNNNAGAGGTTCCTCTTCTGATGGTT
AGCACACGTCTGAACTCTTTCCCTACACGACGCTCTTCCGATCTNNNAGCTCGGAATGCTACTGATC
AGCACACGTCTGAACTCTTTCCCTACACGACGCTCTTCCGATCTNNNAGCTCGGAATGCTACTGATC
AGCACACGTCTGAACTCTTTCCCTACACGACGCTCTTCCGATCTNNNAGCTCGGAATGCTACTGATC
AGCACACGTCTGAACTCTTTCCCTACACGACGCTCTTCCGATCTNNNGGGAGGTACAGAAAGGAGT
AGCACACGTCTGAACTCTTTCCCTACACGACGCTCTTCCGATCTNNNGGGAGGTACAGAAAGGAGT
AGCACACGTCTGAACTCTTTCCCTACACGACGCTCTTCCGATCTNNNGGGAGGTACAGAAAGGAGT
GCACACGTCTGAACTCTTTCCCTACACGACGCTCTTCCGATCTNNNGACATTGCAGTGAGCTGAGATT
GCACACGTCTGAACTCTTTCCCTACACGACGCTCTTCCGATCTNNNGACATTGCAGTGAGCTGAGATT
GCACACGTCTGAACTCTTTCCCTACACGACGCTCTTCCGATCTNNNGACATTGCAGTGAGCTGAGATT
GAGCACACGTCTGAACTCTTTCCCTACACGACGCTCTTCCGATCTNNNAAAGTATGTGGTGCATGTGA
GAGCACACGTCTGAACTCTTTCCCTACACGACGCTCTTCCGATCTNNNAAAGTATGTGGTGCATGTGA
GAGCACACGTCTGAACTCTTTCCCTACACGACGCTCTTCCGATCTNNNAAAGTATGTGGTGCATGTGA
GCACACGTCTGAACTCTTTCCCTACACGACGCTCTTCCGATCTNNNCATCTGTCTGCACAATGTTGGG
GCACACGTCTGAACTCTTTCCCTACACGACGCTCTTCCGATCTNNNCATCTGTCTGCACAATGTTGGG
GCACACGTCTGAACTCTTTCCCTACACGACGCTCTTCCGATCTNNNCATCTGTCTGCACAATGTTGGG
GAGCACACGTCTGAACTCTTTCCCTACACGACGCTCTTCCGATCTNNNCTTGCAGTGAGCTGAGATCA
GAGCACACGTCTGAACTCTTTCCCTACACGACGCTCTTCCGATCTNNNCTTGCAGTGAGCTGAGATCA
GAGCACACGTCTGAACTCTTTCCCTACACGACGCTCTTCCGATCTNNNCTTGCAGTGAGCTGAGATCA
GCACACGTCTGAACTCTTTCCCTACACGACGCTCTTCCGATCTNNNCACGACAGGTGTGAAGAAAAT
GCACACGTCTGAACTCTTTCCCTACACGACGCTCTTCCGATCTNNNCACGACAGGTGTGAAGAAAAT
GCACACGTCTGAACTCTTTCCCTACACGACGCTCTTCCGATCTNNNCACGACAGGTGTGAAGAAAAT
AGCACACGTCTGAACTCTTTCCCTACACGACGCTCTTCCGATCTNNNGCCTCTGGAGTACCTAGTCTTT
AGCACACGTCTGAACTCTTTCCCTACACGACGCTCTTCCGATCTNNNGCCTCTGGAGTACCTAGTCTTT
AGCACACGTCTGAACTCTTTCCCTACACGACGCTCTTCCGATCTNNNGCCTCTGGAGTACCTAGTCTTT
AGCACACGTCTGAACTCTTTCCCTACACGACGCTCTTCCGATCTNNNATGAGCAGACATTTCAAAGCC
AGCACACGTCTGAACTCTTTCCCTACACGACGCTCTTCCGATCTNNNATGAGCAGACATTTCAAAGCC
AGCACACGTCTGAACTCTTTCCCTACACGACGCTCTTCCGATCTNNNATGAGCAGACATTTCAAAGCC
AGCACACGTCTGAACTCTTTCCCTACACGACGCTCTTCCGATCTNNNTACTGAAGCCCACAAGAGGTT
AGCACACGTCTGAACTCTTTCCCTACACGACGCTCTTCCGATCTNNNTACTGAAGCCCACAAGAGGTT
AGCACACGTCTGAACTCTTTCCCTACACGACGCTCTTCCGATCTNNNTACTGAAGCCCACAAGAGGTT
GCACACGTCTGAACTCTTTCCCTACACGACGCTCTTCCGATCTNNNAGACTCCATCAATACCAGGGGT

GCACACGTCTGAACTCTTTCCCTACACGACGCTCTTCCGATCTNNNAGACTCCATCAATACCAGGGGT

GCACACGTCTGAACTCTTTCCCTACACGACGCTCTTCCGATCTNNNAGACTCCATCAATACCAGGGGT

AGAGCACACGTCTGAACTCTTTCCCTACACGACGCTCTTCCGATCTNNNATAAAAATTAGCTGGGTGT

AGAGCACACGTCTGAACTCTTTCCCTACACGACGCTCTTCCGATCTNNNATAAAAATTAGCTGGGTGT

AGAGCACACGTCTGAACTCTTTCCCTACACGACGCTCTTCCGATCTNNNATAAAAATTAGCTGGGTGT

GCACACGTCTGAACTCTTTCCCTACACGACGCTCTTCCGATCTNNNGATCCGAATGGCAAAGACGAGT

GCACACGTCTGAACTCTTTCCCTACACGACGCTCTTCCGATCTNNNGATCCGAATGGCAAAGACGAGT

GCACACGTCTGAACTCTTTCCCTACACGACGCTCTTCCGATCTNNNGATCCGAATGGCAAAGACGAGT

AGCACACGTCTGAACTCTTTCCCTACACGACGCTCTTCCGATCTNNNTAATTCTTTTGGAAAAACTGC

AGCACACGTCTGAACTCTTTCCCTACACGACGCTCTTCCGATCTNNNTAATTCTTTTGGAAAAACTGC

AGCACACGTCTGAACTCTTTCCCTACACGACGCTCTTCCGATCTNNNTAATTCTTTTGGAAAAACTGC

AGCACACGTCTGAACTCTTTCCCTACACGACGCTCTTCCGATCTNNNCTTCCCAAAGAGGAGGTAAG

AGCACACGTCTGAACTCTTTCCCTACACGACGCTCTTCCGATCTNNNCTTCCCAAAGAGGAGGTAAG

AGCACACGTCTGAACTCTTTCCCTACACGACGCTCTTCCGATCTNNNCTTCCCAAAGAGGAGGTAAG

GCACACGTCTGAACTCTTTCCCTACACGACGCTCTTCCGATCTNNNCTCAACAGAACTCCTCTTCTCCC

GCACACGTCTGAACTCTTTCCCTACACGACGCTCTTCCGATCTNNNCTCAACAGAACTCCTCTTCTCCC

GCACACGTCTGAACTCTTTCCCTACACGACGCTCTTCCGATCTNNNCTCAACAGAACTCCTCTTCTCCC

GCACACGTCTGAACTCTTTCCCTACACGACGCTCTTCCGATCTNNNGCAACATGGTGAGACTCCATCT

GCACACGTCTGAACTCTTTCCCTACACGACGCTCTTCCGATCTNNNGCAACATGGTGAGACTCCATCT

GCACACGTCTGAACTCTTTCCCTACACGACGCTCTTCCGATCTNNNGCAACATGGTGAGACTCCATCT

GAGCACACGTCTGAACTCTTTCCCTACACGACGCTCTTCCGATCTNNNGCTACTGACCACTGGACACT

GAGCACACGTCTGAACTCTTTCCCTACACGACGCTCTTCCGATCTNNNGCTACTGACCACTGGACACT

GAGCACACGTCTGAACTCTTTCCCTACACGACGCTCTTCCGATCTNNNGCTACTGACCACTGGACACT

AGCACACGTCTGAACTCTTTCCCTACACGACGCTCTTCCGATCTNNNGTTCATGCCATTCTCTTGCCTC

AGCACACGTCTGAACTCTTTCCCTACACGACGCTCTTCCGATCTNNNGTTCATGCCATTCTCTTGCCTC

AGCACACGTCTGAACTCTTTCCCTACACGACGCTCTTCCGATCTNNNGTTCATGCCATTCTCTTGCCTC

AGCACACGTCTGAACTCTTTCCCTACACGACGCTCTTCCGATCTNNNCTGTGGTAGGAGAATGGTGT

AGCACACGTCTGAACTCTTTCCCTACACGACGCTCTTCCGATCTNNNCTGTGGTAGGAGAATGGTGT

AGCACACGTCTGAACTCTTTCCCTACACGACGCTCTTCCGATCTNNNCTGTGGTAGGAGAATGGTGT

AGCACACGTCTGAACTCTTTCCCTACACGACGCTCTTCCGATCTNNNGTCACTTCTCACTCTGGTCTCTG

AGCACACGTCTGAACTCTTTCCCTACACGACGCTCTTCCGATCTNNNGTCACTTCTCACTCTGGTCTCTG

AGCACACGTCTGAACTCTTTCCCTACACGACGCTCTTCCGATCTNNNGTCACTTCTCACTCTGGTCTCTG

AGCACACGTCTGAACTCTTTCCCTACACGACGCTCTTCCGATCTNNNCACACACACGCACACACATATA

AGCACACGTCTGAACTCTTTCCCTACACGACGCTCTTCCGATCTNNNCACACACACGCACACACATATA

AGCACACGTCTGAACTCTTTCCCTACACGACGCTCTTCCGATCTNNNCACACACACGCACACACATATA

GAGCACACGTCTGAACTCTTTCCCTACACGACGCTCTTCCGATCTNNNCAAGGCAGGAGAATTGCTT

GAGCACACGTCTGAACTCTTTCCCTACACGACGCTCTTCCGATCTNNNCAAGGCAGGAGAATTGCTT

GAGCACACGTCTGAACTCTTTCCCTACACGACGCTCTTCCGATCTNNNCAAGGCAGGAGAATTGCTT

GCACACGTCTGAACTCTTTCCCTACACGACGCTCTTCCGATCTNNNGTAACACACTTTGGAGGCAAGA

GCACACGTCTGAACTCTTTCCCTACACGACGCTCTTCCGATCTNNNGTAACACACTTTGGAGGCAAGA

GCACACGTCTGAACTCTTTCCCTACACGACGCTCTTCCGATCTNNNGTAACACACTTTGGAGGCAAGA

AGCACACGTCTGAACTCTTTCCCTACACGACGCTCTTCCGATCTNNNCAGGAGGTGGAGGTTATGAT

AGCACACGTCTGAACTCTTTCCCTACACGACGCTCTTCCGATCTNNNCAGGAGGTGGAGGTTATGAT

AGCACACGTCTGAACTCTTTCCCTACACGACGCTCTTCCGATCTNNNCAGGAGGTGGAGGTTATGAT

GAGCACACGTCTGAACTCTTTCCCTACACGACGCTCTTCCGATCTNNNCCCAACTAGCTGCAAGAGTTT

GAGCACACGTCTGAACTCTTTCCCTACACGACGCTCTTCCGATCTNNNCCCAACTAGCTGCAAGAGTTT

GAGCACACGTCTGAACTCTTTCCCTACACGACGCTCTTCCGATCTNNNCCCAACTAGCTGCAAGAGTTT

AGCACACGTCTGAACTCTTTCCCTACACGACGCTCTTCCGATCTNNNAGGAGAATTGCTTGAACCCA

AGCACACGTCTGAACTCTTTCCCTACACGACGCTCTTCCGATCTNNNAGGAGAATTGCTTGAACCCA

AGCACACGTCTGAACTCTTTCCCTACACGACGCTCTTCCGATCTNNNAGGAGAATTGCTTGAACCCA

AGAGCACACGTCTGAACTCTTTCCCTACACGACGCTCTTCCGATCTNNNGACACAAGAGCGAAACTA

AGAGCACACGTCTGAACTCTTTCCCTACACGACGCTCTTCCGATCTNNNGACACAAGAGCGAAACTA

AGAGCACACGTCTGAACTCTTTCCCTACACGACGCTCTTCCGATCTNNNGACACAAGAGCGAAACTA

AGCACACGTCTGAACTCTTTCCCTACACGACGCTCTTCCGATCTNNNCTGAGACAGGAGAATCGCTTG

AGCACACGTCTGAACTCTTTCCCTACACGACGCTCTTCCGATCTNNNCTGAGACAGGAGAATCGCTTG

AGCACACGTCTGAACTCTTTCCCTACACGACGCTCTTCCGATCTNNNCTGAGACAGGAGAATCGCTTG

AGCACACGTCTGAACTCTTTCCCTACACGACGCTCTTCCGATCTNNNAAGGGGAAGAAGAGAAGCA

AGCACACGTCTGAACTCTTTCCCTACACGACGCTCTTCCGATCTNNNAAGGGGAAGAAGAGAAGCA

AGCACACGTCTGAACTCTTTCCCTACACGACGCTCTTCCGATCTNNNAAGGGGAAGAAGAGAAGCA

GCACACGTCTGAACTCTTTCCCTACACGACGCTCTTCCGATCTNNNACTGTGATTGTGTGCTTCTCTTT

GCACACGTCTGAACTCTTTCCCTACACGACGCTCTTCCGATCTNNNACTGTGATTGTGTGCTTCTCTTT

GCACACGTCTGAACTCTTTCCCTACACGACGCTCTTCCGATCTNNNACTGTGATTGTGTGCTTCTCTTT

GCACACGTCTGAACTCTTTCCCTACACGACGCTCTTCCGATCTNNNTGCTTTGCTGTATGTCAAACACA

GCACACGTCTGAACTCTTTCCCTACACGACGCTCTTCCGATCTNNNTGCTTTGCTGTATGTCAAACACA

GCACACGTCTGAACTCTTTCCCTACACGACGCTCTTCCGATCTNNNTGCTTTGCTGTATGTCAAACACA

GAGCACACGTCTGAACTCTTTCCCTACACGACGCTCTTCCGATCTNNNTACCTGTAATCCCAGTGCTTT

GAGCACACGTCTGAACTCTTTCCCTACACGACGCTCTTCCGATCTNNNTACCTGTAATCCCAGTGCTTT

GAGCACACGTCTGAACTCTTTCCCTACACGACGCTCTTCCGATCTNNNTACCTGTAATCCCAGTGCTTT

GCACACGTCTGAACTCTTTCCCTACACGACGCTCTTCCGATCTNNNGCCTGTGTTCTTCTTTGTAGCAT

GCACACGTCTGAACTCTTTCCCTACACGACGCTCTTCCGATCTNNNGCCTGTGTTCTTCTTTGTAGCAT

GCACACGTCTGAACTCTTTCCCTACACGACGCTCTTCCGATCTNNNGCCTGTGTTCTTCTTTGTAGCAT

GCACACGTCTGAACTCTTTCCCTACACGACGCTCTTCCGATCTNNNGCTCTTTTGATGACCTACTCCCA

GCACACGTCTGAACTCTTTCCCTACACGACGCTCTTCCGATCTNNNGCTCTTTTGATGACCTACTCCCA

GCACACGTCTGAACTCTTTCCCTACACGACGCTCTTCCGATCTNNNGCTCTTTTGATGACCTACTCCCA

AGCACACGTCTGAACTCTTTCCCTACACGACGCTCTTCCGATCTNNNTTTAGTAGAGATGGGGTTTC

AGCACACGTCTGAACTCTTTCCCTACACGACGCTCTTCCGATCTNNNTTTAGTAGAGATGGGGTTTC

AGCACACGTCTGAACTCTTTCCCTACACGACGCTCTTCCGATCTNNNTTTAGTAGAGATGGGGTTTC

GAGCACACGTCTGAACTCTTTCCCTACACGACGCTCTTCCGATCTNNNAAGAGACGCCCTAATGAATC

GAGCACACGTCTGAACTCTTTCCCTACACGACGCTCTTCCGATCTNNNAAGAGACGCCCTAATGAATC

GAGCACACGTCTGAACTCTTTCCCTACACGACGCTCTTCCGATCTNNNAAGAGACGCCCTAATGAATC

AGCACACGTCTGAACTCTTTCCCTACACGACGCTCTTCCGATCTNNNGCCAAGAGACTGGGTGTTGAA

AGCACACGTCTGAACTCTTTCCCTACACGACGCTCTTCCGATCTNNNGCCAAGAGACTGGGTGTTGAA

AGCACACGTCTGAACTCTTTCCCTACACGACGCTCTTCCGATCTNNNGCCAAGAGACTGGGTGTTGAA

GCACACGTCTGAACTCTTTCCCTACACGACGCTCTTCCGATCTNNNCAAGCAGAGGTTGGAAGAGTTT

GCACACGTCTGAACTCTTTCCCTACACGACGCTCTTCCGATCTNNNCAAGCAGAGGTTGGAAGAGTTT

GCACACGTCTGAACTCTTTCCCTACACGACGCTCTTCCGATCTNNNCAAGCAGAGGTTGGAAGAGTTT

GAGCACACGTCTGAACTCTTTCCCTACACGACGCTCTTCCGATCTNNNCACACACACACACACACACATG

GAGCACACGTCTGAACTCTTTCCCTACACGACGCTCTTCCGATCTNNNCACACACACACACACACACATG

GAGCACACGTCTGAACTCTTTCCCTACACGACGCTCTTCCGATCTNNNCACACACACACACACACACATG

GAGCACACGTCTGAACTCTTTCCCTACACGACGCTCTTCCGATCTNNNGAGGCTGCATTGAGCTATGAT

GAGCACACGTCTGAACTCTTTCCCTACACGACGCTCTTCCGATCTNNNGAGGCTGCATTGAGCTATGAT

GAGCACACGTCTGAACTCTTTCCCTACACGACGCTCTTCCGATCTNNNGAGGCTGCATTGAGCTATGAT

GCACACGTCTGAACTCTTTCCCTACACGACGCTCTTCCGATCTNNNTCTCGTTGTCAATCTCTTACTATAC

GCACACGTCTGAACTCTTTCCCTACACGACGCTCTTCCGATCTNNNTCTCGTTGTCAATCTCTTACTATAC

GCACACGTCTGAACTCTTTCCCTACACGACGCTCTTCCGATCTNNNTCTCGTTGTCAATCTCTTACTATAC

AGCACACGTCTGAACTCTTTCCCTACACGACGCTCTTCCGATCTNNNTCATTGAACATGCACAACAGTC

AGCACACGTCTGAACTCTTTCCCTACACGACGCTCTTCCGATCTNNNTCATTGAACATGCACAACAGTC

AGCACACGTCTGAACTCTTTCCCTACACGACGCTCTTCCGATCTNNNTCATTGAACATGCACAACAGTC

GAGCACACGTCTGAACTCTTTCCCTACACGACGCTCTTCCGATCTNNNGAGATTGAGGCACTGAACTC

GAGCACACGTCTGAACTCTTTCCCTACACGACGCTCTTCCGATCTNNNGAGATTGAGGCACTGAACTC

GAGCACACGTCTGAACTCTTTCCCTACACGACGCTCTTCCGATCTNNNGAGATTGAGGCACTGAACTC

AGCACACGTCTGAACTCTTTCCCTACACGACGCTCTTCCGATCTNNNGAAGTGGTTTGACAGCTGCGAA

AGCACACGTCTGAACTCTTTCCCTACACGACGCTCTTCCGATCTNNNGAAGTGGTTTGACAGCTGCGAA

AGCACACGTCTGAACTCTTTCCCTACACGACGCTCTTCCGATCTNNNGAAGTGGTTTGACAGCTGCGAA

GCACACGTCTGAACTCTTTCCCTACACGACGCTCTTCCGATCTNNNTGTTTGCATGGAGAGATTGGTC

GCACACGTCTGAACTCTTTCCCTACACGACGCTCTTCCGATCTNNNTGTTTGCATGGAGAGATTGGTC

GCACACGTCTGAACTCTTTCCCTACACGACGCTCTTCCGATCTNNNTGTTTGCATGGAGAGATTGGTC

AGCACACGTCTGAACTCTTTCCCTACACGACGCTCTTCCGATCTNNNGCTCTTTCAGTTGTGCATTGTCTC

AGCACACGTCTGAACTCTTTCCCTACACGACGCTCTTCCGATCTNNNGCTCTTTCAGTTGTGCATTGTCTC

AGCACACGTCTGAACTCTTTCCCTACACGACGCTCTTCCGATCTNNNGCTCTTTCAGTTGTGCATTGTCTC

GAGCACACGTCTGAACTCTTTCCCTACACGACGCTCTTCCGATCTNNNGAGGAAGGGAAAGAGGCTG

GAGCACACGTCTGAACTCTTTCCCTACACGACGCTCTTCCGATCTNNNGAGGAAGGGAAAGAGGCTG

GAGCACACGTCTGAACTCTTTCCCTACACGACGCTCTTCCGATCTNNNGAGGAAGGGAAAGAGGCTG

AGCACACGTCTGAACTCTTTCCCTACACGACGCTCTTCCGATCTNNNTCCTCTCTCTCCTCTCTCTCTCA

AGCACACGTCTGAACTCTTTCCCTACACGACGCTCTTCCGATCTNNNTCCTCTCTCTCCTCTCTCTCTCA

AGCACACGTCTGAACTCTTTCCCTACACGACGCTCTTCCGATCTNNNTCCTCTCTCTCCTCTCTCTCTCA

GAGCACACGTCTGAACTCTTTCCCTACACGACGCTCTTCCGATCTNNNAACAACAACAAAACTAGCAGG

GAGCACACGTCTGAACTCTTTCCCTACACGACGCTCTTCCGATCTNNNAACAACAACAAAACTAGCAGG

GAGCACACGTCTGAACTCTTTCCCTACACGACGCTCTTCCGATCTNNNAACAACAACAAAACTAGCAGG

AGCACACGTCTGAACTCTTTCCCTACACGACGCTCTTCCGATCTNNNTTTATGAGAAGAGGCTGGAGA

AGCACACGTCTGAACTCTTTCCCTACACGACGCTCTTCCGATCTNNNTTTATGAGAAGAGGCTGGAGA

AGCACACGTCTGAACTCTTTCCCTACACGACGCTCTTCCGATCTNNNTTTATGAGAAGAGGCTGGAGA

GAGCACACGTCTGAACTCTTTCCCTACACGACGCTCTTCCGATCTNNNGACTTGAAAATCGCTGAACCC

GAGCACACGTCTGAACTCTTTCCCTACACGACGCTCTTCCGATCTNNNGACTTGAAAATCGCTGAACCC

GAGCACACGTCTGAACTCTTTCCCTACACGACGCTCTTCCGATCTNNNGACTTGAAAATCGCTGAACCC

GCACACGTCTGAACTCTTTCCCTACACGACGCTCTTCCGATCTNNNACCAGATAACTTGGTGCTCCATTC

GCACACGTCTGAACTCTTTCCCTACACGACGCTCTTCCGATCTNNNACCAGATAACTTGGTGCTCCATTC

GCACACGTCTGAACTCTTTCCCTACACGACGCTCTTCCGATCTNNNACCAGATAACTTGGTGCTCCATTC

GCACACGTCTGAACTCTTTCCCTACACGACGCTCTTCCGATCTNNNGGATTCTTGTTCAAAAGTGGCCA

GCACACGTCTGAACTCTTTCCCTACACGACGCTCTTCCGATCTNNNGGATTCTTGTTCAAAAGTGGCCA
GCACACGTCTGAACTCTTTCCCTACACGACGCTCTTCCGATCTNNNGGATTCTTGTTCAAAAGTGGCCA
GAGCACACGTCTGAACTCTTTCCCTACACGACGCTCTTCCGATCTNNNAGGAGGCTGAGACAGGAGAA
GAGCACACGTCTGAACTCTTTCCCTACACGACGCTCTTCCGATCTNNNAGGAGGCTGAGACAGGAGAA
GAGCACACGTCTGAACTCTTTCCCTACACGACGCTCTTCCGATCTNNNAGGAGGCTGAGACAGGAGAA
GAGCACACGTCTGAACTCTTTCCCTACACGACGCTCTTCCGATCTNNNACAAACCTGTGGAGTGTGGA
GAGCACACGTCTGAACTCTTTCCCTACACGACGCTCTTCCGATCTNNNACAAACCTGTGGAGTGTGGA
GAGCACACGTCTGAACTCTTTCCCTACACGACGCTCTTCCGATCTNNNACAAACCTGTGGAGTGTGGA
GAGCACACGTCTGAACTCTTTCCCTACACGACGCTCTTCCGATCTNNNCAGCAAGGAGAAAACTGTTC
GAGCACACGTCTGAACTCTTTCCCTACACGACGCTCTTCCGATCTNNNCAGCAAGGAGAAAACTGTTC
GAGCACACGTCTGAACTCTTTCCCTACACGACGCTCTTCCGATCTNNNCAGCAAGGAGAAAACTGTTC
GAGCACACGTCTGAACTCTTTCCCTACACGACGCTCTTCCGATCTNNNGAGAATGGCGTGAATCCAGGA
GAGCACACGTCTGAACTCTTTCCCTACACGACGCTCTTCCGATCTNNNGAGAATGGCGTGAATCCAGGA
GAGCACACGTCTGAACTCTTTCCCTACACGACGCTCTTCCGATCTNNNGAGAATGGCGTGAATCCAGGA
AGCACACGTCTGAACTCTTTCCCTACACGACGCTCTTCCGATCTNNNGCACATGACATCTCACCACAGTA
AGCACACGTCTGAACTCTTTCCCTACACGACGCTCTTCCGATCTNNNGCACATGACATCTCACCACAGTA
AGCACACGTCTGAACTCTTTCCCTACACGACGCTCTTCCGATCTNNNGCACATGACATCTCACCACAGTA
GAGCACACGTCTGAACTCTTTCCCTACACGACGCTCTTCCGATCTNNNGATTGTAAGGCAAGAACTCAG
GAGCACACGTCTGAACTCTTTCCCTACACGACGCTCTTCCGATCTNNNGATTGTAAGGCAAGAACTCAG
GAGCACACGTCTGAACTCTTTCCCTACACGACGCTCTTCCGATCTNNNGATTGTAAGGCAAGAACTCAG
GCACACGTCTGAACTCTTTCCCTACACGACGCTCTTCCGATCTNNNGGCCAAGCTAGTGATTGATGTGA
GCACACGTCTGAACTCTTTCCCTACACGACGCTCTTCCGATCTNNNGGCCAAGCTAGTGATTGATGTGA
GCACACGTCTGAACTCTTTCCCTACACGACGCTCTTCCGATCTNNNGGCCAAGCTAGTGATTGATGTGA
AGCACACGTCTGAACTCTTTCCCTACACGACGCTCTTCCGATCTNNNCTGTAGTCCCAGCTACTCAGGA
AGCACACGTCTGAACTCTTTCCCTACACGACGCTCTTCCGATCTNNNCTGTAGTCCCAGCTACTCAGGA
AGCACACGTCTGAACTCTTTCCCTACACGACGCTCTTCCGATCTNNNCTGTAGTCCCAGCTACTCAGGA
GAGCACACGTCTGAACTCTTTCCCTACACGACGCTCTTCCGATCTNNNGAGAATGGCATGAACCTGGG
GAGCACACGTCTGAACTCTTTCCCTACACGACGCTCTTCCGATCTNNNGAGAATGGCATGAACCTGGG
GAGCACACGTCTGAACTCTTTCCCTACACGACGCTCTTCCGATCTNNNGAGAATGGCATGAACCTGGG
AGAGCACACGTCTGAACTCTTTCCCTACACGACGCTCTTCCGATCTNNNTGCTGCAACATGGATAAACC
AGAGCACACGTCTGAACTCTTTCCCTACACGACGCTCTTCCGATCTNNNTGCTGCAACATGGATAAACC
AGAGCACACGTCTGAACTCTTTCCCTACACGACGCTCTTCCGATCTNNNTGCTGCAACATGGATAAACC
AGAGCACACGTCTGAACTCTTTCCCTACACGACGCTCTTCCGATCTNNNTGTCTGTACCTCCTATCACCAC
AGAGCACACGTCTGAACTCTTTCCCTACACGACGCTCTTCCGATCTNNNTGTCTGTACCTCCTATCACCAC
AGAGCACACGTCTGAACTCTTTCCCTACACGACGCTCTTCCGATCTNNNTGTCTGTACCTCCTATCACCAC
AGCACACGTCTGAACTCTTTCCCTACACGACGCTCTTCCGATCTNNNTGGGATGGCAAAGACTAAGAG
AGCACACGTCTGAACTCTTTCCCTACACGACGCTCTTCCGATCTNNNTGGGATGGCAAAGACTAAGAG
AGCACACGTCTGAACTCTTTCCCTACACGACGCTCTTCCGATCTNNNTGGGATGGCAAAGACTAAGAG
AGCACACGTCTGAACTCTTTCCCTACACGACGCTCTTCCGATCTNNNCAATCCTCCCACTTCAGCCTCCAT
AGCACACGTCTGAACTCTTTCCCTACACGACGCTCTTCCGATCTNNNCAATCCTCCCACTTCAGCCTCCAT
AGCACACGTCTGAACTCTTTCCCTACACGACGCTCTTCCGATCTNNNCAATCCTCCCACTTCAGCCTCCAT
GCACACGTCTGAACTCTTTCCCTACACGACGCTCTTCCGATCTNNNCTTTCCCAACAAAAGCCCTTTCCCA
GCACACGTCTGAACTCTTTCCCTACACGACGCTCTTCCGATCTNNNCTTTCCCAACAAAAGCCCTTTCCCA

GCACACGTCTGAACTCTTTCCCTACACGACGCTCTTCCGATCTNNNCTTTCCCAACAAAAGCCCTTTCCC
AGCACACGTCTGAACTCTTTCCCTACACGACGCTCTTCCGATCTNNNATTGCAGTGAAATGGGATCAT
AGCACACGTCTGAACTCTTTCCCTACACGACGCTCTTCCGATCTNNNATTGCAGTGAAATGGGATCAT
AGCACACGTCTGAACTCTTTCCCTACACGACGCTCTTCCGATCTNNNATTGCAGTGAAATGGGATCAT
GCACACGTCTGAACTCTTTCCCTACACGACGCTCTTCCGATCTNNNTGTAGTCCTAGCTACTCAGGAG
GCACACGTCTGAACTCTTTCCCTACACGACGCTCTTCCGATCTNNNTGTAGTCCTAGCTACTCAGGAG
GCACACGTCTGAACTCTTTCCCTACACGACGCTCTTCCGATCTNNNTGTAGTCCTAGCTACTCAGGAG
GCACACGTCTGAACTCTTTCCCTACACGACGCTCTTCCGATCTNNNCACGGTTCCCACACCCTTAGCAT
GCACACGTCTGAACTCTTTCCCTACACGACGCTCTTCCGATCTNNNCACGGTTCCCACACCCTTAGCAT
GCACACGTCTGAACTCTTTCCCTACACGACGCTCTTCCGATCTNNNCACGGTTCCCACACCCTTAGCAT
GCACACGTCTGAACTCTTTCCCTACACGACGCTCTTCCGATCTNNNAGGTTATCTGGACTTGGAGGG
GCACACGTCTGAACTCTTTCCCTACACGACGCTCTTCCGATCTNNNAGGTTATCTGGACTTGGAGGG
GCACACGTCTGAACTCTTTCCCTACACGACGCTCTTCCGATCTNNNAGGTTATCTGGACTTGGAGGG
AGCACACGTCTGAACTCTTTCCCTACACGACGCTCTTCCGATCTNNNTCAAATCCACCTCCCTTTCACAT
AGCACACGTCTGAACTCTTTCCCTACACGACGCTCTTCCGATCTNNNTCAAATCCACCTCCCTTTCACAT
AGCACACGTCTGAACTCTTTCCCTACACGACGCTCTTCCGATCTNNNTCAAATCCACCTCCCTTTCACAT
GAGCACACGTCTGAACTCTTTCCCTACACGACGCTCTTCCGATCTNNNTCCTTTGCAAGCCTCTCAATA
GAGCACACGTCTGAACTCTTTCCCTACACGACGCTCTTCCGATCTNNNTCCTTTGCAAGCCTCTCAATA
GAGCACACGTCTGAACTCTTTCCCTACACGACGCTCTTCCGATCTNNNTCCTTTGCAAGCCTCTCAATA
GCACACGTCTGAACTCTTTCCCTACACGACGCTCTTCCGATCTNNNCTGGACAGGCAGTCTTTTGATTG
GCACACGTCTGAACTCTTTCCCTACACGACGCTCTTCCGATCTNNNCTGGACAGGCAGTCTTTTGATTG
GCACACGTCTGAACTCTTTCCCTACACGACGCTCTTCCGATCTNNNCTGGACAGGCAGTCTTTTGATTG
GAGCACACGTCTGAACTCTTTCCCTACACGACGCTCTTCCGATCTNNNAAAAATTAGATGGGCGTGG
GAGCACACGTCTGAACTCTTTCCCTACACGACGCTCTTCCGATCTNNNAAAAATTAGATGGGCGTGG
GAGCACACGTCTGAACTCTTTCCCTACACGACGCTCTTCCGATCTNNNAAAAATTAGATGGGCGTGG
GAGCACACGTCTGAACTCTTTCCCTACACGACGCTCTTCCGATCTNNNAGCAGTCATTTATTGCAAAG
GAGCACACGTCTGAACTCTTTCCCTACACGACGCTCTTCCGATCTNNNAGCAGTCATTTATTGCAAAG
GAGCACACGTCTGAACTCTTTCCCTACACGACGCTCTTCCGATCTNNNAGCAGTCATTTATTGCAAAG
AGAGCACACGTCTGAACTCTTTCCCTACACGACGCTCTTCCGATCTNNNACAAAAATTAGCCAAGGA
AGAGCACACGTCTGAACTCTTTCCCTACACGACGCTCTTCCGATCTNNNACAAAAATTAGCCAAGGA
AGAGCACACGTCTGAACTCTTTCCCTACACGACGCTCTTCCGATCTNNNACAAAAATTAGCCAAGGA
GAGCACACGTCTGAACTCTTTCCCTACACGACGCTCTTCCGATCTNNNCGTGCATGAATGTGTTTGTG
GAGCACACGTCTGAACTCTTTCCCTACACGACGCTCTTCCGATCTNNNCGTGCATGAATGTGTTTGTG
GAGCACACGTCTGAACTCTTTCCCTACACGACGCTCTTCCGATCTNNNCGTGCATGAATGTGTTTGTG
GCACACGTCTGAACTCTTTCCCTACACGACGCTCTTCCGATCTNNNGCAAAAGACCATGAAAGGCTCT
GCACACGTCTGAACTCTTTCCCTACACGACGCTCTTCCGATCTNNNGCAAAAGACCATGAAAGGCTCT
GCACACGTCTGAACTCTTTCCCTACACGACGCTCTTCCGATCTNNNGCAAAAGACCATGAAAGGCTCT
AGCACACGTCTGAACTCTTTCCCTACACGACGCTCTTCCGATCTNNNACAAAAATTAGCCAGGCGTG
AGCACACGTCTGAACTCTTTCCCTACACGACGCTCTTCCGATCTNNNACAAAAATTAGCCAGGCGTG
AGCACACGTCTGAACTCTTTCCCTACACGACGCTCTTCCGATCTNNNACAAAAATTAGCCAGGCGTG
CACACGTCTGAACTCTTTCCCTACACGACGCTCTTCCGATCTNNNTGAGTTTATTTTGAAGGTTAACTC
CACACGTCTGAACTCTTTCCCTACACGACGCTCTTCCGATCTNNNTGAGTTTATTTTGAAGGTTAACTC
CACACGTCTGAACTCTTTCCCTACACGACGCTCTTCCGATCTNNNTGAGTTTATTTTGAAGGTTAACTC

GAGCACACGTCTGAACTCTTTCCCTACACGACGCTCTTCCGATCTNNNGGTTGAGTGGGAAAAAAGG
GAGCACACGTCTGAACTCTTTCCCTACACGACGCTCTTCCGATCTNNNGGTTGAGTGGGAAAAAAGG
GAGCACACGTCTGAACTCTTTCCCTACACGACGCTCTTCCGATCTNNNGGTTGAGTGGGAAAAAAGG
GCACACGTCTGAACTCTTTCCCTACACGACGCTCTTCCGATCTNNNATCAAGGGAAGAAATAAAGGAT
GCACACGTCTGAACTCTTTCCCTACACGACGCTCTTCCGATCTNNNATCAAGGGAAGAAATAAAGGAT
GCACACGTCTGAACTCTTTCCCTACACGACGCTCTTCCGATCTNNNATCAAGGGAAGAAATAAAGGAT
GAGCACACGTCTGAACTCTTTCCCTACACGACGCTCTTCCGATCTNNNCAGAGGTTGTAGTGAGCCAA
GAGCACACGTCTGAACTCTTTCCCTACACGACGCTCTTCCGATCTNNNCAGAGGTTGTAGTGAGCCAA
GAGCACACGTCTGAACTCTTTCCCTACACGACGCTCTTCCGATCTNNNCAGAGGTTGTAGTGAGCCAA
AGCACACGTCTGAACTCTTTCCCTACACGACGCTCTTCCGATCTNNNGAGTTCACACAAGCCGTTCACT
AGCACACGTCTGAACTCTTTCCCTACACGACGCTCTTCCGATCTNNNGAGTTCACACAAGCCGTTCACT
AGCACACGTCTGAACTCTTTCCCTACACGACGCTCTTCCGATCTNNNGAGTTCACACAAGCCGTTCACT
AGAGCACACGTCTGAACTCTTTCCCTACACGACGCTCTTCCGATCTNNNGGAACAGAAAACCAAACACC
AGAGCACACGTCTGAACTCTTTCCCTACACGACGCTCTTCCGATCTNNNGGAACAGAAAACCAAACACC
AGAGCACACGTCTGAACTCTTTCCCTACACGACGCTCTTCCGATCTNNNGGAACAGAAAACCAAACACC
AGCACACGTCTGAACTCTTTCCCTACACGACGCTCTTCCGATCTNNNGAGGCAGGAAAATCACTTGAA
AGCACACGTCTGAACTCTTTCCCTACACGACGCTCTTCCGATCTNNNGAGGCAGGAAAATCACTTGAA
AGCACACGTCTGAACTCTTTCCCTACACGACGCTCTTCCGATCTNNNGAGGCAGGAAAATCACTTGAA
GAGCACACGTCTGAACTCTTTCCCTACACGACGCTCTTCCGATCTNNNAAATACAACCCATCACACACGT
GAGCACACGTCTGAACTCTTTCCCTACACGACGCTCTTCCGATCTNNNAAATACAACCCATCACACACGT
GAGCACACGTCTGAACTCTTTCCCTACACGACGCTCTTCCGATCTNNNAAATACAACCCATCACACACGT
AGCACACGTCTGAACTCTTTCCCTACACGACGCTCTTCCGATCTNNNCGGGTAGGAAGGCAGGCTAG
AGCACACGTCTGAACTCTTTCCCTACACGACGCTCTTCCGATCTNNNCGGGTAGGAAGGCAGGCTAG
AGCACACGTCTGAACTCTTTCCCTACACGACGCTCTTCCGATCTNNNCGGGTAGGAAGGCAGGCTAG
AGAGCACACGTCTGAACTCTTTCCCTACACGACGCTCTTCCGATCTNNNCTGTAATCCCAGCTACACAGG
AGAGCACACGTCTGAACTCTTTCCCTACACGACGCTCTTCCGATCTNNNCTGTAATCCCAGCTACACAGG
AGAGCACACGTCTGAACTCTTTCCCTACACGACGCTCTTCCGATCTNNNCTGTAATCCCAGCTACACAGG
GAGCACACGTCTGAACTCTTTCCCTACACGACGCTCTTCCGATCTNNNGCAGAAGAATCACTTGAACC
GAGCACACGTCTGAACTCTTTCCCTACACGACGCTCTTCCGATCTNNNGCAGAAGAATCACTTGAACC
GAGCACACGTCTGAACTCTTTCCCTACACGACGCTCTTCCGATCTNNNGCAGAAGAATCACTTGAACC
AGCACACGTCTGAACTCTTTCCCTACACGACGCTCTTCCGATCTNNNCTGAGGCAGGAGAATCACTTGA
AGCACACGTCTGAACTCTTTCCCTACACGACGCTCTTCCGATCTNNNCTGAGGCAGGAGAATCACTTGA
AGCACACGTCTGAACTCTTTCCCTACACGACGCTCTTCCGATCTNNNCTGAGGCAGGAGAATCACTTGA
AGAGCACACGTCTGAACTCTTTCCCTACACGACGCTCTTCCGATCTNNNGCTTGCAGTGAGCTGAGAT
AGAGCACACGTCTGAACTCTTTCCCTACACGACGCTCTTCCGATCTNNNGCTTGCAGTGAGCTGAGAT
AGAGCACACGTCTGAACTCTTTCCCTACACGACGCTCTTCCGATCTNNNGCTTGCAGTGAGCTGAGAT
GCACACGTCTGAACTCTTTCCCTACACGACGCTCTTCCGATCTNNNGAGAGAGGAGGGGAACAGGAT
GCACACGTCTGAACTCTTTCCCTACACGACGCTCTTCCGATCTNNNGAGAGAGGAGGGGAACAGGAT
GCACACGTCTGAACTCTTTCCCTACACGACGCTCTTCCGATCTNNNGAGAGAGGAGGGGAACAGGAT
GCACACGTCTGAACTCTTTCCCTACACGACGCTCTTCCGATCTNNNCGCCTCCTTTTCGTAACACTGTTC
GCACACGTCTGAACTCTTTCCCTACACGACGCTCTTCCGATCTNNNCGCCTCCTTTTCGTAACACTGTTC
GCACACGTCTGAACTCTTTCCCTACACGACGCTCTTCCGATCTNNNCGCCTCCTTTTCGTAACACTGTTC
AAGAGCACACGTCTGAACTCTTTCCCTACACGACGCTCTTCCGATCTNNNACTCATCAAGTTTTGGGAA

AGAGCACACGTCTGAACTCTTTCCCTACACGACGCTCTTCCGATCTNNNACTCATCAAGTTTTGGGAA
AGAGCACACGTCTGAACTCTTTCCCTACACGACGCTCTTCCGATCTNNNACTCATCAAGTTTTGGGAA
GCACACGTCTGAACTCTTTCCCTACACGACGCTCTTCCGATCTNNNCGATGCACTGATTTCTGTGTGAA
GCACACGTCTGAACTCTTTCCCTACACGACGCTCTTCCGATCTNNNCGATGCACTGATTTCTGTGTGAA
GCACACGTCTGAACTCTTTCCCTACACGACGCTCTTCCGATCTNNNCGATGCACTGATTTCTGTGTGAA
AGAGCACACGTCTGAACTCTTTCCCTACACGACGCTCTTCCGATCTNNNACTGTCTCTTAGGGTCTGTG
AGAGCACACGTCTGAACTCTTTCCCTACACGACGCTCTTCCGATCTNNNACTGTCTCTTAGGGTCTGTG
AGAGCACACGTCTGAACTCTTTCCCTACACGACGCTCTTCCGATCTNNNACTGTCTCTTAGGGTCTGTG
AGCACACGTCTGAACTCTTTCCCTACACGACGCTCTTCCGATCTNNNGCCAGTTATCCCAGCACCATTTA
AGCACACGTCTGAACTCTTTCCCTACACGACGCTCTTCCGATCTNNNGCCAGTTATCCCAGCACCATTTA
AGCACACGTCTGAACTCTTTCCCTACACGACGCTCTTCCGATCTNNNGCCAGTTATCCCAGCACCATTTA
AGCACACGTCTGAACTCTTTCCCTACACGACGCTCTTCCGATCTNNNAGGCAGGAGAATCACTTGAA
AGCACACGTCTGAACTCTTTCCCTACACGACGCTCTTCCGATCTNNNAGGCAGGAGAATCACTTGAA
AGCACACGTCTGAACTCTTTCCCTACACGACGCTCTTCCGATCTNNNAGGCAGGAGAATCACTTGAA
AGCACACGTCTGAACTCTTTCCCTACACGACGCTCTTCCGATCTNNNTCCTAATGTTTTCCCTCCCCTTG
AGCACACGTCTGAACTCTTTCCCTACACGACGCTCTTCCGATCTNNNTCCTAATGTTTTCCCTCCCCTTG
AGCACACGTCTGAACTCTTTCCCTACACGACGCTCTTCCGATCTNNNTCCTAATGTTTTCCCTCCCCTTG
AGCACACGTCTGAACTCTTTCCCTACACGACGCTCTTCCGATCTNNNAAAGTGTCTTCTCATAGAGGGA
AGCACACGTCTGAACTCTTTCCCTACACGACGCTCTTCCGATCTNNNAAAGTGTCTTCTCATAGAGGGA
AGCACACGTCTGAACTCTTTCCCTACACGACGCTCTTCCGATCTNNNAAAGTGTCTTCTCATAGAGGGA
GCACACGTCTGAACTCTTTCCCTACACGACGCTCTTCCGATCTNNNCCATCCTTAGTCCCAGAAATGGC
GCACACGTCTGAACTCTTTCCCTACACGACGCTCTTCCGATCTNNNCCATCCTTAGTCCCAGAAATGGC
GCACACGTCTGAACTCTTTCCCTACACGACGCTCTTCCGATCTNNNCCATCCTTAGTCCCAGAAATGGC
GCACACGTCTGAACTCTTTCCCTACACGACGCTCTTCCGATCTNNNCTTTCTCCTGTTTATGGGTTTCAT
GCACACGTCTGAACTCTTTCCCTACACGACGCTCTTCCGATCTNNNCTTTCTCCTGTTTATGGGTTTCAT
GCACACGTCTGAACTCTTTCCCTACACGACGCTCTTCCGATCTNNNCTTTCTCCTGTTTATGGGTTTCAT
AGCACACGTCTGAACTCTTTCCCTACACGACGCTCTTCCGATCTNNNCCAGTTTTTGGCTGCTGTGAAT
AGCACACGTCTGAACTCTTTCCCTACACGACGCTCTTCCGATCTNNNCCAGTTTTTGGCTGCTGTGAAT
AGCACACGTCTGAACTCTTTCCCTACACGACGCTCTTCCGATCTNNNCCAGTTTTTGGCTGCTGTGAAT
AGCACACGTCTGAACTCTTTCCCTACACGACGCTCTTCCGATCTNNNTCTGTAGTCCCAGCTATTCAGGA
AGCACACGTCTGAACTCTTTCCCTACACGACGCTCTTCCGATCTNNNTCTGTAGTCCCAGCTATTCAGGA
AGCACACGTCTGAACTCTTTCCCTACACGACGCTCTTCCGATCTNNNTCTGTAGTCCCAGCTATTCAGGA
AGCACACGTCTGAACTCTTTCCCTACACGACGCTCTTCCGATCTNNNTGCCATTTTCCTTCTCTGCCTAT
AGCACACGTCTGAACTCTTTCCCTACACGACGCTCTTCCGATCTNNNTGCCATTTTCCTTCTCTGCCTAT
AGCACACGTCTGAACTCTTTCCCTACACGACGCTCTTCCGATCTNNNTGCCATTTTCCTTCTCTGCCTAT
ACACGTCTGAACTCTTTCCCTACACGACGCTCTTCCGATCTNNNCTAGAGGGAGAAATGGCAGTCCA
ACACGTCTGAACTCTTTCCCTACACGACGCTCTTCCGATCTNNNCTAGAGGGAGAAATGGCAGTCCA
ACACGTCTGAACTCTTTCCCTACACGACGCTCTTCCGATCTNNNCTAGAGGGAGAAATGGCAGTCCA
AGCACACGTCTGAACTCTTTCCCTACACGACGCTCTTCCGATCTNNNAGTCTTGGGAGGGGTTTTTCT
AGCACACGTCTGAACTCTTTCCCTACACGACGCTCTTCCGATCTNNNAGTCTTGGGAGGGGTTTTTCT
AGCACACGTCTGAACTCTTTCCCTACACGACGCTCTTCCGATCTNNNAGTCTTGGGAGGGGTTTTTCT
GAGCACACGTCTGAACTCTTTCCCTACACGACGCTCTTCCGATCTNNNTGGGATGAGAGAACACAAAC
GAGCACACGTCTGAACTCTTTCCCTACACGACGCTCTTCCGATCTNNNTGGGATGAGAGAACACAAAC

AGCACACGTCTGAACTCTTTCCCTACACGACGCTCTTCCGATCTNNNTGGGATGAGAGAACACAAAC
AGCACACGTCTGAACTCTTTCCCTACACGACGCTCTTCCGATCTNNNGAAACTGTGAAGATTCCTGTG
AGCACACGTCTGAACTCTTTCCCTACACGACGCTCTTCCGATCTNNNGAAACTGTGAAGATTCCTGTG
AGCACACGTCTGAACTCTTTCCCTACACGACGCTCTTCCGATCTNNNGAAACTGTGAAGATTCCTGTG
AGCACACGTCTGAACTCTTTCCCTACACGACGCTCTTCCGATCTNNNAGAGGAGGCTTAGAGTTGG
AGCACACGTCTGAACTCTTTCCCTACACGACGCTCTTCCGATCTNNNAGAGGAGGCTTAGAGTTGG
AGCACACGTCTGAACTCTTTCCCTACACGACGCTCTTCCGATCTNNNAGAGGAGGCTTAGAGTTGG
AGCACACGTCTGAACTCTTTCCCTACACGACGCTCTTCCGATCTNNNTGATTGAGCCAAGGTCATAC
AGCACACGTCTGAACTCTTTCCCTACACGACGCTCTTCCGATCTNNNTGATTGAGCCAAGGTCATAC
AGCACACGTCTGAACTCTTTCCCTACACGACGCTCTTCCGATCTNNNTGATTGAGCCAAGGTCATAC
AGCACACGTCTGAACTCTTTCCCTACACGACGCTCTTCCGATCTNNNCAAAGGCTCAGGAAACAGACG
AGCACACGTCTGAACTCTTTCCCTACACGACGCTCTTCCGATCTNNNCAAAGGCTCAGGAAACAGACG
AGCACACGTCTGAACTCTTTCCCTACACGACGCTCTTCCGATCTNNNCAAAGGCTCAGGAAACAGACG
GCACACGTCTGAACTCTTTCCCTACACGACGCTCTTCCGATCTNNNAGCTAGTTCCAGAAAGATGGT
GCACACGTCTGAACTCTTTCCCTACACGACGCTCTTCCGATCTNNNAGCTAGTTCCAGAAAGATGGT
GCACACGTCTGAACTCTTTCCCTACACGACGCTCTTCCGATCTNNNAGCTAGTTCCAGAAAGATGGT
GCACACGTCTGAACTCTTTCCCTACACGACGCTCTTCCGATCTNNNGCCCTCTCTGAGTTCTCTAAACC
GCACACGTCTGAACTCTTTCCCTACACGACGCTCTTCCGATCTNNNGCCCTCTCTGAGTTCTCTAAACC
GCACACGTCTGAACTCTTTCCCTACACGACGCTCTTCCGATCTNNNGCCCTCTCTGAGTTCTCTAAACC
GCACACGTCTGAACTCTTTCCCTACACGACGCTCTTCCGATCTNNNCTTATGCCCTCCCTCTCCGATCAT
GCACACGTCTGAACTCTTTCCCTACACGACGCTCTTCCGATCTNNNCTTATGCCCTCCCTCTCCGATCAT
GCACACGTCTGAACTCTTTCCCTACACGACGCTCTTCCGATCTNNNCTTATGCCCTCCCTCTCCGATCAT
ACACGTCTGAACTCTTTCCCTACACGACGCTCTTCCGATCTNNNTCCTACATCTGCTGGCATTCATTCAT
ACACGTCTGAACTCTTTCCCTACACGACGCTCTTCCGATCTNNNTCCTACATCTGCTGGCATTCATTCAT
ACACGTCTGAACTCTTTCCCTACACGACGCTCTTCCGATCTNNNTCCTACATCTGCTGGCATTCATTCAT
GAGCACACGTCTGAACTCTTTCCCTACACGACGCTCTTCCGATCTNNNAAACTAAGGAGTTGTGCGT
GAGCACACGTCTGAACTCTTTCCCTACACGACGCTCTTCCGATCTNNNAAACTAAGGAGTTGTGCGT
GAGCACACGTCTGAACTCTTTCCCTACACGACGCTCTTCCGATCTNNNAAACTAAGGAGTTGTGCGT
AGCACACGTCTGAACTCTTTCCCTACACGACGCTCTTCCGATCTNNNACACCTGTAATCCCAGCACTTT
AGCACACGTCTGAACTCTTTCCCTACACGACGCTCTTCCGATCTNNNACACCTGTAATCCCAGCACTTT
AGCACACGTCTGAACTCTTTCCCTACACGACGCTCTTCCGATCTNNNACACCTGTAATCCCAGCACTTT
AGAGCACACGTCTGAACTCTTTCCCTACACGACGCTCTTCCGATCTNNNGCCTGTAATCTCAGCTACTA
AGAGCACACGTCTGAACTCTTTCCCTACACGACGCTCTTCCGATCTNNNGCCTGTAATCTCAGCTACTA
AGAGCACACGTCTGAACTCTTTCCCTACACGACGCTCTTCCGATCTNNNGCCTGTAATCTCAGCTACTA
AGCACACGTCTGAACTCTTTCCCTACACGACGCTCTTCCGATCTNNNGGGGCCCAAGATGGTATATTA
AGCACACGTCTGAACTCTTTCCCTACACGACGCTCTTCCGATCTNNNGGGGCCCAAGATGGTATATTA
AGCACACGTCTGAACTCTTTCCCTACACGACGCTCTTCCGATCTNNNGGGGCCCAAGATGGTATATTA
AGCACACGTCTGAACTCTTTCCCTACACGACGCTCTTCCGATCTNNNAGTAGCAGACGTGGTTGTTCT
AGCACACGTCTGAACTCTTTCCCTACACGACGCTCTTCCGATCTNNNAGTAGCAGACGTGGTTGTTCT
AGCACACGTCTGAACTCTTTCCCTACACGACGCTCTTCCGATCTNNNAGTAGCAGACGTGGTTGTTCT
GCACACGTCTGAACTCTTTCCCTACACGACGCTCTTCCGATCTNNNAATTAGCCAGAGATGATGGTG
GCACACGTCTGAACTCTTTCCCTACACGACGCTCTTCCGATCTNNNAATTAGCCAGAGATGATGGTG
GCACACGTCTGAACTCTTTCCCTACACGACGCTCTTCCGATCTNNNAATTAGCCAGAGATGATGGTG

AGCACACGTCTGAACTCTTTCCCTACACGACGCTCTTCCGATCTNNNAGGTTTCAGTGAGTCGAGATC

AGCACACGTCTGAACTCTTTCCCTACACGACGCTCTTCCGATCTNNNAGGTTTCAGTGAGTCGAGATC

AGCACACGTCTGAACTCTTTCCCTACACGACGCTCTTCCGATCTNNNAGGTTTCAGTGAGTCGAGATC

GCACACGTCTGAACTCTTTCCCTACACGACGCTCTTCCGATCTNNNCCAAAGTTCTGGAATTACAGGC

GCACACGTCTGAACTCTTTCCCTACACGACGCTCTTCCGATCTNNNCCAAAGTTCTGGAATTACAGGC

GCACACGTCTGAACTCTTTCCCTACACGACGCTCTTCCGATCTNNNCCAAAGTTCTGGAATTACAGGC

GAGCACACGTCTGAACTCTTTCCCTACACGACGCTCTTCCGATCTNNNCACAGGACAATGAATCTCTG

GAGCACACGTCTGAACTCTTTCCCTACACGACGCTCTTCCGATCTNNNCACAGGACAATGAATCTCTG

GAGCACACGTCTGAACTCTTTCCCTACACGACGCTCTTCCGATCTNNNCACAGGACAATGAATCTCTG

GAGCACACGTCTGAACTCTTTCCCTACACGACGCTCTTCCGATCTNNNAGGAAAAGTCCAAACCCATC

GAGCACACGTCTGAACTCTTTCCCTACACGACGCTCTTCCGATCTNNNAGGAAAAGTCCAAACCCATC

GAGCACACGTCTGAACTCTTTCCCTACACGACGCTCTTCCGATCTNNNAGGAAAAGTCCAAACCCATC

AGCACACGTCTGAACTCTTTCCCTACACGACGCTCTTCCGATCTNNNCCTTCTCTTTGTTTGGGTTGGG

AGCACACGTCTGAACTCTTTCCCTACACGACGCTCTTCCGATCTNNNCCTTCTCTTTGTTTGGGTTGGG

AGCACACGTCTGAACTCTTTCCCTACACGACGCTCTTCCGATCTNNNCCTTCTCTTTGTTTGGGTTGGG

AGCACACGTCTGAACTCTTTCCCTACACGACGCTCTTCCGATCTNNNCCAGGTTCAAGTGATTCTCCTG

AGCACACGTCTGAACTCTTTCCCTACACGACGCTCTTCCGATCTNNNCCAGGTTCAAGTGATTCTCCTG

AGCACACGTCTGAACTCTTTCCCTACACGACGCTCTTCCGATCTNNNCCAGGTTCAAGTGATTCTCCTG

AGCACACGTCTGAACTCTTTCCCTACACGACGCTCTTCCGATCTNNNTCAGTGTTATCGCTACTCTCTG

AGCACACGTCTGAACTCTTTCCCTACACGACGCTCTTCCGATCTNNNTCAGTGTTATCGCTACTCTCTG

AGCACACGTCTGAACTCTTTCCCTACACGACGCTCTTCCGATCTNNNTCAGTGTTATCGCTACTCTCTG

GAGCACACGTCTGAACTCTTTCCCTACACGACGCTCTTCCGATCTNNNTGAGCTGAGATCATACCATTG

GAGCACACGTCTGAACTCTTTCCCTACACGACGCTCTTCCGATCTNNNTGAGCTGAGATCATACCATTG

GAGCACACGTCTGAACTCTTTCCCTACACGACGCTCTTCCGATCTNNNTGAGCTGAGATCATACCATTG

AGAGCACACGTCTGAACTCTTTCCCTACACGACGCTCTTCCGATCTNNNATTTGTCTGGGTATGGGG

AGAGCACACGTCTGAACTCTTTCCCTACACGACGCTCTTCCGATCTNNNATTTGTCTGGGTATGGGG

AGAGCACACGTCTGAACTCTTTCCCTACACGACGCTCTTCCGATCTNNNATTTGTCTGGGTATGGGG

GAGCACACGTCTGAACTCTTTCCCTACACGACGCTCTTCCGATCTNNNGAGCTGGGAAGGTGTTTGG

GAGCACACGTCTGAACTCTTTCCCTACACGACGCTCTTCCGATCTNNNGAGCTGGGAAGGTGTTTGG

GAGCACACGTCTGAACTCTTTCCCTACACGACGCTCTTCCGATCTNNNGAGCTGGGAAGGTGTTTGG

AGCACACGTCTGAACTCTTTCCCTACACGACGCTCTTCCGATCTNNNTGCCTTGTGAAGAGTTTTGCTT

AGCACACGTCTGAACTCTTTCCCTACACGACGCTCTTCCGATCTNNNTGCCTTGTGAAGAGTTTTGCTT

AGCACACGTCTGAACTCTTTCCCTACACGACGCTCTTCCGATCTNNNTGCCTTGTGAAGAGTTTTGCTT

GAGCACACGTCTGAACTCTTTCCCTACACGACGCTCTTCCGATCTNNNGGAGGTTGCAGTGAGTCATG

GAGCACACGTCTGAACTCTTTCCCTACACGACGCTCTTCCGATCTNNNGGAGGTTGCAGTGAGTCATG

GAGCACACGTCTGAACTCTTTCCCTACACGACGCTCTTCCGATCTNNNGGAGGTTGCAGTGAGTCATG

GAGCACACGTCTGAACTCTTTCCCTACACGACGCTCTTCCGATCTNNNGATATGTGAGTTTGCCTGTG

GAGCACACGTCTGAACTCTTTCCCTACACGACGCTCTTCCGATCTNNNGATATGTGAGTTTGCCTGTG

GAGCACACGTCTGAACTCTTTCCCTACACGACGCTCTTCCGATCTNNNGATATGTGAGTTTGCCTGTG

GAGCACACGTCTGAACTCTTTCCCTACACGACGCTCTTCCGATCTNNNTCCATCCATGCCCAGAATCTTT

GAGCACACGTCTGAACTCTTTCCCTACACGACGCTCTTCCGATCTNNNTCCATCCATGCCCAGAATCTTT

GAGCACACGTCTGAACTCTTTCCCTACACGACGCTCTTCCGATCTNNNTCCATCCATGCCCAGAATCTTT

GAAGAGCACACGTCTGAACTCTTTCCCTACACGACGCTCTTCCGATCTNNNTAACAGTGAAAGGAATG

AAGAGCACACGTCTGAACTCTTTCCCTACACGACGCTCTTCCGATCTNNNTAACAGTGAAAGGAATG

AAGAGCACACGTCTGAACTCTTTCCCTACACGACGCTCTTCCGATCTNNNTAACAGTGAAAGGAATG

GAGCACACGTCTGAACTCTTTCCCTACACGACGCTCTTCCGATCTNNNTGACCAATCCCTGACAATTGC

GAGCACACGTCTGAACTCTTTCCCTACACGACGCTCTTCCGATCTNNNTGACCAATCCCTGACAATTGC

GAGCACACGTCTGAACTCTTTCCCTACACGACGCTCTTCCGATCTNNNTGACCAATCCCTGACAATTGC

GCACACGTCTGAACTCTTTCCCTACACGACGCTCTTCCGATCTNNNCTGTGAGCTGAGATCGTACCATT

GCACACGTCTGAACTCTTTCCCTACACGACGCTCTTCCGATCTNNNCTGTGAGCTGAGATCGTACCATT

GCACACGTCTGAACTCTTTCCCTACACGACGCTCTTCCGATCTNNNCTGTGAGCTGAGATCGTACCATT

AGCACACGTCTGAACTCTTTCCCTACACGACGCTCTTCCGATCTNNNTCTGAAAGTGGGAAAGGAAT

AGCACACGTCTGAACTCTTTCCCTACACGACGCTCTTCCGATCTNNNTCTGAAAGTGGGAAAGGAAT

AGCACACGTCTGAACTCTTTCCCTACACGACGCTCTTCCGATCTNNNTCTGAAAGTGGGAAAGGAAT

AGCACACGTCTGAACTCTTTCCCTACACGACGCTCTTCCGATCTNNNAAGTTTCCGTTGCATCTTGGT

AGCACACGTCTGAACTCTTTCCCTACACGACGCTCTTCCGATCTNNNAAGTTTCCGTTGCATCTTGGT

AGCACACGTCTGAACTCTTTCCCTACACGACGCTCTTCCGATCTNNNAAGTTTCCGTTGCATCTTGGT

GCACACGTCTGAACTCTTTCCCTACACGACGCTCTTCCGATCTNNNTAGTCCTAGCTACTTGGAAGGC

GCACACGTCTGAACTCTTTCCCTACACGACGCTCTTCCGATCTNNNTAGTCCTAGCTACTTGGAAGGC

GCACACGTCTGAACTCTTTCCCTACACGACGCTCTTCCGATCTNNNTAGTCCTAGCTACTTGGAAGGC

AGCACACGTCTGAACTCTTTCCCTACACGACGCTCTTCCGATCTNNNTTGGCAGAATAGTGTGAGTAG

AGCACACGTCTGAACTCTTTCCCTACACGACGCTCTTCCGATCTNNNTTGGCAGAATAGTGTGAGTAG

AGCACACGTCTGAACTCTTTCCCTACACGACGCTCTTCCGATCTNNNTTGGCAGAATAGTGTGAGTAG

GCACACGTCTGAACTCTTTCCCTACACGACGCTCTTCCGATCTNNNAGACAGAAAGCTCCACTCTCTT

GCACACGTCTGAACTCTTTCCCTACACGACGCTCTTCCGATCTNNNAGACAGAAAGCTCCACTCTCTT

GCACACGTCTGAACTCTTTCCCTACACGACGCTCTTCCGATCTNNNAGACAGAAAGCTCCACTCTCTT

GCACACGTCTGAACTCTTTCCCTACACGACGCTCTTCCGATCTNNNGGCAAGAGAGTGAGACTCCATT

GCACACGTCTGAACTCTTTCCCTACACGACGCTCTTCCGATCTNNNGGCAAGAGAGTGAGACTCCATT

GCACACGTCTGAACTCTTTCCCTACACGACGCTCTTCCGATCTNNNGGCAAGAGAGTGAGACTCCATT

GCACACGTCTGAACTCTTTCCCTACACGACGCTCTTCCGATCTNNNCTCCAGTGCCTAGCACATCTCTT

GCACACGTCTGAACTCTTTCCCTACACGACGCTCTTCCGATCTNNNCTCCAGTGCCTAGCACATCTCTT

GCACACGTCTGAACTCTTTCCCTACACGACGCTCTTCCGATCTNNNCTCCAGTGCCTAGCACATCTCTT

AGCACACGTCTGAACTCTTTCCCTACACGACGCTCTTCCGATCTNNNTGGTGCCTTTTTCAGAAGTTC

AGCACACGTCTGAACTCTTTCCCTACACGACGCTCTTCCGATCTNNNTGGTGCCTTTTTCAGAAGTTC

AGCACACGTCTGAACTCTTTCCCTACACGACGCTCTTCCGATCTNNNTGGTGCCTTTTTCAGAAGTTC

GAGCACACGTCTGAACTCTTTCCCTACACGACGCTCTTCCGATCTNNNTTTCAAGGAAGAGCAACCTCT

GAGCACACGTCTGAACTCTTTCCCTACACGACGCTCTTCCGATCTNNNTTTCAAGGAAGAGCAACCTCT

GAGCACACGTCTGAACTCTTTCCCTACACGACGCTCTTCCGATCTNNNTTTCAAGGAAGAGCAACCTCT

GAGCACACGTCTGAACTCTTTCCCTACACGACGCTCTTCCGATCTNNNTTCCCCTTTTCTTGACCATCCA

GAGCACACGTCTGAACTCTTTCCCTACACGACGCTCTTCCGATCTNNNTTCCCCTTTTCTTGACCATCCA

GAGCACACGTCTGAACTCTTTCCCTACACGACGCTCTTCCGATCTNNNTTCCCCTTTTCTTGACCATCCA

GCACACGTCTGAACTCTTTCCCTACACGACGCTCTTCCGATCTNNNTGGAAACATGCACAATGATGCT

GCACACGTCTGAACTCTTTCCCTACACGACGCTCTTCCGATCTNNNTGGAAACATGCACAATGATGCT

GCACACGTCTGAACTCTTTCCCTACACGACGCTCTTCCGATCTNNNTGGAAACATGCACAATGATGCT

GCACACGTCTGAACTCTTTCCCTACACGACGCTCTTCCGATCTNNNAACCTGGGCAACAAGAGTGAA

GCACACGTCTGAACTCTTTCCCTACACGACGCTCTTCCGATCTNNNAACCTGGGCAACAAGAGTGAA

AGCACACGTCTGAACTCTTTCCCTACACGACGCTCTTCCGATCTNNNAACCTGGGCAACAAGAGTGAA
GCACACGTCTGAACTCTTTCCCTACACGACGCTCTTCCGATCTNNNAAGGTCTGTGTTCTCTGTGAGA
GCACACGTCTGAACTCTTTCCCTACACGACGCTCTTCCGATCTNNNAAGGTCTGTGTTCTCTGTGAGA
GCACACGTCTGAACTCTTTCCCTACACGACGCTCTTCCGATCTNNNAAGGTCTGTGTTCTCTGTGAGA
GAGCACACGTCTGAACTCTTTCCCTACACGACGCTCTTCCGATCTNNNGCTGATCTGCTCCTTGATGT
GAGCACACGTCTGAACTCTTTCCCTACACGACGCTCTTCCGATCTNNNGCTGATCTGCTCCTTGATGT
GAGCACACGTCTGAACTCTTTCCCTACACGACGCTCTTCCGATCTNNNGCTGATCTGCTCCTTGATGT
AGCACACGTCTGAACTCTTTCCCTACACGACGCTCTTCCGATCTNNNGAGGCTGAGGCAGGAGAAT
AGCACACGTCTGAACTCTTTCCCTACACGACGCTCTTCCGATCTNNNGAGGCTGAGGCAGGAGAAT
AGCACACGTCTGAACTCTTTCCCTACACGACGCTCTTCCGATCTNNNGAGGCTGAGGCAGGAGAAT
AGCACACGTCTGAACTCTTTCCCTACACGACGCTCTTCCGATCTNNNACCTGTAGTCCCAGCTACTCGC
AGCACACGTCTGAACTCTTTCCCTACACGACGCTCTTCCGATCTNNNACCTGTAGTCCCAGCTACTCGC
AGCACACGTCTGAACTCTTTCCCTACACGACGCTCTTCCGATCTNNNACCTGTAGTCCCAGCTACTCGC
GAGCACACGTCTGAACTCTTTCCCTACACGACGCTCTTCCGATCTNNNATCATACCACCAGATCTGTTG
GAGCACACGTCTGAACTCTTTCCCTACACGACGCTCTTCCGATCTNNNATCATACCACCAGATCTGTTG
GAGCACACGTCTGAACTCTTTCCCTACACGACGCTCTTCCGATCTNNNATCATACCACCAGATCTGTTG
AGCACACGTCTGAACTCTTTCCCTACACGACGCTCTTCCGATCTNNNTGATAAACCTCTTCATTGCCCC
AGCACACGTCTGAACTCTTTCCCTACACGACGCTCTTCCGATCTNNNTGATAAACCTCTTCATTGCCCC
AGCACACGTCTGAACTCTTTCCCTACACGACGCTCTTCCGATCTNNNTGATAAACCTCTTCATTGCCCC
AGCACACGTCTGAACTCTTTCCCTACACGACGCTCTTCCGATCTNNNCTCCATCCCACTATGCTCTTC
AGCACACGTCTGAACTCTTTCCCTACACGACGCTCTTCCGATCTNNNCTCCATCCCACTATGCTCTTC
AGCACACGTCTGAACTCTTTCCCTACACGACGCTCTTCCGATCTNNNCTCCATCCCACTATGCTCTTC
AGCACACGTCTGAACTCTTTCCCTACACGACGCTCTTCCGATCTNNNTCTGGGTAGCAGTATGGCAAT
AGCACACGTCTGAACTCTTTCCCTACACGACGCTCTTCCGATCTNNNTCTGGGTAGCAGTATGGCAAT
AGCACACGTCTGAACTCTTTCCCTACACGACGCTCTTCCGATCTNNNTCTGGGTAGCAGTATGGCAAT
AGCACACGTCTGAACTCTTTCCCTACACGACGCTCTTCCGATCTNNNCCGCAACCTGTTTTTTGTTTTG
AGCACACGTCTGAACTCTTTCCCTACACGACGCTCTTCCGATCTNNNCCGCAACCTGTTTTTTGTTTTG
AGCACACGTCTGAACTCTTTCCCTACACGACGCTCTTCCGATCTNNNCCGCAACCTGTTTTTTGTTTTG
GCACACGTCTGAACTCTTTCCCTACACGACGCTCTTCCGATCTNNNCCGTCTGAGAAGTGAGGAGCC
GCACACGTCTGAACTCTTTCCCTACACGACGCTCTTCCGATCTNNNCCGTCTGAGAAGTGAGGAGCC
GCACACGTCTGAACTCTTTCCCTACACGACGCTCTTCCGATCTNNNCCGTCTGAGAAGTGAGGAGCC
CACACGTCTGAACTCTTTCCCTACACGACGCTCTTCCGATCTNNNGGCTGATCTTGACAAACATGCAT
CACACGTCTGAACTCTTTCCCTACACGACGCTCTTCCGATCTNNNGGCTGATCTTGACAAACATGCAT
CACACGTCTGAACTCTTTCCCTACACGACGCTCTTCCGATCTNNNGGCTGATCTTGACAAACATGCAT
AGCACACGTCTGAACTCTTTCCCTACACGACGCTCTTCCGATCTNNNTTCAACCTGCAGTGAGCTATG
AGCACACGTCTGAACTCTTTCCCTACACGACGCTCTTCCGATCTNNNTTCAACCTGCAGTGAGCTATG
AGCACACGTCTGAACTCTTTCCCTACACGACGCTCTTCCGATCTNNNTTCAACCTGCAGTGAGCTATG
GAGCACACGTCTGAACTCTTTCCCTACACGACGCTCTTCCGATCTNNNGTGCCTGTAATCCCAGCTACT
GAGCACACGTCTGAACTCTTTCCCTACACGACGCTCTTCCGATCTNNNGTGCCTGTAATCCCAGCTACT
GAGCACACGTCTGAACTCTTTCCCTACACGACGCTCTTCCGATCTNNNGTGCCTGTAATCCCAGCTACT
CACGTCTGAACTCTTTCCCTACACGACGCTCTTCCGATCTNNNCCTAGATGCCCACAATTTTTGCACAT
CACGTCTGAACTCTTTCCCTACACGACGCTCTTCCGATCTNNNCCTAGATGCCCACAATTTTTGCACAT
CACGTCTGAACTCTTTCCCTACACGACGCTCTTCCGATCTNNNCCTAGATGCCCACAATTTTTGCACAT

GCACACGTCTGAACTCTTTCCCTACACGACGCTCTTCCGATCTNNNGTGGGTGTATGGTGGTATCACA

GCACACGTCTGAACTCTTTCCCTACACGACGCTCTTCCGATCTNNNGTGGGTGTATGGTGGTATCACA

GCACACGTCTGAACTCTTTCCCTACACGACGCTCTTCCGATCTNNNGTGGGTGTATGGTGGTATCACA

GAGCACACGTCTGAACTCTTTCCCTACACGACGCTCTTCCGATCTNNNTGGGGGCTTATTTTGGAGA

GAGCACACGTCTGAACTCTTTCCCTACACGACGCTCTTCCGATCTNNNTGGGGGCTTATTTTGGAGA

GAGCACACGTCTGAACTCTTTCCCTACACGACGCTCTTCCGATCTNNNTGGGGGCTTATTTTGGAGA

AGCACACGTCTGAACTCTTTCCCTACACGACGCTCTTCCGATCTNNNACACGACATACACATAGCACCT

AGCACACGTCTGAACTCTTTCCCTACACGACGCTCTTCCGATCTNNNACACGACATACACATAGCACCT

AGCACACGTCTGAACTCTTTCCCTACACGACGCTCTTCCGATCTNNNACACGACATACACATAGCACCT

GCACACGTCTGAACTCTTTCCCTACACGACGCTCTTCCGATCTNNNATGCCTCAGGGACTTTTCTCTTG

GCACACGTCTGAACTCTTTCCCTACACGACGCTCTTCCGATCTNNNATGCCTCAGGGACTTTTCTCTTG

GCACACGTCTGAACTCTTTCCCTACACGACGCTCTTCCGATCTNNNATGCCTCAGGGACTTTTCTCTTG

AGCACACGTCTGAACTCTTTCCCTACACGACGCTCTTCCGATCTNNNGCCTGGCCAATATTCTGTGTGT

AGCACACGTCTGAACTCTTTCCCTACACGACGCTCTTCCGATCTNNNGCCTGGCCAATATTCTGTGTGT

AGCACACGTCTGAACTCTTTCCCTACACGACGCTCTTCCGATCTNNNGCCTGGCCAATATTCTGTGTGT

GAGCACACGTCTGAACTCTTTCCCTACACGACGCTCTTCCGATCTNNNGCTGAGTATTTCCTGTGGACC

GAGCACACGTCTGAACTCTTTCCCTACACGACGCTCTTCCGATCTNNNGCTGAGTATTTCCTGTGGACC

GAGCACACGTCTGAACTCTTTCCCTACACGACGCTCTTCCGATCTNNNGCTGAGTATTTCCTGTGGACC

GAGCACACGTCTGAACTCTTTCCCTACACGACGCTCTTCCGATCTNNNAAACCCAGGTGAGACTAATT

GAGCACACGTCTGAACTCTTTCCCTACACGACGCTCTTCCGATCTNNNAAACCCAGGTGAGACTAATT

GAGCACACGTCTGAACTCTTTCCCTACACGACGCTCTTCCGATCTNNNAAACCCAGGTGAGACTAATT

AGCACACGTCTGAACTCTTTCCCTACACGACGCTCTTCCGATCTNNNGTTGCAGTGAGCCAAGATCAT

AGCACACGTCTGAACTCTTTCCCTACACGACGCTCTTCCGATCTNNNGTTGCAGTGAGCCAAGATCAT

AGCACACGTCTGAACTCTTTCCCTACACGACGCTCTTCCGATCTNNNGTTGCAGTGAGCCAAGATCAT

AGCACACGTCTGAACTCTTTCCCTACACGACGCTCTTCCGATCTNNNATGCAACCATTTGTCTCTCTCCT

AGCACACGTCTGAACTCTTTCCCTACACGACGCTCTTCCGATCTNNNATGCAACCATTTGTCTCTCTCCT

AGCACACGTCTGAACTCTTTCCCTACACGACGCTCTTCCGATCTNNNATGCAACCATTTGTCTCTCTCCT

GAGCACACGTCTGAACTCTTTCCCTACACGACGCTCTTCCGATCTNNNAAGAAGGAACAGAGGGCTTT

GAGCACACGTCTGAACTCTTTCCCTACACGACGCTCTTCCGATCTNNNAAGAAGGAACAGAGGGCTTT

GAGCACACGTCTGAACTCTTTCCCTACACGACGCTCTTCCGATCTNNNAAGAAGGAACAGAGGGCTTT

AGCACACGTCTGAACTCTTTCCCTACACGACGCTCTTCCGATCTNNNTTTCTCTGCAGGTGCCAAAAAA

AGCACACGTCTGAACTCTTTCCCTACACGACGCTCTTCCGATCTNNNTTTCTCTGCAGGTGCCAAAAAA

AGCACACGTCTGAACTCTTTCCCTACACGACGCTCTTCCGATCTNNNTTTCTCTGCAGGTGCCAAAAAA

GAGCACACGTCTGAACTCTTTCCCTACACGACGCTCTTCCGATCTNNNCTGTGTGCGTTAGTTTTCACC

GAGCACACGTCTGAACTCTTTCCCTACACGACGCTCTTCCGATCTNNNCTGTGTGCGTTAGTTTTCACC

GAGCACACGTCTGAACTCTTTCCCTACACGACGCTCTTCCGATCTNNNCTGTGTGCGTTAGTTTTCACC

AGCACACGTCTGAACTCTTTCCCTACACGACGCTCTTCCGATCTNNNTGTTGAATGAAAGTGGACGTG

AGCACACGTCTGAACTCTTTCCCTACACGACGCTCTTCCGATCTNNNTGTTGAATGAAAGTGGACGTG

AGCACACGTCTGAACTCTTTCCCTACACGACGCTCTTCCGATCTNNNTGTTGAATGAAAGTGGACGTG

CACACGTCTGAACTCTTTCCCTACACGACGCTCTTCCGATCTNNNTCATCCTCTTTTTCGGTTGTTGCCA

CACACGTCTGAACTCTTTCCCTACACGACGCTCTTCCGATCTNNNTCATCCTCTTTTTCGGTTGTTGCCA

CACACGTCTGAACTCTTTCCCTACACGACGCTCTTCCGATCTNNNTCATCCTCTTTTTCGGTTGTTGCCA

AGCACACGTCTGAACTCTTTCCCTACACGACGCTCTTCCGATCTNNNGATGCAAACAGACTTAGCGTC

AGCACACGTCTGAACTCTTTCCCTACACGACGCTCTTCCGATCTNNNGATGCAAACAGACTTAGCGTC
AGCACACGTCTGAACTCTTTCCCTACACGACGCTCTTCCGATCTNNNGATGCAAACAGACTTAGCGTC
AGCACACGTCTGAACTCTTTCCCTACACGACGCTCTTCCGATCTNNNCAAGGTCTTCTGCTGCTACAAT
AGCACACGTCTGAACTCTTTCCCTACACGACGCTCTTCCGATCTNNNCAAGGTCTTCTGCTGCTACAAT
AGCACACGTCTGAACTCTTTCCCTACACGACGCTCTTCCGATCTNNNCAAGGTCTTCTGCTGCTACAAT
AGCACACGTCTGAACTCTTTCCCTACACGACGCTCTTCCGATCTNNNTGTTGCCATCCAGATTCTGACT
AGCACACGTCTGAACTCTTTCCCTACACGACGCTCTTCCGATCTNNNTGTTGCCATCCAGATTCTGACT
AGCACACGTCTGAACTCTTTCCCTACACGACGCTCTTCCGATCTNNNTGTTGCCATCCAGATTCTGACT
CACACGTCTGAACTCTTTCCCTACACGACGCTCTTCCGATCTNNNAACCTGAACAGAATTCTTCACCAT
CACACGTCTGAACTCTTTCCCTACACGACGCTCTTCCGATCTNNNAACCTGAACAGAATTCTTCACCAT
CACACGTCTGAACTCTTTCCCTACACGACGCTCTTCCGATCTNNNAACCTGAACAGAATTCTTCACCAT
AGCACACGTCTGAACTCTTTCCCTACACGACGCTCTTCCGATCTNNNCACACACATACACACACACACAT
AGCACACGTCTGAACTCTTTCCCTACACGACGCTCTTCCGATCTNNNCACACACATACACACACACACAT
AGCACACGTCTGAACTCTTTCCCTACACGACGCTCTTCCGATCTNNNCACACACATACACACACACACAT
GAGCACACGTCTGAACTCTTTCCCTACACGACGCTCTTCCGATCTNNNGAGGTTGCAGTGAGCTGAAA
GAGCACACGTCTGAACTCTTTCCCTACACGACGCTCTTCCGATCTNNNGAGGTTGCAGTGAGCTGAAA
GAGCACACGTCTGAACTCTTTCCCTACACGACGCTCTTCCGATCTNNNGAGGTTGCAGTGAGCTGAAA
AGCACACGTCTGAACTCTTTCCCTACACGACGCTCTTCCGATCTNNNCTTTTGTATGTCTGTTGGCCG
AGCACACGTCTGAACTCTTTCCCTACACGACGCTCTTCCGATCTNNNCTTTTGTATGTCTGTTGGCCG
AGCACACGTCTGAACTCTTTCCCTACACGACGCTCTTCCGATCTNNNCTTTTGTATGTCTGTTGGCCG
AGCACACGTCTGAACTCTTTCCCTACACGACGCTCTTCCGATCTNNNACAATCCAGTTGATCAGAAAGC
AGCACACGTCTGAACTCTTTCCCTACACGACGCTCTTCCGATCTNNNACAATCCAGTTGATCAGAAAGC
AGCACACGTCTGAACTCTTTCCCTACACGACGCTCTTCCGATCTNNNACAATCCAGTTGATCAGAAAGC
AGCACACGTCTGAACTCTTTCCCTACACGACGCTCTTCCGATCTNNNCTCTTGGTACATCAGAGGTGC
AGCACACGTCTGAACTCTTTCCCTACACGACGCTCTTCCGATCTNNNCTCTTGGTACATCAGAGGTGC
AGCACACGTCTGAACTCTTTCCCTACACGACGCTCTTCCGATCTNNNCTCTTGGTACATCAGAGGTGC
AGCACACGTCTGAACTCTTTCCCTACACGACGCTCTTCCGATCTNNNGTGGCCAGTTTCAGCTTGTATT
AGCACACGTCTGAACTCTTTCCCTACACGACGCTCTTCCGATCTNNNGTGGCCAGTTTCAGCTTGTATT
AGCACACGTCTGAACTCTTTCCCTACACGACGCTCTTCCGATCTNNNGTGGCCAGTTTCAGCTTGTATT
GCACACGTCTGAACTCTTTCCCTACACGACGCTCTTCCGATCTNNNGGCGACAAGAGTGAAATTCCAT
GCACACGTCTGAACTCTTTCCCTACACGACGCTCTTCCGATCTNNNGGCGACAAGAGTGAAATTCCAT
GCACACGTCTGAACTCTTTCCCTACACGACGCTCTTCCGATCTNNNGGCGACAAGAGTGAAATTCCAT
AGCACACGTCTGAACTCTTTCCCTACACGACGCTCTTCCGATCTNNNTCAGAGGAAGGTAGGGCAAA
AGCACACGTCTGAACTCTTTCCCTACACGACGCTCTTCCGATCTNNNTCAGAGGAAGGTAGGGCAAA
AGCACACGTCTGAACTCTTTCCCTACACGACGCTCTTCCGATCTNNNTCAGAGGAAGGTAGGGCAAA
AGCACACGTCTGAACTCTTTCCCTACACGACGCTCTTCCGATCTNNNAAGCCTTCTGCCACAAAAATGA
AGCACACGTCTGAACTCTTTCCCTACACGACGCTCTTCCGATCTNNNAAGCCTTCTGCCACAAAAATGA
AGCACACGTCTGAACTCTTTCCCTACACGACGCTCTTCCGATCTNNNAAGCCTTCTGCCACAAAAATGA
GAGCACACGTCTGAACTCTTTCCCTACACGACGCTCTTCCGATCTNNNTTAGCTTTAGTCACTGAGAAA
GAGCACACGTCTGAACTCTTTCCCTACACGACGCTCTTCCGATCTNNNTTAGCTTTAGTCACTGAGAAA
GAGCACACGTCTGAACTCTTTCCCTACACGACGCTCTTCCGATCTNNNTTAGCTTTAGTCACTGAGAAA
GAGCACACGTCTGAACTCTTTCCCTACACGACGCTCTTCCGATCTNNNCTGCTGACATCCTTGACCTTGA
GAGCACACGTCTGAACTCTTTCCCTACACGACGCTCTTCCGATCTNNNCTGCTGACATCCTTGACCTTGA

GAGCACACGTCTGAACTCTTTCCCTACACGACGCTCTTCCGATCTNNNCTGCTGACATCCTTGACCTTGA
GAGCACACGTCTGAACTCTTTCCCTACACGACGCTCTTCCGATCTNNNTGAGGCAGGAGAATGGTATG
GAGCACACGTCTGAACTCTTTCCCTACACGACGCTCTTCCGATCTNNNTGAGGCAGGAGAATGGTATG
GAGCACACGTCTGAACTCTTTCCCTACACGACGCTCTTCCGATCTNNNTGAGGCAGGAGAATGGTATG
GAGCACACGTCTGAACTCTTTCCCTACACGACGCTCTTCCGATCTNNNGAGGTTACAGTGAGCCGAGAT
GAGCACACGTCTGAACTCTTTCCCTACACGACGCTCTTCCGATCTNNNGAGGTTACAGTGAGCCGAGAT
GAGCACACGTCTGAACTCTTTCCCTACACGACGCTCTTCCGATCTNNNGAGGTTACAGTGAGCCGAGAT
CACACGTCTGAACTCTTTCCCTACACGACGCTCTTCCGATCTNNNGAGCGAGACTCTGTCACAAAAAACA
CACACGTCTGAACTCTTTCCCTACACGACGCTCTTCCGATCTNNNGAGCGAGACTCTGTCACAAAAAACA
CACACGTCTGAACTCTTTCCCTACACGACGCTCTTCCGATCTNNNGAGCGAGACTCTGTCACAAAAAACA
AGCACACGTCTGAACTCTTTCCCTACACGACGCTCTTCCGATCTNNNCTGGGTCTGTAACAGAGTGAG
AGCACACGTCTGAACTCTTTCCCTACACGACGCTCTTCCGATCTNNNCTGGGTCTGTAACAGAGTGAG
AGCACACGTCTGAACTCTTTCCCTACACGACGCTCTTCCGATCTNNNCTGGGTCTGTAACAGAGTGAG
GCACACGTCTGAACTCTTTCCCTACACGACGCTCTTCCGATCTNNNGTGATTCTCAGCTTAAGCATCATC
GCACACGTCTGAACTCTTTCCCTACACGACGCTCTTCCGATCTNNNGTGATTCTCAGCTTAAGCATCATC
GCACACGTCTGAACTCTTTCCCTACACGACGCTCTTCCGATCTNNNGTGATTCTCAGCTTAAGCATCATC
AGAGCACACGTCTGAACTCTTTCCCTACACGACGCTCTTCCGATCTNNNAGGGTAAGCTGGAAAAGAA
AGAGCACACGTCTGAACTCTTTCCCTACACGACGCTCTTCCGATCTNNNAGGGTAAGCTGGAAAAGAA
AGAGCACACGTCTGAACTCTTTCCCTACACGACGCTCTTCCGATCTNNNAGGGTAAGCTGGAAAAGAA
GAAGAGCACACGTCTGAACTCTTTCCCTACACGACGCTCTTCCGATCTNNNCCCTCCTGGTGTTCGGAA
GAAGAGCACACGTCTGAACTCTTTCCCTACACGACGCTCTTCCGATCTNNNCCCTCCTGGTGTTCGGAA
GAAGAGCACACGTCTGAACTCTTTCCCTACACGACGCTCTTCCGATCTNNNCCCTCCTGGTGTTCGGAA
AGCACACGTCTGAACTCTTTCCCTACACGACGCTCTTCCGATCTNNNCTGAGGCAGGAGAATCACTTGA
AGCACACGTCTGAACTCTTTCCCTACACGACGCTCTTCCGATCTNNNCTGAGGCAGGAGAATCACTTGA
AGCACACGTCTGAACTCTTTCCCTACACGACGCTCTTCCGATCTNNNCTGAGGCAGGAGAATCACTTGA
GCACACGTCTGAACTCTTTCCCTACACGACGCTCTTCCGATCTNNNTTGTCAGAGGCACTAGTGATGTC
GCACACGTCTGAACTCTTTCCCTACACGACGCTCTTCCGATCTNNNTTGTCAGAGGCACTAGTGATGTC
GCACACGTCTGAACTCTTTCCCTACACGACGCTCTTCCGATCTNNNTTGTCAGAGGCACTAGTGATGTC
GAGCACACGTCTGAACTCTTTCCCTACACGACGCTCTTCCGATCTNNNCTGTAGTCCCAGCTACTTCGGA
GAGCACACGTCTGAACTCTTTCCCTACACGACGCTCTTCCGATCTNNNCTGTAGTCCCAGCTACTTCGGA
GAGCACACGTCTGAACTCTTTCCCTACACGACGCTCTTCCGATCTNNNCTGTAGTCCCAGCTACTTCGGA
AGCACACGTCTGAACTCTTTCCCTACACGACGCTCTTCCGATCTNNNTGTGTGAATCTTGTGATTGTGT
AGCACACGTCTGAACTCTTTCCCTACACGACGCTCTTCCGATCTNNNTGTGTGAATCTTGTGATTGTGT
AGCACACGTCTGAACTCTTTCCCTACACGACGCTCTTCCGATCTNNNTGTGTGAATCTTGTGATTGTGT
AGAGCACACGTCTGAACTCTTTCCCTACACGACGCTCTTCCGATCTNNNTACCCATCTCTGACTTACATG
AGAGCACACGTCTGAACTCTTTCCCTACACGACGCTCTTCCGATCTNNNTACCCATCTCTGACTTACATG
AGAGCACACGTCTGAACTCTTTCCCTACACGACGCTCTTCCGATCTNNNTACCCATCTCTGACTTACATG
AGCACACGTCTGAACTCTTTCCCTACACGACGCTCTTCCGATCTNNNTTGTGCTGTAGAAGGCCTAAGT
AGCACACGTCTGAACTCTTTCCCTACACGACGCTCTTCCGATCTNNNTTGTGCTGTAGAAGGCCTAAGT
AGCACACGTCTGAACTCTTTCCCTACACGACGCTCTTCCGATCTNNNTTGTGCTGTAGAAGGCCTAAGT
AGCACACGTCTGAACTCTTTCCCTACACGACGCTCTTCCGATCTNNNTCAGGAGGTTGAGGTAGGAG
AGCACACGTCTGAACTCTTTCCCTACACGACGCTCTTCCGATCTNNNTCAGGAGGTTGAGGTAGGAG
AGCACACGTCTGAACTCTTTCCCTACACGACGCTCTTCCGATCTNNNTCAGGAGGTTGAGGTAGGAG

GCACACGTCTGAACTCTTTCCCTACACGACGCTCTTCCGATCTNNNCACTGCTGAGTGGGCAATATAG
GCACACGTCTGAACTCTTTCCCTACACGACGCTCTTCCGATCTNNNCACTGCTGAGTGGGCAATATAG
GCACACGTCTGAACTCTTTCCCTACACGACGCTCTTCCGATCTNNNCACTGCTGAGTGGGCAATATAG
GAGCACACGTCTGAACTCTTTCCCTACACGACGCTCTTCCGATCTNNNACCATACCTCCAAGTCATCAA
GAGCACACGTCTGAACTCTTTCCCTACACGACGCTCTTCCGATCTNNNACCATACCTCCAAGTCATCAA
GAGCACACGTCTGAACTCTTTCCCTACACGACGCTCTTCCGATCTNNNACCATACCTCCAAGTCATCAA
CACACGTCTGAACTCTTTCCCTACACGACGCTCTTCCGATCTNNNTTGAAGGTTTGTGGCAAAGTTGT
CACACGTCTGAACTCTTTCCCTACACGACGCTCTTCCGATCTNNNTTGAAGGTTTGTGGCAAAGTTGT
CACACGTCTGAACTCTTTCCCTACACGACGCTCTTCCGATCTNNNTTGAAGGTTTGTGGCAAAGTTGT
GCACACGTCTGAACTCTTTCCCTACACGACGCTCTTCCGATCTNNNAATGGGATCATATATATGCTTG
GCACACGTCTGAACTCTTTCCCTACACGACGCTCTTCCGATCTNNNAATGGGATCATATATATGCTTG
GCACACGTCTGAACTCTTTCCCTACACGACGCTCTTCCGATCTNNNAATGGGATCATATATATGCTTG
AGCACACGTCTGAACTCTTTCCCTACACGACGCTCTTCCGATCTNNNTCCAGGAGCCCTTTCTTTCTTTC
AGCACACGTCTGAACTCTTTCCCTACACGACGCTCTTCCGATCTNNNTCCAGGAGCCCTTTCTTTCTTTC
AGCACACGTCTGAACTCTTTCCCTACACGACGCTCTTCCGATCTNNNTCCAGGAGCCCTTTCTTTCTTTC
GCACACGTCTGAACTCTTTCCCTACACGACGCTCTTCCGATCTNNNCTTATGTACCTCAATATCATCCTC
GCACACGTCTGAACTCTTTCCCTACACGACGCTCTTCCGATCTNNNCTTATGTACCTCAATATCATCCTC
GCACACGTCTGAACTCTTTCCCTACACGACGCTCTTCCGATCTNNNCTTATGTACCTCAATATCATCCTC
AGCACACGTCTGAACTCTTTCCCTACACGACGCTCTTCCGATCTNNNGACATTGGCAAATATCCCTTCG
AGCACACGTCTGAACTCTTTCCCTACACGACGCTCTTCCGATCTNNNGACATTGGCAAATATCCCTTCG
AGCACACGTCTGAACTCTTTCCCTACACGACGCTCTTCCGATCTNNNGACATTGGCAAATATCCCTTCG
AGAGCACACGTCTGAACTCTTTCCCTACACGACGCTCTTCCGATCTNNNGTGTTCCATAGCGACACTTT
AGAGCACACGTCTGAACTCTTTCCCTACACGACGCTCTTCCGATCTNNNGTGTTCCATAGCGACACTTT
AGAGCACACGTCTGAACTCTTTCCCTACACGACGCTCTTCCGATCTNNNGTGTTCCATAGCGACACTTT
AGCACACGTCTGAACTCTTTCCCTACACGACGCTCTTCCGATCTNNNGTCCATCCCAATCTCTACCAACA
AGCACACGTCTGAACTCTTTCCCTACACGACGCTCTTCCGATCTNNNGTCCATCCCAATCTCTACCAACA
AGCACACGTCTGAACTCTTTCCCTACACGACGCTCTTCCGATCTNNNGTCCATCCCAATCTCTACCAACA
AGCACACGTCTGAACTCTTTCCCTACACGACGCTCTTCCGATCTNNNCTCTGACAGTCTGACGGTCTTT
AGCACACGTCTGAACTCTTTCCCTACACGACGCTCTTCCGATCTNNNCTCTGACAGTCTGACGGTCTTT
AGCACACGTCTGAACTCTTTCCCTACACGACGCTCTTCCGATCTNNNCTCTGACAGTCTGACGGTCTTT
AGCACACGTCTGAACTCTTTCCCTACACGACGCTCTTCCGATCTNNNCTGGAGACTTAGGCAGGAGA
AGCACACGTCTGAACTCTTTCCCTACACGACGCTCTTCCGATCTNNNCTGGAGACTTAGGCAGGAGA
AGCACACGTCTGAACTCTTTCCCTACACGACGCTCTTCCGATCTNNNCTGGAGACTTAGGCAGGAGA
AGCACACGTCTGAACTCTTTCCCTACACGACGCTCTTCCGATCTNNNTCCACCAGCAAAAAGATTAGC
AGCACACGTCTGAACTCTTTCCCTACACGACGCTCTTCCGATCTNNNTCCACCAGCAAAAAGATTAGC
AGCACACGTCTGAACTCTTTCCCTACACGACGCTCTTCCGATCTNNNTCCACCAGCAAAAAGATTAGC
GAGCACACGTCTGAACTCTTTCCCTACACGACGCTCTTCCGATCTNNNAGCTCAGCTGGTACAACATCT
GAGCACACGTCTGAACTCTTTCCCTACACGACGCTCTTCCGATCTNNNAGCTCAGCTGGTACAACATCT
GAGCACACGTCTGAACTCTTTCCCTACACGACGCTCTTCCGATCTNNNAGCTCAGCTGGTACAACATCT
GAGCACACGTCTGAACTCTTTCCCTACACGACGCTCTTCCGATCTNNNTGGCTTCCTACACAATGGATG
GAGCACACGTCTGAACTCTTTCCCTACACGACGCTCTTCCGATCTNNNTGGCTTCCTACACAATGGATG
GAGCACACGTCTGAACTCTTTCCCTACACGACGCTCTTCCGATCTNNNTGGCTTCCTACACAATGGATG
GAGCACACGTCTGAACTCTTTCCCTACACGACGCTCTTCCGATCTNNNTTCTGTTGTGTCTGAGTCACC

GAGCACACGTCTGAACTCTTTCCCTACACGACGCTCTTCCGATCTNNNTTCTGTTGTGTCTGAGTCACCA
GAGCACACGTCTGAACTCTTTCCCTACACGACGCTCTTCCGATCTNNNTTCTGTTGTGTCTGAGTCACCA
AGAGCACACGTCTGAACTCTTTCCCTACACGACGCTCTTCCGATCTNNNCCCACTAACTCGTCATCTAGCA
AGAGCACACGTCTGAACTCTTTCCCTACACGACGCTCTTCCGATCTNNNCCCACTAACTCGTCATCTAGCA
AGAGCACACGTCTGAACTCTTTCCCTACACGACGCTCTTCCGATCTNNNCCCACTAACTCGTCATCTAGCA
AGCACACGTCTGAACTCTTTCCCTACACGACGCTCTTCCGATCTNNNGTGGGGAGGACTTTGTCTTACA
AGCACACGTCTGAACTCTTTCCCTACACGACGCTCTTCCGATCTNNNGTGGGGAGGACTTTGTCTTACA
AGCACACGTCTGAACTCTTTCCCTACACGACGCTCTTCCGATCTNNNGTGGGGAGGACTTTGTCTTACA
AGCACACGTCTGAACTCTTTCCCTACACGACGCTCTTCCGATCTNNNCTGAGGCAGGAGAATTGCTTG
AGCACACGTCTGAACTCTTTCCCTACACGACGCTCTTCCGATCTNNNCTGAGGCAGGAGAATTGCTTG
AGCACACGTCTGAACTCTTTCCCTACACGACGCTCTTCCGATCTNNNCTGAGGCAGGAGAATTGCTTG
GCACACGTCTGAACTCTTTCCCTACACGACGCTCTTCCGATCTNNNCTCAGAACGCCCAGTAAGAGTAT
GCACACGTCTGAACTCTTTCCCTACACGACGCTCTTCCGATCTNNNCTCAGAACGCCCAGTAAGAGTAT
GCACACGTCTGAACTCTTTCCCTACACGACGCTCTTCCGATCTNNNCTCAGAACGCCCAGTAAGAGTAT
GAGCACACGTCTGAACTCTTTCCCTACACGACGCTCTTCCGATCTNNNGAGTGCTTTGAGACCCATAAG
GAGCACACGTCTGAACTCTTTCCCTACACGACGCTCTTCCGATCTNNNGAGTGCTTTGAGACCCATAAG
GAGCACACGTCTGAACTCTTTCCCTACACGACGCTCTTCCGATCTNNNGAGTGCTTTGAGACCCATAAG
AGCACACGTCTGAACTCTTTCCCTACACGACGCTCTTCCGATCTNNNTGGAAGGCTGTGCATATGTGT
AGCACACGTCTGAACTCTTTCCCTACACGACGCTCTTCCGATCTNNNTGGAAGGCTGTGCATATGTGT
AGCACACGTCTGAACTCTTTCCCTACACGACGCTCTTCCGATCTNNNTGGAAGGCTGTGCATATGTGT
AGCACACGTCTGAACTCTTTCCCTACACGACGCTCTTCCGATCTNNNTAAACCAGTTTCCATGTGGGTC
AGCACACGTCTGAACTCTTTCCCTACACGACGCTCTTCCGATCTNNNTAAACCAGTTTCCATGTGGGTC
AGCACACGTCTGAACTCTTTCCCTACACGACGCTCTTCCGATCTNNNTAAACCAGTTTCCATGTGGGTC
GAGCACACGTCTGAACTCTTTCCCTACACGACGCTCTTCCGATCTNNNCCAAAGTGCTGGGATTACAAG
GAGCACACGTCTGAACTCTTTCCCTACACGACGCTCTTCCGATCTNNNCCAAAGTGCTGGGATTACAAG
GAGCACACGTCTGAACTCTTTCCCTACACGACGCTCTTCCGATCTNNNCCAAAGTGCTGGGATTACAAG
AGCACACGTCTGAACTCTTTCCCTACACGACGCTCTTCCGATCTNNNTCTTTGGTTTTCAACTGTTGGAC
AGCACACGTCTGAACTCTTTCCCTACACGACGCTCTTCCGATCTNNNTCTTTGGTTTTCAACTGTTGGAC
AGCACACGTCTGAACTCTTTCCCTACACGACGCTCTTCCGATCTNNNTCTTTGGTTTTCAACTGTTGGAC
AGCACACGTCTGAACTCTTTCCCTACACGACGCTCTTCCGATCTNNNGTGGTAGGGAGGGAGTAGGC
AGCACACGTCTGAACTCTTTCCCTACACGACGCTCTTCCGATCTNNNGTGGTAGGGAGGGAGTAGGC
AGCACACGTCTGAACTCTTTCCCTACACGACGCTCTTCCGATCTNNNGTGGTAGGGAGGGAGTAGGC
GCACACGTCTGAACTCTTTCCCTACACGACGCTCTTCCGATCTNNNAGGGGAAACCAATGAAGAGTTG
GCACACGTCTGAACTCTTTCCCTACACGACGCTCTTCCGATCTNNNAGGGGAAACCAATGAAGAGTTG
GCACACGTCTGAACTCTTTCCCTACACGACGCTCTTCCGATCTNNNAGGGGAAACCAATGAAGAGTTG
AGCACACGTCTGAACTCTTTCCCTACACGACGCTCTTCCGATCTNNNTGCACCATTTTACCATCCTACCAC
AGCACACGTCTGAACTCTTTCCCTACACGACGCTCTTCCGATCTNNNTGCACCATTTTACCATCCTACCAC
AGCACACGTCTGAACTCTTTCCCTACACGACGCTCTTCCGATCTNNNTGCACCATTTTACCATCCTACCAC
GAGCACACGTCTGAACTCTTTCCCTACACGACGCTCTTCCGATCTNNNCCAAGGATCCTGCCATTTCAAA
GAGCACACGTCTGAACTCTTTCCCTACACGACGCTCTTCCGATCTNNNCCAAGGATCCTGCCATTTCAAA
GAGCACACGTCTGAACTCTTTCCCTACACGACGCTCTTCCGATCTNNNCCAAGGATCCTGCCATTTCAAA
GAGCACACGTCTGAACTCTTTCCCTACACGACGCTCTTCCGATCTNNNGCTACTTGGGAAATGGAGGT
GAGCACACGTCTGAACTCTTTCCCTACACGACGCTCTTCCGATCTNNNGCTACTTGGGAAATGGAGGT

GAGCACACGTCTGAACTCTTTCCCTACACGACGCTCTTCCGATCTNNNGCTACTTGGGAAATGGAGGT

AGAGCACACGTCTGAACTCTTTCCCTACACGACGCTCTTCCGATCTNNNGTGGAGAATCAACACCCTCT

AGAGCACACGTCTGAACTCTTTCCCTACACGACGCTCTTCCGATCTNNNGTGGAGAATCAACACCCTCT

AGAGCACACGTCTGAACTCTTTCCCTACACGACGCTCTTCCGATCTNNNGTGGAGAATCAACACCCTCT

AGCACACGTCTGAACTCTTTCCCTACACGACGCTCTTCCGATCTNNNCTGTAGTCCCAGCTACTCGAGA

AGCACACGTCTGAACTCTTTCCCTACACGACGCTCTTCCGATCTNNNCTGTAGTCCCAGCTACTCGAGA

AGCACACGTCTGAACTCTTTCCCTACACGACGCTCTTCCGATCTNNNCTGTAGTCCCAGCTACTCGAGA

AGCACACGTCTGAACTCTTTCCCTACACGACGCTCTTCCGATCTNNNCCACGCCTGGCTAATTTTTGTAC

AGCACACGTCTGAACTCTTTCCCTACACGACGCTCTTCCGATCTNNNCCACGCCTGGCTAATTTTTGTAC

AGCACACGTCTGAACTCTTTCCCTACACGACGCTCTTCCGATCTNNNCCACGCCTGGCTAATTTTTGTAC

AGCACACGTCTGAACTCTTTCCCTACACGACGCTCTTCCGATCTNNNAAGGAAGGAAAGAAGGGAAG

AGCACACGTCTGAACTCTTTCCCTACACGACGCTCTTCCGATCTNNNAAGGAAGGAAAGAAGGGAAG

AGCACACGTCTGAACTCTTTCCCTACACGACGCTCTTCCGATCTNNNAAGGAAGGAAAGAAGGGAAG

ACACGTCTGAACTCTTTCCCTACACGACGCTCTTCCGATCTNNNAGTAATTAGAACGGAAGGGAACAT

ACACGTCTGAACTCTTTCCCTACACGACGCTCTTCCGATCTNNNAGTAATTAGAACGGAAGGGAACAT

ACACGTCTGAACTCTTTCCCTACACGACGCTCTTCCGATCTNNNAGTAATTAGAACGGAAGGGAACAT

GAGCACACGTCTGAACTCTTTCCCTACACGACGCTCTTCCGATCTNNNCTCACCAGGAAAGTATGTGG

GAGCACACGTCTGAACTCTTTCCCTACACGACGCTCTTCCGATCTNNNCTCACCAGGAAAGTATGTGG

GAGCACACGTCTGAACTCTTTCCCTACACGACGCTCTTCCGATCTNNNCTCACCAGGAAAGTATGTGG

GCACACGTCTGAACTCTTTCCCTACACGACGCTCTTCCGATCTNNNGGAGGCAGTTTTTCAGGATTTCC

GCACACGTCTGAACTCTTTCCCTACACGACGCTCTTCCGATCTNNNGGAGGCAGTTTTTCAGGATTTCC

GCACACGTCTGAACTCTTTCCCTACACGACGCTCTTCCGATCTNNNGGAGGCAGTTTTTCAGGATTTCC

GCACACGTCTGAACTCTTTCCCTACACGACGCTCTTCCGATCTNNNGCCCATCCCAAAATAAAGACCA

GCACACGTCTGAACTCTTTCCCTACACGACGCTCTTCCGATCTNNNGCCCATCCCAAAATAAAGACCA

GCACACGTCTGAACTCTTTCCCTACACGACGCTCTTCCGATCTNNNGCCCATCCCAAAATAAAGACCA

AGCACACGTCTGAACTCTTTCCCTACACGACGCTCTTCCGATCTNNNCTGTAGTCCCAGCTACTCAGGAG

AGCACACGTCTGAACTCTTTCCCTACACGACGCTCTTCCGATCTNNNCTGTAGTCCCAGCTACTCAGGAG

AGCACACGTCTGAACTCTTTCCCTACACGACGCTCTTCCGATCTNNNCTGTAGTCCCAGCTACTCAGGAG

AGAGCACACGTCTGAACTCTTTCCCTACACGACGCTCTTCCGATCTNNNTACCATGCGAGTAAGTGCCC

AGAGCACACGTCTGAACTCTTTCCCTACACGACGCTCTTCCGATCTNNNTACCATGCGAGTAAGTGCCC

AGAGCACACGTCTGAACTCTTTCCCTACACGACGCTCTTCCGATCTNNNTACCATGCGAGTAAGTGCCC

GAGCACACGTCTGAACTCTTTCCCTACACGACGCTCTTCCGATCTNNNCTGAGGCAGGAGAATAGCTTC

GAGCACACGTCTGAACTCTTTCCCTACACGACGCTCTTCCGATCTNNNCTGAGGCAGGAGAATAGCTTC

GAGCACACGTCTGAACTCTTTCCCTACACGACGCTCTTCCGATCTNNNCTGAGGCAGGAGAATAGCTTC

GAGCACACGTCTGAACTCTTTCCCTACACGACGCTCTTCCGATCTNNNCTGTAATCCCAGCTACTCAGGA

GAGCACACGTCTGAACTCTTTCCCTACACGACGCTCTTCCGATCTNNNCTGTAATCCCAGCTACTCAGGA

GAGCACACGTCTGAACTCTTTCCCTACACGACGCTCTTCCGATCTNNNCTGTAATCCCAGCTACTCAGGA

GAGCACACGTCTGAACTCTTTCCCTACACGACGCTCTTCCGATCTNNNGAACTCCCTGGATGTCTGATC

GAGCACACGTCTGAACTCTTTCCCTACACGACGCTCTTCCGATCTNNNGAACTCCCTGGATGTCTGATC

GAGCACACGTCTGAACTCTTTCCCTACACGACGCTCTTCCGATCTNNNGAACTCCCTGGATGTCTGATC

AGAGCACACGTCTGAACTCTTTCCCTACACGACGCTCTTCCGATCTNNNGGAGTTGGGTGATGCATTC

AGAGCACACGTCTGAACTCTTTCCCTACACGACGCTCTTCCGATCTNNNGGAGTTGGGTGATGCATTC

AGAGCACACGTCTGAACTCTTTCCCTACACGACGCTCTTCCGATCTNNNGGAGTTGGGTGATGCATTC

AGCACACGTCTGAACTCTTTCCCTACACGACGCTCTTCCGATCTNNNGCTGAGGCAGGAGAGAGAATA
AGCACACGTCTGAACTCTTTCCCTACACGACGCTCTTCCGATCTNNNGCTGAGGCAGGAGAGAGAATA
AGCACACGTCTGAACTCTTTCCCTACACGACGCTCTTCCGATCTNNNGCTGAGGCAGGAGAGAGAATA
AGAGCACACGTCTGAACTCTTTCCCTACACGACGCTCTTCCGATCTNNNGGAGCTGTTTCTACTTCCTAG
AGAGCACACGTCTGAACTCTTTCCCTACACGACGCTCTTCCGATCTNNNGGAGCTGTTTCTACTTCCTAG
AGAGCACACGTCTGAACTCTTTCCCTACACGACGCTCTTCCGATCTNNNGGAGCTGTTTCTACTTCCTAG
AGCACACGTCTGAACTCTTTCCCTACACGACGCTCTTCCGATCTNNNGGCAAAAGAGTGAGACTGCATC
AGCACACGTCTGAACTCTTTCCCTACACGACGCTCTTCCGATCTNNNGGCAAAAGAGTGAGACTGCATC
AGCACACGTCTGAACTCTTTCCCTACACGACGCTCTTCCGATCTNNNGGCAAAAGAGTGAGACTGCATC
AGCACACGTCTGAACTCTTTCCCTACACGACGCTCTTCCGATCTNNNGGCTTCTTACATTCTAGCCTGG
AGCACACGTCTGAACTCTTTCCCTACACGACGCTCTTCCGATCTNNNGGCTTCTTACATTCTAGCCTGG
AGCACACGTCTGAACTCTTTCCCTACACGACGCTCTTCCGATCTNNNGGCTTCTTACATTCTAGCCTGG
GAGCACACGTCTGAACTCTTTCCCTACACGACGCTCTTCCGATCTNNNGATAACTTGAGCCCAGGAGTT
GAGCACACGTCTGAACTCTTTCCCTACACGACGCTCTTCCGATCTNNNGATAACTTGAGCCCAGGAGTT
GAGCACACGTCTGAACTCTTTCCCTACACGACGCTCTTCCGATCTNNNGATAACTTGAGCCCAGGAGTT
GAGCACACGTCTGAACTCTTTCCCTACACGACGCTCTTCCGATCTNNNGACTGAGGCTGAGGTGTAATA
GAGCACACGTCTGAACTCTTTCCCTACACGACGCTCTTCCGATCTNNNGACTGAGGCTGAGGTGTAATA
GAGCACACGTCTGAACTCTTTCCCTACACGACGCTCTTCCGATCTNNNGACTGAGGCTGAGGTGTAATA
GAGCACACGTCTGAACTCTTTCCCTACACGACGCTCTTCCGATCTNNNTAGTAGTTAATGGTATGGATG
GAGCACACGTCTGAACTCTTTCCCTACACGACGCTCTTCCGATCTNNNTAGTAGTTAATGGTATGGATG
GAGCACACGTCTGAACTCTTTCCCTACACGACGCTCTTCCGATCTNNNTAGTAGTTAATGGTATGGATG
AGAGCACACGTCTGAACTCTTTCCCTACACGACGCTCTTCCGATCTNNNCCACGGAATTGCCTTTGAAAC
AGAGCACACGTCTGAACTCTTTCCCTACACGACGCTCTTCCGATCTNNNCCACGGAATTGCCTTTGAAAC
AGAGCACACGTCTGAACTCTTTCCCTACACGACGCTCTTCCGATCTNNNCCACGGAATTGCCTTTGAAAC
AGCACACGTCTGAACTCTTTCCCTACACGACGCTCTTCCGATCTNNNTTAGGATGGGAGGATGGTGGA
AGCACACGTCTGAACTCTTTCCCTACACGACGCTCTTCCGATCTNNNTTAGGATGGGAGGATGGTGGA
AGCACACGTCTGAACTCTTTCCCTACACGACGCTCTTCCGATCTNNNTTAGGATGGGAGGATGGTGGA
GAGCACACGTCTGAACTCTTTCCCTACACGACGCTCTTCCGATCTNNNATGCTGCTAATCCAAGGAGTG
GAGCACACGTCTGAACTCTTTCCCTACACGACGCTCTTCCGATCTNNNATGCTGCTAATCCAAGGAGTG
GAGCACACGTCTGAACTCTTTCCCTACACGACGCTCTTCCGATCTNNNATGCTGCTAATCCAAGGAGTG
GAGCACACGTCTGAACTCTTTCCCTACACGACGCTCTTCCGATCTNNNTGGGTGACAGAACAAGACTC
GAGCACACGTCTGAACTCTTTCCCTACACGACGCTCTTCCGATCTNNNTGGGTGACAGAACAAGACTC
GAGCACACGTCTGAACTCTTTCCCTACACGACGCTCTTCCGATCTNNNTGGGTGACAGAACAAGACTC
GAGCACACGTCTGAACTCTTTCCCTACACGACGCTCTTCCGATCTNNNTTGACAGAGTCAAGCACTTTCC
GAGCACACGTCTGAACTCTTTCCCTACACGACGCTCTTCCGATCTNNNTTGACAGAGTCAAGCACTTTCC
GAGCACACGTCTGAACTCTTTCCCTACACGACGCTCTTCCGATCTNNNTTGACAGAGTCAAGCACTTTCC
GCACACGTCTGAACTCTTTCCCTACACGACGCTCTTCCGATCTNNNGGAGAGCTAGTGTGATCTTCTGG
GCACACGTCTGAACTCTTTCCCTACACGACGCTCTTCCGATCTNNNGGAGAGCTAGTGTGATCTTCTGG
GCACACGTCTGAACTCTTTCCCTACACGACGCTCTTCCGATCTNNNGGAGAGCTAGTGTGATCTTCTGG
AAGAGCACACGTCTGAACTCTTTCCCTACACGACGCTCTTCCGATCTNNNACATTGTACTTTCAGGTGA
AAGAGCACACGTCTGAACTCTTTCCCTACACGACGCTCTTCCGATCTNNNACATTGTACTTTCAGGTGA
AAGAGCACACGTCTGAACTCTTTCCCTACACGACGCTCTTCCGATCTNNNACATTGTACTTTCAGGTGA
AGCACACGTCTGAACTCTTTCCCTACACGACGCTCTTCCGATCTNNNCAGACTAGTCTTCTTGCCACAGT

AGCACACGTCTGAACTCTTTCCCTACACGACGCTCTTCCGATCTNNNCAGACTAGTCTTCTTGCCACAGT
AGCACACGTCTGAACTCTTTCCCTACACGACGCTCTTCCGATCTNNNCAGACTAGTCTTCTTGCCACAGT
GAGCACACGTCTGAACTCTTTCCCTACACGACGCTCTTCCGATCTNNNCCACGCCCAGCTAATTTTTGT
GAGCACACGTCTGAACTCTTTCCCTACACGACGCTCTTCCGATCTNNNCCACGCCCAGCTAATTTTTGT
GAGCACACGTCTGAACTCTTTCCCTACACGACGCTCTTCCGATCTNNNCCACGCCCAGCTAATTTTTGT
AGAGCACACGTCTGAACTCTTTCCCTACACGACGCTCTTCCGATCTNNNCTGGCTGCAGGAGTCTGGT
AGAGCACACGTCTGAACTCTTTCCCTACACGACGCTCTTCCGATCTNNNCTGGCTGCAGGAGTCTGGT
AGAGCACACGTCTGAACTCTTTCCCTACACGACGCTCTTCCGATCTNNNCTGGCTGCAGGAGTCTGGT
AGCACACGTCTGAACTCTTTCCCTACACGACGCTCTTCCGATCTNNNTCCTACTCTCCCTCACAATCCTC
AGCACACGTCTGAACTCTTTCCCTACACGACGCTCTTCCGATCTNNNTCCTACTCTCCCTCACAATCCTC
AGCACACGTCTGAACTCTTTCCCTACACGACGCTCTTCCGATCTNNNTCCTACTCTCCCTCACAATCCTC
AGAGCACACGTCTGAACTCTTTCCCTACACGACGCTCTTCCGATCTNNNCCGTACAATCTATTCTGTGAA
AGAGCACACGTCTGAACTCTTTCCCTACACGACGCTCTTCCGATCTNNNCCGTACAATCTATTCTGTGAA
AGAGCACACGTCTGAACTCTTTCCCTACACGACGCTCTTCCGATCTNNNCCGTACAATCTATTCTGTGAA
GAGCACACGTCTGAACTCTTTCCCTACACGACGCTCTTCCGATCTNNNGAGCTTACAGTGACCCGAGAT
GAGCACACGTCTGAACTCTTTCCCTACACGACGCTCTTCCGATCTNNNGAGCTTACAGTGACCCGAGAT
GAGCACACGTCTGAACTCTTTCCCTACACGACGCTCTTCCGATCTNNNGAGCTTACAGTGACCCGAGAT
AGCACACGTCTGAACTCTTTCCCTACACGACGCTCTTCCGATCTNNNTCTTGTTTTTGCTCCTACTTGCCCA
AGCACACGTCTGAACTCTTTCCCTACACGACGCTCTTCCGATCTNNNTCTTGTTTTTGCTCCTACTTGCCCA
AGCACACGTCTGAACTCTTTCCCTACACGACGCTCTTCCGATCTNNNTCTTGTTTTTGCTCCTACTTGCCCA
GAGCACACGTCTGAACTCTTTCCCTACACGACGCTCTTCCGATCTNNNGGATCCCCAAACAAATGTGTCT
GAGCACACGTCTGAACTCTTTCCCTACACGACGCTCTTCCGATCTNNNGGATCCCCAAACAAATGTGTCT
GAGCACACGTCTGAACTCTTTCCCTACACGACGCTCTTCCGATCTNNNGGATCCCCAAACAAATGTGTCT
AGCACACGTCTGAACTCTTTCCCTACACGACGCTCTTCCGATCTNNNGGCAGGCACACTTGATGTAACC
AGCACACGTCTGAACTCTTTCCCTACACGACGCTCTTCCGATCTNNNGGCAGGCACACTTGATGTAACC
AGCACACGTCTGAACTCTTTCCCTACACGACGCTCTTCCGATCTNNNGGCAGGCACACTTGATGTAACC
AGCACACGTCTGAACTCTTTCCCTACACGACGCTCTTCCGATCTNNNGTCACCTCACTCAGAATAGCCAT
AGCACACGTCTGAACTCTTTCCCTACACGACGCTCTTCCGATCTNNNGTCACCTCACTCAGAATAGCCAT
AGCACACGTCTGAACTCTTTCCCTACACGACGCTCTTCCGATCTNNNGTCACCTCACTCAGAATAGCCAT
AGCACACGTCTGAACTCTTTCCCTACACGACGCTCTTCCGATCTNNNCTGTAGTCCCAGCTACTCAGGA
AGCACACGTCTGAACTCTTTCCCTACACGACGCTCTTCCGATCTNNNCTGTAGTCCCAGCTACTCAGGA
AGCACACGTCTGAACTCTTTCCCTACACGACGCTCTTCCGATCTNNNCTGTAGTCCCAGCTACTCAGGA
AGCACACGTCTGAACTCTTTCCCTACACGACGCTCTTCCGATCTNNNTACTAGACAGTCCTCAAACATA
AGCACACGTCTGAACTCTTTCCCTACACGACGCTCTTCCGATCTNNNTACTAGACAGTCCTCAAACATA
AGCACACGTCTGAACTCTTTCCCTACACGACGCTCTTCCGATCTNNNTACTAGACAGTCCTCAAACATA
GAGCACACGTCTGAACTCTTTCCCTACACGACGCTCTTCCGATCTNNNGAGAATCACCTGAACCTGAGA
GAGCACACGTCTGAACTCTTTCCCTACACGACGCTCTTCCGATCTNNNGAGAATCACCTGAACCTGAGA
GAGCACACGTCTGAACTCTTTCCCTACACGACGCTCTTCCGATCTNNNGAGAATCACCTGAACCTGAGA
AGCACACGTCTGAACTCTTTCCCTACACGACGCTCTTCCGATCTNNNAACAAAGAAAGAAGTGCTCCA
AGCACACGTCTGAACTCTTTCCCTACACGACGCTCTTCCGATCTNNNAACAAAGAAAGAAGTGCTCCA
AGCACACGTCTGAACTCTTTCCCTACACGACGCTCTTCCGATCTNNNAACAAAGAAAGAAGTGCTCCA
AGCACACGTCTGAACTCTTTCCCTACACGACGCTCTTCCGATCTNNNGATTACAGGCACAAACCATTG
AGCACACGTCTGAACTCTTTCCCTACACGACGCTCTTCCGATCTNNNGATTACAGGCACAAACCATTG

AGCACACGTCTGAACTCTTTCCCTACACGACGCTCTTCCGATCTNNNGATTACAGGCACAAACCATTG
AGCACACGTCTGAACTCTTTCCCTACACGACGCTCTTCCGATCTNNNGCTGTTCTCGTGATAGTGAGT
AGCACACGTCTGAACTCTTTCCCTACACGACGCTCTTCCGATCTNNNGCTGTTCTCGTGATAGTGAGT
AGCACACGTCTGAACTCTTTCCCTACACGACGCTCTTCCGATCTNNNGCTGTTCTCGTGATAGTGAGT
AGCACACGTCTGAACTCTTTCCCTACACGACGCTCTTCCGATCTNNNCTCCTGACCTCAAGTTATCCAC
AGCACACGTCTGAACTCTTTCCCTACACGACGCTCTTCCGATCTNNNCTCCTGACCTCAAGTTATCCAC
AGCACACGTCTGAACTCTTTCCCTACACGACGCTCTTCCGATCTNNNCTCCTGACCTCAAGTTATCCAC
GCACACGTCTGAACTCTTTCCCTACACGACGCTCTTCCGATCTNNNACCATCAAACCCAACCCTAATCA
GCACACGTCTGAACTCTTTCCCTACACGACGCTCTTCCGATCTNNNACCATCAAACCCAACCCTAATCA
GCACACGTCTGAACTCTTTCCCTACACGACGCTCTTCCGATCTNNNACCATCAAACCCAACCCTAATCA
AGCACACGTCTGAACTCTTTCCCTACACGACGCTCTTCCGATCTNNNTACCAACCAACTGAATTCTGGG
AGCACACGTCTGAACTCTTTCCCTACACGACGCTCTTCCGATCTNNNTACCAACCAACTGAATTCTGGG
AGCACACGTCTGAACTCTTTCCCTACACGACGCTCTTCCGATCTNNNTACCAACCAACTGAATTCTGGG
AGCACACGTCTGAACTCTTTCCCTACACGACGCTCTTCCGATCTNNNATCACCTTCCCTATGACACATC
AGCACACGTCTGAACTCTTTCCCTACACGACGCTCTTCCGATCTNNNATCACCTTCCCTATGACACATC
AGCACACGTCTGAACTCTTTCCCTACACGACGCTCTTCCGATCTNNNATCACCTTCCCTATGACACATC
AGCACACGTCTGAACTCTTTCCCTACACGACGCTCTTCCGATCTNNNTTGGTCTCAAGCTACATGGTT
AGCACACGTCTGAACTCTTTCCCTACACGACGCTCTTCCGATCTNNNTTGGTCTCAAGCTACATGGTT
AGCACACGTCTGAACTCTTTCCCTACACGACGCTCTTCCGATCTNNNTTGGTCTCAAGCTACATGGTT
GCACACGTCTGAACTCTTTCCCTACACGACGCTCTTCCGATCTNNNCTTCGATGACTACCAGGGCAGC
GCACACGTCTGAACTCTTTCCCTACACGACGCTCTTCCGATCTNNNCTTCGATGACTACCAGGGCAGC
GCACACGTCTGAACTCTTTCCCTACACGACGCTCTTCCGATCTNNNCTTCGATGACTACCAGGGCAGC
AGCACACGTCTGAACTCTTTCCCTACACGACGCTCTTCCGATCTNNNACAAGACCAGATGAACTCACA
AGCACACGTCTGAACTCTTTCCCTACACGACGCTCTTCCGATCTNNNACAAGACCAGATGAACTCACA
AGCACACGTCTGAACTCTTTCCCTACACGACGCTCTTCCGATCTNNNACAAGACCAGATGAACTCACA
GCACACGTCTGAACTCTTTCCCTACACGACGCTCTTCCGATCTNNNACCAGCTCTTCTTTGTATGTGGT
GCACACGTCTGAACTCTTTCCCTACACGACGCTCTTCCGATCTNNNACCAGCTCTTCTTTGTATGTGGT
GCACACGTCTGAACTCTTTCCCTACACGACGCTCTTCCGATCTNNNACCAGCTCTTCTTTGTATGTGGT
GCACACGTCTGAACTCTTTCCCTACACGACGCTCTTCCGATCTNNNTGCCTCTTGACATTTGTGTTTGT
GCACACGTCTGAACTCTTTCCCTACACGACGCTCTTCCGATCTNNNTGCCTCTTGACATTTGTGTTTGT
GCACACGTCTGAACTCTTTCCCTACACGACGCTCTTCCGATCTNNNTGCCTCTTGACATTTGTGTTTGT
AGCACACGTCTGAACTCTTTCCCTACACGACGCTCTTCCGATCTNNNCCTGAGTAACTGGAACTACAG
AGCACACGTCTGAACTCTTTCCCTACACGACGCTCTTCCGATCTNNNCCTGAGTAACTGGAACTACAG
AGCACACGTCTGAACTCTTTCCCTACACGACGCTCTTCCGATCTNNNCCTGAGTAACTGGAACTACAG
ACACGTCTGAACTCTTTCCCTACACGACGCTCTTCCGATCTNNNCACATCTGACCCAAATGCCATTTCAT
ACACGTCTGAACTCTTTCCCTACACGACGCTCTTCCGATCTNNNCACATCTGACCCAAATGCCATTTCAT
ACACGTCTGAACTCTTTCCCTACACGACGCTCTTCCGATCTNNNCACATCTGACCCAAATGCCATTTCAT
AGCACACGTCTGAACTCTTTCCCTACACGACGCTCTTCCGATCTNNNCATGTGTTCATGTGTGGGATG
AGCACACGTCTGAACTCTTTCCCTACACGACGCTCTTCCGATCTNNNCATGTGTTCATGTGTGGGATG
AGCACACGTCTGAACTCTTTCCCTACACGACGCTCTTCCGATCTNNNCATGTGTTCATGTGTGGGATG
AGCACACGTCTGAACTCTTTCCCTACACGACGCTCTTCCGATCTNNNCTGAGGCAGAAGAAGCACTT
AGCACACGTCTGAACTCTTTCCCTACACGACGCTCTTCCGATCTNNNCTGAGGCAGAAGAAGCACTT
AGCACACGTCTGAACTCTTTCCCTACACGACGCTCTTCCGATCTNNNCTGAGGCAGAAGAAGCACTT

GCACACGTCTGAACTCTTTCCCTACACGACGCTCTTCCGATCTNNNATTTCTGACAACCAAAGGGAAG
GCACACGTCTGAACTCTTTCCCTACACGACGCTCTTCCGATCTNNNATTTCTGACAACCAAAGGGAAG
GCACACGTCTGAACTCTTTCCCTACACGACGCTCTTCCGATCTNNNATTTCTGACAACCAAAGGGAAG
CACACGTCTGAACTCTTTCCCTACACGACGCTCTTCCGATCTNNNGGCTGAAATGCGTCTGTAAACAC
CACACGTCTGAACTCTTTCCCTACACGACGCTCTTCCGATCTNNNGGCTGAAATGCGTCTGTAAACAC
CACACGTCTGAACTCTTTCCCTACACGACGCTCTTCCGATCTNNNGGCTGAAATGCGTCTGTAAACAC
GCACACGTCTGAACTCTTTCCCTACACGACGCTCTTCCGATCTNNNACCAATGACATTTTCCACAGAAC
GCACACGTCTGAACTCTTTCCCTACACGACGCTCTTCCGATCTNNNACCAATGACATTTTCCACAGAAC
GCACACGTCTGAACTCTTTCCCTACACGACGCTCTTCCGATCTNNNACCAATGACATTTTCCACAGAAC
AGCACACGTCTGAACTCTTTCCCTACACGACGCTCTTCCGATCTNNNGGAGCCATAAAGCTACACCTC
AGCACACGTCTGAACTCTTTCCCTACACGACGCTCTTCCGATCTNNNGGAGCCATAAAGCTACACCTC
AGCACACGTCTGAACTCTTTCCCTACACGACGCTCTTCCGATCTNNNGGAGCCATAAAGCTACACCTC
AGCACACGTCTGAACTCTTTCCCTACACGACGCTCTTCCGATCTNNNTTAAATATGAGATTTACCAAG
AGCACACGTCTGAACTCTTTCCCTACACGACGCTCTTCCGATCTNNNTTAAATATGAGATTTACCAAG
AGCACACGTCTGAACTCTTTCCCTACACGACGCTCTTCCGATCTNNNTTAAATATGAGATTTACCAAG
AGCACACGTCTGAACTCTTTCCCTACACGACGCTCTTCCGATCTNNNTTTGCCTCGTTCTCTCTCCTTAC
AGCACACGTCTGAACTCTTTCCCTACACGACGCTCTTCCGATCTNNNTTTGCCTCGTTCTCTCTCCTTAC
AGCACACGTCTGAACTCTTTCCCTACACGACGCTCTTCCGATCTNNNTTTGCCTCGTTCTCTCTCCTTAC
GCACACGTCTGAACTCTTTCCCTACACGACGCTCTTCCGATCTNNNCACAGACCTCAGAGGAATCACT
GCACACGTCTGAACTCTTTCCCTACACGACGCTCTTCCGATCTNNNCACAGACCTCAGAGGAATCACT
GCACACGTCTGAACTCTTTCCCTACACGACGCTCTTCCGATCTNNNCACAGACCTCAGAGGAATCACT
GAAGAGCACACGTCTGAACTCTTTCCCTACACGACGCTCTTCCGATCTNNNATTTTTACCCACCTCGCA
GAAGAGCACACGTCTGAACTCTTTCCCTACACGACGCTCTTCCGATCTNNNATTTTTACCCACCTCGCA
GAAGAGCACACGTCTGAACTCTTTCCCTACACGACGCTCTTCCGATCTNNNATTTTTACCCACCTCGCA
GAGCACACGTCTGAACTCTTTCCCTACACGACGCTCTTCCGATCTNNNCCTGGGCAACAAGAAAGAAA
GAGCACACGTCTGAACTCTTTCCCTACACGACGCTCTTCCGATCTNNNCCTGGGCAACAAGAAAGAAA
GAGCACACGTCTGAACTCTTTCCCTACACGACGCTCTTCCGATCTNNNCCTGGGCAACAAGAAAGAAA
GAGCACACGTCTGAACTCTTTCCCTACACGACGCTCTTCCGATCTNNNTTTACATATCATCTCGTCCAAT
GAGCACACGTCTGAACTCTTTCCCTACACGACGCTCTTCCGATCTNNNTTTACATATCATCTCGTCCAAT
GAGCACACGTCTGAACTCTTTCCCTACACGACGCTCTTCCGATCTNNNTTTACATATCATCTCGTCCAAT
GAGCACACGTCTGAACTCTTTCCCTACACGACGCTCTTCCGATCTNNNTTACTCATGACGTCGAAGGC
GAGCACACGTCTGAACTCTTTCCCTACACGACGCTCTTCCGATCTNNNTTACTCATGACGTCGAAGGC
GAGCACACGTCTGAACTCTTTCCCTACACGACGCTCTTCCGATCTNNNTTACTCATGACGTCGAAGGC
AGCACACGTCTGAACTCTTTCCCTACACGACGCTCTTCCGATCTNNNAATTGGTAAAGGGTAGGAGCT
AGCACACGTCTGAACTCTTTCCCTACACGACGCTCTTCCGATCTNNNAATTGGTAAAGGGTAGGAGCT
AGCACACGTCTGAACTCTTTCCCTACACGACGCTCTTCCGATCTNNNAATTGGTAAAGGGTAGGAGCT
GCACACGTCTGAACTCTTTCCCTACACGACGCTCTTCCGATCTNNNATGATTACACCATCACAAGGCCT
GCACACGTCTGAACTCTTTCCCTACACGACGCTCTTCCGATCTNNNATGATTACACCATCACAAGGCCT
GCACACGTCTGAACTCTTTCCCTACACGACGCTCTTCCGATCTNNNATGATTACACCATCACAAGGCCT
GAGCACACGTCTGAACTCTTTCCCTACACGACGCTCTTCCGATCTNNNGTAATCCCAGCTACTCAAGA
GAGCACACGTCTGAACTCTTTCCCTACACGACGCTCTTCCGATCTNNNGTAATCCCAGCTACTCAAGA
GAGCACACGTCTGAACTCTTTCCCTACACGACGCTCTTCCGATCTNNNGTAATCCCAGCTACTCAAGA
GAGCACACGTCTGAACTCTTTCCCTACACGACGCTCTTCCGATCTNNNGCTGACCCTTCTTCCATTTGT

GAGCACACGTCTGAACTCTTTCCCTACACGACGCTCTTCCGATCTNNNGCTGACCCTTCTTCCATTTGTT

GAGCACACGTCTGAACTCTTTCCCTACACGACGCTCTTCCGATCTNNNGCTGACCCTTCTTCCATTTGTT

GAGCACACGTCTGAACTCTTTCCCTACACGACGCTCTTCCGATCTNNNACTTCCTTCTGTGTGCCATGT

GAGCACACGTCTGAACTCTTTCCCTACACGACGCTCTTCCGATCTNNNACTTCCTTCTGTGTGCCATGT

GAGCACACGTCTGAACTCTTTCCCTACACGACGCTCTTCCGATCTNNNACTTCCTTCTGTGTGCCATGT

GAGCACACGTCTGAACTCTTTCCCTACACGACGCTCTTCCGATCTNNNGAGGCAGGAGAATGGTGTG

GAGCACACGTCTGAACTCTTTCCCTACACGACGCTCTTCCGATCTNNNGAGGCAGGAGAATGGTGTG

GAGCACACGTCTGAACTCTTTCCCTACACGACGCTCTTCCGATCTNNNGAGGCAGGAGAATGGTGTG

AGCACACGTCTGAACTCTTTCCCTACACGACGCTCTTCCGATCTNNNGGATGTCTTTCATTTCTTCAG

AGCACACGTCTGAACTCTTTCCCTACACGACGCTCTTCCGATCTNNNGGATGTCTTTCATTTCTTCAG

AGCACACGTCTGAACTCTTTCCCTACACGACGCTCTTCCGATCTNNNGGATGTCTTTCATTTCTTCAG

GAGCACACGTCTGAACTCTTTCCCTACACGACGCTCTTCCGATCTNNNTTGAGGCAGGAGAATTGCTT

GAGCACACGTCTGAACTCTTTCCCTACACGACGCTCTTCCGATCTNNNTTGAGGCAGGAGAATTGCTT

GAGCACACGTCTGAACTCTTTCCCTACACGACGCTCTTCCGATCTNNNTTGAGGCAGGAGAATTGCTT

GCACACGTCTGAACTCTTTCCCTACACGACGCTCTTCCGATCTNNNGCAAAACCCTGTCTCTGCAAATA

GCACACGTCTGAACTCTTTCCCTACACGACGCTCTTCCGATCTNNNGCAAAACCCTGTCTCTGCAAATA

GCACACGTCTGAACTCTTTCCCTACACGACGCTCTTCCGATCTNNNGCAAAACCCTGTCTCTGCAAATA

AGAGCACACGTCTGAACTCTTTCCCTACACGACGCTCTTCCGATCTNNNAAGATCTGATGGAGGTGG

AGAGCACACGTCTGAACTCTTTCCCTACACGACGCTCTTCCGATCTNNNAAGATCTGATGGAGGTGG

AGAGCACACGTCTGAACTCTTTCCCTACACGACGCTCTTCCGATCTNNNAAGATCTGATGGAGGTGG

AGCACACGTCTGAACTCTTTCCCTACACGACGCTCTTCCGATCTNNNCCGGACTACAACTCCCAGAATG

AGCACACGTCTGAACTCTTTCCCTACACGACGCTCTTCCGATCTNNNCCGGACTACAACTCCCAGAATG

AGCACACGTCTGAACTCTTTCCCTACACGACGCTCTTCCGATCTNNNCCGGACTACAACTCCCAGAATG

AGCACACGTCTGAACTCTTTCCCTACACGACGCTCTTCCGATCTNNNGAGGTTGCAGTGAGCCAAGAT

AGCACACGTCTGAACTCTTTCCCTACACGACGCTCTTCCGATCTNNNGAGGTTGCAGTGAGCCAAGAT

AGCACACGTCTGAACTCTTTCCCTACACGACGCTCTTCCGATCTNNNGAGGTTGCAGTGAGCCAAGAT

GAGCACACGTCTGAACTCTTTCCCTACACGACGCTCTTCCGATCTNNNGATCAGGAGTTCGATAACAG

GAGCACACGTCTGAACTCTTTCCCTACACGACGCTCTTCCGATCTNNNGATCAGGAGTTCGATAACAG

GAGCACACGTCTGAACTCTTTCCCTACACGACGCTCTTCCGATCTNNNGATCAGGAGTTCGATAACAG

GAGCACACGTCTGAACTCTTTCCCTACACGACGCTCTTCCGATCTNNNCTGTAGTCCCAGCTACTCAGG

GAGCACACGTCTGAACTCTTTCCCTACACGACGCTCTTCCGATCTNNNCTGTAGTCCCAGCTACTCAGG

GAGCACACGTCTGAACTCTTTCCCTACACGACGCTCTTCCGATCTNNNCTGTAGTCCCAGCTACTCAGG

GCACACGTCTGAACTCTTTCCCTACACGACGCTCTTCCGATCTNNNGGGTTCAAGTGATTCTCCTACCT

GCACACGTCTGAACTCTTTCCCTACACGACGCTCTTCCGATCTNNNGGGTTCAAGTGATTCTCCTACCT

GCACACGTCTGAACTCTTTCCCTACACGACGCTCTTCCGATCTNNNGGGTTCAAGTGATTCTCCTACCT

GAAGAGCACACGTCTGAACTCTTTCCCTACACGACGCTCTTCCGATCTNNNTTCGGCTTCCTGGGATT

GAAGAGCACACGTCTGAACTCTTTCCCTACACGACGCTCTTCCGATCTNNNTTCGGCTTCCTGGGATT

GAAGAGCACACGTCTGAACTCTTTCCCTACACGACGCTCTTCCGATCTNNNTTCGGCTTCCTGGGATT

GAGCACACGTCTGAACTCTTTCCCTACACGACGCTCTTCCGATCTNNNGGAGGCTTGACACTAGGAG

GAGCACACGTCTGAACTCTTTCCCTACACGACGCTCTTCCGATCTNNNGGAGGCTTGACACTAGGAG

GAGCACACGTCTGAACTCTTTCCCTACACGACGCTCTTCCGATCTNNNGGAGGCTTGACACTAGGAG

AGCACACGTCTGAACTCTTTCCCTACACGACGCTCTTCCGATCTNNNCTTCTTCACGTCCGAATCCCTAA

AGCACACGTCTGAACTCTTTCCCTACACGACGCTCTTCCGATCTNNNCTTCTTCACGTCCGAATCCCTAA

AGCACACGTCTGAACTCTTTCCCTACACGACGCTCTTCCGATCTNNNCTTCTTCACGTCCGAATCCCTAA
AGCACACGTCTGAACTCTTTCCCTACACGACGCTCTTCCGATCTNNNCCCCGTGATTTGTCTCCAAAAT
AGCACACGTCTGAACTCTTTCCCTACACGACGCTCTTCCGATCTNNNCCCCGTGATTTGTCTCCAAAAT
AGCACACGTCTGAACTCTTTCCCTACACGACGCTCTTCCGATCTNNNCCCCGTGATTTGTCTCCAAAAT
AGAGCACACGTCTGAACTCTTTCCCTACACGACGCTCTTCCGATCTNNNAGCCAATCTTCATTCTGCTT
AGAGCACACGTCTGAACTCTTTCCCTACACGACGCTCTTCCGATCTNNNAGCCAATCTTCATTCTGCTT
AGAGCACACGTCTGAACTCTTTCCCTACACGACGCTCTTCCGATCTNNNAGCCAATCTTCATTCTGCTT
AGCACACGTCTGAACTCTTTCCCTACACGACGCTCTTCCGATCTNNNTCTACTGCCTACACAGATCACA
AGCACACGTCTGAACTCTTTCCCTACACGACGCTCTTCCGATCTNNNTCTACTGCCTACACAGATCACA
AGCACACGTCTGAACTCTTTCCCTACACGACGCTCTTCCGATCTNNNTCTACTGCCTACACAGATCACA
AGCACACGTCTGAACTCTTTCCCTACACGACGCTCTTCCGATCTNNNTTTGTGTTTCAGGTGCCATAAC
AGCACACGTCTGAACTCTTTCCCTACACGACGCTCTTCCGATCTNNNTTTGTGTTTCAGGTGCCATAAC
AGCACACGTCTGAACTCTTTCCCTACACGACGCTCTTCCGATCTNNNTTTGTGTTTCAGGTGCCATAAC
AGCACACGTCTGAACTCTTTCCCTACACGACGCTCTTCCGATCTNNNTTTCCTTATTGTTGGTGGAGCC
AGCACACGTCTGAACTCTTTCCCTACACGACGCTCTTCCGATCTNNNTTTCCTTATTGTTGGTGGAGCC
AGCACACGTCTGAACTCTTTCCCTACACGACGCTCTTCCGATCTNNNTTTCCTTATTGTTGGTGGAGCC
AGCACACGTCTGAACTCTTTCCCTACACGACGCTCTTCCGATCTNNNCTTGAAACTGGGAGGTGGAG
AGCACACGTCTGAACTCTTTCCCTACACGACGCTCTTCCGATCTNNNCTTGAAACTGGGAGGTGGAG
AGCACACGTCTGAACTCTTTCCCTACACGACGCTCTTCCGATCTNNNCTTGAAACTGGGAGGTGGAG
GCACACGTCTGAACTCTTTCCCTACACGACGCTCTTCCGATCTNNNTTGACGTTGAAGGGAAAGAGA
GCACACGTCTGAACTCTTTCCCTACACGACGCTCTTCCGATCTNNNTTGACGTTGAAGGGAAAGAGA
GCACACGTCTGAACTCTTTCCCTACACGACGCTCTTCCGATCTNNNTTGACGTTGAAGGGAAAGAGA
AGCACACGTCTGAACTCTTTCCCTACACGACGCTCTTCCGATCTNNNCTCCCAAGTAGCTGAGACTACA
AGCACACGTCTGAACTCTTTCCCTACACGACGCTCTTCCGATCTNNNCTCCCAAGTAGCTGAGACTACA
AGCACACGTCTGAACTCTTTCCCTACACGACGCTCTTCCGATCTNNNCTCCCAAGTAGCTGAGACTACA
CACACGTCTGAACTCTTTCCCTACACGACGCTCTTCCGATCTNNNGCATCAGATAGAAGGTAATGTAT
CACACGTCTGAACTCTTTCCCTACACGACGCTCTTCCGATCTNNNGCATCAGATAGAAGGTAATGTAT
CACACGTCTGAACTCTTTCCCTACACGACGCTCTTCCGATCTNNNGCATCAGATAGAAGGTAATGTAT
CACACGTCTGAACTCTTTCCCTACACGACGCTCTTCCGATCTNNNGTTATTTGGGAGTCTGAGGTGGA
CACACGTCTGAACTCTTTCCCTACACGACGCTCTTCCGATCTNNNGTTATTTGGGAGTCTGAGGTGGA
CACACGTCTGAACTCTTTCCCTACACGACGCTCTTCCGATCTNNNGTTATTTGGGAGTCTGAGGTGGA
AGCACACGTCTGAACTCTTTCCCTACACGACGCTCTTCCGATCTNNNGTACCTGGTCCCAATTTCGAG
AGCACACGTCTGAACTCTTTCCCTACACGACGCTCTTCCGATCTNNNGTACCTGGTCCCAATTTCGAG
AGCACACGTCTGAACTCTTTCCCTACACGACGCTCTTCCGATCTNNNGTACCTGGTCCCAATTTCGAG
AGCACACGTCTGAACTCTTTCCCTACACGACGCTCTTCCGATCTNNNTGGACACGCATTTGGAGATAC
AGCACACGTCTGAACTCTTTCCCTACACGACGCTCTTCCGATCTNNNTGGACACGCATTTGGAGATAC
AGCACACGTCTGAACTCTTTCCCTACACGACGCTCTTCCGATCTNNNTGGACACGCATTTGGAGATAC
AGAGCACACGTCTGAACTCTTTCCCTACACGACGCTCTTCCGATCTNNNACCCAAAGTATAGCACTTGC
AGAGCACACGTCTGAACTCTTTCCCTACACGACGCTCTTCCGATCTNNNACCCAAAGTATAGCACTTGC
AGAGCACACGTCTGAACTCTTTCCCTACACGACGCTCTTCCGATCTNNNACCCAAAGTATAGCACTTGC
AGCACACGTCTGAACTCTTTCCCTACACGACGCTCTTCCGATCTNNNAGTTCTTCCTCTCTTCTCCCTG
AGCACACGTCTGAACTCTTTCCCTACACGACGCTCTTCCGATCTNNNAGTTCTTCCTCTCTTCTCCCTG
AGCACACGTCTGAACTCTTTCCCTACACGACGCTCTTCCGATCTNNNAGTTCTTCCTCTCTTCTCCCTG

GCACACGTCTGAACTCTTTCCCTACACGACGCTCTTCCGATCTNNNAGTCTCTTCATCTCACTACCCCTCA
GCACACGTCTGAACTCTTTCCCTACACGACGCTCTTCCGATCTNNNAGTCTCTTCATCTCACTACCCCTCA
GCACACGTCTGAACTCTTTCCCTACACGACGCTCTTCCGATCTNNNAGTCTCTTCATCTCACTACCCCTCA
GAGCACACGTCTGAACTCTTTCCCTACACGACGCTCTTCCGATCTNNNTACCTGTAATCCCAGCTACTTG
GAGCACACGTCTGAACTCTTTCCCTACACGACGCTCTTCCGATCTNNNTACCTGTAATCCCAGCTACTTG
GAGCACACGTCTGAACTCTTTCCCTACACGACGCTCTTCCGATCTNNNTACCTGTAATCCCAGCTACTTG
GCACACGTCTGAACTCTTTCCCTACACGACGCTCTTCCGATCTNNNTAAGTGTGTGATGGGGAAGAGT
GCACACGTCTGAACTCTTTCCCTACACGACGCTCTTCCGATCTNNNTAAGTGTGTGATGGGGAAGAGT
GCACACGTCTGAACTCTTTCCCTACACGACGCTCTTCCGATCTNNNTAAGTGTGTGATGGGGAAGAGT
GAGCACACGTCTGAACTCTTTCCCTACACGACGCTCTTCCGATCTNNNTGTGTGACTGTAATCCCAGCTA
GAGCACACGTCTGAACTCTTTCCCTACACGACGCTCTTCCGATCTNNNTGTGTGACTGTAATCCCAGCTA
GAGCACACGTCTGAACTCTTTCCCTACACGACGCTCTTCCGATCTNNNTGTGTGACTGTAATCCCAGCTA
GCACACGTCTGAACTCTTTCCCTACACGACGCTCTTCCGATCTNNNCCCCCCAAAGTGTTAGGATTACA
GCACACGTCTGAACTCTTTCCCTACACGACGCTCTTCCGATCTNNNCCCCCCAAAGTGTTAGGATTACA
GCACACGTCTGAACTCTTTCCCTACACGACGCTCTTCCGATCTNNNCCCCCCAAAGTGTTAGGATTACA
CACGTCTGAACTCTTTCCCTACACGACGCTCTTCCGATCTNNNCACACACACATATACACAGTGCGCAT
CACGTCTGAACTCTTTCCCTACACGACGCTCTTCCGATCTNNNCACACACACATATACACAGTGCGCAT
CACGTCTGAACTCTTTCCCTACACGACGCTCTTCCGATCTNNNCACACACACATATACACAGTGCGCAT
GCACACGTCTGAACTCTTTCCCTACACGACGCTCTTCCGATCTNNNAATGAAGCTGGTTGGGTTTGAT
GCACACGTCTGAACTCTTTCCCTACACGACGCTCTTCCGATCTNNNAATGAAGCTGGTTGGGTTTGAT
GCACACGTCTGAACTCTTTCCCTACACGACGCTCTTCCGATCTNNNAATGAAGCTGGTTGGGTTTGAT
AGCACACGTCTGAACTCTTTCCCTACACGACGCTCTTCCGATCTNNNGAACCTTTGCAACCACAATGAG
AGCACACGTCTGAACTCTTTCCCTACACGACGCTCTTCCGATCTNNNGAACCTTTGCAACCACAATGAG
AGCACACGTCTGAACTCTTTCCCTACACGACGCTCTTCCGATCTNNNGAACCTTTGCAACCACAATGAG
AGCACACGTCTGAACTCTTTCCCTACACGACGCTCTTCCGATCTNNNAGTCATCTCAGTCCTGGATGGT
AGCACACGTCTGAACTCTTTCCCTACACGACGCTCTTCCGATCTNNNAGTCATCTCAGTCCTGGATGGT
AGCACACGTCTGAACTCTTTCCCTACACGACGCTCTTCCGATCTNNNAGTCATCTCAGTCCTGGATGGT
AGCACACGTCTGAACTCTTTCCCTACACGACGCTCTTCCGATCTNNNTGAGAGGTTTTTGAGATGGAG
AGCACACGTCTGAACTCTTTCCCTACACGACGCTCTTCCGATCTNNNTGAGAGGTTTTTGAGATGGAG
AGCACACGTCTGAACTCTTTCCCTACACGACGCTCTTCCGATCTNNNTGAGAGGTTTTTGAGATGGAG
GCACACGTCTGAACTCTTTCCCTACACGACGCTCTTCCGATCTNNNCTTCATCTGTATTTATAGCCACTCA
GCACACGTCTGAACTCTTTCCCTACACGACGCTCTTCCGATCTNNNCTTCATCTGTATTTATAGCCACTCA
GCACACGTCTGAACTCTTTCCCTACACGACGCTCTTCCGATCTNNNCTTCATCTGTATTTATAGCCACTCA
AGAGCACACGTCTGAACTCTTTCCCTACACGACGCTCTTCCGATCTNNNACCTGGAAACTCTCCCTAGT
AGAGCACACGTCTGAACTCTTTCCCTACACGACGCTCTTCCGATCTNNNACCTGGAAACTCTCCCTAGT
AGAGCACACGTCTGAACTCTTTCCCTACACGACGCTCTTCCGATCTNNNACCTGGAAACTCTCCCTAGT
AGCACACGTCTGAACTCTTTCCCTACACGACGCTCTTCCGATCTNNNAGGCATGAAGAAAAAAAAAAA
AGCACACGTCTGAACTCTTTCCCTACACGACGCTCTTCCGATCTNNNAGGCATGAAGAAAAAAAAAAA
AGCACACGTCTGAACTCTTTCCCTACACGACGCTCTTCCGATCTNNNAGGCATGAAGAAAAAAAAAAA
GCACACGTCTGAACTCTTTCCCTACACGACGCTCTTCCGATCTNNNAGGTTGCAGTGAGTTGAGATCA
GCACACGTCTGAACTCTTTCCCTACACGACGCTCTTCCGATCTNNNAGGTTGCAGTGAGTTGAGATCA
GCACACGTCTGAACTCTTTCCCTACACGACGCTCTTCCGATCTNNNAGGTTGCAGTGAGTTGAGATCA
GAGCACACGTCTGAACTCTTTCCCTACACGACGCTCTTCCGATCTNNNAAAAATGGCCTTGAGAGACT

GAGCACACGTCTGAACTCTTTCCCTACACGACGCTCTTCCGATCTNNNAAAAATGGCCTTGAGAGACT
GAGCACACGTCTGAACTCTTTCCCTACACGACGCTCTTCCGATCTNNNAAAAATGGCCTTGAGAGACT
AGCACACGTCTGAACTCTTTCCCTACACGACGCTCTTCCGATCTNNNACTTTGCTTGGTTCTCCAGAGA
AGCACACGTCTGAACTCTTTCCCTACACGACGCTCTTCCGATCTNNNACTTTGCTTGGTTCTCCAGAGA
AGCACACGTCTGAACTCTTTCCCTACACGACGCTCTTCCGATCTNNNACTTTGCTTGGTTCTCCAGAGA
GAGCACACGTCTGAACTCTTTCCCTACACGACGCTCTTCCGATCTNNNCCATCTCTCTGTTCTCCTTGGA
GAGCACACGTCTGAACTCTTTCCCTACACGACGCTCTTCCGATCTNNNCCATCTCTCTGTTCTCCTTGGA
GAGCACACGTCTGAACTCTTTCCCTACACGACGCTCTTCCGATCTNNNCCATCTCTCTGTTCTCCTTGGA
AGCACACGTCTGAACTCTTTCCCTACACGACGCTCTTCCGATCTNNNTCTCTCTCACACACACACAAACA
AGCACACGTCTGAACTCTTTCCCTACACGACGCTCTTCCGATCTNNNTCTCTCTCACACACACACAAACA
AGCACACGTCTGAACTCTTTCCCTACACGACGCTCTTCCGATCTNNNTCTCTCTCACACACACACAAACA
GAGCACACGTCTGAACTCTTTCCCTACACGACGCTCTTCCGATCTNNNACAGGAGAATGTCATGAACC
GAGCACACGTCTGAACTCTTTCCCTACACGACGCTCTTCCGATCTNNNACAGGAGAATGTCATGAACC
GAGCACACGTCTGAACTCTTTCCCTACACGACGCTCTTCCGATCTNNNACAGGAGAATGTCATGAACC
GAGCACACGTCTGAACTCTTTCCCTACACGACGCTCTTCCGATCTNNNCGTATCTGTAGTCCCAGCTAC
GAGCACACGTCTGAACTCTTTCCCTACACGACGCTCTTCCGATCTNNNCGTATCTGTAGTCCCAGCTAC
GAGCACACGTCTGAACTCTTTCCCTACACGACGCTCTTCCGATCTNNNCGTATCTGTAGTCCCAGCTAC
GCACACGTCTGAACTCTTTCCCTACACGACGCTCTTCCGATCTNNNTCATGAGAACAGCAAGGAGGA
GCACACGTCTGAACTCTTTCCCTACACGACGCTCTTCCGATCTNNNTCATGAGAACAGCAAGGAGGA
GCACACGTCTGAACTCTTTCCCTACACGACGCTCTTCCGATCTNNNTCATGAGAACAGCAAGGAGGA
AGCACACGTCTGAACTCTTTCCCTACACGACGCTCTTCCGATCTNNNGGAATCACTTGAACCTGGGAG
AGCACACGTCTGAACTCTTTCCCTACACGACGCTCTTCCGATCTNNNGGAATCACTTGAACCTGGGAG
AGCACACGTCTGAACTCTTTCCCTACACGACGCTCTTCCGATCTNNNGGAATCACTTGAACCTGGGAG
GCACACGTCTGAACTCTTTCCCTACACGACGCTCTTCCGATCTNNNTGTCATTGATTTCTGCTCAGTCC
GCACACGTCTGAACTCTTTCCCTACACGACGCTCTTCCGATCTNNNTGTCATTGATTTCTGCTCAGTCC
GCACACGTCTGAACTCTTTCCCTACACGACGCTCTTCCGATCTNNNTGTCATTGATTTCTGCTCAGTCC
GAGCACACGTCTGAACTCTTTCCCTACACGACGCTCTTCCGATCTNNNTCGTTTTGTTTTCCCAGAGCT
GAGCACACGTCTGAACTCTTTCCCTACACGACGCTCTTCCGATCTNNNTCGTTTTGTTTTCCCAGAGCT
GAGCACACGTCTGAACTCTTTCCCTACACGACGCTCTTCCGATCTNNNTCGTTTTGTTTTCCCAGAGCT
GAGCACACGTCTGAACTCTTTCCCTACACGACGCTCTTCCGATCTNNNCTGAGGCAGGAGAATCACTT
GAGCACACGTCTGAACTCTTTCCCTACACGACGCTCTTCCGATCTNNNCTGAGGCAGGAGAATCACTT
GAGCACACGTCTGAACTCTTTCCCTACACGACGCTCTTCCGATCTNNNCTGAGGCAGGAGAATCACTT
AGCACACGTCTGAACTCTTTCCCTACACGACGCTCTTCCGATCTNNNACTGTTGCAGGTGATCAAAACA
AGCACACGTCTGAACTCTTTCCCTACACGACGCTCTTCCGATCTNNNACTGTTGCAGGTGATCAAAACA
AGCACACGTCTGAACTCTTTCCCTACACGACGCTCTTCCGATCTNNNACTGTTGCAGGTGATCAAAACA
GAGCACACGTCTGAACTCTTTCCCTACACGACGCTCTTCCGATCTNNNCTCATGCCTATAATCCCAGCAC
GAGCACACGTCTGAACTCTTTCCCTACACGACGCTCTTCCGATCTNNNCTCATGCCTATAATCCCAGCAC
GAGCACACGTCTGAACTCTTTCCCTACACGACGCTCTTCCGATCTNNNCTCATGCCTATAATCCCAGCAC
AGCACACGTCTGAACTCTTTCCCTACACGACGCTCTTCCGATCTNNNACAAAATTTACAAGTGGTGGC
AGCACACGTCTGAACTCTTTCCCTACACGACGCTCTTCCGATCTNNNACAAAATTTACAAGTGGTGGC
AGCACACGTCTGAACTCTTTCCCTACACGACGCTCTTCCGATCTNNNACAAAATTTACAAGTGGTGGC
AGCACACGTCTGAACTCTTTCCCTACACGACGCTCTTCCGATCTNNNTTTTAGCTCCCTCTTATGTGGG
AGCACACGTCTGAACTCTTTCCCTACACGACGCTCTTCCGATCTNNNTTTTAGCTCCCTCTTATGTGGG

AGCACACGTCTGAACTCTTTCCCTACACGACGCTCTTCCGATCTNNNTTTTAGCTCCCTCTTATGTGGG
AGCACACGTCTGAACTCTTTCCCTACACGACGCTCTTCCGATCTNNNACTACTTCTACTACTACTACACA
AGCACACGTCTGAACTCTTTCCCTACACGACGCTCTTCCGATCTNNNACTACTTCTACTACTACTACACA
AGCACACGTCTGAACTCTTTCCCTACACGACGCTCTTCCGATCTNNNACTACTTCTACTACTACTACACA
AGCACACGTCTGAACTCTTTCCCTACACGACGCTCTTCCGATCTNNNTTGAATAGCTGAGACCACTGA
AGCACACGTCTGAACTCTTTCCCTACACGACGCTCTTCCGATCTNNNTTGAATAGCTGAGACCACTGA
AGCACACGTCTGAACTCTTTCCCTACACGACGCTCTTCCGATCTNNNTTGAATAGCTGAGACCACTGA
GCACACGTCTGAACTCTTTCCCTACACGACGCTCTTCCGATCTNNNTCTTCTTGTTGCAGATAAGGCCT
GCACACGTCTGAACTCTTTCCCTACACGACGCTCTTCCGATCTNNNTCTTCTTGTTGCAGATAAGGCCT
GCACACGTCTGAACTCTTTCCCTACACGACGCTCTTCCGATCTNNNTCTTCTTGTTGCAGATAAGGCCT
GAGCACACGTCTGAACTCTTTCCCTACACGACGCTCTTCCGATCTNNNCAGAGGTTGTAGTGAGCCAAG
GAGCACACGTCTGAACTCTTTCCCTACACGACGCTCTTCCGATCTNNNCAGAGGTTGTAGTGAGCCAAG
GAGCACACGTCTGAACTCTTTCCCTACACGACGCTCTTCCGATCTNNNCAGAGGTTGTAGTGAGCCAAG
AGCACACGTCTGAACTCTTTCCCTACACGACGCTCTTCCGATCTNNNTTCCTTCTCTCTGTTCCCTCACCCA
AGCACACGTCTGAACTCTTTCCCTACACGACGCTCTTCCGATCTNNNTTCCTTCTCTCTGTTCCCTCACCCA
AGCACACGTCTGAACTCTTTCCCTACACGACGCTCTTCCGATCTNNNTTCCTTCTCTCTGTTCCCTCACCCA
AGCACACGTCTGAACTCTTTCCCTACACGACGCTCTTCCGATCTNNNTGGGTCTTTGAGGAATTTCCAC
AGCACACGTCTGAACTCTTTCCCTACACGACGCTCTTCCGATCTNNNTGGGTCTTTGAGGAATTTCCAC
AGCACACGTCTGAACTCTTTCCCTACACGACGCTCTTCCGATCTNNNTGGGTCTTTGAGGAATTTCCAC
GAGCACACGTCTGAACTCTTTCCCTACACGACGCTCTTCCGATCTNNNTGGAAACCTGGTGCATGATTT
GAGCACACGTCTGAACTCTTTCCCTACACGACGCTCTTCCGATCTNNNTGGAAACCTGGTGCATGATTT
GAGCACACGTCTGAACTCTTTCCCTACACGACGCTCTTCCGATCTNNNTGGAAACCTGGTGCATGATTT
GAGCACACGTCTGAACTCTTTCCCTACACGACGCTCTTCCGATCTNNNTATCAGCTGCTAGAATGCAAG
GAGCACACGTCTGAACTCTTTCCCTACACGACGCTCTTCCGATCTNNNTATCAGCTGCTAGAATGCAAG
GAGCACACGTCTGAACTCTTTCCCTACACGACGCTCTTCCGATCTNNNTATCAGCTGCTAGAATGCAAG
AGCACACGTCTGAACTCTTTCCCTACACGACGCTCTTCCGATCTNNNTCCTTCCTTCCTTTCCTTCCTTCCA
AGCACACGTCTGAACTCTTTCCCTACACGACGCTCTTCCGATCTNNNTCCTTCCTTCCTTTCCTTCCTTCCA
AGCACACGTCTGAACTCTTTCCCTACACGACGCTCTTCCGATCTNNNTCCTTCCTTCCTTTCCTTCCTTCCA
AGCACACGTCTGAACTCTTTCCCTACACGACGCTCTTCCGATCTNNNGGCCTATCACAATGCTGTAGGT
AGCACACGTCTGAACTCTTTCCCTACACGACGCTCTTCCGATCTNNNGGCCTATCACAATGCTGTAGGT
AGCACACGTCTGAACTCTTTCCCTACACGACGCTCTTCCGATCTNNNGGCCTATCACAATGCTGTAGGT
AGCACACGTCTGAACTCTTTCCCTACACGACGCTCTTCCGATCTNNNAAATATCTTACCTGCAGTAATCT
AGCACACGTCTGAACTCTTTCCCTACACGACGCTCTTCCGATCTNNNAAATATCTTACCTGCAGTAATCT
AGCACACGTCTGAACTCTTTCCCTACACGACGCTCTTCCGATCTNNNAAATATCTTACCTGCAGTAATCT
GAGCACACGTCTGAACTCTTTCCCTACACGACGCTCTTCCGATCTNNNGAATCCACAGTGTGATGGATG
GAGCACACGTCTGAACTCTTTCCCTACACGACGCTCTTCCGATCTNNNGAATCCACAGTGTGATGGATG
GAGCACACGTCTGAACTCTTTCCCTACACGACGCTCTTCCGATCTNNNGAATCCACAGTGTGATGGATG
CACACGTCTGAACTCTTTCCCTACACGACGCTCTTCCGATCTNNNACCTTCTCTCTCTCTCTCTCTCTCAT
CACACGTCTGAACTCTTTCCCTACACGACGCTCTTCCGATCTNNNACCTTCTCTCTCTCTCTCTCTCTCAT
CACACGTCTGAACTCTTTCCCTACACGACGCTCTTCCGATCTNNNACCTTCTCTCTCTCTCTCTCTCTCAT
AGCACACGTCTGAACTCTTTCCCTACACGACGCTCTTCCGATCTNNNGAGGGGTTGGGGGTATGATTA
AGCACACGTCTGAACTCTTTCCCTACACGACGCTCTTCCGATCTNNNGAGGGGTTGGGGGTATGATTA
AGCACACGTCTGAACTCTTTCCCTACACGACGCTCTTCCGATCTNNNGAGGGGTTGGGGGTATGATTA

AGCACACGTCTGAACTCTTTCCCTACACGACGCTCTTCCGATCTNNNGAGGTTACAGTGAGCCAAGATC

AGCACACGTCTGAACTCTTTCCCTACACGACGCTCTTCCGATCTNNNGAGGTTACAGTGAGCCAAGATC

AGCACACGTCTGAACTCTTTCCCTACACGACGCTCTTCCGATCTNNNGAGGTTACAGTGAGCCAAGATC

GAGCACACGTCTGAACTCTTTCCCTACACGACGCTCTTCCGATCTNNNCATAGGGAGCAAGAGAGGTC

GAGCACACGTCTGAACTCTTTCCCTACACGACGCTCTTCCGATCTNNNCATAGGGAGCAAGAGAGGTC

GAGCACACGTCTGAACTCTTTCCCTACACGACGCTCTTCCGATCTNNNCATAGGGAGCAAGAGAGGTC

CACACGTCTGAACTCTTTCCCTACACGACGCTCTTCCGATCTNNNCTCCTCCCTTCTTTCTTCCTTCCCATC

CACACGTCTGAACTCTTTCCCTACACGACGCTCTTCCGATCTNNNCTCCTCCCTTCTTTCTTCCTTCCCATC

CACACGTCTGAACTCTTTCCCTACACGACGCTCTTCCGATCTNNNCTCCTCCCTTCTTTCTTCCTTCCCATC

GCACACGTCTGAACTCTTTCCCTACACGACGCTCTTCCGATCTNNNCGCAGAAGGACATCAAAAAAGGT

GCACACGTCTGAACTCTTTCCCTACACGACGCTCTTCCGATCTNNNCGCAGAAGGACATCAAAAAAGGT

GCACACGTCTGAACTCTTTCCCTACACGACGCTCTTCCGATCTNNNCGCAGAAGGACATCAAAAAAGGT

GAGCACACGTCTGAACTCTTTCCCTACACGACGCTCTTCCGATCTNNNAGGAAAAGTGTCTCCTTTCCA

GAGCACACGTCTGAACTCTTTCCCTACACGACGCTCTTCCGATCTNNNAGGAAAAGTGTCTCCTTTCCA

GAGCACACGTCTGAACTCTTTCCCTACACGACGCTCTTCCGATCTNNNAGGAAAAGTGTCTCCTTTCCA

GAGCACACGTCTGAACTCTTTCCCTACACGACGCTCTTCCGATCTNNNCACAGAAGTTGCAGTGAATG

GAGCACACGTCTGAACTCTTTCCCTACACGACGCTCTTCCGATCTNNNCACAGAAGTTGCAGTGAATG

GAGCACACGTCTGAACTCTTTCCCTACACGACGCTCTTCCGATCTNNNCACAGAAGTTGCAGTGAATG

GAGCACACGTCTGAACTCTTTCCCTACACGACGCTCTTCCGATCTNNNGAGGCAACAGAATCACTTGAA

GAGCACACGTCTGAACTCTTTCCCTACACGACGCTCTTCCGATCTNNNGAGGCAACAGAATCACTTGAA

GAGCACACGTCTGAACTCTTTCCCTACACGACGCTCTTCCGATCTNNNGAGGCAACAGAATCACTTGAA

GAGCACACGTCTGAACTCTTTCCCTACACGACGCTCTTCCGATCTNNNATGTGAGGGTGTGTGGTGTC

GAGCACACGTCTGAACTCTTTCCCTACACGACGCTCTTCCGATCTNNNATGTGAGGGTGTGTGGTGTC

GAGCACACGTCTGAACTCTTTCCCTACACGACGCTCTTCCGATCTNNNATGTGAGGGTGTGTGGTGTC

AGCACACGTCTGAACTCTTTCCCTACACGACGCTCTTCCGATCTNNNCCTGGTGCATGCTACCTACATA

AGCACACGTCTGAACTCTTTCCCTACACGACGCTCTTCCGATCTNNNCCTGGTGCATGCTACCTACATA

AGCACACGTCTGAACTCTTTCCCTACACGACGCTCTTCCGATCTNNNCCTGGTGCATGCTACCTACATA

AGCACACGTCTGAACTCTTTCCCTACACGACGCTCTTCCGATCTNNNCTCCCAAGACTGAACCATGAAG

AGCACACGTCTGAACTCTTTCCCTACACGACGCTCTTCCGATCTNNNCTCCCAAGACTGAACCATGAAG

AGCACACGTCTGAACTCTTTCCCTACACGACGCTCTTCCGATCTNNNCTCCCAAGACTGAACCATGAAG

AGCACACGTCTGAACTCTTTCCCTACACGACGCTCTTCCGATCTNNNTACAGGTGTCTTCATTCTGCGAA

AGCACACGTCTGAACTCTTTCCCTACACGACGCTCTTCCGATCTNNNTACAGGTGTCTTCATTCTGCGAA

AGCACACGTCTGAACTCTTTCCCTACACGACGCTCTTCCGATCTNNNTACAGGTGTCTTCATTCTGCGAA

AGCACACGTCTGAACTCTTTCCCTACACGACGCTCTTCCGATCTNNNGGAGGCTGAGGCAGGAAAAT

AGCACACGTCTGAACTCTTTCCCTACACGACGCTCTTCCGATCTNNNGGAGGCTGAGGCAGGAAAAT

AGCACACGTCTGAACTCTTTCCCTACACGACGCTCTTCCGATCTNNNGGAGGCTGAGGCAGGAAAAT

GAGCACACGTCTGAACTCTTTCCCTACACGACGCTCTTCCGATCTNNNGACCCTTTTGTTCAGCTTCAGA

GAGCACACGTCTGAACTCTTTCCCTACACGACGCTCTTCCGATCTNNNGACCCTTTTGTTCAGCTTCAGA

GAGCACACGTCTGAACTCTTTCCCTACACGACGCTCTTCCGATCTNNNGACCCTTTTGTTCAGCTTCAGA

GCACACGTCTGAACTCTTTCCCTACACGACGCTCTTCCGATCTNNNGCCTGTAATCCCAGCTACTCGCAT

GCACACGTCTGAACTCTTTCCCTACACGACGCTCTTCCGATCTNNNGCCTGTAATCCCAGCTACTCGCAT

GCACACGTCTGAACTCTTTCCCTACACGACGCTCTTCCGATCTNNNGCCTGTAATCCCAGCTACTCGCAT

AGAGCACACGTCTGAACTCTTTCCCTACACGACGCTCTTCCGATCTNNNGGGTAATGCAAACAGTTGG

AGAGCACACGTCTGAACTCTTTCCCTACACGACGCTCTTCCGATCTNNNGGGTAATGCAAACAGTTGG
AGAGCACACGTCTGAACTCTTTCCCTACACGACGCTCTTCCGATCTNNNGGGTAATGCAAACAGTTGG
AGCACACGTCTGAACTCTTTCCCTACACGACGCTCTTCCGATCTNNNTTGAGCCTAATCTCTTTGTGGG
AGCACACGTCTGAACTCTTTCCCTACACGACGCTCTTCCGATCTNNNTTGAGCCTAATCTCTTTGTGGG
AGCACACGTCTGAACTCTTTCCCTACACGACGCTCTTCCGATCTNNNTTGAGCCTAATCTCTTTGTGGG
GGAAGAGCACACGTCTGAACTCTTTCCCTACACGACGCTCTTCCGATCTNNNATCCACTTGAGGAGCCA
GGAAGAGCACACGTCTGAACTCTTTCCCTACACGACGCTCTTCCGATCTNNNATCCACTTGAGGAGCCA
GGAAGAGCACACGTCTGAACTCTTTCCCTACACGACGCTCTTCCGATCTNNNATCCACTTGAGGAGCCA
GAGCACACGTCTGAACTCTTTCCCTACACGACGCTCTTCCGATCTNNNTCTTTTCCTCCAACTCCACGTA
GAGCACACGTCTGAACTCTTTCCCTACACGACGCTCTTCCGATCTNNNTCTTTTCCTCCAACTCCACGTA
GAGCACACGTCTGAACTCTTTCCCTACACGACGCTCTTCCGATCTNNNTCTTTTCCTCCAACTCCACGTA
AGCACACGTCTGAACTCTTTCCCTACACGACGCTCTTCCGATCTNNNCAAAGATTGCAGTGAGCCAAGA
AGCACACGTCTGAACTCTTTCCCTACACGACGCTCTTCCGATCTNNNCAAAGATTGCAGTGAGCCAAGA
AGCACACGTCTGAACTCTTTCCCTACACGACGCTCTTCCGATCTNNNCAAAGATTGCAGTGAGCCAAGA
GAGCACACGTCTGAACTCTTTCCCTACACGACGCTCTTCCGATCTNNNCCATTCAGCCAGTCTAGGTCTT
GAGCACACGTCTGAACTCTTTCCCTACACGACGCTCTTCCGATCTNNNCCATTCAGCCAGTCTAGGTCTT
GAGCACACGTCTGAACTCTTTCCCTACACGACGCTCTTCCGATCTNNNCCATTCAGCCAGTCTAGGTCTT
AGCACACGTCTGAACTCTTTCCCTACACGACGCTCTTCCGATCTNNNAAGAACCGGCACTTTTCCTCTAT
AGCACACGTCTGAACTCTTTCCCTACACGACGCTCTTCCGATCTNNNAAGAACCGGCACTTTTCCTCTAT
AGCACACGTCTGAACTCTTTCCCTACACGACGCTCTTCCGATCTNNNAAGAACCGGCACTTTTCCTCTAT
AGCACACGTCTGAACTCTTTCCCTACACGACGCTCTTCCGATCTNNNCATAGAAGCCAAAGGGAAGTTG
AGCACACGTCTGAACTCTTTCCCTACACGACGCTCTTCCGATCTNNNCATAGAAGCCAAAGGGAAGTTG
AGCACACGTCTGAACTCTTTCCCTACACGACGCTCTTCCGATCTNNNCATAGAAGCCAAAGGGAAGTTG
AGCACACGTCTGAACTCTTTCCCTACACGACGCTCTTCCGATCTNNNGGAGAATCACTTGAACCCAGGA
AGCACACGTCTGAACTCTTTCCCTACACGACGCTCTTCCGATCTNNNGGAGAATCACTTGAACCCAGGA
AGCACACGTCTGAACTCTTTCCCTACACGACGCTCTTCCGATCTNNNGGAGAATCACTTGAACCCAGGA
AGAGCACACGTCTGAACTCTTTCCCTACACGACGCTCTTCCGATCTNNNGATGCCTGTAATCCCAGCTA
AGAGCACACGTCTGAACTCTTTCCCTACACGACGCTCTTCCGATCTNNNGATGCCTGTAATCCCAGCTA
AGAGCACACGTCTGAACTCTTTCCCTACACGACGCTCTTCCGATCTNNNGATGCCTGTAATCCCAGCTA
AGCACACGTCTGAACTCTTTCCCTACACGACGCTCTTCCGATCTNNNCGTCCTGATGATTTTTATTTTTGC
AGCACACGTCTGAACTCTTTCCCTACACGACGCTCTTCCGATCTNNNCGTCCTGATGATTTTTATTTTTGC
AGCACACGTCTGAACTCTTTCCCTACACGACGCTCTTCCGATCTNNNCGTCCTGATGATTTTTATTTTTGC
AGAGCACACGTCTGAACTCTTTCCCTACACGACGCTCTTCCGATCTNNNGCCTGTAGTCCCAGCTACTTG
AGAGCACACGTCTGAACTCTTTCCCTACACGACGCTCTTCCGATCTNNNGCCTGTAGTCCCAGCTACTTG
AGAGCACACGTCTGAACTCTTTCCCTACACGACGCTCTTCCGATCTNNNGCCTGTAGTCCCAGCTACTTG
GAGCACACGTCTGAACTCTTTCCCTACACGACGCTCTTCCGATCTNNNCTAGTGTCAGTAATGGGCTG
GAGCACACGTCTGAACTCTTTCCCTACACGACGCTCTTCCGATCTNNNCTAGTGTCAGTAATGGGCTG
GAGCACACGTCTGAACTCTTTCCCTACACGACGCTCTTCCGATCTNNNCTAGTGTCAGTAATGGGCTG
AGCACACGTCTGAACTCTTTCCCTACACGACGCTCTTCCGATCTNNNAAACACTATGGAGTGTTCTGGG
AGCACACGTCTGAACTCTTTCCCTACACGACGCTCTTCCGATCTNNNAAACACTATGGAGTGTTCTGGG
AGCACACGTCTGAACTCTTTCCCTACACGACGCTCTTCCGATCTNNNAAACACTATGGAGTGTTCTGGG
GAGCACACGTCTGAACTCTTTCCCTACACGACGCTCTTCCGATCTNNNAGACAGGAGAATTGCTTGAA
GAGCACACGTCTGAACTCTTTCCCTACACGACGCTCTTCCGATCTNNNAGACAGGAGAATTGCTTGAA

GAGCACACGTCTGAACTCTTTCCCTACACGACGCTCTTCCGATCTNNNAGACAGGAGAATTGCTTGAA
GAGCACACGTCTGAACTCTTTCCCTACACGACGCTCTTCCGATCTNNNTGGTAAGGGCGAAGACTGA
GAGCACACGTCTGAACTCTTTCCCTACACGACGCTCTTCCGATCTNNNTGGTAAGGGCGAAGACTGA
GAGCACACGTCTGAACTCTTTCCCTACACGACGCTCTTCCGATCTNNNTGGTAAGGGCGAAGACTGA
AGCACACGTCTGAACTCTTTCCCTACACGACGCTCTTCCGATCTNNNCACAAAACAAACAAAAGGCCA
AGCACACGTCTGAACTCTTTCCCTACACGACGCTCTTCCGATCTNNNCACAAAACAAACAAAAGGCCA
AGCACACGTCTGAACTCTTTCCCTACACGACGCTCTTCCGATCTNNNCACAAAACAAACAAAAGGCCA
ACACGTCTGAACTCTTTCCCTACACGACGCTCTTCCGATCTNNNGTTCTAGGGGAAATGAATGTGCAT
ACACGTCTGAACTCTTTCCCTACACGACGCTCTTCCGATCTNNNGTTCTAGGGGAAATGAATGTGCAT
ACACGTCTGAACTCTTTCCCTACACGACGCTCTTCCGATCTNNNGTTCTAGGGGAAATGAATGTGCAT
GCACACGTCTGAACTCTTTCCCTACACGACGCTCTTCCGATCTNNNGCAGGAGAATTGCTTGAACCT
GCACACGTCTGAACTCTTTCCCTACACGACGCTCTTCCGATCTNNNGCAGGAGAATTGCTTGAACCT
GCACACGTCTGAACTCTTTCCCTACACGACGCTCTTCCGATCTNNNGCAGGAGAATTGCTTGAACCT
AGCACACGTCTGAACTCTTTCCCTACACGACGCTCTTCCGATCTNNNGACCCAGCTGGCAAATATCCTT
AGCACACGTCTGAACTCTTTCCCTACACGACGCTCTTCCGATCTNNNGACCCAGCTGGCAAATATCCTT
AGCACACGTCTGAACTCTTTCCCTACACGACGCTCTTCCGATCTNNNGACCCAGCTGGCAAATATCCTT
AGAGCACACGTCTGAACTCTTTCCCTACACGACGCTCTTCCGATCTNNNACTTGAGCAGCCCGGAAGT
AGAGCACACGTCTGAACTCTTTCCCTACACGACGCTCTTCCGATCTNNNACTTGAGCAGCCCGGAAGT
AGAGCACACGTCTGAACTCTTTCCCTACACGACGCTCTTCCGATCTNNNACTTGAGCAGCCCGGAAGT
GCACACGTCTGAACTCTTTCCCTACACGACGCTCTTCCGATCTNNNCTCAGAAGCATCCTCAGCTGGC
GCACACGTCTGAACTCTTTCCCTACACGACGCTCTTCCGATCTNNNCTCAGAAGCATCCTCAGCTGGC
GCACACGTCTGAACTCTTTCCCTACACGACGCTCTTCCGATCTNNNCTCAGAAGCATCCTCAGCTGGC
GCACACGTCTGAACTCTTTCCCTACACGACGCTCTTCCGATCTNNNGAAAGGTGGAAAGAAATGGGT
GCACACGTCTGAACTCTTTCCCTACACGACGCTCTTCCGATCTNNNGAAAGGTGGAAAGAAATGGGT
GCACACGTCTGAACTCTTTCCCTACACGACGCTCTTCCGATCTNNNGAAAGGTGGAAAGAAATGGGT
GCACACGTCTGAACTCTTTCCCTACACGACGCTCTTCCGATCTNNNATGCTGGTGTTACAGGTGTTA
GCACACGTCTGAACTCTTTCCCTACACGACGCTCTTCCGATCTNNNATGCTGGTGTTACAGGTGTTA
GCACACGTCTGAACTCTTTCCCTACACGACGCTCTTCCGATCTNNNATGCTGGTGTTACAGGTGTTA
AGCACACGTCTGAACTCTTTCCCTACACGACGCTCTTCCGATCTNNNTATTTGTGTGTGTGTGTGT
AGCACACGTCTGAACTCTTTCCCTACACGACGCTCTTCCGATCTNNNTATTTGTGTGTGTGTGTGT
AGCACACGTCTGAACTCTTTCCCTACACGACGCTCTTCCGATCTNNNTATTTGTGTGTGTGTGTGT
AGCACACGTCTGAACTCTTTCCCTACACGACGCTCTTCCGATCTNNNTGAGGCAGGAGAATCACTTGT
AGCACACGTCTGAACTCTTTCCCTACACGACGCTCTTCCGATCTNNNTGAGGCAGGAGAATCACTTGT
AGCACACGTCTGAACTCTTTCCCTACACGACGCTCTTCCGATCTNNNTGAGGCAGGAGAATCACTTGT
AGCACACGTCTGAACTCTTTCCCTACACGACGCTCTTCCGATCTNNNAAGAAATGCTGGAACCTAGG
AGCACACGTCTGAACTCTTTCCCTACACGACGCTCTTCCGATCTNNNAAGAAATGCTGGAACCTAGG
AGCACACGTCTGAACTCTTTCCCTACACGACGCTCTTCCGATCTNNNAAGAAATGCTGGAACCTAGG
GAGCACACGTCTGAACTCTTTCCCTACACGACGCTCTTCCGATCTNNNTACTCATGAGGCTGAGGTGG
GAGCACACGTCTGAACTCTTTCCCTACACGACGCTCTTCCGATCTNNNTACTCATGAGGCTGAGGTGG
GAGCACACGTCTGAACTCTTTCCCTACACGACGCTCTTCCGATCTNNNTACTCATGAGGCTGAGGTGG
GCACACGTCTGAACTCTTTCCCTACACGACGCTCTTCCGATCTNNNTTCATGGGCATTTGCATGGTTT
GCACACGTCTGAACTCTTTCCCTACACGACGCTCTTCCGATCTNNNTTCATGGGCATTTGCATGGTTT
GCACACGTCTGAACTCTTTCCCTACACGACGCTCTTCCGATCTNNNTTCATGGGCATTTGCATGGTTT

GAGCACACGTCTGAACTCTTTCCCTACACGACGCTCTTCCGATCTNNNGCTAAGGCAGGAGAATCACT
GAGCACACGTCTGAACTCTTTCCCTACACGACGCTCTTCCGATCTNNNGCTAAGGCAGGAGAATCACT
GAGCACACGTCTGAACTCTTTCCCTACACGACGCTCTTCCGATCTNNNGCTAAGGCAGGAGAATCACT
GAGCACACGTCTGAACTCTTTCCCTACACGACGCTCTTCCGATCTNNNGCATGATAAACAGCTGGCATC
GAGCACACGTCTGAACTCTTTCCCTACACGACGCTCTTCCGATCTNNNGCATGATAAACAGCTGGCATC
GAGCACACGTCTGAACTCTTTCCCTACACGACGCTCTTCCGATCTNNNGCATGATAAACAGCTGGCATC
AGCACACGTCTGAACTCTTTCCCTACACGACGCTCTTCCGATCTNNNATGCCTGTAATCCAGCATTTTG
AGCACACGTCTGAACTCTTTCCCTACACGACGCTCTTCCGATCTNNNATGCCTGTAATCCAGCATTTTG
AGCACACGTCTGAACTCTTTCCCTACACGACGCTCTTCCGATCTNNNATGCCTGTAATCCAGCATTTTG
AGAGCACACGTCTGAACTCTTTCCCTACACGACGCTCTTCCGATCTNNNCACCAATCCTTGATCATGCCA
AGAGCACACGTCTGAACTCTTTCCCTACACGACGCTCTTCCGATCTNNNCACCAATCCTTGATCATGCCA
AGAGCACACGTCTGAACTCTTTCCCTACACGACGCTCTTCCGATCTNNNCACCAATCCTTGATCATGCCA
GAGCACACGTCTGAACTCTTTCCCTACACGACGCTCTTCCGATCTNNNCCAATCAAAAGACACAGAGTG
GAGCACACGTCTGAACTCTTTCCCTACACGACGCTCTTCCGATCTNNNCCAATCAAAAGACACAGAGTG
GAGCACACGTCTGAACTCTTTCCCTACACGACGCTCTTCCGATCTNNNCCAATCAAAAGACACAGAGTG
ACACGTCTGAACTCTTTCCCTACACGACGCTCTTCCGATCTNNNAGAGGTCACACTAGTACCTTCATCG
ACACGTCTGAACTCTTTCCCTACACGACGCTCTTCCGATCTNNNAGAGGTCACACTAGTACCTTCATCG
ACACGTCTGAACTCTTTCCCTACACGACGCTCTTCCGATCTNNNAGAGGTCACACTAGTACCTTCATCG
GAGCACACGTCTGAACTCTTTCCCTACACGACGCTCTTCCGATCTNNNTCACCCTCAAAGTAGCACATA
GAGCACACGTCTGAACTCTTTCCCTACACGACGCTCTTCCGATCTNNNTCACCCTCAAAGTAGCACATA
GAGCACACGTCTGAACTCTTTCCCTACACGACGCTCTTCCGATCTNNNTCACCCTCAAAGTAGCACATA
AGAGCACACGTCTGAACTCTTTCCCTACACGACGCTCTTCCGATCTNNNGTAAATAGTCTCAGCTCCTC
AGAGCACACGTCTGAACTCTTTCCCTACACGACGCTCTTCCGATCTNNNGTAAATAGTCTCAGCTCCTC
AGAGCACACGTCTGAACTCTTTCCCTACACGACGCTCTTCCGATCTNNNGTAAATAGTCTCAGCTCCTC
AGCACACGTCTGAACTCTTTCCCTACACGACGCTCTTCCGATCTNNNGTGGATGGATGACTAGAGTGA
AGCACACGTCTGAACTCTTTCCCTACACGACGCTCTTCCGATCTNNNGTGGATGGATGACTAGAGTGA
AGCACACGTCTGAACTCTTTCCCTACACGACGCTCTTCCGATCTNNNGTGGATGGATGACTAGAGTGA
GAGCACACGTCTGAACTCTTTCCCTACACGACGCTCTTCCGATCTNNNAGATCTTGAGGGGAGTGGGA
GAGCACACGTCTGAACTCTTTCCCTACACGACGCTCTTCCGATCTNNNAGATCTTGAGGGGAGTGGGA
GAGCACACGTCTGAACTCTTTCCCTACACGACGCTCTTCCGATCTNNNAGATCTTGAGGGGAGTGGGA
AGCACACGTCTGAACTCTTTCCCTACACGACGCTCTTCCGATCTNNNCAGAGCAAGACCCTGTCTCTAAG
AGCACACGTCTGAACTCTTTCCCTACACGACGCTCTTCCGATCTNNNCAGAGCAAGACCCTGTCTCTAAG
AGCACACGTCTGAACTCTTTCCCTACACGACGCTCTTCCGATCTNNNCAGAGCAAGACCCTGTCTCTAAG
AGCACACGTCTGAACTCTTTCCCTACACGACGCTCTTCCGATCTNNNGAATGACGGTTTCCAGCTTCAT
AGCACACGTCTGAACTCTTTCCCTACACGACGCTCTTCCGATCTNNNGAATGACGGTTTCCAGCTTCAT
AGCACACGTCTGAACTCTTTCCCTACACGACGCTCTTCCGATCTNNNGAATGACGGTTTCCAGCTTCAT
AGCACACGTCTGAACTCTTTCCCTACACGACGCTCTTCCGATCTNNNCCCTAGGGCCTATGCTCAATAA
AGCACACGTCTGAACTCTTTCCCTACACGACGCTCTTCCGATCTNNNCCCTAGGGCCTATGCTCAATAA
AGCACACGTCTGAACTCTTTCCCTACACGACGCTCTTCCGATCTNNNCCCTAGGGCCTATGCTCAATAA
ACACGTCTGAACTCTTTCCCTACACGACGCTCTTCCGATCTNNNAGAGGCCAGTTGCTTACAAACTACAT
ACACGTCTGAACTCTTTCCCTACACGACGCTCTTCCGATCTNNNAGAGGCCAGTTGCTTACAAACTACAT
ACACGTCTGAACTCTTTCCCTACACGACGCTCTTCCGATCTNNNAGAGGCCAGTTGCTTACAAACTACAT
GAGCACACGTCTGAACTCTTTCCCTACACGACGCTCTTCCGATCTNNNAAGTTGCAGTAAGCCAAGAT

GAGCACACGTCTGAACTCTTTCCCTACACGACGCTCTTCCGATCTNNNAAGTTGCAGTAAGCCAAGAT
GAGCACACGTCTGAACTCTTTCCCTACACGACGCTCTTCCGATCTNNNAAGTTGCAGTAAGCCAAGAT
AGCACACGTCTGAACTCTTTCCCTACACGACGCTCTTCCGATCTNNNGAAAGCTTTGCAAGAGTATGC
AGCACACGTCTGAACTCTTTCCCTACACGACGCTCTTCCGATCTNNNGAAAGCTTTGCAAGAGTATGC
AGCACACGTCTGAACTCTTTCCCTACACGACGCTCTTCCGATCTNNNGAAAGCTTTGCAAGAGTATGC
AGCACACGTCTGAACTCTTTCCCTACACGACGCTCTTCCGATCTNNNATCATCACTACCTCCTTACCCCA
AGCACACGTCTGAACTCTTTCCCTACACGACGCTCTTCCGATCTNNNATCATCACTACCTCCTTACCCCA
AGCACACGTCTGAACTCTTTCCCTACACGACGCTCTTCCGATCTNNNATCATCACTACCTCCTTACCCCA
AGCACACGTCTGAACTCTTTCCCTACACGACGCTCTTCCGATCTNNNGGATCCTTTCAGCTCTGATTCC
AGCACACGTCTGAACTCTTTCCCTACACGACGCTCTTCCGATCTNNNGGATCCTTTCAGCTCTGATTCC
AGCACACGTCTGAACTCTTTCCCTACACGACGCTCTTCCGATCTNNNGGATCCTTTCAGCTCTGATTCC
GAGCACACGTCTGAACTCTTTCCCTACACGACGCTCTTCCGATCTNNNATCATGAGGTCAGGAGATC
GAGCACACGTCTGAACTCTTTCCCTACACGACGCTCTTCCGATCTNNNATCATGAGGTCAGGAGATC
GAGCACACGTCTGAACTCTTTCCCTACACGACGCTCTTCCGATCTNNNATCATGAGGTCAGGAGATC
GAGCACACGTCTGAACTCTTTCCCTACACGACGCTCTTCCGATCTNNNCTCCCAAGTAGCTGGTACTAC
GAGCACACGTCTGAACTCTTTCCCTACACGACGCTCTTCCGATCTNNNCTCCCAAGTAGCTGGTACTAC
GAGCACACGTCTGAACTCTTTCCCTACACGACGCTCTTCCGATCTNNNCTCCCAAGTAGCTGGTACTAC
GCACACGTCTGAACTCTTTCCCTACACGACGCTCTTCCGATCTNNNTTCCTCCAGTTGCTACTCTGTGAC
GCACACGTCTGAACTCTTTCCCTACACGACGCTCTTCCGATCTNNNTTCCTCCAGTTGCTACTCTGTGAC
GCACACGTCTGAACTCTTTCCCTACACGACGCTCTTCCGATCTNNNTTCCTCCAGTTGCTACTCTGTGAC
GAGCACACGTCTGAACTCTTTCCCTACACGACGCTCTTCCGATCTNNNTGCAGAGAAAGAGTGAGAG
GAGCACACGTCTGAACTCTTTCCCTACACGACGCTCTTCCGATCTNNNTGCAGAGAAAGAGTGAGAG
GAGCACACGTCTGAACTCTTTCCCTACACGACGCTCTTCCGATCTNNNTGCAGAGAAAGAGTGAGAG
AGCACACGTCTGAACTCTTTCCCTACACGACGCTCTTCCGATCTNNNCTTTCTTTTACTGGACTGGGG
AGCACACGTCTGAACTCTTTCCCTACACGACGCTCTTCCGATCTNNNCTTTCTTTTACTGGACTGGGG
AGCACACGTCTGAACTCTTTCCCTACACGACGCTCTTCCGATCTNNNCTTTCTTTTACTGGACTGGGG
AGCACACGTCTGAACTCTTTCCCTACACGACGCTCTTCCGATCTNNNTTAGAGAAGTTTAAGAGGCA
AGCACACGTCTGAACTCTTTCCCTACACGACGCTCTTCCGATCTNNNTTAGAGAAGTTTAAGAGGCA
AGCACACGTCTGAACTCTTTCCCTACACGACGCTCTTCCGATCTNNNTTAGAGAAGTTTAAGAGGCA
ACACGTCTGAACTCTTTCCCTACACGACGCTCTTCCGATCTNNNATATATGTGCCATCTTTCTACATCAT
ACACGTCTGAACTCTTTCCCTACACGACGCTCTTCCGATCTNNNATATATGTGCCATCTTTCTACATCAT
ACACGTCTGAACTCTTTCCCTACACGACGCTCTTCCGATCTNNNATATATGTGCCATCTTTCTACATCAT
AGCACACGTCTGAACTCTTTCCCTACACGACGCTCTTCCGATCTNNNGTACTGAGAGTCTCCTTGGGTT
AGCACACGTCTGAACTCTTTCCCTACACGACGCTCTTCCGATCTNNNGTACTGAGAGTCTCCTTGGGTT
AGCACACGTCTGAACTCTTTCCCTACACGACGCTCTTCCGATCTNNNGTACTGAGAGTCTCCTTGGGTT
AGCACACGTCTGAACTCTTTCCCTACACGACGCTCTTCCGATCTNNNAGGACATAGTGGGTGTCAACA
AGCACACGTCTGAACTCTTTCCCTACACGACGCTCTTCCGATCTNNNAGGACATAGTGGGTGTCAACA
AGCACACGTCTGAACTCTTTCCCTACACGACGCTCTTCCGATCTNNNAGGACATAGTGGGTGTCAACA
AGCACACGTCTGAACTCTTTCCCTACACGACGCTCTTCCGATCTNNNGATGGGACTGTGACACTCTAC
AGCACACGTCTGAACTCTTTCCCTACACGACGCTCTTCCGATCTNNNGATGGGACTGTGACACTCTAC
AGCACACGTCTGAACTCTTTCCCTACACGACGCTCTTCCGATCTNNNGATGGGACTGTGACACTCTAC
GCACACGTCTGAACTCTTTCCCTACACGACGCTCTTCCGATCTNNNCTGAACCCGACTGGCTATCATTT
GCACACGTCTGAACTCTTTCCCTACACGACGCTCTTCCGATCTNNNCTGAACCCGACTGGCTATCATTT

GCACACGTCTGAACTCTTTCCCTACACGACGCTCTTCCGATCTNNNCTGAACCCGACTGGCTATCATTT
AGCACACGTCTGAACTCTTTCCCTACACGACGCTCTTCCGATCTNNNCTGTAGTCCCAGCTACTTGGA
AGCACACGTCTGAACTCTTTCCCTACACGACGCTCTTCCGATCTNNNCTGTAGTCCCAGCTACTTGGA
AGCACACGTCTGAACTCTTTCCCTACACGACGCTCTTCCGATCTNNNCTGTAGTCCCAGCTACTTGGA
CACGTCTGAACTCTTTCCCTACACGACGCTCTTCCGATCTNNNTGGGAATCAAGATCAGTTGGTGACA
CACGTCTGAACTCTTTCCCTACACGACGCTCTTCCGATCTNNNTGGGAATCAAGATCAGTTGGTGACA
CACGTCTGAACTCTTTCCCTACACGACGCTCTTCCGATCTNNNTGGGAATCAAGATCAGTTGGTGACA
GAGCACACGTCTGAACTCTTTCCCTACACGACGCTCTTCCGATCTNNNAAAACGAAAATAAGGAGAG
GAGCACACGTCTGAACTCTTTCCCTACACGACGCTCTTCCGATCTNNNAAAACGAAAATAAGGAGAG
GAGCACACGTCTGAACTCTTTCCCTACACGACGCTCTTCCGATCTNNNAAAACGAAAATAAGGAGAG
AGCACACGTCTGAACTCTTTCCCTACACGACGCTCTTCCGATCTNNNCTTGCAATGAGCTGAGATCGTG
AGCACACGTCTGAACTCTTTCCCTACACGACGCTCTTCCGATCTNNNCTTGCAATGAGCTGAGATCGTG
AGCACACGTCTGAACTCTTTCCCTACACGACGCTCTTCCGATCTNNNCTTGCAATGAGCTGAGATCGTG
AGCACACGTCTGAACTCTTTCCCTACACGACGCTCTTCCGATCTNNNACACCTGTAATCCCAGCTACTT
AGCACACGTCTGAACTCTTTCCCTACACGACGCTCTTCCGATCTNNNACACCTGTAATCCCAGCTACTT
AGCACACGTCTGAACTCTTTCCCTACACGACGCTCTTCCGATCTNNNACACCTGTAATCCCAGCTACTT
AGCACACGTCTGAACTCTTTCCCTACACGACGCTCTTCCGATCTNNNACCTGTAGTCCTAGCTACTCAG
AGCACACGTCTGAACTCTTTCCCTACACGACGCTCTTCCGATCTNNNACCTGTAGTCCTAGCTACTCAG
AGCACACGTCTGAACTCTTTCCCTACACGACGCTCTTCCGATCTNNNACCTGTAGTCCTAGCTACTCAG
AGCACACGTCTGAACTCTTTCCCTACACGACGCTCTTCCGATCTNNNGAACGGGAATACCAGAAGGA
AGCACACGTCTGAACTCTTTCCCTACACGACGCTCTTCCGATCTNNNGAACGGGAATACCAGAAGGA
AGCACACGTCTGAACTCTTTCCCTACACGACGCTCTTCCGATCTNNNGAACGGGAATACCAGAAGGA
GAGCACACGTCTGAACTCTTTCCCTACACGACGCTCTTCCGATCTNNNGGGTGTGTTTGTGTAGCA
GAGCACACGTCTGAACTCTTTCCCTACACGACGCTCTTCCGATCTNNNGGGTGTGTTTGTGTAGCA
GAGCACACGTCTGAACTCTTTCCCTACACGACGCTCTTCCGATCTNNNGGGTGTGTTTGTGTAGCA
CACACGTCTGAACTCTTTCCCTACACGACGCTCTTCCGATCTNNNTCCATTTTATTTGAGTTAGTGAGT
CACACGTCTGAACTCTTTCCCTACACGACGCTCTTCCGATCTNNNTCCATTTTATTTGAGTTAGTGAGT
CACACGTCTGAACTCTTTCCCTACACGACGCTCTTCCGATCTNNNTCCATTTTATTTGAGTTAGTGAGT
AGAGCACACGTCTGAACTCTTTCCCTACACGACGCTCTTCCGATCTNNNATACAAAATTAGCTGGGGG
AGAGCACACGTCTGAACTCTTTCCCTACACGACGCTCTTCCGATCTNNNATACAAAATTAGCTGGGGG
AGAGCACACGTCTGAACTCTTTCCCTACACGACGCTCTTCCGATCTNNNATACAAAATTAGCTGGGGG
GCACACGTCTGAACTCTTTCCCTACACGACGCTCTTCCGATCTNNNTTATGCTAAGGTGCCCATATCCC
GCACACGTCTGAACTCTTTCCCTACACGACGCTCTTCCGATCTNNNTTATGCTAAGGTGCCCATATCCC
GCACACGTCTGAACTCTTTCCCTACACGACGCTCTTCCGATCTNNNTTATGCTAAGGTGCCCATATCCC
CACACGTCTGAACTCTTTCCCTACACGACGCTCTTCCGATCTNNNAACAGTTCATTGTTTCTTTTTTCCA
CACACGTCTGAACTCTTTCCCTACACGACGCTCTTCCGATCTNNNAACAGTTCATTGTTTCTTTTTTCCA
CACACGTCTGAACTCTTTCCCTACACGACGCTCTTCCGATCTNNNAACAGTTCATTGTTTCTTTTTTCCA
GAGCACACGTCTGAACTCTTTCCCTACACGACGCTCTTCCGATCTNNNGAGACAGGAGAATGGCTAG
GAGCACACGTCTGAACTCTTTCCCTACACGACGCTCTTCCGATCTNNNGAGACAGGAGAATGGCTAG
GAGCACACGTCTGAACTCTTTCCCTACACGACGCTCTTCCGATCTNNNGAGACAGGAGAATGGCTAG
AGCACACGTCTGAACTCTTTCCCTACACGACGCTCTTCCGATCTNNNAAGTGTTGGGATTACAGGCA
AGCACACGTCTGAACTCTTTCCCTACACGACGCTCTTCCGATCTNNNAAGTGTTGGGATTACAGGCA
AGCACACGTCTGAACTCTTTCCCTACACGACGCTCTTCCGATCTNNNAAGTGTTGGGATTACAGGCA

GAGCACACGTCTGAACTCTTTCCCTACACGACGCTCTTCCGATCTNNNGGTAGGAATTCCGGCTTGTCA
GAGCACACGTCTGAACTCTTTCCCTACACGACGCTCTTCCGATCTNNNGGTAGGAATTCCGGCTTGTCA
GAGCACACGTCTGAACTCTTTCCCTACACGACGCTCTTCCGATCTNNNGGTAGGAATTCCGGCTTGTCA
GAGCACACGTCTGAACTCTTTCCCTACACGACGCTCTTCCGATCTNNNGAGACATGACCACCTCTTCTAC
GAGCACACGTCTGAACTCTTTCCCTACACGACGCTCTTCCGATCTNNNGAGACATGACCACCTCTTCTAC
GAGCACACGTCTGAACTCTTTCCCTACACGACGCTCTTCCGATCTNNNGAGACATGACCACCTCTTCTAC
AGCACACGTCTGAACTCTTTCCCTACACGACGCTCTTCCGATCTNNNAAAAATTTAGCCAGGTGTGGTC
AGCACACGTCTGAACTCTTTCCCTACACGACGCTCTTCCGATCTNNNAAAAATTTAGCCAGGTGTGGTC
AGCACACGTCTGAACTCTTTCCCTACACGACGCTCTTCCGATCTNNNAAAAATTTAGCCAGGTGTGGTC
GAGCACACGTCTGAACTCTTTCCCTACACGACGCTCTTCCGATCTNNNGGGAGGTTAGCACAGTGAAA
GAGCACACGTCTGAACTCTTTCCCTACACGACGCTCTTCCGATCTNNNGGGAGGTTAGCACAGTGAAA
GAGCACACGTCTGAACTCTTTCCCTACACGACGCTCTTCCGATCTNNNGGGAGGTTAGCACAGTGAAA
AGAGCACACGTCTGAACTCTTTCCCTACACGACGCTCTTCCGATCTNNNAAAGTGCTGGGATTACAGGC
AGAGCACACGTCTGAACTCTTTCCCTACACGACGCTCTTCCGATCTNNNAAAGTGCTGGGATTACAGGC
AGAGCACACGTCTGAACTCTTTCCCTACACGACGCTCTTCCGATCTNNNAAAGTGCTGGGATTACAGGC
GCACACGTCTGAACTCTTTCCCTACACGACGCTCTTCCGATCTNNNTAAAAAGGGCTAGTCGTGCTCATC
GCACACGTCTGAACTCTTTCCCTACACGACGCTCTTCCGATCTNNNTAAAAAGGGCTAGTCGTGCTCATC
GCACACGTCTGAACTCTTTCCCTACACGACGCTCTTCCGATCTNNNTAAAAAGGGCTAGTCGTGCTCATC
CACACGTCTGAACTCTTTCCCTACACGACGCTCTTCCGATCTNNNGATCCAGAAGCAGAATAAGAGCAT
CACACGTCTGAACTCTTTCCCTACACGACGCTCTTCCGATCTNNNGATCCAGAAGCAGAATAAGAGCAT
CACACGTCTGAACTCTTTCCCTACACGACGCTCTTCCGATCTNNNGATCCAGAAGCAGAATAAGAGCAT
GAGCACACGTCTGAACTCTTTCCCTACACGACGCTCTTCCGATCTNNNCAATGAGGATGGTAGGGTGC
GAGCACACGTCTGAACTCTTTCCCTACACGACGCTCTTCCGATCTNNNCAATGAGGATGGTAGGGTGC
GAGCACACGTCTGAACTCTTTCCCTACACGACGCTCTTCCGATCTNNNCAATGAGGATGGTAGGGTGC
AGCACACGTCTGAACTCTTTCCCTACACGACGCTCTTCCGATCTNNNGGCTCACACCTCTAATCCAAGTTC
AGCACACGTCTGAACTCTTTCCCTACACGACGCTCTTCCGATCTNNNGGCTCACACCTCTAATCCAAGTTC
AGCACACGTCTGAACTCTTTCCCTACACGACGCTCTTCCGATCTNNNGGCTCACACCTCTAATCCAAGTTC
AGAGCACACGTCTGAACTCTTTCCCTACACGACGCTCTTCCGATCTNNNCCAAAGGAGTGGTAAAGGCT
AGAGCACACGTCTGAACTCTTTCCCTACACGACGCTCTTCCGATCTNNNCCAAAGGAGTGGTAAAGGCT
AGAGCACACGTCTGAACTCTTTCCCTACACGACGCTCTTCCGATCTNNNCCAAAGGAGTGGTAAAGGCT
AGAGCACACGTCTGAACTCTTTCCCTACACGACGCTCTTCCGATCTNNNTCCCCGTGGGAAATACAGAC
AGAGCACACGTCTGAACTCTTTCCCTACACGACGCTCTTCCGATCTNNNTCCCCGTGGGAAATACAGAC
AGAGCACACGTCTGAACTCTTTCCCTACACGACGCTCTTCCGATCTNNNTCCCCGTGGGAAATACAGAC
AGCACACGTCTGAACTCTTTCCCTACACGACGCTCTTCCGATCTNNNAAGACCCTTCCTTCCAAACACTTC
AGCACACGTCTGAACTCTTTCCCTACACGACGCTCTTCCGATCTNNNAAGACCCTTCCTTCCAAACACTTC
AGCACACGTCTGAACTCTTTCCCTACACGACGCTCTTCCGATCTNNNAAGACCCTTCCTTCCAAACACTTC
GCACACGTCTGAACTCTTTCCCTACACGACGCTCTTCCGATCTNNNGTAAAAACTCTGGGGAAATAAAA
GCACACGTCTGAACTCTTTCCCTACACGACGCTCTTCCGATCTNNNGTAAAAACTCTGGGGAAATAAAA
GCACACGTCTGAACTCTTTCCCTACACGACGCTCTTCCGATCTNNNGTAAAAACTCTGGGGAAATAAAA
AGCACACGTCTGAACTCTTTCCCTACACGACGCTCTTCCGATCTNNNGAGTTGCTTGAACCTGGGAGA
AGCACACGTCTGAACTCTTTCCCTACACGACGCTCTTCCGATCTNNNGAGTTGCTTGAACCTGGGAGA
AGCACACGTCTGAACTCTTTCCCTACACGACGCTCTTCCGATCTNNNGAGTTGCTTGAACCTGGGAGA
GAGCACACGTCTGAACTCTTTCCCTACACGACGCTCTTCCGATCTNNNTGAATGAGGATATCTTCATAG

GAGCACACGTCTGAACTCTTTCCCTACACGACGCTCTTCCGATCTNNNTGAATGAGGATATCTTCATAG
GAGCACACGTCTGAACTCTTTCCCTACACGACGCTCTTCCGATCTNNNTGAATGAGGATATCTTCATAG
GAGCACACGTCTGAACTCTTTCCCTACACGACGCTCTTCCGATCTNNNTTGAATCCCATCCCCATTTTCCT
GAGCACACGTCTGAACTCTTTCCCTACACGACGCTCTTCCGATCTNNNTTGAATCCCATCCCCATTTTCCT
GAGCACACGTCTGAACTCTTTCCCTACACGACGCTCTTCCGATCTNNNTTGAATCCCATCCCCATTTTCCT
GAGCACACGTCTGAACTCTTTCCCTACACGACGCTCTTCCGATCTNNNCTCAACCCCTGTAGTGGATTGA
GAGCACACGTCTGAACTCTTTCCCTACACGACGCTCTTCCGATCTNNNCTCAACCCCTGTAGTGGATTGA
GAGCACACGTCTGAACTCTTTCCCTACACGACGCTCTTCCGATCTNNNCTCAACCCCTGTAGTGGATTGA
AGCACACGTCTGAACTCTTTCCCTACACGACGCTCTTCCGATCTNNNCGATGGAGAAAATTTCAGGCCA
AGCACACGTCTGAACTCTTTCCCTACACGACGCTCTTCCGATCTNNNCGATGGAGAAAATTTCAGGCCA
AGCACACGTCTGAACTCTTTCCCTACACGACGCTCTTCCGATCTNNNCGATGGAGAAAATTTCAGGCCA
GAGCACACGTCTGAACTCTTTCCCTACACGACGCTCTTCCGATCTNNNCTGAATAGCTGGGATTACAGG
GAGCACACGTCTGAACTCTTTCCCTACACGACGCTCTTCCGATCTNNNCTGAATAGCTGGGATTACAGG
GAGCACACGTCTGAACTCTTTCCCTACACGACGCTCTTCCGATCTNNNCTGAATAGCTGGGATTACAGG
GCACACGTCTGAACTCTTTCCCTACACGACGCTCTTCCGATCTNNNGCTTGCAGTGAGCTGAGATCACA
GCACACGTCTGAACTCTTTCCCTACACGACGCTCTTCCGATCTNNNGCTTGCAGTGAGCTGAGATCACA
GCACACGTCTGAACTCTTTCCCTACACGACGCTCTTCCGATCTNNNGCTTGCAGTGAGCTGAGATCACA
AGCACACGTCTGAACTCTTTCCCTACACGACGCTCTTCCGATCTNNNTGGTTGGCCAAGAACTGATAAC
AGCACACGTCTGAACTCTTTCCCTACACGACGCTCTTCCGATCTNNNTGGTTGGCCAAGAACTGATAAC
AGCACACGTCTGAACTCTTTCCCTACACGACGCTCTTCCGATCTNNNTGGTTGGCCAAGAACTGATAAC
AGAGCACACGTCTGAACTCTTTCCCTACACGACGCTCTTCCGATCTNNNGTCACAGGTGTCTGACAACA
AGAGCACACGTCTGAACTCTTTCCCTACACGACGCTCTTCCGATCTNNNGTCACAGGTGTCTGACAACA
AGAGCACACGTCTGAACTCTTTCCCTACACGACGCTCTTCCGATCTNNNGTCACAGGTGTCTGACAACA
AGCACACGTCTGAACTCTTTCCCTACACGACGCTCTTCCGATCTNNNCTTTCCTCACTGAGCACTTCTCT
AGCACACGTCTGAACTCTTTCCCTACACGACGCTCTTCCGATCTNNNCTTTCCTCACTGAGCACTTCTCT
AGCACACGTCTGAACTCTTTCCCTACACGACGCTCTTCCGATCTNNNCTTTCCTCACTGAGCACTTCTCT
AGAGCACACGTCTGAACTCTTTCCCTACACGACGCTCTTCCGATCTNNNACTGTGTCTATATGCCTGGAA
AGAGCACACGTCTGAACTCTTTCCCTACACGACGCTCTTCCGATCTNNNACTGTGTCTATATGCCTGGAA
AGAGCACACGTCTGAACTCTTTCCCTACACGACGCTCTTCCGATCTNNNACTGTGTCTATATGCCTGGAA
AGCACACGTCTGAACTCTTTCCCTACACGACGCTCTTCCGATCTNNNGAGGCAGGAGAATTGCTTGAA
AGCACACGTCTGAACTCTTTCCCTACACGACGCTCTTCCGATCTNNNGAGGCAGGAGAATTGCTTGAA
AGCACACGTCTGAACTCTTTCCCTACACGACGCTCTTCCGATCTNNNGAGGCAGGAGAATTGCTTGAA
AGCACACGTCTGAACTCTTTCCCTACACGACGCTCTTCCGATCTNNNGCCAATCAAGCTTTCATTTTGCT
AGCACACGTCTGAACTCTTTCCCTACACGACGCTCTTCCGATCTNNNGCCAATCAAGCTTTCATTTTGCT
AGCACACGTCTGAACTCTTTCCCTACACGACGCTCTTCCGATCTNNNGCCAATCAAGCTTTCATTTTGCT
GAGCACACGTCTGAACTCTTTCCCTACACGACGCTCTTCCGATCTNNNAATCTCAGCTACTTGGGAGGT
GAGCACACGTCTGAACTCTTTCCCTACACGACGCTCTTCCGATCTNNNAATCTCAGCTACTTGGGAGGT
GAGCACACGTCTGAACTCTTTCCCTACACGACGCTCTTCCGATCTNNNAATCTCAGCTACTTGGGAGGT
AGCACACGTCTGAACTCTTTCCCTACACGACGCTCTTCCGATCTNNNCTGGGACATGGCTTCCACAAAC
AGCACACGTCTGAACTCTTTCCCTACACGACGCTCTTCCGATCTNNNCTGGGACATGGCTTCCACAAAC
AGCACACGTCTGAACTCTTTCCCTACACGACGCTCTTCCGATCTNNNCTGGGACATGGCTTCCACAAAC
GAGCACACGTCTGAACTCTTTCCCTACACGACGCTCTTCCGATCTNNNGGGAGCCATAAGGGAACATTC
GAGCACACGTCTGAACTCTTTCCCTACACGACGCTCTTCCGATCTNNNGGGAGCCATAAGGGAACATTC

GAGCACACGTCTGAACTCTTTCCCTACACGACGCTCTTCCGATCTNNNGGGAGCCATAAGGGAACATT
GAGCACACGTCTGAACTCTTTCCCTACACGACGCTCTTCCGATCTNNNGTGAGTGGTATTCCTTGTTTG
GAGCACACGTCTGAACTCTTTCCCTACACGACGCTCTTCCGATCTNNNGTGAGTGGTATTCCTTGTTTG
GAGCACACGTCTGAACTCTTTCCCTACACGACGCTCTTCCGATCTNNNGTGAGTGGTATTCCTTGTTTG
AGCACACGTCTGAACTCTTTCCCTACACGACGCTCTTCCGATCTNNNTCTTCACACTTGTGTTCTGGGA
AGCACACGTCTGAACTCTTTCCCTACACGACGCTCTTCCGATCTNNNTCTTCACACTTGTGTTCTGGGA
AGCACACGTCTGAACTCTTTCCCTACACGACGCTCTTCCGATCTNNNTCTTCACACTTGTGTTCTGGGA
GAGCACACGTCTGAACTCTTTCCCTACACGACGCTCTTCCGATCTNNNCACAGGAACCAAAACAGCTT
GAGCACACGTCTGAACTCTTTCCCTACACGACGCTCTTCCGATCTNNNCACAGGAACCAAAACAGCTT
GAGCACACGTCTGAACTCTTTCCCTACACGACGCTCTTCCGATCTNNNCACAGGAACCAAAACAGCTT
GCACACGTCTGAACTCTTTCCCTACACGACGCTCTTCCGATCTNNNCACCTGTAATCCAGCTACTTGG
GCACACGTCTGAACTCTTTCCCTACACGACGCTCTTCCGATCTNNNCACCTGTAATCCAGCTACTTGG
GCACACGTCTGAACTCTTTCCCTACACGACGCTCTTCCGATCTNNNCACCTGTAATCCAGCTACTTGG
AGCACACGTCTGAACTCTTTCCCTACACGACGCTCTTCCGATCTNNNTGTGGGGATCCAGGATTTGTA
AGCACACGTCTGAACTCTTTCCCTACACGACGCTCTTCCGATCTNNNTGTGGGGATCCAGGATTTGTA
AGCACACGTCTGAACTCTTTCCCTACACGACGCTCTTCCGATCTNNNTGTGGGGATCCAGGATTTGTA
GCACACGTCTGAACTCTTTCCCTACACGACGCTCTTCCGATCTNNNAAGTAGCTGGGATTATAGGCG
GCACACGTCTGAACTCTTTCCCTACACGACGCTCTTCCGATCTNNNAAGTAGCTGGGATTATAGGCG
GCACACGTCTGAACTCTTTCCCTACACGACGCTCTTCCGATCTNNNAAGTAGCTGGGATTATAGGCG
GAGCACACGTCTGAACTCTTTCCCTACACGACGCTCTTCCGATCTNNNCACAACACCTAAAAAAGCTGG
GAGCACACGTCTGAACTCTTTCCCTACACGACGCTCTTCCGATCTNNNCACAACACCTAAAAAAGCTGG
GAGCACACGTCTGAACTCTTTCCCTACACGACGCTCTTCCGATCTNNNCACAACACCTAAAAAAGCTGG
AGCACACGTCTGAACTCTTTCCCTACACGACGCTCTTCCGATCTNNNTCGTTCCTCTGGAAGTTTTGTC
AGCACACGTCTGAACTCTTTCCCTACACGACGCTCTTCCGATCTNNNTCGTTCCTCTGGAAGTTTTGTC
AGCACACGTCTGAACTCTTTCCCTACACGACGCTCTTCCGATCTNNNTCGTTCCTCTGGAAGTTTTGTC
GCACACGTCTGAACTCTTTCCCTACACGACGCTCTTCCGATCTNNNTTGGTGTAAGGCATGATGGTG
GCACACGTCTGAACTCTTTCCCTACACGACGCTCTTCCGATCTNNNTTGGTGTAAGGCATGATGGTG
GCACACGTCTGAACTCTTTCCCTACACGACGCTCTTCCGATCTNNNTTGGTGTAAGGCATGATGGTG
GCACACGTCTGAACTCTTTCCCTACACGACGCTCTTCCGATCTNNNCACAATAGGGCCAGGTAACAGA
GCACACGTCTGAACTCTTTCCCTACACGACGCTCTTCCGATCTNNNCACAATAGGGCCAGGTAACAGA
GCACACGTCTGAACTCTTTCCCTACACGACGCTCTTCCGATCTNNNCACAATAGGGCCAGGTAACAGA
GAGCACACGTCTGAACTCTTTCCCTACACGACGCTCTTCCGATCTNNNACAAGATAAGGGTAATCACG
GAGCACACGTCTGAACTCTTTCCCTACACGACGCTCTTCCGATCTNNNACAAGATAAGGGTAATCACG
GAGCACACGTCTGAACTCTTTCCCTACACGACGCTCTTCCGATCTNNNACAAGATAAGGGTAATCACG
GAGCACACGTCTGAACTCTTTCCCTACACGACGCTCTTCCGATCTNNNCAGGAGAACCCCTTGAACCTG
GAGCACACGTCTGAACTCTTTCCCTACACGACGCTCTTCCGATCTNNNCAGGAGAACCCCTTGAACCTG
GAGCACACGTCTGAACTCTTTCCCTACACGACGCTCTTCCGATCTNNNCAGGAGAACCCCTTGAACCTG
GAGCACACGTCTGAACTCTTTCCCTACACGACGCTCTTCCGATCTNNNAATGGAGGTGGAAGCTTGC
GAGCACACGTCTGAACTCTTTCCCTACACGACGCTCTTCCGATCTNNNAATGGAGGTGGAAGCTTGC
GAGCACACGTCTGAACTCTTTCCCTACACGACGCTCTTCCGATCTNNNAATGGAGGTGGAAGCTTGC
GAGCACACGTCTGAACTCTTTCCCTACACGACGCTCTTCCGATCTNNNCCCAGGTGAAAATTGAGCCA
GAGCACACGTCTGAACTCTTTCCCTACACGACGCTCTTCCGATCTNNNCCCAGGTGAAAATTGAGCCA
GAGCACACGTCTGAACTCTTTCCCTACACGACGCTCTTCCGATCTNNNCCCAGGTGAAAATTGAGCCA

GAGCACACGTCTGAACTCTTTCCCTACACGACGCTCTTCCGATCTNNNCACGTTCCATGAGCACAGATA
GAGCACACGTCTGAACTCTTTCCCTACACGACGCTCTTCCGATCTNNNCACGTTCCATGAGCACAGATA
GAGCACACGTCTGAACTCTTTCCCTACACGACGCTCTTCCGATCTNNNCACGTTCCATGAGCACAGATA
AGCACACGTCTGAACTCTTTCCCTACACGACGCTCTTCCGATCTNNNTCAGATGTTTCCACAAGGCTCTT
AGCACACGTCTGAACTCTTTCCCTACACGACGCTCTTCCGATCTNNNTCAGATGTTTCCACAAGGCTCTT
AGCACACGTCTGAACTCTTTCCCTACACGACGCTCTTCCGATCTNNNTCAGATGTTTCCACAAGGCTCTT
AGAGCACACGTCTGAACTCTTTCCCTACACGACGCTCTTCCGATCTNNNAAAGGAATTTTGGGCCCAA
AGAGCACACGTCTGAACTCTTTCCCTACACGACGCTCTTCCGATCTNNNAAAGGAATTTTGGGCCCAA
AGAGCACACGTCTGAACTCTTTCCCTACACGACGCTCTTCCGATCTNNNAAAGGAATTTTGGGCCCAA
AGCACACGTCTGAACTCTTTCCCTACACGACGCTCTTCCGATCTNNNAATGGCCTAGAGCTAGACATGT
AGCACACGTCTGAACTCTTTCCCTACACGACGCTCTTCCGATCTNNNAATGGCCTAGAGCTAGACATGT
AGCACACGTCTGAACTCTTTCCCTACACGACGCTCTTCCGATCTNNNAATGGCCTAGAGCTAGACATGT
AGCACACGTCTGAACTCTTTCCCTACACGACGCTCTTCCGATCTNNNATAATCCCAGCTACTTGGAAGG
AGCACACGTCTGAACTCTTTCCCTACACGACGCTCTTCCGATCTNNNATAATCCCAGCTACTTGGAAGG
AGCACACGTCTGAACTCTTTCCCTACACGACGCTCTTCCGATCTNNNATAATCCCAGCTACTTGGAAGG
AGCACACGTCTGAACTCTTTCCCTACACGACGCTCTTCCGATCTNNNGTAGAAGGTGAAGGGGAAGCA
AGCACACGTCTGAACTCTTTCCCTACACGACGCTCTTCCGATCTNNNGTAGAAGGTGAAGGGGAAGCA
AGCACACGTCTGAACTCTTTCCCTACACGACGCTCTTCCGATCTNNNGTAGAAGGTGAAGGGGAAGCA
AGAGCACACGTCTGAACTCTTTCCCTACACGACGCTCTTCCGATCTNNNATCCAAGTCCCTTTTCTCTCAC
AGAGCACACGTCTGAACTCTTTCCCTACACGACGCTCTTCCGATCTNNNATCCAAGTCCCTTTTCTCTCAC
AGAGCACACGTCTGAACTCTTTCCCTACACGACGCTCTTCCGATCTNNNATCCAAGTCCCTTTTCTCTCAC
GCACACGTCTGAACTCTTTCCCTACACGACGCTCTTCCGATCTNNNACTTGGTGTTACTATGTTGCCCAC
GCACACGTCTGAACTCTTTCCCTACACGACGCTCTTCCGATCTNNNACTTGGTGTTACTATGTTGCCCAC
GCACACGTCTGAACTCTTTCCCTACACGACGCTCTTCCGATCTNNNACTTGGTGTTACTATGTTGCCCAC
AGAGCACACGTCTGAACTCTTTCCCTACACGACGCTCTTCCGATCTNNNGGCTCACACTTATAATCCCAG
AGAGCACACGTCTGAACTCTTTCCCTACACGACGCTCTTCCGATCTNNNGGCTCACACTTATAATCCCAG
AGAGCACACGTCTGAACTCTTTCCCTACACGACGCTCTTCCGATCTNNNGGCTCACACTTATAATCCCAG
GCACACGTCTGAACTCTTTCCCTACACGACGCTCTTCCGATCTNNNCCACTATGACCACCTTTCAGTCACA
GCACACGTCTGAACTCTTTCCCTACACGACGCTCTTCCGATCTNNNCCACTATGACCACCTTTCAGTCACA
GCACACGTCTGAACTCTTTCCCTACACGACGCTCTTCCGATCTNNNCCACTATGACCACCTTTCAGTCACA
AGCACACGTCTGAACTCTTTCCCTACACGACGCTCTTCCGATCTNNNCAAAACCCTCTCCTGCTTTCAAG
AGCACACGTCTGAACTCTTTCCCTACACGACGCTCTTCCGATCTNNNCAAAACCCTCTCCTGCTTTCAAG
AGCACACGTCTGAACTCTTTCCCTACACGACGCTCTTCCGATCTNNNCAAAACCCTCTCCTGCTTTCAAG
AGCACACGTCTGAACTCTTTCCCTACACGACGCTCTTCCGATCTNNNTGACTGTCCAAGATCACAGAAG
AGCACACGTCTGAACTCTTTCCCTACACGACGCTCTTCCGATCTNNNTGACTGTCCAAGATCACAGAAG
AGCACACGTCTGAACTCTTTCCCTACACGACGCTCTTCCGATCTNNNTGACTGTCCAAGATCACAGAAG
CACACGTCTGAACTCTTTCCCTACACGACGCTCTTCCGATCTNNNTATAACATTGAAGAACAAGAAGCC
CACACGTCTGAACTCTTTCCCTACACGACGCTCTTCCGATCTNNNTATAACATTGAAGAACAAGAAGCC
CACACGTCTGAACTCTTTCCCTACACGACGCTCTTCCGATCTNNNTATAACATTGAAGAACAAGAAGCC
AGCACACGTCTGAACTCTTTCCCTACACGACGCTCTTCCGATCTNNNCTCTGCATGCAGGGTACTTATCT
AGCACACGTCTGAACTCTTTCCCTACACGACGCTCTTCCGATCTNNNCTCTGCATGCAGGGTACTTATCT
AGCACACGTCTGAACTCTTTCCCTACACGACGCTCTTCCGATCTNNNCTCTGCATGCAGGGTACTTATCT
CACACGTCTGAACTCTTTCCCTACACGACGCTCTTCCGATCTNNNCTGTACATGAGGGTGAGGAGGAC

CACACGTCTGAACTCTTTCCCTACACGACGCTCTTCCGATCTNNNCTGTACATGAGGGTGAGGAGGA
CACACGTCTGAACTCTTTCCCTACACGACGCTCTTCCGATCTNNNCTGTACATGAGGGTGAGGAGGA
GAGCACACGTCTGAACTCTTTCCCTACACGACGCTCTTCCGATCTNNNACACACACACACACACACATAT
GAGCACACGTCTGAACTCTTTCCCTACACGACGCTCTTCCGATCTNNNACACACACACACACACACATAT
GAGCACACGTCTGAACTCTTTCCCTACACGACGCTCTTCCGATCTNNNACACACACACACACACACATAT
GCACACGTCTGAACTCTTTCCCTACACGACGCTCTTCCGATCTNNNAGTTTCTTCTTGATTTTTTTGGAC
GCACACGTCTGAACTCTTTCCCTACACGACGCTCTTCCGATCTNNNAGTTTCTTCTTGATTTTTTTGGAC
GCACACGTCTGAACTCTTTCCCTACACGACGCTCTTCCGATCTNNNAGTTTCTTCTTGATTTTTTTGGAC
GAGCACACGTCTGAACTCTTTCCCTACACGACGCTCTTCCGATCTNNNGCCACCATGCCCAGATAATTTT
GAGCACACGTCTGAACTCTTTCCCTACACGACGCTCTTCCGATCTNNNGCCACCATGCCCAGATAATTTT
GAGCACACGTCTGAACTCTTTCCCTACACGACGCTCTTCCGATCTNNNGCCACCATGCCCAGATAATTTT
GAGCACACGTCTGAACTCTTTCCCTACACGACGCTCTTCCGATCTNNNTCTCTCTCTCTCTCTTTCTTT
GAGCACACGTCTGAACTCTTTCCCTACACGACGCTCTTCCGATCTNNNTCTCTCTCTCTCTCTTTCTTT
GAGCACACGTCTGAACTCTTTCCCTACACGACGCTCTTCCGATCTNNNTCTCTCTCTCTCTCTTTCTTT
GAGCACACGTCTGAACTCTTTCCCTACACGACGCTCTTCCGATCTNNNGGCTCATGTCTGTAATCCCAGA
GAGCACACGTCTGAACTCTTTCCCTACACGACGCTCTTCCGATCTNNNGGCTCATGTCTGTAATCCCAGA
GAGCACACGTCTGAACTCTTTCCCTACACGACGCTCTTCCGATCTNNNGGCTCATGTCTGTAATCCCAGA
GAGCACACGTCTGAACTCTTTCCCTACACGACGCTCTTCCGATCTNNNTGCCTAAATGTTCACAAGCCCA
GAGCACACGTCTGAACTCTTTCCCTACACGACGCTCTTCCGATCTNNNTGCCTAAATGTTCACAAGCCCA
GAGCACACGTCTGAACTCTTTCCCTACACGACGCTCTTCCGATCTNNNTGCCTAAATGTTCACAAGCCCA
GAGCACACGTCTGAACTCTTTCCCTACACGACGCTCTTCCGATCTNNNGAAGTCGCAGTGAGCTAAGA
GAGCACACGTCTGAACTCTTTCCCTACACGACGCTCTTCCGATCTNNNGAAGTCGCAGTGAGCTAAGA
GAGCACACGTCTGAACTCTTTCCCTACACGACGCTCTTCCGATCTNNNGAAGTCGCAGTGAGCTAAGA
GCACACGTCTGAACTCTTTCCCTACACGACGCTCTTCCGATCTNNNGGTGAGTACATCCTCCCCTACTCA
GCACACGTCTGAACTCTTTCCCTACACGACGCTCTTCCGATCTNNNGGTGAGTACATCCTCCCCTACTCA
GCACACGTCTGAACTCTTTCCCTACACGACGCTCTTCCGATCTNNNGGTGAGTACATCCTCCCCTACTCA
AGCACACGTCTGAACTCTTTCCCTACACGACGCTCTTCCGATCTNNNGGAGATTGGGTCAGCCAAGTTA
AGCACACGTCTGAACTCTTTCCCTACACGACGCTCTTCCGATCTNNNGGAGATTGGGTCAGCCAAGTTA
AGCACACGTCTGAACTCTTTCCCTACACGACGCTCTTCCGATCTNNNGGAGATTGGGTCAGCCAAGTTA
AGCACACGTCTGAACTCTTTCCCTACACGACGCTCTTCCGATCTNNNAAGATTCATTTCACTCAACAGA
AGCACACGTCTGAACTCTTTCCCTACACGACGCTCTTCCGATCTNNNAAGATTCATTTCACTCAACAGA
AGCACACGTCTGAACTCTTTCCCTACACGACGCTCTTCCGATCTNNNAAGATTCATTTCACTCAACAGA
GAGCACACGTCTGAACTCTTTCCCTACACGACGCTCTTCCGATCTNNNACCCAGAGATACCCAGAAAA
GAGCACACGTCTGAACTCTTTCCCTACACGACGCTCTTCCGATCTNNNACCCAGAGATACCCAGAAAA
GAGCACACGTCTGAACTCTTTCCCTACACGACGCTCTTCCGATCTNNNACCCAGAGATACCCAGAAAA
AGCACACGTCTGAACTCTTTCCCTACACGACGCTCTTCCGATCTNNNCACCTGTAGTCTCAGCTACTTG
AGCACACGTCTGAACTCTTTCCCTACACGACGCTCTTCCGATCTNNNCACCTGTAGTCTCAGCTACTTG
AGCACACGTCTGAACTCTTTCCCTACACGACGCTCTTCCGATCTNNNCACCTGTAGTCTCAGCTACTTG
AGCACACGTCTGAACTCTTTCCCTACACGACGCTCTTCCGATCTNNNGTTGCAGTGAGCCATGATTACA
AGCACACGTCTGAACTCTTTCCCTACACGACGCTCTTCCGATCTNNNGTTGCAGTGAGCCATGATTACA
AGCACACGTCTGAACTCTTTCCCTACACGACGCTCTTCCGATCTNNNGTTGCAGTGAGCCATGATTACA
AGCACACGTCTGAACTCTTTCCCTACACGACGCTCTTCCGATCTNNNAGCCATCACTGATTTCCAGCTA
AGCACACGTCTGAACTCTTTCCCTACACGACGCTCTTCCGATCTNNNAGCCATCACTGATTTCCAGCTA

AGCACACGTCTGAACTCTTTCCCTACACGACGCTCTTCCGATCTNNNAGCCATCACTGATTTCCAGCTA
GAGCACACGTCTGAACTCTTTCCCTACACGACGCTCTTCCGATCTNNNTCAGGAAGTTGGGAGATA
GAGCACACGTCTGAACTCTTTCCCTACACGACGCTCTTCCGATCTNNNTCAGGAAGTTGGGAGATA
GAGCACACGTCTGAACTCTTTCCCTACACGACGCTCTTCCGATCTNNNTCAGGAAGTTGGGAGATA
AGCACACGTCTGAACTCTTTCCCTACACGACGCTCTTCCGATCTNNNTAGTCTCCGTTTGTTGATCCCT
AGCACACGTCTGAACTCTTTCCCTACACGACGCTCTTCCGATCTNNNTAGTCTCCGTTTGTTGATCCCT
AGCACACGTCTGAACTCTTTCCCTACACGACGCTCTTCCGATCTNNNTAGTCTCCGTTTGTTGATCCCT
AGCACACGTCTGAACTCTTTCCCTACACGACGCTCTTCCGATCTNNNTTTCAGTAGAGGCAGGGTTT
AGCACACGTCTGAACTCTTTCCCTACACGACGCTCTTCCGATCTNNNTTTCAGTAGAGGCAGGGTTT
AGCACACGTCTGAACTCTTTCCCTACACGACGCTCTTCCGATCTNNNTTTCAGTAGAGGCAGGGTTT
AGCACACGTCTGAACTCTTTCCCTACACGACGCTCTTCCGATCTNNNCTGGATGGCAAGAACAGACAA
AGCACACGTCTGAACTCTTTCCCTACACGACGCTCTTCCGATCTNNNCTGGATGGCAAGAACAGACAA
AGCACACGTCTGAACTCTTTCCCTACACGACGCTCTTCCGATCTNNNCTGGATGGCAAGAACAGACAA
AGCACACGTCTGAACTCTTTCCCTACACGACGCTCTTCCGATCTNNNACACCTGTAATGTCAGCTACT
AGCACACGTCTGAACTCTTTCCCTACACGACGCTCTTCCGATCTNNNACACCTGTAATGTCAGCTACT
AGCACACGTCTGAACTCTTTCCCTACACGACGCTCTTCCGATCTNNNACACCTGTAATGTCAGCTACT
AGCACACGTCTGAACTCTTTCCCTACACGACGCTCTTCCGATCTNNNGCGGATTACCTGAGGTTATG
AGCACACGTCTGAACTCTTTCCCTACACGACGCTCTTCCGATCTNNNGCGGATTACCTGAGGTTATG
AGCACACGTCTGAACTCTTTCCCTACACGACGCTCTTCCGATCTNNNGCGGATTACCTGAGGTTATG
AGCACACGTCTGAACTCTTTCCCTACACGACGCTCTTCCGATCTNNNGTAAATAGTCTCAGCTCCTCG
AGCACACGTCTGAACTCTTTCCCTACACGACGCTCTTCCGATCTNNNGTAAATAGTCTCAGCTCCTCG
AGCACACGTCTGAACTCTTTCCCTACACGACGCTCTTCCGATCTNNNGTAAATAGTCTCAGCTCCTCG
GCACACGTCTGAACTCTTTCCCTACACGACGCTCTTCCGATCTNNNTTTGCTTGGGATTCTATTACTTG
GCACACGTCTGAACTCTTTCCCTACACGACGCTCTTCCGATCTNNNTTTGCTTGGGATTCTATTACTTG
GCACACGTCTGAACTCTTTCCCTACACGACGCTCTTCCGATCTNNNTTTGCTTGGGATTCTATTACTTG
AGCACACGTCTGAACTCTTTCCCTACACGACGCTCTTCCGATCTNNNTCATCTCACACCTGTCAGAATG
AGCACACGTCTGAACTCTTTCCCTACACGACGCTCTTCCGATCTNNNTCATCTCACACCTGTCAGAATG
AGCACACGTCTGAACTCTTTCCCTACACGACGCTCTTCCGATCTNNNTCATCTCACACCTGTCAGAATG
GAGCACACGTCTGAACTCTTTCCCTACACGACGCTCTTCCGATCTNNNCCATCCACTCCCAATATGGCA
GAGCACACGTCTGAACTCTTTCCCTACACGACGCTCTTCCGATCTNNNCCATCCACTCCCAATATGGCA
GAGCACACGTCTGAACTCTTTCCCTACACGACGCTCTTCCGATCTNNNCCATCCACTCCCAATATGGCA
GAGCACACGTCTGAACTCTTTCCCTACACGACGCTCTTCCGATCTNNNTCTTTTTGGTCTCAGGTGGT
GAGCACACGTCTGAACTCTTTCCCTACACGACGCTCTTCCGATCTNNNTCTTTTTGGTCTCAGGTGGT
GAGCACACGTCTGAACTCTTTCCCTACACGACGCTCTTCCGATCTNNNTCTTTTTGGTCTCAGGTGGT
AGCACACGTCTGAACTCTTTCCCTACACGACGCTCTTCCGATCTNNNGGGACATCTCCAAATTCCTGCA
AGCACACGTCTGAACTCTTTCCCTACACGACGCTCTTCCGATCTNNNGGGACATCTCCAAATTCCTGCA
AGCACACGTCTGAACTCTTTCCCTACACGACGCTCTTCCGATCTNNNGGGACATCTCCAAATTCCTGCA
ACACGTCTGAACTCTTTCCCTACACGACGCTCTTCCGATCTNNNTTCTGGTAAGCACATTTAATAGACA
ACACGTCTGAACTCTTTCCCTACACGACGCTCTTCCGATCTNNNTTCTGGTAAGCACATTTAATAGACA
ACACGTCTGAACTCTTTCCCTACACGACGCTCTTCCGATCTNNNTTCTGGTAAGCACATTTAATAGACA
GCACACGTCTGAACTCTTTCCCTACACGACGCTCTTCCGATCTNNNCACCTGTAATCCCAGCTACTCAG
GCACACGTCTGAACTCTTTCCCTACACGACGCTCTTCCGATCTNNNCACCTGTAATCCCAGCTACTCAG
GCACACGTCTGAACTCTTTCCCTACACGACGCTCTTCCGATCTNNNCACCTGTAATCCCAGCTACTCAG

GAGCACACGTCTGAACTCTTTCCCTACACGACGCTCTTCCGATCTNNNTTGCCAGCAATGAAGACTTTG
GAGCACACGTCTGAACTCTTTCCCTACACGACGCTCTTCCGATCTNNNTTGCCAGCAATGAAGACTTTG
GAGCACACGTCTGAACTCTTTCCCTACACGACGCTCTTCCGATCTNNNTTGCCAGCAATGAAGACTTTG
ACACGTCTGAACTCTTTCCCTACACGACGCTCTTCCGATCTNNNGCTCCCCACTTCCCAGTAGGCATCGG
ACACGTCTGAACTCTTTCCCTACACGACGCTCTTCCGATCTNNNGCTCCCCACTTCCCAGTAGGCATCGG
ACACGTCTGAACTCTTTCCCTACACGACGCTCTTCCGATCTNNNGCTCCCCACTTCCCAGTAGGCATCGG
GAGCACACGTCTGAACTCTTTCCCTACACGACGCTCTTCCGATCTNNNTGCATTCTGCTTAGGTTTCAAG
GAGCACACGTCTGAACTCTTTCCCTACACGACGCTCTTCCGATCTNNNTGCATTCTGCTTAGGTTTCAAG
GAGCACACGTCTGAACTCTTTCCCTACACGACGCTCTTCCGATCTNNNTGCATTCTGCTTAGGTTTCAAG
AGCACACGTCTGAACTCTTTCCCTACACGACGCTCTTCCGATCTNNNCACCTGTAATCCCAGCATTTTGC
AGCACACGTCTGAACTCTTTCCCTACACGACGCTCTTCCGATCTNNNCACCTGTAATCCCAGCATTTTGC
AGCACACGTCTGAACTCTTTCCCTACACGACGCTCTTCCGATCTNNNCACCTGTAATCCCAGCATTTTGC
GAGCACACGTCTGAACTCTTTCCCTACACGACGCTCTTCCGATCTNNNAGGCCAAGTCAGGAGAATCA
GAGCACACGTCTGAACTCTTTCCCTACACGACGCTCTTCCGATCTNNNAGGCCAAGTCAGGAGAATCA
GAGCACACGTCTGAACTCTTTCCCTACACGACGCTCTTCCGATCTNNNAGGCCAAGTCAGGAGAATCA
AGCACACGTCTGAACTCTTTCCCTACACGACGCTCTTCCGATCTNNNGTCCACGTGCATTCTTGTTACAA
AGCACACGTCTGAACTCTTTCCCTACACGACGCTCTTCCGATCTNNNGTCCACGTGCATTCTTGTTACAA
AGCACACGTCTGAACTCTTTCCCTACACGACGCTCTTCCGATCTNNNGTCCACGTGCATTCTTGTTACAA
GAGCACACGTCTGAACTCTTTCCCTACACGACGCTCTTCCGATCTNNNAAGAGAGAGAGTGAAGGGA
GAGCACACGTCTGAACTCTTTCCCTACACGACGCTCTTCCGATCTNNNAAGAGAGAGAGTGAAGGGA
GAGCACACGTCTGAACTCTTTCCCTACACGACGCTCTTCCGATCTNNNAAGAGAGAGAGTGAAGGGA
GAGCACACGTCTGAACTCTTTCCCTACACGACGCTCTTCCGATCTNNNACAAAACTTCTAATCCCGCCCTC
GAGCACACGTCTGAACTCTTTCCCTACACGACGCTCTTCCGATCTNNNACAAAACTTCTAATCCCGCCCTC
GAGCACACGTCTGAACTCTTTCCCTACACGACGCTCTTCCGATCTNNNACAAAACTTCTAATCCCGCCCTC
AGCACACGTCTGAACTCTTTCCCTACACGACGCTCTTCCGATCTNNNAGATATTCCATGCCAATGAAAT
AGCACACGTCTGAACTCTTTCCCTACACGACGCTCTTCCGATCTNNNAGATATTCCATGCCAATGAAAT
AGCACACGTCTGAACTCTTTCCCTACACGACGCTCTTCCGATCTNNNAGATATTCCATGCCAATGAAAT
GAGCACACGTCTGAACTCTTTCCCTACACGACGCTCTTCCGATCTNNNAGACCTGTCTTCAAGTTCTGAA
GAGCACACGTCTGAACTCTTTCCCTACACGACGCTCTTCCGATCTNNNAGACCTGTCTTCAAGTTCTGAA
GAGCACACGTCTGAACTCTTTCCCTACACGACGCTCTTCCGATCTNNNAGACCTGTCTTCAAGTTCTGAA
GCACACGTCTGAACTCTTTCCCTACACGACGCTCTTCCGATCTNNNTCCACAAACACTCACTGTCAGATCA
GCACACGTCTGAACTCTTTCCCTACACGACGCTCTTCCGATCTNNNTCCACAAACACTCACTGTCAGATCA
GCACACGTCTGAACTCTTTCCCTACACGACGCTCTTCCGATCTNNNTCCACAAACACTCACTGTCAGATCA
GAGCACACGTCTGAACTCTTTCCCTACACGACGCTCTTCCGATCTNNNATTAGCGCTAAATGCCATGCA
GAGCACACGTCTGAACTCTTTCCCTACACGACGCTCTTCCGATCTNNNATTAGCGCTAAATGCCATGCA
GAGCACACGTCTGAACTCTTTCCCTACACGACGCTCTTCCGATCTNNNATTAGCGCTAAATGCCATGCA
GAGCACACGTCTGAACTCTTTCCCTACACGACGCTCTTCCGATCTNNNGATCACTGACAGGCTAAAAGC
GAGCACACGTCTGAACTCTTTCCCTACACGACGCTCTTCCGATCTNNNGATCACTGACAGGCTAAAAGC
GAGCACACGTCTGAACTCTTTCCCTACACGACGCTCTTCCGATCTNNNGATCACTGACAGGCTAAAAGC
GAGCACACGTCTGAACTCTTTCCCTACACGACGCTCTTCCGATCTNNNAAGACAGACTTTGCCATAGTGC
GAGCACACGTCTGAACTCTTTCCCTACACGACGCTCTTCCGATCTNNNAAGACAGACTTTGCCATAGTGC
GAGCACACGTCTGAACTCTTTCCCTACACGACGCTCTTCCGATCTNNNAAGACAGACTTTGCCATAGTGC
AGCACACGTCTGAACTCTTTCCCTACACGACGCTCTTCCGATCTNNNCTGATGCCTATAATCCCAGCACA

AGCACACGTCTGAACTCTTTCCCTACACGACGCTCTTCCGATCTNNNCTGATGCCTATAATCCCAGCAC
AGCACACGTCTGAACTCTTTCCCTACACGACGCTCTTCCGATCTNNNCTGATGCCTATAATCCCAGCAC
GAGCACACGTCTGAACTCTTTCCCTACACGACGCTCTTCCGATCTNNNCCTGGGCTACAAACCTTTACA
GAGCACACGTCTGAACTCTTTCCCTACACGACGCTCTTCCGATCTNNNCCTGGGCTACAAACCTTTACA
GAGCACACGTCTGAACTCTTTCCCTACACGACGCTCTTCCGATCTNNNCCTGGGCTACAAACCTTTACA
AGCACACGTCTGAACTCTTTCCCTACACGACGCTCTTCCGATCTNNNGAGTTTTCAGGCATGAGTGG
AGCACACGTCTGAACTCTTTCCCTACACGACGCTCTTCCGATCTNNNGAGTTTTCAGGCATGAGTGG
AGCACACGTCTGAACTCTTTCCCTACACGACGCTCTTCCGATCTNNNGAGTTTTCAGGCATGAGTGG
ACACGTCTGAACTCTTTCCCTACACGACGCTCTTCCGATCTNNNAGACTCTATCAAGACAGTAAAAAC
ACACGTCTGAACTCTTTCCCTACACGACGCTCTTCCGATCTNNNAGACTCTATCAAGACAGTAAAAAC
ACACGTCTGAACTCTTTCCCTACACGACGCTCTTCCGATCTNNNAGACTCTATCAAGACAGTAAAAAC
GCACACGTCTGAACTCTTTCCCTACACGACGCTCTTCCGATCTNNNAGAAAAAGCAAAGCAGTGTAG
GCACACGTCTGAACTCTTTCCCTACACGACGCTCTTCCGATCTNNNAGAAAAAGCAAAGCAGTGTAG
GCACACGTCTGAACTCTTTCCCTACACGACGCTCTTCCGATCTNNNAGAAAAAGCAAAGCAGTGTAG
AGCACACGTCTGAACTCTTTCCCTACACGACGCTCTTCCGATCTNNNTGTAATCTCAGCACTTTGGGAG
AGCACACGTCTGAACTCTTTCCCTACACGACGCTCTTCCGATCTNNNTGTAATCTCAGCACTTTGGGAG
AGCACACGTCTGAACTCTTTCCCTACACGACGCTCTTCCGATCTNNNTGTAATCTCAGCACTTTGGGAG
AGCACACGTCTGAACTCTTTCCCTACACGACGCTCTTCCGATCTNNNTCTAACAGGCACCAGAAGTCAT
AGCACACGTCTGAACTCTTTCCCTACACGACGCTCTTCCGATCTNNNTCTAACAGGCACCAGAAGTCAT
AGCACACGTCTGAACTCTTTCCCTACACGACGCTCTTCCGATCTNNNTCTAACAGGCACCAGAAGTCAT
GCACACGTCTGAACTCTTTCCCTACACGACGCTCTTCCGATCTNNNGGTGGCAAGTATCTGTAATCCC
GCACACGTCTGAACTCTTTCCCTACACGACGCTCTTCCGATCTNNNGGTGGCAAGTATCTGTAATCCC
GCACACGTCTGAACTCTTTCCCTACACGACGCTCTTCCGATCTNNNGGTGGCAAGTATCTGTAATCCC
AGCACACGTCTGAACTCTTTCCCTACACGACGCTCTTCCGATCTNNNGTGTGTGTGAGAGAGAG
AGCACACGTCTGAACTCTTTCCCTACACGACGCTCTTCCGATCTNNNGTGTGTGTGAGAGAGAG
AGCACACGTCTGAACTCTTTCCCTACACGACGCTCTTCCGATCTNNNGTGTGTGTGAGAGAGAG
AGCACACGTCTGAACTCTTTCCCTACACGACGCTCTTCCGATCTNNNCCCTGGGTTGGTCATATGACA
AGCACACGTCTGAACTCTTTCCCTACACGACGCTCTTCCGATCTNNNCCCTGGGTTGGTCATATGACA
AGCACACGTCTGAACTCTTTCCCTACACGACGCTCTTCCGATCTNNNCCCTGGGTTGGTCATATGACA
GCACACGTCTGAACTCTTTCCCTACACGACGCTCTTCCGATCTNNNAGAATTGCTTGAACCCAGGAGA
GCACACGTCTGAACTCTTTCCCTACACGACGCTCTTCCGATCTNNNAGAATTGCTTGAACCCAGGAGA
GCACACGTCTGAACTCTTTCCCTACACGACGCTCTTCCGATCTNNNAGAATTGCTTGAACCCAGGAGA
GCACACGTCTGAACTCTTTCCCTACACGACGCTCTTCCGATCTNNNCGCAGTCACTAACTTACCCTCAT
GCACACGTCTGAACTCTTTCCCTACACGACGCTCTTCCGATCTNNNCGCAGTCACTAACTTACCCTCAT
GCACACGTCTGAACTCTTTCCCTACACGACGCTCTTCCGATCTNNNCGCAGTCACTAACTTACCCTCAT
GCACACGTCTGAACTCTTTCCCTACACGACGCTCTTCCGATCTNNNCTAGCAGCACAATTAGCACTCAG
GCACACGTCTGAACTCTTTCCCTACACGACGCTCTTCCGATCTNNNCTAGCAGCACAATTAGCACTCAG
GCACACGTCTGAACTCTTTCCCTACACGACGCTCTTCCGATCTNNNCTAGCAGCACAATTAGCACTCAG
GAGCACACGTCTGAACTCTTTCCCTACACGACGCTCTTCCGATCTNNNTGCATTCCAACATTTTTCTGA
GAGCACACGTCTGAACTCTTTCCCTACACGACGCTCTTCCGATCTNNNTGCATTCCAACATTTTTCTGA
GAGCACACGTCTGAACTCTTTCCCTACACGACGCTCTTCCGATCTNNNTGCATTCCAACATTTTTCTGA
AGCACACGTCTGAACTCTTTCCCTACACGACGCTCTTCCGATCTNNNAAATCAACCAGCAGAAGAGCC
AGCACACGTCTGAACTCTTTCCCTACACGACGCTCTTCCGATCTNNNAAATCAACCAGCAGAAGAGCC

AGCACACGTCTGAACTCTTTCCCTACACGACGCTCTTCCGATCTNNNAAATCAACCAGCAGAAGAGCCT
AGAGCACACGTCTGAACTCTTTCCCTACACGACGCTCTTCCGATCTNNNATGACAATCACTTGAACCTG
AGAGCACACGTCTGAACTCTTTCCCTACACGACGCTCTTCCGATCTNNNATGACAATCACTTGAACCTG
AGAGCACACGTCTGAACTCTTTCCCTACACGACGCTCTTCCGATCTNNNATGACAATCACTTGAACCTG
AGCACACGTCTGAACTCTTTCCCTACACGACGCTCTTCCGATCTNNNGATCCACTGAGCTAGACCACTT
AGCACACGTCTGAACTCTTTCCCTACACGACGCTCTTCCGATCTNNNGATCCACTGAGCTAGACCACTT
AGCACACGTCTGAACTCTTTCCCTACACGACGCTCTTCCGATCTNNNGATCCACTGAGCTAGACCACTT
GCACACGTCTGAACTCTTTCCCTACACGACGCTCTTCCGATCTNNNATAAGGAGAGGGAACAGTGGA
GCACACGTCTGAACTCTTTCCCTACACGACGCTCTTCCGATCTNNNATAAGGAGAGGGAACAGTGGA
GCACACGTCTGAACTCTTTCCCTACACGACGCTCTTCCGATCTNNNATAAGGAGAGGGAACAGTGGA
AGCACACGTCTGAACTCTTTCCCTACACGACGCTCTTCCGATCTNNNGGTGACAAAGCAAGACTCTTG
AGCACACGTCTGAACTCTTTCCCTACACGACGCTCTTCCGATCTNNNGGTGACAAAGCAAGACTCTTG
AGCACACGTCTGAACTCTTTCCCTACACGACGCTCTTCCGATCTNNNGGTGACAAAGCAAGACTCTTG
AGCACACGTCTGAACTCTTTCCCTACACGACGCTCTTCCGATCTNNNCTGCAACTGGGTACATTTTGGA
AGCACACGTCTGAACTCTTTCCCTACACGACGCTCTTCCGATCTNNNCTGCAACTGGGTACATTTTGGA
AGCACACGTCTGAACTCTTTCCCTACACGACGCTCTTCCGATCTNNNCTGCAACTGGGTACATTTTGGA
GAGCACACGTCTGAACTCTTTCCCTACACGACGCTCTTCCGATCTNNNTGTAGTACCAGCTACTCGGG
GAGCACACGTCTGAACTCTTTCCCTACACGACGCTCTTCCGATCTNNNTGTAGTACCAGCTACTCGGG
GAGCACACGTCTGAACTCTTTCCCTACACGACGCTCTTCCGATCTNNNTGTAGTACCAGCTACTCGGG
AGCACACGTCTGAACTCTTTCCCTACACGACGCTCTTCCGATCTNNNAGTATCTTTGTGGCGTTCTCTG
AGCACACGTCTGAACTCTTTCCCTACACGACGCTCTTCCGATCTNNNAGTATCTTTGTGGCGTTCTCTG
AGCACACGTCTGAACTCTTTCCCTACACGACGCTCTTCCGATCTNNNAGTATCTTTGTGGCGTTCTCTG
AGCACACGTCTGAACTCTTTCCCTACACGACGCTCTTCCGATCTNNNCAGCTGGATTTGAAGATAGTCC
AGCACACGTCTGAACTCTTTCCCTACACGACGCTCTTCCGATCTNNNCAGCTGGATTTGAAGATAGTCC
AGCACACGTCTGAACTCTTTCCCTACACGACGCTCTTCCGATCTNNNCAGCTGGATTTGAAGATAGTCC
GCACACGTCTGAACTCTTTCCCTACACGACGCTCTTCCGATCTNNNCCCATCTCTGCCATTACAACTCTC
GCACACGTCTGAACTCTTTCCCTACACGACGCTCTTCCGATCTNNNCCCATCTCTGCCATTACAACTCTC
GCACACGTCTGAACTCTTTCCCTACACGACGCTCTTCCGATCTNNNCCCATCTCTGCCATTACAACTCTC
AGCACACGTCTGAACTCTTTCCCTACACGACGCTCTTCCGATCTNNNTGTCCTACTCTGTTCCTGATGGA
AGCACACGTCTGAACTCTTTCCCTACACGACGCTCTTCCGATCTNNNTGTCCTACTCTGTTCCTGATGGA
AGCACACGTCTGAACTCTTTCCCTACACGACGCTCTTCCGATCTNNNTGTCCTACTCTGTTCCTGATGGA
GAGCACACGTCTGAACTCTTTCCCTACACGACGCTCTTCCGATCTNNNGACAATTCCCACCCTACTCTAG
GAGCACACGTCTGAACTCTTTCCCTACACGACGCTCTTCCGATCTNNNGACAATTCCCACCCTACTCTAG
GAGCACACGTCTGAACTCTTTCCCTACACGACGCTCTTCCGATCTNNNGACAATTCCCACCCTACTCTAG
AGCACACGTCTGAACTCTTTCCCTACACGACGCTCTTCCGATCTNNNCTTCAGCTTCTTTCATTGACCCG
AGCACACGTCTGAACTCTTTCCCTACACGACGCTCTTCCGATCTNNNCTTCAGCTTCTTTCATTGACCCG
AGCACACGTCTGAACTCTTTCCCTACACGACGCTCTTCCGATCTNNNCTTCAGCTTCTTTCATTGACCCG
GCACACGTCTGAACTCTTTCCCTACACGACGCTCTTCCGATCTNNNAGCAAACCAGAAAACCCAACTCT
GCACACGTCTGAACTCTTTCCCTACACGACGCTCTTCCGATCTNNNAGCAAACCAGAAAACCCAACTCT
GCACACGTCTGAACTCTTTCCCTACACGACGCTCTTCCGATCTNNNAGCAAACCAGAAAACCCAACTCT
AGCACACGTCTGAACTCTTTCCCTACACGACGCTCTTCCGATCTNNNGTGAGCAGAGATCATGCCACT
AGCACACGTCTGAACTCTTTCCCTACACGACGCTCTTCCGATCTNNNGTGAGCAGAGATCATGCCACT
AGCACACGTCTGAACTCTTTCCCTACACGACGCTCTTCCGATCTNNNGTGAGCAGAGATCATGCCACT

CACACGTCTGAACTCTTTCCCTACACGACGCTCTTCCGATCTNNNACAGTGAAACCCCGTCTCTACTAC
CACACGTCTGAACTCTTTCCCTACACGACGCTCTTCCGATCTNNNACAGTGAAACCCCGTCTCTACTAC
CACACGTCTGAACTCTTTCCCTACACGACGCTCTTCCGATCTNNNACAGTGAAACCCCGTCTCTACTAC
AGCACACGTCTGAACTCTTTCCCTACACGACGCTCTTCCGATCTNNNTCCTGTCCTCCTCTTTTCTCTCTC
AGCACACGTCTGAACTCTTTCCCTACACGACGCTCTTCCGATCTNNNTCCTGTCCTCCTCTTTTCTCTCTC
AGCACACGTCTGAACTCTTTCCCTACACGACGCTCTTCCGATCTNNNTCCTGTCCTCCTCTTTTCTCTCTC
AGAGCACACGTCTGAACTCTTTCCCTACACGACGCTCTTCCGATCTNNNGAAGAGACTCAGGTCAAGA
AGAGCACACGTCTGAACTCTTTCCCTACACGACGCTCTTCCGATCTNNNGAAGAGACTCAGGTCAAGA
AGAGCACACGTCTGAACTCTTTCCCTACACGACGCTCTTCCGATCTNNNGAAGAGACTCAGGTCAAGA
AGAGCACACGTCTGAACTCTTTCCCTACACGACGCTCTTCCGATCTNNNGTGTTCAGATTGAGCCATTG
AGAGCACACGTCTGAACTCTTTCCCTACACGACGCTCTTCCGATCTNNNGTGTTCAGATTGAGCCATTG
AGAGCACACGTCTGAACTCTTTCCCTACACGACGCTCTTCCGATCTNNNGTGTTCAGATTGAGCCATTG
GAGCACACGTCTGAACTCTTTCCCTACACGACGCTCTTCCGATCTNNNTTCTCAGATTCTTGGCTCAAGC
GAGCACACGTCTGAACTCTTTCCCTACACGACGCTCTTCCGATCTNNNTTCTCAGATTCTTGGCTCAAGC
GAGCACACGTCTGAACTCTTTCCCTACACGACGCTCTTCCGATCTNNNTTCTCAGATTCTTGGCTCAAGC
AGCACACGTCTGAACTCTTTCCCTACACGACGCTCTTCCGATCTNNNGTTCAAGTGATTCTCCTCCCTCA
AGCACACGTCTGAACTCTTTCCCTACACGACGCTCTTCCGATCTNNNGTTCAAGTGATTCTCCTCCCTCA
AGCACACGTCTGAACTCTTTCCCTACACGACGCTCTTCCGATCTNNNGTTCAAGTGATTCTCCTCCCTCA
AGCACACGTCTGAACTCTTTCCCTACACGACGCTCTTCCGATCTNNNGGGTGACAGAGTGAGACTACA
AGCACACGTCTGAACTCTTTCCCTACACGACGCTCTTCCGATCTNNNGGGTGACAGAGTGAGACTACA
AGCACACGTCTGAACTCTTTCCCTACACGACGCTCTTCCGATCTNNNGGGTGACAGAGTGAGACTACA
GCACACGTCTGAACTCTTTCCCTACACGACGCTCTTCCGATCTNNNCTCCCTGGTCTGTGTTTGTCTAG
GCACACGTCTGAACTCTTTCCCTACACGACGCTCTTCCGATCTNNNCTCCCTGGTCTGTGTTTGTCTAG
GCACACGTCTGAACTCTTTCCCTACACGACGCTCTTCCGATCTNNNCTCCCTGGTCTGTGTTTGTCTAG
AGCACACGTCTGAACTCTTTCCCTACACGACGCTCTTCCGATCTNNNAGCCAGACTCTTGAAGCTAAAC
AGCACACGTCTGAACTCTTTCCCTACACGACGCTCTTCCGATCTNNNAGCCAGACTCTTGAAGCTAAAC
AGCACACGTCTGAACTCTTTCCCTACACGACGCTCTTCCGATCTNNNAGCCAGACTCTTGAAGCTAAAC
GAGCACACGTCTGAACTCTTTCCCTACACGACGCTCTTCCGATCTNNNGGTGACAGAGAGAGATTCCG
GAGCACACGTCTGAACTCTTTCCCTACACGACGCTCTTCCGATCTNNNGGTGACAGAGAGAGATTCCG
GAGCACACGTCTGAACTCTTTCCCTACACGACGCTCTTCCGATCTNNNGGTGACAGAGAGAGATTCCG
AGCACACGTCTGAACTCTTTCCCTACACGACGCTCTTCCGATCTNNNAAGGAATTGTGTGTGTGTGTG
AGCACACGTCTGAACTCTTTCCCTACACGACGCTCTTCCGATCTNNNAAGGAATTGTGTGTGTGTGTG
AGCACACGTCTGAACTCTTTCCCTACACGACGCTCTTCCGATCTNNNAAGGAATTGTGTGTGTGTGTG
GAGCACACGTCTGAACTCTTTCCCTACACGACGCTCTTCCGATCTNNNGCAGTTTCCAGTCTGACTTTTC
GAGCACACGTCTGAACTCTTTCCCTACACGACGCTCTTCCGATCTNNNGCAGTTTCCAGTCTGACTTTTC
GAGCACACGTCTGAACTCTTTCCCTACACGACGCTCTTCCGATCTNNNGCAGTTTCCAGTCTGACTTTTC
GCACACGTCTGAACTCTTTCCCTACACGACGCTCTTCCGATCTNNNATGGATTGGTGGAATGGTAGTC
GCACACGTCTGAACTCTTTCCCTACACGACGCTCTTCCGATCTNNNATGGATTGGTGGAATGGTAGTC
GCACACGTCTGAACTCTTTCCCTACACGACGCTCTTCCGATCTNNNATGGATTGGTGGAATGGTAGTC
AGCACACGTCTGAACTCTTTCCCTACACGACGCTCTTCCGATCTNNNACAAAAAATTAGCTTGGCGTGC
AGCACACGTCTGAACTCTTTCCCTACACGACGCTCTTCCGATCTNNNACAAAAAATTAGCTTGGCGTGC
AGCACACGTCTGAACTCTTTCCCTACACGACGCTCTTCCGATCTNNNACAAAAAATTAGCTTGGCGTGC
GCACACGTCTGAACTCTTTCCCTACACGACGCTCTTCCGATCTNNNTATCAAGGACTTCTTTTAAGTGAC

CACACGTCTGAACTCTTTCCCTACACGACGCTCTTCCGATCTNNNTATCAAGGACTTCTTTTAAGTGAC

CACACGTCTGAACTCTTTCCCTACACGACGCTCTTCCGATCTNNNTATCAAGGACTTCTTTTAAGTGAC

AGCACACGTCTGAACTCTTTCCCTACACGACGCTCTTCCGATCTNNNGAACCTAGAATTGGGGCTGAG

AGCACACGTCTGAACTCTTTCCCTACACGACGCTCTTCCGATCTNNNGAACCTAGAATTGGGGCTGAG

AGCACACGTCTGAACTCTTTCCCTACACGACGCTCTTCCGATCTNNNGAACCTAGAATTGGGGCTGAG

GCACACGTCTGAACTCTTTCCCTACACGACGCTCTTCCGATCTNNNAAGGTGAGAAGGGAGTTTCAC

GCACACGTCTGAACTCTTTCCCTACACGACGCTCTTCCGATCTNNNAAGGTGAGAAGGGAGTTTCAC

GCACACGTCTGAACTCTTTCCCTACACGACGCTCTTCCGATCTNNNAAGGTGAGAAGGGAGTTTCAC

GAGCACACGTCTGAACTCTTTCCCTACACGACGCTCTTCCGATCTNNNTCTGTGATTTCTTTCAGCAGT

GAGCACACGTCTGAACTCTTTCCCTACACGACGCTCTTCCGATCTNNNTCTGTGATTTCTTTCAGCAGT

GAGCACACGTCTGAACTCTTTCCCTACACGACGCTCTTCCGATCTNNNTCTGTGATTTCTTTCAGCAGT

AGCACACGTCTGAACTCTTTCCCTACACGACGCTCTTCCGATCTNNNCTCCTTCTCTCCTTCCCTCCTTCCA

AGCACACGTCTGAACTCTTTCCCTACACGACGCTCTTCCGATCTNNNCTCCTTCTCTCCTTCCCTCCTTCCA

AGCACACGTCTGAACTCTTTCCCTACACGACGCTCTTCCGATCTNNNCTCCTTCTCTCCTTCCCTCCTTCCA

GCACACGTCTGAACTCTTTCCCTACACGACGCTCTTCCGATCTNNNGGGGGAACATCAGGTAGGGC

GCACACGTCTGAACTCTTTCCCTACACGACGCTCTTCCGATCTNNNGGGGGAACATCAGGTAGGGC

GCACACGTCTGAACTCTTTCCCTACACGACGCTCTTCCGATCTNNNGGGGGAACATCAGGTAGGGC

GCACACGTCTGAACTCTTTCCCTACACGACGCTCTTCCGATCTNNNATTTCTTCAGCATCGCTCCTACCC

GCACACGTCTGAACTCTTTCCCTACACGACGCTCTTCCGATCTNNNATTTCTTCAGCATCGCTCCTACCC

GCACACGTCTGAACTCTTTCCCTACACGACGCTCTTCCGATCTNNNATTTCTTCAGCATCGCTCCTACCC

AGCACACGTCTGAACTCTTTCCCTACACGACGCTCTTCCGATCTNNNGAGAAAATGAGGAAGCAAAA

AGCACACGTCTGAACTCTTTCCCTACACGACGCTCTTCCGATCTNNNGAGAAAATGAGGAAGCAAAA

AGCACACGTCTGAACTCTTTCCCTACACGACGCTCTTCCGATCTNNNGAGAAAATGAGGAAGCAAAA

AGAGCACACGTCTGAACTCTTTCCCTACACGACGCTCTTCCGATCTNNNCACCTCAAGGAGCTAGAGA

AGAGCACACGTCTGAACTCTTTCCCTACACGACGCTCTTCCGATCTNNNCACCTCAAGGAGCTAGAGA

AGAGCACACGTCTGAACTCTTTCCCTACACGACGCTCTTCCGATCTNNNCACCTCAAGGAGCTAGAGA

AGCACACGTCTGAACTCTTTCCCTACACGACGCTCTTCCGATCTNNNGCTCCAAGTGTTTGATGAACT

AGCACACGTCTGAACTCTTTCCCTACACGACGCTCTTCCGATCTNNNGCTCCAAGTGTTTGATGAACT

AGCACACGTCTGAACTCTTTCCCTACACGACGCTCTTCCGATCTNNNGCTCCAAGTGTTTGATGAACT

GAGCACACGTCTGAACTCTTTCCCTACACGACGCTCTTCCGATCTNNNCCAAGTGAGTAATTCAGCACA

GAGCACACGTCTGAACTCTTTCCCTACACGACGCTCTTCCGATCTNNNCCAAGTGAGTAATTCAGCACA

GAGCACACGTCTGAACTCTTTCCCTACACGACGCTCTTCCGATCTNNNCCAAGTGAGTAATTCAGCACA

AGCACACGTCTGAACTCTTTCCCTACACGACGCTCTTCCGATCTNNNGAGAATCGCTTGAACCAGGGA

AGCACACGTCTGAACTCTTTCCCTACACGACGCTCTTCCGATCTNNNGAGAATCGCTTGAACCAGGGA

AGCACACGTCTGAACTCTTTCCCTACACGACGCTCTTCCGATCTNNNGAGAATCGCTTGAACCAGGGA

AGCACACGTCTGAACTCTTTCCCTACACGACGCTCTTCCGATCTNNNGCCACAATTACCAGCGACAAAT

AGCACACGTCTGAACTCTTTCCCTACACGACGCTCTTCCGATCTNNNGCCACAATTACCAGCGACAAAT

AGCACACGTCTGAACTCTTTCCCTACACGACGCTCTTCCGATCTNNNGCCACAATTACCAGCGACAAAT

AGCACACGTCTGAACTCTTTCCCTACACGACGCTCTTCCGATCTNNNCCCAGGAGAGAAAACTAGACT

AGCACACGTCTGAACTCTTTCCCTACACGACGCTCTTCCGATCTNNNCCCAGGAGAGAAAACTAGACT

AGCACACGTCTGAACTCTTTCCCTACACGACGCTCTTCCGATCTNNNCCCAGGAGAGAAAACTAGACT

AGCACACGTCTGAACTCTTTCCCTACACGACGCTCTTCCGATCTNNNGAGATCACACCACTGCACTCCCA

AGCACACGTCTGAACTCTTTCCCTACACGACGCTCTTCCGATCTNNNGAGATCACACCACTGCACTCCCA

AGCACACGTCTGAACTCTTTCCCTACACGACGCTCTTCCGATCTNNNGAGATCACACCACTGCACTCCCA
AGCACACGTCTGAACTCTTTCCCTACACGACGCTCTTCCGATCTNNNAAAAATTAGCCGGGCGTGGTC
AGCACACGTCTGAACTCTTTCCCTACACGACGCTCTTCCGATCTNNNAAAAATTAGCCGGGCGTGGTC
AGCACACGTCTGAACTCTTTCCCTACACGACGCTCTTCCGATCTNNNAAAAATTAGCCGGGCGTGGTC
GAGCACACGTCTGAACTCTTTCCCTACACGACGCTCTTCCGATCTNNNGAGAATGGCATGAACCCAGG
GAGCACACGTCTGAACTCTTTCCCTACACGACGCTCTTCCGATCTNNNGAGAATGGCATGAACCCAGG
GAGCACACGTCTGAACTCTTTCCCTACACGACGCTCTTCCGATCTNNNGAGAATGGCATGAACCCAGG
AGCACACGTCTGAACTCTTTCCCTACACGACGCTCTTCCGATCTNNNCTAGCATCTATTGCTGGCACCAT
AGCACACGTCTGAACTCTTTCCCTACACGACGCTCTTCCGATCTNNNCTAGCATCTATTGCTGGCACCAT
AGCACACGTCTGAACTCTTTCCCTACACGACGCTCTTCCGATCTNNNCTAGCATCTATTGCTGGCACCAT
GAGCACACGTCTGAACTCTTTCCCTACACGACGCTCTTCCGATCTNNNGAGAATCGCTTGAATCCTGGA
GAGCACACGTCTGAACTCTTTCCCTACACGACGCTCTTCCGATCTNNNGAGAATCGCTTGAATCCTGGA
GAGCACACGTCTGAACTCTTTCCCTACACGACGCTCTTCCGATCTNNNGAGAATCGCTTGAATCCTGGA
AGCACACGTCTGAACTCTTTCCCTACACGACGCTCTTCCGATCTNNNATTGTCCATGTCACACGTGTTTC
AGCACACGTCTGAACTCTTTCCCTACACGACGCTCTTCCGATCTNNNATTGTCCATGTCACACGTGTTTC
AGCACACGTCTGAACTCTTTCCCTACACGACGCTCTTCCGATCTNNNATTGTCCATGTCACACGTGTTTC
GAGCACACGTCTGAACTCTTTCCCTACACGACGCTCTTCCGATCTNNNTTTCTCACAGTAGCACGTCATC
GAGCACACGTCTGAACTCTTTCCCTACACGACGCTCTTCCGATCTNNNTTTCTCACAGTAGCACGTCATC
GAGCACACGTCTGAACTCTTTCCCTACACGACGCTCTTCCGATCTNNNTTTCTCACAGTAGCACGTCATC
GCACACGTCTGAACTCTTTCCCTACACGACGCTCTTCCGATCTNNNTAAGTCTCAATTCTCTGCAGCCGC
GCACACGTCTGAACTCTTTCCCTACACGACGCTCTTCCGATCTNNNTAAGTCTCAATTCTCTGCAGCCGC
GCACACGTCTGAACTCTTTCCCTACACGACGCTCTTCCGATCTNNNTAAGTCTCAATTCTCTGCAGCCGC
AGAGCACACGTCTGAACTCTTTCCCTACACGACGCTCTTCCGATCTNNNCTATGGCTCGCATCTTTCACA
AGAGCACACGTCTGAACTCTTTCCCTACACGACGCTCTTCCGATCTNNNCTATGGCTCGCATCTTTCACA
AGAGCACACGTCTGAACTCTTTCCCTACACGACGCTCTTCCGATCTNNNCTATGGCTCGCATCTTTCACA
GAGCACACGTCTGAACTCTTTCCCTACACGACGCTCTTCCGATCTNNNTTCATTCAGGCCCAACTCAGAA
GAGCACACGTCTGAACTCTTTCCCTACACGACGCTCTTCCGATCTNNNTTCATTCAGGCCCAACTCAGAA
GAGCACACGTCTGAACTCTTTCCCTACACGACGCTCTTCCGATCTNNNTTCATTCAGGCCCAACTCAGAA
GAGCACACGTCTGAACTCTTTCCCTACACGACGCTCTTCCGATCTNNNTGAGGGGAATGTTTTGAGGA
GAGCACACGTCTGAACTCTTTCCCTACACGACGCTCTTCCGATCTNNNTGAGGGGAATGTTTTGAGGA
GAGCACACGTCTGAACTCTTTCCCTACACGACGCTCTTCCGATCTNNNTGAGGGGAATGTTTTGAGGA
GAGCACACGTCTGAACTCTTTCCCTACACGACGCTCTTCCGATCTNNNCTGTTGAGAAGAATGCTGCA
GAGCACACGTCTGAACTCTTTCCCTACACGACGCTCTTCCGATCTNNNCTGTTGAGAAGAATGCTGCA
GAGCACACGTCTGAACTCTTTCCCTACACGACGCTCTTCCGATCTNNNCTGTTGAGAAGAATGCTGCA
AGCACACGTCTGAACTCTTTCCCTACACGACGCTCTTCCGATCTNNNCTGTAGTCCCAGCTACTCAGGAG
AGCACACGTCTGAACTCTTTCCCTACACGACGCTCTTCCGATCTNNNCTGTAGTCCCAGCTACTCAGGAG
AGCACACGTCTGAACTCTTTCCCTACACGACGCTCTTCCGATCTNNNCTGTAGTCCCAGCTACTCAGGAG
GAGCACACGTCTGAACTCTTTCCCTACACGACGCTCTTCCGATCTNNNCTGGAACTCCTGACCTCATGA
GAGCACACGTCTGAACTCTTTCCCTACACGACGCTCTTCCGATCTNNNCTGGAACTCCTGACCTCATGA
GAGCACACGTCTGAACTCTTTCCCTACACGACGCTCTTCCGATCTNNNCTGGAACTCCTGACCTCATGA
GAGCACACGTCTGAACTCTTTCCCTACACGACGCTCTTCCGATCTNNNCTGAGGCAGGAGAATCACTTC
GAGCACACGTCTGAACTCTTTCCCTACACGACGCTCTTCCGATCTNNNCTGAGGCAGGAGAATCACTTC
GAGCACACGTCTGAACTCTTTCCCTACACGACGCTCTTCCGATCTNNNCTGAGGCAGGAGAATCACTTC

GAGCACACGTCTGAACTCTTTCCCTACACGACGCTCTTCCGATCTNNNTGTTGTGCCTTGGAGAAGAT

GAGCACACGTCTGAACTCTTTCCCTACACGACGCTCTTCCGATCTNNNTGTTGTGCCTTGGAGAAGAT

GAGCACACGTCTGAACTCTTTCCCTACACGACGCTCTTCCGATCTNNNTGTTGTGCCTTGGAGAAGAT

AGCACACGTCTGAACTCTTTCCCTACACGACGCTCTTCCGATCTNNNTGCATTATTCCCACACATAGCCT

AGCACACGTCTGAACTCTTTCCCTACACGACGCTCTTCCGATCTNNNTGCATTATTCCCACACATAGCCT

AGCACACGTCTGAACTCTTTCCCTACACGACGCTCTTCCGATCTNNNTGCATTATTCCCACACATAGCCT

AGCACACGTCTGAACTCTTTCCCTACACGACGCTCTTCCGATCTNNNGATGACACGCGAATGAAGATG

AGCACACGTCTGAACTCTTTCCCTACACGACGCTCTTCCGATCTNNNGATGACACGCGAATGAAGATG

AGCACACGTCTGAACTCTTTCCCTACACGACGCTCTTCCGATCTNNNGATGACACGCGAATGAAGATG

AAGAGCACACGTCTGAACTCTTTCCCTACACGACGCTCTTCCGATCTNNNCAGACCCGCATCTTGTTGG

AAGAGCACACGTCTGAACTCTTTCCCTACACGACGCTCTTCCGATCTNNNCAGACCCGCATCTTGTTGG

AAGAGCACACGTCTGAACTCTTTCCCTACACGACGCTCTTCCGATCTNNNCAGACCCGCATCTTGTTGG

AGCACACGTCTGAACTCTTTCCCTACACGACGCTCTTCCGATCTNNNAGAATGGCATGAACCCAGGAG

AGCACACGTCTGAACTCTTTCCCTACACGACGCTCTTCCGATCTNNNAGAATGGCATGAACCCAGGAG

AGCACACGTCTGAACTCTTTCCCTACACGACGCTCTTCCGATCTNNNAGAATGGCATGAACCCAGGAG

GAGCACACGTCTGAACTCTTTCCCTACACGACGCTCTTCCGATCTNNNGCCTGTAATCCCAGCTACTCT

GAGCACACGTCTGAACTCTTTCCCTACACGACGCTCTTCCGATCTNNNGCCTGTAATCCCAGCTACTCT

GAGCACACGTCTGAACTCTTTCCCTACACGACGCTCTTCCGATCTNNNGCCTGTAATCCCAGCTACTCT

GAGCACACGTCTGAACTCTTTCCCTACACGACGCTCTTCCGATCTNNNCCAAGGTGGGAGAATAACCT

GAGCACACGTCTGAACTCTTTCCCTACACGACGCTCTTCCGATCTNNNCCAAGGTGGGAGAATAACCT

GAGCACACGTCTGAACTCTTTCCCTACACGACGCTCTTCCGATCTNNNCCAAGGTGGGAGAATAACCT

GAGCACACGTCTGAACTCTTTCCCTACACGACGCTCTTCCGATCTNNNCAGAAGCACTGCCCTAGTTTTT

GAGCACACGTCTGAACTCTTTCCCTACACGACGCTCTTCCGATCTNNNCAGAAGCACTGCCCTAGTTTTT

GAGCACACGTCTGAACTCTTTCCCTACACGACGCTCTTCCGATCTNNNCAGAAGCACTGCCCTAGTTTTT

AGCACACGTCTGAACTCTTTCCCTACACGACGCTCTTCCGATCTNNNGTGAGAAGCCTCTGTTTACCTC

AGCACACGTCTGAACTCTTTCCCTACACGACGCTCTTCCGATCTNNNGTGAGAAGCCTCTGTTTACCTC

AGCACACGTCTGAACTCTTTCCCTACACGACGCTCTTCCGATCTNNNGTGAGAAGCCTCTGTTTACCTC

GAGCACACGTCTGAACTCTTTCCCTACACGACGCTCTTCCGATCTNNNATGAGAGAATGAGAGCCAA

GAGCACACGTCTGAACTCTTTCCCTACACGACGCTCTTCCGATCTNNNATGAGAGAATGAGAGCCAA

GAGCACACGTCTGAACTCTTTCCCTACACGACGCTCTTCCGATCTNNNATGAGAGAATGAGAGCCAA

GAGCACACGTCTGAACTCTTTCCCTACACGACGCTCTTCCGATCTNNNCTGAGGCAGGAGAATTGCTT

GAGCACACGTCTGAACTCTTTCCCTACACGACGCTCTTCCGATCTNNNCTGAGGCAGGAGAATTGCTT

GAGCACACGTCTGAACTCTTTCCCTACACGACGCTCTTCCGATCTNNNCTGAGGCAGGAGAATTGCTT

GAGCACACGTCTGAACTCTTTCCCTACACGACGCTCTTCCGATCTNNNGCACCTGTTGTTTCCTGACTTT

GAGCACACGTCTGAACTCTTTCCCTACACGACGCTCTTCCGATCTNNNGCACCTGTTGTTTCCTGACTTT

GAGCACACGTCTGAACTCTTTCCCTACACGACGCTCTTCCGATCTNNNGCACCTGTTGTTTCCTGACTTT

AGCACACGTCTGAACTCTTTCCCTACACGACGCTCTTCCGATCTNNNACATGGTGAGACCCCTTCTTTAC

AGCACACGTCTGAACTCTTTCCCTACACGACGCTCTTCCGATCTNNNACATGGTGAGACCCCTTCTTTAC

AGCACACGTCTGAACTCTTTCCCTACACGACGCTCTTCCGATCTNNNACATGGTGAGACCCCTTCTTTAC

AGCACACGTCTGAACTCTTTCCCTACACGACGCTCTTCCGATCTNNNCAGACAGTGGGAGTCATGTG

AGCACACGTCTGAACTCTTTCCCTACACGACGCTCTTCCGATCTNNNCAGACAGTGGGAGTCATGTG

AGCACACGTCTGAACTCTTTCCCTACACGACGCTCTTCCGATCTNNNCAGACAGTGGGAGTCATGTG

AGAGCACACGTCTGAACTCTTTCCCTACACGACGCTCTTCCGATCTNNNAACAGCCAACATTGATGAAG

AGAGCACACGTCTGAACTCTTTCCCTACACGACGCTCTTCCGATCTNNNAACAGCCAACATTGATGAAG
AGAGCACACGTCTGAACTCTTTCCCTACACGACGCTCTTCCGATCTNNNAACAGCCAACATTGATGAAG
GAGCACACGTCTGAACTCTTTCCCTACACGACGCTCTTCCGATCTNNNCTGCATGACAGGAGACTCCAT
GAGCACACGTCTGAACTCTTTCCCTACACGACGCTCTTCCGATCTNNNCTGCATGACAGGAGACTCCAT
GAGCACACGTCTGAACTCTTTCCCTACACGACGCTCTTCCGATCTNNNCTGCATGACAGGAGACTCCAT
AGCACACGTCTGAACTCTTTCCCTACACGACGCTCTTCCGATCTNNNGCAGCTGCCATCATATCCAACTA
AGCACACGTCTGAACTCTTTCCCTACACGACGCTCTTCCGATCTNNNGCAGCTGCCATCATATCCAACTA
AGCACACGTCTGAACTCTTTCCCTACACGACGCTCTTCCGATCTNNNGCAGCTGCCATCATATCCAACTA
CACACGTCTGAACTCTTTCCCTACACGACGCTCTTCCGATCTNNNTCTTCCTACCCAGATCCTAACATCG
CACACGTCTGAACTCTTTCCCTACACGACGCTCTTCCGATCTNNNTCTTCCTACCCAGATCCTAACATCG
CACACGTCTGAACTCTTTCCCTACACGACGCTCTTCCGATCTNNNTCTTCCTACCCAGATCCTAACATCG
GAGCACACGTCTGAACTCTTTCCCTACACGACGCTCTTCCGATCTNNNGCAGAAGTGCCACCTTTACTTC
GAGCACACGTCTGAACTCTTTCCCTACACGACGCTCTTCCGATCTNNNGCAGAAGTGCCACCTTTACTTC
GAGCACACGTCTGAACTCTTTCCCTACACGACGCTCTTCCGATCTNNNGCAGAAGTGCCACCTTTACTTC
AGCACACGTCTGAACTCTTTCCCTACACGACGCTCTTCCGATCTNNNAGTCATGGGAAAAGCCTGTAA
AGCACACGTCTGAACTCTTTCCCTACACGACGCTCTTCCGATCTNNNAGTCATGGGAAAAGCCTGTAA
AGCACACGTCTGAACTCTTTCCCTACACGACGCTCTTCCGATCTNNNAGTCATGGGAAAAGCCTGTAA
AGCACACGTCTGAACTCTTTCCCTACACGACGCTCTTCCGATCTNNNTCCTGAGTAGATGGGACTACAG
AGCACACGTCTGAACTCTTTCCCTACACGACGCTCTTCCGATCTNNNTCCTGAGTAGATGGGACTACAG
AGCACACGTCTGAACTCTTTCCCTACACGACGCTCTTCCGATCTNNNTCCTGAGTAGATGGGACTACAG
AGCACACGTCTGAACTCTTTCCCTACACGACGCTCTTCCGATCTNNNAACTGATTCTAGGACCAAGGCA
AGCACACGTCTGAACTCTTTCCCTACACGACGCTCTTCCGATCTNNNAACTGATTCTAGGACCAAGGCA
AGCACACGTCTGAACTCTTTCCCTACACGACGCTCTTCCGATCTNNNAACTGATTCTAGGACCAAGGCA
GAGCACACGTCTGAACTCTTTCCCTACACGACGCTCTTCCGATCTNNNGGCCAAAACGGATGATCTTG
GAGCACACGTCTGAACTCTTTCCCTACACGACGCTCTTCCGATCTNNNGGCCAAAACGGATGATCTTG
GAGCACACGTCTGAACTCTTTCCCTACACGACGCTCTTCCGATCTNNNGGCCAAAACGGATGATCTTG
AGCACACGTCTGAACTCTTTCCCTACACGACGCTCTTCCGATCTNNNTCCTTTGCATTTTCAATGTTGCCA
AGCACACGTCTGAACTCTTTCCCTACACGACGCTCTTCCGATCTNNNTCCTTTGCATTTTCAATGTTGCCA
AGCACACGTCTGAACTCTTTCCCTACACGACGCTCTTCCGATCTNNNTCCTTTGCATTTTCAATGTTGCCA
AGCACACGTCTGAACTCTTTCCCTACACGACGCTCTTCCGATCTNNNAGCAAACCCCTCATCTCTACAAA
AGCACACGTCTGAACTCTTTCCCTACACGACGCTCTTCCGATCTNNNAGCAAACCCCTCATCTCTACAAA
AGCACACGTCTGAACTCTTTCCCTACACGACGCTCTTCCGATCTNNNAGCAAACCCCTCATCTCTACAAA
GAGCACACGTCTGAACTCTTTCCCTACACGACGCTCTTCCGATCTNNNCAAGCCTGGCTAACATTGTGA
GAGCACACGTCTGAACTCTTTCCCTACACGACGCTCTTCCGATCTNNNCAAGCCTGGCTAACATTGTGA
GAGCACACGTCTGAACTCTTTCCCTACACGACGCTCTTCCGATCTNNNCAAGCCTGGCTAACATTGTGA
AGAGCACACGTCTGAACTCTTTCCCTACACGACGCTCTTCCGATCTNNNCTGTAGTTCCAGCTACTTGG
AGAGCACACGTCTGAACTCTTTCCCTACACGACGCTCTTCCGATCTNNNCTGTAGTTCCAGCTACTTGG
AGAGCACACGTCTGAACTCTTTCCCTACACGACGCTCTTCCGATCTNNNCTGTAGTTCCAGCTACTTGG
CACACGTCTGAACTCTTTCCCTACACGACGCTCTTCCGATCTNNNATGTTGGAGACATACCAGACCATC
CACACGTCTGAACTCTTTCCCTACACGACGCTCTTCCGATCTNNNATGTTGGAGACATACCAGACCATC
CACACGTCTGAACTCTTTCCCTACACGACGCTCTTCCGATCTNNNATGTTGGAGACATACCAGACCATC
AGCACACGTCTGAACTCTTTCCCTACACGACGCTCTTCCGATCTNNNAAAAAAGCCTGATTCCTCCTCCT
AGCACACGTCTGAACTCTTTCCCTACACGACGCTCTTCCGATCTNNNAAAAAAGCCTGATTCCTCCTCCT

AGCACACGTCTGAACTCTTTCCCTACACGACGCTCTTCCGATCTNNNAAAAAAGCCTGATTCCTCCTCCT
GAGCACACGTCTGAACTCTTTCCCTACACGACGCTCTTCCGATCTNNNTGATGGTGGTCTGTGAAATCA
GAGCACACGTCTGAACTCTTTCCCTACACGACGCTCTTCCGATCTNNNTGATGGTGGTCTGTGAAATCA
GAGCACACGTCTGAACTCTTTCCCTACACGACGCTCTTCCGATCTNNNTGATGGTGGTCTGTGAAATCA
AGCACACGTCTGAACTCTTTCCCTACACGACGCTCTTCCGATCTNNNAGTAGAATGGTAGAAACACTTC
AGCACACGTCTGAACTCTTTCCCTACACGACGCTCTTCCGATCTNNNAGTAGAATGGTAGAAACACTTC
AGCACACGTCTGAACTCTTTCCCTACACGACGCTCTTCCGATCTNNNAGTAGAATGGTAGAAACACTTC
GCACACGTCTGAACTCTTTCCCTACACGACGCTCTTCCGATCTNNNAGCTCCTACTGTGACATTCTTGGC
GCACACGTCTGAACTCTTTCCCTACACGACGCTCTTCCGATCTNNNAGCTCCTACTGTGACATTCTTGGC
GCACACGTCTGAACTCTTTCCCTACACGACGCTCTTCCGATCTNNNAGCTCCTACTGTGACATTCTTGGC
AGCACACGTCTGAACTCTTTCCCTACACGACGCTCTTCCGATCTNNNAGATTGTGCTACTGTACTCCAG
AGCACACGTCTGAACTCTTTCCCTACACGACGCTCTTCCGATCTNNNAGATTGTGCTACTGTACTCCAG
AGCACACGTCTGAACTCTTTCCCTACACGACGCTCTTCCGATCTNNNAGATTGTGCTACTGTACTCCAG
AGCACACGTCTGAACTCTTTCCCTACACGACGCTCTTCCGATCTNNNTTACCTTTCATCTCCCACCACCTC
AGCACACGTCTGAACTCTTTCCCTACACGACGCTCTTCCGATCTNNNTTACCTTTCATCTCCCACCACCTC
AGCACACGTCTGAACTCTTTCCCTACACGACGCTCTTCCGATCTNNNTTACCTTTCATCTCCCACCACCTC
AGCACACGTCTGAACTCTTTCCCTACACGACGCTCTTCCGATCTNNNCAGACTTCTGGGAGACACTGAA
AGCACACGTCTGAACTCTTTCCCTACACGACGCTCTTCCGATCTNNNCAGACTTCTGGGAGACACTGAA
AGCACACGTCTGAACTCTTTCCCTACACGACGCTCTTCCGATCTNNNCAGACTTCTGGGAGACACTGAA
GAGCACACGTCTGAACTCTTTCCCTACACGACGCTCTTCCGATCTNNNGTCTTCTATCTTCTAGGCCAGC
GAGCACACGTCTGAACTCTTTCCCTACACGACGCTCTTCCGATCTNNNGTCTTCTATCTTCTAGGCCAGC
GAGCACACGTCTGAACTCTTTCCCTACACGACGCTCTTCCGATCTNNNGTCTTCTATCTTCTAGGCCAGC
AGCACACGTCTGAACTCTTTCCCTACACGACGCTCTTCCGATCTNNNTACAGTGAGCTGAGATTGCATC
AGCACACGTCTGAACTCTTTCCCTACACGACGCTCTTCCGATCTNNNTACAGTGAGCTGAGATTGCATC
AGCACACGTCTGAACTCTTTCCCTACACGACGCTCTTCCGATCTNNNTACAGTGAGCTGAGATTGCATC
CACACGTCTGAACTCTTTCCCTACACGACGCTCTTCCGATCTNNNTCCACTTATCAGTCAGGGACAGGCA
CACACGTCTGAACTCTTTCCCTACACGACGCTCTTCCGATCTNNNTCCACTTATCAGTCAGGGACAGGCA
CACACGTCTGAACTCTTTCCCTACACGACGCTCTTCCGATCTNNNTCCACTTATCAGTCAGGGACAGGCA
GAGCACACGTCTGAACTCTTTCCCTACACGACGCTCTTCCGATCTNNNTAGTAGGCGTTGGATCCATTC
GAGCACACGTCTGAACTCTTTCCCTACACGACGCTCTTCCGATCTNNNTAGTAGGCGTTGGATCCATTC
GAGCACACGTCTGAACTCTTTCCCTACACGACGCTCTTCCGATCTNNNTAGTAGGCGTTGGATCCATTC
AGCACACGTCTGAACTCTTTCCCTACACGACGCTCTTCCGATCTNNNTCCCTCTGGATCTTACCCTTCTTC
AGCACACGTCTGAACTCTTTCCCTACACGACGCTCTTCCGATCTNNNTCCCTCTGGATCTTACCCTTCTTC
AGCACACGTCTGAACTCTTTCCCTACACGACGCTCTTCCGATCTNNNTCCCTCTGGATCTTACCCTTCTTC
AGCACACGTCTGAACTCTTTCCCTACACGACGCTCTTCCGATCTNNNAATCCTAGCTACTTTGGAGGCT
AGCACACGTCTGAACTCTTTCCCTACACGACGCTCTTCCGATCTNNNAATCCTAGCTACTTTGGAGGCT
AGCACACGTCTGAACTCTTTCCCTACACGACGCTCTTCCGATCTNNNAATCCTAGCTACTTTGGAGGCT
GAGCACACGTCTGAACTCTTTCCCTACACGACGCTCTTCCGATCTNNNAGATACAAATGGGAGCACCTA
GAGCACACGTCTGAACTCTTTCCCTACACGACGCTCTTCCGATCTNNNAGATACAAATGGGAGCACCTA
GAGCACACGTCTGAACTCTTTCCCTACACGACGCTCTTCCGATCTNNNAGATACAAATGGGAGCACCTA
AGCACACGTCTGAACTCTTTCCCTACACGACGCTCTTCCGATCTNNNACACCTGTAATCCCAGCACTTTC
AGCACACGTCTGAACTCTTTCCCTACACGACGCTCTTCCGATCTNNNACACCTGTAATCCCAGCACTTTC
AGCACACGTCTGAACTCTTTCCCTACACGACGCTCTTCCGATCTNNNACACCTGTAATCCCAGCACTTTC

GCACACGTCTGAACTCTTTCCCTACACGACGCTCTTCCGATCTNNNTGGAGTTTGCAGTGAGCTATGAT

GCACACGTCTGAACTCTTTCCCTACACGACGCTCTTCCGATCTNNNTGGAGTTTGCAGTGAGCTATGAT

GCACACGTCTGAACTCTTTCCCTACACGACGCTCTTCCGATCTNNNTGGAGTTTGCAGTGAGCTATGAT

GAGCACACGTCTGAACTCTTTCCCTACACGACGCTCTTCCGATCTNNNCACACACACACACACACACATA

GAGCACACGTCTGAACTCTTTCCCTACACGACGCTCTTCCGATCTNNNCACACACACACACACACACATA

GAGCACACGTCTGAACTCTTTCCCTACACGACGCTCTTCCGATCTNNNCACACACACACACACACACATA

AGAGCACACGTCTGAACTCTTTCCCTACACGACGCTCTTCCGATCTNNNCTCAGCCTGGAGACTCTGGA

AGAGCACACGTCTGAACTCTTTCCCTACACGACGCTCTTCCGATCTNNNCTCAGCCTGGAGACTCTGGA

AGAGCACACGTCTGAACTCTTTCCCTACACGACGCTCTTCCGATCTNNNCTCAGCCTGGAGACTCTGGA

GAGCACACGTCTGAACTCTTTCCCTACACGACGCTCTTCCGATCTNNNGTGAGTAGGTTGTGGGATCTT

GAGCACACGTCTGAACTCTTTCCCTACACGACGCTCTTCCGATCTNNNGTGAGTAGGTTGTGGGATCTT

GAGCACACGTCTGAACTCTTTCCCTACACGACGCTCTTCCGATCTNNNGTGAGTAGGTTGTGGGATCTT

CACGTCTGAACTCTTTCCCTACACGACGCTCTTCCGATCTNNNTAGTCTCACAGTGCAAGAACCTCCATG

CACGTCTGAACTCTTTCCCTACACGACGCTCTTCCGATCTNNNTAGTCTCACAGTGCAAGAACCTCCATG

CACGTCTGAACTCTTTCCCTACACGACGCTCTTCCGATCTNNNTAGTCTCACAGTGCAAGAACCTCCATG

GAGCACACGTCTGAACTCTTTCCCTACACGACGCTCTTCCGATCTNNNGGAAGCAGGACAGACTAAAA

GAGCACACGTCTGAACTCTTTCCCTACACGACGCTCTTCCGATCTNNNGGAAGCAGGACAGACTAAAA

GAGCACACGTCTGAACTCTTTCCCTACACGACGCTCTTCCGATCTNNNGGAAGCAGGACAGACTAAAA

AGAGCACACGTCTGAACTCTTTCCCTACACGACGCTCTTCCGATCTNNNTGCTTCAGGAGAGTTGCATA

AGAGCACACGTCTGAACTCTTTCCCTACACGACGCTCTTCCGATCTNNNTGCTTCAGGAGAGTTGCATA

AGAGCACACGTCTGAACTCTTTCCCTACACGACGCTCTTCCGATCTNNNTGCTTCAGGAGAGTTGCATA

AGCACACGTCTGAACTCTTTCCCTACACGACGCTCTTCCGATCTNNNAACCCCTGCTTTGTTTTTCTCTGC

AGCACACGTCTGAACTCTTTCCCTACACGACGCTCTTCCGATCTNNNAACCCCTGCTTTGTTTTTCTCTGC

AGCACACGTCTGAACTCTTTCCCTACACGACGCTCTTCCGATCTNNNAACCCCTGCTTTGTTTTTCTCTGC

AGCACACGTCTGAACTCTTTCCCTACACGACGCTCTTCCGATCTNNNGTCCACGTGCATTCTTGTTACAA

AGCACACGTCTGAACTCTTTCCCTACACGACGCTCTTCCGATCTNNNGTCCACGTGCATTCTTGTTACAA

AGCACACGTCTGAACTCTTTCCCTACACGACGCTCTTCCGATCTNNNGTCCACGTGCATTCTTGTTACAA

GAGCACACGTCTGAACTCTTTCCCTACACGACGCTCTTCCGATCTNNNCACCTGTAATCCCAGCTACTCA

GAGCACACGTCTGAACTCTTTCCCTACACGACGCTCTTCCGATCTNNNCACCTGTAATCCCAGCTACTCA

GAGCACACGTCTGAACTCTTTCCCTACACGACGCTCTTCCGATCTNNNCACCTGTAATCCCAGCTACTCA

GCACACGTCTGAACTCTTTCCCTACACGACGCTCTTCCGATCTNNNGGTAGCAGCGAGAAGTGAGGCA

GCACACGTCTGAACTCTTTCCCTACACGACGCTCTTCCGATCTNNNGGTAGCAGCGAGAAGTGAGGCA

GCACACGTCTGAACTCTTTCCCTACACGACGCTCTTCCGATCTNNNGGTAGCAGCGAGAAGTGAGGCA

AGCACACGTCTGAACTCTTTCCCTACACGACGCTCTTCCGATCTNNNAGATCACTGACCACAGATCATCA

AGCACACGTCTGAACTCTTTCCCTACACGACGCTCTTCCGATCTNNNAGATCACTGACCACAGATCATCA

AGCACACGTCTGAACTCTTTCCCTACACGACGCTCTTCCGATCTNNNAGATCACTGACCACAGATCATCA

AGAGCACACGTCTGAACTCTTTCCCTACACGACGCTCTTCCGATCTNNNTGGAAATTCTCTGATGAAGC

AGAGCACACGTCTGAACTCTTTCCCTACACGACGCTCTTCCGATCTNNNTGGAAATTCTCTGATGAAGC

AGAGCACACGTCTGAACTCTTTCCCTACACGACGCTCTTCCGATCTNNNTGGAAATTCTCTGATGAAGC

AGCACACGTCTGAACTCTTTCCCTACACGACGCTCTTCCGATCTNNNTCTGGTCCCATGCATTTTGGATA

AGCACACGTCTGAACTCTTTCCCTACACGACGCTCTTCCGATCTNNNTCTGGTCCCATGCATTTTGGATA

AGCACACGTCTGAACTCTTTCCCTACACGACGCTCTTCCGATCTNNNTCTGGTCCCATGCATTTTGGATA

AGCACACGTCTGAACTCTTTCCCTACACGACGCTCTTCCGATCTNNNGCTACTTGGGAGGCTAATGTAC

AGCACACGTCTGAACTCTTTCCCTACACGACGCTCTTCCGATCTNNNGCTACTTGGGAGGCTAATGTAC
AGCACACGTCTGAACTCTTTCCCTACACGACGCTCTTCCGATCTNNNGCTACTTGGGAGGCTAATGTAC
GAGCACACGTCTGAACTCTTTCCCTACACGACGCTCTTCCGATCTNNNTGGGAAGCAATTTCACACTCA
GAGCACACGTCTGAACTCTTTCCCTACACGACGCTCTTCCGATCTNNNTGGGAAGCAATTTCACACTCA
GAGCACACGTCTGAACTCTTTCCCTACACGACGCTCTTCCGATCTNNNTGGGAAGCAATTTCACACTCA
GCACACGTCTGAACTCTTTCCCTACACGACGCTCTTCCGATCTNNNAGATCCCGGATTTTTGTTTTTTTG
GCACACGTCTGAACTCTTTCCCTACACGACGCTCTTCCGATCTNNNAGATCCCGGATTTTTGTTTTTTTG
GCACACGTCTGAACTCTTTCCCTACACGACGCTCTTCCGATCTNNNAGATCCCGGATTTTTGTTTTTTTG
GAGCACACGTCTGAACTCTTTCCCTACACGACGCTCTTCCGATCTNNNGCCTGTAATCTCAGCTACTCAG
GAGCACACGTCTGAACTCTTTCCCTACACGACGCTCTTCCGATCTNNNGCCTGTAATCTCAGCTACTCAG
GAGCACACGTCTGAACTCTTTCCCTACACGACGCTCTTCCGATCTNNNGCCTGTAATCTCAGCTACTCAG
GAGCACACGTCTGAACTCTTTCCCTACACGACGCTCTTCCGATCTNNNTGGGTTCAAGTGATTCTCCTAC
GAGCACACGTCTGAACTCTTTCCCTACACGACGCTCTTCCGATCTNNNTGGGTTCAAGTGATTCTCCTAC
GAGCACACGTCTGAACTCTTTCCCTACACGACGCTCTTCCGATCTNNNTGGGTTCAAGTGATTCTCCTAC
GCACACGTCTGAACTCTTTCCCTACACGACGCTCTTCCGATCTNNNGCACAACTTGAGAAATCCAGCAA
GCACACGTCTGAACTCTTTCCCTACACGACGCTCTTCCGATCTNNNGCACAACTTGAGAAATCCAGCAA
GCACACGTCTGAACTCTTTCCCTACACGACGCTCTTCCGATCTNNNGCACAACTTGAGAAATCCAGCAA
AGCACACGTCTGAACTCTTTCCCTACACGACGCTCTTCCGATCTNNNCATTGAACTCCTGGATGCAAGT
AGCACACGTCTGAACTCTTTCCCTACACGACGCTCTTCCGATCTNNNCATTGAACTCCTGGATGCAAGT
AGCACACGTCTGAACTCTTTCCCTACACGACGCTCTTCCGATCTNNNCATTGAACTCCTGGATGCAAGT
GCACACGTCTGAACTCTTTCCCTACACGACGCTCTTCCGATCTNNNTGTTTCACTATTGGCACACAGAAC
GCACACGTCTGAACTCTTTCCCTACACGACGCTCTTCCGATCTNNNTGTTTCACTATTGGCACACAGAAC
GCACACGTCTGAACTCTTTCCCTACACGACGCTCTTCCGATCTNNNTGTTTCACTATTGGCACACAGAAC
GAGCACACGTCTGAACTCTTTCCCTACACGACGCTCTTCCGATCTNNNAACCAAGATGTCAATAGGGCC
GAGCACACGTCTGAACTCTTTCCCTACACGACGCTCTTCCGATCTNNNAACCAAGATGTCAATAGGGCC
GAGCACACGTCTGAACTCTTTCCCTACACGACGCTCTTCCGATCTNNNAACCAAGATGTCAATAGGGCC
AGCACACGTCTGAACTCTTTCCCTACACGACGCTCTTCCGATCTNNNAGGTGTAATTTTCTGCTCCCAGT
AGCACACGTCTGAACTCTTTCCCTACACGACGCTCTTCCGATCTNNNAGGTGTAATTTTCTGCTCCCAGT
AGCACACGTCTGAACTCTTTCCCTACACGACGCTCTTCCGATCTNNNAGGTGTAATTTTCTGCTCCCAGT
GAGCACACGTCTGAACTCTTTCCCTACACGACGCTCTTCCGATCTNNNCATTACAAGTCTTGCTGGTGTG
GAGCACACGTCTGAACTCTTTCCCTACACGACGCTCTTCCGATCTNNNCATTACAAGTCTTGCTGGTGTG
GAGCACACGTCTGAACTCTTTCCCTACACGACGCTCTTCCGATCTNNNCATTACAAGTCTTGCTGGTGTG
GCACACGTCTGAACTCTTTCCCTACACGACGCTCTTCCGATCTNNNCCCGACGGATTTTGATCTTTGTTC
GCACACGTCTGAACTCTTTCCCTACACGACGCTCTTCCGATCTNNNCCCGACGGATTTTGATCTTTGTTC
GCACACGTCTGAACTCTTTCCCTACACGACGCTCTTCCGATCTNNNCCCGACGGATTTTGATCTTTGTTC
CACACGTCTGAACTCTTTCCCTACACGACGCTCTTCCGATCTNNNATGATGTCATTGCTCTGGTGACTCA
CACACGTCTGAACTCTTTCCCTACACGACGCTCTTCCGATCTNNNATGATGTCATTGCTCTGGTGACTCA
CACACGTCTGAACTCTTTCCCTACACGACGCTCTTCCGATCTNNNATGATGTCATTGCTCTGGTGACTCA
AGCACACGTCTGAACTCTTTCCCTACACGACGCTCTTCCGATCTNNNCGCAAATCCCATTACTCTGTTCC
AGCACACGTCTGAACTCTTTCCCTACACGACGCTCTTCCGATCTNNNCGCAAATCCCATTACTCTGTTCC
AGCACACGTCTGAACTCTTTCCCTACACGACGCTCTTCCGATCTNNNCGCAAATCCCATTACTCTGTTCC
GAGCACACGTCTGAACTCTTTCCCTACACGACGCTCTTCCGATCTNNNACACCTGTAATCCCAGCACTTTG
GAGCACACGTCTGAACTCTTTCCCTACACGACGCTCTTCCGATCTNNNACACCTGTAATCCCAGCACTTTG

AGCACACGTCTGAACTCTTTCCCTACACGACGCTCTTCCGATCTNNNACACCTGTAATCCCAGCACTTT
GCACACGTCTGAACTCTTTCCCTACACGACGCTCTTCCGATCTNNNTTGATATCCTCGGGAATGAGCAG
GCACACGTCTGAACTCTTTCCCTACACGACGCTCTTCCGATCTNNNTTGATATCCTCGGGAATGAGCAG
GCACACGTCTGAACTCTTTCCCTACACGACGCTCTTCCGATCTNNNTTGATATCCTCGGGAATGAGCAG
AGCACACGTCTGAACTCTTTCCCTACACGACGCTCTTCCGATCTNNNAGGAGCAGAAAGAGAAGAGC
AGCACACGTCTGAACTCTTTCCCTACACGACGCTCTTCCGATCTNNNAGGAGCAGAAAGAGAAGAGC
AGCACACGTCTGAACTCTTTCCCTACACGACGCTCTTCCGATCTNNNAGGAGCAGAAAGAGAAGAGC
AGCACACGTCTGAACTCTTTCCCTACACGACGCTCTTCCGATCTNNNCATTTGAGGTCTGGAGGTTGA
AGCACACGTCTGAACTCTTTCCCTACACGACGCTCTTCCGATCTNNNCATTTGAGGTCTGGAGGTTGA
AGCACACGTCTGAACTCTTTCCCTACACGACGCTCTTCCGATCTNNNCATTTGAGGTCTGGAGGTTGA
AGCACACGTCTGAACTCTTTCCCTACACGACGCTCTTCCGATCTNNNGAACCGCCCTCCTTTCTCAAGCA
AGCACACGTCTGAACTCTTTCCCTACACGACGCTCTTCCGATCTNNNGAACCGCCCTCCTTTCTCAAGCA
AGCACACGTCTGAACTCTTTCCCTACACGACGCTCTTCCGATCTNNNGAACCGCCCTCCTTTCTCAAGCA
CACACGTCTGAACTCTTTCCCTACACGACGCTCTTCCGATCTNNNCACAATCTTGGCTCACTGCAAGCAT
CACACGTCTGAACTCTTTCCCTACACGACGCTCTTCCGATCTNNNCACAATCTTGGCTCACTGCAAGCAT
CACACGTCTGAACTCTTTCCCTACACGACGCTCTTCCGATCTNNNCACAATCTTGGCTCACTGCAAGCAT
GAGCACACGTCTGAACTCTTTCCCTACACGACGCTCTTCCGATCTNNNCTGTAGTCACAGCTACTCAG
GAGCACACGTCTGAACTCTTTCCCTACACGACGCTCTTCCGATCTNNNCTGTAGTCACAGCTACTCAG
GAGCACACGTCTGAACTCTTTCCCTACACGACGCTCTTCCGATCTNNNCTGTAGTCACAGCTACTCAG
AGCACACGTCTGAACTCTTTCCCTACACGACGCTCTTCCGATCTNNNAAAAAATTAGCCTGGCAAGGT
AGCACACGTCTGAACTCTTTCCCTACACGACGCTCTTCCGATCTNNNAAAAAATTAGCCTGGCAAGGT
AGCACACGTCTGAACTCTTTCCCTACACGACGCTCTTCCGATCTNNNAAAAAATTAGCCTGGCAAGGT
AGCACACGTCTGAACTCTTTCCCTACACGACGCTCTTCCGATCTNNNACAGAACAAGACCCTGTCTCAA
AGCACACGTCTGAACTCTTTCCCTACACGACGCTCTTCCGATCTNNNACAGAACAAGACCCTGTCTCAA
AGCACACGTCTGAACTCTTTCCCTACACGACGCTCTTCCGATCTNNNACAGAACAAGACCCTGTCTCAA
AGCACACGTCTGAACTCTTTCCCTACACGACGCTCTTCCGATCTNNNCTGGACAACAAGAGCGAAACT
AGCACACGTCTGAACTCTTTCCCTACACGACGCTCTTCCGATCTNNNCTGGACAACAAGAGCGAAACT
AGCACACGTCTGAACTCTTTCCCTACACGACGCTCTTCCGATCTNNNCTGGACAACAAGAGCGAAACT
AGCACACGTCTGAACTCTTTCCCTACACGACGCTCTTCCGATCTNNNATAGATAGGAGGCCTCTGACGA
AGCACACGTCTGAACTCTTTCCCTACACGACGCTCTTCCGATCTNNNATAGATAGGAGGCCTCTGACGA
AGCACACGTCTGAACTCTTTCCCTACACGACGCTCTTCCGATCTNNNATAGATAGGAGGCCTCTGACGA
AGCACACGTCTGAACTCTTTCCCTACACGACGCTCTTCCGATCTNNNAAGTAGTGTGTGTGGATGTGC
AGCACACGTCTGAACTCTTTCCCTACACGACGCTCTTCCGATCTNNNAAGTAGTGTGTGTGGATGTGC
AGCACACGTCTGAACTCTTTCCCTACACGACGCTCTTCCGATCTNNNAAGTAGTGTGTGTGGATGTGC
AGCACACGTCTGAACTCTTTCCCTACACGACGCTCTTCCGATCTNNNTGTGGGAGAAGAGTGATTTCC
AGCACACGTCTGAACTCTTTCCCTACACGACGCTCTTCCGATCTNNNTGTGGGAGAAGAGTGATTTCC
AGCACACGTCTGAACTCTTTCCCTACACGACGCTCTTCCGATCTNNNTGTGGGAGAAGAGTGATTTCC
GAGCACACGTCTGAACTCTTTCCCTACACGACGCTCTTCCGATCTNNNGTGTAATCCCAGCTACTCAG
GAGCACACGTCTGAACTCTTTCCCTACACGACGCTCTTCCGATCTNNNGTGTAATCCCAGCTACTCAG
GAGCACACGTCTGAACTCTTTCCCTACACGACGCTCTTCCGATCTNNNGTGTAATCCCAGCTACTCAG
AGCACACGTCTGAACTCTTTCCCTACACGACGCTCTTCCGATCTNNNCTCCATGACCAAGAGAGATTG
AGCACACGTCTGAACTCTTTCCCTACACGACGCTCTTCCGATCTNNNCTCCATGACCAAGAGAGATTG
AGCACACGTCTGAACTCTTTCCCTACACGACGCTCTTCCGATCTNNNCTCCATGACCAAGAGAGATTG

AGCACACGTCTGAACTCTTTCCCTACACGACGCTCTTCCGATCTNNNAGACTTTTTCTGAAGGCAGTG
AGCACACGTCTGAACTCTTTCCCTACACGACGCTCTTCCGATCTNNNAGACTTTTTCTGAAGGCAGTG
AGCACACGTCTGAACTCTTTCCCTACACGACGCTCTTCCGATCTNNNAGACTTTTTCTGAAGGCAGTG
CACACGTCTGAACTCTTTCCCTACACGACGCTCTTCCGATCTNNNACTTCTCCAAATTCCATGCTTGCC
CACACGTCTGAACTCTTTCCCTACACGACGCTCTTCCGATCTNNNACTTCTCCAAATTCCATGCTTGCC
CACACGTCTGAACTCTTTCCCTACACGACGCTCTTCCGATCTNNNACTTCTCCAAATTCCATGCTTGCC
GAGCACACGTCTGAACTCTTTCCCTACACGACGCTCTTCCGATCTNNNTTCCTCTGTCTTCTCTCTCTTC
GAGCACACGTCTGAACTCTTTCCCTACACGACGCTCTTCCGATCTNNNTTCCTCTGTCTTCTCTCTCTTC
GAGCACACGTCTGAACTCTTTCCCTACACGACGCTCTTCCGATCTNNNTTCCTCTGTCTTCTCTCTCTTC
AGCACACGTCTGAACTCTTTCCCTACACGACGCTCTTCCGATCTNNNTTTGTGTGTGGCTTCTTTTCG
AGCACACGTCTGAACTCTTTCCCTACACGACGCTCTTCCGATCTNNNTTTGTGTGTGGCTTCTTTTCG
AGCACACGTCTGAACTCTTTCCCTACACGACGCTCTTCCGATCTNNNTTTGTGTGTGGCTTCTTTTCG
AGCACACGTCTGAACTCTTTCCCTACACGACGCTCTTCCGATCTNNNTGCAACCTCTGCTTCCTTTGAA
AGCACACGTCTGAACTCTTTCCCTACACGACGCTCTTCCGATCTNNNTGCAACCTCTGCTTCCTTTGAA
AGCACACGTCTGAACTCTTTCCCTACACGACGCTCTTCCGATCTNNNTGCAACCTCTGCTTCCTTTGAA
GCACACGTCTGAACTCTTTCCCTACACGACGCTCTTCCGATCTNNNTCCATATGCAGAATTTTCCCCAA
GCACACGTCTGAACTCTTTCCCTACACGACGCTCTTCCGATCTNNNTCCATATGCAGAATTTTCCCCAA
GCACACGTCTGAACTCTTTCCCTACACGACGCTCTTCCGATCTNNNTCCATATGCAGAATTTTCCCCAA
AGCACACGTCTGAACTCTTTCCCTACACGACGCTCTTCCGATCTNNNCTCCCAAATGAAAACCATGGC
AGCACACGTCTGAACTCTTTCCCTACACGACGCTCTTCCGATCTNNNCTCCCAAATGAAAACCATGGC
AGCACACGTCTGAACTCTTTCCCTACACGACGCTCTTCCGATCTNNNCTCCCAAATGAAAACCATGGC
GCACACGTCTGAACTCTTTCCCTACACGACGCTCTTCCGATCTNNNTGAAATACTTCTGTCAGCATGC
GCACACGTCTGAACTCTTTCCCTACACGACGCTCTTCCGATCTNNNTGAAATACTTCTGTCAGCATGC
GCACACGTCTGAACTCTTTCCCTACACGACGCTCTTCCGATCTNNNTGAAATACTTCTGTCAGCATGC
ACACGTCTGAACTCTTTCCCTACACGACGCTCTTCCGATCTNNNTTACCTAGGTAATTTTCCATACCCAT
ACACGTCTGAACTCTTTCCCTACACGACGCTCTTCCGATCTNNNTTACCTAGGTAATTTTCCATACCCAT
ACACGTCTGAACTCTTTCCCTACACGACGCTCTTCCGATCTNNNTTACCTAGGTAATTTTCCATACCCAT
GCACACGTCTGAACTCTTTCCCTACACGACGCTCTTCCGATCTNNNCGGTCAACAAACATCAAGTCAG
GCACACGTCTGAACTCTTTCCCTACACGACGCTCTTCCGATCTNNNCGGTCAACAAACATCAAGTCAG
GCACACGTCTGAACTCTTTCCCTACACGACGCTCTTCCGATCTNNNCGGTCAACAAACATCAAGTCAG
GCACACGTCTGAACTCTTTCCCTACACGACGCTCTTCCGATCTNNNACATTGACAAGGAGTATGCAG
GCACACGTCTGAACTCTTTCCCTACACGACGCTCTTCCGATCTNNNACATTGACAAGGAGTATGCAG
GCACACGTCTGAACTCTTTCCCTACACGACGCTCTTCCGATCTNNNACATTGACAAGGAGTATGCAG
CACGTCTGAACTCTTTCCCTACACGACGCTCTTCCGATCTNNNCATTCAGTTGTGTTGAGCAGCAGCAT
CACGTCTGAACTCTTTCCCTACACGACGCTCTTCCGATCTNNNCATTCAGTTGTGTTGAGCAGCAGCAT
CACGTCTGAACTCTTTCCCTACACGACGCTCTTCCGATCTNNNCATTCAGTTGTGTTGAGCAGCAGCAT
AGCACACGTCTGAACTCTTTCCCTACACGACGCTCTTCCGATCTNNNTCCCTTTATGCACTTCCTTGCTG
AGCACACGTCTGAACTCTTTCCCTACACGACGCTCTTCCGATCTNNNTCCCTTTATGCACTTCCTTGCTG
AGCACACGTCTGAACTCTTTCCCTACACGACGCTCTTCCGATCTNNNTCCCTTTATGCACTTCCTTGCTG
GCACACGTCTGAACTCTTTCCCTACACGACGCTCTTCCGATCTNNNGAGAGAGCAAACCTAGGGAAC
GCACACGTCTGAACTCTTTCCCTACACGACGCTCTTCCGATCTNNNGAGAGAGCAAACCTAGGGAAC
GCACACGTCTGAACTCTTTCCCTACACGACGCTCTTCCGATCTNNNGAGAGAGCAAACCTAGGGAAC
GCACACGTCTGAACTCTTTCCCTACACGACGCTCTTCCGATCTNNNGCATTTGGTACTCTCCACCCTTA

AGCACACGTCTGAACTCTTTCCCTACACGACGCTCTTCCGATCTNNNGCATTTGGTACTCTCCACCCTTA
AGCACACGTCTGAACTCTTTCCCTACACGACGCTCTTCCGATCTNNNGCATTTGGTACTCTCCACCCTTA
AGCACACGTCTGAACTCTTTCCCTACACGACGCTCTTCCGATCTNNNAGATGCAAGGGTTTTCTCTGG
AGCACACGTCTGAACTCTTTCCCTACACGACGCTCTTCCGATCTNNNAGATGCAAGGGTTTTCTCTGG
AGCACACGTCTGAACTCTTTCCCTACACGACGCTCTTCCGATCTNNNAGATGCAAGGGTTTTCTCTGG
AGCACACGTCTGAACTCTTTCCCTACACGACGCTCTTCCGATCTNNNTTTTTGAGACAGAGCCATGTG
AGCACACGTCTGAACTCTTTCCCTACACGACGCTCTTCCGATCTNNNTTTTTGAGACAGAGCCATGTG
AGCACACGTCTGAACTCTTTCCCTACACGACGCTCTTCCGATCTNNNTTTTTGAGACAGAGCCATGTG
AGCACACGTCTGAACTCTTTCCCTACACGACGCTCTTCCGATCTNNNTCTTGGTGATCCTACTGACCTC
AGCACACGTCTGAACTCTTTCCCTACACGACGCTCTTCCGATCTNNNTCTTGGTGATCCTACTGACCTC
AGCACACGTCTGAACTCTTTCCCTACACGACGCTCTTCCGATCTNNNTCTTGGTGATCCTACTGACCTC
GAGCACACGTCTGAACTCTTTCCCTACACGACGCTCTTCCGATCTNNNCGAGATCATGCCATTGTACTC
GAGCACACGTCTGAACTCTTTCCCTACACGACGCTCTTCCGATCTNNNCGAGATCATGCCATTGTACTC
GAGCACACGTCTGAACTCTTTCCCTACACGACGCTCTTCCGATCTNNNCGAGATCATGCCATTGTACTC
AGCACACGTCTGAACTCTTTCCCTACACGACGCTCTTCCGATCTNNNCCATTTCGTTTCTGTAGCTGGTC
AGCACACGTCTGAACTCTTTCCCTACACGACGCTCTTCCGATCTNNNCCATTTCGTTTCTGTAGCTGGTC
AGCACACGTCTGAACTCTTTCCCTACACGACGCTCTTCCGATCTNNNCCATTTCGTTTCTGTAGCTGGTC
GCACACGTCTGAACTCTTTCCCTACACGACGCTCTTCCGATCTNNNGCCCCCTTTCTGCCATTAGATTA
GCACACGTCTGAACTCTTTCCCTACACGACGCTCTTCCGATCTNNNGCCCCCTTTCTGCCATTAGATTA
GCACACGTCTGAACTCTTTCCCTACACGACGCTCTTCCGATCTNNNGCCCCCTTTCTGCCATTAGATTA
GCACACGTCTGAACTCTTTCCCTACACGACGCTCTTCCGATCTNNNGTGTTGCAATCAGTGGGCTAAA
GCACACGTCTGAACTCTTTCCCTACACGACGCTCTTCCGATCTNNNGTGTTGCAATCAGTGGGCTAAA
GCACACGTCTGAACTCTTTCCCTACACGACGCTCTTCCGATCTNNNGTGTTGCAATCAGTGGGCTAAA
AGCACACGTCTGAACTCTTTCCCTACACGACGCTCTTCCGATCTNNNACGAAGTTGTTGTGCTGTGTTT
AGCACACGTCTGAACTCTTTCCCTACACGACGCTCTTCCGATCTNNNACGAAGTTGTTGTGCTGTGTTT
AGCACACGTCTGAACTCTTTCCCTACACGACGCTCTTCCGATCTNNNACGAAGTTGTTGTGCTGTGTTT
AGCACACGTCTGAACTCTTTCCCTACACGACGCTCTTCCGATCTNNNGCACCTATAGTCCCAGCTACTT
AGCACACGTCTGAACTCTTTCCCTACACGACGCTCTTCCGATCTNNNGCACCTATAGTCCCAGCTACTT
AGCACACGTCTGAACTCTTTCCCTACACGACGCTCTTCCGATCTNNNGCACCTATAGTCCCAGCTACTT
AGCACACGTCTGAACTCTTTCCCTACACGACGCTCTTCCGATCTNNNTTAGTAGAGATGGGGTTTCAC
AGCACACGTCTGAACTCTTTCCCTACACGACGCTCTTCCGATCTNNNTTAGTAGAGATGGGGTTTCAC
AGCACACGTCTGAACTCTTTCCCTACACGACGCTCTTCCGATCTNNNTTAGTAGAGATGGGGTTTCAC
AGCACACGTCTGAACTCTTTCCCTACACGACGCTCTTCCGATCTNNNCCTGGTAATTCTCATGAAAAAT
AGCACACGTCTGAACTCTTTCCCTACACGACGCTCTTCCGATCTNNNCCTGGTAATTCTCATGAAAAAT
AGCACACGTCTGAACTCTTTCCCTACACGACGCTCTTCCGATCTNNNCCTGGTAATTCTCATGAAAAAT
AGCACACGTCTGAACTCTTTCCCTACACGACGCTCTTCCGATCTNNNCCAGGGTTCAAGCAATTCTCCT
AGCACACGTCTGAACTCTTTCCCTACACGACGCTCTTCCGATCTNNNCCAGGGTTCAAGCAATTCTCCT
AGCACACGTCTGAACTCTTTCCCTACACGACGCTCTTCCGATCTNNNCCAGGGTTCAAGCAATTCTCCT
GAGCACACGTCTGAACTCTTTCCCTACACGACGCTCTTCCGATCTNNNAGGTTGAGGTGAGCCAATAT
GAGCACACGTCTGAACTCTTTCCCTACACGACGCTCTTCCGATCTNNNAGGTTGAGGTGAGCCAATAT
GAGCACACGTCTGAACTCTTTCCCTACACGACGCTCTTCCGATCTNNNAGGTTGAGGTGAGCCAATAT
GAGCACACGTCTGAACTCTTTCCCTACACGACGCTCTTCCGATCTNNNGGAAAGGGTGACTGGATAA
GAGCACACGTCTGAACTCTTTCCCTACACGACGCTCTTCCGATCTNNNGGAAAGGGTGACTGGATAA

GAGCACACGTCTGAACTCTTTCCCTACACGACGCTCTTCCGATCTNNNGGAAAGGGTGACTGGATAA
GCACACGTCTGAACTCTTTCCCTACACGACGCTCTTCCGATCTNNNTGTCCAGAATTAGAAGTACAGT
GCACACGTCTGAACTCTTTCCCTACACGACGCTCTTCCGATCTNNNTGTCCAGAATTAGAAGTACAGT
GCACACGTCTGAACTCTTTCCCTACACGACGCTCTTCCGATCTNNNTGTCCAGAATTAGAAGTACAGT
AGCACACGTCTGAACTCTTTCCCTACACGACGCTCTTCCGATCTNNNGTTCCACCCTTACCAACACTCT
AGCACACGTCTGAACTCTTTCCCTACACGACGCTCTTCCGATCTNNNGTTCCACCCTTACCAACACTCT
AGCACACGTCTGAACTCTTTCCCTACACGACGCTCTTCCGATCTNNNGTTCCACCCTTACCAACACTCT
AGCACACGTCTGAACTCTTTCCCTACACGACGCTCTTCCGATCTNNNCAGGCTGTTCTGTTGACCTGT
AGCACACGTCTGAACTCTTTCCCTACACGACGCTCTTCCGATCTNNNCAGGCTGTTCTGTTGACCTGT
AGCACACGTCTGAACTCTTTCCCTACACGACGCTCTTCCGATCTNNNCAGGCTGTTCTGTTGACCTGT
AGCACACGTCTGAACTCTTTCCCTACACGACGCTCTTCCGATCTNNNGCCAGGAAAGGAAATTGGTG
AGCACACGTCTGAACTCTTTCCCTACACGACGCTCTTCCGATCTNNNGCCAGGAAAGGAAATTGGTG
AGCACACGTCTGAACTCTTTCCCTACACGACGCTCTTCCGATCTNNNGCCAGGAAAGGAAATTGGTG
GAGCACACGTCTGAACTCTTTCCCTACACGACGCTCTTCCGATCTNNNCGCCTGTAATCCCATCTACTG
GAGCACACGTCTGAACTCTTTCCCTACACGACGCTCTTCCGATCTNNNCGCCTGTAATCCCATCTACTG
GAGCACACGTCTGAACTCTTTCCCTACACGACGCTCTTCCGATCTNNNCGCCTGTAATCCCATCTACTG
AGCACACGTCTGAACTCTTTCCCTACACGACGCTCTTCCGATCTNNNGATCATGAATCCGCAGTGAAC
AGCACACGTCTGAACTCTTTCCCTACACGACGCTCTTCCGATCTNNNGATCATGAATCCGCAGTGAAC
AGCACACGTCTGAACTCTTTCCCTACACGACGCTCTTCCGATCTNNNGATCATGAATCCGCAGTGAAC
GCACACGTCTGAACTCTTTCCCTACACGACGCTCTTCCGATCTNNNAAAGAAGGCTAGGTTATGTCGG
GCACACGTCTGAACTCTTTCCCTACACGACGCTCTTCCGATCTNNNAAAGAAGGCTAGGTTATGTCGG
GCACACGTCTGAACTCTTTCCCTACACGACGCTCTTCCGATCTNNNAAAGAAGGCTAGGTTATGTCGG
AGCACACGTCTGAACTCTTTCCCTACACGACGCTCTTCCGATCTNNNCCCAGACACTGGACTATAGCG
AGCACACGTCTGAACTCTTTCCCTACACGACGCTCTTCCGATCTNNNCCCAGACACTGGACTATAGCG
AGCACACGTCTGAACTCTTTCCCTACACGACGCTCTTCCGATCTNNNCCCAGACACTGGACTATAGCG
AGCACACGTCTGAACTCTTTCCCTACACGACGCTCTTCCGATCTNNNGGTGCCTGTAATCCTAGCTAC
AGCACACGTCTGAACTCTTTCCCTACACGACGCTCTTCCGATCTNNNGGTGCCTGTAATCCTAGCTAC
AGCACACGTCTGAACTCTTTCCCTACACGACGCTCTTCCGATCTNNNGGTGCCTGTAATCCTAGCTAC
GAGCACACGTCTGAACTCTTTCCCTACACGACGCTCTTCCGATCTNNNCAGGAGAATCACTTGAACCT
GAGCACACGTCTGAACTCTTTCCCTACACGACGCTCTTCCGATCTNNNCAGGAGAATCACTTGAACCT
GAGCACACGTCTGAACTCTTTCCCTACACGACGCTCTTCCGATCTNNNCAGGAGAATCACTTGAACCT
GCACACGTCTGAACTCTTTCCCTACACGACGCTCTTCCGATCTNNNGAGAAGCCTCTTTTCTACCACC
GCACACGTCTGAACTCTTTCCCTACACGACGCTCTTCCGATCTNNNGAGAAGCCTCTTTTCTACCACC
GCACACGTCTGAACTCTTTCCCTACACGACGCTCTTCCGATCTNNNGAGAAGCCTCTTTTCTACCACC
GAGCACACGTCTGAACTCTTTCCCTACACGACGCTCTTCCGATCTNNNTTGCAGCTAAATTACCATAA
GAGCACACGTCTGAACTCTTTCCCTACACGACGCTCTTCCGATCTNNNTTGCAGCTAAATTACCATAA
GAGCACACGTCTGAACTCTTTCCCTACACGACGCTCTTCCGATCTNNNTTGCAGCTAAATTACCATAA
GCACACGTCTGAACTCTTTCCCTACACGACGCTCTTCCGATCTNNNAAGTGTTGGGATTACAGGCAT
GCACACGTCTGAACTCTTTCCCTACACGACGCTCTTCCGATCTNNNAAGTGTTGGGATTACAGGCAT
GCACACGTCTGAACTCTTTCCCTACACGACGCTCTTCCGATCTNNNAAGTGTTGGGATTACAGGCAT
AGCACACGTCTGAACTCTTTCCCTACACGACGCTCTTCCGATCTNNNCGAAACCCCGTCTCTAAAAAA
AGCACACGTCTGAACTCTTTCCCTACACGACGCTCTTCCGATCTNNNCGAAACCCCGTCTCTAAAAAA
AGCACACGTCTGAACTCTTTCCCTACACGACGCTCTTCCGATCTNNNCGAAACCCCGTCTCTAAAAAA

AGCACACGTCTGAACTCTTTCCCTACACGACGCTCTTCCGATCTNNNCTTTCCTGGGGAGTGAAGAGT

AGCACACGTCTGAACTCTTTCCCTACACGACGCTCTTCCGATCTNNNCTTTCCTGGGGAGTGAAGAGT

AGCACACGTCTGAACTCTTTCCCTACACGACGCTCTTCCGATCTNNNCTTTCCTGGGGAGTGAAGAGT

AGCACACGTCTGAACTCTTTCCCTACACGACGCTCTTCCGATCTNNNGGTCACAGCCCAGATAAACTAC

AGCACACGTCTGAACTCTTTCCCTACACGACGCTCTTCCGATCTNNNGGTCACAGCCCAGATAAACTAC

AGCACACGTCTGAACTCTTTCCCTACACGACGCTCTTCCGATCTNNNGGTCACAGCCCAGATAAACTAC

GAGCACACGTCTGAACTCTTTCCCTACACGACGCTCTTCCGATCTNNNCCCACAACCAGATATCTCCAT

GAGCACACGTCTGAACTCTTTCCCTACACGACGCTCTTCCGATCTNNNCCCACAACCAGATATCTCCAT

GAGCACACGTCTGAACTCTTTCCCTACACGACGCTCTTCCGATCTNNNCCCACAACCAGATATCTCCAT

GAGCACACGTCTGAACTCTTTCCCTACACGACGCTCTTCCGATCTNNNCTGTCATCCCAGCTATTCAGG

GAGCACACGTCTGAACTCTTTCCCTACACGACGCTCTTCCGATCTNNNCTGTCATCCCAGCTATTCAGG

GAGCACACGTCTGAACTCTTTCCCTACACGACGCTCTTCCGATCTNNNCTGTCATCCCAGCTATTCAGG

AGCACACGTCTGAACTCTTTCCCTACACGACGCTCTTCCGATCTNNNTTGTGCCTAGTAGTGAACCTTC

AGCACACGTCTGAACTCTTTCCCTACACGACGCTCTTCCGATCTNNNTTGTGCCTAGTAGTGAACCTTC

AGCACACGTCTGAACTCTTTCCCTACACGACGCTCTTCCGATCTNNNTTGTGCCTAGTAGTGAACCTTC

AGCACACGTCTGAACTCTTTCCCTACACGACGCTCTTCCGATCTNNNTGATTCCACTTTATTAAAAAGC

AGCACACGTCTGAACTCTTTCCCTACACGACGCTCTTCCGATCTNNNTGATTCCACTTTATTAAAAAGC

AGCACACGTCTGAACTCTTTCCCTACACGACGCTCTTCCGATCTNNNTGATTCCACTTTATTAAAAAGC

AGCACACGTCTGAACTCTTTCCCTACACGACGCTCTTCCGATCTNNNTTCATGCCATTCTCCTTCCTCAG

AGCACACGTCTGAACTCTTTCCCTACACGACGCTCTTCCGATCTNNNTTCATGCCATTCTCCTTCCTCAG

AGCACACGTCTGAACTCTTTCCCTACACGACGCTCTTCCGATCTNNNTTCATGCCATTCTCCTTCCTCAG

GAGCACACGTCTGAACTCTTTCCCTACACGACGCTCTTCCGATCTNNNTGAGCCAAGATCACACCATTG

GAGCACACGTCTGAACTCTTTCCCTACACGACGCTCTTCCGATCTNNNTGAGCCAAGATCACACCATTG

GAGCACACGTCTGAACTCTTTCCCTACACGACGCTCTTCCGATCTNNNTGAGCCAAGATCACACCATTG

AGCACACGTCTGAACTCTTTCCCTACACGACGCTCTTCCGATCTNNNGAGGGAGAGAGAGAGAAGG

AGCACACGTCTGAACTCTTTCCCTACACGACGCTCTTCCGATCTNNNGAGGGAGAGAGAGAGAAGG

AGCACACGTCTGAACTCTTTCCCTACACGACGCTCTTCCGATCTNNNGAGGGAGAGAGAGAGAAGG

GCACACGTCTGAACTCTTTCCCTACACGACGCTCTTCCGATCTNNNCTTGCAGTGAGCCGAGATCATC

GCACACGTCTGAACTCTTTCCCTACACGACGCTCTTCCGATCTNNNCTTGCAGTGAGCCGAGATCATC

GCACACGTCTGAACTCTTTCCCTACACGACGCTCTTCCGATCTNNNCTTGCAGTGAGCCGAGATCATC

AGCACACGTCTGAACTCTTTCCCTACACGACGCTCTTCCGATCTNNNTGGCTAAGAAGAGATGTTGG

AGCACACGTCTGAACTCTTTCCCTACACGACGCTCTTCCGATCTNNNTGGCTAAGAAGAGATGTTGG

AGCACACGTCTGAACTCTTTCCCTACACGACGCTCTTCCGATCTNNNTGGCTAAGAAGAGATGTTGG

GCACACGTCTGAACTCTTTCCCTACACGACGCTCTTCCGATCTNNNACCTGGGACACATCAATCACAAC

GCACACGTCTGAACTCTTTCCCTACACGACGCTCTTCCGATCTNNNACCTGGGACACATCAATCACAAC

GCACACGTCTGAACTCTTTCCCTACACGACGCTCTTCCGATCTNNNACCTGGGACACATCAATCACAAC

AGCACACGTCTGAACTCTTTCCCTACACGACGCTCTTCCGATCTNNNGAGATAACGCCACTGCACTCT

AGCACACGTCTGAACTCTTTCCCTACACGACGCTCTTCCGATCTNNNGAGATAACGCCACTGCACTCT

AGCACACGTCTGAACTCTTTCCCTACACGACGCTCTTCCGATCTNNNGAGATAACGCCACTGCACTCT

AGCACACGTCTGAACTCTTTCCCTACACGACGCTCTTCCGATCTNNNAGCTTACAGGCATAGAGGTGT

AGCACACGTCTGAACTCTTTCCCTACACGACGCTCTTCCGATCTNNNAGCTTACAGGCATAGAGGTGT

AGCACACGTCTGAACTCTTTCCCTACACGACGCTCTTCCGATCTNNNAGCTTACAGGCATAGAGGTGT

AGCACACGTCTGAACTCTTTCCCTACACGACGCTCTTCCGATCTNNNGTCATGTGGGGGAAGAAGAC

AGCACACGTCTGAACTCTTTCCCTACACGACGCTCTTCCGATCTNNNGTCATGTGGGGGAAGAAGAC

AGCACACGTCTGAACTCTTTCCCTACACGACGCTCTTCCGATCTNNNGTCATGTGGGGGAAGAAGAC

CACACGTCTGAACTCTTTCCCTACACGACGCTCTTCCGATCTNNNCGGTTGTAGGTTGTCAGAGAGG

CACACGTCTGAACTCTTTCCCTACACGACGCTCTTCCGATCTNNNCGGTTGTAGGTTGTCAGAGAGG

CACACGTCTGAACTCTTTCCCTACACGACGCTCTTCCGATCTNNNCGGTTGTAGGTTGTCAGAGAGG

GAGCACACGTCTGAACTCTTTCCCTACACGACGCTCTTCCGATCTNNNTCAGGCATGCTCAAACAGTA

GAGCACACGTCTGAACTCTTTCCCTACACGACGCTCTTCCGATCTNNNTCAGGCATGCTCAAACAGTA

GAGCACACGTCTGAACTCTTTCCCTACACGACGCTCTTCCGATCTNNNTCAGGCATGCTCAAACAGTA

AGCACACGTCTGAACTCTTTCCCTACACGACGCTCTTCCGATCTNNNGATTACAGGCGTGAGTTACTG

AGCACACGTCTGAACTCTTTCCCTACACGACGCTCTTCCGATCTNNNGATTACAGGCGTGAGTTACTG

AGCACACGTCTGAACTCTTTCCCTACACGACGCTCTTCCGATCTNNNGATTACAGGCGTGAGTTACTG

AGCACACGTCTGAACTCTTTCCCTACACGACGCTCTTCCGATCTNNNACACAGACACACACACACACAT

AGCACACGTCTGAACTCTTTCCCTACACGACGCTCTTCCGATCTNNNACACAGACACACACACACACAT

AGCACACGTCTGAACTCTTTCCCTACACGACGCTCTTCCGATCTNNNACACAGACACACACACACACAT

GAGCACACGTCTGAACTCTTTCCCTACACGACGCTCTTCCGATCTNNNTAGCCTCCATAATCACATGAG

GAGCACACGTCTGAACTCTTTCCCTACACGACGCTCTTCCGATCTNNNTAGCCTCCATAATCACATGAG

GAGCACACGTCTGAACTCTTTCCCTACACGACGCTCTTCCGATCTNNNTAGCCTCCATAATCACATGAG

AGCACACGTCTGAACTCTTTCCCTACACGACGCTCTTCCGATCTNNNATAAAAGGAAGGGGAGACAT

AGCACACGTCTGAACTCTTTCCCTACACGACGCTCTTCCGATCTNNNATAAAAGGAAGGGGAGACAT

AGCACACGTCTGAACTCTTTCCCTACACGACGCTCTTCCGATCTNNNATAAAAGGAAGGGGAGACAT

GCACACGTCTGAACTCTTTCCCTACACGACGCTCTTCCGATCTNNNATGGATGCATGTAGGGAAATGC

GCACACGTCTGAACTCTTTCCCTACACGACGCTCTTCCGATCTNNNATGGATGCATGTAGGGAAATGC

GCACACGTCTGAACTCTTTCCCTACACGACGCTCTTCCGATCTNNNATGGATGCATGTAGGGAAATGC

AGCACACGTCTGAACTCTTTCCCTACACGACGCTCTTCCGATCTNNNAAGTGGTTTCATGTTCACTGCT

AGCACACGTCTGAACTCTTTCCCTACACGACGCTCTTCCGATCTNNNAAGTGGTTTCATGTTCACTGCT

AGCACACGTCTGAACTCTTTCCCTACACGACGCTCTTCCGATCTNNNAAGTGGTTTCATGTTCACTGCT

GCACACGTCTGAACTCTTTCCCTACACGACGCTCTTCCGATCTNNNTGTGCTATGGATTGGTGGAAA

GCACACGTCTGAACTCTTTCCCTACACGACGCTCTTCCGATCTNNNTGTGCTATGGATTGGTGGAAA

GCACACGTCTGAACTCTTTCCCTACACGACGCTCTTCCGATCTNNNTGTGCTATGGATTGGTGGAAA

GAGCACACGTCTGAACTCTTTCCCTACACGACGCTCTTCCGATCTNNNCAATGATGTGTGCCTGTAAT

GAGCACACGTCTGAACTCTTTCCCTACACGACGCTCTTCCGATCTNNNCAATGATGTGTGCCTGTAAT

GAGCACACGTCTGAACTCTTTCCCTACACGACGCTCTTCCGATCTNNNCAATGATGTGTGCCTGTAAT

AGCACACGTCTGAACTCTTTCCCTACACGACGCTCTTCCGATCTNNNAGTATTGCTTGAGCCCAGAAG

AGCACACGTCTGAACTCTTTCCCTACACGACGCTCTTCCGATCTNNNAGTATTGCTTGAGCCCAGAAG

AGCACACGTCTGAACTCTTTCCCTACACGACGCTCTTCCGATCTNNNAGTATTGCTTGAGCCCAGAAG

GAGCACACGTCTGAACTCTTTCCCTACACGACGCTCTTCCGATCTNNNCTTTTCCAAGTGGGTCCCAA

GAGCACACGTCTGAACTCTTTCCCTACACGACGCTCTTCCGATCTNNNCTTTTCCAAGTGGGTCCCAA

GAGCACACGTCTGAACTCTTTCCCTACACGACGCTCTTCCGATCTNNNCTTTTCCAAGTGGGTCCCAA

GAGCACACGTCTGAACTCTTTCCCTACACGACGCTCTTCCGATCTNNNCTATGGGTTTGTCAGAGACA

GAGCACACGTCTGAACTCTTTCCCTACACGACGCTCTTCCGATCTNNNCTATGGGTTTGTCAGAGACA

GAGCACACGTCTGAACTCTTTCCCTACACGACGCTCTTCCGATCTNNNCTATGGGTTTGTCAGAGACA

GCACACGTCTGAACTCTTTCCCTACACGACGCTCTTCCGATCTNNNGGGGAAGGTCTCTCTGGTATT

GCACACGTCTGAACTCTTTCCCTACACGACGCTCTTCCGATCTNNNGGGGAAGGTCTCTCTGGTATT

GCACACGTCTGAACTCTTTCCCTACACGACGCTCTTCCGATCTNNNGGGGAAGGTCTCTCTGGTATT
AGCACACGTCTGAACTCTTTCCCTACACGACGCTCTTCCGATCTNNNTGCACAACAGTTCATTGTTTCT
AGCACACGTCTGAACTCTTTCCCTACACGACGCTCTTCCGATCTNNNTGCACAACAGTTCATTGTTTCT
AGCACACGTCTGAACTCTTTCCCTACACGACGCTCTTCCGATCTNNNTGCACAACAGTTCATTGTTTCT
AGCACACGTCTGAACTCTTTCCCTACACGACGCTCTTCCGATCTNNNGTCTGTGTGGGTGAATGCTTG
AGCACACGTCTGAACTCTTTCCCTACACGACGCTCTTCCGATCTNNNGTCTGTGTGGGTGAATGCTTG
AGCACACGTCTGAACTCTTTCCCTACACGACGCTCTTCCGATCTNNNGTCTGTGTGGGTGAATGCTTG
CACACGTCTGAACTCTTTCCCTACACGACGCTCTTCCGATCTNNNCAACTTCATACCGTCTTCTTCATCG
CACACGTCTGAACTCTTTCCCTACACGACGCTCTTCCGATCTNNNCAACTTCATACCGTCTTCTTCATCG
CACACGTCTGAACTCTTTCCCTACACGACGCTCTTCCGATCTNNNCAACTTCATACCGTCTTCTTCATCG
CACACGTCTGAACTCTTTCCCTACACGACGCTCTTCCGATCTNNNCAGGAGAATTGCTTGAAACTGG
CACACGTCTGAACTCTTTCCCTACACGACGCTCTTCCGATCTNNNCAGGAGAATTGCTTGAAACTGG
CACACGTCTGAACTCTTTCCCTACACGACGCTCTTCCGATCTNNNCAGGAGAATTGCTTGAAACTGG
GCACACGTCTGAACTCTTTCCCTACACGACGCTCTTCCGATCTNNNCTGAGACAGGAGAATCGCTTGA
GCACACGTCTGAACTCTTTCCCTACACGACGCTCTTCCGATCTNNNCTGAGACAGGAGAATCGCTTGA
GCACACGTCTGAACTCTTTCCCTACACGACGCTCTTCCGATCTNNNCTGAGACAGGAGAATCGCTTGA
GCACACGTCTGAACTCTTTCCCTACACGACGCTCTTCCGATCTNNNAGCATTTACCCACAGGAGAAC
GCACACGTCTGAACTCTTTCCCTACACGACGCTCTTCCGATCTNNNAGCATTTACCCACAGGAGAAC
GCACACGTCTGAACTCTTTCCCTACACGACGCTCTTCCGATCTNNNAGCATTTACCCACAGGAGAAC
ACGTCTGAACTCTTTCCCTACACGACGCTCTTCCGATCTNNNCTAGAGAACCTAGGGCCGGAGCATC
ACGTCTGAACTCTTTCCCTACACGACGCTCTTCCGATCTNNNCTAGAGAACCTAGGGCCGGAGCATC
ACGTCTGAACTCTTTCCCTACACGACGCTCTTCCGATCTNNNCTAGAGAACCTAGGGCCGGAGCATC
AGCACACGTCTGAACTCTTTCCCTACACGACGCTCTTCCGATCTNNNTTATTTACTTTCCCTCAGAAGA
AGCACACGTCTGAACTCTTTCCCTACACGACGCTCTTCCGATCTNNNTTATTTACTTTCCCTCAGAAGA
AGCACACGTCTGAACTCTTTCCCTACACGACGCTCTTCCGATCTNNNTTATTTACTTTCCCTCAGAAGA
AGAGCACACGTCTGAACTCTTTCCCTACACGACGCTCTTCCGATCTNNNAGGCCCCAATTAGTAGGCT
AGAGCACACGTCTGAACTCTTTCCCTACACGACGCTCTTCCGATCTNNNAGGCCCCAATTAGTAGGCT
AGAGCACACGTCTGAACTCTTTCCCTACACGACGCTCTTCCGATCTNNNAGGCCCCAATTAGTAGGCT
AGCACACGTCTGAACTCTTTCCCTACACGACGCTCTTCCGATCTNNNTGGGATTTTTTAGCCCTTCAT
AGCACACGTCTGAACTCTTTCCCTACACGACGCTCTTCCGATCTNNNTGGGATTTTTTAGCCCTTCAT
AGCACACGTCTGAACTCTTTCCCTACACGACGCTCTTCCGATCTNNNTGGGATTTTTTAGCCCTTCAT
AGCACACGTCTGAACTCTTTCCCTACACGACGCTCTTCCGATCTNNNACAAACACACACACACACACA
AGCACACGTCTGAACTCTTTCCCTACACGACGCTCTTCCGATCTNNNACAAACACACACACACACACA
AGCACACGTCTGAACTCTTTCCCTACACGACGCTCTTCCGATCTNNNACAAACACACACACACACACA
GCACACGTCTGAACTCTTTCCCTACACGACGCTCTTCCGATCTNNNAGCACTTCACTTCTAACCTCAGG
GCACACGTCTGAACTCTTTCCCTACACGACGCTCTTCCGATCTNNNAGCACTTCACTTCTAACCTCAGG
GCACACGTCTGAACTCTTTCCCTACACGACGCTCTTCCGATCTNNNAGCACTTCACTTCTAACCTCAGG
AGCACACGTCTGAACTCTTTCCCTACACGACGCTCTTCCGATCTNNNCTGAGGCAGGAGAATCACTT
AGCACACGTCTGAACTCTTTCCCTACACGACGCTCTTCCGATCTNNNCTGAGGCAGGAGAATCACTT
AGCACACGTCTGAACTCTTTCCCTACACGACGCTCTTCCGATCTNNNCTGAGGCAGGAGAATCACTT
GCACACGTCTGAACTCTTTCCCTACACGACGCTCTTCCGATCTNNNATGGTCTCTGGTTTTTCTGGTG
GCACACGTCTGAACTCTTTCCCTACACGACGCTCTTCCGATCTNNNATGGTCTCTGGTTTTTCTGGTG
GCACACGTCTGAACTCTTTCCCTACACGACGCTCTTCCGATCTNNNATGGTCTCTGGTTTTTCTGGTG

GAGCACACGTCTGAACTCTTTCCCTACACGACGCTCTTCCGATCTNNNGAGGCTGAAGCAAGAGAAT

GAGCACACGTCTGAACTCTTTCCCTACACGACGCTCTTCCGATCTNNNGAGGCTGAAGCAAGAGAAT

GAGCACACGTCTGAACTCTTTCCCTACACGACGCTCTTCCGATCTNNNGAGGCTGAAGCAAGAGAAT

AGCACACGTCTGAACTCTTTCCCTACACGACGCTCTTCCGATCTNNNGAAGCTTAGCAAGCCCCAAGT

AGCACACGTCTGAACTCTTTCCCTACACGACGCTCTTCCGATCTNNNGAAGCTTAGCAAGCCCCAAGT

AGCACACGTCTGAACTCTTTCCCTACACGACGCTCTTCCGATCTNNNGAAGCTTAGCAAGCCCCAAGT

AGCACACGTCTGAACTCTTTCCCTACACGACGCTCTTCCGATCTNNNATTCCCAGCTATTCAGGTGTCT

AGCACACGTCTGAACTCTTTCCCTACACGACGCTCTTCCGATCTNNNATTCCCAGCTATTCAGGTGTCT

AGCACACGTCTGAACTCTTTCCCTACACGACGCTCTTCCGATCTNNNATTCCCAGCTATTCAGGTGTCT

AGCACACGTCTGAACTCTTTCCCTACACGACGCTCTTCCGATCTNNNCCACCAAGCCCAGCTAATTTTTT

AGCACACGTCTGAACTCTTTCCCTACACGACGCTCTTCCGATCTNNNCCACCAAGCCCAGCTAATTTTTT

AGCACACGTCTGAACTCTTTCCCTACACGACGCTCTTCCGATCTNNNCCACCAAGCCCAGCTAATTTTTT

AAGAGCACACGTCTGAACTCTTTCCCTACACGACGCTCTTCCGATCTNNNGGTTGCTGTGAGCCGAG

AAGAGCACACGTCTGAACTCTTTCCCTACACGACGCTCTTCCGATCTNNNGGTTGCTGTGAGCCGAG

AAGAGCACACGTCTGAACTCTTTCCCTACACGACGCTCTTCCGATCTNNNGGTTGCTGTGAGCCGAG

AGCACACGTCTGAACTCTTTCCCTACACGACGCTCTTCCGATCTNNNCTCCTCATAGTCACTATTGCCC

AGCACACGTCTGAACTCTTTCCCTACACGACGCTCTTCCGATCTNNNCTCCTCATAGTCACTATTGCCC

AGCACACGTCTGAACTCTTTCCCTACACGACGCTCTTCCGATCTNNNCTCCTCATAGTCACTATTGCCC

GCACACGTCTGAACTCTTTCCCTACACGACGCTCTTCCGATCTNNNTGTGCATTTTAATTTCAAAAAG

GCACACGTCTGAACTCTTTCCCTACACGACGCTCTTCCGATCTNNNTGTGCATTTTAATTTCAAAAAG

GCACACGTCTGAACTCTTTCCCTACACGACGCTCTTCCGATCTNNNTGTGCATTTTAATTTCAAAAAG

AGCACACGTCTGAACTCTTTCCCTACACGACGCTCTTCCGATCTNNNACAGCTTCTTCTTTCATGGTGT

AGCACACGTCTGAACTCTTTCCCTACACGACGCTCTTCCGATCTNNNACAGCTTCTTCTTTCATGGTGT

AGCACACGTCTGAACTCTTTCCCTACACGACGCTCTTCCGATCTNNNACAGCTTCTTCTTTCATGGTGT

GAGCACACGTCTGAACTCTTTCCCTACACGACGCTCTTCCGATCTNNNAGGCTGAGGCTAGAGAATC

GAGCACACGTCTGAACTCTTTCCCTACACGACGCTCTTCCGATCTNNNAGGCTGAGGCTAGAGAATC

GAGCACACGTCTGAACTCTTTCCCTACACGACGCTCTTCCGATCTNNNAGGCTGAGGCTAGAGAATC

AGCACACGTCTGAACTCTTTCCCTACACGACGCTCTTCCGATCTNNNTCTGGAGAAGAAGCATGGAG

AGCACACGTCTGAACTCTTTCCCTACACGACGCTCTTCCGATCTNNNTCTGGAGAAGAAGCATGGAG

AGCACACGTCTGAACTCTTTCCCTACACGACGCTCTTCCGATCTNNNTCTGGAGAAGAAGCATGGAG

GAGCACACGTCTGAACTCTTTCCCTACACGACGCTCTTCCGATCTNNNATTCATTTTATTGTGGTATGG

GAGCACACGTCTGAACTCTTTCCCTACACGACGCTCTTCCGATCTNNNATTCATTTTATTGTGGTATGG

GAGCACACGTCTGAACTCTTTCCCTACACGACGCTCTTCCGATCTNNNATTCATTTTATTGTGGTATGG

AGCACACGTCTGAACTCTTTCCCTACACGACGCTCTTCCGATCTNNNTGATACAAAGCTACCCTGCCTC

AGCACACGTCTGAACTCTTTCCCTACACGACGCTCTTCCGATCTNNNTGATACAAAGCTACCCTGCCTC

AGCACACGTCTGAACTCTTTCCCTACACGACGCTCTTCCGATCTNNNTGATACAAAGCTACCCTGCCTC

AGCACACGTCTGAACTCTTTCCCTACACGACGCTCTTCCGATCTNNNCCGGGAGTCGAGATAAATCAG

AGCACACGTCTGAACTCTTTCCCTACACGACGCTCTTCCGATCTNNNCCGGGAGTCGAGATAAATCAG

AGCACACGTCTGAACTCTTTCCCTACACGACGCTCTTCCGATCTNNNCCGGGAGTCGAGATAAATCAG

AGCACACGTCTGAACTCTTTCCCTACACGACGCTCTTCCGATCTNNNCAACTCTGTCTCCTTCCTCTTCC

AGCACACGTCTGAACTCTTTCCCTACACGACGCTCTTCCGATCTNNNCAACTCTGTCTCCTTCCTCTTCC

AGCACACGTCTGAACTCTTTCCCTACACGACGCTCTTCCGATCTNNNCAACTCTGTCTCCTTCCTCTTCC

AGCACACGTCTGAACTCTTTCCCTACACGACGCTCTTCCGATCTNNNGAGCCATTCAAGACACACCTT

GAGCACACGTCTGAACTCTTTCCCTACACGACGCTCTTCCGATCTNNNGAGCCATTCAAGACACACCTTT
GAGCACACGTCTGAACTCTTTCCCTACACGACGCTCTTCCGATCTNNNGAGCCATTCAAGACACACCTTT
AGCACACGTCTGAACTCTTTCCCTACACGACGCTCTTCCGATCTNNNTGTACACACGTAGACACATACT
AGCACACGTCTGAACTCTTTCCCTACACGACGCTCTTCCGATCTNNNTGTACACACGTAGACACATACT
AGCACACGTCTGAACTCTTTCCCTACACGACGCTCTTCCGATCTNNNTGTACACACGTAGACACATACT
AGAGCACACGTCTGAACTCTTTCCCTACACGACGCTCTTCCGATCTNNNTTGAACATTGGAGAGCTAGC
AGAGCACACGTCTGAACTCTTTCCCTACACGACGCTCTTCCGATCTNNNTTGAACATTGGAGAGCTAGC
AGAGCACACGTCTGAACTCTTTCCCTACACGACGCTCTTCCGATCTNNNTTGAACATTGGAGAGCTAGC
AGAGCACACGTCTGAACTCTTTCCCTACACGACGCTCTTCCGATCTNNNTCAACTCCTTTTACAATAGCTC
AGAGCACACGTCTGAACTCTTTCCCTACACGACGCTCTTCCGATCTNNNTCAACTCCTTTTACAATAGCTC
AGAGCACACGTCTGAACTCTTTCCCTACACGACGCTCTTCCGATCTNNNTCAACTCCTTTTACAATAGCTC
GCACACGTCTGAACTCTTTCCCTACACGACGCTCTTCCGATCTNNNCCTTGTCCATGAGAAGCATTGTGC
GCACACGTCTGAACTCTTTCCCTACACGACGCTCTTCCGATCTNNNCCTTGTCCATGAGAAGCATTGTGC
GCACACGTCTGAACTCTTTCCCTACACGACGCTCTTCCGATCTNNNCCTTGTCCATGAGAAGCATTGTGC
GAGCACACGTCTGAACTCTTTCCCTACACGACGCTCTTCCGATCTNNNTGTAGTCCCAGCTACTAGGAA
GAGCACACGTCTGAACTCTTTCCCTACACGACGCTCTTCCGATCTNNNTGTAGTCCCAGCTACTAGGAA
GAGCACACGTCTGAACTCTTTCCCTACACGACGCTCTTCCGATCTNNNTGTAGTCCCAGCTACTAGGAA
GCACACGTCTGAACTCTTTCCCTACACGACGCTCTTCCGATCTNNNCAAATGTGGGCAAGACAAAGCTA
GCACACGTCTGAACTCTTTCCCTACACGACGCTCTTCCGATCTNNNCAAATGTGGGCAAGACAAAGCTA
GCACACGTCTGAACTCTTTCCCTACACGACGCTCTTCCGATCTNNNCAAATGTGGGCAAGACAAAGCTA
AGCACACGTCTGAACTCTTTCCCTACACGACGCTCTTCCGATCTNNNTTTTTTGAGAGCAGTGGCACAA
AGCACACGTCTGAACTCTTTCCCTACACGACGCTCTTCCGATCTNNNTTTTTTGAGAGCAGTGGCACAA
AGCACACGTCTGAACTCTTTCCCTACACGACGCTCTTCCGATCTNNNTTTTTTGAGAGCAGTGGCACAA
GAGCACACGTCTGAACTCTTTCCCTACACGACGCTCTTCCGATCTNNNGAGAATGGTGTGAACCCAGC
GAGCACACGTCTGAACTCTTTCCCTACACGACGCTCTTCCGATCTNNNGAGAATGGTGTGAACCCAGC
GAGCACACGTCTGAACTCTTTCCCTACACGACGCTCTTCCGATCTNNNGAGAATGGTGTGAACCCAGC
AGCACACGTCTGAACTCTTTCCCTACACGACGCTCTTCCGATCTNNNGTAATCCCAGCTACTCAGGAAG
AGCACACGTCTGAACTCTTTCCCTACACGACGCTCTTCCGATCTNNNGTAATCCCAGCTACTCAGGAAG
AGCACACGTCTGAACTCTTTCCCTACACGACGCTCTTCCGATCTNNNGTAATCCCAGCTACTCAGGAAG
GAGCACACGTCTGAACTCTTTCCCTACACGACGCTCTTCCGATCTNNNTGTAATTCCAGCTATTCGGGA
GAGCACACGTCTGAACTCTTTCCCTACACGACGCTCTTCCGATCTNNNTGTAATTCCAGCTATTCGGGA
GAGCACACGTCTGAACTCTTTCCCTACACGACGCTCTTCCGATCTNNNTGTAATTCCAGCTATTCGGGA
AGCACACGTCTGAACTCTTTCCCTACACGACGCTCTTCCGATCTNNNGTCAATCAGGGGTGTGTCTGAC
AGCACACGTCTGAACTCTTTCCCTACACGACGCTCTTCCGATCTNNNGTCAATCAGGGGTGTGTCTGAC
AGCACACGTCTGAACTCTTTCCCTACACGACGCTCTTCCGATCTNNNGTCAATCAGGGGTGTGTCTGAC
AGAGCACACGTCTGAACTCTTTCCCTACACGACGCTCTTCCGATCTNNNAGAGCTACATTGACCAGTCCT
AGAGCACACGTCTGAACTCTTTCCCTACACGACGCTCTTCCGATCTNNNAGAGCTACATTGACCAGTCCT
AGAGCACACGTCTGAACTCTTTCCCTACACGACGCTCTTCCGATCTNNNAGAGCTACATTGACCAGTCCT
AGCACACGTCTGAACTCTTTCCCTACACGACGCTCTTCCGATCTNNNTTTGCCCCTCTTGGATGTATGA
AGCACACGTCTGAACTCTTTCCCTACACGACGCTCTTCCGATCTNNNTTTGCCCCTCTTGGATGTATGA
AGCACACGTCTGAACTCTTTCCCTACACGACGCTCTTCCGATCTNNNTTTGCCCCTCTTGGATGTATGA
GAGCACACGTCTGAACTCTTTCCCTACACGACGCTCTTCCGATCTNNNTCCCGAGGTTTGGCATTACATA
GAGCACACGTCTGAACTCTTTCCCTACACGACGCTCTTCCGATCTNNNTCCCGAGGTTTGGCATTACATA

AGCACACGTCTGAACTCTTTCCCTACACGACGCTCTTCCGATCTNNNTCCCGAGGTTTGGCATTACAT
GCACACGTCTGAACTCTTTCCCTACACGACGCTCTTCCGATCTNNNCTTCTCTGTCATGTCTACCCACCC
GCACACGTCTGAACTCTTTCCCTACACGACGCTCTTCCGATCTNNNCTTCTCTGTCATGTCTACCCACCC
GCACACGTCTGAACTCTTTCCCTACACGACGCTCTTCCGATCTNNNCTTCTCTGTCATGTCTACCCACCC
GCACACGTCTGAACTCTTTCCCTACACGACGCTCTTCCGATCTNNNTGGAGGCATCTCATTCTTTTGG
GCACACGTCTGAACTCTTTCCCTACACGACGCTCTTCCGATCTNNNTGGAGGCATCTCATTCTTTTGG
GCACACGTCTGAACTCTTTCCCTACACGACGCTCTTCCGATCTNNNTGGAGGCATCTCATTCTTTTGG
AGCACACGTCTGAACTCTTTCCCTACACGACGCTCTTCCGATCTNNNAAACTGTACTCCCAAATCCTCC
AGCACACGTCTGAACTCTTTCCCTACACGACGCTCTTCCGATCTNNNAAACTGTACTCCCAAATCCTCC
AGCACACGTCTGAACTCTTTCCCTACACGACGCTCTTCCGATCTNNNAAACTGTACTCCCAAATCCTCC
ACACGTCTGAACTCTTTCCCTACACGACGCTCTTCCGATCTNNNTTCTGGTCTGCATGGCATGAGCAT
ACACGTCTGAACTCTTTCCCTACACGACGCTCTTCCGATCTNNNTTCTGGTCTGCATGGCATGAGCAT
ACACGTCTGAACTCTTTCCCTACACGACGCTCTTCCGATCTNNNTTCTGGTCTGCATGGCATGAGCAT
GCACACGTCTGAACTCTTTCCCTACACGACGCTCTTCCGATCTNNNTGTAGTGAGCCATGATTGCACT
GCACACGTCTGAACTCTTTCCCTACACGACGCTCTTCCGATCTNNNTGTAGTGAGCCATGATTGCACT
GCACACGTCTGAACTCTTTCCCTACACGACGCTCTTCCGATCTNNNTGTAGTGAGCCATGATTGCACT
AGCACACGTCTGAACTCTTTCCCTACACGACGCTCTTCCGATCTNNNCTTGGTGAGGAACAGCTATGA
AGCACACGTCTGAACTCTTTCCCTACACGACGCTCTTCCGATCTNNNCTTGGTGAGGAACAGCTATGA
AGCACACGTCTGAACTCTTTCCCTACACGACGCTCTTCCGATCTNNNCTTGGTGAGGAACAGCTATGA
AGAGCACACGTCTGAACTCTTTCCCTACACGACGCTCTTCCGATCTNNNACCCTGCTATCAGTCTTGAG
AGAGCACACGTCTGAACTCTTTCCCTACACGACGCTCTTCCGATCTNNNACCCTGCTATCAGTCTTGAG
AGAGCACACGTCTGAACTCTTTCCCTACACGACGCTCTTCCGATCTNNNACCCTGCTATCAGTCTTGAG
AGAGCACACGTCTGAACTCTTTCCCTACACGACGCTCTTCCGATCTNNNAGGAGAATTGCTTGATCCT
AGAGCACACGTCTGAACTCTTTCCCTACACGACGCTCTTCCGATCTNNNAGGAGAATTGCTTGATCCT
AGAGCACACGTCTGAACTCTTTCCCTACACGACGCTCTTCCGATCTNNNAGGAGAATTGCTTGATCCT
AGCACACGTCTGAACTCTTTCCCTACACGACGCTCTTCCGATCTNNNCATGCCTGTAATCCCAGCTACT
AGCACACGTCTGAACTCTTTCCCTACACGACGCTCTTCCGATCTNNNCATGCCTGTAATCCCAGCTACT
AGCACACGTCTGAACTCTTTCCCTACACGACGCTCTTCCGATCTNNNCATGCCTGTAATCCCAGCTACT
AGCACACGTCTGAACTCTTTCCCTACACGACGCTCTTCCGATCTNNNCACAAACACACCCACAAACAC
AGCACACGTCTGAACTCTTTCCCTACACGACGCTCTTCCGATCTNNNCACAAACACACCCACAAACAC
AGCACACGTCTGAACTCTTTCCCTACACGACGCTCTTCCGATCTNNNCACAAACACACCCACAAACAC
AGAGCACACGTCTGAACTCTTTCCCTACACGACGCTCTTCCGATCTNNNGAGAATGGTGTGAACCTG
AGAGCACACGTCTGAACTCTTTCCCTACACGACGCTCTTCCGATCTNNNGAGAATGGTGTGAACCTG
AGAGCACACGTCTGAACTCTTTCCCTACACGACGCTCTTCCGATCTNNNGAGAATGGTGTGAACCTG
AGCACACGTCTGAACTCTTTCCCTACACGACGCTCTTCCGATCTNNNGAGCTCAAATCATCACTGCAG
AGCACACGTCTGAACTCTTTCCCTACACGACGCTCTTCCGATCTNNNGAGCTCAAATCATCACTGCAG
AGCACACGTCTGAACTCTTTCCCTACACGACGCTCTTCCGATCTNNNGAGCTCAAATCATCACTGCAG
GCACACGTCTGAACTCTTTCCCTACACGACGCTCTTCCGATCTNNNTGGGTTCTCCTTTCTCTCTCTCTC
GCACACGTCTGAACTCTTTCCCTACACGACGCTCTTCCGATCTNNNTGGGTTCTCCTTTCTCTCTCTCTC
GCACACGTCTGAACTCTTTCCCTACACGACGCTCTTCCGATCTNNNTGGGTTCTCCTTTCTCTCTCTCTC
GCACACGTCTGAACTCTTTCCCTACACGACGCTCTTCCGATCTNNNTGGCATCCCACTATGTTCTCATT
GCACACGTCTGAACTCTTTCCCTACACGACGCTCTTCCGATCTNNNTGGCATCCCACTATGTTCTCATT
GCACACGTCTGAACTCTTTCCCTACACGACGCTCTTCCGATCTNNNTGGCATCCCACTATGTTCTCATT

AGCACACGTCTGAACTCTTTCCCTACACGACGCTCTTCCGATCTNNNTCAACGTCGACCATTCTACAGT
AGCACACGTCTGAACTCTTTCCCTACACGACGCTCTTCCGATCTNNNTCAACGTCGACCATTCTACAGT
AGCACACGTCTGAACTCTTTCCCTACACGACGCTCTTCCGATCTNNNTCAACGTCGACCATTCTACAGT
AGAGCACACGTCTGAACTCTTTCCCTACACGACGCTCTTCCGATCTNNNCTGAGAAATGGAAGACCCT
AGAGCACACGTCTGAACTCTTTCCCTACACGACGCTCTTCCGATCTNNNCTGAGAAATGGAAGACCCT
AGAGCACACGTCTGAACTCTTTCCCTACACGACGCTCTTCCGATCTNNNCTGAGAAATGGAAGACCCT
AGCACACGTCTGAACTCTTTCCCTACACGACGCTCTTCCGATCTNNNACTTGAAGTACCCTCTCCCCAT
AGCACACGTCTGAACTCTTTCCCTACACGACGCTCTTCCGATCTNNNACTTGAAGTACCCTCTCCCCAT
AGCACACGTCTGAACTCTTTCCCTACACGACGCTCTTCCGATCTNNNACTTGAAGTACCCTCTCCCCAT
AGCACACGTCTGAACTCTTTCCCTACACGACGCTCTTCCGATCTNNNCTCCTGGAATCTCTTAGAACCC
AGCACACGTCTGAACTCTTTCCCTACACGACGCTCTTCCGATCTNNNCTCCTGGAATCTCTTAGAACCC
AGCACACGTCTGAACTCTTTCCCTACACGACGCTCTTCCGATCTNNNCTCCTGGAATCTCTTAGAACCC
AGCACACGTCTGAACTCTTTCCCTACACGACGCTCTTCCGATCTNNNAGGCTGAGGAAGGAGAATTG
AGCACACGTCTGAACTCTTTCCCTACACGACGCTCTTCCGATCTNNNAGGCTGAGGAAGGAGAATTG
AGCACACGTCTGAACTCTTTCCCTACACGACGCTCTTCCGATCTNNNAGGCTGAGGAAGGAGAATTG
CACGTCTGAACTCTTTCCCTACACGACGCTCTTCCGATCTNNNTCTGGGGGAAAATGAGAGGTTTTC
CACGTCTGAACTCTTTCCCTACACGACGCTCTTCCGATCTNNNTCTGGGGGAAAATGAGAGGTTTTC
CACGTCTGAACTCTTTCCCTACACGACGCTCTTCCGATCTNNNTCTGGGGGAAAATGAGAGGTTTTC
AGCACACGTCTGAACTCTTTCCCTACACGACGCTCTTCCGATCTNNNCTGCCTCCACGTCATTATCTCTT
AGCACACGTCTGAACTCTTTCCCTACACGACGCTCTTCCGATCTNNNCTGCCTCCACGTCATTATCTCTT
AGCACACGTCTGAACTCTTTCCCTACACGACGCTCTTCCGATCTNNNCTGCCTCCACGTCATTATCTCTT
AGCACACGTCTGAACTCTTTCCCTACACGACGCTCTTCCGATCTNNNTTACAGGGGATGGGAAAGCT
AGCACACGTCTGAACTCTTTCCCTACACGACGCTCTTCCGATCTNNNTTACAGGGGATGGGAAAGCT
AGCACACGTCTGAACTCTTTCCCTACACGACGCTCTTCCGATCTNNNTTACAGGGGATGGGAAAGCT
AGCACACGTCTGAACTCTTTCCCTACACGACGCTCTTCCGATCTNNNTGTACACACGAATGATCCATG
AGCACACGTCTGAACTCTTTCCCTACACGACGCTCTTCCGATCTNNNTGTACACACGAATGATCCATG
AGCACACGTCTGAACTCTTTCCCTACACGACGCTCTTCCGATCTNNNTGTACACACGAATGATCCATG
AGCACACGTCTGAACTCTTTCCCTACACGACGCTCTTCCGATCTNNNTACAATGCCTGTGCCACATTTC
AGCACACGTCTGAACTCTTTCCCTACACGACGCTCTTCCGATCTNNNTACAATGCCTGTGCCACATTTC
AGCACACGTCTGAACTCTTTCCCTACACGACGCTCTTCCGATCTNNNTACAATGCCTGTGCCACATTTC
AGCACACGTCTGAACTCTTTCCCTACACGACGCTCTTCCGATCTNNNAAATTAGCTGGGCATGGTGG
AGCACACGTCTGAACTCTTTCCCTACACGACGCTCTTCCGATCTNNNAAATTAGCTGGGCATGGTGG
AGCACACGTCTGAACTCTTTCCCTACACGACGCTCTTCCGATCTNNNAAATTAGCTGGGCATGGTGG
AGCACACGTCTGAACTCTTTCCCTACACGACGCTCTTCCGATCTNNNGGACAGCTGGATGATTCATCT
AGCACACGTCTGAACTCTTTCCCTACACGACGCTCTTCCGATCTNNNGGACAGCTGGATGATTCATCT
AGCACACGTCTGAACTCTTTCCCTACACGACGCTCTTCCGATCTNNNGGACAGCTGGATGATTCATCT
GCACACGTCTGAACTCTTTCCCTACACGACGCTCTTCCGATCTNNNAAGCAAAGTGAATGTACCTCCC
GCACACGTCTGAACTCTTTCCCTACACGACGCTCTTCCGATCTNNNAAGCAAAGTGAATGTACCTCCC
GCACACGTCTGAACTCTTTCCCTACACGACGCTCTTCCGATCTNNNAAGCAAAGTGAATGTACCTCCC
CACACGTCTGAACTCTTTCCCTACACGACGCTCTTCCGATCTNNNCTTCATCCATTCACCCCCAAGTACA
CACACGTCTGAACTCTTTCCCTACACGACGCTCTTCCGATCTNNNCTTCATCCATTCACCCCCAAGTACA
CACACGTCTGAACTCTTTCCCTACACGACGCTCTTCCGATCTNNNCTTCATCCATTCACCCCCAAGTACA
GCACACGTCTGAACTCTTTCCCTACACGACGCTCTTCCGATCTNNNGCGTGTTAGGCTGAAGTCTCTA

AGCACACGTCTGAACTCTTTCCCTACACGACGCTCTTCCGATCTNNNGCGTGTTAGGCTGAAGTCTCTA
AGCACACGTCTGAACTCTTTCCCTACACGACGCTCTTCCGATCTNNNGCGTGTTAGGCTGAAGTCTCTA
GAGCACACGTCTGAACTCTTTCCCTACACGACGCTCTTCCGATCTNNNCCTTGGCCAAAACATAGCAAA
GAGCACACGTCTGAACTCTTTCCCTACACGACGCTCTTCCGATCTNNNCCTTGGCCAAAACATAGCAAA
GAGCACACGTCTGAACTCTTTCCCTACACGACGCTCTTCCGATCTNNNCCTTGGCCAAAACATAGCAAA
GAGCACACGTCTGAACTCTTTCCCTACACGACGCTCTTCCGATCTNNNAAAAAATTAGCTGGGTGTGG
GAGCACACGTCTGAACTCTTTCCCTACACGACGCTCTTCCGATCTNNNAAAAAATTAGCTGGGTGTGG
GAGCACACGTCTGAACTCTTTCCCTACACGACGCTCTTCCGATCTNNNAAAAAATTAGCTGGGTGTGG
AGCACACGTCTGAACTCTTTCCCTACACGACGCTCTTCCGATCTNNNTGTAGAGATGGGGTTTCACTG
AGCACACGTCTGAACTCTTTCCCTACACGACGCTCTTCCGATCTNNNTGTAGAGATGGGGTTTCACTG
AGCACACGTCTGAACTCTTTCCCTACACGACGCTCTTCCGATCTNNNTGTAGAGATGGGGTTTCACTG
AGCACACGTCTGAACTCTTTCCCTACACGACGCTCTTCCGATCTNNNCACTTTTGCATGTGTGAATCCCT
AGCACACGTCTGAACTCTTTCCCTACACGACGCTCTTCCGATCTNNNCACTTTTGCATGTGTGAATCCCT
AGCACACGTCTGAACTCTTTCCCTACACGACGCTCTTCCGATCTNNNCACTTTTGCATGTGTGAATCCCT
CACACGTCTGAACTCTTTCCCTACACGACGCTCTTCCGATCTNNNTCGCAAAAATCTAAAACTTCTGCT
CACACGTCTGAACTCTTTCCCTACACGACGCTCTTCCGATCTNNNTCGCAAAAATCTAAAACTTCTGCT
CACACGTCTGAACTCTTTCCCTACACGACGCTCTTCCGATCTNNNTCGCAAAAATCTAAAACTTCTGCT
AGCACACGTCTGAACTCTTTCCCTACACGACGCTCTTCCGATCTNNNGTAATCTCAGCTACTCAGGAG
AGCACACGTCTGAACTCTTTCCCTACACGACGCTCTTCCGATCTNNNGTAATCTCAGCTACTCAGGAG
AGCACACGTCTGAACTCTTTCCCTACACGACGCTCTTCCGATCTNNNGTAATCTCAGCTACTCAGGAG
GCACACGTCTGAACTCTTTCCCTACACGACGCTCTTCCGATCTNNNTTTCAGTCAGTAGGTCACTCGTG
GCACACGTCTGAACTCTTTCCCTACACGACGCTCTTCCGATCTNNNTTTCAGTCAGTAGGTCACTCGTG
GCACACGTCTGAACTCTTTCCCTACACGACGCTCTTCCGATCTNNNTTTCAGTCAGTAGGTCACTCGTG
GAGCACACGTCTGAACTCTTTCCCTACACGACGCTCTTCCGATCTNNNGAGAATCCCTTGAACCCAGGA
GAGCACACGTCTGAACTCTTTCCCTACACGACGCTCTTCCGATCTNNNGAGAATCCCTTGAACCCAGGA
GAGCACACGTCTGAACTCTTTCCCTACACGACGCTCTTCCGATCTNNNGAGAATCCCTTGAACCCAGGA
GCACACGTCTGAACTCTTTCCCTACACGACGCTCTTCCGATCTNNNAGGAGCAAGGTACAGGGAAGT
GCACACGTCTGAACTCTTTCCCTACACGACGCTCTTCCGATCTNNNAGGAGCAAGGTACAGGGAAGT
GCACACGTCTGAACTCTTTCCCTACACGACGCTCTTCCGATCTNNNAGGAGCAAGGTACAGGGAAGT
CACACGTCTGAACTCTTTCCCTACACGACGCTCTTCCGATCTNNNAAACACATGTGTCCCCCTTTCTTCAT
CACACGTCTGAACTCTTTCCCTACACGACGCTCTTCCGATCTNNNAAACACATGTGTCCCCCTTTCTTCAT
CACACGTCTGAACTCTTTCCCTACACGACGCTCTTCCGATCTNNNAAACACATGTGTCCCCCTTTCTTCAT
GAGCACACGTCTGAACTCTTTCCCTACACGACGCTCTTCCGATCTNNNTGTTTTCATAACTGCCACTGCT
GAGCACACGTCTGAACTCTTTCCCTACACGACGCTCTTCCGATCTNNNTGTTTTCATAACTGCCACTGCT
GAGCACACGTCTGAACTCTTTCCCTACACGACGCTCTTCCGATCTNNNTGTTTTCATAACTGCCACTGCT
GAGCACACGTCTGAACTCTTTCCCTACACGACGCTCTTCCGATCTNNNATGTATGCATGTAAGCGTGTG
GAGCACACGTCTGAACTCTTTCCCTACACGACGCTCTTCCGATCTNNNATGTATGCATGTAAGCGTGTG
GAGCACACGTCTGAACTCTTTCCCTACACGACGCTCTTCCGATCTNNNATGTATGCATGTAAGCGTGTG
GAGCACACGTCTGAACTCTTTCCCTACACGACGCTCTTCCGATCTNNNGCACCTGTAATCTCAGCTACTC
GAGCACACGTCTGAACTCTTTCCCTACACGACGCTCTTCCGATCTNNNGCACCTGTAATCTCAGCTACTC
GAGCACACGTCTGAACTCTTTCCCTACACGACGCTCTTCCGATCTNNNGCACCTGTAATCTCAGCTACTC
AGCACACGTCTGAACTCTTTCCCTACACGACGCTCTTCCGATCTNNNAGAGTTGGATATCAGTGCATG
AGCACACGTCTGAACTCTTTCCCTACACGACGCTCTTCCGATCTNNNAGAGTTGGATATCAGTGCATG

AGCACACGTCTGAACTCTTTCCCTACACGACGCTCTTCCGATCTNNNAGAGTTGGATATCAGTGCATG
AGCACACGTCTGAACTCTTTCCCTACACGACGCTCTTCCGATCTNNNTCTGCAAAGATAAGCAGCTCT
AGCACACGTCTGAACTCTTTCCCTACACGACGCTCTTCCGATCTNNNTCTGCAAAGATAAGCAGCTCT
AGCACACGTCTGAACTCTTTCCCTACACGACGCTCTTCCGATCTNNNTCTGCAAAGATAAGCAGCTCT
AGCACACGTCTGAACTCTTTCCCTACACGACGCTCTTCCGATCTNNNTTTGAGTTCCTTCTTGGTCTGCT
AGCACACGTCTGAACTCTTTCCCTACACGACGCTCTTCCGATCTNNNTTTGAGTTCCTTCTTGGTCTGCT
AGCACACGTCTGAACTCTTTCCCTACACGACGCTCTTCCGATCTNNNTTTGAGTTCCTTCTTGGTCTGCT
GCACACGTCTGAACTCTTTCCCTACACGACGCTCTTCCGATCTNNNAAGGAGGTACCACTAGTCAACA
GCACACGTCTGAACTCTTTCCCTACACGACGCTCTTCCGATCTNNNAAGGAGGTACCACTAGTCAACA
GCACACGTCTGAACTCTTTCCCTACACGACGCTCTTCCGATCTNNNAAGGAGGTACCACTAGTCAACA
GAGCACACGTCTGAACTCTTTCCCTACACGACGCTCTTCCGATCTNNNATGCCTGTAGTCCCAGCTACT
GAGCACACGTCTGAACTCTTTCCCTACACGACGCTCTTCCGATCTNNNATGCCTGTAGTCCCAGCTACT
GAGCACACGTCTGAACTCTTTCCCTACACGACGCTCTTCCGATCTNNNATGCCTGTAGTCCCAGCTACT
AGCACACGTCTGAACTCTTTCCCTACACGACGCTCTTCCGATCTNNNCTGGCTCTACAATTCCCTTTCCT
AGCACACGTCTGAACTCTTTCCCTACACGACGCTCTTCCGATCTNNNCTGGCTCTACAATTCCCTTTCCT
AGCACACGTCTGAACTCTTTCCCTACACGACGCTCTTCCGATCTNNNCTGGCTCTACAATTCCCTTTCCT
AGAGCACACGTCTGAACTCTTTCCCTACACGACGCTCTTCCGATCTNNNCACCTATCTGCTATAGTGAG
AGAGCACACGTCTGAACTCTTTCCCTACACGACGCTCTTCCGATCTNNNCACCTATCTGCTATAGTGAG
AGAGCACACGTCTGAACTCTTTCCCTACACGACGCTCTTCCGATCTNNNCACCTATCTGCTATAGTGAG
GAGCACACGTCTGAACTCTTTCCCTACACGACGCTCTTCCGATCTNNNGAGGCAGGATAATTGCTTGA
GAGCACACGTCTGAACTCTTTCCCTACACGACGCTCTTCCGATCTNNNGAGGCAGGATAATTGCTTGA
GAGCACACGTCTGAACTCTTTCCCTACACGACGCTCTTCCGATCTNNNGAGGCAGGATAATTGCTTGA
AGCACACGTCTGAACTCTTTCCCTACACGACGCTCTTCCGATCTNNNCAAAACCCTACTGTCTCCACGTA
AGCACACGTCTGAACTCTTTCCCTACACGACGCTCTTCCGATCTNNNCAAAACCCTACTGTCTCCACGTA
AGCACACGTCTGAACTCTTTCCCTACACGACGCTCTTCCGATCTNNNCAAAACCCTACTGTCTCCACGTA
GAGCACACGTCTGAACTCTTTCCCTACACGACGCTCTTCCGATCTNNNAAAGGCCGTTTTGATGTGTTT
GAGCACACGTCTGAACTCTTTCCCTACACGACGCTCTTCCGATCTNNNAAAGGCCGTTTTGATGTGTTT
GAGCACACGTCTGAACTCTTTCCCTACACGACGCTCTTCCGATCTNNNAAAGGCCGTTTTGATGTGTTT
AGCACACGTCTGAACTCTTTCCCTACACGACGCTCTTCCGATCTNNNGTTTGAGTAGCGAATCTGGGTT
AGCACACGTCTGAACTCTTTCCCTACACGACGCTCTTCCGATCTNNNGTTTGAGTAGCGAATCTGGGTT
AGCACACGTCTGAACTCTTTCCCTACACGACGCTCTTCCGATCTNNNGTTTGAGTAGCGAATCTGGGTT
GCACACGTCTGAACTCTTTCCCTACACGACGCTCTTCCGATCTNNNTACACTGTGCAGATTTTAGGCCT
GCACACGTCTGAACTCTTTCCCTACACGACGCTCTTCCGATCTNNNTACACTGTGCAGATTTTAGGCCT
GCACACGTCTGAACTCTTTCCCTACACGACGCTCTTCCGATCTNNNTACACTGTGCAGATTTTAGGCCT
AGCACACGTCTGAACTCTTTCCCTACACGACGCTCTTCCGATCTNNNTGCATTATTTTTTCAGTGCAGA
AGCACACGTCTGAACTCTTTCCCTACACGACGCTCTTCCGATCTNNNTGCATTATTTTTTCAGTGCAGA
AGCACACGTCTGAACTCTTTCCCTACACGACGCTCTTCCGATCTNNNTGCATTATTTTTTCAGTGCAGA
AGCACACGTCTGAACTCTTTCCCTACACGACGCTCTTCCGATCTNNNGAGGCTGAGGCAGGAGAATG
AGCACACGTCTGAACTCTTTCCCTACACGACGCTCTTCCGATCTNNNGAGGCTGAGGCAGGAGAATG
AGCACACGTCTGAACTCTTTCCCTACACGACGCTCTTCCGATCTNNNGAGGCTGAGGCAGGAGAATG
AGAGCACACGTCTGAACTCTTTCCCTACACGACGCTCTTCCGATCTNNNGAGGTTGTAGTGAGCCAA
AGAGCACACGTCTGAACTCTTTCCCTACACGACGCTCTTCCGATCTNNNGAGGTTGTAGTGAGCCAA
AGAGCACACGTCTGAACTCTTTCCCTACACGACGCTCTTCCGATCTNNNGAGGTTGTAGTGAGCCAA

GCACACGTCTGAACTCTTTCCCTACACGACGCTCTTCCGATCTNNNCAGGTGAGGCTTACATACAAGG
GCACACGTCTGAACTCTTTCCCTACACGACGCTCTTCCGATCTNNNCAGGTGAGGCTTACATACAAGG
GCACACGTCTGAACTCTTTCCCTACACGACGCTCTTCCGATCTNNNCAGGTGAGGCTTACATACAAGG
AGCACACGTCTGAACTCTTTCCCTACACGACGCTCTTCCGATCTNNNGTCTTGAGACAGGTGGACACTA
AGCACACGTCTGAACTCTTTCCCTACACGACGCTCTTCCGATCTNNNGTCTTGAGACAGGTGGACACTA
AGCACACGTCTGAACTCTTTCCCTACACGACGCTCTTCCGATCTNNNGTCTTGAGACAGGTGGACACTA
GCACACGTCTGAACTCTTTCCCTACACGACGCTCTTCCGATCTNNNCATGATTATCCTTGGCAGCATGG
GCACACGTCTGAACTCTTTCCCTACACGACGCTCTTCCGATCTNNNCATGATTATCCTTGGCAGCATGG
GCACACGTCTGAACTCTTTCCCTACACGACGCTCTTCCGATCTNNNCATGATTATCCTTGGCAGCATGG
AGCACACGTCTGAACTCTTTCCCTACACGACGCTCTTCCGATCTNNNATGAGACCAGCATCATCCTGAT
AGCACACGTCTGAACTCTTTCCCTACACGACGCTCTTCCGATCTNNNATGAGACCAGCATCATCCTGAT
AGCACACGTCTGAACTCTTTCCCTACACGACGCTCTTCCGATCTNNNATGAGACCAGCATCATCCTGAT
GAGCACACGTCTGAACTCTTTCCCTACACGACGCTCTTCCGATCTNNNGGAGGGGAAAGAAGAGACA
GAGCACACGTCTGAACTCTTTCCCTACACGACGCTCTTCCGATCTNNNGGAGGGGAAAGAAGAGACA
GAGCACACGTCTGAACTCTTTCCCTACACGACGCTCTTCCGATCTNNNGGAGGGGAAAGAAGAGACA
AGCACACGTCTGAACTCTTTCCCTACACGACGCTCTTCCGATCTNNNATTCAGGCAAAGTTAGTCCTGG
AGCACACGTCTGAACTCTTTCCCTACACGACGCTCTTCCGATCTNNNATTCAGGCAAAGTTAGTCCTGG
AGCACACGTCTGAACTCTTTCCCTACACGACGCTCTTCCGATCTNNNATTCAGGCAAAGTTAGTCCTGG
AGCACACGTCTGAACTCTTTCCCTACACGACGCTCTTCCGATCTNNNGATCCCTTTATCCGGGTCTCGCA
AGCACACGTCTGAACTCTTTCCCTACACGACGCTCTTCCGATCTNNNGATCCCTTTATCCGGGTCTCGCA
AGCACACGTCTGAACTCTTTCCCTACACGACGCTCTTCCGATCTNNNGATCCCTTTATCCGGGTCTCGCA
GAGCACACGTCTGAACTCTTTCCCTACACGACGCTCTTCCGATCTNNNCTTATGAAGGCTCCTGTGTCA
GAGCACACGTCTGAACTCTTTCCCTACACGACGCTCTTCCGATCTNNNCTTATGAAGGCTCCTGTGTCA
GAGCACACGTCTGAACTCTTTCCCTACACGACGCTCTTCCGATCTNNNCTTATGAAGGCTCCTGTGTCA
AGCACACGTCTGAACTCTTTCCCTACACGACGCTCTTCCGATCTNNNATACAGTGTGGGATAGCTGCA
AGCACACGTCTGAACTCTTTCCCTACACGACGCTCTTCCGATCTNNNATACAGTGTGGGATAGCTGCA
AGCACACGTCTGAACTCTTTCCCTACACGACGCTCTTCCGATCTNNNATACAGTGTGGGATAGCTGCA
GAGCACACGTCTGAACTCTTTCCCTACACGACGCTCTTCCGATCTNNNCACTGTGTTTGGCTTACCGA
GAGCACACGTCTGAACTCTTTCCCTACACGACGCTCTTCCGATCTNNNCACTGTGTTTGGCTTACCGA
GAGCACACGTCTGAACTCTTTCCCTACACGACGCTCTTCCGATCTNNNCACTGTGTTTGGCTTACCGA
GAGCACACGTCTGAACTCTTTCCCTACACGACGCTCTTCCGATCTNNNTGACTTGAACTTGAGAGGTG
GAGCACACGTCTGAACTCTTTCCCTACACGACGCTCTTCCGATCTNNNTGACTTGAACTTGAGAGGTG
GAGCACACGTCTGAACTCTTTCCCTACACGACGCTCTTCCGATCTNNNTGACTTGAACTTGAGAGGTG
ACACGTCTGAACTCTTTCCCTACACGACGCTCTTCCGATCTNNNATGAATGTTCCATCAAAGGGCATC
ACACGTCTGAACTCTTTCCCTACACGACGCTCTTCCGATCTNNNATGAATGTTCCATCAAAGGGCATC
ACACGTCTGAACTCTTTCCCTACACGACGCTCTTCCGATCTNNNATGAATGTTCCATCAAAGGGCATC
AGAGCACACGTCTGAACTCTTTCCCTACACGACGCTCTTCCGATCTNNNATGTTGCCACCTCTCAGAG
AGAGCACACGTCTGAACTCTTTCCCTACACGACGCTCTTCCGATCTNNNATGTTGCCACCTCTCAGAG
AGAGCACACGTCTGAACTCTTTCCCTACACGACGCTCTTCCGATCTNNNATGTTGCCACCTCTCAGAG
AGCACACGTCTGAACTCTTTCCCTACACGACGCTCTTCCGATCTNNNGGTGACAGAGTGAGAACATG
AGCACACGTCTGAACTCTTTCCCTACACGACGCTCTTCCGATCTNNNGGTGACAGAGTGAGAACATG
AGCACACGTCTGAACTCTTTCCCTACACGACGCTCTTCCGATCTNNNGGTGACAGAGTGAGAACATG
GAGCACACGTCTGAACTCTTTCCCTACACGACGCTCTTCCGATCTNNNGGTAACAGAAGGACGGACTA

GAGCACACGTCTGAACTCTTTCCCTACACGACGCTCTTCCGATCTNNNGGTAACAGAAGGACGGACT
GAGCACACGTCTGAACTCTTTCCCTACACGACGCTCTTCCGATCTNNNGGTAACAGAAGGACGGACT
AGCACACGTCTGAACTCTTTCCCTACACGACGCTCTTCCGATCTNNNTAAGCTTTTGATGTGCTGCT
AGCACACGTCTGAACTCTTTCCCTACACGACGCTCTTCCGATCTNNNTAAGCTTTTGATGTGCTGCT
AGCACACGTCTGAACTCTTTCCCTACACGACGCTCTTCCGATCTNNNTAAGCTTTTGATGTGCTGCT
AGCACACGTCTGAACTCTTTCCCTACACGACGCTCTTCCGATCTNNNTGGCTCCATTTCTGAATCCTTGT
AGCACACGTCTGAACTCTTTCCCTACACGACGCTCTTCCGATCTNNNTGGCTCCATTTCTGAATCCTTGT
AGCACACGTCTGAACTCTTTCCCTACACGACGCTCTTCCGATCTNNNTGGCTCCATTTCTGAATCCTTGT
AGCACACGTCTGAACTCTTTCCCTACACGACGCTCTTCCGATCTNNNCCTACCCCCTGTGATAATCTGT
AGCACACGTCTGAACTCTTTCCCTACACGACGCTCTTCCGATCTNNNCCTACCCCCTGTGATAATCTGT
AGCACACGTCTGAACTCTTTCCCTACACGACGCTCTTCCGATCTNNNCCTACCCCCTGTGATAATCTGT
AGCACACGTCTGAACTCTTTCCCTACACGACGCTCTTCCGATCTNNNGACCTCAGACTCTCTGTTTCCA
AGCACACGTCTGAACTCTTTCCCTACACGACGCTCTTCCGATCTNNNGACCTCAGACTCTCTGTTTCCA
AGCACACGTCTGAACTCTTTCCCTACACGACGCTCTTCCGATCTNNNGACCTCAGACTCTCTGTTTCCA
GAGCACACGTCTGAACTCTTTCCCTACACGACGCTCTTCCGATCTNNNAAGAGTTGGACCTACCTACCT
GAGCACACGTCTGAACTCTTTCCCTACACGACGCTCTTCCGATCTNNNAAGAGTTGGACCTACCTACCT
GAGCACACGTCTGAACTCTTTCCCTACACGACGCTCTTCCGATCTNNNAAGAGTTGGACCTACCTACCT
GCACACGTCTGAACTCTTTCCCTACACGACGCTCTTCCGATCTNNNGGACAGGGAGATTAGAACACA
GCACACGTCTGAACTCTTTCCCTACACGACGCTCTTCCGATCTNNNGGACAGGGAGATTAGAACACA
GCACACGTCTGAACTCTTTCCCTACACGACGCTCTTCCGATCTNNNGGACAGGGAGATTAGAACACA
GAGCACACGTCTGAACTCTTTCCCTACACGACGCTCTTCCGATCTNNNAGACGCATTTTTCTTCACAAC
GAGCACACGTCTGAACTCTTTCCCTACACGACGCTCTTCCGATCTNNNAGACGCATTTTTCTTCACAAC
GAGCACACGTCTGAACTCTTTCCCTACACGACGCTCTTCCGATCTNNNAGACGCATTTTTCTTCACAAC
GCACACGTCTGAACTCTTTCCCTACACGACGCTCTTCCGATCTNNNTCATCAACATCAAGTTTGCCGAC
GCACACGTCTGAACTCTTTCCCTACACGACGCTCTTCCGATCTNNNTCATCAACATCAAGTTTGCCGAC
GCACACGTCTGAACTCTTTCCCTACACGACGCTCTTCCGATCTNNNTCATCAACATCAAGTTTGCCGAC
AGCACACGTCTGAACTCTTTCCCTACACGACGCTCTTCCGATCTNNNGAAAAAAACCCTTCTCGCTCT
AGCACACGTCTGAACTCTTTCCCTACACGACGCTCTTCCGATCTNNNGAAAAAAACCCTTCTCGCTCT
AGCACACGTCTGAACTCTTTCCCTACACGACGCTCTTCCGATCTNNNGAAAAAAACCCTTCTCGCTCT
AGCACACGTCTGAACTCTTTCCCTACACGACGCTCTTCCGATCTNNNTGAGGCAGGAGAATGGTATG
AGCACACGTCTGAACTCTTTCCCTACACGACGCTCTTCCGATCTNNNTGAGGCAGGAGAATGGTATG
AGCACACGTCTGAACTCTTTCCCTACACGACGCTCTTCCGATCTNNNTGAGGCAGGAGAATGGTATG
GCACACGTCTGAACTCTTTCCCTACACGACGCTCTTCCGATCTNNNTCATCTCCTGCAAACACACAAAC
GCACACGTCTGAACTCTTTCCCTACACGACGCTCTTCCGATCTNNNTCATCTCCTGCAAACACACAAAC
GCACACGTCTGAACTCTTTCCCTACACGACGCTCTTCCGATCTNNNTCATCTCCTGCAAACACACAAAC
GCACACGTCTGAACTCTTTCCCTACACGACGCTCTTCCGATCTNNNAACAATGCTGCTATGAACAGTG
GCACACGTCTGAACTCTTTCCCTACACGACGCTCTTCCGATCTNNNAACAATGCTGCTATGAACAGTG
GCACACGTCTGAACTCTTTCCCTACACGACGCTCTTCCGATCTNNNAACAATGCTGCTATGAACAGTG
GAGCACACGTCTGAACTCTTTCCCTACACGACGCTCTTCCGATCTNNNAAACTCCTGACCTCAAGTGAT
GAGCACACGTCTGAACTCTTTCCCTACACGACGCTCTTCCGATCTNNNAAACTCCTGACCTCAAGTGAT
GAGCACACGTCTGAACTCTTTCCCTACACGACGCTCTTCCGATCTNNNAAACTCCTGACCTCAAGTGAT
AGCACACGTCTGAACTCTTTCCCTACACGACGCTCTTCCGATCTNNNGTGACGGGGACTCATAAGGT
AGCACACGTCTGAACTCTTTCCCTACACGACGCTCTTCCGATCTNNNGTGACGGGGACTCATAAGGT

GAGCACACGTCTGAACTCTTTCCCTACACGACGCTCTTCCGATCTNNNGTGACGGGGACTCATAAGGT
AGCACACGTCTGAACTCTTTCCCTACACGACGCTCTTCCGATCTNNNTGAGGCAGGAGAATCACTTGAA
AGCACACGTCTGAACTCTTTCCCTACACGACGCTCTTCCGATCTNNNTGAGGCAGGAGAATCACTTGAA
AGCACACGTCTGAACTCTTTCCCTACACGACGCTCTTCCGATCTNNNTGAGGCAGGAGAATCACTTGAA
AGCACACGTCTGAACTCTTTCCCTACACGACGCTCTTCCGATCTNNNTCAGTCTTTTTCCACCTGTGTCAC
AGCACACGTCTGAACTCTTTCCCTACACGACGCTCTTCCGATCTNNNTCAGTCTTTTTCCACCTGTGTCAC
AGCACACGTCTGAACTCTTTCCCTACACGACGCTCTTCCGATCTNNNTCAGTCTTTTTCCACCTGTGTCAC
AGCACACGTCTGAACTCTTTCCCTACACGACGCTCTTCCGATCTNNNCACCTGTAGTCCCAGCTACTTGC
AGCACACGTCTGAACTCTTTCCCTACACGACGCTCTTCCGATCTNNNCACCTGTAGTCCCAGCTACTTGC
AGCACACGTCTGAACTCTTTCCCTACACGACGCTCTTCCGATCTNNNCACCTGTAGTCCCAGCTACTTGC
GAGCACACGTCTGAACTCTTTCCCTACACGACGCTCTTCCGATCTNNNGGTTCCTGTATTCCCAGCTACT
GAGCACACGTCTGAACTCTTTCCCTACACGACGCTCTTCCGATCTNNNGGTTCCTGTATTCCCAGCTACT
GAGCACACGTCTGAACTCTTTCCCTACACGACGCTCTTCCGATCTNNNGGTTCCTGTATTCCCAGCTACT
GAGCACACGTCTGAACTCTTTCCCTACACGACGCTCTTCCGATCTNNNCCTATGTTCCCCTTTTTGCTGT
GAGCACACGTCTGAACTCTTTCCCTACACGACGCTCTTCCGATCTNNNCCTATGTTCCCCTTTTTGCTGT
GAGCACACGTCTGAACTCTTTCCCTACACGACGCTCTTCCGATCTNNNCCTATGTTCCCCTTTTTGCTGT
AGCACACGTCTGAACTCTTTCCCTACACGACGCTCTTCCGATCTNNNACTCAATAGCAAAAACTCAATCC
AGCACACGTCTGAACTCTTTCCCTACACGACGCTCTTCCGATCTNNNACTCAATAGCAAAAACTCAATCC
AGCACACGTCTGAACTCTTTCCCTACACGACGCTCTTCCGATCTNNNACTCAATAGCAAAAACTCAATCC
AGCACACGTCTGAACTCTTTCCCTACACGACGCTCTTCCGATCTNNNAGTGAGCTATGATTGTGCCAG
AGCACACGTCTGAACTCTTTCCCTACACGACGCTCTTCCGATCTNNNAGTGAGCTATGATTGTGCCAG
AGCACACGTCTGAACTCTTTCCCTACACGACGCTCTTCCGATCTNNNAGTGAGCTATGATTGTGCCAG
AGCACACGTCTGAACTCTTTCCCTACACGACGCTCTTCCGATCTNNNTATTGCTGAGTCCTCTGATTACA
AGCACACGTCTGAACTCTTTCCCTACACGACGCTCTTCCGATCTNNNTATTGCTGAGTCCTCTGATTACA
AGCACACGTCTGAACTCTTTCCCTACACGACGCTCTTCCGATCTNNNTATTGCTGAGTCCTCTGATTACA
GAGCACACGTCTGAACTCTTTCCCTACACGACGCTCTTCCGATCTNNNCAACAGTTTGGGAGAATGAG
GAGCACACGTCTGAACTCTTTCCCTACACGACGCTCTTCCGATCTNNNCAACAGTTTGGGAGAATGAG
GAGCACACGTCTGAACTCTTTCCCTACACGACGCTCTTCCGATCTNNNCAACAGTTTGGGAGAATGAG
AGAGCACACGTCTGAACTCTTTCCCTACACGACGCTCTTCCGATCTNNNAGAGAGAGACAGACAAGGA
AGAGCACACGTCTGAACTCTTTCCCTACACGACGCTCTTCCGATCTNNNAGAGAGAGACAGACAAGGA
AGAGCACACGTCTGAACTCTTTCCCTACACGACGCTCTTCCGATCTNNNAGAGAGAGACAGACAAGGA
GCACACGTCTGAACTCTTTCCCTACACGACGCTCTTCCGATCTNNNTGTGTAGGAATCTTCAGGCTTCC
GCACACGTCTGAACTCTTTCCCTACACGACGCTCTTCCGATCTNNNTGTGTAGGAATCTTCAGGCTTCC
GCACACGTCTGAACTCTTTCCCTACACGACGCTCTTCCGATCTNNNTGTGTAGGAATCTTCAGGCTTCC
GCACACGTCTGAACTCTTTCCCTACACGACGCTCTTCCGATCTNNNCTAGTTCCCAGAGCTTTGCCATAC
GCACACGTCTGAACTCTTTCCCTACACGACGCTCTTCCGATCTNNNCTAGTTCCCAGAGCTTTGCCATAC
GCACACGTCTGAACTCTTTCCCTACACGACGCTCTTCCGATCTNNNCTAGTTCCCAGAGCTTTGCCATAC
AGCACACGTCTGAACTCTTTCCCTACACGACGCTCTTCCGATCTNNNAAGGTCTTTTGAGTGAGAGGCT
AGCACACGTCTGAACTCTTTCCCTACACGACGCTCTTCCGATCTNNNAAGGTCTTTTGAGTGAGAGGCT
AGCACACGTCTGAACTCTTTCCCTACACGACGCTCTTCCGATCTNNNAAGGTCTTTTGAGTGAGAGGCT
GAGCACACGTCTGAACTCTTTCCCTACACGACGCTCTTCCGATCTNNNTGAGGCAGGAGAATCACTTGA
GAGCACACGTCTGAACTCTTTCCCTACACGACGCTCTTCCGATCTNNNTGAGGCAGGAGAATCACTTGA
GAGCACACGTCTGAACTCTTTCCCTACACGACGCTCTTCCGATCTNNNTGAGGCAGGAGAATCACTTGA

AGCACACGTCTGAACTCTTTCCCTACACGACGCTCTTCCGATCTNNNAGACCAAGTCTCTGACTCTGAG

AGCACACGTCTGAACTCTTTCCCTACACGACGCTCTTCCGATCTNNNAGACCAAGTCTCTGACTCTGAG

AGCACACGTCTGAACTCTTTCCCTACACGACGCTCTTCCGATCTNNNAGACCAAGTCTCTGACTCTGAG

GAGCACACGTCTGAACTCTTTCCCTACACGACGCTCTTCCGATCTNNNCTCACCCTACCTGACATAAAC

GAGCACACGTCTGAACTCTTTCCCTACACGACGCTCTTCCGATCTNNNCTCACCCTACCTGACATAAAC

GAGCACACGTCTGAACTCTTTCCCTACACGACGCTCTTCCGATCTNNNCTCACCCTACCTGACATAAAC

AGCACACGTCTGAACTCTTTCCCTACACGACGCTCTTCCGATCTNNNGATTACAGGTGTGAGCTACTG

AGCACACGTCTGAACTCTTTCCCTACACGACGCTCTTCCGATCTNNNGATTACAGGTGTGAGCTACTG

AGCACACGTCTGAACTCTTTCCCTACACGACGCTCTTCCGATCTNNNGATTACAGGTGTGAGCTACTG

AGAGCACACGTCTGAACTCTTTCCCTACACGACGCTCTTCCGATCTNNNTCAGTCAAATAGTTGGCACT

AGAGCACACGTCTGAACTCTTTCCCTACACGACGCTCTTCCGATCTNNNTCAGTCAAATAGTTGGCACT

AGAGCACACGTCTGAACTCTTTCCCTACACGACGCTCTTCCGATCTNNNTCAGTCAAATAGTTGGCACT

AGCACACGTCTGAACTCTTTCCCTACACGACGCTCTTCCGATCTNNNCTGAGGCAGGAGAATCACTTG

AGCACACGTCTGAACTCTTTCCCTACACGACGCTCTTCCGATCTNNNCTGAGGCAGGAGAATCACTTG

AGCACACGTCTGAACTCTTTCCCTACACGACGCTCTTCCGATCTNNNCTGAGGCAGGAGAATCACTTG

AGCACACGTCTGAACTCTTTCCCTACACGACGCTCTTCCGATCTNNNTTGCATTCATAGCTGGGCAGT

AGCACACGTCTGAACTCTTTCCCTACACGACGCTCTTCCGATCTNNNTTGCATTCATAGCTGGGCAGT

AGCACACGTCTGAACTCTTTCCCTACACGACGCTCTTCCGATCTNNNTTGCATTCATAGCTGGGCAGT

AGCACACGTCTGAACTCTTTCCCTACACGACGCTCTTCCGATCTNNNATTTGGTCTGTACTCCCACCTTC

AGCACACGTCTGAACTCTTTCCCTACACGACGCTCTTCCGATCTNNNATTTGGTCTGTACTCCCACCTTC

AGCACACGTCTGAACTCTTTCCCTACACGACGCTCTTCCGATCTNNNATTTGGTCTGTACTCCCACCTTC

AGAGCACACGTCTGAACTCTTTCCCTACACGACGCTCTTCCGATCTNNNCCATAGCACCCTCCAGAAGA

AGAGCACACGTCTGAACTCTTTCCCTACACGACGCTCTTCCGATCTNNNCCATAGCACCCTCCAGAAGA

AGAGCACACGTCTGAACTCTTTCCCTACACGACGCTCTTCCGATCTNNNCCATAGCACCCTCCAGAAGA

GAGCACACGTCTGAACTCTTTCCCTACACGACGCTCTTCCGATCTNNNCTTTTCAGTTCCATCAGCACCA

GAGCACACGTCTGAACTCTTTCCCTACACGACGCTCTTCCGATCTNNNCTTTTCAGTTCCATCAGCACCA

GAGCACACGTCTGAACTCTTTCCCTACACGACGCTCTTCCGATCTNNNCTTTTCAGTTCCATCAGCACCA

GAGCACACGTCTGAACTCTTTCCCTACACGACGCTCTTCCGATCTNNNAATGCTCCATGCACAGGATTA

GAGCACACGTCTGAACTCTTTCCCTACACGACGCTCTTCCGATCTNNNAATGCTCCATGCACAGGATTA

GAGCACACGTCTGAACTCTTTCCCTACACGACGCTCTTCCGATCTNNNAATGCTCCATGCACAGGATTA

GAGCACACGTCTGAACTCTTTCCCTACACGACGCTCTTCCGATCTNNNACCTGTAACTGTGAGATGGC

GAGCACACGTCTGAACTCTTTCCCTACACGACGCTCTTCCGATCTNNNACCTGTAACTGTGAGATGGC

GAGCACACGTCTGAACTCTTTCCCTACACGACGCTCTTCCGATCTNNNACCTGTAACTGTGAGATGGC

GAGCACACGTCTGAACTCTTTCCCTACACGACGCTCTTCCGATCTNNNTTCACTGCATTCTAACCTGGTG

GAGCACACGTCTGAACTCTTTCCCTACACGACGCTCTTCCGATCTNNNTTCACTGCATTCTAACCTGGTG

GAGCACACGTCTGAACTCTTTCCCTACACGACGCTCTTCCGATCTNNNTTCACTGCATTCTAACCTGGTG

GAGCACACGTCTGAACTCTTTCCCTACACGACGCTCTTCCGATCTNNNTCCAGGTAGTTCAGTTTCCTCG

GAGCACACGTCTGAACTCTTTCCCTACACGACGCTCTTCCGATCTNNNTCCAGGTAGTTCAGTTTCCTCG

GAGCACACGTCTGAACTCTTTCCCTACACGACGCTCTTCCGATCTNNNTCCAGGTAGTTCAGTTTCCTCG

AGCACACGTCTGAACTCTTTCCCTACACGACGCTCTTCCGATCTNNNAGCAGCATGATTCACAATTGCA

AGCACACGTCTGAACTCTTTCCCTACACGACGCTCTTCCGATCTNNNAGCAGCATGATTCACAATTGCA

AGCACACGTCTGAACTCTTTCCCTACACGACGCTCTTCCGATCTNNNAGCAGCATGATTCACAATTGCA

GAGCACACGTCTGAACTCTTTCCCTACACGACGCTCTTCCGATCTNNNCAGACACTAACTTTCCAAGCT

GAGCACACGTCTGAACTCTTTCCCTACACGACGCTCTTCCGATCTNNNCAGACACTAACTTTCCAAGCT
GAGCACACGTCTGAACTCTTTCCCTACACGACGCTCTTCCGATCTNNNCAGACACTAACTTTCCAAGCT
GCACACGTCTGAACTCTTTCCCTACACGACGCTCTTCCGATCTNNNCAAAACCACACCCCTTTCAAACC
GCACACGTCTGAACTCTTTCCCTACACGACGCTCTTCCGATCTNNNCAAAACCACACCCCTTTCAAACC
GCACACGTCTGAACTCTTTCCCTACACGACGCTCTTCCGATCTNNNCAAAACCACACCCCTTTCAAACC
GCACACGTCTGAACTCTTTCCCTACACGACGCTCTTCCGATCTNNNTGAATAAATGGGGAGAAGGG
GCACACGTCTGAACTCTTTCCCTACACGACGCTCTTCCGATCTNNNTGAATAAATGGGGAGAAGGG
GCACACGTCTGAACTCTTTCCCTACACGACGCTCTTCCGATCTNNNTGAATAAATGGGGAGAAGGG
GAGCACACGTCTGAACTCTTTCCCTACACGACGCTCTTCCGATCTNNNAACCTGAGCAACAAGAGGG
GAGCACACGTCTGAACTCTTTCCCTACACGACGCTCTTCCGATCTNNNAACCTGAGCAACAAGAGGG
GAGCACACGTCTGAACTCTTTCCCTACACGACGCTCTTCCGATCTNNNAACCTGAGCAACAAGAGGG
GCACACGTCTGAACTCTTTCCCTACACGACGCTCTTCCGATCTNNNTTCCCCTCCACTGTAGAAATTCCC
GCACACGTCTGAACTCTTTCCCTACACGACGCTCTTCCGATCTNNNTTCCCCTCCACTGTAGAAATTCCC
GCACACGTCTGAACTCTTTCCCTACACGACGCTCTTCCGATCTNNNTTCCCCTCCACTGTAGAAATTCCC
AGCACACGTCTGAACTCTTTCCCTACACGACGCTCTTCCGATCTNNNTTTCCAGTTTTGCTTCCACACC
AGCACACGTCTGAACTCTTTCCCTACACGACGCTCTTCCGATCTNNNTTTCCAGTTTTGCTTCCACACC
AGCACACGTCTGAACTCTTTCCCTACACGACGCTCTTCCGATCTNNNTTTCCAGTTTTGCTTCCACACC
GCACACGTCTGAACTCTTTCCCTACACGACGCTCTTCCGATCTNNNCCCCAGTAACATTTGCAAATGA
GCACACGTCTGAACTCTTTCCCTACACGACGCTCTTCCGATCTNNNCCCCAGTAACATTTGCAAATGA
GCACACGTCTGAACTCTTTCCCTACACGACGCTCTTCCGATCTNNNCCCCAGTAACATTTGCAAATGA
CACGTCTGAACTCTTTCCCTACACGACGCTCTTCCGATCTNNNCTGGGGGCCAATATTCAACATTCCAT
CACGTCTGAACTCTTTCCCTACACGACGCTCTTCCGATCTNNNCTGGGGGCCAATATTCAACATTCCAT
CACGTCTGAACTCTTTCCCTACACGACGCTCTTCCGATCTNNNCTGGGGGCCAATATTCAACATTCCAT
GCACACGTCTGAACTCTTTCCCTACACGACGCTCTTCCGATCTNNNTTTTTTTTGGACAGTCTCGCTCCC
GCACACGTCTGAACTCTTTCCCTACACGACGCTCTTCCGATCTNNNTTTTTTTTGGACAGTCTCGCTCCC
GCACACGTCTGAACTCTTTCCCTACACGACGCTCTTCCGATCTNNNTTTTTTTTGGACAGTCTCGCTCCC
GAGCACACGTCTGAACTCTTTCCCTACACGACGCTCTTCCGATCTNNNGTTCAAGCCACAACAAAAGT
GAGCACACGTCTGAACTCTTTCCCTACACGACGCTCTTCCGATCTNNNGTTCAAGCCACAACAAAAGT
GAGCACACGTCTGAACTCTTTCCCTACACGACGCTCTTCCGATCTNNNGTTCAAGCCACAACAAAAGT
AGCACACGTCTGAACTCTTTCCCTACACGACGCTCTTCCGATCTNNNCAGCCTCCTGTATCATCTTCTCC
AGCACACGTCTGAACTCTTTCCCTACACGACGCTCTTCCGATCTNNNCAGCCTCCTGTATCATCTTCTCC
AGCACACGTCTGAACTCTTTCCCTACACGACGCTCTTCCGATCTNNNCAGCCTCCTGTATCATCTTCTCC
AGCACACGTCTGAACTCTTTCCCTACACGACGCTCTTCCGATCTNNNACTCTTCTGTGGATAGTTTCAG
AGCACACGTCTGAACTCTTTCCCTACACGACGCTCTTCCGATCTNNNACTCTTCTGTGGATAGTTTCAG
AGCACACGTCTGAACTCTTTCCCTACACGACGCTCTTCCGATCTNNNACTCTTCTGTGGATAGTTTCAG
GAGCACACGTCTGAACTCTTTCCCTACACGACGCTCTTCCGATCTNNNGTGACAAACTCTCCAACTCA
GAGCACACGTCTGAACTCTTTCCCTACACGACGCTCTTCCGATCTNNNGTGACAAACTCTCCAACTCA
GAGCACACGTCTGAACTCTTTCCCTACACGACGCTCTTCCGATCTNNNGTGACAAACTCTCCAACTCA
GAGCACACGTCTGAACTCTTTCCCTACACGACGCTCTTCCGATCTNNNGAGGACCCTATTGCCTCTTT
GAGCACACGTCTGAACTCTTTCCCTACACGACGCTCTTCCGATCTNNNGAGGACCCTATTGCCTCTTT
GAGCACACGTCTGAACTCTTTCCCTACACGACGCTCTTCCGATCTNNNGAGGACCCTATTGCCTCTTT
AGCACACGTCTGAACTCTTTCCCTACACGACGCTCTTCCGATCTNNNCTTGCAGTGAGCCGAGATCA
AGCACACGTCTGAACTCTTTCCCTACACGACGCTCTTCCGATCTNNNCTTGCAGTGAGCCGAGATCA

GAGCACACGTCTGAACTCTTTCCCTACACGACGCTCTTCCGATCTNNNCTTGCAGTGAGCCGAGATCAT
AGCACACGTCTGAACTCTTTCCCTACACGACGCTCTTCCGATCTNNNAGATCAGCAAGTGACTGTAGT
AGCACACGTCTGAACTCTTTCCCTACACGACGCTCTTCCGATCTNNNAGATCAGCAAGTGACTGTAGT
AGCACACGTCTGAACTCTTTCCCTACACGACGCTCTTCCGATCTNNNAGATCAGCAAGTGACTGTAGT
GAGCACACGTCTGAACTCTTTCCCTACACGACGCTCTTCCGATCTNNNGGTGCCTGTAATCCCAGCTATT
GAGCACACGTCTGAACTCTTTCCCTACACGACGCTCTTCCGATCTNNNGGTGCCTGTAATCCCAGCTATT
GAGCACACGTCTGAACTCTTTCCCTACACGACGCTCTTCCGATCTNNNGGTGCCTGTAATCCCAGCTATT
AGCACACGTCTGAACTCTTTCCCTACACGACGCTCTTCCGATCTNNNCAGCAACAGAATGACACTCGTC
AGCACACGTCTGAACTCTTTCCCTACACGACGCTCTTCCGATCTNNNCAGCAACAGAATGACACTCGTC
AGCACACGTCTGAACTCTTTCCCTACACGACGCTCTTCCGATCTNNNCAGCAACAGAATGACACTCGTC
GCACACGTCTGAACTCTTTCCCTACACGACGCTCTTCCGATCTNNNAAGCACACATACTGCTGACTACAC
GCACACGTCTGAACTCTTTCCCTACACGACGCTCTTCCGATCTNNNAAGCACACATACTGCTGACTACAC
GCACACGTCTGAACTCTTTCCCTACACGACGCTCTTCCGATCTNNNAAGCACACATACTGCTGACTACAC
AGCACACGTCTGAACTCTTTCCCTACACGACGCTCTTCCGATCTNNNTTTACCCTCCCTTCTTTCCTTCCCA
AGCACACGTCTGAACTCTTTCCCTACACGACGCTCTTCCGATCTNNNTTTACCCTCCCTTCTTTCCTTCCCA
AGCACACGTCTGAACTCTTTCCCTACACGACGCTCTTCCGATCTNNNTTTACCCTCCCTTCTTTCCTTCCCA
GAGCACACGTCTGAACTCTTTCCCTACACGACGCTCTTCCGATCTNNNACTCAGTGTTCACATTTTCCCA
GAGCACACGTCTGAACTCTTTCCCTACACGACGCTCTTCCGATCTNNNACTCAGTGTTCACATTTTCCCA
GAGCACACGTCTGAACTCTTTCCCTACACGACGCTCTTCCGATCTNNNACTCAGTGTTCACATTTTCCCA
AAGAGCACACGTCTGAACTCTTTCCCTACACGACGCTCTTCCGATCTNNNGAGGTAGGAGAATGGCGT
AAGAGCACACGTCTGAACTCTTTCCCTACACGACGCTCTTCCGATCTNNNGAGGTAGGAGAATGGCGT
AAGAGCACACGTCTGAACTCTTTCCCTACACGACGCTCTTCCGATCTNNNGAGGTAGGAGAATGGCGT
GAGCACACGTCTGAACTCTTTCCCTACACGACGCTCTTCCGATCTNNNCAACTACACACACATCTGCACA
GAGCACACGTCTGAACTCTTTCCCTACACGACGCTCTTCCGATCTNNNCAACTACACACACATCTGCACA
GAGCACACGTCTGAACTCTTTCCCTACACGACGCTCTTCCGATCTNNNCAACTACACACACATCTGCACA
AGCACACGTCTGAACTCTTTCCCTACACGACGCTCTTCCGATCTNNNAGGAGAATGGCGTAAACCCGCA
AGCACACGTCTGAACTCTTTCCCTACACGACGCTCTTCCGATCTNNNAGGAGAATGGCGTAAACCCGCA
AGCACACGTCTGAACTCTTTCCCTACACGACGCTCTTCCGATCTNNNAGGAGAATGGCGTAAACCCGCA
ACACGTCTGAACTCTTTCCCTACACGACGCTCTTCCGATCTNNNACCACAGTTAGTCTCTTATTGACAAA
ACACGTCTGAACTCTTTCCCTACACGACGCTCTTCCGATCTNNNACCACAGTTAGTCTCTTATTGACAAA
ACACGTCTGAACTCTTTCCCTACACGACGCTCTTCCGATCTNNNACCACAGTTAGTCTCTTATTGACAAA
GCACACGTCTGAACTCTTTCCCTACACGACGCTCTTCCGATCTNNNCAGTTCCATCACACAGCAAACAAC
GCACACGTCTGAACTCTTTCCCTACACGACGCTCTTCCGATCTNNNCAGTTCCATCACACAGCAAACAAC
GCACACGTCTGAACTCTTTCCCTACACGACGCTCTTCCGATCTNNNCAGTTCCATCACACAGCAAACAAC
GCACACGTCTGAACTCTTTCCCTACACGACGCTCTTCCGATCTNNNCACTCTTTCTCCAACACACACACCAT
GCACACGTCTGAACTCTTTCCCTACACGACGCTCTTCCGATCTNNNCACTCTTTCTCCAACACACACACCAT
GCACACGTCTGAACTCTTTCCCTACACGACGCTCTTCCGATCTNNNCACTCTTTCTCCAACACACACACCAT
GAGCACACGTCTGAACTCTTTCCCTACACGACGCTCTTCCGATCTNNNGTGAAGGTTTCTGAGAAAGTA
GAGCACACGTCTGAACTCTTTCCCTACACGACGCTCTTCCGATCTNNNGTGAAGGTTTCTGAGAAAGTA
GAGCACACGTCTGAACTCTTTCCCTACACGACGCTCTTCCGATCTNNNGTGAAGGTTTCTGAGAAAGTA
GAGCACACGTCTGAACTCTTTCCCTACACGACGCTCTTCCGATCTNNNGTGGCCCCAGATGAATTTGTA
GAGCACACGTCTGAACTCTTTCCCTACACGACGCTCTTCCGATCTNNNGTGGCCCCAGATGAATTTGTA
GAGCACACGTCTGAACTCTTTCCCTACACGACGCTCTTCCGATCTNNNGTGGCCCCAGATGAATTTGTA

AAGAGCACACGTCTGAACTCTTTCCCTACACGACGCTCTTCCGATCTNNNACCCTACCTTCCAAGAAAGC
AAGAGCACACGTCTGAACTCTTTCCCTACACGACGCTCTTCCGATCTNNNACCCTACCTTCCAAGAAAGC
AAGAGCACACGTCTGAACTCTTTCCCTACACGACGCTCTTCCGATCTNNNACCCTACCTTCCAAGAAAGC
AGAGCACACGTCTGAACTCTTTCCCTACACGACGCTCTTCCGATCTNNNGTAGTTCCAGATACTGCAGA
AGAGCACACGTCTGAACTCTTTCCCTACACGACGCTCTTCCGATCTNNNGTAGTTCCAGATACTGCAGA
AGAGCACACGTCTGAACTCTTTCCCTACACGACGCTCTTCCGATCTNNNGTAGTTCCAGATACTGCAGA
GAGCACACGTCTGAACTCTTTCCCTACACGACGCTCTTCCGATCTNNNTGTGGGCAAGTGATGGTATA
GAGCACACGTCTGAACTCTTTCCCTACACGACGCTCTTCCGATCTNNNTGTGGGCAAGTGATGGTATA
GAGCACACGTCTGAACTCTTTCCCTACACGACGCTCTTCCGATCTNNNTGTGGGCAAGTGATGGTATA
GAGCACACGTCTGAACTCTTTCCCTACACGACGCTCTTCCGATCTNNNTTTTTTTTGAGACACAGCAAG
GAGCACACGTCTGAACTCTTTCCCTACACGACGCTCTTCCGATCTNNNTTTTTTTTGAGACACAGCAAG
GAGCACACGTCTGAACTCTTTCCCTACACGACGCTCTTCCGATCTNNNTTTTTTTTGAGACACAGCAAG
AGCACACGTCTGAACTCTTTCCCTACACGACGCTCTTCCGATCTNNNGTACAAGTGATTCTCCTGCCTCA
AGCACACGTCTGAACTCTTTCCCTACACGACGCTCTTCCGATCTNNNGTACAAGTGATTCTCCTGCCTCA
AGCACACGTCTGAACTCTTTCCCTACACGACGCTCTTCCGATCTNNNGTACAAGTGATTCTCCTGCCTCA
GAGCACACGTCTGAACTCTTTCCCTACACGACGCTCTTCCGATCTNNNTCTCCAATAAGTGTAGATTTTC
GAGCACACGTCTGAACTCTTTCCCTACACGACGCTCTTCCGATCTNNNTCTCCAATAAGTGTAGATTTTC
GAGCACACGTCTGAACTCTTTCCCTACACGACGCTCTTCCGATCTNNNTCTCCAATAAGTGTAGATTTTC
AGAGCACACGTCTGAACTCTTTCCCTACACGACGCTCTTCCGATCTNNNCAAGCATCCCAGTCAAGAAAA
AGAGCACACGTCTGAACTCTTTCCCTACACGACGCTCTTCCGATCTNNNCAAGCATCCCAGTCAAGAAAA
AGAGCACACGTCTGAACTCTTTCCCTACACGACGCTCTTCCGATCTNNNCAAGCATCCCAGTCAAGAAAA
GAGCACACGTCTGAACTCTTTCCCTACACGACGCTCTTCCGATCTNNNCCTTGCCTCTGTGTTTCTTTGTC
GAGCACACGTCTGAACTCTTTCCCTACACGACGCTCTTCCGATCTNNNCCTTGCCTCTGTGTTTCTTTGTC
GAGCACACGTCTGAACTCTTTCCCTACACGACGCTCTTCCGATCTNNNCCTTGCCTCTGTGTTTCTTTGTC
GAGCACACGTCTGAACTCTTTCCCTACACGACGCTCTTCCGATCTNNNGCTTTGGCCTTTGGATGTATA
GAGCACACGTCTGAACTCTTTCCCTACACGACGCTCTTCCGATCTNNNGCTTTGGCCTTTGGATGTATA
GAGCACACGTCTGAACTCTTTCCCTACACGACGCTCTTCCGATCTNNNGCTTTGGCCTTTGGATGTATA
CACACGTCTGAACTCTTTCCCTACACGACGCTCTTCCGATCTNNNGTCTGACCAGAGAGGACTCATCAC
CACACGTCTGAACTCTTTCCCTACACGACGCTCTTCCGATCTNNNGTCTGACCAGAGAGGACTCATCAC
CACACGTCTGAACTCTTTCCCTACACGACGCTCTTCCGATCTNNNGTCTGACCAGAGAGGACTCATCAC
GAGCACACGTCTGAACTCTTTCCCTACACGACGCTCTTCCGATCTNNNGGCACCTATAATCCCAGCTACT
GAGCACACGTCTGAACTCTTTCCCTACACGACGCTCTTCCGATCTNNNGGCACCTATAATCCCAGCTACT
GAGCACACGTCTGAACTCTTTCCCTACACGACGCTCTTCCGATCTNNNGGCACCTATAATCCCAGCTACT
GCACACGTCTGAACTCTTTCCCTACACGACGCTCTTCCGATCTNNNCTGGGCAACAAGAGCAAAACTCC
GCACACGTCTGAACTCTTTCCCTACACGACGCTCTTCCGATCTNNNCTGGGCAACAAGAGCAAAACTCC
GCACACGTCTGAACTCTTTCCCTACACGACGCTCTTCCGATCTNNNCTGGGCAACAAGAGCAAAACTCC
AGCACACGTCTGAACTCTTTCCCTACACGACGCTCTTCCGATCTNNNGTCCCTCTAAGGTTTTTGCAACC
AGCACACGTCTGAACTCTTTCCCTACACGACGCTCTTCCGATCTNNNGTCCCTCTAAGGTTTTTGCAACC
AGCACACGTCTGAACTCTTTCCCTACACGACGCTCTTCCGATCTNNNGTCCCTCTAAGGTTTTTGCAACC
GAGCACACGTCTGAACTCTTTCCCTACACGACGCTCTTCCGATCTNNNAATGTGAAAACAGAAGGCGTC
GAGCACACGTCTGAACTCTTTCCCTACACGACGCTCTTCCGATCTNNNAATGTGAAAACAGAAGGCGTC
GAGCACACGTCTGAACTCTTTCCCTACACGACGCTCTTCCGATCTNNNAATGTGAAAACAGAAGGCGTC
AGCACACGTCTGAACTCTTTCCCTACACGACGCTCTTCCGATCTNNNTATACACCCACATGCCCAGGTAA

AGCACACGTCTGAACTCTTTCCCTACACGACGCTCTTCCGATCTNNNTATACACCCACATGCCCAGGTA
AGCACACGTCTGAACTCTTTCCCTACACGACGCTCTTCCGATCTNNNTATACACCCACATGCCCAGGTA
AGCACACGTCTGAACTCTTTCCCTACACGACGCTCTTCCGATCTNNNCCCCAAACTGGAAACAACTCA
AGCACACGTCTGAACTCTTTCCCTACACGACGCTCTTCCGATCTNNNCCCCAAACTGGAAACAACTCA
AGCACACGTCTGAACTCTTTCCCTACACGACGCTCTTCCGATCTNNNCCCCAAACTGGAAACAACTCA
GAGCACACGTCTGAACTCTTTCCCTACACGACGCTCTTCCGATCTNNNGCAGGAGAATCAATTGAACC
GAGCACACGTCTGAACTCTTTCCCTACACGACGCTCTTCCGATCTNNNGCAGGAGAATCAATTGAACC
GAGCACACGTCTGAACTCTTTCCCTACACGACGCTCTTCCGATCTNNNGCAGGAGAATCAATTGAACC
AGCACACGTCTGAACTCTTTCCCTACACGACGCTCTTCCGATCTNNNAAGGAATTGTGTGTGTGTGT
AGCACACGTCTGAACTCTTTCCCTACACGACGCTCTTCCGATCTNNNAAGGAATTGTGTGTGTGTGT
AGCACACGTCTGAACTCTTTCCCTACACGACGCTCTTCCGATCTNNNAAGGAATTGTGTGTGTGTGT
AGCACACGTCTGAACTCTTTCCCTACACGACGCTCTTCCGATCTNNNCCACGCCCAGCTAATTTTTGTAT
AGCACACGTCTGAACTCTTTCCCTACACGACGCTCTTCCGATCTNNNCCACGCCCAGCTAATTTTTGTAT
AGCACACGTCTGAACTCTTTCCCTACACGACGCTCTTCCGATCTNNNCCACGCCCAGCTAATTTTTGTAT
AGCACACGTCTGAACTCTTTCCCTACACGACGCTCTTCCGATCTNNNTTCCCTTTCTTTTCCCGAGTTCT
AGCACACGTCTGAACTCTTTCCCTACACGACGCTCTTCCGATCTNNNTTCCCTTTCTTTTCCCGAGTTCT
AGCACACGTCTGAACTCTTTCCCTACACGACGCTCTTCCGATCTNNNTTCCCTTTCTTTTCCCGAGTTCT
AGCACACGTCTGAACTCTTTCCCTACACGACGCTCTTCCGATCTNNNCAGGAGAATGGTGTAAACTG
AGCACACGTCTGAACTCTTTCCCTACACGACGCTCTTCCGATCTNNNCAGGAGAATGGTGTAAACTG
AGCACACGTCTGAACTCTTTCCCTACACGACGCTCTTCCGATCTNNNCAGGAGAATGGTGTAAACTG
AGCACACGTCTGAACTCTTTCCCTACACGACGCTCTTCCGATCTNNNCAGTCCAGACCTCCACTATTTC
AGCACACGTCTGAACTCTTTCCCTACACGACGCTCTTCCGATCTNNNCAGTCCAGACCTCCACTATTTC
AGCACACGTCTGAACTCTTTCCCTACACGACGCTCTTCCGATCTNNNCAGTCCAGACCTCCACTATTTC
GAGCACACGTCTGAACTCTTTCCCTACACGACGCTCTTCCGATCTNNNCTGATGATCTGTCACTGTCTC
GAGCACACGTCTGAACTCTTTCCCTACACGACGCTCTTCCGATCTNNNCTGATGATCTGTCACTGTCTC
GAGCACACGTCTGAACTCTTTCCCTACACGACGCTCTTCCGATCTNNNCTGATGATCTGTCACTGTCTC
GAGCACACGTCTGAACTCTTTCCCTACACGACGCTCTTCCGATCTNNNACCTTGAGCATTCTGCAACAA
GAGCACACGTCTGAACTCTTTCCCTACACGACGCTCTTCCGATCTNNNACCTTGAGCATTCTGCAACAA
GAGCACACGTCTGAACTCTTTCCCTACACGACGCTCTTCCGATCTNNNACCTTGAGCATTCTGCAACAA
GCACACGTCTGAACTCTTTCCCTACACGACGCTCTTCCGATCTNNNCATTTTGGACAAGGGATATTTA
GCACACGTCTGAACTCTTTCCCTACACGACGCTCTTCCGATCTNNNCATTTTGGACAAGGGATATTTA
GCACACGTCTGAACTCTTTCCCTACACGACGCTCTTCCGATCTNNNCATTTTGGACAAGGGATATTTA
GCACACGTCTGAACTCTTTCCCTACACGACGCTCTTCCGATCTNNNCAGTGAGAGGAGAAACCCTTGT
GCACACGTCTGAACTCTTTCCCTACACGACGCTCTTCCGATCTNNNCAGTGAGAGGAGAAACCCTTGT
GCACACGTCTGAACTCTTTCCCTACACGACGCTCTTCCGATCTNNNCAGTGAGAGGAGAAACCCTTGT
GAGCACACGTCTGAACTCTTTCCCTACACGACGCTCTTCCGATCTNNNGAGATAGCGCCACTGCACTC
GAGCACACGTCTGAACTCTTTCCCTACACGACGCTCTTCCGATCTNNNGAGATAGCGCCACTGCACTC
GAGCACACGTCTGAACTCTTTCCCTACACGACGCTCTTCCGATCTNNNGAGATAGCGCCACTGCACTC
AGCACACGTCTGAACTCTTTCCCTACACGACGCTCTTCCGATCTNNNCTTGCGTAAATGTACTAGCGA
AGCACACGTCTGAACTCTTTCCCTACACGACGCTCTTCCGATCTNNNCTTGCGTAAATGTACTAGCGA
AGCACACGTCTGAACTCTTTCCCTACACGACGCTCTTCCGATCTNNNCTTGCGTAAATGTACTAGCGA
GAGCACACGTCTGAACTCTTTCCCTACACGACGCTCTTCCGATCTNNNGAGAGTTTCTTTCCCAAGGCC
GAGCACACGTCTGAACTCTTTCCCTACACGACGCTCTTCCGATCTNNNGAGAGTTTCTTTCCCAAGGCC

GAGCACACGTCTGAACTCTTTCCCTACACGACGCTCTTCCGATCTNNNGAGAGTTTCTTTCCCAAGGCCA
AGCACACGTCTGAACTCTTTCCCTACACGACGCTCTTCCGATCTNNNACCTGCAACACAAAGGAAGAAT
AGCACACGTCTGAACTCTTTCCCTACACGACGCTCTTCCGATCTNNNACCTGCAACACAAAGGAAGAAT
AGCACACGTCTGAACTCTTTCCCTACACGACGCTCTTCCGATCTNNNACCTGCAACACAAAGGAAGAAT
GAGCACACGTCTGAACTCTTTCCCTACACGACGCTCTTCCGATCTNNNTGATGAGAAACTGTCAGCTGT
GAGCACACGTCTGAACTCTTTCCCTACACGACGCTCTTCCGATCTNNNTGATGAGAAACTGTCAGCTGT
GAGCACACGTCTGAACTCTTTCCCTACACGACGCTCTTCCGATCTNNNTGATGAGAAACTGTCAGCTGT
AGCACACGTCTGAACTCTTTCCCTACACGACGCTCTTCCGATCTNNNTGCAAACTCTGGTTCTTTCCATT
AGCACACGTCTGAACTCTTTCCCTACACGACGCTCTTCCGATCTNNNTGCAAACTCTGGTTCTTTCCATT
AGCACACGTCTGAACTCTTTCCCTACACGACGCTCTTCCGATCTNNNTGCAAACTCTGGTTCTTTCCATT
GAGCACACGTCTGAACTCTTTCCCTACACGACGCTCTTCCGATCTNNNACCTCCTTACTCCTGCAAGAAT
GAGCACACGTCTGAACTCTTTCCCTACACGACGCTCTTCCGATCTNNNACCTCCTTACTCCTGCAAGAAT
GAGCACACGTCTGAACTCTTTCCCTACACGACGCTCTTCCGATCTNNNACCTCCTTACTCCTGCAAGAAT
AGCACACGTCTGAACTCTTTCCCTACACGACGCTCTTCCGATCTNNNGGAGGTTGTAGTGAGCCAAGA
AGCACACGTCTGAACTCTTTCCCTACACGACGCTCTTCCGATCTNNNGGAGGTTGTAGTGAGCCAAGA
AGCACACGTCTGAACTCTTTCCCTACACGACGCTCTTCCGATCTNNNGGAGGTTGTAGTGAGCCAAGA
GAGCACACGTCTGAACTCTTTCCCTACACGACGCTCTTCCGATCTNNNCTTTTCCAGTTCGGATGCCTTTT
GAGCACACGTCTGAACTCTTTCCCTACACGACGCTCTTCCGATCTNNNCTTTTCCAGTTCGGATGCCTTTT
GAGCACACGTCTGAACTCTTTCCCTACACGACGCTCTTCCGATCTNNNCTTTTCCAGTTCGGATGCCTTTT
AGCACACGTCTGAACTCTTTCCCTACACGACGCTCTTCCGATCTNNNAAATGAGCCTTCCCATGAGGTA
AGCACACGTCTGAACTCTTTCCCTACACGACGCTCTTCCGATCTNNNAAATGAGCCTTCCCATGAGGTA
AGCACACGTCTGAACTCTTTCCCTACACGACGCTCTTCCGATCTNNNAAATGAGCCTTCCCATGAGGTA
GAGCACACGTCTGAACTCTTTCCCTACACGACGCTCTTCCGATCTNNNACCAAAGGAATGTTAGCCCTT
GAGCACACGTCTGAACTCTTTCCCTACACGACGCTCTTCCGATCTNNNACCAAAGGAATGTTAGCCCTT
GAGCACACGTCTGAACTCTTTCCCTACACGACGCTCTTCCGATCTNNNACCAAAGGAATGTTAGCCCTT
GAGCACACGTCTGAACTCTTTCCCTACACGACGCTCTTCCGATCTNNNCTCTTGCCTAAGTCTCAGGAT
GAGCACACGTCTGAACTCTTTCCCTACACGACGCTCTTCCGATCTNNNCTCTTGCCTAAGTCTCAGGAT
GAGCACACGTCTGAACTCTTTCCCTACACGACGCTCTTCCGATCTNNNCTCTTGCCTAAGTCTCAGGAT
AGCACACGTCTGAACTCTTTCCCTACACGACGCTCTTCCGATCTNNNTAAACTTCGATGTAGCCGACAC
AGCACACGTCTGAACTCTTTCCCTACACGACGCTCTTCCGATCTNNNTAAACTTCGATGTAGCCGACAC
AGCACACGTCTGAACTCTTTCCCTACACGACGCTCTTCCGATCTNNNTAAACTTCGATGTAGCCGACAC
GAGCACACGTCTGAACTCTTTCCCTACACGACGCTCTTCCGATCTNNNTCCAGTGTTTATCAGCTGGTT
GAGCACACGTCTGAACTCTTTCCCTACACGACGCTCTTCCGATCTNNNTCCAGTGTTTATCAGCTGGTT
GAGCACACGTCTGAACTCTTTCCCTACACGACGCTCTTCCGATCTNNNTCCAGTGTTTATCAGCTGGTT
GAGCACACGTCTGAACTCTTTCCCTACACGACGCTCTTCCGATCTNNNAGAATTGCTTGAAACTGGGAA
GAGCACACGTCTGAACTCTTTCCCTACACGACGCTCTTCCGATCTNNNAGAATTGCTTGAAACTGGGAA
GAGCACACGTCTGAACTCTTTCCCTACACGACGCTCTTCCGATCTNNNAGAATTGCTTGAAACTGGGAA
AGCACACGTCTGAACTCTTTCCCTACACGACGCTCTTCCGATCTNNNGCCATATATGACAGATCCACAG
AGCACACGTCTGAACTCTTTCCCTACACGACGCTCTTCCGATCTNNNGCCATATATGACAGATCCACAG
AGCACACGTCTGAACTCTTTCCCTACACGACGCTCTTCCGATCTNNNGCCATATATGACAGATCCACAG
GCACACGTCTGAACTCTTTCCCTACACGACGCTCTTCCGATCTNNNCAAAGTGCTGGTATTGCAGATG
GCACACGTCTGAACTCTTTCCCTACACGACGCTCTTCCGATCTNNNCAAAGTGCTGGTATTGCAGATG
GCACACGTCTGAACTCTTTCCCTACACGACGCTCTTCCGATCTNNNCAAAGTGCTGGTATTGCAGATG

AGCACACGTCTGAACTCTTTCCCTACACGACGCTCTTCCGATCTNNNTACAAAAAATTAGCTGGGCGTG
AGCACACGTCTGAACTCTTTCCCTACACGACGCTCTTCCGATCTNNNTACAAAAAATTAGCTGGGCGTG
AGCACACGTCTGAACTCTTTCCCTACACGACGCTCTTCCGATCTNNNTACAAAAAATTAGCTGGGCGTG
AGCACACGTCTGAACTCTTTCCCTACACGACGCTCTTCCGATCTNNNGCGACAGAGTGAGACTCCATCC
AGCACACGTCTGAACTCTTTCCCTACACGACGCTCTTCCGATCTNNNGCGACAGAGTGAGACTCCATCC
AGCACACGTCTGAACTCTTTCCCTACACGACGCTCTTCCGATCTNNNGCGACAGAGTGAGACTCCATCC
GCACACGTCTGAACTCTTTCCCTACACGACGCTCTTCCGATCTNNNGCCTCCTTAGTAGCTGGGATTAC
GCACACGTCTGAACTCTTTCCCTACACGACGCTCTTCCGATCTNNNGCCTCCTTAGTAGCTGGGATTAC
GCACACGTCTGAACTCTTTCCCTACACGACGCTCTTCCGATCTNNNGCCTCCTTAGTAGCTGGGATTAC
GCACACGTCTGAACTCTTTCCCTACACGACGCTCTTCCGATCTNNNTGTTCCTTCCACGTTGATGTACTC
GCACACGTCTGAACTCTTTCCCTACACGACGCTCTTCCGATCTNNNTGTTCCTTCCACGTTGATGTACTC
GCACACGTCTGAACTCTTTCCCTACACGACGCTCTTCCGATCTNNNTGTTCCTTCCACGTTGATGTACTC
GAGCACACGTCTGAACTCTTTCCCTACACGACGCTCTTCCGATCTNNNTTGGCTAAAACAACATTCTGGT
GAGCACACGTCTGAACTCTTTCCCTACACGACGCTCTTCCGATCTNNNTTGGCTAAAACAACATTCTGGT
GAGCACACGTCTGAACTCTTTCCCTACACGACGCTCTTCCGATCTNNNTTGGCTAAAACAACATTCTGGT
AGCACACGTCTGAACTCTTTCCCTACACGACGCTCTTCCGATCTNNNTCAGGAAACGTACAGTAATGGC
AGCACACGTCTGAACTCTTTCCCTACACGACGCTCTTCCGATCTNNNTCAGGAAACGTACAGTAATGGC
AGCACACGTCTGAACTCTTTCCCTACACGACGCTCTTCCGATCTNNNTCAGGAAACGTACAGTAATGGC
AGCACACGTCTGAACTCTTTCCCTACACGACGCTCTTCCGATCTNNNCTGTAGTCCCAGCTACTCAGAAC
AGCACACGTCTGAACTCTTTCCCTACACGACGCTCTTCCGATCTNNNCTGTAGTCCCAGCTACTCAGAAC
AGCACACGTCTGAACTCTTTCCCTACACGACGCTCTTCCGATCTNNNCTGTAGTCCCAGCTACTCAGAAC
GAGCACACGTCTGAACTCTTTCCCTACACGACGCTCTTCCGATCTNNNGTTCTGAAGTCCTCCAGTGAGA
GAGCACACGTCTGAACTCTTTCCCTACACGACGCTCTTCCGATCTNNNGTTCTGAAGTCCTCCAGTGAGA
GAGCACACGTCTGAACTCTTTCCCTACACGACGCTCTTCCGATCTNNNGTTCTGAAGTCCTCCAGTGAGA
AGAGCACACGTCTGAACTCTTTCCCTACACGACGCTCTTCCGATCTNNNGGGCCAAATGAGGTATTTCC
AGAGCACACGTCTGAACTCTTTCCCTACACGACGCTCTTCCGATCTNNNGGGCCAAATGAGGTATTTCC
AGAGCACACGTCTGAACTCTTTCCCTACACGACGCTCTTCCGATCTNNNGGGCCAAATGAGGTATTTCC
AAGAGCACACGTCTGAACTCTTTCCCTACACGACGCTCTTCCGATCTNNNATTCCATCCAGGTTGCTGTG
AAGAGCACACGTCTGAACTCTTTCCCTACACGACGCTCTTCCGATCTNNNATTCCATCCAGGTTGCTGTG
AAGAGCACACGTCTGAACTCTTTCCCTACACGACGCTCTTCCGATCTNNNATTCCATCCAGGTTGCTGTG
AGAGCACACGTCTGAACTCTTTCCCTACACGACGCTCTTCCGATCTNNNTACACAAAAGGCAGATCCTCC
AGAGCACACGTCTGAACTCTTTCCCTACACGACGCTCTTCCGATCTNNNTACACAAAAGGCAGATCCTCC
AGAGCACACGTCTGAACTCTTTCCCTACACGACGCTCTTCCGATCTNNNTACACAAAAGGCAGATCCTCC
GAGCACACGTCTGAACTCTTTCCCTACACGACGCTCTTCCGATCTNNNATGTGTGTGTGTGTGTGTGTA
GAGCACACGTCTGAACTCTTTCCCTACACGACGCTCTTCCGATCTNNNATGTGTGTGTGTGTGTGTGTA
GAGCACACGTCTGAACTCTTTCCCTACACGACGCTCTTCCGATCTNNNATGTGTGTGTGTGTGTGTGTA
AGCACACGTCTGAACTCTTTCCCTACACGACGCTCTTCCGATCTNNNAATTTCCATGACCACCCAGACAC
AGCACACGTCTGAACTCTTTCCCTACACGACGCTCTTCCGATCTNNNAATTTCCATGACCACCCAGACAC
AGCACACGTCTGAACTCTTTCCCTACACGACGCTCTTCCGATCTNNNAATTTCCATGACCACCCAGACAC
GAGCACACGTCTGAACTCTTTCCCTACACGACGCTCTTCCGATCTNNNAGCAATTTCCGTAAGTGTGGT
GAGCACACGTCTGAACTCTTTCCCTACACGACGCTCTTCCGATCTNNNAGCAATTTCCGTAAGTGTGGT
GAGCACACGTCTGAACTCTTTCCCTACACGACGCTCTTCCGATCTNNNAGCAATTTCCGTAAGTGTGGT
GAGCACACGTCTGAACTCTTTCCCTACACGACGCTCTTCCGATCTNNNGCAATCCAAAGTATGAGAGGT

AGCACACGTCTGAACTCTTTCCCTACACGACGCTCTTCCGATCTNNNGCAATCCAAAGTATGAGAGGT

AGCACACGTCTGAACTCTTTCCCTACACGACGCTCTTCCGATCTNNNGCAATCCAAAGTATGAGAGGT

GAGCACACGTCTGAACTCTTTCCCTACACGACGCTCTTCCGATCTNNNACCTGGTAGTGGGTTTATGTC

GAGCACACGTCTGAACTCTTTCCCTACACGACGCTCTTCCGATCTNNNACCTGGTAGTGGGTTTATGTC

GAGCACACGTCTGAACTCTTTCCCTACACGACGCTCTTCCGATCTNNNACCTGGTAGTGGGTTTATGTC

AGCACACGTCTGAACTCTTTCCCTACACGACGCTCTTCCGATCTNNNACCATGCTTTTTGTTCCTTCCCC

AGCACACGTCTGAACTCTTTCCCTACACGACGCTCTTCCGATCTNNNACCATGCTTTTTGTTCCTTCCCC

AGCACACGTCTGAACTCTTTCCCTACACGACGCTCTTCCGATCTNNNACCATGCTTTTTGTTCCTTCCCC

AGCACACGTCTGAACTCTTTCCCTACACGACGCTCTTCCGATCTNNNATGTGCTGCCTCTAATACCCAAA

AGCACACGTCTGAACTCTTTCCCTACACGACGCTCTTCCGATCTNNNATGTGCTGCCTCTAATACCCAAA

AGCACACGTCTGAACTCTTTCCCTACACGACGCTCTTCCGATCTNNNATGTGCTGCCTCTAATACCCAAA

GAGCACACGTCTGAACTCTTTCCCTACACGACGCTCTTCCGATCTNNNTTGAACTGGCAAGAGAGGAC

GAGCACACGTCTGAACTCTTTCCCTACACGACGCTCTTCCGATCTNNNTTGAACTGGCAAGAGAGGAC

GAGCACACGTCTGAACTCTTTCCCTACACGACGCTCTTCCGATCTNNNTTGAACTGGCAAGAGAGGAC

AGCACACGTCTGAACTCTTTCCCTACACGACGCTCTTCCGATCTNNNCAGTCCCAATGAACAAACATCC

AGCACACGTCTGAACTCTTTCCCTACACGACGCTCTTCCGATCTNNNCAGTCCCAATGAACAAACATCC

AGCACACGTCTGAACTCTTTCCCTACACGACGCTCTTCCGATCTNNNCAGTCCCAATGAACAAACATCC

GCACACGTCTGAACTCTTTCCCTACACGACGCTCTTCCGATCTNNNTTTCGTAGGCCCCGTAATGAAAG

GCACACGTCTGAACTCTTTCCCTACACGACGCTCTTCCGATCTNNNTTTCGTAGGCCCCGTAATGAAAG

GCACACGTCTGAACTCTTTCCCTACACGACGCTCTTCCGATCTNNNTTTCGTAGGCCCCGTAATGAAAG

AGCACACGTCTGAACTCTTTCCCTACACGACGCTCTTCCGATCTNNNACAAGTCCCACAGTGATTTCCTC

AGCACACGTCTGAACTCTTTCCCTACACGACGCTCTTCCGATCTNNNACAAGTCCCACAGTGATTTCCTC

AGCACACGTCTGAACTCTTTCCCTACACGACGCTCTTCCGATCTNNNACAAGTCCCACAGTGATTTCCTC

GAGCACACGTCTGAACTCTTTCCCTACACGACGCTCTTCCGATCTNNNAACGACTGCAGTAAAGGGCA

GAGCACACGTCTGAACTCTTTCCCTACACGACGCTCTTCCGATCTNNNAACGACTGCAGTAAAGGGCA

GAGCACACGTCTGAACTCTTTCCCTACACGACGCTCTTCCGATCTNNNAACGACTGCAGTAAAGGGCA

AGCACACGTCTGAACTCTTTCCCTACACGACGCTCTTCCGATCTNNNGGTGTGAGTGAAATGGAAGA

AGCACACGTCTGAACTCTTTCCCTACACGACGCTCTTCCGATCTNNNGGTGTGAGTGAAATGGAAGA

AGCACACGTCTGAACTCTTTCCCTACACGACGCTCTTCCGATCTNNNGGTGTGAGTGAAATGGAAGA

GAGCACACGTCTGAACTCTTTCCCTACACGACGCTCTTCCGATCTNNNAGAGGATTCTGACTGTTTCAC

GAGCACACGTCTGAACTCTTTCCCTACACGACGCTCTTCCGATCTNNNAGAGGATTCTGACTGTTTCAC

GAGCACACGTCTGAACTCTTTCCCTACACGACGCTCTTCCGATCTNNNAGAGGATTCTGACTGTTTCAC

AGCACACGTCTGAACTCTTTCCCTACACGACGCTCTTCCGATCTNNNCTGTGATTCAGGGAAGCTCTGA

AGCACACGTCTGAACTCTTTCCCTACACGACGCTCTTCCGATCTNNNCTGTGATTCAGGGAAGCTCTGA

AGCACACGTCTGAACTCTTTCCCTACACGACGCTCTTCCGATCTNNNCTGTGATTCAGGGAAGCTCTGA

AGCACACGTCTGAACTCTTTCCCTACACGACGCTCTTCCGATCTNNNCTGTAGTGAACCGAGATCGTAC

AGCACACGTCTGAACTCTTTCCCTACACGACGCTCTTCCGATCTNNNCTGTAGTGAACCGAGATCGTAC

AGCACACGTCTGAACTCTTTCCCTACACGACGCTCTTCCGATCTNNNCTGTAGTGAACCGAGATCGTAC

GCACACGTCTGAACTCTTTCCCTACACGACGCTCTTCCGATCTNNNGAATGGTTTATGTCAACAAAATG

GCACACGTCTGAACTCTTTCCCTACACGACGCTCTTCCGATCTNNNGAATGGTTTATGTCAACAAAATG

GCACACGTCTGAACTCTTTCCCTACACGACGCTCTTCCGATCTNNNGAATGGTTTATGTCAACAAAATG

AGCACACGTCTGAACTCTTTCCCTACACGACGCTCTTCCGATCTNNNAGAATTGCTTGAACCCGGGAG

AGCACACGTCTGAACTCTTTCCCTACACGACGCTCTTCCGATCTNNNAGAATTGCTTGAACCCGGGAG

AGCACACGTCTGAACTCTTTCCCTACACGACGCTCTTCCGATCTNNNAGAATTGCTTGAACCCGGGAG
AGCACACGTCTGAACTCTTTCCCTACACGACGCTCTTCCGATCTNNNGCAGGAGAATTGCTTGAACCA
AGCACACGTCTGAACTCTTTCCCTACACGACGCTCTTCCGATCTNNNGCAGGAGAATTGCTTGAACCA
AGCACACGTCTGAACTCTTTCCCTACACGACGCTCTTCCGATCTNNNGCAGGAGAATTGCTTGAACCA
AGCACACGTCTGAACTCTTTCCCTACACGACGCTCTTCCGATCTNNNGAGGCAGGAGAATGGTGTG
AGCACACGTCTGAACTCTTTCCCTACACGACGCTCTTCCGATCTNNNGAGGCAGGAGAATGGTGTG
AGCACACGTCTGAACTCTTTCCCTACACGACGCTCTTCCGATCTNNNGAGGCAGGAGAATGGTGTG
GAGCACACGTCTGAACTCTTTCCCTACACGACGCTCTTCCGATCTNNNGTGGTTGATTGTCTTGATGC
GAGCACACGTCTGAACTCTTTCCCTACACGACGCTCTTCCGATCTNNNGTGGTTGATTGTCTTGATGC
GAGCACACGTCTGAACTCTTTCCCTACACGACGCTCTTCCGATCTNNNGTGGTTGATTGTCTTGATGC
GAGCACACGTCTGAACTCTTTCCCTACACGACGCTCTTCCGATCTNNNGCACAGGATACAGGATAGG
GAGCACACGTCTGAACTCTTTCCCTACACGACGCTCTTCCGATCTNNNGCACAGGATACAGGATAGG
GAGCACACGTCTGAACTCTTTCCCTACACGACGCTCTTCCGATCTNNNGCACAGGATACAGGATAGG
GCACACGTCTGAACTCTTTCCCTACACGACGCTCTTCCGATCTNNNAGAGACAGTGTCTTACTATGTTT
GCACACGTCTGAACTCTTTCCCTACACGACGCTCTTCCGATCTNNNAGAGACAGTGTCTTACTATGTTT
GCACACGTCTGAACTCTTTCCCTACACGACGCTCTTCCGATCTNNNAGAGACAGTGTCTTACTATGTTT
GAGCACACGTCTGAACTCTTTCCCTACACGACGCTCTTCCGATCTNNNCTGAGACAGGAGAATCGCTT
GAGCACACGTCTGAACTCTTTCCCTACACGACGCTCTTCCGATCTNNNCTGAGACAGGAGAATCGCTT
GAGCACACGTCTGAACTCTTTCCCTACACGACGCTCTTCCGATCTNNNCTGAGACAGGAGAATCGCTT
GAGCACACGTCTGAACTCTTTCCCTACACGACGCTCTTCCGATCTNNNCTTGCAGTAAGCCGAGATCA
GAGCACACGTCTGAACTCTTTCCCTACACGACGCTCTTCCGATCTNNNCTTGCAGTAAGCCGAGATCA
GAGCACACGTCTGAACTCTTTCCCTACACGACGCTCTTCCGATCTNNNCTTGCAGTAAGCCGAGATCA
GCACACGTCTGAACTCTTTCCCTACACGACGCTCTTCCGATCTNNNCGCTCTCTCTCTCTCTCTCTCCA
GCACACGTCTGAACTCTTTCCCTACACGACGCTCTTCCGATCTNNNCGCTCTCTCTCTCTCTCTCTCCA
GCACACGTCTGAACTCTTTCCCTACACGACGCTCTTCCGATCTNNNCGCTCTCTCTCTCTCTCTCTCCA
CACACGTCTGAACTCTTTCCCTACACGACGCTCTTCCGATCTNNNGGACCACTGTAATCCTGAAGACA
CACACGTCTGAACTCTTTCCCTACACGACGCTCTTCCGATCTNNNGGACCACTGTAATCCTGAAGACA
CACACGTCTGAACTCTTTCCCTACACGACGCTCTTCCGATCTNNNGGACCACTGTAATCCTGAAGACA
CACACGTCTGAACTCTTTCCCTACACGACGCTCTTCCGATCTNNNCAGGGTTTCACCAAATTAGCCAGC
CACACGTCTGAACTCTTTCCCTACACGACGCTCTTCCGATCTNNNCAGGGTTTCACCAAATTAGCCAGC
CACACGTCTGAACTCTTTCCCTACACGACGCTCTTCCGATCTNNNCAGGGTTTCACCAAATTAGCCAGC
AGCACACGTCTGAACTCTTTCCCTACACGACGCTCTTCCGATCTNNNTAGGCATGGTGTGTTTACAGT
AGCACACGTCTGAACTCTTTCCCTACACGACGCTCTTCCGATCTNNNTAGGCATGGTGTGTTTACAGT
AGCACACGTCTGAACTCTTTCCCTACACGACGCTCTTCCGATCTNNNTAGGCATGGTGTGTTTACAGT
AGCACACGTCTGAACTCTTTCCCTACACGACGCTCTTCCGATCTNNNAGGAAACAAGAGCTTTTCTCCC
AGCACACGTCTGAACTCTTTCCCTACACGACGCTCTTCCGATCTNNNAGGAAACAAGAGCTTTTCTCCC
AGCACACGTCTGAACTCTTTCCCTACACGACGCTCTTCCGATCTNNNAGGAAACAAGAGCTTTTCTCCC
GAGCACACGTCTGAACTCTTTCCCTACACGACGCTCTTCCGATCTNNNTTCTAGCAGAGACGGGCTTTT
GAGCACACGTCTGAACTCTTTCCCTACACGACGCTCTTCCGATCTNNNTTCTAGCAGAGACGGGCTTTT
GAGCACACGTCTGAACTCTTTCCCTACACGACGCTCTTCCGATCTNNNTTCTAGCAGAGACGGGCTTTT
AGCACACGTCTGAACTCTTTCCCTACACGACGCTCTTCCGATCTNNNTCACAAGGTCAAGAGATCGAG
AGCACACGTCTGAACTCTTTCCCTACACGACGCTCTTCCGATCTNNNTCACAAGGTCAAGAGATCGAG
AGCACACGTCTGAACTCTTTCCCTACACGACGCTCTTCCGATCTNNNTCACAAGGTCAAGAGATCGAG

AGCACACGTCTGAACTCTTTCCCTACACGACGCTCTTCCGATCTNNNAGACAGAAGAATCGCTTGAAC
AGCACACGTCTGAACTCTTTCCCTACACGACGCTCTTCCGATCTNNNAGACAGAAGAATCGCTTGAAC
AGCACACGTCTGAACTCTTTCCCTACACGACGCTCTTCCGATCTNNNAGACAGAAGAATCGCTTGAAC
AGCACACGTCTGAACTCTTTCCCTACACGACGCTCTTCCGATCTNNNGGTAGCTAGTGAGTAGGAG
AGCACACGTCTGAACTCTTTCCCTACACGACGCTCTTCCGATCTNNNGGTAGCTAGTGAGTAGGAG
AGCACACGTCTGAACTCTTTCCCTACACGACGCTCTTCCGATCTNNNGGTAGCTAGTGAGTAGGAG
GCACACGTCTGAACTCTTTCCCTACACGACGCTCTTCCGATCTNNNTCATTGTTTGAGACAGGCTACG
GCACACGTCTGAACTCTTTCCCTACACGACGCTCTTCCGATCTNNNTCATTGTTTGAGACAGGCTACG
GCACACGTCTGAACTCTTTCCCTACACGACGCTCTTCCGATCTNNNTCATTGTTTGAGACAGGCTACG
GCACACGTCTGAACTCTTTCCCTACACGACGCTCTTCCGATCTNNNCCTGCCTTCTCAGTTTCCTAACTC
GCACACGTCTGAACTCTTTCCCTACACGACGCTCTTCCGATCTNNNCCTGCCTTCTCAGTTTCCTAACTC
GCACACGTCTGAACTCTTTCCCTACACGACGCTCTTCCGATCTNNNCCTGCCTTCTCAGTTTCCTAACTC
AGCACACGTCTGAACTCTTTCCCTACACGACGCTCTTCCGATCTNNNTTGAGTTGAAATCATGGTATT
AGCACACGTCTGAACTCTTTCCCTACACGACGCTCTTCCGATCTNNNTTGAGTTGAAATCATGGTATT
AGCACACGTCTGAACTCTTTCCCTACACGACGCTCTTCCGATCTNNNTTGAGTTGAAATCATGGTATT
AGCACACGTCTGAACTCTTTCCCTACACGACGCTCTTCCGATCTNNNCTATTTGGGCTACTCTGGACT
AGCACACGTCTGAACTCTTTCCCTACACGACGCTCTTCCGATCTNNNCTATTTGGGCTACTCTGGACT
AGCACACGTCTGAACTCTTTCCCTACACGACGCTCTTCCGATCTNNNCTATTTGGGCTACTCTGGACT
GAGCACACGTCTGAACTCTTTCCCTACACGACGCTCTTCCGATCTNNNCGAAGGTCACCAACATCAAA
GAGCACACGTCTGAACTCTTTCCCTACACGACGCTCTTCCGATCTNNNCGAAGGTCACCAACATCAAA
GAGCACACGTCTGAACTCTTTCCCTACACGACGCTCTTCCGATCTNNNCGAAGGTCACCAACATCAAA
AGCACACGTCTGAACTCTTTCCCTACACGACGCTCTTCCGATCTNNNTAGTGTCAAAATTCACCACCC
AGCACACGTCTGAACTCTTTCCCTACACGACGCTCTTCCGATCTNNNTAGTGTCAAAATTCACCACCC
AGCACACGTCTGAACTCTTTCCCTACACGACGCTCTTCCGATCTNNNTAGTGTCAAAATTCACCACCC
AGCACACGTCTGAACTCTTTCCCTACACGACGCTCTTCCGATCTNNNTTCCCTCATCTCCCGTGCTTCC
AGCACACGTCTGAACTCTTTCCCTACACGACGCTCTTCCGATCTNNNTTCCCTCATCTCCCGTGCTTCC
AGCACACGTCTGAACTCTTTCCCTACACGACGCTCTTCCGATCTNNNTTCCCTCATCTCCCGTGCTTCC
AGCACACGTCTGAACTCTTTCCCTACACGACGCTCTTCCGATCTNNNGCGGTTCATCTGTATCCTCTGT
AGCACACGTCTGAACTCTTTCCCTACACGACGCTCTTCCGATCTNNNGCGGTTCATCTGTATCCTCTGT
AGCACACGTCTGAACTCTTTCCCTACACGACGCTCTTCCGATCTNNNGCGGTTCATCTGTATCCTCTGT
ACACGTCTGAACTCTTTCCCTACACGACGCTCTTCCGATCTNNNAGAGATTATCCTGGATTGTCTCAT
ACACGTCTGAACTCTTTCCCTACACGACGCTCTTCCGATCTNNNAGAGATTATCCTGGATTGTCTCAT
ACACGTCTGAACTCTTTCCCTACACGACGCTCTTCCGATCTNNNAGAGATTATCCTGGATTGTCTCAT
GCACACGTCTGAACTCTTTCCCTACACGACGCTCTTCCGATCTNNNGATAAGACCTGAAGCAGAGCA
GCACACGTCTGAACTCTTTCCCTACACGACGCTCTTCCGATCTNNNGATAAGACCTGAAGCAGAGCA
GCACACGTCTGAACTCTTTCCCTACACGACGCTCTTCCGATCTNNNGATAAGACCTGAAGCAGAGCA
GCACACGTCTGAACTCTTTCCCTACACGACGCTCTTCCGATCTNNNATGTTGAAACACTTTAGGGGC
GCACACGTCTGAACTCTTTCCCTACACGACGCTCTTCCGATCTNNNATGTTGAAACACTTTAGGGGC
GCACACGTCTGAACTCTTTCCCTACACGACGCTCTTCCGATCTNNNATGTTGAAACACTTTAGGGGC
GAGCACACGTCTGAACTCTTTCCCTACACGACGCTCTTCCGATCTNNNAAACATTGCCAAACATCTCCT
GAGCACACGTCTGAACTCTTTCCCTACACGACGCTCTTCCGATCTNNNAAACATTGCCAAACATCTCCT
GAGCACACGTCTGAACTCTTTCCCTACACGACGCTCTTCCGATCTNNNAAACATTGCCAAACATCTCCT
ACACGTCTGAACTCTTTCCCTACACGACGCTCTTCCGATCTNNNATAAAGGAATGAGTCATCTGTTTC

CACACGTCTGAACTCTTTCCCTACACGACGCTCTTCCGATCTNNNATAAAGGAATGAGTCATCTGTTTCA
CACACGTCTGAACTCTTTCCCTACACGACGCTCTTCCGATCTNNNATAAAGGAATGAGTCATCTGTTTCA
AGCACACGTCTGAACTCTTTCCCTACACGACGCTCTTCCGATCTNNNGGAAGGGTAGGACAAAAGGAG
AGCACACGTCTGAACTCTTTCCCTACACGACGCTCTTCCGATCTNNNGGAAGGGTAGGACAAAAGGAG
AGCACACGTCTGAACTCTTTCCCTACACGACGCTCTTCCGATCTNNNGGAAGGGTAGGACAAAAGGAG
GCACACGTCTGAACTCTTTCCCTACACGACGCTCTTCCGATCTNNNCTTCCTTTACTTGATGCTCTTCATC
GCACACGTCTGAACTCTTTCCCTACACGACGCTCTTCCGATCTNNNCTTCCTTTACTTGATGCTCTTCATC
GCACACGTCTGAACTCTTTCCCTACACGACGCTCTTCCGATCTNNNCTTCCTTTACTTGATGCTCTTCATC
GAGCACACGTCTGAACTCTTTCCCTACACGACGCTCTTCCGATCTNNNACCCAATAACTGGTGATCTAAC
GAGCACACGTCTGAACTCTTTCCCTACACGACGCTCTTCCGATCTNNNACCCAATAACTGGTGATCTAAC
GAGCACACGTCTGAACTCTTTCCCTACACGACGCTCTTCCGATCTNNNACCCAATAACTGGTGATCTAAC
AGAGCACACGTCTGAACTCTTTCCCTACACGACGCTCTTCCGATCTNNNAAGAGTCTGTTAGGAGTAA
AGAGCACACGTCTGAACTCTTTCCCTACACGACGCTCTTCCGATCTNNNAAGAGTCTGTTAGGAGTAA
AGAGCACACGTCTGAACTCTTTCCCTACACGACGCTCTTCCGATCTNNNAAGAGTCTGTTAGGAGTAA
AGCACACGTCTGAACTCTTTCCCTACACGACGCTCTTCCGATCTNNNAGAACAGCGAAAGTCCATAGCA
AGCACACGTCTGAACTCTTTCCCTACACGACGCTCTTCCGATCTNNNAGAACAGCGAAAGTCCATAGCA
AGCACACGTCTGAACTCTTTCCCTACACGACGCTCTTCCGATCTNNNAGAACAGCGAAAGTCCATAGCA
AGCACACGTCTGAACTCTTTCCCTACACGACGCTCTTCCGATCTNNNTAATTGATGATCATCTGATTGAG
AGCACACGTCTGAACTCTTTCCCTACACGACGCTCTTCCGATCTNNNTAATTGATGATCATCTGATTGAG
AGCACACGTCTGAACTCTTTCCCTACACGACGCTCTTCCGATCTNNNTAATTGATGATCATCTGATTGAG
AGCACACGTCTGAACTCTTTCCCTACACGACGCTCTTCCGATCTNNNTCTGAGACTTGGGCACAGATAC
AGCACACGTCTGAACTCTTTCCCTACACGACGCTCTTCCGATCTNNNTCTGAGACTTGGGCACAGATAC
AGCACACGTCTGAACTCTTTCCCTACACGACGCTCTTCCGATCTNNNTCTGAGACTTGGGCACAGATAC
GCACACGTCTGAACTCTTTCCCTACACGACGCTCTTCCGATCTNNNGTGACTTGGAGGTGTTTCTCTCT
GCACACGTCTGAACTCTTTCCCTACACGACGCTCTTCCGATCTNNNGTGACTTGGAGGTGTTTCTCTCT
GCACACGTCTGAACTCTTTCCCTACACGACGCTCTTCCGATCTNNNGTGACTTGGAGGTGTTTCTCTCT
GAGCACACGTCTGAACTCTTTCCCTACACGACGCTCTTCCGATCTNNNGCAACACAGTGAGACCCTATC
GAGCACACGTCTGAACTCTTTCCCTACACGACGCTCTTCCGATCTNNNGCAACACAGTGAGACCCTATC
GAGCACACGTCTGAACTCTTTCCCTACACGACGCTCTTCCGATCTNNNGCAACACAGTGAGACCCTATC
AGCACACGTCTGAACTCTTTCCCTACACGACGCTCTTCCGATCTNNNACCCTTGCTAGCTCTTTCTAACCC
AGCACACGTCTGAACTCTTTCCCTACACGACGCTCTTCCGATCTNNNACCCTTGCTAGCTCTTTCTAACCC
AGCACACGTCTGAACTCTTTCCCTACACGACGCTCTTCCGATCTNNNACCCTTGCTAGCTCTTTCTAACCC
GAGCACACGTCTGAACTCTTTCCCTACACGACGCTCTTCCGATCTNNNTGACCCATGTGCCATTTTGAAA
GAGCACACGTCTGAACTCTTTCCCTACACGACGCTCTTCCGATCTNNNTGACCCATGTGCCATTTTGAAA
GAGCACACGTCTGAACTCTTTCCCTACACGACGCTCTTCCGATCTNNNTGACCCATGTGCCATTTTGAAA
GCACACGTCTGAACTCTTTCCCTACACGACGCTCTTCCGATCTNNNAAAGGAACGAAGGCAAATGGAA
GCACACGTCTGAACTCTTTCCCTACACGACGCTCTTCCGATCTNNNAAAGGAACGAAGGCAAATGGAA
GCACACGTCTGAACTCTTTCCCTACACGACGCTCTTCCGATCTNNNAAAGGAACGAAGGCAAATGGAA
GAGCACACGTCTGAACTCTTTCCCTACACGACGCTCTTCCGATCTNNNGAAGTTGCAGTGAGCTGAGA
GAGCACACGTCTGAACTCTTTCCCTACACGACGCTCTTCCGATCTNNNGAAGTTGCAGTGAGCTGAGA
GAGCACACGTCTGAACTCTTTCCCTACACGACGCTCTTCCGATCTNNNGAAGTTGCAGTGAGCTGAGA
GAGCACACGTCTGAACTCTTTCCCTACACGACGCTCTTCCGATCTNNNGAATGTAGAGGATGAGGAAA
GAGCACACGTCTGAACTCTTTCCCTACACGACGCTCTTCCGATCTNNNGAATGTAGAGGATGAGGAAA

AGCACACGTCTGAACTCTTTCCCTACACGACGCTCTTCCGATCTNNNGAATGTAGAGGATGAGGAA
GCACACGTCTGAACTCTTTCCCTACACGACGCTCTTCCGATCTNNNGGGTAAAAGCTCCATGACATTG
GCACACGTCTGAACTCTTTCCCTACACGACGCTCTTCCGATCTNNNGGGTAAAAGCTCCATGACATTG
GCACACGTCTGAACTCTTTCCCTACACGACGCTCTTCCGATCTNNNGGGTAAAAGCTCCATGACATTG
AGCACACGTCTGAACTCTTTCCCTACACGACGCTCTTCCGATCTNNNCCATTTTGCATTCCCACTAGCA
AGCACACGTCTGAACTCTTTCCCTACACGACGCTCTTCCGATCTNNNCCATTTTGCATTCCCACTAGCA
AGCACACGTCTGAACTCTTTCCCTACACGACGCTCTTCCGATCTNNNCCATTTTGCATTCCCACTAGCA
GCACACGTCTGAACTCTTTCCCTACACGACGCTCTTCCGATCTNNNACCTAGGAATTGACCAAAAAAG
GCACACGTCTGAACTCTTTCCCTACACGACGCTCTTCCGATCTNNNACCTAGGAATTGACCAAAAAAG
GCACACGTCTGAACTCTTTCCCTACACGACGCTCTTCCGATCTNNNACCTAGGAATTGACCAAAAAAG
AGCACACGTCTGAACTCTTTCCCTACACGACGCTCTTCCGATCTNNNAAATGTGAACGGGTCTCCAG
AGCACACGTCTGAACTCTTTCCCTACACGACGCTCTTCCGATCTNNNAAATGTGAACGGGTCTCCAG
AGCACACGTCTGAACTCTTTCCCTACACGACGCTCTTCCGATCTNNNAAATGTGAACGGGTCTCCAG
AGCACACGTCTGAACTCTTTCCCTACACGACGCTCTTCCGATCTNNNAGGTTCAAATGATTCTCCTGC
AGCACACGTCTGAACTCTTTCCCTACACGACGCTCTTCCGATCTNNNAGGTTCAAATGATTCTCCTGC
AGCACACGTCTGAACTCTTTCCCTACACGACGCTCTTCCGATCTNNNAGGTTCAAATGATTCTCCTGC
GAGCACACGTCTGAACTCTTTCCCTACACGACGCTCTTCCGATCTNNNACTATGCTGGCTTCAAGACT
GAGCACACGTCTGAACTCTTTCCCTACACGACGCTCTTCCGATCTNNNACTATGCTGGCTTCAAGACT
GAGCACACGTCTGAACTCTTTCCCTACACGACGCTCTTCCGATCTNNNACTATGCTGGCTTCAAGACT
GAGCACACGTCTGAACTCTTTCCCTACACGACGCTCTTCCGATCTNNNCAGGGTTGAGGTCCTAGATG
GAGCACACGTCTGAACTCTTTCCCTACACGACGCTCTTCCGATCTNNNCAGGGTTGAGGTCCTAGATG
GAGCACACGTCTGAACTCTTTCCCTACACGACGCTCTTCCGATCTNNNCAGGGTTGAGGTCCTAGATG
AGCACACGTCTGAACTCTTTCCCTACACGACGCTCTTCCGATCTNNNTGGATTTTGAGGTTGAACCC
AGCACACGTCTGAACTCTTTCCCTACACGACGCTCTTCCGATCTNNNTGGATTTTGAGGTTGAACCC
AGCACACGTCTGAACTCTTTCCCTACACGACGCTCTTCCGATCTNNNTGGATTTTGAGGTTGAACCC
AGCACACGTCTGAACTCTTTCCCTACACGACGCTCTTCCGATCTNNNAATGCCCCCATTGAGAGTAAG
AGCACACGTCTGAACTCTTTCCCTACACGACGCTCTTCCGATCTNNNAATGCCCCCATTGAGAGTAAG
AGCACACGTCTGAACTCTTTCCCTACACGACGCTCTTCCGATCTNNNAATGCCCCCATTGAGAGTAAG
CACACGTCTGAACTCTTTCCCTACACGACGCTCTTCCGATCTNNNGAGGCTACGGTGAACCAGGATA
CACACGTCTGAACTCTTTCCCTACACGACGCTCTTCCGATCTNNNGAGGCTACGGTGAACCAGGATA
CACACGTCTGAACTCTTTCCCTACACGACGCTCTTCCGATCTNNNGAGGCTACGGTGAACCAGGATA
AGCACACGTCTGAACTCTTTCCCTACACGACGCTCTTCCGATCTNNNGGAGTGGACACATAGGAAA
AGCACACGTCTGAACTCTTTCCCTACACGACGCTCTTCCGATCTNNNGGAGTGGACACATAGGAAA
AGCACACGTCTGAACTCTTTCCCTACACGACGCTCTTCCGATCTNNNGGAGTGGACACATAGGAAA
GCACACGTCTGAACTCTTTCCCTACACGACGCTCTTCCGATCTNNNTGGAACTACATGCATAAGCCAC
GCACACGTCTGAACTCTTTCCCTACACGACGCTCTTCCGATCTNNNTGGAACTACATGCATAAGCCAC
GCACACGTCTGAACTCTTTCCCTACACGACGCTCTTCCGATCTNNNTGGAACTACATGCATAAGCCAC
CACACGTCTGAACTCTTTCCCTACACGACGCTCTTCCGATCTNNNTTTCAAATAAAACTTGTACTGGG
CACACGTCTGAACTCTTTCCCTACACGACGCTCTTCCGATCTNNNTTTCAAATAAAACTTGTACTGGG
CACACGTCTGAACTCTTTCCCTACACGACGCTCTTCCGATCTNNNTTTCAAATAAAACTTGTACTGGG
AGCACACGTCTGAACTCTTTCCCTACACGACGCTCTTCCGATCTNNNTTCTAACGTCCAAGGTCAGGA
AGCACACGTCTGAACTCTTTCCCTACACGACGCTCTTCCGATCTNNNTTCTAACGTCCAAGGTCAGGA
AGCACACGTCTGAACTCTTTCCCTACACGACGCTCTTCCGATCTNNNTTCTAACGTCCAAGGTCAGGA

AGCACACGTCTGAACTCTTTCCCTACACGACGCTCTTCCGATCTNNNGCATTCTTGTAGTCCCAGCTAC
AGCACACGTCTGAACTCTTTCCCTACACGACGCTCTTCCGATCTNNNGCATTCTTGTAGTCCCAGCTAC
AGCACACGTCTGAACTCTTTCCCTACACGACGCTCTTCCGATCTNNNGCATTCTTGTAGTCCCAGCTAC
AGCACACGTCTGAACTCTTTCCCTACACGACGCTCTTCCGATCTNNNGAGCCTGGCTGGATTTTCTTTT
AGCACACGTCTGAACTCTTTCCCTACACGACGCTCTTCCGATCTNNNGAGCCTGGCTGGATTTTCTTTT
AGCACACGTCTGAACTCTTTCCCTACACGACGCTCTTCCGATCTNNNGAGCCTGGCTGGATTTTCTTTT
GCACACGTCTGAACTCTTTCCCTACACGACGCTCTTCCGATCTNNNCTTGCAGTGAGCCAAGATCTCAC
GCACACGTCTGAACTCTTTCCCTACACGACGCTCTTCCGATCTNNNCTTGCAGTGAGCCAAGATCTCAC
GCACACGTCTGAACTCTTTCCCTACACGACGCTCTTCCGATCTNNNCTTGCAGTGAGCCAAGATCTCAC
AGCACACGTCTGAACTCTTTCCCTACACGACGCTCTTCCGATCTNNNCTTATGAGCCAAAAGAAGCCC
AGCACACGTCTGAACTCTTTCCCTACACGACGCTCTTCCGATCTNNNCTTATGAGCCAAAAGAAGCCC
AGCACACGTCTGAACTCTTTCCCTACACGACGCTCTTCCGATCTNNNCTTATGAGCCAAAAGAAGCCC
GCACACGTCTGAACTCTTTCCCTACACGACGCTCTTCCGATCTNNNAGTCTCAACGAAATGGAATGG
GCACACGTCTGAACTCTTTCCCTACACGACGCTCTTCCGATCTNNNAGTCTCAACGAAATGGAATGG
GCACACGTCTGAACTCTTTCCCTACACGACGCTCTTCCGATCTNNNAGTCTCAACGAAATGGAATGG
AAGAGCACACGTCTGAACTCTTTCCCTACACGACGCTCTTCCGATCTNNNAACTCCTGAAGAGGAGG
AAGAGCACACGTCTGAACTCTTTCCCTACACGACGCTCTTCCGATCTNNNAACTCCTGAAGAGGAGG
AAGAGCACACGTCTGAACTCTTTCCCTACACGACGCTCTTCCGATCTNNNAACTCCTGAAGAGGAGG
AGCACACGTCTGAACTCTTTCCCTACACGACGCTCTTCCGATCTNNNGTGCACACACACACACATATT
AGCACACGTCTGAACTCTTTCCCTACACGACGCTCTTCCGATCTNNNGTGCACACACACACACATATT
AGCACACGTCTGAACTCTTTCCCTACACGACGCTCTTCCGATCTNNNGTGCACACACACACACATATT
GCACACGTCTGAACTCTTTCCCTACACGACGCTCTTCCGATCTNNNGAGGCAGGAGAATGGTGTGA
GCACACGTCTGAACTCTTTCCCTACACGACGCTCTTCCGATCTNNNGAGGCAGGAGAATGGTGTGA
GCACACGTCTGAACTCTTTCCCTACACGACGCTCTTCCGATCTNNNGAGGCAGGAGAATGGTGTGA
GCACACGTCTGAACTCTTTCCCTACACGACGCTCTTCCGATCTNNNCCAGATCCATCACACATTCCTCT
GCACACGTCTGAACTCTTTCCCTACACGACGCTCTTCCGATCTNNNCCAGATCCATCACACATTCCTCT
GCACACGTCTGAACTCTTTCCCTACACGACGCTCTTCCGATCTNNNCCAGATCCATCACACATTCCTCT
GCACACGTCTGAACTCTTTCCCTACACGACGCTCTTCCGATCTNNNGCAAGGGGCATTTACATTCACA
GCACACGTCTGAACTCTTTCCCTACACGACGCTCTTCCGATCTNNNGCAAGGGGCATTTACATTCACA
GCACACGTCTGAACTCTTTCCCTACACGACGCTCTTCCGATCTNNNGCAAGGGGCATTTACATTCACA
AGCACACGTCTGAACTCTTTCCCTACACGACGCTCTTCCGATCTNNNGCATCTGCTGTCCTCCCATTATT
AGCACACGTCTGAACTCTTTCCCTACACGACGCTCTTCCGATCTNNNGCATCTGCTGTCCTCCCATTATT
AGCACACGTCTGAACTCTTTCCCTACACGACGCTCTTCCGATCTNNNGCATCTGCTGTCCTCCCATTATT
AGCACACGTCTGAACTCTTTCCCTACACGACGCTCTTCCGATCTNNNGTAATCTCAGCTACTCTTGGG
AGCACACGTCTGAACTCTTTCCCTACACGACGCTCTTCCGATCTNNNGTAATCTCAGCTACTCTTGGG
AGCACACGTCTGAACTCTTTCCCTACACGACGCTCTTCCGATCTNNNGTAATCTCAGCTACTCTTGGG
GAGCACACGTCTGAACTCTTTCCCTACACGACGCTCTTCCGATCTNNNGTCTCAAGGTCTGTTTAGGG
GAGCACACGTCTGAACTCTTTCCCTACACGACGCTCTTCCGATCTNNNGTCTCAAGGTCTGTTTAGGG
GAGCACACGTCTGAACTCTTTCCCTACACGACGCTCTTCCGATCTNNNGTCTCAAGGTCTGTTTAGGG
AGAGCACACGTCTGAACTCTTTCCCTACACGACGCTCTTCCGATCTNNNCACACGTTAGATTGGTGTA
AGAGCACACGTCTGAACTCTTTCCCTACACGACGCTCTTCCGATCTNNNCACACGTTAGATTGGTGTA
AGAGCACACGTCTGAACTCTTTCCCTACACGACGCTCTTCCGATCTNNNCACACGTTAGATTGGTGTA
GCACACGTCTGAACTCTTTCCCTACACGACGCTCTTCCGATCTNNNTGTGGCTGAAAGTGAAGTTCT

AGCACACGTCTGAACTCTTTCCCTACACGACGCTCTTCCGATCTNNNTGTGGCTGAAAGTGAAGTTCTC
AGCACACGTCTGAACTCTTTCCCTACACGACGCTCTTCCGATCTNNNTGTGGCTGAAAGTGAAGTTCTC
GAGCACACGTCTGAACTCTTTCCCTACACGACGCTCTTCCGATCTNNNCCTGGAAAGGCAACCAAACTA
GAGCACACGTCTGAACTCTTTCCCTACACGACGCTCTTCCGATCTNNNCCTGGAAAGGCAACCAAACTA
GAGCACACGTCTGAACTCTTTCCCTACACGACGCTCTTCCGATCTNNNCCTGGAAAGGCAACCAAACTA
GAGCACACGTCTGAACTCTTTCCCTACACGACGCTCTTCCGATCTNNNCAAACCTGTGCTTGGTACATC
GAGCACACGTCTGAACTCTTTCCCTACACGACGCTCTTCCGATCTNNNCAAACCTGTGCTTGGTACATC
GAGCACACGTCTGAACTCTTTCCCTACACGACGCTCTTCCGATCTNNNCAAACCTGTGCTTGGTACATC
AGAGCACACGTCTGAACTCTTTCCCTACACGACGCTCTTCCGATCTNNNTGGTATCTGTGGGCATATGT
AGAGCACACGTCTGAACTCTTTCCCTACACGACGCTCTTCCGATCTNNNTGGTATCTGTGGGCATATGT
AGAGCACACGTCTGAACTCTTTCCCTACACGACGCTCTTCCGATCTNNNTGGTATCTGTGGGCATATGT
GAGCACACGTCTGAACTCTTTCCCTACACGACGCTCTTCCGATCTNNNTCAGCAGTGTTTTGTAGTTCTC
GAGCACACGTCTGAACTCTTTCCCTACACGACGCTCTTCCGATCTNNNTCAGCAGTGTTTTGTAGTTCTC
GAGCACACGTCTGAACTCTTTCCCTACACGACGCTCTTCCGATCTNNNTCAGCAGTGTTTTGTAGTTCTC
AGCACACGTCTGAACTCTTTCCCTACACGACGCTCTTCCGATCTNNNGCAGGAGAATTGCTTGATCTTG
AGCACACGTCTGAACTCTTTCCCTACACGACGCTCTTCCGATCTNNNGCAGGAGAATTGCTTGATCTTG
AGCACACGTCTGAACTCTTTCCCTACACGACGCTCTTCCGATCTNNNGCAGGAGAATTGCTTGATCTTG
AGCACACGTCTGAACTCTTTCCCTACACGACGCTCTTCCGATCTNNNCTCTTCCCCTTGGTTTGTAACAT
AGCACACGTCTGAACTCTTTCCCTACACGACGCTCTTCCGATCTNNNCTCTTCCCCTTGGTTTGTAACAT
AGCACACGTCTGAACTCTTTCCCTACACGACGCTCTTCCGATCTNNNCTCTTCCCCTTGGTTTGTAACAT
GCACACGTCTGAACTCTTTCCCTACACGACGCTCTTCCGATCTNNNAGGTAGTTTCATCTCTATTGCAAT
GCACACGTCTGAACTCTTTCCCTACACGACGCTCTTCCGATCTNNNAGGTAGTTTCATCTCTATTGCAAT
GCACACGTCTGAACTCTTTCCCTACACGACGCTCTTCCGATCTNNNAGGTAGTTTCATCTCTATTGCAAT
AGAGCACACGTCTGAACTCTTTCCCTACACGACGCTCTTCCGATCTNNNCTCATGGCCAGGAAACTTTGA
AGAGCACACGTCTGAACTCTTTCCCTACACGACGCTCTTCCGATCTNNNCTCATGGCCAGGAAACTTTGA
AGAGCACACGTCTGAACTCTTTCCCTACACGACGCTCTTCCGATCTNNNCTCATGGCCAGGAAACTTTGA
AGCACACGTCTGAACTCTTTCCCTACACGACGCTCTTCCGATCTNNNGCTGATGGCCTTTCTAAGCAATC
AGCACACGTCTGAACTCTTTCCCTACACGACGCTCTTCCGATCTNNNGCTGATGGCCTTTCTAAGCAATC
AGCACACGTCTGAACTCTTTCCCTACACGACGCTCTTCCGATCTNNNGCTGATGGCCTTTCTAAGCAATC
AGAGCACACGTCTGAACTCTTTCCCTACACGACGCTCTTCCGATCTNNNATCTAGTTAGTTCCCTCCAGC
AGAGCACACGTCTGAACTCTTTCCCTACACGACGCTCTTCCGATCTNNNATCTAGTTAGTTCCCTCCAGC
AGAGCACACGTCTGAACTCTTTCCCTACACGACGCTCTTCCGATCTNNNATCTAGTTAGTTCCCTCCAGC
AGCACACGTCTGAACTCTTTCCCTACACGACGCTCTTCCGATCTNNNGAGGCAGGAAAATCACTTGAA
AGCACACGTCTGAACTCTTTCCCTACACGACGCTCTTCCGATCTNNNGAGGCAGGAAAATCACTTGAA
AGCACACGTCTGAACTCTTTCCCTACACGACGCTCTTCCGATCTNNNGAGGCAGGAAAATCACTTGAA
AAGAGCACACGTCTGAACTCTTTCCCTACACGACGCTCTTCCGATCTNNNGGAAGAAGGGAGAAAG
AAGAGCACACGTCTGAACTCTTTCCCTACACGACGCTCTTCCGATCTNNNGGAAGAAGGGAGAAAG
AAGAGCACACGTCTGAACTCTTTCCCTACACGACGCTCTTCCGATCTNNNGGAAGAAGGGAGAAAG
GAGCACACGTCTGAACTCTTTCCCTACACGACGCTCTTCCGATCTNNNAAGAATTGCTTGAACCTGGGA
GAGCACACGTCTGAACTCTTTCCCTACACGACGCTCTTCCGATCTNNNAAGAATTGCTTGAACCTGGGA
GAGCACACGTCTGAACTCTTTCCCTACACGACGCTCTTCCGATCTNNNAAGAATTGCTTGAACCTGGGA
GAGCACACGTCTGAACTCTTTCCCTACACGACGCTCTTCCGATCTNNNCGTAAGCCAAAATTGTGCCACT
GAGCACACGTCTGAACTCTTTCCCTACACGACGCTCTTCCGATCTNNNCGTAAGCCAAAATTGTGCCACT

AGCACACGTCTGAACTCTTTCCCTACACGACGCTCTTCCGATCTNNNCGTAAGCCAAAATTGTGCCACT

GAGCACACGTCTGAACTCTTTCCCTACACGACGCTCTTCCGATCTNNNTTATGGTGAGCAGAGATGGT

GAGCACACGTCTGAACTCTTTCCCTACACGACGCTCTTCCGATCTNNNTTATGGTGAGCAGAGATGGT

GAGCACACGTCTGAACTCTTTCCCTACACGACGCTCTTCCGATCTNNNTTATGGTGAGCAGAGATGGT

AGCACACGTCTGAACTCTTTCCCTACACGACGCTCTTCCGATCTNNNCTGTAGTCCCAGCTACTCAAGAG

AGCACACGTCTGAACTCTTTCCCTACACGACGCTCTTCCGATCTNNNCTGTAGTCCCAGCTACTCAAGAG

AGCACACGTCTGAACTCTTTCCCTACACGACGCTCTTCCGATCTNNNCTGTAGTCCCAGCTACTCAAGAG

GCACACGTCTGAACTCTTTCCCTACACGACGCTCTTCCGATCTNNNGAGGCTGAGGCAGGAGAATGCA

GCACACGTCTGAACTCTTTCCCTACACGACGCTCTTCCGATCTNNNGAGGCTGAGGCAGGAGAATGCA

GCACACGTCTGAACTCTTTCCCTACACGACGCTCTTCCGATCTNNNGAGGCTGAGGCAGGAGAATGCA

GAGCACACGTCTGAACTCTTTCCCTACACGACGCTCTTCCGATCTNNNCAGCCTGGACAACATAGCAAA

GAGCACACGTCTGAACTCTTTCCCTACACGACGCTCTTCCGATCTNNNCAGCCTGGACAACATAGCAAA

GAGCACACGTCTGAACTCTTTCCCTACACGACGCTCTTCCGATCTNNNCAGCCTGGACAACATAGCAAA

GCACACGTCTGAACTCTTTCCCTACACGACGCTCTTCCGATCTNNNCCACGTGAGCTGTAGGGAAACCA

GCACACGTCTGAACTCTTTCCCTACACGACGCTCTTCCGATCTNNNCCACGTGAGCTGTAGGGAAACCA

GCACACGTCTGAACTCTTTCCCTACACGACGCTCTTCCGATCTNNNCCACGTGAGCTGTAGGGAAACCA

GCACACGTCTGAACTCTTTCCCTACACGACGCTCTTCCGATCTNNNCATTGGTTCTTCTTTGTGGGAGC

GCACACGTCTGAACTCTTTCCCTACACGACGCTCTTCCGATCTNNNCATTGGTTCTTCTTTGTGGGAGC

GCACACGTCTGAACTCTTTCCCTACACGACGCTCTTCCGATCTNNNCATTGGTTCTTCTTTGTGGGAGC

AGCACACGTCTGAACTCTTTCCCTACACGACGCTCTTCCGATCTNNNGGTTTACAGGGAGAGGCTGTA

AGCACACGTCTGAACTCTTTCCCTACACGACGCTCTTCCGATCTNNNGGTTTACAGGGAGAGGCTGTA

AGCACACGTCTGAACTCTTTCCCTACACGACGCTCTTCCGATCTNNNGGTTTACAGGGAGAGGCTGTA

GAGCACACGTCTGAACTCTTTCCCTACACGACGCTCTTCCGATCTNNNAGGAGGGTCATCAAAATCTGC

GAGCACACGTCTGAACTCTTTCCCTACACGACGCTCTTCCGATCTNNNAGGAGGGTCATCAAAATCTGC

GAGCACACGTCTGAACTCTTTCCCTACACGACGCTCTTCCGATCTNNNAGGAGGGTCATCAAAATCTGC

GCACACGTCTGAACTCTTTCCCTACACGACGCTCTTCCGATCTNNNGAGTATGTGTGTGTGAGGGGA

GCACACGTCTGAACTCTTTCCCTACACGACGCTCTTCCGATCTNNNGAGTATGTGTGTGTGAGGGGA

GCACACGTCTGAACTCTTTCCCTACACGACGCTCTTCCGATCTNNNGAGTATGTGTGTGTGAGGGGA

AGCACACGTCTGAACTCTTTCCCTACACGACGCTCTTCCGATCTNNNCCTGATAGACTGTCTGGCATTG

AGCACACGTCTGAACTCTTTCCCTACACGACGCTCTTCCGATCTNNNCCTGATAGACTGTCTGGCATTG

AGCACACGTCTGAACTCTTTCCCTACACGACGCTCTTCCGATCTNNNCCTGATAGACTGTCTGGCATTG

AGCACACGTCTGAACTCTTTCCCTACACGACGCTCTTCCGATCTNNNGTCATGAGACTGAGACCATCCT

AGCACACGTCTGAACTCTTTCCCTACACGACGCTCTTCCGATCTNNNGTCATGAGACTGAGACCATCCT

AGCACACGTCTGAACTCTTTCCCTACACGACGCTCTTCCGATCTNNNGTCATGAGACTGAGACCATCCT

AGCACACGTCTGAACTCTTTCCCTACACGACGCTCTTCCGATCTNNNCTGTGTTAGCAGCTACTCAGTG

AGCACACGTCTGAACTCTTTCCCTACACGACGCTCTTCCGATCTNNNCTGTGTTAGCAGCTACTCAGTG

AGCACACGTCTGAACTCTTTCCCTACACGACGCTCTTCCGATCTNNNCTGTGTTAGCAGCTACTCAGTG

GCACACGTCTGAACTCTTTCCCTACACGACGCTCTTCCGATCTNNNGACCATTAGCATCACTTAATTTGCA

GCACACGTCTGAACTCTTTCCCTACACGACGCTCTTCCGATCTNNNGACCATTAGCATCACTTAATTTGCA

GCACACGTCTGAACTCTTTCCCTACACGACGCTCTTCCGATCTNNNGACCATTAGCATCACTTAATTTGCA

AGAGCACACGTCTGAACTCTTTCCCTACACGACGCTCTTCCGATCTNNNTACCTCATCAAGTTTGGCAC

AGAGCACACGTCTGAACTCTTTCCCTACACGACGCTCTTCCGATCTNNNTACCTCATCAAGTTTGGCAC

AGAGCACACGTCTGAACTCTTTCCCTACACGACGCTCTTCCGATCTNNNTACCTCATCAAGTTTGGCAC

GAGCACACGTCTGAACTCTTTCCCTACACGACGCTCTTCCGATCTNNNCAGACCCAGACTGTAACCTCA

GAGCACACGTCTGAACTCTTTCCCTACACGACGCTCTTCCGATCTNNNCAGACCCAGACTGTAACCTCA

GAGCACACGTCTGAACTCTTTCCCTACACGACGCTCTTCCGATCTNNNCAGACCCAGACTGTAACCTCA

CACACGTCTGAACTCTTTCCCTACACGACGCTCTTCCGATCTNNNCATAGCACATTTGAGGCATCTGGC

CACACGTCTGAACTCTTTCCCTACACGACGCTCTTCCGATCTNNNCATAGCACATTTGAGGCATCTGGC

CACACGTCTGAACTCTTTCCCTACACGACGCTCTTCCGATCTNNNCATAGCACATTTGAGGCATCTGGC

GAGCACACGTCTGAACTCTTTCCCTACACGACGCTCTTCCGATCTNNNTTGCAGTGAGCCGAGATTGC

GAGCACACGTCTGAACTCTTTCCCTACACGACGCTCTTCCGATCTNNNTTGCAGTGAGCCGAGATTGC

GAGCACACGTCTGAACTCTTTCCCTACACGACGCTCTTCCGATCTNNNTTGCAGTGAGCCGAGATTGC

GAGCACACGTCTGAACTCTTTCCCTACACGACGCTCTTCCGATCTNNNTCACCACATTCTCTTCACTCTG

GAGCACACGTCTGAACTCTTTCCCTACACGACGCTCTTCCGATCTNNNTCACCACATTCTCTTCACTCTG

GAGCACACGTCTGAACTCTTTCCCTACACGACGCTCTTCCGATCTNNNTCACCACATTCTCTTCACTCTG

AGCACACGTCTGAACTCTTTCCCTACACGACGCTCTTCCGATCTNNNTCATCAGTGTCAGGAAAGCAAC

AGCACACGTCTGAACTCTTTCCCTACACGACGCTCTTCCGATCTNNNTCATCAGTGTCAGGAAAGCAAC

AGCACACGTCTGAACTCTTTCCCTACACGACGCTCTTCCGATCTNNNTCATCAGTGTCAGGAAAGCAAC

AGCACACGTCTGAACTCTTTCCCTACACGACGCTCTTCCGATCTNNNTCTCCTCCCCAAAAGTTCCTA

AGCACACGTCTGAACTCTTTCCCTACACGACGCTCTTCCGATCTNNNTCTCCTCCCCAAAAGTTCCTA

AGCACACGTCTGAACTCTTTCCCTACACGACGCTCTTCCGATCTNNNTCTCCTCCCCAAAAGTTCCTA

GAGCACACGTCTGAACTCTTTCCCTACACGACGCTCTTCCGATCTNNNCTGAGGCAGGAGAATCACTT

GAGCACACGTCTGAACTCTTTCCCTACACGACGCTCTTCCGATCTNNNCTGAGGCAGGAGAATCACTT

GAGCACACGTCTGAACTCTTTCCCTACACGACGCTCTTCCGATCTNNNCTGAGGCAGGAGAATCACTT

GAGCACACGTCTGAACTCTTTCCCTACACGACGCTCTTCCGATCTNNNATTTGTGTCATAGTGCAGTG

GAGCACACGTCTGAACTCTTTCCCTACACGACGCTCTTCCGATCTNNNATTTGTGTCATAGTGCAGTG

GAGCACACGTCTGAACTCTTTCCCTACACGACGCTCTTCCGATCTNNNATTTGTGTCATAGTGCAGTG

GAGCACACGTCTGAACTCTTTCCCTACACGACGCTCTTCCGATCTNNNGAGGTAAATTCAAGGAGCCC

GAGCACACGTCTGAACTCTTTCCCTACACGACGCTCTTCCGATCTNNNGAGGTAAATTCAAGGAGCCC

GAGCACACGTCTGAACTCTTTCCCTACACGACGCTCTTCCGATCTNNNGAGGTAAATTCAAGGAGCCC

AGAGCACACGTCTGAACTCTTTCCCTACACGACGCTCTTCCGATCTNNNGGCAGGTGATTTTGAGTTCT

AGAGCACACGTCTGAACTCTTTCCCTACACGACGCTCTTCCGATCTNNNGGCAGGTGATTTTGAGTTCT

AGAGCACACGTCTGAACTCTTTCCCTACACGACGCTCTTCCGATCTNNNGGCAGGTGATTTTGAGTTCT

GAGCACACGTCTGAACTCTTTCCCTACACGACGCTCTTCCGATCTNNNCTGAGGCAGGAGAATTGCTT

GAGCACACGTCTGAACTCTTTCCCTACACGACGCTCTTCCGATCTNNNCTGAGGCAGGAGAATTGCTT

GAGCACACGTCTGAACTCTTTCCCTACACGACGCTCTTCCGATCTNNNCTGAGGCAGGAGAATTGCTT

AGCACACGTCTGAACTCTTTCCCTACACGACGCTCTTCCGATCTNNNAAATATACATGCACCCAACGCA

AGCACACGTCTGAACTCTTTCCCTACACGACGCTCTTCCGATCTNNNAAATATACATGCACCCAACGCA

AGCACACGTCTGAACTCTTTCCCTACACGACGCTCTTCCGATCTNNNAAATATACATGCACCCAACGCA

GAGCACACGTCTGAACTCTTTCCCTACACGACGCTCTTCCGATCTNNNAGACTGTGAGTTCTGTCTTGC

GAGCACACGTCTGAACTCTTTCCCTACACGACGCTCTTCCGATCTNNNAGACTGTGAGTTCTGTCTTGC

GAGCACACGTCTGAACTCTTTCCCTACACGACGCTCTTCCGATCTNNNAGACTGTGAGTTCTGTCTTGC

GCACACGTCTGAACTCTTTCCCTACACGACGCTCTTCCGATCTNNNCACTGTTTTCGGACCTTCCATGCA

GCACACGTCTGAACTCTTTCCCTACACGACGCTCTTCCGATCTNNNCACTGTTTTCGGACCTTCCATGCA

GCACACGTCTGAACTCTTTCCCTACACGACGCTCTTCCGATCTNNNCACTGTTTTCGGACCTTCCATGCA

GAGCACACGTCTGAACTCTTTCCCTACACGACGCTCTTCCGATCTNNNCTGTAATCCCAGCTACTCAGGA

AGCACACGTCTGAACTCTTTCCCTACACGACGCTCTTCCGATCTNNNCTGTAATCCCAGCTACTCAGG
AGCACACGTCTGAACTCTTTCCCTACACGACGCTCTTCCGATCTNNNCTGTAATCCCAGCTACTCAGG
GCACACGTCTGAACTCTTTCCCTACACGACGCTCTTCCGATCTNNNTACTGCCAGAACATCAGAGTCTG
GCACACGTCTGAACTCTTTCCCTACACGACGCTCTTCCGATCTNNNTACTGCCAGAACATCAGAGTCTG
GCACACGTCTGAACTCTTTCCCTACACGACGCTCTTCCGATCTNNNTACTGCCAGAACATCAGAGTCTG
AGAGCACACGTCTGAACTCTTTCCCTACACGACGCTCTTCCGATCTNNNCTGTATGGGAACGTTCACA
AGAGCACACGTCTGAACTCTTTCCCTACACGACGCTCTTCCGATCTNNNCTGTATGGGAACGTTCACA
AGAGCACACGTCTGAACTCTTTCCCTACACGACGCTCTTCCGATCTNNNCTGTATGGGAACGTTCACA
AGCACACGTCTGAACTCTTTCCCTACACGACGCTCTTCCGATCTNNNTCAGTTGTCTTTCCTCACCCATC
AGCACACGTCTGAACTCTTTCCCTACACGACGCTCTTCCGATCTNNNTCAGTTGTCTTTCCTCACCCATC
AGCACACGTCTGAACTCTTTCCCTACACGACGCTCTTCCGATCTNNNTCAGTTGTCTTTCCTCACCCATC
AGCACACGTCTGAACTCTTTCCCTACACGACGCTCTTCCGATCTNNNTACAGAATCATCGTGGTCTTGC
AGCACACGTCTGAACTCTTTCCCTACACGACGCTCTTCCGATCTNNNTACAGAATCATCGTGGTCTTGC
AGCACACGTCTGAACTCTTTCCCTACACGACGCTCTTCCGATCTNNNTACAGAATCATCGTGGTCTTGC
GAGCACACGTCTGAACTCTTTCCCTACACGACGCTCTTCCGATCTNNNACATGCTTGTAATCCCAGCTA
GAGCACACGTCTGAACTCTTTCCCTACACGACGCTCTTCCGATCTNNNACATGCTTGTAATCCCAGCTA
GAGCACACGTCTGAACTCTTTCCCTACACGACGCTCTTCCGATCTNNNACATGCTTGTAATCCCAGCTA
GAGCACACGTCTGAACTCTTTCCCTACACGACGCTCTTCCGATCTNNNTGAAAGAGCAGATAGTTGC
GAGCACACGTCTGAACTCTTTCCCTACACGACGCTCTTCCGATCTNNNTGAAAGAGCAGATAGTTGC
GAGCACACGTCTGAACTCTTTCCCTACACGACGCTCTTCCGATCTNNNTGAAAGAGCAGATAGTTGC
GCACACGTCTGAACTCTTTCCCTACACGACGCTCTTCCGATCTNNNGCACTTGTCTGGTGATGAGTGT
GCACACGTCTGAACTCTTTCCCTACACGACGCTCTTCCGATCTNNNGCACTTGTCTGGTGATGAGTGT
GCACACGTCTGAACTCTTTCCCTACACGACGCTCTTCCGATCTNNNGCACTTGTCTGGTGATGAGTGT
AGAGCACACGTCTGAACTCTTTCCCTACACGACGCTCTTCCGATCTNNNCTAGGAATGGGGAGGTCA
AGAGCACACGTCTGAACTCTTTCCCTACACGACGCTCTTCCGATCTNNNCTAGGAATGGGGAGGTCA
AGAGCACACGTCTGAACTCTTTCCCTACACGACGCTCTTCCGATCTNNNCTAGGAATGGGGAGGTCA
AGCACACGTCTGAACTCTTTCCCTACACGACGCTCTTCCGATCTNNNTGGGAGGTAATTGAATCATG
AGCACACGTCTGAACTCTTTCCCTACACGACGCTCTTCCGATCTNNNTGGGAGGTAATTGAATCATG
AGCACACGTCTGAACTCTTTCCCTACACGACGCTCTTCCGATCTNNNTGGGAGGTAATTGAATCATG
GCACACGTCTGAACTCTTTCCCTACACGACGCTCTTCCGATCTNNNGAGTAGCTGGGATTATAGGCA
GCACACGTCTGAACTCTTTCCCTACACGACGCTCTTCCGATCTNNNGAGTAGCTGGGATTATAGGCA
GCACACGTCTGAACTCTTTCCCTACACGACGCTCTTCCGATCTNNNGAGTAGCTGGGATTATAGGCA
GCACACGTCTGAACTCTTTCCCTACACGACGCTCTTCCGATCTNNNCAGCTGGGGGACACCTAAATGC
GCACACGTCTGAACTCTTTCCCTACACGACGCTCTTCCGATCTNNNCAGCTGGGGGACACCTAAATGC
GCACACGTCTGAACTCTTTCCCTACACGACGCTCTTCCGATCTNNNCAGCTGGGGGACACCTAAATGC
AGCACACGTCTGAACTCTTTCCCTACACGACGCTCTTCCGATCTNNNCCAACAACTGCAGAGCATACAT
AGCACACGTCTGAACTCTTTCCCTACACGACGCTCTTCCGATCTNNNCCAACAACTGCAGAGCATACAT
AGCACACGTCTGAACTCTTTCCCTACACGACGCTCTTCCGATCTNNNCCAACAACTGCAGAGCATACAT
AGCACACGTCTGAACTCTTTCCCTACACGACGCTCTTCCGATCTNNNAGGAGAATGGCGTGAACCTGC
AGCACACGTCTGAACTCTTTCCCTACACGACGCTCTTCCGATCTNNNAGGAGAATGGCGTGAACCTGC
AGCACACGTCTGAACTCTTTCCCTACACGACGCTCTTCCGATCTNNNAGGAGAATGGCGTGAACCTGC
GAGCACACGTCTGAACTCTTTCCCTACACGACGCTCTTCCGATCTNNNGTGTGGTGTGTGTGTGTGT
GAGCACACGTCTGAACTCTTTCCCTACACGACGCTCTTCCGATCTNNNGTGTGGTGTGTGTGTGTGT

AGAGCACACGTCTGAACTCTTTCCCTACACGACGCTCTTCCGATCTNNNGTGTGGTGTGTGTGTGTGT

AGCACACGTCTGAACTCTTTCCCTACACGACGCTCTTCCGATCTNNNCTGACACACCTCACCTTGGAGCA

AGCACACGTCTGAACTCTTTCCCTACACGACGCTCTTCCGATCTNNNCTGACACACCTCACCTTGGAGCA

AGCACACGTCTGAACTCTTTCCCTACACGACGCTCTTCCGATCTNNNCTGACACACCTCACCTTGGAGCA

AGCACACGTCTGAACTCTTTCCCTACACGACGCTCTTCCGATCTNNNCTGCAGATGAGGTGAGGAGAT

AGCACACGTCTGAACTCTTTCCCTACACGACGCTCTTCCGATCTNNNCTGCAGATGAGGTGAGGAGAT

AGCACACGTCTGAACTCTTTCCCTACACGACGCTCTTCCGATCTNNNCTGCAGATGAGGTGAGGAGAT

CACACGTCTGAACTCTTTCCCTACACGACGCTCTTCCGATCTNNNCACCCTCGAGACCTCCTAGGCATCG

CACACGTCTGAACTCTTTCCCTACACGACGCTCTTCCGATCTNNNCACCCTCGAGACCTCCTAGGCATCG

CACACGTCTGAACTCTTTCCCTACACGACGCTCTTCCGATCTNNNCACCCTCGAGACCTCCTAGGCATCG

AGCACACGTCTGAACTCTTTCCCTACACGACGCTCTTCCGATCTNNNCTTGGCTATGGTCTCAAGGAAG

AGCACACGTCTGAACTCTTTCCCTACACGACGCTCTTCCGATCTNNNCTTGGCTATGGTCTCAAGGAAG

AGCACACGTCTGAACTCTTTCCCTACACGACGCTCTTCCGATCTNNNCTTGGCTATGGTCTCAAGGAAG

GAGCACACGTCTGAACTCTTTCCCTACACGACGCTCTTCCGATCTNNNCCCTAAACCGTGGTAACTCTCA

GAGCACACGTCTGAACTCTTTCCCTACACGACGCTCTTCCGATCTNNNCCCTAAACCGTGGTAACTCTCA

GAGCACACGTCTGAACTCTTTCCCTACACGACGCTCTTCCGATCTNNNCCCTAAACCGTGGTAACTCTCA

AGCACACGTCTGAACTCTTTCCCTACACGACGCTCTTCCGATCTNNNTGGGAGGGATTGCCAAGAATT

AGCACACGTCTGAACTCTTTCCCTACACGACGCTCTTCCGATCTNNNTGGGAGGGATTGCCAAGAATT

AGCACACGTCTGAACTCTTTCCCTACACGACGCTCTTCCGATCTNNNTGGGAGGGATTGCCAAGAATT

AGAGCACACGTCTGAACTCTTTCCCTACACGACGCTCTTCCGATCTNNNGAATATAGCCCGTACCCTCAT

AGAGCACACGTCTGAACTCTTTCCCTACACGACGCTCTTCCGATCTNNNGAATATAGCCCGTACCCTCAT

AGAGCACACGTCTGAACTCTTTCCCTACACGACGCTCTTCCGATCTNNNGAATATAGCCCGTACCCTCAT

AGCACACGTCTGAACTCTTTCCCTACACGACGCTCTTCCGATCTNNNCTGACCACTCCTTCTGTGCTATT

AGCACACGTCTGAACTCTTTCCCTACACGACGCTCTTCCGATCTNNNCTGACCACTCCTTCTGTGCTATT

AGCACACGTCTGAACTCTTTCCCTACACGACGCTCTTCCGATCTNNNCTGACCACTCCTTCTGTGCTATT

AGCACACGTCTGAACTCTTTCCCTACACGACGCTCTTCCGATCTNNNGCAGGAGAATGATTGGTGTGA

AGCACACGTCTGAACTCTTTCCCTACACGACGCTCTTCCGATCTNNNGCAGGAGAATGATTGGTGTGA

AGCACACGTCTGAACTCTTTCCCTACACGACGCTCTTCCGATCTNNNGCAGGAGAATGATTGGTGTGA

GAGCACACGTCTGAACTCTTTCCCTACACGACGCTCTTCCGATCTNNNAGTCAAGGTTTGGCCTGTGTA

GAGCACACGTCTGAACTCTTTCCCTACACGACGCTCTTCCGATCTNNNAGTCAAGGTTTGGCCTGTGTA

GAGCACACGTCTGAACTCTTTCCCTACACGACGCTCTTCCGATCTNNNAGTCAAGGTTTGGCCTGTGTA

GAGCACACGTCTGAACTCTTTCCCTACACGACGCTCTTCCGATCTNNNCATTATGCATTAGCCTCCTCCC

GAGCACACGTCTGAACTCTTTCCCTACACGACGCTCTTCCGATCTNNNCATTATGCATTAGCCTCCTCCC

GAGCACACGTCTGAACTCTTTCCCTACACGACGCTCTTCCGATCTNNNCATTATGCATTAGCCTCCTCCC

GAGCACACGTCTGAACTCTTTCCCTACACGACGCTCTTCCGATCTNNNATACAGAGAGACCCTTGTGCA

GAGCACACGTCTGAACTCTTTCCCTACACGACGCTCTTCCGATCTNNNATACAGAGAGACCCTTGTGCA

GAGCACACGTCTGAACTCTTTCCCTACACGACGCTCTTCCGATCTNNNATACAGAGAGACCCTTGTGCA

GCACACGTCTGAACTCTTTCCCTACACGACGCTCTTCCGATCTNNNGTCTCTTCCTAACAAGGGCACTC

GCACACGTCTGAACTCTTTCCCTACACGACGCTCTTCCGATCTNNNGTCTCTTCCTAACAAGGGCACTC

GCACACGTCTGAACTCTTTCCCTACACGACGCTCTTCCGATCTNNNGTCTCTTCCTAACAAGGGCACTC

AGCACACGTCTGAACTCTTTCCCTACACGACGCTCTTCCGATCTNNNAGTGCTGGGATTATAGGCATG

AGCACACGTCTGAACTCTTTCCCTACACGACGCTCTTCCGATCTNNNAGTGCTGGGATTATAGGCATG

AGCACACGTCTGAACTCTTTCCCTACACGACGCTCTTCCGATCTNNNAGTGCTGGGATTATAGGCATG

GAGCACACGTCTGAACTCTTTCCCTACACGACGCTCTTCCGATCTNNNGCAAGAGAATGGCATGAACC
GAGCACACGTCTGAACTCTTTCCCTACACGACGCTCTTCCGATCTNNNGCAAGAGAATGGCATGAACC
GAGCACACGTCTGAACTCTTTCCCTACACGACGCTCTTCCGATCTNNNGCAAGAGAATGGCATGAACC
AGCACACGTCTGAACTCTTTCCCTACACGACGCTCTTCCGATCTNNNAAGCTCTGGCTTATGGGTTTAG
AGCACACGTCTGAACTCTTTCCCTACACGACGCTCTTCCGATCTNNNAAGCTCTGGCTTATGGGTTTAG
AGCACACGTCTGAACTCTTTCCCTACACGACGCTCTTCCGATCTNNNAAGCTCTGGCTTATGGGTTTAG
GAGCACACGTCTGAACTCTTTCCCTACACGACGCTCTTCCGATCTNNNCTAGGCAACAAAGAGAGACTC
GAGCACACGTCTGAACTCTTTCCCTACACGACGCTCTTCCGATCTNNNCTAGGCAACAAAGAGAGACTC
GAGCACACGTCTGAACTCTTTCCCTACACGACGCTCTTCCGATCTNNNCTAGGCAACAAAGAGAGACTC
GAGCACACGTCTGAACTCTTTCCCTACACGACGCTCTTCCGATCTNNNCCATGATGAGGGATTCCATGA
GAGCACACGTCTGAACTCTTTCCCTACACGACGCTCTTCCGATCTNNNCCATGATGAGGGATTCCATGA
GAGCACACGTCTGAACTCTTTCCCTACACGACGCTCTTCCGATCTNNNCCATGATGAGGGATTCCATGA
AGCACACGTCTGAACTCTTTCCCTACACGACGCTCTTCCGATCTNNNGCTGAGACAGGAGAATCACTTG
AGCACACGTCTGAACTCTTTCCCTACACGACGCTCTTCCGATCTNNNGCTGAGACAGGAGAATCACTTG
AGCACACGTCTGAACTCTTTCCCTACACGACGCTCTTCCGATCTNNNGCTGAGACAGGAGAATCACTTG
GAGCACACGTCTGAACTCTTTCCCTACACGACGCTCTTCCGATCTNNNTCAGGAAGTGTGAATCCTTCA
GAGCACACGTCTGAACTCTTTCCCTACACGACGCTCTTCCGATCTNNNTCAGGAAGTGTGAATCCTTCA
GAGCACACGTCTGAACTCTTTCCCTACACGACGCTCTTCCGATCTNNNTCAGGAAGTGTGAATCCTTCA
GAGCACACGTCTGAACTCTTTCCCTACACGACGCTCTTCCGATCTNNNCTTGCAGTGAGCTGAGACACG
GAGCACACGTCTGAACTCTTTCCCTACACGACGCTCTTCCGATCTNNNCTTGCAGTGAGCTGAGACACG
GAGCACACGTCTGAACTCTTTCCCTACACGACGCTCTTCCGATCTNNNCTTGCAGTGAGCTGAGACACG
AGCACACGTCTGAACTCTTTCCCTACACGACGCTCTTCCGATCTNNNCACCTGTAATCCCAGCTACTCGC
AGCACACGTCTGAACTCTTTCCCTACACGACGCTCTTCCGATCTNNNCACCTGTAATCCCAGCTACTCGC
AGCACACGTCTGAACTCTTTCCCTACACGACGCTCTTCCGATCTNNNCACCTGTAATCCCAGCTACTCGC
GAGCACACGTCTGAACTCTTTCCCTACACGACGCTCTTCCGATCTNNNCTGGGCAACAAGAGCAAAACT
GAGCACACGTCTGAACTCTTTCCCTACACGACGCTCTTCCGATCTNNNCTGGGCAACAAGAGCAAAACT
GAGCACACGTCTGAACTCTTTCCCTACACGACGCTCTTCCGATCTNNNCTGGGCAACAAGAGCAAAACT
GAGCACACGTCTGAACTCTTTCCCTACACGACGCTCTTCCGATCTNNNGAAAGCGGACCAGGTAAGGG
GAGCACACGTCTGAACTCTTTCCCTACACGACGCTCTTCCGATCTNNNGAAAGCGGACCAGGTAAGGG
GAGCACACGTCTGAACTCTTTCCCTACACGACGCTCTTCCGATCTNNNGAAAGCGGACCAGGTAAGGG
AGAGCACACGTCTGAACTCTTTCCCTACACGACGCTCTTCCGATCTNNNCCACTAATACAGGCTTGCCCA
AGAGCACACGTCTGAACTCTTTCCCTACACGACGCTCTTCCGATCTNNNCCACTAATACAGGCTTGCCCA
AGAGCACACGTCTGAACTCTTTCCCTACACGACGCTCTTCCGATCTNNNCCACTAATACAGGCTTGCCCA
CACACGTCTGAACTCTTTCCCTACACGACGCTCTTCCGATCTNNNAAGTGTACCTGCAAGTGGGTAATC
CACACGTCTGAACTCTTTCCCTACACGACGCTCTTCCGATCTNNNAAGTGTACCTGCAAGTGGGTAATC
CACACGTCTGAACTCTTTCCCTACACGACGCTCTTCCGATCTNNNAAGTGTACCTGCAAGTGGGTAATC
AGCACACGTCTGAACTCTTTCCCTACACGACGCTCTTCCGATCTNNNGAGAATCACTTGAACCTGGGA
AGCACACGTCTGAACTCTTTCCCTACACGACGCTCTTCCGATCTNNNGAGAATCACTTGAACCTGGGA
AGCACACGTCTGAACTCTTTCCCTACACGACGCTCTTCCGATCTNNNGAGAATCACTTGAACCTGGGA
AGCACACGTCTGAACTCTTTCCCTACACGACGCTCTTCCGATCTNNNTAAAAATTAGCAGAGCGTGGTC
AGCACACGTCTGAACTCTTTCCCTACACGACGCTCTTCCGATCTNNNTAAAAATTAGCAGAGCGTGGTC
AGCACACGTCTGAACTCTTTCCCTACACGACGCTCTTCCGATCTNNNTAAAAATTAGCAGAGCGTGGTC
GAGCACACGTCTGAACTCTTTCCCTACACGACGCTCTTCCGATCTNNNGTGTGTGTGTGTGTATGTC

AGCACACGTCTGAACTCTTTCCCTACACGACGCTCTTCCGATCTNNNGTGTGTGTGTGTGTATGT
AGCACACGTCTGAACTCTTTCCCTACACGACGCTCTTCCGATCTNNNGTGTGTGTGTGTGTATGT
GAGCACACGTCTGAACTCTTTCCCTACACGACGCTCTTCCGATCTNNNCCAGCATGAGTGACAGAGTA
GAGCACACGTCTGAACTCTTTCCCTACACGACGCTCTTCCGATCTNNNCCAGCATGAGTGACAGAGTA
GAGCACACGTCTGAACTCTTTCCCTACACGACGCTCTTCCGATCTNNNCCAGCATGAGTGACAGAGTA
AGCACACGTCTGAACTCTTTCCCTACACGACGCTCTTCCGATCTNNNAACATGGCGAAACCCTGTCTC
AGCACACGTCTGAACTCTTTCCCTACACGACGCTCTTCCGATCTNNNAACATGGCGAAACCCTGTCTC
AGCACACGTCTGAACTCTTTCCCTACACGACGCTCTTCCGATCTNNNAACATGGCGAAACCCTGTCTC
AGCACACGTCTGAACTCTTTCCCTACACGACGCTCTTCCGATCTNNNGGTGGGAAGGAAGTGAGTAT
AGCACACGTCTGAACTCTTTCCCTACACGACGCTCTTCCGATCTNNNGGTGGGAAGGAAGTGAGTAT
AGCACACGTCTGAACTCTTTCCCTACACGACGCTCTTCCGATCTNNNGGTGGGAAGGAAGTGAGTAT
GAGCACACGTCTGAACTCTTTCCCTACACGACGCTCTTCCGATCTNNNTGAGCCACAAGAGTATTTAC
GAGCACACGTCTGAACTCTTTCCCTACACGACGCTCTTCCGATCTNNNTGAGCCACAAGAGTATTTAC
GAGCACACGTCTGAACTCTTTCCCTACACGACGCTCTTCCGATCTNNNTGAGCCACAAGAGTATTTAC
AGCACACGTCTGAACTCTTTCCCTACACGACGCTCTTCCGATCTNNNCACTTTCATTCCTGGCTTGAAG
AGCACACGTCTGAACTCTTTCCCTACACGACGCTCTTCCGATCTNNNCACTTTCATTCCTGGCTTGAAG
AGCACACGTCTGAACTCTTTCCCTACACGACGCTCTTCCGATCTNNNCACTTTCATTCCTGGCTTGAAG
AGCACACGTCTGAACTCTTTCCCTACACGACGCTCTTCCGATCTNNNTAGCCCTGATGACTACCACAG
AGCACACGTCTGAACTCTTTCCCTACACGACGCTCTTCCGATCTNNNTAGCCCTGATGACTACCACAG
AGCACACGTCTGAACTCTTTCCCTACACGACGCTCTTCCGATCTNNNTAGCCCTGATGACTACCACAG
AGCACACGTCTGAACTCTTTCCCTACACGACGCTCTTCCGATCTNNNTTTCCTTCAGCTCTTTCCCTCTC
AGCACACGTCTGAACTCTTTCCCTACACGACGCTCTTCCGATCTNNNTTTCCTTCAGCTCTTTCCCTCTC
AGCACACGTCTGAACTCTTTCCCTACACGACGCTCTTCCGATCTNNNTTTCCTTCAGCTCTTTCCCTCTC
AGCACACGTCTGAACTCTTTCCCTACACGACGCTCTTCCGATCTNNNAGAATCGCTTGAATCAGGGAG
AGCACACGTCTGAACTCTTTCCCTACACGACGCTCTTCCGATCTNNNAGAATCGCTTGAATCAGGGAG
AGCACACGTCTGAACTCTTTCCCTACACGACGCTCTTCCGATCTNNNAGAATCGCTTGAATCAGGGAG
AGCACACGTCTGAACTCTTTCCCTACACGACGCTCTTCCGATCTNNNTTAGTGACTGGGGTATTTCTG
AGCACACGTCTGAACTCTTTCCCTACACGACGCTCTTCCGATCTNNNTTAGTGACTGGGGTATTTCTG
AGCACACGTCTGAACTCTTTCCCTACACGACGCTCTTCCGATCTNNNTTAGTGACTGGGGTATTTCTG
GCACACGTCTGAACTCTTTCCCTACACGACGCTCTTCCGATCTNNNCAGCCTCTCTCATGCTCCTAAATC
GCACACGTCTGAACTCTTTCCCTACACGACGCTCTTCCGATCTNNNCAGCCTCTCTCATGCTCCTAAATC
GCACACGTCTGAACTCTTTCCCTACACGACGCTCTTCCGATCTNNNCAGCCTCTCTCATGCTCCTAAATC
GCACACGTCTGAACTCTTTCCCTACACGACGCTCTTCCGATCTNNNATGCCCAGCCTTCTTTTCTTTTCC
GCACACGTCTGAACTCTTTCCCTACACGACGCTCTTCCGATCTNNNATGCCCAGCCTTCTTTTCTTTTCC
GCACACGTCTGAACTCTTTCCCTACACGACGCTCTTCCGATCTNNNATGCCCAGCCTTCTTTTCTTTTCC
GCACACGTCTGAACTCTTTCCCTACACGACGCTCTTCCGATCTNNNGACAGTTCTACAAAGGAGGCT
GCACACGTCTGAACTCTTTCCCTACACGACGCTCTTCCGATCTNNNGACAGTTCTACAAAGGAGGCT
GCACACGTCTGAACTCTTTCCCTACACGACGCTCTTCCGATCTNNNGACAGTTCTACAAAGGAGGCT
AGCACACGTCTGAACTCTTTCCCTACACGACGCTCTTCCGATCTNNNCTTCAAGTAGGTTTCGTGCTCC
AGCACACGTCTGAACTCTTTCCCTACACGACGCTCTTCCGATCTNNNCTTCAAGTAGGTTTCGTGCTCC
AGCACACGTCTGAACTCTTTCCCTACACGACGCTCTTCCGATCTNNNCTTCAAGTAGGTTTCGTGCTCC
GAGCACACGTCTGAACTCTTTCCCTACACGACGCTCTTCCGATCTNNNGTAATTCTAGCTACTCGGGA
GAGCACACGTCTGAACTCTTTCCCTACACGACGCTCTTCCGATCTNNNGTAATTCTAGCTACTCGGGA

GAGCACACGTCTGAACTCTTTCCCTACACGACGCTCTTCCGATCTNNNGTAATTCTAGCTACTCGGGA
GAGCACACGTCTGAACTCTTTCCCTACACGACGCTCTTCCGATCTNNNTCTGCCTCACAATTGTGTTGA
GAGCACACGTCTGAACTCTTTCCCTACACGACGCTCTTCCGATCTNNNTCTGCCTCACAATTGTGTTGA
GAGCACACGTCTGAACTCTTTCCCTACACGACGCTCTTCCGATCTNNNTCTGCCTCACAATTGTGTTGA
AGCACACGTCTGAACTCTTTCCCTACACGACGCTCTTCCGATCTNNNATGTATCACCCCAATCTGTGCT
AGCACACGTCTGAACTCTTTCCCTACACGACGCTCTTCCGATCTNNNATGTATCACCCCAATCTGTGCT
AGCACACGTCTGAACTCTTTCCCTACACGACGCTCTTCCGATCTNNNATGTATCACCCCAATCTGTGCT
GAGCACACGTCTGAACTCTTTCCCTACACGACGCTCTTCCGATCTNNNCCTGGGAAACATAGTGAGA
GAGCACACGTCTGAACTCTTTCCCTACACGACGCTCTTCCGATCTNNNCCTGGGAAACATAGTGAGA
GAGCACACGTCTGAACTCTTTCCCTACACGACGCTCTTCCGATCTNNNCCTGGGAAACATAGTGAGA
AGCACACGTCTGAACTCTTTCCCTACACGACGCTCTTCCGATCTNNNTTCTTATCTGGGCCTTCCTTTC
AGCACACGTCTGAACTCTTTCCCTACACGACGCTCTTCCGATCTNNNTTCTTATCTGGGCCTTCCTTTC
AGCACACGTCTGAACTCTTTCCCTACACGACGCTCTTCCGATCTNNNTTCTTATCTGGGCCTTCCTTTC
AGCACACGTCTGAACTCTTTCCCTACACGACGCTCTTCCGATCTNNNATCCACTTCAACCATTTTCCAG
AGCACACGTCTGAACTCTTTCCCTACACGACGCTCTTCCGATCTNNNATCCACTTCAACCATTTTCCAG
AGCACACGTCTGAACTCTTTCCCTACACGACGCTCTTCCGATCTNNNATCCACTTCAACCATTTTCCAG
AGCACACGTCTGAACTCTTTCCCTACACGACGCTCTTCCGATCTNNNACTGTTCCCAGATTATCTGCAG
AGCACACGTCTGAACTCTTTCCCTACACGACGCTCTTCCGATCTNNNACTGTTCCCAGATTATCTGCAG
AGCACACGTCTGAACTCTTTCCCTACACGACGCTCTTCCGATCTNNNACTGTTCCCAGATTATCTGCAG
AGCACACGTCTGAACTCTTTCCCTACACGACGCTCTTCCGATCTNNNCCGCAACCTGTTTTTTGTTTTG
AGCACACGTCTGAACTCTTTCCCTACACGACGCTCTTCCGATCTNNNCCGCAACCTGTTTTTTGTTTTG
AGCACACGTCTGAACTCTTTCCCTACACGACGCTCTTCCGATCTNNNCCGCAACCTGTTTTTTGTTTTG
GCACACGTCTGAACTCTTTCCCTACACGACGCTCTTCCGATCTNNNGTAAGAGTTAAAGAGTAAAGC
GCACACGTCTGAACTCTTTCCCTACACGACGCTCTTCCGATCTNNNGTAAGAGTTAAAGAGTAAAGC
GCACACGTCTGAACTCTTTCCCTACACGACGCTCTTCCGATCTNNNGTAAGAGTTAAAGAGTAAAGC
AGCACACGTCTGAACTCTTTCCCTACACGACGCTCTTCCGATCTNNNCTCCTGAGTAGCTGGGATTAC
AGCACACGTCTGAACTCTTTCCCTACACGACGCTCTTCCGATCTNNNCTCCTGAGTAGCTGGGATTAC
AGCACACGTCTGAACTCTTTCCCTACACGACGCTCTTCCGATCTNNNCTCCTGAGTAGCTGGGATTAC
GCACACGTCTGAACTCTTTCCCTACACGACGCTCTTCCGATCTNNNCTCAGCCATTTCAAGACACAAG
GCACACGTCTGAACTCTTTCCCTACACGACGCTCTTCCGATCTNNNCTCAGCCATTTCAAGACACAAG
GCACACGTCTGAACTCTTTCCCTACACGACGCTCTTCCGATCTNNNCTCAGCCATTTCAAGACACAAG
GCACACGTCTGAACTCTTTCCCTACACGACGCTCTTCCGATCTNNNATGCCTGTAATCCCAGTTACTC
GCACACGTCTGAACTCTTTCCCTACACGACGCTCTTCCGATCTNNNATGCCTGTAATCCCAGTTACTC
GCACACGTCTGAACTCTTTCCCTACACGACGCTCTTCCGATCTNNNATGCCTGTAATCCCAGTTACTC
GCACACGTCTGAACTCTTTCCCTACACGACGCTCTTCCGATCTNNNATGTGTGTATGCTAACTTCTGC
GCACACGTCTGAACTCTTTCCCTACACGACGCTCTTCCGATCTNNNATGTGTGTATGCTAACTTCTGC
GCACACGTCTGAACTCTTTCCCTACACGACGCTCTTCCGATCTNNNATGTGTGTATGCTAACTTCTGC
AGCACACGTCTGAACTCTTTCCCTACACGACGCTCTTCCGATCTNNNGAAAAAACAATCCTAGGCCAG
AGCACACGTCTGAACTCTTTCCCTACACGACGCTCTTCCGATCTNNNGAAAAAACAATCCTAGGCCAG
AGCACACGTCTGAACTCTTTCCCTACACGACGCTCTTCCGATCTNNNGAAAAAACAATCCTAGGCCAG
GAGCACACGTCTGAACTCTTTCCCTACACGACGCTCTTCCGATCTNNNCATAGCGTGACTCCTGTCTCT
GAGCACACGTCTGAACTCTTTCCCTACACGACGCTCTTCCGATCTNNNCATAGCGTGACTCCTGTCTCT
GAGCACACGTCTGAACTCTTTCCCTACACGACGCTCTTCCGATCTNNNCATAGCGTGACTCCTGTCTCT

GCACACGTCTGAACTCTTTCCCTACACGACGCTCTTCCGATCTNNNTTTCCTCCCTTCATACACACACAC
GCACACGTCTGAACTCTTTCCCTACACGACGCTCTTCCGATCTNNNTTTCCTCCCTTCATACACACACAC
GCACACGTCTGAACTCTTTCCCTACACGACGCTCTTCCGATCTNNNTTTCCTCCCTTCATACACACACAC
AGCACACGTCTGAACTCTTTCCCTACACGACGCTCTTCCGATCTNNNGCACATCTGTAATCCCAGCTACT
AGCACACGTCTGAACTCTTTCCCTACACGACGCTCTTCCGATCTNNNGCACATCTGTAATCCCAGCTACT
AGCACACGTCTGAACTCTTTCCCTACACGACGCTCTTCCGATCTNNNGCACATCTGTAATCCCAGCTACT
AGAGCACACGTCTGAACTCTTTCCCTACACGACGCTCTTCCGATCTNNNCAATATAGCGAGACCCCGTT
AGAGCACACGTCTGAACTCTTTCCCTACACGACGCTCTTCCGATCTNNNCAATATAGCGAGACCCCGTT
AGAGCACACGTCTGAACTCTTTCCCTACACGACGCTCTTCCGATCTNNNCAATATAGCGAGACCCCGTT
GCACACGTCTGAACTCTTTCCCTACACGACGCTCTTCCGATCTNNNAGCGGGAGATTTTCCAGGTTTTT
GCACACGTCTGAACTCTTTCCCTACACGACGCTCTTCCGATCTNNNAGCGGGAGATTTTCCAGGTTTTT
GCACACGTCTGAACTCTTTCCCTACACGACGCTCTTCCGATCTNNNAGCGGGAGATTTTCCAGGTTTTT
GCACACGTCTGAACTCTTTCCCTACACGACGCTCTTCCGATCTNNNATTCCAAATCCATCCAGTTCCCA
GCACACGTCTGAACTCTTTCCCTACACGACGCTCTTCCGATCTNNNATTCCAAATCCATCCAGTTCCCA
GCACACGTCTGAACTCTTTCCCTACACGACGCTCTTCCGATCTNNNATTCCAAATCCATCCAGTTCCCA
AGCACACGTCTGAACTCTTTCCCTACACGACGCTCTTCCGATCTNNNCAAGAGTTTGAGGTTGCAGTG
AGCACACGTCTGAACTCTTTCCCTACACGACGCTCTTCCGATCTNNNCAAGAGTTTGAGGTTGCAGTG
AGCACACGTCTGAACTCTTTCCCTACACGACGCTCTTCCGATCTNNNCAAGAGTTTGAGGTTGCAGTG
GAGCACACGTCTGAACTCTTTCCCTACACGACGCTCTTCCGATCTNNNAAGATCACACCACTGCACTCT
GAGCACACGTCTGAACTCTTTCCCTACACGACGCTCTTCCGATCTNNNAAGATCACACCACTGCACTCT
GAGCACACGTCTGAACTCTTTCCCTACACGACGCTCTTCCGATCTNNNAAGATCACACCACTGCACTCT
GAGCACACGTCTGAACTCTTTCCCTACACGACGCTCTTCCGATCTNNNTCCTTGTTTGCAGATGCCATA
GAGCACACGTCTGAACTCTTTCCCTACACGACGCTCTTCCGATCTNNNTCCTTGTTTGCAGATGCCATA
GAGCACACGTCTGAACTCTTTCCCTACACGACGCTCTTCCGATCTNNNTCCTTGTTTGCAGATGCCATA
AGCACACGTCTGAACTCTTTCCCTACACGACGCTCTTCCGATCTNNNGGGAGTGAATGGTCCTATCT
AGCACACGTCTGAACTCTTTCCCTACACGACGCTCTTCCGATCTNNNGGGAGTGAATGGTCCTATCT
AGCACACGTCTGAACTCTTTCCCTACACGACGCTCTTCCGATCTNNNGGGAGTGAATGGTCCTATCT
AGCACACGTCTGAACTCTTTCCCTACACGACGCTCTTCCGATCTNNNCCTGGGTAACACAGGGAATTTC
AGCACACGTCTGAACTCTTTCCCTACACGACGCTCTTCCGATCTNNNCCTGGGTAACACAGGGAATTTC
AGCACACGTCTGAACTCTTTCCCTACACGACGCTCTTCCGATCTNNNCCTGGGTAACACAGGGAATTTC
GCACACGTCTGAACTCTTTCCCTACACGACGCTCTTCCGATCTNNNCTTGCAGTTGCTGACTTCTCTTC
GCACACGTCTGAACTCTTTCCCTACACGACGCTCTTCCGATCTNNNCTTGCAGTTGCTGACTTCTCTTC
GCACACGTCTGAACTCTTTCCCTACACGACGCTCTTCCGATCTNNNCTTGCAGTTGCTGACTTCTCTTC
AGCACACGTCTGAACTCTTTCCCTACACGACGCTCTTCCGATCTNNNCATTCAAATTTTACAGGATCTGA
AGCACACGTCTGAACTCTTTCCCTACACGACGCTCTTCCGATCTNNNCATTCAAATTTTACAGGATCTGA
AGCACACGTCTGAACTCTTTCCCTACACGACGCTCTTCCGATCTNNNCATTCAAATTTTACAGGATCTGA
AGCACACGTCTGAACTCTTTCCCTACACGACGCTCTTCCGATCTNNNCAGCTACTTGGGAGGCTAAA
AGCACACGTCTGAACTCTTTCCCTACACGACGCTCTTCCGATCTNNNCAGCTACTTGGGAGGCTAAA
AGCACACGTCTGAACTCTTTCCCTACACGACGCTCTTCCGATCTNNNCAGCTACTTGGGAGGCTAAA
GAGCACACGTCTGAACTCTTTCCCTACACGACGCTCTTCCGATCTNNNGGGAGGGATAGCATTGGGA
GAGCACACGTCTGAACTCTTTCCCTACACGACGCTCTTCCGATCTNNNGGGAGGGATAGCATTGGGA
GAGCACACGTCTGAACTCTTTCCCTACACGACGCTCTTCCGATCTNNNGGGAGGGATAGCATTGGGA
AGCACACGTCTGAACTCTTTCCCTACACGACGCTCTTCCGATCTNNNGGAGGTGGGGAGGCATATGT

AGCACACGTCTGAACTCTTTCCCTACACGACGCTCTTCCGATCTNNNGGAGGTGGGGAGGCATATGT
AGCACACGTCTGAACTCTTTCCCTACACGACGCTCTTCCGATCTNNNGGAGGTGGGGAGGCATATGT
AGCACACGTCTGAACTCTTTCCCTACACGACGCTCTTCCGATCTNNNTTTCATTGTCAGTGTACAGAAA
AGCACACGTCTGAACTCTTTCCCTACACGACGCTCTTCCGATCTNNNTTTCATTGTCAGTGTACAGAAA
AGCACACGTCTGAACTCTTTCCCTACACGACGCTCTTCCGATCTNNNTTTCATTGTCAGTGTACAGAAA
AGCACACGTCTGAACTCTTTCCCTACACGACGCTCTTCCGATCTNNNCATTGCACCATCAGAGTTCACA
AGCACACGTCTGAACTCTTTCCCTACACGACGCTCTTCCGATCTNNNCATTGCACCATCAGAGTTCACA
AGCACACGTCTGAACTCTTTCCCTACACGACGCTCTTCCGATCTNNNCATTGCACCATCAGAGTTCACA
GCACACGTCTGAACTCTTTCCCTACACGACGCTCTTCCGATCTNNNTGGTGCAGGTCACTGTCTATTTG
GCACACGTCTGAACTCTTTCCCTACACGACGCTCTTCCGATCTNNNTGGTGCAGGTCACTGTCTATTTG
GCACACGTCTGAACTCTTTCCCTACACGACGCTCTTCCGATCTNNNTGGTGCAGGTCACTGTCTATTTG
GAGCACACGTCTGAACTCTTTCCCTACACGACGCTCTTCCGATCTNNNAGGAGAAAGAGCTAAAGCA
GAGCACACGTCTGAACTCTTTCCCTACACGACGCTCTTCCGATCTNNNAGGAGAAAGAGCTAAAGCA
GAGCACACGTCTGAACTCTTTCCCTACACGACGCTCTTCCGATCTNNNAGGAGAAAGAGCTAAAGCA
AGCACACGTCTGAACTCTTTCCCTACACGACGCTCTTCCGATCTNNNCTTTGCTTCAGGCTAAGAGCT
AGCACACGTCTGAACTCTTTCCCTACACGACGCTCTTCCGATCTNNNCTTTGCTTCAGGCTAAGAGCT
AGCACACGTCTGAACTCTTTCCCTACACGACGCTCTTCCGATCTNNNCTTTGCTTCAGGCTAAGAGCT
GAGCACACGTCTGAACTCTTTCCCTACACGACGCTCTTCCGATCTNNNTATGCCCACAGTCCAAACTTG
GAGCACACGTCTGAACTCTTTCCCTACACGACGCTCTTCCGATCTNNNTATGCCCACAGTCCAAACTTG
GAGCACACGTCTGAACTCTTTCCCTACACGACGCTCTTCCGATCTNNNTATGCCCACAGTCCAAACTTG
ACACGTCTGAACTCTTTCCCTACACGACGCTCTTCCGATCTNNNCCACGGGTCAAAGACTTCAATTTCA
ACACGTCTGAACTCTTTCCCTACACGACGCTCTTCCGATCTNNNCCACGGGTCAAAGACTTCAATTTCA
ACACGTCTGAACTCTTTCCCTACACGACGCTCTTCCGATCTNNNCCACGGGTCAAAGACTTCAATTTCA
AGCACACGTCTGAACTCTTTCCCTACACGACGCTCTTCCGATCTNNNTAGAGAAGATGATAGAGCAG
AGCACACGTCTGAACTCTTTCCCTACACGACGCTCTTCCGATCTNNNTAGAGAAGATGATAGAGCAG
AGCACACGTCTGAACTCTTTCCCTACACGACGCTCTTCCGATCTNNNTAGAGAAGATGATAGAGCAG
AGCACACGTCTGAACTCTTTCCCTACACGACGCTCTTCCGATCTNNNCACAAAGCGTTTTAAAAAAGA
AGCACACGTCTGAACTCTTTCCCTACACGACGCTCTTCCGATCTNNNCACAAAGCGTTTTAAAAAAGA
AGCACACGTCTGAACTCTTTCCCTACACGACGCTCTTCCGATCTNNNCACAAAGCGTTTTAAAAAAGA
AGCACACGTCTGAACTCTTTCCCTACACGACGCTCTTCCGATCTNNNACAGCTCAGCTCCCTTCTAATTC
AGCACACGTCTGAACTCTTTCCCTACACGACGCTCTTCCGATCTNNNACAGCTCAGCTCCCTTCTAATTC
AGCACACGTCTGAACTCTTTCCCTACACGACGCTCTTCCGATCTNNNACAGCTCAGCTCCCTTCTAATTC
GCACACGTCTGAACTCTTTCCCTACACGACGCTCTTCCGATCTNNNGTCACAAAGCTAGTCAGTTCAGC
GCACACGTCTGAACTCTTTCCCTACACGACGCTCTTCCGATCTNNNGTCACAAAGCTAGTCAGTTCAGC
GCACACGTCTGAACTCTTTCCCTACACGACGCTCTTCCGATCTNNNGTCACAAAGCTAGTCAGTTCAGC
AGCACACGTCTGAACTCTTTCCCTACACGACGCTCTTCCGATCTNNNGCCACATTTCCTAGGCTAGTCTT
AGCACACGTCTGAACTCTTTCCCTACACGACGCTCTTCCGATCTNNNGCCACATTTCCTAGGCTAGTCTT
AGCACACGTCTGAACTCTTTCCCTACACGACGCTCTTCCGATCTNNNGCCACATTTCCTAGGCTAGTCTT
AGCACACGTCTGAACTCTTTCCCTACACGACGCTCTTCCGATCTNNNACAAAAATTAGCCAGGTGTGG
AGCACACGTCTGAACTCTTTCCCTACACGACGCTCTTCCGATCTNNNACAAAAATTAGCCAGGTGTGG
AGCACACGTCTGAACTCTTTCCCTACACGACGCTCTTCCGATCTNNNACAAAAATTAGCCAGGTGTGG
AGCACACGTCTGAACTCTTTCCCTACACGACGCTCTTCCGATCTNNNATTATGTTTACTGGGCAGGTGG
AGCACACGTCTGAACTCTTTCCCTACACGACGCTCTTCCGATCTNNNATTATGTTTACTGGGCAGGTGG

AGCACACGTCTGAACTCTTTCCCTACACGACGCTCTTCCGATCTNNNATTATGTTTACTGGGCAGGTGG
AGCACACGTCTGAACTCTTTCCCTACACGACGCTCTTCCGATCTNNNCCTACTGGGAAGTGAGGACCCC
AGCACACGTCTGAACTCTTTCCCTACACGACGCTCTTCCGATCTNNNCCTACTGGGAAGTGAGGACCCC
AGCACACGTCTGAACTCTTTCCCTACACGACGCTCTTCCGATCTNNNCCTACTGGGAAGTGAGGACCCC
AGCACACGTCTGAACTCTTTCCCTACACGACGCTCTTCCGATCTNNNCATGCTGTAATCCCCACACTTTG
AGCACACGTCTGAACTCTTTCCCTACACGACGCTCTTCCGATCTNNNCATGCTGTAATCCCCACACTTTG
AGCACACGTCTGAACTCTTTCCCTACACGACGCTCTTCCGATCTNNNCATGCTGTAATCCCCACACTTTG
AGAGCACACGTCTGAACTCTTTCCCTACACGACGCTCTTCCGATCTNNNATATCGGGAAAACACATGCA
AGAGCACACGTCTGAACTCTTTCCCTACACGACGCTCTTCCGATCTNNNATATCGGGAAAACACATGCA
AGAGCACACGTCTGAACTCTTTCCCTACACGACGCTCTTCCGATCTNNNATATCGGGAAAACACATGCA
AGCACACGTCTGAACTCTTTCCCTACACGACGCTCTTCCGATCTNNNCAGTGGTAGTCTTCAGCCTCTAC
AGCACACGTCTGAACTCTTTCCCTACACGACGCTCTTCCGATCTNNNCAGTGGTAGTCTTCAGCCTCTAC
AGCACACGTCTGAACTCTTTCCCTACACGACGCTCTTCCGATCTNNNCAGTGGTAGTCTTCAGCCTCTAC
GAGCACACGTCTGAACTCTTTCCCTACACGACGCTCTTCCGATCTNNNTAATCCCAGCTACTCAGGAGA
GAGCACACGTCTGAACTCTTTCCCTACACGACGCTCTTCCGATCTNNNTAATCCCAGCTACTCAGGAGA
GAGCACACGTCTGAACTCTTTCCCTACACGACGCTCTTCCGATCTNNNTAATCCCAGCTACTCAGGAGA
AGAGCACACGTCTGAACTCTTTCCCTACACGACGCTCTTCCGATCTNNNTCCAGGATGAGAAGGAGGA
AGAGCACACGTCTGAACTCTTTCCCTACACGACGCTCTTCCGATCTNNNTCCAGGATGAGAAGGAGGA
AGAGCACACGTCTGAACTCTTTCCCTACACGACGCTCTTCCGATCTNNNTCCAGGATGAGAAGGAGGA
GCACACGTCTGAACTCTTTCCCTACACGACGCTCTTCCGATCTNNNGGCCTGTATCTCCTGACAATTCAC
GCACACGTCTGAACTCTTTCCCTACACGACGCTCTTCCGATCTNNNGGCCTGTATCTCCTGACAATTCAC
GCACACGTCTGAACTCTTTCCCTACACGACGCTCTTCCGATCTNNNGGCCTGTATCTCCTGACAATTCAC
GAGCACACGTCTGAACTCTTTCCCTACACGACGCTCTTCCGATCTNNNCTAAGGCAGGAGAATCGCTTG
GAGCACACGTCTGAACTCTTTCCCTACACGACGCTCTTCCGATCTNNNCTAAGGCAGGAGAATCGCTTG
GAGCACACGTCTGAACTCTTTCCCTACACGACGCTCTTCCGATCTNNNCTAAGGCAGGAGAATCGCTTG
GAGCACACGTCTGAACTCTTTCCCTACACGACGCTCTTCCGATCTNNNTATGTGGTGATGGCAACTCTG
GAGCACACGTCTGAACTCTTTCCCTACACGACGCTCTTCCGATCTNNNTATGTGGTGATGGCAACTCTG
GAGCACACGTCTGAACTCTTTCCCTACACGACGCTCTTCCGATCTNNNTATGTGGTGATGGCAACTCTG
GCACACGTCTGAACTCTTTCCCTACACGACGCTCTTCCGATCTNNNGCTGGATGTGGATGAAAGCATT
GCACACGTCTGAACTCTTTCCCTACACGACGCTCTTCCGATCTNNNGCTGGATGTGGATGAAAGCATT
GCACACGTCTGAACTCTTTCCCTACACGACGCTCTTCCGATCTNNNGCTGGATGTGGATGAAAGCATT
AGCACACGTCTGAACTCTTTCCCTACACGACGCTCTTCCGATCTNNNGCTGGCCAAAGTTCAAATTCTC
AGCACACGTCTGAACTCTTTCCCTACACGACGCTCTTCCGATCTNNNGCTGGCCAAAGTTCAAATTCTC
AGCACACGTCTGAACTCTTTCCCTACACGACGCTCTTCCGATCTNNNGCTGGCCAAAGTTCAAATTCTC
AGCACACGTCTGAACTCTTTCCCTACACGACGCTCTTCCGATCTNNNGGATGACAGGGTGAGACTCAA
AGCACACGTCTGAACTCTTTCCCTACACGACGCTCTTCCGATCTNNNGGATGACAGGGTGAGACTCAA
AGCACACGTCTGAACTCTTTCCCTACACGACGCTCTTCCGATCTNNNGGATGACAGGGTGAGACTCAA
GAGCACACGTCTGAACTCTTTCCCTACACGACGCTCTTCCGATCTNNNGAGATCGCTCCATCATACTCCA
GAGCACACGTCTGAACTCTTTCCCTACACGACGCTCTTCCGATCTNNNGAGATCGCTCCATCATACTCCA
GAGCACACGTCTGAACTCTTTCCCTACACGACGCTCTTCCGATCTNNNGAGATCGCTCCATCATACTCCA
AGCACACGTCTGAACTCTTTCCCTACACGACGCTCTTCCGATCTNNNCATGCCAGGGATGCTTATTCAT
AGCACACGTCTGAACTCTTTCCCTACACGACGCTCTTCCGATCTNNNCATGCCAGGGATGCTTATTCAT
AGCACACGTCTGAACTCTTTCCCTACACGACGCTCTTCCGATCTNNNCATGCCAGGGATGCTTATTCAT

GAGCACACGTCTGAACTCTTTCCCTACACGACGCTCTTCCGATCTNNNAAATCCTCCCAGCCAACTACAA

GAGCACACGTCTGAACTCTTTCCCTACACGACGCTCTTCCGATCTNNNAAATCCTCCCAGCCAACTACAA

GAGCACACGTCTGAACTCTTTCCCTACACGACGCTCTTCCGATCTNNNAAATCCTCCCAGCCAACTACAA

GAGCACACGTCTGAACTCTTTCCCTACACGACGCTCTTCCGATCTNNNCACCTGTAGTCCCAGCTAATT

GAGCACACGTCTGAACTCTTTCCCTACACGACGCTCTTCCGATCTNNNCACCTGTAGTCCCAGCTAATT

GAGCACACGTCTGAACTCTTTCCCTACACGACGCTCTTCCGATCTNNNCACCTGTAGTCCCAGCTAATT

AGAGCACACGTCTGAACTCTTTCCCTACACGACGCTCTTCCGATCTNNNCTGAGGCAGGAGAATCACTT

AGAGCACACGTCTGAACTCTTTCCCTACACGACGCTCTTCCGATCTNNNCTGAGGCAGGAGAATCACTT

AGAGCACACGTCTGAACTCTTTCCCTACACGACGCTCTTCCGATCTNNNCTGAGGCAGGAGAATCACTT

AGCACACGTCTGAACTCTTTCCCTACACGACGCTCTTCCGATCTNNNTACTTGCTTTGGAGACACAGTC

AGCACACGTCTGAACTCTTTCCCTACACGACGCTCTTCCGATCTNNNTACTTGCTTTGGAGACACAGTC

AGCACACGTCTGAACTCTTTCCCTACACGACGCTCTTCCGATCTNNNTACTTGCTTTGGAGACACAGTC

GAGCACACGTCTGAACTCTTTCCCTACACGACGCTCTTCCGATCTNNNGGCACCTGTAATCCTAGCTAC

GAGCACACGTCTGAACTCTTTCCCTACACGACGCTCTTCCGATCTNNNGGCACCTGTAATCCTAGCTAC

GAGCACACGTCTGAACTCTTTCCCTACACGACGCTCTTCCGATCTNNNGGCACCTGTAATCCTAGCTAC

GAGCACACGTCTGAACTCTTTCCCTACACGACGCTCTTCCGATCTNNNCAACAAGAGCATAACCTTGCC

GAGCACACGTCTGAACTCTTTCCCTACACGACGCTCTTCCGATCTNNNCAACAAGAGCATAACCTTGCC

GAGCACACGTCTGAACTCTTTCCCTACACGACGCTCTTCCGATCTNNNCAACAAGAGCATAACCTTGCC

CACGTCTGAACTCTTTCCCTACACGACGCTCTTCCGATCTNNNACAAGGACCCCACCTTCTCCATCGGA

CACGTCTGAACTCTTTCCCTACACGACGCTCTTCCGATCTNNNACAAGGACCCCACCTTCTCCATCGGA

CACGTCTGAACTCTTTCCCTACACGACGCTCTTCCGATCTNNNACAAGGACCCCACCTTCTCCATCGGA

AGCACACGTCTGAACTCTTTCCCTACACGACGCTCTTCCGATCTNNNGGGTAGGTACCTGTTTCTTGCTA

AGCACACGTCTGAACTCTTTCCCTACACGACGCTCTTCCGATCTNNNGGGTAGGTACCTGTTTCTTGCTA

AGCACACGTCTGAACTCTTTCCCTACACGACGCTCTTCCGATCTNNNGGGTAGGTACCTGTTTCTTGCTA

GAGCACACGTCTGAACTCTTTCCCTACACGACGCTCTTCCGATCTNNNGAGAATTGCTTGAACCAGGAC

GAGCACACGTCTGAACTCTTTCCCTACACGACGCTCTTCCGATCTNNNGAGAATTGCTTGAACCAGGAC

GAGCACACGTCTGAACTCTTTCCCTACACGACGCTCTTCCGATCTNNNGAGAATTGCTTGAACCAGGAC

GAGCACACGTCTGAACTCTTTCCCTACACGACGCTCTTCCGATCTNNNCGGATGTAAATTTCGAGTTTTC

GAGCACACGTCTGAACTCTTTCCCTACACGACGCTCTTCCGATCTNNNCGGATGTAAATTTCGAGTTTTC

GAGCACACGTCTGAACTCTTTCCCTACACGACGCTCTTCCGATCTNNNCGGATGTAAATTTCGAGTTTTC

GAGCACACGTCTGAACTCTTTCCCTACACGACGCTCTTCCGATCTNNNAGAGAAGGGGCTGCTGATGA

GAGCACACGTCTGAACTCTTTCCCTACACGACGCTCTTCCGATCTNNNAGAGAAGGGGCTGCTGATGA

GAGCACACGTCTGAACTCTTTCCCTACACGACGCTCTTCCGATCTNNNAGAGAAGGGGCTGCTGATGA

GAGCACACGTCTGAACTCTTTCCCTACACGACGCTCTTCCGATCTNNNTAGTTGGGACACAGGTATGCA

GAGCACACGTCTGAACTCTTTCCCTACACGACGCTCTTCCGATCTNNNTAGTTGGGACACAGGTATGCA

GAGCACACGTCTGAACTCTTTCCCTACACGACGCTCTTCCGATCTNNNTAGTTGGGACACAGGTATGCA

GAGCACACGTCTGAACTCTTTCCCTACACGACGCTCTTCCGATCTNNNTGCCTGTAGTCCCAGCTACTCC

GAGCACACGTCTGAACTCTTTCCCTACACGACGCTCTTCCGATCTNNNTGCCTGTAGTCCCAGCTACTCC

GAGCACACGTCTGAACTCTTTCCCTACACGACGCTCTTCCGATCTNNNTGCCTGTAGTCCCAGCTACTCC

GAGCACACGTCTGAACTCTTTCCCTACACGACGCTCTTCCGATCTNNNACAGAATGGCCAAACTTGCAT

GAGCACACGTCTGAACTCTTTCCCTACACGACGCTCTTCCGATCTNNNACAGAATGGCCAAACTTGCAT

GAGCACACGTCTGAACTCTTTCCCTACACGACGCTCTTCCGATCTNNNACAGAATGGCCAAACTTGCAT

AGCACACGTCTGAACTCTTTCCCTACACGACGCTCTTCCGATCTNNNTGGTGATTCCTTACTGCAACAGT

AGCACACGTCTGAACTCTTTCCCTACACGACGCTCTTCCGATCTNNNTGGTGATTCCTTACTGCAACAG
AGCACACGTCTGAACTCTTTCCCTACACGACGCTCTTCCGATCTNNNTGGTGATTCCTTACTGCAACAG
GCACACGTCTGAACTCTTTCCCTACACGACGCTCTTCCGATCTNNNTACAAAAAATTAGCCGGGCGTG
GCACACGTCTGAACTCTTTCCCTACACGACGCTCTTCCGATCTNNNTACAAAAAATTAGCCGGGCGTG
GCACACGTCTGAACTCTTTCCCTACACGACGCTCTTCCGATCTNNNTACAAAAAATTAGCCGGGCGTG
GAGCACACGTCTGAACTCTTTCCCTACACGACGCTCTTCCGATCTNNNGTAGTCCTGGTGGGGTTCTT
GAGCACACGTCTGAACTCTTTCCCTACACGACGCTCTTCCGATCTNNNGTAGTCCTGGTGGGGTTCTT
GAGCACACGTCTGAACTCTTTCCCTACACGACGCTCTTCCGATCTNNNGTAGTCCTGGTGGGGTTCTT
GAGCACACGTCTGAACTCTTTCCCTACACGACGCTCTTCCGATCTNNNCACCTGTAGTTCCAGCTACT
GAGCACACGTCTGAACTCTTTCCCTACACGACGCTCTTCCGATCTNNNCACCTGTAGTTCCAGCTACT
GAGCACACGTCTGAACTCTTTCCCTACACGACGCTCTTCCGATCTNNNCACCTGTAGTTCCAGCTACT
AGCACACGTCTGAACTCTTTCCCTACACGACGCTCTTCCGATCTNNNAAAGTATGTGTGTTTTCTTACA
AGCACACGTCTGAACTCTTTCCCTACACGACGCTCTTCCGATCTNNNAAAGTATGTGTGTTTTCTTACA
AGCACACGTCTGAACTCTTTCCCTACACGACGCTCTTCCGATCTNNNAAAGTATGTGTGTTTTCTTACA
AGCACACGTCTGAACTCTTTCCCTACACGACGCTCTTCCGATCTNNNCTGAGGCAGGAGAATCACTT
AGCACACGTCTGAACTCTTTCCCTACACGACGCTCTTCCGATCTNNNCTGAGGCAGGAGAATCACTT
AGCACACGTCTGAACTCTTTCCCTACACGACGCTCTTCCGATCTNNNCTGAGGCAGGAGAATCACTT
CACACGTCTGAACTCTTTCCCTACACGACGCTCTTCCGATCTNNNGTTCACTCCTTCCAGCTGCTCCAT
CACACGTCTGAACTCTTTCCCTACACGACGCTCTTCCGATCTNNNGTTCACTCCTTCCAGCTGCTCCAT
CACACGTCTGAACTCTTTCCCTACACGACGCTCTTCCGATCTNNNGTTCACTCCTTCCAGCTGCTCCAT
AGCACACGTCTGAACTCTTTCCCTACACGACGCTCTTCCGATCTNNNCAAGGATGCTCATGTTGGGAA
AGCACACGTCTGAACTCTTTCCCTACACGACGCTCTTCCGATCTNNNCAAGGATGCTCATGTTGGGAA
AGCACACGTCTGAACTCTTTCCCTACACGACGCTCTTCCGATCTNNNCAAGGATGCTCATGTTGGGAA
GCACACGTCTGAACTCTTTCCCTACACGACGCTCTTCCGATCTNNNCTCCACCTCGATCTCATTGGTAG
GCACACGTCTGAACTCTTTCCCTACACGACGCTCTTCCGATCTNNNCTCCACCTCGATCTCATTGGTAG
GCACACGTCTGAACTCTTTCCCTACACGACGCTCTTCCGATCTNNNCTCCACCTCGATCTCATTGGTAG
GCACACGTCTGAACTCTTTCCCTACACGACGCTCTTCCGATCTNNNGAATGCAGGGAAATACAAGGA
GCACACGTCTGAACTCTTTCCCTACACGACGCTCTTCCGATCTNNNGAATGCAGGGAAATACAAGGA
GCACACGTCTGAACTCTTTCCCTACACGACGCTCTTCCGATCTNNNGAATGCAGGGAAATACAAGGA
AGCACACGTCTGAACTCTTTCCCTACACGACGCTCTTCCGATCTNNNTGATTGGCGTCCTTGTAAGA
AGCACACGTCTGAACTCTTTCCCTACACGACGCTCTTCCGATCTNNNTGATTGGCGTCCTTGTAAGA
AGCACACGTCTGAACTCTTTCCCTACACGACGCTCTTCCGATCTNNNTGATTGGCGTCCTTGTAAGA
AGCACACGTCTGAACTCTTTCCCTACACGACGCTCTTCCGATCTNNNTGAGACAGACATTTACACACCC
AGCACACGTCTGAACTCTTTCCCTACACGACGCTCTTCCGATCTNNNTGAGACAGACATTTACACACCC
AGCACACGTCTGAACTCTTTCCCTACACGACGCTCTTCCGATCTNNNTGAGACAGACATTTACACACCC
GCACACGTCTGAACTCTTTCCCTACACGACGCTCTTCCGATCTNNNACTGACTTTTTTCTTTTCTTTTTT
GCACACGTCTGAACTCTTTCCCTACACGACGCTCTTCCGATCTNNNACTGACTTTTTTCTTTTCTTTTTT
GCACACGTCTGAACTCTTTCCCTACACGACGCTCTTCCGATCTNNNACTGACTTTTTTCTTTTCTTTTTT
AGAGCACACGTCTGAACTCTTTCCCTACACGACGCTCTTCCGATCTNNNACATTCCCCTACCATGTTAC
AGAGCACACGTCTGAACTCTTTCCCTACACGACGCTCTTCCGATCTNNNACATTCCCCTACCATGTTAC
AGAGCACACGTCTGAACTCTTTCCCTACACGACGCTCTTCCGATCTNNNACATTCCCCTACCATGTTAC
AGCACACGTCTGAACTCTTTCCCTACACGACGCTCTTCCGATCTNNNATGCCTTTAGTCCCAGCTACTT
AGCACACGTCTGAACTCTTTCCCTACACGACGCTCTTCCGATCTNNNATGCCTTTAGTCCCAGCTACTT

AGCACACGTCTGAACTCTTTCCCTACACGACGCTCTTCCGATCTNNNATGCCTTTAGTCCCAGCTACTT
GAGCACACGTCTGAACTCTTTCCCTACACGACGCTCTTCCGATCTNNNAATCTCAGCTACTTGGGAGGT
GAGCACACGTCTGAACTCTTTCCCTACACGACGCTCTTCCGATCTNNNAATCTCAGCTACTTGGGAGGT
GAGCACACGTCTGAACTCTTTCCCTACACGACGCTCTTCCGATCTNNNAATCTCAGCTACTTGGGAGGT
AGCACACGTCTGAACTCTTTCCCTACACGACGCTCTTCCGATCTNNNCTTTCCCCCACTTTTGTTTGTGT
AGCACACGTCTGAACTCTTTCCCTACACGACGCTCTTCCGATCTNNNCTTTCCCCCACTTTTGTTTGTGT
AGCACACGTCTGAACTCTTTCCCTACACGACGCTCTTCCGATCTNNNCTTTCCCCCACTTTTGTTTGTGT
GCACACGTCTGAACTCTTTCCCTACACGACGCTCTTCCGATCTNNNCTGAGTCATCTCTGTCATCCCAG
GCACACGTCTGAACTCTTTCCCTACACGACGCTCTTCCGATCTNNNCTGAGTCATCTCTGTCATCCCAG
GCACACGTCTGAACTCTTTCCCTACACGACGCTCTTCCGATCTNNNCTGAGTCATCTCTGTCATCCCAG
AGCACACGTCTGAACTCTTTCCCTACACGACGCTCTTCCGATCTNNNCTCCAACGCAGACTGGTTATTT
AGCACACGTCTGAACTCTTTCCCTACACGACGCTCTTCCGATCTNNNCTCCAACGCAGACTGGTTATTT
AGCACACGTCTGAACTCTTTCCCTACACGACGCTCTTCCGATCTNNNCTCCAACGCAGACTGGTTATTT
GCACACGTCTGAACTCTTTCCCTACACGACGCTCTTCCGATCTNNNCCAATGCCCGGCTAATTTTTCTT
GCACACGTCTGAACTCTTTCCCTACACGACGCTCTTCCGATCTNNNCCAATGCCCGGCTAATTTTTCTT
GCACACGTCTGAACTCTTTCCCTACACGACGCTCTTCCGATCTNNNCCAATGCCCGGCTAATTTTTCTT
GCACACGTCTGAACTCTTTCCCTACACGACGCTCTTCCGATCTNNNTGTTGGATTTTATTTGCATTTTCC
GCACACGTCTGAACTCTTTCCCTACACGACGCTCTTCCGATCTNNNTGTTGGATTTTATTTGCATTTTCC
GCACACGTCTGAACTCTTTCCCTACACGACGCTCTTCCGATCTNNNTGTTGGATTTTATTTGCATTTTCC
GAGCACACGTCTGAACTCTTTCCCTACACGACGCTCTTCCGATCTNNNCTGCATCCACTAACTCGTCAT
GAGCACACGTCTGAACTCTTTCCCTACACGACGCTCTTCCGATCTNNNCTGCATCCACTAACTCGTCAT
GAGCACACGTCTGAACTCTTTCCCTACACGACGCTCTTCCGATCTNNNCTGCATCCACTAACTCGTCAT
AGCACACGTCTGAACTCTTTCCCTACACGACGCTCTTCCGATCTNNNTTCCCTCTCCTCTCCTCAAGTAG
AGCACACGTCTGAACTCTTTCCCTACACGACGCTCTTCCGATCTNNNTTCCCTCTCCTCTCCTCAAGTAG
AGCACACGTCTGAACTCTTTCCCTACACGACGCTCTTCCGATCTNNNTTCCCTCTCCTCTCCTCAAGTAG
AGCACACGTCTGAACTCTTTCCCTACACGACGCTCTTCCGATCTNNNCTCCTCTCTTAGCATTCGGTTCT
AGCACACGTCTGAACTCTTTCCCTACACGACGCTCTTCCGATCTNNNCTCCTCTCTTAGCATTCGGTTCT
AGCACACGTCTGAACTCTTTCCCTACACGACGCTCTTCCGATCTNNNCTCCTCTCTTAGCATTCGGTTCT
GAGCACACGTCTGAACTCTTTCCCTACACGACGCTCTTCCGATCTNNNTCTACTACAGTGACACGAAAG
GAGCACACGTCTGAACTCTTTCCCTACACGACGCTCTTCCGATCTNNNTCTACTACAGTGACACGAAAG
GAGCACACGTCTGAACTCTTTCCCTACACGACGCTCTTCCGATCTNNNTCTACTACAGTGACACGAAAG
GAGCACACGTCTGAACTCTTTCCCTACACGACGCTCTTCCGATCTNNNTCATTCCGTCTCTATGAACAGG
GAGCACACGTCTGAACTCTTTCCCTACACGACGCTCTTCCGATCTNNNTCATTCCGTCTCTATGAACAGG
GAGCACACGTCTGAACTCTTTCCCTACACGACGCTCTTCCGATCTNNNTCATTCCGTCTCTATGAACAGG
GAGCACACGTCTGAACTCTTTCCCTACACGACGCTCTTCCGATCTNNNTTACCATGGGACCTGAACTGC
GAGCACACGTCTGAACTCTTTCCCTACACGACGCTCTTCCGATCTNNNTTACCATGGGACCTGAACTGC
GAGCACACGTCTGAACTCTTTCCCTACACGACGCTCTTCCGATCTNNNTTACCATGGGACCTGAACTGC
AGCACACGTCTGAACTCTTTCCCTACACGACGCTCTTCCGATCTNNNTATAGTCCCAGCTACTCGAGAG
AGCACACGTCTGAACTCTTTCCCTACACGACGCTCTTCCGATCTNNNTATAGTCCCAGCTACTCGAGAG
AGCACACGTCTGAACTCTTTCCCTACACGACGCTCTTCCGATCTNNNTATAGTCCCAGCTACTCGAGAG
GAGCACACGTCTGAACTCTTTCCCTACACGACGCTCTTCCGATCTNNNCATAGTGAAACCCTGTCTCTG
GAGCACACGTCTGAACTCTTTCCCTACACGACGCTCTTCCGATCTNNNCATAGTGAAACCCTGTCTCTG
GAGCACACGTCTGAACTCTTTCCCTACACGACGCTCTTCCGATCTNNNCATAGTGAAACCCTGTCTCTG

GAGCACACGTCTGAACTCTTTCCCTACACGACGCTCTTCCGATCTNNNGGATCACTTGAGGTCAGGAG

GAGCACACGTCTGAACTCTTTCCCTACACGACGCTCTTCCGATCTNNNGGATCACTTGAGGTCAGGAG

GAGCACACGTCTGAACTCTTTCCCTACACGACGCTCTTCCGATCTNNNGGATCACTTGAGGTCAGGAG

AGCACACGTCTGAACTCTTTCCCTACACGACGCTCTTCCGATCTNNNGTCGTGATGGATTGAGAAGCA

AGCACACGTCTGAACTCTTTCCCTACACGACGCTCTTCCGATCTNNNGTCGTGATGGATTGAGAAGCA

AGCACACGTCTGAACTCTTTCCCTACACGACGCTCTTCCGATCTNNNGTCGTGATGGATTGAGAAGCA

AGCACACGTCTGAACTCTTTCCCTACACGACGCTCTTCCGATCTNNNTAGATGGAAAATAGGCACCTG

AGCACACGTCTGAACTCTTTCCCTACACGACGCTCTTCCGATCTNNNTAGATGGAAAATAGGCACCTG

AGCACACGTCTGAACTCTTTCCCTACACGACGCTCTTCCGATCTNNNTAGATGGAAAATAGGCACCTG

AGCACACGTCTGAACTCTTTCCCTACACGACGCTCTTCCGATCTNNNCATAGCTCACTACAGCCTCAGTC

AGCACACGTCTGAACTCTTTCCCTACACGACGCTCTTCCGATCTNNNCATAGCTCACTACAGCCTCAGTC

AGCACACGTCTGAACTCTTTCCCTACACGACGCTCTTCCGATCTNNNCATAGCTCACTACAGCCTCAGTC

AGAGCACACGTCTGAACTCTTTCCCTACACGACGCTCTTCCGATCTNNNGGTAGGAGAATCACTTGAAC

AGAGCACACGTCTGAACTCTTTCCCTACACGACGCTCTTCCGATCTNNNGGTAGGAGAATCACTTGAAC

AGAGCACACGTCTGAACTCTTTCCCTACACGACGCTCTTCCGATCTNNNGGTAGGAGAATCACTTGAAC

GCACACGTCTGAACTCTTTCCCTACACGACGCTCTTCCGATCTNNNTAGAACTGCTTTCTGGAGTGTGC

GCACACGTCTGAACTCTTTCCCTACACGACGCTCTTCCGATCTNNNTAGAACTGCTTTCTGGAGTGTGC

GCACACGTCTGAACTCTTTCCCTACACGACGCTCTTCCGATCTNNNTAGAACTGCTTTCTGGAGTGTGC

GAGCACACGTCTGAACTCTTTCCCTACACGACGCTCTTCCGATCTNNNTTGCAAATTTCAAGCCAAGGG

GAGCACACGTCTGAACTCTTTCCCTACACGACGCTCTTCCGATCTNNNTTGCAAATTTCAAGCCAAGGG

GAGCACACGTCTGAACTCTTTCCCTACACGACGCTCTTCCGATCTNNNTTGCAAATTTCAAGCCAAGGG

GAGCACACGTCTGAACTCTTTCCCTACACGACGCTCTTCCGATCTNNNTCTTTCTTTCTTTCCCTCCCTCCC

GAGCACACGTCTGAACTCTTTCCCTACACGACGCTCTTCCGATCTNNNTCTTTCTTTCTTTCCCTCCCTCCC

GAGCACACGTCTGAACTCTTTCCCTACACGACGCTCTTCCGATCTNNNTCTTTCTTTCTTTCCCTCCCTCCC

GAGCACACGTCTGAACTCTTTCCCTACACGACGCTCTTCCGATCTNNNACTCACCACAGTCATCAGCAGC

GAGCACACGTCTGAACTCTTTCCCTACACGACGCTCTTCCGATCTNNNACTCACCACAGTCATCAGCAGC

GAGCACACGTCTGAACTCTTTCCCTACACGACGCTCTTCCGATCTNNNACTCACCACAGTCATCAGCAGC

AGCACACGTCTGAACTCTTTCCCTACACGACGCTCTTCCGATCTNNNCTATCTGAAAATACATTTCCCTC

AGCACACGTCTGAACTCTTTCCCTACACGACGCTCTTCCGATCTNNNCTATCTGAAAATACATTTCCCTC

AGCACACGTCTGAACTCTTTCCCTACACGACGCTCTTCCGATCTNNNCTATCTGAAAATACATTTCCCTC

GAGCACACGTCTGAACTCTTTCCCTACACGACGCTCTTCCGATCTNNNCAGGAGAATTGCTTGAAACCA

GAGCACACGTCTGAACTCTTTCCCTACACGACGCTCTTCCGATCTNNNCAGGAGAATTGCTTGAAACCA

GAGCACACGTCTGAACTCTTTCCCTACACGACGCTCTTCCGATCTNNNCAGGAGAATTGCTTGAAACCA

AGCACACGTCTGAACTCTTTCCCTACACGACGCTCTTCCGATCTNNNACCCACCAAACTGCTAGAAAGA

AGCACACGTCTGAACTCTTTCCCTACACGACGCTCTTCCGATCTNNNACCCACCAAACTGCTAGAAAGA

AGCACACGTCTGAACTCTTTCCCTACACGACGCTCTTCCGATCTNNNACCCACCAAACTGCTAGAAAGA

AGAGCACACGTCTGAACTCTTTCCCTACACGACGCTCTTCCGATCTNNNTTTCCCAGTCAAGGGAAAGG

AGAGCACACGTCTGAACTCTTTCCCTACACGACGCTCTTCCGATCTNNNTTTCCCAGTCAAGGGAAAGG

AGAGCACACGTCTGAACTCTTTCCCTACACGACGCTCTTCCGATCTNNNTTTCCCAGTCAAGGGAAAGG

AGCACACGTCTGAACTCTTTCCCTACACGACGCTCTTCCGATCTNNNTAAAATGGTCCCGGCAATAAAC

AGCACACGTCTGAACTCTTTCCCTACACGACGCTCTTCCGATCTNNNTAAAATGGTCCCGGCAATAAAC

AGCACACGTCTGAACTCTTTCCCTACACGACGCTCTTCCGATCTNNNTAAAATGGTCCCGGCAATAAAC

GCACACGTCTGAACTCTTTCCCTACACGACGCTCTTCCGATCTNNNTGAGCAGGAAGGGAGAATGAAC

GCACACGTCTGAACTCTTTCCCTACACGACGCTCTTCCGATCTNNNTGAGCAGGAAGGGAGAATGAA
GCACACGTCTGAACTCTTTCCCTACACGACGCTCTTCCGATCTNNNTGAGCAGGAAGGGAGAATGAA
GAGCACACGTCTGAACTCTTTCCCTACACGACGCTCTTCCGATCTNNNGAGATCACACCACTACACTCC
GAGCACACGTCTGAACTCTTTCCCTACACGACGCTCTTCCGATCTNNNGAGATCACACCACTACACTCC
GAGCACACGTCTGAACTCTTTCCCTACACGACGCTCTTCCGATCTNNNGAGATCACACCACTACACTCC
GAGCACACGTCTGAACTCTTTCCCTACACGACGCTCTTCCGATCTNNNTGATTGCCCTTCACCTGAAACA
GAGCACACGTCTGAACTCTTTCCCTACACGACGCTCTTCCGATCTNNNTGATTGCCCTTCACCTGAAACA
GAGCACACGTCTGAACTCTTTCCCTACACGACGCTCTTCCGATCTNNNTGATTGCCCTTCACCTGAAACA
AGAGCACACGTCTGAACTCTTTCCCTACACGACGCTCTTCCGATCTNNNCCCAGGATCCACTACAAAAG
AGAGCACACGTCTGAACTCTTTCCCTACACGACGCTCTTCCGATCTNNNCCCAGGATCCACTACAAAAG
AGAGCACACGTCTGAACTCTTTCCCTACACGACGCTCTTCCGATCTNNNCCCAGGATCCACTACAAAAG
AGCACACGTCTGAACTCTTTCCCTACACGACGCTCTTCCGATCTNNNAGAGTACATGGGCTTAGGGCA
AGCACACGTCTGAACTCTTTCCCTACACGACGCTCTTCCGATCTNNNAGAGTACATGGGCTTAGGGCA
AGCACACGTCTGAACTCTTTCCCTACACGACGCTCTTCCGATCTNNNAGAGTACATGGGCTTAGGGCA
GAGCACACGTCTGAACTCTTTCCCTACACGACGCTCTTCCGATCTNNNAGGTTCACCAAGGAGTCACTT
GAGCACACGTCTGAACTCTTTCCCTACACGACGCTCTTCCGATCTNNNAGGTTCACCAAGGAGTCACTT
GAGCACACGTCTGAACTCTTTCCCTACACGACGCTCTTCCGATCTNNNAGGTTCACCAAGGAGTCACTT
AGCACACGTCTGAACTCTTTCCCTACACGACGCTCTTCCGATCTNNNAAAATAGCAGACGTTTCCCCTC
AGCACACGTCTGAACTCTTTCCCTACACGACGCTCTTCCGATCTNNNAAAATAGCAGACGTTTCCCCTC
AGCACACGTCTGAACTCTTTCCCTACACGACGCTCTTCCGATCTNNNAAAATAGCAGACGTTTCCCCTC
GAGCACACGTCTGAACTCTTTCCCTACACGACGCTCTTCCGATCTNNNGTACATTTTTGTGCCCGTGTG
GAGCACACGTCTGAACTCTTTCCCTACACGACGCTCTTCCGATCTNNNGTACATTTTTGTGCCCGTGTG
GAGCACACGTCTGAACTCTTTCCCTACACGACGCTCTTCCGATCTNNNGTACATTTTTGTGCCCGTGTG
GAGCACACGTCTGAACTCTTTCCCTACACGACGCTCTTCCGATCTNNNTTCCTTCCTTCCTTCCTACAAG
GAGCACACGTCTGAACTCTTTCCCTACACGACGCTCTTCCGATCTNNNTTCCTTCCTTCCTTCCTACAAG
GAGCACACGTCTGAACTCTTTCCCTACACGACGCTCTTCCGATCTNNNTTCCTTCCTTCCTTCCTACAAG
GAGCACACGTCTGAACTCTTTCCCTACACGACGCTCTTCCGATCTNNNCTGAGACAGGAGAATGGTG
GAGCACACGTCTGAACTCTTTCCCTACACGACGCTCTTCCGATCTNNNCTGAGACAGGAGAATGGTG
GAGCACACGTCTGAACTCTTTCCCTACACGACGCTCTTCCGATCTNNNCTGAGACAGGAGAATGGTG
GAGCACACGTCTGAACTCTTTCCCTACACGACGCTCTTCCGATCTNNNGAGATCGTGACACTGCACTCC
GAGCACACGTCTGAACTCTTTCCCTACACGACGCTCTTCCGATCTNNNGAGATCGTGACACTGCACTCC
GAGCACACGTCTGAACTCTTTCCCTACACGACGCTCTTCCGATCTNNNGAGATCGTGACACTGCACTCC
AGCACACGTCTGAACTCTTTCCCTACACGACGCTCTTCCGATCTNNNATCACCAAATGGCACCATACGA
AGCACACGTCTGAACTCTTTCCCTACACGACGCTCTTCCGATCTNNNATCACCAAATGGCACCATACGA
AGCACACGTCTGAACTCTTTCCCTACACGACGCTCTTCCGATCTNNNATCACCAAATGGCACCATACGA
GAGCACACGTCTGAACTCTTTCCCTACACGACGCTCTTCCGATCTNNNGCATTTGGTGTAGTCCGTGT
GAGCACACGTCTGAACTCTTTCCCTACACGACGCTCTTCCGATCTNNNGCATTTGGTGTAGTCCGTGT
GAGCACACGTCTGAACTCTTTCCCTACACGACGCTCTTCCGATCTNNNGCATTTGGTGTAGTCCGTGT
GCACACGTCTGAACTCTTTCCCTACACGACGCTCTTCCGATCTNNNTTACCCTACATGGTCTAAAAAGC
GCACACGTCTGAACTCTTTCCCTACACGACGCTCTTCCGATCTNNNTTACCCTACATGGTCTAAAAAGC
GCACACGTCTGAACTCTTTCCCTACACGACGCTCTTCCGATCTNNNTTACCCTACATGGTCTAAAAAGC
GAGCACACGTCTGAACTCTTTCCCTACACGACGCTCTTCCGATCTNNNGCACACACAGAGGTTCCTAAA
GAGCACACGTCTGAACTCTTTCCCTACACGACGCTCTTCCGATCTNNNGCACACACAGAGGTTCCTAAA

GAGCACACGTCTGAACTCTTTCCCTACACGACGCTCTTCCGATCTNNNGCACACACAGAGGTTCCTAAA
GAGCACACGTCTGAACTCTTTCCCTACACGACGCTCTTCCGATCTNNNTTCGGCTTCGAACTGGTGATT
GAGCACACGTCTGAACTCTTTCCCTACACGACGCTCTTCCGATCTNNNTTCGGCTTCGAACTGGTGATT
GAGCACACGTCTGAACTCTTTCCCTACACGACGCTCTTCCGATCTNNNTTCGGCTTCGAACTGGTGATT
AGAGCACACGTCTGAACTCTTTCCCTACACGACGCTCTTCCGATCTNNNGTATGCTCCACAATGCCAAG
AGAGCACACGTCTGAACTCTTTCCCTACACGACGCTCTTCCGATCTNNNGTATGCTCCACAATGCCAAG
AGAGCACACGTCTGAACTCTTTCCCTACACGACGCTCTTCCGATCTNNNGTATGCTCCACAATGCCAAG
GAGCACACGTCTGAACTCTTTCCCTACACGACGCTCTTCCGATCTNNNTTATAGGCACCCAACATGACA
GAGCACACGTCTGAACTCTTTCCCTACACGACGCTCTTCCGATCTNNNTTATAGGCACCCAACATGACA
GAGCACACGTCTGAACTCTTTCCCTACACGACGCTCTTCCGATCTNNNTTATAGGCACCCAACATGACA
GCACACGTCTGAACTCTTTCCCTACACGACGCTCTTCCGATCTNNNTGAATCCTCCTTTTCCACTGACCCA
GCACACGTCTGAACTCTTTCCCTACACGACGCTCTTCCGATCTNNNTGAATCCTCCTTTTCCACTGACCCA
GCACACGTCTGAACTCTTTCCCTACACGACGCTCTTCCGATCTNNNTGAATCCTCCTTTTCCACTGACCCA
AGCACACGTCTGAACTCTTTCCCTACACGACGCTCTTCCGATCTNNNAAAATGTGTGTGTGTGTGTGT
AGCACACGTCTGAACTCTTTCCCTACACGACGCTCTTCCGATCTNNNAAAATGTGTGTGTGTGTGTGT
AGCACACGTCTGAACTCTTTCCCTACACGACGCTCTTCCGATCTNNNAAAATGTGTGTGTGTGTGTGT
GAGCACACGTCTGAACTCTTTCCCTACACGACGCTCTTCCGATCTNNNACAGGCCAACATTCAAATTCA
GAGCACACGTCTGAACTCTTTCCCTACACGACGCTCTTCCGATCTNNNACAGGCCAACATTCAAATTCA
GAGCACACGTCTGAACTCTTTCCCTACACGACGCTCTTCCGATCTNNNACAGGCCAACATTCAAATTCA
AGCACACGTCTGAACTCTTTCCCTACACGACGCTCTTCCGATCTNNNCAAACTCCAACAGGTGATGCTT
AGCACACGTCTGAACTCTTTCCCTACACGACGCTCTTCCGATCTNNNCAAACTCCAACAGGTGATGCTT
AGCACACGTCTGAACTCTTTCCCTACACGACGCTCTTCCGATCTNNNCAAACTCCAACAGGTGATGCTT
GAGCACACGTCTGAACTCTTTCCCTACACGACGCTCTTCCGATCTNNNGTTGCAGTGAACCGAGATCA
GAGCACACGTCTGAACTCTTTCCCTACACGACGCTCTTCCGATCTNNNGTTGCAGTGAACCGAGATCA
GAGCACACGTCTGAACTCTTTCCCTACACGACGCTCTTCCGATCTNNNGTTGCAGTGAACCGAGATCA
GAGCACACGTCTGAACTCTTTCCCTACACGACGCTCTTCCGATCTNNNCTGTAATTCGAGCTACTCCG
GAGCACACGTCTGAACTCTTTCCCTACACGACGCTCTTCCGATCTNNNCTGTAATTCGAGCTACTCCG
GAGCACACGTCTGAACTCTTTCCCTACACGACGCTCTTCCGATCTNNNCTGTAATTCGAGCTACTCCG
AGCACACGTCTGAACTCTTTCCCTACACGACGCTCTTCCGATCTNNNATTCTCTAGAGCACATCAAGG
AGCACACGTCTGAACTCTTTCCCTACACGACGCTCTTCCGATCTNNNATTCTCTAGAGCACATCAAGG
AGCACACGTCTGAACTCTTTCCCTACACGACGCTCTTCCGATCTNNNATTCTCTAGAGCACATCAAGG
GAGCACACGTCTGAACTCTTTCCCTACACGACGCTCTTCCGATCTNNNGGAGAAGGGCATCATGAGAT
GAGCACACGTCTGAACTCTTTCCCTACACGACGCTCTTCCGATCTNNNGGAGAAGGGCATCATGAGAT
GAGCACACGTCTGAACTCTTTCCCTACACGACGCTCTTCCGATCTNNNGGAGAAGGGCATCATGAGAT
AGCACACGTCTGAACTCTTTCCCTACACGACGCTCTTCCGATCTNNNCTGAGGCAGGAGAATCACTTG
AGCACACGTCTGAACTCTTTCCCTACACGACGCTCTTCCGATCTNNNCTGAGGCAGGAGAATCACTTG
AGCACACGTCTGAACTCTTTCCCTACACGACGCTCTTCCGATCTNNNCTGAGGCAGGAGAATCACTTG
AGCACACGTCTGAACTCTTTCCCTACACGACGCTCTTCCGATCTNNNAAGGGCAATCAGCTTTACTCA
AGCACACGTCTGAACTCTTTCCCTACACGACGCTCTTCCGATCTNNNAAGGGCAATCAGCTTTACTCA
AGCACACGTCTGAACTCTTTCCCTACACGACGCTCTTCCGATCTNNNAAGGGCAATCAGCTTTACTCA
AGCACACGTCTGAACTCTTTCCCTACACGACGCTCTTCCGATCTNNNATTGTCCTATGACCCTGCCAAAT
AGCACACGTCTGAACTCTTTCCCTACACGACGCTCTTCCGATCTNNNATTGTCCTATGACCCTGCCAAAT
AGCACACGTCTGAACTCTTTCCCTACACGACGCTCTTCCGATCTNNNATTGTCCTATGACCCTGCCAAAT

AGCACACGTCTGAACTCTTTCCCTACACGACGCTCTTCCGATCTNNNAATCCCAAGTTTGCAATGAGG
AGCACACGTCTGAACTCTTTCCCTACACGACGCTCTTCCGATCTNNNAATCCCAAGTTTGCAATGAGG
AGCACACGTCTGAACTCTTTCCCTACACGACGCTCTTCCGATCTNNNAATCCCAAGTTTGCAATGAGG
GCACACGTCTGAACTCTTTCCCTACACGACGCTCTTCCGATCTNNNTCAAGTGACTTTCCTACCTCAGC
GCACACGTCTGAACTCTTTCCCTACACGACGCTCTTCCGATCTNNNTCAAGTGACTTTCCTACCTCAGC
GCACACGTCTGAACTCTTTCCCTACACGACGCTCTTCCGATCTNNNTCAAGTGACTTTCCTACCTCAGC
GCACACGTCTGAACTCTTTCCCTACACGACGCTCTTCCGATCTNNNCAACAGCCACTAAGTGAAACCT
GCACACGTCTGAACTCTTTCCCTACACGACGCTCTTCCGATCTNNNCAACAGCCACTAAGTGAAACCT
GCACACGTCTGAACTCTTTCCCTACACGACGCTCTTCCGATCTNNNCAACAGCCACTAAGTGAAACCT
CACACGTCTGAACTCTTTCCCTACACGACGCTCTTCCGATCTNNNCAACTTTTGCATGTTTTGTAGACA
CACACGTCTGAACTCTTTCCCTACACGACGCTCTTCCGATCTNNNCAACTTTTGCATGTTTTGTAGACA
CACACGTCTGAACTCTTTCCCTACACGACGCTCTTCCGATCTNNNCAACTTTTGCATGTTTTGTAGACA
AGCACACGTCTGAACTCTTTCCCTACACGACGCTCTTCCGATCTNNNCTGAGGCAGGAGAATCACTTG
AGCACACGTCTGAACTCTTTCCCTACACGACGCTCTTCCGATCTNNNCTGAGGCAGGAGAATCACTTG
AGCACACGTCTGAACTCTTTCCCTACACGACGCTCTTCCGATCTNNNCTGAGGCAGGAGAATCACTTG
CACACGTCTGAACTCTTTCCCTACACGACGCTCTTCCGATCTNNNCTCCTGGGTTCAAGTGATTCTCCC
CACACGTCTGAACTCTTTCCCTACACGACGCTCTTCCGATCTNNNCTCCTGGGTTCAAGTGATTCTCCC
CACACGTCTGAACTCTTTCCCTACACGACGCTCTTCCGATCTNNNCTCCTGGGTTCAAGTGATTCTCCC
GCACACGTCTGAACTCTTTCCCTACACGACGCTCTTCCGATCTNNNCAGAGTTGTTCAACCATCACCAC
GCACACGTCTGAACTCTTTCCCTACACGACGCTCTTCCGATCTNNNCAGAGTTGTTCAACCATCACCAC
GCACACGTCTGAACTCTTTCCCTACACGACGCTCTTCCGATCTNNNCAGAGTTGTTCAACCATCACCAC
AGCACACGTCTGAACTCTTTCCCTACACGACGCTCTTCCGATCTNNNGACTGTAGTCCCAGCTACATG
AGCACACGTCTGAACTCTTTCCCTACACGACGCTCTTCCGATCTNNNGACTGTAGTCCCAGCTACATG
AGCACACGTCTGAACTCTTTCCCTACACGACGCTCTTCCGATCTNNNGACTGTAGTCCCAGCTACATG
GCACACGTCTGAACTCTTTCCCTACACGACGCTCTTCCGATCTNNNACTACAAAGCCAGCACTACAAGA
GCACACGTCTGAACTCTTTCCCTACACGACGCTCTTCCGATCTNNNACTACAAAGCCAGCACTACAAGA
GCACACGTCTGAACTCTTTCCCTACACGACGCTCTTCCGATCTNNNACTACAAAGCCAGCACTACAAGA
GCACACGTCTGAACTCTTTCCCTACACGACGCTCTTCCGATCTNNNACTCCGCTCTCTTTTGATTCCATC
GCACACGTCTGAACTCTTTCCCTACACGACGCTCTTCCGATCTNNNACTCCGCTCTCTTTTGATTCCATC
GCACACGTCTGAACTCTTTCCCTACACGACGCTCTTCCGATCTNNNACTCCGCTCTCTTTTGATTCCATC
AGCACACGTCTGAACTCTTTCCCTACACGACGCTCTTCCGATCTNNNAAAAAATTAGCCGGGTGTGG
AGCACACGTCTGAACTCTTTCCCTACACGACGCTCTTCCGATCTNNNAAAAAATTAGCCGGGTGTGG
AGCACACGTCTGAACTCTTTCCCTACACGACGCTCTTCCGATCTNNNAAAAAATTAGCCGGGTGTGG
GAGCACACGTCTGAACTCTTTCCCTACACGACGCTCTTCCGATCTNNNGAAACCCATTTGAAACCCTTG
GAGCACACGTCTGAACTCTTTCCCTACACGACGCTCTTCCGATCTNNNGAAACCCATTTGAAACCCTTG
GAGCACACGTCTGAACTCTTTCCCTACACGACGCTCTTCCGATCTNNNGAAACCCATTTGAAACCCTTG
GAGCACACGTCTGAACTCTTTCCCTACACGACGCTCTTCCGATCTNNNGTGAATTGAACCATCAGCAT
GAGCACACGTCTGAACTCTTTCCCTACACGACGCTCTTCCGATCTNNNGTGAATTGAACCATCAGCAT
GAGCACACGTCTGAACTCTTTCCCTACACGACGCTCTTCCGATCTNNNGTGAATTGAACCATCAGCAT
AGCACACGTCTGAACTCTTTCCCTACACGACGCTCTTCCGATCTNNNATGGGGAATAGGGACACACT
AGCACACGTCTGAACTCTTTCCCTACACGACGCTCTTCCGATCTNNNATGGGGAATAGGGACACACT
AGCACACGTCTGAACTCTTTCCCTACACGACGCTCTTCCGATCTNNNATGGGGAATAGGGACACACT
GCACACGTCTGAACTCTTTCCCTACACGACGCTCTTCCGATCTNNNTACTGAAGTACTCTGAGACGG

AGCACACGTCTGAACTCTTTCCCTACACGACGCTCTTCCGATCTNNNTACTGAAGTACTCTGAGACGG
AGCACACGTCTGAACTCTTTCCCTACACGACGCTCTTCCGATCTNNNTACTGAAGTACTCTGAGACGG
AGCACACGTCTGAACTCTTTCCCTACACGACGCTCTTCCGATCTNNNGTAGTGTAGTGATCTGGGCTC
AGCACACGTCTGAACTCTTTCCCTACACGACGCTCTTCCGATCTNNNGTAGTGTAGTGATCTGGGCTC
AGCACACGTCTGAACTCTTTCCCTACACGACGCTCTTCCGATCTNNNGTAGTGTAGTGATCTGGGCTC
GAGCACACGTCTGAACTCTTTCCCTACACGACGCTCTTCCGATCTNNNGAGATCGAGCCATTACACTCC
GAGCACACGTCTGAACTCTTTCCCTACACGACGCTCTTCCGATCTNNNGAGATCGAGCCATTACACTCC
GAGCACACGTCTGAACTCTTTCCCTACACGACGCTCTTCCGATCTNNNGAGATCGAGCCATTACACTCC
GAGCACACGTCTGAACTCTTTCCCTACACGACGCTCTTCCGATCTNNNCAATGACCTCATTCCACCAGGT
GAGCACACGTCTGAACTCTTTCCCTACACGACGCTCTTCCGATCTNNNCAATGACCTCATTCCACCAGGT
GAGCACACGTCTGAACTCTTTCCCTACACGACGCTCTTCCGATCTNNNCAATGACCTCATTCCACCAGGT
GAGCACACGTCTGAACTCTTTCCCTACACGACGCTCTTCCGATCTNNNTTTTACCTCCATCAAGGCCCTT
GAGCACACGTCTGAACTCTTTCCCTACACGACGCTCTTCCGATCTNNNTTTTACCTCCATCAAGGCCCTT
GAGCACACGTCTGAACTCTTTCCCTACACGACGCTCTTCCGATCTNNNTTTTACCTCCATCAAGGCCCTT
AGAGCACACGTCTGAACTCTTTCCCTACACGACGCTCTTCCGATCTNNNAGGCAGGATAATCACTTGA
AGAGCACACGTCTGAACTCTTTCCCTACACGACGCTCTTCCGATCTNNNAGGCAGGATAATCACTTGA
AGAGCACACGTCTGAACTCTTTCCCTACACGACGCTCTTCCGATCTNNNAGGCAGGATAATCACTTGA
AGAGCACACGTCTGAACTCTTTCCCTACACGACGCTCTTCCGATCTNNNTGGGGGAGGCTAACCCTAA
AGAGCACACGTCTGAACTCTTTCCCTACACGACGCTCTTCCGATCTNNNTGGGGGAGGCTAACCCTAA
AGAGCACACGTCTGAACTCTTTCCCTACACGACGCTCTTCCGATCTNNNTGGGGGAGGCTAACCCTAA
GAGCACACGTCTGAACTCTTTCCCTACACGACGCTCTTCCGATCTNNNTGCCTGTAATCCCAGCTACTT
GAGCACACGTCTGAACTCTTTCCCTACACGACGCTCTTCCGATCTNNNTGCCTGTAATCCCAGCTACTT
GAGCACACGTCTGAACTCTTTCCCTACACGACGCTCTTCCGATCTNNNTGCCTGTAATCCCAGCTACTT
GAGCACACGTCTGAACTCTTTCCCTACACGACGCTCTTCCGATCTNNNGAGAATGGCTTGAACACAGG
GAGCACACGTCTGAACTCTTTCCCTACACGACGCTCTTCCGATCTNNNGAGAATGGCTTGAACACAGG
GAGCACACGTCTGAACTCTTTCCCTACACGACGCTCTTCCGATCTNNNGAGAATGGCTTGAACACAGG
GAGCACACGTCTGAACTCTTTCCCTACACGACGCTCTTCCGATCTNNNGGAACTGCAAATCAACAACA
GAGCACACGTCTGAACTCTTTCCCTACACGACGCTCTTCCGATCTNNNGGAACTGCAAATCAACAACA
GAGCACACGTCTGAACTCTTTCCCTACACGACGCTCTTCCGATCTNNNGGAACTGCAAATCAACAACA
AGCACACGTCTGAACTCTTTCCCTACACGACGCTCTTCCGATCTNNNCCATCATTCCACCATCTCAGAG
AGCACACGTCTGAACTCTTTCCCTACACGACGCTCTTCCGATCTNNNCCATCATTCCACCATCTCAGAG
AGCACACGTCTGAACTCTTTCCCTACACGACGCTCTTCCGATCTNNNCCATCATTCCACCATCTCAGAG
AGCACACGTCTGAACTCTTTCCCTACACGACGCTCTTCCGATCTNNNTGCACAGAACCATATCCTAAGC
AGCACACGTCTGAACTCTTTCCCTACACGACGCTCTTCCGATCTNNNTGCACAGAACCATATCCTAAGC
AGCACACGTCTGAACTCTTTCCCTACACGACGCTCTTCCGATCTNNNTGCACAGAACCATATCCTAAGC
GCACACGTCTGAACTCTTTCCCTACACGACGCTCTTCCGATCTNNNTTGAAAGCCCAGGGAGACTATT
GCACACGTCTGAACTCTTTCCCTACACGACGCTCTTCCGATCTNNNTTGAAAGCCCAGGGAGACTATT
GCACACGTCTGAACTCTTTCCCTACACGACGCTCTTCCGATCTNNNTTGAAAGCCCAGGGAGACTATT
AGCACACGTCTGAACTCTTTCCCTACACGACGCTCTTCCGATCTNNNCTGGGATCATGGTATCCAGGC
AGCACACGTCTGAACTCTTTCCCTACACGACGCTCTTCCGATCTNNNCTGGGATCATGGTATCCAGGC
AGCACACGTCTGAACTCTTTCCCTACACGACGCTCTTCCGATCTNNNCTGGGATCATGGTATCCAGGC
AGCACACGTCTGAACTCTTTCCCTACACGACGCTCTTCCGATCTNNNTCGTATGTTTGTGTCTTTCTGG
AGCACACGTCTGAACTCTTTCCCTACACGACGCTCTTCCGATCTNNNTCGTATGTTTGTGTCTTTCTGG

AGCACACGTCTGAACTCTTTCCCTACACGACGCTCTTCCGATCTNNNTCGTATGTTTGTGTCTTTCTGG
CACGTCTGAACTCTTTCCCTACACGACGCTCTTCCGATCTNNNGGTGAAGGTACAGACTCATCCAGCA
CACGTCTGAACTCTTTCCCTACACGACGCTCTTCCGATCTNNNGGTGAAGGTACAGACTCATCCAGCA
CACGTCTGAACTCTTTCCCTACACGACGCTCTTCCGATCTNNNGGTGAAGGTACAGACTCATCCAGCA
AGCACACGTCTGAACTCTTTCCCTACACGACGCTCTTCCGATCTNNNTAAGACTTCCTTTACTTTCTCT
AGCACACGTCTGAACTCTTTCCCTACACGACGCTCTTCCGATCTNNNTAAGACTTCCTTTACTTTCTCT
AGCACACGTCTGAACTCTTTCCCTACACGACGCTCTTCCGATCTNNNTAAGACTTCCTTTACTTTCTCT
AGCACACGTCTGAACTCTTTCCCTACACGACGCTCTTCCGATCTNNNTGTTGGGTAATCCTGTGGAAG
AGCACACGTCTGAACTCTTTCCCTACACGACGCTCTTCCGATCTNNNTGTTGGGTAATCCTGTGGAAG
AGCACACGTCTGAACTCTTTCCCTACACGACGCTCTTCCGATCTNNNTGTTGGGTAATCCTGTGGAAG
AGCACACGTCTGAACTCTTTCCCTACACGACGCTCTTCCGATCTNNNAGAGGACGTCAGGATTCACTC
AGCACACGTCTGAACTCTTTCCCTACACGACGCTCTTCCGATCTNNNAGAGGACGTCAGGATTCACTC
AGCACACGTCTGAACTCTTTCCCTACACGACGCTCTTCCGATCTNNNAGAGGACGTCAGGATTCACTC
AGCACACGTCTGAACTCTTTCCCTACACGACGCTCTTCCGATCTNNNAGTTTCATCCATGTTGCTGCAA
AGCACACGTCTGAACTCTTTCCCTACACGACGCTCTTCCGATCTNNNAGTTTCATCCATGTTGCTGCAA
AGCACACGTCTGAACTCTTTCCCTACACGACGCTCTTCCGATCTNNNAGTTTCATCCATGTTGCTGCAA
GCACACGTCTGAACTCTTTCCCTACACGACGCTCTTCCGATCTNNNACCCTGACCTCGAGAATGAATT
GCACACGTCTGAACTCTTTCCCTACACGACGCTCTTCCGATCTNNNACCCTGACCTCGAGAATGAATT
GCACACGTCTGAACTCTTTCCCTACACGACGCTCTTCCGATCTNNNACCCTGACCTCGAGAATGAATT
GCACACGTCTGAACTCTTTCCCTACACGACGCTCTTCCGATCTNNNATCCTGTAACTTGGGAGAGGGT
GCACACGTCTGAACTCTTTCCCTACACGACGCTCTTCCGATCTNNNATCCTGTAACTTGGGAGAGGGT
GCACACGTCTGAACTCTTTCCCTACACGACGCTCTTCCGATCTNNNATCCTGTAACTTGGGAGAGGGT
GCACACGTCTGAACTCTTTCCCTACACGACGCTCTTCCGATCTNNNTGTAATCCCAGCACTTTAGGAG
GCACACGTCTGAACTCTTTCCCTACACGACGCTCTTCCGATCTNNNTGTAATCCCAGCACTTTAGGAG
GCACACGTCTGAACTCTTTCCCTACACGACGCTCTTCCGATCTNNNTGTAATCCCAGCACTTTAGGAG
GAGCACACGTCTGAACTCTTTCCCTACACGACGCTCTTCCGATCTNNNAGCAGCAAGAACAACTTTTG
GAGCACACGTCTGAACTCTTTCCCTACACGACGCTCTTCCGATCTNNNAGCAGCAAGAACAACTTTTG
GAGCACACGTCTGAACTCTTTCCCTACACGACGCTCTTCCGATCTNNNAGCAGCAAGAACAACTTTTG
GCACACGTCTGAACTCTTTCCCTACACGACGCTCTTCCGATCTNNNAATTCTAGAGCCAGAGACAGG
GCACACGTCTGAACTCTTTCCCTACACGACGCTCTTCCGATCTNNNAATTCTAGAGCCAGAGACAGG
GCACACGTCTGAACTCTTTCCCTACACGACGCTCTTCCGATCTNNNAATTCTAGAGCCAGAGACAGG
AGCACACGTCTGAACTCTTTCCCTACACGACGCTCTTCCGATCTNNNTACCACCACTTCATTCTCCTTG
AGCACACGTCTGAACTCTTTCCCTACACGACGCTCTTCCGATCTNNNTACCACCACTTCATTCTCCTTG
AGCACACGTCTGAACTCTTTCCCTACACGACGCTCTTCCGATCTNNNTACCACCACTTCATTCTCCTTG
GCACACGTCTGAACTCTTTCCCTACACGACGCTCTTCCGATCTNNNCTCTCTCACATGGCTTCTTCA
GCACACGTCTGAACTCTTTCCCTACACGACGCTCTTCCGATCTNNNCTCTCTCACATGGCTTCTTCA
GCACACGTCTGAACTCTTTCCCTACACGACGCTCTTCCGATCTNNNCTCTCTCACATGGCTTCTTCA
GCACACGTCTGAACTCTTTCCCTACACGACGCTCTTCCGATCTNNNGTGATGATAGTGGTGGTGG
GCACACGTCTGAACTCTTTCCCTACACGACGCTCTTCCGATCTNNNGTGATGATAGTGGTGGTGG
GCACACGTCTGAACTCTTTCCCTACACGACGCTCTTCCGATCTNNNGTGATGATAGTGGTGGTGG
GAGCACACGTCTGAACTCTTTCCCTACACGACGCTCTTCCGATCTNNNTCTCCTTGGGAATCTCAGTAG
GAGCACACGTCTGAACTCTTTCCCTACACGACGCTCTTCCGATCTNNNTCTCCTTGGGAATCTCAGTAG
GAGCACACGTCTGAACTCTTTCCCTACACGACGCTCTTCCGATCTNNNTCTCCTTGGGAATCTCAGTAG

GAGCACACGTCTGAACTCTTTCCCTACACGACGCTCTTCCGATCTNNNCCCATAGAAGCCGTTATTGG
GAGCACACGTCTGAACTCTTTCCCTACACGACGCTCTTCCGATCTNNNCCCATAGAAGCCGTTATTGG
GAGCACACGTCTGAACTCTTTCCCTACACGACGCTCTTCCGATCTNNNCCCATAGAAGCCGTTATTGG
GCACACGTCTGAACTCTTTCCCTACACGACGCTCTTCCGATCTNNNTCAAGAAGGCATAACAGCATCA
GCACACGTCTGAACTCTTTCCCTACACGACGCTCTTCCGATCTNNNTCAAGAAGGCATAACAGCATCA
GCACACGTCTGAACTCTTTCCCTACACGACGCTCTTCCGATCTNNNTCAAGAAGGCATAACAGCATCA
AGCACACGTCTGAACTCTTTCCCTACACGACGCTCTTCCGATCTNNNTGCTAATGTGGAGTTACAGG
AGCACACGTCTGAACTCTTTCCCTACACGACGCTCTTCCGATCTNNNTGCTAATGTGGAGTTACAGG
AGCACACGTCTGAACTCTTTCCCTACACGACGCTCTTCCGATCTNNNTGCTAATGTGGAGTTACAGG
AGCACACGTCTGAACTCTTTCCCTACACGACGCTCTTCCGATCTNNNTTTCATCATATAGGCCAGGCT
AGCACACGTCTGAACTCTTTCCCTACACGACGCTCTTCCGATCTNNNTTTCATCATATAGGCCAGGCT
AGCACACGTCTGAACTCTTTCCCTACACGACGCTCTTCCGATCTNNNTTTCATCATATAGGCCAGGCT
AGCACACGTCTGAACTCTTTCCCTACACGACGCTCTTCCGATCTNNNTGTTTTGCTGGGAAGGTTGTT
AGCACACGTCTGAACTCTTTCCCTACACGACGCTCTTCCGATCTNNNTGTTTTGCTGGGAAGGTTGTT
AGCACACGTCTGAACTCTTTCCCTACACGACGCTCTTCCGATCTNNNTGTTTTGCTGGGAAGGTTGTT
GAGCACACGTCTGAACTCTTTCCCTACACGACGCTCTTCCGATCTNNNGATCATGAGGTCAGGAGTT
GAGCACACGTCTGAACTCTTTCCCTACACGACGCTCTTCCGATCTNNNGATCATGAGGTCAGGAGTT
GAGCACACGTCTGAACTCTTTCCCTACACGACGCTCTTCCGATCTNNNGATCATGAGGTCAGGAGTT
GAGCACACGTCTGAACTCTTTCCCTACACGACGCTCTTCCGATCTNNNCAGACATGTGTGGAGTGCTT
GAGCACACGTCTGAACTCTTTCCCTACACGACGCTCTTCCGATCTNNNCAGACATGTGTGGAGTGCTT
GAGCACACGTCTGAACTCTTTCCCTACACGACGCTCTTCCGATCTNNNCAGACATGTGTGGAGTGCTT
AGCACACGTCTGAACTCTTTCCCTACACGACGCTCTTCCGATCTNNNTTATTCTATTGCCCAGAAATAG
AGCACACGTCTGAACTCTTTCCCTACACGACGCTCTTCCGATCTNNNTTATTCTATTGCCCAGAAATAG
AGCACACGTCTGAACTCTTTCCCTACACGACGCTCTTCCGATCTNNNTTATTCTATTGCCCAGAAATAG
AGCACACGTCTGAACTCTTTCCCTACACGACGCTCTTCCGATCTNNNGTCAACATGACAAAACCCCATCT
AGCACACGTCTGAACTCTTTCCCTACACGACGCTCTTCCGATCTNNNGTCAACATGACAAAACCCCATCT
AGCACACGTCTGAACTCTTTCCCTACACGACGCTCTTCCGATCTNNNGTCAACATGACAAAACCCCATCT
AGAGCACACGTCTGAACTCTTTCCCTACACGACGCTCTTCCGATCTNNNCTGGCGTAGTGGACTGGAA
AGAGCACACGTCTGAACTCTTTCCCTACACGACGCTCTTCCGATCTNNNCTGGCGTAGTGGACTGGAA
AGAGCACACGTCTGAACTCTTTCCCTACACGACGCTCTTCCGATCTNNNCTGGCGTAGTGGACTGGAA
AGCACACGTCTGAACTCTTTCCCTACACGACGCTCTTCCGATCTNNNAGAGAACTAAAACGCAGTCA
AGCACACGTCTGAACTCTTTCCCTACACGACGCTCTTCCGATCTNNNAGAGAACTAAAACGCAGTCA
AGCACACGTCTGAACTCTTTCCCTACACGACGCTCTTCCGATCTNNNAGAGAACTAAAACGCAGTCA
AGCACACGTCTGAACTCTTTCCCTACACGACGCTCTTCCGATCTNNNCGACAAAACGAGACACTCTGTC
AGCACACGTCTGAACTCTTTCCCTACACGACGCTCTTCCGATCTNNNCGACAAAACGAGACACTCTGTC
AGCACACGTCTGAACTCTTTCCCTACACGACGCTCTTCCGATCTNNNCGACAAAACGAGACACTCTGTC
GCACACGTCTGAACTCTTTCCCTACACGACGCTCTTCCGATCTNNNTGGCTATTTCAAAAAACAGGCA
GCACACGTCTGAACTCTTTCCCTACACGACGCTCTTCCGATCTNNNTGGCTATTTCAAAAAACAGGCA
GCACACGTCTGAACTCTTTCCCTACACGACGCTCTTCCGATCTNNNTGGCTATTTCAAAAAACAGGCA
CACACGTCTGAACTCTTTCCCTACACGACGCTCTTCCGATCTNNNTTCCTACGAAGGTCATGAAACAT
CACACGTCTGAACTCTTTCCCTACACGACGCTCTTCCGATCTNNNTTCCTACGAAGGTCATGAAACAT
CACACGTCTGAACTCTTTCCCTACACGACGCTCTTCCGATCTNNNTTCCTACGAAGGTCATGAAACAT
AGCACACGTCTGAACTCTTTCCCTACACGACGCTCTTCCGATCTNNNCAGGTCATAAGGCTTAGCAGCT

AGCACACGTCTGAACTCTTTCCCTACACGACGCTCTTCCGATCTNNNCAGGTCATAAGGCTTAGCAGCT
AGCACACGTCTGAACTCTTTCCCTACACGACGCTCTTCCGATCTNNNCAGGTCATAAGGCTTAGCAGCT
GAGCACACGTCTGAACTCTTTCCCTACACGACGCTCTTCCGATCTNNNAACCCACAGTTGCAATGGAAA
GAGCACACGTCTGAACTCTTTCCCTACACGACGCTCTTCCGATCTNNNAACCCACAGTTGCAATGGAAA
GAGCACACGTCTGAACTCTTTCCCTACACGACGCTCTTCCGATCTNNNAACCCACAGTTGCAATGGAAA
AGCACACGTCTGAACTCTTTCCCTACACGACGCTCTTCCGATCTNNNAAACCCTATCATCGCCTACTGTC
AGCACACGTCTGAACTCTTTCCCTACACGACGCTCTTCCGATCTNNNAAACCCTATCATCGCCTACTGTC
AGCACACGTCTGAACTCTTTCCCTACACGACGCTCTTCCGATCTNNNAAACCCTATCATCGCCTACTGTC
AGAGCACACGTCTGAACTCTTTCCCTACACGACGCTCTTCCGATCTNNNTCATGTTCCTAGAGCCAGGTA
AGAGCACACGTCTGAACTCTTTCCCTACACGACGCTCTTCCGATCTNNNTCATGTTCCTAGAGCCAGGTA
AGAGCACACGTCTGAACTCTTTCCCTACACGACGCTCTTCCGATCTNNNTCATGTTCCTAGAGCCAGGTA
AGAGCACACGTCTGAACTCTTTCCCTACACGACGCTCTTCCGATCTNNNTATTCAGAAGCTTCAAGGCG
AGAGCACACGTCTGAACTCTTTCCCTACACGACGCTCTTCCGATCTNNNTATTCAGAAGCTTCAAGGCG
AGAGCACACGTCTGAACTCTTTCCCTACACGACGCTCTTCCGATCTNNNTATTCAGAAGCTTCAAGGCG
GAGCACACGTCTGAACTCTTTCCCTACACGACGCTCTTCCGATCTNNNGAACGGAGGCCCAGAGAGAT
GAGCACACGTCTGAACTCTTTCCCTACACGACGCTCTTCCGATCTNNNGAACGGAGGCCCAGAGAGAT
GAGCACACGTCTGAACTCTTTCCCTACACGACGCTCTTCCGATCTNNNGAACGGAGGCCCAGAGAGAT
AGCACACGTCTGAACTCTTTCCCTACACGACGCTCTTCCGATCTNNNGCACTGGTAAAATAAATACTCT
AGCACACGTCTGAACTCTTTCCCTACACGACGCTCTTCCGATCTNNNGCACTGGTAAAATAAATACTCT
AGCACACGTCTGAACTCTTTCCCTACACGACGCTCTTCCGATCTNNNGCACTGGTAAAATAAATACTCT
GAGCACACGTCTGAACTCTTTCCCTACACGACGCTCTTCCGATCTNNNTTTAGTAGAGATGGGGTTTCC
GAGCACACGTCTGAACTCTTTCCCTACACGACGCTCTTCCGATCTNNNTTTAGTAGAGATGGGGTTTCC
GAGCACACGTCTGAACTCTTTCCCTACACGACGCTCTTCCGATCTNNNTTTAGTAGAGATGGGGTTTCC
GAGCACACGTCTGAACTCTTTCCCTACACGACGCTCTTCCGATCTNNNAGATTCATGGGTTTTTGACAG
GAGCACACGTCTGAACTCTTTCCCTACACGACGCTCTTCCGATCTNNNAGATTCATGGGTTTTTGACAG
GAGCACACGTCTGAACTCTTTCCCTACACGACGCTCTTCCGATCTNNNAGATTCATGGGTTTTTGACAG
AGCACACGTCTGAACTCTTTCCCTACACGACGCTCTTCCGATCTNNNTGCAGTTACTTCCTCCACAGAAG
AGCACACGTCTGAACTCTTTCCCTACACGACGCTCTTCCGATCTNNNTGCAGTTACTTCCTCCACAGAAG
AGCACACGTCTGAACTCTTTCCCTACACGACGCTCTTCCGATCTNNNTGCAGTTACTTCCTCCACAGAAG
AGCACACGTCTGAACTCTTTCCCTACACGACGCTCTTCCGATCTNNNAAAAATTAGCCGGGCATGGTGC
AGCACACGTCTGAACTCTTTCCCTACACGACGCTCTTCCGATCTNNNAAAAATTAGCCGGGCATGGTGC
AGCACACGTCTGAACTCTTTCCCTACACGACGCTCTTCCGATCTNNNAAAAATTAGCCGGGCATGGTGC
GCACACGTCTGAACTCTTTCCCTACACGACGCTCTTCCGATCTNNNGGAATCTGCCCTGTGTCGTCCATC
GCACACGTCTGAACTCTTTCCCTACACGACGCTCTTCCGATCTNNNGGAATCTGCCCTGTGTCGTCCATC
GCACACGTCTGAACTCTTTCCCTACACGACGCTCTTCCGATCTNNNGGAATCTGCCCTGTGTCGTCCATC
GAGCACACGTCTGAACTCTTTCCCTACACGACGCTCTTCCGATCTNNNGATGCTACGGACCTTACGTCA
GAGCACACGTCTGAACTCTTTCCCTACACGACGCTCTTCCGATCTNNNGATGCTACGGACCTTACGTCA
GAGCACACGTCTGAACTCTTTCCCTACACGACGCTCTTCCGATCTNNNGATGCTACGGACCTTACGTCA
GCACACGTCTGAACTCTTTCCCTACACGACGCTCTTCCGATCTNNNTCCTCTGCAGTCCAATACTCCTACA
GCACACGTCTGAACTCTTTCCCTACACGACGCTCTTCCGATCTNNNTCCTCTGCAGTCCAATACTCCTACA
GCACACGTCTGAACTCTTTCCCTACACGACGCTCTTCCGATCTNNNTCCTCTGCAGTCCAATACTCCTACA
GAGCACACGTCTGAACTCTTTCCCTACACGACGCTCTTCCGATCTNNNCCTTCTGTCCTGTCAGCTAACAT
GAGCACACGTCTGAACTCTTTCCCTACACGACGCTCTTCCGATCTNNNCCTTCTGTCCTGTCAGCTAACAT

AGCACACGTCTGAACTCTTTCCCTACACGACGCTCTTCCGATCTNNNCCTTCTGTCCTGTCAGCTAACA
ACACGTCTGAACTCTTTCCCTACACGACGCTCTTCCGATCTNNNGTGTGCGTTGTCAGGACTTTGCAT
ACACGTCTGAACTCTTTCCCTACACGACGCTCTTCCGATCTNNNGTGTGCGTTGTCAGGACTTTGCAT
ACACGTCTGAACTCTTTCCCTACACGACGCTCTTCCGATCTNNNGTGTGCGTTGTCAGGACTTTGCAT
GCACACGTCTGAACTCTTTCCCTACACGACGCTCTTCCGATCTNNNTGATTCACATACCATAGGTCCCA
GCACACGTCTGAACTCTTTCCCTACACGACGCTCTTCCGATCTNNNTGATTCACATACCATAGGTCCCA
GCACACGTCTGAACTCTTTCCCTACACGACGCTCTTCCGATCTNNNTGATTCACATACCATAGGTCCCA
GAGCACACGTCTGAACTCTTTCCCTACACGACGCTCTTCCGATCTNNNTATGTGCCTGGCCTTTTTTTC
GAGCACACGTCTGAACTCTTTCCCTACACGACGCTCTTCCGATCTNNNTATGTGCCTGGCCTTTTTTTC
GAGCACACGTCTGAACTCTTTCCCTACACGACGCTCTTCCGATCTNNNTATGTGCCTGGCCTTTTTTTC
AGCACACGTCTGAACTCTTTCCCTACACGACGCTCTTCCGATCTNNNTCAGAGCAGAGAAACTTCAG
AGCACACGTCTGAACTCTTTCCCTACACGACGCTCTTCCGATCTNNNTCAGAGCAGAGAAACTTCAG
AGCACACGTCTGAACTCTTTCCCTACACGACGCTCTTCCGATCTNNNTCAGAGCAGAGAAACTTCAG
AAGAGCACACGTCTGAACTCTTTCCCTACACGACGCTCTTCCGATCTNNNGGTTGCAGTGAGTCGAG
AAGAGCACACGTCTGAACTCTTTCCCTACACGACGCTCTTCCGATCTNNNGGTTGCAGTGAGTCGAG
AAGAGCACACGTCTGAACTCTTTCCCTACACGACGCTCTTCCGATCTNNNGGTTGCAGTGAGTCGAG
CACACGTCTGAACTCTTTCCCTACACGACGCTCTTCCGATCTNNNCATCCTGGCTAACACAGTGAAACC
CACACGTCTGAACTCTTTCCCTACACGACGCTCTTCCGATCTNNNCATCCTGGCTAACACAGTGAAACC
CACACGTCTGAACTCTTTCCCTACACGACGCTCTTCCGATCTNNNCATCCTGGCTAACACAGTGAAACC
AGCACACGTCTGAACTCTTTCCCTACACGACGCTCTTCCGATCTNNNCTACACAAAACACCCTTTGACC
AGCACACGTCTGAACTCTTTCCCTACACGACGCTCTTCCGATCTNNNCTACACAAAACACCCTTTGACC
AGCACACGTCTGAACTCTTTCCCTACACGACGCTCTTCCGATCTNNNCTACACAAAACACCCTTTGACC
AGCACACGTCTGAACTCTTTCCCTACACGACGCTCTTCCGATCTNNNTCTCTCTCTCTCTGTGTGTGT
AGCACACGTCTGAACTCTTTCCCTACACGACGCTCTTCCGATCTNNNTCTCTCTCTCTCTGTGTGTGT
AGCACACGTCTGAACTCTTTCCCTACACGACGCTCTTCCGATCTNNNTCTCTCTCTCTCTGTGTGTGT
GAGCACACGTCTGAACTCTTTCCCTACACGACGCTCTTCCGATCTNNNTTTCTCTCATGTGTCCTTCCCA
GAGCACACGTCTGAACTCTTTCCCTACACGACGCTCTTCCGATCTNNNTTTCTCTCATGTGTCCTTCCCA
GAGCACACGTCTGAACTCTTTCCCTACACGACGCTCTTCCGATCTNNNTTTCTCTCATGTGTCCTTCCCA
AGCACACGTCTGAACTCTTTCCCTACACGACGCTCTTCCGATCTNNNTTGATCATTTGCATGGATGTG
AGCACACGTCTGAACTCTTTCCCTACACGACGCTCTTCCGATCTNNNTTGATCATTTGCATGGATGTG
AGCACACGTCTGAACTCTTTCCCTACACGACGCTCTTCCGATCTNNNTTGATCATTTGCATGGATGTG
AGCACACGTCTGAACTCTTTCCCTACACGACGCTCTTCCGATCTNNNGCTTGTTGGGCACATGAATGT
AGCACACGTCTGAACTCTTTCCCTACACGACGCTCTTCCGATCTNNNGCTTGTTGGGCACATGAATGT
AGCACACGTCTGAACTCTTTCCCTACACGACGCTCTTCCGATCTNNNGCTTGTTGGGCACATGAATGT
GAGCACACGTCTGAACTCTTTCCCTACACGACGCTCTTCCGATCTNNNTTCCTGTAGTCCCAGCTACTC
GAGCACACGTCTGAACTCTTTCCCTACACGACGCTCTTCCGATCTNNNTTCCTGTAGTCCCAGCTACTC
GAGCACACGTCTGAACTCTTTCCCTACACGACGCTCTTCCGATCTNNNTTCCTGTAGTCCCAGCTACTC
AGCACACGTCTGAACTCTTTCCCTACACGACGCTCTTCCGATCTNNNCCAACCAAGGGACAACATAATA
AGCACACGTCTGAACTCTTTCCCTACACGACGCTCTTCCGATCTNNNCCAACCAAGGGACAACATAATA
AGCACACGTCTGAACTCTTTCCCTACACGACGCTCTTCCGATCTNNNCCAACCAAGGGACAACATAATA
AGCACACGTCTGAACTCTTTCCCTACACGACGCTCTTCCGATCTNNNATCATTTGAACTCAGGAGGCA
AGCACACGTCTGAACTCTTTCCCTACACGACGCTCTTCCGATCTNNNATCATTTGAACTCAGGAGGCA
AGCACACGTCTGAACTCTTTCCCTACACGACGCTCTTCCGATCTNNNATCATTTGAACTCAGGAGGCA

AGAGCACACGTCTGAACTCTTTCCCTACACGACGCTCTTCCGATCTNNNGCCTGTAGTCCCAGCTACTT
AGAGCACACGTCTGAACTCTTTCCCTACACGACGCTCTTCCGATCTNNNGCCTGTAGTCCCAGCTACTT
AGAGCACACGTCTGAACTCTTTCCCTACACGACGCTCTTCCGATCTNNNGCCTGTAGTCCCAGCTACTT
AGCACACGTCTGAACTCTTTCCCTACACGACGCTCTTCCGATCTNNNAAGGGACGAGAATCACCTGAA
AGCACACGTCTGAACTCTTTCCCTACACGACGCTCTTCCGATCTNNNAAGGGACGAGAATCACCTGAA
AGCACACGTCTGAACTCTTTCCCTACACGACGCTCTTCCGATCTNNNAAGGGACGAGAATCACCTGAA
AGCACACGTCTGAACTCTTTCCCTACACGACGCTCTTCCGATCTNNNGCTGGAACATTTCTTGTAGTGC
AGCACACGTCTGAACTCTTTCCCTACACGACGCTCTTCCGATCTNNNGCTGGAACATTTCTTGTAGTGC
AGCACACGTCTGAACTCTTTCCCTACACGACGCTCTTCCGATCTNNNGCTGGAACATTTCTTGTAGTGC
GAGCACACGTCTGAACTCTTTCCCTACACGACGCTCTTCCGATCTNNNAATATGGTCATCACGTGGAAG
GAGCACACGTCTGAACTCTTTCCCTACACGACGCTCTTCCGATCTNNNAATATGGTCATCACGTGGAAG
GAGCACACGTCTGAACTCTTTCCCTACACGACGCTCTTCCGATCTNNNAATATGGTCATCACGTGGAAG
AGCACACGTCTGAACTCTTTCCCTACACGACGCTCTTCCGATCTNNNTTTACCATGTCCCACAGCAAAAA
AGCACACGTCTGAACTCTTTCCCTACACGACGCTCTTCCGATCTNNNTTTACCATGTCCCACAGCAAAAA
AGCACACGTCTGAACTCTTTCCCTACACGACGCTCTTCCGATCTNNNTTTACCATGTCCCACAGCAAAAA
AGCACACGTCTGAACTCTTTCCCTACACGACGCTCTTCCGATCTNNNTGCGATCTCGGCTTACTGTAAG
AGCACACGTCTGAACTCTTTCCCTACACGACGCTCTTCCGATCTNNNTGCGATCTCGGCTTACTGTAAG
AGCACACGTCTGAACTCTTTCCCTACACGACGCTCTTCCGATCTNNNTGCGATCTCGGCTTACTGTAAG
AGCACACGTCTGAACTCTTTCCCTACACGACGCTCTTCCGATCTNNNTAGGGAGAGAGATGAGTGCTT
AGCACACGTCTGAACTCTTTCCCTACACGACGCTCTTCCGATCTNNNTAGGGAGAGAGATGAGTGCTT
AGCACACGTCTGAACTCTTTCCCTACACGACGCTCTTCCGATCTNNNTAGGGAGAGAGATGAGTGCTT
AGAGCACACGTCTGAACTCTTTCCCTACACGACGCTCTTCCGATCTNNNCTCAAGCAGAATGTATGCCA
AGAGCACACGTCTGAACTCTTTCCCTACACGACGCTCTTCCGATCTNNNCTCAAGCAGAATGTATGCCA
AGAGCACACGTCTGAACTCTTTCCCTACACGACGCTCTTCCGATCTNNNCTCAAGCAGAATGTATGCCA
AGCACACGTCTGAACTCTTTCCCTACACGACGCTCTTCCGATCTNNNACAATGATGGTCGCTTTCCTCTC
AGCACACGTCTGAACTCTTTCCCTACACGACGCTCTTCCGATCTNNNACAATGATGGTCGCTTTCCTCTC
AGCACACGTCTGAACTCTTTCCCTACACGACGCTCTTCCGATCTNNNACAATGATGGTCGCTTTCCTCTC
AGCACACGTCTGAACTCTTTCCCTACACGACGCTCTTCCGATCTNNNCCATGAGGGAGATTCTGAGGA
AGCACACGTCTGAACTCTTTCCCTACACGACGCTCTTCCGATCTNNNCCATGAGGGAGATTCTGAGGA
AGCACACGTCTGAACTCTTTCCCTACACGACGCTCTTCCGATCTNNNCCATGAGGGAGATTCTGAGGA
AGCACACGTCTGAACTCTTTCCCTACACGACGCTCTTCCGATCTNNNGGAAGGAGGATGGTTGATGTG
AGCACACGTCTGAACTCTTTCCCTACACGACGCTCTTCCGATCTNNNGGAAGGAGGATGGTTGATGTG
AGCACACGTCTGAACTCTTTCCCTACACGACGCTCTTCCGATCTNNNGGAAGGAGGATGGTTGATGTG
AGCACACGTCTGAACTCTTTCCCTACACGACGCTCTTCCGATCTNNNGTTGCAGTGAGCTGAGATTGT
AGCACACGTCTGAACTCTTTCCCTACACGACGCTCTTCCGATCTNNNGTTGCAGTGAGCTGAGATTGT
AGCACACGTCTGAACTCTTTCCCTACACGACGCTCTTCCGATCTNNNGTTGCAGTGAGCTGAGATTGT
AGCACACGTCTGAACTCTTTCCCTACACGACGCTCTTCCGATCTNNNAGGGTCTGTGTGCATTGATGAT
AGCACACGTCTGAACTCTTTCCCTACACGACGCTCTTCCGATCTNNNAGGGTCTGTGTGCATTGATGAT
AGCACACGTCTGAACTCTTTCCCTACACGACGCTCTTCCGATCTNNNAGGGTCTGTGTGCATTGATGAT
AGCACACGTCTGAACTCTTTCCCTACACGACGCTCTTCCGATCTNNNATGATGGGGATTTGAAAGTGC
AGCACACGTCTGAACTCTTTCCCTACACGACGCTCTTCCGATCTNNNATGATGGGGATTTGAAAGTGC
AGCACACGTCTGAACTCTTTCCCTACACGACGCTCTTCCGATCTNNNATGATGGGGATTTGAAAGTGC
GAGCACACGTCTGAACTCTTTCCCTACACGACGCTCTTCCGATCTNNNGAGAGGTGGAGATTGCAGTG

AGCACACGTCTGAACTCTTTCCCTACACGACGCTCTTCCGATCTNNNGAGAGGTGGAGATTGCAGTG
AGCACACGTCTGAACTCTTTCCCTACACGACGCTCTTCCGATCTNNNGAGAGGTGGAGATTGCAGTG
AGAGCACACGTCTGAACTCTTTCCCTACACGACGCTCTTCCGATCTNNNAACACCCAATATGTGCAGGT
AGAGCACACGTCTGAACTCTTTCCCTACACGACGCTCTTCCGATCTNNNAACACCCAATATGTGCAGGT
AGAGCACACGTCTGAACTCTTTCCCTACACGACGCTCTTCCGATCTNNNAACACCCAATATGTGCAGGT
AGCACACGTCTGAACTCTTTCCCTACACGACGCTCTTCCGATCTNNNTTTACATAAGTCGGCTGACCCA
AGCACACGTCTGAACTCTTTCCCTACACGACGCTCTTCCGATCTNNNTTTACATAAGTCGGCTGACCCA
AGCACACGTCTGAACTCTTTCCCTACACGACGCTCTTCCGATCTNNNTTTACATAAGTCGGCTGACCCA
GAGCACACGTCTGAACTCTTTCCCTACACGACGCTCTTCCGATCTNNNAAAAAATTAGCCAGGCGTG
GAGCACACGTCTGAACTCTTTCCCTACACGACGCTCTTCCGATCTNNNAAAAAATTAGCCAGGCGTG
GAGCACACGTCTGAACTCTTTCCCTACACGACGCTCTTCCGATCTNNNAAAAAATTAGCCAGGCGTG
AGCACACGTCTGAACTCTTTCCCTACACGACGCTCTTCCGATCTNNNCATGGCTGCTGTTGCTTTTTTTT
AGCACACGTCTGAACTCTTTCCCTACACGACGCTCTTCCGATCTNNNCATGGCTGCTGTTGCTTTTTTTT
AGCACACGTCTGAACTCTTTCCCTACACGACGCTCTTCCGATCTNNNCATGGCTGCTGTTGCTTTTTTTT
AGCACACGTCTGAACTCTTTCCCTACACGACGCTCTTCCGATCTNNNATTCAGATCATCCCTGGCCAAAT
AGCACACGTCTGAACTCTTTCCCTACACGACGCTCTTCCGATCTNNNATTCAGATCATCCCTGGCCAAAT
AGCACACGTCTGAACTCTTTCCCTACACGACGCTCTTCCGATCTNNNATTCAGATCATCCCTGGCCAAAT
CACACGTCTGAACTCTTTCCCTACACGACGCTCTTCCGATCTNNNCTTTGGTCCCCTTTTCCTTTTCCCAT
CACACGTCTGAACTCTTTCCCTACACGACGCTCTTCCGATCTNNNCTTTGGTCCCCTTTTCCTTTTCCCAT
CACACGTCTGAACTCTTTCCCTACACGACGCTCTTCCGATCTNNNCTTTGGTCCCCTTTTCCTTTTCCCAT
GAGCACACGTCTGAACTCTTTCCCTACACGACGCTCTTCCGATCTNNNGTGATGTACACCTGTAGCCCT
GAGCACACGTCTGAACTCTTTCCCTACACGACGCTCTTCCGATCTNNNGTGATGTACACCTGTAGCCCT
GAGCACACGTCTGAACTCTTTCCCTACACGACGCTCTTCCGATCTNNNGTGATGTACACCTGTAGCCCT
AGCACACGTCTGAACTCTTTCCCTACACGACGCTCTTCCGATCTNNNTAAAGCCCAGTGAGCATATTC
AGCACACGTCTGAACTCTTTCCCTACACGACGCTCTTCCGATCTNNNTAAAGCCCAGTGAGCATATTC
AGCACACGTCTGAACTCTTTCCCTACACGACGCTCTTCCGATCTNNNTAAAGCCCAGTGAGCATATTC
AGCACACGTCTGAACTCTTTCCCTACACGACGCTCTTCCGATCTNNNTTTCTGCCAGAAAGGAAAACG
AGCACACGTCTGAACTCTTTCCCTACACGACGCTCTTCCGATCTNNNTTTCTGCCAGAAAGGAAAACG
AGCACACGTCTGAACTCTTTCCCTACACGACGCTCTTCCGATCTNNNTTTCTGCCAGAAAGGAAAACG
GCACACGTCTGAACTCTTTCCCTACACGACGCTCTTCCGATCTNNNATAGTTGGTGGAGGGGATAAG
GCACACGTCTGAACTCTTTCCCTACACGACGCTCTTCCGATCTNNNATAGTTGGTGGAGGGGATAAG
GCACACGTCTGAACTCTTTCCCTACACGACGCTCTTCCGATCTNNNATAGTTGGTGGAGGGGATAAG
AGCACACGTCTGAACTCTTTCCCTACACGACGCTCTTCCGATCTNNNTTGGGAGCTCCTAGGATAGCA
AGCACACGTCTGAACTCTTTCCCTACACGACGCTCTTCCGATCTNNNTTGGGAGCTCCTAGGATAGCA
AGCACACGTCTGAACTCTTTCCCTACACGACGCTCTTCCGATCTNNNTTGGGAGCTCCTAGGATAGCA
GAGCACACGTCTGAACTCTTTCCCTACACGACGCTCTTCCGATCTNNNGGTCTGTGTGATGTTTTTCCC
GAGCACACGTCTGAACTCTTTCCCTACACGACGCTCTTCCGATCTNNNGGTCTGTGTGATGTTTTTCCC
GAGCACACGTCTGAACTCTTTCCCTACACGACGCTCTTCCGATCTNNNGGTCTGTGTGATGTTTTTCCC
AGCACACGTCTGAACTCTTTCCCTACACGACGCTCTTCCGATCTNNNCTGAAGTACAGTGATGCAATG
AGCACACGTCTGAACTCTTTCCCTACACGACGCTCTTCCGATCTNNNCTGAAGTACAGTGATGCAATG
AGCACACGTCTGAACTCTTTCCCTACACGACGCTCTTCCGATCTNNNCTGAAGTACAGTGATGCAATG
GCACACGTCTGAACTCTTTCCCTACACGACGCTCTTCCGATCTNNNGAACTCCTAGCCTCAAGTGATCC
GCACACGTCTGAACTCTTTCCCTACACGACGCTCTTCCGATCTNNNGAACTCCTAGCCTCAAGTGATCC

GCACACGTCTGAACTCTTTCCCTACACGACGCTCTTCCGATCTNNNGAACTCCTAGCCTCAAGTGATCC
GAGCACACGTCTGAACTCTTTCCCTACACGACGCTCTTCCGATCTNNNACTAAATGCATTTTCCCTTTCC
GAGCACACGTCTGAACTCTTTCCCTACACGACGCTCTTCCGATCTNNNACTAAATGCATTTTCCCTTTCC
GAGCACACGTCTGAACTCTTTCCCTACACGACGCTCTTCCGATCTNNNACTAAATGCATTTTCCCTTTCC
AGCACACGTCTGAACTCTTTCCCTACACGACGCTCTTCCGATCTNNNTGAGGGTTTTGAGTGAGAGT
AGCACACGTCTGAACTCTTTCCCTACACGACGCTCTTCCGATCTNNNTGAGGGTTTTGAGTGAGAGT
AGCACACGTCTGAACTCTTTCCCTACACGACGCTCTTCCGATCTNNNTGAGGGTTTTGAGTGAGAGT
CACACGTCTGAACTCTTTCCCTACACGACGCTCTTCCGATCTNNNGAGTTGTGGCTCCCTTCTCTCCAT
CACACGTCTGAACTCTTTCCCTACACGACGCTCTTCCGATCTNNNGAGTTGTGGCTCCCTTCTCTCCAT
CACACGTCTGAACTCTTTCCCTACACGACGCTCTTCCGATCTNNNGAGTTGTGGCTCCCTTCTCTCCAT
GAGCACACGTCTGAACTCTTTCCCTACACGACGCTCTTCCGATCTNNNACAAACAAGTCACTCACGTTT
GAGCACACGTCTGAACTCTTTCCCTACACGACGCTCTTCCGATCTNNNACAAACAAGTCACTCACGTTT
GAGCACACGTCTGAACTCTTTCCCTACACGACGCTCTTCCGATCTNNNACAAACAAGTCACTCACGTTT
AGCACACGTCTGAACTCTTTCCCTACACGACGCTCTTCCGATCTNNNGCAATTGCTGGGTTGTATGGT
AGCACACGTCTGAACTCTTTCCCTACACGACGCTCTTCCGATCTNNNGCAATTGCTGGGTTGTATGGT
AGCACACGTCTGAACTCTTTCCCTACACGACGCTCTTCCGATCTNNNGCAATTGCTGGGTTGTATGGT
AGCACACGTCTGAACTCTTTCCCTACACGACGCTCTTCCGATCTNNNCTTCTAACAGACAGGACCCTCA
AGCACACGTCTGAACTCTTTCCCTACACGACGCTCTTCCGATCTNNNCTTCTAACAGACAGGACCCTCA
AGCACACGTCTGAACTCTTTCCCTACACGACGCTCTTCCGATCTNNNCTTCTAACAGACAGGACCCTCA
GAGCACACGTCTGAACTCTTTCCCTACACGACGCTCTTCCGATCTNNNGGTGACAGAGCTAGACTGCA
GAGCACACGTCTGAACTCTTTCCCTACACGACGCTCTTCCGATCTNNNGGTGACAGAGCTAGACTGCA
GAGCACACGTCTGAACTCTTTCCCTACACGACGCTCTTCCGATCTNNNGGTGACAGAGCTAGACTGCA
AAGAGCACACGTCTGAACTCTTTCCCTACACGACGCTCTTCCGATCTNNNACTTGGGATTACTTCTCCT
AAGAGCACACGTCTGAACTCTTTCCCTACACGACGCTCTTCCGATCTNNNACTTGGGATTACTTCTCCT
AAGAGCACACGTCTGAACTCTTTCCCTACACGACGCTCTTCCGATCTNNNACTTGGGATTACTTCTCCT
AGCACACGTCTGAACTCTTTCCCTACACGACGCTCTTCCGATCTNNNAGAATCGTCTGATTGCTGGAG
AGCACACGTCTGAACTCTTTCCCTACACGACGCTCTTCCGATCTNNNAGAATCGTCTGATTGCTGGAG
AGCACACGTCTGAACTCTTTCCCTACACGACGCTCTTCCGATCTNNNAGAATCGTCTGATTGCTGGAG
AGCACACGTCTGAACTCTTTCCCTACACGACGCTCTTCCGATCTNNNGAGATGGAGTTTCGCTCTTGT
AGCACACGTCTGAACTCTTTCCCTACACGACGCTCTTCCGATCTNNNGAGATGGAGTTTCGCTCTTGT
AGCACACGTCTGAACTCTTTCCCTACACGACGCTCTTCCGATCTNNNGAGATGGAGTTTCGCTCTTGT
GAGCACACGTCTGAACTCTTTCCCTACACGACGCTCTTCCGATCTNNNGAGATCATGCCACTACACTCC
GAGCACACGTCTGAACTCTTTCCCTACACGACGCTCTTCCGATCTNNNGAGATCATGCCACTACACTCC
GAGCACACGTCTGAACTCTTTCCCTACACGACGCTCTTCCGATCTNNNGAGATCATGCCACTACACTCC
AGCACACGTCTGAACTCTTTCCCTACACGACGCTCTTCCGATCTNNNGGAGGAACTACTAATTGCAC
AGCACACGTCTGAACTCTTTCCCTACACGACGCTCTTCCGATCTNNNGGAGGAACTACTAATTGCAC
AGCACACGTCTGAACTCTTTCCCTACACGACGCTCTTCCGATCTNNNGGAGGAACTACTAATTGCAC
GCACACGTCTGAACTCTTTCCCTACACGACGCTCTTCCGATCTNNNCACCTCGGCTTTCACTGGAGACA
GCACACGTCTGAACTCTTTCCCTACACGACGCTCTTCCGATCTNNNCACCTCGGCTTTCACTGGAGACA
GCACACGTCTGAACTCTTTCCCTACACGACGCTCTTCCGATCTNNNCACCTCGGCTTTCACTGGAGACA
GAGCACACGTCTGAACTCTTTCCCTACACGACGCTCTTCCGATCTNNNTCCCAAACTGATTTGGCCAT
GAGCACACGTCTGAACTCTTTCCCTACACGACGCTCTTCCGATCTNNNTCCCAAACTGATTTGGCCAT
GAGCACACGTCTGAACTCTTTCCCTACACGACGCTCTTCCGATCTNNNTCCCAAACTGATTTGGCCAT

GAGCACACGTCTGAACTCTTTCCCTACACGACGCTCTTCCGATCTNNNGGCTCTACCTGGTGTGCATAT
GAGCACACGTCTGAACTCTTTCCCTACACGACGCTCTTCCGATCTNNNGGCTCTACCTGGTGTGCATAT
GAGCACACGTCTGAACTCTTTCCCTACACGACGCTCTTCCGATCTNNNGGCTCTACCTGGTGTGCATAT
GAGCACACGTCTGAACTCTTTCCCTACACGACGCTCTTCCGATCTNNNATGGCTGTGTATATCTCTGCCT
GAGCACACGTCTGAACTCTTTCCCTACACGACGCTCTTCCGATCTNNNATGGCTGTGTATATCTCTGCCT
GAGCACACGTCTGAACTCTTTCCCTACACGACGCTCTTCCGATCTNNNATGGCTGTGTATATCTCTGCCT
AGCACACGTCTGAACTCTTTCCCTACACGACGCTCTTCCGATCTNNNGAGACACATGCTAGCGCGTCCAT
AGCACACGTCTGAACTCTTTCCCTACACGACGCTCTTCCGATCTNNNGAGACACATGCTAGCGCGTCCAT
AGCACACGTCTGAACTCTTTCCCTACACGACGCTCTTCCGATCTNNNGAGACACATGCTAGCGCGTCCAT
AGCACACGTCTGAACTCTTTCCCTACACGACGCTCTTCCGATCTNNNCAGATCACTTGAGGTCAGGAG
AGCACACGTCTGAACTCTTTCCCTACACGACGCTCTTCCGATCTNNNCAGATCACTTGAGGTCAGGAG
AGCACACGTCTGAACTCTTTCCCTACACGACGCTCTTCCGATCTNNNCAGATCACTTGAGGTCAGGAG
GAGCACACGTCTGAACTCTTTCCCTACACGACGCTCTTCCGATCTNNNGATGTAGAGCGAGAGGAAGA
GAGCACACGTCTGAACTCTTTCCCTACACGACGCTCTTCCGATCTNNNGATGTAGAGCGAGAGGAAGA
GAGCACACGTCTGAACTCTTTCCCTACACGACGCTCTTCCGATCTNNNGATGTAGAGCGAGAGGAAGA
AGCACACGTCTGAACTCTTTCCCTACACGACGCTCTTCCGATCTNNNAGTGAAAGGTTTCTTAGCAAC
AGCACACGTCTGAACTCTTTCCCTACACGACGCTCTTCCGATCTNNNAGTGAAAGGTTTCTTAGCAAC
AGCACACGTCTGAACTCTTTCCCTACACGACGCTCTTCCGATCTNNNAGTGAAAGGTTTCTTAGCAAC
AGCACACGTCTGAACTCTTTCCCTACACGACGCTCTTCCGATCTNNNCTGAGCCCTCCATCCATTTTAGT
AGCACACGTCTGAACTCTTTCCCTACACGACGCTCTTCCGATCTNNNCTGAGCCCTCCATCCATTTTAGT
AGCACACGTCTGAACTCTTTCCCTACACGACGCTCTTCCGATCTNNNCTGAGCCCTCCATCCATTTTAGT
AGCACACGTCTGAACTCTTTCCCTACACGACGCTCTTCCGATCTNNNCGGCTTTTGTCATTCTCCATGTT
AGCACACGTCTGAACTCTTTCCCTACACGACGCTCTTCCGATCTNNNCGGCTTTTGTCATTCTCCATGTT
AGCACACGTCTGAACTCTTTCCCTACACGACGCTCTTCCGATCTNNNCGGCTTTTGTCATTCTCCATGTT
GAGCACACGTCTGAACTCTTTCCCTACACGACGCTCTTCCGATCTNNNTATGAAGATGACACTGGTAAT
GAGCACACGTCTGAACTCTTTCCCTACACGACGCTCTTCCGATCTNNNTATGAAGATGACACTGGTAAT
GAGCACACGTCTGAACTCTTTCCCTACACGACGCTCTTCCGATCTNNNTATGAAGATGACACTGGTAAT
AGCACACGTCTGAACTCTTTCCCTACACGACGCTCTTCCGATCTNNNCGCCTGTAATCCTAGCTACTCT
AGCACACGTCTGAACTCTTTCCCTACACGACGCTCTTCCGATCTNNNCGCCTGTAATCCTAGCTACTCT
AGCACACGTCTGAACTCTTTCCCTACACGACGCTCTTCCGATCTNNNCGCCTGTAATCCTAGCTACTCT
GAGCACACGTCTGAACTCTTTCCCTACACGACGCTCTTCCGATCTNNNCGTCGCTGGGAGGAATTTCA
GAGCACACGTCTGAACTCTTTCCCTACACGACGCTCTTCCGATCTNNNCGTCGCTGGGAGGAATTTCA
GAGCACACGTCTGAACTCTTTCCCTACACGACGCTCTTCCGATCTNNNCGTCGCTGGGAGGAATTTCA
AGCACACGTCTGAACTCTTTCCCTACACGACGCTCTTCCGATCTNNNTGGTGTTACTCCAGACAATGA
AGCACACGTCTGAACTCTTTCCCTACACGACGCTCTTCCGATCTNNNTGGTGTTACTCCAGACAATGA
AGCACACGTCTGAACTCTTTCCCTACACGACGCTCTTCCGATCTNNNTGGTGTTACTCCAGACAATGA
GAGCACACGTCTGAACTCTTTCCCTACACGACGCTCTTCCGATCTNNNCTAAGGCAGGAGAATGGTGT
GAGCACACGTCTGAACTCTTTCCCTACACGACGCTCTTCCGATCTNNNCTAAGGCAGGAGAATGGTGT
GAGCACACGTCTGAACTCTTTCCCTACACGACGCTCTTCCGATCTNNNCTAAGGCAGGAGAATGGTGT
GAGCACACGTCTGAACTCTTTCCCTACACGACGCTCTTCCGATCTNNNGAGGCCCTCTAAACTTGGAGA
GAGCACACGTCTGAACTCTTTCCCTACACGACGCTCTTCCGATCTNNNGAGGCCCTCTAAACTTGGAGA
GAGCACACGTCTGAACTCTTTCCCTACACGACGCTCTTCCGATCTNNNGAGGCCCTCTAAACTTGGAGA
AGCACACGTCTGAACTCTTTCCCTACACGACGCTCTTCCGATCTNNNGAAGCCTGACTACCACCATTGAA

GCACACGTCTGAACTCTTTCCCTACACGACGCTCTTCCGATCTNNNGAAGCCTGACTACCACCATTGAA
GCACACGTCTGAACTCTTTCCCTACACGACGCTCTTCCGATCTNNNGAAGCCTGACTACCACCATTGAA
GAGCACACGTCTGAACTCTTTCCCTACACGACGCTCTTCCGATCTNNNGATGCAGGAGAATTGCTTGAT
GAGCACACGTCTGAACTCTTTCCCTACACGACGCTCTTCCGATCTNNNGATGCAGGAGAATTGCTTGAT
GAGCACACGTCTGAACTCTTTCCCTACACGACGCTCTTCCGATCTNNNGATGCAGGAGAATTGCTTGAT
AGAGCACACGTCTGAACTCTTTCCCTACACGACGCTCTTCCGATCTNNNGAGATAGGGACAGGGTGG
AGAGCACACGTCTGAACTCTTTCCCTACACGACGCTCTTCCGATCTNNNGAGATAGGGACAGGGTGG
AGAGCACACGTCTGAACTCTTTCCCTACACGACGCTCTTCCGATCTNNNGAGATAGGGACAGGGTGG
AGAGCACACGTCTGAACTCTTTCCCTACACGACGCTCTTCCGATCTNNNGAGGCAGGAAAATCACTTGA
AGAGCACACGTCTGAACTCTTTCCCTACACGACGCTCTTCCGATCTNNNGAGGCAGGAAAATCACTTGA
AGAGCACACGTCTGAACTCTTTCCCTACACGACGCTCTTCCGATCTNNNGAGGCAGGAAAATCACTTGA
GCACACGTCTGAACTCTTTCCCTACACGACGCTCTTCCGATCTNNNCTGAGAAAAGCTGGGTTTGGTA
GCACACGTCTGAACTCTTTCCCTACACGACGCTCTTCCGATCTNNNCTGAGAAAAGCTGGGTTTGGTA
GCACACGTCTGAACTCTTTCCCTACACGACGCTCTTCCGATCTNNNCTGAGAAAAGCTGGGTTTGGTA
AGCACACGTCTGAACTCTTTCCCTACACGACGCTCTTCCGATCTNNNCAAGCTCCTATCAGGGTTCACA
AGCACACGTCTGAACTCTTTCCCTACACGACGCTCTTCCGATCTNNNCAAGCTCCTATCAGGGTTCACA
AGCACACGTCTGAACTCTTTCCCTACACGACGCTCTTCCGATCTNNNCAAGCTCCTATCAGGGTTCACA
AGCACACGTCTGAACTCTTTCCCTACACGACGCTCTTCCGATCTNNNGGACAGACTGACACAGATACAC
AGCACACGTCTGAACTCTTTCCCTACACGACGCTCTTCCGATCTNNNGGACAGACTGACACAGATACAC
AGCACACGTCTGAACTCTTTCCCTACACGACGCTCTTCCGATCTNNNGGACAGACTGACACAGATACAC
AGAGCACACGTCTGAACTCTTTCCCTACACGACGCTCTTCCGATCTNNNCACATGCCTATAGTCCCACGT
AGAGCACACGTCTGAACTCTTTCCCTACACGACGCTCTTCCGATCTNNNCACATGCCTATAGTCCCACGT
AGAGCACACGTCTGAACTCTTTCCCTACACGACGCTCTTCCGATCTNNNCACATGCCTATAGTCCCACGT
AGCACACGTCTGAACTCTTTCCCTACACGACGCTCTTCCGATCTNNNTATGTCACCCTCCAGAAACTTGG
AGCACACGTCTGAACTCTTTCCCTACACGACGCTCTTCCGATCTNNNTATGTCACCCTCCAGAAACTTGG
AGCACACGTCTGAACTCTTTCCCTACACGACGCTCTTCCGATCTNNNTATGTCACCCTCCAGAAACTTGG
ACACGTCTGAACTCTTTCCCTACACGACGCTCTTCCGATCTNNNGCTCCCCACTTCCCAGTAGGCATCGG
ACACGTCTGAACTCTTTCCCTACACGACGCTCTTCCGATCTNNNGCTCCCCACTTCCCAGTAGGCATCGG
ACACGTCTGAACTCTTTCCCTACACGACGCTCTTCCGATCTNNNGCTCCCCACTTCCCAGTAGGCATCGG
GAGCACACGTCTGAACTCTTTCCCTACACGACGCTCTTCCGATCTNNNAAGAAATGTTTTTAGTTATGG
GAGCACACGTCTGAACTCTTTCCCTACACGACGCTCTTCCGATCTNNNAAGAAATGTTTTTAGTTATGG
GAGCACACGTCTGAACTCTTTCCCTACACGACGCTCTTCCGATCTNNNAAGAAATGTTTTTAGTTATGG
GAGCACACGTCTGAACTCTTTCCCTACACGACGCTCTTCCGATCTNNNATGGCAAAAGAGACAATCAG
GAGCACACGTCTGAACTCTTTCCCTACACGACGCTCTTCCGATCTNNNATGGCAAAAGAGACAATCAG
GAGCACACGTCTGAACTCTTTCCCTACACGACGCTCTTCCGATCTNNNATGGCAAAAGAGACAATCAG
GAGCACACGTCTGAACTCTTTCCCTACACGACGCTCTTCCGATCTNNNGAGGCAGGAGAATTGCTTGA
GAGCACACGTCTGAACTCTTTCCCTACACGACGCTCTTCCGATCTNNNGAGGCAGGAGAATTGCTTGA
GAGCACACGTCTGAACTCTTTCCCTACACGACGCTCTTCCGATCTNNNGAGGCAGGAGAATTGCTTGA
AGCACACGTCTGAACTCTTTCCCTACACGACGCTCTTCCGATCTNNNTGACTAAATCAGCTAGAGGTTT
AGCACACGTCTGAACTCTTTCCCTACACGACGCTCTTCCGATCTNNNTGACTAAATCAGCTAGAGGTTT
AGCACACGTCTGAACTCTTTCCCTACACGACGCTCTTCCGATCTNNNTGACTAAATCAGCTAGAGGTTT
AGCACACGTCTGAACTCTTTCCCTACACGACGCTCTTCCGATCTNNNTGTAATCTCAGCACTTTGGGAG
AGCACACGTCTGAACTCTTTCCCTACACGACGCTCTTCCGATCTNNNTGTAATCTCAGCACTTTGGGAG

AGCACACGTCTGAACTCTTTCCCTACACGACGCTCTTCCGATCTNNNTGTAATCTCAGCACTTTGGGAG
GAGCACACGTCTGAACTCTTTCCCTACACGACGCTCTTCCGATCTNNNCTGTTGCTGCTACTACTACTG
GAGCACACGTCTGAACTCTTTCCCTACACGACGCTCTTCCGATCTNNNCTGTTGCTGCTACTACTACTG
GAGCACACGTCTGAACTCTTTCCCTACACGACGCTCTTCCGATCTNNNCTGTTGCTGCTACTACTACTG
AGAGCACACGTCTGAACTCTTTCCCTACACGACGCTCTTCCGATCTNNNGTTGTAGTTAGCCGAGATTG
AGAGCACACGTCTGAACTCTTTCCCTACACGACGCTCTTCCGATCTNNNGTTGTAGTTAGCCGAGATTG
AGAGCACACGTCTGAACTCTTTCCCTACACGACGCTCTTCCGATCTNNNGTTGTAGTTAGCCGAGATTG
GAGCACACGTCTGAACTCTTTCCCTACACGACGCTCTTCCGATCTNNNTTCTGTTTGCTTTCAGGAGCT
GAGCACACGTCTGAACTCTTTCCCTACACGACGCTCTTCCGATCTNNNTTCTGTTTGCTTTCAGGAGCT
GAGCACACGTCTGAACTCTTTCCCTACACGACGCTCTTCCGATCTNNNTTCTGTTTGCTTTCAGGAGCT
GAGCACACGTCTGAACTCTTTCCCTACACGACGCTCTTCCGATCTNNNATTCGCCATCACTGTCATCTCT
GAGCACACGTCTGAACTCTTTCCCTACACGACGCTCTTCCGATCTNNNATTCGCCATCACTGTCATCTCT
GAGCACACGTCTGAACTCTTTCCCTACACGACGCTCTTCCGATCTNNNATTCGCCATCACTGTCATCTCT
GAGCACACGTCTGAACTCTTTCCCTACACGACGCTCTTCCGATCTNNNCTAGCTTGAAGGGTATGGAGC
GAGCACACGTCTGAACTCTTTCCCTACACGACGCTCTTCCGATCTNNNCTAGCTTGAAGGGTATGGAGC
GAGCACACGTCTGAACTCTTTCCCTACACGACGCTCTTCCGATCTNNNCTAGCTTGAAGGGTATGGAGC
AGCACACGTCTGAACTCTTTCCCTACACGACGCTCTTCCGATCTNNNTCTCTTCTCTTGTCTCCCTCCATC
AGCACACGTCTGAACTCTTTCCCTACACGACGCTCTTCCGATCTNNNTCTCTTCTCTTGTCTCCCTCCATC
AGCACACGTCTGAACTCTTTCCCTACACGACGCTCTTCCGATCTNNNTCTCTTCTCTTGTCTCCCTCCATC
GAGCACACGTCTGAACTCTTTCCCTACACGACGCTCTTCCGATCTNNNCCCAGTCTCCAGCAGTTCTTTA
GAGCACACGTCTGAACTCTTTCCCTACACGACGCTCTTCCGATCTNNNCCCAGTCTCCAGCAGTTCTTTA
GAGCACACGTCTGAACTCTTTCCCTACACGACGCTCTTCCGATCTNNNCCCAGTCTCCAGCAGTTCTTTA
GAGCACACGTCTGAACTCTTTCCCTACACGACGCTCTTCCGATCTNNNGGTGCCACCACAATAAGGATA
GAGCACACGTCTGAACTCTTTCCCTACACGACGCTCTTCCGATCTNNNGGTGCCACCACAATAAGGATA
GAGCACACGTCTGAACTCTTTCCCTACACGACGCTCTTCCGATCTNNNGGTGCCACCACAATAAGGATA
AGCACACGTCTGAACTCTTTCCCTACACGACGCTCTTCCGATCTNNNACCCTCCTTTCTCTCTCTCTCTCCA
AGCACACGTCTGAACTCTTTCCCTACACGACGCTCTTCCGATCTNNNACCCTCCTTTCTCTCTCTCTCTCCA
AGCACACGTCTGAACTCTTTCCCTACACGACGCTCTTCCGATCTNNNACCCTCCTTTCTCTCTCTCTCTCCA
AGAGCACACGTCTGAACTCTTTCCCTACACGACGCTCTTCCGATCTNNNAGTCAGCGTTAGTATGGAGC
AGAGCACACGTCTGAACTCTTTCCCTACACGACGCTCTTCCGATCTNNNAGTCAGCGTTAGTATGGAGC
AGAGCACACGTCTGAACTCTTTCCCTACACGACGCTCTTCCGATCTNNNAGTCAGCGTTAGTATGGAGC
GAGCACACGTCTGAACTCTTTCCCTACACGACGCTCTTCCGATCTNNNAAGCACTTGGAAAACATTGCA
GAGCACACGTCTGAACTCTTTCCCTACACGACGCTCTTCCGATCTNNNAAGCACTTGGAAAACATTGCA
GAGCACACGTCTGAACTCTTTCCCTACACGACGCTCTTCCGATCTNNNAAGCACTTGGAAAACATTGCA
GAGCACACGTCTGAACTCTTTCCCTACACGACGCTCTTCCGATCTNNNTTTACACTTATCTGGCCACAGCT
GAGCACACGTCTGAACTCTTTCCCTACACGACGCTCTTCCGATCTNNNTTTACACTTATCTGGCCACAGCT
GAGCACACGTCTGAACTCTTTCCCTACACGACGCTCTTCCGATCTNNNTTTACACTTATCTGGCCACAGCT
GAGCACACGTCTGAACTCTTTCCCTACACGACGCTCTTCCGATCTNNNTTCTCTCTCTCTCTCGCTCTCTC
GAGCACACGTCTGAACTCTTTCCCTACACGACGCTCTTCCGATCTNNNTTCTCTCTCTCTCTCGCTCTCTC
GAGCACACGTCTGAACTCTTTCCCTACACGACGCTCTTCCGATCTNNNTTCTCTCTCTCTCTCGCTCTCTC
GAGCACACGTCTGAACTCTTTCCCTACACGACGCTCTTCCGATCTNNNCTATGGAGTAGGGAGGGCAA
GAGCACACGTCTGAACTCTTTCCCTACACGACGCTCTTCCGATCTNNNCTATGGAGTAGGGAGGGCAA
GAGCACACGTCTGAACTCTTTCCCTACACGACGCTCTTCCGATCTNNNCTATGGAGTAGGGAGGGCAA

AGCACACGTCTGAACTCTTTCCCTACACGACGCTCTTCCGATCTNNNGAATGAATTTGGACATGCCCCA
AGCACACGTCTGAACTCTTTCCCTACACGACGCTCTTCCGATCTNNNGAATGAATTTGGACATGCCCCA
AGCACACGTCTGAACTCTTTCCCTACACGACGCTCTTCCGATCTNNNGAATGAATTTGGACATGCCCCA
AGCACACGTCTGAACTCTTTCCCTACACGACGCTCTTCCGATCTNNNCGCAACAAGAATGAGACTCCAT
AGCACACGTCTGAACTCTTTCCCTACACGACGCTCTTCCGATCTNNNCGCAACAAGAATGAGACTCCAT
AGCACACGTCTGAACTCTTTCCCTACACGACGCTCTTCCGATCTNNNCGCAACAAGAATGAGACTCCAT
AGCACACGTCTGAACTCTTTCCCTACACGACGCTCTTCCGATCTNNNGAGAATCGCTTGAACCCAGGAC
AGCACACGTCTGAACTCTTTCCCTACACGACGCTCTTCCGATCTNNNGAGAATCGCTTGAACCCAGGAC
AGCACACGTCTGAACTCTTTCCCTACACGACGCTCTTCCGATCTNNNGAGAATCGCTTGAACCCAGGAC
GAGCACACGTCTGAACTCTTTCCCTACACGACGCTCTTCCGATCTNNNAAGACAATCCTGGCTAACATG
GAGCACACGTCTGAACTCTTTCCCTACACGACGCTCTTCCGATCTNNNAAGACAATCCTGGCTAACATG
GAGCACACGTCTGAACTCTTTCCCTACACGACGCTCTTCCGATCTNNNAAGACAATCCTGGCTAACATG
AGCACACGTCTGAACTCTTTCCCTACACGACGCTCTTCCGATCTNNNGAGGCAGGAAAATCACTTGAAC
AGCACACGTCTGAACTCTTTCCCTACACGACGCTCTTCCGATCTNNNGAGGCAGGAAAATCACTTGAAC
AGCACACGTCTGAACTCTTTCCCTACACGACGCTCTTCCGATCTNNNGAGGCAGGAAAATCACTTGAAC
AGCACACGTCTGAACTCTTTCCCTACACGACGCTCTTCCGATCTNNNTTTATATTGCCTCCTCCTAGCCCC
AGCACACGTCTGAACTCTTTCCCTACACGACGCTCTTCCGATCTNNNTTTATATTGCCTCCTCCTAGCCCC
AGCACACGTCTGAACTCTTTCCCTACACGACGCTCTTCCGATCTNNNTTTATATTGCCTCCTCCTAGCCCC
AGCACACGTCTGAACTCTTTCCCTACACGACGCTCTTCCGATCTNNNGAGGCTGAGGCAGGAGAATGC
AGCACACGTCTGAACTCTTTCCCTACACGACGCTCTTCCGATCTNNNGAGGCTGAGGCAGGAGAATGC
AGCACACGTCTGAACTCTTTCCCTACACGACGCTCTTCCGATCTNNNGAGGCTGAGGCAGGAGAATGC
AGCACACGTCTGAACTCTTTCCCTACACGACGCTCTTCCGATCTNNNAATTAGCCCCTTTCAGTTCACCTC
AGCACACGTCTGAACTCTTTCCCTACACGACGCTCTTCCGATCTNNNAATTAGCCCCTTTCAGTTCACCTC
AGCACACGTCTGAACTCTTTCCCTACACGACGCTCTTCCGATCTNNNAATTAGCCCCTTTCAGTTCACCTC
AGCACACGTCTGAACTCTTTCCCTACACGACGCTCTTCCGATCTNNNGCCCTTCTTTCTTGTTCCCTTCACA
AGCACACGTCTGAACTCTTTCCCTACACGACGCTCTTCCGATCTNNNGCCCTTCTTTCTTGTTCCCTTCACA
AGCACACGTCTGAACTCTTTCCCTACACGACGCTCTTCCGATCTNNNGCCCTTCTTTCTTGTTCCCTTCACA
GAGCACACGTCTGAACTCTTTCCCTACACGACGCTCTTCCGATCTNNNTGACCTCTGGCTCAAGAGATCT
GAGCACACGTCTGAACTCTTTCCCTACACGACGCTCTTCCGATCTNNNTGACCTCTGGCTCAAGAGATCT
GAGCACACGTCTGAACTCTTTCCCTACACGACGCTCTTCCGATCTNNNTGACCTCTGGCTCAAGAGATCT
GAGCACACGTCTGAACTCTTTCCCTACACGACGCTCTTCCGATCTNNNATCCTCCTAGGTACACAACTCC
GAGCACACGTCTGAACTCTTTCCCTACACGACGCTCTTCCGATCTNNNATCCTCCTAGGTACACAACTCC
GAGCACACGTCTGAACTCTTTCCCTACACGACGCTCTTCCGATCTNNNATCCTCCTAGGTACACAACTCC
AGCACACGTCTGAACTCTTTCCCTACACGACGCTCTTCCGATCTNNNCTTTCTGGGAGAACCAAAGGAC
AGCACACGTCTGAACTCTTTCCCTACACGACGCTCTTCCGATCTNNNCTTTCTGGGAGAACCAAAGGAC
AGCACACGTCTGAACTCTTTCCCTACACGACGCTCTTCCGATCTNNNCTTTCTGGGAGAACCAAAGGAC
AGCACACGTCTGAACTCTTTCCCTACACGACGCTCTTCCGATCTNNNGAGGTGGAGCTTGCAGTGAGCA
AGCACACGTCTGAACTCTTTCCCTACACGACGCTCTTCCGATCTNNNGAGGTGGAGCTTGCAGTGAGCA
AGCACACGTCTGAACTCTTTCCCTACACGACGCTCTTCCGATCTNNNGAGGTGGAGCTTGCAGTGAGCA
GAGCACACGTCTGAACTCTTTCCCTACACGACGCTCTTCCGATCTNNNAAAAGACCCAGAATAGCCAAA
GAGCACACGTCTGAACTCTTTCCCTACACGACGCTCTTCCGATCTNNNAAAAGACCCAGAATAGCCAAA
GAGCACACGTCTGAACTCTTTCCCTACACGACGCTCTTCCGATCTNNNAAAAGACCCAGAATAGCCAAA
AGCACACGTCTGAACTCTTTCCCTACACGACGCTCTTCCGATCTNNNTTGTCATCTCCTTGAGAGCAGTC

AGCACACGTCTGAACTCTTTCCCTACACGACGCTCTTCCGATCTNNNTTGTCATCTCCTTGAGAGCAGT
AGCACACGTCTGAACTCTTTCCCTACACGACGCTCTTCCGATCTNNNTTGTCATCTCCTTGAGAGCAGT
GAGCACACGTCTGAACTCTTTCCCTACACGACGCTCTTCCGATCTNNNATCAAGGCATTAGGCAAGTA
GAGCACACGTCTGAACTCTTTCCCTACACGACGCTCTTCCGATCTNNNATCAAGGCATTAGGCAAGTA
GAGCACACGTCTGAACTCTTTCCCTACACGACGCTCTTCCGATCTNNNATCAAGGCATTAGGCAAGTA
GAGCACACGTCTGAACTCTTTCCCTACACGACGCTCTTCCGATCTNNNTCAAAGTGATGGGATTACAG
GAGCACACGTCTGAACTCTTTCCCTACACGACGCTCTTCCGATCTNNNTCAAAGTGATGGGATTACAG
GAGCACACGTCTGAACTCTTTCCCTACACGACGCTCTTCCGATCTNNNTCAAAGTGATGGGATTACAG
GAGCACACGTCTGAACTCTTTCCCTACACGACGCTCTTCCGATCTNNNAACTCACCTGCACTCTCTGTTT
GAGCACACGTCTGAACTCTTTCCCTACACGACGCTCTTCCGATCTNNNAACTCACCTGCACTCTCTGTTT
GAGCACACGTCTGAACTCTTTCCCTACACGACGCTCTTCCGATCTNNNAACTCACCTGCACTCTCTGTTT
AGCACACGTCTGAACTCTTTCCCTACACGACGCTCTTCCGATCTNNNGAGTGACAGAGTGAGACTCCA
AGCACACGTCTGAACTCTTTCCCTACACGACGCTCTTCCGATCTNNNGAGTGACAGAGTGAGACTCCA
AGCACACGTCTGAACTCTTTCCCTACACGACGCTCTTCCGATCTNNNGAGTGACAGAGTGAGACTCCA
GAGCACACGTCTGAACTCTTTCCCTACACGACGCTCTTCCGATCTNNNCATTGGCTCCAAGGAAAGTC
GAGCACACGTCTGAACTCTTTCCCTACACGACGCTCTTCCGATCTNNNCATTGGCTCCAAGGAAAGTC
GAGCACACGTCTGAACTCTTTCCCTACACGACGCTCTTCCGATCTNNNCATTGGCTCCAAGGAAAGTC
GAGCACACGTCTGAACTCTTTCCCTACACGACGCTCTTCCGATCTNNNCCATGGGTGCCAAATCATTAG
GAGCACACGTCTGAACTCTTTCCCTACACGACGCTCTTCCGATCTNNNCCATGGGTGCCAAATCATTAG
GAGCACACGTCTGAACTCTTTCCCTACACGACGCTCTTCCGATCTNNNCCATGGGTGCCAAATCATTAG
GAGCACACGTCTGAACTCTTTCCCTACACGACGCTCTTCCGATCTNNNAGTGAGCTGATATCGCACTA
GAGCACACGTCTGAACTCTTTCCCTACACGACGCTCTTCCGATCTNNNAGTGAGCTGATATCGCACTA
GAGCACACGTCTGAACTCTTTCCCTACACGACGCTCTTCCGATCTNNNAGTGAGCTGATATCGCACTA
AGCACACGTCTGAACTCTTTCCCTACACGACGCTCTTCCGATCTNNNGTGAGCTCCCATCTCAGAATC
AGCACACGTCTGAACTCTTTCCCTACACGACGCTCTTCCGATCTNNNGTGAGCTCCCATCTCAGAATC
AGCACACGTCTGAACTCTTTCCCTACACGACGCTCTTCCGATCTNNNGTGAGCTCCCATCTCAGAATC
GAGCACACGTCTGAACTCTTTCCCTACACGACGCTCTTCCGATCTNNNGCCTGTAGTCCCAGCTACTTG
GAGCACACGTCTGAACTCTTTCCCTACACGACGCTCTTCCGATCTNNNGCCTGTAGTCCCAGCTACTTG
GAGCACACGTCTGAACTCTTTCCCTACACGACGCTCTTCCGATCTNNNGCCTGTAGTCCCAGCTACTTG
GCACACGTCTGAACTCTTTCCCTACACGACGCTCTTCCGATCTNNNCAGAAGAATTGCTTGAACCTGG
GCACACGTCTGAACTCTTTCCCTACACGACGCTCTTCCGATCTNNNCAGAAGAATTGCTTGAACCTGG
GCACACGTCTGAACTCTTTCCCTACACGACGCTCTTCCGATCTNNNCAGAAGAATTGCTTGAACCTGG
AGCACACGTCTGAACTCTTTCCCTACACGACGCTCTTCCGATCTNNNTTGCTAATCCTGTCACCACCATT
AGCACACGTCTGAACTCTTTCCCTACACGACGCTCTTCCGATCTNNNTTGCTAATCCTGTCACCACCATT
AGCACACGTCTGAACTCTTTCCCTACACGACGCTCTTCCGATCTNNNTTGCTAATCCTGTCACCACCATT
GCACACGTCTGAACTCTTTCCCTACACGACGCTCTTCCGATCTNNNTCACATTGACAAGCTGGAAGAG
GCACACGTCTGAACTCTTTCCCTACACGACGCTCTTCCGATCTNNNTCACATTGACAAGCTGGAAGAG
GCACACGTCTGAACTCTTTCCCTACACGACGCTCTTCCGATCTNNNTCACATTGACAAGCTGGAAGAG
AGCACACGTCTGAACTCTTTCCCTACACGACGCTCTTCCGATCTNNNTATCTCAGGGCCTTGTCTTCCT
AGCACACGTCTGAACTCTTTCCCTACACGACGCTCTTCCGATCTNNNTATCTCAGGGCCTTGTCTTCCT
AGCACACGTCTGAACTCTTTCCCTACACGACGCTCTTCCGATCTNNNTATCTCAGGGCCTTGTCTTCCT
GAGCACACGTCTGAACTCTTTCCCTACACGACGCTCTTCCGATCTNNNTGAGCATCATCCGTTATTCAG
GAGCACACGTCTGAACTCTTTCCCTACACGACGCTCTTCCGATCTNNNTGAGCATCATCCGTTATTCAG

GAGCACACGTCTGAACTCTTTCCCTACACGACGCTCTTCCGATCTNNNTGAGCATCATCCGTTATTCAG
AGCACACGTCTGAACTCTTTCCCTACACGACGCTCTTCCGATCTNNNTGTCCAGGCTGATTTCTAACTC
AGCACACGTCTGAACTCTTTCCCTACACGACGCTCTTCCGATCTNNNTGTCCAGGCTGATTTCTAACTC
AGCACACGTCTGAACTCTTTCCCTACACGACGCTCTTCCGATCTNNNTGTCCAGGCTGATTTCTAACTC
GAGCACACGTCTGAACTCTTTCCCTACACGACGCTCTTCCGATCTNNNACCAACCTTTCCTTCCTCTGAT
GAGCACACGTCTGAACTCTTTCCCTACACGACGCTCTTCCGATCTNNNACCAACCTTTCCTTCCTCTGAT
GAGCACACGTCTGAACTCTTTCCCTACACGACGCTCTTCCGATCTNNNACCAACCTTTCCTTCCTCTGAT
AGCACACGTCTGAACTCTTTCCCTACACGACGCTCTTCCGATCTNNNAGCTTGGACTACAGATGTGTG
AGCACACGTCTGAACTCTTTCCCTACACGACGCTCTTCCGATCTNNNAGCTTGGACTACAGATGTGTG
AGCACACGTCTGAACTCTTTCCCTACACGACGCTCTTCCGATCTNNNAGCTTGGACTACAGATGTGTG
GCACACGTCTGAACTCTTTCCCTACACGACGCTCTTCCGATCTNNNTGGACAAAGTTTGGGGGTTTTT
GCACACGTCTGAACTCTTTCCCTACACGACGCTCTTCCGATCTNNNTGGACAAAGTTTGGGGGTTTTT
GCACACGTCTGAACTCTTTCCCTACACGACGCTCTTCCGATCTNNNTGGACAAAGTTTGGGGGTTTTT
AGCACACGTCTGAACTCTTTCCCTACACGACGCTCTTCCGATCTNNNGTTTACACATCATGCCGCCATT
AGCACACGTCTGAACTCTTTCCCTACACGACGCTCTTCCGATCTNNNGTTTACACATCATGCCGCCATT
AGCACACGTCTGAACTCTTTCCCTACACGACGCTCTTCCGATCTNNNGTTTACACATCATGCCGCCATT
AGCACACGTCTGAACTCTTTCCCTACACGACGCTCTTCCGATCTNNNTGGTAGTGGTGAGCAAATGTG
AGCACACGTCTGAACTCTTTCCCTACACGACGCTCTTCCGATCTNNNTGGTAGTGGTGAGCAAATGTG
AGCACACGTCTGAACTCTTTCCCTACACGACGCTCTTCCGATCTNNNTGGTAGTGGTGAGCAAATGTG
AGCACACGTCTGAACTCTTTCCCTACACGACGCTCTTCCGATCTNNNGTGAGCTTTGATTGAGCCATT
AGCACACGTCTGAACTCTTTCCCTACACGACGCTCTTCCGATCTNNNGTGAGCTTTGATTGAGCCATT
AGCACACGTCTGAACTCTTTCCCTACACGACGCTCTTCCGATCTNNNGTGAGCTTTGATTGAGCCATT
GAGCACACGTCTGAACTCTTTCCCTACACGACGCTCTTCCGATCTNNNGAGAGGCTGAGACAGGAGA
GAGCACACGTCTGAACTCTTTCCCTACACGACGCTCTTCCGATCTNNNGAGAGGCTGAGACAGGAGA
GAGCACACGTCTGAACTCTTTCCCTACACGACGCTCTTCCGATCTNNNGAGAGGCTGAGACAGGAGA
GAGCACACGTCTGAACTCTTTCCCTACACGACGCTCTTCCGATCTNNNTGAGCATCAGTAGTCCATGT
GAGCACACGTCTGAACTCTTTCCCTACACGACGCTCTTCCGATCTNNNTGAGCATCAGTAGTCCATGT
GAGCACACGTCTGAACTCTTTCCCTACACGACGCTCTTCCGATCTNNNTGAGCATCAGTAGTCCATGT
GAGCACACGTCTGAACTCTTTCCCTACACGACGCTCTTCCGATCTNNNACTCCACATGGTCATCAACAC
GAGCACACGTCTGAACTCTTTCCCTACACGACGCTCTTCCGATCTNNNACTCCACATGGTCATCAACAC
GAGCACACGTCTGAACTCTTTCCCTACACGACGCTCTTCCGATCTNNNACTCCACATGGTCATCAACAC
AGCACACGTCTGAACTCTTTCCCTACACGACGCTCTTCCGATCTNNNAGCTAAGTGTGGCTATGCTAG
AGCACACGTCTGAACTCTTTCCCTACACGACGCTCTTCCGATCTNNNAGCTAAGTGTGGCTATGCTAG
AGCACACGTCTGAACTCTTTCCCTACACGACGCTCTTCCGATCTNNNAGCTAAGTGTGGCTATGCTAG
AGCACACGTCTGAACTCTTTCCCTACACGACGCTCTTCCGATCTNNNGACAGGATGGTGGTCTTGCTT
AGCACACGTCTGAACTCTTTCCCTACACGACGCTCTTCCGATCTNNNGACAGGATGGTGGTCTTGCTT
AGCACACGTCTGAACTCTTTCCCTACACGACGCTCTTCCGATCTNNNGACAGGATGGTGGTCTTGCTT
GAGCACACGTCTGAACTCTTTCCCTACACGACGCTCTTCCGATCTNNNACAAGTATCTTGCTCTGGGTT
GAGCACACGTCTGAACTCTTTCCCTACACGACGCTCTTCCGATCTNNNACAAGTATCTTGCTCTGGGTT
GAGCACACGTCTGAACTCTTTCCCTACACGACGCTCTTCCGATCTNNNACAAGTATCTTGCTCTGGGTT
AGCACACGTCTGAACTCTTTCCCTACACGACGCTCTTCCGATCTNNNGGAAACCACCACATCTACCTGA
AGCACACGTCTGAACTCTTTCCCTACACGACGCTCTTCCGATCTNNNGGAAACCACCACATCTACCTGA
AGCACACGTCTGAACTCTTTCCCTACACGACGCTCTTCCGATCTNNNGGAAACCACCACATCTACCTGA

AGCACACGTCTGAACTCTTTCCCTACACGACGCTCTTCCGATCTNNNAGAGCCATCTGGGATGTTAG
AGCACACGTCTGAACTCTTTCCCTACACGACGCTCTTCCGATCTNNNAGAGCCATCTGGGATGTTAG
AGCACACGTCTGAACTCTTTCCCTACACGACGCTCTTCCGATCTNNNAGAGCCATCTGGGATGTTAG
AGCACACGTCTGAACTCTTTCCCTACACGACGCTCTTCCGATCTNNNCTGAACCTGAATGTCTGGCAT
AGCACACGTCTGAACTCTTTCCCTACACGACGCTCTTCCGATCTNNNCTGAACCTGAATGTCTGGCAT
AGCACACGTCTGAACTCTTTCCCTACACGACGCTCTTCCGATCTNNNCTGAACCTGAATGTCTGGCAT
AGCACACGTCTGAACTCTTTCCCTACACGACGCTCTTCCGATCTNNNTGGGTGAAAGAGCAAGATTC
AGCACACGTCTGAACTCTTTCCCTACACGACGCTCTTCCGATCTNNNTGGGTGAAAGAGCAAGATTC
AGCACACGTCTGAACTCTTTCCCTACACGACGCTCTTCCGATCTNNNTGGGTGAAAGAGCAAGATTC
GCACACGTCTGAACTCTTTCCCTACACGACGCTCTTCCGATCTNNNATTCACATCATCTTTGTAGGTAT
GCACACGTCTGAACTCTTTCCCTACACGACGCTCTTCCGATCTNNNATTCACATCATCTTTGTAGGTAT
GCACACGTCTGAACTCTTTCCCTACACGACGCTCTTCCGATCTNNNATTCACATCATCTTTGTAGGTAT
AGCACACGTCTGAACTCTTTCCCTACACGACGCTCTTCCGATCTNNNGTAACTCCAAGCCATGCCAAAT
AGCACACGTCTGAACTCTTTCCCTACACGACGCTCTTCCGATCTNNNGTAACTCCAAGCCATGCCAAAT
AGCACACGTCTGAACTCTTTCCCTACACGACGCTCTTCCGATCTNNNGTAACTCCAAGCCATGCCAAAT
ACACGTCTGAACTCTTTCCCTACACGACGCTCTTCCGATCTNNNAGTAACATATCCCGGATGAGCATC
ACACGTCTGAACTCTTTCCCTACACGACGCTCTTCCGATCTNNNAGTAACATATCCCGGATGAGCATC
ACACGTCTGAACTCTTTCCCTACACGACGCTCTTCCGATCTNNNAGTAACATATCCCGGATGAGCATC
AGCACACGTCTGAACTCTTTCCCTACACGACGCTCTTCCGATCTNNNTTGGAGAAGCTTGATCTACCC
AGCACACGTCTGAACTCTTTCCCTACACGACGCTCTTCCGATCTNNNTTGGAGAAGCTTGATCTACCC
AGCACACGTCTGAACTCTTTCCCTACACGACGCTCTTCCGATCTNNNTTGGAGAAGCTTGATCTACCC
AGCACACGTCTGAACTCTTTCCCTACACGACGCTCTTCCGATCTNNNTTCTGGTTGTTTTGTACACTCT
AGCACACGTCTGAACTCTTTCCCTACACGACGCTCTTCCGATCTNNNTTCTGGTTGTTTTGTACACTCT
AGCACACGTCTGAACTCTTTCCCTACACGACGCTCTTCCGATCTNNNTTCTGGTTGTTTTGTACACTCT
AGCACACGTCTGAACTCTTTCCCTACACGACGCTCTTCCGATCTNNNAAAAGAAGAAATTCTTTCCAG
AGCACACGTCTGAACTCTTTCCCTACACGACGCTCTTCCGATCTNNNAAAAGAAGAAATTCTTTCCAG
AGCACACGTCTGAACTCTTTCCCTACACGACGCTCTTCCGATCTNNNAAAAGAAGAAATTCTTTCCAG
GAGCACACGTCTGAACTCTTTCCCTACACGACGCTCTTCCGATCTNNNTTGCCATATCATGTGTGTGTG
GAGCACACGTCTGAACTCTTTCCCTACACGACGCTCTTCCGATCTNNNTTGCCATATCATGTGTGTGTG
GAGCACACGTCTGAACTCTTTCCCTACACGACGCTCTTCCGATCTNNNTTGCCATATCATGTGTGTGTG
GCACACGTCTGAACTCTTTCCCTACACGACGCTCTTCCGATCTNNNCTTTGCCTTTGCCTTCCCTGCATC
GCACACGTCTGAACTCTTTCCCTACACGACGCTCTTCCGATCTNNNCTTTGCCTTTGCCTTCCCTGCATC
GCACACGTCTGAACTCTTTCCCTACACGACGCTCTTCCGATCTNNNCTTTGCCTTTGCCTTCCCTGCATC
GAGCACACGTCTGAACTCTTTCCCTACACGACGCTCTTCCGATCTNNNATGTCAGTACACAGCTAGCT
GAGCACACGTCTGAACTCTTTCCCTACACGACGCTCTTCCGATCTNNNATGTCAGTACACAGCTAGCT
GAGCACACGTCTGAACTCTTTCCCTACACGACGCTCTTCCGATCTNNNATGTCAGTACACAGCTAGCT
GAGCACACGTCTGAACTCTTTCCCTACACGACGCTCTTCCGATCTNNNGAGCCGAGATCATGCCACTG
GAGCACACGTCTGAACTCTTTCCCTACACGACGCTCTTCCGATCTNNNGAGCCGAGATCATGCCACTG
GAGCACACGTCTGAACTCTTTCCCTACACGACGCTCTTCCGATCTNNNGAGCCGAGATCATGCCACTG
AGCACACGTCTGAACTCTTTCCCTACACGACGCTCTTCCGATCTNNNGGCAACATAGTAAGACCCTGT
AGCACACGTCTGAACTCTTTCCCTACACGACGCTCTTCCGATCTNNNGGCAACATAGTAAGACCCTGT
AGCACACGTCTGAACTCTTTCCCTACACGACGCTCTTCCGATCTNNNGGCAACATAGTAAGACCCTGT
GCACACGTCTGAACTCTTTCCCTACACGACGCTCTTCCGATCTNNNCCCCTTTCCTGTTTCAGAGTGTC

AGCACACGTCTGAACTCTTTCCCTACACGACGCTCTTCCGATCTNNNCCCCTTTCCTGTTTCAGAGTGTC
AGCACACGTCTGAACTCTTTCCCTACACGACGCTCTTCCGATCTNNNCCCCTTTCCTGTTTCAGAGTGTC
AGAGCACACGTCTGAACTCTTTCCCTACACGACGCTCTTCCGATCTNNNACGGGTGTATCTAAATGCC
AGAGCACACGTCTGAACTCTTTCCCTACACGACGCTCTTCCGATCTNNNACGGGTGTATCTAAATGCC
AGAGCACACGTCTGAACTCTTTCCCTACACGACGCTCTTCCGATCTNNNACGGGTGTATCTAAATGCC
AGCACACGTCTGAACTCTTTCCCTACACGACGCTCTTCCGATCTNNNCTTTCAAAACATTCCTTTTCTTC
AGCACACGTCTGAACTCTTTCCCTACACGACGCTCTTCCGATCTNNNCTTTCAAAACATTCCTTTTCTTC
AGCACACGTCTGAACTCTTTCCCTACACGACGCTCTTCCGATCTNNNCTTTCAAAACATTCCTTTTCTTC
GAGCACACGTCTGAACTCTTTCCCTACACGACGCTCTTCCGATCTNNNGCCTCCTGTAATCTCAGCTACT
GAGCACACGTCTGAACTCTTTCCCTACACGACGCTCTTCCGATCTNNNGCCTCCTGTAATCTCAGCTACT
GAGCACACGTCTGAACTCTTTCCCTACACGACGCTCTTCCGATCTNNNGCCTCCTGTAATCTCAGCTACT
AGCACACGTCTGAACTCTTTCCCTACACGACGCTCTTCCGATCTNNNCCAAGATTTGGTGCTAGGTGT
AGCACACGTCTGAACTCTTTCCCTACACGACGCTCTTCCGATCTNNNCCAAGATTTGGTGCTAGGTGT
AGCACACGTCTGAACTCTTTCCCTACACGACGCTCTTCCGATCTNNNCCAAGATTTGGTGCTAGGTGT
AGCACACGTCTGAACTCTTTCCCTACACGACGCTCTTCCGATCTNNNTGCATGTCTCCTTATGGGTCTC
AGCACACGTCTGAACTCTTTCCCTACACGACGCTCTTCCGATCTNNNTGCATGTCTCCTTATGGGTCTC
AGCACACGTCTGAACTCTTTCCCTACACGACGCTCTTCCGATCTNNNTGCATGTCTCCTTATGGGTCTC
GAGCACACGTCTGAACTCTTTCCCTACACGACGCTCTTCCGATCTNNNCCCACTGTTATCAAGACATGT
GAGCACACGTCTGAACTCTTTCCCTACACGACGCTCTTCCGATCTNNNCCCACTGTTATCAAGACATGT
GAGCACACGTCTGAACTCTTTCCCTACACGACGCTCTTCCGATCTNNNCCCACTGTTATCAAGACATGT
AGCACACGTCTGAACTCTTTCCCTACACGACGCTCTTCCGATCTNNNCTCTTCTCTCTCTCTCTCCCTCCCA
AGCACACGTCTGAACTCTTTCCCTACACGACGCTCTTCCGATCTNNNCTCTTCTCTCTCTCTCTCCCTCCCA
AGCACACGTCTGAACTCTTTCCCTACACGACGCTCTTCCGATCTNNNCTCTTCTCTCTCTCTCTCCCTCCCA
AGCACACGTCTGAACTCTTTCCCTACACGACGCTCTTCCGATCTNNNCTACACCAACACCATCTTCATGC
AGCACACGTCTGAACTCTTTCCCTACACGACGCTCTTCCGATCTNNNCTACACCAACACCATCTTCATGC
AGCACACGTCTGAACTCTTTCCCTACACGACGCTCTTCCGATCTNNNCTACACCAACACCATCTTCATGC
GAGCACACGTCTGAACTCTTTCCCTACACGACGCTCTTCCGATCTNNNTATCAAGATAGTACAGAGAA
GAGCACACGTCTGAACTCTTTCCCTACACGACGCTCTTCCGATCTNNNTATCAAGATAGTACAGAGAA
GAGCACACGTCTGAACTCTTTCCCTACACGACGCTCTTCCGATCTNNNTATCAAGATAGTACAGAGAA
AGCACACGTCTGAACTCTTTCCCTACACGACGCTCTTCCGATCTNNNCAGATCTAGAGGCTTGAGCAG
AGCACACGTCTGAACTCTTTCCCTACACGACGCTCTTCCGATCTNNNCAGATCTAGAGGCTTGAGCAG
AGCACACGTCTGAACTCTTTCCCTACACGACGCTCTTCCGATCTNNNCAGATCTAGAGGCTTGAGCAG
AGCACACGTCTGAACTCTTTCCCTACACGACGCTCTTCCGATCTNNNTCAATGCCTTTGATAGGGTGG
AGCACACGTCTGAACTCTTTCCCTACACGACGCTCTTCCGATCTNNNTCAATGCCTTTGATAGGGTGG
AGCACACGTCTGAACTCTTTCCCTACACGACGCTCTTCCGATCTNNNTCAATGCCTTTGATAGGGTGG
GAGCACACGTCTGAACTCTTTCCCTACACGACGCTCTTCCGATCTNNNATTCTCTCGCATGTACACACCA
GAGCACACGTCTGAACTCTTTCCCTACACGACGCTCTTCCGATCTNNNATTCTCTCGCATGTACACACCA
GAGCACACGTCTGAACTCTTTCCCTACACGACGCTCTTCCGATCTNNNATTCTCTCGCATGTACACACCA
GAGCACACGTCTGAACTCTTTCCCTACACGACGCTCTTCCGATCTNNNAGCCAAACCATATCAGAAGG
GAGCACACGTCTGAACTCTTTCCCTACACGACGCTCTTCCGATCTNNNAGCCAAACCATATCAGAAGG
GAGCACACGTCTGAACTCTTTCCCTACACGACGCTCTTCCGATCTNNNAGCCAAACCATATCAGAAGG
AGCACACGTCTGAACTCTTTCCCTACACGACGCTCTTCCGATCTNNNTCAGTTCATCCTTCAAGACCTGG
AGCACACGTCTGAACTCTTTCCCTACACGACGCTCTTCCGATCTNNNTCAGTTCATCCTTCAAGACCTGG

AGCACACGTCTGAACTCTTTCCCTACACGACGCTCTTCCGATCTNNNTCAGTTCATCCTTCAAGACCTGG
AGAGCACACGTCTGAACTCTTTCCCTACACGACGCTCTTCCGATCTNNNGAGAATGGTGTGAACCCAGG
AGAGCACACGTCTGAACTCTTTCCCTACACGACGCTCTTCCGATCTNNNGAGAATGGTGTGAACCCAGG
AGAGCACACGTCTGAACTCTTTCCCTACACGACGCTCTTCCGATCTNNNGAGAATGGTGTGAACCCAGG
AGCACACGTCTGAACTCTTTCCCTACACGACGCTCTTCCGATCTNNNGTCCCTTTTTCTCCACAACCTTGCA
AGCACACGTCTGAACTCTTTCCCTACACGACGCTCTTCCGATCTNNNGTCCCTTTTTCTCCACAACCTTGCA
AGCACACGTCTGAACTCTTTCCCTACACGACGCTCTTCCGATCTNNNGTCCCTTTTTCTCCACAACCTTGCA
AGAGCACACGTCTGAACTCTTTCCCTACACGACGCTCTTCCGATCTNNNAATCAAAAGCAGGCACAAAC
AGAGCACACGTCTGAACTCTTTCCCTACACGACGCTCTTCCGATCTNNNAATCAAAAGCAGGCACAAAC
AGAGCACACGTCTGAACTCTTTCCCTACACGACGCTCTTCCGATCTNNNAATCAAAAGCAGGCACAAAC
GAGCACACGTCTGAACTCTTTCCCTACACGACGCTCTTCCGATCTNNNGAAAAGCATGTTCACACACACC
GAGCACACGTCTGAACTCTTTCCCTACACGACGCTCTTCCGATCTNNNGAAAAGCATGTTCACACACACC
GAGCACACGTCTGAACTCTTTCCCTACACGACGCTCTTCCGATCTNNNGAAAAGCATGTTCACACACACC
AGCACACGTCTGAACTCTTTCCCTACACGACGCTCTTCCGATCTNNNAGGAGAATTGCTTGAATCTGG
AGCACACGTCTGAACTCTTTCCCTACACGACGCTCTTCCGATCTNNNAGGAGAATTGCTTGAATCTGG
AGCACACGTCTGAACTCTTTCCCTACACGACGCTCTTCCGATCTNNNAGGAGAATTGCTTGAATCTGG
AGCACACGTCTGAACTCTTTCCCTACACGACGCTCTTCCGATCTNNNGTCTCTCTCTCTCTCTCTCTCCCCA
AGCACACGTCTGAACTCTTTCCCTACACGACGCTCTTCCGATCTNNNGTCTCTCTCTCTCTCTCTCTCCCCA
AGCACACGTCTGAACTCTTTCCCTACACGACGCTCTTCCGATCTNNNGTCTCTCTCTCTCTCTCTCTCCCCA
AGCACACGTCTGAACTCTTTCCCTACACGACGCTCTTCCGATCTNNNTTATGAGGATTTTCAGCCTGGC
AGCACACGTCTGAACTCTTTCCCTACACGACGCTCTTCCGATCTNNNTTATGAGGATTTTCAGCCTGGC
AGCACACGTCTGAACTCTTTCCCTACACGACGCTCTTCCGATCTNNNTTATGAGGATTTTCAGCCTGGC
GAGCACACGTCTGAACTCTTTCCCTACACGACGCTCTTCCGATCTNNNGGCTCATGGGATTGCTTTTGA
GAGCACACGTCTGAACTCTTTCCCTACACGACGCTCTTCCGATCTNNNGGCTCATGGGATTGCTTTTGA
GAGCACACGTCTGAACTCTTTCCCTACACGACGCTCTTCCGATCTNNNGGCTCATGGGATTGCTTTTGA
AGCACACGTCTGAACTCTTTCCCTACACGACGCTCTTCCGATCTNNNCCAACATGGTGAAAACCTGTCTC
AGCACACGTCTGAACTCTTTCCCTACACGACGCTCTTCCGATCTNNNCCAACATGGTGAAAACCTGTCTC
AGCACACGTCTGAACTCTTTCCCTACACGACGCTCTTCCGATCTNNNCCAACATGGTGAAAACCTGTCTC
GAGCACACGTCTGAACTCTTTCCCTACACGACGCTCTTCCGATCTNNNCTACAAGGAATGAGCCACCAT
GAGCACACGTCTGAACTCTTTCCCTACACGACGCTCTTCCGATCTNNNCTACAAGGAATGAGCCACCAT
GAGCACACGTCTGAACTCTTTCCCTACACGACGCTCTTCCGATCTNNNCTACAAGGAATGAGCCACCAT
AGCACACGTCTGAACTCTTTCCCTACACGACGCTCTTCCGATCTNNNATTCTTAGAGCCCAGGAGGTCA
AGCACACGTCTGAACTCTTTCCCTACACGACGCTCTTCCGATCTNNNATTCTTAGAGCCCAGGAGGTCA
AGCACACGTCTGAACTCTTTCCCTACACGACGCTCTTCCGATCTNNNATTCTTAGAGCCCAGGAGGTCA
AGCACACGTCTGAACTCTTTCCCTACACGACGCTCTTCCGATCTNNNCCCCTCAGAACTTGAAGGAACTG
AGCACACGTCTGAACTCTTTCCCTACACGACGCTCTTCCGATCTNNNCCCCTCAGAACTTGAAGGAACTG
AGCACACGTCTGAACTCTTTCCCTACACGACGCTCTTCCGATCTNNNCCCCTCAGAACTTGAAGGAACTG
AGCACACGTCTGAACTCTTTCCCTACACGACGCTCTTCCGATCTNNNGTATATCTGTCAGGCTAGCAGC
AGCACACGTCTGAACTCTTTCCCTACACGACGCTCTTCCGATCTNNNGTATATCTGTCAGGCTAGCAGC
AGCACACGTCTGAACTCTTTCCCTACACGACGCTCTTCCGATCTNNNGTATATCTGTCAGGCTAGCAGC
GAGCACACGTCTGAACTCTTTCCCTACACGACGCTCTTCCGATCTNNNTAGCTATCCTGGCTGAAAGAA
GAGCACACGTCTGAACTCTTTCCCTACACGACGCTCTTCCGATCTNNNTAGCTATCCTGGCTGAAAGAA
GAGCACACGTCTGAACTCTTTCCCTACACGACGCTCTTCCGATCTNNNTAGCTATCCTGGCTGAAAGAA

GAGCACACGTCTGAACTCTTTCCCTACACGACGCTCTTCCGATCTNNNCAGCAACTTGGATGAAGCTA

GAGCACACGTCTGAACTCTTTCCCTACACGACGCTCTTCCGATCTNNNCAGCAACTTGGATGAAGCTA

GAGCACACGTCTGAACTCTTTCCCTACACGACGCTCTTCCGATCTNNNCAGCAACTTGGATGAAGCTA

AGAGCACACGTCTGAACTCTTTCCCTACACGACGCTCTTCCGATCTNNNATGTGTATGACCACTTTGGC

AGAGCACACGTCTGAACTCTTTCCCTACACGACGCTCTTCCGATCTNNNATGTGTATGACCACTTTGGC

AGAGCACACGTCTGAACTCTTTCCCTACACGACGCTCTTCCGATCTNNNATGTGTATGACCACTTTGGC

AGCACACGTCTGAACTCTTTCCCTACACGACGCTCTTCCGATCTNNNAGGGGATTGTGATCAGGTTT

AGCACACGTCTGAACTCTTTCCCTACACGACGCTCTTCCGATCTNNNAGGGGATTGTGATCAGGTTT

AGCACACGTCTGAACTCTTTCCCTACACGACGCTCTTCCGATCTNNNAGGGGATTGTGATCAGGTTT

AGCACACGTCTGAACTCTTTCCCTACACGACGCTCTTCCGATCTNNNCAGACTCTTACCAGCATATATA

AGCACACGTCTGAACTCTTTCCCTACACGACGCTCTTCCGATCTNNNCAGACTCTTACCAGCATATATA

AGCACACGTCTGAACTCTTTCCCTACACGACGCTCTTCCGATCTNNNCAGACTCTTACCAGCATATATA

GCACACGTCTGAACTCTTTCCCTACACGACGCTCTTCCGATCTNNNAGGTTTCAGTGAGCCAAGATCA

GCACACGTCTGAACTCTTTCCCTACACGACGCTCTTCCGATCTNNNAGGTTTCAGTGAGCCAAGATCA

GCACACGTCTGAACTCTTTCCCTACACGACGCTCTTCCGATCTNNNAGGTTTCAGTGAGCCAAGATCA

GAGCACACGTCTGAACTCTTTCCCTACACGACGCTCTTCCGATCTNNNCACTTGAGGTCAGGAGTTCA

GAGCACACGTCTGAACTCTTTCCCTACACGACGCTCTTCCGATCTNNNCACTTGAGGTCAGGAGTTCA

GAGCACACGTCTGAACTCTTTCCCTACACGACGCTCTTCCGATCTNNNCACTTGAGGTCAGGAGTTCA

AGCACACGTCTGAACTCTTTCCCTACACGACGCTCTTCCGATCTNNNTTGTCTCTTCCCCTCTTCCTGAC

AGCACACGTCTGAACTCTTTCCCTACACGACGCTCTTCCGATCTNNNTTGTCTCTTCCCCTCTTCCTGAC

AGCACACGTCTGAACTCTTTCCCTACACGACGCTCTTCCGATCTNNNTTGTCTCTTCCCCTCTTCCTGAC

GCACACGTCTGAACTCTTTCCCTACACGACGCTCTTCCGATCTNNNCCAATGACTGCCTTCTTGCTTTGC

GCACACGTCTGAACTCTTTCCCTACACGACGCTCTTCCGATCTNNNCCAATGACTGCCTTCTTGCTTTGC

GCACACGTCTGAACTCTTTCCCTACACGACGCTCTTCCGATCTNNNCCAATGACTGCCTTCTTGCTTTGC

AGCACACGTCTGAACTCTTTCCCTACACGACGCTCTTCCGATCTNNNTCACTGAAGGGGAATCTGGG

AGCACACGTCTGAACTCTTTCCCTACACGACGCTCTTCCGATCTNNNTCACTGAAGGGGAATCTGGG

AGCACACGTCTGAACTCTTTCCCTACACGACGCTCTTCCGATCTNNNTCACTGAAGGGGAATCTGGG

AGAGCACACGTCTGAACTCTTTCCCTACACGACGCTCTTCCGATCTNNNTACACACATGCACACACCAT

AGAGCACACGTCTGAACTCTTTCCCTACACGACGCTCTTCCGATCTNNNTACACACATGCACACACCAT

AGAGCACACGTCTGAACTCTTTCCCTACACGACGCTCTTCCGATCTNNNTACACACATGCACACACCAT

AGCACACGTCTGAACTCTTTCCCTACACGACGCTCTTCCGATCTNNNATCATTCGCATTCTTACCACCC

AGCACACGTCTGAACTCTTTCCCTACACGACGCTCTTCCGATCTNNNATCATTCGCATTCTTACCACCC

AGCACACGTCTGAACTCTTTCCCTACACGACGCTCTTCCGATCTNNNATCATTCGCATTCTTACCACCC

AGCACACGTCTGAACTCTTTCCCTACACGACGCTCTTCCGATCTNNNTCCAGTCTATCATTGATGGGC

AGCACACGTCTGAACTCTTTCCCTACACGACGCTCTTCCGATCTNNNTCCAGTCTATCATTGATGGGC

AGCACACGTCTGAACTCTTTCCCTACACGACGCTCTTCCGATCTNNNTCCAGTCTATCATTGATGGGC

GAGCACACGTCTGAACTCTTTCCCTACACGACGCTCTTCCGATCTNNNGAAGAAACACACGGAGGAC

GAGCACACGTCTGAACTCTTTCCCTACACGACGCTCTTCCGATCTNNNGAAGAAACACACGGAGGAC

GAGCACACGTCTGAACTCTTTCCCTACACGACGCTCTTCCGATCTNNNGAAGAAACACACGGAGGAC

GCACACGTCTGAACTCTTTCCCTACACGACGCTCTTCCGATCTNNNTGCCTAAGACAGACACTGAACT

GCACACGTCTGAACTCTTTCCCTACACGACGCTCTTCCGATCTNNNTGCCTAAGACAGACACTGAACT

GCACACGTCTGAACTCTTTCCCTACACGACGCTCTTCCGATCTNNNTGCCTAAGACAGACACTGAACT

AGCACACGTCTGAACTCTTTCCCTACACGACGCTCTTCCGATCTNNNTGATTTGCATTGAACACAAAC

GAGCACACGTCTGAACTCTTTCCCTACACGACGCTCTTCCGATCTNNNTGATTTGCATTGAACACAAACT
GAGCACACGTCTGAACTCTTTCCCTACACGACGCTCTTCCGATCTNNNTGATTTGCATTGAACACAAACT
AGAGCACACGTCTGAACTCTTTCCCTACACGACGCTCTTCCGATCTNNNCTTTGAACATCAGCCTCAAAG
AGAGCACACGTCTGAACTCTTTCCCTACACGACGCTCTTCCGATCTNNNCTTTGAACATCAGCCTCAAAG
AGAGCACACGTCTGAACTCTTTCCCTACACGACGCTCTTCCGATCTNNNCTTTGAACATCAGCCTCAAAG
GCACACGTCTGAACTCTTTCCCTACACGACGCTCTTCCGATCTNNNTGTTGACACATCATACTCACTTGA
GCACACGTCTGAACTCTTTCCCTACACGACGCTCTTCCGATCTNNNTGTTGACACATCATACTCACTTGA
GCACACGTCTGAACTCTTTCCCTACACGACGCTCTTCCGATCTNNNTGTTGACACATCATACTCACTTGA
AGCACACGTCTGAACTCTTTCCCTACACGACGCTCTTCCGATCTNNNACTGAGTAGCCATTGATGAGCA
AGCACACGTCTGAACTCTTTCCCTACACGACGCTCTTCCGATCTNNNACTGAGTAGCCATTGATGAGCA
AGCACACGTCTGAACTCTTTCCCTACACGACGCTCTTCCGATCTNNNACTGAGTAGCCATTGATGAGCA
GAGCACACGTCTGAACTCTTTCCCTACACGACGCTCTTCCGATCTNNNCTGAGGCAGGAGAATCACTT
GAGCACACGTCTGAACTCTTTCCCTACACGACGCTCTTCCGATCTNNNCTGAGGCAGGAGAATCACTT
GAGCACACGTCTGAACTCTTTCCCTACACGACGCTCTTCCGATCTNNNCTGAGGCAGGAGAATCACTT
AGAGCACACGTCTGAACTCTTTCCCTACACGACGCTCTTCCGATCTNNNCTGCCTCTCACTGGTGTCTTT
AGAGCACACGTCTGAACTCTTTCCCTACACGACGCTCTTCCGATCTNNNCTGCCTCTCACTGGTGTCTTT
AGAGCACACGTCTGAACTCTTTCCCTACACGACGCTCTTCCGATCTNNNCTGCCTCTCACTGGTGTCTTT
AGCACACGTCTGAACTCTTTCCCTACACGACGCTCTTCCGATCTNNNCTCTCTGGTGTGACAATTGG
AGCACACGTCTGAACTCTTTCCCTACACGACGCTCTTCCGATCTNNNCTCTCTGGTGTGACAATTGG
AGCACACGTCTGAACTCTTTCCCTACACGACGCTCTTCCGATCTNNNCTCTCTGGTGTGACAATTGG
CACACGTCTGAACTCTTTCCCTACACGACGCTCTTCCGATCTNNNTGAGTTTATTTTGAAGGTTAACTCA
CACACGTCTGAACTCTTTCCCTACACGACGCTCTTCCGATCTNNNTGAGTTTATTTTGAAGGTTAACTCA
CACACGTCTGAACTCTTTCCCTACACGACGCTCTTCCGATCTNNNTGAGTTTATTTTGAAGGTTAACTCA
GAGCACACGTCTGAACTCTTTCCCTACACGACGCTCTTCCGATCTNNNAGACAGGAGAATTGCTTGAAG
GAGCACACGTCTGAACTCTTTCCCTACACGACGCTCTTCCGATCTNNNAGACAGGAGAATTGCTTGAAG
GAGCACACGTCTGAACTCTTTCCCTACACGACGCTCTTCCGATCTNNNAGACAGGAGAATTGCTTGAAG
AGCACACGTCTGAACTCTTTCCCTACACGACGCTCTTCCGATCTNNNAGCATCTAGGACTACAAGTGTG
AGCACACGTCTGAACTCTTTCCCTACACGACGCTCTTCCGATCTNNNAGCATCTAGGACTACAAGTGTG
AGCACACGTCTGAACTCTTTCCCTACACGACGCTCTTCCGATCTNNNAGCATCTAGGACTACAAGTGTG
AGCACACGTCTGAACTCTTTCCCTACACGACGCTCTTCCGATCTNNNCTGGGTCCGATAACTGGGCTCC
AGCACACGTCTGAACTCTTTCCCTACACGACGCTCTTCCGATCTNNNCTGGGTCCGATAACTGGGCTCC
AGCACACGTCTGAACTCTTTCCCTACACGACGCTCTTCCGATCTNNNCTGGGTCCGATAACTGGGCTCC
GAGCACACGTCTGAACTCTTTCCCTACACGACGCTCTTCCGATCTNNNAGGAGAATGGCGTGAACCTG
GAGCACACGTCTGAACTCTTTCCCTACACGACGCTCTTCCGATCTNNNAGGAGAATGGCGTGAACCTG
GAGCACACGTCTGAACTCTTTCCCTACACGACGCTCTTCCGATCTNNNAGGAGAATGGCGTGAACCTG
AGCACACGTCTGAACTCTTTCCCTACACGACGCTCTTCCGATCTNNNTCAGATGGTTGTGAAGAAATTG
AGCACACGTCTGAACTCTTTCCCTACACGACGCTCTTCCGATCTNNNTCAGATGGTTGTGAAGAAATTG
AGCACACGTCTGAACTCTTTCCCTACACGACGCTCTTCCGATCTNNNTCAGATGGTTGTGAAGAAATTG
AGCACACGTCTGAACTCTTTCCCTACACGACGCTCTTCCGATCTNNNGCCCTCAAACAAAACACAGAACT
AGCACACGTCTGAACTCTTTCCCTACACGACGCTCTTCCGATCTNNNGCCCTCAAACAAAACACAGAACT
AGCACACGTCTGAACTCTTTCCCTACACGACGCTCTTCCGATCTNNNGCCCTCAAACAAAACACAGAACT
AGCACACGTCTGAACTCTTTCCCTACACGACGCTCTTCCGATCTNNNAGATCTCTTTTGCTGTCTCGGT
AGCACACGTCTGAACTCTTTCCCTACACGACGCTCTTCCGATCTNNNAGATCTCTTTTGCTGTCTCGGT

AGCACACGTCTGAACTCTTTCCCTACACGACGCTCTTCCGATCTNNNAGATCTCTTTTGCTGTCTCGGT
AGCACACGTCTGAACTCTTTCCCTACACGACGCTCTTCCGATCTNNNTCTTCAAATATACTTTGCTTACG
AGCACACGTCTGAACTCTTTCCCTACACGACGCTCTTCCGATCTNNNTCTTCAAATATACTTTGCTTACG
AGCACACGTCTGAACTCTTTCCCTACACGACGCTCTTCCGATCTNNNTCTTCAAATATACTTTGCTTACG
GCACACGTCTGAACTCTTTCCCTACACGACGCTCTTCCGATCTNNNCACCGTATTACTTTGCTGCAGAA
GCACACGTCTGAACTCTTTCCCTACACGACGCTCTTCCGATCTNNNCACCGTATTACTTTGCTGCAGAA
GCACACGTCTGAACTCTTTCCCTACACGACGCTCTTCCGATCTNNNCACCGTATTACTTTGCTGCAGAA
GCACACGTCTGAACTCTTTCCCTACACGACGCTCTTCCGATCTNNNAACGAGCCAAGAAGGAGAGAA
GCACACGTCTGAACTCTTTCCCTACACGACGCTCTTCCGATCTNNNAACGAGCCAAGAAGGAGAGAA
GCACACGTCTGAACTCTTTCCCTACACGACGCTCTTCCGATCTNNNAACGAGCCAAGAAGGAGAGAA
AGCACACGTCTGAACTCTTTCCCTACACGACGCTCTTCCGATCTNNNTTTATTTAGTCCTTCCCCCGG
AGCACACGTCTGAACTCTTTCCCTACACGACGCTCTTCCGATCTNNNTTTATTTAGTCCTTCCCCCGG
AGCACACGTCTGAACTCTTTCCCTACACGACGCTCTTCCGATCTNNNTTTATTTAGTCCTTCCCCCGG
GAGCACACGTCTGAACTCTTTCCCTACACGACGCTCTTCCGATCTNNNAACTGATTCTAGCTGTGAGCC
GAGCACACGTCTGAACTCTTTCCCTACACGACGCTCTTCCGATCTNNNAACTGATTCTAGCTGTGAGCC
GAGCACACGTCTGAACTCTTTCCCTACACGACGCTCTTCCGATCTNNNAACTGATTCTAGCTGTGAGCC
AGCACACGTCTGAACTCTTTCCCTACACGACGCTCTTCCGATCTNNNTATTCCATTCTCCAAGGCCTCTG
AGCACACGTCTGAACTCTTTCCCTACACGACGCTCTTCCGATCTNNNTATTCCATTCTCCAAGGCCTCTG
AGCACACGTCTGAACTCTTTCCCTACACGACGCTCTTCCGATCTNNNTATTCCATTCTCCAAGGCCTCTG
GCACACGTCTGAACTCTTTCCCTACACGACGCTCTTCCGATCTNNNCTGGGTAACAAGAGCGAAACTT
GCACACGTCTGAACTCTTTCCCTACACGACGCTCTTCCGATCTNNNCTGGGTAACAAGAGCGAAACTT
GCACACGTCTGAACTCTTTCCCTACACGACGCTCTTCCGATCTNNNCTGGGTAACAAGAGCGAAACTT
GAGCACACGTCTGAACTCTTTCCCTACACGACGCTCTTCCGATCTNNNCACCTGAAGTCTCAGCTACTC
GAGCACACGTCTGAACTCTTTCCCTACACGACGCTCTTCCGATCTNNNCACCTGAAGTCTCAGCTACTC
GAGCACACGTCTGAACTCTTTCCCTACACGACGCTCTTCCGATCTNNNCACCTGAAGTCTCAGCTACTC
AGCACACGTCTGAACTCTTTCCCTACACGACGCTCTTCCGATCTNNNCTCCAGTGATTTTCCTGCCTCAG
AGCACACGTCTGAACTCTTTCCCTACACGACGCTCTTCCGATCTNNNCTCCAGTGATTTTCCTGCCTCAG
AGCACACGTCTGAACTCTTTCCCTACACGACGCTCTTCCGATCTNNNCTCCAGTGATTTTCCTGCCTCAG
AGCACACGTCTGAACTCTTTCCCTACACGACGCTCTTCCGATCTNNNGACAAAGTCTCTCTCTGTCACC
AGCACACGTCTGAACTCTTTCCCTACACGACGCTCTTCCGATCTNNNGACAAAGTCTCTCTCTGTCACC
AGCACACGTCTGAACTCTTTCCCTACACGACGCTCTTCCGATCTNNNGACAAAGTCTCTCTCTGTCACC
GAGCACACGTCTGAACTCTTTCCCTACACGACGCTCTTCCGATCTNNNGGCCAAATGGAAGTCATTAGA
GAGCACACGTCTGAACTCTTTCCCTACACGACGCTCTTCCGATCTNNNGGCCAAATGGAAGTCATTAGA
GAGCACACGTCTGAACTCTTTCCCTACACGACGCTCTTCCGATCTNNNGGCCAAATGGAAGTCATTAGA
GAGCACACGTCTGAACTCTTTCCCTACACGACGCTCTTCCGATCTNNNAGCAATGACTGTAAAGACTG
GAGCACACGTCTGAACTCTTTCCCTACACGACGCTCTTCCGATCTNNNAGCAATGACTGTAAAGACTG
GAGCACACGTCTGAACTCTTTCCCTACACGACGCTCTTCCGATCTNNNAGCAATGACTGTAAAGACTG
AGCACACGTCTGAACTCTTTCCCTACACGACGCTCTTCCGATCTNNNGTGTGTTGACTTTGTATCCTGC
AGCACACGTCTGAACTCTTTCCCTACACGACGCTCTTCCGATCTNNNGTGTGTTGACTTTGTATCCTGC
AGCACACGTCTGAACTCTTTCCCTACACGACGCTCTTCCGATCTNNNGTGTGTTGACTTTGTATCCTGC
GCACACGTCTGAACTCTTTCCCTACACGACGCTCTTCCGATCTNNNTTCTGCCTCTATTTTGTCTGCCTC
GCACACGTCTGAACTCTTTCCCTACACGACGCTCTTCCGATCTNNNTTCTGCCTCTATTTTGTCTGCCTC
GCACACGTCTGAACTCTTTCCCTACACGACGCTCTTCCGATCTNNNTTCTGCCTCTATTTTGTCTGCCTC

AGCACACGTCTGAACTCTTTCCCTACACGACGCTCTTCCGATCTNNNTCCCCTTCTTGTTTCTTCCTCAACA
AGCACACGTCTGAACTCTTTCCCTACACGACGCTCTTCCGATCTNNNTCCCCTTCTTGTTTCTTCCTCAACA
AGCACACGTCTGAACTCTTTCCCTACACGACGCTCTTCCGATCTNNNTCCCCTTCTTGTTTCTTCCTCAACA
GAGCACACGTCTGAACTCTTTCCCTACACGACGCTCTTCCGATCTNNNTTCAGCTGATCTCATCTGAGTG
GAGCACACGTCTGAACTCTTTCCCTACACGACGCTCTTCCGATCTNNNTTCAGCTGATCTCATCTGAGTG
GAGCACACGTCTGAACTCTTTCCCTACACGACGCTCTTCCGATCTNNNTTCAGCTGATCTCATCTGAGTG
AGCACACGTCTGAACTCTTTCCCTACACGACGCTCTTCCGATCTNNNATGGGAGATCTGACTGTGGTTT
AGCACACGTCTGAACTCTTTCCCTACACGACGCTCTTCCGATCTNNNATGGGAGATCTGACTGTGGTTT
AGCACACGTCTGAACTCTTTCCCTACACGACGCTCTTCCGATCTNNNATGGGAGATCTGACTGTGGTTT
AGCACACGTCTGAACTCTTTCCCTACACGACGCTCTTCCGATCTNNNTACAAAAAATTAGCCGGGTGT
AGCACACGTCTGAACTCTTTCCCTACACGACGCTCTTCCGATCTNNNTACAAAAAATTAGCCGGGTGT
AGCACACGTCTGAACTCTTTCCCTACACGACGCTCTTCCGATCTNNNTACAAAAAATTAGCCGGGTGT
GAGCACACGTCTGAACTCTTTCCCTACACGACGCTCTTCCGATCTNNNGGCTCACACCTGTAATTCCAGT
GAGCACACGTCTGAACTCTTTCCCTACACGACGCTCTTCCGATCTNNNGGCTCACACCTGTAATTCCAGT
GAGCACACGTCTGAACTCTTTCCCTACACGACGCTCTTCCGATCTNNNGGCTCACACCTGTAATTCCAGT
AGAGCACACGTCTGAACTCTTTCCCTACACGACGCTCTTCCGATCTNNNATTTGAAGGTACACAGTGTC
AGAGCACACGTCTGAACTCTTTCCCTACACGACGCTCTTCCGATCTNNNATTTGAAGGTACACAGTGTC
AGAGCACACGTCTGAACTCTTTCCCTACACGACGCTCTTCCGATCTNNNATTTGAAGGTACACAGTGTC
CACACGTCTGAACTCTTTCCCTACACGACGCTCTTCCGATCTNNNTTCCCTTTCCTTGTACATTCTGTCAT
CACACGTCTGAACTCTTTCCCTACACGACGCTCTTCCGATCTNNNTTCCCTTTCCTTGTACATTCTGTCAT
CACACGTCTGAACTCTTTCCCTACACGACGCTCTTCCGATCTNNNTTCCCTTTCCTTGTACATTCTGTCAT
GAGCACACGTCTGAACTCTTTCCCTACACGACGCTCTTCCGATCTNNNCCTTGTCCTTATTTCTCCAGGC
GAGCACACGTCTGAACTCTTTCCCTACACGACGCTCTTCCGATCTNNNCCTTGTCCTTATTTCTCCAGGC
GAGCACACGTCTGAACTCTTTCCCTACACGACGCTCTTCCGATCTNNNCCTTGTCCTTATTTCTCCAGGC
AGCACACGTCTGAACTCTTTCCCTACACGACGCTCTTCCGATCTNNNCTTGCAGTGAGCTGAGATCAT
AGCACACGTCTGAACTCTTTCCCTACACGACGCTCTTCCGATCTNNNCTTGCAGTGAGCTGAGATCAT
AGCACACGTCTGAACTCTTTCCCTACACGACGCTCTTCCGATCTNNNCTTGCAGTGAGCTGAGATCAT
GAGCACACGTCTGAACTCTTTCCCTACACGACGCTCTTCCGATCTNNNCTGAGGCAGGAGAATCACTTG
GAGCACACGTCTGAACTCTTTCCCTACACGACGCTCTTCCGATCTNNNCTGAGGCAGGAGAATCACTTG
GAGCACACGTCTGAACTCTTTCCCTACACGACGCTCTTCCGATCTNNNCTGAGGCAGGAGAATCACTTG
GAGCACACGTCTGAACTCTTTCCCTACACGACGCTCTTCCGATCTNNNGAGAATCTCTTGAGCCCAGTA
GAGCACACGTCTGAACTCTTTCCCTACACGACGCTCTTCCGATCTNNNGAGAATCTCTTGAGCCCAGTA
GAGCACACGTCTGAACTCTTTCCCTACACGACGCTCTTCCGATCTNNNGAGAATCTCTTGAGCCCAGTA
AGCACACGTCTGAACTCTTTCCCTACACGACGCTCTTCCGATCTNNNGAATTGTGGTGGAGGTTGTCA
AGCACACGTCTGAACTCTTTCCCTACACGACGCTCTTCCGATCTNNNGAATTGTGGTGGAGGTTGTCA
AGCACACGTCTGAACTCTTTCCCTACACGACGCTCTTCCGATCTNNNGAATTGTGGTGGAGGTTGTCA
AGAGCACACGTCTGAACTCTTTCCCTACACGACGCTCTTCCGATCTNNNAGGCAGAGAATTGCTTGAAC
AGAGCACACGTCTGAACTCTTTCCCTACACGACGCTCTTCCGATCTNNNAGGCAGAGAATTGCTTGAAC
AGAGCACACGTCTGAACTCTTTCCCTACACGACGCTCTTCCGATCTNNNAGGCAGAGAATTGCTTGAAC
AGCACACGTCTGAACTCTTTCCCTACACGACGCTCTTCCGATCTNNNCACCAGCTGTGCTTATCAGAAAC
AGCACACGTCTGAACTCTTTCCCTACACGACGCTCTTCCGATCTNNNCACCAGCTGTGCTTATCAGAAAC
AGCACACGTCTGAACTCTTTCCCTACACGACGCTCTTCCGATCTNNNCACCAGCTGTGCTTATCAGAAAC
AGCACACGTCTGAACTCTTTCCCTACACGACGCTCTTCCGATCTNNNTGTGAAGAAAGTGTACAGGAC

AGCACACGTCTGAACTCTTTCCCTACACGACGCTCTTCCGATCTNNNTGTGAAGAAAGTGTACAGGA

AGCACACGTCTGAACTCTTTCCCTACACGACGCTCTTCCGATCTNNNTGTGAAGAAAGTGTACAGGA

AGCACACGTCTGAACTCTTTCCCTACACGACGCTCTTCCGATCTNNNGCAACAAATATGGGAGTGCAG

AGCACACGTCTGAACTCTTTCCCTACACGACGCTCTTCCGATCTNNNGCAACAAATATGGGAGTGCAG

AGCACACGTCTGAACTCTTTCCCTACACGACGCTCTTCCGATCTNNNGCAACAAATATGGGAGTGCAG

CACACGTCTGAACTCTTTCCCTACACGACGCTCTTCCGATCTNNNAGAGATTATCCTGGATTGTCTCAT

CACACGTCTGAACTCTTTCCCTACACGACGCTCTTCCGATCTNNNAGAGATTATCCTGGATTGTCTCAT

CACACGTCTGAACTCTTTCCCTACACGACGCTCTTCCGATCTNNNAGAGATTATCCTGGATTGTCTCAT

GAGCACACGTCTGAACTCTTTCCCTACACGACGCTCTTCCGATCTNNNGAGTAGCTGGGATTACAGGA

GAGCACACGTCTGAACTCTTTCCCTACACGACGCTCTTCCGATCTNNNGAGTAGCTGGGATTACAGGA

GAGCACACGTCTGAACTCTTTCCCTACACGACGCTCTTCCGATCTNNNGAGTAGCTGGGATTACAGGA

AGCACACGTCTGAACTCTTTCCCTACACGACGCTCTTCCGATCTNNNTCTCTGAGTAGCTGGGACTACA

AGCACACGTCTGAACTCTTTCCCTACACGACGCTCTTCCGATCTNNNTCTCTGAGTAGCTGGGACTACA

AGCACACGTCTGAACTCTTTCCCTACACGACGCTCTTCCGATCTNNNTCTCTGAGTAGCTGGGACTACA

GAGCACACGTCTGAACTCTTTCCCTACACGACGCTCTTCCGATCTNNNTTTCTAACAAGCTCCCAGATGC

GAGCACACGTCTGAACTCTTTCCCTACACGACGCTCTTCCGATCTNNNTTTCTAACAAGCTCCCAGATGC

GAGCACACGTCTGAACTCTTTCCCTACACGACGCTCTTCCGATCTNNNTTTCTAACAAGCTCCCAGATGC

GAGCACACGTCTGAACTCTTTCCCTACACGACGCTCTTCCGATCTNNNGTAAGGGTGTGTGGATGTGT

GAGCACACGTCTGAACTCTTTCCCTACACGACGCTCTTCCGATCTNNNGTAAGGGTGTGTGGATGTGT

GAGCACACGTCTGAACTCTTTCCCTACACGACGCTCTTCCGATCTNNNGTAAGGGTGTGTGGATGTGT

AGCACACGTCTGAACTCTTTCCCTACACGACGCTCTTCCGATCTNNNTAGAAGCAATTTCTGAAGCAGC

AGCACACGTCTGAACTCTTTCCCTACACGACGCTCTTCCGATCTNNNTAGAAGCAATTTCTGAAGCAGC

AGCACACGTCTGAACTCTTTCCCTACACGACGCTCTTCCGATCTNNNTAGAAGCAATTTCTGAAGCAGC

AGAGCACACGTCTGAACTCTTTCCCTACACGACGCTCTTCCGATCTNNNAGAATGGCTTGAACTTGAGA

AGAGCACACGTCTGAACTCTTTCCCTACACGACGCTCTTCCGATCTNNNAGAATGGCTTGAACTTGAGA

AGAGCACACGTCTGAACTCTTTCCCTACACGACGCTCTTCCGATCTNNNAGAATGGCTTGAACTTGAGA

AGCACACGTCTGAACTCTTTCCCTACACGACGCTCTTCCGATCTNNNTCAACGTCCCTCCTAGAAAACTG

AGCACACGTCTGAACTCTTTCCCTACACGACGCTCTTCCGATCTNNNTCAACGTCCCTCCTAGAAAACTG

AGCACACGTCTGAACTCTTTCCCTACACGACGCTCTTCCGATCTNNNTCAACGTCCCTCCTAGAAAACTG

AGAGCACACGTCTGAACTCTTTCCCTACACGACGCTCTTCCGATCTNNNCATAAGGATGAGGGATGG

AGAGCACACGTCTGAACTCTTTCCCTACACGACGCTCTTCCGATCTNNNCATAAGGATGAGGGATGG

AGAGCACACGTCTGAACTCTTTCCCTACACGACGCTCTTCCGATCTNNNCATAAGGATGAGGGATGG

AGCACACGTCTGAACTCTTTCCCTACACGACGCTCTTCCGATCTNNNTGAGGCAGGAGAATCACTTCA

AGCACACGTCTGAACTCTTTCCCTACACGACGCTCTTCCGATCTNNNTGAGGCAGGAGAATCACTTCA

AGCACACGTCTGAACTCTTTCCCTACACGACGCTCTTCCGATCTNNNTGAGGCAGGAGAATCACTTCA

AGCACACGTCTGAACTCTTTCCCTACACGACGCTCTTCCGATCTNNNAAATGAACAAACCAAGCTCCTC

AGCACACGTCTGAACTCTTTCCCTACACGACGCTCTTCCGATCTNNNAAATGAACAAACCAAGCTCCTC

AGCACACGTCTGAACTCTTTCCCTACACGACGCTCTTCCGATCTNNNAAATGAACAAACCAAGCTCCTC

GAGCACACGTCTGAACTCTTTCCCTACACGACGCTCTTCCGATCTNNNTGTGCAATCATGGTTTACTGC

GAGCACACGTCTGAACTCTTTCCCTACACGACGCTCTTCCGATCTNNNTGTGCAATCATGGTTTACTGC

GAGCACACGTCTGAACTCTTTCCCTACACGACGCTCTTCCGATCTNNNTGTGCAATCATGGTTTACTGC

AGCACACGTCTGAACTCTTTCCCTACACGACGCTCTTCCGATCTNNNAGATCTCTTGAGCCCAGAAGTT

AGCACACGTCTGAACTCTTTCCCTACACGACGCTCTTCCGATCTNNNAGATCTCTTGAGCCCAGAAGTT

AGCACACGTCTGAACTCTTTCCCTACACGACGCTCTTCCGATCTNNNAGATCTCTTGAGCCCAGAAGTT
AGCACACGTCTGAACTCTTTCCCTACACGACGCTCTTCCGATCTNNNGGATTACAGGCATGAGTCACTG
AGCACACGTCTGAACTCTTTCCCTACACGACGCTCTTCCGATCTNNNGGATTACAGGCATGAGTCACTG
AGCACACGTCTGAACTCTTTCCCTACACGACGCTCTTCCGATCTNNNGGATTACAGGCATGAGTCACTG
AGAGCACACGTCTGAACTCTTTCCCTACACGACGCTCTTCCGATCTNNNTGTGCTCAACAGAATGAAGA
AGAGCACACGTCTGAACTCTTTCCCTACACGACGCTCTTCCGATCTNNNTGTGCTCAACAGAATGAAGA
AGAGCACACGTCTGAACTCTTTCCCTACACGACGCTCTTCCGATCTNNNTGTGCTCAACAGAATGAAGA
GAGCACACGTCTGAACTCTTTCCCTACACGACGCTCTTCCGATCTNNNTCTGCTCCCTAAGAGTATGAG
GAGCACACGTCTGAACTCTTTCCCTACACGACGCTCTTCCGATCTNNNTCTGCTCCCTAAGAGTATGAG
GAGCACACGTCTGAACTCTTTCCCTACACGACGCTCTTCCGATCTNNNTCTGCTCCCTAAGAGTATGAG
AGCACACGTCTGAACTCTTTCCCTACACGACGCTCTTCCGATCTNNNCTTGCATTCCTGAGACAAATCCC
AGCACACGTCTGAACTCTTTCCCTACACGACGCTCTTCCGATCTNNNCTTGCATTCCTGAGACAAATCCC
AGCACACGTCTGAACTCTTTCCCTACACGACGCTCTTCCGATCTNNNCTTGCATTCCTGAGACAAATCCC
AGCACACGTCTGAACTCTTTCCCTACACGACGCTCTTCCGATCTNNNAGATTGGAGAGGATTGCTTGA
AGCACACGTCTGAACTCTTTCCCTACACGACGCTCTTCCGATCTNNNAGATTGGAGAGGATTGCTTGA
AGCACACGTCTGAACTCTTTCCCTACACGACGCTCTTCCGATCTNNNAGATTGGAGAGGATTGCTTGA
GAGCACACGTCTGAACTCTTTCCCTACACGACGCTCTTCCGATCTNNNCTGTTTCTGTGTTGTTTCACCC
GAGCACACGTCTGAACTCTTTCCCTACACGACGCTCTTCCGATCTNNNCTGTTTCTGTGTTGTTTCACCC
GAGCACACGTCTGAACTCTTTCCCTACACGACGCTCTTCCGATCTNNNCTGTTTCTGTGTTGTTTCACCC
AGCACACGTCTGAACTCTTTCCCTACACGACGCTCTTCCGATCTNNNCAGAGAATCACTTGAACCCAGG
AGCACACGTCTGAACTCTTTCCCTACACGACGCTCTTCCGATCTNNNCAGAGAATCACTTGAACCCAGG
AGCACACGTCTGAACTCTTTCCCTACACGACGCTCTTCCGATCTNNNCAGAGAATCACTTGAACCCAGG
AGAGCACACGTCTGAACTCTTTCCCTACACGACGCTCTTCCGATCTNNNCCAACATGGTAAAACCCTGTC
AGAGCACACGTCTGAACTCTTTCCCTACACGACGCTCTTCCGATCTNNNCCAACATGGTAAAACCCTGTC
AGAGCACACGTCTGAACTCTTTCCCTACACGACGCTCTTCCGATCTNNNCCAACATGGTAAAACCCTGTC
AGCACACGTCTGAACTCTTTCCCTACACGACGCTCTTCCGATCTNNNCCTCCTGAGTTTGAGCAATTCTC
AGCACACGTCTGAACTCTTTCCCTACACGACGCTCTTCCGATCTNNNCCTCCTGAGTTTGAGCAATTCTC
AGCACACGTCTGAACTCTTTCCCTACACGACGCTCTTCCGATCTNNNCCTCCTGAGTTTGAGCAATTCTC
AGCACACGTCTGAACTCTTTCCCTACACGACGCTCTTCCGATCTNNNCGATGCTCATTTCTCTTCAGCAG
AGCACACGTCTGAACTCTTTCCCTACACGACGCTCTTCCGATCTNNNCGATGCTCATTTCTCTTCAGCAG
AGCACACGTCTGAACTCTTTCCCTACACGACGCTCTTCCGATCTNNNCGATGCTCATTTCTCTTCAGCAG
GCACACGTCTGAACTCTTTCCCTACACGACGCTCTTCCGATCTNNNATGAAATTTGGCTGCTCCCAGACC
GCACACGTCTGAACTCTTTCCCTACACGACGCTCTTCCGATCTNNNATGAAATTTGGCTGCTCCCAGACC
GCACACGTCTGAACTCTTTCCCTACACGACGCTCTTCCGATCTNNNATGAAATTTGGCTGCTCCCAGACC
AGCACACGTCTGAACTCTTTCCCTACACGACGCTCTTCCGATCTNNNATTCTCAGGGAAGGGGATACT
AGCACACGTCTGAACTCTTTCCCTACACGACGCTCTTCCGATCTNNNATTCTCAGGGAAGGGGATACT
AGCACACGTCTGAACTCTTTCCCTACACGACGCTCTTCCGATCTNNNATTCTCAGGGAAGGGGATACT
AGCACACGTCTGAACTCTTTCCCTACACGACGCTCTTCCGATCTNNNTTGAAGAAAACACTGTGACCAG
AGCACACGTCTGAACTCTTTCCCTACACGACGCTCTTCCGATCTNNNTTGAAGAAAACACTGTGACCAG
AGCACACGTCTGAACTCTTTCCCTACACGACGCTCTTCCGATCTNNNTTGAAGAAAACACTGTGACCAG
GAGCACACGTCTGAACTCTTTCCCTACACGACGCTCTTCCGATCTNNNGGTGACAGAGTGAGACTCCAT
GAGCACACGTCTGAACTCTTTCCCTACACGACGCTCTTCCGATCTNNNGGTGACAGAGTGAGACTCCAT
GAGCACACGTCTGAACTCTTTCCCTACACGACGCTCTTCCGATCTNNNGGTGACAGAGTGAGACTCCAT

AGCACACGTCTGAACTCTTTCCCTACACGACGCTCTTCCGATCTNNNCCTGGACACAGTGAAGAACCT
AGCACACGTCTGAACTCTTTCCCTACACGACGCTCTTCCGATCTNNNCCTGGACACAGTGAAGAACCT
AGCACACGTCTGAACTCTTTCCCTACACGACGCTCTTCCGATCTNNNCCTGGACACAGTGAAGAACCT
AGCACACGTCTGAACTCTTTCCCTACACGACGCTCTTCCGATCTNNNGCAGGCAGTTTTACAGATGA
AGCACACGTCTGAACTCTTTCCCTACACGACGCTCTTCCGATCTNNNGCAGGCAGTTTTACAGATGA
AGCACACGTCTGAACTCTTTCCCTACACGACGCTCTTCCGATCTNNNGCAGGCAGTTTTACAGATGA
GCACACGTCTGAACTCTTTCCCTACACGACGCTCTTCCGATCTNNNAGGCTGAATGATGAGATTTCCC
GCACACGTCTGAACTCTTTCCCTACACGACGCTCTTCCGATCTNNNAGGCTGAATGATGAGATTTCCC
GCACACGTCTGAACTCTTTCCCTACACGACGCTCTTCCGATCTNNNAGGCTGAATGATGAGATTTCCC
GCACACGTCTGAACTCTTTCCCTACACGACGCTCTTCCGATCTNNNCTTTCAGGAGCTCCATACACAC
GCACACGTCTGAACTCTTTCCCTACACGACGCTCTTCCGATCTNNNCTTTCAGGAGCTCCATACACAC
GCACACGTCTGAACTCTTTCCCTACACGACGCTCTTCCGATCTNNNCTTTCAGGAGCTCCATACACAC
GCACACGTCTGAACTCTTTCCCTACACGACGCTCTTCCGATCTNNNGGACCCTCTCAATCTGAGTTTT
GCACACGTCTGAACTCTTTCCCTACACGACGCTCTTCCGATCTNNNGGACCCTCTCAATCTGAGTTTT
GCACACGTCTGAACTCTTTCCCTACACGACGCTCTTCCGATCTNNNGGACCCTCTCAATCTGAGTTTT
CACACGTCTGAACTCTTTCCCTACACGACGCTCTTCCGATCTNNNCCAGTTCCTAGCAATTCCTGAGA
CACACGTCTGAACTCTTTCCCTACACGACGCTCTTCCGATCTNNNCCAGTTCCTAGCAATTCCTGAGA
CACACGTCTGAACTCTTTCCCTACACGACGCTCTTCCGATCTNNNCCAGTTCCTAGCAATTCCTGAGA
AGCACACGTCTGAACTCTTTCCCTACACGACGCTCTTCCGATCTNNNCGCCCGGCTTTTCTTTTTTTTT
AGCACACGTCTGAACTCTTTCCCTACACGACGCTCTTCCGATCTNNNCGCCCGGCTTTTCTTTTTTTTT
AGCACACGTCTGAACTCTTTCCCTACACGACGCTCTTCCGATCTNNNCGCCCGGCTTTTCTTTTTTTTT
GCACACGTCTGAACTCTTTCCCTACACGACGCTCTTCCGATCTNNNGGTGACATAGCAAGACTCCGTAT
GCACACGTCTGAACTCTTTCCCTACACGACGCTCTTCCGATCTNNNGGTGACATAGCAAGACTCCGTAT
GCACACGTCTGAACTCTTTCCCTACACGACGCTCTTCCGATCTNNNGGTGACATAGCAAGACTCCGTAT
ACACGTCTGAACTCTTTCCCTACACGACGCTCTTCCGATCTNNNCTAGTGAAAAAAAGAAAAGAAGC
ACACGTCTGAACTCTTTCCCTACACGACGCTCTTCCGATCTNNNCTAGTGAAAAAAAGAAAAGAAGC
ACACGTCTGAACTCTTTCCCTACACGACGCTCTTCCGATCTNNNCTAGTGAAAAAAAGAAAAGAAGC
AGAGCACACGTCTGAACTCTTTCCCTACACGACGCTCTTCCGATCTNNNTGCTGCAGCCTTTCTCATAT
AGAGCACACGTCTGAACTCTTTCCCTACACGACGCTCTTCCGATCTNNNTGCTGCAGCCTTTCTCATAT
AGAGCACACGTCTGAACTCTTTCCCTACACGACGCTCTTCCGATCTNNNTGCTGCAGCCTTTCTCATAT
AGCACACGTCTGAACTCTTTCCCTACACGACGCTCTTCCGATCTNNNTCTCTTCCACAGTGATTCAAGG
AGCACACGTCTGAACTCTTTCCCTACACGACGCTCTTCCGATCTNNNTCTCTTCCACAGTGATTCAAGG
AGCACACGTCTGAACTCTTTCCCTACACGACGCTCTTCCGATCTNNNTCTCTTCCACAGTGATTCAAGG
GAGCACACGTCTGAACTCTTTCCCTACACGACGCTCTTCCGATCTNNNTCATTACAAACACATCCACGC
GAGCACACGTCTGAACTCTTTCCCTACACGACGCTCTTCCGATCTNNNTCATTACAAACACATCCACGC
GAGCACACGTCTGAACTCTTTCCCTACACGACGCTCTTCCGATCTNNNTCATTACAAACACATCCACGC
AGCACACGTCTGAACTCTTTCCCTACACGACGCTCTTCCGATCTNNNTGCAGTGAGTCATGATCAAGT
AGCACACGTCTGAACTCTTTCCCTACACGACGCTCTTCCGATCTNNNTGCAGTGAGTCATGATCAAGT
AGCACACGTCTGAACTCTTTCCCTACACGACGCTCTTCCGATCTNNNTGCAGTGAGTCATGATCAAGT
AGCACACGTCTGAACTCTTTCCCTACACGACGCTCTTCCGATCTNNNAATGGCAAAGTGAGACTCCAT
AGCACACGTCTGAACTCTTTCCCTACACGACGCTCTTCCGATCTNNNAATGGCAAAGTGAGACTCCAT
AGCACACGTCTGAACTCTTTCCCTACACGACGCTCTTCCGATCTNNNAATGGCAAAGTGAGACTCCAT
CACACGTCTGAACTCTTTCCCTACACGACGCTCTTCCGATCTNNNCCACGAAAACCCTAGTGAATCAC

GCACACGTCTGAACTCTTTCCCTACACGACGCTCTTCCGATCTNNNCCACGAAAACCCTAGTGAATCACC
GCACACGTCTGAACTCTTTCCCTACACGACGCTCTTCCGATCTNNNCCACGAAAACCCTAGTGAATCACC
AGCACACGTCTGAACTCTTTCCCTACACGACGCTCTTCCGATCTNNNCTAAGACTACAGGTCACACCAC
AGCACACGTCTGAACTCTTTCCCTACACGACGCTCTTCCGATCTNNNCTAAGACTACAGGTCACACCAC
AGCACACGTCTGAACTCTTTCCCTACACGACGCTCTTCCGATCTNNNCTAAGACTACAGGTCACACCAC
GAGCACACGTCTGAACTCTTTCCCTACACGACGCTCTTCCGATCTNNNGGGTTTGCAATGCATAGGTTC
GAGCACACGTCTGAACTCTTTCCCTACACGACGCTCTTCCGATCTNNNGGGTTTGCAATGCATAGGTTC
GAGCACACGTCTGAACTCTTTCCCTACACGACGCTCTTCCGATCTNNNGGGTTTGCAATGCATAGGTTC
AGCACACGTCTGAACTCTTTCCCTACACGACGCTCTTCCGATCTNNNTGGACATCAGGACTCTGTCACAT
AGCACACGTCTGAACTCTTTCCCTACACGACGCTCTTCCGATCTNNNTGGACATCAGGACTCTGTCACAT
AGCACACGTCTGAACTCTTTCCCTACACGACGCTCTTCCGATCTNNNTGGACATCAGGACTCTGTCACAT
AGCACACGTCTGAACTCTTTCCCTACACGACGCTCTTCCGATCTNNNCCTTTGATTCCTGACCTCCAACAC
AGCACACGTCTGAACTCTTTCCCTACACGACGCTCTTCCGATCTNNNCCTTTGATTCCTGACCTCCAACAC
AGCACACGTCTGAACTCTTTCCCTACACGACGCTCTTCCGATCTNNNCCTTTGATTCCTGACCTCCAACAC
GAGCACACGTCTGAACTCTTTCCCTACACGACGCTCTTCCGATCTNNNATTGTATGCACCTGTAGTCCCA
GAGCACACGTCTGAACTCTTTCCCTACACGACGCTCTTCCGATCTNNNATTGTATGCACCTGTAGTCCCA
GAGCACACGTCTGAACTCTTTCCCTACACGACGCTCTTCCGATCTNNNATTGTATGCACCTGTAGTCCCA
AGCACACGTCTGAACTCTTTCCCTACACGACGCTCTTCCGATCTNNNTTTAGCCACTCACCAACTGAAGC
AGCACACGTCTGAACTCTTTCCCTACACGACGCTCTTCCGATCTNNNTTTAGCCACTCACCAACTGAAGC
AGCACACGTCTGAACTCTTTCCCTACACGACGCTCTTCCGATCTNNNTTTAGCCACTCACCAACTGAAGC
AGCACACGTCTGAACTCTTTCCCTACACGACGCTCTTCCGATCTNNNGAGATCATAAGCCACTGCACTC
AGCACACGTCTGAACTCTTTCCCTACACGACGCTCTTCCGATCTNNNGAGATCATAAGCCACTGCACTC
AGCACACGTCTGAACTCTTTCCCTACACGACGCTCTTCCGATCTNNNGAGATCATAAGCCACTGCACTC
AGCACACGTCTGAACTCTTTCCCTACACGACGCTCTTCCGATCTNNNTTTGGGAAACAAGACCCTTCTGC
AGCACACGTCTGAACTCTTTCCCTACACGACGCTCTTCCGATCTNNNTTTGGGAAACAAGACCCTTCTGC
AGCACACGTCTGAACTCTTTCCCTACACGACGCTCTTCCGATCTNNNTTTGGGAAACAAGACCCTTCTGC
AGCACACGTCTGAACTCTTTCCCTACACGACGCTCTTCCGATCTNNNAAAAAAAAAATTAGCCGGGCG
AGCACACGTCTGAACTCTTTCCCTACACGACGCTCTTCCGATCTNNNAAAAAAAAAATTAGCCGGGCG
AGCACACGTCTGAACTCTTTCCCTACACGACGCTCTTCCGATCTNNNAAAAAAAAAATTAGCCGGGCG
GAGCACACGTCTGAACTCTTTCCCTACACGACGCTCTTCCGATCTNNNTCAGTTCAAGTCCACTGAGGG
GAGCACACGTCTGAACTCTTTCCCTACACGACGCTCTTCCGATCTNNNTCAGTTCAAGTCCACTGAGGG
GAGCACACGTCTGAACTCTTTCCCTACACGACGCTCTTCCGATCTNNNTCAGTTCAAGTCCACTGAGGG
GAGCACACGTCTGAACTCTTTCCCTACACGACGCTCTTCCGATCTNNNAATGTGTACCACTCTGCATGG
GAGCACACGTCTGAACTCTTTCCCTACACGACGCTCTTCCGATCTNNNAATGTGTACCACTCTGCATGG
GAGCACACGTCTGAACTCTTTCCCTACACGACGCTCTTCCGATCTNNNAATGTGTACCACTCTGCATGG
AGAGCACACGTCTGAACTCTTTCCCTACACGACGCTCTTCCGATCTNNNTATGTGTGTGTGTGTGTGTG
AGAGCACACGTCTGAACTCTTTCCCTACACGACGCTCTTCCGATCTNNNTATGTGTGTGTGTGTGTGTG
AGAGCACACGTCTGAACTCTTTCCCTACACGACGCTCTTCCGATCTNNNTATGTGTGTGTGTGTGTGTG
AGCACACGTCTGAACTCTTTCCCTACACGACGCTCTTCCGATCTNNNCCGTCCTCCTTATTTAGTCCCCC
AGCACACGTCTGAACTCTTTCCCTACACGACGCTCTTCCGATCTNNNCCGTCCTCCTTATTTAGTCCCCC
AGCACACGTCTGAACTCTTTCCCTACACGACGCTCTTCCGATCTNNNCCGTCCTCCTTATTTAGTCCCCC
AGAGCACACGTCTGAACTCTTTCCCTACACGACGCTCTTCCGATCTNNNTTTCTATGTTCTTCCTCCTCCC
AGAGCACACGTCTGAACTCTTTCCCTACACGACGCTCTTCCGATCTNNNTTTCTATGTTCTTCCTCCTCCC

AGAGCACACGTCTGAACTCTTTCCCTACACGACGCTCTTCCGATCTNNNTTTCTATGTTCTTCCTCCTCC
AGCACACGTCTGAACTCTTTCCCTACACGACGCTCTTCCGATCTNNNTGAGCAACAGAATGAGTCATC
AGCACACGTCTGAACTCTTTCCCTACACGACGCTCTTCCGATCTNNNTGAGCAACAGAATGAGTCATC
AGCACACGTCTGAACTCTTTCCCTACACGACGCTCTTCCGATCTNNNTGAGCAACAGAATGAGTCATC
AGCACACGTCTGAACTCTTTCCCTACACGACGCTCTTCCGATCTNNNCCAAACTTGGCTTCCTTCTTGA
AGCACACGTCTGAACTCTTTCCCTACACGACGCTCTTCCGATCTNNNCCAAACTTGGCTTCCTTCTTGA
AGCACACGTCTGAACTCTTTCCCTACACGACGCTCTTCCGATCTNNNCCAAACTTGGCTTCCTTCTTGA
CACACGTCTGAACTCTTTCCCTACACGACGCTCTTCCGATCTNNNCATGACACAGTGGCTATGAGGAG
CACACGTCTGAACTCTTTCCCTACACGACGCTCTTCCGATCTNNNCATGACACAGTGGCTATGAGGAG
CACACGTCTGAACTCTTTCCCTACACGACGCTCTTCCGATCTNNNCATGACACAGTGGCTATGAGGAG
GAGCACACGTCTGAACTCTTTCCCTACACGACGCTCTTCCGATCTNNNGCACACCTATGGTTCCAT
GAGCACACGTCTGAACTCTTTCCCTACACGACGCTCTTCCGATCTNNNGCACACCTATGGTTCCAT
GAGCACACGTCTGAACTCTTTCCCTACACGACGCTCTTCCGATCTNNNGCACACCTATGGTTCCAT
GCACACGTCTGAACTCTTTCCCTACACGACGCTCTTCCGATCTNNNTACCTGAATTTCCACGATGACC
GCACACGTCTGAACTCTTTCCCTACACGACGCTCTTCCGATCTNNNTACCTGAATTTCCACGATGACC
GCACACGTCTGAACTCTTTCCCTACACGACGCTCTTCCGATCTNNNTACCTGAATTTCCACGATGACC
AGCACACGTCTGAACTCTTTCCCTACACGACGCTCTTCCGATCTNNNAGACTCAGAGCCAGGATAAAT
AGCACACGTCTGAACTCTTTCCCTACACGACGCTCTTCCGATCTNNNAGACTCAGAGCCAGGATAAAT
AGCACACGTCTGAACTCTTTCCCTACACGACGCTCTTCCGATCTNNNAGACTCAGAGCCAGGATAAAT
GAGCACACGTCTGAACTCTTTCCCTACACGACGCTCTTCCGATCTNNNACAAAAATTAGCCAGACGTG
GAGCACACGTCTGAACTCTTTCCCTACACGACGCTCTTCCGATCTNNNACAAAAATTAGCCAGACGTG
GAGCACACGTCTGAACTCTTTCCCTACACGACGCTCTTCCGATCTNNNACAAAAATTAGCCAGACGTG
AGCACACGTCTGAACTCTTTCCCTACACGACGCTCTTCCGATCTNNNGAGACTTCCTTGTGTCATAATA
AGCACACGTCTGAACTCTTTCCCTACACGACGCTCTTCCGATCTNNNGAGACTTCCTTGTGTCATAATA
AGCACACGTCTGAACTCTTTCCCTACACGACGCTCTTCCGATCTNNNGAGACTTCCTTGTGTCATAATA
AGCACACGTCTGAACTCTTTCCCTACACGACGCTCTTCCGATCTNNNTTCAGGGGTACATATTCAGGA
AGCACACGTCTGAACTCTTTCCCTACACGACGCTCTTCCGATCTNNNTTCAGGGGTACATATTCAGGA
AGCACACGTCTGAACTCTTTCCCTACACGACGCTCTTCCGATCTNNNTTCAGGGGTACATATTCAGGA
AGCACACGTCTGAACTCTTTCCCTACACGACGCTCTTCCGATCTNNNTGGTTCGATGTGACCAGAAGA
AGCACACGTCTGAACTCTTTCCCTACACGACGCTCTTCCGATCTNNNTGGTTCGATGTGACCAGAAGA
AGCACACGTCTGAACTCTTTCCCTACACGACGCTCTTCCGATCTNNNTGGTTCGATGTGACCAGAAGA
AGCACACGTCTGAACTCTTTCCCTACACGACGCTCTTCCGATCTNNNATGAATTGCCTCTACTCCCACA
AGCACACGTCTGAACTCTTTCCCTACACGACGCTCTTCCGATCTNNNATGAATTGCCTCTACTCCCACA
AGCACACGTCTGAACTCTTTCCCTACACGACGCTCTTCCGATCTNNNATGAATTGCCTCTACTCCCACA
AGCACACGTCTGAACTCTTTCCCTACACGACGCTCTTCCGATCTNNNCTGGAGAAGGGGAGAAGAGG
AGCACACGTCTGAACTCTTTCCCTACACGACGCTCTTCCGATCTNNNCTGGAGAAGGGGAGAAGAGG
AGCACACGTCTGAACTCTTTCCCTACACGACGCTCTTCCGATCTNNNCTGGAGAAGGGGAGAAGAGG
GCACACGTCTGAACTCTTTCCCTACACGACGCTCTTCCGATCTNNNAAAAAATTAGCTGGGATGGTG
GCACACGTCTGAACTCTTTCCCTACACGACGCTCTTCCGATCTNNNAAAAAATTAGCTGGGATGGTG
GCACACGTCTGAACTCTTTCCCTACACGACGCTCTTCCGATCTNNNAAAAAATTAGCTGGGATGGTG
AGCACACGTCTGAACTCTTTCCCTACACGACGCTCTTCCGATCTNNNGAGATAACGCAATTGCACTCC
AGCACACGTCTGAACTCTTTCCCTACACGACGCTCTTCCGATCTNNNGAGATAACGCAATTGCACTCC
AGCACACGTCTGAACTCTTTCCCTACACGACGCTCTTCCGATCTNNNGAGATAACGCAATTGCACTCC

AGCACACGTCTGAACTCTTTCCCTACACGACGCTCTTCCGATCTNNNCCTTTCAGAAGGCTAGTGTTCT
AGCACACGTCTGAACTCTTTCCCTACACGACGCTCTTCCGATCTNNNCCTTTCAGAAGGCTAGTGTTCT
AGCACACGTCTGAACTCTTTCCCTACACGACGCTCTTCCGATCTNNNCCTTTCAGAAGGCTAGTGTTCT
AGCACACGTCTGAACTCTTTCCCTACACGACGCTCTTCCGATCTNNNGACTCTGAATTCAGTCACCCCT
AGCACACGTCTGAACTCTTTCCCTACACGACGCTCTTCCGATCTNNNGACTCTGAATTCAGTCACCCCT
AGCACACGTCTGAACTCTTTCCCTACACGACGCTCTTCCGATCTNNNGACTCTGAATTCAGTCACCCCT
AGCACACGTCTGAACTCTTTCCCTACACGACGCTCTTCCGATCTNNNCAGAGAATAGGACCCACTCCCT
AGCACACGTCTGAACTCTTTCCCTACACGACGCTCTTCCGATCTNNNCAGAGAATAGGACCCACTCCCT
AGCACACGTCTGAACTCTTTCCCTACACGACGCTCTTCCGATCTNNNCAGAGAATAGGACCCACTCCCT
AGCACACGTCTGAACTCTTTCCCTACACGACGCTCTTCCGATCTNNNTGTTGGGGTGTTCAAAACATAG
AGCACACGTCTGAACTCTTTCCCTACACGACGCTCTTCCGATCTNNNTGTTGGGGTGTTCAAAACATAG
AGCACACGTCTGAACTCTTTCCCTACACGACGCTCTTCCGATCTNNNTGTTGGGGTGTTCAAAACATAG
AGCACACGTCTGAACTCTTTCCCTACACGACGCTCTTCCGATCTNNNGCACATGGTTGGCATTCAGTAA
AGCACACGTCTGAACTCTTTCCCTACACGACGCTCTTCCGATCTNNNGCACATGGTTGGCATTCAGTAA
AGCACACGTCTGAACTCTTTCCCTACACGACGCTCTTCCGATCTNNNGCACATGGTTGGCATTCAGTAA
GAGCACACGTCTGAACTCTTTCCCTACACGACGCTCTTCCGATCTNNNCTCGCCCATTCCATGAATTTCT
GAGCACACGTCTGAACTCTTTCCCTACACGACGCTCTTCCGATCTNNNCTCGCCCATTCCATGAATTTCT
GAGCACACGTCTGAACTCTTTCCCTACACGACGCTCTTCCGATCTNNNCTCGCCCATTCCATGAATTTCT
AGCACACGTCTGAACTCTTTCCCTACACGACGCTCTTCCGATCTNNNAGGCTTTACAGTTAGTTCTGC
AGCACACGTCTGAACTCTTTCCCTACACGACGCTCTTCCGATCTNNNAGGCTTTACAGTTAGTTCTGC
AGCACACGTCTGAACTCTTTCCCTACACGACGCTCTTCCGATCTNNNAGGCTTTACAGTTAGTTCTGC
AGCACACGTCTGAACTCTTTCCCTACACGACGCTCTTCCGATCTNNNCCATAAGAACAAAGATACATGT
AGCACACGTCTGAACTCTTTCCCTACACGACGCTCTTCCGATCTNNNCCATAAGAACAAAGATACATGT
AGCACACGTCTGAACTCTTTCCCTACACGACGCTCTTCCGATCTNNNCCATAAGAACAAAGATACATGT
GAGCACACGTCTGAACTCTTTCCCTACACGACGCTCTTCCGATCTNNNCTTCAGTAACAACTGCACAGC
GAGCACACGTCTGAACTCTTTCCCTACACGACGCTCTTCCGATCTNNNCTTCAGTAACAACTGCACAGC
GAGCACACGTCTGAACTCTTTCCCTACACGACGCTCTTCCGATCTNNNCTTCAGTAACAACTGCACAGC
GAGCACACGTCTGAACTCTTTCCCTACACGACGCTCTTCCGATCTNNNCTGGGAAGAGAGTGAGGAT
GAGCACACGTCTGAACTCTTTCCCTACACGACGCTCTTCCGATCTNNNCTGGGAAGAGAGTGAGGAT
GAGCACACGTCTGAACTCTTTCCCTACACGACGCTCTTCCGATCTNNNCTGGGAAGAGAGTGAGGAT
GAGCACACGTCTGAACTCTTTCCCTACACGACGCTCTTCCGATCTNNNTAGAAAGTGGTAGTGCCTGC
GAGCACACGTCTGAACTCTTTCCCTACACGACGCTCTTCCGATCTNNNTAGAAAGTGGTAGTGCCTGC
GAGCACACGTCTGAACTCTTTCCCTACACGACGCTCTTCCGATCTNNNTAGAAAGTGGTAGTGCCTGC
AGCACACGTCTGAACTCTTTCCCTACACGACGCTCTTCCGATCTNNNACAGCAAGACCTCATCTGTACT
AGCACACGTCTGAACTCTTTCCCTACACGACGCTCTTCCGATCTNNNACAGCAAGACCTCATCTGTACT
AGCACACGTCTGAACTCTTTCCCTACACGACGCTCTTCCGATCTNNNACAGCAAGACCTCATCTGTACT
GCACACGTCTGAACTCTTTCCCTACACGACGCTCTTCCGATCTNNNCCACACCCAGCTAGTTTTTGTAT
GCACACGTCTGAACTCTTTCCCTACACGACGCTCTTCCGATCTNNNCCACACCCAGCTAGTTTTTGTAT
GCACACGTCTGAACTCTTTCCCTACACGACGCTCTTCCGATCTNNNCCACACCCAGCTAGTTTTTGTAT
AGCACACGTCTGAACTCTTTCCCTACACGACGCTCTTCCGATCTNNNATGACTCATGTCTGCATCTCTG
AGCACACGTCTGAACTCTTTCCCTACACGACGCTCTTCCGATCTNNNATGACTCATGTCTGCATCTCTG
AGCACACGTCTGAACTCTTTCCCTACACGACGCTCTTCCGATCTNNNATGACTCATGTCTGCATCTCTG
GAGCACACGTCTGAACTCTTTCCCTACACGACGCTCTTCCGATCTNNNAAGAATGAAAGCATTGGGA

GAGCACACGTCTGAACTCTTTCCCTACACGACGCTCTTCCGATCTNNNAAGAATGAAAGCATTGGGAC
GAGCACACGTCTGAACTCTTTCCCTACACGACGCTCTTCCGATCTNNNAAGAATGAAAGCATTGGGAC
GCACACGTCTGAACTCTTTCCCTACACGACGCTCTTCCGATCTNNNCTCAGCCTCCCAAAGTACTGGTA
GCACACGTCTGAACTCTTTCCCTACACGACGCTCTTCCGATCTNNNCTCAGCCTCCCAAAGTACTGGTA
GCACACGTCTGAACTCTTTCCCTACACGACGCTCTTCCGATCTNNNCTCAGCCTCCCAAAGTACTGGTA
GAGCACACGTCTGAACTCTTTCCCTACACGACGCTCTTCCGATCTNNNGCTGGGTTTTCCTTGAACTTC
GAGCACACGTCTGAACTCTTTCCCTACACGACGCTCTTCCGATCTNNNGCTGGGTTTTCCTTGAACTTC
GAGCACACGTCTGAACTCTTTCCCTACACGACGCTCTTCCGATCTNNNGCTGGGTTTTCCTTGAACTTC
GAGCACACGTCTGAACTCTTTCCCTACACGACGCTCTTCCGATCTNNNGAAAAGAGAAGAGAGCAGC
GAGCACACGTCTGAACTCTTTCCCTACACGACGCTCTTCCGATCTNNNGAAAAGAGAAGAGAGCAGC
GAGCACACGTCTGAACTCTTTCCCTACACGACGCTCTTCCGATCTNNNGAAAAGAGAAGAGAGCAGC
AGCACACGTCTGAACTCTTTCCCTACACGACGCTCTTCCGATCTNNNGAATAGCTCATCACATGCCACA
AGCACACGTCTGAACTCTTTCCCTACACGACGCTCTTCCGATCTNNNGAATAGCTCATCACATGCCACA
AGCACACGTCTGAACTCTTTCCCTACACGACGCTCTTCCGATCTNNNGAATAGCTCATCACATGCCACA
GCACACGTCTGAACTCTTTCCCTACACGACGCTCTTCCGATCTNNNGACTGTAATCCCAGCTACTCCAG
GCACACGTCTGAACTCTTTCCCTACACGACGCTCTTCCGATCTNNNGACTGTAATCCCAGCTACTCCAG
GCACACGTCTGAACTCTTTCCCTACACGACGCTCTTCCGATCTNNNGACTGTAATCCCAGCTACTCCAG
AGCACACGTCTGAACTCTTTCCCTACACGACGCTCTTCCGATCTNNNCTGTAGTCCCAGCTACTCAGGA
AGCACACGTCTGAACTCTTTCCCTACACGACGCTCTTCCGATCTNNNCTGTAGTCCCAGCTACTCAGGA
AGCACACGTCTGAACTCTTTCCCTACACGACGCTCTTCCGATCTNNNCTGTAGTCCCAGCTACTCAGGA
GAGCACACGTCTGAACTCTTTCCCTACACGACGCTCTTCCGATCTNNNTGCTGGTTTACACACTTGTAT
GAGCACACGTCTGAACTCTTTCCCTACACGACGCTCTTCCGATCTNNNTGCTGGTTTACACACTTGTAT
GAGCACACGTCTGAACTCTTTCCCTACACGACGCTCTTCCGATCTNNNTGCTGGTTTACACACTTGTAT
GAGCACACGTCTGAACTCTTTCCCTACACGACGCTCTTCCGATCTNNNAGCTTGCAGTGAGTCAAAAT
GAGCACACGTCTGAACTCTTTCCCTACACGACGCTCTTCCGATCTNNNAGCTTGCAGTGAGTCAAAAT
GAGCACACGTCTGAACTCTTTCCCTACACGACGCTCTTCCGATCTNNNAGCTTGCAGTGAGTCAAAAT
GCACACGTCTGAACTCTTTCCCTACACGACGCTCTTCCGATCTNNNGATTCCACATAGCAGTGAGACCA
GCACACGTCTGAACTCTTTCCCTACACGACGCTCTTCCGATCTNNNGATTCCACATAGCAGTGAGACCA
GCACACGTCTGAACTCTTTCCCTACACGACGCTCTTCCGATCTNNNGATTCCACATAGCAGTGAGACCA
GAGCACACGTCTGAACTCTTTCCCTACACGACGCTCTTCCGATCTNNNGCCCAGCCAGATGATTCTTTT
GAGCACACGTCTGAACTCTTTCCCTACACGACGCTCTTCCGATCTNNNGCCCAGCCAGATGATTCTTTT
GAGCACACGTCTGAACTCTTTCCCTACACGACGCTCTTCCGATCTNNNGCCCAGCCAGATGATTCTTTT
AGCACACGTCTGAACTCTTTCCCTACACGACGCTCTTCCGATCTNNNCATCACGAGGTTAGGAGATGC
AGCACACGTCTGAACTCTTTCCCTACACGACGCTCTTCCGATCTNNNCATCACGAGGTTAGGAGATGC
AGCACACGTCTGAACTCTTTCCCTACACGACGCTCTTCCGATCTNNNCATCACGAGGTTAGGAGATGC
AGCACACGTCTGAACTCTTTCCCTACACGACGCTCTTCCGATCTNNNTAGCTGGACAGTTGTGTACCAA
AGCACACGTCTGAACTCTTTCCCTACACGACGCTCTTCCGATCTNNNTAGCTGGACAGTTGTGTACCAA
AGCACACGTCTGAACTCTTTCCCTACACGACGCTCTTCCGATCTNNNTAGCTGGACAGTTGTGTACCAA
GAGCACACGTCTGAACTCTTTCCCTACACGACGCTCTTCCGATCTNNNGGAGGTTTTGAGTCAGGAG
GAGCACACGTCTGAACTCTTTCCCTACACGACGCTCTTCCGATCTNNNGGAGGTTTTGAGTCAGGAG
GAGCACACGTCTGAACTCTTTCCCTACACGACGCTCTTCCGATCTNNNGGAGGTTTTGAGTCAGGAG
AGCACACGTCTGAACTCTTTCCCTACACGACGCTCTTCCGATCTNNNTTAGAGTGCAGTGGCGTGATT
AGCACACGTCTGAACTCTTTCCCTACACGACGCTCTTCCGATCTNNNTTAGAGTGCAGTGGCGTGATT

GAGCACACGTCTGAACTCTTTCCCTACACGACGCTCTTCCGATCTNNNTTAGAGTGCAGTGGCGTGATT
GAGCACACGTCTGAACTCTTTCCCTACACGACGCTCTTCCGATCTNNNTGAGACCAGAGTTCAAGATCA
GAGCACACGTCTGAACTCTTTCCCTACACGACGCTCTTCCGATCTNNNTGAGACCAGAGTTCAAGATCA
GAGCACACGTCTGAACTCTTTCCCTACACGACGCTCTTCCGATCTNNNTGAGACCAGAGTTCAAGATCA
AGCACACGTCTGAACTCTTTCCCTACACGACGCTCTTCCGATCTNNNCCACACCTGTTTCTTCTTCTCCTC
AGCACACGTCTGAACTCTTTCCCTACACGACGCTCTTCCGATCTNNNCCACACCTGTTTCTTCTTCTCCTC
AGCACACGTCTGAACTCTTTCCCTACACGACGCTCTTCCGATCTNNNCCACACCTGTTTCTTCTTCTCCTC
GAGCACACGTCTGAACTCTTTCCCTACACGACGCTCTTCCGATCTNNNGTTGCAGTGAGCTGTGATCAT
GAGCACACGTCTGAACTCTTTCCCTACACGACGCTCTTCCGATCTNNNGTTGCAGTGAGCTGTGATCAT
GAGCACACGTCTGAACTCTTTCCCTACACGACGCTCTTCCGATCTNNNGTTGCAGTGAGCTGTGATCAT
GAGCACACGTCTGAACTCTTTCCCTACACGACGCTCTTCCGATCTNNNGATGCTCTCCAGTTTGTTGGA
GAGCACACGTCTGAACTCTTTCCCTACACGACGCTCTTCCGATCTNNNGATGCTCTCCAGTTTGTTGGA
GAGCACACGTCTGAACTCTTTCCCTACACGACGCTCTTCCGATCTNNNGATGCTCTCCAGTTTGTTGGA
GAGCACACGTCTGAACTCTTTCCCTACACGACGCTCTTCCGATCTNNNTTACTTCAGTAGTCCCATGTG
GAGCACACGTCTGAACTCTTTCCCTACACGACGCTCTTCCGATCTNNNTTACTTCAGTAGTCCCATGTG
GAGCACACGTCTGAACTCTTTCCCTACACGACGCTCTTCCGATCTNNNTTACTTCAGTAGTCCCATGTG
GAGCACACGTCTGAACTCTTTCCCTACACGACGCTCTTCCGATCTNNNGAGGAGAAGTTGGTCTTGGT
GAGCACACGTCTGAACTCTTTCCCTACACGACGCTCTTCCGATCTNNNGAGGAGAAGTTGGTCTTGGT
GAGCACACGTCTGAACTCTTTCCCTACACGACGCTCTTCCGATCTNNNGAGGAGAAGTTGGTCTTGGT
CACGTCTGAACTCTTTCCCTACACGACGCTCTTCCGATCTNNNTGACAAGATCAAAGATGAAGGAGG
CACGTCTGAACTCTTTCCCTACACGACGCTCTTCCGATCTNNNTGACAAGATCAAAGATGAAGGAGG
CACGTCTGAACTCTTTCCCTACACGACGCTCTTCCGATCTNNNTGACAAGATCAAAGATGAAGGAGG
ACACGTCTGAACTCTTTCCCTACACGACGCTCTTCCGATCTNNNGAGTAGCTCCCTAGGTTGAGGACAT
ACACGTCTGAACTCTTTCCCTACACGACGCTCTTCCGATCTNNNGAGTAGCTCCCTAGGTTGAGGACAT
ACACGTCTGAACTCTTTCCCTACACGACGCTCTTCCGATCTNNNGAGTAGCTCCCTAGGTTGAGGACAT
AGAGCACACGTCTGAACTCTTTCCCTACACGACGCTCTTCCGATCTNNNACACGGACAACAATTATGC
AGAGCACACGTCTGAACTCTTTCCCTACACGACGCTCTTCCGATCTNNNACACGGACAACAATTATGC
AGAGCACACGTCTGAACTCTTTCCCTACACGACGCTCTTCCGATCTNNNACACGGACAACAATTATGC
AGCACACGTCTGAACTCTTTCCCTACACGACGCTCTTCCGATCTNNNAAAGACTGTGAGTTGTGGTGA
AGCACACGTCTGAACTCTTTCCCTACACGACGCTCTTCCGATCTNNNAAAGACTGTGAGTTGTGGTGA
AGCACACGTCTGAACTCTTTCCCTACACGACGCTCTTCCGATCTNNNAAAGACTGTGAGTTGTGGTGA
AGAGCACACGTCTGAACTCTTTCCCTACACGACGCTCTTCCGATCTNNNTCATGTCACTGCAATCTCCTC
AGAGCACACGTCTGAACTCTTTCCCTACACGACGCTCTTCCGATCTNNNTCATGTCACTGCAATCTCCTC
AGAGCACACGTCTGAACTCTTTCCCTACACGACGCTCTTCCGATCTNNNTCATGTCACTGCAATCTCCTC
AGCACACGTCTGAACTCTTTCCCTACACGACGCTCTTCCGATCTNNNCTTGAACTCCTGACCTCCTGATC
AGCACACGTCTGAACTCTTTCCCTACACGACGCTCTTCCGATCTNNNCTTGAACTCCTGACCTCCTGATC
AGCACACGTCTGAACTCTTTCCCTACACGACGCTCTTCCGATCTNNNCTTGAACTCCTGACCTCCTGATC
AGCACACGTCTGAACTCTTTCCCTACACGACGCTCTTCCGATCTNNNCTGAGATCACACCACTGCACTT
AGCACACGTCTGAACTCTTTCCCTACACGACGCTCTTCCGATCTNNNCTGAGATCACACCACTGCACTT
AGCACACGTCTGAACTCTTTCCCTACACGACGCTCTTCCGATCTNNNCTGAGATCACACCACTGCACTT
GAGCACACGTCTGAACTCTTTCCCTACACGACGCTCTTCCGATCTNNNAACACTCCTCCATCATTTTGGG
GAGCACACGTCTGAACTCTTTCCCTACACGACGCTCTTCCGATCTNNNAACACTCCTCCATCATTTTGGG
GAGCACACGTCTGAACTCTTTCCCTACACGACGCTCTTCCGATCTNNNAACACTCCTCCATCATTTTGGG

AGCACACGTCTGAACTCTTTCCCTACACGACGCTCTTCCGATCTNNNGGACCAACAGTCAGCACTTACA
AGCACACGTCTGAACTCTTTCCCTACACGACGCTCTTCCGATCTNNNGGACCAACAGTCAGCACTTACA
AGCACACGTCTGAACTCTTTCCCTACACGACGCTCTTCCGATCTNNNGGACCAACAGTCAGCACTTACA
AGCACACGTCTGAACTCTTTCCCTACACGACGCTCTTCCGATCTNNNTGGGAAATGCAGGTTTATGAA
AGCACACGTCTGAACTCTTTCCCTACACGACGCTCTTCCGATCTNNNTGGGAAATGCAGGTTTATGAA
AGCACACGTCTGAACTCTTTCCCTACACGACGCTCTTCCGATCTNNNTGGGAAATGCAGGTTTATGAA
AGCACACGTCTGAACTCTTTCCCTACACGACGCTCTTCCGATCTNNNCTGTAGTCCTAGCTACTCGGAG
AGCACACGTCTGAACTCTTTCCCTACACGACGCTCTTCCGATCTNNNCTGTAGTCCTAGCTACTCGGAG
AGCACACGTCTGAACTCTTTCCCTACACGACGCTCTTCCGATCTNNNCTGTAGTCCTAGCTACTCGGAG
GAGCACACGTCTGAACTCTTTCCCTACACGACGCTCTTCCGATCTNNNGAGGCATAGCACCCTGAAGA
GAGCACACGTCTGAACTCTTTCCCTACACGACGCTCTTCCGATCTNNNGAGGCATAGCACCCTGAAGA
GAGCACACGTCTGAACTCTTTCCCTACACGACGCTCTTCCGATCTNNNGAGGCATAGCACCCTGAAGA
CACACGTCTGAACTCTTTCCCTACACGACGCTCTTCCGATCTNNNAAGAAGTCGGTGGACAAGAACAG
CACACGTCTGAACTCTTTCCCTACACGACGCTCTTCCGATCTNNNAAGAAGTCGGTGGACAAGAACAG
CACACGTCTGAACTCTTTCCCTACACGACGCTCTTCCGATCTNNNAAGAAGTCGGTGGACAAGAACAG
GAGCACACGTCTGAACTCTTTCCCTACACGACGCTCTTCCGATCTNNNCAGCAGATAAAAGGCAGTG
GAGCACACGTCTGAACTCTTTCCCTACACGACGCTCTTCCGATCTNNNCAGCAGATAAAAGGCAGTG
GAGCACACGTCTGAACTCTTTCCCTACACGACGCTCTTCCGATCTNNNCAGCAGATAAAAGGCAGTG
GCACACGTCTGAACTCTTTCCCTACACGACGCTCTTCCGATCTNNNATTTTTTGAGACAGTCTCGCCAT
GCACACGTCTGAACTCTTTCCCTACACGACGCTCTTCCGATCTNNNATTTTTTGAGACAGTCTCGCCAT
GCACACGTCTGAACTCTTTCCCTACACGACGCTCTTCCGATCTNNNATTTTTTGAGACAGTCTCGCCAT
AGCACACGTCTGAACTCTTTCCCTACACGACGCTCTTCCGATCTNNNGTGATGCCCTGTTGCAGATATT
AGCACACGTCTGAACTCTTTCCCTACACGACGCTCTTCCGATCTNNNGTGATGCCCTGTTGCAGATATT
AGCACACGTCTGAACTCTTTCCCTACACGACGCTCTTCCGATCTNNNGTGATGCCCTGTTGCAGATATT
AGCACACGTCTGAACTCTTTCCCTACACGACGCTCTTCCGATCTNNNCCCCTCTCTCTTTCTCTGAGGAT
AGCACACGTCTGAACTCTTTCCCTACACGACGCTCTTCCGATCTNNNCCCCTCTCTCTTTCTCTGAGGAT
AGCACACGTCTGAACTCTTTCCCTACACGACGCTCTTCCGATCTNNNCCCCTCTCTCTTTCTCTGAGGAT
AGAGCACACGTCTGAACTCTTTCCCTACACGACGCTCTTCCGATCTNNNAGAGGGAACAGTAATTGAG
AGAGCACACGTCTGAACTCTTTCCCTACACGACGCTCTTCCGATCTNNNAGAGGGAACAGTAATTGAG
AGAGCACACGTCTGAACTCTTTCCCTACACGACGCTCTTCCGATCTNNNAGAGGGAACAGTAATTGAG
AGCACACGTCTGAACTCTTTCCCTACACGACGCTCTTCCGATCTNNNGAGGCAGGAGAATTGCTTGAA
AGCACACGTCTGAACTCTTTCCCTACACGACGCTCTTCCGATCTNNNGAGGCAGGAGAATTGCTTGAA
AGCACACGTCTGAACTCTTTCCCTACACGACGCTCTTCCGATCTNNNGAGGCAGGAGAATTGCTTGAA
AGCACACGTCTGAACTCTTTCCCTACACGACGCTCTTCCGATCTNNNTTCACATCATTCTCCTGCCTCAG
AGCACACGTCTGAACTCTTTCCCTACACGACGCTCTTCCGATCTNNNTTCACATCATTCTCCTGCCTCAG
AGCACACGTCTGAACTCTTTCCCTACACGACGCTCTTCCGATCTNNNTTCACATCATTCTCCTGCCTCAG
GAGCACACGTCTGAACTCTTTCCCTACACGACGCTCTTCCGATCTNNNTAGATATACCCACTCACCCCAC
GAGCACACGTCTGAACTCTTTCCCTACACGACGCTCTTCCGATCTNNNTAGATATACCCACTCACCCCAC
GAGCACACGTCTGAACTCTTTCCCTACACGACGCTCTTCCGATCTNNNTAGATATACCCACTCACCCCAC
AGCACACGTCTGAACTCTTTCCCTACACGACGCTCTTCCGATCTNNNCCATGCCCGGCTAATTTTTTGT
AGCACACGTCTGAACTCTTTCCCTACACGACGCTCTTCCGATCTNNNCCATGCCCGGCTAATTTTTTGT
AGCACACGTCTGAACTCTTTCCCTACACGACGCTCTTCCGATCTNNNCCATGCCCGGCTAATTTTTTGT
AGCACACGTCTGAACTCTTTCCCTACACGACGCTCTTCCGATCTNNNTTTTCATCACCCCAAATACACGC

AGCACACGTCTGAACTCTTTCCCTACACGACGCTCTTCCGATCTNNNTTTTCATCACCCCAAATACACGCC
AGCACACGTCTGAACTCTTTCCCTACACGACGCTCTTCCGATCTNNNTTTTCATCACCCCAAATACACGCC
GAGCACACGTCTGAACTCTTTCCCTACACGACGCTCTTCCGATCTNNNAAACTCTTCGTTTGTGTGCATC
GAGCACACGTCTGAACTCTTTCCCTACACGACGCTCTTCCGATCTNNNAAACTCTTCGTTTGTGTGCATC
GAGCACACGTCTGAACTCTTTCCCTACACGACGCTCTTCCGATCTNNNAAACTCTTCGTTTGTGTGCATC
AGCACACGTCTGAACTCTTTCCCTACACGACGCTCTTCCGATCTNNNAATGCCTACTGGGATAAGCTTC
AGCACACGTCTGAACTCTTTCCCTACACGACGCTCTTCCGATCTNNNAATGCCTACTGGGATAAGCTTC
AGCACACGTCTGAACTCTTTCCCTACACGACGCTCTTCCGATCTNNNAATGCCTACTGGGATAAGCTTC
AGCACACGTCTGAACTCTTTCCCTACACGACGCTCTTCCGATCTNNNTTACTGAGCCCTTTTGTGGACTT
AGCACACGTCTGAACTCTTTCCCTACACGACGCTCTTCCGATCTNNNTTACTGAGCCCTTTTGTGGACTT
AGCACACGTCTGAACTCTTTCCCTACACGACGCTCTTCCGATCTNNNTTACTGAGCCCTTTTGTGGACTT
AGCACACGTCTGAACTCTTTCCCTACACGACGCTCTTCCGATCTNNNGAGATCGCGCCACTGTACTCCA
AGCACACGTCTGAACTCTTTCCCTACACGACGCTCTTCCGATCTNNNGAGATCGCGCCACTGTACTCCA
AGCACACGTCTGAACTCTTTCCCTACACGACGCTCTTCCGATCTNNNGAGATCGCGCCACTGTACTCCA
AGCACACGTCTGAACTCTTTCCCTACACGACGCTCTTCCGATCTNNNGCTGAAGTTACACCAAGTCCTGA
AGCACACGTCTGAACTCTTTCCCTACACGACGCTCTTCCGATCTNNNGCTGAAGTTACACCAAGTCCTGA
AGCACACGTCTGAACTCTTTCCCTACACGACGCTCTTCCGATCTNNNGCTGAAGTTACACCAAGTCCTGA
AGCACACGTCTGAACTCTTTCCCTACACGACGCTCTTCCGATCTNNNAGTTTCCTTTCTAAATACAGCAT
AGCACACGTCTGAACTCTTTCCCTACACGACGCTCTTCCGATCTNNNAGTTTCCTTTCTAAATACAGCAT
AGCACACGTCTGAACTCTTTCCCTACACGACGCTCTTCCGATCTNNNAGTTTCCTTTCTAAATACAGCAT
GAGCACACGTCTGAACTCTTTCCCTACACGACGCTCTTCCGATCTNNNGTGCTTGTAATCCCAGCTACTT
GAGCACACGTCTGAACTCTTTCCCTACACGACGCTCTTCCGATCTNNNGTGCTTGTAATCCCAGCTACTT
GAGCACACGTCTGAACTCTTTCCCTACACGACGCTCTTCCGATCTNNNGTGCTTGTAATCCCAGCTACTT
AGCACACGTCTGAACTCTTTCCCTACACGACGCTCTTCCGATCTNNNGGGAGGCCAAGACAGAAGAAT
AGCACACGTCTGAACTCTTTCCCTACACGACGCTCTTCCGATCTNNNGGGAGGCCAAGACAGAAGAAT
AGCACACGTCTGAACTCTTTCCCTACACGACGCTCTTCCGATCTNNNGGGAGGCCAAGACAGAAGAAT
GAGCACACGTCTGAACTCTTTCCCTACACGACGCTCTTCCGATCTNNNCTGAGTAGCTCGGATTACTGG
GAGCACACGTCTGAACTCTTTCCCTACACGACGCTCTTCCGATCTNNNCTGAGTAGCTCGGATTACTGG
GAGCACACGTCTGAACTCTTTCCCTACACGACGCTCTTCCGATCTNNNCTGAGTAGCTCGGATTACTGG
AAGAGCACACGTCTGAACTCTTTCCCTACACGACGCTCTTCCGATCTNNNCTCAAATGTTGCTTACCGTC
AAGAGCACACGTCTGAACTCTTTCCCTACACGACGCTCTTCCGATCTNNNCTCAAATGTTGCTTACCGTC
AAGAGCACACGTCTGAACTCTTTCCCTACACGACGCTCTTCCGATCTNNNCTCAAATGTTGCTTACCGTC
AGAGCACACGTCTGAACTCTTTCCCTACACGACGCTCTTCCGATCTNNNGAGGCAGAAGAATCACTTGA
AGAGCACACGTCTGAACTCTTTCCCTACACGACGCTCTTCCGATCTNNNGAGGCAGAAGAATCACTTGA
AGAGCACACGTCTGAACTCTTTCCCTACACGACGCTCTTCCGATCTNNNGAGGCAGAAGAATCACTTGA
GAGCACACGTCTGAACTCTTTCCCTACACGACGCTCTTCCGATCTNNNATGCCTGTAATCCCAGCACTTT
GAGCACACGTCTGAACTCTTTCCCTACACGACGCTCTTCCGATCTNNNATGCCTGTAATCCCAGCACTTT
GAGCACACGTCTGAACTCTTTCCCTACACGACGCTCTTCCGATCTNNNATGCCTGTAATCCCAGCACTTT
AGCACACGTCTGAACTCTTTCCCTACACGACGCTCTTCCGATCTNNNCATCTTCACATCAACCATGGAGC
AGCACACGTCTGAACTCTTTCCCTACACGACGCTCTTCCGATCTNNNCATCTTCACATCAACCATGGAGC
AGCACACGTCTGAACTCTTTCCCTACACGACGCTCTTCCGATCTNNNCATCTTCACATCAACCATGGAGC
AGCACACGTCTGAACTCTTTCCCTACACGACGCTCTTCCGATCTNNNGCCTGGGTGACAAAATGAGATT
AGCACACGTCTGAACTCTTTCCCTACACGACGCTCTTCCGATCTNNNGCCTGGGTGACAAAATGAGATT

AGCACACGTCTGAACTCTTTCCCTACACGACGCTCTTCCGATCTNNNGCCTGGGTGACAAAATGAGAT

GCACACGTCTGAACTCTTTCCCTACACGACGCTCTTCCGATCTNNNCCACGAAAACCCTAGTGAATCACC

GCACACGTCTGAACTCTTTCCCTACACGACGCTCTTCCGATCTNNNCCACGAAAACCCTAGTGAATCACC

GCACACGTCTGAACTCTTTCCCTACACGACGCTCTTCCGATCTNNNCCACGAAAACCCTAGTGAATCACC

GCACACGTCTGAACTCTTTCCCTACACGACGCTCTTCCGATCTNNNAAACAGACCCAAGCAGAGTTTCC

GCACACGTCTGAACTCTTTCCCTACACGACGCTCTTCCGATCTNNNAAACAGACCCAAGCAGAGTTTCC

GCACACGTCTGAACTCTTTCCCTACACGACGCTCTTCCGATCTNNNAAACAGACCCAAGCAGAGTTTCC

AGCACACGTCTGAACTCTTTCCCTACACGACGCTCTTCCGATCTNNNGTCTCTGTTTTGCTCCATTGGTC

AGCACACGTCTGAACTCTTTCCCTACACGACGCTCTTCCGATCTNNNGTCTCTGTTTTGCTCCATTGGTC

AGCACACGTCTGAACTCTTTCCCTACACGACGCTCTTCCGATCTNNNGTCTCTGTTTTGCTCCATTGGTC

GCACACGTCTGAACTCTTTCCCTACACGACGCTCTTCCGATCTNNNCACAGGCAAAAATGTACCCCTCTCA

GCACACGTCTGAACTCTTTCCCTACACGACGCTCTTCCGATCTNNNCACAGGCAAAAATGTACCCCTCTCA

GCACACGTCTGAACTCTTTCCCTACACGACGCTCTTCCGATCTNNNCACAGGCAAAAATGTACCCCTCTCA

AGCACACGTCTGAACTCTTTCCCTACACGACGCTCTTCCGATCTNNNCATTTATGTGAGATGTGGTTTT

AGCACACGTCTGAACTCTTTCCCTACACGACGCTCTTCCGATCTNNNCATTTATGTGAGATGTGGTTTT

AGCACACGTCTGAACTCTTTCCCTACACGACGCTCTTCCGATCTNNNCATTTATGTGAGATGTGGTTTT

GAGCACACGTCTGAACTCTTTCCCTACACGACGCTCTTCCGATCTNNNGGTAAGGCAGGAGAATCACTT

GAGCACACGTCTGAACTCTTTCCCTACACGACGCTCTTCCGATCTNNNGGTAAGGCAGGAGAATCACTT

GAGCACACGTCTGAACTCTTTCCCTACACGACGCTCTTCCGATCTNNNGGTAAGGCAGGAGAATCACTT

AGAGCACACGTCTGAACTCTTTCCCTACACGACGCTCTTCCGATCTNNNAACCTTGGAACTGACACCGA

AGAGCACACGTCTGAACTCTTTCCCTACACGACGCTCTTCCGATCTNNNAACCTTGGAACTGACACCGA

AGAGCACACGTCTGAACTCTTTCCCTACACGACGCTCTTCCGATCTNNNAACCTTGGAACTGACACCGA

GCACACGTCTGAACTCTTTCCCTACACGACGCTCTTCCGATCTNNNGTAATAAAACCATTCCCGCCTCCC

GCACACGTCTGAACTCTTTCCCTACACGACGCTCTTCCGATCTNNNGTAATAAAACCATTCCCGCCTCCC

GCACACGTCTGAACTCTTTCCCTACACGACGCTCTTCCGATCTNNNGTAATAAAACCATTCCCGCCTCCC

AGCACACGTCTGAACTCTTTCCCTACACGACGCTCTTCCGATCTNNNCTTGTCTCTACTTCCAGTCCGTCC

AGCACACGTCTGAACTCTTTCCCTACACGACGCTCTTCCGATCTNNNCTTGTCTCTACTTCCAGTCCGTCC

AGCACACGTCTGAACTCTTTCCCTACACGACGCTCTTCCGATCTNNNCTTGTCTCTACTTCCAGTCCGTCC

GAGCACACGTCTGAACTCTTTCCCTACACGACGCTCTTCCGATCTNNNGAACAAGCAGAAGAAAGAGCT

GAGCACACGTCTGAACTCTTTCCCTACACGACGCTCTTCCGATCTNNNGAACAAGCAGAAGAAAGAGCT

GAGCACACGTCTGAACTCTTTCCCTACACGACGCTCTTCCGATCTNNNGAACAAGCAGAAGAAAGAGCT

GAGCACACGTCTGAACTCTTTCCCTACACGACGCTCTTCCGATCTNNNAAGCAGGAGAATCACTTGAAC

GAGCACACGTCTGAACTCTTTCCCTACACGACGCTCTTCCGATCTNNNAAGCAGGAGAATCACTTGAAC

GAGCACACGTCTGAACTCTTTCCCTACACGACGCTCTTCCGATCTNNNAAGCAGGAGAATCACTTGAAC

AGCACACGTCTGAACTCTTTCCCTACACGACGCTCTTCCGATCTNNNTCTGATAAGCAACTGCAGACCT

AGCACACGTCTGAACTCTTTCCCTACACGACGCTCTTCCGATCTNNNTCTGATAAGCAACTGCAGACCT

AGCACACGTCTGAACTCTTTCCCTACACGACGCTCTTCCGATCTNNNTCTGATAAGCAACTGCAGACCT

GAGCACACGTCTGAACTCTTTCCCTACACGACGCTCTTCCGATCTNNNTGCCAGAATGATGCTTGTAAA

GAGCACACGTCTGAACTCTTTCCCTACACGACGCTCTTCCGATCTNNNTGCCAGAATGATGCTTGTAAA

GAGCACACGTCTGAACTCTTTCCCTACACGACGCTCTTCCGATCTNNNTGCCAGAATGATGCTTGTAAA

GAGCACACGTCTGAACTCTTTCCCTACACGACGCTCTTCCGATCTNNNTGAGGAAGGAGAATCGCTTG

GAGCACACGTCTGAACTCTTTCCCTACACGACGCTCTTCCGATCTNNNTGAGGAAGGAGAATCGCTTG

GAGCACACGTCTGAACTCTTTCCCTACACGACGCTCTTCCGATCTNNNTGAGGAAGGAGAATCGCTTG

GAGCACACGTCTGAACTCTTTCCCTACACGACGCTCTTCCGATCTNNNTGGTCTTGAATAAGGTCCGG
GAGCACACGTCTGAACTCTTTCCCTACACGACGCTCTTCCGATCTNNNTGGTCTTGAATAAGGTCCGG
GAGCACACGTCTGAACTCTTTCCCTACACGACGCTCTTCCGATCTNNNTGGTCTTGAATAAGGTCCGG
GCACACGTCTGAACTCTTTCCCTACACGACGCTCTTCCGATCTNNNATGGGATTGAAAACATAGACTT
GCACACGTCTGAACTCTTTCCCTACACGACGCTCTTCCGATCTNNNATGGGATTGAAAACATAGACTT
GCACACGTCTGAACTCTTTCCCTACACGACGCTCTTCCGATCTNNNATGGGATTGAAAACATAGACTT
CACACGTCTGAACTCTTTCCCTACACGACGCTCTTCCGATCTNNNAGGCAGGAGAATTACTTGAACC
CACACGTCTGAACTCTTTCCCTACACGACGCTCTTCCGATCTNNNAGGCAGGAGAATTACTTGAACC
CACACGTCTGAACTCTTTCCCTACACGACGCTCTTCCGATCTNNNAGGCAGGAGAATTACTTGAACC
AGCACACGTCTGAACTCTTTCCCTACACGACGCTCTTCCGATCTNNNTCAGGTTCACTTCAAGACACA
AGCACACGTCTGAACTCTTTCCCTACACGACGCTCTTCCGATCTNNNTCAGGTTCACTTCAAGACACA
AGCACACGTCTGAACTCTTTCCCTACACGACGCTCTTCCGATCTNNNTCAGGTTCACTTCAAGACACA
AGCACACGTCTGAACTCTTTCCCTACACGACGCTCTTCCGATCTNNNGGGAAACTTCCAATGAAGCC
AGCACACGTCTGAACTCTTTCCCTACACGACGCTCTTCCGATCTNNNGGGAAACTTCCAATGAAGCC
AGCACACGTCTGAACTCTTTCCCTACACGACGCTCTTCCGATCTNNNGGGAAACTTCCAATGAAGCC
GCACACGTCTGAACTCTTTCCCTACACGACGCTCTTCCGATCTNNNAAATTAGCTGAATGGCGTGAA
GCACACGTCTGAACTCTTTCCCTACACGACGCTCTTCCGATCTNNNAAATTAGCTGAATGGCGTGAA
GCACACGTCTGAACTCTTTCCCTACACGACGCTCTTCCGATCTNNNAAATTAGCTGAATGGCGTGAA
GAGCACACGTCTGAACTCTTTCCCTACACGACGCTCTTCCGATCTNNNCCCATCGCCCAAATACCTTCT
GAGCACACGTCTGAACTCTTTCCCTACACGACGCTCTTCCGATCTNNNCCCATCGCCCAAATACCTTCT
GAGCACACGTCTGAACTCTTTCCCTACACGACGCTCTTCCGATCTNNNCCCATCGCCCAAATACCTTCT
CACACGTCTGAACTCTTTCCCTACACGACGCTCTTCCGATCTNNNCTCCCACATTGTTGTTTCCAGACCA
CACACGTCTGAACTCTTTCCCTACACGACGCTCTTCCGATCTNNNCTCCCACATTGTTGTTTCCAGACCA
CACACGTCTGAACTCTTTCCCTACACGACGCTCTTCCGATCTNNNCTCCCACATTGTTGTTTCCAGACCA
GCACACGTCTGAACTCTTTCCCTACACGACGCTCTTCCGATCTNNNCAAAATCCACATCCAGGCATCCC
GCACACGTCTGAACTCTTTCCCTACACGACGCTCTTCCGATCTNNNCAAAATCCACATCCAGGCATCCC
GCACACGTCTGAACTCTTTCCCTACACGACGCTCTTCCGATCTNNNCAAAATCCACATCCAGGCATCCC
GCACACGTCTGAACTCTTTCCCTACACGACGCTCTTCCGATCTNNNTGCTCCCTCAAATCAGTGAAAG
GCACACGTCTGAACTCTTTCCCTACACGACGCTCTTCCGATCTNNNTGCTCCCTCAAATCAGTGAAAG
GCACACGTCTGAACTCTTTCCCTACACGACGCTCTTCCGATCTNNNTGCTCCCTCAAATCAGTGAAAG
GCACACGTCTGAACTCTTTCCCTACACGACGCTCTTCCGATCTNNNCACTGGTTTCTGATGTGCTGGC
GCACACGTCTGAACTCTTTCCCTACACGACGCTCTTCCGATCTNNNCACTGGTTTCTGATGTGCTGGC
GCACACGTCTGAACTCTTTCCCTACACGACGCTCTTCCGATCTNNNCACTGGTTTCTGATGTGCTGGC
GAGCACACGTCTGAACTCTTTCCCTACACGACGCTCTTCCGATCTNNNTTAGTAGAGATGGGGTTTC
GAGCACACGTCTGAACTCTTTCCCTACACGACGCTCTTCCGATCTNNNTTAGTAGAGATGGGGTTTC
GAGCACACGTCTGAACTCTTTCCCTACACGACGCTCTTCCGATCTNNNTTAGTAGAGATGGGGTTTC
AGCACACGTCTGAACTCTTTCCCTACACGACGCTCTTCCGATCTNNNTGGAAGGTACTGGAGTTGGG
AGCACACGTCTGAACTCTTTCCCTACACGACGCTCTTCCGATCTNNNTGGAAGGTACTGGAGTTGGG
AGCACACGTCTGAACTCTTTCCCTACACGACGCTCTTCCGATCTNNNTGGAAGGTACTGGAGTTGGG
GCACACGTCTGAACTCTTTCCCTACACGACGCTCTTCCGATCTNNNCAATCTGGGTTCAAGCGATTCT
GCACACGTCTGAACTCTTTCCCTACACGACGCTCTTCCGATCTNNNCAATCTGGGTTCAAGCGATTCT
GCACACGTCTGAACTCTTTCCCTACACGACGCTCTTCCGATCTNNNCAATCTGGGTTCAAGCGATTCT
GCACACGTCTGAACTCTTTCCCTACACGACGCTCTTCCGATCTNNNGAGTAATAGTCTCCAGCCAGGT

GCACACGTCTGAACTCTTTCCCTACACGACGCTCTTCCGATCTNNNGAGTAATAGTCTCCAGCCAGGT
GCACACGTCTGAACTCTTTCCCTACACGACGCTCTTCCGATCTNNNGAGTAATAGTCTCCAGCCAGGT
AGCACACGTCTGAACTCTTTCCCTACACGACGCTCTTCCGATCTNNNTGCCCATATTACTGAGATGGTG
AGCACACGTCTGAACTCTTTCCCTACACGACGCTCTTCCGATCTNNNTGCCCATATTACTGAGATGGTG
AGCACACGTCTGAACTCTTTCCCTACACGACGCTCTTCCGATCTNNNTGCCCATATTACTGAGATGGTG
AGCACACGTCTGAACTCTTTCCCTACACGACGCTCTTCCGATCTNNNATGAGGTTTCACTACATTGGC
AGCACACGTCTGAACTCTTTCCCTACACGACGCTCTTCCGATCTNNNATGAGGTTTCACTACATTGGC
AGCACACGTCTGAACTCTTTCCCTACACGACGCTCTTCCGATCTNNNATGAGGTTTCACTACATTGGC
AGCACACGTCTGAACTCTTTCCCTACACGACGCTCTTCCGATCTNNNGGTCAAACCAAGTAAGATTGT
AGCACACGTCTGAACTCTTTCCCTACACGACGCTCTTCCGATCTNNNGGTCAAACCAAGTAAGATTGT
AGCACACGTCTGAACTCTTTCCCTACACGACGCTCTTCCGATCTNNNGGTCAAACCAAGTAAGATTGT
GAGCACACGTCTGAACTCTTTCCCTACACGACGCTCTTCCGATCTNNNAGAGGTTGTAGTGAGCCGA
GAGCACACGTCTGAACTCTTTCCCTACACGACGCTCTTCCGATCTNNNAGAGGTTGTAGTGAGCCGA
GAGCACACGTCTGAACTCTTTCCCTACACGACGCTCTTCCGATCTNNNAGAGGTTGTAGTGAGCCGA
GAGCACACGTCTGAACTCTTTCCCTACACGACGCTCTTCCGATCTNNNGTGAAACCCCGTCTCTACTAG
GAGCACACGTCTGAACTCTTTCCCTACACGACGCTCTTCCGATCTNNNGTGAAACCCCGTCTCTACTAG
GAGCACACGTCTGAACTCTTTCCCTACACGACGCTCTTCCGATCTNNNGTGAAACCCCGTCTCTACTAG
AGCACACGTCTGAACTCTTTCCCTACACGACGCTCTTCCGATCTNNNACGATCTCAGCTCACTGTAAG
AGCACACGTCTGAACTCTTTCCCTACACGACGCTCTTCCGATCTNNNACGATCTCAGCTCACTGTAAG
AGCACACGTCTGAACTCTTTCCCTACACGACGCTCTTCCGATCTNNNACGATCTCAGCTCACTGTAAG
AGCACACGTCTGAACTCTTTCCCTACACGACGCTCTTCCGATCTNNNATAGTTACTGGTGATGGGGG
AGCACACGTCTGAACTCTTTCCCTACACGACGCTCTTCCGATCTNNNATAGTTACTGGTGATGGGGG
AGCACACGTCTGAACTCTTTCCCTACACGACGCTCTTCCGATCTNNNATAGTTACTGGTGATGGGGG
GCACACGTCTGAACTCTTTCCCTACACGACGCTCTTCCGATCTNNNTCCTAAGTAGCTGGGACTACAG
GCACACGTCTGAACTCTTTCCCTACACGACGCTCTTCCGATCTNNNTCCTAAGTAGCTGGGACTACAG
GCACACGTCTGAACTCTTTCCCTACACGACGCTCTTCCGATCTNNNTCCTAAGTAGCTGGGACTACAG
GCACACGTCTGAACTCTTTCCCTACACGACGCTCTTCCGATCTNNNATTGGCAATGTGTAAGTGTTCC
GCACACGTCTGAACTCTTTCCCTACACGACGCTCTTCCGATCTNNNATTGGCAATGTGTAAGTGTTCC
GCACACGTCTGAACTCTTTCCCTACACGACGCTCTTCCGATCTNNNATTGGCAATGTGTAAGTGTTCC
GCACACGTCTGAACTCTTTCCCTACACGACGCTCTTCCGATCTNNNCGACAGCTACTCCGAAGCCGCAT
GCACACGTCTGAACTCTTTCCCTACACGACGCTCTTCCGATCTNNNCGACAGCTACTCCGAAGCCGCAT
GCACACGTCTGAACTCTTTCCCTACACGACGCTCTTCCGATCTNNNCGACAGCTACTCCGAAGCCGCAT
AGCACACGTCTGAACTCTTTCCCTACACGACGCTCTTCCGATCTNNNTAGTTCCAGCTACTCCAGAGG
AGCACACGTCTGAACTCTTTCCCTACACGACGCTCTTCCGATCTNNNTAGTTCCAGCTACTCCAGAGG
AGCACACGTCTGAACTCTTTCCCTACACGACGCTCTTCCGATCTNNNTAGTTCCAGCTACTCCAGAGG
GAGCACACGTCTGAACTCTTTCCCTACACGACGCTCTTCCGATCTNNNTGCAGATTCCTGTTCTGATAC
GAGCACACGTCTGAACTCTTTCCCTACACGACGCTCTTCCGATCTNNNTGCAGATTCCTGTTCTGATAC
GAGCACACGTCTGAACTCTTTCCCTACACGACGCTCTTCCGATCTNNNTGCAGATTCCTGTTCTGATAC
GAGCACACGTCTGAACTCTTTCCCTACACGACGCTCTTCCGATCTNNNTTGCTCCTTGTCTTGAAGAGC
GAGCACACGTCTGAACTCTTTCCCTACACGACGCTCTTCCGATCTNNNTTGCTCCTTGTCTTGAAGAGC
GAGCACACGTCTGAACTCTTTCCCTACACGACGCTCTTCCGATCTNNNTTGCTCCTTGTCTTGAAGAGC
GAGCACACGTCTGAACTCTTTCCCTACACGACGCTCTTCCGATCTNNNTGGGTGATAGAGTGAGACT
GAGCACACGTCTGAACTCTTTCCCTACACGACGCTCTTCCGATCTNNNTGGGTGATAGAGTGAGACT

GAGCACACGTCTGAACTCTTTCCCTACACGACGCTCTTCCGATCTNNNTGGGTGATAGAGTGAGACT
AGCACACGTCTGAACTCTTTCCCTACACGACGCTCTTCCGATCTNNNACCAACATGATGAAACCCTGT
AGCACACGTCTGAACTCTTTCCCTACACGACGCTCTTCCGATCTNNNACCAACATGATGAAACCCTGT
AGCACACGTCTGAACTCTTTCCCTACACGACGCTCTTCCGATCTNNNACCAACATGATGAAACCCTGT
GAGCACACGTCTGAACTCTTTCCCTACACGACGCTCTTCCGATCTNNNGTAATCCCAGCTACTAAGGA
GAGCACACGTCTGAACTCTTTCCCTACACGACGCTCTTCCGATCTNNNGTAATCCCAGCTACTAAGGA
GAGCACACGTCTGAACTCTTTCCCTACACGACGCTCTTCCGATCTNNNGTAATCCCAGCTACTAAGGA
AGCACACGTCTGAACTCTTTCCCTACACGACGCTCTTCCGATCTNNNACAGTCCCACACCTGTCATTAT
AGCACACGTCTGAACTCTTTCCCTACACGACGCTCTTCCGATCTNNNACAGTCCCACACCTGTCATTAT
AGCACACGTCTGAACTCTTTCCCTACACGACGCTCTTCCGATCTNNNACAGTCCCACACCTGTCATTAT
GCACACGTCTGAACTCTTTCCCTACACGACGCTCTTCCGATCTNNNCTGCTCCCATATCATGCCTTTTG
GCACACGTCTGAACTCTTTCCCTACACGACGCTCTTCCGATCTNNNCTGCTCCCATATCATGCCTTTTG
GCACACGTCTGAACTCTTTCCCTACACGACGCTCTTCCGATCTNNNCTGCTCCCATATCATGCCTTTTG
GCACACGTCTGAACTCTTTCCCTACACGACGCTCTTCCGATCTNNNCGGCAAGGTGATACTCCATCTCC
GCACACGTCTGAACTCTTTCCCTACACGACGCTCTTCCGATCTNNNCGGCAAGGTGATACTCCATCTCC
GCACACGTCTGAACTCTTTCCCTACACGACGCTCTTCCGATCTNNNCGGCAAGGTGATACTCCATCTCC
AGCACACGTCTGAACTCTTTCCCTACACGACGCTCTTCCGATCTNNNAGCAAGTCCCCACTTCTTTTTG
AGCACACGTCTGAACTCTTTCCCTACACGACGCTCTTCCGATCTNNNAGCAAGTCCCCACTTCTTTTTG
AGCACACGTCTGAACTCTTTCCCTACACGACGCTCTTCCGATCTNNNAGCAAGTCCCCACTTCTTTTTG
AAGAGCACACGTCTGAACTCTTTCCCTACACGACGCTCTTCCGATCTNNNGGTGTGATGTTCTTTCTTT
AAGAGCACACGTCTGAACTCTTTCCCTACACGACGCTCTTCCGATCTNNNGGTGTGATGTTCTTTCTTT
AAGAGCACACGTCTGAACTCTTTCCCTACACGACGCTCTTCCGATCTNNNGGTGTGATGTTCTTTCTTT
CACGTCTGAACTCTTTCCCTACACGACGCTCTTCCGATCTNNNAATCCTGGTCCAGTAAAGAGCATCG
CACGTCTGAACTCTTTCCCTACACGACGCTCTTCCGATCTNNNAATCCTGGTCCAGTAAAGAGCATCG
CACGTCTGAACTCTTTCCCTACACGACGCTCTTCCGATCTNNNAATCCTGGTCCAGTAAAGAGCATCG
AGCACACGTCTGAACTCTTTCCCTACACGACGCTCTTCCGATCTNNNTTGTCAAAAGAATACTGGCAG
AGCACACGTCTGAACTCTTTCCCTACACGACGCTCTTCCGATCTNNNTTGTCAAAAGAATACTGGCAG
AGCACACGTCTGAACTCTTTCCCTACACGACGCTCTTCCGATCTNNNTTGTCAAAAGAATACTGGCAG
AGCACACGTCTGAACTCTTTCCCTACACGACGCTCTTCCGATCTNNNGGCTGAGATAGGAGGATCA
AGCACACGTCTGAACTCTTTCCCTACACGACGCTCTTCCGATCTNNNGGCTGAGATAGGAGGATCA
AGCACACGTCTGAACTCTTTCCCTACACGACGCTCTTCCGATCTNNNGGCTGAGATAGGAGGATCA
GAGCACACGTCTGAACTCTTTCCCTACACGACGCTCTTCCGATCTNNNCTTATAACCCTAAGGCAGGC
GAGCACACGTCTGAACTCTTTCCCTACACGACGCTCTTCCGATCTNNNCTTATAACCCTAAGGCAGGC
GAGCACACGTCTGAACTCTTTCCCTACACGACGCTCTTCCGATCTNNNCTTATAACCCTAAGGCAGGC
AGCACACGTCTGAACTCTTTCCCTACACGACGCTCTTCCGATCTNNNAATATCTATGAGTTGGGGCCA
AGCACACGTCTGAACTCTTTCCCTACACGACGCTCTTCCGATCTNNNAATATCTATGAGTTGGGGCCA
AGCACACGTCTGAACTCTTTCCCTACACGACGCTCTTCCGATCTNNNAATATCTATGAGTTGGGGCCA
AGCACACGTCTGAACTCTTTCCCTACACGACGCTCTTCCGATCTNNNTGTATGTGTACAGGTGTGTG
AGCACACGTCTGAACTCTTTCCCTACACGACGCTCTTCCGATCTNNNTGTATGTGTACAGGTGTGTG
AGCACACGTCTGAACTCTTTCCCTACACGACGCTCTTCCGATCTNNNTGTATGTGTACAGGTGTGTG
AGAGCACACGTCTGAACTCTTTCCCTACACGACGCTCTTCCGATCTNNNGTACCCACTAACCAACCCAT
AGAGCACACGTCTGAACTCTTTCCCTACACGACGCTCTTCCGATCTNNNGTACCCACTAACCAACCCAT
AGAGCACACGTCTGAACTCTTTCCCTACACGACGCTCTTCCGATCTNNNGTACCCACTAACCAACCCAT

AGCACACGTCTGAACTCTTTCCCTACACGACGCTCTTCCGATCTNNNGCTTGATATCAGCTCACCACAA
AGCACACGTCTGAACTCTTTCCCTACACGACGCTCTTCCGATCTNNNGCTTGATATCAGCTCACCACAA
AGCACACGTCTGAACTCTTTCCCTACACGACGCTCTTCCGATCTNNNGCTTGATATCAGCTCACCACAA
GAGCACACGTCTGAACTCTTTCCCTACACGACGCTCTTCCGATCTNNNGAGACTGCAGTGAGCTGATA
GAGCACACGTCTGAACTCTTTCCCTACACGACGCTCTTCCGATCTNNNGAGACTGCAGTGAGCTGATA
GAGCACACGTCTGAACTCTTTCCCTACACGACGCTCTTCCGATCTNNNGAGACTGCAGTGAGCTGATA
GAGCACACGTCTGAACTCTTTCCCTACACGACGCTCTTCCGATCTNNNAGAGAGACAGAGAGAGATGG
GAGCACACGTCTGAACTCTTTCCCTACACGACGCTCTTCCGATCTNNNAGAGAGACAGAGAGAGATGG
GAGCACACGTCTGAACTCTTTCCCTACACGACGCTCTTCCGATCTNNNAGAGAGACAGAGAGAGATGG
GCACACGTCTGAACTCTTTCCCTACACGACGCTCTTCCGATCTNNNTACCAACAAAGGTCAAGCTGGAG
GCACACGTCTGAACTCTTTCCCTACACGACGCTCTTCCGATCTNNNTACCAACAAAGGTCAAGCTGGAG
GCACACGTCTGAACTCTTTCCCTACACGACGCTCTTCCGATCTNNNTACCAACAAAGGTCAAGCTGGAG
AGCACACGTCTGAACTCTTTCCCTACACGACGCTCTTCCGATCTNNNTAGATTAGTGCCCTCAGTGGTT
AGCACACGTCTGAACTCTTTCCCTACACGACGCTCTTCCGATCTNNNTAGATTAGTGCCCTCAGTGGTT
AGCACACGTCTGAACTCTTTCCCTACACGACGCTCTTCCGATCTNNNTAGATTAGTGCCCTCAGTGGTT
AGCACACGTCTGAACTCTTTCCCTACACGACGCTCTTCCGATCTNNNCAATCCATCATTGAGTCCTGGCA
AGCACACGTCTGAACTCTTTCCCTACACGACGCTCTTCCGATCTNNNCAATCCATCATTGAGTCCTGGCA
AGCACACGTCTGAACTCTTTCCCTACACGACGCTCTTCCGATCTNNNCAATCCATCATTGAGTCCTGGCA
GAGCACACGTCTGAACTCTTTCCCTACACGACGCTCTTCCGATCTNNNACCTAGCCTACCACACTTACTC
GAGCACACGTCTGAACTCTTTCCCTACACGACGCTCTTCCGATCTNNNACCTAGCCTACCACACTTACTC
GAGCACACGTCTGAACTCTTTCCCTACACGACGCTCTTCCGATCTNNNACCTAGCCTACCACACTTACTC
AGAGCACACGTCTGAACTCTTTCCCTACACGACGCTCTTCCGATCTNNNAAGGAGTATGTTCGTTCTCC
AGAGCACACGTCTGAACTCTTTCCCTACACGACGCTCTTCCGATCTNNNAAGGAGTATGTTCGTTCTCC
AGAGCACACGTCTGAACTCTTTCCCTACACGACGCTCTTCCGATCTNNNAAGGAGTATGTTCGTTCTCC
AGCACACGTCTGAACTCTTTCCCTACACGACGCTCTTCCGATCTNNNCCAAATCTCCATCCCTCAGAGAG
AGCACACGTCTGAACTCTTTCCCTACACGACGCTCTTCCGATCTNNNCCAAATCTCCATCCCTCAGAGAG
AGCACACGTCTGAACTCTTTCCCTACACGACGCTCTTCCGATCTNNNCCAAATCTCCATCCCTCAGAGAG
GAGCACACGTCTGAACTCTTTCCCTACACGACGCTCTTCCGATCTNNNCAGGAGAACTGCTTGAGGAG
GAGCACACGTCTGAACTCTTTCCCTACACGACGCTCTTCCGATCTNNNCAGGAGAACTGCTTGAGGAG
GAGCACACGTCTGAACTCTTTCCCTACACGACGCTCTTCCGATCTNNNCAGGAGAACTGCTTGAGGAG
GAGCACACGTCTGAACTCTTTCCCTACACGACGCTCTTCCGATCTNNNCTGCAGGATGACAGATCGAAA
GAGCACACGTCTGAACTCTTTCCCTACACGACGCTCTTCCGATCTNNNCTGCAGGATGACAGATCGAAA
GAGCACACGTCTGAACTCTTTCCCTACACGACGCTCTTCCGATCTNNNCTGCAGGATGACAGATCGAAA
GAGCACACGTCTGAACTCTTTCCCTACACGACGCTCTTCCGATCTNNNACACATAGTGATACCACCTTG
GAGCACACGTCTGAACTCTTTCCCTACACGACGCTCTTCCGATCTNNNACACATAGTGATACCACCTTG
GAGCACACGTCTGAACTCTTTCCCTACACGACGCTCTTCCGATCTNNNACACATAGTGATACCACCTTG
AGCACACGTCTGAACTCTTTCCCTACACGACGCTCTTCCGATCTNNNTCCCTCTTTGTCTCTTCCTCTCTCA
AGCACACGTCTGAACTCTTTCCCTACACGACGCTCTTCCGATCTNNNTCCCTCTTTGTCTCTTCCTCTCTCA
AGCACACGTCTGAACTCTTTCCCTACACGACGCTCTTCCGATCTNNNTCCCTCTTTGTCTCTTCCTCTCTCA
ACACGTCTGAACTCTTTCCCTACACGACGCTCTTCCGATCTNNNCTCCTGAGTAGCTGGGACTACAGCAT
ACACGTCTGAACTCTTTCCCTACACGACGCTCTTCCGATCTNNNCTCCTGAGTAGCTGGGACTACAGCAT
ACACGTCTGAACTCTTTCCCTACACGACGCTCTTCCGATCTNNNCTCCTGAGTAGCTGGGACTACAGCAT
GCACACGTCTGAACTCTTTCCCTACACGACGCTCTTCCGATCTNNNTTCGTCTCTCTCTTTCTGACTCCCAT

CACACGTCTGAACTCTTTCCCTACACGACGCTCTTCCGATCTNNNTTCGTCTCTCTCTTTCTGACTCCCA
CACACGTCTGAACTCTTTCCCTACACGACGCTCTTCCGATCTNNNTTCGTCTCTCTCTTTCTGACTCCCA
AGCACACGTCTGAACTCTTTCCCTACACGACGCTCTTCCGATCTNNNCATCATCTTTCATCCTCCCACCT
AGCACACGTCTGAACTCTTTCCCTACACGACGCTCTTCCGATCTNNNCATCATCTTTCATCCTCCCACCT
AGCACACGTCTGAACTCTTTCCCTACACGACGCTCTTCCGATCTNNNCATCATCTTTCATCCTCCCACCT
ACACGTCTGAACTCTTTCCCTACACGACGCTCTTCCGATCTNNNCTACGATCAGTCCTACCAATACATC
ACACGTCTGAACTCTTTCCCTACACGACGCTCTTCCGATCTNNNCTACGATCAGTCCTACCAATACATC
ACACGTCTGAACTCTTTCCCTACACGACGCTCTTCCGATCTNNNCTACGATCAGTCCTACCAATACATC
GCACACGTCTGAACTCTTTCCCTACACGACGCTCTTCCGATCTNNNTATTTTTGTGTGTGTGCGTGTG
GCACACGTCTGAACTCTTTCCCTACACGACGCTCTTCCGATCTNNNTATTTTTGTGTGTGTGCGTGTG
GCACACGTCTGAACTCTTTCCCTACACGACGCTCTTCCGATCTNNNTATTTTTGTGTGTGTGCGTGTG
GCACACGTCTGAACTCTTTCCCTACACGACGCTCTTCCGATCTNNNTCCTACCTTGACTTCCCAAAGTG
GCACACGTCTGAACTCTTTCCCTACACGACGCTCTTCCGATCTNNNTCCTACCTTGACTTCCCAAAGTG
GCACACGTCTGAACTCTTTCCCTACACGACGCTCTTCCGATCTNNNTCCTACCTTGACTTCCCAAAGTG
GCACACGTCTGAACTCTTTCCCTACACGACGCTCTTCCGATCTNNNCAAACACAGTGAAACCCCATCT
GCACACGTCTGAACTCTTTCCCTACACGACGCTCTTCCGATCTNNNCAAACACAGTGAAACCCCATCT
GCACACGTCTGAACTCTTTCCCTACACGACGCTCTTCCGATCTNNNCAAACACAGTGAAACCCCATCT
GCACACGTCTGAACTCTTTCCCTACACGACGCTCTTCCGATCTNNNTGCAATTATCTTAACAAGTCTAA
GCACACGTCTGAACTCTTTCCCTACACGACGCTCTTCCGATCTNNNTGCAATTATCTTAACAAGTCTAA
GCACACGTCTGAACTCTTTCCCTACACGACGCTCTTCCGATCTNNNTGCAATTATCTTAACAAGTCTAA
GAGCACACGTCTGAACTCTTTCCCTACACGACGCTCTTCCGATCTNNNGATCAGAGTAGGGATGCAC
GAGCACACGTCTGAACTCTTTCCCTACACGACGCTCTTCCGATCTNNNGATCAGAGTAGGGATGCAC
GAGCACACGTCTGAACTCTTTCCCTACACGACGCTCTTCCGATCTNNNGATCAGAGTAGGGATGCAC
AGAGCACACGTCTGAACTCTTTCCCTACACGACGCTCTTCCGATCTNNNGACAAGGTTTCACTGTGTT
AGAGCACACGTCTGAACTCTTTCCCTACACGACGCTCTTCCGATCTNNNGACAAGGTTTCACTGTGTT
AGAGCACACGTCTGAACTCTTTCCCTACACGACGCTCTTCCGATCTNNNGACAAGGTTTCACTGTGTT
GCACACGTCTGAACTCTTTCCCTACACGACGCTCTTCCGATCTNNNTTCTGAGAGATGTATTTTCTTTT
GCACACGTCTGAACTCTTTCCCTACACGACGCTCTTCCGATCTNNNTTCTGAGAGATGTATTTTCTTTT
GCACACGTCTGAACTCTTTCCCTACACGACGCTCTTCCGATCTNNNTTCTGAGAGATGTATTTTCTTTT
GCACACGTCTGAACTCTTTCCCTACACGACGCTCTTCCGATCTNNNTGTCTAACTCCAAAGCAGGAAC
GCACACGTCTGAACTCTTTCCCTACACGACGCTCTTCCGATCTNNNTGTCTAACTCCAAAGCAGGAAC
GCACACGTCTGAACTCTTTCCCTACACGACGCTCTTCCGATCTNNNTGTCTAACTCCAAAGCAGGAAC
GAGCACACGTCTGAACTCTTTCCCTACACGACGCTCTTCCGATCTNNNCTTGCAGTGAGTAGAGATCC
GAGCACACGTCTGAACTCTTTCCCTACACGACGCTCTTCCGATCTNNNCTTGCAGTGAGTAGAGATCC
GAGCACACGTCTGAACTCTTTCCCTACACGACGCTCTTCCGATCTNNNCTTGCAGTGAGTAGAGATCC
GCACACGTCTGAACTCTTTCCCTACACGACGCTCTTCCGATCTNNNGAAAACCAGAGACCTTTGCTCA
GCACACGTCTGAACTCTTTCCCTACACGACGCTCTTCCGATCTNNNGAAAACCAGAGACCTTTGCTCA
GCACACGTCTGAACTCTTTCCCTACACGACGCTCTTCCGATCTNNNGAAAACCAGAGACCTTTGCTCA
GAGCACACGTCTGAACTCTTTCCCTACACGACGCTCTTCCGATCTNNNGAACATGTTTTTCTCCTTTTT
GAGCACACGTCTGAACTCTTTCCCTACACGACGCTCTTCCGATCTNNNGAACATGTTTTTCTCCTTTTT
GAGCACACGTCTGAACTCTTTCCCTACACGACGCTCTTCCGATCTNNNGAACATGTTTTTCTCCTTTTT
GAGCACACGTCTGAACTCTTTCCCTACACGACGCTCTTCCGATCTNNNTGCAAGTATCCATAGTTTGG
GAGCACACGTCTGAACTCTTTCCCTACACGACGCTCTTCCGATCTNNNTGCAAGTATCCATAGTTTGG

GAGCACACGTCTGAACTCTTTCCCTACACGACGCTCTTCCGATCTNNNTGCAAGTATCCATAGTTTGG
AGCACACGTCTGAACTCTTTCCCTACACGACGCTCTTCCGATCTNNNGACAGAGCGAGACTCCGTCTCA
AGCACACGTCTGAACTCTTTCCCTACACGACGCTCTTCCGATCTNNNGACAGAGCGAGACTCCGTCTCA
AGCACACGTCTGAACTCTTTCCCTACACGACGCTCTTCCGATCTNNNGACAGAGCGAGACTCCGTCTCA
GAGCACACGTCTGAACTCTTTCCCTACACGACGCTCTTCCGATCTNNNAACTCTCCCTGCTGACTTGTC
GAGCACACGTCTGAACTCTTTCCCTACACGACGCTCTTCCGATCTNNNAACTCTCCCTGCTGACTTGTC
GAGCACACGTCTGAACTCTTTCCCTACACGACGCTCTTCCGATCTNNNAACTCTCCCTGCTGACTTGTC
GAGCACACGTCTGAACTCTTTCCCTACACGACGCTCTTCCGATCTNNNTGAGCCTGGGCCATATAAAA
GAGCACACGTCTGAACTCTTTCCCTACACGACGCTCTTCCGATCTNNNTGAGCCTGGGCCATATAAAA
GAGCACACGTCTGAACTCTTTCCCTACACGACGCTCTTCCGATCTNNNTGAGCCTGGGCCATATAAAA
AGCACACGTCTGAACTCTTTCCCTACACGACGCTCTTCCGATCTNNNTGAGGTAGGAGAATCGCTTG
AGCACACGTCTGAACTCTTTCCCTACACGACGCTCTTCCGATCTNNNTGAGGTAGGAGAATCGCTTG
AGCACACGTCTGAACTCTTTCCCTACACGACGCTCTTCCGATCTNNNTGAGGTAGGAGAATCGCTTG
AGCACACGTCTGAACTCTTTCCCTACACGACGCTCTTCCGATCTNNNTGGCAGCATGTGACCAAGTAT
AGCACACGTCTGAACTCTTTCCCTACACGACGCTCTTCCGATCTNNNTGGCAGCATGTGACCAAGTAT
AGCACACGTCTGAACTCTTTCCCTACACGACGCTCTTCCGATCTNNNTGGCAGCATGTGACCAAGTAT
GAGCACACGTCTGAACTCTTTCCCTACACGACGCTCTTCCGATCTNNNTTTGAGTGGCATTGCATCT
GAGCACACGTCTGAACTCTTTCCCTACACGACGCTCTTCCGATCTNNNTTTGAGTGGCATTGCATCT
GAGCACACGTCTGAACTCTTTCCCTACACGACGCTCTTCCGATCTNNNTTTGAGTGGCATTGCATCT
GCACACGTCTGAACTCTTTCCCTACACGACGCTCTTCCGATCTNNNACACCATCACACCTGGCTAATTT
GCACACGTCTGAACTCTTTCCCTACACGACGCTCTTCCGATCTNNNACACCATCACACCTGGCTAATTT
GCACACGTCTGAACTCTTTCCCTACACGACGCTCTTCCGATCTNNNACACCATCACACCTGGCTAATTT
GCACACGTCTGAACTCTTTCCCTACACGACGCTCTTCCGATCTNNNTTCGAACTCCTGAGCTCAATTGA
GCACACGTCTGAACTCTTTCCCTACACGACGCTCTTCCGATCTNNNTTCGAACTCCTGAGCTCAATTGA
GCACACGTCTGAACTCTTTCCCTACACGACGCTCTTCCGATCTNNNTTCGAACTCCTGAGCTCAATTGA
AGCACACGTCTGAACTCTTTCCCTACACGACGCTCTTCCGATCTNNNGCTGGTGTGGCAGAAAGAT
AGCACACGTCTGAACTCTTTCCCTACACGACGCTCTTCCGATCTNNNGCTGGTGTGGCAGAAAGAT
AGCACACGTCTGAACTCTTTCCCTACACGACGCTCTTCCGATCTNNNGCTGGTGTGGCAGAAAGAT
AGCACACGTCTGAACTCTTTCCCTACACGACGCTCTTCCGATCTNNNTCCCTAGGCATAAAAACTCGG
AGCACACGTCTGAACTCTTTCCCTACACGACGCTCTTCCGATCTNNNTCCCTAGGCATAAAAACTCGG
AGCACACGTCTGAACTCTTTCCCTACACGACGCTCTTCCGATCTNNNTCCCTAGGCATAAAAACTCGG
AGCACACGTCTGAACTCTTTCCCTACACGACGCTCTTCCGATCTNNNTGGTTACAGATAGGAGTGTGT
AGCACACGTCTGAACTCTTTCCCTACACGACGCTCTTCCGATCTNNNTGGTTACAGATAGGAGTGTGT
AGCACACGTCTGAACTCTTTCCCTACACGACGCTCTTCCGATCTNNNTGGTTACAGATAGGAGTGTGT
AGCACACGTCTGAACTCTTTCCCTACACGACGCTCTTCCGATCTNNNAATTGCAAGAGTGTGTGATCA
AGCACACGTCTGAACTCTTTCCCTACACGACGCTCTTCCGATCTNNNAATTGCAAGAGTGTGTGATCA
AGCACACGTCTGAACTCTTTCCCTACACGACGCTCTTCCGATCTNNNAATTGCAAGAGTGTGTGATCA
AGCACACGTCTGAACTCTTTCCCTACACGACGCTCTTCCGATCTNNNCCACCTATCCACACATCCTTTC
AGCACACGTCTGAACTCTTTCCCTACACGACGCTCTTCCGATCTNNNCCACCTATCCACACATCCTTTC
AGCACACGTCTGAACTCTTTCCCTACACGACGCTCTTCCGATCTNNNCCACCTATCCACACATCCTTTC
GAGCACACGTCTGAACTCTTTCCCTACACGACGCTCTTCCGATCTNNNGTGAGAATTACTTGTACCCG
GAGCACACGTCTGAACTCTTTCCCTACACGACGCTCTTCCGATCTNNNGTGAGAATTACTTGTACCCG
GAGCACACGTCTGAACTCTTTCCCTACACGACGCTCTTCCGATCTNNNGTGAGAATTACTTGTACCCG

GCACACGTCTGAACTCTTTCCCTACACGACGCTCTTCCGATCTNNNGGACAAGTCTGTTGTCTCGATGA
GCACACGTCTGAACTCTTTCCCTACACGACGCTCTTCCGATCTNNNGGACAAGTCTGTTGTCTCGATGA
GCACACGTCTGAACTCTTTCCCTACACGACGCTCTTCCGATCTNNNGGACAAGTCTGTTGTCTCGATGA
AGCACACGTCTGAACTCTTTCCCTACACGACGCTCTTCCGATCTNNNAATACAAAAAATTAGCCGGGCG
AGCACACGTCTGAACTCTTTCCCTACACGACGCTCTTCCGATCTNNNAATACAAAAAATTAGCCGGGCG
AGCACACGTCTGAACTCTTTCCCTACACGACGCTCTTCCGATCTNNNAATACAAAAAATTAGCCGGGCG
GAGCACACGTCTGAACTCTTTCCCTACACGACGCTCTTCCGATCTNNNGAATCACTTGAACCCGGGAGA
GAGCACACGTCTGAACTCTTTCCCTACACGACGCTCTTCCGATCTNNNGAATCACTTGAACCCGGGAGA
GAGCACACGTCTGAACTCTTTCCCTACACGACGCTCTTCCGATCTNNNGAATCACTTGAACCCGGGAGA
AGCACACGTCTGAACTCTTTCCCTACACGACGCTCTTCCGATCTNNNCCGGATTCAAACGATTCTCCTCTC
AGCACACGTCTGAACTCTTTCCCTACACGACGCTCTTCCGATCTNNNCCGGATTCAAACGATTCTCCTCTC
AGCACACGTCTGAACTCTTTCCCTACACGACGCTCTTCCGATCTNNNCCGGATTCAAACGATTCTCCTCTC
GCACACGTCTGAACTCTTTCCCTACACGACGCTCTTCCGATCTNNNCTTGTTCTAGACAGTGGCCTTCTC
GCACACGTCTGAACTCTTTCCCTACACGACGCTCTTCCGATCTNNNCTTGTTCTAGACAGTGGCCTTCTC
GCACACGTCTGAACTCTTTCCCTACACGACGCTCTTCCGATCTNNNCTTGTTCTAGACAGTGGCCTTCTC
GAGCACACGTCTGAACTCTTTCCCTACACGACGCTCTTCCGATCTNNNTTAGGGAAGTTCAGGGCATTC
GAGCACACGTCTGAACTCTTTCCCTACACGACGCTCTTCCGATCTNNNTTAGGGAAGTTCAGGGCATTC
GAGCACACGTCTGAACTCTTTCCCTACACGACGCTCTTCCGATCTNNNTTAGGGAAGTTCAGGGCATTC
GCACACGTCTGAACTCTTTCCCTACACGACGCTCTTCCGATCTNNNGCAAGTCTTGGAGGCCAATTTTT
GCACACGTCTGAACTCTTTCCCTACACGACGCTCTTCCGATCTNNNGCAAGTCTTGGAGGCCAATTTTT
GCACACGTCTGAACTCTTTCCCTACACGACGCTCTTCCGATCTNNNGCAAGTCTTGGAGGCCAATTTTT
GAGCACACGTCTGAACTCTTTCCCTACACGACGCTCTTCCGATCTNNNGACAAAGTGCCTGAAACTTGT
GAGCACACGTCTGAACTCTTTCCCTACACGACGCTCTTCCGATCTNNNGACAAAGTGCCTGAAACTTGT
GAGCACACGTCTGAACTCTTTCCCTACACGACGCTCTTCCGATCTNNNGACAAAGTGCCTGAAACTTGT
GAGCACACGTCTGAACTCTTTCCCTACACGACGCTCTTCCGATCTNNNTGCACTCATATTCCCCAGAGAT
GAGCACACGTCTGAACTCTTTCCCTACACGACGCTCTTCCGATCTNNNTGCACTCATATTCCCCAGAGAT
GAGCACACGTCTGAACTCTTTCCCTACACGACGCTCTTCCGATCTNNNTGCACTCATATTCCCCAGAGAT
CACACGTCTGAACTCTTTCCCTACACGACGCTCTTCCGATCTNNNCAAGCATGTTCCAGTATTTGCATCG
CACACGTCTGAACTCTTTCCCTACACGACGCTCTTCCGATCTNNNCAAGCATGTTCCAGTATTTGCATCG
CACACGTCTGAACTCTTTCCCTACACGACGCTCTTCCGATCTNNNCAAGCATGTTCCAGTATTTGCATCG
AGCACACGTCTGAACTCTTTCCCTACACGACGCTCTTCCGATCTNNNCACCAGAACGCATTCCTTTCTTCC
AGCACACGTCTGAACTCTTTCCCTACACGACGCTCTTCCGATCTNNNCACCAGAACGCATTCCTTTCTTCC
AGCACACGTCTGAACTCTTTCCCTACACGACGCTCTTCCGATCTNNNCACCAGAACGCATTCCTTTCTTCC
AGAGCACACGTCTGAACTCTTTCCCTACACGACGCTCTTCCGATCTNNNCTGAGGCAGGAGAATCACTT
AGAGCACACGTCTGAACTCTTTCCCTACACGACGCTCTTCCGATCTNNNCTGAGGCAGGAGAATCACTT
AGAGCACACGTCTGAACTCTTTCCCTACACGACGCTCTTCCGATCTNNNCTGAGGCAGGAGAATCACTT
GAGCACACGTCTGAACTCTTTCCCTACACGACGCTCTTCCGATCTNNNTTTCCTCCCTGAGCACCTTTATC
GAGCACACGTCTGAACTCTTTCCCTACACGACGCTCTTCCGATCTNNNTTTCCTCCCTGAGCACCTTTATC
GAGCACACGTCTGAACTCTTTCCCTACACGACGCTCTTCCGATCTNNNTTTCCTCCCTGAGCACCTTTATC
GAGCACACGTCTGAACTCTTTCCCTACACGACGCTCTTCCGATCTNNNCAGGAGGTGGAGATTGTAGT
GAGCACACGTCTGAACTCTTTCCCTACACGACGCTCTTCCGATCTNNNCAGGAGGTGGAGATTGTAGT
GAGCACACGTCTGAACTCTTTCCCTACACGACGCTCTTCCGATCTNNNCAGGAGGTGGAGATTGTAGT
GAGCACACGTCTGAACTCTTTCCCTACACGACGCTCTTCCGATCTNNNATAAGTGGAAGGAGAGGGG

GAGCACACGTCTGAACTCTTTCCCTACACGACGCTCTTCCGATCTNNNATAAGTGGAAGGAGAGGG
GAGCACACGTCTGAACTCTTTCCCTACACGACGCTCTTCCGATCTNNNATAAGTGGAAGGAGAGGG
AGCACACGTCTGAACTCTTTCCCTACACGACGCTCTTCCGATCTNNNCCGAGTAAATAATGGGGGCT
AGCACACGTCTGAACTCTTTCCCTACACGACGCTCTTCCGATCTNNNCCGAGTAAATAATGGGGGCT
AGCACACGTCTGAACTCTTTCCCTACACGACGCTCTTCCGATCTNNNCCGAGTAAATAATGGGGGCT
GAGCACACGTCTGAACTCTTTCCCTACACGACGCTCTTCCGATCTNNNCGCCTATAGTTCCAGCTACTC
GAGCACACGTCTGAACTCTTTCCCTACACGACGCTCTTCCGATCTNNNCGCCTATAGTTCCAGCTACTC
GAGCACACGTCTGAACTCTTTCCCTACACGACGCTCTTCCGATCTNNNCGCCTATAGTTCCAGCTACTC
GCACACGTCTGAACTCTTTCCCTACACGACGCTCTTCCGATCTNNNGGATCACCTGAGGTCAAGAGTT
GCACACGTCTGAACTCTTTCCCTACACGACGCTCTTCCGATCTNNNGGATCACCTGAGGTCAAGAGTT
GCACACGTCTGAACTCTTTCCCTACACGACGCTCTTCCGATCTNNNGGATCACCTGAGGTCAAGAGTT
GCACACGTCTGAACTCTTTCCCTACACGACGCTCTTCCGATCTNNNTGATGGAGTAAGACCCTGACTC
GCACACGTCTGAACTCTTTCCCTACACGACGCTCTTCCGATCTNNNTGATGGAGTAAGACCCTGACTC
GCACACGTCTGAACTCTTTCCCTACACGACGCTCTTCCGATCTNNNTGATGGAGTAAGACCCTGACTC
GAGCACACGTCTGAACTCTTTCCCTACACGACGCTCTTCCGATCTNNNGAGGCTGAGGCAGGAGAAT
GAGCACACGTCTGAACTCTTTCCCTACACGACGCTCTTCCGATCTNNNGAGGCTGAGGCAGGAGAAT
GAGCACACGTCTGAACTCTTTCCCTACACGACGCTCTTCCGATCTNNNGAGGCTGAGGCAGGAGAAT
AGCACACGTCTGAACTCTTTCCCTACACGACGCTCTTCCGATCTNNNTGTCTGTTCATATCCTTTGCCCA
AGCACACGTCTGAACTCTTTCCCTACACGACGCTCTTCCGATCTNNNTGTCTGTTCATATCCTTTGCCCA
AGCACACGTCTGAACTCTTTCCCTACACGACGCTCTTCCGATCTNNNTGTCTGTTCATATCCTTTGCCCA
AGCACACGTCTGAACTCTTTCCCTACACGACGCTCTTCCGATCTNNNAAATCATCTCCAAAAAAAGTCT
AGCACACGTCTGAACTCTTTCCCTACACGACGCTCTTCCGATCTNNNAAATCATCTCCAAAAAAAGTCT
AGCACACGTCTGAACTCTTTCCCTACACGACGCTCTTCCGATCTNNNAAATCATCTCCAAAAAAAGTCT
GCACACGTCTGAACTCTTTCCCTACACGACGCTCTTCCGATCTNNNCCTCAAGGATCCTCTTTCTGACC
GCACACGTCTGAACTCTTTCCCTACACGACGCTCTTCCGATCTNNNCCTCAAGGATCCTCTTTCTGACC
GCACACGTCTGAACTCTTTCCCTACACGACGCTCTTCCGATCTNNNCCTCAAGGATCCTCTTTCTGACC
GAGCACACGTCTGAACTCTTTCCCTACACGACGCTCTTCCGATCTNNNCTTGTCACTTCTCTACCTGTCC
GAGCACACGTCTGAACTCTTTCCCTACACGACGCTCTTCCGATCTNNNCTTGTCACTTCTCTACCTGTCC
GAGCACACGTCTGAACTCTTTCCCTACACGACGCTCTTCCGATCTNNNCTTGTCACTTCTCTACCTGTCC
GCACACGTCTGAACTCTTTCCCTACACGACGCTCTTCCGATCTNNNCCTGTAATCCTAGCACTTTGGG
GCACACGTCTGAACTCTTTCCCTACACGACGCTCTTCCGATCTNNNCCTGTAATCCTAGCACTTTGGG
GCACACGTCTGAACTCTTTCCCTACACGACGCTCTTCCGATCTNNNCCTGTAATCCTAGCACTTTGGG
GAGCACACGTCTGAACTCTTTCCCTACACGACGCTCTTCCGATCTNNNATGCATGTAATCCCAGCACTT
GAGCACACGTCTGAACTCTTTCCCTACACGACGCTCTTCCGATCTNNNATGCATGTAATCCCAGCACTT
GAGCACACGTCTGAACTCTTTCCCTACACGACGCTCTTCCGATCTNNNATGCATGTAATCCCAGCACTT
AGCACACGTCTGAACTCTTTCCCTACACGACGCTCTTCCGATCTNNNTAGGCTCAAAATGAAGGGAT
AGCACACGTCTGAACTCTTTCCCTACACGACGCTCTTCCGATCTNNNTAGGCTCAAAATGAAGGGAT
AGCACACGTCTGAACTCTTTCCCTACACGACGCTCTTCCGATCTNNNTAGGCTCAAAATGAAGGGAT
GAGCACACGTCTGAACTCTTTCCCTACACGACGCTCTTCCGATCTNNNGGATCACTTGAGACCAGGA
GAGCACACGTCTGAACTCTTTCCCTACACGACGCTCTTCCGATCTNNNGGATCACTTGAGACCAGGA
GAGCACACGTCTGAACTCTTTCCCTACACGACGCTCTTCCGATCTNNNGGATCACTTGAGACCAGGA
GCACACGTCTGAACTCTTTCCCTACACGACGCTCTTCCGATCTNNNTTGCTGTTTTTAGACCTGCCATG
GCACACGTCTGAACTCTTTCCCTACACGACGCTCTTCCGATCTNNNTTGCTGTTTTTAGACCTGCCATG

GCACACGTCTGAACTCTTTCCCTACACGACGCTCTTCCGATCTNNNTTGCTGTTTTAGACCTGCCATG
GAGCACACGTCTGAACTCTTTCCCTACACGACGCTCTTCCGATCTNNNAGTTGTAAAAGCAATTTCTTT
GAGCACACGTCTGAACTCTTTCCCTACACGACGCTCTTCCGATCTNNNAGTTGTAAAAGCAATTTCTTT
GAGCACACGTCTGAACTCTTTCCCTACACGACGCTCTTCCGATCTNNNAGTTGTAAAAGCAATTTCTTT
GCACACGTCTGAACTCTTTCCCTACACGACGCTCTTCCGATCTNNNTATGAAATCTGCGCAGGGTCAT
GCACACGTCTGAACTCTTTCCCTACACGACGCTCTTCCGATCTNNNTATGAAATCTGCGCAGGGTCAT
GCACACGTCTGAACTCTTTCCCTACACGACGCTCTTCCGATCTNNNTATGAAATCTGCGCAGGGTCAT
GAGCACACGTCTGAACTCTTTCCCTACACGACGCTCTTCCGATCTNNNTCAGGACTAAAGCAGTGGAA
GAGCACACGTCTGAACTCTTTCCCTACACGACGCTCTTCCGATCTNNNTCAGGACTAAAGCAGTGGAA
GAGCACACGTCTGAACTCTTTCCCTACACGACGCTCTTCCGATCTNNNTCAGGACTAAAGCAGTGGAA
GCACACGTCTGAACTCTTTCCCTACACGACGCTCTTCCGATCTNNNGTGGGATCTCAAGAAGCCAGTA
GCACACGTCTGAACTCTTTCCCTACACGACGCTCTTCCGATCTNNNGTGGGATCTCAAGAAGCCAGTA
GCACACGTCTGAACTCTTTCCCTACACGACGCTCTTCCGATCTNNNGTGGGATCTCAAGAAGCCAGTA
AGAGCACACGTCTGAACTCTTTCCCTACACGACGCTCTTCCGATCTNNNTTCCAGGCTTATTAGGCTTC
AGAGCACACGTCTGAACTCTTTCCCTACACGACGCTCTTCCGATCTNNNTTCCAGGCTTATTAGGCTTC
AGAGCACACGTCTGAACTCTTTCCCTACACGACGCTCTTCCGATCTNNNTTCCAGGCTTATTAGGCTTC
AGAGCACACGTCTGAACTCTTTCCCTACACGACGCTCTTCCGATCTNNNATAAAAGCATTGGTATGGC
AGAGCACACGTCTGAACTCTTTCCCTACACGACGCTCTTCCGATCTNNNATAAAAGCATTGGTATGGC
AGAGCACACGTCTGAACTCTTTCCCTACACGACGCTCTTCCGATCTNNNATAAAAGCATTGGTATGGC
AGCACACGTCTGAACTCTTTCCCTACACGACGCTCTTCCGATCTNNNTAGCACACACTTTTCCACCACAA
AGCACACGTCTGAACTCTTTCCCTACACGACGCTCTTCCGATCTNNNTAGCACACACTTTTCCACCACAA
AGCACACGTCTGAACTCTTTCCCTACACGACGCTCTTCCGATCTNNNTAGCACACACTTTTCCACCACAA
GCACACGTCTGAACTCTTTCCCTACACGACGCTCTTCCGATCTNNNAGAGAGGAGGATGGAAGAGGA
GCACACGTCTGAACTCTTTCCCTACACGACGCTCTTCCGATCTNNNAGAGAGGAGGATGGAAGAGGA
GCACACGTCTGAACTCTTTCCCTACACGACGCTCTTCCGATCTNNNAGAGAGGAGGATGGAAGAGGA
GAGCACACGTCTGAACTCTTTCCCTACACGACGCTCTTCCGATCTNNNACCCCTAGGAAGACTGAAACA
GAGCACACGTCTGAACTCTTTCCCTACACGACGCTCTTCCGATCTNNNACCCCTAGGAAGACTGAAACA
GAGCACACGTCTGAACTCTTTCCCTACACGACGCTCTTCCGATCTNNNACCCCTAGGAAGACTGAAACA
GAGCACACGTCTGAACTCTTTCCCTACACGACGCTCTTCCGATCTNNNTGTGCCCTCAGGTTATTCTAC
GAGCACACGTCTGAACTCTTTCCCTACACGACGCTCTTCCGATCTNNNTGTGCCCTCAGGTTATTCTAC
GAGCACACGTCTGAACTCTTTCCCTACACGACGCTCTTCCGATCTNNNTGTGCCCTCAGGTTATTCTAC
AGCACACGTCTGAACTCTTTCCCTACACGACGCTCTTCCGATCTNNNCTGTAATCCCAGCTACTCAGGA
AGCACACGTCTGAACTCTTTCCCTACACGACGCTCTTCCGATCTNNNCTGTAATCCCAGCTACTCAGGA
AGCACACGTCTGAACTCTTTCCCTACACGACGCTCTTCCGATCTNNNCTGTAATCCCAGCTACTCAGGA
AGCACACGTCTGAACTCTTTCCCTACACGACGCTCTTCCGATCTNNNAACATTGCTTTGCCACTATGCAT
AGCACACGTCTGAACTCTTTCCCTACACGACGCTCTTCCGATCTNNNAACATTGCTTTGCCACTATGCAT
AGCACACGTCTGAACTCTTTCCCTACACGACGCTCTTCCGATCTNNNAACATTGCTTTGCCACTATGCAT
AGCACACGTCTGAACTCTTTCCCTACACGACGCTCTTCCGATCTNNNCTCCTGAGTAGCTGGGACTACA
AGCACACGTCTGAACTCTTTCCCTACACGACGCTCTTCCGATCTNNNCTCCTGAGTAGCTGGGACTACA
AGCACACGTCTGAACTCTTTCCCTACACGACGCTCTTCCGATCTNNNCTCCTGAGTAGCTGGGACTACA
GAGCACACGTCTGAACTCTTTCCCTACACGACGCTCTTCCGATCTNNNCTCTTACAATTTGGGGAAGG
GAGCACACGTCTGAACTCTTTCCCTACACGACGCTCTTCCGATCTNNNCTCTTACAATTTGGGGAAGG
GAGCACACGTCTGAACTCTTTCCCTACACGACGCTCTTCCGATCTNNNCTCTTACAATTTGGGGAAGG

AGCACACGTCTGAACTCTTTCCCTACACGACGCTCTTCCGATCTNNNTGCTATGTGAATTTGGGGGA

AGCACACGTCTGAACTCTTTCCCTACACGACGCTCTTCCGATCTNNNTGCTATGTGAATTTGGGGGA

AGCACACGTCTGAACTCTTTCCCTACACGACGCTCTTCCGATCTNNNTGCTATGTGAATTTGGGGGA

GAGCACACGTCTGAACTCTTTCCCTACACGACGCTCTTCCGATCTNNNGTGTTACAGAGGCAATGAG

GAGCACACGTCTGAACTCTTTCCCTACACGACGCTCTTCCGATCTNNNGTGTTACAGAGGCAATGAG

GAGCACACGTCTGAACTCTTTCCCTACACGACGCTCTTCCGATCTNNNGTGTTACAGAGGCAATGAG

AGAGCACACGTCTGAACTCTTTCCCTACACGACGCTCTTCCGATCTNNNAGAGTTCTCACTTGGTCAT

AGAGCACACGTCTGAACTCTTTCCCTACACGACGCTCTTCCGATCTNNNAGAGTTCTCACTTGGTCAT

AGAGCACACGTCTGAACTCTTTCCCTACACGACGCTCTTCCGATCTNNNAGAGTTCTCACTTGGTCAT

GAGCACACGTCTGAACTCTTTCCCTACACGACGCTCTTCCGATCTNNNTCCAACAAAAAGGTGACCTTG

GAGCACACGTCTGAACTCTTTCCCTACACGACGCTCTTCCGATCTNNNTCCAACAAAAAGGTGACCTTG

GAGCACACGTCTGAACTCTTTCCCTACACGACGCTCTTCCGATCTNNNTCCAACAAAAAGGTGACCTTG

AGAGCACACGTCTGAACTCTTTCCCTACACGACGCTCTTCCGATCTNNNCCTCGAGCATACCACTTTCTT

AGAGCACACGTCTGAACTCTTTCCCTACACGACGCTCTTCCGATCTNNNCCTCGAGCATACCACTTTCTT

AGAGCACACGTCTGAACTCTTTCCCTACACGACGCTCTTCCGATCTNNNCCTCGAGCATACCACTTTCTT

AGCACACGTCTGAACTCTTTCCCTACACGACGCTCTTCCGATCTNNNTGTTCATGTCCTTCACCCACTTT

AGCACACGTCTGAACTCTTTCCCTACACGACGCTCTTCCGATCTNNNTGTTCATGTCCTTCACCCACTTT

AGCACACGTCTGAACTCTTTCCCTACACGACGCTCTTCCGATCTNNNTGTTCATGTCCTTCACCCACTTT

AGAGCACACGTCTGAACTCTTTCCCTACACGACGCTCTTCCGATCTNNNTGAGGTAGGAGAATGGCA

AGAGCACACGTCTGAACTCTTTCCCTACACGACGCTCTTCCGATCTNNNTGAGGTAGGAGAATGGCA

AGAGCACACGTCTGAACTCTTTCCCTACACGACGCTCTTCCGATCTNNNTGAGGTAGGAGAATGGCA

AGCACACGTCTGAACTCTTTCCCTACACGACGCTCTTCCGATCTNNNAATATTGATCTTGCCTTCTTTG

AGCACACGTCTGAACTCTTTCCCTACACGACGCTCTTCCGATCTNNNAATATTGATCTTGCCTTCTTTG

AGCACACGTCTGAACTCTTTCCCTACACGACGCTCTTCCGATCTNNNAATATTGATCTTGCCTTCTTTG

AGCACACGTCTGAACTCTTTCCCTACACGACGCTCTTCCGATCTNNNTTGGCCAAGGAAAACATCATG

AGCACACGTCTGAACTCTTTCCCTACACGACGCTCTTCCGATCTNNNTTGGCCAAGGAAAACATCATG

AGCACACGTCTGAACTCTTTCCCTACACGACGCTCTTCCGATCTNNNTTGGCCAAGGAAAACATCATG

AGCACACGTCTGAACTCTTTCCCTACACGACGCTCTTCCGATCTNNNCCTCCCTTCTGTATTTTCAGCCT

AGCACACGTCTGAACTCTTTCCCTACACGACGCTCTTCCGATCTNNNCCTCCCTTCTGTATTTTCAGCCT

AGCACACGTCTGAACTCTTTCCCTACACGACGCTCTTCCGATCTNNNCCTCCCTTCTGTATTTTCAGCCT

AGCACACGTCTGAACTCTTTCCCTACACGACGCTCTTCCGATCTNNNACAGTGTACAAGGGTTCCCTTT

AGCACACGTCTGAACTCTTTCCCTACACGACGCTCTTCCGATCTNNNACAGTGTACAAGGGTTCCCTTT

AGCACACGTCTGAACTCTTTCCCTACACGACGCTCTTCCGATCTNNNACAGTGTACAAGGGTTCCCTTT

AGCACACGTCTGAACTCTTTCCCTACACGACGCTCTTCCGATCTNNNGAGTTGCTTGAACCTGGGAGA

AGCACACGTCTGAACTCTTTCCCTACACGACGCTCTTCCGATCTNNNGAGTTGCTTGAACCTGGGAGA

AGCACACGTCTGAACTCTTTCCCTACACGACGCTCTTCCGATCTNNNGAGTTGCTTGAACCTGGGAGA

GAGCACACGTCTGAACTCTTTCCCTACACGACGCTCTTCCGATCTNNNGCAGGAAAATCTCTTGAACCC

GAGCACACGTCTGAACTCTTTCCCTACACGACGCTCTTCCGATCTNNNGCAGGAAAATCTCTTGAACCC

GAGCACACGTCTGAACTCTTTCCCTACACGACGCTCTTCCGATCTNNNGCAGGAAAATCTCTTGAACCC

GCACACGTCTGAACTCTTTCCCTACACGACGCTCTTCCGATCTNNNCCCTCACCTCCTGGATAGGGCAT

GCACACGTCTGAACTCTTTCCCTACACGACGCTCTTCCGATCTNNNCCCTCACCTCCTGGATAGGGCAT

GCACACGTCTGAACTCTTTCCCTACACGACGCTCTTCCGATCTNNNCCCTCACCTCCTGGATAGGGCAT

GAGCACACGTCTGAACTCTTTCCCTACACGACGCTCTTCCGATCTNNNCTCAATTTTGGAGAAGCAGC

GAGCACACGTCTGAACTCTTTCCCTACACGACGCTCTTCCGATCTNNNCTCAATTTTGGAGAAGCAGC
GAGCACACGTCTGAACTCTTTCCCTACACGACGCTCTTCCGATCTNNNCTCAATTTTGGAGAAGCAGC
AGCACACGTCTGAACTCTTTCCCTACACGACGCTCTTCCGATCTNNNACCATGCAGAAATCCAAAACCT
AGCACACGTCTGAACTCTTTCCCTACACGACGCTCTTCCGATCTNNNACCATGCAGAAATCCAAAACCT
AGCACACGTCTGAACTCTTTCCCTACACGACGCTCTTCCGATCTNNNACCATGCAGAAATCCAAAACCT
ACACGTCTGAACTCTTTCCCTACACGACGCTCTTCCGATCTNNNTGTGCCACCTTGTTATGTGAGTACA
ACACGTCTGAACTCTTTCCCTACACGACGCTCTTCCGATCTNNNTGTGCCACCTTGTTATGTGAGTACA
ACACGTCTGAACTCTTTCCCTACACGACGCTCTTCCGATCTNNNTGTGCCACCTTGTTATGTGAGTACA
AGCACACGTCTGAACTCTTTCCCTACACGACGCTCTTCCGATCTNNNCACCTGTAATCCCAGCTACTCG
AGCACACGTCTGAACTCTTTCCCTACACGACGCTCTTCCGATCTNNNCACCTGTAATCCCAGCTACTCG
AGCACACGTCTGAACTCTTTCCCTACACGACGCTCTTCCGATCTNNNCACCTGTAATCCCAGCTACTCG
AGCACACGTCTGAACTCTTTCCCTACACGACGCTCTTCCGATCTNNNAGGGTAGCTAAGACCTCACT
AGCACACGTCTGAACTCTTTCCCTACACGACGCTCTTCCGATCTNNNAGGGTAGCTAAGACCTCACT
AGCACACGTCTGAACTCTTTCCCTACACGACGCTCTTCCGATCTNNNAGGGTAGCTAAGACCTCACT
AGCACACGTCTGAACTCTTTCCCTACACGACGCTCTTCCGATCTNNNGGCTCCTCACTTCCCAGTAGG
AGCACACGTCTGAACTCTTTCCCTACACGACGCTCTTCCGATCTNNNGGCTCCTCACTTCCCAGTAGG
AGCACACGTCTGAACTCTTTCCCTACACGACGCTCTTCCGATCTNNNGGCTCCTCACTTCCCAGTAGG
GAGCACACGTCTGAACTCTTTCCCTACACGACGCTCTTCCGATCTNNNCTTGAGAAGACTGGAGAAG
GAGCACACGTCTGAACTCTTTCCCTACACGACGCTCTTCCGATCTNNNCTTGAGAAGACTGGAGAAG
GAGCACACGTCTGAACTCTTTCCCTACACGACGCTCTTCCGATCTNNNCTTGAGAAGACTGGAGAAG
GCACACGTCTGAACTCTTTCCCTACACGACGCTCTTCCGATCTNNNAATCTTATGAATGTCAACAATGT
GCACACGTCTGAACTCTTTCCCTACACGACGCTCTTCCGATCTNNNAATCTTATGAATGTCAACAATGT
GCACACGTCTGAACTCTTTCCCTACACGACGCTCTTCCGATCTNNNAATCTTATGAATGTCAACAATGT
AGCACACGTCTGAACTCTTTCCCTACACGACGCTCTTCCGATCTNNNTAAAGGCTGCTGTGAACTGAG
AGCACACGTCTGAACTCTTTCCCTACACGACGCTCTTCCGATCTNNNTAAAGGCTGCTGTGAACTGAG
AGCACACGTCTGAACTCTTTCCCTACACGACGCTCTTCCGATCTNNNTAAAGGCTGCTGTGAACTGAG
CACACGTCTGAACTCTTTCCCTACACGACGCTCTTCCGATCTNNNCTGTTTGTATTCACCGCTTCACCC
CACACGTCTGAACTCTTTCCCTACACGACGCTCTTCCGATCTNNNCTGTTTGTATTCACCGCTTCACCC
CACACGTCTGAACTCTTTCCCTACACGACGCTCTTCCGATCTNNNCTGTTTGTATTCACCGCTTCACCC
AGCACACGTCTGAACTCTTTCCCTACACGACGCTCTTCCGATCTNNNATCAACTGTTACAGAAATGAAA
AGCACACGTCTGAACTCTTTCCCTACACGACGCTCTTCCGATCTNNNATCAACTGTTACAGAAATGAAA
AGCACACGTCTGAACTCTTTCCCTACACGACGCTCTTCCGATCTNNNATCAACTGTTACAGAAATGAAA
AGAGCACACGTCTGAACTCTTTCCCTACACGACGCTCTTCCGATCTNNNTAGTCCCAACTACTCAGGA
AGAGCACACGTCTGAACTCTTTCCCTACACGACGCTCTTCCGATCTNNNTAGTCCCAACTACTCAGGA
AGAGCACACGTCTGAACTCTTTCCCTACACGACGCTCTTCCGATCTNNNTAGTCCCAACTACTCAGGA
GCACACGTCTGAACTCTTTCCCTACACGACGCTCTTCCGATCTNNNCACTCCTGATGTTCACTCTCCAAC
GCACACGTCTGAACTCTTTCCCTACACGACGCTCTTCCGATCTNNNCACTCCTGATGTTCACTCTCCAAC
GCACACGTCTGAACTCTTTCCCTACACGACGCTCTTCCGATCTNNNCACTCCTGATGTTCACTCTCCAAC
AGCACACGTCTGAACTCTTTCCCTACACGACGCTCTTCCGATCTNNNTTTGCTCATGGCCCCATATTTTC
AGCACACGTCTGAACTCTTTCCCTACACGACGCTCTTCCGATCTNNNTTTGCTCATGGCCCCATATTTTC
AGCACACGTCTGAACTCTTTCCCTACACGACGCTCTTCCGATCTNNNTTTGCTCATGGCCCCATATTTTC
AGAGCACACGTCTGAACTCTTTCCCTACACGACGCTCTTCCGATCTNNNTGTGTGTTCAGCTTTTGTT
AGAGCACACGTCTGAACTCTTTCCCTACACGACGCTCTTCCGATCTNNNTGTGTGTTCAGCTTTTGTT

AAGAGCACACGTCTGAACTCTTTCCCTACACGACGCTCTTCCGATCTNNNTGTGTGTTCAGCTTTTGTT

GAGCACACGTCTGAACTCTTTCCCTACACGACGCTCTTCCGATCTNNNATGACATAACTGCACTCCAAC

GAGCACACGTCTGAACTCTTTCCCTACACGACGCTCTTCCGATCTNNNATGACATAACTGCACTCCAAC

GAGCACACGTCTGAACTCTTTCCCTACACGACGCTCTTCCGATCTNNNATGACATAACTGCACTCCAAC

AGCACACGTCTGAACTCTTTCCCTACACGACGCTCTTCCGATCTNNNTGGGAAATTGTCTAGATTTAC

AGCACACGTCTGAACTCTTTCCCTACACGACGCTCTTCCGATCTNNNTGGGAAATTGTCTAGATTTAC

AGCACACGTCTGAACTCTTTCCCTACACGACGCTCTTCCGATCTNNNTGGGAAATTGTCTAGATTTAC

GAGCACACGTCTGAACTCTTTCCCTACACGACGCTCTTCCGATCTNNNTCCTAGCTACTTAGGAGGCT

GAGCACACGTCTGAACTCTTTCCCTACACGACGCTCTTCCGATCTNNNTCCTAGCTACTTAGGAGGCT

GAGCACACGTCTGAACTCTTTCCCTACACGACGCTCTTCCGATCTNNNTCCTAGCTACTTAGGAGGCT

GAGCACACGTCTGAACTCTTTCCCTACACGACGCTCTTCCGATCTNNNGTTCAAAACCAGCCTGAGCA

GAGCACACGTCTGAACTCTTTCCCTACACGACGCTCTTCCGATCTNNNGTTCAAAACCAGCCTGAGCA

GAGCACACGTCTGAACTCTTTCCCTACACGACGCTCTTCCGATCTNNNGTTCAAAACCAGCCTGAGCA

GCACACGTCTGAACTCTTTCCCTACACGACGCTCTTCCGATCTNNNGAATCCCGTAACAACTGGACTC

GCACACGTCTGAACTCTTTCCCTACACGACGCTCTTCCGATCTNNNGAATCCCGTAACAACTGGACTC

GCACACGTCTGAACTCTTTCCCTACACGACGCTCTTCCGATCTNNNGAATCCCGTAACAACTGGACTC

AGCACACGTCTGAACTCTTTCCCTACACGACGCTCTTCCGATCTNNNGTACAGGAACAGACACATAG

AGCACACGTCTGAACTCTTTCCCTACACGACGCTCTTCCGATCTNNNGTACAGGAACAGACACATAG

AGCACACGTCTGAACTCTTTCCCTACACGACGCTCTTCCGATCTNNNGTACAGGAACAGACACATAG

GAGCACACGTCTGAACTCTTTCCCTACACGACGCTCTTCCGATCTNNNCTGAGGCAGGAGAATCACTT

GAGCACACGTCTGAACTCTTTCCCTACACGACGCTCTTCCGATCTNNNCTGAGGCAGGAGAATCACTT

GAGCACACGTCTGAACTCTTTCCCTACACGACGCTCTTCCGATCTNNNCTGAGGCAGGAGAATCACTT

GCACACGTCTGAACTCTTTCCCTACACGACGCTCTTCCGATCTNNNGAAGCTGAGGCAGGAGAATGG

GCACACGTCTGAACTCTTTCCCTACACGACGCTCTTCCGATCTNNNGAAGCTGAGGCAGGAGAATGG

GCACACGTCTGAACTCTTTCCCTACACGACGCTCTTCCGATCTNNNGAAGCTGAGGCAGGAGAATGG

GCACACGTCTGAACTCTTTCCCTACACGACGCTCTTCCGATCTNNNCACCACAACTGGCGAATTTTTT

GCACACGTCTGAACTCTTTCCCTACACGACGCTCTTCCGATCTNNNCACCACAACTGGCGAATTTTTT

GCACACGTCTGAACTCTTTCCCTACACGACGCTCTTCCGATCTNNNCACCACAACTGGCGAATTTTTT

GAGCACACGTCTGAACTCTTTCCCTACACGACGCTCTTCCGATCTNNNCAAAAAAAAATTAGCCGGGC

GAGCACACGTCTGAACTCTTTCCCTACACGACGCTCTTCCGATCTNNNCAAAAAAAAATTAGCCGGGC

GAGCACACGTCTGAACTCTTTCCCTACACGACGCTCTTCCGATCTNNNCAAAAAAAAATTAGCCGGGC

AGCACACGTCTGAACTCTTTCCCTACACGACGCTCTTCCGATCTNNNGTCACACAATAGCCTAAATATT

AGCACACGTCTGAACTCTTTCCCTACACGACGCTCTTCCGATCTNNNGTCACACAATAGCCTAAATATT

AGCACACGTCTGAACTCTTTCCCTACACGACGCTCTTCCGATCTNNNGTCACACAATAGCCTAAATATT

GCACACGTCTGAACTCTTTCCCTACACGACGCTCTTCCGATCTNNNAAGATTTAATGACCTCAGGTTT

GCACACGTCTGAACTCTTTCCCTACACGACGCTCTTCCGATCTNNNAAGATTTAATGACCTCAGGTTT

GCACACGTCTGAACTCTTTCCCTACACGACGCTCTTCCGATCTNNNAAGATTTAATGACCTCAGGTTT

AGCACACGTCTGAACTCTTTCCCTACACGACGCTCTTCCGATCTNNNTTGATAGGACTGTGAGTGTG

AGCACACGTCTGAACTCTTTCCCTACACGACGCTCTTCCGATCTNNNTTGATAGGACTGTGAGTGTG

AGCACACGTCTGAACTCTTTCCCTACACGACGCTCTTCCGATCTNNNTTGATAGGACTGTGAGTGTG

AGCACACGTCTGAACTCTTTCCCTACACGACGCTCTTCCGATCTNNNTTCAAGTGATTCTCCTTCCCCAG

AGCACACGTCTGAACTCTTTCCCTACACGACGCTCTTCCGATCTNNNTTCAAGTGATTCTCCTTCCCCAG

AGCACACGTCTGAACTCTTTCCCTACACGACGCTCTTCCGATCTNNNTTCAAGTGATTCTCCTTCCCCAG

AGCACACGTCTGAACTCTTTCCCTACACGACGCTCTTCCGATCTNNNGAGTCTTGCTCTGTTGCCCAGC
AGCACACGTCTGAACTCTTTCCCTACACGACGCTCTTCCGATCTNNNGAGTCTTGCTCTGTTGCCCAGC
AGCACACGTCTGAACTCTTTCCCTACACGACGCTCTTCCGATCTNNNGAGTCTTGCTCTGTTGCCCAGC
GCACACGTCTGAACTCTTTCCCTACACGACGCTCTTCCGATCTNNNCTCACCTGTAATCTCAGCACT
GCACACGTCTGAACTCTTTCCCTACACGACGCTCTTCCGATCTNNNCTCACCTGTAATCTCAGCACT
GCACACGTCTGAACTCTTTCCCTACACGACGCTCTTCCGATCTNNNCTCACCTGTAATCTCAGCACT
GAGCACACGTCTGAACTCTTTCCCTACACGACGCTCTTCCGATCTNNNCCAAGCTAAACAACCTGGAA
GAGCACACGTCTGAACTCTTTCCCTACACGACGCTCTTCCGATCTNNNCCAAGCTAAACAACCTGGAA
GAGCACACGTCTGAACTCTTTCCCTACACGACGCTCTTCCGATCTNNNCCAAGCTAAACAACCTGGAA
AGCACACGTCTGAACTCTTTCCCTACACGACGCTCTTCCGATCTNNNTAAAGGAAGCTGCTCTGAAG
AGCACACGTCTGAACTCTTTCCCTACACGACGCTCTTCCGATCTNNNTAAAGGAAGCTGCTCTGAAG
AGCACACGTCTGAACTCTTTCCCTACACGACGCTCTTCCGATCTNNNTAAAGGAAGCTGCTCTGAAG
GAGCACACGTCTGAACTCTTTCCCTACACGACGCTCTTCCGATCTNNNTCCTGTATCCCTCTCAGGCAT
GAGCACACGTCTGAACTCTTTCCCTACACGACGCTCTTCCGATCTNNNTCCTGTATCCCTCTCAGGCAT
GAGCACACGTCTGAACTCTTTCCCTACACGACGCTCTTCCGATCTNNNTCCTGTATCCCTCTCAGGCAT
GCACACGTCTGAACTCTTTCCCTACACGACGCTCTTCCGATCTNNNTTGACATTTGGGATCGCCTCATT
GCACACGTCTGAACTCTTTCCCTACACGACGCTCTTCCGATCTNNNTTGACATTTGGGATCGCCTCATT
GCACACGTCTGAACTCTTTCCCTACACGACGCTCTTCCGATCTNNNTTGACATTTGGGATCGCCTCATT
GCACACGTCTGAACTCTTTCCCTACACGACGCTCTTCCGATCTNNNGAAGCCACTACAAATGGATGTC
GCACACGTCTGAACTCTTTCCCTACACGACGCTCTTCCGATCTNNNGAAGCCACTACAAATGGATGTC
GCACACGTCTGAACTCTTTCCCTACACGACGCTCTTCCGATCTNNNGAAGCCACTACAAATGGATGTC
AGCACACGTCTGAACTCTTTCCCTACACGACGCTCTTCCGATCTNNNCTGTCATATCAGCTACTCGGG
AGCACACGTCTGAACTCTTTCCCTACACGACGCTCTTCCGATCTNNNCTGTCATATCAGCTACTCGGG
AGCACACGTCTGAACTCTTTCCCTACACGACGCTCTTCCGATCTNNNCTGTCATATCAGCTACTCGGG
GAGCACACGTCTGAACTCTTTCCCTACACGACGCTCTTCCGATCTNNNACAAAAAACAGCGGTTCTAT
GAGCACACGTCTGAACTCTTTCCCTACACGACGCTCTTCCGATCTNNNACAAAAAACAGCGGTTCTAT
GAGCACACGTCTGAACTCTTTCCCTACACGACGCTCTTCCGATCTNNNACAAAAAACAGCGGTTCTAT
GCACACGTCTGAACTCTTTCCCTACACGACGCTCTTCCGATCTNNNCAGTTTCTGGTCATGCCCTTGTA
GCACACGTCTGAACTCTTTCCCTACACGACGCTCTTCCGATCTNNNCAGTTTCTGGTCATGCCCTTGTA
GCACACGTCTGAACTCTTTCCCTACACGACGCTCTTCCGATCTNNNCAGTTTCTGGTCATGCCCTTGTA
AGCACACGTCTGAACTCTTTCCCTACACGACGCTCTTCCGATCTNNNATTGTAATAGAATGGAAGGAA
AGCACACGTCTGAACTCTTTCCCTACACGACGCTCTTCCGATCTNNNATTGTAATAGAATGGAAGGAA
AGCACACGTCTGAACTCTTTCCCTACACGACGCTCTTCCGATCTNNNATTGTAATAGAATGGAAGGAA
AGCACACGTCTGAACTCTTTCCCTACACGACGCTCTTCCGATCTNNNCTGGTGAGGACAAAGGTTCC
AGCACACGTCTGAACTCTTTCCCTACACGACGCTCTTCCGATCTNNNCTGGTGAGGACAAAGGTTCC
AGCACACGTCTGAACTCTTTCCCTACACGACGCTCTTCCGATCTNNNCTGGTGAGGACAAAGGTTCC
AGCACACGTCTGAACTCTTTCCCTACACGACGCTCTTCCGATCTNNNGTCAAGAAGCCAACAGTCATGT
AGCACACGTCTGAACTCTTTCCCTACACGACGCTCTTCCGATCTNNNGTCAAGAAGCCAACAGTCATGT
AGCACACGTCTGAACTCTTTCCCTACACGACGCTCTTCCGATCTNNNGTCAAGAAGCCAACAGTCATGT
GCACACGTCTGAACTCTTTCCCTACACGACGCTCTTCCGATCTNNNGAAGTAAGGAGCCCCTCTGCCA
GCACACGTCTGAACTCTTTCCCTACACGACGCTCTTCCGATCTNNNGAAGTAAGGAGCCCCTCTGCCA
GCACACGTCTGAACTCTTTCCCTACACGACGCTCTTCCGATCTNNNGAAGTAAGGAGCCCCTCTGCCA
GAGCACACGTCTGAACTCTTTCCCTACACGACGCTCTTCCGATCTNNNAGTAGGATGGATGGACAG

AGAGCACACGTCTGAACTCTTTCCCTACACGACGCTCTTCCGATCTNNNAGTAGGATGGATGGACAGA

AGAGCACACGTCTGAACTCTTTCCCTACACGACGCTCTTCCGATCTNNNAGTAGGATGGATGGACAGA

AGCACACGTCTGAACTCTTTCCCTACACGACGCTCTTCCGATCTNNNAAGAGGCAAAGGTGGATTTGTT

AGCACACGTCTGAACTCTTTCCCTACACGACGCTCTTCCGATCTNNNAAGAGGCAAAGGTGGATTTGTT

AGCACACGTCTGAACTCTTTCCCTACACGACGCTCTTCCGATCTNNNAAGAGGCAAAGGTGGATTTGTT

AGCACACGTCTGAACTCTTTCCCTACACGACGCTCTTCCGATCTNNNGCAGTTTGCAGATAAACAGGT

AGCACACGTCTGAACTCTTTCCCTACACGACGCTCTTCCGATCTNNNGCAGTTTGCAGATAAACAGGT

AGCACACGTCTGAACTCTTTCCCTACACGACGCTCTTCCGATCTNNNGCAGTTTGCAGATAAACAGGT

GCACACGTCTGAACTCTTTCCCTACACGACGCTCTTCCGATCTNNNTTTGCTGTTGTGTGCCTAATAGC

GCACACGTCTGAACTCTTTCCCTACACGACGCTCTTCCGATCTNNNTTTGCTGTTGTGTGCCTAATAGC

GCACACGTCTGAACTCTTTCCCTACACGACGCTCTTCCGATCTNNNTTTGCTGTTGTGTGCCTAATAGC

AGAGCACACGTCTGAACTCTTTCCCTACACGACGCTCTTCCGATCTNNNCAAGATTGTGCCACTATACT

AGAGCACACGTCTGAACTCTTTCCCTACACGACGCTCTTCCGATCTNNNCAAGATTGTGCCACTATACT

AGAGCACACGTCTGAACTCTTTCCCTACACGACGCTCTTCCGATCTNNNCAAGATTGTGCCACTATACT

GAGCACACGTCTGAACTCTTTCCCTACACGACGCTCTTCCGATCTNNNTAGCACTTTCAGGAGAGCATCA

GAGCACACGTCTGAACTCTTTCCCTACACGACGCTCTTCCGATCTNNNTAGCACTTTCAGGAGAGCATCA

GAGCACACGTCTGAACTCTTTCCCTACACGACGCTCTTCCGATCTNNNTAGCACTTTCAGGAGAGCATCA

ACACGTCTGAACTCTTTCCCTACACGACGCTCTTCCGATCTNNNATATATGTGCCATCTTTCTACATCAT

ACACGTCTGAACTCTTTCCCTACACGACGCTCTTCCGATCTNNNATATATGTGCCATCTTTCTACATCAT

ACACGTCTGAACTCTTTCCCTACACGACGCTCTTCCGATCTNNNATATATGTGCCATCTTTCTACATCAT

AGCACACGTCTGAACTCTTTCCCTACACGACGCTCTTCCGATCTNNNCTAGGTGACAGAGGAAGACTG

AGCACACGTCTGAACTCTTTCCCTACACGACGCTCTTCCGATCTNNNCTAGGTGACAGAGGAAGACTG

AGCACACGTCTGAACTCTTTCCCTACACGACGCTCTTCCGATCTNNNCTAGGTGACAGAGGAAGACTG

GAGCACACGTCTGAACTCTTTCCCTACACGACGCTCTTCCGATCTNNNTCTATGACTTTCCTTCCTCCCCT

GAGCACACGTCTGAACTCTTTCCCTACACGACGCTCTTCCGATCTNNNTCTATGACTTTCCTTCCTCCCCT

GAGCACACGTCTGAACTCTTTCCCTACACGACGCTCTTCCGATCTNNNTCTATGACTTTCCTTCCTCCCCT

AGCACACGTCTGAACTCTTTCCCTACACGACGCTCTTCCGATCTNNNCTTCTAGATTCCCAGGCATGTGT

AGCACACGTCTGAACTCTTTCCCTACACGACGCTCTTCCGATCTNNNCTTCTAGATTCCCAGGCATGTGT

AGCACACGTCTGAACTCTTTCCCTACACGACGCTCTTCCGATCTNNNCTTCTAGATTCCCAGGCATGTGT

AGCACACGTCTGAACTCTTTCCCTACACGACGCTCTTCCGATCTNNNAACTTACCAAGACCTCCTGGGG

AGCACACGTCTGAACTCTTTCCCTACACGACGCTCTTCCGATCTNNNAACTTACCAAGACCTCCTGGGG

AGCACACGTCTGAACTCTTTCCCTACACGACGCTCTTCCGATCTNNNAACTTACCAAGACCTCCTGGGG

AGCACACGTCTGAACTCTTTCCCTACACGACGCTCTTCCGATCTNNNTGCCTGTAATCTCAGCTACTTG

AGCACACGTCTGAACTCTTTCCCTACACGACGCTCTTCCGATCTNNNTGCCTGTAATCTCAGCTACTTG

AGCACACGTCTGAACTCTTTCCCTACACGACGCTCTTCCGATCTNNNTGCCTGTAATCTCAGCTACTTG

AGCACACGTCTGAACTCTTTCCCTACACGACGCTCTTCCGATCTNNNTCAATGACCTCTGGAACTATGG

AGCACACGTCTGAACTCTTTCCCTACACGACGCTCTTCCGATCTNNNTCAATGACCTCTGGAACTATGG

AGCACACGTCTGAACTCTTTCCCTACACGACGCTCTTCCGATCTNNNTCAATGACCTCTGGAACTATGG

GAGCACACGTCTGAACTCTTTCCCTACACGACGCTCTTCCGATCTNNNAGGTTGCAATGAGCTGAGACT

GAGCACACGTCTGAACTCTTTCCCTACACGACGCTCTTCCGATCTNNNAGGTTGCAATGAGCTGAGACT

GAGCACACGTCTGAACTCTTTCCCTACACGACGCTCTTCCGATCTNNNAGGTTGCAATGAGCTGAGACT

GAGCACACGTCTGAACTCTTTCCCTACACGACGCTCTTCCGATCTNNNTGGGTTGAGTAAAAACAGGT

GAGCACACGTCTGAACTCTTTCCCTACACGACGCTCTTCCGATCTNNNTGGGTTGAGTAAAAACAGGT

AGCACACGTCTGAACTCTTTCCCTACACGACGCTCTTCCGATCTNNNTGGGTTGAGTAAAAACAGGT
AGCACACGTCTGAACTCTTTCCCTACACGACGCTCTTCCGATCTNNNAGTCGTTGTGCAAACAGACTCT
AGCACACGTCTGAACTCTTTCCCTACACGACGCTCTTCCGATCTNNNAGTCGTTGTGCAAACAGACTCT
AGCACACGTCTGAACTCTTTCCCTACACGACGCTCTTCCGATCTNNNAGTCGTTGTGCAAACAGACTCT
GCACACGTCTGAACTCTTTCCCTACACGACGCTCTTCCGATCTNNNGCCAAAGCAATGGTACCAGAGT
GCACACGTCTGAACTCTTTCCCTACACGACGCTCTTCCGATCTNNNGCCAAAGCAATGGTACCAGAGT
GCACACGTCTGAACTCTTTCCCTACACGACGCTCTTCCGATCTNNNGCCAAAGCAATGGTACCAGAGT
AGCACACGTCTGAACTCTTTCCCTACACGACGCTCTTCCGATCTNNNCTGCCACACCCAGCTAATTTTTT
AGCACACGTCTGAACTCTTTCCCTACACGACGCTCTTCCGATCTNNNCTGCCACACCCAGCTAATTTTTT
AGCACACGTCTGAACTCTTTCCCTACACGACGCTCTTCCGATCTNNNCTGCCACACCCAGCTAATTTTTT
AGCACACGTCTGAACTCTTTCCCTACACGACGCTCTTCCGATCTNNNGTATTGCTATGTGGGTGAGG
AGCACACGTCTGAACTCTTTCCCTACACGACGCTCTTCCGATCTNNNGTATTGCTATGTGGGTGAGG
AGCACACGTCTGAACTCTTTCCCTACACGACGCTCTTCCGATCTNNNGTATTGCTATGTGGGTGAGG
AGCACACGTCTGAACTCTTTCCCTACACGACGCTCTTCCGATCTNNNGAATCGTTTGAACCTGGGAAG
AGCACACGTCTGAACTCTTTCCCTACACGACGCTCTTCCGATCTNNNGAATCGTTTGAACCTGGGAAG
AGCACACGTCTGAACTCTTTCCCTACACGACGCTCTTCCGATCTNNNGAATCGTTTGAACCTGGGAAG
GAGCACACGTCTGAACTCTTTCCCTACACGACGCTCTTCCGATCTNNNTACAAAAAATTAGCCGGGCA
GAGCACACGTCTGAACTCTTTCCCTACACGACGCTCTTCCGATCTNNNTACAAAAAATTAGCCGGGCA
GAGCACACGTCTGAACTCTTTCCCTACACGACGCTCTTCCGATCTNNNTACAAAAAATTAGCCGGGCA
GCACACGTCTGAACTCTTTCCCTACACGACGCTCTTCCGATCTNNNACTCAAAGAGTTACATGGTACTT
GCACACGTCTGAACTCTTTCCCTACACGACGCTCTTCCGATCTNNNACTCAAAGAGTTACATGGTACTT
GCACACGTCTGAACTCTTTCCCTACACGACGCTCTTCCGATCTNNNACTCAAAGAGTTACATGGTACTT
GAGCACACGTCTGAACTCTTTCCCTACACGACGCTCTTCCGATCTNNNGACAGAGCAAGACTCCGTCT
GAGCACACGTCTGAACTCTTTCCCTACACGACGCTCTTCCGATCTNNNGACAGAGCAAGACTCCGTCT
GAGCACACGTCTGAACTCTTTCCCTACACGACGCTCTTCCGATCTNNNGACAGAGCAAGACTCCGTCT
AGCACACGTCTGAACTCTTTCCCTACACGACGCTCTTCCGATCTNNNTGCAGAGGGTGTGTTCCTATA
AGCACACGTCTGAACTCTTTCCCTACACGACGCTCTTCCGATCTNNNTGCAGAGGGTGTGTTCCTATA
AGCACACGTCTGAACTCTTTCCCTACACGACGCTCTTCCGATCTNNNTGCAGAGGGTGTGTTCCTATA
GAGCACACGTCTGAACTCTTTCCCTACACGACGCTCTTCCGATCTNNNACACCTATCCACAAGTGTCAA
GAGCACACGTCTGAACTCTTTCCCTACACGACGCTCTTCCGATCTNNNACACCTATCCACAAGTGTCAA
GAGCACACGTCTGAACTCTTTCCCTACACGACGCTCTTCCGATCTNNNACACCTATCCACAAGTGTCAA
AGCACACGTCTGAACTCTTTCCCTACACGACGCTCTTCCGATCTNNNGTGACAAAACCAAACTCCGTCT
AGCACACGTCTGAACTCTTTCCCTACACGACGCTCTTCCGATCTNNNGTGACAAAACCAAACTCCGTCT
AGCACACGTCTGAACTCTTTCCCTACACGACGCTCTTCCGATCTNNNGTGACAAAACCAAACTCCGTCT
GCACACGTCTGAACTCTTTCCCTACACGACGCTCTTCCGATCTNNNTCTCAAGCCTGACTGTTTAGGT
GCACACGTCTGAACTCTTTCCCTACACGACGCTCTTCCGATCTNNNTCTCAAGCCTGACTGTTTAGGT
GCACACGTCTGAACTCTTTCCCTACACGACGCTCTTCCGATCTNNNTCTCAAGCCTGACTGTTTAGGT
GAGCACACGTCTGAACTCTTTCCCTACACGACGCTCTTCCGATCTNNNTCATGTTACCCCAGTTAGAAT
GAGCACACGTCTGAACTCTTTCCCTACACGACGCTCTTCCGATCTNNNTCATGTTACCCCAGTTAGAAT
GAGCACACGTCTGAACTCTTTCCCTACACGACGCTCTTCCGATCTNNNTCATGTTACCCCAGTTAGAAT
GAGCACACGTCTGAACTCTTTCCCTACACGACGCTCTTCCGATCTNNNAGGATCATGGTCAGAGTGG
GAGCACACGTCTGAACTCTTTCCCTACACGACGCTCTTCCGATCTNNNAGGATCATGGTCAGAGTGG
GAGCACACGTCTGAACTCTTTCCCTACACGACGCTCTTCCGATCTNNNAGGATCATGGTCAGAGTGG

GAGCACACGTCTGAACTCTTTCCCTACACGACGCTCTTCCGATCTNNNGCCTGTAGTCCCAGCTACTTG
GAGCACACGTCTGAACTCTTTCCCTACACGACGCTCTTCCGATCTNNNGCCTGTAGTCCCAGCTACTTG
GAGCACACGTCTGAACTCTTTCCCTACACGACGCTCTTCCGATCTNNNGCCTGTAGTCCCAGCTACTTG
GAGCACACGTCTGAACTCTTTCCCTACACGACGCTCTTCCGATCTNNNAAGACACTGCCCTGAAAATCA
GAGCACACGTCTGAACTCTTTCCCTACACGACGCTCTTCCGATCTNNNAAGACACTGCCCTGAAAATCA
GAGCACACGTCTGAACTCTTTCCCTACACGACGCTCTTCCGATCTNNNAAGACACTGCCCTGAAAATCA
AGCACACGTCTGAACTCTTTCCCTACACGACGCTCTTCCGATCTNNNGTCTTCAATACCCCACTGACAGT
AGCACACGTCTGAACTCTTTCCCTACACGACGCTCTTCCGATCTNNNGTCTTCAATACCCCACTGACAGT
AGCACACGTCTGAACTCTTTCCCTACACGACGCTCTTCCGATCTNNNGTCTTCAATACCCCACTGACAGT
GCACACGTCTGAACTCTTTCCCTACACGACGCTCTTCCGATCTNNNCCGCAGAAATTTTGATGGGGAGA
GCACACGTCTGAACTCTTTCCCTACACGACGCTCTTCCGATCTNNNCCGCAGAAATTTTGATGGGGAGA
GCACACGTCTGAACTCTTTCCCTACACGACGCTCTTCCGATCTNNNCCGCAGAAATTTTGATGGGGAGA
AGCACACGTCTGAACTCTTTCCCTACACGACGCTCTTCCGATCTNNNGAGGATGTTGCCAATGTGATGC
AGCACACGTCTGAACTCTTTCCCTACACGACGCTCTTCCGATCTNNNGAGGATGTTGCCAATGTGATGC
AGCACACGTCTGAACTCTTTCCCTACACGACGCTCTTCCGATCTNNNGAGGATGTTGCCAATGTGATGC
AGCACACGTCTGAACTCTTTCCCTACACGACGCTCTTCCGATCTNNNCGCTGTTATTTACGCCTTTCCATC
AGCACACGTCTGAACTCTTTCCCTACACGACGCTCTTCCGATCTNNNCGCTGTTATTTACGCCTTTCCATC
AGCACACGTCTGAACTCTTTCCCTACACGACGCTCTTCCGATCTNNNCGCTGTTATTTACGCCTTTCCATC
AGCACACGTCTGAACTCTTTCCCTACACGACGCTCTTCCGATCTNNNGCATTTAGGTTTCCCACATGGT
AGCACACGTCTGAACTCTTTCCCTACACGACGCTCTTCCGATCTNNNGCATTTAGGTTTCCCACATGGT
AGCACACGTCTGAACTCTTTCCCTACACGACGCTCTTCCGATCTNNNGCATTTAGGTTTCCCACATGGT
AGCACACGTCTGAACTCTTTCCCTACACGACGCTCTTCCGATCTNNNCTACCTACTGTGCTAATGAGGCA
AGCACACGTCTGAACTCTTTCCCTACACGACGCTCTTCCGATCTNNNCTACCTACTGTGCTAATGAGGCA
AGCACACGTCTGAACTCTTTCCCTACACGACGCTCTTCCGATCTNNNCTACCTACTGTGCTAATGAGGCA
GAGCACACGTCTGAACTCTTTCCCTACACGACGCTCTTCCGATCTNNNTGGCCCATACATAAAAGGCAA
GAGCACACGTCTGAACTCTTTCCCTACACGACGCTCTTCCGATCTNNNTGGCCCATACATAAAAGGCAA
GAGCACACGTCTGAACTCTTTCCCTACACGACGCTCTTCCGATCTNNNTGGCCCATACATAAAAGGCAA
GAGCACACGTCTGAACTCTTTCCCTACACGACGCTCTTCCGATCTNNNAACCAACCAGCTAACACTGCTA
GAGCACACGTCTGAACTCTTTCCCTACACGACGCTCTTCCGATCTNNNAACCAACCAGCTAACACTGCTA
GAGCACACGTCTGAACTCTTTCCCTACACGACGCTCTTCCGATCTNNNAACCAACCAGCTAACACTGCTA
GAGCACACGTCTGAACTCTTTCCCTACACGACGCTCTTCCGATCTNNNTGATATGACCCTTCGTTTTTTT
GAGCACACGTCTGAACTCTTTCCCTACACGACGCTCTTCCGATCTNNNTGATATGACCCTTCGTTTTTTT
GAGCACACGTCTGAACTCTTTCCCTACACGACGCTCTTCCGATCTNNNTGATATGACCCTTCGTTTTTTT
GAGCACACGTCTGAACTCTTTCCCTACACGACGCTCTTCCGATCTNNNTGAGGTTGAAGACAGGCTTTT
GAGCACACGTCTGAACTCTTTCCCTACACGACGCTCTTCCGATCTNNNTGAGGTTGAAGACAGGCTTTT
GAGCACACGTCTGAACTCTTTCCCTACACGACGCTCTTCCGATCTNNNTGAGGTTGAAGACAGGCTTTT
AGAGCACACGTCTGAACTCTTTCCCTACACGACGCTCTTCCGATCTNNNCTGAGGCAGGAGAATCACTT
AGAGCACACGTCTGAACTCTTTCCCTACACGACGCTCTTCCGATCTNNNCTGAGGCAGGAGAATCACTT
AGAGCACACGTCTGAACTCTTTCCCTACACGACGCTCTTCCGATCTNNNCTGAGGCAGGAGAATCACTT
GAGCACACGTCTGAACTCTTTCCCTACACGACGCTCTTCCGATCTNNNCAGAGTTGCCAGCTCTACAAA
GAGCACACGTCTGAACTCTTTCCCTACACGACGCTCTTCCGATCTNNNCAGAGTTGCCAGCTCTACAAA
GAGCACACGTCTGAACTCTTTCCCTACACGACGCTCTTCCGATCTNNNCAGAGTTGCCAGCTCTACAAA
AGCACACGTCTGAACTCTTTCCCTACACGACGCTCTTCCGATCTNNNTGTCTTAGAACTTCCTCGACACCC

GCACACGTCTGAACTCTTTCCCTACACGACGCTCTTCCGATCTNNNTGTCTTAGAACTTCCTCGACACC
GCACACGTCTGAACTCTTTCCCTACACGACGCTCTTCCGATCTNNNTGTCTTAGAACTTCCTCGACACC
AGCACACGTCTGAACTCTTTCCCTACACGACGCTCTTCCGATCTNNNCCATGGATATGAAGGCCCAAC
AGCACACGTCTGAACTCTTTCCCTACACGACGCTCTTCCGATCTNNNCCATGGATATGAAGGCCCAAC
AGCACACGTCTGAACTCTTTCCCTACACGACGCTCTTCCGATCTNNNCCATGGATATGAAGGCCCAAC
AGAGCACACGTCTGAACTCTTTCCCTACACGACGCTCTTCCGATCTNNNGAAAAGACCAGAAGGAATT
AGAGCACACGTCTGAACTCTTTCCCTACACGACGCTCTTCCGATCTNNNGAAAAGACCAGAAGGAATT
AGAGCACACGTCTGAACTCTTTCCCTACACGACGCTCTTCCGATCTNNNGAAAAGACCAGAAGGAATT
GAGCACACGTCTGAACTCTTTCCCTACACGACGCTCTTCCGATCTNNNTAAGGGGGTCACTTCAAGG
GAGCACACGTCTGAACTCTTTCCCTACACGACGCTCTTCCGATCTNNNTAAGGGGGTCACTTCAAGG
GAGCACACGTCTGAACTCTTTCCCTACACGACGCTCTTCCGATCTNNNTAAGGGGGTCACTTCAAGG
AGAGCACACGTCTGAACTCTTTCCCTACACGACGCTCTTCCGATCTNNNTACGTGAGACTTGGAATTG
AGAGCACACGTCTGAACTCTTTCCCTACACGACGCTCTTCCGATCTNNNTACGTGAGACTTGGAATTG
AGAGCACACGTCTGAACTCTTTCCCTACACGACGCTCTTCCGATCTNNNTACGTGAGACTTGGAATTG
GAGCACACGTCTGAACTCTTTCCCTACACGACGCTCTTCCGATCTNNNATCTTTCTCAGAAGCGAGCTC
GAGCACACGTCTGAACTCTTTCCCTACACGACGCTCTTCCGATCTNNNATCTTTCTCAGAAGCGAGCTC
GAGCACACGTCTGAACTCTTTCCCTACACGACGCTCTTCCGATCTNNNATCTTTCTCAGAAGCGAGCTC
AGCACACGTCTGAACTCTTTCCCTACACGACGCTCTTCCGATCTNNNAGAATCACTTGAACCTGGGAG
AGCACACGTCTGAACTCTTTCCCTACACGACGCTCTTCCGATCTNNNAGAATCACTTGAACCTGGGAG
AGCACACGTCTGAACTCTTTCCCTACACGACGCTCTTCCGATCTNNNAGAATCACTTGAACCTGGGAG
AGCACACGTCTGAACTCTTTCCCTACACGACGCTCTTCCGATCTNNNCCCCATCTTTCCTCTTTCCTTT
AGCACACGTCTGAACTCTTTCCCTACACGACGCTCTTCCGATCTNNNCCCCATCTTTCCTCTTTCCTTT
AGCACACGTCTGAACTCTTTCCCTACACGACGCTCTTCCGATCTNNNCCCCATCTTTCCTCTTTCCTTT
GCACACGTCTGAACTCTTTCCCTACACGACGCTCTTCCGATCTNNNACGATGACCTCCAACATTGCTTT
GCACACGTCTGAACTCTTTCCCTACACGACGCTCTTCCGATCTNNNACGATGACCTCCAACATTGCTTT
GCACACGTCTGAACTCTTTCCCTACACGACGCTCTTCCGATCTNNNACGATGACCTCCAACATTGCTTT
GAGCACACGTCTGAACTCTTTCCCTACACGACGCTCTTCCGATCTNNNATGTAGAAGCTAGTAGGGT
GAGCACACGTCTGAACTCTTTCCCTACACGACGCTCTTCCGATCTNNNATGTAGAAGCTAGTAGGGT
GAGCACACGTCTGAACTCTTTCCCTACACGACGCTCTTCCGATCTNNNATGTAGAAGCTAGTAGGGT
GCACACGTCTGAACTCTTTCCCTACACGACGCTCTTCCGATCTNNNGCTATCCCAGTCTCCATTCTGAG
GCACACGTCTGAACTCTTTCCCTACACGACGCTCTTCCGATCTNNNGCTATCCCAGTCTCCATTCTGAG
GCACACGTCTGAACTCTTTCCCTACACGACGCTCTTCCGATCTNNNGCTATCCCAGTCTCCATTCTGAG
GAGCACACGTCTGAACTCTTTCCCTACACGACGCTCTTCCGATCTNNNTGGTTGAGGATTTCTGTGGG
GAGCACACGTCTGAACTCTTTCCCTACACGACGCTCTTCCGATCTNNNTGGTTGAGGATTTCTGTGGG
GAGCACACGTCTGAACTCTTTCCCTACACGACGCTCTTCCGATCTNNNTGGTTGAGGATTTCTGTGGG
CACACGTCTGAACTCTTTCCCTACACGACGCTCTTCCGATCTNNNGAGGTAGAGTTTGCAGTGAGTC
CACACGTCTGAACTCTTTCCCTACACGACGCTCTTCCGATCTNNNGAGGTAGAGTTTGCAGTGAGTC
CACACGTCTGAACTCTTTCCCTACACGACGCTCTTCCGATCTNNNGAGGTAGAGTTTGCAGTGAGTC
AGCACACGTCTGAACTCTTTCCCTACACGACGCTCTTCCGATCTNNNGAAATGGGAAGGTCAGAGTC
AGCACACGTCTGAACTCTTTCCCTACACGACGCTCTTCCGATCTNNNGAAATGGGAAGGTCAGAGTC
AGCACACGTCTGAACTCTTTCCCTACACGACGCTCTTCCGATCTNNNGAAATGGGAAGGTCAGAGTC
AGCACACGTCTGAACTCTTTCCCTACACGACGCTCTTCCGATCTNNNAACATCTAACCAACAGGAAG
AGCACACGTCTGAACTCTTTCCCTACACGACGCTCTTCCGATCTNNNAACATCTAACCAACAGGAAG

GAGCACACGTCTGAACTCTTTCCCTACACGACGCTCTTCCGATCTNNNAACATCTAACCAACAGGAAGC
AGCACACGTCTGAACTCTTTCCCTACACGACGCTCTTCCGATCTNNNTACAGGTAGCTGCCATCACACT
AGCACACGTCTGAACTCTTTCCCTACACGACGCTCTTCCGATCTNNNTACAGGTAGCTGCCATCACACT
AGCACACGTCTGAACTCTTTCCCTACACGACGCTCTTCCGATCTNNNTACAGGTAGCTGCCATCACACT
AGCACACGTCTGAACTCTTTCCCTACACGACGCTCTTCCGATCTNNNGGGTCCAGAAAGAGAGAGAGA
AGCACACGTCTGAACTCTTTCCCTACACGACGCTCTTCCGATCTNNNGGGTCCAGAAAGAGAGAGAGA
AGCACACGTCTGAACTCTTTCCCTACACGACGCTCTTCCGATCTNNNGGGTCCAGAAAGAGAGAGAGA
GCACACGTCTGAACTCTTTCCCTACACGACGCTCTTCCGATCTNNNCGTCAAACCTCGCCTTCAAATAA
GCACACGTCTGAACTCTTTCCCTACACGACGCTCTTCCGATCTNNNCGTCAAACCTCGCCTTCAAATAA
GCACACGTCTGAACTCTTTCCCTACACGACGCTCTTCCGATCTNNNCGTCAAACCTCGCCTTCAAATAA
AGCACACGTCTGAACTCTTTCCCTACACGACGCTCTTCCGATCTNNNAATCACCTCCTCCAAAGAATCC
AGCACACGTCTGAACTCTTTCCCTACACGACGCTCTTCCGATCTNNNAATCACCTCCTCCAAAGAATCC
AGCACACGTCTGAACTCTTTCCCTACACGACGCTCTTCCGATCTNNNAATCACCTCCTCCAAAGAATCC
AGCACACGTCTGAACTCTTTCCCTACACGACGCTCTTCCGATCTNNNTTTGAAACCCCTCCCTTACCAGA
AGCACACGTCTGAACTCTTTCCCTACACGACGCTCTTCCGATCTNNNTTTGAAACCCCTCCCTTACCAGA
AGCACACGTCTGAACTCTTTCCCTACACGACGCTCTTCCGATCTNNNTTTGAAACCCCTCCCTTACCAGA
AGCACACGTCTGAACTCTTTCCCTACACGACGCTCTTCCGATCTNNNTTACCATATTGCCCATGCTGGT
AGCACACGTCTGAACTCTTTCCCTACACGACGCTCTTCCGATCTNNNTTACCATATTGCCCATGCTGGT
AGCACACGTCTGAACTCTTTCCCTACACGACGCTCTTCCGATCTNNNTTACCATATTGCCCATGCTGGT
AGCACACGTCTGAACTCTTTCCCTACACGACGCTCTTCCGATCTNNNAGCCTGGCCACAAAATTTCTTA
AGCACACGTCTGAACTCTTTCCCTACACGACGCTCTTCCGATCTNNNAGCCTGGCCACAAAATTTCTTA
AGCACACGTCTGAACTCTTTCCCTACACGACGCTCTTCCGATCTNNNAGCCTGGCCACAAAATTTCTTA
CACACGTCTGAACTCTTTCCCTACACGACGCTCTTCCGATCTNNNATTATTCTGAGTTTCTTTCCTTGCAT
CACACGTCTGAACTCTTTCCCTACACGACGCTCTTCCGATCTNNNATTATTCTGAGTTTCTTTCCTTGCAT
CACACGTCTGAACTCTTTCCCTACACGACGCTCTTCCGATCTNNNATTATTCTGAGTTTCTTTCCTTGCAT
AGCACACGTCTGAACTCTTTCCCTACACGACGCTCTTCCGATCTNNNTAAGCATCCAAAGCACAGGAAC
AGCACACGTCTGAACTCTTTCCCTACACGACGCTCTTCCGATCTNNNTAAGCATCCAAAGCACAGGAAC
AGCACACGTCTGAACTCTTTCCCTACACGACGCTCTTCCGATCTNNNTAAGCATCCAAAGCACAGGAAC
GAGCACACGTCTGAACTCTTTCCCTACACGACGCTCTTCCGATCTNNNTTACGTAGTTTGAACTCACGC
GAGCACACGTCTGAACTCTTTCCCTACACGACGCTCTTCCGATCTNNNTTACGTAGTTTGAACTCACGC
GAGCACACGTCTGAACTCTTTCCCTACACGACGCTCTTCCGATCTNNNTTACGTAGTTTGAACTCACGC
GAGCACACGTCTGAACTCTTTCCCTACACGACGCTCTTCCGATCTNNNGCTTACACCTGTAATCCCAACA
GAGCACACGTCTGAACTCTTTCCCTACACGACGCTCTTCCGATCTNNNGCTTACACCTGTAATCCCAACA
GAGCACACGTCTGAACTCTTTCCCTACACGACGCTCTTCCGATCTNNNGCTTACACCTGTAATCCCAACA
GAGCACACGTCTGAACTCTTTCCCTACACGACGCTCTTCCGATCTNNNATGCCCACTGTAGAAGTTCTG
GAGCACACGTCTGAACTCTTTCCCTACACGACGCTCTTCCGATCTNNNATGCCCACTGTAGAAGTTCTG
GAGCACACGTCTGAACTCTTTCCCTACACGACGCTCTTCCGATCTNNNATGCCCACTGTAGAAGTTCTG
AGCACACGTCTGAACTCTTTCCCTACACGACGCTCTTCCGATCTNNNAAAAAAAAGAGTTGCAGCCAA
AGCACACGTCTGAACTCTTTCCCTACACGACGCTCTTCCGATCTNNNAAAAAAAAGAGTTGCAGCCAA
AGCACACGTCTGAACTCTTTCCCTACACGACGCTCTTCCGATCTNNNAAAAAAAAGAGTTGCAGCCAA
GAGCACACGTCTGAACTCTTTCCCTACACGACGCTCTTCCGATCTNNNGCACATACCTGTATTCCCAGCT
GAGCACACGTCTGAACTCTTTCCCTACACGACGCTCTTCCGATCTNNNGCACATACCTGTATTCCCAGCT
GAGCACACGTCTGAACTCTTTCCCTACACGACGCTCTTCCGATCTNNNGCACATACCTGTATTCCCAGCT

GCACACGTCTGAACTCTTTCCCTACACGACGCTCTTCCGATCTNNNTCCTACCATTTTGAGGTGACCTG
GCACACGTCTGAACTCTTTCCCTACACGACGCTCTTCCGATCTNNNTCCTACCATTTTGAGGTGACCTG
GCACACGTCTGAACTCTTTCCCTACACGACGCTCTTCCGATCTNNNTCCTACCATTTTGAGGTGACCTG
AGCACACGTCTGAACTCTTTCCCTACACGACGCTCTTCCGATCTNNNTCAGAATTTGCAACCATCACC
AGCACACGTCTGAACTCTTTCCCTACACGACGCTCTTCCGATCTNNNTCAGAATTTGCAACCATCACC
AGCACACGTCTGAACTCTTTCCCTACACGACGCTCTTCCGATCTNNNTCAGAATTTGCAACCATCACC
GAGCACACGTCTGAACTCTTTCCCTACACGACGCTCTTCCGATCTNNNAATGAGACCATGTATGTCCT
GAGCACACGTCTGAACTCTTTCCCTACACGACGCTCTTCCGATCTNNNAATGAGACCATGTATGTCCT
GAGCACACGTCTGAACTCTTTCCCTACACGACGCTCTTCCGATCTNNNAATGAGACCATGTATGTCCT
CACACGTCTGAACTCTTTCCCTACACGACGCTCTTCCGATCTNNNATGTGAAATGCAAAGAGAAGCC
CACACGTCTGAACTCTTTCCCTACACGACGCTCTTCCGATCTNNNATGTGAAATGCAAAGAGAAGCC
CACACGTCTGAACTCTTTCCCTACACGACGCTCTTCCGATCTNNNATGTGAAATGCAAAGAGAAGCC
GCACACGTCTGAACTCTTTCCCTACACGACGCTCTTCCGATCTNNNTATTGGGATTACAGGTGTGAG
GCACACGTCTGAACTCTTTCCCTACACGACGCTCTTCCGATCTNNNTATTGGGATTACAGGTGTGAG
GCACACGTCTGAACTCTTTCCCTACACGACGCTCTTCCGATCTNNNTATTGGGATTACAGGTGTGAG
AGCACACGTCTGAACTCTTTCCCTACACGACGCTCTTCCGATCTNNNACTGGGTATCTACCTCGAGGA
AGCACACGTCTGAACTCTTTCCCTACACGACGCTCTTCCGATCTNNNACTGGGTATCTACCTCGAGGA
AGCACACGTCTGAACTCTTTCCCTACACGACGCTCTTCCGATCTNNNACTGGGTATCTACCTCGAGGA
AGCACACGTCTGAACTCTTTCCCTACACGACGCTCTTCCGATCTNNNAAGACTGCAATGAGCCACATT
AGCACACGTCTGAACTCTTTCCCTACACGACGCTCTTCCGATCTNNNAAGACTGCAATGAGCCACATT
AGCACACGTCTGAACTCTTTCCCTACACGACGCTCTTCCGATCTNNNAAGACTGCAATGAGCCACATT
GAGCACACGTCTGAACTCTTTCCCTACACGACGCTCTTCCGATCTNNNCTCACCTGAAATCCCAGTA
GAGCACACGTCTGAACTCTTTCCCTACACGACGCTCTTCCGATCTNNNCTCACCTGAAATCCCAGTA
GAGCACACGTCTGAACTCTTTCCCTACACGACGCTCTTCCGATCTNNNCTCACCTGAAATCCCAGTA
AGCACACGTCTGAACTCTTTCCCTACACGACGCTCTTCCGATCTNNNCTCCTTGCCAGTCTCATCTTTCT
AGCACACGTCTGAACTCTTTCCCTACACGACGCTCTTCCGATCTNNNCTCCTTGCCAGTCTCATCTTTCT
AGCACACGTCTGAACTCTTTCCCTACACGACGCTCTTCCGATCTNNNCTCCTTGCCAGTCTCATCTTTCT
AGCACACGTCTGAACTCTTTCCCTACACGACGCTCTTCCGATCTNNNCACCATTGTAGCATCCAAGCAA
AGCACACGTCTGAACTCTTTCCCTACACGACGCTCTTCCGATCTNNNCACCATTGTAGCATCCAAGCAA
AGCACACGTCTGAACTCTTTCCCTACACGACGCTCTTCCGATCTNNNCACCATTGTAGCATCCAAGCAA
AGCACACGTCTGAACTCTTTCCCTACACGACGCTCTTCCGATCTNNNTGTGATTAGGAAGCAGTGAG
AGCACACGTCTGAACTCTTTCCCTACACGACGCTCTTCCGATCTNNNTGTGATTAGGAAGCAGTGAG
AGCACACGTCTGAACTCTTTCCCTACACGACGCTCTTCCGATCTNNNTGTGATTAGGAAGCAGTGAG
GCACACGTCTGAACTCTTTCCCTACACGACGCTCTTCCGATCTNNNATCATTTTCCAAAGCCTCAGAG
GCACACGTCTGAACTCTTTCCCTACACGACGCTCTTCCGATCTNNNATCATTTTCCAAAGCCTCAGAG
GCACACGTCTGAACTCTTTCCCTACACGACGCTCTTCCGATCTNNNATCATTTTCCAAAGCCTCAGAG
GAGCACACGTCTGAACTCTTTCCCTACACGACGCTCTTCCGATCTNNNACAAACAAAAATTAGCCGGG
GAGCACACGTCTGAACTCTTTCCCTACACGACGCTCTTCCGATCTNNNACAAACAAAAATTAGCCGGG
GAGCACACGTCTGAACTCTTTCCCTACACGACGCTCTTCCGATCTNNNACAAACAAAAATTAGCCGGG
GAGCACACGTCTGAACTCTTTCCCTACACGACGCTCTTCCGATCTNNNGGACTGAGACTTTTCCCTCAA
GAGCACACGTCTGAACTCTTTCCCTACACGACGCTCTTCCGATCTNNNGGACTGAGACTTTTCCCTCAA
GAGCACACGTCTGAACTCTTTCCCTACACGACGCTCTTCCGATCTNNNGGACTGAGACTTTTCCCTCAA
GCACACGTCTGAACTCTTTCCCTACACGACGCTCTTCCGATCTNNNCCCATTCCCCTCCTGCTAACAT

AGCACACGTCTGAACTCTTTCCCTACACGACGCTCTTCCGATCTNNNCCATTCCCCCTCCTGCTAACATC

AGCACACGTCTGAACTCTTTCCCTACACGACGCTCTTCCGATCTNNNCCATTCCCCCTCCTGCTAACATC

AGCACACGTCTGAACTCTTTCCCTACACGACGCTCTTCCGATCTNNNGGGCGACAGAGAAAAACTTCAT

AGCACACGTCTGAACTCTTTCCCTACACGACGCTCTTCCGATCTNNNGGGCGACAGAGAAAAACTTCAT

AGCACACGTCTGAACTCTTTCCCTACACGACGCTCTTCCGATCTNNNGGGCGACAGAGAAAAACTTCAT

AGAGCACACGTCTGAACTCTTTCCCTACACGACGCTCTTCCGATCTNNNATGGACACAGGAAGGGAAA

AGAGCACACGTCTGAACTCTTTCCCTACACGACGCTCTTCCGATCTNNNATGGACACAGGAAGGGAAA

AGAGCACACGTCTGAACTCTTTCCCTACACGACGCTCTTCCGATCTNNNATGGACACAGGAAGGGAAA

GCACACGTCTGAACTCTTTCCCTACACGACGCTCTTCCGATCTNNNCTATTACCTTCTGATGACTTCCCAT

GCACACGTCTGAACTCTTTCCCTACACGACGCTCTTCCGATCTNNNCTATTACCTTCTGATGACTTCCCAT

GCACACGTCTGAACTCTTTCCCTACACGACGCTCTTCCGATCTNNNCTATTACCTTCTGATGACTTCCCAT

GCACACGTCTGAACTCTTTCCCTACACGACGCTCTTCCGATCTNNNATCAGCCGCAGCAAGTCGTACAT

GCACACGTCTGAACTCTTTCCCTACACGACGCTCTTCCGATCTNNNATCAGCCGCAGCAAGTCGTACAT

GCACACGTCTGAACTCTTTCCCTACACGACGCTCTTCCGATCTNNNATCAGCCGCAGCAAGTCGTACAT

ACACGTCTGAACTCTTTCCCTACACGACGCTCTTCCGATCTNNNCTGGGATTACAATTGTGAGCCACCA

ACACGTCTGAACTCTTTCCCTACACGACGCTCTTCCGATCTNNNCTGGGATTACAATTGTGAGCCACCA

ACACGTCTGAACTCTTTCCCTACACGACGCTCTTCCGATCTNNNCTGGGATTACAATTGTGAGCCACCA

GAGCACACGTCTGAACTCTTTCCCTACACGACGCTCTTCCGATCTNNNTGTACACTGACCCCAATGTGTA

GAGCACACGTCTGAACTCTTTCCCTACACGACGCTCTTCCGATCTNNNTGTACACTGACCCCAATGTGTA

GAGCACACGTCTGAACTCTTTCCCTACACGACGCTCTTCCGATCTNNNTGTACACTGACCCCAATGTGTA

AGCACACGTCTGAACTCTTTCCCTACACGACGCTCTTCCGATCTNNNCCATGTGGTAACTTTTGCTGGAC

AGCACACGTCTGAACTCTTTCCCTACACGACGCTCTTCCGATCTNNNCCATGTGGTAACTTTTGCTGGAC

AGCACACGTCTGAACTCTTTCCCTACACGACGCTCTTCCGATCTNNNCCATGTGGTAACTTTTGCTGGAC

AAGAGCACACGTCTGAACTCTTTCCCTACACGACGCTCTTCCGATCTNNNACCAAGAGACTGACAGAGA

AAGAGCACACGTCTGAACTCTTTCCCTACACGACGCTCTTCCGATCTNNNACCAAGAGACTGACAGAGA

AAGAGCACACGTCTGAACTCTTTCCCTACACGACGCTCTTCCGATCTNNNACCAAGAGACTGACAGAGA

GCACACGTCTGAACTCTTTCCCTACACGACGCTCTTCCGATCTNNNGTGGCAGAGTAAGACTCCATCTC

GCACACGTCTGAACTCTTTCCCTACACGACGCTCTTCCGATCTNNNGTGGCAGAGTAAGACTCCATCTC

GCACACGTCTGAACTCTTTCCCTACACGACGCTCTTCCGATCTNNNGTGGCAGAGTAAGACTCCATCTC

GAGCACACGTCTGAACTCTTTCCCTACACGACGCTCTTCCGATCTNNNGGATTTGAACCCAGGTCTGTA

GAGCACACGTCTGAACTCTTTCCCTACACGACGCTCTTCCGATCTNNNGGATTTGAACCCAGGTCTGTA

GAGCACACGTCTGAACTCTTTCCCTACACGACGCTCTTCCGATCTNNNGGATTTGAACCCAGGTCTGTA

GAGCACACGTCTGAACTCTTTCCCTACACGACGCTCTTCCGATCTNNNAAGATCTCGCCACTACACTTCA

GAGCACACGTCTGAACTCTTTCCCTACACGACGCTCTTCCGATCTNNNAAGATCTCGCCACTACACTTCA

GAGCACACGTCTGAACTCTTTCCCTACACGACGCTCTTCCGATCTNNNAAGATCTCGCCACTACACTTCA

GAGCACACGTCTGAACTCTTTCCCTACACGACGCTCTTCCGATCTNNNCAGGATTCCTCTCGGTCTTTCT

GAGCACACGTCTGAACTCTTTCCCTACACGACGCTCTTCCGATCTNNNCAGGATTCCTCTCGGTCTTTCT

GAGCACACGTCTGAACTCTTTCCCTACACGACGCTCTTCCGATCTNNNCAGGATTCCTCTCGGTCTTTCT

GAGCACACGTCTGAACTCTTTCCCTACACGACGCTCTTCCGATCTNNNCAAAACCACCTGGAAGATGTC

GAGCACACGTCTGAACTCTTTCCCTACACGACGCTCTTCCGATCTNNNCAAAACCACCTGGAAGATGTC

GAGCACACGTCTGAACTCTTTCCCTACACGACGCTCTTCCGATCTNNNCAAAACCACCTGGAAGATGTC

GAGCACACGTCTGAACTCTTTCCCTACACGACGCTCTTCCGATCTNNNCTGAGGCAGGAGAATCACTTC

GAGCACACGTCTGAACTCTTTCCCTACACGACGCTCTTCCGATCTNNNCTGAGGCAGGAGAATCACTTC

GAGCACACGTCTGAACTCTTTCCCTACACGACGCTCTTCCGATCTNNNCTGAGGCAGGAGAATCACTTG
AGCACACGTCTGAACTCTTTCCCTACACGACGCTCTTCCGATCTNNNTACAAAAAATTAGCCCGGCGTGG
AGCACACGTCTGAACTCTTTCCCTACACGACGCTCTTCCGATCTNNNTACAAAAAATTAGCCCGGCGTGG
AGCACACGTCTGAACTCTTTCCCTACACGACGCTCTTCCGATCTNNNTACAAAAAATTAGCCCGGCGTGG
GAGCACACGTCTGAACTCTTTCCCTACACGACGCTCTTCCGATCTNNNTACAAAAATTAGCGGGGTGTG
GAGCACACGTCTGAACTCTTTCCCTACACGACGCTCTTCCGATCTNNNTACAAAAATTAGCGGGGTGTG
GAGCACACGTCTGAACTCTTTCCCTACACGACGCTCTTCCGATCTNNNTACAAAAATTAGCGGGGTGTG
GAGCACACGTCTGAACTCTTTCCCTACACGACGCTCTTCCGATCTNNNATTTGGGCTGGGAACTGTTCT
GAGCACACGTCTGAACTCTTTCCCTACACGACGCTCTTCCGATCTNNNATTTGGGCTGGGAACTGTTCT
GAGCACACGTCTGAACTCTTTCCCTACACGACGCTCTTCCGATCTNNNATTTGGGCTGGGAACTGTTCT
AGCACACGTCTGAACTCTTTCCCTACACGACGCTCTTCCGATCTNNNAGAGAGGAAAGTGCACATCAA
AGCACACGTCTGAACTCTTTCCCTACACGACGCTCTTCCGATCTNNNAGAGAGGAAAGTGCACATCAA
AGCACACGTCTGAACTCTTTCCCTACACGACGCTCTTCCGATCTNNNAGAGAGGAAAGTGCACATCAA
GAGCACACGTCTGAACTCTTTCCCTACACGACGCTCTTCCGATCTNNNGCCTCTAATCTGCCATCTTGGA
GAGCACACGTCTGAACTCTTTCCCTACACGACGCTCTTCCGATCTNNNGCCTCTAATCTGCCATCTTGGA
GAGCACACGTCTGAACTCTTTCCCTACACGACGCTCTTCCGATCTNNNGCCTCTAATCTGCCATCTTGGA
GAGCACACGTCTGAACTCTTTCCCTACACGACGCTCTTCCGATCTNNNAGTGTCCCAGAGAAACCTTGT
GAGCACACGTCTGAACTCTTTCCCTACACGACGCTCTTCCGATCTNNNAGTGTCCCAGAGAAACCTTGT
GAGCACACGTCTGAACTCTTTCCCTACACGACGCTCTTCCGATCTNNNAGTGTCCCAGAGAAACCTTGT
ACACGTCTGAACTCTTTCCCTACACGACGCTCTTCCGATCTNNNTGTTCTACCTCAGGTATCCGAGACAT
ACACGTCTGAACTCTTTCCCTACACGACGCTCTTCCGATCTNNNTGTTCTACCTCAGGTATCCGAGACAT
ACACGTCTGAACTCTTTCCCTACACGACGCTCTTCCGATCTNNNTGTTCTACCTCAGGTATCCGAGACAT
GAGCACACGTCTGAACTCTTTCCCTACACGACGCTCTTCCGATCTNNNCCCAAGACGCTGACAACATGA
GAGCACACGTCTGAACTCTTTCCCTACACGACGCTCTTCCGATCTNNNCCCAAGACGCTGACAACATGA
GAGCACACGTCTGAACTCTTTCCCTACACGACGCTCTTCCGATCTNNNCCCAAGACGCTGACAACATGA
AGCACACGTCTGAACTCTTTCCCTACACGACGCTCTTCCGATCTNNNACCATGCCCGGCTAATTTTTTTA
AGCACACGTCTGAACTCTTTCCCTACACGACGCTCTTCCGATCTNNNACCATGCCCGGCTAATTTTTTTA
AGCACACGTCTGAACTCTTTCCCTACACGACGCTCTTCCGATCTNNNACCATGCCCGGCTAATTTTTTTA
GAGCACACGTCTGAACTCTTTCCCTACACGACGCTCTTCCGATCTNNNAAAATGACCTTCCTTCCCGTCT
GAGCACACGTCTGAACTCTTTCCCTACACGACGCTCTTCCGATCTNNNAAAATGACCTTCCTTCCCGTCT
GAGCACACGTCTGAACTCTTTCCCTACACGACGCTCTTCCGATCTNNNAAAATGACCTTCCTTCCCGTCT
GCACACGTCTGAACTCTTTCCCTACACGACGCTCTTCCGATCTNNNAGAAGTAAGCTCAGATGACAGC
GCACACGTCTGAACTCTTTCCCTACACGACGCTCTTCCGATCTNNNAGAAGTAAGCTCAGATGACAGC
GCACACGTCTGAACTCTTTCCCTACACGACGCTCTTCCGATCTNNNAGAAGTAAGCTCAGATGACAGC
GAGCACACGTCTGAACTCTTTCCCTACACGACGCTCTTCCGATCTNNNGGATCACTTGAGGTCAGGAG
GAGCACACGTCTGAACTCTTTCCCTACACGACGCTCTTCCGATCTNNNGGATCACTTGAGGTCAGGAG
GAGCACACGTCTGAACTCTTTCCCTACACGACGCTCTTCCGATCTNNNGGATCACTTGAGGTCAGGAG
AGAGCACACGTCTGAACTCTTTCCCTACACGACGCTCTTCCGATCTNNNGCATAGTCTCACTCATAGGTG
AGAGCACACGTCTGAACTCTTTCCCTACACGACGCTCTTCCGATCTNNNGCATAGTCTCACTCATAGGTG
AGAGCACACGTCTGAACTCTTTCCCTACACGACGCTCTTCCGATCTNNNGCATAGTCTCACTCATAGGTG
AGAGCACACGTCTGAACTCTTTCCCTACACGACGCTCTTCCGATCTNNNTCTGGTGTGGTCTGGAAAAT
AGAGCACACGTCTGAACTCTTTCCCTACACGACGCTCTTCCGATCTNNNTCTGGTGTGGTCTGGAAAAT
AGAGCACACGTCTGAACTCTTTCCCTACACGACGCTCTTCCGATCTNNNTCTGGTGTGGTCTGGAAAAT

AGCACACGTCTGAACTCTTTCCCTACACGACGCTCTTCCGATCTNNNACATCACAGGAGACAGGATGA
AGCACACGTCTGAACTCTTTCCCTACACGACGCTCTTCCGATCTNNNACATCACAGGAGACAGGATGA
AGCACACGTCTGAACTCTTTCCCTACACGACGCTCTTCCGATCTNNNACATCACAGGAGACAGGATGA
AGCACACGTCTGAACTCTTTCCCTACACGACGCTCTTCCGATCTNNNTGGGAAAAATCTAGAATGGTT
AGCACACGTCTGAACTCTTTCCCTACACGACGCTCTTCCGATCTNNNTGGGAAAAATCTAGAATGGTT
AGCACACGTCTGAACTCTTTCCCTACACGACGCTCTTCCGATCTNNNTGGGAAAAATCTAGAATGGTT
AGCACACGTCTGAACTCTTTCCCTACACGACGCTCTTCCGATCTNNNGGGGTCGGATTTCATTCTTCTG
AGCACACGTCTGAACTCTTTCCCTACACGACGCTCTTCCGATCTNNNGGGGTCGGATTTCATTCTTCTG
AGCACACGTCTGAACTCTTTCCCTACACGACGCTCTTCCGATCTNNNGGGGTCGGATTTCATTCTTCTG
AGCACACGTCTGAACTCTTTCCCTACACGACGCTCTTCCGATCTNNNATCGCAAACCTTTTTATGCCACA
AGCACACGTCTGAACTCTTTCCCTACACGACGCTCTTCCGATCTNNNATCGCAAACCTTTTTATGCCACA
AGCACACGTCTGAACTCTTTCCCTACACGACGCTCTTCCGATCTNNNATCGCAAACCTTTTTATGCCACA
AGCACACGTCTGAACTCTTTCCCTACACGACGCTCTTCCGATCTNNNTCTGTCAAGGTCTGTCTAGGTT
AGCACACGTCTGAACTCTTTCCCTACACGACGCTCTTCCGATCTNNNTCTGTCAAGGTCTGTCTAGGTT
AGCACACGTCTGAACTCTTTCCCTACACGACGCTCTTCCGATCTNNNTCTGTCAAGGTCTGTCTAGGTT
AGCACACGTCTGAACTCTTTCCCTACACGACGCTCTTCCGATCTNNNCTGTATTCCCCACTGTGACAAGG
AGCACACGTCTGAACTCTTTCCCTACACGACGCTCTTCCGATCTNNNCTGTATTCCCCACTGTGACAAGG
AGCACACGTCTGAACTCTTTCCCTACACGACGCTCTTCCGATCTNNNCTGTATTCCCCACTGTGACAAGG
GCACACGTCTGAACTCTTTCCCTACACGACGCTCTTCCGATCTNNNGAGAGAGACCCCGTCTAAAGAGA
GCACACGTCTGAACTCTTTCCCTACACGACGCTCTTCCGATCTNNNGAGAGAGACCCCGTCTAAAGAGA
GCACACGTCTGAACTCTTTCCCTACACGACGCTCTTCCGATCTNNNGAGAGAGACCCCGTCTAAAGAGA
GAGCACACGTCTGAACTCTTTCCCTACACGACGCTCTTCCGATCTNNNCTCTTCCTTCTGCTAACAACTGC
GAGCACACGTCTGAACTCTTTCCCTACACGACGCTCTTCCGATCTNNNCTCTTCCTTCTGCTAACAACTGC
GAGCACACGTCTGAACTCTTTCCCTACACGACGCTCTTCCGATCTNNNCTCTTCCTTCTGCTAACAACTGC
GCACACGTCTGAACTCTTTCCCTACACGACGCTCTTCCGATCTNNNCTAGCTGGGACTACAAGTGTGAC
GCACACGTCTGAACTCTTTCCCTACACGACGCTCTTCCGATCTNNNCTAGCTGGGACTACAAGTGTGAC
GCACACGTCTGAACTCTTTCCCTACACGACGCTCTTCCGATCTNNNCTAGCTGGGACTACAAGTGTGAC
AGCACACGTCTGAACTCTTTCCCTACACGACGCTCTTCCGATCTNNNGAGAATGGCTTGAACCCAGGAC
AGCACACGTCTGAACTCTTTCCCTACACGACGCTCTTCCGATCTNNNGAGAATGGCTTGAACCCAGGAC
AGCACACGTCTGAACTCTTTCCCTACACGACGCTCTTCCGATCTNNNGAGAATGGCTTGAACCCAGGAC
GCACACGTCTGAACTCTTTCCCTACACGACGCTCTTCCGATCTNNNTTCCCTGCGTACTTCTTTCTACCCAT
GCACACGTCTGAACTCTTTCCCTACACGACGCTCTTCCGATCTNNNTTCCCTGCGTACTTCTTTCTACCCAT
GCACACGTCTGAACTCTTTCCCTACACGACGCTCTTCCGATCTNNNTTCCCTGCGTACTTCTTTCTACCCAT
AGAGCACACGTCTGAACTCTTTCCCTACACGACGCTCTTCCGATCTNNNCCTTACTGGTGTCATGAGCA
AGAGCACACGTCTGAACTCTTTCCCTACACGACGCTCTTCCGATCTNNNCCTTACTGGTGTCATGAGCA
AGAGCACACGTCTGAACTCTTTCCCTACACGACGCTCTTCCGATCTNNNCCTTACTGGTGTCATGAGCA
AGCACACGTCTGAACTCTTTCCCTACACGACGCTCTTCCGATCTNNNCTGCCAACGGATACAGAGATGA
AGCACACGTCTGAACTCTTTCCCTACACGACGCTCTTCCGATCTNNNCTGCCAACGGATACAGAGATGA
AGCACACGTCTGAACTCTTTCCCTACACGACGCTCTTCCGATCTNNNCTGCCAACGGATACAGAGATGA
GAGCACACGTCTGAACTCTTTCCCTACACGACGCTCTTCCGATCTNNNCATACCAGGGATGCAGTAGGT
GAGCACACGTCTGAACTCTTTCCCTACACGACGCTCTTCCGATCTNNNCATACCAGGGATGCAGTAGGT
GAGCACACGTCTGAACTCTTTCCCTACACGACGCTCTTCCGATCTNNNCATACCAGGGATGCAGTAGGT
GCACACGTCTGAACTCTTTCCCTACACGACGCTCTTCCGATCTNNNCTTTTCTCTCCCTTTCCCTCCTCCAT

GCACACGTCTGAACTCTTTCCCTACACGACGCTCTTCCGATCTNNNCTTTCTCTCCCTTTCCCTCCTCCAT
GCACACGTCTGAACTCTTTCCCTACACGACGCTCTTCCGATCTNNNCTTTCTCTCCCTTTCCCTCCTCCAT
GCACACGTCTGAACTCTTTCCCTACACGACGCTCTTCCGATCTNNNGACCACGTTTAGATTCTGAGCAT
GCACACGTCTGAACTCTTTCCCTACACGACGCTCTTCCGATCTNNNGACCACGTTTAGATTCTGAGCAT
GCACACGTCTGAACTCTTTCCCTACACGACGCTCTTCCGATCTNNNGACCACGTTTAGATTCTGAGCAT
AGCACACGTCTGAACTCTTTCCCTACACGACGCTCTTCCGATCTNNNGCAACCTCTACTTACCAGGTTCA
AGCACACGTCTGAACTCTTTCCCTACACGACGCTCTTCCGATCTNNNGCAACCTCTACTTACCAGGTTCA
AGCACACGTCTGAACTCTTTCCCTACACGACGCTCTTCCGATCTNNNGCAACCTCTACTTACCAGGTTCA
GCACACGTCTGAACTCTTTCCCTACACGACGCTCTTCCGATCTNNNTCTTGTCAGGCCACTCTTTTTCTCA
GCACACGTCTGAACTCTTTCCCTACACGACGCTCTTCCGATCTNNNTCTTGTCAGGCCACTCTTTTTCTCA
GCACACGTCTGAACTCTTTCCCTACACGACGCTCTTCCGATCTNNNTCTTGTCAGGCCACTCTTTTTCTCA
GAGCACACGTCTGAACTCTTTCCCTACACGACGCTCTTCCGATCTNNNAAGCTGAAGCAGGAAAATTG
GAGCACACGTCTGAACTCTTTCCCTACACGACGCTCTTCCGATCTNNNAAGCTGAAGCAGGAAAATTG
GAGCACACGTCTGAACTCTTTCCCTACACGACGCTCTTCCGATCTNNNAAGCTGAAGCAGGAAAATTG
GAGCACACGTCTGAACTCTTTCCCTACACGACGCTCTTCCGATCTNNNGTTACGTGGGAGCTAAGCTA
GAGCACACGTCTGAACTCTTTCCCTACACGACGCTCTTCCGATCTNNNGTTACGTGGGAGCTAAGCTA
GAGCACACGTCTGAACTCTTTCCCTACACGACGCTCTTCCGATCTNNNGTTACGTGGGAGCTAAGCTA
GAGCACACGTCTGAACTCTTTCCCTACACGACGCTCTTCCGATCTNNNCATTCTGTCCAACACTCACCATG
GAGCACACGTCTGAACTCTTTCCCTACACGACGCTCTTCCGATCTNNNCATTCTGTCCAACACTCACCATG
GAGCACACGTCTGAACTCTTTCCCTACACGACGCTCTTCCGATCTNNNCATTCTGTCCAACACTCACCATG
CACACGTCTGAACTCTTTCCCTACACGACGCTCTTCCGATCTNNNCCAAGTGACCCTGTGCAGAGCATC
CACACGTCTGAACTCTTTCCCTACACGACGCTCTTCCGATCTNNNCCAAGTGACCCTGTGCAGAGCATC
CACACGTCTGAACTCTTTCCCTACACGACGCTCTTCCGATCTNNNCCAAGTGACCCTGTGCAGAGCATC
AGCACACGTCTGAACTCTTTCCCTACACGACGCTCTTCCGATCTNNNTGAAGTTGTCATCGGTCAGGATG
AGCACACGTCTGAACTCTTTCCCTACACGACGCTCTTCCGATCTNNNTGAAGTTGTCATCGGTCAGGATG
AGCACACGTCTGAACTCTTTCCCTACACGACGCTCTTCCGATCTNNNTGAAGTTGTCATCGGTCAGGATG
GAGCACACGTCTGAACTCTTTCCCTACACGACGCTCTTCCGATCTNNNAAAAATTAGCTGGACATGGTG
GAGCACACGTCTGAACTCTTTCCCTACACGACGCTCTTCCGATCTNNNAAAAATTAGCTGGACATGGTG
GAGCACACGTCTGAACTCTTTCCCTACACGACGCTCTTCCGATCTNNNAAAAATTAGCTGGACATGGTG
GAGCACACGTCTGAACTCTTTCCCTACACGACGCTCTTCCGATCTNNNGAATCACTAGAGCCCACAAGG
GAGCACACGTCTGAACTCTTTCCCTACACGACGCTCTTCCGATCTNNNGAATCACTAGAGCCCACAAGG
GAGCACACGTCTGAACTCTTTCCCTACACGACGCTCTTCCGATCTNNNGAATCACTAGAGCCCACAAGG
GAGCACACGTCTGAACTCTTTCCCTACACGACGCTCTTCCGATCTNNNAGCCTAAACATCACCACTATGC
GAGCACACGTCTGAACTCTTTCCCTACACGACGCTCTTCCGATCTNNNAGCCTAAACATCACCACTATGC
GAGCACACGTCTGAACTCTTTCCCTACACGACGCTCTTCCGATCTNNNAGCCTAAACATCACCACTATGC
AGAGCACACGTCTGAACTCTTTCCCTACACGACGCTCTTCCGATCTNNNGAGGTCTAGAGTTTGAGGCT
AGAGCACACGTCTGAACTCTTTCCCTACACGACGCTCTTCCGATCTNNNGAGGTCTAGAGTTTGAGGCT
AGAGCACACGTCTGAACTCTTTCCCTACACGACGCTCTTCCGATCTNNNGAGGTCTAGAGTTTGAGGCT
GAGCACACGTCTGAACTCTTTCCCTACACGACGCTCTTCCGATCTNNNACAGAGCAAGCTCCTGTTTCAT
GAGCACACGTCTGAACTCTTTCCCTACACGACGCTCTTCCGATCTNNNACAGAGCAAGCTCCTGTTTCAT
GAGCACACGTCTGAACTCTTTCCCTACACGACGCTCTTCCGATCTNNNACAGAGCAAGCTCCTGTTTCAT
GAGCACACGTCTGAACTCTTTCCCTACACGACGCTCTTCCGATCTNNNCAGTGAGGGATGGAAAGAG
GAGCACACGTCTGAACTCTTTCCCTACACGACGCTCTTCCGATCTNNNCAGTGAGGGATGGAAAGAG

GAGCACACGTCTGAACTCTTTCCCTACACGACGCTCTTCCGATCTNNNCAGTGAGGGATGGAAAGAG
AGCACACGTCTGAACTCTTTCCCTACACGACGCTCTTCCGATCTNNNAGCAAAGGGAGTAAAGCTAGA
AGCACACGTCTGAACTCTTTCCCTACACGACGCTCTTCCGATCTNNNAGCAAAGGGAGTAAAGCTAGA
AGCACACGTCTGAACTCTTTCCCTACACGACGCTCTTCCGATCTNNNAGCAAAGGGAGTAAAGCTAGA
GAGCACACGTCTGAACTCTTTCCCTACACGACGCTCTTCCGATCTNNNTACTCAGGGACAAGTGAAAGT
GAGCACACGTCTGAACTCTTTCCCTACACGACGCTCTTCCGATCTNNNTACTCAGGGACAAGTGAAAGT
GAGCACACGTCTGAACTCTTTCCCTACACGACGCTCTTCCGATCTNNNTACTCAGGGACAAGTGAAAGT
AGCACACGTCTGAACTCTTTCCCTACACGACGCTCTTCCGATCTNNNCACCATTGTAGCATCCAAGCAA
AGCACACGTCTGAACTCTTTCCCTACACGACGCTCTTCCGATCTNNNCACCATTGTAGCATCCAAGCAA
AGCACACGTCTGAACTCTTTCCCTACACGACGCTCTTCCGATCTNNNCACCATTGTAGCATCCAAGCAA
GAGCACACGTCTGAACTCTTTCCCTACACGACGCTCTTCCGATCTNNNATAATGATAGTATTCCATTGCC
GAGCACACGTCTGAACTCTTTCCCTACACGACGCTCTTCCGATCTNNNATAATGATAGTATTCCATTGCC
GAGCACACGTCTGAACTCTTTCCCTACACGACGCTCTTCCGATCTNNNATAATGATAGTATTCCATTGCC
AGCACACGTCTGAACTCTTTCCCTACACGACGCTCTTCCGATCTNNNGAGGCTCCTTCTGCACTGTTCAT
AGCACACGTCTGAACTCTTTCCCTACACGACGCTCTTCCGATCTNNNGAGGCTCCTTCTGCACTGTTCAT
AGCACACGTCTGAACTCTTTCCCTACACGACGCTCTTCCGATCTNNNGAGGCTCCTTCTGCACTGTTCAT
AGCACACGTCTGAACTCTTTCCCTACACGACGCTCTTCCGATCTNNNCGCCCGGCTTTTCTTTTTTTTTTC
AGCACACGTCTGAACTCTTTCCCTACACGACGCTCTTCCGATCTNNNCGCCCGGCTTTTCTTTTTTTTTTC
AGCACACGTCTGAACTCTTTCCCTACACGACGCTCTTCCGATCTNNNCGCCCGGCTTTTCTTTTTTTTTTC
GAGCACACGTCTGAACTCTTTCCCTACACGACGCTCTTCCGATCTNNNAGCAAATATAGGTGGTAGCC
GAGCACACGTCTGAACTCTTTCCCTACACGACGCTCTTCCGATCTNNNAGCAAATATAGGTGGTAGCC
GAGCACACGTCTGAACTCTTTCCCTACACGACGCTCTTCCGATCTNNNAGCAAATATAGGTGGTAGCC
AGCACACGTCTGAACTCTTTCCCTACACGACGCTCTTCCGATCTNNNAGATCTGAGAACTGACAGACTG
AGCACACGTCTGAACTCTTTCCCTACACGACGCTCTTCCGATCTNNNAGATCTGAGAACTGACAGACTG
AGCACACGTCTGAACTCTTTCCCTACACGACGCTCTTCCGATCTNNNAGATCTGAGAACTGACAGACTG
AGCACACGTCTGAACTCTTTCCCTACACGACGCTCTTCCGATCTNNNGTTTTCTTCTTGCACGCAAATCCC
AGCACACGTCTGAACTCTTTCCCTACACGACGCTCTTCCGATCTNNNGTTTTCTTCTTGCACGCAAATCCC
AGCACACGTCTGAACTCTTTCCCTACACGACGCTCTTCCGATCTNNNGTTTTCTTCTTGCACGCAAATCCC
AGCACACGTCTGAACTCTTTCCCTACACGACGCTCTTCCGATCTNNNTTTCCTCCACTATTCCCCTTTGGCA
AGCACACGTCTGAACTCTTTCCCTACACGACGCTCTTCCGATCTNNNTTTCCTCCACTATTCCCCTTTGGCA
AGCACACGTCTGAACTCTTTCCCTACACGACGCTCTTCCGATCTNNNTTTCCTCCACTATTCCCCTTTGGCA
AGCACACGTCTGAACTCTTTCCCTACACGACGCTCTTCCGATCTNNNAACACTCGCCACTATACATCCAC
AGCACACGTCTGAACTCTTTCCCTACACGACGCTCTTCCGATCTNNNAACACTCGCCACTATACATCCAC
AGCACACGTCTGAACTCTTTCCCTACACGACGCTCTTCCGATCTNNNAACACTCGCCACTATACATCCAC
GCACACGTCTGAACTCTTTCCCTACACGACGCTCTTCCGATCTNNNTGCCACTGCTTTATCAACTAAGTC
GCACACGTCTGAACTCTTTCCCTACACGACGCTCTTCCGATCTNNNTGCCACTGCTTTATCAACTAAGTC
GCACACGTCTGAACTCTTTCCCTACACGACGCTCTTCCGATCTNNNTGCCACTGCTTTATCAACTAAGTC
GAGCACACGTCTGAACTCTTTCCCTACACGACGCTCTTCCGATCTNNNAAAGACATCCTGCCTAGTTCCC
GAGCACACGTCTGAACTCTTTCCCTACACGACGCTCTTCCGATCTNNNAAAGACATCCTGCCTAGTTCCC
GAGCACACGTCTGAACTCTTTCCCTACACGACGCTCTTCCGATCTNNNAAAGACATCCTGCCTAGTTCCC
AGAGCACACGTCTGAACTCTTTCCCTACACGACGCTCTTCCGATCTNNNTGAGACAGGGAGAATTGCT
AGAGCACACGTCTGAACTCTTTCCCTACACGACGCTCTTCCGATCTNNNTGAGACAGGGAGAATTGCT
AGAGCACACGTCTGAACTCTTTCCCTACACGACGCTCTTCCGATCTNNNTGAGACAGGGAGAATTGCT

CACACGTCTGAACTCTTTCCCTACACGACGCTCTTCCGATCTNNNTGAACAGTGGATGTCGCTTTTTCC
CACACGTCTGAACTCTTTCCCTACACGACGCTCTTCCGATCTNNNTGAACAGTGGATGTCGCTTTTTCC
CACACGTCTGAACTCTTTCCCTACACGACGCTCTTCCGATCTNNNTGAACAGTGGATGTCGCTTTTTCC
AGAGCACACGTCTGAACTCTTTCCCTACACGACGCTCTTCCGATCTNNNAGAATCACTTGAACCCAGGA
AGAGCACACGTCTGAACTCTTTCCCTACACGACGCTCTTCCGATCTNNNAGAATCACTTGAACCCAGGA
AGAGCACACGTCTGAACTCTTTCCCTACACGACGCTCTTCCGATCTNNNAGAATCACTTGAACCCAGGA
AGCACACGTCTGAACTCTTTCCCTACACGACGCTCTTCCGATCTNNNTCTTGGAAGTAACAGAACTTAAT
AGCACACGTCTGAACTCTTTCCCTACACGACGCTCTTCCGATCTNNNTCTTGGAAGTAACAGAACTTAAT
AGCACACGTCTGAACTCTTTCCCTACACGACGCTCTTCCGATCTNNNTCTTGGAAGTAACAGAACTTAAT
GAGCACACGTCTGAACTCTTTCCCTACACGACGCTCTTCCGATCTNNNTAGGTTGCCTGTTCACTATGAC
GAGCACACGTCTGAACTCTTTCCCTACACGACGCTCTTCCGATCTNNNTAGGTTGCCTGTTCACTATGAC
GAGCACACGTCTGAACTCTTTCCCTACACGACGCTCTTCCGATCTNNNTAGGTTGCCTGTTCACTATGAC
GAGCACACGTCTGAACTCTTTCCCTACACGACGCTCTTCCGATCTNNNGCTTGCTTTCTTGTTTGTTAGC
GAGCACACGTCTGAACTCTTTCCCTACACGACGCTCTTCCGATCTNNNGCTTGCTTTCTTGTTTGTTAGC
GAGCACACGTCTGAACTCTTTCCCTACACGACGCTCTTCCGATCTNNNGCTTGCTTTCTTGTTTGTTAGC
GAGCACACGTCTGAACTCTTTCCCTACACGACGCTCTTCCGATCTNNNTCAGACCTTTGCCTTGATCATG
GAGCACACGTCTGAACTCTTTCCCTACACGACGCTCTTCCGATCTNNNTCAGACCTTTGCCTTGATCATG
GAGCACACGTCTGAACTCTTTCCCTACACGACGCTCTTCCGATCTNNNTCAGACCTTTGCCTTGATCATG
GAGCACACGTCTGAACTCTTTCCCTACACGACGCTCTTCCGATCTNNNGAGTTTGCAGTGAGCCGAGAT
GAGCACACGTCTGAACTCTTTCCCTACACGACGCTCTTCCGATCTNNNGAGTTTGCAGTGAGCCGAGAT
GAGCACACGTCTGAACTCTTTCCCTACACGACGCTCTTCCGATCTNNNGAGTTTGCAGTGAGCCGAGAT
GAGCACACGTCTGAACTCTTTCCCTACACGACGCTCTTCCGATCTNNNGAAACCTTGCACATGTATCCCC
GAGCACACGTCTGAACTCTTTCCCTACACGACGCTCTTCCGATCTNNNGAAACCTTGCACATGTATCCCC
GAGCACACGTCTGAACTCTTTCCCTACACGACGCTCTTCCGATCTNNNGAAACCTTGCACATGTATCCCC
GAGCACACGTCTGAACTCTTTCCCTACACGACGCTCTTCCGATCTNNNCTGACAAAGAGCTCATACCCAC
GAGCACACGTCTGAACTCTTTCCCTACACGACGCTCTTCCGATCTNNNCTGACAAAGAGCTCATACCCAC
GAGCACACGTCTGAACTCTTTCCCTACACGACGCTCTTCCGATCTNNNCTGACAAAGAGCTCATACCCAC
GAGCACACGTCTGAACTCTTTCCCTACACGACGCTCTTCCGATCTNNNGAGAGAGATGTAGGCAGGA
GAGCACACGTCTGAACTCTTTCCCTACACGACGCTCTTCCGATCTNNNGAGAGAGATGTAGGCAGGA
GAGCACACGTCTGAACTCTTTCCCTACACGACGCTCTTCCGATCTNNNGAGAGAGATGTAGGCAGGA
GAGCACACGTCTGAACTCTTTCCCTACACGACGCTCTTCCGATCTNNNCTGACCTCGTGATCCAAGAGT
GAGCACACGTCTGAACTCTTTCCCTACACGACGCTCTTCCGATCTNNNCTGACCTCGTGATCCAAGAGT
GAGCACACGTCTGAACTCTTTCCCTACACGACGCTCTTCCGATCTNNNCTGACCTCGTGATCCAAGAGT
GAGCACACGTCTGAACTCTTTCCCTACACGACGCTCTTCCGATCTNNNATTATTTGTTGTTTGATCACTCA
GAGCACACGTCTGAACTCTTTCCCTACACGACGCTCTTCCGATCTNNNATTATTTGTTGTTTGATCACTCA
GAGCACACGTCTGAACTCTTTCCCTACACGACGCTCTTCCGATCTNNNATTATTTGTTGTTTGATCACTCA
CACACGTCTGAACTCTTTCCCTACACGACGCTCTTCCGATCTNNNCTAGAGCTTTTCCTTTTAAGAGACA
CACACGTCTGAACTCTTTCCCTACACGACGCTCTTCCGATCTNNNCTAGAGCTTTTCCTTTTAAGAGACA
CACACGTCTGAACTCTTTCCCTACACGACGCTCTTCCGATCTNNNCTAGAGCTTTTCCTTTTAAGAGACA
AGCACACGTCTGAACTCTTTCCCTACACGACGCTCTTCCGATCTNNNCTGAGGCAGGAGAATCACTTGA
AGCACACGTCTGAACTCTTTCCCTACACGACGCTCTTCCGATCTNNNCTGAGGCAGGAGAATCACTTGA
AGCACACGTCTGAACTCTTTCCCTACACGACGCTCTTCCGATCTNNNCTGAGGCAGGAGAATCACTTGA
AGCACACGTCTGAACTCTTTCCCTACACGACGCTCTTCCGATCTNNNGCCAAAAGCAGTCTAAAGGCAT

AGCACACGTCTGAACTCTTTCCCTACACGACGCTCTTCCGATCTNNNGCCAAAAGCAGTCTAAAGGCAT
AGCACACGTCTGAACTCTTTCCCTACACGACGCTCTTCCGATCTNNNGCCAAAAGCAGTCTAAAGGCAT
AGCACACGTCTGAACTCTTTCCCTACACGACGCTCTTCCGATCTNNNATTGCTTGAGGCTAGGAGTTCA
AGCACACGTCTGAACTCTTTCCCTACACGACGCTCTTCCGATCTNNNATTGCTTGAGGCTAGGAGTTCA
AGCACACGTCTGAACTCTTTCCCTACACGACGCTCTTCCGATCTNNNATTGCTTGAGGCTAGGAGTTCA
AGCACACGTCTGAACTCTTTCCCTACACGACGCTCTTCCGATCTNNNGAAACAGGGCAGGAACGAAAA
AGCACACGTCTGAACTCTTTCCCTACACGACGCTCTTCCGATCTNNNGAAACAGGGCAGGAACGAAAA
AGCACACGTCTGAACTCTTTCCCTACACGACGCTCTTCCGATCTNNNGAAACAGGGCAGGAACGAAAA
GAGCACACGTCTGAACTCTTTCCCTACACGACGCTCTTCCGATCTNNNGCCCTGGAACATGAATCTTTG
GAGCACACGTCTGAACTCTTTCCCTACACGACGCTCTTCCGATCTNNNGCCCTGGAACATGAATCTTTG
GAGCACACGTCTGAACTCTTTCCCTACACGACGCTCTTCCGATCTNNNGCCCTGGAACATGAATCTTTG
GAGCACACGTCTGAACTCTTTCCCTACACGACGCTCTTCCGATCTNNNACAGTTCTCAAGCCTTGGTGC
GAGCACACGTCTGAACTCTTTCCCTACACGACGCTCTTCCGATCTNNNACAGTTCTCAAGCCTTGGTGC
GAGCACACGTCTGAACTCTTTCCCTACACGACGCTCTTCCGATCTNNNACAGTTCTCAAGCCTTGGTGC
AGCACACGTCTGAACTCTTTCCCTACACGACGCTCTTCCGATCTNNNTTCTCCTGTGCTACACATTCACAC
AGCACACGTCTGAACTCTTTCCCTACACGACGCTCTTCCGATCTNNNTTCTCCTGTGCTACACATTCACAC
AGCACACGTCTGAACTCTTTCCCTACACGACGCTCTTCCGATCTNNNTTCTCCTGTGCTACACATTCACAC
GAGCACACGTCTGAACTCTTTCCCTACACGACGCTCTTCCGATCTNNNTCATTCATTCATTCCCCCACAT
GAGCACACGTCTGAACTCTTTCCCTACACGACGCTCTTCCGATCTNNNTCATTCATTCATTCCCCCACAT
GAGCACACGTCTGAACTCTTTCCCTACACGACGCTCTTCCGATCTNNNTCATTCATTCATTCCCCCACAT
AGCACACGTCTGAACTCTTTCCCTACACGACGCTCTTCCGATCTNNNATGTGCTTCAGTAACTCTCTCCC
AGCACACGTCTGAACTCTTTCCCTACACGACGCTCTTCCGATCTNNNATGTGCTTCAGTAACTCTCTCCC
AGCACACGTCTGAACTCTTTCCCTACACGACGCTCTTCCGATCTNNNATGTGCTTCAGTAACTCTCTCCC
CACACGTCTGAACTCTTTCCCTACACGACGCTCTTCCGATCTNNNCTTCTCTTTTCCCTCTCCCCTTCCATCC
CACACGTCTGAACTCTTTCCCTACACGACGCTCTTCCGATCTNNNCTTCTCTTTTCCCTCTCCCCTTCCATCC
CACACGTCTGAACTCTTTCCCTACACGACGCTCTTCCGATCTNNNCTTCTCTTTTCCCTCTCCCCTTCCATCC
AGCACACGTCTGAACTCTTTCCCTACACGACGCTCTTCCGATCTNNNCGCGTGTGTGTATTCCAAATTC
AGCACACGTCTGAACTCTTTCCCTACACGACGCTCTTCCGATCTNNNCGCGTGTGTGTATTCCAAATTC
AGCACACGTCTGAACTCTTTCCCTACACGACGCTCTTCCGATCTNNNCGCGTGTGTGTATTCCAAATTC
AGCACACGTCTGAACTCTTTCCCTACACGACGCTCTTCCGATCTNNNTTGAAGCTGTGCTCCAGAATTTC
AGCACACGTCTGAACTCTTTCCCTACACGACGCTCTTCCGATCTNNNTTGAAGCTGTGCTCCAGAATTTC
AGCACACGTCTGAACTCTTTCCCTACACGACGCTCTTCCGATCTNNNTTGAAGCTGTGCTCCAGAATTTC
AGCACACGTCTGAACTCTTTCCCTACACGACGCTCTTCCGATCTNNNCATTGTTAGGGAGGGAGCAGA
AGCACACGTCTGAACTCTTTCCCTACACGACGCTCTTCCGATCTNNNCATTGTTAGGGAGGGAGCAGA
AGCACACGTCTGAACTCTTTCCCTACACGACGCTCTTCCGATCTNNNCATTGTTAGGGAGGGAGCAGA
GAGCACACGTCTGAACTCTTTCCCTACACGACGCTCTTCCGATCTNNNATGAAGACACAGAAACCATGC
GAGCACACGTCTGAACTCTTTCCCTACACGACGCTCTTCCGATCTNNNATGAAGACACAGAAACCATGC
GAGCACACGTCTGAACTCTTTCCCTACACGACGCTCTTCCGATCTNNNATGAAGACACAGAAACCATGC
GCACACGTCTGAACTCTTTCCCTACACGACGCTCTTCCGATCTNNNCTAGATCGCGCCACTACACTCCAT
GCACACGTCTGAACTCTTTCCCTACACGACGCTCTTCCGATCTNNNCTAGATCGCGCCACTACACTCCAT
GCACACGTCTGAACTCTTTCCCTACACGACGCTCTTCCGATCTNNNCTAGATCGCGCCACTACACTCCAT
GAGCACACGTCTGAACTCTTTCCCTACACGACGCTCTTCCGATCTNNNTGAAGTTTTTGTACTGGCCACA
GAGCACACGTCTGAACTCTTTCCCTACACGACGCTCTTCCGATCTNNNTGAAGTTTTTGTACTGGCCACA

AGCACACGTCTGAACTCTTTCCCTACACGACGCTCTTCCGATCTNNNTGAAGTTTTTGTACTGGCCAC
AGCACACGTCTGAACTCTTTCCCTACACGACGCTCTTCCGATCTNNNAGAACAAGATGGAGACTGCA
AGCACACGTCTGAACTCTTTCCCTACACGACGCTCTTCCGATCTNNNAGAACAAGATGGAGACTGCA
AGCACACGTCTGAACTCTTTCCCTACACGACGCTCTTCCGATCTNNNAGAACAAGATGGAGACTGCA
AGCACACGTCTGAACTCTTTCCCTACACGACGCTCTTCCGATCTNNNCGCAAATTGTAAAAAACCAGC
AGCACACGTCTGAACTCTTTCCCTACACGACGCTCTTCCGATCTNNNCGCAAATTGTAAAAAACCAGC
AGCACACGTCTGAACTCTTTCCCTACACGACGCTCTTCCGATCTNNNCGCAAATTGTAAAAAACCAGC
AGCACACGTCTGAACTCTTTCCCTACACGACGCTCTTCCGATCTNNNTCTATTCTGCACAATCCACTCCC
AGCACACGTCTGAACTCTTTCCCTACACGACGCTCTTCCGATCTNNNTCTATTCTGCACAATCCACTCCC
AGCACACGTCTGAACTCTTTCCCTACACGACGCTCTTCCGATCTNNNTCTATTCTGCACAATCCACTCCC
CACACGTCTGAACTCTTTCCCTACACGACGCTCTTCCGATCTNNNCTAGCTACCCCACCTCTCTCTAGC
CACACGTCTGAACTCTTTCCCTACACGACGCTCTTCCGATCTNNNCTAGCTACCCCACCTCTCTCTAGC
CACACGTCTGAACTCTTTCCCTACACGACGCTCTTCCGATCTNNNCTAGCTACCCCACCTCTCTCTAGC
AGCACACGTCTGAACTCTTTCCCTACACGACGCTCTTCCGATCTNNNCAGTAGCCTTGTTTGGGAAGG
AGCACACGTCTGAACTCTTTCCCTACACGACGCTCTTCCGATCTNNNCAGTAGCCTTGTTTGGGAAGG
AGCACACGTCTGAACTCTTTCCCTACACGACGCTCTTCCGATCTNNNCAGTAGCCTTGTTTGGGAAGG
AGCACACGTCTGAACTCTTTCCCTACACGACGCTCTTCCGATCTNNNATCAGTTGACCTGTTGGCCTAA
AGCACACGTCTGAACTCTTTCCCTACACGACGCTCTTCCGATCTNNNATCAGTTGACCTGTTGGCCTAA
AGCACACGTCTGAACTCTTTCCCTACACGACGCTCTTCCGATCTNNNATCAGTTGACCTGTTGGCCTAA
GCACACGTCTGAACTCTTTCCCTACACGACGCTCTTCCGATCTNNNCTGAGGCAGGAGAATTGCTTG
GCACACGTCTGAACTCTTTCCCTACACGACGCTCTTCCGATCTNNNCTGAGGCAGGAGAATTGCTTG
GCACACGTCTGAACTCTTTCCCTACACGACGCTCTTCCGATCTNNNCTGAGGCAGGAGAATTGCTTG
AGCACACGTCTGAACTCTTTCCCTACACGACGCTCTTCCGATCTNNNCTTGCAGTGAGCTGAGATTGC
AGCACACGTCTGAACTCTTTCCCTACACGACGCTCTTCCGATCTNNNCTTGCAGTGAGCTGAGATTGC
AGCACACGTCTGAACTCTTTCCCTACACGACGCTCTTCCGATCTNNNCTTGCAGTGAGCTGAGATTGC
GAGCACACGTCTGAACTCTTTCCCTACACGACGCTCTTCCGATCTNNNAGGTTACAGTTGGCCAAGAT
GAGCACACGTCTGAACTCTTTCCCTACACGACGCTCTTCCGATCTNNNAGGTTACAGTTGGCCAAGAT
GAGCACACGTCTGAACTCTTTCCCTACACGACGCTCTTCCGATCTNNNAGGTTACAGTTGGCCAAGAT
AGCACACGTCTGAACTCTTTCCCTACACGACGCTCTTCCGATCTNNNCCCGATCCAACCCTTTTCTGAGC
AGCACACGTCTGAACTCTTTCCCTACACGACGCTCTTCCGATCTNNNCCCGATCCAACCCTTTTCTGAGC
AGCACACGTCTGAACTCTTTCCCTACACGACGCTCTTCCGATCTNNNCCCGATCCAACCCTTTTCTGAGC
GAGCACACGTCTGAACTCTTTCCCTACACGACGCTCTTCCGATCTNNNTTGAAAGGCTAGAAAGGAG
GAGCACACGTCTGAACTCTTTCCCTACACGACGCTCTTCCGATCTNNNTTGAAAGGCTAGAAAGGAG
GAGCACACGTCTGAACTCTTTCCCTACACGACGCTCTTCCGATCTNNNTTGAAAGGCTAGAAAGGAG
CACACGTCTGAACTCTTTCCCTACACGACGCTCTTCCGATCTNNNGGTGTCCTTCAGTCTCACTTGATC
CACACGTCTGAACTCTTTCCCTACACGACGCTCTTCCGATCTNNNGGTGTCCTTCAGTCTCACTTGATC
CACACGTCTGAACTCTTTCCCTACACGACGCTCTTCCGATCTNNNGGTGTCCTTCAGTCTCACTTGATC
AGCACACGTCTGAACTCTTTCCCTACACGACGCTCTTCCGATCTNNNCTCAAACAATTCTCCTGCCTCAG
AGCACACGTCTGAACTCTTTCCCTACACGACGCTCTTCCGATCTNNNCTCAAACAATTCTCCTGCCTCAG
AGCACACGTCTGAACTCTTTCCCTACACGACGCTCTTCCGATCTNNNCTCAAACAATTCTCCTGCCTCAG
CACACGTCTGAACTCTTTCCCTACACGACGCTCTTCCGATCTNNNGTACATGGACTAGTGGCCAACTC
CACACGTCTGAACTCTTTCCCTACACGACGCTCTTCCGATCTNNNGTACATGGACTAGTGGCCAACTC
CACACGTCTGAACTCTTTCCCTACACGACGCTCTTCCGATCTNNNGTACATGGACTAGTGGCCAACTC

CACACGTCTGAACTCTTTCCCTACACGACGCTCTTCCGATCTNNNCCCTCTACCCCTTCATCCATCTACA
CACACGTCTGAACTCTTTCCCTACACGACGCTCTTCCGATCTNNNCCCTCTACCCCTTCATCCATCTACA
CACACGTCTGAACTCTTTCCCTACACGACGCTCTTCCGATCTNNNCCCTCTACCCCTTCATCCATCTACA
AGCACACGTCTGAACTCTTTCCCTACACGACGCTCTTCCGATCTNNNGATAGGTGGGGTTAGATGTG
AGCACACGTCTGAACTCTTTCCCTACACGACGCTCTTCCGATCTNNNGATAGGTGGGGTTAGATGTG
AGCACACGTCTGAACTCTTTCCCTACACGACGCTCTTCCGATCTNNNGATAGGTGGGGTTAGATGTG
GCACACGTCTGAACTCTTTCCCTACACGACGCTCTTCCGATCTNNNCACTGGAATTTGGAGGAGCAA
GCACACGTCTGAACTCTTTCCCTACACGACGCTCTTCCGATCTNNNCACTGGAATTTGGAGGAGCAA
GCACACGTCTGAACTCTTTCCCTACACGACGCTCTTCCGATCTNNNCACTGGAATTTGGAGGAGCAA
GAGCACACGTCTGAACTCTTTCCCTACACGACGCTCTTCCGATCTNNNTCCATCCTGATATGTGTGTGT
GAGCACACGTCTGAACTCTTTCCCTACACGACGCTCTTCCGATCTNNNTCCATCCTGATATGTGTGTGT
GAGCACACGTCTGAACTCTTTCCCTACACGACGCTCTTCCGATCTNNNTCCATCCTGATATGTGTGTGT
GAGCACACGTCTGAACTCTTTCCCTACACGACGCTCTTCCGATCTNNNGAGGCTGAGGCAGGAGAAT
GAGCACACGTCTGAACTCTTTCCCTACACGACGCTCTTCCGATCTNNNGAGGCTGAGGCAGGAGAAT
GAGCACACGTCTGAACTCTTTCCCTACACGACGCTCTTCCGATCTNNNGAGGCTGAGGCAGGAGAAT
AGCACACGTCTGAACTCTTTCCCTACACGACGCTCTTCCGATCTNNNCACCACCTGGCAGTATTCTCAT
AGCACACGTCTGAACTCTTTCCCTACACGACGCTCTTCCGATCTNNNCACCACCTGGCAGTATTCTCAT
AGCACACGTCTGAACTCTTTCCCTACACGACGCTCTTCCGATCTNNNCACCACCTGGCAGTATTCTCAT
GCACACGTCTGAACTCTTTCCCTACACGACGCTCTTCCGATCTNNNTTTTCATAACCCATTGGCAGCC
GCACACGTCTGAACTCTTTCCCTACACGACGCTCTTCCGATCTNNNTTTTCATAACCCATTGGCAGCC
GCACACGTCTGAACTCTTTCCCTACACGACGCTCTTCCGATCTNNNTTTTCATAACCCATTGGCAGCC
GCACACGTCTGAACTCTTTCCCTACACGACGCTCTTCCGATCTNNNTGCAGTTGTTACAGATGGACTC
GCACACGTCTGAACTCTTTCCCTACACGACGCTCTTCCGATCTNNNTGCAGTTGTTACAGATGGACTC
GCACACGTCTGAACTCTTTCCCTACACGACGCTCTTCCGATCTNNNTGCAGTTGTTACAGATGGACTC
AGCACACGTCTGAACTCTTTCCCTACACGACGCTCTTCCGATCTNNNATTTTGTTTCTGTTGCTCCTGG
AGCACACGTCTGAACTCTTTCCCTACACGACGCTCTTCCGATCTNNNATTTTGTTTCTGTTGCTCCTGG
AGCACACGTCTGAACTCTTTCCCTACACGACGCTCTTCCGATCTNNNATTTTGTTTCTGTTGCTCCTGG
CACACGTCTGAACTCTTTCCCTACACGACGCTCTTCCGATCTNNNGAAACGTCTAGATGCTCAACCATC
CACACGTCTGAACTCTTTCCCTACACGACGCTCTTCCGATCTNNNGAAACGTCTAGATGCTCAACCATC
CACACGTCTGAACTCTTTCCCTACACGACGCTCTTCCGATCTNNNGAAACGTCTAGATGCTCAACCATC
AGCACACGTCTGAACTCTTTCCCTACACGACGCTCTTCCGATCTNNNTCTGAATTTATCCCACCCCTCA
AGCACACGTCTGAACTCTTTCCCTACACGACGCTCTTCCGATCTNNNTCTGAATTTATCCCACCCCTCA
AGCACACGTCTGAACTCTTTCCCTACACGACGCTCTTCCGATCTNNNTCTGAATTTATCCCACCCCTCA
AGCACACGTCTGAACTCTTTCCCTACACGACGCTCTTCCGATCTNNNGTCATACTTTGGGTGAAACAG
AGCACACGTCTGAACTCTTTCCCTACACGACGCTCTTCCGATCTNNNGTCATACTTTGGGTGAAACAG
AGCACACGTCTGAACTCTTTCCCTACACGACGCTCTTCCGATCTNNNGTCATACTTTGGGTGAAACAG
AGCACACGTCTGAACTCTTTCCCTACACGACGCTCTTCCGATCTNNNGCAGAAGTTGTAGTGAGCTG
AGCACACGTCTGAACTCTTTCCCTACACGACGCTCTTCCGATCTNNNGCAGAAGTTGTAGTGAGCTG
AGCACACGTCTGAACTCTTTCCCTACACGACGCTCTTCCGATCTNNNGCAGAAGTTGTAGTGAGCTG
GCACACGTCTGAACTCTTTCCCTACACGACGCTCTTCCGATCTNNNCAGAGCAAGACCCTGTCTCAAAT
GCACACGTCTGAACTCTTTCCCTACACGACGCTCTTCCGATCTNNNCAGAGCAAGACCCTGTCTCAAAT
GCACACGTCTGAACTCTTTCCCTACACGACGCTCTTCCGATCTNNNCAGAGCAAGACCCTGTCTCAAAT
GCACACGTCTGAACTCTTTCCCTACACGACGCTCTTCCGATCTNNNACAACGCATCATCACTCCAATAC

AGCACACGTCTGAACTCTTTCCCTACACGACGCTCTTCCGATCTNNNACAACGCATCATCACTCCAATAC
AGCACACGTCTGAACTCTTTCCCTACACGACGCTCTTCCGATCTNNNACAACGCATCATCACTCCAATAC
AGCACACGTCTGAACTCTTTCCCTACACGACGCTCTTCCGATCTNNNCAAGATTACGCCACTGCACTCC
AGCACACGTCTGAACTCTTTCCCTACACGACGCTCTTCCGATCTNNNCAAGATTACGCCACTGCACTCC
AGCACACGTCTGAACTCTTTCCCTACACGACGCTCTTCCGATCTNNNCAAGATTACGCCACTGCACTCC
CACACGTCTGAACTCTTTCCCTACACGACGCTCTTCCGATCTNNNCCTTGTGTGCATAGAGATGTTCAC
CACACGTCTGAACTCTTTCCCTACACGACGCTCTTCCGATCTNNNCCTTGTGTGCATAGAGATGTTCAC
CACACGTCTGAACTCTTTCCCTACACGACGCTCTTCCGATCTNNNCCTTGTGTGCATAGAGATGTTCAC
GCACACGTCTGAACTCTTTCCCTACACGACGCTCTTCCGATCTNNNTTACTGGAGAGTTATTTGTACA
GCACACGTCTGAACTCTTTCCCTACACGACGCTCTTCCGATCTNNNTTACTGGAGAGTTATTTGTACA
GCACACGTCTGAACTCTTTCCCTACACGACGCTCTTCCGATCTNNNTTACTGGAGAGTTATTTGTACA
AGCACACGTCTGAACTCTTTCCCTACACGACGCTCTTCCGATCTNNNAGGCCTGACATATGTTTCACTG
AGCACACGTCTGAACTCTTTCCCTACACGACGCTCTTCCGATCTNNNAGGCCTGACATATGTTTCACTG
AGCACACGTCTGAACTCTTTCCCTACACGACGCTCTTCCGATCTNNNAGGCCTGACATATGTTTCACTG
GAGCACACGTCTGAACTCTTTCCCTACACGACGCTCTTCCGATCTNNNAAGACCAACCTGGACAACAT
GAGCACACGTCTGAACTCTTTCCCTACACGACGCTCTTCCGATCTNNNAAGACCAACCTGGACAACAT
GAGCACACGTCTGAACTCTTTCCCTACACGACGCTCTTCCGATCTNNNAAGACCAACCTGGACAACAT
AGCACACGTCTGAACTCTTTCCCTACACGACGCTCTTCCGATCTNNNGTGCAAGTAAAGAGCCTTAGT
AGCACACGTCTGAACTCTTTCCCTACACGACGCTCTTCCGATCTNNNGTGCAAGTAAAGAGCCTTAGT
AGCACACGTCTGAACTCTTTCCCTACACGACGCTCTTCCGATCTNNNGTGCAAGTAAAGAGCCTTAGT
AGCACACGTCTGAACTCTTTCCCTACACGACGCTCTTCCGATCTNNNCTGTAGTCCCAGCTACTCGAG
AGCACACGTCTGAACTCTTTCCCTACACGACGCTCTTCCGATCTNNNCTGTAGTCCCAGCTACTCGAG
AGCACACGTCTGAACTCTTTCCCTACACGACGCTCTTCCGATCTNNNCTGTAGTCCCAGCTACTCGAG
AGCACACGTCTGAACTCTTTCCCTACACGACGCTCTTCCGATCTNNNACAGTGGAATATGATGCAGCT
AGCACACGTCTGAACTCTTTCCCTACACGACGCTCTTCCGATCTNNNACAGTGGAATATGATGCAGCT
AGCACACGTCTGAACTCTTTCCCTACACGACGCTCTTCCGATCTNNNACAGTGGAATATGATGCAGCT
AGCACACGTCTGAACTCTTTCCCTACACGACGCTCTTCCGATCTNNNCCCACCCACAAAAACCTTTTTA
AGCACACGTCTGAACTCTTTCCCTACACGACGCTCTTCCGATCTNNNCCCACCCACAAAAACCTTTTTA
AGCACACGTCTGAACTCTTTCCCTACACGACGCTCTTCCGATCTNNNCCCACCCACAAAAACCTTTTTA
GCACACGTCTGAACTCTTTCCCTACACGACGCTCTTCCGATCTNNNACTGCACCTTTATCAGAATGGCT
GCACACGTCTGAACTCTTTCCCTACACGACGCTCTTCCGATCTNNNACTGCACCTTTATCAGAATGGCT
GCACACGTCTGAACTCTTTCCCTACACGACGCTCTTCCGATCTNNNACTGCACCTTTATCAGAATGGCT
GCACACGTCTGAACTCTTTCCCTACACGACGCTCTTCCGATCTNNNACATGTCTTGTCACTTTAGACCA
GCACACGTCTGAACTCTTTCCCTACACGACGCTCTTCCGATCTNNNACATGTCTTGTCACTTTAGACCA
GCACACGTCTGAACTCTTTCCCTACACGACGCTCTTCCGATCTNNNACATGTCTTGTCACTTTAGACCA
AGCACACGTCTGAACTCTTTCCCTACACGACGCTCTTCCGATCTNNNTGCTTTCTTCCTTTCGTTCGTCA
AGCACACGTCTGAACTCTTTCCCTACACGACGCTCTTCCGATCTNNNTGCTTTCTTCCTTTCGTTCGTCA
AGCACACGTCTGAACTCTTTCCCTACACGACGCTCTTCCGATCTNNNTGCTTTCTTCCTTTCGTTCGTCA
GCACACGTCTGAACTCTTTCCCTACACGACGCTCTTCCGATCTNNNACACCAGTCATGTTTCCTCTCTCC
GCACACGTCTGAACTCTTTCCCTACACGACGCTCTTCCGATCTNNNACACCAGTCATGTTTCCTCTCTCC
GCACACGTCTGAACTCTTTCCCTACACGACGCTCTTCCGATCTNNNACACCAGTCATGTTTCCTCTCTCC
AGCACACGTCTGAACTCTTTCCCTACACGACGCTCTTCCGATCTNNNTAAATGAGTTCCCTGGGTTACC
AGCACACGTCTGAACTCTTTCCCTACACGACGCTCTTCCGATCTNNNTAAATGAGTTCCCTGGGTTACC

AGCACACGTCTGAACTCTTTCCCTACACGACGCTCTTCCGATCTNNNTAAATGAGTTCCCTGGGTTACC
GAGCACACGTCTGAACTCTTTCCCTACACGACGCTCTTCCGATCTNNNGGCAACAGAGTAAGACTCCA
GAGCACACGTCTGAACTCTTTCCCTACACGACGCTCTTCCGATCTNNNGGCAACAGAGTAAGACTCCA
GAGCACACGTCTGAACTCTTTCCCTACACGACGCTCTTCCGATCTNNNGGCAACAGAGTAAGACTCCA
GAGCACACGTCTGAACTCTTTCCCTACACGACGCTCTTCCGATCTNNNCTGAGGCAGGAGAATCACTT
GAGCACACGTCTGAACTCTTTCCCTACACGACGCTCTTCCGATCTNNNCTGAGGCAGGAGAATCACTT
GAGCACACGTCTGAACTCTTTCCCTACACGACGCTCTTCCGATCTNNNCTGAGGCAGGAGAATCACTT
AGCACACGTCTGAACTCTTTCCCTACACGACGCTCTTCCGATCTNNNAAGTGCCCTTGAGACCTAAGA
AGCACACGTCTGAACTCTTTCCCTACACGACGCTCTTCCGATCTNNNAAGTGCCCTTGAGACCTAAGA
AGCACACGTCTGAACTCTTTCCCTACACGACGCTCTTCCGATCTNNNAAGTGCCCTTGAGACCTAAGA
GCACACGTCTGAACTCTTTCCCTACACGACGCTCTTCCGATCTNNNAAATATGCCACTGTTGTCTGTGC
GCACACGTCTGAACTCTTTCCCTACACGACGCTCTTCCGATCTNNNAAATATGCCACTGTTGTCTGTGC
GCACACGTCTGAACTCTTTCCCTACACGACGCTCTTCCGATCTNNNAAATATGCCACTGTTGTCTGTGC
GAGCACACGTCTGAACTCTTTCCCTACACGACGCTCTTCCGATCTNNNTGTTCTGGTTTGCGTTGCATTA
GAGCACACGTCTGAACTCTTTCCCTACACGACGCTCTTCCGATCTNNNTGTTCTGGTTTGCGTTGCATTA
GAGCACACGTCTGAACTCTTTCCCTACACGACGCTCTTCCGATCTNNNTGTTCTGGTTTGCGTTGCATTA
GAGCACACGTCTGAACTCTTTCCCTACACGACGCTCTTCCGATCTNNNCTCAGCCATACCCAATATACCC
GAGCACACGTCTGAACTCTTTCCCTACACGACGCTCTTCCGATCTNNNCTCAGCCATACCCAATATACCC
GAGCACACGTCTGAACTCTTTCCCTACACGACGCTCTTCCGATCTNNNCTCAGCCATACCCAATATACCC
AGCACACGTCTGAACTCTTTCCCTACACGACGCTCTTCCGATCTNNNTTTGGAAGTTAGGCTCTCAGGA
AGCACACGTCTGAACTCTTTCCCTACACGACGCTCTTCCGATCTNNNTTTGGAAGTTAGGCTCTCAGGA
AGCACACGTCTGAACTCTTTCCCTACACGACGCTCTTCCGATCTNNNTTTGGAAGTTAGGCTCTCAGGA
AGCACACGTCTGAACTCTTTCCCTACACGACGCTCTTCCGATCTNNNGGTTGCAGTGAGCTGTGATTG
AGCACACGTCTGAACTCTTTCCCTACACGACGCTCTTCCGATCTNNNGGTTGCAGTGAGCTGTGATTG
AGCACACGTCTGAACTCTTTCCCTACACGACGCTCTTCCGATCTNNNGGTTGCAGTGAGCTGTGATTG
AGCACACGTCTGAACTCTTTCCCTACACGACGCTCTTCCGATCTNNNATCTTTGAATCTGGGAGGCAGA
AGCACACGTCTGAACTCTTTCCCTACACGACGCTCTTCCGATCTNNNATCTTTGAATCTGGGAGGCAGA
AGCACACGTCTGAACTCTTTCCCTACACGACGCTCTTCCGATCTNNNATCTTTGAATCTGGGAGGCAGA
AGCACACGTCTGAACTCTTTCCCTACACGACGCTCTTCCGATCTNNNCCGAGTAGCTGGGACTACAGG
AGCACACGTCTGAACTCTTTCCCTACACGACGCTCTTCCGATCTNNNCCGAGTAGCTGGGACTACAGG
AGCACACGTCTGAACTCTTTCCCTACACGACGCTCTTCCGATCTNNNCCGAGTAGCTGGGACTACAGG
GAGCACACGTCTGAACTCTTTCCCTACACGACGCTCTTCCGATCTNNNTCAATGAATACTTGGCGGACA
GAGCACACGTCTGAACTCTTTCCCTACACGACGCTCTTCCGATCTNNNTCAATGAATACTTGGCGGACA
GAGCACACGTCTGAACTCTTTCCCTACACGACGCTCTTCCGATCTNNNTCAATGAATACTTGGCGGACA
AGCACACGTCTGAACTCTTTCCCTACACGACGCTCTTCCGATCTNNNCAATCTGGGTTCAAGCGATTCT
AGCACACGTCTGAACTCTTTCCCTACACGACGCTCTTCCGATCTNNNCAATCTGGGTTCAAGCGATTCT
AGCACACGTCTGAACTCTTTCCCTACACGACGCTCTTCCGATCTNNNCAATCTGGGTTCAAGCGATTCT
AGAGCACACGTCTGAACTCTTTCCCTACACGACGCTCTTCCGATCTNNNACTAGGGTATCTTGTTGCAG
AGAGCACACGTCTGAACTCTTTCCCTACACGACGCTCTTCCGATCTNNNACTAGGGTATCTTGTTGCAG
AGAGCACACGTCTGAACTCTTTCCCTACACGACGCTCTTCCGATCTNNNACTAGGGTATCTTGTTGCAG
GAGCACACGTCTGAACTCTTTCCCTACACGACGCTCTTCCGATCTNNNCCAGCCTGAGTGACAAAGTTA
GAGCACACGTCTGAACTCTTTCCCTACACGACGCTCTTCCGATCTNNNCCAGCCTGAGTGACAAAGTTA
GAGCACACGTCTGAACTCTTTCCCTACACGACGCTCTTCCGATCTNNNCCAGCCTGAGTGACAAAGTTA

GCACACGTCTGAACTCTTTCCCTACACGACGCTCTTCCGATCTNNNAAGTTCTTTTCAGGGCAATCCG
GCACACGTCTGAACTCTTTCCCTACACGACGCTCTTCCGATCTNNNAAGTTCTTTTCAGGGCAATCCG
GCACACGTCTGAACTCTTTCCCTACACGACGCTCTTCCGATCTNNNAAGTTCTTTTCAGGGCAATCCG
GCACACGTCTGAACTCTTTCCCTACACGACGCTCTTCCGATCTNNNCAGGTGCCCATTTATACGTTTCC
GCACACGTCTGAACTCTTTCCCTACACGACGCTCTTCCGATCTNNNCAGGTGCCCATTTATACGTTTCC
GCACACGTCTGAACTCTTTCCCTACACGACGCTCTTCCGATCTNNNCAGGTGCCCATTTATACGTTTCC
AGCACACGTCTGAACTCTTTCCCTACACGACGCTCTTCCGATCTNNNGAACAGAGGACTAGTTGAGG
AGCACACGTCTGAACTCTTTCCCTACACGACGCTCTTCCGATCTNNNGAACAGAGGACTAGTTGAGG
AGCACACGTCTGAACTCTTTCCCTACACGACGCTCTTCCGATCTNNNGAACAGAGGACTAGTTGAGG
GCACACGTCTGAACTCTTTCCCTACACGACGCTCTTCCGATCTNNNTTTGTCTGCCTCTGTCACACAT
GCACACGTCTGAACTCTTTCCCTACACGACGCTCTTCCGATCTNNNTTTGTCTGCCTCTGTCACACAT
GCACACGTCTGAACTCTTTCCCTACACGACGCTCTTCCGATCTNNNTTTGTCTGCCTCTGTCACACAT
AGCACACGTCTGAACTCTTTCCCTACACGACGCTCTTCCGATCTNNNATTCCTAATGACTTCCCTGAGG
AGCACACGTCTGAACTCTTTCCCTACACGACGCTCTTCCGATCTNNNATTCCTAATGACTTCCCTGAGG
AGCACACGTCTGAACTCTTTCCCTACACGACGCTCTTCCGATCTNNNATTCCTAATGACTTCCCTGAGG
AGCACACGTCTGAACTCTTTCCCTACACGACGCTCTTCCGATCTNNNAAGTAGGAGACACATCAGCAG
AGCACACGTCTGAACTCTTTCCCTACACGACGCTCTTCCGATCTNNNAAGTAGGAGACACATCAGCAG
AGCACACGTCTGAACTCTTTCCCTACACGACGCTCTTCCGATCTNNNAAGTAGGAGACACATCAGCAG
AGCACACGTCTGAACTCTTTCCCTACACGACGCTCTTCCGATCTNNNAGGAGAGAGAGAGGAGAGA
AGCACACGTCTGAACTCTTTCCCTACACGACGCTCTTCCGATCTNNNAGGAGAGAGAGAGGAGAGA
AGCACACGTCTGAACTCTTTCCCTACACGACGCTCTTCCGATCTNNNAGGAGAGAGAGAGGAGAGA
GAGCACACGTCTGAACTCTTTCCCTACACGACGCTCTTCCGATCTNNNCAGGAGATGGAGGCTACAA
GAGCACACGTCTGAACTCTTTCCCTACACGACGCTCTTCCGATCTNNNCAGGAGATGGAGGCTACAA
GAGCACACGTCTGAACTCTTTCCCTACACGACGCTCTTCCGATCTNNNCAGGAGATGGAGGCTACAA
CACACGTCTGAACTCTTTCCCTACACGACGCTCTTCCGATCTNNNAAAGCTTTGAGAATGTTCTTAAA
CACACGTCTGAACTCTTTCCCTACACGACGCTCTTCCGATCTNNNAAAGCTTTGAGAATGTTCTTAAA
CACACGTCTGAACTCTTTCCCTACACGACGCTCTTCCGATCTNNNAAAGCTTTGAGAATGTTCTTAAA
AGCACACGTCTGAACTCTTTCCCTACACGACGCTCTTCCGATCTNNNTCCAAGCAATGTACTCAGTCAC
AGCACACGTCTGAACTCTTTCCCTACACGACGCTCTTCCGATCTNNNTCCAAGCAATGTACTCAGTCAC
AGCACACGTCTGAACTCTTTCCCTACACGACGCTCTTCCGATCTNNNTCCAAGCAATGTACTCAGTCAC
AGAGCACACGTCTGAACTCTTTCCCTACACGACGCTCTTCCGATCTNNNATCCCCCTATTGCCTTTTCT
AGAGCACACGTCTGAACTCTTTCCCTACACGACGCTCTTCCGATCTNNNATCCCCCTATTGCCTTTTCT
AGAGCACACGTCTGAACTCTTTCCCTACACGACGCTCTTCCGATCTNNNATCCCCCTATTGCCTTTTCT
CACACGTCTGAACTCTTTCCCTACACGACGCTCTTCCGATCTNNNGTCGTAGGACAAAGGGCCCCCAT
CACACGTCTGAACTCTTTCCCTACACGACGCTCTTCCGATCTNNNGTCGTAGGACAAAGGGCCCCCAT
CACACGTCTGAACTCTTTCCCTACACGACGCTCTTCCGATCTNNNGTCGTAGGACAAAGGGCCCCCAT
GAGCACACGTCTGAACTCTTTCCCTACACGACGCTCTTCCGATCTNNNTCTCCTCTCCTTCTCCTTCTTCT
GAGCACACGTCTGAACTCTTTCCCTACACGACGCTCTTCCGATCTNNNTCTCCTCTCCTTCTCCTTCTTCT
GAGCACACGTCTGAACTCTTTCCCTACACGACGCTCTTCCGATCTNNNTCTCCTCTCCTTCTCCTTCTTCT
AGCACACGTCTGAACTCTTTCCCTACACGACGCTCTTCCGATCTNNNTCTCTTGCTGTTTTCAGTGTCCT
AGCACACGTCTGAACTCTTTCCCTACACGACGCTCTTCCGATCTNNNTCTCTTGCTGTTTTCAGTGTCCT
AGCACACGTCTGAACTCTTTCCCTACACGACGCTCTTCCGATCTNNNTCTCTTGCTGTTTTCAGTGTCCT
GAGCACACGTCTGAACTCTTTCCCTACACGACGCTCTTCCGATCTNNNTGGTGTACAGGCTAGAAAGC

AGCACACGTCTGAACTCTTTCCCTACACGACGCTCTTCCGATCTNNNTGGTGTACAGGCTAGAAAGC
AGCACACGTCTGAACTCTTTCCCTACACGACGCTCTTCCGATCTNNNTGGTGTACAGGCTAGAAAGC
GAGCACACGTCTGAACTCTTTCCCTACACGACGCTCTTCCGATCTNNNAGTATTGGCAAGAGAAGTG
GAGCACACGTCTGAACTCTTTCCCTACACGACGCTCTTCCGATCTNNNAGTATTGGCAAGAGAAGTG
GAGCACACGTCTGAACTCTTTCCCTACACGACGCTCTTCCGATCTNNNAGTATTGGCAAGAGAAGTG
AGCACACGTCTGAACTCTTTCCCTACACGACGCTCTTCCGATCTNNNGTTTCACCTCCTATCAGACACG
AGCACACGTCTGAACTCTTTCCCTACACGACGCTCTTCCGATCTNNNGTTTCACCTCCTATCAGACACG
AGCACACGTCTGAACTCTTTCCCTACACGACGCTCTTCCGATCTNNNGTTTCACCTCCTATCAGACACG
AGCACACGTCTGAACTCTTTCCCTACACGACGCTCTTCCGATCTNNNAGGTTGCAGTGAGCTGAGAT
AGCACACGTCTGAACTCTTTCCCTACACGACGCTCTTCCGATCTNNNAGGTTGCAGTGAGCTGAGAT
AGCACACGTCTGAACTCTTTCCCTACACGACGCTCTTCCGATCTNNNAGGTTGCAGTGAGCTGAGAT
GCACACGTCTGAACTCTTTCCCTACACGACGCTCTTCCGATCTNNNGCTGTTGTCAAACATCTGGTTC
GCACACGTCTGAACTCTTTCCCTACACGACGCTCTTCCGATCTNNNGCTGTTGTCAAACATCTGGTTC
GCACACGTCTGAACTCTTTCCCTACACGACGCTCTTCCGATCTNNNGCTGTTGTCAAACATCTGGTTC
AGCACACGTCTGAACTCTTTCCCTACACGACGCTCTTCCGATCTNNNAGTTGAGCTACGGAGACAGA
AGCACACGTCTGAACTCTTTCCCTACACGACGCTCTTCCGATCTNNNAGTTGAGCTACGGAGACAGA
AGCACACGTCTGAACTCTTTCCCTACACGACGCTCTTCCGATCTNNNAGTTGAGCTACGGAGACAGA
GCACACGTCTGAACTCTTTCCCTACACGACGCTCTTCCGATCTNNNTCTACACAGAACTCAATGGCCTC
GCACACGTCTGAACTCTTTCCCTACACGACGCTCTTCCGATCTNNNTCTACACAGAACTCAATGGCCTC
GCACACGTCTGAACTCTTTCCCTACACGACGCTCTTCCGATCTNNNTCTACACAGAACTCAATGGCCTC
GCACACGTCTGAACTCTTTCCCTACACGACGCTCTTCCGATCTNNNGGTGAATTACTTTGGTGGCATG
GCACACGTCTGAACTCTTTCCCTACACGACGCTCTTCCGATCTNNNGGTGAATTACTTTGGTGGCATG
GCACACGTCTGAACTCTTTCCCTACACGACGCTCTTCCGATCTNNNGGTGAATTACTTTGGTGGCATG
AGCACACGTCTGAACTCTTTCCCTACACGACGCTCTTCCGATCTNNNTGAGCAGGAATAAAGTGACCT
AGCACACGTCTGAACTCTTTCCCTACACGACGCTCTTCCGATCTNNNTGAGCAGGAATAAAGTGACCT
AGCACACGTCTGAACTCTTTCCCTACACGACGCTCTTCCGATCTNNNTGAGCAGGAATAAAGTGACCT
GCACACGTCTGAACTCTTTCCCTACACGACGCTCTTCCGATCTNNNCTGCTACCACTTTGTAACTTGCC
GCACACGTCTGAACTCTTTCCCTACACGACGCTCTTCCGATCTNNNCTGCTACCACTTTGTAACTTGCC
GCACACGTCTGAACTCTTTCCCTACACGACGCTCTTCCGATCTNNNCTGCTACCACTTTGTAACTTGCC
AGCACACGTCTGAACTCTTTCCCTACACGACGCTCTTCCGATCTNNNCTTCCAGAAGTCTTGGCCAAAT
AGCACACGTCTGAACTCTTTCCCTACACGACGCTCTTCCGATCTNNNCTTCCAGAAGTCTTGGCCAAAT
AGCACACGTCTGAACTCTTTCCCTACACGACGCTCTTCCGATCTNNNCTTCCAGAAGTCTTGGCCAAAT
CACACGTCTGAACTCTTTCCCTACACGACGCTCTTCCGATCTNNNAATCATCTTTGTTGATTTCATCAC
CACACGTCTGAACTCTTTCCCTACACGACGCTCTTCCGATCTNNNAATCATCTTTGTTGATTTCATCAC
CACACGTCTGAACTCTTTCCCTACACGACGCTCTTCCGATCTNNNAATCATCTTTGTTGATTTCATCAC
AGCACACGTCTGAACTCTTTCCCTACACGACGCTCTTCCGATCTNNNGGGTGTGGTGGTGCATAACT
AGCACACGTCTGAACTCTTTCCCTACACGACGCTCTTCCGATCTNNNGGGTGTGGTGGTGCATAACT
AGCACACGTCTGAACTCTTTCCCTACACGACGCTCTTCCGATCTNNNGGGTGTGGTGGTGCATAACT
AGCACACGTCTGAACTCTTTCCCTACACGACGCTCTTCCGATCTNNNTTCAAGGATGCAGTGAGTGAT
AGCACACGTCTGAACTCTTTCCCTACACGACGCTCTTCCGATCTNNNTTCAAGGATGCAGTGAGTGAT
AGCACACGTCTGAACTCTTTCCCTACACGACGCTCTTCCGATCTNNNTTCAAGGATGCAGTGAGTGAT
AGCACACGTCTGAACTCTTTCCCTACACGACGCTCTTCCGATCTNNNGCTCTGATCATGGCAAGTATG
AGCACACGTCTGAACTCTTTCCCTACACGACGCTCTTCCGATCTNNNGCTCTGATCATGGCAAGTATG

AGCACACGTCTGAACTCTTTCCCTACACGACGCTCTTCCGATCTNNNGCTCTGATCATGGCAAGTATG
AGCACACGTCTGAACTCTTTCCCTACACGACGCTCTTCCGATCTNNNTTATGTCATTCCTCGGCATCAG
AGCACACGTCTGAACTCTTTCCCTACACGACGCTCTTCCGATCTNNNTTATGTCATTCCTCGGCATCAG
AGCACACGTCTGAACTCTTTCCCTACACGACGCTCTTCCGATCTNNNTTATGTCATTCCTCGGCATCAG
AGCACACGTCTGAACTCTTTCCCTACACGACGCTCTTCCGATCTNNNTACCAAAAAATTAGCTGGGC
AGCACACGTCTGAACTCTTTCCCTACACGACGCTCTTCCGATCTNNNTACCAAAAAATTAGCTGGGC
AGCACACGTCTGAACTCTTTCCCTACACGACGCTCTTCCGATCTNNNTACCAAAAAATTAGCTGGGC
GCACACGTCTGAACTCTTTCCCTACACGACGCTCTTCCGATCTNNNTTTGGCTGTAGTTTGGTTTCCA
GCACACGTCTGAACTCTTTCCCTACACGACGCTCTTCCGATCTNNNTTTGGCTGTAGTTTGGTTTCCA
GCACACGTCTGAACTCTTTCCCTACACGACGCTCTTCCGATCTNNNTTTGGCTGTAGTTTGGTTTCCA
GAGCACACGTCTGAACTCTTTCCCTACACGACGCTCTTCCGATCTNNNTGAGGCAGGAGAATCACTTG
GAGCACACGTCTGAACTCTTTCCCTACACGACGCTCTTCCGATCTNNNTGAGGCAGGAGAATCACTTG
GAGCACACGTCTGAACTCTTTCCCTACACGACGCTCTTCCGATCTNNNTGAGGCAGGAGAATCACTTG
GAGCACACGTCTGAACTCTTTCCCTACACGACGCTCTTCCGATCTNNNGTTAGGAGTTCTAGACCAGC
GAGCACACGTCTGAACTCTTTCCCTACACGACGCTCTTCCGATCTNNNGTTAGGAGTTCTAGACCAGC
GAGCACACGTCTGAACTCTTTCCCTACACGACGCTCTTCCGATCTNNNGTTAGGAGTTCTAGACCAGC
GAGCACACGTCTGAACTCTTTCCCTACACGACGCTCTTCCGATCTNNNCGGGGCTGAGGAAGAAAGA
GAGCACACGTCTGAACTCTTTCCCTACACGACGCTCTTCCGATCTNNNCGGGGCTGAGGAAGAAAGA
GAGCACACGTCTGAACTCTTTCCCTACACGACGCTCTTCCGATCTNNNCGGGGCTGAGGAAGAAAGA
GCACACGTCTGAACTCTTTCCCTACACGACGCTCTTCCGATCTNNNCTCCTGAGTAGCTGGGACTACA
GCACACGTCTGAACTCTTTCCCTACACGACGCTCTTCCGATCTNNNCTCCTGAGTAGCTGGGACTACA
GCACACGTCTGAACTCTTTCCCTACACGACGCTCTTCCGATCTNNNCTCCTGAGTAGCTGGGACTACA
AGCACACGTCTGAACTCTTTCCCTACACGACGCTCTTCCGATCTNNNAGGAGAATGGCGTGAACCTGC
AGCACACGTCTGAACTCTTTCCCTACACGACGCTCTTCCGATCTNNNAGGAGAATGGCGTGAACCTGC
AGCACACGTCTGAACTCTTTCCCTACACGACGCTCTTCCGATCTNNNAGGAGAATGGCGTGAACCTGC
GCACACGTCTGAACTCTTTCCCTACACGACGCTCTTCCGATCTNNNTGGATTGTATGATTCCCCAGTG
GCACACGTCTGAACTCTTTCCCTACACGACGCTCTTCCGATCTNNNTGGATTGTATGATTCCCCAGTG
GCACACGTCTGAACTCTTTCCCTACACGACGCTCTTCCGATCTNNNTGGATTGTATGATTCCCCAGTG
GAGCACACGTCTGAACTCTTTCCCTACACGACGCTCTTCCGATCTNNNAAAGCACAACACACATGAAC
GAGCACACGTCTGAACTCTTTCCCTACACGACGCTCTTCCGATCTNNNAAAGCACAACACACATGAAC
GAGCACACGTCTGAACTCTTTCCCTACACGACGCTCTTCCGATCTNNNAAAGCACAACACACATGAAC
GAGCACACGTCTGAACTCTTTCCCTACACGACGCTCTTCCGATCTNNNTCGATCTCCTGACCTCATGAT
GAGCACACGTCTGAACTCTTTCCCTACACGACGCTCTTCCGATCTNNNTCGATCTCCTGACCTCATGAT
GAGCACACGTCTGAACTCTTTCCCTACACGACGCTCTTCCGATCTNNNTCGATCTCCTGACCTCATGAT
AGCACACGTCTGAACTCTTTCCCTACACGACGCTCTTCCGATCTNNNTTCCTCCTCTTCATCTTCACTGC
AGCACACGTCTGAACTCTTTCCCTACACGACGCTCTTCCGATCTNNNTTCCTCCTCTTCATCTTCACTGC
AGCACACGTCTGAACTCTTTCCCTACACGACGCTCTTCCGATCTNNNTTCCTCCTCTTCATCTTCACTGC
AGCACACGTCTGAACTCTTTCCCTACACGACGCTCTTCCGATCTNNNTAGTCCAGCTACTCAGGAGGC
AGCACACGTCTGAACTCTTTCCCTACACGACGCTCTTCCGATCTNNNTAGTCCAGCTACTCAGGAGGC
AGCACACGTCTGAACTCTTTCCCTACACGACGCTCTTCCGATCTNNNTAGTCCAGCTACTCAGGAGGC
AGCACACGTCTGAACTCTTTCCCTACACGACGCTCTTCCGATCTNNNATTAGCTTCTCTCTTCTGGCTC
AGCACACGTCTGAACTCTTTCCCTACACGACGCTCTTCCGATCTNNNATTAGCTTCTCTCTTCTGGCTC
AGCACACGTCTGAACTCTTTCCCTACACGACGCTCTTCCGATCTNNNATTAGCTTCTCTCTTCTGGCTC

AGCACACGTCTGAACTCTTTCCCTACACGACGCTCTTCCGATCTNNNCAACTTTCTTGACCAGCTTTCCC
AGCACACGTCTGAACTCTTTCCCTACACGACGCTCTTCCGATCTNNNCAACTTTCTTGACCAGCTTTCCC
AGCACACGTCTGAACTCTTTCCCTACACGACGCTCTTCCGATCTNNNCAACTTTCTTGACCAGCTTTCCC
AGAGCACACGTCTGAACTCTTTCCCTACACGACGCTCTTCCGATCTNNNCAATTGCACCCAGTGTAGGA
AGAGCACACGTCTGAACTCTTTCCCTACACGACGCTCTTCCGATCTNNNCAATTGCACCCAGTGTAGGA
AGAGCACACGTCTGAACTCTTTCCCTACACGACGCTCTTCCGATCTNNNCAATTGCACCCAGTGTAGGA
AGCACACGTCTGAACTCTTTCCCTACACGACGCTCTTCCGATCTNNNTTCCGCTCATCAGATTTTCCAGG
AGCACACGTCTGAACTCTTTCCCTACACGACGCTCTTCCGATCTNNNTTCCGCTCATCAGATTTTCCAGG
AGCACACGTCTGAACTCTTTCCCTACACGACGCTCTTCCGATCTNNNTTCCGCTCATCAGATTTTCCAGG
AGCACACGTCTGAACTCTTTCCCTACACGACGCTCTTCCGATCTNNNGAGGTGACACGACAGTTTTCAA
AGCACACGTCTGAACTCTTTCCCTACACGACGCTCTTCCGATCTNNNGAGGTGACACGACAGTTTTCAA
AGCACACGTCTGAACTCTTTCCCTACACGACGCTCTTCCGATCTNNNGAGGTGACACGACAGTTTTCAA
GAGCACACGTCTGAACTCTTTCCCTACACGACGCTCTTCCGATCTNNNCCTGTGTCCTTTCTCACTTTCCT
GAGCACACGTCTGAACTCTTTCCCTACACGACGCTCTTCCGATCTNNNCCTGTGTCCTTTCTCACTTTCCT
GAGCACACGTCTGAACTCTTTCCCTACACGACGCTCTTCCGATCTNNNCCTGTGTCCTTTCTCACTTTCCT
AGCACACGTCTGAACTCTTTCCCTACACGACGCTCTTCCGATCTNNNTTAGGATGGGAGGATGGTGGA
AGCACACGTCTGAACTCTTTCCCTACACGACGCTCTTCCGATCTNNNTTAGGATGGGAGGATGGTGGA
AGCACACGTCTGAACTCTTTCCCTACACGACGCTCTTCCGATCTNNNTTAGGATGGGAGGATGGTGGA
GAGCACACGTCTGAACTCTTTCCCTACACGACGCTCTTCCGATCTNNNAAAAAATTAGCTGAGCGTGGT
GAGCACACGTCTGAACTCTTTCCCTACACGACGCTCTTCCGATCTNNNAAAAAATTAGCTGAGCGTGGT
GAGCACACGTCTGAACTCTTTCCCTACACGACGCTCTTCCGATCTNNNAAAAAATTAGCTGAGCGTGGT
AGCACACGTCTGAACTCTTTCCCTACACGACGCTCTTCCGATCTNNNGATGGACAAAACAGTGTGTGG
AGCACACGTCTGAACTCTTTCCCTACACGACGCTCTTCCGATCTNNNGATGGACAAAACAGTGTGTGG
AGCACACGTCTGAACTCTTTCCCTACACGACGCTCTTCCGATCTNNNGATGGACAAAACAGTGTGTGG
AGCACACGTCTGAACTCTTTCCCTACACGACGCTCTTCCGATCTNNNCCCAAATTGGAGCTAAACTGAA
AGCACACGTCTGAACTCTTTCCCTACACGACGCTCTTCCGATCTNNNCCCAAATTGGAGCTAAACTGAA
AGCACACGTCTGAACTCTTTCCCTACACGACGCTCTTCCGATCTNNNCCCAAATTGGAGCTAAACTGAA
GAGCACACGTCTGAACTCTTTCCCTACACGACGCTCTTCCGATCTNNNATCTGGGTGACAAGAGCAAG
GAGCACACGTCTGAACTCTTTCCCTACACGACGCTCTTCCGATCTNNNATCTGGGTGACAAGAGCAAG
GAGCACACGTCTGAACTCTTTCCCTACACGACGCTCTTCCGATCTNNNATCTGGGTGACAAGAGCAAG
AGCACACGTCTGAACTCTTTCCCTACACGACGCTCTTCCGATCTNNNGGTGACACTGTACTCAGGTTAC
AGCACACGTCTGAACTCTTTCCCTACACGACGCTCTTCCGATCTNNNGGTGACACTGTACTCAGGTTAC
AGCACACGTCTGAACTCTTTCCCTACACGACGCTCTTCCGATCTNNNGGTGACACTGTACTCAGGTTAC
AGCACACGTCTGAACTCTTTCCCTACACGACGCTCTTCCGATCTNNNGATGGTCTCAATCTCCTGACCTC
AGCACACGTCTGAACTCTTTCCCTACACGACGCTCTTCCGATCTNNNGATGGTCTCAATCTCCTGACCTC
AGCACACGTCTGAACTCTTTCCCTACACGACGCTCTTCCGATCTNNNGATGGTCTCAATCTCCTGACCTC
GAGCACACGTCTGAACTCTTTCCCTACACGACGCTCTTCCGATCTNNNCTGGCCAACATAGTGAAACCT
GAGCACACGTCTGAACTCTTTCCCTACACGACGCTCTTCCGATCTNNNCTGGCCAACATAGTGAAACCT
GAGCACACGTCTGAACTCTTTCCCTACACGACGCTCTTCCGATCTNNNCTGGCCAACATAGTGAAACCT
AGCACACGTCTGAACTCTTTCCCTACACGACGCTCTTCCGATCTNNNTAAACACAAAGTCCTACGCCTG
AGCACACGTCTGAACTCTTTCCCTACACGACGCTCTTCCGATCTNNNTAAACACAAAGTCCTACGCCTG
AGCACACGTCTGAACTCTTTCCCTACACGACGCTCTTCCGATCTNNNTAAACACAAAGTCCTACGCCTG
GAGCACACGTCTGAACTCTTTCCCTACACGACGCTCTTCCGATCTNNNACATACCTGTAGTCCCAGCTA

GAGCACACGTCTGAACTCTTTCCCTACACGACGCTCTTCCGATCTNNNACATACCTGTAGTCCCAGCTA
GAGCACACGTCTGAACTCTTTCCCTACACGACGCTCTTCCGATCTNNNACATACCTGTAGTCCCAGCTA
AGCACACGTCTGAACTCTTTCCCTACACGACGCTCTTCCGATCTNNNATGTGGTGTTGGTTGGTTCT
AGCACACGTCTGAACTCTTTCCCTACACGACGCTCTTCCGATCTNNNATGTGGTGTTGGTTGGTTCT
AGCACACGTCTGAACTCTTTCCCTACACGACGCTCTTCCGATCTNNNATGTGGTGTTGGTTGGTTCT
GAGCACACGTCTGAACTCTTTCCCTACACGACGCTCTTCCGATCTNNNCTTACACTGCTGAAGCGAAA
GAGCACACGTCTGAACTCTTTCCCTACACGACGCTCTTCCGATCTNNNCTTACACTGCTGAAGCGAAA
GAGCACACGTCTGAACTCTTTCCCTACACGACGCTCTTCCGATCTNNNCTTACACTGCTGAAGCGAAA
AGCACACGTCTGAACTCTTTCCCTACACGACGCTCTTCCGATCTNNNAAAAGGTCAGTGGCCTCCAAA
AGCACACGTCTGAACTCTTTCCCTACACGACGCTCTTCCGATCTNNNAAAAGGTCAGTGGCCTCCAAA
AGCACACGTCTGAACTCTTTCCCTACACGACGCTCTTCCGATCTNNNAAAAGGTCAGTGGCCTCCAAA
GCACACGTCTGAACTCTTTCCCTACACGACGCTCTTCCGATCTNNNTGGTCTCCCAAAAAGAGGATCT
GCACACGTCTGAACTCTTTCCCTACACGACGCTCTTCCGATCTNNNTGGTCTCCCAAAAAGAGGATCT
GCACACGTCTGAACTCTTTCCCTACACGACGCTCTTCCGATCTNNNTGGTCTCCCAAAAAGAGGATCT
GCACACGTCTGAACTCTTTCCCTACACGACGCTCTTCCGATCTNNNGCAAAAGGCAGCAGCAAAATT
GCACACGTCTGAACTCTTTCCCTACACGACGCTCTTCCGATCTNNNGCAAAAGGCAGCAGCAAAATT
GCACACGTCTGAACTCTTTCCCTACACGACGCTCTTCCGATCTNNNGCAAAAGGCAGCAGCAAAATT
AGCACACGTCTGAACTCTTTCCCTACACGACGCTCTTCCGATCTNNNCCTTATTTTCCCCCTGGTCCTA
AGCACACGTCTGAACTCTTTCCCTACACGACGCTCTTCCGATCTNNNCCTTATTTTCCCCCTGGTCCTA
AGCACACGTCTGAACTCTTTCCCTACACGACGCTCTTCCGATCTNNNCCTTATTTTCCCCCTGGTCCTA
GAGCACACGTCTGAACTCTTTCCCTACACGACGCTCTTCCGATCTNNNACATGCACACACACACACATG
GAGCACACGTCTGAACTCTTTCCCTACACGACGCTCTTCCGATCTNNNACATGCACACACACACACATG
GAGCACACGTCTGAACTCTTTCCCTACACGACGCTCTTCCGATCTNNNACATGCACACACACACACATG
AGCACACGTCTGAACTCTTTCCCTACACGACGCTCTTCCGATCTNNNGTACTATATGGGGTAGCCACA
AGCACACGTCTGAACTCTTTCCCTACACGACGCTCTTCCGATCTNNNGTACTATATGGGGTAGCCACA
AGCACACGTCTGAACTCTTTCCCTACACGACGCTCTTCCGATCTNNNGTACTATATGGGGTAGCCACA
CACACGTCTGAACTCTTTCCCTACACGACGCTCTTCCGATCTNNNCCCACAGGTTTTAGCTCCCTTTTC
CACACGTCTGAACTCTTTCCCTACACGACGCTCTTCCGATCTNNNCCCACAGGTTTTAGCTCCCTTTTC
CACACGTCTGAACTCTTTCCCTACACGACGCTCTTCCGATCTNNNCCCACAGGTTTTAGCTCCCTTTTC
AGCACACGTCTGAACTCTTTCCCTACACGACGCTCTTCCGATCTNNNTCATCCTGGCTAACATGGTGAA
AGCACACGTCTGAACTCTTTCCCTACACGACGCTCTTCCGATCTNNNTCATCCTGGCTAACATGGTGAA
AGCACACGTCTGAACTCTTTCCCTACACGACGCTCTTCCGATCTNNNTCATCCTGGCTAACATGGTGAA
AGCACACGTCTGAACTCTTTCCCTACACGACGCTCTTCCGATCTNNNTCCGAGGGAATCCTGTATGTTT
AGCACACGTCTGAACTCTTTCCCTACACGACGCTCTTCCGATCTNNNTCCGAGGGAATCCTGTATGTTT
AGCACACGTCTGAACTCTTTCCCTACACGACGCTCTTCCGATCTNNNTCCGAGGGAATCCTGTATGTTT
AGCACACGTCTGAACTCTTTCCCTACACGACGCTCTTCCGATCTNNNTTTAGGAGAGACGGGGTTTC
AGCACACGTCTGAACTCTTTCCCTACACGACGCTCTTCCGATCTNNNTTTAGGAGAGACGGGGTTTC
AGCACACGTCTGAACTCTTTCCCTACACGACGCTCTTCCGATCTNNNTTTAGGAGAGACGGGGTTTC
AGCACACGTCTGAACTCTTTCCCTACACGACGCTCTTCCGATCTNNNCTATGAGGTGTGTCCCTGTGG
AGCACACGTCTGAACTCTTTCCCTACACGACGCTCTTCCGATCTNNNCTATGAGGTGTGTCCCTGTGG
AGCACACGTCTGAACTCTTTCCCTACACGACGCTCTTCCGATCTNNNCTATGAGGTGTGTCCCTGTGG
GAGCACACGTCTGAACTCTTTCCCTACACGACGCTCTTCCGATCTNNNACATACCTGTAGTCCCAGCTA
GAGCACACGTCTGAACTCTTTCCCTACACGACGCTCTTCCGATCTNNNACATACCTGTAGTCCCAGCTA

GAGCACACGTCTGAACTCTTTCCCTACACGACGCTCTTCCGATCTNNNACATACCTGTAGTCCCAGCTA
AGCACACGTCTGAACTCTTTCCCTACACGACGCTCTTCCGATCTNNNCACCTCCACACCTGTATAAATG
AGCACACGTCTGAACTCTTTCCCTACACGACGCTCTTCCGATCTNNNCACCTCCACACCTGTATAAATG
AGCACACGTCTGAACTCTTTCCCTACACGACGCTCTTCCGATCTNNNCACCTCCACACCTGTATAAATG
ACACGTCTGAACTCTTTCCCTACACGACGCTCTTCCGATCTNNNTGACCATTGGTAGTTTCTTGGCTCA
ACACGTCTGAACTCTTTCCCTACACGACGCTCTTCCGATCTNNNTGACCATTGGTAGTTTCTTGGCTCA
ACACGTCTGAACTCTTTCCCTACACGACGCTCTTCCGATCTNNNTGACCATTGGTAGTTTCTTGGCTCA
AGCACACGTCTGAACTCTTTCCCTACACGACGCTCTTCCGATCTNNNTCACTTTCTTTTGCTTGCACCAA
AGCACACGTCTGAACTCTTTCCCTACACGACGCTCTTCCGATCTNNNTCACTTTCTTTTGCTTGCACCAA
AGCACACGTCTGAACTCTTTCCCTACACGACGCTCTTCCGATCTNNNTCACTTTCTTTTGCTTGCACCAA
AGCACACGTCTGAACTCTTTCCCTACACGACGCTCTTCCGATCTNNNTGAGGCAGGAGAATCTCTTGA
AGCACACGTCTGAACTCTTTCCCTACACGACGCTCTTCCGATCTNNNTGAGGCAGGAGAATCTCTTGA
AGCACACGTCTGAACTCTTTCCCTACACGACGCTCTTCCGATCTNNNTGAGGCAGGAGAATCTCTTGA
GCACACGTCTGAACTCTTTCCCTACACGACGCTCTTCCGATCTNNNCAGTGGCCACTTCAGGAAAAAAT
GCACACGTCTGAACTCTTTCCCTACACGACGCTCTTCCGATCTNNNCAGTGGCCACTTCAGGAAAAAAT
GCACACGTCTGAACTCTTTCCCTACACGACGCTCTTCCGATCTNNNCAGTGGCCACTTCAGGAAAAAAT
AGCACACGTCTGAACTCTTTCCCTACACGACGCTCTTCCGATCTNNNGCCTGGGTGAAAAGAGTGAA
AGCACACGTCTGAACTCTTTCCCTACACGACGCTCTTCCGATCTNNNGCCTGGGTGAAAAGAGTGAA
AGCACACGTCTGAACTCTTTCCCTACACGACGCTCTTCCGATCTNNNGCCTGGGTGAAAAGAGTGAA
GCACACGTCTGAACTCTTTCCCTACACGACGCTCTTCCGATCTNNNAGTCCCAGCTACTCAGGAGGCA
GCACACGTCTGAACTCTTTCCCTACACGACGCTCTTCCGATCTNNNAGTCCCAGCTACTCAGGAGGCA
GCACACGTCTGAACTCTTTCCCTACACGACGCTCTTCCGATCTNNNAGTCCCAGCTACTCAGGAGGCA
AGCACACGTCTGAACTCTTTCCCTACACGACGCTCTTCCGATCTNNNGTAATCTCAGCTACTCAGGAG
AGCACACGTCTGAACTCTTTCCCTACACGACGCTCTTCCGATCTNNNGTAATCTCAGCTACTCAGGAG
AGCACACGTCTGAACTCTTTCCCTACACGACGCTCTTCCGATCTNNNGTAATCTCAGCTACTCAGGAG
GCACACGTCTGAACTCTTTCCCTACACGACGCTCTTCCGATCTNNNGTTTCATCCATGTTGTAGCGTGT
GCACACGTCTGAACTCTTTCCCTACACGACGCTCTTCCGATCTNNNGTTTCATCCATGTTGTAGCGTGT
GCACACGTCTGAACTCTTTCCCTACACGACGCTCTTCCGATCTNNNGTTTCATCCATGTTGTAGCGTGT
GCACACGTCTGAACTCTTTCCCTACACGACGCTCTTCCGATCTNNNAGAAAGTGACCAGCCAGTAGAT
GCACACGTCTGAACTCTTTCCCTACACGACGCTCTTCCGATCTNNNAGAAAGTGACCAGCCAGTAGAT
GCACACGTCTGAACTCTTTCCCTACACGACGCTCTTCCGATCTNNNAGAAAGTGACCAGCCAGTAGAT
ACACGTCTGAACTCTTTCCCTACACGACGCTCTTCCGATCTNNNCCTTATGAAAGGCCTGCTTTGACCA
ACACGTCTGAACTCTTTCCCTACACGACGCTCTTCCGATCTNNNCCTTATGAAAGGCCTGCTTTGACCA
ACACGTCTGAACTCTTTCCCTACACGACGCTCTTCCGATCTNNNCCTTATGAAAGGCCTGCTTTGACCA
GCACACGTCTGAACTCTTTCCCTACACGACGCTCTTCCGATCTNNNCAGTTTCGCTAAGACCATTGTGT
GCACACGTCTGAACTCTTTCCCTACACGACGCTCTTCCGATCTNNNCAGTTTCGCTAAGACCATTGTGT
GCACACGTCTGAACTCTTTCCCTACACGACGCTCTTCCGATCTNNNCAGTTTCGCTAAGACCATTGTGT
AGCACACGTCTGAACTCTTTCCCTACACGACGCTCTTCCGATCTNNNTGTTTCTCCTCAGATCCAAGTG
AGCACACGTCTGAACTCTTTCCCTACACGACGCTCTTCCGATCTNNNTGTTTCTCCTCAGATCCAAGTG
AGCACACGTCTGAACTCTTTCCCTACACGACGCTCTTCCGATCTNNNTGTTTCTCCTCAGATCCAAGTG
GAGCACACGTCTGAACTCTTTCCCTACACGACGCTCTTCCGATCTNNNGAGGTTACAGTGAGCCAAGA
GAGCACACGTCTGAACTCTTTCCCTACACGACGCTCTTCCGATCTNNNGAGGTTACAGTGAGCCAAGA
GAGCACACGTCTGAACTCTTTCCCTACACGACGCTCTTCCGATCTNNNGAGGTTACAGTGAGCCAAGA

AGCACACGTCTGAACTCTTTCCCTACACGACGCTCTTCCGATCTNNNGTCCAGACTTCACCACTACACAA
AGCACACGTCTGAACTCTTTCCCTACACGACGCTCTTCCGATCTNNNGTCCAGACTTCACCACTACACAA
AGCACACGTCTGAACTCTTTCCCTACACGACGCTCTTCCGATCTNNNGTCCAGACTTCACCACTACACAA
AGAGCACACGTCTGAACTCTTTCCCTACACGACGCTCTTCCGATCTNNNGACTCCCCTTATGATTATGGC
AGAGCACACGTCTGAACTCTTTCCCTACACGACGCTCTTCCGATCTNNNGACTCCCCTTATGATTATGGC
AGAGCACACGTCTGAACTCTTTCCCTACACGACGCTCTTCCGATCTNNNGACTCCCCTTATGATTATGGC
GAGCACACGTCTGAACTCTTTCCCTACACGACGCTCTTCCGATCTNNNTCCTTCCTGTGCGGAAATAAG
GAGCACACGTCTGAACTCTTTCCCTACACGACGCTCTTCCGATCTNNNTCCTTCCTGTGCGGAAATAAG
GAGCACACGTCTGAACTCTTTCCCTACACGACGCTCTTCCGATCTNNNTCCTTCCTGTGCGGAAATAAG
GCACACGTCTGAACTCTTTCCCTACACGACGCTCTTCCGATCTNNNTCACCATTTGTCTTTGCAGGGAAC
GCACACGTCTGAACTCTTTCCCTACACGACGCTCTTCCGATCTNNNTCACCATTTGTCTTTGCAGGGAAC
GCACACGTCTGAACTCTTTCCCTACACGACGCTCTTCCGATCTNNNTCACCATTTGTCTTTGCAGGGAAC
AGCACACGTCTGAACTCTTTCCCTACACGACGCTCTTCCGATCTNNNAGCCTGTCTGGGTTCTTATGAAC
AGCACACGTCTGAACTCTTTCCCTACACGACGCTCTTCCGATCTNNNAGCCTGTCTGGGTTCTTATGAAC
AGCACACGTCTGAACTCTTTCCCTACACGACGCTCTTCCGATCTNNNAGCCTGTCTGGGTTCTTATGAAC
AGAGCACACGTCTGAACTCTTTCCCTACACGACGCTCTTCCGATCTNNNCAAATTGGAGTGGATATGAT
AGAGCACACGTCTGAACTCTTTCCCTACACGACGCTCTTCCGATCTNNNCAAATTGGAGTGGATATGAT
AGAGCACACGTCTGAACTCTTTCCCTACACGACGCTCTTCCGATCTNNNCAAATTGGAGTGGATATGAT
AGCACACGTCTGAACTCTTTCCCTACACGACGCTCTTCCGATCTNNNTAGCCAAGCAAAGTTCATAGGC
AGCACACGTCTGAACTCTTTCCCTACACGACGCTCTTCCGATCTNNNTAGCCAAGCAAAGTTCATAGGC
AGCACACGTCTGAACTCTTTCCCTACACGACGCTCTTCCGATCTNNNTAGCCAAGCAAAGTTCATAGGC
GAGCACACGTCTGAACTCTTTCCCTACACGACGCTCTTCCGATCTNNNGTGTATGACTCCACAGCCTGTA
GAGCACACGTCTGAACTCTTTCCCTACACGACGCTCTTCCGATCTNNNGTGTATGACTCCACAGCCTGTA
GAGCACACGTCTGAACTCTTTCCCTACACGACGCTCTTCCGATCTNNNGTGTATGACTCCACAGCCTGTA
AGCACACGTCTGAACTCTTTCCCTACACGACGCTCTTCCGATCTNNNGAGAATGGCGTGAACCAGGAA
AGCACACGTCTGAACTCTTTCCCTACACGACGCTCTTCCGATCTNNNGAGAATGGCGTGAACCAGGAA
AGCACACGTCTGAACTCTTTCCCTACACGACGCTCTTCCGATCTNNNGAGAATGGCGTGAACCAGGAA
CACACGTCTGAACTCTTTCCCTACACGACGCTCTTCCGATCTNNNACAGAATGCACACAATAAAGGGTC
CACACGTCTGAACTCTTTCCCTACACGACGCTCTTCCGATCTNNNACAGAATGCACACAATAAAGGGTC
CACACGTCTGAACTCTTTCCCTACACGACGCTCTTCCGATCTNNNACAGAATGCACACAATAAAGGGTC
AGCACACGTCTGAACTCTTTCCCTACACGACGCTCTTCCGATCTNNNAGAAAAAGGCCAGAAGAAAGC
AGCACACGTCTGAACTCTTTCCCTACACGACGCTCTTCCGATCTNNNAGAAAAAGGCCAGAAGAAAGC
AGCACACGTCTGAACTCTTTCCCTACACGACGCTCTTCCGATCTNNNAGAAAAAGGCCAGAAGAAAGC
AGCACACGTCTGAACTCTTTCCCTACACGACGCTCTTCCGATCTNNNCTTACGGAGGATTCCCAGGAAA
AGCACACGTCTGAACTCTTTCCCTACACGACGCTCTTCCGATCTNNNCTTACGGAGGATTCCCAGGAAA
AGCACACGTCTGAACTCTTTCCCTACACGACGCTCTTCCGATCTNNNCTTACGGAGGATTCCCAGGAAA
GCACACGTCTGAACTCTTTCCCTACACGACGCTCTTCCGATCTNNNCTTAGTTACCTTTTTCCATGTTTCAT
GCACACGTCTGAACTCTTTCCCTACACGACGCTCTTCCGATCTNNNCTTAGTTACCTTTTTCCATGTTTCAT
GCACACGTCTGAACTCTTTCCCTACACGACGCTCTTCCGATCTNNNCTTAGTTACCTTTTTCCATGTTTCAT
GAGCACACGTCTGAACTCTTTCCCTACACGACGCTCTTCCGATCTNNNACTGCCATAAAAATGCTCTGT
GAGCACACGTCTGAACTCTTTCCCTACACGACGCTCTTCCGATCTNNNACTGCCATAAAAATGCTCTGT
GAGCACACGTCTGAACTCTTTCCCTACACGACGCTCTTCCGATCTNNNACTGCCATAAAAATGCTCTGT
AGAGCACACGTCTGAACTCTTTCCCTACACGACGCTCTTCCGATCTNNNAAAAATTAGCCTAGCATGGTC

GAGCACACGTCTGAACTCTTTCCCTACACGACGCTCTTCCGATCTNNNAAAAATTAGCCTAGCATGGT
GAGCACACGTCTGAACTCTTTCCCTACACGACGCTCTTCCGATCTNNNAAAAATTAGCCTAGCATGGT
GCACACGTCTGAACTCTTTCCCTACACGACGCTCTTCCGATCTNNNATCACTTGAGCCAGTAGGTGA
GCACACGTCTGAACTCTTTCCCTACACGACGCTCTTCCGATCTNNNATCACTTGAGCCAGTAGGTGA
GCACACGTCTGAACTCTTTCCCTACACGACGCTCTTCCGATCTNNNATCACTTGAGCCAGTAGGTGA
GCACACGTCTGAACTCTTTCCCTACACGACGCTCTTCCGATCTNNNGAGGCAGGAGAATTGCTTGAA
GCACACGTCTGAACTCTTTCCCTACACGACGCTCTTCCGATCTNNNGAGGCAGGAGAATTGCTTGAA
GCACACGTCTGAACTCTTTCCCTACACGACGCTCTTCCGATCTNNNGAGGCAGGAGAATTGCTTGAA
AGCACACGTCTGAACTCTTTCCCTACACGACGCTCTTCCGATCTNNNCTGTAATCCCAGCTACTCAGG
AGCACACGTCTGAACTCTTTCCCTACACGACGCTCTTCCGATCTNNNCTGTAATCCCAGCTACTCAGG
AGCACACGTCTGAACTCTTTCCCTACACGACGCTCTTCCGATCTNNNCTGTAATCCCAGCTACTCAGG
AGCACACGTCTGAACTCTTTCCCTACACGACGCTCTTCCGATCTNNNCAGCAGGGATATAGCATCCTG
AGCACACGTCTGAACTCTTTCCCTACACGACGCTCTTCCGATCTNNNCAGCAGGGATATAGCATCCTG
AGCACACGTCTGAACTCTTTCCCTACACGACGCTCTTCCGATCTNNNCAGCAGGGATATAGCATCCTG
GCACACGTCTGAACTCTTTCCCTACACGACGCTCTTCCGATCTNNNTTTCTTTCTCCCCTCCTCAAGTG
GCACACGTCTGAACTCTTTCCCTACACGACGCTCTTCCGATCTNNNTTTCTTTCTCCCCTCCTCAAGTG
GCACACGTCTGAACTCTTTCCCTACACGACGCTCTTCCGATCTNNNTTTCTTTCTCCCCTCCTCAAGTG
GAGCACACGTCTGAACTCTTTCCCTACACGACGCTCTTCCGATCTNNNCAACTACTTGGGAGGCTGAG
GAGCACACGTCTGAACTCTTTCCCTACACGACGCTCTTCCGATCTNNNCAACTACTTGGGAGGCTGAG
GAGCACACGTCTGAACTCTTTCCCTACACGACGCTCTTCCGATCTNNNCAACTACTTGGGAGGCTGAG
AGCACACGTCTGAACTCTTTCCCTACACGACGCTCTTCCGATCTNNNGATTGCTTGAGGCCAGTAGTT
AGCACACGTCTGAACTCTTTCCCTACACGACGCTCTTCCGATCTNNNGATTGCTTGAGGCCAGTAGTT
AGCACACGTCTGAACTCTTTCCCTACACGACGCTCTTCCGATCTNNNGATTGCTTGAGGCCAGTAGTT
AGAGCACACGTCTGAACTCTTTCCCTACACGACGCTCTTCCGATCTNNNAGGGAGTCAGTAAGAGAC
AGAGCACACGTCTGAACTCTTTCCCTACACGACGCTCTTCCGATCTNNNAGGGAGTCAGTAAGAGAC
AGAGCACACGTCTGAACTCTTTCCCTACACGACGCTCTTCCGATCTNNNAGGGAGTCAGTAAGAGAC
GAGCACACGTCTGAACTCTTTCCCTACACGACGCTCTTCCGATCTNNNCGGCTGGAACACTGAATGA
GAGCACACGTCTGAACTCTTTCCCTACACGACGCTCTTCCGATCTNNNCGGCTGGAACACTGAATGA
GAGCACACGTCTGAACTCTTTCCCTACACGACGCTCTTCCGATCTNNNCGGCTGGAACACTGAATGA
AGCACACGTCTGAACTCTTTCCCTACACGACGCTCTTCCGATCTNNNCAGAGATTGCTCCACTCCACTC
AGCACACGTCTGAACTCTTTCCCTACACGACGCTCTTCCGATCTNNNCAGAGATTGCTCCACTCCACTC
AGCACACGTCTGAACTCTTTCCCTACACGACGCTCTTCCGATCTNNNCAGAGATTGCTCCACTCCACTC
AGCACACGTCTGAACTCTTTCCCTACACGACGCTCTTCCGATCTNNNGTGAAGTATGCCTGAGGAG
AGCACACGTCTGAACTCTTTCCCTACACGACGCTCTTCCGATCTNNNGTGAAGTATGCCTGAGGAG
AGCACACGTCTGAACTCTTTCCCTACACGACGCTCTTCCGATCTNNNGTGAAGTATGCCTGAGGAG
GAGCACACGTCTGAACTCTTTCCCTACACGACGCTCTTCCGATCTNNNTTTGGGAGGTCACAAGTTGT
GAGCACACGTCTGAACTCTTTCCCTACACGACGCTCTTCCGATCTNNNTTTGGGAGGTCACAAGTTGT
GAGCACACGTCTGAACTCTTTCCCTACACGACGCTCTTCCGATCTNNNTTTGGGAGGTCACAAGTTGT
GCACACGTCTGAACTCTTTCCCTACACGACGCTCTTCCGATCTNNNTCTCCTTTCTCCATCACCCAAAG
GCACACGTCTGAACTCTTTCCCTACACGACGCTCTTCCGATCTNNNTCTCCTTTCTCCATCACCCAAAG
GCACACGTCTGAACTCTTTCCCTACACGACGCTCTTCCGATCTNNNTCTCCTTTCTCCATCACCCAAAG
AGCACACGTCTGAACTCTTTCCCTACACGACGCTCTTCCGATCTNNNATTCAGATGATTGGACTATTCA
AGCACACGTCTGAACTCTTTCCCTACACGACGCTCTTCCGATCTNNNATTCAGATGATTGGACTATTCA

AGCACACGTCTGAACTCTTTCCCTACACGACGCTCTTCCGATCTNNNATTCAGATGATTGGACTATTCA
GAGCACACGTCTGAACTCTTTCCCTACACGACGCTCTTCCGATCTNNNCTACTGAGTGGGACCCTTCAC
GAGCACACGTCTGAACTCTTTCCCTACACGACGCTCTTCCGATCTNNNCTACTGAGTGGGACCCTTCAC
GAGCACACGTCTGAACTCTTTCCCTACACGACGCTCTTCCGATCTNNNCTACTGAGTGGGACCCTTCAC
AGCACACGTCTGAACTCTTTCCCTACACGACGCTCTTCCGATCTNNNGGTGACCTAAACAGCGAGTTA
AGCACACGTCTGAACTCTTTCCCTACACGACGCTCTTCCGATCTNNNGGTGACCTAAACAGCGAGTTA
AGCACACGTCTGAACTCTTTCCCTACACGACGCTCTTCCGATCTNNNGGTGACCTAAACAGCGAGTTA
GAGCACACGTCTGAACTCTTTCCCTACACGACGCTCTTCCGATCTNNNCTGCTCCCTTTGGTAAGTCAT
GAGCACACGTCTGAACTCTTTCCCTACACGACGCTCTTCCGATCTNNNCTGCTCCCTTTGGTAAGTCAT
GAGCACACGTCTGAACTCTTTCCCTACACGACGCTCTTCCGATCTNNNCTGCTCCCTTTGGTAAGTCAT
GAGCACACGTCTGAACTCTTTCCCTACACGACGCTCTTCCGATCTNNNAAGGGTGTGTGGACTCTCTG
GAGCACACGTCTGAACTCTTTCCCTACACGACGCTCTTCCGATCTNNNAAGGGTGTGTGGACTCTCTG
GAGCACACGTCTGAACTCTTTCCCTACACGACGCTCTTCCGATCTNNNAAGGGTGTGTGGACTCTCTG
GAGCACACGTCTGAACTCTTTCCCTACACGACGCTCTTCCGATCTNNNTTTTTCAGTTGGAGTCTCGCT
GAGCACACGTCTGAACTCTTTCCCTACACGACGCTCTTCCGATCTNNNTTTTTCAGTTGGAGTCTCGCT
GAGCACACGTCTGAACTCTTTCCCTACACGACGCTCTTCCGATCTNNNTTTTTCAGTTGGAGTCTCGCT
AGCACACGTCTGAACTCTTTCCCTACACGACGCTCTTCCGATCTNNNAAAAAATTAGCTGGCCATGGT
AGCACACGTCTGAACTCTTTCCCTACACGACGCTCTTCCGATCTNNNAAAAAATTAGCTGGCCATGGT
AGCACACGTCTGAACTCTTTCCCTACACGACGCTCTTCCGATCTNNNAAAAAATTAGCTGGCCATGGT
AGCACACGTCTGAACTCTTTCCCTACACGACGCTCTTCCGATCTNNNCTACCAAGACATCACTGGGTCA
AGCACACGTCTGAACTCTTTCCCTACACGACGCTCTTCCGATCTNNNCTACCAAGACATCACTGGGTCA
AGCACACGTCTGAACTCTTTCCCTACACGACGCTCTTCCGATCTNNNCTACCAAGACATCACTGGGTCA
GAGCACACGTCTGAACTCTTTCCCTACACGACGCTCTTCCGATCTNNNTCCCTAACTGTGGATCCTACTG
GAGCACACGTCTGAACTCTTTCCCTACACGACGCTCTTCCGATCTNNNTCCCTAACTGTGGATCCTACTG
GAGCACACGTCTGAACTCTTTCCCTACACGACGCTCTTCCGATCTNNNTCCCTAACTGTGGATCCTACTG
GAGCACACGTCTGAACTCTTTCCCTACACGACGCTCTTCCGATCTNNNCAGAAGGCTGAGGTAGGAG
GAGCACACGTCTGAACTCTTTCCCTACACGACGCTCTTCCGATCTNNNCAGAAGGCTGAGGTAGGAG
GAGCACACGTCTGAACTCTTTCCCTACACGACGCTCTTCCGATCTNNNCAGAAGGCTGAGGTAGGAG
GCACACGTCTGAACTCTTTCCCTACACGACGCTCTTCCGATCTNNNCTGAGGTGAGCCTGTGTTCTCTC
GCACACGTCTGAACTCTTTCCCTACACGACGCTCTTCCGATCTNNNCTGAGGTGAGCCTGTGTTCTCTC
GCACACGTCTGAACTCTTTCCCTACACGACGCTCTTCCGATCTNNNCTGAGGTGAGCCTGTGTTCTCTC
GAGCACACGTCTGAACTCTTTCCCTACACGACGCTCTTCCGATCTNNNCTGTAGTCACAGCTACTCAGG
GAGCACACGTCTGAACTCTTTCCCTACACGACGCTCTTCCGATCTNNNCTGTAGTCACAGCTACTCAGG
GAGCACACGTCTGAACTCTTTCCCTACACGACGCTCTTCCGATCTNNNCTGTAGTCACAGCTACTCAGG
GAGCACACGTCTGAACTCTTTCCCTACACGACGCTCTTCCGATCTNNNCCATTCTCCTACCTCAGTCTCCT
GAGCACACGTCTGAACTCTTTCCCTACACGACGCTCTTCCGATCTNNNCCATTCTCCTACCTCAGTCTCCT
GAGCACACGTCTGAACTCTTTCCCTACACGACGCTCTTCCGATCTNNNCCATTCTCCTACCTCAGTCTCCT
GAGCACACGTCTGAACTCTTTCCCTACACGACGCTCTTCCGATCTNNNAATCTTTAGCAGGGAAGTGCT
GAGCACACGTCTGAACTCTTTCCCTACACGACGCTCTTCCGATCTNNNAATCTTTAGCAGGGAAGTGCT
GAGCACACGTCTGAACTCTTTCCCTACACGACGCTCTTCCGATCTNNNAATCTTTAGCAGGGAAGTGCT
AGCACACGTCTGAACTCTTTCCCTACACGACGCTCTTCCGATCTNNNGCAAAGCTACACCTTTCCTCAT
AGCACACGTCTGAACTCTTTCCCTACACGACGCTCTTCCGATCTNNNGCAAAGCTACACCTTTCCTCAT
AGCACACGTCTGAACTCTTTCCCTACACGACGCTCTTCCGATCTNNNGCAAAGCTACACCTTTCCTCAT

AGAGCACACGTCTGAACTCTTTCCCTACACGACGCTCTTCCGATCTNNNAGAAGTGGCCCAGATGACT
AGAGCACACGTCTGAACTCTTTCCCTACACGACGCTCTTCCGATCTNNNAGAAGTGGCCCAGATGACT
AGAGCACACGTCTGAACTCTTTCCCTACACGACGCTCTTCCGATCTNNNAGAAGTGGCCCAGATGACT
AGCACACGTCTGAACTCTTTCCCTACACGACGCTCTTCCGATCTNNNGCTAACTCCTCTCAGAGTGATG
AGCACACGTCTGAACTCTTTCCCTACACGACGCTCTTCCGATCTNNNGCTAACTCCTCTCAGAGTGATG
AGCACACGTCTGAACTCTTTCCCTACACGACGCTCTTCCGATCTNNNGCTAACTCCTCTCAGAGTGATG
AGCACACGTCTGAACTCTTTCCCTACACGACGCTCTTCCGATCTNNNCTGAGGCAGGAGAATCACTTG
AGCACACGTCTGAACTCTTTCCCTACACGACGCTCTTCCGATCTNNNCTGAGGCAGGAGAATCACTTG
AGCACACGTCTGAACTCTTTCCCTACACGACGCTCTTCCGATCTNNNCTGAGGCAGGAGAATCACTTG
AGCACACGTCTGAACTCTTTCCCTACACGACGCTCTTCCGATCTNNNTACAGGCCCAAGTGCTATTGAAT
AGCACACGTCTGAACTCTTTCCCTACACGACGCTCTTCCGATCTNNNTACAGGCCCAAGTGCTATTGAAT
AGCACACGTCTGAACTCTTTCCCTACACGACGCTCTTCCGATCTNNNTACAGGCCCAAGTGCTATTGAAT
AGCACACGTCTGAACTCTTTCCCTACACGACGCTCTTCCGATCTNNNCTTGCAGTGAGCCGAGATTGCA
AGCACACGTCTGAACTCTTTCCCTACACGACGCTCTTCCGATCTNNNCTTGCAGTGAGCCGAGATTGCA
AGCACACGTCTGAACTCTTTCCCTACACGACGCTCTTCCGATCTNNNCTTGCAGTGAGCCGAGATTGCA
GAGCACACGTCTGAACTCTTTCCCTACACGACGCTCTTCCGATCTNNNGAGGCTGAGGTAGAAGAATT
GAGCACACGTCTGAACTCTTTCCCTACACGACGCTCTTCCGATCTNNNGAGGCTGAGGTAGAAGAATT
GAGCACACGTCTGAACTCTTTCCCTACACGACGCTCTTCCGATCTNNNGAGGCTGAGGTAGAAGAATT
AGCACACGTCTGAACTCTTTCCCTACACGACGCTCTTCCGATCTNNNTAGCTAGGATTACAGTCACGCA
AGCACACGTCTGAACTCTTTCCCTACACGACGCTCTTCCGATCTNNNTAGCTAGGATTACAGTCACGCA
AGCACACGTCTGAACTCTTTCCCTACACGACGCTCTTCCGATCTNNNTAGCTAGGATTACAGTCACGCA
AGCACACGTCTGAACTCTTTCCCTACACGACGCTCTTCCGATCTNNNCACCTGTAATTCCAGCACTTTG
AGCACACGTCTGAACTCTTTCCCTACACGACGCTCTTCCGATCTNNNCACCTGTAATTCCAGCACTTTG
AGCACACGTCTGAACTCTTTCCCTACACGACGCTCTTCCGATCTNNNCACCTGTAATTCCAGCACTTTG
GCACACGTCTGAACTCTTTCCCTACACGACGCTCTTCCGATCTNNNAAAGGTCACATACTCGGCTCTGC
GCACACGTCTGAACTCTTTCCCTACACGACGCTCTTCCGATCTNNNAAAGGTCACATACTCGGCTCTGC
GCACACGTCTGAACTCTTTCCCTACACGACGCTCTTCCGATCTNNNAAAGGTCACATACTCGGCTCTGC
AGCACACGTCTGAACTCTTTCCCTACACGACGCTCTTCCGATCTNNNCCTCAACTTTGCCTCTCTCCTTA
AGCACACGTCTGAACTCTTTCCCTACACGACGCTCTTCCGATCTNNNCCTCAACTTTGCCTCTCTCCTTA
AGCACACGTCTGAACTCTTTCCCTACACGACGCTCTTCCGATCTNNNCCTCAACTTTGCCTCTCTCCTTA
AGAGCACACGTCTGAACTCTTTCCCTACACGACGCTCTTCCGATCTNNNTGGACAAGTAACACCTCTCA
AGAGCACACGTCTGAACTCTTTCCCTACACGACGCTCTTCCGATCTNNNTGGACAAGTAACACCTCTCA
AGAGCACACGTCTGAACTCTTTCCCTACACGACGCTCTTCCGATCTNNNTGGACAAGTAACACCTCTCA
AGCACACGTCTGAACTCTTTCCCTACACGACGCTCTTCCGATCTNNNTCAAAAGCAAGCCAGAGTAGCT
AGCACACGTCTGAACTCTTTCCCTACACGACGCTCTTCCGATCTNNNTCAAAAGCAAGCCAGAGTAGCT
AGCACACGTCTGAACTCTTTCCCTACACGACGCTCTTCCGATCTNNNTCAAAAGCAAGCCAGAGTAGCT
AGCACACGTCTGAACTCTTTCCCTACACGACGCTCTTCCGATCTNNNGAAGGATCACTTGAGTCCAGG
AGCACACGTCTGAACTCTTTCCCTACACGACGCTCTTCCGATCTNNNGAAGGATCACTTGAGTCCAGG
AGCACACGTCTGAACTCTTTCCCTACACGACGCTCTTCCGATCTNNNGAAGGATCACTTGAGTCCAGG
GAGCACACGTCTGAACTCTTTCCCTACACGACGCTCTTCCGATCTNNNTCTTCGATTATCCAGGTGAAC
GAGCACACGTCTGAACTCTTTCCCTACACGACGCTCTTCCGATCTNNNTCTTCGATTATCCAGGTGAAC
GAGCACACGTCTGAACTCTTTCCCTACACGACGCTCTTCCGATCTNNNTCTTCGATTATCCAGGTGAAC
GCACACGTCTGAACTCTTTCCCTACACGACGCTCTTCCGATCTNNNCCCGTTCTGTTTGTAGAGTCTCA

GCACACGTCTGAACTCTTTCCCTACACGACGCTCTTCCGATCTNNNCCCGTTCTGTTTGTAGAGTCTCA
GCACACGTCTGAACTCTTTCCCTACACGACGCTCTTCCGATCTNNNCCCGTTCTGTTTGTAGAGTCTCA
AGCACACGTCTGAACTCTTTCCCTACACGACGCTCTTCCGATCTNNNAGTTTGAGAACGTCTGGCCAA
AGCACACGTCTGAACTCTTTCCCTACACGACGCTCTTCCGATCTNNNAGTTTGAGAACGTCTGGCCAA
AGCACACGTCTGAACTCTTTCCCTACACGACGCTCTTCCGATCTNNNAGTTTGAGAACGTCTGGCCAA
GAGCACACGTCTGAACTCTTTCCCTACACGACGCTCTTCCGATCTNNNGAAAAGTGAATTCCCCACAGT
GAGCACACGTCTGAACTCTTTCCCTACACGACGCTCTTCCGATCTNNNGAAAAGTGAATTCCCCACAGT
GAGCACACGTCTGAACTCTTTCCCTACACGACGCTCTTCCGATCTNNNGAAAAGTGAATTCCCCACAGT
GAGCACACGTCTGAACTCTTTCCCTACACGACGCTCTTCCGATCTNNNAAAAAATTAGCTGGGTGTG
GAGCACACGTCTGAACTCTTTCCCTACACGACGCTCTTCCGATCTNNNAAAAAATTAGCTGGGTGTG
GAGCACACGTCTGAACTCTTTCCCTACACGACGCTCTTCCGATCTNNNAAAAAATTAGCTGGGTGTG
GAGCACACGTCTGAACTCTTTCCCTACACGACGCTCTTCCGATCTNNNGCTGATGTGGGAGGATCAC
GAGCACACGTCTGAACTCTTTCCCTACACGACGCTCTTCCGATCTNNNGCTGATGTGGGAGGATCAC
GAGCACACGTCTGAACTCTTTCCCTACACGACGCTCTTCCGATCTNNNGCTGATGTGGGAGGATCAC
GAGCACACGTCTGAACTCTTTCCCTACACGACGCTCTTCCGATCTNNNTAAGCTTTTTGATGTGCTGCT
GAGCACACGTCTGAACTCTTTCCCTACACGACGCTCTTCCGATCTNNNTAAGCTTTTTGATGTGCTGCT
GAGCACACGTCTGAACTCTTTCCCTACACGACGCTCTTCCGATCTNNNTAAGCTTTTTGATGTGCTGCT
AGCACACGTCTGAACTCTTTCCCTACACGACGCTCTTCCGATCTNNNTGTGTGTATGTGTGTGTCTCC
AGCACACGTCTGAACTCTTTCCCTACACGACGCTCTTCCGATCTNNNTGTGTGTATGTGTGTGTCTCC
AGCACACGTCTGAACTCTTTCCCTACACGACGCTCTTCCGATCTNNNTGTGTGTATGTGTGTGTCTCC
AGCACACGTCTGAACTCTTTCCCTACACGACGCTCTTCCGATCTNNNCAAAGGAGCTGATCGCAAATCA
AGCACACGTCTGAACTCTTTCCCTACACGACGCTCTTCCGATCTNNNCAAAGGAGCTGATCGCAAATCA
AGCACACGTCTGAACTCTTTCCCTACACGACGCTCTTCCGATCTNNNCAAAGGAGCTGATCGCAAATCA
GAGCACACGTCTGAACTCTTTCCCTACACGACGCTCTTCCGATCTNNNGTAAGAGGCCAGACTTCTAG
GAGCACACGTCTGAACTCTTTCCCTACACGACGCTCTTCCGATCTNNNGTAAGAGGCCAGACTTCTAG
GAGCACACGTCTGAACTCTTTCCCTACACGACGCTCTTCCGATCTNNNGTAAGAGGCCAGACTTCTAG
AGCACACGTCTGAACTCTTTCCCTACACGACGCTCTTCCGATCTNNNTGTGTGTGTGTCTCTTCTCCAT
AGCACACGTCTGAACTCTTTCCCTACACGACGCTCTTCCGATCTNNNTGTGTGTGTGTCTCTTCTCCAT
AGCACACGTCTGAACTCTTTCCCTACACGACGCTCTTCCGATCTNNNTGTGTGTGTGTCTCTTCTCCAT
AGCACACGTCTGAACTCTTTCCCTACACGACGCTCTTCCGATCTNNNATACAAAAAAATTAGCCGGGC
AGCACACGTCTGAACTCTTTCCCTACACGACGCTCTTCCGATCTNNNATACAAAAAAATTAGCCGGGC
AGCACACGTCTGAACTCTTTCCCTACACGACGCTCTTCCGATCTNNNATACAAAAAAATTAGCCGGGC
AGCACACGTCTGAACTCTTTCCCTACACGACGCTCTTCCGATCTNNNAGCTCCCTTATCTGTTTGTTGGT
AGCACACGTCTGAACTCTTTCCCTACACGACGCTCTTCCGATCTNNNAGCTCCCTTATCTGTTTGTTGGT
AGCACACGTCTGAACTCTTTCCCTACACGACGCTCTTCCGATCTNNNAGCTCCCTTATCTGTTTGTTGGT
AGCACACGTCTGAACTCTTTCCCTACACGACGCTCTTCCGATCTNNNCTGAGGCAGGAGAATCACTTG
AGCACACGTCTGAACTCTTTCCCTACACGACGCTCTTCCGATCTNNNCTGAGGCAGGAGAATCACTTG
AGCACACGTCTGAACTCTTTCCCTACACGACGCTCTTCCGATCTNNNCTGAGGCAGGAGAATCACTTG
AGCACACGTCTGAACTCTTTCCCTACACGACGCTCTTCCGATCTNNNGTCATCCTAGCACTTTGGGAG
AGCACACGTCTGAACTCTTTCCCTACACGACGCTCTTCCGATCTNNNGTCATCCTAGCACTTTGGGAG
AGCACACGTCTGAACTCTTTCCCTACACGACGCTCTTCCGATCTNNNGTCATCCTAGCACTTTGGGAG
GAGCACACGTCTGAACTCTTTCCCTACACGACGCTCTTCCGATCTNNNACAGAGCTTCAGACTGCTGTT
GAGCACACGTCTGAACTCTTTCCCTACACGACGCTCTTCCGATCTNNNACAGAGCTTCAGACTGCTGTT

GAGCACACGTCTGAACTCTTTCCCTACACGACGCTCTTCCGATCTNNNACAGAGCTTCAGACTGCTGTT
GCACACGTCTGAACTCTTTCCCTACACGACGCTCTTCCGATCTNNNGTTGGGAAAATTATCACTATGA
GCACACGTCTGAACTCTTTCCCTACACGACGCTCTTCCGATCTNNNGTTGGGAAAATTATCACTATGA
GCACACGTCTGAACTCTTTCCCTACACGACGCTCTTCCGATCTNNNGTTGGGAAAATTATCACTATGA
AGCACACGTCTGAACTCTTTCCCTACACGACGCTCTTCCGATCTNNNACCATTCGCCAATGTCCTTGTT
AGCACACGTCTGAACTCTTTCCCTACACGACGCTCTTCCGATCTNNNACCATTCGCCAATGTCCTTGTT
AGCACACGTCTGAACTCTTTCCCTACACGACGCTCTTCCGATCTNNNACCATTCGCCAATGTCCTTGTT
AGCACACGTCTGAACTCTTTCCCTACACGACGCTCTTCCGATCTNNNCAAGCTTGCATTTCTCTCCAGAA
AGCACACGTCTGAACTCTTTCCCTACACGACGCTCTTCCGATCTNNNCAAGCTTGCATTTCTCTCCAGAA
AGCACACGTCTGAACTCTTTCCCTACACGACGCTCTTCCGATCTNNNCAAGCTTGCATTTCTCTCCAGAA
CACGTCTGAACTCTTTCCCTACACGACGCTCTTCCGATCTNNNCAAAGGGCTTTTCAGTTAGTGGGCA
CACGTCTGAACTCTTTCCCTACACGACGCTCTTCCGATCTNNNCAAAGGGCTTTTCAGTTAGTGGGCA
CACGTCTGAACTCTTTCCCTACACGACGCTCTTCCGATCTNNNCAAAGGGCTTTTCAGTTAGTGGGCA
GAGCACACGTCTGAACTCTTTCCCTACACGACGCTCTTCCGATCTNNNATGTAAGCACCCTATTGGTTC
GAGCACACGTCTGAACTCTTTCCCTACACGACGCTCTTCCGATCTNNNATGTAAGCACCCTATTGGTTC
GAGCACACGTCTGAACTCTTTCCCTACACGACGCTCTTCCGATCTNNNATGTAAGCACCCTATTGGTTC
GAGCACACGTCTGAACTCTTTCCCTACACGACGCTCTTCCGATCTNNNCTAAAAGGGTTTTCAGGCTG
GAGCACACGTCTGAACTCTTTCCCTACACGACGCTCTTCCGATCTNNNCTAAAAGGGTTTTCAGGCTG
GAGCACACGTCTGAACTCTTTCCCTACACGACGCTCTTCCGATCTNNNCTAAAAGGGTTTTCAGGCTG
GAGCACACGTCTGAACTCTTTCCCTACACGACGCTCTTCCGATCTNNNGTGCCACCTCAGAATTCAAG
GAGCACACGTCTGAACTCTTTCCCTACACGACGCTCTTCCGATCTNNNGTGCCACCTCAGAATTCAAG
GAGCACACGTCTGAACTCTTTCCCTACACGACGCTCTTCCGATCTNNNGTGCCACCTCAGAATTCAAG
GAGCACACGTCTGAACTCTTTCCCTACACGACGCTCTTCCGATCTNNNGAGATCACACCAATGCACTCC
GAGCACACGTCTGAACTCTTTCCCTACACGACGCTCTTCCGATCTNNNGAGATCACACCAATGCACTCC
GAGCACACGTCTGAACTCTTTCCCTACACGACGCTCTTCCGATCTNNNGAGATCACACCAATGCACTCC
AGCACACGTCTGAACTCTTTCCCTACACGACGCTCTTCCGATCTNNNTCATGAGCCAAATCCTGCAGT
AGCACACGTCTGAACTCTTTCCCTACACGACGCTCTTCCGATCTNNNTCATGAGCCAAATCCTGCAGT
AGCACACGTCTGAACTCTTTCCCTACACGACGCTCTTCCGATCTNNNTCATGAGCCAAATCCTGCAGT
GCACACGTCTGAACTCTTTCCCTACACGACGCTCTTCCGATCTNNNGAACTCCTGACCTCATGTGATCC
GCACACGTCTGAACTCTTTCCCTACACGACGCTCTTCCGATCTNNNGAACTCCTGACCTCATGTGATCC
GCACACGTCTGAACTCTTTCCCTACACGACGCTCTTCCGATCTNNNGAACTCCTGACCTCATGTGATCC
CACACGTCTGAACTCTTTCCCTACACGACGCTCTTCCGATCTNNNCAATTCAATCCATCGCACTCTCCCA
CACACGTCTGAACTCTTTCCCTACACGACGCTCTTCCGATCTNNNCAATTCAATCCATCGCACTCTCCCA
CACACGTCTGAACTCTTTCCCTACACGACGCTCTTCCGATCTNNNCAATTCAATCCATCGCACTCTCCCA
GAGCACACGTCTGAACTCTTTCCCTACACGACGCTCTTCCGATCTNNNCCATGACTCCCCAGACTTCTTA
GAGCACACGTCTGAACTCTTTCCCTACACGACGCTCTTCCGATCTNNNCCATGACTCCCCAGACTTCTTA
GAGCACACGTCTGAACTCTTTCCCTACACGACGCTCTTCCGATCTNNNCCATGACTCCCCAGACTTCTTA
AGCACACGTCTGAACTCTTTCCCTACACGACGCTCTTCCGATCTNNNGAGGATTGATTGAGCCTGGGA
AGCACACGTCTGAACTCTTTCCCTACACGACGCTCTTCCGATCTNNNGAGGATTGATTGAGCCTGGGA
AGCACACGTCTGAACTCTTTCCCTACACGACGCTCTTCCGATCTNNNGAGGATTGATTGAGCCTGGGA
AGCACACGTCTGAACTCTTTCCCTACACGACGCTCTTCCGATCTNNNAGGTTCCAGGATTACAACCTCT
AGCACACGTCTGAACTCTTTCCCTACACGACGCTCTTCCGATCTNNNAGGTTCCAGGATTACAACCTCT
AGCACACGTCTGAACTCTTTCCCTACACGACGCTCTTCCGATCTNNNAGGTTCCAGGATTACAACCTCT

GAGCACACGTCTGAACTCTTTCCCTACACGACGCTCTTCCGATCTNNNCAAAGGAGCCAAGTTCTGTTCA
GAGCACACGTCTGAACTCTTTCCCTACACGACGCTCTTCCGATCTNNNCAAAGGAGCCAAGTTCTGTTCA
GAGCACACGTCTGAACTCTTTCCCTACACGACGCTCTTCCGATCTNNNCAAAGGAGCCAAGTTCTGTTCA
AAGAGCACACGTCTGAACTCTTTCCCTACACGACGCTCTTCCGATCTNNNGGAGGGAGAAAGGAAAG
AAGAGCACACGTCTGAACTCTTTCCCTACACGACGCTCTTCCGATCTNNNGGAGGGAGAAAGGAAAG
AAGAGCACACGTCTGAACTCTTTCCCTACACGACGCTCTTCCGATCTNNNGGAGGGAGAAAGGAAAG
GAGCACACGTCTGAACTCTTTCCCTACACGACGCTCTTCCGATCTNNNTGCATTTTCTACCGTGCTCATT
GAGCACACGTCTGAACTCTTTCCCTACACGACGCTCTTCCGATCTNNNTGCATTTTCTACCGTGCTCATT
GAGCACACGTCTGAACTCTTTCCCTACACGACGCTCTTCCGATCTNNNTGCATTTTCTACCGTGCTCATT
GAGCACACGTCTGAACTCTTTCCCTACACGACGCTCTTCCGATCTNNNGTGAGAGAGAGAGCAAGAGA
GAGCACACGTCTGAACTCTTTCCCTACACGACGCTCTTCCGATCTNNNGTGAGAGAGAGAGCAAGAGA
GAGCACACGTCTGAACTCTTTCCCTACACGACGCTCTTCCGATCTNNNGTGAGAGAGAGAGCAAGAGA
CACACGTCTGAACTCTTTCCCTACACGACGCTCTTCCGATCTNNNAGCTGCTGTGCTCGATTTCTCCATC
CACACGTCTGAACTCTTTCCCTACACGACGCTCTTCCGATCTNNNAGCTGCTGTGCTCGATTTCTCCATC
CACACGTCTGAACTCTTTCCCTACACGACGCTCTTCCGATCTNNNAGCTGCTGTGCTCGATTTCTCCATC
AGCACACGTCTGAACTCTTTCCCTACACGACGCTCTTCCGATCTNNNAGTAGCTGGGACTATAGGTGT
AGCACACGTCTGAACTCTTTCCCTACACGACGCTCTTCCGATCTNNNAGTAGCTGGGACTATAGGTGT
AGCACACGTCTGAACTCTTTCCCTACACGACGCTCTTCCGATCTNNNAGTAGCTGGGACTATAGGTGT
GAGCACACGTCTGAACTCTTTCCCTACACGACGCTCTTCCGATCTNNNGTGATTATACCAACGTAGGGC
GAGCACACGTCTGAACTCTTTCCCTACACGACGCTCTTCCGATCTNNNGTGATTATACCAACGTAGGGC
GAGCACACGTCTGAACTCTTTCCCTACACGACGCTCTTCCGATCTNNNGTGATTATACCAACGTAGGGC
GAGCACACGTCTGAACTCTTTCCCTACACGACGCTCTTCCGATCTNNNCTGAGGCAGGAGAATCACTTG
GAGCACACGTCTGAACTCTTTCCCTACACGACGCTCTTCCGATCTNNNCTGAGGCAGGAGAATCACTTG
GAGCACACGTCTGAACTCTTTCCCTACACGACGCTCTTCCGATCTNNNCTGAGGCAGGAGAATCACTTG
AGCACACGTCTGAACTCTTTCCCTACACGACGCTCTTCCGATCTNNNCTGAGGCAGGAGAATCACTTGA
AGCACACGTCTGAACTCTTTCCCTACACGACGCTCTTCCGATCTNNNCTGAGGCAGGAGAATCACTTGA
AGCACACGTCTGAACTCTTTCCCTACACGACGCTCTTCCGATCTNNNCTGAGGCAGGAGAATCACTTGA
AGCACACGTCTGAACTCTTTCCCTACACGACGCTCTTCCGATCTNNNCAACCTAGGTGACAGAGTGAG
AGCACACGTCTGAACTCTTTCCCTACACGACGCTCTTCCGATCTNNNCAACCTAGGTGACAGAGTGAG
AGCACACGTCTGAACTCTTTCCCTACACGACGCTCTTCCGATCTNNNCAACCTAGGTGACAGAGTGAG
AGCACACGTCTGAACTCTTTCCCTACACGACGCTCTTCCGATCTNNNTCCTAATTCATATCCCTGGCTGC
AGCACACGTCTGAACTCTTTCCCTACACGACGCTCTTCCGATCTNNNTCCTAATTCATATCCCTGGCTGC
AGCACACGTCTGAACTCTTTCCCTACACGACGCTCTTCCGATCTNNNTCCTAATTCATATCCCTGGCTGC
AGCACACGTCTGAACTCTTTCCCTACACGACGCTCTTCCGATCTNNNCTGGCCTACACTCTCTTAGCTGC
AGCACACGTCTGAACTCTTTCCCTACACGACGCTCTTCCGATCTNNNCTGGCCTACACTCTCTTAGCTGC
AGCACACGTCTGAACTCTTTCCCTACACGACGCTCTTCCGATCTNNNCTGGCCTACACTCTCTTAGCTGC
AGCACACGTCTGAACTCTTTCCCTACACGACGCTCTTCCGATCTNNNTAAGATCACACCATTGCACTCCA
AGCACACGTCTGAACTCTTTCCCTACACGACGCTCTTCCGATCTNNNTAAGATCACACCATTGCACTCCA
AGCACACGTCTGAACTCTTTCCCTACACGACGCTCTTCCGATCTNNNTAAGATCACACCATTGCACTCCA
GAGCACACGTCTGAACTCTTTCCCTACACGACGCTCTTCCGATCTNNNTTTTTCAGCTCCTGGCATTCAA
GAGCACACGTCTGAACTCTTTCCCTACACGACGCTCTTCCGATCTNNNTTTTTCAGCTCCTGGCATTCAA
GAGCACACGTCTGAACTCTTTCCCTACACGACGCTCTTCCGATCTNNNTTTTTCAGCTCCTGGCATTCAA
AGCACACGTCTGAACTCTTTCCCTACACGACGCTCTTCCGATCTNNNAGAAACTGGCTCCAAGTGATGA

AGCACACGTCTGAACTCTTTCCCTACACGACGCTCTTCCGATCTNNNAGAAACTGGCTCCAAGTGATGA
AGCACACGTCTGAACTCTTTCCCTACACGACGCTCTTCCGATCTNNNAGAAACTGGCTCCAAGTGATGA
GAGCACACGTCTGAACTCTTTCCCTACACGACGCTCTTCCGATCTNNNCAGACCACAAACCAGGAGACA
GAGCACACGTCTGAACTCTTTCCCTACACGACGCTCTTCCGATCTNNNCAGACCACAAACCAGGAGACA
GAGCACACGTCTGAACTCTTTCCCTACACGACGCTCTTCCGATCTNNNCAGACCACAAACCAGGAGACA
AGCACACGTCTGAACTCTTTCCCTACACGACGCTCTTCCGATCTNNNGGTGACAGAGTGAGACTCCATC
AGCACACGTCTGAACTCTTTCCCTACACGACGCTCTTCCGATCTNNNGGTGACAGAGTGAGACTCCATC
AGCACACGTCTGAACTCTTTCCCTACACGACGCTCTTCCGATCTNNNGGTGACAGAGTGAGACTCCATC
AGAGCACACGTCTGAACTCTTTCCCTACACGACGCTCTTCCGATCTNNNTTAGGGCACCTGCAGTTTGA
AGAGCACACGTCTGAACTCTTTCCCTACACGACGCTCTTCCGATCTNNNTTAGGGCACCTGCAGTTTGA
AGAGCACACGTCTGAACTCTTTCCCTACACGACGCTCTTCCGATCTNNNTTAGGGCACCTGCAGTTTGA
GAGCACACGTCTGAACTCTTTCCCTACACGACGCTCTTCCGATCTNNNACTCCTGAAACAGATGGGAAA
GAGCACACGTCTGAACTCTTTCCCTACACGACGCTCTTCCGATCTNNNACTCCTGAAACAGATGGGAAA
GAGCACACGTCTGAACTCTTTCCCTACACGACGCTCTTCCGATCTNNNACTCCTGAAACAGATGGGAAA
AGCACACGTCTGAACTCTTTCCCTACACGACGCTCTTCCGATCTNNNGGCACAGATGAGACTAGAAAC
AGCACACGTCTGAACTCTTTCCCTACACGACGCTCTTCCGATCTNNNGGCACAGATGAGACTAGAAAC
AGCACACGTCTGAACTCTTTCCCTACACGACGCTCTTCCGATCTNNNGGCACAGATGAGACTAGAAAC
AGCACACGTCTGAACTCTTTCCCTACACGACGCTCTTCCGATCTNNNCTCCCTCGTGGTTCAGACAGCAT
AGCACACGTCTGAACTCTTTCCCTACACGACGCTCTTCCGATCTNNNCTCCCTCGTGGTTCAGACAGCAT
AGCACACGTCTGAACTCTTTCCCTACACGACGCTCTTCCGATCTNNNCTCCCTCGTGGTTCAGACAGCAT
GAGCACACGTCTGAACTCTTTCCCTACACGACGCTCTTCCGATCTNNNCTCTCTCTCTCACACACACAC
GAGCACACGTCTGAACTCTTTCCCTACACGACGCTCTTCCGATCTNNNCTCTCTCTCTCACACACACAC
GAGCACACGTCTGAACTCTTTCCCTACACGACGCTCTTCCGATCTNNNCTCTCTCTCTCACACACACAC
AGAGCACACGTCTGAACTCTTTCCCTACACGACGCTCTTCCGATCTNNNCAGGAGTTTTTGCTGGGTGC
AGAGCACACGTCTGAACTCTTTCCCTACACGACGCTCTTCCGATCTNNNCAGGAGTTTTTGCTGGGTGC
AGAGCACACGTCTGAACTCTTTCCCTACACGACGCTCTTCCGATCTNNNCAGGAGTTTTTGCTGGGTGC
AGAGCACACGTCTGAACTCTTTCCCTACACGACGCTCTTCCGATCTNNNGGGACTTAGAGACCAAAGA
AGAGCACACGTCTGAACTCTTTCCCTACACGACGCTCTTCCGATCTNNNGGGACTTAGAGACCAAAGA
AGAGCACACGTCTGAACTCTTTCCCTACACGACGCTCTTCCGATCTNNNGGGACTTAGAGACCAAAGA
AGAGCACACGTCTGAACTCTTTCCCTACACGACGCTCTTCCGATCTNNNAATCCTAGCTACTCAGGAGG
AGAGCACACGTCTGAACTCTTTCCCTACACGACGCTCTTCCGATCTNNNAATCCTAGCTACTCAGGAGG
AGAGCACACGTCTGAACTCTTTCCCTACACGACGCTCTTCCGATCTNNNAATCCTAGCTACTCAGGAGG
GCACACGTCTGAACTCTTTCCCTACACGACGCTCTTCCGATCTNNNACCTAGTTGTCACCTTTGTATAAC
GCACACGTCTGAACTCTTTCCCTACACGACGCTCTTCCGATCTNNNACCTAGTTGTCACCTTTGTATAAC
GCACACGTCTGAACTCTTTCCCTACACGACGCTCTTCCGATCTNNNACCTAGTTGTCACCTTTGTATAAC
AGCACACGTCTGAACTCTTTCCCTACACGACGCTCTTCCGATCTNNNAGGGAAAGGAGCATGGTTGTT
AGCACACGTCTGAACTCTTTCCCTACACGACGCTCTTCCGATCTNNNAGGGAAAGGAGCATGGTTGTT
AGCACACGTCTGAACTCTTTCCCTACACGACGCTCTTCCGATCTNNNAGGGAAAGGAGCATGGTTGTT
AGCACACGTCTGAACTCTTTCCCTACACGACGCTCTTCCGATCTNNNGGCAGTGGATTTTAGCTTGCTA
AGCACACGTCTGAACTCTTTCCCTACACGACGCTCTTCCGATCTNNNGGCAGTGGATTTTAGCTTGCTA
AGCACACGTCTGAACTCTTTCCCTACACGACGCTCTTCCGATCTNNNGGCAGTGGATTTTAGCTTGCTA
GAGCACACGTCTGAACTCTTTCCCTACACGACGCTCTTCCGATCTNNNTCTTTACACTGCTTTTGTGATG
GAGCACACGTCTGAACTCTTTCCCTACACGACGCTCTTCCGATCTNNNTCTTTACACTGCTTTTGTGATG

GAGCACACGTCTGAACTCTTTCCCTACACGACGCTCTTCCGATCTNNNTCTTTACACTGCTTTTGTGATG
GAGCACACGTCTGAACTCTTTCCCTACACGACGCTCTTCCGATCTNNNAGTAGAGCAAGGTGGTTGAA
GAGCACACGTCTGAACTCTTTCCCTACACGACGCTCTTCCGATCTNNNAGTAGAGCAAGGTGGTTGAA
GAGCACACGTCTGAACTCTTTCCCTACACGACGCTCTTCCGATCTNNNAGTAGAGCAAGGTGGTTGAA
GAGCACACGTCTGAACTCTTTCCCTACACGACGCTCTTCCGATCTNNNAAGAAATGCGACCTTTCCAACA
GAGCACACGTCTGAACTCTTTCCCTACACGACGCTCTTCCGATCTNNNAAGAAATGCGACCTTTCCAACA
GAGCACACGTCTGAACTCTTTCCCTACACGACGCTCTTCCGATCTNNNAAGAAATGCGACCTTTCCAACA
AGCACACGTCTGAACTCTTTCCCTACACGACGCTCTTCCGATCTNNNAGGTTCTGAGTCCAAGTTCCAA
AGCACACGTCTGAACTCTTTCCCTACACGACGCTCTTCCGATCTNNNAGGTTCTGAGTCCAAGTTCCAA
AGCACACGTCTGAACTCTTTCCCTACACGACGCTCTTCCGATCTNNNAGGTTCTGAGTCCAAGTTCCAA
AGCACACGTCTGAACTCTTTCCCTACACGACGCTCTTCCGATCTNNNTGGGTAAGCTTTTTGATGTGCT
AGCACACGTCTGAACTCTTTCCCTACACGACGCTCTTCCGATCTNNNTGGGTAAGCTTTTTGATGTGCT
AGCACACGTCTGAACTCTTTCCCTACACGACGCTCTTCCGATCTNNNTGGGTAAGCTTTTTGATGTGCT
AGCACACGTCTGAACTCTTTCCCTACACGACGCTCTTCCGATCTNNNTCTTTCTCTCTCTCTCTCCCTCCCA
AGCACACGTCTGAACTCTTTCCCTACACGACGCTCTTCCGATCTNNNTCTTTCTCTCTCTCTCTCCCTCCCA
AGCACACGTCTGAACTCTTTCCCTACACGACGCTCTTCCGATCTNNNTCTTTCTCTCTCTCTCTCCCTCCCA
AGAGCACACGTCTGAACTCTTTCCCTACACGACGCTCTTCCGATCTNNNTTATATGTCAGTCTCTGCCTG
AGAGCACACGTCTGAACTCTTTCCCTACACGACGCTCTTCCGATCTNNNTTATATGTCAGTCTCTGCCTG
AGAGCACACGTCTGAACTCTTTCCCTACACGACGCTCTTCCGATCTNNNTTATATGTCAGTCTCTGCCTG
AGAGCACACGTCTGAACTCTTTCCCTACACGACGCTCTTCCGATCTNNNGAGTTCACACCAGTACACTCC
AGAGCACACGTCTGAACTCTTTCCCTACACGACGCTCTTCCGATCTNNNGAGTTCACACCAGTACACTCC
AGAGCACACGTCTGAACTCTTTCCCTACACGACGCTCTTCCGATCTNNNGAGTTCACACCAGTACACTCC
ACACGTCTGAACTCTTTCCCTACACGACGCTCTTCCGATCTNNNCTAGGTGTCAGACGGCTGTCCATCG
ACACGTCTGAACTCTTTCCCTACACGACGCTCTTCCGATCTNNNCTAGGTGTCAGACGGCTGTCCATCG
ACACGTCTGAACTCTTTCCCTACACGACGCTCTTCCGATCTNNNCTAGGTGTCAGACGGCTGTCCATCG
GAGCACACGTCTGAACTCTTTCCCTACACGACGCTCTTCCGATCTNNNTAATGGTCTCCAATTCCATCCG
GAGCACACGTCTGAACTCTTTCCCTACACGACGCTCTTCCGATCTNNNTAATGGTCTCCAATTCCATCCG
GAGCACACGTCTGAACTCTTTCCCTACACGACGCTCTTCCGATCTNNNTAATGGTCTCCAATTCCATCCG
AGAGCACACGTCTGAACTCTTTCCCTACACGACGCTCTTCCGATCTNNNAGATACTTGGGAGGCTGAA
AGAGCACACGTCTGAACTCTTTCCCTACACGACGCTCTTCCGATCTNNNAGATACTTGGGAGGCTGAA
AGAGCACACGTCTGAACTCTTTCCCTACACGACGCTCTTCCGATCTNNNAGATACTTGGGAGGCTGAA
GAGCACACGTCTGAACTCTTTCCCTACACGACGCTCTTCCGATCTNNNAATACAGGTGGAAAAGGCTG
GAGCACACGTCTGAACTCTTTCCCTACACGACGCTCTTCCGATCTNNNAATACAGGTGGAAAAGGCTG
GAGCACACGTCTGAACTCTTTCCCTACACGACGCTCTTCCGATCTNNNAATACAGGTGGAAAAGGCTG
AGCACACGTCTGAACTCTTTCCCTACACGACGCTCTTCCGATCTNNNTGCCTGTAGTCCCAGCTACTCCA
AGCACACGTCTGAACTCTTTCCCTACACGACGCTCTTCCGATCTNNNTGCCTGTAGTCCCAGCTACTCCA
AGCACACGTCTGAACTCTTTCCCTACACGACGCTCTTCCGATCTNNNTGCCTGTAGTCCCAGCTACTCCA
GAGCACACGTCTGAACTCTTTCCCTACACGACGCTCTTCCGATCTNNNTGGCCTCAGAGGATATGTACA
GAGCACACGTCTGAACTCTTTCCCTACACGACGCTCTTCCGATCTNNNTGGCCTCAGAGGATATGTACA
GAGCACACGTCTGAACTCTTTCCCTACACGACGCTCTTCCGATCTNNNTGGCCTCAGAGGATATGTACA
GAGCACACGTCTGAACTCTTTCCCTACACGACGCTCTTCCGATCTNNNAGTGCTCTTATGGTGAAGCAC
GAGCACACGTCTGAACTCTTTCCCTACACGACGCTCTTCCGATCTNNNAGTGCTCTTATGGTGAAGCAC
GAGCACACGTCTGAACTCTTTCCCTACACGACGCTCTTCCGATCTNNNAGTGCTCTTATGGTGAAGCAC

AGAGCACACGTCTGAACTCTTTCCCTACACGACGCTCTTCCGATCTNNNCCCTCTTTGAAGTGAAGACA
AGAGCACACGTCTGAACTCTTTCCCTACACGACGCTCTTCCGATCTNNNCCCTCTTTGAAGTGAAGACA
AGAGCACACGTCTGAACTCTTTCCCTACACGACGCTCTTCCGATCTNNNCCCTCTTTGAAGTGAAGACA
GAGCACACGTCTGAACTCTTTCCCTACACGACGCTCTTCCGATCTNNNCCAACCCCACATTATGGATGGA
GAGCACACGTCTGAACTCTTTCCCTACACGACGCTCTTCCGATCTNNNCCAACCCCACATTATGGATGGA
GAGCACACGTCTGAACTCTTTCCCTACACGACGCTCTTCCGATCTNNNCCAACCCCACATTATGGATGGA
AGAGCACACGTCTGAACTCTTTCCCTACACGACGCTCTTCCGATCTNNNTCCACTTCTCTAGCCTAACCT
AGAGCACACGTCTGAACTCTTTCCCTACACGACGCTCTTCCGATCTNNNTCCACTTCTCTAGCCTAACCT
AGAGCACACGTCTGAACTCTTTCCCTACACGACGCTCTTCCGATCTNNNTCCACTTCTCTAGCCTAACCT
AAGAGCACACGTCTGAACTCTTTCCCTACACGACGCTCTTCCGATCTNNNCCACTTGGAATTGTCAAAG
AAGAGCACACGTCTGAACTCTTTCCCTACACGACGCTCTTCCGATCTNNNCCACTTGGAATTGTCAAAG
AAGAGCACACGTCTGAACTCTTTCCCTACACGACGCTCTTCCGATCTNNNCCACTTGGAATTGTCAAAG
AGCACACGTCTGAACTCTTTCCCTACACGACGCTCTTCCGATCTNNNTGGAAGTCAGAAGGCCTAAGTT
AGCACACGTCTGAACTCTTTCCCTACACGACGCTCTTCCGATCTNNNTGGAAGTCAGAAGGCCTAAGTT
AGCACACGTCTGAACTCTTTCCCTACACGACGCTCTTCCGATCTNNNTGGAAGTCAGAAGGCCTAAGTT
GCACACGTCTGAACTCTTTCCCTACACGACGCTCTTCCGATCTNNNCTGAGGCAGGAGAATCACTTGAA
GCACACGTCTGAACTCTTTCCCTACACGACGCTCTTCCGATCTNNNCTGAGGCAGGAGAATCACTTGAA
GCACACGTCTGAACTCTTTCCCTACACGACGCTCTTCCGATCTNNNCTGAGGCAGGAGAATCACTTGAA
GAGCACACGTCTGAACTCTTTCCCTACACGACGCTCTTCCGATCTNNNACACACACATGCACACACTAT
GAGCACACGTCTGAACTCTTTCCCTACACGACGCTCTTCCGATCTNNNACACACACATGCACACACTAT
GAGCACACGTCTGAACTCTTTCCCTACACGACGCTCTTCCGATCTNNNACACACACATGCACACACTAT
AGCACACGTCTGAACTCTTTCCCTACACGACGCTCTTCCGATCTNNNAGCCACTTGGTTAGGACATAG
AGCACACGTCTGAACTCTTTCCCTACACGACGCTCTTCCGATCTNNNAGCCACTTGGTTAGGACATAG
AGCACACGTCTGAACTCTTTCCCTACACGACGCTCTTCCGATCTNNNAGCCACTTGGTTAGGACATAG
AGCACACGTCTGAACTCTTTCCCTACACGACGCTCTTCCGATCTNNNACAACATGTCGAGAACCAGAAC
AGCACACGTCTGAACTCTTTCCCTACACGACGCTCTTCCGATCTNNNACAACATGTCGAGAACCAGAAC
AGCACACGTCTGAACTCTTTCCCTACACGACGCTCTTCCGATCTNNNACAACATGTCGAGAACCAGAAC
GAGCACACGTCTGAACTCTTTCCCTACACGACGCTCTTCCGATCTNNNGCCTATTTGACATCAAGAGTG
GAGCACACGTCTGAACTCTTTCCCTACACGACGCTCTTCCGATCTNNNGCCTATTTGACATCAAGAGTG
GAGCACACGTCTGAACTCTTTCCCTACACGACGCTCTTCCGATCTNNNGCCTATTTGACATCAAGAGTG
AGAGCACACGTCTGAACTCTTTCCCTACACGACGCTCTTCCGATCTNNNTGAGCTCTGAAATTGAGGCA
AGAGCACACGTCTGAACTCTTTCCCTACACGACGCTCTTCCGATCTNNNTGAGCTCTGAAATTGAGGCA
AGAGCACACGTCTGAACTCTTTCCCTACACGACGCTCTTCCGATCTNNNTGAGCTCTGAAATTGAGGCA
GAGCACACGTCTGAACTCTTTCCCTACACGACGCTCTTCCGATCTNNNACTGAGGTAGGAGGATCACTT
GAGCACACGTCTGAACTCTTTCCCTACACGACGCTCTTCCGATCTNNNACTGAGGTAGGAGGATCACTT
GAGCACACGTCTGAACTCTTTCCCTACACGACGCTCTTCCGATCTNNNACTGAGGTAGGAGGATCACTT
AGCACACGTCTGAACTCTTTCCCTACACGACGCTCTTCCGATCTNNNCTCCAACCATCAGAGGTACAGA
AGCACACGTCTGAACTCTTTCCCTACACGACGCTCTTCCGATCTNNNCTCCAACCATCAGAGGTACAGA
AGCACACGTCTGAACTCTTTCCCTACACGACGCTCTTCCGATCTNNNCTCCAACCATCAGAGGTACAGA
GAGCACACGTCTGAACTCTTTCCCTACACGACGCTCTTCCGATCTNNNGATAGAGCAAGACGGCGTCT
GAGCACACGTCTGAACTCTTTCCCTACACGACGCTCTTCCGATCTNNNGATAGAGCAAGACGGCGTCT
GAGCACACGTCTGAACTCTTTCCCTACACGACGCTCTTCCGATCTNNNGATAGAGCAAGACGGCGTCT
GAGCACACGTCTGAACTCTTTCCCTACACGACGCTCTTCCGATCTNNNAGGTGGATTATGAGGTCAG

GAGCACACGTCTGAACTCTTTCCCTACACGACGCTCTTCCGATCTNNNAGGTGGATTATGAGGTCAG
GAGCACACGTCTGAACTCTTTCCCTACACGACGCTCTTCCGATCTNNNAGGTGGATTATGAGGTCAG
AGCACACGTCTGAACTCTTTCCCTACACGACGCTCTTCCGATCTNNNTGTGTCCAGCCGAATTCACACAT
AGCACACGTCTGAACTCTTTCCCTACACGACGCTCTTCCGATCTNNNTGTGTCCAGCCGAATTCACACAT
AGCACACGTCTGAACTCTTTCCCTACACGACGCTCTTCCGATCTNNNTGTGTCCAGCCGAATTCACACAT
GAGCACACGTCTGAACTCTTTCCCTACACGACGCTCTTCCGATCTNNNCTGTAGTCCCAGCTACTCAGGA
GAGCACACGTCTGAACTCTTTCCCTACACGACGCTCTTCCGATCTNNNCTGTAGTCCCAGCTACTCAGGA
GAGCACACGTCTGAACTCTTTCCCTACACGACGCTCTTCCGATCTNNNCTGTAGTCCCAGCTACTCAGGA
GAGCACACGTCTGAACTCTTTCCCTACACGACGCTCTTCCGATCTNNNCTGAGGTGGGAGAATCACTT
GAGCACACGTCTGAACTCTTTCCCTACACGACGCTCTTCCGATCTNNNCTGAGGTGGGAGAATCACTT
GAGCACACGTCTGAACTCTTTCCCTACACGACGCTCTTCCGATCTNNNCTGAGGTGGGAGAATCACTT
GAGCACACGTCTGAACTCTTTCCCTACACGACGCTCTTCCGATCTNNNGTATTCCAACAGAGGTTCCCA
GAGCACACGTCTGAACTCTTTCCCTACACGACGCTCTTCCGATCTNNNGTATTCCAACAGAGGTTCCCA
GAGCACACGTCTGAACTCTTTCCCTACACGACGCTCTTCCGATCTNNNGTATTCCAACAGAGGTTCCCA
GAGCACACGTCTGAACTCTTTCCCTACACGACGCTCTTCCGATCTNNNCTCTAGGACACAGCCCACGAA
GAGCACACGTCTGAACTCTTTCCCTACACGACGCTCTTCCGATCTNNNCTCTAGGACACAGCCCACGAA
GAGCACACGTCTGAACTCTTTCCCTACACGACGCTCTTCCGATCTNNNCTCTAGGACACAGCCCACGAA
GAGCACACGTCTGAACTCTTTCCCTACACGACGCTCTTCCGATCTNNNTCGGTGTTGGAGAGAAATTC
GAGCACACGTCTGAACTCTTTCCCTACACGACGCTCTTCCGATCTNNNTCGGTGTTGGAGAGAAATTC
GAGCACACGTCTGAACTCTTTCCCTACACGACGCTCTTCCGATCTNNNTCGGTGTTGGAGAGAAATTC
ACACGTCTGAACTCTTTCCCTACACGACGCTCTTCCGATCTNNNCACCATGAGGACTCCCAACAAGCAT
ACACGTCTGAACTCTTTCCCTACACGACGCTCTTCCGATCTNNNCACCATGAGGACTCCCAACAAGCAT
ACACGTCTGAACTCTTTCCCTACACGACGCTCTTCCGATCTNNNCACCATGAGGACTCCCAACAAGCAT
GAGCACACGTCTGAACTCTTTCCCTACACGACGCTCTTCCGATCTNNNACAAGAGAAAATGTAAGCCTC
GAGCACACGTCTGAACTCTTTCCCTACACGACGCTCTTCCGATCTNNNACAAGAGAAAATGTAAGCCTC
GAGCACACGTCTGAACTCTTTCCCTACACGACGCTCTTCCGATCTNNNACAAGAGAAAATGTAAGCCTC
GAGCACACGTCTGAACTCTTTCCCTACACGACGCTCTTCCGATCTNNNCGCAACAAACAGCAAAAACCA
GAGCACACGTCTGAACTCTTTCCCTACACGACGCTCTTCCGATCTNNNCGCAACAAACAGCAAAAACCA
GAGCACACGTCTGAACTCTTTCCCTACACGACGCTCTTCCGATCTNNNCGCAACAAACAGCAAAAACCA
AGAGCACACGTCTGAACTCTTTCCCTACACGACGCTCTTCCGATCTNNNACTCAGCTCACAGGTAGAAT
AGAGCACACGTCTGAACTCTTTCCCTACACGACGCTCTTCCGATCTNNNACTCAGCTCACAGGTAGAAT
AGAGCACACGTCTGAACTCTTTCCCTACACGACGCTCTTCCGATCTNNNACTCAGCTCACAGGTAGAAT
AGCACACGTCTGAACTCTTTCCCTACACGACGCTCTTCCGATCTNNNCTGGACAAGAGAACAAGACTCC
AGCACACGTCTGAACTCTTTCCCTACACGACGCTCTTCCGATCTNNNCTGGACAAGAGAACAAGACTCC
AGCACACGTCTGAACTCTTTCCCTACACGACGCTCTTCCGATCTNNNCTGGACAAGAGAACAAGACTCC
AGCACACGTCTGAACTCTTTCCCTACACGACGCTCTTCCGATCTNNNAATGATGTGAACCCAGGTGTGA
AGCACACGTCTGAACTCTTTCCCTACACGACGCTCTTCCGATCTNNNAATGATGTGAACCCAGGTGTGA
AGCACACGTCTGAACTCTTTCCCTACACGACGCTCTTCCGATCTNNNAATGATGTGAACCCAGGTGTGA
GAGCACACGTCTGAACTCTTTCCCTACACGACGCTCTTCCGATCTNNNTAGCTAGGATTACAGGCACA
GAGCACACGTCTGAACTCTTTCCCTACACGACGCTCTTCCGATCTNNNTAGCTAGGATTACAGGCACA
GAGCACACGTCTGAACTCTTTCCCTACACGACGCTCTTCCGATCTNNNTAGCTAGGATTACAGGCACA
AGCACACGTCTGAACTCTTTCCCTACACGACGCTCTTCCGATCTNNNCAAGGAGTTCTTGGCTTTCCTCA
AGCACACGTCTGAACTCTTTCCCTACACGACGCTCTTCCGATCTNNNCAAGGAGTTCTTGGCTTTCCTCA

AGCACACGTCTGAACTCTTTCCCTACACGACGCTCTTCCGATCTNNNCAAGGAGTTCTTGGCTTTCCTC

GCACACGTCTGAACTCTTTCCCTACACGACGCTCTTCCGATCTNNNTTTCTGAGAAACTGGATTTGCC

GCACACGTCTGAACTCTTTCCCTACACGACGCTCTTCCGATCTNNNTTTCTGAGAAACTGGATTTGCC

GCACACGTCTGAACTCTTTCCCTACACGACGCTCTTCCGATCTNNNTTTCTGAGAAACTGGATTTGCC

AGCACACGTCTGAACTCTTTCCCTACACGACGCTCTTCCGATCTNNNGAAGCAAAGCTGACATTGGA

AGCACACGTCTGAACTCTTTCCCTACACGACGCTCTTCCGATCTNNNGAAGCAAAGCTGACATTGGA

AGCACACGTCTGAACTCTTTCCCTACACGACGCTCTTCCGATCTNNNGAAGCAAAGCTGACATTGGA

CACACGTCTGAACTCTTTCCCTACACGACGCTCTTCCGATCTNNNTCCTCCATGGTCTGGTAAGTGTA

CACACGTCTGAACTCTTTCCCTACACGACGCTCTTCCGATCTNNNTCCTCCATGGTCTGGTAAGTGTA

CACACGTCTGAACTCTTTCCCTACACGACGCTCTTCCGATCTNNNTCCTCCATGGTCTGGTAAGTGTA

ACACGTCTGAACTCTTTCCCTACACGACGCTCTTCCGATCTNNNAGAAAAGTTCCCCTCCCCAAAATCA

ACACGTCTGAACTCTTTCCCTACACGACGCTCTTCCGATCTNNNAGAAAAGTTCCCCTCCCCAAAATCA

ACACGTCTGAACTCTTTCCCTACACGACGCTCTTCCGATCTNNNAGAAAAGTTCCCCTCCCCAAAATCA

AGCACACGTCTGAACTCTTTCCCTACACGACGCTCTTCCGATCTNNNTAAAGTGGAGCCTGAAGTAG

AGCACACGTCTGAACTCTTTCCCTACACGACGCTCTTCCGATCTNNNTAAAGTGGAGCCTGAAGTAG

AGCACACGTCTGAACTCTTTCCCTACACGACGCTCTTCCGATCTNNNTAAAGTGGAGCCTGAAGTAG

AGCACACGTCTGAACTCTTTCCCTACACGACGCTCTTCCGATCTNNNACAAAAATTAGCTGGGTGTGG

AGCACACGTCTGAACTCTTTCCCTACACGACGCTCTTCCGATCTNNNACAAAAATTAGCTGGGTGTGG

AGCACACGTCTGAACTCTTTCCCTACACGACGCTCTTCCGATCTNNNACAAAAATTAGCTGGGTGTGG

AGCACACGTCTGAACTCTTTCCCTACACGACGCTCTTCCGATCTNNNGGTCTGATCACCCCAACGTATT

AGCACACGTCTGAACTCTTTCCCTACACGACGCTCTTCCGATCTNNNGGTCTGATCACCCCAACGTATT

AGCACACGTCTGAACTCTTTCCCTACACGACGCTCTTCCGATCTNNNGGTCTGATCACCCCAACGTATT

GCACACGTCTGAACTCTTTCCCTACACGACGCTCTTCCGATCTNNNTAGGCATTACCAGCATGATGGA

GCACACGTCTGAACTCTTTCCCTACACGACGCTCTTCCGATCTNNNTAGGCATTACCAGCATGATGGA

GCACACGTCTGAACTCTTTCCCTACACGACGCTCTTCCGATCTNNNTAGGCATTACCAGCATGATGGA

GAGCACACGTCTGAACTCTTTCCCTACACGACGCTCTTCCGATCTNNNCATGTCAACCTTTTTCTTTTCT

GAGCACACGTCTGAACTCTTTCCCTACACGACGCTCTTCCGATCTNNNCATGTCAACCTTTTTCTTTTCT

GAGCACACGTCTGAACTCTTTCCCTACACGACGCTCTTCCGATCTNNNCATGTCAACCTTTTTCTTTTCT

AGCACACGTCTGAACTCTTTCCCTACACGACGCTCTTCCGATCTNNNTCATAATTGCATTGACCCGAGT

AGCACACGTCTGAACTCTTTCCCTACACGACGCTCTTCCGATCTNNNTCATAATTGCATTGACCCGAGT

AGCACACGTCTGAACTCTTTCCCTACACGACGCTCTTCCGATCTNNNTCATAATTGCATTGACCCGAGT

GAGCACACGTCTGAACTCTTTCCCTACACGACGCTCTTCCGATCTNNNAAATGAGAATCGCTTGAACC

GAGCACACGTCTGAACTCTTTCCCTACACGACGCTCTTCCGATCTNNNAAATGAGAATCGCTTGAACC

GAGCACACGTCTGAACTCTTTCCCTACACGACGCTCTTCCGATCTNNNAAATGAGAATCGCTTGAACC

GAGCACACGTCTGAACTCTTTCCCTACACGACGCTCTTCCGATCTNNNGGAGGCCTCAGGAAACTTA

GAGCACACGTCTGAACTCTTTCCCTACACGACGCTCTTCCGATCTNNNGGAGGCCTCAGGAAACTTA

GAGCACACGTCTGAACTCTTTCCCTACACGACGCTCTTCCGATCTNNNGGAGGCCTCAGGAAACTTA

GCACACGTCTGAACTCTTTCCCTACACGACGCTCTTCCGATCTNNNTATCTGTCCAGTGCTGAAAGTG

GCACACGTCTGAACTCTTTCCCTACACGACGCTCTTCCGATCTNNNTATCTGTCCAGTGCTGAAAGTG

GCACACGTCTGAACTCTTTCCCTACACGACGCTCTTCCGATCTNNNTATCTGTCCAGTGCTGAAAGTG

GAGCACACGTCTGAACTCTTTCCCTACACGACGCTCTTCCGATCTNNNAGAGGGAACAGTAATTGAG

GAGCACACGTCTGAACTCTTTCCCTACACGACGCTCTTCCGATCTNNNAGAGGGAACAGTAATTGAG

GAGCACACGTCTGAACTCTTTCCCTACACGACGCTCTTCCGATCTNNNAGAGGGAACAGTAATTGAG

AGCACACGTCTGAACTCTTTCCCTACACGACGCTCTTCCGATCTNNNCCTCCACGTAAACACAGTTACCT
AGCACACGTCTGAACTCTTTCCCTACACGACGCTCTTCCGATCTNNNCCTCCACGTAAACACAGTTACCT
AGCACACGTCTGAACTCTTTCCCTACACGACGCTCTTCCGATCTNNNCCTCCACGTAAACACAGTTACCT
AGCACACGTCTGAACTCTTTCCCTACACGACGCTCTTCCGATCTNNNCACCCCAGGACTTTACTGATGAA
AGCACACGTCTGAACTCTTTCCCTACACGACGCTCTTCCGATCTNNNCACCCCAGGACTTTACTGATGAA
AGCACACGTCTGAACTCTTTCCCTACACGACGCTCTTCCGATCTNNNCACCCCAGGACTTTACTGATGAA
AGCACACGTCTGAACTCTTTCCCTACACGACGCTCTTCCGATCTNNNAAAAAATTAGCCAGGCGTGGT
AGCACACGTCTGAACTCTTTCCCTACACGACGCTCTTCCGATCTNNNAAAAAATTAGCCAGGCGTGGT
AGCACACGTCTGAACTCTTTCCCTACACGACGCTCTTCCGATCTNNNAAAAAATTAGCCAGGCGTGGT
GAGCACACGTCTGAACTCTTTCCCTACACGACGCTCTTCCGATCTNNNGCCTGTAATCCCAGTACTTTG
GAGCACACGTCTGAACTCTTTCCCTACACGACGCTCTTCCGATCTNNNGCCTGTAATCCCAGTACTTTG
GAGCACACGTCTGAACTCTTTCCCTACACGACGCTCTTCCGATCTNNNGCCTGTAATCCCAGTACTTTG
AGCACACGTCTGAACTCTTTCCCTACACGACGCTCTTCCGATCTNNNATTAGCCTTGCTAGGGGTCTTTC
AGCACACGTCTGAACTCTTTCCCTACACGACGCTCTTCCGATCTNNNATTAGCCTTGCTAGGGGTCTTTC
AGCACACGTCTGAACTCTTTCCCTACACGACGCTCTTCCGATCTNNNATTAGCCTTGCTAGGGGTCTTTC
GAGCACACGTCTGAACTCTTTCCCTACACGACGCTCTTCCGATCTNNNTGATGACTGGATGAAAGGACA
GAGCACACGTCTGAACTCTTTCCCTACACGACGCTCTTCCGATCTNNNTGATGACTGGATGAAAGGACA
GAGCACACGTCTGAACTCTTTCCCTACACGACGCTCTTCCGATCTNNNTGATGACTGGATGAAAGGACA
GAGCACACGTCTGAACTCTTTCCCTACACGACGCTCTTCCGATCTNNNATAGACCAAAGGGAAAGCGA
GAGCACACGTCTGAACTCTTTCCCTACACGACGCTCTTCCGATCTNNNATAGACCAAAGGGAAAGCGA
GAGCACACGTCTGAACTCTTTCCCTACACGACGCTCTTCCGATCTNNNATAGACCAAAGGGAAAGCGA
AGCACACGTCTGAACTCTTTCCCTACACGACGCTCTTCCGATCTNNNGAATCAAATGGGGGGAGCTT
AGCACACGTCTGAACTCTTTCCCTACACGACGCTCTTCCGATCTNNNGAATCAAATGGGGGGAGCTT
AGCACACGTCTGAACTCTTTCCCTACACGACGCTCTTCCGATCTNNNGAATCAAATGGGGGGAGCTT
GAGCACACGTCTGAACTCTTTCCCTACACGACGCTCTTCCGATCTNNNACATACCACTGAGCACAAAGT
GAGCACACGTCTGAACTCTTTCCCTACACGACGCTCTTCCGATCTNNNACATACCACTGAGCACAAAGT
GAGCACACGTCTGAACTCTTTCCCTACACGACGCTCTTCCGATCTNNNACATACCACTGAGCACAAAGT
AGAGCACACGTCTGAACTCTTTCCCTACACGACGCTCTTCCGATCTNNNAGTATCTGGGATTACAGGTG
AGAGCACACGTCTGAACTCTTTCCCTACACGACGCTCTTCCGATCTNNNAGTATCTGGGATTACAGGTG
AGAGCACACGTCTGAACTCTTTCCCTACACGACGCTCTTCCGATCTNNNAGTATCTGGGATTACAGGTG
ACACGTCTGAACTCTTTCCCTACACGACGCTCTTCCGATCTNNNGATGCGGAAATCAGGATCCACTTCA
ACACGTCTGAACTCTTTCCCTACACGACGCTCTTCCGATCTNNNGATGCGGAAATCAGGATCCACTTCA
ACACGTCTGAACTCTTTCCCTACACGACGCTCTTCCGATCTNNNGATGCGGAAATCAGGATCCACTTCA
AGCACACGTCTGAACTCTTTCCCTACACGACGCTCTTCCGATCTNNNGCAATGTAAGAGATAATTATTC
AGCACACGTCTGAACTCTTTCCCTACACGACGCTCTTCCGATCTNNNGCAATGTAAGAGATAATTATTC
AGCACACGTCTGAACTCTTTCCCTACACGACGCTCTTCCGATCTNNNGCAATGTAAGAGATAATTATTC
AGCACACGTCTGAACTCTTTCCCTACACGACGCTCTTCCGATCTNNNAAGCAAGCAGAGCTCATTAGTC
AGCACACGTCTGAACTCTTTCCCTACACGACGCTCTTCCGATCTNNNAAGCAAGCAGAGCTCATTAGTC
AGCACACGTCTGAACTCTTTCCCTACACGACGCTCTTCCGATCTNNNAAGCAAGCAGAGCTCATTAGTC
AGCACACGTCTGAACTCTTTCCCTACACGACGCTCTTCCGATCTNNNGACATTGCAGTGAGCTGAGATT
AGCACACGTCTGAACTCTTTCCCTACACGACGCTCTTCCGATCTNNNGACATTGCAGTGAGCTGAGATT
AGCACACGTCTGAACTCTTTCCCTACACGACGCTCTTCCGATCTNNNGACATTGCAGTGAGCTGAGATT
AGCACACGTCTGAACTCTTTCCCTACACGACGCTCTTCCGATCTNNNGCGAAGGAACAAAACTCCATC

AGCACACGTCTGAACTCTTTCCCTACACGACGCTCTTCCGATCTNNNGCGAAGGAACAAAACTCCATC
AGCACACGTCTGAACTCTTTCCCTACACGACGCTCTTCCGATCTNNNGCGAAGGAACAAAACTCCATC
GAGCACACGTCTGAACTCTTTCCCTACACGACGCTCTTCCGATCTNNNAGGTTGCAGTGAGATGTGA
GAGCACACGTCTGAACTCTTTCCCTACACGACGCTCTTCCGATCTNNNAGGTTGCAGTGAGATGTGA
GAGCACACGTCTGAACTCTTTCCCTACACGACGCTCTTCCGATCTNNNAGGTTGCAGTGAGATGTGA
GAGCACACGTCTGAACTCTTTCCCTACACGACGCTCTTCCGATCTNNNGATTCAGCAAAGGAGACCGA
GAGCACACGTCTGAACTCTTTCCCTACACGACGCTCTTCCGATCTNNNGATTCAGCAAAGGAGACCGA
GAGCACACGTCTGAACTCTTTCCCTACACGACGCTCTTCCGATCTNNNGATTCAGCAAAGGAGACCGA
AGCACACGTCTGAACTCTTTCCCTACACGACGCTCTTCCGATCTNNNTTGCTATTGAGGGACATTGTG
AGCACACGTCTGAACTCTTTCCCTACACGACGCTCTTCCGATCTNNNTTGCTATTGAGGGACATTGTG
AGCACACGTCTGAACTCTTTCCCTACACGACGCTCTTCCGATCTNNNTTGCTATTGAGGGACATTGTG
GCACACGTCTGAACTCTTTCCCTACACGACGCTCTTCCGATCTNNNACCTCCAGTGCAATGTTGAATT
GCACACGTCTGAACTCTTTCCCTACACGACGCTCTTCCGATCTNNNACCTCCAGTGCAATGTTGAATT
GCACACGTCTGAACTCTTTCCCTACACGACGCTCTTCCGATCTNNNACCTCCAGTGCAATGTTGAATT
AGCACACGTCTGAACTCTTTCCCTACACGACGCTCTTCCGATCTNNNGAAAGTACACTTGGAAAAGGG
AGCACACGTCTGAACTCTTTCCCTACACGACGCTCTTCCGATCTNNNGAAAGTACACTTGGAAAAGGG
AGCACACGTCTGAACTCTTTCCCTACACGACGCTCTTCCGATCTNNNGAAAGTACACTTGGAAAAGGG
AGCACACGTCTGAACTCTTTCCCTACACGACGCTCTTCCGATCTNNNGTACCATTGTACTCTAGCCTGG
AGCACACGTCTGAACTCTTTCCCTACACGACGCTCTTCCGATCTNNNGTACCATTGTACTCTAGCCTGG
AGCACACGTCTGAACTCTTTCCCTACACGACGCTCTTCCGATCTNNNGTACCATTGTACTCTAGCCTGG
AGCACACGTCTGAACTCTTTCCCTACACGACGCTCTTCCGATCTNNNATAATTCTCATGCCTCAGCCTCC
AGCACACGTCTGAACTCTTTCCCTACACGACGCTCTTCCGATCTNNNATAATTCTCATGCCTCAGCCTCC
AGCACACGTCTGAACTCTTTCCCTACACGACGCTCTTCCGATCTNNNATAATTCTCATGCCTCAGCCTCC
GCACACGTCTGAACTCTTTCCCTACACGACGCTCTTCCGATCTNNNGGTGGGAGGATTACTTGAACT
GCACACGTCTGAACTCTTTCCCTACACGACGCTCTTCCGATCTNNNGGTGGGAGGATTACTTGAACT
GCACACGTCTGAACTCTTTCCCTACACGACGCTCTTCCGATCTNNNGGTGGGAGGATTACTTGAACT
AGCACACGTCTGAACTCTTTCCCTACACGACGCTCTTCCGATCTNNNCTGTAATCCCAGCTACTCAGG
AGCACACGTCTGAACTCTTTCCCTACACGACGCTCTTCCGATCTNNNCTGTAATCCCAGCTACTCAGG
AGCACACGTCTGAACTCTTTCCCTACACGACGCTCTTCCGATCTNNNCTGTAATCCCAGCTACTCAGG
GAGCACACGTCTGAACTCTTTCCCTACACGACGCTCTTCCGATCTNNNGATCACGTGAGGTCAGGAG
GAGCACACGTCTGAACTCTTTCCCTACACGACGCTCTTCCGATCTNNNGATCACGTGAGGTCAGGAG
GAGCACACGTCTGAACTCTTTCCCTACACGACGCTCTTCCGATCTNNNGATCACGTGAGGTCAGGAG
AGCACACGTCTGAACTCTTTCCCTACACGACGCTCTTCCGATCTNNNAGCTGGCTACAGAATCCTGAA
AGCACACGTCTGAACTCTTTCCCTACACGACGCTCTTCCGATCTNNNAGCTGGCTACAGAATCCTGAA
AGCACACGTCTGAACTCTTTCCCTACACGACGCTCTTCCGATCTNNNAGCTGGCTACAGAATCCTGAA
AGCACACGTCTGAACTCTTTCCCTACACGACGCTCTTCCGATCTNNNTCTGCCTCCTTCTTGATCTCCTA
AGCACACGTCTGAACTCTTTCCCTACACGACGCTCTTCCGATCTNNNTCTGCCTCCTTCTTGATCTCCTA
AGCACACGTCTGAACTCTTTCCCTACACGACGCTCTTCCGATCTNNNTCTGCCTCCTTCTTGATCTCCTA
GAGCACACGTCTGAACTCTTTCCCTACACGACGCTCTTCCGATCTNNNCTGTAATCCCAGCTACTCGGA
GAGCACACGTCTGAACTCTTTCCCTACACGACGCTCTTCCGATCTNNNCTGTAATCCCAGCTACTCGGA
GAGCACACGTCTGAACTCTTTCCCTACACGACGCTCTTCCGATCTNNNCTGTAATCCCAGCTACTCGGA
AGAGCACACGTCTGAACTCTTTCCCTACACGACGCTCTTCCGATCTNNNGAAGTTGTAGTGAGCCGAG
AGAGCACACGTCTGAACTCTTTCCCTACACGACGCTCTTCCGATCTNNNGAAGTTGTAGTGAGCCGAG

AGAGCACACGTCTGAACTCTTTCCCTACACGACGCTCTTCCGATCTNNNGAAGTTGTAGTGAGCCGAG
GAGCACACGTCTGAACTCTTTCCCTACACGACGCTCTTCCGATCTNNNTCACAGCCTCAAGAAATCCAAG
GAGCACACGTCTGAACTCTTTCCCTACACGACGCTCTTCCGATCTNNNTCACAGCCTCAAGAAATCCAAG
GAGCACACGTCTGAACTCTTTCCCTACACGACGCTCTTCCGATCTNNNTCACAGCCTCAAGAAATCCAAG
AGCACACGTCTGAACTCTTTCCCTACACGACGCTCTTCCGATCTNNNGAATCCAGAGGTCATACAGCTC
AGCACACGTCTGAACTCTTTCCCTACACGACGCTCTTCCGATCTNNNGAATCCAGAGGTCATACAGCTC
AGCACACGTCTGAACTCTTTCCCTACACGACGCTCTTCCGATCTNNNGAATCCAGAGGTCATACAGCTC
CACACGTCTGAACTCTTTCCCTACACGACGCTCTTCCGATCTNNNGTTCAAGAGATTCTCCTGCCTCACA
CACACGTCTGAACTCTTTCCCTACACGACGCTCTTCCGATCTNNNGTTCAAGAGATTCTCCTGCCTCACA
CACACGTCTGAACTCTTTCCCTACACGACGCTCTTCCGATCTNNNGTTCAAGAGATTCTCCTGCCTCACA
AGCACACGTCTGAACTCTTTCCCTACACGACGCTCTTCCGATCTNNNCTTACTTCCCTTCTCCTGCGAATC
AGCACACGTCTGAACTCTTTCCCTACACGACGCTCTTCCGATCTNNNCTTACTTCCCTTCTCCTGCGAATC
AGCACACGTCTGAACTCTTTCCCTACACGACGCTCTTCCGATCTNNNCTTACTTCCCTTCTCCTGCGAATC
AGCACACGTCTGAACTCTTTCCCTACACGACGCTCTTCCGATCTNNNTATTACAGTCGTGAACCACCGT
AGCACACGTCTGAACTCTTTCCCTACACGACGCTCTTCCGATCTNNNTATTACAGTCGTGAACCACCGT
AGCACACGTCTGAACTCTTTCCCTACACGACGCTCTTCCGATCTNNNTATTACAGTCGTGAACCACCGT
GAGCACACGTCTGAACTCTTTCCCTACACGACGCTCTTCCGATCTNNNGGTATCTGGGGAGAAAAGAA
GAGCACACGTCTGAACTCTTTCCCTACACGACGCTCTTCCGATCTNNNGGTATCTGGGGAGAAAAGAA
GAGCACACGTCTGAACTCTTTCCCTACACGACGCTCTTCCGATCTNNNGGTATCTGGGGAGAAAAGAA
GAGCACACGTCTGAACTCTTTCCCTACACGACGCTCTTCCGATCTNNNAGAGGTTGAAATGAGCCGAG
GAGCACACGTCTGAACTCTTTCCCTACACGACGCTCTTCCGATCTNNNAGAGGTTGAAATGAGCCGAG
GAGCACACGTCTGAACTCTTTCCCTACACGACGCTCTTCCGATCTNNNAGAGGTTGAAATGAGCCGAG
GAGCACACGTCTGAACTCTTTCCCTACACGACGCTCTTCCGATCTNNNCCACTCTGTGAACATTTCTACTC
GAGCACACGTCTGAACTCTTTCCCTACACGACGCTCTTCCGATCTNNNCCACTCTGTGAACATTTCTACTC
GAGCACACGTCTGAACTCTTTCCCTACACGACGCTCTTCCGATCTNNNCCACTCTGTGAACATTTCTACTC
AGAGCACACGTCTGAACTCTTTCCCTACACGACGCTCTTCCGATCTNNNTGGCTTTTTTTTTCTCCTGAC
AGAGCACACGTCTGAACTCTTTCCCTACACGACGCTCTTCCGATCTNNNTGGCTTTTTTTTTCTCCTGAC
AGAGCACACGTCTGAACTCTTTCCCTACACGACGCTCTTCCGATCTNNNTGGCTTTTTTTTTCTCCTGAC
AAGAGCACACGTCTGAACTCTTTCCCTACACGACGCTCTTCCGATCTNNNGCTGCAGTGAGCTATGATT
AAGAGCACACGTCTGAACTCTTTCCCTACACGACGCTCTTCCGATCTNNNGCTGCAGTGAGCTATGATT
AAGAGCACACGTCTGAACTCTTTCCCTACACGACGCTCTTCCGATCTNNNGCTGCAGTGAGCTATGATT
AGCACACGTCTGAACTCTTTCCCTACACGACGCTCTTCCGATCTNNNATATTACATTTCAGAGTTCACCT
AGCACACGTCTGAACTCTTTCCCTACACGACGCTCTTCCGATCTNNNATATTACATTTCAGAGTTCACCT
AGCACACGTCTGAACTCTTTCCCTACACGACGCTCTTCCGATCTNNNATATTACATTTCAGAGTTCACCT
GAGCACACGTCTGAACTCTTTCCCTACACGACGCTCTTCCGATCTNNNCTCTCAGCTACTCGGGACTTTA
GAGCACACGTCTGAACTCTTTCCCTACACGACGCTCTTCCGATCTNNNCTCTCAGCTACTCGGGACTTTA
GAGCACACGTCTGAACTCTTTCCCTACACGACGCTCTTCCGATCTNNNCTCTCAGCTACTCGGGACTTTA
AGCACACGTCTGAACTCTTTCCCTACACGACGCTCTTCCGATCTNNNATTTCAGCAGCCTTCTCAGTTTG
AGCACACGTCTGAACTCTTTCCCTACACGACGCTCTTCCGATCTNNNATTTCAGCAGCCTTCTCAGTTTG
AGCACACGTCTGAACTCTTTCCCTACACGACGCTCTTCCGATCTNNNATTTCAGCAGCCTTCTCAGTTTG
AGCACACGTCTGAACTCTTTCCCTACACGACGCTCTTCCGATCTNNNTTGATTGTTTCCTTTGCTGTGCA
AGCACACGTCTGAACTCTTTCCCTACACGACGCTCTTCCGATCTNNNTTGATTGTTTCCTTTGCTGTGCA
AGCACACGTCTGAACTCTTTCCCTACACGACGCTCTTCCGATCTNNNTTGATTGTTTCCTTTGCTGTGCA

AGCACACGTCTGAACTCTTTCCCTACACGACGCTCTTCCGATCTNNNGTCCCTCTCTGTCACTTTCTCTG
AGCACACGTCTGAACTCTTTCCCTACACGACGCTCTTCCGATCTNNNGTCCCTCTCTGTCACTTTCTCTG
AGCACACGTCTGAACTCTTTCCCTACACGACGCTCTTCCGATCTNNNGTCCCTCTCTGTCACTTTCTCTG
GCACACGTCTGAACTCTTTCCCTACACGACGCTCTTCCGATCTNNNGACCACCCCAGGGATTCTAATA
GCACACGTCTGAACTCTTTCCCTACACGACGCTCTTCCGATCTNNNGACCACCCCAGGGATTCTAATA
GCACACGTCTGAACTCTTTCCCTACACGACGCTCTTCCGATCTNNNGACCACCCCAGGGATTCTAATA
GAGCACACGTCTGAACTCTTTCCCTACACGACGCTCTTCCGATCTNNNCCTAGAAACATTCACCCTCCC
GAGCACACGTCTGAACTCTTTCCCTACACGACGCTCTTCCGATCTNNNCCTAGAAACATTCACCCTCCC
GAGCACACGTCTGAACTCTTTCCCTACACGACGCTCTTCCGATCTNNNCCTAGAAACATTCACCCTCCC
AGCACACGTCTGAACTCTTTCCCTACACGACGCTCTTCCGATCTNNNTCCAGAGAAAGATGCAGTGA
AGCACACGTCTGAACTCTTTCCCTACACGACGCTCTTCCGATCTNNNTCCAGAGAAAGATGCAGTGA
AGCACACGTCTGAACTCTTTCCCTACACGACGCTCTTCCGATCTNNNTCCAGAGAAAGATGCAGTGA
GAGCACACGTCTGAACTCTTTCCCTACACGACGCTCTTCCGATCTNNNAACACAGGAACAACCTCTGA
GAGCACACGTCTGAACTCTTTCCCTACACGACGCTCTTCCGATCTNNNAACACAGGAACAACCTCTGA
GAGCACACGTCTGAACTCTTTCCCTACACGACGCTCTTCCGATCTNNNAACACAGGAACAACCTCTGA
AGCACACGTCTGAACTCTTTCCCTACACGACGCTCTTCCGATCTNNNTCCATTTTCCTAAGTGAGACTT
AGCACACGTCTGAACTCTTTCCCTACACGACGCTCTTCCGATCTNNNTCCATTTTCCTAAGTGAGACTT
AGCACACGTCTGAACTCTTTCCCTACACGACGCTCTTCCGATCTNNNTCCATTTTCCTAAGTGAGACTT
AGCACACGTCTGAACTCTTTCCCTACACGACGCTCTTCCGATCTNNNCCAGCACATTCCCAAATGGTTT
AGCACACGTCTGAACTCTTTCCCTACACGACGCTCTTCCGATCTNNNCCAGCACATTCCCAAATGGTTT
AGCACACGTCTGAACTCTTTCCCTACACGACGCTCTTCCGATCTNNNCCAGCACATTCCCAAATGGTTT
GCACACGTCTGAACTCTTTCCCTACACGACGCTCTTCCGATCTNNNCATTGCTACTGGAGTGTCTTTGC
GCACACGTCTGAACTCTTTCCCTACACGACGCTCTTCCGATCTNNNCATTGCTACTGGAGTGTCTTTGC
GCACACGTCTGAACTCTTTCCCTACACGACGCTCTTCCGATCTNNNCATTGCTACTGGAGTGTCTTTGC
GCACACGTCTGAACTCTTTCCCTACACGACGCTCTTCCGATCTNNNTCTAGTTTCTTGAGGTGGGAG
GCACACGTCTGAACTCTTTCCCTACACGACGCTCTTCCGATCTNNNTCTAGTTTCTTGAGGTGGGAG
GCACACGTCTGAACTCTTTCCCTACACGACGCTCTTCCGATCTNNNTCTAGTTTCTTGAGGTGGGAG
CACACGTCTGAACTCTTTCCCTACACGACGCTCTTCCGATCTNNNAGCTTTAAAGTACTGTAGATGTGC
CACACGTCTGAACTCTTTCCCTACACGACGCTCTTCCGATCTNNNAGCTTTAAAGTACTGTAGATGTGC
CACACGTCTGAACTCTTTCCCTACACGACGCTCTTCCGATCTNNNAGCTTTAAAGTACTGTAGATGTGC
CACACGTCTGAACTCTTTCCCTACACGACGCTCTTCCGATCTNNNAAGCCAATTTCCAAAGTGAACCCC
CACACGTCTGAACTCTTTCCCTACACGACGCTCTTCCGATCTNNNAAGCCAATTTCCAAAGTGAACCCC
CACACGTCTGAACTCTTTCCCTACACGACGCTCTTCCGATCTNNNAAGCCAATTTCCAAAGTGAACCCC
AGCACACGTCTGAACTCTTTCCCTACACGACGCTCTTCCGATCTNNNGACGCCTAAGACTCTCCCTCTT
AGCACACGTCTGAACTCTTTCCCTACACGACGCTCTTCCGATCTNNNGACGCCTAAGACTCTCCCTCTT
AGCACACGTCTGAACTCTTTCCCTACACGACGCTCTTCCGATCTNNNGACGCCTAAGACTCTCCCTCTT
GCACACGTCTGAACTCTTTCCCTACACGACGCTCTTCCGATCTNNNCTAAGGCTGTGGCTTGGAGTG
GCACACGTCTGAACTCTTTCCCTACACGACGCTCTTCCGATCTNNNCTAAGGCTGTGGCTTGGAGTG
GCACACGTCTGAACTCTTTCCCTACACGACGCTCTTCCGATCTNNNCTAAGGCTGTGGCTTGGAGTG
AGCACACGTCTGAACTCTTTCCCTACACGACGCTCTTCCGATCTNNNCTGTAGTCCTACCTGCTCTGG
AGCACACGTCTGAACTCTTTCCCTACACGACGCTCTTCCGATCTNNNCTGTAGTCCTACCTGCTCTGG
AGCACACGTCTGAACTCTTTCCCTACACGACGCTCTTCCGATCTNNNCTGTAGTCCTACCTGCTCTGG
GAGCACACGTCTGAACTCTTTCCCTACACGACGCTCTTCCGATCTNNNTACAAAAAATTAGCCGGGCT

GAGCACACGTCTGAACTCTTTCCCTACACGACGCTCTTCCGATCTNNNTACAAAAAATTAGCCGGGCT
GAGCACACGTCTGAACTCTTTCCCTACACGACGCTCTTCCGATCTNNNTACAAAAAATTAGCCGGGCT
AGCACACGTCTGAACTCTTTCCCTACACGACGCTCTTCCGATCTNNNAGAGAAGATGGTTGACAAGG
AGCACACGTCTGAACTCTTTCCCTACACGACGCTCTTCCGATCTNNNAGAGAAGATGGTTGACAAGG
AGCACACGTCTGAACTCTTTCCCTACACGACGCTCTTCCGATCTNNNAGAGAAGATGGTTGACAAGG
GCACACGTCTGAACTCTTTCCCTACACGACGCTCTTCCGATCTNNNAGCTCCTGGGGGTAATCTTGA
GCACACGTCTGAACTCTTTCCCTACACGACGCTCTTCCGATCTNNNAGCTCCTGGGGGTAATCTTGA
GCACACGTCTGAACTCTTTCCCTACACGACGCTCTTCCGATCTNNNAGCTCCTGGGGGTAATCTTGA
GCACACGTCTGAACTCTTTCCCTACACGACGCTCTTCCGATCTNNNGACATATCTGGACTTCCCTTCCC
GCACACGTCTGAACTCTTTCCCTACACGACGCTCTTCCGATCTNNNGACATATCTGGACTTCCCTTCCC
GCACACGTCTGAACTCTTTCCCTACACGACGCTCTTCCGATCTNNNGACATATCTGGACTTCCCTTCCC
GAGCACACGTCTGAACTCTTTCCCTACACGACGCTCTTCCGATCTNNNTATTTAGGATGCTGAGGTGG
GAGCACACGTCTGAACTCTTTCCCTACACGACGCTCTTCCGATCTNNNTATTTAGGATGCTGAGGTGG
GAGCACACGTCTGAACTCTTTCCCTACACGACGCTCTTCCGATCTNNNTATTTAGGATGCTGAGGTGG
GCACACGTCTGAACTCTTTCCCTACACGACGCTCTTCCGATCTNNNCACTCAGAGAAATTCAAGACTG
GCACACGTCTGAACTCTTTCCCTACACGACGCTCTTCCGATCTNNNCACTCAGAGAAATTCAAGACTG
GCACACGTCTGAACTCTTTCCCTACACGACGCTCTTCCGATCTNNNCACTCAGAGAAATTCAAGACTG
AGCACACGTCTGAACTCTTTCCCTACACGACGCTCTTCCGATCTNNNCCTACACTCCTCAAGAAGCAG
AGCACACGTCTGAACTCTTTCCCTACACGACGCTCTTCCGATCTNNNCCTACACTCCTCAAGAAGCAG
AGCACACGTCTGAACTCTTTCCCTACACGACGCTCTTCCGATCTNNNCCTACACTCCTCAAGAAGCAG
AGCACACGTCTGAACTCTTTCCCTACACGACGCTCTTCCGATCTNNNGACTTGAGTGATGCTTGCTCA
AGCACACGTCTGAACTCTTTCCCTACACGACGCTCTTCCGATCTNNNGACTTGAGTGATGCTTGCTCA
AGCACACGTCTGAACTCTTTCCCTACACGACGCTCTTCCGATCTNNNGACTTGAGTGATGCTTGCTCA
AGCACACGTCTGAACTCTTTCCCTACACGACGCTCTTCCGATCTNNNGAGGTTGCAGTAAGCTGAGAT
AGCACACGTCTGAACTCTTTCCCTACACGACGCTCTTCCGATCTNNNGAGGTTGCAGTAAGCTGAGAT
AGCACACGTCTGAACTCTTTCCCTACACGACGCTCTTCCGATCTNNNGAGGTTGCAGTAAGCTGAGAT
AGCACACGTCTGAACTCTTTCCCTACACGACGCTCTTCCGATCTNNNATTGCCCCAAATGTGTCTTCAA
AGCACACGTCTGAACTCTTTCCCTACACGACGCTCTTCCGATCTNNNATTGCCCCAAATGTGTCTTCAA
AGCACACGTCTGAACTCTTTCCCTACACGACGCTCTTCCGATCTNNNATTGCCCCAAATGTGTCTTCAA
GAGCACACGTCTGAACTCTTTCCCTACACGACGCTCTTCCGATCTNNNGTGCTCAGCCAACATCAGTAT
GAGCACACGTCTGAACTCTTTCCCTACACGACGCTCTTCCGATCTNNNGTGCTCAGCCAACATCAGTAT
GAGCACACGTCTGAACTCTTTCCCTACACGACGCTCTTCCGATCTNNNGTGCTCAGCCAACATCAGTAT
GCACACGTCTGAACTCTTTCCCTACACGACGCTCTTCCGATCTNNNCTGGCTTCATTTCCCTCCCTTATC
GCACACGTCTGAACTCTTTCCCTACACGACGCTCTTCCGATCTNNNCTGGCTTCATTTCCCTCCCTTATC
GCACACGTCTGAACTCTTTCCCTACACGACGCTCTTCCGATCTNNNCTGGCTTCATTTCCCTCCCTTATC
AGCACACGTCTGAACTCTTTCCCTACACGACGCTCTTCCGATCTNNNGGGGCCTAATGCAGTACTGTT
AGCACACGTCTGAACTCTTTCCCTACACGACGCTCTTCCGATCTNNNGGGGCCTAATGCAGTACTGTT
AGCACACGTCTGAACTCTTTCCCTACACGACGCTCTTCCGATCTNNNGGGGCCTAATGCAGTACTGTT
GCACACGTCTGAACTCTTTCCCTACACGACGCTCTTCCGATCTNNNGTTTGCCTTTACTTCCCTGTTCCC
GCACACGTCTGAACTCTTTCCCTACACGACGCTCTTCCGATCTNNNGTTTGCCTTTACTTCCCTGTTCCC
GCACACGTCTGAACTCTTTCCCTACACGACGCTCTTCCGATCTNNNGTTTGCCTTTACTTCCCTGTTCCC
GCACACGTCTGAACTCTTTCCCTACACGACGCTCTTCCGATCTNNNCAGATCTGCTCGCACCCTTCCAT
GCACACGTCTGAACTCTTTCCCTACACGACGCTCTTCCGATCTNNNCAGATCTGCTCGCACCCTTCCAT

GCACACGTCTGAACTCTTTCCCTACACGACGCTCTTCCGATCTNNNCAGATCTGCTCGCACCCTTCCAT
AGCACACGTCTGAACTCTTTCCCTACACGACGCTCTTCCGATCTNNNCATGCCAATGGAGGTGCATTTT
AGCACACGTCTGAACTCTTTCCCTACACGACGCTCTTCCGATCTNNNCATGCCAATGGAGGTGCATTTT
AGCACACGTCTGAACTCTTTCCCTACACGACGCTCTTCCGATCTNNNCATGCCAATGGAGGTGCATTTT
GCACACGTCTGAACTCTTTCCCTACACGACGCTCTTCCGATCTNNNATAGCTGGGATTACAGGCATG
GCACACGTCTGAACTCTTTCCCTACACGACGCTCTTCCGATCTNNNATAGCTGGGATTACAGGCATG
GCACACGTCTGAACTCTTTCCCTACACGACGCTCTTCCGATCTNNNATAGCTGGGATTACAGGCATG
AGCACACGTCTGAACTCTTTCCCTACACGACGCTCTTCCGATCTNNNGAGAATCGCTTGAATCCAGGA
AGCACACGTCTGAACTCTTTCCCTACACGACGCTCTTCCGATCTNNNGAGAATCGCTTGAATCCAGGA
AGCACACGTCTGAACTCTTTCCCTACACGACGCTCTTCCGATCTNNNGAGAATCGCTTGAATCCAGGA
AGCACACGTCTGAACTCTTTCCCTACACGACGCTCTTCCGATCTNNNGGAGAATCGCTTGAACCTGG
AGCACACGTCTGAACTCTTTCCCTACACGACGCTCTTCCGATCTNNNGGAGAATCGCTTGAACCTGG
AGCACACGTCTGAACTCTTTCCCTACACGACGCTCTTCCGATCTNNNGGAGAATCGCTTGAACCTGG
GAGCACACGTCTGAACTCTTTCCCTACACGACGCTCTTCCGATCTNNNACTTTCTTGATTCTCCTGAGG
GAGCACACGTCTGAACTCTTTCCCTACACGACGCTCTTCCGATCTNNNACTTTCTTGATTCTCCTGAGG
GAGCACACGTCTGAACTCTTTCCCTACACGACGCTCTTCCGATCTNNNACTTTCTTGATTCTCCTGAGG
AGCACACGTCTGAACTCTTTCCCTACACGACGCTCTTCCGATCTNNNGTCTTCAGACTTGGACTGGA
AGCACACGTCTGAACTCTTTCCCTACACGACGCTCTTCCGATCTNNNGTCTTCAGACTTGGACTGGA
AGCACACGTCTGAACTCTTTCCCTACACGACGCTCTTCCGATCTNNNGTCTTCAGACTTGGACTGGA
AGCACACGTCTGAACTCTTTCCCTACACGACGCTCTTCCGATCTNNNGAATCTCTGGTTTGGGGTTTT
AGCACACGTCTGAACTCTTTCCCTACACGACGCTCTTCCGATCTNNNGAATCTCTGGTTTGGGGTTTT
AGCACACGTCTGAACTCTTTCCCTACACGACGCTCTTCCGATCTNNNGAATCTCTGGTTTGGGGTTTT
AGCACACGTCTGAACTCTTTCCCTACACGACGCTCTTCCGATCTNNNTCAGCATTGAAGTTGCTGGA
AGCACACGTCTGAACTCTTTCCCTACACGACGCTCTTCCGATCTNNNTCAGCATTGAAGTTGCTGGA
AGCACACGTCTGAACTCTTTCCCTACACGACGCTCTTCCGATCTNNNTCAGCATTGAAGTTGCTGGA
GAGCACACGTCTGAACTCTTTCCCTACACGACGCTCTTCCGATCTNNNGAAGGAGAGTGGGAAGAG
GAGCACACGTCTGAACTCTTTCCCTACACGACGCTCTTCCGATCTNNNGAAGGAGAGTGGGAAGAG
GAGCACACGTCTGAACTCTTTCCCTACACGACGCTCTTCCGATCTNNNGAAGGAGAGTGGGAAGAG
AGCACACGTCTGAACTCTTTCCCTACACGACGCTCTTCCGATCTNNNTTTTCATCAACCACGTCTTTGG
AGCACACGTCTGAACTCTTTCCCTACACGACGCTCTTCCGATCTNNNTTTTCATCAACCACGTCTTTGG
AGCACACGTCTGAACTCTTTCCCTACACGACGCTCTTCCGATCTNNNTTTTCATCAACCACGTCTTTGG
ACACGTCTGAACTCTTTCCCTACACGACGCTCTTCCGATCTNNNGAAGTTGAACTCTCCCTGTTTGCCA
ACACGTCTGAACTCTTTCCCTACACGACGCTCTTCCGATCTNNNGAAGTTGAACTCTCCCTGTTTGCCA
ACACGTCTGAACTCTTTCCCTACACGACGCTCTTCCGATCTNNNGAAGTTGAACTCTCCCTGTTTGCCA
CACGTCTGAACTCTTTCCCTACACGACGCTCTTCCGATCTNNNCTCCTGAGTACCTGCACAATACACAT
CACGTCTGAACTCTTTCCCTACACGACGCTCTTCCGATCTNNNCTCCTGAGTACCTGCACAATACACAT
CACGTCTGAACTCTTTCCCTACACGACGCTCTTCCGATCTNNNCTCCTGAGTACCTGCACAATACACAT
AGCACACGTCTGAACTCTTTCCCTACACGACGCTCTTCCGATCTNNNTTGCAGAGCATGGGTAAGAT
AGCACACGTCTGAACTCTTTCCCTACACGACGCTCTTCCGATCTNNNTTGCAGAGCATGGGTAAGAT
AGCACACGTCTGAACTCTTTCCCTACACGACGCTCTTCCGATCTNNNTTGCAGAGCATGGGTAAGAT
CACACGTCTGAACTCTTTCCCTACACGACGCTCTTCCGATCTNNNCTCAGGCTCCAGACCACAACCATC
CACACGTCTGAACTCTTTCCCTACACGACGCTCTTCCGATCTNNNCTCAGGCTCCAGACCACAACCATC
CACACGTCTGAACTCTTTCCCTACACGACGCTCTTCCGATCTNNNCTCAGGCTCCAGACCACAACCATC

AGCACACGTCTGAACTCTTTCCCTACACGACGCTCTTCCGATCTNNNTTGGCCAAAAGTACTGCATGG
AGCACACGTCTGAACTCTTTCCCTACACGACGCTCTTCCGATCTNNNTTGGCCAAAAGTACTGCATGG
AGCACACGTCTGAACTCTTTCCCTACACGACGCTCTTCCGATCTNNNTTGGCCAAAAGTACTGCATGG
AGCACACGTCTGAACTCTTTCCCTACACGACGCTCTTCCGATCTNNNGTCTCAAACAAGACCGAGGAT
AGCACACGTCTGAACTCTTTCCCTACACGACGCTCTTCCGATCTNNNGTCTCAAACAAGACCGAGGAT
AGCACACGTCTGAACTCTTTCCCTACACGACGCTCTTCCGATCTNNNGTCTCAAACAAGACCGAGGAT
AGAGCACACGTCTGAACTCTTTCCCTACACGACGCTCTTCCGATCTNNNGCCTGTAGTCCCAGTTACCT
AGAGCACACGTCTGAACTCTTTCCCTACACGACGCTCTTCCGATCTNNNGCCTGTAGTCCCAGTTACCT
AGAGCACACGTCTGAACTCTTTCCCTACACGACGCTCTTCCGATCTNNNGCCTGTAGTCCCAGTTACCT
GAGCACACGTCTGAACTCTTTCCCTACACGACGCTCTTCCGATCTNNNGAGATAGCGTCACTGCACTCC
GAGCACACGTCTGAACTCTTTCCCTACACGACGCTCTTCCGATCTNNNGAGATAGCGTCACTGCACTCC
GAGCACACGTCTGAACTCTTTCCCTACACGACGCTCTTCCGATCTNNNGAGATAGCGTCACTGCACTCC
AGCACACGTCTGAACTCTTTCCCTACACGACGCTCTTCCGATCTNNNGGTATCTGTCCGGACAACTTTCT
AGCACACGTCTGAACTCTTTCCCTACACGACGCTCTTCCGATCTNNNGGTATCTGTCCGGACAACTTTCT
AGCACACGTCTGAACTCTTTCCCTACACGACGCTCTTCCGATCTNNNGGTATCTGTCCGGACAACTTTCT
AGCACACGTCTGAACTCTTTCCCTACACGACGCTCTTCCGATCTNNNAGTACAAAAACTAGCCATGCGT
AGCACACGTCTGAACTCTTTCCCTACACGACGCTCTTCCGATCTNNNAGTACAAAAACTAGCCATGCGT
AGCACACGTCTGAACTCTTTCCCTACACGACGCTCTTCCGATCTNNNAGTACAAAAACTAGCCATGCGT
GAGCACACGTCTGAACTCTTTCCCTACACGACGCTCTTCCGATCTNNNTCCCCAGCTTTTTGTTTTTG
GAGCACACGTCTGAACTCTTTCCCTACACGACGCTCTTCCGATCTNNNTCCCCAGCTTTTTGTTTTTG
GAGCACACGTCTGAACTCTTTCCCTACACGACGCTCTTCCGATCTNNNTCCCCAGCTTTTTGTTTTTG
AGCACACGTCTGAACTCTTTCCCTACACGACGCTCTTCCGATCTNNNCAGTCATTATCTCATCAACATTT
AGCACACGTCTGAACTCTTTCCCTACACGACGCTCTTCCGATCTNNNCAGTCATTATCTCATCAACATTT
AGCACACGTCTGAACTCTTTCCCTACACGACGCTCTTCCGATCTNNNCAGTCATTATCTCATCAACATTT
GCACACGTCTGAACTCTTTCCCTACACGACGCTCTTCCGATCTNNNCAACCGGAATAGTCTTTCAGCAA
GCACACGTCTGAACTCTTTCCCTACACGACGCTCTTCCGATCTNNNCAACCGGAATAGTCTTTCAGCAA
GCACACGTCTGAACTCTTTCCCTACACGACGCTCTTCCGATCTNNNCAACCGGAATAGTCTTTCAGCAA
GAGCACACGTCTGAACTCTTTCCCTACACGACGCTCTTCCGATCTNNNAGGACAAAAACAATTAGAAG
GAGCACACGTCTGAACTCTTTCCCTACACGACGCTCTTCCGATCTNNNAGGACAAAAACAATTAGAAG
GAGCACACGTCTGAACTCTTTCCCTACACGACGCTCTTCCGATCTNNNAGGACAAAAACAATTAGAAG
GAGCACACGTCTGAACTCTTTCCCTACACGACGCTCTTCCGATCTNNNGGTTGTCAAAAGGGAAGTA
GAGCACACGTCTGAACTCTTTCCCTACACGACGCTCTTCCGATCTNNNGGTTGTCAAAAGGGAAGTA
GAGCACACGTCTGAACTCTTTCCCTACACGACGCTCTTCCGATCTNNNGGTTGTCAAAAGGGAAGTA
AGAGCACACGTCTGAACTCTTTCCCTACACGACGCTCTTCCGATCTNNNCGCCTGTAATCCTAGCTACTC
AGAGCACACGTCTGAACTCTTTCCCTACACGACGCTCTTCCGATCTNNNCGCCTGTAATCCTAGCTACTC
AGAGCACACGTCTGAACTCTTTCCCTACACGACGCTCTTCCGATCTNNNCGCCTGTAATCCTAGCTACTC
AGCACACGTCTGAACTCTTTCCCTACACGACGCTCTTCCGATCTNNNCGCTTATAATCCCAGCACTTTG
AGCACACGTCTGAACTCTTTCCCTACACGACGCTCTTCCGATCTNNNCGCTTATAATCCCAGCACTTTG
AGCACACGTCTGAACTCTTTCCCTACACGACGCTCTTCCGATCTNNNCGCTTATAATCCCAGCACTTTG
AGAGCACACGTCTGAACTCTTTCCCTACACGACGCTCTTCCGATCTNNNTCATAGGCACAAAGTCTCTG
AGAGCACACGTCTGAACTCTTTCCCTACACGACGCTCTTCCGATCTNNNTCATAGGCACAAAGTCTCTG
AGAGCACACGTCTGAACTCTTTCCCTACACGACGCTCTTCCGATCTNNNTCATAGGCACAAAGTCTCTG
AGCACACGTCTGAACTCTTTCCCTACACGACGCTCTTCCGATCTNNNGTTCAAGACTAGCCTGGGTAA

AGCACACGTCTGAACTCTTTCCCTACACGACGCTCTTCCGATCTNNNGTTCAAGACTAGCCTGGGTAA
AGCACACGTCTGAACTCTTTCCCTACACGACGCTCTTCCGATCTNNNGTTCAAGACTAGCCTGGGTAA
AGCACACGTCTGAACTCTTTCCCTACACGACGCTCTTCCGATCTNNNTAGTAGACAAATCCACTTGCCC
AGCACACGTCTGAACTCTTTCCCTACACGACGCTCTTCCGATCTNNNTAGTAGACAAATCCACTTGCCC
AGCACACGTCTGAACTCTTTCCCTACACGACGCTCTTCCGATCTNNNTAGTAGACAAATCCACTTGCCC
AGCACACGTCTGAACTCTTTCCCTACACGACGCTCTTCCGATCTNNNTCAGTGCTTTGATGTAATCCA
AGCACACGTCTGAACTCTTTCCCTACACGACGCTCTTCCGATCTNNNTCAGTGCTTTGATGTAATCCA
AGCACACGTCTGAACTCTTTCCCTACACGACGCTCTTCCGATCTNNNTCAGTGCTTTGATGTAATCCA
AGCACACGTCTGAACTCTTTCCCTACACGACGCTCTTCCGATCTNNNAGGCAGGAGAATCACTTGAAC
AGCACACGTCTGAACTCTTTCCCTACACGACGCTCTTCCGATCTNNNAGGCAGGAGAATCACTTGAAC
AGCACACGTCTGAACTCTTTCCCTACACGACGCTCTTCCGATCTNNNAGGCAGGAGAATCACTTGAAC
GCACACGTCTGAACTCTTTCCCTACACGACGCTCTTCCGATCTNNNACCCTCCTTTCTCTCTCTCTCTCCA
GCACACGTCTGAACTCTTTCCCTACACGACGCTCTTCCGATCTNNNACCCTCCTTTCTCTCTCTCTCTCCA
GCACACGTCTGAACTCTTTCCCTACACGACGCTCTTCCGATCTNNNACCCTCCTTTCTCTCTCTCTCTCCA
AGCACACGTCTGAACTCTTTCCCTACACGACGCTCTTCCGATCTNNNCAAGTTCCTGAGGTAACATGG
AGCACACGTCTGAACTCTTTCCCTACACGACGCTCTTCCGATCTNNNCAAGTTCCTGAGGTAACATGG
AGCACACGTCTGAACTCTTTCCCTACACGACGCTCTTCCGATCTNNNCAAGTTCCTGAGGTAACATGG
GCACACGTCTGAACTCTTTCCCTACACGACGCTCTTCCGATCTNNNTTTCAGGTGACAGAGTGAGACT
GCACACGTCTGAACTCTTTCCCTACACGACGCTCTTCCGATCTNNNTTTCAGGTGACAGAGTGAGACT
GCACACGTCTGAACTCTTTCCCTACACGACGCTCTTCCGATCTNNNTTTCAGGTGACAGAGTGAGACT
GCACACGTCTGAACTCTTTCCCTACACGACGCTCTTCCGATCTNNNACCAGAAATGAAGGTGGAGAC
GCACACGTCTGAACTCTTTCCCTACACGACGCTCTTCCGATCTNNNACCAGAAATGAAGGTGGAGAC
GCACACGTCTGAACTCTTTCCCTACACGACGCTCTTCCGATCTNNNACCAGAAATGAAGGTGGAGAC
AGCACACGTCTGAACTCTTTCCCTACACGACGCTCTTCCGATCTNNNTCCTGGTATCTCCACTGTCATCT
AGCACACGTCTGAACTCTTTCCCTACACGACGCTCTTCCGATCTNNNTCCTGGTATCTCCACTGTCATCT
AGCACACGTCTGAACTCTTTCCCTACACGACGCTCTTCCGATCTNNNTCCTGGTATCTCCACTGTCATCT
AGCACACGTCTGAACTCTTTCCCTACACGACGCTCTTCCGATCTNNNAAGAGTAATGGTCTGAAGAG
AGCACACGTCTGAACTCTTTCCCTACACGACGCTCTTCCGATCTNNNAAGAGTAATGGTCTGAAGAG
AGCACACGTCTGAACTCTTTCCCTACACGACGCTCTTCCGATCTNNNAAGAGTAATGGTCTGAAGAG
GAGCACACGTCTGAACTCTTTCCCTACACGACGCTCTTCCGATCTNNNCACCTGTAATCCCAGCTACTC
GAGCACACGTCTGAACTCTTTCCCTACACGACGCTCTTCCGATCTNNNCACCTGTAATCCCAGCTACTC
GAGCACACGTCTGAACTCTTTCCCTACACGACGCTCTTCCGATCTNNNCACCTGTAATCCCAGCTACTC
GCACACGTCTGAACTCTTTCCCTACACGACGCTCTTCCGATCTNNNATGGTTGGCTAGATTTCCAAGG
GCACACGTCTGAACTCTTTCCCTACACGACGCTCTTCCGATCTNNNATGGTTGGCTAGATTTCCAAGG
GCACACGTCTGAACTCTTTCCCTACACGACGCTCTTCCGATCTNNNATGGTTGGCTAGATTTCCAAGG
GAGCACACGTCTGAACTCTTTCCCTACACGACGCTCTTCCGATCTNNNAAAGAGAGGCTGTTAGACA
GAGCACACGTCTGAACTCTTTCCCTACACGACGCTCTTCCGATCTNNNAAAGAGAGGCTGTTAGACA
GAGCACACGTCTGAACTCTTTCCCTACACGACGCTCTTCCGATCTNNNAAAGAGAGGCTGTTAGACA
AGCACACGTCTGAACTCTTTCCCTACACGACGCTCTTCCGATCTNNNCTGTAGTCCCAGCTACTCAGGA
AGCACACGTCTGAACTCTTTCCCTACACGACGCTCTTCCGATCTNNNCTGTAGTCCCAGCTACTCAGGA
AGCACACGTCTGAACTCTTTCCCTACACGACGCTCTTCCGATCTNNNCTGTAGTCCCAGCTACTCAGGA
AGCACACGTCTGAACTCTTTCCCTACACGACGCTCTTCCGATCTNNNTGTCTAGCTAGTGGCCTATCTG
AGCACACGTCTGAACTCTTTCCCTACACGACGCTCTTCCGATCTNNNTGTCTAGCTAGTGGCCTATCTG

AGCACACGTCTGAACTCTTTCCCTACACGACGCTCTTCCGATCTNNNTGTCTAGCTAGTGGCCTATCTG
AGCACACGTCTGAACTCTTTCCCTACACGACGCTCTTCCGATCTNNNTGAGGCAGGAGAATCTCTTG
AGCACACGTCTGAACTCTTTCCCTACACGACGCTCTTCCGATCTNNNTGAGGCAGGAGAATCTCTTG
AGCACACGTCTGAACTCTTTCCCTACACGACGCTCTTCCGATCTNNNTGAGGCAGGAGAATCTCTTG
AGCACACGTCTGAACTCTTTCCCTACACGACGCTCTTCCGATCTNNNCCAACATGTTGAAACCCCATCT
AGCACACGTCTGAACTCTTTCCCTACACGACGCTCTTCCGATCTNNNCCAACATGTTGAAACCCCATCT
AGCACACGTCTGAACTCTTTCCCTACACGACGCTCTTCCGATCTNNNCCAACATGTTGAAACCCCATCT
GAGCACACGTCTGAACTCTTTCCCTACACGACGCTCTTCCGATCTNNNTTGTAGTCCCAGCTACTCAGG
GAGCACACGTCTGAACTCTTTCCCTACACGACGCTCTTCCGATCTNNNTTGTAGTCCCAGCTACTCAGG
GAGCACACGTCTGAACTCTTTCCCTACACGACGCTCTTCCGATCTNNNTTGTAGTCCCAGCTACTCAGG
AGCACACGTCTGAACTCTTTCCCTACACGACGCTCTTCCGATCTNNNTGGTAGGAATGTTGGTCTGG
AGCACACGTCTGAACTCTTTCCCTACACGACGCTCTTCCGATCTNNNTGGTAGGAATGTTGGTCTGG
AGCACACGTCTGAACTCTTTCCCTACACGACGCTCTTCCGATCTNNNTGGTAGGAATGTTGGTCTGG
CACACGTCTGAACTCTTTCCCTACACGACGCTCTTCCGATCTNNNGTCACCTGTGATTTCTTTCAGCACA
CACACGTCTGAACTCTTTCCCTACACGACGCTCTTCCGATCTNNNGTCACCTGTGATTTCTTTCAGCACA
CACACGTCTGAACTCTTTCCCTACACGACGCTCTTCCGATCTNNNGTCACCTGTGATTTCTTTCAGCACA
AGCACACGTCTGAACTCTTTCCCTACACGACGCTCTTCCGATCTNNNCTGAGGCAGGAGAATCACTTG
AGCACACGTCTGAACTCTTTCCCTACACGACGCTCTTCCGATCTNNNCTGAGGCAGGAGAATCACTTG
AGCACACGTCTGAACTCTTTCCCTACACGACGCTCTTCCGATCTNNNCTGAGGCAGGAGAATCACTTG
AGAGCACACGTCTGAACTCTTTCCCTACACGACGCTCTTCCGATCTNNNCTACTTGCTCCTAAGAAGCA
AGAGCACACGTCTGAACTCTTTCCCTACACGACGCTCTTCCGATCTNNNCTACTTGCTCCTAAGAAGCA
AGAGCACACGTCTGAACTCTTTCCCTACACGACGCTCTTCCGATCTNNNCTACTTGCTCCTAAGAAGCA
GAGCACACGTCTGAACTCTTTCCCTACACGACGCTCTTCCGATCTNNNACTCCAGAGACAATGCTAAG
GAGCACACGTCTGAACTCTTTCCCTACACGACGCTCTTCCGATCTNNNACTCCAGAGACAATGCTAAG
GAGCACACGTCTGAACTCTTTCCCTACACGACGCTCTTCCGATCTNNNACTCCAGAGACAATGCTAAG
GAGCACACGTCTGAACTCTTTCCCTACACGACGCTCTTCCGATCTNNNAACGAAGGTAAGGTTCTTGG
GAGCACACGTCTGAACTCTTTCCCTACACGACGCTCTTCCGATCTNNNAACGAAGGTAAGGTTCTTGG
GAGCACACGTCTGAACTCTTTCCCTACACGACGCTCTTCCGATCTNNNAACGAAGGTAAGGTTCTTGG
AGCACACGTCTGAACTCTTTCCCTACACGACGCTCTTCCGATCTNNNAGTGCCACTGAGTACATTTGT
AGCACACGTCTGAACTCTTTCCCTACACGACGCTCTTCCGATCTNNNAGTGCCACTGAGTACATTTGT
AGCACACGTCTGAACTCTTTCCCTACACGACGCTCTTCCGATCTNNNAGTGCCACTGAGTACATTTGT
GCACACGTCTGAACTCTTTCCCTACACGACGCTCTTCCGATCTNNNTTCTCTTTTTTCCCTCTGTGCCTCA
GCACACGTCTGAACTCTTTCCCTACACGACGCTCTTCCGATCTNNNTTCTCTTTTTTCCCTCTGTGCCTCA
GCACACGTCTGAACTCTTTCCCTACACGACGCTCTTCCGATCTNNNTTCTCTTTTTTCCCTCTGTGCCTCA
GAGCACACGTCTGAACTCTTTCCCTACACGACGCTCTTCCGATCTNNNCTCAAAGTGTTGGGATTACAG
GAGCACACGTCTGAACTCTTTCCCTACACGACGCTCTTCCGATCTNNNCTCAAAGTGTTGGGATTACAG
GAGCACACGTCTGAACTCTTTCCCTACACGACGCTCTTCCGATCTNNNCTCAAAGTGTTGGGATTACAG
AGCACACGTCTGAACTCTTTCCCTACACGACGCTCTTCCGATCTNNNATGGTGGTAGGTGCCTATAAT
AGCACACGTCTGAACTCTTTCCCTACACGACGCTCTTCCGATCTNNNATGGTGGTAGGTGCCTATAAT
AGCACACGTCTGAACTCTTTCCCTACACGACGCTCTTCCGATCTNNNATGGTGGTAGGTGCCTATAAT
AGCACACGTCTGAACTCTTTCCCTACACGACGCTCTTCCGATCTNNNTGTTGTGTGTTGACTTTGGAA
AGCACACGTCTGAACTCTTTCCCTACACGACGCTCTTCCGATCTNNNTGTTGTGTGTTGACTTTGGAA
AGCACACGTCTGAACTCTTTCCCTACACGACGCTCTTCCGATCTNNNTGTTGTGTGTTGACTTTGGAA

AGCACACGTCTGAACTCTTTCCCTACACGACGCTCTTCCGATCTNNNCCCATTCGTATGACTCCAAGC
AGCACACGTCTGAACTCTTTCCCTACACGACGCTCTTCCGATCTNNNCCCATTCGTATGACTCCAAGC
AGCACACGTCTGAACTCTTTCCCTACACGACGCTCTTCCGATCTNNNCCCATTCGTATGACTCCAAGC
AGAGCACACGTCTGAACTCTTTCCCTACACGACGCTCTTCCGATCTNNNAGACCTTTAGATGTAGCTG
AGAGCACACGTCTGAACTCTTTCCCTACACGACGCTCTTCCGATCTNNNAGACCTTTAGATGTAGCTG
AGAGCACACGTCTGAACTCTTTCCCTACACGACGCTCTTCCGATCTNNNAGACCTTTAGATGTAGCTG
AGCACACGTCTGAACTCTTTCCCTACACGACGCTCTTCCGATCTNNNATTGGACTGATGTGACTGAA
AGCACACGTCTGAACTCTTTCCCTACACGACGCTCTTCCGATCTNNNATTGGACTGATGTGACTGAA
AGCACACGTCTGAACTCTTTCCCTACACGACGCTCTTCCGATCTNNNATTGGACTGATGTGACTGAA
GCACACGTCTGAACTCTTTCCCTACACGACGCTCTTCCGATCTNNNGCACAGATGACAGCAGTTACAT
GCACACGTCTGAACTCTTTCCCTACACGACGCTCTTCCGATCTNNNGCACAGATGACAGCAGTTACAT
GCACACGTCTGAACTCTTTCCCTACACGACGCTCTTCCGATCTNNNGCACAGATGACAGCAGTTACAT
AGCACACGTCTGAACTCTTTCCCTACACGACGCTCTTCCGATCTNNNTGTCCTGCCAGTCTTCTTTTTCT
AGCACACGTCTGAACTCTTTCCCTACACGACGCTCTTCCGATCTNNNTGTCCTGCCAGTCTTCTTTTTCT
AGCACACGTCTGAACTCTTTCCCTACACGACGCTCTTCCGATCTNNNTGTCCTGCCAGTCTTCTTTTTCT
AGCACACGTCTGAACTCTTTCCCTACACGACGCTCTTCCGATCTNNNAGTGTCTATTCAGATCCTTTGT
AGCACACGTCTGAACTCTTTCCCTACACGACGCTCTTCCGATCTNNNAGTGTCTATTCAGATCCTTTGT
AGCACACGTCTGAACTCTTTCCCTACACGACGCTCTTCCGATCTNNNAGTGTCTATTCAGATCCTTTGT
AGCACACGTCTGAACTCTTTCCCTACACGACGCTCTTCCGATCTNNNTATGCTTGTTGGCTGCATGTA
AGCACACGTCTGAACTCTTTCCCTACACGACGCTCTTCCGATCTNNNTATGCTTGTTGGCTGCATGTA
AGCACACGTCTGAACTCTTTCCCTACACGACGCTCTTCCGATCTNNNTATGCTTGTTGGCTGCATGTA
GCACACGTCTGAACTCTTTCCCTACACGACGCTCTTCCGATCTNNNACAGACATCCATGAAATACAATA
GCACACGTCTGAACTCTTTCCCTACACGACGCTCTTCCGATCTNNNACAGACATCCATGAAATACAATA
GCACACGTCTGAACTCTTTCCCTACACGACGCTCTTCCGATCTNNNACAGACATCCATGAAATACAATA
GAGCACACGTCTGAACTCTTTCCCTACACGACGCTCTTCCGATCTNNNGAGTACTCAGGGAGTAAACA
GAGCACACGTCTGAACTCTTTCCCTACACGACGCTCTTCCGATCTNNNGAGTACTCAGGGAGTAAACA
GAGCACACGTCTGAACTCTTTCCCTACACGACGCTCTTCCGATCTNNNGAGTACTCAGGGAGTAAACA
GCACACGTCTGAACTCTTTCCCTACACGACGCTCTTCCGATCTNNNGCAGACCACACACACATGCAT
GCACACGTCTGAACTCTTTCCCTACACGACGCTCTTCCGATCTNNNGCAGACCACACACACATGCAT
GCACACGTCTGAACTCTTTCCCTACACGACGCTCTTCCGATCTNNNGCAGACCACACACACATGCAT
GAGCACACGTCTGAACTCTTTCCCTACACGACGCTCTTCCGATCTNNNACCTCCTGAGTAGCTGGGAT
GAGCACACGTCTGAACTCTTTCCCTACACGACGCTCTTCCGATCTNNNACCTCCTGAGTAGCTGGGAT
GAGCACACGTCTGAACTCTTTCCCTACACGACGCTCTTCCGATCTNNNACCTCCTGAGTAGCTGGGAT
ACACGTCTGAACTCTTTCCCTACACGACGCTCTTCCGATCTNNNACCAGTCCTCTTCCTCCATCATCCAT
ACACGTCTGAACTCTTTCCCTACACGACGCTCTTCCGATCTNNNACCAGTCCTCTTCCTCCATCATCCAT
ACACGTCTGAACTCTTTCCCTACACGACGCTCTTCCGATCTNNNACCAGTCCTCTTCCTCCATCATCCAT
AGAGCACACGTCTGAACTCTTTCCCTACACGACGCTCTTCCGATCTNNNGAGAATCCCTTGAACCCAG
AGAGCACACGTCTGAACTCTTTCCCTACACGACGCTCTTCCGATCTNNNGAGAATCCCTTGAACCCAG
AGAGCACACGTCTGAACTCTTTCCCTACACGACGCTCTTCCGATCTNNNGAGAATCCCTTGAACCCAG
CACACGTCTGAACTCTTTCCCTACACGACGCTCTTCCGATCTNNNCCCCAAAATCCAACGTGTCATTCC
CACACGTCTGAACTCTTTCCCTACACGACGCTCTTCCGATCTNNNCCCCAAAATCCAACGTGTCATTCC
CACACGTCTGAACTCTTTCCCTACACGACGCTCTTCCGATCTNNNCCCCAAAATCCAACGTGTCATTCC
AGCACACGTCTGAACTCTTTCCCTACACGACGCTCTTCCGATCTNNNTCGGAAGCCTACACATACGTTA

AGCACACGTCTGAACTCTTTCCCTACACGACGCTCTTCCGATCTNNNTCGGAAGCCTACACATACGTTAT
AGCACACGTCTGAACTCTTTCCCTACACGACGCTCTTCCGATCTNNNTCGGAAGCCTACACATACGTTAT
AGAGCACACGTCTGAACTCTTTCCCTACACGACGCTCTTCCGATCTNNNGCTTTGGTCATGTGAAGTT
AGAGCACACGTCTGAACTCTTTCCCTACACGACGCTCTTCCGATCTNNNGCTTTGGTCATGTGAAGTT
AGAGCACACGTCTGAACTCTTTCCCTACACGACGCTCTTCCGATCTNNNGCTTTGGTCATGTGAAGTT
GAGCACACGTCTGAACTCTTTCCCTACACGACGCTCTTCCGATCTNNNCCAGGCCATGACAAGTAAAACA
GAGCACACGTCTGAACTCTTTCCCTACACGACGCTCTTCCGATCTNNNCCAGGCCATGACAAGTAAAACA
GAGCACACGTCTGAACTCTTTCCCTACACGACGCTCTTCCGATCTNNNCCAGGCCATGACAAGTAAAACA
GCACACGTCTGAACTCTTTCCCTACACGACGCTCTTCCGATCTNNNATGAAGTCGAGCGTGCTGTGCAT
GCACACGTCTGAACTCTTTCCCTACACGACGCTCTTCCGATCTNNNATGAAGTCGAGCGTGCTGTGCAT
GCACACGTCTGAACTCTTTCCCTACACGACGCTCTTCCGATCTNNNATGAAGTCGAGCGTGCTGTGCAT
AGCACACGTCTGAACTCTTTCCCTACACGACGCTCTTCCGATCTNNNCTGAGGCAGGAGAATCACTTGA
AGCACACGTCTGAACTCTTTCCCTACACGACGCTCTTCCGATCTNNNCTGAGGCAGGAGAATCACTTGA
AGCACACGTCTGAACTCTTTCCCTACACGACGCTCTTCCGATCTNNNCTGAGGCAGGAGAATCACTTGA
GAGCACACGTCTGAACTCTTTCCCTACACGACGCTCTTCCGATCTNNNGATGGTCTCAATCTCCTGACCT
GAGCACACGTCTGAACTCTTTCCCTACACGACGCTCTTCCGATCTNNNGATGGTCTCAATCTCCTGACCT
GAGCACACGTCTGAACTCTTTCCCTACACGACGCTCTTCCGATCTNNNGATGGTCTCAATCTCCTGACCT
GAGCACACGTCTGAACTCTTTCCCTACACGACGCTCTTCCGATCTNNNTTCTCTCTCTGTGTCTCTTGAG
GAGCACACGTCTGAACTCTTTCCCTACACGACGCTCTTCCGATCTNNNTTCTCTCTCTGTGTCTCTTGAG
GAGCACACGTCTGAACTCTTTCCCTACACGACGCTCTTCCGATCTNNNTTCTCTCTCTGTGTCTCTTGAG
AGAGCACACGTCTGAACTCTTTCCCTACACGACGCTCTTCCGATCTNNNAGGATCATTTGAAGCCAGGA
AGAGCACACGTCTGAACTCTTTCCCTACACGACGCTCTTCCGATCTNNNAGGATCATTTGAAGCCAGGA
AGAGCACACGTCTGAACTCTTTCCCTACACGACGCTCTTCCGATCTNNNAGGATCATTTGAAGCCAGGA
AGCACACGTCTGAACTCTTTCCCTACACGACGCTCTTCCGATCTNNNGAGAATAAGCAGGTATGGCTGG
AGCACACGTCTGAACTCTTTCCCTACACGACGCTCTTCCGATCTNNNGAGAATAAGCAGGTATGGCTGG
AGCACACGTCTGAACTCTTTCCCTACACGACGCTCTTCCGATCTNNNGAGAATAAGCAGGTATGGCTGG
AGCACACGTCTGAACTCTTTCCCTACACGACGCTCTTCCGATCTNNNCAGATCTTCAACTTACTTTCTTTG
AGCACACGTCTGAACTCTTTCCCTACACGACGCTCTTCCGATCTNNNCAGATCTTCAACTTACTTTCTTTG
AGCACACGTCTGAACTCTTTCCCTACACGACGCTCTTCCGATCTNNNCAGATCTTCAACTTACTTTCTTTG
AGCACACGTCTGAACTCTTTCCCTACACGACGCTCTTCCGATCTNNNCTCCCACCCAAAGTGATGTTAG
AGCACACGTCTGAACTCTTTCCCTACACGACGCTCTTCCGATCTNNNCTCCCACCCAAAGTGATGTTAG
AGCACACGTCTGAACTCTTTCCCTACACGACGCTCTTCCGATCTNNNCTCCCACCCAAAGTGATGTTAG
AGCACACGTCTGAACTCTTTCCCTACACGACGCTCTTCCGATCTNNNTAGTAAACACTCTATCAATTGGT
AGCACACGTCTGAACTCTTTCCCTACACGACGCTCTTCCGATCTNNNTAGTAAACACTCTATCAATTGGT
AGCACACGTCTGAACTCTTTCCCTACACGACGCTCTTCCGATCTNNNTAGTAAACACTCTATCAATTGGT
GAGCACACGTCTGAACTCTTTCCCTACACGACGCTCTTCCGATCTNNNAGATCCCTCCAAAATCAAGTGG
GAGCACACGTCTGAACTCTTTCCCTACACGACGCTCTTCCGATCTNNNAGATCCCTCCAAAATCAAGTGG
GAGCACACGTCTGAACTCTTTCCCTACACGACGCTCTTCCGATCTNNNAGATCCCTCCAAAATCAAGTGG
GAGCACACGTCTGAACTCTTTCCCTACACGACGCTCTTCCGATCTNNNCTGTGGTCTAGACTCTCTCCTCT
GAGCACACGTCTGAACTCTTTCCCTACACGACGCTCTTCCGATCTNNNCTGTGGTCTAGACTCTCTCCTCT
GAGCACACGTCTGAACTCTTTCCCTACACGACGCTCTTCCGATCTNNNCTGTGGTCTAGACTCTCTCCTCT
GAGCACACGTCTGAACTCTTTCCCTACACGACGCTCTTCCGATCTNNNGAGAAGAAGCGCATCACTTCA
GAGCACACGTCTGAACTCTTTCCCTACACGACGCTCTTCCGATCTNNNGAGAAGAAGCGCATCACTTCA

AGCACACGTCTGAACTCTTTCCCTACACGACGCTCTTCCGATCTNNNGAGAAGAAGCGCATCACTTC
AGCACACGTCTGAACTCTTTCCCTACACGACGCTCTTCCGATCTNNNGTGCCTGTAATCCCAGCTACT
AGCACACGTCTGAACTCTTTCCCTACACGACGCTCTTCCGATCTNNNGTGCCTGTAATCCCAGCTACT
AGCACACGTCTGAACTCTTTCCCTACACGACGCTCTTCCGATCTNNNGTGCCTGTAATCCCAGCTACT
GCACACGTCTGAACTCTTTCCCTACACGACGCTCTTCCGATCTNNNAGCAAGAGACAGCAGGATTAG
GCACACGTCTGAACTCTTTCCCTACACGACGCTCTTCCGATCTNNNAGCAAGAGACAGCAGGATTAG
GCACACGTCTGAACTCTTTCCCTACACGACGCTCTTCCGATCTNNNAGCAAGAGACAGCAGGATTAG
AGCACACGTCTGAACTCTTTCCCTACACGACGCTCTTCCGATCTNNNGGGGATCTGCTTGGAGTTTTC
AGCACACGTCTGAACTCTTTCCCTACACGACGCTCTTCCGATCTNNNGGGGATCTGCTTGGAGTTTTC
AGCACACGTCTGAACTCTTTCCCTACACGACGCTCTTCCGATCTNNNGGGGATCTGCTTGGAGTTTTC
AGCACACGTCTGAACTCTTTCCCTACACGACGCTCTTCCGATCTNNNAAAAGAACGCAACTGGTCAT
AGCACACGTCTGAACTCTTTCCCTACACGACGCTCTTCCGATCTNNNAAAAGAACGCAACTGGTCAT
AGCACACGTCTGAACTCTTTCCCTACACGACGCTCTTCCGATCTNNNAAAAGAACGCAACTGGTCAT
GCACACGTCTGAACTCTTTCCCTACACGACGCTCTTCCGATCTNNNTTCTCCATTCTCCATTGTGGTCCC
GCACACGTCTGAACTCTTTCCCTACACGACGCTCTTCCGATCTNNNTTCTCCATTCTCCATTGTGGTCCC
GCACACGTCTGAACTCTTTCCCTACACGACGCTCTTCCGATCTNNNTTCTCCATTCTCCATTGTGGTCCC
GAGCACACGTCTGAACTCTTTCCCTACACGACGCTCTTCCGATCTNNNGGGGTAAACATGACAGCTTT
GAGCACACGTCTGAACTCTTTCCCTACACGACGCTCTTCCGATCTNNNGGGGTAAACATGACAGCTTT
GAGCACACGTCTGAACTCTTTCCCTACACGACGCTCTTCCGATCTNNNGGGGTAAACATGACAGCTTT
CACACGTCTGAACTCTTTCCCTACACGACGCTCTTCCGATCTNNNTACAAGGGGGAGGAATCTGGAT
CACACGTCTGAACTCTTTCCCTACACGACGCTCTTCCGATCTNNNTACAAGGGGGAGGAATCTGGAT
CACACGTCTGAACTCTTTCCCTACACGACGCTCTTCCGATCTNNNTACAAGGGGGAGGAATCTGGAT
AGCACACGTCTGAACTCTTTCCCTACACGACGCTCTTCCGATCTNNNGCCCCTAACTTAGACAGCAAGT
AGCACACGTCTGAACTCTTTCCCTACACGACGCTCTTCCGATCTNNNGCCCCTAACTTAGACAGCAAGT
AGCACACGTCTGAACTCTTTCCCTACACGACGCTCTTCCGATCTNNNGCCCCTAACTTAGACAGCAAGT
GCACACGTCTGAACTCTTTCCCTACACGACGCTCTTCCGATCTNNNAGGTCTGTTGCCTGTTGTGATA
GCACACGTCTGAACTCTTTCCCTACACGACGCTCTTCCGATCTNNNAGGTCTGTTGCCTGTTGTGATA
GCACACGTCTGAACTCTTTCCCTACACGACGCTCTTCCGATCTNNNAGGTCTGTTGCCTGTTGTGATA
GAGCACACGTCTGAACTCTTTCCCTACACGACGCTCTTCCGATCTNNNGAGAATTCCTTGAACCTGGG
GAGCACACGTCTGAACTCTTTCCCTACACGACGCTCTTCCGATCTNNNGAGAATTCCTTGAACCTGGG
GAGCACACGTCTGAACTCTTTCCCTACACGACGCTCTTCCGATCTNNNGAGAATTCCTTGAACCTGGG
GCACACGTCTGAACTCTTTCCCTACACGACGCTCTTCCGATCTNNNAAGTAGCTGGGATTACAGTCA
GCACACGTCTGAACTCTTTCCCTACACGACGCTCTTCCGATCTNNNAAGTAGCTGGGATTACAGTCA
GCACACGTCTGAACTCTTTCCCTACACGACGCTCTTCCGATCTNNNAAGTAGCTGGGATTACAGTCA
AGCACACGTCTGAACTCTTTCCCTACACGACGCTCTTCCGATCTNNNCTCCTGCTCTCTACCATCCTAGA
AGCACACGTCTGAACTCTTTCCCTACACGACGCTCTTCCGATCTNNNCTCCTGCTCTCTACCATCCTAGA
AGCACACGTCTGAACTCTTTCCCTACACGACGCTCTTCCGATCTNNNCTCCTGCTCTCTACCATCCTAGA
GCACACGTCTGAACTCTTTCCCTACACGACGCTCTTCCGATCTNNNCCATTTGCTAAGTGACCTTGAG
GCACACGTCTGAACTCTTTCCCTACACGACGCTCTTCCGATCTNNNCCATTTGCTAAGTGACCTTGAG
GCACACGTCTGAACTCTTTCCCTACACGACGCTCTTCCGATCTNNNCCATTTGCTAAGTGACCTTGAG
GAGCACACGTCTGAACTCTTTCCCTACACGACGCTCTTCCGATCTNNNCAAGCCCGGCTATTTCTTTCT
GAGCACACGTCTGAACTCTTTCCCTACACGACGCTCTTCCGATCTNNNCAAGCCCGGCTATTTCTTTCT
GAGCACACGTCTGAACTCTTTCCCTACACGACGCTCTTCCGATCTNNNCAAGCCCGGCTATTTCTTTCT

GAGCACACGTCTGAACTCTTTCCCTACACGACGCTCTTCCGATCTNNNCTGTAGTTCCAGATACTGCG
GAGCACACGTCTGAACTCTTTCCCTACACGACGCTCTTCCGATCTNNNCTGTAGTTCCAGATACTGCG
GAGCACACGTCTGAACTCTTTCCCTACACGACGCTCTTCCGATCTNNNCTGTAGTTCCAGATACTGCG
GCACACGTCTGAACTCTTTCCCTACACGACGCTCTTCCGATCTNNNTCCTCCTAACAGTTACCACCTGAC
GCACACGTCTGAACTCTTTCCCTACACGACGCTCTTCCGATCTNNNTCCTCCTAACAGTTACCACCTGAC
GCACACGTCTGAACTCTTTCCCTACACGACGCTCTTCCGATCTNNNTCCTCCTAACAGTTACCACCTGAC
GAGCACACGTCTGAACTCTTTCCCTACACGACGCTCTTCCGATCTNNNCCACAGTGTGAATACTCCCTG
GAGCACACGTCTGAACTCTTTCCCTACACGACGCTCTTCCGATCTNNNCCACAGTGTGAATACTCCCTG
GAGCACACGTCTGAACTCTTTCCCTACACGACGCTCTTCCGATCTNNNCCACAGTGTGAATACTCCCTG
AGCACACGTCTGAACTCTTTCCCTACACGACGCTCTTCCGATCTNNNTTTAGTAGAGACGGGGCTTC
AGCACACGTCTGAACTCTTTCCCTACACGACGCTCTTCCGATCTNNNTTTAGTAGAGACGGGGCTTC
AGCACACGTCTGAACTCTTTCCCTACACGACGCTCTTCCGATCTNNNTTTAGTAGAGACGGGGCTTC
AGCACACGTCTGAACTCTTTCCCTACACGACGCTCTTCCGATCTNNNAGAAGTGCATTTGAGGTTTGC
AGCACACGTCTGAACTCTTTCCCTACACGACGCTCTTCCGATCTNNNAGAAGTGCATTTGAGGTTTGC
AGCACACGTCTGAACTCTTTCCCTACACGACGCTCTTCCGATCTNNNAGAAGTGCATTTGAGGTTTGC
AGCACACGTCTGAACTCTTTCCCTACACGACGCTCTTCCGATCTNNNGGTGCCTGTAATCCCAGATACT
AGCACACGTCTGAACTCTTTCCCTACACGACGCTCTTCCGATCTNNNGGTGCCTGTAATCCCAGATACT
AGCACACGTCTGAACTCTTTCCCTACACGACGCTCTTCCGATCTNNNGGTGCCTGTAATCCCAGATACT
GAGCACACGTCTGAACTCTTTCCCTACACGACGCTCTTCCGATCTNNNGGCCACAAAGAAAGAGCAC
GAGCACACGTCTGAACTCTTTCCCTACACGACGCTCTTCCGATCTNNNGGCCACAAAGAAAGAGCAC
GAGCACACGTCTGAACTCTTTCCCTACACGACGCTCTTCCGATCTNNNGGCCACAAAGAAAGAGCAC
AGCACACGTCTGAACTCTTTCCCTACACGACGCTCTTCCGATCTNNNCTCTTGCAAACGAGCCACTTTTT
AGCACACGTCTGAACTCTTTCCCTACACGACGCTCTTCCGATCTNNNCTCTTGCAAACGAGCCACTTTTT
AGCACACGTCTGAACTCTTTCCCTACACGACGCTCTTCCGATCTNNNCTCTTGCAAACGAGCCACTTTTT
GCACACGTCTGAACTCTTTCCCTACACGACGCTCTTCCGATCTNNNAGAGTCTCCAGCTCTGACATCTA
GCACACGTCTGAACTCTTTCCCTACACGACGCTCTTCCGATCTNNNAGAGTCTCCAGCTCTGACATCTA
GCACACGTCTGAACTCTTTCCCTACACGACGCTCTTCCGATCTNNNAGAGTCTCCAGCTCTGACATCTA
AGCACACGTCTGAACTCTTTCCCTACACGACGCTCTTCCGATCTNNNCGACTTTTCCTGCTGAAATGAC
AGCACACGTCTGAACTCTTTCCCTACACGACGCTCTTCCGATCTNNNCGACTTTTCCTGCTGAAATGAC
AGCACACGTCTGAACTCTTTCCCTACACGACGCTCTTCCGATCTNNNCGACTTTTCCTGCTGAAATGAC
AAGAGCACACGTCTGAACTCTTTCCCTACACGACGCTCTTCCGATCTNNNTTCCACCTCTGAGTCTCATG
AAGAGCACACGTCTGAACTCTTTCCCTACACGACGCTCTTCCGATCTNNNTTCCACCTCTGAGTCTCATG
AAGAGCACACGTCTGAACTCTTTCCCTACACGACGCTCTTCCGATCTNNNTTCCACCTCTGAGTCTCATG
GAGCACACGTCTGAACTCTTTCCCTACACGACGCTCTTCCGATCTNNNAAGTGCTGAGATTACAGGCA
GAGCACACGTCTGAACTCTTTCCCTACACGACGCTCTTCCGATCTNNNAAGTGCTGAGATTACAGGCA
GAGCACACGTCTGAACTCTTTCCCTACACGACGCTCTTCCGATCTNNNAAGTGCTGAGATTACAGGCA
GAGCACACGTCTGAACTCTTTCCCTACACGACGCTCTTCCGATCTNNNTTCTTTGAGGTATTTTATGAT
GAGCACACGTCTGAACTCTTTCCCTACACGACGCTCTTCCGATCTNNNTTCTTTGAGGTATTTTATGAT
GAGCACACGTCTGAACTCTTTCCCTACACGACGCTCTTCCGATCTNNNTTCTTTGAGGTATTTTATGAT
CACACGTCTGAACTCTTTCCCTACACGACGCTCTTCCGATCTNNNCGCTGTATTATTTGGGGTCCTCTCA
CACACGTCTGAACTCTTTCCCTACACGACGCTCTTCCGATCTNNNCGCTGTATTATTTGGGGTCCTCTCA
CACACGTCTGAACTCTTTCCCTACACGACGCTCTTCCGATCTNNNCGCTGTATTATTTGGGGTCCTCTCA
GCACACGTCTGAACTCTTTCCCTACACGACGCTCTTCCGATCTNNNGCATCTCAGTTTGGAGGCATTTT

GCACACGTCTGAACTCTTTCCCTACACGACGCTCTTCCGATCTNNNGCATCTCAGTTTGGAGGCATTTT

GCACACGTCTGAACTCTTTCCCTACACGACGCTCTTCCGATCTNNNGCATCTCAGTTTGGAGGCATTTT

AGCACACGTCTGAACTCTTTCCCTACACGACGCTCTTCCGATCTNNNGAAGTGGGTGTATCTTTCTCCC

AGCACACGTCTGAACTCTTTCCCTACACGACGCTCTTCCGATCTNNNGAAGTGGGTGTATCTTTCTCCC

AGCACACGTCTGAACTCTTTCCCTACACGACGCTCTTCCGATCTNNNGAAGTGGGTGTATCTTTCTCCC

GAGCACACGTCTGAACTCTTTCCCTACACGACGCTCTTCCGATCTNNNGGATGGAAGGGAAAAACAG

GAGCACACGTCTGAACTCTTTCCCTACACGACGCTCTTCCGATCTNNNGGATGGAAGGGAAAAACAG

GAGCACACGTCTGAACTCTTTCCCTACACGACGCTCTTCCGATCTNNNGGATGGAAGGGAAAAACAG

GAGCACACGTCTGAACTCTTTCCCTACACGACGCTCTTCCGATCTNNNGGAATGTCCCAAGGCTCAGT

GAGCACACGTCTGAACTCTTTCCCTACACGACGCTCTTCCGATCTNNNGGAATGTCCCAAGGCTCAGT

GAGCACACGTCTGAACTCTTTCCCTACACGACGCTCTTCCGATCTNNNGGAATGTCCCAAGGCTCAGT

AGCACACGTCTGAACTCTTTCCCTACACGACGCTCTTCCGATCTNNNGGTGACATAGCGAGACTCCATC

AGCACACGTCTGAACTCTTTCCCTACACGACGCTCTTCCGATCTNNNGGTGACATAGCGAGACTCCATC

AGCACACGTCTGAACTCTTTCCCTACACGACGCTCTTCCGATCTNNNGGTGACATAGCGAGACTCCATC

AGCACACGTCTGAACTCTTTCCCTACACGACGCTCTTCCGATCTNNNACTGGGAAGAATGGCTAGAAT

AGCACACGTCTGAACTCTTTCCCTACACGACGCTCTTCCGATCTNNNACTGGGAAGAATGGCTAGAAT

AGCACACGTCTGAACTCTTTCCCTACACGACGCTCTTCCGATCTNNNACTGGGAAGAATGGCTAGAAT

AAGAGCACACGTCTGAACTCTTTCCCTACACGACGCTCTTCCGATCTNNNATCTTGGTCAGGAAGACTC

AAGAGCACACGTCTGAACTCTTTCCCTACACGACGCTCTTCCGATCTNNNATCTTGGTCAGGAAGACTC

AAGAGCACACGTCTGAACTCTTTCCCTACACGACGCTCTTCCGATCTNNNATCTTGGTCAGGAAGACTC

GAGCACACGTCTGAACTCTTTCCCTACACGACGCTCTTCCGATCTNNNCAATGCTTTCTTCCAACCATCCA

GAGCACACGTCTGAACTCTTTCCCTACACGACGCTCTTCCGATCTNNNCAATGCTTTCTTCCAACCATCCA

GAGCACACGTCTGAACTCTTTCCCTACACGACGCTCTTCCGATCTNNNCAATGCTTTCTTCCAACCATCCA

AGCACACGTCTGAACTCTTTCCCTACACGACGCTCTTCCGATCTNNNTCCTAACTTTGATGACAGAAAAC

AGCACACGTCTGAACTCTTTCCCTACACGACGCTCTTCCGATCTNNNTCCTAACTTTGATGACAGAAAAC

AGCACACGTCTGAACTCTTTCCCTACACGACGCTCTTCCGATCTNNNTCCTAACTTTGATGACAGAAAAC

GAGCACACGTCTGAACTCTTTCCCTACACGACGCTCTTCCGATCTNNNGTTTAGAACAGAAACTTAGCT

GAGCACACGTCTGAACTCTTTCCCTACACGACGCTCTTCCGATCTNNNGTTTAGAACAGAAACTTAGCT

GAGCACACGTCTGAACTCTTTCCCTACACGACGCTCTTCCGATCTNNNGTTTAGAACAGAAACTTAGCT

GCACACGTCTGAACTCTTTCCCTACACGACGCTCTTCCGATCTNNNTATCATTTCAGGCAAGGTCCCTCCA

GCACACGTCTGAACTCTTTCCCTACACGACGCTCTTCCGATCTNNNTATCATTTCAGGCAAGGTCCCTCCA

GCACACGTCTGAACTCTTTCCCTACACGACGCTCTTCCGATCTNNNTATCATTTCAGGCAAGGTCCCTCCA

AGCACACGTCTGAACTCTTTCCCTACACGACGCTCTTCCGATCTNNNACAATGGTGGTCGCATAAGACT

AGCACACGTCTGAACTCTTTCCCTACACGACGCTCTTCCGATCTNNNACAATGGTGGTCGCATAAGACT

AGCACACGTCTGAACTCTTTCCCTACACGACGCTCTTCCGATCTNNNACAATGGTGGTCGCATAAGACT

GAGCACACGTCTGAACTCTTTCCCTACACGACGCTCTTCCGATCTNNNACCAACCCTACCAGAAGTTTGT

GAGCACACGTCTGAACTCTTTCCCTACACGACGCTCTTCCGATCTNNNACCAACCCTACCAGAAGTTTGT

GAGCACACGTCTGAACTCTTTCCCTACACGACGCTCTTCCGATCTNNNACCAACCCTACCAGAAGTTTGT

AGCACACGTCTGAACTCTTTCCCTACACGACGCTCTTCCGATCTNNNTTCCTCAAAGGCCATTTAGAGGC

AGCACACGTCTGAACTCTTTCCCTACACGACGCTCTTCCGATCTNNNTTCCTCAAAGGCCATTTAGAGGC

AGCACACGTCTGAACTCTTTCCCTACACGACGCTCTTCCGATCTNNNTTCCTCAAAGGCCATTTAGAGGC

AGCACACGTCTGAACTCTTTCCCTACACGACGCTCTTCCGATCTNNNCAAAGGATCGAGGACATGAGC

AGCACACGTCTGAACTCTTTCCCTACACGACGCTCTTCCGATCTNNNCAAAGGATCGAGGACATGAGC

AGCACACGTCTGAACTCTTTCCCTACACGACGCTCTTCCGATCTNNNCAAAGGATCGAGGACATGAGG

AGAGCACACGTCTGAACTCTTTCCCTACACGACGCTCTTCCGATCTNNNACTCCATCTCTCTCTCTCTCTC

AGAGCACACGTCTGAACTCTTTCCCTACACGACGCTCTTCCGATCTNNNACTCCATCTCTCTCTCTCTCTC

AGAGCACACGTCTGAACTCTTTCCCTACACGACGCTCTTCCGATCTNNNACTCCATCTCTCTCTCTCTCTC

GAGCACACGTCTGAACTCTTTCCCTACACGACGCTCTTCCGATCTNNNAGGCAGGAGAATTGCTTGAAC

GAGCACACGTCTGAACTCTTTCCCTACACGACGCTCTTCCGATCTNNNAGGCAGGAGAATTGCTTGAAC

GAGCACACGTCTGAACTCTTTCCCTACACGACGCTCTTCCGATCTNNNAGGCAGGAGAATTGCTTGAAC

AGCACACGTCTGAACTCTTTCCCTACACGACGCTCTTCCGATCTNNNAGCAAAACAGATGGAAGGAAG

AGCACACGTCTGAACTCTTTCCCTACACGACGCTCTTCCGATCTNNNAGCAAAACAGATGGAAGGAAG

AGCACACGTCTGAACTCTTTCCCTACACGACGCTCTTCCGATCTNNNAGCAAAACAGATGGAAGGAAG

AGCACACGTCTGAACTCTTTCCCTACACGACGCTCTTCCGATCTNNNGTGGAGTTCTAGTCGGCCATAA

AGCACACGTCTGAACTCTTTCCCTACACGACGCTCTTCCGATCTNNNGTGGAGTTCTAGTCGGCCATAA

AGCACACGTCTGAACTCTTTCCCTACACGACGCTCTTCCGATCTNNNGTGGAGTTCTAGTCGGCCATAA

AGCACACGTCTGAACTCTTTCCCTACACGACGCTCTTCCGATCTNNNAGCCATCATCCTATTGCTGAACA

AGCACACGTCTGAACTCTTTCCCTACACGACGCTCTTCCGATCTNNNAGCCATCATCCTATTGCTGAACA

AGCACACGTCTGAACTCTTTCCCTACACGACGCTCTTCCGATCTNNNAGCCATCATCCTATTGCTGAACA

GAGCACACGTCTGAACTCTTTCCCTACACGACGCTCTTCCGATCTNNNATGCACATCACAACCACAATG

GAGCACACGTCTGAACTCTTTCCCTACACGACGCTCTTCCGATCTNNNATGCACATCACAACCACAATG

GAGCACACGTCTGAACTCTTTCCCTACACGACGCTCTTCCGATCTNNNATGCACATCACAACCACAATG

ACACGTCTGAACTCTTTCCCTACACGACGCTCTTCCGATCTNNNAATCTTTGCCTGGCTCTATGTCTCATG

ACACGTCTGAACTCTTTCCCTACACGACGCTCTTCCGATCTNNNAATCTTTGCCTGGCTCTATGTCTCATG

ACACGTCTGAACTCTTTCCCTACACGACGCTCTTCCGATCTNNNAATCTTTGCCTGGCTCTATGTCTCATG

AGCACACGTCTGAACTCTTTCCCTACACGACGCTCTTCCGATCTNNNAGTTTCCAGAAGTCAGTACAGC

AGCACACGTCTGAACTCTTTCCCTACACGACGCTCTTCCGATCTNNNAGTTTCCAGAAGTCAGTACAGC

AGCACACGTCTGAACTCTTTCCCTACACGACGCTCTTCCGATCTNNNAGTTTCCAGAAGTCAGTACAGC

GAGCACACGTCTGAACTCTTTCCCTACACGACGCTCTTCCGATCTNNNTCCAAAAAGACAGAGTCCTGA

GAGCACACGTCTGAACTCTTTCCCTACACGACGCTCTTCCGATCTNNNTCCAAAAAGACAGAGTCCTGA

GAGCACACGTCTGAACTCTTTCCCTACACGACGCTCTTCCGATCTNNNTCCAAAAAGACAGAGTCCTGA

AGCACACGTCTGAACTCTTTCCCTACACGACGCTCTTCCGATCTNNNTTTAGTAGAGACGGGATTTCGC

AGCACACGTCTGAACTCTTTCCCTACACGACGCTCTTCCGATCTNNNTTTAGTAGAGACGGGATTTCGC

AGCACACGTCTGAACTCTTTCCCTACACGACGCTCTTCCGATCTNNNTTTAGTAGAGACGGGATTTCGC

GAGCACACGTCTGAACTCTTTCCCTACACGACGCTCTTCCGATCTNNNTGTAATCCTGGTGCTTTGGGA

GAGCACACGTCTGAACTCTTTCCCTACACGACGCTCTTCCGATCTNNNTGTAATCCTGGTGCTTTGGGA

GAGCACACGTCTGAACTCTTTCCCTACACGACGCTCTTCCGATCTNNNTGTAATCCTGGTGCTTTGGGA

GCACACGTCTGAACTCTTTCCCTACACGACGCTCTTCCGATCTNNNGCTCTGAACTCAGGAAACAACAC

GCACACGTCTGAACTCTTTCCCTACACGACGCTCTTCCGATCTNNNGCTCTGAACTCAGGAAACAACAC

GCACACGTCTGAACTCTTTCCCTACACGACGCTCTTCCGATCTNNNGCTCTGAACTCAGGAAACAACAC

AGCACACGTCTGAACTCTTTCCCTACACGACGCTCTTCCGATCTNNNTGACAATGTCAAAAATTGGTCAC

AGCACACGTCTGAACTCTTTCCCTACACGACGCTCTTCCGATCTNNNTGACAATGTCAAAAATTGGTCAC

AGCACACGTCTGAACTCTTTCCCTACACGACGCTCTTCCGATCTNNNTGACAATGTCAAAAATTGGTCAC

AGCACACGTCTGAACTCTTTCCCTACACGACGCTCTTCCGATCTNNNCTTATGCAGACGGACTCTCTACC

AGCACACGTCTGAACTCTTTCCCTACACGACGCTCTTCCGATCTNNNCTTATGCAGACGGACTCTCTACC

AGCACACGTCTGAACTCTTTCCCTACACGACGCTCTTCCGATCTNNNCTTATGCAGACGGACTCTCTACC

GAGCACACGTCTGAACTCTTTCCCTACACGACGCTCTTCCGATCTNNNCTTTTCAAACTTCGCCAGGG
GAGCACACGTCTGAACTCTTTCCCTACACGACGCTCTTCCGATCTNNNCTTTTCAAACTTCGCCAGGG
GAGCACACGTCTGAACTCTTTCCCTACACGACGCTCTTCCGATCTNNNCTTTTCAAACTTCGCCAGGG
AGCACACGTCTGAACTCTTTCCCTACACGACGCTCTTCCGATCTNNNGTGAGCCATGATAGTGCCACT
AGCACACGTCTGAACTCTTTCCCTACACGACGCTCTTCCGATCTNNNGTGAGCCATGATAGTGCCACT
AGCACACGTCTGAACTCTTTCCCTACACGACGCTCTTCCGATCTNNNGTGAGCCATGATAGTGCCACT
GCACACGTCTGAACTCTTTCCCTACACGACGCTCTTCCGATCTNNNGCTGTGACTGGATTCTCTCTCT
GCACACGTCTGAACTCTTTCCCTACACGACGCTCTTCCGATCTNNNGCTGTGACTGGATTCTCTCTCT
GCACACGTCTGAACTCTTTCCCTACACGACGCTCTTCCGATCTNNNGCTGTGACTGGATTCTCTCTCT
GAGCACACGTCTGAACTCTTTCCCTACACGACGCTCTTCCGATCTNNNATGTTGTGACCTCTTTTCCTG
GAGCACACGTCTGAACTCTTTCCCTACACGACGCTCTTCCGATCTNNNATGTTGTGACCTCTTTTCCTG
GAGCACACGTCTGAACTCTTTCCCTACACGACGCTCTTCCGATCTNNNATGTTGTGACCTCTTTTCCTG
GAGCACACGTCTGAACTCTTTCCCTACACGACGCTCTTCCGATCTNNNCTAAGGTGGGAGAATCACTG
GAGCACACGTCTGAACTCTTTCCCTACACGACGCTCTTCCGATCTNNNCTAAGGTGGGAGAATCACTG
GAGCACACGTCTGAACTCTTTCCCTACACGACGCTCTTCCGATCTNNNCTAAGGTGGGAGAATCACTG
AGCACACGTCTGAACTCTTTCCCTACACGACGCTCTTCCGATCTNNNTGCATGGTTTTGAGCTTTATA
AGCACACGTCTGAACTCTTTCCCTACACGACGCTCTTCCGATCTNNNTGCATGGTTTTGAGCTTTATA
AGCACACGTCTGAACTCTTTCCCTACACGACGCTCTTCCGATCTNNNTGCATGGTTTTGAGCTTTATA
AGCACACGTCTGAACTCTTTCCCTACACGACGCTCTTCCGATCTNNNTAGTAGAGACGGGGTTTCTC
AGCACACGTCTGAACTCTTTCCCTACACGACGCTCTTCCGATCTNNNTAGTAGAGACGGGGTTTCTC
AGCACACGTCTGAACTCTTTCCCTACACGACGCTCTTCCGATCTNNNTAGTAGAGACGGGGTTTCTC
GCACACGTCTGAACTCTTTCCCTACACGACGCTCTTCCGATCTNNNGCTTCAGGCATTGTTCAGAGTTT
GCACACGTCTGAACTCTTTCCCTACACGACGCTCTTCCGATCTNNNGCTTCAGGCATTGTTCAGAGTTT
GCACACGTCTGAACTCTTTCCCTACACGACGCTCTTCCGATCTNNNGCTTCAGGCATTGTTCAGAGTTT
GCACACGTCTGAACTCTTTCCCTACACGACGCTCTTCCGATCTNNNCAGGAGGCTGAGGTATGAGAA
GCACACGTCTGAACTCTTTCCCTACACGACGCTCTTCCGATCTNNNCAGGAGGCTGAGGTATGAGAA
GCACACGTCTGAACTCTTTCCCTACACGACGCTCTTCCGATCTNNNCAGGAGGCTGAGGTATGAGAA
AGCACACGTCTGAACTCTTTCCCTACACGACGCTCTTCCGATCTNNNGGATTGTAGGAATTCTGACT
AGCACACGTCTGAACTCTTTCCCTACACGACGCTCTTCCGATCTNNNGGATTGTAGGAATTCTGACT
AGCACACGTCTGAACTCTTTCCCTACACGACGCTCTTCCGATCTNNNGGATTGTAGGAATTCTGACT
GCACACGTCTGAACTCTTTCCCTACACGACGCTCTTCCGATCTNNNCCAATGACAAAAGCGTCTTTCCC
GCACACGTCTGAACTCTTTCCCTACACGACGCTCTTCCGATCTNNNCCAATGACAAAAGCGTCTTTCCC
GCACACGTCTGAACTCTTTCCCTACACGACGCTCTTCCGATCTNNNCCAATGACAAAAGCGTCTTTCCC
AGCACACGTCTGAACTCTTTCCCTACACGACGCTCTTCCGATCTNNNGAATGCCTGTAATCCCAGCTAC
AGCACACGTCTGAACTCTTTCCCTACACGACGCTCTTCCGATCTNNNGAATGCCTGTAATCCCAGCTAC
AGCACACGTCTGAACTCTTTCCCTACACGACGCTCTTCCGATCTNNNGAATGCCTGTAATCCCAGCTAC
AGCACACGTCTGAACTCTTTCCCTACACGACGCTCTTCCGATCTNNNAACATCAGGAAATGGTCTCTC
AGCACACGTCTGAACTCTTTCCCTACACGACGCTCTTCCGATCTNNNAACATCAGGAAATGGTCTCTC
AGCACACGTCTGAACTCTTTCCCTACACGACGCTCTTCCGATCTNNNAACATCAGGAAATGGTCTCTC
AGCACACGTCTGAACTCTTTCCCTACACGACGCTCTTCCGATCTNNNATTTCCTCTTCCCTTTCTGCCTT
AGCACACGTCTGAACTCTTTCCCTACACGACGCTCTTCCGATCTNNNATTTCCTCTTCCCTTTCTGCCTT
AGCACACGTCTGAACTCTTTCCCTACACGACGCTCTTCCGATCTNNNATTTCCTCTTCCCTTTCTGCCTT
GCACACGTCTGAACTCTTTCCCTACACGACGCTCTTCCGATCTNNNTGTGTGACAGAGCGAGACTCCA

GCACACGTCTGAACTCTTTCCCTACACGACGCTCTTCCGATCTNNNTGTGTGACAGAGCGAGACTCCA

GCACACGTCTGAACTCTTTCCCTACACGACGCTCTTCCGATCTNNNTGTGTGACAGAGCGAGACTCCA

GCACACGTCTGAACTCTTTCCCTACACGACGCTCTTCCGATCTNNNAGCAATTACAACTTGGCTGCTTT

GCACACGTCTGAACTCTTTCCCTACACGACGCTCTTCCGATCTNNNAGCAATTACAACTTGGCTGCTTT

GCACACGTCTGAACTCTTTCCCTACACGACGCTCTTCCGATCTNNNAGCAATTACAACTTGGCTGCTTT

AGCACACGTCTGAACTCTTTCCCTACACGACGCTCTTCCGATCTNNNTGGGCAACACAGTAAGATTCTG

AGCACACGTCTGAACTCTTTCCCTACACGACGCTCTTCCGATCTNNNTGGGCAACACAGTAAGATTCTG

AGCACACGTCTGAACTCTTTCCCTACACGACGCTCTTCCGATCTNNNTGGGCAACACAGTAAGATTCTG

GAGCACACGTCTGAACTCTTTCCCTACACGACGCTCTTCCGATCTNNNCCTCAGTAAAGACTCACCTCC

GAGCACACGTCTGAACTCTTTCCCTACACGACGCTCTTCCGATCTNNNCCTCAGTAAAGACTCACCTCC

GAGCACACGTCTGAACTCTTTCCCTACACGACGCTCTTCCGATCTNNNCCTCAGTAAAGACTCACCTCC

GAGCACACGTCTGAACTCTTTCCCTACACGACGCTCTTCCGATCTNNNGAGATGGGACCCTGAACTAA

GAGCACACGTCTGAACTCTTTCCCTACACGACGCTCTTCCGATCTNNNGAGATGGGACCCTGAACTAA

GAGCACACGTCTGAACTCTTTCCCTACACGACGCTCTTCCGATCTNNNGAGATGGGACCCTGAACTAA

AGCACACGTCTGAACTCTTTCCCTACACGACGCTCTTCCGATCTNNNAAAACACCAAAGACCTGGACA

AGCACACGTCTGAACTCTTTCCCTACACGACGCTCTTCCGATCTNNNAAAACACCAAAGACCTGGACA

AGCACACGTCTGAACTCTTTCCCTACACGACGCTCTTCCGATCTNNNAAAACACCAAAGACCTGGACA

GCACACGTCTGAACTCTTTCCCTACACGACGCTCTTCCGATCTNNNGTGAAAAAGGAGGTGGTGTTT

GCACACGTCTGAACTCTTTCCCTACACGACGCTCTTCCGATCTNNNGTGAAAAAGGAGGTGGTGTTT

GCACACGTCTGAACTCTTTCCCTACACGACGCTCTTCCGATCTNNNGTGAAAAAGGAGGTGGTGTTT

AGCACACGTCTGAACTCTTTCCCTACACGACGCTCTTCCGATCTNNNTATTCAGCCCCAAATGCCAAAA

AGCACACGTCTGAACTCTTTCCCTACACGACGCTCTTCCGATCTNNNTATTCAGCCCCAAATGCCAAAA

AGCACACGTCTGAACTCTTTCCCTACACGACGCTCTTCCGATCTNNNTATTCAGCCCCAAATGCCAAAA

AGCACACGTCTGAACTCTTTCCCTACACGACGCTCTTCCGATCTNNNTAAGAGGTAGAACTAGCACCC

AGCACACGTCTGAACTCTTTCCCTACACGACGCTCTTCCGATCTNNNTAAGAGGTAGAACTAGCACCC

AGCACACGTCTGAACTCTTTCCCTACACGACGCTCTTCCGATCTNNNTAAGAGGTAGAACTAGCACCC

AGCACACGTCTGAACTCTTTCCCTACACGACGCTCTTCCGATCTNNNTTCCAACAGTGAACCCCTGTTT

AGCACACGTCTGAACTCTTTCCCTACACGACGCTCTTCCGATCTNNNTTCCAACAGTGAACCCCTGTTT

AGCACACGTCTGAACTCTTTCCCTACACGACGCTCTTCCGATCTNNNTTCCAACAGTGAACCCCTGTTT

AGCACACGTCTGAACTCTTTCCCTACACGACGCTCTTCCGATCTNNNAAAAAATTAGCCAGGTGTGG

AGCACACGTCTGAACTCTTTCCCTACACGACGCTCTTCCGATCTNNNAAAAAATTAGCCAGGTGTGG

AGCACACGTCTGAACTCTTTCCCTACACGACGCTCTTCCGATCTNNNAAAAAATTAGCCAGGTGTGG

AGCACACGTCTGAACTCTTTCCCTACACGACGCTCTTCCGATCTNNNGCATCTTAGAAGTCTGCCTAC

AGCACACGTCTGAACTCTTTCCCTACACGACGCTCTTCCGATCTNNNGCATCTTAGAAGTCTGCCTAC

AGCACACGTCTGAACTCTTTCCCTACACGACGCTCTTCCGATCTNNNGCATCTTAGAAGTCTGCCTAC

AGCACACGTCTGAACTCTTTCCCTACACGACGCTCTTCCGATCTNNNACCATGTCATCCTTCCCTTTCTC

AGCACACGTCTGAACTCTTTCCCTACACGACGCTCTTCCGATCTNNNACCATGTCATCCTTCCCTTTCTC

AGCACACGTCTGAACTCTTTCCCTACACGACGCTCTTCCGATCTNNNACCATGTCATCCTTCCCTTTCTC

AGCACACGTCTGAACTCTTTCCCTACACGACGCTCTTCCGATCTNNNAGATCACACAGAACTTGCTTTG

AGCACACGTCTGAACTCTTTCCCTACACGACGCTCTTCCGATCTNNNAGATCACACAGAACTTGCTTTG

AGCACACGTCTGAACTCTTTCCCTACACGACGCTCTTCCGATCTNNNAGATCACACAGAACTTGCTTTG

GCACACGTCTGAACTCTTTCCCTACACGACGCTCTTCCGATCTNNNCTGTAATTACTGCGGCTCCACTCC

GCACACGTCTGAACTCTTTCCCTACACGACGCTCTTCCGATCTNNNCTGTAATTACTGCGGCTCCACTCC

GCACACGTCTGAACTCTTTCCCTACACGACGCTCTTCCGATCTNNNCTGTAATTACTGCGGCTCCACTCC
GAGCACACGTCTGAACTCTTTCCCTACACGACGCTCTTCCGATCTNNNTATGCCTGTAATCCCAGCTTCT
GAGCACACGTCTGAACTCTTTCCCTACACGACGCTCTTCCGATCTNNNTATGCCTGTAATCCCAGCTTCT
GAGCACACGTCTGAACTCTTTCCCTACACGACGCTCTTCCGATCTNNNTATGCCTGTAATCCCAGCTTCT
GAGCACACGTCTGAACTCTTTCCCTACACGACGCTCTTCCGATCTNNNCCACTTCTTCCCACAGGACCCAT
GAGCACACGTCTGAACTCTTTCCCTACACGACGCTCTTCCGATCTNNNCCACTTCTTCCCACAGGACCCAT
GAGCACACGTCTGAACTCTTTCCCTACACGACGCTCTTCCGATCTNNNCCACTTCTTCCCACAGGACCCAT
GAGCACACGTCTGAACTCTTTCCCTACACGACGCTCTTCCGATCTNNNAAGAGAATGGCGTGAACCCAC
GAGCACACGTCTGAACTCTTTCCCTACACGACGCTCTTCCGATCTNNNAAGAGAATGGCGTGAACCCAC
GAGCACACGTCTGAACTCTTTCCCTACACGACGCTCTTCCGATCTNNNAAGAGAATGGCGTGAACCCAC
GAGCACACGTCTGAACTCTTTCCCTACACGACGCTCTTCCGATCTNNNATCTGTAGTCCTAGCTACTCGG
GAGCACACGTCTGAACTCTTTCCCTACACGACGCTCTTCCGATCTNNNATCTGTAGTCCTAGCTACTCGG
GAGCACACGTCTGAACTCTTTCCCTACACGACGCTCTTCCGATCTNNNATCTGTAGTCCTAGCTACTCGG
GAGCACACGTCTGAACTCTTTCCCTACACGACGCTCTTCCGATCTNNNTCAGAACCTCACACTTACACAC
GAGCACACGTCTGAACTCTTTCCCTACACGACGCTCTTCCGATCTNNNTCAGAACCTCACACTTACACAC
GAGCACACGTCTGAACTCTTTCCCTACACGACGCTCTTCCGATCTNNNTCAGAACCTCACACTTACACAC
GAGCACACGTCTGAACTCTTTCCCTACACGACGCTCTTCCGATCTNNNTCTAGCTTCAGGTCTTTGCACA
GAGCACACGTCTGAACTCTTTCCCTACACGACGCTCTTCCGATCTNNNTCTAGCTTCAGGTCTTTGCACA
GAGCACACGTCTGAACTCTTTCCCTACACGACGCTCTTCCGATCTNNNTCTAGCTTCAGGTCTTTGCACA
AGCACACGTCTGAACTCTTTCCCTACACGACGCTCTTCCGATCTNNNAGCATACTGTGTGTCCACTTGAT
AGCACACGTCTGAACTCTTTCCCTACACGACGCTCTTCCGATCTNNNAGCATACTGTGTGTCCACTTGAT
AGCACACGTCTGAACTCTTTCCCTACACGACGCTCTTCCGATCTNNNAGCATACTGTGTGTCCACTTGAT
GCACACGTCTGAACTCTTTCCCTACACGACGCTCTTCCGATCTNNNTGTGGTGTTTGGTTTTCTGTTCCC
GCACACGTCTGAACTCTTTCCCTACACGACGCTCTTCCGATCTNNNTGTGGTGTTTGGTTTTCTGTTCCC
GCACACGTCTGAACTCTTTCCCTACACGACGCTCTTCCGATCTNNNTGTGGTGTTTGGTTTTCTGTTCCC
GCACACGTCTGAACTCTTTCCCTACACGACGCTCTTCCGATCTNNNCCTTAAAATAGAATGTGGACCTAC
GCACACGTCTGAACTCTTTCCCTACACGACGCTCTTCCGATCTNNNCCTTAAAATAGAATGTGGACCTAC
GCACACGTCTGAACTCTTTCCCTACACGACGCTCTTCCGATCTNNNCCTTAAAATAGAATGTGGACCTAC
AGCACACGTCTGAACTCTTTCCCTACACGACGCTCTTCCGATCTNNNGAGGCTGAGGCAGGAGAATG
AGCACACGTCTGAACTCTTTCCCTACACGACGCTCTTCCGATCTNNNGAGGCTGAGGCAGGAGAATG
AGCACACGTCTGAACTCTTTCCCTACACGACGCTCTTCCGATCTNNNGAGGCTGAGGCAGGAGAATG
AGCACACGTCTGAACTCTTTCCCTACACGACGCTCTTCCGATCTNNNGACTCACCTTGGGAGAGAACTT
AGCACACGTCTGAACTCTTTCCCTACACGACGCTCTTCCGATCTNNNGACTCACCTTGGGAGAGAACTT
AGCACACGTCTGAACTCTTTCCCTACACGACGCTCTTCCGATCTNNNGACTCACCTTGGGAGAGAACTT
AGCACACGTCTGAACTCTTTCCCTACACGACGCTCTTCCGATCTNNNGTCACCCTGACTCACTGATTCATC
AGCACACGTCTGAACTCTTTCCCTACACGACGCTCTTCCGATCTNNNGTCACCCTGACTCACTGATTCATC
AGCACACGTCTGAACTCTTTCCCTACACGACGCTCTTCCGATCTNNNGTCACCCTGACTCACTGATTCATC
GAGCACACGTCTGAACTCTTTCCCTACACGACGCTCTTCCGATCTNNNAATGCTGTATTCCCAGAGACAC
GAGCACACGTCTGAACTCTTTCCCTACACGACGCTCTTCCGATCTNNNAATGCTGTATTCCCAGAGACAC
GAGCACACGTCTGAACTCTTTCCCTACACGACGCTCTTCCGATCTNNNAATGCTGTATTCCCAGAGACAC
AGAGCACACGTCTGAACTCTTTCCCTACACGACGCTCTTCCGATCTNNNGTACCTGTAATCTCAGCTACC
AGAGCACACGTCTGAACTCTTTCCCTACACGACGCTCTTCCGATCTNNNGTACCTGTAATCTCAGCTACC
AGAGCACACGTCTGAACTCTTTCCCTACACGACGCTCTTCCGATCTNNNGTACCTGTAATCTCAGCTACC

AGCACACGTCTGAACTCTTTCCCTACACGACGCTCTTCCGATCTNNNAACAGGAAATATGGCTTCCAC
AGCACACGTCTGAACTCTTTCCCTACACGACGCTCTTCCGATCTNNNAACAGGAAATATGGCTTCCAC
AGCACACGTCTGAACTCTTTCCCTACACGACGCTCTTCCGATCTNNNAACAGGAAATATGGCTTCCAC
GAGCACACGTCTGAACTCTTTCCCTACACGACGCTCTTCCGATCTNNNCCCTTCCCAGTTACGAGAATC
GAGCACACGTCTGAACTCTTTCCCTACACGACGCTCTTCCGATCTNNNCCCTTCCCAGTTACGAGAATC
GAGCACACGTCTGAACTCTTTCCCTACACGACGCTCTTCCGATCTNNNCCCTTCCCAGTTACGAGAATC
CACACGTCTGAACTCTTTCCCTACACGACGCTCTTCCGATCTNNNTAAAAATGTGGTTTCTTTAGTCCCA
CACACGTCTGAACTCTTTCCCTACACGACGCTCTTCCGATCTNNNTAAAAATGTGGTTTCTTTAGTCCCA
CACACGTCTGAACTCTTTCCCTACACGACGCTCTTCCGATCTNNNTAAAAATGTGGTTTCTTTAGTCCCA
AGCACACGTCTGAACTCTTTCCCTACACGACGCTCTTCCGATCTNNNTTGCTGTTCTCCCTTTACATCCT
AGCACACGTCTGAACTCTTTCCCTACACGACGCTCTTCCGATCTNNNTTGCTGTTCTCCCTTTACATCCT
AGCACACGTCTGAACTCTTTCCCTACACGACGCTCTTCCGATCTNNNTTGCTGTTCTCCCTTTACATCCT
GCACACGTCTGAACTCTTTCCCTACACGACGCTCTTCCGATCTNNNATGACTTCTTTGTGCATGTTGG
GCACACGTCTGAACTCTTTCCCTACACGACGCTCTTCCGATCTNNNATGACTTCTTTGTGCATGTTGG
GCACACGTCTGAACTCTTTCCCTACACGACGCTCTTCCGATCTNNNATGACTTCTTTGTGCATGTTGG
GCACACGTCTGAACTCTTTCCCTACACGACGCTCTTCCGATCTNNNAGAGAGAGAGAGAGAGAAAG
GCACACGTCTGAACTCTTTCCCTACACGACGCTCTTCCGATCTNNNAGAGAGAGAGAGAGAGAAAG
GCACACGTCTGAACTCTTTCCCTACACGACGCTCTTCCGATCTNNNAGAGAGAGAGAGAGAGAAAG
GCACACGTCTGAACTCTTTCCCTACACGACGCTCTTCCGATCTNNNGGGTGTAGAATTTTAGGCTGGG
GCACACGTCTGAACTCTTTCCCTACACGACGCTCTTCCGATCTNNNGGGTGTAGAATTTTAGGCTGGG
GCACACGTCTGAACTCTTTCCCTACACGACGCTCTTCCGATCTNNNGGGTGTAGAATTTTAGGCTGGG
GCACACGTCTGAACTCTTTCCCTACACGACGCTCTTCCGATCTNNNATACTTTTACTTCTTTACTGGGT
GCACACGTCTGAACTCTTTCCCTACACGACGCTCTTCCGATCTNNNATACTTTTACTTCTTTACTGGGT
GCACACGTCTGAACTCTTTCCCTACACGACGCTCTTCCGATCTNNNATACTTTTACTTCTTTACTGGGT
AGCACACGTCTGAACTCTTTCCCTACACGACGCTCTTCCGATCTNNNTAATTCATCTGCCTCAGCCTCTC
AGCACACGTCTGAACTCTTTCCCTACACGACGCTCTTCCGATCTNNNTAATTCATCTGCCTCAGCCTCTC
AGCACACGTCTGAACTCTTTCCCTACACGACGCTCTTCCGATCTNNNTAATTCATCTGCCTCAGCCTCTC
AGCACACGTCTGAACTCTTTCCCTACACGACGCTCTTCCGATCTNNNTCCTGGGCATGACATGAGAAA
AGCACACGTCTGAACTCTTTCCCTACACGACGCTCTTCCGATCTNNNTCCTGGGCATGACATGAGAAA
AGCACACGTCTGAACTCTTTCCCTACACGACGCTCTTCCGATCTNNNTCCTGGGCATGACATGAGAAA
GCACACGTCTGAACTCTTTCCCTACACGACGCTCTTCCGATCTNNNGGTAGAAAGAGTGTGCTGGGTA
GCACACGTCTGAACTCTTTCCCTACACGACGCTCTTCCGATCTNNNGGTAGAAAGAGTGTGCTGGGTA
GCACACGTCTGAACTCTTTCCCTACACGACGCTCTTCCGATCTNNNGGTAGAAAGAGTGTGCTGGGTA
CACACGTCTGAACTCTTTCCCTACACGACGCTCTTCCGATCTNNNACCCAATCTAAAGTTGCAACCGAC
CACACGTCTGAACTCTTTCCCTACACGACGCTCTTCCGATCTNNNACCCAATCTAAAGTTGCAACCGAC
CACACGTCTGAACTCTTTCCCTACACGACGCTCTTCCGATCTNNNACCCAATCTAAAGTTGCAACCGAC
GCACACGTCTGAACTCTTTCCCTACACGACGCTCTTCCGATCTNNNTCTTTCTGTGTCGAATTTTATAGC
GCACACGTCTGAACTCTTTCCCTACACGACGCTCTTCCGATCTNNNTCTTTCTGTGTCGAATTTTATAGC
GCACACGTCTGAACTCTTTCCCTACACGACGCTCTTCCGATCTNNNTCTTTCTGTGTCGAATTTTATAGC
AGCACACGTCTGAACTCTTTCCCTACACGACGCTCTTCCGATCTNNNACATGGTCAGGAAAACTTCCCA
AGCACACGTCTGAACTCTTTCCCTACACGACGCTCTTCCGATCTNNNACATGGTCAGGAAAACTTCCCA
AGCACACGTCTGAACTCTTTCCCTACACGACGCTCTTCCGATCTNNNACATGGTCAGGAAAACTTCCCA
AGCACACGTCTGAACTCTTTCCCTACACGACGCTCTTCCGATCTNNNAAGTAGAGTTGCAGCTCCTTT

AGCACACGTCTGAACTCTTTCCCTACACGACGCTCTTCCGATCTNNNAAGTAGAGTTGCAGCTCCTTT

AGCACACGTCTGAACTCTTTCCCTACACGACGCTCTTCCGATCTNNNAAGTAGAGTTGCAGCTCCTTT

GAGCACACGTCTGAACTCTTTCCCTACACGACGCTCTTCCGATCTNNNGAGAATTGCTTGAACCCAGG

GAGCACACGTCTGAACTCTTTCCCTACACGACGCTCTTCCGATCTNNNGAGAATTGCTTGAACCCAGG

GAGCACACGTCTGAACTCTTTCCCTACACGACGCTCTTCCGATCTNNNGAGAATTGCTTGAACCCAGG

GAGCACACGTCTGAACTCTTTCCCTACACGACGCTCTTCCGATCTNNNCATGGAATCAACCTAAGTGT

GAGCACACGTCTGAACTCTTTCCCTACACGACGCTCTTCCGATCTNNNCATGGAATCAACCTAAGTGT

GAGCACACGTCTGAACTCTTTCCCTACACGACGCTCTTCCGATCTNNNCATGGAATCAACCTAAGTGT

ACACGTCTGAACTCTTTCCCTACACGACGCTCTTCCGATCTNNNCTCAGCTCCCACACCGTACGCATCG

ACACGTCTGAACTCTTTCCCTACACGACGCTCTTCCGATCTNNNCTCAGCTCCCACACCGTACGCATCG

ACACGTCTGAACTCTTTCCCTACACGACGCTCTTCCGATCTNNNCTCAGCTCCCACACCGTACGCATCG

AGCACACGTCTGAACTCTTTCCCTACACGACGCTCTTCCGATCTNNNCAAAGAACACCCTTCCCCTTCT

AGCACACGTCTGAACTCTTTCCCTACACGACGCTCTTCCGATCTNNNCAAAGAACACCCTTCCCCTTCT

AGCACACGTCTGAACTCTTTCCCTACACGACGCTCTTCCGATCTNNNCAAAGAACACCCTTCCCCTTCT

AGCACACGTCTGAACTCTTTCCCTACACGACGCTCTTCCGATCTNNNGCCTATAATCCCAGCTACTA

AGCACACGTCTGAACTCTTTCCCTACACGACGCTCTTCCGATCTNNNGCCTATAATCCCAGCTACTA

AGCACACGTCTGAACTCTTTCCCTACACGACGCTCTTCCGATCTNNNGCCTATAATCCCAGCTACTA

GCACACGTCTGAACTCTTTCCCTACACGACGCTCTTCCGATCTNNNGAAGGCCTGGAGGTATGAAAC

GCACACGTCTGAACTCTTTCCCTACACGACGCTCTTCCGATCTNNNGAAGGCCTGGAGGTATGAAAC

GCACACGTCTGAACTCTTTCCCTACACGACGCTCTTCCGATCTNNNGAAGGCCTGGAGGTATGAAAC

GAGCACACGTCTGAACTCTTTCCCTACACGACGCTCTTCCGATCTNNNCTGGGACTACAGGTGTGCAC

GAGCACACGTCTGAACTCTTTCCCTACACGACGCTCTTCCGATCTNNNCTGGGACTACAGGTGTGCAC

GAGCACACGTCTGAACTCTTTCCCTACACGACGCTCTTCCGATCTNNNCTGGGACTACAGGTGTGCAC

GAGCACACGTCTGAACTCTTTCCCTACACGACGCTCTTCCGATCTNNNTAAGTGATCATATTTACCACC

GAGCACACGTCTGAACTCTTTCCCTACACGACGCTCTTCCGATCTNNNTAAGTGATCATATTTACCACC

GAGCACACGTCTGAACTCTTTCCCTACACGACGCTCTTCCGATCTNNNTAAGTGATCATATTTACCACC

GAGCACACGTCTGAACTCTTTCCCTACACGACGCTCTTCCGATCTNNNCATCCCATCTCAAAAAAAAA

GAGCACACGTCTGAACTCTTTCCCTACACGACGCTCTTCCGATCTNNNCATCCCATCTCAAAAAAAAA

GAGCACACGTCTGAACTCTTTCCCTACACGACGCTCTTCCGATCTNNNCATCCCATCTCAAAAAAAAA

AGCACACGTCTGAACTCTTTCCCTACACGACGCTCTTCCGATCTNNNATACAGAATTAGCCGGGCATG

AGCACACGTCTGAACTCTTTCCCTACACGACGCTCTTCCGATCTNNNATACAGAATTAGCCGGGCATG

AGCACACGTCTGAACTCTTTCCCTACACGACGCTCTTCCGATCTNNNATACAGAATTAGCCGGGCATG

GAGCACACGTCTGAACTCTTTCCCTACACGACGCTCTTCCGATCTNNNGGAAGAACTCACACTGGGGA

GAGCACACGTCTGAACTCTTTCCCTACACGACGCTCTTCCGATCTNNNGGAAGAACTCACACTGGGGA

GAGCACACGTCTGAACTCTTTCCCTACACGACGCTCTTCCGATCTNNNGGAAGAACTCACACTGGGGA

AGCACACGTCTGAACTCTTTCCCTACACGACGCTCTTCCGATCTNNNACAGTGGTGCAAATTCAGCTT

AGCACACGTCTGAACTCTTTCCCTACACGACGCTCTTCCGATCTNNNACAGTGGTGCAAATTCAGCTT

AGCACACGTCTGAACTCTTTCCCTACACGACGCTCTTCCGATCTNNNACAGTGGTGCAAATTCAGCTT

GAGCACACGTCTGAACTCTTTCCCTACACGACGCTCTTCCGATCTNNNGTCTGCAATCTGAAAGCAACA

GAGCACACGTCTGAACTCTTTCCCTACACGACGCTCTTCCGATCTNNNGTCTGCAATCTGAAAGCAACA

GAGCACACGTCTGAACTCTTTCCCTACACGACGCTCTTCCGATCTNNNGTCTGCAATCTGAAAGCAACA

AGCACACGTCTGAACTCTTTCCCTACACGACGCTCTTCCGATCTNNNAGTGTCAGTTCATGTTGCCTCT

AGCACACGTCTGAACTCTTTCCCTACACGACGCTCTTCCGATCTNNNAGTGTCAGTTCATGTTGCCTCT

AGCACACGTCTGAACTCTTTCCCTACACGACGCTCTTCCGATCTNNNAGTGTCAGTTCATGTTGCCTCTC
AGCACACGTCTGAACTCTTTCCCTACACGACGCTCTTCCGATCTNNNGGTACCCCTCTCCAATTATTTGT
AGCACACGTCTGAACTCTTTCCCTACACGACGCTCTTCCGATCTNNNGGTACCCCTCTCCAATTATTTGT
AGCACACGTCTGAACTCTTTCCCTACACGACGCTCTTCCGATCTNNNGGTACCCCTCTCCAATTATTTGT
GAGCACACGTCTGAACTCTTTCCCTACACGACGCTCTTCCGATCTNNNGTTCAAGCAATTCTCATGCCTC
GAGCACACGTCTGAACTCTTTCCCTACACGACGCTCTTCCGATCTNNNGTTCAAGCAATTCTCATGCCTC
GAGCACACGTCTGAACTCTTTCCCTACACGACGCTCTTCCGATCTNNNGTTCAAGCAATTCTCATGCCTC
AGCACACGTCTGAACTCTTTCCCTACACGACGCTCTTCCGATCTNNNCTATATTGGTCAGGCTGGTCTCC
AGCACACGTCTGAACTCTTTCCCTACACGACGCTCTTCCGATCTNNNCTATATTGGTCAGGCTGGTCTCC
AGCACACGTCTGAACTCTTTCCCTACACGACGCTCTTCCGATCTNNNCTATATTGGTCAGGCTGGTCTCC
GAGCACACGTCTGAACTCTTTCCCTACACGACGCTCTTCCGATCTNNNCCTGTAGTCCCAGCTACTCTAC
GAGCACACGTCTGAACTCTTTCCCTACACGACGCTCTTCCGATCTNNNCCTGTAGTCCCAGCTACTCTAC
GAGCACACGTCTGAACTCTTTCCCTACACGACGCTCTTCCGATCTNNNCCTGTAGTCCCAGCTACTCTAC
AGCACACGTCTGAACTCTTTCCCTACACGACGCTCTTCCGATCTNNNAATAGAAGGCATGGGAAGATC
AGCACACGTCTGAACTCTTTCCCTACACGACGCTCTTCCGATCTNNNAATAGAAGGCATGGGAAGATC
AGCACACGTCTGAACTCTTTCCCTACACGACGCTCTTCCGATCTNNNAATAGAAGGCATGGGAAGATC
GAGCACACGTCTGAACTCTTTCCCTACACGACGCTCTTCCGATCTNNNTTCACAGTTTCTCAGACTTGG
GAGCACACGTCTGAACTCTTTCCCTACACGACGCTCTTCCGATCTNNNTTCACAGTTTCTCAGACTTGG
GAGCACACGTCTGAACTCTTTCCCTACACGACGCTCTTCCGATCTNNNTTCACAGTTTCTCAGACTTGG
AGCACACGTCTGAACTCTTTCCCTACACGACGCTCTTCCGATCTNNNCTGTAGTCCCAGCTACTCAGGA
AGCACACGTCTGAACTCTTTCCCTACACGACGCTCTTCCGATCTNNNCTGTAGTCCCAGCTACTCAGGA
AGCACACGTCTGAACTCTTTCCCTACACGACGCTCTTCCGATCTNNNCTGTAGTCCCAGCTACTCAGGA
AGCACACGTCTGAACTCTTTCCCTACACGACGCTCTTCCGATCTNNNGCTCATGCCTGTAATCTCATCAC
AGCACACGTCTGAACTCTTTCCCTACACGACGCTCTTCCGATCTNNNGCTCATGCCTGTAATCTCATCAC
AGCACACGTCTGAACTCTTTCCCTACACGACGCTCTTCCGATCTNNNGCTCATGCCTGTAATCTCATCAC
AGCACACGTCTGAACTCTTTCCCTACACGACGCTCTTCCGATCTNNNTGGACATCATCATTTTACCCAGA
AGCACACGTCTGAACTCTTTCCCTACACGACGCTCTTCCGATCTNNNTGGACATCATCATTTTACCCAGA
AGCACACGTCTGAACTCTTTCCCTACACGACGCTCTTCCGATCTNNNTGGACATCATCATTTTACCCAGA
GAGCACACGTCTGAACTCTTTCCCTACACGACGCTCTTCCGATCTNNNTCTTTGGATTTCCAGGTGCATT
GAGCACACGTCTGAACTCTTTCCCTACACGACGCTCTTCCGATCTNNNTCTTTGGATTTCCAGGTGCATT
GAGCACACGTCTGAACTCTTTCCCTACACGACGCTCTTCCGATCTNNNTCTTTGGATTTCCAGGTGCATT
GAGCACACGTCTGAACTCTTTCCCTACACGACGCTCTTCCGATCTNNNTCCAAAAACACCCATCAGTCTC
GAGCACACGTCTGAACTCTTTCCCTACACGACGCTCTTCCGATCTNNNTCCAAAAACACCCATCAGTCTC
GAGCACACGTCTGAACTCTTTCCCTACACGACGCTCTTCCGATCTNNNTCCAAAAACACCCATCAGTCTC
AGCACACGTCTGAACTCTTTCCCTACACGACGCTCTTCCGATCTNNNACTGAAATGCCCCTAATGGATCA
AGCACACGTCTGAACTCTTTCCCTACACGACGCTCTTCCGATCTNNNACTGAAATGCCCCTAATGGATCA
AGCACACGTCTGAACTCTTTCCCTACACGACGCTCTTCCGATCTNNNACTGAAATGCCCCTAATGGATCA
AGCACACGTCTGAACTCTTTCCCTACACGACGCTCTTCCGATCTNNNGAGGCTCCAGAATGATTTACCC
AGCACACGTCTGAACTCTTTCCCTACACGACGCTCTTCCGATCTNNNGAGGCTCCAGAATGATTTACCC
AGCACACGTCTGAACTCTTTCCCTACACGACGCTCTTCCGATCTNNNGAGGCTCCAGAATGATTTACCC
AGCACACGTCTGAACTCTTTCCCTACACGACGCTCTTCCGATCTNNNCCCTGCACATTCTGGGATAATC
AGCACACGTCTGAACTCTTTCCCTACACGACGCTCTTCCGATCTNNNCCCTGCACATTCTGGGATAATC
AGCACACGTCTGAACTCTTTCCCTACACGACGCTCTTCCGATCTNNNCCCTGCACATTCTGGGATAATC

AGCACACGTCTGAACTCTTTCCCTACACGACGCTCTTCCGATCTNNNACTGGAATACAAACAAGTTGG
AGCACACGTCTGAACTCTTTCCCTACACGACGCTCTTCCGATCTNNNACTGGAATACAAACAAGTTGG
AGCACACGTCTGAACTCTTTCCCTACACGACGCTCTTCCGATCTNNNACTGGAATACAAACAAGTTGG
AGCACACGTCTGAACTCTTTCCCTACACGACGCTCTTCCGATCTNNNCTCCTTCCTTCCTTCCTTCCTTCCA
AGCACACGTCTGAACTCTTTCCCTACACGACGCTCTTCCGATCTNNNCTCCTTCCTTCCTTCCTTCCTTCCA
AGCACACGTCTGAACTCTTTCCCTACACGACGCTCTTCCGATCTNNNCTCCTTCCTTCCTTCCTTCCTTCCA
AGCACACGTCTGAACTCTTTCCCTACACGACGCTCTTCCGATCTNNNTCTTAGGCGTCTGAACACTTCTC
AGCACACGTCTGAACTCTTTCCCTACACGACGCTCTTCCGATCTNNNTCTTAGGCGTCTGAACACTTCTC
AGCACACGTCTGAACTCTTTCCCTACACGACGCTCTTCCGATCTNNNTCTTAGGCGTCTGAACACTTCTC
AGCACACGTCTGAACTCTTTCCCTACACGACGCTCTTCCGATCTNNNTGAGCCAAGATTGTGCCACTCA
AGCACACGTCTGAACTCTTTCCCTACACGACGCTCTTCCGATCTNNNTGAGCCAAGATTGTGCCACTCA
AGCACACGTCTGAACTCTTTCCCTACACGACGCTCTTCCGATCTNNNTGAGCCAAGATTGTGCCACTCA
AGCACACGTCTGAACTCTTTCCCTACACGACGCTCTTCCGATCTNNNAACTCTCACTCACAGCATCACATC
AGCACACGTCTGAACTCTTTCCCTACACGACGCTCTTCCGATCTNNNAACTCTCACTCACAGCATCACATC
AGCACACGTCTGAACTCTTTCCCTACACGACGCTCTTCCGATCTNNNAACTCTCACTCACAGCATCACATC
GAGCACACGTCTGAACTCTTTCCCTACACGACGCTCTTCCGATCTNNNCTCTCCCACAAGAACCCTATGG
GAGCACACGTCTGAACTCTTTCCCTACACGACGCTCTTCCGATCTNNNCTCTCCCACAAGAACCCTATGG
GAGCACACGTCTGAACTCTTTCCCTACACGACGCTCTTCCGATCTNNNCTCTCCCACAAGAACCCTATGG
AGCACACGTCTGAACTCTTTCCCTACACGACGCTCTTCCGATCTNNNATTTGTAAGGCGTGACTCCTGAT
AGCACACGTCTGAACTCTTTCCCTACACGACGCTCTTCCGATCTNNNATTTGTAAGGCGTGACTCCTGAT
AGCACACGTCTGAACTCTTTCCCTACACGACGCTCTTCCGATCTNNNATTTGTAAGGCGTGACTCCTGAT
ACACGTCTGAACTCTTTCCCTACACGACGCTCTTCCGATCTNNNATGCACATGAAGCCTCTAGGTAGCAT
ACACGTCTGAACTCTTTCCCTACACGACGCTCTTCCGATCTNNNATGCACATGAAGCCTCTAGGTAGCAT
ACACGTCTGAACTCTTTCCCTACACGACGCTCTTCCGATCTNNNATGCACATGAAGCCTCTAGGTAGCAT
GAGCACACGTCTGAACTCTTTCCCTACACGACGCTCTTCCGATCTNNNCTTGCAATGAGCTGAGATGGT
GAGCACACGTCTGAACTCTTTCCCTACACGACGCTCTTCCGATCTNNNCTTGCAATGAGCTGAGATGGT
GAGCACACGTCTGAACTCTTTCCCTACACGACGCTCTTCCGATCTNNNCTTGCAATGAGCTGAGATGGT
GAGCACACGTCTGAACTCTTTCCCTACACGACGCTCTTCCGATCTNNNGCAGAGGTTATAGTGAGCCAA
GAGCACACGTCTGAACTCTTTCCCTACACGACGCTCTTCCGATCTNNNGCAGAGGTTATAGTGAGCCAA
GAGCACACGTCTGAACTCTTTCCCTACACGACGCTCTTCCGATCTNNNGCAGAGGTTATAGTGAGCCAA
GAGCACACGTCTGAACTCTTTCCCTACACGACGCTCTTCCGATCTNNNGTACCTGTAGTCCCAGCTATTC
GAGCACACGTCTGAACTCTTTCCCTACACGACGCTCTTCCGATCTNNNGTACCTGTAGTCCCAGCTATTC
GAGCACACGTCTGAACTCTTTCCCTACACGACGCTCTTCCGATCTNNNGTACCTGTAGTCCCAGCTATTC
AGCACACGTCTGAACTCTTTCCCTACACGACGCTCTTCCGATCTNNNACCTTTCCTTGCCTCTTTCCTAGC
AGCACACGTCTGAACTCTTTCCCTACACGACGCTCTTCCGATCTNNNACCTTTCCTTGCCTCTTTCCTAGC
AGCACACGTCTGAACTCTTTCCCTACACGACGCTCTTCCGATCTNNNACCTTTCCTTGCCTCTTTCCTAGC
GAGCACACGTCTGAACTCTTTCCCTACACGACGCTCTTCCGATCTNNNGTAATTCCAGCACTTTGGTAG
GAGCACACGTCTGAACTCTTTCCCTACACGACGCTCTTCCGATCTNNNGTAATTCCAGCACTTTGGTAG
GAGCACACGTCTGAACTCTTTCCCTACACGACGCTCTTCCGATCTNNNGTAATTCCAGCACTTTGGTAG
AGAGCACACGTCTGAACTCTTTCCCTACACGACGCTCTTCCGATCTNNNGAGAAGGCCCTGCTTTCAGA
AGAGCACACGTCTGAACTCTTTCCCTACACGACGCTCTTCCGATCTNNNGAGAAGGCCCTGCTTTCAGA
AGAGCACACGTCTGAACTCTTTCCCTACACGACGCTCTTCCGATCTNNNGAGAAGGCCCTGCTTTCAGA
AGAGCACACGTCTGAACTCTTTCCCTACACGACGCTCTTCCGATCTNNNTGCTCAAAATCATCGCCTTCA

AGAGCACACGTCTGAACTCTTTCCCTACACGACGCTCTTCCGATCTNNNTGCTCAAAATCATCGCCTTCA
AGAGCACACGTCTGAACTCTTTCCCTACACGACGCTCTTCCGATCTNNNTGCTCAAAATCATCGCCTTCA
AGCACACGTCTGAACTCTTTCCCTACACGACGCTCTTCCGATCTNNNATCCAGAATTTCTTCCCCCAAAT
AGCACACGTCTGAACTCTTTCCCTACACGACGCTCTTCCGATCTNNNATCCAGAATTTCTTCCCCCAAAT
AGCACACGTCTGAACTCTTTCCCTACACGACGCTCTTCCGATCTNNNATCCAGAATTTCTTCCCCCAAAT
GAGCACACGTCTGAACTCTTTCCCTACACGACGCTCTTCCGATCTNNNTGAAGGTTGCAGTGAGTTGA
GAGCACACGTCTGAACTCTTTCCCTACACGACGCTCTTCCGATCTNNNTGAAGGTTGCAGTGAGTTGA
GAGCACACGTCTGAACTCTTTCCCTACACGACGCTCTTCCGATCTNNNTGAAGGTTGCAGTGAGTTGA
AGCACACGTCTGAACTCTTTCCCTACACGACGCTCTTCCGATCTNNNGCTCAACAATGATACACACACG
AGCACACGTCTGAACTCTTTCCCTACACGACGCTCTTCCGATCTNNNGCTCAACAATGATACACACACG
AGCACACGTCTGAACTCTTTCCCTACACGACGCTCTTCCGATCTNNNGCTCAACAATGATACACACACG
GAGCACACGTCTGAACTCTTTCCCTACACGACGCTCTTCCGATCTNNNTAGCTCTGCATCAATGGATA
GAGCACACGTCTGAACTCTTTCCCTACACGACGCTCTTCCGATCTNNNTAGCTCTGCATCAATGGATA
GAGCACACGTCTGAACTCTTTCCCTACACGACGCTCTTCCGATCTNNNTAGCTCTGCATCAATGGATA
GAGCACACGTCTGAACTCTTTCCCTACACGACGCTCTTCCGATCTNNNGACTGGGCACACACTAGGATC
GAGCACACGTCTGAACTCTTTCCCTACACGACGCTCTTCCGATCTNNNGACTGGGCACACACTAGGATC
GAGCACACGTCTGAACTCTTTCCCTACACGACGCTCTTCCGATCTNNNGACTGGGCACACACTAGGATC
GAGCACACGTCTGAACTCTTTCCCTACACGACGCTCTTCCGATCTNNNTCATGAGGCTGAAGCAAGAG
GAGCACACGTCTGAACTCTTTCCCTACACGACGCTCTTCCGATCTNNNTCATGAGGCTGAAGCAAGAG
GAGCACACGTCTGAACTCTTTCCCTACACGACGCTCTTCCGATCTNNNTCATGAGGCTGAAGCAAGAG
CACACGTCTGAACTCTTTCCCTACACGACGCTCTTCCGATCTNNNCATTCCCTTGGTGGCAAATCAATCA
CACACGTCTGAACTCTTTCCCTACACGACGCTCTTCCGATCTNNNCATTCCCTTGGTGGCAAATCAATCA
CACACGTCTGAACTCTTTCCCTACACGACGCTCTTCCGATCTNNNCATTCCCTTGGTGGCAAATCAATCA
AGCACACGTCTGAACTCTTTCCCTACACGACGCTCTTCCGATCTNNNCACCCGGCACTATAAACGTTTAT
AGCACACGTCTGAACTCTTTCCCTACACGACGCTCTTCCGATCTNNNCACCCGGCACTATAAACGTTTAT
AGCACACGTCTGAACTCTTTCCCTACACGACGCTCTTCCGATCTNNNCACCCGGCACTATAAACGTTTAT
CACGTCTGAACTCTTTCCCTACACGACGCTCTTCCGATCTNNNTTATAGATGCGAATCCTTTCCCATCGG
CACGTCTGAACTCTTTCCCTACACGACGCTCTTCCGATCTNNNTTATAGATGCGAATCCTTTCCCATCGG
CACGTCTGAACTCTTTCCCTACACGACGCTCTTCCGATCTNNNTTATAGATGCGAATCCTTTCCCATCGG
GAGCACACGTCTGAACTCTTTCCCTACACGACGCTCTTCCGATCTNNNAATACAAAAAATTAGCCGGGC
GAGCACACGTCTGAACTCTTTCCCTACACGACGCTCTTCCGATCTNNNAATACAAAAAATTAGCCGGGC
GAGCACACGTCTGAACTCTTTCCCTACACGACGCTCTTCCGATCTNNNAATACAAAAAATTAGCCGGGC
AGCACACGTCTGAACTCTTTCCCTACACGACGCTCTTCCGATCTNNNGTGCCCGGCCTATTCCTATATTT
AGCACACGTCTGAACTCTTTCCCTACACGACGCTCTTCCGATCTNNNGTGCCCGGCCTATTCCTATATTT
AGCACACGTCTGAACTCTTTCCCTACACGACGCTCTTCCGATCTNNNGTGCCCGGCCTATTCCTATATTT
GAGCACACGTCTGAACTCTTTCCCTACACGACGCTCTTCCGATCTNNNACAAGTCAGGGGTTACTGTGT
GAGCACACGTCTGAACTCTTTCCCTACACGACGCTCTTCCGATCTNNNACAAGTCAGGGGTTACTGTGT
GAGCACACGTCTGAACTCTTTCCCTACACGACGCTCTTCCGATCTNNNACAAGTCAGGGGTTACTGTGT
AGCACACGTCTGAACTCTTTCCCTACACGACGCTCTTCCGATCTNNNTATGCCTGTAATCCCAGCACTTT
AGCACACGTCTGAACTCTTTCCCTACACGACGCTCTTCCGATCTNNNTATGCCTGTAATCCCAGCACTTT
AGCACACGTCTGAACTCTTTCCCTACACGACGCTCTTCCGATCTNNNTATGCCTGTAATCCCAGCACTTT
GAGCACACGTCTGAACTCTTTCCCTACACGACGCTCTTCCGATCTNNNCATGCACATACACACACACACA
GAGCACACGTCTGAACTCTTTCCCTACACGACGCTCTTCCGATCTNNNCATGCACATACACACACACACA

AGCACACGTCTGAACTCTTTCCCTACACGACGCTCTTCCGATCTNNNCATGCACATACACACACACAC
CACACGTCTGAACTCTTTCCCTACACGACGCTCTTCCGATCTNNNCTCCCTCACTGGTGACATTCTACCA
CACACGTCTGAACTCTTTCCCTACACGACGCTCTTCCGATCTNNNCTCCCTCACTGGTGACATTCTACCA
CACACGTCTGAACTCTTTCCCTACACGACGCTCTTCCGATCTNNNCTCCCTCACTGGTGACATTCTACCA
GCACACGTCTGAACTCTTTCCCTACACGACGCTCTTCCGATCTNNNATTTCCCACCGCCTCTCTTTGCAT
GCACACGTCTGAACTCTTTCCCTACACGACGCTCTTCCGATCTNNNATTTCCCACCGCCTCTCTTTGCAT
GCACACGTCTGAACTCTTTCCCTACACGACGCTCTTCCGATCTNNNATTTCCCACCGCCTCTCTTTGCAT
AGCACACGTCTGAACTCTTTCCCTACACGACGCTCTTCCGATCTNNNAGGCCAAATCAGGAGGATCAT
AGCACACGTCTGAACTCTTTCCCTACACGACGCTCTTCCGATCTNNNAGGCCAAATCAGGAGGATCAT
AGCACACGTCTGAACTCTTTCCCTACACGACGCTCTTCCGATCTNNNAGGCCAAATCAGGAGGATCAT
ACACGTCTGAACTCTTTCCCTACACGACGCTCTTCCGATCTNNNACAAAGGGTGAGCATGTATCCTTC
ACACGTCTGAACTCTTTCCCTACACGACGCTCTTCCGATCTNNNACAAAGGGTGAGCATGTATCCTTC
ACACGTCTGAACTCTTTCCCTACACGACGCTCTTCCGATCTNNNACAAAGGGTGAGCATGTATCCTTC
CACACGTCTGAACTCTTTCCCTACACGACGCTCTTCCGATCTNNNAACTGGCAAGTCTGAAGGTACTT
CACACGTCTGAACTCTTTCCCTACACGACGCTCTTCCGATCTNNNAACTGGCAAGTCTGAAGGTACTT
CACACGTCTGAACTCTTTCCCTACACGACGCTCTTCCGATCTNNNAACTGGCAAGTCTGAAGGTACTT
GCACACGTCTGAACTCTTTCCCTACACGACGCTCTTCCGATCTNNNCAAAGGGGTAAAAAAAAAAAG
GCACACGTCTGAACTCTTTCCCTACACGACGCTCTTCCGATCTNNNCAAAGGGGTAAAAAAAAAAAG
GCACACGTCTGAACTCTTTCCCTACACGACGCTCTTCCGATCTNNNCAAAGGGGTAAAAAAAAAAAG
AGCACACGTCTGAACTCTTTCCCTACACGACGCTCTTCCGATCTNNNTGGACACACAGGTCTTCTTCA
AGCACACGTCTGAACTCTTTCCCTACACGACGCTCTTCCGATCTNNNTGGACACACAGGTCTTCTTCA
AGCACACGTCTGAACTCTTTCCCTACACGACGCTCTTCCGATCTNNNTGGACACACAGGTCTTCTTCA
AGCACACGTCTGAACTCTTTCCCTACACGACGCTCTTCCGATCTNNNTGCATGTATCAAGAGTTCCCTC
AGCACACGTCTGAACTCTTTCCCTACACGACGCTCTTCCGATCTNNNTGCATGTATCAAGAGTTCCCTC
AGCACACGTCTGAACTCTTTCCCTACACGACGCTCTTCCGATCTNNNTGCATGTATCAAGAGTTCCCTC
GAGCACACGTCTGAACTCTTTCCCTACACGACGCTCTTCCGATCTNNNGTGAGCCAAGATTTTGCCAC
GAGCACACGTCTGAACTCTTTCCCTACACGACGCTCTTCCGATCTNNNGTGAGCCAAGATTTTGCCAC
GAGCACACGTCTGAACTCTTTCCCTACACGACGCTCTTCCGATCTNNNGTGAGCCAAGATTTTGCCAC
GCACACGTCTGAACTCTTTCCCTACACGACGCTCTTCCGATCTNNNTAATGTCTCCCATCACCCCTAGA
GCACACGTCTGAACTCTTTCCCTACACGACGCTCTTCCGATCTNNNTAATGTCTCCCATCACCCCTAGA
GCACACGTCTGAACTCTTTCCCTACACGACGCTCTTCCGATCTNNNTAATGTCTCCCATCACCCCTAGA
AGCACACGTCTGAACTCTTTCCCTACACGACGCTCTTCCGATCTNNNCTGAGGCAGGAGAATCACTTG
AGCACACGTCTGAACTCTTTCCCTACACGACGCTCTTCCGATCTNNNCTGAGGCAGGAGAATCACTTG
AGCACACGTCTGAACTCTTTCCCTACACGACGCTCTTCCGATCTNNNCTGAGGCAGGAGAATCACTTG
AGCACACGTCTGAACTCTTTCCCTACACGACGCTCTTCCGATCTNNNTGAGGTAGGAGAATCGCTTG
AGCACACGTCTGAACTCTTTCCCTACACGACGCTCTTCCGATCTNNNTGAGGTAGGAGAATCGCTTG
AGCACACGTCTGAACTCTTTCCCTACACGACGCTCTTCCGATCTNNNTGAGGTAGGAGAATCGCTTG
AGCACACGTCTGAACTCTTTCCCTACACGACGCTCTTCCGATCTNNNCCAGCAATTCTATTCCCAGGT
AGCACACGTCTGAACTCTTTCCCTACACGACGCTCTTCCGATCTNNNCCAGCAATTCTATTCCCAGGT
AGCACACGTCTGAACTCTTTCCCTACACGACGCTCTTCCGATCTNNNCCAGCAATTCTATTCCCAGGT
AGCACACGTCTGAACTCTTTCCCTACACGACGCTCTTCCGATCTNNNCACCCCTACTCCTTTTTCTCCA
AGCACACGTCTGAACTCTTTCCCTACACGACGCTCTTCCGATCTNNNCACCCCTACTCCTTTTTCTCCA
AGCACACGTCTGAACTCTTTCCCTACACGACGCTCTTCCGATCTNNNCACCCCTACTCCTTTTTCTCCA

GCACACGTCTGAACTCTTTCCCTACACGACGCTCTTCCGATCTNNNATATGTGTGTGTGTGTGTGTGT
GCACACGTCTGAACTCTTTCCCTACACGACGCTCTTCCGATCTNNNATATGTGTGTGTGTGTGTGTGT
GCACACGTCTGAACTCTTTCCCTACACGACGCTCTTCCGATCTNNNATATGTGTGTGTGTGTGTGTGT
AGAGCACACGTCTGAACTCTTTCCCTACACGACGCTCTTCCGATCTNNNTGTACACACAGACACATACA
AGAGCACACGTCTGAACTCTTTCCCTACACGACGCTCTTCCGATCTNNNTGTACACACAGACACATACA
AGAGCACACGTCTGAACTCTTTCCCTACACGACGCTCTTCCGATCTNNNTGTACACACAGACACATACA
AGCACACGTCTGAACTCTTTCCCTACACGACGCTCTTCCGATCTNNNAGTGCTAGGATTACAGGCGTA
AGCACACGTCTGAACTCTTTCCCTACACGACGCTCTTCCGATCTNNNAGTGCTAGGATTACAGGCGTA
AGCACACGTCTGAACTCTTTCCCTACACGACGCTCTTCCGATCTNNNAGTGCTAGGATTACAGGCGTA
GCACACGTCTGAACTCTTTCCCTACACGACGCTCTTCCGATCTNNNCATACACCCCTCCCTTTTTGAGAC
GCACACGTCTGAACTCTTTCCCTACACGACGCTCTTCCGATCTNNNCATACACCCCTCCCTTTTTGAGAC
GCACACGTCTGAACTCTTTCCCTACACGACGCTCTTCCGATCTNNNCATACACCCCTCCCTTTTTGAGAC
GCACACGTCTGAACTCTTTCCCTACACGACGCTCTTCCGATCTNNNCCTCTTGAAAAAAGACTGAGGG
GCACACGTCTGAACTCTTTCCCTACACGACGCTCTTCCGATCTNNNCCTCTTGAAAAAAGACTGAGGG
GCACACGTCTGAACTCTTTCCCTACACGACGCTCTTCCGATCTNNNCCTCTTGAAAAAAGACTGAGGG
GAGCACACGTCTGAACTCTTTCCCTACACGACGCTCTTCCGATCTNNNTGGCTCACACTTGTAATTCCAG
GAGCACACGTCTGAACTCTTTCCCTACACGACGCTCTTCCGATCTNNNTGGCTCACACTTGTAATTCCAG
GAGCACACGTCTGAACTCTTTCCCTACACGACGCTCTTCCGATCTNNNTGGCTCACACTTGTAATTCCAG
AGAGCACACGTCTGAACTCTTTCCCTACACGACGCTCTTCCGATCTNNNAGAGAGTGTTCCAGAGGAA
AGAGCACACGTCTGAACTCTTTCCCTACACGACGCTCTTCCGATCTNNNAGAGAGTGTTCCAGAGGAA
AGAGCACACGTCTGAACTCTTTCCCTACACGACGCTCTTCCGATCTNNNAGAGAGTGTTCCAGAGGAA
AGCACACGTCTGAACTCTTTCCCTACACGACGCTCTTCCGATCTNNNAAGGATCGCTTGAGTCTGGAA
AGCACACGTCTGAACTCTTTCCCTACACGACGCTCTTCCGATCTNNNAAGGATCGCTTGAGTCTGGAA
AGCACACGTCTGAACTCTTTCCCTACACGACGCTCTTCCGATCTNNNAAGGATCGCTTGAGTCTGGAA
GCACACGTCTGAACTCTTTCCCTACACGACGCTCTTCCGATCTNNNCAACGAACTATCATTCCTGACGCC
GCACACGTCTGAACTCTTTCCCTACACGACGCTCTTCCGATCTNNNCAACGAACTATCATTCCTGACGCC
GCACACGTCTGAACTCTTTCCCTACACGACGCTCTTCCGATCTNNNCAACGAACTATCATTCCTGACGCC
AGCACACGTCTGAACTCTTTCCCTACACGACGCTCTTCCGATCTNNNTGAACCATGCTAAACAGAGAGC
AGCACACGTCTGAACTCTTTCCCTACACGACGCTCTTCCGATCTNNNTGAACCATGCTAAACAGAGAGC
AGCACACGTCTGAACTCTTTCCCTACACGACGCTCTTCCGATCTNNNTGAACCATGCTAAACAGAGAGC
GCACACGTCTGAACTCTTTCCCTACACGACGCTCTTCCGATCTNNNCCACAATAGCTTTCTTCTCCCCACA
GCACACGTCTGAACTCTTTCCCTACACGACGCTCTTCCGATCTNNNCCACAATAGCTTTCTTCTCCCCACA
GCACACGTCTGAACTCTTTCCCTACACGACGCTCTTCCGATCTNNNCCACAATAGCTTTCTTCTCCCCACA
AGCACACGTCTGAACTCTTTCCCTACACGACGCTCTTCCGATCTNNNCGAATAAATCCCACGTGGTCAT
AGCACACGTCTGAACTCTTTCCCTACACGACGCTCTTCCGATCTNNNCGAATAAATCCCACGTGGTCAT
AGCACACGTCTGAACTCTTTCCCTACACGACGCTCTTCCGATCTNNNCGAATAAATCCCACGTGGTCAT
GAGCACACGTCTGAACTCTTTCCCTACACGACGCTCTTCCGATCTNNNTGGAGATTTCAGTGAGCCAAG
GAGCACACGTCTGAACTCTTTCCCTACACGACGCTCTTCCGATCTNNNTGGAGATTTCAGTGAGCCAAG
GAGCACACGTCTGAACTCTTTCCCTACACGACGCTCTTCCGATCTNNNTGGAGATTTCAGTGAGCCAAG
GAGCACACGTCTGAACTCTTTCCCTACACGACGCTCTTCCGATCTNNNCCCTTTTCGTTCTTATCTCCCCA
GAGCACACGTCTGAACTCTTTCCCTACACGACGCTCTTCCGATCTNNNCCCTTTTCGTTCTTATCTCCCCA
GAGCACACGTCTGAACTCTTTCCCTACACGACGCTCTTCCGATCTNNNCCCTTTTCGTTCTTATCTCCCCA
GAGCACACGTCTGAACTCTTTCCCTACACGACGCTCTTCCGATCTNNNAGAAGCCTTGTTCTTTCCTGAG

AGCACACGTCTGAACTCTTTCCCTACACGACGCTCTTCCGATCTNNNAGAAGCCTTGTTCTTTCCTGAG
AGCACACGTCTGAACTCTTTCCCTACACGACGCTCTTCCGATCTNNNAGAAGCCTTGTTCTTTCCTGAG
GAGCACACGTCTGAACTCTTTCCCTACACGACGCTCTTCCGATCTNNNAAAAGAAAAAAAACCTCGCC
GAGCACACGTCTGAACTCTTTCCCTACACGACGCTCTTCCGATCTNNNAAAAGAAAAAAAACCTCGCC
GAGCACACGTCTGAACTCTTTCCCTACACGACGCTCTTCCGATCTNNNAAAAGAAAAAAAACCTCGCC
GCACACGTCTGAACTCTTTCCCTACACGACGCTCTTCCGATCTNNNTCAGATTGGCTGCTGATTCTGAT
GCACACGTCTGAACTCTTTCCCTACACGACGCTCTTCCGATCTNNNTCAGATTGGCTGCTGATTCTGAT
GCACACGTCTGAACTCTTTCCCTACACGACGCTCTTCCGATCTNNNTCAGATTGGCTGCTGATTCTGAT
AGCACACGTCTGAACTCTTTCCCTACACGACGCTCTTCCGATCTNNNTGAGATCACATGGTCACAGGA
AGCACACGTCTGAACTCTTTCCCTACACGACGCTCTTCCGATCTNNNTGAGATCACATGGTCACAGGA
AGCACACGTCTGAACTCTTTCCCTACACGACGCTCTTCCGATCTNNNTGAGATCACATGGTCACAGGA
AGCACACGTCTGAACTCTTTCCCTACACGACGCTCTTCCGATCTNNNCACTGGACACACACAGTACAA
AGCACACGTCTGAACTCTTTCCCTACACGACGCTCTTCCGATCTNNNCACTGGACACACACAGTACAA
AGCACACGTCTGAACTCTTTCCCTACACGACGCTCTTCCGATCTNNNCACTGGACACACACAGTACAA
GAGCACACGTCTGAACTCTTTCCCTACACGACGCTCTTCCGATCTNNNAAAAAATTAGCTGGGTGTG
GAGCACACGTCTGAACTCTTTCCCTACACGACGCTCTTCCGATCTNNNAAAAAATTAGCTGGGTGTG
GAGCACACGTCTGAACTCTTTCCCTACACGACGCTCTTCCGATCTNNNAAAAAATTAGCTGGGTGTG
AGCACACGTCTGAACTCTTTCCCTACACGACGCTCTTCCGATCTNNNGGAACAGAACAAGACCCTGTC
AGCACACGTCTGAACTCTTTCCCTACACGACGCTCTTCCGATCTNNNGGAACAGAACAAGACCCTGTC
AGCACACGTCTGAACTCTTTCCCTACACGACGCTCTTCCGATCTNNNGGAACAGAACAAGACCCTGTC
AGAGCACACGTCTGAACTCTTTCCCTACACGACGCTCTTCCGATCTNNNGGTTAGGAAGCTGCTGCCA
AGAGCACACGTCTGAACTCTTTCCCTACACGACGCTCTTCCGATCTNNNGGTTAGGAAGCTGCTGCCA
AGAGCACACGTCTGAACTCTTTCCCTACACGACGCTCTTCCGATCTNNNGGTTAGGAAGCTGCTGCCA
AGCACACGTCTGAACTCTTTCCCTACACGACGCTCTTCCGATCTNNNTGAAAGAGGTGGAGGTTGTA
AGCACACGTCTGAACTCTTTCCCTACACGACGCTCTTCCGATCTNNNTGAAAGAGGTGGAGGTTGTA
AGCACACGTCTGAACTCTTTCCCTACACGACGCTCTTCCGATCTNNNTGAAAGAGGTGGAGGTTGTA
GAGCACACGTCTGAACTCTTTCCCTACACGACGCTCTTCCGATCTNNNAGATGAGGATTGGCAGGAG
GAGCACACGTCTGAACTCTTTCCCTACACGACGCTCTTCCGATCTNNNAGATGAGGATTGGCAGGAG
GAGCACACGTCTGAACTCTTTCCCTACACGACGCTCTTCCGATCTNNNAGATGAGGATTGGCAGGAG
GAGCACACGTCTGAACTCTTTCCCTACACGACGCTCTTCCGATCTNNNCTGTAATCCCAGCTACTCAGG
GAGCACACGTCTGAACTCTTTCCCTACACGACGCTCTTCCGATCTNNNCTGTAATCCCAGCTACTCAGG
GAGCACACGTCTGAACTCTTTCCCTACACGACGCTCTTCCGATCTNNNCTGTAATCCCAGCTACTCAGG
AGCACACGTCTGAACTCTTTCCCTACACGACGCTCTTCCGATCTNNNTGTGGTGGCTAGAAACTAGGA
AGCACACGTCTGAACTCTTTCCCTACACGACGCTCTTCCGATCTNNNTGTGGTGGCTAGAAACTAGGA
AGCACACGTCTGAACTCTTTCCCTACACGACGCTCTTCCGATCTNNNTGTGGTGGCTAGAAACTAGGA
AGCACACGTCTGAACTCTTTCCCTACACGACGCTCTTCCGATCTNNNACTGTACTCTAGCCTGGTGAG
AGCACACGTCTGAACTCTTTCCCTACACGACGCTCTTCCGATCTNNNACTGTACTCTAGCCTGGTGAG
AGCACACGTCTGAACTCTTTCCCTACACGACGCTCTTCCGATCTNNNACTGTACTCTAGCCTGGTGAG
AGAGCACACGTCTGAACTCTTTCCCTACACGACGCTCTTCCGATCTNNNGTAAATAGTCTCAGCTCCTC
AGAGCACACGTCTGAACTCTTTCCCTACACGACGCTCTTCCGATCTNNNGTAAATAGTCTCAGCTCCTC
AGAGCACACGTCTGAACTCTTTCCCTACACGACGCTCTTCCGATCTNNNGTAAATAGTCTCAGCTCCTC
GAGCACACGTCTGAACTCTTTCCCTACACGACGCTCTTCCGATCTNNNAGGCTCCTCTTTCTGATCCTC
GAGCACACGTCTGAACTCTTTCCCTACACGACGCTCTTCCGATCTNNNAGGCTCCTCTTTCTGATCCTC

GAGCACACGTCTGAACTCTTTCCCTACACGACGCTCTTCCGATCTNNNAGGCTCCTCTTTCTGATCCTC
GCACACGTCTGAACTCTTTCCCTACACGACGCTCTTCCGATCTNNNATATGGGGTTTCAACATGTTGG
GCACACGTCTGAACTCTTTCCCTACACGACGCTCTTCCGATCTNNNATATGGGGTTTCAACATGTTGG
GCACACGTCTGAACTCTTTCCCTACACGACGCTCTTCCGATCTNNNATATGGGGTTTCAACATGTTGG
AGCACACGTCTGAACTCTTTCCCTACACGACGCTCTTCCGATCTNNNGAGGATCATTTGAGTCCAGGA
AGCACACGTCTGAACTCTTTCCCTACACGACGCTCTTCCGATCTNNNGAGGATCATTTGAGTCCAGGA
AGCACACGTCTGAACTCTTTCCCTACACGACGCTCTTCCGATCTNNNGAGGATCATTTGAGTCCAGGA
AGCACACGTCTGAACTCTTTCCCTACACGACGCTCTTCCGATCTNNNGAAGTGGAGCTTGCAATGAG
AGCACACGTCTGAACTCTTTCCCTACACGACGCTCTTCCGATCTNNNGAAGTGGAGCTTGCAATGAG
AGCACACGTCTGAACTCTTTCCCTACACGACGCTCTTCCGATCTNNNGAAGTGGAGCTTGCAATGAG
GAGCACACGTCTGAACTCTTTCCCTACACGACGCTCTTCCGATCTNNNCTGAGGCAGGAGAATCACTT
GAGCACACGTCTGAACTCTTTCCCTACACGACGCTCTTCCGATCTNNNCTGAGGCAGGAGAATCACTT
GAGCACACGTCTGAACTCTTTCCCTACACGACGCTCTTCCGATCTNNNCTGAGGCAGGAGAATCACTT
GCACACGTCTGAACTCTTTCCCTACACGACGCTCTTCCGATCTNNNCTCTTGACCTTGTGATCTGTCCA
GCACACGTCTGAACTCTTTCCCTACACGACGCTCTTCCGATCTNNNCTCTTGACCTTGTGATCTGTCCA
GCACACGTCTGAACTCTTTCCCTACACGACGCTCTTCCGATCTNNNCTCTTGACCTTGTGATCTGTCCA
AGCACACGTCTGAACTCTTTCCCTACACGACGCTCTTCCGATCTNNNGGAGAAACACAATTCTGAAG
AGCACACGTCTGAACTCTTTCCCTACACGACGCTCTTCCGATCTNNNGGAGAAACACAATTCTGAAG
AGCACACGTCTGAACTCTTTCCCTACACGACGCTCTTCCGATCTNNNGGAGAAACACAATTCTGAAG
AGCACACGTCTGAACTCTTTCCCTACACGACGCTCTTCCGATCTNNNGAGGTGAACTAACCAACTCTC
AGCACACGTCTGAACTCTTTCCCTACACGACGCTCTTCCGATCTNNNGAGGTGAACTAACCAACTCTC
AGCACACGTCTGAACTCTTTCCCTACACGACGCTCTTCCGATCTNNNGAGGTGAACTAACCAACTCTC
GCACACGTCTGAACTCTTTCCCTACACGACGCTCTTCCGATCTNNNCTCTGCAATTTCCCTGGAACTGAC
GCACACGTCTGAACTCTTTCCCTACACGACGCTCTTCCGATCTNNNCTCTGCAATTTCCCTGGAACTGAC
GCACACGTCTGAACTCTTTCCCTACACGACGCTCTTCCGATCTNNNCTCTGCAATTTCCCTGGAACTGAC
AGCACACGTCTGAACTCTTTCCCTACACGACGCTCTTCCGATCTNNNAGCCTTGAATTCTTCTTTTTCAT
AGCACACGTCTGAACTCTTTCCCTACACGACGCTCTTCCGATCTNNNAGCCTTGAATTCTTCTTTTTCAT
AGCACACGTCTGAACTCTTTCCCTACACGACGCTCTTCCGATCTNNNAGCCTTGAATTCTTCTTTTTCAT
AGCACACGTCTGAACTCTTTCCCTACACGACGCTCTTCCGATCTNNNCACAAACAGATTGCATTAT
AGCACACGTCTGAACTCTTTCCCTACACGACGCTCTTCCGATCTNNNCACAAACAGATTGCATTAT
AGCACACGTCTGAACTCTTTCCCTACACGACGCTCTTCCGATCTNNNCACAAACAGATTGCATTAT
CACACGTCTGAACTCTTTCCCTACACGACGCTCTTCCGATCTNNNGTGAAGTTTACACCATGTGTTAG
CACACGTCTGAACTCTTTCCCTACACGACGCTCTTCCGATCTNNNGTGAAGTTTACACCATGTGTTAG
CACACGTCTGAACTCTTTCCCTACACGACGCTCTTCCGATCTNNNGTGAAGTTTACACCATGTGTTAG
GCACACGTCTGAACTCTTTCCCTACACGACGCTCTTCCGATCTNNNTTTACCACTAGAAACTAACAGTT
GCACACGTCTGAACTCTTTCCCTACACGACGCTCTTCCGATCTNNNTTTACCACTAGAAACTAACAGTT
GCACACGTCTGAACTCTTTCCCTACACGACGCTCTTCCGATCTNNNTTTACCACTAGAAACTAACAGTT
AGAGCACACGTCTGAACTCTTTCCCTACACGACGCTCTTCCGATCTNNNAGGATCATAGAGGTCCTGT
AGAGCACACGTCTGAACTCTTTCCCTACACGACGCTCTTCCGATCTNNNAGGATCATAGAGGTCCTGT
AGAGCACACGTCTGAACTCTTTCCCTACACGACGCTCTTCCGATCTNNNAGGATCATAGAGGTCCTGT
AGCACACGTCTGAACTCTTTCCCTACACGACGCTCTTCCGATCTNNNGTGTGTGTTCCTTTAGGCAGA
AGCACACGTCTGAACTCTTTCCCTACACGACGCTCTTCCGATCTNNNGTGTGTGTTCCTTTAGGCAGA
AGCACACGTCTGAACTCTTTCCCTACACGACGCTCTTCCGATCTNNNGTGTGTGTTCCTTTAGGCAGA

GCACACGTCTGAACTCTTTCCCTACACGACGCTCTTCCGATCTNNNATACACAAGCCCCTTCACAATCCCA
GCACACGTCTGAACTCTTTCCCTACACGACGCTCTTCCGATCTNNNATACACAAGCCCCTTCACAATCCCA
GCACACGTCTGAACTCTTTCCCTACACGACGCTCTTCCGATCTNNNATACACAAGCCCCTTCACAATCCCA
AGCACACGTCTGAACTCTTTCCCTACACGACGCTCTTCCGATCTNNNGGAAGACTCACTTGCTCCAAAA
AGCACACGTCTGAACTCTTTCCCTACACGACGCTCTTCCGATCTNNNGGAAGACTCACTTGCTCCAAAA
AGCACACGTCTGAACTCTTTCCCTACACGACGCTCTTCCGATCTNNNGGAAGACTCACTTGCTCCAAAA
GAGCACACGTCTGAACTCTTTCCCTACACGACGCTCTTCCGATCTNNNCTGCAGTGAGCTGTGATTTAC
GAGCACACGTCTGAACTCTTTCCCTACACGACGCTCTTCCGATCTNNNCTGCAGTGAGCTGTGATTTAC
GAGCACACGTCTGAACTCTTTCCCTACACGACGCTCTTCCGATCTNNNCTGCAGTGAGCTGTGATTTAC
AGCACACGTCTGAACTCTTTCCCTACACGACGCTCTTCCGATCTNNNTGTTCACTCATTAAATTTTGTCT
AGCACACGTCTGAACTCTTTCCCTACACGACGCTCTTCCGATCTNNNTGTTCACTCATTAAATTTTGTCT
AGCACACGTCTGAACTCTTTCCCTACACGACGCTCTTCCGATCTNNNTGTTCACTCATTAAATTTTGTCT
GAGCACACGTCTGAACTCTTTCCCTACACGACGCTCTTCCGATCTNNNGGCATGTAACAGTCAGGGTT
GAGCACACGTCTGAACTCTTTCCCTACACGACGCTCTTCCGATCTNNNGGCATGTAACAGTCAGGGTT
GAGCACACGTCTGAACTCTTTCCCTACACGACGCTCTTCCGATCTNNNGGCATGTAACAGTCAGGGTT
GAGCACACGTCTGAACTCTTTCCCTACACGACGCTCTTCCGATCTNNNTCTTGTACCCATCAGTCTTTGG
GAGCACACGTCTGAACTCTTTCCCTACACGACGCTCTTCCGATCTNNNTCTTGTACCCATCAGTCTTTGG
GAGCACACGTCTGAACTCTTTCCCTACACGACGCTCTTCCGATCTNNNTCTTGTACCCATCAGTCTTTGG
ACACGTCTGAACTCTTTCCCTACACGACGCTCTTCCGATCTNNNTCTACACTTTAGCGATTCCCACCCATC
ACACGTCTGAACTCTTTCCCTACACGACGCTCTTCCGATCTNNNTCTACACTTTAGCGATTCCCACCCATC
ACACGTCTGAACTCTTTCCCTACACGACGCTCTTCCGATCTNNNTCTACACTTTAGCGATTCCCACCCATC
GAGCACACGTCTGAACTCTTTCCCTACACGACGCTCTTCCGATCTNNNCAATAGGTATGAGACGGACCA
GAGCACACGTCTGAACTCTTTCCCTACACGACGCTCTTCCGATCTNNNCAATAGGTATGAGACGGACCA
GAGCACACGTCTGAACTCTTTCCCTACACGACGCTCTTCCGATCTNNNCAATAGGTATGAGACGGACCA
AGCACACGTCTGAACTCTTTCCCTACACGACGCTCTTCCGATCTNNNCCCAAATGATGAAGTGATGCAC
AGCACACGTCTGAACTCTTTCCCTACACGACGCTCTTCCGATCTNNNCCCAAATGATGAAGTGATGCAC
AGCACACGTCTGAACTCTTTCCCTACACGACGCTCTTCCGATCTNNNCCCAAATGATGAAGTGATGCAC
GAGCACACGTCTGAACTCTTTCCCTACACGACGCTCTTCCGATCTNNNGCATGCCAACTACTGTTTTCAC
GAGCACACGTCTGAACTCTTTCCCTACACGACGCTCTTCCGATCTNNNGCATGCCAACTACTGTTTTCAC
GAGCACACGTCTGAACTCTTTCCCTACACGACGCTCTTCCGATCTNNNGCATGCCAACTACTGTTTTCAC
AGAGCACACGTCTGAACTCTTTCCCTACACGACGCTCTTCCGATCTNNNGTTGAGGTGGGAGGATCAC
AGAGCACACGTCTGAACTCTTTCCCTACACGACGCTCTTCCGATCTNNNGTTGAGGTGGGAGGATCAC
AGAGCACACGTCTGAACTCTTTCCCTACACGACGCTCTTCCGATCTNNNGTTGAGGTGGGAGGATCAC
GCACACGTCTGAACTCTTTCCCTACACGACGCTCTTCCGATCTNNNAAGGATGAGATTGCCCCTCTTCTC
GCACACGTCTGAACTCTTTCCCTACACGACGCTCTTCCGATCTNNNAAGGATGAGATTGCCCCTCTTCTC
GCACACGTCTGAACTCTTTCCCTACACGACGCTCTTCCGATCTNNNAAGGATGAGATTGCCCCTCTTCTC
GAGCACACGTCTGAACTCTTTCCCTACACGACGCTCTTCCGATCTNNNCCAGGTTCATCCATTGTTGCAA
GAGCACACGTCTGAACTCTTTCCCTACACGACGCTCTTCCGATCTNNNCCAGGTTCATCCATTGTTGCAA
GAGCACACGTCTGAACTCTTTCCCTACACGACGCTCTTCCGATCTNNNCCAGGTTCATCCATTGTTGCAA
AGCACACGTCTGAACTCTTTCCCTACACGACGCTCTTCCGATCTNNNATGCAACAACCATGAGAGATCA
AGCACACGTCTGAACTCTTTCCCTACACGACGCTCTTCCGATCTNNNATGCAACAACCATGAGAGATCA
AGCACACGTCTGAACTCTTTCCCTACACGACGCTCTTCCGATCTNNNATGCAACAACCATGAGAGATCA
GAGCACACGTCTGAACTCTTTCCCTACACGACGCTCTTCCGATCTNNNGAGCTCTGACAAAAGGAGGAC

AGCACACGTCTGAACTCTTTCCCTACACGACGCTCTTCCGATCTNNNGAGCTCTGACAAAAGGAGGA
AGCACACGTCTGAACTCTTTCCCTACACGACGCTCTTCCGATCTNNNGAGCTCTGACAAAAGGAGGA
AGCACACGTCTGAACTCTTTCCCTACACGACGCTCTTCCGATCTNNNCAGAAACAGCACTGGGGAATT
AGCACACGTCTGAACTCTTTCCCTACACGACGCTCTTCCGATCTNNNCAGAAACAGCACTGGGGAATT
AGCACACGTCTGAACTCTTTCCCTACACGACGCTCTTCCGATCTNNNCAGAAACAGCACTGGGGAATT
AGCACACGTCTGAACTCTTTCCCTACACGACGCTCTTCCGATCTNNNCATAAACACACACGGCTTCCA
AGCACACGTCTGAACTCTTTCCCTACACGACGCTCTTCCGATCTNNNCATAAACACACACGGCTTCCA
AGCACACGTCTGAACTCTTTCCCTACACGACGCTCTTCCGATCTNNNCATAAACACACACGGCTTCCA
GCACACGTCTGAACTCTTTCCCTACACGACGCTCTTCCGATCTNNNAATACAAACATTAGCCAGGCGT
GCACACGTCTGAACTCTTTCCCTACACGACGCTCTTCCGATCTNNNAATACAAACATTAGCCAGGCGT
GCACACGTCTGAACTCTTTCCCTACACGACGCTCTTCCGATCTNNNAATACAAACATTAGCCAGGCGT
GAGCACACGTCTGAACTCTTTCCCTACACGACGCTCTTCCGATCTNNNGTGGTTGCTTTGAAGAGTG
GAGCACACGTCTGAACTCTTTCCCTACACGACGCTCTTCCGATCTNNNGTGGTTGCTTTGAAGAGTG
GAGCACACGTCTGAACTCTTTCCCTACACGACGCTCTTCCGATCTNNNGTGGTTGCTTTGAAGAGTG
ACACGTCTGAACTCTTTCCCTACACGACGCTCTTCCGATCTNNNGGCTCCTCACTTCCCAGTAGGCATC
ACACGTCTGAACTCTTTCCCTACACGACGCTCTTCCGATCTNNNGGCTCCTCACTTCCCAGTAGGCATC
ACACGTCTGAACTCTTTCCCTACACGACGCTCTTCCGATCTNNNGGCTCCTCACTTCCCAGTAGGCATC
GCACACGTCTGAACTCTTTCCCTACACGACGCTCTTCCGATCTNNNCCTGGTATGACGATGAATTTGG
GCACACGTCTGAACTCTTTCCCTACACGACGCTCTTCCGATCTNNNCCTGGTATGACGATGAATTTGG
GCACACGTCTGAACTCTTTCCCTACACGACGCTCTTCCGATCTNNNCCTGGTATGACGATGAATTTGG
GAGCACACGTCTGAACTCTTTCCCTACACGACGCTCTTCCGATCTNNNGAGCCTGTAATCCCAGCTACT
GAGCACACGTCTGAACTCTTTCCCTACACGACGCTCTTCCGATCTNNNGAGCCTGTAATCCCAGCTACT
GAGCACACGTCTGAACTCTTTCCCTACACGACGCTCTTCCGATCTNNNGAGCCTGTAATCCCAGCTACT
AGCACACGTCTGAACTCTTTCCCTACACGACGCTCTTCCGATCTNNNGCCTATAATCCCAGCTACTCA
AGCACACGTCTGAACTCTTTCCCTACACGACGCTCTTCCGATCTNNNGCCTATAATCCCAGCTACTCA
AGCACACGTCTGAACTCTTTCCCTACACGACGCTCTTCCGATCTNNNGCCTATAATCCCAGCTACTCA
AGCACACGTCTGAACTCTTTCCCTACACGACGCTCTTCCGATCTNNNAGCGTCCTCTTTCCATCTTTCAT
AGCACACGTCTGAACTCTTTCCCTACACGACGCTCTTCCGATCTNNNAGCGTCCTCTTTCCATCTTTCAT
AGCACACGTCTGAACTCTTTCCCTACACGACGCTCTTCCGATCTNNNAGCGTCCTCTTTCCATCTTTCAT
GCACACGTCTGAACTCTTTCCCTACACGACGCTCTTCCGATCTNNNGGAATCTGCCCTGTGTCGTCCAT
GCACACGTCTGAACTCTTTCCCTACACGACGCTCTTCCGATCTNNNGGAATCTGCCCTGTGTCGTCCAT
GCACACGTCTGAACTCTTTCCCTACACGACGCTCTTCCGATCTNNNGGAATCTGCCCTGTGTCGTCCAT
GCACACGTCTGAACTCTTTCCCTACACGACGCTCTTCCGATCTNNNAACAAAAAGACACCTGCACTCA
GCACACGTCTGAACTCTTTCCCTACACGACGCTCTTCCGATCTNNNAACAAAAAGACACCTGCACTCA
GCACACGTCTGAACTCTTTCCCTACACGACGCTCTTCCGATCTNNNAACAAAAAGACACCTGCACTCA
GCACACGTCTGAACTCTTTCCCTACACGACGCTCTTCCGATCTNNNTCACCTTCCTTTCCTCTCTCTCTCA
GCACACGTCTGAACTCTTTCCCTACACGACGCTCTTCCGATCTNNNTCACCTTCCTTTCCTCTCTCTCTCA
GCACACGTCTGAACTCTTTCCCTACACGACGCTCTTCCGATCTNNNTCACCTTCCTTTCCTCTCTCTCTCA
AGCACACGTCTGAACTCTTTCCCTACACGACGCTCTTCCGATCTNNNCTGTAGTCCCAGCTACTCGAGA
AGCACACGTCTGAACTCTTTCCCTACACGACGCTCTTCCGATCTNNNCTGTAGTCCCAGCTACTCGAGA
AGCACACGTCTGAACTCTTTCCCTACACGACGCTCTTCCGATCTNNNCTGTAGTCCCAGCTACTCGAGA
AGCACACGTCTGAACTCTTTCCCTACACGACGCTCTTCCGATCTNNNGAGCAGAGATCACACCATCCT
AGCACACGTCTGAACTCTTTCCCTACACGACGCTCTTCCGATCTNNNGAGCAGAGATCACACCATCCT

AGCACACGTCTGAACTCTTTCCCTACACGACGCTCTTCCGATCTNNNGAGCAGAGATCACACCATCCT
GCACACGTCTGAACTCTTTCCCTACACGACGCTCTTCCGATCTNNNGGCTCGAATAATTTCTTGCACGA
GCACACGTCTGAACTCTTTCCCTACACGACGCTCTTCCGATCTNNNGGCTCGAATAATTTCTTGCACGA
GCACACGTCTGAACTCTTTCCCTACACGACGCTCTTCCGATCTNNNGGCTCGAATAATTTCTTGCACGA
AGCACACGTCTGAACTCTTTCCCTACACGACGCTCTTCCGATCTNNNTCATTATGAGGAGTTGTGGAG
AGCACACGTCTGAACTCTTTCCCTACACGACGCTCTTCCGATCTNNNTCATTATGAGGAGTTGTGGAG
AGCACACGTCTGAACTCTTTCCCTACACGACGCTCTTCCGATCTNNNTCATTATGAGGAGTTGTGGAG
AGCACACGTCTGAACTCTTTCCCTACACGACGCTCTTCCGATCTNNNCCTTTACCCCAAACAAACCCTCC
AGCACACGTCTGAACTCTTTCCCTACACGACGCTCTTCCGATCTNNNCCTTTACCCCAAACAAACCCTCC
AGCACACGTCTGAACTCTTTCCCTACACGACGCTCTTCCGATCTNNNCCTTTACCCCAAACAAACCCTCC
AGCACACGTCTGAACTCTTTCCCTACACGACGCTCTTCCGATCTNNNGGCGAAACCTCGTCTCTACTAA
AGCACACGTCTGAACTCTTTCCCTACACGACGCTCTTCCGATCTNNNGGCGAAACCTCGTCTCTACTAA
AGCACACGTCTGAACTCTTTCCCTACACGACGCTCTTCCGATCTNNNGGCGAAACCTCGTCTCTACTAA
GCACACGTCTGAACTCTTTCCCTACACGACGCTCTTCCGATCTNNNTCATTCAGTGGGATTCCTTCCTG
GCACACGTCTGAACTCTTTCCCTACACGACGCTCTTCCGATCTNNNTCATTCAGTGGGATTCCTTCCTG
GCACACGTCTGAACTCTTTCCCTACACGACGCTCTTCCGATCTNNNTCATTCAGTGGGATTCCTTCCTG
AGCACACGTCTGAACTCTTTCCCTACACGACGCTCTTCCGATCTNNNATTACAAAAAATTAGCCGGGC
AGCACACGTCTGAACTCTTTCCCTACACGACGCTCTTCCGATCTNNNATTACAAAAAATTAGCCGGGC
AGCACACGTCTGAACTCTTTCCCTACACGACGCTCTTCCGATCTNNNATTACAAAAAATTAGCCGGGC
AGCACACGTCTGAACTCTTTCCCTACACGACGCTCTTCCGATCTNNNTAGCTGGGATTACAGACACACA
AGCACACGTCTGAACTCTTTCCCTACACGACGCTCTTCCGATCTNNNTAGCTGGGATTACAGACACACA
AGCACACGTCTGAACTCTTTCCCTACACGACGCTCTTCCGATCTNNNTAGCTGGGATTACAGACACACA
GAGCACACGTCTGAACTCTTTCCCTACACGACGCTCTTCCGATCTNNNCACCTGTAATCCCAGCTACTCA
GAGCACACGTCTGAACTCTTTCCCTACACGACGCTCTTCCGATCTNNNCACCTGTAATCCCAGCTACTCA
GAGCACACGTCTGAACTCTTTCCCTACACGACGCTCTTCCGATCTNNNCACCTGTAATCCCAGCTACTCA
AGAGCACACGTCTGAACTCTTTCCCTACACGACGCTCTTCCGATCTNNNCAGGAAATTCAGTTGGAGGA
AGAGCACACGTCTGAACTCTTTCCCTACACGACGCTCTTCCGATCTNNNCAGGAAATTCAGTTGGAGGA
AGAGCACACGTCTGAACTCTTTCCCTACACGACGCTCTTCCGATCTNNNCAGGAAATTCAGTTGGAGGA
AGCACACGTCTGAACTCTTTCCCTACACGACGCTCTTCCGATCTNNNTATGAGGATGGGTGTGGGAGA
AGCACACGTCTGAACTCTTTCCCTACACGACGCTCTTCCGATCTNNNTATGAGGATGGGTGTGGGAGA
AGCACACGTCTGAACTCTTTCCCTACACGACGCTCTTCCGATCTNNNTATGAGGATGGGTGTGGGAGA
GCACACGTCTGAACTCTTTCCCTACACGACGCTCTTCCGATCTNNNTAAACGTTACTGCAGGTTCCTGA
GCACACGTCTGAACTCTTTCCCTACACGACGCTCTTCCGATCTNNNTAAACGTTACTGCAGGTTCCTGA
GCACACGTCTGAACTCTTTCCCTACACGACGCTCTTCCGATCTNNNTAAACGTTACTGCAGGTTCCTGA
AGCACACGTCTGAACTCTTTCCCTACACGACGCTCTTCCGATCTNNNGTTTCCCCATAGTCCGCTTTTG
AGCACACGTCTGAACTCTTTCCCTACACGACGCTCTTCCGATCTNNNGTTTCCCCATAGTCCGCTTTTG
AGCACACGTCTGAACTCTTTCCCTACACGACGCTCTTCCGATCTNNNGTTTCCCCATAGTCCGCTTTTG
GAGCACACGTCTGAACTCTTTCCCTACACGACGCTCTTCCGATCTNNNCAGAGCTTTCAGTGAGCCTAG
GAGCACACGTCTGAACTCTTTCCCTACACGACGCTCTTCCGATCTNNNCAGAGCTTTCAGTGAGCCTAG
GAGCACACGTCTGAACTCTTTCCCTACACGACGCTCTTCCGATCTNNNCAGAGCTTTCAGTGAGCCTAG
GCACACGTCTGAACTCTTTCCCTACACGACGCTCTTCCGATCTNNNCCAGGAGTTCAAGGTTACAGTG
GCACACGTCTGAACTCTTTCCCTACACGACGCTCTTCCGATCTNNNCCAGGAGTTCAAGGTTACAGTG
GCACACGTCTGAACTCTTTCCCTACACGACGCTCTTCCGATCTNNNCCAGGAGTTCAAGGTTACAGTG

AGCACACGTCTGAACTCTTTCCCTACACGACGCTCTTCCGATCTNNNATGTTTACAGCCACACAATTGC
AGCACACGTCTGAACTCTTTCCCTACACGACGCTCTTCCGATCTNNNATGTTTACAGCCACACAATTGC
AGCACACGTCTGAACTCTTTCCCTACACGACGCTCTTCCGATCTNNNATGTTTACAGCCACACAATTGC
ACACGTCTGAACTCTTTCCCTACACGACGCTCTTCCGATCTNNNGGGATCCAAGACTGATTCACCATC
ACACGTCTGAACTCTTTCCCTACACGACGCTCTTCCGATCTNNNGGGATCCAAGACTGATTCACCATC
ACACGTCTGAACTCTTTCCCTACACGACGCTCTTCCGATCTNNNGGGATCCAAGACTGATTCACCATC
AGCACACGTCTGAACTCTTTCCCTACACGACGCTCTTCCGATCTNNNCTCACACCTGTAATCCCAACTC
AGCACACGTCTGAACTCTTTCCCTACACGACGCTCTTCCGATCTNNNCTCACACCTGTAATCCCAACTC
AGCACACGTCTGAACTCTTTCCCTACACGACGCTCTTCCGATCTNNNCTCACACCTGTAATCCCAACTC
AGCACACGTCTGAACTCTTTCCCTACACGACGCTCTTCCGATCTNNNTAGGCAAGTCCAAGTGCTAA
AGCACACGTCTGAACTCTTTCCCTACACGACGCTCTTCCGATCTNNNTAGGCAAGTCCAAGTGCTAA
AGCACACGTCTGAACTCTTTCCCTACACGACGCTCTTCCGATCTNNNTAGGCAAGTCCAAGTGCTAA
GAGCACACGTCTGAACTCTTTCCCTACACGACGCTCTTCCGATCTNNNCCTCTTCACATCCTCACCTGTT
GAGCACACGTCTGAACTCTTTCCCTACACGACGCTCTTCCGATCTNNNCCTCTTCACATCCTCACCTGTT
GAGCACACGTCTGAACTCTTTCCCTACACGACGCTCTTCCGATCTNNNCCTCTTCACATCCTCACCTGTT
GAGCACACGTCTGAACTCTTTCCCTACACGACGCTCTTCCGATCTNNNCAGAGGTCGTGCCATTGTCT
GAGCACACGTCTGAACTCTTTCCCTACACGACGCTCTTCCGATCTNNNCAGAGGTCGTGCCATTGTCT
GAGCACACGTCTGAACTCTTTCCCTACACGACGCTCTTCCGATCTNNNCAGAGGTCGTGCCATTGTCT
AGCACACGTCTGAACTCTTTCCCTACACGACGCTCTTCCGATCTNNNCAAGGCTCATAGTGAGGAGG
AGCACACGTCTGAACTCTTTCCCTACACGACGCTCTTCCGATCTNNNCAAGGCTCATAGTGAGGAGG
AGCACACGTCTGAACTCTTTCCCTACACGACGCTCTTCCGATCTNNNCAAGGCTCATAGTGAGGAGG
AGCACACGTCTGAACTCTTTCCCTACACGACGCTCTTCCGATCTNNNAATACAAAAAATTAGCCGGG
AGCACACGTCTGAACTCTTTCCCTACACGACGCTCTTCCGATCTNNNAATACAAAAAATTAGCCGGG
AGCACACGTCTGAACTCTTTCCCTACACGACGCTCTTCCGATCTNNNAATACAAAAAATTAGCCGGG
AGCACACGTCTGAACTCTTTCCCTACACGACGCTCTTCCGATCTNNNGGCCTGAATTCCTAGATGAG
AGCACACGTCTGAACTCTTTCCCTACACGACGCTCTTCCGATCTNNNGGCCTGAATTCCTAGATGAG
AGCACACGTCTGAACTCTTTCCCTACACGACGCTCTTCCGATCTNNNGGCCTGAATTCCTAGATGAG
AGCACACGTCTGAACTCTTTCCCTACACGACGCTCTTCCGATCTNNNTCTTCTCTATGGGAATCACCGG
AGCACACGTCTGAACTCTTTCCCTACACGACGCTCTTCCGATCTNNNTCTTCTCTATGGGAATCACCGG
AGCACACGTCTGAACTCTTTCCCTACACGACGCTCTTCCGATCTNNNTCTTCTCTATGGGAATCACCGG
AGCACACGTCTGAACTCTTTCCCTACACGACGCTCTTCCGATCTNNNCTTGAACCAGGGAGTTAGAG
AGCACACGTCTGAACTCTTTCCCTACACGACGCTCTTCCGATCTNNNCTTGAACCAGGGAGTTAGAG
AGCACACGTCTGAACTCTTTCCCTACACGACGCTCTTCCGATCTNNNCTTGAACCAGGGAGTTAGAG
GCACACGTCTGAACTCTTTCCCTACACGACGCTCTTCCGATCTNNNGCATGTTCTTGGCAACTTTGTC
GCACACGTCTGAACTCTTTCCCTACACGACGCTCTTCCGATCTNNNGCATGTTCTTGGCAACTTTGTC
GCACACGTCTGAACTCTTTCCCTACACGACGCTCTTCCGATCTNNNGCATGTTCTTGGCAACTTTGTC
GCACACGTCTGAACTCTTTCCCTACACGACGCTCTTCCGATCTNNNTAGAACAAGGGCAGGTGGTTT
GCACACGTCTGAACTCTTTCCCTACACGACGCTCTTCCGATCTNNNTAGAACAAGGGCAGGTGGTTT
GCACACGTCTGAACTCTTTCCCTACACGACGCTCTTCCGATCTNNNTAGAACAAGGGCAGGTGGTTT
GAGCACACGTCTGAACTCTTTCCCTACACGACGCTCTTCCGATCTNNNTCACAACTTAAGCTGGTGTC
GAGCACACGTCTGAACTCTTTCCCTACACGACGCTCTTCCGATCTNNNTCACAACTTAAGCTGGTGTC
GAGCACACGTCTGAACTCTTTCCCTACACGACGCTCTTCCGATCTNNNTCACAACTTAAGCTGGTGTC
AGCACACGTCTGAACTCTTTCCCTACACGACGCTCTTCCGATCTNNNGAGTTTGCAGTGAGCCGAGA

GAGCACACGTCTGAACTCTTTCCCTACACGACGCTCTTCCGATCTNNNGAGTTTGCAGTGAGCCGAGA
GAGCACACGTCTGAACTCTTTCCCTACACGACGCTCTTCCGATCTNNNGAGTTTGCAGTGAGCCGAGA
GAGCACACGTCTGAACTCTTTCCCTACACGACGCTCTTCCGATCTNNNCTTCAATCAACAGAGACCATG
GAGCACACGTCTGAACTCTTTCCCTACACGACGCTCTTCCGATCTNNNCTTCAATCAACAGAGACCATG
GAGCACACGTCTGAACTCTTTCCCTACACGACGCTCTTCCGATCTNNNCTTCAATCAACAGAGACCATG
GAGCACACGTCTGAACTCTTTCCCTACACGACGCTCTTCCGATCTNNNGAAAGATGGACACAGCAAC
GAGCACACGTCTGAACTCTTTCCCTACACGACGCTCTTCCGATCTNNNGAAAGATGGACACAGCAAC
GAGCACACGTCTGAACTCTTTCCCTACACGACGCTCTTCCGATCTNNNGAAAGATGGACACAGCAAC
AGCACACGTCTGAACTCTTTCCCTACACGACGCTCTTCCGATCTNNNGTTGTGATGAGCCGAGATTGT
AGCACACGTCTGAACTCTTTCCCTACACGACGCTCTTCCGATCTNNNGTTGTGATGAGCCGAGATTGT
AGCACACGTCTGAACTCTTTCCCTACACGACGCTCTTCCGATCTNNNGTTGTGATGAGCCGAGATTGT
GCACACGTCTGAACTCTTTCCCTACACGACGCTCTTCCGATCTNNNCCTTAGCTCATTTCCGCCTATCCC
GCACACGTCTGAACTCTTTCCCTACACGACGCTCTTCCGATCTNNNCCTTAGCTCATTTCCGCCTATCCC
GCACACGTCTGAACTCTTTCCCTACACGACGCTCTTCCGATCTNNNCCTTAGCTCATTTCCGCCTATCCC
GCACACGTCTGAACTCTTTCCCTACACGACGCTCTTCCGATCTNNNAAATGTCTCTCTCTCTGTGTGCA
GCACACGTCTGAACTCTTTCCCTACACGACGCTCTTCCGATCTNNNAAATGTCTCTCTCTCTGTGTGCA
GCACACGTCTGAACTCTTTCCCTACACGACGCTCTTCCGATCTNNNAAATGTCTCTCTCTCTGTGTGCA
AAGAGCACACGTCTGAACTCTTTCCCTACACGACGCTCTTCCGATCTNNNCCAATCTACATTCTGGCCA
AAGAGCACACGTCTGAACTCTTTCCCTACACGACGCTCTTCCGATCTNNNCCAATCTACATTCTGGCCA
AAGAGCACACGTCTGAACTCTTTCCCTACACGACGCTCTTCCGATCTNNNCCAATCTACATTCTGGCCA
GAGCACACGTCTGAACTCTTTCCCTACACGACGCTCTTCCGATCTNNNTGCACACAACATTCACCAAAA
GAGCACACGTCTGAACTCTTTCCCTACACGACGCTCTTCCGATCTNNNTGCACACAACATTCACCAAAA
GAGCACACGTCTGAACTCTTTCCCTACACGACGCTCTTCCGATCTNNNTGCACACAACATTCACCAAAA
AGCACACGTCTGAACTCTTTCCCTACACGACGCTCTTCCGATCTNNNCTGGGTTCAAGCCAATTCTCTT
AGCACACGTCTGAACTCTTTCCCTACACGACGCTCTTCCGATCTNNNCTGGGTTCAAGCCAATTCTCTT
AGCACACGTCTGAACTCTTTCCCTACACGACGCTCTTCCGATCTNNNCTGGGTTCAAGCCAATTCTCTT
AGCACACGTCTGAACTCTTTCCCTACACGACGCTCTTCCGATCTNNNCCTGGGACATGATGAGTCATAG
AGCACACGTCTGAACTCTTTCCCTACACGACGCTCTTCCGATCTNNNCCTGGGACATGATGAGTCATAG
AGCACACGTCTGAACTCTTTCCCTACACGACGCTCTTCCGATCTNNNCCTGGGACATGATGAGTCATAG
GCACACGTCTGAACTCTTTCCCTACACGACGCTCTTCCGATCTNNNGGCTAGTCTTGAACTGCTGATCT
GCACACGTCTGAACTCTTTCCCTACACGACGCTCTTCCGATCTNNNGGCTAGTCTTGAACTGCTGATCT
GCACACGTCTGAACTCTTTCCCTACACGACGCTCTTCCGATCTNNNGGCTAGTCTTGAACTGCTGATCT
GCACACGTCTGAACTCTTTCCCTACACGACGCTCTTCCGATCTNNNTGTGAGTGAGTGTGTGTGTATG
GCACACGTCTGAACTCTTTCCCTACACGACGCTCTTCCGATCTNNNTGTGAGTGAGTGTGTGTGTATG
GCACACGTCTGAACTCTTTCCCTACACGACGCTCTTCCGATCTNNNTGTGAGTGAGTGTGTGTGTATG
AGCACACGTCTGAACTCTTTCCCTACACGACGCTCTTCCGATCTNNNCTGATCATTGCCATCCAGACCTA
AGCACACGTCTGAACTCTTTCCCTACACGACGCTCTTCCGATCTNNNCTGATCATTGCCATCCAGACCTA
AGCACACGTCTGAACTCTTTCCCTACACGACGCTCTTCCGATCTNNNCTGATCATTGCCATCCAGACCTA
AGCACACGTCTGAACTCTTTCCCTACACGACGCTCTTCCGATCTNNNCCAAGTATCTGGGACTACAGGC
AGCACACGTCTGAACTCTTTCCCTACACGACGCTCTTCCGATCTNNNCCAAGTATCTGGGACTACAGGC
AGCACACGTCTGAACTCTTTCCCTACACGACGCTCTTCCGATCTNNNCCAAGTATCTGGGACTACAGGC
AGCACACGTCTGAACTCTTTCCCTACACGACGCTCTTCCGATCTNNNGGATACTATATGCTGTTGCCC
AGCACACGTCTGAACTCTTTCCCTACACGACGCTCTTCCGATCTNNNGGATACTATATGCTGTTGCCC

AGCACACGTCTGAACTCTTTCCCTACACGACGCTCTTCCGATCTNNNGGGATACTATATGCTGTTGCCC
AGCACACGTCTGAACTCTTTCCCTACACGACGCTCTTCCGATCTNNNTTCATGACTTGTATTTGATGAT
AGCACACGTCTGAACTCTTTCCCTACACGACGCTCTTCCGATCTNNNTTCATGACTTGTATTTGATGAT
AGCACACGTCTGAACTCTTTCCCTACACGACGCTCTTCCGATCTNNNTTCATGACTTGTATTTGATGAT
GCACACGTCTGAACTCTTTCCCTACACGACGCTCTTCCGATCTNNNGAGGAAGATCTGCAAGTGCTGTA
GCACACGTCTGAACTCTTTCCCTACACGACGCTCTTCCGATCTNNNGAGGAAGATCTGCAAGTGCTGTA
GCACACGTCTGAACTCTTTCCCTACACGACGCTCTTCCGATCTNNNGAGGAAGATCTGCAAGTGCTGTA
GAGCACACGTCTGAACTCTTTCCCTACACGACGCTCTTCCGATCTNNNGCCATGTCAGAAACAAATGAC
GAGCACACGTCTGAACTCTTTCCCTACACGACGCTCTTCCGATCTNNNGCCATGTCAGAAACAAATGAC
GAGCACACGTCTGAACTCTTTCCCTACACGACGCTCTTCCGATCTNNNGCCATGTCAGAAACAAATGAC
AGCACACGTCTGAACTCTTTCCCTACACGACGCTCTTCCGATCTNNNTCCCCGTGTCCTCAAATTCATTA
AGCACACGTCTGAACTCTTTCCCTACACGACGCTCTTCCGATCTNNNTCCCCGTGTCCTCAAATTCATTA
AGCACACGTCTGAACTCTTTCCCTACACGACGCTCTTCCGATCTNNNTCCCCGTGTCCTCAAATTCATTA
AGCACACGTCTGAACTCTTTCCCTACACGACGCTCTTCCGATCTNNNCCAAAATCCAAAGTGCTGGGAA
AGCACACGTCTGAACTCTTTCCCTACACGACGCTCTTCCGATCTNNNCCAAAATCCAAAGTGCTGGGAA
AGCACACGTCTGAACTCTTTCCCTACACGACGCTCTTCCGATCTNNNCCAAAATCCAAAGTGCTGGGAA
GAGCACACGTCTGAACTCTTTCCCTACACGACGCTCTTCCGATCTNNNTTCTGTGTAGCTGGGATTACA
GAGCACACGTCTGAACTCTTTCCCTACACGACGCTCTTCCGATCTNNNTTCTGTGTAGCTGGGATTACA
GAGCACACGTCTGAACTCTTTCCCTACACGACGCTCTTCCGATCTNNNTTCTGTGTAGCTGGGATTACA
AGCACACGTCTGAACTCTTTCCCTACACGACGCTCTTCCGATCTNNNAAATACAGGTCTGCACCAGTAG
AGCACACGTCTGAACTCTTTCCCTACACGACGCTCTTCCGATCTNNNAAATACAGGTCTGCACCAGTAG
AGCACACGTCTGAACTCTTTCCCTACACGACGCTCTTCCGATCTNNNAAATACAGGTCTGCACCAGTAG
GCACACGTCTGAACTCTTTCCCTACACGACGCTCTTCCGATCTNNNTATTAGACTGGCTGAGGTCCATGC
GCACACGTCTGAACTCTTTCCCTACACGACGCTCTTCCGATCTNNNTATTAGACTGGCTGAGGTCCATGC
GCACACGTCTGAACTCTTTCCCTACACGACGCTCTTCCGATCTNNNTATTAGACTGGCTGAGGTCCATGC
AGAGCACACGTCTGAACTCTTTCCCTACACGACGCTCTTCCGATCTNNNCAAGGAGGCATACATCTTGCA
AGAGCACACGTCTGAACTCTTTCCCTACACGACGCTCTTCCGATCTNNNCAAGGAGGCATACATCTTGCA
AGAGCACACGTCTGAACTCTTTCCCTACACGACGCTCTTCCGATCTNNNCAAGGAGGCATACATCTTGCA
ACACGTCTGAACTCTTTCCCTACACGACGCTCTTCCGATCTNNNGTAGACACTTAGGCTGCTTCCATCAT
ACACGTCTGAACTCTTTCCCTACACGACGCTCTTCCGATCTNNNGTAGACACTTAGGCTGCTTCCATCAT
ACACGTCTGAACTCTTTCCCTACACGACGCTCTTCCGATCTNNNGTAGACACTTAGGCTGCTTCCATCAT
GCACACGTCTGAACTCTTTCCCTACACGACGCTCTTCCGATCTNNNGAAGCATTCTGTCCACTGAACAG
GCACACGTCTGAACTCTTTCCCTACACGACGCTCTTCCGATCTNNNGAAGCATTCTGTCCACTGAACAG
GCACACGTCTGAACTCTTTCCCTACACGACGCTCTTCCGATCTNNNGAAGCATTCTGTCCACTGAACAG
AGCACACGTCTGAACTCTTTCCCTACACGACGCTCTTCCGATCTNNNCAAAACATTAGCTGGGTGTGGT
AGCACACGTCTGAACTCTTTCCCTACACGACGCTCTTCCGATCTNNNCAAAACATTAGCTGGGTGTGGT
AGCACACGTCTGAACTCTTTCCCTACACGACGCTCTTCCGATCTNNNCAAAACATTAGCTGGGTGTGGT
GAGCACACGTCTGAACTCTTTCCCTACACGACGCTCTTCCGATCTNNNCGTATAGCACCATCTCCCATCT
GAGCACACGTCTGAACTCTTTCCCTACACGACGCTCTTCCGATCTNNNCGTATAGCACCATCTCCCATCT
GAGCACACGTCTGAACTCTTTCCCTACACGACGCTCTTCCGATCTNNNCGTATAGCACCATCTCCCATCT
AGCACACGTCTGAACTCTTTCCCTACACGACGCTCTTCCGATCTNNNGTGTCTCTCCATTGTCCCCATAA
AGCACACGTCTGAACTCTTTCCCTACACGACGCTCTTCCGATCTNNNGTGTCTCTCCATTGTCCCCATAA
AGCACACGTCTGAACTCTTTCCCTACACGACGCTCTTCCGATCTNNNGTGTCTCTCCATTGTCCCCATAA

AGAGCACACGTCTGAACTCTTTCCCTACACGACGCTCTTCCGATCTNNNAGGTTTCATGAAGCTGCCAA
AGAGCACACGTCTGAACTCTTTCCCTACACGACGCTCTTCCGATCTNNNAGGTTTCATGAAGCTGCCAA
AGAGCACACGTCTGAACTCTTTCCCTACACGACGCTCTTCCGATCTNNNAGGTTTCATGAAGCTGCCAA
AGAGCACACGTCTGAACTCTTTCCCTACACGACGCTCTTCCGATCTNNNTTCTTCATGAAGACCTCACAG
AGAGCACACGTCTGAACTCTTTCCCTACACGACGCTCTTCCGATCTNNNTTCTTCATGAAGACCTCACAG
AGAGCACACGTCTGAACTCTTTCCCTACACGACGCTCTTCCGATCTNNNTTCTTCATGAAGACCTCACAG
AGAGCACACGTCTGAACTCTTTCCCTACACGACGCTCTTCCGATCTNNNTGATGGAGACAAACCATAG
AGAGCACACGTCTGAACTCTTTCCCTACACGACGCTCTTCCGATCTNNNTGATGGAGACAAACCATAG
AGAGCACACGTCTGAACTCTTTCCCTACACGACGCTCTTCCGATCTNNNTGATGGAGACAAACCATAG
GAGCACACGTCTGAACTCTTTCCCTACACGACGCTCTTCCGATCTNNNCTGAGGCAGGAGAATCACTTG
GAGCACACGTCTGAACTCTTTCCCTACACGACGCTCTTCCGATCTNNNCTGAGGCAGGAGAATCACTTG
GAGCACACGTCTGAACTCTTTCCCTACACGACGCTCTTCCGATCTNNNCTGAGGCAGGAGAATCACTTG
AGCACACGTCTGAACTCTTTCCCTACACGACGCTCTTCCGATCTNNNTTTGAGACAGTCTCGCTCTCTC
AGCACACGTCTGAACTCTTTCCCTACACGACGCTCTTCCGATCTNNNTTTGAGACAGTCTCGCTCTCTC
AGCACACGTCTGAACTCTTTCCCTACACGACGCTCTTCCGATCTNNNTTTGAGACAGTCTCGCTCTCTC
GAGCACACGTCTGAACTCTTTCCCTACACGACGCTCTTCCGATCTNNNCACCTTGGAGAGTGGGAAAGA
GAGCACACGTCTGAACTCTTTCCCTACACGACGCTCTTCCGATCTNNNCACCTTGGAGAGTGGGAAAGA
GAGCACACGTCTGAACTCTTTCCCTACACGACGCTCTTCCGATCTNNNCACCTTGGAGAGTGGGAAAGA
AGCACACGTCTGAACTCTTTCCCTACACGACGCTCTTCCGATCTNNNAGGCTTATTTGCTCTCTCATGTT
AGCACACGTCTGAACTCTTTCCCTACACGACGCTCTTCCGATCTNNNAGGCTTATTTGCTCTCTCATGTT
AGCACACGTCTGAACTCTTTCCCTACACGACGCTCTTCCGATCTNNNAGGCTTATTTGCTCTCTCATGTT
AGCACACGTCTGAACTCTTTCCCTACACGACGCTCTTCCGATCTNNNGAGGCAGGAAAATCACTTGAA
AGCACACGTCTGAACTCTTTCCCTACACGACGCTCTTCCGATCTNNNGAGGCAGGAAAATCACTTGAA
AGCACACGTCTGAACTCTTTCCCTACACGACGCTCTTCCGATCTNNNGAGGCAGGAAAATCACTTGAA
GCACACGTCTGAACTCTTTCCCTACACGACGCTCTTCCGATCTNNNATGGTCCAGTTTAGGAGAGGAG
GCACACGTCTGAACTCTTTCCCTACACGACGCTCTTCCGATCTNNNATGGTCCAGTTTAGGAGAGGAG
GCACACGTCTGAACTCTTTCCCTACACGACGCTCTTCCGATCTNNNATGGTCCAGTTTAGGAGAGGAG
AGCACACGTCTGAACTCTTTCCCTACACGACGCTCTTCCGATCTNNNGGTGACAGAGCAAGATTCCAT
AGCACACGTCTGAACTCTTTCCCTACACGACGCTCTTCCGATCTNNNGGTGACAGAGCAAGATTCCAT
AGCACACGTCTGAACTCTTTCCCTACACGACGCTCTTCCGATCTNNNGGTGACAGAGCAAGATTCCAT
AAGAGCACACGTCTGAACTCTTTCCCTACACGACGCTCTTCCGATCTNNNTGAGTCCATCAGCCAGTTTT
AAGAGCACACGTCTGAACTCTTTCCCTACACGACGCTCTTCCGATCTNNNTGAGTCCATCAGCCAGTTTT
AAGAGCACACGTCTGAACTCTTTCCCTACACGACGCTCTTCCGATCTNNNTGAGTCCATCAGCCAGTTTT
GAGCACACGTCTGAACTCTTTCCCTACACGACGCTCTTCCGATCTNNNGTATTATGATCAGGACTCACA
GAGCACACGTCTGAACTCTTTCCCTACACGACGCTCTTCCGATCTNNNGTATTATGATCAGGACTCACA
GAGCACACGTCTGAACTCTTTCCCTACACGACGCTCTTCCGATCTNNNGTATTATGATCAGGACTCACA
AGCACACGTCTGAACTCTTTCCCTACACGACGCTCTTCCGATCTNNNGATAGCCATGTGCCCTCAATTT
AGCACACGTCTGAACTCTTTCCCTACACGACGCTCTTCCGATCTNNNGATAGCCATGTGCCCTCAATTT
AGCACACGTCTGAACTCTTTCCCTACACGACGCTCTTCCGATCTNNNGATAGCCATGTGCCCTCAATTT
CACACGTCTGAACTCTTTCCCTACACGACGCTCTTCCGATCTNNNGGCTCCTCCAATGTCTTCTCTTTCAT
CACACGTCTGAACTCTTTCCCTACACGACGCTCTTCCGATCTNNNGGCTCCTCCAATGTCTTCTCTTTCAT
CACACGTCTGAACTCTTTCCCTACACGACGCTCTTCCGATCTNNNGGCTCCTCCAATGTCTTCTCTTTCAT
AGCACACGTCTGAACTCTTTCCCTACACGACGCTCTTCCGATCTNNNAGGTCCCTCAAGGGAGAAATT

AGCACACGTCTGAACTCTTTCCCTACACGACGCTCTTCCGATCTNNNAGGTCCCTCAAGGGAGAAATTG
AGCACACGTCTGAACTCTTTCCCTACACGACGCTCTTCCGATCTNNNAGGTCCCTCAAGGGAGAAATTG
CACACGTCTGAACTCTTTCCCTACACGACGCTCTTCCGATCTNNNAGTCAAAGTGGAAAGAGGTCACA
CACACGTCTGAACTCTTTCCCTACACGACGCTCTTCCGATCTNNNAGTCAAAGTGGAAAGAGGTCACA
CACACGTCTGAACTCTTTCCCTACACGACGCTCTTCCGATCTNNNAGTCAAAGTGGAAAGAGGTCACA
GAGCACACGTCTGAACTCTTTCCCTACACGACGCTCTTCCGATCTNNNACCTGTAATCCTAGCACTTTGG
GAGCACACGTCTGAACTCTTTCCCTACACGACGCTCTTCCGATCTNNNACCTGTAATCCTAGCACTTTGG
GAGCACACGTCTGAACTCTTTCCCTACACGACGCTCTTCCGATCTNNNACCTGTAATCCTAGCACTTTGG
GAGCACACGTCTGAACTCTTTCCCTACACGACGCTCTTCCGATCTNNNTGAGGCAGGAGAATCTCTTGA
GAGCACACGTCTGAACTCTTTCCCTACACGACGCTCTTCCGATCTNNNTGAGGCAGGAGAATCTCTTGA
GAGCACACGTCTGAACTCTTTCCCTACACGACGCTCTTCCGATCTNNNTGAGGCAGGAGAATCTCTTGA
AGCACACGTCTGAACTCTTTCCCTACACGACGCTCTTCCGATCTNNNAATCTCCAAGTTTCTGTTGCAGC
AGCACACGTCTGAACTCTTTCCCTACACGACGCTCTTCCGATCTNNNAATCTCCAAGTTTCTGTTGCAGC
AGCACACGTCTGAACTCTTTCCCTACACGACGCTCTTCCGATCTNNNAATCTCCAAGTTTCTGTTGCAGC
AGAGCACACGTCTGAACTCTTTCCCTACACGACGCTCTTCCGATCTNNNAGAACTGAGGTTTGGGAATC
AGAGCACACGTCTGAACTCTTTCCCTACACGACGCTCTTCCGATCTNNNAGAACTGAGGTTTGGGAATC
AGAGCACACGTCTGAACTCTTTCCCTACACGACGCTCTTCCGATCTNNNAGAACTGAGGTTTGGGAATC
AGCACACGTCTGAACTCTTTCCCTACACGACGCTCTTCCGATCTNNNTGCCTGTAATCCCAGCTAATCAC
AGCACACGTCTGAACTCTTTCCCTACACGACGCTCTTCCGATCTNNNTGCCTGTAATCCCAGCTAATCAC
AGCACACGTCTGAACTCTTTCCCTACACGACGCTCTTCCGATCTNNNTGCCTGTAATCCCAGCTAATCAC
GAGCACACGTCTGAACTCTTTCCCTACACGACGCTCTTCCGATCTNNNAGGTGAGTTGACAAATAGCCT
GAGCACACGTCTGAACTCTTTCCCTACACGACGCTCTTCCGATCTNNNAGGTGAGTTGACAAATAGCCT
GAGCACACGTCTGAACTCTTTCCCTACACGACGCTCTTCCGATCTNNNAGGTGAGTTGACAAATAGCCT
GAGCACACGTCTGAACTCTTTCCCTACACGACGCTCTTCCGATCTNNNCTGGGTGACAAGAGCAAAAC
GAGCACACGTCTGAACTCTTTCCCTACACGACGCTCTTCCGATCTNNNCTGGGTGACAAGAGCAAAAC
GAGCACACGTCTGAACTCTTTCCCTACACGACGCTCTTCCGATCTNNNCTGGGTGACAAGAGCAAAAC
AGAGCACACGTCTGAACTCTTTCCCTACACGACGCTCTTCCGATCTNNNTCAATCACGTACTGTCTGCAG
AGAGCACACGTCTGAACTCTTTCCCTACACGACGCTCTTCCGATCTNNNTCAATCACGTACTGTCTGCAG
AGAGCACACGTCTGAACTCTTTCCCTACACGACGCTCTTCCGATCTNNNTCAATCACGTACTGTCTGCAG
AGCACACGTCTGAACTCTTTCCCTACACGACGCTCTTCCGATCTNNNTCAACATGAGCTCTCTCTCCCAA
AGCACACGTCTGAACTCTTTCCCTACACGACGCTCTTCCGATCTNNNTCAACATGAGCTCTCTCTCCCAA
AGCACACGTCTGAACTCTTTCCCTACACGACGCTCTTCCGATCTNNNTCAACATGAGCTCTCTCTCCCAA
GAGCACACGTCTGAACTCTTTCCCTACACGACGCTCTTCCGATCTNNNTCCAGCAGAAATTGAGAGTGT
GAGCACACGTCTGAACTCTTTCCCTACACGACGCTCTTCCGATCTNNNTCCAGCAGAAATTGAGAGTGT
GAGCACACGTCTGAACTCTTTCCCTACACGACGCTCTTCCGATCTNNNTCCAGCAGAAATTGAGAGTGT
AGCACACGTCTGAACTCTTTCCCTACACGACGCTCTTCCGATCTNNNGCTTACATGTGCTAGGGTAGCT
AGCACACGTCTGAACTCTTTCCCTACACGACGCTCTTCCGATCTNNNGCTTACATGTGCTAGGGTAGCT
AGCACACGTCTGAACTCTTTCCCTACACGACGCTCTTCCGATCTNNNGCTTACATGTGCTAGGGTAGCT
AGCACACGTCTGAACTCTTTCCCTACACGACGCTCTTCCGATCTNNNCGTCTTCTGGATGTGGAGGAA
AGCACACGTCTGAACTCTTTCCCTACACGACGCTCTTCCGATCTNNNCGTCTTCTGGATGTGGAGGAA
AGCACACGTCTGAACTCTTTCCCTACACGACGCTCTTCCGATCTNNNCGTCTTCTGGATGTGGAGGAA
GAGCACACGTCTGAACTCTTTCCCTACACGACGCTCTTCCGATCTNNNCATAGCAAACTGAAACCTCTGC
GAGCACACGTCTGAACTCTTTCCCTACACGACGCTCTTCCGATCTNNNCATAGCAAACTGAAACCTCTGC

AGCACACGTCTGAACTCTTTCCCTACACGACGCTCTTCCGATCTNNNCATAGCAAACTGAAACCTCTG
AGCACACGTCTGAACTCTTTCCCTACACGACGCTCTTCCGATCTNNNTTACTGTAACCTCAAACTCCCG
AGCACACGTCTGAACTCTTTCCCTACACGACGCTCTTCCGATCTNNNTTACTGTAACCTCAAACTCCCG
AGCACACGTCTGAACTCTTTCCCTACACGACGCTCTTCCGATCTNNNTTACTGTAACCTCAAACTCCCG
GCACACGTCTGAACTCTTTCCCTACACGACGCTCTTCCGATCTNNNACTGGATGATATGGTTTGCCTG
GCACACGTCTGAACTCTTTCCCTACACGACGCTCTTCCGATCTNNNACTGGATGATATGGTTTGCCTG
GCACACGTCTGAACTCTTTCCCTACACGACGCTCTTCCGATCTNNNACTGGATGATATGGTTTGCCTG
GAGCACACGTCTGAACTCTTTCCCTACACGACGCTCTTCCGATCTNNNTGGGATTCAATAGTCAGGCA
GAGCACACGTCTGAACTCTTTCCCTACACGACGCTCTTCCGATCTNNNTGGGATTCAATAGTCAGGCA
GAGCACACGTCTGAACTCTTTCCCTACACGACGCTCTTCCGATCTNNNTGGGATTCAATAGTCAGGCA
CACACGTCTGAACTCTTTCCCTACACGACGCTCTTCCGATCTNNNGGATTACAGGTGTGAGCTACCAT
CACACGTCTGAACTCTTTCCCTACACGACGCTCTTCCGATCTNNNGGATTACAGGTGTGAGCTACCAT
CACACGTCTGAACTCTTTCCCTACACGACGCTCTTCCGATCTNNNGGATTACAGGTGTGAGCTACCAT
AGCACACGTCTGAACTCTTTCCCTACACGACGCTCTTCCGATCTNNNGAGAATGGTGTGAACCCAGG
AGCACACGTCTGAACTCTTTCCCTACACGACGCTCTTCCGATCTNNNGAGAATGGTGTGAACCCAGG
AGCACACGTCTGAACTCTTTCCCTACACGACGCTCTTCCGATCTNNNGAGAATGGTGTGAACCCAGG
GAGCACACGTCTGAACTCTTTCCCTACACGACGCTCTTCCGATCTNNNGCAAGCTCAAACTCCATAAAG
GAGCACACGTCTGAACTCTTTCCCTACACGACGCTCTTCCGATCTNNNGCAAGCTCAAACTCCATAAAG
GAGCACACGTCTGAACTCTTTCCCTACACGACGCTCTTCCGATCTNNNGCAAGCTCAAACTCCATAAAG
GAGCACACGTCTGAACTCTTTCCCTACACGACGCTCTTCCGATCTNNNTGTGCTGTATGAATTCAAGC
GAGCACACGTCTGAACTCTTTCCCTACACGACGCTCTTCCGATCTNNNTGTGCTGTATGAATTCAAGC
GAGCACACGTCTGAACTCTTTCCCTACACGACGCTCTTCCGATCTNNNTGTGCTGTATGAATTCAAGC
GCACACGTCTGAACTCTTTCCCTACACGACGCTCTTCCGATCTNNNAAAGTGCTGGGATTACAGGTG
GCACACGTCTGAACTCTTTCCCTACACGACGCTCTTCCGATCTNNNAAAGTGCTGGGATTACAGGTG
GCACACGTCTGAACTCTTTCCCTACACGACGCTCTTCCGATCTNNNAAAGTGCTGGGATTACAGGTG
GCACACGTCTGAACTCTTTCCCTACACGACGCTCTTCCGATCTNNNTTATGTAGCTGGTCCTGCTTCAG
GCACACGTCTGAACTCTTTCCCTACACGACGCTCTTCCGATCTNNNTTATGTAGCTGGTCCTGCTTCAG
GCACACGTCTGAACTCTTTCCCTACACGACGCTCTTCCGATCTNNNTTATGTAGCTGGTCCTGCTTCAG
AGCACACGTCTGAACTCTTTCCCTACACGACGCTCTTCCGATCTNNNAAAGTGCTGGGATTACAGGT
AGCACACGTCTGAACTCTTTCCCTACACGACGCTCTTCCGATCTNNNAAAGTGCTGGGATTACAGGT
AGCACACGTCTGAACTCTTTCCCTACACGACGCTCTTCCGATCTNNNAAAGTGCTGGGATTACAGGT
AGCACACGTCTGAACTCTTTCCCTACACGACGCTCTTCCGATCTNNNTCTACAGACATACACCGTCATG
AGCACACGTCTGAACTCTTTCCCTACACGACGCTCTTCCGATCTNNNTCTACAGACATACACCGTCATG
AGCACACGTCTGAACTCTTTCCCTACACGACGCTCTTCCGATCTNNNTCTACAGACATACACCGTCATG
AGCACACGTCTGAACTCTTTCCCTACACGACGCTCTTCCGATCTNNNAGTAACCATTGCAGAATTGGC
AGCACACGTCTGAACTCTTTCCCTACACGACGCTCTTCCGATCTNNNAGTAACCATTGCAGAATTGGC
AGCACACGTCTGAACTCTTTCCCTACACGACGCTCTTCCGATCTNNNAGTAACCATTGCAGAATTGGC
GAGCACACGTCTGAACTCTTTCCCTACACGACGCTCTTCCGATCTNNNACAGCAACAACATCTCCACTT
GAGCACACGTCTGAACTCTTTCCCTACACGACGCTCTTCCGATCTNNNACAGCAACAACATCTCCACTT
GAGCACACGTCTGAACTCTTTCCCTACACGACGCTCTTCCGATCTNNNACAGCAACAACATCTCCACTT
CACGTCTGAACTCTTTCCCTACACGACGCTCTTCCGATCTNNNAGTTTTGGTATGCTGATTGGCTTCA
CACGTCTGAACTCTTTCCCTACACGACGCTCTTCCGATCTNNNAGTTTTGGTATGCTGATTGGCTTCA
CACGTCTGAACTCTTTCCCTACACGACGCTCTTCCGATCTNNNAGTTTTGGTATGCTGATTGGCTTCA

AGCACACGTCTGAACTCTTTCCCTACACGACGCTCTTCCGATCTNNNAAACTACTGGGATTACAGGCGT

AGCACACGTCTGAACTCTTTCCCTACACGACGCTCTTCCGATCTNNNAAACTACTGGGATTACAGGCGT

AGCACACGTCTGAACTCTTTCCCTACACGACGCTCTTCCGATCTNNNAAACTACTGGGATTACAGGCGT

GAGCACACGTCTGAACTCTTTCCCTACACGACGCTCTTCCGATCTNNNAGCCCTTTACCCAAGATAGTG

GAGCACACGTCTGAACTCTTTCCCTACACGACGCTCTTCCGATCTNNNAGCCCTTTACCCAAGATAGTG

GAGCACACGTCTGAACTCTTTCCCTACACGACGCTCTTCCGATCTNNNAGCCCTTTACCCAAGATAGTG

AGCACACGTCTGAACTCTTTCCCTACACGACGCTCTTCCGATCTNNNATGTGTGTGTGTGTGTGTGTA

AGCACACGTCTGAACTCTTTCCCTACACGACGCTCTTCCGATCTNNNATGTGTGTGTGTGTGTGTGTA

AGCACACGTCTGAACTCTTTCCCTACACGACGCTCTTCCGATCTNNNATGTGTGTGTGTGTGTGTGTA

GAGCACACGTCTGAACTCTTTCCCTACACGACGCTCTTCCGATCTNNNAAACACAAAAGATCATCAGGG

GAGCACACGTCTGAACTCTTTCCCTACACGACGCTCTTCCGATCTNNNAAACACAAAAGATCATCAGGG

GAGCACACGTCTGAACTCTTTCCCTACACGACGCTCTTCCGATCTNNNAAACACAAAAGATCATCAGGG

AGCACACGTCTGAACTCTTTCCCTACACGACGCTCTTCCGATCTNNNGCAACAATAGCGAAACTCCATC

AGCACACGTCTGAACTCTTTCCCTACACGACGCTCTTCCGATCTNNNGCAACAATAGCGAAACTCCATC

AGCACACGTCTGAACTCTTTCCCTACACGACGCTCTTCCGATCTNNNGCAACAATAGCGAAACTCCATC

GAGCACACGTCTGAACTCTTTCCCTACACGACGCTCTTCCGATCTNNNAGGCAGTAGAATCACTTGAA

GAGCACACGTCTGAACTCTTTCCCTACACGACGCTCTTCCGATCTNNNAGGCAGTAGAATCACTTGAA

GAGCACACGTCTGAACTCTTTCCCTACACGACGCTCTTCCGATCTNNNAGGCAGTAGAATCACTTGAA

GAGCACACGTCTGAACTCTTTCCCTACACGACGCTCTTCCGATCTNNNGAAAACATTTGGCTGTGTCTG

GAGCACACGTCTGAACTCTTTCCCTACACGACGCTCTTCCGATCTNNNGAAAACATTTGGCTGTGTCTG

GAGCACACGTCTGAACTCTTTCCCTACACGACGCTCTTCCGATCTNNNGAAAACATTTGGCTGTGTCTG

GAGCACACGTCTGAACTCTTTCCCTACACGACGCTCTTCCGATCTNNNGCCTGGCCTTTTTCTAATGTCT

GAGCACACGTCTGAACTCTTTCCCTACACGACGCTCTTCCGATCTNNNGCCTGGCCTTTTTCTAATGTCT

GAGCACACGTCTGAACTCTTTCCCTACACGACGCTCTTCCGATCTNNNGCCTGGCCTTTTTCTAATGTCT

AGAGCACACGTCTGAACTCTTTCCCTACACGACGCTCTTCCGATCTNNNGCTCTAGTGGGGAGAGTAC

AGAGCACACGTCTGAACTCTTTCCCTACACGACGCTCTTCCGATCTNNNGCTCTAGTGGGGAGAGTAC

AGAGCACACGTCTGAACTCTTTCCCTACACGACGCTCTTCCGATCTNNNGCTCTAGTGGGGAGAGTAC

AGAGCACACGTCTGAACTCTTTCCCTACACGACGCTCTTCCGATCTNNNTGGACCTGTAGCACTATTCC

AGAGCACACGTCTGAACTCTTTCCCTACACGACGCTCTTCCGATCTNNNTGGACCTGTAGCACTATTCC

AGAGCACACGTCTGAACTCTTTCCCTACACGACGCTCTTCCGATCTNNNTGGACCTGTAGCACTATTCC

AGCACACGTCTGAACTCTTTCCCTACACGACGCTCTTCCGATCTNNNGAGGGTGAAGGCACTGTTCAG

AGCACACGTCTGAACTCTTTCCCTACACGACGCTCTTCCGATCTNNNGAGGGTGAAGGCACTGTTCAG

AGCACACGTCTGAACTCTTTCCCTACACGACGCTCTTCCGATCTNNNGAGGGTGAAGGCACTGTTCAG

AGCACACGTCTGAACTCTTTCCCTACACGACGCTCTTCCGATCTNNNAAAATGTGTGTGTGTGTGTGT

AGCACACGTCTGAACTCTTTCCCTACACGACGCTCTTCCGATCTNNNAAAATGTGTGTGTGTGTGTGT

AGCACACGTCTGAACTCTTTCCCTACACGACGCTCTTCCGATCTNNNAAAATGTGTGTGTGTGTGTGT

AGCACACGTCTGAACTCTTTCCCTACACGACGCTCTTCCGATCTNNNAAGGCCCAAAGGGTACATATT

AGCACACGTCTGAACTCTTTCCCTACACGACGCTCTTCCGATCTNNNAAGGCCCAAAGGGTACATATT

AGCACACGTCTGAACTCTTTCCCTACACGACGCTCTTCCGATCTNNNAAGGCCCAAAGGGTACATATT

GAGCACACGTCTGAACTCTTTCCCTACACGACGCTCTTCCGATCTNNNGGTTCAAGTGATTCTCCTGT

GAGCACACGTCTGAACTCTTTCCCTACACGACGCTCTTCCGATCTNNNGGTTCAAGTGATTCTCCTGT

GAGCACACGTCTGAACTCTTTCCCTACACGACGCTCTTCCGATCTNNNGGTTCAAGTGATTCTCCTGT

GAGCACACGTCTGAACTCTTTCCCTACACGACGCTCTTCCGATCTNNNGATCTTGGCTCACTGCAAGCTC

GAGCACACGTCTGAACTCTTTCCCTACACGACGCTCTTCCGATCTNNNGATCTTGGCTCACTGCAAGCTC
GAGCACACGTCTGAACTCTTTCCCTACACGACGCTCTTCCGATCTNNNGATCTTGGCTCACTGCAAGCTC
GAGCACACGTCTGAACTCTTTCCCTACACGACGCTCTTCCGATCTNNNGCAAAGTCAAGAACCGACAG
GAGCACACGTCTGAACTCTTTCCCTACACGACGCTCTTCCGATCTNNNGCAAAGTCAAGAACCGACAG
GAGCACACGTCTGAACTCTTTCCCTACACGACGCTCTTCCGATCTNNNGCAAAGTCAAGAACCGACAG
CACACGTCTGAACTCTTTCCCTACACGACGCTCTTCCGATCTNNNTCATATCAAGAGCCGTGTGTGTGC
CACACGTCTGAACTCTTTCCCTACACGACGCTCTTCCGATCTNNNTCATATCAAGAGCCGTGTGTGTGC
CACACGTCTGAACTCTTTCCCTACACGACGCTCTTCCGATCTNNNTCATATCAAGAGCCGTGTGTGTGC
GCACACGTCTGAACTCTTTCCCTACACGACGCTCTTCCGATCTNNNGTTCCTGTAGTCCCAGCTACTTGC
GCACACGTCTGAACTCTTTCCCTACACGACGCTCTTCCGATCTNNNGTTCCTGTAGTCCCAGCTACTTGC
GCACACGTCTGAACTCTTTCCCTACACGACGCTCTTCCGATCTNNNGTTCCTGTAGTCCCAGCTACTTGC
GAGCACACGTCTGAACTCTTTCCCTACACGACGCTCTTCCGATCTNNNGAAGTTGCAGTGAGCCAAGAT
GAGCACACGTCTGAACTCTTTCCCTACACGACGCTCTTCCGATCTNNNGAAGTTGCAGTGAGCCAAGAT
GAGCACACGTCTGAACTCTTTCCCTACACGACGCTCTTCCGATCTNNNGAAGTTGCAGTGAGCCAAGAT
GAGCACACGTCTGAACTCTTTCCCTACACGACGCTCTTCCGATCTNNNAGACAGGAGAATTGCTTGAA
GAGCACACGTCTGAACTCTTTCCCTACACGACGCTCTTCCGATCTNNNAGACAGGAGAATTGCTTGAA
GAGCACACGTCTGAACTCTTTCCCTACACGACGCTCTTCCGATCTNNNAGACAGGAGAATTGCTTGAA
GAGCACACGTCTGAACTCTTTCCCTACACGACGCTCTTCCGATCTNNNCAAAAATTAGCTGGGTGTGG
GAGCACACGTCTGAACTCTTTCCCTACACGACGCTCTTCCGATCTNNNCAAAAATTAGCTGGGTGTGG
GAGCACACGTCTGAACTCTTTCCCTACACGACGCTCTTCCGATCTNNNCAAAAATTAGCTGGGTGTGG
GAGCACACGTCTGAACTCTTTCCCTACACGACGCTCTTCCGATCTNNNACAAAACCAGAACCAAGAGGA
GAGCACACGTCTGAACTCTTTCCCTACACGACGCTCTTCCGATCTNNNACAAAACCAGAACCAAGAGGA
GAGCACACGTCTGAACTCTTTCCCTACACGACGCTCTTCCGATCTNNNACAAAACCAGAACCAAGAGGA
GAGCACACGTCTGAACTCTTTCCCTACACGACGCTCTTCCGATCTNNNAGGCATCTCCCATTTTTCTCACT
GAGCACACGTCTGAACTCTTTCCCTACACGACGCTCTTCCGATCTNNNAGGCATCTCCCATTTTTCTCACT
GAGCACACGTCTGAACTCTTTCCCTACACGACGCTCTTCCGATCTNNNAGGCATCTCCCATTTTTCTCACT
GCACACGTCTGAACTCTTTCCCTACACGACGCTCTTCCGATCTNNNTCTGAGAAGTGAGGAGCCTCTCCA
GCACACGTCTGAACTCTTTCCCTACACGACGCTCTTCCGATCTNNNTCTGAGAAGTGAGGAGCCTCTCCA
GCACACGTCTGAACTCTTTCCCTACACGACGCTCTTCCGATCTNNNTCTGAGAAGTGAGGAGCCTCTCCA
GAGCACACGTCTGAACTCTTTCCCTACACGACGCTCTTCCGATCTNNNGCCTGGGTGATAGAATGAGA
GAGCACACGTCTGAACTCTTTCCCTACACGACGCTCTTCCGATCTNNNGCCTGGGTGATAGAATGAGA
GAGCACACGTCTGAACTCTTTCCCTACACGACGCTCTTCCGATCTNNNGCCTGGGTGATAGAATGAGA
GAGCACACGTCTGAACTCTTTCCCTACACGACGCTCTTCCGATCTNNNAATTACCCCGAGACAAAAACAC
GAGCACACGTCTGAACTCTTTCCCTACACGACGCTCTTCCGATCTNNNAATTACCCCGAGACAAAAACAC
GAGCACACGTCTGAACTCTTTCCCTACACGACGCTCTTCCGATCTNNNAATTACCCCGAGACAAAAACAC
AGAGCACACGTCTGAACTCTTTCCCTACACGACGCTCTTCCGATCTNNNACTTCTGCAGTATTTGGGGC
AGAGCACACGTCTGAACTCTTTCCCTACACGACGCTCTTCCGATCTNNNACTTCTGCAGTATTTGGGGC
AGAGCACACGTCTGAACTCTTTCCCTACACGACGCTCTTCCGATCTNNNACTTCTGCAGTATTTGGGGC
AGCACACGTCTGAACTCTTTCCCTACACGACGCTCTTCCGATCTNNNAAGAGGTAGAGTCCAGTGAGG
AGCACACGTCTGAACTCTTTCCCTACACGACGCTCTTCCGATCTNNNAAGAGGTAGAGTCCAGTGAGG
AGCACACGTCTGAACTCTTTCCCTACACGACGCTCTTCCGATCTNNNAAGAGGTAGAGTCCAGTGAGG
GAGCACACGTCTGAACTCTTTCCCTACACGACGCTCTTCCGATCTNNNAACCATTGCTTTAGGTCAGGA
GAGCACACGTCTGAACTCTTTCCCTACACGACGCTCTTCCGATCTNNNAACCATTGCTTTAGGTCAGGA

GAGCACACGTCTGAACTCTTTCCCTACACGACGCTCTTCCGATCTNNNAACCATTGCTTTAGGTCAGGA

GAGCACACGTCTGAACTCTTTCCCTACACGACGCTCTTCCGATCTNNNCTCCAGCTTCAAGACAAGGACT

GAGCACACGTCTGAACTCTTTCCCTACACGACGCTCTTCCGATCTNNNCTCCAGCTTCAAGACAAGGACT

GAGCACACGTCTGAACTCTTTCCCTACACGACGCTCTTCCGATCTNNNCTCCAGCTTCAAGACAAGGACT

GAGCACACGTCTGAACTCTTTCCCTACACGACGCTCTTCCGATCTNNNGAGATTGTGGTGAGCGGAGA

GAGCACACGTCTGAACTCTTTCCCTACACGACGCTCTTCCGATCTNNNGAGATTGTGGTGAGCGGAGA

GAGCACACGTCTGAACTCTTTCCCTACACGACGCTCTTCCGATCTNNNGAGATTGTGGTGAGCGGAGA

GAGCACACGTCTGAACTCTTTCCCTACACGACGCTCTTCCGATCTNNNGCTTGTCACAATAGGGCTGTT

GAGCACACGTCTGAACTCTTTCCCTACACGACGCTCTTCCGATCTNNNGCTTGTCACAATAGGGCTGTT

GAGCACACGTCTGAACTCTTTCCCTACACGACGCTCTTCCGATCTNNNGCTTGTCACAATAGGGCTGTT

GCACACGTCTGAACTCTTTCCCTACACGACGCTCTTCCGATCTNNNAGCTCCTTGTTACCTGAGTCCATCA

GCACACGTCTGAACTCTTTCCCTACACGACGCTCTTCCGATCTNNNAGCTCCTTGTTACCTGAGTCCATCA

GCACACGTCTGAACTCTTTCCCTACACGACGCTCTTCCGATCTNNNAGCTCCTTGTTACCTGAGTCCATCA

AGCACACGTCTGAACTCTTTCCCTACACGACGCTCTTCCGATCTNNNCTAGCATCCCTGGCTCGAAACCA

AGCACACGTCTGAACTCTTTCCCTACACGACGCTCTTCCGATCTNNNCTAGCATCCCTGGCTCGAAACCA

AGCACACGTCTGAACTCTTTCCCTACACGACGCTCTTCCGATCTNNNCTAGCATCCCTGGCTCGAAACCA

GAGCACACGTCTGAACTCTTTCCCTACACGACGCTCTTCCGATCTNNNGCAAGAGAATCACTTGAACCC

GAGCACACGTCTGAACTCTTTCCCTACACGACGCTCTTCCGATCTNNNGCAAGAGAATCACTTGAACCC

GAGCACACGTCTGAACTCTTTCCCTACACGACGCTCTTCCGATCTNNNGCAAGAGAATCACTTGAACCC

AGAGCACACGTCTGAACTCTTTCCCTACACGACGCTCTTCCGATCTNNNTTACATGGTGGAAGCCTCTTC

AGAGCACACGTCTGAACTCTTTCCCTACACGACGCTCTTCCGATCTNNNTTACATGGTGGAAGCCTCTTC

AGAGCACACGTCTGAACTCTTTCCCTACACGACGCTCTTCCGATCTNNNTTACATGGTGGAAGCCTCTTC

GAGCACACGTCTGAACTCTTTCCCTACACGACGCTCTTCCGATCTNNNCTGAAGCAGGAGAATGGTGT

GAGCACACGTCTGAACTCTTTCCCTACACGACGCTCTTCCGATCTNNNCTGAAGCAGGAGAATGGTGT

GAGCACACGTCTGAACTCTTTCCCTACACGACGCTCTTCCGATCTNNNCTGAAGCAGGAGAATGGTGT

GCACACGTCTGAACTCTTTCCCTACACGACGCTCTTCCGATCTNNNCGAGGCCATAAAAAGGATGTGT

GCACACGTCTGAACTCTTTCCCTACACGACGCTCTTCCGATCTNNNCGAGGCCATAAAAAGGATGTGT

GCACACGTCTGAACTCTTTCCCTACACGACGCTCTTCCGATCTNNNCGAGGCCATAAAAAGGATGTGT

AGCACACGTCTGAACTCTTTCCCTACACGACGCTCTTCCGATCTNNNAATTTCCTTCCTTCCTTCCTCCCCA

AGCACACGTCTGAACTCTTTCCCTACACGACGCTCTTCCGATCTNNNAATTTCCTTCCTTCCTTCCTCCCCA

AGCACACGTCTGAACTCTTTCCCTACACGACGCTCTTCCGATCTNNNAATTTCCTTCCTTCCTTCCTCCCCA

AGCACACGTCTGAACTCTTTCCCTACACGACGCTCTTCCGATCTNNNGAGGTTACAGTGAGCCAAGAT

AGCACACGTCTGAACTCTTTCCCTACACGACGCTCTTCCGATCTNNNGAGGTTACAGTGAGCCAAGAT

AGCACACGTCTGAACTCTTTCCCTACACGACGCTCTTCCGATCTNNNGAGGTTACAGTGAGCCAAGAT

GAGCACACGTCTGAACTCTTTCCCTACACGACGCTCTTCCGATCTNNNTGCCTCCTGATGAAAAAACAC

GAGCACACGTCTGAACTCTTTCCCTACACGACGCTCTTCCGATCTNNNTGCCTCCTGATGAAAAAACAC

GAGCACACGTCTGAACTCTTTCCCTACACGACGCTCTTCCGATCTNNNTGCCTCCTGATGAAAAAACAC

AGCACACGTCTGAACTCTTTCCCTACACGACGCTCTTCCGATCTNNNCTGGATCTAAAGATTATGCAGA

AGCACACGTCTGAACTCTTTCCCTACACGACGCTCTTCCGATCTNNNCTGGATCTAAAGATTATGCAGA

AGCACACGTCTGAACTCTTTCCCTACACGACGCTCTTCCGATCTNNNCTGGATCTAAAGATTATGCAGA

AGCACACGTCTGAACTCTTTCCCTACACGACGCTCTTCCGATCTNNNTGCTACACCCCTCATTCTACCTA

AGCACACGTCTGAACTCTTTCCCTACACGACGCTCTTCCGATCTNNNTGCTACACCCCTCATTCTACCTA

AGCACACGTCTGAACTCTTTCCCTACACGACGCTCTTCCGATCTNNNTGCTACACCCCTCATTCTACCTA

AGCACACGTCTGAACTCTTTCCCTACACGACGCTCTTCCGATCTNNNCTCACCTCTGTCTTGGTAGCTG
AGCACACGTCTGAACTCTTTCCCTACACGACGCTCTTCCGATCTNNNCTCACCTCTGTCTTGGTAGCTG
AGCACACGTCTGAACTCTTTCCCTACACGACGCTCTTCCGATCTNNNCTCACCTCTGTCTTGGTAGCTG
AGCACACGTCTGAACTCTTTCCCTACACGACGCTCTTCCGATCTNNNGGTTCAAGTGATTATGCTGCCT
AGCACACGTCTGAACTCTTTCCCTACACGACGCTCTTCCGATCTNNNGGTTCAAGTGATTATGCTGCCT
AGCACACGTCTGAACTCTTTCCCTACACGACGCTCTTCCGATCTNNNGGTTCAAGTGATTATGCTGCCT
GAGCACACGTCTGAACTCTTTCCCTACACGACGCTCTTCCGATCTNNNCTCCCAGTGGTAGCCTAGTCT
GAGCACACGTCTGAACTCTTTCCCTACACGACGCTCTTCCGATCTNNNCTCCCAGTGGTAGCCTAGTCT
GAGCACACGTCTGAACTCTTTCCCTACACGACGCTCTTCCGATCTNNNCTCCCAGTGGTAGCCTAGTCT
AAGAGCACACGTCTGAACTCTTTCCCTACACGACGCTCTTCCGATCTNNNGTGAGCTGAGATCATGCC
AAGAGCACACGTCTGAACTCTTTCCCTACACGACGCTCTTCCGATCTNNNGTGAGCTGAGATCATGCC
AAGAGCACACGTCTGAACTCTTTCCCTACACGACGCTCTTCCGATCTNNNGTGAGCTGAGATCATGCC
GAGCACACGTCTGAACTCTTTCCCTACACGACGCTCTTCCGATCTNNNCCCTGAGAACCTCTAACATCAC
GAGCACACGTCTGAACTCTTTCCCTACACGACGCTCTTCCGATCTNNNCCCTGAGAACCTCTAACATCAC
GAGCACACGTCTGAACTCTTTCCCTACACGACGCTCTTCCGATCTNNNCCCTGAGAACCTCTAACATCAC
GAGCACACGTCTGAACTCTTTCCCTACACGACGCTCTTCCGATCTNNNTAGTCCCAGCTACTCAGGAGA
GAGCACACGTCTGAACTCTTTCCCTACACGACGCTCTTCCGATCTNNNTAGTCCCAGCTACTCAGGAGA
GAGCACACGTCTGAACTCTTTCCCTACACGACGCTCTTCCGATCTNNNTAGTCCCAGCTACTCAGGAGA
AGAGCACACGTCTGAACTCTTTCCCTACACGACGCTCTTCCGATCTNNNCCCATCTACTTGGAAGACTG
AGAGCACACGTCTGAACTCTTTCCCTACACGACGCTCTTCCGATCTNNNCCCATCTACTTGGAAGACTG
AGAGCACACGTCTGAACTCTTTCCCTACACGACGCTCTTCCGATCTNNNCCCATCTACTTGGAAGACTG
AGAGCACACGTCTGAACTCTTTCCCTACACGACGCTCTTCCGATCTNNNCCTGAGTGAAGAAGCCAAG
AGAGCACACGTCTGAACTCTTTCCCTACACGACGCTCTTCCGATCTNNNCCTGAGTGAAGAAGCCAAG
AGAGCACACGTCTGAACTCTTTCCCTACACGACGCTCTTCCGATCTNNNCCTGAGTGAAGAAGCCAAG
CACACGTCTGAACTCTTTCCCTACACGACGCTCTTCCGATCTNNNTTTATTTGTAATTTTTGCTGCTGCAT
CACACGTCTGAACTCTTTCCCTACACGACGCTCTTCCGATCTNNNTTTATTTGTAATTTTTGCTGCTGCAT
CACACGTCTGAACTCTTTCCCTACACGACGCTCTTCCGATCTNNNTTTATTTGTAATTTTTGCTGCTGCAT
AGCACACGTCTGAACTCTTTCCCTACACGACGCTCTTCCGATCTNNNGAGTGACTCCTGCTTCAACCTT
AGCACACGTCTGAACTCTTTCCCTACACGACGCTCTTCCGATCTNNNGAGTGACTCCTGCTTCAACCTT
AGCACACGTCTGAACTCTTTCCCTACACGACGCTCTTCCGATCTNNNGAGTGACTCCTGCTTCAACCTT
AGCACACGTCTGAACTCTTTCCCTACACGACGCTCTTCCGATCTNNNCTGGACAGGTAGGGTTGGGG
AGCACACGTCTGAACTCTTTCCCTACACGACGCTCTTCCGATCTNNNCTGGACAGGTAGGGTTGGGG
AGCACACGTCTGAACTCTTTCCCTACACGACGCTCTTCCGATCTNNNCTGGACAGGTAGGGTTGGGG
GAGCACACGTCTGAACTCTTTCCCTACACGACGCTCTTCCGATCTNNNACTCAAGATTGGCCCATTTAC
GAGCACACGTCTGAACTCTTTCCCTACACGACGCTCTTCCGATCTNNNACTCAAGATTGGCCCATTTAC
GAGCACACGTCTGAACTCTTTCCCTACACGACGCTCTTCCGATCTNNNACTCAAGATTGGCCCATTTAC
GAGCACACGTCTGAACTCTTTCCCTACACGACGCTCTTCCGATCTNNNTGTCATCACTCTGACCACAGC
GAGCACACGTCTGAACTCTTTCCCTACACGACGCTCTTCCGATCTNNNTGTCATCACTCTGACCACAGC
GAGCACACGTCTGAACTCTTTCCCTACACGACGCTCTTCCGATCTNNNTGTCATCACTCTGACCACAGC
AGCACACGTCTGAACTCTTTCCCTACACGACGCTCTTCCGATCTNNNTTCCACTTGGGTCTTCAGATAG
AGCACACGTCTGAACTCTTTCCCTACACGACGCTCTTCCGATCTNNNTTCCACTTGGGTCTTCAGATAG
AGCACACGTCTGAACTCTTTCCCTACACGACGCTCTTCCGATCTNNNTTCCACTTGGGTCTTCAGATAG
AGCACACGTCTGAACTCTTTCCCTACACGACGCTCTTCCGATCTNNNTTTGATCACAATGGGAAAATT

AGCACACGTCTGAACTCTTTCCCTACACGACGCTCTTCCGATCTNNNTTTGATCACAATGGGAAAATT
AGCACACGTCTGAACTCTTTCCCTACACGACGCTCTTCCGATCTNNNTTTGATCACAATGGGAAAATT
GCACACGTCTGAACTCTTTCCCTACACGACGCTCTTCCGATCTNNNGGTACTTCTGTCACTGCAGAGA
GCACACGTCTGAACTCTTTCCCTACACGACGCTCTTCCGATCTNNNGGTACTTCTGTCACTGCAGAGA
GCACACGTCTGAACTCTTTCCCTACACGACGCTCTTCCGATCTNNNGGTACTTCTGTCACTGCAGAGA
GAGCACACGTCTGAACTCTTTCCCTACACGACGCTCTTCCGATCTNNNCTGGCCAACATGATGAAGAC
GAGCACACGTCTGAACTCTTTCCCTACACGACGCTCTTCCGATCTNNNCTGGCCAACATGATGAAGAC
GAGCACACGTCTGAACTCTTTCCCTACACGACGCTCTTCCGATCTNNNCTGGCCAACATGATGAAGAC
AGCACACGTCTGAACTCTTTCCCTACACGACGCTCTTCCGATCTNNNGCCTGTAGTCCCAGATACTTG
AGCACACGTCTGAACTCTTTCCCTACACGACGCTCTTCCGATCTNNNGCCTGTAGTCCCAGATACTTG
AGCACACGTCTGAACTCTTTCCCTACACGACGCTCTTCCGATCTNNNGCCTGTAGTCCCAGATACTTG
GAGCACACGTCTGAACTCTTTCCCTACACGACGCTCTTCCGATCTNNNAGATTGCTTGAGTCCAGGA
GAGCACACGTCTGAACTCTTTCCCTACACGACGCTCTTCCGATCTNNNAGATTGCTTGAGTCCAGGA
GAGCACACGTCTGAACTCTTTCCCTACACGACGCTCTTCCGATCTNNNAGATTGCTTGAGTCCAGGA
GCACACGTCTGAACTCTTTCCCTACACGACGCTCTTCCGATCTNNNCCAGCTACCAAGACAGGAGAAT
GCACACGTCTGAACTCTTTCCCTACACGACGCTCTTCCGATCTNNNCCAGCTACCAAGACAGGAGAAT
GCACACGTCTGAACTCTTTCCCTACACGACGCTCTTCCGATCTNNNCCAGCTACCAAGACAGGAGAAT
GAGCACACGTCTGAACTCTTTCCCTACACGACGCTCTTCCGATCTNNNTGACAAGAGTGAAGCTCCTT
GAGCACACGTCTGAACTCTTTCCCTACACGACGCTCTTCCGATCTNNNTGACAAGAGTGAAGCTCCTT
GAGCACACGTCTGAACTCTTTCCCTACACGACGCTCTTCCGATCTNNNTGACAAGAGTGAAGCTCCTT
AGCACACGTCTGAACTCTTTCCCTACACGACGCTCTTCCGATCTNNNCACGCCCAGCTAATTTTTGTAG
AGCACACGTCTGAACTCTTTCCCTACACGACGCTCTTCCGATCTNNNCACGCCCAGCTAATTTTTGTAG
AGCACACGTCTGAACTCTTTCCCTACACGACGCTCTTCCGATCTNNNCACGCCCAGCTAATTTTTGTAG
GAGCACACGTCTGAACTCTTTCCCTACACGACGCTCTTCCGATCTNNNGGAGTTTACTGTGAGCCGA
GAGCACACGTCTGAACTCTTTCCCTACACGACGCTCTTCCGATCTNNNGGAGTTTACTGTGAGCCGA
GAGCACACGTCTGAACTCTTTCCCTACACGACGCTCTTCCGATCTNNNGGAGTTTACTGTGAGCCGA
GCACACGTCTGAACTCTTTCCCTACACGACGCTCTTCCGATCTNNNCTGAGGCAGGAGAATCACTTG
GCACACGTCTGAACTCTTTCCCTACACGACGCTCTTCCGATCTNNNCTGAGGCAGGAGAATCACTTG
GCACACGTCTGAACTCTTTCCCTACACGACGCTCTTCCGATCTNNNCTGAGGCAGGAGAATCACTTG
GCACACGTCTGAACTCTTTCCCTACACGACGCTCTTCCGATCTNNNAAATTAGTTGGGCGTGGTAAT
GCACACGTCTGAACTCTTTCCCTACACGACGCTCTTCCGATCTNNNAAATTAGTTGGGCGTGGTAAT
GCACACGTCTGAACTCTTTCCCTACACGACGCTCTTCCGATCTNNNAAATTAGTTGGGCGTGGTAAT
GAGCACACGTCTGAACTCTTTCCCTACACGACGCTCTTCCGATCTNNNCCAAGATCGTACCACTGCACT
GAGCACACGTCTGAACTCTTTCCCTACACGACGCTCTTCCGATCTNNNCCAAGATCGTACCACTGCACT
GAGCACACGTCTGAACTCTTTCCCTACACGACGCTCTTCCGATCTNNNCCAAGATCGTACCACTGCACT
GCACACGTCTGAACTCTTTCCCTACACGACGCTCTTCCGATCTNNNATTCTTTTTGTCCCTTCCCTCCCC
GCACACGTCTGAACTCTTTCCCTACACGACGCTCTTCCGATCTNNNATTCTTTTTGTCCCTTCCCTCCCC
GCACACGTCTGAACTCTTTCCCTACACGACGCTCTTCCGATCTNNNATTCTTTTTGTCCCTTCCCTCCCC
AGCACACGTCTGAACTCTTTCCCTACACGACGCTCTTCCGATCTNNNAGTATAGGTATGTGCCACCAT
AGCACACGTCTGAACTCTTTCCCTACACGACGCTCTTCCGATCTNNNAGTATAGGTATGTGCCACCAT
AGCACACGTCTGAACTCTTTCCCTACACGACGCTCTTCCGATCTNNNAGTATAGGTATGTGCCACCAT
GCACACGTCTGAACTCTTTCCCTACACGACGCTCTTCCGATCTNNNCATGTGGGAATGCATGCCTAAT
GCACACGTCTGAACTCTTTCCCTACACGACGCTCTTCCGATCTNNNCATGTGGGAATGCATGCCTAAT

GCACACGTCTGAACTCTTTCCCTACACGACGCTCTTCCGATCTNNNCATGTGGGAATGCATGCCTAATA

GAGCACACGTCTGAACTCTTTCCCTACACGACGCTCTTCCGATCTNNNCCTGTAGTCCCAGCTACTCCAG

GAGCACACGTCTGAACTCTTTCCCTACACGACGCTCTTCCGATCTNNNCCTGTAGTCCCAGCTACTCCAG

GAGCACACGTCTGAACTCTTTCCCTACACGACGCTCTTCCGATCTNNNCCTGTAGTCCCAGCTACTCCAG

ACACGTCTGAACTCTTTCCCTACACGACGCTCTTCCGATCTNNNCTGTGAATTTCGATGCTGGAGACCA

ACACGTCTGAACTCTTTCCCTACACGACGCTCTTCCGATCTNNNCTGTGAATTTCGATGCTGGAGACCA

ACACGTCTGAACTCTTTCCCTACACGACGCTCTTCCGATCTNNNCTGTGAATTTCGATGCTGGAGACCA

CACACGTCTGAACTCTTTCCCTACACGACGCTCTTCCGATCTNNNAGGTTCACCTCATCCATTTACTTCTC

CACACGTCTGAACTCTTTCCCTACACGACGCTCTTCCGATCTNNNAGGTTCACCTCATCCATTTACTTCTC

CACACGTCTGAACTCTTTCCCTACACGACGCTCTTCCGATCTNNNAGGTTCACCTCATCCATTTACTTCTC

AGCACACGTCTGAACTCTTTCCCTACACGACGCTCTTCCGATCTNNNAGTCGAGATATTGACATCAATG

AGCACACGTCTGAACTCTTTCCCTACACGACGCTCTTCCGATCTNNNAGTCGAGATATTGACATCAATG

AGCACACGTCTGAACTCTTTCCCTACACGACGCTCTTCCGATCTNNNAGTCGAGATATTGACATCAATG

GAGCACACGTCTGAACTCTTTCCCTACACGACGCTCTTCCGATCTNNNGTGCCTGTAATCCCAGTTACTC

GAGCACACGTCTGAACTCTTTCCCTACACGACGCTCTTCCGATCTNNNGTGCCTGTAATCCCAGTTACTC

GAGCACACGTCTGAACTCTTTCCCTACACGACGCTCTTCCGATCTNNNGTGCCTGTAATCCCAGTTACTC

GAGCACACGTCTGAACTCTTTCCCTACACGACGCTCTTCCGATCTNNNGAAAGCTGAGGCATGAGAATC

GAGCACACGTCTGAACTCTTTCCCTACACGACGCTCTTCCGATCTNNNGAAAGCTGAGGCATGAGAATC

GAGCACACGTCTGAACTCTTTCCCTACACGACGCTCTTCCGATCTNNNGAAAGCTGAGGCATGAGAATC

AGCACACGTCTGAACTCTTTCCCTACACGACGCTCTTCCGATCTNNNTGTCTTTCTGTGCCTGGCTTATTC

AGCACACGTCTGAACTCTTTCCCTACACGACGCTCTTCCGATCTNNNTGTCTTTCTGTGCCTGGCTTATTC

AGCACACGTCTGAACTCTTTCCCTACACGACGCTCTTCCGATCTNNNTGTCTTTCTGTGCCTGGCTTATTC

AGCACACGTCTGAACTCTTTCCCTACACGACGCTCTTCCGATCTNNNCCACCTTTGCGTTTCTAACTCTG

AGCACACGTCTGAACTCTTTCCCTACACGACGCTCTTCCGATCTNNNCCACCTTTGCGTTTCTAACTCTG

AGCACACGTCTGAACTCTTTCCCTACACGACGCTCTTCCGATCTNNNCCACCTTTGCGTTTCTAACTCTG

GCACACGTCTGAACTCTTTCCCTACACGACGCTCTTCCGATCTNNNTGTCTTACTTGCTCTGATCTGGCC

GCACACGTCTGAACTCTTTCCCTACACGACGCTCTTCCGATCTNNNTGTCTTACTTGCTCTGATCTGGCC

GCACACGTCTGAACTCTTTCCCTACACGACGCTCTTCCGATCTNNNTGTCTTACTTGCTCTGATCTGGCC

AGCACACGTCTGAACTCTTTCCCTACACGACGCTCTTCCGATCTNNNAGATGAGCGTTACGATTTACCGT

AGCACACGTCTGAACTCTTTCCCTACACGACGCTCTTCCGATCTNNNAGATGAGCGTTACGATTTACCGT

AGCACACGTCTGAACTCTTTCCCTACACGACGCTCTTCCGATCTNNNAGATGAGCGTTACGATTTACCGT

GAGCACACGTCTGAACTCTTTCCCTACACGACGCTCTTCCGATCTNNNGAGACTGAGGCAGGAGAATC

GAGCACACGTCTGAACTCTTTCCCTACACGACGCTCTTCCGATCTNNNGAGACTGAGGCAGGAGAATC

GAGCACACGTCTGAACTCTTTCCCTACACGACGCTCTTCCGATCTNNNGAGACTGAGGCAGGAGAATC

GAGCACACGTCTGAACTCTTTCCCTACACGACGCTCTTCCGATCTNNNTATGATAGCTTCCCACCGTTTC

GAGCACACGTCTGAACTCTTTCCCTACACGACGCTCTTCCGATCTNNNTATGATAGCTTCCCACCGTTTC

GAGCACACGTCTGAACTCTTTCCCTACACGACGCTCTTCCGATCTNNNTATGATAGCTTCCCACCGTTTC

AGCACACGTCTGAACTCTTTCCCTACACGACGCTCTTCCGATCTNNNATCTTGACATCATGATCTGCCCCC

AGCACACGTCTGAACTCTTTCCCTACACGACGCTCTTCCGATCTNNNATCTTGACATCATGATCTGCCCCC

AGCACACGTCTGAACTCTTTCCCTACACGACGCTCTTCCGATCTNNNATCTTGACATCATGATCTGCCCCC

CACACGTCTGAACTCTTTCCCTACACGACGCTCTTCCGATCTNNNGTACCACAGTTTCTTCTGTAACATCC

CACACGTCTGAACTCTTTCCCTACACGACGCTCTTCCGATCTNNNGTACCACAGTTTCTTCTGTAACATCC

CACACGTCTGAACTCTTTCCCTACACGACGCTCTTCCGATCTNNNGTACCACAGTTTCTTCTGTAACATCC

AGCACACGTCTGAACTCTTTCCCTACACGACGCTCTTCCGATCTNNNCTGAGGCAGGAGAATCGTTTG
AGCACACGTCTGAACTCTTTCCCTACACGACGCTCTTCCGATCTNNNCTGAGGCAGGAGAATCGTTTG
AGCACACGTCTGAACTCTTTCCCTACACGACGCTCTTCCGATCTNNNCTGAGGCAGGAGAATCGTTTG
GAGCACACGTCTGAACTCTTTCCCTACACGACGCTCTTCCGATCTNNNAGATGGTGGAAATATGTGTG
GAGCACACGTCTGAACTCTTTCCCTACACGACGCTCTTCCGATCTNNNAGATGGTGGAAATATGTGTG
GAGCACACGTCTGAACTCTTTCCCTACACGACGCTCTTCCGATCTNNNAGATGGTGGAAATATGTGTG
GAGCACACGTCTGAACTCTTTCCCTACACGACGCTCTTCCGATCTNNNCCCAAGGCCTCCAAATAGTCT
GAGCACACGTCTGAACTCTTTCCCTACACGACGCTCTTCCGATCTNNNCCCAAGGCCTCCAAATAGTCT
GAGCACACGTCTGAACTCTTTCCCTACACGACGCTCTTCCGATCTNNNCCCAAGGCCTCCAAATAGTCT
GAGCACACGTCTGAACTCTTTCCCTACACGACGCTCTTCCGATCTNNNAGTGAGCCATGATCATGTCAC
GAGCACACGTCTGAACTCTTTCCCTACACGACGCTCTTCCGATCTNNNAGTGAGCCATGATCATGTCAC
GAGCACACGTCTGAACTCTTTCCCTACACGACGCTCTTCCGATCTNNNAGTGAGCCATGATCATGTCAC
AGCACACGTCTGAACTCTTTCCCTACACGACGCTCTTCCGATCTNNNCTGTCTCTGTCTTTCCTTCCCTC
AGCACACGTCTGAACTCTTTCCCTACACGACGCTCTTCCGATCTNNNCTGTCTCTGTCTTTCCTTCCCTC
AGCACACGTCTGAACTCTTTCCCTACACGACGCTCTTCCGATCTNNNCTGTCTCTGTCTTTCCTTCCCTC
AGCACACGTCTGAACTCTTTCCCTACACGACGCTCTTCCGATCTNNNCGCCTGGCTAATTTGTGTGTGC
AGCACACGTCTGAACTCTTTCCCTACACGACGCTCTTCCGATCTNNNCGCCTGGCTAATTTGTGTGTGC
AGCACACGTCTGAACTCTTTCCCTACACGACGCTCTTCCGATCTNNNCGCCTGGCTAATTTGTGTGTGC
AGCACACGTCTGAACTCTTTCCCTACACGACGCTCTTCCGATCTNNNGGATTGAGTCCACAGGGAGAA
AGCACACGTCTGAACTCTTTCCCTACACGACGCTCTTCCGATCTNNNGGATTGAGTCCACAGGGAGAA
AGCACACGTCTGAACTCTTTCCCTACACGACGCTCTTCCGATCTNNNGGATTGAGTCCACAGGGAGAA
GCACACGTCTGAACTCTTTCCCTACACGACGCTCTTCCGATCTNNNATGAGAGCAAAGGGAGAATTCG
GCACACGTCTGAACTCTTTCCCTACACGACGCTCTTCCGATCTNNNATGAGAGCAAAGGGAGAATTCG
GCACACGTCTGAACTCTTTCCCTACACGACGCTCTTCCGATCTNNNATGAGAGCAAAGGGAGAATTCG
AGCACACGTCTGAACTCTTTCCCTACACGACGCTCTTCCGATCTNNNTCTCCCTCTCTCTCTCTCTCTCA
AGCACACGTCTGAACTCTTTCCCTACACGACGCTCTTCCGATCTNNNTCTCCCTCTCTCTCTCTCTCTCA
AGCACACGTCTGAACTCTTTCCCTACACGACGCTCTTCCGATCTNNNTCTCCCTCTCTCTCTCTCTCTCA
GCACACGTCTGAACTCTTTCCCTACACGACGCTCTTCCGATCTNNNCAGCTAATCGAGAACTGGAGAG
GCACACGTCTGAACTCTTTCCCTACACGACGCTCTTCCGATCTNNNCAGCTAATCGAGAACTGGAGAG
GCACACGTCTGAACTCTTTCCCTACACGACGCTCTTCCGATCTNNNCAGCTAATCGAGAACTGGAGAG
AGCACACGTCTGAACTCTTTCCCTACACGACGCTCTTCCGATCTNNNGTCCCTCTCTGTCACTTTCTCTG
AGCACACGTCTGAACTCTTTCCCTACACGACGCTCTTCCGATCTNNNGTCCCTCTCTGTCACTTTCTCTG
AGCACACGTCTGAACTCTTTCCCTACACGACGCTCTTCCGATCTNNNGTCCCTCTCTGTCACTTTCTCTG
AGCACACGTCTGAACTCTTTCCCTACACGACGCTCTTCCGATCTNNNTTGAGGCAGGAGAATTGCTTG
AGCACACGTCTGAACTCTTTCCCTACACGACGCTCTTCCGATCTNNNTTGAGGCAGGAGAATTGCTTG
AGCACACGTCTGAACTCTTTCCCTACACGACGCTCTTCCGATCTNNNTTGAGGCAGGAGAATTGCTTG
AGCACACGTCTGAACTCTTTCCCTACACGACGCTCTTCCGATCTNNNGGAACTTCCAAATAATGTGTTT
AGCACACGTCTGAACTCTTTCCCTACACGACGCTCTTCCGATCTNNNGGAACTTCCAAATAATGTGTTT
AGCACACGTCTGAACTCTTTCCCTACACGACGCTCTTCCGATCTNNNGGAACTTCCAAATAATGTGTTT
AGCACACGTCTGAACTCTTTCCCTACACGACGCTCTTCCGATCTNNNAGAAATGAAGTGGAAATATGA
AGCACACGTCTGAACTCTTTCCCTACACGACGCTCTTCCGATCTNNNAGAAATGAAGTGGAAATATGA
AGCACACGTCTGAACTCTTTCCCTACACGACGCTCTTCCGATCTNNNAGAAATGAAGTGGAAATATGA
AGCACACGTCTGAACTCTTTCCCTACACGACGCTCTTCCGATCTNNNGCTGAAGCAGGAGAAATGTTT

AGCACACGTCTGAACTCTTTCCCTACACGACGCTCTTCCGATCTNNNGCTGAAGCAGGAGAAATGTTT

AGCACACGTCTGAACTCTTTCCCTACACGACGCTCTTCCGATCTNNNGCTGAAGCAGGAGAAATGTTT

AGCACACGTCTGAACTCTTTCCCTACACGACGCTCTTCCGATCTNNNGAAAGAAAGAGTTGAGCAAG

AGCACACGTCTGAACTCTTTCCCTACACGACGCTCTTCCGATCTNNNGAAAGAAAGAGTTGAGCAAG

AGCACACGTCTGAACTCTTTCCCTACACGACGCTCTTCCGATCTNNNGAAAGAAAGAGTTGAGCAAG

AGCACACGTCTGAACTCTTTCCCTACACGACGCTCTTCCGATCTNNNGAGACAGGAGAATTGCTTGAA

AGCACACGTCTGAACTCTTTCCCTACACGACGCTCTTCCGATCTNNNGAGACAGGAGAATTGCTTGAA

AGCACACGTCTGAACTCTTTCCCTACACGACGCTCTTCCGATCTNNNGAGACAGGAGAATTGCTTGAA

GAGCACACGTCTGAACTCTTTCCCTACACGACGCTCTTCCGATCTNNNTGTTGTGGTGATTCTATCCAG

GAGCACACGTCTGAACTCTTTCCCTACACGACGCTCTTCCGATCTNNNTGTTGTGGTGATTCTATCCAG

GAGCACACGTCTGAACTCTTTCCCTACACGACGCTCTTCCGATCTNNNTGTTGTGGTGATTCTATCCAG

GAGCACACGTCTGAACTCTTTCCCTACACGACGCTCTTCCGATCTNNNAGATGGGGAAGGGTTTGCTT

GAGCACACGTCTGAACTCTTTCCCTACACGACGCTCTTCCGATCTNNNAGATGGGGAAGGGTTTGCTT

GAGCACACGTCTGAACTCTTTCCCTACACGACGCTCTTCCGATCTNNNAGATGGGGAAGGGTTTGCTT

AGCACACGTCTGAACTCTTTCCCTACACGACGCTCTTCCGATCTNNNACACTCTTGTAATCCCAGCAACT

AGCACACGTCTGAACTCTTTCCCTACACGACGCTCTTCCGATCTNNNACACTCTTGTAATCCCAGCAACT

AGCACACGTCTGAACTCTTTCCCTACACGACGCTCTTCCGATCTNNNACACTCTTGTAATCCCAGCAACT

AGCACACGTCTGAACTCTTTCCCTACACGACGCTCTTCCGATCTNNNATCTACAGGAGGGTAAAGAGG

AGCACACGTCTGAACTCTTTCCCTACACGACGCTCTTCCGATCTNNNATCTACAGGAGGGTAAAGAGG

AGCACACGTCTGAACTCTTTCCCTACACGACGCTCTTCCGATCTNNNATCTACAGGAGGGTAAAGAGG

AGCACACGTCTGAACTCTTTCCCTACACGACGCTCTTCCGATCTNNNTAGAGCTGCTTTGGTGTCTTCA

AGCACACGTCTGAACTCTTTCCCTACACGACGCTCTTCCGATCTNNNTAGAGCTGCTTTGGTGTCTTCA

AGCACACGTCTGAACTCTTTCCCTACACGACGCTCTTCCGATCTNNNTAGAGCTGCTTTGGTGTCTTCA

GAGCACACGTCTGAACTCTTTCCCTACACGACGCTCTTCCGATCTNNNGTACCCGTAATCTCAGCTACTC

GAGCACACGTCTGAACTCTTTCCCTACACGACGCTCTTCCGATCTNNNGTACCCGTAATCTCAGCTACTC

GAGCACACGTCTGAACTCTTTCCCTACACGACGCTCTTCCGATCTNNNGTACCCGTAATCTCAGCTACTC

GCACACGTCTGAACTCTTTCCCTACACGACGCTCTTCCGATCTNNNATAGTCACTTACCACCATGCCTGC

GCACACGTCTGAACTCTTTCCCTACACGACGCTCTTCCGATCTNNNATAGTCACTTACCACCATGCCTGC

GCACACGTCTGAACTCTTTCCCTACACGACGCTCTTCCGATCTNNNATAGTCACTTACCACCATGCCTGC

AAGAGCACACGTCTGAACTCTTTCCCTACACGACGCTCTTCCGATCTNNNTGAGCCATGATCATGCCAC

AAGAGCACACGTCTGAACTCTTTCCCTACACGACGCTCTTCCGATCTNNNTGAGCCATGATCATGCCAC

AAGAGCACACGTCTGAACTCTTTCCCTACACGACGCTCTTCCGATCTNNNTGAGCCATGATCATGCCAC

AGCACACGTCTGAACTCTTTCCCTACACGACGCTCTTCCGATCTNNNGGACAGTAAGTGGAGCCTTCTA

AGCACACGTCTGAACTCTTTCCCTACACGACGCTCTTCCGATCTNNNGGACAGTAAGTGGAGCCTTCTA

AGCACACGTCTGAACTCTTTCCCTACACGACGCTCTTCCGATCTNNNGGACAGTAAGTGGAGCCTTCTA

AGCACACGTCTGAACTCTTTCCCTACACGACGCTCTTCCGATCTNNNTTCAGTTGGGACGTAATAAGG

AGCACACGTCTGAACTCTTTCCCTACACGACGCTCTTCCGATCTNNNTTCAGTTGGGACGTAATAAGG

AGCACACGTCTGAACTCTTTCCCTACACGACGCTCTTCCGATCTNNNTTCAGTTGGGACGTAATAAGG

GAGCACACGTCTGAACTCTTTCCCTACACGACGCTCTTCCGATCTNNNTAAGAAATTGGTGGCTAGAAT

GAGCACACGTCTGAACTCTTTCCCTACACGACGCTCTTCCGATCTNNNTAAGAAATTGGTGGCTAGAAT

GAGCACACGTCTGAACTCTTTCCCTACACGACGCTCTTCCGATCTNNNTAAGAAATTGGTGGCTAGAAT

GAAGAGCACACGTCTGAACTCTTTCCCTACACGACGCTCTTCCGATCTNNNCACTGCATGTCTTTGTGCA

GAAGAGCACACGTCTGAACTCTTTCCCTACACGACGCTCTTCCGATCTNNNCACTGCATGTCTTTGTGCA

AAGAGCACACGTCTGAACTCTTTCCCTACACGACGCTCTTCCGATCTNNNCACTGCATGTCTTTGTGC
CACACGTCTGAACTCTTTCCCTACACGACGCTCTTCCGATCTNNNACTCCATTCTAGGCAACAGAGTG
CACACGTCTGAACTCTTTCCCTACACGACGCTCTTCCGATCTNNNACTCCATTCTAGGCAACAGAGTG
CACACGTCTGAACTCTTTCCCTACACGACGCTCTTCCGATCTNNNACTCCATTCTAGGCAACAGAGTG
GCACACGTCTGAACTCTTTCCCTACACGACGCTCTTCCGATCTNNNCTTCAGCCTTCCAAAGTGTTGAG
GCACACGTCTGAACTCTTTCCCTACACGACGCTCTTCCGATCTNNNCTTCAGCCTTCCAAAGTGTTGAG
GCACACGTCTGAACTCTTTCCCTACACGACGCTCTTCCGATCTNNNCTTCAGCCTTCCAAAGTGTTGAG
GCACACGTCTGAACTCTTTCCCTACACGACGCTCTTCCGATCTNNNGTTCAAGCAATTCTCTGCCTCAG
GCACACGTCTGAACTCTTTCCCTACACGACGCTCTTCCGATCTNNNGTTCAAGCAATTCTCTGCCTCAG
GCACACGTCTGAACTCTTTCCCTACACGACGCTCTTCCGATCTNNNGTTCAAGCAATTCTCTGCCTCAG
GCACACGTCTGAACTCTTTCCCTACACGACGCTCTTCCGATCTNNNGTTGGTGAGTTTAGGTGAAAG
GCACACGTCTGAACTCTTTCCCTACACGACGCTCTTCCGATCTNNNGTTGGTGAGTTTAGGTGAAAG
GCACACGTCTGAACTCTTTCCCTACACGACGCTCTTCCGATCTNNNGTTGGTGAGTTTAGGTGAAAG
AGCACACGTCTGAACTCTTTCCCTACACGACGCTCTTCCGATCTNNNTGGTCTTGAACTACTGACCTCA
AGCACACGTCTGAACTCTTTCCCTACACGACGCTCTTCCGATCTNNNTGGTCTTGAACTACTGACCTCA
AGCACACGTCTGAACTCTTTCCCTACACGACGCTCTTCCGATCTNNNTGGTCTTGAACTACTGACCTCA
AGCACACGTCTGAACTCTTTCCCTACACGACGCTCTTCCGATCTNNNCTCTGTTCCCCTCAGGATTGAT
AGCACACGTCTGAACTCTTTCCCTACACGACGCTCTTCCGATCTNNNCTCTGTTCCCCTCAGGATTGAT
AGCACACGTCTGAACTCTTTCCCTACACGACGCTCTTCCGATCTNNNCTCTGTTCCCCTCAGGATTGAT
GCACACGTCTGAACTCTTTCCCTACACGACGCTCTTCCGATCTNNNTGCGTGTTTCAGTGGACTGTAT
GCACACGTCTGAACTCTTTCCCTACACGACGCTCTTCCGATCTNNNTGCGTGTTTCAGTGGACTGTAT
GCACACGTCTGAACTCTTTCCCTACACGACGCTCTTCCGATCTNNNTGCGTGTTTCAGTGGACTGTAT
GAGCACACGTCTGAACTCTTTCCCTACACGACGCTCTTCCGATCTNNNCAAGCATTGGATGTTGTTCCA
GAGCACACGTCTGAACTCTTTCCCTACACGACGCTCTTCCGATCTNNNCAAGCATTGGATGTTGTTCCA
GAGCACACGTCTGAACTCTTTCCCTACACGACGCTCTTCCGATCTNNNCAAGCATTGGATGTTGTTCCA
GAGCACACGTCTGAACTCTTTCCCTACACGACGCTCTTCCGATCTNNNGAGGCTGAGGCAGAAGAAT
GAGCACACGTCTGAACTCTTTCCCTACACGACGCTCTTCCGATCTNNNGAGGCTGAGGCAGAAGAAT
GAGCACACGTCTGAACTCTTTCCCTACACGACGCTCTTCCGATCTNNNGAGGCTGAGGCAGAAGAAT
AGCACACGTCTGAACTCTTTCCCTACACGACGCTCTTCCGATCTNNNCTTCTCACCTTAGGCGATCATC
AGCACACGTCTGAACTCTTTCCCTACACGACGCTCTTCCGATCTNNNCTTCTCACCTTAGGCGATCATC
AGCACACGTCTGAACTCTTTCCCTACACGACGCTCTTCCGATCTNNNCTTCTCACCTTAGGCGATCATC
AGCACACGTCTGAACTCTTTCCCTACACGACGCTCTTCCGATCTNNNTAATTTTAGGAGTAGGATTGA
AGCACACGTCTGAACTCTTTCCCTACACGACGCTCTTCCGATCTNNNTAATTTTAGGAGTAGGATTGA
AGCACACGTCTGAACTCTTTCCCTACACGACGCTCTTCCGATCTNNNTAATTTTAGGAGTAGGATTGA
AGCACACGTCTGAACTCTTTCCCTACACGACGCTCTTCCGATCTNNNTTCTGGCTGAGGTCCTAAATG
AGCACACGTCTGAACTCTTTCCCTACACGACGCTCTTCCGATCTNNNTTCTGGCTGAGGTCCTAAATG
AGCACACGTCTGAACTCTTTCCCTACACGACGCTCTTCCGATCTNNNTTCTGGCTGAGGTCCTAAATG
GCACACGTCTGAACTCTTTCCCTACACGACGCTCTTCCGATCTNNNTTGACATTTGTTCAGCTCACAC
GCACACGTCTGAACTCTTTCCCTACACGACGCTCTTCCGATCTNNNTTGACATTTGTTCAGCTCACAC
GCACACGTCTGAACTCTTTCCCTACACGACGCTCTTCCGATCTNNNTTGACATTTGTTCAGCTCACAC
GAGCACACGTCTGAACTCTTTCCCTACACGACGCTCTTCCGATCTNNNTTGTCCCACATGGTCTAAGT
GAGCACACGTCTGAACTCTTTCCCTACACGACGCTCTTCCGATCTNNNTTGTCCCACATGGTCTAAGT
GAGCACACGTCTGAACTCTTTCCCTACACGACGCTCTTCCGATCTNNNTTGTCCCACATGGTCTAAGT

GAGCACACGTCTGAACTCTTTCCCTACACGACGCTCTTCCGATCTNNNAGCAGGTGTATGTGTGCATG
GAGCACACGTCTGAACTCTTTCCCTACACGACGCTCTTCCGATCTNNNAGCAGGTGTATGTGTGCATG
GAGCACACGTCTGAACTCTTTCCCTACACGACGCTCTTCCGATCTNNNAGCAGGTGTATGTGTGCATG
AGAGCACACGTCTGAACTCTTTCCCTACACGACGCTCTTCCGATCTNNNGTACAGCACCCTAGTTCAGAT
AGAGCACACGTCTGAACTCTTTCCCTACACGACGCTCTTCCGATCTNNNGTACAGCACCCTAGTTCAGAT
AGAGCACACGTCTGAACTCTTTCCCTACACGACGCTCTTCCGATCTNNNGTACAGCACCCTAGTTCAGAT
GAGCACACGTCTGAACTCTTTCCCTACACGACGCTCTTCCGATCTNNNTGTCACTCTAGGGAAAGGGTT
GAGCACACGTCTGAACTCTTTCCCTACACGACGCTCTTCCGATCTNNNTGTCACTCTAGGGAAAGGGTT
GAGCACACGTCTGAACTCTTTCCCTACACGACGCTCTTCCGATCTNNNTGTCACTCTAGGGAAAGGGTT
GAGCACACGTCTGAACTCTTTCCCTACACGACGCTCTTCCGATCTNNNGATGCAGTCACTGAACATGCT
GAGCACACGTCTGAACTCTTTCCCTACACGACGCTCTTCCGATCTNNNGATGCAGTCACTGAACATGCT
GAGCACACGTCTGAACTCTTTCCCTACACGACGCTCTTCCGATCTNNNGATGCAGTCACTGAACATGCT
AGCACACGTCTGAACTCTTTCCCTACACGACGCTCTTCCGATCTNNNATAGAAGGTCCGGAAACAATGC
AGCACACGTCTGAACTCTTTCCCTACACGACGCTCTTCCGATCTNNNATAGAAGGTCCGGAAACAATGC
AGCACACGTCTGAACTCTTTCCCTACACGACGCTCTTCCGATCTNNNATAGAAGGTCCGGAAACAATGC
AGCACACGTCTGAACTCTTTCCCTACACGACGCTCTTCCGATCTNNNGTGAATTTGTTCCTGGCTTCTCC
AGCACACGTCTGAACTCTTTCCCTACACGACGCTCTTCCGATCTNNNGTGAATTTGTTCCTGGCTTCTCC
AGCACACGTCTGAACTCTTTCCCTACACGACGCTCTTCCGATCTNNNGTGAATTTGTTCCTGGCTTCTCC
GAGCACACGTCTGAACTCTTTCCCTACACGACGCTCTTCCGATCTNNNGCAAGAGAATCACTTGAACCC
GAGCACACGTCTGAACTCTTTCCCTACACGACGCTCTTCCGATCTNNNGCAAGAGAATCACTTGAACCC
GAGCACACGTCTGAACTCTTTCCCTACACGACGCTCTTCCGATCTNNNGCAAGAGAATCACTTGAACCC
AGCACACGTCTGAACTCTTTCCCTACACGACGCTCTTCCGATCTNNNTCAGCCTCCTGAGTAGCTATGAT
AGCACACGTCTGAACTCTTTCCCTACACGACGCTCTTCCGATCTNNNTCAGCCTCCTGAGTAGCTATGAT
AGCACACGTCTGAACTCTTTCCCTACACGACGCTCTTCCGATCTNNNTCAGCCTCCTGAGTAGCTATGAT
GAGCACACGTCTGAACTCTTTCCCTACACGACGCTCTTCCGATCTNNNCTGTAATCCCAGCTACTCAGGA
GAGCACACGTCTGAACTCTTTCCCTACACGACGCTCTTCCGATCTNNNCTGTAATCCCAGCTACTCAGGA
GAGCACACGTCTGAACTCTTTCCCTACACGACGCTCTTCCGATCTNNNCTGTAATCCCAGCTACTCAGGA
CACACGTCTGAACTCTTTCCCTACACGACGCTCTTCCGATCTNNNAGGTTCACCTCATCCATTTACTTCCA
CACACGTCTGAACTCTTTCCCTACACGACGCTCTTCCGATCTNNNAGGTTCACCTCATCCATTTACTTCCA
CACACGTCTGAACTCTTTCCCTACACGACGCTCTTCCGATCTNNNAGGTTCACCTCATCCATTTACTTCCA
CACACGTCTGAACTCTTTCCCTACACGACGCTCTTCCGATCTNNNATCCACGCTGCACTTGAATTCTCCA
CACACGTCTGAACTCTTTCCCTACACGACGCTCTTCCGATCTNNNATCCACGCTGCACTTGAATTCTCCA
CACACGTCTGAACTCTTTCCCTACACGACGCTCTTCCGATCTNNNATCCACGCTGCACTTGAATTCTCCA
AGCACACGTCTGAACTCTTTCCCTACACGACGCTCTTCCGATCTNNNTGTGAAGGTTGTAAGGACATTT
AGCACACGTCTGAACTCTTTCCCTACACGACGCTCTTCCGATCTNNNTGTGAAGGTTGTAAGGACATTT
AGCACACGTCTGAACTCTTTCCCTACACGACGCTCTTCCGATCTNNNTGTGAAGGTTGTAAGGACATTT
GAGCACACGTCTGAACTCTTTCCCTACACGACGCTCTTCCGATCTNNNGAGGCTGAGGCATAAGAATCA
GAGCACACGTCTGAACTCTTTCCCTACACGACGCTCTTCCGATCTNNNGAGGCTGAGGCATAAGAATCA
GAGCACACGTCTGAACTCTTTCCCTACACGACGCTCTTCCGATCTNNNGAGGCTGAGGCATAAGAATCA
GCACACGTCTGAACTCTTTCCCTACACGACGCTCTTCCGATCTNNNAAATTAGTGGCCATGATCATGCC
GCACACGTCTGAACTCTTTCCCTACACGACGCTCTTCCGATCTNNNAAATTAGTGGCCATGATCATGCC
GCACACGTCTGAACTCTTTCCCTACACGACGCTCTTCCGATCTNNNAAATTAGTGGCCATGATCATGCC
AGCACACGTCTGAACTCTTTCCCTACACGACGCTCTTCCGATCTNNNCCAGCTCTATTCTTATAAACTTC

AGCACACGTCTGAACTCTTTCCCTACACGACGCTCTTCCGATCTNNNCCAGCTCTATTCTTATAAACTTC
AGCACACGTCTGAACTCTTTCCCTACACGACGCTCTTCCGATCTNNNCCAGCTCTATTCTTATAAACTTC
AGCACACGTCTGAACTCTTTCCCTACACGACGCTCTTCCGATCTNNNTTGAACTCCTGACCTCATGATC
AGCACACGTCTGAACTCTTTCCCTACACGACGCTCTTCCGATCTNNNTTGAACTCCTGACCTCATGATC
AGCACACGTCTGAACTCTTTCCCTACACGACGCTCTTCCGATCTNNNTTGAACTCCTGACCTCATGATC
GAGCACACGTCTGAACTCTTTCCCTACACGACGCTCTTCCGATCTNNNCGTGATAAGTCCCGTCTCAGG
GAGCACACGTCTGAACTCTTTCCCTACACGACGCTCTTCCGATCTNNNCGTGATAAGTCCCGTCTCAGG
GAGCACACGTCTGAACTCTTTCCCTACACGACGCTCTTCCGATCTNNNCGTGATAAGTCCCGTCTCAGG
GAGCACACGTCTGAACTCTTTCCCTACACGACGCTCTTCCGATCTNNNGAGGCAGGAGAATTTTGTG
GAGCACACGTCTGAACTCTTTCCCTACACGACGCTCTTCCGATCTNNNGAGGCAGGAGAATTTTGTG
GAGCACACGTCTGAACTCTTTCCCTACACGACGCTCTTCCGATCTNNNGAGGCAGGAGAATTTTGTG
AGCACACGTCTGAACTCTTTCCCTACACGACGCTCTTCCGATCTNNNTGGGTAGAGAAGGGCATATG
AGCACACGTCTGAACTCTTTCCCTACACGACGCTCTTCCGATCTNNNTGGGTAGAGAAGGGCATATG
AGCACACGTCTGAACTCTTTCCCTACACGACGCTCTTCCGATCTNNNTGGGTAGAGAAGGGCATATG
GAGCACACGTCTGAACTCTTTCCCTACACGACGCTCTTCCGATCTNNNATTTGGCAATATCATCGAAG
GAGCACACGTCTGAACTCTTTCCCTACACGACGCTCTTCCGATCTNNNATTTGGCAATATCATCGAAG
GAGCACACGTCTGAACTCTTTCCCTACACGACGCTCTTCCGATCTNNNATTTGGCAATATCATCGAAG
AGCACACGTCTGAACTCTTTCCCTACACGACGCTCTTCCGATCTNNNAAGTGCTGCATTGTACAGTCAT
AGCACACGTCTGAACTCTTTCCCTACACGACGCTCTTCCGATCTNNNAAGTGCTGCATTGTACAGTCAT
AGCACACGTCTGAACTCTTTCCCTACACGACGCTCTTCCGATCTNNNAAGTGCTGCATTGTACAGTCAT
GCACACGTCTGAACTCTTTCCCTACACGACGCTCTTCCGATCTNNNTGATAGATAGGTGGAGTCTGCC
GCACACGTCTGAACTCTTTCCCTACACGACGCTCTTCCGATCTNNNTGATAGATAGGTGGAGTCTGCC
GCACACGTCTGAACTCTTTCCCTACACGACGCTCTTCCGATCTNNNTGATAGATAGGTGGAGTCTGCC
GCACACGTCTGAACTCTTTCCCTACACGACGCTCTTCCGATCTNNNTCCATGCCTCTCTCTCTCTCTATC
GCACACGTCTGAACTCTTTCCCTACACGACGCTCTTCCGATCTNNNTCCATGCCTCTCTCTCTCTCTATC
GCACACGTCTGAACTCTTTCCCTACACGACGCTCTTCCGATCTNNNTCCATGCCTCTCTCTCTCTCTATC
GCACACGTCTGAACTCTTTCCCTACACGACGCTCTTCCGATCTNNNTATGGGGCACAAACACTACACA
GCACACGTCTGAACTCTTTCCCTACACGACGCTCTTCCGATCTNNNTATGGGGCACAAACACTACACA
GCACACGTCTGAACTCTTTCCCTACACGACGCTCTTCCGATCTNNNTATGGGGCACAAACACTACACA
AGCACACGTCTGAACTCTTTCCCTACACGACGCTCTTCCGATCTNNNTTTGTGTCAGTGTTCATCGTGG
AGCACACGTCTGAACTCTTTCCCTACACGACGCTCTTCCGATCTNNNTTTGTGTCAGTGTTCATCGTGG
AGCACACGTCTGAACTCTTTCCCTACACGACGCTCTTCCGATCTNNNTTTGTGTCAGTGTTCATCGTGG
AGCACACGTCTGAACTCTTTCCCTACACGACGCTCTTCCGATCTNNNAAACTGCCAAATTGAAGGAGC
AGCACACGTCTGAACTCTTTCCCTACACGACGCTCTTCCGATCTNNNAAACTGCCAAATTGAAGGAGC
AGCACACGTCTGAACTCTTTCCCTACACGACGCTCTTCCGATCTNNNAAACTGCCAAATTGAAGGAGC
AGCACACGTCTGAACTCTTTCCCTACACGACGCTCTTCCGATCTNNNTCTGTTGGTTCTAAGCCCAAGT
AGCACACGTCTGAACTCTTTCCCTACACGACGCTCTTCCGATCTNNNTCTGTTGGTTCTAAGCCCAAGT
AGCACACGTCTGAACTCTTTCCCTACACGACGCTCTTCCGATCTNNNTCTGTTGGTTCTAAGCCCAAGT
GAGCACACGTCTGAACTCTTTCCCTACACGACGCTCTTCCGATCTNNNGCAGTGTTTTGTGTGTGCTTT
GAGCACACGTCTGAACTCTTTCCCTACACGACGCTCTTCCGATCTNNNGCAGTGTTTTGTGTGTGCTTT
GAGCACACGTCTGAACTCTTTCCCTACACGACGCTCTTCCGATCTNNNGCAGTGTTTTGTGTGTGCTTT
AGCACACGTCTGAACTCTTTCCCTACACGACGCTCTTCCGATCTNNNGGGAGAATCACTTGAACCCA
AGCACACGTCTGAACTCTTTCCCTACACGACGCTCTTCCGATCTNNNGGGAGAATCACTTGAACCCA

GAGCACACGTCTGAACTCTTTCCCTACACGACGCTCTTCCGATCTNNNGGGAGAATCACTTGAACCCAG
GCACACGTCTGAACTCTTTCCCTACACGACGCTCTTCCGATCTNNNCTAAATAGGCTGTGAATGGGGG
GCACACGTCTGAACTCTTTCCCTACACGACGCTCTTCCGATCTNNNCTAAATAGGCTGTGAATGGGGG
GCACACGTCTGAACTCTTTCCCTACACGACGCTCTTCCGATCTNNNCTAAATAGGCTGTGAATGGGGG
GAGCACACGTCTGAACTCTTTCCCTACACGACGCTCTTCCGATCTNNNAATCTTACAGATAGCAGCCCTC
GAGCACACGTCTGAACTCTTTCCCTACACGACGCTCTTCCGATCTNNNAATCTTACAGATAGCAGCCCTC
GAGCACACGTCTGAACTCTTTCCCTACACGACGCTCTTCCGATCTNNNAATCTTACAGATAGCAGCCCTC
GAGCACACGTCTGAACTCTTTCCCTACACGACGCTCTTCCGATCTNNNGACAGAGCTAGACTCCGTCTC
GAGCACACGTCTGAACTCTTTCCCTACACGACGCTCTTCCGATCTNNNGACAGAGCTAGACTCCGTCTC
GAGCACACGTCTGAACTCTTTCCCTACACGACGCTCTTCCGATCTNNNGACAGAGCTAGACTCCGTCTC
AGCACACGTCTGAACTCTTTCCCTACACGACGCTCTTCCGATCTNNNGATCTCAGCTCACTACAACCTCC
AGCACACGTCTGAACTCTTTCCCTACACGACGCTCTTCCGATCTNNNGATCTCAGCTCACTACAACCTCC
AGCACACGTCTGAACTCTTTCCCTACACGACGCTCTTCCGATCTNNNGATCTCAGCTCACTACAACCTCC
GAGCACACGTCTGAACTCTTTCCCTACACGACGCTCTTCCGATCTNNNCAGTCTGAGCAAGAGAGTGA
GAGCACACGTCTGAACTCTTTCCCTACACGACGCTCTTCCGATCTNNNCAGTCTGAGCAAGAGAGTGA
GAGCACACGTCTGAACTCTTTCCCTACACGACGCTCTTCCGATCTNNNCAGTCTGAGCAAGAGAGTGA
AGCACACGTCTGAACTCTTTCCCTACACGACGCTCTTCCGATCTNNNCAAACACGATGATCCCTTTTGCT
AGCACACGTCTGAACTCTTTCCCTACACGACGCTCTTCCGATCTNNNCAAACACGATGATCCCTTTTGCT
AGCACACGTCTGAACTCTTTCCCTACACGACGCTCTTCCGATCTNNNCAAACACGATGATCCCTTTTGCT
GAGCACACGTCTGAACTCTTTCCCTACACGACGCTCTTCCGATCTNNNAGAGGTGGTGATTGAAACTA
GAGCACACGTCTGAACTCTTTCCCTACACGACGCTCTTCCGATCTNNNAGAGGTGGTGATTGAAACTA
GAGCACACGTCTGAACTCTTTCCCTACACGACGCTCTTCCGATCTNNNAGAGGTGGTGATTGAAACTA
GAGCACACGTCTGAACTCTTTCCCTACACGACGCTCTTCCGATCTNNNCTTCTTGGCTTTAAGTAAGTA
GAGCACACGTCTGAACTCTTTCCCTACACGACGCTCTTCCGATCTNNNCTTCTTGGCTTTAAGTAAGTA
GAGCACACGTCTGAACTCTTTCCCTACACGACGCTCTTCCGATCTNNNCTTCTTGGCTTTAAGTAAGTA
GCACACGTCTGAACTCTTTCCCTACACGACGCTCTTCCGATCTNNNAGAGGAACTACAGGATAAGGGC
GCACACGTCTGAACTCTTTCCCTACACGACGCTCTTCCGATCTNNNAGAGGAACTACAGGATAAGGGC
GCACACGTCTGAACTCTTTCCCTACACGACGCTCTTCCGATCTNNNAGAGGAACTACAGGATAAGGGC
AAGAGCACACGTCTGAACTCTTTCCCTACACGACGCTCTTCCGATCTNNNGCCAAGATCATGCCATTGT
AAGAGCACACGTCTGAACTCTTTCCCTACACGACGCTCTTCCGATCTNNNGCCAAGATCATGCCATTGT
AAGAGCACACGTCTGAACTCTTTCCCTACACGACGCTCTTCCGATCTNNNGCCAAGATCATGCCATTGT
AGCACACGTCTGAACTCTTTCCCTACACGACGCTCTTCCGATCTNNNACATTTGATACAAACTGGCAGCC
AGCACACGTCTGAACTCTTTCCCTACACGACGCTCTTCCGATCTNNNACATTTGATACAAACTGGCAGCC
AGCACACGTCTGAACTCTTTCCCTACACGACGCTCTTCCGATCTNNNACATTTGATACAAACTGGCAGCC
AGAGCACACGTCTGAACTCTTTCCCTACACGACGCTCTTCCGATCTNNNTCTTCTGACAGCTTGCTTCAG
AGAGCACACGTCTGAACTCTTTCCCTACACGACGCTCTTCCGATCTNNNTCTTCTGACAGCTTGCTTCAG
AGAGCACACGTCTGAACTCTTTCCCTACACGACGCTCTTCCGATCTNNNTCTTCTGACAGCTTGCTTCAG
AGCACACGTCTGAACTCTTTCCCTACACGACGCTCTTCCGATCTNNNTGGTAGGAATTCTCACTCACAGC
AGCACACGTCTGAACTCTTTCCCTACACGACGCTCTTCCGATCTNNNTGGTAGGAATTCTCACTCACAGC
AGCACACGTCTGAACTCTTTCCCTACACGACGCTCTTCCGATCTNNNTGGTAGGAATTCTCACTCACAGC
GAGCACACGTCTGAACTCTTTCCCTACACGACGCTCTTCCGATCTNNNTATTCCATGGTGTCTGTGTGTC
GAGCACACGTCTGAACTCTTTCCCTACACGACGCTCTTCCGATCTNNNTATTCCATGGTGTCTGTGTGTC
GAGCACACGTCTGAACTCTTTCCCTACACGACGCTCTTCCGATCTNNNTATTCCATGGTGTCTGTGTGTC

AGCACACGTCTGAACTCTTTCCCTACACGACGCTCTTCCGATCTNNNAAGTTCAAGACCAGACTGCTCA
AGCACACGTCTGAACTCTTTCCCTACACGACGCTCTTCCGATCTNNNAAGTTCAAGACCAGACTGCTCA
AGCACACGTCTGAACTCTTTCCCTACACGACGCTCTTCCGATCTNNNAAGTTCAAGACCAGACTGCTCA
AGCACACGTCTGAACTCTTTCCCTACACGACGCTCTTCCGATCTNNNTCTGTGTGTGTGTGTGTGTGTT
AGCACACGTCTGAACTCTTTCCCTACACGACGCTCTTCCGATCTNNNTCTGTGTGTGTGTGTGTGTGTT
AGCACACGTCTGAACTCTTTCCCTACACGACGCTCTTCCGATCTNNNTCTGTGTGTGTGTGTGTGTGTT
GAGCACACGTCTGAACTCTTTCCCTACACGACGCTCTTCCGATCTNNNTCAGACAACACAATTTGGCAT
GAGCACACGTCTGAACTCTTTCCCTACACGACGCTCTTCCGATCTNNNTCAGACAACACAATTTGGCAT
GAGCACACGTCTGAACTCTTTCCCTACACGACGCTCTTCCGATCTNNNTCAGACAACACAATTTGGCAT
GAGCACACGTCTGAACTCTTTCCCTACACGACGCTCTTCCGATCTNNNAATACAGCACAGTGATGGGT
GAGCACACGTCTGAACTCTTTCCCTACACGACGCTCTTCCGATCTNNNAATACAGCACAGTGATGGGT
GAGCACACGTCTGAACTCTTTCCCTACACGACGCTCTTCCGATCTNNNAATACAGCACAGTGATGGGT
GAGCACACGTCTGAACTCTTTCCCTACACGACGCTCTTCCGATCTNNNACATTGACCTGTGCCACATACT
GAGCACACGTCTGAACTCTTTCCCTACACGACGCTCTTCCGATCTNNNACATTGACCTGTGCCACATACT
GAGCACACGTCTGAACTCTTTCCCTACACGACGCTCTTCCGATCTNNNACATTGACCTGTGCCACATACT
GAGCACACGTCTGAACTCTTTCCCTACACGACGCTCTTCCGATCTNNNTACTTCGATGTGTCAGCCTGA
GAGCACACGTCTGAACTCTTTCCCTACACGACGCTCTTCCGATCTNNNTACTTCGATGTGTCAGCCTGA
GAGCACACGTCTGAACTCTTTCCCTACACGACGCTCTTCCGATCTNNNTACTTCGATGTGTCAGCCTGA
GCACACGTCTGAACTCTTTCCCTACACGACGCTCTTCCGATCTNNNTTTGTCCTCCAAAGACAGAGACC
GCACACGTCTGAACTCTTTCCCTACACGACGCTCTTCCGATCTNNNTTTGTCCTCCAAAGACAGAGACC
GCACACGTCTGAACTCTTTCCCTACACGACGCTCTTCCGATCTNNNTTTGTCCTCCAAAGACAGAGACC
GAGCACACGTCTGAACTCTTTCCCTACACGACGCTCTTCCGATCTNNNGTTGTTGTTCTTTGTAGGCCC
GAGCACACGTCTGAACTCTTTCCCTACACGACGCTCTTCCGATCTNNNGTTGTTGTTCTTTGTAGGCCC
GAGCACACGTCTGAACTCTTTCCCTACACGACGCTCTTCCGATCTNNNGTTGTTGTTCTTTGTAGGCCC
CACGTCTGAACTCTTTCCCTACACGACGCTCTTCCGATCTNNNTCATAGCTATTGTGTGGCTTTCTCATC
CACGTCTGAACTCTTTCCCTACACGACGCTCTTCCGATCTNNNTCATAGCTATTGTGTGGCTTTCTCATC
CACGTCTGAACTCTTTCCCTACACGACGCTCTTCCGATCTNNNTCATAGCTATTGTGTGGCTTTCTCATC
GAGCACACGTCTGAACTCTTTCCCTACACGACGCTCTTCCGATCTNNNACTCAAAGGCTGGAGGTAAG
GAGCACACGTCTGAACTCTTTCCCTACACGACGCTCTTCCGATCTNNNACTCAAAGGCTGGAGGTAAG
GAGCACACGTCTGAACTCTTTCCCTACACGACGCTCTTCCGATCTNNNACTCAAAGGCTGGAGGTAAG
AGCACACGTCTGAACTCTTTCCCTACACGACGCTCTTCCGATCTNNNGGATCATGATGTCAGGAGTTC
AGCACACGTCTGAACTCTTTCCCTACACGACGCTCTTCCGATCTNNNGGATCATGATGTCAGGAGTTC
AGCACACGTCTGAACTCTTTCCCTACACGACGCTCTTCCGATCTNNNGGATCATGATGTCAGGAGTTC
AGCACACGTCTGAACTCTTTCCCTACACGACGCTCTTCCGATCTNNNCTGACCAGCTAGATCCCACTTCT
AGCACACGTCTGAACTCTTTCCCTACACGACGCTCTTCCGATCTNNNCTGACCAGCTAGATCCCACTTCT
AGCACACGTCTGAACTCTTTCCCTACACGACGCTCTTCCGATCTNNNCTGACCAGCTAGATCCCACTTCT
GAGCACACGTCTGAACTCTTTCCCTACACGACGCTCTTCCGATCTNNNAGCTCCTCCAAGATTCTCTCTCT
GAGCACACGTCTGAACTCTTTCCCTACACGACGCTCTTCCGATCTNNNAGCTCCTCCAAGATTCTCTCTCT
GAGCACACGTCTGAACTCTTTCCCTACACGACGCTCTTCCGATCTNNNAGCTCCTCCAAGATTCTCTCTCT
AGCACACGTCTGAACTCTTTCCCTACACGACGCTCTTCCGATCTNNNTTCAGAAATAGCAGGCTTGTGG
AGCACACGTCTGAACTCTTTCCCTACACGACGCTCTTCCGATCTNNNTTCAGAAATAGCAGGCTTGTGG
AGCACACGTCTGAACTCTTTCCCTACACGACGCTCTTCCGATCTNNNTTCAGAAATAGCAGGCTTGTGG
GAGCACACGTCTGAACTCTTTCCCTACACGACGCTCTTCCGATCTNNNTATGGAGAAGCACACATGAA

GAGCACACGTCTGAACTCTTTCCCTACACGACGCTCTTCCGATCTNNNTATGGAGAAGCACACATGAA
GAGCACACGTCTGAACTCTTTCCCTACACGACGCTCTTCCGATCTNNNTATGGAGAAGCACACATGAA
AGCACACGTCTGAACTCTTTCCCTACACGACGCTCTTCCGATCTNNNCAGGTGACAGTGAGCTATGAT
AGCACACGTCTGAACTCTTTCCCTACACGACGCTCTTCCGATCTNNNCAGGTGACAGTGAGCTATGAT
AGCACACGTCTGAACTCTTTCCCTACACGACGCTCTTCCGATCTNNNCAGGTGACAGTGAGCTATGAT
CACACGTCTGAACTCTTTCCCTACACGACGCTCTTCCGATCTNNNGGCTCCTCACTTCCCAGTAGGCAT
CACACGTCTGAACTCTTTCCCTACACGACGCTCTTCCGATCTNNNGGCTCCTCACTTCCCAGTAGGCAT
CACACGTCTGAACTCTTTCCCTACACGACGCTCTTCCGATCTNNNGGCTCCTCACTTCCCAGTAGGCAT
AGCACACGTCTGAACTCTTTCCCTACACGACGCTCTTCCGATCTNNNGGGATGCTGAATCAGGAGAA
AGCACACGTCTGAACTCTTTCCCTACACGACGCTCTTCCGATCTNNNGGGATGCTGAATCAGGAGAA
AGCACACGTCTGAACTCTTTCCCTACACGACGCTCTTCCGATCTNNNGGGATGCTGAATCAGGAGAA
GAGCACACGTCTGAACTCTTTCCCTACACGACGCTCTTCCGATCTNNNTGAGGCAGGAGAATCTCTT
GAGCACACGTCTGAACTCTTTCCCTACACGACGCTCTTCCGATCTNNNTGAGGCAGGAGAATCTCTT
GAGCACACGTCTGAACTCTTTCCCTACACGACGCTCTTCCGATCTNNNTGAGGCAGGAGAATCTCTT
GCACACGTCTGAACTCTTTCCCTACACGACGCTCTTCCGATCTNNNGAGGAAGAACTGGCGAAGGG
GCACACGTCTGAACTCTTTCCCTACACGACGCTCTTCCGATCTNNNGAGGAAGAACTGGCGAAGGG
GCACACGTCTGAACTCTTTCCCTACACGACGCTCTTCCGATCTNNNGAGGAAGAACTGGCGAAGGG
GCACACGTCTGAACTCTTTCCCTACACGACGCTCTTCCGATCTNNNATTTCAGGACTTGCTACATTTTCC
GCACACGTCTGAACTCTTTCCCTACACGACGCTCTTCCGATCTNNNATTTCAGGACTTGCTACATTTTCC
GCACACGTCTGAACTCTTTCCCTACACGACGCTCTTCCGATCTNNNATTTCAGGACTTGCTACATTTTCC
GCACACGTCTGAACTCTTTCCCTACACGACGCTCTTCCGATCTNNNATCATGAGCCTGGTAGGTTGAA
GCACACGTCTGAACTCTTTCCCTACACGACGCTCTTCCGATCTNNNATCATGAGCCTGGTAGGTTGAA
GCACACGTCTGAACTCTTTCCCTACACGACGCTCTTCCGATCTNNNATCATGAGCCTGGTAGGTTGAA
GCACACGTCTGAACTCTTTCCCTACACGACGCTCTTCCGATCTNNNGCATTCCAATTTTCACCTCCAGC
GCACACGTCTGAACTCTTTCCCTACACGACGCTCTTCCGATCTNNNGCATTCCAATTTTCACCTCCAGC
GCACACGTCTGAACTCTTTCCCTACACGACGCTCTTCCGATCTNNNGCATTCCAATTTTCACCTCCAGC
GAGCACACGTCTGAACTCTTTCCCTACACGACGCTCTTCCGATCTNNNCTGAGGCAGGAGAATTGCTT
GAGCACACGTCTGAACTCTTTCCCTACACGACGCTCTTCCGATCTNNNCTGAGGCAGGAGAATTGCTT
GAGCACACGTCTGAACTCTTTCCCTACACGACGCTCTTCCGATCTNNNCTGAGGCAGGAGAATTGCTT
GCACACGTCTGAACTCTTTCCCTACACGACGCTCTTCCGATCTNNNCTGATTGCTTCGAAGAGCATGT
GCACACGTCTGAACTCTTTCCCTACACGACGCTCTTCCGATCTNNNCTGATTGCTTCGAAGAGCATGT
GCACACGTCTGAACTCTTTCCCTACACGACGCTCTTCCGATCTNNNCTGATTGCTTCGAAGAGCATGT
AGCACACGTCTGAACTCTTTCCCTACACGACGCTCTTCCGATCTNNNAACCAGACCAAATACCCATAC
AGCACACGTCTGAACTCTTTCCCTACACGACGCTCTTCCGATCTNNNAACCAGACCAAATACCCATAC
AGCACACGTCTGAACTCTTTCCCTACACGACGCTCTTCCGATCTNNNAACCAGACCAAATACCCATAC
AGCACACGTCTGAACTCTTTCCCTACACGACGCTCTTCCGATCTNNNCAGCTACAGATGGTACAGGTG
AGCACACGTCTGAACTCTTTCCCTACACGACGCTCTTCCGATCTNNNCAGCTACAGATGGTACAGGTG
AGCACACGTCTGAACTCTTTCCCTACACGACGCTCTTCCGATCTNNNCAGCTACAGATGGTACAGGTG
GAGCACACGTCTGAACTCTTTCCCTACACGACGCTCTTCCGATCTNNNGAAAGCAGAATGGCAGACA
GAGCACACGTCTGAACTCTTTCCCTACACGACGCTCTTCCGATCTNNNGAAAGCAGAATGGCAGACA
GAGCACACGTCTGAACTCTTTCCCTACACGACGCTCTTCCGATCTNNNGAAAGCAGAATGGCAGACA
AGCACACGTCTGAACTCTTTCCCTACACGACGCTCTTCCGATCTNNNGTAATCCCAGCTATACAGGAG
AGCACACGTCTGAACTCTTTCCCTACACGACGCTCTTCCGATCTNNNGTAATCCCAGCTATACAGGAG

GAGCACACGTCTGAACTCTTTCCCTACACGACGCTCTTCCGATCTNNNGTAATCCCAGCTATACAGGAG
GAGCACACGTCTGAACTCTTTCCCTACACGACGCTCTTCCGATCTNNNGAGAATGGATCCTGGTCCCTT
GAGCACACGTCTGAACTCTTTCCCTACACGACGCTCTTCCGATCTNNNGAGAATGGATCCTGGTCCCTT
GAGCACACGTCTGAACTCTTTCCCTACACGACGCTCTTCCGATCTNNNGAGAATGGATCCTGGTCCCTT
AGCACACGTCTGAACTCTTTCCCTACACGACGCTCTTCCGATCTNNNAGTGGCAAGTACACTCAAAAGA
AGCACACGTCTGAACTCTTTCCCTACACGACGCTCTTCCGATCTNNNAGTGGCAAGTACACTCAAAAGA
AGCACACGTCTGAACTCTTTCCCTACACGACGCTCTTCCGATCTNNNAGTGGCAAGTACACTCAAAAGA
GAGCACACGTCTGAACTCTTTCCCTACACGACGCTCTTCCGATCTNNNAAAGGATGTGTCCGTCTGGTC
GAGCACACGTCTGAACTCTTTCCCTACACGACGCTCTTCCGATCTNNNAAAGGATGTGTCCGTCTGGTC
GAGCACACGTCTGAACTCTTTCCCTACACGACGCTCTTCCGATCTNNNAAAGGATGTGTCCGTCTGGTC
AGAGCACACGTCTGAACTCTTTCCCTACACGACGCTCTTCCGATCTNNNTGCAGAACTCCAAACACTAG
AGAGCACACGTCTGAACTCTTTCCCTACACGACGCTCTTCCGATCTNNNTGCAGAACTCCAAACACTAG
AGAGCACACGTCTGAACTCTTTCCCTACACGACGCTCTTCCGATCTNNNTGCAGAACTCCAAACACTAG
GAGCACACGTCTGAACTCTTTCCCTACACGACGCTCTTCCGATCTNNNACCTCCTTGTACATTGTTTTGC
GAGCACACGTCTGAACTCTTTCCCTACACGACGCTCTTCCGATCTNNNACCTCCTTGTACATTGTTTTGC
GAGCACACGTCTGAACTCTTTCCCTACACGACGCTCTTCCGATCTNNNACCTCCTTGTACATTGTTTTGC
AGCACACGTCTGAACTCTTTCCCTACACGACGCTCTTCCGATCTNNNCTTCCCAGGTTCAAGCAATCTTC
AGCACACGTCTGAACTCTTTCCCTACACGACGCTCTTCCGATCTNNNCTTCCCAGGTTCAAGCAATCTTC
AGCACACGTCTGAACTCTTTCCCTACACGACGCTCTTCCGATCTNNNCTTCCCAGGTTCAAGCAATCTTC
AGAGCACACGTCTGAACTCTTTCCCTACACGACGCTCTTCCGATCTNNNCTGTAATCCCAGCTACTCGG
AGAGCACACGTCTGAACTCTTTCCCTACACGACGCTCTTCCGATCTNNNCTGTAATCCCAGCTACTCGG
AGAGCACACGTCTGAACTCTTTCCCTACACGACGCTCTTCCGATCTNNNCTGTAATCCCAGCTACTCGG
CACACGTCTGAACTCTTTCCCTACACGACGCTCTTCCGATCTNNNCACAACGACCACATCCTCATAGACA
CACACGTCTGAACTCTTTCCCTACACGACGCTCTTCCGATCTNNNCACAACGACCACATCCTCATAGACA
CACACGTCTGAACTCTTTCCCTACACGACGCTCTTCCGATCTNNNCACAACGACCACATCCTCATAGACA
AGCACACGTCTGAACTCTTTCCCTACACGACGCTCTTCCGATCTNNNATACAGGTGGGTTGTCTTGAG
AGCACACGTCTGAACTCTTTCCCTACACGACGCTCTTCCGATCTNNNATACAGGTGGGTTGTCTTGAG
AGCACACGTCTGAACTCTTTCCCTACACGACGCTCTTCCGATCTNNNATACAGGTGGGTTGTCTTGAG
AGCACACGTCTGAACTCTTTCCCTACACGACGCTCTTCCGATCTNNNCCAAGCACTCATGAATCTAAGC
AGCACACGTCTGAACTCTTTCCCTACACGACGCTCTTCCGATCTNNNCCAAGCACTCATGAATCTAAGC
AGCACACGTCTGAACTCTTTCCCTACACGACGCTCTTCCGATCTNNNCCAAGCACTCATGAATCTAAGC
AGCACACGTCTGAACTCTTTCCCTACACGACGCTCTTCCGATCTNNNGGTGACAGAGTGAGTCTTCAA
AGCACACGTCTGAACTCTTTCCCTACACGACGCTCTTCCGATCTNNNGGTGACAGAGTGAGTCTTCAA
AGCACACGTCTGAACTCTTTCCCTACACGACGCTCTTCCGATCTNNNGGTGACAGAGTGAGTCTTCAA
AGCACACGTCTGAACTCTTTCCCTACACGACGCTCTTCCGATCTNNNAGGCCAACATGATCTATCCATCC
AGCACACGTCTGAACTCTTTCCCTACACGACGCTCTTCCGATCTNNNAGGCCAACATGATCTATCCATCC
AGCACACGTCTGAACTCTTTCCCTACACGACGCTCTTCCGATCTNNNAGGCCAACATGATCTATCCATCC
AGCACACGTCTGAACTCTTTCCCTACACGACGCTCTTCCGATCTNNNCAGAGGAAATGAGGAAATGGG
AGCACACGTCTGAACTCTTTCCCTACACGACGCTCTTCCGATCTNNNCAGAGGAAATGAGGAAATGGG
AGCACACGTCTGAACTCTTTCCCTACACGACGCTCTTCCGATCTNNNCAGAGGAAATGAGGAAATGGG
AGCACACGTCTGAACTCTTTCCCTACACGACGCTCTTCCGATCTNNNAGCATCTAGGTTGTTTCCTCTG
AGCACACGTCTGAACTCTTTCCCTACACGACGCTCTTCCGATCTNNNAGCATCTAGGTTGTTTCCTCTG
AGCACACGTCTGAACTCTTTCCCTACACGACGCTCTTCCGATCTNNNAGCATCTAGGTTGTTTCCTCTG

AGCACACGTCTGAACTCTTTCCCTACACGACGCTCTTCCGATCTNNNTGAGACAGAAATTGAGGAGTG

AGCACACGTCTGAACTCTTTCCCTACACGACGCTCTTCCGATCTNNNTGAGACAGAAATTGAGGAGTG

AGCACACGTCTGAACTCTTTCCCTACACGACGCTCTTCCGATCTNNNTGAGACAGAAATTGAGGAGTG

GCACACGTCTGAACTCTTTCCCTACACGACGCTCTTCCGATCTNNNTGCCATTTCCTTTATTTCCAGGTCA

GCACACGTCTGAACTCTTTCCCTACACGACGCTCTTCCGATCTNNNTGCCATTTCCTTTATTTCCAGGTCA

GCACACGTCTGAACTCTTTCCCTACACGACGCTCTTCCGATCTNNNTGCCATTTCCTTTATTTCCAGGTCA

AGCACACGTCTGAACTCTTTCCCTACACGACGCTCTTCCGATCTNNNGTGGTTCCAAGGAAATGCAATG

AGCACACGTCTGAACTCTTTCCCTACACGACGCTCTTCCGATCTNNNGTGGTTCCAAGGAAATGCAATG

AGCACACGTCTGAACTCTTTCCCTACACGACGCTCTTCCGATCTNNNGTGGTTCCAAGGAAATGCAATG

AGCACACGTCTGAACTCTTTCCCTACACGACGCTCTTCCGATCTNNNGGTGAGGACAACTATTCCGATC

AGCACACGTCTGAACTCTTTCCCTACACGACGCTCTTCCGATCTNNNGGTGAGGACAACTATTCCGATC

AGCACACGTCTGAACTCTTTCCCTACACGACGCTCTTCCGATCTNNNGGTGAGGACAACTATTCCGATC

GAGCACACGTCTGAACTCTTTCCCTACACGACGCTCTTCCGATCTNNNGGTCTTCTGCTTTATCCCTGTC

GAGCACACGTCTGAACTCTTTCCCTACACGACGCTCTTCCGATCTNNNGGTCTTCTGCTTTATCCCTGTC

GAGCACACGTCTGAACTCTTTCCCTACACGACGCTCTTCCGATCTNNNGGTCTTCTGCTTTATCCCTGTC

AGCACACGTCTGAACTCTTTCCCTACACGACGCTCTTCCGATCTNNNTCCTTCTTTCTCTGTCTTTCTCCTC

AGCACACGTCTGAACTCTTTCCCTACACGACGCTCTTCCGATCTNNNTCCTTCTTTCTCTGTCTTTCTCCTC

AGCACACGTCTGAACTCTTTCCCTACACGACGCTCTTCCGATCTNNNTCCTTCTTTCTCTGTCTTTCTCCTC

AGAGCACACGTCTGAACTCTTTCCCTACACGACGCTCTTCCGATCTNNNGAGAAGAAGCAAGCATTCA

AGAGCACACGTCTGAACTCTTTCCCTACACGACGCTCTTCCGATCTNNNGAGAAGAAGCAAGCATTCA

AGAGCACACGTCTGAACTCTTTCCCTACACGACGCTCTTCCGATCTNNNGAGAAGAAGCAAGCATTCA

AGAGCACACGTCTGAACTCTTTCCCTACACGACGCTCTTCCGATCTNNNGAGGCTGAGGCAGGAGAAT

AGAGCACACGTCTGAACTCTTTCCCTACACGACGCTCTTCCGATCTNNNGAGGCTGAGGCAGGAGAAT

AGAGCACACGTCTGAACTCTTTCCCTACACGACGCTCTTCCGATCTNNNGAGGCTGAGGCAGGAGAAT

AGAGCACACGTCTGAACTCTTTCCCTACACGACGCTCTTCCGATCTNNNCACTCAGATTCCTAAACCCGA

AGAGCACACGTCTGAACTCTTTCCCTACACGACGCTCTTCCGATCTNNNCACTCAGATTCCTAAACCCGA

AGAGCACACGTCTGAACTCTTTCCCTACACGACGCTCTTCCGATCTNNNCACTCAGATTCCTAAACCCGA

AGCACACGTCTGAACTCTTTCCCTACACGACGCTCTTCCGATCTNNNTCAGGTACAAGCAGTCAGTTCTC

AGCACACGTCTGAACTCTTTCCCTACACGACGCTCTTCCGATCTNNNTCAGGTACAAGCAGTCAGTTCTC

AGCACACGTCTGAACTCTTTCCCTACACGACGCTCTTCCGATCTNNNTCAGGTACAAGCAGTCAGTTCTC

GAGCACACGTCTGAACTCTTTCCCTACACGACGCTCTTCCGATCTNNNCTGGAATAGCTGTAGGAACG

GAGCACACGTCTGAACTCTTTCCCTACACGACGCTCTTCCGATCTNNNCTGGAATAGCTGTAGGAACG

GAGCACACGTCTGAACTCTTTCCCTACACGACGCTCTTCCGATCTNNNCTGGAATAGCTGTAGGAACG

AAGAGCACACGTCTGAACTCTTTCCCTACACGACGCTCTTCCGATCTNNNGGCAGGAGAATCACTTGAA

AAGAGCACACGTCTGAACTCTTTCCCTACACGACGCTCTTCCGATCTNNNGGCAGGAGAATCACTTGAA

AAGAGCACACGTCTGAACTCTTTCCCTACACGACGCTCTTCCGATCTNNNGGCAGGAGAATCACTTGAA

GAGCACACGTCTGAACTCTTTCCCTACACGACGCTCTTCCGATCTNNNCACACCTGTAATCCCAGCTACTC

GAGCACACGTCTGAACTCTTTCCCTACACGACGCTCTTCCGATCTNNNCACACCTGTAATCCCAGCTACTC

GAGCACACGTCTGAACTCTTTCCCTACACGACGCTCTTCCGATCTNNNCACACCTGTAATCCCAGCTACTC

GCACACGTCTGAACTCTTTCCCTACACGACGCTCTTCCGATCTNNNTTCAAGCTATTCTCCTGCCTCAGC

GCACACGTCTGAACTCTTTCCCTACACGACGCTCTTCCGATCTNNNTTCAAGCTATTCTCCTGCCTCAGC

GCACACGTCTGAACTCTTTCCCTACACGACGCTCTTCCGATCTNNNTTCAAGCTATTCTCCTGCCTCAGC

GAGCACACGTCTGAACTCTTTCCCTACACGACGCTCTTCCGATCTNNNGGAGCTAAGCTATGAGGATG

AGCACACGTCTGAACTCTTTCCCTACACGACGCTCTTCCGATCTNNNGGAGCTAAGCTATGAGGATG
AGCACACGTCTGAACTCTTTCCCTACACGACGCTCTTCCGATCTNNNGGAGCTAAGCTATGAGGATG
GCACACGTCTGAACTCTTTCCCTACACGACGCTCTTCCGATCTNNNTTTCAATCATAAGGTCGGCCCAG
GCACACGTCTGAACTCTTTCCCTACACGACGCTCTTCCGATCTNNNTTTCAATCATAAGGTCGGCCCAG
GCACACGTCTGAACTCTTTCCCTACACGACGCTCTTCCGATCTNNNTTTCAATCATAAGGTCGGCCCAG
GCACACGTCTGAACTCTTTCCCTACACGACGCTCTTCCGATCTNNNCAGAAATGAAGACCCAAACAGG
GCACACGTCTGAACTCTTTCCCTACACGACGCTCTTCCGATCTNNNCAGAAATGAAGACCCAAACAGG
GCACACGTCTGAACTCTTTCCCTACACGACGCTCTTCCGATCTNNNCAGAAATGAAGACCCAAACAGG
GAGCACACGTCTGAACTCTTTCCCTACACGACGCTCTTCCGATCTNNNTTTGCTTCACATTCTTGACAG
GAGCACACGTCTGAACTCTTTCCCTACACGACGCTCTTCCGATCTNNNTTTGCTTCACATTCTTGACAG
GAGCACACGTCTGAACTCTTTCCCTACACGACGCTCTTCCGATCTNNNTTTGCTTCACATTCTTGACAG
GCACACGTCTGAACTCTTTCCCTACACGACGCTCTTCCGATCTNNNACCTCCAAACCTCCTTTACCTCAC
GCACACGTCTGAACTCTTTCCCTACACGACGCTCTTCCGATCTNNNACCTCCAAACCTCCTTTACCTCAC
GCACACGTCTGAACTCTTTCCCTACACGACGCTCTTCCGATCTNNNACCTCCAAACCTCCTTTACCTCAC
CACACGTCTGAACTCTTTCCCTACACGACGCTCTTCCGATCTNNNCTGATAAGGCAGCCTGTAAACAC
CACACGTCTGAACTCTTTCCCTACACGACGCTCTTCCGATCTNNNCTGATAAGGCAGCCTGTAAACAC
CACACGTCTGAACTCTTTCCCTACACGACGCTCTTCCGATCTNNNCTGATAAGGCAGCCTGTAAACAC
GCACACGTCTGAACTCTTTCCCTACACGACGCTCTTCCGATCTNNNTGAAGTGAGCTGATATTGCACC
GCACACGTCTGAACTCTTTCCCTACACGACGCTCTTCCGATCTNNNTGAAGTGAGCTGATATTGCACC
GCACACGTCTGAACTCTTTCCCTACACGACGCTCTTCCGATCTNNNTGAAGTGAGCTGATATTGCACC
AGAGCACACGTCTGAACTCTTTCCCTACACGACGCTCTTCCGATCTNNNCTCAAAGTGGTCCACTCTCA
AGAGCACACGTCTGAACTCTTTCCCTACACGACGCTCTTCCGATCTNNNCTCAAAGTGGTCCACTCTCA
AGAGCACACGTCTGAACTCTTTCCCTACACGACGCTCTTCCGATCTNNNCTCAAAGTGGTCCACTCTCA
GCACACGTCTGAACTCTTTCCCTACACGACGCTCTTCCGATCTNNNATCCCAAGGAAAATGTCGTCTG
GCACACGTCTGAACTCTTTCCCTACACGACGCTCTTCCGATCTNNNATCCCAAGGAAAATGTCGTCTG
GCACACGTCTGAACTCTTTCCCTACACGACGCTCTTCCGATCTNNNATCCCAAGGAAAATGTCGTCTG
ACACGTCTGAACTCTTTCCCTACACGACGCTCTTCCGATCTNNNAAAGTCCTGTAGCTCATAGTCATCG
ACACGTCTGAACTCTTTCCCTACACGACGCTCTTCCGATCTNNNAAAGTCCTGTAGCTCATAGTCATCG
ACACGTCTGAACTCTTTCCCTACACGACGCTCTTCCGATCTNNNAAAGTCCTGTAGCTCATAGTCATCG
AGCACACGTCTGAACTCTTTCCCTACACGACGCTCTTCCGATCTNNNAGGACAGAACATGGAAAGCCT
AGCACACGTCTGAACTCTTTCCCTACACGACGCTCTTCCGATCTNNNAGGACAGAACATGGAAAGCCT
AGCACACGTCTGAACTCTTTCCCTACACGACGCTCTTCCGATCTNNNAGGACAGAACATGGAAAGCCT
AGCACACGTCTGAACTCTTTCCCTACACGACGCTCTTCCGATCTNNNCTTATCAGGCAACAGAAGGCA
AGCACACGTCTGAACTCTTTCCCTACACGACGCTCTTCCGATCTNNNCTTATCAGGCAACAGAAGGCA
AGCACACGTCTGAACTCTTTCCCTACACGACGCTCTTCCGATCTNNNCTTATCAGGCAACAGAAGGCA
GCACACGTCTGAACTCTTTCCCTACACGACGCTCTTCCGATCTNNNCTGGGGCCAACGAAAAGAGACA
GCACACGTCTGAACTCTTTCCCTACACGACGCTCTTCCGATCTNNNCTGGGGCCAACGAAAAGAGACA
GCACACGTCTGAACTCTTTCCCTACACGACGCTCTTCCGATCTNNNCTGGGGCCAACGAAAAGAGACA
CACACGTCTGAACTCTTTCCCTACACGACGCTCTTCCGATCTNNNGCCTCACCTGATCATAGTACCTGC
CACACGTCTGAACTCTTTCCCTACACGACGCTCTTCCGATCTNNNGCCTCACCTGATCATAGTACCTGC
CACACGTCTGAACTCTTTCCCTACACGACGCTCTTCCGATCTNNNGCCTCACCTGATCATAGTACCTGC
GCACACGTCTGAACTCTTTCCCTACACGACGCTCTTCCGATCTNNNGTTCCACACGCTTTGTTTCCTAA
GCACACGTCTGAACTCTTTCCCTACACGACGCTCTTCCGATCTNNNGTTCCACACGCTTTGTTTCCTAA

AGCACACGTCTGAACTCTTTCCCTACACGACGCTCTTCCGATCTNNNGTTCCACACGCTTTGTTTCCTAA
AGCACACGTCTGAACTCTTTCCCTACACGACGCTCTTCCGATCTNNNACTTGGGTATCTGGCAGACAAT
AGCACACGTCTGAACTCTTTCCCTACACGACGCTCTTCCGATCTNNNACTTGGGTATCTGGCAGACAAT
AGCACACGTCTGAACTCTTTCCCTACACGACGCTCTTCCGATCTNNNACTTGGGTATCTGGCAGACAAT
AGCACACGTCTGAACTCTTTCCCTACACGACGCTCTTCCGATCTNNNTTCTTACATGGACACAGGAAG
AGCACACGTCTGAACTCTTTCCCTACACGACGCTCTTCCGATCTNNNTTCTTACATGGACACAGGAAG
AGCACACGTCTGAACTCTTTCCCTACACGACGCTCTTCCGATCTNNNTTCTTACATGGACACAGGAAG
AGAGCACACGTCTGAACTCTTTCCCTACACGACGCTCTTCCGATCTNNNCAGCTTAAGTAACTGGCTTG
AGAGCACACGTCTGAACTCTTTCCCTACACGACGCTCTTCCGATCTNNNCAGCTTAAGTAACTGGCTTG
AGAGCACACGTCTGAACTCTTTCCCTACACGACGCTCTTCCGATCTNNNCAGCTTAAGTAACTGGCTTG
GAGCACACGTCTGAACTCTTTCCCTACACGACGCTCTTCCGATCTNNNCTGAGGCAGGAGAATCACTT
GAGCACACGTCTGAACTCTTTCCCTACACGACGCTCTTCCGATCTNNNCTGAGGCAGGAGAATCACTT
GAGCACACGTCTGAACTCTTTCCCTACACGACGCTCTTCCGATCTNNNCTGAGGCAGGAGAATCACTT
GAGCACACGTCTGAACTCTTTCCCTACACGACGCTCTTCCGATCTNNNGTAACAGGCACCTATAGTCCC
GAGCACACGTCTGAACTCTTTCCCTACACGACGCTCTTCCGATCTNNNGTAACAGGCACCTATAGTCCC
GAGCACACGTCTGAACTCTTTCCCTACACGACGCTCTTCCGATCTNNNGTAACAGGCACCTATAGTCCC
AGAGCACACGTCTGAACTCTTTCCCTACACGACGCTCTTCCGATCTNNNCATCCTGGCTAACATGGTGA
AGAGCACACGTCTGAACTCTTTCCCTACACGACGCTCTTCCGATCTNNNCATCCTGGCTAACATGGTGA
AGAGCACACGTCTGAACTCTTTCCCTACACGACGCTCTTCCGATCTNNNCATCCTGGCTAACATGGTGA
AGCACACGTCTGAACTCTTTCCCTACACGACGCTCTTCCGATCTNNNGACAGGACAGGACACTTGGG
AGCACACGTCTGAACTCTTTCCCTACACGACGCTCTTCCGATCTNNNGACAGGACAGGACACTTGGG
AGCACACGTCTGAACTCTTTCCCTACACGACGCTCTTCCGATCTNNNGACAGGACAGGACACTTGGG
GCACACGTCTGAACTCTTTCCCTACACGACGCTCTTCCGATCTNNNTCTATTACATGCTTAGGGAAATT
GCACACGTCTGAACTCTTTCCCTACACGACGCTCTTCCGATCTNNNTCTATTACATGCTTAGGGAAATT
GCACACGTCTGAACTCTTTCCCTACACGACGCTCTTCCGATCTNNNTCTATTACATGCTTAGGGAAATT
AGAGCACACGTCTGAACTCTTTCCCTACACGACGCTCTTCCGATCTNNNTTTTTCTTTGTTGTTGCAGCT
AGAGCACACGTCTGAACTCTTTCCCTACACGACGCTCTTCCGATCTNNNTTTTTCTTTGTTGTTGCAGCT
AGAGCACACGTCTGAACTCTTTCCCTACACGACGCTCTTCCGATCTNNNTTTTTCTTTGTTGTTGCAGCT
AGCACACGTCTGAACTCTTTCCCTACACGACGCTCTTCCGATCTNNNGAATGCAGTGGTGTGATCATA
AGCACACGTCTGAACTCTTTCCCTACACGACGCTCTTCCGATCTNNNGAATGCAGTGGTGTGATCATA
AGCACACGTCTGAACTCTTTCCCTACACGACGCTCTTCCGATCTNNNGAATGCAGTGGTGTGATCATA
AGCACACGTCTGAACTCTTTCCCTACACGACGCTCTTCCGATCTNNNAAGATCGCACTACTGCAATCCAG
AGCACACGTCTGAACTCTTTCCCTACACGACGCTCTTCCGATCTNNNAAGATCGCACTACTGCAATCCAG
AGCACACGTCTGAACTCTTTCCCTACACGACGCTCTTCCGATCTNNNAAGATCGCACTACTGCAATCCAG
GAGCACACGTCTGAACTCTTTCCCTACACGACGCTCTTCCGATCTNNNCCTGGCTCATGGATGATGACA
GAGCACACGTCTGAACTCTTTCCCTACACGACGCTCTTCCGATCTNNNCCTGGCTCATGGATGATGACA
GAGCACACGTCTGAACTCTTTCCCTACACGACGCTCTTCCGATCTNNNCCTGGCTCATGGATGATGACA
GCACACGTCTGAACTCTTTCCCTACACGACGCTCTTCCGATCTNNNGTAACAGACGACTTCACGTTTCGC
GCACACGTCTGAACTCTTTCCCTACACGACGCTCTTCCGATCTNNNGTAACAGACGACTTCACGTTTCGC
GCACACGTCTGAACTCTTTCCCTACACGACGCTCTTCCGATCTNNNGTAACAGACGACTTCACGTTTCGC
AGCACACGTCTGAACTCTTTCCCTACACGACGCTCTTCCGATCTNNNAGACATCCATGCTCATACACCAG
AGCACACGTCTGAACTCTTTCCCTACACGACGCTCTTCCGATCTNNNAGACATCCATGCTCATACACCAG
AGCACACGTCTGAACTCTTTCCCTACACGACGCTCTTCCGATCTNNNAGACATCCATGCTCATACACCAG

GAGCACACGTCTGAACTCTTTCCCTACACGACGCTCTTCCGATCTNNNCAGGCAACAATGTGAAACTCCA
GAGCACACGTCTGAACTCTTTCCCTACACGACGCTCTTCCGATCTNNNCAGGCAACAATGTGAAACTCCA
GAGCACACGTCTGAACTCTTTCCCTACACGACGCTCTTCCGATCTNNNCAGGCAACAATGTGAAACTCCA
AGAGCACACGTCTGAACTCTTTCCCTACACGACGCTCTTCCGATCTNNNGTTGCAGTGAGCTGTGATC
AGAGCACACGTCTGAACTCTTTCCCTACACGACGCTCTTCCGATCTNNNGTTGCAGTGAGCTGTGATC
AGAGCACACGTCTGAACTCTTTCCCTACACGACGCTCTTCCGATCTNNNGTTGCAGTGAGCTGTGATC
CACGTCTGAACTCTTTCCCTACACGACGCTCTTCCGATCTNNNGAGGAGGAGAGACTGTGAGTGGCAT
CACGTCTGAACTCTTTCCCTACACGACGCTCTTCCGATCTNNNGAGGAGGAGAGACTGTGAGTGGCAT
CACGTCTGAACTCTTTCCCTACACGACGCTCTTCCGATCTNNNGAGGAGGAGAGACTGTGAGTGGCAT
AGCACACGTCTGAACTCTTTCCCTACACGACGCTCTTCCGATCTNNNAGACCCACATACGAGACACAATT
AGCACACGTCTGAACTCTTTCCCTACACGACGCTCTTCCGATCTNNNAGACCCACATACGAGACACAATT
AGCACACGTCTGAACTCTTTCCCTACACGACGCTCTTCCGATCTNNNAGACCCACATACGAGACACAATT
ACACGTCTGAACTCTTTCCCTACACGACGCTCTTCCGATCTNNNAACACATCACCACCTGGACATCCATC
ACACGTCTGAACTCTTTCCCTACACGACGCTCTTCCGATCTNNNAACACATCACCACCTGGACATCCATC
ACACGTCTGAACTCTTTCCCTACACGACGCTCTTCCGATCTNNNAACACATCACCACCTGGACATCCATC
GCACACGTCTGAACTCTTTCCCTACACGACGCTCTTCCGATCTNNNTGTTATGACGATTACACGAGCCA
GCACACGTCTGAACTCTTTCCCTACACGACGCTCTTCCGATCTNNNTGTTATGACGATTACACGAGCCA
GCACACGTCTGAACTCTTTCCCTACACGACGCTCTTCCGATCTNNNTGTTATGACGATTACACGAGCCA
GAGCACACGTCTGAACTCTTTCCCTACACGACGCTCTTCCGATCTNNNGAGGTAACACCACTGCACTCTA
GAGCACACGTCTGAACTCTTTCCCTACACGACGCTCTTCCGATCTNNNGAGGTAACACCACTGCACTCTA
GAGCACACGTCTGAACTCTTTCCCTACACGACGCTCTTCCGATCTNNNGAGGTAACACCACTGCACTCTA
AGCACACGTCTGAACTCTTTCCCTACACGACGCTCTTCCGATCTNNNAGACACCCCAGTTTAGCAGTTA
AGCACACGTCTGAACTCTTTCCCTACACGACGCTCTTCCGATCTNNNAGACACCCCAGTTTAGCAGTTA
AGCACACGTCTGAACTCTTTCCCTACACGACGCTCTTCCGATCTNNNAGACACCCCAGTTTAGCAGTTA
GAGCACACGTCTGAACTCTTTCCCTACACGACGCTCTTCCGATCTNNNGGGAGTCTAGTATGTTGGTGT
GAGCACACGTCTGAACTCTTTCCCTACACGACGCTCTTCCGATCTNNNGGGAGTCTAGTATGTTGGTGT
GAGCACACGTCTGAACTCTTTCCCTACACGACGCTCTTCCGATCTNNNGGGAGTCTAGTATGTTGGTGT
AGCACACGTCTGAACTCTTTCCCTACACGACGCTCTTCCGATCTNNNGGGGAGGATGGTACTGGATCT
AGCACACGTCTGAACTCTTTCCCTACACGACGCTCTTCCGATCTNNNGGGGAGGATGGTACTGGATCT
AGCACACGTCTGAACTCTTTCCCTACACGACGCTCTTCCGATCTNNNGGGGAGGATGGTACTGGATCT
AGCACACGTCTGAACTCTTTCCCTACACGACGCTCTTCCGATCTNNNACTGAAGGTTGGAGCCATGTAA
AGCACACGTCTGAACTCTTTCCCTACACGACGCTCTTCCGATCTNNNACTGAAGGTTGGAGCCATGTAA
AGCACACGTCTGAACTCTTTCCCTACACGACGCTCTTCCGATCTNNNACTGAAGGTTGGAGCCATGTAA
AGCACACGTCTGAACTCTTTCCCTACACGACGCTCTTCCGATCTNNNCTTCCTCTAACACTACCCTCTGCCA
AGCACACGTCTGAACTCTTTCCCTACACGACGCTCTTCCGATCTNNNCTTCCTCTAACACTACCCTCTGCCA
AGCACACGTCTGAACTCTTTCCCTACACGACGCTCTTCCGATCTNNNCTTCCTCTAACACTACCCTCTGCCA
AGAGCACACGTCTGAACTCTTTCCCTACACGACGCTCTTCCGATCTNNNACAATACAGGGGACAAGAG
AGAGCACACGTCTGAACTCTTTCCCTACACGACGCTCTTCCGATCTNNNACAATACAGGGGACAAGAG
AGAGCACACGTCTGAACTCTTTCCCTACACGACGCTCTTCCGATCTNNNACAATACAGGGGACAAGAG
AGCACACGTCTGAACTCTTTCCCTACACGACGCTCTTCCGATCTNNNGAGGCAGGAGAATTGCTTGAA
AGCACACGTCTGAACTCTTTCCCTACACGACGCTCTTCCGATCTNNNGAGGCAGGAGAATTGCTTGAA
AGCACACGTCTGAACTCTTTCCCTACACGACGCTCTTCCGATCTNNNGAGGCAGGAGAATTGCTTGAA
GAGCACACGTCTGAACTCTTTCCCTACACGACGCTCTTCCGATCTNNNAATACCTGAGGTGGAACAGGT

AGCACACGTCTGAACTCTTTCCCTACACGACGCTCTTCCGATCTNNNAATACCTGAGGTGGAACAGG
AGCACACGTCTGAACTCTTTCCCTACACGACGCTCTTCCGATCTNNNAATACCTGAGGTGGAACAGG
AGCACACGTCTGAACTCTTTCCCTACACGACGCTCTTCCGATCTNNNTAATACTAGCACATTGGGAGG
AGCACACGTCTGAACTCTTTCCCTACACGACGCTCTTCCGATCTNNNTAATACTAGCACATTGGGAGG
AGCACACGTCTGAACTCTTTCCCTACACGACGCTCTTCCGATCTNNNTAATACTAGCACATTGGGAGG
AGCACACGTCTGAACTCTTTCCCTACACGACGCTCTTCCGATCTNNNTAATACTAGCACATTGGGAGG
AGCACACGTCTGAACTCTTTCCCTACACGACGCTCTTCCGATCTNNNTAATACTAGCACATTGGGAGG
AGCACACGTCTGAACTCTTTCCCTACACGACGCTCTTCCGATCTNNNTAATACTAGCACATTGGGAGG
GCACACGTCTGAACTCTTTCCCTACACGACGCTCTTCCGATCTNNNATCACAGCAGCTAGTTTCCTCAT
GCACACGTCTGAACTCTTTCCCTACACGACGCTCTTCCGATCTNNNATCACAGCAGCTAGTTTCCTCAT
GCACACGTCTGAACTCTTTCCCTACACGACGCTCTTCCGATCTNNNATCACAGCAGCTAGTTTCCTCAT
GCACACGTCTGAACTCTTTCCCTACACGACGCTCTTCCGATCTNNNATCACAGCAGCTAGTTTCCTCAT
GCACACGTCTGAACTCTTTCCCTACACGACGCTCTTCCGATCTNNNATCACAGCAGCTAGTTTCCTCAT
GCACACGTCTGAACTCTTTCCCTACACGACGCTCTTCCGATCTNNNATCACAGCAGCTAGTTTCCTCAT
GAGCACACGTCTGAACTCTTTCCCTACACGACGCTCTTCCGATCTNNNTCTGAGTAGCAGGCAAGAAG
GAGCACACGTCTGAACTCTTTCCCTACACGACGCTCTTCCGATCTNNNTCTGAGTAGCAGGCAAGAAG
GAGCACACGTCTGAACTCTTTCCCTACACGACGCTCTTCCGATCTNNNTCTGAGTAGCAGGCAAGAAG
GAGCACACGTCTGAACTCTTTCCCTACACGACGCTCTTCCGATCTNNNTCTGAGTAGCAGGCAAGAAG
GAGCACACGTCTGAACTCTTTCCCTACACGACGCTCTTCCGATCTNNNTCTGAGTAGCAGGCAAGAAG
GAGCACACGTCTGAACTCTTTCCCTACACGACGCTCTTCCGATCTNNNTCTGAGTAGCAGGCAAGAAG
AAGAGCACACGTCTGAACTCTTTCCCTACACGACGCTCTTCCGATCTNNNGTACTTGTCCTCCATGTGT
AAGAGCACACGTCTGAACTCTTTCCCTACACGACGCTCTTCCGATCTNNNGTACTTGTCCTCCATGTGT
AAGAGCACACGTCTGAACTCTTTCCCTACACGACGCTCTTCCGATCTNNNGTACTTGTCCTCCATGTGT
AAGAGCACACGTCTGAACTCTTTCCCTACACGACGCTCTTCCGATCTNNNGTACTTGTCCTCCATGTGT
AAGAGCACACGTCTGAACTCTTTCCCTACACGACGCTCTTCCGATCTNNNGTACTTGTCCTCCATGTGT
AAGAGCACACGTCTGAACTCTTTCCCTACACGACGCTCTTCCGATCTNNNGTACTTGTCCTCCATGTGT
AGCACACGTCTGAACTCTTTCCCTACACGACGCTCTTCCGATCTNNNCCAAAACAACATGGTACTGGC
AGCACACGTCTGAACTCTTTCCCTACACGACGCTCTTCCGATCTNNNCCAAAACAACATGGTACTGGC
AGCACACGTCTGAACTCTTTCCCTACACGACGCTCTTCCGATCTNNNCCAAAACAACATGGTACTGGC
AGCACACGTCTGAACTCTTTCCCTACACGACGCTCTTCCGATCTNNNCCAAAACAACATGGTACTGGC
AGCACACGTCTGAACTCTTTCCCTACACGACGCTCTTCCGATCTNNNCCAAAACAACATGGTACTGGC
AGCACACGTCTGAACTCTTTCCCTACACGACGCTCTTCCGATCTNNNCCAAAACAACATGGTACTGGC
GAGCACACGTCTGAACTCTTTCCCTACACGACGCTCTTCCGATCTNNNCTTCTCTCTAACCCCCAATGC
GAGCACACGTCTGAACTCTTTCCCTACACGACGCTCTTCCGATCTNNNCTTCTCTCTAACCCCCAATGC
GAGCACACGTCTGAACTCTTTCCCTACACGACGCTCTTCCGATCTNNNCTTCTCTCTAACCCCCAATGC
GAGCACACGTCTGAACTCTTTCCCTACACGACGCTCTTCCGATCTNNNCTTCTCTCTAACCCCCAATGC
GAGCACACGTCTGAACTCTTTCCCTACACGACGCTCTTCCGATCTNNNCTTCTCTCTAACCCCCAATGC
GAGCACACGTCTGAACTCTTTCCCTACACGACGCTCTTCCGATCTNNNCTTCTCTCTAACCCCCAATGC
GAGCACACGTCTGAACTCTTTCCCTACACGACGCTCTTCCGATCTNNNATGGTTCCACATGAAAGCCA
GAGCACACGTCTGAACTCTTTCCCTACACGACGCTCTTCCGATCTNNNATGGTTCCACATGAAAGCCA
GAGCACACGTCTGAACTCTTTCCCTACACGACGCTCTTCCGATCTNNNATGGTTCCACATGAAAGCCA
GAGCACACGTCTGAACTCTTTCCCTACACGACGCTCTTCCGATCTNNNATGGTTCCACATGAAAGCCA
GAGCACACGTCTGAACTCTTTCCCTACACGACGCTCTTCCGATCTNNNATGGTTCCACATGAAAGCCA

GAGCACACGTCTGAACTCTTTCCCTACACGACGCTCTTCCGATCTNNNATGGTTCCACATGAAAGCCA
AGCACACGTCTGAACTCTTTCCCTACACGACGCTCTTCCGATCTNNNAGCTAGGATGAAAACCCTGTG
AGCACACGTCTGAACTCTTTCCCTACACGACGCTCTTCCGATCTNNNAGCTAGGATGAAAACCCTGTG
AGCACACGTCTGAACTCTTTCCCTACACGACGCTCTTCCGATCTNNNAGCTAGGATGAAAACCCTGTG
AGCACACGTCTGAACTCTTTCCCTACACGACGCTCTTCCGATCTNNNAGCTAGGATGAAAACCCTGTG
AGCACACGTCTGAACTCTTTCCCTACACGACGCTCTTCCGATCTNNNAGCTAGGATGAAAACCCTGTG
AGCACACGTCTGAACTCTTTCCCTACACGACGCTCTTCCGATCTNNNAGCTAGGATGAAAACCCTGTG
AGCACACGTCTGAACTCTTTCCCTACACGACGCTCTTCCGATCTNNNCGACAGAACGAGACTCCGTCC
AGCACACGTCTGAACTCTTTCCCTACACGACGCTCTTCCGATCTNNNCGACAGAACGAGACTCCGTCC
AGCACACGTCTGAACTCTTTCCCTACACGACGCTCTTCCGATCTNNNCGACAGAACGAGACTCCGTCC
AGCACACGTCTGAACTCTTTCCCTACACGACGCTCTTCCGATCTNNNCGACAGAACGAGACTCCGTCC
AGCACACGTCTGAACTCTTTCCCTACACGACGCTCTTCCGATCTNNNCGACAGAACGAGACTCCGTCC
AGCACACGTCTGAACTCTTTCCCTACACGACGCTCTTCCGATCTNNNCGACAGAACGAGACTCCGTCC
AGCACACGTCTGAACTCTTTCCCTACACGACGCTCTTCCGATCTNNNTTTCTTCCCATTTCTCAGCTCGA
AGCACACGTCTGAACTCTTTCCCTACACGACGCTCTTCCGATCTNNNTTTCTTCCCATTTCTCAGCTCGA
AGCACACGTCTGAACTCTTTCCCTACACGACGCTCTTCCGATCTNNNTTTCTTCCCATTTCTCAGCTCGA
AGCACACGTCTGAACTCTTTCCCTACACGACGCTCTTCCGATCTNNNTTTCTTCCCATTTCTCAGCTCGA
AGCACACGTCTGAACTCTTTCCCTACACGACGCTCTTCCGATCTNNNTTTCTTCCCATTTCTCAGCTCGA
AGCACACGTCTGAACTCTTTCCCTACACGACGCTCTTCCGATCTNNNTTTCTTCCCATTTCTCAGCTCGA
GAGCACACGTCTGAACTCTTTCCCTACACGACGCTCTTCCGATCTNNNCAGAAAAGTGACACCCCAAAG
GAGCACACGTCTGAACTCTTTCCCTACACGACGCTCTTCCGATCTNNNCAGAAAAGTGACACCCCAAAG
GAGCACACGTCTGAACTCTTTCCCTACACGACGCTCTTCCGATCTNNNCAGAAAAGTGACACCCCAAAG
GAGCACACGTCTGAACTCTTTCCCTACACGACGCTCTTCCGATCTNNNCAGAAAAGTGACACCCCAAAG
GAGCACACGTCTGAACTCTTTCCCTACACGACGCTCTTCCGATCTNNNCAGAAAAGTGACACCCCAAAG
GAGCACACGTCTGAACTCTTTCCCTACACGACGCTCTTCCGATCTNNNCAGAAAAGTGACACCCCAAAG
GAGCACACGTCTGAACTCTTTCCCTACACGACGCTCTTCCGATCTNNNCTGGAGCTCAGGTGAACTTAT
GAGCACACGTCTGAACTCTTTCCCTACACGACGCTCTTCCGATCTNNNCTGGAGCTCAGGTGAACTTAT
GAGCACACGTCTGAACTCTTTCCCTACACGACGCTCTTCCGATCTNNNCTGGAGCTCAGGTGAACTTAT
GAGCACACGTCTGAACTCTTTCCCTACACGACGCTCTTCCGATCTNNNCTGGAGCTCAGGTGAACTTAT
GAGCACACGTCTGAACTCTTTCCCTACACGACGCTCTTCCGATCTNNNCTGGAGCTCAGGTGAACTTAT
GAGCACACGTCTGAACTCTTTCCCTACACGACGCTCTTCCGATCTNNNCTGGAGCTCAGGTGAACTTAT
AGCACACGTCTGAACTCTTTCCCTACACGACGCTCTTCCGATCTNNNATTACAAAAAATTAGCCGGGC
AGCACACGTCTGAACTCTTTCCCTACACGACGCTCTTCCGATCTNNNATTACAAAAAATTAGCCGGGC
AGCACACGTCTGAACTCTTTCCCTACACGACGCTCTTCCGATCTNNNATTACAAAAAATTAGCCGGGC
AGCACACGTCTGAACTCTTTCCCTACACGACGCTCTTCCGATCTNNNATTACAAAAAATTAGCCGGGC
AGCACACGTCTGAACTCTTTCCCTACACGACGCTCTTCCGATCTNNNATTACAAAAAATTAGCCGGGC
AGCACACGTCTGAACTCTTTCCCTACACGACGCTCTTCCGATCTNNNATTACAAAAAATTAGCCGGGC
GCACACGTCTGAACTCTTTCCCTACACGACGCTCTTCCGATCTNNNCCAGTGGGATATTATCTAACTTT
GCACACGTCTGAACTCTTTCCCTACACGACGCTCTTCCGATCTNNNCCAGTGGGATATTATCTAACTTT
GCACACGTCTGAACTCTTTCCCTACACGACGCTCTTCCGATCTNNNCCAGTGGGATATTATCTAACTTT
GCACACGTCTGAACTCTTTCCCTACACGACGCTCTTCCGATCTNNNCCAGTGGGATATTATCTAACTTT
GCACACGTCTGAACTCTTTCCCTACACGACGCTCTTCCGATCTNNNCCAGTGGGATATTATCTAACTTT
GCACACGTCTGAACTCTTTCCCTACACGACGCTCTTCCGATCTNNNCCAGTGGGATATTATCTAACTTT

AGCACACGTCTGAACTCTTTCCCTACACGACGCTCTTCCGATCTNNNCTTCCACACTCCCAAACTCATCC

AGCACACGTCTGAACTCTTTCCCTACACGACGCTCTTCCGATCTNNNCTTCCACACTCCCAAACTCATCC

AGCACACGTCTGAACTCTTTCCCTACACGACGCTCTTCCGATCTNNNCTTCCACACTCCCAAACTCATCC

AGCACACGTCTGAACTCTTTCCCTACACGACGCTCTTCCGATCTNNNCTTCCACACTCCCAAACTCATCC

AGCACACGTCTGAACTCTTTCCCTACACGACGCTCTTCCGATCTNNNCTTCCACACTCCCAAACTCATCC

AGCACACGTCTGAACTCTTTCCCTACACGACGCTCTTCCGATCTNNNCTTCCACACTCCCAAACTCATCC

GAGCACACGTCTGAACTCTTTCCCTACACGACGCTCTTCCGATCTNNNGATGCGATCAACTGGAAGAA

GAGCACACGTCTGAACTCTTTCCCTACACGACGCTCTTCCGATCTNNNGATGCGATCAACTGGAAGAA

GAGCACACGTCTGAACTCTTTCCCTACACGACGCTCTTCCGATCTNNNGATGCGATCAACTGGAAGAA

GAGCACACGTCTGAACTCTTTCCCTACACGACGCTCTTCCGATCTNNNGATGCGATCAACTGGAAGAA

GAGCACACGTCTGAACTCTTTCCCTACACGACGCTCTTCCGATCTNNNGATGCGATCAACTGGAAGAA

GAGCACACGTCTGAACTCTTTCCCTACACGACGCTCTTCCGATCTNNNGATGCGATCAACTGGAAGAA

AGCACACGTCTGAACTCTTTCCCTACACGACGCTCTTCCGATCTNNNCAGGAAAATCGCTTGAACCCA

AGCACACGTCTGAACTCTTTCCCTACACGACGCTCTTCCGATCTNNNCAGGAAAATCGCTTGAACCCA

AGCACACGTCTGAACTCTTTCCCTACACGACGCTCTTCCGATCTNNNCAGGAAAATCGCTTGAACCCA

AGCACACGTCTGAACTCTTTCCCTACACGACGCTCTTCCGATCTNNNCAGGAAAATCGCTTGAACCCA

AGCACACGTCTGAACTCTTTCCCTACACGACGCTCTTCCGATCTNNNCAGGAAAATCGCTTGAACCCA

AGCACACGTCTGAACTCTTTCCCTACACGACGCTCTTCCGATCTNNNCAGGAAAATCGCTTGAACCCA

GAGCACACGTCTGAACTCTTTCCCTACACGACGCTCTTCCGATCTNNNGAGGTAACAGGAAAAGGGG

GAGCACACGTCTGAACTCTTTCCCTACACGACGCTCTTCCGATCTNNNGAGGTAACAGGAAAAGGGG

GAGCACACGTCTGAACTCTTTCCCTACACGACGCTCTTCCGATCTNNNGAGGTAACAGGAAAAGGGG

GAGCACACGTCTGAACTCTTTCCCTACACGACGCTCTTCCGATCTNNNGAGGTAACAGGAAAAGGGG

GAGCACACGTCTGAACTCTTTCCCTACACGACGCTCTTCCGATCTNNNGAGGTAACAGGAAAAGGGG

GAGCACACGTCTGAACTCTTTCCCTACACGACGCTCTTCCGATCTNNNGAGGTAACAGGAAAAGGGG

AGCACACGTCTGAACTCTTTCCCTACACGACGCTCTTCCGATCTNNNTCTCTAGTGATGTGACTCCCAG

AGCACACGTCTGAACTCTTTCCCTACACGACGCTCTTCCGATCTNNNTCTCTAGTGATGTGACTCCCAG

AGCACACGTCTGAACTCTTTCCCTACACGACGCTCTTCCGATCTNNNTCTCTAGTGATGTGACTCCCAG

AGCACACGTCTGAACTCTTTCCCTACACGACGCTCTTCCGATCTNNNTCTCTAGTGATGTGACTCCCAG

AGCACACGTCTGAACTCTTTCCCTACACGACGCTCTTCCGATCTNNNTCTCTAGTGATGTGACTCCCAG

AGCACACGTCTGAACTCTTTCCCTACACGACGCTCTTCCGATCTNNNTCTCTAGTGATGTGACTCCCAG

GCACACGTCTGAACTCTTTCCCTACACGACGCTCTTCCGATCTNNNTCCATGCTTCTCTCTCTCTCTCA

GCACACGTCTGAACTCTTTCCCTACACGACGCTCTTCCGATCTNNNTCCATGCTTCTCTCTCTCTCTCA

GCACACGTCTGAACTCTTTCCCTACACGACGCTCTTCCGATCTNNNTCCATGCTTCTCTCTCTCTCTCA

GCACACGTCTGAACTCTTTCCCTACACGACGCTCTTCCGATCTNNNTCCATGCTTCTCTCTCTCTCTCA

GCACACGTCTGAACTCTTTCCCTACACGACGCTCTTCCGATCTNNNTCCATGCTTCTCTCTCTCTCTCA

GCACACGTCTGAACTCTTTCCCTACACGACGCTCTTCCGATCTNNNTCCATGCTTCTCTCTCTCTCTCA

AGCACACGTCTGAACTCTTTCCCTACACGACGCTCTTCCGATCTNNNGTGGCAAGGGATGTTTGTTAC

AGCACACGTCTGAACTCTTTCCCTACACGACGCTCTTCCGATCTNNNGTGGCAAGGGATGTTTGTTAC

AGCACACGTCTGAACTCTTTCCCTACACGACGCTCTTCCGATCTNNNGTGGCAAGGGATGTTTGTTAC

AGCACACGTCTGAACTCTTTCCCTACACGACGCTCTTCCGATCTNNNGTGGCAAGGGATGTTTGTTAC

AGCACACGTCTGAACTCTTTCCCTACACGACGCTCTTCCGATCTNNNGTGGCAAGGGATGTTTGTTAC

AGCACACGTCTGAACTCTTTCCCTACACGACGCTCTTCCGATCTNNNGTGGCAAGGGATGTTTGTTAC

AGCACACGTCTGAACTCTTTCCCTACACGACGCTCTTCCGATCTNNNCTGTAGTCCCAGCTACTCAGGA

AGCACACGTCTGAACTCTTTCCCTACACGACGCTCTTCCGATCTNNNCTGTAGTCCCAGCTACTCAGGA
AGCACACGTCTGAACTCTTTCCCTACACGACGCTCTTCCGATCTNNNCTGTAGTCCCAGCTACTCAGGA
AGCACACGTCTGAACTCTTTCCCTACACGACGCTCTTCCGATCTNNNCTGTAGTCCCAGCTACTCAGGA
AGCACACGTCTGAACTCTTTCCCTACACGACGCTCTTCCGATCTNNNCTGTAGTCCCAGCTACTCAGGA
AGCACACGTCTGAACTCTTTCCCTACACGACGCTCTTCCGATCTNNNCTGTAGTCCCAGCTACTCAGGA
AGCACACGTCTGAACTCTTTCCCTACACGACGCTCTTCCGATCTNNNTCGAGAAGCTGAGTGAACATCA
AGCACACGTCTGAACTCTTTCCCTACACGACGCTCTTCCGATCTNNNTCGAGAAGCTGAGTGAACATCA
AGCACACGTCTGAACTCTTTCCCTACACGACGCTCTTCCGATCTNNNTCGAGAAGCTGAGTGAACATCA
AGCACACGTCTGAACTCTTTCCCTACACGACGCTCTTCCGATCTNNNTCGAGAAGCTGAGTGAACATCA
AGCACACGTCTGAACTCTTTCCCTACACGACGCTCTTCCGATCTNNNTCGAGAAGCTGAGTGAACATCA
AGCACACGTCTGAACTCTTTCCCTACACGACGCTCTTCCGATCTNNNTCGAGAAGCTGAGTGAACATCA
GAGCACACGTCTGAACTCTTTCCCTACACGACGCTCTTCCGATCTNNNTAGGAATAGCCAGGCCAGAA
GAGCACACGTCTGAACTCTTTCCCTACACGACGCTCTTCCGATCTNNNTAGGAATAGCCAGGCCAGAA
GAGCACACGTCTGAACTCTTTCCCTACACGACGCTCTTCCGATCTNNNTAGGAATAGCCAGGCCAGAA
GAGCACACGTCTGAACTCTTTCCCTACACGACGCTCTTCCGATCTNNNTAGGAATAGCCAGGCCAGAA
GAGCACACGTCTGAACTCTTTCCCTACACGACGCTCTTCCGATCTNNNTAGGAATAGCCAGGCCAGAA
GAGCACACGTCTGAACTCTTTCCCTACACGACGCTCTTCCGATCTNNNTAGGAATAGCCAGGCCAGAA
AGCACACGTCTGAACTCTTTCCCTACACGACGCTCTTCCGATCTNNNCCATGCCCGGCTAATTTTTCTTT
AGCACACGTCTGAACTCTTTCCCTACACGACGCTCTTCCGATCTNNNCCATGCCCGGCTAATTTTTCTTT
AGCACACGTCTGAACTCTTTCCCTACACGACGCTCTTCCGATCTNNNCCATGCCCGGCTAATTTTTCTTT
AGCACACGTCTGAACTCTTTCCCTACACGACGCTCTTCCGATCTNNNCCATGCCCGGCTAATTTTTCTTT
AGCACACGTCTGAACTCTTTCCCTACACGACGCTCTTCCGATCTNNNCCATGCCCGGCTAATTTTTCTTT
AGCACACGTCTGAACTCTTTCCCTACACGACGCTCTTCCGATCTNNNCCATGCCCGGCTAATTTTTCTTT
AGCACACGTCTGAACTCTTTCCCTACACGACGCTCTTCCGATCTNNNCTCTCTTCCTCTCTCTCTCCCCA
AGCACACGTCTGAACTCTTTCCCTACACGACGCTCTTCCGATCTNNNCTCTCTTCCTCTCTCTCTCCCCA
AGCACACGTCTGAACTCTTTCCCTACACGACGCTCTTCCGATCTNNNCTCTCTTCCTCTCTCTCTCCCCA
AGCACACGTCTGAACTCTTTCCCTACACGACGCTCTTCCGATCTNNNCTCTCTTCCTCTCTCTCTCCCCA
AGCACACGTCTGAACTCTTTCCCTACACGACGCTCTTCCGATCTNNNCTCTCTTCCTCTCTCTCTCCCCA
AGCACACGTCTGAACTCTTTCCCTACACGACGCTCTTCCGATCTNNNCTCTCTTCCTCTCTCTCTCCCCA
AGCACACGTCTGAACTCTTTCCCTACACGACGCTCTTCCGATCTNNNGAGAGAGAGACAGACCCAGTA
AGCACACGTCTGAACTCTTTCCCTACACGACGCTCTTCCGATCTNNNGAGAGAGAGACAGACCCAGTA
AGCACACGTCTGAACTCTTTCCCTACACGACGCTCTTCCGATCTNNNGAGAGAGAGACAGACCCAGTA
AGCACACGTCTGAACTCTTTCCCTACACGACGCTCTTCCGATCTNNNGAGAGAGAGACAGACCCAGTA
AGCACACGTCTGAACTCTTTCCCTACACGACGCTCTTCCGATCTNNNGAGAGAGAGACAGACCCAGTA
AGCACACGTCTGAACTCTTTCCCTACACGACGCTCTTCCGATCTNNNGAGAGAGAGACAGACCCAGTA
GAGCACACGTCTGAACTCTTTCCCTACACGACGCTCTTCCGATCTNNNAGAACATTCTTGTCACCCCAGA
GAGCACACGTCTGAACTCTTTCCCTACACGACGCTCTTCCGATCTNNNAGAACATTCTTGTCACCCCAGA
GAGCACACGTCTGAACTCTTTCCCTACACGACGCTCTTCCGATCTNNNAGAACATTCTTGTCACCCCAGA
GAGCACACGTCTGAACTCTTTCCCTACACGACGCTCTTCCGATCTNNNAGAACATTCTTGTCACCCCAGA
GAGCACACGTCTGAACTCTTTCCCTACACGACGCTCTTCCGATCTNNNAGAACATTCTTGTCACCCCAGA
GAGCACACGTCTGAACTCTTTCCCTACACGACGCTCTTCCGATCTNNNAGAACATTCTTGTCACCCCAGA
GCACACGTCTGAACTCTTTCCCTACACGACGCTCTTCCGATCTNNNGAAGTGTCCACCCTACTCCTCTTC
GCACACGTCTGAACTCTTTCCCTACACGACGCTCTTCCGATCTNNNGAAGTGTCCACCCTACTCCTCTTC

GCACACGTCTGAACTCTTTCCCTACACGACGCTCTTCCGATCTNNNGAAGTGTCCACCCTACTCCTCTTC

GCACACGTCTGAACTCTTTCCCTACACGACGCTCTTCCGATCTNNNGAAGTGTCCACCCTACTCCTCTTC

GCACACGTCTGAACTCTTTCCCTACACGACGCTCTTCCGATCTNNNGAAGTGTCCACCCTACTCCTCTTC

GCACACGTCTGAACTCTTTCCCTACACGACGCTCTTCCGATCTNNNGAAGTGTCCACCCTACTCCTCTTC

GAGCACACGTCTGAACTCTTTCCCTACACGACGCTCTTCCGATCTNNNGATTCTGGACAGACAGACAGA

GAGCACACGTCTGAACTCTTTCCCTACACGACGCTCTTCCGATCTNNNGATTCTGGACAGACAGACAGA

GAGCACACGTCTGAACTCTTTCCCTACACGACGCTCTTCCGATCTNNNGATTCTGGACAGACAGACAGA

GAGCACACGTCTGAACTCTTTCCCTACACGACGCTCTTCCGATCTNNNGATTCTGGACAGACAGACAGA

GAGCACACGTCTGAACTCTTTCCCTACACGACGCTCTTCCGATCTNNNGATTCTGGACAGACAGACAGA

GAGCACACGTCTGAACTCTTTCCCTACACGACGCTCTTCCGATCTNNNGATTCTGGACAGACAGACAGA

GCACACGTCTGAACTCTTTCCCTACACGACGCTCTTCCGATCTNNNGACACCTCCCCTCTGATCTCATTCA

GCACACGTCTGAACTCTTTCCCTACACGACGCTCTTCCGATCTNNNGACACCTCCCCTCTGATCTCATTCA

GCACACGTCTGAACTCTTTCCCTACACGACGCTCTTCCGATCTNNNGACACCTCCCCTCTGATCTCATTCA

GCACACGTCTGAACTCTTTCCCTACACGACGCTCTTCCGATCTNNNGACACCTCCCCTCTGATCTCATTCA

GCACACGTCTGAACTCTTTCCCTACACGACGCTCTTCCGATCTNNNGACACCTCCCCTCTGATCTCATTCA

GCACACGTCTGAACTCTTTCCCTACACGACGCTCTTCCGATCTNNNGACACCTCCCCTCTGATCTCATTCA

GAGCACACGTCTGAACTCTTTCCCTACACGACGCTCTTCCGATCTNNNCAGCAATGTAGCATATCCTGT

GAGCACACGTCTGAACTCTTTCCCTACACGACGCTCTTCCGATCTNNNCAGCAATGTAGCATATCCTGT

GAGCACACGTCTGAACTCTTTCCCTACACGACGCTCTTCCGATCTNNNCAGCAATGTAGCATATCCTGT

GAGCACACGTCTGAACTCTTTCCCTACACGACGCTCTTCCGATCTNNNCAGCAATGTAGCATATCCTGT

GAGCACACGTCTGAACTCTTTCCCTACACGACGCTCTTCCGATCTNNNCAGCAATGTAGCATATCCTGT

GAGCACACGTCTGAACTCTTTCCCTACACGACGCTCTTCCGATCTNNNCAGCAATGTAGCATATCCTGT

AGCACACGTCTGAACTCTTTCCCTACACGACGCTCTTCCGATCTNNNAAAGCTGGAATTCAGAGCGCA

AGCACACGTCTGAACTCTTTCCCTACACGACGCTCTTCCGATCTNNNAAAGCTGGAATTCAGAGCGCA

AGCACACGTCTGAACTCTTTCCCTACACGACGCTCTTCCGATCTNNNAAAGCTGGAATTCAGAGCGCA

AGCACACGTCTGAACTCTTTCCCTACACGACGCTCTTCCGATCTNNNAAAGCTGGAATTCAGAGCGCA

AGCACACGTCTGAACTCTTTCCCTACACGACGCTCTTCCGATCTNNNAAAGCTGGAATTCAGAGCGCA

AGCACACGTCTGAACTCTTTCCCTACACGACGCTCTTCCGATCTNNNAAAGCTGGAATTCAGAGCGCA

AGAGCACACGTCTGAACTCTTTCCCTACACGACGCTCTTCCGATCTNNNGCCTAATCCCAAGTGTCCTTT

AGAGCACACGTCTGAACTCTTTCCCTACACGACGCTCTTCCGATCTNNNGCCTAATCCCAAGTGTCCTTT

AGAGCACACGTCTGAACTCTTTCCCTACACGACGCTCTTCCGATCTNNNGCCTAATCCCAAGTGTCCTTT

AGAGCACACGTCTGAACTCTTTCCCTACACGACGCTCTTCCGATCTNNNGCCTAATCCCAAGTGTCCTTT

AGAGCACACGTCTGAACTCTTTCCCTACACGACGCTCTTCCGATCTNNNGCCTAATCCCAAGTGTCCTTT

AGAGCACACGTCTGAACTCTTTCCCTACACGACGCTCTTCCGATCTNNNGCCTAATCCCAAGTGTCCTTT

AGCACACGTCTGAACTCTTTCCCTACACGACGCTCTTCCGATCTNNNCATACATATAGGGCCCCTGCTCT

AGCACACGTCTGAACTCTTTCCCTACACGACGCTCTTCCGATCTNNNCATACATATAGGGCCCCTGCTCT

AGCACACGTCTGAACTCTTTCCCTACACGACGCTCTTCCGATCTNNNCATACATATAGGGCCCCTGCTCT

AGCACACGTCTGAACTCTTTCCCTACACGACGCTCTTCCGATCTNNNCATACATATAGGGCCCCTGCTCT

AGCACACGTCTGAACTCTTTCCCTACACGACGCTCTTCCGATCTNNNCATACATATAGGGCCCCTGCTCT

AGCACACGTCTGAACTCTTTCCCTACACGACGCTCTTCCGATCTNNNCATACATATAGGGCCCCTGCTCT

CACACGTCTGAACTCTTTCCCTACACGACGCTCTTCCGATCTNNNGCCTCATGGACCCACAAGATAAACA

CACACGTCTGAACTCTTTCCCTACACGACGCTCTTCCGATCTNNNGCCTCATGGACCCACAAGATAAACA

CACACGTCTGAACTCTTTCCCTACACGACGCTCTTCCGATCTNNNGCCTCATGGACCCACAAGATAAACA

CACACGTCTGAACTCTTTCCCTACACGACGCTCTTCCGATCTNNNGCCTCATGGACCCACAAGATAAACA
CACACGTCTGAACTCTTTCCCTACACGACGCTCTTCCGATCTNNNGCCTCATGGACCCACAAGATAAACA
CACACGTCTGAACTCTTTCCCTACACGACGCTCTTCCGATCTNNNGCCTCATGGACCCACAAGATAAACA
CACACGTCTGAACTCTTTCCCTACACGACGCTCTTCCGATCTNNNAATCCCCATATCCCACTATCCTCAT
CACACGTCTGAACTCTTTCCCTACACGACGCTCTTCCGATCTNNNAATCCCCATATCCCACTATCCTCAT
CACACGTCTGAACTCTTTCCCTACACGACGCTCTTCCGATCTNNNAATCCCCATATCCCACTATCCTCAT
CACACGTCTGAACTCTTTCCCTACACGACGCTCTTCCGATCTNNNAATCCCCATATCCCACTATCCTCAT
CACACGTCTGAACTCTTTCCCTACACGACGCTCTTCCGATCTNNNAATCCCCATATCCCACTATCCTCAT
CACACGTCTGAACTCTTTCCCTACACGACGCTCTTCCGATCTNNNAATCCCCATATCCCACTATCCTCAT
AGAGCACACGTCTGAACTCTTTCCCTACACGACGCTCTTCCGATCTNNNGCAACATGATAAAGCCCCAT
AGAGCACACGTCTGAACTCTTTCCCTACACGACGCTCTTCCGATCTNNNGCAACATGATAAAGCCCCAT
AGAGCACACGTCTGAACTCTTTCCCTACACGACGCTCTTCCGATCTNNNGCAACATGATAAAGCCCCAT
AGAGCACACGTCTGAACTCTTTCCCTACACGACGCTCTTCCGATCTNNNGCAACATGATAAAGCCCCAT
AGAGCACACGTCTGAACTCTTTCCCTACACGACGCTCTTCCGATCTNNNGCAACATGATAAAGCCCCAT
AGAGCACACGTCTGAACTCTTTCCCTACACGACGCTCTTCCGATCTNNNGCAACATGATAAAGCCCCAT
AGCACACGTCTGAACTCTTTCCCTACACGACGCTCTTCCGATCTNNNGTGTGTGCTGGATTTCCACATG
AGCACACGTCTGAACTCTTTCCCTACACGACGCTCTTCCGATCTNNNGTGTGTGCTGGATTTCCACATG
AGCACACGTCTGAACTCTTTCCCTACACGACGCTCTTCCGATCTNNNGTGTGTGCTGGATTTCCACATG
AGCACACGTCTGAACTCTTTCCCTACACGACGCTCTTCCGATCTNNNGTGTGTGCTGGATTTCCACATG
AGCACACGTCTGAACTCTTTCCCTACACGACGCTCTTCCGATCTNNNGTGTGTGCTGGATTTCCACATG
AGCACACGTCTGAACTCTTTCCCTACACGACGCTCTTCCGATCTNNNGTGTGTGCTGGATTTCCACATG
GAGCACACGTCTGAACTCTTTCCCTACACGACGCTCTTCCGATCTNNNGAGAATCCCTTGAACCCAGGA
GAGCACACGTCTGAACTCTTTCCCTACACGACGCTCTTCCGATCTNNNGAGAATCCCTTGAACCCAGGA
GAGCACACGTCTGAACTCTTTCCCTACACGACGCTCTTCCGATCTNNNGAGAATCCCTTGAACCCAGGA
GAGCACACGTCTGAACTCTTTCCCTACACGACGCTCTTCCGATCTNNNGAGAATCCCTTGAACCCAGGA
GAGCACACGTCTGAACTCTTTCCCTACACGACGCTCTTCCGATCTNNNGAGAATCCCTTGAACCCAGGA
GAGCACACGTCTGAACTCTTTCCCTACACGACGCTCTTCCGATCTNNNGAGAATCCCTTGAACCCAGGA
GAGCACACGTCTGAACTCTTTCCCTACACGACGCTCTTCCGATCTNNNTGAGGTCAGATAAGGGTGAC
GAGCACACGTCTGAACTCTTTCCCTACACGACGCTCTTCCGATCTNNNTGAGGTCAGATAAGGGTGAC
GAGCACACGTCTGAACTCTTTCCCTACACGACGCTCTTCCGATCTNNNTGAGGTCAGATAAGGGTGAC
GAGCACACGTCTGAACTCTTTCCCTACACGACGCTCTTCCGATCTNNNTGAGGTCAGATAAGGGTGAC
GAGCACACGTCTGAACTCTTTCCCTACACGACGCTCTTCCGATCTNNNTGAGGTCAGATAAGGGTGAC
GAGCACACGTCTGAACTCTTTCCCTACACGACGCTCTTCCGATCTNNNTGAGGTCAGATAAGGGTGAC
AGCACACGTCTGAACTCTTTCCCTACACGACGCTCTTCCGATCTNNNATTTCCTTGGGAGGCCCTTTATT
AGCACACGTCTGAACTCTTTCCCTACACGACGCTCTTCCGATCTNNNATTTCCTTGGGAGGCCCTTTATT
AGCACACGTCTGAACTCTTTCCCTACACGACGCTCTTCCGATCTNNNATTTCCTTGGGAGGCCCTTTATT
AGCACACGTCTGAACTCTTTCCCTACACGACGCTCTTCCGATCTNNNATTTCCTTGGGAGGCCCTTTATT
AGCACACGTCTGAACTCTTTCCCTACACGACGCTCTTCCGATCTNNNATTTCCTTGGGAGGCCCTTTATT
AGCACACGTCTGAACTCTTTCCCTACACGACGCTCTTCCGATCTNNNATTTCCTTGGGAGGCCCTTTATT
GAGCACACGTCTGAACTCTTTCCCTACACGACGCTCTTCCGATCTNNNCCTGGGCAACAAGAGTGAAA
GAGCACACGTCTGAACTCTTTCCCTACACGACGCTCTTCCGATCTNNNCCTGGGCAACAAGAGTGAAA
GAGCACACGTCTGAACTCTTTCCCTACACGACGCTCTTCCGATCTNNNCCTGGGCAACAAGAGTGAAA
GAGCACACGTCTGAACTCTTTCCCTACACGACGCTCTTCCGATCTNNNCCTGGGCAACAAGAGTGAAA

GAGCACACGTCTGAACTCTTTCCCTACACGACGCTCTTCCGATCTNNNCCTGGGCAACAAGAGTGAAA
GAGCACACGTCTGAACTCTTTCCCTACACGACGCTCTTCCGATCTNNNCCTGGGCAACAAGAGTGAAA
GAGCACACGTCTGAACTCTTTCCCTACACGACGCTCTTCCGATCTNNNCTGTTCCATTGTATCACCCCAG
GAGCACACGTCTGAACTCTTTCCCTACACGACGCTCTTCCGATCTNNNCTGTTCCATTGTATCACCCCAG
GAGCACACGTCTGAACTCTTTCCCTACACGACGCTCTTCCGATCTNNNCTGTTCCATTGTATCACCCCAG
GAGCACACGTCTGAACTCTTTCCCTACACGACGCTCTTCCGATCTNNNCTGTTCCATTGTATCACCCCAG
GAGCACACGTCTGAACTCTTTCCCTACACGACGCTCTTCCGATCTNNNCTGTTCCATTGTATCACCCCAG
GAGCACACGTCTGAACTCTTTCCCTACACGACGCTCTTCCGATCTNNNCTGTTCCATTGTATCACCCCAG
GCACACGTCTGAACTCTTTCCCTACACGACGCTCTTCCGATCTNNNAGAAAAGGCAAAAGAATTGTTCC
GCACACGTCTGAACTCTTTCCCTACACGACGCTCTTCCGATCTNNNAGAAAAGGCAAAAGAATTGTTCC
GCACACGTCTGAACTCTTTCCCTACACGACGCTCTTCCGATCTNNNAGAAAAGGCAAAAGAATTGTTCC
GCACACGTCTGAACTCTTTCCCTACACGACGCTCTTCCGATCTNNNAGAAAAGGCAAAAGAATTGTTCC
GCACACGTCTGAACTCTTTCCCTACACGACGCTCTTCCGATCTNNNAGAAAAGGCAAAAGAATTGTTCC
GCACACGTCTGAACTCTTTCCCTACACGACGCTCTTCCGATCTNNNAGAAAAGGCAAAAGAATTGTTCC
GAGCACACGTCTGAACTCTTTCCCTACACGACGCTCTTCCGATCTNNNAAGGCACAGGCAACAAAAGA
GAGCACACGTCTGAACTCTTTCCCTACACGACGCTCTTCCGATCTNNNAAGGCACAGGCAACAAAAGA
GAGCACACGTCTGAACTCTTTCCCTACACGACGCTCTTCCGATCTNNNAAGGCACAGGCAACAAAAGA
GAGCACACGTCTGAACTCTTTCCCTACACGACGCTCTTCCGATCTNNNAAGGCACAGGCAACAAAAGA
GAGCACACGTCTGAACTCTTTCCCTACACGACGCTCTTCCGATCTNNNAAGGCACAGGCAACAAAAGA
GAGCACACGTCTGAACTCTTTCCCTACACGACGCTCTTCCGATCTNNNAAGGCACAGGCAACAAAAGA
AGCACACGTCTGAACTCTTTCCCTACACGACGCTCTTCCGATCTNNNGAGGGAGAGGGAGAGCTCTTC
AGCACACGTCTGAACTCTTTCCCTACACGACGCTCTTCCGATCTNNNGAGGGAGAGGGAGAGCTCTTC
AGCACACGTCTGAACTCTTTCCCTACACGACGCTCTTCCGATCTNNNGAGGGAGAGGGAGAGCTCTTC
AGCACACGTCTGAACTCTTTCCCTACACGACGCTCTTCCGATCTNNNGAGGGAGAGGGAGAGCTCTTC
AGCACACGTCTGAACTCTTTCCCTACACGACGCTCTTCCGATCTNNNGAGGGAGAGGGAGAGCTCTTC
AGCACACGTCTGAACTCTTTCCCTACACGACGCTCTTCCGATCTNNNGAGGGAGAGGGAGAGCTCTTC
GAGCACACGTCTGAACTCTTTCCCTACACGACGCTCTTCCGATCTNNNTGAGAATCACTTGAACCCAGG
GAGCACACGTCTGAACTCTTTCCCTACACGACGCTCTTCCGATCTNNNTGAGAATCACTTGAACCCAGG
GAGCACACGTCTGAACTCTTTCCCTACACGACGCTCTTCCGATCTNNNTGAGAATCACTTGAACCCAGG
GAGCACACGTCTGAACTCTTTCCCTACACGACGCTCTTCCGATCTNNNTGAGAATCACTTGAACCCAGG
GAGCACACGTCTGAACTCTTTCCCTACACGACGCTCTTCCGATCTNNNTGAGAATCACTTGAACCCAGG
GAGCACACGTCTGAACTCTTTCCCTACACGACGCTCTTCCGATCTNNNTGAGAATCACTTGAACCCAGG
GAGCACACGTCTGAACTCTTTCCCTACACGACGCTCTTCCGATCTNNNATTAGGGGGTCTGTGCCTACT
GAGCACACGTCTGAACTCTTTCCCTACACGACGCTCTTCCGATCTNNNATTAGGGGGTCTGTGCCTACT
GAGCACACGTCTGAACTCTTTCCCTACACGACGCTCTTCCGATCTNNNATTAGGGGGTCTGTGCCTACT
GAGCACACGTCTGAACTCTTTCCCTACACGACGCTCTTCCGATCTNNNATTAGGGGGTCTGTGCCTACT
GAGCACACGTCTGAACTCTTTCCCTACACGACGCTCTTCCGATCTNNNATTAGGGGGTCTGTGCCTACT
GAGCACACGTCTGAACTCTTTCCCTACACGACGCTCTTCCGATCTNNNATTAGGGGGTCTGTGCCTACT
GAGCACACGTCTGAACTCTTTCCCTACACGACGCTCTTCCGATCTNNNAGAGGATTCTGACTGTTTCAC
GAGCACACGTCTGAACTCTTTCCCTACACGACGCTCTTCCGATCTNNNAGAGGATTCTGACTGTTTCAC
GAGCACACGTCTGAACTCTTTCCCTACACGACGCTCTTCCGATCTNNNAGAGGATTCTGACTGTTTCAC
GAGCACACGTCTGAACTCTTTCCCTACACGACGCTCTTCCGATCTNNNAGAGGATTCTGACTGTTTCAC
GAGCACACGTCTGAACTCTTTCCCTACACGACGCTCTTCCGATCTNNNAGAGGATTCTGACTGTTTCAC

AGCACACGTCTGAACTCTTTCCCTACACGACGCTCTTCCGATCTNNNAGAGGATTCTGACTGTTTCAC
GCACACGTCTGAACTCTTTCCCTACACGACGCTCTTCCGATCTNNNACTGTGCTTTCCCATTACATTGC
GCACACGTCTGAACTCTTTCCCTACACGACGCTCTTCCGATCTNNNACTGTGCTTTCCCATTACATTGC
GCACACGTCTGAACTCTTTCCCTACACGACGCTCTTCCGATCTNNNACTGTGCTTTCCCATTACATTGC
GCACACGTCTGAACTCTTTCCCTACACGACGCTCTTCCGATCTNNNACTGTGCTTTCCCATTACATTGC
GCACACGTCTGAACTCTTTCCCTACACGACGCTCTTCCGATCTNNNACTGTGCTTTCCCATTACATTGC
GCACACGTCTGAACTCTTTCCCTACACGACGCTCTTCCGATCTNNNACTGTGCTTTCCCATTACATTGC
AGCACACGTCTGAACTCTTTCCCTACACGACGCTCTTCCGATCTNNNAGGAGCTGGTACCATTTCTTCT
AGCACACGTCTGAACTCTTTCCCTACACGACGCTCTTCCGATCTNNNAGGAGCTGGTACCATTTCTTCT
AGCACACGTCTGAACTCTTTCCCTACACGACGCTCTTCCGATCTNNNAGGAGCTGGTACCATTTCTTCT
AGCACACGTCTGAACTCTTTCCCTACACGACGCTCTTCCGATCTNNNAGGAGCTGGTACCATTTCTTCT
AGCACACGTCTGAACTCTTTCCCTACACGACGCTCTTCCGATCTNNNAGGAGCTGGTACCATTTCTTCT
AGCACACGTCTGAACTCTTTCCCTACACGACGCTCTTCCGATCTNNNAGGAGCTGGTACCATTTCTTCT
AGCACACGTCTGAACTCTTTCCCTACACGACGCTCTTCCGATCTNNNGGTCTTTGGTGGAAAAGGGA
AGCACACGTCTGAACTCTTTCCCTACACGACGCTCTTCCGATCTNNNGGTCTTTGGTGGAAAAGGGA
AGCACACGTCTGAACTCTTTCCCTACACGACGCTCTTCCGATCTNNNGGTCTTTGGTGGAAAAGGGA
AGCACACGTCTGAACTCTTTCCCTACACGACGCTCTTCCGATCTNNNGGTCTTTGGTGGAAAAGGGA
AGCACACGTCTGAACTCTTTCCCTACACGACGCTCTTCCGATCTNNNGGTCTTTGGTGGAAAAGGGA
AGCACACGTCTGAACTCTTTCCCTACACGACGCTCTTCCGATCTNNNGGTCTTTGGTGGAAAAGGGA
AGCACACGTCTGAACTCTTTCCCTACACGACGCTCTTCCGATCTNNNGAAAATAGACTTGGCTGCTTG
AGCACACGTCTGAACTCTTTCCCTACACGACGCTCTTCCGATCTNNNGAAAATAGACTTGGCTGCTTG
AGCACACGTCTGAACTCTTTCCCTACACGACGCTCTTCCGATCTNNNGAAAATAGACTTGGCTGCTTG
AGCACACGTCTGAACTCTTTCCCTACACGACGCTCTTCCGATCTNNNGAAAATAGACTTGGCTGCTTG
AGCACACGTCTGAACTCTTTCCCTACACGACGCTCTTCCGATCTNNNGAAAATAGACTTGGCTGCTTG
AGCACACGTCTGAACTCTTTCCCTACACGACGCTCTTCCGATCTNNNGAAAATAGACTTGGCTGCTTG
GAGCACACGTCTGAACTCTTTCCCTACACGACGCTCTTCCGATCTNNNCGCCCGGCTAATTATTTTGTG
GAGCACACGTCTGAACTCTTTCCCTACACGACGCTCTTCCGATCTNNNCGCCCGGCTAATTATTTTGTG
GAGCACACGTCTGAACTCTTTCCCTACACGACGCTCTTCCGATCTNNNCGCCCGGCTAATTATTTTGTG
GAGCACACGTCTGAACTCTTTCCCTACACGACGCTCTTCCGATCTNNNCGCCCGGCTAATTATTTTGTG
GAGCACACGTCTGAACTCTTTCCCTACACGACGCTCTTCCGATCTNNNCGCCCGGCTAATTATTTTGTG
GAGCACACGTCTGAACTCTTTCCCTACACGACGCTCTTCCGATCTNNNCGCCCGGCTAATTATTTTGTG
CACACGTCTGAACTCTTTCCCTACACGACGCTCTTCCGATCTNNNCATCCTGGCTAACACAGTGAAACC
CACACGTCTGAACTCTTTCCCTACACGACGCTCTTCCGATCTNNNCATCCTGGCTAACACAGTGAAACC
CACACGTCTGAACTCTTTCCCTACACGACGCTCTTCCGATCTNNNCATCCTGGCTAACACAGTGAAACC
CACACGTCTGAACTCTTTCCCTACACGACGCTCTTCCGATCTNNNCATCCTGGCTAACACAGTGAAACC
CACACGTCTGAACTCTTTCCCTACACGACGCTCTTCCGATCTNNNCATCCTGGCTAACACAGTGAAACC
CACACGTCTGAACTCTTTCCCTACACGACGCTCTTCCGATCTNNNCATCCTGGCTAACACAGTGAAACC
AGCACACGTCTGAACTCTTTCCCTACACGACGCTCTTCCGATCTNNNTCAAAAGGTAGAGGAGGAG
AGCACACGTCTGAACTCTTTCCCTACACGACGCTCTTCCGATCTNNNTCAAAAGGTAGAGGAGGAG
AGCACACGTCTGAACTCTTTCCCTACACGACGCTCTTCCGATCTNNNTCAAAAGGTAGAGGAGGAG
AGCACACGTCTGAACTCTTTCCCTACACGACGCTCTTCCGATCTNNNTCAAAAGGTAGAGGAGGAG
AGCACACGTCTGAACTCTTTCCCTACACGACGCTCTTCCGATCTNNNTCAAAAGGTAGAGGAGGAG
AGCACACGTCTGAACTCTTTCCCTACACGACGCTCTTCCGATCTNNNTCAAAAGGTAGAGGAGGAG

GCACACGTCTGAACTCTTTCCCTACACGACGCTCTTCCGATCTNNNCTGTCACTGAGGGGATTGGCCA
GCACACGTCTGAACTCTTTCCCTACACGACGCTCTTCCGATCTNNNCTGTCACTGAGGGGATTGGCCA
GCACACGTCTGAACTCTTTCCCTACACGACGCTCTTCCGATCTNNNCTGTCACTGAGGGGATTGGCCA
GCACACGTCTGAACTCTTTCCCTACACGACGCTCTTCCGATCTNNNCTGTCACTGAGGGGATTGGCCA
GCACACGTCTGAACTCTTTCCCTACACGACGCTCTTCCGATCTNNNCTGTCACTGAGGGGATTGGCCA
GCACACGTCTGAACTCTTTCCCTACACGACGCTCTTCCGATCTNNNCTGTCACTGAGGGGATTGGCCA
GAGCACACGTCTGAACTCTTTCCCTACACGACGCTCTTCCGATCTNNNGAGGCTGAGGCAGGAAAAT
GAGCACACGTCTGAACTCTTTCCCTACACGACGCTCTTCCGATCTNNNGAGGCTGAGGCAGGAAAAT
GAGCACACGTCTGAACTCTTTCCCTACACGACGCTCTTCCGATCTNNNGAGGCTGAGGCAGGAAAAT
GAGCACACGTCTGAACTCTTTCCCTACACGACGCTCTTCCGATCTNNNGAGGCTGAGGCAGGAAAAT
GAGCACACGTCTGAACTCTTTCCCTACACGACGCTCTTCCGATCTNNNGAGGCTGAGGCAGGAAAAT
GAGCACACGTCTGAACTCTTTCCCTACACGACGCTCTTCCGATCTNNNGAGGCTGAGGCAGGAAAAT
GAGCACACGTCTGAACTCTTTCCCTACACGACGCTCTTCCGATCTNNNTGGAGAGATTGGGTGTTTGT
GAGCACACGTCTGAACTCTTTCCCTACACGACGCTCTTCCGATCTNNNTGGAGAGATTGGGTGTTTGT
GAGCACACGTCTGAACTCTTTCCCTACACGACGCTCTTCCGATCTNNNTGGAGAGATTGGGTGTTTGT
GAGCACACGTCTGAACTCTTTCCCTACACGACGCTCTTCCGATCTNNNTGGAGAGATTGGGTGTTTGT
GAGCACACGTCTGAACTCTTTCCCTACACGACGCTCTTCCGATCTNNNTGGAGAGATTGGGTGTTTGT
GAGCACACGTCTGAACTCTTTCCCTACACGACGCTCTTCCGATCTNNNTGGAGAGATTGGGTGTTTGT
GCACACGTCTGAACTCTTTCCCTACACGACGCTCTTCCGATCTNNNAGACACAGCGCCCTTTCTTCCAT
GCACACGTCTGAACTCTTTCCCTACACGACGCTCTTCCGATCTNNNAGACACAGCGCCCTTTCTTCCAT
GCACACGTCTGAACTCTTTCCCTACACGACGCTCTTCCGATCTNNNAGACACAGCGCCCTTTCTTCCAT
GCACACGTCTGAACTCTTTCCCTACACGACGCTCTTCCGATCTNNNAGACACAGCGCCCTTTCTTCCAT
GCACACGTCTGAACTCTTTCCCTACACGACGCTCTTCCGATCTNNNAGACACAGCGCCCTTTCTTCCAT
GCACACGTCTGAACTCTTTCCCTACACGACGCTCTTCCGATCTNNNAGACACAGCGCCCTTTCTTCCAT
AGCACACGTCTGAACTCTTTCCCTACACGACGCTCTTCCGATCTNNNGTGGATCACTTGAAGTCAGG
AGCACACGTCTGAACTCTTTCCCTACACGACGCTCTTCCGATCTNNNGTGGATCACTTGAAGTCAGG
AGCACACGTCTGAACTCTTTCCCTACACGACGCTCTTCCGATCTNNNGTGGATCACTTGAAGTCAGG
AGCACACGTCTGAACTCTTTCCCTACACGACGCTCTTCCGATCTNNNGTGGATCACTTGAAGTCAGG
AGCACACGTCTGAACTCTTTCCCTACACGACGCTCTTCCGATCTNNNGTGGATCACTTGAAGTCAGG
AGCACACGTCTGAACTCTTTCCCTACACGACGCTCTTCCGATCTNNNGTGGATCACTTGAAGTCAGG
AGCACACGTCTGAACTCTTTCCCTACACGACGCTCTTCCGATCTNNNGAGGGGCTCAAGAGCAGATT
AGCACACGTCTGAACTCTTTCCCTACACGACGCTCTTCCGATCTNNNGAGGGGCTCAAGAGCAGATT
AGCACACGTCTGAACTCTTTCCCTACACGACGCTCTTCCGATCTNNNGAGGGGCTCAAGAGCAGATT
AGCACACGTCTGAACTCTTTCCCTACACGACGCTCTTCCGATCTNNNGAGGGGCTCAAGAGCAGATT
AGCACACGTCTGAACTCTTTCCCTACACGACGCTCTTCCGATCTNNNGAGGGGCTCAAGAGCAGATT
AGCACACGTCTGAACTCTTTCCCTACACGACGCTCTTCCGATCTNNNGAGGGGCTCAAGAGCAGATT
GCACACGTCTGAACTCTTTCCCTACACGACGCTCTTCCGATCTNNNGAACTGACTCCCTCCCTTTTCTCC
GCACACGTCTGAACTCTTTCCCTACACGACGCTCTTCCGATCTNNNGAACTGACTCCCTCCCTTTTCTCC
GCACACGTCTGAACTCTTTCCCTACACGACGCTCTTCCGATCTNNNGAACTGACTCCCTCCCTTTTCTCC
GCACACGTCTGAACTCTTTCCCTACACGACGCTCTTCCGATCTNNNGAACTGACTCCCTCCCTTTTCTCC
GCACACGTCTGAACTCTTTCCCTACACGACGCTCTTCCGATCTNNNGAACTGACTCCCTCCCTTTTCTCC
GCACACGTCTGAACTCTTTCCCTACACGACGCTCTTCCGATCTNNNGAACTGACTCCCTCCCTTTTCTCC
GCACACGTCTGAACTCTTTCCCTACACGACGCTCTTCCGATCTNNNAATGCATAAAACCCCTGGGAG

AGCACACGTCTGAACTCTTTCCCTACACGACGCTCTTCCGATCTNNNAATGCATAAAACCCCTGGGAGA
AGCACACGTCTGAACTCTTTCCCTACACGACGCTCTTCCGATCTNNNAATGCATAAAACCCCTGGGAGA
AGCACACGTCTGAACTCTTTCCCTACACGACGCTCTTCCGATCTNNNAATGCATAAAACCCCTGGGAGA
AGCACACGTCTGAACTCTTTCCCTACACGACGCTCTTCCGATCTNNNAATGCATAAAACCCCTGGGAGA
AGCACACGTCTGAACTCTTTCCCTACACGACGCTCTTCCGATCTNNNAATGCATAAAACCCCTGGGAGA
GCACACGTCTGAACTCTTTCCCTACACGACGCTCTTCCGATCTNNNCTTGGCATTAGCTTGGTGTTTCCC
GCACACGTCTGAACTCTTTCCCTACACGACGCTCTTCCGATCTNNNCTTGGCATTAGCTTGGTGTTTCCC
GCACACGTCTGAACTCTTTCCCTACACGACGCTCTTCCGATCTNNNCTTGGCATTAGCTTGGTGTTTCCC
GCACACGTCTGAACTCTTTCCCTACACGACGCTCTTCCGATCTNNNCTTGGCATTAGCTTGGTGTTTCCC
GCACACGTCTGAACTCTTTCCCTACACGACGCTCTTCCGATCTNNNCTTGGCATTAGCTTGGTGTTTCCC
GCACACGTCTGAACTCTTTCCCTACACGACGCTCTTCCGATCTNNNCTTGGCATTAGCTTGGTGTTTCCC
AGCACACGTCTGAACTCTTTCCCTACACGACGCTCTTCCGATCTNNNCTGTAATCCCAGCTACTCGAGAG
AGCACACGTCTGAACTCTTTCCCTACACGACGCTCTTCCGATCTNNNCTGTAATCCCAGCTACTCGAGAG
AGCACACGTCTGAACTCTTTCCCTACACGACGCTCTTCCGATCTNNNCTGTAATCCCAGCTACTCGAGAG
AGCACACGTCTGAACTCTTTCCCTACACGACGCTCTTCCGATCTNNNCTGTAATCCCAGCTACTCGAGAG
AGCACACGTCTGAACTCTTTCCCTACACGACGCTCTTCCGATCTNNNCTGTAATCCCAGCTACTCGAGAG
AGCACACGTCTGAACTCTTTCCCTACACGACGCTCTTCCGATCTNNNCTGTAATCCCAGCTACTCGAGAG
GCACACGTCTGAACTCTTTCCCTACACGACGCTCTTCCGATCTNNNGAGATATGTCTAGGTCAGGCCAT
GCACACGTCTGAACTCTTTCCCTACACGACGCTCTTCCGATCTNNNGAGATATGTCTAGGTCAGGCCAT
GCACACGTCTGAACTCTTTCCCTACACGACGCTCTTCCGATCTNNNGAGATATGTCTAGGTCAGGCCAT
GCACACGTCTGAACTCTTTCCCTACACGACGCTCTTCCGATCTNNNGAGATATGTCTAGGTCAGGCCAT
GCACACGTCTGAACTCTTTCCCTACACGACGCTCTTCCGATCTNNNGAGATATGTCTAGGTCAGGCCAT
GCACACGTCTGAACTCTTTCCCTACACGACGCTCTTCCGATCTNNNGAGATATGTCTAGGTCAGGCCAT
GAGCACACGTCTGAACTCTTTCCCTACACGACGCTCTTCCGATCTNNNCTGAGGCAGGAGAATCACTT
GAGCACACGTCTGAACTCTTTCCCTACACGACGCTCTTCCGATCTNNNCTGAGGCAGGAGAATCACTT
GAGCACACGTCTGAACTCTTTCCCTACACGACGCTCTTCCGATCTNNNCTGAGGCAGGAGAATCACTT
GAGCACACGTCTGAACTCTTTCCCTACACGACGCTCTTCCGATCTNNNCTGAGGCAGGAGAATCACTT
GAGCACACGTCTGAACTCTTTCCCTACACGACGCTCTTCCGATCTNNNCTGAGGCAGGAGAATCACTT
GAGCACACGTCTGAACTCTTTCCCTACACGACGCTCTTCCGATCTNNNCTGAGGCAGGAGAATCACTT
AGAGCACACGTCTGAACTCTTTCCCTACACGACGCTCTTCCGATCTNNNAGAGCTTGCAGTGAGATGA
AGAGCACACGTCTGAACTCTTTCCCTACACGACGCTCTTCCGATCTNNNAGAGCTTGCAGTGAGATGA
AGAGCACACGTCTGAACTCTTTCCCTACACGACGCTCTTCCGATCTNNNAGAGCTTGCAGTGAGATGA
AGAGCACACGTCTGAACTCTTTCCCTACACGACGCTCTTCCGATCTNNNAGAGCTTGCAGTGAGATGA
AGAGCACACGTCTGAACTCTTTCCCTACACGACGCTCTTCCGATCTNNNAGAGCTTGCAGTGAGATGA
AGAGCACACGTCTGAACTCTTTCCCTACACGACGCTCTTCCGATCTNNNAGAGCTTGCAGTGAGATGA
GAGCACACGTCTGAACTCTTTCCCTACACGACGCTCTTCCGATCTNNNTGCACCAAACTAATATTTCCAC
GAGCACACGTCTGAACTCTTTCCCTACACGACGCTCTTCCGATCTNNNTGCACCAAACTAATATTTCCAC
GAGCACACGTCTGAACTCTTTCCCTACACGACGCTCTTCCGATCTNNNTGCACCAAACTAATATTTCCAC
GAGCACACGTCTGAACTCTTTCCCTACACGACGCTCTTCCGATCTNNNTGCACCAAACTAATATTTCCAC
GAGCACACGTCTGAACTCTTTCCCTACACGACGCTCTTCCGATCTNNNTGCACCAAACTAATATTTCCAC
GAGCACACGTCTGAACTCTTTCCCTACACGACGCTCTTCCGATCTNNNTGCACCAAACTAATATTTCCAC
GAGCACACGTCTGAACTCTTTCCCTACACGACGCTCTTCCGATCTNNNGTTGGGTCTTGTGGGCTAAT
GAGCACACGTCTGAACTCTTTCCCTACACGACGCTCTTCCGATCTNNNGTTGGGTCTTGTGGGCTAAT

GAGCACACGTCTGAACTCTTTCCCTACACGACGCTCTTCCGATCTNNNGTTGGGTCTTGTGGGCTAAT

GAGCACACGTCTGAACTCTTTCCCTACACGACGCTCTTCCGATCTNNNGTTGGGTCTTGTGGGCTAAT

GAGCACACGTCTGAACTCTTTCCCTACACGACGCTCTTCCGATCTNNNGTTGGGTCTTGTGGGCTAAT

GAGCACACGTCTGAACTCTTTCCCTACACGACGCTCTTCCGATCTNNNGTTGGGTCTTGTGGGCTAAT

AGCACACGTCTGAACTCTTTCCCTACACGACGCTCTTCCGATCTNNNATTGACCTGGGAGTCCCTGAAT

AGCACACGTCTGAACTCTTTCCCTACACGACGCTCTTCCGATCTNNNATTGACCTGGGAGTCCCTGAAT

AGCACACGTCTGAACTCTTTCCCTACACGACGCTCTTCCGATCTNNNATTGACCTGGGAGTCCCTGAAT

AGCACACGTCTGAACTCTTTCCCTACACGACGCTCTTCCGATCTNNNATTGACCTGGGAGTCCCTGAAT

AGCACACGTCTGAACTCTTTCCCTACACGACGCTCTTCCGATCTNNNATTGACCTGGGAGTCCCTGAAT

AGCACACGTCTGAACTCTTTCCCTACACGACGCTCTTCCGATCTNNNATTGACCTGGGAGTCCCTGAAT

GAGCACACGTCTGAACTCTTTCCCTACACGACGCTCTTCCGATCTNNNCAGGAGGCAGAAGTTACAGT

GAGCACACGTCTGAACTCTTTCCCTACACGACGCTCTTCCGATCTNNNCAGGAGGCAGAAGTTACAGT

GAGCACACGTCTGAACTCTTTCCCTACACGACGCTCTTCCGATCTNNNCAGGAGGCAGAAGTTACAGT

GAGCACACGTCTGAACTCTTTCCCTACACGACGCTCTTCCGATCTNNNCAGGAGGCAGAAGTTACAGT

GAGCACACGTCTGAACTCTTTCCCTACACGACGCTCTTCCGATCTNNNCAGGAGGCAGAAGTTACAGT

GAGCACACGTCTGAACTCTTTCCCTACACGACGCTCTTCCGATCTNNNCAGGAGGCAGAAGTTACAGT

AGCACACGTCTGAACTCTTTCCCTACACGACGCTCTTCCGATCTNNNCGGTTTTCAAGGCAAACAACAT

AGCACACGTCTGAACTCTTTCCCTACACGACGCTCTTCCGATCTNNNCGGTTTTCAAGGCAAACAACAT

AGCACACGTCTGAACTCTTTCCCTACACGACGCTCTTCCGATCTNNNCGGTTTTCAAGGCAAACAACAT

AGCACACGTCTGAACTCTTTCCCTACACGACGCTCTTCCGATCTNNNCGGTTTTCAAGGCAAACAACAT

AGCACACGTCTGAACTCTTTCCCTACACGACGCTCTTCCGATCTNNNCGGTTTTCAAGGCAAACAACAT

AGCACACGTCTGAACTCTTTCCCTACACGACGCTCTTCCGATCTNNNCGGTTTTCAAGGCAAACAACAT

GAGCACACGTCTGAACTCTTTCCCTACACGACGCTCTTCCGATCTNNNTAGTCCCAGCTACTCAGGAGA

GAGCACACGTCTGAACTCTTTCCCTACACGACGCTCTTCCGATCTNNNTAGTCCCAGCTACTCAGGAGA

GAGCACACGTCTGAACTCTTTCCCTACACGACGCTCTTCCGATCTNNNTAGTCCCAGCTACTCAGGAGA

GAGCACACGTCTGAACTCTTTCCCTACACGACGCTCTTCCGATCTNNNTAGTCCCAGCTACTCAGGAGA

GAGCACACGTCTGAACTCTTTCCCTACACGACGCTCTTCCGATCTNNNTAGTCCCAGCTACTCAGGAGA

GAGCACACGTCTGAACTCTTTCCCTACACGACGCTCTTCCGATCTNNNTAGTCCCAGCTACTCAGGAGA

AGCACACGTCTGAACTCTTTCCCTACACGACGCTCTTCCGATCTNNNAGAAATAGGTCAGCAGGGAACA

AGCACACGTCTGAACTCTTTCCCTACACGACGCTCTTCCGATCTNNNAGAAATAGGTCAGCAGGGAACA

AGCACACGTCTGAACTCTTTCCCTACACGACGCTCTTCCGATCTNNNAGAAATAGGTCAGCAGGGAACA

AGCACACGTCTGAACTCTTTCCCTACACGACGCTCTTCCGATCTNNNAGAAATAGGTCAGCAGGGAACA

AGCACACGTCTGAACTCTTTCCCTACACGACGCTCTTCCGATCTNNNAGAAATAGGTCAGCAGGGAACA

AGCACACGTCTGAACTCTTTCCCTACACGACGCTCTTCCGATCTNNNAGAAATAGGTCAGCAGGGAACA

AGAGCACACGTCTGAACTCTTTCCCTACACGACGCTCTTCCGATCTNNNAATTTGCTCAGTTGCAGGGA

AGAGCACACGTCTGAACTCTTTCCCTACACGACGCTCTTCCGATCTNNNAATTTGCTCAGTTGCAGGGA

AGAGCACACGTCTGAACTCTTTCCCTACACGACGCTCTTCCGATCTNNNAATTTGCTCAGTTGCAGGGA

AGAGCACACGTCTGAACTCTTTCCCTACACGACGCTCTTCCGATCTNNNAATTTGCTCAGTTGCAGGGA

AGAGCACACGTCTGAACTCTTTCCCTACACGACGCTCTTCCGATCTNNNAATTTGCTCAGTTGCAGGGA

AGAGCACACGTCTGAACTCTTTCCCTACACGACGCTCTTCCGATCTNNNAATTTGCTCAGTTGCAGGGA

GCACACGTCTGAACTCTTTCCCTACACGACGCTCTTCCGATCTNNNGAGGGACTTTGTGGGAGGTAAT

GCACACGTCTGAACTCTTTCCCTACACGACGCTCTTCCGATCTNNNGAGGGACTTTGTGGGAGGTAAT

GCACACGTCTGAACTCTTTCCCTACACGACGCTCTTCCGATCTNNNGAGGGACTTTGTGGGAGGTAAT

GCACACGTCTGAACTCTTTCCCTACACGACGCTCTTCCGATCTNNNGAGGGACTTTGTGGGAGGTAA
GCACACGTCTGAACTCTTTCCCTACACGACGCTCTTCCGATCTNNNGAGGGACTTTGTGGGAGGTAA
GCACACGTCTGAACTCTTTCCCTACACGACGCTCTTCCGATCTNNNGAGGGACTTTGTGGGAGGTAA
GCACACGTCTGAACTCTTTCCCTACACGACGCTCTTCCGATCTNNNGAGTAGTTGGGATTACAGGCT
GCACACGTCTGAACTCTTTCCCTACACGACGCTCTTCCGATCTNNNGAGTAGTTGGGATTACAGGCT
GCACACGTCTGAACTCTTTCCCTACACGACGCTCTTCCGATCTNNNGAGTAGTTGGGATTACAGGCT
GCACACGTCTGAACTCTTTCCCTACACGACGCTCTTCCGATCTNNNGAGTAGTTGGGATTACAGGCT
GCACACGTCTGAACTCTTTCCCTACACGACGCTCTTCCGATCTNNNGAGTAGTTGGGATTACAGGCT
GCACACGTCTGAACTCTTTCCCTACACGACGCTCTTCCGATCTNNNGAGTAGTTGGGATTACAGGCT
GAGCACACGTCTGAACTCTTTCCCTACACGACGCTCTTCCGATCTNNNTCACACCTGTAATTCCAGCAC
GAGCACACGTCTGAACTCTTTCCCTACACGACGCTCTTCCGATCTNNNTCACACCTGTAATTCCAGCAC
GAGCACACGTCTGAACTCTTTCCCTACACGACGCTCTTCCGATCTNNNTCACACCTGTAATTCCAGCAC
GAGCACACGTCTGAACTCTTTCCCTACACGACGCTCTTCCGATCTNNNTCACACCTGTAATTCCAGCAC
GAGCACACGTCTGAACTCTTTCCCTACACGACGCTCTTCCGATCTNNNTCACACCTGTAATTCCAGCAC
GAGCACACGTCTGAACTCTTTCCCTACACGACGCTCTTCCGATCTNNNTCACACCTGTAATTCCAGCAC
GAGCACACGTCTGAACTCTTTCCCTACACGACGCTCTTCCGATCTNNNTCTACCTCCCTCTGAACCTCTT
GAGCACACGTCTGAACTCTTTCCCTACACGACGCTCTTCCGATCTNNNTCTACCTCCCTCTGAACCTCTT
GAGCACACGTCTGAACTCTTTCCCTACACGACGCTCTTCCGATCTNNNTCTACCTCCCTCTGAACCTCTT
GAGCACACGTCTGAACTCTTTCCCTACACGACGCTCTTCCGATCTNNNTCTACCTCCCTCTGAACCTCTT
GAGCACACGTCTGAACTCTTTCCCTACACGACGCTCTTCCGATCTNNNTCTACCTCCCTCTGAACCTCTT
GAGCACACGTCTGAACTCTTTCCCTACACGACGCTCTTCCGATCTNNNTCTACCTCCCTCTGAACCTCTT
GAGCACACGTCTGAACTCTTTCCCTACACGACGCTCTTCCGATCTNNNGGTTTCCCAGAGTTGGTCAG
GAGCACACGTCTGAACTCTTTCCCTACACGACGCTCTTCCGATCTNNNGGTTTCCCAGAGTTGGTCAG
GAGCACACGTCTGAACTCTTTCCCTACACGACGCTCTTCCGATCTNNNGGTTTCCCAGAGTTGGTCAG
GAGCACACGTCTGAACTCTTTCCCTACACGACGCTCTTCCGATCTNNNGGTTTCCCAGAGTTGGTCAG
GAGCACACGTCTGAACTCTTTCCCTACACGACGCTCTTCCGATCTNNNGGTTTCCCAGAGTTGGTCAG
GAGCACACGTCTGAACTCTTTCCCTACACGACGCTCTTCCGATCTNNNGGTTTCCCAGAGTTGGTCAG
AGCACACGTCTGAACTCTTTCCCTACACGACGCTCTTCCGATCTNNNACAAAGACACCTGGCCTAAA
AGCACACGTCTGAACTCTTTCCCTACACGACGCTCTTCCGATCTNNNACAAAGACACCTGGCCTAAA
AGCACACGTCTGAACTCTTTCCCTACACGACGCTCTTCCGATCTNNNACAAAGACACCTGGCCTAAA
AGCACACGTCTGAACTCTTTCCCTACACGACGCTCTTCCGATCTNNNACAAAGACACCTGGCCTAAA
AGCACACGTCTGAACTCTTTCCCTACACGACGCTCTTCCGATCTNNNACAAAGACACCTGGCCTAAA
AGCACACGTCTGAACTCTTTCCCTACACGACGCTCTTCCGATCTNNNACAAAGACACCTGGCCTAAA
GAGCACACGTCTGAACTCTTTCCCTACACGACGCTCTTCCGATCTNNNGTGAGCTGAGATCATGCCAC
GAGCACACGTCTGAACTCTTTCCCTACACGACGCTCTTCCGATCTNNNGTGAGCTGAGATCATGCCAC
GAGCACACGTCTGAACTCTTTCCCTACACGACGCTCTTCCGATCTNNNGTGAGCTGAGATCATGCCAC
GAGCACACGTCTGAACTCTTTCCCTACACGACGCTCTTCCGATCTNNNGTGAGCTGAGATCATGCCAC
GAGCACACGTCTGAACTCTTTCCCTACACGACGCTCTTCCGATCTNNNGTGAGCTGAGATCATGCCAC
GAGCACACGTCTGAACTCTTTCCCTACACGACGCTCTTCCGATCTNNNGTGAGCTGAGATCATGCCAC
GAGCACACGTCTGAACTCTTTCCCTACACGACGCTCTTCCGATCTNNNTGAGTTTCAAAATGTGGCAA
GAGCACACGTCTGAACTCTTTCCCTACACGACGCTCTTCCGATCTNNNTGAGTTTCAAAATGTGGCAA
GAGCACACGTCTGAACTCTTTCCCTACACGACGCTCTTCCGATCTNNNTGAGTTTCAAAATGTGGCAA
GAGCACACGTCTGAACTCTTTCCCTACACGACGCTCTTCCGATCTNNNTGAGTTTCAAAATGTGGCAA

GAGCACACGTCTGAACTCTTTCCCTACACGACGCTCTTCCGATCTNNNTGAGTTTCAAAATGTGGCAA
GAGCACACGTCTGAACTCTTTCCCTACACGACGCTCTTCCGATCTNNNTGAGTTTCAAAATGTGGCAA
GCACACGTCTGAACTCTTTCCCTACACGACGCTCTTCCGATCTNNNCCCAAAAAAACCCATTCTGTCCT
GCACACGTCTGAACTCTTTCCCTACACGACGCTCTTCCGATCTNNNCCCAAAAAAACCCATTCTGTCCT
GCACACGTCTGAACTCTTTCCCTACACGACGCTCTTCCGATCTNNNCCCAAAAAAACCCATTCTGTCCT
GCACACGTCTGAACTCTTTCCCTACACGACGCTCTTCCGATCTNNNCCCAAAAAAACCCATTCTGTCCT
GCACACGTCTGAACTCTTTCCCTACACGACGCTCTTCCGATCTNNNCCCAAAAAAACCCATTCTGTCCT
GCACACGTCTGAACTCTTTCCCTACACGACGCTCTTCCGATCTNNNCCCAAAAAAACCCATTCTGTCCT
AGCACACGTCTGAACTCTTTCCCTACACGACGCTCTTCCGATCTNNNCTATCCCTCCACTAGCCTCCTC
AGCACACGTCTGAACTCTTTCCCTACACGACGCTCTTCCGATCTNNNCTATCCCTCCACTAGCCTCCTC
AGCACACGTCTGAACTCTTTCCCTACACGACGCTCTTCCGATCTNNNCTATCCCTCCACTAGCCTCCTC
AGCACACGTCTGAACTCTTTCCCTACACGACGCTCTTCCGATCTNNNCTATCCCTCCACTAGCCTCCTC
AGCACACGTCTGAACTCTTTCCCTACACGACGCTCTTCCGATCTNNNCTATCCCTCCACTAGCCTCCTC
AGCACACGTCTGAACTCTTTCCCTACACGACGCTCTTCCGATCTNNNCTATCCCTCCACTAGCCTCCTC
GCACACGTCTGAACTCTTTCCCTACACGACGCTCTTCCGATCTNNNAGCATCATATTGGGCAGTATAC
GCACACGTCTGAACTCTTTCCCTACACGACGCTCTTCCGATCTNNNAGCATCATATTGGGCAGTATAC
GCACACGTCTGAACTCTTTCCCTACACGACGCTCTTCCGATCTNNNAGCATCATATTGGGCAGTATAC
GCACACGTCTGAACTCTTTCCCTACACGACGCTCTTCCGATCTNNNAGCATCATATTGGGCAGTATAC
GCACACGTCTGAACTCTTTCCCTACACGACGCTCTTCCGATCTNNNAGCATCATATTGGGCAGTATAC
GCACACGTCTGAACTCTTTCCCTACACGACGCTCTTCCGATCTNNNAGCATCATATTGGGCAGTATAC
AGCACACGTCTGAACTCTTTCCCTACACGACGCTCTTCCGATCTNNNGGCATCTTTTGAGTGGGAACT
AGCACACGTCTGAACTCTTTCCCTACACGACGCTCTTCCGATCTNNNGGCATCTTTTGAGTGGGAACT
AGCACACGTCTGAACTCTTTCCCTACACGACGCTCTTCCGATCTNNNGGCATCTTTTGAGTGGGAACT
AGCACACGTCTGAACTCTTTCCCTACACGACGCTCTTCCGATCTNNNGGCATCTTTTGAGTGGGAACT
AGCACACGTCTGAACTCTTTCCCTACACGACGCTCTTCCGATCTNNNGGCATCTTTTGAGTGGGAACT
AGCACACGTCTGAACTCTTTCCCTACACGACGCTCTTCCGATCTNNNGGCATCTTTTGAGTGGGAACT
GAGCACACGTCTGAACTCTTTCCCTACACGACGCTCTTCCGATCTNNNTGGCTCCAGTGTTGTTTTTTT
GAGCACACGTCTGAACTCTTTCCCTACACGACGCTCTTCCGATCTNNNTGGCTCCAGTGTTGTTTTTTT
GAGCACACGTCTGAACTCTTTCCCTACACGACGCTCTTCCGATCTNNNTGGCTCCAGTGTTGTTTTTTT
GAGCACACGTCTGAACTCTTTCCCTACACGACGCTCTTCCGATCTNNNTGGCTCCAGTGTTGTTTTTTT
GAGCACACGTCTGAACTCTTTCCCTACACGACGCTCTTCCGATCTNNNTGGCTCCAGTGTTGTTTTTTT
GAGCACACGTCTGAACTCTTTCCCTACACGACGCTCTTCCGATCTNNNTGGCTCCAGTGTTGTTTTTTT
CACACGTCTGAACTCTTTCCCTACACGACGCTCTTCCGATCTNNNAAAGAAAAAGCTGTGACAGGGT
CACACGTCTGAACTCTTTCCCTACACGACGCTCTTCCGATCTNNNAAAGAAAAAGCTGTGACAGGGT
CACACGTCTGAACTCTTTCCCTACACGACGCTCTTCCGATCTNNNAAAGAAAAAGCTGTGACAGGGT
CACACGTCTGAACTCTTTCCCTACACGACGCTCTTCCGATCTNNNAAAGAAAAAGCTGTGACAGGGT
CACACGTCTGAACTCTTTCCCTACACGACGCTCTTCCGATCTNNNAAAGAAAAAGCTGTGACAGGGT
CACACGTCTGAACTCTTTCCCTACACGACGCTCTTCCGATCTNNNAAAGAAAAAGCTGTGACAGGGT
AGCACACGTCTGAACTCTTTCCCTACACGACGCTCTTCCGATCTNNNCCACCCCTCCATAAATGAGTT
AGCACACGTCTGAACTCTTTCCCTACACGACGCTCTTCCGATCTNNNCCACCCCTCCATAAATGAGTT
AGCACACGTCTGAACTCTTTCCCTACACGACGCTCTTCCGATCTNNNCCACCCCTCCATAAATGAGTT
AGCACACGTCTGAACTCTTTCCCTACACGACGCTCTTCCGATCTNNNCCACCCCTCCATAAATGAGTT
AGCACACGTCTGAACTCTTTCCCTACACGACGCTCTTCCGATCTNNNCCACCCCTCCATAAATGAGTT

AGCACACGTCTGAACTCTTTCCCTACACGACGCTCTTCCGATCTNNNCCCACCCCTCCATAAATGAGTT
GCACACGTCTGAACTCTTTCCCTACACGACGCTCTTCCGATCTNNNAGAGACACTGAAGAGGGTAGG
GCACACGTCTGAACTCTTTCCCTACACGACGCTCTTCCGATCTNNNAGAGACACTGAAGAGGGTAGG
GCACACGTCTGAACTCTTTCCCTACACGACGCTCTTCCGATCTNNNAGAGACACTGAAGAGGGTAGG
GCACACGTCTGAACTCTTTCCCTACACGACGCTCTTCCGATCTNNNAGAGACACTGAAGAGGGTAGG
GCACACGTCTGAACTCTTTCCCTACACGACGCTCTTCCGATCTNNNAGAGACACTGAAGAGGGTAGG
GCACACGTCTGAACTCTTTCCCTACACGACGCTCTTCCGATCTNNNAGAGACACTGAAGAGGGTAGG
AGCACACGTCTGAACTCTTTCCCTACACGACGCTCTTCCGATCTNNNCAGATCTATAAACCCAGCCCCT
AGCACACGTCTGAACTCTTTCCCTACACGACGCTCTTCCGATCTNNNCAGATCTATAAACCCAGCCCCT
AGCACACGTCTGAACTCTTTCCCTACACGACGCTCTTCCGATCTNNNCAGATCTATAAACCCAGCCCCT
AGCACACGTCTGAACTCTTTCCCTACACGACGCTCTTCCGATCTNNNCAGATCTATAAACCCAGCCCCT
AGCACACGTCTGAACTCTTTCCCTACACGACGCTCTTCCGATCTNNNCAGATCTATAAACCCAGCCCCT
AGCACACGTCTGAACTCTTTCCCTACACGACGCTCTTCCGATCTNNNCAGATCTATAAACCCAGCCCCT
AGCACACGTCTGAACTCTTTCCCTACACGACGCTCTTCCGATCTNNNATCATGAGGTCAGGAGATGG
AGCACACGTCTGAACTCTTTCCCTACACGACGCTCTTCCGATCTNNNATCATGAGGTCAGGAGATGG
AGCACACGTCTGAACTCTTTCCCTACACGACGCTCTTCCGATCTNNNATCATGAGGTCAGGAGATGG
AGCACACGTCTGAACTCTTTCCCTACACGACGCTCTTCCGATCTNNNATCATGAGGTCAGGAGATGG
AGCACACGTCTGAACTCTTTCCCTACACGACGCTCTTCCGATCTNNNATCATGAGGTCAGGAGATGG
AGCACACGTCTGAACTCTTTCCCTACACGACGCTCTTCCGATCTNNNATCATGAGGTCAGGAGATGG
AGCACACGTCTGAACTCTTTCCCTACACGACGCTCTTCCGATCTNNNAAAGAGCAGCAGCCAGAGATT
AGCACACGTCTGAACTCTTTCCCTACACGACGCTCTTCCGATCTNNNAAAGAGCAGCAGCCAGAGATT
AGCACACGTCTGAACTCTTTCCCTACACGACGCTCTTCCGATCTNNNAAAGAGCAGCAGCCAGAGATT
AGCACACGTCTGAACTCTTTCCCTACACGACGCTCTTCCGATCTNNNAAAGAGCAGCAGCCAGAGATT
AGCACACGTCTGAACTCTTTCCCTACACGACGCTCTTCCGATCTNNNAAAGAGCAGCAGCCAGAGATT
AGCACACGTCTGAACTCTTTCCCTACACGACGCTCTTCCGATCTNNNAAAGAGCAGCAGCCAGAGATT
ACACGTCTGAACTCTTTCCCTACACGACGCTCTTCCGATCTNNNTTCCTTTCATTCGAGGAGAGACGC
ACACGTCTGAACTCTTTCCCTACACGACGCTCTTCCGATCTNNNTTCCTTTCATTCGAGGAGAGACGC
ACACGTCTGAACTCTTTCCCTACACGACGCTCTTCCGATCTNNNTTCCTTTCATTCGAGGAGAGACGC
ACACGTCTGAACTCTTTCCCTACACGACGCTCTTCCGATCTNNNTTCCTTTCATTCGAGGAGAGACGC
ACACGTCTGAACTCTTTCCCTACACGACGCTCTTCCGATCTNNNTTCCTTTCATTCGAGGAGAGACGC
ACACGTCTGAACTCTTTCCCTACACGACGCTCTTCCGATCTNNNTTCCTTTCATTCGAGGAGAGACGC
GCACACGTCTGAACTCTTTCCCTACACGACGCTCTTCCGATCTNNNGGATTGTTTGAGCCTGGGAAA
GCACACGTCTGAACTCTTTCCCTACACGACGCTCTTCCGATCTNNNGGATTGTTTGAGCCTGGGAAA
GCACACGTCTGAACTCTTTCCCTACACGACGCTCTTCCGATCTNNNGGATTGTTTGAGCCTGGGAAA
GCACACGTCTGAACTCTTTCCCTACACGACGCTCTTCCGATCTNNNGGATTGTTTGAGCCTGGGAAA
GCACACGTCTGAACTCTTTCCCTACACGACGCTCTTCCGATCTNNNGGATTGTTTGAGCCTGGGAAA
GCACACGTCTGAACTCTTTCCCTACACGACGCTCTTCCGATCTNNNGGATTGTTTGAGCCTGGGAAA
GCACACGTCTGAACTCTTTCCCTACACGACGCTCTTCCGATCTNNNAGCACTGGGTTATAGATGCCA
GCACACGTCTGAACTCTTTCCCTACACGACGCTCTTCCGATCTNNNAGCACTGGGTTATAGATGCCA
GCACACGTCTGAACTCTTTCCCTACACGACGCTCTTCCGATCTNNNAGCACTGGGTTATAGATGCCA
GCACACGTCTGAACTCTTTCCCTACACGACGCTCTTCCGATCTNNNAGCACTGGGTTATAGATGCCA
GCACACGTCTGAACTCTTTCCCTACACGACGCTCTTCCGATCTNNNAGCACTGGGTTATAGATGCCA
GCACACGTCTGAACTCTTTCCCTACACGACGCTCTTCCGATCTNNNAGCACTGGGTTATAGATGCCA

GCACACGTCTGAACTCTTTCCCTACACGACGCTCTTCCGATCTNNNAATCAGGGCAATACTGGGTTCA

GCACACGTCTGAACTCTTTCCCTACACGACGCTCTTCCGATCTNNNAATCAGGGCAATACTGGGTTCA

GCACACGTCTGAACTCTTTCCCTACACGACGCTCTTCCGATCTNNNAATCAGGGCAATACTGGGTTCA

GCACACGTCTGAACTCTTTCCCTACACGACGCTCTTCCGATCTNNNAATCAGGGCAATACTGGGTTCA

GCACACGTCTGAACTCTTTCCCTACACGACGCTCTTCCGATCTNNNAATCAGGGCAATACTGGGTTCA

GCACACGTCTGAACTCTTTCCCTACACGACGCTCTTCCGATCTNNNAATCAGGGCAATACTGGGTTCA

ACACGTCTGAACTCTTTCCCTACACGACGCTCTTCCGATCTNNNCTTCTTCTCCATTGTCTCCTCATCGG

ACACGTCTGAACTCTTTCCCTACACGACGCTCTTCCGATCTNNNCTTCTTCTCCATTGTCTCCTCATCGG

ACACGTCTGAACTCTTTCCCTACACGACGCTCTTCCGATCTNNNCTTCTTCTCCATTGTCTCCTCATCGG

ACACGTCTGAACTCTTTCCCTACACGACGCTCTTCCGATCTNNNCTTCTTCTCCATTGTCTCCTCATCGG

ACACGTCTGAACTCTTTCCCTACACGACGCTCTTCCGATCTNNNCTTCTTCTCCATTGTCTCCTCATCGG

ACACGTCTGAACTCTTTCCCTACACGACGCTCTTCCGATCTNNNCTTCTTCTCCATTGTCTCCTCATCGG

AGCACACGTCTGAACTCTTTCCCTACACGACGCTCTTCCGATCTNNNATCTCGTCTTACTGCAACCTCT

AGCACACGTCTGAACTCTTTCCCTACACGACGCTCTTCCGATCTNNNATCTCGTCTTACTGCAACCTCT

AGCACACGTCTGAACTCTTTCCCTACACGACGCTCTTCCGATCTNNNATCTCGTCTTACTGCAACCTCT

AGCACACGTCTGAACTCTTTCCCTACACGACGCTCTTCCGATCTNNNATCTCGTCTTACTGCAACCTCT

AGCACACGTCTGAACTCTTTCCCTACACGACGCTCTTCCGATCTNNNATCTCGTCTTACTGCAACCTCT

AGCACACGTCTGAACTCTTTCCCTACACGACGCTCTTCCGATCTNNNATCTCGTCTTACTGCAACCTCT

AGCACACGTCTGAACTCTTTCCCTACACGACGCTCTTCCGATCTNNNGCTTTCCCTGTGTAAAGAGCA

AGCACACGTCTGAACTCTTTCCCTACACGACGCTCTTCCGATCTNNNGCTTTCCCTGTGTAAAGAGCA

AGCACACGTCTGAACTCTTTCCCTACACGACGCTCTTCCGATCTNNNGCTTTCCCTGTGTAAAGAGCA

AGCACACGTCTGAACTCTTTCCCTACACGACGCTCTTCCGATCTNNNGCTTTCCCTGTGTAAAGAGCA

AGCACACGTCTGAACTCTTTCCCTACACGACGCTCTTCCGATCTNNNGCTTTCCCTGTGTAAAGAGCA

AGCACACGTCTGAACTCTTTCCCTACACGACGCTCTTCCGATCTNNNGCTTTCCCTGTGTAAAGAGCA

AGCACACGTCTGAACTCTTTCCCTACACGACGCTCTTCCGATCTNNNACTCCAGACCTATGAGATGGT

AGCACACGTCTGAACTCTTTCCCTACACGACGCTCTTCCGATCTNNNACTCCAGACCTATGAGATGGT

AGCACACGTCTGAACTCTTTCCCTACACGACGCTCTTCCGATCTNNNACTCCAGACCTATGAGATGGT

AGCACACGTCTGAACTCTTTCCCTACACGACGCTCTTCCGATCTNNNACTCCAGACCTATGAGATGGT

AGCACACGTCTGAACTCTTTCCCTACACGACGCTCTTCCGATCTNNNACTCCAGACCTATGAGATGGT

AGCACACGTCTGAACTCTTTCCCTACACGACGCTCTTCCGATCTNNNACTCCAGACCTATGAGATGGT

GCACACGTCTGAACTCTTTCCCTACACGACGCTCTTCCGATCTNNNCTCGCTTTGAGAGACAGGTAT

GCACACGTCTGAACTCTTTCCCTACACGACGCTCTTCCGATCTNNNCTCGCTTTGAGAGACAGGTAT

GCACACGTCTGAACTCTTTCCCTACACGACGCTCTTCCGATCTNNNCTCGCTTTGAGAGACAGGTAT

GCACACGTCTGAACTCTTTCCCTACACGACGCTCTTCCGATCTNNNCTCGCTTTGAGAGACAGGTAT

GCACACGTCTGAACTCTTTCCCTACACGACGCTCTTCCGATCTNNNCTCGCTTTGAGAGACAGGTAT

GCACACGTCTGAACTCTTTCCCTACACGACGCTCTTCCGATCTNNNCTCGCTTTGAGAGACAGGTAT

AGCACACGTCTGAACTCTTTCCCTACACGACGCTCTTCCGATCTNNNTGACCTATGCTGCAGTTTTGT

AGCACACGTCTGAACTCTTTCCCTACACGACGCTCTTCCGATCTNNNTGACCTATGCTGCAGTTTTGT

AGCACACGTCTGAACTCTTTCCCTACACGACGCTCTTCCGATCTNNNTGACCTATGCTGCAGTTTTGT

AGCACACGTCTGAACTCTTTCCCTACACGACGCTCTTCCGATCTNNNTGACCTATGCTGCAGTTTTGT

AGCACACGTCTGAACTCTTTCCCTACACGACGCTCTTCCGATCTNNNTGACCTATGCTGCAGTTTTGT

AGCACACGTCTGAACTCTTTCCCTACACGACGCTCTTCCGATCTNNNTGACCTATGCTGCAGTTTTGT

GCACACGTCTGAACTCTTTCCCTACACGACGCTCTTCCGATCTNNNGAGGCTGAGGCACTAGAATTTC

GCACACGTCTGAACTCTTTCCCTACACGACGCTCTTCCGATCTNNNGAGGCTGAGGCACTAGAATTTC
GCACACGTCTGAACTCTTTCCCTACACGACGCTCTTCCGATCTNNNGAGGCTGAGGCACTAGAATTTC
GCACACGTCTGAACTCTTTCCCTACACGACGCTCTTCCGATCTNNNGAGGCTGAGGCACTAGAATTTC
GCACACGTCTGAACTCTTTCCCTACACGACGCTCTTCCGATCTNNNGAGGCTGAGGCACTAGAATTTC
GCACACGTCTGAACTCTTTCCCTACACGACGCTCTTCCGATCTNNNGAGGCTGAGGCACTAGAATTTC
GAGCACACGTCTGAACTCTTTCCCTACACGACGCTCTTCCGATCTNNNATCACTAGGTCAGGAGTTCG
GAGCACACGTCTGAACTCTTTCCCTACACGACGCTCTTCCGATCTNNNATCACTAGGTCAGGAGTTCG
GAGCACACGTCTGAACTCTTTCCCTACACGACGCTCTTCCGATCTNNNATCACTAGGTCAGGAGTTCG
GAGCACACGTCTGAACTCTTTCCCTACACGACGCTCTTCCGATCTNNNATCACTAGGTCAGGAGTTCG
GAGCACACGTCTGAACTCTTTCCCTACACGACGCTCTTCCGATCTNNNATCACTAGGTCAGGAGTTCG
GAGCACACGTCTGAACTCTTTCCCTACACGACGCTCTTCCGATCTNNNATCACTAGGTCAGGAGTTCG
AGCACACGTCTGAACTCTTTCCCTACACGACGCTCTTCCGATCTNNNATGCTTGTAATCCCAGCACTTT
AGCACACGTCTGAACTCTTTCCCTACACGACGCTCTTCCGATCTNNNATGCTTGTAATCCCAGCACTTT
AGCACACGTCTGAACTCTTTCCCTACACGACGCTCTTCCGATCTNNNATGCTTGTAATCCCAGCACTTT
AGCACACGTCTGAACTCTTTCCCTACACGACGCTCTTCCGATCTNNNATGCTTGTAATCCCAGCACTTT
AGCACACGTCTGAACTCTTTCCCTACACGACGCTCTTCCGATCTNNNATGCTTGTAATCCCAGCACTTT
AGCACACGTCTGAACTCTTTCCCTACACGACGCTCTTCCGATCTNNNATGCTTGTAATCCCAGCACTTT
GAGCACACGTCTGAACTCTTTCCCTACACGACGCTCTTCCGATCTNNNTGCAGTAGGGTCACATATCA
GAGCACACGTCTGAACTCTTTCCCTACACGACGCTCTTCCGATCTNNNTGCAGTAGGGTCACATATCA
GAGCACACGTCTGAACTCTTTCCCTACACGACGCTCTTCCGATCTNNNTGCAGTAGGGTCACATATCA
GAGCACACGTCTGAACTCTTTCCCTACACGACGCTCTTCCGATCTNNNTGCAGTAGGGTCACATATCA
GAGCACACGTCTGAACTCTTTCCCTACACGACGCTCTTCCGATCTNNNTGCAGTAGGGTCACATATCA
GAGCACACGTCTGAACTCTTTCCCTACACGACGCTCTTCCGATCTNNNTGCAGTAGGGTCACATATCA
GCACACGTCTGAACTCTTTCCCTACACGACGCTCTTCCGATCTNNNGTAAGAGTTAAAGAGTAAAGC
GCACACGTCTGAACTCTTTCCCTACACGACGCTCTTCCGATCTNNNGTAAGAGTTAAAGAGTAAAGC
GCACACGTCTGAACTCTTTCCCTACACGACGCTCTTCCGATCTNNNGTAAGAGTTAAAGAGTAAAGC
GCACACGTCTGAACTCTTTCCCTACACGACGCTCTTCCGATCTNNNGTAAGAGTTAAAGAGTAAAGC
GCACACGTCTGAACTCTTTCCCTACACGACGCTCTTCCGATCTNNNGTAAGAGTTAAAGAGTAAAGC
GCACACGTCTGAACTCTTTCCCTACACGACGCTCTTCCGATCTNNNGTAAGAGTTAAAGAGTAAAGC
GCACACGTCTGAACTCTTTCCCTACACGACGCTCTTCCGATCTNNNAATGGGTTCTTGTCATACGACC
GCACACGTCTGAACTCTTTCCCTACACGACGCTCTTCCGATCTNNNAATGGGTTCTTGTCATACGACC
GCACACGTCTGAACTCTTTCCCTACACGACGCTCTTCCGATCTNNNAATGGGTTCTTGTCATACGACC
GCACACGTCTGAACTCTTTCCCTACACGACGCTCTTCCGATCTNNNAATGGGTTCTTGTCATACGACC
GCACACGTCTGAACTCTTTCCCTACACGACGCTCTTCCGATCTNNNAATGGGTTCTTGTCATACGACC
GCACACGTCTGAACTCTTTCCCTACACGACGCTCTTCCGATCTNNNAATGGGTTCTTGTCATACGACC
AGCACACGTCTGAACTCTTTCCCTACACGACGCTCTTCCGATCTNNNCAGTATCGTGTCACTGTGCTCC
AGCACACGTCTGAACTCTTTCCCTACACGACGCTCTTCCGATCTNNNCAGTATCGTGTCACTGTGCTCC
AGCACACGTCTGAACTCTTTCCCTACACGACGCTCTTCCGATCTNNNCAGTATCGTGTCACTGTGCTCC
AGCACACGTCTGAACTCTTTCCCTACACGACGCTCTTCCGATCTNNNCAGTATCGTGTCACTGTGCTCC
AGCACACGTCTGAACTCTTTCCCTACACGACGCTCTTCCGATCTNNNCAGTATCGTGTCACTGTGCTCC
AGCACACGTCTGAACTCTTTCCCTACACGACGCTCTTCCGATCTNNNCAGTATCGTGTCACTGTGCTCC
AGCACACGTCTGAACTCTTTCCCTACACGACGCTCTTCCGATCTNNNAGGCTGGTATCAAAAATGTGC
AGCACACGTCTGAACTCTTTCCCTACACGACGCTCTTCCGATCTNNNAGGCTGGTATCAAAAATGTGC

AGCACACGTCTGAACTCTTTCCCTACACGACGCTCTTCCGATCTNNNAGGCTGGTATCAAAAATGTGC
AGCACACGTCTGAACTCTTTCCCTACACGACGCTCTTCCGATCTNNNAGGCTGGTATCAAAAATGTGC
AGCACACGTCTGAACTCTTTCCCTACACGACGCTCTTCCGATCTNNNAGGCTGGTATCAAAAATGTGC
AGCACACGTCTGAACTCTTTCCCTACACGACGCTCTTCCGATCTNNNAGGCTGGTATCAAAAATGTGC
GCACACGTCTGAACTCTTTCCCTACACGACGCTCTTCCGATCTNNNGATTATTTGAGCCCCGGAAGTT
GCACACGTCTGAACTCTTTCCCTACACGACGCTCTTCCGATCTNNNGATTATTTGAGCCCCGGAAGTT
GCACACGTCTGAACTCTTTCCCTACACGACGCTCTTCCGATCTNNNGATTATTTGAGCCCCGGAAGTT
GCACACGTCTGAACTCTTTCCCTACACGACGCTCTTCCGATCTNNNGATTATTTGAGCCCCGGAAGTT
GCACACGTCTGAACTCTTTCCCTACACGACGCTCTTCCGATCTNNNGATTATTTGAGCCCCGGAAGTT
GCACACGTCTGAACTCTTTCCCTACACGACGCTCTTCCGATCTNNNGATTATTTGAGCCCCGGAAGTT
GCACACGTCTGAACTCTTTCCCTACACGACGCTCTTCCGATCTNNNTTTCCCTGGAGAATTGGTCACA
GCACACGTCTGAACTCTTTCCCTACACGACGCTCTTCCGATCTNNNTTTCCCTGGAGAATTGGTCACA
GCACACGTCTGAACTCTTTCCCTACACGACGCTCTTCCGATCTNNNTTTCCCTGGAGAATTGGTCACA
GCACACGTCTGAACTCTTTCCCTACACGACGCTCTTCCGATCTNNNTTTCCCTGGAGAATTGGTCACA
GCACACGTCTGAACTCTTTCCCTACACGACGCTCTTCCGATCTNNNTTTCCCTGGAGAATTGGTCACA
GCACACGTCTGAACTCTTTCCCTACACGACGCTCTTCCGATCTNNNTTTCCCTGGAGAATTGGTCACA
GAGCACACGTCTGAACTCTTTCCCTACACGACGCTCTTCCGATCTNNNGAGGCAGGAAAATCACTTG
GAGCACACGTCTGAACTCTTTCCCTACACGACGCTCTTCCGATCTNNNGAGGCAGGAAAATCACTTG
GAGCACACGTCTGAACTCTTTCCCTACACGACGCTCTTCCGATCTNNNGAGGCAGGAAAATCACTTG
GAGCACACGTCTGAACTCTTTCCCTACACGACGCTCTTCCGATCTNNNGAGGCAGGAAAATCACTTG
GAGCACACGTCTGAACTCTTTCCCTACACGACGCTCTTCCGATCTNNNGAGGCAGGAAAATCACTTG
GAGCACACGTCTGAACTCTTTCCCTACACGACGCTCTTCCGATCTNNNGAGGCAGGAAAATCACTTG
GCACACGTCTGAACTCTTTCCCTACACGACGCTCTTCCGATCTNNNAAAAAACACACACACACATG
GCACACGTCTGAACTCTTTCCCTACACGACGCTCTTCCGATCTNNNAAAAAACACACACACACATG
GCACACGTCTGAACTCTTTCCCTACACGACGCTCTTCCGATCTNNNAAAAAACACACACACACATG
GCACACGTCTGAACTCTTTCCCTACACGACGCTCTTCCGATCTNNNAAAAAACACACACACACATG
GCACACGTCTGAACTCTTTCCCTACACGACGCTCTTCCGATCTNNNAAAAAACACACACACACATG
GCACACGTCTGAACTCTTTCCCTACACGACGCTCTTCCGATCTNNNAAAAAACACACACACACATG
AGCACACGTCTGAACTCTTTCCCTACACGACGCTCTTCCGATCTNNNGACTACAGAGTGAGACCCTGT
AGCACACGTCTGAACTCTTTCCCTACACGACGCTCTTCCGATCTNNNGACTACAGAGTGAGACCCTGT
AGCACACGTCTGAACTCTTTCCCTACACGACGCTCTTCCGATCTNNNGACTACAGAGTGAGACCCTGT
AGCACACGTCTGAACTCTTTCCCTACACGACGCTCTTCCGATCTNNNGACTACAGAGTGAGACCCTGT
AGCACACGTCTGAACTCTTTCCCTACACGACGCTCTTCCGATCTNNNGACTACAGAGTGAGACCCTGT
AGCACACGTCTGAACTCTTTCCCTACACGACGCTCTTCCGATCTNNNGACTACAGAGTGAGACCCTGT
CACACGTCTGAACTCTTTCCCTACACGACGCTCTTCCGATCTNNNTTAAACAGGACTCACAAACGCATC
CACACGTCTGAACTCTTTCCCTACACGACGCTCTTCCGATCTNNNTTAAACAGGACTCACAAACGCATC
CACACGTCTGAACTCTTTCCCTACACGACGCTCTTCCGATCTNNNTTAAACAGGACTCACAAACGCATC
CACACGTCTGAACTCTTTCCCTACACGACGCTCTTCCGATCTNNNTTAAACAGGACTCACAAACGCATC
CACACGTCTGAACTCTTTCCCTACACGACGCTCTTCCGATCTNNNTTAAACAGGACTCACAAACGCATC
CACACGTCTGAACTCTTTCCCTACACGACGCTCTTCCGATCTNNNTTAAACAGGACTCACAAACGCATC
GAGCACACGTCTGAACTCTTTCCCTACACGACGCTCTTCCGATCTNNNATTTTTTTTACAAGCCGTGG
GAGCACACGTCTGAACTCTTTCCCTACACGACGCTCTTCCGATCTNNNATTTTTTTTACAAGCCGTGG
GAGCACACGTCTGAACTCTTTCCCTACACGACGCTCTTCCGATCTNNNATTTTTTTTACAAGCCGTGG

GAGCACACGTCTGAACTCTTTCCCTACACGACGCTCTTCCGATCTNNNATTTTTTTTACAAGCCGTGG
GAGCACACGTCTGAACTCTTTCCCTACACGACGCTCTTCCGATCTNNNATTTTTTTTACAAGCCGTGG
GAGCACACGTCTGAACTCTTTCCCTACACGACGCTCTTCCGATCTNNNATTTTTTTTACAAGCCGTGG
GCACACGTCTGAACTCTTTCCCTACACGACGCTCTTCCGATCTNNNACACCATACCACATGAATATCTG
GCACACGTCTGAACTCTTTCCCTACACGACGCTCTTCCGATCTNNNACACCATACCACATGAATATCTG
GCACACGTCTGAACTCTTTCCCTACACGACGCTCTTCCGATCTNNNACACCATACCACATGAATATCTG
GCACACGTCTGAACTCTTTCCCTACACGACGCTCTTCCGATCTNNNACACCATACCACATGAATATCTG
GCACACGTCTGAACTCTTTCCCTACACGACGCTCTTCCGATCTNNNACACCATACCACATGAATATCTG
GCACACGTCTGAACTCTTTCCCTACACGACGCTCTTCCGATCTNNNACACCATACCACATGAATATCTG
AGCACACGTCTGAACTCTTTCCCTACACGACGCTCTTCCGATCTNNNACAAGTAGGGAGGTGATGA
AGCACACGTCTGAACTCTTTCCCTACACGACGCTCTTCCGATCTNNNACAAGTAGGGAGGTGATGA
AGCACACGTCTGAACTCTTTCCCTACACGACGCTCTTCCGATCTNNNACAAGTAGGGAGGTGATGA
AGCACACGTCTGAACTCTTTCCCTACACGACGCTCTTCCGATCTNNNACAAGTAGGGAGGTGATGA
AGCACACGTCTGAACTCTTTCCCTACACGACGCTCTTCCGATCTNNNACAAGTAGGGAGGTGATGA
AGCACACGTCTGAACTCTTTCCCTACACGACGCTCTTCCGATCTNNNACAAGTAGGGAGGTGATGA
GAGCACACGTCTGAACTCTTTCCCTACACGACGCTCTTCCGATCTNNNCCCTGTCTCGACAAAACAAA
GAGCACACGTCTGAACTCTTTCCCTACACGACGCTCTTCCGATCTNNNCCCTGTCTCGACAAAACAAA
GAGCACACGTCTGAACTCTTTCCCTACACGACGCTCTTCCGATCTNNNCCCTGTCTCGACAAAACAAA
GAGCACACGTCTGAACTCTTTCCCTACACGACGCTCTTCCGATCTNNNCCCTGTCTCGACAAAACAAA
GAGCACACGTCTGAACTCTTTCCCTACACGACGCTCTTCCGATCTNNNCCCTGTCTCGACAAAACAAA
GAGCACACGTCTGAACTCTTTCCCTACACGACGCTCTTCCGATCTNNNCCCTGTCTCGACAAAACAAA
AGCACACGTCTGAACTCTTTCCCTACACGACGCTCTTCCGATCTNNNAGTAACATTTAAACCTGTAAA
AGCACACGTCTGAACTCTTTCCCTACACGACGCTCTTCCGATCTNNNAGTAACATTTAAACCTGTAAA
AGCACACGTCTGAACTCTTTCCCTACACGACGCTCTTCCGATCTNNNAGTAACATTTAAACCTGTAAA
AGCACACGTCTGAACTCTTTCCCTACACGACGCTCTTCCGATCTNNNAGTAACATTTAAACCTGTAAA
AGCACACGTCTGAACTCTTTCCCTACACGACGCTCTTCCGATCTNNNAGTAACATTTAAACCTGTAAA
AGCACACGTCTGAACTCTTTCCCTACACGACGCTCTTCCGATCTNNNAGTAACATTTAAACCTGTAAA
AGCACACGTCTGAACTCTTTCCCTACACGACGCTCTTCCGATCTNNNGACAGTATGGCCAGATTGGTA
AGCACACGTCTGAACTCTTTCCCTACACGACGCTCTTCCGATCTNNNGACAGTATGGCCAGATTGGTA
AGCACACGTCTGAACTCTTTCCCTACACGACGCTCTTCCGATCTNNNGACAGTATGGCCAGATTGGTA
AGCACACGTCTGAACTCTTTCCCTACACGACGCTCTTCCGATCTNNNGACAGTATGGCCAGATTGGTA
AGCACACGTCTGAACTCTTTCCCTACACGACGCTCTTCCGATCTNNNGACAGTATGGCCAGATTGGTA
AGCACACGTCTGAACTCTTTCCCTACACGACGCTCTTCCGATCTNNNGACAGTATGGCCAGATTGGTA
AGCACACGTCTGAACTCTTTCCCTACACGACGCTCTTCCGATCTNNNGCTGGATCTTACATACCCTGA
AGCACACGTCTGAACTCTTTCCCTACACGACGCTCTTCCGATCTNNNGCTGGATCTTACATACCCTGA
AGCACACGTCTGAACTCTTTCCCTACACGACGCTCTTCCGATCTNNNGCTGGATCTTACATACCCTGA
AGCACACGTCTGAACTCTTTCCCTACACGACGCTCTTCCGATCTNNNGCTGGATCTTACATACCCTGA
AGCACACGTCTGAACTCTTTCCCTACACGACGCTCTTCCGATCTNNNGCTGGATCTTACATACCCTGA
AGCACACGTCTGAACTCTTTCCCTACACGACGCTCTTCCGATCTNNNGCTGGATCTTACATACCCTGA
AGCACACGTCTGAACTCTTTCCCTACACGACGCTCTTCCGATCTNNNCTTCCTACTCTGTCCTCACAAG
AGCACACGTCTGAACTCTTTCCCTACACGACGCTCTTCCGATCTNNNCTTCCTACTCTGTCCTCACAAG
AGCACACGTCTGAACTCTTTCCCTACACGACGCTCTTCCGATCTNNNCTTCCTACTCTGTCCTCACAAG
AGCACACGTCTGAACTCTTTCCCTACACGACGCTCTTCCGATCTNNNCTTCCTACTCTGTCCTCACAAG

AGCACACGTCTGAACTCTTTCCCTACACGACGCTCTTCCGATCTNNNCTTCCTACTCTGTCCTCACAAG
AGCACACGTCTGAACTCTTTCCCTACACGACGCTCTTCCGATCTNNNCTTCCTACTCTGTCCTCACAAG
AGCACACGTCTGAACTCTTTCCCTACACGACGCTCTTCCGATCTNNNAGGTTTAGTGTTACAAACTCAG
AGCACACGTCTGAACTCTTTCCCTACACGACGCTCTTCCGATCTNNNAGGTTTAGTGTTACAAACTCAG
AGCACACGTCTGAACTCTTTCCCTACACGACGCTCTTCCGATCTNNNAGGTTTAGTGTTACAAACTCAG
AGCACACGTCTGAACTCTTTCCCTACACGACGCTCTTCCGATCTNNNAGGTTTAGTGTTACAAACTCAG
AGCACACGTCTGAACTCTTTCCCTACACGACGCTCTTCCGATCTNNNAGGTTTAGTGTTACAAACTCAG
AGCACACGTCTGAACTCTTTCCCTACACGACGCTCTTCCGATCTNNNAGGTTTAGTGTTACAAACTCAG
AGCACACGTCTGAACTCTTTCCCTACACGACGCTCTTCCGATCTNNNCTATAGTCCAGCCTTGGTGACA
AGCACACGTCTGAACTCTTTCCCTACACGACGCTCTTCCGATCTNNNCTATAGTCCAGCCTTGGTGACA
AGCACACGTCTGAACTCTTTCCCTACACGACGCTCTTCCGATCTNNNCTATAGTCCAGCCTTGGTGACA
AGCACACGTCTGAACTCTTTCCCTACACGACGCTCTTCCGATCTNNNCTATAGTCCAGCCTTGGTGACA
AGCACACGTCTGAACTCTTTCCCTACACGACGCTCTTCCGATCTNNNCTATAGTCCAGCCTTGGTGACA
AGCACACGTCTGAACTCTTTCCCTACACGACGCTCTTCCGATCTNNNCTATAGTCCAGCCTTGGTGACA
AGCACACGTCTGAACTCTTTCCCTACACGACGCTCTTCCGATCTNNNCCCAAATGATGAAGTGATGCA
AGCACACGTCTGAACTCTTTCCCTACACGACGCTCTTCCGATCTNNNCCCAAATGATGAAGTGATGCA
AGCACACGTCTGAACTCTTTCCCTACACGACGCTCTTCCGATCTNNNCCCAAATGATGAAGTGATGCA
AGCACACGTCTGAACTCTTTCCCTACACGACGCTCTTCCGATCTNNNCCCAAATGATGAAGTGATGCA
AGCACACGTCTGAACTCTTTCCCTACACGACGCTCTTCCGATCTNNNCCCAAATGATGAAGTGATGCA
AGCACACGTCTGAACTCTTTCCCTACACGACGCTCTTCCGATCTNNNCCCAAATGATGAAGTGATGCA
AGCACACGTCTGAACTCTTTCCCTACACGACGCTCTTCCGATCTNNNTTCACTGCATTCTAACCTGGT
AGCACACGTCTGAACTCTTTCCCTACACGACGCTCTTCCGATCTNNNTTCACTGCATTCTAACCTGGT
AGCACACGTCTGAACTCTTTCCCTACACGACGCTCTTCCGATCTNNNTTCACTGCATTCTAACCTGGT
AGCACACGTCTGAACTCTTTCCCTACACGACGCTCTTCCGATCTNNNTTCACTGCATTCTAACCTGGT
AGCACACGTCTGAACTCTTTCCCTACACGACGCTCTTCCGATCTNNNTTCACTGCATTCTAACCTGGT
AGCACACGTCTGAACTCTTTCCCTACACGACGCTCTTCCGATCTNNNTTCACTGCATTCTAACCTGGT
AGCACACGTCTGAACTCTTTCCCTACACGACGCTCTTCCGATCTNNNTTTCACCTTAGCCGTACAGGA
AGCACACGTCTGAACTCTTTCCCTACACGACGCTCTTCCGATCTNNNTTTCACCTTAGCCGTACAGGA
AGCACACGTCTGAACTCTTTCCCTACACGACGCTCTTCCGATCTNNNTTTCACCTTAGCCGTACAGGA
AGCACACGTCTGAACTCTTTCCCTACACGACGCTCTTCCGATCTNNNTTTCACCTTAGCCGTACAGGA
AGCACACGTCTGAACTCTTTCCCTACACGACGCTCTTCCGATCTNNNTTTCACCTTAGCCGTACAGGA
AGCACACGTCTGAACTCTTTCCCTACACGACGCTCTTCCGATCTNNNTTTCACCTTAGCCGTACAGGA
AGAGCACACGTCTGAACTCTTTCCCTACACGACGCTCTTCCGATCTNNNAATCTGCCTGCTTTTGTTCA
AGAGCACACGTCTGAACTCTTTCCCTACACGACGCTCTTCCGATCTNNNAATCTGCCTGCTTTTGTTCA
AGAGCACACGTCTGAACTCTTTCCCTACACGACGCTCTTCCGATCTNNNAATCTGCCTGCTTTTGTTCA
AGAGCACACGTCTGAACTCTTTCCCTACACGACGCTCTTCCGATCTNNNAATCTGCCTGCTTTTGTTCA
AGAGCACACGTCTGAACTCTTTCCCTACACGACGCTCTTCCGATCTNNNAATCTGCCTGCTTTTGTTCA
AGAGCACACGTCTGAACTCTTTCCCTACACGACGCTCTTCCGATCTNNNAATCTGCCTGCTTTTGTTCA
AGCACACGTCTGAACTCTTTCCCTACACGACGCTCTTCCGATCTNNNAATTTTGGCTCACTGCAACCTC
AGCACACGTCTGAACTCTTTCCCTACACGACGCTCTTCCGATCTNNNAATTTTGGCTCACTGCAACCTC
AGCACACGTCTGAACTCTTTCCCTACACGACGCTCTTCCGATCTNNNAATTTTGGCTCACTGCAACCTC
AGCACACGTCTGAACTCTTTCCCTACACGACGCTCTTCCGATCTNNNAATTTTGGCTCACTGCAACCTC
AGCACACGTCTGAACTCTTTCCCTACACGACGCTCTTCCGATCTNNNAATTTTGGCTCACTGCAACCTC

AGCACACGTCTGAACTCTTTCCCTACACGACGCTCTTCCGATCTNNNAATTTTGGCTCACTGCAACCTCC
GCACACGTCTGAACTCTTTCCCTACACGACGCTCTTCCGATCTNNNCCTTTCTTTCTTCCTCCCTCCTGC
GCACACGTCTGAACTCTTTCCCTACACGACGCTCTTCCGATCTNNNCCTTTCTTTCTTCCTCCCTCCTGC
GCACACGTCTGAACTCTTTCCCTACACGACGCTCTTCCGATCTNNNCCTTTCTTTCTTCCTCCCTCCTGC
GCACACGTCTGAACTCTTTCCCTACACGACGCTCTTCCGATCTNNNCCTTTCTTTCTTCCTCCCTCCTGC
GCACACGTCTGAACTCTTTCCCTACACGACGCTCTTCCGATCTNNNCCTTTCTTTCTTCCTCCCTCCTGC
GCACACGTCTGAACTCTTTCCCTACACGACGCTCTTCCGATCTNNNCCTTTCTTTCTTCCTCCCTCCTGC
AGCACACGTCTGAACTCTTTCCCTACACGACGCTCTTCCGATCTNNNGAGAACTCACCAAACTGCTTTC
AGCACACGTCTGAACTCTTTCCCTACACGACGCTCTTCCGATCTNNNGAGAACTCACCAAACTGCTTTC
AGCACACGTCTGAACTCTTTCCCTACACGACGCTCTTCCGATCTNNNGAGAACTCACCAAACTGCTTTC
AGCACACGTCTGAACTCTTTCCCTACACGACGCTCTTCCGATCTNNNGAGAACTCACCAAACTGCTTTC
AGCACACGTCTGAACTCTTTCCCTACACGACGCTCTTCCGATCTNNNGAGAACTCACCAAACTGCTTTC
AGCACACGTCTGAACTCTTTCCCTACACGACGCTCTTCCGATCTNNNGAGAACTCACCAAACTGCTTTC
CACACGTCTGAACTCTTTCCCTACACGACGCTCTTCCGATCTNNNGATTCCCATCCGTCCTTAGGTTGC
CACACGTCTGAACTCTTTCCCTACACGACGCTCTTCCGATCTNNNGATTCCCATCCGTCCTTAGGTTGC
CACACGTCTGAACTCTTTCCCTACACGACGCTCTTCCGATCTNNNGATTCCCATCCGTCCTTAGGTTGC
CACACGTCTGAACTCTTTCCCTACACGACGCTCTTCCGATCTNNNGATTCCCATCCGTCCTTAGGTTGC
CACACGTCTGAACTCTTTCCCTACACGACGCTCTTCCGATCTNNNGATTCCCATCCGTCCTTAGGTTGC
CACACGTCTGAACTCTTTCCCTACACGACGCTCTTCCGATCTNNNGATTCCCATCCGTCCTTAGGTTGC
GAGCACACGTCTGAACTCTTTCCCTACACGACGCTCTTCCGATCTNNNTAACAACAGAGCAAGACTCC
GAGCACACGTCTGAACTCTTTCCCTACACGACGCTCTTCCGATCTNNNTAACAACAGAGCAAGACTCC
GAGCACACGTCTGAACTCTTTCCCTACACGACGCTCTTCCGATCTNNNTAACAACAGAGCAAGACTCC
GAGCACACGTCTGAACTCTTTCCCTACACGACGCTCTTCCGATCTNNNTAACAACAGAGCAAGACTCC
GAGCACACGTCTGAACTCTTTCCCTACACGACGCTCTTCCGATCTNNNTAACAACAGAGCAAGACTCC
GAGCACACGTCTGAACTCTTTCCCTACACGACGCTCTTCCGATCTNNNTAACAACAGAGCAAGACTCC
AGCACACGTCTGAACTCTTTCCCTACACGACGCTCTTCCGATCTNNNTTACCCAGATGTGCTCACTTGA
AGCACACGTCTGAACTCTTTCCCTACACGACGCTCTTCCGATCTNNNTTACCCAGATGTGCTCACTTGA
AGCACACGTCTGAACTCTTTCCCTACACGACGCTCTTCCGATCTNNNTTACCCAGATGTGCTCACTTGA
AGCACACGTCTGAACTCTTTCCCTACACGACGCTCTTCCGATCTNNNTTACCCAGATGTGCTCACTTGA
AGCACACGTCTGAACTCTTTCCCTACACGACGCTCTTCCGATCTNNNTTACCCAGATGTGCTCACTTGA
AGCACACGTCTGAACTCTTTCCCTACACGACGCTCTTCCGATCTNNNTTACCCAGATGTGCTCACTTGA
AGCACACGTCTGAACTCTTTCCCTACACGACGCTCTTCCGATCTNNNACCAGTACCATGCTGTTTTTCT
AGCACACGTCTGAACTCTTTCCCTACACGACGCTCTTCCGATCTNNNACCAGTACCATGCTGTTTTTCT
AGCACACGTCTGAACTCTTTCCCTACACGACGCTCTTCCGATCTNNNACCAGTACCATGCTGTTTTTCT
AGCACACGTCTGAACTCTTTCCCTACACGACGCTCTTCCGATCTNNNACCAGTACCATGCTGTTTTTCT
AGCACACGTCTGAACTCTTTCCCTACACGACGCTCTTCCGATCTNNNACCAGTACCATGCTGTTTTTCT
AGCACACGTCTGAACTCTTTCCCTACACGACGCTCTTCCGATCTNNNACCAGTACCATGCTGTTTTTCT
GCACACGTCTGAACTCTTTCCCTACACGACGCTCTTCCGATCTNNNGCTGATAGAGCCAGACTTTGTC
GCACACGTCTGAACTCTTTCCCTACACGACGCTCTTCCGATCTNNNGCTGATAGAGCCAGACTTTGTC
GCACACGTCTGAACTCTTTCCCTACACGACGCTCTTCCGATCTNNNGCTGATAGAGCCAGACTTTGTC
GCACACGTCTGAACTCTTTCCCTACACGACGCTCTTCCGATCTNNNGCTGATAGAGCCAGACTTTGTC
GCACACGTCTGAACTCTTTCCCTACACGACGCTCTTCCGATCTNNNGCTGATAGAGCCAGACTTTGTC
GCACACGTCTGAACTCTTTCCCTACACGACGCTCTTCCGATCTNNNGCTGATAGAGCCAGACTTTGTC

AGCACACGTCTGAACTCTTTCCCTACACGACGCTCTTCCGATCTNNNAACGCCAAAGTGTTGAAAGAAG
AGCACACGTCTGAACTCTTTCCCTACACGACGCTCTTCCGATCTNNNAACGCCAAAGTGTTGAAAGAAG
AGCACACGTCTGAACTCTTTCCCTACACGACGCTCTTCCGATCTNNNAACGCCAAAGTGTTGAAAGAAG
AGCACACGTCTGAACTCTTTCCCTACACGACGCTCTTCCGATCTNNNAACGCCAAAGTGTTGAAAGAAG
AGCACACGTCTGAACTCTTTCCCTACACGACGCTCTTCCGATCTNNNAACGCCAAAGTGTTGAAAGAAG
AGCACACGTCTGAACTCTTTCCCTACACGACGCTCTTCCGATCTNNNAACGCCAAAGTGTTGAAAGAAG
CACACGTCTGAACTCTTTCCCTACACGACGCTCTTCCGATCTNNNATTGATCAGGATTTATTCACAGACA
CACACGTCTGAACTCTTTCCCTACACGACGCTCTTCCGATCTNNNATTGATCAGGATTTATTCACAGACA
CACACGTCTGAACTCTTTCCCTACACGACGCTCTTCCGATCTNNNATTGATCAGGATTTATTCACAGACA
CACACGTCTGAACTCTTTCCCTACACGACGCTCTTCCGATCTNNNATTGATCAGGATTTATTCACAGACA
CACACGTCTGAACTCTTTCCCTACACGACGCTCTTCCGATCTNNNATTGATCAGGATTTATTCACAGACA
CACACGTCTGAACTCTTTCCCTACACGACGCTCTTCCGATCTNNNATTGATCAGGATTTATTCACAGACA
AGCACACGTCTGAACTCTTTCCCTACACGACGCTCTTCCGATCTNNNTCCTGAGTAGCTGAGACTACAG
AGCACACGTCTGAACTCTTTCCCTACACGACGCTCTTCCGATCTNNNTCCTGAGTAGCTGAGACTACAG
AGCACACGTCTGAACTCTTTCCCTACACGACGCTCTTCCGATCTNNNTCCTGAGTAGCTGAGACTACAG
AGCACACGTCTGAACTCTTTCCCTACACGACGCTCTTCCGATCTNNNTCCTGAGTAGCTGAGACTACAG
AGCACACGTCTGAACTCTTTCCCTACACGACGCTCTTCCGATCTNNNTCCTGAGTAGCTGAGACTACAG
AGCACACGTCTGAACTCTTTCCCTACACGACGCTCTTCCGATCTNNNTCCTGAGTAGCTGAGACTACAG
AGCACACGTCTGAACTCTTTCCCTACACGACGCTCTTCCGATCTNNNTGTAATCCCAGCTACTCAGGAG
AGCACACGTCTGAACTCTTTCCCTACACGACGCTCTTCCGATCTNNNTGTAATCCCAGCTACTCAGGAG
AGCACACGTCTGAACTCTTTCCCTACACGACGCTCTTCCGATCTNNNTGTAATCCCAGCTACTCAGGAG
AGCACACGTCTGAACTCTTTCCCTACACGACGCTCTTCCGATCTNNNTGTAATCCCAGCTACTCAGGAG
AGCACACGTCTGAACTCTTTCCCTACACGACGCTCTTCCGATCTNNNTGTAATCCCAGCTACTCAGGAG
AGCACACGTCTGAACTCTTTCCCTACACGACGCTCTTCCGATCTNNNTGTAATCCCAGCTACTCAGGAG
GAGCACACGTCTGAACTCTTTCCCTACACGACGCTCTTCCGATCTNNNCACTGCACCATCTCCAGAGTTA
GAGCACACGTCTGAACTCTTTCCCTACACGACGCTCTTCCGATCTNNNCACTGCACCATCTCCAGAGTTA
GAGCACACGTCTGAACTCTTTCCCTACACGACGCTCTTCCGATCTNNNCACTGCACCATCTCCAGAGTTA
GAGCACACGTCTGAACTCTTTCCCTACACGACGCTCTTCCGATCTNNNCACTGCACCATCTCCAGAGTTA
GAGCACACGTCTGAACTCTTTCCCTACACGACGCTCTTCCGATCTNNNCACTGCACCATCTCCAGAGTTA
GAGCACACGTCTGAACTCTTTCCCTACACGACGCTCTTCCGATCTNNNCACTGCACCATCTCCAGAGTTA
AGCACACGTCTGAACTCTTTCCCTACACGACGCTCTTCCGATCTNNNGAAGGGGGGAGAAGAAAGTA
AGCACACGTCTGAACTCTTTCCCTACACGACGCTCTTCCGATCTNNNGAAGGGGGGAGAAGAAAGTA
AGCACACGTCTGAACTCTTTCCCTACACGACGCTCTTCCGATCTNNNGAAGGGGGGAGAAGAAAGTA
AGCACACGTCTGAACTCTTTCCCTACACGACGCTCTTCCGATCTNNNGAAGGGGGGAGAAGAAAGTA
AGCACACGTCTGAACTCTTTCCCTACACGACGCTCTTCCGATCTNNNGAAGGGGGGAGAAGAAAGTA
AGCACACGTCTGAACTCTTTCCCTACACGACGCTCTTCCGATCTNNNGAAGGGGGGAGAAGAAAGTA
GAGCACACGTCTGAACTCTTTCCCTACACGACGCTCTTCCGATCTNNNGGGCTACAGAGTGAGATGTTC
GAGCACACGTCTGAACTCTTTCCCTACACGACGCTCTTCCGATCTNNNGGGCTACAGAGTGAGATGTTC
GAGCACACGTCTGAACTCTTTCCCTACACGACGCTCTTCCGATCTNNNGGGCTACAGAGTGAGATGTTC
GAGCACACGTCTGAACTCTTTCCCTACACGACGCTCTTCCGATCTNNNGGGCTACAGAGTGAGATGTTC
GAGCACACGTCTGAACTCTTTCCCTACACGACGCTCTTCCGATCTNNNGGGCTACAGAGTGAGATGTTC
GAGCACACGTCTGAACTCTTTCCCTACACGACGCTCTTCCGATCTNNNGGGCTACAGAGTGAGATGTTC
AGCACACGTCTGAACTCTTTCCCTACACGACGCTCTTCCGATCTNNNTGGGTGTAGGAATGACACTAG

AGCACACGTCTGAACTCTTTCCCTACACGACGCTCTTCCGATCTNNNTGGGTGTAGGAATGACACTAG
AGCACACGTCTGAACTCTTTCCCTACACGACGCTCTTCCGATCTNNNTGGGTGTAGGAATGACACTAG
AGCACACGTCTGAACTCTTTCCCTACACGACGCTCTTCCGATCTNNNTGGGTGTAGGAATGACACTAG
AGCACACGTCTGAACTCTTTCCCTACACGACGCTCTTCCGATCTNNNTGGGTGTAGGAATGACACTAG
AGCACACGTCTGAACTCTTTCCCTACACGACGCTCTTCCGATCTNNNTGGGTGTAGGAATGACACTAG
AGCACACGTCTGAACTCTTTCCCTACACGACGCTCTTCCGATCTNNNAATACTATTGTCCCAGGTGAC
AGCACACGTCTGAACTCTTTCCCTACACGACGCTCTTCCGATCTNNNAATACTATTGTCCCAGGTGAC
AGCACACGTCTGAACTCTTTCCCTACACGACGCTCTTCCGATCTNNNAATACTATTGTCCCAGGTGAC
AGCACACGTCTGAACTCTTTCCCTACACGACGCTCTTCCGATCTNNNAATACTATTGTCCCAGGTGAC
AGCACACGTCTGAACTCTTTCCCTACACGACGCTCTTCCGATCTNNNAATACTATTGTCCCAGGTGAC
AGCACACGTCTGAACTCTTTCCCTACACGACGCTCTTCCGATCTNNNAATACTATTGTCCCAGGTGAC
GCACACGTCTGAACTCTTTCCCTACACGACGCTCTTCCGATCTNNNCTGTAATCCCAGCTACTCAGGAG
GCACACGTCTGAACTCTTTCCCTACACGACGCTCTTCCGATCTNNNCTGTAATCCCAGCTACTCAGGAG
GCACACGTCTGAACTCTTTCCCTACACGACGCTCTTCCGATCTNNNCTGTAATCCCAGCTACTCAGGAG
GCACACGTCTGAACTCTTTCCCTACACGACGCTCTTCCGATCTNNNCTGTAATCCCAGCTACTCAGGAG
GCACACGTCTGAACTCTTTCCCTACACGACGCTCTTCCGATCTNNNCTGTAATCCCAGCTACTCAGGAG
GCACACGTCTGAACTCTTTCCCTACACGACGCTCTTCCGATCTNNNCTGTAATCCCAGCTACTCAGGAG
GAGCACACGTCTGAACTCTTTCCCTACACGACGCTCTTCCGATCTNNNTCACTCCATGTTTTCCCTTTTCC
GAGCACACGTCTGAACTCTTTCCCTACACGACGCTCTTCCGATCTNNNTCACTCCATGTTTTCCCTTTTCC
GAGCACACGTCTGAACTCTTTCCCTACACGACGCTCTTCCGATCTNNNTCACTCCATGTTTTCCCTTTTCC
GAGCACACGTCTGAACTCTTTCCCTACACGACGCTCTTCCGATCTNNNTCACTCCATGTTTTCCCTTTTCC
GAGCACACGTCTGAACTCTTTCCCTACACGACGCTCTTCCGATCTNNNTCACTCCATGTTTTCCCTTTTCC
GAGCACACGTCTGAACTCTTTCCCTACACGACGCTCTTCCGATCTNNNTCACTCCATGTTTTCCCTTTTCC
AGCACACGTCTGAACTCTTTCCCTACACGACGCTCTTCCGATCTNNNTCCCAATGAAAGTAAAGTACAA
AGCACACGTCTGAACTCTTTCCCTACACGACGCTCTTCCGATCTNNNTCCCAATGAAAGTAAAGTACAA
AGCACACGTCTGAACTCTTTCCCTACACGACGCTCTTCCGATCTNNNTCCCAATGAAAGTAAAGTACAA
AGCACACGTCTGAACTCTTTCCCTACACGACGCTCTTCCGATCTNNNTCCCAATGAAAGTAAAGTACAA
AGCACACGTCTGAACTCTTTCCCTACACGACGCTCTTCCGATCTNNNTCCCAATGAAAGTAAAGTACAA
AGCACACGTCTGAACTCTTTCCCTACACGACGCTCTTCCGATCTNNNTCCCAATGAAAGTAAAGTACAA
AGCACACGTCTGAACTCTTTCCCTACACGACGCTCTTCCGATCTNNNTCTGTGACTACCTATGGTTTTGT
AGCACACGTCTGAACTCTTTCCCTACACGACGCTCTTCCGATCTNNNTCTGTGACTACCTATGGTTTTGT
AGCACACGTCTGAACTCTTTCCCTACACGACGCTCTTCCGATCTNNNTCTGTGACTACCTATGGTTTTGT
AGCACACGTCTGAACTCTTTCCCTACACGACGCTCTTCCGATCTNNNTCTGTGACTACCTATGGTTTTGT
AGCACACGTCTGAACTCTTTCCCTACACGACGCTCTTCCGATCTNNNTCTGTGACTACCTATGGTTTTGT
AGCACACGTCTGAACTCTTTCCCTACACGACGCTCTTCCGATCTNNNTCTGTGACTACCTATGGTTTTGT
AGCACACGTCTGAACTCTTTCCCTACACGACGCTCTTCCGATCTNNNTCTTCTAATGCACCTCGCTTTCT
AGCACACGTCTGAACTCTTTCCCTACACGACGCTCTTCCGATCTNNNTCTTCTAATGCACCTCGCTTTCT
AGCACACGTCTGAACTCTTTCCCTACACGACGCTCTTCCGATCTNNNTCTTCTAATGCACCTCGCTTTCT
AGCACACGTCTGAACTCTTTCCCTACACGACGCTCTTCCGATCTNNNTCTTCTAATGCACCTCGCTTTCT
AGCACACGTCTGAACTCTTTCCCTACACGACGCTCTTCCGATCTNNNTCTTCTAATGCACCTCGCTTTCT
AGCACACGTCTGAACTCTTTCCCTACACGACGCTCTTCCGATCTNNNTCTTCTAATGCACCTCGCTTTCT
AGCACACGTCTGAACTCTTTCCCTACACGACGCTCTTCCGATCTNNNAAGAATTTCACCTTGTTCCCAA
AGCACACGTCTGAACTCTTTCCCTACACGACGCTCTTCCGATCTNNNAAGAATTTCACCTTGTTCCCAA

AGCACACGTCTGAACTCTTTCCCTACACGACGCTCTTCCGATCTNNNAAGAATTTCACCTTGTTCCCAA
AGCACACGTCTGAACTCTTTCCCTACACGACGCTCTTCCGATCTNNNAAGAATTTCACCTTGTTCCCAA
AGCACACGTCTGAACTCTTTCCCTACACGACGCTCTTCCGATCTNNNAAGAATTTCACCTTGTTCCCAA
AGCACACGTCTGAACTCTTTCCCTACACGACGCTCTTCCGATCTNNNAAGAATTTCACCTTGTTCCCAA
GAGCACACGTCTGAACTCTTTCCCTACACGACGCTCTTCCGATCTNNNGAGGCTGAGACAGGAGAATC
GAGCACACGTCTGAACTCTTTCCCTACACGACGCTCTTCCGATCTNNNGAGGCTGAGACAGGAGAATC
GAGCACACGTCTGAACTCTTTCCCTACACGACGCTCTTCCGATCTNNNGAGGCTGAGACAGGAGAATC
GAGCACACGTCTGAACTCTTTCCCTACACGACGCTCTTCCGATCTNNNGAGGCTGAGACAGGAGAATC
GAGCACACGTCTGAACTCTTTCCCTACACGACGCTCTTCCGATCTNNNGAGGCTGAGACAGGAGAATC
GAGCACACGTCTGAACTCTTTCCCTACACGACGCTCTTCCGATCTNNNGAGGCTGAGACAGGAGAATC
AGCACACGTCTGAACTCTTTCCCTACACGACGCTCTTCCGATCTNNNAGTGAGCAGAGATTGTACCAC
AGCACACGTCTGAACTCTTTCCCTACACGACGCTCTTCCGATCTNNNAGTGAGCAGAGATTGTACCAC
AGCACACGTCTGAACTCTTTCCCTACACGACGCTCTTCCGATCTNNNAGTGAGCAGAGATTGTACCAC
AGCACACGTCTGAACTCTTTCCCTACACGACGCTCTTCCGATCTNNNAGTGAGCAGAGATTGTACCAC
AGCACACGTCTGAACTCTTTCCCTACACGACGCTCTTCCGATCTNNNAGTGAGCAGAGATTGTACCAC
AGCACACGTCTGAACTCTTTCCCTACACGACGCTCTTCCGATCTNNNAGTGAGCAGAGATTGTACCAC
GAGCACACGTCTGAACTCTTTCCCTACACGACGCTCTTCCGATCTNNNGCTAATCTAGCGAGAATGTG
GAGCACACGTCTGAACTCTTTCCCTACACGACGCTCTTCCGATCTNNNGCTAATCTAGCGAGAATGTG
GAGCACACGTCTGAACTCTTTCCCTACACGACGCTCTTCCGATCTNNNGCTAATCTAGCGAGAATGTG
GAGCACACGTCTGAACTCTTTCCCTACACGACGCTCTTCCGATCTNNNGCTAATCTAGCGAGAATGTG
GAGCACACGTCTGAACTCTTTCCCTACACGACGCTCTTCCGATCTNNNGCTAATCTAGCGAGAATGTG
GAGCACACGTCTGAACTCTTTCCCTACACGACGCTCTTCCGATCTNNNGCTAATCTAGCGAGAATGTG
CACGTCTGAACTCTTTCCCTACACGACGCTCTTCCGATCTNNNTAAAAAATAAGGCTATTGGATGCCA
CACGTCTGAACTCTTTCCCTACACGACGCTCTTCCGATCTNNNTAAAAAATAAGGCTATTGGATGCCA
CACGTCTGAACTCTTTCCCTACACGACGCTCTTCCGATCTNNNTAAAAAATAAGGCTATTGGATGCCA
CACGTCTGAACTCTTTCCCTACACGACGCTCTTCCGATCTNNNTAAAAAATAAGGCTATTGGATGCCA
CACGTCTGAACTCTTTCCCTACACGACGCTCTTCCGATCTNNNTAAAAAATAAGGCTATTGGATGCCA
CACGTCTGAACTCTTTCCCTACACGACGCTCTTCCGATCTNNNTAAAAAATAAGGCTATTGGATGCCA
GAGCACACGTCTGAACTCTTTCCCTACACGACGCTCTTCCGATCTNNNAGACTGACACTTTAGGGAG
GAGCACACGTCTGAACTCTTTCCCTACACGACGCTCTTCCGATCTNNNAGACTGACACTTTAGGGAG
GAGCACACGTCTGAACTCTTTCCCTACACGACGCTCTTCCGATCTNNNAGACTGACACTTTAGGGAG
GAGCACACGTCTGAACTCTTTCCCTACACGACGCTCTTCCGATCTNNNAGACTGACACTTTAGGGAG
GAGCACACGTCTGAACTCTTTCCCTACACGACGCTCTTCCGATCTNNNAGACTGACACTTTAGGGAG
GAGCACACGTCTGAACTCTTTCCCTACACGACGCTCTTCCGATCTNNNAGACTGACACTTTAGGGAG
GCACACGTCTGAACTCTTTCCCTACACGACGCTCTTCCGATCTNNNAGGGTTCCAATTTCTCCACCTAC
GCACACGTCTGAACTCTTTCCCTACACGACGCTCTTCCGATCTNNNAGGGTTCCAATTTCTCCACCTAC
GCACACGTCTGAACTCTTTCCCTACACGACGCTCTTCCGATCTNNNAGGGTTCCAATTTCTCCACCTAC
GCACACGTCTGAACTCTTTCCCTACACGACGCTCTTCCGATCTNNNAGGGTTCCAATTTCTCCACCTAC
GCACACGTCTGAACTCTTTCCCTACACGACGCTCTTCCGATCTNNNAGGGTTCCAATTTCTCCACCTAC
GCACACGTCTGAACTCTTTCCCTACACGACGCTCTTCCGATCTNNNAGGGTTCCAATTTCTCCACCTAC
CACACGTCTGAACTCTTTCCCTACACGACGCTCTTCCGATCTNNNTTCATCCATCCATGTACCCACTCCA
CACACGTCTGAACTCTTTCCCTACACGACGCTCTTCCGATCTNNNTTCATCCATCCATGTACCCACTCCA
CACACGTCTGAACTCTTTCCCTACACGACGCTCTTCCGATCTNNNTTCATCCATCCATGTACCCACTCCA

GCACACGTCTGAACTCTTTCCCTACACGACGCTCTTCCGATCTNNNTTCATCCATCCATGTACCCACTCCA
GCACACGTCTGAACTCTTTCCCTACACGACGCTCTTCCGATCTNNNTTCATCCATCCATGTACCCACTCCA
GCACACGTCTGAACTCTTTCCCTACACGACGCTCTTCCGATCTNNNTTCATCCATCCATGTACCCACTCCA
GAGCACACGTCTGAACTCTTTCCCTACACGACGCTCTTCCGATCTNNNGAGGTTGCATTGAGGTGAGA
GAGCACACGTCTGAACTCTTTCCCTACACGACGCTCTTCCGATCTNNNGAGGTTGCATTGAGGTGAGA
GAGCACACGTCTGAACTCTTTCCCTACACGACGCTCTTCCGATCTNNNGAGGTTGCATTGAGGTGAGA
GAGCACACGTCTGAACTCTTTCCCTACACGACGCTCTTCCGATCTNNNGAGGTTGCATTGAGGTGAGA
GAGCACACGTCTGAACTCTTTCCCTACACGACGCTCTTCCGATCTNNNGAGGTTGCATTGAGGTGAGA
GAGCACACGTCTGAACTCTTTCCCTACACGACGCTCTTCCGATCTNNNGAGGTTGCATTGAGGTGAGA
AGCACACGTCTGAACTCTTTCCCTACACGACGCTCTTCCGATCTNNNCAAAATGCAAAGCTTTGTGGAG
AGCACACGTCTGAACTCTTTCCCTACACGACGCTCTTCCGATCTNNNCAAAATGCAAAGCTTTGTGGAG
AGCACACGTCTGAACTCTTTCCCTACACGACGCTCTTCCGATCTNNNCAAAATGCAAAGCTTTGTGGAG
AGCACACGTCTGAACTCTTTCCCTACACGACGCTCTTCCGATCTNNNCAAAATGCAAAGCTTTGTGGAG
AGCACACGTCTGAACTCTTTCCCTACACGACGCTCTTCCGATCTNNNCAAAATGCAAAGCTTTGTGGAG
AGCACACGTCTGAACTCTTTCCCTACACGACGCTCTTCCGATCTNNNCAAAATGCAAAGCTTTGTGGAG
AGCACACGTCTGAACTCTTTCCCTACACGACGCTCTTCCGATCTNNNCATCTGTAGTCCCAGCTACTCAG
AGCACACGTCTGAACTCTTTCCCTACACGACGCTCTTCCGATCTNNNCATCTGTAGTCCCAGCTACTCAG
AGCACACGTCTGAACTCTTTCCCTACACGACGCTCTTCCGATCTNNNCATCTGTAGTCCCAGCTACTCAG
AGCACACGTCTGAACTCTTTCCCTACACGACGCTCTTCCGATCTNNNCATCTGTAGTCCCAGCTACTCAG
AGCACACGTCTGAACTCTTTCCCTACACGACGCTCTTCCGATCTNNNCATCTGTAGTCCCAGCTACTCAG
AGCACACGTCTGAACTCTTTCCCTACACGACGCTCTTCCGATCTNNNCATCTGTAGTCCCAGCTACTCAG
AGCACACGTCTGAACTCTTTCCCTACACGACGCTCTTCCGATCTNNNGCATAATGTTGAGGTTTGGGCT
AGCACACGTCTGAACTCTTTCCCTACACGACGCTCTTCCGATCTNNNGCATAATGTTGAGGTTTGGGCT
AGCACACGTCTGAACTCTTTCCCTACACGACGCTCTTCCGATCTNNNGCATAATGTTGAGGTTTGGGCT
AGCACACGTCTGAACTCTTTCCCTACACGACGCTCTTCCGATCTNNNGCATAATGTTGAGGTTTGGGCT
AGCACACGTCTGAACTCTTTCCCTACACGACGCTCTTCCGATCTNNNGCATAATGTTGAGGTTTGGGCT
AGCACACGTCTGAACTCTTTCCCTACACGACGCTCTTCCGATCTNNNGCATAATGTTGAGGTTTGGGCT
AGCACACGTCTGAACTCTTTCCCTACACGACGCTCTTCCGATCTNNNTATACCAAGAGGGAAACAAGCC
AGCACACGTCTGAACTCTTTCCCTACACGACGCTCTTCCGATCTNNNTATACCAAGAGGGAAACAAGCC
AGCACACGTCTGAACTCTTTCCCTACACGACGCTCTTCCGATCTNNNTATACCAAGAGGGAAACAAGCC
AGCACACGTCTGAACTCTTTCCCTACACGACGCTCTTCCGATCTNNNTATACCAAGAGGGAAACAAGCC
AGCACACGTCTGAACTCTTTCCCTACACGACGCTCTTCCGATCTNNNTATACCAAGAGGGAAACAAGCC
AGCACACGTCTGAACTCTTTCCCTACACGACGCTCTTCCGATCTNNNTATACCAAGAGGGAAACAAGCC
GAGCACACGTCTGAACTCTTTCCCTACACGACGCTCTTCCGATCTNNNTTAGTGTTCATAGCCAGCCAAG
GAGCACACGTCTGAACTCTTTCCCTACACGACGCTCTTCCGATCTNNNTTAGTGTTCATAGCCAGCCAAG
GAGCACACGTCTGAACTCTTTCCCTACACGACGCTCTTCCGATCTNNNTTAGTGTTCATAGCCAGCCAAG
GAGCACACGTCTGAACTCTTTCCCTACACGACGCTCTTCCGATCTNNNTTAGTGTTCATAGCCAGCCAAG
GAGCACACGTCTGAACTCTTTCCCTACACGACGCTCTTCCGATCTNNNTTAGTGTTCATAGCCAGCCAAG
GAGCACACGTCTGAACTCTTTCCCTACACGACGCTCTTCCGATCTNNNTTAGTGTTCATAGCCAGCCAAG
GAGCACACGTCTGAACTCTTTCCCTACACGACGCTCTTCCGATCTNNNTTTTTGAGACAGGGTCTTACCC
GAGCACACGTCTGAACTCTTTCCCTACACGACGCTCTTCCGATCTNNNTTTTTGAGACAGGGTCTTACCC
GAGCACACGTCTGAACTCTTTCCCTACACGACGCTCTTCCGATCTNNNTTTTTGAGACAGGGTCTTACCC
GAGCACACGTCTGAACTCTTTCCCTACACGACGCTCTTCCGATCTNNNTTTTTGAGACAGGGTCTTACCC

AGCACACGTCTGAACTCTTTCCCTACACGACGCTCTTCCGATCTNNNTTTTTGAGACAGGGTCTTACC
AGCACACGTCTGAACTCTTTCCCTACACGACGCTCTTCCGATCTNNNTTTTTGAGACAGGGTCTTACC
AGCACACGTCTGAACTCTTTCCCTACACGACGCTCTTCCGATCTNNNATGAGCAAATTCCTTCACACTG
AGCACACGTCTGAACTCTTTCCCTACACGACGCTCTTCCGATCTNNNATGAGCAAATTCCTTCACACTG
AGCACACGTCTGAACTCTTTCCCTACACGACGCTCTTCCGATCTNNNATGAGCAAATTCCTTCACACTG
AGCACACGTCTGAACTCTTTCCCTACACGACGCTCTTCCGATCTNNNATGAGCAAATTCCTTCACACTG
AGCACACGTCTGAACTCTTTCCCTACACGACGCTCTTCCGATCTNNNATGAGCAAATTCCTTCACACTG
AGCACACGTCTGAACTCTTTCCCTACACGACGCTCTTCCGATCTNNNATGAGCAAATTCCTTCACACTG
AGCACACGTCTGAACTCTTTCCCTACACGACGCTCTTCCGATCTNNNATAATCTACAATTATCCAGGCT
AGCACACGTCTGAACTCTTTCCCTACACGACGCTCTTCCGATCTNNNATAATCTACAATTATCCAGGCT
AGCACACGTCTGAACTCTTTCCCTACACGACGCTCTTCCGATCTNNNATAATCTACAATTATCCAGGCT
AGCACACGTCTGAACTCTTTCCCTACACGACGCTCTTCCGATCTNNNATAATCTACAATTATCCAGGCT
AGCACACGTCTGAACTCTTTCCCTACACGACGCTCTTCCGATCTNNNATAATCTACAATTATCCAGGCT
AGCACACGTCTGAACTCTTTCCCTACACGACGCTCTTCCGATCTNNNATAATCTACAATTATCCAGGCT
AGCACACGTCTGAACTCTTTCCCTACACGACGCTCTTCCGATCTNNNGAGGGAAGGAGAAAGGAAG
AGCACACGTCTGAACTCTTTCCCTACACGACGCTCTTCCGATCTNNNGAGGGAAGGAGAAAGGAAG
AGCACACGTCTGAACTCTTTCCCTACACGACGCTCTTCCGATCTNNNGAGGGAAGGAGAAAGGAAG
AGCACACGTCTGAACTCTTTCCCTACACGACGCTCTTCCGATCTNNNGAGGGAAGGAGAAAGGAAG
AGCACACGTCTGAACTCTTTCCCTACACGACGCTCTTCCGATCTNNNGAGGGAAGGAGAAAGGAAG
AGCACACGTCTGAACTCTTTCCCTACACGACGCTCTTCCGATCTNNNGAGGGAAGGAGAAAGGAAG
GCACACGTCTGAACTCTTTCCCTACACGACGCTCTTCCGATCTNNNCCATGGAGGGTAAGGGTACTT
GCACACGTCTGAACTCTTTCCCTACACGACGCTCTTCCGATCTNNNCCATGGAGGGTAAGGGTACTT
GCACACGTCTGAACTCTTTCCCTACACGACGCTCTTCCGATCTNNNCCATGGAGGGTAAGGGTACTT
GCACACGTCTGAACTCTTTCCCTACACGACGCTCTTCCGATCTNNNCCATGGAGGGTAAGGGTACTT
GCACACGTCTGAACTCTTTCCCTACACGACGCTCTTCCGATCTNNNCCATGGAGGGTAAGGGTACTT
GCACACGTCTGAACTCTTTCCCTACACGACGCTCTTCCGATCTNNNCCATGGAGGGTAAGGGTACTT
AGCACACGTCTGAACTCTTTCCCTACACGACGCTCTTCCGATCTNNNGAGTTCTCACATATCTACGCCC
AGCACACGTCTGAACTCTTTCCCTACACGACGCTCTTCCGATCTNNNGAGTTCTCACATATCTACGCCC
AGCACACGTCTGAACTCTTTCCCTACACGACGCTCTTCCGATCTNNNGAGTTCTCACATATCTACGCCC
AGCACACGTCTGAACTCTTTCCCTACACGACGCTCTTCCGATCTNNNGAGTTCTCACATATCTACGCCC
AGCACACGTCTGAACTCTTTCCCTACACGACGCTCTTCCGATCTNNNGAGTTCTCACATATCTACGCCC
AGCACACGTCTGAACTCTTTCCCTACACGACGCTCTTCCGATCTNNNGAGTTCTCACATATCTACGCCC
AGCACACGTCTGAACTCTTTCCCTACACGACGCTCTTCCGATCTNNNCTGTAATCCCAGCTACTCAGG
AGCACACGTCTGAACTCTTTCCCTACACGACGCTCTTCCGATCTNNNCTGTAATCCCAGCTACTCAGG
AGCACACGTCTGAACTCTTTCCCTACACGACGCTCTTCCGATCTNNNCTGTAATCCCAGCTACTCAGG
AGCACACGTCTGAACTCTTTCCCTACACGACGCTCTTCCGATCTNNNCTGTAATCCCAGCTACTCAGG
AGCACACGTCTGAACTCTTTCCCTACACGACGCTCTTCCGATCTNNNCTGTAATCCCAGCTACTCAGG
AGCACACGTCTGAACTCTTTCCCTACACGACGCTCTTCCGATCTNNNCTGTAATCCCAGCTACTCAGG
AGCACACGTCTGAACTCTTTCCCTACACGACGCTCTTCCGATCTNNNGAGGTCAGAAGTTCAAAACCA
AGCACACGTCTGAACTCTTTCCCTACACGACGCTCTTCCGATCTNNNGAGGTCAGAAGTTCAAAACCA
AGCACACGTCTGAACTCTTTCCCTACACGACGCTCTTCCGATCTNNNGAGGTCAGAAGTTCAAAACCA
AGCACACGTCTGAACTCTTTCCCTACACGACGCTCTTCCGATCTNNNGAGGTCAGAAGTTCAAAACCA
AGCACACGTCTGAACTCTTTCCCTACACGACGCTCTTCCGATCTNNNGAGGTCAGAAGTTCAAAACCA

AGCACACGTCTGAACTCTTTCCCTACACGACGCTCTTCCGATCTNNNGAGGTCAGAAGTTCAAAACCA
AGCACACGTCTGAACTCTTTCCCTACACGACGCTCTTCCGATCTNNNTGTTTGTTGGCCTGAACATTTG
AGCACACGTCTGAACTCTTTCCCTACACGACGCTCTTCCGATCTNNNTGTTTGTTGGCCTGAACATTTG
AGCACACGTCTGAACTCTTTCCCTACACGACGCTCTTCCGATCTNNNTGTTTGTTGGCCTGAACATTTG
AGCACACGTCTGAACTCTTTCCCTACACGACGCTCTTCCGATCTNNNTGTTTGTTGGCCTGAACATTTG
AGCACACGTCTGAACTCTTTCCCTACACGACGCTCTTCCGATCTNNNTGTTTGTTGGCCTGAACATTTG
AGCACACGTCTGAACTCTTTCCCTACACGACGCTCTTCCGATCTNNNTGTTTGTTGGCCTGAACATTTG
AGCACACGTCTGAACTCTTTCCCTACACGACGCTCTTCCGATCTNNNAAAGAATATCAGAGCAGGGCT
AGCACACGTCTGAACTCTTTCCCTACACGACGCTCTTCCGATCTNNNAAAGAATATCAGAGCAGGGCT
AGCACACGTCTGAACTCTTTCCCTACACGACGCTCTTCCGATCTNNNAAAGAATATCAGAGCAGGGCT
AGCACACGTCTGAACTCTTTCCCTACACGACGCTCTTCCGATCTNNNAAAGAATATCAGAGCAGGGCT
AGCACACGTCTGAACTCTTTCCCTACACGACGCTCTTCCGATCTNNNAAAGAATATCAGAGCAGGGCT
AGCACACGTCTGAACTCTTTCCCTACACGACGCTCTTCCGATCTNNNAAAGAATATCAGAGCAGGGCT
AGCACACGTCTGAACTCTTTCCCTACACGACGCTCTTCCGATCTNNNCCTCCTCAAGGGTTTAGGATG
AGCACACGTCTGAACTCTTTCCCTACACGACGCTCTTCCGATCTNNNCCTCCTCAAGGGTTTAGGATG
AGCACACGTCTGAACTCTTTCCCTACACGACGCTCTTCCGATCTNNNCCTCCTCAAGGGTTTAGGATG
AGCACACGTCTGAACTCTTTCCCTACACGACGCTCTTCCGATCTNNNCCTCCTCAAGGGTTTAGGATG
AGCACACGTCTGAACTCTTTCCCTACACGACGCTCTTCCGATCTNNNCCTCCTCAAGGGTTTAGGATG
AGCACACGTCTGAACTCTTTCCCTACACGACGCTCTTCCGATCTNNNCCTCCTCAAGGGTTTAGGATG
AGCACACGTCTGAACTCTTTCCCTACACGACGCTCTTCCGATCTNNNCTGAGGCAGGAGAATAGCTT
AGCACACGTCTGAACTCTTTCCCTACACGACGCTCTTCCGATCTNNNCTGAGGCAGGAGAATAGCTT
AGCACACGTCTGAACTCTTTCCCTACACGACGCTCTTCCGATCTNNNCTGAGGCAGGAGAATAGCTT
AGCACACGTCTGAACTCTTTCCCTACACGACGCTCTTCCGATCTNNNCTGAGGCAGGAGAATAGCTT
AGCACACGTCTGAACTCTTTCCCTACACGACGCTCTTCCGATCTNNNCTGAGGCAGGAGAATAGCTT
AGCACACGTCTGAACTCTTTCCCTACACGACGCTCTTCCGATCTNNNCTGAGGCAGGAGAATAGCTT
AGCACACGTCTGAACTCTTTCCCTACACGACGCTCTTCCGATCTNNNCAAAAAGACACTGTGAGGGCT
AGCACACGTCTGAACTCTTTCCCTACACGACGCTCTTCCGATCTNNNCAAAAAGACACTGTGAGGGCT
AGCACACGTCTGAACTCTTTCCCTACACGACGCTCTTCCGATCTNNNCAAAAAGACACTGTGAGGGCT
AGCACACGTCTGAACTCTTTCCCTACACGACGCTCTTCCGATCTNNNCAAAAAGACACTGTGAGGGCT
AGCACACGTCTGAACTCTTTCCCTACACGACGCTCTTCCGATCTNNNCAAAAAGACACTGTGAGGGCT
AGCACACGTCTGAACTCTTTCCCTACACGACGCTCTTCCGATCTNNNCAAAAAGACACTGTGAGGGCT
AGAGCACACGTCTGAACTCTTTCCCTACACGACGCTCTTCCGATCTNNNAAAAAAAAATTAGCCGGGCA
AGAGCACACGTCTGAACTCTTTCCCTACACGACGCTCTTCCGATCTNNNAAAAAAAAATTAGCCGGGCA
AGAGCACACGTCTGAACTCTTTCCCTACACGACGCTCTTCCGATCTNNNAAAAAAAAATTAGCCGGGCA
AGAGCACACGTCTGAACTCTTTCCCTACACGACGCTCTTCCGATCTNNNAAAAAAAAATTAGCCGGGCA
AGAGCACACGTCTGAACTCTTTCCCTACACGACGCTCTTCCGATCTNNNAAAAAAAAATTAGCCGGGCA
AGAGCACACGTCTGAACTCTTTCCCTACACGACGCTCTTCCGATCTNNNAAAAAAAAATTAGCCGGGCA
AGCACACGTCTGAACTCTTTCCCTACACGACGCTCTTCCGATCTNNNCTTGTCCCTGACATTCTCAGTGA
AGCACACGTCTGAACTCTTTCCCTACACGACGCTCTTCCGATCTNNNCTTGTCCCTGACATTCTCAGTGA
AGCACACGTCTGAACTCTTTCCCTACACGACGCTCTTCCGATCTNNNCTTGTCCCTGACATTCTCAGTGA
AGCACACGTCTGAACTCTTTCCCTACACGACGCTCTTCCGATCTNNNCTTGTCCCTGACATTCTCAGTGA
AGCACACGTCTGAACTCTTTCCCTACACGACGCTCTTCCGATCTNNNCTTGTCCCTGACATTCTCAGTGA
AGCACACGTCTGAACTCTTTCCCTACACGACGCTCTTCCGATCTNNNCTTGTCCCTGACATTCTCAGTGA

GCACACGTCTGAACTCTTTCCCTACACGACGCTCTTCCGATCTNNNGCCTCACCTGATCATAGTACCTG

GCACACGTCTGAACTCTTTCCCTACACGACGCTCTTCCGATCTNNNGCCTCACCTGATCATAGTACCTG

GCACACGTCTGAACTCTTTCCCTACACGACGCTCTTCCGATCTNNNGCCTCACCTGATCATAGTACCTG

GCACACGTCTGAACTCTTTCCCTACACGACGCTCTTCCGATCTNNNGCCTCACCTGATCATAGTACCTG

GCACACGTCTGAACTCTTTCCCTACACGACGCTCTTCCGATCTNNNGCCTCACCTGATCATAGTACCTG

GCACACGTCTGAACTCTTTCCCTACACGACGCTCTTCCGATCTNNNGCCTCACCTGATCATAGTACCTG

GCACACGTCTGAACTCTTTCCCTACACGACGCTCTTCCGATCTNNNCAGCCTAGGCAAAAATGGTGAA

GCACACGTCTGAACTCTTTCCCTACACGACGCTCTTCCGATCTNNNCAGCCTAGGCAAAAATGGTGAA

GCACACGTCTGAACTCTTTCCCTACACGACGCTCTTCCGATCTNNNCAGCCTAGGCAAAAATGGTGAA

GCACACGTCTGAACTCTTTCCCTACACGACGCTCTTCCGATCTNNNCAGCCTAGGCAAAAATGGTGAA

GCACACGTCTGAACTCTTTCCCTACACGACGCTCTTCCGATCTNNNCAGCCTAGGCAAAAATGGTGAA

GCACACGTCTGAACTCTTTCCCTACACGACGCTCTTCCGATCTNNNCAGCCTAGGCAAAAATGGTGAA

GAGCACACGTCTGAACTCTTTCCCTACACGACGCTCTTCCGATCTNNNCTGTAATCCCAGCTACTCAGG

GAGCACACGTCTGAACTCTTTCCCTACACGACGCTCTTCCGATCTNNNCTGTAATCCCAGCTACTCAGG

GAGCACACGTCTGAACTCTTTCCCTACACGACGCTCTTCCGATCTNNNCTGTAATCCCAGCTACTCAGG

GAGCACACGTCTGAACTCTTTCCCTACACGACGCTCTTCCGATCTNNNCTGTAATCCCAGCTACTCAGG

GAGCACACGTCTGAACTCTTTCCCTACACGACGCTCTTCCGATCTNNNCTGTAATCCCAGCTACTCAGG

GAGCACACGTCTGAACTCTTTCCCTACACGACGCTCTTCCGATCTNNNCTGTAATCCCAGCTACTCAGG

AGCACACGTCTGAACTCTTTCCCTACACGACGCTCTTCCGATCTNNNTGCTATGTTACCCAGGCTGATT

AGCACACGTCTGAACTCTTTCCCTACACGACGCTCTTCCGATCTNNNTGCTATGTTACCCAGGCTGATT

AGCACACGTCTGAACTCTTTCCCTACACGACGCTCTTCCGATCTNNNTGCTATGTTACCCAGGCTGATT

AGCACACGTCTGAACTCTTTCCCTACACGACGCTCTTCCGATCTNNNTGCTATGTTACCCAGGCTGATT

AGCACACGTCTGAACTCTTTCCCTACACGACGCTCTTCCGATCTNNNTGCTATGTTACCCAGGCTGATT

AGCACACGTCTGAACTCTTTCCCTACACGACGCTCTTCCGATCTNNNTGCTATGTTACCCAGGCTGATT

AGCACACGTCTGAACTCTTTCCCTACACGACGCTCTTCCGATCTNNNGAGAATTGCTTGAACCCAGGA

AGCACACGTCTGAACTCTTTCCCTACACGACGCTCTTCCGATCTNNNGAGAATTGCTTGAACCCAGGA

AGCACACGTCTGAACTCTTTCCCTACACGACGCTCTTCCGATCTNNNGAGAATTGCTTGAACCCAGGA

AGCACACGTCTGAACTCTTTCCCTACACGACGCTCTTCCGATCTNNNGAGAATTGCTTGAACCCAGGA

AGCACACGTCTGAACTCTTTCCCTACACGACGCTCTTCCGATCTNNNGAGAATTGCTTGAACCCAGGA

AGCACACGTCTGAACTCTTTCCCTACACGACGCTCTTCCGATCTNNNGAGAATTGCTTGAACCCAGGA

AGCACACGTCTGAACTCTTTCCCTACACGACGCTCTTCCGATCTNNNCTGAGGCAGGAAAATTGCTTG

AGCACACGTCTGAACTCTTTCCCTACACGACGCTCTTCCGATCTNNNCTGAGGCAGGAAAATTGCTTG

AGCACACGTCTGAACTCTTTCCCTACACGACGCTCTTCCGATCTNNNCTGAGGCAGGAAAATTGCTTG

AGCACACGTCTGAACTCTTTCCCTACACGACGCTCTTCCGATCTNNNCTGAGGCAGGAAAATTGCTTG

AGCACACGTCTGAACTCTTTCCCTACACGACGCTCTTCCGATCTNNNCTGAGGCAGGAAAATTGCTTG

AGCACACGTCTGAACTCTTTCCCTACACGACGCTCTTCCGATCTNNNCTGAGGCAGGAAAATTGCTTG

AGCACACGTCTGAACTCTTTCCCTACACGACGCTCTTCCGATCTNNNATGGCTAATTCTAGGACTTGG

AGCACACGTCTGAACTCTTTCCCTACACGACGCTCTTCCGATCTNNNATGGCTAATTCTAGGACTTGG

AGCACACGTCTGAACTCTTTCCCTACACGACGCTCTTCCGATCTNNNATGGCTAATTCTAGGACTTGG

AGCACACGTCTGAACTCTTTCCCTACACGACGCTCTTCCGATCTNNNATGGCTAATTCTAGGACTTGG

AGCACACGTCTGAACTCTTTCCCTACACGACGCTCTTCCGATCTNNNATGGCTAATTCTAGGACTTGG

AGCACACGTCTGAACTCTTTCCCTACACGACGCTCTTCCGATCTNNNATGGCTAATTCTAGGACTTGG

GAGCACACGTCTGAACTCTTTCCCTACACGACGCTCTTCCGATCTNNNATCTTCTCTGTCCCTATCCGAC

GAGCACACGTCTGAACTCTTTCCCTACACGACGCTCTTCCGATCTNNNATCTTCTCTGTCCCTATCCGAC
GAGCACACGTCTGAACTCTTTCCCTACACGACGCTCTTCCGATCTNNNATCTTCTCTGTCCCTATCCGAC
GAGCACACGTCTGAACTCTTTCCCTACACGACGCTCTTCCGATCTNNNATCTTCTCTGTCCCTATCCGAC
GAGCACACGTCTGAACTCTTTCCCTACACGACGCTCTTCCGATCTNNNATCTTCTCTGTCCCTATCCGAC
GAGCACACGTCTGAACTCTTTCCCTACACGACGCTCTTCCGATCTNNNATCTTCTCTGTCCCTATCCGAC
GAGCACACGTCTGAACTCTTTCCCTACACGACGCTCTTCCGATCTNNNCTCTTTTCCCTCCTCCTCACTTC
GAGCACACGTCTGAACTCTTTCCCTACACGACGCTCTTCCGATCTNNNCTCTTTTCCCTCCTCCTCACTTC
GAGCACACGTCTGAACTCTTTCCCTACACGACGCTCTTCCGATCTNNNCTCTTTTCCCTCCTCCTCACTTC
GAGCACACGTCTGAACTCTTTCCCTACACGACGCTCTTCCGATCTNNNCTCTTTTCCCTCCTCCTCACTTC
GAGCACACGTCTGAACTCTTTCCCTACACGACGCTCTTCCGATCTNNNCTCTTTTCCCTCCTCCTCACTTC
GAGCACACGTCTGAACTCTTTCCCTACACGACGCTCTTCCGATCTNNNCTCTTTTCCCTCCTCCTCACTTC
GAGCACACGTCTGAACTCTTTCCCTACACGACGCTCTTCCGATCTNNNGCTACTCCTAATTTGGGGGG
GAGCACACGTCTGAACTCTTTCCCTACACGACGCTCTTCCGATCTNNNGCTACTCCTAATTTGGGGGG
GAGCACACGTCTGAACTCTTTCCCTACACGACGCTCTTCCGATCTNNNGCTACTCCTAATTTGGGGGG
GAGCACACGTCTGAACTCTTTCCCTACACGACGCTCTTCCGATCTNNNGCTACTCCTAATTTGGGGGG
GAGCACACGTCTGAACTCTTTCCCTACACGACGCTCTTCCGATCTNNNGCTACTCCTAATTTGGGGGG
GAGCACACGTCTGAACTCTTTCCCTACACGACGCTCTTCCGATCTNNNGCTACTCCTAATTTGGGGGG
GCACACGTCTGAACTCTTTCCCTACACGACGCTCTTCCGATCTNNNGAGGTTGCAGTGAGCTGAGATC
GCACACGTCTGAACTCTTTCCCTACACGACGCTCTTCCGATCTNNNGAGGTTGCAGTGAGCTGAGATC
GCACACGTCTGAACTCTTTCCCTACACGACGCTCTTCCGATCTNNNGAGGTTGCAGTGAGCTGAGATC
GCACACGTCTGAACTCTTTCCCTACACGACGCTCTTCCGATCTNNNGAGGTTGCAGTGAGCTGAGATC
GCACACGTCTGAACTCTTTCCCTACACGACGCTCTTCCGATCTNNNGAGGTTGCAGTGAGCTGAGATC
GCACACGTCTGAACTCTTTCCCTACACGACGCTCTTCCGATCTNNNGAGGTTGCAGTGAGCTGAGATC
GCACACGTCTGAACTCTTTCCCTACACGACGCTCTTCCGATCTNNNTGGCATTCTTTTACATGTTTGCA
GCACACGTCTGAACTCTTTCCCTACACGACGCTCTTCCGATCTNNNTGGCATTCTTTTACATGTTTGCA
GCACACGTCTGAACTCTTTCCCTACACGACGCTCTTCCGATCTNNNTGGCATTCTTTTACATGTTTGCA
GCACACGTCTGAACTCTTTCCCTACACGACGCTCTTCCGATCTNNNTGGCATTCTTTTACATGTTTGCA
GCACACGTCTGAACTCTTTCCCTACACGACGCTCTTCCGATCTNNNTGGCATTCTTTTACATGTTTGCA
GCACACGTCTGAACTCTTTCCCTACACGACGCTCTTCCGATCTNNNTGGCATTCTTTTACATGTTTGCA
GAGCACACGTCTGAACTCTTTCCCTACACGACGCTCTTCCGATCTNNNCTGTAGTCCCAGCTACTCAG
GAGCACACGTCTGAACTCTTTCCCTACACGACGCTCTTCCGATCTNNNCTGTAGTCCCAGCTACTCAG
GAGCACACGTCTGAACTCTTTCCCTACACGACGCTCTTCCGATCTNNNCTGTAGTCCCAGCTACTCAG
GAGCACACGTCTGAACTCTTTCCCTACACGACGCTCTTCCGATCTNNNCTGTAGTCCCAGCTACTCAG
GAGCACACGTCTGAACTCTTTCCCTACACGACGCTCTTCCGATCTNNNCTGTAGTCCCAGCTACTCAG
GAGCACACGTCTGAACTCTTTCCCTACACGACGCTCTTCCGATCTNNNCTGTAGTCCCAGCTACTCAG
AGCACACGTCTGAACTCTTTCCCTACACGACGCTCTTCCGATCTNNNTTCAGGCTTCTTCTTGCTGAGA
AGCACACGTCTGAACTCTTTCCCTACACGACGCTCTTCCGATCTNNNTTCAGGCTTCTTCTTGCTGAGA
AGCACACGTCTGAACTCTTTCCCTACACGACGCTCTTCCGATCTNNNTTCAGGCTTCTTCTTGCTGAGA
AGCACACGTCTGAACTCTTTCCCTACACGACGCTCTTCCGATCTNNNTTCAGGCTTCTTCTTGCTGAGA
AGCACACGTCTGAACTCTTTCCCTACACGACGCTCTTCCGATCTNNNTTCAGGCTTCTTCTTGCTGAGA
AGCACACGTCTGAACTCTTTCCCTACACGACGCTCTTCCGATCTNNNTTCAGGCTTCTTCTTGCTGAGA
AGCACACGTCTGAACTCTTTCCCTACACGACGCTCTTCCGATCTNNNAGAAAGGCTGAACAGAAGG
AGCACACGTCTGAACTCTTTCCCTACACGACGCTCTTCCGATCTNNNAGAAAGGCTGAACAGAAGG

GAGCACACGTCTGAACTCTTTCCCTACACGACGCTCTTCCGATCTNNNAGAAAGGCTGAACAGAAGGA
GAGCACACGTCTGAACTCTTTCCCTACACGACGCTCTTCCGATCTNNNAGAAAGGCTGAACAGAAGGA
GAGCACACGTCTGAACTCTTTCCCTACACGACGCTCTTCCGATCTNNNAGAAAGGCTGAACAGAAGGA
GAGCACACGTCTGAACTCTTTCCCTACACGACGCTCTTCCGATCTNNNAGAAAGGCTGAACAGAAGGA
GAGCACACGTCTGAACTCTTTCCCTACACGACGCTCTTCCGATCTNNNTTCACCTACTGTAATCTAGTTT
GAGCACACGTCTGAACTCTTTCCCTACACGACGCTCTTCCGATCTNNNTTCACCTACTGTAATCTAGTTT
GAGCACACGTCTGAACTCTTTCCCTACACGACGCTCTTCCGATCTNNNTTCACCTACTGTAATCTAGTTT
GAGCACACGTCTGAACTCTTTCCCTACACGACGCTCTTCCGATCTNNNTTCACCTACTGTAATCTAGTTT
GAGCACACGTCTGAACTCTTTCCCTACACGACGCTCTTCCGATCTNNNTTCACCTACTGTAATCTAGTTT
GAGCACACGTCTGAACTCTTTCCCTACACGACGCTCTTCCGATCTNNNTTCACCTACTGTAATCTAGTTT
AGAGCACACGTCTGAACTCTTTCCCTACACGACGCTCTTCCGATCTNNNGGATCAGGTATAGCTGCTG
AGAGCACACGTCTGAACTCTTTCCCTACACGACGCTCTTCCGATCTNNNGGATCAGGTATAGCTGCTG
AGAGCACACGTCTGAACTCTTTCCCTACACGACGCTCTTCCGATCTNNNGGATCAGGTATAGCTGCTG
AGAGCACACGTCTGAACTCTTTCCCTACACGACGCTCTTCCGATCTNNNGGATCAGGTATAGCTGCTG
AGAGCACACGTCTGAACTCTTTCCCTACACGACGCTCTTCCGATCTNNNGGATCAGGTATAGCTGCTG
AGAGCACACGTCTGAACTCTTTCCCTACACGACGCTCTTCCGATCTNNNGGATCAGGTATAGCTGCTG
GAAGAGCACACGTCTGAACTCTTTCCCTACACGACGCTCTTCCGATCTNNNAATTGAGCTCCTCGGACA
GAAGAGCACACGTCTGAACTCTTTCCCTACACGACGCTCTTCCGATCTNNNAATTGAGCTCCTCGGACA
GAAGAGCACACGTCTGAACTCTTTCCCTACACGACGCTCTTCCGATCTNNNAATTGAGCTCCTCGGACA
GAAGAGCACACGTCTGAACTCTTTCCCTACACGACGCTCTTCCGATCTNNNAATTGAGCTCCTCGGACA
GAAGAGCACACGTCTGAACTCTTTCCCTACACGACGCTCTTCCGATCTNNNAATTGAGCTCCTCGGACA
GAAGAGCACACGTCTGAACTCTTTCCCTACACGACGCTCTTCCGATCTNNNAATTGAGCTCCTCGGACA
AGCACACGTCTGAACTCTTTCCCTACACGACGCTCTTCCGATCTNNNCTAGCATAGGAGCAAGGAAGGA
AGCACACGTCTGAACTCTTTCCCTACACGACGCTCTTCCGATCTNNNCTAGCATAGGAGCAAGGAAGGA
AGCACACGTCTGAACTCTTTCCCTACACGACGCTCTTCCGATCTNNNCTAGCATAGGAGCAAGGAAGGA
AGCACACGTCTGAACTCTTTCCCTACACGACGCTCTTCCGATCTNNNCTAGCATAGGAGCAAGGAAGGA
AGCACACGTCTGAACTCTTTCCCTACACGACGCTCTTCCGATCTNNNCTAGCATAGGAGCAAGGAAGGA
AGCACACGTCTGAACTCTTTCCCTACACGACGCTCTTCCGATCTNNNCTAGCATAGGAGCAAGGAAGGA
AGCACACGTCTGAACTCTTTCCCTACACGACGCTCTTCCGATCTNNNGGAGGTTGTAGTGAGCCAAGA
AGCACACGTCTGAACTCTTTCCCTACACGACGCTCTTCCGATCTNNNGGAGGTTGTAGTGAGCCAAGA
AGCACACGTCTGAACTCTTTCCCTACACGACGCTCTTCCGATCTNNNGGAGGTTGTAGTGAGCCAAGA
AGCACACGTCTGAACTCTTTCCCTACACGACGCTCTTCCGATCTNNNGGAGGTTGTAGTGAGCCAAGA
AGCACACGTCTGAACTCTTTCCCTACACGACGCTCTTCCGATCTNNNGGAGGTTGTAGTGAGCCAAGA
AGCACACGTCTGAACTCTTTCCCTACACGACGCTCTTCCGATCTNNNGGAGGTTGTAGTGAGCCAAGA
GAGCACACGTCTGAACTCTTTCCCTACACGACGCTCTTCCGATCTNNNCTGTAATCCCAGCTACTCAGG
GAGCACACGTCTGAACTCTTTCCCTACACGACGCTCTTCCGATCTNNNCTGTAATCCCAGCTACTCAGG
GAGCACACGTCTGAACTCTTTCCCTACACGACGCTCTTCCGATCTNNNCTGTAATCCCAGCTACTCAGG
GAGCACACGTCTGAACTCTTTCCCTACACGACGCTCTTCCGATCTNNNCTGTAATCCCAGCTACTCAGG
GAGCACACGTCTGAACTCTTTCCCTACACGACGCTCTTCCGATCTNNNCTGTAATCCCAGCTACTCAGG
GAGCACACGTCTGAACTCTTTCCCTACACGACGCTCTTCCGATCTNNNCTGTAATCCCAGCTACTCAGG
GAGCACACGTCTGAACTCTTTCCCTACACGACGCTCTTCCGATCTNNNAGGCATTATGTGGGTGCCAA
GAGCACACGTCTGAACTCTTTCCCTACACGACGCTCTTCCGATCTNNNAGGCATTATGTGGGTGCCAA
GAGCACACGTCTGAACTCTTTCCCTACACGACGCTCTTCCGATCTNNNAGGCATTATGTGGGTGCCAA

GAGCACACGTCTGAACTCTTTCCCTACACGACGCTCTTCCGATCTNNNAGGCATTATGTGGGTGCCAA
GAGCACACGTCTGAACTCTTTCCCTACACGACGCTCTTCCGATCTNNNAGGCATTATGTGGGTGCCAA
GAGCACACGTCTGAACTCTTTCCCTACACGACGCTCTTCCGATCTNNNAGGCATTATGTGGGTGCCAA
AGCACACGTCTGAACTCTTTCCCTACACGACGCTCTTCCGATCTNNNTCTCTCTCTCTCTGTGTGTGT
AGCACACGTCTGAACTCTTTCCCTACACGACGCTCTTCCGATCTNNNTCTCTCTCTCTCTGTGTGTGT
AGCACACGTCTGAACTCTTTCCCTACACGACGCTCTTCCGATCTNNNTCTCTCTCTCTCTGTGTGTGT
AGCACACGTCTGAACTCTTTCCCTACACGACGCTCTTCCGATCTNNNTCTCTCTCTCTCTGTGTGTGT
AGCACACGTCTGAACTCTTTCCCTACACGACGCTCTTCCGATCTNNNTCTCTCTCTCTCTGTGTGTGT
AGCACACGTCTGAACTCTTTCCCTACACGACGCTCTTCCGATCTNNNTCTCTCTCTCTCTGTGTGTGT
GAGCACACGTCTGAACTCTTTCCCTACACGACGCTCTTCCGATCTNNNTACCCAGTCTCCTGTAGCATCT
GAGCACACGTCTGAACTCTTTCCCTACACGACGCTCTTCCGATCTNNNTACCCAGTCTCCTGTAGCATCT
GAGCACACGTCTGAACTCTTTCCCTACACGACGCTCTTCCGATCTNNNTACCCAGTCTCCTGTAGCATCT
GAGCACACGTCTGAACTCTTTCCCTACACGACGCTCTTCCGATCTNNNTACCCAGTCTCCTGTAGCATCT
GAGCACACGTCTGAACTCTTTCCCTACACGACGCTCTTCCGATCTNNNTACCCAGTCTCCTGTAGCATCT
GAGCACACGTCTGAACTCTTTCCCTACACGACGCTCTTCCGATCTNNNTACCCAGTCTCCTGTAGCATCT
AGCACACGTCTGAACTCTTTCCCTACACGACGCTCTTCCGATCTNNNAACAGGTTAATTCCCAATTTAA
AGCACACGTCTGAACTCTTTCCCTACACGACGCTCTTCCGATCTNNNAACAGGTTAATTCCCAATTTAA
AGCACACGTCTGAACTCTTTCCCTACACGACGCTCTTCCGATCTNNNAACAGGTTAATTCCCAATTTAA
AGCACACGTCTGAACTCTTTCCCTACACGACGCTCTTCCGATCTNNNAACAGGTTAATTCCCAATTTAA
AGCACACGTCTGAACTCTTTCCCTACACGACGCTCTTCCGATCTNNNAACAGGTTAATTCCCAATTTAA
AGCACACGTCTGAACTCTTTCCCTACACGACGCTCTTCCGATCTNNNAACAGGTTAATTCCCAATTTAA
GCACACGTCTGAACTCTTTCCCTACACGACGCTCTTCCGATCTNNNGGAGAAAAAACAATGTGGAGG
GCACACGTCTGAACTCTTTCCCTACACGACGCTCTTCCGATCTNNNGGAGAAAAAACAATGTGGAGG
GCACACGTCTGAACTCTTTCCCTACACGACGCTCTTCCGATCTNNNGGAGAAAAAACAATGTGGAGG
GCACACGTCTGAACTCTTTCCCTACACGACGCTCTTCCGATCTNNNGGAGAAAAAACAATGTGGAGG
GCACACGTCTGAACTCTTTCCCTACACGACGCTCTTCCGATCTNNNGGAGAAAAAACAATGTGGAGG
GCACACGTCTGAACTCTTTCCCTACACGACGCTCTTCCGATCTNNNGGAGAAAAAACAATGTGGAGG
AGCACACGTCTGAACTCTTTCCCTACACGACGCTCTTCCGATCTNNNCACACTGTTCTGCATCTTGCTTT
AGCACACGTCTGAACTCTTTCCCTACACGACGCTCTTCCGATCTNNNCACACTGTTCTGCATCTTGCTTT
AGCACACGTCTGAACTCTTTCCCTACACGACGCTCTTCCGATCTNNNCACACTGTTCTGCATCTTGCTTT
AGCACACGTCTGAACTCTTTCCCTACACGACGCTCTTCCGATCTNNNCACACTGTTCTGCATCTTGCTTT
AGCACACGTCTGAACTCTTTCCCTACACGACGCTCTTCCGATCTNNNCACACTGTTCTGCATCTTGCTTT
AGCACACGTCTGAACTCTTTCCCTACACGACGCTCTTCCGATCTNNNCACACTGTTCTGCATCTTGCTTT
GAGCACACGTCTGAACTCTTTCCCTACACGACGCTCTTCCGATCTNNNTCTAGTTGATCACCTTTGGCA
GAGCACACGTCTGAACTCTTTCCCTACACGACGCTCTTCCGATCTNNNTCTAGTTGATCACCTTTGGCA
GAGCACACGTCTGAACTCTTTCCCTACACGACGCTCTTCCGATCTNNNTCTAGTTGATCACCTTTGGCA
GAGCACACGTCTGAACTCTTTCCCTACACGACGCTCTTCCGATCTNNNTCTAGTTGATCACCTTTGGCA
GAGCACACGTCTGAACTCTTTCCCTACACGACGCTCTTCCGATCTNNNTCTAGTTGATCACCTTTGGCA
GAGCACACGTCTGAACTCTTTCCCTACACGACGCTCTTCCGATCTNNNTCTAGTTGATCACCTTTGGCA
GAGCACACGTCTGAACTCTTTCCCTACACGACGCTCTTCCGATCTNNNCATTTTTTGTCCCTGACTACGG
GAGCACACGTCTGAACTCTTTCCCTACACGACGCTCTTCCGATCTNNNCATTTTTTGTCCCTGACTACGG
GAGCACACGTCTGAACTCTTTCCCTACACGACGCTCTTCCGATCTNNNCATTTTTTGTCCCTGACTACGG
GAGCACACGTCTGAACTCTTTCCCTACACGACGCTCTTCCGATCTNNNCATTTTTTGTCCCTGACTACGG

GAGCACACGTCTGAACTCTTTCCCTACACGACGCTCTTCCGATCTNNNCATTTTTGTCCCTGACTACGG
GAGCACACGTCTGAACTCTTTCCCTACACGACGCTCTTCCGATCTNNNCATTTTTGTCCCTGACTACGG
GCACACGTCTGAACTCTTTCCCTACACGACGCTCTTCCGATCTNNNAGATTCATAGCATACCCCTCCCCC
GCACACGTCTGAACTCTTTCCCTACACGACGCTCTTCCGATCTNNNAGATTCATAGCATACCCCTCCCCC
GCACACGTCTGAACTCTTTCCCTACACGACGCTCTTCCGATCTNNNAGATTCATAGCATACCCCTCCCCC
GCACACGTCTGAACTCTTTCCCTACACGACGCTCTTCCGATCTNNNAGATTCATAGCATACCCCTCCCCC
GCACACGTCTGAACTCTTTCCCTACACGACGCTCTTCCGATCTNNNAGATTCATAGCATACCCCTCCCCC
GCACACGTCTGAACTCTTTCCCTACACGACGCTCTTCCGATCTNNNAGATTCATAGCATACCCCTCCCCC
AGCACACGTCTGAACTCTTTCCCTACACGACGCTCTTCCGATCTNNNGAAAGCAAATGTGTAACCATC
AGCACACGTCTGAACTCTTTCCCTACACGACGCTCTTCCGATCTNNNGAAAGCAAATGTGTAACCATC
AGCACACGTCTGAACTCTTTCCCTACACGACGCTCTTCCGATCTNNNGAAAGCAAATGTGTAACCATC
AGCACACGTCTGAACTCTTTCCCTACACGACGCTCTTCCGATCTNNNGAAAGCAAATGTGTAACCATC
AGCACACGTCTGAACTCTTTCCCTACACGACGCTCTTCCGATCTNNNGAAAGCAAATGTGTAACCATC
AGCACACGTCTGAACTCTTTCCCTACACGACGCTCTTCCGATCTNNNGAAAGCAAATGTGTAACCATC
AGCACACGTCTGAACTCTTTCCCTACACGACGCTCTTCCGATCTNNNGAGATTTCTCTCCTTTACCCCTT
AGCACACGTCTGAACTCTTTCCCTACACGACGCTCTTCCGATCTNNNGAGATTTCTCTCCTTTACCCCTT
AGCACACGTCTGAACTCTTTCCCTACACGACGCTCTTCCGATCTNNNGAGATTTCTCTCCTTTACCCCTT
AGCACACGTCTGAACTCTTTCCCTACACGACGCTCTTCCGATCTNNNGAGATTTCTCTCCTTTACCCCTT
AGCACACGTCTGAACTCTTTCCCTACACGACGCTCTTCCGATCTNNNGAGATTTCTCTCCTTTACCCCTT
AGCACACGTCTGAACTCTTTCCCTACACGACGCTCTTCCGATCTNNNGAGATTTCTCTCCTTTACCCCTT
AGCACACGTCTGAACTCTTTCCCTACACGACGCTCTTCCGATCTNNNGGCTGGGGATGGAGTTGATAT
AGCACACGTCTGAACTCTTTCCCTACACGACGCTCTTCCGATCTNNNGGCTGGGGATGGAGTTGATAT
AGCACACGTCTGAACTCTTTCCCTACACGACGCTCTTCCGATCTNNNGGCTGGGGATGGAGTTGATAT
AGCACACGTCTGAACTCTTTCCCTACACGACGCTCTTCCGATCTNNNGGCTGGGGATGGAGTTGATAT
AGCACACGTCTGAACTCTTTCCCTACACGACGCTCTTCCGATCTNNNGGCTGGGGATGGAGTTGATAT
AGCACACGTCTGAACTCTTTCCCTACACGACGCTCTTCCGATCTNNNGGCTGGGGATGGAGTTGATAT
GCACACGTCTGAACTCTTTCCCTACACGACGCTCTTCCGATCTNNNAGAATCTTGCCCAAATCCTCCAAC
GCACACGTCTGAACTCTTTCCCTACACGACGCTCTTCCGATCTNNNAGAATCTTGCCCAAATCCTCCAAC
GCACACGTCTGAACTCTTTCCCTACACGACGCTCTTCCGATCTNNNAGAATCTTGCCCAAATCCTCCAAC
GCACACGTCTGAACTCTTTCCCTACACGACGCTCTTCCGATCTNNNAGAATCTTGCCCAAATCCTCCAAC
GCACACGTCTGAACTCTTTCCCTACACGACGCTCTTCCGATCTNNNAGAATCTTGCCCAAATCCTCCAAC
GCACACGTCTGAACTCTTTCCCTACACGACGCTCTTCCGATCTNNNAGAATCTTGCCCAAATCCTCCAAC
AGCACACGTCTGAACTCTTTCCCTACACGACGCTCTTCCGATCTNNNAGCCTCCTGAGTAGCTAGAAT
AGCACACGTCTGAACTCTTTCCCTACACGACGCTCTTCCGATCTNNNAGCCTCCTGAGTAGCTAGAAT
AGCACACGTCTGAACTCTTTCCCTACACGACGCTCTTCCGATCTNNNAGCCTCCTGAGTAGCTAGAAT
AGCACACGTCTGAACTCTTTCCCTACACGACGCTCTTCCGATCTNNNAGCCTCCTGAGTAGCTAGAAT
AGCACACGTCTGAACTCTTTCCCTACACGACGCTCTTCCGATCTNNNAGCCTCCTGAGTAGCTAGAAT
AGCACACGTCTGAACTCTTTCCCTACACGACGCTCTTCCGATCTNNNAGCCTCCTGAGTAGCTAGAAT
GAGCACACGTCTGAACTCTTTCCCTACACGACGCTCTTCCGATCTNNNCTGACGTAGGAGAATCGCATG
GAGCACACGTCTGAACTCTTTCCCTACACGACGCTCTTCCGATCTNNNCTGACGTAGGAGAATCGCATG
GAGCACACGTCTGAACTCTTTCCCTACACGACGCTCTTCCGATCTNNNCTGACGTAGGAGAATCGCATG
GAGCACACGTCTGAACTCTTTCCCTACACGACGCTCTTCCGATCTNNNCTGACGTAGGAGAATCGCATG
GAGCACACGTCTGAACTCTTTCCCTACACGACGCTCTTCCGATCTNNNCTGACGTAGGAGAATCGCATG

GAGCACACGTCTGAACTCTTTCCCTACACGACGCTCTTCCGATCTNNNCTGACGTAGGAGAATCGCATG

AGCACACGTCTGAACTCTTTCCCTACACGACGCTCTTCCGATCTNNNCTTTCCAATGGAGCGATCTGGA

AGCACACGTCTGAACTCTTTCCCTACACGACGCTCTTCCGATCTNNNCTTTCCAATGGAGCGATCTGGA

AGCACACGTCTGAACTCTTTCCCTACACGACGCTCTTCCGATCTNNNCTTTCCAATGGAGCGATCTGGA

AGCACACGTCTGAACTCTTTCCCTACACGACGCTCTTCCGATCTNNNCTTTCCAATGGAGCGATCTGGA

AGCACACGTCTGAACTCTTTCCCTACACGACGCTCTTCCGATCTNNNCTTTCCAATGGAGCGATCTGGA

AGCACACGTCTGAACTCTTTCCCTACACGACGCTCTTCCGATCTNNNCTTTCCAATGGAGCGATCTGGA

GAGCACACGTCTGAACTCTTTCCCTACACGACGCTCTTCCGATCTNNNCCACGTGTGGCTATTTTTTCTG

GAGCACACGTCTGAACTCTTTCCCTACACGACGCTCTTCCGATCTNNNCCACGTGTGGCTATTTTTTCTG

GAGCACACGTCTGAACTCTTTCCCTACACGACGCTCTTCCGATCTNNNCCACGTGTGGCTATTTTTTCTG

GAGCACACGTCTGAACTCTTTCCCTACACGACGCTCTTCCGATCTNNNCCACGTGTGGCTATTTTTTCTG

GAGCACACGTCTGAACTCTTTCCCTACACGACGCTCTTCCGATCTNNNCCACGTGTGGCTATTTTTTCTG

GAGCACACGTCTGAACTCTTTCCCTACACGACGCTCTTCCGATCTNNNCCACGTGTGGCTATTTTTTCTG

AGCACACGTCTGAACTCTTTCCCTACACGACGCTCTTCCGATCTNNNATCTAGGGTCTCTTATGGGCCT

AGCACACGTCTGAACTCTTTCCCTACACGACGCTCTTCCGATCTNNNATCTAGGGTCTCTTATGGGCCT

AGCACACGTCTGAACTCTTTCCCTACACGACGCTCTTCCGATCTNNNATCTAGGGTCTCTTATGGGCCT

AGCACACGTCTGAACTCTTTCCCTACACGACGCTCTTCCGATCTNNNATCTAGGGTCTCTTATGGGCCT

AGCACACGTCTGAACTCTTTCCCTACACGACGCTCTTCCGATCTNNNATCTAGGGTCTCTTATGGGCCT

AGCACACGTCTGAACTCTTTCCCTACACGACGCTCTTCCGATCTNNNATCTAGGGTCTCTTATGGGCCT

AGCACACGTCTGAACTCTTTCCCTACACGACGCTCTTCCGATCTNNNTAGAAAAAAATTAGCCGGGCGT

AGCACACGTCTGAACTCTTTCCCTACACGACGCTCTTCCGATCTNNNTAGAAAAAAATTAGCCGGGCGT

AGCACACGTCTGAACTCTTTCCCTACACGACGCTCTTCCGATCTNNNTAGAAAAAAATTAGCCGGGCGT

AGCACACGTCTGAACTCTTTCCCTACACGACGCTCTTCCGATCTNNNTAGAAAAAAATTAGCCGGGCGT

AGCACACGTCTGAACTCTTTCCCTACACGACGCTCTTCCGATCTNNNTAGAAAAAAATTAGCCGGGCGT

AGCACACGTCTGAACTCTTTCCCTACACGACGCTCTTCCGATCTNNNTAGAAAAAAATTAGCCGGGCGT

AGCACACGTCTGAACTCTTTCCCTACACGACGCTCTTCCGATCTNNNGTTTCTGCAAACATAGTCGGAG

AGCACACGTCTGAACTCTTTCCCTACACGACGCTCTTCCGATCTNNNGTTTCTGCAAACATAGTCGGAG

AGCACACGTCTGAACTCTTTCCCTACACGACGCTCTTCCGATCTNNNGTTTCTGCAAACATAGTCGGAG

AGCACACGTCTGAACTCTTTCCCTACACGACGCTCTTCCGATCTNNNGTTTCTGCAAACATAGTCGGAG

AGCACACGTCTGAACTCTTTCCCTACACGACGCTCTTCCGATCTNNNGTTTCTGCAAACATAGTCGGAG

AGCACACGTCTGAACTCTTTCCCTACACGACGCTCTTCCGATCTNNNGTTTCTGCAAACATAGTCGGAG

AGCACACGTCTGAACTCTTTCCCTACACGACGCTCTTCCGATCTNNNAGCTAACAGGTGTGTGTGTATG

AGCACACGTCTGAACTCTTTCCCTACACGACGCTCTTCCGATCTNNNAGCTAACAGGTGTGTGTGTATG

AGCACACGTCTGAACTCTTTCCCTACACGACGCTCTTCCGATCTNNNAGCTAACAGGTGTGTGTGTATG

AGCACACGTCTGAACTCTTTCCCTACACGACGCTCTTCCGATCTNNNAGCTAACAGGTGTGTGTGTATG

AGCACACGTCTGAACTCTTTCCCTACACGACGCTCTTCCGATCTNNNAGCTAACAGGTGTGTGTGTATG

AGCACACGTCTGAACTCTTTCCCTACACGACGCTCTTCCGATCTNNNAGCTAACAGGTGTGTGTGTATG

GAGCACACGTCTGAACTCTTTCCCTACACGACGCTCTTCCGATCTNNNGGAGACCTACCAGTAGACATG

GAGCACACGTCTGAACTCTTTCCCTACACGACGCTCTTCCGATCTNNNGGAGACCTACCAGTAGACATG

GAGCACACGTCTGAACTCTTTCCCTACACGACGCTCTTCCGATCTNNNGGAGACCTACCAGTAGACATG

GAGCACACGTCTGAACTCTTTCCCTACACGACGCTCTTCCGATCTNNNGGAGACCTACCAGTAGACATG

GAGCACACGTCTGAACTCTTTCCCTACACGACGCTCTTCCGATCTNNNGGAGACCTACCAGTAGACATG

GAGCACACGTCTGAACTCTTTCCCTACACGACGCTCTTCCGATCTNNNGGAGACCTACCAGTAGACATG

CACGTCTGAACTCTTTCCCTACACGACGCTCTTCCGATCTNNNTAGTCTCACAGTGCAAGAACCTCCAT
CACGTCTGAACTCTTTCCCTACACGACGCTCTTCCGATCTNNNTAGTCTCACAGTGCAAGAACCTCCAT
CACGTCTGAACTCTTTCCCTACACGACGCTCTTCCGATCTNNNTAGTCTCACAGTGCAAGAACCTCCAT
CACGTCTGAACTCTTTCCCTACACGACGCTCTTCCGATCTNNNTAGTCTCACAGTGCAAGAACCTCCAT
CACGTCTGAACTCTTTCCCTACACGACGCTCTTCCGATCTNNNTAGTCTCACAGTGCAAGAACCTCCAT
CACGTCTGAACTCTTTCCCTACACGACGCTCTTCCGATCTNNNTAGTCTCACAGTGCAAGAACCTCCAT
GAGCACACGTCTGAACTCTTTCCCTACACGACGCTCTTCCGATCTNNNTATACATTGCATTCCCTCTGG
GAGCACACGTCTGAACTCTTTCCCTACACGACGCTCTTCCGATCTNNNTATACATTGCATTCCCTCTGG
GAGCACACGTCTGAACTCTTTCCCTACACGACGCTCTTCCGATCTNNNTATACATTGCATTCCCTCTGG
GAGCACACGTCTGAACTCTTTCCCTACACGACGCTCTTCCGATCTNNNTATACATTGCATTCCCTCTGG
GAGCACACGTCTGAACTCTTTCCCTACACGACGCTCTTCCGATCTNNNTATACATTGCATTCCCTCTGG
GAGCACACGTCTGAACTCTTTCCCTACACGACGCTCTTCCGATCTNNNTATACATTGCATTCCCTCTGG
CACGTCTGAACTCTTTCCCTACACGACGCTCTTCCGATCTNNNAGTGGAAGAGTCAGAACACACAGC
CACGTCTGAACTCTTTCCCTACACGACGCTCTTCCGATCTNNNAGTGGAAGAGTCAGAACACACAGC
CACGTCTGAACTCTTTCCCTACACGACGCTCTTCCGATCTNNNAGTGGAAGAGTCAGAACACACAGC
CACGTCTGAACTCTTTCCCTACACGACGCTCTTCCGATCTNNNAGTGGAAGAGTCAGAACACACAGC
CACGTCTGAACTCTTTCCCTACACGACGCTCTTCCGATCTNNNAGTGGAAGAGTCAGAACACACAGC
CACGTCTGAACTCTTTCCCTACACGACGCTCTTCCGATCTNNNAGTGGAAGAGTCAGAACACACAGC
AGCACACGTCTGAACTCTTTCCCTACACGACGCTCTTCCGATCTNNNTTGTCAACAAGCGAAGAAAAC
AGCACACGTCTGAACTCTTTCCCTACACGACGCTCTTCCGATCTNNNTTGTCAACAAGCGAAGAAAAC
AGCACACGTCTGAACTCTTTCCCTACACGACGCTCTTCCGATCTNNNTTGTCAACAAGCGAAGAAAAC
AGCACACGTCTGAACTCTTTCCCTACACGACGCTCTTCCGATCTNNNTTGTCAACAAGCGAAGAAAAC
AGCACACGTCTGAACTCTTTCCCTACACGACGCTCTTCCGATCTNNNTTGTCAACAAGCGAAGAAAAC
AGCACACGTCTGAACTCTTTCCCTACACGACGCTCTTCCGATCTNNNTTGTCAACAAGCGAAGAAAAC
AGCACACGTCTGAACTCTTTCCCTACACGACGCTCTTCCGATCTNNNCTGTAATGGATTGGTGGCAA
AGCACACGTCTGAACTCTTTCCCTACACGACGCTCTTCCGATCTNNNCTGTAATGGATTGGTGGCAA
AGCACACGTCTGAACTCTTTCCCTACACGACGCTCTTCCGATCTNNNCTGTAATGGATTGGTGGCAA
AGCACACGTCTGAACTCTTTCCCTACACGACGCTCTTCCGATCTNNNCTGTAATGGATTGGTGGCAA
AGCACACGTCTGAACTCTTTCCCTACACGACGCTCTTCCGATCTNNNCTGTAATGGATTGGTGGCAA
AGCACACGTCTGAACTCTTTCCCTACACGACGCTCTTCCGATCTNNNCTGTAATGGATTGGTGGCAA
AGCACACGTCTGAACTCTTTCCCTACACGACGCTCTTCCGATCTNNNAACTCCTGGCCTCAAGTAATCT
AGCACACGTCTGAACTCTTTCCCTACACGACGCTCTTCCGATCTNNNAACTCCTGGCCTCAAGTAATCT
AGCACACGTCTGAACTCTTTCCCTACACGACGCTCTTCCGATCTNNNAACTCCTGGCCTCAAGTAATCT
AGCACACGTCTGAACTCTTTCCCTACACGACGCTCTTCCGATCTNNNAACTCCTGGCCTCAAGTAATCT
AGCACACGTCTGAACTCTTTCCCTACACGACGCTCTTCCGATCTNNNAACTCCTGGCCTCAAGTAATCT
AGCACACGTCTGAACTCTTTCCCTACACGACGCTCTTCCGATCTNNNAACTCCTGGCCTCAAGTAATCT
GAGCACACGTCTGAACTCTTTCCCTACACGACGCTCTTCCGATCTNNNGGAGAGAGAGAGAGAGGA
GAGCACACGTCTGAACTCTTTCCCTACACGACGCTCTTCCGATCTNNNGGAGAGAGAGAGAGAGGA
GAGCACACGTCTGAACTCTTTCCCTACACGACGCTCTTCCGATCTNNNGGAGAGAGAGAGAGAGGA
GAGCACACGTCTGAACTCTTTCCCTACACGACGCTCTTCCGATCTNNNGGAGAGAGAGAGAGAGGA
GAGCACACGTCTGAACTCTTTCCCTACACGACGCTCTTCCGATCTNNNGGAGAGAGAGAGAGAGGA
GAGCACACGTCTGAACTCTTTCCCTACACGACGCTCTTCCGATCTNNNGGAGAGAGAGAGAGAGGA
AGCACACGTCTGAACTCTTTCCCTACACGACGCTCTTCCGATCTNNNCTGCAGTAAGTCAAGATTGTG

GAGCACACGTCTGAACTCTTTCCCTACACGACGCTCTTCCGATCTNNNCTGCAGTAAGTCAAGATTGTG
GAGCACACGTCTGAACTCTTTCCCTACACGACGCTCTTCCGATCTNNNCTGCAGTAAGTCAAGATTGTG
GAGCACACGTCTGAACTCTTTCCCTACACGACGCTCTTCCGATCTNNNCTGCAGTAAGTCAAGATTGTG
GAGCACACGTCTGAACTCTTTCCCTACACGACGCTCTTCCGATCTNNNCTGCAGTAAGTCAAGATTGTG
GAGCACACGTCTGAACTCTTTCCCTACACGACGCTCTTCCGATCTNNNCTGCAGTAAGTCAAGATTGTG
AGCACACGTCTGAACTCTTTCCCTACACGACGCTCTTCCGATCTNNNCCTCTCTTAGGAGCAGCTTTC
AGCACACGTCTGAACTCTTTCCCTACACGACGCTCTTCCGATCTNNNCCTCTCTTAGGAGCAGCTTTC
AGCACACGTCTGAACTCTTTCCCTACACGACGCTCTTCCGATCTNNNCCTCTCTTAGGAGCAGCTTTC
AGCACACGTCTGAACTCTTTCCCTACACGACGCTCTTCCGATCTNNNCCTCTCTTAGGAGCAGCTTTC
AGCACACGTCTGAACTCTTTCCCTACACGACGCTCTTCCGATCTNNNCCTCTCTTAGGAGCAGCTTTC
AGCACACGTCTGAACTCTTTCCCTACACGACGCTCTTCCGATCTNNNCCTCTCTTAGGAGCAGCTTTC
AGCACACGTCTGAACTCTTTCCCTACACGACGCTCTTCCGATCTNNNTGTGAAACAGTGAGATTCCGT
AGCACACGTCTGAACTCTTTCCCTACACGACGCTCTTCCGATCTNNNTGTGAAACAGTGAGATTCCGT
AGCACACGTCTGAACTCTTTCCCTACACGACGCTCTTCCGATCTNNNTGTGAAACAGTGAGATTCCGT
AGCACACGTCTGAACTCTTTCCCTACACGACGCTCTTCCGATCTNNNTGTGAAACAGTGAGATTCCGT
AGCACACGTCTGAACTCTTTCCCTACACGACGCTCTTCCGATCTNNNTGTGAAACAGTGAGATTCCGT
AGCACACGTCTGAACTCTTTCCCTACACGACGCTCTTCCGATCTNNNTGTGAAACAGTGAGATTCCGT
AGCACACGTCTGAACTCTTTCCCTACACGACGCTCTTCCGATCTNNNTCTACCGCCTTGTGATACCATTT
AGCACACGTCTGAACTCTTTCCCTACACGACGCTCTTCCGATCTNNNTCTACCGCCTTGTGATACCATTT
AGCACACGTCTGAACTCTTTCCCTACACGACGCTCTTCCGATCTNNNTCTACCGCCTTGTGATACCATTT
AGCACACGTCTGAACTCTTTCCCTACACGACGCTCTTCCGATCTNNNTCTACCGCCTTGTGATACCATTT
AGCACACGTCTGAACTCTTTCCCTACACGACGCTCTTCCGATCTNNNTCTACCGCCTTGTGATACCATTT
AGCACACGTCTGAACTCTTTCCCTACACGACGCTCTTCCGATCTNNNTCTACCGCCTTGTGATACCATTT
AGAGCACACGTCTGAACTCTTTCCCTACACGACGCTCTTCCGATCTNNNCTCACTCCATCCTGTCCATTG
AGAGCACACGTCTGAACTCTTTCCCTACACGACGCTCTTCCGATCTNNNCTCACTCCATCCTGTCCATTG
AGAGCACACGTCTGAACTCTTTCCCTACACGACGCTCTTCCGATCTNNNCTCACTCCATCCTGTCCATTG
AGAGCACACGTCTGAACTCTTTCCCTACACGACGCTCTTCCGATCTNNNCTCACTCCATCCTGTCCATTG
AGAGCACACGTCTGAACTCTTTCCCTACACGACGCTCTTCCGATCTNNNCTCACTCCATCCTGTCCATTG
AGAGCACACGTCTGAACTCTTTCCCTACACGACGCTCTTCCGATCTNNNCTCACTCCATCCTGTCCATTG
GAGCACACGTCTGAACTCTTTCCCTACACGACGCTCTTCCGATCTNNNCTGAGGCAAGAGGATCACTT
GAGCACACGTCTGAACTCTTTCCCTACACGACGCTCTTCCGATCTNNNCTGAGGCAAGAGGATCACTT
GAGCACACGTCTGAACTCTTTCCCTACACGACGCTCTTCCGATCTNNNCTGAGGCAAGAGGATCACTT
GAGCACACGTCTGAACTCTTTCCCTACACGACGCTCTTCCGATCTNNNCTGAGGCAAGAGGATCACTT
GAGCACACGTCTGAACTCTTTCCCTACACGACGCTCTTCCGATCTNNNCTGAGGCAAGAGGATCACTT
GAGCACACGTCTGAACTCTTTCCCTACACGACGCTCTTCCGATCTNNNCTGAGGCAAGAGGATCACTT
GAGCACACGTCTGAACTCTTTCCCTACACGACGCTCTTCCGATCTNNNCTGTAGTCCCAGCTACTCGGG
GAGCACACGTCTGAACTCTTTCCCTACACGACGCTCTTCCGATCTNNNCTGTAGTCCCAGCTACTCGGG
GAGCACACGTCTGAACTCTTTCCCTACACGACGCTCTTCCGATCTNNNCTGTAGTCCCAGCTACTCGGG
GAGCACACGTCTGAACTCTTTCCCTACACGACGCTCTTCCGATCTNNNCTGTAGTCCCAGCTACTCGGG
GAGCACACGTCTGAACTCTTTCCCTACACGACGCTCTTCCGATCTNNNCTGTAGTCCCAGCTACTCGGG
GAGCACACGTCTGAACTCTTTCCCTACACGACGCTCTTCCGATCTNNNCTGTAGTCCCAGCTACTCGGG
CACACGTCTGAACTCTTTCCCTACACGACGCTCTTCCGATCTNNNTAGTCTCACAGTGCAAGAACCTCCA
CACACGTCTGAACTCTTTCCCTACACGACGCTCTTCCGATCTNNNTAGTCTCACAGTGCAAGAACCTCCA

ACACGTCTGAACTCTTTCCCTACACGACGCTCTTCCGATCTNNNTAGTCTCACAGTGCAAGAACCTCCA
ACACGTCTGAACTCTTTCCCTACACGACGCTCTTCCGATCTNNNTAGTCTCACAGTGCAAGAACCTCCA
ACACGTCTGAACTCTTTCCCTACACGACGCTCTTCCGATCTNNNTAGTCTCACAGTGCAAGAACCTCCA
ACACGTCTGAACTCTTTCCCTACACGACGCTCTTCCGATCTNNNTAGTCTCACAGTGCAAGAACCTCCA
GCACACGTCTGAACTCTTTCCCTACACGACGCTCTTCCGATCTNNNCTCTTACAAAGAAGGGCTGGACT
GCACACGTCTGAACTCTTTCCCTACACGACGCTCTTCCGATCTNNNCTCTTACAAAGAAGGGCTGGACT
GCACACGTCTGAACTCTTTCCCTACACGACGCTCTTCCGATCTNNNCTCTTACAAAGAAGGGCTGGACT
GCACACGTCTGAACTCTTTCCCTACACGACGCTCTTCCGATCTNNNCTCTTACAAAGAAGGGCTGGACT
GCACACGTCTGAACTCTTTCCCTACACGACGCTCTTCCGATCTNNNCTCTTACAAAGAAGGGCTGGACT
GCACACGTCTGAACTCTTTCCCTACACGACGCTCTTCCGATCTNNNCTCTTACAAAGAAGGGCTGGACT
AGCACACGTCTGAACTCTTTCCCTACACGACGCTCTTCCGATCTNNNGCAAGTATCTGAGAGATATGG
AGCACACGTCTGAACTCTTTCCCTACACGACGCTCTTCCGATCTNNNGCAAGTATCTGAGAGATATGG
AGCACACGTCTGAACTCTTTCCCTACACGACGCTCTTCCGATCTNNNGCAAGTATCTGAGAGATATGG
AGCACACGTCTGAACTCTTTCCCTACACGACGCTCTTCCGATCTNNNGCAAGTATCTGAGAGATATGG
AGCACACGTCTGAACTCTTTCCCTACACGACGCTCTTCCGATCTNNNGCAAGTATCTGAGAGATATGG
AGCACACGTCTGAACTCTTTCCCTACACGACGCTCTTCCGATCTNNNGCAAGTATCTGAGAGATATGG
GCACACGTCTGAACTCTTTCCCTACACGACGCTCTTCCGATCTNNNTGATTTGCCTCTAAGAGCCCAT
GCACACGTCTGAACTCTTTCCCTACACGACGCTCTTCCGATCTNNNTGATTTGCCTCTAAGAGCCCAT
GCACACGTCTGAACTCTTTCCCTACACGACGCTCTTCCGATCTNNNTGATTTGCCTCTAAGAGCCCAT
GCACACGTCTGAACTCTTTCCCTACACGACGCTCTTCCGATCTNNNTGATTTGCCTCTAAGAGCCCAT
GCACACGTCTGAACTCTTTCCCTACACGACGCTCTTCCGATCTNNNTGATTTGCCTCTAAGAGCCCAT
GCACACGTCTGAACTCTTTCCCTACACGACGCTCTTCCGATCTNNNTGATTTGCCTCTAAGAGCCCAT
AGCACACGTCTGAACTCTTTCCCTACACGACGCTCTTCCGATCTNNNTGAACAGCTTCTCCTCATCTTCC
AGCACACGTCTGAACTCTTTCCCTACACGACGCTCTTCCGATCTNNNTGAACAGCTTCTCCTCATCTTCC
AGCACACGTCTGAACTCTTTCCCTACACGACGCTCTTCCGATCTNNNTGAACAGCTTCTCCTCATCTTCC
AGCACACGTCTGAACTCTTTCCCTACACGACGCTCTTCCGATCTNNNTGAACAGCTTCTCCTCATCTTCC
AGCACACGTCTGAACTCTTTCCCTACACGACGCTCTTCCGATCTNNNTGAACAGCTTCTCCTCATCTTCC
AGCACACGTCTGAACTCTTTCCCTACACGACGCTCTTCCGATCTNNNTGAACAGCTTCTCCTCATCTTCC
AGCACACGTCTGAACTCTTTCCCTACACGACGCTCTTCCGATCTNNNTCCCTTACTCCCTTTGGGTCTT
AGCACACGTCTGAACTCTTTCCCTACACGACGCTCTTCCGATCTNNNTCCCTTACTCCCTTTGGGTCTT
AGCACACGTCTGAACTCTTTCCCTACACGACGCTCTTCCGATCTNNNTCCCTTACTCCCTTTGGGTCTT
AGCACACGTCTGAACTCTTTCCCTACACGACGCTCTTCCGATCTNNNTCCCTTACTCCCTTTGGGTCTT
AGCACACGTCTGAACTCTTTCCCTACACGACGCTCTTCCGATCTNNNTCCCTTACTCCCTTTGGGTCTT
AGCACACGTCTGAACTCTTTCCCTACACGACGCTCTTCCGATCTNNNTCCCTTACTCCCTTTGGGTCTT
AGCACACGTCTGAACTCTTTCCCTACACGACGCTCTTCCGATCTNNNCCCTGTACTATTACTCAACCGC
AGCACACGTCTGAACTCTTTCCCTACACGACGCTCTTCCGATCTNNNCCCTGTACTATTACTCAACCGC
AGCACACGTCTGAACTCTTTCCCTACACGACGCTCTTCCGATCTNNNCCCTGTACTATTACTCAACCGC
AGCACACGTCTGAACTCTTTCCCTACACGACGCTCTTCCGATCTNNNCCCTGTACTATTACTCAACCGC
AGCACACGTCTGAACTCTTTCCCTACACGACGCTCTTCCGATCTNNNCCCTGTACTATTACTCAACCGC
AGCACACGTCTGAACTCTTTCCCTACACGACGCTCTTCCGATCTNNNCCCTGTACTATTACTCAACCGC
GAGCACACGTCTGAACTCTTTCCCTACACGACGCTCTTCCGATCTNNNTTAGGCTTTACAAATCATATC
GAGCACACGTCTGAACTCTTTCCCTACACGACGCTCTTCCGATCTNNNTTAGGCTTTACAAATCATATC
GAGCACACGTCTGAACTCTTTCCCTACACGACGCTCTTCCGATCTNNNTTAGGCTTTACAAATCATATC

GAGCACACGTCTGAACTCTTTCCCTACACGACGCTCTTCCGATCTNNNTTAGGCTTTACAAATCATATC
GAGCACACGTCTGAACTCTTTCCCTACACGACGCTCTTCCGATCTNNNTTAGGCTTTACAAATCATATC
GAGCACACGTCTGAACTCTTTCCCTACACGACGCTCTTCCGATCTNNNTTAGGCTTTACAAATCATATC
AGCACACGTCTGAACTCTTTCCCTACACGACGCTCTTCCGATCTNNNAAAAAATTAGCAGGACGTGG
AGCACACGTCTGAACTCTTTCCCTACACGACGCTCTTCCGATCTNNNAAAAAATTAGCAGGACGTGG
AGCACACGTCTGAACTCTTTCCCTACACGACGCTCTTCCGATCTNNNAAAAAATTAGCAGGACGTGG
AGCACACGTCTGAACTCTTTCCCTACACGACGCTCTTCCGATCTNNNAAAAAATTAGCAGGACGTGG
AGCACACGTCTGAACTCTTTCCCTACACGACGCTCTTCCGATCTNNNAAAAAATTAGCAGGACGTGG
AGCACACGTCTGAACTCTTTCCCTACACGACGCTCTTCCGATCTNNNAAAAAATTAGCAGGACGTGG
GAGCACACGTCTGAACTCTTTCCCTACACGACGCTCTTCCGATCTNNNGCAGGGGGGATATGAATG
GAGCACACGTCTGAACTCTTTCCCTACACGACGCTCTTCCGATCTNNNGCAGGGGGGATATGAATG
GAGCACACGTCTGAACTCTTTCCCTACACGACGCTCTTCCGATCTNNNGCAGGGGGGATATGAATG
GAGCACACGTCTGAACTCTTTCCCTACACGACGCTCTTCCGATCTNNNGCAGGGGGGATATGAATG
GAGCACACGTCTGAACTCTTTCCCTACACGACGCTCTTCCGATCTNNNGCAGGGGGGATATGAATG
GAGCACACGTCTGAACTCTTTCCCTACACGACGCTCTTCCGATCTNNNGCAGGGGGGATATGAATG
AGCACACGTCTGAACTCTTTCCCTACACGACGCTCTTCCGATCTNNNGTCTCCGAGGGGAAAAAAAG
AGCACACGTCTGAACTCTTTCCCTACACGACGCTCTTCCGATCTNNNGTCTCCGAGGGGAAAAAAAG
AGCACACGTCTGAACTCTTTCCCTACACGACGCTCTTCCGATCTNNNGTCTCCGAGGGGAAAAAAAG
AGCACACGTCTGAACTCTTTCCCTACACGACGCTCTTCCGATCTNNNGTCTCCGAGGGGAAAAAAAG
AGCACACGTCTGAACTCTTTCCCTACACGACGCTCTTCCGATCTNNNGTCTCCGAGGGGAAAAAAAG
AGCACACGTCTGAACTCTTTCCCTACACGACGCTCTTCCGATCTNNNGTCTCCGAGGGGAAAAAAAG
CACGTCTGAACTCTTTCCCTACACGACGCTCTTCCGATCTNNNGAACTCCTGACCTCAAGTGATCCCAT
CACGTCTGAACTCTTTCCCTACACGACGCTCTTCCGATCTNNNGAACTCCTGACCTCAAGTGATCCCAT
CACGTCTGAACTCTTTCCCTACACGACGCTCTTCCGATCTNNNGAACTCCTGACCTCAAGTGATCCCAT
CACGTCTGAACTCTTTCCCTACACGACGCTCTTCCGATCTNNNGAACTCCTGACCTCAAGTGATCCCAT
CACGTCTGAACTCTTTCCCTACACGACGCTCTTCCGATCTNNNGAACTCCTGACCTCAAGTGATCCCAT
CACGTCTGAACTCTTTCCCTACACGACGCTCTTCCGATCTNNNGAACTCCTGACCTCAAGTGATCCCAT
AGCACACGTCTGAACTCTTTCCCTACACGACGCTCTTCCGATCTNNNTTCTTTCAGTATGTTACAGAAG
AGCACACGTCTGAACTCTTTCCCTACACGACGCTCTTCCGATCTNNNTTCTTTCAGTATGTTACAGAAG
AGCACACGTCTGAACTCTTTCCCTACACGACGCTCTTCCGATCTNNNTTCTTTCAGTATGTTACAGAAG
AGCACACGTCTGAACTCTTTCCCTACACGACGCTCTTCCGATCTNNNTTCTTTCAGTATGTTACAGAAG
AGCACACGTCTGAACTCTTTCCCTACACGACGCTCTTCCGATCTNNNTTCTTTCAGTATGTTACAGAAG
AGCACACGTCTGAACTCTTTCCCTACACGACGCTCTTCCGATCTNNNTTCTTTCAGTATGTTACAGAAG
AGCACACGTCTGAACTCTTTCCCTACACGACGCTCTTCCGATCTNNNCTGAGGCAGGAGAATCACTTG
AGCACACGTCTGAACTCTTTCCCTACACGACGCTCTTCCGATCTNNNCTGAGGCAGGAGAATCACTTG
AGCACACGTCTGAACTCTTTCCCTACACGACGCTCTTCCGATCTNNNCTGAGGCAGGAGAATCACTTG
AGCACACGTCTGAACTCTTTCCCTACACGACGCTCTTCCGATCTNNNCTGAGGCAGGAGAATCACTTG
AGCACACGTCTGAACTCTTTCCCTACACGACGCTCTTCCGATCTNNNCTGAGGCAGGAGAATCACTTG
AGCACACGTCTGAACTCTTTCCCTACACGACGCTCTTCCGATCTNNNCTGAGGCAGGAGAATCACTTG
AGCACACGTCTGAACTCTTTCCCTACACGACGCTCTTCCGATCTNNNACAACTCATTCTTCCAACAAACA
AGCACACGTCTGAACTCTTTCCCTACACGACGCTCTTCCGATCTNNNACAACTCATTCTTCCAACAAACA
AGCACACGTCTGAACTCTTTCCCTACACGACGCTCTTCCGATCTNNNACAACTCATTCTTCCAACAAACA
AGCACACGTCTGAACTCTTTCCCTACACGACGCTCTTCCGATCTNNNACAACTCATTCTTCCAACAAACA

AGCACACGTCTGAACTCTTTCCCTACACGACGCTCTTCCGATCTNNNACAACTCATTCTTCCAACAAACA
AGCACACGTCTGAACTCTTTCCCTACACGACGCTCTTCCGATCTNNNACAACTCATTCTTCCAACAAACA
AGCACACGTCTGAACTCTTTCCCTACACGACGCTCTTCCGATCTNNNTCAGAAACCCAGAAGGATGTC
AGCACACGTCTGAACTCTTTCCCTACACGACGCTCTTCCGATCTNNNTCAGAAACCCAGAAGGATGTC
AGCACACGTCTGAACTCTTTCCCTACACGACGCTCTTCCGATCTNNNTCAGAAACCCAGAAGGATGTC
AGCACACGTCTGAACTCTTTCCCTACACGACGCTCTTCCGATCTNNNTCAGAAACCCAGAAGGATGTC
AGCACACGTCTGAACTCTTTCCCTACACGACGCTCTTCCGATCTNNNTCAGAAACCCAGAAGGATGTC
AGCACACGTCTGAACTCTTTCCCTACACGACGCTCTTCCGATCTNNNTCAGAAACCCAGAAGGATGTC
GCACACGTCTGAACTCTTTCCCTACACGACGCTCTTCCGATCTNNNATGTCTGCACATCCTGTACTTGT
GCACACGTCTGAACTCTTTCCCTACACGACGCTCTTCCGATCTNNNATGTCTGCACATCCTGTACTTGT
GCACACGTCTGAACTCTTTCCCTACACGACGCTCTTCCGATCTNNNATGTCTGCACATCCTGTACTTGT
GCACACGTCTGAACTCTTTCCCTACACGACGCTCTTCCGATCTNNNATGTCTGCACATCCTGTACTTGT
GCACACGTCTGAACTCTTTCCCTACACGACGCTCTTCCGATCTNNNATGTCTGCACATCCTGTACTTGT
GCACACGTCTGAACTCTTTCCCTACACGACGCTCTTCCGATCTNNNATGTCTGCACATCCTGTACTTGT
GCACACGTCTGAACTCTTTCCCTACACGACGCTCTTCCGATCTNNNGCTAAGCTGCAAACTACATAGC
GCACACGTCTGAACTCTTTCCCTACACGACGCTCTTCCGATCTNNNGCTAAGCTGCAAACTACATAGC
GCACACGTCTGAACTCTTTCCCTACACGACGCTCTTCCGATCTNNNGCTAAGCTGCAAACTACATAGC
GCACACGTCTGAACTCTTTCCCTACACGACGCTCTTCCGATCTNNNGCTAAGCTGCAAACTACATAGC
GCACACGTCTGAACTCTTTCCCTACACGACGCTCTTCCGATCTNNNGCTAAGCTGCAAACTACATAGC
GCACACGTCTGAACTCTTTCCCTACACGACGCTCTTCCGATCTNNNGCTAAGCTGCAAACTACATAGC
GAGCACACGTCTGAACTCTTTCCCTACACGACGCTCTTCCGATCTNNNTGAGGCAGAAGAATTGCTT
GAGCACACGTCTGAACTCTTTCCCTACACGACGCTCTTCCGATCTNNNTGAGGCAGAAGAATTGCTT
GAGCACACGTCTGAACTCTTTCCCTACACGACGCTCTTCCGATCTNNNTGAGGCAGAAGAATTGCTT
GAGCACACGTCTGAACTCTTTCCCTACACGACGCTCTTCCGATCTNNNTGAGGCAGAAGAATTGCTT
GAGCACACGTCTGAACTCTTTCCCTACACGACGCTCTTCCGATCTNNNTGAGGCAGAAGAATTGCTT
GAGCACACGTCTGAACTCTTTCCCTACACGACGCTCTTCCGATCTNNNTGAGGCAGAAGAATTGCTT
GAGCACACGTCTGAACTCTTTCCCTACACGACGCTCTTCCGATCTNNNTGAGGCAGAAGAATTGCTT
GAGCACACGTCTGAACTCTTTCCCTACACGACGCTCTTCCGATCTNNNTGAGGCAGAAGAATTGCTT
GAGCACACGTCTGAACTCTTTCCCTACACGACGCTCTTCCGATCTNNNTGAGGCAGAAGAATTGCTT
AGCACACGTCTGAACTCTTTCCCTACACGACGCTCTTCCGATCTNNNGCCTGTAGTCCCAGCTACTTG
AGCACACGTCTGAACTCTTTCCCTACACGACGCTCTTCCGATCTNNNGCCTGTAGTCCCAGCTACTTG
AGCACACGTCTGAACTCTTTCCCTACACGACGCTCTTCCGATCTNNNGCCTGTAGTCCCAGCTACTTG
AGCACACGTCTGAACTCTTTCCCTACACGACGCTCTTCCGATCTNNNGCCTGTAGTCCCAGCTACTTG
AGCACACGTCTGAACTCTTTCCCTACACGACGCTCTTCCGATCTNNNGCCTGTAGTCCCAGCTACTTG
AGCACACGTCTGAACTCTTTCCCTACACGACGCTCTTCCGATCTNNNGCCTGTAGTCCCAGCTACTTG
AGCACACGTCTGAACTCTTTCCCTACACGACGCTCTTCCGATCTNNNGCCTGTAGTCCCAGCTACTTG
AGCACACGTCTGAACTCTTTCCCTACACGACGCTCTTCCGATCTNNNGCCTGTAGTCCCAGCTACTTG
AGCACACGTCTGAACTCTTTCCCTACACGACGCTCTTCCGATCTNNNGCCTGTAGTCCCAGCTACTTG
GAGCACACGTCTGAACTCTTTCCCTACACGACGCTCTTCCGATCTNNNATCTCACACTATTGCCCAGAC
GAGCACACGTCTGAACTCTTTCCCTACACGACGCTCTTCCGATCTNNNATCTCACACTATTGCCCAGAC
GAGCACACGTCTGAACTCTTTCCCTACACGACGCTCTTCCGATCTNNNATCTCACACTATTGCCCAGAC
GAGCACACGTCTGAACTCTTTCCCTACACGACGCTCTTCCGATCTNNNATCTCACACTATTGCCCAGAC
GAGCACACGTCTGAACTCTTTCCCTACACGACGCTCTTCCGATCTNNNATCTCACACTATTGCCCAGAC

GAGCACACGTCTGAACTCTTTCCCTACACGACGCTCTTCCGATCTNNNATCTCACACTATTGCCCAGAC
GAGCACACGTCTGAACTCTTTCCCTACACGACGCTCTTCCGATCTNNNATCTCACACTATTGCCCAGAC
GAGCACACGTCTGAACTCTTTCCCTACACGACGCTCTTCCGATCTNNNATCTCACACTATTGCCCAGAC
GAGCACACGTCTGAACTCTTTCCCTACACGACGCTCTTCCGATCTNNNATCTCACACTATTGCCCAGAC
GCACACGTCTGAACTCTTTCCCTACACGACGCTCTTCCGATCTNNNCTTTCAAGCAGCTGGGATTACT
GCACACGTCTGAACTCTTTCCCTACACGACGCTCTTCCGATCTNNNCTTTCAAGCAGCTGGGATTACT
GCACACGTCTGAACTCTTTCCCTACACGACGCTCTTCCGATCTNNNCTTTCAAGCAGCTGGGATTACT
GCACACGTCTGAACTCTTTCCCTACACGACGCTCTTCCGATCTNNNCTTTCAAGCAGCTGGGATTACT
GCACACGTCTGAACTCTTTCCCTACACGACGCTCTTCCGATCTNNNCTTTCAAGCAGCTGGGATTACT
GCACACGTCTGAACTCTTTCCCTACACGACGCTCTTCCGATCTNNNCTTTCAAGCAGCTGGGATTACT
GCACACGTCTGAACTCTTTCCCTACACGACGCTCTTCCGATCTNNNCTTTCAAGCAGCTGGGATTACT
GCACACGTCTGAACTCTTTCCCTACACGACGCTCTTCCGATCTNNNCTTTCAAGCAGCTGGGATTACT
GCACACGTCTGAACTCTTTCCCTACACGACGCTCTTCCGATCTNNNCTTTCAAGCAGCTGGGATTACT
GCACACGTCTGAACTCTTTCCCTACACGACGCTCTTCCGATCTNNNTCCACGGTGGATGACCTAATTTA
GCACACGTCTGAACTCTTTCCCTACACGACGCTCTTCCGATCTNNNTCCACGGTGGATGACCTAATTTA
GCACACGTCTGAACTCTTTCCCTACACGACGCTCTTCCGATCTNNNTCCACGGTGGATGACCTAATTTA
GCACACGTCTGAACTCTTTCCCTACACGACGCTCTTCCGATCTNNNTCCACGGTGGATGACCTAATTTA
GCACACGTCTGAACTCTTTCCCTACACGACGCTCTTCCGATCTNNNTCCACGGTGGATGACCTAATTTA
GCACACGTCTGAACTCTTTCCCTACACGACGCTCTTCCGATCTNNNTCCACGGTGGATGACCTAATTTA
GCACACGTCTGAACTCTTTCCCTACACGACGCTCTTCCGATCTNNNTCCACGGTGGATGACCTAATTTA
GCACACGTCTGAACTCTTTCCCTACACGACGCTCTTCCGATCTNNNTCCACGGTGGATGACCTAATTTA
GCACACGTCTGAACTCTTTCCCTACACGACGCTCTTCCGATCTNNNTCCACGGTGGATGACCTAATTTA
GAGCACACGTCTGAACTCTTTCCCTACACGACGCTCTTCCGATCTNNNGAATCTTCAACCTCTGCCAGC
GAGCACACGTCTGAACTCTTTCCCTACACGACGCTCTTCCGATCTNNNGAATCTTCAACCTCTGCCAGC
GAGCACACGTCTGAACTCTTTCCCTACACGACGCTCTTCCGATCTNNNGAATCTTCAACCTCTGCCAGC
GAGCACACGTCTGAACTCTTTCCCTACACGACGCTCTTCCGATCTNNNGAATCTTCAACCTCTGCCAGC
GAGCACACGTCTGAACTCTTTCCCTACACGACGCTCTTCCGATCTNNNGAATCTTCAACCTCTGCCAGC
GAGCACACGTCTGAACTCTTTCCCTACACGACGCTCTTCCGATCTNNNGAATCTTCAACCTCTGCCAGC
GAGCACACGTCTGAACTCTTTCCCTACACGACGCTCTTCCGATCTNNNGAATCTTCAACCTCTGCCAGC
GAGCACACGTCTGAACTCTTTCCCTACACGACGCTCTTCCGATCTNNNGAATCTTCAACCTCTGCCAGC
GAGCACACGTCTGAACTCTTTCCCTACACGACGCTCTTCCGATCTNNNGAATCTTCAACCTCTGCCAGC
GAGCACACGTCTGAACTCTTTCCCTACACGACGCTCTTCCGATCTNNNGGAGCATTTGCTTTAGGGAC
GAGCACACGTCTGAACTCTTTCCCTACACGACGCTCTTCCGATCTNNNGGAGCATTTGCTTTAGGGAC
GAGCACACGTCTGAACTCTTTCCCTACACGACGCTCTTCCGATCTNNNGGAGCATTTGCTTTAGGGAC
GAGCACACGTCTGAACTCTTTCCCTACACGACGCTCTTCCGATCTNNNGGAGCATTTGCTTTAGGGAC
GAGCACACGTCTGAACTCTTTCCCTACACGACGCTCTTCCGATCTNNNGGAGCATTTGCTTTAGGGAC
GAGCACACGTCTGAACTCTTTCCCTACACGACGCTCTTCCGATCTNNNGGAGCATTTGCTTTAGGGAC
GAGCACACGTCTGAACTCTTTCCCTACACGACGCTCTTCCGATCTNNNGGAGCATTTGCTTTAGGGAC
GAGCACACGTCTGAACTCTTTCCCTACACGACGCTCTTCCGATCTNNNGGAGCATTTGCTTTAGGGAC
GAGCACACGTCTGAACTCTTTCCCTACACGACGCTCTTCCGATCTNNNGGAGCATTTGCTTTAGGGAC
GAGCACACGTCTGAACTCTTTCCCTACACGACGCTCTTCCGATCTNNNGGGTATGATGATGCTATTTT
GAGCACACGTCTGAACTCTTTCCCTACACGACGCTCTTCCGATCTNNNGGGTATGATGATGCTATTTT
GAGCACACGTCTGAACTCTTTCCCTACACGACGCTCTTCCGATCTNNNGGGTATGATGATGCTATTTT

AGAGCACACGTCTGAACTCTTTCCCTACACGACGCTCTTCCGATCTNNNGGGTATGATGATGCTATTTT
AGAGCACACGTCTGAACTCTTTCCCTACACGACGCTCTTCCGATCTNNNGGGTATGATGATGCTATTTT
AGAGCACACGTCTGAACTCTTTCCCTACACGACGCTCTTCCGATCTNNNGGGTATGATGATGCTATTTT
AGAGCACACGTCTGAACTCTTTCCCTACACGACGCTCTTCCGATCTNNNGGGTATGATGATGCTATTTT
AGAGCACACGTCTGAACTCTTTCCCTACACGACGCTCTTCCGATCTNNNGGGTATGATGATGCTATTTT
AGAGCACACGTCTGAACTCTTTCCCTACACGACGCTCTTCCGATCTNNNGGGTATGATGATGCTATTTT
AGAGCACACGTCTGAACTCTTTCCCTACACGACGCTCTTCCGATCTNNNCTTGAACATGGGTGGGGTTT
AGAGCACACGTCTGAACTCTTTCCCTACACGACGCTCTTCCGATCTNNNCTTGAACATGGGTGGGGTTT
AGAGCACACGTCTGAACTCTTTCCCTACACGACGCTCTTCCGATCTNNNCTTGAACATGGGTGGGGTTT
AGAGCACACGTCTGAACTCTTTCCCTACACGACGCTCTTCCGATCTNNNCTTGAACATGGGTGGGGTTT
AGAGCACACGTCTGAACTCTTTCCCTACACGACGCTCTTCCGATCTNNNCTTGAACATGGGTGGGGTTT
AGAGCACACGTCTGAACTCTTTCCCTACACGACGCTCTTCCGATCTNNNCTTGAACATGGGTGGGGTTT
AGAGCACACGTCTGAACTCTTTCCCTACACGACGCTCTTCCGATCTNNNCTTGAACATGGGTGGGGTTT
AGAGCACACGTCTGAACTCTTTCCCTACACGACGCTCTTCCGATCTNNNCTTGAACATGGGTGGGGTTT
AGAGCACACGTCTGAACTCTTTCCCTACACGACGCTCTTCCGATCTNNNCTTGAACATGGGTGGGGTTT
AGCACACGTCTGAACTCTTTCCCTACACGACGCTCTTCCGATCTNNNACAAATCTTCCTGGCATTCCTGT
AGCACACGTCTGAACTCTTTCCCTACACGACGCTCTTCCGATCTNNNACAAATCTTCCTGGCATTCCTGT
AGCACACGTCTGAACTCTTTCCCTACACGACGCTCTTCCGATCTNNNACAAATCTTCCTGGCATTCCTGT
AGCACACGTCTGAACTCTTTCCCTACACGACGCTCTTCCGATCTNNNACAAATCTTCCTGGCATTCCTGT
AGCACACGTCTGAACTCTTTCCCTACACGACGCTCTTCCGATCTNNNACAAATCTTCCTGGCATTCCTGT
AGCACACGTCTGAACTCTTTCCCTACACGACGCTCTTCCGATCTNNNACAAATCTTCCTGGCATTCCTGT
AGCACACGTCTGAACTCTTTCCCTACACGACGCTCTTCCGATCTNNNACAAATCTTCCTGGCATTCCTGT
AGCACACGTCTGAACTCTTTCCCTACACGACGCTCTTCCGATCTNNNACAAATCTTCCTGGCATTCCTGT
AGCACACGTCTGAACTCTTTCCCTACACGACGCTCTTCCGATCTNNNACAAATCTTCCTGGCATTCCTGT
GAGCACACGTCTGAACTCTTTCCCTACACGACGCTCTTCCGATCTNNNCTGAGATGGAGGAATCGCTTC
GAGCACACGTCTGAACTCTTTCCCTACACGACGCTCTTCCGATCTNNNCTGAGATGGAGGAATCGCTTC
GAGCACACGTCTGAACTCTTTCCCTACACGACGCTCTTCCGATCTNNNCTGAGATGGAGGAATCGCTTC
GAGCACACGTCTGAACTCTTTCCCTACACGACGCTCTTCCGATCTNNNCTGAGATGGAGGAATCGCTTC
GAGCACACGTCTGAACTCTTTCCCTACACGACGCTCTTCCGATCTNNNCTGAGATGGAGGAATCGCTTC
GAGCACACGTCTGAACTCTTTCCCTACACGACGCTCTTCCGATCTNNNCTGAGATGGAGGAATCGCTTC
GAGCACACGTCTGAACTCTTTCCCTACACGACGCTCTTCCGATCTNNNCTGAGATGGAGGAATCGCTTC
GAGCACACGTCTGAACTCTTTCCCTACACGACGCTCTTCCGATCTNNNCTGAGATGGAGGAATCGCTTC
GAGCACACGTCTGAACTCTTTCCCTACACGACGCTCTTCCGATCTNNNCTGAGATGGAGGAATCGCTTC
AGCACACGTCTGAACTCTTTCCCTACACGACGCTCTTCCGATCTNNNCATGCCTCTAATCCCAGCTACTC
AGCACACGTCTGAACTCTTTCCCTACACGACGCTCTTCCGATCTNNNCATGCCTCTAATCCCAGCTACTC
AGCACACGTCTGAACTCTTTCCCTACACGACGCTCTTCCGATCTNNNCATGCCTCTAATCCCAGCTACTC
AGCACACGTCTGAACTCTTTCCCTACACGACGCTCTTCCGATCTNNNCATGCCTCTAATCCCAGCTACTC
AGCACACGTCTGAACTCTTTCCCTACACGACGCTCTTCCGATCTNNNCATGCCTCTAATCCCAGCTACTC
AGCACACGTCTGAACTCTTTCCCTACACGACGCTCTTCCGATCTNNNCATGCCTCTAATCCCAGCTACTC
AGCACACGTCTGAACTCTTTCCCTACACGACGCTCTTCCGATCTNNNCATGCCTCTAATCCCAGCTACTC
AGCACACGTCTGAACTCTTTCCCTACACGACGCTCTTCCGATCTNNNCATGCCTCTAATCCCAGCTACTC
AGCACACGTCTGAACTCTTTCCCTACACGACGCTCTTCCGATCTNNNCATGCCTCTAATCCCAGCTACTC
AGCACACGTCTGAACTCTTTCCCTACACGACGCTCTTCCGATCTNNNGACTTACATAGAAGGCCTGGG

AGCACACGTCTGAACTCTTTCCCTACACGACGCTCTTCCGATCTNNNGACTTACATAGAAGGCCTGGG
AGCACACGTCTGAACTCTTTCCCTACACGACGCTCTTCCGATCTNNNGACTTACATAGAAGGCCTGGG
AGCACACGTCTGAACTCTTTCCCTACACGACGCTCTTCCGATCTNNNGACTTACATAGAAGGCCTGGG
AGCACACGTCTGAACTCTTTCCCTACACGACGCTCTTCCGATCTNNNGACTTACATAGAAGGCCTGGG
AGCACACGTCTGAACTCTTTCCCTACACGACGCTCTTCCGATCTNNNGACTTACATAGAAGGCCTGGG
AGCACACGTCTGAACTCTTTCCCTACACGACGCTCTTCCGATCTNNNGACTTACATAGAAGGCCTGGG
AGCACACGTCTGAACTCTTTCCCTACACGACGCTCTTCCGATCTNNNGACTTACATAGAAGGCCTGGG
AGCACACGTCTGAACTCTTTCCCTACACGACGCTCTTCCGATCTNNNGACTTACATAGAAGGCCTGGG
AGCACACGTCTGAACTCTTTCCCTACACGACGCTCTTCCGATCTNNNCCCAAAGTCCTTTGTGTCATTCC
AGCACACGTCTGAACTCTTTCCCTACACGACGCTCTTCCGATCTNNNCCCAAAGTCCTTTGTGTCATTCC
AGCACACGTCTGAACTCTTTCCCTACACGACGCTCTTCCGATCTNNNCCCAAAGTCCTTTGTGTCATTCC
AGCACACGTCTGAACTCTTTCCCTACACGACGCTCTTCCGATCTNNNCCCAAAGTCCTTTGTGTCATTCC
AGCACACGTCTGAACTCTTTCCCTACACGACGCTCTTCCGATCTNNNCCCAAAGTCCTTTGTGTCATTCC
AGCACACGTCTGAACTCTTTCCCTACACGACGCTCTTCCGATCTNNNCCCAAAGTCCTTTGTGTCATTCC
AGCACACGTCTGAACTCTTTCCCTACACGACGCTCTTCCGATCTNNNCCCAAAGTCCTTTGTGTCATTCC
AGCACACGTCTGAACTCTTTCCCTACACGACGCTCTTCCGATCTNNNCCCAAAGTCCTTTGTGTCATTCC
AGCACACGTCTGAACTCTTTCCCTACACGACGCTCTTCCGATCTNNNCCCAAAGTCCTTTGTGTCATTCC
AGCACACGTCTGAACTCTTTCCCTACACGACGCTCTTCCGATCTNNNAGCTGAGGAAACTTACCTACCA
AGCACACGTCTGAACTCTTTCCCTACACGACGCTCTTCCGATCTNNNAGCTGAGGAAACTTACCTACCA
AGCACACGTCTGAACTCTTTCCCTACACGACGCTCTTCCGATCTNNNAGCTGAGGAAACTTACCTACCA
AGCACACGTCTGAACTCTTTCCCTACACGACGCTCTTCCGATCTNNNAGCTGAGGAAACTTACCTACCA
AGCACACGTCTGAACTCTTTCCCTACACGACGCTCTTCCGATCTNNNAGCTGAGGAAACTTACCTACCA
AGCACACGTCTGAACTCTTTCCCTACACGACGCTCTTCCGATCTNNNAGCTGAGGAAACTTACCTACCA
AGCACACGTCTGAACTCTTTCCCTACACGACGCTCTTCCGATCTNNNAGCTGAGGAAACTTACCTACCA
AGCACACGTCTGAACTCTTTCCCTACACGACGCTCTTCCGATCTNNNAGCTGAGGAAACTTACCTACCA
AGCACACGTCTGAACTCTTTCCCTACACGACGCTCTTCCGATCTNNNAGCTGAGGAAACTTACCTACCA
GAGCACACGTCTGAACTCTTTCCCTACACGACGCTCTTCCGATCTNNNCTATGGAGTTGGAAGCACCTT
GAGCACACGTCTGAACTCTTTCCCTACACGACGCTCTTCCGATCTNNNCTATGGAGTTGGAAGCACCTT
GAGCACACGTCTGAACTCTTTCCCTACACGACGCTCTTCCGATCTNNNCTATGGAGTTGGAAGCACCTT
GAGCACACGTCTGAACTCTTTCCCTACACGACGCTCTTCCGATCTNNNCTATGGAGTTGGAAGCACCTT
GAGCACACGTCTGAACTCTTTCCCTACACGACGCTCTTCCGATCTNNNCTATGGAGTTGGAAGCACCTT
GAGCACACGTCTGAACTCTTTCCCTACACGACGCTCTTCCGATCTNNNCTATGGAGTTGGAAGCACCTT
GAGCACACGTCTGAACTCTTTCCCTACACGACGCTCTTCCGATCTNNNCTATGGAGTTGGAAGCACCTT
GAGCACACGTCTGAACTCTTTCCCTACACGACGCTCTTCCGATCTNNNCTATGGAGTTGGAAGCACCTT
GAGCACACGTCTGAACTCTTTCCCTACACGACGCTCTTCCGATCTNNNCTATGGAGTTGGAAGCACCTT
GAAGAGCACACGTCTGAACTCTTTCCCTACACGACGCTCTTCCGATCTNNNGTTGTCTCTAATTGCAGCC
GAAGAGCACACGTCTGAACTCTTTCCCTACACGACGCTCTTCCGATCTNNNGTTGTCTCTAATTGCAGCC
GAAGAGCACACGTCTGAACTCTTTCCCTACACGACGCTCTTCCGATCTNNNGTTGTCTCTAATTGCAGCC
GAAGAGCACACGTCTGAACTCTTTCCCTACACGACGCTCTTCCGATCTNNNGTTGTCTCTAATTGCAGCC
GAAGAGCACACGTCTGAACTCTTTCCCTACACGACGCTCTTCCGATCTNNNGTTGTCTCTAATTGCAGCC
GAAGAGCACACGTCTGAACTCTTTCCCTACACGACGCTCTTCCGATCTNNNGTTGTCTCTAATTGCAGCC
GAAGAGCACACGTCTGAACTCTTTCCCTACACGACGCTCTTCCGATCTNNNGTTGTCTCTAATTGCAGCC
GAAGAGCACACGTCTGAACTCTTTCCCTACACGACGCTCTTCCGATCTNNNGTTGTCTCTAATTGCAGCC

GAAGAGCACACGTCTGAACTCTTTCCCTACACGACGCTCTTCCGATCTNNNGTTGTCTCTAATTGCAGC
AGCACACGTCTGAACTCTTTCCCTACACGACGCTCTTCCGATCTNNNCTGTAGTCCCAGCTACTCAGGA
AGCACACGTCTGAACTCTTTCCCTACACGACGCTCTTCCGATCTNNNCTGTAGTCCCAGCTACTCAGGA
AGCACACGTCTGAACTCTTTCCCTACACGACGCTCTTCCGATCTNNNCTGTAGTCCCAGCTACTCAGGA
AGCACACGTCTGAACTCTTTCCCTACACGACGCTCTTCCGATCTNNNCTGTAGTCCCAGCTACTCAGGA
AGCACACGTCTGAACTCTTTCCCTACACGACGCTCTTCCGATCTNNNCTGTAGTCCCAGCTACTCAGGA
AGCACACGTCTGAACTCTTTCCCTACACGACGCTCTTCCGATCTNNNCTGTAGTCCCAGCTACTCAGGA
AGCACACGTCTGAACTCTTTCCCTACACGACGCTCTTCCGATCTNNNCTGTAGTCCCAGCTACTCAGGA
AGCACACGTCTGAACTCTTTCCCTACACGACGCTCTTCCGATCTNNNCTGTAGTCCCAGCTACTCAGGA
AGCACACGTCTGAACTCTTTCCCTACACGACGCTCTTCCGATCTNNNCTGTAGTCCCAGCTACTCAGGA
AGCACACGTCTGAACTCTTTCCCTACACGACGCTCTTCCGATCTNNNGGGATTGGTGGTTATATCCCGT
AGCACACGTCTGAACTCTTTCCCTACACGACGCTCTTCCGATCTNNNGGGATTGGTGGTTATATCCCGT
AGCACACGTCTGAACTCTTTCCCTACACGACGCTCTTCCGATCTNNNGGGATTGGTGGTTATATCCCGT
AGCACACGTCTGAACTCTTTCCCTACACGACGCTCTTCCGATCTNNNGGGATTGGTGGTTATATCCCGT
AGCACACGTCTGAACTCTTTCCCTACACGACGCTCTTCCGATCTNNNGGGATTGGTGGTTATATCCCGT
AGCACACGTCTGAACTCTTTCCCTACACGACGCTCTTCCGATCTNNNGGGATTGGTGGTTATATCCCGT
AGCACACGTCTGAACTCTTTCCCTACACGACGCTCTTCCGATCTNNNGGGATTGGTGGTTATATCCCGT
AGCACACGTCTGAACTCTTTCCCTACACGACGCTCTTCCGATCTNNNGGGATTGGTGGTTATATCCCGT
AGCACACGTCTGAACTCTTTCCCTACACGACGCTCTTCCGATCTNNNGGGATTGGTGGTTATATCCCGT
AGCACACGTCTGAACTCTTTCCCTACACGACGCTCTTCCGATCTNNNGAAATGGGGTTTCTCCATGTTG
AGCACACGTCTGAACTCTTTCCCTACACGACGCTCTTCCGATCTNNNGAAATGGGGTTTCTCCATGTTG
AGCACACGTCTGAACTCTTTCCCTACACGACGCTCTTCCGATCTNNNGAAATGGGGTTTCTCCATGTTG
AGCACACGTCTGAACTCTTTCCCTACACGACGCTCTTCCGATCTNNNGAAATGGGGTTTCTCCATGTTG
AGCACACGTCTGAACTCTTTCCCTACACGACGCTCTTCCGATCTNNNGAAATGGGGTTTCTCCATGTTG
AGCACACGTCTGAACTCTTTCCCTACACGACGCTCTTCCGATCTNNNGAAATGGGGTTTCTCCATGTTG
AGCACACGTCTGAACTCTTTCCCTACACGACGCTCTTCCGATCTNNNGAAATGGGGTTTCTCCATGTTG
AGCACACGTCTGAACTCTTTCCCTACACGACGCTCTTCCGATCTNNNGAAATGGGGTTTCTCCATGTTG
AGCACACGTCTGAACTCTTTCCCTACACGACGCTCTTCCGATCTNNNGAAATGGGGTTTCTCCATGTTG
GAGCACACGTCTGAACTCTTTCCCTACACGACGCTCTTCCGATCTNNNTTTGTTTGTAAGGAAGGGGA
GAGCACACGTCTGAACTCTTTCCCTACACGACGCTCTTCCGATCTNNNTTTGTTTGTAAGGAAGGGGA
GAGCACACGTCTGAACTCTTTCCCTACACGACGCTCTTCCGATCTNNNTTTGTTTGTAAGGAAGGGGA
GAGCACACGTCTGAACTCTTTCCCTACACGACGCTCTTCCGATCTNNNTTTGTTTGTAAGGAAGGGGA
GAGCACACGTCTGAACTCTTTCCCTACACGACGCTCTTCCGATCTNNNTTTGTTTGTAAGGAAGGGGA
GAGCACACGTCTGAACTCTTTCCCTACACGACGCTCTTCCGATCTNNNTTTGTTTGTAAGGAAGGGGA
GAGCACACGTCTGAACTCTTTCCCTACACGACGCTCTTCCGATCTNNNTTTGTTTGTAAGGAAGGGGA
GAGCACACGTCTGAACTCTTTCCCTACACGACGCTCTTCCGATCTNNNTTTGTTTGTAAGGAAGGGGA
GAGCACACGTCTGAACTCTTTCCCTACACGACGCTCTTCCGATCTNNNTTTGTTTGTAAGGAAGGGGA
GAGCACACGTCTGAACTCTTTCCCTACACGACGCTCTTCCGATCTNNNACTCCTGTAATCCCAGCACTTT
GAGCACACGTCTGAACTCTTTCCCTACACGACGCTCTTCCGATCTNNNACTCCTGTAATCCCAGCACTTT
GAGCACACGTCTGAACTCTTTCCCTACACGACGCTCTTCCGATCTNNNACTCCTGTAATCCCAGCACTTT
GAGCACACGTCTGAACTCTTTCCCTACACGACGCTCTTCCGATCTNNNACTCCTGTAATCCCAGCACTTT
GAGCACACGTCTGAACTCTTTCCCTACACGACGCTCTTCCGATCTNNNACTCCTGTAATCCCAGCACTTT
GAGCACACGTCTGAACTCTTTCCCTACACGACGCTCTTCCGATCTNNNACTCCTGTAATCCCAGCACTTT

GAGCACACGTCTGAACTCTTTCCCTACACGACGCTCTTCCGATCTNNNACTCCTGTAATCCCAGCACTTT

GAGCACACGTCTGAACTCTTTCCCTACACGACGCTCTTCCGATCTNNNACTCCTGTAATCCCAGCACTTT

GAGCACACGTCTGAACTCTTTCCCTACACGACGCTCTTCCGATCTNNNACTCCTGTAATCCCAGCACTTT

AGCACACGTCTGAACTCTTTCCCTACACGACGCTCTTCCGATCTNNNCATTCATCATGATTGCTTCCCCC

AGCACACGTCTGAACTCTTTCCCTACACGACGCTCTTCCGATCTNNNCATTCATCATGATTGCTTCCCCC

AGCACACGTCTGAACTCTTTCCCTACACGACGCTCTTCCGATCTNNNCATTCATCATGATTGCTTCCCCC

AGCACACGTCTGAACTCTTTCCCTACACGACGCTCTTCCGATCTNNNCATTCATCATGATTGCTTCCCCC

AGCACACGTCTGAACTCTTTCCCTACACGACGCTCTTCCGATCTNNNCATTCATCATGATTGCTTCCCCC

AGCACACGTCTGAACTCTTTCCCTACACGACGCTCTTCCGATCTNNNCATTCATCATGATTGCTTCCCCC

AGCACACGTCTGAACTCTTTCCCTACACGACGCTCTTCCGATCTNNNCATTCATCATGATTGCTTCCCCC

AGCACACGTCTGAACTCTTTCCCTACACGACGCTCTTCCGATCTNNNCATTCATCATGATTGCTTCCCCC

AGCACACGTCTGAACTCTTTCCCTACACGACGCTCTTCCGATCTNNNCATTCATCATGATTGCTTCCCCC

GAGCACACGTCTGAACTCTTTCCCTACACGACGCTCTTCCGATCTNNNACGCCTTCTCTCCACATATGTT

GAGCACACGTCTGAACTCTTTCCCTACACGACGCTCTTCCGATCTNNNACGCCTTCTCTCCACATATGTT

GAGCACACGTCTGAACTCTTTCCCTACACGACGCTCTTCCGATCTNNNACGCCTTCTCTCCACATATGTT

GAGCACACGTCTGAACTCTTTCCCTACACGACGCTCTTCCGATCTNNNACGCCTTCTCTCCACATATGTT

GAGCACACGTCTGAACTCTTTCCCTACACGACGCTCTTCCGATCTNNNACGCCTTCTCTCCACATATGTT

GAGCACACGTCTGAACTCTTTCCCTACACGACGCTCTTCCGATCTNNNACGCCTTCTCTCCACATATGTT

GAGCACACGTCTGAACTCTTTCCCTACACGACGCTCTTCCGATCTNNNACGCCTTCTCTCCACATATGTT

GAGCACACGTCTGAACTCTTTCCCTACACGACGCTCTTCCGATCTNNNACGCCTTCTCTCCACATATGTT

GAGCACACGTCTGAACTCTTTCCCTACACGACGCTCTTCCGATCTNNNACGCCTTCTCTCCACATATGTT

GAGCACACGTCTGAACTCTTTCCCTACACGACGCTCTTCCGATCTNNNTGACTTCCCATTCGGTTTACTG

GAGCACACGTCTGAACTCTTTCCCTACACGACGCTCTTCCGATCTNNNTGACTTCCCATTCGGTTTACTG

GAGCACACGTCTGAACTCTTTCCCTACACGACGCTCTTCCGATCTNNNTGACTTCCCATTCGGTTTACTG

GAGCACACGTCTGAACTCTTTCCCTACACGACGCTCTTCCGATCTNNNTGACTTCCCATTCGGTTTACTG

GAGCACACGTCTGAACTCTTTCCCTACACGACGCTCTTCCGATCTNNNTGACTTCCCATTCGGTTTACTG

GAGCACACGTCTGAACTCTTTCCCTACACGACGCTCTTCCGATCTNNNTGACTTCCCATTCGGTTTACTG

GAGCACACGTCTGAACTCTTTCCCTACACGACGCTCTTCCGATCTNNNTGACTTCCCATTCGGTTTACTG

GAGCACACGTCTGAACTCTTTCCCTACACGACGCTCTTCCGATCTNNNTGACTTCCCATTCGGTTTACTG

GAGCACACGTCTGAACTCTTTCCCTACACGACGCTCTTCCGATCTNNNTGACTTCCCATTCGGTTTACTG

AGCACACGTCTGAACTCTTTCCCTACACGACGCTCTTCCGATCTNNNAGTGCTGGGATTATAGGCATG

AGCACACGTCTGAACTCTTTCCCTACACGACGCTCTTCCGATCTNNNAGTGCTGGGATTATAGGCATG

AGCACACGTCTGAACTCTTTCCCTACACGACGCTCTTCCGATCTNNNAGTGCTGGGATTATAGGCATG

AGCACACGTCTGAACTCTTTCCCTACACGACGCTCTTCCGATCTNNNAGTGCTGGGATTATAGGCATG

AGCACACGTCTGAACTCTTTCCCTACACGACGCTCTTCCGATCTNNNAGTGCTGGGATTATAGGCATG

AGCACACGTCTGAACTCTTTCCCTACACGACGCTCTTCCGATCTNNNAGTGCTGGGATTATAGGCATG

AGCACACGTCTGAACTCTTTCCCTACACGACGCTCTTCCGATCTNNNAGTGCTGGGATTATAGGCATG

AGCACACGTCTGAACTCTTTCCCTACACGACGCTCTTCCGATCTNNNAGTGCTGGGATTATAGGCATG

AGCACACGTCTGAACTCTTTCCCTACACGACGCTCTTCCGATCTNNNAGTGCTGGGATTATAGGCATG

GAGCACACGTCTGAACTCTTTCCCTACACGACGCTCTTCCGATCTNNNGTTTGAAGTGTACCAGTTGCA

GAGCACACGTCTGAACTCTTTCCCTACACGACGCTCTTCCGATCTNNNGTTTGAAGTGTACCAGTTGCA

GAGCACACGTCTGAACTCTTTCCCTACACGACGCTCTTCCGATCTNNNGTTTGAAGTGTACCAGTTGCA

GAGCACACGTCTGAACTCTTTCCCTACACGACGCTCTTCCGATCTNNNGTTTGAAGTGTACCAGTTGCA

GAGCACACGTCTGAACTCTTTCCCTACACGACGCTCTTCCGATCTNNNGTTTGAAGTGTACCAGTTGCA
GAGCACACGTCTGAACTCTTTCCCTACACGACGCTCTTCCGATCTNNNGTTTGAAGTGTACCAGTTGCA
GAGCACACGTCTGAACTCTTTCCCTACACGACGCTCTTCCGATCTNNNGTTTGAAGTGTACCAGTTGCA
GAGCACACGTCTGAACTCTTTCCCTACACGACGCTCTTCCGATCTNNNGTTTGAAGTGTACCAGTTGCA
GAGCACACGTCTGAACTCTTTCCCTACACGACGCTCTTCCGATCTNNNGTTTGAAGTGTACCAGTTGCA
GAGCACACGTCTGAACTCTTTCCCTACACGACGCTCTTCCGATCTNNNAATCAGGTTGAGAGCATGAGA
GAGCACACGTCTGAACTCTTTCCCTACACGACGCTCTTCCGATCTNNNAATCAGGTTGAGAGCATGAGA
GAGCACACGTCTGAACTCTTTCCCTACACGACGCTCTTCCGATCTNNNAATCAGGTTGAGAGCATGAGA
GAGCACACGTCTGAACTCTTTCCCTACACGACGCTCTTCCGATCTNNNAATCAGGTTGAGAGCATGAGA
GAGCACACGTCTGAACTCTTTCCCTACACGACGCTCTTCCGATCTNNNAATCAGGTTGAGAGCATGAGA
GAGCACACGTCTGAACTCTTTCCCTACACGACGCTCTTCCGATCTNNNAATCAGGTTGAGAGCATGAGA
GAGCACACGTCTGAACTCTTTCCCTACACGACGCTCTTCCGATCTNNNAATCAGGTTGAGAGCATGAGA
GAGCACACGTCTGAACTCTTTCCCTACACGACGCTCTTCCGATCTNNNAATCAGGTTGAGAGCATGAGA
GAGCACACGTCTGAACTCTTTCCCTACACGACGCTCTTCCGATCTNNNAATCAGGTTGAGAGCATGAGA
AGAGCACACGTCTGAACTCTTTCCCTACACGACGCTCTTCCGATCTNNNTATAGTCCAGCACCTCCAGTA
AGAGCACACGTCTGAACTCTTTCCCTACACGACGCTCTTCCGATCTNNNTATAGTCCAGCACCTCCAGTA
AGAGCACACGTCTGAACTCTTTCCCTACACGACGCTCTTCCGATCTNNNTATAGTCCAGCACCTCCAGTA
AGAGCACACGTCTGAACTCTTTCCCTACACGACGCTCTTCCGATCTNNNTATAGTCCAGCACCTCCAGTA
AGAGCACACGTCTGAACTCTTTCCCTACACGACGCTCTTCCGATCTNNNTATAGTCCAGCACCTCCAGTA
AGAGCACACGTCTGAACTCTTTCCCTACACGACGCTCTTCCGATCTNNNTATAGTCCAGCACCTCCAGTA
AGAGCACACGTCTGAACTCTTTCCCTACACGACGCTCTTCCGATCTNNNTATAGTCCAGCACCTCCAGTA
AGAGCACACGTCTGAACTCTTTCCCTACACGACGCTCTTCCGATCTNNNTATAGTCCAGCACCTCCAGTA
AGAGCACACGTCTGAACTCTTTCCCTACACGACGCTCTTCCGATCTNNNTATAGTCCAGCACCTCCAGTA
GAGCACACGTCTGAACTCTTTCCCTACACGACGCTCTTCCGATCTNNNACCCTCAATGACTCTTGAAGG
GAGCACACGTCTGAACTCTTTCCCTACACGACGCTCTTCCGATCTNNNACCCTCAATGACTCTTGAAGG
GAGCACACGTCTGAACTCTTTCCCTACACGACGCTCTTCCGATCTNNNACCCTCAATGACTCTTGAAGG
GAGCACACGTCTGAACTCTTTCCCTACACGACGCTCTTCCGATCTNNNACCCTCAATGACTCTTGAAGG
GAGCACACGTCTGAACTCTTTCCCTACACGACGCTCTTCCGATCTNNNACCCTCAATGACTCTTGAAGG
GAGCACACGTCTGAACTCTTTCCCTACACGACGCTCTTCCGATCTNNNACCCTCAATGACTCTTGAAGG
GAGCACACGTCTGAACTCTTTCCCTACACGACGCTCTTCCGATCTNNNACCCTCAATGACTCTTGAAGG
GAGCACACGTCTGAACTCTTTCCCTACACGACGCTCTTCCGATCTNNNACCCTCAATGACTCTTGAAGG
GAGCACACGTCTGAACTCTTTCCCTACACGACGCTCTTCCGATCTNNNACCCTCAATGACTCTTGAAGG
GAGCACACGTCTGAACTCTTTCCCTACACGACGCTCTTCCGATCTNNNGTTTTGATAAGACAGCTGGGA
GAGCACACGTCTGAACTCTTTCCCTACACGACGCTCTTCCGATCTNNNGTTTTGATAAGACAGCTGGGA
GAGCACACGTCTGAACTCTTTCCCTACACGACGCTCTTCCGATCTNNNGTTTTGATAAGACAGCTGGGA
GAGCACACGTCTGAACTCTTTCCCTACACGACGCTCTTCCGATCTNNNGTTTTGATAAGACAGCTGGGA
GAGCACACGTCTGAACTCTTTCCCTACACGACGCTCTTCCGATCTNNNGTTTTGATAAGACAGCTGGGA
GAGCACACGTCTGAACTCTTTCCCTACACGACGCTCTTCCGATCTNNNGTTTTGATAAGACAGCTGGGA
GAGCACACGTCTGAACTCTTTCCCTACACGACGCTCTTCCGATCTNNNGTTTTGATAAGACAGCTGGGA
GAGCACACGTCTGAACTCTTTCCCTACACGACGCTCTTCCGATCTNNNGTTTTGATAAGACAGCTGGGA
GAGCACACGTCTGAACTCTTTCCCTACACGACGCTCTTCCGATCTNNNGTTTTGATAAGACAGCTGGGA
GAGCACACGTCTGAACTCTTTCCCTACACGACGCTCTTCCGATCTNNNTCCTCTAGCCAAAGGACCAGA
GAGCACACGTCTGAACTCTTTCCCTACACGACGCTCTTCCGATCTNNNTCCTCTAGCCAAAGGACCAGA

GAGCACACGTCTGAACTCTTTCCCTACACGACGCTCTTCCGATCTNNNTCCTCTAGCCAAAGGACCAGA
GAGCACACGTCTGAACTCTTTCCCTACACGACGCTCTTCCGATCTNNNTCCTCTAGCCAAAGGACCAGA
GAGCACACGTCTGAACTCTTTCCCTACACGACGCTCTTCCGATCTNNNTCCTCTAGCCAAAGGACCAGA
GAGCACACGTCTGAACTCTTTCCCTACACGACGCTCTTCCGATCTNNNTCCTCTAGCCAAAGGACCAGA
GAGCACACGTCTGAACTCTTTCCCTACACGACGCTCTTCCGATCTNNNTCCTCTAGCCAAAGGACCAGA
GAGCACACGTCTGAACTCTTTCCCTACACGACGCTCTTCCGATCTNNNTCCTCTAGCCAAAGGACCAGA
GAGCACACGTCTGAACTCTTTCCCTACACGACGCTCTTCCGATCTNNNTCCTCTAGCCAAAGGACCAGA
AGCACACGTCTGAACTCTTTCCCTACACGACGCTCTTCCGATCTNNNCTTGGGAGCTGAGACATGAGA
AGCACACGTCTGAACTCTTTCCCTACACGACGCTCTTCCGATCTNNNCTTGGGAGCTGAGACATGAGA
AGCACACGTCTGAACTCTTTCCCTACACGACGCTCTTCCGATCTNNNCTTGGGAGCTGAGACATGAGA
AGCACACGTCTGAACTCTTTCCCTACACGACGCTCTTCCGATCTNNNCTTGGGAGCTGAGACATGAGA
AGCACACGTCTGAACTCTTTCCCTACACGACGCTCTTCCGATCTNNNCTTGGGAGCTGAGACATGAGA
AGCACACGTCTGAACTCTTTCCCTACACGACGCTCTTCCGATCTNNNCTTGGGAGCTGAGACATGAGA
AGCACACGTCTGAACTCTTTCCCTACACGACGCTCTTCCGATCTNNNCTTGGGAGCTGAGACATGAGA
AGCACACGTCTGAACTCTTTCCCTACACGACGCTCTTCCGATCTNNNCTTGGGAGCTGAGACATGAGA
AGCACACGTCTGAACTCTTTCCCTACACGACGCTCTTCCGATCTNNNCTTGGGAGCTGAGACATGAGA
GAGCACACGTCTGAACTCTTTCCCTACACGACGCTCTTCCGATCTNNNATGCCTGTAATCCCAGCTACTT
GAGCACACGTCTGAACTCTTTCCCTACACGACGCTCTTCCGATCTNNNATGCCTGTAATCCCAGCTACTT
GAGCACACGTCTGAACTCTTTCCCTACACGACGCTCTTCCGATCTNNNATGCCTGTAATCCCAGCTACTT
GAGCACACGTCTGAACTCTTTCCCTACACGACGCTCTTCCGATCTNNNATGCCTGTAATCCCAGCTACTT
GAGCACACGTCTGAACTCTTTCCCTACACGACGCTCTTCCGATCTNNNATGCCTGTAATCCCAGCTACTT
GAGCACACGTCTGAACTCTTTCCCTACACGACGCTCTTCCGATCTNNNATGCCTGTAATCCCAGCTACTT
GAGCACACGTCTGAACTCTTTCCCTACACGACGCTCTTCCGATCTNNNATGCCTGTAATCCCAGCTACTT
GAGCACACGTCTGAACTCTTTCCCTACACGACGCTCTTCCGATCTNNNATGCCTGTAATCCCAGCTACTT
GAGCACACGTCTGAACTCTTTCCCTACACGACGCTCTTCCGATCTNNNATGCCTGTAATCCCAGCTACTT
GCACACGTCTGAACTCTTTCCCTACACGACGCTCTTCCGATCTNNNTCAAGTCTTCTCTTGCCATGACAC
GCACACGTCTGAACTCTTTCCCTACACGACGCTCTTCCGATCTNNNTCAAGTCTTCTCTTGCCATGACAC
GCACACGTCTGAACTCTTTCCCTACACGACGCTCTTCCGATCTNNNTCAAGTCTTCTCTTGCCATGACAC
GCACACGTCTGAACTCTTTCCCTACACGACGCTCTTCCGATCTNNNTCAAGTCTTCTCTTGCCATGACAC
GCACACGTCTGAACTCTTTCCCTACACGACGCTCTTCCGATCTNNNTCAAGTCTTCTCTTGCCATGACAC
GCACACGTCTGAACTCTTTCCCTACACGACGCTCTTCCGATCTNNNTCAAGTCTTCTCTTGCCATGACAC
GCACACGTCTGAACTCTTTCCCTACACGACGCTCTTCCGATCTNNNTCAAGTCTTCTCTTGCCATGACAC
GCACACGTCTGAACTCTTTCCCTACACGACGCTCTTCCGATCTNNNTCAAGTCTTCTCTTGCCATGACAC
GCACACGTCTGAACTCTTTCCCTACACGACGCTCTTCCGATCTNNNTCAAGTCTTCTCTTGCCATGACAC
AGCACACGTCTGAACTCTTTCCCTACACGACGCTCTTCCGATCTNNNTCCCCAGAGAGTCAGTAGAAAG
AGCACACGTCTGAACTCTTTCCCTACACGACGCTCTTCCGATCTNNNTCCCCAGAGAGTCAGTAGAAAG
AGCACACGTCTGAACTCTTTCCCTACACGACGCTCTTCCGATCTNNNTCCCCAGAGAGTCAGTAGAAAG
AGCACACGTCTGAACTCTTTCCCTACACGACGCTCTTCCGATCTNNNTCCCCAGAGAGTCAGTAGAAAG
AGCACACGTCTGAACTCTTTCCCTACACGACGCTCTTCCGATCTNNNTCCCCAGAGAGTCAGTAGAAAG
AGCACACGTCTGAACTCTTTCCCTACACGACGCTCTTCCGATCTNNNTCCCCAGAGAGTCAGTAGAAAG
AGCACACGTCTGAACTCTTTCCCTACACGACGCTCTTCCGATCTNNNTCCCCAGAGAGTCAGTAGAAAG
AGCACACGTCTGAACTCTTTCCCTACACGACGCTCTTCCGATCTNNNTCCCCAGAGAGTCAGTAGAAAG
AGCACACGTCTGAACTCTTTCCCTACACGACGCTCTTCCGATCTNNNTCCCCAGAGAGTCAGTAGAAAG

AGCACACGTCTGAACTCTTTCCCTACACGACGCTCTTCCGATCTNNNCAAACAGCAGCCCTACAGAAG
AGCACACGTCTGAACTCTTTCCCTACACGACGCTCTTCCGATCTNNNCAAACAGCAGCCCTACAGAAG
AGCACACGTCTGAACTCTTTCCCTACACGACGCTCTTCCGATCTNNNCAAACAGCAGCCCTACAGAAG
AGCACACGTCTGAACTCTTTCCCTACACGACGCTCTTCCGATCTNNNCAAACAGCAGCCCTACAGAAG
AGCACACGTCTGAACTCTTTCCCTACACGACGCTCTTCCGATCTNNNCAAACAGCAGCCCTACAGAAG
AGCACACGTCTGAACTCTTTCCCTACACGACGCTCTTCCGATCTNNNCAAACAGCAGCCCTACAGAAG
AGCACACGTCTGAACTCTTTCCCTACACGACGCTCTTCCGATCTNNNCAAACAGCAGCCCTACAGAAG
AGCACACGTCTGAACTCTTTCCCTACACGACGCTCTTCCGATCTNNNCAAACAGCAGCCCTACAGAAG
AGCACACGTCTGAACTCTTTCCCTACACGACGCTCTTCCGATCTNNNCAAACAGCAGCCCTACAGAAG
GCACACGTCTGAACTCTTTCCCTACACGACGCTCTTCCGATCTNNNGTATTCCCAGTTACTCAGGAGG
GCACACGTCTGAACTCTTTCCCTACACGACGCTCTTCCGATCTNNNGTATTCCCAGTTACTCAGGAGG
GCACACGTCTGAACTCTTTCCCTACACGACGCTCTTCCGATCTNNNGTATTCCCAGTTACTCAGGAGG
GCACACGTCTGAACTCTTTCCCTACACGACGCTCTTCCGATCTNNNGTATTCCCAGTTACTCAGGAGG
GCACACGTCTGAACTCTTTCCCTACACGACGCTCTTCCGATCTNNNGTATTCCCAGTTACTCAGGAGG
GCACACGTCTGAACTCTTTCCCTACACGACGCTCTTCCGATCTNNNGTATTCCCAGTTACTCAGGAGG
GCACACGTCTGAACTCTTTCCCTACACGACGCTCTTCCGATCTNNNGTATTCCCAGTTACTCAGGAGG
GCACACGTCTGAACTCTTTCCCTACACGACGCTCTTCCGATCTNNNGTATTCCCAGTTACTCAGGAGG
GCACACGTCTGAACTCTTTCCCTACACGACGCTCTTCCGATCTNNNGTATTCCCAGTTACTCAGGAGG
GCACACGTCTGAACTCTTTCCCTACACGACGCTCTTCCGATCTNNNATGGGAAATGTTCTGGCTGTA
GCACACGTCTGAACTCTTTCCCTACACGACGCTCTTCCGATCTNNNATGGGAAATGTTCTGGCTGTA
GCACACGTCTGAACTCTTTCCCTACACGACGCTCTTCCGATCTNNNATGGGAAATGTTCTGGCTGTA
GCACACGTCTGAACTCTTTCCCTACACGACGCTCTTCCGATCTNNNATGGGAAATGTTCTGGCTGTA
GCACACGTCTGAACTCTTTCCCTACACGACGCTCTTCCGATCTNNNATGGGAAATGTTCTGGCTGTA
GCACACGTCTGAACTCTTTCCCTACACGACGCTCTTCCGATCTNNNATGGGAAATGTTCTGGCTGTA
GCACACGTCTGAACTCTTTCCCTACACGACGCTCTTCCGATCTNNNATGGGAAATGTTCTGGCTGTA
GCACACGTCTGAACTCTTTCCCTACACGACGCTCTTCCGATCTNNNATGGGAAATGTTCTGGCTGTA
GCACACGTCTGAACTCTTTCCCTACACGACGCTCTTCCGATCTNNNATGGGAAATGTTCTGGCTGTA
AGCACACGTCTGAACTCTTTCCCTACACGACGCTCTTCCGATCTNNNGACTTCTGCTCCAAAGGTTCC
AGCACACGTCTGAACTCTTTCCCTACACGACGCTCTTCCGATCTNNNGACTTCTGCTCCAAAGGTTCC
AGCACACGTCTGAACTCTTTCCCTACACGACGCTCTTCCGATCTNNNGACTTCTGCTCCAAAGGTTCC
AGCACACGTCTGAACTCTTTCCCTACACGACGCTCTTCCGATCTNNNGACTTCTGCTCCAAAGGTTCC
AGCACACGTCTGAACTCTTTCCCTACACGACGCTCTTCCGATCTNNNGACTTCTGCTCCAAAGGTTCC
AGCACACGTCTGAACTCTTTCCCTACACGACGCTCTTCCGATCTNNNGACTTCTGCTCCAAAGGTTCC
AGCACACGTCTGAACTCTTTCCCTACACGACGCTCTTCCGATCTNNNGACTTCTGCTCCAAAGGTTCC
AGCACACGTCTGAACTCTTTCCCTACACGACGCTCTTCCGATCTNNNGACTTCTGCTCCAAAGGTTCC
AGCACACGTCTGAACTCTTTCCCTACACGACGCTCTTCCGATCTNNNGACTTCTGCTCCAAAGGTTCC
GCACACGTCTGAACTCTTTCCCTACACGACGCTCTTCCGATCTNNNCCCTGTTCCGGATAATTCACGTA
GCACACGTCTGAACTCTTTCCCTACACGACGCTCTTCCGATCTNNNCCCTGTTCCGGATAATTCACGTA
GCACACGTCTGAACTCTTTCCCTACACGACGCTCTTCCGATCTNNNCCCTGTTCCGGATAATTCACGTA
GCACACGTCTGAACTCTTTCCCTACACGACGCTCTTCCGATCTNNNCCCTGTTCCGGATAATTCACGTA
GCACACGTCTGAACTCTTTCCCTACACGACGCTCTTCCGATCTNNNCCCTGTTCCGGATAATTCACGTA
GCACACGTCTGAACTCTTTCCCTACACGACGCTCTTCCGATCTNNNCCCTGTTCCGGATAATTCACGTA
GCACACGTCTGAACTCTTTCCCTACACGACGCTCTTCCGATCTNNNCCCTGTTCCGGATAATTCACGTA

GCACACGTCTGAACTCTTTCCCTACACGACGCTCTTCCGATCTNNNCCCTGTTCCGGATAATTCACGTA
GCACACGTCTGAACTCTTTCCCTACACGACGCTCTTCCGATCTNNNCCCTGTTCCGGATAATTCACGTA
AGCACACGTCTGAACTCTTTCCCTACACGACGCTCTTCCGATCTNNNCCCTACATTCTGGAGAACTAGG
AGCACACGTCTGAACTCTTTCCCTACACGACGCTCTTCCGATCTNNNCCCTACATTCTGGAGAACTAGG
AGCACACGTCTGAACTCTTTCCCTACACGACGCTCTTCCGATCTNNNCCCTACATTCTGGAGAACTAGG
AGCACACGTCTGAACTCTTTCCCTACACGACGCTCTTCCGATCTNNNCCCTACATTCTGGAGAACTAGG
AGCACACGTCTGAACTCTTTCCCTACACGACGCTCTTCCGATCTNNNCCCTACATTCTGGAGAACTAGG
AGCACACGTCTGAACTCTTTCCCTACACGACGCTCTTCCGATCTNNNCCCTACATTCTGGAGAACTAGG
AGCACACGTCTGAACTCTTTCCCTACACGACGCTCTTCCGATCTNNNCCCTACATTCTGGAGAACTAGG
AGCACACGTCTGAACTCTTTCCCTACACGACGCTCTTCCGATCTNNNCCCTACATTCTGGAGAACTAGG
AGCACACGTCTGAACTCTTTCCCTACACGACGCTCTTCCGATCTNNNCCCTACATTCTGGAGAACTAGG
GCACACGTCTGAACTCTTTCCCTACACGACGCTCTTCCGATCTNNNGAGAATCCTTTGAACCTGACAGG
GCACACGTCTGAACTCTTTCCCTACACGACGCTCTTCCGATCTNNNGAGAATCCTTTGAACCTGACAGG
GCACACGTCTGAACTCTTTCCCTACACGACGCTCTTCCGATCTNNNGAGAATCCTTTGAACCTGACAGG
GCACACGTCTGAACTCTTTCCCTACACGACGCTCTTCCGATCTNNNGAGAATCCTTTGAACCTGACAGG
GCACACGTCTGAACTCTTTCCCTACACGACGCTCTTCCGATCTNNNGAGAATCCTTTGAACCTGACAGG
GCACACGTCTGAACTCTTTCCCTACACGACGCTCTTCCGATCTNNNGAGAATCCTTTGAACCTGACAGG
GCACACGTCTGAACTCTTTCCCTACACGACGCTCTTCCGATCTNNNGAGAATCCTTTGAACCTGACAGG
GCACACGTCTGAACTCTTTCCCTACACGACGCTCTTCCGATCTNNNGAGAATCCTTTGAACCTGACAGG
GCACACGTCTGAACTCTTTCCCTACACGACGCTCTTCCGATCTNNNGAGAATCCTTTGAACCTGACAGG
GAGCACACGTCTGAACTCTTTCCCTACACGACGCTCTTCCGATCTNNNGAGATTGAGGCACTGAACTC
GAGCACACGTCTGAACTCTTTCCCTACACGACGCTCTTCCGATCTNNNGAGATTGAGGCACTGAACTC
GAGCACACGTCTGAACTCTTTCCCTACACGACGCTCTTCCGATCTNNNGAGATTGAGGCACTGAACTC
GAGCACACGTCTGAACTCTTTCCCTACACGACGCTCTTCCGATCTNNNGAGATTGAGGCACTGAACTC
GAGCACACGTCTGAACTCTTTCCCTACACGACGCTCTTCCGATCTNNNGAGATTGAGGCACTGAACTC
GAGCACACGTCTGAACTCTTTCCCTACACGACGCTCTTCCGATCTNNNGAGATTGAGGCACTGAACTC
GAGCACACGTCTGAACTCTTTCCCTACACGACGCTCTTCCGATCTNNNGAGATTGAGGCACTGAACTC
GAGCACACGTCTGAACTCTTTCCCTACACGACGCTCTTCCGATCTNNNGAGATTGAGGCACTGAACTC
GAGCACACGTCTGAACTCTTTCCCTACACGACGCTCTTCCGATCTNNNGAGATTGAGGCACTGAACTC
AGCACACGTCTGAACTCTTTCCCTACACGACGCTCTTCCGATCTNNNACTCCACAGTAACACAAGTTGT
AGCACACGTCTGAACTCTTTCCCTACACGACGCTCTTCCGATCTNNNACTCCACAGTAACACAAGTTGT
AGCACACGTCTGAACTCTTTCCCTACACGACGCTCTTCCGATCTNNNACTCCACAGTAACACAAGTTGT
AGCACACGTCTGAACTCTTTCCCTACACGACGCTCTTCCGATCTNNNACTCCACAGTAACACAAGTTGT
AGCACACGTCTGAACTCTTTCCCTACACGACGCTCTTCCGATCTNNNACTCCACAGTAACACAAGTTGT
AGCACACGTCTGAACTCTTTCCCTACACGACGCTCTTCCGATCTNNNACTCCACAGTAACACAAGTTGT
AGCACACGTCTGAACTCTTTCCCTACACGACGCTCTTCCGATCTNNNACTCCACAGTAACACAAGTTGT
AGCACACGTCTGAACTCTTTCCCTACACGACGCTCTTCCGATCTNNNACTCCACAGTAACACAAGTTGT
AGCACACGTCTGAACTCTTTCCCTACACGACGCTCTTCCGATCTNNNACTCCACAGTAACACAAGTTGT
GAGCACACGTCTGAACTCTTTCCCTACACGACGCTCTTCCGATCTNNNAGAATCACTTGAAACCGGGA
GAGCACACGTCTGAACTCTTTCCCTACACGACGCTCTTCCGATCTNNNAGAATCACTTGAAACCGGGA
GAGCACACGTCTGAACTCTTTCCCTACACGACGCTCTTCCGATCTNNNAGAATCACTTGAAACCGGGA
GAGCACACGTCTGAACTCTTTCCCTACACGACGCTCTTCCGATCTNNNAGAATCACTTGAAACCGGGA
GAGCACACGTCTGAACTCTTTCCCTACACGACGCTCTTCCGATCTNNNAGAATCACTTGAAACCGGGA

GAGCACACGTCTGAACTCTTTCCCTACACGACGCTCTTCCGATCTNNNAGAATCACTTGAAACCGGGA
GAGCACACGTCTGAACTCTTTCCCTACACGACGCTCTTCCGATCTNNNAGAATCACTTGAAACCGGGA
GAGCACACGTCTGAACTCTTTCCCTACACGACGCTCTTCCGATCTNNNAGAATCACTTGAAACCGGGA
GAGCACACGTCTGAACTCTTTCCCTACACGACGCTCTTCCGATCTNNNAGAATCACTTGAAACCGGGA
AGCACACGTCTGAACTCTTTCCCTACACGACGCTCTTCCGATCTNNNGTGGCTTTCATTCCCCTATAT
AGCACACGTCTGAACTCTTTCCCTACACGACGCTCTTCCGATCTNNNGTGGCTTTCATTCCCCTATAT
AGCACACGTCTGAACTCTTTCCCTACACGACGCTCTTCCGATCTNNNGTGGCTTTCATTCCCCTATAT
AGCACACGTCTGAACTCTTTCCCTACACGACGCTCTTCCGATCTNNNGTGGCTTTCATTCCCCTATAT
AGCACACGTCTGAACTCTTTCCCTACACGACGCTCTTCCGATCTNNNGTGGCTTTCATTCCCCTATAT
AGCACACGTCTGAACTCTTTCCCTACACGACGCTCTTCCGATCTNNNGTGGCTTTCATTCCCCTATAT
AGCACACGTCTGAACTCTTTCCCTACACGACGCTCTTCCGATCTNNNGTGGCTTTCATTCCCCTATAT
AGCACACGTCTGAACTCTTTCCCTACACGACGCTCTTCCGATCTNNNGTGGCTTTCATTCCCCTATAT
AGCACACGTCTGAACTCTTTCCCTACACGACGCTCTTCCGATCTNNNGTGGCTTTCATTCCCCTATAT
AGCACACGTCTGAACTCTTTCCCTACACGACGCTCTTCCGATCTNNNAGCTTACAGTGAGCCAAGATCA
AGCACACGTCTGAACTCTTTCCCTACACGACGCTCTTCCGATCTNNNAGCTTACAGTGAGCCAAGATCA
AGCACACGTCTGAACTCTTTCCCTACACGACGCTCTTCCGATCTNNNAGCTTACAGTGAGCCAAGATCA
AGCACACGTCTGAACTCTTTCCCTACACGACGCTCTTCCGATCTNNNAGCTTACAGTGAGCCAAGATCA
AGCACACGTCTGAACTCTTTCCCTACACGACGCTCTTCCGATCTNNNAGCTTACAGTGAGCCAAGATCA
AGCACACGTCTGAACTCTTTCCCTACACGACGCTCTTCCGATCTNNNAGCTTACAGTGAGCCAAGATCA
AGCACACGTCTGAACTCTTTCCCTACACGACGCTCTTCCGATCTNNNAGCTTACAGTGAGCCAAGATCA
AGCACACGTCTGAACTCTTTCCCTACACGACGCTCTTCCGATCTNNNAGCTTACAGTGAGCCAAGATCA
AGCACACGTCTGAACTCTTTCCCTACACGACGCTCTTCCGATCTNNNAGCTTACAGTGAGCCAAGATCA
GAGCACACGTCTGAACTCTTTCCCTACACGACGCTCTTCCGATCTNNNGCCTGTAATCCCAGATACTCA
GAGCACACGTCTGAACTCTTTCCCTACACGACGCTCTTCCGATCTNNNGCCTGTAATCCCAGATACTCA
GAGCACACGTCTGAACTCTTTCCCTACACGACGCTCTTCCGATCTNNNGCCTGTAATCCCAGATACTCA
GAGCACACGTCTGAACTCTTTCCCTACACGACGCTCTTCCGATCTNNNGCCTGTAATCCCAGATACTCA
GAGCACACGTCTGAACTCTTTCCCTACACGACGCTCTTCCGATCTNNNGCCTGTAATCCCAGATACTCA
GAGCACACGTCTGAACTCTTTCCCTACACGACGCTCTTCCGATCTNNNGCCTGTAATCCCAGATACTCA
GAGCACACGTCTGAACTCTTTCCCTACACGACGCTCTTCCGATCTNNNGCCTGTAATCCCAGATACTCA
GAGCACACGTCTGAACTCTTTCCCTACACGACGCTCTTCCGATCTNNNGCCTGTAATCCCAGATACTCA
GAGCACACGTCTGAACTCTTTCCCTACACGACGCTCTTCCGATCTNNNGCCTGTAATCCCAGATACTCA
AGCACACGTCTGAACTCTTTCCCTACACGACGCTCTTCCGATCTNNNGAGAATCGCTTGAACCCTGGA
AGCACACGTCTGAACTCTTTCCCTACACGACGCTCTTCCGATCTNNNGAGAATCGCTTGAACCCTGGA
AGCACACGTCTGAACTCTTTCCCTACACGACGCTCTTCCGATCTNNNGAGAATCGCTTGAACCCTGGA
AGCACACGTCTGAACTCTTTCCCTACACGACGCTCTTCCGATCTNNNGAGAATCGCTTGAACCCTGGA
AGCACACGTCTGAACTCTTTCCCTACACGACGCTCTTCCGATCTNNNGAGAATCGCTTGAACCCTGGA
AGCACACGTCTGAACTCTTTCCCTACACGACGCTCTTCCGATCTNNNGAGAATCGCTTGAACCCTGGA
AGCACACGTCTGAACTCTTTCCCTACACGACGCTCTTCCGATCTNNNGAGAATCGCTTGAACCCTGGA
AGCACACGTCTGAACTCTTTCCCTACACGACGCTCTTCCGATCTNNNGAGAATCGCTTGAACCCTGGA
AGCACACGTCTGAACTCTTTCCCTACACGACGCTCTTCCGATCTNNNGAGAATCGCTTGAACCCTGGA
GAGCACACGTCTGAACTCTTTCCCTACACGACGCTCTTCCGATCTNNNACCATGCCAGGCTAGTTTTTG
GAGCACACGTCTGAACTCTTTCCCTACACGACGCTCTTCCGATCTNNNACCATGCCAGGCTAGTTTTTG
GAGCACACGTCTGAACTCTTTCCCTACACGACGCTCTTCCGATCTNNNACCATGCCAGGCTAGTTTTTG

GAGCACACGTCTGAACTCTTTCCCTACACGACGCTCTTCCGATCTNNNACCATGCCAGGCTAGTTTTG
GAGCACACGTCTGAACTCTTTCCCTACACGACGCTCTTCCGATCTNNNACCATGCCAGGCTAGTTTTG
GAGCACACGTCTGAACTCTTTCCCTACACGACGCTCTTCCGATCTNNNACCATGCCAGGCTAGTTTTG
GAGCACACGTCTGAACTCTTTCCCTACACGACGCTCTTCCGATCTNNNACCATGCCAGGCTAGTTTTG
GAGCACACGTCTGAACTCTTTCCCTACACGACGCTCTTCCGATCTNNNACCATGCCAGGCTAGTTTTG
GAGCACACGTCTGAACTCTTTCCCTACACGACGCTCTTCCGATCTNNNACCATGCCAGGCTAGTTTTG
GAGCACACGTCTGAACTCTTTCCCTACACGACGCTCTTCCGATCTNNNTGGAGTTTCACTCTGTTGCCTA
GAGCACACGTCTGAACTCTTTCCCTACACGACGCTCTTCCGATCTNNNTGGAGTTTCACTCTGTTGCCTA
GAGCACACGTCTGAACTCTTTCCCTACACGACGCTCTTCCGATCTNNNTGGAGTTTCACTCTGTTGCCTA
GAGCACACGTCTGAACTCTTTCCCTACACGACGCTCTTCCGATCTNNNTGGAGTTTCACTCTGTTGCCTA
GAGCACACGTCTGAACTCTTTCCCTACACGACGCTCTTCCGATCTNNNTGGAGTTTCACTCTGTTGCCTA
GAGCACACGTCTGAACTCTTTCCCTACACGACGCTCTTCCGATCTNNNTGGAGTTTCACTCTGTTGCCTA
GAGCACACGTCTGAACTCTTTCCCTACACGACGCTCTTCCGATCTNNNTGGAGTTTCACTCTGTTGCCTA
GAGCACACGTCTGAACTCTTTCCCTACACGACGCTCTTCCGATCTNNNTGGAGTTTCACTCTGTTGCCTA
GAGCACACGTCTGAACTCTTTCCCTACACGACGCTCTTCCGATCTNNNTGGAGTTTCACTCTGTTGCCTA
AGCACACGTCTGAACTCTTTCCCTACACGACGCTCTTCCGATCTNNNATTCCATCCAGGTTGCTGAGAA
AGCACACGTCTGAACTCTTTCCCTACACGACGCTCTTCCGATCTNNNATTCCATCCAGGTTGCTGAGAA
AGCACACGTCTGAACTCTTTCCCTACACGACGCTCTTCCGATCTNNNATTCCATCCAGGTTGCTGAGAA
AGCACACGTCTGAACTCTTTCCCTACACGACGCTCTTCCGATCTNNNATTCCATCCAGGTTGCTGAGAA
GAGCACACGTCTGAACTCTTTCCCTACACGACGCTCTTCCGATCTNNNAGTGAGGCATGATGGTAGGC
GAGCACACGTCTGAACTCTTTCCCTACACGACGCTCTTCCGATCTNNNAGTGAGGCATGATGGTAGGC
GAGCACACGTCTGAACTCTTTCCCTACACGACGCTCTTCCGATCTNNNAGTGAGGCATGATGGTAGGC
GAGCACACGTCTGAACTCTTTCCCTACACGACGCTCTTCCGATCTNNNAGTGAGGCATGATGGTAGGC
AGCACACGTCTGAACTCTTTCCCTACACGACGCTCTTCCGATCTNNNTTTTAGGTGGCCTTTTCATCAT
AGCACACGTCTGAACTCTTTCCCTACACGACGCTCTTCCGATCTNNNTTTTAGGTGGCCTTTTCATCAT
AGCACACGTCTGAACTCTTTCCCTACACGACGCTCTTCCGATCTNNNTTTTAGGTGGCCTTTTCATCAT
AGCACACGTCTGAACTCTTTCCCTACACGACGCTCTTCCGATCTNNNTTTTAGGTGGCCTTTTCATCAT
GAGCACACGTCTGAACTCTTTCCCTACACGACGCTCTTCCGATCTNNNTTGCTGGAAGAGTATAAGCCT
GAGCACACGTCTGAACTCTTTCCCTACACGACGCTCTTCCGATCTNNNTTGCTGGAAGAGTATAAGCCT
GAGCACACGTCTGAACTCTTTCCCTACACGACGCTCTTCCGATCTNNNTTGCTGGAAGAGTATAAGCCT
GAGCACACGTCTGAACTCTTTCCCTACACGACGCTCTTCCGATCTNNNTTGCTGGAAGAGTATAAGCCT
GAGCACACGTCTGAACTCTTTCCCTACACGACGCTCTTCCGATCTNNNAAGGCCAGCATTACTCTGATA
GAGCACACGTCTGAACTCTTTCCCTACACGACGCTCTTCCGATCTNNNAAGGCCAGCATTACTCTGATA
GAGCACACGTCTGAACTCTTTCCCTACACGACGCTCTTCCGATCTNNNAAGGCCAGCATTACTCTGATA
GAGCACACGTCTGAACTCTTTCCCTACACGACGCTCTTCCGATCTNNNAAGGCCAGCATTACTCTGATA
AGCACACGTCTGAACTCTTTCCCTACACGACGCTCTTCCGATCTNNNTGTTCCCAAAGTTCTAGCTCCTCC
AGCACACGTCTGAACTCTTTCCCTACACGACGCTCTTCCGATCTNNNTGTTCCCAAAGTTCTAGCTCCTCC
AGCACACGTCTGAACTCTTTCCCTACACGACGCTCTTCCGATCTNNNTGTTCCCAAAGTTCTAGCTCCTCC
AGCACACGTCTGAACTCTTTCCCTACACGACGCTCTTCCGATCTNNNTGTTCCCAAAGTTCTAGCTCCTCC
AGAGCACACGTCTGAACTCTTTCCCTACACGACGCTCTTCCGATCTNNNGAGCCTGTAATCCCAGCTACT
AGAGCACACGTCTGAACTCTTTCCCTACACGACGCTCTTCCGATCTNNNGAGCCTGTAATCCCAGCTACT
AGAGCACACGTCTGAACTCTTTCCCTACACGACGCTCTTCCGATCTNNNGAGCCTGTAATCCCAGCTACT
AGAGCACACGTCTGAACTCTTTCCCTACACGACGCTCTTCCGATCTNNNGAGCCTGTAATCCCAGCTACT

GAGCACACGTCTGAACTCTTTCCCTACACGACGCTCTTCCGATCTNNNCTAGGCAAACGAGTGAGACT
GAGCACACGTCTGAACTCTTTCCCTACACGACGCTCTTCCGATCTNNNCTAGGCAAACGAGTGAGACT
GAGCACACGTCTGAACTCTTTCCCTACACGACGCTCTTCCGATCTNNNCTAGGCAAACGAGTGAGACT
GAGCACACGTCTGAACTCTTTCCCTACACGACGCTCTTCCGATCTNNNCTAGGCAAACGAGTGAGACT
AGCACACGTCTGAACTCTTTCCCTACACGACGCTCTTCCGATCTNNNAAAAAATTAGCTGGGCGTGGT
AGCACACGTCTGAACTCTTTCCCTACACGACGCTCTTCCGATCTNNNAAAAAATTAGCTGGGCGTGGT
AGCACACGTCTGAACTCTTTCCCTACACGACGCTCTTCCGATCTNNNAAAAAATTAGCTGGGCGTGGT
AGCACACGTCTGAACTCTTTCCCTACACGACGCTCTTCCGATCTNNNAAAAAATTAGCTGGGCGTGGT
GCACACGTCTGAACTCTTTCCCTACACGACGCTCTTCCGATCTNNNGGCAGCTTGTTTCTTATTGGAT
GCACACGTCTGAACTCTTTCCCTACACGACGCTCTTCCGATCTNNNGGCAGCTTGTTTCTTATTGGAT
GCACACGTCTGAACTCTTTCCCTACACGACGCTCTTCCGATCTNNNGGCAGCTTGTTTCTTATTGGAT
GCACACGTCTGAACTCTTTCCCTACACGACGCTCTTCCGATCTNNNGGCAGCTTGTTTCTTATTGGAT
GCACACGTCTGAACTCTTTCCCTACACGACGCTCTTCCGATCTNNNGGTGGTTCATGCCTGTAATCCTA
GCACACGTCTGAACTCTTTCCCTACACGACGCTCTTCCGATCTNNNGGTGGTTCATGCCTGTAATCCTA
GCACACGTCTGAACTCTTTCCCTACACGACGCTCTTCCGATCTNNNGGTGGTTCATGCCTGTAATCCTA
GCACACGTCTGAACTCTTTCCCTACACGACGCTCTTCCGATCTNNNGGTGGTTCATGCCTGTAATCCTA
AGCACACGTCTGAACTCTTTCCCTACACGACGCTCTTCCGATCTNNNTGCTCTAGGATTCCACACTTCTG
AGCACACGTCTGAACTCTTTCCCTACACGACGCTCTTCCGATCTNNNTGCTCTAGGATTCCACACTTCTG
AGCACACGTCTGAACTCTTTCCCTACACGACGCTCTTCCGATCTNNNTGCTCTAGGATTCCACACTTCTG
AGCACACGTCTGAACTCTTTCCCTACACGACGCTCTTCCGATCTNNNTGCTCTAGGATTCCACACTTCTG
GCACACGTCTGAACTCTTTCCCTACACGACGCTCTTCCGATCTNNNGAGCTCAGAAATTGACTCACACA
GCACACGTCTGAACTCTTTCCCTACACGACGCTCTTCCGATCTNNNGAGCTCAGAAATTGACTCACACA
GCACACGTCTGAACTCTTTCCCTACACGACGCTCTTCCGATCTNNNGAGCTCAGAAATTGACTCACACA
GCACACGTCTGAACTCTTTCCCTACACGACGCTCTTCCGATCTNNNGAGCTCAGAAATTGACTCACACA
GAGCACACGTCTGAACTCTTTCCCTACACGACGCTCTTCCGATCTNNNGCAATTCCTGCCAACTAAACAA
GAGCACACGTCTGAACTCTTTCCCTACACGACGCTCTTCCGATCTNNNGCAATTCCTGCCAACTAAACAA
GAGCACACGTCTGAACTCTTTCCCTACACGACGCTCTTCCGATCTNNNGCAATTCCTGCCAACTAAACAA
GAGCACACGTCTGAACTCTTTCCCTACACGACGCTCTTCCGATCTNNNGCAATTCCTGCCAACTAAACAA
AGCACACGTCTGAACTCTTTCCCTACACGACGCTCTTCCGATCTNNNCATTTACACACTGTGTCCCACAC
AGCACACGTCTGAACTCTTTCCCTACACGACGCTCTTCCGATCTNNNCATTTACACACTGTGTCCCACAC
AGCACACGTCTGAACTCTTTCCCTACACGACGCTCTTCCGATCTNNNCATTTACACACTGTGTCCCACAC
AGCACACGTCTGAACTCTTTCCCTACACGACGCTCTTCCGATCTNNNCATTTACACACTGTGTCCCACAC
AGCACACGTCTGAACTCTTTCCCTACACGACGCTCTTCCGATCTNNNGTGATCTGGGAATTTCAGGAG
AGCACACGTCTGAACTCTTTCCCTACACGACGCTCTTCCGATCTNNNGTGATCTGGGAATTTCAGGAG
AGCACACGTCTGAACTCTTTCCCTACACGACGCTCTTCCGATCTNNNGTGATCTGGGAATTTCAGGAG
AGCACACGTCTGAACTCTTTCCCTACACGACGCTCTTCCGATCTNNNGTGATCTGGGAATTTCAGGAG
AGCACACGTCTGAACTCTTTCCCTACACGACGCTCTTCCGATCTNNNGAGGTTGCAGTGAGCTGAGAT
AGCACACGTCTGAACTCTTTCCCTACACGACGCTCTTCCGATCTNNNGAGGTTGCAGTGAGCTGAGAT
AGCACACGTCTGAACTCTTTCCCTACACGACGCTCTTCCGATCTNNNGAGGTTGCAGTGAGCTGAGAT
AGCACACGTCTGAACTCTTTCCCTACACGACGCTCTTCCGATCTNNNGAGGTTGCAGTGAGCTGAGAT
GAGCACACGTCTGAACTCTTTCCCTACACGACGCTCTTCCGATCTNNNTCTTCAGAACTTTCAGAGAAA
GAGCACACGTCTGAACTCTTTCCCTACACGACGCTCTTCCGATCTNNNTCTTCAGAACTTTCAGAGAAA
GAGCACACGTCTGAACTCTTTCCCTACACGACGCTCTTCCGATCTNNNTCTTCAGAACTTTCAGAGAAA

GAGCACACGTCTGAACTCTTTCCCTACACGACGCTCTTCCGATCTNNNTCTTCAGAACTTTCAGAGAAA

GCACACGTCTGAACTCTTTCCCTACACGACGCTCTTCCGATCTNNNACCTACGTCTAGCTACCTACCCA

GCACACGTCTGAACTCTTTCCCTACACGACGCTCTTCCGATCTNNNACCTACGTCTAGCTACCTACCCA

GCACACGTCTGAACTCTTTCCCTACACGACGCTCTTCCGATCTNNNACCTACGTCTAGCTACCTACCCA

GCACACGTCTGAACTCTTTCCCTACACGACGCTCTTCCGATCTNNNACCTACGTCTAGCTACCTACCCA

GCACACGTCTGAACTCTTTCCCTACACGACGCTCTTCCGATCTNNNGGGGAGTGTATTAGACAGACA

GCACACGTCTGAACTCTTTCCCTACACGACGCTCTTCCGATCTNNNGGGGAGTGTATTAGACAGACA

GCACACGTCTGAACTCTTTCCCTACACGACGCTCTTCCGATCTNNNGGGGAGTGTATTAGACAGACA

GCACACGTCTGAACTCTTTCCCTACACGACGCTCTTCCGATCTNNNGGGGAGTGTATTAGACAGACA

AGCACACGTCTGAACTCTTTCCCTACACGACGCTCTTCCGATCTNNNGAACCAGGGAGTCGGAGATT

AGCACACGTCTGAACTCTTTCCCTACACGACGCTCTTCCGATCTNNNGAACCAGGGAGTCGGAGATT

AGCACACGTCTGAACTCTTTCCCTACACGACGCTCTTCCGATCTNNNGAACCAGGGAGTCGGAGATT

AGCACACGTCTGAACTCTTTCCCTACACGACGCTCTTCCGATCTNNNGAACCAGGGAGTCGGAGATT

AGCACACGTCTGAACTCTTTCCCTACACGACGCTCTTCCGATCTNNNTCCCCATTAGCTTTTCACCTAA

AGCACACGTCTGAACTCTTTCCCTACACGACGCTCTTCCGATCTNNNTCCCCATTAGCTTTTCACCTAA

AGCACACGTCTGAACTCTTTCCCTACACGACGCTCTTCCGATCTNNNTCCCCATTAGCTTTTCACCTAA

AGCACACGTCTGAACTCTTTCCCTACACGACGCTCTTCCGATCTNNNTCCCCATTAGCTTTTCACCTAA

AGCACACGTCTGAACTCTTTCCCTACACGACGCTCTTCCGATCTNNNTGAGCCAAGATCGTTCCACTGC

AGCACACGTCTGAACTCTTTCCCTACACGACGCTCTTCCGATCTNNNTGAGCCAAGATCGTTCCACTGC

AGCACACGTCTGAACTCTTTCCCTACACGACGCTCTTCCGATCTNNNTGAGCCAAGATCGTTCCACTGC

AGCACACGTCTGAACTCTTTCCCTACACGACGCTCTTCCGATCTNNNTGAGCCAAGATCGTTCCACTGC

GAGCACACGTCTGAACTCTTTCCCTACACGACGCTCTTCCGATCTNNNGCCTTCTTTGTCTGTTCCATG

GAGCACACGTCTGAACTCTTTCCCTACACGACGCTCTTCCGATCTNNNGCCTTCTTTGTCTGTTCCATG

GAGCACACGTCTGAACTCTTTCCCTACACGACGCTCTTCCGATCTNNNGCCTTCTTTGTCTGTTCCATG

GAGCACACGTCTGAACTCTTTCCCTACACGACGCTCTTCCGATCTNNNGCCTTCTTTGTCTGTTCCATG

CACACGTCTGAACTCTTTCCCTACACGACGCTCTTCCGATCTNNNGCTGAACTTGCGTGGCTAATATTC

CACACGTCTGAACTCTTTCCCTACACGACGCTCTTCCGATCTNNNGCTGAACTTGCGTGGCTAATATTC

CACACGTCTGAACTCTTTCCCTACACGACGCTCTTCCGATCTNNNGCTGAACTTGCGTGGCTAATATTC

CACACGTCTGAACTCTTTCCCTACACGACGCTCTTCCGATCTNNNGCTGAACTTGCGTGGCTAATATTC

AGCACACGTCTGAACTCTTTCCCTACACGACGCTCTTCCGATCTNNNAGACAGGAGAATCACTTGAA

AGCACACGTCTGAACTCTTTCCCTACACGACGCTCTTCCGATCTNNNAGACAGGAGAATCACTTGAA

AGCACACGTCTGAACTCTTTCCCTACACGACGCTCTTCCGATCTNNNAGACAGGAGAATCACTTGAA

AGCACACGTCTGAACTCTTTCCCTACACGACGCTCTTCCGATCTNNNAGACAGGAGAATCACTTGAA

GCACACGTCTGAACTCTTTCCCTACACGACGCTCTTCCGATCTNNNTGTATGCTTGTAGTCCCAGCTA

GCACACGTCTGAACTCTTTCCCTACACGACGCTCTTCCGATCTNNNTGTATGCTTGTAGTCCCAGCTA

GCACACGTCTGAACTCTTTCCCTACACGACGCTCTTCCGATCTNNNTGTATGCTTGTAGTCCCAGCTA

GCACACGTCTGAACTCTTTCCCTACACGACGCTCTTCCGATCTNNNTGTATGCTTGTAGTCCCAGCTA

GAGCACACGTCTGAACTCTTTCCCTACACGACGCTCTTCCGATCTNNNAGGGAACATAACAGGTGGG

GAGCACACGTCTGAACTCTTTCCCTACACGACGCTCTTCCGATCTNNNAGGGAACATAACAGGTGGG

GAGCACACGTCTGAACTCTTTCCCTACACGACGCTCTTCCGATCTNNNAGGGAACATAACAGGTGGG

GAGCACACGTCTGAACTCTTTCCCTACACGACGCTCTTCCGATCTNNNAGGGAACATAACAGGTGGG

GAGCACACGTCTGAACTCTTTCCCTACACGACGCTCTTCCGATCTNNNTGATGAGATTATGAGTGATT

GAGCACACGTCTGAACTCTTTCCCTACACGACGCTCTTCCGATCTNNNTGATGAGATTATGAGTGATT

GAGCACACGTCTGAACTCTTTCCCTACACGACGCTCTTCCGATCTNNNTGATGAGATTATGAGTGATT
GAGCACACGTCTGAACTCTTTCCCTACACGACGCTCTTCCGATCTNNNTGATGAGATTATGAGTGATT
CACACGTCTGAACTCTTTCCCTACACGACGCTCTTCCGATCTNNNTGGTGTGATGATTATATCTGCCAC
CACACGTCTGAACTCTTTCCCTACACGACGCTCTTCCGATCTNNNTGGTGTGATGATTATATCTGCCAC
CACACGTCTGAACTCTTTCCCTACACGACGCTCTTCCGATCTNNNTGGTGTGATGATTATATCTGCCAC
CACACGTCTGAACTCTTTCCCTACACGACGCTCTTCCGATCTNNNTGGTGTGATGATTATATCTGCCAC
GCACACGTCTGAACTCTTTCCCTACACGACGCTCTTCCGATCTNNNCCTGATGTCAATTTCTTCAGGGG
GCACACGTCTGAACTCTTTCCCTACACGACGCTCTTCCGATCTNNNCCTGATGTCAATTTCTTCAGGGG
GCACACGTCTGAACTCTTTCCCTACACGACGCTCTTCCGATCTNNNCCTGATGTCAATTTCTTCAGGGG
GCACACGTCTGAACTCTTTCCCTACACGACGCTCTTCCGATCTNNNCCTGATGTCAATTTCTTCAGGGG
AGCACACGTCTGAACTCTTTCCCTACACGACGCTCTTCCGATCTNNNAGTATGGCATCTCCTCATCTCCT
AGCACACGTCTGAACTCTTTCCCTACACGACGCTCTTCCGATCTNNNAGTATGGCATCTCCTCATCTCCT
AGCACACGTCTGAACTCTTTCCCTACACGACGCTCTTCCGATCTNNNAGTATGGCATCTCCTCATCTCCT
AGCACACGTCTGAACTCTTTCCCTACACGACGCTCTTCCGATCTNNNAGTATGGCATCTCCTCATCTCCT
GAGCACACGTCTGAACTCTTTCCCTACACGACGCTCTTCCGATCTNNNTGACAGAGACTCATTCCACAG
GAGCACACGTCTGAACTCTTTCCCTACACGACGCTCTTCCGATCTNNNTGACAGAGACTCATTCCACAG
GAGCACACGTCTGAACTCTTTCCCTACACGACGCTCTTCCGATCTNNNTGACAGAGACTCATTCCACAG
GAGCACACGTCTGAACTCTTTCCCTACACGACGCTCTTCCGATCTNNNTGACAGAGACTCATTCCACAG
GCACACGTCTGAACTCTTTCCCTACACGACGCTCTTCCGATCTNNNTCCACTTTGAAGGCCATTGTGTA
GCACACGTCTGAACTCTTTCCCTACACGACGCTCTTCCGATCTNNNTCCACTTTGAAGGCCATTGTGTA
GCACACGTCTGAACTCTTTCCCTACACGACGCTCTTCCGATCTNNNTCCACTTTGAAGGCCATTGTGTA
GCACACGTCTGAACTCTTTCCCTACACGACGCTCTTCCGATCTNNNTCCACTTTGAAGGCCATTGTGTA
GCACACGTCTGAACTCTTTCCCTACACGACGCTCTTCCGATCTNNNTGTGTGACAGAGCGAGACTCCA
GCACACGTCTGAACTCTTTCCCTACACGACGCTCTTCCGATCTNNNTGTGTGACAGAGCGAGACTCCA
GCACACGTCTGAACTCTTTCCCTACACGACGCTCTTCCGATCTNNNTGTGTGACAGAGCGAGACTCCA
GCACACGTCTGAACTCTTTCCCTACACGACGCTCTTCCGATCTNNNTGTGTGACAGAGCGAGACTCCA
AGCACACGTCTGAACTCTTTCCCTACACGACGCTCTTCCGATCTNNNTTCAAGCTATTCTCCTGCCTCA
AGCACACGTCTGAACTCTTTCCCTACACGACGCTCTTCCGATCTNNNTTCAAGCTATTCTCCTGCCTCA
AGCACACGTCTGAACTCTTTCCCTACACGACGCTCTTCCGATCTNNNTTCAAGCTATTCTCCTGCCTCA
AGCACACGTCTGAACTCTTTCCCTACACGACGCTCTTCCGATCTNNNTTCAAGCTATTCTCCTGCCTCA
AGCACACGTCTGAACTCTTTCCCTACACGACGCTCTTCCGATCTNNNCATGGGCTTACTTCTTGGCTTTT
AGCACACGTCTGAACTCTTTCCCTACACGACGCTCTTCCGATCTNNNCATGGGCTTACTTCTTGGCTTTT
AGCACACGTCTGAACTCTTTCCCTACACGACGCTCTTCCGATCTNNNCATGGGCTTACTTCTTGGCTTTT
AGCACACGTCTGAACTCTTTCCCTACACGACGCTCTTCCGATCTNNNCATGGGCTTACTTCTTGGCTTTT
AGCACACGTCTGAACTCTTTCCCTACACGACGCTCTTCCGATCTNNNAATCGGAGAAGTCACAGTAGC
AGCACACGTCTGAACTCTTTCCCTACACGACGCTCTTCCGATCTNNNAATCGGAGAAGTCACAGTAGC
AGCACACGTCTGAACTCTTTCCCTACACGACGCTCTTCCGATCTNNNAATCGGAGAAGTCACAGTAGC
AGCACACGTCTGAACTCTTTCCCTACACGACGCTCTTCCGATCTNNNAATCGGAGAAGTCACAGTAGC
AGCACACGTCTGAACTCTTTCCCTACACGACGCTCTTCCGATCTNNNCCAAAGATGGTTCAGCATACAC
AGCACACGTCTGAACTCTTTCCCTACACGACGCTCTTCCGATCTNNNCCAAAGATGGTTCAGCATACAC
AGCACACGTCTGAACTCTTTCCCTACACGACGCTCTTCCGATCTNNNCCAAAGATGGTTCAGCATACAC
AGCACACGTCTGAACTCTTTCCCTACACGACGCTCTTCCGATCTNNNCCAAAGATGGTTCAGCATACAC
AGCACACGTCTGAACTCTTTCCCTACACGACGCTCTTCCGATCTNNNGGGTCAGAGAAGCCATATTTCC

AGCACACGTCTGAACTCTTTCCCTACACGACGCTCTTCCGATCTNNNGGGTCAGAGAAGCCATATTTCC
AGCACACGTCTGAACTCTTTCCCTACACGACGCTCTTCCGATCTNNNGGGTCAGAGAAGCCATATTTCC
AGCACACGTCTGAACTCTTTCCCTACACGACGCTCTTCCGATCTNNNGGGTCAGAGAAGCCATATTTCC
AGCACACGTCTGAACTCTTTCCCTACACGACGCTCTTCCGATCTNNNTGGTTGGGAGAGATTTTTTCCT
AGCACACGTCTGAACTCTTTCCCTACACGACGCTCTTCCGATCTNNNTGGTTGGGAGAGATTTTTTCCT
AGCACACGTCTGAACTCTTTCCCTACACGACGCTCTTCCGATCTNNNTGGTTGGGAGAGATTTTTTCCT
AGCACACGTCTGAACTCTTTCCCTACACGACGCTCTTCCGATCTNNNTGGTTGGGAGAGATTTTTTCCT
AGAGCACACGTCTGAACTCTTTCCCTACACGACGCTCTTCCGATCTNNNGACTCATGCCTATAATCCCAG
AGAGCACACGTCTGAACTCTTTCCCTACACGACGCTCTTCCGATCTNNNGACTCATGCCTATAATCCCAG
AGAGCACACGTCTGAACTCTTTCCCTACACGACGCTCTTCCGATCTNNNGACTCATGCCTATAATCCCAG
AGAGCACACGTCTGAACTCTTTCCCTACACGACGCTCTTCCGATCTNNNGACTCATGCCTATAATCCCAG
AGCACACGTCTGAACTCTTTCCCTACACGACGCTCTTCCGATCTNNNTTATCCACCTTTCCCACAACACA
AGCACACGTCTGAACTCTTTCCCTACACGACGCTCTTCCGATCTNNNTTATCCACCTTTCCCACAACACA
AGCACACGTCTGAACTCTTTCCCTACACGACGCTCTTCCGATCTNNNTTATCCACCTTTCCCACAACACA
AGCACACGTCTGAACTCTTTCCCTACACGACGCTCTTCCGATCTNNNTTATCCACCTTTCCCACAACACA
GAGCACACGTCTGAACTCTTTCCCTACACGACGCTCTTCCGATCTNNNCTCTTCCTCTGTTCTGCAGCAAT
GAGCACACGTCTGAACTCTTTCCCTACACGACGCTCTTCCGATCTNNNCTCTTCCTCTGTTCTGCAGCAAT
GAGCACACGTCTGAACTCTTTCCCTACACGACGCTCTTCCGATCTNNNCTCTTCCTCTGTTCTGCAGCAAT
GAGCACACGTCTGAACTCTTTCCCTACACGACGCTCTTCCGATCTNNNCTCTTCCTCTGTTCTGCAGCAAT
GAGCACACGTCTGAACTCTTTCCCTACACGACGCTCTTCCGATCTNNNTTTCCTGGTTTCCTTTTTGCCTC
GAGCACACGTCTGAACTCTTTCCCTACACGACGCTCTTCCGATCTNNNTTTCCTGGTTTCCTTTTTGCCTC
GAGCACACGTCTGAACTCTTTCCCTACACGACGCTCTTCCGATCTNNNTTTCCTGGTTTCCTTTTTGCCTC
GAGCACACGTCTGAACTCTTTCCCTACACGACGCTCTTCCGATCTNNNTTTCCTGGTTTCCTTTTTGCCTC
AGCACACGTCTGAACTCTTTCCCTACACGACGCTCTTCCGATCTNNNAGCAAGTCCTCTTGATACCAGAT
AGCACACGTCTGAACTCTTTCCCTACACGACGCTCTTCCGATCTNNNAGCAAGTCCTCTTGATACCAGAT
AGCACACGTCTGAACTCTTTCCCTACACGACGCTCTTCCGATCTNNNAGCAAGTCCTCTTGATACCAGAT
AGCACACGTCTGAACTCTTTCCCTACACGACGCTCTTCCGATCTNNNAGCAAGTCCTCTTGATACCAGAT
GAGCACACGTCTGAACTCTTTCCCTACACGACGCTCTTCCGATCTNNNCCATGCTCATCACCTTCATAGG
GAGCACACGTCTGAACTCTTTCCCTACACGACGCTCTTCCGATCTNNNCCATGCTCATCACCTTCATAGG
GAGCACACGTCTGAACTCTTTCCCTACACGACGCTCTTCCGATCTNNNCCATGCTCATCACCTTCATAGG
GAGCACACGTCTGAACTCTTTCCCTACACGACGCTCTTCCGATCTNNNCCATGCTCATCACCTTCATAGG
CACACGTCTGAACTCTTTCCCTACACGACGCTCTTCCGATCTNNNGGGTATAATTATGATCTCAGACACA
CACACGTCTGAACTCTTTCCCTACACGACGCTCTTCCGATCTNNNGGGTATAATTATGATCTCAGACACA
CACACGTCTGAACTCTTTCCCTACACGACGCTCTTCCGATCTNNNGGGTATAATTATGATCTCAGACACA
CACACGTCTGAACTCTTTCCCTACACGACGCTCTTCCGATCTNNNGGGTATAATTATGATCTCAGACACA
AGCACACGTCTGAACTCTTTCCCTACACGACGCTCTTCCGATCTNNNAAAAAAAGCCCAAGACCAGACA
AGCACACGTCTGAACTCTTTCCCTACACGACGCTCTTCCGATCTNNNAAAAAAAGCCCAAGACCAGACA
AGCACACGTCTGAACTCTTTCCCTACACGACGCTCTTCCGATCTNNNAAAAAAAGCCCAAGACCAGACA
AGCACACGTCTGAACTCTTTCCCTACACGACGCTCTTCCGATCTNNNAAAAAAAGCCCAAGACCAGACA
AGCACACGTCTGAACTCTTTCCCTACACGACGCTCTTCCGATCTNNNAGTGCTATGTTTCACTAAACCCC
AGCACACGTCTGAACTCTTTCCCTACACGACGCTCTTCCGATCTNNNAGTGCTATGTTTCACTAAACCCC
AGCACACGTCTGAACTCTTTCCCTACACGACGCTCTTCCGATCTNNNAGTGCTATGTTTCACTAAACCCC
AGCACACGTCTGAACTCTTTCCCTACACGACGCTCTTCCGATCTNNNAGTGCTATGTTTCACTAAACCCC

GAGCACACGTCTGAACTCTTTCCCTACACGACGCTCTTCCGATCTNNNCAGGAGAATTGCTTGAACCA
GAGCACACGTCTGAACTCTTTCCCTACACGACGCTCTTCCGATCTNNNCAGGAGAATTGCTTGAACCA
GAGCACACGTCTGAACTCTTTCCCTACACGACGCTCTTCCGATCTNNNCAGGAGAATTGCTTGAACCA
GAGCACACGTCTGAACTCTTTCCCTACACGACGCTCTTCCGATCTNNNCAGGAGAATTGCTTGAACCA
AGCACACGTCTGAACTCTTTCCCTACACGACGCTCTTCCGATCTNNNCTGTAGTCCCAGCTACTCAGG
AGCACACGTCTGAACTCTTTCCCTACACGACGCTCTTCCGATCTNNNCTGTAGTCCCAGCTACTCAGG
AGCACACGTCTGAACTCTTTCCCTACACGACGCTCTTCCGATCTNNNCTGTAGTCCCAGCTACTCAGG
AGCACACGTCTGAACTCTTTCCCTACACGACGCTCTTCCGATCTNNNCTGTAGTCCCAGCTACTCAGG
AGAGCACACGTCTGAACTCTTTCCCTACACGACGCTCTTCCGATCTNNNACTTGAAATTGGAAGGTGG
AGAGCACACGTCTGAACTCTTTCCCTACACGACGCTCTTCCGATCTNNNACTTGAAATTGGAAGGTGG
AGAGCACACGTCTGAACTCTTTCCCTACACGACGCTCTTCCGATCTNNNACTTGAAATTGGAAGGTGG
AGAGCACACGTCTGAACTCTTTCCCTACACGACGCTCTTCCGATCTNNNACTTGAAATTGGAAGGTGG
GCACACGTCTGAACTCTTTCCCTACACGACGCTCTTCCGATCTNNNTGTGGCAAAATATTCAACCTCAC
GCACACGTCTGAACTCTTTCCCTACACGACGCTCTTCCGATCTNNNTGTGGCAAAATATTCAACCTCAC
GCACACGTCTGAACTCTTTCCCTACACGACGCTCTTCCGATCTNNNTGTGGCAAAATATTCAACCTCAC
GCACACGTCTGAACTCTTTCCCTACACGACGCTCTTCCGATCTNNNTGTGGCAAAATATTCAACCTCAC
AGCACACGTCTGAACTCTTTCCCTACACGACGCTCTTCCGATCTNNNCCTCAGCATCTCAGACCAACAT
AGCACACGTCTGAACTCTTTCCCTACACGACGCTCTTCCGATCTNNNCCTCAGCATCTCAGACCAACAT
AGCACACGTCTGAACTCTTTCCCTACACGACGCTCTTCCGATCTNNNCCTCAGCATCTCAGACCAACAT
AGCACACGTCTGAACTCTTTCCCTACACGACGCTCTTCCGATCTNNNCCTCAGCATCTCAGACCAACAT
AGCACACGTCTGAACTCTTTCCCTACACGACGCTCTTCCGATCTNNNTTGAACAAAATAGGATCGCCC
AGCACACGTCTGAACTCTTTCCCTACACGACGCTCTTCCGATCTNNNTTGAACAAAATAGGATCGCCC
AGCACACGTCTGAACTCTTTCCCTACACGACGCTCTTCCGATCTNNNTTGAACAAAATAGGATCGCCC
AGCACACGTCTGAACTCTTTCCCTACACGACGCTCTTCCGATCTNNNTTGAACAAAATAGGATCGCCC
CACACGTCTGAACTCTTTCCCTACACGACGCTCTTCCGATCTNNNGACTCCAGGTTGTCCATCTGCCAT
CACACGTCTGAACTCTTTCCCTACACGACGCTCTTCCGATCTNNNGACTCCAGGTTGTCCATCTGCCAT
CACACGTCTGAACTCTTTCCCTACACGACGCTCTTCCGATCTNNNGACTCCAGGTTGTCCATCTGCCAT
CACACGTCTGAACTCTTTCCCTACACGACGCTCTTCCGATCTNNNGACTCCAGGTTGTCCATCTGCCAT
GCACACGTCTGAACTCTTTCCCTACACGACGCTCTTCCGATCTNNNAGAAGAATGGTGTGAACCCAG
GCACACGTCTGAACTCTTTCCCTACACGACGCTCTTCCGATCTNNNAGAAGAATGGTGTGAACCCAG
GCACACGTCTGAACTCTTTCCCTACACGACGCTCTTCCGATCTNNNAGAAGAATGGTGTGAACCCAG
GCACACGTCTGAACTCTTTCCCTACACGACGCTCTTCCGATCTNNNAGAAGAATGGTGTGAACCCAG
AGCACACGTCTGAACTCTTTCCCTACACGACGCTCTTCCGATCTNNNATACCTGTGGTCCCAGCTACTT
AGCACACGTCTGAACTCTTTCCCTACACGACGCTCTTCCGATCTNNNATACCTGTGGTCCCAGCTACTT
AGCACACGTCTGAACTCTTTCCCTACACGACGCTCTTCCGATCTNNNATACCTGTGGTCCCAGCTACTT
AGCACACGTCTGAACTCTTTCCCTACACGACGCTCTTCCGATCTNNNATACCTGTGGTCCCAGCTACTT
GAGCACACGTCTGAACTCTTTCCCTACACGACGCTCTTCCGATCTNNNTGAGGAGGAGGTTGTAG
GAGCACACGTCTGAACTCTTTCCCTACACGACGCTCTTCCGATCTNNNTGAGGAGGAGGTTGTAG
GAGCACACGTCTGAACTCTTTCCCTACACGACGCTCTTCCGATCTNNNTGAGGAGGAGGTTGTAG
GAGCACACGTCTGAACTCTTTCCCTACACGACGCTCTTCCGATCTNNNTGAGGAGGAGGTTGTAG
AGCACACGTCTGAACTCTTTCCCTACACGACGCTCTTCCGATCTNNNTCTATCATTGACGGGCATGTAG
AGCACACGTCTGAACTCTTTCCCTACACGACGCTCTTCCGATCTNNNTCTATCATTGACGGGCATGTAG
AGCACACGTCTGAACTCTTTCCCTACACGACGCTCTTCCGATCTNNNTCTATCATTGACGGGCATGTAG

AGCACACGTCTGAACTCTTTCCCTACACGACGCTCTTCCGATCTNNNTCTATCATTGACGGGCATGTAG
GAGCACACGTCTGAACTCTTTCCCTACACGACGCTCTTCCGATCTNNNAACATTTTGCTGCCCAACCAAT
GAGCACACGTCTGAACTCTTTCCCTACACGACGCTCTTCCGATCTNNNAACATTTTGCTGCCCAACCAAT
GAGCACACGTCTGAACTCTTTCCCTACACGACGCTCTTCCGATCTNNNAACATTTTGCTGCCCAACCAAT
GAGCACACGTCTGAACTCTTTCCCTACACGACGCTCTTCCGATCTNNNAACATTTTGCTGCCCAACCAAT
AGCACACGTCTGAACTCTTTCCCTACACGACGCTCTTCCGATCTNNNTTCCGGTAGAATTCAGCTGTGA
AGCACACGTCTGAACTCTTTCCCTACACGACGCTCTTCCGATCTNNNTTCCGGTAGAATTCAGCTGTGA
AGCACACGTCTGAACTCTTTCCCTACACGACGCTCTTCCGATCTNNNTTCCGGTAGAATTCAGCTGTGA
AGCACACGTCTGAACTCTTTCCCTACACGACGCTCTTCCGATCTNNNTTCCGGTAGAATTCAGCTGTGA
AGCACACGTCTGAACTCTTTCCCTACACGACGCTCTTCCGATCTNNNAGCAACCAGATGTCCCTCAATA
AGCACACGTCTGAACTCTTTCCCTACACGACGCTCTTCCGATCTNNNAGCAACCAGATGTCCCTCAATA
AGCACACGTCTGAACTCTTTCCCTACACGACGCTCTTCCGATCTNNNAGCAACCAGATGTCCCTCAATA
AGCACACGTCTGAACTCTTTCCCTACACGACGCTCTTCCGATCTNNNAGCAACCAGATGTCCCTCAATA
GCACACGTCTGAACTCTTTCCCTACACGACGCTCTTCCGATCTNNNCATCTCACGATGTCAGAGAGGACC
GCACACGTCTGAACTCTTTCCCTACACGACGCTCTTCCGATCTNNNCATCTCACGATGTCAGAGAGGACC
GCACACGTCTGAACTCTTTCCCTACACGACGCTCTTCCGATCTNNNCATCTCACGATGTCAGAGAGGACC
GCACACGTCTGAACTCTTTCCCTACACGACGCTCTTCCGATCTNNNCATCTCACGATGTCAGAGAGGACC
AGCACACGTCTGAACTCTTTCCCTACACGACGCTCTTCCGATCTNNNTTTAGAGGGAAAACGGTGATGC
AGCACACGTCTGAACTCTTTCCCTACACGACGCTCTTCCGATCTNNNTTTAGAGGGAAAACGGTGATGC
AGCACACGTCTGAACTCTTTCCCTACACGACGCTCTTCCGATCTNNNTTTAGAGGGAAAACGGTGATGC
AGCACACGTCTGAACTCTTTCCCTACACGACGCTCTTCCGATCTNNNTTTAGAGGGAAAACGGTGATGC
AGCACACGTCTGAACTCTTTCCCTACACGACGCTCTTCCGATCTNNNGAAACGCATGTCTTCTGTCACAA
AGCACACGTCTGAACTCTTTCCCTACACGACGCTCTTCCGATCTNNNGAAACGCATGTCTTCTGTCACAA
AGCACACGTCTGAACTCTTTCCCTACACGACGCTCTTCCGATCTNNNGAAACGCATGTCTTCTGTCACAA
AGCACACGTCTGAACTCTTTCCCTACACGACGCTCTTCCGATCTNNNGAAACGCATGTCTTCTGTCACAA
AGCACACGTCTGAACTCTTTCCCTACACGACGCTCTTCCGATCTNNNATCAGGATAACCCTCCTCCCTTGC
AGCACACGTCTGAACTCTTTCCCTACACGACGCTCTTCCGATCTNNNATCAGGATAACCCTCCTCCCTTGC
AGCACACGTCTGAACTCTTTCCCTACACGACGCTCTTCCGATCTNNNATCAGGATAACCCTCCTCCCTTGC
AGCACACGTCTGAACTCTTTCCCTACACGACGCTCTTCCGATCTNNNATCAGGATAACCCTCCTCCCTTGC
AGCACACGTCTGAACTCTTTCCCTACACGACGCTCTTCCGATCTNNNAGGGTGAACACATCAGATGTC
AGCACACGTCTGAACTCTTTCCCTACACGACGCTCTTCCGATCTNNNAGGGTGAACACATCAGATGTC
AGCACACGTCTGAACTCTTTCCCTACACGACGCTCTTCCGATCTNNNAGGGTGAACACATCAGATGTC
AGCACACGTCTGAACTCTTTCCCTACACGACGCTCTTCCGATCTNNNAGGGTGAACACATCAGATGTC
AGAGCACACGTCTGAACTCTTTCCCTACACGACGCTCTTCCGATCTNNNGGGCTCCAGCATATTATAGCA
AGAGCACACGTCTGAACTCTTTCCCTACACGACGCTCTTCCGATCTNNNGGGCTCCAGCATATTATAGCA
AGAGCACACGTCTGAACTCTTTCCCTACACGACGCTCTTCCGATCTNNNGGGCTCCAGCATATTATAGCA
AGAGCACACGTCTGAACTCTTTCCCTACACGACGCTCTTCCGATCTNNNGGGCTCCAGCATATTATAGCA
AGAGCACACGTCTGAACTCTTTCCCTACACGACGCTCTTCCGATCTNNNAGAACAGGTAAGTACGGGA
AGAGCACACGTCTGAACTCTTTCCCTACACGACGCTCTTCCGATCTNNNAGAACAGGTAAGTACGGGA
AGAGCACACGTCTGAACTCTTTCCCTACACGACGCTCTTCCGATCTNNNAGAACAGGTAAGTACGGGA
AGAGCACACGTCTGAACTCTTTCCCTACACGACGCTCTTCCGATCTNNNAGAACAGGTAAGTACGGGA
AGCACACGTCTGAACTCTTTCCCTACACGACGCTCTTCCGATCTNNNTTCCAAGTAGCTGGGTTTACAG
AGCACACGTCTGAACTCTTTCCCTACACGACGCTCTTCCGATCTNNNTTCCAAGTAGCTGGGTTTACAG

AGCACACGTCTGAACTCTTTCCCTACACGACGCTCTTCCGATCTNNNTTCCAAGTAGCTGGGTTTACAG
AGCACACGTCTGAACTCTTTCCCTACACGACGCTCTTCCGATCTNNNTTCCAAGTAGCTGGGTTTACAG
AGCACACGTCTGAACTCTTTCCCTACACGACGCTCTTCCGATCTNNNAGGAGAATTGCTGCTTGAACT
AGCACACGTCTGAACTCTTTCCCTACACGACGCTCTTCCGATCTNNNAGGAGAATTGCTGCTTGAACT
AGCACACGTCTGAACTCTTTCCCTACACGACGCTCTTCCGATCTNNNAGGAGAATTGCTGCTTGAACT
AGCACACGTCTGAACTCTTTCCCTACACGACGCTCTTCCGATCTNNNAGGAGAATTGCTGCTTGAACT
GAGCACACGTCTGAACTCTTTCCCTACACGACGCTCTTCCGATCTNNNATCCACTGCTTTAGATTGGGA
GAGCACACGTCTGAACTCTTTCCCTACACGACGCTCTTCCGATCTNNNATCCACTGCTTTAGATTGGGA
GAGCACACGTCTGAACTCTTTCCCTACACGACGCTCTTCCGATCTNNNATCCACTGCTTTAGATTGGGA
GAGCACACGTCTGAACTCTTTCCCTACACGACGCTCTTCCGATCTNNNATCCACTGCTTTAGATTGGGA
GAGCACACGTCTGAACTCTTTCCCTACACGACGCTCTTCCGATCTNNNGCAACAAGAATGAAACTCCAG
GAGCACACGTCTGAACTCTTTCCCTACACGACGCTCTTCCGATCTNNNGCAACAAGAATGAAACTCCAG
GAGCACACGTCTGAACTCTTTCCCTACACGACGCTCTTCCGATCTNNNGCAACAAGAATGAAACTCCAG
GAGCACACGTCTGAACTCTTTCCCTACACGACGCTCTTCCGATCTNNNGCAACAAGAATGAAACTCCAG
GCACACGTCTGAACTCTTTCCCTACACGACGCTCTTCCGATCTNNNGCTCATGTCTGCCTAACTACCTA
GCACACGTCTGAACTCTTTCCCTACACGACGCTCTTCCGATCTNNNGCTCATGTCTGCCTAACTACCTA
GCACACGTCTGAACTCTTTCCCTACACGACGCTCTTCCGATCTNNNGCTCATGTCTGCCTAACTACCTA
GCACACGTCTGAACTCTTTCCCTACACGACGCTCTTCCGATCTNNNGCTCATGTCTGCCTAACTACCTA
GAGCACACGTCTGAACTCTTTCCCTACACGACGCTCTTCCGATCTNNNGGTAAAGTGCTGGCTCTAAC
GAGCACACGTCTGAACTCTTTCCCTACACGACGCTCTTCCGATCTNNNGGTAAAGTGCTGGCTCTAAC
GAGCACACGTCTGAACTCTTTCCCTACACGACGCTCTTCCGATCTNNNGGTAAAGTGCTGGCTCTAAC
GAGCACACGTCTGAACTCTTTCCCTACACGACGCTCTTCCGATCTNNNGGTAAAGTGCTGGCTCTAAC
AGCACACGTCTGAACTCTTTCCCTACACGACGCTCTTCCGATCTNNNTTTTGGGTAGCGGTCCTTTATC
AGCACACGTCTGAACTCTTTCCCTACACGACGCTCTTCCGATCTNNNTTTTGGGTAGCGGTCCTTTATC
AGCACACGTCTGAACTCTTTCCCTACACGACGCTCTTCCGATCTNNNTTTTGGGTAGCGGTCCTTTATC
AGCACACGTCTGAACTCTTTCCCTACACGACGCTCTTCCGATCTNNNTTTTGGGTAGCGGTCCTTTATC
AGCACACGTCTGAACTCTTTCCCTACACGACGCTCTTCCGATCTNNNTCATATGTTTGTTGGCCATTTGT
AGCACACGTCTGAACTCTTTCCCTACACGACGCTCTTCCGATCTNNNTCATATGTTTGTTGGCCATTTGT
AGCACACGTCTGAACTCTTTCCCTACACGACGCTCTTCCGATCTNNNTCATATGTTTGTTGGCCATTTGT
AGCACACGTCTGAACTCTTTCCCTACACGACGCTCTTCCGATCTNNNTCATATGTTTGTTGGCCATTTGT
GAGCACACGTCTGAACTCTTTCCCTACACGACGCTCTTCCGATCTNNNTTTGCAGTGAGCCAATATTGT
GAGCACACGTCTGAACTCTTTCCCTACACGACGCTCTTCCGATCTNNNTTTGCAGTGAGCCAATATTGT
GAGCACACGTCTGAACTCTTTCCCTACACGACGCTCTTCCGATCTNNNTTTGCAGTGAGCCAATATTGT
GAGCACACGTCTGAACTCTTTCCCTACACGACGCTCTTCCGATCTNNNTTTGCAGTGAGCCAATATTGT
GAGCACACGTCTGAACTCTTTCCCTACACGACGCTCTTCCGATCTNNNTTCAACAAATTAGGCAGTGG
GAGCACACGTCTGAACTCTTTCCCTACACGACGCTCTTCCGATCTNNNTTCAACAAATTAGGCAGTGG
GAGCACACGTCTGAACTCTTTCCCTACACGACGCTCTTCCGATCTNNNTTCAACAAATTAGGCAGTGG
GAGCACACGTCTGAACTCTTTCCCTACACGACGCTCTTCCGATCTNNNTTCAACAAATTAGGCAGTGG
AGAGCACACGTCTGAACTCTTTCCCTACACGACGCTCTTCCGATCTNNNCAGTTTGCTCACTTCTTTGG
AGAGCACACGTCTGAACTCTTTCCCTACACGACGCTCTTCCGATCTNNNCAGTTTGCTCACTTCTTTGG
AGAGCACACGTCTGAACTCTTTCCCTACACGACGCTCTTCCGATCTNNNCAGTTTGCTCACTTCTTTGG
AGAGCACACGTCTGAACTCTTTCCCTACACGACGCTCTTCCGATCTNNNCAGTTTGCTCACTTCTTTGG
AGCACACGTCTGAACTCTTTCCCTACACGACGCTCTTCCGATCTNNNCACTCCTTCCCCATCAATACTGC

AGCACACGTCTGAACTCTTTCCCTACACGACGCTCTTCCGATCTNNNCACTCCTTCCCCATCAATACTGC
AGCACACGTCTGAACTCTTTCCCTACACGACGCTCTTCCGATCTNNNCACTCCTTCCCCATCAATACTGC
AGCACACGTCTGAACTCTTTCCCTACACGACGCTCTTCCGATCTNNNCACTCCTTCCCCATCAATACTGC
AGCACACGTCTGAACTCTTTCCCTACACGACGCTCTTCCGATCTNNNCTCATGCCTGTAATCCTAGCACT
AGCACACGTCTGAACTCTTTCCCTACACGACGCTCTTCCGATCTNNNCTCATGCCTGTAATCCTAGCACT
AGCACACGTCTGAACTCTTTCCCTACACGACGCTCTTCCGATCTNNNCTCATGCCTGTAATCCTAGCACT
AGCACACGTCTGAACTCTTTCCCTACACGACGCTCTTCCGATCTNNNCTCATGCCTGTAATCCTAGCACT
AGCACACGTCTGAACTCTTTCCCTACACGACGCTCTTCCGATCTNNNGAATGACGTGAACCCAGGAG
AGCACACGTCTGAACTCTTTCCCTACACGACGCTCTTCCGATCTNNNGAATGACGTGAACCCAGGAG
AGCACACGTCTGAACTCTTTCCCTACACGACGCTCTTCCGATCTNNNGAATGACGTGAACCCAGGAG
AGCACACGTCTGAACTCTTTCCCTACACGACGCTCTTCCGATCTNNNGAATGACGTGAACCCAGGAG
GCACACGTCTGAACTCTTTCCCTACACGACGCTCTTCCGATCTNNNAGATTTCTGTTTGGGCTTGTGT
GCACACGTCTGAACTCTTTCCCTACACGACGCTCTTCCGATCTNNNAGATTTCTGTTTGGGCTTGTGT
GCACACGTCTGAACTCTTTCCCTACACGACGCTCTTCCGATCTNNNAGATTTCTGTTTGGGCTTGTGT
GCACACGTCTGAACTCTTTCCCTACACGACGCTCTTCCGATCTNNNAGATTTCTGTTTGGGCTTGTGT
AGAGCACACGTCTGAACTCTTTCCCTACACGACGCTCTTCCGATCTNNNTGAGGAACCATGGACTTTT
AGAGCACACGTCTGAACTCTTTCCCTACACGACGCTCTTCCGATCTNNNTGAGGAACCATGGACTTTT
AGAGCACACGTCTGAACTCTTTCCCTACACGACGCTCTTCCGATCTNNNTGAGGAACCATGGACTTTT
AGAGCACACGTCTGAACTCTTTCCCTACACGACGCTCTTCCGATCTNNNTGAGGAACCATGGACTTTT
AGCACACGTCTGAACTCTTTCCCTACACGACGCTCTTCCGATCTNNNATGGTCCTGATGACTCACTGT
AGCACACGTCTGAACTCTTTCCCTACACGACGCTCTTCCGATCTNNNATGGTCCTGATGACTCACTGT
AGCACACGTCTGAACTCTTTCCCTACACGACGCTCTTCCGATCTNNNATGGTCCTGATGACTCACTGT
AGCACACGTCTGAACTCTTTCCCTACACGACGCTCTTCCGATCTNNNATGGTCCTGATGACTCACTGT
AGCACACGTCTGAACTCTTTCCCTACACGACGCTCTTCCGATCTNNNGGAGAATGGCGTAAACCTGG
AGCACACGTCTGAACTCTTTCCCTACACGACGCTCTTCCGATCTNNNGGAGAATGGCGTAAACCTGG
AGCACACGTCTGAACTCTTTCCCTACACGACGCTCTTCCGATCTNNNGGAGAATGGCGTAAACCTGG
AGCACACGTCTGAACTCTTTCCCTACACGACGCTCTTCCGATCTNNNGGAGAATGGCGTAAACCTGG
GCACACGTCTGAACTCTTTCCCTACACGACGCTCTTCCGATCTNNNAACCCAGCACCTCATGTCTAATG
GCACACGTCTGAACTCTTTCCCTACACGACGCTCTTCCGATCTNNNAACCCAGCACCTCATGTCTAATG
GCACACGTCTGAACTCTTTCCCTACACGACGCTCTTCCGATCTNNNAACCCAGCACCTCATGTCTAATG
GCACACGTCTGAACTCTTTCCCTACACGACGCTCTTCCGATCTNNNAACCCAGCACCTCATGTCTAATG
AGCACACGTCTGAACTCTTTCCCTACACGACGCTCTTCCGATCTNNNTAGGCGATACATGCTGCAGTA
AGCACACGTCTGAACTCTTTCCCTACACGACGCTCTTCCGATCTNNNTAGGCGATACATGCTGCAGTA
AGCACACGTCTGAACTCTTTCCCTACACGACGCTCTTCCGATCTNNNTAGGCGATACATGCTGCAGTA
AGCACACGTCTGAACTCTTTCCCTACACGACGCTCTTCCGATCTNNNTAGGCGATACATGCTGCAGTA
AGCACACGTCTGAACTCTTTCCCTACACGACGCTCTTCCGATCTNNNTGAGAATCACTTGAACCTGGG
AGCACACGTCTGAACTCTTTCCCTACACGACGCTCTTCCGATCTNNNTGAGAATCACTTGAACCTGGG
AGCACACGTCTGAACTCTTTCCCTACACGACGCTCTTCCGATCTNNNTGAGAATCACTTGAACCTGGG
AGCACACGTCTGAACTCTTTCCCTACACGACGCTCTTCCGATCTNNNTGAGAATCACTTGAACCTGGG
AGCACACGTCTGAACTCTTTCCCTACACGACGCTCTTCCGATCTNNNCACCTGTAATCCCAGCTACTCG
AGCACACGTCTGAACTCTTTCCCTACACGACGCTCTTCCGATCTNNNCACCTGTAATCCCAGCTACTCG
AGCACACGTCTGAACTCTTTCCCTACACGACGCTCTTCCGATCTNNNCACCTGTAATCCCAGCTACTCG
AGCACACGTCTGAACTCTTTCCCTACACGACGCTCTTCCGATCTNNNCACCTGTAATCCCAGCTACTCG

AGAGCACACGTCTGAACTCTTTCCCTACACGACGCTCTTCCGATCTNNNAGTGGTTTCCACAAGAAGGC
AGAGCACACGTCTGAACTCTTTCCCTACACGACGCTCTTCCGATCTNNNAGTGGTTTCCACAAGAAGGC
AGAGCACACGTCTGAACTCTTTCCCTACACGACGCTCTTCCGATCTNNNAGTGGTTTCCACAAGAAGGC
AGAGCACACGTCTGAACTCTTTCCCTACACGACGCTCTTCCGATCTNNNAGTGGTTTCCACAAGAAGGC
GAGCACACGTCTGAACTCTTTCCCTACACGACGCTCTTCCGATCTNNNAATACAAAAATTAGCCGGGCG
GAGCACACGTCTGAACTCTTTCCCTACACGACGCTCTTCCGATCTNNNAATACAAAAATTAGCCGGGCG
GAGCACACGTCTGAACTCTTTCCCTACACGACGCTCTTCCGATCTNNNAATACAAAAATTAGCCGGGCG
GAGCACACGTCTGAACTCTTTCCCTACACGACGCTCTTCCGATCTNNNAATACAAAAATTAGCCGGGCG
GAGCACACGTCTGAACTCTTTCCCTACACGACGCTCTTCCGATCTNNNATCATCTGGAGCAGTTTCTTG
GAGCACACGTCTGAACTCTTTCCCTACACGACGCTCTTCCGATCTNNNATCATCTGGAGCAGTTTCTTG
GAGCACACGTCTGAACTCTTTCCCTACACGACGCTCTTCCGATCTNNNATCATCTGGAGCAGTTTCTTG
GAGCACACGTCTGAACTCTTTCCCTACACGACGCTCTTCCGATCTNNNATCATCTGGAGCAGTTTCTTG
AGCACACGTCTGAACTCTTTCCCTACACGACGCTCTTCCGATCTNNNAAACACAGCATCATTTCAGACC
AGCACACGTCTGAACTCTTTCCCTACACGACGCTCTTCCGATCTNNNAAACACAGCATCATTTCAGACC
AGCACACGTCTGAACTCTTTCCCTACACGACGCTCTTCCGATCTNNNAAACACAGCATCATTTCAGACC
AGCACACGTCTGAACTCTTTCCCTACACGACGCTCTTCCGATCTNNNAAACACAGCATCATTTCAGACC
AGAGCACACGTCTGAACTCTTTCCCTACACGACGCTCTTCCGATCTNNNCTTGCAGTGAGATGAGATTC
AGAGCACACGTCTGAACTCTTTCCCTACACGACGCTCTTCCGATCTNNNCTTGCAGTGAGATGAGATTC
AGAGCACACGTCTGAACTCTTTCCCTACACGACGCTCTTCCGATCTNNNCTTGCAGTGAGATGAGATTC
AGAGCACACGTCTGAACTCTTTCCCTACACGACGCTCTTCCGATCTNNNCTTGCAGTGAGATGAGATTC
GAGCACACGTCTGAACTCTTTCCCTACACGACGCTCTTCCGATCTNNNCCGGCCGCTTTTTTTTTTTTTT
GAGCACACGTCTGAACTCTTTCCCTACACGACGCTCTTCCGATCTNNNCCGGCCGCTTTTTTTTTTTTTT
GAGCACACGTCTGAACTCTTTCCCTACACGACGCTCTTCCGATCTNNNCCGGCCGCTTTTTTTTTTTTTT
GAGCACACGTCTGAACTCTTTCCCTACACGACGCTCTTCCGATCTNNNCCGGCCGCTTTTTTTTTTTTTT
GAGCACACGTCTGAACTCTTTCCCTACACGACGCTCTTCCGATCTNNNCTGTATTCTGCTAACAGGAGG
GAGCACACGTCTGAACTCTTTCCCTACACGACGCTCTTCCGATCTNNNCTGTATTCTGCTAACAGGAGG
GAGCACACGTCTGAACTCTTTCCCTACACGACGCTCTTCCGATCTNNNCTGTATTCTGCTAACAGGAGG
GAGCACACGTCTGAACTCTTTCCCTACACGACGCTCTTCCGATCTNNNCTGTATTCTGCTAACAGGAGG
AGCACACGTCTGAACTCTTTCCCTACACGACGCTCTTCCGATCTNNNACAACTCTAGCACTTACCACCAG
AGCACACGTCTGAACTCTTTCCCTACACGACGCTCTTCCGATCTNNNACAACTCTAGCACTTACCACCAG
AGCACACGTCTGAACTCTTTCCCTACACGACGCTCTTCCGATCTNNNACAACTCTAGCACTTACCACCAG
AGCACACGTCTGAACTCTTTCCCTACACGACGCTCTTCCGATCTNNNACAACTCTAGCACTTACCACCAG
AGCACACGTCTGAACTCTTTCCCTACACGACGCTCTTCCGATCTNNNGTGAAATACTCCACGTTGCACAC
AGCACACGTCTGAACTCTTTCCCTACACGACGCTCTTCCGATCTNNNGTGAAATACTCCACGTTGCACAC
AGCACACGTCTGAACTCTTTCCCTACACGACGCTCTTCCGATCTNNNGTGAAATACTCCACGTTGCACAC
AGCACACGTCTGAACTCTTTCCCTACACGACGCTCTTCCGATCTNNNGTGAAATACTCCACGTTGCACAC
AGCACACGTCTGAACTCTTTCCCTACACGACGCTCTTCCGATCTNNNCTGTAGTTGAAGCTGTGTGTTG
AGCACACGTCTGAACTCTTTCCCTACACGACGCTCTTCCGATCTNNNCTGTAGTTGAAGCTGTGTGTTG
AGCACACGTCTGAACTCTTTCCCTACACGACGCTCTTCCGATCTNNNCTGTAGTTGAAGCTGTGTGTTG
AGCACACGTCTGAACTCTTTCCCTACACGACGCTCTTCCGATCTNNNCTGTAGTTGAAGCTGTGTGTTG
GCACACGTCTGAACTCTTTCCCTACACGACGCTCTTCCGATCTNNNTCATCCCTCCTCCTACCCATAAACAT
GCACACGTCTGAACTCTTTCCCTACACGACGCTCTTCCGATCTNNNTCATCCCTCCTCCTACCCATAAACAT
GCACACGTCTGAACTCTTTCCCTACACGACGCTCTTCCGATCTNNNTCATCCCTCCTCCTACCCATAAACAT

GCACACGTCTGAACTCTTTCCCTACACGACGCTCTTCCGATCTNNNTCATCCCTCCTCCTACCCATAAACAT
GAGCACACGTCTGAACTCTTTCCCTACACGACGCTCTTCCGATCTNNNAAGTGTTGGGATTACAGGTG
GAGCACACGTCTGAACTCTTTCCCTACACGACGCTCTTCCGATCTNNNAAGTGTTGGGATTACAGGTG
GAGCACACGTCTGAACTCTTTCCCTACACGACGCTCTTCCGATCTNNNAAGTGTTGGGATTACAGGTG
GAGCACACGTCTGAACTCTTTCCCTACACGACGCTCTTCCGATCTNNNAAGTGTTGGGATTACAGGTG
GAGCACACGTCTGAACTCTTTCCCTACACGACGCTCTTCCGATCTNNNCTCATGCCTGTAATCCTAGCAC
GAGCACACGTCTGAACTCTTTCCCTACACGACGCTCTTCCGATCTNNNCTCATGCCTGTAATCCTAGCAC
GAGCACACGTCTGAACTCTTTCCCTACACGACGCTCTTCCGATCTNNNCTCATGCCTGTAATCCTAGCAC
GAGCACACGTCTGAACTCTTTCCCTACACGACGCTCTTCCGATCTNNNCTCATGCCTGTAATCCTAGCAC
AGCACACGTCTGAACTCTTTCCCTACACGACGCTCTTCCGATCTNNNAATGCAATGATTGTCAACCTGG
AGCACACGTCTGAACTCTTTCCCTACACGACGCTCTTCCGATCTNNNAATGCAATGATTGTCAACCTGG
AGCACACGTCTGAACTCTTTCCCTACACGACGCTCTTCCGATCTNNNAATGCAATGATTGTCAACCTGG
AGCACACGTCTGAACTCTTTCCCTACACGACGCTCTTCCGATCTNNNAATGCAATGATTGTCAACCTGG
AGAGCACACGTCTGAACTCTTTCCCTACACGACGCTCTTCCGATCTNNNCGGATCACAAGGTCAGAAGA
AGAGCACACGTCTGAACTCTTTCCCTACACGACGCTCTTCCGATCTNNNCGGATCACAAGGTCAGAAGA
AGAGCACACGTCTGAACTCTTTCCCTACACGACGCTCTTCCGATCTNNNCGGATCACAAGGTCAGAAGA
AGAGCACACGTCTGAACTCTTTCCCTACACGACGCTCTTCCGATCTNNNCGGATCACAAGGTCAGAAGA
CACACGTCTGAACTCTTTCCCTACACGACGCTCTTCCGATCTNNNCTCTAGGAAGGGAGGAAGAGAGC
CACACGTCTGAACTCTTTCCCTACACGACGCTCTTCCGATCTNNNCTCTAGGAAGGGAGGAAGAGAGC
CACACGTCTGAACTCTTTCCCTACACGACGCTCTTCCGATCTNNNCTCTAGGAAGGGAGGAAGAGAGC
CACACGTCTGAACTCTTTCCCTACACGACGCTCTTCCGATCTNNNCTCTAGGAAGGGAGGAAGAGAGC
AGCACACGTCTGAACTCTTTCCCTACACGACGCTCTTCCGATCTNNNCTCTCTAGCTATCCCCTCTCCA
AGCACACGTCTGAACTCTTTCCCTACACGACGCTCTTCCGATCTNNNCTCTCTAGCTATCCCCTCTCCA
AGCACACGTCTGAACTCTTTCCCTACACGACGCTCTTCCGATCTNNNCTCTCTAGCTATCCCCTCTCCA
AGCACACGTCTGAACTCTTTCCCTACACGACGCTCTTCCGATCTNNNCTCTCTAGCTATCCCCTCTCCA
GCACACGTCTGAACTCTTTCCCTACACGACGCTCTTCCGATCTNNNCCTTCCTGGAGTTCCTTGTTGTAC
GCACACGTCTGAACTCTTTCCCTACACGACGCTCTTCCGATCTNNNCCTTCCTGGAGTTCCTTGTTGTAC
GCACACGTCTGAACTCTTTCCCTACACGACGCTCTTCCGATCTNNNCCTTCCTGGAGTTCCTTGTTGTAC
GCACACGTCTGAACTCTTTCCCTACACGACGCTCTTCCGATCTNNNCCTTCCTGGAGTTCCTTGTTGTAC
GAGCACACGTCTGAACTCTTTCCCTACACGACGCTCTTCCGATCTNNNGTTTTAGATCCGGGTTGAGGA
GAGCACACGTCTGAACTCTTTCCCTACACGACGCTCTTCCGATCTNNNGTTTTAGATCCGGGTTGAGGA
GAGCACACGTCTGAACTCTTTCCCTACACGACGCTCTTCCGATCTNNNGTTTTAGATCCGGGTTGAGGA
GAGCACACGTCTGAACTCTTTCCCTACACGACGCTCTTCCGATCTNNNGTTTTAGATCCGGGTTGAGGA
GAGCACACGTCTGAACTCTTTCCCTACACGACGCTCTTCCGATCTNNNCTTGTGCCAGTTTTCAAAGGG
GAGCACACGTCTGAACTCTTTCCCTACACGACGCTCTTCCGATCTNNNCTTGTGCCAGTTTTCAAAGGG
GAGCACACGTCTGAACTCTTTCCCTACACGACGCTCTTCCGATCTNNNCTTGTGCCAGTTTTCAAAGGG
GAGCACACGTCTGAACTCTTTCCCTACACGACGCTCTTCCGATCTNNNCTTGTGCCAGTTTTCAAAGGG
AGAGCACACGTCTGAACTCTTTCCCTACACGACGCTCTTCCGATCTNNNGAGCTGAGGTCACAGCAAC
AGAGCACACGTCTGAACTCTTTCCCTACACGACGCTCTTCCGATCTNNNGAGCTGAGGTCACAGCAAC
AGAGCACACGTCTGAACTCTTTCCCTACACGACGCTCTTCCGATCTNNNGAGCTGAGGTCACAGCAAC
AGAGCACACGTCTGAACTCTTTCCCTACACGACGCTCTTCCGATCTNNNGAGCTGAGGTCACAGCAAC
GCACACGTCTGAACTCTTTCCCTACACGACGCTCTTCCGATCTNNNCACCTGTAATCCCAGCTACTCAGCA
GCACACGTCTGAACTCTTTCCCTACACGACGCTCTTCCGATCTNNNCACCTGTAATCCCAGCTACTCAGCA

GCACACGTCTGAACTCTTTCCCTACACGACGCTCTTCCGATCTNNNCACCTGTAATCCCAGCTACTCAGCA
GCACACGTCTGAACTCTTTCCCTACACGACGCTCTTCCGATCTNNNCACCTGTAATCCCAGCTACTCAGCA
AGCACACGTCTGAACTCTTTCCCTACACGACGCTCTTCCGATCTNNNGGAGAACCTCCTGTGCCATATG
AGCACACGTCTGAACTCTTTCCCTACACGACGCTCTTCCGATCTNNNGGAGAACCTCCTGTGCCATATG
AGCACACGTCTGAACTCTTTCCCTACACGACGCTCTTCCGATCTNNNGGAGAACCTCCTGTGCCATATG
AGCACACGTCTGAACTCTTTCCCTACACGACGCTCTTCCGATCTNNNGGAGAACCTCCTGTGCCATATG
AGCACACGTCTGAACTCTTTCCCTACACGACGCTCTTCCGATCTNNNCAGCCTCAAGTAAGTCTCCACA
AGCACACGTCTGAACTCTTTCCCTACACGACGCTCTTCCGATCTNNNCAGCCTCAAGTAAGTCTCCACA
AGCACACGTCTGAACTCTTTCCCTACACGACGCTCTTCCGATCTNNNCAGCCTCAAGTAAGTCTCCACA
AGCACACGTCTGAACTCTTTCCCTACACGACGCTCTTCCGATCTNNNCAGCCTCAAGTAAGTCTCCACA
AGCACACGTCTGAACTCTTTCCCTACACGACGCTCTTCCGATCTNNNTCCAAAGCTACATTTTAATGGCA
AGCACACGTCTGAACTCTTTCCCTACACGACGCTCTTCCGATCTNNNTCCAAAGCTACATTTTAATGGCA
AGCACACGTCTGAACTCTTTCCCTACACGACGCTCTTCCGATCTNNNTCCAAAGCTACATTTTAATGGCA
AGCACACGTCTGAACTCTTTCCCTACACGACGCTCTTCCGATCTNNNTCCAAAGCTACATTTTAATGGCA
GAGCACACGTCTGAACTCTTTCCCTACACGACGCTCTTCCGATCTNNNTGTCCTAACATAAACCCAGACC
GAGCACACGTCTGAACTCTTTCCCTACACGACGCTCTTCCGATCTNNNTGTCCTAACATAAACCCAGACC
GAGCACACGTCTGAACTCTTTCCCTACACGACGCTCTTCCGATCTNNNTGTCCTAACATAAACCCAGACC
GAGCACACGTCTGAACTCTTTCCCTACACGACGCTCTTCCGATCTNNNTGTCCTAACATAAACCCAGACC
GAGCACACGTCTGAACTCTTTCCCTACACGACGCTCTTCCGATCTNNNATCAGCTTGACCAGGATCTCAT
GAGCACACGTCTGAACTCTTTCCCTACACGACGCTCTTCCGATCTNNNATCAGCTTGACCAGGATCTCAT
GAGCACACGTCTGAACTCTTTCCCTACACGACGCTCTTCCGATCTNNNATCAGCTTGACCAGGATCTCAT
GAGCACACGTCTGAACTCTTTCCCTACACGACGCTCTTCCGATCTNNNATCAGCTTGACCAGGATCTCAT
GCACACGTCTGAACTCTTTCCCTACACGACGCTCTTCCGATCTNNNGTCCAGTTGAAAACTTGATTCCC
GCACACGTCTGAACTCTTTCCCTACACGACGCTCTTCCGATCTNNNGTCCAGTTGAAAACTTGATTCCC
GCACACGTCTGAACTCTTTCCCTACACGACGCTCTTCCGATCTNNNGTCCAGTTGAAAACTTGATTCCC
GCACACGTCTGAACTCTTTCCCTACACGACGCTCTTCCGATCTNNNGTCCAGTTGAAAACTTGATTCCC
AGCACACGTCTGAACTCTTTCCCTACACGACGCTCTTCCGATCTNNNCTCAGGTGATCTTCCCACTTCAC
AGCACACGTCTGAACTCTTTCCCTACACGACGCTCTTCCGATCTNNNCTCAGGTGATCTTCCCACTTCAC
AGCACACGTCTGAACTCTTTCCCTACACGACGCTCTTCCGATCTNNNCTCAGGTGATCTTCCCACTTCAC
AGCACACGTCTGAACTCTTTCCCTACACGACGCTCTTCCGATCTNNNCTCAGGTGATCTTCCCACTTCAC
AGCACACGTCTGAACTCTTTCCCTACACGACGCTCTTCCGATCTNNNATTTGAAAAATTGCTTCAATCTG
AGCACACGTCTGAACTCTTTCCCTACACGACGCTCTTCCGATCTNNNATTTGAAAAATTGCTTCAATCTG
AGCACACGTCTGAACTCTTTCCCTACACGACGCTCTTCCGATCTNNNATTTGAAAAATTGCTTCAATCTG
AGCACACGTCTGAACTCTTTCCCTACACGACGCTCTTCCGATCTNNNATTTGAAAAATTGCTTCAATCTG
GAGCACACGTCTGAACTCTTTCCCTACACGACGCTCTTCCGATCTNNNTAGAAGGGGAGAAGTTGTTG
GAGCACACGTCTGAACTCTTTCCCTACACGACGCTCTTCCGATCTNNNTAGAAGGGGAGAAGTTGTTG
GAGCACACGTCTGAACTCTTTCCCTACACGACGCTCTTCCGATCTNNNTAGAAGGGGAGAAGTTGTTG
GAGCACACGTCTGAACTCTTTCCCTACACGACGCTCTTCCGATCTNNNTAGAAGGGGAGAAGTTGTTG
GAGCACACGTCTGAACTCTTTCCCTACACGACGCTCTTCCGATCTNNNAGCTTGCAGTGAGCCTAGATC
GAGCACACGTCTGAACTCTTTCCCTACACGACGCTCTTCCGATCTNNNAGCTTGCAGTGAGCCTAGATC
GAGCACACGTCTGAACTCTTTCCCTACACGACGCTCTTCCGATCTNNNAGCTTGCAGTGAGCCTAGATC
GAGCACACGTCTGAACTCTTTCCCTACACGACGCTCTTCCGATCTNNNAGCTTGCAGTGAGCCTAGATC
AGAGCACACGTCTGAACTCTTTCCCTACACGACGCTCTTCCGATCTNNNCAATGCTCTCTGCTCCATGAT

GAGCACACGTCTGAACTCTTTCCCTACACGACGCTCTTCCGATCTNNNCAATGCTCTCTGCTCCATGAT
GAGCACACGTCTGAACTCTTTCCCTACACGACGCTCTTCCGATCTNNNCAATGCTCTCTGCTCCATGAT
GAGCACACGTCTGAACTCTTTCCCTACACGACGCTCTTCCGATCTNNNCAATGCTCTCTGCTCCATGAT
CACACGTCTGAACTCTTTCCCTACACGACGCTCTTCCGATCTNNNGGAATCTGCCCTGTGTCGTCCAT
CACACGTCTGAACTCTTTCCCTACACGACGCTCTTCCGATCTNNNGGAATCTGCCCTGTGTCGTCCAT
CACACGTCTGAACTCTTTCCCTACACGACGCTCTTCCGATCTNNNGGAATCTGCCCTGTGTCGTCCAT
CACACGTCTGAACTCTTTCCCTACACGACGCTCTTCCGATCTNNNGGAATCTGCCCTGTGTCGTCCAT
GAGCACACGTCTGAACTCTTTCCCTACACGACGCTCTTCCGATCTNNNGAGGCAGAAGAATCGCTTGA
GAGCACACGTCTGAACTCTTTCCCTACACGACGCTCTTCCGATCTNNNGAGGCAGAAGAATCGCTTGA
GAGCACACGTCTGAACTCTTTCCCTACACGACGCTCTTCCGATCTNNNGAGGCAGAAGAATCGCTTGA
GAGCACACGTCTGAACTCTTTCCCTACACGACGCTCTTCCGATCTNNNGAGGCAGAAGAATCGCTTGA
GCACACGTCTGAACTCTTTCCCTACACGACGCTCTTCCGATCTNNNGTATAATATCTGGAAGGTACTG
GCACACGTCTGAACTCTTTCCCTACACGACGCTCTTCCGATCTNNNGTATAATATCTGGAAGGTACTG
GCACACGTCTGAACTCTTTCCCTACACGACGCTCTTCCGATCTNNNGTATAATATCTGGAAGGTACTG
GCACACGTCTGAACTCTTTCCCTACACGACGCTCTTCCGATCTNNNGTATAATATCTGGAAGGTACTG
GAGCACACGTCTGAACTCTTTCCCTACACGACGCTCTTCCGATCTNNNTATGAGTGATGATGACCCAT
GAGCACACGTCTGAACTCTTTCCCTACACGACGCTCTTCCGATCTNNNTATGAGTGATGATGACCCAT
GAGCACACGTCTGAACTCTTTCCCTACACGACGCTCTTCCGATCTNNNTATGAGTGATGATGACCCAT
GAGCACACGTCTGAACTCTTTCCCTACACGACGCTCTTCCGATCTNNNTATGAGTGATGATGACCCAT
AGCACACGTCTGAACTCTTTCCCTACACGACGCTCTTCCGATCTNNNCCTACAGCCATGGCAATATTG
AGCACACGTCTGAACTCTTTCCCTACACGACGCTCTTCCGATCTNNNCCTACAGCCATGGCAATATTG
AGCACACGTCTGAACTCTTTCCCTACACGACGCTCTTCCGATCTNNNCCTACAGCCATGGCAATATTG
AGCACACGTCTGAACTCTTTCCCTACACGACGCTCTTCCGATCTNNNCCTACAGCCATGGCAATATTG
GAGCACACGTCTGAACTCTTTCCCTACACGACGCTCTTCCGATCTNNNATACTTGAGCCTGGGAGTTT
GAGCACACGTCTGAACTCTTTCCCTACACGACGCTCTTCCGATCTNNNATACTTGAGCCTGGGAGTTT
GAGCACACGTCTGAACTCTTTCCCTACACGACGCTCTTCCGATCTNNNATACTTGAGCCTGGGAGTTT
GAGCACACGTCTGAACTCTTTCCCTACACGACGCTCTTCCGATCTNNNATACTTGAGCCTGGGAGTTT
GCACACGTCTGAACTCTTTCCCTACACGACGCTCTTCCGATCTNNNGTCTTGGAGATGTGGCATAGG
GCACACGTCTGAACTCTTTCCCTACACGACGCTCTTCCGATCTNNNGTCTTGGAGATGTGGCATAGG
GCACACGTCTGAACTCTTTCCCTACACGACGCTCTTCCGATCTNNNGTCTTGGAGATGTGGCATAGG
GCACACGTCTGAACTCTTTCCCTACACGACGCTCTTCCGATCTNNNGTCTTGGAGATGTGGCATAGG
AGCACACGTCTGAACTCTTTCCCTACACGACGCTCTTCCGATCTNNNCCTCCAAACTCCTTTACCCAAG
AGCACACGTCTGAACTCTTTCCCTACACGACGCTCTTCCGATCTNNNCCTCCAAACTCCTTTACCCAAG
AGCACACGTCTGAACTCTTTCCCTACACGACGCTCTTCCGATCTNNNCCTCCAAACTCCTTTACCCAAG
AGCACACGTCTGAACTCTTTCCCTACACGACGCTCTTCCGATCTNNNCCTCCAAACTCCTTTACCCAAG
AGCACACGTCTGAACTCTTTCCCTACACGACGCTCTTCCGATCTNNNGCTGACTTTCACGACACTTGAA
AGCACACGTCTGAACTCTTTCCCTACACGACGCTCTTCCGATCTNNNGCTGACTTTCACGACACTTGAA
AGCACACGTCTGAACTCTTTCCCTACACGACGCTCTTCCGATCTNNNGCTGACTTTCACGACACTTGAA
AGCACACGTCTGAACTCTTTCCCTACACGACGCTCTTCCGATCTNNNGCTGACTTTCACGACACTTGAA
AGCACACGTCTGAACTCTTTCCCTACACGACGCTCTTCCGATCTNNNGGCAGAAAAAAGTGCTTGA
AGCACACGTCTGAACTCTTTCCCTACACGACGCTCTTCCGATCTNNNGGCAGAAAAAAGTGCTTGA
AGCACACGTCTGAACTCTTTCCCTACACGACGCTCTTCCGATCTNNNGGCAGAAAAAAGTGCTTGA
AGCACACGTCTGAACTCTTTCCCTACACGACGCTCTTCCGATCTNNNGGCAGAAAAAAGTGCTTGA

AGCACACGTCTGAACTCTTTCCCTACACGACGCTCTTCCGATCTNNNCAGTTCAAAAACATTCACAGCCC
AGCACACGTCTGAACTCTTTCCCTACACGACGCTCTTCCGATCTNNNCAGTTCAAAAACATTCACAGCCC
AGCACACGTCTGAACTCTTTCCCTACACGACGCTCTTCCGATCTNNNCAGTTCAAAAACATTCACAGCCC
AGCACACGTCTGAACTCTTTCCCTACACGACGCTCTTCCGATCTNNNCAGTTCAAAAACATTCACAGCCC
GAGCACACGTCTGAACTCTTTCCCTACACGACGCTCTTCCGATCTNNNTCACAAGGTCAGGAGTTTGAG
GAGCACACGTCTGAACTCTTTCCCTACACGACGCTCTTCCGATCTNNNTCACAAGGTCAGGAGTTTGAG
GAGCACACGTCTGAACTCTTTCCCTACACGACGCTCTTCCGATCTNNNTCACAAGGTCAGGAGTTTGAG
GAGCACACGTCTGAACTCTTTCCCTACACGACGCTCTTCCGATCTNNNTCACAAGGTCAGGAGTTTGAG
GAGCACACGTCTGAACTCTTTCCCTACACGACGCTCTTCCGATCTNNNCTGAGGCAGGAGAATCACTT
GAGCACACGTCTGAACTCTTTCCCTACACGACGCTCTTCCGATCTNNNCTGAGGCAGGAGAATCACTT
GAGCACACGTCTGAACTCTTTCCCTACACGACGCTCTTCCGATCTNNNCTGAGGCAGGAGAATCACTT
GAGCACACGTCTGAACTCTTTCCCTACACGACGCTCTTCCGATCTNNNCTGAGGCAGGAGAATCACTT
GAGCACACGTCTGAACTCTTTCCCTACACGACGCTCTTCCGATCTNNNATATTGGTCAGGCTGGTCTTG
GAGCACACGTCTGAACTCTTTCCCTACACGACGCTCTTCCGATCTNNNATATTGGTCAGGCTGGTCTTG
GAGCACACGTCTGAACTCTTTCCCTACACGACGCTCTTCCGATCTNNNATATTGGTCAGGCTGGTCTTG
GAGCACACGTCTGAACTCTTTCCCTACACGACGCTCTTCCGATCTNNNATATTGGTCAGGCTGGTCTTG
AGCACACGTCTGAACTCTTTCCCTACACGACGCTCTTCCGATCTNNNGAAATCCAGTCCGGCTTTTCCAG
AGCACACGTCTGAACTCTTTCCCTACACGACGCTCTTCCGATCTNNNGAAATCCAGTCCGGCTTTTCCAG
AGCACACGTCTGAACTCTTTCCCTACACGACGCTCTTCCGATCTNNNGAAATCCAGTCCGGCTTTTCCAG
AGCACACGTCTGAACTCTTTCCCTACACGACGCTCTTCCGATCTNNNGAAATCCAGTCCGGCTTTTCCAG
AGCACACGTCTGAACTCTTTCCCTACACGACGCTCTTCCGATCTNNNTGTAATTCCAGCTACTTGGGAC
AGCACACGTCTGAACTCTTTCCCTACACGACGCTCTTCCGATCTNNNTGTAATTCCAGCTACTTGGGAC
AGCACACGTCTGAACTCTTTCCCTACACGACGCTCTTCCGATCTNNNTGTAATTCCAGCTACTTGGGAC
AGCACACGTCTGAACTCTTTCCCTACACGACGCTCTTCCGATCTNNNTGTAATTCCAGCTACTTGGGAC
GAGCACACGTCTGAACTCTTTCCCTACACGACGCTCTTCCGATCTNNNCTGCTGAGCATGGACTAAGA
GAGCACACGTCTGAACTCTTTCCCTACACGACGCTCTTCCGATCTNNNCTGCTGAGCATGGACTAAGA
GAGCACACGTCTGAACTCTTTCCCTACACGACGCTCTTCCGATCTNNNCTGCTGAGCATGGACTAAGA
GAGCACACGTCTGAACTCTTTCCCTACACGACGCTCTTCCGATCTNNNCTGCTGAGCATGGACTAAGA
GCACACGTCTGAACTCTTTCCCTACACGACGCTCTTCCGATCTNNNAGGAATATTCGAGAAATGGTGT
GCACACGTCTGAACTCTTTCCCTACACGACGCTCTTCCGATCTNNNAGGAATATTCGAGAAATGGTGT
GCACACGTCTGAACTCTTTCCCTACACGACGCTCTTCCGATCTNNNAGGAATATTCGAGAAATGGTGT
GCACACGTCTGAACTCTTTCCCTACACGACGCTCTTCCGATCTNNNAGGAATATTCGAGAAATGGTGT
AGCACACGTCTGAACTCTTTCCCTACACGACGCTCTTCCGATCTNNNCAGAGCCAAATTCTACCAGAGG
AGCACACGTCTGAACTCTTTCCCTACACGACGCTCTTCCGATCTNNNCAGAGCCAAATTCTACCAGAGG
AGCACACGTCTGAACTCTTTCCCTACACGACGCTCTTCCGATCTNNNCAGAGCCAAATTCTACCAGAGG
AGCACACGTCTGAACTCTTTCCCTACACGACGCTCTTCCGATCTNNNCAGAGCCAAATTCTACCAGAGG
GCACACGTCTGAACTCTTTCCCTACACGACGCTCTTCCGATCTNNNACTCTTACTTAGGGAAAGAAAAT
GCACACGTCTGAACTCTTTCCCTACACGACGCTCTTCCGATCTNNNACTCTTACTTAGGGAAAGAAAAT
GCACACGTCTGAACTCTTTCCCTACACGACGCTCTTCCGATCTNNNACTCTTACTTAGGGAAAGAAAAT
GCACACGTCTGAACTCTTTCCCTACACGACGCTCTTCCGATCTNNNACTCTTACTTAGGGAAAGAAAAT
AGCACACGTCTGAACTCTTTCCCTACACGACGCTCTTCCGATCTNNNAAGATCGGGAGCAAAGTTGAC
AGCACACGTCTGAACTCTTTCCCTACACGACGCTCTTCCGATCTNNNAAGATCGGGAGCAAAGTTGAC
AGCACACGTCTGAACTCTTTCCCTACACGACGCTCTTCCGATCTNNNAAGATCGGGAGCAAAGTTGAC

AGCACACGTCTGAACTCTTTCCCTACACGACGCTCTTCCGATCTNNNAAGATCGGGAGCAAAGTTGA
AGCACACGTCTGAACTCTTTCCCTACACGACGCTCTTCCGATCTNNNTCCGTTTCTAAAGCACAGGAA
AGCACACGTCTGAACTCTTTCCCTACACGACGCTCTTCCGATCTNNNTCCGTTTCTAAAGCACAGGAA
AGCACACGTCTGAACTCTTTCCCTACACGACGCTCTTCCGATCTNNNTCCGTTTCTAAAGCACAGGAA
AGCACACGTCTGAACTCTTTCCCTACACGACGCTCTTCCGATCTNNNTCCGTTTCTAAAGCACAGGAA
AGCACACGTCTGAACTCTTTCCCTACACGACGCTCTTCCGATCTNNNTTTCCGGCTCTTTAGTATGGG
AGCACACGTCTGAACTCTTTCCCTACACGACGCTCTTCCGATCTNNNTTTCCGGCTCTTTAGTATGGG
AGCACACGTCTGAACTCTTTCCCTACACGACGCTCTTCCGATCTNNNTTTCCGGCTCTTTAGTATGGG
AGCACACGTCTGAACTCTTTCCCTACACGACGCTCTTCCGATCTNNNTTTCCGGCTCTTTAGTATGGG
GAGCACACGTCTGAACTCTTTCCCTACACGACGCTCTTCCGATCTNNNCCTGGACCATTTTCTCCAGCT
GAGCACACGTCTGAACTCTTTCCCTACACGACGCTCTTCCGATCTNNNCCTGGACCATTTTCTCCAGCT
GAGCACACGTCTGAACTCTTTCCCTACACGACGCTCTTCCGATCTNNNCCTGGACCATTTTCTCCAGCT
GAGCACACGTCTGAACTCTTTCCCTACACGACGCTCTTCCGATCTNNNCCTGGACCATTTTCTCCAGCT
GAGCACACGTCTGAACTCTTTCCCTACACGACGCTCTTCCGATCTNNNATGAGCAGCCAACCTTAGTT
GAGCACACGTCTGAACTCTTTCCCTACACGACGCTCTTCCGATCTNNNATGAGCAGCCAACCTTAGTT
GAGCACACGTCTGAACTCTTTCCCTACACGACGCTCTTCCGATCTNNNATGAGCAGCCAACCTTAGTT
GAGCACACGTCTGAACTCTTTCCCTACACGACGCTCTTCCGATCTNNNATGAGCAGCCAACCTTAGTT
AGCACACGTCTGAACTCTTTCCCTACACGACGCTCTTCCGATCTNNNCAGATGGACAGATAGACAGA
AGCACACGTCTGAACTCTTTCCCTACACGACGCTCTTCCGATCTNNNCAGATGGACAGATAGACAGA
AGCACACGTCTGAACTCTTTCCCTACACGACGCTCTTCCGATCTNNNCAGATGGACAGATAGACAGA
AGCACACGTCTGAACTCTTTCCCTACACGACGCTCTTCCGATCTNNNCAGATGGACAGATAGACAGA
AGCACACGTCTGAACTCTTTCCCTACACGACGCTCTTCCGATCTNNNCTTAGCAAAGCAGTTTGCCTCT
AGCACACGTCTGAACTCTTTCCCTACACGACGCTCTTCCGATCTNNNCTTAGCAAAGCAGTTTGCCTCT
AGCACACGTCTGAACTCTTTCCCTACACGACGCTCTTCCGATCTNNNCTTAGCAAAGCAGTTTGCCTCT
AGCACACGTCTGAACTCTTTCCCTACACGACGCTCTTCCGATCTNNNCTTAGCAAAGCAGTTTGCCTCT
AGCACACGTCTGAACTCTTTCCCTACACGACGCTCTTCCGATCTNNNCATAGGCTGTCTCTCCTCATTC
AGCACACGTCTGAACTCTTTCCCTACACGACGCTCTTCCGATCTNNNCATAGGCTGTCTCTCCTCATTC
AGCACACGTCTGAACTCTTTCCCTACACGACGCTCTTCCGATCTNNNCATAGGCTGTCTCTCCTCATTC
AGCACACGTCTGAACTCTTTCCCTACACGACGCTCTTCCGATCTNNNCATAGGCTGTCTCTCCTCATTC
CACACGTCTGAACTCTTTCCCTACACGACGCTCTTCCGATCTNNNGCTGACGCAACTTTATCCCAGTA
CACACGTCTGAACTCTTTCCCTACACGACGCTCTTCCGATCTNNNGCTGACGCAACTTTATCCCAGTA
CACACGTCTGAACTCTTTCCCTACACGACGCTCTTCCGATCTNNNGCTGACGCAACTTTATCCCAGTA
CACACGTCTGAACTCTTTCCCTACACGACGCTCTTCCGATCTNNNGCTGACGCAACTTTATCCCAGTA
AGCACACGTCTGAACTCTTTCCCTACACGACGCTCTTCCGATCTNNNAATATGCAAATTCCACACCAGG
AGCACACGTCTGAACTCTTTCCCTACACGACGCTCTTCCGATCTNNNAATATGCAAATTCCACACCAGG
AGCACACGTCTGAACTCTTTCCCTACACGACGCTCTTCCGATCTNNNAATATGCAAATTCCACACCAGG
AGCACACGTCTGAACTCTTTCCCTACACGACGCTCTTCCGATCTNNNAATATGCAAATTCCACACCAGG
AGCACACGTCTGAACTCTTTCCCTACACGACGCTCTTCCGATCTNNNGGATCCATTTCTAGGTCTGCC
AGCACACGTCTGAACTCTTTCCCTACACGACGCTCTTCCGATCTNNNGGATCCATTTCTAGGTCTGCC
AGCACACGTCTGAACTCTTTCCCTACACGACGCTCTTCCGATCTNNNGGATCCATTTCTAGGTCTGCC
AGCACACGTCTGAACTCTTTCCCTACACGACGCTCTTCCGATCTNNNGGATCCATTTCTAGGTCTGCC
AGCACACGTCTGAACTCTTTCCCTACACGACGCTCTTCCGATCTNNNGAGGCTGAGGCAGGAGAATG
AGCACACGTCTGAACTCTTTCCCTACACGACGCTCTTCCGATCTNNNGAGGCTGAGGCAGGAGAATG

GAGCACACGTCTGAACTCTTTCCCTACACGACGCTCTTCCGATCTNNNGAGGCTGAGGCAGGAGAATG

GAGCACACGTCTGAACTCTTTCCCTACACGACGCTCTTCCGATCTNNNGAGGCTGAGGCAGGAGAATG

AGCACACGTCTGAACTCTTTCCCTACACGACGCTCTTCCGATCTNNNTCCTACTGAGATTGCTCTGCCTT

AGCACACGTCTGAACTCTTTCCCTACACGACGCTCTTCCGATCTNNNTCCTACTGAGATTGCTCTGCCTT

AGCACACGTCTGAACTCTTTCCCTACACGACGCTCTTCCGATCTNNNTCCTACTGAGATTGCTCTGCCTT

AGCACACGTCTGAACTCTTTCCCTACACGACGCTCTTCCGATCTNNNTCCTACTGAGATTGCTCTGCCTT

AGCACACGTCTGAACTCTTTCCCTACACGACGCTCTTCCGATCTNNNTGAGGTCCTTCTCTCTCTCTCC

AGCACACGTCTGAACTCTTTCCCTACACGACGCTCTTCCGATCTNNNTGAGGTCCTTCTCTCTCTCTCC

AGCACACGTCTGAACTCTTTCCCTACACGACGCTCTTCCGATCTNNNTGAGGTCCTTCTCTCTCTCTCC

AGCACACGTCTGAACTCTTTCCCTACACGACGCTCTTCCGATCTNNNTGAGGTCCTTCTCTCTCTCTCC

GCACACGTCTGAACTCTTTCCCTACACGACGCTCTTCCGATCTNNNTTCTGAAACTGAAGAGTCAAACAT

GCACACGTCTGAACTCTTTCCCTACACGACGCTCTTCCGATCTNNNTTCTGAAACTGAAGAGTCAAACAT

GCACACGTCTGAACTCTTTCCCTACACGACGCTCTTCCGATCTNNNTTCTGAAACTGAAGAGTCAAACAT

GCACACGTCTGAACTCTTTCCCTACACGACGCTCTTCCGATCTNNNTTCTGAAACTGAAGAGTCAAACAT

AGAGCACACGTCTGAACTCTTTCCCTACACGACGCTCTTCCGATCTNNNAGAATTGCTTGAATCAGGGA

AGAGCACACGTCTGAACTCTTTCCCTACACGACGCTCTTCCGATCTNNNAGAATTGCTTGAATCAGGGA

AGAGCACACGTCTGAACTCTTTCCCTACACGACGCTCTTCCGATCTNNNAGAATTGCTTGAATCAGGGA

AGAGCACACGTCTGAACTCTTTCCCTACACGACGCTCTTCCGATCTNNNAGAATTGCTTGAATCAGGGA

GAGCACACGTCTGAACTCTTTCCCTACACGACGCTCTTCCGATCTNNNGGATCACTTGAGGTCAGGAGT

GAGCACACGTCTGAACTCTTTCCCTACACGACGCTCTTCCGATCTNNNGGATCACTTGAGGTCAGGAGT

GAGCACACGTCTGAACTCTTTCCCTACACGACGCTCTTCCGATCTNNNGGATCACTTGAGGTCAGGAGT

GAGCACACGTCTGAACTCTTTCCCTACACGACGCTCTTCCGATCTNNNGGATCACTTGAGGTCAGGAGT

GAGCACACGTCTGAACTCTTTCCCTACACGACGCTCTTCCGATCTNNNTTGAGCTTTCATAACCTGCTGC

GAGCACACGTCTGAACTCTTTCCCTACACGACGCTCTTCCGATCTNNNTTGAGCTTTCATAACCTGCTGC

GAGCACACGTCTGAACTCTTTCCCTACACGACGCTCTTCCGATCTNNNTTGAGCTTTCATAACCTGCTGC

GAGCACACGTCTGAACTCTTTCCCTACACGACGCTCTTCCGATCTNNNTTGAGCTTTCATAACCTGCTGC

AGAGCACACGTCTGAACTCTTTCCCTACACGACGCTCTTCCGATCTNNNTGGAAACAAGGTAAAGGTG

AGAGCACACGTCTGAACTCTTTCCCTACACGACGCTCTTCCGATCTNNNTGGAAACAAGGTAAAGGTG

AGAGCACACGTCTGAACTCTTTCCCTACACGACGCTCTTCCGATCTNNNTGGAAACAAGGTAAAGGTG

AGAGCACACGTCTGAACTCTTTCCCTACACGACGCTCTTCCGATCTNNNTGGAAACAAGGTAAAGGTG

AGCACACGTCTGAACTCTTTCCCTACACGACGCTCTTCCGATCTNNNACCTTGGAGGGTAGTGCTTTTA

AGCACACGTCTGAACTCTTTCCCTACACGACGCTCTTCCGATCTNNNACCTTGGAGGGTAGTGCTTTTA

AGCACACGTCTGAACTCTTTCCCTACACGACGCTCTTCCGATCTNNNACCTTGGAGGGTAGTGCTTTTA

AGCACACGTCTGAACTCTTTCCCTACACGACGCTCTTCCGATCTNNNACCTTGGAGGGTAGTGCTTTTA

AGCACACGTCTGAACTCTTTCCCTACACGACGCTCTTCCGATCTNNNACTGTGGTCTATAGATCCCTGG

AGCACACGTCTGAACTCTTTCCCTACACGACGCTCTTCCGATCTNNNACTGTGGTCTATAGATCCCTGG

AGCACACGTCTGAACTCTTTCCCTACACGACGCTCTTCCGATCTNNNACTGTGGTCTATAGATCCCTGG

AGCACACGTCTGAACTCTTTCCCTACACGACGCTCTTCCGATCTNNNACTGTGGTCTATAGATCCCTGG

GCACACGTCTGAACTCTTTCCCTACACGACGCTCTTCCGATCTNNNCACTGAGGTGCATCACTCTTTTCC

GCACACGTCTGAACTCTTTCCCTACACGACGCTCTTCCGATCTNNNCACTGAGGTGCATCACTCTTTTCC

GCACACGTCTGAACTCTTTCCCTACACGACGCTCTTCCGATCTNNNCACTGAGGTGCATCACTCTTTTCC

GCACACGTCTGAACTCTTTCCCTACACGACGCTCTTCCGATCTNNNCACTGAGGTGCATCACTCTTTTCC

GAGCACACGTCTGAACTCTTTCCCTACACGACGCTCTTCCGATCTNNNCTGCTATTATACTCCAGCCTGG

GAGCACACGTCTGAACTCTTTCCCTACACGACGCTCTTCCGATCTNNNCTGCTATTATACTCCAGCCTGG
GAGCACACGTCTGAACTCTTTCCCTACACGACGCTCTTCCGATCTNNNCTGCTATTATACTCCAGCCTGG
GAGCACACGTCTGAACTCTTTCCCTACACGACGCTCTTCCGATCTNNNCTGCTATTATACTCCAGCCTGG
GAGCACACGTCTGAACTCTTTCCCTACACGACGCTCTTCCGATCTNNNTCCTGAGTAGCTGGGATTACA
GAGCACACGTCTGAACTCTTTCCCTACACGACGCTCTTCCGATCTNNNTCCTGAGTAGCTGGGATTACA
GAGCACACGTCTGAACTCTTTCCCTACACGACGCTCTTCCGATCTNNNTCCTGAGTAGCTGGGATTACA
GAGCACACGTCTGAACTCTTTCCCTACACGACGCTCTTCCGATCTNNNTCCTGAGTAGCTGGGATTACA
GAGCACACGTCTGAACTCTTTCCCTACACGACGCTCTTCCGATCTNNNTGTGAGCCTAACTGGAGTTTCT
GAGCACACGTCTGAACTCTTTCCCTACACGACGCTCTTCCGATCTNNNTGTGAGCCTAACTGGAGTTTCT
GAGCACACGTCTGAACTCTTTCCCTACACGACGCTCTTCCGATCTNNNTGTGAGCCTAACTGGAGTTTCT
GAGCACACGTCTGAACTCTTTCCCTACACGACGCTCTTCCGATCTNNNTGTGAGCCTAACTGGAGTTTCT
GAGCACACGTCTGAACTCTTTCCCTACACGACGCTCTTCCGATCTNNNGCCATTTGTGTGGAGCCATATA
GAGCACACGTCTGAACTCTTTCCCTACACGACGCTCTTCCGATCTNNNGCCATTTGTGTGGAGCCATATA
GAGCACACGTCTGAACTCTTTCCCTACACGACGCTCTTCCGATCTNNNGCCATTTGTGTGGAGCCATATA
GAGCACACGTCTGAACTCTTTCCCTACACGACGCTCTTCCGATCTNNNGCCATTTGTGTGGAGCCATATA
GAGCACACGTCTGAACTCTTTCCCTACACGACGCTCTTCCGATCTNNNCAGTGAAGAGAAGAGGAAG
GAGCACACGTCTGAACTCTTTCCCTACACGACGCTCTTCCGATCTNNNCAGTGAAGAGAAGAGGAAG
GAGCACACGTCTGAACTCTTTCCCTACACGACGCTCTTCCGATCTNNNCAGTGAAGAGAAGAGGAAG
GAGCACACGTCTGAACTCTTTCCCTACACGACGCTCTTCCGATCTNNNCAGTGAAGAGAAGAGGAAG
GAGCACACGTCTGAACTCTTTCCCTACACGACGCTCTTCCGATCTNNNAATAAGAGTGCTGAGTGGG
GAGCACACGTCTGAACTCTTTCCCTACACGACGCTCTTCCGATCTNNNAATAAGAGTGCTGAGTGGG
GAGCACACGTCTGAACTCTTTCCCTACACGACGCTCTTCCGATCTNNNAATAAGAGTGCTGAGTGGG
GAGCACACGTCTGAACTCTTTCCCTACACGACGCTCTTCCGATCTNNNAATAAGAGTGCTGAGTGGG
GAGCACACGTCTGAACTCTTTCCCTACACGACGCTCTTCCGATCTNNNATCCTCATACACAAAACAGCCA
GAGCACACGTCTGAACTCTTTCCCTACACGACGCTCTTCCGATCTNNNATCCTCATACACAAAACAGCCA
GAGCACACGTCTGAACTCTTTCCCTACACGACGCTCTTCCGATCTNNNATCCTCATACACAAAACAGCCA
GAGCACACGTCTGAACTCTTTCCCTACACGACGCTCTTCCGATCTNNNATCCTCATACACAAAACAGCCA
GAGCACACGTCTGAACTCTTTCCCTACACGACGCTCTTCCGATCTNNNGCTTAGTAGTATTCCATGGTGT
GAGCACACGTCTGAACTCTTTCCCTACACGACGCTCTTCCGATCTNNNGCTTAGTAGTATTCCATGGTGT
GAGCACACGTCTGAACTCTTTCCCTACACGACGCTCTTCCGATCTNNNGCTTAGTAGTATTCCATGGTGT
GAGCACACGTCTGAACTCTTTCCCTACACGACGCTCTTCCGATCTNNNGCTTAGTAGTATTCCATGGTGT
AGAGCACACGTCTGAACTCTTTCCCTACACGACGCTCTTCCGATCTNNNAAAAAATTAGCCGGGTGTG
AGAGCACACGTCTGAACTCTTTCCCTACACGACGCTCTTCCGATCTNNNAAAAAATTAGCCGGGTGTG
AGAGCACACGTCTGAACTCTTTCCCTACACGACGCTCTTCCGATCTNNNAAAAAATTAGCCGGGTGTG
AGAGCACACGTCTGAACTCTTTCCCTACACGACGCTCTTCCGATCTNNNAAAAAATTAGCCGGGTGTG
GCACACGTCTGAACTCTTTCCCTACACGACGCTCTTCCGATCTNNNGATTCCACATAGCAGTGAGACCA
GCACACGTCTGAACTCTTTCCCTACACGACGCTCTTCCGATCTNNNGATTCCACATAGCAGTGAGACCA
GCACACGTCTGAACTCTTTCCCTACACGACGCTCTTCCGATCTNNNGATTCCACATAGCAGTGAGACCA
GCACACGTCTGAACTCTTTCCCTACACGACGCTCTTCCGATCTNNNGATTCCACATAGCAGTGAGACCA
GAGCACACGTCTGAACTCTTTCCCTACACGACGCTCTTCCGATCTNNNGCAAAGCCAAACAAGTTGTAA
GAGCACACGTCTGAACTCTTTCCCTACACGACGCTCTTCCGATCTNNNGCAAAGCCAAACAAGTTGTAA
GAGCACACGTCTGAACTCTTTCCCTACACGACGCTCTTCCGATCTNNNGCAAAGCCAAACAAGTTGTAA
GAGCACACGTCTGAACTCTTTCCCTACACGACGCTCTTCCGATCTNNNGCAAAGCCAAACAAGTTGTAA

AGCACACGTCTGAACTCTTTCCCTACACGACGCTCTTCCGATCTNNNAAGGGGGCAAGTACTACATCAA
AGCACACGTCTGAACTCTTTCCCTACACGACGCTCTTCCGATCTNNNAAGGGGGCAAGTACTACATCAA
AGCACACGTCTGAACTCTTTCCCTACACGACGCTCTTCCGATCTNNNAAGGGGGCAAGTACTACATCAA
AGCACACGTCTGAACTCTTTCCCTACACGACGCTCTTCCGATCTNNNAAGGGGGCAAGTACTACATCAA
CACACGTCTGAACTCTTTCCCTACACGACGCTCTTCCGATCTNNNGTAACAATTGATGTGAGGCCTCG
CACACGTCTGAACTCTTTCCCTACACGACGCTCTTCCGATCTNNNGTAACAATTGATGTGAGGCCTCG
CACACGTCTGAACTCTTTCCCTACACGACGCTCTTCCGATCTNNNGTAACAATTGATGTGAGGCCTCG
CACACGTCTGAACTCTTTCCCTACACGACGCTCTTCCGATCTNNNGTAACAATTGATGTGAGGCCTCG
AGCACACGTCTGAACTCTTTCCCTACACGACGCTCTTCCGATCTNNNGAGAATCACTTGAACCTGGGA
AGCACACGTCTGAACTCTTTCCCTACACGACGCTCTTCCGATCTNNNGAGAATCACTTGAACCTGGGA
AGCACACGTCTGAACTCTTTCCCTACACGACGCTCTTCCGATCTNNNGAGAATCACTTGAACCTGGGA
AGCACACGTCTGAACTCTTTCCCTACACGACGCTCTTCCGATCTNNNGAGAATCACTTGAACCTGGGA
GAGCACACGTCTGAACTCTTTCCCTACACGACGCTCTTCCGATCTNNNGGACCACACTTTGAGAACTGT
GAGCACACGTCTGAACTCTTTCCCTACACGACGCTCTTCCGATCTNNNGGACCACACTTTGAGAACTGT
GAGCACACGTCTGAACTCTTTCCCTACACGACGCTCTTCCGATCTNNNGGACCACACTTTGAGAACTGT
GAGCACACGTCTGAACTCTTTCCCTACACGACGCTCTTCCGATCTNNNGGACCACACTTTGAGAACTGT
AGCACACGTCTGAACTCTTTCCCTACACGACGCTCTTCCGATCTNNNTCCCCATATAAGTTCTTAATTT
AGCACACGTCTGAACTCTTTCCCTACACGACGCTCTTCCGATCTNNNTCCCCATATAAGTTCTTAATTT
AGCACACGTCTGAACTCTTTCCCTACACGACGCTCTTCCGATCTNNNTCCCCATATAAGTTCTTAATTT
AGCACACGTCTGAACTCTTTCCCTACACGACGCTCTTCCGATCTNNNTCCCCATATAAGTTCTTAATTT
AGCACACGTCTGAACTCTTTCCCTACACGACGCTCTTCCGATCTNNNTGAGCTGAGATTGCACCACTG
AGCACACGTCTGAACTCTTTCCCTACACGACGCTCTTCCGATCTNNNTGAGCTGAGATTGCACCACTG
AGCACACGTCTGAACTCTTTCCCTACACGACGCTCTTCCGATCTNNNTGAGCTGAGATTGCACCACTG
AGCACACGTCTGAACTCTTTCCCTACACGACGCTCTTCCGATCTNNNTGAGCTGAGATTGCACCACTG
AGCACACGTCTGAACTCTTTCCCTACACGACGCTCTTCCGATCTNNNCACTTGTAGTCCCAGCTACTCG
AGCACACGTCTGAACTCTTTCCCTACACGACGCTCTTCCGATCTNNNCACTTGTAGTCCCAGCTACTCG
AGCACACGTCTGAACTCTTTCCCTACACGACGCTCTTCCGATCTNNNCACTTGTAGTCCCAGCTACTCG
AGCACACGTCTGAACTCTTTCCCTACACGACGCTCTTCCGATCTNNNCACTTGTAGTCCCAGCTACTCG
GCACACGTCTGAACTCTTTCCCTACACGACGCTCTTCCGATCTNNNAAAGATCAGGTGGATGGGTACA
GCACACGTCTGAACTCTTTCCCTACACGACGCTCTTCCGATCTNNNAAAGATCAGGTGGATGGGTACA
GCACACGTCTGAACTCTTTCCCTACACGACGCTCTTCCGATCTNNNAAAGATCAGGTGGATGGGTACA
GCACACGTCTGAACTCTTTCCCTACACGACGCTCTTCCGATCTNNNAAAGATCAGGTGGATGGGTACA
AGCACACGTCTGAACTCTTTCCCTACACGACGCTCTTCCGATCTNNNCCTCAGTGACAAGAGTGAGAC
AGCACACGTCTGAACTCTTTCCCTACACGACGCTCTTCCGATCTNNNCCTCAGTGACAAGAGTGAGAC
AGCACACGTCTGAACTCTTTCCCTACACGACGCTCTTCCGATCTNNNCCTCAGTGACAAGAGTGAGAC
AGCACACGTCTGAACTCTTTCCCTACACGACGCTCTTCCGATCTNNNCCTCAGTGACAAGAGTGAGAC
GCACACGTCTGAACTCTTTCCCTACACGACGCTCTTCCGATCTNNNGATCACGAAGTCAGGAGCTTGA
GCACACGTCTGAACTCTTTCCCTACACGACGCTCTTCCGATCTNNNGATCACGAAGTCAGGAGCTTGA
GCACACGTCTGAACTCTTTCCCTACACGACGCTCTTCCGATCTNNNGATCACGAAGTCAGGAGCTTGA
GCACACGTCTGAACTCTTTCCCTACACGACGCTCTTCCGATCTNNNGATCACGAAGTCAGGAGCTTGA
GCACACGTCTGAACTCTTTCCCTACACGACGCTCTTCCGATCTNNNTACCTCAGCCTCCCAAAAATGTTC
GCACACGTCTGAACTCTTTCCCTACACGACGCTCTTCCGATCTNNNTACCTCAGCCTCCCAAAAATGTTC
GCACACGTCTGAACTCTTTCCCTACACGACGCTCTTCCGATCTNNNTACCTCAGCCTCCCAAAAATGTTC

GCACACGTCTGAACTCTTTCCCTACACGACGCTCTTCCGATCTNNNTACCTCAGCCTCCCAAAAATGTTC

AGCACACGTCTGAACTCTTTCCCTACACGACGCTCTTCCGATCTNNNGACTTGCCTCATGTGGACATTT

AGCACACGTCTGAACTCTTTCCCTACACGACGCTCTTCCGATCTNNNGACTTGCCTCATGTGGACATTT

AGCACACGTCTGAACTCTTTCCCTACACGACGCTCTTCCGATCTNNNGACTTGCCTCATGTGGACATTT

AGCACACGTCTGAACTCTTTCCCTACACGACGCTCTTCCGATCTNNNGACTTGCCTCATGTGGACATTT

AGCACACGTCTGAACTCTTTCCCTACACGACGCTCTTCCGATCTNNNGTGAAAATGAGTGACTGCAGCT

AGCACACGTCTGAACTCTTTCCCTACACGACGCTCTTCCGATCTNNNGTGAAAATGAGTGACTGCAGCT

AGCACACGTCTGAACTCTTTCCCTACACGACGCTCTTCCGATCTNNNGTGAAAATGAGTGACTGCAGCT

AGCACACGTCTGAACTCTTTCCCTACACGACGCTCTTCCGATCTNNNGTGAAAATGAGTGACTGCAGCT

AGCACACGTCTGAACTCTTTCCCTACACGACGCTCTTCCGATCTNNNAGTTCTAGTGTGACCTTGCTCTC

AGCACACGTCTGAACTCTTTCCCTACACGACGCTCTTCCGATCTNNNAGTTCTAGTGTGACCTTGCTCTC

AGCACACGTCTGAACTCTTTCCCTACACGACGCTCTTCCGATCTNNNAGTTCTAGTGTGACCTTGCTCTC

AGCACACGTCTGAACTCTTTCCCTACACGACGCTCTTCCGATCTNNNAGTTCTAGTGTGACCTTGCTCTC

AGCACACGTCTGAACTCTTTCCCTACACGACGCTCTTCCGATCTNNNTATAGTCCCAGCTACTCGAGAG

AGCACACGTCTGAACTCTTTCCCTACACGACGCTCTTCCGATCTNNNTATAGTCCCAGCTACTCGAGAG

AGCACACGTCTGAACTCTTTCCCTACACGACGCTCTTCCGATCTNNNTATAGTCCCAGCTACTCGAGAG

AGCACACGTCTGAACTCTTTCCCTACACGACGCTCTTCCGATCTNNNTATAGTCCCAGCTACTCGAGAG

AGCACACGTCTGAACTCTTTCCCTACACGACGCTCTTCCGATCTNNNTCACTGAGTGGGTAGAGAAAA

AGCACACGTCTGAACTCTTTCCCTACACGACGCTCTTCCGATCTNNNTCACTGAGTGGGTAGAGAAAA

AGCACACGTCTGAACTCTTTCCCTACACGACGCTCTTCCGATCTNNNTCACTGAGTGGGTAGAGAAAA

AGCACACGTCTGAACTCTTTCCCTACACGACGCTCTTCCGATCTNNNTCACTGAGTGGGTAGAGAAAA

AGCACACGTCTGAACTCTTTCCCTACACGACGCTCTTCCGATCTNNNCTGAGGTGAGAGGATTGCTTG

AGCACACGTCTGAACTCTTTCCCTACACGACGCTCTTCCGATCTNNNCTGAGGTGAGAGGATTGCTTG

AGCACACGTCTGAACTCTTTCCCTACACGACGCTCTTCCGATCTNNNCTGAGGTGAGAGGATTGCTTG

AGCACACGTCTGAACTCTTTCCCTACACGACGCTCTTCCGATCTNNNCTGAGGTGAGAGGATTGCTTG

AGCACACGTCTGAACTCTTTCCCTACACGACGCTCTTCCGATCTNNNGTGACCTTACCTGACACTTTCCTC

AGCACACGTCTGAACTCTTTCCCTACACGACGCTCTTCCGATCTNNNGTGACCTTACCTGACACTTTCCTC

AGCACACGTCTGAACTCTTTCCCTACACGACGCTCTTCCGATCTNNNGTGACCTTACCTGACACTTTCCTC

AGCACACGTCTGAACTCTTTCCCTACACGACGCTCTTCCGATCTNNNGTGACCTTACCTGACACTTTCCTC

GAGCACACGTCTGAACTCTTTCCCTACACGACGCTCTTCCGATCTNNNCTAAGGTGGGAGGATTGCTT

GAGCACACGTCTGAACTCTTTCCCTACACGACGCTCTTCCGATCTNNNCTAAGGTGGGAGGATTGCTT

GAGCACACGTCTGAACTCTTTCCCTACACGACGCTCTTCCGATCTNNNCTAAGGTGGGAGGATTGCTT

GAGCACACGTCTGAACTCTTTCCCTACACGACGCTCTTCCGATCTNNNCTAAGGTGGGAGGATTGCTT

AGCACACGTCTGAACTCTTTCCCTACACGACGCTCTTCCGATCTNNNCCACCATAGGAAGGACAAATTG

AGCACACGTCTGAACTCTTTCCCTACACGACGCTCTTCCGATCTNNNCCACCATAGGAAGGACAAATTG

AGCACACGTCTGAACTCTTTCCCTACACGACGCTCTTCCGATCTNNNCCACCATAGGAAGGACAAATTG

AGCACACGTCTGAACTCTTTCCCTACACGACGCTCTTCCGATCTNNNCCACCATAGGAAGGACAAATTG

AGCACACGTCTGAACTCTTTCCCTACACGACGCTCTTCCGATCTNNNTCTTATGACCCAGTTCTCCCCAG

AGCACACGTCTGAACTCTTTCCCTACACGACGCTCTTCCGATCTNNNTCTTATGACCCAGTTCTCCCCAG

AGCACACGTCTGAACTCTTTCCCTACACGACGCTCTTCCGATCTNNNTCTTATGACCCAGTTCTCCCCAG

AGCACACGTCTGAACTCTTTCCCTACACGACGCTCTTCCGATCTNNNTCTTATGACCCAGTTCTCCCCAG

AGCACACGTCTGAACTCTTTCCCTACACGACGCTCTTCCGATCTNNNAGTCCACAACATGCTGAGGATA

AGCACACGTCTGAACTCTTTCCCTACACGACGCTCTTCCGATCTNNNAGTCCACAACATGCTGAGGATA

GCACACGTCTGAACTCTTTCCCTACACGACGCTCTTCCGATCTNNNAGTCCACAACATGCTGAGGATA

GCACACGTCTGAACTCTTTCCCTACACGACGCTCTTCCGATCTNNNAGTCCACAACATGCTGAGGATA

AGCACACGTCTGAACTCTTTCCCTACACGACGCTCTTCCGATCTNNNCTTACAGAGAGCCGAGATCAC

AGCACACGTCTGAACTCTTTCCCTACACGACGCTCTTCCGATCTNNNCTTACAGAGAGCCGAGATCAC

AGCACACGTCTGAACTCTTTCCCTACACGACGCTCTTCCGATCTNNNCTTACAGAGAGCCGAGATCAC

AGCACACGTCTGAACTCTTTCCCTACACGACGCTCTTCCGATCTNNNCTTACAGAGAGCCGAGATCAC

GAGCACACGTCTGAACTCTTTCCCTACACGACGCTCTTCCGATCTNNNAAAATGAAAAAGGGCTGGA

GAGCACACGTCTGAACTCTTTCCCTACACGACGCTCTTCCGATCTNNNAAAATGAAAAAGGGCTGGA

GAGCACACGTCTGAACTCTTTCCCTACACGACGCTCTTCCGATCTNNNAAAATGAAAAAGGGCTGGA

GAGCACACGTCTGAACTCTTTCCCTACACGACGCTCTTCCGATCTNNNAAAATGAAAAAGGGCTGGA

GAGCACACGTCTGAACTCTTTCCCTACACGACGCTCTTCCGATCTNNNCACTTGAGGTCAGGAGTTCA

GAGCACACGTCTGAACTCTTTCCCTACACGACGCTCTTCCGATCTNNNCACTTGAGGTCAGGAGTTCA

GAGCACACGTCTGAACTCTTTCCCTACACGACGCTCTTCCGATCTNNNCACTTGAGGTCAGGAGTTCA

GAGCACACGTCTGAACTCTTTCCCTACACGACGCTCTTCCGATCTNNNCACTTGAGGTCAGGAGTTCA

CACGTCTGAACTCTTTCCCTACACGACGCTCTTCCGATCTNNNTGTCTTACATTCTTGTATTCCAGCACA

CACGTCTGAACTCTTTCCCTACACGACGCTCTTCCGATCTNNNTGTCTTACATTCTTGTATTCCAGCACA

CACGTCTGAACTCTTTCCCTACACGACGCTCTTCCGATCTNNNTGTCTTACATTCTTGTATTCCAGCACA

CACGTCTGAACTCTTTCCCTACACGACGCTCTTCCGATCTNNNTGTCTTACATTCTTGTATTCCAGCACA

GAGCACACGTCTGAACTCTTTCCCTACACGACGCTCTTCCGATCTNNNTCATGTCCCACAACCTAACTC

GAGCACACGTCTGAACTCTTTCCCTACACGACGCTCTTCCGATCTNNNTCATGTCCCACAACCTAACTC

GAGCACACGTCTGAACTCTTTCCCTACACGACGCTCTTCCGATCTNNNTCATGTCCCACAACCTAACTC

GAGCACACGTCTGAACTCTTTCCCTACACGACGCTCTTCCGATCTNNNTCATGTCCCACAACCTAACTC

GAGCACACGTCTGAACTCTTTCCCTACACGACGCTCTTCCGATCTNNNCCTGTAAAATCCCAGCAGTTT

GAGCACACGTCTGAACTCTTTCCCTACACGACGCTCTTCCGATCTNNNCCTGTAAAATCCCAGCAGTTT

GAGCACACGTCTGAACTCTTTCCCTACACGACGCTCTTCCGATCTNNNCCTGTAAAATCCCAGCAGTTT

GAGCACACGTCTGAACTCTTTCCCTACACGACGCTCTTCCGATCTNNNCCTGTAAAATCCCAGCAGTTT

GCACACGTCTGAACTCTTTCCCTACACGACGCTCTTCCGATCTNNNTCCTACTATACTGGAGACACTGC

GCACACGTCTGAACTCTTTCCCTACACGACGCTCTTCCGATCTNNNTCCTACTATACTGGAGACACTGC

GCACACGTCTGAACTCTTTCCCTACACGACGCTCTTCCGATCTNNNTCCTACTATACTGGAGACACTGC

GCACACGTCTGAACTCTTTCCCTACACGACGCTCTTCCGATCTNNNTCCTACTATACTGGAGACACTGC

AGCACACGTCTGAACTCTTTCCCTACACGACGCTCTTCCGATCTNNNAAAATTAGTTGGGTGCAATG

AGCACACGTCTGAACTCTTTCCCTACACGACGCTCTTCCGATCTNNNAAAATTAGTTGGGTGCAATG

AGCACACGTCTGAACTCTTTCCCTACACGACGCTCTTCCGATCTNNNAAAATTAGTTGGGTGCAATG

AGCACACGTCTGAACTCTTTCCCTACACGACGCTCTTCCGATCTNNNAAAATTAGTTGGGTGCAATG

ACGTCTGAACTCTTTCCCTACACGACGCTCTTCCGATCTNNNGTCTCCTCCCTTCTCACCCCCATCGGAC

ACGTCTGAACTCTTTCCCTACACGACGCTCTTCCGATCTNNNGTCTCCTCCCTTCTCACCCCCATCGGAC

ACGTCTGAACTCTTTCCCTACACGACGCTCTTCCGATCTNNNGTCTCCTCCCTTCTCACCCCCATCGGAC

ACGTCTGAACTCTTTCCCTACACGACGCTCTTCCGATCTNNNGTCTCCTCCCTTCTCACCCCCATCGGAC

GCACACGTCTGAACTCTTTCCCTACACGACGCTCTTCCGATCTNNNTCACATTTTTTTGAACAATTCAC

GCACACGTCTGAACTCTTTCCCTACACGACGCTCTTCCGATCTNNNTCACATTTTTTTGAACAATTCAC

GCACACGTCTGAACTCTTTCCCTACACGACGCTCTTCCGATCTNNNTCACATTTTTTTGAACAATTCAC

GCACACGTCTGAACTCTTTCCCTACACGACGCTCTTCCGATCTNNNTCACATTTTTTTGAACAATTCAC

AGCACACGTCTGAACTCTTTCCCTACACGACGCTCTTCCGATCTNNNCCTCAAGATACCTTCCCACCTCA

AGCACACGTCTGAACTCTTTCCCTACACGACGCTCTTCCGATCTNNNCCTCAAGATACCTTCCCACCTCA
AGCACACGTCTGAACTCTTTCCCTACACGACGCTCTTCCGATCTNNNCCTCAAGATACCTTCCCACCTCA
AGCACACGTCTGAACTCTTTCCCTACACGACGCTCTTCCGATCTNNNCCTCAAGATACCTTCCCACCTCA
AGCACACGTCTGAACTCTTTCCCTACACGACGCTCTTCCGATCTNNNATGTCTCATTGAGAATGGCCTC
AGCACACGTCTGAACTCTTTCCCTACACGACGCTCTTCCGATCTNNNATGTCTCATTGAGAATGGCCTC
AGCACACGTCTGAACTCTTTCCCTACACGACGCTCTTCCGATCTNNNATGTCTCATTGAGAATGGCCTC
AGCACACGTCTGAACTCTTTCCCTACACGACGCTCTTCCGATCTNNNATGTCTCATTGAGAATGGCCTC
AGCACACGTCTGAACTCTTTCCCTACACGACGCTCTTCCGATCTNNNTGACTGTCAGTCCCATATTCAC
AGCACACGTCTGAACTCTTTCCCTACACGACGCTCTTCCGATCTNNNTGACTGTCAGTCCCATATTCAC
AGCACACGTCTGAACTCTTTCCCTACACGACGCTCTTCCGATCTNNNTGACTGTCAGTCCCATATTCAC
AGCACACGTCTGAACTCTTTCCCTACACGACGCTCTTCCGATCTNNNTGACTGTCAGTCCCATATTCAC
GAGCACACGTCTGAACTCTTTCCCTACACGACGCTCTTCCGATCTNNNTTTAGTCTGCAGATGTGGAA
GAGCACACGTCTGAACTCTTTCCCTACACGACGCTCTTCCGATCTNNNTTTAGTCTGCAGATGTGGAA
GAGCACACGTCTGAACTCTTTCCCTACACGACGCTCTTCCGATCTNNNTTTAGTCTGCAGATGTGGAA
GAGCACACGTCTGAACTCTTTCCCTACACGACGCTCTTCCGATCTNNNTTTAGTCTGCAGATGTGGAA
AGCACACGTCTGAACTCTTTCCCTACACGACGCTCTTCCGATCTNNNCCCAAAATGTTGCCACAGCTTT
AGCACACGTCTGAACTCTTTCCCTACACGACGCTCTTCCGATCTNNNCCCAAAATGTTGCCACAGCTTT
AGCACACGTCTGAACTCTTTCCCTACACGACGCTCTTCCGATCTNNNCCCAAAATGTTGCCACAGCTTT
AGCACACGTCTGAACTCTTTCCCTACACGACGCTCTTCCGATCTNNNCCCAAAATGTTGCCACAGCTTT
GAGCACACGTCTGAACTCTTTCCCTACACGACGCTCTTCCGATCTNNNGTTTGCCATGACTCGGATTTC
GAGCACACGTCTGAACTCTTTCCCTACACGACGCTCTTCCGATCTNNNGTTTGCCATGACTCGGATTTC
GAGCACACGTCTGAACTCTTTCCCTACACGACGCTCTTCCGATCTNNNGTTTGCCATGACTCGGATTTC
GAGCACACGTCTGAACTCTTTCCCTACACGACGCTCTTCCGATCTNNNGTTTGCCATGACTCGGATTTC
GCACACGTCTGAACTCTTTCCCTACACGACGCTCTTCCGATCTNNNTCCACTAGGAGGAAACAGAAA
GCACACGTCTGAACTCTTTCCCTACACGACGCTCTTCCGATCTNNNTCCACTAGGAGGAAACAGAAA
GCACACGTCTGAACTCTTTCCCTACACGACGCTCTTCCGATCTNNNTCCACTAGGAGGAAACAGAAA
GCACACGTCTGAACTCTTTCCCTACACGACGCTCTTCCGATCTNNNTCCACTAGGAGGAAACAGAAA
GAAGAGCACACGTCTGAACTCTTTCCCTACACGACGCTCTTCCGATCTNNNGAGGCAGGAGAATGGT
GAAGAGCACACGTCTGAACTCTTTCCCTACACGACGCTCTTCCGATCTNNNGAGGCAGGAGAATGGT
GAAGAGCACACGTCTGAACTCTTTCCCTACACGACGCTCTTCCGATCTNNNGAGGCAGGAGAATGGT
GAAGAGCACACGTCTGAACTCTTTCCCTACACGACGCTCTTCCGATCTNNNGAGGCAGGAGAATGGT
GCACACGTCTGAACTCTTTCCCTACACGACGCTCTTCCGATCTNNNCACTGTGCTTGGTCTGTTTTCTT
GCACACGTCTGAACTCTTTCCCTACACGACGCTCTTCCGATCTNNNCACTGTGCTTGGTCTGTTTTCTT
GCACACGTCTGAACTCTTTCCCTACACGACGCTCTTCCGATCTNNNCACTGTGCTTGGTCTGTTTTCTT
GCACACGTCTGAACTCTTTCCCTACACGACGCTCTTCCGATCTNNNCACTGTGCTTGGTCTGTTTTCTT
GAGCACACGTCTGAACTCTTTCCCTACACGACGCTCTTCCGATCTNNNTATTCCCGATGACCCTGTCTT
GAGCACACGTCTGAACTCTTTCCCTACACGACGCTCTTCCGATCTNNNTATTCCCGATGACCCTGTCTT
GAGCACACGTCTGAACTCTTTCCCTACACGACGCTCTTCCGATCTNNNTATTCCCGATGACCCTGTCTT
GAGCACACGTCTGAACTCTTTCCCTACACGACGCTCTTCCGATCTNNNTATTCCCGATGACCCTGTCTT
GCACACGTCTGAACTCTTTCCCTACACGACGCTCTTCCGATCTNNNTGGTCCCTTCTTTCAGCGTGTAT
GCACACGTCTGAACTCTTTCCCTACACGACGCTCTTCCGATCTNNNTGGTCCCTTCTTTCAGCGTGTAT
GCACACGTCTGAACTCTTTCCCTACACGACGCTCTTCCGATCTNNNTGGTCCCTTCTTTCAGCGTGTAT
GCACACGTCTGAACTCTTTCCCTACACGACGCTCTTCCGATCTNNNTGGTCCCTTCTTTCAGCGTGTAT

AGCACACGTCTGAACTCTTTCCCTACACGACGCTCTTCCGATCTNNNTTTCACTTTGTCCTTGTCTGCCA
AGCACACGTCTGAACTCTTTCCCTACACGACGCTCTTCCGATCTNNNTTTCACTTTGTCCTTGTCTGCCA
AGCACACGTCTGAACTCTTTCCCTACACGACGCTCTTCCGATCTNNNTTTCACTTTGTCCTTGTCTGCCA
AGCACACGTCTGAACTCTTTCCCTACACGACGCTCTTCCGATCTNNNTTTCACTTTGTCCTTGTCTGCCA
AGCACACGTCTGAACTCTTTCCCTACACGACGCTCTTCCGATCTNNNGCATCAGCAGAAGTCAGACTA
AGCACACGTCTGAACTCTTTCCCTACACGACGCTCTTCCGATCTNNNGCATCAGCAGAAGTCAGACTA
AGCACACGTCTGAACTCTTTCCCTACACGACGCTCTTCCGATCTNNNGCATCAGCAGAAGTCAGACTA
AGCACACGTCTGAACTCTTTCCCTACACGACGCTCTTCCGATCTNNNGCATCAGCAGAAGTCAGACTA
AGCACACGTCTGAACTCTTTCCCTACACGACGCTCTTCCGATCTNNNCCGAGAAGAAGTGGCTAATCT
AGCACACGTCTGAACTCTTTCCCTACACGACGCTCTTCCGATCTNNNCCGAGAAGAAGTGGCTAATCT
AGCACACGTCTGAACTCTTTCCCTACACGACGCTCTTCCGATCTNNNCCGAGAAGAAGTGGCTAATCT
AGCACACGTCTGAACTCTTTCCCTACACGACGCTCTTCCGATCTNNNCCGAGAAGAAGTGGCTAATCT
AGCACACGTCTGAACTCTTTCCCTACACGACGCTCTTCCGATCTNNNCTTGCAGTGAGCTGAGATCGT
AGCACACGTCTGAACTCTTTCCCTACACGACGCTCTTCCGATCTNNNCTTGCAGTGAGCTGAGATCGT
AGCACACGTCTGAACTCTTTCCCTACACGACGCTCTTCCGATCTNNNCTTGCAGTGAGCTGAGATCGT
AGCACACGTCTGAACTCTTTCCCTACACGACGCTCTTCCGATCTNNNCTTGCAGTGAGCTGAGATCGT
AGCACACGTCTGAACTCTTTCCCTACACGACGCTCTTCCGATCTNNNGAGGAGGAGAGAGAGAAGA
AGCACACGTCTGAACTCTTTCCCTACACGACGCTCTTCCGATCTNNNGAGGAGGAGAGAGAGAAGA
AGCACACGTCTGAACTCTTTCCCTACACGACGCTCTTCCGATCTNNNGAGGAGGAGAGAGAGAAGA
AGCACACGTCTGAACTCTTTCCCTACACGACGCTCTTCCGATCTNNNGAGGAGGAGAGAGAGAAGA
GCACACGTCTGAACTCTTTCCCTACACGACGCTCTTCCGATCTNNNGCTAAACCAAGAACACAAACCCA
GCACACGTCTGAACTCTTTCCCTACACGACGCTCTTCCGATCTNNNGCTAAACCAAGAACACAAACCCA
GCACACGTCTGAACTCTTTCCCTACACGACGCTCTTCCGATCTNNNGCTAAACCAAGAACACAAACCCA
GCACACGTCTGAACTCTTTCCCTACACGACGCTCTTCCGATCTNNNGCTAAACCAAGAACACAAACCCA
AGCACACGTCTGAACTCTTTCCCTACACGACGCTCTTCCGATCTNNNCCTGCAGAGCTGTAAACCAAT
AGCACACGTCTGAACTCTTTCCCTACACGACGCTCTTCCGATCTNNNCCTGCAGAGCTGTAAACCAAT
AGCACACGTCTGAACTCTTTCCCTACACGACGCTCTTCCGATCTNNNCCTGCAGAGCTGTAAACCAAT
AGCACACGTCTGAACTCTTTCCCTACACGACGCTCTTCCGATCTNNNCCTGCAGAGCTGTAAACCAAT
GCACACGTCTGAACTCTTTCCCTACACGACGCTCTTCCGATCTNNNAAGTTGGTTGTACTCTCCCCATC
GCACACGTCTGAACTCTTTCCCTACACGACGCTCTTCCGATCTNNNAAGTTGGTTGTACTCTCCCCATC
GCACACGTCTGAACTCTTTCCCTACACGACGCTCTTCCGATCTNNNAAGTTGGTTGTACTCTCCCCATC
GCACACGTCTGAACTCTTTCCCTACACGACGCTCTTCCGATCTNNNAAGTTGGTTGTACTCTCCCCATC
GCACACGTCTGAACTCTTTCCCTACACGACGCTCTTCCGATCTNNNGAATGCCAAAGGGGGAGTTTT
GCACACGTCTGAACTCTTTCCCTACACGACGCTCTTCCGATCTNNNGAATGCCAAAGGGGGAGTTTT
GCACACGTCTGAACTCTTTCCCTACACGACGCTCTTCCGATCTNNNGAATGCCAAAGGGGGAGTTTT
GCACACGTCTGAACTCTTTCCCTACACGACGCTCTTCCGATCTNNNGAATGCCAAAGGGGGAGTTTT
AGCACACGTCTGAACTCTTTCCCTACACGACGCTCTTCCGATCTNNNACTCTTTCTCCAGGTTGGACAA
AGCACACGTCTGAACTCTTTCCCTACACGACGCTCTTCCGATCTNNNACTCTTTCTCCAGGTTGGACAA
AGCACACGTCTGAACTCTTTCCCTACACGACGCTCTTCCGATCTNNNACTCTTTCTCCAGGTTGGACAA
AGCACACGTCTGAACTCTTTCCCTACACGACGCTCTTCCGATCTNNNACTCTTTCTCCAGGTTGGACAA
AGCACACGTCTGAACTCTTTCCCTACACGACGCTCTTCCGATCTNNNTCTTTTCTTTGCAACCTCACCA
AGCACACGTCTGAACTCTTTCCCTACACGACGCTCTTCCGATCTNNNTCTTTTCTTTGCAACCTCACCA
AGCACACGTCTGAACTCTTTCCCTACACGACGCTCTTCCGATCTNNNTCTTTTCTTTGCAACCTCACCA

GAGCACACGTCTGAACTCTTTCCCTACACGACGCTCTTCCGATCTNNNTCTTTTCTTTGCAACCTCACCAG

GAGCACACGTCTGAACTCTTTCCCTACACGACGCTCTTCCGATCTNNNAGGGGAACACCAAAATCTCAG

GAGCACACGTCTGAACTCTTTCCCTACACGACGCTCTTCCGATCTNNNAGGGGAACACCAAAATCTCAG

GAGCACACGTCTGAACTCTTTCCCTACACGACGCTCTTCCGATCTNNNAGGGGAACACCAAAATCTCAG

GAGCACACGTCTGAACTCTTTCCCTACACGACGCTCTTCCGATCTNNNAGGGGAACACCAAAATCTCAG

AGCACACGTCTGAACTCTTTCCCTACACGACGCTCTTCCGATCTNNNCTGTCACTGTTTTGTGTGTGTG

AGCACACGTCTGAACTCTTTCCCTACACGACGCTCTTCCGATCTNNNCTGTCACTGTTTTGTGTGTGTG

AGCACACGTCTGAACTCTTTCCCTACACGACGCTCTTCCGATCTNNNCTGTCACTGTTTTGTGTGTGTG

AGCACACGTCTGAACTCTTTCCCTACACGACGCTCTTCCGATCTNNNCTGTCACTGTTTTGTGTGTGTG

AGCACACGTCTGAACTCTTTCCCTACACGACGCTCTTCCGATCTNNNTGAGTTGACAGAGTGAGACCTT

AGCACACGTCTGAACTCTTTCCCTACACGACGCTCTTCCGATCTNNNTGAGTTGACAGAGTGAGACCTT

AGCACACGTCTGAACTCTTTCCCTACACGACGCTCTTCCGATCTNNNTGAGTTGACAGAGTGAGACCTT

AGCACACGTCTGAACTCTTTCCCTACACGACGCTCTTCCGATCTNNNTGAGTTGACAGAGTGAGACCTT

GAGCACACGTCTGAACTCTTTCCCTACACGACGCTCTTCCGATCTNNNTCTGATTTCTCCACTTGCTCAGT

GAGCACACGTCTGAACTCTTTCCCTACACGACGCTCTTCCGATCTNNNTCTGATTTCTCCACTTGCTCAGT

GAGCACACGTCTGAACTCTTTCCCTACACGACGCTCTTCCGATCTNNNTCTGATTTCTCCACTTGCTCAGT

GAGCACACGTCTGAACTCTTTCCCTACACGACGCTCTTCCGATCTNNNTCTGATTTCTCCACTTGCTCAGT

GCACACGTCTGAACTCTTTCCCTACACGACGCTCTTCCGATCTNNNTTTCTGTTGGAAAGCTGCTCTGC

GCACACGTCTGAACTCTTTCCCTACACGACGCTCTTCCGATCTNNNTTTCTGTTGGAAAGCTGCTCTGC

GCACACGTCTGAACTCTTTCCCTACACGACGCTCTTCCGATCTNNNTTTCTGTTGGAAAGCTGCTCTGC

GCACACGTCTGAACTCTTTCCCTACACGACGCTCTTCCGATCTNNNTTTCTGTTGGAAAGCTGCTCTGC

GAGCACACGTCTGAACTCTTTCCCTACACGACGCTCTTCCGATCTNNNTCTGCCCATGTTCAGATCATTC

GAGCACACGTCTGAACTCTTTCCCTACACGACGCTCTTCCGATCTNNNTCTGCCCATGTTCAGATCATTC

GAGCACACGTCTGAACTCTTTCCCTACACGACGCTCTTCCGATCTNNNTCTGCCCATGTTCAGATCATTC

GAGCACACGTCTGAACTCTTTCCCTACACGACGCTCTTCCGATCTNNNTCTGCCCATGTTCAGATCATTC

AGCACACGTCTGAACTCTTTCCCTACACGACGCTCTTCCGATCTNNNGAGGGAGAGGAATGGAGGGA

AGCACACGTCTGAACTCTTTCCCTACACGACGCTCTTCCGATCTNNNGAGGGAGAGGAATGGAGGGA

AGCACACGTCTGAACTCTTTCCCTACACGACGCTCTTCCGATCTNNNGAGGGAGAGGAATGGAGGGA

AGCACACGTCTGAACTCTTTCCCTACACGACGCTCTTCCGATCTNNNGAGGGAGAGGAATGGAGGGA

GAGCACACGTCTGAACTCTTTCCCTACACGACGCTCTTCCGATCTNNNGCTAAGGCAGGAGAATCACTT

GAGCACACGTCTGAACTCTTTCCCTACACGACGCTCTTCCGATCTNNNGCTAAGGCAGGAGAATCACTT

GAGCACACGTCTGAACTCTTTCCCTACACGACGCTCTTCCGATCTNNNGCTAAGGCAGGAGAATCACTT

GAGCACACGTCTGAACTCTTTCCCTACACGACGCTCTTCCGATCTNNNGCTAAGGCAGGAGAATCACTT

GAGCACACGTCTGAACTCTTTCCCTACACGACGCTCTTCCGATCTNNNTGTGGTATCCTTTCAGAGTTCA

GAGCACACGTCTGAACTCTTTCCCTACACGACGCTCTTCCGATCTNNNTGTGGTATCCTTTCAGAGTTCA

GAGCACACGTCTGAACTCTTTCCCTACACGACGCTCTTCCGATCTNNNTGTGGTATCCTTTCAGAGTTCA

GAGCACACGTCTGAACTCTTTCCCTACACGACGCTCTTCCGATCTNNNTGTGGTATCCTTTCAGAGTTCA

GAGCACACGTCTGAACTCTTTCCCTACACGACGCTCTTCCGATCTNNNAAGGTGTGAAAAACGTGGGA

GAGCACACGTCTGAACTCTTTCCCTACACGACGCTCTTCCGATCTNNNAAGGTGTGAAAAACGTGGGA

GAGCACACGTCTGAACTCTTTCCCTACACGACGCTCTTCCGATCTNNNAAGGTGTGAAAAACGTGGGA

GAGCACACGTCTGAACTCTTTCCCTACACGACGCTCTTCCGATCTNNNAAGGTGTGAAAAACGTGGGA

CACACGTCTGAACTCTTTCCCTACACGACGCTCTTCCGATCTNNNCCATTGGAAGCCACAATATGACACA

CACACGTCTGAACTCTTTCCCTACACGACGCTCTTCCGATCTNNNCCATTGGAAGCCACAATATGACACA

CACACGTCTGAACTCTTTCCCTACACGACGCTCTTCCGATCTNNNCCATTGGAAGCCACAATATGACACA
CACACGTCTGAACTCTTTCCCTACACGACGCTCTTCCGATCTNNNCCATTGGAAGCCACAATATGACACA
GCACACGTCTGAACTCTTTCCCTACACGACGCTCTTCCGATCTNNNATGGCCAATTAGTTGAACATGG
GCACACGTCTGAACTCTTTCCCTACACGACGCTCTTCCGATCTNNNATGGCCAATTAGTTGAACATGG
GCACACGTCTGAACTCTTTCCCTACACGACGCTCTTCCGATCTNNNATGGCCAATTAGTTGAACATGG
GCACACGTCTGAACTCTTTCCCTACACGACGCTCTTCCGATCTNNNATGGCCAATTAGTTGAACATGG
AGCACACGTCTGAACTCTTTCCCTACACGACGCTCTTCCGATCTNNNCAGGATGCTCCCAAGTACTGAT
AGCACACGTCTGAACTCTTTCCCTACACGACGCTCTTCCGATCTNNNCAGGATGCTCCCAAGTACTGAT
AGCACACGTCTGAACTCTTTCCCTACACGACGCTCTTCCGATCTNNNCAGGATGCTCCCAAGTACTGAT
AGCACACGTCTGAACTCTTTCCCTACACGACGCTCTTCCGATCTNNNCAGGATGCTCCCAAGTACTGAT
AGAGCACACGTCTGAACTCTTTCCCTACACGACGCTCTTCCGATCTNNNTGTAGTACAATACCAGGCA
AGAGCACACGTCTGAACTCTTTCCCTACACGACGCTCTTCCGATCTNNNTGTAGTACAATACCAGGCA
AGAGCACACGTCTGAACTCTTTCCCTACACGACGCTCTTCCGATCTNNNTGTAGTACAATACCAGGCA
AGAGCACACGTCTGAACTCTTTCCCTACACGACGCTCTTCCGATCTNNNTGTAGTACAATACCAGGCA
AGCACACGTCTGAACTCTTTCCCTACACGACGCTCTTCCGATCTNNNTGAGGCAGGAGAATGACTTGA
AGCACACGTCTGAACTCTTTCCCTACACGACGCTCTTCCGATCTNNNTGAGGCAGGAGAATGACTTGA
AGCACACGTCTGAACTCTTTCCCTACACGACGCTCTTCCGATCTNNNTGAGGCAGGAGAATGACTTGA
AGCACACGTCTGAACTCTTTCCCTACACGACGCTCTTCCGATCTNNNTGAGGCAGGAGAATGACTTGA
AGCACACGTCTGAACTCTTTCCCTACACGACGCTCTTCCGATCTNNNTGGGAAACAGAGCAAGATCTT
AGCACACGTCTGAACTCTTTCCCTACACGACGCTCTTCCGATCTNNNTGGGAAACAGAGCAAGATCTT
AGCACACGTCTGAACTCTTTCCCTACACGACGCTCTTCCGATCTNNNTGGGAAACAGAGCAAGATCTT
AGCACACGTCTGAACTCTTTCCCTACACGACGCTCTTCCGATCTNNNTGGGAAACAGAGCAAGATCTT
CACACGTCTGAACTCTTTCCCTACACGACGCTCTTCCGATCTNNNGCCCCAGGTATTTCTTTAGAGCTCA
CACACGTCTGAACTCTTTCCCTACACGACGCTCTTCCGATCTNNNGCCCCAGGTATTTCTTTAGAGCTCA
CACACGTCTGAACTCTTTCCCTACACGACGCTCTTCCGATCTNNNGCCCCAGGTATTTCTTTAGAGCTCA
CACACGTCTGAACTCTTTCCCTACACGACGCTCTTCCGATCTNNNGCCCCAGGTATTTCTTTAGAGCTCA
AGCACACGTCTGAACTCTTTCCCTACACGACGCTCTTCCGATCTNNNGAGACAGGAGAATTGCTTGAA
AGCACACGTCTGAACTCTTTCCCTACACGACGCTCTTCCGATCTNNNGAGACAGGAGAATTGCTTGAA
AGCACACGTCTGAACTCTTTCCCTACACGACGCTCTTCCGATCTNNNGAGACAGGAGAATTGCTTGAA
AGCACACGTCTGAACTCTTTCCCTACACGACGCTCTTCCGATCTNNNGAGACAGGAGAATTGCTTGAA
AGCACACGTCTGAACTCTTTCCCTACACGACGCTCTTCCGATCTNNNTGGGTCTTTGAGGAATTTCCAC
AGCACACGTCTGAACTCTTTCCCTACACGACGCTCTTCCGATCTNNNTGGGTCTTTGAGGAATTTCCAC
AGCACACGTCTGAACTCTTTCCCTACACGACGCTCTTCCGATCTNNNTGGGTCTTTGAGGAATTTCCAC
AGCACACGTCTGAACTCTTTCCCTACACGACGCTCTTCCGATCTNNNTGGGTCTTTGAGGAATTTCCAC
AGCACACGTCTGAACTCTTTCCCTACACGACGCTCTTCCGATCTNNNAGGTCAGGAGATTTAGGGGTC
AGCACACGTCTGAACTCTTTCCCTACACGACGCTCTTCCGATCTNNNAGGTCAGGAGATTTAGGGGTC
AGCACACGTCTGAACTCTTTCCCTACACGACGCTCTTCCGATCTNNNAGGTCAGGAGATTTAGGGGTC
AGCACACGTCTGAACTCTTTCCCTACACGACGCTCTTCCGATCTNNNAGGTCAGGAGATTTAGGGGTC
GAGCACACGTCTGAACTCTTTCCCTACACGACGCTCTTCCGATCTNNNGGGATAGCATTGGGAGATAT
GAGCACACGTCTGAACTCTTTCCCTACACGACGCTCTTCCGATCTNNNGGGATAGCATTGGGAGATAT
GAGCACACGTCTGAACTCTTTCCCTACACGACGCTCTTCCGATCTNNNGGGATAGCATTGGGAGATAT
GAGCACACGTCTGAACTCTTTCCCTACACGACGCTCTTCCGATCTNNNGGGATAGCATTGGGAGATAT
GCACACGTCTGAACTCTTTCCCTACACGACGCTCTTCCGATCTNNNGTGAGGTTAGTGGTTACATCCCT

GCACACGTCTGAACTCTTTCCCTACACGACGCTCTTCCGATCTNNNGTGAGGTTAGTGGTTACATCCCT
GCACACGTCTGAACTCTTTCCCTACACGACGCTCTTCCGATCTNNNGTGAGGTTAGTGGTTACATCCCT
GCACACGTCTGAACTCTTTCCCTACACGACGCTCTTCCGATCTNNNGTGAGGTTAGTGGTTACATCCCT
AGCACACGTCTGAACTCTTTCCCTACACGACGCTCTTCCGATCTNNNGAGGTTGCAATGAGCTGAGATT
AGCACACGTCTGAACTCTTTCCCTACACGACGCTCTTCCGATCTNNNGAGGTTGCAATGAGCTGAGATT
AGCACACGTCTGAACTCTTTCCCTACACGACGCTCTTCCGATCTNNNGAGGTTGCAATGAGCTGAGATT
AGCACACGTCTGAACTCTTTCCCTACACGACGCTCTTCCGATCTNNNGAGGTTGCAATGAGCTGAGATT
GAGCACACGTCTGAACTCTTTCCCTACACGACGCTCTTCCGATCTNNNAGGAGAACTTTTCCCTTCTGT
GAGCACACGTCTGAACTCTTTCCCTACACGACGCTCTTCCGATCTNNNAGGAGAACTTTTCCCTTCTGT
GAGCACACGTCTGAACTCTTTCCCTACACGACGCTCTTCCGATCTNNNAGGAGAACTTTTCCCTTCTGT
GAGCACACGTCTGAACTCTTTCCCTACACGACGCTCTTCCGATCTNNNAGGAGAACTTTTCCCTTCTGT
GAGCACACGTCTGAACTCTTTCCCTACACGACGCTCTTCCGATCTNNNACGCCTGTAATCCCAGTACTTT
GAGCACACGTCTGAACTCTTTCCCTACACGACGCTCTTCCGATCTNNNACGCCTGTAATCCCAGTACTTT
GAGCACACGTCTGAACTCTTTCCCTACACGACGCTCTTCCGATCTNNNACGCCTGTAATCCCAGTACTTT
GAGCACACGTCTGAACTCTTTCCCTACACGACGCTCTTCCGATCTNNNACGCCTGTAATCCCAGTACTTT
AGCACACGTCTGAACTCTTTCCCTACACGACGCTCTTCCGATCTNNNTGCAGAATTCACAGAATCCTTGT
AGCACACGTCTGAACTCTTTCCCTACACGACGCTCTTCCGATCTNNNTGCAGAATTCACAGAATCCTTGT
AGCACACGTCTGAACTCTTTCCCTACACGACGCTCTTCCGATCTNNNTGCAGAATTCACAGAATCCTTGT
AGCACACGTCTGAACTCTTTCCCTACACGACGCTCTTCCGATCTNNNTGCAGAATTCACAGAATCCTTGT
GAGCACACGTCTGAACTCTTTCCCTACACGACGCTCTTCCGATCTNNNACGTGGAGATGGCACTGAATA
GAGCACACGTCTGAACTCTTTCCCTACACGACGCTCTTCCGATCTNNNACGTGGAGATGGCACTGAATA
GAGCACACGTCTGAACTCTTTCCCTACACGACGCTCTTCCGATCTNNNACGTGGAGATGGCACTGAATA
GAGCACACGTCTGAACTCTTTCCCTACACGACGCTCTTCCGATCTNNNACGTGGAGATGGCACTGAATA
AGCACACGTCTGAACTCTTTCCCTACACGACGCTCTTCCGATCTNNNTCCTCATAGCTTAGCTCCCACTT
AGCACACGTCTGAACTCTTTCCCTACACGACGCTCTTCCGATCTNNNTCCTCATAGCTTAGCTCCCACTT
AGCACACGTCTGAACTCTTTCCCTACACGACGCTCTTCCGATCTNNNTCCTCATAGCTTAGCTCCCACTT
AGCACACGTCTGAACTCTTTCCCTACACGACGCTCTTCCGATCTNNNTCCTCATAGCTTAGCTCCCACTT
AGCACACGTCTGAACTCTTTCCCTACACGACGCTCTTCCGATCTNNNTCCACGAGTTAGAGGAGTGTA
AGCACACGTCTGAACTCTTTCCCTACACGACGCTCTTCCGATCTNNNTCCACGAGTTAGAGGAGTGTA
AGCACACGTCTGAACTCTTTCCCTACACGACGCTCTTCCGATCTNNNTCCACGAGTTAGAGGAGTGTA
AGCACACGTCTGAACTCTTTCCCTACACGACGCTCTTCCGATCTNNNTCCACGAGTTAGAGGAGTGTA
AGCACACGTCTGAACTCTTTCCCTACACGACGCTCTTCCGATCTNNNTCACTTAGCTTTTCCCTAAAATA
AGCACACGTCTGAACTCTTTCCCTACACGACGCTCTTCCGATCTNNNTCACTTAGCTTTTCCCTAAAATA
AGCACACGTCTGAACTCTTTCCCTACACGACGCTCTTCCGATCTNNNTCACTTAGCTTTTCCCTAAAATA
AGCACACGTCTGAACTCTTTCCCTACACGACGCTCTTCCGATCTNNNTCACTTAGCTTTTCCCTAAAATA
GAGCACACGTCTGAACTCTTTCCCTACACGACGCTCTTCCGATCTNNNGCCTATGCTGTTCTTTGGAAT
GAGCACACGTCTGAACTCTTTCCCTACACGACGCTCTTCCGATCTNNNGCCTATGCTGTTCTTTGGAAT
GAGCACACGTCTGAACTCTTTCCCTACACGACGCTCTTCCGATCTNNNGCCTATGCTGTTCTTTGGAAT
GAGCACACGTCTGAACTCTTTCCCTACACGACGCTCTTCCGATCTNNNGCCTATGCTGTTCTTTGGAAT
AGCACACGTCTGAACTCTTTCCCTACACGACGCTCTTCCGATCTNNNGATACGGTGTCAATGCAACAA
AGCACACGTCTGAACTCTTTCCCTACACGACGCTCTTCCGATCTNNNGATACGGTGTCAATGCAACAA
AGCACACGTCTGAACTCTTTCCCTACACGACGCTCTTCCGATCTNNNGATACGGTGTCAATGCAACAA
AGCACACGTCTGAACTCTTTCCCTACACGACGCTCTTCCGATCTNNNGATACGGTGTCAATGCAACAA

AGCACACGTCTGAACTCTTTCCCTACACGACGCTCTTCCGATCTNNNAAAAAGGAATGCTTACCAGTG
AGCACACGTCTGAACTCTTTCCCTACACGACGCTCTTCCGATCTNNNAAAAAGGAATGCTTACCAGTG
AGCACACGTCTGAACTCTTTCCCTACACGACGCTCTTCCGATCTNNNAAAAAGGAATGCTTACCAGTG
AGCACACGTCTGAACTCTTTCCCTACACGACGCTCTTCCGATCTNNNAAAAAGGAATGCTTACCAGTG
AGCACACGTCTGAACTCTTTCCCTACACGACGCTCTTCCGATCTNNNCGTTTGTGGAAACTGCTCTACT
AGCACACGTCTGAACTCTTTCCCTACACGACGCTCTTCCGATCTNNNCGTTTGTGGAAACTGCTCTACT
AGCACACGTCTGAACTCTTTCCCTACACGACGCTCTTCCGATCTNNNCGTTTGTGGAAACTGCTCTACT
AGCACACGTCTGAACTCTTTCCCTACACGACGCTCTTCCGATCTNNNCGTTTGTGGAAACTGCTCTACT
AGCACACGTCTGAACTCTTTCCCTACACGACGCTCTTCCGATCTNNNAGGACCTGATTTCCTTACTGCC
AGCACACGTCTGAACTCTTTCCCTACACGACGCTCTTCCGATCTNNNAGGACCTGATTTCCTTACTGCC
AGCACACGTCTGAACTCTTTCCCTACACGACGCTCTTCCGATCTNNNAGGACCTGATTTCCTTACTGCC
AGCACACGTCTGAACTCTTTCCCTACACGACGCTCTTCCGATCTNNNAGGACCTGATTTCCTTACTGCC
AGCACACGTCTGAACTCTTTCCCTACACGACGCTCTTCCGATCTNNNTGTGGATCATGAATCTGCAGT
AGCACACGTCTGAACTCTTTCCCTACACGACGCTCTTCCGATCTNNNTGTGGATCATGAATCTGCAGT
AGCACACGTCTGAACTCTTTCCCTACACGACGCTCTTCCGATCTNNNTGTGGATCATGAATCTGCAGT
AGCACACGTCTGAACTCTTTCCCTACACGACGCTCTTCCGATCTNNNTGTGGATCATGAATCTGCAGT
GAGCACACGTCTGAACTCTTTCCCTACACGACGCTCTTCCGATCTNNNGGAGCTGGAGGCCATTATT
GAGCACACGTCTGAACTCTTTCCCTACACGACGCTCTTCCGATCTNNNGGAGCTGGAGGCCATTATT
GAGCACACGTCTGAACTCTTTCCCTACACGACGCTCTTCCGATCTNNNGGAGCTGGAGGCCATTATT
GAGCACACGTCTGAACTCTTTCCCTACACGACGCTCTTCCGATCTNNNGGAGCTGGAGGCCATTATT
AGCACACGTCTGAACTCTTTCCCTACACGACGCTCTTCCGATCTNNNAAAAATTAGCTGGGAATGGT
AGCACACGTCTGAACTCTTTCCCTACACGACGCTCTTCCGATCTNNNAAAAATTAGCTGGGAATGGT
AGCACACGTCTGAACTCTTTCCCTACACGACGCTCTTCCGATCTNNNAAAAATTAGCTGGGAATGGT
AGCACACGTCTGAACTCTTTCCCTACACGACGCTCTTCCGATCTNNNAAAAATTAGCTGGGAATGGT
AGAGCACACGTCTGAACTCTTTCCCTACACGACGCTCTTCCGATCTNNNTCAGTCTAAAGTGTGTGTC
AGAGCACACGTCTGAACTCTTTCCCTACACGACGCTCTTCCGATCTNNNTCAGTCTAAAGTGTGTGTC
AGAGCACACGTCTGAACTCTTTCCCTACACGACGCTCTTCCGATCTNNNTCAGTCTAAAGTGTGTGTC
AGAGCACACGTCTGAACTCTTTCCCTACACGACGCTCTTCCGATCTNNNTCAGTCTAAAGTGTGTGTC
GCACACGTCTGAACTCTTTCCCTACACGACGCTCTTCCGATCTNNNCTCCTGAGTAGCTGGGATTACA
GCACACGTCTGAACTCTTTCCCTACACGACGCTCTTCCGATCTNNNCTCCTGAGTAGCTGGGATTACA
GCACACGTCTGAACTCTTTCCCTACACGACGCTCTTCCGATCTNNNCTCCTGAGTAGCTGGGATTACA
GCACACGTCTGAACTCTTTCCCTACACGACGCTCTTCCGATCTNNNCTCCTGAGTAGCTGGGATTACA
GCACACGTCTGAACTCTTTCCCTACACGACGCTCTTCCGATCTNNNTGGAGGGAAGAATCTCAGAGCT
GCACACGTCTGAACTCTTTCCCTACACGACGCTCTTCCGATCTNNNTGGAGGGAAGAATCTCAGAGCT
GCACACGTCTGAACTCTTTCCCTACACGACGCTCTTCCGATCTNNNTGGAGGGAAGAATCTCAGAGCT
GCACACGTCTGAACTCTTTCCCTACACGACGCTCTTCCGATCTNNNTGGAGGGAAGAATCTCAGAGCT
GAGCACACGTCTGAACTCTTTCCCTACACGACGCTCTTCCGATCTNNNCTCCTGTAGTCCCAGCTACTC
GAGCACACGTCTGAACTCTTTCCCTACACGACGCTCTTCCGATCTNNNCTCCTGTAGTCCCAGCTACTC
GAGCACACGTCTGAACTCTTTCCCTACACGACGCTCTTCCGATCTNNNCTCCTGTAGTCCCAGCTACTC
GAGCACACGTCTGAACTCTTTCCCTACACGACGCTCTTCCGATCTNNNCTCCTGTAGTCCCAGCTACTC
AGCACACGTCTGAACTCTTTCCCTACACGACGCTCTTCCGATCTNNNGCAACAGATTTTTCTGGGATC
AGCACACGTCTGAACTCTTTCCCTACACGACGCTCTTCCGATCTNNNGCAACAGATTTTTCTGGGATC
AGCACACGTCTGAACTCTTTCCCTACACGACGCTCTTCCGATCTNNNGCAACAGATTTTTCTGGGATC

AGCACACGTCTGAACTCTTTCCCTACACGACGCTCTTCCGATCTNNNGCAACAGATTTTTCTGGGATCC
GAGCACACGTCTGAACTCTTTCCCTACACGACGCTCTTCCGATCTNNNTACCTGTAATCCCAGCTATTCG
GAGCACACGTCTGAACTCTTTCCCTACACGACGCTCTTCCGATCTNNNTACCTGTAATCCCAGCTATTCG
GAGCACACGTCTGAACTCTTTCCCTACACGACGCTCTTCCGATCTNNNTACCTGTAATCCCAGCTATTCG
GAGCACACGTCTGAACTCTTTCCCTACACGACGCTCTTCCGATCTNNNTACCTGTAATCCCAGCTATTCG
AGAGCACACGTCTGAACTCTTTCCCTACACGACGCTCTTCCGATCTNNNAAAAACTATCTGTGGCAGTG
AGAGCACACGTCTGAACTCTTTCCCTACACGACGCTCTTCCGATCTNNNAAAAACTATCTGTGGCAGTG
AGAGCACACGTCTGAACTCTTTCCCTACACGACGCTCTTCCGATCTNNNAAAAACTATCTGTGGCAGTG
AGAGCACACGTCTGAACTCTTTCCCTACACGACGCTCTTCCGATCTNNNAAAAACTATCTGTGGCAGTG
GAGCACACGTCTGAACTCTTTCCCTACACGACGCTCTTCCGATCTNNNGCACTCATATGTTCATTGCAGC
GAGCACACGTCTGAACTCTTTCCCTACACGACGCTCTTCCGATCTNNNGCACTCATATGTTCATTGCAGC
GAGCACACGTCTGAACTCTTTCCCTACACGACGCTCTTCCGATCTNNNGCACTCATATGTTCATTGCAGC
GAGCACACGTCTGAACTCTTTCCCTACACGACGCTCTTCCGATCTNNNGCACTCATATGTTCATTGCAGC
CACACGTCTGAACTCTTTCCCTACACGACGCTCTTCCGATCTNNNCGAGTAACTGGGACAACAGGTGC
CACACGTCTGAACTCTTTCCCTACACGACGCTCTTCCGATCTNNNCGAGTAACTGGGACAACAGGTGC
CACACGTCTGAACTCTTTCCCTACACGACGCTCTTCCGATCTNNNCGAGTAACTGGGACAACAGGTGC
CACACGTCTGAACTCTTTCCCTACACGACGCTCTTCCGATCTNNNCGAGTAACTGGGACAACAGGTGC
GAGCACACGTCTGAACTCTTTCCCTACACGACGCTCTTCCGATCTNNNCTATCTCTCCCCTTCTGCCATTC
GAGCACACGTCTGAACTCTTTCCCTACACGACGCTCTTCCGATCTNNNCTATCTCTCCCCTTCTGCCATTC
GAGCACACGTCTGAACTCTTTCCCTACACGACGCTCTTCCGATCTNNNCTATCTCTCCCCTTCTGCCATTC
GAGCACACGTCTGAACTCTTTCCCTACACGACGCTCTTCCGATCTNNNCTATCTCTCCCCTTCTGCCATTC
AGCACACGTCTGAACTCTTTCCCTACACGACGCTCTTCCGATCTNNNGGCTACAATATGGACCAACCTGA
AGCACACGTCTGAACTCTTTCCCTACACGACGCTCTTCCGATCTNNNGGCTACAATATGGACCAACCTGA
AGCACACGTCTGAACTCTTTCCCTACACGACGCTCTTCCGATCTNNNGGCTACAATATGGACCAACCTGA
AGCACACGTCTGAACTCTTTCCCTACACGACGCTCTTCCGATCTNNNGGCTACAATATGGACCAACCTGA
AGCACACGTCTGAACTCTTTCCCTACACGACGCTCTTCCGATCTNNNTTGAAACTTGCTGGCAGTTATG
AGCACACGTCTGAACTCTTTCCCTACACGACGCTCTTCCGATCTNNNTTGAAACTTGCTGGCAGTTATG
AGCACACGTCTGAACTCTTTCCCTACACGACGCTCTTCCGATCTNNNTTGAAACTTGCTGGCAGTTATG
AGCACACGTCTGAACTCTTTCCCTACACGACGCTCTTCCGATCTNNNTTGAAACTTGCTGGCAGTTATG
AGCACACGTCTGAACTCTTTCCCTACACGACGCTCTTCCGATCTNNNAGAATCATTTGAACCCGGGAGA
AGCACACGTCTGAACTCTTTCCCTACACGACGCTCTTCCGATCTNNNAGAATCATTTGAACCCGGGAGA
AGCACACGTCTGAACTCTTTCCCTACACGACGCTCTTCCGATCTNNNAGAATCATTTGAACCCGGGAGA
AGCACACGTCTGAACTCTTTCCCTACACGACGCTCTTCCGATCTNNNAGAATCATTTGAACCCGGGAGA
GAGCACACGTCTGAACTCTTTCCCTACACGACGCTCTTCCGATCTNNNCTTCTGGAGAGGCAAGAACTC
GAGCACACGTCTGAACTCTTTCCCTACACGACGCTCTTCCGATCTNNNCTTCTGGAGAGGCAAGAACTC
GAGCACACGTCTGAACTCTTTCCCTACACGACGCTCTTCCGATCTNNNCTTCTGGAGAGGCAAGAACTC
GAGCACACGTCTGAACTCTTTCCCTACACGACGCTCTTCCGATCTNNNCTTCTGGAGAGGCAAGAACTC
AGCACACGTCTGAACTCTTTCCCTACACGACGCTCTTCCGATCTNNNCTCCCACTTACCCCTTTCAATCAC
AGCACACGTCTGAACTCTTTCCCTACACGACGCTCTTCCGATCTNNNCTCCCACTTACCCCTTTCAATCAC
AGCACACGTCTGAACTCTTTCCCTACACGACGCTCTTCCGATCTNNNCTCCCACTTACCCCTTTCAATCAC
AGCACACGTCTGAACTCTTTCCCTACACGACGCTCTTCCGATCTNNNCTCCCACTTACCCCTTTCAATCAC
AGCACACGTCTGAACTCTTTCCCTACACGACGCTCTTCCGATCTNNNTAAGCTTTTTGAAGGCAAGGAG
AGCACACGTCTGAACTCTTTCCCTACACGACGCTCTTCCGATCTNNNTAAGCTTTTTGAAGGCAAGGAG

GCACACGTCTGAACTCTTTCCCTACACGACGCTCTTCCGATCTNNNTAAGCTTTTGAAGGCAAGGAG
GCACACGTCTGAACTCTTTCCCTACACGACGCTCTTCCGATCTNNNTAAGCTTTTGAAGGCAAGGAG
GCACACGTCTGAACTCTTTCCCTACACGACGCTCTTCCGATCTNNNTGTCTGAGAGTAGGCCGTAAG
GCACACGTCTGAACTCTTTCCCTACACGACGCTCTTCCGATCTNNNTGTCTGAGAGTAGGCCGTAAG
GCACACGTCTGAACTCTTTCCCTACACGACGCTCTTCCGATCTNNNTGTCTGAGAGTAGGCCGTAAG
GCACACGTCTGAACTCTTTCCCTACACGACGCTCTTCCGATCTNNNTGTCTGAGAGTAGGCCGTAAG
AGCACACGTCTGAACTCTTTCCCTACACGACGCTCTTCCGATCTNNNAGGTTCAAGCAATTCTCCTGT
AGCACACGTCTGAACTCTTTCCCTACACGACGCTCTTCCGATCTNNNAGGTTCAAGCAATTCTCCTGT
AGCACACGTCTGAACTCTTTCCCTACACGACGCTCTTCCGATCTNNNAGGTTCAAGCAATTCTCCTGT
AGCACACGTCTGAACTCTTTCCCTACACGACGCTCTTCCGATCTNNNAGGTTCAAGCAATTCTCCTGT
AGAGCACACGTCTGAACTCTTTCCCTACACGACGCTCTTCCGATCTNNNTGGTCCATGCAGAGTATAT
AGAGCACACGTCTGAACTCTTTCCCTACACGACGCTCTTCCGATCTNNNTGGTCCATGCAGAGTATAT
AGAGCACACGTCTGAACTCTTTCCCTACACGACGCTCTTCCGATCTNNNTGGTCCATGCAGAGTATAT
AGAGCACACGTCTGAACTCTTTCCCTACACGACGCTCTTCCGATCTNNNTGGTCCATGCAGAGTATAT
AGCACACGTCTGAACTCTTTCCCTACACGACGCTCTTCCGATCTNNNCCTCAGAACTTTGCAGAGTCTC
AGCACACGTCTGAACTCTTTCCCTACACGACGCTCTTCCGATCTNNNCCTCAGAACTTTGCAGAGTCTC
AGCACACGTCTGAACTCTTTCCCTACACGACGCTCTTCCGATCTNNNCCTCAGAACTTTGCAGAGTCTC
AGCACACGTCTGAACTCTTTCCCTACACGACGCTCTTCCGATCTNNNCCTCAGAACTTTGCAGAGTCTC
GAGCACACGTCTGAACTCTTTCCCTACACGACGCTCTTCCGATCTNNNCTGTCTGACAGCTTGAAGAG
GAGCACACGTCTGAACTCTTTCCCTACACGACGCTCTTCCGATCTNNNCTGTCTGACAGCTTGAAGAG
GAGCACACGTCTGAACTCTTTCCCTACACGACGCTCTTCCGATCTNNNCTGTCTGACAGCTTGAAGAG
GAGCACACGTCTGAACTCTTTCCCTACACGACGCTCTTCCGATCTNNNCTGTCTGACAGCTTGAAGAG
AGCACACGTCTGAACTCTTTCCCTACACGACGCTCTTCCGATCTNNNGAGAATGGCATGAACCTGGGA
AGCACACGTCTGAACTCTTTCCCTACACGACGCTCTTCCGATCTNNNGAGAATGGCATGAACCTGGGA
AGCACACGTCTGAACTCTTTCCCTACACGACGCTCTTCCGATCTNNNGAGAATGGCATGAACCTGGGA
AGCACACGTCTGAACTCTTTCCCTACACGACGCTCTTCCGATCTNNNGAGAATGGCATGAACCTGGGA
GAGCACACGTCTGAACTCTTTCCCTACACGACGCTCTTCCGATCTNNNTTGGACCAAAGGAAACCTA
GAGCACACGTCTGAACTCTTTCCCTACACGACGCTCTTCCGATCTNNNTTGGACCAAAGGAAACCTA
GAGCACACGTCTGAACTCTTTCCCTACACGACGCTCTTCCGATCTNNNTTGGACCAAAGGAAACCTA
GAGCACACGTCTGAACTCTTTCCCTACACGACGCTCTTCCGATCTNNNTTGGACCAAAGGAAACCTA
AGCACACGTCTGAACTCTTTCCCTACACGACGCTCTTCCGATCTNNNAAATTACTTGAGCCCAGGAGG
AGCACACGTCTGAACTCTTTCCCTACACGACGCTCTTCCGATCTNNNAAATTACTTGAGCCCAGGAGG
AGCACACGTCTGAACTCTTTCCCTACACGACGCTCTTCCGATCTNNNAAATTACTTGAGCCCAGGAGG
AGCACACGTCTGAACTCTTTCCCTACACGACGCTCTTCCGATCTNNNAAATTACTTGAGCCCAGGAGG
AGCACACGTCTGAACTCTTTCCCTACACGACGCTCTTCCGATCTNNNTCCAGGTTTAGGAGGGGTCT
AGCACACGTCTGAACTCTTTCCCTACACGACGCTCTTCCGATCTNNNTCCAGGTTTAGGAGGGGTCT
AGCACACGTCTGAACTCTTTCCCTACACGACGCTCTTCCGATCTNNNTCCAGGTTTAGGAGGGGTCT
AGCACACGTCTGAACTCTTTCCCTACACGACGCTCTTCCGATCTNNNTCCAGGTTTAGGAGGGGTCT
AGAGCACACGTCTGAACTCTTTCCCTACACGACGCTCTTCCGATCTNNNTCATCTGCCGACTCTCAACT
AGAGCACACGTCTGAACTCTTTCCCTACACGACGCTCTTCCGATCTNNNTCATCTGCCGACTCTCAACT
AGAGCACACGTCTGAACTCTTTCCCTACACGACGCTCTTCCGATCTNNNTCATCTGCCGACTCTCAACT
AGAGCACACGTCTGAACTCTTTCCCTACACGACGCTCTTCCGATCTNNNTCATCTGCCGACTCTCAACT
AGCACACGTCTGAACTCTTTCCCTACACGACGCTCTTCCGATCTNNNTCCAGGAGGATCTTCTCATCTC

AGCACACGTCTGAACTCTTTCCCTACACGACGCTCTTCCGATCTNNNTCCAGGAGGATCTTCTCATCTC
AGCACACGTCTGAACTCTTTCCCTACACGACGCTCTTCCGATCTNNNTCCAGGAGGATCTTCTCATCTC
AGCACACGTCTGAACTCTTTCCCTACACGACGCTCTTCCGATCTNNNTCCAGGAGGATCTTCTCATCTC
GAGCACACGTCTGAACTCTTTCCCTACACGACGCTCTTCCGATCTNNNCACGCTCAACAATTATCCACC
GAGCACACGTCTGAACTCTTTCCCTACACGACGCTCTTCCGATCTNNNCACGCTCAACAATTATCCACC
GAGCACACGTCTGAACTCTTTCCCTACACGACGCTCTTCCGATCTNNNCACGCTCAACAATTATCCACC
GAGCACACGTCTGAACTCTTTCCCTACACGACGCTCTTCCGATCTNNNCACGCTCAACAATTATCCACC
AGCACACGTCTGAACTCTTTCCCTACACGACGCTCTTCCGATCTNNNGGTTCAGATTTCACTGCCTCAT
AGCACACGTCTGAACTCTTTCCCTACACGACGCTCTTCCGATCTNNNGGTTCAGATTTCACTGCCTCAT
AGCACACGTCTGAACTCTTTCCCTACACGACGCTCTTCCGATCTNNNGGTTCAGATTTCACTGCCTCAT
AGCACACGTCTGAACTCTTTCCCTACACGACGCTCTTCCGATCTNNNGGTTCAGATTTCACTGCCTCAT
AGCACACGTCTGAACTCTTTCCCTACACGACGCTCTTCCGATCTNNNTAGTCCCAACTACTTGAGAGG
AGCACACGTCTGAACTCTTTCCCTACACGACGCTCTTCCGATCTNNNTAGTCCCAACTACTTGAGAGG
AGCACACGTCTGAACTCTTTCCCTACACGACGCTCTTCCGATCTNNNTAGTCCCAACTACTTGAGAGG
AGCACACGTCTGAACTCTTTCCCTACACGACGCTCTTCCGATCTNNNTAGTCCCAACTACTTGAGAGG
GCACACGTCTGAACTCTTTCCCTACACGACGCTCTTCCGATCTNNNTGGTTTCTGATCAGTACCTTGG
GCACACGTCTGAACTCTTTCCCTACACGACGCTCTTCCGATCTNNNTGGTTTCTGATCAGTACCTTGG
GCACACGTCTGAACTCTTTCCCTACACGACGCTCTTCCGATCTNNNTGGTTTCTGATCAGTACCTTGG
GCACACGTCTGAACTCTTTCCCTACACGACGCTCTTCCGATCTNNNTGGTTTCTGATCAGTACCTTGG
GAGCACACGTCTGAACTCTTTCCCTACACGACGCTCTTCCGATCTNNNAGCCATCTCATCAGCCATTTT
GAGCACACGTCTGAACTCTTTCCCTACACGACGCTCTTCCGATCTNNNAGCCATCTCATCAGCCATTTT
GAGCACACGTCTGAACTCTTTCCCTACACGACGCTCTTCCGATCTNNNAGCCATCTCATCAGCCATTTT
GAGCACACGTCTGAACTCTTTCCCTACACGACGCTCTTCCGATCTNNNAGCCATCTCATCAGCCATTTT
AGCACACGTCTGAACTCTTTCCCTACACGACGCTCTTCCGATCTNNNAGACAACCTCAACTAACTCCAG
AGCACACGTCTGAACTCTTTCCCTACACGACGCTCTTCCGATCTNNNAGACAACCTCAACTAACTCCAG
AGCACACGTCTGAACTCTTTCCCTACACGACGCTCTTCCGATCTNNNAGACAACCTCAACTAACTCCAG
AGCACACGTCTGAACTCTTTCCCTACACGACGCTCTTCCGATCTNNNAGACAACCTCAACTAACTCCAG
GCACACGTCTGAACTCTTTCCCTACACGACGCTCTTCCGATCTNNNTTTCCAGAACAGTCATGCAACA
GCACACGTCTGAACTCTTTCCCTACACGACGCTCTTCCGATCTNNNTTTCCAGAACAGTCATGCAACA
GCACACGTCTGAACTCTTTCCCTACACGACGCTCTTCCGATCTNNNTTTCCAGAACAGTCATGCAACA
GCACACGTCTGAACTCTTTCCCTACACGACGCTCTTCCGATCTNNNTTTCCAGAACAGTCATGCAACA
AGCACACGTCTGAACTCTTTCCCTACACGACGCTCTTCCGATCTNNNCTAGAAGAGAACCAGGGCCAA
AGCACACGTCTGAACTCTTTCCCTACACGACGCTCTTCCGATCTNNNCTAGAAGAGAACCAGGGCCAA
AGCACACGTCTGAACTCTTTCCCTACACGACGCTCTTCCGATCTNNNCTAGAAGAGAACCAGGGCCAA
AGCACACGTCTGAACTCTTTCCCTACACGACGCTCTTCCGATCTNNNCTAGAAGAGAACCAGGGCCAA
AGCACACGTCTGAACTCTTTCCCTACACGACGCTCTTCCGATCTNNNATCCCAGCACCATTTGTTGAAT
AGCACACGTCTGAACTCTTTCCCTACACGACGCTCTTCCGATCTNNNATCCCAGCACCATTTGTTGAAT
AGCACACGTCTGAACTCTTTCCCTACACGACGCTCTTCCGATCTNNNATCCCAGCACCATTTGTTGAAT
AGCACACGTCTGAACTCTTTCCCTACACGACGCTCTTCCGATCTNNNATCCCAGCACCATTTGTTGAAT
GAGCACACGTCTGAACTCTTTCCCTACACGACGCTCTTCCGATCTNNNCATCTTTCAGGAGCAAGATC
GAGCACACGTCTGAACTCTTTCCCTACACGACGCTCTTCCGATCTNNNCATCTTTCAGGAGCAAGATC
GAGCACACGTCTGAACTCTTTCCCTACACGACGCTCTTCCGATCTNNNCATCTTTCAGGAGCAAGATC
GAGCACACGTCTGAACTCTTTCCCTACACGACGCTCTTCCGATCTNNNCATCTTTCAGGAGCAAGATC

AGCACACGTCTGAACTCTTTCCCTACACGACGCTCTTCCGATCTNNNTGTAGTCCCAGCTACTCAGGT
AGCACACGTCTGAACTCTTTCCCTACACGACGCTCTTCCGATCTNNNTGTAGTCCCAGCTACTCAGGT
AGCACACGTCTGAACTCTTTCCCTACACGACGCTCTTCCGATCTNNNTGTAGTCCCAGCTACTCAGGT
AGCACACGTCTGAACTCTTTCCCTACACGACGCTCTTCCGATCTNNNTGTAGTCCCAGCTACTCAGGT
AGCACACGTCTGAACTCTTTCCCTACACGACGCTCTTCCGATCTNNNAGTGGTCAAAAGCATGGACT
AGCACACGTCTGAACTCTTTCCCTACACGACGCTCTTCCGATCTNNNAGTGGTCAAAAGCATGGACT
AGCACACGTCTGAACTCTTTCCCTACACGACGCTCTTCCGATCTNNNAGTGGTCAAAAGCATGGACT
AGCACACGTCTGAACTCTTTCCCTACACGACGCTCTTCCGATCTNNNAGTGGTCAAAAGCATGGACT
GCACACGTCTGAACTCTTTCCCTACACGACGCTCTTCCGATCTNNNACAAAAATTAGCCAGGCCTGTT
GCACACGTCTGAACTCTTTCCCTACACGACGCTCTTCCGATCTNNNACAAAAATTAGCCAGGCCTGTT
GCACACGTCTGAACTCTTTCCCTACACGACGCTCTTCCGATCTNNNACAAAAATTAGCCAGGCCTGTT
GCACACGTCTGAACTCTTTCCCTACACGACGCTCTTCCGATCTNNNACAAAAATTAGCCAGGCCTGTT
AGCACACGTCTGAACTCTTTCCCTACACGACGCTCTTCCGATCTNNNACCATCTCACACCAGTCAGAAT
AGCACACGTCTGAACTCTTTCCCTACACGACGCTCTTCCGATCTNNNACCATCTCACACCAGTCAGAAT
AGCACACGTCTGAACTCTTTCCCTACACGACGCTCTTCCGATCTNNNACCATCTCACACCAGTCAGAAT
AGCACACGTCTGAACTCTTTCCCTACACGACGCTCTTCCGATCTNNNACCATCTCACACCAGTCAGAAT
GCACACGTCTGAACTCTTTCCCTACACGACGCTCTTCCGATCTNNNGAGGCAGGAGAATGGTGTGA
GCACACGTCTGAACTCTTTCCCTACACGACGCTCTTCCGATCTNNNGAGGCAGGAGAATGGTGTGA
GCACACGTCTGAACTCTTTCCCTACACGACGCTCTTCCGATCTNNNGAGGCAGGAGAATGGTGTGA
GCACACGTCTGAACTCTTTCCCTACACGACGCTCTTCCGATCTNNNGAGGCAGGAGAATGGTGTGA
AGCACACGTCTGAACTCTTTCCCTACACGACGCTCTTCCGATCTNNNTTTTGGTAGAGACAGGGTTTT
AGCACACGTCTGAACTCTTTCCCTACACGACGCTCTTCCGATCTNNNTTTTGGTAGAGACAGGGTTTT
AGCACACGTCTGAACTCTTTCCCTACACGACGCTCTTCCGATCTNNNTTTTGGTAGAGACAGGGTTTT
AGCACACGTCTGAACTCTTTCCCTACACGACGCTCTTCCGATCTNNNTTTTGGTAGAGACAGGGTTTT
AGAGCACACGTCTGAACTCTTTCCCTACACGACGCTCTTCCGATCTNNNACACACACACACACACACAA
AGAGCACACGTCTGAACTCTTTCCCTACACGACGCTCTTCCGATCTNNNACACACACACACACACACAA
AGAGCACACGTCTGAACTCTTTCCCTACACGACGCTCTTCCGATCTNNNACACACACACACACACACAA
AGAGCACACGTCTGAACTCTTTCCCTACACGACGCTCTTCCGATCTNNNACACACACACACACACACAA
AGCACACGTCTGAACTCTTTCCCTACACGACGCTCTTCCGATCTNNNGATTCCTGGCTGGGGTCTGTC
AGCACACGTCTGAACTCTTTCCCTACACGACGCTCTTCCGATCTNNNGATTCCTGGCTGGGGTCTGTC
AGCACACGTCTGAACTCTTTCCCTACACGACGCTCTTCCGATCTNNNGATTCCTGGCTGGGGTCTGTC
AGCACACGTCTGAACTCTTTCCCTACACGACGCTCTTCCGATCTNNNGATTCCTGGCTGGGGTCTGTC
AGCACACGTCTGAACTCTTTCCCTACACGACGCTCTTCCGATCTNNNGTTGCAGTGAGCCAAGATTGT
AGCACACGTCTGAACTCTTTCCCTACACGACGCTCTTCCGATCTNNNGTTGCAGTGAGCCAAGATTGT
AGCACACGTCTGAACTCTTTCCCTACACGACGCTCTTCCGATCTNNNGTTGCAGTGAGCCAAGATTGT
AGCACACGTCTGAACTCTTTCCCTACACGACGCTCTTCCGATCTNNNGTTGCAGTGAGCCAAGATTGT
GCACACGTCTGAACTCTTTCCCTACACGACGCTCTTCCGATCTNNNGCTGAGACAGGAGAATCACTT
GCACACGTCTGAACTCTTTCCCTACACGACGCTCTTCCGATCTNNNGCTGAGACAGGAGAATCACTT
GCACACGTCTGAACTCTTTCCCTACACGACGCTCTTCCGATCTNNNGCTGAGACAGGAGAATCACTT
GCACACGTCTGAACTCTTTCCCTACACGACGCTCTTCCGATCTNNNGCTGAGACAGGAGAATCACTT
GAGCACACGTCTGAACTCTTTCCCTACACGACGCTCTTCCGATCTNNNGTGGATCACCTATGGTCAAG
GAGCACACGTCTGAACTCTTTCCCTACACGACGCTCTTCCGATCTNNNGTGGATCACCTATGGTCAAG
GAGCACACGTCTGAACTCTTTCCCTACACGACGCTCTTCCGATCTNNNGTGGATCACCTATGGTCAAG

GAGCACACGTCTGAACTCTTTCCCTACACGACGCTCTTCCGATCTNNNGTGGATCACCTATGGTCAAG
GCACACGTCTGAACTCTTTCCCTACACGACGCTCTTCCGATCTNNNGGTGACAAAGTGAGACTCAGTC
GCACACGTCTGAACTCTTTCCCTACACGACGCTCTTCCGATCTNNNGGTGACAAAGTGAGACTCAGTC
GCACACGTCTGAACTCTTTCCCTACACGACGCTCTTCCGATCTNNNGGTGACAAAGTGAGACTCAGTC
GCACACGTCTGAACTCTTTCCCTACACGACGCTCTTCCGATCTNNNGGTGACAAAGTGAGACTCAGTC
AGCACACGTCTGAACTCTTTCCCTACACGACGCTCTTCCGATCTNNNGAGGTTGCAGTGAGCTAAGAT
AGCACACGTCTGAACTCTTTCCCTACACGACGCTCTTCCGATCTNNNGAGGTTGCAGTGAGCTAAGAT
AGCACACGTCTGAACTCTTTCCCTACACGACGCTCTTCCGATCTNNNGAGGTTGCAGTGAGCTAAGAT
AGCACACGTCTGAACTCTTTCCCTACACGACGCTCTTCCGATCTNNNGAGGTTGCAGTGAGCTAAGAT
AGCACACGTCTGAACTCTTTCCCTACACGACGCTCTTCCGATCTNNNCCCAACTAAGCATGGTGATTTC
AGCACACGTCTGAACTCTTTCCCTACACGACGCTCTTCCGATCTNNNCCCAACTAAGCATGGTGATTTC
AGCACACGTCTGAACTCTTTCCCTACACGACGCTCTTCCGATCTNNNCCCAACTAAGCATGGTGATTTC
AGCACACGTCTGAACTCTTTCCCTACACGACGCTCTTCCGATCTNNNCCCAACTAAGCATGGTGATTTC
GCACACGTCTGAACTCTTTCCCTACACGACGCTCTTCCGATCTNNNCTTTCTTGGGGTGCTGAGTCTAT
GCACACGTCTGAACTCTTTCCCTACACGACGCTCTTCCGATCTNNNCTTTCTTGGGGTGCTGAGTCTAT
GCACACGTCTGAACTCTTTCCCTACACGACGCTCTTCCGATCTNNNCTTTCTTGGGGTGCTGAGTCTAT
GCACACGTCTGAACTCTTTCCCTACACGACGCTCTTCCGATCTNNNCTTTCTTGGGGTGCTGAGTCTAT
CACACGTCTGAACTCTTTCCCTACACGACGCTCTTCCGATCTNNNCTCTTCTGTTCTCCCCTCCCTTCCAT
CACACGTCTGAACTCTTTCCCTACACGACGCTCTTCCGATCTNNNCTCTTCTGTTCTCCCCTCCCTTCCAT
CACACGTCTGAACTCTTTCCCTACACGACGCTCTTCCGATCTNNNCTCTTCTGTTCTCCCCTCCCTTCCAT
CACACGTCTGAACTCTTTCCCTACACGACGCTCTTCCGATCTNNNCTCTTCTGTTCTCCCCTCCCTTCCAT
AGAGCACACGTCTGAACTCTTTCCCTACACGACGCTCTTCCGATCTNNNCTAATTCAGGCAGGGCAAA
AGAGCACACGTCTGAACTCTTTCCCTACACGACGCTCTTCCGATCTNNNCTAATTCAGGCAGGGCAAA
AGAGCACACGTCTGAACTCTTTCCCTACACGACGCTCTTCCGATCTNNNCTAATTCAGGCAGGGCAAA
AGAGCACACGTCTGAACTCTTTCCCTACACGACGCTCTTCCGATCTNNNCTAATTCAGGCAGGGCAAA
GAGCACACGTCTGAACTCTTTCCCTACACGACGCTCTTCCGATCTNNNCATGATCTCGGCTCACTGTAA
GAGCACACGTCTGAACTCTTTCCCTACACGACGCTCTTCCGATCTNNNCATGATCTCGGCTCACTGTAA
GAGCACACGTCTGAACTCTTTCCCTACACGACGCTCTTCCGATCTNNNCATGATCTCGGCTCACTGTAA
GAGCACACGTCTGAACTCTTTCCCTACACGACGCTCTTCCGATCTNNNCATGATCTCGGCTCACTGTAA
AGCACACGTCTGAACTCTTTCCCTACACGACGCTCTTCCGATCTNNNATTCCACTTGGTCATGGTGTG
AGCACACGTCTGAACTCTTTCCCTACACGACGCTCTTCCGATCTNNNATTCCACTTGGTCATGGTGTG
AGCACACGTCTGAACTCTTTCCCTACACGACGCTCTTCCGATCTNNNATTCCACTTGGTCATGGTGTG
AGCACACGTCTGAACTCTTTCCCTACACGACGCTCTTCCGATCTNNNATTCCACTTGGTCATGGTGTG
AGAGCACACGTCTGAACTCTTTCCCTACACGACGCTCTTCCGATCTNNNAGAATCACTTGAACCTGGG
AGAGCACACGTCTGAACTCTTTCCCTACACGACGCTCTTCCGATCTNNNAGAATCACTTGAACCTGGG
AGAGCACACGTCTGAACTCTTTCCCTACACGACGCTCTTCCGATCTNNNAGAATCACTTGAACCTGGG
AGAGCACACGTCTGAACTCTTTCCCTACACGACGCTCTTCCGATCTNNNAGAATCACTTGAACCTGGG
AGAGCACACGTCTGAACTCTTTCCCTACACGACGCTCTTCCGATCTNNNCGCTGATCTGTTTTGTTTGT
AGAGCACACGTCTGAACTCTTTCCCTACACGACGCTCTTCCGATCTNNNCGCTGATCTGTTTTGTTTGT
AGAGCACACGTCTGAACTCTTTCCCTACACGACGCTCTTCCGATCTNNNCGCTGATCTGTTTTGTTTGT
AGAGCACACGTCTGAACTCTTTCCCTACACGACGCTCTTCCGATCTNNNCGCTGATCTGTTTTGTTTGT
GCACACGTCTGAACTCTTTCCCTACACGACGCTCTTCCGATCTNNNCTCCACCAGGTCTGTGATGTTTT
GCACACGTCTGAACTCTTTCCCTACACGACGCTCTTCCGATCTNNNCTCCACCAGGTCTGTGATGTTTT

AGCACACGTCTGAACTCTTTCCCTACACGACGCTCTTCCGATCTNNNCTCCACCAGGTCTGTGATGTTTT
AGCACACGTCTGAACTCTTTCCCTACACGACGCTCTTCCGATCTNNNCTCCACCAGGTCTGTGATGTTTT
AGCACACGTCTGAACTCTTTCCCTACACGACGCTCTTCCGATCTNNNTGAGAGCAGAGACATCTTTCCT
AGCACACGTCTGAACTCTTTCCCTACACGACGCTCTTCCGATCTNNNTGAGAGCAGAGACATCTTTCCT
AGCACACGTCTGAACTCTTTCCCTACACGACGCTCTTCCGATCTNNNTGAGAGCAGAGACATCTTTCCT
AGCACACGTCTGAACTCTTTCCCTACACGACGCTCTTCCGATCTNNNTGAGAGCAGAGACATCTTTCCT
AGCACACGTCTGAACTCTTTCCCTACACGACGCTCTTCCGATCTNNNGTTCAAGAGATTCTCCTGCCTCA
AGCACACGTCTGAACTCTTTCCCTACACGACGCTCTTCCGATCTNNNGTTCAAGAGATTCTCCTGCCTCA
AGCACACGTCTGAACTCTTTCCCTACACGACGCTCTTCCGATCTNNNGTTCAAGAGATTCTCCTGCCTCA
AGCACACGTCTGAACTCTTTCCCTACACGACGCTCTTCCGATCTNNNGTTCAAGAGATTCTCCTGCCTCA
GAGCACACGTCTGAACTCTTTCCCTACACGACGCTCTTCCGATCTNNNCACCTGTACTCCCAGCTATTCA
GAGCACACGTCTGAACTCTTTCCCTACACGACGCTCTTCCGATCTNNNCACCTGTACTCCCAGCTATTCA
GAGCACACGTCTGAACTCTTTCCCTACACGACGCTCTTCCGATCTNNNCACCTGTACTCCCAGCTATTCA
GAGCACACGTCTGAACTCTTTCCCTACACGACGCTCTTCCGATCTNNNCACCTGTACTCCCAGCTATTCA
GAGCACACGTCTGAACTCTTTCCCTACACGACGCTCTTCCGATCTNNNAAAATTAGCTGGGCATAGTC
GAGCACACGTCTGAACTCTTTCCCTACACGACGCTCTTCCGATCTNNNAAAATTAGCTGGGCATAGTC
GAGCACACGTCTGAACTCTTTCCCTACACGACGCTCTTCCGATCTNNNAAAATTAGCTGGGCATAGTC
GAGCACACGTCTGAACTCTTTCCCTACACGACGCTCTTCCGATCTNNNAAAATTAGCTGGGCATAGTC
GAGCACACGTCTGAACTCTTTCCCTACACGACGCTCTTCCGATCTNNNTAGGAGAATCGTTTGAACCTG
GAGCACACGTCTGAACTCTTTCCCTACACGACGCTCTTCCGATCTNNNTAGGAGAATCGTTTGAACCTG
GAGCACACGTCTGAACTCTTTCCCTACACGACGCTCTTCCGATCTNNNTAGGAGAATCGTTTGAACCTG
GAGCACACGTCTGAACTCTTTCCCTACACGACGCTCTTCCGATCTNNNTAGGAGAATCGTTTGAACCTG
GAGCACACGTCTGAACTCTTTCCCTACACGACGCTCTTCCGATCTNNNTGCTGATCCAGAGAAATGAA
GAGCACACGTCTGAACTCTTTCCCTACACGACGCTCTTCCGATCTNNNTGCTGATCCAGAGAAATGAA
GAGCACACGTCTGAACTCTTTCCCTACACGACGCTCTTCCGATCTNNNTGCTGATCCAGAGAAATGAA
GAGCACACGTCTGAACTCTTTCCCTACACGACGCTCTTCCGATCTNNNTGCTGATCCAGAGAAATGAA
AGAGCACACGTCTGAACTCTTTCCCTACACGACGCTCTTCCGATCTNNNAACTTGCCAGACACTACATT
AGAGCACACGTCTGAACTCTTTCCCTACACGACGCTCTTCCGATCTNNNAACTTGCCAGACACTACATT
AGAGCACACGTCTGAACTCTTTCCCTACACGACGCTCTTCCGATCTNNNAACTTGCCAGACACTACATT
AGAGCACACGTCTGAACTCTTTCCCTACACGACGCTCTTCCGATCTNNNAACTTGCCAGACACTACATT
GAGCACACGTCTGAACTCTTTCCCTACACGACGCTCTTCCGATCTNNNCTCAAACAGGCGCAATTCAAT
GAGCACACGTCTGAACTCTTTCCCTACACGACGCTCTTCCGATCTNNNCTCAAACAGGCGCAATTCAAT
GAGCACACGTCTGAACTCTTTCCCTACACGACGCTCTTCCGATCTNNNCTCAAACAGGCGCAATTCAAT
GAGCACACGTCTGAACTCTTTCCCTACACGACGCTCTTCCGATCTNNNCTCAAACAGGCGCAATTCAAT
AGCACACGTCTGAACTCTTTCCCTACACGACGCTCTTCCGATCTNNNTTTTTGCTATGTTTCCCAGGCT
AGCACACGTCTGAACTCTTTCCCTACACGACGCTCTTCCGATCTNNNTTTTTGCTATGTTTCCCAGGCT
AGCACACGTCTGAACTCTTTCCCTACACGACGCTCTTCCGATCTNNNTTTTTGCTATGTTTCCCAGGCT
AGCACACGTCTGAACTCTTTCCCTACACGACGCTCTTCCGATCTNNNTTTTTGCTATGTTTCCCAGGCT
AGAGCACACGTCTGAACTCTTTCCCTACACGACGCTCTTCCGATCTNNNTCAGGCTTCCTAAGATGCA
AGAGCACACGTCTGAACTCTTTCCCTACACGACGCTCTTCCGATCTNNNTCAGGCTTCCTAAGATGCA
AGAGCACACGTCTGAACTCTTTCCCTACACGACGCTCTTCCGATCTNNNTCAGGCTTCCTAAGATGCA
AGAGCACACGTCTGAACTCTTTCCCTACACGACGCTCTTCCGATCTNNNTCAGGCTTCCTAAGATGCA
AGAGCACACGTCTGAACTCTTTCCCTACACGACGCTCTTCCGATCTNNNGATCTCAAACTTCTCAGGCT

GAGCACACGTCTGAACTCTTTCCCTACACGACGCTCTTCCGATCTNNNGATCTCAAACTTCTCAGGCT
GAGCACACGTCTGAACTCTTTCCCTACACGACGCTCTTCCGATCTNNNGATCTCAAACTTCTCAGGCT
GAGCACACGTCTGAACTCTTTCCCTACACGACGCTCTTCCGATCTNNNGATCTCAAACTTCTCAGGCT
GAGCACACGTCTGAACTCTTTCCCTACACGACGCTCTTCCGATCTNNNAGACTACCCAGAAAAACCCAT
GAGCACACGTCTGAACTCTTTCCCTACACGACGCTCTTCCGATCTNNNAGACTACCCAGAAAAACCCAT
GAGCACACGTCTGAACTCTTTCCCTACACGACGCTCTTCCGATCTNNNAGACTACCCAGAAAAACCCAT
GAGCACACGTCTGAACTCTTTCCCTACACGACGCTCTTCCGATCTNNNAGACTACCCAGAAAAACCCAT
AGCACACGTCTGAACTCTTTCCCTACACGACGCTCTTCCGATCTNNNGCAGGAGAATTGCTTGAACCT
AGCACACGTCTGAACTCTTTCCCTACACGACGCTCTTCCGATCTNNNGCAGGAGAATTGCTTGAACCT
AGCACACGTCTGAACTCTTTCCCTACACGACGCTCTTCCGATCTNNNGCAGGAGAATTGCTTGAACCT
AGCACACGTCTGAACTCTTTCCCTACACGACGCTCTTCCGATCTNNNGCAGGAGAATTGCTTGAACCT
AGCACACGTCTGAACTCTTTCCCTACACGACGCTCTTCCGATCTNNNAGGAGAATGGCGTGAACCTG
AGCACACGTCTGAACTCTTTCCCTACACGACGCTCTTCCGATCTNNNAGGAGAATGGCGTGAACCTG
AGCACACGTCTGAACTCTTTCCCTACACGACGCTCTTCCGATCTNNNAGGAGAATGGCGTGAACCTG
AGCACACGTCTGAACTCTTTCCCTACACGACGCTCTTCCGATCTNNNAGGAGAATGGCGTGAACCTG
GAGCACACGTCTGAACTCTTTCCCTACACGACGCTCTTCCGATCTNNNAGCAGGAGAATCACTTGAA
GAGCACACGTCTGAACTCTTTCCCTACACGACGCTCTTCCGATCTNNNAGCAGGAGAATCACTTGAA
GAGCACACGTCTGAACTCTTTCCCTACACGACGCTCTTCCGATCTNNNAGCAGGAGAATCACTTGAA
GAGCACACGTCTGAACTCTTTCCCTACACGACGCTCTTCCGATCTNNNAGCAGGAGAATCACTTGAA
GCACACGTCTGAACTCTTTCCCTACACGACGCTCTTCCGATCTNNNGAGCTATTATCCCCACATGCTCA
GCACACGTCTGAACTCTTTCCCTACACGACGCTCTTCCGATCTNNNGAGCTATTATCCCCACATGCTCA
GCACACGTCTGAACTCTTTCCCTACACGACGCTCTTCCGATCTNNNGAGCTATTATCCCCACATGCTCA
GCACACGTCTGAACTCTTTCCCTACACGACGCTCTTCCGATCTNNNGAGCTATTATCCCCACATGCTCA
GAGCACACGTCTGAACTCTTTCCCTACACGACGCTCTTCCGATCTNNNTATGTGCCTGTAATCCCAGCT
GAGCACACGTCTGAACTCTTTCCCTACACGACGCTCTTCCGATCTNNNTATGTGCCTGTAATCCCAGCT
GAGCACACGTCTGAACTCTTTCCCTACACGACGCTCTTCCGATCTNNNTATGTGCCTGTAATCCCAGCT
GAGCACACGTCTGAACTCTTTCCCTACACGACGCTCTTCCGATCTNNNTATGTGCCTGTAATCCCAGCT
AGCACACGTCTGAACTCTTTCCCTACACGACGCTCTTCCGATCTNNNGCAAGAGAATCACTTGAACA
AGCACACGTCTGAACTCTTTCCCTACACGACGCTCTTCCGATCTNNNGCAAGAGAATCACTTGAACA
AGCACACGTCTGAACTCTTTCCCTACACGACGCTCTTCCGATCTNNNGCAAGAGAATCACTTGAACA
AGCACACGTCTGAACTCTTTCCCTACACGACGCTCTTCCGATCTNNNGCAAGAGAATCACTTGAACA
GAGCACACGTCTGAACTCTTTCCCTACACGACGCTCTTCCGATCTNNNGAGGTGGAGGTTGCAGTGA
GAGCACACGTCTGAACTCTTTCCCTACACGACGCTCTTCCGATCTNNNGAGGTGGAGGTTGCAGTGA
GAGCACACGTCTGAACTCTTTCCCTACACGACGCTCTTCCGATCTNNNGAGGTGGAGGTTGCAGTGA
GAGCACACGTCTGAACTCTTTCCCTACACGACGCTCTTCCGATCTNNNGAGGTGGAGGTTGCAGTGA
AGCACACGTCTGAACTCTTTCCCTACACGACGCTCTTCCGATCTNNNTCTATGTATTGTAATTCTAGC
AGCACACGTCTGAACTCTTTCCCTACACGACGCTCTTCCGATCTNNNTCTATGTATTGTAATTCTAGC
AGCACACGTCTGAACTCTTTCCCTACACGACGCTCTTCCGATCTNNNTCTATGTATTGTAATTCTAGC
AGCACACGTCTGAACTCTTTCCCTACACGACGCTCTTCCGATCTNNNTCTATGTATTGTAATTCTAGC
CACACGTCTGAACTCTTTCCCTACACGACGCTCTTCCGATCTNNNCAAAAGGTGTGAGTTTGAAAAT
CACACGTCTGAACTCTTTCCCTACACGACGCTCTTCCGATCTNNNCAAAAGGTGTGAGTTTGAAAAT
CACACGTCTGAACTCTTTCCCTACACGACGCTCTTCCGATCTNNNCAAAAGGTGTGAGTTTGAAAAT
CACACGTCTGAACTCTTTCCCTACACGACGCTCTTCCGATCTNNNCAAAAGGTGTGAGTTTGAAAAT

AGCACACGTCTGAACTCTTTCCCTACACGACGCTCTTCCGATCTNNNTGCAGATTCTGATTGGGTAGGT
AGCACACGTCTGAACTCTTTCCCTACACGACGCTCTTCCGATCTNNNTGCAGATTCTGATTGGGTAGGT
AGCACACGTCTGAACTCTTTCCCTACACGACGCTCTTCCGATCTNNNTGCAGATTCTGATTGGGTAGGT
AGCACACGTCTGAACTCTTTCCCTACACGACGCTCTTCCGATCTNNNTGCAGATTCTGATTGGGTAGGT
AGCACACGTCTGAACTCTTTCCCTACACGACGCTCTTCCGATCTNNNCTCCCAGGTCCTTCTGTCCTTCCA
AGCACACGTCTGAACTCTTTCCCTACACGACGCTCTTCCGATCTNNNCTCCCAGGTCCTTCTGTCCTTCCA
AGCACACGTCTGAACTCTTTCCCTACACGACGCTCTTCCGATCTNNNCTCCCAGGTCCTTCTGTCCTTCCA
AGCACACGTCTGAACTCTTTCCCTACACGACGCTCTTCCGATCTNNNCTCCCAGGTCCTTCTGTCCTTCCA
AGAGCACACGTCTGAACTCTTTCCCTACACGACGCTCTTCCGATCTNNNTGGACAGATCATGGTTTCAA
AGAGCACACGTCTGAACTCTTTCCCTACACGACGCTCTTCCGATCTNNNTGGACAGATCATGGTTTCAA
AGAGCACACGTCTGAACTCTTTCCCTACACGACGCTCTTCCGATCTNNNTGGACAGATCATGGTTTCAA
AGAGCACACGTCTGAACTCTTTCCCTACACGACGCTCTTCCGATCTNNNTGGACAGATCATGGTTTCAA
GAGCACACGTCTGAACTCTTTCCCTACACGACGCTCTTCCGATCTNNNTTTGGAAAGCCCACAGTACAA
GAGCACACGTCTGAACTCTTTCCCTACACGACGCTCTTCCGATCTNNNTTTGGAAAGCCCACAGTACAA
GAGCACACGTCTGAACTCTTTCCCTACACGACGCTCTTCCGATCTNNNTTTGGAAAGCCCACAGTACAA
GAGCACACGTCTGAACTCTTTCCCTACACGACGCTCTTCCGATCTNNNTTTGGAAAGCCCACAGTACAA
GCACACGTCTGAACTCTTTCCCTACACGACGCTCTTCCGATCTNNNTAGAGTGTAGATGCATCAAAATC
GCACACGTCTGAACTCTTTCCCTACACGACGCTCTTCCGATCTNNNTAGAGTGTAGATGCATCAAAATC
GCACACGTCTGAACTCTTTCCCTACACGACGCTCTTCCGATCTNNNTAGAGTGTAGATGCATCAAAATC
GCACACGTCTGAACTCTTTCCCTACACGACGCTCTTCCGATCTNNNTAGAGTGTAGATGCATCAAAATC
GAGCACACGTCTGAACTCTTTCCCTACACGACGCTCTTCCGATCTNNNAAAATTAGCCAGGCATGATGC
GAGCACACGTCTGAACTCTTTCCCTACACGACGCTCTTCCGATCTNNNAAAATTAGCCAGGCATGATGC
GAGCACACGTCTGAACTCTTTCCCTACACGACGCTCTTCCGATCTNNNAAAATTAGCCAGGCATGATGC
GAGCACACGTCTGAACTCTTTCCCTACACGACGCTCTTCCGATCTNNNAAAATTAGCCAGGCATGATGC
AGCACACGTCTGAACTCTTTCCCTACACGACGCTCTTCCGATCTNNNAAAGGATTCCAGAACCTTCCTCC
AGCACACGTCTGAACTCTTTCCCTACACGACGCTCTTCCGATCTNNNAAAGGATTCCAGAACCTTCCTCC
AGCACACGTCTGAACTCTTTCCCTACACGACGCTCTTCCGATCTNNNAAAGGATTCCAGAACCTTCCTCC
AGCACACGTCTGAACTCTTTCCCTACACGACGCTCTTCCGATCTNNNAAAGGATTCCAGAACCTTCCTCC
GAGCACACGTCTGAACTCTTTCCCTACACGACGCTCTTCCGATCTNNNAAATATTGAGGGTCTGGCACT
GAGCACACGTCTGAACTCTTTCCCTACACGACGCTCTTCCGATCTNNNAAATATTGAGGGTCTGGCACT
GAGCACACGTCTGAACTCTTTCCCTACACGACGCTCTTCCGATCTNNNAAATATTGAGGGTCTGGCACT
GAGCACACGTCTGAACTCTTTCCCTACACGACGCTCTTCCGATCTNNNAAATATTGAGGGTCTGGCACT
GAGCACACGTCTGAACTCTTTCCCTACACGACGCTCTTCCGATCTNNNGCTAAAGAAGTGGCCTCAAAC
GAGCACACGTCTGAACTCTTTCCCTACACGACGCTCTTCCGATCTNNNGCTAAAGAAGTGGCCTCAAAC
GAGCACACGTCTGAACTCTTTCCCTACACGACGCTCTTCCGATCTNNNGCTAAAGAAGTGGCCTCAAAC
GAGCACACGTCTGAACTCTTTCCCTACACGACGCTCTTCCGATCTNNNGCTAAAGAAGTGGCCTCAAAC
AGCACACGTCTGAACTCTTTCCCTACACGACGCTCTTCCGATCTNNNTCTCCTAGGAGAGAAAAGAGG
AGCACACGTCTGAACTCTTTCCCTACACGACGCTCTTCCGATCTNNNTCTCCTAGGAGAGAAAAGAGG
AGCACACGTCTGAACTCTTTCCCTACACGACGCTCTTCCGATCTNNNTCTCCTAGGAGAGAAAAGAGG
AGCACACGTCTGAACTCTTTCCCTACACGACGCTCTTCCGATCTNNNTCTCCTAGGAGAGAAAAGAGG
AGCACACGTCTGAACTCTTTCCCTACACGACGCTCTTCCGATCTNNNAGGCCTGTAATCCCAGTACTTTG
AGCACACGTCTGAACTCTTTCCCTACACGACGCTCTTCCGATCTNNNAGGCCTGTAATCCCAGTACTTTG
AGCACACGTCTGAACTCTTTCCCTACACGACGCTCTTCCGATCTNNNAGGCCTGTAATCCCAGTACTTTG

GCACACGTCTGAACTCTTTCCCTACACGACGCTCTTCCGATCTNNNAGGCCTGTAATCCCAGTACTTTG
AGCACACGTCTGAACTCTTTCCCTACACGACGCTCTTCCGATCTNNNAGCCTTGTACATGCTGATAGCT
AGCACACGTCTGAACTCTTTCCCTACACGACGCTCTTCCGATCTNNNAGCCTTGTACATGCTGATAGCT
AGCACACGTCTGAACTCTTTCCCTACACGACGCTCTTCCGATCTNNNAGCCTTGTACATGCTGATAGCT
AGCACACGTCTGAACTCTTTCCCTACACGACGCTCTTCCGATCTNNNAGCCTTGTACATGCTGATAGCT
AGCACACGTCTGAACTCTTTCCCTACACGACGCTCTTCCGATCTNNNCAGGAGCATCACTTGAACCCAA
AGCACACGTCTGAACTCTTTCCCTACACGACGCTCTTCCGATCTNNNCAGGAGCATCACTTGAACCCAA
AGCACACGTCTGAACTCTTTCCCTACACGACGCTCTTCCGATCTNNNCAGGAGCATCACTTGAACCCAA
AGCACACGTCTGAACTCTTTCCCTACACGACGCTCTTCCGATCTNNNCAGGAGCATCACTTGAACCCAA
AGCACACGTCTGAACTCTTTCCCTACACGACGCTCTTCCGATCTNNNATTTCAGGTAGAGGAAGCCAC
AGCACACGTCTGAACTCTTTCCCTACACGACGCTCTTCCGATCTNNNATTTCAGGTAGAGGAAGCCAC
AGCACACGTCTGAACTCTTTCCCTACACGACGCTCTTCCGATCTNNNATTTCAGGTAGAGGAAGCCAC
AGCACACGTCTGAACTCTTTCCCTACACGACGCTCTTCCGATCTNNNATTTCAGGTAGAGGAAGCCAC
AGCACACGTCTGAACTCTTTCCCTACACGACGCTCTTCCGATCTNNNATGGCAAACACCCACCTTTACA
AGCACACGTCTGAACTCTTTCCCTACACGACGCTCTTCCGATCTNNNATGGCAAACACCCACCTTTACA
AGCACACGTCTGAACTCTTTCCCTACACGACGCTCTTCCGATCTNNNATGGCAAACACCCACCTTTACA
AGCACACGTCTGAACTCTTTCCCTACACGACGCTCTTCCGATCTNNNATGGCAAACACCCACCTTTACA
GAGCACACGTCTGAACTCTTTCCCTACACGACGCTCTTCCGATCTNNNACTTCTAGGGGCTTTTCCAACT
GAGCACACGTCTGAACTCTTTCCCTACACGACGCTCTTCCGATCTNNNACTTCTAGGGGCTTTTCCAACT
GAGCACACGTCTGAACTCTTTCCCTACACGACGCTCTTCCGATCTNNNACTTCTAGGGGCTTTTCCAACT
GAGCACACGTCTGAACTCTTTCCCTACACGACGCTCTTCCGATCTNNNACTTCTAGGGGCTTTTCCAACT
AGCACACGTCTGAACTCTTTCCCTACACGACGCTCTTCCGATCTNNNATAGTGGTGCATGCCTGTAGT
AGCACACGTCTGAACTCTTTCCCTACACGACGCTCTTCCGATCTNNNATAGTGGTGCATGCCTGTAGT
AGCACACGTCTGAACTCTTTCCCTACACGACGCTCTTCCGATCTNNNATAGTGGTGCATGCCTGTAGT
AGCACACGTCTGAACTCTTTCCCTACACGACGCTCTTCCGATCTNNNATAGTGGTGCATGCCTGTAGT
GAGCACACGTCTGAACTCTTTCCCTACACGACGCTCTTCCGATCTNNNCCTGTAATCCCAAATACTCCGG
GAGCACACGTCTGAACTCTTTCCCTACACGACGCTCTTCCGATCTNNNCCTGTAATCCCAAATACTCCGG
GAGCACACGTCTGAACTCTTTCCCTACACGACGCTCTTCCGATCTNNNCCTGTAATCCCAAATACTCCGG
GAGCACACGTCTGAACTCTTTCCCTACACGACGCTCTTCCGATCTNNNCCTGTAATCCCAAATACTCCGG
GAGCACACGTCTGAACTCTTTCCCTACACGACGCTCTTCCGATCTNNNCGCTTCTAATTTCACCAGTAGC
GAGCACACGTCTGAACTCTTTCCCTACACGACGCTCTTCCGATCTNNNCGCTTCTAATTTCACCAGTAGC
GAGCACACGTCTGAACTCTTTCCCTACACGACGCTCTTCCGATCTNNNCGCTTCTAATTTCACCAGTAGC
GAGCACACGTCTGAACTCTTTCCCTACACGACGCTCTTCCGATCTNNNCGCTTCTAATTTCACCAGTAGC
AGCACACGTCTGAACTCTTTCCCTACACGACGCTCTTCCGATCTNNNCAGGTGGTCCTTCTCGAGTTTT
AGCACACGTCTGAACTCTTTCCCTACACGACGCTCTTCCGATCTNNNCAGGTGGTCCTTCTCGAGTTTT
AGCACACGTCTGAACTCTTTCCCTACACGACGCTCTTCCGATCTNNNCAGGTGGTCCTTCTCGAGTTTT
AGCACACGTCTGAACTCTTTCCCTACACGACGCTCTTCCGATCTNNNCAGGTGGTCCTTCTCGAGTTTT
GAGCACACGTCTGAACTCTTTCCCTACACGACGCTCTTCCGATCTNNNGAGATTGGGCCACTGCACTC
GAGCACACGTCTGAACTCTTTCCCTACACGACGCTCTTCCGATCTNNNGAGATTGGGCCACTGCACTC
GAGCACACGTCTGAACTCTTTCCCTACACGACGCTCTTCCGATCTNNNGAGATTGGGCCACTGCACTC
GAGCACACGTCTGAACTCTTTCCCTACACGACGCTCTTCCGATCTNNNGAGATTGGGCCACTGCACTC
AGAGCACACGTCTGAACTCTTTCCCTACACGACGCTCTTCCGATCTNNNTTCACCAAATCCACTTCCCTA
AGAGCACACGTCTGAACTCTTTCCCTACACGACGCTCTTCCGATCTNNNTTCACCAAATCCACTTCCCTA

GAGCACACGTCTGAACTCTTTCCCTACACGACGCTCTTCCGATCTNNNTTCACCAAATCCACTTCCCTA
GAGCACACGTCTGAACTCTTTCCCTACACGACGCTCTTCCGATCTNNNTTCACCAAATCCACTTCCCTA
GCACACGTCTGAACTCTTTCCCTACACGACGCTCTTCCGATCTNNNCTTAGCTACAGGCAGTTTCCCA
GCACACGTCTGAACTCTTTCCCTACACGACGCTCTTCCGATCTNNNCTTAGCTACAGGCAGTTTCCCA
GCACACGTCTGAACTCTTTCCCTACACGACGCTCTTCCGATCTNNNCTTAGCTACAGGCAGTTTCCCA
GCACACGTCTGAACTCTTTCCCTACACGACGCTCTTCCGATCTNNNCTTAGCTACAGGCAGTTTCCCA
CACACGTCTGAACTCTTTCCCTACACGACGCTCTTCCGATCTNNNAGGTTGGAATCAGTTGGAGAGT
CACACGTCTGAACTCTTTCCCTACACGACGCTCTTCCGATCTNNNAGGTTGGAATCAGTTGGAGAGT
CACACGTCTGAACTCTTTCCCTACACGACGCTCTTCCGATCTNNNAGGTTGGAATCAGTTGGAGAGT
CACACGTCTGAACTCTTTCCCTACACGACGCTCTTCCGATCTNNNAGGTTGGAATCAGTTGGAGAGT
GAGCACACGTCTGAACTCTTTCCCTACACGACGCTCTTCCGATCTNNNATCTCTGACTTCAAGGAGCTG
GAGCACACGTCTGAACTCTTTCCCTACACGACGCTCTTCCGATCTNNNATCTCTGACTTCAAGGAGCTG
GAGCACACGTCTGAACTCTTTCCCTACACGACGCTCTTCCGATCTNNNATCTCTGACTTCAAGGAGCTG
GAGCACACGTCTGAACTCTTTCCCTACACGACGCTCTTCCGATCTNNNATCTCTGACTTCAAGGAGCTG
GAGCACACGTCTGAACTCTTTCCCTACACGACGCTCTTCCGATCTNNNAACTTTCATGACAGGAGGT
GAGCACACGTCTGAACTCTTTCCCTACACGACGCTCTTCCGATCTNNNAACTTTCATGACAGGAGGT
GAGCACACGTCTGAACTCTTTCCCTACACGACGCTCTTCCGATCTNNNAACTTTCATGACAGGAGGT
GAGCACACGTCTGAACTCTTTCCCTACACGACGCTCTTCCGATCTNNNAACTTTCATGACAGGAGGT
AGCACACGTCTGAACTCTTTCCCTACACGACGCTCTTCCGATCTNNNGCAGAAACAACGAATTTCCTCC
AGCACACGTCTGAACTCTTTCCCTACACGACGCTCTTCCGATCTNNNGCAGAAACAACGAATTTCCTCC
AGCACACGTCTGAACTCTTTCCCTACACGACGCTCTTCCGATCTNNNGCAGAAACAACGAATTTCCTCC
AGCACACGTCTGAACTCTTTCCCTACACGACGCTCTTCCGATCTNNNGCAGAAACAACGAATTTCCTCC
AGCACACGTCTGAACTCTTTCCCTACACGACGCTCTTCCGATCTNNNCTGAGGCAGAGAATTGCTTG
AGCACACGTCTGAACTCTTTCCCTACACGACGCTCTTCCGATCTNNNCTGAGGCAGAGAATTGCTTG
AGCACACGTCTGAACTCTTTCCCTACACGACGCTCTTCCGATCTNNNCTGAGGCAGAGAATTGCTTG
AGCACACGTCTGAACTCTTTCCCTACACGACGCTCTTCCGATCTNNNCTGAGGCAGAGAATTGCTTG
AGCACACGTCTGAACTCTTTCCCTACACGACGCTCTTCCGATCTNNNCTTGCAGTAAGCCGAGATTGC
AGCACACGTCTGAACTCTTTCCCTACACGACGCTCTTCCGATCTNNNCTTGCAGTAAGCCGAGATTGC
AGCACACGTCTGAACTCTTTCCCTACACGACGCTCTTCCGATCTNNNCTTGCAGTAAGCCGAGATTGC
AGCACACGTCTGAACTCTTTCCCTACACGACGCTCTTCCGATCTNNNCTTGCAGTAAGCCGAGATTGC
AGCACACGTCTGAACTCTTTCCCTACACGACGCTCTTCCGATCTNNNTCAGTCTCCTGAGTAGCTGAG
AGCACACGTCTGAACTCTTTCCCTACACGACGCTCTTCCGATCTNNNTCAGTCTCCTGAGTAGCTGAG
AGCACACGTCTGAACTCTTTCCCTACACGACGCTCTTCCGATCTNNNTCAGTCTCCTGAGTAGCTGAG
AGCACACGTCTGAACTCTTTCCCTACACGACGCTCTTCCGATCTNNNTCAGTCTCCTGAGTAGCTGAG
GCACACGTCTGAACTCTTTCCCTACACGACGCTCTTCCGATCTNNNCTTTTCATTCGTTTCACTTGACAT
GCACACGTCTGAACTCTTTCCCTACACGACGCTCTTCCGATCTNNNCTTTTCATTCGTTTCACTTGACAT
GCACACGTCTGAACTCTTTCCCTACACGACGCTCTTCCGATCTNNNCTTTTCATTCGTTTCACTTGACAT
GCACACGTCTGAACTCTTTCCCTACACGACGCTCTTCCGATCTNNNCTTTTCATTCGTTTCACTTGACAT
GCACACGTCTGAACTCTTTCCCTACACGACGCTCTTCCGATCTNNNAAAGCTACCACATCAGTCTGAC
GCACACGTCTGAACTCTTTCCCTACACGACGCTCTTCCGATCTNNNAAAGCTACCACATCAGTCTGAC
GCACACGTCTGAACTCTTTCCCTACACGACGCTCTTCCGATCTNNNAAAGCTACCACATCAGTCTGAC
GCACACGTCTGAACTCTTTCCCTACACGACGCTCTTCCGATCTNNNAAAGCTACCACATCAGTCTGAC
GCACACGTCTGAACTCTTTCCCTACACGACGCTCTTCCGATCTNNNAACAGTTCTCCGTCTCCAACTTT

AGCACACGTCTGAACTCTTTCCCTACACGACGCTCTTCCGATCTNNNAACAGTTCTCCGTCTCCAACTTT
AGCACACGTCTGAACTCTTTCCCTACACGACGCTCTTCCGATCTNNNAACAGTTCTCCGTCTCCAACTTT
AGCACACGTCTGAACTCTTTCCCTACACGACGCTCTTCCGATCTNNNAACAGTTCTCCGTCTCCAACTTT
GAGCACACGTCTGAACTCTTTCCCTACACGACGCTCTTCCGATCTNNNCAGGAAGAGGAGGAGTTTT
GAGCACACGTCTGAACTCTTTCCCTACACGACGCTCTTCCGATCTNNNCAGGAAGAGGAGGAGTTTT
GAGCACACGTCTGAACTCTTTCCCTACACGACGCTCTTCCGATCTNNNCAGGAAGAGGAGGAGTTTT
GAGCACACGTCTGAACTCTTTCCCTACACGACGCTCTTCCGATCTNNNCAGGAAGAGGAGGAGTTTT
AAGAGCACACGTCTGAACTCTTTCCCTACACGACGCTCTTCCGATCTNNNTGAGAATCACTTGAACCTG
AAGAGCACACGTCTGAACTCTTTCCCTACACGACGCTCTTCCGATCTNNNTGAGAATCACTTGAACCTG
AAGAGCACACGTCTGAACTCTTTCCCTACACGACGCTCTTCCGATCTNNNTGAGAATCACTTGAACCTG
AAGAGCACACGTCTGAACTCTTTCCCTACACGACGCTCTTCCGATCTNNNTGAGAATCACTTGAACCTG
GAGCACACGTCTGAACTCTTTCCCTACACGACGCTCTTCCGATCTNNNTTCTGAGAGACCTGGCCATTC
GAGCACACGTCTGAACTCTTTCCCTACACGACGCTCTTCCGATCTNNNTTCTGAGAGACCTGGCCATTC
GAGCACACGTCTGAACTCTTTCCCTACACGACGCTCTTCCGATCTNNNTTCTGAGAGACCTGGCCATTC
GAGCACACGTCTGAACTCTTTCCCTACACGACGCTCTTCCGATCTNNNTTCTGAGAGACCTGGCCATTC
GAGCACACGTCTGAACTCTTTCCCTACACGACGCTCTTCCGATCTNNNAAAAAAAAATTGCCAGGCGT
GAGCACACGTCTGAACTCTTTCCCTACACGACGCTCTTCCGATCTNNNAAAAAAAAATTGCCAGGCGT
GAGCACACGTCTGAACTCTTTCCCTACACGACGCTCTTCCGATCTNNNAAAAAAAAATTGCCAGGCGT
GAGCACACGTCTGAACTCTTTCCCTACACGACGCTCTTCCGATCTNNNAAAAAAAAATTGCCAGGCGT
AGCACACGTCTGAACTCTTTCCCTACACGACGCTCTTCCGATCTNNNGCAATTGGGTTGGTTTCAAGT
AGCACACGTCTGAACTCTTTCCCTACACGACGCTCTTCCGATCTNNNGCAATTGGGTTGGTTTCAAGT
AGCACACGTCTGAACTCTTTCCCTACACGACGCTCTTCCGATCTNNNGCAATTGGGTTGGTTTCAAGT
AGCACACGTCTGAACTCTTTCCCTACACGACGCTCTTCCGATCTNNNGCAATTGGGTTGGTTTCAAGT
CACGTCTGAACTCTTTCCCTACACGACGCTCTTCCGATCTNNNCCTCAGCCTTGTGTCTTTGTTTTCAT
CACGTCTGAACTCTTTCCCTACACGACGCTCTTCCGATCTNNNCCTCAGCCTTGTGTCTTTGTTTTCAT
CACGTCTGAACTCTTTCCCTACACGACGCTCTTCCGATCTNNNCCTCAGCCTTGTGTCTTTGTTTTCAT
CACGTCTGAACTCTTTCCCTACACGACGCTCTTCCGATCTNNNCCTCAGCCTTGTGTCTTTGTTTTCAT
GAGCACACGTCTGAACTCTTTCCCTACACGACGCTCTTCCGATCTNNNTGGGGAAGAATCTGACAGC
GAGCACACGTCTGAACTCTTTCCCTACACGACGCTCTTCCGATCTNNNTGGGGAAGAATCTGACAGC
GAGCACACGTCTGAACTCTTTCCCTACACGACGCTCTTCCGATCTNNNTGGGGAAGAATCTGACAGC
GAGCACACGTCTGAACTCTTTCCCTACACGACGCTCTTCCGATCTNNNTGGGGAAGAATCTGACAGC
AGCACACGTCTGAACTCTTTCCCTACACGACGCTCTTCCGATCTNNNTAACTGAAGATATTCCAACTCA
AGCACACGTCTGAACTCTTTCCCTACACGACGCTCTTCCGATCTNNNTAACTGAAGATATTCCAACTCA
AGCACACGTCTGAACTCTTTCCCTACACGACGCTCTTCCGATCTNNNTAACTGAAGATATTCCAACTCA
AGCACACGTCTGAACTCTTTCCCTACACGACGCTCTTCCGATCTNNNTAACTGAAGATATTCCAACTCA
AGCACACGTCTGAACTCTTTCCCTACACGACGCTCTTCCGATCTNNNCTGAGGCAGGAGAATTGCTT
AGCACACGTCTGAACTCTTTCCCTACACGACGCTCTTCCGATCTNNNCTGAGGCAGGAGAATTGCTT
AGCACACGTCTGAACTCTTTCCCTACACGACGCTCTTCCGATCTNNNCTGAGGCAGGAGAATTGCTT
AGCACACGTCTGAACTCTTTCCCTACACGACGCTCTTCCGATCTNNNCTGAGGCAGGAGAATTGCTT
GCACACGTCTGAACTCTTTCCCTACACGACGCTCTTCCGATCTNNNAAAGGTCTACTCAGTGGCAAGA
GCACACGTCTGAACTCTTTCCCTACACGACGCTCTTCCGATCTNNNAAAGGTCTACTCAGTGGCAAGA
GCACACGTCTGAACTCTTTCCCTACACGACGCTCTTCCGATCTNNNAAAGGTCTACTCAGTGGCAAGA
GCACACGTCTGAACTCTTTCCCTACACGACGCTCTTCCGATCTNNNAAAGGTCTACTCAGTGGCAAGA

ACACGTCTGAACTCTTTCCCTACACGACGCTCTTCCGATCTNNNGTTCAAATTCTGGTTGAAAGATCAT
ACACGTCTGAACTCTTTCCCTACACGACGCTCTTCCGATCTNNNGTTCAAATTCTGGTTGAAAGATCAT
ACACGTCTGAACTCTTTCCCTACACGACGCTCTTCCGATCTNNNGTTCAAATTCTGGTTGAAAGATCAT
ACACGTCTGAACTCTTTCCCTACACGACGCTCTTCCGATCTNNNGTTCAAATTCTGGTTGAAAGATCAT
AGCACACGTCTGAACTCTTTCCCTACACGACGCTCTTCCGATCTNNNCACCTGTAATCCCAGCAGTTTT
AGCACACGTCTGAACTCTTTCCCTACACGACGCTCTTCCGATCTNNNCACCTGTAATCCCAGCAGTTTT
AGCACACGTCTGAACTCTTTCCCTACACGACGCTCTTCCGATCTNNNCACCTGTAATCCCAGCAGTTTT
AGCACACGTCTGAACTCTTTCCCTACACGACGCTCTTCCGATCTNNNCACCTGTAATCCCAGCAGTTTT
AGCACACGTCTGAACTCTTTCCCTACACGACGCTCTTCCGATCTNNNGCCTGTAGTCCCAGCTACTTGC
AGCACACGTCTGAACTCTTTCCCTACACGACGCTCTTCCGATCTNNNGCCTGTAGTCCCAGCTACTTGC
AGCACACGTCTGAACTCTTTCCCTACACGACGCTCTTCCGATCTNNNGCCTGTAGTCCCAGCTACTTGC
AGCACACGTCTGAACTCTTTCCCTACACGACGCTCTTCCGATCTNNNGCCTGTAGTCCCAGCTACTTGC
AGCACACGTCTGAACTCTTTCCCTACACGACGCTCTTCCGATCTNNNAAAGCCCAAAAATTAGGGAGG
AGCACACGTCTGAACTCTTTCCCTACACGACGCTCTTCCGATCTNNNAAAGCCCAAAAATTAGGGAGG
AGCACACGTCTGAACTCTTTCCCTACACGACGCTCTTCCGATCTNNNAAAGCCCAAAAATTAGGGAGG
AGCACACGTCTGAACTCTTTCCCTACACGACGCTCTTCCGATCTNNNAAAGCCCAAAAATTAGGGAGG
AGCACACGTCTGAACTCTTTCCCTACACGACGCTCTTCCGATCTNNNAGAATTGCTTGAACCCAGGAG
AGCACACGTCTGAACTCTTTCCCTACACGACGCTCTTCCGATCTNNNAGAATTGCTTGAACCCAGGAG
AGCACACGTCTGAACTCTTTCCCTACACGACGCTCTTCCGATCTNNNAGAATTGCTTGAACCCAGGAG
AGCACACGTCTGAACTCTTTCCCTACACGACGCTCTTCCGATCTNNNAGAATTGCTTGAACCCAGGAG
AGCACACGTCTGAACTCTTTCCCTACACGACGCTCTTCCGATCTNNNTCCCATGATCTCTCTCTCTCTC
AGCACACGTCTGAACTCTTTCCCTACACGACGCTCTTCCGATCTNNNTCCCATGATCTCTCTCTCTCTC
AGCACACGTCTGAACTCTTTCCCTACACGACGCTCTTCCGATCTNNNTCCCATGATCTCTCTCTCTCTC
AGCACACGTCTGAACTCTTTCCCTACACGACGCTCTTCCGATCTNNNTCCCATGATCTCTCTCTCTCTC
GAGCACACGTCTGAACTCTTTCCCTACACGACGCTCTTCCGATCTNNNGCAACATAGTGAGACCCCATC
GAGCACACGTCTGAACTCTTTCCCTACACGACGCTCTTCCGATCTNNNGCAACATAGTGAGACCCCATC
GAGCACACGTCTGAACTCTTTCCCTACACGACGCTCTTCCGATCTNNNGCAACATAGTGAGACCCCATC
GAGCACACGTCTGAACTCTTTCCCTACACGACGCTCTTCCGATCTNNNGCAACATAGTGAGACCCCATC
GAGCACACGTCTGAACTCTTTCCCTACACGACGCTCTTCCGATCTNNNATAAACGGGCAGAAGACAGT
GAGCACACGTCTGAACTCTTTCCCTACACGACGCTCTTCCGATCTNNNATAAACGGGCAGAAGACAGT
GAGCACACGTCTGAACTCTTTCCCTACACGACGCTCTTCCGATCTNNNATAAACGGGCAGAAGACAGT
GAGCACACGTCTGAACTCTTTCCCTACACGACGCTCTTCCGATCTNNNATAAACGGGCAGAAGACAGT
AGCACACGTCTGAACTCTTTCCCTACACGACGCTCTTCCGATCTNNNCTCAAGCAATCTTCCCACTTCAG
AGCACACGTCTGAACTCTTTCCCTACACGACGCTCTTCCGATCTNNNCTCAAGCAATCTTCCCACTTCAG
AGCACACGTCTGAACTCTTTCCCTACACGACGCTCTTCCGATCTNNNCTCAAGCAATCTTCCCACTTCAG
AGCACACGTCTGAACTCTTTCCCTACACGACGCTCTTCCGATCTNNNCTCAAGCAATCTTCCCACTTCAG
AGCACACGTCTGAACTCTTTCCCTACACGACGCTCTTCCGATCTNNNTTAGCCATGATGAAAGTCCAT
AGCACACGTCTGAACTCTTTCCCTACACGACGCTCTTCCGATCTNNNTTAGCCATGATGAAAGTCCAT
AGCACACGTCTGAACTCTTTCCCTACACGACGCTCTTCCGATCTNNNTTAGCCATGATGAAAGTCCAT
AGCACACGTCTGAACTCTTTCCCTACACGACGCTCTTCCGATCTNNNTTAGCCATGATGAAAGTCCAT
AGCACACGTCTGAACTCTTTCCCTACACGACGCTCTTCCGATCTNNNCTGTGTGTCTGTGTCCTCATCT
AGCACACGTCTGAACTCTTTCCCTACACGACGCTCTTCCGATCTNNNCTGTGTGTCTGTGTCCTCATCT
AGCACACGTCTGAACTCTTTCCCTACACGACGCTCTTCCGATCTNNNCTGTGTGTCTGTGTCCTCATCT

AGCACACGTCTGAACTCTTTCCCTACACGACGCTCTTCCGATCTNNNCTGTGTGTCTGTGTCCTCATCT
CACACGTCTGAACTCTTTCCCTACACGACGCTCTTCCGATCTNNNCATTAGCTGGAGAAAGTGGAGG
CACACGTCTGAACTCTTTCCCTACACGACGCTCTTCCGATCTNNNCATTAGCTGGAGAAAGTGGAGG
CACACGTCTGAACTCTTTCCCTACACGACGCTCTTCCGATCTNNNCATTAGCTGGAGAAAGTGGAGG
CACACGTCTGAACTCTTTCCCTACACGACGCTCTTCCGATCTNNNCATTAGCTGGAGAAAGTGGAGG
AGCACACGTCTGAACTCTTTCCCTACACGACGCTCTTCCGATCTNNNTGAAGTTTTTACCCTGAGCCTC
AGCACACGTCTGAACTCTTTCCCTACACGACGCTCTTCCGATCTNNNTGAAGTTTTTACCCTGAGCCTC
AGCACACGTCTGAACTCTTTCCCTACACGACGCTCTTCCGATCTNNNTGAAGTTTTTACCCTGAGCCTC
AGCACACGTCTGAACTCTTTCCCTACACGACGCTCTTCCGATCTNNNTGAAGTTTTTACCCTGAGCCTC
AGCACACGTCTGAACTCTTTCCCTACACGACGCTCTTCCGATCTNNNTCTGTTTTGTGAGCAGTTCTCC
AGCACACGTCTGAACTCTTTCCCTACACGACGCTCTTCCGATCTNNNTCTGTTTTGTGAGCAGTTCTCC
AGCACACGTCTGAACTCTTTCCCTACACGACGCTCTTCCGATCTNNNTCTGTTTTGTGAGCAGTTCTCC
AGCACACGTCTGAACTCTTTCCCTACACGACGCTCTTCCGATCTNNNTCTGTTTTGTGAGCAGTTCTCC
AGCACACGTCTGAACTCTTTCCCTACACGACGCTCTTCCGATCTNNNCACCCATCAACCCATCATCTACA
AGCACACGTCTGAACTCTTTCCCTACACGACGCTCTTCCGATCTNNNCACCCATCAACCCATCATCTACA
AGCACACGTCTGAACTCTTTCCCTACACGACGCTCTTCCGATCTNNNCACCCATCAACCCATCATCTACA
AGCACACGTCTGAACTCTTTCCCTACACGACGCTCTTCCGATCTNNNCACCCATCAACCCATCATCTACA
GCACACGTCTGAACTCTTTCCCTACACGACGCTCTTCCGATCTNNNAAGCAAAAACTGGCCATTTGAG
GCACACGTCTGAACTCTTTCCCTACACGACGCTCTTCCGATCTNNNAAGCAAAAACTGGCCATTTGAG
GCACACGTCTGAACTCTTTCCCTACACGACGCTCTTCCGATCTNNNAAGCAAAAACTGGCCATTTGAG
GCACACGTCTGAACTCTTTCCCTACACGACGCTCTTCCGATCTNNNAAGCAAAAACTGGCCATTTGAG
AGCACACGTCTGAACTCTTTCCCTACACGACGCTCTTCCGATCTNNNGCAGCAATCTGTCTTTTCTTGT
AGCACACGTCTGAACTCTTTCCCTACACGACGCTCTTCCGATCTNNNGCAGCAATCTGTCTTTTCTTGT
AGCACACGTCTGAACTCTTTCCCTACACGACGCTCTTCCGATCTNNNGCAGCAATCTGTCTTTTCTTGT
AGCACACGTCTGAACTCTTTCCCTACACGACGCTCTTCCGATCTNNNGCAGCAATCTGTCTTTTCTTGT
GCACACGTCTGAACTCTTTCCCTACACGACGCTCTTCCGATCTNNNAAATGACATGCTCAGGGCCATT
GCACACGTCTGAACTCTTTCCCTACACGACGCTCTTCCGATCTNNNAAATGACATGCTCAGGGCCATT
GCACACGTCTGAACTCTTTCCCTACACGACGCTCTTCCGATCTNNNAAATGACATGCTCAGGGCCATT
GCACACGTCTGAACTCTTTCCCTACACGACGCTCTTCCGATCTNNNAAATGACATGCTCAGGGCCATT
AGCACACGTCTGAACTCTTTCCCTACACGACGCTCTTCCGATCTNNNGGGTATGGCCAGATTACTATT
AGCACACGTCTGAACTCTTTCCCTACACGACGCTCTTCCGATCTNNNGGGTATGGCCAGATTACTATT
AGCACACGTCTGAACTCTTTCCCTACACGACGCTCTTCCGATCTNNNGGGTATGGCCAGATTACTATT
AGCACACGTCTGAACTCTTTCCCTACACGACGCTCTTCCGATCTNNNGGGTATGGCCAGATTACTATT
AGCACACGTCTGAACTCTTTCCCTACACGACGCTCTTCCGATCTNNNTTGTACACAAATCACCAAAAGC
AGCACACGTCTGAACTCTTTCCCTACACGACGCTCTTCCGATCTNNNTTGTACACAAATCACCAAAAGC
AGCACACGTCTGAACTCTTTCCCTACACGACGCTCTTCCGATCTNNNTTGTACACAAATCACCAAAAGC
AGCACACGTCTGAACTCTTTCCCTACACGACGCTCTTCCGATCTNNNTTGTACACAAATCACCAAAAGC
GAGCACACGTCTGAACTCTTTCCCTACACGACGCTCTTCCGATCTNNNAAGTCACCTAAGCTTGCTACT
GAGCACACGTCTGAACTCTTTCCCTACACGACGCTCTTCCGATCTNNNAAGTCACCTAAGCTTGCTACT
GAGCACACGTCTGAACTCTTTCCCTACACGACGCTCTTCCGATCTNNNAAGTCACCTAAGCTTGCTACT
GAGCACACGTCTGAACTCTTTCCCTACACGACGCTCTTCCGATCTNNNAAGTCACCTAAGCTTGCTACT
GCACACGTCTGAACTCTTTCCCTACACGACGCTCTTCCGATCTNNNCCTATAGTCCCAGCTACTCGGGC
GCACACGTCTGAACTCTTTCCCTACACGACGCTCTTCCGATCTNNNCCTATAGTCCCAGCTACTCGGGC

GCACACGTCTGAACTCTTTCCCTACACGACGCTCTTCCGATCTNNNCCTATAGTCCCAGCTACTCGGG
GCACACGTCTGAACTCTTTCCCTACACGACGCTCTTCCGATCTNNNCCTATAGTCCCAGCTACTCGGG
AGCACACGTCTGAACTCTTTCCCTACACGACGCTCTTCCGATCTNNNCAGCTCTCATCTAGGCACTGTA
AGCACACGTCTGAACTCTTTCCCTACACGACGCTCTTCCGATCTNNNCAGCTCTCATCTAGGCACTGTA
AGCACACGTCTGAACTCTTTCCCTACACGACGCTCTTCCGATCTNNNCAGCTCTCATCTAGGCACTGTA
AGCACACGTCTGAACTCTTTCCCTACACGACGCTCTTCCGATCTNNNCAGCTCTCATCTAGGCACTGTA
AGCACACGTCTGAACTCTTTCCCTACACGACGCTCTTCCGATCTNNNGTACTGCGGAGTGACCTCACC
AGCACACGTCTGAACTCTTTCCCTACACGACGCTCTTCCGATCTNNNGTACTGCGGAGTGACCTCACC
AGCACACGTCTGAACTCTTTCCCTACACGACGCTCTTCCGATCTNNNGTACTGCGGAGTGACCTCACC
AGCACACGTCTGAACTCTTTCCCTACACGACGCTCTTCCGATCTNNNGTACTGCGGAGTGACCTCACC
GAGCACACGTCTGAACTCTTTCCCTACACGACGCTCTTCCGATCTNNNACATCCATTGAACACCAAGT
GAGCACACGTCTGAACTCTTTCCCTACACGACGCTCTTCCGATCTNNNACATCCATTGAACACCAAGT
GAGCACACGTCTGAACTCTTTCCCTACACGACGCTCTTCCGATCTNNNACATCCATTGAACACCAAGT
GAGCACACGTCTGAACTCTTTCCCTACACGACGCTCTTCCGATCTNNNACATCCATTGAACACCAAGT
AGCACACGTCTGAACTCTTTCCCTACACGACGCTCTTCCGATCTNNNAAAGGCAAGTAAACAATCTT
AGCACACGTCTGAACTCTTTCCCTACACGACGCTCTTCCGATCTNNNAAAGGCAAGTAAACAATCTT
AGCACACGTCTGAACTCTTTCCCTACACGACGCTCTTCCGATCTNNNAAAGGCAAGTAAACAATCTT
AGCACACGTCTGAACTCTTTCCCTACACGACGCTCTTCCGATCTNNNAAAGGCAAGTAAACAATCTT
AGCACACGTCTGAACTCTTTCCCTACACGACGCTCTTCCGATCTNNNCTGAGGCAGGAGAATGATGT
AGCACACGTCTGAACTCTTTCCCTACACGACGCTCTTCCGATCTNNNCTGAGGCAGGAGAATGATGT
AGCACACGTCTGAACTCTTTCCCTACACGACGCTCTTCCGATCTNNNCTGAGGCAGGAGAATGATGT
AGCACACGTCTGAACTCTTTCCCTACACGACGCTCTTCCGATCTNNNCTGAGGCAGGAGAATGATGT
GAGCACACGTCTGAACTCTTTCCCTACACGACGCTCTTCCGATCTNNNGCATGCCTGTAGTCCCAACTA
GAGCACACGTCTGAACTCTTTCCCTACACGACGCTCTTCCGATCTNNNGCATGCCTGTAGTCCCAACTA
GAGCACACGTCTGAACTCTTTCCCTACACGACGCTCTTCCGATCTNNNGCATGCCTGTAGTCCCAACTA
GAGCACACGTCTGAACTCTTTCCCTACACGACGCTCTTCCGATCTNNNGCATGCCTGTAGTCCCAACTA
AGCACACGTCTGAACTCTTTCCCTACACGACGCTCTTCCGATCTNNNAGGAGGAAAAAGAGTTCAAG
AGCACACGTCTGAACTCTTTCCCTACACGACGCTCTTCCGATCTNNNAGGAGGAAAAAGAGTTCAAG
AGCACACGTCTGAACTCTTTCCCTACACGACGCTCTTCCGATCTNNNAGGAGGAAAAAGAGTTCAAG
AGCACACGTCTGAACTCTTTCCCTACACGACGCTCTTCCGATCTNNNAGGAGGAAAAAGAGTTCAAG
AGCACACGTCTGAACTCTTTCCCTACACGACGCTCTTCCGATCTNNNTTATAGAGCAGGTTATAAGA
AGCACACGTCTGAACTCTTTCCCTACACGACGCTCTTCCGATCTNNNTTATAGAGCAGGTTATAAGA
AGCACACGTCTGAACTCTTTCCCTACACGACGCTCTTCCGATCTNNNTTATAGAGCAGGTTATAAGA
AGCACACGTCTGAACTCTTTCCCTACACGACGCTCTTCCGATCTNNNTTATAGAGCAGGTTATAAGA
GCACACGTCTGAACTCTTTCCCTACACGACGCTCTTCCGATCTNNNGGAGTTGGGGCTTCAACATATG
GCACACGTCTGAACTCTTTCCCTACACGACGCTCTTCCGATCTNNNGGAGTTGGGGCTTCAACATATG
GCACACGTCTGAACTCTTTCCCTACACGACGCTCTTCCGATCTNNNGGAGTTGGGGCTTCAACATATG
GCACACGTCTGAACTCTTTCCCTACACGACGCTCTTCCGATCTNNNGGAGTTGGGGCTTCAACATATG
GAGCACACGTCTGAACTCTTTCCCTACACGACGCTCTTCCGATCTNNNGCTCTTCTAGGGTGTTTGTC
GAGCACACGTCTGAACTCTTTCCCTACACGACGCTCTTCCGATCTNNNGCTCTTCTAGGGTGTTTGTC
GAGCACACGTCTGAACTCTTTCCCTACACGACGCTCTTCCGATCTNNNGCTCTTCTAGGGTGTTTGTC
GAGCACACGTCTGAACTCTTTCCCTACACGACGCTCTTCCGATCTNNNGCTCTTCTAGGGTGTTTGTC
GAGCACACGTCTGAACTCTTTCCCTACACGACGCTCTTCCGATCTNNNTTCAAAAATACACCTGCACAC

GAGCACACGTCTGAACTCTTTCCCTACACGACGCTCTTCCGATCTNNNTTCAAAAATACACCTGCACAC

GAGCACACGTCTGAACTCTTTCCCTACACGACGCTCTTCCGATCTNNNTTCAAAAATACACCTGCACAC

GAGCACACGTCTGAACTCTTTCCCTACACGACGCTCTTCCGATCTNNNTTCAAAAATACACCTGCACAC

GAGCACACGTCTGAACTCTTTCCCTACACGACGCTCTTCCGATCTNNNACCACAGAGTGAACATGGT

GAGCACACGTCTGAACTCTTTCCCTACACGACGCTCTTCCGATCTNNNACCACAGAGTGAACATGGT

GAGCACACGTCTGAACTCTTTCCCTACACGACGCTCTTCCGATCTNNNACCACAGAGTGAACATGGT

GAGCACACGTCTGAACTCTTTCCCTACACGACGCTCTTCCGATCTNNNACCACAGAGTGAACATGGT

GCACACGTCTGAACTCTTTCCCTACACGACGCTCTTCCGATCTNNNCTCATGCCTATAATCCCAGCACT

GCACACGTCTGAACTCTTTCCCTACACGACGCTCTTCCGATCTNNNCTCATGCCTATAATCCCAGCACT

GCACACGTCTGAACTCTTTCCCTACACGACGCTCTTCCGATCTNNNCTCATGCCTATAATCCCAGCACT

GCACACGTCTGAACTCTTTCCCTACACGACGCTCTTCCGATCTNNNCTCATGCCTATAATCCCAGCACT

AGCACACGTCTGAACTCTTTCCCTACACGACGCTCTTCCGATCTNNNGTTAGGGGCAAAGTCAGGAC

AGCACACGTCTGAACTCTTTCCCTACACGACGCTCTTCCGATCTNNNGTTAGGGGCAAAGTCAGGAC

AGCACACGTCTGAACTCTTTCCCTACACGACGCTCTTCCGATCTNNNGTTAGGGGCAAAGTCAGGAC

AGCACACGTCTGAACTCTTTCCCTACACGACGCTCTTCCGATCTNNNGTTAGGGGCAAAGTCAGGAC

GCACACGTCTGAACTCTTTCCCTACACGACGCTCTTCCGATCTNNNTACTGGGAAGTGAGGAGCCCC

GCACACGTCTGAACTCTTTCCCTACACGACGCTCTTCCGATCTNNNTACTGGGAAGTGAGGAGCCCC

GCACACGTCTGAACTCTTTCCCTACACGACGCTCTTCCGATCTNNNTACTGGGAAGTGAGGAGCCCC

GCACACGTCTGAACTCTTTCCCTACACGACGCTCTTCCGATCTNNNTACTGGGAAGTGAGGAGCCCC

GCACACGTCTGAACTCTTTCCCTACACGACGCTCTTCCGATCTNNNAAACTGGGAGGAGAAACACATC

GCACACGTCTGAACTCTTTCCCTACACGACGCTCTTCCGATCTNNNAAACTGGGAGGAGAAACACATC

GCACACGTCTGAACTCTTTCCCTACACGACGCTCTTCCGATCTNNNAAACTGGGAGGAGAAACACATC

GCACACGTCTGAACTCTTTCCCTACACGACGCTCTTCCGATCTNNNAAACTGGGAGGAGAAACACATC

GCACACGTCTGAACTCTTTCCCTACACGACGCTCTTCCGATCTNNNTTGGGGTTGTCTTTTCACTTCCT

GCACACGTCTGAACTCTTTCCCTACACGACGCTCTTCCGATCTNNNTTGGGGTTGTCTTTTCACTTCCT

GCACACGTCTGAACTCTTTCCCTACACGACGCTCTTCCGATCTNNNTTGGGGTTGTCTTTTCACTTCCT

GCACACGTCTGAACTCTTTCCCTACACGACGCTCTTCCGATCTNNNTTGGGGTTGTCTTTTCACTTCCT

CACGTCTGAACTCTTTCCCTACACGACGCTCTTCCGATCTNNNAATTCCCAGGTGATGCACTTTTGCAT

CACGTCTGAACTCTTTCCCTACACGACGCTCTTCCGATCTNNNAATTCCCAGGTGATGCACTTTTGCAT

CACGTCTGAACTCTTTCCCTACACGACGCTCTTCCGATCTNNNAATTCCCAGGTGATGCACTTTTGCAT

CACGTCTGAACTCTTTCCCTACACGACGCTCTTCCGATCTNNNAATTCCCAGGTGATGCACTTTTGCAT

GCACACGTCTGAACTCTTTCCCTACACGACGCTCTTCCGATCTNNNCCCAGGTTCAAGCAATTCTCATG

GCACACGTCTGAACTCTTTCCCTACACGACGCTCTTCCGATCTNNNCCCAGGTTCAAGCAATTCTCATG

GCACACGTCTGAACTCTTTCCCTACACGACGCTCTTCCGATCTNNNCCCAGGTTCAAGCAATTCTCATG

GCACACGTCTGAACTCTTTCCCTACACGACGCTCTTCCGATCTNNNCCCAGGTTCAAGCAATTCTCATG

GAGCACACGTCTGAACTCTTTCCCTACACGACGCTCTTCCGATCTNNNATCTAAGGAAAGTGCGATG

GAGCACACGTCTGAACTCTTTCCCTACACGACGCTCTTCCGATCTNNNATCTAAGGAAAGTGCGATG

GAGCACACGTCTGAACTCTTTCCCTACACGACGCTCTTCCGATCTNNNATCTAAGGAAAGTGCGATG

GAGCACACGTCTGAACTCTTTCCCTACACGACGCTCTTCCGATCTNNNATCTAAGGAAAGTGCGATG

AGCACACGTCTGAACTCTTTCCCTACACGACGCTCTTCCGATCTNNNGGTAGTTTGTTCAGGGAACC

AGCACACGTCTGAACTCTTTCCCTACACGACGCTCTTCCGATCTNNNGGTAGTTTGTTCAGGGAACC

AGCACACGTCTGAACTCTTTCCCTACACGACGCTCTTCCGATCTNNNGGTAGTTTGTTCAGGGAACC

AGCACACGTCTGAACTCTTTCCCTACACGACGCTCTTCCGATCTNNNGGTAGTTTGTTCAGGGAACC

GCACACGTCTGAACTCTTTCCCTACACGACGCTCTTCCGATCTNNNGTTTGCAACAGCTCTCTGGTAA
GCACACGTCTGAACTCTTTCCCTACACGACGCTCTTCCGATCTNNNGTTTGCAACAGCTCTCTGGTAA
GCACACGTCTGAACTCTTTCCCTACACGACGCTCTTCCGATCTNNNGTTTGCAACAGCTCTCTGGTAA
GCACACGTCTGAACTCTTTCCCTACACGACGCTCTTCCGATCTNNNGTTTGCAACAGCTCTCTGGTAA
AGCACACGTCTGAACTCTTTCCCTACACGACGCTCTTCCGATCTNNNGCCAAAGAGTATTGAAGAGG
AGCACACGTCTGAACTCTTTCCCTACACGACGCTCTTCCGATCTNNNGCCAAAGAGTATTGAAGAGG
AGCACACGTCTGAACTCTTTCCCTACACGACGCTCTTCCGATCTNNNGCCAAAGAGTATTGAAGAGG
AGCACACGTCTGAACTCTTTCCCTACACGACGCTCTTCCGATCTNNNGCCAAAGAGTATTGAAGAGG
AGCACACGTCTGAACTCTTTCCCTACACGACGCTCTTCCGATCTNNNATTCATTTCAACTTGGTACATTT
AGCACACGTCTGAACTCTTTCCCTACACGACGCTCTTCCGATCTNNNATTCATTTCAACTTGGTACATTT
AGCACACGTCTGAACTCTTTCCCTACACGACGCTCTTCCGATCTNNNATTCATTTCAACTTGGTACATTT
AGCACACGTCTGAACTCTTTCCCTACACGACGCTCTTCCGATCTNNNATTCATTTCAACTTGGTACATTT
AGCACACGTCTGAACTCTTTCCCTACACGACGCTCTTCCGATCTNNNACTACTTCAAGGGCTAGGAAT
AGCACACGTCTGAACTCTTTCCCTACACGACGCTCTTCCGATCTNNNACTACTTCAAGGGCTAGGAAT
AGCACACGTCTGAACTCTTTCCCTACACGACGCTCTTCCGATCTNNNACTACTTCAAGGGCTAGGAAT
AGCACACGTCTGAACTCTTTCCCTACACGACGCTCTTCCGATCTNNNACTACTTCAAGGGCTAGGAAT
AGCACACGTCTGAACTCTTTCCCTACACGACGCTCTTCCGATCTNNNCAAACAAACACACACACACACA
AGCACACGTCTGAACTCTTTCCCTACACGACGCTCTTCCGATCTNNNCAAACAAACACACACACACACA
AGCACACGTCTGAACTCTTTCCCTACACGACGCTCTTCCGATCTNNNCAAACAAACACACACACACACA
AGCACACGTCTGAACTCTTTCCCTACACGACGCTCTTCCGATCTNNNCAAACAAACACACACACACACA
AGCACACGTCTGAACTCTTTCCCTACACGACGCTCTTCCGATCTNNNGAGGTTGCAGTGAGCTGAGA
AGCACACGTCTGAACTCTTTCCCTACACGACGCTCTTCCGATCTNNNGAGGTTGCAGTGAGCTGAGA
AGCACACGTCTGAACTCTTTCCCTACACGACGCTCTTCCGATCTNNNGAGGTTGCAGTGAGCTGAGA
AGCACACGTCTGAACTCTTTCCCTACACGACGCTCTTCCGATCTNNNGAGGTTGCAGTGAGCTGAGA
GAGCACACGTCTGAACTCTTTCCCTACACGACGCTCTTCCGATCTNNNTGGAAGGTGAACCGGAAAA
GAGCACACGTCTGAACTCTTTCCCTACACGACGCTCTTCCGATCTNNNTGGAAGGTGAACCGGAAAA
GAGCACACGTCTGAACTCTTTCCCTACACGACGCTCTTCCGATCTNNNTGGAAGGTGAACCGGAAAA
GAGCACACGTCTGAACTCTTTCCCTACACGACGCTCTTCCGATCTNNNTGGAAGGTGAACCGGAAAA
AGCACACGTCTGAACTCTTTCCCTACACGACGCTCTTCCGATCTNNNTGCCCTGTTGCATTAGTTCTTC
AGCACACGTCTGAACTCTTTCCCTACACGACGCTCTTCCGATCTNNNTGCCCTGTTGCATTAGTTCTTC
AGCACACGTCTGAACTCTTTCCCTACACGACGCTCTTCCGATCTNNNTGCCCTGTTGCATTAGTTCTTC
AGCACACGTCTGAACTCTTTCCCTACACGACGCTCTTCCGATCTNNNTGCCCTGTTGCATTAGTTCTTC
AGCACACGTCTGAACTCTTTCCCTACACGACGCTCTTCCGATCTNNNTATACAAATAATGCCACGGTG
AGCACACGTCTGAACTCTTTCCCTACACGACGCTCTTCCGATCTNNNTATACAAATAATGCCACGGTG
AGCACACGTCTGAACTCTTTCCCTACACGACGCTCTTCCGATCTNNNTATACAAATAATGCCACGGTG
AGCACACGTCTGAACTCTTTCCCTACACGACGCTCTTCCGATCTNNNTATACAAATAATGCCACGGTG
AGCACACGTCTGAACTCTTTCCCTACACGACGCTCTTCCGATCTNNNGGAAGAGGAGTAAGCAGTCC
AGCACACGTCTGAACTCTTTCCCTACACGACGCTCTTCCGATCTNNNGGAAGAGGAGTAAGCAGTCC
AGCACACGTCTGAACTCTTTCCCTACACGACGCTCTTCCGATCTNNNGGAAGAGGAGTAAGCAGTCC
AGCACACGTCTGAACTCTTTCCCTACACGACGCTCTTCCGATCTNNNGGAAGAGGAGTAAGCAGTCC
AGCACACGTCTGAACTCTTTCCCTACACGACGCTCTTCCGATCTNNNGCAATGAGAGTGAGACTCCA
AGCACACGTCTGAACTCTTTCCCTACACGACGCTCTTCCGATCTNNNGCAATGAGAGTGAGACTCCA
AGCACACGTCTGAACTCTTTCCCTACACGACGCTCTTCCGATCTNNNGCAATGAGAGTGAGACTCCA

GAGCACACGTCTGAACTCTTTCCCTACACGACGCTCTTCCGATCTNNNGCAATGAGAGTGAGACTCCAT
AGCACACGTCTGAACTCTTTCCCTACACGACGCTCTTCCGATCTNNNGGTGACAGAGAGAGACTCCAT
AGCACACGTCTGAACTCTTTCCCTACACGACGCTCTTCCGATCTNNNGGTGACAGAGAGAGACTCCAT
AGCACACGTCTGAACTCTTTCCCTACACGACGCTCTTCCGATCTNNNGGTGACAGAGAGAGACTCCAT
AGCACACGTCTGAACTCTTTCCCTACACGACGCTCTTCCGATCTNNNGGTGACAGAGAGAGACTCCAT
AGCACACGTCTGAACTCTTTCCCTACACGACGCTCTTCCGATCTNNNAGGATGCTTTCTGTCTCCTCCAT
AGCACACGTCTGAACTCTTTCCCTACACGACGCTCTTCCGATCTNNNAGGATGCTTTCTGTCTCCTCCAT
AGCACACGTCTGAACTCTTTCCCTACACGACGCTCTTCCGATCTNNNAGGATGCTTTCTGTCTCCTCCAT
AGCACACGTCTGAACTCTTTCCCTACACGACGCTCTTCCGATCTNNNAGGATGCTTTCTGTCTCCTCCAT
GAGCACACGTCTGAACTCTTTCCCTACACGACGCTCTTCCGATCTNNNAAATTTAGCCAGGCGTGGTTG
GAGCACACGTCTGAACTCTTTCCCTACACGACGCTCTTCCGATCTNNNAAATTTAGCCAGGCGTGGTTG
GAGCACACGTCTGAACTCTTTCCCTACACGACGCTCTTCCGATCTNNNAAATTTAGCCAGGCGTGGTTG
GAGCACACGTCTGAACTCTTTCCCTACACGACGCTCTTCCGATCTNNNAAATTTAGCCAGGCGTGGTTG
GAGCACACGTCTGAACTCTTTCCCTACACGACGCTCTTCCGATCTNNNTGAGGAGAGAGGAAAGAGA
GAGCACACGTCTGAACTCTTTCCCTACACGACGCTCTTCCGATCTNNNTGAGGAGAGAGGAAAGAGA
GAGCACACGTCTGAACTCTTTCCCTACACGACGCTCTTCCGATCTNNNTGAGGAGAGAGGAAAGAGA
GAGCACACGTCTGAACTCTTTCCCTACACGACGCTCTTCCGATCTNNNTGAGGAGAGAGGAAAGAGA
AGCACACGTCTGAACTCTTTCCCTACACGACGCTCTTCCGATCTNNNCAAGAGAATTGCTTGAACCCAG
AGCACACGTCTGAACTCTTTCCCTACACGACGCTCTTCCGATCTNNNCAAGAGAATTGCTTGAACCCAG
AGCACACGTCTGAACTCTTTCCCTACACGACGCTCTTCCGATCTNNNCAAGAGAATTGCTTGAACCCAG
AGCACACGTCTGAACTCTTTCCCTACACGACGCTCTTCCGATCTNNNCAAGAGAATTGCTTGAACCCAG
AGCACACGTCTGAACTCTTTCCCTACACGACGCTCTTCCGATCTNNNAAATGCAAACACAGATGCCTA
AGCACACGTCTGAACTCTTTCCCTACACGACGCTCTTCCGATCTNNNAAATGCAAACACAGATGCCTA
AGCACACGTCTGAACTCTTTCCCTACACGACGCTCTTCCGATCTNNNAAATGCAAACACAGATGCCTA
AGCACACGTCTGAACTCTTTCCCTACACGACGCTCTTCCGATCTNNNAAATGCAAACACAGATGCCTA
GCACACGTCTGAACTCTTTCCCTACACGACGCTCTTCCGATCTNNNATAGATAGGCCAGCGAGACAAA
GCACACGTCTGAACTCTTTCCCTACACGACGCTCTTCCGATCTNNNATAGATAGGCCAGCGAGACAAA
GCACACGTCTGAACTCTTTCCCTACACGACGCTCTTCCGATCTNNNATAGATAGGCCAGCGAGACAAA
GCACACGTCTGAACTCTTTCCCTACACGACGCTCTTCCGATCTNNNATAGATAGGCCAGCGAGACAAA
AGCACACGTCTGAACTCTTTCCCTACACGACGCTCTTCCGATCTNNNTTTGGTCTGAAGCTGTGTTGTT
AGCACACGTCTGAACTCTTTCCCTACACGACGCTCTTCCGATCTNNNTTTGGTCTGAAGCTGTGTTGTT
AGCACACGTCTGAACTCTTTCCCTACACGACGCTCTTCCGATCTNNNTTTGGTCTGAAGCTGTGTTGTT
AGCACACGTCTGAACTCTTTCCCTACACGACGCTCTTCCGATCTNNNTTTGGTCTGAAGCTGTGTTGTT
AGCACACGTCTGAACTCTTTCCCTACACGACGCTCTTCCGATCTNNNGCTCCTCTTTGGTAGAATTTGG
AGCACACGTCTGAACTCTTTCCCTACACGACGCTCTTCCGATCTNNNGCTCCTCTTTGGTAGAATTTGG
AGCACACGTCTGAACTCTTTCCCTACACGACGCTCTTCCGATCTNNNGCTCCTCTTTGGTAGAATTTGG
AGCACACGTCTGAACTCTTTCCCTACACGACGCTCTTCCGATCTNNNGCTCCTCTTTGGTAGAATTTGG
GCACACGTCTGAACTCTTTCCCTACACGACGCTCTTCCGATCTNNNTTCTTCCTTCTGTTTGCTTTGGACA
GCACACGTCTGAACTCTTTCCCTACACGACGCTCTTCCGATCTNNNTTCTTCCTTCTGTTTGCTTTGGACA
GCACACGTCTGAACTCTTTCCCTACACGACGCTCTTCCGATCTNNNTTCTTCCTTCTGTTTGCTTTGGACA
GCACACGTCTGAACTCTTTCCCTACACGACGCTCTTCCGATCTNNNTTCTTCCTTCTGTTTGCTTTGGACA
GAGCACACGTCTGAACTCTTTCCCTACACGACGCTCTTCCGATCTNNNCTGGCCAAGGGATTTTGTGAA
GAGCACACGTCTGAACTCTTTCCCTACACGACGCTCTTCCGATCTNNNCTGGCCAAGGGATTTTGTGAA

GAGCACACGTCTGAACTCTTTCCCTACACGACGCTCTTCCGATCTNNNCTGGCCAAGGGATTTTGTGAA
GAGCACACGTCTGAACTCTTTCCCTACACGACGCTCTTCCGATCTNNNCTGGCCAAGGGATTTTGTGAA
AGCACACGTCTGAACTCTTTCCCTACACGACGCTCTTCCGATCTNNNTGCTTTCATTAGCACAGAGCAT
AGCACACGTCTGAACTCTTTCCCTACACGACGCTCTTCCGATCTNNNTGCTTTCATTAGCACAGAGCAT
AGCACACGTCTGAACTCTTTCCCTACACGACGCTCTTCCGATCTNNNTGCTTTCATTAGCACAGAGCAT
AGCACACGTCTGAACTCTTTCCCTACACGACGCTCTTCCGATCTNNNTGCTTTCATTAGCACAGAGCAT
AGCACACGTCTGAACTCTTTCCCTACACGACGCTCTTCCGATCTNNNAGTGCTGCAATCTAAGCTCACT
AGCACACGTCTGAACTCTTTCCCTACACGACGCTCTTCCGATCTNNNAGTGCTGCAATCTAAGCTCACT
AGCACACGTCTGAACTCTTTCCCTACACGACGCTCTTCCGATCTNNNAGTGCTGCAATCTAAGCTCACT
AGCACACGTCTGAACTCTTTCCCTACACGACGCTCTTCCGATCTNNNAGTGCTGCAATCTAAGCTCACT
AGCACACGTCTGAACTCTTTCCCTACACGACGCTCTTCCGATCTNNNACATTTTGCAACACTTACAATGC
AGCACACGTCTGAACTCTTTCCCTACACGACGCTCTTCCGATCTNNNACATTTTGCAACACTTACAATGC
AGCACACGTCTGAACTCTTTCCCTACACGACGCTCTTCCGATCTNNNACATTTTGCAACACTTACAATGC
AGCACACGTCTGAACTCTTTCCCTACACGACGCTCTTCCGATCTNNNACATTTTGCAACACTTACAATGC
AGCACACGTCTGAACTCTTTCCCTACACGACGCTCTTCCGATCTNNNAACTCATGTTCACTTTCCCTGGT
AGCACACGTCTGAACTCTTTCCCTACACGACGCTCTTCCGATCTNNNAACTCATGTTCACTTTCCCTGGT
AGCACACGTCTGAACTCTTTCCCTACACGACGCTCTTCCGATCTNNNAACTCATGTTCACTTTCCCTGGT
AGCACACGTCTGAACTCTTTCCCTACACGACGCTCTTCCGATCTNNNAACTCATGTTCACTTTCCCTGGT
GAGCACACGTCTGAACTCTTTCCCTACACGACGCTCTTCCGATCTNNNCAATGGAAGGAAGAACTTGT
GAGCACACGTCTGAACTCTTTCCCTACACGACGCTCTTCCGATCTNNNCAATGGAAGGAAGAACTTGT
GAGCACACGTCTGAACTCTTTCCCTACACGACGCTCTTCCGATCTNNNCAATGGAAGGAAGAACTTGT
GAGCACACGTCTGAACTCTTTCCCTACACGACGCTCTTCCGATCTNNNCAATGGAAGGAAGAACTTGT
GCACACGTCTGAACTCTTTCCCTACACGACGCTCTTCCGATCTNNNAAACTCCTGACCTCAAGTGATCC
GCACACGTCTGAACTCTTTCCCTACACGACGCTCTTCCGATCTNNNAAACTCCTGACCTCAAGTGATCC
GCACACGTCTGAACTCTTTCCCTACACGACGCTCTTCCGATCTNNNAAACTCCTGACCTCAAGTGATCC
GCACACGTCTGAACTCTTTCCCTACACGACGCTCTTCCGATCTNNNAAACTCCTGACCTCAAGTGATCC
GAGCACACGTCTGAACTCTTTCCCTACACGACGCTCTTCCGATCTNNNGGTTCATCCATGTTGTATAGC
GAGCACACGTCTGAACTCTTTCCCTACACGACGCTCTTCCGATCTNNNGGTTCATCCATGTTGTATAGC
GAGCACACGTCTGAACTCTTTCCCTACACGACGCTCTTCCGATCTNNNGGTTCATCCATGTTGTATAGC
GAGCACACGTCTGAACTCTTTCCCTACACGACGCTCTTCCGATCTNNNGGTTCATCCATGTTGTATAGC
AGCACACGTCTGAACTCTTTCCCTACACGACGCTCTTCCGATCTNNNATGTCCTTTTCATCTTCTGCAGC
AGCACACGTCTGAACTCTTTCCCTACACGACGCTCTTCCGATCTNNNATGTCCTTTTCATCTTCTGCAGC
AGCACACGTCTGAACTCTTTCCCTACACGACGCTCTTCCGATCTNNNATGTCCTTTTCATCTTCTGCAGC
AGCACACGTCTGAACTCTTTCCCTACACGACGCTCTTCCGATCTNNNATGTCCTTTTCATCTTCTGCAGC
AGAGCACACGTCTGAACTCTTTCCCTACACGACGCTCTTCCGATCTNNNGCAGAGGTGGAAGGATTAC
AGAGCACACGTCTGAACTCTTTCCCTACACGACGCTCTTCCGATCTNNNGCAGAGGTGGAAGGATTAC
AGAGCACACGTCTGAACTCTTTCCCTACACGACGCTCTTCCGATCTNNNGCAGAGGTGGAAGGATTAC
AGAGCACACGTCTGAACTCTTTCCCTACACGACGCTCTTCCGATCTNNNGCAGAGGTGGAAGGATTAC
AGCACACGTCTGAACTCTTTCCCTACACGACGCTCTTCCGATCTNNNTGCCTTGGAATTTGGTTTGTGAT
AGCACACGTCTGAACTCTTTCCCTACACGACGCTCTTCCGATCTNNNTGCCTTGGAATTTGGTTTGTGAT
AGCACACGTCTGAACTCTTTCCCTACACGACGCTCTTCCGATCTNNNTGCCTTGGAATTTGGTTTGTGAT
AGCACACGTCTGAACTCTTTCCCTACACGACGCTCTTCCGATCTNNNTGCCTTGGAATTTGGTTTGTGAT
GAGCACACGTCTGAACTCTTTCCCTACACGACGCTCTTCCGATCTNNNTTGGAGGAGGGGGTAAAAA

GAGCACACGTCTGAACTCTTTCCCTACACGACGCTCTTCCGATCTNNNTTGGAGGAGGGGGTAAAAA
GAGCACACGTCTGAACTCTTTCCCTACACGACGCTCTTCCGATCTNNNTTGGAGGAGGGGGTAAAAA
GAGCACACGTCTGAACTCTTTCCCTACACGACGCTCTTCCGATCTNNNTTGGAGGAGGGGGTAAAAA
GAGCACACGTCTGAACTCTTTCCCTACACGACGCTCTTCCGATCTNNNTCCCTTGGTTTCCTTGAACTCA
GAGCACACGTCTGAACTCTTTCCCTACACGACGCTCTTCCGATCTNNNTCCCTTGGTTTCCTTGAACTCA
GAGCACACGTCTGAACTCTTTCCCTACACGACGCTCTTCCGATCTNNNTCCCTTGGTTTCCTTGAACTCA
GAGCACACGTCTGAACTCTTTCCCTACACGACGCTCTTCCGATCTNNNTCCCTTGGTTTCCTTGAACTCA
GAGCACACGTCTGAACTCTTTCCCTACACGACGCTCTTCCGATCTNNNTCCTCAAGGTTCATCCATGTTC
GAGCACACGTCTGAACTCTTTCCCTACACGACGCTCTTCCGATCTNNNTCCTCAAGGTTCATCCATGTTC
GAGCACACGTCTGAACTCTTTCCCTACACGACGCTCTTCCGATCTNNNTCCTCAAGGTTCATCCATGTTC
GAGCACACGTCTGAACTCTTTCCCTACACGACGCTCTTCCGATCTNNNTCCTCAAGGTTCATCCATGTTC
CACACGTCTGAACTCTTTCCCTACACGACGCTCTTCCGATCTNNNCTACGAGACCTTCATCAGTGACGCA
CACACGTCTGAACTCTTTCCCTACACGACGCTCTTCCGATCTNNNCTACGAGACCTTCATCAGTGACGCA
CACACGTCTGAACTCTTTCCCTACACGACGCTCTTCCGATCTNNNCTACGAGACCTTCATCAGTGACGCA
CACACGTCTGAACTCTTTCCCTACACGACGCTCTTCCGATCTNNNCTACGAGACCTTCATCAGTGACGCA
GAGCACACGTCTGAACTCTTTCCCTACACGACGCTCTTCCGATCTNNNAGGAACATTTCTCTCATGCCAC
GAGCACACGTCTGAACTCTTTCCCTACACGACGCTCTTCCGATCTNNNAGGAACATTTCTCTCATGCCAC
GAGCACACGTCTGAACTCTTTCCCTACACGACGCTCTTCCGATCTNNNAGGAACATTTCTCTCATGCCAC
GAGCACACGTCTGAACTCTTTCCCTACACGACGCTCTTCCGATCTNNNAGGAACATTTCTCTCATGCCAC
AGCACACGTCTGAACTCTTTCCCTACACGACGCTCTTCCGATCTNNNCCGAGCAACAAGACCAAAAATC
AGCACACGTCTGAACTCTTTCCCTACACGACGCTCTTCCGATCTNNNCCGAGCAACAAGACCAAAAATC
AGCACACGTCTGAACTCTTTCCCTACACGACGCTCTTCCGATCTNNNCCGAGCAACAAGACCAAAAATC
AGCACACGTCTGAACTCTTTCCCTACACGACGCTCTTCCGATCTNNNCCGAGCAACAAGACCAAAAATC
AGCACACGTCTGAACTCTTTCCCTACACGACGCTCTTCCGATCTNNNAAGATCAACCTGGGAAGCGTT
AGCACACGTCTGAACTCTTTCCCTACACGACGCTCTTCCGATCTNNNAAGATCAACCTGGGAAGCGTT
AGCACACGTCTGAACTCTTTCCCTACACGACGCTCTTCCGATCTNNNAAGATCAACCTGGGAAGCGTT
AGCACACGTCTGAACTCTTTCCCTACACGACGCTCTTCCGATCTNNNAAGATCAACCTGGGAAGCGTT
AGCACACGTCTGAACTCTTTCCCTACACGACGCTCTTCCGATCTNNNTCCGACTTCCACTTGCAATACAG
AGCACACGTCTGAACTCTTTCCCTACACGACGCTCTTCCGATCTNNNTCCGACTTCCACTTGCAATACAG
AGCACACGTCTGAACTCTTTCCCTACACGACGCTCTTCCGATCTNNNTCCGACTTCCACTTGCAATACAG
AGCACACGTCTGAACTCTTTCCCTACACGACGCTCTTCCGATCTNNNTCCGACTTCCACTTGCAATACAG
GAGCACACGTCTGAACTCTTTCCCTACACGACGCTCTTCCGATCTNNNACACCTATAGTCCCAGCTACTC
GAGCACACGTCTGAACTCTTTCCCTACACGACGCTCTTCCGATCTNNNACACCTATAGTCCCAGCTACTC
GAGCACACGTCTGAACTCTTTCCCTACACGACGCTCTTCCGATCTNNNACACCTATAGTCCCAGCTACTC
GAGCACACGTCTGAACTCTTTCCCTACACGACGCTCTTCCGATCTNNNACACCTATAGTCCCAGCTACTC
AGCACACGTCTGAACTCTTTCCCTACACGACGCTCTTCCGATCTNNNGCTGGTAAAGTTCAGGTCATG
AGCACACGTCTGAACTCTTTCCCTACACGACGCTCTTCCGATCTNNNGCTGGTAAAGTTCAGGTCATG
AGCACACGTCTGAACTCTTTCCCTACACGACGCTCTTCCGATCTNNNGCTGGTAAAGTTCAGGTCATG
AGCACACGTCTGAACTCTTTCCCTACACGACGCTCTTCCGATCTNNNGCTGGTAAAGTTCAGGTCATG
AAGAGCACACGTCTGAACTCTTTCCCTACACGACGCTCTTCCGATCTNNNGCAAAAAATATGGAACCAC
AAGAGCACACGTCTGAACTCTTTCCCTACACGACGCTCTTCCGATCTNNNGCAAAAAATATGGAACCAC
AAGAGCACACGTCTGAACTCTTTCCCTACACGACGCTCTTCCGATCTNNNGCAAAAAATATGGAACCAC
AAGAGCACACGTCTGAACTCTTTCCCTACACGACGCTCTTCCGATCTNNNGCAAAAAATATGGAACCAC

AGAGCACACGTCTGAACTCTTTCCCTACACGACGCTCTTCCGATCTNNNTCACTATGTTGGTGAGGGAT
AGAGCACACGTCTGAACTCTTTCCCTACACGACGCTCTTCCGATCTNNNTCACTATGTTGGTGAGGGAT
AGAGCACACGTCTGAACTCTTTCCCTACACGACGCTCTTCCGATCTNNNTCACTATGTTGGTGAGGGAT
AGAGCACACGTCTGAACTCTTTCCCTACACGACGCTCTTCCGATCTNNNTCACTATGTTGGTGAGGGAT
AGAGCACACGTCTGAACTCTTTCCCTACACGACGCTCTTCCGATCTNNNGAGAATCACTTGAACCCAGG
AGAGCACACGTCTGAACTCTTTCCCTACACGACGCTCTTCCGATCTNNNGAGAATCACTTGAACCCAGG
AGAGCACACGTCTGAACTCTTTCCCTACACGACGCTCTTCCGATCTNNNGAGAATCACTTGAACCCAGG
AGAGCACACGTCTGAACTCTTTCCCTACACGACGCTCTTCCGATCTNNNGAGAATCACTTGAACCCAGG
AGCACACGTCTGAACTCTTTCCCTACACGACGCTCTTCCGATCTNNNTGGAATGCTTTGAACGAATGAA
AGCACACGTCTGAACTCTTTCCCTACACGACGCTCTTCCGATCTNNNTGGAATGCTTTGAACGAATGAA
AGCACACGTCTGAACTCTTTCCCTACACGACGCTCTTCCGATCTNNNTGGAATGCTTTGAACGAATGAA
AGCACACGTCTGAACTCTTTCCCTACACGACGCTCTTCCGATCTNNNTGGAATGCTTTGAACGAATGAA
GAGCACACGTCTGAACTCTTTCCCTACACGACGCTCTTCCGATCTNNNCAATACTTTGGGAAGCCAAGG
GAGCACACGTCTGAACTCTTTCCCTACACGACGCTCTTCCGATCTNNNCAATACTTTGGGAAGCCAAGG
GAGCACACGTCTGAACTCTTTCCCTACACGACGCTCTTCCGATCTNNNCAATACTTTGGGAAGCCAAGG
GAGCACACGTCTGAACTCTTTCCCTACACGACGCTCTTCCGATCTNNNCAATACTTTGGGAAGCCAAGG
CACACGTCTGAACTCTTTCCCTACACGACGCTCTTCCGATCTNNNAGCTAAGGACTTACTACTGCCATCAT
CACACGTCTGAACTCTTTCCCTACACGACGCTCTTCCGATCTNNNAGCTAAGGACTTACTACTGCCATCAT
CACACGTCTGAACTCTTTCCCTACACGACGCTCTTCCGATCTNNNAGCTAAGGACTTACTACTGCCATCAT
CACACGTCTGAACTCTTTCCCTACACGACGCTCTTCCGATCTNNNAGCTAAGGACTTACTACTGCCATCAT
AGCACACGTCTGAACTCTTTCCCTACACGACGCTCTTCCGATCTNNNTTCAAGTGATTCTCCTGCCTCAG
AGCACACGTCTGAACTCTTTCCCTACACGACGCTCTTCCGATCTNNNTTCAAGTGATTCTCCTGCCTCAG
AGCACACGTCTGAACTCTTTCCCTACACGACGCTCTTCCGATCTNNNTTCAAGTGATTCTCCTGCCTCAG
AGCACACGTCTGAACTCTTTCCCTACACGACGCTCTTCCGATCTNNNTTCAAGTGATTCTCCTGCCTCAG
GCACACGTCTGAACTCTTTCCCTACACGACGCTCTTCCGATCTNNNGAAAACTGGCACGTCTTACCAAA
GCACACGTCTGAACTCTTTCCCTACACGACGCTCTTCCGATCTNNNGAAAACTGGCACGTCTTACCAAA
GCACACGTCTGAACTCTTTCCCTACACGACGCTCTTCCGATCTNNNGAAAACTGGCACGTCTTACCAAA
GCACACGTCTGAACTCTTTCCCTACACGACGCTCTTCCGATCTNNNGAAAACTGGCACGTCTTACCAAA
AGAGCACACGTCTGAACTCTTTCCCTACACGACGCTCTTCCGATCTNNNATTTGGGGAATGAGAGGAA
AGAGCACACGTCTGAACTCTTTCCCTACACGACGCTCTTCCGATCTNNNATTTGGGGAATGAGAGGAA
AGAGCACACGTCTGAACTCTTTCCCTACACGACGCTCTTCCGATCTNNNATTTGGGGAATGAGAGGAA
AGAGCACACGTCTGAACTCTTTCCCTACACGACGCTCTTCCGATCTNNNATTTGGGGAATGAGAGGAA
AGCACACGTCTGAACTCTTTCCCTACACGACGCTCTTCCGATCTNNNAAGGATACCAATAAACGGCACC
AGCACACGTCTGAACTCTTTCCCTACACGACGCTCTTCCGATCTNNNAAGGATACCAATAAACGGCACC
AGCACACGTCTGAACTCTTTCCCTACACGACGCTCTTCCGATCTNNNAAGGATACCAATAAACGGCACC
AGCACACGTCTGAACTCTTTCCCTACACGACGCTCTTCCGATCTNNNAAGGATACCAATAAACGGCACC
GAGCACACGTCTGAACTCTTTCCCTACACGACGCTCTTCCGATCTNNNCAAACCTGTGCTTGGTACATC
GAGCACACGTCTGAACTCTTTCCCTACACGACGCTCTTCCGATCTNNNCAAACCTGTGCTTGGTACATC
GAGCACACGTCTGAACTCTTTCCCTACACGACGCTCTTCCGATCTNNNCAAACCTGTGCTTGGTACATC
GAGCACACGTCTGAACTCTTTCCCTACACGACGCTCTTCCGATCTNNNCAAACCTGTGCTTGGTACATC
AGAGCACACGTCTGAACTCTTTCCCTACACGACGCTCTTCCGATCTNNNCATCGGGTCAGTCATAGAGC
AGAGCACACGTCTGAACTCTTTCCCTACACGACGCTCTTCCGATCTNNNCATCGGGTCAGTCATAGAGC
AGAGCACACGTCTGAACTCTTTCCCTACACGACGCTCTTCCGATCTNNNCATCGGGTCAGTCATAGAGC

GAGCACACGTCTGAACTCTTTCCCTACACGACGCTCTTCCGATCTNNNCATCGGGTCAGTCATAGAGC
AGCACACGTCTGAACTCTTTCCCTACACGACGCTCTTCCGATCTNNNTTCAAGGACTTTTTGGCATCTG
AGCACACGTCTGAACTCTTTCCCTACACGACGCTCTTCCGATCTNNNTTCAAGGACTTTTTGGCATCTG
AGCACACGTCTGAACTCTTTCCCTACACGACGCTCTTCCGATCTNNNTTCAAGGACTTTTTGGCATCTG
AGCACACGTCTGAACTCTTTCCCTACACGACGCTCTTCCGATCTNNNTTCAAGGACTTTTTGGCATCTG
ACACGTCTGAACTCTTTCCCTACACGACGCTCTTCCGATCTNNNTCATATCATTAGGTACACAGAAACA
ACACGTCTGAACTCTTTCCCTACACGACGCTCTTCCGATCTNNNTCATATCATTAGGTACACAGAAACA
ACACGTCTGAACTCTTTCCCTACACGACGCTCTTCCGATCTNNNTCATATCATTAGGTACACAGAAACA
ACACGTCTGAACTCTTTCCCTACACGACGCTCTTCCGATCTNNNTCATATCATTAGGTACACAGAAACA
AGCACACGTCTGAACTCTTTCCCTACACGACGCTCTTCCGATCTNNNGGTCTTGAACTCCTGACCTCAG
AGCACACGTCTGAACTCTTTCCCTACACGACGCTCTTCCGATCTNNNGGTCTTGAACTCCTGACCTCAG
AGCACACGTCTGAACTCTTTCCCTACACGACGCTCTTCCGATCTNNNGGTCTTGAACTCCTGACCTCAG
AGCACACGTCTGAACTCTTTCCCTACACGACGCTCTTCCGATCTNNNGGTCTTGAACTCCTGACCTCAG
GCACACGTCTGAACTCTTTCCCTACACGACGCTCTTCCGATCTNNNGGAGTTGCTCTTCCTTCTCCTTTC
GCACACGTCTGAACTCTTTCCCTACACGACGCTCTTCCGATCTNNNGGAGTTGCTCTTCCTTCTCCTTTC
GCACACGTCTGAACTCTTTCCCTACACGACGCTCTTCCGATCTNNNGGAGTTGCTCTTCCTTCTCCTTTC
GCACACGTCTGAACTCTTTCCCTACACGACGCTCTTCCGATCTNNNGGAGTTGCTCTTCCTTCTCCTTTC
AGCACACGTCTGAACTCTTTCCCTACACGACGCTCTTCCGATCTNNNCACTTGGATCCCGTACCTGACCA
AGCACACGTCTGAACTCTTTCCCTACACGACGCTCTTCCGATCTNNNCACTTGGATCCCGTACCTGACCA
AGCACACGTCTGAACTCTTTCCCTACACGACGCTCTTCCGATCTNNNCACTTGGATCCCGTACCTGACCA
AGCACACGTCTGAACTCTTTCCCTACACGACGCTCTTCCGATCTNNNCACTTGGATCCCGTACCTGACCA
AGCACACGTCTGAACTCTTTCCCTACACGACGCTCTTCCGATCTNNNCGTGTGAATGAAAAGGATGTG
AGCACACGTCTGAACTCTTTCCCTACACGACGCTCTTCCGATCTNNNCGTGTGAATGAAAAGGATGTG
AGCACACGTCTGAACTCTTTCCCTACACGACGCTCTTCCGATCTNNNCGTGTGAATGAAAAGGATGTG
AGCACACGTCTGAACTCTTTCCCTACACGACGCTCTTCCGATCTNNNCGTGTGAATGAAAAGGATGTG
AGCACACGTCTGAACTCTTTCCCTACACGACGCTCTTCCGATCTNNNCACCTGTAATCCCAGCTACTCAG
AGCACACGTCTGAACTCTTTCCCTACACGACGCTCTTCCGATCTNNNCACCTGTAATCCCAGCTACTCAG
AGCACACGTCTGAACTCTTTCCCTACACGACGCTCTTCCGATCTNNNCACCTGTAATCCCAGCTACTCAG
AGCACACGTCTGAACTCTTTCCCTACACGACGCTCTTCCGATCTNNNCACCTGTAATCCCAGCTACTCAG
GAGCACACGTCTGAACTCTTTCCCTACACGACGCTCTTCCGATCTNNNTAGGGAGACTGAGGCAGGAA
GAGCACACGTCTGAACTCTTTCCCTACACGACGCTCTTCCGATCTNNNTAGGGAGACTGAGGCAGGAA
GAGCACACGTCTGAACTCTTTCCCTACACGACGCTCTTCCGATCTNNNTAGGGAGACTGAGGCAGGAA
GAGCACACGTCTGAACTCTTTCCCTACACGACGCTCTTCCGATCTNNNTAGGGAGACTGAGGCAGGAA
AAGAGCACACGTCTGAACTCTTTCCCTACACGACGCTCTTCCGATCTNNNGAAGATGCAGTGAGCCAC
AAGAGCACACGTCTGAACTCTTTCCCTACACGACGCTCTTCCGATCTNNNGAAGATGCAGTGAGCCAC
AAGAGCACACGTCTGAACTCTTTCCCTACACGACGCTCTTCCGATCTNNNGAAGATGCAGTGAGCCAC
AAGAGCACACGTCTGAACTCTTTCCCTACACGACGCTCTTCCGATCTNNNGAAGATGCAGTGAGCCAC
GAGCACACGTCTGAACTCTTTCCCTACACGACGCTCTTCCGATCTNNNCCATCAACTCCAAAGTCTAAGG
GAGCACACGTCTGAACTCTTTCCCTACACGACGCTCTTCCGATCTNNNCCATCAACTCCAAAGTCTAAGG
GAGCACACGTCTGAACTCTTTCCCTACACGACGCTCTTCCGATCTNNNCCATCAACTCCAAAGTCTAAGG
GAGCACACGTCTGAACTCTTTCCCTACACGACGCTCTTCCGATCTNNNCCATCAACTCCAAAGTCTAAGG
GAGCACACGTCTGAACTCTTTCCCTACACGACGCTCTTCCGATCTNNNAATAAAAAAGTCAAGGGTGC
GAGCACACGTCTGAACTCTTTCCCTACACGACGCTCTTCCGATCTNNNAATAAAAAAGTCAAGGGTGC

GAGCACACGTCTGAACTCTTTCCCTACACGACGCTCTTCCGATCTNNNAATAAAAAAGTCAAGGGTGC
GAGCACACGTCTGAACTCTTTCCCTACACGACGCTCTTCCGATCTNNNAATAAAAAAGTCAAGGGTGC
AGCACACGTCTGAACTCTTTCCCTACACGACGCTCTTCCGATCTNNNGTGTGCCTGTAACCTTAGCTACT
AGCACACGTCTGAACTCTTTCCCTACACGACGCTCTTCCGATCTNNNGTGTGCCTGTAACCTTAGCTACT
AGCACACGTCTGAACTCTTTCCCTACACGACGCTCTTCCGATCTNNNGTGTGCCTGTAACCTTAGCTACT
AGCACACGTCTGAACTCTTTCCCTACACGACGCTCTTCCGATCTNNNGTGTGCCTGTAACCTTAGCTACT
GAGCACACGTCTGAACTCTTTCCCTACACGACGCTCTTCCGATCTNNNTTTTGCATTTGCTTCTGTTGT
GAGCACACGTCTGAACTCTTTCCCTACACGACGCTCTTCCGATCTNNNTTTTGCATTTGCTTCTGTTGT
GAGCACACGTCTGAACTCTTTCCCTACACGACGCTCTTCCGATCTNNNTTTTGCATTTGCTTCTGTTGT
GAGCACACGTCTGAACTCTTTCCCTACACGACGCTCTTCCGATCTNNNTTTTGCATTTGCTTCTGTTGT
GAGCACACGTCTGAACTCTTTCCCTACACGACGCTCTTCCGATCTNNNATCACTTGAAACCTTGAGGCA
GAGCACACGTCTGAACTCTTTCCCTACACGACGCTCTTCCGATCTNNNATCACTTGAAACCTTGAGGCA
GAGCACACGTCTGAACTCTTTCCCTACACGACGCTCTTCCGATCTNNNATCACTTGAAACCTTGAGGCA
GAGCACACGTCTGAACTCTTTCCCTACACGACGCTCTTCCGATCTNNNATCACTTGAAACCTTGAGGCA
GCACACGTCTGAACTCTTTCCCTACACGACGCTCTTCCGATCTNNNGATGAAACGTATGCAGATGGCAT
GCACACGTCTGAACTCTTTCCCTACACGACGCTCTTCCGATCTNNNGATGAAACGTATGCAGATGGCAT
GCACACGTCTGAACTCTTTCCCTACACGACGCTCTTCCGATCTNNNGATGAAACGTATGCAGATGGCAT
GCACACGTCTGAACTCTTTCCCTACACGACGCTCTTCCGATCTNNNGATGAAACGTATGCAGATGGCAT
AGCACACGTCTGAACTCTTTCCCTACACGACGCTCTTCCGATCTNNNGCAATCCCTATCAAAATCCCAGC
AGCACACGTCTGAACTCTTTCCCTACACGACGCTCTTCCGATCTNNNGCAATCCCTATCAAAATCCCAGC
AGCACACGTCTGAACTCTTTCCCTACACGACGCTCTTCCGATCTNNNGCAATCCCTATCAAAATCCCAGC
AGCACACGTCTGAACTCTTTCCCTACACGACGCTCTTCCGATCTNNNGCAATCCCTATCAAAATCCCAGC
GAGCACACGTCTGAACTCTTTCCCTACACGACGCTCTTCCGATCTNNNGAGGATGTAGGGATCTGGG
GAGCACACGTCTGAACTCTTTCCCTACACGACGCTCTTCCGATCTNNNGAGGATGTAGGGATCTGGG
GAGCACACGTCTGAACTCTTTCCCTACACGACGCTCTTCCGATCTNNNGAGGATGTAGGGATCTGGG
GAGCACACGTCTGAACTCTTTCCCTACACGACGCTCTTCCGATCTNNNGAGGATGTAGGGATCTGGG
AGCACACGTCTGAACTCTTTCCCTACACGACGCTCTTCCGATCTNNNCTCCTGAGTAGCTGGGACTACA
AGCACACGTCTGAACTCTTTCCCTACACGACGCTCTTCCGATCTNNNCTCCTGAGTAGCTGGGACTACA
AGCACACGTCTGAACTCTTTCCCTACACGACGCTCTTCCGATCTNNNCTCCTGAGTAGCTGGGACTACA
AGCACACGTCTGAACTCTTTCCCTACACGACGCTCTTCCGATCTNNNCTCCTGAGTAGCTGGGACTACA
AGCACACGTCTGAACTCTTTCCCTACACGACGCTCTTCCGATCTNNNCTGAGGCAGGAGAATCACTTGA
AGCACACGTCTGAACTCTTTCCCTACACGACGCTCTTCCGATCTNNNCTGAGGCAGGAGAATCACTTGA
AGCACACGTCTGAACTCTTTCCCTACACGACGCTCTTCCGATCTNNNCTGAGGCAGGAGAATCACTTGA
AGCACACGTCTGAACTCTTTCCCTACACGACGCTCTTCCGATCTNNNCTGAGGCAGGAGAATCACTTGA
GCACACGTCTGAACTCTTTCCCTACACGACGCTCTTCCGATCTNNNGTGGTATTGCATTGTGGTTTCGA
GCACACGTCTGAACTCTTTCCCTACACGACGCTCTTCCGATCTNNNGTGGTATTGCATTGTGGTTTCGA
GCACACGTCTGAACTCTTTCCCTACACGACGCTCTTCCGATCTNNNGTGGTATTGCATTGTGGTTTCGA
GCACACGTCTGAACTCTTTCCCTACACGACGCTCTTCCGATCTNNNGTGGTATTGCATTGTGGTTTCGA
AGCACACGTCTGAACTCTTTCCCTACACGACGCTCTTCCGATCTNNNTGGGTGTTTGCATAGCAGAATT
AGCACACGTCTGAACTCTTTCCCTACACGACGCTCTTCCGATCTNNNTGGGTGTTTGCATAGCAGAATT
AGCACACGTCTGAACTCTTTCCCTACACGACGCTCTTCCGATCTNNNTGGGTGTTTGCATAGCAGAATT
AGCACACGTCTGAACTCTTTCCCTACACGACGCTCTTCCGATCTNNNTGGGTGTTTGCATAGCAGAATT
AGCACACGTCTGAACTCTTTCCCTACACGACGCTCTTCCGATCTNNNACAGCTGTACCAATAGGAAAG

AGCACACGTCTGAACTCTTTCCCTACACGACGCTCTTCCGATCTNNNACAGCTGTACCAATAGGAAAG
AGCACACGTCTGAACTCTTTCCCTACACGACGCTCTTCCGATCTNNNACAGCTGTACCAATAGGAAAG
AGCACACGTCTGAACTCTTTCCCTACACGACGCTCTTCCGATCTNNNACAGCTGTACCAATAGGAAAG
GAGCACACGTCTGAACTCTTTCCCTACACGACGCTCTTCCGATCTNNNTGTCCCTGTTTCTCTCTCTC
GAGCACACGTCTGAACTCTTTCCCTACACGACGCTCTTCCGATCTNNNTGTCCCTGTTTCTCTCTCTC
GAGCACACGTCTGAACTCTTTCCCTACACGACGCTCTTCCGATCTNNNTGTCCCTGTTTCTCTCTCTC
GAGCACACGTCTGAACTCTTTCCCTACACGACGCTCTTCCGATCTNNNTGTCCCTGTTTCTCTCTCTC
GAGCACACGTCTGAACTCTTTCCCTACACGACGCTCTTCCGATCTNNNGCATTGTTTTTGCTGCTGTT
GAGCACACGTCTGAACTCTTTCCCTACACGACGCTCTTCCGATCTNNNGCATTGTTTTTGCTGCTGTT
GAGCACACGTCTGAACTCTTTCCCTACACGACGCTCTTCCGATCTNNNGCATTGTTTTTGCTGCTGTT
GAGCACACGTCTGAACTCTTTCCCTACACGACGCTCTTCCGATCTNNNGCATTGTTTTTGCTGCTGTT
AGCACACGTCTGAACTCTTTCCCTACACGACGCTCTTCCGATCTNNNGAGATCCTGCTACTGCACTCTA
AGCACACGTCTGAACTCTTTCCCTACACGACGCTCTTCCGATCTNNNGAGATCCTGCTACTGCACTCTA
AGCACACGTCTGAACTCTTTCCCTACACGACGCTCTTCCGATCTNNNGAGATCCTGCTACTGCACTCTA
AGCACACGTCTGAACTCTTTCCCTACACGACGCTCTTCCGATCTNNNGAGATCCTGCTACTGCACTCTA
GAGCACACGTCTGAACTCTTTCCCTACACGACGCTCTTCCGATCTNNNCAGATCATGAGGTCAGGAG
GAGCACACGTCTGAACTCTTTCCCTACACGACGCTCTTCCGATCTNNNCAGATCATGAGGTCAGGAG
GAGCACACGTCTGAACTCTTTCCCTACACGACGCTCTTCCGATCTNNNCAGATCATGAGGTCAGGAG
GAGCACACGTCTGAACTCTTTCCCTACACGACGCTCTTCCGATCTNNNCAGATCATGAGGTCAGGAG
CACACGTCTGAACTCTTTCCCTACACGACGCTCTTCCGATCTNNNATGCAAAAATGGGATGATGTGCT
CACACGTCTGAACTCTTTCCCTACACGACGCTCTTCCGATCTNNNATGCAAAAATGGGATGATGTGCT
CACACGTCTGAACTCTTTCCCTACACGACGCTCTTCCGATCTNNNATGCAAAAATGGGATGATGTGCT
CACACGTCTGAACTCTTTCCCTACACGACGCTCTTCCGATCTNNNATGCAAAAATGGGATGATGTGCT
GCACACGTCTGAACTCTTTCCCTACACGACGCTCTTCCGATCTNNNCAGACAAGAGTAGCCCACACAG
GCACACGTCTGAACTCTTTCCCTACACGACGCTCTTCCGATCTNNNCAGACAAGAGTAGCCCACACAG
GCACACGTCTGAACTCTTTCCCTACACGACGCTCTTCCGATCTNNNCAGACAAGAGTAGCCCACACAG
GCACACGTCTGAACTCTTTCCCTACACGACGCTCTTCCGATCTNNNCAGACAAGAGTAGCCCACACAG
AGCACACGTCTGAACTCTTTCCCTACACGACGCTCTTCCGATCTNNNGCATATGCCTGTATTCCCAGCT
AGCACACGTCTGAACTCTTTCCCTACACGACGCTCTTCCGATCTNNNGCATATGCCTGTATTCCCAGCT
AGCACACGTCTGAACTCTTTCCCTACACGACGCTCTTCCGATCTNNNGCATATGCCTGTATTCCCAGCT
AGCACACGTCTGAACTCTTTCCCTACACGACGCTCTTCCGATCTNNNGCATATGCCTGTATTCCCAGCT
AGCACACGTCTGAACTCTTTCCCTACACGACGCTCTTCCGATCTNNNTGGGCTTCCAAGTGGATACTA
AGCACACGTCTGAACTCTTTCCCTACACGACGCTCTTCCGATCTNNNTGGGCTTCCAAGTGGATACTA
AGCACACGTCTGAACTCTTTCCCTACACGACGCTCTTCCGATCTNNNTGGGCTTCCAAGTGGATACTA
AGCACACGTCTGAACTCTTTCCCTACACGACGCTCTTCCGATCTNNNTGGGCTTCCAAGTGGATACTA
GCACACGTCTGAACTCTTTCCCTACACGACGCTCTTCCGATCTNNNGAATTCACACCATTGCACTCCAG
GCACACGTCTGAACTCTTTCCCTACACGACGCTCTTCCGATCTNNNGAATTCACACCATTGCACTCCAG
GCACACGTCTGAACTCTTTCCCTACACGACGCTCTTCCGATCTNNNGAATTCACACCATTGCACTCCAG
GCACACGTCTGAACTCTTTCCCTACACGACGCTCTTCCGATCTNNNGAATTCACACCATTGCACTCCAG
CACACGTCTGAACTCTTTCCCTACACGACGCTCTTCCGATCTNNNTCCAAAGCCTTTCTTCCTCTCTTCA
CACACGTCTGAACTCTTTCCCTACACGACGCTCTTCCGATCTNNNTCCAAAGCCTTTCTTCCTCTCTTCA
CACACGTCTGAACTCTTTCCCTACACGACGCTCTTCCGATCTNNNTCCAAAGCCTTTCTTCCTCTCTTCA
CACACGTCTGAACTCTTTCCCTACACGACGCTCTTCCGATCTNNNTCCAAAGCCTTTCTTCCTCTCTTCA

GAGCACACGTCTGAACTCTTTCCCTACACGACGCTCTTCCGATCTNNNTCTGTTCCCATACATCTTCCTG
GAGCACACGTCTGAACTCTTTCCCTACACGACGCTCTTCCGATCTNNNTCTGTTCCCATACATCTTCCTG
GAGCACACGTCTGAACTCTTTCCCTACACGACGCTCTTCCGATCTNNNTCTGTTCCCATACATCTTCCTG
GAGCACACGTCTGAACTCTTTCCCTACACGACGCTCTTCCGATCTNNNTCTGTTCCCATACATCTTCCTG
ACACGTCTGAACTCTTTCCCTACACGACGCTCTTCCGATCTNNNTAGAAAAAGTTAAACCAAACAGGC
ACACGTCTGAACTCTTTCCCTACACGACGCTCTTCCGATCTNNNTAGAAAAAGTTAAACCAAACAGGC
ACACGTCTGAACTCTTTCCCTACACGACGCTCTTCCGATCTNNNTAGAAAAAGTTAAACCAAACAGGC
ACACGTCTGAACTCTTTCCCTACACGACGCTCTTCCGATCTNNNTAGAAAAAGTTAAACCAAACAGGC
GAGCACACGTCTGAACTCTTTCCCTACACGACGCTCTTCCGATCTNNNAGGGCAAGAAATCATTCCAG
GAGCACACGTCTGAACTCTTTCCCTACACGACGCTCTTCCGATCTNNNAGGGCAAGAAATCATTCCAG
GAGCACACGTCTGAACTCTTTCCCTACACGACGCTCTTCCGATCTNNNAGGGCAAGAAATCATTCCAG
GAGCACACGTCTGAACTCTTTCCCTACACGACGCTCTTCCGATCTNNNAGGGCAAGAAATCATTCCAG
AGCACACGTCTGAACTCTTTCCCTACACGACGCTCTTCCGATCTNNNTAGAGACACGGTTTCACCATG
AGCACACGTCTGAACTCTTTCCCTACACGACGCTCTTCCGATCTNNNTAGAGACACGGTTTCACCATG
AGCACACGTCTGAACTCTTTCCCTACACGACGCTCTTCCGATCTNNNTAGAGACACGGTTTCACCATG
AGCACACGTCTGAACTCTTTCCCTACACGACGCTCTTCCGATCTNNNTAGAGACACGGTTTCACCATG
GCACACGTCTGAACTCTTTCCCTACACGACGCTCTTCCGATCTNNNTATGGGAAATGTGTGTGCATGT
GCACACGTCTGAACTCTTTCCCTACACGACGCTCTTCCGATCTNNNTATGGGAAATGTGTGTGCATGT
GCACACGTCTGAACTCTTTCCCTACACGACGCTCTTCCGATCTNNNTATGGGAAATGTGTGTGCATGT
GCACACGTCTGAACTCTTTCCCTACACGACGCTCTTCCGATCTNNNTATGGGAAATGTGTGTGCATGT
AGCACACGTCTGAACTCTTTCCCTACACGACGCTCTTCCGATCTNNNGCCCTAGTTGACATATGCTTCG
AGCACACGTCTGAACTCTTTCCCTACACGACGCTCTTCCGATCTNNNGCCCTAGTTGACATATGCTTCG
AGCACACGTCTGAACTCTTTCCCTACACGACGCTCTTCCGATCTNNNGCCCTAGTTGACATATGCTTCG
AGCACACGTCTGAACTCTTTCCCTACACGACGCTCTTCCGATCTNNNGCCCTAGTTGACATATGCTTCG
ACACGTCTGAACTCTTTCCCTACACGACGCTCTTCCGATCTNNNGGAAGTTTTCTGAAGATCAAAGCA
ACACGTCTGAACTCTTTCCCTACACGACGCTCTTCCGATCTNNNGGAAGTTTTCTGAAGATCAAAGCA
ACACGTCTGAACTCTTTCCCTACACGACGCTCTTCCGATCTNNNGGAAGTTTTCTGAAGATCAAAGCA
ACACGTCTGAACTCTTTCCCTACACGACGCTCTTCCGATCTNNNGGAAGTTTTCTGAAGATCAAAGCA
GAGCACACGTCTGAACTCTTTCCCTACACGACGCTCTTCCGATCTNNNTAAGGTAAGGCTGTCCTTCT
GAGCACACGTCTGAACTCTTTCCCTACACGACGCTCTTCCGATCTNNNTAAGGTAAGGCTGTCCTTCT
GAGCACACGTCTGAACTCTTTCCCTACACGACGCTCTTCCGATCTNNNTAAGGTAAGGCTGTCCTTCT
GAGCACACGTCTGAACTCTTTCCCTACACGACGCTCTTCCGATCTNNNTAAGGTAAGGCTGTCCTTCT
AGCACACGTCTGAACTCTTTCCCTACACGACGCTCTTCCGATCTNNNCTGAATTCACCCCCATTCGTTT
AGCACACGTCTGAACTCTTTCCCTACACGACGCTCTTCCGATCTNNNCTGAATTCACCCCCATTCGTTT
AGCACACGTCTGAACTCTTTCCCTACACGACGCTCTTCCGATCTNNNCTGAATTCACCCCCATTCGTTT
AGCACACGTCTGAACTCTTTCCCTACACGACGCTCTTCCGATCTNNNCTGAATTCACCCCCATTCGTTT
AGCACACGTCTGAACTCTTTCCCTACACGACGCTCTTCCGATCTNNNACAACTGTAATCCCAGCTACT
AGCACACGTCTGAACTCTTTCCCTACACGACGCTCTTCCGATCTNNNACAACTGTAATCCCAGCTACT
AGCACACGTCTGAACTCTTTCCCTACACGACGCTCTTCCGATCTNNNACAACTGTAATCCCAGCTACT
AGCACACGTCTGAACTCTTTCCCTACACGACGCTCTTCCGATCTNNNACAACTGTAATCCCAGCTACT
GCACACGTCTGAACTCTTTCCCTACACGACGCTCTTCCGATCTNNNGATTCAAGCATTTCTCCTGCCTCC
GCACACGTCTGAACTCTTTCCCTACACGACGCTCTTCCGATCTNNNGATTCAAGCATTTCTCCTGCCTCC
GCACACGTCTGAACTCTTTCCCTACACGACGCTCTTCCGATCTNNNGATTCAAGCATTTCTCCTGCCTCC

GCACACGTCTGAACTCTTTCCCTACACGACGCTCTTCCGATCTNNNGATTCAAGCATTTCTCCTGCCTCC
AGAGCACACGTCTGAACTCTTTCCCTACACGACGCTCTTCCGATCTNNNATACAAAAATTAGCTGGGCG
AGAGCACACGTCTGAACTCTTTCCCTACACGACGCTCTTCCGATCTNNNATACAAAAATTAGCTGGGCG
AGAGCACACGTCTGAACTCTTTCCCTACACGACGCTCTTCCGATCTNNNATACAAAAATTAGCTGGGCG
AGAGCACACGTCTGAACTCTTTCCCTACACGACGCTCTTCCGATCTNNNATACAAAAATTAGCTGGGCG
AGCACACGTCTGAACTCTTTCCCTACACGACGCTCTTCCGATCTNNNGCCATTGTGAATAATGCTGCTCT
AGCACACGTCTGAACTCTTTCCCTACACGACGCTCTTCCGATCTNNNGCCATTGTGAATAATGCTGCTCT
AGCACACGTCTGAACTCTTTCCCTACACGACGCTCTTCCGATCTNNNGCCATTGTGAATAATGCTGCTCT
AGCACACGTCTGAACTCTTTCCCTACACGACGCTCTTCCGATCTNNNGCCATTGTGAATAATGCTGCTCT
GAGCACACGTCTGAACTCTTTCCCTACACGACGCTCTTCCGATCTNNNCAGCCTCCGAAAATGTGGAAA
GAGCACACGTCTGAACTCTTTCCCTACACGACGCTCTTCCGATCTNNNCAGCCTCCGAAAATGTGGAAA
GAGCACACGTCTGAACTCTTTCCCTACACGACGCTCTTCCGATCTNNNCAGCCTCCGAAAATGTGGAAA
GAGCACACGTCTGAACTCTTTCCCTACACGACGCTCTTCCGATCTNNNCAGCCTCCGAAAATGTGGAAA
AGAGCACACGTCTGAACTCTTTCCCTACACGACGCTCTTCCGATCTNNNAGAATCACTTGAACCTGGGA
AGAGCACACGTCTGAACTCTTTCCCTACACGACGCTCTTCCGATCTNNNAGAATCACTTGAACCTGGGA
AGAGCACACGTCTGAACTCTTTCCCTACACGACGCTCTTCCGATCTNNNAGAATCACTTGAACCTGGGA
AGAGCACACGTCTGAACTCTTTCCCTACACGACGCTCTTCCGATCTNNNAGAATCACTTGAACCTGGGA
GAGCACACGTCTGAACTCTTTCCCTACACGACGCTCTTCCGATCTNNNTTTATCAATAATCTTGGAGGCT
GAGCACACGTCTGAACTCTTTCCCTACACGACGCTCTTCCGATCTNNNTTTATCAATAATCTTGGAGGCT
GAGCACACGTCTGAACTCTTTCCCTACACGACGCTCTTCCGATCTNNNTTTATCAATAATCTTGGAGGCT
GAGCACACGTCTGAACTCTTTCCCTACACGACGCTCTTCCGATCTNNNTTTATCAATAATCTTGGAGGCT
AGCACACGTCTGAACTCTTTCCCTACACGACGCTCTTCCGATCTNNNTGCTTTTCCTTTCTGAAGATGCT
AGCACACGTCTGAACTCTTTCCCTACACGACGCTCTTCCGATCTNNNTGCTTTTCCTTTCTGAAGATGCT
AGCACACGTCTGAACTCTTTCCCTACACGACGCTCTTCCGATCTNNNTGCTTTTCCTTTCTGAAGATGCT
AGCACACGTCTGAACTCTTTCCCTACACGACGCTCTTCCGATCTNNNTGCTTTTCCTTTCTGAAGATGCT
GAGCACACGTCTGAACTCTTTCCCTACACGACGCTCTTCCGATCTNNNGAGTGAAATCCTCACGCTTCCT
GAGCACACGTCTGAACTCTTTCCCTACACGACGCTCTTCCGATCTNNNGAGTGAAATCCTCACGCTTCCT
GAGCACACGTCTGAACTCTTTCCCTACACGACGCTCTTCCGATCTNNNGAGTGAAATCCTCACGCTTCCT
GAGCACACGTCTGAACTCTTTCCCTACACGACGCTCTTCCGATCTNNNGAGTGAAATCCTCACGCTTCCT
GCACACGTCTGAACTCTTTCCCTACACGACGCTCTTCCGATCTNNNATCCTGATGTACCCCTGCAAAATC
GCACACGTCTGAACTCTTTCCCTACACGACGCTCTTCCGATCTNNNATCCTGATGTACCCCTGCAAAATC
GCACACGTCTGAACTCTTTCCCTACACGACGCTCTTCCGATCTNNNATCCTGATGTACCCCTGCAAAATC
GCACACGTCTGAACTCTTTCCCTACACGACGCTCTTCCGATCTNNNATCCTGATGTACCCCTGCAAAATC
GAGCACACGTCTGAACTCTTTCCCTACACGACGCTCTTCCGATCTNNNTCATGCAAATGAATCCCCTGGA
GAGCACACGTCTGAACTCTTTCCCTACACGACGCTCTTCCGATCTNNNTCATGCAAATGAATCCCCTGGA
GAGCACACGTCTGAACTCTTTCCCTACACGACGCTCTTCCGATCTNNNTCATGCAAATGAATCCCCTGGA
GAGCACACGTCTGAACTCTTTCCCTACACGACGCTCTTCCGATCTNNNTCATGCAAATGAATCCCCTGGA
AGCACACGTCTGAACTCTTTCCCTACACGACGCTCTTCCGATCTNNNCCCTGATGATGCCTTGTGTTTTT
AGCACACGTCTGAACTCTTTCCCTACACGACGCTCTTCCGATCTNNNCCCTGATGATGCCTTGTGTTTTT
AGCACACGTCTGAACTCTTTCCCTACACGACGCTCTTCCGATCTNNNCCCTGATGATGCCTTGTGTTTTT
AGCACACGTCTGAACTCTTTCCCTACACGACGCTCTTCCGATCTNNNCCCTGATGATGCCTTGTGTTTTT
AAGAGCACACGTCTGAACTCTTTCCCTACACGACGCTCTTCCGATCTNNNCTCCAGCCTGGGTGACGAA
AAGAGCACACGTCTGAACTCTTTCCCTACACGACGCTCTTCCGATCTNNNCTCCAGCCTGGGTGACGAA

AGAGCACACGTCTGAACTCTTTCCCTACACGACGCTCTTCCGATCTNNNCTCCAGCCTGGGTGACGAA
AGAGCACACGTCTGAACTCTTTCCCTACACGACGCTCTTCCGATCTNNNCTCCAGCCTGGGTGACGAA
AGCACACGTCTGAACTCTTTCCCTACACGACGCTCTTCCGATCTNNNTCCCTATCATAGCCTAAGAAGC
AGCACACGTCTGAACTCTTTCCCTACACGACGCTCTTCCGATCTNNNTCCCTATCATAGCCTAAGAAGC
AGCACACGTCTGAACTCTTTCCCTACACGACGCTCTTCCGATCTNNNTCCCTATCATAGCCTAAGAAGC
AGCACACGTCTGAACTCTTTCCCTACACGACGCTCTTCCGATCTNNNTCCCTATCATAGCCTAAGAAGC
GCACACGTCTGAACTCTTTCCCTACACGACGCTCTTCCGATCTNNNCCCAAACAAAAGAGGCCAAATG
GCACACGTCTGAACTCTTTCCCTACACGACGCTCTTCCGATCTNNNCCCAAACAAAAGAGGCCAAATG
GCACACGTCTGAACTCTTTCCCTACACGACGCTCTTCCGATCTNNNCCCAAACAAAAGAGGCCAAATG
GCACACGTCTGAACTCTTTCCCTACACGACGCTCTTCCGATCTNNNCCCAAACAAAAGAGGCCAAATG
AGCACACGTCTGAACTCTTTCCCTACACGACGCTCTTCCGATCTNNNCAGGACTGTCAAGCAGAGAAT
AGCACACGTCTGAACTCTTTCCCTACACGACGCTCTTCCGATCTNNNCAGGACTGTCAAGCAGAGAAT
AGCACACGTCTGAACTCTTTCCCTACACGACGCTCTTCCGATCTNNNCAGGACTGTCAAGCAGAGAAT
AGCACACGTCTGAACTCTTTCCCTACACGACGCTCTTCCGATCTNNNCAGGACTGTCAAGCAGAGAAT
GAGCACACGTCTGAACTCTTTCCCTACACGACGCTCTTCCGATCTNNNCAAGTCCTGTAGTGAAGTT
GAGCACACGTCTGAACTCTTTCCCTACACGACGCTCTTCCGATCTNNNCAAGTCCTGTAGTGAAGTT
GAGCACACGTCTGAACTCTTTCCCTACACGACGCTCTTCCGATCTNNNCAAGTCCTGTAGTGAAGTT
GAGCACACGTCTGAACTCTTTCCCTACACGACGCTCTTCCGATCTNNNCAAGTCCTGTAGTGAAGTT
GCACACGTCTGAACTCTTTCCCTACACGACGCTCTTCCGATCTNNNATGCTAGATCTACAATGGCCCA
GCACACGTCTGAACTCTTTCCCTACACGACGCTCTTCCGATCTNNNATGCTAGATCTACAATGGCCCA
GCACACGTCTGAACTCTTTCCCTACACGACGCTCTTCCGATCTNNNATGCTAGATCTACAATGGCCCA
GCACACGTCTGAACTCTTTCCCTACACGACGCTCTTCCGATCTNNNATGCTAGATCTACAATGGCCCA
GCACACGTCTGAACTCTTTCCCTACACGACGCTCTTCCGATCTNNNCTCCCTGGCTCTCTTGCTTTTTCC
GCACACGTCTGAACTCTTTCCCTACACGACGCTCTTCCGATCTNNNCTCCCTGGCTCTCTTGCTTTTTCC
GCACACGTCTGAACTCTTTCCCTACACGACGCTCTTCCGATCTNNNCTCCCTGGCTCTCTTGCTTTTTCC
GCACACGTCTGAACTCTTTCCCTACACGACGCTCTTCCGATCTNNNCTCCCTGGCTCTCTTGCTTTTTCC
GCACACGTCTGAACTCTTTCCCTACACGACGCTCTTCCGATCTNNNCTTGCTGTGAGCCGAGATCACA
GCACACGTCTGAACTCTTTCCCTACACGACGCTCTTCCGATCTNNNCTTGCTGTGAGCCGAGATCACA
GCACACGTCTGAACTCTTTCCCTACACGACGCTCTTCCGATCTNNNCTTGCTGTGAGCCGAGATCACA
GCACACGTCTGAACTCTTTCCCTACACGACGCTCTTCCGATCTNNNCTTGCTGTGAGCCGAGATCACA
GAGCACACGTCTGAACTCTTTCCCTACACGACGCTCTTCCGATCTNNNAGGAGAAATGCTTGAACCT
GAGCACACGTCTGAACTCTTTCCCTACACGACGCTCTTCCGATCTNNNAGGAGAAATGCTTGAACCT
GAGCACACGTCTGAACTCTTTCCCTACACGACGCTCTTCCGATCTNNNAGGAGAAATGCTTGAACCT
GAGCACACGTCTGAACTCTTTCCCTACACGACGCTCTTCCGATCTNNNAGGAGAAATGCTTGAACCT
GCACACGTCTGAACTCTTTCCCTACACGACGCTCTTCCGATCTNNNATCACAAAGGTGCAAGAGCTA
GCACACGTCTGAACTCTTTCCCTACACGACGCTCTTCCGATCTNNNATCACAAAGGTGCAAGAGCTA
GCACACGTCTGAACTCTTTCCCTACACGACGCTCTTCCGATCTNNNATCACAAAGGTGCAAGAGCTA
GCACACGTCTGAACTCTTTCCCTACACGACGCTCTTCCGATCTNNNATCACAAAGGTGCAAGAGCTA
GAGCACACGTCTGAACTCTTTCCCTACACGACGCTCTTCCGATCTNNNAGGGTTGAGAAGAACATTG
GAGCACACGTCTGAACTCTTTCCCTACACGACGCTCTTCCGATCTNNNAGGGTTGAGAAGAACATTG
GAGCACACGTCTGAACTCTTTCCCTACACGACGCTCTTCCGATCTNNNAGGGTTGAGAAGAACATTG
GAGCACACGTCTGAACTCTTTCCCTACACGACGCTCTTCCGATCTNNNAGGGTTGAGAAGAACATTG
AGCACACGTCTGAACTCTTTCCCTACACGACGCTCTTCCGATCTNNNCTGTAATCCCAAGCTACTCGGG

AGCACACGTCTGAACTCTTTCCCTACACGACGCTCTTCCGATCTNNNCTGTAATCCCAAGCTACTCGGG
AGCACACGTCTGAACTCTTTCCCTACACGACGCTCTTCCGATCTNNNCTGTAATCCCAAGCTACTCGGG
AGCACACGTCTGAACTCTTTCCCTACACGACGCTCTTCCGATCTNNNCTGTAATCCCAAGCTACTCGGG
AGCACACGTCTGAACTCTTTCCCTACACGACGCTCTTCCGATCTNNNGAGGCTGAGGCAGGAGAAT
AGCACACGTCTGAACTCTTTCCCTACACGACGCTCTTCCGATCTNNNGAGGCTGAGGCAGGAGAAT
AGCACACGTCTGAACTCTTTCCCTACACGACGCTCTTCCGATCTNNNGAGGCTGAGGCAGGAGAAT
AGCACACGTCTGAACTCTTTCCCTACACGACGCTCTTCCGATCTNNNGAGGCTGAGGCAGGAGAAT
GAGCACACGTCTGAACTCTTTCCCTACACGACGCTCTTCCGATCTNNNGTACCTGTGGATTTGGGAG
GAGCACACGTCTGAACTCTTTCCCTACACGACGCTCTTCCGATCTNNNGTACCTGTGGATTTGGGAG
GAGCACACGTCTGAACTCTTTCCCTACACGACGCTCTTCCGATCTNNNGTACCTGTGGATTTGGGAG
GAGCACACGTCTGAACTCTTTCCCTACACGACGCTCTTCCGATCTNNNGTACCTGTGGATTTGGGAG
AGCACACGTCTGAACTCTTTCCCTACACGACGCTCTTCCGATCTNNNGAAGATGCTGAGTGGACATTT
AGCACACGTCTGAACTCTTTCCCTACACGACGCTCTTCCGATCTNNNGAAGATGCTGAGTGGACATTT
AGCACACGTCTGAACTCTTTCCCTACACGACGCTCTTCCGATCTNNNGAAGATGCTGAGTGGACATTT
AGCACACGTCTGAACTCTTTCCCTACACGACGCTCTTCCGATCTNNNGAAGATGCTGAGTGGACATTT
GCACACGTCTGAACTCTTTCCCTACACGACGCTCTTCCGATCTNNNAGAAAGAGAATGCACACCTAG
GCACACGTCTGAACTCTTTCCCTACACGACGCTCTTCCGATCTNNNAGAAAGAGAATGCACACCTAG
GCACACGTCTGAACTCTTTCCCTACACGACGCTCTTCCGATCTNNNAGAAAGAGAATGCACACCTAG
GCACACGTCTGAACTCTTTCCCTACACGACGCTCTTCCGATCTNNNAGAAAGAGAATGCACACCTAG
GCACACGTCTGAACTCTTTCCCTACACGACGCTCTTCCGATCTNNNCCCCTGCCCCCATATTTATTTTC
GCACACGTCTGAACTCTTTCCCTACACGACGCTCTTCCGATCTNNNCCCCTGCCCCCATATTTATTTTC
GCACACGTCTGAACTCTTTCCCTACACGACGCTCTTCCGATCTNNNCCCCTGCCCCCATATTTATTTTC
GCACACGTCTGAACTCTTTCCCTACACGACGCTCTTCCGATCTNNNCCCCTGCCCCCATATTTATTTTC
ACACGTCTGAACTCTTTCCCTACACGACGCTCTTCCGATCTNNNGACTGAGATTCCACTGCTCGGCAT
ACACGTCTGAACTCTTTCCCTACACGACGCTCTTCCGATCTNNNGACTGAGATTCCACTGCTCGGCAT
ACACGTCTGAACTCTTTCCCTACACGACGCTCTTCCGATCTNNNGACTGAGATTCCACTGCTCGGCAT
ACACGTCTGAACTCTTTCCCTACACGACGCTCTTCCGATCTNNNGACTGAGATTCCACTGCTCGGCAT
AGCACACGTCTGAACTCTTTCCCTACACGACGCTCTTCCGATCTNNNGGACAAATCCATCCATTATGAC
AGCACACGTCTGAACTCTTTCCCTACACGACGCTCTTCCGATCTNNNGGACAAATCCATCCATTATGAC
AGCACACGTCTGAACTCTTTCCCTACACGACGCTCTTCCGATCTNNNGGACAAATCCATCCATTATGAC
AGCACACGTCTGAACTCTTTCCCTACACGACGCTCTTCCGATCTNNNGGACAAATCCATCCATTATGAC
GCACACGTCTGAACTCTTTCCCTACACGACGCTCTTCCGATCTNNNTGTTGACAGAGTGAGACTCCAT
GCACACGTCTGAACTCTTTCCCTACACGACGCTCTTCCGATCTNNNTGTTGACAGAGTGAGACTCCAT
GCACACGTCTGAACTCTTTCCCTACACGACGCTCTTCCGATCTNNNTGTTGACAGAGTGAGACTCCAT
GCACACGTCTGAACTCTTTCCCTACACGACGCTCTTCCGATCTNNNTGTTGACAGAGTGAGACTCCAT
CACACGTCTGAACTCTTTCCCTACACGACGCTCTTCCGATCTNNNCTTCCTTGTCCCCACATTTGAACC
CACACGTCTGAACTCTTTCCCTACACGACGCTCTTCCGATCTNNNCTTCCTTGTCCCCACATTTGAACC
CACACGTCTGAACTCTTTCCCTACACGACGCTCTTCCGATCTNNNCTTCCTTGTCCCCACATTTGAACC
CACACGTCTGAACTCTTTCCCTACACGACGCTCTTCCGATCTNNNCTTCCTTGTCCCCACATTTGAACC
GCACACGTCTGAACTCTTTCCCTACACGACGCTCTTCCGATCTNNNGATGGTTTCCAGCTTCATCCATG
GCACACGTCTGAACTCTTTCCCTACACGACGCTCTTCCGATCTNNNGATGGTTTCCAGCTTCATCCATG
GCACACGTCTGAACTCTTTCCCTACACGACGCTCTTCCGATCTNNNGATGGTTTCCAGCTTCATCCATG
GCACACGTCTGAACTCTTTCCCTACACGACGCTCTTCCGATCTNNNGATGGTTTCCAGCTTCATCCATG

AGCACACGTCTGAACTCTTTCCCTACACGACGCTCTTCCGATCTNNNCAGAAGAATTGCTTGAACCCAG
AGCACACGTCTGAACTCTTTCCCTACACGACGCTCTTCCGATCTNNNCAGAAGAATTGCTTGAACCCAG
AGCACACGTCTGAACTCTTTCCCTACACGACGCTCTTCCGATCTNNNCAGAAGAATTGCTTGAACCCAG
AGCACACGTCTGAACTCTTTCCCTACACGACGCTCTTCCGATCTNNNCAGAAGAATTGCTTGAACCCAG
CACACGTCTGAACTCTTTCCCTACACGACGCTCTTCCGATCTNNNACATGGCAAACAGGAGACCTAGCA
CACACGTCTGAACTCTTTCCCTACACGACGCTCTTCCGATCTNNNACATGGCAAACAGGAGACCTAGCA
CACACGTCTGAACTCTTTCCCTACACGACGCTCTTCCGATCTNNNACATGGCAAACAGGAGACCTAGCA
CACACGTCTGAACTCTTTCCCTACACGACGCTCTTCCGATCTNNNACATGGCAAACAGGAGACCTAGCA
GAGCACACGTCTGAACTCTTTCCCTACACGACGCTCTTCCGATCTNNNGTGTCTGTCTGTCTGTCTGTCT
GAGCACACGTCTGAACTCTTTCCCTACACGACGCTCTTCCGATCTNNNGTGTCTGTCTGTCTGTCTGTCT
GAGCACACGTCTGAACTCTTTCCCTACACGACGCTCTTCCGATCTNNNGTGTCTGTCTGTCTGTCTGTCT
GAGCACACGTCTGAACTCTTTCCCTACACGACGCTCTTCCGATCTNNNGTGTCTGTCTGTCTGTCTGTCT
AGCACACGTCTGAACTCTTTCCCTACACGACGCTCTTCCGATCTNNNATTGAGAGAGACAGAGACAGG
AGCACACGTCTGAACTCTTTCCCTACACGACGCTCTTCCGATCTNNNATTGAGAGAGACAGAGACAGG
AGCACACGTCTGAACTCTTTCCCTACACGACGCTCTTCCGATCTNNNATTGAGAGAGACAGAGACAGG
AGCACACGTCTGAACTCTTTCCCTACACGACGCTCTTCCGATCTNNNATTGAGAGAGACAGAGACAGG
GAGCACACGTCTGAACTCTTTCCCTACACGACGCTCTTCCGATCTNNNTAGGCATGAACCATCATACCCA
GAGCACACGTCTGAACTCTTTCCCTACACGACGCTCTTCCGATCTNNNTAGGCATGAACCATCATACCCA
GAGCACACGTCTGAACTCTTTCCCTACACGACGCTCTTCCGATCTNNNTAGGCATGAACCATCATACCCA
GAGCACACGTCTGAACTCTTTCCCTACACGACGCTCTTCCGATCTNNNTAGGCATGAACCATCATACCCA
CACACGTCTGAACTCTTTCCCTACACGACGCTCTTCCGATCTNNNTTGTAAATTCAACTCATTCTTTTCCA
CACACGTCTGAACTCTTTCCCTACACGACGCTCTTCCGATCTNNNTTGTAAATTCAACTCATTCTTTTCCA
CACACGTCTGAACTCTTTCCCTACACGACGCTCTTCCGATCTNNNTTGTAAATTCAACTCATTCTTTTCCA
CACACGTCTGAACTCTTTCCCTACACGACGCTCTTCCGATCTNNNTTGTAAATTCAACTCATTCTTTTCCA
AGCACACGTCTGAACTCTTTCCCTACACGACGCTCTTCCGATCTNNNAATATTGTAACCGCCAGCCCTTT
AGCACACGTCTGAACTCTTTCCCTACACGACGCTCTTCCGATCTNNNAATATTGTAACCGCCAGCCCTTT
AGCACACGTCTGAACTCTTTCCCTACACGACGCTCTTCCGATCTNNNAATATTGTAACCGCCAGCCCTTT
AGCACACGTCTGAACTCTTTCCCTACACGACGCTCTTCCGATCTNNNAATATTGTAACCGCCAGCCCTTT
AGCACACGTCTGAACTCTTTCCCTACACGACGCTCTTCCGATCTNNNCGAGGCAGGAGAATTGATTGA
AGCACACGTCTGAACTCTTTCCCTACACGACGCTCTTCCGATCTNNNCGAGGCAGGAGAATTGATTGA
AGCACACGTCTGAACTCTTTCCCTACACGACGCTCTTCCGATCTNNNCGAGGCAGGAGAATTGATTGA
AGCACACGTCTGAACTCTTTCCCTACACGACGCTCTTCCGATCTNNNCGAGGCAGGAGAATTGATTGA
GCACACGTCTGAACTCTTTCCCTACACGACGCTCTTCCGATCTNNNTTGAGGAGGGGAGAAGTTACTCT
GCACACGTCTGAACTCTTTCCCTACACGACGCTCTTCCGATCTNNNTTGAGGAGGGGAGAAGTTACTCT
GCACACGTCTGAACTCTTTCCCTACACGACGCTCTTCCGATCTNNNTTGAGGAGGGGAGAAGTTACTCT
GCACACGTCTGAACTCTTTCCCTACACGACGCTCTTCCGATCTNNNTTGAGGAGGGGAGAAGTTACTCT
GAGCACACGTCTGAACTCTTTCCCTACACGACGCTCTTCCGATCTNNNTTTTTCTGTGTGTGTGTGTGTC
GAGCACACGTCTGAACTCTTTCCCTACACGACGCTCTTCCGATCTNNNTTTTTCTGTGTGTGTGTGTGTC
GAGCACACGTCTGAACTCTTTCCCTACACGACGCTCTTCCGATCTNNNTTTTTCTGTGTGTGTGTGTGTC
GAGCACACGTCTGAACTCTTTCCCTACACGACGCTCTTCCGATCTNNNTTTTTCTGTGTGTGTGTGTGTC
AGCACACGTCTGAACTCTTTCCCTACACGACGCTCTTCCGATCTNNNTCAATGCAACTTGTTTTATGTAC
AGCACACGTCTGAACTCTTTCCCTACACGACGCTCTTCCGATCTNNNTCAATGCAACTTGTTTTATGTAC
AGCACACGTCTGAACTCTTTCCCTACACGACGCTCTTCCGATCTNNNTCAATGCAACTTGTTTTATGTAC

AGCACACGTCTGAACTCTTTCCCTACACGACGCTCTTCCGATCTNNNTCAATGCAACTTGTTTTATGTA
AGAGCACACGTCTGAACTCTTTCCCTACACGACGCTCTTCCGATCTNNNAGGAAGAGAGATGGGTCCT
AGAGCACACGTCTGAACTCTTTCCCTACACGACGCTCTTCCGATCTNNNAGGAAGAGAGATGGGTCCT
AGAGCACACGTCTGAACTCTTTCCCTACACGACGCTCTTCCGATCTNNNAGGAAGAGAGATGGGTCCT
AGAGCACACGTCTGAACTCTTTCCCTACACGACGCTCTTCCGATCTNNNAGGAAGAGAGATGGGTCCT
GAGCACACGTCTGAACTCTTTCCCTACACGACGCTCTTCCGATCTNNNAGTCCCAGAAGAATTGAGCA
GAGCACACGTCTGAACTCTTTCCCTACACGACGCTCTTCCGATCTNNNAGTCCCAGAAGAATTGAGCA
GAGCACACGTCTGAACTCTTTCCCTACACGACGCTCTTCCGATCTNNNAGTCCCAGAAGAATTGAGCA
GAGCACACGTCTGAACTCTTTCCCTACACGACGCTCTTCCGATCTNNNAGTCCCAGAAGAATTGAGCA
AGAGCACACGTCTGAACTCTTTCCCTACACGACGCTCTTCCGATCTNNNTAAAAAGACGCACGATTCAT
AGAGCACACGTCTGAACTCTTTCCCTACACGACGCTCTTCCGATCTNNNTAAAAAGACGCACGATTCAT
AGAGCACACGTCTGAACTCTTTCCCTACACGACGCTCTTCCGATCTNNNTAAAAAGACGCACGATTCAT
AGAGCACACGTCTGAACTCTTTCCCTACACGACGCTCTTCCGATCTNNNTAAAAAGACGCACGATTCAT
AGCACACGTCTGAACTCTTTCCCTACACGACGCTCTTCCGATCTNNNTGTGAGCATGTTTTCGAATACG
AGCACACGTCTGAACTCTTTCCCTACACGACGCTCTTCCGATCTNNNTGTGAGCATGTTTTCGAATACG
AGCACACGTCTGAACTCTTTCCCTACACGACGCTCTTCCGATCTNNNTGTGAGCATGTTTTCGAATACG
AGCACACGTCTGAACTCTTTCCCTACACGACGCTCTTCCGATCTNNNTGTGAGCATGTTTTCGAATACG
CACACGTCTGAACTCTTTCCCTACACGACGCTCTTCCGATCTNNNGAACGGATCGTCTCCTCGTTGCATC
CACACGTCTGAACTCTTTCCCTACACGACGCTCTTCCGATCTNNNGAACGGATCGTCTCCTCGTTGCATC
CACACGTCTGAACTCTTTCCCTACACGACGCTCTTCCGATCTNNNGAACGGATCGTCTCCTCGTTGCATC
CACACGTCTGAACTCTTTCCCTACACGACGCTCTTCCGATCTNNNGAACGGATCGTCTCCTCGTTGCATC
AGCACACGTCTGAACTCTTTCCCTACACGACGCTCTTCCGATCTNNNTGGAGGAGAAAAGCTTGTTACA
AGCACACGTCTGAACTCTTTCCCTACACGACGCTCTTCCGATCTNNNTGGAGGAGAAAAGCTTGTTACA
AGCACACGTCTGAACTCTTTCCCTACACGACGCTCTTCCGATCTNNNTGGAGGAGAAAAGCTTGTTACA
AGCACACGTCTGAACTCTTTCCCTACACGACGCTCTTCCGATCTNNNTGGAGGAGAAAAGCTTGTTACA
AGCACACGTCTGAACTCTTTCCCTACACGACGCTCTTCCGATCTNNNGCCTAAGATCTCTTGAGGGCAA
AGCACACGTCTGAACTCTTTCCCTACACGACGCTCTTCCGATCTNNNGCCTAAGATCTCTTGAGGGCAA
AGCACACGTCTGAACTCTTTCCCTACACGACGCTCTTCCGATCTNNNGCCTAAGATCTCTTGAGGGCAA
AGCACACGTCTGAACTCTTTCCCTACACGACGCTCTTCCGATCTNNNGCCTAAGATCTCTTGAGGGCAA
AGCACACGTCTGAACTCTTTCCCTACACGACGCTCTTCCGATCTNNNAGCTGTCATCTCTGCAGATTTCA
AGCACACGTCTGAACTCTTTCCCTACACGACGCTCTTCCGATCTNNNAGCTGTCATCTCTGCAGATTTCA
AGCACACGTCTGAACTCTTTCCCTACACGACGCTCTTCCGATCTNNNAGCTGTCATCTCTGCAGATTTCA
AGCACACGTCTGAACTCTTTCCCTACACGACGCTCTTCCGATCTNNNAGCTGTCATCTCTGCAGATTTCA
AGCACACGTCTGAACTCTTTCCCTACACGACGCTCTTCCGATCTNNNCTGAGGCAGGAGAATCAGTTG
AGCACACGTCTGAACTCTTTCCCTACACGACGCTCTTCCGATCTNNNCTGAGGCAGGAGAATCAGTTG
AGCACACGTCTGAACTCTTTCCCTACACGACGCTCTTCCGATCTNNNCTGAGGCAGGAGAATCAGTTG
AGCACACGTCTGAACTCTTTCCCTACACGACGCTCTTCCGATCTNNNCTGAGGCAGGAGAATCAGTTG
GCACACGTCTGAACTCTTTCCCTACACGACGCTCTTCCGATCTNNNACTACATGTGAGTCTTCCCCTTG
GCACACGTCTGAACTCTTTCCCTACACGACGCTCTTCCGATCTNNNACTACATGTGAGTCTTCCCCTTG
GCACACGTCTGAACTCTTTCCCTACACGACGCTCTTCCGATCTNNNACTACATGTGAGTCTTCCCCTTG
GCACACGTCTGAACTCTTTCCCTACACGACGCTCTTCCGATCTNNNACTACATGTGAGTCTTCCCCTTG
GAGCACACGTCTGAACTCTTTCCCTACACGACGCTCTTCCGATCTNNNCGGTAGATCATGGGACTTCTC
GAGCACACGTCTGAACTCTTTCCCTACACGACGCTCTTCCGATCTNNNCGGTAGATCATGGGACTTCTC

GAGCACACGTCTGAACTCTTTCCCTACACGACGCTCTTCCGATCTNNNCGGTAGATCATGGGACTTCTC
GAGCACACGTCTGAACTCTTTCCCTACACGACGCTCTTCCGATCTNNNCGGTAGATCATGGGACTTCTC
AGCACACGTCTGAACTCTTTCCCTACACGACGCTCTTCCGATCTNNNGAAACTAGGTGTGTATTGTTTG
AGCACACGTCTGAACTCTTTCCCTACACGACGCTCTTCCGATCTNNNGAAACTAGGTGTGTATTGTTTG
AGCACACGTCTGAACTCTTTCCCTACACGACGCTCTTCCGATCTNNNGAAACTAGGTGTGTATTGTTTG
AGCACACGTCTGAACTCTTTCCCTACACGACGCTCTTCCGATCTNNNGAAACTAGGTGTGTATTGTTTG
AGCACACGTCTGAACTCTTTCCCTACACGACGCTCTTCCGATCTNNNCCCGAGTAGCTAGGATTACAAG
AGCACACGTCTGAACTCTTTCCCTACACGACGCTCTTCCGATCTNNNCCCGAGTAGCTAGGATTACAAG
AGCACACGTCTGAACTCTTTCCCTACACGACGCTCTTCCGATCTNNNCCCGAGTAGCTAGGATTACAAG
AGCACACGTCTGAACTCTTTCCCTACACGACGCTCTTCCGATCTNNNCCCGAGTAGCTAGGATTACAAG
AGAGCACACGTCTGAACTCTTTCCCTACACGACGCTCTTCCGATCTNNNTCCTACTGAGATTGCTCTGCC
AGAGCACACGTCTGAACTCTTTCCCTACACGACGCTCTTCCGATCTNNNTCCTACTGAGATTGCTCTGCC
AGAGCACACGTCTGAACTCTTTCCCTACACGACGCTCTTCCGATCTNNNTCCTACTGAGATTGCTCTGCC
AGAGCACACGTCTGAACTCTTTCCCTACACGACGCTCTTCCGATCTNNNTCCTACTGAGATTGCTCTGCC
GAGCACACGTCTGAACTCTTTCCCTACACGACGCTCTTCCGATCTNNNCCCATATCTGAAGATAAATTT
GAGCACACGTCTGAACTCTTTCCCTACACGACGCTCTTCCGATCTNNNCCCATATCTGAAGATAAATTT
GAGCACACGTCTGAACTCTTTCCCTACACGACGCTCTTCCGATCTNNNCCCATATCTGAAGATAAATTT
GAGCACACGTCTGAACTCTTTCCCTACACGACGCTCTTCCGATCTNNNCCCATATCTGAAGATAAATTT
AGCACACGTCTGAACTCTTTCCCTACACGACGCTCTTCCGATCTNNNTGGCCAATACTTGACATTTGTG
AGCACACGTCTGAACTCTTTCCCTACACGACGCTCTTCCGATCTNNNTGGCCAATACTTGACATTTGTG
AGCACACGTCTGAACTCTTTCCCTACACGACGCTCTTCCGATCTNNNTGGCCAATACTTGACATTTGTG
AGCACACGTCTGAACTCTTTCCCTACACGACGCTCTTCCGATCTNNNTGGCCAATACTTGACATTTGTG
GAGCACACGTCTGAACTCTTTCCCTACACGACGCTCTTCCGATCTNNNCCTGGATTGGAGAGGAGAAC
GAGCACACGTCTGAACTCTTTCCCTACACGACGCTCTTCCGATCTNNNCCTGGATTGGAGAGGAGAAC
GAGCACACGTCTGAACTCTTTCCCTACACGACGCTCTTCCGATCTNNNCCTGGATTGGAGAGGAGAAC
GAGCACACGTCTGAACTCTTTCCCTACACGACGCTCTTCCGATCTNNNCCTGGATTGGAGAGGAGAAC
AGCACACGTCTGAACTCTTTCCCTACACGACGCTCTTCCGATCTNNNGTCTTCATTCTTATCATCTTCACCA
AGCACACGTCTGAACTCTTTCCCTACACGACGCTCTTCCGATCTNNNGTCTTCATTCTTATCATCTTCACCA
AGCACACGTCTGAACTCTTTCCCTACACGACGCTCTTCCGATCTNNNGTCTTCATTCTTATCATCTTCACCA
AGCACACGTCTGAACTCTTTCCCTACACGACGCTCTTCCGATCTNNNGTCTTCATTCTTATCATCTTCACCA
AGCACACGTCTGAACTCTTTCCCTACACGACGCTCTTCCGATCTNNNACCACGCCCAGCTAAATTTTTTT
AGCACACGTCTGAACTCTTTCCCTACACGACGCTCTTCCGATCTNNNACCACGCCCAGCTAAATTTTTTT
AGCACACGTCTGAACTCTTTCCCTACACGACGCTCTTCCGATCTNNNACCACGCCCAGCTAAATTTTTTT
AGCACACGTCTGAACTCTTTCCCTACACGACGCTCTTCCGATCTNNNACCACGCCCAGCTAAATTTTTTT
GAGCACACGTCTGAACTCTTTCCCTACACGACGCTCTTCCGATCTNNNAACTGCGTTCTCCCACAGAAAA
GAGCACACGTCTGAACTCTTTCCCTACACGACGCTCTTCCGATCTNNNAACTGCGTTCTCCCACAGAAAA
GAGCACACGTCTGAACTCTTTCCCTACACGACGCTCTTCCGATCTNNNAACTGCGTTCTCCCACAGAAAA
GAGCACACGTCTGAACTCTTTCCCTACACGACGCTCTTCCGATCTNNNAACTGCGTTCTCCCACAGAAAA
AGCACACGTCTGAACTCTTTCCCTACACGACGCTCTTCCGATCTNNNCCAGATGCTCCATTCTCACCTAA
AGCACACGTCTGAACTCTTTCCCTACACGACGCTCTTCCGATCTNNNCCAGATGCTCCATTCTCACCTAA
AGCACACGTCTGAACTCTTTCCCTACACGACGCTCTTCCGATCTNNNCCAGATGCTCCATTCTCACCTAA
AGCACACGTCTGAACTCTTTCCCTACACGACGCTCTTCCGATCTNNNCCAGATGCTCCATTCTCACCTAA
GAGCACACGTCTGAACTCTTTCCCTACACGACGCTCTTCCGATCTNNNGAAGATGGATATGACCAGTG

AGCACACGTCTGAACTCTTTCCCTACACGACGCTCTTCCGATCTNNNGAAGATGGATATGACCAGTG

AGCACACGTCTGAACTCTTTCCCTACACGACGCTCTTCCGATCTNNNGAAGATGGATATGACCAGTG

AGCACACGTCTGAACTCTTTCCCTACACGACGCTCTTCCGATCTNNNGAAGATGGATATGACCAGTG

GAGCACACGTCTGAACTCTTTCCCTACACGACGCTCTTCCGATCTNNNAGTGAGCTGAGATTGTACCA

GAGCACACGTCTGAACTCTTTCCCTACACGACGCTCTTCCGATCTNNNAGTGAGCTGAGATTGTACCA

GAGCACACGTCTGAACTCTTTCCCTACACGACGCTCTTCCGATCTNNNAGTGAGCTGAGATTGTACCA

GAGCACACGTCTGAACTCTTTCCCTACACGACGCTCTTCCGATCTNNNAGTGAGCTGAGATTGTACCA

AGCACACGTCTGAACTCTTTCCCTACACGACGCTCTTCCGATCTNNNCTGTAATCCCAGCTACTCAGGA

AGCACACGTCTGAACTCTTTCCCTACACGACGCTCTTCCGATCTNNNCTGTAATCCCAGCTACTCAGGA

AGCACACGTCTGAACTCTTTCCCTACACGACGCTCTTCCGATCTNNNCTGTAATCCCAGCTACTCAGGA

AGCACACGTCTGAACTCTTTCCCTACACGACGCTCTTCCGATCTNNNCTGTAATCCCAGCTACTCAGGA

GAGCACACGTCTGAACTCTTTCCCTACACGACGCTCTTCCGATCTNNNGAACCATGGGAAGTGAGA

GAGCACACGTCTGAACTCTTTCCCTACACGACGCTCTTCCGATCTNNNGAACCATGGGAAGTGAGA

GAGCACACGTCTGAACTCTTTCCCTACACGACGCTCTTCCGATCTNNNGAACCATGGGAAGTGAGA

GAGCACACGTCTGAACTCTTTCCCTACACGACGCTCTTCCGATCTNNNGAACCATGGGAAGTGAGA

GCACACGTCTGAACTCTTTCCCTACACGACGCTCTTCCGATCTNNNGCATAAGTTCATCCATGTTGTCA

GCACACGTCTGAACTCTTTCCCTACACGACGCTCTTCCGATCTNNNGCATAAGTTCATCCATGTTGTCA

GCACACGTCTGAACTCTTTCCCTACACGACGCTCTTCCGATCTNNNGCATAAGTTCATCCATGTTGTCA

GCACACGTCTGAACTCTTTCCCTACACGACGCTCTTCCGATCTNNNGCATAAGTTCATCCATGTTGTCA

GAGCACACGTCTGAACTCTTTCCCTACACGACGCTCTTCCGATCTNNNCCACCATTTTCCCTGTAGATCC

GAGCACACGTCTGAACTCTTTCCCTACACGACGCTCTTCCGATCTNNNCCACCATTTTCCCTGTAGATCC

GAGCACACGTCTGAACTCTTTCCCTACACGACGCTCTTCCGATCTNNNCCACCATTTTCCCTGTAGATCC

GAGCACACGTCTGAACTCTTTCCCTACACGACGCTCTTCCGATCTNNNCCACCATTTTCCCTGTAGATCC

AGCACACGTCTGAACTCTTTCCCTACACGACGCTCTTCCGATCTNNNTTATCCTTCCAATGTGGGCAGA

AGCACACGTCTGAACTCTTTCCCTACACGACGCTCTTCCGATCTNNNTTATCCTTCCAATGTGGGCAGA

AGCACACGTCTGAACTCTTTCCCTACACGACGCTCTTCCGATCTNNNTTATCCTTCCAATGTGGGCAGA

AGCACACGTCTGAACTCTTTCCCTACACGACGCTCTTCCGATCTNNNTTATCCTTCCAATGTGGGCAGA

GCACACGTCTGAACTCTTTCCCTACACGACGCTCTTCCGATCTNNNAGCTGAGAAAGGAAGCAGACA

GCACACGTCTGAACTCTTTCCCTACACGACGCTCTTCCGATCTNNNAGCTGAGAAAGGAAGCAGACA

GCACACGTCTGAACTCTTTCCCTACACGACGCTCTTCCGATCTNNNAGCTGAGAAAGGAAGCAGACA

GCACACGTCTGAACTCTTTCCCTACACGACGCTCTTCCGATCTNNNAGCTGAGAAAGGAAGCAGACA

AGCACACGTCTGAACTCTTTCCCTACACGACGCTCTTCCGATCTNNNTGCAGAGAAACCTACCATTTGT

AGCACACGTCTGAACTCTTTCCCTACACGACGCTCTTCCGATCTNNNTGCAGAGAAACCTACCATTTGT

AGCACACGTCTGAACTCTTTCCCTACACGACGCTCTTCCGATCTNNNTGCAGAGAAACCTACCATTTGT

AGCACACGTCTGAACTCTTTCCCTACACGACGCTCTTCCGATCTNNNTGCAGAGAAACCTACCATTTGT

AGCACACGTCTGAACTCTTTCCCTACACGACGCTCTTCCGATCTNNNTCTATCTGCCCATGAACACAAC

AGCACACGTCTGAACTCTTTCCCTACACGACGCTCTTCCGATCTNNNTCTATCTGCCCATGAACACAAC

AGCACACGTCTGAACTCTTTCCCTACACGACGCTCTTCCGATCTNNNTCTATCTGCCCATGAACACAAC

AGCACACGTCTGAACTCTTTCCCTACACGACGCTCTTCCGATCTNNNTCTATCTGCCCATGAACACAAC

GCACACGTCTGAACTCTTTCCCTACACGACGCTCTTCCGATCTNNNCGCCCGGCTAATTCTTGTATTTT

GCACACGTCTGAACTCTTTCCCTACACGACGCTCTTCCGATCTNNNCGCCCGGCTAATTCTTGTATTTT

GCACACGTCTGAACTCTTTCCCTACACGACGCTCTTCCGATCTNNNCGCCCGGCTAATTCTTGTATTTT

GCACACGTCTGAACTCTTTCCCTACACGACGCTCTTCCGATCTNNNCGCCCGGCTAATTCTTGTATTTT

AGCACACGTCTGAACTCTTTCCCTACACGACGCTCTTCCGATCTNNNTCTGCCTCCTAGTTTCAAGCAAT
AGCACACGTCTGAACTCTTTCCCTACACGACGCTCTTCCGATCTNNNTCTGCCTCCTAGTTTCAAGCAAT
AGCACACGTCTGAACTCTTTCCCTACACGACGCTCTTCCGATCTNNNTCTGCCTCCTAGTTTCAAGCAAT
AGCACACGTCTGAACTCTTTCCCTACACGACGCTCTTCCGATCTNNNTCTGCCTCCTAGTTTCAAGCAAT
AGCACACGTCTGAACTCTTTCCCTACACGACGCTCTTCCGATCTNNNTGTTCTTACTCCCTTGCTTCCAA
AGCACACGTCTGAACTCTTTCCCTACACGACGCTCTTCCGATCTNNNTGTTCTTACTCCCTTGCTTCCAA
AGCACACGTCTGAACTCTTTCCCTACACGACGCTCTTCCGATCTNNNTGTTCTTACTCCCTTGCTTCCAA
AGCACACGTCTGAACTCTTTCCCTACACGACGCTCTTCCGATCTNNNTGTTCTTACTCCCTTGCTTCCAA
AGCACACGTCTGAACTCTTTCCCTACACGACGCTCTTCCGATCTNNNGCCACCCTTTATTGAACACATCC
AGCACACGTCTGAACTCTTTCCCTACACGACGCTCTTCCGATCTNNNGCCACCCTTTATTGAACACATCC
AGCACACGTCTGAACTCTTTCCCTACACGACGCTCTTCCGATCTNNNGCCACCCTTTATTGAACACATCC
AGCACACGTCTGAACTCTTTCCCTACACGACGCTCTTCCGATCTNNNGCCACCCTTTATTGAACACATCC
GAGCACACGTCTGAACTCTTTCCCTACACGACGCTCTTCCGATCTNNNGTGTTTGAAGGGGAGGAAG
GAGCACACGTCTGAACTCTTTCCCTACACGACGCTCTTCCGATCTNNNGTGTTTGAAGGGGAGGAAG
GAGCACACGTCTGAACTCTTTCCCTACACGACGCTCTTCCGATCTNNNGTGTTTGAAGGGGAGGAAG
GAGCACACGTCTGAACTCTTTCCCTACACGACGCTCTTCCGATCTNNNGTGTTTGAAGGGGAGGAAG
GAGCACACGTCTGAACTCTTTCCCTACACGACGCTCTTCCGATCTNNNCTGTATTCCCAGCTACTCAGAG
GAGCACACGTCTGAACTCTTTCCCTACACGACGCTCTTCCGATCTNNNCTGTATTCCCAGCTACTCAGAG
GAGCACACGTCTGAACTCTTTCCCTACACGACGCTCTTCCGATCTNNNCTGTATTCCCAGCTACTCAGAG
GAGCACACGTCTGAACTCTTTCCCTACACGACGCTCTTCCGATCTNNNCTGTATTCCCAGCTACTCAGAG
CACACGTCTGAACTCTTTCCCTACACGACGCTCTTCCGATCTNNNCAAAGAAAAAAAACAAGGCCGGG
CACACGTCTGAACTCTTTCCCTACACGACGCTCTTCCGATCTNNNCAAAGAAAAAAAACAAGGCCGGG
CACACGTCTGAACTCTTTCCCTACACGACGCTCTTCCGATCTNNNCAAAGAAAAAAAACAAGGCCGGG
CACACGTCTGAACTCTTTCCCTACACGACGCTCTTCCGATCTNNNCAAAGAAAAAAAACAAGGCCGGG
GAGCACACGTCTGAACTCTTTCCCTACACGACGCTCTTCCGATCTNNNTCCATTCTCACTTGTACCACAG
GAGCACACGTCTGAACTCTTTCCCTACACGACGCTCTTCCGATCTNNNTCCATTCTCACTTGTACCACAG
GAGCACACGTCTGAACTCTTTCCCTACACGACGCTCTTCCGATCTNNNTCCATTCTCACTTGTACCACAG
GAGCACACGTCTGAACTCTTTCCCTACACGACGCTCTTCCGATCTNNNTCCATTCTCACTTGTACCACAG
AGCACACGTCTGAACTCTTTCCCTACACGACGCTCTTCCGATCTNNNCCTTGATTAGGCTCAGGTTCGG
AGCACACGTCTGAACTCTTTCCCTACACGACGCTCTTCCGATCTNNNCCTTGATTAGGCTCAGGTTCGG
AGCACACGTCTGAACTCTTTCCCTACACGACGCTCTTCCGATCTNNNCCTTGATTAGGCTCAGGTTCGG
AGCACACGTCTGAACTCTTTCCCTACACGACGCTCTTCCGATCTNNNCCTTGATTAGGCTCAGGTTCGG
GAGCACACGTCTGAACTCTTTCCCTACACGACGCTCTTCCGATCTNNNTGTAGACAGTCATGTACCCAC
GAGCACACGTCTGAACTCTTTCCCTACACGACGCTCTTCCGATCTNNNTGTAGACAGTCATGTACCCAC
GAGCACACGTCTGAACTCTTTCCCTACACGACGCTCTTCCGATCTNNNTGTAGACAGTCATGTACCCAC
GAGCACACGTCTGAACTCTTTCCCTACACGACGCTCTTCCGATCTNNNTGTAGACAGTCATGTACCCAC
AGCACACGTCTGAACTCTTTCCCTACACGACGCTCTTCCGATCTNNNTGGATAACTTTATGCAGTGCAA
AGCACACGTCTGAACTCTTTCCCTACACGACGCTCTTCCGATCTNNNTGGATAACTTTATGCAGTGCAA
AGCACACGTCTGAACTCTTTCCCTACACGACGCTCTTCCGATCTNNNTGGATAACTTTATGCAGTGCAA
AGCACACGTCTGAACTCTTTCCCTACACGACGCTCTTCCGATCTNNNTGGATAACTTTATGCAGTGCAA
AGAGCACACGTCTGAACTCTTTCCCTACACGACGCTCTTCCGATCTNNNGAGGAGAAGAGAGGAGA
AGAGCACACGTCTGAACTCTTTCCCTACACGACGCTCTTCCGATCTNNNGAGGAGAAGAGAGGAGA
AGAGCACACGTCTGAACTCTTTCCCTACACGACGCTCTTCCGATCTNNNGAGGAGAAGAGAGGAGA

AGAGCACACGTCTGAACTCTTTCCCTACACGACGCTCTTCCGATCTNNNGAGGAGAAGAGAGGAGA
GAGCACACGTCTGAACTCTTTCCCTACACGACGCTCTTCCGATCTNNNGACCTCCTGACTGAGACACTC
GAGCACACGTCTGAACTCTTTCCCTACACGACGCTCTTCCGATCTNNNGACCTCCTGACTGAGACACTC
GAGCACACGTCTGAACTCTTTCCCTACACGACGCTCTTCCGATCTNNNGACCTCCTGACTGAGACACTC
GAGCACACGTCTGAACTCTTTCCCTACACGACGCTCTTCCGATCTNNNGACCTCCTGACTGAGACACTC
AGCACACGTCTGAACTCTTTCCCTACACGACGCTCTTCCGATCTNNNACAAAAATTAGCCCAGTGTGG
AGCACACGTCTGAACTCTTTCCCTACACGACGCTCTTCCGATCTNNNACAAAAATTAGCCCAGTGTGG
AGCACACGTCTGAACTCTTTCCCTACACGACGCTCTTCCGATCTNNNACAAAAATTAGCCCAGTGTGG
AGCACACGTCTGAACTCTTTCCCTACACGACGCTCTTCCGATCTNNNACAAAAATTAGCCCAGTGTGG
GAGCACACGTCTGAACTCTTTCCCTACACGACGCTCTTCCGATCTNNNTGAAAGAGCAGGCATCCCTA
GAGCACACGTCTGAACTCTTTCCCTACACGACGCTCTTCCGATCTNNNTGAAAGAGCAGGCATCCCTA
GAGCACACGTCTGAACTCTTTCCCTACACGACGCTCTTCCGATCTNNNTGAAAGAGCAGGCATCCCTA
GAGCACACGTCTGAACTCTTTCCCTACACGACGCTCTTCCGATCTNNNTGAAAGAGCAGGCATCCCTA
AGCACACGTCTGAACTCTTTCCCTACACGACGCTCTTCCGATCTNNNGTACAGGGATTACAGGAGTG
AGCACACGTCTGAACTCTTTCCCTACACGACGCTCTTCCGATCTNNNGTACAGGGATTACAGGAGTG
AGCACACGTCTGAACTCTTTCCCTACACGACGCTCTTCCGATCTNNNGTACAGGGATTACAGGAGTG
AGCACACGTCTGAACTCTTTCCCTACACGACGCTCTTCCGATCTNNNGTACAGGGATTACAGGAGTG
GAGCACACGTCTGAACTCTTTCCCTACACGACGCTCTTCCGATCTNNNATAGCTCACTGCAGCCATAAC
GAGCACACGTCTGAACTCTTTCCCTACACGACGCTCTTCCGATCTNNNATAGCTCACTGCAGCCATAAC
GAGCACACGTCTGAACTCTTTCCCTACACGACGCTCTTCCGATCTNNNATAGCTCACTGCAGCCATAAC
GAGCACACGTCTGAACTCTTTCCCTACACGACGCTCTTCCGATCTNNNATAGCTCACTGCAGCCATAAC
GCACACGTCTGAACTCTTTCCCTACACGACGCTCTTCCGATCTNNNAGGAATGCAGGTATCTCTTTGA
GCACACGTCTGAACTCTTTCCCTACACGACGCTCTTCCGATCTNNNAGGAATGCAGGTATCTCTTTGA
GCACACGTCTGAACTCTTTCCCTACACGACGCTCTTCCGATCTNNNAGGAATGCAGGTATCTCTTTGA
GCACACGTCTGAACTCTTTCCCTACACGACGCTCTTCCGATCTNNNAGGAATGCAGGTATCTCTTTGA
AGCACACGTCTGAACTCTTTCCCTACACGACGCTCTTCCGATCTNNNAGAGCCTCGTGGTTTGTTTTTT
AGCACACGTCTGAACTCTTTCCCTACACGACGCTCTTCCGATCTNNNAGAGCCTCGTGGTTTGTTTTTT
AGCACACGTCTGAACTCTTTCCCTACACGACGCTCTTCCGATCTNNNAGAGCCTCGTGGTTTGTTTTTT
AGCACACGTCTGAACTCTTTCCCTACACGACGCTCTTCCGATCTNNNAGAGCCTCGTGGTTTGTTTTTT
AGCACACGTCTGAACTCTTTCCCTACACGACGCTCTTCCGATCTNNNGCCAAGATCATGCCATTGTACT
AGCACACGTCTGAACTCTTTCCCTACACGACGCTCTTCCGATCTNNNGCCAAGATCATGCCATTGTACT
AGCACACGTCTGAACTCTTTCCCTACACGACGCTCTTCCGATCTNNNGCCAAGATCATGCCATTGTACT
AGCACACGTCTGAACTCTTTCCCTACACGACGCTCTTCCGATCTNNNGCCAAGATCATGCCATTGTACT
AGCACACGTCTGAACTCTTTCCCTACACGACGCTCTTCCGATCTNNNTAATGACTTCCAAAGCTGGGTC
AGCACACGTCTGAACTCTTTCCCTACACGACGCTCTTCCGATCTNNNTAATGACTTCCAAAGCTGGGTC
AGCACACGTCTGAACTCTTTCCCTACACGACGCTCTTCCGATCTNNNTAATGACTTCCAAAGCTGGGTC
AGCACACGTCTGAACTCTTTCCCTACACGACGCTCTTCCGATCTNNNTAATGACTTCCAAAGCTGGGTC
AGCACACGTCTGAACTCTTTCCCTACACGACGCTCTTCCGATCTNNNTGCTTTTCTGAGGGACTATCAC
AGCACACGTCTGAACTCTTTCCCTACACGACGCTCTTCCGATCTNNNTGCTTTTCTGAGGGACTATCAC
AGCACACGTCTGAACTCTTTCCCTACACGACGCTCTTCCGATCTNNNTGCTTTTCTGAGGGACTATCAC
AGCACACGTCTGAACTCTTTCCCTACACGACGCTCTTCCGATCTNNNTGCTTTTCTGAGGGACTATCAC
GAGCACACGTCTGAACTCTTTCCCTACACGACGCTCTTCCGATCTNNNACCACCATGCCCAGCTAATATT
GAGCACACGTCTGAACTCTTTCCCTACACGACGCTCTTCCGATCTNNNACCACCATGCCCAGCTAATATT

GAGCACACGTCTGAACTCTTTCCCTACACGACGCTCTTCCGATCTNNNACCACCATGCCCAGCTAATATT
GAGCACACGTCTGAACTCTTTCCCTACACGACGCTCTTCCGATCTNNNACCACCATGCCCAGCTAATATT
GAGCACACGTCTGAACTCTTTCCCTACACGACGCTCTTCCGATCTNNNGCAATAGAGTGAGGTCCTGT
GAGCACACGTCTGAACTCTTTCCCTACACGACGCTCTTCCGATCTNNNGCAATAGAGTGAGGTCCTGT
GAGCACACGTCTGAACTCTTTCCCTACACGACGCTCTTCCGATCTNNNGCAATAGAGTGAGGTCCTGT
GAGCACACGTCTGAACTCTTTCCCTACACGACGCTCTTCCGATCTNNNGCAATAGAGTGAGGTCCTGT
AGCACACGTCTGAACTCTTTCCCTACACGACGCTCTTCCGATCTNNNTCTCATCCTATTATTGCGCTGCT
AGCACACGTCTGAACTCTTTCCCTACACGACGCTCTTCCGATCTNNNTCTCATCCTATTATTGCGCTGCT
AGCACACGTCTGAACTCTTTCCCTACACGACGCTCTTCCGATCTNNNTCTCATCCTATTATTGCGCTGCT
AGCACACGTCTGAACTCTTTCCCTACACGACGCTCTTCCGATCTNNNTCTCATCCTATTATTGCGCTGCT
GCACACGTCTGAACTCTTTCCCTACACGACGCTCTTCCGATCTNNNTTCGTTTTCCATCACTTTGTGCACA
GCACACGTCTGAACTCTTTCCCTACACGACGCTCTTCCGATCTNNNTTCGTTTTCCATCACTTTGTGCACA
GCACACGTCTGAACTCTTTCCCTACACGACGCTCTTCCGATCTNNNTTCGTTTTCCATCACTTTGTGCACA
GCACACGTCTGAACTCTTTCCCTACACGACGCTCTTCCGATCTNNNTTCGTTTTCCATCACTTTGTGCACA
AGCACACGTCTGAACTCTTTCCCTACACGACGCTCTTCCGATCTNNNAAAAGAATGTGCTTCAGAGGG
AGCACACGTCTGAACTCTTTCCCTACACGACGCTCTTCCGATCTNNNAAAAGAATGTGCTTCAGAGGG
AGCACACGTCTGAACTCTTTCCCTACACGACGCTCTTCCGATCTNNNAAAAGAATGTGCTTCAGAGGG
AGCACACGTCTGAACTCTTTCCCTACACGACGCTCTTCCGATCTNNNAAAAGAATGTGCTTCAGAGGG
GAGCACACGTCTGAACTCTTTCCCTACACGACGCTCTTCCGATCTNNNCAAATCTTCTTCTTCCGAGCAG
GAGCACACGTCTGAACTCTTTCCCTACACGACGCTCTTCCGATCTNNNCAAATCTTCTTCTTCCGAGCAG
GAGCACACGTCTGAACTCTTTCCCTACACGACGCTCTTCCGATCTNNNCAAATCTTCTTCTTCCGAGCAG
GAGCACACGTCTGAACTCTTTCCCTACACGACGCTCTTCCGATCTNNNCAAATCTTCTTCTTCCGAGCAG
GAGCACACGTCTGAACTCTTTCCCTACACGACGCTCTTCCGATCTNNNACCTGTGGCTACCTCTGTTTTA
GAGCACACGTCTGAACTCTTTCCCTACACGACGCTCTTCCGATCTNNNACCTGTGGCTACCTCTGTTTTA
GAGCACACGTCTGAACTCTTTCCCTACACGACGCTCTTCCGATCTNNNACCTGTGGCTACCTCTGTTTTA
GAGCACACGTCTGAACTCTTTCCCTACACGACGCTCTTCCGATCTNNNACCTGTGGCTACCTCTGTTTTA
AGCACACGTCTGAACTCTTTCCCTACACGACGCTCTTCCGATCTNNNGAGGCTGAGGCAGGAGAATG
AGCACACGTCTGAACTCTTTCCCTACACGACGCTCTTCCGATCTNNNGAGGCTGAGGCAGGAGAATG
AGCACACGTCTGAACTCTTTCCCTACACGACGCTCTTCCGATCTNNNGAGGCTGAGGCAGGAGAATG
AGCACACGTCTGAACTCTTTCCCTACACGACGCTCTTCCGATCTNNNGAGGCTGAGGCAGGAGAATG
AGCACACGTCTGAACTCTTTCCCTACACGACGCTCTTCCGATCTNNNATTCATTTCAACTTGGTACATTT
AGCACACGTCTGAACTCTTTCCCTACACGACGCTCTTCCGATCTNNNATTCATTTCAACTTGGTACATTT
AGCACACGTCTGAACTCTTTCCCTACACGACGCTCTTCCGATCTNNNATTCATTTCAACTTGGTACATTT
AGCACACGTCTGAACTCTTTCCCTACACGACGCTCTTCCGATCTNNNATTCATTTCAACTTGGTACATTT
ACACGTCTGAACTCTTTCCCTACACGACGCTCTTCCGATCTNNNAACTCCTGATCTTGTGATCCACCCAT
ACACGTCTGAACTCTTTCCCTACACGACGCTCTTCCGATCTNNNAACTCCTGATCTTGTGATCCACCCAT
ACACGTCTGAACTCTTTCCCTACACGACGCTCTTCCGATCTNNNAACTCCTGATCTTGTGATCCACCCAT
ACACGTCTGAACTCTTTCCCTACACGACGCTCTTCCGATCTNNNAACTCCTGATCTTGTGATCCACCCAT
AGAGCACACGTCTGAACTCTTTCCCTACACGACGCTCTTCCGATCTNNNGAATCACTTGAGCTGGGGA
AGAGCACACGTCTGAACTCTTTCCCTACACGACGCTCTTCCGATCTNNNGAATCACTTGAGCTGGGGA
AGAGCACACGTCTGAACTCTTTCCCTACACGACGCTCTTCCGATCTNNNGAATCACTTGAGCTGGGGA
AGAGCACACGTCTGAACTCTTTCCCTACACGACGCTCTTCCGATCTNNNGAATCACTTGAGCTGGGGA
AGCACACGTCTGAACTCTTTCCCTACACGACGCTCTTCCGATCTNNNTTGTTAGTGTTTTGGGCAGATG

AGCACACGTCTGAACTCTTTCCCTACACGACGCTCTTCCGATCTNNNTTGTTAGTGTTTTGGGCAGATG
AGCACACGTCTGAACTCTTTCCCTACACGACGCTCTTCCGATCTNNNTTGTTAGTGTTTTGGGCAGATG
AGCACACGTCTGAACTCTTTCCCTACACGACGCTCTTCCGATCTNNNTTGTTAGTGTTTTGGGCAGATG
GAGCACACGTCTGAACTCTTTCCCTACACGACGCTCTTCCGATCTNNNTGAGGCAGAAGAATGGTTTG
GAGCACACGTCTGAACTCTTTCCCTACACGACGCTCTTCCGATCTNNNTGAGGCAGAAGAATGGTTTG
GAGCACACGTCTGAACTCTTTCCCTACACGACGCTCTTCCGATCTNNNTGAGGCAGAAGAATGGTTTG
GAGCACACGTCTGAACTCTTTCCCTACACGACGCTCTTCCGATCTNNNTGAGGCAGAAGAATGGTTTG
GAGCACACGTCTGAACTCTTTCCCTACACGACGCTCTTCCGATCTNNNGTAGATGGGGGAGGATACA
GAGCACACGTCTGAACTCTTTCCCTACACGACGCTCTTCCGATCTNNNGTAGATGGGGGAGGATACA
GAGCACACGTCTGAACTCTTTCCCTACACGACGCTCTTCCGATCTNNNGTAGATGGGGGAGGATACA
GAGCACACGTCTGAACTCTTTCCCTACACGACGCTCTTCCGATCTNNNGTAGATGGGGGAGGATACA
GAGCACACGTCTGAACTCTTTCCCTACACGACGCTCTTCCGATCTNNNCTATCAGAACTCAACCTCACCCC
GAGCACACGTCTGAACTCTTTCCCTACACGACGCTCTTCCGATCTNNNCTATCAGAACTCAACCTCACCCC
GAGCACACGTCTGAACTCTTTCCCTACACGACGCTCTTCCGATCTNNNCTATCAGAACTCAACCTCACCCC
GAGCACACGTCTGAACTCTTTCCCTACACGACGCTCTTCCGATCTNNNCTATCAGAACTCAACCTCACCCC
AGAGCACACGTCTGAACTCTTTCCCTACACGACGCTCTTCCGATCTNNNAATTCTTTCCCAGTCCACGTA
AGAGCACACGTCTGAACTCTTTCCCTACACGACGCTCTTCCGATCTNNNAATTCTTTCCCAGTCCACGTA
AGAGCACACGTCTGAACTCTTTCCCTACACGACGCTCTTCCGATCTNNNAATTCTTTCCCAGTCCACGTA
AGAGCACACGTCTGAACTCTTTCCCTACACGACGCTCTTCCGATCTNNNAATTCTTTCCCAGTCCACGTA
AGCACACGTCTGAACTCTTTCCCTACACGACGCTCTTCCGATCTNNNATCATGTTTTAGTTGGAGGCGG
AGCACACGTCTGAACTCTTTCCCTACACGACGCTCTTCCGATCTNNNATCATGTTTTAGTTGGAGGCGG
AGCACACGTCTGAACTCTTTCCCTACACGACGCTCTTCCGATCTNNNATCATGTTTTAGTTGGAGGCGG
AGCACACGTCTGAACTCTTTCCCTACACGACGCTCTTCCGATCTNNNATCATGTTTTAGTTGGAGGCGG
GAGCACACGTCTGAACTCTTTCCCTACACGACGCTCTTCCGATCTNNNGCGATATTGAGCAGTGTCCCA
GAGCACACGTCTGAACTCTTTCCCTACACGACGCTCTTCCGATCTNNNGCGATATTGAGCAGTGTCCCA
GAGCACACGTCTGAACTCTTTCCCTACACGACGCTCTTCCGATCTNNNGCGATATTGAGCAGTGTCCCA
GAGCACACGTCTGAACTCTTTCCCTACACGACGCTCTTCCGATCTNNNGCGATATTGAGCAGTGTCCCA
AGCACACGTCTGAACTCTTTCCCTACACGACGCTCTTCCGATCTNNNATTTAACAACAAGCATGAAAAA
AGCACACGTCTGAACTCTTTCCCTACACGACGCTCTTCCGATCTNNNATTTAACAACAAGCATGAAAAA
AGCACACGTCTGAACTCTTTCCCTACACGACGCTCTTCCGATCTNNNATTTAACAACAAGCATGAAAAA
AGCACACGTCTGAACTCTTTCCCTACACGACGCTCTTCCGATCTNNNATTTAACAACAAGCATGAAAAA
AGCACACGTCTGAACTCTTTCCCTACACGACGCTCTTCCGATCTNNNAGCTTTATATGTGGAGGTGGG
AGCACACGTCTGAACTCTTTCCCTACACGACGCTCTTCCGATCTNNNAGCTTTATATGTGGAGGTGGG
AGCACACGTCTGAACTCTTTCCCTACACGACGCTCTTCCGATCTNNNAGCTTTATATGTGGAGGTGGG
AGCACACGTCTGAACTCTTTCCCTACACGACGCTCTTCCGATCTNNNAGCTTTATATGTGGAGGTGGG
GCACACGTCTGAACTCTTTCCCTACACGACGCTCTTCCGATCTNNNTCCATACTCTTTGGACTGTAGGTC
GCACACGTCTGAACTCTTTCCCTACACGACGCTCTTCCGATCTNNNTCCATACTCTTTGGACTGTAGGTC
GCACACGTCTGAACTCTTTCCCTACACGACGCTCTTCCGATCTNNNTCCATACTCTTTGGACTGTAGGTC
GCACACGTCTGAACTCTTTCCCTACACGACGCTCTTCCGATCTNNNTCCATACTCTTTGGACTGTAGGTC
AGCACACGTCTGAACTCTTTCCCTACACGACGCTCTTCCGATCTNNNTAGCACCTCTGTTCTCTTTCGTC
AGCACACGTCTGAACTCTTTCCCTACACGACGCTCTTCCGATCTNNNTAGCACCTCTGTTCTCTTTCGTC
AGCACACGTCTGAACTCTTTCCCTACACGACGCTCTTCCGATCTNNNTAGCACCTCTGTTCTCTTTCGTC
AGCACACGTCTGAACTCTTTCCCTACACGACGCTCTTCCGATCTNNNTAGCACCTCTGTTCTCTTTCGTC

CACACGTCTGAACTCTTTCCCTACACGACGCTCTTCCGATCTNNNCACAACGACCACATCCTCATAGACA
CACACGTCTGAACTCTTTCCCTACACGACGCTCTTCCGATCTNNNCACAACGACCACATCCTCATAGACA
CACACGTCTGAACTCTTTCCCTACACGACGCTCTTCCGATCTNNNCACAACGACCACATCCTCATAGACA
CACACGTCTGAACTCTTTCCCTACACGACGCTCTTCCGATCTNNNCACAACGACCACATCCTCATAGACA
GAGCACACGTCTGAACTCTTTCCCTACACGACGCTCTTCCGATCTNNNGAGGTTGCAGTGAGCTGAGA
GAGCACACGTCTGAACTCTTTCCCTACACGACGCTCTTCCGATCTNNNGAGGTTGCAGTGAGCTGAGA
GAGCACACGTCTGAACTCTTTCCCTACACGACGCTCTTCCGATCTNNNGAGGTTGCAGTGAGCTGAGA
GAGCACACGTCTGAACTCTTTCCCTACACGACGCTCTTCCGATCTNNNGAGGTTGCAGTGAGCTGAGA
GCACACGTCTGAACTCTTTCCCTACACGACGCTCTTCCGATCTNNNTCACCTATATGTCATGACAGATGA
GCACACGTCTGAACTCTTTCCCTACACGACGCTCTTCCGATCTNNNTCACCTATATGTCATGACAGATGA
GCACACGTCTGAACTCTTTCCCTACACGACGCTCTTCCGATCTNNNTCACCTATATGTCATGACAGATGA
GCACACGTCTGAACTCTTTCCCTACACGACGCTCTTCCGATCTNNNTCACCTATATGTCATGACAGATGA
AGCACACGTCTGAACTCTTTCCCTACACGACGCTCTTCCGATCTNNNCACCTGTAGTCCCAGCAACTCCA
AGCACACGTCTGAACTCTTTCCCTACACGACGCTCTTCCGATCTNNNCACCTGTAGTCCCAGCAACTCCA
AGCACACGTCTGAACTCTTTCCCTACACGACGCTCTTCCGATCTNNNCACCTGTAGTCCCAGCAACTCCA
AGCACACGTCTGAACTCTTTCCCTACACGACGCTCTTCCGATCTNNNCACCTGTAGTCCCAGCAACTCCA
AGCACACGTCTGAACTCTTTCCCTACACGACGCTCTTCCGATCTNNNCTCACACTCTGGACTTAGAGAG
AGCACACGTCTGAACTCTTTCCCTACACGACGCTCTTCCGATCTNNNCTCACACTCTGGACTTAGAGAG
AGCACACGTCTGAACTCTTTCCCTACACGACGCTCTTCCGATCTNNNCTCACACTCTGGACTTAGAGAG
AGCACACGTCTGAACTCTTTCCCTACACGACGCTCTTCCGATCTNNNCTCACACTCTGGACTTAGAGAG
GAGCACACGTCTGAACTCTTTCCCTACACGACGCTCTTCCGATCTNNNTAGGAGACTGAGGCAGGAGA
GAGCACACGTCTGAACTCTTTCCCTACACGACGCTCTTCCGATCTNNNTAGGAGACTGAGGCAGGAGA
GAGCACACGTCTGAACTCTTTCCCTACACGACGCTCTTCCGATCTNNNTAGGAGACTGAGGCAGGAGA
GAGCACACGTCTGAACTCTTTCCCTACACGACGCTCTTCCGATCTNNNTAGGAGACTGAGGCAGGAGA
AGCACACGTCTGAACTCTTTCCCTACACGACGCTCTTCCGATCTNNNTGAACAGGCTTTGTCTATGTGG
AGCACACGTCTGAACTCTTTCCCTACACGACGCTCTTCCGATCTNNNTGAACAGGCTTTGTCTATGTGG
AGCACACGTCTGAACTCTTTCCCTACACGACGCTCTTCCGATCTNNNTGAACAGGCTTTGTCTATGTGG
AGCACACGTCTGAACTCTTTCCCTACACGACGCTCTTCCGATCTNNNTGAACAGGCTTTGTCTATGTGG
AGCACACGTCTGAACTCTTTCCCTACACGACGCTCTTCCGATCTNNNAAATTCCTGTCAGTCCTCCAAG
AGCACACGTCTGAACTCTTTCCCTACACGACGCTCTTCCGATCTNNNAAATTCCTGTCAGTCCTCCAAG
AGCACACGTCTGAACTCTTTCCCTACACGACGCTCTTCCGATCTNNNAAATTCCTGTCAGTCCTCCAAG
AGCACACGTCTGAACTCTTTCCCTACACGACGCTCTTCCGATCTNNNAAATTCCTGTCAGTCCTCCAAG
AGCACACGTCTGAACTCTTTCCCTACACGACGCTCTTCCGATCTNNNGCAACAGACGGAGACTCTGTAT
AGCACACGTCTGAACTCTTTCCCTACACGACGCTCTTCCGATCTNNNGCAACAGACGGAGACTCTGTAT
AGCACACGTCTGAACTCTTTCCCTACACGACGCTCTTCCGATCTNNNGCAACAGACGGAGACTCTGTAT
AGCACACGTCTGAACTCTTTCCCTACACGACGCTCTTCCGATCTNNNGCAACAGACGGAGACTCTGTAT
ACACGTCTGAACTCTTTCCCTACACGACGCTCTTCCGATCTNNNCTCTCAATGAGCTGTTGGGTACACA
ACACGTCTGAACTCTTTCCCTACACGACGCTCTTCCGATCTNNNCTCTCAATGAGCTGTTGGGTACACA
ACACGTCTGAACTCTTTCCCTACACGACGCTCTTCCGATCTNNNCTCTCAATGAGCTGTTGGGTACACA
ACACGTCTGAACTCTTTCCCTACACGACGCTCTTCCGATCTNNNCTCTCAATGAGCTGTTGGGTACACA
GAGCACACGTCTGAACTCTTTCCCTACACGACGCTCTTCCGATCTNNNATGGGAAAATTCTCTCCCTCTT
GAGCACACGTCTGAACTCTTTCCCTACACGACGCTCTTCCGATCTNNNATGGGAAAATTCTCTCCCTCTT
GAGCACACGTCTGAACTCTTTCCCTACACGACGCTCTTCCGATCTNNNATGGGAAAATTCTCTCCCTCTT

AGCACACGTCTGAACTCTTTCCCTACACGACGCTCTTCCGATCTNNNATGGGAAAATTCTCTCCCTCTT
CACGTCTGAACTCTTTCCCTACACGACGCTCTTCCGATCTNNNGCAAACTCAGAACCTTTGTGTGTCAT
CACGTCTGAACTCTTTCCCTACACGACGCTCTTCCGATCTNNNGCAAACTCAGAACCTTTGTGTGTCAT
CACGTCTGAACTCTTTCCCTACACGACGCTCTTCCGATCTNNNGCAAACTCAGAACCTTTGTGTGTCAT
CACGTCTGAACTCTTTCCCTACACGACGCTCTTCCGATCTNNNGCAAACTCAGAACCTTTGTGTGTCAT
GCACACGTCTGAACTCTTTCCCTACACGACGCTCTTCCGATCTNNNGTAATCCCAGCTACTTGAGAGG
GCACACGTCTGAACTCTTTCCCTACACGACGCTCTTCCGATCTNNNGTAATCCCAGCTACTTGAGAGG
GCACACGTCTGAACTCTTTCCCTACACGACGCTCTTCCGATCTNNNGTAATCCCAGCTACTTGAGAGG
GCACACGTCTGAACTCTTTCCCTACACGACGCTCTTCCGATCTNNNGTAATCCCAGCTACTTGAGAGG
GCACACGTCTGAACTCTTTCCCTACACGACGCTCTTCCGATCTNNNGGCTTCTAAAGATGAGCCATCC
GCACACGTCTGAACTCTTTCCCTACACGACGCTCTTCCGATCTNNNGGCTTCTAAAGATGAGCCATCC
GCACACGTCTGAACTCTTTCCCTACACGACGCTCTTCCGATCTNNNGGCTTCTAAAGATGAGCCATCC
GCACACGTCTGAACTCTTTCCCTACACGACGCTCTTCCGATCTNNNGGCTTCTAAAGATGAGCCATCC
AGCACACGTCTGAACTCTTTCCCTACACGACGCTCTTCCGATCTNNNTGAGGCAGGAGAATGGTATG
AGCACACGTCTGAACTCTTTCCCTACACGACGCTCTTCCGATCTNNNTGAGGCAGGAGAATGGTATG
AGCACACGTCTGAACTCTTTCCCTACACGACGCTCTTCCGATCTNNNTGAGGCAGGAGAATGGTATG
AGCACACGTCTGAACTCTTTCCCTACACGACGCTCTTCCGATCTNNNTGAGGCAGGAGAATGGTATG
GCACACGTCTGAACTCTTTCCCTACACGACGCTCTTCCGATCTNNNGCCAAGATCATGCCATTGTACT
GCACACGTCTGAACTCTTTCCCTACACGACGCTCTTCCGATCTNNNGCCAAGATCATGCCATTGTACT
GCACACGTCTGAACTCTTTCCCTACACGACGCTCTTCCGATCTNNNGCCAAGATCATGCCATTGTACT
GCACACGTCTGAACTCTTTCCCTACACGACGCTCTTCCGATCTNNNGCCAAGATCATGCCATTGTACT
AGCACACGTCTGAACTCTTTCCCTACACGACGCTCTTCCGATCTNNNTCTACTATAGCATCTCCAAAGG
AGCACACGTCTGAACTCTTTCCCTACACGACGCTCTTCCGATCTNNNTCTACTATAGCATCTCCAAAGG
AGCACACGTCTGAACTCTTTCCCTACACGACGCTCTTCCGATCTNNNTCTACTATAGCATCTCCAAAGG
AGCACACGTCTGAACTCTTTCCCTACACGACGCTCTTCCGATCTNNNTCTACTATAGCATCTCCAAAGG
GCACACGTCTGAACTCTTTCCCTACACGACGCTCTTCCGATCTNNNTGAGCACCTACTAGAAGCC
GCACACGTCTGAACTCTTTCCCTACACGACGCTCTTCCGATCTNNNTGAGCACCTACTAGAAGCC
GCACACGTCTGAACTCTTTCCCTACACGACGCTCTTCCGATCTNNNTGAGCACCTACTAGAAGCC
GCACACGTCTGAACTCTTTCCCTACACGACGCTCTTCCGATCTNNNTGAGCACCTACTAGAAGCC
AGCACACGTCTGAACTCTTTCCCTACACGACGCTCTTCCGATCTNNNTGTAAGCAATCTTCCCTCCTCA
AGCACACGTCTGAACTCTTTCCCTACACGACGCTCTTCCGATCTNNNTGTAAGCAATCTTCCCTCCTCA
AGCACACGTCTGAACTCTTTCCCTACACGACGCTCTTCCGATCTNNNTGTAAGCAATCTTCCCTCCTCA
AGCACACGTCTGAACTCTTTCCCTACACGACGCTCTTCCGATCTNNNTGTAAGCAATCTTCCCTCCTCA
AGCACACGTCTGAACTCTTTCCCTACACGACGCTCTTCCGATCTNNNATGTGCCTGTAATTCCAGCTAC
AGCACACGTCTGAACTCTTTCCCTACACGACGCTCTTCCGATCTNNNATGTGCCTGTAATTCCAGCTAC
AGCACACGTCTGAACTCTTTCCCTACACGACGCTCTTCCGATCTNNNATGTGCCTGTAATTCCAGCTAC
AGCACACGTCTGAACTCTTTCCCTACACGACGCTCTTCCGATCTNNNATGTGCCTGTAATTCCAGCTAC
GAGCACACGTCTGAACTCTTTCCCTACACGACGCTCTTCCGATCTNNNCCTCCTTGTCATTGATCTTTG
GAGCACACGTCTGAACTCTTTCCCTACACGACGCTCTTCCGATCTNNNCCTCCTTGTCATTGATCTTTG
GAGCACACGTCTGAACTCTTTCCCTACACGACGCTCTTCCGATCTNNNCCTCCTTGTCATTGATCTTTG
GAGCACACGTCTGAACTCTTTCCCTACACGACGCTCTTCCGATCTNNNCCTCCTTGTCATTGATCTTTG
AGCACACGTCTGAACTCTTTCCCTACACGACGCTCTTCCGATCTNNNTGTAATTCCAGCTACTTGGGA
AGCACACGTCTGAACTCTTTCCCTACACGACGCTCTTCCGATCTNNNTGTAATTCCAGCTACTTGGGA

AGCACACGTCTGAACTCTTTCCCTACACGACGCTCTTCCGATCTNNNTGTAATTCCAGCTACTTGGGA
AGCACACGTCTGAACTCTTTCCCTACACGACGCTCTTCCGATCTNNNTGTAATTCCAGCTACTTGGGA
AGCACACGTCTGAACTCTTTCCCTACACGACGCTCTTCCGATCTNNNTAACTCTGTCTTGTCGGCTGAA
AGCACACGTCTGAACTCTTTCCCTACACGACGCTCTTCCGATCTNNNTAACTCTGTCTTGTCGGCTGAA
AGCACACGTCTGAACTCTTTCCCTACACGACGCTCTTCCGATCTNNNTAACTCTGTCTTGTCGGCTGAA
AGCACACGTCTGAACTCTTTCCCTACACGACGCTCTTCCGATCTNNNTAACTCTGTCTTGTCGGCTGAA
CACGTCTGAACTCTTTCCCTACACGACGCTCTTCCGATCTNNNGGCTTTTTTGGTTGGTAGGCTACAT
CACGTCTGAACTCTTTCCCTACACGACGCTCTTCCGATCTNNNGGCTTTTTTGGTTGGTAGGCTACAT
CACGTCTGAACTCTTTCCCTACACGACGCTCTTCCGATCTNNNGGCTTTTTTGGTTGGTAGGCTACAT
CACGTCTGAACTCTTTCCCTACACGACGCTCTTCCGATCTNNNGGCTTTTTTGGTTGGTAGGCTACAT
AGCACACGTCTGAACTCTTTCCCTACACGACGCTCTTCCGATCTNNNGTGATGACTTCTCACTTCAGCA
AGCACACGTCTGAACTCTTTCCCTACACGACGCTCTTCCGATCTNNNGTGATGACTTCTCACTTCAGCA
AGCACACGTCTGAACTCTTTCCCTACACGACGCTCTTCCGATCTNNNGTGATGACTTCTCACTTCAGCA
AGCACACGTCTGAACTCTTTCCCTACACGACGCTCTTCCGATCTNNNGTGATGACTTCTCACTTCAGCA
GCACACGTCTGAACTCTTTCCCTACACGACGCTCTTCCGATCTNNNGAGTTGAAGACTCCAGCCTCCTC
GCACACGTCTGAACTCTTTCCCTACACGACGCTCTTCCGATCTNNNGAGTTGAAGACTCCAGCCTCCTC
GCACACGTCTGAACTCTTTCCCTACACGACGCTCTTCCGATCTNNNGAGTTGAAGACTCCAGCCTCCTC
GCACACGTCTGAACTCTTTCCCTACACGACGCTCTTCCGATCTNNNGAGTTGAAGACTCCAGCCTCCTC
GAGCACACGTCTGAACTCTTTCCCTACACGACGCTCTTCCGATCTNNNTGGCCCCACAATGAGATATG
GAGCACACGTCTGAACTCTTTCCCTACACGACGCTCTTCCGATCTNNNTGGCCCCACAATGAGATATG
GAGCACACGTCTGAACTCTTTCCCTACACGACGCTCTTCCGATCTNNNTGGCCCCACAATGAGATATG
GAGCACACGTCTGAACTCTTTCCCTACACGACGCTCTTCCGATCTNNNTGGCCCCACAATGAGATATG
AGCACACGTCTGAACTCTTTCCCTACACGACGCTCTTCCGATCTNNNTACGGAGATTAGCATTGTCCCT
AGCACACGTCTGAACTCTTTCCCTACACGACGCTCTTCCGATCTNNNTACGGAGATTAGCATTGTCCCT
AGCACACGTCTGAACTCTTTCCCTACACGACGCTCTTCCGATCTNNNTACGGAGATTAGCATTGTCCCT
AGCACACGTCTGAACTCTTTCCCTACACGACGCTCTTCCGATCTNNNTACGGAGATTAGCATTGTCCCT
GCACACGTCTGAACTCTTTCCCTACACGACGCTCTTCCGATCTNNNCTCCCATTCTGTAGGTTGTCTGT
GCACACGTCTGAACTCTTTCCCTACACGACGCTCTTCCGATCTNNNCTCCCATTCTGTAGGTTGTCTGT
GCACACGTCTGAACTCTTTCCCTACACGACGCTCTTCCGATCTNNNCTCCCATTCTGTAGGTTGTCTGT
GCACACGTCTGAACTCTTTCCCTACACGACGCTCTTCCGATCTNNNCTCCCATTCTGTAGGTTGTCTGT
AGAGCACACGTCTGAACTCTTTCCCTACACGACGCTCTTCCGATCTNNNACAAAATTAGTTGGGCATG
AGAGCACACGTCTGAACTCTTTCCCTACACGACGCTCTTCCGATCTNNNACAAAATTAGTTGGGCATG
AGAGCACACGTCTGAACTCTTTCCCTACACGACGCTCTTCCGATCTNNNACAAAATTAGTTGGGCATG
AGAGCACACGTCTGAACTCTTTCCCTACACGACGCTCTTCCGATCTNNNACAAAATTAGTTGGGCATG
GAGCACACGTCTGAACTCTTTCCCTACACGACGCTCTTCCGATCTNNNCAAAAGATTAGCCAGGTGTG
GAGCACACGTCTGAACTCTTTCCCTACACGACGCTCTTCCGATCTNNNCAAAAGATTAGCCAGGTGTG
GAGCACACGTCTGAACTCTTTCCCTACACGACGCTCTTCCGATCTNNNCAAAAGATTAGCCAGGTGTG
GAGCACACGTCTGAACTCTTTCCCTACACGACGCTCTTCCGATCTNNNCAAAAGATTAGCCAGGTGTG
GCACACGTCTGAACTCTTTCCCTACACGACGCTCTTCCGATCTNNNGATCCACCACCACACTTAGCTAA
GCACACGTCTGAACTCTTTCCCTACACGACGCTCTTCCGATCTNNNGATCCACCACCACACTTAGCTAA
GCACACGTCTGAACTCTTTCCCTACACGACGCTCTTCCGATCTNNNGATCCACCACCACACTTAGCTAA
GCACACGTCTGAACTCTTTCCCTACACGACGCTCTTCCGATCTNNNGATCCACCACCACACTTAGCTAA
AGCACACGTCTGAACTCTTTCCCTACACGACGCTCTTCCGATCTNNNGGAAACTGATTAGCACAGAA

GAGCACACGTCTGAACTCTTTCCCTACACGACGCTCTTCCGATCTNNNGGAAACTGATTAGCACAGAAG
GAGCACACGTCTGAACTCTTTCCCTACACGACGCTCTTCCGATCTNNNGGAAACTGATTAGCACAGAAG
GAGCACACGTCTGAACTCTTTCCCTACACGACGCTCTTCCGATCTNNNGGAAACTGATTAGCACAGAAG
AGAGCACACGTCTGAACTCTTTCCCTACACGACGCTCTTCCGATCTNNNTATGTCTTTCAGGTCTGCCTC
AGAGCACACGTCTGAACTCTTTCCCTACACGACGCTCTTCCGATCTNNNTATGTCTTTCAGGTCTGCCTC
AGAGCACACGTCTGAACTCTTTCCCTACACGACGCTCTTCCGATCTNNNTATGTCTTTCAGGTCTGCCTC
AGAGCACACGTCTGAACTCTTTCCCTACACGACGCTCTTCCGATCTNNNTATGTCTTTCAGGTCTGCCTC
GAGCACACGTCTGAACTCTTTCCCTACACGACGCTCTTCCGATCTNNNCCTGGCCAACATGATGAAATC
GAGCACACGTCTGAACTCTTTCCCTACACGACGCTCTTCCGATCTNNNCCTGGCCAACATGATGAAATC
GAGCACACGTCTGAACTCTTTCCCTACACGACGCTCTTCCGATCTNNNCCTGGCCAACATGATGAAATC
GAGCACACGTCTGAACTCTTTCCCTACACGACGCTCTTCCGATCTNNNCCTGGCCAACATGATGAAATC
AGCACACGTCTGAACTCTTTCCCTACACGACGCTCTTCCGATCTNNNTGTGTCTGTGTGTGTGTGTGTA
AGCACACGTCTGAACTCTTTCCCTACACGACGCTCTTCCGATCTNNNTGTGTCTGTGTGTGTGTGTGTA
AGCACACGTCTGAACTCTTTCCCTACACGACGCTCTTCCGATCTNNNTGTGTCTGTGTGTGTGTGTGTA
AGCACACGTCTGAACTCTTTCCCTACACGACGCTCTTCCGATCTNNNTGTGTCTGTGTGTGTGTGTGTA
AGCACACGTCTGAACTCTTTCCCTACACGACGCTCTTCCGATCTNNNATTCTCCTTAAAATTCATACCAG
AGCACACGTCTGAACTCTTTCCCTACACGACGCTCTTCCGATCTNNNATTCTCCTTAAAATTCATACCAG
AGCACACGTCTGAACTCTTTCCCTACACGACGCTCTTCCGATCTNNNATTCTCCTTAAAATTCATACCAG
AGCACACGTCTGAACTCTTTCCCTACACGACGCTCTTCCGATCTNNNATTCTCCTTAAAATTCATACCAG
AGCACACGTCTGAACTCTTTCCCTACACGACGCTCTTCCGATCTNNNTGTAGTCCCAGCTACTTAGGAG
AGCACACGTCTGAACTCTTTCCCTACACGACGCTCTTCCGATCTNNNTGTAGTCCCAGCTACTTAGGAG
AGCACACGTCTGAACTCTTTCCCTACACGACGCTCTTCCGATCTNNNTGTAGTCCCAGCTACTTAGGAG
AGCACACGTCTGAACTCTTTCCCTACACGACGCTCTTCCGATCTNNNTGTAGTCCCAGCTACTTAGGAG
CACACGTCTGAACTCTTTCCCTACACGACGCTCTTCCGATCTNNNTAGGTGCGGTAGTCCAGGAAGCAT
CACACGTCTGAACTCTTTCCCTACACGACGCTCTTCCGATCTNNNTAGGTGCGGTAGTCCAGGAAGCAT
CACACGTCTGAACTCTTTCCCTACACGACGCTCTTCCGATCTNNNTAGGTGCGGTAGTCCAGGAAGCAT
CACACGTCTGAACTCTTTCCCTACACGACGCTCTTCCGATCTNNNTAGGTGCGGTAGTCCAGGAAGCAT
AGCACACGTCTGAACTCTTTCCCTACACGACGCTCTTCCGATCTNNNGAAGTTGAGAACACAGTGACAG
AGCACACGTCTGAACTCTTTCCCTACACGACGCTCTTCCGATCTNNNGAAGTTGAGAACACAGTGACAG
AGCACACGTCTGAACTCTTTCCCTACACGACGCTCTTCCGATCTNNNGAAGTTGAGAACACAGTGACAG
AGCACACGTCTGAACTCTTTCCCTACACGACGCTCTTCCGATCTNNNGAAGTTGAGAACACAGTGACAG
GAGCACACGTCTGAACTCTTTCCCTACACGACGCTCTTCCGATCTNNNAGCTGTAGTGTGCAAGATGAT
GAGCACACGTCTGAACTCTTTCCCTACACGACGCTCTTCCGATCTNNNAGCTGTAGTGTGCAAGATGAT
GAGCACACGTCTGAACTCTTTCCCTACACGACGCTCTTCCGATCTNNNAGCTGTAGTGTGCAAGATGAT
GAGCACACGTCTGAACTCTTTCCCTACACGACGCTCTTCCGATCTNNNAGCTGTAGTGTGCAAGATGAT
AGCACACGTCTGAACTCTTTCCCTACACGACGCTCTTCCGATCTNNNGAGGTTGCAGTGAGCTGAGAT
AGCACACGTCTGAACTCTTTCCCTACACGACGCTCTTCCGATCTNNNGAGGTTGCAGTGAGCTGAGAT
AGCACACGTCTGAACTCTTTCCCTACACGACGCTCTTCCGATCTNNNGAGGTTGCAGTGAGCTGAGAT
AGCACACGTCTGAACTCTTTCCCTACACGACGCTCTTCCGATCTNNNGAGGTTGCAGTGAGCTGAGAT
GCACACGTCTGAACTCTTTCCCTACACGACGCTCTTCCGATCTNNNTTTTACTCCTCTAACCCCTGCAGCA
GCACACGTCTGAACTCTTTCCCTACACGACGCTCTTCCGATCTNNNTTTTACTCCTCTAACCCCTGCAGCA
GCACACGTCTGAACTCTTTCCCTACACGACGCTCTTCCGATCTNNNTTTTACTCCTCTAACCCCTGCAGCA
GCACACGTCTGAACTCTTTCCCTACACGACGCTCTTCCGATCTNNNTTTTACTCCTCTAACCCCTGCAGCA

AGCACACGTCTGAACTCTTTCCCTACACGACGCTCTTCCGATCTNNNGAATCTCTGGGCCAAGAGGTAA

AGCACACGTCTGAACTCTTTCCCTACACGACGCTCTTCCGATCTNNNGAATCTCTGGGCCAAGAGGTAA

AGCACACGTCTGAACTCTTTCCCTACACGACGCTCTTCCGATCTNNNGAATCTCTGGGCCAAGAGGTAA

AGCACACGTCTGAACTCTTTCCCTACACGACGCTCTTCCGATCTNNNGAATCTCTGGGCCAAGAGGTAA

AGCACACGTCTGAACTCTTTCCCTACACGACGCTCTTCCGATCTNNNAAACCATGCAATGTTTCTGGTCC

AGCACACGTCTGAACTCTTTCCCTACACGACGCTCTTCCGATCTNNNAAACCATGCAATGTTTCTGGTCC

AGCACACGTCTGAACTCTTTCCCTACACGACGCTCTTCCGATCTNNNAAACCATGCAATGTTTCTGGTCC

AGCACACGTCTGAACTCTTTCCCTACACGACGCTCTTCCGATCTNNNAAACCATGCAATGTTTCTGGTCC

CACACGTCTGAACTCTTTCCCTACACGACGCTCTTCCGATCTNNNCTGTGAACCTAGACATCCCTGTGCA

CACACGTCTGAACTCTTTCCCTACACGACGCTCTTCCGATCTNNNCTGTGAACCTAGACATCCCTGTGCA

CACACGTCTGAACTCTTTCCCTACACGACGCTCTTCCGATCTNNNCTGTGAACCTAGACATCCCTGTGCA

CACACGTCTGAACTCTTTCCCTACACGACGCTCTTCCGATCTNNNCTGTGAACCTAGACATCCCTGTGCA

AGAGCACACGTCTGAACTCTTTCCCTACACGACGCTCTTCCGATCTNNNCAGTATACGTCTGTAGTCCCA

AGAGCACACGTCTGAACTCTTTCCCTACACGACGCTCTTCCGATCTNNNCAGTATACGTCTGTAGTCCCA

AGAGCACACGTCTGAACTCTTTCCCTACACGACGCTCTTCCGATCTNNNCAGTATACGTCTGTAGTCCCA

AGAGCACACGTCTGAACTCTTTCCCTACACGACGCTCTTCCGATCTNNNCAGTATACGTCTGTAGTCCCA

GCACACGTCTGAACTCTTTCCCTACACGACGCTCTTCCGATCTNNNTGTTGCCTATGCTTTGGTCCTATC

GCACACGTCTGAACTCTTTCCCTACACGACGCTCTTCCGATCTNNNTGTTGCCTATGCTTTGGTCCTATC

GCACACGTCTGAACTCTTTCCCTACACGACGCTCTTCCGATCTNNNTGTTGCCTATGCTTTGGTCCTATC

GCACACGTCTGAACTCTTTCCCTACACGACGCTCTTCCGATCTNNNTGTTGCCTATGCTTTGGTCCTATC

GAGCACACGTCTGAACTCTTTCCCTACACGACGCTCTTCCGATCTNNNACTTGTTGTCACATAAATTGGA

GAGCACACGTCTGAACTCTTTCCCTACACGACGCTCTTCCGATCTNNNACTTGTTGTCACATAAATTGGA

GAGCACACGTCTGAACTCTTTCCCTACACGACGCTCTTCCGATCTNNNACTTGTTGTCACATAAATTGGA

GAGCACACGTCTGAACTCTTTCCCTACACGACGCTCTTCCGATCTNNNACTTGTTGTCACATAAATTGGA

GAGCACACGTCTGAACTCTTTCCCTACACGACGCTCTTCCGATCTNNNGAGGACCTACGAACAGTGACA

GAGCACACGTCTGAACTCTTTCCCTACACGACGCTCTTCCGATCTNNNGAGGACCTACGAACAGTGACA

GAGCACACGTCTGAACTCTTTCCCTACACGACGCTCTTCCGATCTNNNGAGGACCTACGAACAGTGACA

GAGCACACGTCTGAACTCTTTCCCTACACGACGCTCTTCCGATCTNNNGAGGACCTACGAACAGTGACA

GAGCACACGTCTGAACTCTTTCCCTACACGACGCTCTTCCGATCTNNNTTTTGTGTGTCTAGAGCGTG

GAGCACACGTCTGAACTCTTTCCCTACACGACGCTCTTCCGATCTNNNTTTTGTGTGTCTAGAGCGTG

GAGCACACGTCTGAACTCTTTCCCTACACGACGCTCTTCCGATCTNNNTTTTGTGTGTCTAGAGCGTG

GAGCACACGTCTGAACTCTTTCCCTACACGACGCTCTTCCGATCTNNNTTTTGTGTGTCTAGAGCGTG

GAGCACACGTCTGAACTCTTTCCCTACACGACGCTCTTCCGATCTNNNCACCTTAGCCTCCTGATTAGCT

GAGCACACGTCTGAACTCTTTCCCTACACGACGCTCTTCCGATCTNNNCACCTTAGCCTCCTGATTAGCT

GAGCACACGTCTGAACTCTTTCCCTACACGACGCTCTTCCGATCTNNNCACCTTAGCCTCCTGATTAGCT

GAGCACACGTCTGAACTCTTTCCCTACACGACGCTCTTCCGATCTNNNCACCTTAGCCTCCTGATTAGCT

AGCACACGTCTGAACTCTTTCCCTACACGACGCTCTTCCGATCTNNNGGTAAGCCAAAAAGCTGCACAT

AGCACACGTCTGAACTCTTTCCCTACACGACGCTCTTCCGATCTNNNGGTAAGCCAAAAAGCTGCACAT

AGCACACGTCTGAACTCTTTCCCTACACGACGCTCTTCCGATCTNNNGGTAAGCCAAAAAGCTGCACAT

AGCACACGTCTGAACTCTTTCCCTACACGACGCTCTTCCGATCTNNNGGTAAGCCAAAAAGCTGCACAT

AGCACACGTCTGAACTCTTTCCCTACACGACGCTCTTCCGATCTNNNGCTGAGCAAAAAATCACTAGGC

AGCACACGTCTGAACTCTTTCCCTACACGACGCTCTTCCGATCTNNNGCTGAGCAAAAAATCACTAGGC

AGCACACGTCTGAACTCTTTCCCTACACGACGCTCTTCCGATCTNNNGCTGAGCAAAAAATCACTAGGC

GCACACGTCTGAACTCTTTCCCTACACGACGCTCTTCCGATCTNNNGCTGAGCAAAAAATCACTAGGC
AGCACACGTCTGAACTCTTTCCCTACACGACGCTCTTCCGATCTNNNTCCTCCCTTTCTGACATCTCCAT
AGCACACGTCTGAACTCTTTCCCTACACGACGCTCTTCCGATCTNNNTCCTCCCTTTCTGACATCTCCAT
AGCACACGTCTGAACTCTTTCCCTACACGACGCTCTTCCGATCTNNNTCCTCCCTTTCTGACATCTCCAT
AGCACACGTCTGAACTCTTTCCCTACACGACGCTCTTCCGATCTNNNTCCTCCCTTTCTGACATCTCCAT
AGCACACGTCTGAACTCTTTCCCTACACGACGCTCTTCCGATCTNNNGCTGAAGCAGGAGAAATGTTT
AGCACACGTCTGAACTCTTTCCCTACACGACGCTCTTCCGATCTNNNGCTGAAGCAGGAGAAATGTTT
AGCACACGTCTGAACTCTTTCCCTACACGACGCTCTTCCGATCTNNNGCTGAAGCAGGAGAAATGTTT
AGCACACGTCTGAACTCTTTCCCTACACGACGCTCTTCCGATCTNNNGCTGAAGCAGGAGAAATGTTT
GAGCACACGTCTGAACTCTTTCCCTACACGACGCTCTTCCGATCTNNNTGATTACTGGAAGCGCACTG
GAGCACACGTCTGAACTCTTTCCCTACACGACGCTCTTCCGATCTNNNTGATTACTGGAAGCGCACTG
GAGCACACGTCTGAACTCTTTCCCTACACGACGCTCTTCCGATCTNNNTGATTACTGGAAGCGCACTG
GAGCACACGTCTGAACTCTTTCCCTACACGACGCTCTTCCGATCTNNNTGATTACTGGAAGCGCACTG
GCACACGTCTGAACTCTTTCCCTACACGACGCTCTTCCGATCTNNNAAGAACAGACCATGAGCAGAGG
GCACACGTCTGAACTCTTTCCCTACACGACGCTCTTCCGATCTNNNAAGAACAGACCATGAGCAGAGG
GCACACGTCTGAACTCTTTCCCTACACGACGCTCTTCCGATCTNNNAAGAACAGACCATGAGCAGAGG
GCACACGTCTGAACTCTTTCCCTACACGACGCTCTTCCGATCTNNNAAGAACAGACCATGAGCAGAGG
GAGCACACGTCTGAACTCTTTCCCTACACGACGCTCTTCCGATCTNNNATTTCTATCCTTCCCCAAGCA
GAGCACACGTCTGAACTCTTTCCCTACACGACGCTCTTCCGATCTNNNATTTCTATCCTTCCCCAAGCA
GAGCACACGTCTGAACTCTTTCCCTACACGACGCTCTTCCGATCTNNNATTTCTATCCTTCCCCAAGCA
GAGCACACGTCTGAACTCTTTCCCTACACGACGCTCTTCCGATCTNNNATTTCTATCCTTCCCCAAGCA
AGCACACGTCTGAACTCTTTCCCTACACGACGCTCTTCCGATCTNNNGGAGGTAGAGGGTGCTCTCT
AGCACACGTCTGAACTCTTTCCCTACACGACGCTCTTCCGATCTNNNGGAGGTAGAGGGTGCTCTCT
AGCACACGTCTGAACTCTTTCCCTACACGACGCTCTTCCGATCTNNNGGAGGTAGAGGGTGCTCTCT
AGCACACGTCTGAACTCTTTCCCTACACGACGCTCTTCCGATCTNNNGGAGGTAGAGGGTGCTCTCT
GAGCACACGTCTGAACTCTTTCCCTACACGACGCTCTTCCGATCTNNNTCCTAGGTAGGAGTTTTGG
GAGCACACGTCTGAACTCTTTCCCTACACGACGCTCTTCCGATCTNNNTCCTAGGTAGGAGTTTTGG
GAGCACACGTCTGAACTCTTTCCCTACACGACGCTCTTCCGATCTNNNTCCTAGGTAGGAGTTTTGG
GAGCACACGTCTGAACTCTTTCCCTACACGACGCTCTTCCGATCTNNNTCCTAGGTAGGAGTTTTGG
GAGCACACGTCTGAACTCTTTCCCTACACGACGCTCTTCCGATCTNNNACATGCAAAAGAGAGTGTT
GAGCACACGTCTGAACTCTTTCCCTACACGACGCTCTTCCGATCTNNNACATGCAAAAGAGAGTGTT
GAGCACACGTCTGAACTCTTTCCCTACACGACGCTCTTCCGATCTNNNACATGCAAAAGAGAGTGTT
GAGCACACGTCTGAACTCTTTCCCTACACGACGCTCTTCCGATCTNNNACATGCAAAAGAGAGTGTT
GAGCACACGTCTGAACTCTTTCCCTACACGACGCTCTTCCGATCTNNNACACCTGGCTGAAACTCACTA
GAGCACACGTCTGAACTCTTTCCCTACACGACGCTCTTCCGATCTNNNACACCTGGCTGAAACTCACTA
GAGCACACGTCTGAACTCTTTCCCTACACGACGCTCTTCCGATCTNNNACACCTGGCTGAAACTCACTA
GAGCACACGTCTGAACTCTTTCCCTACACGACGCTCTTCCGATCTNNNACACCTGGCTGAAACTCACTA
AGCACACGTCTGAACTCTTTCCCTACACGACGCTCTTCCGATCTNNNTCCTAACTTCAGTGTTGTCCCC
AGCACACGTCTGAACTCTTTCCCTACACGACGCTCTTCCGATCTNNNTCCTAACTTCAGTGTTGTCCCC
AGCACACGTCTGAACTCTTTCCCTACACGACGCTCTTCCGATCTNNNTCCTAACTTCAGTGTTGTCCCC
AGCACACGTCTGAACTCTTTCCCTACACGACGCTCTTCCGATCTNNNTCCTAACTTCAGTGTTGTCCCC
GCACACGTCTGAACTCTTTCCCTACACGACGCTCTTCCGATCTNNNTGAGGCAGGAGAATAGGTTGA
GCACACGTCTGAACTCTTTCCCTACACGACGCTCTTCCGATCTNNNTGAGGCAGGAGAATAGGTTGA

GCACACGTCTGAACTCTTTCCCTACACGACGCTCTTCCGATCTNNNTGAGGCAGGAGAATAGGTTGA
GCACACGTCTGAACTCTTTCCCTACACGACGCTCTTCCGATCTNNNTGAGGCAGGAGAATAGGTTGA
AGCACACGTCTGAACTCTTTCCCTACACGACGCTCTTCCGATCTNNNTGGAGCAAGACTCTGTCTCAA
AGCACACGTCTGAACTCTTTCCCTACACGACGCTCTTCCGATCTNNNTGGAGCAAGACTCTGTCTCAA
AGCACACGTCTGAACTCTTTCCCTACACGACGCTCTTCCGATCTNNNTGGAGCAAGACTCTGTCTCAA
AGCACACGTCTGAACTCTTTCCCTACACGACGCTCTTCCGATCTNNNTGGAGCAAGACTCTGTCTCAA
AGCACACGTCTGAACTCTTTCCCTACACGACGCTCTTCCGATCTNNNAGCTGACATATAAAGCAAGGC
AGCACACGTCTGAACTCTTTCCCTACACGACGCTCTTCCGATCTNNNAGCTGACATATAAAGCAAGGC
AGCACACGTCTGAACTCTTTCCCTACACGACGCTCTTCCGATCTNNNAGCTGACATATAAAGCAAGGC
AGCACACGTCTGAACTCTTTCCCTACACGACGCTCTTCCGATCTNNNAGCTGACATATAAAGCAAGGC
AGCACACGTCTGAACTCTTTCCCTACACGACGCTCTTCCGATCTNNNCCACCAGGTGACCTATGTTCT
AGCACACGTCTGAACTCTTTCCCTACACGACGCTCTTCCGATCTNNNCCACCAGGTGACCTATGTTCT
AGCACACGTCTGAACTCTTTCCCTACACGACGCTCTTCCGATCTNNNCCACCAGGTGACCTATGTTCT
AGCACACGTCTGAACTCTTTCCCTACACGACGCTCTTCCGATCTNNNCCACCAGGTGACCTATGTTCT
GCACACGTCTGAACTCTTTCCCTACACGACGCTCTTCCGATCTNNNTCTTGGAAGACCTGGAGTTTTG
GCACACGTCTGAACTCTTTCCCTACACGACGCTCTTCCGATCTNNNTCTTGGAAGACCTGGAGTTTTG
GCACACGTCTGAACTCTTTCCCTACACGACGCTCTTCCGATCTNNNTCTTGGAAGACCTGGAGTTTTG
GCACACGTCTGAACTCTTTCCCTACACGACGCTCTTCCGATCTNNNTCTTGGAAGACCTGGAGTTTTG
AGCACACGTCTGAACTCTTTCCCTACACGACGCTCTTCCGATCTNNNTCTGTACTAGACTGCACACCA
AGCACACGTCTGAACTCTTTCCCTACACGACGCTCTTCCGATCTNNNTCTGTACTAGACTGCACACCA
AGCACACGTCTGAACTCTTTCCCTACACGACGCTCTTCCGATCTNNNTCTGTACTAGACTGCACACCA
AGCACACGTCTGAACTCTTTCCCTACACGACGCTCTTCCGATCTNNNTCTGTACTAGACTGCACACCA
GAGCACACGTCTGAACTCTTTCCCTACACGACGCTCTTCCGATCTNNNATACCTGTGGTCCCAGATACT
GAGCACACGTCTGAACTCTTTCCCTACACGACGCTCTTCCGATCTNNNATACCTGTGGTCCCAGATACT
GAGCACACGTCTGAACTCTTTCCCTACACGACGCTCTTCCGATCTNNNATACCTGTGGTCCCAGATACT
GAGCACACGTCTGAACTCTTTCCCTACACGACGCTCTTCCGATCTNNNATACCTGTGGTCCCAGATACT
GAGCACACGTCTGAACTCTTTCCCTACACGACGCTCTTCCGATCTNNNGTTGAGAAACCCTGGGTTAG
GAGCACACGTCTGAACTCTTTCCCTACACGACGCTCTTCCGATCTNNNGTTGAGAAACCCTGGGTTAG
GAGCACACGTCTGAACTCTTTCCCTACACGACGCTCTTCCGATCTNNNGTTGAGAAACCCTGGGTTAG
GAGCACACGTCTGAACTCTTTCCCTACACGACGCTCTTCCGATCTNNNGTTGAGAAACCCTGGGTTAG
AGCACACGTCTGAACTCTTTCCCTACACGACGCTCTTCCGATCTNNNACCTTCCATTGCCCTCTTTACTT
AGCACACGTCTGAACTCTTTCCCTACACGACGCTCTTCCGATCTNNNACCTTCCATTGCCCTCTTTACTT
AGCACACGTCTGAACTCTTTCCCTACACGACGCTCTTCCGATCTNNNACCTTCCATTGCCCTCTTTACTT
AGCACACGTCTGAACTCTTTCCCTACACGACGCTCTTCCGATCTNNNACCTTCCATTGCCCTCTTTACTT
AGCACACGTCTGAACTCTTTCCCTACACGACGCTCTTCCGATCTNNNGATCTCCTGACCTCATGATCCA
AGCACACGTCTGAACTCTTTCCCTACACGACGCTCTTCCGATCTNNNGATCTCCTGACCTCATGATCCA
AGCACACGTCTGAACTCTTTCCCTACACGACGCTCTTCCGATCTNNNGATCTCCTGACCTCATGATCCA
AGCACACGTCTGAACTCTTTCCCTACACGACGCTCTTCCGATCTNNNGATCTCCTGACCTCATGATCCA
GAGCACACGTCTGAACTCTTTCCCTACACGACGCTCTTCCGATCTNNNAGTAAAAATTGTAGTCCCAG
GAGCACACGTCTGAACTCTTTCCCTACACGACGCTCTTCCGATCTNNNAGTAAAAATTGTAGTCCCAG
GAGCACACGTCTGAACTCTTTCCCTACACGACGCTCTTCCGATCTNNNAGTAAAAATTGTAGTCCCAG
GAGCACACGTCTGAACTCTTTCCCTACACGACGCTCTTCCGATCTNNNAGTAAAAATTGTAGTCCCAG
AGCACACGTCTGAACTCTTTCCCTACACGACGCTCTTCCGATCTNNNTTGGTGTGTGTGTATAGTCCC

GAGCACACGTCTGAACTCTTTCCCTACACGACGCTCTTCCGATCTNNNTTGGTGTGTGTGTATAGTCC
GAGCACACGTCTGAACTCTTTCCCTACACGACGCTCTTCCGATCTNNNTTGGTGTGTGTGTATAGTCC
GAGCACACGTCTGAACTCTTTCCCTACACGACGCTCTTCCGATCTNNNTTGGTGTGTGTGTATAGTCC
AGCACACGTCTGAACTCTTTCCCTACACGACGCTCTTCCGATCTNNNGTAATCCCAGCTACTCAGGAAG
AGCACACGTCTGAACTCTTTCCCTACACGACGCTCTTCCGATCTNNNGTAATCCCAGCTACTCAGGAAG
AGCACACGTCTGAACTCTTTCCCTACACGACGCTCTTCCGATCTNNNGTAATCCCAGCTACTCAGGAAG
AGCACACGTCTGAACTCTTTCCCTACACGACGCTCTTCCGATCTNNNGTAATCCCAGCTACTCAGGAAG
AGCACACGTCTGAACTCTTTCCCTACACGACGCTCTTCCGATCTNNNATGTCTACTTACTGGGTTGCCT
AGCACACGTCTGAACTCTTTCCCTACACGACGCTCTTCCGATCTNNNATGTCTACTTACTGGGTTGCCT
AGCACACGTCTGAACTCTTTCCCTACACGACGCTCTTCCGATCTNNNATGTCTACTTACTGGGTTGCCT
AGCACACGTCTGAACTCTTTCCCTACACGACGCTCTTCCGATCTNNNATGTCTACTTACTGGGTTGCCT
GAGCACACGTCTGAACTCTTTCCCTACACGACGCTCTTCCGATCTNNNGAGGCTCTGAACTGGGTTAG
GAGCACACGTCTGAACTCTTTCCCTACACGACGCTCTTCCGATCTNNNGAGGCTCTGAACTGGGTTAG
GAGCACACGTCTGAACTCTTTCCCTACACGACGCTCTTCCGATCTNNNGAGGCTCTGAACTGGGTTAG
GAGCACACGTCTGAACTCTTTCCCTACACGACGCTCTTCCGATCTNNNGAGGCTCTGAACTGGGTTAG
AGCACACGTCTGAACTCTTTCCCTACACGACGCTCTTCCGATCTNNNCTGGACAACATGGTAAAACCC
AGCACACGTCTGAACTCTTTCCCTACACGACGCTCTTCCGATCTNNNCTGGACAACATGGTAAAACCC
AGCACACGTCTGAACTCTTTCCCTACACGACGCTCTTCCGATCTNNNCTGGACAACATGGTAAAACCC
AGCACACGTCTGAACTCTTTCCCTACACGACGCTCTTCCGATCTNNNCTGGACAACATGGTAAAACCC
GCACACGTCTGAACTCTTTCCCTACACGACGCTCTTCCGATCTNNNGAAAGGAGCCATGATGTGAGA
GCACACGTCTGAACTCTTTCCCTACACGACGCTCTTCCGATCTNNNGAAAGGAGCCATGATGTGAGA
GCACACGTCTGAACTCTTTCCCTACACGACGCTCTTCCGATCTNNNGAAAGGAGCCATGATGTGAGA
GCACACGTCTGAACTCTTTCCCTACACGACGCTCTTCCGATCTNNNGAAAGGAGCCATGATGTGAGA
GCACACGTCTGAACTCTTTCCCTACACGACGCTCTTCCGATCTNNNCAGAAGTCCAAAACGGGAAGTT
GCACACGTCTGAACTCTTTCCCTACACGACGCTCTTCCGATCTNNNCAGAAGTCCAAAACGGGAAGTT
GCACACGTCTGAACTCTTTCCCTACACGACGCTCTTCCGATCTNNNCAGAAGTCCAAAACGGGAAGTT
GCACACGTCTGAACTCTTTCCCTACACGACGCTCTTCCGATCTNNNCAGAAGTCCAAAACGGGAAGTT
AGCACACGTCTGAACTCTTTCCCTACACGACGCTCTTCCGATCTNNNGAGAATCGCTTGAACTCAGGA
AGCACACGTCTGAACTCTTTCCCTACACGACGCTCTTCCGATCTNNNGAGAATCGCTTGAACTCAGGA
AGCACACGTCTGAACTCTTTCCCTACACGACGCTCTTCCGATCTNNNGAGAATCGCTTGAACTCAGGA
AGCACACGTCTGAACTCTTTCCCTACACGACGCTCTTCCGATCTNNNGAGAATCGCTTGAACTCAGGA
GAGCACACGTCTGAACTCTTTCCCTACACGACGCTCTTCCGATCTNNNTTCTGATTGCTGTCCCTCACT
GAGCACACGTCTGAACTCTTTCCCTACACGACGCTCTTCCGATCTNNNTTCTGATTGCTGTCCCTCACT
GAGCACACGTCTGAACTCTTTCCCTACACGACGCTCTTCCGATCTNNNTTCTGATTGCTGTCCCTCACT
GAGCACACGTCTGAACTCTTTCCCTACACGACGCTCTTCCGATCTNNNTTCTGATTGCTGTCCCTCACT
CACACGTCTGAACTCTTTCCCTACACGACGCTCTTCCGATCTNNNCTGATAGATTCCAGAGAGATTGC
CACACGTCTGAACTCTTTCCCTACACGACGCTCTTCCGATCTNNNCTGATAGATTCCAGAGAGATTGC
CACACGTCTGAACTCTTTCCCTACACGACGCTCTTCCGATCTNNNCTGATAGATTCCAGAGAGATTGC
CACACGTCTGAACTCTTTCCCTACACGACGCTCTTCCGATCTNNNCTGATAGATTCCAGAGAGATTGC
CACACGTCTGAACTCTTTCCCTACACGACGCTCTTCCGATCTNNNACTCTTGTCTTCATTCTGCCACAC
CACACGTCTGAACTCTTTCCCTACACGACGCTCTTCCGATCTNNNACTCTTGTCTTCATTCTGCCACAC
CACACGTCTGAACTCTTTCCCTACACGACGCTCTTCCGATCTNNNACTCTTGTCTTCATTCTGCCACAC
CACACGTCTGAACTCTTTCCCTACACGACGCTCTTCCGATCTNNNACTCTTGTCTTCATTCTGCCACAC

AGCACACGTCTGAACTCTTTCCCTACACGACGCTCTTCCGATCTNNNCCTATGATTTCATCTCTGCCCCA
AGCACACGTCTGAACTCTTTCCCTACACGACGCTCTTCCGATCTNNNCCTATGATTTCATCTCTGCCCCA
AGCACACGTCTGAACTCTTTCCCTACACGACGCTCTTCCGATCTNNNCCTATGATTTCATCTCTGCCCCA
AGCACACGTCTGAACTCTTTCCCTACACGACGCTCTTCCGATCTNNNCCTATGATTTCATCTCTGCCCCA
AGCACACGTCTGAACTCTTTCCCTACACGACGCTCTTCCGATCTNNNCTGTAGCATGCTATGATTGTG
AGCACACGTCTGAACTCTTTCCCTACACGACGCTCTTCCGATCTNNNCTGTAGCATGCTATGATTGTG
AGCACACGTCTGAACTCTTTCCCTACACGACGCTCTTCCGATCTNNNCTGTAGCATGCTATGATTGTG
AGCACACGTCTGAACTCTTTCCCTACACGACGCTCTTCCGATCTNNNCTGTAGCATGCTATGATTGTG
AGCACACGTCTGAACTCTTTCCCTACACGACGCTCTTCCGATCTNNNATGTTGCTGCCTGATCATTCTTG
AGCACACGTCTGAACTCTTTCCCTACACGACGCTCTTCCGATCTNNNATGTTGCTGCCTGATCATTCTTG
AGCACACGTCTGAACTCTTTCCCTACACGACGCTCTTCCGATCTNNNATGTTGCTGCCTGATCATTCTTG
AGCACACGTCTGAACTCTTTCCCTACACGACGCTCTTCCGATCTNNNATGTTGCTGCCTGATCATTCTTG
GAGCACACGTCTGAACTCTTTCCCTACACGACGCTCTTCCGATCTNNNAAAGCCCAAGCATATCACCATT
GAGCACACGTCTGAACTCTTTCCCTACACGACGCTCTTCCGATCTNNNAAAGCCCAAGCATATCACCATT
GAGCACACGTCTGAACTCTTTCCCTACACGACGCTCTTCCGATCTNNNAAAGCCCAAGCATATCACCATT
GAGCACACGTCTGAACTCTTTCCCTACACGACGCTCTTCCGATCTNNNAAAGCCCAAGCATATCACCATT
AGCACACGTCTGAACTCTTTCCCTACACGACGCTCTTCCGATCTNNNAAAATGCTGGGATTACAGGCA
AGCACACGTCTGAACTCTTTCCCTACACGACGCTCTTCCGATCTNNNAAAATGCTGGGATTACAGGCA
AGCACACGTCTGAACTCTTTCCCTACACGACGCTCTTCCGATCTNNNAAAATGCTGGGATTACAGGCA
AGCACACGTCTGAACTCTTTCCCTACACGACGCTCTTCCGATCTNNNAAAATGCTGGGATTACAGGCA
GAGCACACGTCTGAACTCTTTCCCTACACGACGCTCTTCCGATCTNNNGGAAAGGGCTTTTGTTGGGA
GAGCACACGTCTGAACTCTTTCCCTACACGACGCTCTTCCGATCTNNNGGAAAGGGCTTTTGTTGGGA
GAGCACACGTCTGAACTCTTTCCCTACACGACGCTCTTCCGATCTNNNGGAAAGGGCTTTTGTTGGGA
GAGCACACGTCTGAACTCTTTCCCTACACGACGCTCTTCCGATCTNNNGGAAAGGGCTTTTGTTGGGA
GCACACGTCTGAACTCTTTCCCTACACGACGCTCTTCCGATCTNNNACTTTGGTGGCATTTGAGAAGC
GCACACGTCTGAACTCTTTCCCTACACGACGCTCTTCCGATCTNNNACTTTGGTGGCATTTGAGAAGC
GCACACGTCTGAACTCTTTCCCTACACGACGCTCTTCCGATCTNNNACTTTGGTGGCATTTGAGAAGC
GCACACGTCTGAACTCTTTCCCTACACGACGCTCTTCCGATCTNNNACTTTGGTGGCATTTGAGAAGC
AGCACACGTCTGAACTCTTTCCCTACACGACGCTCTTCCGATCTNNNTGTAGGCCTTGGAATTAGAAG
AGCACACGTCTGAACTCTTTCCCTACACGACGCTCTTCCGATCTNNNTGTAGGCCTTGGAATTAGAAG
AGCACACGTCTGAACTCTTTCCCTACACGACGCTCTTCCGATCTNNNTGTAGGCCTTGGAATTAGAAG
AGCACACGTCTGAACTCTTTCCCTACACGACGCTCTTCCGATCTNNNTGTAGGCCTTGGAATTAGAAG
AGCACACGTCTGAACTCTTTCCCTACACGACGCTCTTCCGATCTNNNAAGGTCGTTTTCACCTGTATGTC
AGCACACGTCTGAACTCTTTCCCTACACGACGCTCTTCCGATCTNNNAAGGTCGTTTTCACCTGTATGTC
AGCACACGTCTGAACTCTTTCCCTACACGACGCTCTTCCGATCTNNNAAGGTCGTTTTCACCTGTATGTC
AGCACACGTCTGAACTCTTTCCCTACACGACGCTCTTCCGATCTNNNAAGGTCGTTTTCACCTGTATGTC
AGCACACGTCTGAACTCTTTCCCTACACGACGCTCTTCCGATCTNNNAGACCAAAGGAACAGAGTAG
AGCACACGTCTGAACTCTTTCCCTACACGACGCTCTTCCGATCTNNNAGACCAAAGGAACAGAGTAG
AGCACACGTCTGAACTCTTTCCCTACACGACGCTCTTCCGATCTNNNAGACCAAAGGAACAGAGTAG
AGCACACGTCTGAACTCTTTCCCTACACGACGCTCTTCCGATCTNNNAGACCAAAGGAACAGAGTAG
CACACGTCTGAACTCTTTCCCTACACGACGCTCTTCCGATCTNNNTCATCCCCTTTCACACACAAAGACA
CACACGTCTGAACTCTTTCCCTACACGACGCTCTTCCGATCTNNNTCATCCCCTTTCACACACAAAGACA
CACACGTCTGAACTCTTTCCCTACACGACGCTCTTCCGATCTNNNTCATCCCCTTTCACACACAAAGACA

CACACGTCTGAACTCTTTCCCTACACGACGCTCTTCCGATCTNNNTCATCCCCTTTCACACACAAAGACAT
AGCACACGTCTGAACTCTTTCCCTACACGACGCTCTTCCGATCTNNNAAAAGTAGATAGGCATGGTGG
AGCACACGTCTGAACTCTTTCCCTACACGACGCTCTTCCGATCTNNNAAAAGTAGATAGGCATGGTGG
AGCACACGTCTGAACTCTTTCCCTACACGACGCTCTTCCGATCTNNNAAAAGTAGATAGGCATGGTGG
AGCACACGTCTGAACTCTTTCCCTACACGACGCTCTTCCGATCTNNNAAAAGTAGATAGGCATGGTGG
AGCACACGTCTGAACTCTTTCCCTACACGACGCTCTTCCGATCTNNNATGAGCTGATATCATGCCATTG
AGCACACGTCTGAACTCTTTCCCTACACGACGCTCTTCCGATCTNNNATGAGCTGATATCATGCCATTG
AGCACACGTCTGAACTCTTTCCCTACACGACGCTCTTCCGATCTNNNATGAGCTGATATCATGCCATTG
AGCACACGTCTGAACTCTTTCCCTACACGACGCTCTTCCGATCTNNNATGAGCTGATATCATGCCATTG
AGCACACGTCTGAACTCTTTCCCTACACGACGCTCTTCCGATCTNNNTCTAAGCTTTTTTTCGGCAGTGG
AGCACACGTCTGAACTCTTTCCCTACACGACGCTCTTCCGATCTNNNTCTAAGCTTTTTTTCGGCAGTGG
AGCACACGTCTGAACTCTTTCCCTACACGACGCTCTTCCGATCTNNNTCTAAGCTTTTTTTCGGCAGTGG
AGCACACGTCTGAACTCTTTCCCTACACGACGCTCTTCCGATCTNNNTCTAAGCTTTTTTTCGGCAGTGG
AGCACACGTCTGAACTCTTTCCCTACACGACGCTCTTCCGATCTNNNATCTGGGATTACATGCACCTAC
AGCACACGTCTGAACTCTTTCCCTACACGACGCTCTTCCGATCTNNNATCTGGGATTACATGCACCTAC
AGCACACGTCTGAACTCTTTCCCTACACGACGCTCTTCCGATCTNNNATCTGGGATTACATGCACCTAC
AGCACACGTCTGAACTCTTTCCCTACACGACGCTCTTCCGATCTNNNATCTGGGATTACATGCACCTAC
AGCACACGTCTGAACTCTTTCCCTACACGACGCTCTTCCGATCTNNNTAGATAGAACACTGGAACACAG
AGCACACGTCTGAACTCTTTCCCTACACGACGCTCTTCCGATCTNNNTAGATAGAACACTGGAACACAG
AGCACACGTCTGAACTCTTTCCCTACACGACGCTCTTCCGATCTNNNTAGATAGAACACTGGAACACAG
AGCACACGTCTGAACTCTTTCCCTACACGACGCTCTTCCGATCTNNNTAGATAGAACACTGGAACACAG
GCACACGTCTGAACTCTTTCCCTACACGACGCTCTTCCGATCTNNNCTGTAGATGTCACCTCCTAAGCCC
GCACACGTCTGAACTCTTTCCCTACACGACGCTCTTCCGATCTNNNCTGTAGATGTCACCTCCTAAGCCC
GCACACGTCTGAACTCTTTCCCTACACGACGCTCTTCCGATCTNNNCTGTAGATGTCACCTCCTAAGCCC
GCACACGTCTGAACTCTTTCCCTACACGACGCTCTTCCGATCTNNNCTGTAGATGTCACCTCCTAAGCCC
GAGCACACGTCTGAACTCTTTCCCTACACGACGCTCTTCCGATCTNNNCTGTAGTCCCAGCTACTCAGGA
GAGCACACGTCTGAACTCTTTCCCTACACGACGCTCTTCCGATCTNNNCTGTAGTCCCAGCTACTCAGGA
GAGCACACGTCTGAACTCTTTCCCTACACGACGCTCTTCCGATCTNNNCTGTAGTCCCAGCTACTCAGGA
GAGCACACGTCTGAACTCTTTCCCTACACGACGCTCTTCCGATCTNNNCTGTAGTCCCAGCTACTCAGGA
GAGCACACGTCTGAACTCTTTCCCTACACGACGCTCTTCCGATCTNNNTTCGTGATAGTGGTTCGTTGC
GAGCACACGTCTGAACTCTTTCCCTACACGACGCTCTTCCGATCTNNNTTCGTGATAGTGGTTCGTTGC
GAGCACACGTCTGAACTCTTTCCCTACACGACGCTCTTCCGATCTNNNTTCGTGATAGTGGTTCGTTGC
GAGCACACGTCTGAACTCTTTCCCTACACGACGCTCTTCCGATCTNNNTTCGTGATAGTGGTTCGTTGC
GAGCACACGTCTGAACTCTTTCCCTACACGACGCTCTTCCGATCTNNNGGAGCCTGTGGAGATGCTATA
GAGCACACGTCTGAACTCTTTCCCTACACGACGCTCTTCCGATCTNNNGGAGCCTGTGGAGATGCTATA
GAGCACACGTCTGAACTCTTTCCCTACACGACGCTCTTCCGATCTNNNGGAGCCTGTGGAGATGCTATA
GAGCACACGTCTGAACTCTTTCCCTACACGACGCTCTTCCGATCTNNNGGAGCCTGTGGAGATGCTATA
AGAGCACACGTCTGAACTCTTTCCCTACACGACGCTCTTCCGATCTNNNATGCTACCTTTTTAGTGCATG
AGAGCACACGTCTGAACTCTTTCCCTACACGACGCTCTTCCGATCTNNNATGCTACCTTTTTAGTGCATG
AGAGCACACGTCTGAACTCTTTCCCTACACGACGCTCTTCCGATCTNNNATGCTACCTTTTTAGTGCATG
AGAGCACACGTCTGAACTCTTTCCCTACACGACGCTCTTCCGATCTNNNATGCTACCTTTTTAGTGCATG
AGAGCACACGTCTGAACTCTTTCCCTACACGACGCTCTTCCGATCTNNNCTCTCCTCGATTTGCACGGTC
AGAGCACACGTCTGAACTCTTTCCCTACACGACGCTCTTCCGATCTNNNCTCTCCTCGATTTGCACGGTC

GAGCACACGTCTGAACTCTTTCCCTACACGACGCTCTTCCGATCTNNNCTCTCCTCGATTTGCACGGTC
GAGCACACGTCTGAACTCTTTCCCTACACGACGCTCTTCCGATCTNNNCTCTCCTCGATTTGCACGGTC
GCACACGTCTGAACTCTTTCCCTACACGACGCTCTTCCGATCTNNNTGCTTGCTTGAGCTTTCTACCTA
GCACACGTCTGAACTCTTTCCCTACACGACGCTCTTCCGATCTNNNTGCTTGCTTGAGCTTTCTACCTA
GCACACGTCTGAACTCTTTCCCTACACGACGCTCTTCCGATCTNNNTGCTTGCTTGAGCTTTCTACCTA
GCACACGTCTGAACTCTTTCCCTACACGACGCTCTTCCGATCTNNNTGCTTGCTTGAGCTTTCTACCTA
AGCACACGTCTGAACTCTTTCCCTACACGACGCTCTTCCGATCTNNNAGAGGTATGTGTATGTGTGCC
AGCACACGTCTGAACTCTTTCCCTACACGACGCTCTTCCGATCTNNNAGAGGTATGTGTATGTGTGCC
AGCACACGTCTGAACTCTTTCCCTACACGACGCTCTTCCGATCTNNNAGAGGTATGTGTATGTGTGCC
AGCACACGTCTGAACTCTTTCCCTACACGACGCTCTTCCGATCTNNNAGAGGTATGTGTATGTGTGCC
GAGCACACGTCTGAACTCTTTCCCTACACGACGCTCTTCCGATCTNNNGCTGAGGAAGGTGGATAACT
GAGCACACGTCTGAACTCTTTCCCTACACGACGCTCTTCCGATCTNNNGCTGAGGAAGGTGGATAACT
GAGCACACGTCTGAACTCTTTCCCTACACGACGCTCTTCCGATCTNNNGCTGAGGAAGGTGGATAACT
GAGCACACGTCTGAACTCTTTCCCTACACGACGCTCTTCCGATCTNNNGCTGAGGAAGGTGGATAACT
GCACACGTCTGAACTCTTTCCCTACACGACGCTCTTCCGATCTNNNAAAAATTAGCCGGGTGTGTTG
GCACACGTCTGAACTCTTTCCCTACACGACGCTCTTCCGATCTNNNAAAAATTAGCCGGGTGTGTTG
GCACACGTCTGAACTCTTTCCCTACACGACGCTCTTCCGATCTNNNAAAAATTAGCCGGGTGTGTTG
GCACACGTCTGAACTCTTTCCCTACACGACGCTCTTCCGATCTNNNAAAAATTAGCCGGGTGTGTTG
AGCACACGTCTGAACTCTTTCCCTACACGACGCTCTTCCGATCTNNNCACAATGGAATATCATTCAACT
AGCACACGTCTGAACTCTTTCCCTACACGACGCTCTTCCGATCTNNNCACAATGGAATATCATTCAACT
AGCACACGTCTGAACTCTTTCCCTACACGACGCTCTTCCGATCTNNNCACAATGGAATATCATTCAACT
AGCACACGTCTGAACTCTTTCCCTACACGACGCTCTTCCGATCTNNNCACAATGGAATATCATTCAACT
GCACACGTCTGAACTCTTTCCCTACACGACGCTCTTCCGATCTNNNTCTCTAACAGTGCCTTACATGCAC
GCACACGTCTGAACTCTTTCCCTACACGACGCTCTTCCGATCTNNNTCTCTAACAGTGCCTTACATGCAC
GCACACGTCTGAACTCTTTCCCTACACGACGCTCTTCCGATCTNNNTCTCTAACAGTGCCTTACATGCAC
GCACACGTCTGAACTCTTTCCCTACACGACGCTCTTCCGATCTNNNTCTCTAACAGTGCCTTACATGCAC
AGCACACGTCTGAACTCTTTCCCTACACGACGCTCTTCCGATCTNNNGAGAAAGCAAAGGAATTCTTG
AGCACACGTCTGAACTCTTTCCCTACACGACGCTCTTCCGATCTNNNGAGAAAGCAAAGGAATTCTTG
AGCACACGTCTGAACTCTTTCCCTACACGACGCTCTTCCGATCTNNNGAGAAAGCAAAGGAATTCTTG
AGCACACGTCTGAACTCTTTCCCTACACGACGCTCTTCCGATCTNNNGAGAAAGCAAAGGAATTCTTG
GCACACGTCTGAACTCTTTCCCTACACGACGCTCTTCCGATCTNNNACCTGATTCTTTACTCCCAACCCC
GCACACGTCTGAACTCTTTCCCTACACGACGCTCTTCCGATCTNNNACCTGATTCTTTACTCCCAACCCC
GCACACGTCTGAACTCTTTCCCTACACGACGCTCTTCCGATCTNNNACCTGATTCTTTACTCCCAACCCC
GCACACGTCTGAACTCTTTCCCTACACGACGCTCTTCCGATCTNNNACCTGATTCTTTACTCCCAACCCC
GAGCACACGTCTGAACTCTTTCCCTACACGACGCTCTTCCGATCTNNNTTAAGGGAGCAAAGAAGGC
GAGCACACGTCTGAACTCTTTCCCTACACGACGCTCTTCCGATCTNNNTTAAGGGAGCAAAGAAGGC
GAGCACACGTCTGAACTCTTTCCCTACACGACGCTCTTCCGATCTNNNTTAAGGGAGCAAAGAAGGC
GAGCACACGTCTGAACTCTTTCCCTACACGACGCTCTTCCGATCTNNNTTAAGGGAGCAAAGAAGGC
CACACGTCTGAACTCTTTCCCTACACGACGCTCTTCCGATCTNNNACGTGGATTGCCTTTACTTTGGTC
CACACGTCTGAACTCTTTCCCTACACGACGCTCTTCCGATCTNNNACGTGGATTGCCTTTACTTTGGTC
CACACGTCTGAACTCTTTCCCTACACGACGCTCTTCCGATCTNNNACGTGGATTGCCTTTACTTTGGTC
CACACGTCTGAACTCTTTCCCTACACGACGCTCTTCCGATCTNNNACGTGGATTGCCTTTACTTTGGTC
AGCACACGTCTGAACTCTTTCCCTACACGACGCTCTTCCGATCTNNNCTAAGGCAGGAGAATCGCTTG

AGCACACGTCTGAACTCTTTCCCTACACGACGCTCTTCCGATCTNNNCTAAGGCAGGAGAATCGCTTG
AGCACACGTCTGAACTCTTTCCCTACACGACGCTCTTCCGATCTNNNCTAAGGCAGGAGAATCGCTTG
AGCACACGTCTGAACTCTTTCCCTACACGACGCTCTTCCGATCTNNNCTAAGGCAGGAGAATCGCTTG
GAGCACACGTCTGAACTCTTTCCCTACACGACGCTCTTCCGATCTNNNCTGTAGTCCCAGCTACTCAGGA
GAGCACACGTCTGAACTCTTTCCCTACACGACGCTCTTCCGATCTNNNCTGTAGTCCCAGCTACTCAGGA
GAGCACACGTCTGAACTCTTTCCCTACACGACGCTCTTCCGATCTNNNCTGTAGTCCCAGCTACTCAGGA
GAGCACACGTCTGAACTCTTTCCCTACACGACGCTCTTCCGATCTNNNCTGTAGTCCCAGCTACTCAGGA
AGCACACGTCTGAACTCTTTCCCTACACGACGCTCTTCCGATCTNNNTGGACAATAGGCATTATTCTAG
AGCACACGTCTGAACTCTTTCCCTACACGACGCTCTTCCGATCTNNNTGGACAATAGGCATTATTCTAG
AGCACACGTCTGAACTCTTTCCCTACACGACGCTCTTCCGATCTNNNTGGACAATAGGCATTATTCTAG
AGCACACGTCTGAACTCTTTCCCTACACGACGCTCTTCCGATCTNNNTGGACAATAGGCATTATTCTAG
GCACACGTCTGAACTCTTTCCCTACACGACGCTCTTCCGATCTNNNAGCAAAGAAGGCCAAGAGAGTT
GCACACGTCTGAACTCTTTCCCTACACGACGCTCTTCCGATCTNNNAGCAAAGAAGGCCAAGAGAGTT
GCACACGTCTGAACTCTTTCCCTACACGACGCTCTTCCGATCTNNNAGCAAAGAAGGCCAAGAGAGTT
GCACACGTCTGAACTCTTTCCCTACACGACGCTCTTCCGATCTNNNAGCAAAGAAGGCCAAGAGAGTT
AGAGCACACGTCTGAACTCTTTCCCTACACGACGCTCTTCCGATCTNNNTCCCAGGATATGCTTAGGAG
AGAGCACACGTCTGAACTCTTTCCCTACACGACGCTCTTCCGATCTNNNTCCCAGGATATGCTTAGGAG
AGAGCACACGTCTGAACTCTTTCCCTACACGACGCTCTTCCGATCTNNNTCCCAGGATATGCTTAGGAG
AGAGCACACGTCTGAACTCTTTCCCTACACGACGCTCTTCCGATCTNNNTCCCAGGATATGCTTAGGAG
GAGCACACGTCTGAACTCTTTCCCTACACGACGCTCTTCCGATCTNNNCTGAGGCAGGAGAATCACTT
GAGCACACGTCTGAACTCTTTCCCTACACGACGCTCTTCCGATCTNNNCTGAGGCAGGAGAATCACTT
GAGCACACGTCTGAACTCTTTCCCTACACGACGCTCTTCCGATCTNNNCTGAGGCAGGAGAATCACTT
GAGCACACGTCTGAACTCTTTCCCTACACGACGCTCTTCCGATCTNNNCTGAGGCAGGAGAATCACTT
GAGCACACGTCTGAACTCTTTCCCTACACGACGCTCTTCCGATCTNNNCGTAATCCCGGAACATCTTTC
GAGCACACGTCTGAACTCTTTCCCTACACGACGCTCTTCCGATCTNNNCGTAATCCCGGAACATCTTTC
GAGCACACGTCTGAACTCTTTCCCTACACGACGCTCTTCCGATCTNNNCGTAATCCCGGAACATCTTTC
GAGCACACGTCTGAACTCTTTCCCTACACGACGCTCTTCCGATCTNNNCGTAATCCCGGAACATCTTTC
GAGCACACGTCTGAACTCTTTCCCTACACGACGCTCTTCCGATCTNNNTCCTCAAAACTTAGGATACCC
GAGCACACGTCTGAACTCTTTCCCTACACGACGCTCTTCCGATCTNNNTCCTCAAAACTTAGGATACCC
GAGCACACGTCTGAACTCTTTCCCTACACGACGCTCTTCCGATCTNNNTCCTCAAAACTTAGGATACCC
GAGCACACGTCTGAACTCTTTCCCTACACGACGCTCTTCCGATCTNNNTCCTCAAAACTTAGGATACCC
AGCACACGTCTGAACTCTTTCCCTACACGACGCTCTTCCGATCTNNNAGATGATTATAGGCGTGCATT
AGCACACGTCTGAACTCTTTCCCTACACGACGCTCTTCCGATCTNNNAGATGATTATAGGCGTGCATT
AGCACACGTCTGAACTCTTTCCCTACACGACGCTCTTCCGATCTNNNAGATGATTATAGGCGTGCATT
AGCACACGTCTGAACTCTTTCCCTACACGACGCTCTTCCGATCTNNNAGATGATTATAGGCGTGCATT
AGCACACGTCTGAACTCTTTCCCTACACGACGCTCTTCCGATCTNNNACAAAAAACAGATTCTGCTACCA
AGCACACGTCTGAACTCTTTCCCTACACGACGCTCTTCCGATCTNNNACAAAAAACAGATTCTGCTACCA
AGCACACGTCTGAACTCTTTCCCTACACGACGCTCTTCCGATCTNNNACAAAAAACAGATTCTGCTACCA
AGCACACGTCTGAACTCTTTCCCTACACGACGCTCTTCCGATCTNNNACAAAAAACAGATTCTGCTACCA
AGCACACGTCTGAACTCTTTCCCTACACGACGCTCTTCCGATCTNNNCCTTCCTTTCTCTCTCCTTCCACCA
AGCACACGTCTGAACTCTTTCCCTACACGACGCTCTTCCGATCTNNNCCTTCCTTTCTCTCTCCTTCCACCA
AGCACACGTCTGAACTCTTTCCCTACACGACGCTCTTCCGATCTNNNCCTTCCTTTCTCTCTCCTTCCACCA
AGCACACGTCTGAACTCTTTCCCTACACGACGCTCTTCCGATCTNNNCCTTCCTTTCTCTCTCCTTCCACCA

AGCACACGTCTGAACTCTTTCCCTACACGACGCTCTTCCGATCTNNNGTTTAGGCGTGGTCTTGAAAAG
AGCACACGTCTGAACTCTTTCCCTACACGACGCTCTTCCGATCTNNNGTTTAGGCGTGGTCTTGAAAAG
AGCACACGTCTGAACTCTTTCCCTACACGACGCTCTTCCGATCTNNNGTTTAGGCGTGGTCTTGAAAAG
AGCACACGTCTGAACTCTTTCCCTACACGACGCTCTTCCGATCTNNNGTTTAGGCGTGGTCTTGAAAAG
GCACACGTCTGAACTCTTTCCCTACACGACGCTCTTCCGATCTNNNTTGGGTTTGGTTTGCTCTTGTTTC
GCACACGTCTGAACTCTTTCCCTACACGACGCTCTTCCGATCTNNNTTGGGTTTGGTTTGCTCTTGTTTC
GCACACGTCTGAACTCTTTCCCTACACGACGCTCTTCCGATCTNNNTTGGGTTTGGTTTGCTCTTGTTTC
GCACACGTCTGAACTCTTTCCCTACACGACGCTCTTCCGATCTNNNTTGGGTTTGGTTTGCTCTTGTTTC
GAGCACACGTCTGAACTCTTTCCCTACACGACGCTCTTCCGATCTNNNGAGATCACACCACTGCACTCTA
GAGCACACGTCTGAACTCTTTCCCTACACGACGCTCTTCCGATCTNNNGAGATCACACCACTGCACTCTA
GAGCACACGTCTGAACTCTTTCCCTACACGACGCTCTTCCGATCTNNNGAGATCACACCACTGCACTCTA
GAGCACACGTCTGAACTCTTTCCCTACACGACGCTCTTCCGATCTNNNGAGATCACACCACTGCACTCTA
AGCACACGTCTGAACTCTTTCCCTACACGACGCTCTTCCGATCTNNNATCCCCTGGGGTTCTTTTCTAAC
AGCACACGTCTGAACTCTTTCCCTACACGACGCTCTTCCGATCTNNNATCCCCTGGGGTTCTTTTCTAAC
AGCACACGTCTGAACTCTTTCCCTACACGACGCTCTTCCGATCTNNNATCCCCTGGGGTTCTTTTCTAAC
AGCACACGTCTGAACTCTTTCCCTACACGACGCTCTTCCGATCTNNNATCCCCTGGGGTTCTTTTCTAAC
AGCACACGTCTGAACTCTTTCCCTACACGACGCTCTTCCGATCTNNNAACTGGTTCCTCTTTGTCCTAGG
AGCACACGTCTGAACTCTTTCCCTACACGACGCTCTTCCGATCTNNNAACTGGTTCCTCTTTGTCCTAGG
AGCACACGTCTGAACTCTTTCCCTACACGACGCTCTTCCGATCTNNNAACTGGTTCCTCTTTGTCCTAGG
AGCACACGTCTGAACTCTTTCCCTACACGACGCTCTTCCGATCTNNNAACTGGTTCCTCTTTGTCCTAGG
AGCACACGTCTGAACTCTTTCCCTACACGACGCTCTTCCGATCTNNNACCTGTAATCTCAGCTACTTGG
AGCACACGTCTGAACTCTTTCCCTACACGACGCTCTTCCGATCTNNNACCTGTAATCTCAGCTACTTGG
AGCACACGTCTGAACTCTTTCCCTACACGACGCTCTTCCGATCTNNNACCTGTAATCTCAGCTACTTGG
AGCACACGTCTGAACTCTTTCCCTACACGACGCTCTTCCGATCTNNNACCTGTAATCTCAGCTACTTGG
AGCACACGTCTGAACTCTTTCCCTACACGACGCTCTTCCGATCTNNNGGAGCTAAGCTATGAGGACACA
AGCACACGTCTGAACTCTTTCCCTACACGACGCTCTTCCGATCTNNNGGAGCTAAGCTATGAGGACACA
AGCACACGTCTGAACTCTTTCCCTACACGACGCTCTTCCGATCTNNNGGAGCTAAGCTATGAGGACACA
AGCACACGTCTGAACTCTTTCCCTACACGACGCTCTTCCGATCTNNNGGAGCTAAGCTATGAGGACACA
GCACACGTCTGAACTCTTTCCCTACACGACGCTCTTCCGATCTNNNATCCTGGCTAACACGGTGAAACCA
GCACACGTCTGAACTCTTTCCCTACACGACGCTCTTCCGATCTNNNATCCTGGCTAACACGGTGAAACCA
GCACACGTCTGAACTCTTTCCCTACACGACGCTCTTCCGATCTNNNATCCTGGCTAACACGGTGAAACCA
GCACACGTCTGAACTCTTTCCCTACACGACGCTCTTCCGATCTNNNATCCTGGCTAACACGGTGAAACCA
GAGCACACGTCTGAACTCTTTCCCTACACGACGCTCTTCCGATCTNNNGTGTCTGTAGTCCCAGCTACTT
GAGCACACGTCTGAACTCTTTCCCTACACGACGCTCTTCCGATCTNNNGTGTCTGTAGTCCCAGCTACTT
GAGCACACGTCTGAACTCTTTCCCTACACGACGCTCTTCCGATCTNNNGTGTCTGTAGTCCCAGCTACTT
GAGCACACGTCTGAACTCTTTCCCTACACGACGCTCTTCCGATCTNNNGTGTCTGTAGTCCCAGCTACTT
AGCACACGTCTGAACTCTTTCCCTACACGACGCTCTTCCGATCTNNNTTAGGGATGGTGGATAGGAGC
AGCACACGTCTGAACTCTTTCCCTACACGACGCTCTTCCGATCTNNNTTAGGGATGGTGGATAGGAGC
AGCACACGTCTGAACTCTTTCCCTACACGACGCTCTTCCGATCTNNNTTAGGGATGGTGGATAGGAGC
AGCACACGTCTGAACTCTTTCCCTACACGACGCTCTTCCGATCTNNNTTAGGGATGGTGGATAGGAGC
AGCACACGTCTGAACTCTTTCCCTACACGACGCTCTTCCGATCTNNNAGGAGAATTGTTTGAACCCAGC
AGCACACGTCTGAACTCTTTCCCTACACGACGCTCTTCCGATCTNNNAGGAGAATTGTTTGAACCCAGC
AGCACACGTCTGAACTCTTTCCCTACACGACGCTCTTCCGATCTNNNAGGAGAATTGTTTGAACCCAGC

AGCACACGTCTGAACTCTTTCCCTACACGACGCTCTTCCGATCTNNNAGGAGAATTGTTTGAACCCAG
AGCACACGTCTGAACTCTTTCCCTACACGACGCTCTTCCGATCTNNNTGTAATCCAGCACTTTGGTAG
AGCACACGTCTGAACTCTTTCCCTACACGACGCTCTTCCGATCTNNNTGTAATCCAGCACTTTGGTAG
AGCACACGTCTGAACTCTTTCCCTACACGACGCTCTTCCGATCTNNNTGTAATCCAGCACTTTGGTAG
AGCACACGTCTGAACTCTTTCCCTACACGACGCTCTTCCGATCTNNNTGTAATCCAGCACTTTGGTAG
GAGCACACGTCTGAACTCTTTCCCTACACGACGCTCTTCCGATCTNNNGTCTCTCCATAAGGGCATCTA
GAGCACACGTCTGAACTCTTTCCCTACACGACGCTCTTCCGATCTNNNGTCTCTCCATAAGGGCATCTA
GAGCACACGTCTGAACTCTTTCCCTACACGACGCTCTTCCGATCTNNNGTCTCTCCATAAGGGCATCTA
GAGCACACGTCTGAACTCTTTCCCTACACGACGCTCTTCCGATCTNNNGTCTCTCCATAAGGGCATCTA
GAGCACACGTCTGAACTCTTTCCCTACACGACGCTCTTCCGATCTNNNCTGAGGCAAGAGAACAGTAG
GAGCACACGTCTGAACTCTTTCCCTACACGACGCTCTTCCGATCTNNNCTGAGGCAAGAGAACAGTAG
GAGCACACGTCTGAACTCTTTCCCTACACGACGCTCTTCCGATCTNNNCTGAGGCAAGAGAACAGTAG
GAGCACACGTCTGAACTCTTTCCCTACACGACGCTCTTCCGATCTNNNCTGAGGCAAGAGAACAGTAG
GCACACGTCTGAACTCTTTCCCTACACGACGCTCTTCCGATCTNNNTTGAAGGCTTGATATGGTTTGG
GCACACGTCTGAACTCTTTCCCTACACGACGCTCTTCCGATCTNNNTTGAAGGCTTGATATGGTTTGG
GCACACGTCTGAACTCTTTCCCTACACGACGCTCTTCCGATCTNNNTTGAAGGCTTGATATGGTTTGG
GCACACGTCTGAACTCTTTCCCTACACGACGCTCTTCCGATCTNNNTTGAAGGCTTGATATGGTTTGG
GCACACGTCTGAACTCTTTCCCTACACGACGCTCTTCCGATCTNNNAGTACTGATGAAGATCACCACG
GCACACGTCTGAACTCTTTCCCTACACGACGCTCTTCCGATCTNNNAGTACTGATGAAGATCACCACG
GCACACGTCTGAACTCTTTCCCTACACGACGCTCTTCCGATCTNNNAGTACTGATGAAGATCACCACG
GCACACGTCTGAACTCTTTCCCTACACGACGCTCTTCCGATCTNNNAGTACTGATGAAGATCACCACG
AGCACACGTCTGAACTCTTTCCCTACACGACGCTCTTCCGATCTNNNCAATAGCATAGAGGCAGAAGG
AGCACACGTCTGAACTCTTTCCCTACACGACGCTCTTCCGATCTNNNCAATAGCATAGAGGCAGAAGG
AGCACACGTCTGAACTCTTTCCCTACACGACGCTCTTCCGATCTNNNCAATAGCATAGAGGCAGAAGG
AGCACACGTCTGAACTCTTTCCCTACACGACGCTCTTCCGATCTNNNCAATAGCATAGAGGCAGAAGG
GCACACGTCTGAACTCTTTCCCTACACGACGCTCTTCCGATCTNNNCTCTTTCAGCTCTGAGTCCACATC
GCACACGTCTGAACTCTTTCCCTACACGACGCTCTTCCGATCTNNNCTCTTTCAGCTCTGAGTCCACATC
GCACACGTCTGAACTCTTTCCCTACACGACGCTCTTCCGATCTNNNCTCTTTCAGCTCTGAGTCCACATC
GCACACGTCTGAACTCTTTCCCTACACGACGCTCTTCCGATCTNNNCTCTTTCAGCTCTGAGTCCACATC
GCACACGTCTGAACTCTTTCCCTACACGACGCTCTTCCGATCTNNNCCCTCAAACACTGAAAACAAGCT
GCACACGTCTGAACTCTTTCCCTACACGACGCTCTTCCGATCTNNNCCCTCAAACACTGAAAACAAGCT
GCACACGTCTGAACTCTTTCCCTACACGACGCTCTTCCGATCTNNNCCCTCAAACACTGAAAACAAGCT
GCACACGTCTGAACTCTTTCCCTACACGACGCTCTTCCGATCTNNNCCCTCAAACACTGAAAACAAGCT
CACACGTCTGAACTCTTTCCCTACACGACGCTCTTCCGATCTNNNCTCGATCTCCTGACTTCGTGATCC
CACACGTCTGAACTCTTTCCCTACACGACGCTCTTCCGATCTNNNCTCGATCTCCTGACTTCGTGATCC
CACACGTCTGAACTCTTTCCCTACACGACGCTCTTCCGATCTNNNCTCGATCTCCTGACTTCGTGATCC
CACACGTCTGAACTCTTTCCCTACACGACGCTCTTCCGATCTNNNCTCGATCTCCTGACTTCGTGATCC
GAGCACACGTCTGAACTCTTTCCCTACACGACGCTCTTCCGATCTNNNAAGGAAGACGAGGAATTCA
GAGCACACGTCTGAACTCTTTCCCTACACGACGCTCTTCCGATCTNNNAAGGAAGACGAGGAATTCA
GAGCACACGTCTGAACTCTTTCCCTACACGACGCTCTTCCGATCTNNNAAGGAAGACGAGGAATTCA
GAGCACACGTCTGAACTCTTTCCCTACACGACGCTCTTCCGATCTNNNAAGGAAGACGAGGAATTCA
AGCACACGTCTGAACTCTTTCCCTACACGACGCTCTTCCGATCTNNNTGGAGCATAAGCACACCTATT
AGCACACGTCTGAACTCTTTCCCTACACGACGCTCTTCCGATCTNNNTGGAGCATAAGCACACCTATT

AGCACACGTCTGAACTCTTTCCCTACACGACGCTCTTCCGATCTNNNTGGAGCATAAGCACACCTATTC
AGCACACGTCTGAACTCTTTCCCTACACGACGCTCTTCCGATCTNNNTGGAGCATAAGCACACCTATTC
GAGCACACGTCTGAACTCTTTCCCTACACGACGCTCTTCCGATCTNNNAAGTACAGTGCTGGTCTGTA
GAGCACACGTCTGAACTCTTTCCCTACACGACGCTCTTCCGATCTNNNAAGTACAGTGCTGGTCTGTA
GAGCACACGTCTGAACTCTTTCCCTACACGACGCTCTTCCGATCTNNNAAGTACAGTGCTGGTCTGTA
GAGCACACGTCTGAACTCTTTCCCTACACGACGCTCTTCCGATCTNNNAAGTACAGTGCTGGTCTGTA
GAGCACACGTCTGAACTCTTTCCCTACACGACGCTCTTCCGATCTNNNCTGAGATGAGAGGATGGCT
GAGCACACGTCTGAACTCTTTCCCTACACGACGCTCTTCCGATCTNNNCTGAGATGAGAGGATGGCT
GAGCACACGTCTGAACTCTTTCCCTACACGACGCTCTTCCGATCTNNNCTGAGATGAGAGGATGGCT
GAGCACACGTCTGAACTCTTTCCCTACACGACGCTCTTCCGATCTNNNCTGAGATGAGAGGATGGCT
GAGCACACGTCTGAACTCTTTCCCTACACGACGCTCTTCCGATCTNNNTTTGGTGACTTCCGGTCTTA
GAGCACACGTCTGAACTCTTTCCCTACACGACGCTCTTCCGATCTNNNTTTGGTGACTTCCGGTCTTA
GAGCACACGTCTGAACTCTTTCCCTACACGACGCTCTTCCGATCTNNNTTTGGTGACTTCCGGTCTTA
GAGCACACGTCTGAACTCTTTCCCTACACGACGCTCTTCCGATCTNNNTTTGGTGACTTCCGGTCTTA
AGCACACGTCTGAACTCTTTCCCTACACGACGCTCTTCCGATCTNNNGAAATGAAAACCTTAGACCGG
AGCACACGTCTGAACTCTTTCCCTACACGACGCTCTTCCGATCTNNNGAAATGAAAACCTTAGACCGG
AGCACACGTCTGAACTCTTTCCCTACACGACGCTCTTCCGATCTNNNGAAATGAAAACCTTAGACCGG
AGCACACGTCTGAACTCTTTCCCTACACGACGCTCTTCCGATCTNNNGAAATGAAAACCTTAGACCGG
AGCACACGTCTGAACTCTTTCCCTACACGACGCTCTTCCGATCTNNNATCCAGCCAGAACAGAACTTTC
AGCACACGTCTGAACTCTTTCCCTACACGACGCTCTTCCGATCTNNNATCCAGCCAGAACAGAACTTTC
AGCACACGTCTGAACTCTTTCCCTACACGACGCTCTTCCGATCTNNNATCCAGCCAGAACAGAACTTTC
AGCACACGTCTGAACTCTTTCCCTACACGACGCTCTTCCGATCTNNNATCCAGCCAGAACAGAACTTTC
GAGCACACGTCTGAACTCTTTCCCTACACGACGCTCTTCCGATCTNNNCTACTCTCTGCAGACAAGACC
GAGCACACGTCTGAACTCTTTCCCTACACGACGCTCTTCCGATCTNNNCTACTCTCTGCAGACAAGACC
GAGCACACGTCTGAACTCTTTCCCTACACGACGCTCTTCCGATCTNNNCTACTCTCTGCAGACAAGACC
GAGCACACGTCTGAACTCTTTCCCTACACGACGCTCTTCCGATCTNNNCTACTCTCTGCAGACAAGACC
AGCACACGTCTGAACTCTTTCCCTACACGACGCTCTTCCGATCTNNNTGTATACATGTGGGCATCAGG
AGCACACGTCTGAACTCTTTCCCTACACGACGCTCTTCCGATCTNNNTGTATACATGTGGGCATCAGG
AGCACACGTCTGAACTCTTTCCCTACACGACGCTCTTCCGATCTNNNTGTATACATGTGGGCATCAGG
AGCACACGTCTGAACTCTTTCCCTACACGACGCTCTTCCGATCTNNNTGTATACATGTGGGCATCAGG
AGCACACGTCTGAACTCTTTCCCTACACGACGCTCTTCCGATCTNNNCAAATGTGACACAGTGAAGGC
AGCACACGTCTGAACTCTTTCCCTACACGACGCTCTTCCGATCTNNNCAAATGTGACACAGTGAAGGC
AGCACACGTCTGAACTCTTTCCCTACACGACGCTCTTCCGATCTNNNCAAATGTGACACAGTGAAGGC
AGCACACGTCTGAACTCTTTCCCTACACGACGCTCTTCCGATCTNNNCAAATGTGACACAGTGAAGGC
GAGCACACGTCTGAACTCTTTCCCTACACGACGCTCTTCCGATCTNNNGAATGCTGTCAACTGAGACT
GAGCACACGTCTGAACTCTTTCCCTACACGACGCTCTTCCGATCTNNNGAATGCTGTCAACTGAGACT
GAGCACACGTCTGAACTCTTTCCCTACACGACGCTCTTCCGATCTNNNGAATGCTGTCAACTGAGACT
GAGCACACGTCTGAACTCTTTCCCTACACGACGCTCTTCCGATCTNNNGAATGCTGTCAACTGAGACT
AGCACACGTCTGAACTCTTTCCCTACACGACGCTCTTCCGATCTNNNACACAAAAGAGATGCAATGTC
AGCACACGTCTGAACTCTTTCCCTACACGACGCTCTTCCGATCTNNNACACAAAAGAGATGCAATGTC
AGCACACGTCTGAACTCTTTCCCTACACGACGCTCTTCCGATCTNNNACACAAAAGAGATGCAATGTC
AGCACACGTCTGAACTCTTTCCCTACACGACGCTCTTCCGATCTNNNACACAAAAGAGATGCAATGTC
GCACACGTCTGAACTCTTTCCCTACACGACGCTCTTCCGATCTNNNTCTTTTTCCCTTCCTTCCCTTCCCAT

GCACACGTCTGAACTCTTTCCCTACACGACGCTCTTCCGATCTNNNTCTTTTCCCTTCCTTCCCTTCCCAT
GCACACGTCTGAACTCTTTCCCTACACGACGCTCTTCCGATCTNNNTCTTTTCCCTTCCTTCCCTTCCCAT
GCACACGTCTGAACTCTTTCCCTACACGACGCTCTTCCGATCTNNNTCTTTTCCCTTCCTTCCCTTCCCAT
GAGCACACGTCTGAACTCTTTCCCTACACGACGCTCTTCCGATCTNNNTTTGAAGGTCAAGTCTGCAGC
GAGCACACGTCTGAACTCTTTCCCTACACGACGCTCTTCCGATCTNNNTTTGAAGGTCAAGTCTGCAGC
GAGCACACGTCTGAACTCTTTCCCTACACGACGCTCTTCCGATCTNNNTTTGAAGGTCAAGTCTGCAGC
GAGCACACGTCTGAACTCTTTCCCTACACGACGCTCTTCCGATCTNNNTTTGAAGGTCAAGTCTGCAGC
AGCACACGTCTGAACTCTTTCCCTACACGACGCTCTTCCGATCTNNNTAGTGTGAATTTTCCCAGGCTG
AGCACACGTCTGAACTCTTTCCCTACACGACGCTCTTCCGATCTNNNTAGTGTGAATTTTCCCAGGCTG
AGCACACGTCTGAACTCTTTCCCTACACGACGCTCTTCCGATCTNNNTAGTGTGAATTTTCCCAGGCTG
AGCACACGTCTGAACTCTTTCCCTACACGACGCTCTTCCGATCTNNNTAGTGTGAATTTTCCCAGGCTG
AGAGCACACGTCTGAACTCTTTCCCTACACGACGCTCTTCCGATCTNNNTCAGTCCCAATGAGATGAACT
AGAGCACACGTCTGAACTCTTTCCCTACACGACGCTCTTCCGATCTNNNTCAGTCCCAATGAGATGAACT
AGAGCACACGTCTGAACTCTTTCCCTACACGACGCTCTTCCGATCTNNNTCAGTCCCAATGAGATGAACT
AGAGCACACGTCTGAACTCTTTCCCTACACGACGCTCTTCCGATCTNNNTCAGTCCCAATGAGATGAACT
AGCACACGTCTGAACTCTTTCCCTACACGACGCTCTTCCGATCTNNNTTTCCCAGTGCACTATAGTGGTC
AGCACACGTCTGAACTCTTTCCCTACACGACGCTCTTCCGATCTNNNTTTCCCAGTGCACTATAGTGGTC
AGCACACGTCTGAACTCTTTCCCTACACGACGCTCTTCCGATCTNNNTTTCCCAGTGCACTATAGTGGTC
AGCACACGTCTGAACTCTTTCCCTACACGACGCTCTTCCGATCTNNNTTTCCCAGTGCACTATAGTGGTC
AGCACACGTCTGAACTCTTTCCCTACACGACGCTCTTCCGATCTNNNTGGCAGACTCAACTTCTTAGCAT
AGCACACGTCTGAACTCTTTCCCTACACGACGCTCTTCCGATCTNNNTGGCAGACTCAACTTCTTAGCAT
AGCACACGTCTGAACTCTTTCCCTACACGACGCTCTTCCGATCTNNNTGGCAGACTCAACTTCTTAGCAT
AGCACACGTCTGAACTCTTTCCCTACACGACGCTCTTCCGATCTNNNTGGCAGACTCAACTTCTTAGCAT
AGCACACGTCTGAACTCTTTCCCTACACGACGCTCTTCCGATCTNNNAAAAAAAATTAGCCAGGCGTGC
AGCACACGTCTGAACTCTTTCCCTACACGACGCTCTTCCGATCTNNNAAAAAAAATTAGCCAGGCGTGC
AGCACACGTCTGAACTCTTTCCCTACACGACGCTCTTCCGATCTNNNAAAAAAAATTAGCCAGGCGTGC
AGCACACGTCTGAACTCTTTCCCTACACGACGCTCTTCCGATCTNNNAAAAAAAATTAGCCAGGCGTGC
GAGCACACGTCTGAACTCTTTCCCTACACGACGCTCTTCCGATCTNNNAAAAGGGATGGACAGAATCAC
GAGCACACGTCTGAACTCTTTCCCTACACGACGCTCTTCCGATCTNNNAAAAGGGATGGACAGAATCAC
GAGCACACGTCTGAACTCTTTCCCTACACGACGCTCTTCCGATCTNNNAAAAGGGATGGACAGAATCAC
GAGCACACGTCTGAACTCTTTCCCTACACGACGCTCTTCCGATCTNNNAAAAGGGATGGACAGAATCAC
GAGCACACGTCTGAACTCTTTCCCTACACGACGCTCTTCCGATCTNNNCTGGATGACAAGAGCGAAAC
GAGCACACGTCTGAACTCTTTCCCTACACGACGCTCTTCCGATCTNNNCTGGATGACAAGAGCGAAAC
GAGCACACGTCTGAACTCTTTCCCTACACGACGCTCTTCCGATCTNNNCTGGATGACAAGAGCGAAAC
GAGCACACGTCTGAACTCTTTCCCTACACGACGCTCTTCCGATCTNNNCTGGATGACAAGAGCGAAAC
GAGCACACGTCTGAACTCTTTCCCTACACGACGCTCTTCCGATCTNNNGGTAGCACCAGTAGTTGAGTG
GAGCACACGTCTGAACTCTTTCCCTACACGACGCTCTTCCGATCTNNNGGTAGCACCAGTAGTTGAGTG
GAGCACACGTCTGAACTCTTTCCCTACACGACGCTCTTCCGATCTNNNGGTAGCACCAGTAGTTGAGTG
GAGCACACGTCTGAACTCTTTCCCTACACGACGCTCTTCCGATCTNNNGGTAGCACCAGTAGTTGAGTG
AGCACACGTCTGAACTCTTTCCCTACACGACGCTCTTCCGATCTNNNGCAGAGAATTTCTTGAACCCGA
AGCACACGTCTGAACTCTTTCCCTACACGACGCTCTTCCGATCTNNNGCAGAGAATTTCTTGAACCCGA
AGCACACGTCTGAACTCTTTCCCTACACGACGCTCTTCCGATCTNNNGCAGAGAATTTCTTGAACCCGA
AGCACACGTCTGAACTCTTTCCCTACACGACGCTCTTCCGATCTNNNGCAGAGAATTTCTTGAACCCGA

GCACACGTCTGAACTCTTTCCCTACACGACGCTCTTCCGATCTNNNGTTATGTGCTCCTCTACCTTCCCC
GCACACGTCTGAACTCTTTCCCTACACGACGCTCTTCCGATCTNNNGTTATGTGCTCCTCTACCTTCCCC
GCACACGTCTGAACTCTTTCCCTACACGACGCTCTTCCGATCTNNNGTTATGTGCTCCTCTACCTTCCCC
GCACACGTCTGAACTCTTTCCCTACACGACGCTCTTCCGATCTNNNGTTATGTGCTCCTCTACCTTCCCC
GAGCACACGTCTGAACTCTTTCCCTACACGACGCTCTTCCGATCTNNNGAAAGGAAAATGCCAGGGC
GAGCACACGTCTGAACTCTTTCCCTACACGACGCTCTTCCGATCTNNNGAAAGGAAAATGCCAGGGC
GAGCACACGTCTGAACTCTTTCCCTACACGACGCTCTTCCGATCTNNNGAAAGGAAAATGCCAGGGC
GAGCACACGTCTGAACTCTTTCCCTACACGACGCTCTTCCGATCTNNNGAAAGGAAAATGCCAGGGC
AGCACACGTCTGAACTCTTTCCCTACACGACGCTCTTCCGATCTNNNACAGAGAGTGACTGATAAAAG
AGCACACGTCTGAACTCTTTCCCTACACGACGCTCTTCCGATCTNNNACAGAGAGTGACTGATAAAAG
AGCACACGTCTGAACTCTTTCCCTACACGACGCTCTTCCGATCTNNNACAGAGAGTGACTGATAAAAG
AGCACACGTCTGAACTCTTTCCCTACACGACGCTCTTCCGATCTNNNACAGAGAGTGACTGATAAAAG
AGAGCACACGTCTGAACTCTTTCCCTACACGACGCTCTTCCGATCTNNNGAAGAAGACCCTGTCCCTC
AGAGCACACGTCTGAACTCTTTCCCTACACGACGCTCTTCCGATCTNNNGAAGAAGACCCTGTCCCTC
AGAGCACACGTCTGAACTCTTTCCCTACACGACGCTCTTCCGATCTNNNGAAGAAGACCCTGTCCCTC
AGAGCACACGTCTGAACTCTTTCCCTACACGACGCTCTTCCGATCTNNNGAAGAAGACCCTGTCCCTC
GAGCACACGTCTGAACTCTTTCCCTACACGACGCTCTTCCGATCTNNNGAGATTCAGGGGTGGTACTT
GAGCACACGTCTGAACTCTTTCCCTACACGACGCTCTTCCGATCTNNNGAGATTCAGGGGTGGTACTT
GAGCACACGTCTGAACTCTTTCCCTACACGACGCTCTTCCGATCTNNNGAGATTCAGGGGTGGTACTT
GAGCACACGTCTGAACTCTTTCCCTACACGACGCTCTTCCGATCTNNNGAGATTCAGGGGTGGTACTT
GAGCACACGTCTGAACTCTTTCCCTACACGACGCTCTTCCGATCTNNNTGTGAGTTTGGGCATTGACA
GAGCACACGTCTGAACTCTTTCCCTACACGACGCTCTTCCGATCTNNNTGTGAGTTTGGGCATTGACA
GAGCACACGTCTGAACTCTTTCCCTACACGACGCTCTTCCGATCTNNNTGTGAGTTTGGGCATTGACA
GAGCACACGTCTGAACTCTTTCCCTACACGACGCTCTTCCGATCTNNNTGTGAGTTTGGGCATTGACA
AGCACACGTCTGAACTCTTTCCCTACACGACGCTCTTCCGATCTNNNGGGAGGGTGATATGGTTTAC
AGCACACGTCTGAACTCTTTCCCTACACGACGCTCTTCCGATCTNNNGGGAGGGTGATATGGTTTAC
AGCACACGTCTGAACTCTTTCCCTACACGACGCTCTTCCGATCTNNNGGGAGGGTGATATGGTTTAC
AGCACACGTCTGAACTCTTTCCCTACACGACGCTCTTCCGATCTNNNGGGAGGGTGATATGGTTTAC
GCACACGTCTGAACTCTTTCCCTACACGACGCTCTTCCGATCTNNNGGATGAGGATTGCTTGAAAAC
GCACACGTCTGAACTCTTTCCCTACACGACGCTCTTCCGATCTNNNGGATGAGGATTGCTTGAAAAC
GCACACGTCTGAACTCTTTCCCTACACGACGCTCTTCCGATCTNNNGGATGAGGATTGCTTGAAAAC
GCACACGTCTGAACTCTTTCCCTACACGACGCTCTTCCGATCTNNNGGATGAGGATTGCTTGAAAAC
GCACACGTCTGAACTCTTTCCCTACACGACGCTCTTCCGATCTNNNACAGAACCAGAATTTAAGGTG
GCACACGTCTGAACTCTTTCCCTACACGACGCTCTTCCGATCTNNNACAGAACCAGAATTTAAGGTG
GCACACGTCTGAACTCTTTCCCTACACGACGCTCTTCCGATCTNNNACAGAACCAGAATTTAAGGTG
GCACACGTCTGAACTCTTTCCCTACACGACGCTCTTCCGATCTNNNACAGAACCAGAATTTAAGGTG
GCACACGTCTGAACTCTTTCCCTACACGACGCTCTTCCGATCTNNNAATGAGATCTTCGTATTCCGGG
GCACACGTCTGAACTCTTTCCCTACACGACGCTCTTCCGATCTNNNAATGAGATCTTCGTATTCCGGG
GCACACGTCTGAACTCTTTCCCTACACGACGCTCTTCCGATCTNNNAATGAGATCTTCGTATTCCGGG
GCACACGTCTGAACTCTTTCCCTACACGACGCTCTTCCGATCTNNNAATGAGATCTTCGTATTCCGGG
GCACACGTCTGAACTCTTTCCCTACACGACGCTCTTCCGATCTNNNATACCTATATTGTGCTTTAAGGT
GCACACGTCTGAACTCTTTCCCTACACGACGCTCTTCCGATCTNNNATACCTATATTGTGCTTTAAGGT
GCACACGTCTGAACTCTTTCCCTACACGACGCTCTTCCGATCTNNNATACCTATATTGTGCTTTAAGGT

GCACACGTCTGAACTCTTTCCCTACACGACGCTCTTCCGATCTNNNATACCTATATTGTGCTTTAAGGT
CACACGTCTGAACTCTTTCCCTACACGACGCTCTTCCGATCTNNNTGCTGAATTGACTTCTTTGTCATT
CACACGTCTGAACTCTTTCCCTACACGACGCTCTTCCGATCTNNNTGCTGAATTGACTTCTTTGTCATT
CACACGTCTGAACTCTTTCCCTACACGACGCTCTTCCGATCTNNNTGCTGAATTGACTTCTTTGTCATT
CACACGTCTGAACTCTTTCCCTACACGACGCTCTTCCGATCTNNNTGCTGAATTGACTTCTTTGTCATT
GCACACGTCTGAACTCTTTCCCTACACGACGCTCTTCCGATCTNNNCATGCAGTTTGGACTAACAGGT
GCACACGTCTGAACTCTTTCCCTACACGACGCTCTTCCGATCTNNNCATGCAGTTTGGACTAACAGGT
GCACACGTCTGAACTCTTTCCCTACACGACGCTCTTCCGATCTNNNCATGCAGTTTGGACTAACAGGT
GCACACGTCTGAACTCTTTCCCTACACGACGCTCTTCCGATCTNNNCATGCAGTTTGGACTAACAGGT
GCACACGTCTGAACTCTTTCCCTACACGACGCTCTTCCGATCTNNNAAGTAGCTGAGACTACAGGCAT
GCACACGTCTGAACTCTTTCCCTACACGACGCTCTTCCGATCTNNNAAGTAGCTGAGACTACAGGCAT
GCACACGTCTGAACTCTTTCCCTACACGACGCTCTTCCGATCTNNNAAGTAGCTGAGACTACAGGCAT
GCACACGTCTGAACTCTTTCCCTACACGACGCTCTTCCGATCTNNNAAGTAGCTGAGACTACAGGCAT
AGCACACGTCTGAACTCTTTCCCTACACGACGCTCTTCCGATCTNNNGTCCTCTCAGCTTTTCGGTCCC
AGCACACGTCTGAACTCTTTCCCTACACGACGCTCTTCCGATCTNNNGTCCTCTCAGCTTTTCGGTCCC
AGCACACGTCTGAACTCTTTCCCTACACGACGCTCTTCCGATCTNNNGTCCTCTCAGCTTTTCGGTCCC
AGCACACGTCTGAACTCTTTCCCTACACGACGCTCTTCCGATCTNNNGTCCTCTCAGCTTTTCGGTCCC
GCACACGTCTGAACTCTTTCCCTACACGACGCTCTTCCGATCTNNNGTTTTGACTTTGTGAACCCCAA
GCACACGTCTGAACTCTTTCCCTACACGACGCTCTTCCGATCTNNNGTTTTGACTTTGTGAACCCCAA
GCACACGTCTGAACTCTTTCCCTACACGACGCTCTTCCGATCTNNNGTTTTGACTTTGTGAACCCCAA
GCACACGTCTGAACTCTTTCCCTACACGACGCTCTTCCGATCTNNNGTTTTGACTTTGTGAACCCCAA
GCACACGTCTGAACTCTTTCCCTACACGACGCTCTTCCGATCTNNNACCCTATTCCTACCAAATGCACT
GCACACGTCTGAACTCTTTCCCTACACGACGCTCTTCCGATCTNNNACCCTATTCCTACCAAATGCACT
GCACACGTCTGAACTCTTTCCCTACACGACGCTCTTCCGATCTNNNACCCTATTCCTACCAAATGCACT
GCACACGTCTGAACTCTTTCCCTACACGACGCTCTTCCGATCTNNNACCCTATTCCTACCAAATGCACT
AGCACACGTCTGAACTCTTTCCCTACACGACGCTCTTCCGATCTNNNATGTGATTGTGTGTGTGTGT
AGCACACGTCTGAACTCTTTCCCTACACGACGCTCTTCCGATCTNNNATGTGATTGTGTGTGTGTGT
AGCACACGTCTGAACTCTTTCCCTACACGACGCTCTTCCGATCTNNNATGTGATTGTGTGTGTGTGT
AGCACACGTCTGAACTCTTTCCCTACACGACGCTCTTCCGATCTNNNATGTGATTGTGTGTGTGTGT
GAGCACACGTCTGAACTCTTTCCCTACACGACGCTCTTCCGATCTNNNTCAGGAGAATCACTTGAACC
GAGCACACGTCTGAACTCTTTCCCTACACGACGCTCTTCCGATCTNNNTCAGGAGAATCACTTGAACC
GAGCACACGTCTGAACTCTTTCCCTACACGACGCTCTTCCGATCTNNNTCAGGAGAATCACTTGAACC
GAGCACACGTCTGAACTCTTTCCCTACACGACGCTCTTCCGATCTNNNTCAGGAGAATCACTTGAACC
GCACACGTCTGAACTCTTTCCCTACACGACGCTCTTCCGATCTNNNACAAATCCACTGGGCTATACAC
GCACACGTCTGAACTCTTTCCCTACACGACGCTCTTCCGATCTNNNACAAATCCACTGGGCTATACAC
GCACACGTCTGAACTCTTTCCCTACACGACGCTCTTCCGATCTNNNACAAATCCACTGGGCTATACAC
GCACACGTCTGAACTCTTTCCCTACACGACGCTCTTCCGATCTNNNACAAATCCACTGGGCTATACAC
GAGCACACGTCTGAACTCTTTCCCTACACGACGCTCTTCCGATCTNNNAATGTTTGTGTTTTAGCCTG
GAGCACACGTCTGAACTCTTTCCCTACACGACGCTCTTCCGATCTNNNAATGTTTGTGTTTTAGCCTG
GAGCACACGTCTGAACTCTTTCCCTACACGACGCTCTTCCGATCTNNNAATGTTTGTGTTTTAGCCTG
GAGCACACGTCTGAACTCTTTCCCTACACGACGCTCTTCCGATCTNNNAATGTTTGTGTTTTAGCCTG
GCACACGTCTGAACTCTTTCCCTACACGACGCTCTTCCGATCTNNNTAAGGGGGGAAAGCAAGCATT
GCACACGTCTGAACTCTTTCCCTACACGACGCTCTTCCGATCTNNNTAAGGGGGGAAAGCAAGCATT

AGCACACGTCTGAACTCTTTCCCTACACGACGCTCTTCCGATCTNNNTAAGGGGGGAAAGCAAGCATTA

AGCACACGTCTGAACTCTTTCCCTACACGACGCTCTTCCGATCTNNNTAAGGGGGGAAAGCAAGCATTA

AGCACACGTCTGAACTCTTTCCCTACACGACGCTCTTCCGATCTNNNGCTGTAGCAGTTTCTAAAAATAA

AGCACACGTCTGAACTCTTTCCCTACACGACGCTCTTCCGATCTNNNGCTGTAGCAGTTTCTAAAAATAA

AGCACACGTCTGAACTCTTTCCCTACACGACGCTCTTCCGATCTNNNGCTGTAGCAGTTTCTAAAAATAA

AGCACACGTCTGAACTCTTTCCCTACACGACGCTCTTCCGATCTNNNGCTGTAGCAGTTTCTAAAAATAA

AGCACACGTCTGAACTCTTTCCCTACACGACGCTCTTCCGATCTNNNTGTGTTGTTCTCTGTGCCTACTT

AGCACACGTCTGAACTCTTTCCCTACACGACGCTCTTCCGATCTNNNTGTGTTGTTCTCTGTGCCTACTT

AGCACACGTCTGAACTCTTTCCCTACACGACGCTCTTCCGATCTNNNTGTGTTGTTCTCTGTGCCTACTT

AGCACACGTCTGAACTCTTTCCCTACACGACGCTCTTCCGATCTNNNTGTGTTGTTCTCTGTGCCTACTT

GAGCACACGTCTGAACTCTTTCCCTACACGACGCTCTTCCGATCTNNNATGCCTGTAATCCCAGTTACTC

GAGCACACGTCTGAACTCTTTCCCTACACGACGCTCTTCCGATCTNNNATGCCTGTAATCCCAGTTACTC

GAGCACACGTCTGAACTCTTTCCCTACACGACGCTCTTCCGATCTNNNATGCCTGTAATCCCAGTTACTC

GAGCACACGTCTGAACTCTTTCCCTACACGACGCTCTTCCGATCTNNNATGCCTGTAATCCCAGTTACTC

GAGCACACGTCTGAACTCTTTCCCTACACGACGCTCTTCCGATCTNNNGAGCTTGTAGTGAGCCGAGA

GAGCACACGTCTGAACTCTTTCCCTACACGACGCTCTTCCGATCTNNNGAGCTTGTAGTGAGCCGAGA

GAGCACACGTCTGAACTCTTTCCCTACACGACGCTCTTCCGATCTNNNGAGCTTGTAGTGAGCCGAGA

GAGCACACGTCTGAACTCTTTCCCTACACGACGCTCTTCCGATCTNNNGAGCTTGTAGTGAGCCGAGA

AGCACACGTCTGAACTCTTTCCCTACACGACGCTCTTCCGATCTNNNCTGGGTTCAGAATCTGTATACCA

AGCACACGTCTGAACTCTTTCCCTACACGACGCTCTTCCGATCTNNNCTGGGTTCAGAATCTGTATACCA

AGCACACGTCTGAACTCTTTCCCTACACGACGCTCTTCCGATCTNNNCTGGGTTCAGAATCTGTATACCA

AGCACACGTCTGAACTCTTTCCCTACACGACGCTCTTCCGATCTNNNCTGGGTTCAGAATCTGTATACCA

AGCACACGTCTGAACTCTTTCCCTACACGACGCTCTTCCGATCTNNNTGAGGCCAGCTTAGCTAACATA

AGCACACGTCTGAACTCTTTCCCTACACGACGCTCTTCCGATCTNNNTGAGGCCAGCTTAGCTAACATA

AGCACACGTCTGAACTCTTTCCCTACACGACGCTCTTCCGATCTNNNTGAGGCCAGCTTAGCTAACATA

AGCACACGTCTGAACTCTTTCCCTACACGACGCTCTTCCGATCTNNNTGAGGCCAGCTTAGCTAACATA

GCACACGTCTGAACTCTTTCCCTACACGACGCTCTTCCGATCTNNNGAATGTGCCCGATCTCATCAAAG

GCACACGTCTGAACTCTTTCCCTACACGACGCTCTTCCGATCTNNNGAATGTGCCCGATCTCATCAAAG

GCACACGTCTGAACTCTTTCCCTACACGACGCTCTTCCGATCTNNNGAATGTGCCCGATCTCATCAAAG

GCACACGTCTGAACTCTTTCCCTACACGACGCTCTTCCGATCTNNNGAATGTGCCCGATCTCATCAAAG

GAGCACACGTCTGAACTCTTTCCCTACACGACGCTCTTCCGATCTNNNGAGATCACACCACTGCTCTCCC

GAGCACACGTCTGAACTCTTTCCCTACACGACGCTCTTCCGATCTNNNGAGATCACACCACTGCTCTCCC

GAGCACACGTCTGAACTCTTTCCCTACACGACGCTCTTCCGATCTNNNGAGATCACACCACTGCTCTCCC

GAGCACACGTCTGAACTCTTTCCCTACACGACGCTCTTCCGATCTNNNGAGATCACACCACTGCTCTCCC

GAGCACACGTCTGAACTCTTTCCCTACACGACGCTCTTCCGATCTNNNATGCAAATCAAGCTACTACCCC

GAGCACACGTCTGAACTCTTTCCCTACACGACGCTCTTCCGATCTNNNATGCAAATCAAGCTACTACCCC

GAGCACACGTCTGAACTCTTTCCCTACACGACGCTCTTCCGATCTNNNATGCAAATCAAGCTACTACCCC

GAGCACACGTCTGAACTCTTTCCCTACACGACGCTCTTCCGATCTNNNATGCAAATCAAGCTACTACCCC

GAGCACACGTCTGAACTCTTTCCCTACACGACGCTCTTCCGATCTNNNATGATCTTGGCTCACTGCAACT

GAGCACACGTCTGAACTCTTTCCCTACACGACGCTCTTCCGATCTNNNATGATCTTGGCTCACTGCAACT

GAGCACACGTCTGAACTCTTTCCCTACACGACGCTCTTCCGATCTNNNATGATCTTGGCTCACTGCAACT

GAGCACACGTCTGAACTCTTTCCCTACACGACGCTCTTCCGATCTNNNATGATCTTGGCTCACTGCAACT

AGAGCACACGTCTGAACTCTTTCCCTACACGACGCTCTTCCGATCTNNNGGCCACAAAGAAAGAGCACA

GAGCACACGTCTGAACTCTTTCCCTACACGACGCTCTTCCGATCTNNNGGCCACAAAGAAAGAGCAC
GAGCACACGTCTGAACTCTTTCCCTACACGACGCTCTTCCGATCTNNNGGCCACAAAGAAAGAGCAC
GAGCACACGTCTGAACTCTTTCCCTACACGACGCTCTTCCGATCTNNNGGCCACAAAGAAAGAGCAC
AGCACACGTCTGAACTCTTTCCCTACACGACGCTCTTCCGATCTNNNACAGTTCATTCTCTAAGTGAT
AGCACACGTCTGAACTCTTTCCCTACACGACGCTCTTCCGATCTNNNACAGTTCATTCTCTAAGTGAT
AGCACACGTCTGAACTCTTTCCCTACACGACGCTCTTCCGATCTNNNACAGTTCATTCTCTAAGTGAT
AGCACACGTCTGAACTCTTTCCCTACACGACGCTCTTCCGATCTNNNACAGTTCATTCTCTAAGTGAT
AGCACACGTCTGAACTCTTTCCCTACACGACGCTCTTCCGATCTNNNAAATCAAAGATGTTCGGATTC
AGCACACGTCTGAACTCTTTCCCTACACGACGCTCTTCCGATCTNNNAAATCAAAGATGTTCGGATTC
AGCACACGTCTGAACTCTTTCCCTACACGACGCTCTTCCGATCTNNNAAATCAAAGATGTTCGGATTC
AGCACACGTCTGAACTCTTTCCCTACACGACGCTCTTCCGATCTNNNAAATCAAAGATGTTCGGATTC
GAGCACACGTCTGAACTCTTTCCCTACACGACGCTCTTCCGATCTNNNGACAGAGCAAGACTCCGTCT
GAGCACACGTCTGAACTCTTTCCCTACACGACGCTCTTCCGATCTNNNGACAGAGCAAGACTCCGTCT
GAGCACACGTCTGAACTCTTTCCCTACACGACGCTCTTCCGATCTNNNGACAGAGCAAGACTCCGTCT
GAGCACACGTCTGAACTCTTTCCCTACACGACGCTCTTCCGATCTNNNGACAGAGCAAGACTCCGTCT
AGCACACGTCTGAACTCTTTCCCTACACGACGCTCTTCCGATCTNNNGGGAGAAAGAGGGAGACTCC
AGCACACGTCTGAACTCTTTCCCTACACGACGCTCTTCCGATCTNNNGGGAGAAAGAGGGAGACTCC
AGCACACGTCTGAACTCTTTCCCTACACGACGCTCTTCCGATCTNNNGGGAGAAAGAGGGAGACTCC
AGCACACGTCTGAACTCTTTCCCTACACGACGCTCTTCCGATCTNNNGGGAGAAAGAGGGAGACTCC
AGCACACGTCTGAACTCTTTCCCTACACGACGCTCTTCCGATCTNNNAGAAGGACACAAATGACAAA
AGCACACGTCTGAACTCTTTCCCTACACGACGCTCTTCCGATCTNNNAGAAGGACACAAATGACAAA
AGCACACGTCTGAACTCTTTCCCTACACGACGCTCTTCCGATCTNNNAGAAGGACACAAATGACAAA
AGCACACGTCTGAACTCTTTCCCTACACGACGCTCTTCCGATCTNNNAGAAGGACACAAATGACAAA
AGCACACGTCTGAACTCTTTCCCTACACGACGCTCTTCCGATCTNNNAGCAAGTCCCTTTCACCTATGA
AGCACACGTCTGAACTCTTTCCCTACACGACGCTCTTCCGATCTNNNAGCAAGTCCCTTTCACCTATGA
AGCACACGTCTGAACTCTTTCCCTACACGACGCTCTTCCGATCTNNNAGCAAGTCCCTTTCACCTATGA
AGCACACGTCTGAACTCTTTCCCTACACGACGCTCTTCCGATCTNNNAGCAAGTCCCTTTCACCTATGA
AGCACACGTCTGAACTCTTTCCCTACACGACGCTCTTCCGATCTNNNGCAAGATGCTAGTGTGTGAT
AGCACACGTCTGAACTCTTTCCCTACACGACGCTCTTCCGATCTNNNGCAAGATGCTAGTGTGTGAT
AGCACACGTCTGAACTCTTTCCCTACACGACGCTCTTCCGATCTNNNGCAAGATGCTAGTGTGTGAT
AGCACACGTCTGAACTCTTTCCCTACACGACGCTCTTCCGATCTNNNGCAAGATGCTAGTGTGTGAT
AGCACACGTCTGAACTCTTTCCCTACACGACGCTCTTCCGATCTNNNCTGAGGCAGGAAAATTGCTTG
AGCACACGTCTGAACTCTTTCCCTACACGACGCTCTTCCGATCTNNNCTGAGGCAGGAAAATTGCTTG
AGCACACGTCTGAACTCTTTCCCTACACGACGCTCTTCCGATCTNNNCTGAGGCAGGAAAATTGCTTG
AGCACACGTCTGAACTCTTTCCCTACACGACGCTCTTCCGATCTNNNCTGAGGCAGGAAAATTGCTTG
AGCACACGTCTGAACTCTTTCCCTACACGACGCTCTTCCGATCTNNNTGGCCTACTTTTGGGGATATC
AGCACACGTCTGAACTCTTTCCCTACACGACGCTCTTCCGATCTNNNTGGCCTACTTTTGGGGATATC
AGCACACGTCTGAACTCTTTCCCTACACGACGCTCTTCCGATCTNNNTGGCCTACTTTTGGGGATATC
AGCACACGTCTGAACTCTTTCCCTACACGACGCTCTTCCGATCTNNNTGGCCTACTTTTGGGGATATC
AGCACACGTCTGAACTCTTTCCCTACACGACGCTCTTCCGATCTNNNTACTTCTGGCAGCTTCAGCAA
AGCACACGTCTGAACTCTTTCCCTACACGACGCTCTTCCGATCTNNNTACTTCTGGCAGCTTCAGCAA
AGCACACGTCTGAACTCTTTCCCTACACGACGCTCTTCCGATCTNNNTACTTCTGGCAGCTTCAGCAA
AGCACACGTCTGAACTCTTTCCCTACACGACGCTCTTCCGATCTNNNTACTTCTGGCAGCTTCAGCAA

AGCACACGTCTGAACTCTTTCCCTACACGACGCTCTTCCGATCTNNNCATCTTTGGCATTGACAGGTT
AGCACACGTCTGAACTCTTTCCCTACACGACGCTCTTCCGATCTNNNCATCTTTGGCATTGACAGGTT
AGCACACGTCTGAACTCTTTCCCTACACGACGCTCTTCCGATCTNNNCATCTTTGGCATTGACAGGTT
AGCACACGTCTGAACTCTTTCCCTACACGACGCTCTTCCGATCTNNNCATCTTTGGCATTGACAGGTT
AGCACACGTCTGAACTCTTTCCCTACACGACGCTCTTCCGATCTNNNCAGGGGCCAGAAAGTTATTAG
AGCACACGTCTGAACTCTTTCCCTACACGACGCTCTTCCGATCTNNNCAGGGGCCAGAAAGTTATTAG
AGCACACGTCTGAACTCTTTCCCTACACGACGCTCTTCCGATCTNNNCAGGGGCCAGAAAGTTATTAG
AGCACACGTCTGAACTCTTTCCCTACACGACGCTCTTCCGATCTNNNCAGGGGCCAGAAAGTTATTAG
GCACACGTCTGAACTCTTTCCCTACACGACGCTCTTCCGATCTNNNAGGGCAACATAGTGAGACATT
GCACACGTCTGAACTCTTTCCCTACACGACGCTCTTCCGATCTNNNAGGGCAACATAGTGAGACATT
GCACACGTCTGAACTCTTTCCCTACACGACGCTCTTCCGATCTNNNAGGGCAACATAGTGAGACATT
GCACACGTCTGAACTCTTTCCCTACACGACGCTCTTCCGATCTNNNAGGGCAACATAGTGAGACATT
CACACGTCTGAACTCTTTCCCTACACGACGCTCTTCCGATCTNNNGGTTTCCACTGAAAAGTCTGCTG
CACACGTCTGAACTCTTTCCCTACACGACGCTCTTCCGATCTNNNGGTTTCCACTGAAAAGTCTGCTG
CACACGTCTGAACTCTTTCCCTACACGACGCTCTTCCGATCTNNNGGTTTCCACTGAAAAGTCTGCTG
CACACGTCTGAACTCTTTCCCTACACGACGCTCTTCCGATCTNNNGGTTTCCACTGAAAAGTCTGCTG
GAGCACACGTCTGAACTCTTTCCCTACACGACGCTCTTCCGATCTNNNTTTCTGGCCATGTTTGAGAG
GAGCACACGTCTGAACTCTTTCCCTACACGACGCTCTTCCGATCTNNNTTTCTGGCCATGTTTGAGAG
GAGCACACGTCTGAACTCTTTCCCTACACGACGCTCTTCCGATCTNNNTTTCTGGCCATGTTTGAGAG
GAGCACACGTCTGAACTCTTTCCCTACACGACGCTCTTCCGATCTNNNTTTCTGGCCATGTTTGAGAG
AGCACACGTCTGAACTCTTTCCCTACACGACGCTCTTCCGATCTNNNGAGGCTGAGGCAGGAGAATG
AGCACACGTCTGAACTCTTTCCCTACACGACGCTCTTCCGATCTNNNGAGGCTGAGGCAGGAGAATG
AGCACACGTCTGAACTCTTTCCCTACACGACGCTCTTCCGATCTNNNGAGGCTGAGGCAGGAGAATG
AGCACACGTCTGAACTCTTTCCCTACACGACGCTCTTCCGATCTNNNGAGGCTGAGGCAGGAGAATG
GAGCACACGTCTGAACTCTTTCCCTACACGACGCTCTTCCGATCTNNNCAACTCGAAGACCAGGAGAT
GAGCACACGTCTGAACTCTTTCCCTACACGACGCTCTTCCGATCTNNNCAACTCGAAGACCAGGAGAT
GAGCACACGTCTGAACTCTTTCCCTACACGACGCTCTTCCGATCTNNNCAACTCGAAGACCAGGAGAT
GAGCACACGTCTGAACTCTTTCCCTACACGACGCTCTTCCGATCTNNNCAACTCGAAGACCAGGAGAT
ACACGTCTGAACTCTTTCCCTACACGACGCTCTTCCGATCTNNNTCCTTCCTTCCCTTCTTTCTCTTCATC
ACACGTCTGAACTCTTTCCCTACACGACGCTCTTCCGATCTNNNTCCTTCCTTCCCTTCTTTCTCTTCATC
ACACGTCTGAACTCTTTCCCTACACGACGCTCTTCCGATCTNNNTCCTTCCTTCCCTTCTTTCTCTTCATC
ACACGTCTGAACTCTTTCCCTACACGACGCTCTTCCGATCTNNNTCCTTCCTTCCCTTCTTTCTCTTCATC
GAGCACACGTCTGAACTCTTTCCCTACACGACGCTCTTCCGATCTNNNGCTGAAGGAAAAAAAAACTG
GAGCACACGTCTGAACTCTTTCCCTACACGACGCTCTTCCGATCTNNNGCTGAAGGAAAAAAAAACTG
GAGCACACGTCTGAACTCTTTCCCTACACGACGCTCTTCCGATCTNNNGCTGAAGGAAAAAAAAACTG
GAGCACACGTCTGAACTCTTTCCCTACACGACGCTCTTCCGATCTNNNGCTGAAGGAAAAAAAAACTG
GCACACGTCTGAACTCTTTCCCTACACGACGCTCTTCCGATCTNNNTGGTTTTGGTATCAGGGAACA
GCACACGTCTGAACTCTTTCCCTACACGACGCTCTTCCGATCTNNNTGGTTTTGGTATCAGGGAACA
GCACACGTCTGAACTCTTTCCCTACACGACGCTCTTCCGATCTNNNTGGTTTTGGTATCAGGGAACA
GCACACGTCTGAACTCTTTCCCTACACGACGCTCTTCCGATCTNNNTGGTTTTGGTATCAGGGAACA
AGCACACGTCTGAACTCTTTCCCTACACGACGCTCTTCCGATCTNNNTGGTGCAACAGACATAGTGAT
AGCACACGTCTGAACTCTTTCCCTACACGACGCTCTTCCGATCTNNNTGGTGCAACAGACATAGTGAT
AGCACACGTCTGAACTCTTTCCCTACACGACGCTCTTCCGATCTNNNTGGTGCAACAGACATAGTGAT

AGCACACGTCTGAACTCTTTCCCTACACGACGCTCTTCCGATCTNNNTGGTGCAACAGACATAGTGAT
AGCACACGTCTGAACTCTTTCCCTACACGACGCTCTTCCGATCTNNNCCAGCCTGGGAAATAGAGTG
AGCACACGTCTGAACTCTTTCCCTACACGACGCTCTTCCGATCTNNNCCAGCCTGGGAAATAGAGTG
AGCACACGTCTGAACTCTTTCCCTACACGACGCTCTTCCGATCTNNNCCAGCCTGGGAAATAGAGTG
AGCACACGTCTGAACTCTTTCCCTACACGACGCTCTTCCGATCTNNNCCAGCCTGGGAAATAGAGTG
GCACACGTCTGAACTCTTTCCCTACACGACGCTCTTCCGATCTNNNCCATCCATGTGTATTGTGTGTG
GCACACGTCTGAACTCTTTCCCTACACGACGCTCTTCCGATCTNNNCCATCCATGTGTATTGTGTGTG
GCACACGTCTGAACTCTTTCCCTACACGACGCTCTTCCGATCTNNNCCATCCATGTGTATTGTGTGTG
GCACACGTCTGAACTCTTTCCCTACACGACGCTCTTCCGATCTNNNCCATCCATGTGTATTGTGTGTG
GCACACGTCTGAACTCTTTCCCTACACGACGCTCTTCCGATCTNNNCTGGGATTCTGTTCTGAGGTTC
GCACACGTCTGAACTCTTTCCCTACACGACGCTCTTCCGATCTNNNCTGGGATTCTGTTCTGAGGTTC
GCACACGTCTGAACTCTTTCCCTACACGACGCTCTTCCGATCTNNNCTGGGATTCTGTTCTGAGGTTC
GCACACGTCTGAACTCTTTCCCTACACGACGCTCTTCCGATCTNNNCTGGGATTCTGTTCTGAGGTTC
AGCACACGTCTGAACTCTTTCCCTACACGACGCTCTTCCGATCTNNNGAGAATTGCTTGAACCCAGGA
AGCACACGTCTGAACTCTTTCCCTACACGACGCTCTTCCGATCTNNNGAGAATTGCTTGAACCCAGGA
AGCACACGTCTGAACTCTTTCCCTACACGACGCTCTTCCGATCTNNNGAGAATTGCTTGAACCCAGGA
AGCACACGTCTGAACTCTTTCCCTACACGACGCTCTTCCGATCTNNNGAGAATTGCTTGAACCCAGGA
GAGCACACGTCTGAACTCTTTCCCTACACGACGCTCTTCCGATCTNNNCATTGCATTGATATGATGGAT
GAGCACACGTCTGAACTCTTTCCCTACACGACGCTCTTCCGATCTNNNCATTGCATTGATATGATGGAT
GAGCACACGTCTGAACTCTTTCCCTACACGACGCTCTTCCGATCTNNNCATTGCATTGATATGATGGAT
GAGCACACGTCTGAACTCTTTCCCTACACGACGCTCTTCCGATCTNNNCATTGCATTGATATGATGGAT
GCACACGTCTGAACTCTTTCCCTACACGACGCTCTTCCGATCTNNNAGACAGATCATGACTCCAGATG
GCACACGTCTGAACTCTTTCCCTACACGACGCTCTTCCGATCTNNNAGACAGATCATGACTCCAGATG
GCACACGTCTGAACTCTTTCCCTACACGACGCTCTTCCGATCTNNNAGACAGATCATGACTCCAGATG
GCACACGTCTGAACTCTTTCCCTACACGACGCTCTTCCGATCTNNNAGACAGATCATGACTCCAGATG
GCACACGTCTGAACTCTTTCCCTACACGACGCTCTTCCGATCTNNNCTTGAAGGAACCCCCATGGAGC
GCACACGTCTGAACTCTTTCCCTACACGACGCTCTTCCGATCTNNNCTTGAAGGAACCCCCATGGAGC
GCACACGTCTGAACTCTTTCCCTACACGACGCTCTTCCGATCTNNNCTTGAAGGAACCCCCATGGAGC
GCACACGTCTGAACTCTTTCCCTACACGACGCTCTTCCGATCTNNNCTTGAAGGAACCCCCATGGAGC
GAGCACACGTCTGAACTCTTTCCCTACACGACGCTCTTCCGATCTNNNGTACCTGTAATCTCAGCTAC
GAGCACACGTCTGAACTCTTTCCCTACACGACGCTCTTCCGATCTNNNGTACCTGTAATCTCAGCTAC
GAGCACACGTCTGAACTCTTTCCCTACACGACGCTCTTCCGATCTNNNGTACCTGTAATCTCAGCTAC
GAGCACACGTCTGAACTCTTTCCCTACACGACGCTCTTCCGATCTNNNGTACCTGTAATCTCAGCTAC
GCACACGTCTGAACTCTTTCCCTACACGACGCTCTTCCGATCTNNNAGACTCTGTGGAAAGTATGCT
GCACACGTCTGAACTCTTTCCCTACACGACGCTCTTCCGATCTNNNAGACTCTGTGGAAAGTATGCT
GCACACGTCTGAACTCTTTCCCTACACGACGCTCTTCCGATCTNNNAGACTCTGTGGAAAGTATGCT
GCACACGTCTGAACTCTTTCCCTACACGACGCTCTTCCGATCTNNNAGACTCTGTGGAAAGTATGCT
GCACACGTCTGAACTCTTTCCCTACACGACGCTCTTCCGATCTNNNTCCCTCTTTTTGCTCTCTGCTAG
GCACACGTCTGAACTCTTTCCCTACACGACGCTCTTCCGATCTNNNTCCCTCTTTTTGCTCTCTGCTAG
GCACACGTCTGAACTCTTTCCCTACACGACGCTCTTCCGATCTNNNTCCCTCTTTTTGCTCTCTGCTAG
GCACACGTCTGAACTCTTTCCCTACACGACGCTCTTCCGATCTNNNTCCCTCTTTTTGCTCTCTGCTAG
GCACACGTCTGAACTCTTTCCCTACACGACGCTCTTCCGATCTNNNGTCAGTGTATGTAGTAACCATA
GCACACGTCTGAACTCTTTCCCTACACGACGCTCTTCCGATCTNNNGTCAGTGTATGTAGTAACCATA

GCACACGTCTGAACTCTTTCCCTACACGACGCTCTTCCGATCTNNNGTCAGTGTATGTAGTAACCATA
GCACACGTCTGAACTCTTTCCCTACACGACGCTCTTCCGATCTNNNGTCAGTGTATGTAGTAACCATA
AGCACACGTCTGAACTCTTTCCCTACACGACGCTCTTCCGATCTNNNGCTCATGCATGTAATCCCAACA
AGCACACGTCTGAACTCTTTCCCTACACGACGCTCTTCCGATCTNNNGCTCATGCATGTAATCCCAACA
AGCACACGTCTGAACTCTTTCCCTACACGACGCTCTTCCGATCTNNNGCTCATGCATGTAATCCCAACA
AGCACACGTCTGAACTCTTTCCCTACACGACGCTCTTCCGATCTNNNGCTCATGCATGTAATCCCAACA
GCACACGTCTGAACTCTTTCCCTACACGACGCTCTTCCGATCTNNNGCTACCTTGCATCTGGTTGTTTT
GCACACGTCTGAACTCTTTCCCTACACGACGCTCTTCCGATCTNNNGCTACCTTGCATCTGGTTGTTTT
GCACACGTCTGAACTCTTTCCCTACACGACGCTCTTCCGATCTNNNGCTACCTTGCATCTGGTTGTTTT
GCACACGTCTGAACTCTTTCCCTACACGACGCTCTTCCGATCTNNNGCTACCTTGCATCTGGTTGTTTT
AGCACACGTCTGAACTCTTTCCCTACACGACGCTCTTCCGATCTNNNAAACCATTACAGTAGCCAGTT
AGCACACGTCTGAACTCTTTCCCTACACGACGCTCTTCCGATCTNNNAAACCATTACAGTAGCCAGTT
AGCACACGTCTGAACTCTTTCCCTACACGACGCTCTTCCGATCTNNNAAACCATTACAGTAGCCAGTT
AGCACACGTCTGAACTCTTTCCCTACACGACGCTCTTCCGATCTNNNAAACCATTACAGTAGCCAGTT
CACACGTCTGAACTCTTTCCCTACACGACGCTCTTCCGATCTNNNAGTCCCTACCCTAACAGGATTCAC
CACACGTCTGAACTCTTTCCCTACACGACGCTCTTCCGATCTNNNAGTCCCTACCCTAACAGGATTCAC
CACACGTCTGAACTCTTTCCCTACACGACGCTCTTCCGATCTNNNAGTCCCTACCCTAACAGGATTCAC
CACACGTCTGAACTCTTTCCCTACACGACGCTCTTCCGATCTNNNAGTCCCTACCCTAACAGGATTCAC
GAGCACACGTCTGAACTCTTTCCCTACACGACGCTCTTCCGATCTNNNGAGTTCACACCAGTACACTCC
GAGCACACGTCTGAACTCTTTCCCTACACGACGCTCTTCCGATCTNNNGAGTTCACACCAGTACACTCC
GAGCACACGTCTGAACTCTTTCCCTACACGACGCTCTTCCGATCTNNNGAGTTCACACCAGTACACTCC
GAGCACACGTCTGAACTCTTTCCCTACACGACGCTCTTCCGATCTNNNGAGTTCACACCAGTACACTCC
GAGCACACGTCTGAACTCTTTCCCTACACGACGCTCTTCCGATCTNNNTGATCATTAGATCTGGGGT
GAGCACACGTCTGAACTCTTTCCCTACACGACGCTCTTCCGATCTNNNTGATCATTAGATCTGGGGT
GAGCACACGTCTGAACTCTTTCCCTACACGACGCTCTTCCGATCTNNNTGATCATTAGATCTGGGGT
GAGCACACGTCTGAACTCTTTCCCTACACGACGCTCTTCCGATCTNNNTGATCATTAGATCTGGGGT
AGCACACGTCTGAACTCTTTCCCTACACGACGCTCTTCCGATCTNNNATCCTGGGAAGTTGATGCTGT
AGCACACGTCTGAACTCTTTCCCTACACGACGCTCTTCCGATCTNNNATCCTGGGAAGTTGATGCTGT
AGCACACGTCTGAACTCTTTCCCTACACGACGCTCTTCCGATCTNNNATCCTGGGAAGTTGATGCTGT
AGCACACGTCTGAACTCTTTCCCTACACGACGCTCTTCCGATCTNNNATCCTGGGAAGTTGATGCTGT
AGCACACGTCTGAACTCTTTCCCTACACGACGCTCTTCCGATCTNNNAACGGACAGAAGTTGGAAAA
AGCACACGTCTGAACTCTTTCCCTACACGACGCTCTTCCGATCTNNNAACGGACAGAAGTTGGAAAA
AGCACACGTCTGAACTCTTTCCCTACACGACGCTCTTCCGATCTNNNAACGGACAGAAGTTGGAAAA
AGCACACGTCTGAACTCTTTCCCTACACGACGCTCTTCCGATCTNNNAACGGACAGAAGTTGGAAAA
AGCACACGTCTGAACTCTTTCCCTACACGACGCTCTTCCGATCTNNNCTGTAATCCCAGCTACTCGAG
AGCACACGTCTGAACTCTTTCCCTACACGACGCTCTTCCGATCTNNNCTGTAATCCCAGCTACTCGAG
AGCACACGTCTGAACTCTTTCCCTACACGACGCTCTTCCGATCTNNNCTGTAATCCCAGCTACTCGAG
AGCACACGTCTGAACTCTTTCCCTACACGACGCTCTTCCGATCTNNNCTGTAATCCCAGCTACTCGAG
AGCACACGTCTGAACTCTTTCCCTACACGACGCTCTTCCGATCTNNNCCTGTTGCTCTGGGACTGATTA
AGCACACGTCTGAACTCTTTCCCTACACGACGCTCTTCCGATCTNNNCCTGTTGCTCTGGGACTGATTA
AGCACACGTCTGAACTCTTTCCCTACACGACGCTCTTCCGATCTNNNCCTGTTGCTCTGGGACTGATTA
AGCACACGTCTGAACTCTTTCCCTACACGACGCTCTTCCGATCTNNNCCTGTTGCTCTGGGACTGATTA
CACACGTCTGAACTCTTTCCCTACACGACGCTCTTCCGATCTNNNAATGACTACAAAAAACTGTCAAT

GCACACGTCTGAACTCTTTCCCTACACGACGCTCTTCCGATCTNNNAATGACTACAAAAAACTGTCAAT

GCACACGTCTGAACTCTTTCCCTACACGACGCTCTTCCGATCTNNNAATGACTACAAAAAACTGTCAAT

GCACACGTCTGAACTCTTTCCCTACACGACGCTCTTCCGATCTNNNAATGACTACAAAAAACTGTCAAT

GAGCACACGTCTGAACTCTTTCCCTACACGACGCTCTTCCGATCTNNNTCTGGTCTCTTCCTTGACACA

GAGCACACGTCTGAACTCTTTCCCTACACGACGCTCTTCCGATCTNNNTCTGGTCTCTTCCTTGACACA

GAGCACACGTCTGAACTCTTTCCCTACACGACGCTCTTCCGATCTNNNTCTGGTCTCTTCCTTGACACA

GAGCACACGTCTGAACTCTTTCCCTACACGACGCTCTTCCGATCTNNNTCTGGTCTCTTCCTTGACACA

AGCACACGTCTGAACTCTTTCCCTACACGACGCTCTTCCGATCTNNNAGACATTCCAAAGGCTGATCA

AGCACACGTCTGAACTCTTTCCCTACACGACGCTCTTCCGATCTNNNAGACATTCCAAAGGCTGATCA

AGCACACGTCTGAACTCTTTCCCTACACGACGCTCTTCCGATCTNNNAGACATTCCAAAGGCTGATCA

AGCACACGTCTGAACTCTTTCCCTACACGACGCTCTTCCGATCTNNNAGACATTCCAAAGGCTGATCA

AGCACACGTCTGAACTCTTTCCCTACACGACGCTCTTCCGATCTNNNGAGGCAGGAGAATTGCTTGA

AGCACACGTCTGAACTCTTTCCCTACACGACGCTCTTCCGATCTNNNGAGGCAGGAGAATTGCTTGA

AGCACACGTCTGAACTCTTTCCCTACACGACGCTCTTCCGATCTNNNGAGGCAGGAGAATTGCTTGA

AGCACACGTCTGAACTCTTTCCCTACACGACGCTCTTCCGATCTNNNGAGGCAGGAGAATTGCTTGA

GAGCACACGTCTGAACTCTTTCCCTACACGACGCTCTTCCGATCTNNNCAGTTACCACCTGTGTCCCAA

GAGCACACGTCTGAACTCTTTCCCTACACGACGCTCTTCCGATCTNNNCAGTTACCACCTGTGTCCCAA

GAGCACACGTCTGAACTCTTTCCCTACACGACGCTCTTCCGATCTNNNCAGTTACCACCTGTGTCCCAA

GAGCACACGTCTGAACTCTTTCCCTACACGACGCTCTTCCGATCTNNNCAGTTACCACCTGTGTCCCAA

GAGCACACGTCTGAACTCTTTCCCTACACGACGCTCTTCCGATCTNNNCACGCCCGGCTAATTTTTGT

GAGCACACGTCTGAACTCTTTCCCTACACGACGCTCTTCCGATCTNNNCACGCCCGGCTAATTTTTGT

GAGCACACGTCTGAACTCTTTCCCTACACGACGCTCTTCCGATCTNNNCACGCCCGGCTAATTTTTGT

GAGCACACGTCTGAACTCTTTCCCTACACGACGCTCTTCCGATCTNNNCACGCCCGGCTAATTTTTGT

AGCACACGTCTGAACTCTTTCCCTACACGACGCTCTTCCGATCTNNNAGGGCAGACTCCATCTCAAAAA

AGCACACGTCTGAACTCTTTCCCTACACGACGCTCTTCCGATCTNNNAGGGCAGACTCCATCTCAAAAA

AGCACACGTCTGAACTCTTTCCCTACACGACGCTCTTCCGATCTNNNAGGGCAGACTCCATCTCAAAAA

AGCACACGTCTGAACTCTTTCCCTACACGACGCTCTTCCGATCTNNNAGGGCAGACTCCATCTCAAAAA

GAGCACACGTCTGAACTCTTTCCCTACACGACGCTCTTCCGATCTNNNCTGTGATCCCAGCTACTTGAG

GAGCACACGTCTGAACTCTTTCCCTACACGACGCTCTTCCGATCTNNNCTGTGATCCCAGCTACTTGAG

GAGCACACGTCTGAACTCTTTCCCTACACGACGCTCTTCCGATCTNNNCTGTGATCCCAGCTACTTGAG

GAGCACACGTCTGAACTCTTTCCCTACACGACGCTCTTCCGATCTNNNCTGTGATCCCAGCTACTTGAG

AGCACACGTCTGAACTCTTTCCCTACACGACGCTCTTCCGATCTNNNGATCGCTTGAGCCTAGGAAGT

AGCACACGTCTGAACTCTTTCCCTACACGACGCTCTTCCGATCTNNNGATCGCTTGAGCCTAGGAAGT

AGCACACGTCTGAACTCTTTCCCTACACGACGCTCTTCCGATCTNNNGATCGCTTGAGCCTAGGAAGT

AGCACACGTCTGAACTCTTTCCCTACACGACGCTCTTCCGATCTNNNGATCGCTTGAGCCTAGGAAGT

GAGCACACGTCTGAACTCTTTCCCTACACGACGCTCTTCCGATCTNNNTCTCGATCTCCTGACCTCATGA

GAGCACACGTCTGAACTCTTTCCCTACACGACGCTCTTCCGATCTNNNTCTCGATCTCCTGACCTCATGA

GAGCACACGTCTGAACTCTTTCCCTACACGACGCTCTTCCGATCTNNNTCTCGATCTCCTGACCTCATGA

GAGCACACGTCTGAACTCTTTCCCTACACGACGCTCTTCCGATCTNNNTCTCGATCTCCTGACCTCATGA

GAGCACACGTCTGAACTCTTTCCCTACACGACGCTCTTCCGATCTNNNGTAAATAGTCTCAGCTCCTCG

GAGCACACGTCTGAACTCTTTCCCTACACGACGCTCTTCCGATCTNNNGTAAATAGTCTCAGCTCCTCG

GAGCACACGTCTGAACTCTTTCCCTACACGACGCTCTTCCGATCTNNNGTAAATAGTCTCAGCTCCTCG

GAGCACACGTCTGAACTCTTTCCCTACACGACGCTCTTCCGATCTNNNGTAAATAGTCTCAGCTCCTCG

AGAGCACACGTCTGAACTCTTTCCCTACACGACGCTCTTCCGATCTNNNAGAATCACTTGAACCCAGGA
AGAGCACACGTCTGAACTCTTTCCCTACACGACGCTCTTCCGATCTNNNAGAATCACTTGAACCCAGGA
AGAGCACACGTCTGAACTCTTTCCCTACACGACGCTCTTCCGATCTNNNAGAATCACTTGAACCCAGGA
AGAGCACACGTCTGAACTCTTTCCCTACACGACGCTCTTCCGATCTNNNAGAATCACTTGAACCCAGGA
AGCACACGTCTGAACTCTTTCCCTACACGACGCTCTTCCGATCTNNNCACAGCCGAGAAAGAAATTCC
AGCACACGTCTGAACTCTTTCCCTACACGACGCTCTTCCGATCTNNNCACAGCCGAGAAAGAAATTCC
AGCACACGTCTGAACTCTTTCCCTACACGACGCTCTTCCGATCTNNNCACAGCCGAGAAAGAAATTCC
AGCACACGTCTGAACTCTTTCCCTACACGACGCTCTTCCGATCTNNNCACAGCCGAGAAAGAAATTCC
AGCACACGTCTGAACTCTTTCCCTACACGACGCTCTTCCGATCTNNNCCAAAATCCAAAGTGCTGGGAA
AGCACACGTCTGAACTCTTTCCCTACACGACGCTCTTCCGATCTNNNCCAAAATCCAAAGTGCTGGGAA
AGCACACGTCTGAACTCTTTCCCTACACGACGCTCTTCCGATCTNNNCCAAAATCCAAAGTGCTGGGAA
AGCACACGTCTGAACTCTTTCCCTACACGACGCTCTTCCGATCTNNNCCAAAATCCAAAGTGCTGGGAA
AGCACACGTCTGAACTCTTTCCCTACACGACGCTCTTCCGATCTNNNTTAGCATCCCAAGCAGAAAGGA
AGCACACGTCTGAACTCTTTCCCTACACGACGCTCTTCCGATCTNNNTTAGCATCCCAAGCAGAAAGGA
AGCACACGTCTGAACTCTTTCCCTACACGACGCTCTTCCGATCTNNNTTAGCATCCCAAGCAGAAAGGA
AGCACACGTCTGAACTCTTTCCCTACACGACGCTCTTCCGATCTNNNTTAGCATCCCAAGCAGAAAGGA
AGCACACGTCTGAACTCTTTCCCTACACGACGCTCTTCCGATCTNNNGAGAATCACTTGAACCCAGGA
AGCACACGTCTGAACTCTTTCCCTACACGACGCTCTTCCGATCTNNNGAGAATCACTTGAACCCAGGA
AGCACACGTCTGAACTCTTTCCCTACACGACGCTCTTCCGATCTNNNGAGAATCACTTGAACCCAGGA
AGCACACGTCTGAACTCTTTCCCTACACGACGCTCTTCCGATCTNNNGAGAATCACTTGAACCCAGGA
GAGCACACGTCTGAACTCTTTCCCTACACGACGCTCTTCCGATCTNNNGTTTGTGGAGAGAGGGAGA
GAGCACACGTCTGAACTCTTTCCCTACACGACGCTCTTCCGATCTNNNGTTTGTGGAGAGAGGGAGA
GAGCACACGTCTGAACTCTTTCCCTACACGACGCTCTTCCGATCTNNNGTTTGTGGAGAGAGGGAGA
GAGCACACGTCTGAACTCTTTCCCTACACGACGCTCTTCCGATCTNNNGTTTGTGGAGAGAGGGAGA
GCACACGTCTGAACTCTTTCCCTACACGACGCTCTTCCGATCTNNNCGTCCTCAACCTTAGCAAAACACC
GCACACGTCTGAACTCTTTCCCTACACGACGCTCTTCCGATCTNNNCGTCCTCAACCTTAGCAAAACACC
GCACACGTCTGAACTCTTTCCCTACACGACGCTCTTCCGATCTNNNCGTCCTCAACCTTAGCAAAACACC
GCACACGTCTGAACTCTTTCCCTACACGACGCTCTTCCGATCTNNNCGTCCTCAACCTTAGCAAAACACC
GAGCACACGTCTGAACTCTTTCCCTACACGACGCTCTTCCGATCTNNNTGTAATCCCCACTACTCAGGA
GAGCACACGTCTGAACTCTTTCCCTACACGACGCTCTTCCGATCTNNNTGTAATCCCCACTACTCAGGA
GAGCACACGTCTGAACTCTTTCCCTACACGACGCTCTTCCGATCTNNNTGTAATCCCCACTACTCAGGA
GAGCACACGTCTGAACTCTTTCCCTACACGACGCTCTTCCGATCTNNNTGTAATCCCCACTACTCAGGA
GAGCACACGTCTGAACTCTTTCCCTACACGACGCTCTTCCGATCTNNNCTTGAGCCCAGGAAGTCGAG
GAGCACACGTCTGAACTCTTTCCCTACACGACGCTCTTCCGATCTNNNCTTGAGCCCAGGAAGTCGAG
GAGCACACGTCTGAACTCTTTCCCTACACGACGCTCTTCCGATCTNNNCTTGAGCCCAGGAAGTCGAG
GAGCACACGTCTGAACTCTTTCCCTACACGACGCTCTTCCGATCTNNNCTTGAGCCCAGGAAGTCGAG
GAGCACACGTCTGAACTCTTTCCCTACACGACGCTCTTCCGATCTNNNTTGAGGGTGATTCCTGCACAT
GAGCACACGTCTGAACTCTTTCCCTACACGACGCTCTTCCGATCTNNNTTGAGGGTGATTCCTGCACAT
GAGCACACGTCTGAACTCTTTCCCTACACGACGCTCTTCCGATCTNNNTTGAGGGTGATTCCTGCACAT
GAGCACACGTCTGAACTCTTTCCCTACACGACGCTCTTCCGATCTNNNTTGAGGGTGATTCCTGCACAT
ACACGTCTGAACTCTTTCCCTACACGACGCTCTTCCGATCTNNNCCCTTCTTGATGCAGAAAATGGTCA
ACACGTCTGAACTCTTTCCCTACACGACGCTCTTCCGATCTNNNCCCTTCTTGATGCAGAAAATGGTCA
ACACGTCTGAACTCTTTCCCTACACGACGCTCTTCCGATCTNNNCCCTTCTTGATGCAGAAAATGGTCA

ACACGTCTGAACTCTTTCCCTACACGACGCTCTTCCGATCTNNNCCCTTCTTGATGCAGAAAATGGTCA

AGCACACGTCTGAACTCTTTCCCTACACGACGCTCTTCCGATCTNNNCCTTGGCTCCATACTCTAAACCA

AGCACACGTCTGAACTCTTTCCCTACACGACGCTCTTCCGATCTNNNCCTTGGCTCCATACTCTAAACCA

AGCACACGTCTGAACTCTTTCCCTACACGACGCTCTTCCGATCTNNNCCTTGGCTCCATACTCTAAACCA

AGCACACGTCTGAACTCTTTCCCTACACGACGCTCTTCCGATCTNNNCCTTGGCTCCATACTCTAAACCA

GAGCACACGTCTGAACTCTTTCCCTACACGACGCTCTTCCGATCTNNNCACCAAGATCTCCATCCTGGA

GAGCACACGTCTGAACTCTTTCCCTACACGACGCTCTTCCGATCTNNNCACCAAGATCTCCATCCTGGA

GAGCACACGTCTGAACTCTTTCCCTACACGACGCTCTTCCGATCTNNNCACCAAGATCTCCATCCTGGA

GAGCACACGTCTGAACTCTTTCCCTACACGACGCTCTTCCGATCTNNNCACCAAGATCTCCATCCTGGA

GAGCACACGTCTGAACTCTTTCCCTACACGACGCTCTTCCGATCTNNNGGTCATTCCAGCAATTCCCATT

GAGCACACGTCTGAACTCTTTCCCTACACGACGCTCTTCCGATCTNNNGGTCATTCCAGCAATTCCCATT

GAGCACACGTCTGAACTCTTTCCCTACACGACGCTCTTCCGATCTNNNGGTCATTCCAGCAATTCCCATT

GAGCACACGTCTGAACTCTTTCCCTACACGACGCTCTTCCGATCTNNNGGTCATTCCAGCAATTCCCATT

AGCACACGTCTGAACTCTTTCCCTACACGACGCTCTTCCGATCTNNNGCAGGGATGATCTTTCCTGTG

AGCACACGTCTGAACTCTTTCCCTACACGACGCTCTTCCGATCTNNNGCAGGGATGATCTTTCCTGTG

AGCACACGTCTGAACTCTTTCCCTACACGACGCTCTTCCGATCTNNNGCAGGGATGATCTTTCCTGTG

AGCACACGTCTGAACTCTTTCCCTACACGACGCTCTTCCGATCTNNNGCAGGGATGATCTTTCCTGTG

GCACACGTCTGAACTCTTTCCCTACACGACGCTCTTCCGATCTNNNGTGCCTGCAAATTCATACATGGT

GCACACGTCTGAACTCTTTCCCTACACGACGCTCTTCCGATCTNNNGTGCCTGCAAATTCATACATGGT

GCACACGTCTGAACTCTTTCCCTACACGACGCTCTTCCGATCTNNNGTGCCTGCAAATTCATACATGGT

GCACACGTCTGAACTCTTTCCCTACACGACGCTCTTCCGATCTNNNGTGCCTGCAAATTCATACATGGT

AGCACACGTCTGAACTCTTTCCCTACACGACGCTCTTCCGATCTNNNTTTAGACTGTGGTCCCAATCAC

AGCACACGTCTGAACTCTTTCCCTACACGACGCTCTTCCGATCTNNNTTTAGACTGTGGTCCCAATCAC

AGCACACGTCTGAACTCTTTCCCTACACGACGCTCTTCCGATCTNNNTTTAGACTGTGGTCCCAATCAC

AGCACACGTCTGAACTCTTTCCCTACACGACGCTCTTCCGATCTNNNTTTAGACTGTGGTCCCAATCAC

GCACACGTCTGAACTCTTTCCCTACACGACGCTCTTCCGATCTNNNGCAGCAAGATAGGTAGTAAGGC

GCACACGTCTGAACTCTTTCCCTACACGACGCTCTTCCGATCTNNNGCAGCAAGATAGGTAGTAAGGC

GCACACGTCTGAACTCTTTCCCTACACGACGCTCTTCCGATCTNNNGCAGCAAGATAGGTAGTAAGGC

GCACACGTCTGAACTCTTTCCCTACACGACGCTCTTCCGATCTNNNGCAGCAAGATAGGTAGTAAGGC

GAGCACACGTCTGAACTCTTTCCCTACACGACGCTCTTCCGATCTNNNCACTTGAGGTCAGGAGTTCA

GAGCACACGTCTGAACTCTTTCCCTACACGACGCTCTTCCGATCTNNNCACTTGAGGTCAGGAGTTCA

GAGCACACGTCTGAACTCTTTCCCTACACGACGCTCTTCCGATCTNNNCACTTGAGGTCAGGAGTTCA

GAGCACACGTCTGAACTCTTTCCCTACACGACGCTCTTCCGATCTNNNCACTTGAGGTCAGGAGTTCA

GAGCACACGTCTGAACTCTTTCCCTACACGACGCTCTTCCGATCTNNNGTGAGTCAGAGTGTACAGAG

GAGCACACGTCTGAACTCTTTCCCTACACGACGCTCTTCCGATCTNNNGTGAGTCAGAGTGTACAGAG

GAGCACACGTCTGAACTCTTTCCCTACACGACGCTCTTCCGATCTNNNGTGAGTCAGAGTGTACAGAG

GAGCACACGTCTGAACTCTTTCCCTACACGACGCTCTTCCGATCTNNNGTGAGTCAGAGTGTACAGAG

AGCACACGTCTGAACTCTTTCCCTACACGACGCTCTTCCGATCTNNNTTGCCTGATCTTTTCTCCTCTGG

AGCACACGTCTGAACTCTTTCCCTACACGACGCTCTTCCGATCTNNNTTGCCTGATCTTTTCTCCTCTGG

AGCACACGTCTGAACTCTTTCCCTACACGACGCTCTTCCGATCTNNNTTGCCTGATCTTTTCTCCTCTGG

AGCACACGTCTGAACTCTTTCCCTACACGACGCTCTTCCGATCTNNNTTGCCTGATCTTTTCTCCTCTGG

GCACACGTCTGAACTCTTTCCCTACACGACGCTCTTCCGATCTNNNCAGGTTTAGAAGGGGTGTCTGG

GCACACGTCTGAACTCTTTCCCTACACGACGCTCTTCCGATCTNNNCAGGTTTAGAAGGGGTGTCTGG

GCACACGTCTGAACTCTTTCCCTACACGACGCTCTTCCGATCTNNNCAGGTTTAGAAGGGGTGTCTGG
GCACACGTCTGAACTCTTTCCCTACACGACGCTCTTCCGATCTNNNCAGGTTTAGAAGGGGTGTCTGG
GAGCACACGTCTGAACTCTTTCCCTACACGACGCTCTTCCGATCTNNNCCCCAAACCCATACCCTTCTATC
GAGCACACGTCTGAACTCTTTCCCTACACGACGCTCTTCCGATCTNNNCCCCAAACCCATACCCTTCTATC
GAGCACACGTCTGAACTCTTTCCCTACACGACGCTCTTCCGATCTNNNCCCCAAACCCATACCCTTCTATC
GAGCACACGTCTGAACTCTTTCCCTACACGACGCTCTTCCGATCTNNNCCCCAAACCCATACCCTTCTATC
AAGAGCACACGTCTGAACTCTTTCCCTACACGACGCTCTTCCGATCTNNNCTGTACAAATGCAGGTTGC
AAGAGCACACGTCTGAACTCTTTCCCTACACGACGCTCTTCCGATCTNNNCTGTACAAATGCAGGTTGC
AAGAGCACACGTCTGAACTCTTTCCCTACACGACGCTCTTCCGATCTNNNCTGTACAAATGCAGGTTGC
AAGAGCACACGTCTGAACTCTTTCCCTACACGACGCTCTTCCGATCTNNNCTGTACAAATGCAGGTTGC
AGAGCACACGTCTGAACTCTTTCCCTACACGACGCTCTTCCGATCTNNNAAAGGGGACATTACCACTGA
AGAGCACACGTCTGAACTCTTTCCCTACACGACGCTCTTCCGATCTNNNAAAGGGGACATTACCACTGA
AGAGCACACGTCTGAACTCTTTCCCTACACGACGCTCTTCCGATCTNNNAAAGGGGACATTACCACTGA
AGAGCACACGTCTGAACTCTTTCCCTACACGACGCTCTTCCGATCTNNNAAAGGGGACATTACCACTGA
AGAGCACACGTCTGAACTCTTTCCCTACACGACGCTCTTCCGATCTNNNTTCCATCTCATCTGCTTGGTA
AGAGCACACGTCTGAACTCTTTCCCTACACGACGCTCTTCCGATCTNNNTTCCATCTCATCTGCTTGGTA
AGAGCACACGTCTGAACTCTTTCCCTACACGACGCTCTTCCGATCTNNNTTCCATCTCATCTGCTTGGTA
AGAGCACACGTCTGAACTCTTTCCCTACACGACGCTCTTCCGATCTNNNTTCCATCTCATCTGCTTGGTA
AGCACACGTCTGAACTCTTTCCCTACACGACGCTCTTCCGATCTNNNGGTGACAAGGTGAGACTCTTT
AGCACACGTCTGAACTCTTTCCCTACACGACGCTCTTCCGATCTNNNGGTGACAAGGTGAGACTCTTT
AGCACACGTCTGAACTCTTTCCCTACACGACGCTCTTCCGATCTNNNGGTGACAAGGTGAGACTCTTT
AGCACACGTCTGAACTCTTTCCCTACACGACGCTCTTCCGATCTNNNGGTGACAAGGTGAGACTCTTT
AGCACACGTCTGAACTCTTTCCCTACACGACGCTCTTCCGATCTNNNGCTGAGTTTACTACCTGAACACC
AGCACACGTCTGAACTCTTTCCCTACACGACGCTCTTCCGATCTNNNGCTGAGTTTACTACCTGAACACC
AGCACACGTCTGAACTCTTTCCCTACACGACGCTCTTCCGATCTNNNGCTGAGTTTACTACCTGAACACC
AGCACACGTCTGAACTCTTTCCCTACACGACGCTCTTCCGATCTNNNGCTGAGTTTACTACCTGAACACC
AGAGCACACGTCTGAACTCTTTCCCTACACGACGCTCTTCCGATCTNNNGATGCTAGAGGAGGAGGCA
AGAGCACACGTCTGAACTCTTTCCCTACACGACGCTCTTCCGATCTNNNGATGCTAGAGGAGGAGGCA
AGAGCACACGTCTGAACTCTTTCCCTACACGACGCTCTTCCGATCTNNNGATGCTAGAGGAGGAGGCA
AGAGCACACGTCTGAACTCTTTCCCTACACGACGCTCTTCCGATCTNNNGATGCTAGAGGAGGAGGCA
AGCACACGTCTGAACTCTTTCCCTACACGACGCTCTTCCGATCTNNNGAGATACTGGCTCATCATACTAC
AGCACACGTCTGAACTCTTTCCCTACACGACGCTCTTCCGATCTNNNGAGATACTGGCTCATCATACTAC
AGCACACGTCTGAACTCTTTCCCTACACGACGCTCTTCCGATCTNNNGAGATACTGGCTCATCATACTAC
AGCACACGTCTGAACTCTTTCCCTACACGACGCTCTTCCGATCTNNNGAGATACTGGCTCATCATACTAC
AGCACACGTCTGAACTCTTTCCCTACACGACGCTCTTCCGATCTNNNTAGGATAGCCAGTAGTAGCCTC
AGCACACGTCTGAACTCTTTCCCTACACGACGCTCTTCCGATCTNNNTAGGATAGCCAGTAGTAGCCTC
AGCACACGTCTGAACTCTTTCCCTACACGACGCTCTTCCGATCTNNNTAGGATAGCCAGTAGTAGCCTC
AGCACACGTCTGAACTCTTTCCCTACACGACGCTCTTCCGATCTNNNTAGGATAGCCAGTAGTAGCCTC
AGCACACGTCTGAACTCTTTCCCTACACGACGCTCTTCCGATCTNNNTAGCTGGCAAATAAACCTGGTC
AGCACACGTCTGAACTCTTTCCCTACACGACGCTCTTCCGATCTNNNTAGCTGGCAAATAAACCTGGTC
AGCACACGTCTGAACTCTTTCCCTACACGACGCTCTTCCGATCTNNNTAGCTGGCAAATAAACCTGGTC
AGCACACGTCTGAACTCTTTCCCTACACGACGCTCTTCCGATCTNNNTAGCTGGCAAATAAACCTGGTC
GAGCACACGTCTGAACTCTTTCCCTACACGACGCTCTTCCGATCTNNNGAGTGCTGCTTCACATCCTACA

GAGCACACGTCTGAACTCTTTCCCTACACGACGCTCTTCCGATCTNNNGAGTGCTGCTTCACATCCTACA
GAGCACACGTCTGAACTCTTTCCCTACACGACGCTCTTCCGATCTNNNGAGTGCTGCTTCACATCCTACA
GAGCACACGTCTGAACTCTTTCCCTACACGACGCTCTTCCGATCTNNNGAGTGCTGCTTCACATCCTACA
AGCACACGTCTGAACTCTTTCCCTACACGACGCTCTTCCGATCTNNNGGGAGGTATCATGCAATCAT
AGCACACGTCTGAACTCTTTCCCTACACGACGCTCTTCCGATCTNNNGGGAGGTATCATGCAATCAT
AGCACACGTCTGAACTCTTTCCCTACACGACGCTCTTCCGATCTNNNGGGAGGTATCATGCAATCAT
AGCACACGTCTGAACTCTTTCCCTACACGACGCTCTTCCGATCTNNNGGGAGGTATCATGCAATCAT
GCACACGTCTGAACTCTTTCCCTACACGACGCTCTTCCGATCTNNNCACTGCCTAAATCATCCCTTCCTCA
GCACACGTCTGAACTCTTTCCCTACACGACGCTCTTCCGATCTNNNCACTGCCTAAATCATCCCTTCCTCA
GCACACGTCTGAACTCTTTCCCTACACGACGCTCTTCCGATCTNNNCACTGCCTAAATCATCCCTTCCTCA
GCACACGTCTGAACTCTTTCCCTACACGACGCTCTTCCGATCTNNNCACTGCCTAAATCATCCCTTCCTCA
GAGCACACGTCTGAACTCTTTCCCTACACGACGCTCTTCCGATCTNNNCTCTATCTCCAGACCTTGTGAG
GAGCACACGTCTGAACTCTTTCCCTACACGACGCTCTTCCGATCTNNNCTCTATCTCCAGACCTTGTGAG
GAGCACACGTCTGAACTCTTTCCCTACACGACGCTCTTCCGATCTNNNCTCTATCTCCAGACCTTGTGAG
GAGCACACGTCTGAACTCTTTCCCTACACGACGCTCTTCCGATCTNNNCTCTATCTCCAGACCTTGTGAG
AGCACACGTCTGAACTCTTTCCCTACACGACGCTCTTCCGATCTNNNCAATTCCAGTGCAGTGTATGTG
AGCACACGTCTGAACTCTTTCCCTACACGACGCTCTTCCGATCTNNNCAATTCCAGTGCAGTGTATGTG
AGCACACGTCTGAACTCTTTCCCTACACGACGCTCTTCCGATCTNNNCAATTCCAGTGCAGTGTATGTG
AGCACACGTCTGAACTCTTTCCCTACACGACGCTCTTCCGATCTNNNCAATTCCAGTGCAGTGTATGTG
GAGCACACGTCTGAACTCTTTCCCTACACGACGCTCTTCCGATCTNNNGAAATAGGGTCTTTGCAGATG
GAGCACACGTCTGAACTCTTTCCCTACACGACGCTCTTCCGATCTNNNGAAATAGGGTCTTTGCAGATG
GAGCACACGTCTGAACTCTTTCCCTACACGACGCTCTTCCGATCTNNNGAAATAGGGTCTTTGCAGATG
GAGCACACGTCTGAACTCTTTCCCTACACGACGCTCTTCCGATCTNNNGAAATAGGGTCTTTGCAGATG
GAGCACACGTCTGAACTCTTTCCCTACACGACGCTCTTCCGATCTNNNAAAGTGCTGGGATTACAGGC
GAGCACACGTCTGAACTCTTTCCCTACACGACGCTCTTCCGATCTNNNAAAGTGCTGGGATTACAGGC
GAGCACACGTCTGAACTCTTTCCCTACACGACGCTCTTCCGATCTNNNAAAGTGCTGGGATTACAGGC
GAGCACACGTCTGAACTCTTTCCCTACACGACGCTCTTCCGATCTNNNAAAGTGCTGGGATTACAGGC
GAGCACACGTCTGAACTCTTTCCCTACACGACGCTCTTCCGATCTNNNCTCGATCTCTTGACCTCGTGAT
GAGCACACGTCTGAACTCTTTCCCTACACGACGCTCTTCCGATCTNNNCTCGATCTCTTGACCTCGTGAT
GAGCACACGTCTGAACTCTTTCCCTACACGACGCTCTTCCGATCTNNNCTCGATCTCTTGACCTCGTGAT
GAGCACACGTCTGAACTCTTTCCCTACACGACGCTCTTCCGATCTNNNCTCGATCTCTTGACCTCGTGAT
AGCACACGTCTGAACTCTTTCCCTACACGACGCTCTTCCGATCTNNNCCCTTAGCCACTGAGTTTGTCTT
AGCACACGTCTGAACTCTTTCCCTACACGACGCTCTTCCGATCTNNNCCCTTAGCCACTGAGTTTGTCTT
AGCACACGTCTGAACTCTTTCCCTACACGACGCTCTTCCGATCTNNNCCCTTAGCCACTGAGTTTGTCTT
AGCACACGTCTGAACTCTTTCCCTACACGACGCTCTTCCGATCTNNNCCCTTAGCCACTGAGTTTGTCTT
AGCACACGTCTGAACTCTTTCCCTACACGACGCTCTTCCGATCTNNNCTGAGACAGGAGATCGCTTGA
AGCACACGTCTGAACTCTTTCCCTACACGACGCTCTTCCGATCTNNNCTGAGACAGGAGATCGCTTGA
AGCACACGTCTGAACTCTTTCCCTACACGACGCTCTTCCGATCTNNNCTGAGACAGGAGATCGCTTGA
AGCACACGTCTGAACTCTTTCCCTACACGACGCTCTTCCGATCTNNNCTGAGACAGGAGATCGCTTGA
AGCACACGTCTGAACTCTTTCCCTACACGACGCTCTTCCGATCTNNNTGCCAGTAATTCCAGCATTTTG
AGCACACGTCTGAACTCTTTCCCTACACGACGCTCTTCCGATCTNNNTGCCAGTAATTCCAGCATTTTG
AGCACACGTCTGAACTCTTTCCCTACACGACGCTCTTCCGATCTNNNTGCCAGTAATTCCAGCATTTTG
AGCACACGTCTGAACTCTTTCCCTACACGACGCTCTTCCGATCTNNNTGCCAGTAATTCCAGCATTTTG

AGCACACGTCTGAACTCTTTCCCTACACGACGCTCTTCCGATCTNNNACAGGACTGCCTAACTTATCAG
AGCACACGTCTGAACTCTTTCCCTACACGACGCTCTTCCGATCTNNNACAGGACTGCCTAACTTATCAG
AGCACACGTCTGAACTCTTTCCCTACACGACGCTCTTCCGATCTNNNACAGGACTGCCTAACTTATCAG
AGCACACGTCTGAACTCTTTCCCTACACGACGCTCTTCCGATCTNNNACAGGACTGCCTAACTTATCAG
AGCACACGTCTGAACTCTTTCCCTACACGACGCTCTTCCGATCTNNNTCAGTATGCACGTCTTCACTTC
AGCACACGTCTGAACTCTTTCCCTACACGACGCTCTTCCGATCTNNNTCAGTATGCACGTCTTCACTTC
AGCACACGTCTGAACTCTTTCCCTACACGACGCTCTTCCGATCTNNNTCAGTATGCACGTCTTCACTTC
AGCACACGTCTGAACTCTTTCCCTACACGACGCTCTTCCGATCTNNNTCAGTATGCACGTCTTCACTTC
GCACACGTCTGAACTCTTTCCCTACACGACGCTCTTCCGATCTNNNACGATTCTCTGATCTCCTGCTTCC
GCACACGTCTGAACTCTTTCCCTACACGACGCTCTTCCGATCTNNNACGATTCTCTGATCTCCTGCTTCC
GCACACGTCTGAACTCTTTCCCTACACGACGCTCTTCCGATCTNNNACGATTCTCTGATCTCCTGCTTCC
GCACACGTCTGAACTCTTTCCCTACACGACGCTCTTCCGATCTNNNACGATTCTCTGATCTCCTGCTTCC
AGCACACGTCTGAACTCTTTCCCTACACGACGCTCTTCCGATCTNNNCTGTAATCCCAGCTACTCAGGA
AGCACACGTCTGAACTCTTTCCCTACACGACGCTCTTCCGATCTNNNCTGTAATCCCAGCTACTCAGGA
AGCACACGTCTGAACTCTTTCCCTACACGACGCTCTTCCGATCTNNNCTGTAATCCCAGCTACTCAGGA
AGCACACGTCTGAACTCTTTCCCTACACGACGCTCTTCCGATCTNNNCTGTAATCCCAGCTACTCAGGA
AGCACACGTCTGAACTCTTTCCCTACACGACGCTCTTCCGATCTNNNAGTTGATGATGTTTGTGAAGC
AGCACACGTCTGAACTCTTTCCCTACACGACGCTCTTCCGATCTNNNAGTTGATGATGTTTGTGAAGC
AGCACACGTCTGAACTCTTTCCCTACACGACGCTCTTCCGATCTNNNAGTTGATGATGTTTGTGAAGC
AGCACACGTCTGAACTCTTTCCCTACACGACGCTCTTCCGATCTNNNAGTTGATGATGTTTGTGAAGC
AGCACACGTCTGAACTCTTTCCCTACACGACGCTCTTCCGATCTNNNAGTGACCTTAGCAGTGTCCAA
AGCACACGTCTGAACTCTTTCCCTACACGACGCTCTTCCGATCTNNNAGTGACCTTAGCAGTGTCCAA
AGCACACGTCTGAACTCTTTCCCTACACGACGCTCTTCCGATCTNNNAGTGACCTTAGCAGTGTCCAA
AGCACACGTCTGAACTCTTTCCCTACACGACGCTCTTCCGATCTNNNAGTGACCTTAGCAGTGTCCAA
GCACACGTCTGAACTCTTTCCCTACACGACGCTCTTCCGATCTNNNAAACCATATTTAGCCAGTAGTCTC
GCACACGTCTGAACTCTTTCCCTACACGACGCTCTTCCGATCTNNNAAACCATATTTAGCCAGTAGTCTC
GCACACGTCTGAACTCTTTCCCTACACGACGCTCTTCCGATCTNNNAAACCATATTTAGCCAGTAGTCTC
GCACACGTCTGAACTCTTTCCCTACACGACGCTCTTCCGATCTNNNAAACCATATTTAGCCAGTAGTCTC
GAGCACACGTCTGAACTCTTTCCCTACACGACGCTCTTCCGATCTNNNCAATCGCCTTTTCCAAGAACTG
GAGCACACGTCTGAACTCTTTCCCTACACGACGCTCTTCCGATCTNNNCAATCGCCTTTTCCAAGAACTG
GAGCACACGTCTGAACTCTTTCCCTACACGACGCTCTTCCGATCTNNNCAATCGCCTTTTCCAAGAACTG
GAGCACACGTCTGAACTCTTTCCCTACACGACGCTCTTCCGATCTNNNCAATCGCCTTTTCCAAGAACTG
AGCACACGTCTGAACTCTTTCCCTACACGACGCTCTTCCGATCTNNNAGACCTAAGACAACAGGAACA
AGCACACGTCTGAACTCTTTCCCTACACGACGCTCTTCCGATCTNNNAGACCTAAGACAACAGGAACA
AGCACACGTCTGAACTCTTTCCCTACACGACGCTCTTCCGATCTNNNAGACCTAAGACAACAGGAACA
AGCACACGTCTGAACTCTTTCCCTACACGACGCTCTTCCGATCTNNNAGACCTAAGACAACAGGAACA
GAGCACACGTCTGAACTCTTTCCCTACACGACGCTCTTCCGATCTNNNCAACAAGCTGTGATCACACC
GAGCACACGTCTGAACTCTTTCCCTACACGACGCTCTTCCGATCTNNNCAACAAGCTGTGATCACACC
GAGCACACGTCTGAACTCTTTCCCTACACGACGCTCTTCCGATCTNNNCAACAAGCTGTGATCACACC
GAGCACACGTCTGAACTCTTTCCCTACACGACGCTCTTCCGATCTNNNCAACAAGCTGTGATCACACC
AGCACACGTCTGAACTCTTTCCCTACACGACGCTCTTCCGATCTNNNTCAGAAGGAAACCACCTACTTC
AGCACACGTCTGAACTCTTTCCCTACACGACGCTCTTCCGATCTNNNTCAGAAGGAAACCACCTACTTC
AGCACACGTCTGAACTCTTTCCCTACACGACGCTCTTCCGATCTNNNTCAGAAGGAAACCACCTACTTC

AGCACACGTCTGAACTCTTTCCCTACACGACGCTCTTCCGATCTNNNTCAGAAGGAAACCACCTACTTC
AGCACACGTCTGAACTCTTTCCCTACACGACGCTCTTCCGATCTNNNGGTGTGAATGCTGTATCCACA
AGCACACGTCTGAACTCTTTCCCTACACGACGCTCTTCCGATCTNNNGGTGTGAATGCTGTATCCACA
AGCACACGTCTGAACTCTTTCCCTACACGACGCTCTTCCGATCTNNNGGTGTGAATGCTGTATCCACA
AGCACACGTCTGAACTCTTTCCCTACACGACGCTCTTCCGATCTNNNGGTGTGAATGCTGTATCCACA
AGCACACGTCTGAACTCTTTCCCTACACGACGCTCTTCCGATCTNNNCTGAGGCAGGAGAATCACTTG
AGCACACGTCTGAACTCTTTCCCTACACGACGCTCTTCCGATCTNNNCTGAGGCAGGAGAATCACTTG
AGCACACGTCTGAACTCTTTCCCTACACGACGCTCTTCCGATCTNNNCTGAGGCAGGAGAATCACTTG
AGCACACGTCTGAACTCTTTCCCTACACGACGCTCTTCCGATCTNNNCTGAGGCAGGAGAATCACTTG
GAGCACACGTCTGAACTCTTTCCCTACACGACGCTCTTCCGATCTNNNAAGGAACATAGGCATCTGG
GAGCACACGTCTGAACTCTTTCCCTACACGACGCTCTTCCGATCTNNNAAGGAACATAGGCATCTGG
GAGCACACGTCTGAACTCTTTCCCTACACGACGCTCTTCCGATCTNNNAAGGAACATAGGCATCTGG
GAGCACACGTCTGAACTCTTTCCCTACACGACGCTCTTCCGATCTNNNAAGGAACATAGGCATCTGG
GCACACGTCTGAACTCTTTCCCTACACGACGCTCTTCCGATCTNNNAAGAAAGGCATCCAGAATTGG
GCACACGTCTGAACTCTTTCCCTACACGACGCTCTTCCGATCTNNNAAGAAAGGCATCCAGAATTGG
GCACACGTCTGAACTCTTTCCCTACACGACGCTCTTCCGATCTNNNAAGAAAGGCATCCAGAATTGG
GCACACGTCTGAACTCTTTCCCTACACGACGCTCTTCCGATCTNNNAAGAAAGGCATCCAGAATTGG
AGCACACGTCTGAACTCTTTCCCTACACGACGCTCTTCCGATCTNNNAAATCTCCAATCCCCTGACAGT
AGCACACGTCTGAACTCTTTCCCTACACGACGCTCTTCCGATCTNNNAAATCTCCAATCCCCTGACAGT
AGCACACGTCTGAACTCTTTCCCTACACGACGCTCTTCCGATCTNNNAAATCTCCAATCCCCTGACAGT
AGCACACGTCTGAACTCTTTCCCTACACGACGCTCTTCCGATCTNNNAAATCTCCAATCCCCTGACAGT
GAGCACACGTCTGAACTCTTTCCCTACACGACGCTCTTCCGATCTNNNCCACCTTCCATGTTCAAGCAAT
GAGCACACGTCTGAACTCTTTCCCTACACGACGCTCTTCCGATCTNNNCCACCTTCCATGTTCAAGCAAT
GAGCACACGTCTGAACTCTTTCCCTACACGACGCTCTTCCGATCTNNNCCACCTTCCATGTTCAAGCAAT
GAGCACACGTCTGAACTCTTTCCCTACACGACGCTCTTCCGATCTNNNCCACCTTCCATGTTCAAGCAAT
GAGCACACGTCTGAACTCTTTCCCTACACGACGCTCTTCCGATCTNNNTGGGAGCCTTACTGTGAAAAT
GAGCACACGTCTGAACTCTTTCCCTACACGACGCTCTTCCGATCTNNNTGGGAGCCTTACTGTGAAAAT
GAGCACACGTCTGAACTCTTTCCCTACACGACGCTCTTCCGATCTNNNTGGGAGCCTTACTGTGAAAAT
GAGCACACGTCTGAACTCTTTCCCTACACGACGCTCTTCCGATCTNNNTGGGAGCCTTACTGTGAAAAT
GCACACGTCTGAACTCTTTCCCTACACGACGCTCTTCCGATCTNNNTGAGAATCACTTGAACCAGGGA
GCACACGTCTGAACTCTTTCCCTACACGACGCTCTTCCGATCTNNNTGAGAATCACTTGAACCAGGGA
GCACACGTCTGAACTCTTTCCCTACACGACGCTCTTCCGATCTNNNTGAGAATCACTTGAACCAGGGA
GCACACGTCTGAACTCTTTCCCTACACGACGCTCTTCCGATCTNNNTGAGAATCACTTGAACCAGGGA
AGCACACGTCTGAACTCTTTCCCTACACGACGCTCTTCCGATCTNNNCTTTCCAAACTCTCTCCGCTCTCC
AGCACACGTCTGAACTCTTTCCCTACACGACGCTCTTCCGATCTNNNCTTTCCAAACTCTCTCCGCTCTCC
AGCACACGTCTGAACTCTTTCCCTACACGACGCTCTTCCGATCTNNNCTTTCCAAACTCTCTCCGCTCTCC
AGCACACGTCTGAACTCTTTCCCTACACGACGCTCTTCCGATCTNNNCTTTCCAAACTCTCTCCGCTCTCC
GAGCACACGTCTGAACTCTTTCCCTACACGACGCTCTTCCGATCTNNNCACACTACACTTCTGTGGGAA
GAGCACACGTCTGAACTCTTTCCCTACACGACGCTCTTCCGATCTNNNCACACTACACTTCTGTGGGAA
GAGCACACGTCTGAACTCTTTCCCTACACGACGCTCTTCCGATCTNNNCACACTACACTTCTGTGGGAA
GAGCACACGTCTGAACTCTTTCCCTACACGACGCTCTTCCGATCTNNNCACACTACACTTCTGTGGGAA
GAGCACACGTCTGAACTCTTTCCCTACACGACGCTCTTCCGATCTNNNTATAGTGAGGCCCTGTCTGT
GAGCACACGTCTGAACTCTTTCCCTACACGACGCTCTTCCGATCTNNNTATAGTGAGGCCCTGTCTGT

GAGCACACGTCTGAACTCTTTCCCTACACGACGCTCTTCCGATCTNNNTATAGTGAGGCCCTGTCTGTA

GAGCACACGTCTGAACTCTTTCCCTACACGACGCTCTTCCGATCTNNNTATAGTGAGGCCCTGTCTGTA

GCACACGTCTGAACTCTTTCCCTACACGACGCTCTTCCGATCTNNNAAGATCTGGGCAACACAGTCAGC

GCACACGTCTGAACTCTTTCCCTACACGACGCTCTTCCGATCTNNNAAGATCTGGGCAACACAGTCAGC

GCACACGTCTGAACTCTTTCCCTACACGACGCTCTTCCGATCTNNNAAGATCTGGGCAACACAGTCAGC

GCACACGTCTGAACTCTTTCCCTACACGACGCTCTTCCGATCTNNNAAGATCTGGGCAACACAGTCAGC

GCACACGTCTGAACTCTTTCCCTACACGACGCTCTTCCGATCTNNNAGACTTACAGGAACTGCAGCCTA

GCACACGTCTGAACTCTTTCCCTACACGACGCTCTTCCGATCTNNNAGACTTACAGGAACTGCAGCCTA

GCACACGTCTGAACTCTTTCCCTACACGACGCTCTTCCGATCTNNNAGACTTACAGGAACTGCAGCCTA

GCACACGTCTGAACTCTTTCCCTACACGACGCTCTTCCGATCTNNNAGACTTACAGGAACTGCAGCCTA

AGAGCACACGTCTGAACTCTTTCCCTACACGACGCTCTTCCGATCTNNNGAGAATCACTTGAACCCGGG

AGAGCACACGTCTGAACTCTTTCCCTACACGACGCTCTTCCGATCTNNNGAGAATCACTTGAACCCGGG

AGAGCACACGTCTGAACTCTTTCCCTACACGACGCTCTTCCGATCTNNNGAGAATCACTTGAACCCGGG

AGAGCACACGTCTGAACTCTTTCCCTACACGACGCTCTTCCGATCTNNNGAGAATCACTTGAACCCGGG

AGCACACGTCTGAACTCTTTCCCTACACGACGCTCTTCCGATCTNNNCCTCAGCCTCCCAAATAGCTAAA

AGCACACGTCTGAACTCTTTCCCTACACGACGCTCTTCCGATCTNNNCCTCAGCCTCCCAAATAGCTAAA

AGCACACGTCTGAACTCTTTCCCTACACGACGCTCTTCCGATCTNNNCCTCAGCCTCCCAAATAGCTAAA

AGCACACGTCTGAACTCTTTCCCTACACGACGCTCTTCCGATCTNNNCCTCAGCCTCCCAAATAGCTAAA

GAGCACACGTCTGAACTCTTTCCCTACACGACGCTCTTCCGATCTNNNTGCCATGTAGAAAAGGCAGTT

GAGCACACGTCTGAACTCTTTCCCTACACGACGCTCTTCCGATCTNNNTGCCATGTAGAAAAGGCAGTT

GAGCACACGTCTGAACTCTTTCCCTACACGACGCTCTTCCGATCTNNNTGCCATGTAGAAAAGGCAGTT

GAGCACACGTCTGAACTCTTTCCCTACACGACGCTCTTCCGATCTNNNTGCCATGTAGAAAAGGCAGTT

AGCACACGTCTGAACTCTTTCCCTACACGACGCTCTTCCGATCTNNNCCCCTGACTTTACGGCCACGCATG

AGCACACGTCTGAACTCTTTCCCTACACGACGCTCTTCCGATCTNNNCCCCTGACTTTACGGCCACGCATG

AGCACACGTCTGAACTCTTTCCCTACACGACGCTCTTCCGATCTNNNCCCCTGACTTTACGGCCACGCATG

AGCACACGTCTGAACTCTTTCCCTACACGACGCTCTTCCGATCTNNNCCCCTGACTTTACGGCCACGCATG

GAGCACACGTCTGAACTCTTTCCCTACACGACGCTCTTCCGATCTNNNAACTCTGAGGTTCATGCAATCC

GAGCACACGTCTGAACTCTTTCCCTACACGACGCTCTTCCGATCTNNNAACTCTGAGGTTCATGCAATCC

GAGCACACGTCTGAACTCTTTCCCTACACGACGCTCTTCCGATCTNNNAACTCTGAGGTTCATGCAATCC

GAGCACACGTCTGAACTCTTTCCCTACACGACGCTCTTCCGATCTNNNAACTCTGAGGTTCATGCAATCC

AGCACACGTCTGAACTCTTTCCCTACACGACGCTCTTCCGATCTNNNAAAATTTAGGATGGTTTTCTTG

AGCACACGTCTGAACTCTTTCCCTACACGACGCTCTTCCGATCTNNNAAAATTTAGGATGGTTTTCTTG

AGCACACGTCTGAACTCTTTCCCTACACGACGCTCTTCCGATCTNNNAAAATTTAGGATGGTTTTCTTG

AGCACACGTCTGAACTCTTTCCCTACACGACGCTCTTCCGATCTNNNAAAATTTAGGATGGTTTTCTTG

AGCACACGTCTGAACTCTTTCCCTACACGACGCTCTTCCGATCTNNNTTTAGTCAGGTTCCTCTGAGTCC

AGCACACGTCTGAACTCTTTCCCTACACGACGCTCTTCCGATCTNNNTTTAGTCAGGTTCCTCTGAGTCC

AGCACACGTCTGAACTCTTTCCCTACACGACGCTCTTCCGATCTNNNTTTAGTCAGGTTCCTCTGAGTCC

AGCACACGTCTGAACTCTTTCCCTACACGACGCTCTTCCGATCTNNNTTTAGTCAGGTTCCTCTGAGTCC

GAGCACACGTCTGAACTCTTTCCCTACACGACGCTCTTCCGATCTNNNGTGAAACCCCGTCTCTACAAAA

GAGCACACGTCTGAACTCTTTCCCTACACGACGCTCTTCCGATCTNNNGTGAAACCCCGTCTCTACAAAA

GAGCACACGTCTGAACTCTTTCCCTACACGACGCTCTTCCGATCTNNNGTGAAACCCCGTCTCTACAAAA

GAGCACACGTCTGAACTCTTTCCCTACACGACGCTCTTCCGATCTNNNGTGAAACCCCGTCTCTACAAAA

AGCACACGTCTGAACTCTTTCCCTACACGACGCTCTTCCGATCTNNNGTAAATTCTGCCCAGTCCAAAC

AGCACACGTCTGAACTCTTTCCCTACACGACGCTCTTCCGATCTNNNGTAAATTCTGCCCAGTCCAAAC
AGCACACGTCTGAACTCTTTCCCTACACGACGCTCTTCCGATCTNNNGTAAATTCTGCCCAGTCCAAAC
AGCACACGTCTGAACTCTTTCCCTACACGACGCTCTTCCGATCTNNNGTAAATTCTGCCCAGTCCAAAC
GAGCACACGTCTGAACTCTTTCCCTACACGACGCTCTTCCGATCTNNNGTGAGCTGAGATTGCACCATT
GAGCACACGTCTGAACTCTTTCCCTACACGACGCTCTTCCGATCTNNNGTGAGCTGAGATTGCACCATT
GAGCACACGTCTGAACTCTTTCCCTACACGACGCTCTTCCGATCTNNNGTGAGCTGAGATTGCACCATT
GAGCACACGTCTGAACTCTTTCCCTACACGACGCTCTTCCGATCTNNNGTGAGCTGAGATTGCACCATT
AGCACACGTCTGAACTCTTTCCCTACACGACGCTCTTCCGATCTNNNAAGCCTATAGTCCCAGCTACTT
AGCACACGTCTGAACTCTTTCCCTACACGACGCTCTTCCGATCTNNNAAGCCTATAGTCCCAGCTACTT
AGCACACGTCTGAACTCTTTCCCTACACGACGCTCTTCCGATCTNNNAAGCCTATAGTCCCAGCTACTT
AGCACACGTCTGAACTCTTTCCCTACACGACGCTCTTCCGATCTNNNAAGCCTATAGTCCCAGCTACTT
AGCACACGTCTGAACTCTTTCCCTACACGACGCTCTTCCGATCTNNNAGAACACACAGCAGATCATCCT
AGCACACGTCTGAACTCTTTCCCTACACGACGCTCTTCCGATCTNNNAGAACACACAGCAGATCATCCT
AGCACACGTCTGAACTCTTTCCCTACACGACGCTCTTCCGATCTNNNAGAACACACAGCAGATCATCCT
AGCACACGTCTGAACTCTTTCCCTACACGACGCTCTTCCGATCTNNNAGAACACACAGCAGATCATCCT
AAGAGCACACGTCTGAACTCTTTCCCTACACGACGCTCTTCCGATCTNNNGGACATCAAACCCCTTCTCT
AAGAGCACACGTCTGAACTCTTTCCCTACACGACGCTCTTCCGATCTNNNGGACATCAAACCCCTTCTCT
AAGAGCACACGTCTGAACTCTTTCCCTACACGACGCTCTTCCGATCTNNNGGACATCAAACCCCTTCTCT
AAGAGCACACGTCTGAACTCTTTCCCTACACGACGCTCTTCCGATCTNNNGGACATCAAACCCCTTCTCT
AGCACACGTCTGAACTCTTTCCCTACACGACGCTCTTCCGATCTNNNTGCCTTGTAGCCACAGATAACAT
AGCACACGTCTGAACTCTTTCCCTACACGACGCTCTTCCGATCTNNNTGCCTTGTAGCCACAGATAACAT
AGCACACGTCTGAACTCTTTCCCTACACGACGCTCTTCCGATCTNNNTGCCTTGTAGCCACAGATAACAT
AGCACACGTCTGAACTCTTTCCCTACACGACGCTCTTCCGATCTNNNTGCCTTGTAGCCACAGATAACAT
AGCACACGTCTGAACTCTTTCCCTACACGACGCTCTTCCGATCTNNNTCTACCTGCCTAGTGGACTTTCC
AGCACACGTCTGAACTCTTTCCCTACACGACGCTCTTCCGATCTNNNTCTACCTGCCTAGTGGACTTTCC
AGCACACGTCTGAACTCTTTCCCTACACGACGCTCTTCCGATCTNNNTCTACCTGCCTAGTGGACTTTCC
AGCACACGTCTGAACTCTTTCCCTACACGACGCTCTTCCGATCTNNNTCTACCTGCCTAGTGGACTTTCC
GCACACGTCTGAACTCTTTCCCTACACGACGCTCTTCCGATCTNNNAAAACCACTCCAGTAAAGTAGCC
GCACACGTCTGAACTCTTTCCCTACACGACGCTCTTCCGATCTNNNAAAACCACTCCAGTAAAGTAGCC
GCACACGTCTGAACTCTTTCCCTACACGACGCTCTTCCGATCTNNNAAAACCACTCCAGTAAAGTAGCC
GCACACGTCTGAACTCTTTCCCTACACGACGCTCTTCCGATCTNNNAAAACCACTCCAGTAAAGTAGCC
GAGCACACGTCTGAACTCTTTCCCTACACGACGCTCTTCCGATCTNNNCACCCGGCTTGTATCGAAAAT
GAGCACACGTCTGAACTCTTTCCCTACACGACGCTCTTCCGATCTNNNCACCCGGCTTGTATCGAAAAT
GAGCACACGTCTGAACTCTTTCCCTACACGACGCTCTTCCGATCTNNNCACCCGGCTTGTATCGAAAAT
GAGCACACGTCTGAACTCTTTCCCTACACGACGCTCTTCCGATCTNNNCACCCGGCTTGTATCGAAAAT
GAGCACACGTCTGAACTCTTTCCCTACACGACGCTCTTCCGATCTNNNGTGACCCTTAGTTCCCAACTAG
GAGCACACGTCTGAACTCTTTCCCTACACGACGCTCTTCCGATCTNNNGTGACCCTTAGTTCCCAACTAG
GAGCACACGTCTGAACTCTTTCCCTACACGACGCTCTTCCGATCTNNNGTGACCCTTAGTTCCCAACTAG
GAGCACACGTCTGAACTCTTTCCCTACACGACGCTCTTCCGATCTNNNGTGACCCTTAGTTCCCAACTAG
GCACACGTCTGAACTCTTTCCCTACACGACGCTCTTCCGATCTNNNAGGCATAAGCAGCAACTGACAAT
GCACACGTCTGAACTCTTTCCCTACACGACGCTCTTCCGATCTNNNAGGCATAAGCAGCAACTGACAAT
GCACACGTCTGAACTCTTTCCCTACACGACGCTCTTCCGATCTNNNAGGCATAAGCAGCAACTGACAAT
GCACACGTCTGAACTCTTTCCCTACACGACGCTCTTCCGATCTNNNAGGCATAAGCAGCAACTGACAAT

AGCACACGTCTGAACTCTTTCCCTACACGACGCTCTTCCGATCTNNNAGAACAAATGTTGAGGCATCA
AGCACACGTCTGAACTCTTTCCCTACACGACGCTCTTCCGATCTNNNAGAACAAATGTTGAGGCATCA
AGCACACGTCTGAACTCTTTCCCTACACGACGCTCTTCCGATCTNNNAGAACAAATGTTGAGGCATCA
AGCACACGTCTGAACTCTTTCCCTACACGACGCTCTTCCGATCTNNNAGAACAAATGTTGAGGCATCA
AGAGCACACGTCTGAACTCTTTCCCTACACGACGCTCTTCCGATCTNNNAATTCCTGGACACATACACC
AGAGCACACGTCTGAACTCTTTCCCTACACGACGCTCTTCCGATCTNNNAATTCCTGGACACATACACC
AGAGCACACGTCTGAACTCTTTCCCTACACGACGCTCTTCCGATCTNNNAATTCCTGGACACATACACC
AGAGCACACGTCTGAACTCTTTCCCTACACGACGCTCTTCCGATCTNNNAATTCCTGGACACATACACC
AGCACACGTCTGAACTCTTTCCCTACACGACGCTCTTCCGATCTNNNAGTCACACTCTCCTAAGCAACT
AGCACACGTCTGAACTCTTTCCCTACACGACGCTCTTCCGATCTNNNAGTCACACTCTCCTAAGCAACT
AGCACACGTCTGAACTCTTTCCCTACACGACGCTCTTCCGATCTNNNAGTCACACTCTCCTAAGCAACT
AGCACACGTCTGAACTCTTTCCCTACACGACGCTCTTCCGATCTNNNAGTCACACTCTCCTAAGCAACT
GAGCACACGTCTGAACTCTTTCCCTACACGACGCTCTTCCGATCTNNNCCCAGCCTGGTCAATAATAGT
GAGCACACGTCTGAACTCTTTCCCTACACGACGCTCTTCCGATCTNNNCCCAGCCTGGTCAATAATAGT
GAGCACACGTCTGAACTCTTTCCCTACACGACGCTCTTCCGATCTNNNCCCAGCCTGGTCAATAATAGT
GAGCACACGTCTGAACTCTTTCCCTACACGACGCTCTTCCGATCTNNNCCCAGCCTGGTCAATAATAGT
CACACGTCTGAACTCTTTCCCTACACGACGCTCTTCCGATCTNNNTTTGAATTTATGGTATTCTGTGCC
CACACGTCTGAACTCTTTCCCTACACGACGCTCTTCCGATCTNNNTTTGAATTTATGGTATTCTGTGCC
CACACGTCTGAACTCTTTCCCTACACGACGCTCTTCCGATCTNNNTTTGAATTTATGGTATTCTGTGCC
CACACGTCTGAACTCTTTCCCTACACGACGCTCTTCCGATCTNNNTTTGAATTTATGGTATTCTGTGCC
AGCACACGTCTGAACTCTTTCCCTACACGACGCTCTTCCGATCTNNNTATACAACGACTCTTGAGCCCA
AGCACACGTCTGAACTCTTTCCCTACACGACGCTCTTCCGATCTNNNTATACAACGACTCTTGAGCCCA
AGCACACGTCTGAACTCTTTCCCTACACGACGCTCTTCCGATCTNNNTATACAACGACTCTTGAGCCCA
AGCACACGTCTGAACTCTTTCCCTACACGACGCTCTTCCGATCTNNNTATACAACGACTCTTGAGCCCA
GAGCACACGTCTGAACTCTTTCCCTACACGACGCTCTTCCGATCTNNNAAAGAGTAATCTCCAGGTTG
GAGCACACGTCTGAACTCTTTCCCTACACGACGCTCTTCCGATCTNNNAAAGAGTAATCTCCAGGTTG
GAGCACACGTCTGAACTCTTTCCCTACACGACGCTCTTCCGATCTNNNAAAGAGTAATCTCCAGGTTG
GAGCACACGTCTGAACTCTTTCCCTACACGACGCTCTTCCGATCTNNNAAAGAGTAATCTCCAGGTTG
GCACACGTCTGAACTCTTTCCCTACACGACGCTCTTCCGATCTNNNGTGGATCACTTGAACTCAGGAG
GCACACGTCTGAACTCTTTCCCTACACGACGCTCTTCCGATCTNNNGTGGATCACTTGAACTCAGGAG
GCACACGTCTGAACTCTTTCCCTACACGACGCTCTTCCGATCTNNNGTGGATCACTTGAACTCAGGAG
GCACACGTCTGAACTCTTTCCCTACACGACGCTCTTCCGATCTNNNGTGGATCACTTGAACTCAGGAG
AGCACACGTCTGAACTCTTTCCCTACACGACGCTCTTCCGATCTNNNATCCTGATGTACCCCTGCAAAA
AGCACACGTCTGAACTCTTTCCCTACACGACGCTCTTCCGATCTNNNATCCTGATGTACCCCTGCAAAA
AGCACACGTCTGAACTCTTTCCCTACACGACGCTCTTCCGATCTNNNATCCTGATGTACCCCTGCAAAA
AGCACACGTCTGAACTCTTTCCCTACACGACGCTCTTCCGATCTNNNATCCTGATGTACCCCTGCAAAA
GCACACGTCTGAACTCTTTCCCTACACGACGCTCTTCCGATCTNNNCATGAGAATCGCTTGAACCAGTG
GCACACGTCTGAACTCTTTCCCTACACGACGCTCTTCCGATCTNNNCATGAGAATCGCTTGAACCAGTG
GCACACGTCTGAACTCTTTCCCTACACGACGCTCTTCCGATCTNNNCATGAGAATCGCTTGAACCAGTG
GCACACGTCTGAACTCTTTCCCTACACGACGCTCTTCCGATCTNNNCATGAGAATCGCTTGAACCAGTG
GAGCACACGTCTGAACTCTTTCCCTACACGACGCTCTTCCGATCTNNNGGTCCTTACTTGGTTAGATGC
GAGCACACGTCTGAACTCTTTCCCTACACGACGCTCTTCCGATCTNNNGGTCCTTACTTGGTTAGATGC
GAGCACACGTCTGAACTCTTTCCCTACACGACGCTCTTCCGATCTNNNGGTCCTTACTTGGTTAGATGC

AGCACACGTCTGAACTCTTTCCCTACACGACGCTCTTCCGATCTNNNGGTCCTTACTTGGTTAGATGC

GCACACGTCTGAACTCTTTCCCTACACGACGCTCTTCCGATCTNNNAGTAAGGTCTTTTCAGCATTTG

GCACACGTCTGAACTCTTTCCCTACACGACGCTCTTCCGATCTNNNAGTAAGGTCTTTTCAGCATTTG

GCACACGTCTGAACTCTTTCCCTACACGACGCTCTTCCGATCTNNNAGTAAGGTCTTTTCAGCATTTG

GCACACGTCTGAACTCTTTCCCTACACGACGCTCTTCCGATCTNNNAGTAAGGTCTTTTCAGCATTTG

AGCACACGTCTGAACTCTTTCCCTACACGACGCTCTTCCGATCTNNNTACTGGGAAGTGAGGAGCCC

AGCACACGTCTGAACTCTTTCCCTACACGACGCTCTTCCGATCTNNNTACTGGGAAGTGAGGAGCCC

AGCACACGTCTGAACTCTTTCCCTACACGACGCTCTTCCGATCTNNNTACTGGGAAGTGAGGAGCCC

AGCACACGTCTGAACTCTTTCCCTACACGACGCTCTTCCGATCTNNNTACTGGGAAGTGAGGAGCCC

AGCACACGTCTGAACTCTTTCCCTACACGACGCTCTTCCGATCTNNNTGGCCTGACCTTGTTTTTGTTT

AGCACACGTCTGAACTCTTTCCCTACACGACGCTCTTCCGATCTNNNTGGCCTGACCTTGTTTTTGTTT

AGCACACGTCTGAACTCTTTCCCTACACGACGCTCTTCCGATCTNNNTGGCCTGACCTTGTTTTTGTTT

AGCACACGTCTGAACTCTTTCCCTACACGACGCTCTTCCGATCTNNNTGGCCTGACCTTGTTTTTGTTT

CACACGTCTGAACTCTTTCCCTACACGACGCTCTTCCGATCTNNNTGAAGTCTTTTAGTTGAAATGCG

CACACGTCTGAACTCTTTCCCTACACGACGCTCTTCCGATCTNNNTGAAGTCTTTTAGTTGAAATGCG

CACACGTCTGAACTCTTTCCCTACACGACGCTCTTCCGATCTNNNTGAAGTCTTTTAGTTGAAATGCG

CACACGTCTGAACTCTTTCCCTACACGACGCTCTTCCGATCTNNNTGAAGTCTTTTAGTTGAAATGCG

GCACACGTCTGAACTCTTTCCCTACACGACGCTCTTCCGATCTNNNTCATTCCCTTTTGTGGCTGAGT

GCACACGTCTGAACTCTTTCCCTACACGACGCTCTTCCGATCTNNNTCATTCCCTTTTGTGGCTGAGT

GCACACGTCTGAACTCTTTCCCTACACGACGCTCTTCCGATCTNNNTCATTCCCTTTTGTGGCTGAGT

GCACACGTCTGAACTCTTTCCCTACACGACGCTCTTCCGATCTNNNTCATTCCCTTTTGTGGCTGAGT

AGCACACGTCTGAACTCTTTCCCTACACGACGCTCTTCCGATCTNNNAGAGCCCTACTTTGGGCTACA

AGCACACGTCTGAACTCTTTCCCTACACGACGCTCTTCCGATCTNNNAGAGCCCTACTTTGGGCTACA

AGCACACGTCTGAACTCTTTCCCTACACGACGCTCTTCCGATCTNNNAGAGCCCTACTTTGGGCTACA

AGCACACGTCTGAACTCTTTCCCTACACGACGCTCTTCCGATCTNNNAGAGCCCTACTTTGGGCTACA

GCACACGTCTGAACTCTTTCCCTACACGACGCTCTTCCGATCTNNNGGAATCTGCCCTGTGTCGTCCAT

GCACACGTCTGAACTCTTTCCCTACACGACGCTCTTCCGATCTNNNGGAATCTGCCCTGTGTCGTCCAT

GCACACGTCTGAACTCTTTCCCTACACGACGCTCTTCCGATCTNNNGGAATCTGCCCTGTGTCGTCCAT

GCACACGTCTGAACTCTTTCCCTACACGACGCTCTTCCGATCTNNNGGAATCTGCCCTGTGTCGTCCAT

CACGTCTGAACTCTTTCCCTACACGACGCTCTTCCGATCTNNNGAGTTAGATATGGAATTGGGTAGC

CACGTCTGAACTCTTTCCCTACACGACGCTCTTCCGATCTNNNGAGTTAGATATGGAATTGGGTAGC

CACGTCTGAACTCTTTCCCTACACGACGCTCTTCCGATCTNNNGAGTTAGATATGGAATTGGGTAGC

CACGTCTGAACTCTTTCCCTACACGACGCTCTTCCGATCTNNNGAGTTAGATATGGAATTGGGTAGC

GCACACGTCTGAACTCTTTCCCTACACGACGCTCTTCCGATCTNNNATACAGAGGCTACATTCGTGGT

GCACACGTCTGAACTCTTTCCCTACACGACGCTCTTCCGATCTNNNATACAGAGGCTACATTCGTGGT

GCACACGTCTGAACTCTTTCCCTACACGACGCTCTTCCGATCTNNNATACAGAGGCTACATTCGTGGT

GCACACGTCTGAACTCTTTCCCTACACGACGCTCTTCCGATCTNNNATACAGAGGCTACATTCGTGGT

GAGCACACGTCTGAACTCTTTCCCTACACGACGCTCTTCCGATCTNNNCGGATCATGAGGTCAAGAG

GAGCACACGTCTGAACTCTTTCCCTACACGACGCTCTTCCGATCTNNNCGGATCATGAGGTCAAGAG

GAGCACACGTCTGAACTCTTTCCCTACACGACGCTCTTCCGATCTNNNCGGATCATGAGGTCAAGAG

GAGCACACGTCTGAACTCTTTCCCTACACGACGCTCTTCCGATCTNNNCGGATCATGAGGTCAAGAG

AGCACACGTCTGAACTCTTTCCCTACACGACGCTCTTCCGATCTNNNGCTAGCAGAGGGTGTACTAT

AGCACACGTCTGAACTCTTTCCCTACACGACGCTCTTCCGATCTNNNGCTAGCAGAGGGTGTACTAT

GAGCACACGTCTGAACTCTTTCCCTACACGACGCTCTTCCGATCTNNNGCTAGCAGAGGGTGTACTATA
GAGCACACGTCTGAACTCTTTCCCTACACGACGCTCTTCCGATCTNNNGCTAGCAGAGGGTGTACTATA
AGCACACGTCTGAACTCTTTCCCTACACGACGCTCTTCCGATCTNNNTAACTCCTTCTGAATCTCCAGCC
AGCACACGTCTGAACTCTTTCCCTACACGACGCTCTTCCGATCTNNNTAACTCCTTCTGAATCTCCAGCC
AGCACACGTCTGAACTCTTTCCCTACACGACGCTCTTCCGATCTNNNTAACTCCTTCTGAATCTCCAGCC
AGCACACGTCTGAACTCTTTCCCTACACGACGCTCTTCCGATCTNNNTAACTCCTTCTGAATCTCCAGCC
GCACACGTCTGAACTCTTTCCCTACACGACGCTCTTCCGATCTNNNCAGACTCCACGTGTGTCTTTTTCC
GCACACGTCTGAACTCTTTCCCTACACGACGCTCTTCCGATCTNNNCAGACTCCACGTGTGTCTTTTTCC
GCACACGTCTGAACTCTTTCCCTACACGACGCTCTTCCGATCTNNNCAGACTCCACGTGTGTCTTTTTCC
GCACACGTCTGAACTCTTTCCCTACACGACGCTCTTCCGATCTNNNCAGACTCCACGTGTGTCTTTTTCC
GCACACGTCTGAACTCTTTCCCTACACGACGCTCTTCCGATCTNNNAGCTTTTGGTGAAGATCCAGGA
GCACACGTCTGAACTCTTTCCCTACACGACGCTCTTCCGATCTNNNAGCTTTTGGTGAAGATCCAGGA
GCACACGTCTGAACTCTTTCCCTACACGACGCTCTTCCGATCTNNNAGCTTTTGGTGAAGATCCAGGA
GCACACGTCTGAACTCTTTCCCTACACGACGCTCTTCCGATCTNNNAGCTTTTGGTGAAGATCCAGGA
AGCACACGTCTGAACTCTTTCCCTACACGACGCTCTTCCGATCTNNNCTGTAATCCCAGCTACTCAGGA
AGCACACGTCTGAACTCTTTCCCTACACGACGCTCTTCCGATCTNNNCTGTAATCCCAGCTACTCAGGA
AGCACACGTCTGAACTCTTTCCCTACACGACGCTCTTCCGATCTNNNCTGTAATCCCAGCTACTCAGGA
AGCACACGTCTGAACTCTTTCCCTACACGACGCTCTTCCGATCTNNNCTGTAATCCCAGCTACTCAGGA
GCACACGTCTGAACTCTTTCCCTACACGACGCTCTTCCGATCTNNNTAGATCGTCTGAGGTCAGGAGT
GCACACGTCTGAACTCTTTCCCTACACGACGCTCTTCCGATCTNNNTAGATCGTCTGAGGTCAGGAGT
GCACACGTCTGAACTCTTTCCCTACACGACGCTCTTCCGATCTNNNTAGATCGTCTGAGGTCAGGAGT
GCACACGTCTGAACTCTTTCCCTACACGACGCTCTTCCGATCTNNNTAGATCGTCTGAGGTCAGGAGT
AGCACACGTCTGAACTCTTTCCCTACACGACGCTCTTCCGATCTNNNTCTGGTGTTTTTTGTTTGTTTG
AGCACACGTCTGAACTCTTTCCCTACACGACGCTCTTCCGATCTNNNTCTGGTGTTTTTTGTTTGTTTG
AGCACACGTCTGAACTCTTTCCCTACACGACGCTCTTCCGATCTNNNTCTGGTGTTTTTTGTTTGTTTG
AGCACACGTCTGAACTCTTTCCCTACACGACGCTCTTCCGATCTNNNTCTGGTGTTTTTTGTTTGTTTG
GAGCACACGTCTGAACTCTTTCCCTACACGACGCTCTTCCGATCTNNNGGGGTGATGTGTGTACCAAA
GAGCACACGTCTGAACTCTTTCCCTACACGACGCTCTTCCGATCTNNNGGGGTGATGTGTGTACCAAA
GAGCACACGTCTGAACTCTTTCCCTACACGACGCTCTTCCGATCTNNNGGGGTGATGTGTGTACCAAA
GAGCACACGTCTGAACTCTTTCCCTACACGACGCTCTTCCGATCTNNNGGGGTGATGTGTGTACCAAA
GAGCACACGTCTGAACTCTTTCCCTACACGACGCTCTTCCGATCTNNNAACAAGAAGGACGAAGATCT
GAGCACACGTCTGAACTCTTTCCCTACACGACGCTCTTCCGATCTNNNAACAAGAAGGACGAAGATCT
GAGCACACGTCTGAACTCTTTCCCTACACGACGCTCTTCCGATCTNNNAACAAGAAGGACGAAGATCT
GAGCACACGTCTGAACTCTTTCCCTACACGACGCTCTTCCGATCTNNNAACAAGAAGGACGAAGATCT
ACACGTCTGAACTCTTTCCCTACACGACGCTCTTCCGATCTNNNCTCCTTGGTGAGGTCTCCCTACATCC
ACACGTCTGAACTCTTTCCCTACACGACGCTCTTCCGATCTNNNCTCCTTGGTGAGGTCTCCCTACATCC
ACACGTCTGAACTCTTTCCCTACACGACGCTCTTCCGATCTNNNCTCCTTGGTGAGGTCTCCCTACATCC
ACACGTCTGAACTCTTTCCCTACACGACGCTCTTCCGATCTNNNCTCCTTGGTGAGGTCTCCCTACATCC
GAGCACACGTCTGAACTCTTTCCCTACACGACGCTCTTCCGATCTNNNAGTGTGGTAACCTATTGCTGC
GAGCACACGTCTGAACTCTTTCCCTACACGACGCTCTTCCGATCTNNNAGTGTGGTAACCTATTGCTGC
GAGCACACGTCTGAACTCTTTCCCTACACGACGCTCTTCCGATCTNNNAGTGTGGTAACCTATTGCTGC
GAGCACACGTCTGAACTCTTTCCCTACACGACGCTCTTCCGATCTNNNAGTGTGGTAACCTATTGCTGC
AGCACACGTCTGAACTCTTTCCCTACACGACGCTCTTCCGATCTNNNTGGCTATTGTGGCTGTCATCTA

AGCACACGTCTGAACTCTTTCCCTACACGACGCTCTTCCGATCTNNNTGGCTATTGTGGCTGTCATCT
AGCACACGTCTGAACTCTTTCCCTACACGACGCTCTTCCGATCTNNNTGGCTATTGTGGCTGTCATCT
AGCACACGTCTGAACTCTTTCCCTACACGACGCTCTTCCGATCTNNNTGGCTATTGTGGCTGTCATCT
GCACACGTCTGAACTCTTTCCCTACACGACGCTCTTCCGATCTNNNTCTTTCTTTCCTTCCTCCCTTCTCA
GCACACGTCTGAACTCTTTCCCTACACGACGCTCTTCCGATCTNNNTCTTTCTTTCCTTCCTCCCTTCTCA
GCACACGTCTGAACTCTTTCCCTACACGACGCTCTTCCGATCTNNNTCTTTCTTTCCTTCCTCCCTTCTCA
GCACACGTCTGAACTCTTTCCCTACACGACGCTCTTCCGATCTNNNTCTTTCTTTCCTTCCTCCCTTCTCA
AGCACACGTCTGAACTCTTTCCCTACACGACGCTCTTCCGATCTNNNTACCTCTTCTGCTCCCTTTTTCC
AGCACACGTCTGAACTCTTTCCCTACACGACGCTCTTCCGATCTNNNTACCTCTTCTGCTCCCTTTTTCC
AGCACACGTCTGAACTCTTTCCCTACACGACGCTCTTCCGATCTNNNTACCTCTTCTGCTCCCTTTTTCC
AGCACACGTCTGAACTCTTTCCCTACACGACGCTCTTCCGATCTNNNTACCTCTTCTGCTCCCTTTTTCC
GAGCACACGTCTGAACTCTTTCCCTACACGACGCTCTTCCGATCTNNNGTGTGAAAATGGAAGAGG
GAGCACACGTCTGAACTCTTTCCCTACACGACGCTCTTCCGATCTNNNGTGTGAAAATGGAAGAGG
GAGCACACGTCTGAACTCTTTCCCTACACGACGCTCTTCCGATCTNNNGTGTGAAAATGGAAGAGG
GAGCACACGTCTGAACTCTTTCCCTACACGACGCTCTTCCGATCTNNNGTGTGAAAATGGAAGAGG
GCACACGTCTGAACTCTTTCCCTACACGACGCTCTTCCGATCTNNNTATGGACCATTCTAGCCACACA
GCACACGTCTGAACTCTTTCCCTACACGACGCTCTTCCGATCTNNNTATGGACCATTCTAGCCACACA
GCACACGTCTGAACTCTTTCCCTACACGACGCTCTTCCGATCTNNNTATGGACCATTCTAGCCACACA
GCACACGTCTGAACTCTTTCCCTACACGACGCTCTTCCGATCTNNNTATGGACCATTCTAGCCACACA
GCACACGTCTGAACTCTTTCCCTACACGACGCTCTTCCGATCTNNNAGGAAGCTGAGACAGGAGAAT
GCACACGTCTGAACTCTTTCCCTACACGACGCTCTTCCGATCTNNNAGGAAGCTGAGACAGGAGAAT
GCACACGTCTGAACTCTTTCCCTACACGACGCTCTTCCGATCTNNNAGGAAGCTGAGACAGGAGAAT
GCACACGTCTGAACTCTTTCCCTACACGACGCTCTTCCGATCTNNNAGGAAGCTGAGACAGGAGAAT
GCACACGTCTGAACTCTTTCCCTACACGACGCTCTTCCGATCTNNNCACTTGCTTTGAACACCAGTGT
GCACACGTCTGAACTCTTTCCCTACACGACGCTCTTCCGATCTNNNCACTTGCTTTGAACACCAGTGT
GCACACGTCTGAACTCTTTCCCTACACGACGCTCTTCCGATCTNNNCACTTGCTTTGAACACCAGTGT
GCACACGTCTGAACTCTTTCCCTACACGACGCTCTTCCGATCTNNNCACTTGCTTTGAACACCAGTGT
GCACACGTCTGAACTCTTTCCCTACACGACGCTCTTCCGATCTNNNCCTGGCCAGAACTTCCAATACTA
GCACACGTCTGAACTCTTTCCCTACACGACGCTCTTCCGATCTNNNCCTGGCCAGAACTTCCAATACTA
GCACACGTCTGAACTCTTTCCCTACACGACGCTCTTCCGATCTNNNCCTGGCCAGAACTTCCAATACTA
GCACACGTCTGAACTCTTTCCCTACACGACGCTCTTCCGATCTNNNCCTGGCCAGAACTTCCAATACTA
AGCACACGTCTGAACTCTTTCCCTACACGACGCTCTTCCGATCTNNNGACCCTTACAAACACTCGTTGC
AGCACACGTCTGAACTCTTTCCCTACACGACGCTCTTCCGATCTNNNGACCCTTACAAACACTCGTTGC
AGCACACGTCTGAACTCTTTCCCTACACGACGCTCTTCCGATCTNNNGACCCTTACAAACACTCGTTGC
AGCACACGTCTGAACTCTTTCCCTACACGACGCTCTTCCGATCTNNNGACCCTTACAAACACTCGTTGC
AGCACACGTCTGAACTCTTTCCCTACACGACGCTCTTCCGATCTNNNGAAGCTTTAGGCACTGGTGGT
AGCACACGTCTGAACTCTTTCCCTACACGACGCTCTTCCGATCTNNNGAAGCTTTAGGCACTGGTGGT
AGCACACGTCTGAACTCTTTCCCTACACGACGCTCTTCCGATCTNNNGAAGCTTTAGGCACTGGTGGT
AGCACACGTCTGAACTCTTTCCCTACACGACGCTCTTCCGATCTNNNGAAGCTTTAGGCACTGGTGGT
GAGCACACGTCTGAACTCTTTCCCTACACGACGCTCTTCCGATCTNNNCACCTGTAATCCCAGCTACTC
GAGCACACGTCTGAACTCTTTCCCTACACGACGCTCTTCCGATCTNNNCACCTGTAATCCCAGCTACTC
GAGCACACGTCTGAACTCTTTCCCTACACGACGCTCTTCCGATCTNNNCACCTGTAATCCCAGCTACTC
GAGCACACGTCTGAACTCTTTCCCTACACGACGCTCTTCCGATCTNNNCACCTGTAATCCCAGCTACTC

GCACACGTCTGAACTCTTTCCCTACACGACGCTCTTCCGATCTNNNTTTTGTTGCTGCTTCAAAGACCA

GCACACGTCTGAACTCTTTCCCTACACGACGCTCTTCCGATCTNNNTTTTGTTGCTGCTTCAAAGACCA

GCACACGTCTGAACTCTTTCCCTACACGACGCTCTTCCGATCTNNNTTTTGTTGCTGCTTCAAAGACCA

GCACACGTCTGAACTCTTTCCCTACACGACGCTCTTCCGATCTNNNTTTTGTTGCTGCTTCAAAGACCA

AGCACACGTCTGAACTCTTTCCCTACACGACGCTCTTCCGATCTNNNGGGAGGTGTGAGGTAAAAGT

AGCACACGTCTGAACTCTTTCCCTACACGACGCTCTTCCGATCTNNNGGGAGGTGTGAGGTAAAAGT

AGCACACGTCTGAACTCTTTCCCTACACGACGCTCTTCCGATCTNNNGGGAGGTGTGAGGTAAAAGT

AGCACACGTCTGAACTCTTTCCCTACACGACGCTCTTCCGATCTNNNGGGAGGTGTGAGGTAAAAGT

AGCACACGTCTGAACTCTTTCCCTACACGACGCTCTTCCGATCTNNNCCTCCTCTCCATCAAAGTCTGA

AGCACACGTCTGAACTCTTTCCCTACACGACGCTCTTCCGATCTNNNCCTCCTCTCCATCAAAGTCTGA

AGCACACGTCTGAACTCTTTCCCTACACGACGCTCTTCCGATCTNNNCCTCCTCTCCATCAAAGTCTGA

AGCACACGTCTGAACTCTTTCCCTACACGACGCTCTTCCGATCTNNNCCTCCTCTCCATCAAAGTCTGA

CACACGTCTGAACTCTTTCCCTACACGACGCTCTTCCGATCTNNNCACTCACAGCGAAAACCAGAAATC

CACACGTCTGAACTCTTTCCCTACACGACGCTCTTCCGATCTNNNCACTCACAGCGAAAACCAGAAATC

CACACGTCTGAACTCTTTCCCTACACGACGCTCTTCCGATCTNNNCACTCACAGCGAAAACCAGAAATC

CACACGTCTGAACTCTTTCCCTACACGACGCTCTTCCGATCTNNNCACTCACAGCGAAAACCAGAAATC

CACACGTCTGAACTCTTTCCCTACACGACGCTCTTCCGATCTNNNCTTTGGAGTCCTTGCTACCAGCCA

CACACGTCTGAACTCTTTCCCTACACGACGCTCTTCCGATCTNNNCTTTGGAGTCCTTGCTACCAGCCA

CACACGTCTGAACTCTTTCCCTACACGACGCTCTTCCGATCTNNNCTTTGGAGTCCTTGCTACCAGCCA

CACACGTCTGAACTCTTTCCCTACACGACGCTCTTCCGATCTNNNCTTTGGAGTCCTTGCTACCAGCCA

GAGCACACGTCTGAACTCTTTCCCTACACGACGCTCTTCCGATCTNNNGAGATCACGTCACTGCATTCC

GAGCACACGTCTGAACTCTTTCCCTACACGACGCTCTTCCGATCTNNNGAGATCACGTCACTGCATTCC

GAGCACACGTCTGAACTCTTTCCCTACACGACGCTCTTCCGATCTNNNGAGATCACGTCACTGCATTCC

GAGCACACGTCTGAACTCTTTCCCTACACGACGCTCTTCCGATCTNNNGAGATCACGTCACTGCATTCC

AGCACACGTCTGAACTCTTTCCCTACACGACGCTCTTCCGATCTNNNAGTGCTGGAGAGTAAAAGGA

AGCACACGTCTGAACTCTTTCCCTACACGACGCTCTTCCGATCTNNNAGTGCTGGAGAGTAAAAGGA

AGCACACGTCTGAACTCTTTCCCTACACGACGCTCTTCCGATCTNNNAGTGCTGGAGAGTAAAAGGA

AGCACACGTCTGAACTCTTTCCCTACACGACGCTCTTCCGATCTNNNAGTGCTGGAGAGTAAAAGGA

AGAGCACACGTCTGAACTCTTTCCCTACACGACGCTCTTCCGATCTNNNCACCAACTGGACAAAGCCA

AGAGCACACGTCTGAACTCTTTCCCTACACGACGCTCTTCCGATCTNNNCACCAACTGGACAAAGCCA

AGAGCACACGTCTGAACTCTTTCCCTACACGACGCTCTTCCGATCTNNNCACCAACTGGACAAAGCCA

AGAGCACACGTCTGAACTCTTTCCCTACACGACGCTCTTCCGATCTNNNCACCAACTGGACAAAGCCA

GAGCACACGTCTGAACTCTTTCCCTACACGACGCTCTTCCGATCTNNNGAGGCAGGAGAATTGTTTGA

GAGCACACGTCTGAACTCTTTCCCTACACGACGCTCTTCCGATCTNNNGAGGCAGGAGAATTGTTTGA

GAGCACACGTCTGAACTCTTTCCCTACACGACGCTCTTCCGATCTNNNGAGGCAGGAGAATTGTTTGA

GAGCACACGTCTGAACTCTTTCCCTACACGACGCTCTTCCGATCTNNNGAGGCAGGAGAATTGTTTGA

AGCACACGTCTGAACTCTTTCCCTACACGACGCTCTTCCGATCTNNNTAGCTTTCCAAGTAGAAGCCA

AGCACACGTCTGAACTCTTTCCCTACACGACGCTCTTCCGATCTNNNTAGCTTTCCAAGTAGAAGCCA

AGCACACGTCTGAACTCTTTCCCTACACGACGCTCTTCCGATCTNNNTAGCTTTCCAAGTAGAAGCCA

AGCACACGTCTGAACTCTTTCCCTACACGACGCTCTTCCGATCTNNNTAGCTTTCCAAGTAGAAGCCA

GAGCACACGTCTGAACTCTTTCCCTACACGACGCTCTTCCGATCTNNNCCCTGCTATAGTCAAGAACCCT

GAGCACACGTCTGAACTCTTTCCCTACACGACGCTCTTCCGATCTNNNCCCTGCTATAGTCAAGAACCCT

GAGCACACGTCTGAACTCTTTCCCTACACGACGCTCTTCCGATCTNNNCCCTGCTATAGTCAAGAACCCT

GAGCACACGTCTGAACTCTTTCCCTACACGACGCTCTTCCGATCTNNNCCCTGCTATAGTCAAGAACCCT
AGCACACGTCTGAACTCTTTCCCTACACGACGCTCTTCCGATCTNNNAGGTGAGAGAGGAAGACAGAA
AGCACACGTCTGAACTCTTTCCCTACACGACGCTCTTCCGATCTNNNAGGTGAGAGAGGAAGACAGAA
AGCACACGTCTGAACTCTTTCCCTACACGACGCTCTTCCGATCTNNNAGGTGAGAGAGGAAGACAGAA
AGCACACGTCTGAACTCTTTCCCTACACGACGCTCTTCCGATCTNNNAGGTGAGAGAGGAAGACAGAA
GAGCACACGTCTGAACTCTTTCCCTACACGACGCTCTTCCGATCTNNNTATTTGAAAACGTTACCTGAAA
GAGCACACGTCTGAACTCTTTCCCTACACGACGCTCTTCCGATCTNNNTATTTGAAAACGTTACCTGAAA
GAGCACACGTCTGAACTCTTTCCCTACACGACGCTCTTCCGATCTNNNTATTTGAAAACGTTACCTGAAA
GAGCACACGTCTGAACTCTTTCCCTACACGACGCTCTTCCGATCTNNNTATTTGAAAACGTTACCTGAAA
GAGCACACGTCTGAACTCTTTCCCTACACGACGCTCTTCCGATCTNNNTCACCTTCATTTTGTTTTATGG
GAGCACACGTCTGAACTCTTTCCCTACACGACGCTCTTCCGATCTNNNTCACCTTCATTTTGTTTTATGG
GAGCACACGTCTGAACTCTTTCCCTACACGACGCTCTTCCGATCTNNNTCACCTTCATTTTGTTTTATGG
GAGCACACGTCTGAACTCTTTCCCTACACGACGCTCTTCCGATCTNNNTCACCTTCATTTTGTTTTATGG
AGCACACGTCTGAACTCTTTCCCTACACGACGCTCTTCCGATCTNNNTGGGTCCATATGTAAACATTTC
AGCACACGTCTGAACTCTTTCCCTACACGACGCTCTTCCGATCTNNNTGGGTCCATATGTAAACATTTC
AGCACACGTCTGAACTCTTTCCCTACACGACGCTCTTCCGATCTNNNTGGGTCCATATGTAAACATTTC
AGCACACGTCTGAACTCTTTCCCTACACGACGCTCTTCCGATCTNNNTGGGTCCATATGTAAACATTTC
GAGCACACGTCTGAACTCTTTCCCTACACGACGCTCTTCCGATCTNNNGATCACTTGAGCCCAGGATGT
GAGCACACGTCTGAACTCTTTCCCTACACGACGCTCTTCCGATCTNNNGATCACTTGAGCCCAGGATGT
GAGCACACGTCTGAACTCTTTCCCTACACGACGCTCTTCCGATCTNNNGATCACTTGAGCCCAGGATGT
GAGCACACGTCTGAACTCTTTCCCTACACGACGCTCTTCCGATCTNNNGATCACTTGAGCCCAGGATGT
GCACACGTCTGAACTCTTTCCCTACACGACGCTCTTCCGATCTNNNAACAATCCTTCTTGGCATATGGA
GCACACGTCTGAACTCTTTCCCTACACGACGCTCTTCCGATCTNNNAACAATCCTTCTTGGCATATGGA
GCACACGTCTGAACTCTTTCCCTACACGACGCTCTTCCGATCTNNNAACAATCCTTCTTGGCATATGGA
GCACACGTCTGAACTCTTTCCCTACACGACGCTCTTCCGATCTNNNAACAATCCTTCTTGGCATATGGA
GAGCACACGTCTGAACTCTTTCCCTACACGACGCTCTTCCGATCTNNNCAGGCAGGTCAATTCCTTGTTT
GAGCACACGTCTGAACTCTTTCCCTACACGACGCTCTTCCGATCTNNNCAGGCAGGTCAATTCCTTGTTT
GAGCACACGTCTGAACTCTTTCCCTACACGACGCTCTTCCGATCTNNNCAGGCAGGTCAATTCCTTGTTT
GAGCACACGTCTGAACTCTTTCCCTACACGACGCTCTTCCGATCTNNNCAGGCAGGTCAATTCCTTGTTT
GCACACGTCTGAACTCTTTCCCTACACGACGCTCTTCCGATCTNNNTTATCTCCAAAGAGTGTGCCTCCC
GCACACGTCTGAACTCTTTCCCTACACGACGCTCTTCCGATCTNNNTTATCTCCAAAGAGTGTGCCTCCC
GCACACGTCTGAACTCTTTCCCTACACGACGCTCTTCCGATCTNNNTTATCTCCAAAGAGTGTGCCTCCC
GCACACGTCTGAACTCTTTCCCTACACGACGCTCTTCCGATCTNNNTTATCTCCAAAGAGTGTGCCTCCC
GCACACGTCTGAACTCTTTCCCTACACGACGCTCTTCCGATCTNNNCCTTTCCTCCTTCCCCCTGGCATCG
GCACACGTCTGAACTCTTTCCCTACACGACGCTCTTCCGATCTNNNCCTTTCCTCCTTCCCCCTGGCATCG
GCACACGTCTGAACTCTTTCCCTACACGACGCTCTTCCGATCTNNNCCTTTCCTCCTTCCCCCTGGCATCG
GCACACGTCTGAACTCTTTCCCTACACGACGCTCTTCCGATCTNNNCCTTTCCTCCTTCCCCCTGGCATCG
AGCACACGTCTGAACTCTTTCCCTACACGACGCTCTTCCGATCTNNNTCACTCATTCTGGAAAAGCTCCT
AGCACACGTCTGAACTCTTTCCCTACACGACGCTCTTCCGATCTNNNTCACTCATTCTGGAAAAGCTCCT
AGCACACGTCTGAACTCTTTCCCTACACGACGCTCTTCCGATCTNNNTCACTCATTCTGGAAAAGCTCCT
AGCACACGTCTGAACTCTTTCCCTACACGACGCTCTTCCGATCTNNNTCACTCATTCTGGAAAAGCTCCT
GAGCACACGTCTGAACTCTTTCCCTACACGACGCTCTTCCGATCTNNNGTGGCTCTGTTCCCAAAAAGA
GAGCACACGTCTGAACTCTTTCCCTACACGACGCTCTTCCGATCTNNNGTGGCTCTGTTCCCAAAAAGA

GAGCACACGTCTGAACTCTTTCCCTACACGACGCTCTTCCGATCTNNNGTGGCTCTGTTCCCAAAAAG
GAGCACACGTCTGAACTCTTTCCCTACACGACGCTCTTCCGATCTNNNGTGGCTCTGTTCCCAAAAAG
AGCACACGTCTGAACTCTTTCCCTACACGACGCTCTTCCGATCTNNNCATGCCTGTAATCCCAGCTACT
AGCACACGTCTGAACTCTTTCCCTACACGACGCTCTTCCGATCTNNNCATGCCTGTAATCCCAGCTACT
AGCACACGTCTGAACTCTTTCCCTACACGACGCTCTTCCGATCTNNNCATGCCTGTAATCCCAGCTACT
AGCACACGTCTGAACTCTTTCCCTACACGACGCTCTTCCGATCTNNNCATGCCTGTAATCCCAGCTACT
GAGCACACGTCTGAACTCTTTCCCTACACGACGCTCTTCCGATCTNNNAGAGAGGTTTAGGTGTCTG
GAGCACACGTCTGAACTCTTTCCCTACACGACGCTCTTCCGATCTNNNAGAGAGGTTTAGGTGTCTG
GAGCACACGTCTGAACTCTTTCCCTACACGACGCTCTTCCGATCTNNNAGAGAGGTTTAGGTGTCTG
GAGCACACGTCTGAACTCTTTCCCTACACGACGCTCTTCCGATCTNNNAGAGAGGTTTAGGTGTCTG
AGCACACGTCTGAACTCTTTCCCTACACGACGCTCTTCCGATCTNNNCAGATCACGCCACTGTACTACA
AGCACACGTCTGAACTCTTTCCCTACACGACGCTCTTCCGATCTNNNCAGATCACGCCACTGTACTACA
AGCACACGTCTGAACTCTTTCCCTACACGACGCTCTTCCGATCTNNNCAGATCACGCCACTGTACTACA
AGCACACGTCTGAACTCTTTCCCTACACGACGCTCTTCCGATCTNNNCAGATCACGCCACTGTACTACA
GCACACGTCTGAACTCTTTCCCTACACGACGCTCTTCCGATCTNNNGAGAATCGCTTGAACCCAGGAG
GCACACGTCTGAACTCTTTCCCTACACGACGCTCTTCCGATCTNNNGAGAATCGCTTGAACCCAGGAG
GCACACGTCTGAACTCTTTCCCTACACGACGCTCTTCCGATCTNNNGAGAATCGCTTGAACCCAGGAG
GCACACGTCTGAACTCTTTCCCTACACGACGCTCTTCCGATCTNNNGAGAATCGCTTGAACCCAGGAG
AGCACACGTCTGAACTCTTTCCCTACACGACGCTCTTCCGATCTNNNCATGCCTGTAATCCCAGCTACT
AGCACACGTCTGAACTCTTTCCCTACACGACGCTCTTCCGATCTNNNCATGCCTGTAATCCCAGCTACT
AGCACACGTCTGAACTCTTTCCCTACACGACGCTCTTCCGATCTNNNCATGCCTGTAATCCCAGCTACT
AGCACACGTCTGAACTCTTTCCCTACACGACGCTCTTCCGATCTNNNCATGCCTGTAATCCCAGCTACT
AGCACACGTCTGAACTCTTTCCCTACACGACGCTCTTCCGATCTNNNTTAGTAGAGATGGGGTTTCAC
AGCACACGTCTGAACTCTTTCCCTACACGACGCTCTTCCGATCTNNNTTAGTAGAGATGGGGTTTCAC
AGCACACGTCTGAACTCTTTCCCTACACGACGCTCTTCCGATCTNNNTTAGTAGAGATGGGGTTTCAC
AGCACACGTCTGAACTCTTTCCCTACACGACGCTCTTCCGATCTNNNTTAGTAGAGATGGGGTTTCAC
GAGCACACGTCTGAACTCTTTCCCTACACGACGCTCTTCCGATCTNNNTGAGGGCCTATCGATATATA
GAGCACACGTCTGAACTCTTTCCCTACACGACGCTCTTCCGATCTNNNTGAGGGCCTATCGATATATA
GAGCACACGTCTGAACTCTTTCCCTACACGACGCTCTTCCGATCTNNNTGAGGGCCTATCGATATATA
GAGCACACGTCTGAACTCTTTCCCTACACGACGCTCTTCCGATCTNNNTGAGGGCCTATCGATATATA
GAGCACACGTCTGAACTCTTTCCCTACACGACGCTCTTCCGATCTNNNTGCTTCTTCATAGGTCAGCTG
GAGCACACGTCTGAACTCTTTCCCTACACGACGCTCTTCCGATCTNNNTGCTTCTTCATAGGTCAGCTG
GAGCACACGTCTGAACTCTTTCCCTACACGACGCTCTTCCGATCTNNNTGCTTCTTCATAGGTCAGCTG
GAGCACACGTCTGAACTCTTTCCCTACACGACGCTCTTCCGATCTNNNTGCTTCTTCATAGGTCAGCTG
AGCACACGTCTGAACTCTTTCCCTACACGACGCTCTTCCGATCTNNNCTCTTCTAGGGGGAGAGGTC
AGCACACGTCTGAACTCTTTCCCTACACGACGCTCTTCCGATCTNNNCTCTTCTAGGGGGAGAGGTC
AGCACACGTCTGAACTCTTTCCCTACACGACGCTCTTCCGATCTNNNCTCTTCTAGGGGGAGAGGTC
AGCACACGTCTGAACTCTTTCCCTACACGACGCTCTTCCGATCTNNNCTCTTCTAGGGGGAGAGGTC
GAGCACACGTCTGAACTCTTTCCCTACACGACGCTCTTCCGATCTNNNTATACACACACACACACACAC
GAGCACACGTCTGAACTCTTTCCCTACACGACGCTCTTCCGATCTNNNTATACACACACACACACACAC
GAGCACACGTCTGAACTCTTTCCCTACACGACGCTCTTCCGATCTNNNTATACACACACACACACACAC
GAGCACACGTCTGAACTCTTTCCCTACACGACGCTCTTCCGATCTNNNTATACACACACACACACACAC
AGCACACGTCTGAACTCTTTCCCTACACGACGCTCTTCCGATCTNNNTGTGATCCCAGCTATTCAGGA

AGCACACGTCTGAACTCTTTCCCTACACGACGCTCTTCCGATCTNNNTGTGATCCCAGCTATTCAGGA
AGCACACGTCTGAACTCTTTCCCTACACGACGCTCTTCCGATCTNNNTGTGATCCCAGCTATTCAGGA
AGCACACGTCTGAACTCTTTCCCTACACGACGCTCTTCCGATCTNNNTGTGATCCCAGCTATTCAGGA
AGCACACGTCTGAACTCTTTCCCTACACGACGCTCTTCCGATCTNNNCAGGTTCTTCCTCGACTCAGAA
AGCACACGTCTGAACTCTTTCCCTACACGACGCTCTTCCGATCTNNNCAGGTTCTTCCTCGACTCAGAA
AGCACACGTCTGAACTCTTTCCCTACACGACGCTCTTCCGATCTNNNCAGGTTCTTCCTCGACTCAGAA
AGCACACGTCTGAACTCTTTCCCTACACGACGCTCTTCCGATCTNNNCAGGTTCTTCCTCGACTCAGAA
AGCACACGTCTGAACTCTTTCCCTACACGACGCTCTTCCGATCTNNNAGAGCTTGCAGTGAGTGGAT
AGCACACGTCTGAACTCTTTCCCTACACGACGCTCTTCCGATCTNNNAGAGCTTGCAGTGAGTGGAT
AGCACACGTCTGAACTCTTTCCCTACACGACGCTCTTCCGATCTNNNAGAGCTTGCAGTGAGTGGAT
AGCACACGTCTGAACTCTTTCCCTACACGACGCTCTTCCGATCTNNNAGAGCTTGCAGTGAGTGGAT
GCACACGTCTGAACTCTTTCCCTACACGACGCTCTTCCGATCTNNNGGTGAGGTAGATTTCCGGAAT
GCACACGTCTGAACTCTTTCCCTACACGACGCTCTTCCGATCTNNNGGTGAGGTAGATTTCCGGAAT
GCACACGTCTGAACTCTTTCCCTACACGACGCTCTTCCGATCTNNNGGTGAGGTAGATTTCCGGAAT
GCACACGTCTGAACTCTTTCCCTACACGACGCTCTTCCGATCTNNNGGTGAGGTAGATTTCCGGAAT
GCACACGTCTGAACTCTTTCCCTACACGACGCTCTTCCGATCTNNNGATGTAGTCCCAGCTACTCAGG
GCACACGTCTGAACTCTTTCCCTACACGACGCTCTTCCGATCTNNNGATGTAGTCCCAGCTACTCAGG
GCACACGTCTGAACTCTTTCCCTACACGACGCTCTTCCGATCTNNNGATGTAGTCCCAGCTACTCAGG
GCACACGTCTGAACTCTTTCCCTACACGACGCTCTTCCGATCTNNNGATGTAGTCCCAGCTACTCAGG
AGCACACGTCTGAACTCTTTCCCTACACGACGCTCTTCCGATCTNNNCTGTAGTTCCAGCTACTCGGGA
AGCACACGTCTGAACTCTTTCCCTACACGACGCTCTTCCGATCTNNNCTGTAGTTCCAGCTACTCGGGA
AGCACACGTCTGAACTCTTTCCCTACACGACGCTCTTCCGATCTNNNCTGTAGTTCCAGCTACTCGGGA
AGCACACGTCTGAACTCTTTCCCTACACGACGCTCTTCCGATCTNNNCTGTAGTTCCAGCTACTCGGGA
GAGCACACGTCTGAACTCTTTCCCTACACGACGCTCTTCCGATCTNNNTGACAGTAGCTTGGGGAATA
GAGCACACGTCTGAACTCTTTCCCTACACGACGCTCTTCCGATCTNNNTGACAGTAGCTTGGGGAATA
GAGCACACGTCTGAACTCTTTCCCTACACGACGCTCTTCCGATCTNNNTGACAGTAGCTTGGGGAATA
GAGCACACGTCTGAACTCTTTCCCTACACGACGCTCTTCCGATCTNNNTGACAGTAGCTTGGGGAATA
GCACACGTCTGAACTCTTTCCCTACACGACGCTCTTCCGATCTNNNTAAACTTCGATGTAGCCGACACA
GCACACGTCTGAACTCTTTCCCTACACGACGCTCTTCCGATCTNNNTAAACTTCGATGTAGCCGACACA
GCACACGTCTGAACTCTTTCCCTACACGACGCTCTTCCGATCTNNNTAAACTTCGATGTAGCCGACACA
GCACACGTCTGAACTCTTTCCCTACACGACGCTCTTCCGATCTNNNTAAACTTCGATGTAGCCGACACA
AGCACACGTCTGAACTCTTTCCCTACACGACGCTCTTCCGATCTNNNCTGTAATCCCAGCTACTCAGGA
AGCACACGTCTGAACTCTTTCCCTACACGACGCTCTTCCGATCTNNNCTGTAATCCCAGCTACTCAGGA
AGCACACGTCTGAACTCTTTCCCTACACGACGCTCTTCCGATCTNNNCTGTAATCCCAGCTACTCAGGA
AGCACACGTCTGAACTCTTTCCCTACACGACGCTCTTCCGATCTNNNCTGTAATCCCAGCTACTCAGGA
GAGCACACGTCTGAACTCTTTCCCTACACGACGCTCTTCCGATCTNNNCTTGCAGTGAGCTGAGATTG
GAGCACACGTCTGAACTCTTTCCCTACACGACGCTCTTCCGATCTNNNCTTGCAGTGAGCTGAGATTG
GAGCACACGTCTGAACTCTTTCCCTACACGACGCTCTTCCGATCTNNNCTTGCAGTGAGCTGAGATTG
GAGCACACGTCTGAACTCTTTCCCTACACGACGCTCTTCCGATCTNNNCTTGCAGTGAGCTGAGATTG
AGCACACGTCTGAACTCTTTCCCTACACGACGCTCTTCCGATCTNNNCATACATGCACCCATTCATTCC
AGCACACGTCTGAACTCTTTCCCTACACGACGCTCTTCCGATCTNNNCATACATGCACCCATTCATTCC
AGCACACGTCTGAACTCTTTCCCTACACGACGCTCTTCCGATCTNNNCATACATGCACCCATTCATTCC
AGCACACGTCTGAACTCTTTCCCTACACGACGCTCTTCCGATCTNNNCATACATGCACCCATTCATTCC

AGAGCACACGTCTGAACTCTTTCCCTACACGACGCTCTTCCGATCTNNNTGGGGGACAGAATGAGA
AGAGCACACGTCTGAACTCTTTCCCTACACGACGCTCTTCCGATCTNNNTGGGGGACAGAATGAGA
AGAGCACACGTCTGAACTCTTTCCCTACACGACGCTCTTCCGATCTNNNTGGGGGACAGAATGAGA
AGAGCACACGTCTGAACTCTTTCCCTACACGACGCTCTTCCGATCTNNNTGGGGGACAGAATGAGA
AGCACACGTCTGAACTCTTTCCCTACACGACGCTCTTCCGATCTNNNGTGCCTGTAATCCCAGCTACTT
AGCACACGTCTGAACTCTTTCCCTACACGACGCTCTTCCGATCTNNNGTGCCTGTAATCCCAGCTACTT
AGCACACGTCTGAACTCTTTCCCTACACGACGCTCTTCCGATCTNNNGTGCCTGTAATCCCAGCTACTT
AGCACACGTCTGAACTCTTTCCCTACACGACGCTCTTCCGATCTNNNGTGCCTGTAATCCCAGCTACTT
AGCACACGTCTGAACTCTTTCCCTACACGACGCTCTTCCGATCTNNNCCAGGTTGGTCTCGAACTTCTA
AGCACACGTCTGAACTCTTTCCCTACACGACGCTCTTCCGATCTNNNCCAGGTTGGTCTCGAACTTCTA
AGCACACGTCTGAACTCTTTCCCTACACGACGCTCTTCCGATCTNNNCCAGGTTGGTCTCGAACTTCTA
AGCACACGTCTGAACTCTTTCCCTACACGACGCTCTTCCGATCTNNNCCAGGTTGGTCTCGAACTTCTA
GAGCACACGTCTGAACTCTTTCCCTACACGACGCTCTTCCGATCTNNNCCGGCTAACACAATGAAATC
GAGCACACGTCTGAACTCTTTCCCTACACGACGCTCTTCCGATCTNNNCCGGCTAACACAATGAAATC
GAGCACACGTCTGAACTCTTTCCCTACACGACGCTCTTCCGATCTNNNCCGGCTAACACAATGAAATC
GAGCACACGTCTGAACTCTTTCCCTACACGACGCTCTTCCGATCTNNNCCGGCTAACACAATGAAATC
AGCACACGTCTGAACTCTTTCCCTACACGACGCTCTTCCGATCTNNNACATCTTGTGAGGGTTGTTCC
AGCACACGTCTGAACTCTTTCCCTACACGACGCTCTTCCGATCTNNNACATCTTGTGAGGGTTGTTCC
AGCACACGTCTGAACTCTTTCCCTACACGACGCTCTTCCGATCTNNNACATCTTGTGAGGGTTGTTCC
AGCACACGTCTGAACTCTTTCCCTACACGACGCTCTTCCGATCTNNNACATCTTGTGAGGGTTGTTCC
CACACGTCTGAACTCTTTCCCTACACGACGCTCTTCCGATCTNNNGAGGTTGCAGTGAGTGGAGATC
CACACGTCTGAACTCTTTCCCTACACGACGCTCTTCCGATCTNNNGAGGTTGCAGTGAGTGGAGATC
CACACGTCTGAACTCTTTCCCTACACGACGCTCTTCCGATCTNNNGAGGTTGCAGTGAGTGGAGATC
CACACGTCTGAACTCTTTCCCTACACGACGCTCTTCCGATCTNNNGAGGTTGCAGTGAGTGGAGATC
AGCACACGTCTGAACTCTTTCCCTACACGACGCTCTTCCGATCTNNNTCTCGAGTAGCTGGGACTATA
AGCACACGTCTGAACTCTTTCCCTACACGACGCTCTTCCGATCTNNNTCTCGAGTAGCTGGGACTATA
AGCACACGTCTGAACTCTTTCCCTACACGACGCTCTTCCGATCTNNNTCTCGAGTAGCTGGGACTATA
AGCACACGTCTGAACTCTTTCCCTACACGACGCTCTTCCGATCTNNNTCTCGAGTAGCTGGGACTATA
AGCACACGTCTGAACTCTTTCCCTACACGACGCTCTTCCGATCTNNNATAACGCTATCTGGTAGGGCT
AGCACACGTCTGAACTCTTTCCCTACACGACGCTCTTCCGATCTNNNATAACGCTATCTGGTAGGGCT
AGCACACGTCTGAACTCTTTCCCTACACGACGCTCTTCCGATCTNNNATAACGCTATCTGGTAGGGCT
AGCACACGTCTGAACTCTTTCCCTACACGACGCTCTTCCGATCTNNNATAACGCTATCTGGTAGGGCT
AGCACACGTCTGAACTCTTTCCCTACACGACGCTCTTCCGATCTNNNAAAAAATTAGCCAGGTGTGG
AGCACACGTCTGAACTCTTTCCCTACACGACGCTCTTCCGATCTNNNAAAAAATTAGCCAGGTGTGG
AGCACACGTCTGAACTCTTTCCCTACACGACGCTCTTCCGATCTNNNAAAAAATTAGCCAGGTGTGG
AGCACACGTCTGAACTCTTTCCCTACACGACGCTCTTCCGATCTNNNAAAAAATTAGCCAGGTGTGG
AGCACACGTCTGAACTCTTTCCCTACACGACGCTCTTCCGATCTNNNTTGAGATGGAGTCTTGCTCTG
AGCACACGTCTGAACTCTTTCCCTACACGACGCTCTTCCGATCTNNNTTGAGATGGAGTCTTGCTCTG
AGCACACGTCTGAACTCTTTCCCTACACGACGCTCTTCCGATCTNNNTTGAGATGGAGTCTTGCTCTG
AGCACACGTCTGAACTCTTTCCCTACACGACGCTCTTCCGATCTNNNTTGAGATGGAGTCTTGCTCTG
AGCACACGTCTGAACTCTTTCCCTACACGACGCTCTTCCGATCTNNNCGAGTGATGGCATGTGACTT
AGCACACGTCTGAACTCTTTCCCTACACGACGCTCTTCCGATCTNNNCGAGTGATGGCATGTGACTT
AGCACACGTCTGAACTCTTTCCCTACACGACGCTCTTCCGATCTNNNCGAGTGATGGCATGTGACTT

AGCACACGTCTGAACTCTTTCCCTACACGACGCTCTTCCGATCTNNNCGAGTGATGGCATGTGACTT
GAGCACACGTCTGAACTCTTTCCCTACACGACGCTCTTCCGATCTNNNGCTTGTGACTAGGGATTTGT
GAGCACACGTCTGAACTCTTTCCCTACACGACGCTCTTCCGATCTNNNGCTTGTGACTAGGGATTTGT
GAGCACACGTCTGAACTCTTTCCCTACACGACGCTCTTCCGATCTNNNGCTTGTGACTAGGGATTTGT
GAGCACACGTCTGAACTCTTTCCCTACACGACGCTCTTCCGATCTNNNGCTTGTGACTAGGGATTTGT
AGCACACGTCTGAACTCTTTCCCTACACGACGCTCTTCCGATCTNNNCAAAAATTAGCTGGACCTGGT
AGCACACGTCTGAACTCTTTCCCTACACGACGCTCTTCCGATCTNNNCAAAAATTAGCTGGACCTGGT
AGCACACGTCTGAACTCTTTCCCTACACGACGCTCTTCCGATCTNNNCAAAAATTAGCTGGACCTGGT
AGCACACGTCTGAACTCTTTCCCTACACGACGCTCTTCCGATCTNNNCAAAAATTAGCTGGACCTGGT
AGCACACGTCTGAACTCTTTCCCTACACGACGCTCTTCCGATCTNNNGATGGTCTCTTCAGAACCCTTG
AGCACACGTCTGAACTCTTTCCCTACACGACGCTCTTCCGATCTNNNGATGGTCTCTTCAGAACCCTTG
AGCACACGTCTGAACTCTTTCCCTACACGACGCTCTTCCGATCTNNNGATGGTCTCTTCAGAACCCTTG
AGCACACGTCTGAACTCTTTCCCTACACGACGCTCTTCCGATCTNNNGATGGTCTCTTCAGAACCCTTG
GAGCACACGTCTGAACTCTTTCCCTACACGACGCTCTTCCGATCTNNNGTCAAGAGATCGAGACCATC
GAGCACACGTCTGAACTCTTTCCCTACACGACGCTCTTCCGATCTNNNGTCAAGAGATCGAGACCATC
GAGCACACGTCTGAACTCTTTCCCTACACGACGCTCTTCCGATCTNNNGTCAAGAGATCGAGACCATC
GAGCACACGTCTGAACTCTTTCCCTACACGACGCTCTTCCGATCTNNNGTCAAGAGATCGAGACCATC
AGCACACGTCTGAACTCTTTCCCTACACGACGCTCTTCCGATCTNNNCTGCCTATGCCCCCTCTAGTCC
AGCACACGTCTGAACTCTTTCCCTACACGACGCTCTTCCGATCTNNNCTGCCTATGCCCCCTCTAGTCC
AGCACACGTCTGAACTCTTTCCCTACACGACGCTCTTCCGATCTNNNCTGCCTATGCCCCCTCTAGTCC
AGCACACGTCTGAACTCTTTCCCTACACGACGCTCTTCCGATCTNNNCTGCCTATGCCCCCTCTAGTCC
GCACACGTCTGAACTCTTTCCCTACACGACGCTCTTCCGATCTNNNATCATGACACCCAAGTCTCCAAG
GCACACGTCTGAACTCTTTCCCTACACGACGCTCTTCCGATCTNNNATCATGACACCCAAGTCTCCAAG
GCACACGTCTGAACTCTTTCCCTACACGACGCTCTTCCGATCTNNNATCATGACACCCAAGTCTCCAAG
GCACACGTCTGAACTCTTTCCCTACACGACGCTCTTCCGATCTNNNATCATGACACCCAAGTCTCCAAG
AGCACACGTCTGAACTCTTTCCCTACACGACGCTCTTCCGATCTNNNGCAGCATTATTCTCAACAGCC
AGCACACGTCTGAACTCTTTCCCTACACGACGCTCTTCCGATCTNNNGCAGCATTATTCTCAACAGCC
AGCACACGTCTGAACTCTTTCCCTACACGACGCTCTTCCGATCTNNNGCAGCATTATTCTCAACAGCC
AGCACACGTCTGAACTCTTTCCCTACACGACGCTCTTCCGATCTNNNGCAGCATTATTCTCAACAGCC
GCACACGTCTGAACTCTTTCCCTACACGACGCTCTTCCGATCTNNNGGTGGTTGTGGTGACCTGATA
GCACACGTCTGAACTCTTTCCCTACACGACGCTCTTCCGATCTNNNGGTGGTTGTGGTGACCTGATA
GCACACGTCTGAACTCTTTCCCTACACGACGCTCTTCCGATCTNNNGGTGGTTGTGGTGACCTGATA
GCACACGTCTGAACTCTTTCCCTACACGACGCTCTTCCGATCTNNNGGTGGTTGTGGTGACCTGATA
CACACGTCTGAACTCTTTCCCTACACGACGCTCTTCCGATCTNNNCTGGCTTCTCCTGTAGGACACCA
CACACGTCTGAACTCTTTCCCTACACGACGCTCTTCCGATCTNNNCTGGCTTCTCCTGTAGGACACCA
CACACGTCTGAACTCTTTCCCTACACGACGCTCTTCCGATCTNNNCTGGCTTCTCCTGTAGGACACCA
CACACGTCTGAACTCTTTCCCTACACGACGCTCTTCCGATCTNNNCTGGCTTCTCCTGTAGGACACCA
AGCACACGTCTGAACTCTTTCCCTACACGACGCTCTTCCGATCTNNNAAAAAATTAGCCAGGCGTGG
AGCACACGTCTGAACTCTTTCCCTACACGACGCTCTTCCGATCTNNNAAAAAATTAGCCAGGCGTGG
AGCACACGTCTGAACTCTTTCCCTACACGACGCTCTTCCGATCTNNNAAAAAATTAGCCAGGCGTGG
AGCACACGTCTGAACTCTTTCCCTACACGACGCTCTTCCGATCTNNNAAAAAATTAGCCAGGCGTGG
GAGCACACGTCTGAACTCTTTCCCTACACGACGCTCTTCCGATCTNNNGTAGGAAAAGCAATGACAG
GAGCACACGTCTGAACTCTTTCCCTACACGACGCTCTTCCGATCTNNNGTAGGAAAAGCAATGACAG

GAGCACACGTCTGAACTCTTTCCCTACACGACGCTCTTCCGATCTNNNGTAGGAAAAGCAATGACAG
GAGCACACGTCTGAACTCTTTCCCTACACGACGCTCTTCCGATCTNNNGTAGGAAAAGCAATGACAG
AGCACACGTCTGAACTCTTTCCCTACACGACGCTCTTCCGATCTNNNATGTTATGTCTCTGGGTTCCAC
AGCACACGTCTGAACTCTTTCCCTACACGACGCTCTTCCGATCTNNNATGTTATGTCTCTGGGTTCCAC
AGCACACGTCTGAACTCTTTCCCTACACGACGCTCTTCCGATCTNNNATGTTATGTCTCTGGGTTCCAC
AGCACACGTCTGAACTCTTTCCCTACACGACGCTCTTCCGATCTNNNATGTTATGTCTCTGGGTTCCAC
CACACGTCTGAACTCTTTCCCTACACGACGCTCTTCCGATCTNNNAAAATCACAGCATTGTAAGAAAT
CACACGTCTGAACTCTTTCCCTACACGACGCTCTTCCGATCTNNNAAAATCACAGCATTGTAAGAAAT
CACACGTCTGAACTCTTTCCCTACACGACGCTCTTCCGATCTNNNAAAATCACAGCATTGTAAGAAAT
CACACGTCTGAACTCTTTCCCTACACGACGCTCTTCCGATCTNNNAAAATCACAGCATTGTAAGAAAT
GAGCACACGTCTGAACTCTTTCCCTACACGACGCTCTTCCGATCTNNNGGATGACCAGAATGAGAGA
GAGCACACGTCTGAACTCTTTCCCTACACGACGCTCTTCCGATCTNNNGGATGACCAGAATGAGAGA
GAGCACACGTCTGAACTCTTTCCCTACACGACGCTCTTCCGATCTNNNGGATGACCAGAATGAGAGA
GAGCACACGTCTGAACTCTTTCCCTACACGACGCTCTTCCGATCTNNNGGATGACCAGAATGAGAGA
GAGCACACGTCTGAACTCTTTCCCTACACGACGCTCTTCCGATCTNNNTGATCATTCTCCACAAACTGG
GAGCACACGTCTGAACTCTTTCCCTACACGACGCTCTTCCGATCTNNNTGATCATTCTCCACAAACTGG
GAGCACACGTCTGAACTCTTTCCCTACACGACGCTCTTCCGATCTNNNTGATCATTCTCCACAAACTGG
GAGCACACGTCTGAACTCTTTCCCTACACGACGCTCTTCCGATCTNNNTGATCATTCTCCACAAACTGG
AGCACACGTCTGAACTCTTTCCCTACACGACGCTCTTCCGATCTNNNCGTTGATTTACAGACCCATAC
AGCACACGTCTGAACTCTTTCCCTACACGACGCTCTTCCGATCTNNNCGTTGATTTACAGACCCATAC
AGCACACGTCTGAACTCTTTCCCTACACGACGCTCTTCCGATCTNNNCGTTGATTTACAGACCCATAC
AGCACACGTCTGAACTCTTTCCCTACACGACGCTCTTCCGATCTNNNCGTTGATTTACAGACCCATAC
AGCACACGTCTGAACTCTTTCCCTACACGACGCTCTTCCGATCTNNNCAACAAGGAGAGAGATGCTGT
AGCACACGTCTGAACTCTTTCCCTACACGACGCTCTTCCGATCTNNNCAACAAGGAGAGAGATGCTGT
AGCACACGTCTGAACTCTTTCCCTACACGACGCTCTTCCGATCTNNNCAACAAGGAGAGAGATGCTGT
AGCACACGTCTGAACTCTTTCCCTACACGACGCTCTTCCGATCTNNNCAACAAGGAGAGAGATGCTGT
GAGCACACGTCTGAACTCTTTCCCTACACGACGCTCTTCCGATCTNNNTAACATTCTCTGTCCTTGGGC
GAGCACACGTCTGAACTCTTTCCCTACACGACGCTCTTCCGATCTNNNTAACATTCTCTGTCCTTGGGC
GAGCACACGTCTGAACTCTTTCCCTACACGACGCTCTTCCGATCTNNNTAACATTCTCTGTCCTTGGGC
GAGCACACGTCTGAACTCTTTCCCTACACGACGCTCTTCCGATCTNNNTAACATTCTCTGTCCTTGGGC
GCACACGTCTGAACTCTTTCCCTACACGACGCTCTTCCGATCTNNNTGATCCATGATCGGGCCATTGT
GCACACGTCTGAACTCTTTCCCTACACGACGCTCTTCCGATCTNNNTGATCCATGATCGGGCCATTGT
GCACACGTCTGAACTCTTTCCCTACACGACGCTCTTCCGATCTNNNTGATCCATGATCGGGCCATTGT
GCACACGTCTGAACTCTTTCCCTACACGACGCTCTTCCGATCTNNNTGATCCATGATCGGGCCATTGT
GCACACGTCTGAACTCTTTCCCTACACGACGCTCTTCCGATCTNNNTCACAGACATGATTTGACCCAGT
GCACACGTCTGAACTCTTTCCCTACACGACGCTCTTCCGATCTNNNTCACAGACATGATTTGACCCAGT
GCACACGTCTGAACTCTTTCCCTACACGACGCTCTTCCGATCTNNNTCACAGACATGATTTGACCCAGT
GCACACGTCTGAACTCTTTCCCTACACGACGCTCTTCCGATCTNNNTCACAGACATGATTTGACCCAGT
GCACACGTCTGAACTCTTTCCCTACACGACGCTCTTCCGATCTNNNAACTTGTTAAAATTCACAAAGTG
GCACACGTCTGAACTCTTTCCCTACACGACGCTCTTCCGATCTNNNAACTTGTTAAAATTCACAAAGTG
GCACACGTCTGAACTCTTTCCCTACACGACGCTCTTCCGATCTNNNAACTTGTTAAAATTCACAAAGTG
GCACACGTCTGAACTCTTTCCCTACACGACGCTCTTCCGATCTNNNAACTTGTTAAAATTCACAAAGTG
GAGCACACGTCTGAACTCTTTCCCTACACGACGCTCTTCCGATCTNNNACTCAAGTGATCTACCTGCCTT

GAGCACACGTCTGAACTCTTTCCCTACACGACGCTCTTCCGATCTNNNACTCAAGTGATCTACCTGCCTT
GAGCACACGTCTGAACTCTTTCCCTACACGACGCTCTTCCGATCTNNNACTCAAGTGATCTACCTGCCTT
GAGCACACGTCTGAACTCTTTCCCTACACGACGCTCTTCCGATCTNNNACTCAAGTGATCTACCTGCCTT
GAGCACACGTCTGAACTCTTTCCCTACACGACGCTCTTCCGATCTNNNTCCTTTCCTCTCTTCCCTTCTCT
GAGCACACGTCTGAACTCTTTCCCTACACGACGCTCTTCCGATCTNNNTCCTTTCCTCTCTTCCCTTCTCT
GAGCACACGTCTGAACTCTTTCCCTACACGACGCTCTTCCGATCTNNNTCCTTTCCTCTCTTCCCTTCTCT
GAGCACACGTCTGAACTCTTTCCCTACACGACGCTCTTCCGATCTNNNTCCTTTCCTCTCTTCCCTTCTCT
GAGCACACGTCTGAACTCTTTCCCTACACGACGCTCTTCCGATCTNNNTATCTATGTAGTGGGGAGCC
GAGCACACGTCTGAACTCTTTCCCTACACGACGCTCTTCCGATCTNNNTATCTATGTAGTGGGGAGCC
GAGCACACGTCTGAACTCTTTCCCTACACGACGCTCTTCCGATCTNNNTATCTATGTAGTGGGGAGCC
GAGCACACGTCTGAACTCTTTCCCTACACGACGCTCTTCCGATCTNNNTATCTATGTAGTGGGGAGCC
AGCACACGTCTGAACTCTTTCCCTACACGACGCTCTTCCGATCTNNNCCATCCTGCCCTACTACATCACG
AGCACACGTCTGAACTCTTTCCCTACACGACGCTCTTCCGATCTNNNCCATCCTGCCCTACTACATCACG
AGCACACGTCTGAACTCTTTCCCTACACGACGCTCTTCCGATCTNNNCCATCCTGCCCTACTACATCACG
AGCACACGTCTGAACTCTTTCCCTACACGACGCTCTTCCGATCTNNNCCATCCTGCCCTACTACATCACG
AGCACACGTCTGAACTCTTTCCCTACACGACGCTCTTCCGATCTNNNATCAAGAGTGCATATTGGCAA
AGCACACGTCTGAACTCTTTCCCTACACGACGCTCTTCCGATCTNNNATCAAGAGTGCATATTGGCAA
AGCACACGTCTGAACTCTTTCCCTACACGACGCTCTTCCGATCTNNNATCAAGAGTGCATATTGGCAA
AGCACACGTCTGAACTCTTTCCCTACACGACGCTCTTCCGATCTNNNATCAAGAGTGCATATTGGCAA
GAGCACACGTCTGAACTCTTTCCCTACACGACGCTCTTCCGATCTNNNCTCACTGTGACCTGGGATTA
GAGCACACGTCTGAACTCTTTCCCTACACGACGCTCTTCCGATCTNNNCTCACTGTGACCTGGGATTA
GAGCACACGTCTGAACTCTTTCCCTACACGACGCTCTTCCGATCTNNNCTCACTGTGACCTGGGATTA
GAGCACACGTCTGAACTCTTTCCCTACACGACGCTCTTCCGATCTNNNCTCACTGTGACCTGGGATTA
GAGCACACGTCTGAACTCTTTCCCTACACGACGCTCTTCCGATCTNNNTTAGAATTCCAGGATGGTGTG
GAGCACACGTCTGAACTCTTTCCCTACACGACGCTCTTCCGATCTNNNTTAGAATTCCAGGATGGTGTG
GAGCACACGTCTGAACTCTTTCCCTACACGACGCTCTTCCGATCTNNNTTAGAATTCCAGGATGGTGTG
GAGCACACGTCTGAACTCTTTCCCTACACGACGCTCTTCCGATCTNNNTTAGAATTCCAGGATGGTGTG
AGCACACGTCTGAACTCTTTCCCTACACGACGCTCTTCCGATCTNNNCATCGCCCTTCCTCTTTACCCCAT
AGCACACGTCTGAACTCTTTCCCTACACGACGCTCTTCCGATCTNNNCATCGCCCTTCCTCTTTACCCCAT
AGCACACGTCTGAACTCTTTCCCTACACGACGCTCTTCCGATCTNNNCATCGCCCTTCCTCTTTACCCCAT
AGCACACGTCTGAACTCTTTCCCTACACGACGCTCTTCCGATCTNNNCATCGCCCTTCCTCTTTACCCCAT
ACACGTCTGAACTCTTTCCCTACACGACGCTCTTCCGATCTNNNGAGGTTGAAACTCGGGTTCCAGCA
ACACGTCTGAACTCTTTCCCTACACGACGCTCTTCCGATCTNNNGAGGTTGAAACTCGGGTTCCAGCA
ACACGTCTGAACTCTTTCCCTACACGACGCTCTTCCGATCTNNNGAGGTTGAAACTCGGGTTCCAGCA
ACACGTCTGAACTCTTTCCCTACACGACGCTCTTCCGATCTNNNGAGGTTGAAACTCGGGTTCCAGCA
AGAGCACACGTCTGAACTCTTTCCCTACACGACGCTCTTCCGATCTNNNAACAATATGGCAGTGAGTT
AGAGCACACGTCTGAACTCTTTCCCTACACGACGCTCTTCCGATCTNNNAACAATATGGCAGTGAGTT
AGAGCACACGTCTGAACTCTTTCCCTACACGACGCTCTTCCGATCTNNNAACAATATGGCAGTGAGTT
AGAGCACACGTCTGAACTCTTTCCCTACACGACGCTCTTCCGATCTNNNAACAATATGGCAGTGAGTT
GAGCACACGTCTGAACTCTTTCCCTACACGACGCTCTTCCGATCTNNNTATAATCCCAGCTACTTGGAG
GAGCACACGTCTGAACTCTTTCCCTACACGACGCTCTTCCGATCTNNNTATAATCCCAGCTACTTGGAG
GAGCACACGTCTGAACTCTTTCCCTACACGACGCTCTTCCGATCTNNNTATAATCCCAGCTACTTGGAG
GAGCACACGTCTGAACTCTTTCCCTACACGACGCTCTTCCGATCTNNNTATAATCCCAGCTACTTGGAG

AGCACACGTCTGAACTCTTTCCCTACACGACGCTCTTCCGATCTNNNGATGAAATTTTAGAGAAAAGG
AGCACACGTCTGAACTCTTTCCCTACACGACGCTCTTCCGATCTNNNGATGAAATTTTAGAGAAAAGG
AGCACACGTCTGAACTCTTTCCCTACACGACGCTCTTCCGATCTNNNGATGAAATTTTAGAGAAAAGG
AGCACACGTCTGAACTCTTTCCCTACACGACGCTCTTCCGATCTNNNGATGAAATTTTAGAGAAAAGG
AGCACACGTCTGAACTCTTTCCCTACACGACGCTCTTCCGATCTNNNTTTTACTTACTTCCGCTCCCACC
AGCACACGTCTGAACTCTTTCCCTACACGACGCTCTTCCGATCTNNNTTTTACTTACTTCCGCTCCCACC
AGCACACGTCTGAACTCTTTCCCTACACGACGCTCTTCCGATCTNNNTTTTACTTACTTCCGCTCCCACC
AGCACACGTCTGAACTCTTTCCCTACACGACGCTCTTCCGATCTNNNTTTTACTTACTTCCGCTCCCACC
GAGCACACGTCTGAACTCTTTCCCTACACGACGCTCTTCCGATCTNNNAAAAATCTTGGACACACGGAC
GAGCACACGTCTGAACTCTTTCCCTACACGACGCTCTTCCGATCTNNNAAAAATCTTGGACACACGGAC
GAGCACACGTCTGAACTCTTTCCCTACACGACGCTCTTCCGATCTNNNAAAAATCTTGGACACACGGAC
GAGCACACGTCTGAACTCTTTCCCTACACGACGCTCTTCCGATCTNNNAAAAATCTTGGACACACGGAC
GAGCACACGTCTGAACTCTTTCCCTACACGACGCTCTTCCGATCTNNNCCTCCAGACTTTTATTCCACCCA
GAGCACACGTCTGAACTCTTTCCCTACACGACGCTCTTCCGATCTNNNCCTCCAGACTTTTATTCCACCCA
GAGCACACGTCTGAACTCTTTCCCTACACGACGCTCTTCCGATCTNNNCCTCCAGACTTTTATTCCACCCA
GAGCACACGTCTGAACTCTTTCCCTACACGACGCTCTTCCGATCTNNNCCTCCAGACTTTTATTCCACCCA
GAGCACACGTCTGAACTCTTTCCCTACACGACGCTCTTCCGATCTNNNGAGTAGCTGGGACTACAGTTG
GAGCACACGTCTGAACTCTTTCCCTACACGACGCTCTTCCGATCTNNNGAGTAGCTGGGACTACAGTTG
GAGCACACGTCTGAACTCTTTCCCTACACGACGCTCTTCCGATCTNNNGAGTAGCTGGGACTACAGTTG
GAGCACACGTCTGAACTCTTTCCCTACACGACGCTCTTCCGATCTNNNGAGTAGCTGGGACTACAGTTG
GAGCACACGTCTGAACTCTTTCCCTACACGACGCTCTTCCGATCTNNNAAGGCAAAAATTAGTGGGTG
GAGCACACGTCTGAACTCTTTCCCTACACGACGCTCTTCCGATCTNNNAAGGCAAAAATTAGTGGGTG
GAGCACACGTCTGAACTCTTTCCCTACACGACGCTCTTCCGATCTNNNAAGGCAAAAATTAGTGGGTG
GAGCACACGTCTGAACTCTTTCCCTACACGACGCTCTTCCGATCTNNNAAGGCAAAAATTAGTGGGTG
GAGCACACGTCTGAACTCTTTCCCTACACGACGCTCTTCCGATCTNNNCTGTAGTTCCAGCTTCTCAGGA
GAGCACACGTCTGAACTCTTTCCCTACACGACGCTCTTCCGATCTNNNCTGTAGTTCCAGCTTCTCAGGA
GAGCACACGTCTGAACTCTTTCCCTACACGACGCTCTTCCGATCTNNNCTGTAGTTCCAGCTTCTCAGGA
GAGCACACGTCTGAACTCTTTCCCTACACGACGCTCTTCCGATCTNNNCTGTAGTTCCAGCTTCTCAGGA
AGAGCACACGTCTGAACTCTTTCCCTACACGACGCTCTTCCGATCTNNNGATATGGACATCTATTGGGG
AGAGCACACGTCTGAACTCTTTCCCTACACGACGCTCTTCCGATCTNNNGATATGGACATCTATTGGGG
AGAGCACACGTCTGAACTCTTTCCCTACACGACGCTCTTCCGATCTNNNGATATGGACATCTATTGGGG
AGAGCACACGTCTGAACTCTTTCCCTACACGACGCTCTTCCGATCTNNNGATATGGACATCTATTGGGG
AGCACACGTCTGAACTCTTTCCCTACACGACGCTCTTCCGATCTNNNAGAAAGTGACCAGCCAGTAGAT
AGCACACGTCTGAACTCTTTCCCTACACGACGCTCTTCCGATCTNNNAGAAAGTGACCAGCCAGTAGAT
AGCACACGTCTGAACTCTTTCCCTACACGACGCTCTTCCGATCTNNNAGAAAGTGACCAGCCAGTAGAT
AGCACACGTCTGAACTCTTTCCCTACACGACGCTCTTCCGATCTNNNAGAAAGTGACCAGCCAGTAGAT
GAGCACACGTCTGAACTCTTTCCCTACACGACGCTCTTCCGATCTNNNCGCTTTGTGGAATTATCGTTG
GAGCACACGTCTGAACTCTTTCCCTACACGACGCTCTTCCGATCTNNNCGCTTTGTGGAATTATCGTTG
GAGCACACGTCTGAACTCTTTCCCTACACGACGCTCTTCCGATCTNNNCGCTTTGTGGAATTATCGTTG
GAGCACACGTCTGAACTCTTTCCCTACACGACGCTCTTCCGATCTNNNCGCTTTGTGGAATTATCGTTG
AGCACACGTCTGAACTCTTTCCCTACACGACGCTCTTCCGATCTNNNAGGGGAAGCTGATAAACACTTG
AGCACACGTCTGAACTCTTTCCCTACACGACGCTCTTCCGATCTNNNAGGGGAAGCTGATAAACACTTG
AGCACACGTCTGAACTCTTTCCCTACACGACGCTCTTCCGATCTNNNAGGGGAAGCTGATAAACACTTG

AGCACACGTCTGAACTCTTTCCCTACACGACGCTCTTCCGATCTNNNAGGGGAAGCTGATAAACACTT
AGCACACGTCTGAACTCTTTCCCTACACGACGCTCTTCCGATCTNNNACCCGGCCTTTTTTCTTTCTTT
AGCACACGTCTGAACTCTTTCCCTACACGACGCTCTTCCGATCTNNNACCCGGCCTTTTTTCTTTCTTT
AGCACACGTCTGAACTCTTTCCCTACACGACGCTCTTCCGATCTNNNACCCGGCCTTTTTTCTTTCTTT
AGCACACGTCTGAACTCTTTCCCTACACGACGCTCTTCCGATCTNNNACCCGGCCTTTTTTCTTTCTTT
AGCACACGTCTGAACTCTTTCCCTACACGACGCTCTTCCGATCTNNNCCATACTGGCCTCTCTCACTTAC
AGCACACGTCTGAACTCTTTCCCTACACGACGCTCTTCCGATCTNNNCCATACTGGCCTCTCTCACTTAC
AGCACACGTCTGAACTCTTTCCCTACACGACGCTCTTCCGATCTNNNCCATACTGGCCTCTCTCACTTAC
AGCACACGTCTGAACTCTTTCCCTACACGACGCTCTTCCGATCTNNNCCATACTGGCCTCTCTCACTTAC
AGCACACGTCTGAACTCTTTCCCTACACGACGCTCTTCCGATCTNNNGACAGATTCGCTTTGTCTCAAC
AGCACACGTCTGAACTCTTTCCCTACACGACGCTCTTCCGATCTNNNGACAGATTCGCTTTGTCTCAAC
AGCACACGTCTGAACTCTTTCCCTACACGACGCTCTTCCGATCTNNNGACAGATTCGCTTTGTCTCAAC
AGCACACGTCTGAACTCTTTCCCTACACGACGCTCTTCCGATCTNNNGACAGATTCGCTTTGTCTCAAC
GAGCACACGTCTGAACTCTTTCCCTACACGACGCTCTTCCGATCTNNNCTGTTGCTGTCAGCTGCATA
GAGCACACGTCTGAACTCTTTCCCTACACGACGCTCTTCCGATCTNNNCTGTTGCTGTCAGCTGCATA
GAGCACACGTCTGAACTCTTTCCCTACACGACGCTCTTCCGATCTNNNCTGTTGCTGTCAGCTGCATA
GAGCACACGTCTGAACTCTTTCCCTACACGACGCTCTTCCGATCTNNNCTGTTGCTGTCAGCTGCATA
GAGCACACGTCTGAACTCTTTCCCTACACGACGCTCTTCCGATCTNNNGTGCCCAAAGACTATCGGAA
GAGCACACGTCTGAACTCTTTCCCTACACGACGCTCTTCCGATCTNNNGTGCCCAAAGACTATCGGAA
GAGCACACGTCTGAACTCTTTCCCTACACGACGCTCTTCCGATCTNNNGTGCCCAAAGACTATCGGAA
GAGCACACGTCTGAACTCTTTCCCTACACGACGCTCTTCCGATCTNNNGTGCCCAAAGACTATCGGAA
GAGCACACGTCTGAACTCTTTCCCTACACGACGCTCTTCCGATCTNNNACAATAGGGAACTTTGGAGG
GAGCACACGTCTGAACTCTTTCCCTACACGACGCTCTTCCGATCTNNNACAATAGGGAACTTTGGAGG
GAGCACACGTCTGAACTCTTTCCCTACACGACGCTCTTCCGATCTNNNACAATAGGGAACTTTGGAGG
GAGCACACGTCTGAACTCTTTCCCTACACGACGCTCTTCCGATCTNNNACAATAGGGAACTTTGGAGG
AGCACACGTCTGAACTCTTTCCCTACACGACGCTCTTCCGATCTNNNGGGCAACATAGTGAAACCACAT
AGCACACGTCTGAACTCTTTCCCTACACGACGCTCTTCCGATCTNNNGGGCAACATAGTGAAACCACAT
AGCACACGTCTGAACTCTTTCCCTACACGACGCTCTTCCGATCTNNNGGGCAACATAGTGAAACCACAT
AGCACACGTCTGAACTCTTTCCCTACACGACGCTCTTCCGATCTNNNGGGCAACATAGTGAAACCACAT
AGCACACGTCTGAACTCTTTCCCTACACGACGCTCTTCCGATCTNNNTGGATGGGTTTCATGGTTACA
AGCACACGTCTGAACTCTTTCCCTACACGACGCTCTTCCGATCTNNNTGGATGGGTTTCATGGTTACA
AGCACACGTCTGAACTCTTTCCCTACACGACGCTCTTCCGATCTNNNTGGATGGGTTTCATGGTTACA
AGCACACGTCTGAACTCTTTCCCTACACGACGCTCTTCCGATCTNNNTGGATGGGTTTCATGGTTACA
GAGCACACGTCTGAACTCTTTCCCTACACGACGCTCTTCCGATCTNNNGAGTTCAGGAGATCGAGACC
GAGCACACGTCTGAACTCTTTCCCTACACGACGCTCTTCCGATCTNNNGAGTTCAGGAGATCGAGACC
GAGCACACGTCTGAACTCTTTCCCTACACGACGCTCTTCCGATCTNNNGAGTTCAGGAGATCGAGACC
GAGCACACGTCTGAACTCTTTCCCTACACGACGCTCTTCCGATCTNNNGAGTTCAGGAGATCGAGACC
AGCACACGTCTGAACTCTTTCCCTACACGACGCTCTTCCGATCTNNNCTATATTGGTCAGGCTGGTCTC
AGCACACGTCTGAACTCTTTCCCTACACGACGCTCTTCCGATCTNNNCTATATTGGTCAGGCTGGTCTC
AGCACACGTCTGAACTCTTTCCCTACACGACGCTCTTCCGATCTNNNCTATATTGGTCAGGCTGGTCTC
AGCACACGTCTGAACTCTTTCCCTACACGACGCTCTTCCGATCTNNNCTATATTGGTCAGGCTGGTCTC
AGCACACGTCTGAACTCTTTCCCTACACGACGCTCTTCCGATCTNNNCAGAGATCGCACCACTGTACTC
AGCACACGTCTGAACTCTTTCCCTACACGACGCTCTTCCGATCTNNNCAGAGATCGCACCACTGTACTC

AGCACACGTCTGAACTCTTTCCCTACACGACGCTCTTCCGATCTNNNCAGAGATCGCACCACTGTACTC
AGCACACGTCTGAACTCTTTCCCTACACGACGCTCTTCCGATCTNNNCAGAGATCGCACCACTGTACTC
GAGCACACGTCTGAACTCTTTCCCTACACGACGCTCTTCCGATCTNNNCGGTGAAAGTCCGTCTCTACT
GAGCACACGTCTGAACTCTTTCCCTACACGACGCTCTTCCGATCTNNNCGGTGAAAGTCCGTCTCTACT
GAGCACACGTCTGAACTCTTTCCCTACACGACGCTCTTCCGATCTNNNCGGTGAAAGTCCGTCTCTACT
GAGCACACGTCTGAACTCTTTCCCTACACGACGCTCTTCCGATCTNNNCGGTGAAAGTCCGTCTCTACT
GCACACGTCTGAACTCTTTCCCTACACGACGCTCTTCCGATCTNNNCTGTTTGGCTCAAAATTAGATGC
GCACACGTCTGAACTCTTTCCCTACACGACGCTCTTCCGATCTNNNCTGTTTGGCTCAAAATTAGATGC
GCACACGTCTGAACTCTTTCCCTACACGACGCTCTTCCGATCTNNNCTGTTTGGCTCAAAATTAGATGC
GCACACGTCTGAACTCTTTCCCTACACGACGCTCTTCCGATCTNNNCTGTTTGGCTCAAAATTAGATGC
GAGCACACGTCTGAACTCTTTCCCTACACGACGCTCTTCCGATCTNNNTTGCAATGAACAACCACATAC
GAGCACACGTCTGAACTCTTTCCCTACACGACGCTCTTCCGATCTNNNTTGCAATGAACAACCACATAC
GAGCACACGTCTGAACTCTTTCCCTACACGACGCTCTTCCGATCTNNNTTGCAATGAACAACCACATAC
GAGCACACGTCTGAACTCTTTCCCTACACGACGCTCTTCCGATCTNNNTTGCAATGAACAACCACATAC
GAGCACACGTCTGAACTCTTTCCCTACACGACGCTCTTCCGATCTNNNAAATGAACAAGAGAACGCAA
GAGCACACGTCTGAACTCTTTCCCTACACGACGCTCTTCCGATCTNNNAAATGAACAAGAGAACGCAA
GAGCACACGTCTGAACTCTTTCCCTACACGACGCTCTTCCGATCTNNNAAATGAACAAGAGAACGCAA
GAGCACACGTCTGAACTCTTTCCCTACACGACGCTCTTCCGATCTNNNAAATGAACAAGAGAACGCAA
GAGCACACGTCTGAACTCTTTCCCTACACGACGCTCTTCCGATCTNNNATTTGCCAAATTTTGATCCCGG
GAGCACACGTCTGAACTCTTTCCCTACACGACGCTCTTCCGATCTNNNATTTGCCAAATTTTGATCCCGG
GAGCACACGTCTGAACTCTTTCCCTACACGACGCTCTTCCGATCTNNNATTTGCCAAATTTTGATCCCGG
GAGCACACGTCTGAACTCTTTCCCTACACGACGCTCTTCCGATCTNNNATTTGCCAAATTTTGATCCCGG
AGCACACGTCTGAACTCTTTCCCTACACGACGCTCTTCCGATCTNNNCAAGACCTAGCTGATGGTACCA
AGCACACGTCTGAACTCTTTCCCTACACGACGCTCTTCCGATCTNNNCAAGACCTAGCTGATGGTACCA
AGCACACGTCTGAACTCTTTCCCTACACGACGCTCTTCCGATCTNNNCAAGACCTAGCTGATGGTACCA
AGCACACGTCTGAACTCTTTCCCTACACGACGCTCTTCCGATCTNNNCAAGACCTAGCTGATGGTACCA
GAGCACACGTCTGAACTCTTTCCCTACACGACGCTCTTCCGATCTNNNGTTGTTGTTCTTTGTAGGCCC
GAGCACACGTCTGAACTCTTTCCCTACACGACGCTCTTCCGATCTNNNGTTGTTGTTCTTTGTAGGCCC
GAGCACACGTCTGAACTCTTTCCCTACACGACGCTCTTCCGATCTNNNGTTGTTGTTCTTTGTAGGCCC
GAGCACACGTCTGAACTCTTTCCCTACACGACGCTCTTCCGATCTNNNGTTGTTGTTCTTTGTAGGCCC
GAGCACACGTCTGAACTCTTTCCCTACACGACGCTCTTCCGATCTNNNCTGGGGACTTAGGGATGTAA
GAGCACACGTCTGAACTCTTTCCCTACACGACGCTCTTCCGATCTNNNCTGGGGACTTAGGGATGTAA
GAGCACACGTCTGAACTCTTTCCCTACACGACGCTCTTCCGATCTNNNCTGGGGACTTAGGGATGTAA
GAGCACACGTCTGAACTCTTTCCCTACACGACGCTCTTCCGATCTNNNCTGGGGACTTAGGGATGTAA
GCACACGTCTGAACTCTTTCCCTACACGACGCTCTTCCGATCTNNNCACGCCACTACACCTAGGTATTTC
GCACACGTCTGAACTCTTTCCCTACACGACGCTCTTCCGATCTNNNCACGCCACTACACCTAGGTATTTC
GCACACGTCTGAACTCTTTCCCTACACGACGCTCTTCCGATCTNNNCACGCCACTACACCTAGGTATTTC
GCACACGTCTGAACTCTTTCCCTACACGACGCTCTTCCGATCTNNNCACGCCACTACACCTAGGTATTTC
AGCACACGTCTGAACTCTTTCCCTACACGACGCTCTTCCGATCTNNNCTTATTTAAGAAATTGCCACAC
AGCACACGTCTGAACTCTTTCCCTACACGACGCTCTTCCGATCTNNNCTTATTTAAGAAATTGCCACAC
AGCACACGTCTGAACTCTTTCCCTACACGACGCTCTTCCGATCTNNNCTTATTTAAGAAATTGCCACAC
AGCACACGTCTGAACTCTTTCCCTACACGACGCTCTTCCGATCTNNNCTTATTTAAGAAATTGCCACAC
GCACACGTCTGAACTCTTTCCCTACACGACGCTCTTCCGATCTNNNTGCAAGTTTTCCACTAAAGGCACC

GCACACGTCTGAACTCTTTCCCTACACGACGCTCTTCCGATCTNNNTGCAAGTTTTCCACTAAAGGCACC
GCACACGTCTGAACTCTTTCCCTACACGACGCTCTTCCGATCTNNNTGCAAGTTTTCCACTAAAGGCACC
GCACACGTCTGAACTCTTTCCCTACACGACGCTCTTCCGATCTNNNTGCAAGTTTTCCACTAAAGGCACC
GCACACGTCTGAACTCTTTCCCTACACGACGCTCTTCCGATCTNNNATCATTTCTCTTCTGTCCCCCTGCA
GCACACGTCTGAACTCTTTCCCTACACGACGCTCTTCCGATCTNNNATCATTTCTCTTCTGTCCCCCTGCA
GCACACGTCTGAACTCTTTCCCTACACGACGCTCTTCCGATCTNNNATCATTTCTCTTCTGTCCCCCTGCA
GCACACGTCTGAACTCTTTCCCTACACGACGCTCTTCCGATCTNNNATCATTTCTCTTCTGTCCCCCTGCA
GCACACGTCTGAACTCTTTCCCTACACGACGCTCTTCCGATCTNNNCCACGAAAACCCTAGTGAATCACC
GCACACGTCTGAACTCTTTCCCTACACGACGCTCTTCCGATCTNNNCCACGAAAACCCTAGTGAATCACC
GCACACGTCTGAACTCTTTCCCTACACGACGCTCTTCCGATCTNNNCCACGAAAACCCTAGTGAATCACC
GCACACGTCTGAACTCTTTCCCTACACGACGCTCTTCCGATCTNNNCCACGAAAACCCTAGTGAATCACC
AGCACACGTCTGAACTCTTTCCCTACACGACGCTCTTCCGATCTNNNTGCCATATTTCAAGCACCACTTCC
AGCACACGTCTGAACTCTTTCCCTACACGACGCTCTTCCGATCTNNNTGCCATATTTCAAGCACCACTTCC
AGCACACGTCTGAACTCTTTCCCTACACGACGCTCTTCCGATCTNNNTGCCATATTTCAAGCACCACTTCC
AGCACACGTCTGAACTCTTTCCCTACACGACGCTCTTCCGATCTNNNTGCCATATTTCAAGCACCACTTCC
AGCACACGTCTGAACTCTTTCCCTACACGACGCTCTTCCGATCTNNNAAGTCTGAGATCAAGGTGTCAG
AGCACACGTCTGAACTCTTTCCCTACACGACGCTCTTCCGATCTNNNAAGTCTGAGATCAAGGTGTCAG
AGCACACGTCTGAACTCTTTCCCTACACGACGCTCTTCCGATCTNNNAAGTCTGAGATCAAGGTGTCAG
AGCACACGTCTGAACTCTTTCCCTACACGACGCTCTTCCGATCTNNNAAGTCTGAGATCAAGGTGTCAG
AGCACACGTCTGAACTCTTTCCCTACACGACGCTCTTCCGATCTNNNGTGAGTGTGTGTGTATGTG
AGCACACGTCTGAACTCTTTCCCTACACGACGCTCTTCCGATCTNNNGTGAGTGTGTGTGTATGTG
AGCACACGTCTGAACTCTTTCCCTACACGACGCTCTTCCGATCTNNNGTGAGTGTGTGTGTATGTG
AGCACACGTCTGAACTCTTTCCCTACACGACGCTCTTCCGATCTNNNGTGAGTGTGTGTGTATGTG
AGCACACGTCTGAACTCTTTCCCTACACGACGCTCTTCCGATCTNNNGGGAGATTTTCTTCCTGGGAGA
AGCACACGTCTGAACTCTTTCCCTACACGACGCTCTTCCGATCTNNNGGGAGATTTTCTTCCTGGGAGA
AGCACACGTCTGAACTCTTTCCCTACACGACGCTCTTCCGATCTNNNGGGAGATTTTCTTCCTGGGAGA
AGCACACGTCTGAACTCTTTCCCTACACGACGCTCTTCCGATCTNNNGGGAGATTTTCTTCCTGGGAGA
AGCACACGTCTGAACTCTTTCCCTACACGACGCTCTTCCGATCTNNNTTGAACCCAGAAGTAGGAGGTT
AGCACACGTCTGAACTCTTTCCCTACACGACGCTCTTCCGATCTNNNTTGAACCCAGAAGTAGGAGGTT
AGCACACGTCTGAACTCTTTCCCTACACGACGCTCTTCCGATCTNNNTTGAACCCAGAAGTAGGAGGTT
AGCACACGTCTGAACTCTTTCCCTACACGACGCTCTTCCGATCTNNNTTGAACCCAGAAGTAGGAGGTT
AAGAGCACACGTCTGAACTCTTTCCCTACACGACGCTCTTCCGATCTNNNTCAAGGTCAACCTACAGTG
AAGAGCACACGTCTGAACTCTTTCCCTACACGACGCTCTTCCGATCTNNNTCAAGGTCAACCTACAGTG
AAGAGCACACGTCTGAACTCTTTCCCTACACGACGCTCTTCCGATCTNNNTCAAGGTCAACCTACAGTG
AAGAGCACACGTCTGAACTCTTTCCCTACACGACGCTCTTCCGATCTNNNTCAAGGTCAACCTACAGTG
GAGCACACGTCTGAACTCTTTCCCTACACGACGCTCTTCCGATCTNNNGATATGAGGGGTGGAAGGG
GAGCACACGTCTGAACTCTTTCCCTACACGACGCTCTTCCGATCTNNNGATATGAGGGGTGGAAGGG
GAGCACACGTCTGAACTCTTTCCCTACACGACGCTCTTCCGATCTNNNGATATGAGGGGTGGAAGGG
GAGCACACGTCTGAACTCTTTCCCTACACGACGCTCTTCCGATCTNNNGATATGAGGGGTGGAAGGG
GAGCACACGTCTGAACTCTTTCCCTACACGACGCTCTTCCGATCTNNNCATTTTTAGTTGCCACAGCTGC
GAGCACACGTCTGAACTCTTTCCCTACACGACGCTCTTCCGATCTNNNCATTTTTAGTTGCCACAGCTGC
GAGCACACGTCTGAACTCTTTCCCTACACGACGCTCTTCCGATCTNNNCATTTTTAGTTGCCACAGCTGC
GAGCACACGTCTGAACTCTTTCCCTACACGACGCTCTTCCGATCTNNNCATTTTTAGTTGCCACAGCTGC

GCACACGTCTGAACTCTTTCCCTACACGACGCTCTTCCGATCTNNNGGCCCTATAGAGGTGAACCATTT
GCACACGTCTGAACTCTTTCCCTACACGACGCTCTTCCGATCTNNNGGCCCTATAGAGGTGAACCATTT
GCACACGTCTGAACTCTTTCCCTACACGACGCTCTTCCGATCTNNNGGCCCTATAGAGGTGAACCATTT
GCACACGTCTGAACTCTTTCCCTACACGACGCTCTTCCGATCTNNNGGCCCTATAGAGGTGAACCATTT
GCACACGTCTGAACTCTTTCCCTACACGACGCTCTTCCGATCTNNNGAGGTTGCAGTGAGCTGAGAT
GCACACGTCTGAACTCTTTCCCTACACGACGCTCTTCCGATCTNNNGAGGTTGCAGTGAGCTGAGAT
GCACACGTCTGAACTCTTTCCCTACACGACGCTCTTCCGATCTNNNGAGGTTGCAGTGAGCTGAGAT
GCACACGTCTGAACTCTTTCCCTACACGACGCTCTTCCGATCTNNNGAGGTTGCAGTGAGCTGAGAT
AGAGCACACGTCTGAACTCTTTCCCTACACGACGCTCTTCCGATCTNNNTCCGAATACTGGGATTAGA
AGAGCACACGTCTGAACTCTTTCCCTACACGACGCTCTTCCGATCTNNNTCCGAATACTGGGATTAGA
AGAGCACACGTCTGAACTCTTTCCCTACACGACGCTCTTCCGATCTNNNTCCGAATACTGGGATTAGA
AGAGCACACGTCTGAACTCTTTCCCTACACGACGCTCTTCCGATCTNNNTCCGAATACTGGGATTAGA
AGCACACGTCTGAACTCTTTCCCTACACGACGCTCTTCCGATCTNNNCTGAGGCAGGAGAATTGGTT
AGCACACGTCTGAACTCTTTCCCTACACGACGCTCTTCCGATCTNNNCTGAGGCAGGAGAATTGGTT
AGCACACGTCTGAACTCTTTCCCTACACGACGCTCTTCCGATCTNNNCTGAGGCAGGAGAATTGGTT
AGCACACGTCTGAACTCTTTCCCTACACGACGCTCTTCCGATCTNNNCTGAGGCAGGAGAATTGGTT
GCACACGTCTGAACTCTTTCCCTACACGACGCTCTTCCGATCTNNNCTTTCCTGATCTAATTGTTCATA
GCACACGTCTGAACTCTTTCCCTACACGACGCTCTTCCGATCTNNNCTTTCCTGATCTAATTGTTCATA
GCACACGTCTGAACTCTTTCCCTACACGACGCTCTTCCGATCTNNNCTTTCCTGATCTAATTGTTCATA
GCACACGTCTGAACTCTTTCCCTACACGACGCTCTTCCGATCTNNNCTTTCCTGATCTAATTGTTCATA
GAGCACACGTCTGAACTCTTTCCCTACACGACGCTCTTCCGATCTNNNAGGAAAAGCACCTCATTTTC
GAGCACACGTCTGAACTCTTTCCCTACACGACGCTCTTCCGATCTNNNAGGAAAAGCACCTCATTTTC
GAGCACACGTCTGAACTCTTTCCCTACACGACGCTCTTCCGATCTNNNAGGAAAAGCACCTCATTTTC
GAGCACACGTCTGAACTCTTTCCCTACACGACGCTCTTCCGATCTNNNAGGAAAAGCACCTCATTTTC
AGCACACGTCTGAACTCTTTCCCTACACGACGCTCTTCCGATCTNNNGTTCCAGTGAGCAGAGATCTC
AGCACACGTCTGAACTCTTTCCCTACACGACGCTCTTCCGATCTNNNGTTCCAGTGAGCAGAGATCTC
AGCACACGTCTGAACTCTTTCCCTACACGACGCTCTTCCGATCTNNNGTTCCAGTGAGCAGAGATCTC
AGCACACGTCTGAACTCTTTCCCTACACGACGCTCTTCCGATCTNNNGTTCCAGTGAGCAGAGATCTC
GCACACGTCTGAACTCTTTCCCTACACGACGCTCTTCCGATCTNNNACCAAAAACCACTTGTACTCCCA
GCACACGTCTGAACTCTTTCCCTACACGACGCTCTTCCGATCTNNNACCAAAAACCACTTGTACTCCCA
GCACACGTCTGAACTCTTTCCCTACACGACGCTCTTCCGATCTNNNACCAAAAACCACTTGTACTCCCA
GCACACGTCTGAACTCTTTCCCTACACGACGCTCTTCCGATCTNNNACCAAAAACCACTTGTACTCCCA
AGCACACGTCTGAACTCTTTCCCTACACGACGCTCTTCCGATCTNNNGGAGGATATCTCTCTGCCTGTT
AGCACACGTCTGAACTCTTTCCCTACACGACGCTCTTCCGATCTNNNGGAGGATATCTCTCTGCCTGTT
AGCACACGTCTGAACTCTTTCCCTACACGACGCTCTTCCGATCTNNNGGAGGATATCTCTCTGCCTGTT
AGCACACGTCTGAACTCTTTCCCTACACGACGCTCTTCCGATCTNNNGGAGGATATCTCTCTGCCTGTT
AGCACACGTCTGAACTCTTTCCCTACACGACGCTCTTCCGATCTNNNAAAATAAAATGTGGAGCCCC
AGCACACGTCTGAACTCTTTCCCTACACGACGCTCTTCCGATCTNNNAAAATAAAATGTGGAGCCCC
AGCACACGTCTGAACTCTTTCCCTACACGACGCTCTTCCGATCTNNNAAAATAAAATGTGGAGCCCC
AGCACACGTCTGAACTCTTTCCCTACACGACGCTCTTCCGATCTNNNAAAATAAAATGTGGAGCCCC
GAGCACACGTCTGAACTCTTTCCCTACACGACGCTCTTCCGATCTNNNGTTCTGTCTCACTAGGGTTCC
GAGCACACGTCTGAACTCTTTCCCTACACGACGCTCTTCCGATCTNNNGTTCTGTCTCACTAGGGTTCC
GAGCACACGTCTGAACTCTTTCCCTACACGACGCTCTTCCGATCTNNNGTTCTGTCTCACTAGGGTTCC

GAGCACACGTCTGAACTCTTTCCCTACACGACGCTCTTCCGATCTNNNGTTCTGTCTCACTAGGGTTCC
GAGCACACGTCTGAACTCTTTCCCTACACGACGCTCTTCCGATCTNNNAGTCTGTCCCTGGATCTAAG
GAGCACACGTCTGAACTCTTTCCCTACACGACGCTCTTCCGATCTNNNAGTCTGTCCCTGGATCTAAG
GAGCACACGTCTGAACTCTTTCCCTACACGACGCTCTTCCGATCTNNNAGTCTGTCCCTGGATCTAAG
GAGCACACGTCTGAACTCTTTCCCTACACGACGCTCTTCCGATCTNNNAGTCTGTCCCTGGATCTAAG
GCACACGTCTGAACTCTTTCCCTACACGACGCTCTTCCGATCTNNNCTTCTTCACTGTACCTGCAGGAT
GCACACGTCTGAACTCTTTCCCTACACGACGCTCTTCCGATCTNNNCTTCTTCACTGTACCTGCAGGAT
GCACACGTCTGAACTCTTTCCCTACACGACGCTCTTCCGATCTNNNCTTCTTCACTGTACCTGCAGGAT
GCACACGTCTGAACTCTTTCCCTACACGACGCTCTTCCGATCTNNNCTTCTTCACTGTACCTGCAGGAT
AGCACACGTCTGAACTCTTTCCCTACACGACGCTCTTCCGATCTNNNAATCACATGAACCTAGGAGGC
AGCACACGTCTGAACTCTTTCCCTACACGACGCTCTTCCGATCTNNNAATCACATGAACCTAGGAGGC
AGCACACGTCTGAACTCTTTCCCTACACGACGCTCTTCCGATCTNNNAATCACATGAACCTAGGAGGC
AGCACACGTCTGAACTCTTTCCCTACACGACGCTCTTCCGATCTNNNAATCACATGAACCTAGGAGGC
AGCACACGTCTGAACTCTTTCCCTACACGACGCTCTTCCGATCTNNNACTGAATGCCCCACAATGCTAT
AGCACACGTCTGAACTCTTTCCCTACACGACGCTCTTCCGATCTNNNACTGAATGCCCCACAATGCTAT
AGCACACGTCTGAACTCTTTCCCTACACGACGCTCTTCCGATCTNNNACTGAATGCCCCACAATGCTAT
AGCACACGTCTGAACTCTTTCCCTACACGACGCTCTTCCGATCTNNNACTGAATGCCCCACAATGCTAT
GAGCACACGTCTGAACTCTTTCCCTACACGACGCTCTTCCGATCTNNNTTAGAGAGACCAGTTGTGCC
GAGCACACGTCTGAACTCTTTCCCTACACGACGCTCTTCCGATCTNNNTTAGAGAGACCAGTTGTGCC
GAGCACACGTCTGAACTCTTTCCCTACACGACGCTCTTCCGATCTNNNTTAGAGAGACCAGTTGTGCC
GAGCACACGTCTGAACTCTTTCCCTACACGACGCTCTTCCGATCTNNNTTAGAGAGACCAGTTGTGCC
GCACACGTCTGAACTCTTTCCCTACACGACGCTCTTCCGATCTNNNGAGGGGCAACTGTATAACTGG
GCACACGTCTGAACTCTTTCCCTACACGACGCTCTTCCGATCTNNNGAGGGGCAACTGTATAACTGG
GCACACGTCTGAACTCTTTCCCTACACGACGCTCTTCCGATCTNNNGAGGGGCAACTGTATAACTGG
GCACACGTCTGAACTCTTTCCCTACACGACGCTCTTCCGATCTNNNGAGGGGCAACTGTATAACTGG
GCACACGTCTGAACTCTTTCCCTACACGACGCTCTTCCGATCTNNNTACAAAAAATTAGCTGGGCGTG
GCACACGTCTGAACTCTTTCCCTACACGACGCTCTTCCGATCTNNNTACAAAAAATTAGCTGGGCGTG
GCACACGTCTGAACTCTTTCCCTACACGACGCTCTTCCGATCTNNNTACAAAAAATTAGCTGGGCGTG
GCACACGTCTGAACTCTTTCCCTACACGACGCTCTTCCGATCTNNNTACAAAAAATTAGCTGGGCGTG
AGCACACGTCTGAACTCTTTCCCTACACGACGCTCTTCCGATCTNNNTAGAGAGAGAGAGAGAGGG
AGCACACGTCTGAACTCTTTCCCTACACGACGCTCTTCCGATCTNNNTAGAGAGAGAGAGAGAGGG
AGCACACGTCTGAACTCTTTCCCTACACGACGCTCTTCCGATCTNNNTAGAGAGAGAGAGAGAGGG
AGCACACGTCTGAACTCTTTCCCTACACGACGCTCTTCCGATCTNNNTAGAGAGAGAGAGAGAGGG
AGCACACGTCTGAACTCTTTCCCTACACGACGCTCTTCCGATCTNNNTCCAATGACTTTTTCCTCCCCTT
AGCACACGTCTGAACTCTTTCCCTACACGACGCTCTTCCGATCTNNNTCCAATGACTTTTTCCTCCCCTT
AGCACACGTCTGAACTCTTTCCCTACACGACGCTCTTCCGATCTNNNTCCAATGACTTTTTCCTCCCCTT
AGCACACGTCTGAACTCTTTCCCTACACGACGCTCTTCCGATCTNNNTCCAATGACTTTTTCCTCCCCTT
GAGCACACGTCTGAACTCTTTCCCTACACGACGCTCTTCCGATCTNNNCCCTCCACACCTACAAACTGT
GAGCACACGTCTGAACTCTTTCCCTACACGACGCTCTTCCGATCTNNNCCCTCCACACCTACAAACTGT
GAGCACACGTCTGAACTCTTTCCCTACACGACGCTCTTCCGATCTNNNCCCTCCACACCTACAAACTGT
GAGCACACGTCTGAACTCTTTCCCTACACGACGCTCTTCCGATCTNNNCCCTCCACACCTACAAACTGT
ACACGTCTGAACTCTTTCCCTACACGACGCTCTTCCGATCTNNNCTTTGTTTTGTGCTGAAAGTCGCCA
ACACGTCTGAACTCTTTCCCTACACGACGCTCTTCCGATCTNNNCTTTGTTTTGTGCTGAAAGTCGCCA

CACACGTCTGAACTCTTTCCCTACACGACGCTCTTCCGATCTNNNCTTTGTTTTGTGCTGAAAGTCGCCA
CACACGTCTGAACTCTTTCCCTACACGACGCTCTTCCGATCTNNNCTTTGTTTTGTGCTGAAAGTCGCCA
AGCACACGTCTGAACTCTTTCCCTACACGACGCTCTTCCGATCTNNNGAGCCTAACACTCTCCAACCTAC
AGCACACGTCTGAACTCTTTCCCTACACGACGCTCTTCCGATCTNNNGAGCCTAACACTCTCCAACCTAC
AGCACACGTCTGAACTCTTTCCCTACACGACGCTCTTCCGATCTNNNGAGCCTAACACTCTCCAACCTAC
AGCACACGTCTGAACTCTTTCCCTACACGACGCTCTTCCGATCTNNNGAGCCTAACACTCTCCAACCTAC
GAGCACACGTCTGAACTCTTTCCCTACACGACGCTCTTCCGATCTNNNCCCCACTCCTTTCAGTGAAATTC
GAGCACACGTCTGAACTCTTTCCCTACACGACGCTCTTCCGATCTNNNCCCCACTCCTTTCAGTGAAATTC
GAGCACACGTCTGAACTCTTTCCCTACACGACGCTCTTCCGATCTNNNCCCCACTCCTTTCAGTGAAATTC
GAGCACACGTCTGAACTCTTTCCCTACACGACGCTCTTCCGATCTNNNCCCCACTCCTTTCAGTGAAATTC
AGAGCACACGTCTGAACTCTTTCCCTACACGACGCTCTTCCGATCTNNNCAATGGTGTCACAGAAAGAG
AGAGCACACGTCTGAACTCTTTCCCTACACGACGCTCTTCCGATCTNNNCAATGGTGTCACAGAAAGAG
AGAGCACACGTCTGAACTCTTTCCCTACACGACGCTCTTCCGATCTNNNCAATGGTGTCACAGAAAGAG
AGAGCACACGTCTGAACTCTTTCCCTACACGACGCTCTTCCGATCTNNNCAATGGTGTCACAGAAAGAG
GAGCACACGTCTGAACTCTTTCCCTACACGACGCTCTTCCGATCTNNNCCTGACCCCACTCTTTTCTTTCT
GAGCACACGTCTGAACTCTTTCCCTACACGACGCTCTTCCGATCTNNNCCTGACCCCACTCTTTTCTTTCT
GAGCACACGTCTGAACTCTTTCCCTACACGACGCTCTTCCGATCTNNNCCTGACCCCACTCTTTTCTTTCT
GAGCACACGTCTGAACTCTTTCCCTACACGACGCTCTTCCGATCTNNNCCTGACCCCACTCTTTTCTTTCT
AGCACACGTCTGAACTCTTTCCCTACACGACGCTCTTCCGATCTNNNATTTGCATCTGGTAAGAGCCTC
AGCACACGTCTGAACTCTTTCCCTACACGACGCTCTTCCGATCTNNNATTTGCATCTGGTAAGAGCCTC
AGCACACGTCTGAACTCTTTCCCTACACGACGCTCTTCCGATCTNNNATTTGCATCTGGTAAGAGCCTC
AGCACACGTCTGAACTCTTTCCCTACACGACGCTCTTCCGATCTNNNATTTGCATCTGGTAAGAGCCTC
AGCACACGTCTGAACTCTTTCCCTACACGACGCTCTTCCGATCTNNNGTATGGCTTCTGGGTCATTGCAT
AGCACACGTCTGAACTCTTTCCCTACACGACGCTCTTCCGATCTNNNGTATGGCTTCTGGGTCATTGCAT
AGCACACGTCTGAACTCTTTCCCTACACGACGCTCTTCCGATCTNNNGTATGGCTTCTGGGTCATTGCAT
AGCACACGTCTGAACTCTTTCCCTACACGACGCTCTTCCGATCTNNNGTATGGCTTCTGGGTCATTGCAT
GAGCACACGTCTGAACTCTTTCCCTACACGACGCTCTTCCGATCTNNNATTTCAGCAAGATGAACAGCC
GAGCACACGTCTGAACTCTTTCCCTACACGACGCTCTTCCGATCTNNNATTTCAGCAAGATGAACAGCC
GAGCACACGTCTGAACTCTTTCCCTACACGACGCTCTTCCGATCTNNNATTTCAGCAAGATGAACAGCC
GAGCACACGTCTGAACTCTTTCCCTACACGACGCTCTTCCGATCTNNNATTTCAGCAAGATGAACAGCC
AGCACACGTCTGAACTCTTTCCCTACACGACGCTCTTCCGATCTNNNCTGCCTGTGAATGTTGTGATCTG
AGCACACGTCTGAACTCTTTCCCTACACGACGCTCTTCCGATCTNNNCTGCCTGTGAATGTTGTGATCTG
AGCACACGTCTGAACTCTTTCCCTACACGACGCTCTTCCGATCTNNNCTGCCTGTGAATGTTGTGATCTG
AGCACACGTCTGAACTCTTTCCCTACACGACGCTCTTCCGATCTNNNCTGCCTGTGAATGTTGTGATCTG
AGCACACGTCTGAACTCTTTCCCTACACGACGCTCTTCCGATCTNNNTGAGACTCATGTTTTTACCCCTTC
AGCACACGTCTGAACTCTTTCCCTACACGACGCTCTTCCGATCTNNNTGAGACTCATGTTTTTACCCCTTC
AGCACACGTCTGAACTCTTTCCCTACACGACGCTCTTCCGATCTNNNTGAGACTCATGTTTTTACCCCTTC
AGCACACGTCTGAACTCTTTCCCTACACGACGCTCTTCCGATCTNNNTGAGACTCATGTTTTTACCCCTTC
CACACGTCTGAACTCTTTCCCTACACGACGCTCTTCCGATCTNNNCTCTACTTCCGGAGCCTGGAGCATC
CACACGTCTGAACTCTTTCCCTACACGACGCTCTTCCGATCTNNNCTCTACTTCCGGAGCCTGGAGCATC
CACACGTCTGAACTCTTTCCCTACACGACGCTCTTCCGATCTNNNCTCTACTTCCGGAGCCTGGAGCATC
CACACGTCTGAACTCTTTCCCTACACGACGCTCTTCCGATCTNNNCTCTACTTCCGGAGCCTGGAGCATC
GCACACGTCTGAACTCTTTCCCTACACGACGCTCTTCCGATCTNNNAATGACATTCTCCAGAGTCCCACC

GCACACGTCTGAACTCTTTCCCTACACGACGCTCTTCCGATCTNNNAATGACATTCTCCAGAGTCCCAC
GCACACGTCTGAACTCTTTCCCTACACGACGCTCTTCCGATCTNNNAATGACATTCTCCAGAGTCCCAC
GCACACGTCTGAACTCTTTCCCTACACGACGCTCTTCCGATCTNNNAATGACATTCTCCAGAGTCCCAC
AGCACACGTCTGAACTCTTTCCCTACACGACGCTCTTCCGATCTNNNCATCCCCTTAAATTGGATTAAA
AGCACACGTCTGAACTCTTTCCCTACACGACGCTCTTCCGATCTNNNCATCCCCTTAAATTGGATTAAA
AGCACACGTCTGAACTCTTTCCCTACACGACGCTCTTCCGATCTNNNCATCCCCTTAAATTGGATTAAA
AGCACACGTCTGAACTCTTTCCCTACACGACGCTCTTCCGATCTNNNCATCCCCTTAAATTGGATTAAA
AGAGCACACGTCTGAACTCTTTCCCTACACGACGCTCTTCCGATCTNNNCCTGGTTGTAATCCCAGCTA
AGAGCACACGTCTGAACTCTTTCCCTACACGACGCTCTTCCGATCTNNNCCTGGTTGTAATCCCAGCTA
AGAGCACACGTCTGAACTCTTTCCCTACACGACGCTCTTCCGATCTNNNCCTGGTTGTAATCCCAGCTA
AGAGCACACGTCTGAACTCTTTCCCTACACGACGCTCTTCCGATCTNNNCCTGGTTGTAATCCCAGCTA
GAGCACACGTCTGAACTCTTTCCCTACACGACGCTCTTCCGATCTNNNTAAGCTGAGATCATGTCACTG
GAGCACACGTCTGAACTCTTTCCCTACACGACGCTCTTCCGATCTNNNTAAGCTGAGATCATGTCACTG
GAGCACACGTCTGAACTCTTTCCCTACACGACGCTCTTCCGATCTNNNTAAGCTGAGATCATGTCACTG
GAGCACACGTCTGAACTCTTTCCCTACACGACGCTCTTCCGATCTNNNTAAGCTGAGATCATGTCACTG
AGCACACGTCTGAACTCTTTCCCTACACGACGCTCTTCCGATCTNNNTAATAATCCCCACGAGTCAAGG
AGCACACGTCTGAACTCTTTCCCTACACGACGCTCTTCCGATCTNNNTAATAATCCCCACGAGTCAAGG
AGCACACGTCTGAACTCTTTCCCTACACGACGCTCTTCCGATCTNNNTAATAATCCCCACGAGTCAAGG
AGCACACGTCTGAACTCTTTCCCTACACGACGCTCTTCCGATCTNNNTAATAATCCCCACGAGTCAAGG
AGCACACGTCTGAACTCTTTCCCTACACGACGCTCTTCCGATCTNNNATATTACATTTCAGAGTTCACC
AGCACACGTCTGAACTCTTTCCCTACACGACGCTCTTCCGATCTNNNATATTACATTTCAGAGTTCACC
AGCACACGTCTGAACTCTTTCCCTACACGACGCTCTTCCGATCTNNNATATTACATTTCAGAGTTCACC
AGCACACGTCTGAACTCTTTCCCTACACGACGCTCTTCCGATCTNNNATATTACATTTCAGAGTTCACC
AGCACACGTCTGAACTCTTTCCCTACACGACGCTCTTCCGATCTNNNGAGTTGTGGTGACAGAGATGT
AGCACACGTCTGAACTCTTTCCCTACACGACGCTCTTCCGATCTNNNGAGTTGTGGTGACAGAGATGT
AGCACACGTCTGAACTCTTTCCCTACACGACGCTCTTCCGATCTNNNGAGTTGTGGTGACAGAGATGT
AGCACACGTCTGAACTCTTTCCCTACACGACGCTCTTCCGATCTNNNGAGTTGTGGTGACAGAGATGT
AGCACACGTCTGAACTCTTTCCCTACACGACGCTCTTCCGATCTNNNTAAGGCAGGAGAATTGCTTGA
AGCACACGTCTGAACTCTTTCCCTACACGACGCTCTTCCGATCTNNNTAAGGCAGGAGAATTGCTTGA
AGCACACGTCTGAACTCTTTCCCTACACGACGCTCTTCCGATCTNNNTAAGGCAGGAGAATTGCTTGA
AGCACACGTCTGAACTCTTTCCCTACACGACGCTCTTCCGATCTNNNTAAGGCAGGAGAATTGCTTGA
AGCACACGTCTGAACTCTTTCCCTACACGACGCTCTTCCGATCTNNNAGCTGAGACAGGAGAATCACT
AGCACACGTCTGAACTCTTTCCCTACACGACGCTCTTCCGATCTNNNAGCTGAGACAGGAGAATCACT
AGCACACGTCTGAACTCTTTCCCTACACGACGCTCTTCCGATCTNNNAGCTGAGACAGGAGAATCACT
AGCACACGTCTGAACTCTTTCCCTACACGACGCTCTTCCGATCTNNNAGCTGAGACAGGAGAATCACT
AGCACACGTCTGAACTCTTTCCCTACACGACGCTCTTCCGATCTNNNCACTACTCCGAGCCGAAAATAC
AGCACACGTCTGAACTCTTTCCCTACACGACGCTCTTCCGATCTNNNCACTACTCCGAGCCGAAAATAC
AGCACACGTCTGAACTCTTTCCCTACACGACGCTCTTCCGATCTNNNCACTACTCCGAGCCGAAAATAC
AGCACACGTCTGAACTCTTTCCCTACACGACGCTCTTCCGATCTNNNCACTACTCCGAGCCGAAAATAC
AGCACACGTCTGAACTCTTTCCCTACACGACGCTCTTCCGATCTNNNAGAGTTGGGAGAAAGGGATG
AGCACACGTCTGAACTCTTTCCCTACACGACGCTCTTCCGATCTNNNAGAGTTGGGAGAAAGGGATG
AGCACACGTCTGAACTCTTTCCCTACACGACGCTCTTCCGATCTNNNAGAGTTGGGAGAAAGGGATG
AGCACACGTCTGAACTCTTTCCCTACACGACGCTCTTCCGATCTNNNAGAGTTGGGAGAAAGGGATG

AGCACACGTCTGAACTCTTTCCCTACACGACGCTCTTCCGATCTNNNGTTCTCTCTCTCACCCTCTCT
AGCACACGTCTGAACTCTTTCCCTACACGACGCTCTTCCGATCTNNNGTTCTCTCTCTCACCCTCTCT
AGCACACGTCTGAACTCTTTCCCTACACGACGCTCTTCCGATCTNNNGTTCTCTCTCTCACCCTCTCT
AGCACACGTCTGAACTCTTTCCCTACACGACGCTCTTCCGATCTNNNGTTCTCTCTCTCACCCTCTCT
AGCACACGTCTGAACTCTTTCCCTACACGACGCTCTTCCGATCTNNNCCATGCCCGGCTAATTTTTCTA
AGCACACGTCTGAACTCTTTCCCTACACGACGCTCTTCCGATCTNNNCCATGCCCGGCTAATTTTTCTA
AGCACACGTCTGAACTCTTTCCCTACACGACGCTCTTCCGATCTNNNCCATGCCCGGCTAATTTTTCTA
AGCACACGTCTGAACTCTTTCCCTACACGACGCTCTTCCGATCTNNNCCATGCCCGGCTAATTTTTCTA
GAGCACACGTCTGAACTCTTTCCCTACACGACGCTCTTCCGATCTNNNTTTTGCGGAGGTCAGATGTT
GAGCACACGTCTGAACTCTTTCCCTACACGACGCTCTTCCGATCTNNNTTTTGCGGAGGTCAGATGTT
GAGCACACGTCTGAACTCTTTCCCTACACGACGCTCTTCCGATCTNNNTTTTGCGGAGGTCAGATGTT
GAGCACACGTCTGAACTCTTTCCCTACACGACGCTCTTCCGATCTNNNTTTTGCGGAGGTCAGATGTT
GCACACGTCTGAACTCTTTCCCTACACGACGCTCTTCCGATCTNNNGAGAATGGCATGAACCCAGGA
GCACACGTCTGAACTCTTTCCCTACACGACGCTCTTCCGATCTNNNGAGAATGGCATGAACCCAGGA
GCACACGTCTGAACTCTTTCCCTACACGACGCTCTTCCGATCTNNNGAGAATGGCATGAACCCAGGA
GCACACGTCTGAACTCTTTCCCTACACGACGCTCTTCCGATCTNNNGAGAATGGCATGAACCCAGGA
GAGCACACGTCTGAACTCTTTCCCTACACGACGCTCTTCCGATCTNNNCCTGTTTGATGTGGAAGCCTT
GAGCACACGTCTGAACTCTTTCCCTACACGACGCTCTTCCGATCTNNNCCTGTTTGATGTGGAAGCCTT
GAGCACACGTCTGAACTCTTTCCCTACACGACGCTCTTCCGATCTNNNCCTGTTTGATGTGGAAGCCTT
GAGCACACGTCTGAACTCTTTCCCTACACGACGCTCTTCCGATCTNNNCCTGTTTGATGTGGAAGCCTT
AGCACACGTCTGAACTCTTTCCCTACACGACGCTCTTCCGATCTNNNGTGGTAATTTCTGCAGCTGAAC
AGCACACGTCTGAACTCTTTCCCTACACGACGCTCTTCCGATCTNNNGTGGTAATTTCTGCAGCTGAAC
AGCACACGTCTGAACTCTTTCCCTACACGACGCTCTTCCGATCTNNNGTGGTAATTTCTGCAGCTGAAC
AGCACACGTCTGAACTCTTTCCCTACACGACGCTCTTCCGATCTNNNGTGGTAATTTCTGCAGCTGAAC
AGCACACGTCTGAACTCTTTCCCTACACGACGCTCTTCCGATCTNNNTTGCAACAAATGTCACCAACAG
AGCACACGTCTGAACTCTTTCCCTACACGACGCTCTTCCGATCTNNNTTGCAACAAATGTCACCAACAG
AGCACACGTCTGAACTCTTTCCCTACACGACGCTCTTCCGATCTNNNTTGCAACAAATGTCACCAACAG
AGCACACGTCTGAACTCTTTCCCTACACGACGCTCTTCCGATCTNNNTTGCAACAAATGTCACCAACAG
GAGCACACGTCTGAACTCTTTCCCTACACGACGCTCTTCCGATCTNNNTGGATTGTGGTCATGGTTAC
GAGCACACGTCTGAACTCTTTCCCTACACGACGCTCTTCCGATCTNNNTGGATTGTGGTCATGGTTAC
GAGCACACGTCTGAACTCTTTCCCTACACGACGCTCTTCCGATCTNNNTGGATTGTGGTCATGGTTAC
GAGCACACGTCTGAACTCTTTCCCTACACGACGCTCTTCCGATCTNNNTGGATTGTGGTCATGGTTAC
AGCACACGTCTGAACTCTTTCCCTACACGACGCTCTTCCGATCTNNNACACAGCCATGTCCACTCATTT
AGCACACGTCTGAACTCTTTCCCTACACGACGCTCTTCCGATCTNNNACACAGCCATGTCCACTCATTT
AGCACACGTCTGAACTCTTTCCCTACACGACGCTCTTCCGATCTNNNACACAGCCATGTCCACTCATTT
AGCACACGTCTGAACTCTTTCCCTACACGACGCTCTTCCGATCTNNNACACAGCCATGTCCACTCATTT
AGCACACGTCTGAACTCTTTCCCTACACGACGCTCTTCCGATCTNNNTCTCAAAGGTCAGATCTCTGCA
AGCACACGTCTGAACTCTTTCCCTACACGACGCTCTTCCGATCTNNNTCTCAAAGGTCAGATCTCTGCA
AGCACACGTCTGAACTCTTTCCCTACACGACGCTCTTCCGATCTNNNTCTCAAAGGTCAGATCTCTGCA
AGCACACGTCTGAACTCTTTCCCTACACGACGCTCTTCCGATCTNNNTCTCAAAGGTCAGATCTCTGCA
GCACACGTCTGAACTCTTTCCCTACACGACGCTCTTCCGATCTNNNTCAGATCAAGGTTCCCATTTCCA
GCACACGTCTGAACTCTTTCCCTACACGACGCTCTTCCGATCTNNNTCAGATCAAGGTTCCCATTTCCA
GCACACGTCTGAACTCTTTCCCTACACGACGCTCTTCCGATCTNNNTCAGATCAAGGTTCCCATTTCCA

AGCACACGTCTGAACTCTTTCCCTACACGACGCTCTTCCGATCTNNNTCAGATCAAGGTTCCCATTTCCA
AGAGCACACGTCTGAACTCTTTCCCTACACGACGCTCTTCCGATCTNNNGGTCTACCTGATGCATGATC
AGAGCACACGTCTGAACTCTTTCCCTACACGACGCTCTTCCGATCTNNNGGTCTACCTGATGCATGATC
AGAGCACACGTCTGAACTCTTTCCCTACACGACGCTCTTCCGATCTNNNGGTCTACCTGATGCATGATC
AGAGCACACGTCTGAACTCTTTCCCTACACGACGCTCTTCCGATCTNNNGGTCTACCTGATGCATGATC
AGCACACGTCTGAACTCTTTCCCTACACGACGCTCTTCCGATCTNNNCAGGAGAATCACTTGAACCAGC
AGCACACGTCTGAACTCTTTCCCTACACGACGCTCTTCCGATCTNNNCAGGAGAATCACTTGAACCAGC
AGCACACGTCTGAACTCTTTCCCTACACGACGCTCTTCCGATCTNNNCAGGAGAATCACTTGAACCAGC
AGCACACGTCTGAACTCTTTCCCTACACGACGCTCTTCCGATCTNNNCAGGAGAATCACTTGAACCAGC
GAGCACACGTCTGAACTCTTTCCCTACACGACGCTCTTCCGATCTNNNATCATGAGGTCAGGAGATCG
GAGCACACGTCTGAACTCTTTCCCTACACGACGCTCTTCCGATCTNNNATCATGAGGTCAGGAGATCG
GAGCACACGTCTGAACTCTTTCCCTACACGACGCTCTTCCGATCTNNNATCATGAGGTCAGGAGATCG
GAGCACACGTCTGAACTCTTTCCCTACACGACGCTCTTCCGATCTNNNATCATGAGGTCAGGAGATCG
AGCACACGTCTGAACTCTTTCCCTACACGACGCTCTTCCGATCTNNNTGAGAAGGACAACTGACCTGAA
AGCACACGTCTGAACTCTTTCCCTACACGACGCTCTTCCGATCTNNNTGAGAAGGACAACTGACCTGAA
AGCACACGTCTGAACTCTTTCCCTACACGACGCTCTTCCGATCTNNNTGAGAAGGACAACTGACCTGAA
AGCACACGTCTGAACTCTTTCCCTACACGACGCTCTTCCGATCTNNNTGAGAAGGACAACTGACCTGAA
GAGCACACGTCTGAACTCTTTCCCTACACGACGCTCTTCCGATCTNNNAGGAGTTCTCTAAACCCAAGC
GAGCACACGTCTGAACTCTTTCCCTACACGACGCTCTTCCGATCTNNNAGGAGTTCTCTAAACCCAAGC
GAGCACACGTCTGAACTCTTTCCCTACACGACGCTCTTCCGATCTNNNAGGAGTTCTCTAAACCCAAGC
GAGCACACGTCTGAACTCTTTCCCTACACGACGCTCTTCCGATCTNNNAGGAGTTCTCTAAACCCAAGC
GAGCACACGTCTGAACTCTTTCCCTACACGACGCTCTTCCGATCTNNNTCTTCAAGGAGCACGAGATGA
GAGCACACGTCTGAACTCTTTCCCTACACGACGCTCTTCCGATCTNNNTCTTCAAGGAGCACGAGATGA
GAGCACACGTCTGAACTCTTTCCCTACACGACGCTCTTCCGATCTNNNTCTTCAAGGAGCACGAGATGA
GAGCACACGTCTGAACTCTTTCCCTACACGACGCTCTTCCGATCTNNNTCTTCAAGGAGCACGAGATGA
CACACGTCTGAACTCTTTCCCTACACGACGCTCTTCCGATCTNNNCCCTCAAGCCATACAGAGATTGTCA
CACACGTCTGAACTCTTTCCCTACACGACGCTCTTCCGATCTNNNCCCTCAAGCCATACAGAGATTGTCA
CACACGTCTGAACTCTTTCCCTACACGACGCTCTTCCGATCTNNNCCCTCAAGCCATACAGAGATTGTCA
CACACGTCTGAACTCTTTCCCTACACGACGCTCTTCCGATCTNNNCCCTCAAGCCATACAGAGATTGTCA
AGCACACGTCTGAACTCTTTCCCTACACGACGCTCTTCCGATCTNNNAAGACTGAGAGTGAGGTGGAC
AGCACACGTCTGAACTCTTTCCCTACACGACGCTCTTCCGATCTNNNAAGACTGAGAGTGAGGTGGAC
AGCACACGTCTGAACTCTTTCCCTACACGACGCTCTTCCGATCTNNNAAGACTGAGAGTGAGGTGGAC
AGCACACGTCTGAACTCTTTCCCTACACGACGCTCTTCCGATCTNNNAAGACTGAGAGTGAGGTGGAC
AGAGCACACGTCTGAACTCTTTCCCTACACGACGCTCTTCCGATCTNNNGGAGGTTACAGTGGGTCAA
AGAGCACACGTCTGAACTCTTTCCCTACACGACGCTCTTCCGATCTNNNGGAGGTTACAGTGGGTCAA
AGAGCACACGTCTGAACTCTTTCCCTACACGACGCTCTTCCGATCTNNNGGAGGTTACAGTGGGTCAA
AGAGCACACGTCTGAACTCTTTCCCTACACGACGCTCTTCCGATCTNNNGGAGGTTACAGTGGGTCAA
AGCACACGTCTGAACTCTTTCCCTACACGACGCTCTTCCGATCTNNNCTGGAGTACAGTGATCTCAGCT
AGCACACGTCTGAACTCTTTCCCTACACGACGCTCTTCCGATCTNNNCTGGAGTACAGTGATCTCAGCT
AGCACACGTCTGAACTCTTTCCCTACACGACGCTCTTCCGATCTNNNCTGGAGTACAGTGATCTCAGCT
AGCACACGTCTGAACTCTTTCCCTACACGACGCTCTTCCGATCTNNNCTGGAGTACAGTGATCTCAGCT
AGAGCACACGTCTGAACTCTTTCCCTACACGACGCTCTTCCGATCTNNNAGCAATAGACTCCGTCTCCAA
AGAGCACACGTCTGAACTCTTTCCCTACACGACGCTCTTCCGATCTNNNAGCAATAGACTCCGTCTCCAA

AGAGCACACGTCTGAACTCTTTCCCTACACGACGCTCTTCCGATCTNNNAGCAATAGACTCCGTCTCCA
AGAGCACACGTCTGAACTCTTTCCCTACACGACGCTCTTCCGATCTNNNAGCAATAGACTCCGTCTCCA
GCACACGTCTGAACTCTTTCCCTACACGACGCTCTTCCGATCTNNNGAAGGGATCCAGTTTCAGCTTTT
GCACACGTCTGAACTCTTTCCCTACACGACGCTCTTCCGATCTNNNGAAGGGATCCAGTTTCAGCTTTT
GCACACGTCTGAACTCTTTCCCTACACGACGCTCTTCCGATCTNNNGAAGGGATCCAGTTTCAGCTTTT
GCACACGTCTGAACTCTTTCCCTACACGACGCTCTTCCGATCTNNNGAAGGGATCCAGTTTCAGCTTTT
AGCACACGTCTGAACTCTTTCCCTACACGACGCTCTTCCGATCTNNNCTGAGGCAGGAGAATCACTTGA
AGCACACGTCTGAACTCTTTCCCTACACGACGCTCTTCCGATCTNNNCTGAGGCAGGAGAATCACTTGA
AGCACACGTCTGAACTCTTTCCCTACACGACGCTCTTCCGATCTNNNCTGAGGCAGGAGAATCACTTGA
AGCACACGTCTGAACTCTTTCCCTACACGACGCTCTTCCGATCTNNNCTGAGGCAGGAGAATCACTTGA
AGCACACGTCTGAACTCTTTCCCTACACGACGCTCTTCCGATCTNNNAAGGAGTCTTCAGCCCTAGAAG
AGCACACGTCTGAACTCTTTCCCTACACGACGCTCTTCCGATCTNNNAAGGAGTCTTCAGCCCTAGAAG
AGCACACGTCTGAACTCTTTCCCTACACGACGCTCTTCCGATCTNNNAAGGAGTCTTCAGCCCTAGAAG
AGCACACGTCTGAACTCTTTCCCTACACGACGCTCTTCCGATCTNNNAAGGAGTCTTCAGCCCTAGAAG
GAGCACACGTCTGAACTCTTTCCCTACACGACGCTCTTCCGATCTNNNGGAGGTTGCAATGAACCAAG
GAGCACACGTCTGAACTCTTTCCCTACACGACGCTCTTCCGATCTNNNGGAGGTTGCAATGAACCAAG
GAGCACACGTCTGAACTCTTTCCCTACACGACGCTCTTCCGATCTNNNGGAGGTTGCAATGAACCAAG
GAGCACACGTCTGAACTCTTTCCCTACACGACGCTCTTCCGATCTNNNGGAGGTTGCAATGAACCAAG
AGCACACGTCTGAACTCTTTCCCTACACGACGCTCTTCCGATCTNNNACCCTGTCTGAGAAGTGAGGAG
AGCACACGTCTGAACTCTTTCCCTACACGACGCTCTTCCGATCTNNNACCCTGTCTGAGAAGTGAGGAG
AGCACACGTCTGAACTCTTTCCCTACACGACGCTCTTCCGATCTNNNACCCTGTCTGAGAAGTGAGGAG
AGCACACGTCTGAACTCTTTCCCTACACGACGCTCTTCCGATCTNNNACCCTGTCTGAGAAGTGAGGAG
AGCACACGTCTGAACTCTTTCCCTACACGACGCTCTTCCGATCTNNNACTTTGCCATGGAAAGAAGA
AGCACACGTCTGAACTCTTTCCCTACACGACGCTCTTCCGATCTNNNACTTTGCCATGGAAAGAAGA
AGCACACGTCTGAACTCTTTCCCTACACGACGCTCTTCCGATCTNNNACTTTGCCATGGAAAGAAGA
AGCACACGTCTGAACTCTTTCCCTACACGACGCTCTTCCGATCTNNNACTTTGCCATGGAAAGAAGA
AGAGCACACGTCTGAACTCTTTCCCTACACGACGCTCTTCCGATCTNNNGCAGGAGAATTTCTTGAATC
AGAGCACACGTCTGAACTCTTTCCCTACACGACGCTCTTCCGATCTNNNGCAGGAGAATTTCTTGAATC
AGAGCACACGTCTGAACTCTTTCCCTACACGACGCTCTTCCGATCTNNNGCAGGAGAATTTCTTGAATC
AGAGCACACGTCTGAACTCTTTCCCTACACGACGCTCTTCCGATCTNNNGCAGGAGAATTTCTTGAATC
GAGCACACGTCTGAACTCTTTCCCTACACGACGCTCTTCCGATCTNNNGATTTTGCTCTCTGCTGTTCCA
GAGCACACGTCTGAACTCTTTCCCTACACGACGCTCTTCCGATCTNNNGATTTTGCTCTCTGCTGTTCCA
GAGCACACGTCTGAACTCTTTCCCTACACGACGCTCTTCCGATCTNNNGATTTTGCTCTCTGCTGTTCCA
GAGCACACGTCTGAACTCTTTCCCTACACGACGCTCTTCCGATCTNNNGATTTTGCTCTCTGCTGTTCCA
GAGCACACGTCTGAACTCTTTCCCTACACGACGCTCTTCCGATCTNNNATCCTTACCAGCAGACAGAAC
GAGCACACGTCTGAACTCTTTCCCTACACGACGCTCTTCCGATCTNNNATCCTTACCAGCAGACAGAAC
GAGCACACGTCTGAACTCTTTCCCTACACGACGCTCTTCCGATCTNNNATCCTTACCAGCAGACAGAAC
GAGCACACGTCTGAACTCTTTCCCTACACGACGCTCTTCCGATCTNNNATCCTTACCAGCAGACAGAAC
GAGCACACGTCTGAACTCTTTCCCTACACGACGCTCTTCCGATCTNNNGAGCCTGTAATCCCAGCTACTC
GAGCACACGTCTGAACTCTTTCCCTACACGACGCTCTTCCGATCTNNNGAGCCTGTAATCCCAGCTACTC
GAGCACACGTCTGAACTCTTTCCCTACACGACGCTCTTCCGATCTNNNGAGCCTGTAATCCCAGCTACTC
GAGCACACGTCTGAACTCTTTCCCTACACGACGCTCTTCCGATCTNNNGAGCCTGTAATCCCAGCTACTC
AGCACACGTCTGAACTCTTTCCCTACACGACGCTCTTCCGATCTNNNTGGGCGTGGTTTTTCTTTTCTT

AGCACACGTCTGAACTCTTTCCCTACACGACGCTCTTCCGATCTNNNTGGGCGTGGTTTTTCTTTTCTT
AGCACACGTCTGAACTCTTTCCCTACACGACGCTCTTCCGATCTNNNTGGGCGTGGTTTTTCTTTTCTT
AGCACACGTCTGAACTCTTTCCCTACACGACGCTCTTCCGATCTNNNTGGGCGTGGTTTTTCTTTTCTT
GCACACGTCTGAACTCTTTCCCTACACGACGCTCTTCCGATCTNNNTGAAGTAATGTCCCCAGTCTGT
GCACACGTCTGAACTCTTTCCCTACACGACGCTCTTCCGATCTNNNTGAAGTAATGTCCCCAGTCTGT
GCACACGTCTGAACTCTTTCCCTACACGACGCTCTTCCGATCTNNNTGAAGTAATGTCCCCAGTCTGT
GCACACGTCTGAACTCTTTCCCTACACGACGCTCTTCCGATCTNNNTGAAGTAATGTCCCCAGTCTGT
AGAGCACACGTCTGAACTCTTTCCCTACACGACGCTCTTCCGATCTNNNGAAGGTGCAATTGTGTGTA
AGAGCACACGTCTGAACTCTTTCCCTACACGACGCTCTTCCGATCTNNNGAAGGTGCAATTGTGTGTA
AGAGCACACGTCTGAACTCTTTCCCTACACGACGCTCTTCCGATCTNNNGAAGGTGCAATTGTGTGTA
AGAGCACACGTCTGAACTCTTTCCCTACACGACGCTCTTCCGATCTNNNGAAGGTGCAATTGTGTGTA
AGCACACGTCTGAACTCTTTCCCTACACGACGCTCTTCCGATCTNNNAATCTTCCACCTCCACGTTCAAT
AGCACACGTCTGAACTCTTTCCCTACACGACGCTCTTCCGATCTNNNAATCTTCCACCTCCACGTTCAAT
AGCACACGTCTGAACTCTTTCCCTACACGACGCTCTTCCGATCTNNNAATCTTCCACCTCCACGTTCAAT
AGCACACGTCTGAACTCTTTCCCTACACGACGCTCTTCCGATCTNNNAATCTTCCACCTCCACGTTCAAT
GAGCACACGTCTGAACTCTTTCCCTACACGACGCTCTTCCGATCTNNNCTGAGATTTCTGGAGACACA
GAGCACACGTCTGAACTCTTTCCCTACACGACGCTCTTCCGATCTNNNCTGAGATTTCTGGAGACACA
GAGCACACGTCTGAACTCTTTCCCTACACGACGCTCTTCCGATCTNNNCTGAGATTTCTGGAGACACA
GAGCACACGTCTGAACTCTTTCCCTACACGACGCTCTTCCGATCTNNNCTGAGATTTCTGGAGACACA
AGCACACGTCTGAACTCTTTCCCTACACGACGCTCTTCCGATCTNNNTTTCATTAGCAGGTCCTGAGG
AGCACACGTCTGAACTCTTTCCCTACACGACGCTCTTCCGATCTNNNTTTCATTAGCAGGTCCTGAGG
AGCACACGTCTGAACTCTTTCCCTACACGACGCTCTTCCGATCTNNNTTTCATTAGCAGGTCCTGAGG
AGCACACGTCTGAACTCTTTCCCTACACGACGCTCTTCCGATCTNNNTTTCATTAGCAGGTCCTGAGG
GAGCACACGTCTGAACTCTTTCCCTACACGACGCTCTTCCGATCTNNNCTGCATTGAATGAACCAAGTC
GAGCACACGTCTGAACTCTTTCCCTACACGACGCTCTTCCGATCTNNNCTGCATTGAATGAACCAAGTC
GAGCACACGTCTGAACTCTTTCCCTACACGACGCTCTTCCGATCTNNNCTGCATTGAATGAACCAAGTC
GAGCACACGTCTGAACTCTTTCCCTACACGACGCTCTTCCGATCTNNNCTGCATTGAATGAACCAAGTC
AGCACACGTCTGAACTCTTTCCCTACACGACGCTCTTCCGATCTNNNGGTGACAGAGTGATATCCGTC
AGCACACGTCTGAACTCTTTCCCTACACGACGCTCTTCCGATCTNNNGGTGACAGAGTGATATCCGTC
AGCACACGTCTGAACTCTTTCCCTACACGACGCTCTTCCGATCTNNNGGTGACAGAGTGATATCCGTC
AGCACACGTCTGAACTCTTTCCCTACACGACGCTCTTCCGATCTNNNGGTGACAGAGTGATATCCGTC
AGCACACGTCTGAACTCTTTCCCTACACGACGCTCTTCCGATCTNNNTGTTATGTCCTCTGTCTCTCTCT
AGCACACGTCTGAACTCTTTCCCTACACGACGCTCTTCCGATCTNNNTGTTATGTCCTCTGTCTCTCTCT
AGCACACGTCTGAACTCTTTCCCTACACGACGCTCTTCCGATCTNNNTGTTATGTCCTCTGTCTCTCTCT
AGCACACGTCTGAACTCTTTCCCTACACGACGCTCTTCCGATCTNNNTGTTATGTCCTCTGTCTCTCTCT
GAGCACACGTCTGAACTCTTTCCCTACACGACGCTCTTCCGATCTNNNGGCACCAAACAAGTCTCAAA
GAGCACACGTCTGAACTCTTTCCCTACACGACGCTCTTCCGATCTNNNGGCACCAAACAAGTCTCAAA
GAGCACACGTCTGAACTCTTTCCCTACACGACGCTCTTCCGATCTNNNGGCACCAAACAAGTCTCAAA
GAGCACACGTCTGAACTCTTTCCCTACACGACGCTCTTCCGATCTNNNGGCACCAAACAAGTCTCAAA
AGCACACGTCTGAACTCTTTCCCTACACGACGCTCTTCCGATCTNNNCAGCAGTTTGACAATGATGTG
AGCACACGTCTGAACTCTTTCCCTACACGACGCTCTTCCGATCTNNNCAGCAGTTTGACAATGATGTG
AGCACACGTCTGAACTCTTTCCCTACACGACGCTCTTCCGATCTNNNCAGCAGTTTGACAATGATGTG
AGCACACGTCTGAACTCTTTCCCTACACGACGCTCTTCCGATCTNNNCAGCAGTTTGACAATGATGTG

AGCACACGTCTGAACTCTTTCCCTACACGACGCTCTTCCGATCTNNNTGCTGCCACACCTGTCTATATTT

AGCACACGTCTGAACTCTTTCCCTACACGACGCTCTTCCGATCTNNNTGCTGCCACACCTGTCTATATTT

AGCACACGTCTGAACTCTTTCCCTACACGACGCTCTTCCGATCTNNNTGCTGCCACACCTGTCTATATTT

AGCACACGTCTGAACTCTTTCCCTACACGACGCTCTTCCGATCTNNNTGCTGCCACACCTGTCTATATTT

AGCACACGTCTGAACTCTTTCCCTACACGACGCTCTTCCGATCTNNNTTGCACAAGATGACAGACCAG

AGCACACGTCTGAACTCTTTCCCTACACGACGCTCTTCCGATCTNNNTTGCACAAGATGACAGACCAG

AGCACACGTCTGAACTCTTTCCCTACACGACGCTCTTCCGATCTNNNTTGCACAAGATGACAGACCAG

AGCACACGTCTGAACTCTTTCCCTACACGACGCTCTTCCGATCTNNNTTGCACAAGATGACAGACCAG

GAGCACACGTCTGAACTCTTTCCCTACACGACGCTCTTCCGATCTNNNGAGAATTGCTTGAATGCAGG

GAGCACACGTCTGAACTCTTTCCCTACACGACGCTCTTCCGATCTNNNGAGAATTGCTTGAATGCAGG

GAGCACACGTCTGAACTCTTTCCCTACACGACGCTCTTCCGATCTNNNGAGAATTGCTTGAATGCAGG

GAGCACACGTCTGAACTCTTTCCCTACACGACGCTCTTCCGATCTNNNGAGAATTGCTTGAATGCAGG

AGCACACGTCTGAACTCTTTCCCTACACGACGCTCTTCCGATCTNNNACCCCTCAAATGCTTTTCCCAT

AGCACACGTCTGAACTCTTTCCCTACACGACGCTCTTCCGATCTNNNACCCCTCAAATGCTTTTCCCAT

AGCACACGTCTGAACTCTTTCCCTACACGACGCTCTTCCGATCTNNNACCCCTCAAATGCTTTTCCCAT

AGCACACGTCTGAACTCTTTCCCTACACGACGCTCTTCCGATCTNNNACCCCTCAAATGCTTTTCCCAT

AGCACACGTCTGAACTCTTTCCCTACACGACGCTCTTCCGATCTNNNACTCATTCTACTTCCTGGCCTTG

AGCACACGTCTGAACTCTTTCCCTACACGACGCTCTTCCGATCTNNNACTCATTCTACTTCCTGGCCTTG

AGCACACGTCTGAACTCTTTCCCTACACGACGCTCTTCCGATCTNNNACTCATTCTACTTCCTGGCCTTG

AGCACACGTCTGAACTCTTTCCCTACACGACGCTCTTCCGATCTNNNACTCATTCTACTTCCTGGCCTTG

AGCACACGTCTGAACTCTTTCCCTACACGACGCTCTTCCGATCTNNNGAGAATGTCATGAACCTGGGA

AGCACACGTCTGAACTCTTTCCCTACACGACGCTCTTCCGATCTNNNGAGAATGTCATGAACCTGGGA

AGCACACGTCTGAACTCTTTCCCTACACGACGCTCTTCCGATCTNNNGAGAATGTCATGAACCTGGGA

AGCACACGTCTGAACTCTTTCCCTACACGACGCTCTTCCGATCTNNNGAGAATGTCATGAACCTGGGA

GCACACGTCTGAACTCTTTCCCTACACGACGCTCTTCCGATCTNNNTCTCCTCATTGCCTTAGAACACCC

GCACACGTCTGAACTCTTTCCCTACACGACGCTCTTCCGATCTNNNTCTCCTCATTGCCTTAGAACACCC

GCACACGTCTGAACTCTTTCCCTACACGACGCTCTTCCGATCTNNNTCTCCTCATTGCCTTAGAACACCC

GCACACGTCTGAACTCTTTCCCTACACGACGCTCTTCCGATCTNNNTCTCCTCATTGCCTTAGAACACCC

AGAGCACACGTCTGAACTCTTTCCCTACACGACGCTCTTCCGATCTNNNTCTCTCTTCAGCTTTTGTGC

AGAGCACACGTCTGAACTCTTTCCCTACACGACGCTCTTCCGATCTNNNTCTCTCTTCAGCTTTTGTGC

AGAGCACACGTCTGAACTCTTTCCCTACACGACGCTCTTCCGATCTNNNTCTCTCTTCAGCTTTTGTGC

AGAGCACACGTCTGAACTCTTTCCCTACACGACGCTCTTCCGATCTNNNTCTCTCTTCAGCTTTTGTGC

AGCACACGTCTGAACTCTTTCCCTACACGACGCTCTTCCGATCTNNNCTTTCCCCTGTCTCTCCCAACCAT

AGCACACGTCTGAACTCTTTCCCTACACGACGCTCTTCCGATCTNNNCTTTCCCCTGTCTCTCCCAACCAT

AGCACACGTCTGAACTCTTTCCCTACACGACGCTCTTCCGATCTNNNCTTTCCCCTGTCTCTCCCAACCAT

AGCACACGTCTGAACTCTTTCCCTACACGACGCTCTTCCGATCTNNNCTTTCCCCTGTCTCTCCCAACCAT

GCACACGTCTGAACTCTTTCCCTACACGACGCTCTTCCGATCTNNNCTGAGGCAGGAGAATCACTTGA

GCACACGTCTGAACTCTTTCCCTACACGACGCTCTTCCGATCTNNNCTGAGGCAGGAGAATCACTTGA

GCACACGTCTGAACTCTTTCCCTACACGACGCTCTTCCGATCTNNNCTGAGGCAGGAGAATCACTTGA

GCACACGTCTGAACTCTTTCCCTACACGACGCTCTTCCGATCTNNNCTGAGGCAGGAGAATCACTTGA

AGCACACGTCTGAACTCTTTCCCTACACGACGCTCTTCCGATCTNNNGCATGCTCAAGGATCAAAGACT

AGCACACGTCTGAACTCTTTCCCTACACGACGCTCTTCCGATCTNNNGCATGCTCAAGGATCAAAGACT

AGCACACGTCTGAACTCTTTCCCTACACGACGCTCTTCCGATCTNNNGCATGCTCAAGGATCAAAGACT

AGCACACGTCTGAACTCTTTCCCTACACGACGCTCTTCCGATCTNNNGCATGCTCAAGGATCAAAGACT
AGAGCACACGTCTGAACTCTTTCCCTACACGACGCTCTTCCGATCTNNNTCCTAAGGTGCCTCTTGCTT
AGAGCACACGTCTGAACTCTTTCCCTACACGACGCTCTTCCGATCTNNNTCCTAAGGTGCCTCTTGCTT
AGAGCACACGTCTGAACTCTTTCCCTACACGACGCTCTTCCGATCTNNNTCCTAAGGTGCCTCTTGCTT
AGAGCACACGTCTGAACTCTTTCCCTACACGACGCTCTTCCGATCTNNNTCCTAAGGTGCCTCTTGCTT
GAGCACACGTCTGAACTCTTTCCCTACACGACGCTCTTCCGATCTNNNCCATTTTGAACTCCCCATGTGA
GAGCACACGTCTGAACTCTTTCCCTACACGACGCTCTTCCGATCTNNNCCATTTTGAACTCCCCATGTGA
GAGCACACGTCTGAACTCTTTCCCTACACGACGCTCTTCCGATCTNNNCCATTTTGAACTCCCCATGTGA
GAGCACACGTCTGAACTCTTTCCCTACACGACGCTCTTCCGATCTNNNCCATTTTGAACTCCCCATGTGA
GAGCACACGTCTGAACTCTTTCCCTACACGACGCTCTTCCGATCTNNNATTCTTTACTGCTCTGCTACCCC
GAGCACACGTCTGAACTCTTTCCCTACACGACGCTCTTCCGATCTNNNATTCTTTACTGCTCTGCTACCCC
GAGCACACGTCTGAACTCTTTCCCTACACGACGCTCTTCCGATCTNNNATTCTTTACTGCTCTGCTACCCC
GAGCACACGTCTGAACTCTTTCCCTACACGACGCTCTTCCGATCTNNNATTCTTTACTGCTCTGCTACCCC
AGCACACGTCTGAACTCTTTCCCTACACGACGCTCTTCCGATCTNNNCCTGTGGTAGGCACTAAGGAA
AGCACACGTCTGAACTCTTTCCCTACACGACGCTCTTCCGATCTNNNCCTGTGGTAGGCACTAAGGAA
AGCACACGTCTGAACTCTTTCCCTACACGACGCTCTTCCGATCTNNNCCTGTGGTAGGCACTAAGGAA
AGCACACGTCTGAACTCTTTCCCTACACGACGCTCTTCCGATCTNNNCCTGTGGTAGGCACTAAGGAA
GAGCACACGTCTGAACTCTTTCCCTACACGACGCTCTTCCGATCTNNNCATGCCTGAACCTTCTACTGAC
GAGCACACGTCTGAACTCTTTCCCTACACGACGCTCTTCCGATCTNNNCATGCCTGAACCTTCTACTGAC
GAGCACACGTCTGAACTCTTTCCCTACACGACGCTCTTCCGATCTNNNCATGCCTGAACCTTCTACTGAC
GAGCACACGTCTGAACTCTTTCCCTACACGACGCTCTTCCGATCTNNNCATGCCTGAACCTTCTACTGAC
GAGCACACGTCTGAACTCTTTCCCTACACGACGCTCTTCCGATCTNNNCTAGATGTCTCCGGTGGCAAA
GAGCACACGTCTGAACTCTTTCCCTACACGACGCTCTTCCGATCTNNNCTAGATGTCTCCGGTGGCAAA
GAGCACACGTCTGAACTCTTTCCCTACACGACGCTCTTCCGATCTNNNCTAGATGTCTCCGGTGGCAAA
GAGCACACGTCTGAACTCTTTCCCTACACGACGCTCTTCCGATCTNNNCTAGATGTCTCCGGTGGCAAA
AGCACACGTCTGAACTCTTTCCCTACACGACGCTCTTCCGATCTNNNCTAATGCCACCTAGGACTCCAT
AGCACACGTCTGAACTCTTTCCCTACACGACGCTCTTCCGATCTNNNCTAATGCCACCTAGGACTCCAT
AGCACACGTCTGAACTCTTTCCCTACACGACGCTCTTCCGATCTNNNCTAATGCCACCTAGGACTCCAT
AGCACACGTCTGAACTCTTTCCCTACACGACGCTCTTCCGATCTNNNCTAATGCCACCTAGGACTCCAT
AGAGCACACGTCTGAACTCTTTCCCTACACGACGCTCTTCCGATCTNNNAATCTCCTGACCTTGTGATC
AGAGCACACGTCTGAACTCTTTCCCTACACGACGCTCTTCCGATCTNNNAATCTCCTGACCTTGTGATC
AGAGCACACGTCTGAACTCTTTCCCTACACGACGCTCTTCCGATCTNNNAATCTCCTGACCTTGTGATC
AGAGCACACGTCTGAACTCTTTCCCTACACGACGCTCTTCCGATCTNNNAATCTCCTGACCTTGTGATC
GCACACGTCTGAACTCTTTCCCTACACGACGCTCTTCCGATCTNNNCATGTGGTTCTCTCAGTCTGGAA
GCACACGTCTGAACTCTTTCCCTACACGACGCTCTTCCGATCTNNNCATGTGGTTCTCTCAGTCTGGAA
GCACACGTCTGAACTCTTTCCCTACACGACGCTCTTCCGATCTNNNCATGTGGTTCTCTCAGTCTGGAA
GCACACGTCTGAACTCTTTCCCTACACGACGCTCTTCCGATCTNNNCATGTGGTTCTCTCAGTCTGGAA
AGCACACGTCTGAACTCTTTCCCTACACGACGCTCTTCCGATCTNNNCGCTCATCACTTGATACTTGTGT
AGCACACGTCTGAACTCTTTCCCTACACGACGCTCTTCCGATCTNNNCGCTCATCACTTGATACTTGTGT
AGCACACGTCTGAACTCTTTCCCTACACGACGCTCTTCCGATCTNNNCGCTCATCACTTGATACTTGTGT
AGCACACGTCTGAACTCTTTCCCTACACGACGCTCTTCCGATCTNNNCGCTCATCACTTGATACTTGTGT
GAGCACACGTCTGAACTCTTTCCCTACACGACGCTCTTCCGATCTNNNTCGATCTCCTGACTTTGTGATC
GAGCACACGTCTGAACTCTTTCCCTACACGACGCTCTTCCGATCTNNNTCGATCTCCTGACTTTGTGATC

GAGCACACGTCTGAACTCTTTCCCTACACGACGCTCTTCCGATCTNNNTCGATCTCCTGACTTTGTGATC
GAGCACACGTCTGAACTCTTTCCCTACACGACGCTCTTCCGATCTNNNTCGATCTCCTGACTTTGTGATC
AGCACACGTCTGAACTCTTTCCCTACACGACGCTCTTCCGATCTNNNCAGGATGAACTGCTTGAATGTC
AGCACACGTCTGAACTCTTTCCCTACACGACGCTCTTCCGATCTNNNCAGGATGAACTGCTTGAATGTC
AGCACACGTCTGAACTCTTTCCCTACACGACGCTCTTCCGATCTNNNCAGGATGAACTGCTTGAATGTC
AGCACACGTCTGAACTCTTTCCCTACACGACGCTCTTCCGATCTNNNCAGGATGAACTGCTTGAATGTC
GAGCACACGTCTGAACTCTTTCCCTACACGACGCTCTTCCGATCTNNNGCCTGGCAATCAGAGCTCTTT
GAGCACACGTCTGAACTCTTTCCCTACACGACGCTCTTCCGATCTNNNGCCTGGCAATCAGAGCTCTTT
GAGCACACGTCTGAACTCTTTCCCTACACGACGCTCTTCCGATCTNNNGCCTGGCAATCAGAGCTCTTT
GAGCACACGTCTGAACTCTTTCCCTACACGACGCTCTTCCGATCTNNNGCCTGGCAATCAGAGCTCTTT
AGCACACGTCTGAACTCTTTCCCTACACGACGCTCTTCCGATCTNNNAGGTTGAAGTAATTCTCGTGCC
AGCACACGTCTGAACTCTTTCCCTACACGACGCTCTTCCGATCTNNNAGGTTGAAGTAATTCTCGTGCC
AGCACACGTCTGAACTCTTTCCCTACACGACGCTCTTCCGATCTNNNAGGTTGAAGTAATTCTCGTGCC
AGCACACGTCTGAACTCTTTCCCTACACGACGCTCTTCCGATCTNNNAGGTTGAAGTAATTCTCGTGCC
AGCACACGTCTGAACTCTTTCCCTACACGACGCTCTTCCGATCTNNNGAATCATTGTCCAGAGCTGCTTC
AGCACACGTCTGAACTCTTTCCCTACACGACGCTCTTCCGATCTNNNGAATCATTGTCCAGAGCTGCTTC
AGCACACGTCTGAACTCTTTCCCTACACGACGCTCTTCCGATCTNNNGAATCATTGTCCAGAGCTGCTTC
AGCACACGTCTGAACTCTTTCCCTACACGACGCTCTTCCGATCTNNNGAATCATTGTCCAGAGCTGCTTC
GAGCACACGTCTGAACTCTTTCCCTACACGACGCTCTTCCGATCTNNNCTCTGGTGATGGATCTGATCA
GAGCACACGTCTGAACTCTTTCCCTACACGACGCTCTTCCGATCTNNNCTCTGGTGATGGATCTGATCA
GAGCACACGTCTGAACTCTTTCCCTACACGACGCTCTTCCGATCTNNNCTCTGGTGATGGATCTGATCA
GAGCACACGTCTGAACTCTTTCCCTACACGACGCTCTTCCGATCTNNNCTCTGGTGATGGATCTGATCA
GAGCACACGTCTGAACTCTTTCCCTACACGACGCTCTTCCGATCTNNNCTAAGCCATAAAATGCCAGCA
GAGCACACGTCTGAACTCTTTCCCTACACGACGCTCTTCCGATCTNNNCTAAGCCATAAAATGCCAGCA
GAGCACACGTCTGAACTCTTTCCCTACACGACGCTCTTCCGATCTNNNCTAAGCCATAAAATGCCAGCA
GAGCACACGTCTGAACTCTTTCCCTACACGACGCTCTTCCGATCTNNNCTAAGCCATAAAATGCCAGCA
AGAGCACACGTCTGAACTCTTTCCCTACACGACGCTCTTCCGATCTNNNAAAGCCATCCTGAAGGTTTGT
AGAGCACACGTCTGAACTCTTTCCCTACACGACGCTCTTCCGATCTNNNAAAGCCATCCTGAAGGTTTGT
AGAGCACACGTCTGAACTCTTTCCCTACACGACGCTCTTCCGATCTNNNAAAGCCATCCTGAAGGTTTGT
AGAGCACACGTCTGAACTCTTTCCCTACACGACGCTCTTCCGATCTNNNAAAGCCATCCTGAAGGTTTGT
AGAGCACACGTCTGAACTCTTTCCCTACACGACGCTCTTCCGATCTNNNAGACTGACATTTGATTCGCTC
AGAGCACACGTCTGAACTCTTTCCCTACACGACGCTCTTCCGATCTNNNAGACTGACATTTGATTCGCTC
AGAGCACACGTCTGAACTCTTTCCCTACACGACGCTCTTCCGATCTNNNAGACTGACATTTGATTCGCTC
AGAGCACACGTCTGAACTCTTTCCCTACACGACGCTCTTCCGATCTNNNAGACTGACATTTGATTCGCTC
AGAGCACACGTCTGAACTCTTTCCCTACACGACGCTCTTCCGATCTNNNATCCTTGTCCACGTAGTTTTC
AGAGCACACGTCTGAACTCTTTCCCTACACGACGCTCTTCCGATCTNNNATCCTTGTCCACGTAGTTTTC
AGAGCACACGTCTGAACTCTTTCCCTACACGACGCTCTTCCGATCTNNNATCCTTGTCCACGTAGTTTTC
AGAGCACACGTCTGAACTCTTTCCCTACACGACGCTCTTCCGATCTNNNATCCTTGTCCACGTAGTTTTC
AGAGCACACGTCTGAACTCTTTCCCTACACGACGCTCTTCCGATCTNNNAGCAAAGGATTAATTCCAAA
AGAGCACACGTCTGAACTCTTTCCCTACACGACGCTCTTCCGATCTNNNAGCAAAGGATTAATTCCAAA
AGAGCACACGTCTGAACTCTTTCCCTACACGACGCTCTTCCGATCTNNNAGCAAAGGATTAATTCCAAA
AGAGCACACGTCTGAACTCTTTCCCTACACGACGCTCTTCCGATCTNNNAGCAAAGGATTAATTCCAAA
AGAGCACACGTCTGAACTCTTTCCCTACACGACGCTCTTCCGATCTNNNTATGCCTGTAGTCCCAGCTA

AGAGCACACGTCTGAACTCTTTCCCTACACGACGCTCTTCCGATCTNNNTATGCCTGTAGTCCCAGCTA

AGAGCACACGTCTGAACTCTTTCCCTACACGACGCTCTTCCGATCTNNNTATGCCTGTAGTCCCAGCTA

AGAGCACACGTCTGAACTCTTTCCCTACACGACGCTCTTCCGATCTNNNTATGCCTGTAGTCCCAGCTA

GAGCACACGTCTGAACTCTTTCCCTACACGACGCTCTTCCGATCTNNNCTGCTCTCACCTCTGTTTCTCTG

GAGCACACGTCTGAACTCTTTCCCTACACGACGCTCTTCCGATCTNNNCTGCTCTCACCTCTGTTTCTCTG

GAGCACACGTCTGAACTCTTTCCCTACACGACGCTCTTCCGATCTNNNCTGCTCTCACCTCTGTTTCTCTG

GAGCACACGTCTGAACTCTTTCCCTACACGACGCTCTTCCGATCTNNNCTGCTCTCACCTCTGTTTCTCTG

AGCACACGTCTGAACTCTTTCCCTACACGACGCTCTTCCGATCTNNNTTTGTGAGATGACCTCGTGTGAT

AGCACACGTCTGAACTCTTTCCCTACACGACGCTCTTCCGATCTNNNTTTGTGAGATGACCTCGTGTGAT

AGCACACGTCTGAACTCTTTCCCTACACGACGCTCTTCCGATCTNNNTTTGTGAGATGACCTCGTGTGAT

AGCACACGTCTGAACTCTTTCCCTACACGACGCTCTTCCGATCTNNNTTTGTGAGATGACCTCGTGTGAT

GAGCACACGTCTGAACTCTTTCCCTACACGACGCTCTTCCGATCTNNNTTGCCTTTGTTTTCCTCCTCCTC

GAGCACACGTCTGAACTCTTTCCCTACACGACGCTCTTCCGATCTNNNTTGCCTTTGTTTTCCTCCTCCTC

GAGCACACGTCTGAACTCTTTCCCTACACGACGCTCTTCCGATCTNNNTTGCCTTTGTTTTCCTCCTCCTC

GAGCACACGTCTGAACTCTTTCCCTACACGACGCTCTTCCGATCTNNNTTGCCTTTGTTTTCCTCCTCCTC

GAGCACACGTCTGAACTCTTTCCCTACACGACGCTCTTCCGATCTNNNGCAAATACAATTCCTGACCTGG

GAGCACACGTCTGAACTCTTTCCCTACACGACGCTCTTCCGATCTNNNGCAAATACAATTCCTGACCTGG

GAGCACACGTCTGAACTCTTTCCCTACACGACGCTCTTCCGATCTNNNGCAAATACAATTCCTGACCTGG

GAGCACACGTCTGAACTCTTTCCCTACACGACGCTCTTCCGATCTNNNGCAAATACAATTCCTGACCTGG

AGCACACGTCTGAACTCTTTCCCTACACGACGCTCTTCCGATCTNNNAAGACTGGGTCCCAAAGAGAC

AGCACACGTCTGAACTCTTTCCCTACACGACGCTCTTCCGATCTNNNAAGACTGGGTCCCAAAGAGAC

AGCACACGTCTGAACTCTTTCCCTACACGACGCTCTTCCGATCTNNNAAGACTGGGTCCCAAAGAGAC

AGCACACGTCTGAACTCTTTCCCTACACGACGCTCTTCCGATCTNNNAAGACTGGGTCCCAAAGAGAC

GAGCACACGTCTGAACTCTTTCCCTACACGACGCTCTTCCGATCTNNNGCATAGCCTGAACCAGTTTTGA

GAGCACACGTCTGAACTCTTTCCCTACACGACGCTCTTCCGATCTNNNGCATAGCCTGAACCAGTTTTGA

GAGCACACGTCTGAACTCTTTCCCTACACGACGCTCTTCCGATCTNNNGCATAGCCTGAACCAGTTTTGA

GAGCACACGTCTGAACTCTTTCCCTACACGACGCTCTTCCGATCTNNNGCATAGCCTGAACCAGTTTTGA

AGCACACGTCTGAACTCTTTCCCTACACGACGCTCTTCCGATCTNNNACACCAAGGAGTGTCTCTCTCT

AGCACACGTCTGAACTCTTTCCCTACACGACGCTCTTCCGATCTNNNACACCAAGGAGTGTCTCTCTCT

AGCACACGTCTGAACTCTTTCCCTACACGACGCTCTTCCGATCTNNNACACCAAGGAGTGTCTCTCTCT

AGCACACGTCTGAACTCTTTCCCTACACGACGCTCTTCCGATCTNNNACACCAAGGAGTGTCTCTCTCT

AGAGCACACGTCTGAACTCTTTCCCTACACGACGCTCTTCCGATCTNNNAGTGAGAGGTCCCCATGTGT

AGAGCACACGTCTGAACTCTTTCCCTACACGACGCTCTTCCGATCTNNNAGTGAGAGGTCCCCATGTGT

AGAGCACACGTCTGAACTCTTTCCCTACACGACGCTCTTCCGATCTNNNAGTGAGAGGTCCCCATGTGT

AGAGCACACGTCTGAACTCTTTCCCTACACGACGCTCTTCCGATCTNNNAGTGAGAGGTCCCCATGTGT

CACACGTCTGAACTCTTTCCCTACACGACGCTCTTCCGATCTNNNTCTGCGTGCCAAAATAGTCTTTGCA

CACACGTCTGAACTCTTTCCCTACACGACGCTCTTCCGATCTNNNTCTGCGTGCCAAAATAGTCTTTGCA

CACACGTCTGAACTCTTTCCCTACACGACGCTCTTCCGATCTNNNTCTGCGTGCCAAAATAGTCTTTGCA

CACACGTCTGAACTCTTTCCCTACACGACGCTCTTCCGATCTNNNTCTGCGTGCCAAAATAGTCTTTGCA

AGCACACGTCTGAACTCTTTCCCTACACGACGCTCTTCCGATCTNNNTGGGTATATGTGTGCATGTCTC

AGCACACGTCTGAACTCTTTCCCTACACGACGCTCTTCCGATCTNNNTGGGTATATGTGTGCATGTCTC

AGCACACGTCTGAACTCTTTCCCTACACGACGCTCTTCCGATCTNNNTGGGTATATGTGTGCATGTCTC

AGCACACGTCTGAACTCTTTCCCTACACGACGCTCTTCCGATCTNNNTGGGTATATGTGTGCATGTCTC

GAGCACACGTCTGAACTCTTTCCCTACACGACGCTCTTCCGATCTNNNGGAGAATCACTTGAACCTGG

GAGCACACGTCTGAACTCTTTCCCTACACGACGCTCTTCCGATCTNNNGGAGAATCACTTGAACCTGG

GAGCACACGTCTGAACTCTTTCCCTACACGACGCTCTTCCGATCTNNNGGAGAATCACTTGAACCTGG

GAGCACACGTCTGAACTCTTTCCCTACACGACGCTCTTCCGATCTNNNGGAGAATCACTTGAACCTGG

GAGCACACGTCTGAACTCTTTCCCTACACGACGCTCTTCCGATCTNNNGAGATTGCAGTGAGCCAAG

GAGCACACGTCTGAACTCTTTCCCTACACGACGCTCTTCCGATCTNNNGAGATTGCAGTGAGCCAAG

GAGCACACGTCTGAACTCTTTCCCTACACGACGCTCTTCCGATCTNNNGAGATTGCAGTGAGCCAAG

GAGCACACGTCTGAACTCTTTCCCTACACGACGCTCTTCCGATCTNNNGAGATTGCAGTGAGCCAAG

GCACACGTCTGAACTCTTTCCCTACACGACGCTCTTCCGATCTNNNGAGTTGAGGAGTGGGGCGAC

GCACACGTCTGAACTCTTTCCCTACACGACGCTCTTCCGATCTNNNGAGTTGAGGAGTGGGGCGAC

GCACACGTCTGAACTCTTTCCCTACACGACGCTCTTCCGATCTNNNGAGTTGAGGAGTGGGGCGAC

GCACACGTCTGAACTCTTTCCCTACACGACGCTCTTCCGATCTNNNGAGTTGAGGAGTGGGGCGAC

AGCACACGTCTGAACTCTTTCCCTACACGACGCTCTTCCGATCTNNNAGAATTGAGGAGCTAATGCCT

AGCACACGTCTGAACTCTTTCCCTACACGACGCTCTTCCGATCTNNNAGAATTGAGGAGCTAATGCCT

AGCACACGTCTGAACTCTTTCCCTACACGACGCTCTTCCGATCTNNNAGAATTGAGGAGCTAATGCCT

AGCACACGTCTGAACTCTTTCCCTACACGACGCTCTTCCGATCTNNNAGAATTGAGGAGCTAATGCCT

AGCACACGTCTGAACTCTTTCCCTACACGACGCTCTTCCGATCTNNNTTCTCAGCAGCTGTCCTAGAAT

AGCACACGTCTGAACTCTTTCCCTACACGACGCTCTTCCGATCTNNNTTCTCAGCAGCTGTCCTAGAAT

AGCACACGTCTGAACTCTTTCCCTACACGACGCTCTTCCGATCTNNNTTCTCAGCAGCTGTCCTAGAAT

AGCACACGTCTGAACTCTTTCCCTACACGACGCTCTTCCGATCTNNNTTCTCAGCAGCTGTCCTAGAAT

GAGCACACGTCTGAACTCTTTCCCTACACGACGCTCTTCCGATCTNNNAGTGCTGGGATTACAAAGA

GAGCACACGTCTGAACTCTTTCCCTACACGACGCTCTTCCGATCTNNNAGTGCTGGGATTACAAAGA

GAGCACACGTCTGAACTCTTTCCCTACACGACGCTCTTCCGATCTNNNAGTGCTGGGATTACAAAGA

GAGCACACGTCTGAACTCTTTCCCTACACGACGCTCTTCCGATCTNNNAGTGCTGGGATTACAAAGA

CACACGTCTGAACTCTTTCCCTACACGACGCTCTTCCGATCTNNNATTTCCAGAGTGCCCATTGTGTG

CACACGTCTGAACTCTTTCCCTACACGACGCTCTTCCGATCTNNNATTTCCAGAGTGCCCATTGTGTG

CACACGTCTGAACTCTTTCCCTACACGACGCTCTTCCGATCTNNNATTTCCAGAGTGCCCATTGTGTG

CACACGTCTGAACTCTTTCCCTACACGACGCTCTTCCGATCTNNNATTTCCAGAGTGCCCATTGTGTG

AGCACACGTCTGAACTCTTTCCCTACACGACGCTCTTCCGATCTNNNGCGGATTCTAATACATAGGG

AGCACACGTCTGAACTCTTTCCCTACACGACGCTCTTCCGATCTNNNGCGGATTCTAATACATAGGG

AGCACACGTCTGAACTCTTTCCCTACACGACGCTCTTCCGATCTNNNGCGGATTCTAATACATAGGG

AGCACACGTCTGAACTCTTTCCCTACACGACGCTCTTCCGATCTNNNGCGGATTCTAATACATAGGG

GCACACGTCTGAACTCTTTCCCTACACGACGCTCTTCCGATCTNNNAAGTTTTTCAGTTTTGGGGTG

GCACACGTCTGAACTCTTTCCCTACACGACGCTCTTCCGATCTNNNAAGTTTTTCAGTTTTGGGGTG

GCACACGTCTGAACTCTTTCCCTACACGACGCTCTTCCGATCTNNNAAGTTTTTCAGTTTTGGGGTG

GCACACGTCTGAACTCTTTCCCTACACGACGCTCTTCCGATCTNNNAAGTTTTTCAGTTTTGGGGTG

AGCACACGTCTGAACTCTTTCCCTACACGACGCTCTTCCGATCTNNNGAGAATATGGCAGAGGAGAT

AGCACACGTCTGAACTCTTTCCCTACACGACGCTCTTCCGATCTNNNGAGAATATGGCAGAGGAGAT

AGCACACGTCTGAACTCTTTCCCTACACGACGCTCTTCCGATCTNNNGAGAATATGGCAGAGGAGAT

AGCACACGTCTGAACTCTTTCCCTACACGACGCTCTTCCGATCTNNNGAGAATATGGCAGAGGAGAT

GCACACGTCTGAACTCTTTCCCTACACGACGCTCTTCCGATCTNNNGGATATTCCCAAGTTCCAGGAG

GCACACGTCTGAACTCTTTCCCTACACGACGCTCTTCCGATCTNNNGGATATTCCCAAGTTCCAGGAG

GCACACGTCTGAACTCTTTCCCTACACGACGCTCTTCCGATCTNNNGGATATTCCCAAGTTCCAGGAG

GCACACGTCTGAACTCTTTCCCTACACGACGCTCTTCCGATCTNNNGGATATTCCCAAGTTCCAGGAGG
AGCACACGTCTGAACTCTTTCCCTACACGACGCTCTTCCGATCTNNNGAGGCAAGAGAATCACTTGAA
AGCACACGTCTGAACTCTTTCCCTACACGACGCTCTTCCGATCTNNNGAGGCAAGAGAATCACTTGAA
AGCACACGTCTGAACTCTTTCCCTACACGACGCTCTTCCGATCTNNNGAGGCAAGAGAATCACTTGAA
AGCACACGTCTGAACTCTTTCCCTACACGACGCTCTTCCGATCTNNNGAGGCAAGAGAATCACTTGAA
GAGCACACGTCTGAACTCTTTCCCTACACGACGCTCTTCCGATCTNNNAATTGCTTGAACCTAGGAGAC
GAGCACACGTCTGAACTCTTTCCCTACACGACGCTCTTCCGATCTNNNAATTGCTTGAACCTAGGAGAC
GAGCACACGTCTGAACTCTTTCCCTACACGACGCTCTTCCGATCTNNNAATTGCTTGAACCTAGGAGAC
GAGCACACGTCTGAACTCTTTCCCTACACGACGCTCTTCCGATCTNNNAATTGCTTGAACCTAGGAGAC
GAGCACACGTCTGAACTCTTTCCCTACACGACGCTCTTCCGATCTNNNTCTGGTAGTATGACACCTACA
GAGCACACGTCTGAACTCTTTCCCTACACGACGCTCTTCCGATCTNNNTCTGGTAGTATGACACCTACA
GAGCACACGTCTGAACTCTTTCCCTACACGACGCTCTTCCGATCTNNNTCTGGTAGTATGACACCTACA
GAGCACACGTCTGAACTCTTTCCCTACACGACGCTCTTCCGATCTNNNTCTGGTAGTATGACACCTACA
AGCACACGTCTGAACTCTTTCCCTACACGACGCTCTTCCGATCTNNNGTAGATCACGAGGTCAGGAG
AGCACACGTCTGAACTCTTTCCCTACACGACGCTCTTCCGATCTNNNGTAGATCACGAGGTCAGGAG
AGCACACGTCTGAACTCTTTCCCTACACGACGCTCTTCCGATCTNNNGTAGATCACGAGGTCAGGAG
AGCACACGTCTGAACTCTTTCCCTACACGACGCTCTTCCGATCTNNNGTAGATCACGAGGTCAGGAG
GCACACGTCTGAACTCTTTCCCTACACGACGCTCTTCCGATCTNNNCCAGATCAAAGACAACATCGGT
GCACACGTCTGAACTCTTTCCCTACACGACGCTCTTCCGATCTNNNCCAGATCAAAGACAACATCGGT
GCACACGTCTGAACTCTTTCCCTACACGACGCTCTTCCGATCTNNNCCAGATCAAAGACAACATCGGT
GCACACGTCTGAACTCTTTCCCTACACGACGCTCTTCCGATCTNNNCCAGATCAAAGACAACATCGGT
AGCACACGTCTGAACTCTTTCCCTACACGACGCTCTTCCGATCTNNNACAACACATGGGGATTATGGG
AGCACACGTCTGAACTCTTTCCCTACACGACGCTCTTCCGATCTNNNACAACACATGGGGATTATGGG
AGCACACGTCTGAACTCTTTCCCTACACGACGCTCTTCCGATCTNNNACAACACATGGGGATTATGGG
AGCACACGTCTGAACTCTTTCCCTACACGACGCTCTTCCGATCTNNNACAACACATGGGGATTATGGG
GAGCACACGTCTGAACTCTTTCCCTACACGACGCTCTTCCGATCTNNNAGGAACTGGTATGACAGAT
GAGCACACGTCTGAACTCTTTCCCTACACGACGCTCTTCCGATCTNNNAGGAACTGGTATGACAGAT
GAGCACACGTCTGAACTCTTTCCCTACACGACGCTCTTCCGATCTNNNAGGAACTGGTATGACAGAT
GAGCACACGTCTGAACTCTTTCCCTACACGACGCTCTTCCGATCTNNNAGGAACTGGTATGACAGAT
AGCACACGTCTGAACTCTTTCCCTACACGACGCTCTTCCGATCTNNNAGTACGTGTTCACTGTTTTCTGT
AGCACACGTCTGAACTCTTTCCCTACACGACGCTCTTCCGATCTNNNAGTACGTGTTCACTGTTTTCTGT
AGCACACGTCTGAACTCTTTCCCTACACGACGCTCTTCCGATCTNNNAGTACGTGTTCACTGTTTTCTGT
AGCACACGTCTGAACTCTTTCCCTACACGACGCTCTTCCGATCTNNNAGTACGTGTTCACTGTTTTCTGT
GAGCACACGTCTGAACTCTTTCCCTACACGACGCTCTTCCGATCTNNNGCCTGAGCACAAGTAACATG
GAGCACACGTCTGAACTCTTTCCCTACACGACGCTCTTCCGATCTNNNGCCTGAGCACAAGTAACATG
GAGCACACGTCTGAACTCTTTCCCTACACGACGCTCTTCCGATCTNNNGCCTGAGCACAAGTAACATG
GAGCACACGTCTGAACTCTTTCCCTACACGACGCTCTTCCGATCTNNNGCCTGAGCACAAGTAACATG
AGCACACGTCTGAACTCTTTCCCTACACGACGCTCTTCCGATCTNNNTACTCGGCAAAGTCTCCTTTGT
AGCACACGTCTGAACTCTTTCCCTACACGACGCTCTTCCGATCTNNNTACTCGGCAAAGTCTCCTTTGT
AGCACACGTCTGAACTCTTTCCCTACACGACGCTCTTCCGATCTNNNTACTCGGCAAAGTCTCCTTTGT
AGCACACGTCTGAACTCTTTCCCTACACGACGCTCTTCCGATCTNNNTACTCGGCAAAGTCTCCTTTGT
AGCACACGTCTGAACTCTTTCCCTACACGACGCTCTTCCGATCTNNNCACACATAGGTTCTTGGCCAA
AGCACACGTCTGAACTCTTTCCCTACACGACGCTCTTCCGATCTNNNCACACATAGGTTCTTGGCCAA

AGCACACGTCTGAACTCTTTCCCTACACGACGCTCTTCCGATCTNNNCACACATAGGTTCTTGGCCAA
AGCACACGTCTGAACTCTTTCCCTACACGACGCTCTTCCGATCTNNNCACACATAGGTTCTTGGCCAA
GAGCACACGTCTGAACTCTTTCCCTACACGACGCTCTTCCGATCTNNNCGTCTCTCACCTGTGTCTTCCC
GAGCACACGTCTGAACTCTTTCCCTACACGACGCTCTTCCGATCTNNNCGTCTCTCACCTGTGTCTTCCC
GAGCACACGTCTGAACTCTTTCCCTACACGACGCTCTTCCGATCTNNNCGTCTCTCACCTGTGTCTTCCC
GAGCACACGTCTGAACTCTTTCCCTACACGACGCTCTTCCGATCTNNNCGTCTCTCACCTGTGTCTTCCC
GCACACGTCTGAACTCTTTCCCTACACGACGCTCTTCCGATCTNNNACTGCCTCTAACCCAATCTCTTTC
GCACACGTCTGAACTCTTTCCCTACACGACGCTCTTCCGATCTNNNACTGCCTCTAACCCAATCTCTTTC
GCACACGTCTGAACTCTTTCCCTACACGACGCTCTTCCGATCTNNNACTGCCTCTAACCCAATCTCTTTC
GCACACGTCTGAACTCTTTCCCTACACGACGCTCTTCCGATCTNNNACTGCCTCTAACCCAATCTCTTTC
GCACACGTCTGAACTCTTTCCCTACACGACGCTCTTCCGATCTNNNACTGCCTCTAACCCAATCTCTTTC
GCACACGTCTGAACTCTTTCCCTACACGACGCTCTTCCGATCTNNNACTGCCTCTAACCCAATCTCTTTC
GCACACGTCTGAACTCTTTCCCTACACGACGCTCTTCCGATCTNNNACTGCCTCTAACCCAATCTCTTTC
GCACACGTCTGAACTCTTTCCCTACACGACGCTCTTCCGATCTNNNACTGCCTCTAACCCAATCTCTTTC
GCACACGTCTGAACTCTTTCCCTACACGACGCTCTTCCGATCTNNNACTGCCTCTAACCCAATCTCTTTC
GCACACGTCTGAACTCTTTCCCTACACGACGCTCTTCCGATCTNNNACTGCCTCTAACCCAATCTCTTTC
GAGCACACGTCTGAACTCTTTCCCTACACGACGCTCTTCCGATCTNNNGGAGGCTGAAGTAGGAGAA
GAGCACACGTCTGAACTCTTTCCCTACACGACGCTCTTCCGATCTNNNGGAGGCTGAAGTAGGAGAA
GAGCACACGTCTGAACTCTTTCCCTACACGACGCTCTTCCGATCTNNNGGAGGCTGAAGTAGGAGAA
GAGCACACGTCTGAACTCTTTCCCTACACGACGCTCTTCCGATCTNNNGGAGGCTGAAGTAGGAGAA
GAGCACACGTCTGAACTCTTTCCCTACACGACGCTCTTCCGATCTNNNGGAGGCTGAAGTAGGAGAA
GAGCACACGTCTGAACTCTTTCCCTACACGACGCTCTTCCGATCTNNNGGAGGCTGAAGTAGGAGAA
GAGCACACGTCTGAACTCTTTCCCTACACGACGCTCTTCCGATCTNNNGGAGGCTGAAGTAGGAGAA
GAGCACACGTCTGAACTCTTTCCCTACACGACGCTCTTCCGATCTNNNGGAGGCTGAAGTAGGAGAA
GAGCACACGTCTGAACTCTTTCCCTACACGACGCTCTTCCGATCTNNNGGAGGCTGAAGTAGGAGAA
GAGCACACGTCTGAACTCTTTCCCTACACGACGCTCTTCCGATCTNNNGGAGGCTGAAGTAGGAGAA
GCACACGTCTGAACTCTTTCCCTACACGACGCTCTTCCGATCTNNNTTCCTTACTTCTCCAGGATCTGCC
GCACACGTCTGAACTCTTTCCCTACACGACGCTCTTCCGATCTNNNTTCCTTACTTCTCCAGGATCTGCC
GCACACGTCTGAACTCTTTCCCTACACGACGCTCTTCCGATCTNNNTTCCTTACTTCTCCAGGATCTGCC
GCACACGTCTGAACTCTTTCCCTACACGACGCTCTTCCGATCTNNNTTCCTTACTTCTCCAGGATCTGCC
GCACACGTCTGAACTCTTTCCCTACACGACGCTCTTCCGATCTNNNTTCCTTACTTCTCCAGGATCTGCC
GCACACGTCTGAACTCTTTCCCTACACGACGCTCTTCCGATCTNNNTTCCTTACTTCTCCAGGATCTGCC
GCACACGTCTGAACTCTTTCCCTACACGACGCTCTTCCGATCTNNNTTCCTTACTTCTCCAGGATCTGCC
GCACACGTCTGAACTCTTTCCCTACACGACGCTCTTCCGATCTNNNTTCCTTACTTCTCCAGGATCTGCC
GCACACGTCTGAACTCTTTCCCTACACGACGCTCTTCCGATCTNNNTTCCTTACTTCTCCAGGATCTGCC
GCACACGTCTGAACTCTTTCCCTACACGACGCTCTTCCGATCTNNNTTCCTTACTTCTCCAGGATCTGCC
GAGCACACGTCTGAACTCTTTCCCTACACGACGCTCTTCCGATCTNNNAATACCAAAATTAGCTGGGCG
GAGCACACGTCTGAACTCTTTCCCTACACGACGCTCTTCCGATCTNNNAATACCAAAATTAGCTGGGCG
GAGCACACGTCTGAACTCTTTCCCTACACGACGCTCTTCCGATCTNNNAATACCAAAATTAGCTGGGCG
GAGCACACGTCTGAACTCTTTCCCTACACGACGCTCTTCCGATCTNNNAATACCAAAATTAGCTGGGCG
GAGCACACGTCTGAACTCTTTCCCTACACGACGCTCTTCCGATCTNNNAATACCAAAATTAGCTGGGCG
GAGCACACGTCTGAACTCTTTCCCTACACGACGCTCTTCCGATCTNNNAATACCAAAATTAGCTGGGCG
GAGCACACGTCTGAACTCTTTCCCTACACGACGCTCTTCCGATCTNNNAATACCAAAATTAGCTGGGCG

GAGCACACGTCTGAACTCTTTCCCTACACGACGCTCTTCCGATCTNNNAATACCAAAATTAGCTGGGCG
GAGCACACGTCTGAACTCTTTCCCTACACGACGCTCTTCCGATCTNNNAATACCAAAATTAGCTGGGCG
GAGCACACGTCTGAACTCTTTCCCTACACGACGCTCTTCCGATCTNNNAATACCAAAATTAGCTGGGCG
AGCACACGTCTGAACTCTTTCCCTACACGACGCTCTTCCGATCTNNNCATGGATCAAGGTATCCCGGAT
AGCACACGTCTGAACTCTTTCCCTACACGACGCTCTTCCGATCTNNNCATGGATCAAGGTATCCCGGAT
AGCACACGTCTGAACTCTTTCCCTACACGACGCTCTTCCGATCTNNNCATGGATCAAGGTATCCCGGAT
AGCACACGTCTGAACTCTTTCCCTACACGACGCTCTTCCGATCTNNNCATGGATCAAGGTATCCCGGAT
AGCACACGTCTGAACTCTTTCCCTACACGACGCTCTTCCGATCTNNNCATGGATCAAGGTATCCCGGAT
AGCACACGTCTGAACTCTTTCCCTACACGACGCTCTTCCGATCTNNNCATGGATCAAGGTATCCCGGAT
AGCACACGTCTGAACTCTTTCCCTACACGACGCTCTTCCGATCTNNNCATGGATCAAGGTATCCCGGAT
AGCACACGTCTGAACTCTTTCCCTACACGACGCTCTTCCGATCTNNNCATGGATCAAGGTATCCCGGAT
AGCACACGTCTGAACTCTTTCCCTACACGACGCTCTTCCGATCTNNNCATGGATCAAGGTATCCCGGAT
AGCACACGTCTGAACTCTTTCCCTACACGACGCTCTTCCGATCTNNNCATGGATCAAGGTATCCCGGAT
GAGCACACGTCTGAACTCTTTCCCTACACGACGCTCTTCCGATCTNNNCTTCAGCCTCCCAAGTAGCAT
GAGCACACGTCTGAACTCTTTCCCTACACGACGCTCTTCCGATCTNNNCTTCAGCCTCCCAAGTAGCAT
GAGCACACGTCTGAACTCTTTCCCTACACGACGCTCTTCCGATCTNNNCTTCAGCCTCCCAAGTAGCAT
GAGCACACGTCTGAACTCTTTCCCTACACGACGCTCTTCCGATCTNNNCTTCAGCCTCCCAAGTAGCAT
GAGCACACGTCTGAACTCTTTCCCTACACGACGCTCTTCCGATCTNNNCTTCAGCCTCCCAAGTAGCAT
GAGCACACGTCTGAACTCTTTCCCTACACGACGCTCTTCCGATCTNNNCTTCAGCCTCCCAAGTAGCAT
GAGCACACGTCTGAACTCTTTCCCTACACGACGCTCTTCCGATCTNNNCTTCAGCCTCCCAAGTAGCAT
GAGCACACGTCTGAACTCTTTCCCTACACGACGCTCTTCCGATCTNNNCTTCAGCCTCCCAAGTAGCAT
GAGCACACGTCTGAACTCTTTCCCTACACGACGCTCTTCCGATCTNNNCTTCAGCCTCCCAAGTAGCAT
GAGCACACGTCTGAACTCTTTCCCTACACGACGCTCTTCCGATCTNNNCTTCAGCCTCCCAAGTAGCAT
AGCACACGTCTGAACTCTTTCCCTACACGACGCTCTTCCGATCTNNNTGGCACTAGTGATGTGTACTTCA
AGCACACGTCTGAACTCTTTCCCTACACGACGCTCTTCCGATCTNNNTGGCACTAGTGATGTGTACTTCA
AGCACACGTCTGAACTCTTTCCCTACACGACGCTCTTCCGATCTNNNTGGCACTAGTGATGTGTACTTCA
AGCACACGTCTGAACTCTTTCCCTACACGACGCTCTTCCGATCTNNNTGGCACTAGTGATGTGTACTTCA
AGCACACGTCTGAACTCTTTCCCTACACGACGCTCTTCCGATCTNNNTGGCACTAGTGATGTGTACTTCA
AGCACACGTCTGAACTCTTTCCCTACACGACGCTCTTCCGATCTNNNTGGCACTAGTGATGTGTACTTCA
AGCACACGTCTGAACTCTTTCCCTACACGACGCTCTTCCGATCTNNNTGGCACTAGTGATGTGTACTTCA
AGCACACGTCTGAACTCTTTCCCTACACGACGCTCTTCCGATCTNNNTGGCACTAGTGATGTGTACTTCA
AGCACACGTCTGAACTCTTTCCCTACACGACGCTCTTCCGATCTNNNTGGCACTAGTGATGTGTACTTCA
AGCACACGTCTGAACTCTTTCCCTACACGACGCTCTTCCGATCTNNNTGGCACTAGTGATGTGTACTTCA
GAGCACACGTCTGAACTCTTTCCCTACACGACGCTCTTCCGATCTNNNTCCAGAATACTCCATGGATGT
GAGCACACGTCTGAACTCTTTCCCTACACGACGCTCTTCCGATCTNNNTCCAGAATACTCCATGGATGT
GAGCACACGTCTGAACTCTTTCCCTACACGACGCTCTTCCGATCTNNNTCCAGAATACTCCATGGATGT
GAGCACACGTCTGAACTCTTTCCCTACACGACGCTCTTCCGATCTNNNTCCAGAATACTCCATGGATGT
GAGCACACGTCTGAACTCTTTCCCTACACGACGCTCTTCCGATCTNNNTCCAGAATACTCCATGGATGT
GAGCACACGTCTGAACTCTTTCCCTACACGACGCTCTTCCGATCTNNNTCCAGAATACTCCATGGATGT
GAGCACACGTCTGAACTCTTTCCCTACACGACGCTCTTCCGATCTNNNTCCAGAATACTCCATGGATGT
GAGCACACGTCTGAACTCTTTCCCTACACGACGCTCTTCCGATCTNNNTCCAGAATACTCCATGGATGT
GAGCACACGTCTGAACTCTTTCCCTACACGACGCTCTTCCGATCTNNNTCCAGAATACTCCATGGATGT
GAGCACACGTCTGAACTCTTTCCCTACACGACGCTCTTCCGATCTNNNTCCAGAATACTCCATGGATGT

AGCACACGTCTGAACTCTTTCCCTACACGACGCTCTTCCGATCTNNNATGTACCTGTAGTCCCAGCTA
AGCACACGTCTGAACTCTTTCCCTACACGACGCTCTTCCGATCTNNNATGTACCTGTAGTCCCAGCTA
AGCACACGTCTGAACTCTTTCCCTACACGACGCTCTTCCGATCTNNNATGTACCTGTAGTCCCAGCTA
AGCACACGTCTGAACTCTTTCCCTACACGACGCTCTTCCGATCTNNNATGTACCTGTAGTCCCAGCTA
AGCACACGTCTGAACTCTTTCCCTACACGACGCTCTTCCGATCTNNNATGTACCTGTAGTCCCAGCTA
AGCACACGTCTGAACTCTTTCCCTACACGACGCTCTTCCGATCTNNNATGTACCTGTAGTCCCAGCTA
AGCACACGTCTGAACTCTTTCCCTACACGACGCTCTTCCGATCTNNNATGTACCTGTAGTCCCAGCTA
AGCACACGTCTGAACTCTTTCCCTACACGACGCTCTTCCGATCTNNNATGTACCTGTAGTCCCAGCTA
AGCACACGTCTGAACTCTTTCCCTACACGACGCTCTTCCGATCTNNNATGTACCTGTAGTCCCAGCTA
AGCACACGTCTGAACTCTTTCCCTACACGACGCTCTTCCGATCTNNNATGTACCTGTAGTCCCAGCTA
AGCACACGTCTGAACTCTTTCCCTACACGACGCTCTTCCGATCTNNNATGGGGATCACATGACAGGA
AGCACACGTCTGAACTCTTTCCCTACACGACGCTCTTCCGATCTNNNATGGGGATCACATGACAGGA
AGCACACGTCTGAACTCTTTCCCTACACGACGCTCTTCCGATCTNNNATGGGGATCACATGACAGGA
AGCACACGTCTGAACTCTTTCCCTACACGACGCTCTTCCGATCTNNNATGGGGATCACATGACAGGA
AGCACACGTCTGAACTCTTTCCCTACACGACGCTCTTCCGATCTNNNATGGGGATCACATGACAGGA
AGCACACGTCTGAACTCTTTCCCTACACGACGCTCTTCCGATCTNNNATGGGGATCACATGACAGGA
AGCACACGTCTGAACTCTTTCCCTACACGACGCTCTTCCGATCTNNNATGGGGATCACATGACAGGA
AGCACACGTCTGAACTCTTTCCCTACACGACGCTCTTCCGATCTNNNATGGGGATCACATGACAGGA
AGCACACGTCTGAACTCTTTCCCTACACGACGCTCTTCCGATCTNNNATGGGGATCACATGACAGGA
AGCACACGTCTGAACTCTTTCCCTACACGACGCTCTTCCGATCTNNNATGGGGATCACATGACAGGA
AGCACACGTCTGAACTCTTTCCCTACACGACGCTCTTCCGATCTNNNGAGAATCGCTTGAATCCAGG
AGCACACGTCTGAACTCTTTCCCTACACGACGCTCTTCCGATCTNNNGAGAATCGCTTGAATCCAGG
AGCACACGTCTGAACTCTTTCCCTACACGACGCTCTTCCGATCTNNNGAGAATCGCTTGAATCCAGG
AGCACACGTCTGAACTCTTTCCCTACACGACGCTCTTCCGATCTNNNGAGAATCGCTTGAATCCAGG
AGCACACGTCTGAACTCTTTCCCTACACGACGCTCTTCCGATCTNNNGAGAATCGCTTGAATCCAGG
AGCACACGTCTGAACTCTTTCCCTACACGACGCTCTTCCGATCTNNNGAGAATCGCTTGAATCCAGG
AGCACACGTCTGAACTCTTTCCCTACACGACGCTCTTCCGATCTNNNGAGAATCGCTTGAATCCAGG
AGCACACGTCTGAACTCTTTCCCTACACGACGCTCTTCCGATCTNNNGAGAATCGCTTGAATCCAGG
AGCACACGTCTGAACTCTTTCCCTACACGACGCTCTTCCGATCTNNNGAGAATCGCTTGAATCCAGG
AGCACACGTCTGAACTCTTTCCCTACACGACGCTCTTCCGATCTNNNGAGAATCGCTTGAATCCAGG
AGCACACGTCTGAACTCTTTCCCTACACGACGCTCTTCCGATCTNNNTCTATGCTCCAGAACAGCTCTT
AGCACACGTCTGAACTCTTTCCCTACACGACGCTCTTCCGATCTNNNTCTATGCTCCAGAACAGCTCTT
AGCACACGTCTGAACTCTTTCCCTACACGACGCTCTTCCGATCTNNNTCTATGCTCCAGAACAGCTCTT
AGCACACGTCTGAACTCTTTCCCTACACGACGCTCTTCCGATCTNNNTCTATGCTCCAGAACAGCTCTT
AGCACACGTCTGAACTCTTTCCCTACACGACGCTCTTCCGATCTNNNTCTATGCTCCAGAACAGCTCTT
AGCACACGTCTGAACTCTTTCCCTACACGACGCTCTTCCGATCTNNNTCTATGCTCCAGAACAGCTCTT
AGCACACGTCTGAACTCTTTCCCTACACGACGCTCTTCCGATCTNNNTCTATGCTCCAGAACAGCTCTT
AGCACACGTCTGAACTCTTTCCCTACACGACGCTCTTCCGATCTNNNTCTATGCTCCAGAACAGCTCTT
AGCACACGTCTGAACTCTTTCCCTACACGACGCTCTTCCGATCTNNNTCTATGCTCCAGAACAGCTCTT
AGCACACGTCTGAACTCTTTCCCTACACGACGCTCTTCCGATCTNNNTCTATGCTCCAGAACAGCTCTT
CACACGTCTGAACTCTTTCCCTACACGACGCTCTTCCGATCTNNNTCTTGAACTCCTGACCTCAAGTGC
CACACGTCTGAACTCTTTCCCTACACGACGCTCTTCCGATCTNNNTCTTGAACTCCTGACCTCAAGTGC
CACACGTCTGAACTCTTTCCCTACACGACGCTCTTCCGATCTNNNTCTTGAACTCCTGACCTCAAGTGC

GCACACGTCTGAACTCTTTCCCTACACGACGCTCTTCCGATCTNNNTCTTGAACTCCTGACCTCAAGTGCA
GCACACGTCTGAACTCTTTCCCTACACGACGCTCTTCCGATCTNNNTCTTGAACTCCTGACCTCAAGTGCA
GCACACGTCTGAACTCTTTCCCTACACGACGCTCTTCCGATCTNNNTCTTGAACTCCTGACCTCAAGTGCA
GCACACGTCTGAACTCTTTCCCTACACGACGCTCTTCCGATCTNNNTCTTGAACTCCTGACCTCAAGTGCA
GCACACGTCTGAACTCTTTCCCTACACGACGCTCTTCCGATCTNNNTCTTGAACTCCTGACCTCAAGTGCA
GCACACGTCTGAACTCTTTCCCTACACGACGCTCTTCCGATCTNNNTCTTGAACTCCTGACCTCAAGTGCA
GCACACGTCTGAACTCTTTCCCTACACGACGCTCTTCCGATCTNNNTCTTGAACTCCTGACCTCAAGTGCA
AGCACACGTCTGAACTCTTTCCCTACACGACGCTCTTCCGATCTNNNTCCTGTAGTCTGAGCTACTTGG
AGCACACGTCTGAACTCTTTCCCTACACGACGCTCTTCCGATCTNNNTCCTGTAGTCTGAGCTACTTGG
AGCACACGTCTGAACTCTTTCCCTACACGACGCTCTTCCGATCTNNNTCCTGTAGTCTGAGCTACTTGG
AGCACACGTCTGAACTCTTTCCCTACACGACGCTCTTCCGATCTNNNTCCTGTAGTCTGAGCTACTTGG
AGCACACGTCTGAACTCTTTCCCTACACGACGCTCTTCCGATCTNNNTCCTGTAGTCTGAGCTACTTGG
AGCACACGTCTGAACTCTTTCCCTACACGACGCTCTTCCGATCTNNNTCCTGTAGTCTGAGCTACTTGG
AGCACACGTCTGAACTCTTTCCCTACACGACGCTCTTCCGATCTNNNTCCTGTAGTCTGAGCTACTTGG
AGCACACGTCTGAACTCTTTCCCTACACGACGCTCTTCCGATCTNNNTCCTGTAGTCTGAGCTACTTGG
AGCACACGTCTGAACTCTTTCCCTACACGACGCTCTTCCGATCTNNNTCCTGTAGTCTGAGCTACTTGG
AGCACACGTCTGAACTCTTTCCCTACACGACGCTCTTCCGATCTNNNTCCTGTAGTCTGAGCTACTTGG
AGCACACGTCTGAACTCTTTCCCTACACGACGCTCTTCCGATCTNNNAAATGCAAGGACACCTAGCCAT
AGCACACGTCTGAACTCTTTCCCTACACGACGCTCTTCCGATCTNNNAAATGCAAGGACACCTAGCCAT
AGCACACGTCTGAACTCTTTCCCTACACGACGCTCTTCCGATCTNNNAAATGCAAGGACACCTAGCCAT
AGCACACGTCTGAACTCTTTCCCTACACGACGCTCTTCCGATCTNNNAAATGCAAGGACACCTAGCCAT
AGCACACGTCTGAACTCTTTCCCTACACGACGCTCTTCCGATCTNNNAAATGCAAGGACACCTAGCCAT
AGCACACGTCTGAACTCTTTCCCTACACGACGCTCTTCCGATCTNNNAAATGCAAGGACACCTAGCCAT
AGCACACGTCTGAACTCTTTCCCTACACGACGCTCTTCCGATCTNNNAAATGCAAGGACACCTAGCCAT
AGCACACGTCTGAACTCTTTCCCTACACGACGCTCTTCCGATCTNNNAAATGCAAGGACACCTAGCCAT
AGCACACGTCTGAACTCTTTCCCTACACGACGCTCTTCCGATCTNNNAAATGCAAGGACACCTAGCCAT
AGCACACGTCTGAACTCTTTCCCTACACGACGCTCTTCCGATCTNNNAAATGCAAGGACACCTAGCCAT
AAGAGCACACGTCTGAACTCTTTCCCTACACGACGCTCTTCCGATCTNNNAGGAGAATTGCTTGAACTC
AAGAGCACACGTCTGAACTCTTTCCCTACACGACGCTCTTCCGATCTNNNAGGAGAATTGCTTGAACTC
AAGAGCACACGTCTGAACTCTTTCCCTACACGACGCTCTTCCGATCTNNNAGGAGAATTGCTTGAACTC
AAGAGCACACGTCTGAACTCTTTCCCTACACGACGCTCTTCCGATCTNNNAGGAGAATTGCTTGAACTC
AAGAGCACACGTCTGAACTCTTTCCCTACACGACGCTCTTCCGATCTNNNAGGAGAATTGCTTGAACTC
AAGAGCACACGTCTGAACTCTTTCCCTACACGACGCTCTTCCGATCTNNNAGGAGAATTGCTTGAACTC
AAGAGCACACGTCTGAACTCTTTCCCTACACGACGCTCTTCCGATCTNNNAGGAGAATTGCTTGAACTC
AAGAGCACACGTCTGAACTCTTTCCCTACACGACGCTCTTCCGATCTNNNAGGAGAATTGCTTGAACTC
AAGAGCACACGTCTGAACTCTTTCCCTACACGACGCTCTTCCGATCTNNNAGGAGAATTGCTTGAACTC
AAGAGCACACGTCTGAACTCTTTCCCTACACGACGCTCTTCCGATCTNNNAGGAGAATTGCTTGAACTC
AGCACACGTCTGAACTCTTTCCCTACACGACGCTCTTCCGATCTNNNTCACTGAGGTGACAGAAACTGT
AGCACACGTCTGAACTCTTTCCCTACACGACGCTCTTCCGATCTNNNTCACTGAGGTGACAGAAACTGT
AGCACACGTCTGAACTCTTTCCCTACACGACGCTCTTCCGATCTNNNTCACTGAGGTGACAGAAACTGT
AGCACACGTCTGAACTCTTTCCCTACACGACGCTCTTCCGATCTNNNTCACTGAGGTGACAGAAACTGT
AGCACACGTCTGAACTCTTTCCCTACACGACGCTCTTCCGATCTNNNTCACTGAGGTGACAGAAACTGT
AGCACACGTCTGAACTCTTTCCCTACACGACGCTCTTCCGATCTNNNTCACTGAGGTGACAGAAACTGT

AGCACACGTCTGAACTCTTTCCCTACACGACGCTCTTCCGATCTNNNTCACTGAGGTGACAGAAACTGT
AGCACACGTCTGAACTCTTTCCCTACACGACGCTCTTCCGATCTNNNTCACTGAGGTGACAGAAACTGT
AGCACACGTCTGAACTCTTTCCCTACACGACGCTCTTCCGATCTNNNTCACTGAGGTGACAGAAACTGT
AGCACACGTCTGAACTCTTTCCCTACACGACGCTCTTCCGATCTNNNTCACTGAGGTGACAGAAACTGT
AGCACACGTCTGAACTCTTTCCCTACACGACGCTCTTCCGATCTNNNTGTGTACAGACTCTCCAGGAAA
AGCACACGTCTGAACTCTTTCCCTACACGACGCTCTTCCGATCTNNNTGTGTACAGACTCTCCAGGAAA
AGCACACGTCTGAACTCTTTCCCTACACGACGCTCTTCCGATCTNNNTGTGTACAGACTCTCCAGGAAA
AGCACACGTCTGAACTCTTTCCCTACACGACGCTCTTCCGATCTNNNTGTGTACAGACTCTCCAGGAAA
AGCACACGTCTGAACTCTTTCCCTACACGACGCTCTTCCGATCTNNNTGTGTACAGACTCTCCAGGAAA
AGCACACGTCTGAACTCTTTCCCTACACGACGCTCTTCCGATCTNNNTGTGTACAGACTCTCCAGGAAA
AGCACACGTCTGAACTCTTTCCCTACACGACGCTCTTCCGATCTNNNTGTGTACAGACTCTCCAGGAAA
AGCACACGTCTGAACTCTTTCCCTACACGACGCTCTTCCGATCTNNNTGTGTACAGACTCTCCAGGAAA
AGCACACGTCTGAACTCTTTCCCTACACGACGCTCTTCCGATCTNNNTGTGTACAGACTCTCCAGGAAA
AGCACACGTCTGAACTCTTTCCCTACACGACGCTCTTCCGATCTNNNTGTGTACAGACTCTCCAGGAAA
AGAGCACACGTCTGAACTCTTTCCCTACACGACGCTCTTCCGATCTNNNCTTCTCTCTAACCCCCAATGCC
AGAGCACACGTCTGAACTCTTTCCCTACACGACGCTCTTCCGATCTNNNCTTCTCTCTAACCCCCAATGCC
AGAGCACACGTCTGAACTCTTTCCCTACACGACGCTCTTCCGATCTNNNCTTCTCTCTAACCCCCAATGCC
AGAGCACACGTCTGAACTCTTTCCCTACACGACGCTCTTCCGATCTNNNCTTCTCTCTAACCCCCAATGCC
AGAGCACACGTCTGAACTCTTTCCCTACACGACGCTCTTCCGATCTNNNCTTCTCTCTAACCCCCAATGCC
AGAGCACACGTCTGAACTCTTTCCCTACACGACGCTCTTCCGATCTNNNCTTCTCTCTAACCCCCAATGCC
AGAGCACACGTCTGAACTCTTTCCCTACACGACGCTCTTCCGATCTNNNCTTCTCTCTAACCCCCAATGCC
AGAGCACACGTCTGAACTCTTTCCCTACACGACGCTCTTCCGATCTNNNCTTCTCTCTAACCCCCAATGCC
AGAGCACACGTCTGAACTCTTTCCCTACACGACGCTCTTCCGATCTNNNCTTCTCTCTAACCCCCAATGCC
AGAGCACACGTCTGAACTCTTTCCCTACACGACGCTCTTCCGATCTNNNCTTCTCTCTAACCCCCAATGCC
AGCACACGTCTGAACTCTTTCCCTACACGACGCTCTTCCGATCTNNNGCAAAACCCTGTCTCTGCAAATA
AGCACACGTCTGAACTCTTTCCCTACACGACGCTCTTCCGATCTNNNGCAAAACCCTGTCTCTGCAAATA
AGCACACGTCTGAACTCTTTCCCTACACGACGCTCTTCCGATCTNNNGCAAAACCCTGTCTCTGCAAATA
AGCACACGTCTGAACTCTTTCCCTACACGACGCTCTTCCGATCTNNNGCAAAACCCTGTCTCTGCAAATA
AGCACACGTCTGAACTCTTTCCCTACACGACGCTCTTCCGATCTNNNGCAAAACCCTGTCTCTGCAAATA
AGCACACGTCTGAACTCTTTCCCTACACGACGCTCTTCCGATCTNNNGCAAAACCCTGTCTCTGCAAATA
AGCACACGTCTGAACTCTTTCCCTACACGACGCTCTTCCGATCTNNNGCAAAACCCTGTCTCTGCAAATA
AGCACACGTCTGAACTCTTTCCCTACACGACGCTCTTCCGATCTNNNGCAAAACCCTGTCTCTGCAAATA
AGCACACGTCTGAACTCTTTCCCTACACGACGCTCTTCCGATCTNNNGCAAAACCCTGTCTCTGCAAATA
AGCACACGTCTGAACTCTTTCCCTACACGACGCTCTTCCGATCTNNNGCAAAACCCTGTCTCTGCAAATA
AGCACACGTCTGAACTCTTTCCCTACACGACGCTCTTCCGATCTNNNTAGCTGGGATTATAGGTGCATG
AGCACACGTCTGAACTCTTTCCCTACACGACGCTCTTCCGATCTNNNTAGCTGGGATTATAGGTGCATG
AGCACACGTCTGAACTCTTTCCCTACACGACGCTCTTCCGATCTNNNTAGCTGGGATTATAGGTGCATG
AGCACACGTCTGAACTCTTTCCCTACACGACGCTCTTCCGATCTNNNTAGCTGGGATTATAGGTGCATG
AGCACACGTCTGAACTCTTTCCCTACACGACGCTCTTCCGATCTNNNTAGCTGGGATTATAGGTGCATG
AGCACACGTCTGAACTCTTTCCCTACACGACGCTCTTCCGATCTNNNTAGCTGGGATTATAGGTGCATG
AGCACACGTCTGAACTCTTTCCCTACACGACGCTCTTCCGATCTNNNTAGCTGGGATTATAGGTGCATG
AGCACACGTCTGAACTCTTTCCCTACACGACGCTCTTCCGATCTNNNTAGCTGGGATTATAGGTGCATG
AGCACACGTCTGAACTCTTTCCCTACACGACGCTCTTCCGATCTNNNTAGCTGGGATTATAGGTGCATG

AGCACACGTCTGAACTCTTTCCCTACACGACGCTCTTCCGATCTNNNTAGCTGGGATTATAGGTGCATG
CACACGTCTGAACTCTTTCCCTACACGACGCTCTTCCGATCTNNNTTAGAGTAGATTATATACGACCAA
CACACGTCTGAACTCTTTCCCTACACGACGCTCTTCCGATCTNNNTTAGAGTAGATTATATACGACCAA
CACACGTCTGAACTCTTTCCCTACACGACGCTCTTCCGATCTNNNTTAGAGTAGATTATATACGACCAA
CACACGTCTGAACTCTTTCCCTACACGACGCTCTTCCGATCTNNNTTAGAGTAGATTATATACGACCAA
CACACGTCTGAACTCTTTCCCTACACGACGCTCTTCCGATCTNNNTTAGAGTAGATTATATACGACCAA
CACACGTCTGAACTCTTTCCCTACACGACGCTCTTCCGATCTNNNTTAGAGTAGATTATATACGACCAA
CACACGTCTGAACTCTTTCCCTACACGACGCTCTTCCGATCTNNNTTAGAGTAGATTATATACGACCAA
CACACGTCTGAACTCTTTCCCTACACGACGCTCTTCCGATCTNNNTTAGAGTAGATTATATACGACCAA
CACACGTCTGAACTCTTTCCCTACACGACGCTCTTCCGATCTNNNTTAGAGTAGATTATATACGACCAA
CACACGTCTGAACTCTTTCCCTACACGACGCTCTTCCGATCTNNNTTAGAGTAGATTATATACGACCAA
GCACACGTCTGAACTCTTTCCCTACACGACGCTCTTCCGATCTNNNGCTCTTTTCCTGGCTTGAAGATG
GCACACGTCTGAACTCTTTCCCTACACGACGCTCTTCCGATCTNNNGCTCTTTTCCTGGCTTGAAGATG
GCACACGTCTGAACTCTTTCCCTACACGACGCTCTTCCGATCTNNNGCTCTTTTCCTGGCTTGAAGATG
GCACACGTCTGAACTCTTTCCCTACACGACGCTCTTCCGATCTNNNGCTCTTTTCCTGGCTTGAAGATG
GCACACGTCTGAACTCTTTCCCTACACGACGCTCTTCCGATCTNNNGCTCTTTTCCTGGCTTGAAGATG
GCACACGTCTGAACTCTTTCCCTACACGACGCTCTTCCGATCTNNNGCTCTTTTCCTGGCTTGAAGATG
GCACACGTCTGAACTCTTTCCCTACACGACGCTCTTCCGATCTNNNGCTCTTTTCCTGGCTTGAAGATG
GCACACGTCTGAACTCTTTCCCTACACGACGCTCTTCCGATCTNNNGCTCTTTTCCTGGCTTGAAGATG
GCACACGTCTGAACTCTTTCCCTACACGACGCTCTTCCGATCTNNNGCTCTTTTCCTGGCTTGAAGATG
GCACACGTCTGAACTCTTTCCCTACACGACGCTCTTCCGATCTNNNGCTCTTTTCCTGGCTTGAAGATG
GAGCACACGTCTGAACTCTTTCCCTACACGACGCTCTTCCGATCTNNNTCATTTGAACCCAGGAGACAG
GAGCACACGTCTGAACTCTTTCCCTACACGACGCTCTTCCGATCTNNNTCATTTGAACCCAGGAGACAG
GAGCACACGTCTGAACTCTTTCCCTACACGACGCTCTTCCGATCTNNNTCATTTGAACCCAGGAGACAG
GAGCACACGTCTGAACTCTTTCCCTACACGACGCTCTTCCGATCTNNNTCATTTGAACCCAGGAGACAG
GAGCACACGTCTGAACTCTTTCCCTACACGACGCTCTTCCGATCTNNNTCATTTGAACCCAGGAGACAG
GAGCACACGTCTGAACTCTTTCCCTACACGACGCTCTTCCGATCTNNNTCATTTGAACCCAGGAGACAG
GAGCACACGTCTGAACTCTTTCCCTACACGACGCTCTTCCGATCTNNNTCATTTGAACCCAGGAGACAG
GAGCACACGTCTGAACTCTTTCCCTACACGACGCTCTTCCGATCTNNNTCATTTGAACCCAGGAGACAG
GAGCACACGTCTGAACTCTTTCCCTACACGACGCTCTTCCGATCTNNNTCATTTGAACCCAGGAGACAG
GAGCACACGTCTGAACTCTTTCCCTACACGACGCTCTTCCGATCTNNNTCATTTGAACCCAGGAGACAG
GAGCACACGTCTGAACTCTTTCCCTACACGACGCTCTTCCGATCTNNNAACAGAGCTCGCTTAGATGGA
GAGCACACGTCTGAACTCTTTCCCTACACGACGCTCTTCCGATCTNNNAACAGAGCTCGCTTAGATGGA
GAGCACACGTCTGAACTCTTTCCCTACACGACGCTCTTCCGATCTNNNAACAGAGCTCGCTTAGATGGA
GAGCACACGTCTGAACTCTTTCCCTACACGACGCTCTTCCGATCTNNNAACAGAGCTCGCTTAGATGGA
GAGCACACGTCTGAACTCTTTCCCTACACGACGCTCTTCCGATCTNNNAACAGAGCTCGCTTAGATGGA
GAGCACACGTCTGAACTCTTTCCCTACACGACGCTCTTCCGATCTNNNAACAGAGCTCGCTTAGATGGA
GAGCACACGTCTGAACTCTTTCCCTACACGACGCTCTTCCGATCTNNNAACAGAGCTCGCTTAGATGGA
GAGCACACGTCTGAACTCTTTCCCTACACGACGCTCTTCCGATCTNNNAACAGAGCTCGCTTAGATGGA
GAGCACACGTCTGAACTCTTTCCCTACACGACGCTCTTCCGATCTNNNAACAGAGCTCGCTTAGATGGA
GAGCACACGTCTGAACTCTTTCCCTACACGACGCTCTTCCGATCTNNNAACAGAGCTCGCTTAGATGGA
AGCACACGTCTGAACTCTTTCCCTACACGACGCTCTTCCGATCTNNNGGCTGAGATAGGAGGATCACT
AGCACACGTCTGAACTCTTTCCCTACACGACGCTCTTCCGATCTNNNGGCTGAGATAGGAGGATCACT

AGCACACGTCTGAACTCTTTCCCTACACGACGCTCTTCCGATCTNNNGGCTGAGATAGGAGGATCACT
AGCACACGTCTGAACTCTTTCCCTACACGACGCTCTTCCGATCTNNNGGCTGAGATAGGAGGATCACT
AGCACACGTCTGAACTCTTTCCCTACACGACGCTCTTCCGATCTNNNGGCTGAGATAGGAGGATCACT
AGCACACGTCTGAACTCTTTCCCTACACGACGCTCTTCCGATCTNNNGGCTGAGATAGGAGGATCACT
AGCACACGTCTGAACTCTTTCCCTACACGACGCTCTTCCGATCTNNNGGCTGAGATAGGAGGATCACT
AGCACACGTCTGAACTCTTTCCCTACACGACGCTCTTCCGATCTNNNGGCTGAGATAGGAGGATCACT
AGCACACGTCTGAACTCTTTCCCTACACGACGCTCTTCCGATCTNNNGGCTGAGATAGGAGGATCACT
AGCACACGTCTGAACTCTTTCCCTACACGACGCTCTTCCGATCTNNNGGCTGAGATAGGAGGATCACT
AGAGCACACGTCTGAACTCTTTCCCTACACGACGCTCTTCCGATCTNNNGAGAAAACCAAGATTGCTTA
AGAGCACACGTCTGAACTCTTTCCCTACACGACGCTCTTCCGATCTNNNGAGAAAACCAAGATTGCTTA
AGAGCACACGTCTGAACTCTTTCCCTACACGACGCTCTTCCGATCTNNNGAGAAAACCAAGATTGCTTA
AGAGCACACGTCTGAACTCTTTCCCTACACGACGCTCTTCCGATCTNNNGAGAAAACCAAGATTGCTTA
AGAGCACACGTCTGAACTCTTTCCCTACACGACGCTCTTCCGATCTNNNGAGAAAACCAAGATTGCTTA
AGAGCACACGTCTGAACTCTTTCCCTACACGACGCTCTTCCGATCTNNNGAGAAAACCAAGATTGCTTA
AGAGCACACGTCTGAACTCTTTCCCTACACGACGCTCTTCCGATCTNNNGAGAAAACCAAGATTGCTTA
AGAGCACACGTCTGAACTCTTTCCCTACACGACGCTCTTCCGATCTNNNGAGAAAACCAAGATTGCTTA
AGAGCACACGTCTGAACTCTTTCCCTACACGACGCTCTTCCGATCTNNNGAGAAAACCAAGATTGCTTA
AGAGCACACGTCTGAACTCTTTCCCTACACGACGCTCTTCCGATCTNNNGAGAAAACCAAGATTGCTTA
GCACACGTCTGAACTCTTTCCCTACACGACGCTCTTCCGATCTNNNTTTGACTCAGAGATGCCATGTGT
GCACACGTCTGAACTCTTTCCCTACACGACGCTCTTCCGATCTNNNTTTGACTCAGAGATGCCATGTGT
GCACACGTCTGAACTCTTTCCCTACACGACGCTCTTCCGATCTNNNTTTGACTCAGAGATGCCATGTGT
GCACACGTCTGAACTCTTTCCCTACACGACGCTCTTCCGATCTNNNTTTGACTCAGAGATGCCATGTGT
GCACACGTCTGAACTCTTTCCCTACACGACGCTCTTCCGATCTNNNTTTGACTCAGAGATGCCATGTGT
GCACACGTCTGAACTCTTTCCCTACACGACGCTCTTCCGATCTNNNTTTGACTCAGAGATGCCATGTGT
GCACACGTCTGAACTCTTTCCCTACACGACGCTCTTCCGATCTNNNTTTGACTCAGAGATGCCATGTGT
GCACACGTCTGAACTCTTTCCCTACACGACGCTCTTCCGATCTNNNTTTGACTCAGAGATGCCATGTGT
GCACACGTCTGAACTCTTTCCCTACACGACGCTCTTCCGATCTNNNTTTGACTCAGAGATGCCATGTGT
GCACACGTCTGAACTCTTTCCCTACACGACGCTCTTCCGATCTNNNTTTGACTCAGAGATGCCATGTGT
AGCACACGTCTGAACTCTTTCCCTACACGACGCTCTTCCGATCTNNNATTCGACCCAGTAATCCTGCTAC
AGCACACGTCTGAACTCTTTCCCTACACGACGCTCTTCCGATCTNNNATTCGACCCAGTAATCCTGCTAC
AGCACACGTCTGAACTCTTTCCCTACACGACGCTCTTCCGATCTNNNATTCGACCCAGTAATCCTGCTAC
AGCACACGTCTGAACTCTTTCCCTACACGACGCTCTTCCGATCTNNNATTCGACCCAGTAATCCTGCTAC
AGCACACGTCTGAACTCTTTCCCTACACGACGCTCTTCCGATCTNNNATTCGACCCAGTAATCCTGCTAC
AGCACACGTCTGAACTCTTTCCCTACACGACGCTCTTCCGATCTNNNATTCGACCCAGTAATCCTGCTAC
AGCACACGTCTGAACTCTTTCCCTACACGACGCTCTTCCGATCTNNNATTCGACCCAGTAATCCTGCTAC
AGCACACGTCTGAACTCTTTCCCTACACGACGCTCTTCCGATCTNNNATTCGACCCAGTAATCCTGCTAC
AGCACACGTCTGAACTCTTTCCCTACACGACGCTCTTCCGATCTNNNATTCGACCCAGTAATCCTGCTAC
AGCACACGTCTGAACTCTTTCCCTACACGACGCTCTTCCGATCTNNNATTCGACCCAGTAATCCTGCTAC
AGCACACGTCTGAACTCTTTCCCTACACGACGCTCTTCCGATCTNNNCTTCAGGTAGATTAAGGTATCTT
AGCACACGTCTGAACTCTTTCCCTACACGACGCTCTTCCGATCTNNNCTTCAGGTAGATTAAGGTATCTT
AGCACACGTCTGAACTCTTTCCCTACACGACGCTCTTCCGATCTNNNCTTCAGGTAGATTAAGGTATCTT
AGCACACGTCTGAACTCTTTCCCTACACGACGCTCTTCCGATCTNNNCTTCAGGTAGATTAAGGTATCTT
AGCACACGTCTGAACTCTTTCCCTACACGACGCTCTTCCGATCTNNNCTTCAGGTAGATTAAGGTATCTT

GCACACGTCTGAACTCTTTCCCTACACGACGCTCTTCCGATCTNNNCTTCAGGTAGATTAAGGTATCTT
GCACACGTCTGAACTCTTTCCCTACACGACGCTCTTCCGATCTNNNCTTCAGGTAGATTAAGGTATCTT
GCACACGTCTGAACTCTTTCCCTACACGACGCTCTTCCGATCTNNNCTTCAGGTAGATTAAGGTATCTT
GCACACGTCTGAACTCTTTCCCTACACGACGCTCTTCCGATCTNNNCTTCAGGTAGATTAAGGTATCTT
GCACACGTCTGAACTCTTTCCCTACACGACGCTCTTCCGATCTNNNCTTCAGGTAGATTAAGGTATCTT
CACACGTCTGAACTCTTTCCCTACACGACGCTCTTCCGATCTNNNAAGGACTGAAGACCTGTAAGAGC
CACACGTCTGAACTCTTTCCCTACACGACGCTCTTCCGATCTNNNAAGGACTGAAGACCTGTAAGAGC
CACACGTCTGAACTCTTTCCCTACACGACGCTCTTCCGATCTNNNAAGGACTGAAGACCTGTAAGAGC
CACACGTCTGAACTCTTTCCCTACACGACGCTCTTCCGATCTNNNAAGGACTGAAGACCTGTAAGAGC
CACACGTCTGAACTCTTTCCCTACACGACGCTCTTCCGATCTNNNAAGGACTGAAGACCTGTAAGAGC
CACACGTCTGAACTCTTTCCCTACACGACGCTCTTCCGATCTNNNAAGGACTGAAGACCTGTAAGAGC
CACACGTCTGAACTCTTTCCCTACACGACGCTCTTCCGATCTNNNAAGGACTGAAGACCTGTAAGAGC
CACACGTCTGAACTCTTTCCCTACACGACGCTCTTCCGATCTNNNAAGGACTGAAGACCTGTAAGAGC
CACACGTCTGAACTCTTTCCCTACACGACGCTCTTCCGATCTNNNAAGGACTGAAGACCTGTAAGAGC
CACACGTCTGAACTCTTTCCCTACACGACGCTCTTCCGATCTNNNAAGGACTGAAGACCTGTAAGAGC
GAGCACACGTCTGAACTCTTTCCCTACACGACGCTCTTCCGATCTNNNCAAAAGGCAGGTCACCTACAA
GAGCACACGTCTGAACTCTTTCCCTACACGACGCTCTTCCGATCTNNNCAAAAGGCAGGTCACCTACAA
GAGCACACGTCTGAACTCTTTCCCTACACGACGCTCTTCCGATCTNNNCAAAAGGCAGGTCACCTACAA
GAGCACACGTCTGAACTCTTTCCCTACACGACGCTCTTCCGATCTNNNCAAAAGGCAGGTCACCTACAA
GAGCACACGTCTGAACTCTTTCCCTACACGACGCTCTTCCGATCTNNNCAAAAGGCAGGTCACCTACAA
GAGCACACGTCTGAACTCTTTCCCTACACGACGCTCTTCCGATCTNNNCAAAAGGCAGGTCACCTACAA
GAGCACACGTCTGAACTCTTTCCCTACACGACGCTCTTCCGATCTNNNCAAAAGGCAGGTCACCTACAA
GAGCACACGTCTGAACTCTTTCCCTACACGACGCTCTTCCGATCTNNNCAAAAGGCAGGTCACCTACAA
GAGCACACGTCTGAACTCTTTCCCTACACGACGCTCTTCCGATCTNNNCAAAAGGCAGGTCACCTACAA
GAGCACACGTCTGAACTCTTTCCCTACACGACGCTCTTCCGATCTNNNCAAAAGGCAGGTCACCTACAA
GCACACGTCTGAACTCTTTCCCTACACGACGCTCTTCCGATCTNNNCATGTCTGCGCATTTCTAGACTG
GCACACGTCTGAACTCTTTCCCTACACGACGCTCTTCCGATCTNNNCATGTCTGCGCATTTCTAGACTG
GCACACGTCTGAACTCTTTCCCTACACGACGCTCTTCCGATCTNNNCATGTCTGCGCATTTCTAGACTG
GCACACGTCTGAACTCTTTCCCTACACGACGCTCTTCCGATCTNNNCATGTCTGCGCATTTCTAGACTG
GCACACGTCTGAACTCTTTCCCTACACGACGCTCTTCCGATCTNNNCATGTCTGCGCATTTCTAGACTG
GCACACGTCTGAACTCTTTCCCTACACGACGCTCTTCCGATCTNNNCATGTCTGCGCATTTCTAGACTG
GCACACGTCTGAACTCTTTCCCTACACGACGCTCTTCCGATCTNNNCATGTCTGCGCATTTCTAGACTG
GCACACGTCTGAACTCTTTCCCTACACGACGCTCTTCCGATCTNNNCATGTCTGCGCATTTCTAGACTG
GCACACGTCTGAACTCTTTCCCTACACGACGCTCTTCCGATCTNNNCATGTCTGCGCATTTCTAGACTG
GCACACGTCTGAACTCTTTCCCTACACGACGCTCTTCCGATCTNNNCATGTCTGCGCATTTCTAGACTG
GAGCACACGTCTGAACTCTTTCCCTACACGACGCTCTTCCGATCTNNNGTGCAAGATCAGAGCCAAAT
GAGCACACGTCTGAACTCTTTCCCTACACGACGCTCTTCCGATCTNNNGTGCAAGATCAGAGCCAAAT
GAGCACACGTCTGAACTCTTTCCCTACACGACGCTCTTCCGATCTNNNGTGCAAGATCAGAGCCAAAT
GAGCACACGTCTGAACTCTTTCCCTACACGACGCTCTTCCGATCTNNNGTGCAAGATCAGAGCCAAAT
GAGCACACGTCTGAACTCTTTCCCTACACGACGCTCTTCCGATCTNNNGTGCAAGATCAGAGCCAAAT
GAGCACACGTCTGAACTCTTTCCCTACACGACGCTCTTCCGATCTNNNGTGCAAGATCAGAGCCAAAT
GAGCACACGTCTGAACTCTTTCCCTACACGACGCTCTTCCGATCTNNNGTGCAAGATCAGAGCCAAAT
GAGCACACGTCTGAACTCTTTCCCTACACGACGCTCTTCCGATCTNNNGTGCAAGATCAGAGCCAAAT

GAGCACACGTCTGAACTCTTTCCCTACACGACGCTCTTCCGATCTNNNGTGCAAGATCAGAGCCAAAT
GAGCACACGTCTGAACTCTTTCCCTACACGACGCTCTTCCGATCTNNNGTGCAAGATCAGAGCCAAAT
AGCACACGTCTGAACTCTTTCCCTACACGACGCTCTTCCGATCTNNNTGATCTCAGGCACATTCTGACT
AGCACACGTCTGAACTCTTTCCCTACACGACGCTCTTCCGATCTNNNTGATCTCAGGCACATTCTGACT
AGCACACGTCTGAACTCTTTCCCTACACGACGCTCTTCCGATCTNNNAGGCAAAAGAATCGCTTGAAC
AGCACACGTCTGAACTCTTTCCCTACACGACGCTCTTCCGATCTNNNAGGCAAAAGAATCGCTTGAAC
GAGCACACGTCTGAACTCTTTCCCTACACGACGCTCTTCCGATCTNNNCAGAAGGTTGCAGTGAGTC
GAGCACACGTCTGAACTCTTTCCCTACACGACGCTCTTCCGATCTNNNCAGAAGGTTGCAGTGAGTC
AGCACACGTCTGAACTCTTTCCCTACACGACGCTCTTCCGATCTNNNGGGAGTTACCTCTCTGCAAAC
AGCACACGTCTGAACTCTTTCCCTACACGACGCTCTTCCGATCTNNNGGGAGTTACCTCTCTGCAAAC
CACACGTCTGAACTCTTTCCCTACACGACGCTCTTCCGATCTNNNGTGAAAAACAAATCATCATCATCC
CACACGTCTGAACTCTTTCCCTACACGACGCTCTTCCGATCTNNNGTGAAAAACAAATCATCATCATCC
GAGCACACGTCTGAACTCTTTCCCTACACGACGCTCTTCCGATCTNNNGTGGATGCTGAGAAATTGG
GAGCACACGTCTGAACTCTTTCCCTACACGACGCTCTTCCGATCTNNNGTGGATGCTGAGAAATTGG
GAGCACACGTCTGAACTCTTTCCCTACACGACGCTCTTCCGATCTNNNTGCTCACCTGTAATCTCAGCT
GAGCACACGTCTGAACTCTTTCCCTACACGACGCTCTTCCGATCTNNNTGCTCACCTGTAATCTCAGCT
GAGCACACGTCTGAACTCTTTCCCTACACGACGCTCTTCCGATCTNNNACCTACTTGATCATGGTGGAT
GAGCACACGTCTGAACTCTTTCCCTACACGACGCTCTTCCGATCTNNNACCTACTTGATCATGGTGGAT
AGCACACGTCTGAACTCTTTCCCTACACGACGCTCTTCCGATCTNNNTGGACAACAGAGAAAGACCCT
AGCACACGTCTGAACTCTTTCCCTACACGACGCTCTTCCGATCTNNNTGGACAACAGAGAAAGACCCT
GCACACGTCTGAACTCTTTCCCTACACGACGCTCTTCCGATCTNNNCTGTAGTCCCAGCTACTCAGGAG
GCACACGTCTGAACTCTTTCCCTACACGACGCTCTTCCGATCTNNNCTGTAGTCCCAGCTACTCAGGAG
AAGAGCACACGTCTGAACTCTTTCCCTACACGACGCTCTTCCGATCTNNNGGCGTTTCAGCTAAACCC
AAGAGCACACGTCTGAACTCTTTCCCTACACGACGCTCTTCCGATCTNNNGGCGTTTCAGCTAAACCC
AGCACACGTCTGAACTCTTTCCCTACACGACGCTCTTCCGATCTNNNCAGGTCCTTAATGACAGCAGA
AGCACACGTCTGAACTCTTTCCCTACACGACGCTCTTCCGATCTNNNCAGGTCCTTAATGACAGCAGA
AGCACACGTCTGAACTCTTTCCCTACACGACGCTCTTCCGATCTNNNTAACCTGTGAGCTGATCCTTG
AGCACACGTCTGAACTCTTTCCCTACACGACGCTCTTCCGATCTNNNTAACCTGTGAGCTGATCCTTG
AGCACACGTCTGAACTCTTTCCCTACACGACGCTCTTCCGATCTNNNAACAATTATGTGTGGATAAGA
AGCACACGTCTGAACTCTTTCCCTACACGACGCTCTTCCGATCTNNNAACAATTATGTGTGGATAAGA
AGCACACGTCTGAACTCTTTCCCTACACGACGCTCTTCCGATCTNNNGGAGAAGAGAAGTAAGCTGC
AGCACACGTCTGAACTCTTTCCCTACACGACGCTCTTCCGATCTNNNGGAGAAGAGAAGTAAGCTGC
GAGCACACGTCTGAACTCTTTCCCTACACGACGCTCTTCCGATCTNNNCTTGAAAACAATGTGCGTCCT
GAGCACACGTCTGAACTCTTTCCCTACACGACGCTCTTCCGATCTNNNCTTGAAAACAATGTGCGTCCT
AGCACACGTCTGAACTCTTTCCCTACACGACGCTCTTCCGATCTNNNCCGTCTCAAATATAAGGGGC
AGCACACGTCTGAACTCTTTCCCTACACGACGCTCTTCCGATCTNNNCCGTCTCAAATATAAGGGGC
AGCACACGTCTGAACTCTTTCCCTACACGACGCTCTTCCGATCTNNNAAGGCAGGAGAATCACTTGA
AGCACACGTCTGAACTCTTTCCCTACACGACGCTCTTCCGATCTNNNAAGGCAGGAGAATCACTTGA
AGCACACGTCTGAACTCTTTCCCTACACGACGCTCTTCCGATCTNNNTAAACCGCCAGTGGGAATGTA
AGCACACGTCTGAACTCTTTCCCTACACGACGCTCTTCCGATCTNNNTAAACCGCCAGTGGGAATGTA
AGCACACGTCTGAACTCTTTCCCTACACGACGCTCTTCCGATCTNNNTCTTTTGCACAGGAAGCTAGC
AGCACACGTCTGAACTCTTTCCCTACACGACGCTCTTCCGATCTNNNTCTTTTGCACAGGAAGCTAGC
GCACACGTCTGAACTCTTTCCCTACACGACGCTCTTCCGATCTNNNTATAGTCCCAGCTACTCAGGAG

GCACACGTCTGAACTCTTTCCCTACACGACGCTCTTCCGATCTNNNTATAGTCCCAGCTACTCAGGAG
GAGCACACGTCTGAACTCTTTCCCTACACGACGCTCTTCCGATCTNNNCACCTGTAATCCCAGCTACTTG
GAGCACACGTCTGAACTCTTTCCCTACACGACGCTCTTCCGATCTNNNCACCTGTAATCCCAGCTACTTG
GCACACGTCTGAACTCTTTCCCTACACGACGCTCTTCCGATCTNNNCAAGCAGGTTTTGGACAATAGC
GCACACGTCTGAACTCTTTCCCTACACGACGCTCTTCCGATCTNNNCAAGCAGGTTTTGGACAATAGC
GAGCACACGTCTGAACTCTTTCCCTACACGACGCTCTTCCGATCTNNNTAGGTGAGGGAATGAGACTC
GAGCACACGTCTGAACTCTTTCCCTACACGACGCTCTTCCGATCTNNNTAGGTGAGGGAATGAGACTC
AGCACACGTCTGAACTCTTTCCCTACACGACGCTCTTCCGATCTNNNTGGATGACAGAAGGAGACTCT
AGCACACGTCTGAACTCTTTCCCTACACGACGCTCTTCCGATCTNNNTGGATGACAGAAGGAGACTCT
AGCACACGTCTGAACTCTTTCCCTACACGACGCTCTTCCGATCTNNNAGCCCTTTAGCCTCTGTTTACAA
AGCACACGTCTGAACTCTTTCCCTACACGACGCTCTTCCGATCTNNNAGCCCTTTAGCCTCTGTTTACAA
GAGCACACGTCTGAACTCTTTCCCTACACGACGCTCTTCCGATCTNNNCCTCACAGTCTGGCACTTCAT
GAGCACACGTCTGAACTCTTTCCCTACACGACGCTCTTCCGATCTNNNCCTCACAGTCTGGCACTTCAT
AGCACACGTCTGAACTCTTTCCCTACACGACGCTCTTCCGATCTNNNGATCCCAGGAGTATGACTTCA
AGCACACGTCTGAACTCTTTCCCTACACGACGCTCTTCCGATCTNNNGATCCCAGGAGTATGACTTCA
AAGAGCACACGTCTGAACTCTTTCCCTACACGACGCTCTTCCGATCTNNNACCTACACCTCAGCAGTTA
AAGAGCACACGTCTGAACTCTTTCCCTACACGACGCTCTTCCGATCTNNNACCTACACCTCAGCAGTTA
GAGCACACGTCTGAACTCTTTCCCTACACGACGCTCTTCCGATCTNNNTAACTGTGGGTTTTCTCCTCC
GAGCACACGTCTGAACTCTTTCCCTACACGACGCTCTTCCGATCTNNNTAACTGTGGGTTTTCTCCTCC
GAGCACACGTCTGAACTCTTTCCCTACACGACGCTCTTCCGATCTNNNCTCAAGCCCCAACAATTCTGAA
GAGCACACGTCTGAACTCTTTCCCTACACGACGCTCTTCCGATCTNNNCTCAAGCCCCAACAATTCTGAA
AGCACACGTCTGAACTCTTTCCCTACACGACGCTCTTCCGATCTNNNTCCTCCACTTTATAGGGGTCACT
AGCACACGTCTGAACTCTTTCCCTACACGACGCTCTTCCGATCTNNNTCCTCCACTTTATAGGGGTCACT
GAGCACACGTCTGAACTCTTTCCCTACACGACGCTCTTCCGATCTNNNGAAGTCTTACAGGCCAGGAG
GAGCACACGTCTGAACTCTTTCCCTACACGACGCTCTTCCGATCTNNNGAAGTCTTACAGGCCAGGAG
GCACACGTCTGAACTCTTTCCCTACACGACGCTCTTCCGATCTNNNAGGATAATACACATCACAGTAA
GCACACGTCTGAACTCTTTCCCTACACGACGCTCTTCCGATCTNNNAGGATAATACACATCACAGTAA
AGCACACGTCTGAACTCTTTCCCTACACGACGCTCTTCCGATCTNNNTGAACTCCTGAGCTCAAGTGAT
AGCACACGTCTGAACTCTTTCCCTACACGACGCTCTTCCGATCTNNNTGAACTCCTGAGCTCAAGTGAT
AGAGCACACGTCTGAACTCTTTCCCTACACGACGCTCTTCCGATCTNNNCCCTGGGTCTAGTTTCCCTTT
AGAGCACACGTCTGAACTCTTTCCCTACACGACGCTCTTCCGATCTNNNCCCTGGGTCTAGTTTCCCTTT
CACACGTCTGAACTCTTTCCCTACACGACGCTCTTCCGATCTNNNTTCTAAGGTTTCCTTGGCGTTCTC
CACACGTCTGAACTCTTTCCCTACACGACGCTCTTCCGATCTNNNTTCTAAGGTTTCCTTGGCGTTCTC
AGAGCACACGTCTGAACTCTTTCCCTACACGACGCTCTTCCGATCTNNNATCATCTGAGTCTGGGAGG
AGAGCACACGTCTGAACTCTTTCCCTACACGACGCTCTTCCGATCTNNNATCATCTGAGTCTGGGAGG
AGCACACGTCTGAACTCTTTCCCTACACGACGCTCTTCCGATCTNNNGCAGTGTCTCCTCATTACAGTG
AGCACACGTCTGAACTCTTTCCCTACACGACGCTCTTCCGATCTNNNGCAGTGTCTCCTCATTACAGTG
GAGCACACGTCTGAACTCTTTCCCTACACGACGCTCTTCCGATCTNNNTGTGTGTGTATATTGTGTGTG
GAGCACACGTCTGAACTCTTTCCCTACACGACGCTCTTCCGATCTNNNTGTGTGTGTATATTGTGTGTG
GAGCACACGTCTGAACTCTTTCCCTACACGACGCTCTTCCGATCTNNNTACAAAAATTAGCTGGGCGTG
GAGCACACGTCTGAACTCTTTCCCTACACGACGCTCTTCCGATCTNNNTACAAAAATTAGCTGGGCGTG
AAGAGCACACGTCTGAACTCTTTCCCTACACGACGCTCTTCCGATCTNNNAGGAAGGAAGGAACAGA
AAGAGCACACGTCTGAACTCTTTCCCTACACGACGCTCTTCCGATCTNNNAGGAAGGAAGGAACAGA

AGCACACGTCTGAACTCTTTCCCTACACGACGCTCTTCCGATCTNNNTACCCTGGACTCCTAGACATCT
AGCACACGTCTGAACTCTTTCCCTACACGACGCTCTTCCGATCTNNNTACCCTGGACTCCTAGACATCT
AGCACACGTCTGAACTCTTTCCCTACACGACGCTCTTCCGATCTNNNTGAGGCAGGAGAATCTCTTGA
AGCACACGTCTGAACTCTTTCCCTACACGACGCTCTTCCGATCTNNNTGAGGCAGGAGAATCTCTTGA
CACACGTCTGAACTCTTTCCCTACACGACGCTCTTCCGATCTNNNTGACATGGAAAATTTTGATAATCC
CACACGTCTGAACTCTTTCCCTACACGACGCTCTTCCGATCTNNNTGACATGGAAAATTTTGATAATCC
GCACACGTCTGAACTCTTTCCCTACACGACGCTCTTCCGATCTNNNAGGCTCTCTGAGAGTAGAAAG
GCACACGTCTGAACTCTTTCCCTACACGACGCTCTTCCGATCTNNNAGGCTCTCTGAGAGTAGAAAG
AGCACACGTCTGAACTCTTTCCCTACACGACGCTCTTCCGATCTNNNCACAGCATGGATGTTCTGTAAT
AGCACACGTCTGAACTCTTTCCCTACACGACGCTCTTCCGATCTNNNCACAGCATGGATGTTCTGTAAT
AGCACACGTCTGAACTCTTTCCCTACACGACGCTCTTCCGATCTNNNCTGCTGGGTTCTCTTATGACAC
AGCACACGTCTGAACTCTTTCCCTACACGACGCTCTTCCGATCTNNNCTGCTGGGTTCTCTTATGACAC
AGCACACGTCTGAACTCTTTCCCTACACGACGCTCTTCCGATCTNNNCAGGTTCAAGCAATTCTCCTGC
AGCACACGTCTGAACTCTTTCCCTACACGACGCTCTTCCGATCTNNNCAGGTTCAAGCAATTCTCCTGC
AGCACACGTCTGAACTCTTTCCCTACACGACGCTCTTCCGATCTNNNGTCTGAGCTCACCTAACACTGA
AGCACACGTCTGAACTCTTTCCCTACACGACGCTCTTCCGATCTNNNGTCTGAGCTCACCTAACACTGA
GAGCACACGTCTGAACTCTTTCCCTACACGACGCTCTTCCGATCTNNNTGCAGCCTATCATTGATGGAC
GAGCACACGTCTGAACTCTTTCCCTACACGACGCTCTTCCGATCTNNNTGCAGCCTATCATTGATGGAC
AGCACACGTCTGAACTCTTTCCCTACACGACGCTCTTCCGATCTNNNTTCTCACTTTGCGTCATGGTTT
AGCACACGTCTGAACTCTTTCCCTACACGACGCTCTTCCGATCTNNNTTCTCACTTTGCGTCATGGTTT
AGCACACGTCTGAACTCTTTCCCTACACGACGCTCTTCCGATCTNNNGAGAATCACTGGAACTCGGGA
AGCACACGTCTGAACTCTTTCCCTACACGACGCTCTTCCGATCTNNNGAGAATCACTGGAACTCGGGA
GAGCACACGTCTGAACTCTTTCCCTACACGACGCTCTTCCGATCTNNNGAGGGGAGTGAATCAGATC
GAGCACACGTCTGAACTCTTTCCCTACACGACGCTCTTCCGATCTNNNGAGGGGAGTGAATCAGATC
AGCACACGTCTGAACTCTTTCCCTACACGACGCTCTTCCGATCTNNNAGGAGGCTGAGTAGGATAAT
AGCACACGTCTGAACTCTTTCCCTACACGACGCTCTTCCGATCTNNNAGGAGGCTGAGTAGGATAAT
GAGCACACGTCTGAACTCTTTCCCTACACGACGCTCTTCCGATCTNNNGTGCCTGTAATCCCAGCTACT
GAGCACACGTCTGAACTCTTTCCCTACACGACGCTCTTCCGATCTNNNGTGCCTGTAATCCCAGCTACT
GCACACGTCTGAACTCTTTCCCTACACGACGCTCTTCCGATCTNNNTCTCTTTTCATTTCCATCCCCTCTC
GCACACGTCTGAACTCTTTCCCTACACGACGCTCTTCCGATCTNNNTCTCTTTTCATTTCCATCCCCTCTC
AGAGCACACGTCTGAACTCTTTCCCTACACGACGCTCTTCCGATCTNNNGCAAATCCTGAAAAGAGAC
AGAGCACACGTCTGAACTCTTTCCCTACACGACGCTCTTCCGATCTNNNGCAAATCCTGAAAAGAGAC
AGCACACGTCTGAACTCTTTCCCTACACGACGCTCTTCCGATCTNNNCACAATGTCACCTGAGCAAGT
AGCACACGTCTGAACTCTTTCCCTACACGACGCTCTTCCGATCTNNNCACAATGTCACCTGAGCAAGT
AGCACACGTCTGAACTCTTTCCCTACACGACGCTCTTCCGATCTNNNTGTTCCATTGGTCTAGGTGTCT
AGCACACGTCTGAACTCTTTCCCTACACGACGCTCTTCCGATCTNNNTGTTCCATTGGTCTAGGTGTCT
GAGCACACGTCTGAACTCTTTCCCTACACGACGCTCTTCCGATCTNNNAAAGCCACTCTCATAATGAG
GAGCACACGTCTGAACTCTTTCCCTACACGACGCTCTTCCGATCTNNNAAAGCCACTCTCATAATGAG
GAGCACACGTCTGAACTCTTTCCCTACACGACGCTCTTCCGATCTNNNTAGGAGAATCTAGTGCCCGAT
GAGCACACGTCTGAACTCTTTCCCTACACGACGCTCTTCCGATCTNNNTAGGAGAATCTAGTGCCCGAT
GAGCACACGTCTGAACTCTTTCCCTACACGACGCTCTTCCGATCTNNNCTTCTACTAATGAATGGTAGT
GAGCACACGTCTGAACTCTTTCCCTACACGACGCTCTTCCGATCTNNNCTTCTACTAATGAATGGTAGT
CACACGTCTGAACTCTTTCCCTACACGACGCTCTTCCGATCTNNNCTTCTGCTTCTCGCTCTTGAGCCAT

CACACGTCTGAACTCTTTCCCTACACGACGCTCTTCCGATCTNNNCTTCTGCTTCTCGCTCTTGAGCCAT
CACACGTCTGAACTCTTTCCCTACACGACGCTCTTCCGATCTNNNGATGGAGTTTCACTCTTGTTGCCC
CACACGTCTGAACTCTTTCCCTACACGACGCTCTTCCGATCTNNNGATGGAGTTTCACTCTTGTTGCCC
GAGCACACGTCTGAACTCTTTCCCTACACGACGCTCTTCCGATCTNNNGTATGTTACAGGCTGCCTAGA
GAGCACACGTCTGAACTCTTTCCCTACACGACGCTCTTCCGATCTNNNGTATGTTACAGGCTGCCTAGA
AAGAGCACACGTCTGAACTCTTTCCCTACACGACGCTCTTCCGATCTNNNTTCCGCATTTAAGCGAATCA
AAGAGCACACGTCTGAACTCTTTCCCTACACGACGCTCTTCCGATCTNNNTTCCGCATTTAAGCGAATCA
AGCACACGTCTGAACTCTTTCCCTACACGACGCTCTTCCGATCTNNNAGTGAGCTGAGATCATGCCATT
AGCACACGTCTGAACTCTTTCCCTACACGACGCTCTTCCGATCTNNNAGTGAGCTGAGATCATGCCATT
GAGCACACGTCTGAACTCTTTCCCTACACGACGCTCTTCCGATCTNNNTATCTTCTCAAAGCAAGGGTG
GAGCACACGTCTGAACTCTTTCCCTACACGACGCTCTTCCGATCTNNNTATCTTCTCAAAGCAAGGGTG
AGCACACGTCTGAACTCTTTCCCTACACGACGCTCTTCCGATCTNNNTCCTCATAGCAAGATGACGTGA
AGCACACGTCTGAACTCTTTCCCTACACGACGCTCTTCCGATCTNNNTCCTCATAGCAAGATGACGTGA
GAGCACACGTCTGAACTCTTTCCCTACACGACGCTCTTCCGATCTNNNTCAACATGCAAAAATCAGTAG
GAGCACACGTCTGAACTCTTTCCCTACACGACGCTCTTCCGATCTNNNTCAACATGCAAAAATCAGTAG
AGAGCACACGTCTGAACTCTTTCCCTACACGACGCTCTTCCGATCTNNNTGGGATTTCAGAAGAAGGA
AGAGCACACGTCTGAACTCTTTCCCTACACGACGCTCTTCCGATCTNNNTGGGATTTCAGAAGAAGGA
GAGCACACGTCTGAACTCTTTCCCTACACGACGCTCTTCCGATCTNNNTGTTTGTTTCTCTCTCTGCCTG
GAGCACACGTCTGAACTCTTTCCCTACACGACGCTCTTCCGATCTNNNTGTTTGTTTCTCTCTCTGCCTG
AAGAGCACACGTCTGAACTCTTTCCCTACACGACGCTCTTCCGATCTNNNAGGCGGGAGGGTCACATA
AAGAGCACACGTCTGAACTCTTTCCCTACACGACGCTCTTCCGATCTNNNAGGCGGGAGGGTCACATA
AGCACACGTCTGAACTCTTTCCCTACACGACGCTCTTCCGATCTNNNCTGAGGCAGGAGAATTGCTTG
AGCACACGTCTGAACTCTTTCCCTACACGACGCTCTTCCGATCTNNNCTGAGGCAGGAGAATTGCTTG
GAGCACACGTCTGAACTCTTTCCCTACACGACGCTCTTCCGATCTNNNGGACTCTACATTGTCCTGATGC
GAGCACACGTCTGAACTCTTTCCCTACACGACGCTCTTCCGATCTNNNGGACTCTACATTGTCCTGATGC
GAGCACACGTCTGAACTCTTTCCCTACACGACGCTCTTCCGATCTNNNAAGGTTTTTTTTTTGGGGGG
GAGCACACGTCTGAACTCTTTCCCTACACGACGCTCTTCCGATCTNNNAAGGTTTTTTTTTTGGGGGG
AGCACACGTCTGAACTCTTTCCCTACACGACGCTCTTCCGATCTNNNTCTCTGTCTATGTCTGTGTGCAC
AGCACACGTCTGAACTCTTTCCCTACACGACGCTCTTCCGATCTNNNTCTCTGTCTATGTCTGTGTGCAC
GAGCACACGTCTGAACTCTTTCCCTACACGACGCTCTTCCGATCTNNNGAGACCAAAAGAACAACCCTC
GAGCACACGTCTGAACTCTTTCCCTACACGACGCTCTTCCGATCTNNNGAGACCAAAAGAACAACCCTC
GAGCACACGTCTGAACTCTTTCCCTACACGACGCTCTTCCGATCTNNNACACAGGGATCTTTGCAGTGA
GAGCACACGTCTGAACTCTTTCCCTACACGACGCTCTTCCGATCTNNNACACAGGGATCTTTGCAGTGA
GCACACGTCTGAACTCTTTCCCTACACGACGCTCTTCCGATCTNNNAGGCATCAATCCAAATGACTTGC
GCACACGTCTGAACTCTTTCCCTACACGACGCTCTTCCGATCTNNNAGGCATCAATCCAAATGACTTGC
GAGCACACGTCTGAACTCTTTCCCTACACGACGCTCTTCCGATCTNNNAAACAACCCCTGCAATCCAAA
GAGCACACGTCTGAACTCTTTCCCTACACGACGCTCTTCCGATCTNNNAAACAACCCCTGCAATCCAAA
AGCACACGTCTGAACTCTTTCCCTACACGACGCTCTTCCGATCTNNNCTCTCTGTGTACTCACTGGCTTT
AGCACACGTCTGAACTCTTTCCCTACACGACGCTCTTCCGATCTNNNCTCTCTGTGTACTCACTGGCTTT
GCACACGTCTGAACTCTTTCCCTACACGACGCTCTTCCGATCTNNNTGTTCCAGTTTGTGCACTAACAA
GCACACGTCTGAACTCTTTCCCTACACGACGCTCTTCCGATCTNNNTGTTCCAGTTTGTGCACTAACAA
GCACACGTCTGAACTCTTTCCCTACACGACGCTCTTCCGATCTNNNACAAAAGATGTTTCAGGGAGCT
GCACACGTCTGAACTCTTTCCCTACACGACGCTCTTCCGATCTNNNACAAAAGATGTTTCAGGGAGCT

AGCACACGTCTGAACTCTTTCCCTACACGACGCTCTTCCGATCTNNNTGCATGACACAAGTTCCAGTTCA

AGCACACGTCTGAACTCTTTCCCTACACGACGCTCTTCCGATCTNNNTGCATGACACAAGTTCCAGTTCA

AGCACACGTCTGAACTCTTTCCCTACACGACGCTCTTCCGATCTNNNACACCTGTAGTCCCAGCTACTAA

AGCACACGTCTGAACTCTTTCCCTACACGACGCTCTTCCGATCTNNNACACCTGTAGTCCCAGCTACTAA

AGAGCACACGTCTGAACTCTTTCCCTACACGACGCTCTTCCGATCTNNNCAGTTGTTCCCAGCAATGTCA

AGAGCACACGTCTGAACTCTTTCCCTACACGACGCTCTTCCGATCTNNNCAGTTGTTCCCAGCAATGTCA

GAGCACACGTCTGAACTCTTTCCCTACACGACGCTCTTCCGATCTNNNCGGAACATCTGTGCTAATCAAC

GAGCACACGTCTGAACTCTTTCCCTACACGACGCTCTTCCGATCTNNNCGGAACATCTGTGCTAATCAAC

GAGCACACGTCTGAACTCTTTCCCTACACGACGCTCTTCCGATCTNNNACTGCAGAATTTTCATGGGTT

GAGCACACGTCTGAACTCTTTCCCTACACGACGCTCTTCCGATCTNNNACTGCAGAATTTTCATGGGTT

AGCACACGTCTGAACTCTTTCCCTACACGACGCTCTTCCGATCTNNNCCCCTCTCTCTTTCTCTGAGGATC

AGCACACGTCTGAACTCTTTCCCTACACGACGCTCTTCCGATCTNNNCCCCTCTCTCTTTCTCTGAGGATC

AGCACACGTCTGAACTCTTTCCCTACACGACGCTCTTCCGATCTNNNAACAACCAAGTCCGAGACTGGC

AGCACACGTCTGAACTCTTTCCCTACACGACGCTCTTCCGATCTNNNAACAACCAAGTCCGAGACTGGC

GAGCACACGTCTGAACTCTTTCCCTACACGACGCTCTTCCGATCTNNNTTCCCTGATCTCCAGACTGAGT

GAGCACACGTCTGAACTCTTTCCCTACACGACGCTCTTCCGATCTNNNTTCCCTGATCTCCAGACTGAGT

AGCACACGTCTGAACTCTTTCCCTACACGACGCTCTTCCGATCTNNNTGCCTGCATAGTACATGTTCAGT

AGCACACGTCTGAACTCTTTCCCTACACGACGCTCTTCCGATCTNNNTGCCTGCATAGTACATGTTCAGT

GAGCACACGTCTGAACTCTTTCCCTACACGACGCTCTTCCGATCTNNNCATAGCACCACTGAGGCCATGC

GAGCACACGTCTGAACTCTTTCCCTACACGACGCTCTTCCGATCTNNNCATAGCACCACTGAGGCCATGC

AGAGCACACGTCTGAACTCTTTCCCTACACGACGCTCTTCCGATCTNNNGAGAGGGAGAGAGAGAGA

AGAGCACACGTCTGAACTCTTTCCCTACACGACGCTCTTCCGATCTNNNGAGAGGGAGAGAGAGAGA

GCACACGTCTGAACTCTTTCCCTACACGACGCTCTTCCGATCTNNNCACACTGCTCGGACCAAGAACATC

GCACACGTCTGAACTCTTTCCCTACACGACGCTCTTCCGATCTNNNCACACTGCTCGGACCAAGAACATC

GAGCACACGTCTGAACTCTTTCCCTACACGACGCTCTTCCGATCTNNNTCCTTCCTTCCTTCCTTCCTTTCC

GAGCACACGTCTGAACTCTTTCCCTACACGACGCTCTTCCGATCTNNNTCCTTCCTTCCTTCCTTCCTTTCC

AGCACACGTCTGAACTCTTTCCCTACACGACGCTCTTCCGATCTNNNTAATCTCAGCTACTCAGGAGGCT

AGCACACGTCTGAACTCTTTCCCTACACGACGCTCTTCCGATCTNNNTAATCTCAGCTACTCAGGAGGCT

AGCACACGTCTGAACTCTTTCCCTACACGACGCTCTTCCGATCTNNNGGTCTAAGGTCTCAGCCTCTCTA

AGCACACGTCTGAACTCTTTCCCTACACGACGCTCTTCCGATCTNNNGGTCTAAGGTCTCAGCCTCTCTA

AGCACACGTCTGAACTCTTTCCCTACACGACGCTCTTCCGATCTNNNGGGTGGAAGTAAACACAAAAAC

AGCACACGTCTGAACTCTTTCCCTACACGACGCTCTTCCGATCTNNNGGGTGGAAGTAAACACAAAAAC

GAGCACACGTCTGAACTCTTTCCCTACACGACGCTCTTCCGATCTNNNAGCTACCATAACCAAGTGTGAT

GAGCACACGTCTGAACTCTTTCCCTACACGACGCTCTTCCGATCTNNNAGCTACCATAACCAAGTGTGAT

AGCACACGTCTGAACTCTTTCCCTACACGACGCTCTTCCGATCTNNNCTGGGTTGTAGAATGAGACTG

AGCACACGTCTGAACTCTTTCCCTACACGACGCTCTTCCGATCTNNNCTGGGTTGTAGAATGAGACTG

GCACACGTCTGAACTCTTTCCCTACACGACGCTCTTCCGATCTNNNGAAGGTTGCAGTGAGCCGAAAT

GCACACGTCTGAACTCTTTCCCTACACGACGCTCTTCCGATCTNNNGAAGGTTGCAGTGAGCCGAAAT

GAGCACACGTCTGAACTCTTTCCCTACACGACGCTCTTCCGATCTNNNTTGTGAGTGTGTTGAGACTTC

GAGCACACGTCTGAACTCTTTCCCTACACGACGCTCTTCCGATCTNNNTTGTGAGTGTGTTGAGACTTC

AGCACACGTCTGAACTCTTTCCCTACACGACGCTCTTCCGATCTNNNGTAGCGGCTAAGAGAAGGAAC

AGCACACGTCTGAACTCTTTCCCTACACGACGCTCTTCCGATCTNNNGTAGCGGCTAAGAGAAGGAAC

GAGCACACGTCTGAACTCTTTCCCTACACGACGCTCTTCCGATCTNNNTCCAATGCCCTGGGTAAAGTAA

AGCACACGTCTGAACTCTTTCCCTACACGACGCTCTTCCGATCTNNNTCCAATGCCCTGGGTAAAGTA
CACACGTCTGAACTCTTTCCCTACACGACGCTCTTCCGATCTNNNCTGAGAAGGGCTGATCTTTGCTC
CACACGTCTGAACTCTTTCCCTACACGACGCTCTTCCGATCTNNNCTGAGAAGGGCTGATCTTTGCTC
AGCACACGTCTGAACTCTTTCCCTACACGACGCTCTTCCGATCTNNNACAGAAGCTGAAGACTTGCT
AGCACACGTCTGAACTCTTTCCCTACACGACGCTCTTCCGATCTNNNACAGAAGCTGAAGACTTGCT
ACACGTCTGAACTCTTTCCCTACACGACGCTCTTCCGATCTNNNGCAACAGAGTGAGACCCTGTCCAT
ACACGTCTGAACTCTTTCCCTACACGACGCTCTTCCGATCTNNNGCAACAGAGTGAGACCCTGTCCAT
GAGCACACGTCTGAACTCTTTCCCTACACGACGCTCTTCCGATCTNNNGTAATCCTAGCTACTTGGGA
GAGCACACGTCTGAACTCTTTCCCTACACGACGCTCTTCCGATCTNNNGTAATCCTAGCTACTTGGGA
GCACACGTCTGAACTCTTTCCCTACACGACGCTCTTCCGATCTNNNGTTCACCATTCTCCTCCCTCCA
GCACACGTCTGAACTCTTTCCCTACACGACGCTCTTCCGATCTNNNGTTCACCATTCTCCTCCCTCCA
GCACACGTCTGAACTCTTTCCCTACACGACGCTCTTCCGATCTNNNACAAGGAAATTGGTAATCTGAG
GCACACGTCTGAACTCTTTCCCTACACGACGCTCTTCCGATCTNNNACAAGGAAATTGGTAATCTGAG
AGCACACGTCTGAACTCTTTCCCTACACGACGCTCTTCCGATCTNNNCAGTTACTTAGGAAGCTGAGG
AGCACACGTCTGAACTCTTTCCCTACACGACGCTCTTCCGATCTNNNCAGTTACTTAGGAAGCTGAGG
AGCACACGTCTGAACTCTTTCCCTACACGACGCTCTTCCGATCTNNNTGCACCTATAGTCCCAGCTACT
AGCACACGTCTGAACTCTTTCCCTACACGACGCTCTTCCGATCTNNNTGCACCTATAGTCCCAGCTACT
AGCACACGTCTGAACTCTTTCCCTACACGACGCTCTTCCGATCTNNNACCTACAGCTTTTTGTATGCCCT
AGCACACGTCTGAACTCTTTCCCTACACGACGCTCTTCCGATCTNNNACCTACAGCTTTTTGTATGCCCT
GCACACGTCTGAACTCTTTCCCTACACGACGCTCTTCCGATCTNNNGCACCACATTTGCCTTCGTATAA
GCACACGTCTGAACTCTTTCCCTACACGACGCTCTTCCGATCTNNNGCACCACATTTGCCTTCGTATAA
AGCACACGTCTGAACTCTTTCCCTACACGACGCTCTTCCGATCTNNNAAGACGTAGATTTTTCTGGAA
AGCACACGTCTGAACTCTTTCCCTACACGACGCTCTTCCGATCTNNNAAGACGTAGATTTTTCTGGAA
AGCACACGTCTGAACTCTTTCCCTACACGACGCTCTTCCGATCTNNNTCCCAAAGTGCTGAGATTACA
AGCACACGTCTGAACTCTTTCCCTACACGACGCTCTTCCGATCTNNNTCCCAAAGTGCTGAGATTACA
GAGCACACGTCTGAACTCTTTCCCTACACGACGCTCTTCCGATCTNNNTACTGCGTTGTAAAGCACTG
GAGCACACGTCTGAACTCTTTCCCTACACGACGCTCTTCCGATCTNNNTACTGCGTTGTAAAGCACTG
GAGCACACGTCTGAACTCTTTCCCTACACGACGCTCTTCCGATCTNNNCTGAGGCAGGAGAATCACTT
GAGCACACGTCTGAACTCTTTCCCTACACGACGCTCTTCCGATCTNNNCTGAGGCAGGAGAATCACTT
AGAGCACACGTCTGAACTCTTTCCCTACACGACGCTCTTCCGATCTNNNTACAGACACACACACACAA
AGAGCACACGTCTGAACTCTTTCCCTACACGACGCTCTTCCGATCTNNNTACAGACACACACACACAA
AGCACACGTCTGAACTCTTTCCCTACACGACGCTCTTCCGATCTNNNGCAGAGAAAAACAGGTGAGT
AGCACACGTCTGAACTCTTTCCCTACACGACGCTCTTCCGATCTNNNGCAGAGAAAAACAGGTGAGT
AGCACACGTCTGAACTCTTTCCCTACACGACGCTCTTCCGATCTNNNAAGAGGACCTACACAGTTCCA
AGCACACGTCTGAACTCTTTCCCTACACGACGCTCTTCCGATCTNNNAAGAGGACCTACACAGTTCCA
GCACACGTCTGAACTCTTTCCCTACACGACGCTCTTCCGATCTNNNAAGTAGCTGGGATTACAGGCA
GCACACGTCTGAACTCTTTCCCTACACGACGCTCTTCCGATCTNNNAAGTAGCTGGGATTACAGGCA
AGCACACGTCTGAACTCTTTCCCTACACGACGCTCTTCCGATCTNNNGCACCAGGCAAGGTTAGATA
AGCACACGTCTGAACTCTTTCCCTACACGACGCTCTTCCGATCTNNNGCACCAGGCAAGGTTAGATA
GCACACGTCTGAACTCTTTCCCTACACGACGCTCTTCCGATCTNNNTCCTCAAGGTTCATTCGTGTTG
GCACACGTCTGAACTCTTTCCCTACACGACGCTCTTCCGATCTNNNTCCTCAAGGTTCATTCGTGTTG
GAGCACACGTCTGAACTCTTTCCCTACACGACGCTCTTCCGATCTNNNGATAAGGGCTAATCTGGGG
GAGCACACGTCTGAACTCTTTCCCTACACGACGCTCTTCCGATCTNNNGATAAGGGCTAATCTGGGG

AGCACACGTCTGAACTCTTTCCCTACACGACGCTCTTCCGATCTNNNCTGAGGCAGGAGAATCACTTG

AGCACACGTCTGAACTCTTTCCCTACACGACGCTCTTCCGATCTNNNCTGAGGCAGGAGAATCACTTG

ACACGTCTGAACTCTTTCCCTACACGACGCTCTTCCGATCTNNNGAGACCCGGAATGCGCAGGGCATC

ACACGTCTGAACTCTTTCCCTACACGACGCTCTTCCGATCTNNNGAGACCCGGAATGCGCAGGGCATC

GCACACGTCTGAACTCTTTCCCTACACGACGCTCTTCCGATCTNNNGCTGGGAGGTGGATAACTAGG

GCACACGTCTGAACTCTTTCCCTACACGACGCTCTTCCGATCTNNNGCTGGGAGGTGGATAACTAGG

GAGCACACGTCTGAACTCTTTCCCTACACGACGCTCTTCCGATCTNNNGAAGGGAAGGGGAAGGAA

GAGCACACGTCTGAACTCTTTCCCTACACGACGCTCTTCCGATCTNNNGAAGGGAAGGGGAAGGAA

GAGCACACGTCTGAACTCTTTCCCTACACGACGCTCTTCCGATCTNNNACTCATGGTGAGGACAAAGG

GAGCACACGTCTGAACTCTTTCCCTACACGACGCTCTTCCGATCTNNNACTCATGGTGAGGACAAAGG

AGAGCACACGTCTGAACTCTTTCCCTACACGACGCTCTTCCGATCTNNNGAAACAACAGCCTTACAGG

AGAGCACACGTCTGAACTCTTTCCCTACACGACGCTCTTCCGATCTNNNGAAACAACAGCCTTACAGG

AGCACACGTCTGAACTCTTTCCCTACACGACGCTCTTCCGATCTNNNCTCGTGCCTGAATAGACAGATC

AGCACACGTCTGAACTCTTTCCCTACACGACGCTCTTCCGATCTNNNCTCGTGCCTGAATAGACAGATC

GAGCACACGTCTGAACTCTTTCCCTACACGACGCTCTTCCGATCTNNNGGAACTGTGAGTCCACTAAA

GAGCACACGTCTGAACTCTTTCCCTACACGACGCTCTTCCGATCTNNNGGAACTGTGAGTCCACTAAA

AGCACACGTCTGAACTCTTTCCCTACACGACGCTCTTCCGATCTNNNAATTTAGAGCCGTCCTTGTCCAG

AGCACACGTCTGAACTCTTTCCCTACACGACGCTCTTCCGATCTNNNAATTTAGAGCCGTCCTTGTCCAG

GCACACGTCTGAACTCTTTCCCTACACGACGCTCTTCCGATCTNNNACTCGCCGATGATAACTTTGTCC

GCACACGTCTGAACTCTTTCCCTACACGACGCTCTTCCGATCTNNNACTCGCCGATGATAACTTTGTCC

GAGCACACGTCTGAACTCTTTCCCTACACGACGCTCTTCCGATCTNNNCGTGGTTGTAGCTGCTATTGA

GAGCACACGTCTGAACTCTTTCCCTACACGACGCTCTTCCGATCTNNNCGTGGTTGTAGCTGCTATTGA

AGCACACGTCTGAACTCTTTCCCTACACGACGCTCTTCCGATCTNNNTCTGCAGTGCAAAAGACATTCA

AGCACACGTCTGAACTCTTTCCCTACACGACGCTCTTCCGATCTNNNTCTGCAGTGCAAAAGACATTCA

GAGCACACGTCTGAACTCTTTCCCTACACGACGCTCTTCCGATCTNNNCGAGGTCAGACTTGTTGTGG

GAGCACACGTCTGAACTCTTTCCCTACACGACGCTCTTCCGATCTNNNCGAGGTCAGACTTGTTGTGG

AGAGCACACGTCTGAACTCTTTCCCTACACGACGCTCTTCCGATCTNNNATACTGGTGTTTTGCGAGTC

AGAGCACACGTCTGAACTCTTTCCCTACACGACGCTCTTCCGATCTNNNATACTGGTGTTTTGCGAGTC

GAGCACACGTCTGAACTCTTTCCCTACACGACGCTCTTCCGATCTNNNCTTTATTGCTCAGGCTGGTCT

GAGCACACGTCTGAACTCTTTCCCTACACGACGCTCTTCCGATCTNNNCTTTATTGCTCAGGCTGGTCT

GCACACGTCTGAACTCTTTCCCTACACGACGCTCTTCCGATCTNNNACACTTTTCACATGGTAGGTGCC

GCACACGTCTGAACTCTTTCCCTACACGACGCTCTTCCGATCTNNNACACTTTTCACATGGTAGGTGCC

GAGCACACGTCTGAACTCTTTCCCTACACGACGCTCTTCCGATCTNNNTGCGATTGCCATCTCTACCAA

GAGCACACGTCTGAACTCTTTCCCTACACGACGCTCTTCCGATCTNNNTGCGATTGCCATCTCTACCAA

GAGCACACGTCTGAACTCTTTCCCTACACGACGCTCTTCCGATCTNNNCTGGCTAGCAAATTGGCATT

GAGCACACGTCTGAACTCTTTCCCTACACGACGCTCTTCCGATCTNNNCTGGCTAGCAAATTGGCATT

AGAGCACACGTCTGAACTCTTTCCCTACACGACGCTCTTCCGATCTNNNACTCAGCAAGCTAAGATAG

AGAGCACACGTCTGAACTCTTTCCCTACACGACGCTCTTCCGATCTNNNACTCAGCAAGCTAAGATAG

AGCACACGTCTGAACTCTTTCCCTACACGACGCTCTTCCGATCTNNNGCACTACTGCTCATGTAGTTGG

AGCACACGTCTGAACTCTTTCCCTACACGACGCTCTTCCGATCTNNNGCACTACTGCTCATGTAGTTGG

AGCACACGTCTGAACTCTTTCCCTACACGACGCTCTTCCGATCTNNNACTCAAAAGCCCCTCTTAGTCT

AGCACACGTCTGAACTCTTTCCCTACACGACGCTCTTCCGATCTNNNACTCAAAAGCCCCTCTTAGTCT

AGCACACGTCTGAACTCTTTCCCTACACGACGCTCTTCCGATCTNNNATGGACTACCATGCCTGGCTTC

AGCACACGTCTGAACTCTTTCCCTACACGACGCTCTTCCGATCTNNNATGGACTACCATGCCTGGCTTCC

AGCACACGTCTGAACTCTTTCCCTACACGACGCTCTTCCGATCTNNNCAATCCTCTCCCGTCCTCTTCCATC

AGCACACGTCTGAACTCTTTCCCTACACGACGCTCTTCCGATCTNNNCAATCCTCTCCCGTCCTCTTCCATC

AGAGCACACGTCTGAACTCTTTCCCTACACGACGCTCTTCCGATCTNNNTCTGTGAGGTGAATGACGTC

AGAGCACACGTCTGAACTCTTTCCCTACACGACGCTCTTCCGATCTNNNTCTGTGAGGTGAATGACGTC

AGAGCACACGTCTGAACTCTTTCCCTACACGACGCTCTTCCGATCTNNNGTGACACAGCCAAACAATCAA

AGAGCACACGTCTGAACTCTTTCCCTACACGACGCTCTTCCGATCTNNNGTGACACAGCCAAACAATCAA

GAGCACACGTCTGAACTCTTTCCCTACACGACGCTCTTCCGATCTNNNGCCCTTTATCGTGAGTAACTTC

GAGCACACGTCTGAACTCTTTCCCTACACGACGCTCTTCCGATCTNNNGCCCTTTATCGTGAGTAACTTC

AGCACACGTCTGAACTCTTTCCCTACACGACGCTCTTCCGATCTNNNCCCAGCCAGAACTTCCCTTTTA

AGCACACGTCTGAACTCTTTCCCTACACGACGCTCTTCCGATCTNNNCCCAGCCAGAACTTCCCTTTTA

AGCACACGTCTGAACTCTTTCCCTACACGACGCTCTTCCGATCTNNNTTCTAAAGCTCAGAGTTCCCACCC

AGCACACGTCTGAACTCTTTCCCTACACGACGCTCTTCCGATCTNNNTTCTAAAGCTCAGAGTTCCCACCC

AGCACACGTCTGAACTCTTTCCCTACACGACGCTCTTCCGATCTNNNCCCTTTGGAGAAGGTTTGGGA

AGCACACGTCTGAACTCTTTCCCTACACGACGCTCTTCCGATCTNNNCCCTTTGGAGAAGGTTTGGGA

AGAGCACACGTCTGAACTCTTTCCCTACACGACGCTCTTCCGATCTNNNGGACATGAGGAGAATGTGC

AGAGCACACGTCTGAACTCTTTCCCTACACGACGCTCTTCCGATCTNNNGGACATGAGGAGAATGTGC

AGCACACGTCTGAACTCTTTCCCTACACGACGCTCTTCCGATCTNNNTTGCCCTGAACATAACATTCTGC

AGCACACGTCTGAACTCTTTCCCTACACGACGCTCTTCCGATCTNNNTTGCCCTGAACATAACATTCTGC

GCACACGTCTGAACTCTTTCCCTACACGACGCTCTTCCGATCTNNNTTCTTCCAAATCCCTCGTTGTCTCA

GCACACGTCTGAACTCTTTCCCTACACGACGCTCTTCCGATCTNNNTTCTTCCAAATCCCTCGTTGTCTCA

GCACACGTCTGAACTCTTTCCCTACACGACGCTCTTCCGATCTNNNCTGTTTCTCCCTTGTCTCTCCTCCAT

GCACACGTCTGAACTCTTTCCCTACACGACGCTCTTCCGATCTNNNCTGTTTCTCCCTTGTCTCTCCTCCAT

GAGCACACGTCTGAACTCTTTCCCTACACGACGCTCTTCCGATCTNNNATTGCTTGAGTCCAGGAGTTTC

GAGCACACGTCTGAACTCTTTCCCTACACGACGCTCTTCCGATCTNNNATTGCTTGAGTCCAGGAGTTTC

AGCACACGTCTGAACTCTTTCCCTACACGACGCTCTTCCGATCTNNNAGGACTTCTTTCAACTCCACCTG

AGCACACGTCTGAACTCTTTCCCTACACGACGCTCTTCCGATCTNNNAGGACTTCTTTCAACTCCACCTG

GAGCACACGTCTGAACTCTTTCCCTACACGACGCTCTTCCGATCTNNNGGGCAGTTTGCTTCCCTAATTT

GAGCACACGTCTGAACTCTTTCCCTACACGACGCTCTTCCGATCTNNNGGGCAGTTTGCTTCCCTAATTT

AGCACACGTCTGAACTCTTTCCCTACACGACGCTCTTCCGATCTNNNGTAAACTGAGATCATGCCACTG

AGCACACGTCTGAACTCTTTCCCTACACGACGCTCTTCCGATCTNNNGTAAACTGAGATCATGCCACTG

AGAGCACACGTCTGAACTCTTTCCCTACACGACGCTCTTCCGATCTNNNGCATCTGTAGTCCCAGCTACT

AGAGCACACGTCTGAACTCTTTCCCTACACGACGCTCTTCCGATCTNNNGCATCTGTAGTCCCAGCTACT

AGAGCACACGTCTGAACTCTTTCCCTACACGACGCTCTTCCGATCTNNNACCGTTAGGATGCATAGGAC

AGAGCACACGTCTGAACTCTTTCCCTACACGACGCTCTTCCGATCTNNNACCGTTAGGATGCATAGGAC

AGCACACGTCTGAACTCTTTCCCTACACGACGCTCTTCCGATCTNNNCACCACGCCATGCTAATTTTTTTC

AGCACACGTCTGAACTCTTTCCCTACACGACGCTCTTCCGATCTNNNCACCACGCCATGCTAATTTTTTTC

AGCACACGTCTGAACTCTTTCCCTACACGACGCTCTTCCGATCTNNNCGTGAGACCAGGAAGGGAAATA

AGCACACGTCTGAACTCTTTCCCTACACGACGCTCTTCCGATCTNNNCGTGAGACCAGGAAGGGAAATA

AGCACACGTCTGAACTCTTTCCCTACACGACGCTCTTCCGATCTNNNAGGAAGGAGAATTGCTTGAAC

AGCACACGTCTGAACTCTTTCCCTACACGACGCTCTTCCGATCTNNNAGGAAGGAGAATTGCTTGAAC

GAGCACACGTCTGAACTCTTTCCCTACACGACGCTCTTCCGATCTNNNCCAAATAGCTGGGACTACAGG

GAGCACACGTCTGAACTCTTTCCCTACACGACGCTCTTCCGATCTNNNCCAAATAGCTGGGACTACAGG

AGCACACGTCTGAACTCTTTCCCTACACGACGCTCTTCCGATCTNNNAAAAAATTAGCCGGGTGTGGT
AGCACACGTCTGAACTCTTTCCCTACACGACGCTCTTCCGATCTNNNAAAAAATTAGCCGGGTGTGGT
GAGCACACGTCTGAACTCTTTCCCTACACGACGCTCTTCCGATCTNNNCCACACCTGGCCTGATTATTCT
GAGCACACGTCTGAACTCTTTCCCTACACGACGCTCTTCCGATCTNNNCCACACCTGGCCTGATTATTCT
AGAGCACACGTCTGAACTCTTTCCCTACACGACGCTCTTCCGATCTNNNGTGATTCCATTTATGACAGG
AGAGCACACGTCTGAACTCTTTCCCTACACGACGCTCTTCCGATCTNNNGTGATTCCATTTATGACAGG
AGCACACGTCTGAACTCTTTCCCTACACGACGCTCTTCCGATCTNNNGAAAACTTCACCGAAGCGATAG
AGCACACGTCTGAACTCTTTCCCTACACGACGCTCTTCCGATCTNNNGAAAACTTCACCGAAGCGATAG
AGCACACGTCTGAACTCTTTCCCTACACGACGCTCTTCCGATCTNNNGCTGAAGGAGAACAGGAGAT
AGCACACGTCTGAACTCTTTCCCTACACGACGCTCTTCCGATCTNNNGCTGAAGGAGAACAGGAGAT
AGCACACGTCTGAACTCTTTCCCTACACGACGCTCTTCCGATCTNNNCAAGTAGAGGTCAGTCTCATG
AGCACACGTCTGAACTCTTTCCCTACACGACGCTCTTCCGATCTNNNCAAGTAGAGGTCAGTCTCATG
GAGCACACGTCTGAACTCTTTCCCTACACGACGCTCTTCCGATCTNNNCCTGACTTGAGAAAGGGATCT
GAGCACACGTCTGAACTCTTTCCCTACACGACGCTCTTCCGATCTNNNCCTGACTTGAGAAAGGGATCT
AGAGCACACGTCTGAACTCTTTCCCTACACGACGCTCTTCCGATCTNNNAAGCAATTCTCTGTCTCAGCC
AGAGCACACGTCTGAACTCTTTCCCTACACGACGCTCTTCCGATCTNNNAAGCAATTCTCTGTCTCAGCC
AGCACACGTCTGAACTCTTTCCCTACACGACGCTCTTCCGATCTNNNGCTACCTCACAAGAGTGTTGAG
AGCACACGTCTGAACTCTTTCCCTACACGACGCTCTTCCGATCTNNNGCTACCTCACAAGAGTGTTGAG
AGCACACGTCTGAACTCTTTCCCTACACGACGCTCTTCCGATCTNNNTTGGATCATGCAGCAAAATAGC
AGCACACGTCTGAACTCTTTCCCTACACGACGCTCTTCCGATCTNNNTTGGATCATGCAGCAAAATAGC
GAGCACACGTCTGAACTCTTTCCCTACACGACGCTCTTCCGATCTNNNCCGGCCTTTACACATTTTGAAG
GAGCACACGTCTGAACTCTTTCCCTACACGACGCTCTTCCGATCTNNNCCGGCCTTTACACATTTTGAAG
AGCACACGTCTGAACTCTTTCCCTACACGACGCTCTTCCGATCTNNNCTGTTTTTCCATTACGACCCTGC
AGCACACGTCTGAACTCTTTCCCTACACGACGCTCTTCCGATCTNNNCTGTTTTTCCATTACGACCCTGC
CACACGTCTGAACTCTTTCCCTACACGACGCTCTTCCGATCTNNNTTCTGGAAAAGCTGGAAAAGCAT
CACACGTCTGAACTCTTTCCCTACACGACGCTCTTCCGATCTNNNTTCTGGAAAAGCTGGAAAAGCAT
AGCACACGTCTGAACTCTTTCCCTACACGACGCTCTTCCGATCTNNNTCTGAATCAAGGAAGCCAAGGA
AGCACACGTCTGAACTCTTTCCCTACACGACGCTCTTCCGATCTNNNTCTGAATCAAGGAAGCCAAGGA
AGCACACGTCTGAACTCTTTCCCTACACGACGCTCTTCCGATCTNNNTCTCAGCTTCACCCTCATCATTCC
AGCACACGTCTGAACTCTTTCCCTACACGACGCTCTTCCGATCTNNNTCTCAGCTTCACCCTCATCATTCC
AGCACACGTCTGAACTCTTTCCCTACACGACGCTCTTCCGATCTNNNCGATGGATGCCAAGTGTCCTAT
AGCACACGTCTGAACTCTTTCCCTACACGACGCTCTTCCGATCTNNNCGATGGATGCCAAGTGTCCTAT
AGCACACGTCTGAACTCTTTCCCTACACGACGCTCTTCCGATCTNNNGTGAGGACACTAAAAGCCAAT
AGCACACGTCTGAACTCTTTCCCTACACGACGCTCTTCCGATCTNNNGTGAGGACACTAAAAGCCAAT
GAGCACACGTCTGAACTCTTTCCCTACACGACGCTCTTCCGATCTNNNTTGTCTCAGACAGGACACAGA
GAGCACACGTCTGAACTCTTTCCCTACACGACGCTCTTCCGATCTNNNTTGTCTCAGACAGGACACAGA
GAGCACACGTCTGAACTCTTTCCCTACACGACGCTCTTCCGATCTNNNCCTGAGCAAATGAGCAGGTAT
GAGCACACGTCTGAACTCTTTCCCTACACGACGCTCTTCCGATCTNNNCCTGAGCAAATGAGCAGGTAT
AGCACACGTCTGAACTCTTTCCCTACACGACGCTCTTCCGATCTNNNTTATCTAGGGCCTCTCTCTGGT
AGCACACGTCTGAACTCTTTCCCTACACGACGCTCTTCCGATCTNNNTTATCTAGGGCCTCTCTCTGGT
AGAGCACACGTCTGAACTCTTTCCCTACACGACGCTCTTCCGATCTNNNAGGAGAATGGCGTGAACCT
AGAGCACACGTCTGAACTCTTTCCCTACACGACGCTCTTCCGATCTNNNAGGAGAATGGCGTGAACCT
AGCACACGTCTGAACTCTTTCCCTACACGACGCTCTTCCGATCTNNNGGCTGAGATAGGAGGATCACT

AGCACACGTCTGAACTCTTTCCCTACACGACGCTCTTCCGATCTNNNGGCTGAGATAGGAGGATCACT

AGCACACGTCTGAACTCTTTCCCTACACGACGCTCTTCCGATCTNNNTTCTCCTATGACCCTTTCTGCAC

AGCACACGTCTGAACTCTTTCCCTACACGACGCTCTTCCGATCTNNNTTCTCCTATGACCCTTTCTGCAC

GAGCACACGTCTGAACTCTTTCCCTACACGACGCTCTTCCGATCTNNNCGTTGTATTTGTTACAGCAGC

GAGCACACGTCTGAACTCTTTCCCTACACGACGCTCTTCCGATCTNNNCGTTGTATTTGTTACAGCAGC

CACACGTCTGAACTCTTTCCCTACACGACGCTCTTCCGATCTNNNCGGGATGCAATTCCTTGGGAGCAT

CACACGTCTGAACTCTTTCCCTACACGACGCTCTTCCGATCTNNNCGGGATGCAATTCCTTGGGAGCAT

GAGCACACGTCTGAACTCTTTCCCTACACGACGCTCTTCCGATCTNNNTCCCAATCCAATCCCAATTCCA

GAGCACACGTCTGAACTCTTTCCCTACACGACGCTCTTCCGATCTNNNTCCCAATCCAATCCCAATTCCA

GAGCACACGTCTGAACTCTTTCCCTACACGACGCTCTTCCGATCTNNNAGCCAAACCAAAGTCACCTAG

GAGCACACGTCTGAACTCTTTCCCTACACGACGCTCTTCCGATCTNNNAGCCAAACCAAAGTCACCTAG

GCACACGTCTGAACTCTTTCCCTACACGACGCTCTTCCGATCTNNNTCTGAGGGTCTCAAAATGTAGG

GCACACGTCTGAACTCTTTCCCTACACGACGCTCTTCCGATCTNNNTCTGAGGGTCTCAAAATGTAGG

AGCACACGTCTGAACTCTTTCCCTACACGACGCTCTTCCGATCTNNNGGGGAACAGGTGGTATTTGGT

AGCACACGTCTGAACTCTTTCCCTACACGACGCTCTTCCGATCTNNNGGGGAACAGGTGGTATTTGGT

AGCACACGTCTGAACTCTTTCCCTACACGACGCTCTTCCGATCTNNNGTGACTTTGGGTGGCAGTATTT

AGCACACGTCTGAACTCTTTCCCTACACGACGCTCTTCCGATCTNNNGTGACTTTGGGTGGCAGTATTT

AGCACACGTCTGAACTCTTTCCCTACACGACGCTCTTCCGATCTNNNAAACTCTGACTCCCAGGTTCAA

AGCACACGTCTGAACTCTTTCCCTACACGACGCTCTTCCGATCTNNNAAACTCTGACTCCCAGGTTCAA

AGCACACGTCTGAACTCTTTCCCTACACGACGCTCTTCCGATCTNNNACTTCAAAGCCCTCGTCCATATT

AGCACACGTCTGAACTCTTTCCCTACACGACGCTCTTCCGATCTNNNACTTCAAAGCCCTCGTCCATATT

GAGCACACGTCTGAACTCTTTCCCTACACGACGCTCTTCCGATCTNNNCAGCCTGGACAACAAGAGAGA

GAGCACACGTCTGAACTCTTTCCCTACACGACGCTCTTCCGATCTNNNCAGCCTGGACAACAAGAGAGA

GAGCACACGTCTGAACTCTTTCCCTACACGACGCTCTTCCGATCTNNNTTTAGATGGAGTCTCGCTCTCA

GAGCACACGTCTGAACTCTTTCCCTACACGACGCTCTTCCGATCTNNNTTTAGATGGAGTCTCGCTCTCA

AGCACACGTCTGAACTCTTTCCCTACACGACGCTCTTCCGATCTNNNTGTAATCCTAGCACTTTGGGAG

AGCACACGTCTGAACTCTTTCCCTACACGACGCTCTTCCGATCTNNNTGTAATCCTAGCACTTTGGGAG

GCACACGTCTGAACTCTTTCCCTACACGACGCTCTTCCGATCTNNNAAAGCCATCCACTATAAAATTTT

GCACACGTCTGAACTCTTTCCCTACACGACGCTCTTCCGATCTNNNAAAGCCATCCACTATAAAATTTT

AGAGCACACGTCTGAACTCTTTCCCTACACGACGCTCTTCCGATCTNNNGCATGCCTGTAATCCTAGCTA

AGAGCACACGTCTGAACTCTTTCCCTACACGACGCTCTTCCGATCTNNNGCATGCCTGTAATCCTAGCTA

AGCACACGTCTGAACTCTTTCCCTACACGACGCTCTTCCGATCTNNNCAAGTCCTTGACCTAACCTAGC

AGCACACGTCTGAACTCTTTCCCTACACGACGCTCTTCCGATCTNNNCAAGTCCTTGACCTAACCTAGC

GAGCACACGTCTGAACTCTTTCCCTACACGACGCTCTTCCGATCTNNNAGTGAGCAGAGATTGTACCAC

GAGCACACGTCTGAACTCTTTCCCTACACGACGCTCTTCCGATCTNNNAGTGAGCAGAGATTGTACCAC

GAGCACACGTCTGAACTCTTTCCCTACACGACGCTCTTCCGATCTNNNTGCAGCTTTGATGTTTGAACA

GAGCACACGTCTGAACTCTTTCCCTACACGACGCTCTTCCGATCTNNNTGCAGCTTTGATGTTTGAACA

GCACACGTCTGAACTCTTTCCCTACACGACGCTCTTCCGATCTNNNTTGTGAGGAGATTTCTGAGTGC

GCACACGTCTGAACTCTTTCCCTACACGACGCTCTTCCGATCTNNNTTGTGAGGAGATTTCTGAGTGC

GAGCACACGTCTGAACTCTTTCCCTACACGACGCTCTTCCGATCTNNNGGATCACTTGAGGTCAGGAG

GAGCACACGTCTGAACTCTTTCCCTACACGACGCTCTTCCGATCTNNNGGATCACTTGAGGTCAGGAG

AGCACACGTCTGAACTCTTTCCCTACACGACGCTCTTCCGATCTNNNGAGACAGGAGAATTGCTTGAA

AGCACACGTCTGAACTCTTTCCCTACACGACGCTCTTCCGATCTNNNGAGACAGGAGAATTGCTTGAA

AGCACACGTCTGAACTCTTTCCCTACACGACGCTCTTCCGATCTNNNTTTCAGCGAGTCGTGGTCATAG
AGCACACGTCTGAACTCTTTCCCTACACGACGCTCTTCCGATCTNNNTTTCAGCGAGTCGTGGTCATAG
GCACACGTCTGAACTCTTTCCCTACACGACGCTCTTCCGATCTNNNATTGTTTCTGCAGGGAGATGACT
GCACACGTCTGAACTCTTTCCCTACACGACGCTCTTCCGATCTNNNATTGTTTCTGCAGGGAGATGACT
AGCACACGTCTGAACTCTTTCCCTACACGACGCTCTTCCGATCTNNNTTCTCAACCAGGAGGGATTTT
AGCACACGTCTGAACTCTTTCCCTACACGACGCTCTTCCGATCTNNNTTCTCAACCAGGAGGGATTTT
AGCACACGTCTGAACTCTTTCCCTACACGACGCTCTTCCGATCTNNNGACACTAAGCCCACCTACAACAG
AGCACACGTCTGAACTCTTTCCCTACACGACGCTCTTCCGATCTNNNGACACTAAGCCCACCTACAACAG
GAGCACACGTCTGAACTCTTTCCCTACACGACGCTCTTCCGATCTNNNCCAGTTTGCTGCAACGTCTGCA
GAGCACACGTCTGAACTCTTTCCCTACACGACGCTCTTCCGATCTNNNCCAGTTTGCTGCAACGTCTGCA
AAGAGCACACGTCTGAACTCTTTCCCTACACGACGCTCTTCCGATCTNNNGCCTGTAATCCCAGATACT
AAGAGCACACGTCTGAACTCTTTCCCTACACGACGCTCTTCCGATCTNNNGCCTGTAATCCCAGATACT
AGCACACGTCTGAACTCTTTCCCTACACGACGCTCTTCCGATCTNNNTCATGCCTCCCAAGATTGAATCA
AGCACACGTCTGAACTCTTTCCCTACACGACGCTCTTCCGATCTNNNTCATGCCTCCCAAGATTGAATCA
AGCACACGTCTGAACTCTTTCCCTACACGACGCTCTTCCGATCTNNNTCTCACGAAGAAAGAGAACCCT
AGCACACGTCTGAACTCTTTCCCTACACGACGCTCTTCCGATCTNNNTCTCACGAAGAAAGAGAACCCT
GAGCACACGTCTGAACTCTTTCCCTACACGACGCTCTTCCGATCTNNNGGGAAGAGACTGACAAAGGG
GAGCACACGTCTGAACTCTTTCCCTACACGACGCTCTTCCGATCTNNNGGGAAGAGACTGACAAAGGG
AGCACACGTCTGAACTCTTTCCCTACACGACGCTCTTCCGATCTNNNGCCTGGTGTGACATAAGCTGAA
AGCACACGTCTGAACTCTTTCCCTACACGACGCTCTTCCGATCTNNNGCCTGGTGTGACATAAGCTGAA
AAGAGCACACGTCTGAACTCTTTCCCTACACGACGCTCTTCCGATCTNNNAGTGCTGGGATTACAGAT
AAGAGCACACGTCTGAACTCTTTCCCTACACGACGCTCTTCCGATCTNNNAGTGCTGGGATTACAGAT
GCACACGTCTGAACTCTTTCCCTACACGACGCTCTTCCGATCTNNNTTGCAGCTCATAGACCTTATCCC
GCACACGTCTGAACTCTTTCCCTACACGACGCTCTTCCGATCTNNNTTGCAGCTCATAGACCTTATCCC
AGCACACGTCTGAACTCTTTCCCTACACGACGCTCTTCCGATCTNNNAGGCATGGCAGTTCTACTACAA
AGCACACGTCTGAACTCTTTCCCTACACGACGCTCTTCCGATCTNNNAGGCATGGCAGTTCTACTACAA
AGCACACGTCTGAACTCTTTCCCTACACGACGCTCTTCCGATCTNNNCTGCAATCCTAGCTACTCAGGA
AGCACACGTCTGAACTCTTTCCCTACACGACGCTCTTCCGATCTNNNCTGCAATCCTAGCTACTCAGGA
GAGCACACGTCTGAACTCTTTCCCTACACGACGCTCTTCCGATCTNNNGGTTGCAGTGAGCCAAGATTA
GAGCACACGTCTGAACTCTTTCCCTACACGACGCTCTTCCGATCTNNNGGTTGCAGTGAGCCAAGATTA
AGCACACGTCTGAACTCTTTCCCTACACGACGCTCTTCCGATCTNNNAATCATCACAGCTGCTCAGGAG
AGCACACGTCTGAACTCTTTCCCTACACGACGCTCTTCCGATCTNNNAATCATCACAGCTGCTCAGGAG
GAGCACACGTCTGAACTCTTTCCCTACACGACGCTCTTCCGATCTNNNCCTGGTCACTAACTGCTTGGA
GAGCACACGTCTGAACTCTTTCCCTACACGACGCTCTTCCGATCTNNNCCTGGTCACTAACTGCTTGGA
GAGCACACGTCTGAACTCTTTCCCTACACGACGCTCTTCCGATCTNNNAGGCCTGTAATTCCAGCATTT
GAGCACACGTCTGAACTCTTTCCCTACACGACGCTCTTCCGATCTNNNAGGCCTGTAATTCCAGCATTT
GAGCACACGTCTGAACTCTTTCCCTACACGACGCTCTTCCGATCTNNNGGCTGCAGCTGATAAATGTG
GAGCACACGTCTGAACTCTTTCCCTACACGACGCTCTTCCGATCTNNNGGCTGCAGCTGATAAATGTG
GAGCACACGTCTGAACTCTTTCCCTACACGACGCTCTTCCGATCTNNNGGAGTTTGTGGCTGCAATGA
GAGCACACGTCTGAACTCTTTCCCTACACGACGCTCTTCCGATCTNNNGGAGTTTGTGGCTGCAATGA
ACACGTCTGAACTCTTTCCCTACACGACGCTCTTCCGATCTNNNCTCCTCCCCTCTCATCCCCCCATCGGA
ACACGTCTGAACTCTTTCCCTACACGACGCTCTTCCGATCTNNNCTCCTCCCCTCTCATCCCCCCATCGGA
AAGAGCACACGTCTGAACTCTTTCCCTACACGACGCTCTTCCGATCTNNNTAGAGTTGATATGCACACA

AAGAGCACACGTCTGAACTCTTTCCCTACACGACGCTCTTCCGATCTNNNTAGAGTTGATATGCACACA
AGCACACGTCTGAACTCTTTCCCTACACGACGCTCTTCCGATCTNNNCAGAGTCTGACCTCCCACTTTAC
AGCACACGTCTGAACTCTTTCCCTACACGACGCTCTTCCGATCTNNNCAGAGTCTGACCTCCCACTTTAC
GAGCACACGTCTGAACTCTTTCCCTACACGACGCTCTTCCGATCTNNNGAGTAAAATGAAGGAGCACC
GAGCACACGTCTGAACTCTTTCCCTACACGACGCTCTTCCGATCTNNNGAGTAAAATGAAGGAGCACC
AGCACACGTCTGAACTCTTTCCCTACACGACGCTCTTCCGATCTNNNCTAGGCAGACTACACTCCCTCTT
AGCACACGTCTGAACTCTTTCCCTACACGACGCTCTTCCGATCTNNNCTAGGCAGACTACACTCCCTCTT
AGCACACGTCTGAACTCTTTCCCTACACGACGCTCTTCCGATCTNNNACAAAGGTTCTCCCACTGGAAA
AGCACACGTCTGAACTCTTTCCCTACACGACGCTCTTCCGATCTNNNACAAAGGTTCTCCCACTGGAAA
AGCACACGTCTGAACTCTTTCCCTACACGACGCTCTTCCGATCTNNNGCAAGAATTTCAAGCCAGTGGA
AGCACACGTCTGAACTCTTTCCCTACACGACGCTCTTCCGATCTNNNGCAAGAATTTCAAGCCAGTGGA
AGCACACGTCTGAACTCTTTCCCTACACGACGCTCTTCCGATCTNNNGAGGCAAGAGAATCGCTTGAAT
AGCACACGTCTGAACTCTTTCCCTACACGACGCTCTTCCGATCTNNNGAGGCAAGAGAATCGCTTGAAT
GAGCACACGTCTGAACTCTTTCCCTACACGACGCTCTTCCGATCTNNNGGTGCCTATAATCCCAGCTACT
GAGCACACGTCTGAACTCTTTCCCTACACGACGCTCTTCCGATCTNNNGGTGCCTATAATCCCAGCTACT
GAGCACACGTCTGAACTCTTTCCCTACACGACGCTCTTCCGATCTNNNAGACATGCAAAAATTCAATCC
GAGCACACGTCTGAACTCTTTCCCTACACGACGCTCTTCCGATCTNNNAGACATGCAAAAATTCAATCC
AGCACACGTCTGAACTCTTTCCCTACACGACGCTCTTCCGATCTNNNGACCCACTCTGCACAGTATCAG
AGCACACGTCTGAACTCTTTCCCTACACGACGCTCTTCCGATCTNNNGACCCACTCTGCACAGTATCAG
AGCACACGTCTGAACTCTTTCCCTACACGACGCTCTTCCGATCTNNNTCCTCTCATCTCAGTCTCCAGAG
AGCACACGTCTGAACTCTTTCCCTACACGACGCTCTTCCGATCTNNNTCCTCTCATCTCAGTCTCCAGAG
GAGCACACGTCTGAACTCTTTCCCTACACGACGCTCTTCCGATCTNNNTTACCTGCCCTCAACTGATTCT
GAGCACACGTCTGAACTCTTTCCCTACACGACGCTCTTCCGATCTNNNTTACCTGCCCTCAACTGATTCT
GCACACGTCTGAACTCTTTCCCTACACGACGCTCTTCCGATCTNNNACAGATAAGTGACATCGTGCAGT
GCACACGTCTGAACTCTTTCCCTACACGACGCTCTTCCGATCTNNNACAGATAAGTGACATCGTGCAGT
GAGCACACGTCTGAACTCTTTCCCTACACGACGCTCTTCCGATCTNNNCTGTAGTCCCAGCTACTCAGAA
GAGCACACGTCTGAACTCTTTCCCTACACGACGCTCTTCCGATCTNNNCTGTAGTCCCAGCTACTCAGAA
AGCACACGTCTGAACTCTTTCCCTACACGACGCTCTTCCGATCTNNNGTAGTCTCAGCTATTCAGGAGG
AGCACACGTCTGAACTCTTTCCCTACACGACGCTCTTCCGATCTNNNGTAGTCTCAGCTATTCAGGAGG
AGAGCACACGTCTGAACTCTTTCCCTACACGACGCTCTTCCGATCTNNNTGCTCAAGAACCTTAAGGAG
AGAGCACACGTCTGAACTCTTTCCCTACACGACGCTCTTCCGATCTNNNTGCTCAAGAACCTTAAGGAG
GAGCACACGTCTGAACTCTTTCCCTACACGACGCTCTTCCGATCTNNNCAAGACCTACTGATCTGTTGC
GAGCACACGTCTGAACTCTTTCCCTACACGACGCTCTTCCGATCTNNNCAAGACCTACTGATCTGTTGC
CACACGTCTGAACTCTTTCCCTACACGACGCTCTTCCGATCTNNNCCAAGACAACGCACAATACCTTTCAT
CACACGTCTGAACTCTTTCCCTACACGACGCTCTTCCGATCTNNNCCAAGACAACGCACAATACCTTTCAT
GAGCACACGTCTGAACTCTTTCCCTACACGACGCTCTTCCGATCTNNNGTTCTACCCCTTGCTCGCAGCA
GAGCACACGTCTGAACTCTTTCCCTACACGACGCTCTTCCGATCTNNNGTTCTACCCCTTGCTCGCAGCA
AGCACACGTCTGAACTCTTTCCCTACACGACGCTCTTCCGATCTNNNTGACATCTTTTAGGGCATCCACA
AGCACACGTCTGAACTCTTTCCCTACACGACGCTCTTCCGATCTNNNTGACATCTTTTAGGGCATCCACA
GCACACGTCTGAACTCTTTCCCTACACGACGCTCTTCCGATCTNNNTCTCTCTCTTTCTTCCCGTCTCAT
GCACACGTCTGAACTCTTTCCCTACACGACGCTCTTCCGATCTNNNTCTCTCTCTTTCTTCCCGTCTCAT
GAGCACACGTCTGAACTCTTTCCCTACACGACGCTCTTCCGATCTNNNTCTTGGTTGGCACCAGTGTTTA
GAGCACACGTCTGAACTCTTTCCCTACACGACGCTCTTCCGATCTNNNTCTTGGTTGGCACCAGTGTTTA

GAGCACACGTCTGAACTCTTTCCCTACACGACGCTCTTCCGATCTNNNCAGAACTGCTGGAAAGACAA
GAGCACACGTCTGAACTCTTTCCCTACACGACGCTCTTCCGATCTNNNCAGAACTGCTGGAAAGACAA
AGAGCACACGTCTGAACTCTTTCCCTACACGACGCTCTTCCGATCTNNNGAGGTTGCAGTAAGTCGAG
AGAGCACACGTCTGAACTCTTTCCCTACACGACGCTCTTCCGATCTNNNGAGGTTGCAGTAAGTCGAG
AGAGCACACGTCTGAACTCTTTCCCTACACGACGCTCTTCCGATCTNNNTACATGCAAAGGAGACAGT
AGAGCACACGTCTGAACTCTTTCCCTACACGACGCTCTTCCGATCTNNNTACATGCAAAGGAGACAGT
AGCACACGTCTGAACTCTTTCCCTACACGACGCTCTTCCGATCTNNNACTTTTGGTTTTGGTTCACATGG
AGCACACGTCTGAACTCTTTCCCTACACGACGCTCTTCCGATCTNNNACTTTTGGTTTTGGTTCACATGG
AGCACACGTCTGAACTCTTTCCCTACACGACGCTCTTCCGATCTNNNGAGACAGGAGAATTGCTTGAA
AGCACACGTCTGAACTCTTTCCCTACACGACGCTCTTCCGATCTNNNGAGACAGGAGAATTGCTTGAA
AGCACACGTCTGAACTCTTTCCCTACACGACGCTCTTCCGATCTNNNACCATAAAACGGCTTATATGTT
AGCACACGTCTGAACTCTTTCCCTACACGACGCTCTTCCGATCTNNNACCATAAAACGGCTTATATGTT
AGCACACGTCTGAACTCTTTCCCTACACGACGCTCTTCCGATCTNNNTGCCTGTAATCCCAGCTATTCA
AGCACACGTCTGAACTCTTTCCCTACACGACGCTCTTCCGATCTNNNTGCCTGTAATCCCAGCTATTCA
AGCACACGTCTGAACTCTTTCCCTACACGACGCTCTTCCGATCTNNNTCCCAATGTTTCAGGAACTGAC
AGCACACGTCTGAACTCTTTCCCTACACGACGCTCTTCCGATCTNNNTCCCAATGTTTCAGGAACTGAC
GAGCACACGTCTGAACTCTTTCCCTACACGACGCTCTTCCGATCTNNNGAAGGAAAACTAGTTTGCGA
GAGCACACGTCTGAACTCTTTCCCTACACGACGCTCTTCCGATCTNNNGAAGGAAAACTAGTTTGCGA
AGCACACGTCTGAACTCTTTCCCTACACGACGCTCTTCCGATCTNNNGAGGTTGTGGTGAGCTGAGAT
AGCACACGTCTGAACTCTTTCCCTACACGACGCTCTTCCGATCTNNNGAGGTTGTGGTGAGCTGAGAT
GAGCACACGTCTGAACTCTTTCCCTACACGACGCTCTTCCGATCTNNNGACCCCGTCTCTGCATTTTACA
GAGCACACGTCTGAACTCTTTCCCTACACGACGCTCTTCCGATCTNNNGACCCCGTCTCTGCATTTTACA
AGAGCACACGTCTGAACTCTTTCCCTACACGACGCTCTTCCGATCTNNNTGCTGAAGGAAAAGAGAAA
AGAGCACACGTCTGAACTCTTTCCCTACACGACGCTCTTCCGATCTNNNTGCTGAAGGAAAAGAGAAA
AAGAGCACACGTCTGAACTCTTTCCCTACACGACGCTCTTCCGATCTNNNGTGCCTGTAATCCCAGCTA
AAGAGCACACGTCTGAACTCTTTCCCTACACGACGCTCTTCCGATCTNNNGTGCCTGTAATCCCAGCTA
AGCACACGTCTGAACTCTTTCCCTACACGACGCTCTTCCGATCTNNNGCAGTCAGGGAGGTTTCATATC
AGCACACGTCTGAACTCTTTCCCTACACGACGCTCTTCCGATCTNNNGCAGTCAGGGAGGTTTCATATC
GAGCACACGTCTGAACTCTTTCCCTACACGACGCTCTTCCGATCTNNNGAGGTTGCAGTGAGCTGAGA
GAGCACACGTCTGAACTCTTTCCCTACACGACGCTCTTCCGATCTNNNGAGGTTGCAGTGAGCTGAGA
GAGCACACGTCTGAACTCTTTCCCTACACGACGCTCTTCCGATCTNNNTCATGTTACCCCAGTTAGAAT
GAGCACACGTCTGAACTCTTTCCCTACACGACGCTCTTCCGATCTNNNTCATGTTACCCCAGTTAGAAT
GAGCACACGTCTGAACTCTTTCCCTACACGACGCTCTTCCGATCTNNNCTAGCTACTCTGGAGACAAG
GAGCACACGTCTGAACTCTTTCCCTACACGACGCTCTTCCGATCTNNNCTAGCTACTCTGGAGACAAG
AGCACACGTCTGAACTCTTTCCCTACACGACGCTCTTCCGATCTNNNCCATAATCATGAGAGCCAGGT
AGCACACGTCTGAACTCTTTCCCTACACGACGCTCTTCCGATCTNNNCCATAATCATGAGAGCCAGGT
GAGCACACGTCTGAACTCTTTCCCTACACGACGCTCTTCCGATCTNNNGAGAATGGCATGAATCCAGG
GAGCACACGTCTGAACTCTTTCCCTACACGACGCTCTTCCGATCTNNNGAGAATGGCATGAATCCAGG
GAGCACACGTCTGAACTCTTTCCCTACACGACGCTCTTCCGATCTNNNAACCTAAGTGCCCATCAACTG
GAGCACACGTCTGAACTCTTTCCCTACACGACGCTCTTCCGATCTNNNAACCTAAGTGCCCATCAACTG
AGCACACGTCTGAACTCTTTCCCTACACGACGCTCTTCCGATCTNNNTCATCTTTTTTCCTGCCCCAAAC
AGCACACGTCTGAACTCTTTCCCTACACGACGCTCTTCCGATCTNNNTCATCTTTTTTCCTGCCCCAAAC
AGCACACGTCTGAACTCTTTCCCTACACGACGCTCTTCCGATCTNNNAATACAAAATTAGTCCGGCGTG

AGCACACGTCTGAACTCTTTCCCTACACGACGCTCTTCCGATCTNNNAATACAAAATTAGTCCGGCGTG
GAGCACACGTCTGAACTCTTTCCCTACACGACGCTCTTCCGATCTNNNCTTGCAGTGAGCCGAGATTGT
GAGCACACGTCTGAACTCTTTCCCTACACGACGCTCTTCCGATCTNNNCTTGCAGTGAGCCGAGATTGT
AGCACACGTCTGAACTCTTTCCCTACACGACGCTCTTCCGATCTNNNTATGAGGACGCAAAGGCATAA
AGCACACGTCTGAACTCTTTCCCTACACGACGCTCTTCCGATCTNNNTATGAGGACGCAAAGGCATAA
AGCACACGTCTGAACTCTTTCCCTACACGACGCTCTTCCGATCTNNNGGGGTACAGGTGGTATTTGGT
AGCACACGTCTGAACTCTTTCCCTACACGACGCTCTTCCGATCTNNNGGGGTACAGGTGGTATTTGGT
GAGCACACGTCTGAACTCTTTCCCTACACGACGCTCTTCCGATCTNNNCCAAACTGAGGTTCCAGAGCTA
GAGCACACGTCTGAACTCTTTCCCTACACGACGCTCTTCCGATCTNNNCCAAACTGAGGTTCCAGAGCTA
AGCACACGTCTGAACTCTTTCCCTACACGACGCTCTTCCGATCTNNNTAGAATGACAGCCTTTTTCAGCA
AGCACACGTCTGAACTCTTTCCCTACACGACGCTCTTCCGATCTNNNTAGAATGACAGCCTTTTTCAGCA
AGCACACGTCTGAACTCTTTCCCTACACGACGCTCTTCCGATCTNNNACACATACCCTAAGACCCAGAA
AGCACACGTCTGAACTCTTTCCCTACACGACGCTCTTCCGATCTNNNACACATACCCTAAGACCCAGAA
AGCACACGTCTGAACTCTTTCCCTACACGACGCTCTTCCGATCTNNNAACTTCCACCATCACTCCCAAAT
AGCACACGTCTGAACTCTTTCCCTACACGACGCTCTTCCGATCTNNNAACTTCCACCATCACTCCCAAAT
AGCACACGTCTGAACTCTTTCCCTACACGACGCTCTTCCGATCTNNNCATAGCATGCATGAAATGGGT
AGCACACGTCTGAACTCTTTCCCTACACGACGCTCTTCCGATCTNNNCATAGCATGCATGAAATGGGT
AGCACACGTCTGAACTCTTTCCCTACACGACGCTCTTCCGATCTNNNCAAAGCACAGTGTTATGCAGG
AGCACACGTCTGAACTCTTTCCCTACACGACGCTCTTCCGATCTNNNCAAAGCACAGTGTTATGCAGG
AGCACACGTCTGAACTCTTTCCCTACACGACGCTCTTCCGATCTNNNACACAGTGAAACCCCATCTCTA
AGCACACGTCTGAACTCTTTCCCTACACGACGCTCTTCCGATCTNNNACACAGTGAAACCCCATCTCTA
AGCACACGTCTGAACTCTTTCCCTACACGACGCTCTTCCGATCTNNNCATCCTGCCTAACATGGTGAAAC
AGCACACGTCTGAACTCTTTCCCTACACGACGCTCTTCCGATCTNNNCATCCTGCCTAACATGGTGAAAC
AAGAGCACACGTCTGAACTCTTTCCCTACACGACGCTCTTCCGATCTNNNTCTCTGGCTTCACTCTCTCT
AAGAGCACACGTCTGAACTCTTTCCCTACACGACGCTCTTCCGATCTNNNTCTCTGGCTTCACTCTCTCT
GAGCACACGTCTGAACTCTTTCCCTACACGACGCTCTTCCGATCTNNNGTGTCCTCAGATTTCCCAGTGT
GAGCACACGTCTGAACTCTTTCCCTACACGACGCTCTTCCGATCTNNNGTGTCCTCAGATTTCCCAGTGT
GAGCACACGTCTGAACTCTTTCCCTACACGACGCTCTTCCGATCTNNNAATCAGCTGTAAATCCGTCTG
GAGCACACGTCTGAACTCTTTCCCTACACGACGCTCTTCCGATCTNNNAATCAGCTGTAAATCCGTCTG
AGCACACGTCTGAACTCTTTCCCTACACGACGCTCTTCCGATCTNNNAGATTGCCAGAGCTCATGAAAG
AGCACACGTCTGAACTCTTTCCCTACACGACGCTCTTCCGATCTNNNAGATTGCCAGAGCTCATGAAAG
AGCACACGTCTGAACTCTTTCCCTACACGACGCTCTTCCGATCTNNNAAAACAGATTAGTGGTTGCCA
AGCACACGTCTGAACTCTTTCCCTACACGACGCTCTTCCGATCTNNNAAAACAGATTAGTGGTTGCCA
CACACGTCTGAACTCTTTCCCTACACGACGCTCTTCCGATCTNNNAGGTCTTTTGTTCACTGGATGGTC
CACACGTCTGAACTCTTTCCCTACACGACGCTCTTCCGATCTNNNAGGTCTTTTGTTCACTGGATGGTC
GCACACGTCTGAACTCTTTCCCTACACGACGCTCTTCCGATCTNNNACCCTCCACCAGCAAAAAGATAA
GCACACGTCTGAACTCTTTCCCTACACGACGCTCTTCCGATCTNNNACCCTCCACCAGCAAAAAGATAA
AGCACACGTCTGAACTCTTTCCCTACACGACGCTCTTCCGATCTNNNCTCAAGTGATTGACCTGCTTCAG
AGCACACGTCTGAACTCTTTCCCTACACGACGCTCTTCCGATCTNNNCTCAAGTGATTGACCTGCTTCAG
AGCACACGTCTGAACTCTTTCCCTACACGACGCTCTTCCGATCTNNNCGCTTTCACTCACTATCCACCAT
AGCACACGTCTGAACTCTTTCCCTACACGACGCTCTTCCGATCTNNNCGCTTTCACTCACTATCCACCAT
GCACACGTCTGAACTCTTTCCCTACACGACGCTCTTCCGATCTNNNATCCCCATCACACTTGAACTCAACA
GCACACGTCTGAACTCTTTCCCTACACGACGCTCTTCCGATCTNNNATCCCCATCACACTTGAACTCAACA

AGCACACGTCTGAACTCTTTCCCTACACGACGCTCTTCCGATCTNNNCTGATGGTTTCAAAGTGGCAG
AGCACACGTCTGAACTCTTTCCCTACACGACGCTCTTCCGATCTNNNCTGATGGTTTCAAAGTGGCAG
AGCACACGTCTGAACTCTTTCCCTACACGACGCTCTTCCGATCTNNNGATCTCAGCTCACTGTAACCTC
AGCACACGTCTGAACTCTTTCCCTACACGACGCTCTTCCGATCTNNNGATCTCAGCTCACTGTAACCTC
GCACACGTCTGAACTCTTTCCCTACACGACGCTCTTCCGATCTNNNTGGCGAAACTCTGTCAAAACAA
GCACACGTCTGAACTCTTTCCCTACACGACGCTCTTCCGATCTNNNTGGCGAAACTCTGTCAAAACAA
AGCACACGTCTGAACTCTTTCCCTACACGACGCTCTTCCGATCTNNNGGTTCTCCAGTCATCTTGGATG
AGCACACGTCTGAACTCTTTCCCTACACGACGCTCTTCCGATCTNNNGGTTCTCCAGTCATCTTGGATG
AGCACACGTCTGAACTCTTTCCCTACACGACGCTCTTCCGATCTNNNCTCCTCCTTCCTTTCTCTCTC
AGCACACGTCTGAACTCTTTCCCTACACGACGCTCTTCCGATCTNNNCTCCTCCTTCCTTTCTCTCTC
AGCACACGTCTGAACTCTTTCCCTACACGACGCTCTTCCGATCTNNNATCAGAGACCTACAGGCACAA
AGCACACGTCTGAACTCTTTCCCTACACGACGCTCTTCCGATCTNNNATCAGAGACCTACAGGCACAA
AGCACACGTCTGAACTCTTTCCCTACACGACGCTCTTCCGATCTNNNAGCAATGGAATATCCGCCTGTA
AGCACACGTCTGAACTCTTTCCCTACACGACGCTCTTCCGATCTNNNAGCAATGGAATATCCGCCTGTA
AGAGCACACGTCTGAACTCTTTCCCTACACGACGCTCTTCCGATCTNNNAGGGAGAAGAGGGAGAA
AGAGCACACGTCTGAACTCTTTCCCTACACGACGCTCTTCCGATCTNNNAGGGAGAAGAGGGAGAA
GCACACGTCTGAACTCTTTCCCTACACGACGCTCTTCCGATCTNNNATTGTGCGCAATCATGTCTCATT
GCACACGTCTGAACTCTTTCCCTACACGACGCTCTTCCGATCTNNNATTGTGCGCAATCATGTCTCATT
GAGCACACGTCTGAACTCTTTCCCTACACGACGCTCTTCCGATCTNNNGCAGCAAAATAGCTGCATTC
GAGCACACGTCTGAACTCTTTCCCTACACGACGCTCTTCCGATCTNNNGCAGCAAAATAGCTGCATTC
GCACACGTCTGAACTCTTTCCCTACACGACGCTCTTCCGATCTNNNACCATGCCTAGTTGATTTTAAA
GCACACGTCTGAACTCTTTCCCTACACGACGCTCTTCCGATCTNNNACCATGCCTAGTTGATTTTAAA
AGAGCACACGTCTGAACTCTTTCCCTACACGACGCTCTTCCGATCTNNNATGCTAGTGGTTGATGGGA
AGAGCACACGTCTGAACTCTTTCCCTACACGACGCTCTTCCGATCTNNNATGCTAGTGGTTGATGGGA
CACACGTCTGAACTCTTTCCCTACACGACGCTCTTCCGATCTNNNTCTGGAACGGAGTCTGATTTTGG
CACACGTCTGAACTCTTTCCCTACACGACGCTCTTCCGATCTNNNTCTGGAACGGAGTCTGATTTTGG
GCACACGTCTGAACTCTTTCCCTACACGACGCTCTTCCGATCTNNNTGTTTCCAGACCCTAGCACAAAT
GCACACGTCTGAACTCTTTCCCTACACGACGCTCTTCCGATCTNNNTGTTTCCAGACCCTAGCACAAAT
AGCACACGTCTGAACTCTTTCCCTACACGACGCTCTTCCGATCTNNNTCTCTTTGTGGCTTTGATTTGC
AGCACACGTCTGAACTCTTTCCCTACACGACGCTCTTCCGATCTNNNTCTCTTTGTGGCTTTGATTTGC
AGCACACGTCTGAACTCTTTCCCTACACGACGCTCTTCCGATCTNNNCTACAGGAGTGTACCACCATGT
AGCACACGTCTGAACTCTTTCCCTACACGACGCTCTTCCGATCTNNNCTACAGGAGTGTACCACCATGT
AGCACACGTCTGAACTCTTTCCCTACACGACGCTCTTCCGATCTNNNACCATGCCCCTATTGTGAGAAA
AGCACACGTCTGAACTCTTTCCCTACACGACGCTCTTCCGATCTNNNACCATGCCCCTATTGTGAGAAA
GCACACGTCTGAACTCTTTCCCTACACGACGCTCTTCCGATCTNNNTGTAGTCATAGCCATCCTAACCTC
GCACACGTCTGAACTCTTTCCCTACACGACGCTCTTCCGATCTNNNTGTAGTCATAGCCATCCTAACCTC
GCACACGTCTGAACTCTTTCCCTACACGACGCTCTTCCGATCTNNNGCTAAGGCAGGAGAATCACTTG
GCACACGTCTGAACTCTTTCCCTACACGACGCTCTTCCGATCTNNNGCTAAGGCAGGAGAATCACTTG
AAGAGCACACGTCTGAACTCTTTCCCTACACGACGCTCTTCCGATCTNNNTAGCTCTTCTAGACACCAAC
AAGAGCACACGTCTGAACTCTTTCCCTACACGACGCTCTTCCGATCTNNNTAGCTCTTCTAGACACCAAC
AGCACACGTCTGAACTCTTTCCCTACACGACGCTCTTCCGATCTNNNAGTGGGACAAGATATGAAGG
AGCACACGTCTGAACTCTTTCCCTACACGACGCTCTTCCGATCTNNNAGTGGGACAAGATATGAAGG
AGCACACGTCTGAACTCTTTCCCTACACGACGCTCTTCCGATCTNNNTTCACCAGAAAACACACACACAC

AGCACACGTCTGAACTCTTTCCCTACACGACGCTCTTCCGATCTNNNTTCACCAGAAAACACACACACAC
AGAGCACACGTCTGAACTCTTTCCCTACACGACGCTCTTCCGATCTNNNAAAAAAATAGCAGGGCATG
AGAGCACACGTCTGAACTCTTTCCCTACACGACGCTCTTCCGATCTNNNAAAAAAATAGCAGGGCATG
GAGCACACGTCTGAACTCTTTCCCTACACGACGCTCTTCCGATCTNNNGCAGAGACACCTGACTGTTAG
GAGCACACGTCTGAACTCTTTCCCTACACGACGCTCTTCCGATCTNNNGCAGAGACACCTGACTGTTAG
GAGCACACGTCTGAACTCTTTCCCTACACGACGCTCTTCCGATCTNNNGATAGGCTGAGGCAGGAGAA
GAGCACACGTCTGAACTCTTTCCCTACACGACGCTCTTCCGATCTNNNGATAGGCTGAGGCAGGAGAA
AGCACACGTCTGAACTCTTTCCCTACACGACGCTCTTCCGATCTNNNCCATGTTGGTCAGGTTAGTCTCA
AGCACACGTCTGAACTCTTTCCCTACACGACGCTCTTCCGATCTNNNCCATGTTGGTCAGGTTAGTCTCA
GAGCACACGTCTGAACTCTTTCCCTACACGACGCTCTTCCGATCTNNNCAAGTGTAGCAATGGCACTTA
GAGCACACGTCTGAACTCTTTCCCTACACGACGCTCTTCCGATCTNNNCAAGTGTAGCAATGGCACTTA
GAGCACACGTCTGAACTCTTTCCCTACACGACGCTCTTCCGATCTNNNGGCAAAGGATAAATGCTTGA
GAGCACACGTCTGAACTCTTTCCCTACACGACGCTCTTCCGATCTNNNGGCAAAGGATAAATGCTTGA
AGCACACGTCTGAACTCTTTCCCTACACGACGCTCTTCCGATCTNNNAAACATTCATGAAACATTTGTT
AGCACACGTCTGAACTCTTTCCCTACACGACGCTCTTCCGATCTNNNAAACATTCATGAAACATTTGTT
GCACACGTCTGAACTCTTTCCCTACACGACGCTCTTCCGATCTNNNATTAATACCAGGATGCAACTGCA
GCACACGTCTGAACTCTTTCCCTACACGACGCTCTTCCGATCTNNNATTAATACCAGGATGCAACTGCA
AGCACACGTCTGAACTCTTTCCCTACACGACGCTCTTCCGATCTNNNAAGATCAAGCCACTTCAGTCCA
AGCACACGTCTGAACTCTTTCCCTACACGACGCTCTTCCGATCTNNNAAGATCAAGCCACTTCAGTCCA
GAGCACACGTCTGAACTCTTTCCCTACACGACGCTCTTCCGATCTNNNTGCAGGCATCCATTTCCACTATA
GAGCACACGTCTGAACTCTTTCCCTACACGACGCTCTTCCGATCTNNNTGCAGGCATCCATTTCCACTATA
GAGCACACGTCTGAACTCTTTCCCTACACGACGCTCTTCCGATCTNNNGAGGCTGAGACAGGAGAAT
GAGCACACGTCTGAACTCTTTCCCTACACGACGCTCTTCCGATCTNNNGAGGCTGAGACAGGAGAAT
AGCACACGTCTGAACTCTTTCCCTACACGACGCTCTTCCGATCTNNNGCTAGGATGCCATAACCTACAC
AGCACACGTCTGAACTCTTTCCCTACACGACGCTCTTCCGATCTNNNGCTAGGATGCCATAACCTACAC
AGAGCACACGTCTGAACTCTTTCCCTACACGACGCTCTTCCGATCTNNNAAGCCTTCATTCTCATGTTGG
AGAGCACACGTCTGAACTCTTTCCCTACACGACGCTCTTCCGATCTNNNAAGCCTTCATTCTCATGTTGG
AGCACACGTCTGAACTCTTTCCCTACACGACGCTCTTCCGATCTNNNGCAGGCCTACAAAATGTTGTCTT
AGCACACGTCTGAACTCTTTCCCTACACGACGCTCTTCCGATCTNNNGCAGGCCTACAAAATGTTGTCTT
GAGCACACGTCTGAACTCTTTCCCTACACGACGCTCTTCCGATCTNNNCAGGGATACTTGTGTCACTCCA
GAGCACACGTCTGAACTCTTTCCCTACACGACGCTCTTCCGATCTNNNCAGGGATACTTGTGTCACTCCA
AGCACACGTCTGAACTCTTTCCCTACACGACGCTCTTCCGATCTNNNCACAGAGCTAGCTTCCTTCAGA
AGCACACGTCTGAACTCTTTCCCTACACGACGCTCTTCCGATCTNNNCACAGAGCTAGCTTCCTTCAGA
GCACACGTCTGAACTCTTTCCCTACACGACGCTCTTCCGATCTNNNAAATACACAAAATTAGCCGGGCG
GCACACGTCTGAACTCTTTCCCTACACGACGCTCTTCCGATCTNNNAAATACACAAAATTAGCCGGGCG
GAGCACACGTCTGAACTCTTTCCCTACACGACGCTCTTCCGATCTNNNCATGGAACTGAATGAGCAAAC
GAGCACACGTCTGAACTCTTTCCCTACACGACGCTCTTCCGATCTNNNCATGGAACTGAATGAGCAAAC
AGCACACGTCTGAACTCTTTCCCTACACGACGCTCTTCCGATCTNNNTCTGTAATCCCAGCTACTCAGGA
AGCACACGTCTGAACTCTTTCCCTACACGACGCTCTTCCGATCTNNNTCTGTAATCCCAGCTACTCAGGA
AGCACACGTCTGAACTCTTTCCCTACACGACGCTCTTCCGATCTNNNGAGGAGGAACAAAAACAGCCT
AGCACACGTCTGAACTCTTTCCCTACACGACGCTCTTCCGATCTNNNGAGGAGGAACAAAAACAGCCT
GAGCACACGTCTGAACTCTTTCCCTACACGACGCTCTTCCGATCTNNNACTGGCTCAATTTCAGAACTTG
GAGCACACGTCTGAACTCTTTCCCTACACGACGCTCTTCCGATCTNNNACTGGCTCAATTTCAGAACTTG

AGCACACGTCTGAACTCTTTCCCTACACGACGCTCTTCCGATCTNNNCAAAAAATTAGCCGGGCGTAG
AGCACACGTCTGAACTCTTTCCCTACACGACGCTCTTCCGATCTNNNCAAAAAATTAGCCGGGCGTAG
GCACACGTCTGAACTCTTTCCCTACACGACGCTCTTCCGATCTNNNATGAGTTCTGCTGCTTCTTCAGA
GCACACGTCTGAACTCTTTCCCTACACGACGCTCTTCCGATCTNNNATGAGTTCTGCTGCTTCTTCAGA
AGCACACGTCTGAACTCTTTCCCTACACGACGCTCTTCCGATCTNNNTACAAAAATTAGCCTGGCGTG
AGCACACGTCTGAACTCTTTCCCTACACGACGCTCTTCCGATCTNNNTACAAAAATTAGCCTGGCGTG
CACGTCTGAACTCTTTCCCTACACGACGCTCTTCCGATCTNNNCAGTTCACCTCCCTATCCCTTCCATCG
CACGTCTGAACTCTTTCCCTACACGACGCTCTTCCGATCTNNNCAGTTCACCTCCCTATCCCTTCCATCG
GAGCACACGTCTGAACTCTTTCCCTACACGACGCTCTTCCGATCTNNNACCTGGTAGGTGCTCCATAA
GAGCACACGTCTGAACTCTTTCCCTACACGACGCTCTTCCGATCTNNNACCTGGTAGGTGCTCCATAA
AGCACACGTCTGAACTCTTTCCCTACACGACGCTCTTCCGATCTNNNAGCTAAGATTTGGCAGTAGAG
AGCACACGTCTGAACTCTTTCCCTACACGACGCTCTTCCGATCTNNNAGCTAAGATTTGGCAGTAGAG
AGCACACGTCTGAACTCTTTCCCTACACGACGCTCTTCCGATCTNNNCTGAGGCAGGAGAATCACTT
AGCACACGTCTGAACTCTTTCCCTACACGACGCTCTTCCGATCTNNNCTGAGGCAGGAGAATCACTT
GAGCACACGTCTGAACTCTTTCCCTACACGACGCTCTTCCGATCTNNNAGAAGCTCAGAAGGCTTTGC
GAGCACACGTCTGAACTCTTTCCCTACACGACGCTCTTCCGATCTNNNAGAAGCTCAGAAGGCTTTGC
AGCACACGTCTGAACTCTTTCCCTACACGACGCTCTTCCGATCTNNNGTGGGAGTAACCTTGGCATAT
AGCACACGTCTGAACTCTTTCCCTACACGACGCTCTTCCGATCTNNNGTGGGAGTAACCTTGGCATAT
AGCACACGTCTGAACTCTTTCCCTACACGACGCTCTTCCGATCTNNNTTCACTGTTAGTCCTGGAAGTC
AGCACACGTCTGAACTCTTTCCCTACACGACGCTCTTCCGATCTNNNTTCACTGTTAGTCCTGGAAGTC
AGCACACGTCTGAACTCTTTCCCTACACGACGCTCTTCCGATCTNNNCTCAGGAGGATTAGGGAGGA
AGCACACGTCTGAACTCTTTCCCTACACGACGCTCTTCCGATCTNNNCTCAGGAGGATTAGGGAGGA
AGCACACGTCTGAACTCTTTCCCTACACGACGCTCTTCCGATCTNNNATGCCAAAAATGCAAAGTCCT
AGCACACGTCTGAACTCTTTCCCTACACGACGCTCTTCCGATCTNNNATGCCAAAAATGCAAAGTCCT
GAGCACACGTCTGAACTCTTTCCCTACACGACGCTCTTCCGATCTNNNAATACAAAAAATTAGCCGGG
GAGCACACGTCTGAACTCTTTCCCTACACGACGCTCTTCCGATCTNNNAATACAAAAAATTAGCCGGG
AGCACACGTCTGAACTCTTTCCCTACACGACGCTCTTCCGATCTNNNCAACAAAAGTGATCCCTAGCTG
AGCACACGTCTGAACTCTTTCCCTACACGACGCTCTTCCGATCTNNNCAACAAAAGTGATCCCTAGCTG
GCACACGTCTGAACTCTTTCCCTACACGACGCTCTTCCGATCTNNNGGGCTCCTAAATATCCCTTTGCA
GCACACGTCTGAACTCTTTCCCTACACGACGCTCTTCCGATCTNNNGGGCTCCTAAATATCCCTTTGCA
AGCACACGTCTGAACTCTTTCCCTACACGACGCTCTTCCGATCTNNNTGGTTCCTTCTGGTGCTTTTACT
AGCACACGTCTGAACTCTTTCCCTACACGACGCTCTTCCGATCTNNNTGGTTCCTTCTGGTGCTTTTACT
AGCACACGTCTGAACTCTTTCCCTACACGACGCTCTTCCGATCTNNNTCCAATGGAGCAAGTGTCTTTT
AGCACACGTCTGAACTCTTTCCCTACACGACGCTCTTCCGATCTNNNTCCAATGGAGCAAGTGTCTTTT
CACACGTCTGAACTCTTTCCCTACACGACGCTCTTCCGATCTNNNCAGAGTTTGTGCCCAGTCAAAAT
CACACGTCTGAACTCTTTCCCTACACGACGCTCTTCCGATCTNNNCAGAGTTTGTGCCCAGTCAAAAT
CACACGTCTGAACTCTTTCCCTACACGACGCTCTTCCGATCTNNNCTAAAAATACAAAAATTAGCTGG
CACACGTCTGAACTCTTTCCCTACACGACGCTCTTCCGATCTNNNCTAAAAATACAAAAATTAGCTGG
GCACACGTCTGAACTCTTTCCCTACACGACGCTCTTCCGATCTNNNAGACATGGAAAGGGAGTCATCC
GCACACGTCTGAACTCTTTCCCTACACGACGCTCTTCCGATCTNNNAGACATGGAAAGGGAGTCATCC
CACACGTCTGAACTCTTTCCCTACACGACGCTCTTCCGATCTNNNGTCCGATCTGCTTAGTGTCAAGA
CACACGTCTGAACTCTTTCCCTACACGACGCTCTTCCGATCTNNNGTCCGATCTGCTTAGTGTCAAGA
GAAGAGCACACGTCTGAACTCTTTCCCTACACGACGCTCTTCCGATCTNNNACACACATTCACAGTGG

GAAGAGCACACGTCTGAACTCTTTCCCTACACGACGCTCTTCCGATCTNNNACACACATTCACAGTGG
AGCACACGTCTGAACTCTTTCCCTACACGACGCTCTTCCGATCTNNNATATCTGAGGATTCCCCAACC
AGCACACGTCTGAACTCTTTCCCTACACGACGCTCTTCCGATCTNNNATATCTGAGGATTCCCCAACC
AGCACACGTCTGAACTCTTTCCCTACACGACGCTCTTCCGATCTNNNCTTTCCTGGCTGATGTGCATT
AGCACACGTCTGAACTCTTTCCCTACACGACGCTCTTCCGATCTNNNCTTTCCTGGCTGATGTGCATT
GAGCACACGTCTGAACTCTTTCCCTACACGACGCTCTTCCGATCTNNNCTGTAGTCACAGCTACTCAG
GAGCACACGTCTGAACTCTTTCCCTACACGACGCTCTTCCGATCTNNNCTGTAGTCACAGCTACTCAG
GCACACGTCTGAACTCTTTCCCTACACGACGCTCTTCCGATCTNNNTGATATGTGAGGCTTTTTCCT
GCACACGTCTGAACTCTTTCCCTACACGACGCTCTTCCGATCTNNNTGATATGTGAGGCTTTTTCCT
GAGCACACGTCTGAACTCTTTCCCTACACGACGCTCTTCCGATCTNNNTGAGGAAAAGATGGGTGAC
GAGCACACGTCTGAACTCTTTCCCTACACGACGCTCTTCCGATCTNNNTGAGGAAAAGATGGGTGAC
GCACACGTCTGAACTCTTTCCCTACACGACGCTCTTCCGATCTNNNGCCACCAGAACTCTTACTGCTA
GCACACGTCTGAACTCTTTCCCTACACGACGCTCTTCCGATCTNNNGCCACCAGAACTCTTACTGCTA
AGCACACGTCTGAACTCTTTCCCTACACGACGCTCTTCCGATCTNNNCAGAGTGTGTGCTGGAGGAA
AGCACACGTCTGAACTCTTTCCCTACACGACGCTCTTCCGATCTNNNCAGAGTGTGTGCTGGAGGAA
GAGCACACGTCTGAACTCTTTCCCTACACGACGCTCTTCCGATCTNNNGAAAAGGGAGAGAAACCAA
GAGCACACGTCTGAACTCTTTCCCTACACGACGCTCTTCCGATCTNNNGAAAAGGGAGAGAAACCAA
GCACACGTCTGAACTCTTTCCCTACACGACGCTCTTCCGATCTNNNCAAAAATTACCCTGGTGTGGTG
GCACACGTCTGAACTCTTTCCCTACACGACGCTCTTCCGATCTNNNCAAAAATTACCCTGGTGTGGTG
GCACACGTCTGAACTCTTTCCCTACACGACGCTCTTCCGATCTNNNTAGTTATGGATTATGTGAGAAG
GCACACGTCTGAACTCTTTCCCTACACGACGCTCTTCCGATCTNNNTAGTTATGGATTATGTGAGAAG
AGCACACGTCTGAACTCTTTCCCTACACGACGCTCTTCCGATCTNNNTAATCAGCACTTCTGTCTGGCA
AGCACACGTCTGAACTCTTTCCCTACACGACGCTCTTCCGATCTNNNTAATCAGCACTTCTGTCTGGCA
AGCACACGTCTGAACTCTTTCCCTACACGACGCTCTTCCGATCTNNNAGCTACTTGGGAGGTTGAAGT
AGCACACGTCTGAACTCTTTCCCTACACGACGCTCTTCCGATCTNNNAGCTACTTGGGAGGTTGAAGT
GAGCACACGTCTGAACTCTTTCCCTACACGACGCTCTTCCGATCTNNNCTACTCTGGAGAAACAGCGT
GAGCACACGTCTGAACTCTTTCCCTACACGACGCTCTTCCGATCTNNNCTACTCTGGAGAAACAGCGT
AGCACACGTCTGAACTCTTTCCCTACACGACGCTCTTCCGATCTNNNTCCACGAGACATCATCAGTGT
AGCACACGTCTGAACTCTTTCCCTACACGACGCTCTTCCGATCTNNNTCCACGAGACATCATCAGTGT
AGCACACGTCTGAACTCTTTCCCTACACGACGCTCTTCCGATCTNNNGGACAAAACTCCTTCTGCTTGA
AGCACACGTCTGAACTCTTTCCCTACACGACGCTCTTCCGATCTNNNGGACAAAACTCCTTCTGCTTGA
GCACACGTCTGAACTCTTTCCCTACACGACGCTCTTCCGATCTNNNTGCTTATTCACTCTGGGGATTCG
GCACACGTCTGAACTCTTTCCCTACACGACGCTCTTCCGATCTNNNTGCTTATTCACTCTGGGGATTCG
GAGCACACGTCTGAACTCTTTCCCTACACGACGCTCTTCCGATCTNNNAACACGAACCCTTTTGTGAAC
GAGCACACGTCTGAACTCTTTCCCTACACGACGCTCTTCCGATCTNNNAACACGAACCCTTTTGTGAAC
AGCACACGTCTGAACTCTTTCCCTACACGACGCTCTTCCGATCTNNNATGGAAAGCTTTTCCTGACCT
AGCACACGTCTGAACTCTTTCCCTACACGACGCTCTTCCGATCTNNNATGGAAAGCTTTTCCTGACCT
GAGCACACGTCTGAACTCTTTCCCTACACGACGCTCTTCCGATCTNNNTGAGGCCAGGAGTTTGAGA
GAGCACACGTCTGAACTCTTTCCCTACACGACGCTCTTCCGATCTNNNTGAGGCCAGGAGTTTGAGA
CACACGTCTGAACTCTTTCCCTACACGACGCTCTTCCGATCTNNNCTCACTCCTGGACTAGAGCAAAG
CACACGTCTGAACTCTTTCCCTACACGACGCTCTTCCGATCTNNNCTCACTCCTGGACTAGAGCAAAG
GCACACGTCTGAACTCTTTCCCTACACGACGCTCTTCCGATCTNNNGAACTGTGATCATTCTGCTGCAC
GCACACGTCTGAACTCTTTCCCTACACGACGCTCTTCCGATCTNNNGAACTGTGATCATTCTGCTGCAC

AGCACACGTCTGAACTCTTTCCCTACACGACGCTCTTCCGATCTNNNAGACAGAGCCTAAACATCCCC
AGCACACGTCTGAACTCTTTCCCTACACGACGCTCTTCCGATCTNNNAGACAGAGCCTAAACATCCCC
GCACACGTCTGAACTCTTTCCCTACACGACGCTCTTCCGATCTNNNTGTCTGGAACTCATTCTTCATCC
GCACACGTCTGAACTCTTTCCCTACACGACGCTCTTCCGATCTNNNTGTCTGGAACTCATTCTTCATCC
AGCACACGTCTGAACTCTTTCCCTACACGACGCTCTTCCGATCTNNNAGATGGAATTTCGCTCTTGTT
AGCACACGTCTGAACTCTTTCCCTACACGACGCTCTTCCGATCTNNNAGATGGAATTTCGCTCTTGTT
CACACGTCTGAACTCTTTCCCTACACGACGCTCTTCCGATCTNNNTTTTCATAAGGTTATAGCTCAGC
CACACGTCTGAACTCTTTCCCTACACGACGCTCTTCCGATCTNNNTTTTCATAAGGTTATAGCTCAGC
GCACACGTCTGAACTCTTTCCCTACACGACGCTCTTCCGATCTNNNGACTGGGTGGTTGATTTCAAT
GCACACGTCTGAACTCTTTCCCTACACGACGCTCTTCCGATCTNNNGACTGGGTGGTTGATTTCAAT
GCACACGTCTGAACTCTTTCCCTACACGACGCTCTTCCGATCTNNNGAACGATAAAGGGGTACATGG
GCACACGTCTGAACTCTTTCCCTACACGACGCTCTTCCGATCTNNNGAACGATAAAGGGGTACATGG
AGCACACGTCTGAACTCTTTCCCTACACGACGCTCTTCCGATCTNNNCTCTCTGGTGTGACAATTGG
AGCACACGTCTGAACTCTTTCCCTACACGACGCTCTTCCGATCTNNNCTCTCTGGTGTGACAATTGG
AGCACACGTCTGAACTCTTTCCCTACACGACGCTCTTCCGATCTNNNATATATAAGCTGGGTGTGGT
AGCACACGTCTGAACTCTTTCCCTACACGACGCTCTTCCGATCTNNNATATATAAGCTGGGTGTGGT
AGCACACGTCTGAACTCTTTCCCTACACGACGCTCTTCCGATCTNNNCTGAGGCAGGAGAATCACTT
AGCACACGTCTGAACTCTTTCCCTACACGACGCTCTTCCGATCTNNNCTGAGGCAGGAGAATCACTT
GAGCACACGTCTGAACTCTTTCCCTACACGACGCTCTTCCGATCTNNNATTTCCCCATCTTCTCACCAAC
GAGCACACGTCTGAACTCTTTCCCTACACGACGCTCTTCCGATCTNNNATTTCCCCATCTTCTCACCAAC
GCACACGTCTGAACTCTTTCCCTACACGACGCTCTTCCGATCTNNNGCTGAGCCTGTTCTTTTCTTCTGC
GCACACGTCTGAACTCTTTCCCTACACGACGCTCTTCCGATCTNNNGCTGAGCCTGTTCTTTTCTTCTGC
GCACACGTCTGAACTCTTTCCCTACACGACGCTCTTCCGATCTNNNTGGTCTCAAACTCCTGACTTCAG
GCACACGTCTGAACTCTTTCCCTACACGACGCTCTTCCGATCTNNNTGGTCTCAAACTCCTGACTTCAG
AGCACACGTCTGAACTCTTTCCCTACACGACGCTCTTCCGATCTNNNGTTTGGATCTTCACAGCGCAG
AGCACACGTCTGAACTCTTTCCCTACACGACGCTCTTCCGATCTNNNGTTTGGATCTTCACAGCGCAG
AGCACACGTCTGAACTCTTTCCCTACACGACGCTCTTCCGATCTNNNGAGGTTGCAGTGAGCTGAGA
AGCACACGTCTGAACTCTTTCCCTACACGACGCTCTTCCGATCTNNNGAGGTTGCAGTGAGCTGAGA
GAGCACACGTCTGAACTCTTTCCCTACACGACGCTCTTCCGATCTNNNCGGAACCATTCGGAACAGAG
GAGCACACGTCTGAACTCTTTCCCTACACGACGCTCTTCCGATCTNNNCGGAACCATTCGGAACAGAG
AGCACACGTCTGAACTCTTTCCCTACACGACGCTCTTCCGATCTNNNGCAGGTAGAACCAGGTCTAA
AGCACACGTCTGAACTCTTTCCCTACACGACGCTCTTCCGATCTNNNGCAGGTAGAACCAGGTCTAA
AGCACACGTCTGAACTCTTTCCCTACACGACGCTCTTCCGATCTNNNTAAGGCAGCAGAATCACTTGA
AGCACACGTCTGAACTCTTTCCCTACACGACGCTCTTCCGATCTNNNTAAGGCAGCAGAATCACTTGA
AGCACACGTCTGAACTCTTTCCCTACACGACGCTCTTCCGATCTNNNAATTCAGATGGCCAACAAACCT
AGCACACGTCTGAACTCTTTCCCTACACGACGCTCTTCCGATCTNNNAATTCAGATGGCCAACAAACCT
AGCACACGTCTGAACTCTTTCCCTACACGACGCTCTTCCGATCTNNNGCTTGAGTCCAGGAGTTAGA
AGCACACGTCTGAACTCTTTCCCTACACGACGCTCTTCCGATCTNNNGCTTGAGTCCAGGAGTTAGA
CACACGTCTGAACTCTTTCCCTACACGACGCTCTTCCGATCTNNNGGCAAGAGGATTCAAATAAAGG
CACACGTCTGAACTCTTTCCCTACACGACGCTCTTCCGATCTNNNGGCAAGAGGATTCAAATAAAGG
GAGCACACGTCTGAACTCTTTCCCTACACGACGCTCTTCCGATCTNNNCATCACCACCACTCAGAGTCC
GAGCACACGTCTGAACTCTTTCCCTACACGACGCTCTTCCGATCTNNNCATCACCACCACTCAGAGTCC
GCACACGTCTGAACTCTTTCCCTACACGACGCTCTTCCGATCTNNNTTTCCTTTCATTTCCCTCCCTCCC

AGCACACGTCTGAACTCTTTCCCTACACGACGCTCTTCCGATCTNNNTTTCCTTTCATTTCCCTCCCTCCCA
AGAGCACACGTCTGAACTCTTTCCCTACACGACGCTCTTCCGATCTNNNCTTTGTAGTTGCCTTCGTGGA
AGAGCACACGTCTGAACTCTTTCCCTACACGACGCTCTTCCGATCTNNNCTTTGTAGTTGCCTTCGTGGA
GAGCACACGTCTGAACTCTTTCCCTACACGACGCTCTTCCGATCTNNNGAGGCAGGAAAATGACGTGA
GAGCACACGTCTGAACTCTTTCCCTACACGACGCTCTTCCGATCTNNNGAGGCAGGAAAATGACGTGA
GAGCACACGTCTGAACTCTTTCCCTACACGACGCTCTTCCGATCTNNNGGATGCTAAAAGGCCCATTCTT
GAGCACACGTCTGAACTCTTTCCCTACACGACGCTCTTCCGATCTNNNGGATGCTAAAAGGCCCATTCTT
GAGCACACGTCTGAACTCTTTCCCTACACGACGCTCTTCCGATCTNNNCAGCATGATTTCGAACTGACCT
GAGCACACGTCTGAACTCTTTCCCTACACGACGCTCTTCCGATCTNNNCAGCATGATTTCGAACTGACCT
AGAGCACACGTCTGAACTCTTTCCCTACACGACGCTCTTCCGATCTNNNTGCCTTGGTGGTCTTGAATT
AGAGCACACGTCTGAACTCTTTCCCTACACGACGCTCTTCCGATCTNNNTGCCTTGGTGGTCTTGAATT
AGCACACGTCTGAACTCTTTCCCTACACGACGCTCTTCCGATCTNNNCAGCACTTCCTAGGAGTTAGTG
AGCACACGTCTGAACTCTTTCCCTACACGACGCTCTTCCGATCTNNNCAGCACTTCCTAGGAGTTAGTG
GAGCACACGTCTGAACTCTTTCCCTACACGACGCTCTTCCGATCTNNNATGGATTGAGCCAGATGCCAT
GAGCACACGTCTGAACTCTTTCCCTACACGACGCTCTTCCGATCTNNNATGGATTGAGCCAGATGCCAT
GAGCACACGTCTGAACTCTTTCCCTACACGACGCTCTTCCGATCTNNNGTGTTTCTGTGGAGTCAGTG
GAGCACACGTCTGAACTCTTTCCCTACACGACGCTCTTCCGATCTNNNGTGTTTCTGTGGAGTCAGTG
GCACACGTCTGAACTCTTTCCCTACACGACGCTCTTCCGATCTNNNTCTTCCCCATAAGTCCCCAAAAGC
GCACACGTCTGAACTCTTTCCCTACACGACGCTCTTCCGATCTNNNTCTTCCCCATAAGTCCCCAAAAGC
AGCACACGTCTGAACTCTTTCCCTACACGACGCTCTTCCGATCTNNNGAGGCAGGATAATTGCTGAAAC
AGCACACGTCTGAACTCTTTCCCTACACGACGCTCTTCCGATCTNNNGAGGCAGGATAATTGCTGAAAC
AGCACACGTCTGAACTCTTTCCCTACACGACGCTCTTCCGATCTNNNGGTGACAGAGTGAGACTCCATC
AGCACACGTCTGAACTCTTTCCCTACACGACGCTCTTCCGATCTNNNGGTGACAGAGTGAGACTCCATC
ACACGTCTGAACTCTTTCCCTACACGACGCTCTTCCGATCTNNNTTCTCAATGAGCTGTTGGGCACATC
ACACGTCTGAACTCTTTCCCTACACGACGCTCTTCCGATCTNNNTTCTCAATGAGCTGTTGGGCACATC
AGCACACGTCTGAACTCTTTCCCTACACGACGCTCTTCCGATCTNNNAAGGCCACTTCCCTAAATATGG
AGCACACGTCTGAACTCTTTCCCTACACGACGCTCTTCCGATCTNNNAAGGCCACTTCCCTAAATATGG
AGCACACGTCTGAACTCTTTCCCTACACGACGCTCTTCCGATCTNNNGCTGGAGGAAAATTAGATGCCA
AGCACACGTCTGAACTCTTTCCCTACACGACGCTCTTCCGATCTNNNGCTGGAGGAAAATTAGATGCCA
GAGCACACGTCTGAACTCTTTCCCTACACGACGCTCTTCCGATCTNNNATGGAGTTTTGCTCTTGTTGCT
GAGCACACGTCTGAACTCTTTCCCTACACGACGCTCTTCCGATCTNNNATGGAGTTTTGCTCTTGTTGCT
AGAGCACACGTCTGAACTCTTTCCCTACACGACGCTCTTCCGATCTNNNTCAATGGTTCCTGTTGGCTC
AGAGCACACGTCTGAACTCTTTCCCTACACGACGCTCTTCCGATCTNNNTCAATGGTTCCTGTTGGCTC
AGCACACGTCTGAACTCTTTCCCTACACGACGCTCTTCCGATCTNNNAATGCCTAAAGGTCTCCAACTCC
AGCACACGTCTGAACTCTTTCCCTACACGACGCTCTTCCGATCTNNNAATGCCTAAAGGTCTCCAACTCC
AGCACACGTCTGAACTCTTTCCCTACACGACGCTCTTCCGATCTNNNGGATCAGTGCTGGATGTTTGTT
AGCACACGTCTGAACTCTTTCCCTACACGACGCTCTTCCGATCTNNNGGATCAGTGCTGGATGTTTGTT
GAGCACACGTCTGAACTCTTTCCCTACACGACGCTCTTCCGATCTNNNGACTGGGTAAATTCTGAGCCT
GAGCACACGTCTGAACTCTTTCCCTACACGACGCTCTTCCGATCTNNNGACTGGGTAAATTCTGAGCCT
AGCACACGTCTGAACTCTTTCCCTACACGACGCTCTTCCGATCTNNNCCACAACCGGCTAATTTTTGTGT
AGCACACGTCTGAACTCTTTCCCTACACGACGCTCTTCCGATCTNNNCCACAACCGGCTAATTTTTGTGT
AGCACACGTCTGAACTCTTTCCCTACACGACGCTCTTCCGATCTNNNCTCATCTCTTCTCTTTTCCTCCAT
AGCACACGTCTGAACTCTTTCCCTACACGACGCTCTTCCGATCTNNNCTCATCTCTTCTCTTTTCCTCCAT

GAGCACACGTCTGAACTCTTTCCCTACACGACGCTCTTCCGATCTNNNAGCAACTCTGTCTTTATCGAGG

GAGCACACGTCTGAACTCTTTCCCTACACGACGCTCTTCCGATCTNNNAGCAACTCTGTCTTTATCGAGG

AGCACACGTCTGAACTCTTTCCCTACACGACGCTCTTCCGATCTNNNCCAAAATCCAAAGTGCTGGGAA

AGCACACGTCTGAACTCTTTCCCTACACGACGCTCTTCCGATCTNNNCCAAAATCCAAAGTGCTGGGAA

AGCACACGTCTGAACTCTTTCCCTACACGACGCTCTTCCGATCTNNNCTCTGTGTTGAAGAGGTAGCAG

AGCACACGTCTGAACTCTTTCCCTACACGACGCTCTTCCGATCTNNNCTCTGTGTTGAAGAGGTAGCAG

GCACACGTCTGAACTCTTTCCCTACACGACGCTCTTCCGATCTNNNAACTGGGGAGCTGCCTAAAAATA

GCACACGTCTGAACTCTTTCCCTACACGACGCTCTTCCGATCTNNNAACTGGGGAGCTGCCTAAAAATA

AGCACACGTCTGAACTCTTTCCCTACACGACGCTCTTCCGATCTNNNAGAATTGGCTCATGTTGTTGGT

AGCACACGTCTGAACTCTTTCCCTACACGACGCTCTTCCGATCTNNNAGAATTGGCTCATGTTGTTGGT

AGAGCACACGTCTGAACTCTTTCCCTACACGACGCTCTTCCGATCTNNNGTTGCAGTGAGCTGAGATCA

AGAGCACACGTCTGAACTCTTTCCCTACACGACGCTCTTCCGATCTNNNGTTGCAGTGAGCTGAGATCA

GAGCACACGTCTGAACTCTTTCCCTACACGACGCTCTTCCGATCTNNNTCCGCTACACTTGAATCGATGA

GAGCACACGTCTGAACTCTTTCCCTACACGACGCTCTTCCGATCTNNNTCCGCTACACTTGAATCGATGA

AGCACACGTCTGAACTCTTTCCCTACACGACGCTCTTCCGATCTNNNGAGGTTGCAGTGAACCAAGATT

AGCACACGTCTGAACTCTTTCCCTACACGACGCTCTTCCGATCTNNNGAGGTTGCAGTGAACCAAGATT

GAGCACACGTCTGAACTCTTTCCCTACACGACGCTCTTCCGATCTNNNAACAGTTCTGTCTTTCTGGGG

GAGCACACGTCTGAACTCTTTCCCTACACGACGCTCTTCCGATCTNNNAACAGTTCTGTCTTTCTGGGG

AGCACACGTCTGAACTCTTTCCCTACACGACGCTCTTCCGATCTNNNATTGGTCAGATCTCGAACTCCTG

AGCACACGTCTGAACTCTTTCCCTACACGACGCTCTTCCGATCTNNNATTGGTCAGATCTCGAACTCCTG

CACACGTCTGAACTCTTTCCCTACACGACGCTCTTCCGATCTNNNGTCTCAAACTCCTCACCTGAAGTCAT

CACACGTCTGAACTCTTTCCCTACACGACGCTCTTCCGATCTNNNGTCTCAAACTCCTCACCTGAAGTCAT

GAGCACACGTCTGAACTCTTTCCCTACACGACGCTCTTCCGATCTNNNCTGTGTCCTGCTGTTCTCTGTA

GAGCACACGTCTGAACTCTTTCCCTACACGACGCTCTTCCGATCTNNNCTGTGTCCTGCTGTTCTCTGTA

GAGCACACGTCTGAACTCTTTCCCTACACGACGCTCTTCCGATCTNNNGCAAAGATGAGGAGTGGAAA

GAGCACACGTCTGAACTCTTTCCCTACACGACGCTCTTCCGATCTNNNGCAAAGATGAGGAGTGGAAA

GCACACGTCTGAACTCTTTCCCTACACGACGCTCTTCCGATCTNNNGATCATAGCTCACTACAGCCTCCC

GCACACGTCTGAACTCTTTCCCTACACGACGCTCTTCCGATCTNNNGATCATAGCTCACTACAGCCTCCC

AGCACACGTCTGAACTCTTTCCCTACACGACGCTCTTCCGATCTNNNCAAAATAATTAGCTGGGCGTG

AGCACACGTCTGAACTCTTTCCCTACACGACGCTCTTCCGATCTNNNCAAAATAATTAGCTGGGCGTG

AGCACACGTCTGAACTCTTTCCCTACACGACGCTCTTCCGATCTNNNTCACTCAGGAATTCAGGCACTT

AGCACACGTCTGAACTCTTTCCCTACACGACGCTCTTCCGATCTNNNTCACTCAGGAATTCAGGCACTT

AGCACACGTCTGAACTCTTTCCCTACACGACGCTCTTCCGATCTNNNAAGCTTACATCTTCCTCAGAGCC

AGCACACGTCTGAACTCTTTCCCTACACGACGCTCTTCCGATCTNNNAAGCTTACATCTTCCTCAGAGCC

AGCACACGTCTGAACTCTTTCCCTACACGACGCTCTTCCGATCTNNNGCAAAGTGGCTAAAATGAAAC

AGCACACGTCTGAACTCTTTCCCTACACGACGCTCTTCCGATCTNNNGCAAAGTGGCTAAAATGAAAC

GAGCACACGTCTGAACTCTTTCCCTACACGACGCTCTTCCGATCTNNNCTGAGACAGGAGAATCGCTTG

GAGCACACGTCTGAACTCTTTCCCTACACGACGCTCTTCCGATCTNNNCTGAGACAGGAGAATCGCTTG

GCACACGTCTGAACTCTTTCCCTACACGACGCTCTTCCGATCTNNNAGGATAAGTCTCTGGGTTAGCCT

GCACACGTCTGAACTCTTTCCCTACACGACGCTCTTCCGATCTNNNAGGATAAGTCTCTGGGTTAGCCT

AGCACACGTCTGAACTCTTTCCCTACACGACGCTCTTCCGATCTNNNTCTCTCACACACACACACAAA

AGCACACGTCTGAACTCTTTCCCTACACGACGCTCTTCCGATCTNNNTCTCTCACACACACACACAAA

GAGCACACGTCTGAACTCTTTCCCTACACGACGCTCTTCCGATCTNNNCAAGGAATGGCCAAGCTAGAG

GAGCACACGTCTGAACTCTTTCCCTACACGACGCTCTTCCGATCTNNNCAAGGAATGGCCAAGCTAGAC
GAGCACACGTCTGAACTCTTTCCCTACACGACGCTCTTCCGATCTNNNATCACGGCTGTTCAAAGTTGA
GAGCACACGTCTGAACTCTTTCCCTACACGACGCTCTTCCGATCTNNNATCACGGCTGTTCAAAGTTGA
AGCACACGTCTGAACTCTTTCCCTACACGACGCTCTTCCGATCTNNNTTTCCTAGCAAAGCCAGACTCT
AGCACACGTCTGAACTCTTTCCCTACACGACGCTCTTCCGATCTNNNTTTCCTAGCAAAGCCAGACTCT
GAGCACACGTCTGAACTCTTTCCCTACACGACGCTCTTCCGATCTNNNGCCTCCCAAGTAGCTAGGATT
GAGCACACGTCTGAACTCTTTCCCTACACGACGCTCTTCCGATCTNNNGCCTCCCAAGTAGCTAGGATT
GAGCACACGTCTGAACTCTTTCCCTACACGACGCTCTTCCGATCTNNNGCAGGAGAATCGTTTGAACC
GAGCACACGTCTGAACTCTTTCCCTACACGACGCTCTTCCGATCTNNNGCAGGAGAATCGTTTGAACC
AGAGCACACGTCTGAACTCTTTCCCTACACGACGCTCTTCCGATCTNNNCTAGGCTCCCAGGAATCTGA
AGAGCACACGTCTGAACTCTTTCCCTACACGACGCTCTTCCGATCTNNNCTAGGCTCCCAGGAATCTGA
GAGCACACGTCTGAACTCTTTCCCTACACGACGCTCTTCCGATCTNNNATTGCAAAACCTCGTCCCTAC
GAGCACACGTCTGAACTCTTTCCCTACACGACGCTCTTCCGATCTNNNATTGCAAAACCTCGTCCCTAC
GAGCACACGTCTGAACTCTTTCCCTACACGACGCTCTTCCGATCTNNNTTGGAGTTGCAGTTGATGTGT
GAGCACACGTCTGAACTCTTTCCCTACACGACGCTCTTCCGATCTNNNTTGGAGTTGCAGTTGATGTGT
GAGCACACGTCTGAACTCTTTCCCTACACGACGCTCTTCCGATCTNNNTTTTTCCACCTCAGTCTCCCAAA
GAGCACACGTCTGAACTCTTTCCCTACACGACGCTCTTCCGATCTNNNTTTTTCCACCTCAGTCTCCCAAA
GAGCACACGTCTGAACTCTTTCCCTACACGACGCTCTTCCGATCTNNNCAAAAAAAATTAGCCGGGCGT
GAGCACACGTCTGAACTCTTTCCCTACACGACGCTCTTCCGATCTNNNCAAAAAAAATTAGCCGGGCGT
AGCACACGTCTGAACTCTTTCCCTACACGACGCTCTTCCGATCTNNNCCCAAGTCCCACTGTCAGTAAAT
AGCACACGTCTGAACTCTTTCCCTACACGACGCTCTTCCGATCTNNNCCCAAGTCCCACTGTCAGTAAAT
CACACGTCTGAACTCTTTCCCTACACGACGCTCTTCCGATCTNNNCTGGCCCCCTACTATGTTCTCTACA
CACACGTCTGAACTCTTTCCCTACACGACGCTCTTCCGATCTNNNCTGGCCCCCTACTATGTTCTCTACA
GAGCACACGTCTGAACTCTTTCCCTACACGACGCTCTTCCGATCTNNNTTATCCTATCCTGTAGTGGGG
GAGCACACGTCTGAACTCTTTCCCTACACGACGCTCTTCCGATCTNNNTTATCCTATCCTGTAGTGGGG
AGAGCACACGTCTGAACTCTTTCCCTACACGACGCTCTTCCGATCTNNNCCCAGAAGCTTCTTTGGTAGA
AGAGCACACGTCTGAACTCTTTCCCTACACGACGCTCTTCCGATCTNNNCCCAGAAGCTTCTTTGGTAGA
AGCACACGTCTGAACTCTTTCCCTACACGACGCTCTTCCGATCTNNNGAGAATCGCTTGAACTTGGGAG
AGCACACGTCTGAACTCTTTCCCTACACGACGCTCTTCCGATCTNNNGAGAATCGCTTGAACTTGGGAG
AGCACACGTCTGAACTCTTTCCCTACACGACGCTCTTCCGATCTNNNTCTGGCACTGACACAAATGTTT
AGCACACGTCTGAACTCTTTCCCTACACGACGCTCTTCCGATCTNNNTCTGGCACTGACACAAATGTTT
AGAGCACACGTCTGAACTCTTTCCCTACACGACGCTCTTCCGATCTNNNGAAAATTGCTTGAACTCGGG
AGAGCACACGTCTGAACTCTTTCCCTACACGACGCTCTTCCGATCTNNNGAAAATTGCTTGAACTCGGG
AGCACACGTCTGAACTCTTTCCCTACACGACGCTCTTCCGATCTNNNCCACTTGCACTTTTCACAACACA
AGCACACGTCTGAACTCTTTCCCTACACGACGCTCTTCCGATCTNNNCCACTTGCACTTTTCACAACACA
AGCACACGTCTGAACTCTTTCCCTACACGACGCTCTTCCGATCTNNNCATTCCCCTCTTACTGTGTTCCAC
AGCACACGTCTGAACTCTTTCCCTACACGACGCTCTTCCGATCTNNNCATTCCCCTCTTACTGTGTTCCAC
GAGCACACGTCTGAACTCTTTCCCTACACGACGCTCTTCCGATCTNNNCCAATGCAGCCAGCTAATTCA
GAGCACACGTCTGAACTCTTTCCCTACACGACGCTCTTCCGATCTNNNCCAATGCAGCCAGCTAATTCA
AGAGCACACGTCTGAACTCTTTCCCTACACGACGCTCTTCCGATCTNNNATGAACTACCATGAGCCGTC
AGAGCACACGTCTGAACTCTTTCCCTACACGACGCTCTTCCGATCTNNNATGAACTACCATGAGCCGTC
GAGCACACGTCTGAACTCTTTCCCTACACGACGCTCTTCCGATCTNNNTGCCAGGCACTAATTTTCACAA
GAGCACACGTCTGAACTCTTTCCCTACACGACGCTCTTCCGATCTNNNTGCCAGGCACTAATTTTCACAA

AGCACACGTCTGAACTCTTTCCCTACACGACGCTCTTCCGATCTNNNACTGATAAAATCCCAGCATCACA

AGCACACGTCTGAACTCTTTCCCTACACGACGCTCTTCCGATCTNNNACTGATAAAATCCCAGCATCACA

GAGCACACGTCTGAACTCTTTCCCTACACGACGCTCTTCCGATCTNNNAGTATACCTACAATGCTGTGC

GAGCACACGTCTGAACTCTTTCCCTACACGACGCTCTTCCGATCTNNNAGTATACCTACAATGCTGTGC

AGCACACGTCTGAACTCTTTCCCTACACGACGCTCTTCCGATCTNNNGAGTTTCCACCATGCATGCACCA

AGCACACGTCTGAACTCTTTCCCTACACGACGCTCTTCCGATCTNNNGAGTTTCCACCATGCATGCACCA

AGCACACGTCTGAACTCTTTCCCTACACGACGCTCTTCCGATCTNNNTGAAAGTTTTGGCAGTGGTTGT

AGCACACGTCTGAACTCTTTCCCTACACGACGCTCTTCCGATCTNNNTGAAAGTTTTGGCAGTGGTTGT

AGCACACGTCTGAACTCTTTCCCTACACGACGCTCTTCCGATCTNNNGCATGTCTGTAATCCCAGCTACT

AGCACACGTCTGAACTCTTTCCCTACACGACGCTCTTCCGATCTNNNGCATGTCTGTAATCCCAGCTACT

GAGCACACGTCTGAACTCTTTCCCTACACGACGCTCTTCCGATCTNNNCAACATGGCAAAACTCCATCTC

GAGCACACGTCTGAACTCTTTCCCTACACGACGCTCTTCCGATCTNNNCAACATGGCAAAACTCCATCTC

AGCACACGTCTGAACTCTTTCCCTACACGACGCTCTTCCGATCTNNNCATACACATAGACCCATGGAGCA

AGCACACGTCTGAACTCTTTCCCTACACGACGCTCTTCCGATCTNNNCATACACATAGACCCATGGAGCA

AGCACACGTCTGAACTCTTTCCCTACACGACGCTCTTCCGATCTNNNCTATCTGAAAATACATTTCCCTC

AGCACACGTCTGAACTCTTTCCCTACACGACGCTCTTCCGATCTNNNCTATCTGAAAATACATTTCCCTC

AGCACACGTCTGAACTCTTTCCCTACACGACGCTCTTCCGATCTNNNCCTGCTTAGGATCTTACCTGTGT

AGCACACGTCTGAACTCTTTCCCTACACGACGCTCTTCCGATCTNNNCCTGCTTAGGATCTTACCTGTGT

AGCACACGTCTGAACTCTTTCCCTACACGACGCTCTTCCGATCTNNNGGAGAGAGATGGAGACCTTGTT

AGCACACGTCTGAACTCTTTCCCTACACGACGCTCTTCCGATCTNNNGGAGAGAGATGGAGACCTTGTT

GAGCACACGTCTGAACTCTTTCCCTACACGACGCTCTTCCGATCTNNNTCCTTTCTCTTGGCTGACCTTA

GAGCACACGTCTGAACTCTTTCCCTACACGACGCTCTTCCGATCTNNNTCCTTTCTCTTGGCTGACCTTA

AGCACACGTCTGAACTCTTTCCCTACACGACGCTCTTCCGATCTNNNCTTATAGGTAAGCCCTTGTCCCC

AGCACACGTCTGAACTCTTTCCCTACACGACGCTCTTCCGATCTNNNCTTATAGGTAAGCCCTTGTCCCC

AGAGCACACGTCTGAACTCTTTCCCTACACGACGCTCTTCCGATCTNNNAGTCGCTTGTATTGGACAGA

AGAGCACACGTCTGAACTCTTTCCCTACACGACGCTCTTCCGATCTNNNAGTCGCTTGTATTGGACAGA

CACACGTCTGAACTCTTTCCCTACACGACGCTCTTCCGATCTNNNAAAAAAAAATTAGCTGGGCGTGGT

CACACGTCTGAACTCTTTCCCTACACGACGCTCTTCCGATCTNNNAAAAAAAAATTAGCTGGGCGTGGT

AGAGCACACGTCTGAACTCTTTCCCTACACGACGCTCTTCCGATCTNNNTCAGCTAGCAATCATCACGAT

AGAGCACACGTCTGAACTCTTTCCCTACACGACGCTCTTCCGATCTNNNTCAGCTAGCAATCATCACGAT

GAGCACACGTCTGAACTCTTTCCCTACACGACGCTCTTCCGATCTNNNAGTGGCAGGACAATACGTAAT

GAGCACACGTCTGAACTCTTTCCCTACACGACGCTCTTCCGATCTNNNAGTGGCAGGACAATACGTAAT

GAGCACACGTCTGAACTCTTTCCCTACACGACGCTCTTCCGATCTNNNAGCAGGAGAATAGCTTGAAT

GAGCACACGTCTGAACTCTTTCCCTACACGACGCTCTTCCGATCTNNNAGCAGGAGAATAGCTTGAAT

GAGCACACGTCTGAACTCTTTCCCTACACGACGCTCTTCCGATCTNNNGGGAGGGAAAGGACACAAA

GAGCACACGTCTGAACTCTTTCCCTACACGACGCTCTTCCGATCTNNNGGGAGGGAAAGGACACAAA

AGAGCACACGTCTGAACTCTTTCCCTACACGACGCTCTTCCGATCTNNNCATGTCAGATTCTGTGGCTAC

AGAGCACACGTCTGAACTCTTTCCCTACACGACGCTCTTCCGATCTNNNCATGTCAGATTCTGTGGCTAC

GAGCACACGTCTGAACTCTTTCCCTACACGACGCTCTTCCGATCTNNNCTGACTGGCACTGATTACTCTG

GAGCACACGTCTGAACTCTTTCCCTACACGACGCTCTTCCGATCTNNNCTGACTGGCACTGATTACTCTG

AGAGCACACGTCTGAACTCTTTCCCTACACGACGCTCTTCCGATCTNNNAGAGATGCCATACTGCTTAG

AGAGCACACGTCTGAACTCTTTCCCTACACGACGCTCTTCCGATCTNNNAGAGATGCCATACTGCTTAG

AGCACACGTCTGAACTCTTTCCCTACACGACGCTCTTCCGATCTNNNATCCTCTCCAGCATCTGTTGTTTC

GCACACGTCTGAACTCTTTCCCTACACGACGCTCTTCCGATCTNNNATCCTCTCCAGCATCTGTTGTTT
GAGCACACGTCTGAACTCTTTCCCTACACGACGCTCTTCCGATCTNNNAGGTAAGATTTGGGTGAGG
GAGCACACGTCTGAACTCTTTCCCTACACGACGCTCTTCCGATCTNNNAGGTAAGATTTGGGTGAGG
AGCACACGTCTGAACTCTTTCCCTACACGACGCTCTTCCGATCTNNNCAGTATGATTGGCCCAGCTTTT
AGCACACGTCTGAACTCTTTCCCTACACGACGCTCTTCCGATCTNNNCAGTATGATTGGCCCAGCTTTT
AGCACACGTCTGAACTCTTTCCCTACACGACGCTCTTCCGATCTNNNATCCTTTTGGGGCTGCTTTTAT
AGCACACGTCTGAACTCTTTCCCTACACGACGCTCTTCCGATCTNNNATCCTTTTGGGGCTGCTTTTAT
GAGCACACGTCTGAACTCTTTCCCTACACGACGCTCTTCCGATCTNNNATACAAAAATTAGCCGGGCAT
GAGCACACGTCTGAACTCTTTCCCTACACGACGCTCTTCCGATCTNNNATACAAAAATTAGCCGGGCAT
GCACACGTCTGAACTCTTTCCCTACACGACGCTCTTCCGATCTNNNTCTCTCTTTTGGAATGGGTGTGT
GCACACGTCTGAACTCTTTCCCTACACGACGCTCTTCCGATCTNNNTCTCTCTTTTGGAATGGGTGTGT
AGCACACGTCTGAACTCTTTCCCTACACGACGCTCTTCCGATCTNNNGAGAAAGGGATCCCAGCAAAC
AGCACACGTCTGAACTCTTTCCCTACACGACGCTCTTCCGATCTNNNGAGAAAGGGATCCCAGCAAAC
GAGCACACGTCTGAACTCTTTCCCTACACGACGCTCTTCCGATCTNNNTCATGGCAGAAATAGGATGT
GAGCACACGTCTGAACTCTTTCCCTACACGACGCTCTTCCGATCTNNNTCATGGCAGAAATAGGATGT
AGCACACGTCTGAACTCTTTCCCTACACGACGCTCTTCCGATCTNNNCTTTAGAGTCTCCAACACCAGC
AGCACACGTCTGAACTCTTTCCCTACACGACGCTCTTCCGATCTNNNCTTTAGAGTCTCCAACACCAGC
AGCACACGTCTGAACTCTTTCCCTACACGACGCTCTTCCGATCTNNNATGGTCTTCAGTTGCCTATGCT
AGCACACGTCTGAACTCTTTCCCTACACGACGCTCTTCCGATCTNNNATGGTCTTCAGTTGCCTATGCT
GAGCACACGTCTGAACTCTTTCCCTACACGACGCTCTTCCGATCTNNNTTTGAGGGCAAAAGAAGTTT
GAGCACACGTCTGAACTCTTTCCCTACACGACGCTCTTCCGATCTNNNTTTGAGGGCAAAAGAAGTTT
GAGCACACGTCTGAACTCTTTCCCTACACGACGCTCTTCCGATCTNNNCTGGCAAGAAAAACATCTCCA
GAGCACACGTCTGAACTCTTTCCCTACACGACGCTCTTCCGATCTNNNCTGGCAAGAAAAACATCTCCA
GCACACGTCTGAACTCTTTCCCTACACGACGCTCTTCCGATCTNNNGTTCAGTACACACACACACACAG
GCACACGTCTGAACTCTTTCCCTACACGACGCTCTTCCGATCTNNNGTTCAGTACACACACACACACAG
GAGCACACGTCTGAACTCTTTCCCTACACGACGCTCTTCCGATCTNNNCCAGGTACAGTCTTGAAGCAG
GAGCACACGTCTGAACTCTTTCCCTACACGACGCTCTTCCGATCTNNNCCAGGTACAGTCTTGAAGCAG
AGCACACGTCTGAACTCTTTCCCTACACGACGCTCTTCCGATCTNNNAAAGCATGAAGAGTCCCAGAA
AGCACACGTCTGAACTCTTTCCCTACACGACGCTCTTCCGATCTNNNAAAGCATGAAGAGTCCCAGAA
AGCACACGTCTGAACTCTTTCCCTACACGACGCTCTTCCGATCTNNNATCTACACCCTACTGCTGACCAA
AGCACACGTCTGAACTCTTTCCCTACACGACGCTCTTCCGATCTNNNATCTACACCCTACTGCTGACCAA
AGCACACGTCTGAACTCTTTCCCTACACGACGCTCTTCCGATCTNNNGAAAGCACCAGGCTTCAAAATC
AGCACACGTCTGAACTCTTTCCCTACACGACGCTCTTCCGATCTNNNGAAAGCACCAGGCTTCAAAATC
AGCACACGTCTGAACTCTTTCCCTACACGACGCTCTTCCGATCTNNNTTCACCTTTGGAGGATACTAG
AGCACACGTCTGAACTCTTTCCCTACACGACGCTCTTCCGATCTNNNTTCACCTTTGGAGGATACTAG
AGCACACGTCTGAACTCTTTCCCTACACGACGCTCTTCCGATCTNNNTGTGGGGCATTTGTTCATAGA
AGCACACGTCTGAACTCTTTCCCTACACGACGCTCTTCCGATCTNNNTGTGGGGCATTTGTTCATAGA
AGCACACGTCTGAACTCTTTCCCTACACGACGCTCTTCCGATCTNNNCAAACAGTTGAGGAGTAGCAC
AGCACACGTCTGAACTCTTTCCCTACACGACGCTCTTCCGATCTNNNCAAACAGTTGAGGAGTAGCAC
AGCACACGTCTGAACTCTTTCCCTACACGACGCTCTTCCGATCTNNNGAGCTTGCAGTGAGTCAAGAT
AGCACACGTCTGAACTCTTTCCCTACACGACGCTCTTCCGATCTNNNGAGCTTGCAGTGAGTCAAGAT
GAGCACACGTCTGAACTCTTTCCCTACACGACGCTCTTCCGATCTNNNCTGTGAAAGCTGCAGAGATG
GAGCACACGTCTGAACTCTTTCCCTACACGACGCTCTTCCGATCTNNNCTGTGAAAGCTGCAGAGATG

AGCACACGTCTGAACTCTTTCCCTACACGACGCTCTTCCGATCTNNNATGTGGAAGATTGCAGGACTT
AGCACACGTCTGAACTCTTTCCCTACACGACGCTCTTCCGATCTNNNATGTGGAAGATTGCAGGACTT
AGCACACGTCTGAACTCTTTCCCTACACGACGCTCTTCCGATCTNNNCCCAACCACACATGAAATTTCT
AGCACACGTCTGAACTCTTTCCCTACACGACGCTCTTCCGATCTNNNCCCAACCACACATGAAATTTCT
GAGCACACGTCTGAACTCTTTCCCTACACGACGCTCTTCCGATCTNNNTGATGGGAGGAACCAATGA
GAGCACACGTCTGAACTCTTTCCCTACACGACGCTCTTCCGATCTNNNTGATGGGAGGAACCAATGA
AGCACACGTCTGAACTCTTTCCCTACACGACGCTCTTCCGATCTNNNAGGTAAGCTCCGAAAACCTCA
AGCACACGTCTGAACTCTTTCCCTACACGACGCTCTTCCGATCTNNNAGGTAAGCTCCGAAAACCTCA
AGCACACGTCTGAACTCTTTCCCTACACGACGCTCTTCCGATCTNNNAGACGCATGGAAGAAGAAGA
AGCACACGTCTGAACTCTTTCCCTACACGACGCTCTTCCGATCTNNNAGACGCATGGAAGAAGAAGA
AGCACACGTCTGAACTCTTTCCCTACACGACGCTCTTCCGATCTNNNAAGGAGAAGGAGAGGAAGA
AGCACACGTCTGAACTCTTTCCCTACACGACGCTCTTCCGATCTNNNAAGGAGAAGGAGAGGAAGA
GAGCACACGTCTGAACTCTTTCCCTACACGACGCTCTTCCGATCTNNNCAATGTACAGTCTTTGCCACG
GAGCACACGTCTGAACTCTTTCCCTACACGACGCTCTTCCGATCTNNNCAATGTACAGTCTTTGCCACG
GCACACGTCTGAACTCTTTCCCTACACGACGCTCTTCCGATCTNNNATTTGAAGGCCAGTTTGTGTG
GCACACGTCTGAACTCTTTCCCTACACGACGCTCTTCCGATCTNNNATTTGAAGGCCAGTTTGTGTG
AGAGCACACGTCTGAACTCTTTCCCTACACGACGCTCTTCCGATCTNNNCTGTAATCCCAGCTACTCAG
AGAGCACACGTCTGAACTCTTTCCCTACACGACGCTCTTCCGATCTNNNCTGTAATCCCAGCTACTCAG
AGCACACGTCTGAACTCTTTCCCTACACGACGCTCTTCCGATCTNNNGGCAACATGGTGAAATCCCAT
AGCACACGTCTGAACTCTTTCCCTACACGACGCTCTTCCGATCTNNNGGCAACATGGTGAAATCCCAT
AGAGCACACGTCTGAACTCTTTCCCTACACGACGCTCTTCCGATCTNNNTTGGTATGGCTTCTTGCTC
AGAGCACACGTCTGAACTCTTTCCCTACACGACGCTCTTCCGATCTNNNTTGGTATGGCTTCTTGCTC
GAGCACACGTCTGAACTCTTTCCCTACACGACGCTCTTCCGATCTNNNGTCAGGAGTTTCAGACCAGC
GAGCACACGTCTGAACTCTTTCCCTACACGACGCTCTTCCGATCTNNNGTCAGGAGTTTCAGACCAGC
AGCACACGTCTGAACTCTTTCCCTACACGACGCTCTTCCGATCTNNNCTCAGCCTACCAAAGTGCTAG
AGCACACGTCTGAACTCTTTCCCTACACGACGCTCTTCCGATCTNNNCTCAGCCTACCAAAGTGCTAG
AGAGCACACGTCTGAACTCTTTCCCTACACGACGCTCTTCCGATCTNNNGAGATGGTGGCAGTACAG
AGAGCACACGTCTGAACTCTTTCCCTACACGACGCTCTTCCGATCTNNNGAGATGGTGGCAGTACAG
GAGCACACGTCTGAACTCTTTCCCTACACGACGCTCTTCCGATCTNNNGCAGGAGAATCATTTGAACC
GAGCACACGTCTGAACTCTTTCCCTACACGACGCTCTTCCGATCTNNNGCAGGAGAATCATTTGAACC
AGCACACGTCTGAACTCTTTCCCTACACGACGCTCTTCCGATCTNNNCCTGCCTCATCCTCTGTTTTACT
AGCACACGTCTGAACTCTTTCCCTACACGACGCTCTTCCGATCTNNNCCTGCCTCATCCTCTGTTTTACT
AGCACACGTCTGAACTCTTTCCCTACACGACGCTCTTCCGATCTNNNCTAGATAGCACCACTGCAGTC
AGCACACGTCTGAACTCTTTCCCTACACGACGCTCTTCCGATCTNNNCTAGATAGCACCACTGCAGTC
GCACACGTCTGAACTCTTTCCCTACACGACGCTCTTCCGATCTNNNGGAATCTGCCCTGTGTCGTCCAT
GCACACGTCTGAACTCTTTCCCTACACGACGCTCTTCCGATCTNNNGGAATCTGCCCTGTGTCGTCCAT
AGCACACGTCTGAACTCTTTCCCTACACGACGCTCTTCCGATCTNNNCTGAGGCAGGAGAATCACTT
AGCACACGTCTGAACTCTTTCCCTACACGACGCTCTTCCGATCTNNNCTGAGGCAGGAGAATCACTT
AGCACACGTCTGAACTCTTTCCCTACACGACGCTCTTCCGATCTNNNCAAGCCCAAATCCAGCATTCC
AGCACACGTCTGAACTCTTTCCCTACACGACGCTCTTCCGATCTNNNCAAGCCCAAATCCAGCATTCC
GAGCACACGTCTGAACTCTTTCCCTACACGACGCTCTTCCGATCTNNNAACTCTTGCTTTATTTGGGTG
GAGCACACGTCTGAACTCTTTCCCTACACGACGCTCTTCCGATCTNNNAACTCTTGCTTTATTTGGGTG
AGCACACGTCTGAACTCTTTCCCTACACGACGCTCTTCCGATCTNNNGAGAATCCTTTGAACCCAGG

GAGCACACGTCTGAACTCTTTCCCTACACGACGCTCTTCCGATCTNNNGAGAATCCTTTGAACCCAGGA
GCACACGTCTGAACTCTTTCCCTACACGACGCTCTTCCGATCTNNNGCTAAAGTAAAAAAACAAGGTTT
GCACACGTCTGAACTCTTTCCCTACACGACGCTCTTCCGATCTNNNGCTAAAGTAAAAAAACAAGGTTT
AGCACACGTCTGAACTCTTTCCCTACACGACGCTCTTCCGATCTNNNGGGAAACAGAGAGAGACTCCA
AGCACACGTCTGAACTCTTTCCCTACACGACGCTCTTCCGATCTNNNGGGAAACAGAGAGAGACTCCA
AGCACACGTCTGAACTCTTTCCCTACACGACGCTCTTCCGATCTNNNAAAATTTAGCCAGGCGTGGTA
AGCACACGTCTGAACTCTTTCCCTACACGACGCTCTTCCGATCTNNNAAAATTTAGCCAGGCGTGGTA
GAGCACACGTCTGAACTCTTTCCCTACACGACGCTCTTCCGATCTNNNTGGATGTTTCATGGCATCCTC
GAGCACACGTCTGAACTCTTTCCCTACACGACGCTCTTCCGATCTNNNTGGATGTTTCATGGCATCCTC
AGCACACGTCTGAACTCTTTCCCTACACGACGCTCTTCCGATCTNNNTGGTTGGAATTTTTATGGTCGG
AGCACACGTCTGAACTCTTTCCCTACACGACGCTCTTCCGATCTNNNTGGTTGGAATTTTTATGGTCGG
GAGCACACGTCTGAACTCTTTCCCTACACGACGCTCTTCCGATCTNNNGGATCACTTGAACTTGGGAGC
GAGCACACGTCTGAACTCTTTCCCTACACGACGCTCTTCCGATCTNNNGGATCACTTGAACTTGGGAGC
AGCACACGTCTGAACTCTTTCCCTACACGACGCTCTTCCGATCTNNNACACCTGTAATTCCAGCTACTCG
AGCACACGTCTGAACTCTTTCCCTACACGACGCTCTTCCGATCTNNNACACCTGTAATTCCAGCTACTCG
AGCACACGTCTGAACTCTTTCCCTACACGACGCTCTTCCGATCTNNNTTTCACTTGCCTGTGTATCACCTC
AGCACACGTCTGAACTCTTTCCCTACACGACGCTCTTCCGATCTNNNTTTCACTTGCCTGTGTATCACCTC
AGCACACGTCTGAACTCTTTCCCTACACGACGCTCTTCCGATCTNNNATGTTCCAAATACTGCATGGGA
AGCACACGTCTGAACTCTTTCCCTACACGACGCTCTTCCGATCTNNNATGTTCCAAATACTGCATGGGA
GAGCACACGTCTGAACTCTTTCCCTACACGACGCTCTTCCGATCTNNNACTGCTCAGTCCTTCACTCATTT
GAGCACACGTCTGAACTCTTTCCCTACACGACGCTCTTCCGATCTNNNACTGCTCAGTCCTTCACTCATTT
GAGCACACGTCTGAACTCTTTCCCTACACGACGCTCTTCCGATCTNNNTCCTGTTTCTCCCACTGTACCAT
GAGCACACGTCTGAACTCTTTCCCTACACGACGCTCTTCCGATCTNNNTCCTGTTTCTCCCACTGTACCAT
AGCACACGTCTGAACTCTTTCCCTACACGACGCTCTTCCGATCTNNNTGCTTTATGGTGGTGGTCTAAA
AGCACACGTCTGAACTCTTTCCCTACACGACGCTCTTCCGATCTNNNTGCTTTATGGTGGTGGTCTAAA
GAGCACACGTCTGAACTCTTTCCCTACACGACGCTCTTCCGATCTNNNCCAGTTGCAGCAACAAATCCT
GAGCACACGTCTGAACTCTTTCCCTACACGACGCTCTTCCGATCTNNNCCAGTTGCAGCAACAAATCCT
GAGCACACGTCTGAACTCTTTCCCTACACGACGCTCTTCCGATCTNNNGTGTGCTCAGATATGTTGGAT
GAGCACACGTCTGAACTCTTTCCCTACACGACGCTCTTCCGATCTNNNGTGTGCTCAGATATGTTGGAT
AGAGCACACGTCTGAACTCTTTCCCTACACGACGCTCTTCCGATCTNNNACAGTGAGGCATTTTGTAAA
AGAGCACACGTCTGAACTCTTTCCCTACACGACGCTCTTCCGATCTNNNACAGTGAGGCATTTTGTAAA
GCACACGTCTGAACTCTTTCCCTACACGACGCTCTTCCGATCTNNNTCCATATACCTGAACATCGCCAGC
GCACACGTCTGAACTCTTTCCCTACACGACGCTCTTCCGATCTNNNTCCATATACCTGAACATCGCCAGC
GCACACGTCTGAACTCTTTCCCTACACGACGCTCTTCCGATCTNNNTATACCTCTCGTGCATGAAGCCTC
GCACACGTCTGAACTCTTTCCCTACACGACGCTCTTCCGATCTNNNTATACCTCTCGTGCATGAAGCCTC
AGCACACGTCTGAACTCTTTCCCTACACGACGCTCTTCCGATCTNNNCCTGTCTCTCTGGTTCTGATGAC
AGCACACGTCTGAACTCTTTCCCTACACGACGCTCTTCCGATCTNNNCCTGTCTCTCTGGTTCTGATGAC
AGCACACGTCTGAACTCTTTCCCTACACGACGCTCTTCCGATCTNNNGACTACAGGCACCTGATACCATC
AGCACACGTCTGAACTCTTTCCCTACACGACGCTCTTCCGATCTNNNGACTACAGGCACCTGATACCATC
AGCACACGTCTGAACTCTTTCCCTACACGACGCTCTTCCGATCTNNNGCTTTTATGTTCAGCCCCATCCT
AGCACACGTCTGAACTCTTTCCCTACACGACGCTCTTCCGATCTNNNGCTTTTATGTTCAGCCCCATCCT
AGCACACGTCTGAACTCTTTCCCTACACGACGCTCTTCCGATCTNNNGTCTTTTCTGAAGAAGTTGTTTC
AGCACACGTCTGAACTCTTTCCCTACACGACGCTCTTCCGATCTNNNGTCTTTTCTGAAGAAGTTGTTTC

GAGCACACGTCTGAACTCTTTCCCTACACGACGCTCTTCCGATCTNNNAGGAAGCTGAGACTGGAGA
GAGCACACGTCTGAACTCTTTCCCTACACGACGCTCTTCCGATCTNNNAGGAAGCTGAGACTGGAGA
AGCACACGTCTGAACTCTTTCCCTACACGACGCTCTTCCGATCTNNNATAACAATTGTGTGTGTGTGC
AGCACACGTCTGAACTCTTTCCCTACACGACGCTCTTCCGATCTNNNATAACAATTGTGTGTGTGTGC
GAGCACACGTCTGAACTCTTTCCCTACACGACGCTCTTCCGATCTNNNAGATTCCTTCCTTCCTTCGTTC
GAGCACACGTCTGAACTCTTTCCCTACACGACGCTCTTCCGATCTNNNAGATTCCTTCCTTCCTTCGTTC
AGCACACGTCTGAACTCTTTCCCTACACGACGCTCTTCCGATCTNNNACGGACAAATTCCTGGACACA
AGCACACGTCTGAACTCTTTCCCTACACGACGCTCTTCCGATCTNNNACGGACAAATTCCTGGACACA
AGCACACGTCTGAACTCTTTCCCTACACGACGCTCTTCCGATCTNNNGCAAAGCATTCAAGATGTGAC
AGCACACGTCTGAACTCTTTCCCTACACGACGCTCTTCCGATCTNNNGCAAAGCATTCAAGATGTGAC
GCACACGTCTGAACTCTTTCCCTACACGACGCTCTTCCGATCTNNNCTTGGTGAACCTTAGTACCTCAT
GCACACGTCTGAACTCTTTCCCTACACGACGCTCTTCCGATCTNNNCTTGGTGAACCTTAGTACCTCAT
AGCACACGTCTGAACTCTTTCCCTACACGACGCTCTTCCGATCTNNNAGAACACTCTTGCTCTGTGTCA
AGCACACGTCTGAACTCTTTCCCTACACGACGCTCTTCCGATCTNNNAGAACACTCTTGCTCTGTGTCA
GAGCACACGTCTGAACTCTTTCCCTACACGACGCTCTTCCGATCTNNNTTGGTTTTGTTTTGTTGTGG
GAGCACACGTCTGAACTCTTTCCCTACACGACGCTCTTCCGATCTNNNTTGGTTTTGTTTTGTTGTGG
AGCACACGTCTGAACTCTTTCCCTACACGACGCTCTTCCGATCTNNNTGTCATGGGGTTTGTTGTACA
AGCACACGTCTGAACTCTTTCCCTACACGACGCTCTTCCGATCTNNNTGTCATGGGGTTTGTTGTACA
AGCACACGTCTGAACTCTTTCCCTACACGACGCTCTTCCGATCTNNNGAGAGACTGCTCCTCTGTGCC
AGCACACGTCTGAACTCTTTCCCTACACGACGCTCTTCCGATCTNNNGAGAGACTGCTCCTCTGTGCC
GAGCACACGTCTGAACTCTTTCCCTACACGACGCTCTTCCGATCTNNNAGAATTGCTTGAATCTGGGA
GAGCACACGTCTGAACTCTTTCCCTACACGACGCTCTTCCGATCTNNNAGAATTGCTTGAATCTGGGA
AGCACACGTCTGAACTCTTTCCCTACACGACGCTCTTCCGATCTNNNTGTTGAGTGAAAGAAGCCAGT
AGCACACGTCTGAACTCTTTCCCTACACGACGCTCTTCCGATCTNNNTGTTGAGTGAAAGAAGCCAGT
AGCACACGTCTGAACTCTTTCCCTACACGACGCTCTTCCGATCTNNNTGACTCCACCATACTACTTCCA
AGCACACGTCTGAACTCTTTCCCTACACGACGCTCTTCCGATCTNNNTGACTCCACCATACTACTTCCA
AGCACACGTCTGAACTCTTTCCCTACACGACGCTCTTCCGATCTNNNTTTATCCATGTCCCTGCAAAGG
AGCACACGTCTGAACTCTTTCCCTACACGACGCTCTTCCGATCTNNNTTTATCCATGTCCCTGCAAAGG
GAGCACACGTCTGAACTCTTTCCCTACACGACGCTCTTCCGATCTNNNAGAAGTAAAATGAGAGGGG
GAGCACACGTCTGAACTCTTTCCCTACACGACGCTCTTCCGATCTNNNAGAAGTAAAATGAGAGGGG
GAGCACACGTCTGAACTCTTTCCCTACACGACGCTCTTCCGATCTNNNAAAACAGTGAAATGCTGTCC
GAGCACACGTCTGAACTCTTTCCCTACACGACGCTCTTCCGATCTNNNAAAACAGTGAAATGCTGTCC
AGCACACGTCTGAACTCTTTCCCTACACGACGCTCTTCCGATCTNNNTGGCCTACAGGTCTTTGCATAT
AGCACACGTCTGAACTCTTTCCCTACACGACGCTCTTCCGATCTNNNTGGCCTACAGGTCTTTGCATAT
GCACACGTCTGAACTCTTTCCCTACACGACGCTCTTCCGATCTNNNATGGAGTCTGTCTGTCTGTTCTG
GCACACGTCTGAACTCTTTCCCTACACGACGCTCTTCCGATCTNNNATGGAGTCTGTCTGTCTGTTCTG
GAGCACACGTCTGAACTCTTTCCCTACACGACGCTCTTCCGATCTNNNAGAGAGGAGAGAGGGAAA
GAGCACACGTCTGAACTCTTTCCCTACACGACGCTCTTCCGATCTNNNAGAGAGGAGAGAGGGAAA
GAGCACACGTCTGAACTCTTTCCCTACACGACGCTCTTCCGATCTNNNGTGCCAGGAAAGATGAGACA
GAGCACACGTCTGAACTCTTTCCCTACACGACGCTCTTCCGATCTNNNGTGCCAGGAAAGATGAGACA
GAGCACACGTCTGAACTCTTTCCCTACACGACGCTCTTCCGATCTNNNCGTGGAATCTCGGGGTCTAT
GAGCACACGTCTGAACTCTTTCCCTACACGACGCTCTTCCGATCTNNNCGTGGAATCTCGGGGTCTAT
GCACACGTCTGAACTCTTTCCCTACACGACGCTCTTCCGATCTNNNAAAAAATCTGCTCTGCTCTGGTG

GCACACGTCTGAACTCTTTCCCTACACGACGCTCTTCCGATCTNNNAAAAAATCTGCTCTGCTCTGGT
AGCACACGTCTGAACTCTTTCCCTACACGACGCTCTTCCGATCTNNNTGATGCACACCTGTAATTCCAG
AGCACACGTCTGAACTCTTTCCCTACACGACGCTCTTCCGATCTNNNTGATGCACACCTGTAATTCCAG
GCACACGTCTGAACTCTTTCCCTACACGACGCTCTTCCGATCTNNNGATGAGTGTGTACGTCGTCGAG
GCACACGTCTGAACTCTTTCCCTACACGACGCTCTTCCGATCTNNNGATGAGTGTGTACGTCGTCGAG
AGCACACGTCTGAACTCTTTCCCTACACGACGCTCTTCCGATCTNNNGCCCAGATTGTAGATTTGTGA
AGCACACGTCTGAACTCTTTCCCTACACGACGCTCTTCCGATCTNNNGCCCAGATTGTAGATTTGTGA
AGCACACGTCTGAACTCTTTCCCTACACGACGCTCTTCCGATCTNNNAGATTGCAGTGAGCTGAGAT
AGCACACGTCTGAACTCTTTCCCTACACGACGCTCTTCCGATCTNNNAGATTGCAGTGAGCTGAGAT
GAGCACACGTCTGAACTCTTTCCCTACACGACGCTCTTCCGATCTNNNCCTGGGAGAAACAGATGGT
GAGCACACGTCTGAACTCTTTCCCTACACGACGCTCTTCCGATCTNNNCCTGGGAGAAACAGATGGT
AGCACACGTCTGAACTCTTTCCCTACACGACGCTCTTCCGATCTNNNATCACGCAGATGAAGAGAGG
AGCACACGTCTGAACTCTTTCCCTACACGACGCTCTTCCGATCTNNNATCACGCAGATGAAGAGAGG
GCACACGTCTGAACTCTTTCCCTACACGACGCTCTTCCGATCTNNNCAGATGGTATTTTAAGATTCCT
GCACACGTCTGAACTCTTTCCCTACACGACGCTCTTCCGATCTNNNCAGATGGTATTTTAAGATTCCT
GAGCACACGTCTGAACTCTTTCCCTACACGACGCTCTTCCGATCTNNNTCTCCTTTCAGGGTGGCAAG
GAGCACACGTCTGAACTCTTTCCCTACACGACGCTCTTCCGATCTNNNTCTCCTTTCAGGGTGGCAAG
AGCACACGTCTGAACTCTTTCCCTACACGACGCTCTTCCGATCTNNNCTGAGGCAGGAGAATCACTTG
AGCACACGTCTGAACTCTTTCCCTACACGACGCTCTTCCGATCTNNNCTGAGGCAGGAGAATCACTTG
GCACACGTCTGAACTCTTTCCCTACACGACGCTCTTCCGATCTNNNTTGGTGACATGTTGGCTTGATT
GCACACGTCTGAACTCTTTCCCTACACGACGCTCTTCCGATCTNNNTTGGTGACATGTTGGCTTGATT
GAGCACACGTCTGAACTCTTTCCCTACACGACGCTCTTCCGATCTNNNGCTGTCTGCATCTTGGTAAA
GAGCACACGTCTGAACTCTTTCCCTACACGACGCTCTTCCGATCTNNNGCTGTCTGCATCTTGGTAAA
AGCACACGTCTGAACTCTTTCCCTACACGACGCTCTTCCGATCTNNNAGGATTACAACTCTCCTGCTTT
AGCACACGTCTGAACTCTTTCCCTACACGACGCTCTTCCGATCTNNNAGGATTACAACTCTCCTGCTTT
AGCACACGTCTGAACTCTTTCCCTACACGACGCTCTTCCGATCTNNNAATGTCTGGCTTACTGTCAAC
AGCACACGTCTGAACTCTTTCCCTACACGACGCTCTTCCGATCTNNNAATGTCTGGCTTACTGTCAAC
AGCACACGTCTGAACTCTTTCCCTACACGACGCTCTTCCGATCTNNNATTACAAGTGCTCACCACCACT
AGCACACGTCTGAACTCTTTCCCTACACGACGCTCTTCCGATCTNNNATTACAAGTGCTCACCACCACT
AGCACACGTCTGAACTCTTTCCCTACACGACGCTCTTCCGATCTNNNAACAGAACAGAGCCCTCAGAA
AGCACACGTCTGAACTCTTTCCCTACACGACGCTCTTCCGATCTNNNAACAGAACAGAGCCCTCAGAA
AGCACACGTCTGAACTCTTTCCCTACACGACGCTCTTCCGATCTNNNAGACATGGACACAGTACATCCA
AGCACACGTCTGAACTCTTTCCCTACACGACGCTCTTCCGATCTNNNAGACATGGACACAGTACATCCA
AGCACACGTCTGAACTCTTTCCCTACACGACGCTCTTCCGATCTNNNGATTCTAAGGGGTGAGCATCC
AGCACACGTCTGAACTCTTTCCCTACACGACGCTCTTCCGATCTNNNGATTCTAAGGGGTGAGCATCC
GCACACGTCTGAACTCTTTCCCTACACGACGCTCTTCCGATCTNNNACCACAAAGCAGATAATAAGCC
GCACACGTCTGAACTCTTTCCCTACACGACGCTCTTCCGATCTNNNACCACAAAGCAGATAATAAGCC
AGCACACGTCTGAACTCTTTCCCTACACGACGCTCTTCCGATCTNNNTGGCACTATCACAGATTACTG
AGCACACGTCTGAACTCTTTCCCTACACGACGCTCTTCCGATCTNNNTGGCACTATCACAGATTACTG
AGCACACGTCTGAACTCTTTCCCTACACGACGCTCTTCCGATCTNNNTTACTGAATGTTAGCTGGTCCC
AGCACACGTCTGAACTCTTTCCCTACACGACGCTCTTCCGATCTNNNTTACTGAATGTTAGCTGGTCCC
GAGCACACGTCTGAACTCTTTCCCTACACGACGCTCTTCCGATCTNNNTGTACCTACTTTCCCTCTGTCC
GAGCACACGTCTGAACTCTTTCCCTACACGACGCTCTTCCGATCTNNNTGTACCTACTTTCCCTCTGTCC

AGAGCACACGTCTGAACTCTTTCCCTACACGACGCTCTTCCGATCTNNNGACAGACAATGCAGTAAGCT
AGAGCACACGTCTGAACTCTTTCCCTACACGACGCTCTTCCGATCTNNNGACAGACAATGCAGTAAGCT
AAGAGCACACGTCTGAACTCTTTCCCTACACGACGCTCTTCCGATCTNNNGTAGAGGCAGAGAGAACA
AAGAGCACACGTCTGAACTCTTTCCCTACACGACGCTCTTCCGATCTNNNGTAGAGGCAGAGAGAACA
GAGCACACGTCTGAACTCTTTCCCTACACGACGCTCTTCCGATCTNNNGTTTAGAATGTCCTTGTGTCG
GAGCACACGTCTGAACTCTTTCCCTACACGACGCTCTTCCGATCTNNNGTTTAGAATGTCCTTGTGTCG
GAGCACACGTCTGAACTCTTTCCCTACACGACGCTCTTCCGATCTNNNGACCCAAGTACAGACACAGAT
GAGCACACGTCTGAACTCTTTCCCTACACGACGCTCTTCCGATCTNNNGACCCAAGTACAGACACAGAT
AGCACACGTCTGAACTCTTTCCCTACACGACGCTCTTCCGATCTNNNTGTATGTGTTCCTGACCTCACTG
AGCACACGTCTGAACTCTTTCCCTACACGACGCTCTTCCGATCTNNNTGTATGTGTTCCTGACCTCACTG
AGAGCACACGTCTGAACTCTTTCCCTACACGACGCTCTTCCGATCTNNNGATCACTTGAGCTCAGGAGT
AGAGCACACGTCTGAACTCTTTCCCTACACGACGCTCTTCCGATCTNNNGATCACTTGAGCTCAGGAGT
GAGCACACGTCTGAACTCTTTCCCTACACGACGCTCTTCCGATCTNNNCAGAGTGAGACTGACACACA
GAGCACACGTCTGAACTCTTTCCCTACACGACGCTCTTCCGATCTNNNCAGAGTGAGACTGACACACA
AGCACACGTCTGAACTCTTTCCCTACACGACGCTCTTCCGATCTNNNTATCTTGCTTTCAGTCCCCATG
AGCACACGTCTGAACTCTTTCCCTACACGACGCTCTTCCGATCTNNNTATCTTGCTTTCAGTCCCCATG
AGCACACGTCTGAACTCTTTCCCTACACGACGCTCTTCCGATCTNNNCTCCAGGCAGTAAGAAAAACGT
AGCACACGTCTGAACTCTTTCCCTACACGACGCTCTTCCGATCTNNNCTCCAGGCAGTAAGAAAAACGT
CACACGTCTGAACTCTTTCCCTACACGACGCTCTTCCGATCTNNNAATGAGCTGTTGAGTACACCTCCC
CACACGTCTGAACTCTTTCCCTACACGACGCTCTTCCGATCTNNNAATGAGCTGTTGAGTACACCTCCC
GAGCACACGTCTGAACTCTTTCCCTACACGACGCTCTTCCGATCTNNNAAGCTGAAGCAGGAAAATTG
GAGCACACGTCTGAACTCTTTCCCTACACGACGCTCTTCCGATCTNNNAAGCTGAAGCAGGAAAATTG
AAGAGCACACGTCTGAACTCTTTCCCTACACGACGCTCTTCCGATCTNNNAATCCCAGCTACTTGAGAG
AAGAGCACACGTCTGAACTCTTTCCCTACACGACGCTCTTCCGATCTNNNAATCCCAGCTACTTGAGAG
AGCACACGTCTGAACTCTTTCCCTACACGACGCTCTTCCGATCTNNNTTGTTGAGATTTGCCTTGTGGA
AGCACACGTCTGAACTCTTTCCCTACACGACGCTCTTCCGATCTNNNTTGTTGAGATTTGCCTTGTGGA
GAGCACACGTCTGAACTCTTTCCCTACACGACGCTCTTCCGATCTNNNCGACAAGTGCGAAACTCCATC
GAGCACACGTCTGAACTCTTTCCCTACACGACGCTCTTCCGATCTNNNCGACAAGTGCGAAACTCCATC
GCACACGTCTGAACTCTTTCCCTACACGACGCTCTTCCGATCTNNNTTGCTCAATTTTTCATTTGGTGTT
GCACACGTCTGAACTCTTTCCCTACACGACGCTCTTCCGATCTNNNTTGCTCAATTTTTCATTTGGTGTT
GAGCACACGTCTGAACTCTTTCCCTACACGACGCTCTTCCGATCTNNNGATGCTGTCTGAGTCTGTTCT
GAGCACACGTCTGAACTCTTTCCCTACACGACGCTCTTCCGATCTNNNGATGCTGTCTGAGTCTGTTCT
AGCACACGTCTGAACTCTTTCCCTACACGACGCTCTTCCGATCTNNNAGGTCCACAAGCATCTAACAGA
AGCACACGTCTGAACTCTTTCCCTACACGACGCTCTTCCGATCTNNNAGGTCCACAAGCATCTAACAGA
GAGCACACGTCTGAACTCTTTCCCTACACGACGCTCTTCCGATCTNNNAGACCTATAATCTGTTCCCCCT
GAGCACACGTCTGAACTCTTTCCCTACACGACGCTCTTCCGATCTNNNAGACCTATAATCTGTTCCCCCT
GCACACGTCTGAACTCTTTCCCTACACGACGCTCTTCCGATCTNNNCTTCACATAAAGGAACATGGTTCA
GCACACGTCTGAACTCTTTCCCTACACGACGCTCTTCCGATCTNNNCTTCACATAAAGGAACATGGTTCA
GAGCACACGTCTGAACTCTTTCCCTACACGACGCTCTTCCGATCTNNNTGGACGTTGCTGGTGTATAGA
GAGCACACGTCTGAACTCTTTCCCTACACGACGCTCTTCCGATCTNNNTGGACGTTGCTGGTGTATAGA
AGCACACGTCTGAACTCTTTCCCTACACGACGCTCTTCCGATCTNNNACTTTCTGCTACCACACAAATGC
AGCACACGTCTGAACTCTTTCCCTACACGACGCTCTTCCGATCTNNNACTTTCTGCTACCACACAAATGC
AGAGCACACGTCTGAACTCTTTCCCTACACGACGCTCTTCCGATCTNNNTGGGTACACTGAACAAACCA

AGAGCACACGTCTGAACTCTTTCCCTACACGACGCTCTTCCGATCTNNNTGGGTACACTGAACAAACCA
GAGCACACGTCTGAACTCTTTCCCTACACGACGCTCTTCCGATCTNNNCACCTCAATCTCCTGAGTAGCT
GAGCACACGTCTGAACTCTTTCCCTACACGACGCTCTTCCGATCTNNNCACCTCAATCTCCTGAGTAGCT
GAGCACACGTCTGAACTCTTTCCCTACACGACGCTCTTCCGATCTNNNACGCCTGTAATCCCAGCTATTT
GAGCACACGTCTGAACTCTTTCCCTACACGACGCTCTTCCGATCTNNNACGCCTGTAATCCCAGCTATTT
GAGCACACGTCTGAACTCTTTCCCTACACGACGCTCTTCCGATCTNNNCAAATCCAAGAGCAAGGCAGA
GAGCACACGTCTGAACTCTTTCCCTACACGACGCTCTTCCGATCTNNNCAAATCCAAGAGCAAGGCAGA
GAGCACACGTCTGAACTCTTTCCCTACACGACGCTCTTCCGATCTNNNTGAGTATGAGACCACCTCCTTC
GAGCACACGTCTGAACTCTTTCCCTACACGACGCTCTTCCGATCTNNNTGAGTATGAGACCACCTCCTTC
GAGCACACGTCTGAACTCTTTCCCTACACGACGCTCTTCCGATCTNNNTGGATAGCACTCATAATGTGG
GAGCACACGTCTGAACTCTTTCCCTACACGACGCTCTTCCGATCTNNNTGGATAGCACTCATAATGTGG
GAGCACACGTCTGAACTCTTTCCCTACACGACGCTCTTCCGATCTNNNGAGCTTGCAGTGAGCCAAGA
GAGCACACGTCTGAACTCTTTCCCTACACGACGCTCTTCCGATCTNNNGAGCTTGCAGTGAGCCAAGA
AGAGCACACGTCTGAACTCTTTCCCTACACGACGCTCTTCCGATCTNNNAAATAATCCTCCAGGCTCAG
AGAGCACACGTCTGAACTCTTTCCCTACACGACGCTCTTCCGATCTNNNAAATAATCCTCCAGGCTCAG
AGCACACGTCTGAACTCTTTCCCTACACGACGCTCTTCCGATCTNNNGAGTCTTGCTCTGTTGCCCAGC
AGCACACGTCTGAACTCTTTCCCTACACGACGCTCTTCCGATCTNNNGAGTCTTGCTCTGTTGCCCAGC
AGAGCACACGTCTGAACTCTTTCCCTACACGACGCTCTTCCGATCTNNNAGAGCTAGACCCTGTCTCTGA
AGAGCACACGTCTGAACTCTTTCCCTACACGACGCTCTTCCGATCTNNNAGAGCTAGACCCTGTCTCTGA
GAGCACACGTCTGAACTCTTTCCCTACACGACGCTCTTCCGATCTNNNTCAGGTGATTCTGTCAGCCTCT
GAGCACACGTCTGAACTCTTTCCCTACACGACGCTCTTCCGATCTNNNTCAGGTGATTCTGTCAGCCTCT
GAGCACACGTCTGAACTCTTTCCCTACACGACGCTCTTCCGATCTNNNTCTCCAGTGAGACCAGCAGATA
GAGCACACGTCTGAACTCTTTCCCTACACGACGCTCTTCCGATCTNNNTCTCCAGTGAGACCAGCAGATA
GAGCACACGTCTGAACTCTTTCCCTACACGACGCTCTTCCGATCTNNNTCACCATTGCACTCCAATCTA
GAGCACACGTCTGAACTCTTTCCCTACACGACGCTCTTCCGATCTNNNTCACCATTGCACTCCAATCTA
AGCACACGTCTGAACTCTTTCCCTACACGACGCTCTTCCGATCTNNNTCATACATTAAGCCGCAAGCAGT
AGCACACGTCTGAACTCTTTCCCTACACGACGCTCTTCCGATCTNNNTCATACATTAAGCCGCAAGCAGT
GCACACGTCTGAACTCTTTCCCTACACGACGCTCTTCCGATCTNNNCAGGAAACTTACAATCGTGGCAC
GCACACGTCTGAACTCTTTCCCTACACGACGCTCTTCCGATCTNNNCAGGAAACTTACAATCGTGGCAC
GCACACGTCTGAACTCTTTCCCTACACGACGCTCTTCCGATCTNNNAGGCTGATCTGTGAGGAAATGG
GCACACGTCTGAACTCTTTCCCTACACGACGCTCTTCCGATCTNNNAGGCTGATCTGTGAGGAAATGG
GAGCACACGTCTGAACTCTTTCCCTACACGACGCTCTTCCGATCTNNNTATTGTTCAGACCTCAGCTCCT
GAGCACACGTCTGAACTCTTTCCCTACACGACGCTCTTCCGATCTNNNTATTGTTCAGACCTCAGCTCCT
GAGCACACGTCTGAACTCTTTCCCTACACGACGCTCTTCCGATCTNNNTGGGCAACAGAATAAGACTGT
GAGCACACGTCTGAACTCTTTCCCTACACGACGCTCTTCCGATCTNNNTGGGCAACAGAATAAGACTGT
GAGCACACGTCTGAACTCTTTCCCTACACGACGCTCTTCCGATCTNNNCTCTGTCTCTGTCTCTCTCTCCA
GAGCACACGTCTGAACTCTTTCCCTACACGACGCTCTTCCGATCTNNNCTCTGTCTCTGTCTCTCTCTCCA
GAGCACACGTCTGAACTCTTTCCCTACACGACGCTCTTCCGATCTNNNATCAGAATTAAGACTAGAGAT
GAGCACACGTCTGAACTCTTTCCCTACACGACGCTCTTCCGATCTNNNATCAGAATTAAGACTAGAGAT
GAGCACACGTCTGAACTCTTTCCCTACACGACGCTCTTCCGATCTNNNATTCTACCCACATAAAGCTGCCA
GAGCACACGTCTGAACTCTTTCCCTACACGACGCTCTTCCGATCTNNNATTCTACCCACATAAAGCTGCCA
AGCACACGTCTGAACTCTTTCCCTACACGACGCTCTTCCGATCTNNNGACTAGAGGGAAGAAGAAGTG
AGCACACGTCTGAACTCTTTCCCTACACGACGCTCTTCCGATCTNNNGACTAGAGGGAAGAAGAAGTG

CACACGTCTGAACTCTTTCCCTACACGACGCTCTTCCGATCTNNNGAGATGACAAAAATGAGACACC
CACACGTCTGAACTCTTTCCCTACACGACGCTCTTCCGATCTNNNGAGATGACAAAAATGAGACACC
GCACACGTCTGAACTCTTTCCCTACACGACGCTCTTCCGATCTNNNATCAAACCCCTGGACCAACTAA
GCACACGTCTGAACTCTTTCCCTACACGACGCTCTTCCGATCTNNNATCAAACCCCTGGACCAACTAA
AGCACACGTCTGAACTCTTTCCCTACACGACGCTCTTCCGATCTNNNACCAGGACTCAAACTCAAATCC
AGCACACGTCTGAACTCTTTCCCTACACGACGCTCTTCCGATCTNNNACCAGGACTCAAACTCAAATCC
CACACGTCTGAACTCTTTCCCTACACGACGCTCTTCCGATCTNNNCTTTGCTGCTGTCCACTATGATTC
CACACGTCTGAACTCTTTCCCTACACGACGCTCTTCCGATCTNNNCTTTGCTGCTGTCCACTATGATTC
AGCACACGTCTGAACTCTTTCCCTACACGACGCTCTTCCGATCTNNNCTGTGCTGCTGGCTTTCTATTA
AGCACACGTCTGAACTCTTTCCCTACACGACGCTCTTCCGATCTNNNCTGTGCTGCTGGCTTTCTATTA
GAGCACACGTCTGAACTCTTTCCCTACACGACGCTCTTCCGATCTNNNGAGGCAGGAGAATCGCTTG
GAGCACACGTCTGAACTCTTTCCCTACACGACGCTCTTCCGATCTNNNGAGGCAGGAGAATCGCTTG
AGCACACGTCTGAACTCTTTCCCTACACGACGCTCTTCCGATCTNNNTTTCCCATTCCTGTTTTGCCTTG
AGCACACGTCTGAACTCTTTCCCTACACGACGCTCTTCCGATCTNNNTTTCCCATTCCTGTTTTGCCTTG
AGCACACGTCTGAACTCTTTCCCTACACGACGCTCTTCCGATCTNNNTGCTGCAAATGACAGAATTTC
AGCACACGTCTGAACTCTTTCCCTACACGACGCTCTTCCGATCTNNNTGCTGCAAATGACAGAATTTC
GAGCACACGTCTGAACTCTTTCCCTACACGACGCTCTTCCGATCTNNNGGAACTTGCAGTGAGCTGA
GAGCACACGTCTGAACTCTTTCCCTACACGACGCTCTTCCGATCTNNNGGAACTTGCAGTGAGCTGA
AGCACACGTCTGAACTCTTTCCCTACACGACGCTCTTCCGATCTNNNTGAATCTATGGATTGCTTTGG
AGCACACGTCTGAACTCTTTCCCTACACGACGCTCTTCCGATCTNNNTGAATCTATGGATTGCTTTGG
AGCACACGTCTGAACTCTTTCCCTACACGACGCTCTTCCGATCTNNNTTGTGGTGACCTGAGATTGTA
AGCACACGTCTGAACTCTTTCCCTACACGACGCTCTTCCGATCTNNNTTGTGGTGACCTGAGATTGTA
AGCACACGTCTGAACTCTTTCCCTACACGACGCTCTTCCGATCTNNNCTGGAGAGAGGGCAAAGATT
AGCACACGTCTGAACTCTTTCCCTACACGACGCTCTTCCGATCTNNNCTGGAGAGAGGGCAAAGATT
GCACACGTCTGAACTCTTTCCCTACACGACGCTCTTCCGATCTNNNAAAATAGAAAACAATGAATGG
GCACACGTCTGAACTCTTTCCCTACACGACGCTCTTCCGATCTNNNAAAATAGAAAACAATGAATGG
GAGCACACGTCTGAACTCTTTCCCTACACGACGCTCTTCCGATCTNNNTGAACTCCTTTAGAAATAGCA
GAGCACACGTCTGAACTCTTTCCCTACACGACGCTCTTCCGATCTNNNTGAACTCCTTTAGAAATAGCA
CACACGTCTGAACTCTTTCCCTACACGACGCTCTTCCGATCTNNNCTACTGGGAAGTGAGGAGCCCC
CACACGTCTGAACTCTTTCCCTACACGACGCTCTTCCGATCTNNNCTACTGGGAAGTGAGGAGCCCC
AGCACACGTCTGAACTCTTTCCCTACACGACGCTCTTCCGATCTNNNGTAATCCTAGCTACTTGGGAG
AGCACACGTCTGAACTCTTTCCCTACACGACGCTCTTCCGATCTNNNGTAATCCTAGCTACTTGGGAG
CACACGTCTGAACTCTTTCCCTACACGACGCTCTTCCGATCTNNNTGAGCCTTTGTAAGAGGGCAAT
CACACGTCTGAACTCTTTCCCTACACGACGCTCTTCCGATCTNNNTGAGCCTTTGTAAGAGGGCAAT
GAGCACACGTCTGAACTCTTTCCCTACACGACGCTCTTCCGATCTNNNAAGCTGGAGAGGGAAGGG
GAGCACACGTCTGAACTCTTTCCCTACACGACGCTCTTCCGATCTNNNAAGCTGGAGAGGGAAGGG
AGCACACGTCTGAACTCTTTCCCTACACGACGCTCTTCCGATCTNNNAGGACAATTGCTTGAACCTGA
AGCACACGTCTGAACTCTTTCCCTACACGACGCTCTTCCGATCTNNNAGGACAATTGCTTGAACCTGA
AGCACACGTCTGAACTCTTTCCCTACACGACGCTCTTCCGATCTNNNAAATCATGCACATCACTGACCA
AGCACACGTCTGAACTCTTTCCCTACACGACGCTCTTCCGATCTNNNAAATCATGCACATCACTGACCA
GCACACGTCTGAACTCTTTCCCTACACGACGCTCTTCCGATCTNNNATACCTGTAGTCCCAGCTTCTTG
GCACACGTCTGAACTCTTTCCCTACACGACGCTCTTCCGATCTNNNATACCTGTAGTCCCAGCTTCTTG
AGCACACGTCTGAACTCTTTCCCTACACGACGCTCTTCCGATCTNNNCTGAAATAAGGTGAGGATGC

AGCACACGTCTGAACTCTTTCCCTACACGACGCTCTTCCGATCTNNNCTGAAATAAGGTGAGGATGC
GCACACGTCTGAACTCTTTCCCTACACGACGCTCTTCCGATCTNNNAGGCAGAAGACCTACTTTTCAG
GCACACGTCTGAACTCTTTCCCTACACGACGCTCTTCCGATCTNNNAGGCAGAAGACCTACTTTTCAG
AGCACACGTCTGAACTCTTTCCCTACACGACGCTCTTCCGATCTNNNTCTGAGAAAGACGGAAAACT
AGCACACGTCTGAACTCTTTCCCTACACGACGCTCTTCCGATCTNNNTCTGAGAAAGACGGAAAACT
AGCACACGTCTGAACTCTTTCCCTACACGACGCTCTTCCGATCTNNNCCTTTCCAAGATGACGGGTTG
AGCACACGTCTGAACTCTTTCCCTACACGACGCTCTTCCGATCTNNNCCTTTCCAAGATGACGGGTTG
AGCACACGTCTGAACTCTTTCCCTACACGACGCTCTTCCGATCTNNNGAGAAGCAAGAAGAAACCAA
AGCACACGTCTGAACTCTTTCCCTACACGACGCTCTTCCGATCTNNNGAGAAGCAAGAAGAAACCAA
AGCACACGTCTGAACTCTTTCCCTACACGACGCTCTTCCGATCTNNNTGGGTGACAGAGCAAGATTCT
AGCACACGTCTGAACTCTTTCCCTACACGACGCTCTTCCGATCTNNNTGGGTGACAGAGCAAGATTCT
AGCACACGTCTGAACTCTTTCCCTACACGACGCTCTTCCGATCTNNNGAAAGTAAGTCCTCACCTCGTC
AGCACACGTCTGAACTCTTTCCCTACACGACGCTCTTCCGATCTNNNGAAAGTAAGTCCTCACCTCGTC
AGCACACGTCTGAACTCTTTCCCTACACGACGCTCTTCCGATCTNNNGCTGGATTTCTGGCTTTTCTGA
AGCACACGTCTGAACTCTTTCCCTACACGACGCTCTTCCGATCTNNNGCTGGATTTCTGGCTTTTCTGA
GAGCACACGTCTGAACTCTTTCCCTACACGACGCTCTTCCGATCTNNNGTAATCCCGGCTACTCAGGA
GAGCACACGTCTGAACTCTTTCCCTACACGACGCTCTTCCGATCTNNNGTAATCCCGGCTACTCAGGA
AGCACACGTCTGAACTCTTTCCCTACACGACGCTCTTCCGATCTNNNGGAGCAATTGGCTGACTTCTA
AGCACACGTCTGAACTCTTTCCCTACACGACGCTCTTCCGATCTNNNGGAGCAATTGGCTGACTTCTA
GCACACGTCTGAACTCTTTCCCTACACGACGCTCTTCCGATCTNNNGTACTCCAGCCTGGGTAAGAAA
GCACACGTCTGAACTCTTTCCCTACACGACGCTCTTCCGATCTNNNGTACTCCAGCCTGGGTAAGAAA
GCACACGTCTGAACTCTTTCCCTACACGACGCTCTTCCGATCTNNNTTCTCCCAGGGGAACATACTGT
GCACACGTCTGAACTCTTTCCCTACACGACGCTCTTCCGATCTNNNTTCTCCCAGGGGAACATACTGT
AGCACACGTCTGAACTCTTTCCCTACACGACGCTCTTCCGATCTNNNAGAATAGTTTGAACCTGGGAG
AGCACACGTCTGAACTCTTTCCCTACACGACGCTCTTCCGATCTNNNAGAATAGTTTGAACCTGGGAG
AGCACACGTCTGAACTCTTTCCCTACACGACGCTCTTCCGATCTNNNGCATTTCAGGGATAAAGCCTA
AGCACACGTCTGAACTCTTTCCCTACACGACGCTCTTCCGATCTNNNGCATTTCAGGGATAAAGCCTA
AGCACACGTCTGAACTCTTTCCCTACACGACGCTCTTCCGATCTNNNCATGCTGCTGTGCTCAACTAAT
AGCACACGTCTGAACTCTTTCCCTACACGACGCTCTTCCGATCTNNNCATGCTGCTGTGCTCAACTAAT
AGCACACGTCTGAACTCTTTCCCTACACGACGCTCTTCCGATCTNNNAGAAGGCAAATGGTATAGGC
AGCACACGTCTGAACTCTTTCCCTACACGACGCTCTTCCGATCTNNNAGAAGGCAAATGGTATAGGC
AGCACACGTCTGAACTCTTTCCCTACACGACGCTCTTCCGATCTNNNGTCACCTCTTTCTTCTTTTGGC
AGCACACGTCTGAACTCTTTCCCTACACGACGCTCTTCCGATCTNNNGTCACCTCTTTCTTCTTTTGGC
GCACACGTCTGAACTCTTTCCCTACACGACGCTCTTCCGATCTNNNGCCCTTTATCAGCATAACACCAC
GCACACGTCTGAACTCTTTCCCTACACGACGCTCTTCCGATCTNNNGCCCTTTATCAGCATAACACCAC
AGCACACGTCTGAACTCTTTCCCTACACGACGCTCTTCCGATCTNNNACTGTGTTAGTGGAATGAGTG
AGCACACGTCTGAACTCTTTCCCTACACGACGCTCTTCCGATCTNNNACTGTGTTAGTGGAATGAGTG
AGCACACGTCTGAACTCTTTCCCTACACGACGCTCTTCCGATCTNNNACGTCCCAGATATACTCTACAA
AGCACACGTCTGAACTCTTTCCCTACACGACGCTCTTCCGATCTNNNACGTCCCAGATATACTCTACAA
AGCACACGTCTGAACTCTTTCCCTACACGACGCTCTTCCGATCTNNNAGTGAGATGCCTTTTCAGTAG
AGCACACGTCTGAACTCTTTCCCTACACGACGCTCTTCCGATCTNNNAGTGAGATGCCTTTTCAGTAG
GAGCACACGTCTGAACTCTTTCCCTACACGACGCTCTTCCGATCTNNNAATGGTGTGTTTTCATCTGC
GAGCACACGTCTGAACTCTTTCCCTACACGACGCTCTTCCGATCTNNNAATGGTGTGTTTTCATCTGC

AGCACACGTCTGAACTCTTTCCCTACACGACGCTCTTCCGATCTNNNAGTGCTGGGATTATAGGCATG

AGCACACGTCTGAACTCTTTCCCTACACGACGCTCTTCCGATCTNNNAGTGCTGGGATTATAGGCATG

GAGCACACGTCTGAACTCTTTCCCTACACGACGCTCTTCCGATCTNNNATGGCATGATATGTGGCCTTT

GAGCACACGTCTGAACTCTTTCCCTACACGACGCTCTTCCGATCTNNNATGGCATGATATGTGGCCTTT

AGCACACGTCTGAACTCTTTCCCTACACGACGCTCTTCCGATCTNNNGGATTGCCTGAGGTAAGGAGT

AGCACACGTCTGAACTCTTTCCCTACACGACGCTCTTCCGATCTNNNGGATTGCCTGAGGTAAGGAGT

ACACGTCTGAACTCTTTCCCTACACGACGCTCTTCCGATCTNNNTGGTATTGAGGTTATGCCAAGCTCA

ACACGTCTGAACTCTTTCCCTACACGACGCTCTTCCGATCTNNNTGGTATTGAGGTTATGCCAAGCTCA

GAGCACACGTCTGAACTCTTTCCCTACACGACGCTCTTCCGATCTNNNGAGAGTTTCGCTGCAAAAACA

GAGCACACGTCTGAACTCTTTCCCTACACGACGCTCTTCCGATCTNNNGAGAGTTTCGCTGCAAAAACA

AGCACACGTCTGAACTCTTTCCCTACACGACGCTCTTCCGATCTNNNAAAGTGGATGGAGCATACACA

AGCACACGTCTGAACTCTTTCCCTACACGACGCTCTTCCGATCTNNNAAAGTGGATGGAGCATACACA

AGCACACGTCTGAACTCTTTCCCTACACGACGCTCTTCCGATCTNNNCTTTATTCACCCCGTTTGCCAAAC

AGCACACGTCTGAACTCTTTCCCTACACGACGCTCTTCCGATCTNNNCTTTATTCACCCCGTTTGCCAAAC

AGCACACGTCTGAACTCTTTCCCTACACGACGCTCTTCCGATCTNNNTGTCAGAACTTTGTCACCTGTCT

AGCACACGTCTGAACTCTTTCCCTACACGACGCTCTTCCGATCTNNNTGTCAGAACTTTGTCACCTGTCT

AGCACACGTCTGAACTCTTTCCCTACACGACGCTCTTCCGATCTNNNGCCATGATGATCCCTGATTCTCT

AGCACACGTCTGAACTCTTTCCCTACACGACGCTCTTCCGATCTNNNGCCATGATGATCCCTGATTCTCT

AGCACACGTCTGAACTCTTTCCCTACACGACGCTCTTCCGATCTNNNTCCCTGGGGACTTTAGGAATTG

AGCACACGTCTGAACTCTTTCCCTACACGACGCTCTTCCGATCTNNNTCCCTGGGGACTTTAGGAATTG

AGCACACGTCTGAACTCTTTCCCTACACGACGCTCTTCCGATCTNNNGTTTCAGTGAGCCAAGATCGTG

AGCACACGTCTGAACTCTTTCCCTACACGACGCTCTTCCGATCTNNNGTTTCAGTGAGCCAAGATCGTG

AGCACACGTCTGAACTCTTTCCCTACACGACGCTCTTCCGATCTNNNGAGGCAGGAGAATGGTGTGA

AGCACACGTCTGAACTCTTTCCCTACACGACGCTCTTCCGATCTNNNGAGGCAGGAGAATGGTGTGA

GAGCACACGTCTGAACTCTTTCCCTACACGACGCTCTTCCGATCTNNNTGAGGCAGAGAATGATGTGA

GAGCACACGTCTGAACTCTTTCCCTACACGACGCTCTTCCGATCTNNNTGAGGCAGAGAATGATGTGA

AGCACACGTCTGAACTCTTTCCCTACACGACGCTCTTCCGATCTNNNTAGAAAGAGGAGACAGGTGTT

AGCACACGTCTGAACTCTTTCCCTACACGACGCTCTTCCGATCTNNNTAGAAAGAGGAGACAGGTGTT

GCACACGTCTGAACTCTTTCCCTACACGACGCTCTTCCGATCTNNNCAGGGTTGGGCAGAGGTCAGCA

GCACACGTCTGAACTCTTTCCCTACACGACGCTCTTCCGATCTNNNCAGGGTTGGGCAGAGGTCAGCA

GCACACGTCTGAACTCTTTCCCTACACGACGCTCTTCCGATCTNNNGCCTCAGTTGGAAATGCAGAAAT

GCACACGTCTGAACTCTTTCCCTACACGACGCTCTTCCGATCTNNNGCCTCAGTTGGAAATGCAGAAAT

GAGCACACGTCTGAACTCTTTCCCTACACGACGCTCTTCCGATCTNNNGCACATCACAAAGTTTCCTGTC

GAGCACACGTCTGAACTCTTTCCCTACACGACGCTCTTCCGATCTNNNGCACATCACAAAGTTTCCTGTC

AGCACACGTCTGAACTCTTTCCCTACACGACGCTCTTCCGATCTNNNTCCTTTGCTTTCTCCATGCAAAC

AGCACACGTCTGAACTCTTTCCCTACACGACGCTCTTCCGATCTNNNTCCTTTGCTTTCTCCATGCAAAC

AGCACACGTCTGAACTCTTTCCCTACACGACGCTCTTCCGATCTNNNTGTTTGCCAGACAGGATTTCCTA

AGCACACGTCTGAACTCTTTCCCTACACGACGCTCTTCCGATCTNNNTGTTTGCCAGACAGGATTTCCTA

AGCACACGTCTGAACTCTTTCCCTACACGACGCTCTTCCGATCTNNNTGACAGATTTGCTTTGAAATGC

AGCACACGTCTGAACTCTTTCCCTACACGACGCTCTTCCGATCTNNNTGACAGATTTGCTTTGAAATGC

GAGCACACGTCTGAACTCTTTCCCTACACGACGCTCTTCCGATCTNNNTCTTACCCACAGCAGCCATAAA

GAGCACACGTCTGAACTCTTTCCCTACACGACGCTCTTCCGATCTNNNTCTTACCCACAGCAGCCATAAA

GAGCACACGTCTGAACTCTTTCCCTACACGACGCTCTTCCGATCTNNNTCTCTCCCTCTCTCTCTCTTTCTC

GAGCACACGTCTGAACTCTTTCCCTACACGACGCTCTTCCGATCTNNNTCTCTCCCTCTCTCTCTCTTTCTC
AGAGCACACGTCTGAACTCTTTCCCTACACGACGCTCTTCCGATCTNNNCTGAGGCAGGAGAATCACTT
AGAGCACACGTCTGAACTCTTTCCCTACACGACGCTCTTCCGATCTNNNCTGAGGCAGGAGAATCACTT
AGCACACGTCTGAACTCTTTCCCTACACGACGCTCTTCCGATCTNNNGGCATTACCACTGATCTCACAGA
AGCACACGTCTGAACTCTTTCCCTACACGACGCTCTTCCGATCTNNNGGCATTACCACTGATCTCACAGA
GAGCACACGTCTGAACTCTTTCCCTACACGACGCTCTTCCGATCTNNNCAGTAGCTGGGATTACAGGCA
GAGCACACGTCTGAACTCTTTCCCTACACGACGCTCTTCCGATCTNNNCAGTAGCTGGGATTACAGGCA
AGAGCACACGTCTGAACTCTTTCCCTACACGACGCTCTTCCGATCTNNNAAATATGGATACCTGCATCC
AGAGCACACGTCTGAACTCTTTCCCTACACGACGCTCTTCCGATCTNNNAAATATGGATACCTGCATCC
AGCACACGTCTGAACTCTTTCCCTACACGACGCTCTTCCGATCTNNNAGTCACCATGTTGTGTTACCAAA
AGCACACGTCTGAACTCTTTCCCTACACGACGCTCTTCCGATCTNNNAGTCACCATGTTGTGTTACCAAA
GCACACGTCTGAACTCTTTCCCTACACGACGCTCTTCCGATCTNNNCTCATCTGGAGGTCAACTTTCCAC
GCACACGTCTGAACTCTTTCCCTACACGACGCTCTTCCGATCTNNNCTCATCTGGAGGTCAACTTTCCAC
GAGCACACGTCTGAACTCTTTCCCTACACGACGCTCTTCCGATCTNNNTCTCTGACCAAGCTCACAAAAC
GAGCACACGTCTGAACTCTTTCCCTACACGACGCTCTTCCGATCTNNNTCTCTGACCAAGCTCACAAAAC
GAGCACACGTCTGAACTCTTTCCCTACACGACGCTCTTCCGATCTNNNCCTCCTGAGTAGCTAGGATCAC
GAGCACACGTCTGAACTCTTTCCCTACACGACGCTCTTCCGATCTNNNCCTCCTGAGTAGCTAGGATCAC
CACACGTCTGAACTCTTTCCCTACACGACGCTCTTCCGATCTNNNGTTTCTGTGGTTCTCATTCGGTCCA
CACACGTCTGAACTCTTTCCCTACACGACGCTCTTCCGATCTNNNGTTTCTGTGGTTCTCATTCGGTCCA
GAGCACACGTCTGAACTCTTTCCCTACACGACGCTCTTCCGATCTNNNCTGAGGCAGAGAATTGCTTGA
GAGCACACGTCTGAACTCTTTCCCTACACGACGCTCTTCCGATCTNNNCTGAGGCAGAGAATTGCTTGA
AGCACACGTCTGAACTCTTTCCCTACACGACGCTCTTCCGATCTNNNGCAACAGAATGAGACTCCGTCT
AGCACACGTCTGAACTCTTTCCCTACACGACGCTCTTCCGATCTNNNGCAACAGAATGAGACTCCGTCT
AAGAGCACACGTCTGAACTCTTTCCCTACACGACGCTCTTCCGATCTNNNTGAATCTTTTCCTTACTCTTG
AAGAGCACACGTCTGAACTCTTTCCCTACACGACGCTCTTCCGATCTNNNTGAATCTTTTCCTTACTCTTG
AGCACACGTCTGAACTCTTTCCCTACACGACGCTCTTCCGATCTNNNCGGGGAAAGAAAGTCAAAGGT
AGCACACGTCTGAACTCTTTCCCTACACGACGCTCTTCCGATCTNNNCGGGGAAAGAAAGTCAAAGGT
AGCACACGTCTGAACTCTTTCCCTACACGACGCTCTTCCGATCTNNNTTAGGATGGGAGGATGGTGGA
AGCACACGTCTGAACTCTTTCCCTACACGACGCTCTTCCGATCTNNNTTAGGATGGGAGGATGGTGGA
GAGCACACGTCTGAACTCTTTCCCTACACGACGCTCTTCCGATCTNNNGTAATCCCAGCTACTCAAGAG
GAGCACACGTCTGAACTCTTTCCCTACACGACGCTCTTCCGATCTNNNGTAATCCCAGCTACTCAAGAG
GAGCACACGTCTGAACTCTTTCCCTACACGACGCTCTTCCGATCTNNNCTTACTCTTTCCATGCACCATG
GAGCACACGTCTGAACTCTTTCCCTACACGACGCTCTTCCGATCTNNNCTTACTCTTTCCATGCACCATG
GAGCACACGTCTGAACTCTTTCCCTACACGACGCTCTTCCGATCTNNNACATTCTGAGATCTGGTGAGC
GAGCACACGTCTGAACTCTTTCCCTACACGACGCTCTTCCGATCTNNNACATTCTGAGATCTGGTGAGC
GAGCACACGTCTGAACTCTTTCCCTACACGACGCTCTTCCGATCTNNNTGTAGTCCCAGCTACTCAGGA
GAGCACACGTCTGAACTCTTTCCCTACACGACGCTCTTCCGATCTNNNTGTAGTCCCAGCTACTCAGGA
AGCACACGTCTGAACTCTTTCCCTACACGACGCTCTTCCGATCTNNNAATACAAAAAATTAGCCGGGCG
AGCACACGTCTGAACTCTTTCCCTACACGACGCTCTTCCGATCTNNNAATACAAAAAATTAGCCGGGCG
GAGCACACGTCTGAACTCTTTCCCTACACGACGCTCTTCCGATCTNNNGAGGCTGAGGCAGGAGAATC
GAGCACACGTCTGAACTCTTTCCCTACACGACGCTCTTCCGATCTNNNGAGGCTGAGGCAGGAGAATC
GAGCACACGTCTGAACTCTTTCCCTACACGACGCTCTTCCGATCTNNNCATTTACTCACGGTGGAGGT
GAGCACACGTCTGAACTCTTTCCCTACACGACGCTCTTCCGATCTNNNCATTTACTCACGGTGGAGGT

AGCACACGTCTGAACTCTTTCCCTACACGACGCTCTTCCGATCTNNNTGTAGTAGTAGGCCAAGATCC
AGCACACGTCTGAACTCTTTCCCTACACGACGCTCTTCCGATCTNNNTGTAGTAGTAGGCCAAGATCC
AGCACACGTCTGAACTCTTTCCCTACACGACGCTCTTCCGATCTNNNCAAGTCCCAAAAACCTAACCCA
AGCACACGTCTGAACTCTTTCCCTACACGACGCTCTTCCGATCTNNNCAAGTCCCAAAAACCTAACCCA
GAGCACACGTCTGAACTCTTTCCCTACACGACGCTCTTCCGATCTNNNAGGAGAATGGCGTGAACCTG
GAGCACACGTCTGAACTCTTTCCCTACACGACGCTCTTCCGATCTNNNAGGAGAATGGCGTGAACCTG
AGCACACGTCTGAACTCTTTCCCTACACGACGCTCTTCCGATCTNNNGAAACCAAAGTGACAACATCCG
AGCACACGTCTGAACTCTTTCCCTACACGACGCTCTTCCGATCTNNNGAAACCAAAGTGACAACATCCG
AGCACACGTCTGAACTCTTTCCCTACACGACGCTCTTCCGATCTNNNTGCACCTGGCCAGTTATATCATT
AGCACACGTCTGAACTCTTTCCCTACACGACGCTCTTCCGATCTNNNTGCACCTGGCCAGTTATATCATT
GAGCACACGTCTGAACTCTTTCCCTACACGACGCTCTTCCGATCTNNNCTGTAGTCCCAGCTACTCCAG
GAGCACACGTCTGAACTCTTTCCCTACACGACGCTCTTCCGATCTNNNCTGTAGTCCCAGCTACTCCAG
AAGAGCACACGTCTGAACTCTTTCCCTACACGACGCTCTTCCGATCTNNNGAGCAAGACTCCATCTGAA
AAGAGCACACGTCTGAACTCTTTCCCTACACGACGCTCTTCCGATCTNNNGAGCAAGACTCCATCTGAA
GAGCACACGTCTGAACTCTTTCCCTACACGACGCTCTTCCGATCTNNNAGGCAAGAAGTAGAAATGAC
GAGCACACGTCTGAACTCTTTCCCTACACGACGCTCTTCCGATCTNNNAGGCAAGAAGTAGAAATGAC
AGCACACGTCTGAACTCTTTCCCTACACGACGCTCTTCCGATCTNNNAGTTGCATAGTAGGCAGTTGT
AGCACACGTCTGAACTCTTTCCCTACACGACGCTCTTCCGATCTNNNAGTTGCATAGTAGGCAGTTGT
GAGCACACGTCTGAACTCTTTCCCTACACGACGCTCTTCCGATCTNNNCACAATGTGCACATGTACCCTA
GAGCACACGTCTGAACTCTTTCCCTACACGACGCTCTTCCGATCTNNNCACAATGTGCACATGTACCCTA
GAGCACACGTCTGAACTCTTTCCCTACACGACGCTCTTCCGATCTNNNGAAGGTTGCAGTGAGCTGAA
GAGCACACGTCTGAACTCTTTCCCTACACGACGCTCTTCCGATCTNNNGAAGGTTGCAGTGAGCTGAA
CACACGTCTGAACTCTTTCCCTACACGACGCTCTTCCGATCTNNNCTGGAGCTCGTGGTCATGTTCCAT
CACACGTCTGAACTCTTTCCCTACACGACGCTCTTCCGATCTNNNCTGGAGCTCGTGGTCATGTTCCAT
GCACACGTCTGAACTCTTTCCCTACACGACGCTCTTCCGATCTNNNCTCCTGAAGGCCAAAGAAACTT
GCACACGTCTGAACTCTTTCCCTACACGACGCTCTTCCGATCTNNNCTCCTGAAGGCCAAAGAAACTT
GAGCACACGTCTGAACTCTTTCCCTACACGACGCTCTTCCGATCTNNNGCTCTATCCTTCTGGCTCCTAA
GAGCACACGTCTGAACTCTTTCCCTACACGACGCTCTTCCGATCTNNNGCTCTATCCTTCTGGCTCCTAA
GAGCACACGTCTGAACTCTTTCCCTACACGACGCTCTTCCGATCTNNNATCACGACAAAAGTTGGCAC
GAGCACACGTCTGAACTCTTTCCCTACACGACGCTCTTCCGATCTNNNATCACGACAAAAGTTGGCAC
AGCACACGTCTGAACTCTTTCCCTACACGACGCTCTTCCGATCTNNNGAACCTGCCTTGCCCCATAAAT
AGCACACGTCTGAACTCTTTCCCTACACGACGCTCTTCCGATCTNNNGAACCTGCCTTGCCCCATAAAT
AGCACACGTCTGAACTCTTTCCCTACACGACGCTCTTCCGATCTNNNTTCTGGTTTTCAGCACATTGTG
AGCACACGTCTGAACTCTTTCCCTACACGACGCTCTTCCGATCTNNNTTCTGGTTTTCAGCACATTGTG
AAGAGCACACGTCTGAACTCTTTCCCTACACGACGCTCTTCCGATCTNNNCTGAGGCAGGAGAATCAC
AAGAGCACACGTCTGAACTCTTTCCCTACACGACGCTCTTCCGATCTNNNCTGAGGCAGGAGAATCAC
AGCACACGTCTGAACTCTTTCCCTACACGACGCTCTTCCGATCTNNNGGTGACACAATAAGACCCTGTC
AGCACACGTCTGAACTCTTTCCCTACACGACGCTCTTCCGATCTNNNGGTGACACAATAAGACCCTGTC
GAGCACACGTCTGAACTCTTTCCCTACACGACGCTCTTCCGATCTNNNGATGGAGTCTCACTCTGTCAT
GAGCACACGTCTGAACTCTTTCCCTACACGACGCTCTTCCGATCTNNNGATGGAGTCTCACTCTGTCAT
CACACGTCTGAACTCTTTCCCTACACGACGCTCTTCCGATCTNNNGTCTTTGAGACTGAAGAGAACTTC
CACACGTCTGAACTCTTTCCCTACACGACGCTCTTCCGATCTNNNGTCTTTGAGACTGAAGAGAACTTC
GAGCACACGTCTGAACTCTTTCCCTACACGACGCTCTTCCGATCTNNNGAAAAGGATTCACCCACTTT

GAGCACACGTCTGAACTCTTTCCCTACACGACGCTCTTCCGATCTNNNGAAAAGGATTCACCCACTTT
AGCACACGTCTGAACTCTTTCCCTACACGACGCTCTTCCGATCTNNNCCCCTAAGGTTATTCCAGATTCT
AGCACACGTCTGAACTCTTTCCCTACACGACGCTCTTCCGATCTNNNCCCCTAAGGTTATTCCAGATTCT
AGCACACGTCTGAACTCTTTCCCTACACGACGCTCTTCCGATCTNNNTTATAATCCAGCGCTCAGTGAC
AGCACACGTCTGAACTCTTTCCCTACACGACGCTCTTCCGATCTNNNTTATAATCCAGCGCTCAGTGAC
AGCACACGTCTGAACTCTTTCCCTACACGACGCTCTTCCGATCTNNNCTAAGCTATGAGGACGCAAA
AGCACACGTCTGAACTCTTTCCCTACACGACGCTCTTCCGATCTNNNCTAAGCTATGAGGACGCAAA
ACACGTCTGAACTCTTTCCCTACACGACGCTCTTCCGATCTNNNGATGCCCGTCCTCGTCAATATGCAT
ACACGTCTGAACTCTTTCCCTACACGACGCTCTTCCGATCTNNNGATGCCCGTCCTCGTCAATATGCAT
AGCACACGTCTGAACTCTTTCCCTACACGACGCTCTTCCGATCTNNNTTCTAGGGAAGGCATGAGCAA
AGCACACGTCTGAACTCTTTCCCTACACGACGCTCTTCCGATCTNNNTTCTAGGGAAGGCATGAGCAA
AGCACACGTCTGAACTCTTTCCCTACACGACGCTCTTCCGATCTNNNCCCCTGGAAGATAATGCTTGA
AGCACACGTCTGAACTCTTTCCCTACACGACGCTCTTCCGATCTNNNCCCCTGGAAGATAATGCTTGA
GAGCACACGTCTGAACTCTTTCCCTACACGACGCTCTTCCGATCTNNNGCATCCTGTACCTCAAACCTAC
GAGCACACGTCTGAACTCTTTCCCTACACGACGCTCTTCCGATCTNNNGCATCCTGTACCTCAAACCTAC
GCACACGTCTGAACTCTTTCCCTACACGACGCTCTTCCGATCTNNNAGCAAATCCCAGAAACCATTTCA
GCACACGTCTGAACTCTTTCCCTACACGACGCTCTTCCGATCTNNNAGCAAATCCCAGAAACCATTTCA
AGCACACGTCTGAACTCTTTCCCTACACGACGCTCTTCCGATCTNNNGCCCTAATCTGAGGTACCTTCT
AGCACACGTCTGAACTCTTTCCCTACACGACGCTCTTCCGATCTNNNGCCCTAATCTGAGGTACCTTCT
CACACGTCTGAACTCTTTCCCTACACGACGCTCTTCCGATCTNNNCTGAAATCCCAGCTACTCAGGAG
CACACGTCTGAACTCTTTCCCTACACGACGCTCTTCCGATCTNNNCTGAAATCCCAGCTACTCAGGAG
CACACGTCTGAACTCTTTCCCTACACGACGCTCTTCCGATCTNNNTATTACTATGAGTCAAGCCACTAC
CACACGTCTGAACTCTTTCCCTACACGACGCTCTTCCGATCTNNNTATTACTATGAGTCAAGCCACTAC
ACACGTCTGAACTCTTTCCCTACACGACGCTCTTCCGATCTNNNCATCACTGTCTTCACAAACCCATCG
ACACGTCTGAACTCTTTCCCTACACGACGCTCTTCCGATCTNNNCATCACTGTCTTCACAAACCCATCG
AGCACACGTCTGAACTCTTTCCCTACACGACGCTCTTCCGATCTNNNAGACAGGAGAATCACTTGAAC
AGCACACGTCTGAACTCTTTCCCTACACGACGCTCTTCCGATCTNNNAGACAGGAGAATCACTTGAAC
GCACACGTCTGAACTCTTTCCCTACACGACGCTCTTCCGATCTNNNTGGTTTGCTCTTGCTTCTCTAGT
GCACACGTCTGAACTCTTTCCCTACACGACGCTCTTCCGATCTNNNTGGTTTGCTCTTGCTTCTCTAGT
AGCACACGTCTGAACTCTTTCCCTACACGACGCTCTTCCGATCTNNNCTGAGGCAGGAGAATCACTTG
AGCACACGTCTGAACTCTTTCCCTACACGACGCTCTTCCGATCTNNNCTGAGGCAGGAGAATCACTTG
GAGCACACGTCTGAACTCTTTCCCTACACGACGCTCTTCCGATCTNNNGAGATCTTTGCGGTGATGG
GAGCACACGTCTGAACTCTTTCCCTACACGACGCTCTTCCGATCTNNNGAGATCTTTGCGGTGATGG
GCACACGTCTGAACTCTTTCCCTACACGACGCTCTTCCGATCTNNNGCGTATAAGCTGACCTAGACCAT
GCACACGTCTGAACTCTTTCCCTACACGACGCTCTTCCGATCTNNNGCGTATAAGCTGACCTAGACCAT
AGCACACGTCTGAACTCTTTCCCTACACGACGCTCTTCCGATCTNNNCTTCTGAGATCTCTCTCCTTGG
AGCACACGTCTGAACTCTTTCCCTACACGACGCTCTTCCGATCTNNNCTTCTGAGATCTCTCTCCTTGG
AGCACACGTCTGAACTCTTTCCCTACACGACGCTCTTCCGATCTNNNTGGGGACACAGAGATCCAATCT
AGCACACGTCTGAACTCTTTCCCTACACGACGCTCTTCCGATCTNNNTGGGGACACAGAGATCCAATCT
GAGCACACGTCTGAACTCTTTCCCTACACGACGCTCTTCCGATCTNNNAAAAATTCACCGCAGTGCAT
GAGCACACGTCTGAACTCTTTCCCTACACGACGCTCTTCCGATCTNNNAAAAATTCACCGCAGTGCAT
AGCACACGTCTGAACTCTTTCCCTACACGACGCTCTTCCGATCTNNNCTGTAATCCCAGCTACTCAGGA
AGCACACGTCTGAACTCTTTCCCTACACGACGCTCTTCCGATCTNNNCTGTAATCCCAGCTACTCAGGA

AGCACACGTCTGAACTCTTTCCCTACACGACGCTCTTCCGATCTNNNATAACTGGTAACCACAGGGA

AGCACACGTCTGAACTCTTTCCCTACACGACGCTCTTCCGATCTNNNATAACTGGTAACCACAGGGA

GCACACGTCTGAACTCTTTCCCTACACGACGCTCTTCCGATCTNNNTCAGGTTTTCTCATCTGTGTGG

GCACACGTCTGAACTCTTTCCCTACACGACGCTCTTCCGATCTNNNTCAGGTTTTCTCATCTGTGTGG

AGCACACGTCTGAACTCTTTCCCTACACGACGCTCTTCCGATCTNNNACAGAGTGAGTGAGAGTCTG

AGCACACGTCTGAACTCTTTCCCTACACGACGCTCTTCCGATCTNNNACAGAGTGAGTGAGAGTCTG

GAGCACACGTCTGAACTCTTTCCCTACACGACGCTCTTCCGATCTNNNTATCCTGGCTAACACGGTGT

GAGCACACGTCTGAACTCTTTCCCTACACGACGCTCTTCCGATCTNNNTATCCTGGCTAACACGGTGT

GCACACGTCTGAACTCTTTCCCTACACGACGCTCTTCCGATCTNNNCTGAAGGAGGATGAAACAGTG

GCACACGTCTGAACTCTTTCCCTACACGACGCTCTTCCGATCTNNNCTGAAGGAGGATGAAACAGTG

GCACACGTCTGAACTCTTTCCCTACACGACGCTCTTCCGATCTNNNTTCCTTCCTTCCTTCCTTCCTTCCA

GCACACGTCTGAACTCTTTCCCTACACGACGCTCTTCCGATCTNNNTTCCTTCCTTCCTTCCTTCCTTCCA

AGCACACGTCTGAACTCTTTCCCTACACGACGCTCTTCCGATCTNNNAAATTGGGGCAGGACTATGA

AGCACACGTCTGAACTCTTTCCCTACACGACGCTCTTCCGATCTNNNAAATTGGGGCAGGACTATGA

GCACACGTCTGAACTCTTTCCCTACACGACGCTCTTCCGATCTNNNGATTCCAAGTTCAGTGAAGTG

GCACACGTCTGAACTCTTTCCCTACACGACGCTCTTCCGATCTNNNGATTCCAAGTTCAGTGAAGTG

AGCACACGTCTGAACTCTTTCCCTACACGACGCTCTTCCGATCTNNNGAGTTTGGAAGTTTGAGACCA

AGCACACGTCTGAACTCTTTCCCTACACGACGCTCTTCCGATCTNNNGAGTTTGGAAGTTTGAGACCA

AGCACACGTCTGAACTCTTTCCCTACACGACGCTCTTCCGATCTNNNCCATCAGTGAAAAGGACACA

AGCACACGTCTGAACTCTTTCCCTACACGACGCTCTTCCGATCTNNNCCATCAGTGAAAAGGACACA

AGCACACGTCTGAACTCTTTCCCTACACGACGCTCTTCCGATCTNNNGACAAGGTCTCACTCTGTAAC

AGCACACGTCTGAACTCTTTCCCTACACGACGCTCTTCCGATCTNNNGACAAGGTCTCACTCTGTAAC

GAGCACACGTCTGAACTCTTTCCCTACACGACGCTCTTCCGATCTNNNGCTAAGGCAGGAGAATCACT

GAGCACACGTCTGAACTCTTTCCCTACACGACGCTCTTCCGATCTNNNGCTAAGGCAGGAGAATCACT

AGCACACGTCTGAACTCTTTCCCTACACGACGCTCTTCCGATCTNNNCTGTAGTCCCAGCTACTCAGGA

AGCACACGTCTGAACTCTTTCCCTACACGACGCTCTTCCGATCTNNNCTGTAGTCCCAGCTACTCAGGA

AGCACACGTCTGAACTCTTTCCCTACACGACGCTCTTCCGATCTNNNGCTTTGCTGTGAGAAGTACC

AGCACACGTCTGAACTCTTTCCCTACACGACGCTCTTCCGATCTNNNGCTTTGCTGTGAGAAGTACC

GCACACGTCTGAACTCTTTCCCTACACGACGCTCTTCCGATCTNNNTCCCTCCTGGAAATCTAGAGTC

GCACACGTCTGAACTCTTTCCCTACACGACGCTCTTCCGATCTNNNTCCCTCCTGGAAATCTAGAGTC

AGCACACGTCTGAACTCTTTCCCTACACGACGCTCTTCCGATCTNNNGGGTTTCTACAGCCAGTTCCTG

AGCACACGTCTGAACTCTTTCCCTACACGACGCTCTTCCGATCTNNNGGGTTTCTACAGCCAGTTCCTG

AGCACACGTCTGAACTCTTTCCCTACACGACGCTCTTCCGATCTNNNGATGTCTGCATGCCCATGTTT

AGCACACGTCTGAACTCTTTCCCTACACGACGCTCTTCCGATCTNNNGATGTCTGCATGCCCATGTTT

AGCACACGTCTGAACTCTTTCCCTACACGACGCTCTTCCGATCTNNNGGGTATAGCTGCAGAGGCTT

AGCACACGTCTGAACTCTTTCCCTACACGACGCTCTTCCGATCTNNNGGGTATAGCTGCAGAGGCTT

GAGCACACGTCTGAACTCTTTCCCTACACGACGCTCTTCCGATCTNNNAGTGAGCTGAGATTGTACCA

GAGCACACGTCTGAACTCTTTCCCTACACGACGCTCTTCCGATCTNNNAGTGAGCTGAGATTGTACCA

GCACACGTCTGAACTCTTTCCCTACACGACGCTCTTCCGATCTNNNAAAACATGGCAAAACCCCATCT

GCACACGTCTGAACTCTTTCCCTACACGACGCTCTTCCGATCTNNNAAAACATGGCAAAACCCCATCT

GCACACGTCTGAACTCTTTCCCTACACGACGCTCTTCCGATCTNNNATTTGCCAGTTGACCTTATGCAC

GCACACGTCTGAACTCTTTCCCTACACGACGCTCTTCCGATCTNNNATTTGCCAGTTGACCTTATGCAC

CACACGTCTGAACTCTTTCCCTACACGACGCTCTTCCGATCTNNNGTAGGGGCTTCGGACTTGAGCA

CACACGTCTGAACTCTTTCCCTACACGACGCTCTTCCGATCTNNNGTAGGGGCTTCGGACTTGAGCAT
AGCACACGTCTGAACTCTTTCCCTACACGACGCTCTTCCGATCTNNNCCATCCTACTGGTGTAAGGTG
AGCACACGTCTGAACTCTTTCCCTACACGACGCTCTTCCGATCTNNNCCATCCTACTGGTGTAAGGTG
AGCACACGTCTGAACTCTTTCCCTACACGACGCTCTTCCGATCTNNNTTCCTTTTCCCACTATGTCCCA
AGCACACGTCTGAACTCTTTCCCTACACGACGCTCTTCCGATCTNNNTTCCTTTTCCCACTATGTCCCA
GCACACGTCTGAACTCTTTCCCTACACGACGCTCTTCCGATCTNNNAAAAGGAAGAGTAGCTCCCGA
GCACACGTCTGAACTCTTTCCCTACACGACGCTCTTCCGATCTNNNAAAAGGAAGAGTAGCTCCCGA
ACACGTCTGAACTCTTTCCCTACACGACGCTCTTCCGATCTNNNAACACTGTCTGGGTTTGATGCCCAT
ACACGTCTGAACTCTTTCCCTACACGACGCTCTTCCGATCTNNNAACACTGTCTGGGTTTGATGCCCAT
ACACGTCTGAACTCTTTCCCTACACGACGCTCTTCCGATCTNNNGGCCCTCTTCAAGTTTTCTTTCTCAT
ACACGTCTGAACTCTTTCCCTACACGACGCTCTTCCGATCTNNNGGCCCTCTTCAAGTTTTCTTTCTCAT
AGCACACGTCTGAACTCTTTCCCTACACGACGCTCTTCCGATCTNNNAGGCACATTGGGTAACAAGG
AGCACACGTCTGAACTCTTTCCCTACACGACGCTCTTCCGATCTNNNAGGCACATTGGGTAACAAGG
GCACACGTCTGAACTCTTTCCCTACACGACGCTCTTCCGATCTNNNAGTCCTCCATAAATTATGTAACC
GCACACGTCTGAACTCTTTCCCTACACGACGCTCTTCCGATCTNNNAGTCCTCCATAAATTATGTAACC
CACACGTCTGAACTCTTTCCCTACACGACGCTCTTCCGATCTNNNCTTACCTTCCTCCCCCTCCTCTTCAT
CACACGTCTGAACTCTTTCCCTACACGACGCTCTTCCGATCTNNNCTTACCTTCCTCCCCCTCCTCTTCAT
AAGAGCACACGTCTGAACTCTTTCCCTACACGACGCTCTTCCGATCTNNNGTATTGGTGGATTTATGG
AAGAGCACACGTCTGAACTCTTTCCCTACACGACGCTCTTCCGATCTNNNGTATTGGTGGATTTATGG
AGCACACGTCTGAACTCTTTCCCTACACGACGCTCTTCCGATCTNNNATAAAACAATTAGTCGGGCGT
AGCACACGTCTGAACTCTTTCCCTACACGACGCTCTTCCGATCTNNNATAAAACAATTAGTCGGGCGT
AGCACACGTCTGAACTCTTTCCCTACACGACGCTCTTCCGATCTNNNAATAATGCCTTTTCCCCCCTCTT
AGCACACGTCTGAACTCTTTCCCTACACGACGCTCTTCCGATCTNNNAATAATGCCTTTTCCCCCCTCTT
GCACACGTCTGAACTCTTTCCCTACACGACGCTCTTCCGATCTNNNAATACATGCACACACACAGTTG
GCACACGTCTGAACTCTTTCCCTACACGACGCTCTTCCGATCTNNNAATACATGCACACACACAGTTG
GAGCACACGTCTGAACTCTTTCCCTACACGACGCTCTTCCGATCTNNNGCGACAGAGTGAGACTCCAT
GAGCACACGTCTGAACTCTTTCCCTACACGACGCTCTTCCGATCTNNNGCGACAGAGTGAGACTCCAT
AGCACACGTCTGAACTCTTTCCCTACACGACGCTCTTCCGATCTNNNCGTGAGACAAGATCATGCCAT
AGCACACGTCTGAACTCTTTCCCTACACGACGCTCTTCCGATCTNNNCGTGAGACAAGATCATGCCAT
AGCACACGTCTGAACTCTTTCCCTACACGACGCTCTTCCGATCTNNNGCAGGAGAATCAATTGAACCA
AGCACACGTCTGAACTCTTTCCCTACACGACGCTCTTCCGATCTNNNGCAGGAGAATCAATTGAACCA
AGCACACGTCTGAACTCTTTCCCTACACGACGCTCTTCCGATCTNNNTAGCGAAGGAAGATGCAAAC
AGCACACGTCTGAACTCTTTCCCTACACGACGCTCTTCCGATCTNNNTAGCGAAGGAAGATGCAAAC
GCACACGTCTGAACTCTTTCCCTACACGACGCTCTTCCGATCTNNNCCCTTCCAGATGCTTCCTTTACT
GCACACGTCTGAACTCTTTCCCTACACGACGCTCTTCCGATCTNNNCCCTTCCAGATGCTTCCTTTACT
CACACGTCTGAACTCTTTCCCTACACGACGCTCTTCCGATCTNNNTTTTTCTTACCCCTTGGGCTTCCCA
CACACGTCTGAACTCTTTCCCTACACGACGCTCTTCCGATCTNNNTTTTTCTTACCCCTTGGGCTTCCCA
GAGCACACGTCTGAACTCTTTCCCTACACGACGCTCTTCCGATCTNNNGGATTTGAAAGCGAAACTGG
GAGCACACGTCTGAACTCTTTCCCTACACGACGCTCTTCCGATCTNNNGGATTTGAAAGCGAAACTGG
AGCACACGTCTGAACTCTTTCCCTACACGACGCTCTTCCGATCTNNNAGTGTTTGTTCGTGTTCTTTGC
AGCACACGTCTGAACTCTTTCCCTACACGACGCTCTTCCGATCTNNNAGTGTTTGTTCGTGTTCTTTGC
ACACGTCTGAACTCTTTCCCTACACGACGCTCTTCCGATCTNNNCTTCCCTGGCCATGTCTTTAGAGCA
ACACGTCTGAACTCTTTCCCTACACGACGCTCTTCCGATCTNNNCTTCCCTGGCCATGTCTTTAGAGCA

GAGCACACGTCTGAACTCTTTCCCTACACGACGCTCTTCCGATCTNNNTGATCAGGGCCTTGAAAGAT
GAGCACACGTCTGAACTCTTTCCCTACACGACGCTCTTCCGATCTNNNTGATCAGGGCCTTGAAAGAT
AGCACACGTCTGAACTCTTTCCCTACACGACGCTCTTCCGATCTNNNACGTGGTGACTTTCAGTTCTCA
AGCACACGTCTGAACTCTTTCCCTACACGACGCTCTTCCGATCTNNNACGTGGTGACTTTCAGTTCTCA
AGCACACGTCTGAACTCTTTCCCTACACGACGCTCTTCCGATCTNNNGAGAATTGCTTGAACCTGGG
AGCACACGTCTGAACTCTTTCCCTACACGACGCTCTTCCGATCTNNNGAGAATTGCTTGAACCTGGG
GAGCACACGTCTGAACTCTTTCCCTACACGACGCTCTTCCGATCTNNNGAGGTACCCCACTTGATTCT
GAGCACACGTCTGAACTCTTTCCCTACACGACGCTCTTCCGATCTNNNGAGGTACCCCACTTGATTCT
AGCACACGTCTGAACTCTTTCCCTACACGACGCTCTTCCGATCTNNNAATCACTTGAACCCAGGAGG
AGCACACGTCTGAACTCTTTCCCTACACGACGCTCTTCCGATCTNNNAATCACTTGAACCCAGGAGG
GAGCACACGTCTGAACTCTTTCCCTACACGACGCTCTTCCGATCTNNNTTAGGAGCTCGTTCCCCTGAT
GAGCACACGTCTGAACTCTTTCCCTACACGACGCTCTTCCGATCTNNNTTAGGAGCTCGTTCCCCTGAT
AGCACACGTCTGAACTCTTTCCCTACACGACGCTCTTCCGATCTNNNCCCTGCAAACACTTCTGTGATCT
AGCACACGTCTGAACTCTTTCCCTACACGACGCTCTTCCGATCTNNNCCCTGCAAACACTTCTGTGATCT
GCACACGTCTGAACTCTTTCCCTACACGACGCTCTTCCGATCTNNNTTATGTTGAGGGACACATGGCT
GCACACGTCTGAACTCTTTCCCTACACGACGCTCTTCCGATCTNNNTTATGTTGAGGGACACATGGCT
AGCACACGTCTGAACTCTTTCCCTACACGACGCTCTTCCGATCTNNNCCATGAACTGTTCTTTATGGCC
AGCACACGTCTGAACTCTTTCCCTACACGACGCTCTTCCGATCTNNNCCATGAACTGTTCTTTATGGCC
AGCACACGTCTGAACTCTTTCCCTACACGACGCTCTTCCGATCTNNNAGAGATGGGGTTTCACCGTAT
AGCACACGTCTGAACTCTTTCCCTACACGACGCTCTTCCGATCTNNNAGAGATGGGGTTTCACCGTAT
AGAGCACACGTCTGAACTCTTTCCCTACACGACGCTCTTCCGATCTNNNAAAAAATTAGCTGGGCATG
AGAGCACACGTCTGAACTCTTTCCCTACACGACGCTCTTCCGATCTNNNAAAAAATTAGCTGGGCATG
AGCACACGTCTGAACTCTTTCCCTACACGACGCTCTTCCGATCTNNNGAGGCAAGAGAATGGGTGTG
AGCACACGTCTGAACTCTTTCCCTACACGACGCTCTTCCGATCTNNNGAGGCAAGAGAATGGGTGTG
AGAGCACACGTCTGAACTCTTTCCCTACACGACGCTCTTCCGATCTNNNCTGTAGTCCCAGCTACTCAG
AGAGCACACGTCTGAACTCTTTCCCTACACGACGCTCTTCCGATCTNNNCTGTAGTCCCAGCTACTCAG
AGCACACGTCTGAACTCTTTCCCTACACGACGCTCTTCCGATCTNNNGCAATCCTAGCACTTTGGGAG
AGCACACGTCTGAACTCTTTCCCTACACGACGCTCTTCCGATCTNNNGCAATCCTAGCACTTTGGGAG
GAGCACACGTCTGAACTCTTTCCCTACACGACGCTCTTCCGATCTNNNTCATTCATTCATTCCCCCACA
GAGCACACGTCTGAACTCTTTCCCTACACGACGCTCTTCCGATCTNNNTCATTCATTCATTCCCCCACA
AGCACACGTCTGAACTCTTTCCCTACACGACGCTCTTCCGATCTNNNACACTAGAACAGGGGTTGGAA
AGCACACGTCTGAACTCTTTCCCTACACGACGCTCTTCCGATCTNNNACACTAGAACAGGGGTTGGAA
GCACACGTCTGAACTCTTTCCCTACACGACGCTCTTCCGATCTNNNGAGCAAGACTCCGTCTCAAATAA
GCACACGTCTGAACTCTTTCCCTACACGACGCTCTTCCGATCTNNNGAGCAAGACTCCGTCTCAAATAA
AGCACACGTCTGAACTCTTTCCCTACACGACGCTCTTCCGATCTNNNCAGGGATGTTGGCTTGAAGT
AGCACACGTCTGAACTCTTTCCCTACACGACGCTCTTCCGATCTNNNCAGGGATGTTGGCTTGAAGT
GAGCACACGTCTGAACTCTTTCCCTACACGACGCTCTTCCGATCTNNNCAAGAGAATGGCTTGAAGC
GAGCACACGTCTGAACTCTTTCCCTACACGACGCTCTTCCGATCTNNNCAAGAGAATGGCTTGAAGC
AGCACACGTCTGAACTCTTTCCCTACACGACGCTCTTCCGATCTNNNCTGTAATCCCAGCTACTCAGGA
AGCACACGTCTGAACTCTTTCCCTACACGACGCTCTTCCGATCTNNNCTGTAATCCCAGCTACTCAGGA
AGCACACGTCTGAACTCTTTCCCTACACGACGCTCTTCCGATCTNNNGCCCTATTACCTGGACCTAATC
AGCACACGTCTGAACTCTTTCCCTACACGACGCTCTTCCGATCTNNNGCCCTATTACCTGGACCTAATC
AGCACACGTCTGAACTCTTTCCCTACACGACGCTCTTCCGATCTNNNTCTTTTGCCTATCAGCATGCCT

AGCACACGTCTGAACTCTTTCCCTACACGACGCTCTTCCGATCTNNNTCTTTTGCCTATCAGCATGCCT
AGCACACGTCTGAACTCTTTCCCTACACGACGCTCTTCCGATCTNNNAGAGGTGGAGGGAGAAAATA
AGCACACGTCTGAACTCTTTCCCTACACGACGCTCTTCCGATCTNNNAGAGGTGGAGGGAGAAAATA
AGCACACGTCTGAACTCTTTCCCTACACGACGCTCTTCCGATCTNNNGATGAAAAAGAACCAGTGCAC
AGCACACGTCTGAACTCTTTCCCTACACGACGCTCTTCCGATCTNNNGATGAAAAAGAACCAGTGCAC
GAGCACACGTCTGAACTCTTTCCCTACACGACGCTCTTCCGATCTNNNGCAGAGAGGGGAGAGAAAG
GAGCACACGTCTGAACTCTTTCCCTACACGACGCTCTTCCGATCTNNNGCAGAGAGGGGAGAGAAAG
AGCACACGTCTGAACTCTTTCCCTACACGACGCTCTTCCGATCTNNNCCGAGATATAAAGGAGCCAG
AGCACACGTCTGAACTCTTTCCCTACACGACGCTCTTCCGATCTNNNCCGAGATATAAAGGAGCCAG
AGCACACGTCTGAACTCTTTCCCTACACGACGCTCTTCCGATCTNNNGGTGTGATTGGGCTCAAAGG
AGCACACGTCTGAACTCTTTCCCTACACGACGCTCTTCCGATCTNNNGGTGTGATTGGGCTCAAAGG
CACACGTCTGAACTCTTTCCCTACACGACGCTCTTCCGATCTNNNCGCTCAGCTGTTCTTTCTTTCTTCAT
CACACGTCTGAACTCTTTCCCTACACGACGCTCTTCCGATCTNNNCGCTCAGCTGTTCTTTCTTTCTTCAT
AGCACACGTCTGAACTCTTTCCCTACACGACGCTCTTCCGATCTNNNAGATAAGGTCTCACCGTGTTAG
AGCACACGTCTGAACTCTTTCCCTACACGACGCTCTTCCGATCTNNNAGATAAGGTCTCACCGTGTTAG
AGCACACGTCTGAACTCTTTCCCTACACGACGCTCTTCCGATCTNNNGGGTGATGAAGTTCCCCACTA
AGCACACGTCTGAACTCTTTCCCTACACGACGCTCTTCCGATCTNNNGGGTGATGAAGTTCCCCACTA
AGCACACGTCTGAACTCTTTCCCTACACGACGCTCTTCCGATCTNNNGCTGGAGTACTGTGACATGAT
AGCACACGTCTGAACTCTTTCCCTACACGACGCTCTTCCGATCTNNNGCTGGAGTACTGTGACATGAT
GCACACGTCTGAACTCTTTCCCTACACGACGCTCTTCCGATCTNNNTCAACACTATCCACCTTGTCCTGC
GCACACGTCTGAACTCTTTCCCTACACGACGCTCTTCCGATCTNNNTCAACACTATCCACCTTGTCCTGC
GAGCACACGTCTGAACTCTTTCCCTACACGACGCTCTTCCGATCTNNNAACCTGCCAAACAAATCAATG
GAGCACACGTCTGAACTCTTTCCCTACACGACGCTCTTCCGATCTNNNAACCTGCCAAACAAATCAATG
AGCACACGTCTGAACTCTTTCCCTACACGACGCTCTTCCGATCTNNNGATTGAGTGCCAGCTTTTTCAC
AGCACACGTCTGAACTCTTTCCCTACACGACGCTCTTCCGATCTNNNGATTGAGTGCCAGCTTTTTCAC
GAGCACACGTCTGAACTCTTTCCCTACACGACGCTCTTCCGATCTNNNACAAAGACGCCTCTAATACCA
GAGCACACGTCTGAACTCTTTCCCTACACGACGCTCTTCCGATCTNNNACAAAGACGCCTCTAATACCA
AGCACACGTCTGAACTCTTTCCCTACACGACGCTCTTCCGATCTNNNAAGCACACAGTGACTGTTGAAT
AGCACACGTCTGAACTCTTTCCCTACACGACGCTCTTCCGATCTNNNAAGCACACAGTGACTGTTGAAT
GAGCACACGTCTGAACTCTTTCCCTACACGACGCTCTTCCGATCTNNNTGAAGGTTGCAGTAAGCTGA
GAGCACACGTCTGAACTCTTTCCCTACACGACGCTCTTCCGATCTNNNTGAAGGTTGCAGTAAGCTGA
AGCACACGTCTGAACTCTTTCCCTACACGACGCTCTTCCGATCTNNNGGATGATCATAAAGCACGGT
AGCACACGTCTGAACTCTTTCCCTACACGACGCTCTTCCGATCTNNNGGATGATCATAAAGCACGGT
GAGCACACGTCTGAACTCTTTCCCTACACGACGCTCTTCCGATCTNNNAGCACATTCATGAATTAGGG
GAGCACACGTCTGAACTCTTTCCCTACACGACGCTCTTCCGATCTNNNAGCACATTCATGAATTAGGG
GAGCACACGTCTGAACTCTTTCCCTACACGACGCTCTTCCGATCTNNNGAGGCAGGAGAATAGCGTG
GAGCACACGTCTGAACTCTTTCCCTACACGACGCTCTTCCGATCTNNNGAGGCAGGAGAATAGCGTG
AGCACACGTCTGAACTCTTTCCCTACACGACGCTCTTCCGATCTNNNGAAAATGCATCCGCTCACAAA
AGCACACGTCTGAACTCTTTCCCTACACGACGCTCTTCCGATCTNNNGAAAATGCATCCGCTCACAAA
AGCACACGTCTGAACTCTTTCCCTACACGACGCTCTTCCGATCTNNNTTGAAGGAATTTGGGAAGAG
AGCACACGTCTGAACTCTTTCCCTACACGACGCTCTTCCGATCTNNNTTGAAGGAATTTGGGAAGAG
ACACGTCTGAACTCTTTCCCTACACGACGCTCTTCCGATCTNNNGAAGATACTTACTGTAGTTCTCACA
ACACGTCTGAACTCTTTCCCTACACGACGCTCTTCCGATCTNNNGAAGATACTTACTGTAGTTCTCACA

GAGCACACGTCTGAACTCTTTCCCTACACGACGCTCTTCCGATCTNNNACACACAGAGACCCTTGAGT

GAGCACACGTCTGAACTCTTTCCCTACACGACGCTCTTCCGATCTNNNACACACAGAGACCCTTGAGT

GAGCACACGTCTGAACTCTTTCCCTACACGACGCTCTTCCGATCTNNNTCACATGTAATCCCACCATCTT

GAGCACACGTCTGAACTCTTTCCCTACACGACGCTCTTCCGATCTNNNTCACATGTAATCCCACCATCTT

CACACGTCTGAACTCTTTCCCTACACGACGCTCTTCCGATCTNNNCGATTTTCATGGTTACCAGGCTGC

CACACGTCTGAACTCTTTCCCTACACGACGCTCTTCCGATCTNNNCGATTTTCATGGTTACCAGGCTGC

GAGCACACGTCTGAACTCTTTCCCTACACGACGCTCTTCCGATCTNNNAACTGCATTCTCTTCTCCCAGT

GAGCACACGTCTGAACTCTTTCCCTACACGACGCTCTTCCGATCTNNNAACTGCATTCTCTTCTCCCAGT

AGCACACGTCTGAACTCTTTCCCTACACGACGCTCTTCCGATCTNNNCTGTTCCTATTCGGCCATCTTCT

AGCACACGTCTGAACTCTTTCCCTACACGACGCTCTTCCGATCTNNNCTGTTCCTATTCGGCCATCTTCT

AGCACACGTCTGAACTCTTTCCCTACACGACGCTCTTCCGATCTNNNCTGTAATCCCAGCTACTCAGG

AGCACACGTCTGAACTCTTTCCCTACACGACGCTCTTCCGATCTNNNCTGTAATCCCAGCTACTCAGG

GCACACGTCTGAACTCTTTCCCTACACGACGCTCTTCCGATCTNNNGATAACAAGAAGGAGCCAGAC

GCACACGTCTGAACTCTTTCCCTACACGACGCTCTTCCGATCTNNNGATAACAAGAAGGAGCCAGAC

GAGCACACGTCTGAACTCTTTCCCTACACGACGCTCTTCCGATCTNNNTTGGTTTTGGTTCATGATACG

GAGCACACGTCTGAACTCTTTCCCTACACGACGCTCTTCCGATCTNNNTTGGTTTTGGTTCATGATACG

GAGCACACGTCTGAACTCTTTCCCTACACGACGCTCTTCCGATCTNNNCTGAGTCATTGTGGCAGAAA

GAGCACACGTCTGAACTCTTTCCCTACACGACGCTCTTCCGATCTNNNCTGAGTCATTGTGGCAGAAA

AGCACACGTCTGAACTCTTTCCCTACACGACGCTCTTCCGATCTNNNCTTGAACTCTTGACCTCAGGTG

AGCACACGTCTGAACTCTTTCCCTACACGACGCTCTTCCGATCTNNNCTTGAACTCTTGACCTCAGGTG

GAGCACACGTCTGAACTCTTTCCCTACACGACGCTCTTCCGATCTNNNTGAGTCTATGTGTGTGCATG

GAGCACACGTCTGAACTCTTTCCCTACACGACGCTCTTCCGATCTNNNTGAGTCTATGTGTGTGCATG

GCACACGTCTGAACTCTTTCCCTACACGACGCTCTTCCGATCTNNNGCCATTTGCCAGAGGTAAGAACT

GCACACGTCTGAACTCTTTCCCTACACGACGCTCTTCCGATCTNNNGCCATTTGCCAGAGGTAAGAACT

AGCACACGTCTGAACTCTTTCCCTACACGACGCTCTTCCGATCTNNNTGCTATGAGGACTCAACAGCA

AGCACACGTCTGAACTCTTTCCCTACACGACGCTCTTCCGATCTNNNTGCTATGAGGACTCAACAGCA

GCACACGTCTGAACTCTTTCCCTACACGACGCTCTTCCGATCTNNNTTAGGATGGGAGGATGGTGGA

GCACACGTCTGAACTCTTTCCCTACACGACGCTCTTCCGATCTNNNTTAGGATGGGAGGATGGTGGA

AGCACACGTCTGAACTCTTTCCCTACACGACGCTCTTCCGATCTNNNTTCAAACGATCCTCCTGCCTTA

AGCACACGTCTGAACTCTTTCCCTACACGACGCTCTTCCGATCTNNNTTCAAACGATCCTCCTGCCTTA

AGCACACGTCTGAACTCTTTCCCTACACGACGCTCTTCCGATCTNNNGATCAAGGAGAGACATGAGTG

AGCACACGTCTGAACTCTTTCCCTACACGACGCTCTTCCGATCTNNNGATCAAGGAGAGACATGAGTG

GCACACGTCTGAACTCTTTCCCTACACGACGCTCTTCCGATCTNNNCCATCAGTCACGTTTCTGCATTTC

GCACACGTCTGAACTCTTTCCCTACACGACGCTCTTCCGATCTNNNCCATCAGTCACGTTTCTGCATTTC

GAGCACACGTCTGAACTCTTTCCCTACACGACGCTCTTCCGATCTNNNTTGGGAGAACATGAAACACA

GAGCACACGTCTGAACTCTTTCCCTACACGACGCTCTTCCGATCTNNNTTGGGAGAACATGAAACACA

AGCACACGTCTGAACTCTTTCCCTACACGACGCTCTTCCGATCTNNNGGTGGCAGTAAGTTCTCTGAA

AGCACACGTCTGAACTCTTTCCCTACACGACGCTCTTCCGATCTNNNGGTGGCAGTAAGTTCTCTGAA

GAGCACACGTCTGAACTCTTTCCCTACACGACGCTCTTCCGATCTNNNCTACACCACACAACCCTGAACT

GAGCACACGTCTGAACTCTTTCCCTACACGACGCTCTTCCGATCTNNNCTACACCACACAACCCTGAACT

AGCACACGTCTGAACTCTTTCCCTACACGACGCTCTTCCGATCTNNNTGAGGCAGGAGAATCTCTTGA

AGCACACGTCTGAACTCTTTCCCTACACGACGCTCTTCCGATCTNNNTGAGGCAGGAGAATCTCTTGA

GCACACGTCTGAACTCTTTCCCTACACGACGCTCTTCCGATCTNNNACTCACTTTCCACAAAGGTACCA

AGCACACGTCTGAACTCTTTCCCTACACGACGCTCTTCCGATCTNNNACTCACTTTCCACAAAGGTACCA
AGCACACGTCTGAACTCTTTCCCTACACGACGCTCTTCCGATCTNNNGAACGTTCAGGTTCATCAGCTC
AGCACACGTCTGAACTCTTTCCCTACACGACGCTCTTCCGATCTNNNGAACGTTCAGGTTCATCAGCTC
GAGCACACGTCTGAACTCTTTCCCTACACGACGCTCTTCCGATCTNNNCCTGTTTTCAGCCTATGTGCAA
GAGCACACGTCTGAACTCTTTCCCTACACGACGCTCTTCCGATCTNNNCCTGTTTTCAGCCTATGTGCAA
GAGCACACGTCTGAACTCTTTCCCTACACGACGCTCTTCCGATCTNNNCATGCCTGTAATCCCAGCTACT
GAGCACACGTCTGAACTCTTTCCCTACACGACGCTCTTCCGATCTNNNCATGCCTGTAATCCCAGCTACT
AGAGCACACGTCTGAACTCTTTCCCTACACGACGCTCTTCCGATCTNNNAGCTGCATGTTTATTTGCCC
AGAGCACACGTCTGAACTCTTTCCCTACACGACGCTCTTCCGATCTNNNAGCTGCATGTTTATTTGCCC
AGCACACGTCTGAACTCTTTCCCTACACGACGCTCTTCCGATCTNNNCTGTAGTTCCAGCTACATGGGA
AGCACACGTCTGAACTCTTTCCCTACACGACGCTCTTCCGATCTNNNCTGTAGTTCCAGCTACATGGGA
GAGCACACGTCTGAACTCTTTCCCTACACGACGCTCTTCCGATCTNNNCTGAGGCAGGAAAATTGCTT
GAGCACACGTCTGAACTCTTTCCCTACACGACGCTCTTCCGATCTNNNCTGAGGCAGGAAAATTGCTT
GAGCACACGTCTGAACTCTTTCCCTACACGACGCTCTTCCGATCTNNNAGGTACTCCCGGCCCTAGTCCA
GAGCACACGTCTGAACTCTTTCCCTACACGACGCTCTTCCGATCTNNNAGGTACTCCCGGCCCTAGTCCA
GAGCACACGTCTGAACTCTTTCCCTACACGACGCTCTTCCGATCTNNNAAGAAGTGGAGGTTGCAGTG
GAGCACACGTCTGAACTCTTTCCCTACACGACGCTCTTCCGATCTNNNAAGAAGTGGAGGTTGCAGTG
AGCACACGTCTGAACTCTTTCCCTACACGACGCTCTTCCGATCTNNNGCCTGTAGTCCTAAGCTACTTGG
AGCACACGTCTGAACTCTTTCCCTACACGACGCTCTTCCGATCTNNNGCCTGTAGTCCTAAGCTACTTGG
AGCACACGTCTGAACTCTTTCCCTACACGACGCTCTTCCGATCTNNNCAGTGACCTATGACTGGACTAC
AGCACACGTCTGAACTCTTTCCCTACACGACGCTCTTCCGATCTNNNCAGTGACCTATGACTGGACTAC
GAGCACACGTCTGAACTCTTTCCCTACACGACGCTCTTCCGATCTNNNTCAGGGATTTGATTTCTTCCTG
GAGCACACGTCTGAACTCTTTCCCTACACGACGCTCTTCCGATCTNNNTCAGGGATTTGATTTCTTCCTG
GAGCACACGTCTGAACTCTTTCCCTACACGACGCTCTTCCGATCTNNNTACCGTTGCTACAACAACAGCA
GAGCACACGTCTGAACTCTTTCCCTACACGACGCTCTTCCGATCTNNNTACCGTTGCTACAACAACAGCA
AGCACACGTCTGAACTCTTTCCCTACACGACGCTCTTCCGATCTNNNGGCATAGCATCACCAAGTTGGA
AGCACACGTCTGAACTCTTTCCCTACACGACGCTCTTCCGATCTNNNGGCATAGCATCACCAAGTTGGA
GCACACGTCTGAACTCTTTCCCTACACGACGCTCTTCCGATCTNNNGATTACAGAGGTAACCTGCCACA
GCACACGTCTGAACTCTTTCCCTACACGACGCTCTTCCGATCTNNNGATTACAGAGGTAACCTGCCACA
AAGAGCACACGTCTGAACTCTTTCCCTACACGACGCTCTTCCGATCTNNNAAAGGGGCTGTTTGATTGA
AAGAGCACACGTCTGAACTCTTTCCCTACACGACGCTCTTCCGATCTNNNAAAGGGGCTGTTTGATTGA
AAGAGCACACGTCTGAACTCTTTCCCTACACGACGCTCTTCCGATCTNNNCTAGGGGCTGACAATTCCA
AAGAGCACACGTCTGAACTCTTTCCCTACACGACGCTCTTCCGATCTNNNCTAGGGGCTGACAATTCCA
GAGCACACGTCTGAACTCTTTCCCTACACGACGCTCTTCCGATCTNNNGATGCAATGAAAGGCCATAG
GAGCACACGTCTGAACTCTTTCCCTACACGACGCTCTTCCGATCTNNNGATGCAATGAAAGGCCATAG
GAGCACACGTCTGAACTCTTTCCCTACACGACGCTCTTCCGATCTNNNCTTCCTTTCCTGTCCTTTGCCTA
GAGCACACGTCTGAACTCTTTCCCTACACGACGCTCTTCCGATCTNNNCTTCCTTTCCTGTCCTTTGCCTA
GAGCACACGTCTGAACTCTTTCCCTACACGACGCTCTTCCGATCTNNNAATTTTATTCAGTGGGGGCAT
GAGCACACGTCTGAACTCTTTCCCTACACGACGCTCTTCCGATCTNNNAATTTTATTCAGTGGGGGCAT
CACACGTCTGAACTCTTTCCCTACACGACGCTCTTCCGATCTNNNTCTTTTGAGTAAAGTTTTCTTGTCAT
CACACGTCTGAACTCTTTCCCTACACGACGCTCTTCCGATCTNNNTCTTTTGAGTAAAGTTTTCTTGTCAT
GAGCACACGTCTGAACTCTTTCCCTACACGACGCTCTTCCGATCTNNNGTGCTTGTGTCATTCCACCAAT
GAGCACACGTCTGAACTCTTTCCCTACACGACGCTCTTCCGATCTNNNGTGCTTGTGTCATTCCACCAAT

GAGCACACGTCTGAACTCTTTCCCTACACGACGCTCTTCCGATCTNNNGTACCTGTAGTCCCAGCTACTT

GAGCACACGTCTGAACTCTTTCCCTACACGACGCTCTTCCGATCTNNNGTACCTGTAGTCCCAGCTACTT

AGCACACGTCTGAACTCTTTCCCTACACGACGCTCTTCCGATCTNNNCTGCCAGAACTTTCATGCTTCAA

AGCACACGTCTGAACTCTTTCCCTACACGACGCTCTTCCGATCTNNNCTGCCAGAACTTTCATGCTTCAA

GCACACGTCTGAACTCTTTCCCTACACGACGCTCTTCCGATCTNNNACCCGTAACACAAACAGGTTCAT

GCACACGTCTGAACTCTTTCCCTACACGACGCTCTTCCGATCTNNNACCCGTAACACAAACAGGTTCAT

GAGCACACGTCTGAACTCTTTCCCTACACGACGCTCTTCCGATCTNNNTAGAAGTATCTCCTTCCCACCC

GAGCACACGTCTGAACTCTTTCCCTACACGACGCTCTTCCGATCTNNNTAGAAGTATCTCCTTCCCACCC

AGCACACGTCTGAACTCTTTCCCTACACGACGCTCTTCCGATCTNNNGCCTGTAGTCCCAGCTACTTGCA

AGCACACGTCTGAACTCTTTCCCTACACGACGCTCTTCCGATCTNNNGCCTGTAGTCCCAGCTACTTGCA

GCACACGTCTGAACTCTTTCCCTACACGACGCTCTTCCGATCTNNNTGAAGTTGCCTATGACTCAGCTT

GCACACGTCTGAACTCTTTCCCTACACGACGCTCTTCCGATCTNNNTGAAGTTGCCTATGACTCAGCTT

AGAGCACACGTCTGAACTCTTTCCCTACACGACGCTCTTCCGATCTNNNTCTGCCTTCTAGGACATTCA

AGAGCACACGTCTGAACTCTTTCCCTACACGACGCTCTTCCGATCTNNNTCTGCCTTCTAGGACATTCA

AGCACACGTCTGAACTCTTTCCCTACACGACGCTCTTCCGATCTNNNCTCCTGAGTAGCTGGGATTACA

AGCACACGTCTGAACTCTTTCCCTACACGACGCTCTTCCGATCTNNNCTCCTGAGTAGCTGGGATTACA

ACACGTCTGAACTCTTTCCCTACACGACGCTCTTCCGATCTNNNTAGTCATAATGGCTGTCAAAACATCG

ACACGTCTGAACTCTTTCCCTACACGACGCTCTTCCGATCTNNNTAGTCATAATGGCTGTCAAAACATCG

GAGCACACGTCTGAACTCTTTCCCTACACGACGCTCTTCCGATCTNNNAGCCTGGGTAATAGAGTGAG

GAGCACACGTCTGAACTCTTTCCCTACACGACGCTCTTCCGATCTNNNAGCCTGGGTAATAGAGTGAG

GAGCACACGTCTGAACTCTTTCCCTACACGACGCTCTTCCGATCTNNNGCCAGTAAGAGGGTCAGTTA

GAGCACACGTCTGAACTCTTTCCCTACACGACGCTCTTCCGATCTNNNGCCAGTAAGAGGGTCAGTTA

ACACGTCTGAACTCTTTCCCTACACGACGCTCTTCCGATCTNNNAGTCTGAATTAGCTTGGACCTGGCA

ACACGTCTGAACTCTTTCCCTACACGACGCTCTTCCGATCTNNNAGTCTGAATTAGCTTGGACCTGGCA

GAGCACACGTCTGAACTCTTTCCCTACACGACGCTCTTCCGATCTNNNGTGGATCACTTGAGGTCAGG

GAGCACACGTCTGAACTCTTTCCCTACACGACGCTCTTCCGATCTNNNGTGGATCACTTGAGGTCAGG

GAGCACACGTCTGAACTCTTTCCCTACACGACGCTCTTCCGATCTNNNCTGAGGCAGGAGAATTGCTT

GAGCACACGTCTGAACTCTTTCCCTACACGACGCTCTTCCGATCTNNNCTGAGGCAGGAGAATTGCTT

GAGCACACGTCTGAACTCTTTCCCTACACGACGCTCTTCCGATCTNNNGGTTCTAGGACTGGATCAGG

GAGCACACGTCTGAACTCTTTCCCTACACGACGCTCTTCCGATCTNNNGGTTCTAGGACTGGATCAGG

AGCACACGTCTGAACTCTTTCCCTACACGACGCTCTTCCGATCTNNNGAGATCAAGACACTGGACTCCA

AGCACACGTCTGAACTCTTTCCCTACACGACGCTCTTCCGATCTNNNGAGATCAAGACACTGGACTCCA

GCACACGTCTGAACTCTTTCCCTACACGACGCTCTTCCGATCTNNNAAAATGTCCTGGGTTGTTTCTGC

GCACACGTCTGAACTCTTTCCCTACACGACGCTCTTCCGATCTNNNAAAATGTCCTGGGTTGTTTCTGC

AGCACACGTCTGAACTCTTTCCCTACACGACGCTCTTCCGATCTNNNGTGAGTGCAGGGAACTTCTCT

AGCACACGTCTGAACTCTTTCCCTACACGACGCTCTTCCGATCTNNNGTGAGTGCAGGGAACTTCTCT

GAGCACACGTCTGAACTCTTTCCCTACACGACGCTCTTCCGATCTNNNTGTCAGTTACTGCCAATAAAC

GAGCACACGTCTGAACTCTTTCCCTACACGACGCTCTTCCGATCTNNNTGTCAGTTACTGCCAATAAAC

GAGCACACGTCTGAACTCTTTCCCTACACGACGCTCTTCCGATCTNNNTCTCCTATCACTGCGAGATTGT

GAGCACACGTCTGAACTCTTTCCCTACACGACGCTCTTCCGATCTNNNTCTCCTATCACTGCGAGATTGT

AGCACACGTCTGAACTCTTTCCCTACACGACGCTCTTCCGATCTNNNTGTGGATTTTGGTAAGTGCCTT

AGCACACGTCTGAACTCTTTCCCTACACGACGCTCTTCCGATCTNNNTGTGGATTTTGGTAAGTGCCTT

AGCACACGTCTGAACTCTTTCCCTACACGACGCTCTTCCGATCTNNNGCACCTGATATCTAGTAGCCGA

AGCACACGTCTGAACTCTTTCCCTACACGACGCTCTTCCGATCTNNNGCACCTGATATCTAGTAGCCGA
GAGCACACGTCTGAACTCTTTCCCTACACGACGCTCTTCCGATCTNNNGAGGCAGGAGAATTGCTTGA
GAGCACACGTCTGAACTCTTTCCCTACACGACGCTCTTCCGATCTNNNGAGGCAGGAGAATTGCTTGA
CACACGTCTGAACTCTTTCCCTACACGACGCTCTTCCGATCTNNNCCTTCTCCAGTATGACCTCATCCCA
CACACGTCTGAACTCTTTCCCTACACGACGCTCTTCCGATCTNNNCCTTCTCCAGTATGACCTCATCCCA
GCACACGTCTGAACTCTTTCCCTACACGACGCTCTTCCGATCTNNNGGACGTTGTAGTAGGACAAAGG
GCACACGTCTGAACTCTTTCCCTACACGACGCTCTTCCGATCTNNNGGACGTTGTAGTAGGACAAAGG
GCACACGTCTGAACTCTTTCCCTACACGACGCTCTTCCGATCTNNNGAGAATGGCATGAACCCGAGAG
GCACACGTCTGAACTCTTTCCCTACACGACGCTCTTCCGATCTNNNGAGAATGGCATGAACCCGAGAG
AGCACACGTCTGAACTCTTTCCCTACACGACGCTCTTCCGATCTNNNCTGGTGATGAAACAGCCCATTT
AGCACACGTCTGAACTCTTTCCCTACACGACGCTCTTCCGATCTNNNCTGGTGATGAAACAGCCCATTT
AGCACACGTCTGAACTCTTTCCCTACACGACGCTCTTCCGATCTNNNATGTTCTCCATGTACCAGGCTT
AGCACACGTCTGAACTCTTTCCCTACACGACGCTCTTCCGATCTNNNATGTTCTCCATGTACCAGGCTT
GAGCACACGTCTGAACTCTTTCCCTACACGACGCTCTTCCGATCTNNNCCTTTTCTGAGCCTTTGTTGCT
GAGCACACGTCTGAACTCTTTCCCTACACGACGCTCTTCCGATCTNNNCCTTTTCTGAGCCTTTGTTGCT
AGAGCACACGTCTGAACTCTTTCCCTACACGACGCTCTTCCGATCTNNNCCTTTCTCGTTCTCTCTCCAGT
AGAGCACACGTCTGAACTCTTTCCCTACACGACGCTCTTCCGATCTNNNCCTTTCTCGTTCTCTCTCCAGT
AGCACACGTCTGAACTCTTTCCCTACACGACGCTCTTCCGATCTNNNTTCTGTGTGCACCCTCAAGAAT
AGCACACGTCTGAACTCTTTCCCTACACGACGCTCTTCCGATCTNNNTTCTGTGTGCACCCTCAAGAAT
GAGCACACGTCTGAACTCTTTCCCTACACGACGCTCTTCCGATCTNNNAGTAGTTCTCAACCTTGGCTTG
GAGCACACGTCTGAACTCTTTCCCTACACGACGCTCTTCCGATCTNNNAGTAGTTCTCAACCTTGGCTTG
CACACGTCTGAACTCTTTCCCTACACGACGCTCTTCCGATCTNNNCTCTTTGCTGCCTAGAGTCATGACA
CACACGTCTGAACTCTTTCCCTACACGACGCTCTTCCGATCTNNNCTCTTTGCTGCCTAGAGTCATGACA
AGCACACGTCTGAACTCTTTCCCTACACGACGCTCTTCCGATCTNNNTGCCTGTATTTGTGTGACACTC
AGCACACGTCTGAACTCTTTCCCTACACGACGCTCTTCCGATCTNNNTGCCTGTATTTGTGTGACACTC
AGCACACGTCTGAACTCTTTCCCTACACGACGCTCTTCCGATCTNNNGCGAATTGATCTAAAACAGATA
AGCACACGTCTGAACTCTTTCCCTACACGACGCTCTTCCGATCTNNNGCGAATTGATCTAAAACAGATA
GCACACGTCTGAACTCTTTCCCTACACGACGCTCTTCCGATCTNNNTACATCACTGAAGTTCATATCATC
GCACACGTCTGAACTCTTTCCCTACACGACGCTCTTCCGATCTNNNTACATCACTGAAGTTCATATCATC
AGAGCACACGTCTGAACTCTTTCCCTACACGACGCTCTTCCGATCTNNNCCATTTGGGAGAGTGTCTCA
AGAGCACACGTCTGAACTCTTTCCCTACACGACGCTCTTCCGATCTNNNCCATTTGGGAGAGTGTCTCA
AGCACACGTCTGAACTCTTTCCCTACACGACGCTCTTCCGATCTNNNATCTGTGTCTGAAAAGTGGCTC
AGCACACGTCTGAACTCTTTCCCTACACGACGCTCTTCCGATCTNNNATCTGTGTCTGAAAAGTGGCTC
AGAGCACACGTCTGAACTCTTTCCCTACACGACGCTCTTCCGATCTNNNGCTGAGATCATGTCATTGCA
AGAGCACACGTCTGAACTCTTTCCCTACACGACGCTCTTCCGATCTNNNGCTGAGATCATGTCATTGCA
GAGCACACGTCTGAACTCTTTCCCTACACGACGCTCTTCCGATCTNNNGAACCATTCGGAACAGAGAG
GAGCACACGTCTGAACTCTTTCCCTACACGACGCTCTTCCGATCTNNNGAACCATTCGGAACAGAGAG
AGAGCACACGTCTGAACTCTTTCCCTACACGACGCTCTTCCGATCTNNNGAAGGACAGGTCAGTGGAC
AGAGCACACGTCTGAACTCTTTCCCTACACGACGCTCTTCCGATCTNNNGAAGGACAGGTCAGTGGAC
AGCACACGTCTGAACTCTTTCCCTACACGACGCTCTTCCGATCTNNNAAAAATTAGCTGGGCATGGTG
AGCACACGTCTGAACTCTTTCCCTACACGACGCTCTTCCGATCTNNNAAAAATTAGCTGGGCATGGTG
AGCACACGTCTGAACTCTTTCCCTACACGACGCTCTTCCGATCTNNNCCTATAGTCCCAGCTACTCGGGC
AGCACACGTCTGAACTCTTTCCCTACACGACGCTCTTCCGATCTNNNCCTATAGTCCCAGCTACTCGGGC

AGCACACGTCTGAACTCTTTCCCTACACGACGCTCTTCCGATCTNNNCATTTTCCTTTTCCCTGGCTCTGC

AGCACACGTCTGAACTCTTTCCCTACACGACGCTCTTCCGATCTNNNCATTTTCCTTTTCCCTGGCTCTGC

GAGCACACGTCTGAACTCTTTCCCTACACGACGCTCTTCCGATCTNNNTTGGTGGAGACTTCTATTCTG

GAGCACACGTCTGAACTCTTTCCCTACACGACGCTCTTCCGATCTNNNTTGGTGGAGACTTCTATTCTG

GAGCACACGTCTGAACTCTTTCCCTACACGACGCTCTTCCGATCTNNNGGTGACACAGTGAGATTCCAT

GAGCACACGTCTGAACTCTTTCCCTACACGACGCTCTTCCGATCTNNNGGTGACACAGTGAGATTCCAT

AGCACACGTCTGAACTCTTTCCCTACACGACGCTCTTCCGATCTNNNTCCTAATGCTATCCCTCCCCTAG

AGCACACGTCTGAACTCTTTCCCTACACGACGCTCTTCCGATCTNNNTCCTAATGCTATCCCTCCCCTAG

AGAGCACACGTCTGAACTCTTTCCCTACACGACGCTCTTCCGATCTNNNTCACTTGAACTAAGGAGGC

AGAGCACACGTCTGAACTCTTTCCCTACACGACGCTCTTCCGATCTNNNTCACTTGAACTAAGGAGGC

AGCACACGTCTGAACTCTTTCCCTACACGACGCTCTTCCGATCTNNNTGAACTCCTGACCTTGTGATCT

AGCACACGTCTGAACTCTTTCCCTACACGACGCTCTTCCGATCTNNNTGAACTCCTGACCTTGTGATCT

AGCACACGTCTGAACTCTTTCCCTACACGACGCTCTTCCGATCTNNNCTTCACGTATAAATGGGATGG

AGCACACGTCTGAACTCTTTCCCTACACGACGCTCTTCCGATCTNNNCTTCACGTATAAATGGGATGG

AGCACACGTCTGAACTCTTTCCCTACACGACGCTCTTCCGATCTNNNGGTCAAGCACCAAAAACTGAA

AGCACACGTCTGAACTCTTTCCCTACACGACGCTCTTCCGATCTNNNGGTCAAGCACCAAAAACTGAA

ACACGTCTGAACTCTTTCCCTACACGACGCTCTTCCGATCTNNNGCAATTGGATGCCATAGTTCAGACAT

ACACGTCTGAACTCTTTCCCTACACGACGCTCTTCCGATCTNNNGCAATTGGATGCCATAGTTCAGACAT

AGCACACGTCTGAACTCTTTCCCTACACGACGCTCTTCCGATCTNNNGTGTGGGCACATTTTTTTTTCC

AGCACACGTCTGAACTCTTTCCCTACACGACGCTCTTCCGATCTNNNGTGTGGGCACATTTTTTTTTCC

GAGCACACGTCTGAACTCTTTCCCTACACGACGCTCTTCCGATCTNNNTCTCCTAGTGATACTCAGGCAC

GAGCACACGTCTGAACTCTTTCCCTACACGACGCTCTTCCGATCTNNNTCTCCTAGTGATACTCAGGCAC

AGCACACGTCTGAACTCTTTCCCTACACGACGCTCTTCCGATCTNNNTTCGTTTTCCTAGCACCTGTCATC

AGCACACGTCTGAACTCTTTCCCTACACGACGCTCTTCCGATCTNNNTTCGTTTTCCTAGCACCTGTCATC

AGCACACGTCTGAACTCTTTCCCTACACGACGCTCTTCCGATCTNNNTGAAACGGAAATACCCTTTGCC

AGCACACGTCTGAACTCTTTCCCTACACGACGCTCTTCCGATCTNNNTGAAACGGAAATACCCTTTGCC

GAGCACACGTCTGAACTCTTTCCCTACACGACGCTCTTCCGATCTNNNTCTCTAACTGTAAGCATACCTG

GAGCACACGTCTGAACTCTTTCCCTACACGACGCTCTTCCGATCTNNNTCTCTAACTGTAAGCATACCTG

AGCACACGTCTGAACTCTTTCCCTACACGACGCTCTTCCGATCTNNNATGAAAACCAGCCTATCCCAACT

AGCACACGTCTGAACTCTTTCCCTACACGACGCTCTTCCGATCTNNNATGAAAACCAGCCTATCCCAACT

AGCACACGTCTGAACTCTTTCCCTACACGACGCTCTTCCGATCTNNNTTCAGCAATGGATCTCACCAAGA

AGCACACGTCTGAACTCTTTCCCTACACGACGCTCTTCCGATCTNNNTTCAGCAATGGATCTCACCAAGA

GAGCACACGTCTGAACTCTTTCCCTACACGACGCTCTTCCGATCTNNNTGCCACTGTAGGCTTGAGATA

GAGCACACGTCTGAACTCTTTCCCTACACGACGCTCTTCCGATCTNNNTGCCACTGTAGGCTTGAGATA

GAGCACACGTCTGAACTCTTTCCCTACACGACGCTCTTCCGATCTNNNCCAAGCCCAGCTAATTTGGTT

GAGCACACGTCTGAACTCTTTCCCTACACGACGCTCTTCCGATCTNNNCCAAGCCCAGCTAATTTGGTT

GAGCACACGTCTGAACTCTTTCCCTACACGACGCTCTTCCGATCTNNNAGTAGCCAGGATTATAGCTG

GAGCACACGTCTGAACTCTTTCCCTACACGACGCTCTTCCGATCTNNNAGTAGCCAGGATTATAGCTG

AGCACACGTCTGAACTCTTTCCCTACACGACGCTCTTCCGATCTNNNGCTCGGAGAAGTAGAGCGCAC

AGCACACGTCTGAACTCTTTCCCTACACGACGCTCTTCCGATCTNNNGCTCGGAGAAGTAGAGCGCAC

GAGCACACGTCTGAACTCTTTCCCTACACGACGCTCTTCCGATCTNNNCTATCCCTGCTTGTCTCTGTCA

GAGCACACGTCTGAACTCTTTCCCTACACGACGCTCTTCCGATCTNNNCTATCCCTGCTTGTCTCTGTCA

GAGCACACGTCTGAACTCTTTCCCTACACGACGCTCTTCCGATCTNNNTTGAGGTCAGGAGTTCAAAAC

AGCACACGTCTGAACTCTTTCCCTACACGACGCTCTTCCGATCTNNNTTGAGGTCAGGAGTTCAAAA
AGCACACGTCTGAACTCTTTCCCTACACGACGCTCTTCCGATCTNNNACATCTCTAGTCCCAGCTACTC
AGCACACGTCTGAACTCTTTCCCTACACGACGCTCTTCCGATCTNNNACATCTCTAGTCCCAGCTACTC
AGCACACGTCTGAACTCTTTCCCTACACGACGCTCTTCCGATCTNNNAGTGAAAATCTGGAGAGCAC
AGCACACGTCTGAACTCTTTCCCTACACGACGCTCTTCCGATCTNNNAGTGAAAATCTGGAGAGCAC
AGCACACGTCTGAACTCTTTCCCTACACGACGCTCTTCCGATCTNNNCTTCTCACCTTAGGCGATCATC
AGCACACGTCTGAACTCTTTCCCTACACGACGCTCTTCCGATCTNNNCTTCTCACCTTAGGCGATCATC
AGCACACGTCTGAACTCTTTCCCTACACGACGCTCTTCCGATCTNNNAGCATGTTTCTGTAGGTGGTC
AGCACACGTCTGAACTCTTTCCCTACACGACGCTCTTCCGATCTNNNAGCATGTTTCTGTAGGTGGTC
AGCACACGTCTGAACTCTTTCCCTACACGACGCTCTTCCGATCTNNNCCTGTGGAACTGCAGTACCC
AGCACACGTCTGAACTCTTTCCCTACACGACGCTCTTCCGATCTNNNCCTGTGGAACTGCAGTACCC
GCACACGTCTGAACTCTTTCCCTACACGACGCTCTTCCGATCTNNNGGTTTCTCGCATGTTGTTCTCAT
GCACACGTCTGAACTCTTTCCCTACACGACGCTCTTCCGATCTNNNGGTTTCTCGCATGTTGTTCTCAT
AGCACACGTCTGAACTCTTTCCCTACACGACGCTCTTCCGATCTNNNCATCCCTCATTTCCCCTCAAAG
AGCACACGTCTGAACTCTTTCCCTACACGACGCTCTTCCGATCTNNNCATCCCTCATTTCCCCTCAAAG
CACACGTCTGAACTCTTTCCCTACACGACGCTCTTCCGATCTNNNATGGGCTGTGTTTCCATTTGTTTC
CACACGTCTGAACTCTTTCCCTACACGACGCTCTTCCGATCTNNNATGGGCTGTGTTTCCATTTGTTTC
GAGCACACGTCTGAACTCTTTCCCTACACGACGCTCTTCCGATCTNNNAGAAAGAGGAGGAGGAGA
GAGCACACGTCTGAACTCTTTCCCTACACGACGCTCTTCCGATCTNNNAGAAAGAGGAGGAGGAGA
AGCACACGTCTGAACTCTTTCCCTACACGACGCTCTTCCGATCTNNNCATTGTGCATATGGCCTCAAGA
AGCACACGTCTGAACTCTTTCCCTACACGACGCTCTTCCGATCTNNNCATTGTGCATATGGCCTCAAGA
AGCACACGTCTGAACTCTTTCCCTACACGACGCTCTTCCGATCTNNNGAGGGTGGGCCTTATGAAAT
AGCACACGTCTGAACTCTTTCCCTACACGACGCTCTTCCGATCTNNNGAGGGTGGGCCTTATGAAAT
AGCACACGTCTGAACTCTTTCCCTACACGACGCTCTTCCGATCTNNNTTCTAGTGCCAAAATGCTGTG
AGCACACGTCTGAACTCTTTCCCTACACGACGCTCTTCCGATCTNNNTTCTAGTGCCAAAATGCTGTG
AGCACACGTCTGAACTCTTTCCCTACACGACGCTCTTCCGATCTNNNCTGTAATCCCAGCTACTCAGGA
AGCACACGTCTGAACTCTTTCCCTACACGACGCTCTTCCGATCTNNNCTGTAATCCCAGCTACTCAGGA
GCACACGTCTGAACTCTTTCCCTACACGACGCTCTTCCGATCTNNNTTCAAGGGATTCTTCTGCCTCAG
GCACACGTCTGAACTCTTTCCCTACACGACGCTCTTCCGATCTNNNTTCAAGGGATTCTTCTGCCTCAG
GCACACGTCTGAACTCTTTCCCTACACGACGCTCTTCCGATCTNNNTCAAGGTGTCAAATCCCACGTTT
GCACACGTCTGAACTCTTTCCCTACACGACGCTCTTCCGATCTNNNTCAAGGTGTCAAATCCCACGTTT
AGCACACGTCTGAACTCTTTCCCTACACGACGCTCTTCCGATCTNNNAGACACAAGAGTCTTAGGTTT
AGCACACGTCTGAACTCTTTCCCTACACGACGCTCTTCCGATCTNNNAGACACAAGAGTCTTAGGTTT
AGCACACGTCTGAACTCTTTCCCTACACGACGCTCTTCCGATCTNNNGTTCCTTTTGAGAACAATTCT
AGCACACGTCTGAACTCTTTCCCTACACGACGCTCTTCCGATCTNNNGTTCCTTTTGAGAACAATTCT
GAGCACACGTCTGAACTCTTTCCCTACACGACGCTCTTCCGATCTNNNAGCAAACCAACCACAAAACT
GAGCACACGTCTGAACTCTTTCCCTACACGACGCTCTTCCGATCTNNNAGCAAACCAACCACAAAACT
GCACACGTCTGAACTCTTTCCCTACACGACGCTCTTCCGATCTNNNAATAAACACGCTTGGCTAAGAG
GCACACGTCTGAACTCTTTCCCTACACGACGCTCTTCCGATCTNNNAATAAACACGCTTGGCTAAGAG
GCACACGTCTGAACTCTTTCCCTACACGACGCTCTTCCGATCTNNNGTTCTGTGGATTTGACAAAGGT
GCACACGTCTGAACTCTTTCCCTACACGACGCTCTTCCGATCTNNNGTTCTGTGGATTTGACAAAGGT
GAGCACACGTCTGAACTCTTTCCCTACACGACGCTCTTCCGATCTNNNTTGCGCACGTATGTTTATAG
GAGCACACGTCTGAACTCTTTCCCTACACGACGCTCTTCCGATCTNNNTTGCGCACGTATGTTTATAG

AGCACACGTCTGAACTCTTTCCCTACACGACGCTCTTCCGATCTNNNCCGTTTTCATGACAACCTCTAGC

AGCACACGTCTGAACTCTTTCCCTACACGACGCTCTTCCGATCTNNNCCGTTTTCATGACAACCTCTAGC

GCACACGTCTGAACTCTTTCCCTACACGACGCTCTTCCGATCTNNNCCTGAAACACCCTTCTCTTCACTCA

GCACACGTCTGAACTCTTTCCCTACACGACGCTCTTCCGATCTNNNCCTGAAACACCCTTCTCTTCACTCA

AGCACACGTCTGAACTCTTTCCCTACACGACGCTCTTCCGATCTNNNAAAAATTGGGAACCAGCTTGG

AGCACACGTCTGAACTCTTTCCCTACACGACGCTCTTCCGATCTNNNAAAAATTGGGAACCAGCTTGG

AGCACACGTCTGAACTCTTTCCCTACACGACGCTCTTCCGATCTNNNATTGGGACTGATCAGGGAAACA

AGCACACGTCTGAACTCTTTCCCTACACGACGCTCTTCCGATCTNNNATTGGGACTGATCAGGGAAACA

AGCACACGTCTGAACTCTTTCCCTACACGACGCTCTTCCGATCTNNNAGGAAGCCATAGAGAATGGGC

AGCACACGTCTGAACTCTTTCCCTACACGACGCTCTTCCGATCTNNNAGGAAGCCATAGAGAATGGGC

AGAGCACACGTCTGAACTCTTTCCCTACACGACGCTCTTCCGATCTNNNTCATTTGGTGCTGATGCCAT

AGAGCACACGTCTGAACTCTTTCCCTACACGACGCTCTTCCGATCTNNNTCATTTGGTGCTGATGCCAT

AGCACACGTCTGAACTCTTTCCCTACACGACGCTCTTCCGATCTNNNTTTGCCTGTTGATTTTGCATCCTC

AGCACACGTCTGAACTCTTTCCCTACACGACGCTCTTCCGATCTNNNTTTGCCTGTTGATTTTGCATCCTC

GAGCACACGTCTGAACTCTTTCCCTACACGACGCTCTTCCGATCTNNNCACAAGTAGCTGGGACTACAG

GAGCACACGTCTGAACTCTTTCCCTACACGACGCTCTTCCGATCTNNNCACAAGTAGCTGGGACTACAG

AGCACACGTCTGAACTCTTTCCCTACACGACGCTCTTCCGATCTNNNGATCTCTGAGATTTTGGTGCACC

AGCACACGTCTGAACTCTTTCCCTACACGACGCTCTTCCGATCTNNNGATCTCTGAGATTTTGGTGCACC

GAGCACACGTCTGAACTCTTTCCCTACACGACGCTCTTCCGATCTNNNAATACAAAAAATTAGCCGGGC

GAGCACACGTCTGAACTCTTTCCCTACACGACGCTCTTCCGATCTNNNAATACAAAAAATTAGCCGGGC

AAGAGCACACGTCTGAACTCTTTCCCTACACGACGCTCTTCCGATCTNNNTCAGAGCTGAAATGAAGC

AAGAGCACACGTCTGAACTCTTTCCCTACACGACGCTCTTCCGATCTNNNTCAGAGCTGAAATGAAGC

AGCACACGTCTGAACTCTTTCCCTACACGACGCTCTTCCGATCTNNNAGTGCTCTTAGAGTGAATCGGT

AGCACACGTCTGAACTCTTTCCCTACACGACGCTCTTCCGATCTNNNAGTGCTCTTAGAGTGAATCGGT

GAGCACACGTCTGAACTCTTTCCCTACACGACGCTCTTCCGATCTNNNAAGAGGAAGCAGGGAGAAA

GAGCACACGTCTGAACTCTTTCCCTACACGACGCTCTTCCGATCTNNNAAGAGGAAGCAGGGAGAAA

GAGCACACGTCTGAACTCTTTCCCTACACGACGCTCTTCCGATCTNNNCACCTCCATAGCTACTGCAAGA

GAGCACACGTCTGAACTCTTTCCCTACACGACGCTCTTCCGATCTNNNCACCTCCATAGCTACTGCAAGA

GAGCACACGTCTGAACTCTTTCCCTACACGACGCTCTTCCGATCTNNNTTAGAGAGGGGGGTGAAGA

GAGCACACGTCTGAACTCTTTCCCTACACGACGCTCTTCCGATCTNNNTTAGAGAGGGGGGTGAAGA

GAGCACACGTCTGAACTCTTTCCCTACACGACGCTCTTCCGATCTNNNCTGAGGCAGGAGAATCACTTG

GAGCACACGTCTGAACTCTTTCCCTACACGACGCTCTTCCGATCTNNNCTGAGGCAGGAGAATCACTTG

GAGCACACGTCTGAACTCTTTCCCTACACGACGCTCTTCCGATCTNNNACAAAAATTAGCTGGGCATGC

GAGCACACGTCTGAACTCTTTCCCTACACGACGCTCTTCCGATCTNNNACAAAAATTAGCTGGGCATGC

AGCACACGTCTGAACTCTTTCCCTACACGACGCTCTTCCGATCTNNNGATCTCCGCTCACTACAAACTCCC

AGCACACGTCTGAACTCTTTCCCTACACGACGCTCTTCCGATCTNNNGATCTCCGCTCACTACAAACTCCC

AGCACACGTCTGAACTCTTTCCCTACACGACGCTCTTCCGATCTNNNAGTTGCAATTTCACTTTCCTGGG

AGCACACGTCTGAACTCTTTCCCTACACGACGCTCTTCCGATCTNNNAGTTGCAATTTCACTTTCCTGGG

AGCACACGTCTGAACTCTTTCCCTACACGACGCTCTTCCGATCTNNNTGTCTGCATCTTGCTATTGGACT

AGCACACGTCTGAACTCTTTCCCTACACGACGCTCTTCCGATCTNNNTGTCTGCATCTTGCTATTGGACT

GAGCACACGTCTGAACTCTTTCCCTACACGACGCTCTTCCGATCTNNNCATGTTCTTCAACACGACGGAA

GAGCACACGTCTGAACTCTTTCCCTACACGACGCTCTTCCGATCTNNNCATGTTCTTCAACACGACGGAA

AGCACACGTCTGAACTCTTTCCCTACACGACGCTCTTCCGATCTNNNTTCTCTGATCCTCCAGACTTCCTC

GCACACGTCTGAACTCTTTCCCTACACGACGCTCTTCCGATCTNNNTTCTCTGATCCTCCAGACTTCCTC

GAGCACACGTCTGAACTCTTTCCCTACACGACGCTCTTCCGATCTNNNGAGAATTGCTTGAACCTGGG

GAGCACACGTCTGAACTCTTTCCCTACACGACGCTCTTCCGATCTNNNGAGAATTGCTTGAACCTGGG

GAGCACACGTCTGAACTCTTTCCCTACACGACGCTCTTCCGATCTNNNATAAAAATTAGCTGGGCGTG

GAGCACACGTCTGAACTCTTTCCCTACACGACGCTCTTCCGATCTNNNATAAAAATTAGCTGGGCGTG

AGCACACGTCTGAACTCTTTCCCTACACGACGCTCTTCCGATCTNNNCCTCTCCCCAGAAGACCTAGTA

AGCACACGTCTGAACTCTTTCCCTACACGACGCTCTTCCGATCTNNNCCTCTCCCCAGAAGACCTAGTA

GCACACGTCTGAACTCTTTCCCTACACGACGCTCTTCCGATCTNNNCTTTGGTGCTGCTGCTGCTCCA

GCACACGTCTGAACTCTTTCCCTACACGACGCTCTTCCGATCTNNNCTTTGGTGCTGCTGCTGCTCCA

GCACACGTCTGAACTCTTTCCCTACACGACGCTCTTCCGATCTNNNAGCTCTAATTTCACTGGCTGTGT

GCACACGTCTGAACTCTTTCCCTACACGACGCTCTTCCGATCTNNNAGCTCTAATTTCACTGGCTGTGT

AGCACACGTCTGAACTCTTTCCCTACACGACGCTCTTCCGATCTNNNCATACTGAGAATCTAGGCCACC

AGCACACGTCTGAACTCTTTCCCTACACGACGCTCTTCCGATCTNNNCATACTGAGAATCTAGGCCACC

GAGCACACGTCTGAACTCTTTCCCTACACGACGCTCTTCCGATCTNNNCTCCTCTTGATTTCGGGTGTC

GAGCACACGTCTGAACTCTTTCCCTACACGACGCTCTTCCGATCTNNNCTCCTCTTGATTTCGGGTGTC

GAGCACACGTCTGAACTCTTTCCCTACACGACGCTCTTCCGATCTNNNCTGAGGCAAGAGAATTGCTG

GAGCACACGTCTGAACTCTTTCCCTACACGACGCTCTTCCGATCTNNNCTGAGGCAAGAGAATTGCTG

AGCACACGTCTGAACTCTTTCCCTACACGACGCTCTTCCGATCTNNNCAGAGGAAGTGGGAAGTGAT

AGCACACGTCTGAACTCTTTCCCTACACGACGCTCTTCCGATCTNNNCAGAGGAAGTGGGAAGTGAT

GAGCACACGTCTGAACTCTTTCCCTACACGACGCTCTTCCGATCTNNNAAAAGACACATCACACAATCG

GAGCACACGTCTGAACTCTTTCCCTACACGACGCTCTTCCGATCTNNNAAAAGACACATCACACAATCG

GCACACGTCTGAACTCTTTCCCTACACGACGCTCTTCCGATCTNNNCACACTGTCTTGGTTGTGGTTTT

GCACACGTCTGAACTCTTTCCCTACACGACGCTCTTCCGATCTNNNCACACTGTCTTGGTTGTGGTTTT

GCACACGTCTGAACTCTTTCCCTACACGACGCTCTTCCGATCTNNNGGAATCTGCCCTGTGTCGTCCAT

GCACACGTCTGAACTCTTTCCCTACACGACGCTCTTCCGATCTNNNGGAATCTGCCCTGTGTCGTCCAT

AGCACACGTCTGAACTCTTTCCCTACACGACGCTCTTCCGATCTNNNATACAAAAATTAGCTGGGCGT

AGCACACGTCTGAACTCTTTCCCTACACGACGCTCTTCCGATCTNNNATACAAAAATTAGCTGGGCGT

AGCACACGTCTGAACTCTTTCCCTACACGACGCTCTTCCGATCTNNNCTGAGGCAGGAGAATCACTTG

AGCACACGTCTGAACTCTTTCCCTACACGACGCTCTTCCGATCTNNNCTGAGGCAGGAGAATCACTTG

GAGCACACGTCTGAACTCTTTCCCTACACGACGCTCTTCCGATCTNNNACAACACATGGGGATTATGG

GAGCACACGTCTGAACTCTTTCCCTACACGACGCTCTTCCGATCTNNNACAACACATGGGGATTATGG

GAGCACACGTCTGAACTCTTTCCCTACACGACGCTCTTCCGATCTNNNTTGGGAACTATCTTTAGGGG

GAGCACACGTCTGAACTCTTTCCCTACACGACGCTCTTCCGATCTNNNTTGGGAACTATCTTTAGGGG

AGCACACGTCTGAACTCTTTCCCTACACGACGCTCTTCCGATCTNNNTGTGCCAATCAGTAATGCACAG

AGCACACGTCTGAACTCTTTCCCTACACGACGCTCTTCCGATCTNNNTGTGCCAATCAGTAATGCACAG

AGCACACGTCTGAACTCTTTCCCTACACGACGCTCTTCCGATCTNNNAAAATATCGGCTCTGTGTTTCA

AGCACACGTCTGAACTCTTTCCCTACACGACGCTCTTCCGATCTNNNAAAATATCGGCTCTGTGTTTCA

AGCACACGTCTGAACTCTTTCCCTACACGACGCTCTTCCGATCTNNNGTGCCTGAACTGGGATGTCA

AGCACACGTCTGAACTCTTTCCCTACACGACGCTCTTCCGATCTNNNGTGCCTGAACTGGGATGTCA

CACACGTCTGAACTCTTTCCCTACACGACGCTCTTCCGATCTNNNCTCCCTTCTCTCTACCCAAATCCAT

CACACGTCTGAACTCTTTCCCTACACGACGCTCTTCCGATCTNNNCTCCCTTCTCTCTACCCAAATCCAT

AGCACACGTCTGAACTCTTTCCCTACACGACGCTCTTCCGATCTNNNCCGGTAACTGAGCAAACACTA

AGCACACGTCTGAACTCTTTCCCTACACGACGCTCTTCCGATCTNNNCCGGTAACTGAGCAAACACTA

AGCACACGTCTGAACTCTTTCCCTACACGACGCTCTTCCGATCTNNNATACAAAAAAATTAGCCGGGC
AGCACACGTCTGAACTCTTTCCCTACACGACGCTCTTCCGATCTNNNATACAAAAAAATTAGCCGGGC
AGCACACGTCTGAACTCTTTCCCTACACGACGCTCTTCCGATCTNNNTAAAGTGGGAAGATGGCTTG
AGCACACGTCTGAACTCTTTCCCTACACGACGCTCTTCCGATCTNNNTAAAGTGGGAAGATGGCTTG
AGCACACGTCTGAACTCTTTCCCTACACGACGCTCTTCCGATCTNNNTTTGCTTTTCCATTAGATCAGC
AGCACACGTCTGAACTCTTTCCCTACACGACGCTCTTCCGATCTNNNTTTGCTTTTCCATTAGATCAGC
AGCACACGTCTGAACTCTTTCCCTACACGACGCTCTTCCGATCTNNNCACCTGTAATCCCAGCTACTTG
AGCACACGTCTGAACTCTTTCCCTACACGACGCTCTTCCGATCTNNNCACCTGTAATCCCAGCTACTTG
CACACGTCTGAACTCTTTCCCTACACGACGCTCTTCCGATCTNNNAATTTTTGTATTTGTATCGGCGGC
CACACGTCTGAACTCTTTCCCTACACGACGCTCTTCCGATCTNNNAATTTTTGTATTTGTATCGGCGGC
GAGCACACGTCTGAACTCTTTCCCTACACGACGCTCTTCCGATCTNNNCTGGGAGAGAGAGAGAGA
GAGCACACGTCTGAACTCTTTCCCTACACGACGCTCTTCCGATCTNNNCTGGGAGAGAGAGAGAGA
GCACACGTCTGAACTCTTTCCCTACACGACGCTCTTCCGATCTNNNGTTCTCCTGCTTCAGCCTCTCCAT
GCACACGTCTGAACTCTTTCCCTACACGACGCTCTTCCGATCTNNNGTTCTCCTGCTTCAGCCTCTCCAT
AGCACACGTCTGAACTCTTTCCCTACACGACGCTCTTCCGATCTNNNGAGTAGATTGGATGGAGCTGA
AGCACACGTCTGAACTCTTTCCCTACACGACGCTCTTCCGATCTNNNGAGTAGATTGGATGGAGCTGA
GAGCACACGTCTGAACTCTTTCCCTACACGACGCTCTTCCGATCTNNNCTAGCCTACAGAACAGTGAG
GAGCACACGTCTGAACTCTTTCCCTACACGACGCTCTTCCGATCTNNNCTAGCCTACAGAACAGTGAG
GCACACGTCTGAACTCTTTCCCTACACGACGCTCTTCCGATCTNNNTCACTTGCCATCAAGCTGATTCT
GCACACGTCTGAACTCTTTCCCTACACGACGCTCTTCCGATCTNNNTCACTTGCCATCAAGCTGATTCT
AGCACACGTCTGAACTCTTTCCCTACACGACGCTCTTCCGATCTNNNTGATATCTACATTTGTTTCTGCA
AGCACACGTCTGAACTCTTTCCCTACACGACGCTCTTCCGATCTNNNTGATATCTACATTTGTTTCTGCA
AGCACACGTCTGAACTCTTTCCCTACACGACGCTCTTCCGATCTNNNGCTACCATGCCCAGCTAATTTTT
AGCACACGTCTGAACTCTTTCCCTACACGACGCTCTTCCGATCTNNNGCTACCATGCCCAGCTAATTTTT
AGCACACGTCTGAACTCTTTCCCTACACGACGCTCTTCCGATCTNNNTCTGGGTGGACTAGTGAACTG
AGCACACGTCTGAACTCTTTCCCTACACGACGCTCTTCCGATCTNNNTCTGGGTGGACTAGTGAACTG
GAGCACACGTCTGAACTCTTTCCCTACACGACGCTCTTCCGATCTNNNTTCCACGGTGTATATGTACCA
GAGCACACGTCTGAACTCTTTCCCTACACGACGCTCTTCCGATCTNNNTTCCACGGTGTATATGTACCA
AGCACACGTCTGAACTCTTTCCCTACACGACGCTCTTCCGATCTNNNGCGACAGAGTAAGACTCCATC
AGCACACGTCTGAACTCTTTCCCTACACGACGCTCTTCCGATCTNNNGCGACAGAGTAAGACTCCATC
AAGAGCACACGTCTGAACTCTTTCCCTACACGACGCTCTTCCGATCTNNNCTGAGGCAGGAGAATCAC
AAGAGCACACGTCTGAACTCTTTCCCTACACGACGCTCTTCCGATCTNNNCTGAGGCAGGAGAATCAC
GAGCACACGTCTGAACTCTTTCCCTACACGACGCTCTTCCGATCTNNNGGATAGACAACACCACATACG
GAGCACACGTCTGAACTCTTTCCCTACACGACGCTCTTCCGATCTNNNGGATAGACAACACCACATACG
AGCACACGTCTGAACTCTTTCCCTACACGACGCTCTTCCGATCTNNNGAAGTGTCTGTTCACGTCCTTT
AGCACACGTCTGAACTCTTTCCCTACACGACGCTCTTCCGATCTNNNGAAGTGTCTGTTCACGTCCTTT
AGCACACGTCTGAACTCTTTCCCTACACGACGCTCTTCCGATCTNNNGCCTGTAGTCCCAGCTACTTGC
AGCACACGTCTGAACTCTTTCCCTACACGACGCTCTTCCGATCTNNNGCCTGTAGTCCCAGCTACTTGC
GAGCACACGTCTGAACTCTTTCCCTACACGACGCTCTTCCGATCTNNNCTTTCAGGGGGAGGTTGAA
GAGCACACGTCTGAACTCTTTCCCTACACGACGCTCTTCCGATCTNNNCTTTCAGGGGGAGGTTGAA
GCACACGTCTGAACTCTTTCCCTACACGACGCTCTTCCGATCTNNNCGGTGGAAAATGGAGGAAGTA
GCACACGTCTGAACTCTTTCCCTACACGACGCTCTTCCGATCTNNNCGGTGGAAAATGGAGGAAGTA
AGAGCACACGTCTGAACTCTTTCCCTACACGACGCTCTTCCGATCTNNNCCCGGCTCCAAGCATCTTTT

AGAGCACACGTCTGAACTCTTTCCCTACACGACGCTCTTCCGATCTNNNCCCGGCTCCAAGCATCTTTT
CACGTCTGAACTCTTTCCCTACACGACGCTCTTCCGATCTNNNACTGGGAGAAATCTGAGAAAGCTCA
CACGTCTGAACTCTTTCCCTACACGACGCTCTTCCGATCTNNNACTGGGAGAAATCTGAGAAAGCTCA
AGCACACGTCTGAACTCTTTCCCTACACGACGCTCTTCCGATCTNNNGGATGCAACTAAGAAGGGAAG
AGCACACGTCTGAACTCTTTCCCTACACGACGCTCTTCCGATCTNNNGGATGCAACTAAGAAGGGAAG
GAGCACACGTCTGAACTCTTTCCCTACACGACGCTCTTCCGATCTNNNCTGTAATGAGCCATGACTGAG
GAGCACACGTCTGAACTCTTTCCCTACACGACGCTCTTCCGATCTNNNCTGTAATGAGCCATGACTGAG
AGCACACGTCTGAACTCTTTCCCTACACGACGCTCTTCCGATCTNNNATCAGAGACTAGGATTGCAAC
AGCACACGTCTGAACTCTTTCCCTACACGACGCTCTTCCGATCTNNNATCAGAGACTAGGATTGCAAC
GCACACGTCTGAACTCTTTCCCTACACGACGCTCTTCCGATCTNNNATCTTCTGACCTCATGATCCACCC
GCACACGTCTGAACTCTTTCCCTACACGACGCTCTTCCGATCTNNNATCTTCTGACCTCATGATCCACCC
GAGCACACGTCTGAACTCTTTCCCTACACGACGCTCTTCCGATCTNNNGATGAGCTGAGATCATGCCA
GAGCACACGTCTGAACTCTTTCCCTACACGACGCTCTTCCGATCTNNNGATGAGCTGAGATCATGCCA
GCACACGTCTGAACTCTTTCCCTACACGACGCTCTTCCGATCTNNNCAGTATGGTCGATGATGCTGGT
GCACACGTCTGAACTCTTTCCCTACACGACGCTCTTCCGATCTNNNCAGTATGGTCGATGATGCTGGT
GCACACGTCTGAACTCTTTCCCTACACGACGCTCTTCCGATCTNNNTTACTGTCCCTGGTTCTCTCCATC
GCACACGTCTGAACTCTTTCCCTACACGACGCTCTTCCGATCTNNNTTACTGTCCCTGGTTCTCTCCATC
AGCACACGTCTGAACTCTTTCCCTACACGACGCTCTTCCGATCTNNNAGACACTAAAACCAATTTTGAA
AGCACACGTCTGAACTCTTTCCCTACACGACGCTCTTCCGATCTNNNAGACACTAAAACCAATTTTGAA
AGCACACGTCTGAACTCTTTCCCTACACGACGCTCTTCCGATCTNNNCATTCCTAGAAGACACCATCCCC
AGCACACGTCTGAACTCTTTCCCTACACGACGCTCTTCCGATCTNNNCATTCCTAGAAGACACCATCCCC
AGAGCACACGTCTGAACTCTTTCCCTACACGACGCTCTTCCGATCTNNNCTTGGGAAGCTGAGGTAG
AGAGCACACGTCTGAACTCTTTCCCTACACGACGCTCTTCCGATCTNNNCTTGGGAAGCTGAGGTAG
GCACACGTCTGAACTCTTTCCCTACACGACGCTCTTCCGATCTNNNCCACGCCTGGCTAATTTTTTGTA
GCACACGTCTGAACTCTTTCCCTACACGACGCTCTTCCGATCTNNNCCACGCCTGGCTAATTTTTTGTA
AGCACACGTCTGAACTCTTTCCCTACACGACGCTCTTCCGATCTNNNCCATTGTTCGGTCTATTCAGCA
AGCACACGTCTGAACTCTTTCCCTACACGACGCTCTTCCGATCTNNNCCATTGTTCGGTCTATTCAGCA
AGCACACGTCTGAACTCTTTCCCTACACGACGCTCTTCCGATCTNNNAAAATATTGGGCAACAGAGCG
AGCACACGTCTGAACTCTTTCCCTACACGACGCTCTTCCGATCTNNNAAAATATTGGGCAACAGAGCG
CACACGTCTGAACTCTTTCCCTACACGACGCTCTTCCGATCTNNNCTGTAGTCCCAGCTACTCAGGATC
CACACGTCTGAACTCTTTCCCTACACGACGCTCTTCCGATCTNNNCTGTAGTCCCAGCTACTCAGGATC
GAGCACACGTCTGAACTCTTTCCCTACACGACGCTCTTCCGATCTNNNAACCCACAGTTGCAATGGAAA
GAGCACACGTCTGAACTCTTTCCCTACACGACGCTCTTCCGATCTNNNAACCCACAGTTGCAATGGAAA
GAGCACACGTCTGAACTCTTTCCCTACACGACGCTCTTCCGATCTNNNAGCAAGACGCCATCTCAAAAA
GAGCACACGTCTGAACTCTTTCCCTACACGACGCTCTTCCGATCTNNNAGCAAGACGCCATCTCAAAAA
GAGCACACGTCTGAACTCTTTCCCTACACGACGCTCTTCCGATCTNNNAGAATCGCTTGAACCAAGGA
GAGCACACGTCTGAACTCTTTCCCTACACGACGCTCTTCCGATCTNNNAGAATCGCTTGAACCAAGGA
GAGCACACGTCTGAACTCTTTCCCTACACGACGCTCTTCCGATCTNNNATGCTTGTAATCCCAGCACTT
GAGCACACGTCTGAACTCTTTCCCTACACGACGCTCTTCCGATCTNNNATGCTTGTAATCCCAGCACTT
AGCACACGTCTGAACTCTTTCCCTACACGACGCTCTTCCGATCTNNNTCTGTAGGAGCAACTGGAGAG
AGCACACGTCTGAACTCTTTCCCTACACGACGCTCTTCCGATCTNNNTCTGTAGGAGCAACTGGAGAG
GAGCACACGTCTGAACTCTTTCCCTACACGACGCTCTTCCGATCTNNNTGTAGTCCTAGCTACTCAGGA
GAGCACACGTCTGAACTCTTTCCCTACACGACGCTCTTCCGATCTNNNTGTAGTCCTAGCTACTCAGGA

GAGCACACGTCTGAACTCTTTCCCTACACGACGCTCTTCCGATCTNNNTTGTGAAAGTAGCTACACTCCC
GAGCACACGTCTGAACTCTTTCCCTACACGACGCTCTTCCGATCTNNNTTGTGAAAGTAGCTACACTCCC
AGCACACGTCTGAACTCTTTCCCTACACGACGCTCTTCCGATCTNNNGCAGGAGAATCATTTGAACCCA
AGCACACGTCTGAACTCTTTCCCTACACGACGCTCTTCCGATCTNNNGCAGGAGAATCATTTGAACCCA
GAGCACACGTCTGAACTCTTTCCCTACACGACGCTCTTCCGATCTNNNTGCCTGTAATCTCAGCTACTTC
GAGCACACGTCTGAACTCTTTCCCTACACGACGCTCTTCCGATCTNNNTGCCTGTAATCTCAGCTACTTC
AGCACACGTCTGAACTCTTTCCCTACACGACGCTCTTCCGATCTNNNAGCAAAAATGATGTGTTCCCAG
AGCACACGTCTGAACTCTTTCCCTACACGACGCTCTTCCGATCTNNNAGCAAAAATGATGTGTTCCCAG
GAGCACACGTCTGAACTCTTTCCCTACACGACGCTCTTCCGATCTNNNAACTCAGGAGGTAGATTGCA
GAGCACACGTCTGAACTCTTTCCCTACACGACGCTCTTCCGATCTNNNAACTCAGGAGGTAGATTGCA
AGCACACGTCTGAACTCTTTCCCTACACGACGCTCTTCCGATCTNNNATCTGTCCCGTATCGTAGTTGTC
AGCACACGTCTGAACTCTTTCCCTACACGACGCTCTTCCGATCTNNNATCTGTCCCGTATCGTAGTTGTC
AGCACACGTCTGAACTCTTTCCCTACACGACGCTCTTCCGATCTNNNTAGGCAATAGAGAGAGACTCC
AGCACACGTCTGAACTCTTTCCCTACACGACGCTCTTCCGATCTNNNTAGGCAATAGAGAGAGACTCC
AGCACACGTCTGAACTCTTTCCCTACACGACGCTCTTCCGATCTNNNTCTCAAAGCTGCATCTCAAATGC
AGCACACGTCTGAACTCTTTCCCTACACGACGCTCTTCCGATCTNNNTCTCAAAGCTGCATCTCAAATGC
GAGCACACGTCTGAACTCTTTCCCTACACGACGCTCTTCCGATCTNNNTTTTGGATGCCGATATGGTTT
GAGCACACGTCTGAACTCTTTCCCTACACGACGCTCTTCCGATCTNNNTTTTGGATGCCGATATGGTTT
GAGCACACGTCTGAACTCTTTCCCTACACGACGCTCTTCCGATCTNNNTGAGGCAGGAGAATCTCTTGA
GAGCACACGTCTGAACTCTTTCCCTACACGACGCTCTTCCGATCTNNNTGAGGCAGGAGAATCTCTTGA
AGCACACGTCTGAACTCTTTCCCTACACGACGCTCTTCCGATCTNNNCACCTAGCTACCCTGGACTACCCA
AGCACACGTCTGAACTCTTTCCCTACACGACGCTCTTCCGATCTNNNCACCTAGCTACCCTGGACTACCCA
AGCACACGTCTGAACTCTTTCCCTACACGACGCTCTTCCGATCTNNNAGTGTGGACATGAAAGAGAGT
AGCACACGTCTGAACTCTTTCCCTACACGACGCTCTTCCGATCTNNNAGTGTGGACATGAAAGAGAGT
GAGCACACGTCTGAACTCTTTCCCTACACGACGCTCTTCCGATCTNNNCTGGCTTGCTTTCTTGCTTTCTT
GAGCACACGTCTGAACTCTTTCCCTACACGACGCTCTTCCGATCTNNNCTGGCTTGCTTTCTTGCTTTCTT
AGCACACGTCTGAACTCTTTCCCTACACGACGCTCTTCCGATCTNNNGATTTGATGGGGATCAGTGCCT
AGCACACGTCTGAACTCTTTCCCTACACGACGCTCTTCCGATCTNNNGATTTGATGGGGATCAGTGCCT
CACACGTCTGAACTCTTTCCCTACACGACGCTCTTCCGATCTNNNGATGCCCGTCCTCGTCAATATGCATC
CACACGTCTGAACTCTTTCCCTACACGACGCTCTTCCGATCTNNNGATGCCCGTCCTCGTCAATATGCATC
GCACACGTCTGAACTCTTTCCCTACACGACGCTCTTCCGATCTNNNCATGAATGCAGAACTCACAGCTA
GCACACGTCTGAACTCTTTCCCTACACGACGCTCTTCCGATCTNNNCATGAATGCAGAACTCACAGCTA
AGCACACGTCTGAACTCTTTCCCTACACGACGCTCTTCCGATCTNNNTTTTCCCTTTTCGTGTTCTGCAGC
AGCACACGTCTGAACTCTTTCCCTACACGACGCTCTTCCGATCTNNNTTTTCCCTTTTCGTGTTCTGCAGC
AGAGCACACGTCTGAACTCTTTCCCTACACGACGCTCTTCCGATCTNNNGTCTTTGTCCCTGTACCTGTC
AGAGCACACGTCTGAACTCTTTCCCTACACGACGCTCTTCCGATCTNNNGTCTTTGTCCCTGTACCTGTC
GCACACGTCTGAACTCTTTCCCTACACGACGCTCTTCCGATCTNNNAAACTGGACCCCTTCTTCTTTGTC
GCACACGTCTGAACTCTTTCCCTACACGACGCTCTTCCGATCTNNNAAACTGGACCCCTTCTTCTTTGTC
GAGCACACGTCTGAACTCTTTCCCTACACGACGCTCTTCCGATCTNNNCAAGACTGCACCACTGTACTCC
GAGCACACGTCTGAACTCTTTCCCTACACGACGCTCTTCCGATCTNNNCAAGACTGCACCACTGTACTCC
AGCACACGTCTGAACTCTTTCCCTACACGACGCTCTTCCGATCTNNNAGATCTTTGGTAAGGCAGGTCT
AGCACACGTCTGAACTCTTTCCCTACACGACGCTCTTCCGATCTNNNAGATCTTTGGTAAGGCAGGTCT
AGAGCACACGTCTGAACTCTTTCCCTACACGACGCTCTTCCGATCTNNNGGAGTTCGCAGAGTGGAA

AGAGCACACGTCTGAACTCTTTCCCTACACGACGCTCTTCCGATCTNNNGGAGTTCGCAGAGTGGAA
GCACACGTCTGAACTCTTTCCCTACACGACGCTCTTCCGATCTNNNGGAATGGAAGCTCTGTTGAAA
GCACACGTCTGAACTCTTTCCCTACACGACGCTCTTCCGATCTNNNGGAATGGAAGCTCTGTTGAAA
AGCACACGTCTGAACTCTTTCCCTACACGACGCTCTTCCGATCTNNNCTGTAGTCCCAGCTACCCAGA
AGCACACGTCTGAACTCTTTCCCTACACGACGCTCTTCCGATCTNNNCTGTAGTCCCAGCTACCCAGA
AGCACACGTCTGAACTCTTTCCCTACACGACGCTCTTCCGATCTNNNCAATCCAACAGCTTCACAGAT
AGCACACGTCTGAACTCTTTCCCTACACGACGCTCTTCCGATCTNNNCAATCCAACAGCTTCACAGAT
AGCACACGTCTGAACTCTTTCCCTACACGACGCTCTTCCGATCTNNNGGTGACAGAGTGAGACTCCAT
AGCACACGTCTGAACTCTTTCCCTACACGACGCTCTTCCGATCTNNNGGTGACAGAGTGAGACTCCAT
AGCACACGTCTGAACTCTTTCCCTACACGACGCTCTTCCGATCTNNNTTTGGGACTCACACTCCTAGAG
AGCACACGTCTGAACTCTTTCCCTACACGACGCTCTTCCGATCTNNNTTTGGGACTCACACTCCTAGAG
AGCACACGTCTGAACTCTTTCCCTACACGACGCTCTTCCGATCTNNNGCATGGACAACAAGAGCAAAA
AGCACACGTCTGAACTCTTTCCCTACACGACGCTCTTCCGATCTNNNGCATGGACAACAAGAGCAAAA
AGCACACGTCTGAACTCTTTCCCTACACGACGCTCTTCCGATCTNNNCTGAGTATCTGGGAGCACAGG
AGCACACGTCTGAACTCTTTCCCTACACGACGCTCTTCCGATCTNNNCTGAGTATCTGGGAGCACAGG
AGCACACGTCTGAACTCTTTCCCTACACGACGCTCTTCCGATCTNNNGAGTTTCGCTTTGGAGGATGT
AGCACACGTCTGAACTCTTTCCCTACACGACGCTCTTCCGATCTNNNGAGTTTCGCTTTGGAGGATGT
GCACACGTCTGAACTCTTTCCCTACACGACGCTCTTCCGATCTNNNACAAGGGAGAAATCCAAAACCT
GCACACGTCTGAACTCTTTCCCTACACGACGCTCTTCCGATCTNNNACAAGGGAGAAATCCAAAACCT
GAGCACACGTCTGAACTCTTTCCCTACACGACGCTCTTCCGATCTNNNTAGTTCCAAGCTATTCAGGAG
GAGCACACGTCTGAACTCTTTCCCTACACGACGCTCTTCCGATCTNNNTAGTTCCAAGCTATTCAGGAG
CACACGTCTGAACTCTTTCCCTACACGACGCTCTTCCGATCTNNNACTTTGTGAGTTCAGGTCCGGC
CACACGTCTGAACTCTTTCCCTACACGACGCTCTTCCGATCTNNNACTTTGTGAGTTCAGGTCCGGC
AGCACACGTCTGAACTCTTTCCCTACACGACGCTCTTCCGATCTNNNGGCTGTGTATGTCCATAGT
AGCACACGTCTGAACTCTTTCCCTACACGACGCTCTTCCGATCTNNNGGCTGTGTATGTCCATAGT
GCACACGTCTGAACTCTTTCCCTACACGACGCTCTTCCGATCTNNNGGGCCAGTATGTATTTGGGGA
GCACACGTCTGAACTCTTTCCCTACACGACGCTCTTCCGATCTNNNGGGCCAGTATGTATTTGGGGA
AGCACACGTCTGAACTCTTTCCCTACACGACGCTCTTCCGATCTNNNGCCTGTAGTCCCAGCTACTTG
AGCACACGTCTGAACTCTTTCCCTACACGACGCTCTTCCGATCTNNNGCCTGTAGTCCCAGCTACTTG
AGCACACGTCTGAACTCTTTCCCTACACGACGCTCTTCCGATCTNNNGGCCCCGGGAATTATTTCTTT
AGCACACGTCTGAACTCTTTCCCTACACGACGCTCTTCCGATCTNNNGGCCCCGGGAATTATTTCTTT
GCACACGTCTGAACTCTTTCCCTACACGACGCTCTTCCGATCTNNNTGACAGAGCAAGACCTTGTCTC
GCACACGTCTGAACTCTTTCCCTACACGACGCTCTTCCGATCTNNNTGACAGAGCAAGACCTTGTCTC
AGCACACGTCTGAACTCTTTCCCTACACGACGCTCTTCCGATCTNNNCATGGTGGCATGTGTCTGTAA
AGCACACGTCTGAACTCTTTCCCTACACGACGCTCTTCCGATCTNNNCATGGTGGCATGTGTCTGTAA
AGCACACGTCTGAACTCTTTCCCTACACGACGCTCTTCCGATCTNNNGGTTCAAGGGATTCTCCTCTCT
AGCACACGTCTGAACTCTTTCCCTACACGACGCTCTTCCGATCTNNNGGTTCAAGGGATTCTCCTCTCT
AGCACACGTCTGAACTCTTTCCCTACACGACGCTCTTCCGATCTNNNCAGTTTGCAGTGAGCTGAGAT
AGCACACGTCTGAACTCTTTCCCTACACGACGCTCTTCCGATCTNNNCAGTTTGCAGTGAGCTGAGAT
GAGCACACGTCTGAACTCTTTCCCTACACGACGCTCTTCCGATCTNNNTGAGTAACAGAGCAAGACTG
GAGCACACGTCTGAACTCTTTCCCTACACGACGCTCTTCCGATCTNNNTGAGTAACAGAGCAAGACTG
AGCACACGTCTGAACTCTTTCCCTACACGACGCTCTTCCGATCTNNNTTTCTAGAGAGTCTTCAGCCA
AGCACACGTCTGAACTCTTTCCCTACACGACGCTCTTCCGATCTNNNTTTCTAGAGAGTCTTCAGCCA

GCACACGTCTGAACTCTTTCCCTACACGACGCTCTTCCGATCTNNNACGCCTCTTAGACCTCAACACTAC

GCACACGTCTGAACTCTTTCCCTACACGACGCTCTTCCGATCTNNNACGCCTCTTAGACCTCAACACTAC

GCACACGTCTGAACTCTTTCCCTACACGACGCTCTTCCGATCTNNNTTGACCCATGGAGAACAGAGAA

GCACACGTCTGAACTCTTTCCCTACACGACGCTCTTCCGATCTNNNTTGACCCATGGAGAACAGAGAA

GAGCACACGTCTGAACTCTTTCCCTACACGACGCTCTTCCGATCTNNNTTTACAACATTGGGGGCATT

GAGCACACGTCTGAACTCTTTCCCTACACGACGCTCTTCCGATCTNNNTTTACAACATTGGGGGCATT

AGAGCACACGTCTGAACTCTTTCCCTACACGACGCTCTTCCGATCTNNNTGCCACCTGCTAATTGTAGA

AGAGCACACGTCTGAACTCTTTCCCTACACGACGCTCTTCCGATCTNNNTGCCACCTGCTAATTGTAGA

GAGCACACGTCTGAACTCTTTCCCTACACGACGCTCTTCCGATCTNNNAAATCTCTCCCTTCCTCCCTGTC

GAGCACACGTCTGAACTCTTTCCCTACACGACGCTCTTCCGATCTNNNAAATCTCTCCCTTCCTCCCTGTC

AGCACACGTCTGAACTCTTTCCCTACACGACGCTCTTCCGATCTNNNAGCTGAGATTACAGGTTCACAC

AGCACACGTCTGAACTCTTTCCCTACACGACGCTCTTCCGATCTNNNAGCTGAGATTACAGGTTCACAC

GAGCACACGTCTGAACTCTTTCCCTACACGACGCTCTTCCGATCTNNNGCTAGGGGACTGGGATAGA

GAGCACACGTCTGAACTCTTTCCCTACACGACGCTCTTCCGATCTNNNGCTAGGGGACTGGGATAGA

AGCACACGTCTGAACTCTTTCCCTACACGACGCTCTTCCGATCTNNNTATGGGTTGGTAGAGGAACAG

AGCACACGTCTGAACTCTTTCCCTACACGACGCTCTTCCGATCTNNNTATGGGTTGGTAGAGGAACAG

GCACACGTCTGAACTCTTTCCCTACACGACGCTCTTCCGATCTNNNCACGCCTATAATCCCAGCTACTCC

GCACACGTCTGAACTCTTTCCCTACACGACGCTCTTCCGATCTNNNCACGCCTATAATCCCAGCTACTCC

CACGTCTGAACTCTTTCCCTACACGACGCTCTTCCGATCTNNNCATGTTCAAGACCTTCCTGCTTGCATC

CACGTCTGAACTCTTTCCCTACACGACGCTCTTCCGATCTNNNCATGTTCAAGACCTTCCTGCTTGCATC

GAGCACACGTCTGAACTCTTTCCCTACACGACGCTCTTCCGATCTNNNCCCTAGTAGCTGGGACTACAG

GAGCACACGTCTGAACTCTTTCCCTACACGACGCTCTTCCGATCTNNNCCCTAGTAGCTGGGACTACAG

GAGCACACGTCTGAACTCTTTCCCTACACGACGCTCTTCCGATCTNNNGAGGCCAAATGACAGTTCTC

GAGCACACGTCTGAACTCTTTCCCTACACGACGCTCTTCCGATCTNNNGAGGCCAAATGACAGTTCTC

GAGCACACGTCTGAACTCTTTCCCTACACGACGCTCTTCCGATCTNNNTGTGATATTGCACCTCTGTTA

GAGCACACGTCTGAACTCTTTCCCTACACGACGCTCTTCCGATCTNNNTGTGATATTGCACCTCTGTTA

GAGCACACGTCTGAACTCTTTCCCTACACGACGCTCTTCCGATCTNNNAATGGCAAAGTGAGACTCCAT

GAGCACACGTCTGAACTCTTTCCCTACACGACGCTCTTCCGATCTNNNAATGGCAAAGTGAGACTCCAT

AGCACACGTCTGAACTCTTTCCCTACACGACGCTCTTCCGATCTNNNGCCTGGGTGAAAAGAGTGAA

AGCACACGTCTGAACTCTTTCCCTACACGACGCTCTTCCGATCTNNNGCCTGGGTGAAAAGAGTGAA

GAGCACACGTCTGAACTCTTTCCCTACACGACGCTCTTCCGATCTNNNAAGATATTTCTGGGCTGGGT

GAGCACACGTCTGAACTCTTTCCCTACACGACGCTCTTCCGATCTNNNAAGATATTTCTGGGCTGGGT

GAGCACACGTCTGAACTCTTTCCCTACACGACGCTCTTCCGATCTNNNGCATTCAGCTTCAAGGAACCA

GAGCACACGTCTGAACTCTTTCCCTACACGACGCTCTTCCGATCTNNNGCATTCAGCTTCAAGGAACCA

AGCACACGTCTGAACTCTTTCCCTACACGACGCTCTTCCGATCTNNNATACCCGTTGGCCATTTCTATGT

AGCACACGTCTGAACTCTTTCCCTACACGACGCTCTTCCGATCTNNNATACCCGTTGGCCATTTCTATGT

GAGCACACGTCTGAACTCTTTCCCTACACGACGCTCTTCCGATCTNNNTAATCCCAACACTTTAGGAGG

GAGCACACGTCTGAACTCTTTCCCTACACGACGCTCTTCCGATCTNNNTAATCCCAACACTTTAGGAGG

AGAGCACACGTCTGAACTCTTTCCCTACACGACGCTCTTCCGATCTNNNCTGATGTAGGAGGATCGCT

AGAGCACACGTCTGAACTCTTTCCCTACACGACGCTCTTCCGATCTNNNCTGATGTAGGAGGATCGCT

GAGCACACGTCTGAACTCTTTCCCTACACGACGCTCTTCCGATCTNNNAGGCTGACATCTAGCTACATG

GAGCACACGTCTGAACTCTTTCCCTACACGACGCTCTTCCGATCTNNNAGGCTGACATCTAGCTACATG

CACACGTCTGAACTCTTTCCCTACACGACGCTCTTCCGATCTNNNAAGGCATATGGAGATGTAGTTACA

CACACGTCTGAACTCTTTCCCTACACGACGCTCTTCCGATCTNNNAAGGCATATGGAGATGTAGTTACA
AGCACACGTCTGAACTCTTTCCCTACACGACGCTCTTCCGATCTNNNTCTGTTACAGACTTGCCAGATGT
AGCACACGTCTGAACTCTTTCCCTACACGACGCTCTTCCGATCTNNNTCTGTTACAGACTTGCCAGATGT
GCACACGTCTGAACTCTTTCCCTACACGACGCTCTTCCGATCTNNNCTCCAGGTCCTTATCTGTGTCACC
GCACACGTCTGAACTCTTTCCCTACACGACGCTCTTCCGATCTNNNCTCCAGGTCCTTATCTGTGTCACC
GAGCACACGTCTGAACTCTTTCCCTACACGACGCTCTTCCGATCTNNNAAGGCAAGTACCTCCTCTCTCT
GAGCACACGTCTGAACTCTTTCCCTACACGACGCTCTTCCGATCTNNNAAGGCAAGTACCTCCTCTCTCT
AGCACACGTCTGAACTCTTTCCCTACACGACGCTCTTCCGATCTNNNACTACAGCACCATCCTTTCTGAA
AGCACACGTCTGAACTCTTTCCCTACACGACGCTCTTCCGATCTNNNACTACAGCACCATCCTTTCTGAA
GAGCACACGTCTGAACTCTTTCCCTACACGACGCTCTTCCGATCTNNNGAATGGCTTGAACCCAGGATT
GAGCACACGTCTGAACTCTTTCCCTACACGACGCTCTTCCGATCTNNNGAATGGCTTGAACCCAGGATT
GCACACGTCTGAACTCTTTCCCTACACGACGCTCTTCCGATCTNNNCAAACAAAAGAACCTGGCAGGAG
GCACACGTCTGAACTCTTTCCCTACACGACGCTCTTCCGATCTNNNCAAACAAAAGAACCTGGCAGGAG
AGCACACGTCTGAACTCTTTCCCTACACGACGCTCTTCCGATCTNNNAGTCTTACCTTTCCTGCCACTTT
AGCACACGTCTGAACTCTTTCCCTACACGACGCTCTTCCGATCTNNNAGTCTTACCTTTCCTGCCACTTT
AGAGCACACGTCTGAACTCTTTCCCTACACGACGCTCTTCCGATCTNNNGCACTCAGGGTCTTAGTGAA
AGAGCACACGTCTGAACTCTTTCCCTACACGACGCTCTTCCGATCTNNNGCACTCAGGGTCTTAGTGAA
GAGCACACGTCTGAACTCTTTCCCTACACGACGCTCTTCCGATCTNNNCCTCAGCTAACAGCACTGAACA
GAGCACACGTCTGAACTCTTTCCCTACACGACGCTCTTCCGATCTNNNCCTCAGCTAACAGCACTGAACA
GAGCACACGTCTGAACTCTTTCCCTACACGACGCTCTTCCGATCTNNNTTAGTCCCTGGCAACTACTGT
GAGCACACGTCTGAACTCTTTCCCTACACGACGCTCTTCCGATCTNNNTTAGTCCCTGGCAACTACTGT
GAGCACACGTCTGAACTCTTTCCCTACACGACGCTCTTCCGATCTNNNAGTGAGCTTAGATCGCACCAT
GAGCACACGTCTGAACTCTTTCCCTACACGACGCTCTTCCGATCTNNNAGTGAGCTTAGATCGCACCAT
AGCACACGTCTGAACTCTTTCCCTACACGACGCTCTTCCGATCTNNNTTTTATGTGCTTATTGGCCGCA
AGCACACGTCTGAACTCTTTCCCTACACGACGCTCTTCCGATCTNNNTTTTATGTGCTTATTGGCCGCA
AGCACACGTCTGAACTCTTTCCCTACACGACGCTCTTCCGATCTNNNGAGGTTTCAGTGAGCCAAGATT
AGCACACGTCTGAACTCTTTCCCTACACGACGCTCTTCCGATCTNNNGAGGTTTCAGTGAGCCAAGATT
AGCACACGTCTGAACTCTTTCCCTACACGACGCTCTTCCGATCTNNNCTACTTCCCTTCTTGCAGGCATTC
AGCACACGTCTGAACTCTTTCCCTACACGACGCTCTTCCGATCTNNNCTACTTCCCTTCTTGCAGGCATTC
AGCACACGTCTGAACTCTTTCCCTACACGACGCTCTTCCGATCTNNNACAAGTCACAGTACAGTAGGCTT
AGCACACGTCTGAACTCTTTCCCTACACGACGCTCTTCCGATCTNNNACAAGTCACAGTACAGTAGGCTT
AGCACACGTCTGAACTCTTTCCCTACACGACGCTCTTCCGATCTNNNGCTTCTCTATCCAGCATAGGTCC
AGCACACGTCTGAACTCTTTCCCTACACGACGCTCTTCCGATCTNNNGCTTCTCTATCCAGCATAGGTCC
GAGCACACGTCTGAACTCTTTCCCTACACGACGCTCTTCCGATCTNNNTGAGGTCAGGAGTTTGAGAA
GAGCACACGTCTGAACTCTTTCCCTACACGACGCTCTTCCGATCTNNNTGAGGTCAGGAGTTTGAGAA
GAGCACACGTCTGAACTCTTTCCCTACACGACGCTCTTCCGATCTNNNCATGCTTCTCCTTTGTACGTGT
GAGCACACGTCTGAACTCTTTCCCTACACGACGCTCTTCCGATCTNNNCATGCTTCTCCTTTGTACGTGT
GAGCACACGTCTGAACTCTTTCCCTACACGACGCTCTTCCGATCTNNNGTATCTTGCCCTGTGCCATTTT
GAGCACACGTCTGAACTCTTTCCCTACACGACGCTCTTCCGATCTNNNGTATCTTGCCCTGTGCCATTTT
GAGCACACGTCTGAACTCTTTCCCTACACGACGCTCTTCCGATCTNNNGAGAATCTCTTGAACCTGGGA
GAGCACACGTCTGAACTCTTTCCCTACACGACGCTCTTCCGATCTNNNGAGAATCTCTTGAACCTGGGA
GAGCACACGTCTGAACTCTTTCCCTACACGACGCTCTTCCGATCTNNNCAGTCTCATGAATTCTGACAA
GAGCACACGTCTGAACTCTTTCCCTACACGACGCTCTTCCGATCTNNNCAGTCTCATGAATTCTGACAA

AGCACACGTCTGAACTCTTTCCCTACACGACGCTCTTCCGATCTNNNGACTTCCCAAGCCTCCACAATAA

AGCACACGTCTGAACTCTTTCCCTACACGACGCTCTTCCGATCTNNNGACTTCCCAAGCCTCCACAATAA

AGAGCACACGTCTGAACTCTTTCCCTACACGACGCTCTTCCGATCTNNNGAAAAGAGAAGAGAGCAGC

AGAGCACACGTCTGAACTCTTTCCCTACACGACGCTCTTCCGATCTNNNGAAAAGAGAAGAGAGCAGC

GAGCACACGTCTGAACTCTTTCCCTACACGACGCTCTTCCGATCTNNNGTTAGGAGTTCTAGACCAGC

GAGCACACGTCTGAACTCTTTCCCTACACGACGCTCTTCCGATCTNNNGTTAGGAGTTCTAGACCAGC

AAGAGCACACGTCTGAACTCTTTCCCTACACGACGCTCTTCCGATCTNNNTGGAAAACCTACAGTGAA

AAGAGCACACGTCTGAACTCTTTCCCTACACGACGCTCTTCCGATCTNNNTGGAAAACCTACAGTGAA

GCACACGTCTGAACTCTTTCCCTACACGACGCTCTTCCGATCTNNNGCTTCGGGACTTGTTTCTCCTAA

GCACACGTCTGAACTCTTTCCCTACACGACGCTCTTCCGATCTNNNGCTTCGGGACTTGTTTCTCCTAA

GAGCACACGTCTGAACTCTTTCCCTACACGACGCTCTTCCGATCTNNNTGGCCATCTTCCAAGTGTACT

GAGCACACGTCTGAACTCTTTCCCTACACGACGCTCTTCCGATCTNNNTGGCCATCTTCCAAGTGTACT

AGCACACGTCTGAACTCTTTCCCTACACGACGCTCTTCCGATCTNNNTTCTTCCAGTCAGAAACCACCAA

AGCACACGTCTGAACTCTTTCCCTACACGACGCTCTTCCGATCTNNNTTCTTCCAGTCAGAAACCACCAA

AGCACACGTCTGAACTCTTTCCCTACACGACGCTCTTCCGATCTNNNGTTAAGATCTGGGAATCGACCA

AGCACACGTCTGAACTCTTTCCCTACACGACGCTCTTCCGATCTNNNGTTAAGATCTGGGAATCGACCA

AGAGCACACGTCTGAACTCTTTCCCTACACGACGCTCTTCCGATCTNNNGAAACAACAGCCTTACAGG

AGAGCACACGTCTGAACTCTTTCCCTACACGACGCTCTTCCGATCTNNNGAAACAACAGCCTTACAGG

AGAGCACACGTCTGAACTCTTTCCCTACACGACGCTCTTCCGATCTNNNGTGAGCTGAGATCATGCTAC

AGAGCACACGTCTGAACTCTTTCCCTACACGACGCTCTTCCGATCTNNNGTGAGCTGAGATCATGCTAC

GAGCACACGTCTGAACTCTTTCCCTACACGACGCTCTTCCGATCTNNNGCTCATGCTTATAATCCCAGC

GAGCACACGTCTGAACTCTTTCCCTACACGACGCTCTTCCGATCTNNNGCTCATGCTTATAATCCCAGC

GCACACGTCTGAACTCTTTCCCTACACGACGCTCTTCCGATCTNNNCCAGCTGTTTGCATCTTTTCTCACA

GCACACGTCTGAACTCTTTCCCTACACGACGCTCTTCCGATCTNNNCCAGCTGTTTGCATCTTTTCTCACA

AAGAGCACACGTCTGAACTCTTTCCCTACACGACGCTCTTCCGATCTNNNCTTGTAGTGAGCTGAGAT

AAGAGCACACGTCTGAACTCTTTCCCTACACGACGCTCTTCCGATCTNNNCTTGTAGTGAGCTGAGAT

AGAGCACACGTCTGAACTCTTTCCCTACACGACGCTCTTCCGATCTNNNAGAAGAATCGCTTGAACCCA

AGAGCACACGTCTGAACTCTTTCCCTACACGACGCTCTTCCGATCTNNNAGAAGAATCGCTTGAACCCA

GAGCACACGTCTGAACTCTTTCCCTACACGACGCTCTTCCGATCTNNNGGCAGGAGAATCACTTGAACC

GAGCACACGTCTGAACTCTTTCCCTACACGACGCTCTTCCGATCTNNNGGCAGGAGAATCACTTGAACC

GAGCACACGTCTGAACTCTTTCCCTACACGACGCTCTTCCGATCTNNNTTTTCATGAATTGGTGTGGG

GAGCACACGTCTGAACTCTTTCCCTACACGACGCTCTTCCGATCTNNNTTTTCATGAATTGGTGTGGG

GAGCACACGTCTGAACTCTTTCCCTACACGACGCTCTTCCGATCTNNNAAAAAATTAGCCAGGTGTGG

GAGCACACGTCTGAACTCTTTCCCTACACGACGCTCTTCCGATCTNNNAAAAAATTAGCCAGGTGTGG

AGCACACGTCTGAACTCTTTCCCTACACGACGCTCTTCCGATCTNNNGGCATTACAAGCATTGACCTCAC

AGCACACGTCTGAACTCTTTCCCTACACGACGCTCTTCCGATCTNNNGGCATTACAAGCATTGACCTCAC

AGCACACGTCTGAACTCTTTCCCTACACGACGCTCTTCCGATCTNNNTTTCCTTGGTTCTCTTCCCCAAAC

AGCACACGTCTGAACTCTTTCCCTACACGACGCTCTTCCGATCTNNNTTTCCTTGGTTCTCTTCCCCAAAC

AAGAGCACACGTCTGAACTCTTTCCCTACACGACGCTCTTCCGATCTNNNTGTGGTAGTTGCACACAAT

AAGAGCACACGTCTGAACTCTTTCCCTACACGACGCTCTTCCGATCTNNNTGTGGTAGTTGCACACAAT

AGCACACGTCTGAACTCTTTCCCTACACGACGCTCTTCCGATCTNNNAATAACAGTGGTGACATCGGAC

AGCACACGTCTGAACTCTTTCCCTACACGACGCTCTTCCGATCTNNNAATAACAGTGGTGACATCGGAC

AGCACACGTCTGAACTCTTTCCCTACACGACGCTCTTCCGATCTNNNGCAGCAACTCTGTGTTATCTTCA

AGCACACGTCTGAACTCTTTCCCTACACGACGCTCTTCCGATCTNNNGCAGCAACTCTGTGTTATCTTCA

AGCACACGTCTGAACTCTTTCCCTACACGACGCTCTTCCGATCTNNNGTGACAGAGTGAGACTCCTTCA

AGCACACGTCTGAACTCTTTCCCTACACGACGCTCTTCCGATCTNNNGTGACAGAGTGAGACTCCTTCA

CACACGTCTGAACTCTTTCCCTACACGACGCTCTTCCGATCTNNNGGTTCATGGGATTAGAAGAATTCA

CACACGTCTGAACTCTTTCCCTACACGACGCTCTTCCGATCTNNNGGTTCATGGGATTAGAAGAATTCA

AGCACACGTCTGAACTCTTTCCCTACACGACGCTCTTCCGATCTNNNAGGGAAGCAAGGCACAACTTAT

AGCACACGTCTGAACTCTTTCCCTACACGACGCTCTTCCGATCTNNNAGGGAAGCAAGGCACAACTTAT

AGCACACGTCTGAACTCTTTCCCTACACGACGCTCTTCCGATCTNNNATTGAGCAGAGATCGTGCCATT

AGCACACGTCTGAACTCTTTCCCTACACGACGCTCTTCCGATCTNNNATTGAGCAGAGATCGTGCCATT

GAGCACACGTCTGAACTCTTTCCCTACACGACGCTCTTCCGATCTNNNGGACTCCTCCCTAACTCATTCTC

GAGCACACGTCTGAACTCTTTCCCTACACGACGCTCTTCCGATCTNNNGGACTCCTCCCTAACTCATTCTC

GAGCACACGTCTGAACTCTTTCCCTACACGACGCTCTTCCGATCTNNNAGTTCTGGGATTACAGGTGTC

GAGCACACGTCTGAACTCTTTCCCTACACGACGCTCTTCCGATCTNNNAGTTCTGGGATTACAGGTGTC

GAGCACACGTCTGAACTCTTTCCCTACACGACGCTCTTCCGATCTNNNCTGTCTCCGTCTGTCTTATGCT

GAGCACACGTCTGAACTCTTTCCCTACACGACGCTCTTCCGATCTNNNCTGTCTCCGTCTGTCTTATGCT

AGCACACGTCTGAACTCTTTCCCTACACGACGCTCTTCCGATCTNNNCACGGCTTTCCTGCTTTGATAAA

AGCACACGTCTGAACTCTTTCCCTACACGACGCTCTTCCGATCTNNNCACGGCTTTCCTGCTTTGATAAA

GAGCACACGTCTGAACTCTTTCCCTACACGACGCTCTTCCGATCTNNNATTTGCAAACCCAAGGAAAGA

GAGCACACGTCTGAACTCTTTCCCTACACGACGCTCTTCCGATCTNNNATTTGCAAACCCAAGGAAAGA

GAGCACACGTCTGAACTCTTTCCCTACACGACGCTCTTCCGATCTNNNCAGGGTCGCTGATCTCTTCATA

GAGCACACGTCTGAACTCTTTCCCTACACGACGCTCTTCCGATCTNNNCAGGGTCGCTGATCTCTTCATA

AGCACACGTCTGAACTCTTTCCCTACACGACGCTCTTCCGATCTNNNAATGAATGCCCAGAACCTGTTT

AGCACACGTCTGAACTCTTTCCCTACACGACGCTCTTCCGATCTNNNAATGAATGCCCAGAACCTGTTT

AGAGCACACGTCTGAACTCTTTCCCTACACGACGCTCTTCCGATCTNNNGGCATGAATAAGGAACACTG

AGAGCACACGTCTGAACTCTTTCCCTACACGACGCTCTTCCGATCTNNNGGCATGAATAAGGAACACTG

GAGCACACGTCTGAACTCTTTCCCTACACGACGCTCTTCCGATCTNNNGCAAAGTGCAATAAAACAAGC

GAGCACACGTCTGAACTCTTTCCCTACACGACGCTCTTCCGATCTNNNGCAAAGTGCAATAAAACAAGC

AGCACACGTCTGAACTCTTTCCCTACACGACGCTCTTCCGATCTNNNGATCACATCACTGTACTCCAGCT

AGCACACGTCTGAACTCTTTCCCTACACGACGCTCTTCCGATCTNNNGATCACATCACTGTACTCCAGCT

GCACACGTCTGAACTCTTTCCCTACACGACGCTCTTCCGATCTNNNCTGTTTCTCCCTTGTCTCTCCTCCAT

GCACACGTCTGAACTCTTTCCCTACACGACGCTCTTCCGATCTNNNCTGTTTCTCCCTTGTCTCTCCTCCAT

AGCACACGTCTGAACTCTTTCCCTACACGACGCTCTTCCGATCTNNNGGAATGCAGTGACATGATCAT

AGCACACGTCTGAACTCTTTCCCTACACGACGCTCTTCCGATCTNNNGGAATGCAGTGACATGATCAT

GAGCACACGTCTGAACTCTTTCCCTACACGACGCTCTTCCGATCTNNNACACACTTCTTTGTTTTCTTTTG

GAGCACACGTCTGAACTCTTTCCCTACACGACGCTCTTCCGATCTNNNACACACTTCTTTGTTTTCTTTTG

GAGCACACGTCTGAACTCTTTCCCTACACGACGCTCTTCCGATCTNNNCAAAAAGTTTCAGCAGGGGTC

GAGCACACGTCTGAACTCTTTCCCTACACGACGCTCTTCCGATCTNNNCAAAAAGTTTCAGCAGGGGTC

AGCACACGTCTGAACTCTTTCCCTACACGACGCTCTTCCGATCTNNNGCATGCATTTACAGGTGAAGTC

AGCACACGTCTGAACTCTTTCCCTACACGACGCTCTTCCGATCTNNNGCATGCATTTACAGGTGAAGTC

GCACACGTCTGAACTCTTTCCCTACACGACGCTCTTCCGATCTNNNTCCTTCTGAGGTTCCAAACAACAC

GCACACGTCTGAACTCTTTCCCTACACGACGCTCTTCCGATCTNNNTCCTTCTGAGGTTCCAAACAACAC

GCACACGTCTGAACTCTTTCCCTACACGACGCTCTTCCGATCTNNNCTTCAATAACCTGCGCGGAGACA

GCACACGTCTGAACTCTTTCCCTACACGACGCTCTTCCGATCTNNNCTTCAATAACCTGCGCGGAGACA

AGCACACGTCTGAACTCTTTCCCTACACGACGCTCTTCCGATCTNNNAGTTGGGATTACAGGTGCACA
AGCACACGTCTGAACTCTTTCCCTACACGACGCTCTTCCGATCTNNNAGTTGGGATTACAGGTGCACA
AGCACACGTCTGAACTCTTTCCCTACACGACGCTCTTCCGATCTNNNCAGCTTCCATAATAATGTGAAC
AGCACACGTCTGAACTCTTTCCCTACACGACGCTCTTCCGATCTNNNCAGCTTCCATAATAATGTGAAC
GCACACGTCTGAACTCTTTCCCTACACGACGCTCTTCCGATCTNNNCTCATGAGGCTCAGCTGTCTTT
GCACACGTCTGAACTCTTTCCCTACACGACGCTCTTCCGATCTNNNCTCATGAGGCTCAGCTGTCTTT
ACACGTCTGAACTCTTTCCCTACACGACGCTCTTCCGATCTNNNGATGAGACGGGAGGTGACAAGCA
ACACGTCTGAACTCTTTCCCTACACGACGCTCTTCCGATCTNNNGATGAGACGGGAGGTGACAAGCA
AGCACACGTCTGAACTCTTTCCCTACACGACGCTCTTCCGATCTNNNGTTTCCAGATTGAATGCAGTC
AGCACACGTCTGAACTCTTTCCCTACACGACGCTCTTCCGATCTNNNGTTTCCAGATTGAATGCAGTC
GAGCACACGTCTGAACTCTTTCCCTACACGACGCTCTTCCGATCTNNNGTAGCAGCTTCACTGCCTAGA
GAGCACACGTCTGAACTCTTTCCCTACACGACGCTCTTCCGATCTNNNGTAGCAGCTTCACTGCCTAGA
GAGCACACGTCTGAACTCTTTCCCTACACGACGCTCTTCCGATCTNNNCTCACCTCGCTCATCTCCGCC
GAGCACACGTCTGAACTCTTTCCCTACACGACGCTCTTCCGATCTNNNCTCACCTCGCTCATCTCCGCC
GCACACGTCTGAACTCTTTCCCTACACGACGCTCTTCCGATCTNNNTAGTACAAGCACTATGGGGAAC
GCACACGTCTGAACTCTTTCCCTACACGACGCTCTTCCGATCTNNNTAGTACAAGCACTATGGGGAAC
AGCACACGTCTGAACTCTTTCCCTACACGACGCTCTTCCGATCTNNNTCTCCTGTCCATTTTTCTTGGGA
AGCACACGTCTGAACTCTTTCCCTACACGACGCTCTTCCGATCTNNNTCTCCTGTCCATTTTTCTTGGGA
AGCACACGTCTGAACTCTTTCCCTACACGACGCTCTTCCGATCTNNNCCCAATAACTAGAGCCCACTTT
AGCACACGTCTGAACTCTTTCCCTACACGACGCTCTTCCGATCTNNNCCCAATAACTAGAGCCCACTTT
GAGCACACGTCTGAACTCTTTCCCTACACGACGCTCTTCCGATCTNNNGAAGAGTCTGGTCAAGCTGG
GAGCACACGTCTGAACTCTTTCCCTACACGACGCTCTTCCGATCTNNNGAAGAGTCTGGTCAAGCTGG
AAGAGCACACGTCTGAACTCTTTCCCTACACGACGCTCTTCCGATCTNNNGCTTCTGTTAGTGACCAA
AAGAGCACACGTCTGAACTCTTTCCCTACACGACGCTCTTCCGATCTNNNGCTTCTGTTAGTGACCAA
AGCACACGTCTGAACTCTTTCCCTACACGACGCTCTTCCGATCTNNNGATCATACCACTGTACTCCAGC
AGCACACGTCTGAACTCTTTCCCTACACGACGCTCTTCCGATCTNNNGATCATACCACTGTACTCCAGC
GAGCACACGTCTGAACTCTTTCCCTACACGACGCTCTTCCGATCTNNNACTTTATCGCTTCTCAGCCTTC
GAGCACACGTCTGAACTCTTTCCCTACACGACGCTCTTCCGATCTNNNACTTTATCGCTTCTCAGCCTTC
GCACACGTCTGAACTCTTTCCCTACACGACGCTCTTCCGATCTNNNGGAAGTGAGGAGCGTCTCTGC
GCACACGTCTGAACTCTTTCCCTACACGACGCTCTTCCGATCTNNNGGAAGTGAGGAGCGTCTCTGC
AGCACACGTCTGAACTCTTTCCCTACACGACGCTCTTCCGATCTNNNCTGAGGAAAGAATGGCTTGA
AGCACACGTCTGAACTCTTTCCCTACACGACGCTCTTCCGATCTNNNCTGAGGAAAGAATGGCTTGA
GCACACGTCTGAACTCTTTCCCTACACGACGCTCTTCCGATCTNNNAGTGAGACCCTGCCTCTACTAA
GCACACGTCTGAACTCTTTCCCTACACGACGCTCTTCCGATCTNNNAGTGAGACCCTGCCTCTACTAA
AGCACACGTCTGAACTCTTTCCCTACACGACGCTCTTCCGATCTNNNCCCAAATTTGTGTCTTTCCCACA
AGCACACGTCTGAACTCTTTCCCTACACGACGCTCTTCCGATCTNNNCCCAAATTTGTGTCTTTCCCACA
GAGCACACGTCTGAACTCTTTCCCTACACGACGCTCTTCCGATCTNNNCAAAGGACCAGAAACTTCCA
GAGCACACGTCTGAACTCTTTCCCTACACGACGCTCTTCCGATCTNNNCAAAGGACCAGAAACTTCCA
GCACACGTCTGAACTCTTTCCCTACACGACGCTCTTCCGATCTNNNCACTGATGATGAGGAGGAGGA
GCACACGTCTGAACTCTTTCCCTACACGACGCTCTTCCGATCTNNNCACTGATGATGAGGAGGAGGA
GCACACGTCTGAACTCTTTCCCTACACGACGCTCTTCCGATCTNNNTATATTCCCTGTTGCTCAAGCGA
GCACACGTCTGAACTCTTTCCCTACACGACGCTCTTCCGATCTNNNTATATTCCCTGTTGCTCAAGCGA
AGCACACGTCTGAACTCTTTCCCTACACGACGCTCTTCCGATCTNNNACAGTTCAGCTCAGTGACAAG

AGCACACGTCTGAACTCTTTCCCTACACGACGCTCTTCCGATCTNNNACAGTTCAGCTCAGTGACAAG
CACACGTCTGAACTCTTTCCCTACACGACGCTCTTCCGATCTNNNATCCTTTAGCCCTCCCTAGAAGTC
CACACGTCTGAACTCTTTCCCTACACGACGCTCTTCCGATCTNNNATCCTTTAGCCCTCCCTAGAAGTC
GAGCACACGTCTGAACTCTTTCCCTACACGACGCTCTTCCGATCTNNNTTTACATTTAGGCCCAGTGA
GAGCACACGTCTGAACTCTTTCCCTACACGACGCTCTTCCGATCTNNNTTTACATTTAGGCCCAGTGA
AGCACACGTCTGAACTCTTTCCCTACACGACGCTCTTCCGATCTNNNGCTTGGTTTTCATCCATGTGC
AGCACACGTCTGAACTCTTTCCCTACACGACGCTCTTCCGATCTNNNGCTTGGTTTTCATCCATGTGC
AGCACACGTCTGAACTCTTTCCCTACACGACGCTCTTCCGATCTNNNATGATGACGCACACCTATAGT
AGCACACGTCTGAACTCTTTCCCTACACGACGCTCTTCCGATCTNNNATGATGACGCACACCTATAGT
GAGCACACGTCTGAACTCTTTCCCTACACGACGCTCTTCCGATCTNNNGCCTGTAGTCCCAGCTACTT
GAGCACACGTCTGAACTCTTTCCCTACACGACGCTCTTCCGATCTNNNGCCTGTAGTCCCAGCTACTT
ACACGTCTGAACTCTTTCCCTACACGACGCTCTTCCGATCTNNNGACCAATGTCTGTCGTGTCGCCAT
ACACGTCTGAACTCTTTCCCTACACGACGCTCTTCCGATCTNNNGACCAATGTCTGTCGTGTCGCCAT
GAGCACACGTCTGAACTCTTTCCCTACACGACGCTCTTCCGATCTNNNGAAAAGTCAGACACTTCAGG
GAGCACACGTCTGAACTCTTTCCCTACACGACGCTCTTCCGATCTNNNGAAAAGTCAGACACTTCAGG
AGCACACGTCTGAACTCTTTCCCTACACGACGCTCTTCCGATCTNNNCATTCCTTAGCAAGTTTGGCCA
AGCACACGTCTGAACTCTTTCCCTACACGACGCTCTTCCGATCTNNNCATTCCTTAGCAAGTTTGGCCA
AGCACACGTCTGAACTCTTTCCCTACACGACGCTCTTCCGATCTNNNTACAGATTCCTTGGCCATAACC
AGCACACGTCTGAACTCTTTCCCTACACGACGCTCTTCCGATCTNNNTACAGATTCCTTGGCCATAACC
GAGCACACGTCTGAACTCTTTCCCTACACGACGCTCTTCCGATCTNNNGGGCCTCCATTTGTTTTCTT
GAGCACACGTCTGAACTCTTTCCCTACACGACGCTCTTCCGATCTNNNGGGCCTCCATTTGTTTTCTT
GCACACGTCTGAACTCTTTCCCTACACGACGCTCTTCCGATCTNNNGATAAGGGGGGCAATAACTAG
GCACACGTCTGAACTCTTTCCCTACACGACGCTCTTCCGATCTNNNGATAAGGGGGGCAATAACTAG
AGCACACGTCTGAACTCTTTCCCTACACGACGCTCTTCCGATCTNNNGGAAAAAAAAAAGGCATTGC
AGCACACGTCTGAACTCTTTCCCTACACGACGCTCTTCCGATCTNNNGGAAAAAAAAAAGGCATTGC
AGCACACGTCTGAACTCTTTCCCTACACGACGCTCTTCCGATCTNNNGAGAACCACTTGAACCCAGGA
AGCACACGTCTGAACTCTTTCCCTACACGACGCTCTTCCGATCTNNNGAGAACCACTTGAACCCAGGA
GAGCACACGTCTGAACTCTTTCCCTACACGACGCTCTTCCGATCTNNNGGTAGGATTAGGGAGGCAA
GAGCACACGTCTGAACTCTTTCCCTACACGACGCTCTTCCGATCTNNNGGTAGGATTAGGGAGGCAA
AGCACACGTCTGAACTCTTTCCCTACACGACGCTCTTCCGATCTNNNACCCCTATAATCCCAGCACTTT
AGCACACGTCTGAACTCTTTCCCTACACGACGCTCTTCCGATCTNNNACCCCTATAATCCCAGCACTTT
AGCACACGTCTGAACTCTTTCCCTACACGACGCTCTTCCGATCTNNNGCAGCCCACAAGAATGAGAG
AGCACACGTCTGAACTCTTTCCCTACACGACGCTCTTCCGATCTNNNGCAGCCCACAAGAATGAGAG
AGCACACGTCTGAACTCTTTCCCTACACGACGCTCTTCCGATCTNNNCGAGGCAGATTAGTATGAACC
AGCACACGTCTGAACTCTTTCCCTACACGACGCTCTTCCGATCTNNNCGAGGCAGATTAGTATGAACC
AGCACACGTCTGAACTCTTTCCCTACACGACGCTCTTCCGATCTNNNGAGACAGGAGAATTGCTTGA
AGCACACGTCTGAACTCTTTCCCTACACGACGCTCTTCCGATCTNNNGAGACAGGAGAATTGCTTGA
GCACACGTCTGAACTCTTTCCCTACACGACGCTCTTCCGATCTNNNCTTCTTAAAATACTGCTCACCTG
GCACACGTCTGAACTCTTTCCCTACACGACGCTCTTCCGATCTNNNCTTCTTAAAATACTGCTCACCTG
GCACACGTCTGAACTCTTTCCCTACACGACGCTCTTCCGATCTNNNGAGGATCTATTCACTCTCTGGG
GCACACGTCTGAACTCTTTCCCTACACGACGCTCTTCCGATCTNNNGAGGATCTATTCACTCTCTGGG
AGCACACGTCTGAACTCTTTCCCTACACGACGCTCTTCCGATCTNNNCACTTCCTAAGAGGCACCCCCAT
AGCACACGTCTGAACTCTTTCCCTACACGACGCTCTTCCGATCTNNNCACTTCCTAAGAGGCACCCCCAT

GAGCACACGTCTGAACTCTTTCCCTACACGACGCTCTTCCGATCTNNNCCAGTTGATGTTAGGATTCCA
GAGCACACGTCTGAACTCTTTCCCTACACGACGCTCTTCCGATCTNNNCCAGTTGATGTTAGGATTCCA
AGCACACGTCTGAACTCTTTCCCTACACGACGCTCTTCCGATCTNNNTCATGGTTTCTCCAAGCTCTCA
AGCACACGTCTGAACTCTTTCCCTACACGACGCTCTTCCGATCTNNNTCATGGTTTCTCCAAGCTCTCA
GAGCACACGTCTGAACTCTTTCCCTACACGACGCTCTTCCGATCTNNNTGGAATGGGGGAGGATAGG
GAGCACACGTCTGAACTCTTTCCCTACACGACGCTCTTCCGATCTNNNTGGAATGGGGGAGGATAGG
GAGCACACGTCTGAACTCTTTCCCTACACGACGCTCTTCCGATCTNNNCCAGCAACTTTACCCAGACTGT
GAGCACACGTCTGAACTCTTTCCCTACACGACGCTCTTCCGATCTNNNCCAGCAACTTTACCCAGACTGT
GAGCACACGTCTGAACTCTTTCCCTACACGACGCTCTTCCGATCTNNNCACAGAAAGCTCTCCAGGTCT
GAGCACACGTCTGAACTCTTTCCCTACACGACGCTCTTCCGATCTNNNCACAGAAAGCTCTCCAGGTCT
GAGCACACGTCTGAACTCTTTCCCTACACGACGCTCTTCCGATCTNNNCTTGCAGTGAGCCAAGATGG
GAGCACACGTCTGAACTCTTTCCCTACACGACGCTCTTCCGATCTNNNCTTGCAGTGAGCCAAGATGG
AGCACACGTCTGAACTCTTTCCCTACACGACGCTCTTCCGATCTNNNCACACACACACACACACATACA
AGCACACGTCTGAACTCTTTCCCTACACGACGCTCTTCCGATCTNNNCACACACACACACACACATACA
AGCACACGTCTGAACTCTTTCCCTACACGACGCTCTTCCGATCTNNNAATGATCTGCCTCAACACTTCC
AGCACACGTCTGAACTCTTTCCCTACACGACGCTCTTCCGATCTNNNAATGATCTGCCTCAACACTTCC
AGCACACGTCTGAACTCTTTCCCTACACGACGCTCTTCCGATCTNNNCTGTAGTCCCAGCTACTCAGGA
AGCACACGTCTGAACTCTTTCCCTACACGACGCTCTTCCGATCTNNNCTGTAGTCCCAGCTACTCAGGA
AGCACACGTCTGAACTCTTTCCCTACACGACGCTCTTCCGATCTNNNTTACGGATACATTTTTGTGGGC
AGCACACGTCTGAACTCTTTCCCTACACGACGCTCTTCCGATCTNNNTTACGGATACATTTTTGTGGGC
AGAGCACACGTCTGAACTCTTTCCCTACACGACGCTCTTCCGATCTNNNTGAGGCAGGAGAATAGTG
AGAGCACACGTCTGAACTCTTTCCCTACACGACGCTCTTCCGATCTNNNTGAGGCAGGAGAATAGTG
AGCACACGTCTGAACTCTTTCCCTACACGACGCTCTTCCGATCTNNNAGTAAACCAGTGAAAGTTCCCC
AGCACACGTCTGAACTCTTTCCCTACACGACGCTCTTCCGATCTNNNAGTAAACCAGTGAAAGTTCCCC
GAGCACACGTCTGAACTCTTTCCCTACACGACGCTCTTCCGATCTNNNCAGAATGTAGCCTTTTCCGAT
GAGCACACGTCTGAACTCTTTCCCTACACGACGCTCTTCCGATCTNNNCAGAATGTAGCCTTTTCCGAT
AGCACACGTCTGAACTCTTTCCCTACACGACGCTCTTCCGATCTNNNCTCAAGTGATTGACCTGCTTCA
AGCACACGTCTGAACTCTTTCCCTACACGACGCTCTTCCGATCTNNNCTCAAGTGATTGACCTGCTTCA
GAGCACACGTCTGAACTCTTTCCCTACACGACGCTCTTCCGATCTNNNGAAAACCTGTTGTAACCCTCA
GAGCACACGTCTGAACTCTTTCCCTACACGACGCTCTTCCGATCTNNNGAAAACCTGTTGTAACCCTCA
AGCACACGTCTGAACTCTTTCCCTACACGACGCTCTTCCGATCTNNNTTTTTCCCCACTTTCTCACCTCAC
AGCACACGTCTGAACTCTTTCCCTACACGACGCTCTTCCGATCTNNNTTTTTCCCCACTTTCTCACCTCAC
AGCACACGTCTGAACTCTTTCCCTACACGACGCTCTTCCGATCTNNNTGAAAATTCCACCACTGTGTGA
AGCACACGTCTGAACTCTTTCCCTACACGACGCTCTTCCGATCTNNNTGAAAATTCCACCACTGTGTGA
AGCACACGTCTGAACTCTTTCCCTACACGACGCTCTTCCGATCTNNNGCAAACCAAAGATCAGTAAGAG
AGCACACGTCTGAACTCTTTCCCTACACGACGCTCTTCCGATCTNNNGCAAACCAAAGATCAGTAAGAG
GCACACGTCTGAACTCTTTCCCTACACGACGCTCTTCCGATCTNNNTAGTCCCAGCTACTCAGGAGTCC
GCACACGTCTGAACTCTTTCCCTACACGACGCTCTTCCGATCTNNNTAGTCCCAGCTACTCAGGAGTCC
AGCACACGTCTGAACTCTTTCCCTACACGACGCTCTTCCGATCTNNNGGCTCCAGGTCAGAGTCTAAAT
AGCACACGTCTGAACTCTTTCCCTACACGACGCTCTTCCGATCTNNNGGCTCCAGGTCAGAGTCTAAAT
GAGCACACGTCTGAACTCTTTCCCTACACGACGCTCTTCCGATCTNNNCCTGAACCCCTAGACTGCAAT
GAGCACACGTCTGAACTCTTTCCCTACACGACGCTCTTCCGATCTNNNCCTGAACCCCTAGACTGCAAT
GAGCACACGTCTGAACTCTTTCCCTACACGACGCTCTTCCGATCTNNNCTGTAGTCCCAGCTACTCAGGA

AGCACACGTCTGAACTCTTTCCCTACACGACGCTCTTCCGATCTNNNCTGTAGTCCCAGCTACTCAGG
AGCACACGTCTGAACTCTTTCCCTACACGACGCTCTTCCGATCTNNNGAAGTCCCTCTATGTGTGTGTG
AGCACACGTCTGAACTCTTTCCCTACACGACGCTCTTCCGATCTNNNGAAGTCCCTCTATGTGTGTGTG
GCACACGTCTGAACTCTTTCCCTACACGACGCTCTTCCGATCTNNNCCTCGCCCCTACTTTTTTCTTTCC
GCACACGTCTGAACTCTTTCCCTACACGACGCTCTTCCGATCTNNNCCTCGCCCCTACTTTTTTCTTTCC
AGCACACGTCTGAACTCTTTCCCTACACGACGCTCTTCCGATCTNNNGGCAGAATAATTGCTTGAACC
AGCACACGTCTGAACTCTTTCCCTACACGACGCTCTTCCGATCTNNNGGCAGAATAATTGCTTGAACC
GAGCACACGTCTGAACTCTTTCCCTACACGACGCTCTTCCGATCTNNNGGCTGGCCACCAAAGAGTAT
GAGCACACGTCTGAACTCTTTCCCTACACGACGCTCTTCCGATCTNNNGGCTGGCCACCAAAGAGTAT
AGCACACGTCTGAACTCTTTCCCTACACGACGCTCTTCCGATCTNNNTTGCCTGTAATCCCAGCATTTT
AGCACACGTCTGAACTCTTTCCCTACACGACGCTCTTCCGATCTNNNTTGCCTGTAATCCCAGCATTTT
AGCACACGTCTGAACTCTTTCCCTACACGACGCTCTTCCGATCTNNNCATCCCTGTTTTCCAGATGAGG
AGCACACGTCTGAACTCTTTCCCTACACGACGCTCTTCCGATCTNNNCATCCCTGTTTTCCAGATGAGG
AGCACACGTCTGAACTCTTTCCCTACACGACGCTCTTCCGATCTNNNTGTTCCACTGGTCTATGTGTCT
AGCACACGTCTGAACTCTTTCCCTACACGACGCTCTTCCGATCTNNNTGTTCCACTGGTCTATGTGTCT
GCACACGTCTGAACTCTTTCCCTACACGACGCTCTTCCGATCTNNNAGATCGAGACCATCCTGTGAATG
GCACACGTCTGAACTCTTTCCCTACACGACGCTCTTCCGATCTNNNAGATCGAGACCATCCTGTGAATG
GAGCACACGTCTGAACTCTTTCCCTACACGACGCTCTTCCGATCTNNNTATCAGACCCTTTCGACACAC
GAGCACACGTCTGAACTCTTTCCCTACACGACGCTCTTCCGATCTNNNTATCAGACCCTTTCGACACAC
AGCACACGTCTGAACTCTTTCCCTACACGACGCTCTTCCGATCTNNNTATTTGTTTTGGGACGCTGCA
AGCACACGTCTGAACTCTTTCCCTACACGACGCTCTTCCGATCTNNNTATTTGTTTTGGGACGCTGCA
AGCACACGTCTGAACTCTTTCCCTACACGACGCTCTTCCGATCTNNNGGGGACACAAAGCAAAACCAT
AGCACACGTCTGAACTCTTTCCCTACACGACGCTCTTCCGATCTNNNGGGGACACAAAGCAAAACCAT
GAGCACACGTCTGAACTCTTTCCCTACACGACGCTCTTCCGATCTNNNCAAAATGAGACCCATGCCTTC
GAGCACACGTCTGAACTCTTTCCCTACACGACGCTCTTCCGATCTNNNCAAAATGAGACCCATGCCTTC
AGCACACGTCTGAACTCTTTCCCTACACGACGCTCTTCCGATCTNNNTTCCTCCTTTTTTCCCCAGTGAA
AGCACACGTCTGAACTCTTTCCCTACACGACGCTCTTCCGATCTNNNTTCCTCCTTTTTTCCCCAGTGAA
GAGCACACGTCTGAACTCTTTCCCTACACGACGCTCTTCCGATCTNNNGGAATGGTGTGAACCCAAGA
GAGCACACGTCTGAACTCTTTCCCTACACGACGCTCTTCCGATCTNNNGGAATGGTGTGAACCCAAGA
AGCACACGTCTGAACTCTTTCCCTACACGACGCTCTTCCGATCTNNNTTCCTCCCAAGAAACATTTGCT
AGCACACGTCTGAACTCTTTCCCTACACGACGCTCTTCCGATCTNNNTTCCTCCCAAGAAACATTTGCT
AGCACACGTCTGAACTCTTTCCCTACACGACGCTCTTCCGATCTNNNGCTGCTGGTCATCTTCACTTTA
AGCACACGTCTGAACTCTTTCCCTACACGACGCTCTTCCGATCTNNNGCTGCTGGTCATCTTCACTTTA
AGAGCACACGTCTGAACTCTTTCCCTACACGACGCTCTTCCGATCTNNNATGGTATCAGGAATGGACT
AGAGCACACGTCTGAACTCTTTCCCTACACGACGCTCTTCCGATCTNNNATGGTATCAGGAATGGACT
GCACACGTCTGAACTCTTTCCCTACACGACGCTCTTCCGATCTNNNTAGACTCTGCCTCAGTTTCCTTG
GCACACGTCTGAACTCTTTCCCTACACGACGCTCTTCCGATCTNNNTAGACTCTGCCTCAGTTTCCTTG
GCACACGTCTGAACTCTTTCCCTACACGACGCTCTTCCGATCTNNNGAACTGCATAGGAATACACGCA
GCACACGTCTGAACTCTTTCCCTACACGACGCTCTTCCGATCTNNNGAACTGCATAGGAATACACGCA
GAGCACACGTCTGAACTCTTTCCCTACACGACGCTCTTCCGATCTNNNGGAGGTTTCAGTGAGCCTAG
GAGCACACGTCTGAACTCTTTCCCTACACGACGCTCTTCCGATCTNNNGGAGGTTTCAGTGAGCCTAG
GCACACGTCTGAACTCTTTCCCTACACGACGCTCTTCCGATCTNNNGTAAGAGCCGATCCAGACAAT
GCACACGTCTGAACTCTTTCCCTACACGACGCTCTTCCGATCTNNNGTAAGAGCCGATCCAGACAAT

GCACACGTCTGAACTCTTTCCCTACACGACGCTCTTCCGATCTNNNAACAGATTCTCCCTTGTACAGGT
GCACACGTCTGAACTCTTTCCCTACACGACGCTCTTCCGATCTNNNAACAGATTCTCCCTTGTACAGGT
AGCACACGTCTGAACTCTTTCCCTACACGACGCTCTTCCGATCTNNNCTGTAATCCCAGCTACTCAGGA
AGCACACGTCTGAACTCTTTCCCTACACGACGCTCTTCCGATCTNNNCTGTAATCCCAGCTACTCAGGA
AGCACACGTCTGAACTCTTTCCCTACACGACGCTCTTCCGATCTNNNACCTGTAATTCTAGCACTTGG
AGCACACGTCTGAACTCTTTCCCTACACGACGCTCTTCCGATCTNNNACCTGTAATTCTAGCACTTGG
AGCACACGTCTGAACTCTTTCCCTACACGACGCTCTTCCGATCTNNNGAAACCCTACAAGCCAGAAGA
AGCACACGTCTGAACTCTTTCCCTACACGACGCTCTTCCGATCTNNNGAAACCCTACAAGCCAGAAGA
GCACACGTCTGAACTCTTTCCCTACACGACGCTCTTCCGATCTNNNGCCAGGGTTTCTTTGCTTTCTTT
GCACACGTCTGAACTCTTTCCCTACACGACGCTCTTCCGATCTNNNGCCAGGGTTTCTTTGCTTTCTTT
AGCACACGTCTGAACTCTTTCCCTACACGACGCTCTTCCGATCTNNNACATACACACACACACCCAGTA
AGCACACGTCTGAACTCTTTCCCTACACGACGCTCTTCCGATCTNNNACATACACACACACACCCAGTA
AGCACACGTCTGAACTCTTTCCCTACACGACGCTCTTCCGATCTNNNCAGGAGGCAGAGGTTACAGCC
AGCACACGTCTGAACTCTTTCCCTACACGACGCTCTTCCGATCTNNNCAGGAGGCAGAGGTTACAGCC
GAGCACACGTCTGAACTCTTTCCCTACACGACGCTCTTCCGATCTNNNTCCCCACTCCAAGTGTTTATT
GAGCACACGTCTGAACTCTTTCCCTACACGACGCTCTTCCGATCTNNNTCCCCACTCCAAGTGTTTATT
GCACACGTCTGAACTCTTTCCCTACACGACGCTCTTCCGATCTNNNTCGAGACTTCTTTTTCTAGTGGA
GCACACGTCTGAACTCTTTCCCTACACGACGCTCTTCCGATCTNNNTCGAGACTTCTTTTTCTAGTGGA
GAGCACACGTCTGAACTCTTTCCCTACACGACGCTCTTCCGATCTNNNACATCCCATCCTAATGACCAC
GAGCACACGTCTGAACTCTTTCCCTACACGACGCTCTTCCGATCTNNNACATCCCATCCTAATGACCAC
GAGCACACGTCTGAACTCTTTCCCTACACGACGCTCTTCCGATCTNNNTTCAAGTGATTCTCCTGCCTC
GAGCACACGTCTGAACTCTTTCCCTACACGACGCTCTTCCGATCTNNNTTCAAGTGATTCTCCTGCCTC
GAGCACACGTCTGAACTCTTTCCCTACACGACGCTCTTCCGATCTNNNGCTCATGTCTGTAATCCCAG
GAGCACACGTCTGAACTCTTTCCCTACACGACGCTCTTCCGATCTNNNGCTCATGTCTGTAATCCCAG
AGCACACGTCTGAACTCTTTCCCTACACGACGCTCTTCCGATCTNNNATCGCACCACTGCATTATAGT
AGCACACGTCTGAACTCTTTCCCTACACGACGCTCTTCCGATCTNNNATCGCACCACTGCATTATAGT
AGCACACGTCTGAACTCTTTCCCTACACGACGCTCTTCCGATCTNNNAAAGTCGAAACCCCATGTCTA
AGCACACGTCTGAACTCTTTCCCTACACGACGCTCTTCCGATCTNNNAAAGTCGAAACCCCATGTCTA
GCACACGTCTGAACTCTTTCCCTACACGACGCTCTTCCGATCTNNNGAGAATCGCATGAACTGGAGA
GCACACGTCTGAACTCTTTCCCTACACGACGCTCTTCCGATCTNNNGAGAATCGCATGAACTGGAGA
GCACACGTCTGAACTCTTTCCCTACACGACGCTCTTCCGATCTNNNTCATAAGAATCAGCTCAAAAAA
GCACACGTCTGAACTCTTTCCCTACACGACGCTCTTCCGATCTNNNTCATAAGAATCAGCTCAAAAAA
AGAGCACACGTCTGAACTCTTTCCCTACACGACGCTCTTCCGATCTNNNAGTGTACTGGCTTCAAGT
AGAGCACACGTCTGAACTCTTTCCCTACACGACGCTCTTCCGATCTNNNAGTGTACTGGCTTCAAGT
AGCACACGTCTGAACTCTTTCCCTACACGACGCTCTTCCGATCTNNNACAAAAGTGTTGAATATTGTG
AGCACACGTCTGAACTCTTTCCCTACACGACGCTCTTCCGATCTNNNACAAAAGTGTTGAATATTGTG
GCACACGTCTGAACTCTTTCCCTACACGACGCTCTTCCGATCTNNNACCACACCTGGATGCTTTTTTTT
GCACACGTCTGAACTCTTTCCCTACACGACGCTCTTCCGATCTNNNACCACACCTGGATGCTTTTTTTT
GCACACGTCTGAACTCTTTCCCTACACGACGCTCTTCCGATCTNNNAGTTTTGGCCAAGGAAACATCTG
GCACACGTCTGAACTCTTTCCCTACACGACGCTCTTCCGATCTNNNAGTTTTGGCCAAGGAAACATCTG
GAGCACACGTCTGAACTCTTTCCCTACACGACGCTCTTCCGATCTNNNGCAAGGTCATTGTTCTTAGC
GAGCACACGTCTGAACTCTTTCCCTACACGACGCTCTTCCGATCTNNNGCAAGGTCATTGTTCTTAGC
AGCACACGTCTGAACTCTTTCCCTACACGACGCTCTTCCGATCTNNNTTTTTTCTTCTGCAGCACACTC

AGCACACGTCTGAACTCTTTCCCTACACGACGCTCTTCCGATCTNNNTTTTTCTTCTGCAGCACACTCC
GCACACGTCTGAACTCTTTCCCTACACGACGCTCTTCCGATCTNNNAAGAAGTTGTGTTCCTGAGAGC
GCACACGTCTGAACTCTTTCCCTACACGACGCTCTTCCGATCTNNNAAGAAGTTGTGTTCCTGAGAGC
AGCACACGTCTGAACTCTTTCCCTACACGACGCTCTTCCGATCTNNNGTAGCTATCCAGGGGTTATT
AGCACACGTCTGAACTCTTTCCCTACACGACGCTCTTCCGATCTNNNGTAGCTATCCAGGGGTTATT
GCACACGTCTGAACTCTTTCCCTACACGACGCTCTTCCGATCTNNNTGTGTACAAGTTTCACATTTTAC
GCACACGTCTGAACTCTTTCCCTACACGACGCTCTTCCGATCTNNNTGTGTACAAGTTTCACATTTTAC
GCACACGTCTGAACTCTTTCCCTACACGACGCTCTTCCGATCTNNNACTTCATGCTTTATCCCAGGCTAC
GCACACGTCTGAACTCTTTCCCTACACGACGCTCTTCCGATCTNNNACTTCATGCTTTATCCCAGGCTAC
AGCACACGTCTGAACTCTTTCCCTACACGACGCTCTTCCGATCTNNNATCCAGGTAGTAAGTAGTGG
AGCACACGTCTGAACTCTTTCCCTACACGACGCTCTTCCGATCTNNNATCCAGGTAGTAAGTAGTGG
GAGCACACGTCTGAACTCTTTCCCTACACGACGCTCTTCCGATCTNNNCATGTGACTTCACCTGGATG
GAGCACACGTCTGAACTCTTTCCCTACACGACGCTCTTCCGATCTNNNCATGTGACTTCACCTGGATG
GAGCACACGTCTGAACTCTTTCCCTACACGACGCTCTTCCGATCTNNNACAAGGAAGGAATGTTAGG
GAGCACACGTCTGAACTCTTTCCCTACACGACGCTCTTCCGATCTNNNACAAGGAAGGAATGTTAGG
AGCACACGTCTGAACTCTTTCCCTACACGACGCTCTTCCGATCTNNNAGAGGCAGTGCAGCTCATTAAT
AGCACACGTCTGAACTCTTTCCCTACACGACGCTCTTCCGATCTNNNAGAGGCAGTGCAGCTCATTAAT
GAGCACACGTCTGAACTCTTTCCCTACACGACGCTCTTCCGATCTNNNAAAAATATGAGTGCAAGCCA
GAGCACACGTCTGAACTCTTTCCCTACACGACGCTCTTCCGATCTNNNAAAAATATGAGTGCAAGCCA
AGCACACGTCTGAACTCTTTCCCTACACGACGCTCTTCCGATCTNNNCTTTTTCCTTTGGGTTTCCTCAT
AGCACACGTCTGAACTCTTTCCCTACACGACGCTCTTCCGATCTNNNCTTTTTCCTTTGGGTTTCCTCAT
GAGCACACGTCTGAACTCTTTCCCTACACGACGCTCTTCCGATCTNNNGAATGGCTGAACAAACCCTG
GAGCACACGTCTGAACTCTTTCCCTACACGACGCTCTTCCGATCTNNNGAATGGCTGAACAAACCCTG
GAGCACACGTCTGAACTCTTTCCCTACACGACGCTCTTCCGATCTNNNGCCATGGAGAATGTAGGGG
GAGCACACGTCTGAACTCTTTCCCTACACGACGCTCTTCCGATCTNNNGCCATGGAGAATGTAGGGG
AGCACACGTCTGAACTCTTTCCCTACACGACGCTCTTCCGATCTNNNCTCATGAATAAATGCCTGTTGC
AGCACACGTCTGAACTCTTTCCCTACACGACGCTCTTCCGATCTNNNCTCATGAATAAATGCCTGTTGC
GCACACGTCTGAACTCTTTCCCTACACGACGCTCTTCCGATCTNNNGATCCCCCTTAGAGAAGCTCAT
GCACACGTCTGAACTCTTTCCCTACACGACGCTCTTCCGATCTNNNGATCCCCCTTAGAGAAGCTCAT
GAGCACACGTCTGAACTCTTTCCCTACACGACGCTCTTCCGATCTNNNTTATAGCATGGTGTGGCCTCA
GAGCACACGTCTGAACTCTTTCCCTACACGACGCTCTTCCGATCTNNNTTATAGCATGGTGTGGCCTCA
AGCACACGTCTGAACTCTTTCCCTACACGACGCTCTTCCGATCTNNNACATCTGGTTATGCAGTTGGG
AGCACACGTCTGAACTCTTTCCCTACACGACGCTCTTCCGATCTNNNACATCTGGTTATGCAGTTGGG
AGCACACGTCTGAACTCTTTCCCTACACGACGCTCTTCCGATCTNNNAAACAGGCCAGGAGAGAAAG
AGCACACGTCTGAACTCTTTCCCTACACGACGCTCTTCCGATCTNNNAAACAGGCCAGGAGAGAAAG
GCACACGTCTGAACTCTTTCCCTACACGACGCTCTTCCGATCTNNNGGAGACATCCACTCTTATTCCCC
GCACACGTCTGAACTCTTTCCCTACACGACGCTCTTCCGATCTNNNGGAGACATCCACTCTTATTCCCC
AGCACACGTCTGAACTCTTTCCCTACACGACGCTCTTCCGATCTNNNCACCTGGTCACAGTATGCAAT
AGCACACGTCTGAACTCTTTCCCTACACGACGCTCTTCCGATCTNNNCACCTGGTCACAGTATGCAAT
GCACACGTCTGAACTCTTTCCCTACACGACGCTCTTCCGATCTNNNTTTTCCTGCCTTGTTGCATGATT
GCACACGTCTGAACTCTTTCCCTACACGACGCTCTTCCGATCTNNNTTTTCCTGCCTTGTTGCATGATT
GCACACGTCTGAACTCTTTCCCTACACGACGCTCTTCCGATCTNNNGCTCTCGTAATACAAAAGCAGC
GCACACGTCTGAACTCTTTCCCTACACGACGCTCTTCCGATCTNNNGCTCTCGTAATACAAAAGCAGC

AGCACACGTCTGAACTCTTTCCCTACACGACGCTCTTCCGATCTNNNCTCCCAATTGTAAGAGTTGTGG
AGCACACGTCTGAACTCTTTCCCTACACGACGCTCTTCCGATCTNNNCTCCCAATTGTAAGAGTTGTGG
GCACACGTCTGAACTCTTTCCCTACACGACGCTCTTCCGATCTNNNAGGGGTTTCTGTTCTACTTTTGT
GCACACGTCTGAACTCTTTCCCTACACGACGCTCTTCCGATCTNNNAGGGGTTTCTGTTCTACTTTTGT
GAGCACACGTCTGAACTCTTTCCCTACACGACGCTCTTCCGATCTNNNAAGGCAAGTACCTCCTCTCTCT
GAGCACACGTCTGAACTCTTTCCCTACACGACGCTCTTCCGATCTNNNAAGGCAAGTACCTCCTCTCTCT
AGCACACGTCTGAACTCTTTCCCTACACGACGCTCTTCCGATCTNNNTAAGGAAAGGAGAGCCGTTTG
AGCACACGTCTGAACTCTTTCCCTACACGACGCTCTTCCGATCTNNNTAAGGAAAGGAGAGCCGTTTG
AGCACACGTCTGAACTCTTTCCCTACACGACGCTCTTCCGATCTNNNTATAGTCCCAGCTACTCAGGAG
AGCACACGTCTGAACTCTTTCCCTACACGACGCTCTTCCGATCTNNNTATAGTCCCAGCTACTCAGGAG
AGCACACGTCTGAACTCTTTCCCTACACGACGCTCTTCCGATCTNNNGTTTACAGTGAGCTCGAGATTG
AGCACACGTCTGAACTCTTTCCCTACACGACGCTCTTCCGATCTNNNGTTTACAGTGAGCTCGAGATTG
GAGCACACGTCTGAACTCTTTCCCTACACGACGCTCTTCCGATCTNNNCTGTAATCCCAGCACTTTGGAA
GAGCACACGTCTGAACTCTTTCCCTACACGACGCTCTTCCGATCTNNNCTGTAATCCCAGCACTTTGGAA
AGCACACGTCTGAACTCTTTCCCTACACGACGCTCTTCCGATCTNNNCTGCAGATCTGTTTTTCCACGTT
AGCACACGTCTGAACTCTTTCCCTACACGACGCTCTTCCGATCTNNNCTGCAGATCTGTTTTTCCACGTT
AGCACACGTCTGAACTCTTTCCCTACACGACGCTCTTCCGATCTNNNTCAGCATGCAAATGATGTTTGC
AGCACACGTCTGAACTCTTTCCCTACACGACGCTCTTCCGATCTNNNTCAGCATGCAAATGATGTTTGC
AGCACACGTCTGAACTCTTTCCCTACACGACGCTCTTCCGATCTNNNCTGTAATCCCAGCTCCTCAGAAG
AGCACACGTCTGAACTCTTTCCCTACACGACGCTCTTCCGATCTNNNCTGTAATCCCAGCTCCTCAGAAG
AGAGCACACGTCTGAACTCTTTCCCTACACGACGCTCTTCCGATCTNNNTCTGGTGTGGTCTGGAAAAT
AGAGCACACGTCTGAACTCTTTCCCTACACGACGCTCTTCCGATCTNNNTCTGGTGTGGTCTGGAAAAT
GAGCACACGTCTGAACTCTTTCCCTACACGACGCTCTTCCGATCTNNNGTAGTACCAGCTATTCAGGA
GAGCACACGTCTGAACTCTTTCCCTACACGACGCTCTTCCGATCTNNNGTAGTACCAGCTATTCAGGA
GAGCACACGTCTGAACTCTTTCCCTACACGACGCTCTTCCGATCTNNNGGAGGAAGAAGTTCAAGGC
GAGCACACGTCTGAACTCTTTCCCTACACGACGCTCTTCCGATCTNNNGGAGGAAGAAGTTCAAGGC
AGCACACGTCTGAACTCTTTCCCTACACGACGCTCTTCCGATCTNNNAAAAATAGCGTGTGTGTGAGG
AGCACACGTCTGAACTCTTTCCCTACACGACGCTCTTCCGATCTNNNAAAAATAGCGTGTGTGTGAGG
AGCACACGTCTGAACTCTTTCCCTACACGACGCTCTTCCGATCTNNNCTGTCCTTCACTTCCAGGAACTT
AGCACACGTCTGAACTCTTTCCCTACACGACGCTCTTCCGATCTNNNCTGTCCTTCACTTCCAGGAACTT
GAGCACACGTCTGAACTCTTTCCCTACACGACGCTCTTCCGATCTNNNGTATCTATGCTTTCTAGACAGA
GAGCACACGTCTGAACTCTTTCCCTACACGACGCTCTTCCGATCTNNNGTATCTATGCTTTCTAGACAGA
GAGCACACGTCTGAACTCTTTCCCTACACGACGCTCTTCCGATCTNNNAAGAGAATCACTTGAACCCGG
GAGCACACGTCTGAACTCTTTCCCTACACGACGCTCTTCCGATCTNNNAAGAGAATCACTTGAACCCGG
GAGCACACGTCTGAACTCTTTCCCTACACGACGCTCTTCCGATCTNNNCTGTAATCCCAGACACTCAGG
GAGCACACGTCTGAACTCTTTCCCTACACGACGCTCTTCCGATCTNNNCTGTAATCCCAGACACTCAGG
GCACACGTCTGAACTCTTTCCCTACACGACGCTCTTCCGATCTNNNATAGTCACTTACCACCATGCCTGC
GCACACGTCTGAACTCTTTCCCTACACGACGCTCTTCCGATCTNNNATAGTCACTTACCACCATGCCTGC
GAGCACACGTCTGAACTCTTTCCCTACACGACGCTCTTCCGATCTNNNACAAGAGTGAAACACCCTTGT
GAGCACACGTCTGAACTCTTTCCCTACACGACGCTCTTCCGATCTNNNACAAGAGTGAAACACCCTTGT
AGCACACGTCTGAACTCTTTCCCTACACGACGCTCTTCCGATCTNNNTGATTAGGCACACATCTGGGTA
AGCACACGTCTGAACTCTTTCCCTACACGACGCTCTTCCGATCTNNNTGATTAGGCACACATCTGGGTA
GAGCACACGTCTGAACTCTTTCCCTACACGACGCTCTTCCGATCTNNNAGAATCGCTTGAACCTGGGAG

GAGCACACGTCTGAACTCTTTCCCTACACGACGCTCTTCCGATCTNNNAGAATCGCTTGAACCTGGGAG
AGCACACGTCTGAACTCTTTCCCTACACGACGCTCTTCCGATCTNNNGAAACAATTTCTCCCCAAGAGC
AGCACACGTCTGAACTCTTTCCCTACACGACGCTCTTCCGATCTNNNGAAACAATTTCTCCCCAAGAGC
AGAGCACACGTCTGAACTCTTTCCCTACACGACGCTCTTCCGATCTNNNAGGAAGGTGAAGTGAACAA
AGAGCACACGTCTGAACTCTTTCCCTACACGACGCTCTTCCGATCTNNNAGGAAGGTGAAGTGAACAA
AGCACACGTCTGAACTCTTTCCCTACACGACGCTCTTCCGATCTNNNCGCTACCACACCTGGATAATTTT
AGCACACGTCTGAACTCTTTCCCTACACGACGCTCTTCCGATCTNNNCGCTACCACACCTGGATAATTTT
AGCACACGTCTGAACTCTTTCCCTACACGACGCTCTTCCGATCTNNNAAGCTGTCTCAGGAGCAAACTC
AGCACACGTCTGAACTCTTTCCCTACACGACGCTCTTCCGATCTNNNAAGCTGTCTCAGGAGCAAACTC
AGCACACGTCTGAACTCTTTCCCTACACGACGCTCTTCCGATCTNNNCAAGTAGTAATTGATGGAGAA
AGCACACGTCTGAACTCTTTCCCTACACGACGCTCTTCCGATCTNNNCAAGTAGTAATTGATGGAGAA
AGCACACGTCTGAACTCTTTCCCTACACGACGCTCTTCCGATCTNNNTATCTGTTTTGGTACCAGTGCC
AGCACACGTCTGAACTCTTTCCCTACACGACGCTCTTCCGATCTNNNTATCTGTTTTGGTACCAGTGCC
AGCACACGTCTGAACTCTTTCCCTACACGACGCTCTTCCGATCTNNNAACTTCTCCATCTGTCTCTTGGC
AGCACACGTCTGAACTCTTTCCCTACACGACGCTCTTCCGATCTNNNAACTTCTCCATCTGTCTCTTGGC
AGCACACGTCTGAACTCTTTCCCTACACGACGCTCTTCCGATCTNNNGAAGCAGGAGAATCGCTTGAA
AGCACACGTCTGAACTCTTTCCCTACACGACGCTCTTCCGATCTNNNGAAGCAGGAGAATCGCTTGAA
GCACACGTCTGAACTCTTTCCCTACACGACGCTCTTCCGATCTNNNCATTCTGTTGGAGCTGCTTTTGGC
GCACACGTCTGAACTCTTTCCCTACACGACGCTCTTCCGATCTNNNCATTCTGTTGGAGCTGCTTTTGGC
GAGCACACGTCTGAACTCTTTCCCTACACGACGCTCTTCCGATCTNNNGATAGGGAAGGAAGTCTGGC
GAGCACACGTCTGAACTCTTTCCCTACACGACGCTCTTCCGATCTNNNGATAGGGAAGGAAGTCTGGC
GCACACGTCTGAACTCTTTCCCTACACGACGCTCTTCCGATCTNNNCTAAGTCAACATCTGGGTCCTCCC
GCACACGTCTGAACTCTTTCCCTACACGACGCTCTTCCGATCTNNNCTAAGTCAACATCTGGGTCCTCCC
AGCACACGTCTGAACTCTTTCCCTACACGACGCTCTTCCGATCTNNNTAAAAAAATTAGCTGGGCGTG
AGCACACGTCTGAACTCTTTCCCTACACGACGCTCTTCCGATCTNNNTAAAAAAATTAGCTGGGCGTG
GAGCACACGTCTGAACTCTTTCCCTACACGACGCTCTTCCGATCTNNNAATCCCAACTACTCAAGAGGCT
GAGCACACGTCTGAACTCTTTCCCTACACGACGCTCTTCCGATCTNNNAATCCCAACTACTCAAGAGGCT
AGAGCACACGTCTGAACTCTTTCCCTACACGACGCTCTTCCGATCTNNNTGTGGTTCATGTTTTTGTGT
AGAGCACACGTCTGAACTCTTTCCCTACACGACGCTCTTCCGATCTNNNTGTGGTTCATGTTTTTGTGT
GAGCACACGTCTGAACTCTTTCCCTACACGACGCTCTTCCGATCTNNNGAGGCTAAGGCAAGAGAATC
GAGCACACGTCTGAACTCTTTCCCTACACGACGCTCTTCCGATCTNNNGAGGCTAAGGCAAGAGAATC
GAGCACACGTCTGAACTCTTTCCCTACACGACGCTCTTCCGATCTNNNGTTTTGGTCCTTTTTTGTGGGC
GAGCACACGTCTGAACTCTTTCCCTACACGACGCTCTTCCGATCTNNNGTTTTGGTCCTTTTTTGTGGGC
AGCACACGTCTGAACTCTTTCCCTACACGACGCTCTTCCGATCTNNNATTGGAAGGGTATGGTGACTCA
AGCACACGTCTGAACTCTTTCCCTACACGACGCTCTTCCGATCTNNNATTGGAAGGGTATGGTGACTCA
GAGCACACGTCTGAACTCTTTCCCTACACGACGCTCTTCCGATCTNNNCAACTCTCTAGCTGCTAAGGGA
GAGCACACGTCTGAACTCTTTCCCTACACGACGCTCTTCCGATCTNNNCAACTCTCTAGCTGCTAAGGGA
AGCACACGTCTGAACTCTTTCCCTACACGACGCTCTTCCGATCTNNNTGCATGGTAGGGATACTCAGA
AGCACACGTCTGAACTCTTTCCCTACACGACGCTCTTCCGATCTNNNTGCATGGTAGGGATACTCAGA
AGCACACGTCTGAACTCTTTCCCTACACGACGCTCTTCCGATCTNNNGTGGATCATGAGGTCAGGAGT
AGCACACGTCTGAACTCTTTCCCTACACGACGCTCTTCCGATCTNNNGTGGATCATGAGGTCAGGAGT
AGCACACGTCTGAACTCTTTCCCTACACGACGCTCTTCCGATCTNNNGAGAATCACTTGAATCCGGGAC
AGCACACGTCTGAACTCTTTCCCTACACGACGCTCTTCCGATCTNNNGAGAATCACTTGAATCCGGGAC

AGCACACGTCTGAACTCTTTCCCTACACGACGCTCTTCCGATCTNNNACTTACGTAAATGTACGCAGCC

AGCACACGTCTGAACTCTTTCCCTACACGACGCTCTTCCGATCTNNNACTTACGTAAATGTACGCAGCC

AGCACACGTCTGAACTCTTTCCCTACACGACGCTCTTCCGATCTNNNTTTCATGGAATAGCTGCACGT

AGCACACGTCTGAACTCTTTCCCTACACGACGCTCTTCCGATCTNNNTTTCATGGAATAGCTGCACGT

GCACACGTCTGAACTCTTTCCCTACACGACGCTCTTCCGATCTNNNGGTGTGGTAACCTGGGATACAA

GCACACGTCTGAACTCTTTCCCTACACGACGCTCTTCCGATCTNNNGGTGTGGTAACCTGGGATACAA

GAGCACACGTCTGAACTCTTTCCCTACACGACGCTCTTCCGATCTNNNCAGACTTCTAAGGCTTGGTTG

GAGCACACGTCTGAACTCTTTCCCTACACGACGCTCTTCCGATCTNNNCAGACTTCTAAGGCTTGGTTG

GCACACGTCTGAACTCTTTCCCTACACGACGCTCTTCCGATCTNNNGTCTGCTCTCTTTTTCAGGTAAC

GCACACGTCTGAACTCTTTCCCTACACGACGCTCTTCCGATCTNNNGTCTGCTCTCTTTTTCAGGTAAC

GAGCACACGTCTGAACTCTTTCCCTACACGACGCTCTTCCGATCTNNNTGCTCTCACTCTCACTTCCTTT

GAGCACACGTCTGAACTCTTTCCCTACACGACGCTCTTCCGATCTNNNTGCTCTCACTCTCACTTCCTTT

GAGCACACGTCTGAACTCTTTCCCTACACGACGCTCTTCCGATCTNNNTGGGGAGTAGGTGCAAAAG

GAGCACACGTCTGAACTCTTTCCCTACACGACGCTCTTCCGATCTNNNTGGGGAGTAGGTGCAAAAG

GCACACGTCTGAACTCTTTCCCTACACGACGCTCTTCCGATCTNNNTTGGTGAACTCTTGGAAAGCAA

GCACACGTCTGAACTCTTTCCCTACACGACGCTCTTCCGATCTNNNTTGGTGAACTCTTGGAAAGCAA

GAGCACACGTCTGAACTCTTTCCCTACACGACGCTCTTCCGATCTNNNGCAGTGGTAAAATGCCCTATG

GAGCACACGTCTGAACTCTTTCCCTACACGACGCTCTTCCGATCTNNNGCAGTGGTAAAATGCCCTATG

AGCACACGTCTGAACTCTTTCCCTACACGACGCTCTTCCGATCTNNNAGGCTGAGACACAAGAATTCC

AGCACACGTCTGAACTCTTTCCCTACACGACGCTCTTCCGATCTNNNAGGCTGAGACACAAGAATTCC

GAGCACACGTCTGAACTCTTTCCCTACACGACGCTCTTCCGATCTNNNCAGTCAAGTGGAAAGCTGTC

GAGCACACGTCTGAACTCTTTCCCTACACGACGCTCTTCCGATCTNNNCAGTCAAGTGGAAAGCTGTC

AGCACACGTCTGAACTCTTTCCCTACACGACGCTCTTCCGATCTNNNCTCCAATCCTCTTTCTCCCTTCC

AGCACACGTCTGAACTCTTTCCCTACACGACGCTCTTCCGATCTNNNCTCCAATCCTCTTTCTCCCTTCC

GAGCACACGTCTGAACTCTTTCCCTACACGACGCTCTTCCGATCTNNNGGACTACAGAGGGAGACTCC

GAGCACACGTCTGAACTCTTTCCCTACACGACGCTCTTCCGATCTNNNGGACTACAGAGGGAGACTCC

GCACACGTCTGAACTCTTTCCCTACACGACGCTCTTCCGATCTNNNTTCCACCATTTTGGGGTACAGAT

GCACACGTCTGAACTCTTTCCCTACACGACGCTCTTCCGATCTNNNTTCCACCATTTTGGGGTACAGAT

AGCACACGTCTGAACTCTTTCCCTACACGACGCTCTTCCGATCTNNNGTAAGTTCAGCCATGAGCCATT

AGCACACGTCTGAACTCTTTCCCTACACGACGCTCTTCCGATCTNNNGTAAGTTCAGCCATGAGCCATT

GCACACGTCTGAACTCTTTCCCTACACGACGCTCTTCCGATCTNNNCTTGCAGTGAGCCGAGATTGTC

GCACACGTCTGAACTCTTTCCCTACACGACGCTCTTCCGATCTNNNCTTGCAGTGAGCCGAGATTGTC

AGCACACGTCTGAACTCTTTCCCTACACGACGCTCTTCCGATCTNNNTGAGGGAGGAGAATCACTTGA

AGCACACGTCTGAACTCTTTCCCTACACGACGCTCTTCCGATCTNNNTGAGGGAGGAGAATCACTTGA

AGAGCACACGTCTGAACTCTTTCCCTACACGACGCTCTTCCGATCTNNNCAGAAGCTGGTGAAGTGG

AGAGCACACGTCTGAACTCTTTCCCTACACGACGCTCTTCCGATCTNNNCAGAAGCTGGTGAAGTGG

AGCACACGTCTGAACTCTTTCCCTACACGACGCTCTTCCGATCTNNNTAGTGATTCAGATAGTAGTTG

AGCACACGTCTGAACTCTTTCCCTACACGACGCTCTTCCGATCTNNNTAGTGATTCAGATAGTAGTTG

AGCACACGTCTGAACTCTTTCCCTACACGACGCTCTTCCGATCTNNNTGATAAAACCTCAGCTTGACCC

AGCACACGTCTGAACTCTTTCCCTACACGACGCTCTTCCGATCTNNNTGATAAAACCTCAGCTTGACCC

GAGCACACGTCTGAACTCTTTCCCTACACGACGCTCTTCCGATCTNNNAGTGTCCACCAGTGAATGGTA

GAGCACACGTCTGAACTCTTTCCCTACACGACGCTCTTCCGATCTNNNAGTGTCCACCAGTGAATGGTA

AGCACACGTCTGAACTCTTTCCCTACACGACGCTCTTCCGATCTNNNATAGTTCACATGCTTGACCAGG

AGCACACGTCTGAACTCTTTCCCTACACGACGCTCTTCCGATCTNNNATAGTTCACATGCTTGACCAGG

GAGCACACGTCTGAACTCTTTCCCTACACGACGCTCTTCCGATCTNNNGTTGTGGTAAGCCTAGATTGT

GAGCACACGTCTGAACTCTTTCCCTACACGACGCTCTTCCGATCTNNNGTTGTGGTAAGCCTAGATTGT

AGCACACGTCTGAACTCTTTCCCTACACGACGCTCTTCCGATCTNNNCCAGGCCTGGCTAATTTTTTGTA

AGCACACGTCTGAACTCTTTCCCTACACGACGCTCTTCCGATCTNNNCCAGGCCTGGCTAATTTTTTGTA

GCACACGTCTGAACTCTTTCCCTACACGACGCTCTTCCGATCTNNNCTCCCTTCCTTCCTTCCTGTTTTCA

GCACACGTCTGAACTCTTTCCCTACACGACGCTCTTCCGATCTNNNCTCCCTTCCTTCCTTCCTGTTTTCA

GAGCACACGTCTGAACTCTTTCCCTACACGACGCTCTTCCGATCTNNNATATTTCCCCCATGGCTTTCAGT

GAGCACACGTCTGAACTCTTTCCCTACACGACGCTCTTCCGATCTNNNATATTTCCCCCATGGCTTTCAGT

GAGCACACGTCTGAACTCTTTCCCTACACGACGCTCTTCCGATCTNNNAGATGATTGTAGGTGTGCAG

GAGCACACGTCTGAACTCTTTCCCTACACGACGCTCTTCCGATCTNNNAGATGATTGTAGGTGTGCAG

GCACACGTCTGAACTCTTTCCCTACACGACGCTCTTCCGATCTNNNGGTGTAAACACTGAAACCACGTC

GCACACGTCTGAACTCTTTCCCTACACGACGCTCTTCCGATCTNNNGGTGTAAACACTGAAACCACGTC

CACGTCTGAACTCTTTCCCTACACGACGCTCTTCCGATCTNNNGGTGGTGGTTGTACAAGTCTGAACA

CACGTCTGAACTCTTTCCCTACACGACGCTCTTCCGATCTNNNGGTGGTGGTTGTACAAGTCTGAACA

GAGCACACGTCTGAACTCTTTCCCTACACGACGCTCTTCCGATCTNNNTCATGTGTTTCTCTCTGCCTTGT

GAGCACACGTCTGAACTCTTTCCCTACACGACGCTCTTCCGATCTNNNTCATGTGTTTCTCTCTGCCTTGT

AGCACACGTCTGAACTCTTTCCCTACACGACGCTCTTCCGATCTNNNAGAGACTCTTGTCCTCTTCCCTTC

AGCACACGTCTGAACTCTTTCCCTACACGACGCTCTTCCGATCTNNNAGAGACTCTTGTCCTCTTCCCTTC

AGCACACGTCTGAACTCTTTCCCTACACGACGCTCTTCCGATCTNNNTTATAAGCAGTGCCAAAAATAA

AGCACACGTCTGAACTCTTTCCCTACACGACGCTCTTCCGATCTNNNTTATAAGCAGTGCCAAAAATAA

ACACGTCTGAACTCTTTCCCTACACGACGCTCTTCCGATCTNNNCAACTTCTACAACTTCAGGAGCATCG

ACACGTCTGAACTCTTTCCCTACACGACGCTCTTCCGATCTNNNCAACTTCTACAACTTCAGGAGCATCG

AGCACACGTCTGAACTCTTTCCCTACACGACGCTCTTCCGATCTNNNTTCCCTTTTCTCCATGTCCTCACCA

AGCACACGTCTGAACTCTTTCCCTACACGACGCTCTTCCGATCTNNNTTCCCTTTTCTCCATGTCCTCACCA

GAGCACACGTCTGAACTCTTTCCCTACACGACGCTCTTCCGATCTNNNGCTTTCATGAACTTGCCCAGAA

GAGCACACGTCTGAACTCTTTCCCTACACGACGCTCTTCCGATCTNNNGCTTTCATGAACTTGCCCAGAA

GAGCACACGTCTGAACTCTTTCCCTACACGACGCTCTTCCGATCTNNNAGAAGAACTAGGACCAGAAAC

GAGCACACGTCTGAACTCTTTCCCTACACGACGCTCTTCCGATCTNNNAGAAGAACTAGGACCAGAAAC

GCACACGTCTGAACTCTTTCCCTACACGACGCTCTTCCGATCTNNNTCTTTCATTTTCCCTCTCCCCTGCAT

GCACACGTCTGAACTCTTTCCCTACACGACGCTCTTCCGATCTNNNTCTTTCATTTTCCCTCTCCCCTGCAT

GCACACGTCTGAACTCTTTCCCTACACGACGCTCTTCCGATCTNNNCAGGATAGGCTGTATCCCTGACA

GCACACGTCTGAACTCTTTCCCTACACGACGCTCTTCCGATCTNNNCAGGATAGGCTGTATCCCTGACA

GAGCACACGTCTGAACTCTTTCCCTACACGACGCTCTTCCGATCTNNNCTCAGAACTAACAGGCAAGGG

GAGCACACGTCTGAACTCTTTCCCTACACGACGCTCTTCCGATCTNNNCTCAGAACTAACAGGCAAGGG

GAGCACACGTCTGAACTCTTTCCCTACACGACGCTCTTCCGATCTNNNTTGAGAGTAGTAAACTGGGC

GAGCACACGTCTGAACTCTTTCCCTACACGACGCTCTTCCGATCTNNNTTGAGAGTAGTAAACTGGGC

GCACACGTCTGAACTCTTTCCCTACACGACGCTCTTCCGATCTNNNTCAGGGATATTGGCCTGTAATTT

GCACACGTCTGAACTCTTTCCCTACACGACGCTCTTCCGATCTNNNTCAGGGATATTGGCCTGTAATTT

AGCACACGTCTGAACTCTTTCCCTACACGACGCTCTTCCGATCTNNNGGTCAAAAGGTGAAGCAACTCT

AGCACACGTCTGAACTCTTTCCCTACACGACGCTCTTCCGATCTNNNGGTCAAAAGGTGAAGCAACTCT

GAGCACACGTCTGAACTCTTTCCCTACACGACGCTCTTCCGATCTNNNCAACAACAGGATGCCAGAAG

GAGCACACGTCTGAACTCTTTCCCTACACGACGCTCTTCCGATCTNNNCAACAACAGGATGCCAGAAG

AGCACACGTCTGAACTCTTTCCCTACACGACGCTCTTCCGATCTNNNGAGATGGTTCCTCTGACTTCTC

AGCACACGTCTGAACTCTTTCCCTACACGACGCTCTTCCGATCTNNNGAGATGGTTCCTCTGACTTCTC

AGCACACGTCTGAACTCTTTCCCTACACGACGCTCTTCCGATCTNNNTGTGGGACATACAGAACTGA

AGCACACGTCTGAACTCTTTCCCTACACGACGCTCTTCCGATCTNNNTGTGGGACATACAGAACTGA

AGCACACGTCTGAACTCTTTCCCTACACGACGCTCTTCCGATCTNNNTCAGTGCCAAATCTTCTGTCTG

AGCACACGTCTGAACTCTTTCCCTACACGACGCTCTTCCGATCTNNNTCAGTGCCAAATCTTCTGTCTG

GAGCACACGTCTGAACTCTTTCCCTACACGACGCTCTTCCGATCTNNNCTATCTCCTGACCTTGTGATC

GAGCACACGTCTGAACTCTTTCCCTACACGACGCTCTTCCGATCTNNNCTATCTCCTGACCTTGTGATC

AGCACACGTCTGAACTCTTTCCCTACACGACGCTCTTCCGATCTNNNGACAGACTAGGGGTTTGGAT

AGCACACGTCTGAACTCTTTCCCTACACGACGCTCTTCCGATCTNNNGACAGACTAGGGGTTTGGAT

GAGCACACGTCTGAACTCTTTCCCTACACGACGCTCTTCCGATCTNNNACAAAAATTAGCCAGGCACA

GAGCACACGTCTGAACTCTTTCCCTACACGACGCTCTTCCGATCTNNNACAAAAATTAGCCAGGCACA

AGCACACGTCTGAACTCTTTCCCTACACGACGCTCTTCCGATCTNNNCTACTTCTTCCACTTCCTCTACAT

AGCACACGTCTGAACTCTTTCCCTACACGACGCTCTTCCGATCTNNNCTACTTCTTCCACTTCCTCTACAT

AGCACACGTCTGAACTCTTTCCCTACACGACGCTCTTCCGATCTNNNGCCAGATGGACTGCAGTATGT

AGCACACGTCTGAACTCTTTCCCTACACGACGCTCTTCCGATCTNNNGCCAGATGGACTGCAGTATGT

AGCACACGTCTGAACTCTTTCCCTACACGACGCTCTTCCGATCTNNNTGGAACATTTCTCTTGATTGCG

AGCACACGTCTGAACTCTTTCCCTACACGACGCTCTTCCGATCTNNNTGGAACATTTCTCTTGATTGCG

GCACACGTCTGAACTCTTTCCCTACACGACGCTCTTCCGATCTNNNGCATCATTCATGGCATGGATAG

GCACACGTCTGAACTCTTTCCCTACACGACGCTCTTCCGATCTNNNGCATCATTCATGGCATGGATAG

AGCACACGTCTGAACTCTTTCCCTACACGACGCTCTTCCGATCTNNNTTCATCATTGTGAACACACCAG

AGCACACGTCTGAACTCTTTCCCTACACGACGCTCTTCCGATCTNNNTTCATCATTGTGAACACACCAG

AGAGCACACGTCTGAACTCTTTCCCTACACGACGCTCTTCCGATCTNNNCCTTCAGAAAGCTCCAATCA

AGAGCACACGTCTGAACTCTTTCCCTACACGACGCTCTTCCGATCTNNNCCTTCAGAAAGCTCCAATCA

GAGCACACGTCTGAACTCTTTCCCTACACGACGCTCTTCCGATCTNNNCTGGAGTTGGGGAGACTTC

GAGCACACGTCTGAACTCTTTCCCTACACGACGCTCTTCCGATCTNNNCTGGAGTTGGGGAGACTTC

AGCACACGTCTGAACTCTTTCCCTACACGACGCTCTTCCGATCTNNNTTCCAGAGTTGACAGGGAGAA

AGCACACGTCTGAACTCTTTCCCTACACGACGCTCTTCCGATCTNNNTTCCAGAGTTGACAGGGAGAA

AGCACACGTCTGAACTCTTTCCCTACACGACGCTCTTCCGATCTNNNGGGTTTTACAAATATAAGTCA

AGCACACGTCTGAACTCTTTCCCTACACGACGCTCTTCCGATCTNNNGGGTTTTACAAATATAAGTCA

GCACACGTCTGAACTCTTTCCCTACACGACGCTCTTCCGATCTNNNTCAGACAGAGGCGTCTATAAGT

GCACACGTCTGAACTCTTTCCCTACACGACGCTCTTCCGATCTNNNTCAGACAGAGGCGTCTATAAGT

GCACACGTCTGAACTCTTTCCCTACACGACGCTCTTCCGATCTNNNTCCAGGATTCTTATGGTTTGGG

GCACACGTCTGAACTCTTTCCCTACACGACGCTCTTCCGATCTNNNTCCAGGATTCTTATGGTTTGGG

AGCACACGTCTGAACTCTTTCCCTACACGACGCTCTTCCGATCTNNNCACCAGGGAATCTGAATAACC

AGCACACGTCTGAACTCTTTCCCTACACGACGCTCTTCCGATCTNNNCACCAGGGAATCTGAATAACC

AGCACACGTCTGAACTCTTTCCCTACACGACGCTCTTCCGATCTNNNCCAGCTCCTCTTTGTATCTCTGG

AGCACACGTCTGAACTCTTTCCCTACACGACGCTCTTCCGATCTNNNCCAGCTCCTCTTTGTATCTCTGG

GCACACGTCTGAACTCTTTCCCTACACGACGCTCTTCCGATCTNNNACCTCTGTTGTAACACCTTTCCTC

GCACACGTCTGAACTCTTTCCCTACACGACGCTCTTCCGATCTNNNACCTCTGTTGTAACACCTTTCCTC

GAGCACACGTCTGAACTCTTTCCCTACACGACGCTCTTCCGATCTNNNTGCTTTCTCCCCTAGCTGAAT

GAGCACACGTCTGAACTCTTTCCCTACACGACGCTCTTCCGATCTNNNTGCTTTCTCCCCTAGCTGAAT

AGCACACGTCTGAACTCTTTCCCTACACGACGCTCTTCCGATCTNNNGATCACGAGGTCAGGAGATC

AGCACACGTCTGAACTCTTTCCCTACACGACGCTCTTCCGATCTNNNGATCACGAGGTCAGGAGATC
AGCACACGTCTGAACTCTTTCCCTACACGACGCTCTTCCGATCTNNNTGAGGCAGGAGAATCTCTTGA
AGCACACGTCTGAACTCTTTCCCTACACGACGCTCTTCCGATCTNNNTGAGGCAGGAGAATCTCTTGA
GCACACGTCTGAACTCTTTCCCTACACGACGCTCTTCCGATCTNNNTTTCCCCGATGGCTAATTATGT
GCACACGTCTGAACTCTTTCCCTACACGACGCTCTTCCGATCTNNNTTTCCCCGATGGCTAATTATGT
AGCACACGTCTGAACTCTTTCCCTACACGACGCTCTTCCGATCTNNNGCTCTGCACATATCTGCCTAAC
AGCACACGTCTGAACTCTTTCCCTACACGACGCTCTTCCGATCTNNNGCTCTGCACATATCTGCCTAAC
GCACACGTCTGAACTCTTTCCCTACACGACGCTCTTCCGATCTNNNGAACAACGACCAGGACAAGAAG
GCACACGTCTGAACTCTTTCCCTACACGACGCTCTTCCGATCTNNNGAACAACGACCAGGACAAGAAG
AGCACACGTCTGAACTCTTTCCCTACACGACGCTCTTCCGATCTNNNAAATGCCCACCAATCAATGAG
AGCACACGTCTGAACTCTTTCCCTACACGACGCTCTTCCGATCTNNNAAATGCCCACCAATCAATGAG
AGCACACGTCTGAACTCTTTCCCTACACGACGCTCTTCCGATCTNNNGTGATCTCGGTTCACTACAACC
AGCACACGTCTGAACTCTTTCCCTACACGACGCTCTTCCGATCTNNNGTGATCTCGGTTCACTACAACC
AGCACACGTCTGAACTCTTTCCCTACACGACGCTCTTCCGATCTNNNGTAGAACTGCCTGAACCTGG
AGCACACGTCTGAACTCTTTCCCTACACGACGCTCTTCCGATCTNNNGTAGAACTGCCTGAACCTGG
AGCACACGTCTGAACTCTTTCCCTACACGACGCTCTTCCGATCTNNNGAAATGTGTGCGTGCCTCCTC
AGCACACGTCTGAACTCTTTCCCTACACGACGCTCTTCCGATCTNNNGAAATGTGTGCGTGCCTCCTC
GCACACGTCTGAACTCTTTCCCTACACGACGCTCTTCCGATCTNNNTATACAGGCCAGGTACATAGG
GCACACGTCTGAACTCTTTCCCTACACGACGCTCTTCCGATCTNNNTATACAGGCCAGGTACATAGG
AGCACACGTCTGAACTCTTTCCCTACACGACGCTCTTCCGATCTNNNGAGAGAGCTCATGAAGGACAC
AGCACACGTCTGAACTCTTTCCCTACACGACGCTCTTCCGATCTNNNGAGAGAGCTCATGAAGGACAC
GAGCACACGTCTGAACTCTTTCCCTACACGACGCTCTTCCGATCTNNNCAGGCAGTTTAAATCTCGTT
GAGCACACGTCTGAACTCTTTCCCTACACGACGCTCTTCCGATCTNNNCAGGCAGTTTAAATCTCGTT
AGAGCACACGTCTGAACTCTTTCCCTACACGACGCTCTTCCGATCTNNNTCTGGTTGTTCTCTCCATTTC
AGAGCACACGTCTGAACTCTTTCCCTACACGACGCTCTTCCGATCTNNNTCTGGTTGTTCTCTCCATTTC
GCACACGTCTGAACTCTTTCCCTACACGACGCTCTTCCGATCTNNNATCACTTGAAACCAGAAGGCAG
GCACACGTCTGAACTCTTTCCCTACACGACGCTCTTCCGATCTNNNATCACTTGAAACCAGAAGGCAG
GCACACGTCTGAACTCTTTCCCTACACGACGCTCTTCCGATCTNNNCCAAACTCTGCAGCTTTGAAGA
GCACACGTCTGAACTCTTTCCCTACACGACGCTCTTCCGATCTNNNCCAAACTCTGCAGCTTTGAAGA
GCACACGTCTGAACTCTTTCCCTACACGACGCTCTTCCGATCTNNNATGATGGGGTTGAGATGACAC
GCACACGTCTGAACTCTTTCCCTACACGACGCTCTTCCGATCTNNNATGATGGGGTTGAGATGACAC
GAGCACACGTCTGAACTCTTTCCCTACACGACGCTCTTCCGATCTNNNCAGTAATCCCAGCTACTCAG
GAGCACACGTCTGAACTCTTTCCCTACACGACGCTCTTCCGATCTNNNCAGTAATCCCAGCTACTCAG
AGCACACGTCTGAACTCTTTCCCTACACGACGCTCTTCCGATCTNNNCACCATGCCCAGCTAATTTTTGT
AGCACACGTCTGAACTCTTTCCCTACACGACGCTCTTCCGATCTNNNCACCATGCCCAGCTAATTTTTGT
AGCACACGTCTGAACTCTTTCCCTACACGACGCTCTTCCGATCTNNNTGGCACTGTCCACCCTTATTTA
AGCACACGTCTGAACTCTTTCCCTACACGACGCTCTTCCGATCTNNNTGGCACTGTCCACCCTTATTTA
GAGCACACGTCTGAACTCTTTCCCTACACGACGCTCTTCCGATCTNNNCCAACATGTTGAAACCCCATC
GAGCACACGTCTGAACTCTTTCCCTACACGACGCTCTTCCGATCTNNNCCAACATGTTGAAACCCCATC
AGCACACGTCTGAACTCTTTCCCTACACGACGCTCTTCCGATCTNNNGAGTCTCACTGCCAACTTTTTC
AGCACACGTCTGAACTCTTTCCCTACACGACGCTCTTCCGATCTNNNGAGTCTCACTGCCAACTTTTTC
AGCACACGTCTGAACTCTTTCCCTACACGACGCTCTTCCGATCTNNNCATTCCTACCAGTGTTGCACAA
AGCACACGTCTGAACTCTTTCCCTACACGACGCTCTTCCGATCTNNNCATTCCTACCAGTGTTGCACAA

CACGTCTGAACTCTTTCCCTACACGACGCTCTTCCGATCTNNNATGTGCTGTTCAAAAGGGCTTTTCAT

CACGTCTGAACTCTTTCCCTACACGACGCTCTTCCGATCTNNNATGTGCTGTTCAAAAGGGCTTTTCAT

GAGCACACGTCTGAACTCTTTCCCTACACGACGCTCTTCCGATCTNNNCTTGTAACTCGGGACCCTTG

GAGCACACGTCTGAACTCTTTCCCTACACGACGCTCTTCCGATCTNNNCTTGTAACTCGGGACCCTTG

AGCACACGTCTGAACTCTTTCCCTACACGACGCTCTTCCGATCTNNNCATGTCAGCTCATGCCTGTAA

AGCACACGTCTGAACTCTTTCCCTACACGACGCTCTTCCGATCTNNNCATGTCAGCTCATGCCTGTAA

GAGCACACGTCTGAACTCTTTCCCTACACGACGCTCTTCCGATCTNNNGCACATGCATGTTTATAGCAG

GAGCACACGTCTGAACTCTTTCCCTACACGACGCTCTTCCGATCTNNNGCACATGCATGTTTATAGCAG

GAGCACACGTCTGAACTCTTTCCCTACACGACGCTCTTCCGATCTNNNAAATAACTGAAGCCAGACGT

GAGCACACGTCTGAACTCTTTCCCTACACGACGCTCTTCCGATCTNNNAAATAACTGAAGCCAGACGT

AGCACACGTCTGAACTCTTTCCCTACACGACGCTCTTCCGATCTNNNGAGAATCGCTTGAATCCAGGA

AGCACACGTCTGAACTCTTTCCCTACACGACGCTCTTCCGATCTNNNGAGAATCGCTTGAATCCAGGA

GAGCACACGTCTGAACTCTTTCCCTACACGACGCTCTTCCGATCTNNNGGTCAGTACCAAAGAATGAG

GAGCACACGTCTGAACTCTTTCCCTACACGACGCTCTTCCGATCTNNNGGTCAGTACCAAAGAATGAG

GCACACGTCTGAACTCTTTCCCTACACGACGCTCTTCCGATCTNNNTGGCATTTATTTGGCTTGAGGT

GCACACGTCTGAACTCTTTCCCTACACGACGCTCTTCCGATCTNNNTGGCATTTATTTGGCTTGAGGT

GCACACGTCTGAACTCTTTCCCTACACGACGCTCTTCCGATCTNNNCACCTGGCCACAATTTACTCTTT

GCACACGTCTGAACTCTTTCCCTACACGACGCTCTTCCGATCTNNNCACCTGGCCACAATTTACTCTTT

GAGCACACGTCTGAACTCTTTCCCTACACGACGCTCTTCCGATCTNNNAGTAAGCTGTGATCACACCA

GAGCACACGTCTGAACTCTTTCCCTACACGACGCTCTTCCGATCTNNNAGTAAGCTGTGATCACACCA

AGCACACGTCTGAACTCTTTCCCTACACGACGCTCTTCCGATCTNNNGGGCCTTGGATGAATATTTTG

AGCACACGTCTGAACTCTTTCCCTACACGACGCTCTTCCGATCTNNNGGGCCTTGGATGAATATTTTG

GCACACGTCTGAACTCTTTCCCTACACGACGCTCTTCCGATCTNNNTTCTTTTATATACTGTGCGCGCG

GCACACGTCTGAACTCTTTCCCTACACGACGCTCTTCCGATCTNNNTTCTTTTATATACTGTGCGCGCG

GCACACGTCTGAACTCTTTCCCTACACGACGCTCTTCCGATCTNNNGGATTCCTCCTGTCTCTCTCTCT

GCACACGTCTGAACTCTTTCCCTACACGACGCTCTTCCGATCTNNNGGATTCCTCCTGTCTCTCTCTCT

GAGCACACGTCTGAACTCTTTCCCTACACGACGCTCTTCCGATCTNNNATTCATTTTATTGTGGTATGG

GAGCACACGTCTGAACTCTTTCCCTACACGACGCTCTTCCGATCTNNNATTCATTTTATTGTGGTATGG

AGAGCACACGTCTGAACTCTTTCCCTACACGACGCTCTTCCGATCTNNNCAACACACAGTGGTTGAGA

AGAGCACACGTCTGAACTCTTTCCCTACACGACGCTCTTCCGATCTNNNCAACACACAGTGGTTGAGA

AGCACACGTCTGAACTCTTTCCCTACACGACGCTCTTCCGATCTNNNTTGCTTCTGTAAGAGATGCCAC

AGCACACGTCTGAACTCTTTCCCTACACGACGCTCTTCCGATCTNNNTTGCTTCTGTAAGAGATGCCAC

CACACGTCTGAACTCTTTCCCTACACGACGCTCTTCCGATCTNNNTGTAGTCCCAGCTACTCAGGAGC

CACACGTCTGAACTCTTTCCCTACACGACGCTCTTCCGATCTNNNTGTAGTCCCAGCTACTCAGGAGC

GCACACGTCTGAACTCTTTCCCTACACGACGCTCTTCCGATCTNNNGATCAAACCAGCCTTCAGAGCT

GCACACGTCTGAACTCTTTCCCTACACGACGCTCTTCCGATCTNNNGATCAAACCAGCCTTCAGAGCT

GAGCACACGTCTGAACTCTTTCCCTACACGACGCTCTTCCGATCTNNNCACGGGCCCTCTAGAGAAAA

GAGCACACGTCTGAACTCTTTCCCTACACGACGCTCTTCCGATCTNNNCACGGGCCCTCTAGAGAAAA

GAGCACACGTCTGAACTCTTTCCCTACACGACGCTCTTCCGATCTNNNAGATCACCAGAGGTCAGGAG

GAGCACACGTCTGAACTCTTTCCCTACACGACGCTCTTCCGATCTNNNAGATCACCAGAGGTCAGGAG

GCACACGTCTGAACTCTTTCCCTACACGACGCTCTTCCGATCTNNNTCTTTCTCAGCATTGCCAACCTT

GCACACGTCTGAACTCTTTCCCTACACGACGCTCTTCCGATCTNNNTCTTTCTCAGCATTGCCAACCTT

GCACACGTCTGAACTCTTTCCCTACACGACGCTCTTCCGATCTNNNGACAAGGTTTGATTTCTGGCTC

AGCACACGTCTGAACTCTTTCCCTACACGACGCTCTTCCGATCTNNNGACAAGGTTTGATTTCTGGCTC
AGCACACGTCTGAACTCTTTCCCTACACGACGCTCTTCCGATCTNNNACTACAGCTACTTAGGCAGGA
AGCACACGTCTGAACTCTTTCCCTACACGACGCTCTTCCGATCTNNNACTACAGCTACTTAGGCAGGA
GAGCACACGTCTGAACTCTTTCCCTACACGACGCTCTTCCGATCTNNNCCAAGACTGTGGGGTTTTCTA
GAGCACACGTCTGAACTCTTTCCCTACACGACGCTCTTCCGATCTNNNCCAAGACTGTGGGGTTTTCTA
AGCACACGTCTGAACTCTTTCCCTACACGACGCTCTTCCGATCTNNNGTGCGGAGAACACCTTGAGAG
AGCACACGTCTGAACTCTTTCCCTACACGACGCTCTTCCGATCTNNNGTGCGGAGAACACCTTGAGAG
AGAGCACACGTCTGAACTCTTTCCCTACACGACGCTCTTCCGATCTNNNAGGTGACTGATAGCAAAAG
AGAGCACACGTCTGAACTCTTTCCCTACACGACGCTCTTCCGATCTNNNAGGTGACTGATAGCAAAAG
AGAGCACACGTCTGAACTCTTTCCCTACACGACGCTCTTCCGATCTNNNCAAGACGGGCTACCCAGAAT
AGAGCACACGTCTGAACTCTTTCCCTACACGACGCTCTTCCGATCTNNNCAAGACGGGCTACCCAGAAT
AGCACACGTCTGAACTCTTTCCCTACACGACGCTCTTCCGATCTNNNATGCCAACACACCCAGCTAACAT
AGCACACGTCTGAACTCTTTCCCTACACGACGCTCTTCCGATCTNNNATGCCAACACACCCAGCTAACAT
GAGCACACGTCTGAACTCTTTCCCTACACGACGCTCTTCCGATCTNNNCTGAGGCAGGAGAATCACTTG
GAGCACACGTCTGAACTCTTTCCCTACACGACGCTCTTCCGATCTNNNCTGAGGCAGGAGAATCACTTG
GAGCACACGTCTGAACTCTTTCCCTACACGACGCTCTTCCGATCTNNNGAACAGAAGTGGGATTTGTG
GAGCACACGTCTGAACTCTTTCCCTACACGACGCTCTTCCGATCTNNNGAACAGAAGTGGGATTTGTG
AGCACACGTCTGAACTCTTTCCCTACACGACGCTCTTCCGATCTNNNAAGGCAAAATGAGGGCAAAAT
AGCACACGTCTGAACTCTTTCCCTACACGACGCTCTTCCGATCTNNNAAGGCAAAATGAGGGCAAAAT
GAGCACACGTCTGAACTCTTTCCCTACACGACGCTCTTCCGATCTNNNCAGAGGCTGAGGTGAGAGA
GAGCACACGTCTGAACTCTTTCCCTACACGACGCTCTTCCGATCTNNNCAGAGGCTGAGGTGAGAGA
AGCACACGTCTGAACTCTTTCCCTACACGACGCTCTTCCGATCTNNNGAGAATCACTCGAACCAGGGA
AGCACACGTCTGAACTCTTTCCCTACACGACGCTCTTCCGATCTNNNGAGAATCACTCGAACCAGGGA
AGCACACGTCTGAACTCTTTCCCTACACGACGCTCTTCCGATCTNNNCATCCTAAACTCCGCTTTCAGTG
AGCACACGTCTGAACTCTTTCCCTACACGACGCTCTTCCGATCTNNNCATCCTAAACTCCGCTTTCAGTG
AGCACACGTCTGAACTCTTTCCCTACACGACGCTCTTCCGATCTNNNAGTCTCATACAGGTAAGCATTG
AGCACACGTCTGAACTCTTTCCCTACACGACGCTCTTCCGATCTNNNAGTCTCATACAGGTAAGCATTG
AGCACACGTCTGAACTCTTTCCCTACACGACGCTCTTCCGATCTNNNTTAATGTACAGAGTTTTTGGAT
AGCACACGTCTGAACTCTTTCCCTACACGACGCTCTTCCGATCTNNNTTAATGTACAGAGTTTTTGGAT
GCACACGTCTGAACTCTTTCCCTACACGACGCTCTTCCGATCTNNNCTGAAATCCCCACTACTCAGGAG
GCACACGTCTGAACTCTTTCCCTACACGACGCTCTTCCGATCTNNNCTGAAATCCCCACTACTCAGGAG
GAGCACACGTCTGAACTCTTTCCCTACACGACGCTCTTCCGATCTNNNAAAAAAAAAAAGAGGGAGG
GAGCACACGTCTGAACTCTTTCCCTACACGACGCTCTTCCGATCTNNNAAAAAAAAAAAGAGGGAGG
AGAGCACACGTCTGAACTCTTTCCCTACACGACGCTCTTCCGATCTNNNGCAAAGACACACATAGGCTC
AGAGCACACGTCTGAACTCTTTCCCTACACGACGCTCTTCCGATCTNNNGCAAAGACACACATAGGCTC
AGAGCACACGTCTGAACTCTTTCCCTACACGACGCTCTTCCGATCTNNNACCTATAGGAAGGGTCACC
AGAGCACACGTCTGAACTCTTTCCCTACACGACGCTCTTCCGATCTNNNACCTATAGGAAGGGTCACC
GAGCACACGTCTGAACTCTTTCCCTACACGACGCTCTTCCGATCTNNNTTGCAAGCTCCCTCTGAAAAG
GAGCACACGTCTGAACTCTTTCCCTACACGACGCTCTTCCGATCTNNNTTGCAAGCTCCCTCTGAAAAG
AGCACACGTCTGAACTCTTTCCCTACACGACGCTCTTCCGATCTNNNATATTACATTTCAGAGTTCACC
AGCACACGTCTGAACTCTTTCCCTACACGACGCTCTTCCGATCTNNNATATTACATTTCAGAGTTCACC
GAGCACACGTCTGAACTCTTTCCCTACACGACGCTCTTCCGATCTNNNCCATGCACCAACCTACAAGAG
GAGCACACGTCTGAACTCTTTCCCTACACGACGCTCTTCCGATCTNNNCCATGCACCAACCTACAAGAG

AGCACACGTCTGAACTCTTTCCCTACACGACGCTCTTCCGATCTNNNATGACAGAAGCGAGCAAGAAC
AGCACACGTCTGAACTCTTTCCCTACACGACGCTCTTCCGATCTNNNATGACAGAAGCGAGCAAGAAC
AGCACACGTCTGAACTCTTTCCCTACACGACGCTCTTCCGATCTNNNTGTAATCCCAGCTACTCAGGAG
AGCACACGTCTGAACTCTTTCCCTACACGACGCTCTTCCGATCTNNNTGTAATCCCAGCTACTCAGGAG
GAGCACACGTCTGAACTCTTTCCCTACACGACGCTCTTCCGATCTNNNTGAGGCAGGAGAATCACTTG
GAGCACACGTCTGAACTCTTTCCCTACACGACGCTCTTCCGATCTNNNTGAGGCAGGAGAATCACTTG
GAGCACACGTCTGAACTCTTTCCCTACACGACGCTCTTCCGATCTNNNAGTGAACTACGATTTTGCCAC
GAGCACACGTCTGAACTCTTTCCCTACACGACGCTCTTCCGATCTNNNAGTGAACTACGATTTTGCCAC
AGCACACGTCTGAACTCTTTCCCTACACGACGCTCTTCCGATCTNNNGCACTGAGGCAATTGTTCTTCA
AGCACACGTCTGAACTCTTTCCCTACACGACGCTCTTCCGATCTNNNGCACTGAGGCAATTGTTCTTCA
GAGCACACGTCTGAACTCTTTCCCTACACGACGCTCTTCCGATCTNNNGGTGCCTATAATCCCAGCTAC
GAGCACACGTCTGAACTCTTTCCCTACACGACGCTCTTCCGATCTNNNGGTGCCTATAATCCCAGCTAC
GAGCACACGTCTGAACTCTTTCCCTACACGACGCTCTTCCGATCTNNNGATGCCACGTCCATTTGCTTC
GAGCACACGTCTGAACTCTTTCCCTACACGACGCTCTTCCGATCTNNNGATGCCACGTCCATTTGCTTC
CACACGTCTGAACTCTTTCCCTACACGACGCTCTTCCGATCTNNNGAGGGGAAGGAGGAGGAGGAA
CACACGTCTGAACTCTTTCCCTACACGACGCTCTTCCGATCTNNNGAGGGGAAGGAGGAGGAGGAA
AGCACACGTCTGAACTCTTTCCCTACACGACGCTCTTCCGATCTNNNGGCAAGAAGAGTGAAATTCCG
AGCACACGTCTGAACTCTTTCCCTACACGACGCTCTTCCGATCTNNNGGCAAGAAGAGTGAAATTCCG
AGCACACGTCTGAACTCTTTCCCTACACGACGCTCTTCCGATCTNNNGATCAAATTGGTCCCAATGGCT
AGCACACGTCTGAACTCTTTCCCTACACGACGCTCTTCCGATCTNNNGATCAAATTGGTCCCAATGGCT
GCACACGTCTGAACTCTTTCCCTACACGACGCTCTTCCGATCTNNNAAGGTAAAAGAAATCATTGAGT
GCACACGTCTGAACTCTTTCCCTACACGACGCTCTTCCGATCTNNNAAGGTAAAAGAAATCATTGAGT
ACACGTCTGAACTCTTTCCCTACACGACGCTCTTCCGATCTNNNTTTCAATGAGCTGTTGGGTACACCA
ACACGTCTGAACTCTTTCCCTACACGACGCTCTTCCGATCTNNNTTTCAATGAGCTGTTGGGTACACCA
CACACGTCTGAACTCTTTCCCTACACGACGCTCTTCCGATCTNNNAAGCCCTAACCTCCAATGTGATTC
CACACGTCTGAACTCTTTCCCTACACGACGCTCTTCCGATCTNNNAAGCCCTAACCTCCAATGTGATTC
AGCACACGTCTGAACTCTTTCCCTACACGACGCTCTTCCGATCTNNNTTGTGACTGACAGAAAAAGGT
AGCACACGTCTGAACTCTTTCCCTACACGACGCTCTTCCGATCTNNNTTGTGACTGACAGAAAAAGGT
AGCACACGTCTGAACTCTTTCCCTACACGACGCTCTTCCGATCTNNNGACAAGACTGCTAGTGTCTCTC
AGCACACGTCTGAACTCTTTCCCTACACGACGCTCTTCCGATCTNNNGACAAGACTGCTAGTGTCTCTC
AGCACACGTCTGAACTCTTTCCCTACACGACGCTCTTCCGATCTNNNCATCTGCAATCTTGGTGCTTTG
AGCACACGTCTGAACTCTTTCCCTACACGACGCTCTTCCGATCTNNNCATCTGCAATCTTGGTGCTTTG
GAGCACACGTCTGAACTCTTTCCCTACACGACGCTCTTCCGATCTNNNCTCTCAAGAAATAGGTGTGGC
GAGCACACGTCTGAACTCTTTCCCTACACGACGCTCTTCCGATCTNNNCTCTCAAGAAATAGGTGTGGC
GCACACGTCTGAACTCTTTCCCTACACGACGCTCTTCCGATCTNNNCAAGATGAAGCTCTAACCCCAAC
GCACACGTCTGAACTCTTTCCCTACACGACGCTCTTCCGATCTNNNCAAGATGAAGCTCTAACCCCAAC
AGAGCACACGTCTGAACTCTTTCCCTACACGACGCTCTTCCGATCTNNNTCAAGACTATCCGGGCTAAC
AGAGCACACGTCTGAACTCTTTCCCTACACGACGCTCTTCCGATCTNNNTCAAGACTATCCGGGCTAAC
GAGCACACGTCTGAACTCTTTCCCTACACGACGCTCTTCCGATCTNNNGAATCGCTTGAATCCAGGAGG
GAGCACACGTCTGAACTCTTTCCCTACACGACGCTCTTCCGATCTNNNGAATCGCTTGAATCCAGGAGG
GAGCACACGTCTGAACTCTTTCCCTACACGACGCTCTTCCGATCTNNNCCTCTTTGTCTCTCTGCCTCTTT
GAGCACACGTCTGAACTCTTTCCCTACACGACGCTCTTCCGATCTNNNCCTCTTTGTCTCTCTGCCTCTTT
AGCACACGTCTGAACTCTTTCCCTACACGACGCTCTTCCGATCTNNNGAAACAAATGGAATGCGACAT

AGCACACGTCTGAACTCTTTCCCTACACGACGCTCTTCCGATCTNNNGAAACAAATGGAATGCGACAT
AGCACACGTCTGAACTCTTTCCCTACACGACGCTCTTCCGATCTNNNGGTCACAGTGAGCTGAGATTG
AGCACACGTCTGAACTCTTTCCCTACACGACGCTCTTCCGATCTNNNGGTCACAGTGAGCTGAGATTG
AGCACACGTCTGAACTCTTTCCCTACACGACGCTCTTCCGATCTNNNTGGTCTCTGATTTGTAACCTCGT
AGCACACGTCTGAACTCTTTCCCTACACGACGCTCTTCCGATCTNNNTGGTCTCTGATTTGTAACCTCGT
GAGCACACGTCTGAACTCTTTCCCTACACGACGCTCTTCCGATCTNNNGAAGAAAGGGAAGAGAGGG
GAGCACACGTCTGAACTCTTTCCCTACACGACGCTCTTCCGATCTNNNGAAGAAAGGGAAGAGAGGG
AGCACACGTCTGAACTCTTTCCCTACACGACGCTCTTCCGATCTNNNCTGTAATCCCAGCTACTCAGGA
AGCACACGTCTGAACTCTTTCCCTACACGACGCTCTTCCGATCTNNNCTGTAATCCCAGCTACTCAGGA
GAGCACACGTCTGAACTCTTTCCCTACACGACGCTCTTCCGATCTNNNGGGAAGTCCATGTGAATCA
GAGCACACGTCTGAACTCTTTCCCTACACGACGCTCTTCCGATCTNNNGGGAAGTCCATGTGAATCA
ACACGTCTGAACTCTTTCCCTACACGACGCTCTTCCGATCTNNNATCTTCTGTGAAATGTTTAGGACAT
ACACGTCTGAACTCTTTCCCTACACGACGCTCTTCCGATCTNNNATCTTCTGTGAAATGTTTAGGACAT
GAGCACACGTCTGAACTCTTTCCCTACACGACGCTCTTCCGATCTNNNGATCACACTACTACACTCCAGC
GAGCACACGTCTGAACTCTTTCCCTACACGACGCTCTTCCGATCTNNNGATCACACTACTACACTCCAGC
GAGCACACGTCTGAACTCTTTCCCTACACGACGCTCTTCCGATCTNNNGGAGGATGAAGCAGGAGA
GAGCACACGTCTGAACTCTTTCCCTACACGACGCTCTTCCGATCTNNNGGAGGATGAAGCAGGAGA
GAGCACACGTCTGAACTCTTTCCCTACACGACGCTCTTCCGATCTNNNCTTGTGGTGAGCTGAAATCG
GAGCACACGTCTGAACTCTTTCCCTACACGACGCTCTTCCGATCTNNNCTTGTGGTGAGCTGAAATCG
GAGCACACGTCTGAACTCTTTCCCTACACGACGCTCTTCCGATCTNNNTTTCTATCCCTGTCTGTTCCTG
GAGCACACGTCTGAACTCTTTCCCTACACGACGCTCTTCCGATCTNNNTTTCTATCCCTGTCTGTTCCTG
AGCACACGTCTGAACTCTTTCCCTACACGACGCTCTTCCGATCTNNNGGAGTTGTACTCTGCTCAATCC
AGCACACGTCTGAACTCTTTCCCTACACGACGCTCTTCCGATCTNNNGGAGTTGTACTCTGCTCAATCC
GCACACGTCTGAACTCTTTCCCTACACGACGCTCTTCCGATCTNNNAGACAAGAGAGTAGCCAGCTGA
GCACACGTCTGAACTCTTTCCCTACACGACGCTCTTCCGATCTNNNAGACAAGAGAGTAGCCAGCTGA
GAGCACACGTCTGAACTCTTTCCCTACACGACGCTCTTCCGATCTNNNCCTCCAGAGAACCCTAATACA
GAGCACACGTCTGAACTCTTTCCCTACACGACGCTCTTCCGATCTNNNCCTCCAGAGAACCCTAATACA
AGCACACGTCTGAACTCTTTCCCTACACGACGCTCTTCCGATCTNNNCTCTGACTCAAGCCAAATCCT
AGCACACGTCTGAACTCTTTCCCTACACGACGCTCTTCCGATCTNNNCTCTGACTCAAGCCAAATCCT
GAGCACACGTCTGAACTCTTTCCCTACACGACGCTCTTCCGATCTNNNCGGCAAAAAGGTTATATTTAT
GAGCACACGTCTGAACTCTTTCCCTACACGACGCTCTTCCGATCTNNNCGGCAAAAAGGTTATATTTAT
GAGCACACGTCTGAACTCTTTCCCTACACGACGCTCTTCCGATCTNNNTACTGCAACAGAACACTATGC
GAGCACACGTCTGAACTCTTTCCCTACACGACGCTCTTCCGATCTNNNTACTGCAACAGAACACTATGC
GAGCACACGTCTGAACTCTTTCCCTACACGACGCTCTTCCGATCTNNNTGTGCAGACCAAGGAATTGA
GAGCACACGTCTGAACTCTTTCCCTACACGACGCTCTTCCGATCTNNNTGTGCAGACCAAGGAATTGA
AGCACACGTCTGAACTCTTTCCCTACACGACGCTCTTCCGATCTNNNCCCAATGTACCTCCTTTCTTCCA
AGCACACGTCTGAACTCTTTCCCTACACGACGCTCTTCCGATCTNNNCCCAATGTACCTCCTTTCTTCCA
AGCACACGTCTGAACTCTTTCCCTACACGACGCTCTTCCGATCTNNNTGTGTTTCTGTCAGAGAGTTGC
AGCACACGTCTGAACTCTTTCCCTACACGACGCTCTTCCGATCTNNNTGTGTTTCTGTCAGAGAGTTGC
GAGCACACGTCTGAACTCTTTCCCTACACGACGCTCTTCCGATCTNNNGGGTAGAAAAAATGAATACT
GAGCACACGTCTGAACTCTTTCCCTACACGACGCTCTTCCGATCTNNNGGGTAGAAAAAATGAATACT
GAGCACACGTCTGAACTCTTTCCCTACACGACGCTCTTCCGATCTNNNAAAACTTAGCCAGGCATAGT
GAGCACACGTCTGAACTCTTTCCCTACACGACGCTCTTCCGATCTNNNAAAACTTAGCCAGGCATAGT

AGCACACGTCTGAACTCTTTCCCTACACGACGCTCTTCCGATCTNNNGAGCTCAGAGGAAAGAACTC
AGCACACGTCTGAACTCTTTCCCTACACGACGCTCTTCCGATCTNNNGAGCTCAGAGGAAAGAACTC
GCACACGTCTGAACTCTTTCCCTACACGACGCTCTTCCGATCTNNNTGTTCTTCTCCTCTGTTTGTCCTC
GCACACGTCTGAACTCTTTCCCTACACGACGCTCTTCCGATCTNNNTGTTCTTCTCCTCTGTTTGTCCTC
AGCACACGTCTGAACTCTTTCCCTACACGACGCTCTTCCGATCTNNNTGGCGCTGCTATAATATATGG
AGCACACGTCTGAACTCTTTCCCTACACGACGCTCTTCCGATCTNNNTGGCGCTGCTATAATATATGG
GAGCACACGTCTGAACTCTTTCCCTACACGACGCTCTTCCGATCTNNNGAAGTCTCTCCCACATCAAG
GAGCACACGTCTGAACTCTTTCCCTACACGACGCTCTTCCGATCTNNNGAAGTCTCTCCCACATCAAG
GCACACGTCTGAACTCTTTCCCTACACGACGCTCTTCCGATCTNNNACGGAATCACTCACACATTCTG
GCACACGTCTGAACTCTTTCCCTACACGACGCTCTTCCGATCTNNNACGGAATCACTCACACATTCTG
GCACACGTCTGAACTCTTTCCCTACACGACGCTCTTCCGATCTNNNGGTGAAAAAGAAGGGAAGAC
GCACACGTCTGAACTCTTTCCCTACACGACGCTCTTCCGATCTNNNGGTGAAAAAGAAGGGAAGAC
GCACACGTCTGAACTCTTTCCCTACACGACGCTCTTCCGATCTNNNCACCTTTCAGTGCCTCCAAGAAT
GCACACGTCTGAACTCTTTCCCTACACGACGCTCTTCCGATCTNNNCACCTTTCAGTGCCTCCAAGAAT
GCACACGTCTGAACTCTTTCCCTACACGACGCTCTTCCGATCTNNNGCCACCACATCTGACCCTTATTTC
GCACACGTCTGAACTCTTTCCCTACACGACGCTCTTCCGATCTNNNGCCACCACATCTGACCCTTATTTC
GCACACGTCTGAACTCTTTCCCTACACGACGCTCTTCCGATCTNNNCCAACTACTCACATGTCTCCACA
GCACACGTCTGAACTCTTTCCCTACACGACGCTCTTCCGATCTNNNCCAACTACTCACATGTCTCCACA
AGCACACGTCTGAACTCTTTCCCTACACGACGCTCTTCCGATCTNNNGTGCATGTAATCCCAGCTACT
AGCACACGTCTGAACTCTTTCCCTACACGACGCTCTTCCGATCTNNNGTGCATGTAATCCCAGCTACT
AGAGCACACGTCTGAACTCTTTCCCTACACGACGCTCTTCCGATCTNNNTGCATCTGTAATCCCAGCT
AGAGCACACGTCTGAACTCTTTCCCTACACGACGCTCTTCCGATCTNNNTGCATCTGTAATCCCAGCT
GCACACGTCTGAACTCTTTCCCTACACGACGCTCTTCCGATCTNNNGTCTGAATATGGAACCGAGGTG
GCACACGTCTGAACTCTTTCCCTACACGACGCTCTTCCGATCTNNNGTCTGAATATGGAACCGAGGTG
CACACGTCTGAACTCTTTCCCTACACGACGCTCTTCCGATCTNNNGCTGAGGAGTGAAATTTGAACC
CACACGTCTGAACTCTTTCCCTACACGACGCTCTTCCGATCTNNNGCTGAGGAGTGAAATTTGAACC
GAGCACACGTCTGAACTCTTTCCCTACACGACGCTCTTCCGATCTNNNCCATGCTGGTGTCAAACCTCT
GAGCACACGTCTGAACTCTTTCCCTACACGACGCTCTTCCGATCTNNNCCATGCTGGTGTCAAACCTCT
AGCACACGTCTGAACTCTTTCCCTACACGACGCTCTTCCGATCTNNNGGGGGGTGAAAGTTGAAAA
AGCACACGTCTGAACTCTTTCCCTACACGACGCTCTTCCGATCTNNNGGGGGGTGAAAGTTGAAAA
CACACGTCTGAACTCTTTCCCTACACGACGCTCTTCCGATCTNNNCAGGAGAATTGCTTGAACTTGG
CACACGTCTGAACTCTTTCCCTACACGACGCTCTTCCGATCTNNNCAGGAGAATTGCTTGAACTTGG
AGAGCACACGTCTGAACTCTTTCCCTACACGACGCTCTTCCGATCTNNNGCAGTGCTCTTTATTCGAAG
AGAGCACACGTCTGAACTCTTTCCCTACACGACGCTCTTCCGATCTNNNGCAGTGCTCTTTATTCGAAG
AGCACACGTCTGAACTCTTTCCCTACACGACGCTCTTCCGATCTNNNGATCTCCTTTTTCACTGGCGAA
AGCACACGTCTGAACTCTTTCCCTACACGACGCTCTTCCGATCTNNNGATCTCCTTTTTCACTGGCGAA
GCACACGTCTGAACTCTTTCCCTACACGACGCTCTTCCGATCTNNNGTTGTGGTGACCTGAGATCGT
GCACACGTCTGAACTCTTTCCCTACACGACGCTCTTCCGATCTNNNGTTGTGGTGACCTGAGATCGT
AGCACACGTCTGAACTCTTTCCCTACACGACGCTCTTCCGATCTNNNTTATACCCCTGCTTCATTCCAC
AGCACACGTCTGAACTCTTTCCCTACACGACGCTCTTCCGATCTNNNTTATACCCCTGCTTCATTCCAC
GAGCACACGTCTGAACTCTTTCCCTACACGACGCTCTTCCGATCTNNNTATTTTTCTTCCCTTCCCTCCC
GAGCACACGTCTGAACTCTTTCCCTACACGACGCTCTTCCGATCTNNNTATTTTTCTTCCCTTCCCTCCC
AGCACACGTCTGAACTCTTTCCCTACACGACGCTCTTCCGATCTNNNATCAATCCATCCTAAGGCTTGC

AGCACACGTCTGAACTCTTTCCCTACACGACGCTCTTCCGATCTNNNATCAATCCATCCTAAGGCTTGC
GAGCACACGTCTGAACTCTTTCCCTACACGACGCTCTTCCGATCTNNNCCTGGCCAGAAAGGGAAAAA
GAGCACACGTCTGAACTCTTTCCCTACACGACGCTCTTCCGATCTNNNCCTGGCCAGAAAGGGAAAAA
AGCACACGTCTGAACTCTTTCCCTACACGACGCTCTTCCGATCTNNNCCTTTGGACTGAATCTCATGAG
AGCACACGTCTGAACTCTTTCCCTACACGACGCTCTTCCGATCTNNNCCTTTGGACTGAATCTCATGAG
GAGCACACGTCTGAACTCTTTCCCTACACGACGCTCTTCCGATCTNNNGCTTCTTTTCCCTCAGCTTTCAT
GAGCACACGTCTGAACTCTTTCCCTACACGACGCTCTTCCGATCTNNNGCTTCTTTTCCCTCAGCTTTCAT
GAGCACACGTCTGAACTCTTTCCCTACACGACGCTCTTCCGATCTNNNGAGGGGCATTGACATAAGCA
GAGCACACGTCTGAACTCTTTCCCTACACGACGCTCTTCCGATCTNNNGAGGGGCATTGACATAAGCA
ACACGTCTGAACTCTTTCCCTACACGACGCTCTTCCGATCTNNNCTGGGATTACAATTGTGAGCCATCG
ACACGTCTGAACTCTTTCCCTACACGACGCTCTTCCGATCTNNNCTGGGATTACAATTGTGAGCCATCG
GAGCACACGTCTGAACTCTTTCCCTACACGACGCTCTTCCGATCTNNNCTTCACAAAATGCAGACCAGG
GAGCACACGTCTGAACTCTTTCCCTACACGACGCTCTTCCGATCTNNNCTTCACAAAATGCAGACCAGG
AGCACACGTCTGAACTCTTTCCCTACACGACGCTCTTCCGATCTNNNGAGATCACGCCATTGCACTCTA
AGCACACGTCTGAACTCTTTCCCTACACGACGCTCTTCCGATCTNNNGAGATCACGCCATTGCACTCTA
GCACACGTCTGAACTCTTTCCCTACACGACGCTCTTCCGATCTNNNCCTTATATTGCCCTCTCACCCTCC
GCACACGTCTGAACTCTTTCCCTACACGACGCTCTTCCGATCTNNNCCTTATATTGCCCTCTCACCCTCC
GAGCACACGTCTGAACTCTTTCCCTACACGACGCTCTTCCGATCTNNNCAGGGACATTGGCCTGAAATT
GAGCACACGTCTGAACTCTTTCCCTACACGACGCTCTTCCGATCTNNNCAGGGACATTGGCCTGAAATT
GAGCACACGTCTGAACTCTTTCCCTACACGACGCTCTTCCGATCTNNNTTCATCCGGAATCTATCGGTT
GAGCACACGTCTGAACTCTTTCCCTACACGACGCTCTTCCGATCTNNNTTCATCCGGAATCTATCGGTT
GCACACGTCTGAACTCTTTCCCTACACGACGCTCTTCCGATCTNNNTGGGTAGGGCAAAAGCAAAAT
GCACACGTCTGAACTCTTTCCCTACACGACGCTCTTCCGATCTNNNTGGGTAGGGCAAAAGCAAAAT
AGCACACGTCTGAACTCTTTCCCTACACGACGCTCTTCCGATCTNNNGGCGAGATCCTGACTCAAAAAA
AGCACACGTCTGAACTCTTTCCCTACACGACGCTCTTCCGATCTNNNGGCGAGATCCTGACTCAAAAAA
GAGCACACGTCTGAACTCTTTCCCTACACGACGCTCTTCCGATCTNNNATCATGCCATTATACTCCAGC
GAGCACACGTCTGAACTCTTTCCCTACACGACGCTCTTCCGATCTNNNATCATGCCATTATACTCCAGC
GAGCACACGTCTGAACTCTTTCCCTACACGACGCTCTTCCGATCTNNNCACCTGTAATCCCAGCTACTCT
GAGCACACGTCTGAACTCTTTCCCTACACGACGCTCTTCCGATCTNNNCACCTGTAATCCCAGCTACTCT
GAGCACACGTCTGAACTCTTTCCCTACACGACGCTCTTCCGATCTNNNAGCCCCCAGAGAATCTGGTAA
GAGCACACGTCTGAACTCTTTCCCTACACGACGCTCTTCCGATCTNNNAGCCCCCAGAGAATCTGGTAA
GAGCACACGTCTGAACTCTTTCCCTACACGACGCTCTTCCGATCTNNNGCCCTCATTTCCATCAGAGAG
GAGCACACGTCTGAACTCTTTCCCTACACGACGCTCTTCCGATCTNNNGCCCTCATTTCCATCAGAGAG
GAGCACACGTCTGAACTCTTTCCCTACACGACGCTCTTCCGATCTNNNAGGGTCTCCTGTAATACACCT
GAGCACACGTCTGAACTCTTTCCCTACACGACGCTCTTCCGATCTNNNAGGGTCTCCTGTAATACACCT
GAGCACACGTCTGAACTCTTTCCCTACACGACGCTCTTCCGATCTNNNAATGAGCTGAGTTGCATTTCC
GAGCACACGTCTGAACTCTTTCCCTACACGACGCTCTTCCGATCTNNNAATGAGCTGAGTTGCATTTCC
AGCACACGTCTGAACTCTTTCCCTACACGACGCTCTTCCGATCTNNNGCAAAGGAGACCCCTCCCATTC
AGCACACGTCTGAACTCTTTCCCTACACGACGCTCTTCCGATCTNNNGCAAAGGAGACCCCTCCCATTC
GAGCACACGTCTGAACTCTTTCCCTACACGACGCTCTTCCGATCTNNNTCCGGTCCTAAAGCTTGAAAC
GAGCACACGTCTGAACTCTTTCCCTACACGACGCTCTTCCGATCTNNNTCCGGTCCTAAAGCTTGAAAC
GCACACGTCTGAACTCTTTCCCTACACGACGCTCTTCCGATCTNNNAGACTGAAGGTATCTTTGGCCA
GCACACGTCTGAACTCTTTCCCTACACGACGCTCTTCCGATCTNNNAGACTGAAGGTATCTTTGGCCA

GCACACGTCTGAACTCTTTCCCTACACGACGCTCTTCCGATCTNNNGTTCTCTGAAGCTCCCCTTACAA
GCACACGTCTGAACTCTTTCCCTACACGACGCTCTTCCGATCTNNNGTTCTCTGAAGCTCCCCTTACAA
GAGCACACGTCTGAACTCTTTCCCTACACGACGCTCTTCCGATCTNNNAGCCCAGACGTCTCCTCTAAT
GAGCACACGTCTGAACTCTTTCCCTACACGACGCTCTTCCGATCTNNNAGCCCAGACGTCTCCTCTAAT
AGCACACGTCTGAACTCTTTCCCTACACGACGCTCTTCCGATCTNNNAAAAAAATTAGCCGGGTGTG
AGCACACGTCTGAACTCTTTCCCTACACGACGCTCTTCCGATCTNNNAAAAAAATTAGCCGGGTGTG
GAGCACACGTCTGAACTCTTTCCCTACACGACGCTCTTCCGATCTNNNAGGAGTTTGTTTGTTCCTTTG
GAGCACACGTCTGAACTCTTTCCCTACACGACGCTCTTCCGATCTNNNAGGAGTTTGTTTGTTCCTTTG
GAGCACACGTCTGAACTCTTTCCCTACACGACGCTCTTCCGATCTNNNACACATAGCTCAAACATCCTG
GAGCACACGTCTGAACTCTTTCCCTACACGACGCTCTTCCGATCTNNNACACATAGCTCAAACATCCTG
CACACGTCTGAACTCTTTCCCTACACGACGCTCTTCCGATCTNNNACAATCAGTGAGTAAATAGACAG
CACACGTCTGAACTCTTTCCCTACACGACGCTCTTCCGATCTNNNACAATCAGTGAGTAAATAGACAG
GCACACGTCTGAACTCTTTCCCTACACGACGCTCTTCCGATCTNNNCCACACTTCTTTGCAATCTCCTGC
GCACACGTCTGAACTCTTTCCCTACACGACGCTCTTCCGATCTNNNCCACACTTCTTTGCAATCTCCTGC
CACGTCTGAACTCTTTCCCTACACGACGCTCTTCCGATCTNNNTGGAAAACCTTCGTTACATGACCCAT
CACGTCTGAACTCTTTCCCTACACGACGCTCTTCCGATCTNNNTGGAAAACCTTCGTTACATGACCCAT
GCACACGTCTGAACTCTTTCCCTACACGACGCTCTTCCGATCTNNNCCGGTTAGCTCAGTTGGTAAGA
GCACACGTCTGAACTCTTTCCCTACACGACGCTCTTCCGATCTNNNCCGGTTAGCTCAGTTGGTAAGA
AGCACACGTCTGAACTCTTTCCCTACACGACGCTCTTCCGATCTNNNTCTCCAGAAGAAATCAAGCCA
AGCACACGTCTGAACTCTTTCCCTACACGACGCTCTTCCGATCTNNNTCTCCAGAAGAAATCAAGCCA
AGCACACGTCTGAACTCTTTCCCTACACGACGCTCTTCCGATCTNNNCCCGAGGACTGGAGGAGAAG
AGCACACGTCTGAACTCTTTCCCTACACGACGCTCTTCCGATCTNNNCCCGAGGACTGGAGGAGAAG
GAGCACACGTCTGAACTCTTTCCCTACACGACGCTCTTCCGATCTNNNATCAAATGCCAGCAAGTGTT
GAGCACACGTCTGAACTCTTTCCCTACACGACGCTCTTCCGATCTNNNATCAAATGCCAGCAAGTGTT
GAGCACACGTCTGAACTCTTTCCCTACACGACGCTCTTCCGATCTNNNAGAGAATCGCTTCACAGGAG
GAGCACACGTCTGAACTCTTTCCCTACACGACGCTCTTCCGATCTNNNAGAGAATCGCTTCACAGGAG
AGCACACGTCTGAACTCTTTCCCTACACGACGCTCTTCCGATCTNNNGAGTCTCCTGAAATGCAGCTT
AGCACACGTCTGAACTCTTTCCCTACACGACGCTCTTCCGATCTNNNGAGTCTCCTGAAATGCAGCTT
AGCACACGTCTGAACTCTTTCCCTACACGACGCTCTTCCGATCTNNNATTTTGAGAGCTTTTGGGAAG
AGCACACGTCTGAACTCTTTCCCTACACGACGCTCTTCCGATCTNNNATTTTGAGAGCTTTTGGGAAG
AGAGCACACGTCTGAACTCTTTCCCTACACGACGCTCTTCCGATCTNNNGGAGAGAAGTGAACCAGT
AGAGCACACGTCTGAACTCTTTCCCTACACGACGCTCTTCCGATCTNNNGGAGAGAAGTGAACCAGT
GAGCACACGTCTGAACTCTTTCCCTACACGACGCTCTTCCGATCTNNNTGAGGCAGGAGAATCTCTTG
GAGCACACGTCTGAACTCTTTCCCTACACGACGCTCTTCCGATCTNNNTGAGGCAGGAGAATCTCTTG
AGCACACGTCTGAACTCTTTCCCTACACGACGCTCTTCCGATCTNNNTCCCCACAAAAGACATGAACTC
AGCACACGTCTGAACTCTTTCCCTACACGACGCTCTTCCGATCTNNNTCCCCACAAAAGACATGAACTC
AGCACACGTCTGAACTCTTTCCCTACACGACGCTCTTCCGATCTNNNTGGTTGTATGTCCACAGTCAA
AGCACACGTCTGAACTCTTTCCCTACACGACGCTCTTCCGATCTNNNTGGTTGTATGTCCACAGTCAA
AGAGCACACGTCTGAACTCTTTCCCTACACGACGCTCTTCCGATCTNNNGAGAATGGCATGAACCTGG
AGAGCACACGTCTGAACTCTTTCCCTACACGACGCTCTTCCGATCTNNNGAGAATGGCATGAACCTGG
AGCACACGTCTGAACTCTTTCCCTACACGACGCTCTTCCGATCTNNNCAAGATGGATAGGACTGGGG
AGCACACGTCTGAACTCTTTCCCTACACGACGCTCTTCCGATCTNNNCAAGATGGATAGGACTGGGG
AGAGCACACGTCTGAACTCTTTCCCTACACGACGCTCTTCCGATCTNNNTGGAGATGTCAGAAAAGC

AGAGCACACGTCTGAACTCTTTCCCTACACGACGCTCTTCCGATCTNNNTGGAGATGTCAGAAAAGCA
AGCACACGTCTGAACTCTTTCCCTACACGACGCTCTTCCGATCTNNNTGCGTGTGTAATCTCAGCTACTT
AGCACACGTCTGAACTCTTTCCCTACACGACGCTCTTCCGATCTNNNTGCGTGTGTAATCTCAGCTACTT
AGCACACGTCTGAACTCTTTCCCTACACGACGCTCTTCCGATCTNNNCCTTTCTCACCTCAGAATCTGCT
AGCACACGTCTGAACTCTTTCCCTACACGACGCTCTTCCGATCTNNNCCTTTCTCACCTCAGAATCTGCT
GAGCACACGTCTGAACTCTTTCCCTACACGACGCTCTTCCGATCTNNNCAGAACTGAAGTGTGGGAGG
GAGCACACGTCTGAACTCTTTCCCTACACGACGCTCTTCCGATCTNNNCAGAACTGAAGTGTGGGAGG
AGCACACGTCTGAACTCTTTCCCTACACGACGCTCTTCCGATCTNNNCTCTTCGTCGTTTACCAGCTCAA
AGCACACGTCTGAACTCTTTCCCTACACGACGCTCTTCCGATCTNNNCTCTTCGTCGTTTACCAGCTCAA
AGCACACGTCTGAACTCTTTCCCTACACGACGCTCTTCCGATCTNNNCTGGGTGATAAAGTGAGACTC
AGCACACGTCTGAACTCTTTCCCTACACGACGCTCTTCCGATCTNNNCTGGGTGATAAAGTGAGACTC
GCACACGTCTGAACTCTTTCCCTACACGACGCTCTTCCGATCTNNNTGAGCTTCCATCTGTTTCTTCCCC
GCACACGTCTGAACTCTTTCCCTACACGACGCTCTTCCGATCTNNNTGAGCTTCCATCTGTTTCTTCCCC
GAGCACACGTCTGAACTCTTTCCCTACACGACGCTCTTCCGATCTNNNGTTAGGAAAAGGGACAGAG
GAGCACACGTCTGAACTCTTTCCCTACACGACGCTCTTCCGATCTNNNGTTAGGAAAAGGGACAGAG
GAGCACACGTCTGAACTCTTTCCCTACACGACGCTCTTCCGATCTNNNATCTTGGCACTTCTGGAAGCAT
GAGCACACGTCTGAACTCTTTCCCTACACGACGCTCTTCCGATCTNNNATCTTGGCACTTCTGGAAGCAT
GAGCACACGTCTGAACTCTTTCCCTACACGACGCTCTTCCGATCTNNNCTTTGTCATCAGAGGCGGAA
GAGCACACGTCTGAACTCTTTCCCTACACGACGCTCTTCCGATCTNNNCTTTGTCATCAGAGGCGGAA
AGAGCACACGTCTGAACTCTTTCCCTACACGACGCTCTTCCGATCTNNNCTGTAATCCCAGCTACTCAG
AGAGCACACGTCTGAACTCTTTCCCTACACGACGCTCTTCCGATCTNNNCTGTAATCCCAGCTACTCAG
AGCACACGTCTGAACTCTTTCCCTACACGACGCTCTTCCGATCTNNNACTTTGTGTGTAGGTATGTGTT
AGCACACGTCTGAACTCTTTCCCTACACGACGCTCTTCCGATCTNNNACTTTGTGTGTAGGTATGTGTT
GAGCACACGTCTGAACTCTTTCCCTACACGACGCTCTTCCGATCTNNNTTTCTACCTTATTTACTTGCA
GAGCACACGTCTGAACTCTTTCCCTACACGACGCTCTTCCGATCTNNNTTTCTACCTTATTTACTTGCA
GAGCACACGTCTGAACTCTTTCCCTACACGACGCTCTTCCGATCTNNNCTCCCGAGTAGCTGAGATTAC
GAGCACACGTCTGAACTCTTTCCCTACACGACGCTCTTCCGATCTNNNCTCCCGAGTAGCTGAGATTAC
AGAGCACACGTCTGAACTCTTTCCCTACACGACGCTCTTCCGATCTNNNACAGTGGTAAACAAAACCGT
AGAGCACACGTCTGAACTCTTTCCCTACACGACGCTCTTCCGATCTNNNACAGTGGTAAACAAAACCGT
GAGCACACGTCTGAACTCTTTCCCTACACGACGCTCTTCCGATCTNNNAGTACCTTTTGGAACCCAGAT
GAGCACACGTCTGAACTCTTTCCCTACACGACGCTCTTCCGATCTNNNAGTACCTTTTGGAACCCAGAT
GAGCACACGTCTGAACTCTTTCCCTACACGACGCTCTTCCGATCTNNNGCTCTAGAAGGAAGGCAGAG
GAGCACACGTCTGAACTCTTTCCCTACACGACGCTCTTCCGATCTNNNGCTCTAGAAGGAAGGCAGAG
CACACGTCTGAACTCTTTCCCTACACGACGCTCTTCCGATCTNNNAGAAAAGTTCCCCTCCCCAAAATCA
CACACGTCTGAACTCTTTCCCTACACGACGCTCTTCCGATCTNNNAGAAAAGTTCCCCTCCCCAAAATCA
GAGCACACGTCTGAACTCTTTCCCTACACGACGCTCTTCCGATCTNNNAGTGGGTAGTGAGGATCCTT
GAGCACACGTCTGAACTCTTTCCCTACACGACGCTCTTCCGATCTNNNAGTGGGTAGTGAGGATCCTT
GAGCACACGTCTGAACTCTTTCCCTACACGACGCTCTTCCGATCTNNNTGAGGTAGGAGAATCGCTTG
GAGCACACGTCTGAACTCTTTCCCTACACGACGCTCTTCCGATCTNNNTGAGGTAGGAGAATCGCTTG
AGCACACGTCTGAACTCTTTCCCTACACGACGCTCTTCCGATCTNNNGACAGCGGTTCTCGAGACTAG
AGCACACGTCTGAACTCTTTCCCTACACGACGCTCTTCCGATCTNNNGACAGCGGTTCTCGAGACTAG
AGCACACGTCTGAACTCTTTCCCTACACGACGCTCTTCCGATCTNNNGAGCTTGTAGTGAGCCGAGAT
AGCACACGTCTGAACTCTTTCCCTACACGACGCTCTTCCGATCTNNNGAGCTTGTAGTGAGCCGAGAT

GAGCACACGTCTGAACTCTTTCCCTACACGACGCTCTTCCGATCTNNNGGAAGGGTGCACAAACTAAG

GAGCACACGTCTGAACTCTTTCCCTACACGACGCTCTTCCGATCTNNNGGAAGGGTGCACAAACTAAG

AGCACACGTCTGAACTCTTTCCCTACACGACGCTCTTCCGATCTNNNTGGTGTATCAGACAGGGAAACA

AGCACACGTCTGAACTCTTTCCCTACACGACGCTCTTCCGATCTNNNTGGTGTATCAGACAGGGAAACA

GAGCACACGTCTGAACTCTTTCCCTACACGACGCTCTTCCGATCTNNNAATTCACCTGCACACATGAACT

GAGCACACGTCTGAACTCTTTCCCTACACGACGCTCTTCCGATCTNNNAATTCACCTGCACACATGAACT

AGCACACGTCTGAACTCTTTCCCTACACGACGCTCTTCCGATCTNNNTTGACTTCCACCTCCCAGGTTTA

AGCACACGTCTGAACTCTTTCCCTACACGACGCTCTTCCGATCTNNNTTGACTTCCACCTCCCAGGTTTA

GCACACGTCTGAACTCTTTCCCTACACGACGCTCTTCCGATCTNNNATATGGGGTCTCACTCTGCTGTC

GCACACGTCTGAACTCTTTCCCTACACGACGCTCTTCCGATCTNNNATATGGGGTCTCACTCTGCTGTC

AGCACACGTCTGAACTCTTTCCCTACACGACGCTCTTCCGATCTNNNGCAATTTTTGACTGGATGGCGA

AGCACACGTCTGAACTCTTTCCCTACACGACGCTCTTCCGATCTNNNGCAATTTTTGACTGGATGGCGA

AGAGCACACGTCTGAACTCTTTCCCTACACGACGCTCTTCCGATCTNNNCTCACAAGAAGCCTCTCCTGT

AGAGCACACGTCTGAACTCTTTCCCTACACGACGCTCTTCCGATCTNNNCTCACAAGAAGCCTCTCCTGT

GCACACGTCTGAACTCTTTCCCTACACGACGCTCTTCCGATCTNNNCCATGAAGCAAGCTGTCAGAAAC

GCACACGTCTGAACTCTTTCCCTACACGACGCTCTTCCGATCTNNNCCATGAAGCAAGCTGTCAGAAAC

GAGCACACGTCTGAACTCTTTCCCTACACGACGCTCTTCCGATCTNNNCCTCAGAGTGGTTCTCTCTCT

GAGCACACGTCTGAACTCTTTCCCTACACGACGCTCTTCCGATCTNNNCCTCAGAGTGGTTCTCTCTCT

GAGCACACGTCTGAACTCTTTCCCTACACGACGCTCTTCCGATCTNNNAGCTTTTCAAAACCCCGACTTT

GAGCACACGTCTGAACTCTTTCCCTACACGACGCTCTTCCGATCTNNNAGCTTTTCAAAACCCCGACTTT

AGCACACGTCTGAACTCTTTCCCTACACGACGCTCTTCCGATCTNNNGTGTTAGGAAACCTTGCCTTTG

AGCACACGTCTGAACTCTTTCCCTACACGACGCTCTTCCGATCTNNNGTGTTAGGAAACCTTGCCTTTG

GAGCACACGTCTGAACTCTTTCCCTACACGACGCTCTTCCGATCTNNNCCACAGAATCCCCAGCACACCA

GAGCACACGTCTGAACTCTTTCCCTACACGACGCTCTTCCGATCTNNNCCACAGAATCCCCAGCACACCA

GAGCACACGTCTGAACTCTTTCCCTACACGACGCTCTTCCGATCTNNNCGACAGAGAGAGACTCCGTCT

GAGCACACGTCTGAACTCTTTCCCTACACGACGCTCTTCCGATCTNNNCGACAGAGAGAGACTCCGTCT

GAGCACACGTCTGAACTCTTTCCCTACACGACGCTCTTCCGATCTNNNCTGAAGTGGGAGGATCTCTTC

GAGCACACGTCTGAACTCTTTCCCTACACGACGCTCTTCCGATCTNNNCTGAAGTGGGAGGATCTCTTC

GAGCACACGTCTGAACTCTTTCCCTACACGACGCTCTTCCGATCTNNNGACCCAGGTATGCAAGAAGTG

GAGCACACGTCTGAACTCTTTCCCTACACGACGCTCTTCCGATCTNNNGACCCAGGTATGCAAGAAGTG

AGCACACGTCTGAACTCTTTCCCTACACGACGCTCTTCCGATCTNNNAAAATCAAACTGTAGTTGTTTTC

AGCACACGTCTGAACTCTTTCCCTACACGACGCTCTTCCGATCTNNNAAAATCAAACTGTAGTTGTTTTC

GAGCACACGTCTGAACTCTTTCCCTACACGACGCTCTTCCGATCTNNNTAAATGCTAGAGAACTGGAGC

GAGCACACGTCTGAACTCTTTCCCTACACGACGCTCTTCCGATCTNNNTAAATGCTAGAGAACTGGAGC

GAGCACACGTCTGAACTCTTTCCCTACACGACGCTCTTCCGATCTNNNACTGCCTGCTGATAACTGAGA

GAGCACACGTCTGAACTCTTTCCCTACACGACGCTCTTCCGATCTNNNACTGCCTGCTGATAACTGAGA

GAGCACACGTCTGAACTCTTTCCCTACACGACGCTCTTCCGATCTNNNCTCCCCACTCTAGGAGTCTGTA

GAGCACACGTCTGAACTCTTTCCCTACACGACGCTCTTCCGATCTNNNCTCCCCACTCTAGGAGTCTGTA

AGCACACGTCTGAACTCTTTCCCTACACGACGCTCTTCCGATCTNNNAGCTCTACCCTGAAGAAAATGC

AGCACACGTCTGAACTCTTTCCCTACACGACGCTCTTCCGATCTNNNAGCTCTACCCTGAAGAAAATGC

AGCACACGTCTGAACTCTTTCCCTACACGACGCTCTTCCGATCTNNNTGGTATTGCCTAAGTCCCAACAC

AGCACACGTCTGAACTCTTTCCCTACACGACGCTCTTCCGATCTNNNTGGTATTGCCTAAGTCCCAACAC

GAGCACACGTCTGAACTCTTTCCCTACACGACGCTCTTCCGATCTNNNTGGGGTTTTCAACATAATGGC

GAGCACACGTCTGAACTCTTTCCCTACACGACGCTCTTCCGATCTNNNTGGGGTTTTCAACATAATGGC

AGCACACGTCTGAACTCTTTCCCTACACGACGCTCTTCCGATCTNNNGAAATTATCTCCTGCTCATAGTA

AGCACACGTCTGAACTCTTTCCCTACACGACGCTCTTCCGATCTNNNGAAATTATCTCCTGCTCATAGTA

AGCACACGTCTGAACTCTTTCCCTACACGACGCTCTTCCGATCTNNNGTTGTAGTGAACCAAGATCGCA

AGCACACGTCTGAACTCTTTCCCTACACGACGCTCTTCCGATCTNNNGTTGTAGTGAACCAAGATCGCA

GAGCACACGTCTGAACTCTTTCCCTACACGACGCTCTTCCGATCTNNNGGCAAATACTACAAATTGGGC

GAGCACACGTCTGAACTCTTTCCCTACACGACGCTCTTCCGATCTNNNGGCAAATACTACAAATTGGGC

GAGCACACGTCTGAACTCTTTCCCTACACGACGCTCTTCCGATCTNNNAAAATTGCTTCAATCTGGGAG

GAGCACACGTCTGAACTCTTTCCCTACACGACGCTCTTCCGATCTNNNAAAATTGCTTCAATCTGGGAG

AGCACACGTCTGAACTCTTTCCCTACACGACGCTCTTCCGATCTNNNGCAGCTAAGGAATCCAGCAAA

AGCACACGTCTGAACTCTTTCCCTACACGACGCTCTTCCGATCTNNNGCAGCTAAGGAATCCAGCAAA

AGCACACGTCTGAACTCTTTCCCTACACGACGCTCTTCCGATCTNNNCAGCAAGCACCATTTACTTCCTT

AGCACACGTCTGAACTCTTTCCCTACACGACGCTCTTCCGATCTNNNCAGCAAGCACCATTTACTTCCTT

AGCACACGTCTGAACTCTTTCCCTACACGACGCTCTTCCGATCTNNNCTGAGGCAGGAGAATCACTTGA

AGCACACGTCTGAACTCTTTCCCTACACGACGCTCTTCCGATCTNNNCTGAGGCAGGAGAATCACTTGA

GAGCACACGTCTGAACTCTTTCCCTACACGACGCTCTTCCGATCTNNNCAAGTAGCTAGGACTATAGG

GAGCACACGTCTGAACTCTTTCCCTACACGACGCTCTTCCGATCTNNNCAAGTAGCTAGGACTATAGG

AGAGCACACGTCTGAACTCTTTCCCTACACGACGCTCTTCCGATCTNNNTAAGAGAGCTCTGGATAGCC

AGAGCACACGTCTGAACTCTTTCCCTACACGACGCTCTTCCGATCTNNNTAAGAGAGCTCTGGATAGCC

GAGCACACGTCTGAACTCTTTCCCTACACGACGCTCTTCCGATCTNNNTCTCTCCTTCTGTCTCCTCTG

GAGCACACGTCTGAACTCTTTCCCTACACGACGCTCTTCCGATCTNNNTCTCTCCTTCTGTCTCCTCTG

AGCACACGTCTGAACTCTTTCCCTACACGACGCTCTTCCGATCTNNNCTGTGAGCCATGATTACACCAG

AGCACACGTCTGAACTCTTTCCCTACACGACGCTCTTCCGATCTNNNCTGTGAGCCATGATTACACCAG

ACACGTCTGAACTCTTTCCCTACACGACGCTCTTCCGATCTNNNAGAGTGGTAAAAACAAGGCCTCACA

ACACGTCTGAACTCTTTCCCTACACGACGCTCTTCCGATCTNNNAGAGTGGTAAAAACAAGGCCTCACA

GCACACGTCTGAACTCTTTCCCTACACGACGCTCTTCCGATCTNNNTTTCCCTTGAAAACCAGCACAAGC

GCACACGTCTGAACTCTTTCCCTACACGACGCTCTTCCGATCTNNNTTTCCCTTGAAAACCAGCACAAGC

GAGCACACGTCTGAACTCTTTCCCTACACGACGCTCTTCCGATCTNNNAGACAAACTTCTGAGGTGATG

GAGCACACGTCTGAACTCTTTCCCTACACGACGCTCTTCCGATCTNNNAGACAAACTTCTGAGGTGATG

GAGCACACGTCTGAACTCTTTCCCTACACGACGCTCTTCCGATCTNNNAGAGGAGAAAAGGGGAGGC

GAGCACACGTCTGAACTCTTTCCCTACACGACGCTCTTCCGATCTNNNAGAGGAGAAAAGGGGAGGC

AGCACACGTCTGAACTCTTTCCCTACACGACGCTCTTCCGATCTNNNCACCTGTAATCCCAGCTACTCAG

AGCACACGTCTGAACTCTTTCCCTACACGACGCTCTTCCGATCTNNNCACCTGTAATCCCAGCTACTCAG

AGAGCACACGTCTGAACTCTTTCCCTACACGACGCTCTTCCGATCTNNNACTATCTACTTCCTTAGCCTG

AGAGCACACGTCTGAACTCTTTCCCTACACGACGCTCTTCCGATCTNNNACTATCTACTTCCTTAGCCTG

GAGCACACGTCTGAACTCTTTCCCTACACGACGCTCTTCCGATCTNNNACATTCAGCTTAAAACTTACTG

GAGCACACGTCTGAACTCTTTCCCTACACGACGCTCTTCCGATCTNNNACATTCAGCTTAAAACTTACTG

GAGCACACGTCTGAACTCTTTCCCTACACGACGCTCTTCCGATCTNNNTAGTTGTGACTACAGGTGTGC

GAGCACACGTCTGAACTCTTTCCCTACACGACGCTCTTCCGATCTNNNTAGTTGTGACTACAGGTGTGC

GAGCACACGTCTGAACTCTTTCCCTACACGACGCTCTTCCGATCTNNNACATATGTATGCATGTGCCAT

GAGCACACGTCTGAACTCTTTCCCTACACGACGCTCTTCCGATCTNNNACATATGTATGCATGTGCCAT

GCACACGTCTGAACTCTTTCCCTACACGACGCTCTTCCGATCTNNNAGAAGAGCTGATTTTGGTCCCATC

GCACACGTCTGAACTCTTTCCCTACACGACGCTCTTCCGATCTNNNAGAAGAGCTGATTTTGGTCCCATC

AGCACACGTCTGAACTCTTTCCCTACACGACGCTCTTCCGATCTNNNAGTGCTGAGATTACAGGTGTGA
AGCACACGTCTGAACTCTTTCCCTACACGACGCTCTTCCGATCTNNNAGTGCTGAGATTACAGGTGTGA
AGCACACGTCTGAACTCTTTCCCTACACGACGCTCTTCCGATCTNNNGCCTGTAGTCCCAGCTACTTGCA
AGCACACGTCTGAACTCTTTCCCTACACGACGCTCTTCCGATCTNNNGCCTGTAGTCCCAGCTACTTGCA
GAGCACACGTCTGAACTCTTTCCCTACACGACGCTCTTCCGATCTNNNGAGACATGAGGGGAGATATG
GAGCACACGTCTGAACTCTTTCCCTACACGACGCTCTTCCGATCTNNNGAGACATGAGGGGAGATATG
AGCACACGTCTGAACTCTTTCCCTACACGACGCTCTTCCGATCTNNNATCCTAGGTGACAGAGTGAGA
AGCACACGTCTGAACTCTTTCCCTACACGACGCTCTTCCGATCTNNNATCCTAGGTGACAGAGTGAGA
AGCACACGTCTGAACTCTTTCCCTACACGACGCTCTTCCGATCTNNNGTTTCCAATTCCCACCTCCTTTGC
AGCACACGTCTGAACTCTTTCCCTACACGACGCTCTTCCGATCTNNNGTTTCCAATTCCCACCTCCTTTGC
GAGCACACGTCTGAACTCTTTCCCTACACGACGCTCTTCCGATCTNNNGGGAAAGATCTCTCCCTCTCTC
GAGCACACGTCTGAACTCTTTCCCTACACGACGCTCTTCCGATCTNNNGGGAAAGATCTCTCCCTCTCTC
GAGCACACGTCTGAACTCTTTCCCTACACGACGCTCTTCCGATCTNNNAGACCCCTTCTCAGAACAATGT
GAGCACACGTCTGAACTCTTTCCCTACACGACGCTCTTCCGATCTNNNAGACCCCTTCTCAGAACAATGT
AGCACACGTCTGAACTCTTTCCCTACACGACGCTCTTCCGATCTNNNCTCACACCCTACATAAACACCCA
AGCACACGTCTGAACTCTTTCCCTACACGACGCTCTTCCGATCTNNNCTCACACCCTACATAAACACCCA
GAGCACACGTCTGAACTCTTTCCCTACACGACGCTCTTCCGATCTNNNAAGACCAATACATCTTTATCCA
GAGCACACGTCTGAACTCTTTCCCTACACGACGCTCTTCCGATCTNNNAAGACCAATACATCTTTATCCA
AGCACACGTCTGAACTCTTTCCCTACACGACGCTCTTCCGATCTNNNCCAACACCTGAGTAGCTGGACT
AGCACACGTCTGAACTCTTTCCCTACACGACGCTCTTCCGATCTNNNCCAACACCTGAGTAGCTGGACT
AGCACACGTCTGAACTCTTTCCCTACACGACGCTCTTCCGATCTNNNCTGTTGAGACCCACTTCTCTTCTC
AGCACACGTCTGAACTCTTTCCCTACACGACGCTCTTCCGATCTNNNCTGTTGAGACCCACTTCTCTTCTC
AGCACACGTCTGAACTCTTTCCCTACACGACGCTCTTCCGATCTNNNTGAACTAGAGAAGCAAGAGCAA
AGCACACGTCTGAACTCTTTCCCTACACGACGCTCTTCCGATCTNNNTGAACTAGAGAAGCAAGAGCAA
GAGCACACGTCTGAACTCTTTCCCTACACGACGCTCTTCCGATCTNNNGAGAATCGCTTGAATCCTGGA
GAGCACACGTCTGAACTCTTTCCCTACACGACGCTCTTCCGATCTNNNGAGAATCGCTTGAATCCTGGA
CACACGTCTGAACTCTTTCCCTACACGACGCTCTTCCGATCTNNNAACCTCATTCTGTTTTGTTCTCATCG
CACACGTCTGAACTCTTTCCCTACACGACGCTCTTCCGATCTNNNAACCTCATTCTGTTTTGTTCTCATCG
GAGCACACGTCTGAACTCTTTCCCTACACGACGCTCTTCCGATCTNNNTGGTCTCAAACTTCTGAGCTCA
GAGCACACGTCTGAACTCTTTCCCTACACGACGCTCTTCCGATCTNNNTGGTCTCAAACTTCTGAGCTCA
AGCACACGTCTGAACTCTTTCCCTACACGACGCTCTTCCGATCTNNNTTTTCTCACAGCTCTGGACATCA
AGCACACGTCTGAACTCTTTCCCTACACGACGCTCTTCCGATCTNNNTTTTCTCACAGCTCTGGACATCA
GCACACGTCTGAACTCTTTCCCTACACGACGCTCTTCCGATCTNNNTTAGCTTCTGAAATGCCACTCCTC
GCACACGTCTGAACTCTTTCCCTACACGACGCTCTTCCGATCTNNNTTAGCTTCTGAAATGCCACTCCTC
GAGCACACGTCTGAACTCTTTCCCTACACGACGCTCTTCCGATCTNNNAGAGCAAGAGATAGGAGTCT
GAGCACACGTCTGAACTCTTTCCCTACACGACGCTCTTCCGATCTNNNAGAGCAAGAGATAGGAGTCT
CACACGTCTGAACTCTTTCCCTACACGACGCTCTTCCGATCTNNNATAAAGAATGCTTTTAGGCCGGG
CACACGTCTGAACTCTTTCCCTACACGACGCTCTTCCGATCTNNNATAAAGAATGCTTTTAGGCCGGG
AGCACACGTCTGAACTCTTTCCCTACACGACGCTCTTCCGATCTNNNGAAGCAGGACAATCACTTGAAC
AGCACACGTCTGAACTCTTTCCCTACACGACGCTCTTCCGATCTNNNGAAGCAGGACAATCACTTGAAC
GAGCACACGTCTGAACTCTTTCCCTACACGACGCTCTTCCGATCTNNNATGGGGAGAGGAAGAGGTGA
GAGCACACGTCTGAACTCTTTCCCTACACGACGCTCTTCCGATCTNNNATGGGGAGAGGAAGAGGTGA
GCACACGTCTGAACTCTTTCCCTACACGACGCTCTTCCGATCTNNNCCACAGATGCTCCTTCCATTTTCCA

GCACACGTCTGAACTCTTTCCCTACACGACGCTCTTCCGATCTNNNCCACAGATGCTCCTTCCATTTTCCA
AGAGCACACGTCTGAACTCTTTCCCTACACGACGCTCTTCCGATCTNNNCAATGCACACAAGACTCCTCA
AGAGCACACGTCTGAACTCTTTCCCTACACGACGCTCTTCCGATCTNNNCAATGCACACAAGACTCCTCA
CACACGTCTGAACTCTTTCCCTACACGACGCTCTTCCGATCTNNNCACAACGACCACATCCTCATAGACA
CACACGTCTGAACTCTTTCCCTACACGACGCTCTTCCGATCTNNNCACAACGACCACATCCTCATAGACA
GAGCACACGTCTGAACTCTTTCCCTACACGACGCTCTTCCGATCTNNNATGGTAGGAGAGATTCTTCCC
GAGCACACGTCTGAACTCTTTCCCTACACGACGCTCTTCCGATCTNNNATGGTAGGAGAGATTCTTCCC
GAGCACACGTCTGAACTCTTTCCCTACACGACGCTCTTCCGATCTNNNAGAACTGGAACTTCACTACAC
GAGCACACGTCTGAACTCTTTCCCTACACGACGCTCTTCCGATCTNNNAGAACTGGAACTTCACTACAC
AGCACACGTCTGAACTCTTTCCCTACACGACGCTCTTCCGATCTNNNGGTTCAATCAATTCTCCTGCCTCC
AGCACACGTCTGAACTCTTTCCCTACACGACGCTCTTCCGATCTNNNGGTTCAATCAATTCTCCTGCCTCC
AGCACACGTCTGAACTCTTTCCCTACACGACGCTCTTCCGATCTNNNGAGAATGGCTTGAACCTGGGA
AGCACACGTCTGAACTCTTTCCCTACACGACGCTCTTCCGATCTNNNGAGAATGGCTTGAACCTGGGA
AGCACACGTCTGAACTCTTTCCCTACACGACGCTCTTCCGATCTNNNAGCAGATGGGAAGTGATGAGA
AGCACACGTCTGAACTCTTTCCCTACACGACGCTCTTCCGATCTNNNAGCAGATGGGAAGTGATGAGA
GAGCACACGTCTGAACTCTTTCCCTACACGACGCTCTTCCGATCTNNNAGAAGAATCGCTTGAATCCAC
GAGCACACGTCTGAACTCTTTCCCTACACGACGCTCTTCCGATCTNNNAGAAGAATCGCTTGAATCCAC
AGCACACGTCTGAACTCTTTCCCTACACGACGCTCTTCCGATCTNNNTGTGTGTGTGTATGTGTGTCTC
AGCACACGTCTGAACTCTTTCCCTACACGACGCTCTTCCGATCTNNNTGTGTGTGTGTATGTGTGTCTC
AGCACACGTCTGAACTCTTTCCCTACACGACGCTCTTCCGATCTNNNTGTTATCCAATTGCCAACAGCTC
AGCACACGTCTGAACTCTTTCCCTACACGACGCTCTTCCGATCTNNNTGTTATCCAATTGCCAACAGCTC
GAGCACACGTCTGAACTCTTTCCCTACACGACGCTCTTCCGATCTNNNTGGAGTTCAATAGAGCGTTCT
GAGCACACGTCTGAACTCTTTCCCTACACGACGCTCTTCCGATCTNNNTGGAGTTCAATAGAGCGTTCT
GCACACGTCTGAACTCTTTCCCTACACGACGCTCTTCCGATCTNNNCAGGCAAGTCAACAAATTACCCCC
GCACACGTCTGAACTCTTTCCCTACACGACGCTCTTCCGATCTNNNCAGGCAAGTCAACAAATTACCCCC
GAGCACACGTCTGAACTCTTTCCCTACACGACGCTCTTCCGATCTNNNAAAGATACACATAGCCAGAAA
GAGCACACGTCTGAACTCTTTCCCTACACGACGCTCTTCCGATCTNNNAAAGATACACATAGCCAGAAA
GAGCACACGTCTGAACTCTTTCCCTACACGACGCTCTTCCGATCTNNNCCTTATCACTACTTCCAGAAGC
GAGCACACGTCTGAACTCTTTCCCTACACGACGCTCTTCCGATCTNNNCCTTATCACTACTTCCAGAAGC
GAGCACACGTCTGAACTCTTTCCCTACACGACGCTCTTCCGATCTNNNAAAAAAATTAGCCGGGCATTG
GAGCACACGTCTGAACTCTTTCCCTACACGACGCTCTTCCGATCTNNNAAAAAAATTAGCCGGGCATTG
AGCACACGTCTGAACTCTTTCCCTACACGACGCTCTTCCGATCTNNNATTATCCCATCTTGCTCTGCCTCCA
AGCACACGTCTGAACTCTTTCCCTACACGACGCTCTTCCGATCTNNNATTATCCCATCTTGCTCTGCCTCCA
GAGCACACGTCTGAACTCTTTCCCTACACGACGCTCTTCCGATCTNNNTGACTGCATCCAGAAAATCCAC
GAGCACACGTCTGAACTCTTTCCCTACACGACGCTCTTCCGATCTNNNTGACTGCATCCAGAAAATCCAC
GAGCACACGTCTGAACTCTTTCCCTACACGACGCTCTTCCGATCTNNNCGGCCTGGAAAAGCATTTCTAA
GAGCACACGTCTGAACTCTTTCCCTACACGACGCTCTTCCGATCTNNNCGGCCTGGAAAAGCATTTCTAA
AGCACACGTCTGAACTCTTTCCCTACACGACGCTCTTCCGATCTNNNAGGCAGAAGAATCACTTGAAC
AGCACACGTCTGAACTCTTTCCCTACACGACGCTCTTCCGATCTNNNAGGCAGAAGAATCACTTGAAC
ACACGTCTGAACTCTTTCCCTACACGACGCTCTTCCGATCTNNNCCTTTCCCCCAGTTTTCTAGACCCATC
ACACGTCTGAACTCTTTCCCTACACGACGCTCTTCCGATCTNNNCCTTTCCCCCAGTTTTCTAGACCCATC
AGCACACGTCTGAACTCTTTCCCTACACGACGCTCTTCCGATCTNNNTGGCATTCTGGGGAAAATTTGA
AGCACACGTCTGAACTCTTTCCCTACACGACGCTCTTCCGATCTNNNTGGCATTCTGGGGAAAATTTGA

AGCACACGTCTGAACTCTTTCCCTACACGACGCTCTTCCGATCTNNNTGCTTACCCTTGCTCCAATTACT
AGCACACGTCTGAACTCTTTCCCTACACGACGCTCTTCCGATCTNNNTGCTTACCCTTGCTCCAATTACT
AGCACACGTCTGAACTCTTTCCCTACACGACGCTCTTCCGATCTNNNCTGAGGGTCTTGGGATATATG
AGCACACGTCTGAACTCTTTCCCTACACGACGCTCTTCCGATCTNNNCTGAGGGTCTTGGGATATATG
GAGCACACGTCTGAACTCTTTCCCTACACGACGCTCTTCCGATCTNNNAACAAGGCAAGACCCCATCTC
GAGCACACGTCTGAACTCTTTCCCTACACGACGCTCTTCCGATCTNNNAACAAGGCAAGACCCCATCTC
AGCACACGTCTGAACTCTTTCCCTACACGACGCTCTTCCGATCTNNNCTTGGGGTAATGTGCAGCAAT
AGCACACGTCTGAACTCTTTCCCTACACGACGCTCTTCCGATCTNNNCTTGGGGTAATGTGCAGCAAT
AGCACACGTCTGAACTCTTTCCCTACACGACGCTCTTCCGATCTNNNCTCCTCTGTAAAATGGGGTAG
AGCACACGTCTGAACTCTTTCCCTACACGACGCTCTTCCGATCTNNNCTCCTCTGTAAAATGGGGTAG
AGCACACGTCTGAACTCTTTCCCTACACGACGCTCTTCCGATCTNNNGCTGCATCTATGTGACTCAGTT
AGCACACGTCTGAACTCTTTCCCTACACGACGCTCTTCCGATCTNNNGCTGCATCTATGTGACTCAGTT
AGCACACGTCTGAACTCTTTCCCTACACGACGCTCTTCCGATCTNNNATCGCTGTTCATCAAGGATGC
AGCACACGTCTGAACTCTTTCCCTACACGACGCTCTTCCGATCTNNNATCGCTGTTCATCAAGGATGC
GAGCACACGTCTGAACTCTTTCCCTACACGACGCTCTTCCGATCTNNNTCTCTCTCTCTCTCTTCCTCT
GAGCACACGTCTGAACTCTTTCCCTACACGACGCTCTTCCGATCTNNNTCTCTCTCTCTCTCTTCCTCT
AGAGCACACGTCTGAACTCTTTCCCTACACGACGCTCTTCCGATCTNNNGATTGCATCACAGCCTCAAC
AGAGCACACGTCTGAACTCTTTCCCTACACGACGCTCTTCCGATCTNNNGATTGCATCACAGCCTCAAC
AGCACACGTCTGAACTCTTTCCCTACACGACGCTCTTCCGATCTNNNTACTGGGAAGTGAGGAGCTC
AGCACACGTCTGAACTCTTTCCCTACACGACGCTCTTCCGATCTNNNTACTGGGAAGTGAGGAGCTC
GCACACGTCTGAACTCTTTCCCTACACGACGCTCTTCCGATCTNNNCTAATATGGGCAATAGGGGAGG
GCACACGTCTGAACTCTTTCCCTACACGACGCTCTTCCGATCTNNNCTAATATGGGCAATAGGGGAGG
AGCACACGTCTGAACTCTTTCCCTACACGACGCTCTTCCGATCTNNNAAGGTTGTAGTGAGCTGAGA
AGCACACGTCTGAACTCTTTCCCTACACGACGCTCTTCCGATCTNNNAAGGTTGTAGTGAGCTGAGA
GCACACGTCTGAACTCTTTCCCTACACGACGCTCTTCCGATCTNNNGGGAAATTTCTACGTGGTTGG
GCACACGTCTGAACTCTTTCCCTACACGACGCTCTTCCGATCTNNNGGGAAATTTCTACGTGGTTGG
AGCACACGTCTGAACTCTTTCCCTACACGACGCTCTTCCGATCTNNNCCTAAACAACTGGGGATCCTC
AGCACACGTCTGAACTCTTTCCCTACACGACGCTCTTCCGATCTNNNCCTAAACAACTGGGGATCCTC
AGCACACGTCTGAACTCTTTCCCTACACGACGCTCTTCCGATCTNNNGCCTGTAGTCCCAGCTACTTGC
AGCACACGTCTGAACTCTTTCCCTACACGACGCTCTTCCGATCTNNNGCCTGTAGTCCCAGCTACTTGC
GAGCACACGTCTGAACTCTTTCCCTACACGACGCTCTTCCGATCTNNNTGTAGTCCCAGCTACTCAGG
GAGCACACGTCTGAACTCTTTCCCTACACGACGCTCTTCCGATCTNNNTGTAGTCCCAGCTACTCAGG
GAGCACACGTCTGAACTCTTTCCCTACACGACGCTCTTCCGATCTNNNGTGAAGGCAGTGAGATGTG
GAGCACACGTCTGAACTCTTTCCCTACACGACGCTCTTCCGATCTNNNGTGAAGGCAGTGAGATGTG
AGAGCACACGTCTGAACTCTTTCCCTACACGACGCTCTTCCGATCTNNNAGGAGAATCACTGGAACCT
AGAGCACACGTCTGAACTCTTTCCCTACACGACGCTCTTCCGATCTNNNAGGAGAATCACTGGAACCT
GAGCACACGTCTGAACTCTTTCCCTACACGACGCTCTTCCGATCTNNNCAACTTTCCTCCATGACCAGG
GAGCACACGTCTGAACTCTTTCCCTACACGACGCTCTTCCGATCTNNNCAACTTTCCTCCATGACCAGG
AGCACACGTCTGAACTCTTTCCCTACACGACGCTCTTCCGATCTNNNCATGAGCCAAGGAAGAACCAA
AGCACACGTCTGAACTCTTTCCCTACACGACGCTCTTCCGATCTNNNCATGAGCCAAGGAAGAACCAA
AGCACACGTCTGAACTCTTTCCCTACACGACGCTCTTCCGATCTNNNGACTAAGAGGGGAAACTCAG
AGCACACGTCTGAACTCTTTCCCTACACGACGCTCTTCCGATCTNNNGACTAAGAGGGGAAACTCAG
GAGCACACGTCTGAACTCTTTCCCTACACGACGCTCTTCCGATCTNNNTGCCCATAATACCAGCTACTT

GAGCACACGTCTGAACTCTTTCCCTACACGACGCTCTTCCGATCTNNNTGCCCATAATACCAGCTACTT
AGCACACGTCTGAACTCTTTCCCTACACGACGCTCTTCCGATCTNNNGAAGCTGGTTATCAGTTCTGG
AGCACACGTCTGAACTCTTTCCCTACACGACGCTCTTCCGATCTNNNGAAGCTGGTTATCAGTTCTGG
AGCACACGTCTGAACTCTTTCCCTACACGACGCTCTTCCGATCTNNNAGATCAGCAAGTGACTGTAGT
AGCACACGTCTGAACTCTTTCCCTACACGACGCTCTTCCGATCTNNNAGATCAGCAAGTGACTGTAGT
AGCACACGTCTGAACTCTTTCCCTACACGACGCTCTTCCGATCTNNNGCATGTTGCTTTTGTACCGTC
AGCACACGTCTGAACTCTTTCCCTACACGACGCTCTTCCGATCTNNNGCATGTTGCTTTTGTACCGTC
GCACACGTCTGAACTCTTTCCCTACACGACGCTCTTCCGATCTNNNGTAAAGTATGTGTGTGTGTGT
GCACACGTCTGAACTCTTTCCCTACACGACGCTCTTCCGATCTNNNGTAAAGTATGTGTGTGTGTGT
AGCACACGTCTGAACTCTTTCCCTACACGACGCTCTTCCGATCTNNNTACACAGACACAGACATGCAC
AGCACACGTCTGAACTCTTTCCCTACACGACGCTCTTCCGATCTNNNTACACAGACACAGACATGCAC
CACACGTCTGAACTCTTTCCCTACACGACGCTCTTCCGATCTNNNGGAGCTTTTCTACCCAGAACTCAC
CACACGTCTGAACTCTTTCCCTACACGACGCTCTTCCGATCTNNNGGAGCTTTTCTACCCAGAACTCAC
GCACACGTCTGAACTCTTTCCCTACACGACGCTCTTCCGATCTNNNTGCCTTTTTCTCCTGGATGAGTT
GCACACGTCTGAACTCTTTCCCTACACGACGCTCTTCCGATCTNNNTGCCTTTTTCTCCTGGATGAGTT
GAGCACACGTCTGAACTCTTTCCCTACACGACGCTCTTCCGATCTNNNCCTCCAGACTTCTGAGCAATG
GAGCACACGTCTGAACTCTTTCCCTACACGACGCTCTTCCGATCTNNNCCTCCAGACTTCTGAGCAATG
AGCACACGTCTGAACTCTTTCCCTACACGACGCTCTTCCGATCTNNNCAAGCTGAGCTAGCAAGGGAT
AGCACACGTCTGAACTCTTTCCCTACACGACGCTCTTCCGATCTNNNCAAGCTGAGCTAGCAAGGGAT
AGCACACGTCTGAACTCTTTCCCTACACGACGCTCTTCCGATCTNNNCTCCTATCCTCCTCTGCATCCTA
AGCACACGTCTGAACTCTTTCCCTACACGACGCTCTTCCGATCTNNNCTCCTATCCTCCTCTGCATCCTA
GCACACGTCTGAACTCTTTCCCTACACGACGCTCTTCCGATCTNNNAGCTCAGAATGTAATCTTCTTTC
GCACACGTCTGAACTCTTTCCCTACACGACGCTCTTCCGATCTNNNAGCTCAGAATGTAATCTTCTTTC
AGAGCACACGTCTGAACTCTTTCCCTACACGACGCTCTTCCGATCTNNNACCACCTTGTTTTTCTCAAT
AGAGCACACGTCTGAACTCTTTCCCTACACGACGCTCTTCCGATCTNNNACCACCTTGTTTTTCTCAAT
AGCACACGTCTGAACTCTTTCCCTACACGACGCTCTTCCGATCTNNNCGATGAAGTCGTAGAAGCTG
AGCACACGTCTGAACTCTTTCCCTACACGACGCTCTTCCGATCTNNNCGATGAAGTCGTAGAAGCTG
AGCACACGTCTGAACTCTTTCCCTACACGACGCTCTTCCGATCTNNNTGTAAGAACTTTGCAGTGTTG
AGCACACGTCTGAACTCTTTCCCTACACGACGCTCTTCCGATCTNNNTGTAAGAACTTTGCAGTGTTG
GCACACGTCTGAACTCTTTCCCTACACGACGCTCTTCCGATCTNNNAGCCATGAAAGGAATGGAGTA
GCACACGTCTGAACTCTTTCCCTACACGACGCTCTTCCGATCTNNNAGCCATGAAAGGAATGGAGTA
GCACACGTCTGAACTCTTTCCCTACACGACGCTCTTCCGATCTNNNCAGACCACCCTCTGTTTTTGTAA
GCACACGTCTGAACTCTTTCCCTACACGACGCTCTTCCGATCTNNNCAGACCACCCTCTGTTTTTGTAA
AGCACACGTCTGAACTCTTTCCCTACACGACGCTCTTCCGATCTNNNCTCCTCCCCTCACCCCTGTTCAT
AGCACACGTCTGAACTCTTTCCCTACACGACGCTCTTCCGATCTNNNCTCCTCCCCTCACCCCTGTTCAT
GCACACGTCTGAACTCTTTCCCTACACGACGCTCTTCCGATCTNNNGGTGTGACCATTGCTGAAAACT
GCACACGTCTGAACTCTTTCCCTACACGACGCTCTTCCGATCTNNNGGTGTGACCATTGCTGAAAACT
AGCACACGTCTGAACTCTTTCCCTACACGACGCTCTTCCGATCTNNNAGCCAAGATCACACCATTGTA
AGCACACGTCTGAACTCTTTCCCTACACGACGCTCTTCCGATCTNNNAGCCAAGATCACACCATTGTA
GCACACGTCTGAACTCTTTCCCTACACGACGCTCTTCCGATCTNNNCTCCTAGATGCACCATGCTGTAT
GCACACGTCTGAACTCTTTCCCTACACGACGCTCTTCCGATCTNNNCTCCTAGATGCACCATGCTGTAT
GCACACGTCTGAACTCTTTCCCTACACGACGCTCTTCCGATCTNNNCCTGACCCATTTCTACAACCCAG
GCACACGTCTGAACTCTTTCCCTACACGACGCTCTTCCGATCTNNNCCTGACCCATTTCTACAACCCAG

AGCACACGTCTGAACTCTTTCCCTACACGACGCTCTTCCGATCTNNNGAATGACATCAGGATCCTGG
AGCACACGTCTGAACTCTTTCCCTACACGACGCTCTTCCGATCTNNNGAATGACATCAGGATCCTGG
GCACACGTCTGAACTCTTTCCCTACACGACGCTCTTCCGATCTNNNCTGTTTCCCCTTCTCCCTAAGACC
GCACACGTCTGAACTCTTTCCCTACACGACGCTCTTCCGATCTNNNCTGTTTCCCCTTCTCCCTAAGACC
AGCACACGTCTGAACTCTTTCCCTACACGACGCTCTTCCGATCTNNNGCAGGCTTACATGCCTGTTTGA
AGCACACGTCTGAACTCTTTCCCTACACGACGCTCTTCCGATCTNNNGCAGGCTTACATGCCTGTTTGA
GCACACGTCTGAACTCTTTCCCTACACGACGCTCTTCCGATCTNNNTTTTCAAAGGTGATGGTGGTG
GCACACGTCTGAACTCTTTCCCTACACGACGCTCTTCCGATCTNNNTTTTCAAAGGTGATGGTGGTG
AGCACACGTCTGAACTCTTTCCCTACACGACGCTCTTCCGATCTNNNCTTGGTTACTGGCCACCTGAT
AGCACACGTCTGAACTCTTTCCCTACACGACGCTCTTCCGATCTNNNCTTGGTTACTGGCCACCTGAT
AGCACACGTCTGAACTCTTTCCCTACACGACGCTCTTCCGATCTNNNTTTAGGATATTTTTGAGTGTC
AGCACACGTCTGAACTCTTTCCCTACACGACGCTCTTCCGATCTNNNTTTAGGATATTTTTGAGTGTC
GAGCACACGTCTGAACTCTTTCCCTACACGACGCTCTTCCGATCTNNNATACCAGGGGCTGAAGGATG
GAGCACACGTCTGAACTCTTTCCCTACACGACGCTCTTCCGATCTNNNATACCAGGGGCTGAAGGATG
GAGCACACGTCTGAACTCTTTCCCTACACGACGCTCTTCCGATCTNNNCTCGATAGCTCTTGCATGATG
GAGCACACGTCTGAACTCTTTCCCTACACGACGCTCTTCCGATCTNNNCTCGATAGCTCTTGCATGATG
GAGCACACGTCTGAACTCTTTCCCTACACGACGCTCTTCCGATCTNNNAGACCACCAAATTCTCCTGAA
GAGCACACGTCTGAACTCTTTCCCTACACGACGCTCTTCCGATCTNNNAGACCACCAAATTCTCCTGAA
GCACACGTCTGAACTCTTTCCCTACACGACGCTCTTCCGATCTNNNGAGAATTGCTTGAACCTGGGAA
GCACACGTCTGAACTCTTTCCCTACACGACGCTCTTCCGATCTNNNGAGAATTGCTTGAACCTGGGAA
AGCACACGTCTGAACTCTTTCCCTACACGACGCTCTTCCGATCTNNNACACCAGTTACTGCATCCACTTA
AGCACACGTCTGAACTCTTTCCCTACACGACGCTCTTCCGATCTNNNACACCAGTTACTGCATCCACTTA
AGCACACGTCTGAACTCTTTCCCTACACGACGCTCTTCCGATCTNNNCTGAGGGAGGGGAATCATTTG
AGCACACGTCTGAACTCTTTCCCTACACGACGCTCTTCCGATCTNNNCTGAGGGAGGGGAATCATTTG
GCACACGTCTGAACTCTTTCCCTACACGACGCTCTTCCGATCTNNNCCTGGCTTCTTCCCATTTCCTTTC
GCACACGTCTGAACTCTTTCCCTACACGACGCTCTTCCGATCTNNNCCTGGCTTCTTCCCATTTCCTTTC
AGCACACGTCTGAACTCTTTCCCTACACGACGCTCTTCCGATCTNNNATCACAGCCCTCTGGAAAAAG
AGCACACGTCTGAACTCTTTCCCTACACGACGCTCTTCCGATCTNNNATCACAGCCCTCTGGAAAAAG
GCACACGTCTGAACTCTTTCCCTACACGACGCTCTTCCGATCTNNNCCATAAATGACAGTCAAGCCAC
GCACACGTCTGAACTCTTTCCCTACACGACGCTCTTCCGATCTNNNCCATAAATGACAGTCAAGCCAC
AGCACACGTCTGAACTCTTTCCCTACACGACGCTCTTCCGATCTNNNAATCATTAATTATTGCACAATG
AGCACACGTCTGAACTCTTTCCCTACACGACGCTCTTCCGATCTNNNAATCATTAATTATTGCACAATG
AGCACACGTCTGAACTCTTTCCCTACACGACGCTCTTCCGATCTNNNGTTTTTCAACCTTTGTCCCCCTC
AGCACACGTCTGAACTCTTTCCCTACACGACGCTCTTCCGATCTNNNGTTTTTCAACCTTTGTCCCCCTC
AGCACACGTCTGAACTCTTTCCCTACACGACGCTCTTCCGATCTNNNCTACCTCAACAATTCTCCTGCCT
AGCACACGTCTGAACTCTTTCCCTACACGACGCTCTTCCGATCTNNNCTACCTCAACAATTCTCCTGCCT
AGCACACGTCTGAACTCTTTCCCTACACGACGCTCTTCCGATCTNNNCTGCCTTGGCTATCAGATGGA
AGCACACGTCTGAACTCTTTCCCTACACGACGCTCTTCCGATCTNNNCTGCCTTGGCTATCAGATGGA
AGCACACGTCTGAACTCTTTCCCTACACGACGCTCTTCCGATCTNNNGTTATGGAGGGACCGGTGTGC
AGCACACGTCTGAACTCTTTCCCTACACGACGCTCTTCCGATCTNNNGTTATGGAGGGACCGGTGTGC
GAGCACACGTCTGAACTCTTTCCCTACACGACGCTCTTCCGATCTNNNTACAAAATTAGCTGGGCATG
GAGCACACGTCTGAACTCTTTCCCTACACGACGCTCTTCCGATCTNNNTACAAAATTAGCTGGGCATG
CACGTCTGAACTCTTTCCCTACACGACGCTCTTCCGATCTNNNTACTCGGTTTTGCTGCCTTCTATCATC

CACGTCTGAACTCTTTCCCTACACGACGCTCTTCCGATCTNNNTACTCGGTTTTGCTGCCTTCTATCATC
GCACACGTCTGAACTCTTTCCCTACACGACGCTCTTCCGATCTNNNTCCTGACTACCTGCTAACCACTA
GCACACGTCTGAACTCTTTCCCTACACGACGCTCTTCCGATCTNNNTCCTGACTACCTGCTAACCACTA
GCACACGTCTGAACTCTTTCCCTACACGACGCTCTTCCGATCTNNNGCTTCAGGCATTGTTCAGAGTTT
GCACACGTCTGAACTCTTTCCCTACACGACGCTCTTCCGATCTNNNGCTTCAGGCATTGTTCAGAGTTT
AGCACACGTCTGAACTCTTTCCCTACACGACGCTCTTCCGATCTNNNGTATTTTCTTAGCTTTCCTGTA
AGCACACGTCTGAACTCTTTCCCTACACGACGCTCTTCCGATCTNNNGTATTTTCTTAGCTTTCCTGTA
ACACGTCTGAACTCTTTCCCTACACGACGCTCTTCCGATCTNNNGAAGAAGACATTGTTGAAGTGCAT
ACACGTCTGAACTCTTTCCCTACACGACGCTCTTCCGATCTNNNGAAGAAGACATTGTTGAAGTGCAT
GCACACGTCTGAACTCTTTCCCTACACGACGCTCTTCCGATCTNNNTGGGGTACATATTTAGCAGTG
GCACACGTCTGAACTCTTTCCCTACACGACGCTCTTCCGATCTNNNTGGGGTACATATTTAGCAGTG
CACACGTCTGAACTCTTTCCCTACACGACGCTCTTCCGATCTNNNTCCATAATGGTGAGAGGTGACAG
CACACGTCTGAACTCTTTCCCTACACGACGCTCTTCCGATCTNNNTCCATAATGGTGAGAGGTGACAG
AGCACACGTCTGAACTCTTTCCCTACACGACGCTCTTCCGATCTNNNGTTCCATAACCTGTTTATAACA
AGCACACGTCTGAACTCTTTCCCTACACGACGCTCTTCCGATCTNNNGTTCCATAACCTGTTTATAACA
ACACGTCTGAACTCTTTCCCTACACGACGCTCTTCCGATCTNNNGTTCTTCCACTGCTTACTACCATCGG
ACACGTCTGAACTCTTTCCCTACACGACGCTCTTCCGATCTNNNGTTCTTCCACTGCTTACTACCATCGG
AGCACACGTCTGAACTCTTTCCCTACACGACGCTCTTCCGATCTNNNCACCATGCCCAGCTAATTTTCTT
AGCACACGTCTGAACTCTTTCCCTACACGACGCTCTTCCGATCTNNNCACCATGCCCAGCTAATTTTCTT
GCACACGTCTGAACTCTTTCCCTACACGACGCTCTTCCGATCTNNNATATTGTTGGGCCAGTCTCTCCT
GCACACGTCTGAACTCTTTCCCTACACGACGCTCTTCCGATCTNNNATATTGTTGGGCCAGTCTCTCCT
GAGCACACGTCTGAACTCTTTCCCTACACGACGCTCTTCCGATCTNNNAACAGATGTGTGTATGTGTG
GAGCACACGTCTGAACTCTTTCCCTACACGACGCTCTTCCGATCTNNNAACAGATGTGTGTATGTGTG
AGCACACGTCTGAACTCTTTCCCTACACGACGCTCTTCCGATCTNNNGGGAATCTGATTGGACTTCGA
AGCACACGTCTGAACTCTTTCCCTACACGACGCTCTTCCGATCTNNNGGGAATCTGATTGGACTTCGA
AGCACACGTCTGAACTCTTTCCCTACACGACGCTCTTCCGATCTNNNTTTTTTTGCACCAGTAGCATGCT
AGCACACGTCTGAACTCTTTCCCTACACGACGCTCTTCCGATCTNNNTTTTTTTGCACCAGTAGCATGCT
AGCACACGTCTGAACTCTTTCCCTACACGACGCTCTTCCGATCTNNNTACGAAAAAAATTAGCTGGGC
AGCACACGTCTGAACTCTTTCCCTACACGACGCTCTTCCGATCTNNNTACGAAAAAAATTAGCTGGGC
CACACGTCTGAACTCTTTCCCTACACGACGCTCTTCCGATCTNNNAGTTATTCAGGGGGTTGAAGTGG
CACACGTCTGAACTCTTTCCCTACACGACGCTCTTCCGATCTNNNAGTTATTCAGGGGGTTGAAGTGG
AGCACACGTCTGAACTCTTTCCCTACACGACGCTCTTCCGATCTNNNATCAAGCAATTCCTCGCCAAA
AGCACACGTCTGAACTCTTTCCCTACACGACGCTCTTCCGATCTNNNATCAAGCAATTCCTCGCCAAA
AGCACACGTCTGAACTCTTTCCCTACACGACGCTCTTCCGATCTNNNCTCTAGGGGACAAGAACCACAT
AGCACACGTCTGAACTCTTTCCCTACACGACGCTCTTCCGATCTNNNCTCTAGGGGACAAGAACCACAT
AGCACACGTCTGAACTCTTTCCCTACACGACGCTCTTCCGATCTNNNGAGTGTCCAGAGAATAGCTTC
AGCACACGTCTGAACTCTTTCCCTACACGACGCTCTTCCGATCTNNNGAGTGTCCAGAGAATAGCTTC
GCACACGTCTGAACTCTTTCCCTACACGACGCTCTTCCGATCTNNNAAAATGCTGGGATTACACGCAT
GCACACGTCTGAACTCTTTCCCTACACGACGCTCTTCCGATCTNNNAAAATGCTGGGATTACACGCAT
AGCACACGTCTGAACTCTTTCCCTACACGACGCTCTTCCGATCTNNNTTAGGCTTCTGTTCTGATGAGC
AGCACACGTCTGAACTCTTTCCCTACACGACGCTCTTCCGATCTNNNTTAGGCTTCTGTTCTGATGAGC
AGCACACGTCTGAACTCTTTCCCTACACGACGCTCTTCCGATCTNNNTCCTGCCTCTGCATTCTCATTTC
AGCACACGTCTGAACTCTTTCCCTACACGACGCTCTTCCGATCTNNNTCCTGCCTCTGCATTCTCATTTC

AGCACACGTCTGAACTCTTTCCCTACACGACGCTCTTCCGATCTNNNAATATCTACACTTTGGGAGGCC
AGCACACGTCTGAACTCTTTCCCTACACGACGCTCTTCCGATCTNNNAATATCTACACTTTGGGAGGCC
AGCACACGTCTGAACTCTTTCCCTACACGACGCTCTTCCGATCTNNNGCTGATGCTCCAATTCTGTCATT
AGCACACGTCTGAACTCTTTCCCTACACGACGCTCTTCCGATCTNNNGCTGATGCTCCAATTCTGTCATT
GAGCACACGTCTGAACTCTTTCCCTACACGACGCTCTTCCGATCTNNNACATGGCAAAACCCTGTCTTT
GAGCACACGTCTGAACTCTTTCCCTACACGACGCTCTTCCGATCTNNNACATGGCAAAACCCTGTCTTT
GAGCACACGTCTGAACTCTTTCCCTACACGACGCTCTTCCGATCTNNNACCCAGACCCAATAAAGGAAA
GAGCACACGTCTGAACTCTTTCCCTACACGACGCTCTTCCGATCTNNNACCCAGACCCAATAAAGGAAA
GAGCACACGTCTGAACTCTTTCCCTACACGACGCTCTTCCGATCTNNNACAGACAAGCAAATGTTGACA
GAGCACACGTCTGAACTCTTTCCCTACACGACGCTCTTCCGATCTNNNACAGACAAGCAAATGTTGACA
AGCACACGTCTGAACTCTTTCCCTACACGACGCTCTTCCGATCTNNNTGTCCCCAAAATCATCCTCACTTC
AGCACACGTCTGAACTCTTTCCCTACACGACGCTCTTCCGATCTNNNTGTCCCCAAAATCATCCTCACTTC
GAGCACACGTCTGAACTCTTTCCCTACACGACGCTCTTCCGATCTNNNAAGCAGACCAGGAAGATACTC
GAGCACACGTCTGAACTCTTTCCCTACACGACGCTCTTCCGATCTNNNAAGCAGACCAGGAAGATACTC
AGCACACGTCTGAACTCTTTCCCTACACGACGCTCTTCCGATCTNNNATCGCTTGAGTTCAGGTGATCA
AGCACACGTCTGAACTCTTTCCCTACACGACGCTCTTCCGATCTNNNATCGCTTGAGTTCAGGTGATCA
GAGCACACGTCTGAACTCTTTCCCTACACGACGCTCTTCCGATCTNNNGGCGACTATAATCCCAGCTACT
GAGCACACGTCTGAACTCTTTCCCTACACGACGCTCTTCCGATCTNNNGGCGACTATAATCCCAGCTACT
GAGCACACGTCTGAACTCTTTCCCTACACGACGCTCTTCCGATCTNNNAGACAGGAGAATCACTTGAAC
GAGCACACGTCTGAACTCTTTCCCTACACGACGCTCTTCCGATCTNNNAGACAGGAGAATCACTTGAAC
GAGCACACGTCTGAACTCTTTCCCTACACGACGCTCTTCCGATCTNNNCCTCCAAGCCTGGCTAATCTTA
GAGCACACGTCTGAACTCTTTCCCTACACGACGCTCTTCCGATCTNNNCCTCCAAGCCTGGCTAATCTTA
GAGCACACGTCTGAACTCTTTCCCTACACGACGCTCTTCCGATCTNNNCTACCTTGATGGCCTGGAACT
GAGCACACGTCTGAACTCTTTCCCTACACGACGCTCTTCCGATCTNNNCTACCTTGATGGCCTGGAACT
GAGCACACGTCTGAACTCTTTCCCTACACGACGCTCTTCCGATCTNNNTGCGGCCTTGCTAAACTTTTTA
GAGCACACGTCTGAACTCTTTCCCTACACGACGCTCTTCCGATCTNNNTGCGGCCTTGCTAAACTTTTTA
GAGCACACGTCTGAACTCTTTCCCTACACGACGCTCTTCCGATCTNNNAGCTCCCTTCTGTGTAGGAAC
GAGCACACGTCTGAACTCTTTCCCTACACGACGCTCTTCCGATCTNNNAGCTCCCTTCTGTGTAGGAAC
GAGCACACGTCTGAACTCTTTCCCTACACGACGCTCTTCCGATCTNNNTGCATCTAGTGAGATAAACGT
GAGCACACGTCTGAACTCTTTCCCTACACGACGCTCTTCCGATCTNNNTGCATCTAGTGAGATAAACGT
AGCACACGTCTGAACTCTTTCCCTACACGACGCTCTTCCGATCTNNNGGTCATCAAAAGCAACGTTCTCA
AGCACACGTCTGAACTCTTTCCCTACACGACGCTCTTCCGATCTNNNGGTCATCAAAAGCAACGTTCTCA
AGCACACGTCTGAACTCTTTCCCTACACGACGCTCTTCCGATCTNNNGCTTCTCTCTACCCTGAACACAA
AGCACACGTCTGAACTCTTTCCCTACACGACGCTCTTCCGATCTNNNGCTTCTCTCTACCCTGAACACAA
GCACACGTCTGAACTCTTTCCCTACACGACGCTCTTCCGATCTNNNTGTAGAAGCAGAAGTTGTGGAG
GCACACGTCTGAACTCTTTCCCTACACGACGCTCTTCCGATCTNNNTGTAGAAGCAGAAGTTGTGGAG
GCACACGTCTGAACTCTTTCCCTACACGACGCTCTTCCGATCTNNNAGACGTTCAACAATGGCTCTACT
GCACACGTCTGAACTCTTTCCCTACACGACGCTCTTCCGATCTNNNAGACGTTCAACAATGGCTCTACT
GCACACGTCTGAACTCTTTCCCTACACGACGCTCTTCCGATCTNNNATTTCCAGCCTCTTGCAAAACTCCA
GCACACGTCTGAACTCTTTCCCTACACGACGCTCTTCCGATCTNNNATTTCCAGCCTCTTGCAAAACTCCA
AGCACACGTCTGAACTCTTTCCCTACACGACGCTCTTCCGATCTNNNGTTGCAGTGAGCTGAGATTGTG
AGCACACGTCTGAACTCTTTCCCTACACGACGCTCTTCCGATCTNNNGTTGCAGTGAGCTGAGATTGTG
AGCACACGTCTGAACTCTTTCCCTACACGACGCTCTTCCGATCTNNNACAAAAATTAGCCAGGCTTGGT

AGCACACGTCTGAACTCTTTCCCTACACGACGCTCTTCCGATCTNNNACAAAAATTAGCCAGGCTTGGT
AAGAGCACACGTCTGAACTCTTTCCCTACACGACGCTCTTCCGATCTNNNAAAAATTAGCTGGTCTTG
AAGAGCACACGTCTGAACTCTTTCCCTACACGACGCTCTTCCGATCTNNNAAAAATTAGCTGGTCTTG
GAGCACACGTCTGAACTCTTTCCCTACACGACGCTCTTCCGATCTNNNGGCTCTGTCTCAAGAAAACAA
GAGCACACGTCTGAACTCTTTCCCTACACGACGCTCTTCCGATCTNNNGGCTCTGTCTCAAGAAAACAA
AGAGCACACGTCTGAACTCTTTCCCTACACGACGCTCTTCCGATCTNNNCATGCAGTGAGCCAAGATTG
AGAGCACACGTCTGAACTCTTTCCCTACACGACGCTCTTCCGATCTNNNCATGCAGTGAGCCAAGATTG
AGCACACGTCTGAACTCTTTCCCTACACGACGCTCTTCCGATCTNNNGAGGCAACAGAATCACTTGAA
AGCACACGTCTGAACTCTTTCCCTACACGACGCTCTTCCGATCTNNNGAGGCAACAGAATCACTTGAA
GCACACGTCTGAACTCTTTCCCTACACGACGCTCTTCCGATCTNNNTCGACTATAGCTGTTATGTTAATC
GCACACGTCTGAACTCTTTCCCTACACGACGCTCTTCCGATCTNNNTCGACTATAGCTGTTATGTTAATC
GAGCACACGTCTGAACTCTTTCCCTACACGACGCTCTTCCGATCTNNNTTCTGCTGGATATGGTTGGCT
GAGCACACGTCTGAACTCTTTCCCTACACGACGCTCTTCCGATCTNNNTTCTGCTGGATATGGTTGGCT
AGCACACGTCTGAACTCTTTCCCTACACGACGCTCTTCCGATCTNNNCTGCCTCAGGTCTTTCTTGCTCA
AGCACACGTCTGAACTCTTTCCCTACACGACGCTCTTCCGATCTNNNCTGCCTCAGGTCTTTCTTGCTCA
GAGCACACGTCTGAACTCTTTCCCTACACGACGCTCTTCCGATCTNNNTTCTATGAGTCCAGCATTCTCT
GAGCACACGTCTGAACTCTTTCCCTACACGACGCTCTTCCGATCTNNNTTCTATGAGTCCAGCATTCTCT
AGCACACGTCTGAACTCTTTCCCTACACGACGCTCTTCCGATCTNNNTTCTCAGCTAGGGACAAACGA
AGCACACGTCTGAACTCTTTCCCTACACGACGCTCTTCCGATCTNNNTTCTCAGCTAGGGACAAACGA
AGCACACGTCTGAACTCTTTCCCTACACGACGCTCTTCCGATCTNNNCTGTAATCCAAGCTACTCGGG
AGCACACGTCTGAACTCTTTCCCTACACGACGCTCTTCCGATCTNNNCTGTAATCCAAGCTACTCGGG
AGCACACGTCTGAACTCTTTCCCTACACGACGCTCTTCCGATCTNNNTGATGGTAGAGTAGAAGCAG
AGCACACGTCTGAACTCTTTCCCTACACGACGCTCTTCCGATCTNNNTGATGGTAGAGTAGAAGCAG
AGAGCACACGTCTGAACTCTTTCCCTACACGACGCTCTTCCGATCTNNNCATGCCTGTAATCCCAGCTA
AGAGCACACGTCTGAACTCTTTCCCTACACGACGCTCTTCCGATCTNNNCATGCCTGTAATCCCAGCTA
GAGCACACGTCTGAACTCTTTCCCTACACGACGCTCTTCCGATCTNNNTGAATGGAGAGAGAAGCAA
GAGCACACGTCTGAACTCTTTCCCTACACGACGCTCTTCCGATCTNNNTGAATGGAGAGAGAAGCAA
GAGCACACGTCTGAACTCTTTCCCTACACGACGCTCTTCCGATCTNNNAGTCAATGAATCCAGGAGAG
GAGCACACGTCTGAACTCTTTCCCTACACGACGCTCTTCCGATCTNNNAGTCAATGAATCCAGGAGAG
AGCACACGTCTGAACTCTTTCCCTACACGACGCTCTTCCGATCTNNNGCTGGAATCTGAACTCATTGCT
AGCACACGTCTGAACTCTTTCCCTACACGACGCTCTTCCGATCTNNNGCTGGAATCTGAACTCATTGCT
GCACACGTCTGAACTCTTTCCCTACACGACGCTCTTCCGATCTNNNCCTGAGCAACAGAGTGATACTCCC
GCACACGTCTGAACTCTTTCCCTACACGACGCTCTTCCGATCTNNNCCTGAGCAACAGAGTGATACTCCC
AGCACACGTCTGAACTCTTTCCCTACACGACGCTCTTCCGATCTNNNTTTCATAATGCAAGCAGAAGC
AGCACACGTCTGAACTCTTTCCCTACACGACGCTCTTCCGATCTNNNTTTCATAATGCAAGCAGAAGC
AGCACACGTCTGAACTCTTTCCCTACACGACGCTCTTCCGATCTNNNTGGAAATGTAAGGGACCAGAA
AGCACACGTCTGAACTCTTTCCCTACACGACGCTCTTCCGATCTNNNTGGAAATGTAAGGGACCAGAA
AGCACACGTCTGAACTCTTTCCCTACACGACGCTCTTCCGATCTNNNAAAAAAAAGAGAAGCAGCAGC
AGCACACGTCTGAACTCTTTCCCTACACGACGCTCTTCCGATCTNNNAAAAAAAAGAGAAGCAGCAGC
AGCACACGTCTGAACTCTTTCCCTACACGACGCTCTTCCGATCTNNNCATCCCTGAAGTTGCTAGTCACT
AGCACACGTCTGAACTCTTTCCCTACACGACGCTCTTCCGATCTNNNCATCCCTGAAGTTGCTAGTCACT
AGCACACGTCTGAACTCTTTCCCTACACGACGCTCTTCCGATCTNNNCTTTAGTCCATGAGGTCAAGG
AGCACACGTCTGAACTCTTTCCCTACACGACGCTCTTCCGATCTNNNCTTTAGTCCATGAGGTCAAGG

GAGCACACGTCTGAACTCTTTCCCTACACGACGCTCTTCCGATCTNNNGGAGACTGAGGCAGAAGAAT

GAGCACACGTCTGAACTCTTTCCCTACACGACGCTCTTCCGATCTNNNGGAGACTGAGGCAGAAGAAT

AGCACACGTCTGAACTCTTTCCCTACACGACGCTCTTCCGATCTNNNCTTGTCTCCTGAGCTCAAGTGA

AGCACACGTCTGAACTCTTTCCCTACACGACGCTCTTCCGATCTNNNCTTGTCTCCTGAGCTCAAGTGA

GAGCACACGTCTGAACTCTTTCCCTACACGACGCTCTTCCGATCTNNNTAGCTGGGACTACAAGCATGT

GAGCACACGTCTGAACTCTTTCCCTACACGACGCTCTTCCGATCTNNNTAGCTGGGACTACAAGCATGT

GCACACGTCTGAACTCTTTCCCTACACGACGCTCTTCCGATCTNNNCAGGAGTAAGTCACTGTTCCTGG

GCACACGTCTGAACTCTTTCCCTACACGACGCTCTTCCGATCTNNNCAGGAGTAAGTCACTGTTCCTGG

GAGCACACGTCTGAACTCTTTCCCTACACGACGCTCTTCCGATCTNNNTGTGGAGATCTTTACCTCCCTCA

GAGCACACGTCTGAACTCTTTCCCTACACGACGCTCTTCCGATCTNNNTGTGGAGATCTTTACCTCCCTCA

ACACGTCTGAACTCTTTCCCTACACGACGCTCTTCCGATCTNNNTAACTTTCAGCTGTTAGGATTCCATC

ACACGTCTGAACTCTTTCCCTACACGACGCTCTTCCGATCTNNNTAACTTTCAGCTGTTAGGATTCCATC

GAGCACACGTCTGAACTCTTTCCCTACACGACGCTCTTCCGATCTNNNGCCGGAGTAACAACAGCAAA

GAGCACACGTCTGAACTCTTTCCCTACACGACGCTCTTCCGATCTNNNGCCGGAGTAACAACAGCAAA

AGCACACGTCTGAACTCTTTCCCTACACGACGCTCTTCCGATCTNNNTGTGTTCAATGTATGTGAGCTCT

AGCACACGTCTGAACTCTTTCCCTACACGACGCTCTTCCGATCTNNNTGTGTTCAATGTATGTGAGCTCT

GAGCACACGTCTGAACTCTTTCCCTACACGACGCTCTTCCGATCTNNNTGCCTGTAATTCCAGCTACTCA

GAGCACACGTCTGAACTCTTTCCCTACACGACGCTCTTCCGATCTNNNTGCCTGTAATTCCAGCTACTCA

AGCACACGTCTGAACTCTTTCCCTACACGACGCTCTTCCGATCTNNNGCTTTGAAATCACTGGCCCATA

AGCACACGTCTGAACTCTTTCCCTACACGACGCTCTTCCGATCTNNNGCTTTGAAATCACTGGCCCATA

GAGCACACGTCTGAACTCTTTCCCTACACGACGCTCTTCCGATCTNNNCCATTCTGTGGGCTGTCTTTCT

GAGCACACGTCTGAACTCTTTCCCTACACGACGCTCTTCCGATCTNNNCCATTCTGTGGGCTGTCTTTCT

AGCACACGTCTGAACTCTTTCCCTACACGACGCTCTTCCGATCTNNNATACAAAAATTAGCTGGGCGT

AGCACACGTCTGAACTCTTTCCCTACACGACGCTCTTCCGATCTNNNATACAAAAATTAGCTGGGCGT

GAGCACACGTCTGAACTCTTTCCCTACACGACGCTCTTCCGATCTNNNAGTAGCACAATCATGGCTGAA

GAGCACACGTCTGAACTCTTTCCCTACACGACGCTCTTCCGATCTNNNAGTAGCACAATCATGGCTGAA

AGAGCACACGTCTGAACTCTTTCCCTACACGACGCTCTTCCGATCTNNNTGCCTTCTCTCTGTCTCTGTCT

AGAGCACACGTCTGAACTCTTTCCCTACACGACGCTCTTCCGATCTNNNTGCCTTCTCTCTGTCTCTGTCT

AGCACACGTCTGAACTCTTTCCCTACACGACGCTCTTCCGATCTNNNCATAGCAAGACCCCCATCTCTA

AGCACACGTCTGAACTCTTTCCCTACACGACGCTCTTCCGATCTNNNCATAGCAAGACCCCCATCTCTA

AGCACACGTCTGAACTCTTTCCCTACACGACGCTCTTCCGATCTNNNGAGGTTGTAGTGAGCCGAGAT

AGCACACGTCTGAACTCTTTCCCTACACGACGCTCTTCCGATCTNNNGAGGTTGTAGTGAGCCGAGAT

GAGCACACGTCTGAACTCTTTCCCTACACGACGCTCTTCCGATCTNNNCAATATTGTGAGAGAAGAAG

GAGCACACGTCTGAACTCTTTCCCTACACGACGCTCTTCCGATCTNNNCAATATTGTGAGAGAAGAAG

GAGCACACGTCTGAACTCTTTCCCTACACGACGCTCTTCCGATCTNNNCTTAGTGTCCAGTTTCCACCCA

GAGCACACGTCTGAACTCTTTCCCTACACGACGCTCTTCCGATCTNNNCTTAGTGTCCAGTTTCCACCCA

GAGCACACGTCTGAACTCTTTCCCTACACGACGCTCTTCCGATCTNNNGTCAAAGCCAAGAGGTCCTGA

GAGCACACGTCTGAACTCTTTCCCTACACGACGCTCTTCCGATCTNNNGTCAAAGCCAAGAGGTCCTGA

AGCACACGTCTGAACTCTTTCCCTACACGACGCTCTTCCGATCTNNNAGCCCTGTGAAAATTCAACCCTA

AGCACACGTCTGAACTCTTTCCCTACACGACGCTCTTCCGATCTNNNAGCCCTGTGAAAATTCAACCCTA

AGCACACGTCTGAACTCTTTCCCTACACGACGCTCTTCCGATCTNNNTGCTGCCTCATCTGTCTCTAAT

AGCACACGTCTGAACTCTTTCCCTACACGACGCTCTTCCGATCTNNNTGCTGCCTCATCTGTCTCTAAT

GCACACGTCTGAACTCTTTCCCTACACGACGCTCTTCCGATCTNNNCTGCAGAGAAGAAGGACTGGA

GCACACGTCTGAACTCTTTCCCTACACGACGCTCTTCCGATCTNNNCTGCAGAGAAGAAGGACTGGA
AGCACACGTCTGAACTCTTTCCCTACACGACGCTCTTCCGATCTNNNGCTTGTAAAATCGCTGATCCT
AGCACACGTCTGAACTCTTTCCCTACACGACGCTCTTCCGATCTNNNGCTTGTAAAATCGCTGATCCT
AGCACACGTCTGAACTCTTTCCCTACACGACGCTCTTCCGATCTNNNTTCAGTCACATCTTCAGGCTCT
AGCACACGTCTGAACTCTTTCCCTACACGACGCTCTTCCGATCTNNNTTCAGTCACATCTTCAGGCTCT
AGCACACGTCTGAACTCTTTCCCTACACGACGCTCTTCCGATCTNNNACAATCCAGGCTAGATCATTG
AGCACACGTCTGAACTCTTTCCCTACACGACGCTCTTCCGATCTNNNACAATCCAGGCTAGATCATTG
CACACGTCTGAACTCTTTCCCTACACGACGCTCTTCCGATCTNNNCTGCAGTTTGGATAAGGCAGGA
CACACGTCTGAACTCTTTCCCTACACGACGCTCTTCCGATCTNNNCTGCAGTTTGGATAAGGCAGGA
GCACACGTCTGAACTCTTTCCCTACACGACGCTCTTCCGATCTNNNTCCACAACTTTTAAGTCATGTTT
GCACACGTCTGAACTCTTTCCCTACACGACGCTCTTCCGATCTNNNTCCACAACTTTTAAGTCATGTTT
GCACACGTCTGAACTCTTTCCCTACACGACGCTCTTCCGATCTNNNTCACACTCCATCTTTTGACTCCAC
GCACACGTCTGAACTCTTTCCCTACACGACGCTCTTCCGATCTNNNTCACACTCCATCTTTTGACTCCAC
GCACACGTCTGAACTCTTTCCCTACACGACGCTCTTCCGATCTNNNACACACAATGCCACACAAAGAT
GCACACGTCTGAACTCTTTCCCTACACGACGCTCTTCCGATCTNNNACACACAATGCCACACAAAGAT
GCACACGTCTGAACTCTTTCCCTACACGACGCTCTTCCGATCTNNNAGAGAGCTCTTCATGCAGAAA
GCACACGTCTGAACTCTTTCCCTACACGACGCTCTTCCGATCTNNNAGAGAGCTCTTCATGCAGAAA
AGCACACGTCTGAACTCTTTCCCTACACGACGCTCTTCCGATCTNNNAGAAAAGCACATCCGTGAAA
AGCACACGTCTGAACTCTTTCCCTACACGACGCTCTTCCGATCTNNNAGAAAAGCACATCCGTGAAA
GCACACGTCTGAACTCTTTCCCTACACGACGCTCTTCCGATCTNNNCCTGAGCAAATGAGCAGGTAT
GCACACGTCTGAACTCTTTCCCTACACGACGCTCTTCCGATCTNNNCCTGAGCAAATGAGCAGGTAT
AGCACACGTCTGAACTCTTTCCCTACACGACGCTCTTCCGATCTNNNAACTTTCTTTCTAACACTGTTTT
AGCACACGTCTGAACTCTTTCCCTACACGACGCTCTTCCGATCTNNNAACTTTCTTTCTAACACTGTTTT
GAGCACACGTCTGAACTCTTTCCCTACACGACGCTCTTCCGATCTNNNTCCTGCCTTCCTTCCTTTCTTTT
GAGCACACGTCTGAACTCTTTCCCTACACGACGCTCTTCCGATCTNNNTCCTGCCTTCCTTCCTTTCTTTT
AGCACACGTCTGAACTCTTTCCCTACACGACGCTCTTCCGATCTNNNGAGGTTGCAGTGAGCTGAGAT
AGCACACGTCTGAACTCTTTCCCTACACGACGCTCTTCCGATCTNNNGAGGTTGCAGTGAGCTGAGAT
AGCACACGTCTGAACTCTTTCCCTACACGACGCTCTTCCGATCTNNNAGGAGGAGAGGAGGAAGAA
AGCACACGTCTGAACTCTTTCCCTACACGACGCTCTTCCGATCTNNNAGGAGGAGAGGAGGAAGAA
AGCACACGTCTGAACTCTTTCCCTACACGACGCTCTTCCGATCTNNNTAGGGATGGAATATGCCTGCT
AGCACACGTCTGAACTCTTTCCCTACACGACGCTCTTCCGATCTNNNTAGGGATGGAATATGCCTGCT
GAGCACACGTCTGAACTCTTTCCCTACACGACGCTCTTCCGATCTNNNAGAGATTGCAGTGAGCTGA
GAGCACACGTCTGAACTCTTTCCCTACACGACGCTCTTCCGATCTNNNAGAGATTGCAGTGAGCTGA
AGCACACGTCTGAACTCTTTCCCTACACGACGCTCTTCCGATCTNNNATCACCTTCTGATGGCTAAGAC
AGCACACGTCTGAACTCTTTCCCTACACGACGCTCTTCCGATCTNNNATCACCTTCTGATGGCTAAGAC
AGCACACGTCTGAACTCTTTCCCTACACGACGCTCTTCCGATCTNNNCCTCATGTTTGTGCCATTTCACA
AGCACACGTCTGAACTCTTTCCCTACACGACGCTCTTCCGATCTNNNCCTCATGTTTGTGCCATTTCACA
GCACACGTCTGAACTCTTTCCCTACACGACGCTCTTCCGATCTNNNGATGCCTGTAATCCCAGCTACTT
GCACACGTCTGAACTCTTTCCCTACACGACGCTCTTCCGATCTNNNGATGCCTGTAATCCCAGCTACTT
GAGCACACGTCTGAACTCTTTCCCTACACGACGCTCTTCCGATCTNNNACGCCTATAATGCCAGCTCTT
GAGCACACGTCTGAACTCTTTCCCTACACGACGCTCTTCCGATCTNNNACGCCTATAATGCCAGCTCTT
GAGCACACGTCTGAACTCTTTCCCTACACGACGCTCTTCCGATCTNNNTTCCTCCTCACACTAGACAAG
GAGCACACGTCTGAACTCTTTCCCTACACGACGCTCTTCCGATCTNNNTTCCTCCTCACACTAGACAAG

GCACACGTCTGAACTCTTTCCCTACACGACGCTCTTCCGATCTNNNATTGGTCTATATCTTTGTTTTGG
GCACACGTCTGAACTCTTTCCCTACACGACGCTCTTCCGATCTNNNATTGGTCTATATCTTTGTTTTGG
GAGCACACGTCTGAACTCTTTCCCTACACGACGCTCTTCCGATCTNNNAGTGGAGCTATATGGGTCCT
GAGCACACGTCTGAACTCTTTCCCTACACGACGCTCTTCCGATCTNNNAGTGGAGCTATATGGGTCCT
AGCACACGTCTGAACTCTTTCCCTACACGACGCTCTTCCGATCTNNNCATGCCTGTAATCCCAGCTACT
AGCACACGTCTGAACTCTTTCCCTACACGACGCTCTTCCGATCTNNNCATGCCTGTAATCCCAGCTACT
AGCACACGTCTGAACTCTTTCCCTACACGACGCTCTTCCGATCTNNNTTCCCTAGAGCTATAAATGGT
AGCACACGTCTGAACTCTTTCCCTACACGACGCTCTTCCGATCTNNNTTCCCTAGAGCTATAAATGGT
AGAGCACACGTCTGAACTCTTTCCCTACACGACGCTCTTCCGATCTNNNGCTGAGATCATGTCATTGCA
AGAGCACACGTCTGAACTCTTTCCCTACACGACGCTCTTCCGATCTNNNGCTGAGATCATGTCATTGCA
GCACACGTCTGAACTCTTTCCCTACACGACGCTCTTCCGATCTNNNCCTGTGAGTTGAATGCACAAGT
GCACACGTCTGAACTCTTTCCCTACACGACGCTCTTCCGATCTNNNCCTGTGAGTTGAATGCACAAGT
AGCACACGTCTGAACTCTTTCCCTACACGACGCTCTTCCGATCTNNNCTGTAAATACAAAGCTTTCAAA
AGCACACGTCTGAACTCTTTCCCTACACGACGCTCTTCCGATCTNNNCTGTAAATACAAAGCTTTCAAA
GCACACGTCTGAACTCTTTCCCTACACGACGCTCTTCCGATCTNNNCTGAGGCAGGAGAATCACTTGA
GCACACGTCTGAACTCTTTCCCTACACGACGCTCTTCCGATCTNNNCTGAGGCAGGAGAATCACTTGA
AGCACACGTCTGAACTCTTTCCCTACACGACGCTCTTCCGATCTNNNCACCAAGAACCATAAACCGAG
AGCACACGTCTGAACTCTTTCCCTACACGACGCTCTTCCGATCTNNNCACCAAGAACCATAAACCGAG
AGCACACGTCTGAACTCTTTCCCTACACGACGCTCTTCCGATCTNNNCAGGATGAACTGCTTGAATGT
AGCACACGTCTGAACTCTTTCCCTACACGACGCTCTTCCGATCTNNNCAGGATGAACTGCTTGAATGT
AGCACACGTCTGAACTCTTTCCCTACACGACGCTCTTCCGATCTNNNTCAAAGATCACTTATCGCCCACA
AGCACACGTCTGAACTCTTTCCCTACACGACGCTCTTCCGATCTNNNTCAAAGATCACTTATCGCCCACA
AGCACACGTCTGAACTCTTTCCCTACACGACGCTCTTCCGATCTNNNGTCAAGACACTGTAGGCACTG
AGCACACGTCTGAACTCTTTCCCTACACGACGCTCTTCCGATCTNNNGTCAAGACACTGTAGGCACTG
AGCACACGTCTGAACTCTTTCCCTACACGACGCTCTTCCGATCTNNNATGTGTGTGTGTTTGTCTGTG
AGCACACGTCTGAACTCTTTCCCTACACGACGCTCTTCCGATCTNNNATGTGTGTGTGTTTGTCTGTG
AGCACACGTCTGAACTCTTTCCCTACACGACGCTCTTCCGATCTNNNGAGCCTGTGGTCCTAGCTACTA
AGCACACGTCTGAACTCTTTCCCTACACGACGCTCTTCCGATCTNNNGAGCCTGTGGTCCTAGCTACTA
GAGCACACGTCTGAACTCTTTCCCTACACGACGCTCTTCCGATCTNNNTTAACCGAAAGGTGGCAACA
GAGCACACGTCTGAACTCTTTCCCTACACGACGCTCTTCCGATCTNNNTTAACCGAAAGGTGGCAACA
AGCACACGTCTGAACTCTTTCCCTACACGACGCTCTTCCGATCTNNNGATGGTCCTCTGCAAGAAGG
AGCACACGTCTGAACTCTTTCCCTACACGACGCTCTTCCGATCTNNNGATGGTCCTCTGCAAGAAGG
GAGCACACGTCTGAACTCTTTCCCTACACGACGCTCTTCCGATCTNNNATAATCCTAGCACTTTGGGA
GAGCACACGTCTGAACTCTTTCCCTACACGACGCTCTTCCGATCTNNNATAATCCTAGCACTTTGGGA
AAGAGCACACGTCTGAACTCTTTCCCTACACGACGCTCTTCCGATCTNNNACCTGTAGTCCCAGCTACT
AAGAGCACACGTCTGAACTCTTTCCCTACACGACGCTCTTCCGATCTNNNACCTGTAGTCCCAGCTACT
GAGCACACGTCTGAACTCTTTCCCTACACGACGCTCTTCCGATCTNNNGAGGCTGAGGCAGGAGAAT
GAGCACACGTCTGAACTCTTTCCCTACACGACGCTCTTCCGATCTNNNGAGGCTGAGGCAGGAGAAT
GCACACGTCTGAACTCTTTCCCTACACGACGCTCTTCCGATCTNNNGTTCCCCTGAGACCTACTGAAT
GCACACGTCTGAACTCTTTCCCTACACGACGCTCTTCCGATCTNNNGTTCCCCTGAGACCTACTGAAT
AGCACACGTCTGAACTCTTTCCCTACACGACGCTCTTCCGATCTNNNACAATCTTCCACACCATTCCTG
AGCACACGTCTGAACTCTTTCCCTACACGACGCTCTTCCGATCTNNNACAATCTTCCACACCATTCCTG
GAGCACACGTCTGAACTCTTTCCCTACACGACGCTCTTCCGATCTNNNGTAGGACCTGCATGGGATA

GAGCACACGTCTGAACTCTTTCCCTACACGACGCTCTTCCGATCTNNNGTAGGACCTGCATGGGATA
AGCACACGTCTGAACTCTTTCCCTACACGACGCTCTTCCGATCTNNNAAAAGGGCCTTCAGAGCTGATT
AGCACACGTCTGAACTCTTTCCCTACACGACGCTCTTCCGATCTNNNAAAAGGGCCTTCAGAGCTGATT
AGCACACGTCTGAACTCTTTCCCTACACGACGCTCTTCCGATCTNNNACCTCCTTTCCGAACTAAACTGT
AGCACACGTCTGAACTCTTTCCCTACACGACGCTCTTCCGATCTNNNACCTCCTTTCCGAACTAAACTGT
AGCACACGTCTGAACTCTTTCCCTACACGACGCTCTTCCGATCTNNNATTCAGAGACAGCATCCTGGA
AGCACACGTCTGAACTCTTTCCCTACACGACGCTCTTCCGATCTNNNATTCAGAGACAGCATCCTGGA
ACACGTCTGAACTCTTTCCCTACACGACGCTCTTCCGATCTNNNCTTCTCGCCACTCTTACTAGGGTCAT
ACACGTCTGAACTCTTTCCCTACACGACGCTCTTCCGATCTNNNCTTCTCGCCACTCTTACTAGGGTCAT
GAGCACACGTCTGAACTCTTTCCCTACACGACGCTCTTCCGATCTNNNATCAACCCAACAAAGCCTACCA
GAGCACACGTCTGAACTCTTTCCCTACACGACGCTCTTCCGATCTNNNATCAACCCAACAAAGCCTACCA
GAGCACACGTCTGAACTCTTTCCCTACACGACGCTCTTCCGATCTNNNTCTGATTTGTCAGCATTCTTG
GAGCACACGTCTGAACTCTTTCCCTACACGACGCTCTTCCGATCTNNNTCTGATTTGTCAGCATTCTTG
GCACACGTCTGAACTCTTTCCCTACACGACGCTCTTCCGATCTNNNTCTCAAACATGGTTGTGCACAAA
GCACACGTCTGAACTCTTTCCCTACACGACGCTCTTCCGATCTNNNTCTCAAACATGGTTGTGCACAAA
GAGCACACGTCTGAACTCTTTCCCTACACGACGCTCTTCCGATCTNNNTTGAAATTGCCATTCATCCAG
GAGCACACGTCTGAACTCTTTCCCTACACGACGCTCTTCCGATCTNNNTTGAAATTGCCATTCATCCAG
AAGAGCACACGTCTGAACTCTTTCCCTACACGACGCTCTTCCGATCTNNNGAACACCTGTAGTCCCAGT
AAGAGCACACGTCTGAACTCTTTCCCTACACGACGCTCTTCCGATCTNNNGAACACCTGTAGTCCCAGT
AGCACACGTCTGAACTCTTTCCCTACACGACGCTCTTCCGATCTNNNGTAGACGATACAGGAATGGGC
AGCACACGTCTGAACTCTTTCCCTACACGACGCTCTTCCGATCTNNNGTAGACGATACAGGAATGGGC
GAGCACACGTCTGAACTCTTTCCCTACACGACGCTCTTCCGATCTNNNAGCCCGAGCAAATATTGTGA
GAGCACACGTCTGAACTCTTTCCCTACACGACGCTCTTCCGATCTNNNAGCCCGAGCAAATATTGTGA
AGCACACGTCTGAACTCTTTCCCTACACGACGCTCTTCCGATCTNNNCTGAGGCAGGAGAATCGTTGA
AGCACACGTCTGAACTCTTTCCCTACACGACGCTCTTCCGATCTNNNCTGAGGCAGGAGAATCGTTGA
AGAGCACACGTCTGAACTCTTTCCCTACACGACGCTCTTCCGATCTNNNGCTCAGTCCCTTGAGGCAAT
AGAGCACACGTCTGAACTCTTTCCCTACACGACGCTCTTCCGATCTNNNGCTCAGTCCCTTGAGGCAAT
GCACACGTCTGAACTCTTTCCCTACACGACGCTCTTCCGATCTNNNCCTCTCTCAAAGCTTCCCTCTTTC
GCACACGTCTGAACTCTTTCCCTACACGACGCTCTTCCGATCTNNNCCTCTCTCAAAGCTTCCCTCTTTC
GAGCACACGTCTGAACTCTTTCCCTACACGACGCTCTTCCGATCTNNNGAGGTTGCAGTGAGCTGAGA
GAGCACACGTCTGAACTCTTTCCCTACACGACGCTCTTCCGATCTNNNGAGGTTGCAGTGAGCTGAGA
AGCACACGTCTGAACTCTTTCCCTACACGACGCTCTTCCGATCTNNNCCATTTCCTGGGTTCAAGCAATT
AGCACACGTCTGAACTCTTTCCCTACACGACGCTCTTCCGATCTNNNCCATTTCCTGGGTTCAAGCAATT
AGCACACGTCTGAACTCTTTCCCTACACGACGCTCTTCCGATCTNNNCTGAAATCACTTGTGATCTTGCA
AGCACACGTCTGAACTCTTTCCCTACACGACGCTCTTCCGATCTNNNCTGAAATCACTTGTGATCTTGCA
GCACACGTCTGAACTCTTTCCCTACACGACGCTCTTCCGATCTNNNCATAGGTCTTTCATCCCTTAATAC
GCACACGTCTGAACTCTTTCCCTACACGACGCTCTTCCGATCTNNNCATAGGTCTTTCATCCCTTAATAC
AGCACACGTCTGAACTCTTTCCCTACACGACGCTCTTCCGATCTNNNGTTGTGCACATGTACCCTAGAA
AGCACACGTCTGAACTCTTTCCCTACACGACGCTCTTCCGATCTNNNGTTGTGCACATGTACCCTAGAA
AGCACACGTCTGAACTCTTTCCCTACACGACGCTCTTCCGATCTNNNCTGTTTCAGCCTTTGGGATTCA
AGCACACGTCTGAACTCTTTCCCTACACGACGCTCTTCCGATCTNNNCTGTTTCAGCCTTTGGGATTCA
AGCACACGTCTGAACTCTTTCCCTACACGACGCTCTTCCGATCTNNNATGATGGTGGGTGACTGTAAT
AGCACACGTCTGAACTCTTTCCCTACACGACGCTCTTCCGATCTNNNATGATGGTGGGTGACTGTAAT

AGCACACGTCTGAACTCTTTCCCTACACGACGCTCTTCCGATCTNNNCTGGGAAATAGAGATCCTGC
AGCACACGTCTGAACTCTTTCCCTACACGACGCTCTTCCGATCTNNNCTGGGAAATAGAGATCCTGC
GCACACGTCTGAACTCTTTCCCTACACGACGCTCTTCCGATCTNNNCCATCACATCCAGCTAGCACTTA
GCACACGTCTGAACTCTTTCCCTACACGACGCTCTTCCGATCTNNNCCATCACATCCAGCTAGCACTTA
AGCACACGTCTGAACTCTTTCCCTACACGACGCTCTTCCGATCTNNNTTAGGAAGATGCAGCTGTTT
AGCACACGTCTGAACTCTTTCCCTACACGACGCTCTTCCGATCTNNNTTAGGAAGATGCAGCTGTTT
AGCACACGTCTGAACTCTTTCCCTACACGACGCTCTTCCGATCTNNNCTGAGGCAGGAGAATCACTTG
AGCACACGTCTGAACTCTTTCCCTACACGACGCTCTTCCGATCTNNNCTGAGGCAGGAGAATCACTTG
AGCACACGTCTGAACTCTTTCCCTACACGACGCTCTTCCGATCTNNNATGATTTGGGCCATGAATTTG
AGCACACGTCTGAACTCTTTCCCTACACGACGCTCTTCCGATCTNNNATGATTTGGGCCATGAATTTG
GAGCACACGTCTGAACTCTTTCCCTACACGACGCTCTTCCGATCTNNNACATGAGCAAGACTCTGTCTC
GAGCACACGTCTGAACTCTTTCCCTACACGACGCTCTTCCGATCTNNNACATGAGCAAGACTCTGTCTC
AGAGCACACGTCTGAACTCTTTCCCTACACGACGCTCTTCCGATCTNNNGATTGAAAAGGATGCTGG
AGAGCACACGTCTGAACTCTTTCCCTACACGACGCTCTTCCGATCTNNNGATTGAAAAGGATGCTGG
CACACGTCTGAACTCTTTCCCTACACGACGCTCTTCCGATCTNNNAGTTCATGATTCAAACTGCACTC
CACACGTCTGAACTCTTTCCCTACACGACGCTCTTCCGATCTNNNAGTTCATGATTCAAACTGCACTC
GCACACGTCTGAACTCTTTCCCTACACGACGCTCTTCCGATCTNNNATTTCACGAGTCACCCATCACAG
GCACACGTCTGAACTCTTTCCCTACACGACGCTCTTCCGATCTNNNATTTCACGAGTCACCCATCACAG
AGCACACGTCTGAACTCTTTCCCTACACGACGCTCTTCCGATCTNNNACAAAAATTAGGCAGGCATGG
AGCACACGTCTGAACTCTTTCCCTACACGACGCTCTTCCGATCTNNNACAAAAATTAGGCAGGCATGG
CACACGTCTGAACTCTTTCCCTACACGACGCTCTTCCGATCTNNNGTAAAATGCTTGGTTCAGTGCCCC
CACACGTCTGAACTCTTTCCCTACACGACGCTCTTCCGATCTNNNGTAAAATGCTTGGTTCAGTGCCCC
GCACACGTCTGAACTCTTTCCCTACACGACGCTCTTCCGATCTNNNATAATGCCAGCTACTTAGGAGG
GCACACGTCTGAACTCTTTCCCTACACGACGCTCTTCCGATCTNNNATAATGCCAGCTACTTAGGAGG
GAGCACACGTCTGAACTCTTTCCCTACACGACGCTCTTCCGATCTNNNGGTGTGGTAGTGTATGCCT
GAGCACACGTCTGAACTCTTTCCCTACACGACGCTCTTCCGATCTNNNGGTGTGGTAGTGTATGCCT
GCACACGTCTGAACTCTTTCCCTACACGACGCTCTTCCGATCTNNNGCCGCTTCTTCTTCTTCTTCTTCC
GCACACGTCTGAACTCTTTCCCTACACGACGCTCTTCCGATCTNNNGCCGCTTCTTCTTCTTCTTCTTCC
GCACACGTCTGAACTCTTTCCCTACACGACGCTCTTCCGATCTNNNCATCCCCCTATGTAGAATCTCAT
GCACACGTCTGAACTCTTTCCCTACACGACGCTCTTCCGATCTNNNCATCCCCCTATGTAGAATCTCAT
AGCACACGTCTGAACTCTTTCCCTACACGACGCTCTTCCGATCTNNNTATAGTCCCAGCTACTCAGGA
AGCACACGTCTGAACTCTTTCCCTACACGACGCTCTTCCGATCTNNNTATAGTCCCAGCTACTCAGGA
AGCACACGTCTGAACTCTTTCCCTACACGACGCTCTTCCGATCTNNNACGAGGTCAATTCAGCAAGAG
AGCACACGTCTGAACTCTTTCCCTACACGACGCTCTTCCGATCTNNNACGAGGTCAATTCAGCAAGAG
GAGCACACGTCTGAACTCTTTCCCTACACGACGCTCTTCCGATCTNNNGAGTACAGTCAGTCCTCCAC
GAGCACACGTCTGAACTCTTTCCCTACACGACGCTCTTCCGATCTNNNGAGTACAGTCAGTCCTCCAC
AGCACACGTCTGAACTCTTTCCCTACACGACGCTCTTCCGATCTNNNTGGCAAGAACACAACATGAGA
AGCACACGTCTGAACTCTTTCCCTACACGACGCTCTTCCGATCTNNNTGGCAAGAACACAACATGAGA
GAGCACACGTCTGAACTCTTTCCCTACACGACGCTCTTCCGATCTNNNCTGTTGCAAGACTGGAATTC
GAGCACACGTCTGAACTCTTTCCCTACACGACGCTCTTCCGATCTNNNCTGTTGCAAGACTGGAATTC
AGCACACGTCTGAACTCTTTCCCTACACGACGCTCTTCCGATCTNNNGTGTTGGGATTACAGGCATG
AGCACACGTCTGAACTCTTTCCCTACACGACGCTCTTCCGATCTNNNGTGTTGGGATTACAGGCATG
AGCACACGTCTGAACTCTTTCCCTACACGACGCTCTTCCGATCTNNNGTAGAGAAGGGGTTTCACCAT

AGCACACGTCTGAACTCTTTCCCTACACGACGCTCTTCCGATCTNNNGTAGAGAAGGGGTTTCACCAT
AGCACACGTCTGAACTCTTTCCCTACACGACGCTCTTCCGATCTNNNATCATGAGGTCAGGAGATCG
AGCACACGTCTGAACTCTTTCCCTACACGACGCTCTTCCGATCTNNNATCATGAGGTCAGGAGATCG
AGCACACGTCTGAACTCTTTCCCTACACGACGCTCTTCCGATCTNNNTGCTCATGGATGGGTAGAAT
AGCACACGTCTGAACTCTTTCCCTACACGACGCTCTTCCGATCTNNNTGCTCATGGATGGGTAGAAT
GCACACGTCTGAACTCTTTCCCTACACGACGCTCTTCCGATCTNNNGTCTTTCTTCATGTCCTTTGCCCC
GCACACGTCTGAACTCTTTCCCTACACGACGCTCTTCCGATCTNNNGTCTTTCTTCATGTCCTTTGCCCC
AGCACACGTCTGAACTCTTTCCCTACACGACGCTCTTCCGATCTNNNTCACACACGTCACTGCTGTAAT
AGCACACGTCTGAACTCTTTCCCTACACGACGCTCTTCCGATCTNNNTCACACACGTCACTGCTGTAAT
GCACACGTCTGAACTCTTTCCCTACACGACGCTCTTCCGATCTNNNCCCTTACCATATTAATGTTGAAT
GCACACGTCTGAACTCTTTCCCTACACGACGCTCTTCCGATCTNNNCCCTTACCATATTAATGTTGAAT
GCACACGTCTGAACTCTTTCCCTACACGACGCTCTTCCGATCTNNNCTCCTGAGTAGCTGGGACTACA
GCACACGTCTGAACTCTTTCCCTACACGACGCTCTTCCGATCTNNNCTCCTGAGTAGCTGGGACTACA
GCACACGTCTGAACTCTTTCCCTACACGACGCTCTTCCGATCTNNNAGGGTTTCTCCATGTTGGTCAG
GCACACGTCTGAACTCTTTCCCTACACGACGCTCTTCCGATCTNNNAGGGTTTCTCCATGTTGGTCAG
AGCACACGTCTGAACTCTTTCCCTACACGACGCTCTTCCGATCTNNNATTCAGCAGCTGCCTCTTTTTT
AGCACACGTCTGAACTCTTTCCCTACACGACGCTCTTCCGATCTNNNATTCAGCAGCTGCCTCTTTTTT
CACACGTCTGAACTCTTTCCCTACACGACGCTCTTCCGATCTNNNGATCACAAGGTCAGGAGATCGAG
CACACGTCTGAACTCTTTCCCTACACGACGCTCTTCCGATCTNNNGATCACAAGGTCAGGAGATCGAG
GCACACGTCTGAACTCTTTCCCTACACGACGCTCTTCCGATCTNNNATGCCATAAAGTCCTGAGTGGTT
GCACACGTCTGAACTCTTTCCCTACACGACGCTCTTCCGATCTNNNATGCCATAAAGTCCTGAGTGGTT
AGCACACGTCTGAACTCTTTCCCTACACGACGCTCTTCCGATCTNNNCTTTCCTGCCCTCTCCTATCTTC
AGCACACGTCTGAACTCTTTCCCTACACGACGCTCTTCCGATCTNNNCTTTCCTGCCCTCTCCTATCTTC
GAGCACACGTCTGAACTCTTTCCCTACACGACGCTCTTCCGATCTNNNCTTGAGTCCAGGAGATCAAA
GAGCACACGTCTGAACTCTTTCCCTACACGACGCTCTTCCGATCTNNNCTTGAGTCCAGGAGATCAAA
AGCACACGTCTGAACTCTTTCCCTACACGACGCTCTTCCGATCTNNNTGGATGAGAGAGCACAGTCA
AGCACACGTCTGAACTCTTTCCCTACACGACGCTCTTCCGATCTNNNTGGATGAGAGAGCACAGTCA
AGCACACGTCTGAACTCTTTCCCTACACGACGCTCTTCCGATCTNNNGGGAGAATCTGTTTCCTTGTC
AGCACACGTCTGAACTCTTTCCCTACACGACGCTCTTCCGATCTNNNGGGAGAATCTGTTTCCTTGTC
AGCACACGTCTGAACTCTTTCCCTACACGACGCTCTTCCGATCTNNNGAAGCCTTATGTGTGTGGTGT
AGCACACGTCTGAACTCTTTCCCTACACGACGCTCTTCCGATCTNNNGAAGCCTTATGTGTGTGGTGT
AGCACACGTCTGAACTCTTTCCCTACACGACGCTCTTCCGATCTNNNCTCCTGAGTAGCTGGGACTACA
AGCACACGTCTGAACTCTTTCCCTACACGACGCTCTTCCGATCTNNNCTCCTGAGTAGCTGGGACTACA
AGAGCACACGTCTGAACTCTTTCCCTACACGACGCTCTTCCGATCTNNNTGGTATTCCCACCCAAAGTA
AGAGCACACGTCTGAACTCTTTCCCTACACGACGCTCTTCCGATCTNNNTGGTATTCCCACCCAAAGTA
AGCACACGTCTGAACTCTTTCCCTACACGACGCTCTTCCGATCTNNNTGAGGCAGGAGAATCACTTGA
AGCACACGTCTGAACTCTTTCCCTACACGACGCTCTTCCGATCTNNNTGAGGCAGGAGAATCACTTGA
AGCACACGTCTGAACTCTTTCCCTACACGACGCTCTTCCGATCTNNNTTAGGGAAGTTCAGGGCATT
AGCACACGTCTGAACTCTTTCCCTACACGACGCTCTTCCGATCTNNNTTAGGGAAGTTCAGGGCATT
AGCACACGTCTGAACTCTTTCCCTACACGACGCTCTTCCGATCTNNNGAGATTGGGTTGTCACTGTGT
AGCACACGTCTGAACTCTTTCCCTACACGACGCTCTTCCGATCTNNNGAGATTGGGTTGTCACTGTGT
AGCACACGTCTGAACTCTTTCCCTACACGACGCTCTTCCGATCTNNNTCAAATACCAGTTCCCTTCCTC
AGCACACGTCTGAACTCTTTCCCTACACGACGCTCTTCCGATCTNNNTCAAATACCAGTTCCCTTCCTC

AGCACACGTCTGAACTCTTTCCCTACACGACGCTCTTCCGATCTNNNGATGCCCTGATTTAGAACCAA
AGCACACGTCTGAACTCTTTCCCTACACGACGCTCTTCCGATCTNNNGATGCCCTGATTTAGAACCAA
AGCACACGTCTGAACTCTTTCCCTACACGACGCTCTTCCGATCTNNNATTGCAGTGAGCCAAGATAAT
AGCACACGTCTGAACTCTTTCCCTACACGACGCTCTTCCGATCTNNNATTGCAGTGAGCCAAGATAAT
GAGCACACGTCTGAACTCTTTCCCTACACGACGCTCTTCCGATCTNNNCTGGTCTTTTCCTTTTCTGCCC
GAGCACACGTCTGAACTCTTTCCCTACACGACGCTCTTCCGATCTNNNCTGGTCTTTTCCTTTTCTGCCC
AGCACACGTCTGAACTCTTTCCCTACACGACGCTCTTCCGATCTNNNAGTCTGGCTACTCTCTTTCAAG
AGCACACGTCTGAACTCTTTCCCTACACGACGCTCTTCCGATCTNNNAGTCTGGCTACTCTCTTTCAAG
GCACACGTCTGAACTCTTTCCCTACACGACGCTCTTCCGATCTNNNTCAGTCCAGTGATGTTTGCACTA
GCACACGTCTGAACTCTTTCCCTACACGACGCTCTTCCGATCTNNNTCAGTCCAGTGATGTTTGCACTA
AGCACACGTCTGAACTCTTTCCCTACACGACGCTCTTCCGATCTNNNCACCTGTAATCCCAGCTACTTG
AGCACACGTCTGAACTCTTTCCCTACACGACGCTCTTCCGATCTNNNCACCTGTAATCCCAGCTACTTG
GCACACGTCTGAACTCTTTCCCTACACGACGCTCTTCCGATCTNNNTATCAGGAGGAGTGCTTGTTCA
GCACACGTCTGAACTCTTTCCCTACACGACGCTCTTCCGATCTNNNTATCAGGAGGAGTGCTTGTTCA
GAGCACACGTCTGAACTCTTTCCCTACACGACGCTCTTCCGATCTNNNCATACGCACACAAACAACACA
GAGCACACGTCTGAACTCTTTCCCTACACGACGCTCTTCCGATCTNNNCATACGCACACAAACAACACA
AGCACACGTCTGAACTCTTTCCCTACACGACGCTCTTCCGATCTNNNACGGATCAGATAAGCAAGTGT
AGCACACGTCTGAACTCTTTCCCTACACGACGCTCTTCCGATCTNNNACGGATCAGATAAGCAAGTGT
AGCACACGTCTGAACTCTTTCCCTACACGACGCTCTTCCGATCTNNNTCCAGTTTTTCCACAAATGCC
AGCACACGTCTGAACTCTTTCCCTACACGACGCTCTTCCGATCTNNNTCCAGTTTTTCCACAAATGCC
GCACACGTCTGAACTCTTTCCCTACACGACGCTCTTCCGATCTNNNCATGGTTGTTGCCAAAGACCAT
GCACACGTCTGAACTCTTTCCCTACACGACGCTCTTCCGATCTNNNCATGGTTGTTGCCAAAGACCAT
GAGCACACGTCTGAACTCTTTCCCTACACGACGCTCTTCCGATCTNNNTCCTGTGTCCATGTGTTCTCAT
GAGCACACGTCTGAACTCTTTCCCTACACGACGCTCTTCCGATCTNNNTCCTGTGTCCATGTGTTCTCAT
AGCACACGTCTGAACTCTTTCCCTACACGACGCTCTTCCGATCTNNNAGGCCTCCAAAATTAGTCATG
AGCACACGTCTGAACTCTTTCCCTACACGACGCTCTTCCGATCTNNNAGGCCTCCAAAATTAGTCATG
AGCACACGTCTGAACTCTTTCCCTACACGACGCTCTTCCGATCTNNNTACAAGGGAATAGGCAGGTG
AGCACACGTCTGAACTCTTTCCCTACACGACGCTCTTCCGATCTNNNTACAAGGGAATAGGCAGGTG
GCACACGTCTGAACTCTTTCCCTACACGACGCTCTTCCGATCTNNNCTCTGTCCTCACAGAGTTCAAGC
GCACACGTCTGAACTCTTTCCCTACACGACGCTCTTCCGATCTNNNCTCTGTCCTCACAGAGTTCAAGC
AGCACACGTCTGAACTCTTTCCCTACACGACGCTCTTCCGATCTNNNATAATTTACCCCCAGAAGCTC
AGCACACGTCTGAACTCTTTCCCTACACGACGCTCTTCCGATCTNNNATAATTTACCCCCAGAAGCTC
GCACACGTCTGAACTCTTTCCCTACACGACGCTCTTCCGATCTNNNGGCCCAGACAGACAGATATACA
GCACACGTCTGAACTCTTTCCCTACACGACGCTCTTCCGATCTNNNGGCCCAGACAGACAGATATACA
AGCACACGTCTGAACTCTTTCCCTACACGACGCTCTTCCGATCTNNNTTTCAGTAGAGACGGGGTTTC
AGCACACGTCTGAACTCTTTCCCTACACGACGCTCTTCCGATCTNNNTTTCAGTAGAGACGGGGTTTC
GCACACGTCTGAACTCTTTCCCTACACGACGCTCTTCCGATCTNNNTAGGGACCAACAGGACTTCAAT
GCACACGTCTGAACTCTTTCCCTACACGACGCTCTTCCGATCTNNNTAGGGACCAACAGGACTTCAAT
GAGCACACGTCTGAACTCTTTCCCTACACGACGCTCTTCCGATCTNNNGCTGAGAGAGGACATCACTT
GAGCACACGTCTGAACTCTTTCCCTACACGACGCTCTTCCGATCTNNNGCTGAGAGAGGACATCACTT
GCACACGTCTGAACTCTTTCCCTACACGACGCTCTTCCGATCTNNNTCCTGTTTTCAATTCTTTGGGGT
GCACACGTCTGAACTCTTTCCCTACACGACGCTCTTCCGATCTNNNTCCTGTTTTCAATTCTTTGGGGT
GAGCACACGTCTGAACTCTTTCCCTACACGACGCTCTTCCGATCTNNNCAATGACTATCTGAAGCCAAG

AGCACACGTCTGAACTCTTTCCCTACACGACGCTCTTCCGATCTNNNCAATGACTATCTGAAGCCAAG
GAGCACACGTCTGAACTCTTTCCCTACACGACGCTCTTCCGATCTNNNTGGTCTCAAAGTGATCCTCC
GAGCACACGTCTGAACTCTTTCCCTACACGACGCTCTTCCGATCTNNNTGGTCTCAAAGTGATCCTCC
CACACGTCTGAACTCTTTCCCTACACGACGCTCTTCCGATCTNNNCCCTTGGAGAAGTGGACTCTTAG
CACACGTCTGAACTCTTTCCCTACACGACGCTCTTCCGATCTNNNCCCTTGGAGAAGTGGACTCTTAG
AGCACACGTCTGAACTCTTTCCCTACACGACGCTCTTCCGATCTNNNCTGAGGCAGGAGAATCACTTG
AGCACACGTCTGAACTCTTTCCCTACACGACGCTCTTCCGATCTNNNCTGAGGCAGGAGAATCACTTG
GCACACGTCTGAACTCTTTCCCTACACGACGCTCTTCCGATCTNNNTTTCTGGTTTGGACTCTGGACA
GCACACGTCTGAACTCTTTCCCTACACGACGCTCTTCCGATCTNNNTTTCTGGTTTGGACTCTGGACA
GAGCACACGTCTGAACTCTTTCCCTACACGACGCTCTTCCGATCTNNNTCCTTTCCTTTCCTTTCCTTTCC
GAGCACACGTCTGAACTCTTTCCCTACACGACGCTCTTCCGATCTNNNTCCTTTCCTTTCCTTTCCTTTCC
GAGCACACGTCTGAACTCTTTCCCTACACGACGCTCTTCCGATCTNNNCAGACTCATTGAGACAACCT
GAGCACACGTCTGAACTCTTTCCCTACACGACGCTCTTCCGATCTNNNCAGACTCATTGAGACAACCT
GAGCACACGTCTGAACTCTTTCCCTACACGACGCTCTTCCGATCTNNNCCTGTAGTCCCAGCTACTTG
GAGCACACGTCTGAACTCTTTCCCTACACGACGCTCTTCCGATCTNNNCCTGTAGTCCCAGCTACTTG
AGCACACGTCTGAACTCTTTCCCTACACGACGCTCTTCCGATCTNNNCAGCAGTGGTGTTCTCTAGAC
AGCACACGTCTGAACTCTTTCCCTACACGACGCTCTTCCGATCTNNNCAGCAGTGGTGTTCTCTAGAC
CACGTCTGAACTCTTTCCCTACACGACGCTCTTCCGATCTNNNTATTTGGATGACAGAAACACTTTCAT
CACGTCTGAACTCTTTCCCTACACGACGCTCTTCCGATCTNNNTATTTGGATGACAGAAACACTTTCAT
AGCACACGTCTGAACTCTTTCCCTACACGACGCTCTTCCGATCTNNNCTAGTCAAGGTAGCGAGACAC
AGCACACGTCTGAACTCTTTCCCTACACGACGCTCTTCCGATCTNNNCTAGTCAAGGTAGCGAGACAC
AGCACACGTCTGAACTCTTTCCCTACACGACGCTCTTCCGATCTNNNGCATACCTGTAATCCCAGCTA
AGCACACGTCTGAACTCTTTCCCTACACGACGCTCTTCCGATCTNNNGCATACCTGTAATCCCAGCTA
CACACGTCTGAACTCTTTCCCTACACGACGCTCTTCCGATCTNNNTGAGAATCACAATCACTGCACAT
CACACGTCTGAACTCTTTCCCTACACGACGCTCTTCCGATCTNNNTGAGAATCACAATCACTGCACAT
GCACACGTCTGAACTCTTTCCCTACACGACGCTCTTCCGATCTNNNGAATGAAACCCACCACCTCTTTG
GCACACGTCTGAACTCTTTCCCTACACGACGCTCTTCCGATCTNNNGAATGAAACCCACCACCTCTTTG
AGCACACGTCTGAACTCTTTCCCTACACGACGCTCTTCCGATCTNNNGTGGGTATGCACCTTATTTCT
AGCACACGTCTGAACTCTTTCCCTACACGACGCTCTTCCGATCTNNNGTGGGTATGCACCTTATTTCT
AGCACACGTCTGAACTCTTTCCCTACACGACGCTCTTCCGATCTNNNAGGCTTACAAATGGCAAAGG
AGCACACGTCTGAACTCTTTCCCTACACGACGCTCTTCCGATCTNNNAGGCTTACAAATGGCAAAGG
CACACGTCTGAACTCTTTCCCTACACGACGCTCTTCCGATCTNNNTGGCCATTCATTTTCTTCCTCTTCA
CACACGTCTGAACTCTTTCCCTACACGACGCTCTTCCGATCTNNNTGGCCATTCATTTTCTTCCTCTTCA
AGCACACGTCTGAACTCTTTCCCTACACGACGCTCTTCCGATCTNNNTGCACTCACTACTTGAAAACC
AGCACACGTCTGAACTCTTTCCCTACACGACGCTCTTCCGATCTNNNTGCACTCACTACTTGAAAACC
AGCACACGTCTGAACTCTTTCCCTACACGACGCTCTTCCGATCTNNNGAGATCGCGCATTACACTCCA
AGCACACGTCTGAACTCTTTCCCTACACGACGCTCTTCCGATCTNNNGAGATCGCGCATTACACTCCA
GCACACGTCTGAACTCTTTCCCTACACGACGCTCTTCCGATCTNNNCCTGGCTGCTGAACTCTCTTATT
GCACACGTCTGAACTCTTTCCCTACACGACGCTCTTCCGATCTNNNCCTGGCTGCTGAACTCTCTTATT
GCACACGTCTGAACTCTTTCCCTACACGACGCTCTTCCGATCTNNNTGGCTATTATCCCAGGTTCTTG
GCACACGTCTGAACTCTTTCCCTACACGACGCTCTTCCGATCTNNNTGGCTATTATCCCAGGTTCTTG
AGCACACGTCTGAACTCTTTCCCTACACGACGCTCTTCCGATCTNNNCTGAGGCAGGAGAATCACTTG
AGCACACGTCTGAACTCTTTCCCTACACGACGCTCTTCCGATCTNNNCTGAGGCAGGAGAATCACTTG

AGCACACGTCTGAACTCTTTCCCTACACGACGCTCTTCCGATCTNNNTGAGCTCTGATTATACCACTGC
AGCACACGTCTGAACTCTTTCCCTACACGACGCTCTTCCGATCTNNNTGAGCTCTGATTATACCACTGC
GCACACGTCTGAACTCTTTCCCTACACGACGCTCTTCCGATCTNNNTTACTTTGCAGTCTTCAGGAGGG
GCACACGTCTGAACTCTTTCCCTACACGACGCTCTTCCGATCTNNNTTACTTTGCAGTCTTCAGGAGGG
GAGCACACGTCTGAACTCTTTCCCTACACGACGCTCTTCCGATCTNNNAATGGGGTCTCTTGTCTGTGT
GAGCACACGTCTGAACTCTTTCCCTACACGACGCTCTTCCGATCTNNNAATGGGGTCTCTTGTCTGTGT
AGCACACGTCTGAACTCTTTCCCTACACGACGCTCTTCCGATCTNNNGAGAGAGAGAGAGAAGACCCA
AGCACACGTCTGAACTCTTTCCCTACACGACGCTCTTCCGATCTNNNGAGAGAGAGAGAGAAGACCCA
GAGCACACGTCTGAACTCTTTCCCTACACGACGCTCTTCCGATCTNNNTTCTTCTTCCTTGATAGCCTCAT
GAGCACACGTCTGAACTCTTTCCCTACACGACGCTCTTCCGATCTNNNTTCTTCTTCCTTGATAGCCTCAT
AGCACACGTCTGAACTCTTTCCCTACACGACGCTCTTCCGATCTNNNCCCTTGTTGTGAAAAAGCAACC
AGCACACGTCTGAACTCTTTCCCTACACGACGCTCTTCCGATCTNNNCCCTTGTTGTGAAAAAGCAACC
GAGCACACGTCTGAACTCTTTCCCTACACGACGCTCTTCCGATCTNNNAGCTGGTTATTCTACTGTTGG
GAGCACACGTCTGAACTCTTTCCCTACACGACGCTCTTCCGATCTNNNAGCTGGTTATTCTACTGTTGG
AGCACACGTCTGAACTCTTTCCCTACACGACGCTCTTCCGATCTNNNCAGTCCTGACAGGTTCCTCATTA
AGCACACGTCTGAACTCTTTCCCTACACGACGCTCTTCCGATCTNNNCAGTCCTGACAGGTTCCTCATTA
AGAGCACACGTCTGAACTCTTTCCCTACACGACGCTCTTCCGATCTNNNTGAGGCAGGAGAATCACTTG
AGAGCACACGTCTGAACTCTTTCCCTACACGACGCTCTTCCGATCTNNNTGAGGCAGGAGAATCACTTG
GAGCACACGTCTGAACTCTTTCCCTACACGACGCTCTTCCGATCTNNNGAGGTCCCATCAGGTGCTATA
GAGCACACGTCTGAACTCTTTCCCTACACGACGCTCTTCCGATCTNNNGAGGTCCCATCAGGTGCTATA
GAGCACACGTCTGAACTCTTTCCCTACACGACGCTCTTCCGATCTNNNGCAAACTGAAGCAAATATCAC
GAGCACACGTCTGAACTCTTTCCCTACACGACGCTCTTCCGATCTNNNGCAAACTGAAGCAAATATCAC
GCACACGTCTGAACTCTTTCCCTACACGACGCTCTTCCGATCTNNNTATCATCTCGTCCAATAATAGAAC
GCACACGTCTGAACTCTTTCCCTACACGACGCTCTTCCGATCTNNNTATCATCTCGTCCAATAATAGAAC
AGCACACGTCTGAACTCTTTCCCTACACGACGCTCTTCCGATCTNNNGATCGATAAGCTGAAGGAGGA
AGCACACGTCTGAACTCTTTCCCTACACGACGCTCTTCCGATCTNNNGATCGATAAGCTGAAGGAGGA
GAGCACACGTCTGAACTCTTTCCCTACACGACGCTCTTCCGATCTNNNTTGAGCTTAGGAGGCAGATG
GAGCACACGTCTGAACTCTTTCCCTACACGACGCTCTTCCGATCTNNNTTGAGCTTAGGAGGCAGATG
GAGCACACGTCTGAACTCTTTCCCTACACGACGCTCTTCCGATCTNNNTGTAATCCCAGCTACTCAGGAG
GAGCACACGTCTGAACTCTTTCCCTACACGACGCTCTTCCGATCTNNNTGTAATCCCAGCTACTCAGGAG
GCACACGTCTGAACTCTTTCCCTACACGACGCTCTTCCGATCTNNNCTCCTGAGAAGCTGGGATTACAG
GCACACGTCTGAACTCTTTCCCTACACGACGCTCTTCCGATCTNNNCTCCTGAGAAGCTGGGATTACAG
AGAGCACACGTCTGAACTCTTTCCCTACACGACGCTCTTCCGATCTNNNGCATATGTGACTGGACGTG
AGAGCACACGTCTGAACTCTTTCCCTACACGACGCTCTTCCGATCTNNNGCATATGTGACTGGACGTG
GAGCACACGTCTGAACTCTTTCCCTACACGACGCTCTTCCGATCTNNNCACCATGACCGGCTAATTTTC
GAGCACACGTCTGAACTCTTTCCCTACACGACGCTCTTCCGATCTNNNCACCATGACCGGCTAATTTTC
AGCACACGTCTGAACTCTTTCCCTACACGACGCTCTTCCGATCTNNNTGGAGACAGGAGAATTGCTTG
AGCACACGTCTGAACTCTTTCCCTACACGACGCTCTTCCGATCTNNNTGGAGACAGGAGAATTGCTTG
GAGCACACGTCTGAACTCTTTCCCTACACGACGCTCTTCCGATCTNNNCTATGGTCGTTATTTTCTCATTT
GAGCACACGTCTGAACTCTTTCCCTACACGACGCTCTTCCGATCTNNNCTATGGTCGTTATTTTCTCATTT
AGAGCACACGTCTGAACTCTTTCCCTACACGACGCTCTTCCGATCTNNNTTGAGCTACTCAGGAGGCTG
AGAGCACACGTCTGAACTCTTTCCCTACACGACGCTCTTCCGATCTNNNTTGAGCTACTCAGGAGGCTG
AGAGCACACGTCTGAACTCTTTCCCTACACGACGCTCTTCCGATCTNNNTTGCTGTTGTCATGTGTGTTT

GAGCACACGTCTGAACTCTTTCCCTACACGACGCTCTTCCGATCTNNNTTGCTGTTGTCATGTGTGTTT
AGAGCACACGTCTGAACTCTTTCCCTACACGACGCTCTTCCGATCTNNNAGGCCTGGAATTACAGTTG
AGAGCACACGTCTGAACTCTTTCCCTACACGACGCTCTTCCGATCTNNNAGGCCTGGAATTACAGTTG
GAGCACACGTCTGAACTCTTTCCCTACACGACGCTCTTCCGATCTNNNGAGGCTGAGGCAGGAGAAT
GAGCACACGTCTGAACTCTTTCCCTACACGACGCTCTTCCGATCTNNNGAGGCTGAGGCAGGAGAAT
GCACACGTCTGAACTCTTTCCCTACACGACGCTCTTCCGATCTNNNCTTGAACACTACCAAGCCAGTTG
GCACACGTCTGAACTCTTTCCCTACACGACGCTCTTCCGATCTNNNCTTGAACACTACCAAGCCAGTTG
GCACACGTCTGAACTCTTTCCCTACACGACGCTCTTCCGATCTNNNAGACTCTGTCTCCCAAACAAACA
GCACACGTCTGAACTCTTTCCCTACACGACGCTCTTCCGATCTNNNAGACTCTGTCTCCCAAACAAACA
GCACACGTCTGAACTCTTTCCCTACACGACGCTCTTCCGATCTNNNTAGGATGGGATCAGGGTCTGAG
GCACACGTCTGAACTCTTTCCCTACACGACGCTCTTCCGATCTNNNTAGGATGGGATCAGGGTCTGAG
AGCACACGTCTGAACTCTTTCCCTACACGACGCTCTTCCGATCTNNNACAAAAAATTAGCCGGGCATT
AGCACACGTCTGAACTCTTTCCCTACACGACGCTCTTCCGATCTNNNACAAAAAATTAGCCGGGCATT
AGCACACGTCTGAACTCTTTCCCTACACGACGCTCTTCCGATCTNNNAAGTACTGAACAGTCCTTTGTG
AGCACACGTCTGAACTCTTTCCCTACACGACGCTCTTCCGATCTNNNAAGTACTGAACAGTCCTTTGTG
AGCACACGTCTGAACTCTTTCCCTACACGACGCTCTTCCGATCTNNNGGTATGGTTTGGATCTGTGTCC
AGCACACGTCTGAACTCTTTCCCTACACGACGCTCTTCCGATCTNNNGGTATGGTTTGGATCTGTGTCC
AGCACACGTCTGAACTCTTTCCCTACACGACGCTCTTCCGATCTNNNATCAGGCATTTGAGCTCTTTGC
AGCACACGTCTGAACTCTTTCCCTACACGACGCTCTTCCGATCTNNNATCAGGCATTTGAGCTCTTTGC
GAGCACACGTCTGAACTCTTTCCCTACACGACGCTCTTCCGATCTNNNATACAAAAATTAGCTGGGCG
GAGCACACGTCTGAACTCTTTCCCTACACGACGCTCTTCCGATCTNNNATACAAAAATTAGCTGGGCG
AGCACACGTCTGAACTCTTTCCCTACACGACGCTCTTCCGATCTNNNGGTCTCAAACTTCTGAGCCAAA
AGCACACGTCTGAACTCTTTCCCTACACGACGCTCTTCCGATCTNNNGGTCTCAAACTTCTGAGCCAAA
GAGCACACGTCTGAACTCTTTCCCTACACGACGCTCTTCCGATCTNNNTCTAGACAGAGGAGGATGAA
GAGCACACGTCTGAACTCTTTCCCTACACGACGCTCTTCCGATCTNNNTCTAGACAGAGGAGGATGAA
GCACACGTCTGAACTCTTTCCCTACACGACGCTCTTCCGATCTNNNTACAAAAAATTAGCCGGGCATG
GCACACGTCTGAACTCTTTCCCTACACGACGCTCTTCCGATCTNNNTACAAAAAATTAGCCGGGCATG
AGCACACGTCTGAACTCTTTCCCTACACGACGCTCTTCCGATCTNNNATACCAAAAGCAGACAAAGG
AGCACACGTCTGAACTCTTTCCCTACACGACGCTCTTCCGATCTNNNATACCAAAAGCAGACAAAGG
GAGCACACGTCTGAACTCTTTCCCTACACGACGCTCTTCCGATCTNNNGAGCAAGCCCAGAACTTGAA
GAGCACACGTCTGAACTCTTTCCCTACACGACGCTCTTCCGATCTNNNGAGCAAGCCCAGAACTTGAA
AGCACACGTCTGAACTCTTTCCCTACACGACGCTCTTCCGATCTNNNCCTGGTCAGGCTACTTTCTTCTT
AGCACACGTCTGAACTCTTTCCCTACACGACGCTCTTCCGATCTNNNCCTGGTCAGGCTACTTTCTTCTT
GAGCACACGTCTGAACTCTTTCCCTACACGACGCTCTTCCGATCTNNNTCATGCCTGTAATCGTAGCAC
GAGCACACGTCTGAACTCTTTCCCTACACGACGCTCTTCCGATCTNNNTCATGCCTGTAATCGTAGCAC
GAGCACACGTCTGAACTCTTTCCCTACACGACGCTCTTCCGATCTNNNAGACAGTCTTGCTATGTTGAC
GAGCACACGTCTGAACTCTTTCCCTACACGACGCTCTTCCGATCTNNNAGACAGTCTTGCTATGTTGAC
GAGCACACGTCTGAACTCTTTCCCTACACGACGCTCTTCCGATCTNNNAAGAATCACTTGAACCTGGG
GAGCACACGTCTGAACTCTTTCCCTACACGACGCTCTTCCGATCTNNNAAGAATCACTTGAACCTGGG
AGCACACGTCTGAACTCTTTCCCTACACGACGCTCTTCCGATCTNNNGGATGATCCGCATTGGTCTGA
AGCACACGTCTGAACTCTTTCCCTACACGACGCTCTTCCGATCTNNNGGATGATCCGCATTGGTCTGA
AGCACACGTCTGAACTCTTTCCCTACACGACGCTCTTCCGATCTNNNGATTCTTTGGGTGTCCAGTCTC
AGCACACGTCTGAACTCTTTCCCTACACGACGCTCTTCCGATCTNNNGATTCTTTGGGTGTCCAGTCTC

AGCACACGTCTGAACTCTTTCCCTACACGACGCTCTTCCGATCTNNNCCATCTTGGTTTTGGTGGGTTTT
AGCACACGTCTGAACTCTTTCCCTACACGACGCTCTTCCGATCTNNNCCATCTTGGTTTTGGTGGGTTTT
AGCACACGTCTGAACTCTTTCCCTACACGACGCTCTTCCGATCTNNNATAGCGAAACCCTGCCTCTACTA
AGCACACGTCTGAACTCTTTCCCTACACGACGCTCTTCCGATCTNNNATAGCGAAACCCTGCCTCTACTA
AGCACACGTCTGAACTCTTTCCCTACACGACGCTCTTCCGATCTNNNTAAATATCAGCTCTGGAATAACA
AGCACACGTCTGAACTCTTTCCCTACACGACGCTCTTCCGATCTNNNTAAATATCAGCTCTGGAATAACA
AGAGCACACGTCTGAACTCTTTCCCTACACGACGCTCTTCCGATCTNNNTGACTTCTGATAAGGGCTGC
AGAGCACACGTCTGAACTCTTTCCCTACACGACGCTCTTCCGATCTNNNTGACTTCTGATAAGGGCTGC
AGCACACGTCTGAACTCTTTCCCTACACGACGCTCTTCCGATCTNNNACAAAATCTCCTACCGCTTTTTGT
AGCACACGTCTGAACTCTTTCCCTACACGACGCTCTTCCGATCTNNNACAAAATCTCCTACCGCTTTTTGT
AGCACACGTCTGAACTCTTTCCCTACACGACGCTCTTCCGATCTNNNACATCTGCACTTTCCAATAGGGT
AGCACACGTCTGAACTCTTTCCCTACACGACGCTCTTCCGATCTNNNACATCTGCACTTTCCAATAGGGT
GAGCACACGTCTGAACTCTTTCCCTACACGACGCTCTTCCGATCTNNNGACACACAGAGGAAAGAAAT
GAGCACACGTCTGAACTCTTTCCCTACACGACGCTCTTCCGATCTNNNGACACACAGAGGAAAGAAAT
GAGCACACGTCTGAACTCTTTCCCTACACGACGCTCTTCCGATCTNNNATCATGGCTGTAGTGCTCAGA
GAGCACACGTCTGAACTCTTTCCCTACACGACGCTCTTCCGATCTNNNATCATGGCTGTAGTGCTCAGA
AGCACACGTCTGAACTCTTTCCCTACACGACGCTCTTCCGATCTNNNGGTTCAAGTGATCCTCCTGTCT
AGCACACGTCTGAACTCTTTCCCTACACGACGCTCTTCCGATCTNNNGGTTCAAGTGATCCTCCTGTCT
AGCACACGTCTGAACTCTTTCCCTACACGACGCTCTTCCGATCTNNNACCTCCATACTGCATTTCTGTGT
AGCACACGTCTGAACTCTTTCCCTACACGACGCTCTTCCGATCTNNNACCTCCATACTGCATTTCTGTGT
GAGCACACGTCTGAACTCTTTCCCTACACGACGCTCTTCCGATCTNNNTAATTCTCCCCATTTTCCAGGCT
GAGCACACGTCTGAACTCTTTCCCTACACGACGCTCTTCCGATCTNNNTAATTCTCCCCATTTTCCAGGCT
AGAGCACACGTCTGAACTCTTTCCCTACACGACGCTCTTCCGATCTNNNCTTAGCATCTCCCAGACTGTC
AGAGCACACGTCTGAACTCTTTCCCTACACGACGCTCTTCCGATCTNNNCTTAGCATCTCCCAGACTGTC
AGCACACGTCTGAACTCTTTCCCTACACGACGCTCTTCCGATCTNNNAAGCTCAAGTACCTCCAGAAAG
AGCACACGTCTGAACTCTTTCCCTACACGACGCTCTTCCGATCTNNNAAGCTCAAGTACCTCCAGAAAG
GAGCACACGTCTGAACTCTTTCCCTACACGACGCTCTTCCGATCTNNNCACCAGCCACTTTTCACTGTTA
GAGCACACGTCTGAACTCTTTCCCTACACGACGCTCTTCCGATCTNNNCACCAGCCACTTTTCACTGTTA
AGCACACGTCTGAACTCTTTCCCTACACGACGCTCTTCCGATCTNNNGCTGGGTAGAGTGGCTTTTACTT
AGCACACGTCTGAACTCTTTCCCTACACGACGCTCTTCCGATCTNNNGCTGGGTAGAGTGGCTTTTACTT
AGCACACGTCTGAACTCTTTCCCTACACGACGCTCTTCCGATCTNNNCCCAAATTAAACGTAAATGTTTT
AGCACACGTCTGAACTCTTTCCCTACACGACGCTCTTCCGATCTNNNCCCAAATTAAACGTAAATGTTTT
GAGCACACGTCTGAACTCTTTCCCTACACGACGCTCTTCCGATCTNNNCTCCGAACAAAATACCCCTCTC
GAGCACACGTCTGAACTCTTTCCCTACACGACGCTCTTCCGATCTNNNCTCCGAACAAAATACCCCTCTC
GAGCACACGTCTGAACTCTTTCCCTACACGACGCTCTTCCGATCTNNNAGCAGTGACTGAGCCAACAG
GAGCACACGTCTGAACTCTTTCCCTACACGACGCTCTTCCGATCTNNNAGCAGTGACTGAGCCAACAG
AGCACACGTCTGAACTCTTTCCCTACACGACGCTCTTCCGATCTNNNGTCTGAGGGGACTTTGAGTTT
AGCACACGTCTGAACTCTTTCCCTACACGACGCTCTTCCGATCTNNNGTCTGAGGGGACTTTGAGTTT
AGCACACGTCTGAACTCTTTCCCTACACGACGCTCTTCCGATCTNNNTTGGTAGAACTGGGGTTTTGT
AGCACACGTCTGAACTCTTTCCCTACACGACGCTCTTCCGATCTNNNTTGGTAGAACTGGGGTTTTGT
AGAGCACACGTCTGAACTCTTTCCCTACACGACGCTCTTCCGATCTNNNTGATATCCTTGGGCAAGATC
AGAGCACACGTCTGAACTCTTTCCCTACACGACGCTCTTCCGATCTNNNTGATATCCTTGGGCAAGATC
AGAGCACACGTCTGAACTCTTTCCCTACACGACGCTCTTCCGATCTNNNCCCTTTTGCCTAACAGTGGG

AGAGCACACGTCTGAACTCTTTCCCTACACGACGCTCTTCCGATCTNNNCCCTTTTGCCTAACAGTGGG
AGCACACGTCTGAACTCTTTCCCTACACGACGCTCTTCCGATCTNNNTTCATGGTAAAAACCCTGCACT
AGCACACGTCTGAACTCTTTCCCTACACGACGCTCTTCCGATCTNNNTTCATGGTAAAAACCCTGCACT
GAGCACACGTCTGAACTCTTTCCCTACACGACGCTCTTCCGATCTNNNCAGAAGTTGTAGTGAGCCCAG
GAGCACACGTCTGAACTCTTTCCCTACACGACGCTCTTCCGATCTNNNCAGAAGTTGTAGTGAGCCCAG
AGCACACGTCTGAACTCTTTCCCTACACGACGCTCTTCCGATCTNNNTGCAAAGGACATGAACTCATCCT
AGCACACGTCTGAACTCTTTCCCTACACGACGCTCTTCCGATCTNNNTGCAAAGGACATGAACTCATCCT
GAGCACACGTCTGAACTCTTTCCCTACACGACGCTCTTCCGATCTNNNCAGGAGAATCACTTGAACCCA
GAGCACACGTCTGAACTCTTTCCCTACACGACGCTCTTCCGATCTNNNCAGGAGAATCACTTGAACCCA
AGCACACGTCTGAACTCTTTCCCTACACGACGCTCTTCCGATCTNNNTATACACACCAACAGGCACAGA
AGCACACGTCTGAACTCTTTCCCTACACGACGCTCTTCCGATCTNNNTATACACACCAACAGGCACAGA
GAGCACACGTCTGAACTCTTTCCCTACACGACGCTCTTCCGATCTNNNCTGGGAAACAAGAGTGAAAAC
GAGCACACGTCTGAACTCTTTCCCTACACGACGCTCTTCCGATCTNNNCTGGGAAACAAGAGTGAAAAC
AGAGCACACGTCTGAACTCTTTCCCTACACGACGCTCTTCCGATCTNNNAAGAGGGTGACATTCTGCCT
AGAGCACACGTCTGAACTCTTTCCCTACACGACGCTCTTCCGATCTNNNAAGAGGGTGACATTCTGCCT
GAGCACACGTCTGAACTCTTTCCCTACACGACGCTCTTCCGATCTNNNCCAGAAAGCCCAAGAACATCA
GAGCACACGTCTGAACTCTTTCCCTACACGACGCTCTTCCGATCTNNNCCAGAAAGCCCAAGAACATCA
AGCACACGTCTGAACTCTTTCCCTACACGACGCTCTTCCGATCTNNNTGTAGTCCCAGTTACTCAGGAG
AGCACACGTCTGAACTCTTTCCCTACACGACGCTCTTCCGATCTNNNTGTAGTCCCAGTTACTCAGGAG
GCACACGTCTGAACTCTTTCCCTACACGACGCTCTTCCGATCTNNNTCCCATTCTTTCCTCTCACAAGCA
GCACACGTCTGAACTCTTTCCCTACACGACGCTCTTCCGATCTNNNTCCCATTCTTTCCTCTCACAAGCA
GAGCACACGTCTGAACTCTTTCCCTACACGACGCTCTTCCGATCTNNNGAGGCAGGAGAATCGCTTGA
GAGCACACGTCTGAACTCTTTCCCTACACGACGCTCTTCCGATCTNNNGAGGCAGGAGAATCGCTTGA
ACACGTCTGAACTCTTTCCCTACACGACGCTCTTCCGATCTNNNTCAGAAAACGAGGATGAGGATGGCA
ACACGTCTGAACTCTTTCCCTACACGACGCTCTTCCGATCTNNNTCAGAAAACGAGGATGAGGATGGCA
GAGCACACGTCTGAACTCTTTCCCTACACGACGCTCTTCCGATCTNNNTGCTTACTAAGGAGACATTGG
GAGCACACGTCTGAACTCTTTCCCTACACGACGCTCTTCCGATCTNNNTGCTTACTAAGGAGACATTGG
GAGCACACGTCTGAACTCTTTCCCTACACGACGCTCTTCCGATCTNNNTGGTATTGCCCTATAATAATTC
GAGCACACGTCTGAACTCTTTCCCTACACGACGCTCTTCCGATCTNNNTGGTATTGCCCTATAATAATTC
AAGAGCACACGTCTGAACTCTTTCCCTACACGACGCTCTTCCGATCTNNNGGGCAGGCAAACAGATAT
AAGAGCACACGTCTGAACTCTTTCCCTACACGACGCTCTTCCGATCTNNNGGGCAGGCAAACAGATAT
GAGCACACGTCTGAACTCTTTCCCTACACGACGCTCTTCCGATCTNNNGAGAAAGAGAGCAGTTCCAGT
GAGCACACGTCTGAACTCTTTCCCTACACGACGCTCTTCCGATCTNNNGAGAAAGAGAGCAGTTCCAGT
GAGCACACGTCTGAACTCTTTCCCTACACGACGCTCTTCCGATCTNNNTGCTCGCTAACACTTCTCATTG
GAGCACACGTCTGAACTCTTTCCCTACACGACGCTCTTCCGATCTNNNTGCTCGCTAACACTTCTCATTG
AGCACACGTCTGAACTCTTTCCCTACACGACGCTCTTCCGATCTNNNGATCCACTGAGCTAGACCACTT
AGCACACGTCTGAACTCTTTCCCTACACGACGCTCTTCCGATCTNNNGATCCACTGAGCTAGACCACTT
AGCACACGTCTGAACTCTTTCCCTACACGACGCTCTTCCGATCTNNNACCCAACCGTCACTCTTGTATTTC
AGCACACGTCTGAACTCTTTCCCTACACGACGCTCTTCCGATCTNNNACCCAACCGTCACTCTTGTATTTC
AGCACACGTCTGAACTCTTTCCCTACACGACGCTCTTCCGATCTNNNTAGGGGTAGGAGCTACTTGAGT
AGCACACGTCTGAACTCTTTCCCTACACGACGCTCTTCCGATCTNNNTAGGGGTAGGAGCTACTTGAGT
GAGCACACGTCTGAACTCTTTCCCTACACGACGCTCTTCCGATCTNNNCCCATTTTCTTTTGGTGACGCT
GAGCACACGTCTGAACTCTTTCCCTACACGACGCTCTTCCGATCTNNNCCCATTTTCTTTTGGTGACGCT

GAGCACACGTCTGAACTCTTTCCCTACACGACGCTCTTCCGATCTNNNTGATCCCTGGAGGCTAAACAA
GAGCACACGTCTGAACTCTTTCCCTACACGACGCTCTTCCGATCTNNNTGATCCCTGGAGGCTAAACAA
AGCACACGTCTGAACTCTTTCCCTACACGACGCTCTTCCGATCTNNNTGCTACTTTATACCCGTCTTCCC
AGCACACGTCTGAACTCTTTCCCTACACGACGCTCTTCCGATCTNNNTGCTACTTTATACCCGTCTTCCC
GCACACGTCTGAACTCTTTCCCTACACGACGCTCTTCCGATCTNNNAACAGGAGTTAAAAGACGTGGG
GCACACGTCTGAACTCTTTCCCTACACGACGCTCTTCCGATCTNNNAACAGGAGTTAAAAGACGTGGG
GCACACGTCTGAACTCTTTCCCTACACGACGCTCTTCCGATCTNNNGAGGTTGCAGTAAGCCAAGATT
GCACACGTCTGAACTCTTTCCCTACACGACGCTCTTCCGATCTNNNGAGGTTGCAGTAAGCCAAGATT
GAGCACACGTCTGAACTCTTTCCCTACACGACGCTCTTCCGATCTNNNCTGCAGGTCTAGGGGTGATA
GAGCACACGTCTGAACTCTTTCCCTACACGACGCTCTTCCGATCTNNNCTGCAGGTCTAGGGGTGATA
AGCACACGTCTGAACTCTTTCCCTACACGACGCTCTTCCGATCTNNNGAGAATGGCTTGAATCCGGGA
AGCACACGTCTGAACTCTTTCCCTACACGACGCTCTTCCGATCTNNNGAGAATGGCTTGAATCCGGGA
AGCACACGTCTGAACTCTTTCCCTACACGACGCTCTTCCGATCTNNNTCACAAGATACAGGGAGCATTC
AGCACACGTCTGAACTCTTTCCCTACACGACGCTCTTCCGATCTNNNTCACAAGATACAGGGAGCATTC
GAGCACACGTCTGAACTCTTTCCCTACACGACGCTCTTCCGATCTNNNCAAGAGGGAACTGGGATGAA
GAGCACACGTCTGAACTCTTTCCCTACACGACGCTCTTCCGATCTNNNCAAGAGGGAACTGGGATGAA
GAGCACACGTCTGAACTCTTTCCCTACACGACGCTCTTCCGATCTNNNCTATCACTTGAGCCCAGGAGAT
GAGCACACGTCTGAACTCTTTCCCTACACGACGCTCTTCCGATCTNNNCTATCACTTGAGCCCAGGAGAT
GCACACGTCTGAACTCTTTCCCTACACGACGCTCTTCCGATCTNNNGTGACCCCAGATCATACCATTGT
GCACACGTCTGAACTCTTTCCCTACACGACGCTCTTCCGATCTNNNGTGACCCCAGATCATACCATTGT
AGCACACGTCTGAACTCTTTCCCTACACGACGCTCTTCCGATCTNNNAGCGTTTCTCTCTCTCTCTCTC
AGCACACGTCTGAACTCTTTCCCTACACGACGCTCTTCCGATCTNNNAGCGTTTCTCTCTCTCTCTCTC
GAGCACACGTCTGAACTCTTTCCCTACACGACGCTCTTCCGATCTNNNGTTGAGACGGGAATTGCTTG
GAGCACACGTCTGAACTCTTTCCCTACACGACGCTCTTCCGATCTNNNGTTGAGACGGGAATTGCTTG
AGCACACGTCTGAACTCTTTCCCTACACGACGCTCTTCCGATCTNNNAGCCGCTCCATGAGAAACTTAAT
AGCACACGTCTGAACTCTTTCCCTACACGACGCTCTTCCGATCTNNNAGCCGCTCCATGAGAAACTTAAT
GAGCACACGTCTGAACTCTTTCCCTACACGACGCTCTTCCGATCTNNNAAAAAGGTGTGAATGAAGCCT
GAGCACACGTCTGAACTCTTTCCCTACACGACGCTCTTCCGATCTNNNAAAAAGGTGTGAATGAAGCCT
GAGCACACGTCTGAACTCTTTCCCTACACGACGCTCTTCCGATCTNNNCGGAGGTTACAATGAGCTGA
GAGCACACGTCTGAACTCTTTCCCTACACGACGCTCTTCCGATCTNNNCGGAGGTTACAATGAGCTGA
AGCACACGTCTGAACTCTTTCCCTACACGACGCTCTTCCGATCTNNNTGACAGTGGCAGGAAATATCA
AGCACACGTCTGAACTCTTTCCCTACACGACGCTCTTCCGATCTNNNTGACAGTGGCAGGAAATATCA
AGCACACGTCTGAACTCTTTCCCTACACGACGCTCTTCCGATCTNNNGTACCACTAAACTCCAGTCAGCA
AGCACACGTCTGAACTCTTTCCCTACACGACGCTCTTCCGATCTNNNGTACCACTAAACTCCAGTCAGCA
AGCACACGTCTGAACTCTTTCCCTACACGACGCTCTTCCGATCTNNNTCCTTCTCCCACACAGATATCCTC
AGCACACGTCTGAACTCTTTCCCTACACGACGCTCTTCCGATCTNNNTCCTTCTCCCACACAGATATCCTC
GAGCACACGTCTGAACTCTTTCCCTACACGACGCTCTTCCGATCTNNNAGGTTCAAGTGATTCTTCTGCC
GAGCACACGTCTGAACTCTTTCCCTACACGACGCTCTTCCGATCTNNNAGGTTCAAGTGATTCTTCTGCC
AGCACACGTCTGAACTCTTTCCCTACACGACGCTCTTCCGATCTNNNGAAAAACCCCAAAGCTTCAGGA
AGCACACGTCTGAACTCTTTCCCTACACGACGCTCTTCCGATCTNNNGAAAAACCCCAAAGCTTCAGGA
GAGCACACGTCTGAACTCTTTCCCTACACGACGCTCTTCCGATCTNNNTGCTTTCATGAATATTTTCCCC
GAGCACACGTCTGAACTCTTTCCCTACACGACGCTCTTCCGATCTNNNTGCTTTCATGAATATTTTCCCC
AGCACACGTCTGAACTCTTTCCCTACACGACGCTCTTCCGATCTNNNACTGGGATGTTTTCCCCTCTGTA

AGCACACGTCTGAACTCTTTCCCTACACGACGCTCTTCCGATCTNNNACTGGGATGTTTTCCCCTCTGT

GAGCACACGTCTGAACTCTTTCCCTACACGACGCTCTTCCGATCTNNNTATGGAGAAACCCCGTCTCTA

GAGCACACGTCTGAACTCTTTCCCTACACGACGCTCTTCCGATCTNNNTATGGAGAAACCCCGTCTCTA

GAGCACACGTCTGAACTCTTTCCCTACACGACGCTCTTCCGATCTNNNTTCTCACATTGTCAGAGCAGG

GAGCACACGTCTGAACTCTTTCCCTACACGACGCTCTTCCGATCTNNNTTCTCACATTGTCAGAGCAGG

GAGCACACGTCTGAACTCTTTCCCTACACGACGCTCTTCCGATCTNNNAGCTCAGTTGGAAGCTCAAGT

GAGCACACGTCTGAACTCTTTCCCTACACGACGCTCTTCCGATCTNNNAGCTCAGTTGGAAGCTCAAGT

AGAGCACACGTCTGAACTCTTTCCCTACACGACGCTCTTCCGATCTNNNCTGTAATCCCAGCTACTCGGT

AGAGCACACGTCTGAACTCTTTCCCTACACGACGCTCTTCCGATCTNNNCTGTAATCCCAGCTACTCGGT

CACACGTCTGAACTCTTTCCCTACACGACGCTCTTCCGATCTNNNTCACATTCTGAAGCTGATAGGCTC

CACACGTCTGAACTCTTTCCCTACACGACGCTCTTCCGATCTNNNTCACATTCTGAAGCTGATAGGCTC

AGCACACGTCTGAACTCTTTCCCTACACGACGCTCTTCCGATCTNNNCCAGGGAAACAGAGACTCCATC

AGCACACGTCTGAACTCTTTCCCTACACGACGCTCTTCCGATCTNNNCCAGGGAAACAGAGACTCCATC

GCACACGTCTGAACTCTTTCCCTACACGACGCTCTTCCGATCTNNNTTGCAATGTGTGGTAGAATCCCT

GCACACGTCTGAACTCTTTCCCTACACGACGCTCTTCCGATCTNNNTTGCAATGTGTGGTAGAATCCCT

AGCACACGTCTGAACTCTTTCCCTACACGACGCTCTTCCGATCTNNNCTGGTTATGAGACCTTCCCAAC

AGCACACGTCTGAACTCTTTCCCTACACGACGCTCTTCCGATCTNNNCTGGTTATGAGACCTTCCCAAC

GAGCACACGTCTGAACTCTTTCCCTACACGACGCTCTTCCGATCTNNNGAAGGGAATCACAACACAGA

GAGCACACGTCTGAACTCTTTCCCTACACGACGCTCTTCCGATCTNNNGAAGGGAATCACAACACAGA

AGAGCACACGTCTGAACTCTTTCCCTACACGACGCTCTTCCGATCTNNNATACCTGAATCCAGTGGAGG

AGAGCACACGTCTGAACTCTTTCCCTACACGACGCTCTTCCGATCTNNNATACCTGAATCCAGTGGAGG

AGCACACGTCTGAACTCTTTCCCTACACGACGCTCTTCCGATCTNNNACAGTGCTACAATAGACAGTGC

AGCACACGTCTGAACTCTTTCCCTACACGACGCTCTTCCGATCTNNNACAGTGCTACAATAGACAGTGC

AGCACACGTCTGAACTCTTTCCCTACACGACGCTCTTCCGATCTNNNCCTCCTGGGTTCAAGTGATTCTC

AGCACACGTCTGAACTCTTTCCCTACACGACGCTCTTCCGATCTNNNCCTCCTGGGTTCAAGTGATTCTC

AGCACACGTCTGAACTCTTTCCCTACACGACGCTCTTCCGATCTNNNTGGAGGATTCACCCACTCTTTT

AGCACACGTCTGAACTCTTTCCCTACACGACGCTCTTCCGATCTNNNTGGAGGATTCACCCACTCTTTT

AGCACACGTCTGAACTCTTTCCCTACACGACGCTCTTCCGATCTNNNGAGGTTGCATTGAGCCGAAAT

AGCACACGTCTGAACTCTTTCCCTACACGACGCTCTTCCGATCTNNNGAGGTTGCATTGAGCCGAAAT

AGCACACGTCTGAACTCTTTCCCTACACGACGCTCTTCCGATCTNNNGGTGACAGAGAAAGATTCCGT

AGCACACGTCTGAACTCTTTCCCTACACGACGCTCTTCCGATCTNNNGGTGACAGAGAAAGATTCCGT

GCACACGTCTGAACTCTTTCCCTACACGACGCTCTTCCGATCTNNNCTAGGATTTCCAAGTTGCAGGGG

GCACACGTCTGAACTCTTTCCCTACACGACGCTCTTCCGATCTNNNCTAGGATTTCCAAGTTGCAGGGG

GAGCACACGTCTGAACTCTTTCCCTACACGACGCTCTTCCGATCTNNNCATATTTGAGCAGGACTCCCA

GAGCACACGTCTGAACTCTTTCCCTACACGACGCTCTTCCGATCTNNNCATATTTGAGCAGGACTCCCA

AGCACACGTCTGAACTCTTTCCCTACACGACGCTCTTCCGATCTNNNCTCCTGATTAGCTGGGACTACA

AGCACACGTCTGAACTCTTTCCCTACACGACGCTCTTCCGATCTNNNCTCCTGATTAGCTGGGACTACA

AGAGCACACGTCTGAACTCTTTCCCTACACGACGCTCTTCCGATCTNNNGAGGTGGGAGAATCACTT

AGAGCACACGTCTGAACTCTTTCCCTACACGACGCTCTTCCGATCTNNNGAGGTGGGAGAATCACTT

AGCACACGTCTGAACTCTTTCCCTACACGACGCTCTTCCGATCTNNNAAAAAATTAGCTGGGTGTGGT

AGCACACGTCTGAACTCTTTCCCTACACGACGCTCTTCCGATCTNNNAAAAAATTAGCTGGGTGTGGT

AGCACACGTCTGAACTCTTTCCCTACACGACGCTCTTCCGATCTNNNAATGCTCTACATGCAGGCTTAC

AGCACACGTCTGAACTCTTTCCCTACACGACGCTCTTCCGATCTNNNAATGCTCTACATGCAGGCTTAC

GCACACGTCTGAACTCTTTCCCTACACGACGCTCTTCCGATCTNNNCCTTTTCAGTCTCCTCATCTGCAC
GCACACGTCTGAACTCTTTCCCTACACGACGCTCTTCCGATCTNNNCCTTTTCAGTCTCCTCATCTGCAC
GAGCACACGTCTGAACTCTTTCCCTACACGACGCTCTTCCGATCTNNNGTATATTGCATACCTGATCCA
GAGCACACGTCTGAACTCTTTCCCTACACGACGCTCTTCCGATCTNNNGTATATTGCATACCTGATCCA
GAGCACACGTCTGAACTCTTTCCCTACACGACGCTCTTCCGATCTNNNTTGCCCAAGGTACTGCTGATT
GAGCACACGTCTGAACTCTTTCCCTACACGACGCTCTTCCGATCTNNNTTGCCCAAGGTACTGCTGATT
AGCACACGTCTGAACTCTTTCCCTACACGACGCTCTTCCGATCTNNNGAAGGCTGAGGCAGGAGAAT
AGCACACGTCTGAACTCTTTCCCTACACGACGCTCTTCCGATCTNNNGAAGGCTGAGGCAGGAGAAT
GCACACGTCTGAACTCTTTCCCTACACGACGCTCTTCCGATCTNNNGGAGGAGGAGGAACAGAACA
GCACACGTCTGAACTCTTTCCCTACACGACGCTCTTCCGATCTNNNGGAGGAGGAGGAACAGAACA
GAGCACACGTCTGAACTCTTTCCCTACACGACGCTCTTCCGATCTNNNATATACTCTGTGGCAGGCAT
GAGCACACGTCTGAACTCTTTCCCTACACGACGCTCTTCCGATCTNNNATATACTCTGTGGCAGGCAT
AGCACACGTCTGAACTCTTTCCCTACACGACGCTCTTCCGATCTNNNGGCCACAGAAGAAACAAAAA
AGCACACGTCTGAACTCTTTCCCTACACGACGCTCTTCCGATCTNNNGGCCACAGAAGAAACAAAAA
GAGCACACGTCTGAACTCTTTCCCTACACGACGCTCTTCCGATCTNNNGGCCACAAGTCCTTTAGGAG
GAGCACACGTCTGAACTCTTTCCCTACACGACGCTCTTCCGATCTNNNGGCCACAAGTCCTTTAGGAG
GCACACGTCTGAACTCTTTCCCTACACGACGCTCTTCCGATCTNNNTTTGATGCTGCTTTCCCAACTTC
GCACACGTCTGAACTCTTTCCCTACACGACGCTCTTCCGATCTNNNTTTGATGCTGCTTTCCCAACTTC
GAGCACACGTCTGAACTCTTTCCCTACACGACGCTCTTCCGATCTNNNTAAAGAGTCAAGACCCATCCG
GAGCACACGTCTGAACTCTTTCCCTACACGACGCTCTTCCGATCTNNNTAAAGAGTCAAGACCCATCCG
GCACACGTCTGAACTCTTTCCCTACACGACGCTCTTCCGATCTNNNGGACAAAAAATCTTAAGTGTACT
GCACACGTCTGAACTCTTTCCCTACACGACGCTCTTCCGATCTNNNGGACAAAAAATCTTAAGTGTACT
AGAGCACACGTCTGAACTCTTTCCCTACACGACGCTCTTCCGATCTNNNTAGTGCCCCACCATAGAAGA
AGAGCACACGTCTGAACTCTTTCCCTACACGACGCTCTTCCGATCTNNNTAGTGCCCCACCATAGAAGA
AGAGCACACGTCTGAACTCTTTCCCTACACGACGCTCTTCCGATCTNNNGCCTGAGAATTGCTTGAAA
AGAGCACACGTCTGAACTCTTTCCCTACACGACGCTCTTCCGATCTNNNGCCTGAGAATTGCTTGAAA
GAGCACACGTCTGAACTCTTTCCCTACACGACGCTCTTCCGATCTNNNTTTAGTAGAGACGGGGTTTC
GAGCACACGTCTGAACTCTTTCCCTACACGACGCTCTTCCGATCTNNNTTTAGTAGAGACGGGGTTTC
AGCACACGTCTGAACTCTTTCCCTACACGACGCTCTTCCGATCTNNNTGAGTTGAGAGGATCACTTGA
AGCACACGTCTGAACTCTTTCCCTACACGACGCTCTTCCGATCTNNNTGAGTTGAGAGGATCACTTGA
GCACACGTCTGAACTCTTTCCCTACACGACGCTCTTCCGATCTNNNTTTCCTCTCTGCTGTCTGATGAC
GCACACGTCTGAACTCTTTCCCTACACGACGCTCTTCCGATCTNNNTTTCCTCTCTGCTGTCTGATGAC
GCACACGTCTGAACTCTTTCCCTACACGACGCTCTTCCGATCTNNNGTAATCCCAGCTACTCAAGGAG
GCACACGTCTGAACTCTTTCCCTACACGACGCTCTTCCGATCTNNNGTAATCCCAGCTACTCAAGGAG
GAGCACACGTCTGAACTCTTTCCCTACACGACGCTCTTCCGATCTNNNAGGAGCTCCCTCTTTATATGG
GAGCACACGTCTGAACTCTTTCCCTACACGACGCTCTTCCGATCTNNNAGGAGCTCCCTCTTTATATGG
GCACACGTCTGAACTCTTTCCCTACACGACGCTCTTCCGATCTNNNCTGGGTTGAAGTGATTCTCTTG
GCACACGTCTGAACTCTTTCCCTACACGACGCTCTTCCGATCTNNNCTGGGTTGAAGTGATTCTCTTG
AGAGCACACGTCTGAACTCTTTCCCTACACGACGCTCTTCCGATCTNNNGGAGAATCACCTGAACATG
AGAGCACACGTCTGAACTCTTTCCCTACACGACGCTCTTCCGATCTNNNGGAGAATCACCTGAACATG
GCACACGTCTGAACTCTTTCCCTACACGACGCTCTTCCGATCTNNNCGATCTGTGTTCTGACATTTCCCC
GCACACGTCTGAACTCTTTCCCTACACGACGCTCTTCCGATCTNNNCGATCTGTGTTCTGACATTTCCCC
AGCACACGTCTGAACTCTTTCCCTACACGACGCTCTTCCGATCTNNNGCTTACATGTGCTAGGGTAGC

AGCACACGTCTGAACTCTTTCCCTACACGACGCTCTTCCGATCTNNNGCTTACATGTGCTAGGGTAGC
AGCACACGTCTGAACTCTTTCCCTACACGACGCTCTTCCGATCTNNNTTTTCAGGCTCTCATGTTTCTT
AGCACACGTCTGAACTCTTTCCCTACACGACGCTCTTCCGATCTNNNTTTTCAGGCTCTCATGTTTCTT
GCACACGTCTGAACTCTTTCCCTACACGACGCTCTTCCGATCTNNNAATAATGGGCTCCAACTCCATCC
GCACACGTCTGAACTCTTTCCCTACACGACGCTCTTCCGATCTNNNAATAATGGGCTCCAACTCCATCC
AGCACACGTCTGAACTCTTTCCCTACACGACGCTCTTCCGATCTNNNCCACCCCCCAAAAAAAAGAA
AGCACACGTCTGAACTCTTTCCCTACACGACGCTCTTCCGATCTNNNCCACCCCCCAAAAAAAAGAA
GAGCACACGTCTGAACTCTTTCCCTACACGACGCTCTTCCGATCTNNNAAAATACAAAATTAGCCGG
GAGCACACGTCTGAACTCTTTCCCTACACGACGCTCTTCCGATCTNNNAAAATACAAAATTAGCCGG
GAGCACACGTCTGAACTCTTTCCCTACACGACGCTCTTCCGATCTNNNTTGCTTCCCTCTTTCCCTCTTAC
GAGCACACGTCTGAACTCTTTCCCTACACGACGCTCTTCCGATCTNNNTTGCTTCCCTCTTTCCCTCTTAC
GCACACGTCTGAACTCTTTCCCTACACGACGCTCTTCCGATCTNNNATACTCTTTCTGACTGATGACTA
GCACACGTCTGAACTCTTTCCCTACACGACGCTCTTCCGATCTNNNATACTCTTTCTGACTGATGACTA
GAGCACACGTCTGAACTCTTTCCCTACACGACGCTCTTCCGATCTNNNGTGTAATCCCAGCTACTCAG
GAGCACACGTCTGAACTCTTTCCCTACACGACGCTCTTCCGATCTNNNGTGTAATCCCAGCTACTCAG
GCACACGTCTGAACTCTTTCCCTACACGACGCTCTTCCGATCTNNNACCACGTCCAGCCAAAAAAAAA
GCACACGTCTGAACTCTTTCCCTACACGACGCTCTTCCGATCTNNNACCACGTCCAGCCAAAAAAAAA
CACACGTCTGAACTCTTTCCCTACACGACGCTCTTCCGATCTNNNAGGGCCTTCTCTTTGACTTCTTCC
CACACGTCTGAACTCTTTCCCTACACGACGCTCTTCCGATCTNNNAGGGCCTTCTCTTTGACTTCTTCC
AGCACACGTCTGAACTCTTTCCCTACACGACGCTCTTCCGATCTNNNTTTCATTGTCAGTGTACAGAAA
AGCACACGTCTGAACTCTTTCCCTACACGACGCTCTTCCGATCTNNNTTTCATTGTCAGTGTACAGAAA
AGCACACGTCTGAACTCTTTCCCTACACGACGCTCTTCCGATCTNNNGCCAACATAGGGAAACATGGT
AGCACACGTCTGAACTCTTTCCCTACACGACGCTCTTCCGATCTNNNGCCAACATAGGGAAACATGGT
AGCACACGTCTGAACTCTTTCCCTACACGACGCTCTTCCGATCTNNNCTGGCTCAAGTTCAGTCAGACT
AGCACACGTCTGAACTCTTTCCCTACACGACGCTCTTCCGATCTNNNCTGGCTCAAGTTCAGTCAGACT
GAGCACACGTCTGAACTCTTTCCCTACACGACGCTCTTCCGATCTNNNTGCTCACATCTGTAATCCCAA
GAGCACACGTCTGAACTCTTTCCCTACACGACGCTCTTCCGATCTNNNTGCTCACATCTGTAATCCCAA
GAGCACACGTCTGAACTCTTTCCCTACACGACGCTCTTCCGATCTNNNGGGGTTCTGAGAGCTCCAA
GAGCACACGTCTGAACTCTTTCCCTACACGACGCTCTTCCGATCTNNNGGGGTTCTGAGAGCTCCAA
GAGCACACGTCTGAACTCTTTCCCTACACGACGCTCTTCCGATCTNNNATAATCTCAGCTACTTGGGA
GAGCACACGTCTGAACTCTTTCCCTACACGACGCTCTTCCGATCTNNNATAATCTCAGCTACTTGGGA
AGCACACGTCTGAACTCTTTCCCTACACGACGCTCTTCCGATCTNNNGCCCTTTGGATTTAGGCAGTTT
AGCACACGTCTGAACTCTTTCCCTACACGACGCTCTTCCGATCTNNNGCCCTTTGGATTTAGGCAGTTT
GAGCACACGTCTGAACTCTTTCCCTACACGACGCTCTTCCGATCTNNNTGGGGAATACATGACGTTGA
GAGCACACGTCTGAACTCTTTCCCTACACGACGCTCTTCCGATCTNNNTGGGGAATACATGACGTTGA
AGCACACGTCTGAACTCTTTCCCTACACGACGCTCTTCCGATCTNNNTTCTAGCAGGCCTTCTTGTACT
AGCACACGTCTGAACTCTTTCCCTACACGACGCTCTTCCGATCTNNNTTCTAGCAGGCCTTCTTGTACT
GCACACGTCTGAACTCTTTCCCTACACGACGCTCTTCCGATCTNNNTGCTGTGCAGAAGCTTTTTAGT
GCACACGTCTGAACTCTTTCCCTACACGACGCTCTTCCGATCTNNNTGCTGTGCAGAAGCTTTTTAGT
CACACGTCTGAACTCTTTCCCTACACGACGCTCTTCCGATCTNNNGAAGCAGCTTTCAGATCTGGTGA
CACACGTCTGAACTCTTTCCCTACACGACGCTCTTCCGATCTNNNGAAGCAGCTTTCAGATCTGGTGA
AGCACACGTCTGAACTCTTTCCCTACACGACGCTCTTCCGATCTNNNAACATGGCAAAACCCCATCTCT
AGCACACGTCTGAACTCTTTCCCTACACGACGCTCTTCCGATCTNNNAACATGGCAAAACCCCATCTCT

AGCACACGTCTGAACTCTTTCCCTACACGACGCTCTTCCGATCTNNNAGGATATTCTTCCTGCTTAATGC
AGCACACGTCTGAACTCTTTCCCTACACGACGCTCTTCCGATCTNNNAGGATATTCTTCCTGCTTAATGC
AGCACACGTCTGAACTCTTTCCCTACACGACGCTCTTCCGATCTNNNCCAGCGAGTCTTTCTCCAGTAAA
AGCACACGTCTGAACTCTTTCCCTACACGACGCTCTTCCGATCTNNNCCAGCGAGTCTTTCTCCAGTAAA
GAGCACACGTCTGAACTCTTTCCCTACACGACGCTCTTCCGATCTNNNAGAGGATCATTTGAACCCAGCA
GAGCACACGTCTGAACTCTTTCCCTACACGACGCTCTTCCGATCTNNNAGAGGATCATTTGAACCCAGCA
AGAGCACACGTCTGAACTCTTTCCCTACACGACGCTCTTCCGATCTNNNAAAAAATTAGCCAGGTGTGG
AGAGCACACGTCTGAACTCTTTCCCTACACGACGCTCTTCCGATCTNNNAAAAAATTAGCCAGGTGTGG
GAGCACACGTCTGAACTCTTTCCCTACACGACGCTCTTCCGATCTNNNTTGGATTTGTGTCTCTGCATC
GAGCACACGTCTGAACTCTTTCCCTACACGACGCTCTTCCGATCTNNNTTGGATTTGTGTCTCTGCATC
GAGCACACGTCTGAACTCTTTCCCTACACGACGCTCTTCCGATCTNNNAACTGCTGTTCCAGGTCTTCTT
GAGCACACGTCTGAACTCTTTCCCTACACGACGCTCTTCCGATCTNNNAACTGCTGTTCCAGGTCTTCTT
GAGCACACGTCTGAACTCTTTCCCTACACGACGCTCTTCCGATCTNNNAAAGCAGGAGCAAGAGAGAG
GAGCACACGTCTGAACTCTTTCCCTACACGACGCTCTTCCGATCTNNNAAAGCAGGAGCAAGAGAGAG
AGAGCACACGTCTGAACTCTTTCCCTACACGACGCTCTTCCGATCTNNNCAGGCAACAGAACAAGACTC
AGAGCACACGTCTGAACTCTTTCCCTACACGACGCTCTTCCGATCTNNNCAGGCAACAGAACAAGACTC
AGCACACGTCTGAACTCTTTCCCTACACGACGCTCTTCCGATCTNNNTCATATGCTTATCAGCCACGTGT
AGCACACGTCTGAACTCTTTCCCTACACGACGCTCTTCCGATCTNNNTCATATGCTTATCAGCCACGTGT
AGCACACGTCTGAACTCTTTCCCTACACGACGCTCTTCCGATCTNNNCTTGGTGGCATGTGACTGTAAT
AGCACACGTCTGAACTCTTTCCCTACACGACGCTCTTCCGATCTNNNCTTGGTGGCATGTGACTGTAAT
AGCACACGTCTGAACTCTTTCCCTACACGACGCTCTTCCGATCTNNNTGTCCTTTTTGTAGAGACACGG
AGCACACGTCTGAACTCTTTCCCTACACGACGCTCTTCCGATCTNNNTGTCCTTTTTGTAGAGACACGG
AGCACACGTCTGAACTCTTTCCCTACACGACGCTCTTCCGATCTNNNGAGTTTGCAGTGAGCCAAGATT
AGCACACGTCTGAACTCTTTCCCTACACGACGCTCTTCCGATCTNNNGAGTTTGCAGTGAGCCAAGATT
AAGAGCACACGTCTGAACTCTTTCCCTACACGACGCTCTTCCGATCTNNNTCAGCAAGGTCTTTATAAC
AAGAGCACACGTCTGAACTCTTTCCCTACACGACGCTCTTCCGATCTNNNTCAGCAAGGTCTTTATAAC
GAGCACACGTCTGAACTCTTTCCCTACACGACGCTCTTCCGATCTNNNACTCATTTAGGTCTCGGTTTCC
GAGCACACGTCTGAACTCTTTCCCTACACGACGCTCTTCCGATCTNNNACTCATTTAGGTCTCGGTTTCC
AGCACACGTCTGAACTCTTTCCCTACACGACGCTCTTCCGATCTNNNAGAAACTCCTACAATACAGGGCC
AGCACACGTCTGAACTCTTTCCCTACACGACGCTCTTCCGATCTNNNAGAAACTCCTACAATACAGGGCC
GAGCACACGTCTGAACTCTTTCCCTACACGACGCTCTTCCGATCTNNNCACCTGTACTCCCAGCTATTCA
GAGCACACGTCTGAACTCTTTCCCTACACGACGCTCTTCCGATCTNNNCACCTGTACTCCCAGCTATTCA
GAGCACACGTCTGAACTCTTTCCCTACACGACGCTCTTCCGATCTNNNGGCTTATACCTGTAATCCCTGC
GAGCACACGTCTGAACTCTTTCCCTACACGACGCTCTTCCGATCTNNNGGCTTATACCTGTAATCCCTGC
AGCACACGTCTGAACTCTTTCCCTACACGACGCTCTTCCGATCTNNNCTGAGACAGGAGAATGGCTTG
AGCACACGTCTGAACTCTTTCCCTACACGACGCTCTTCCGATCTNNNCTGAGACAGGAGAATGGCTTG
GAGCACACGTCTGAACTCTTTCCCTACACGACGCTCTTCCGATCTNNNGAGGTGGGAGGATCACTTGA
GAGCACACGTCTGAACTCTTTCCCTACACGACGCTCTTCCGATCTNNNGAGGTGGGAGGATCACTTGA
GAGCACACGTCTGAACTCTTTCCCTACACGACGCTCTTCCGATCTNNNATACAAAAATTGGCTGGGTGT
GAGCACACGTCTGAACTCTTTCCCTACACGACGCTCTTCCGATCTNNNATACAAAAATTGGCTGGGTGT
GAGCACACGTCTGAACTCTTTCCCTACACGACGCTCTTCCGATCTNNNACTGCATGGTAGAAGGAAGC
GAGCACACGTCTGAACTCTTTCCCTACACGACGCTCTTCCGATCTNNNACTGCATGGTAGAAGGAAGC
AGAGCACACGTCTGAACTCTTTCCCTACACGACGCTCTTCCGATCTNNNGCGCAACTGTAATCTCAGCTA

GAGCACACGTCTGAACTCTTTCCCTACACGACGCTCTTCCGATCTNNNGCGCAACTGTAATCTCAGCT
GCACACGTCTGAACTCTTTCCCTACACGACGCTCTTCCGATCTNNNTAAGGGGAGCCTGTAAGAGGA
GCACACGTCTGAACTCTTTCCCTACACGACGCTCTTCCGATCTNNNTAAGGGGAGCCTGTAAGAGGA
AGCACACGTCTGAACTCTTTCCCTACACGACGCTCTTCCGATCTNNNTGCCTGTAATCCCAGATACTTG
AGCACACGTCTGAACTCTTTCCCTACACGACGCTCTTCCGATCTNNNTGCCTGTAATCCCAGATACTTG
CACACGTCTGAACTCTTTCCCTACACGACGCTCTTCCGATCTNNNACCCAGAATACCCCTGACTACATC
CACACGTCTGAACTCTTTCCCTACACGACGCTCTTCCGATCTNNNACCCAGAATACCCCTGACTACATC
GCACACGTCTGAACTCTTTCCCTACACGACGCTCTTCCGATCTNNNCTTTGTCACATCCTAGCGCTTTC
GCACACGTCTGAACTCTTTCCCTACACGACGCTCTTCCGATCTNNNCTTTGTCACATCCTAGCGCTTTC
AGAGCACACGTCTGAACTCTTTCCCTACACGACGCTCTTCCGATCTNNNGTCTGTGCTTGGTTTTCTTC
AGAGCACACGTCTGAACTCTTTCCCTACACGACGCTCTTCCGATCTNNNGTCTGTGCTTGGTTTTCTTC
GCACACGTCTGAACTCTTTCCCTACACGACGCTCTTCCGATCTNNNTTCATAGCTTCCCTTTCCCATCCC
GCACACGTCTGAACTCTTTCCCTACACGACGCTCTTCCGATCTNNNTTCATAGCTTCCCTTTCCCATCCC
CACACGTCTGAACTCTTTCCCTACACGACGCTCTTCCGATCTNNNTTAATGTAAACGGTTATCTTTCCC
CACACGTCTGAACTCTTTCCCTACACGACGCTCTTCCGATCTNNNTTAATGTAAACGGTTATCTTTCCC
GAGCACACGTCTGAACTCTTTCCCTACACGACGCTCTTCCGATCTNNNGCTGTAAGATCACTGCCTATG
GAGCACACGTCTGAACTCTTTCCCTACACGACGCTCTTCCGATCTNNNGCTGTAAGATCACTGCCTATG
AAGAGCACACGTCTGAACTCTTTCCCTACACGACGCTCTTCCGATCTNNNGCGCCTGGCTGAAAATGA
AAGAGCACACGTCTGAACTCTTTCCCTACACGACGCTCTTCCGATCTNNNGCGCCTGGCTGAAAATGA
GAGCACACGTCTGAACTCTTTCCCTACACGACGCTCTTCCGATCTNNNTTACACATTCTGAGACCTGGC
GAGCACACGTCTGAACTCTTTCCCTACACGACGCTCTTCCGATCTNNNTTACACATTCTGAGACCTGGC
AGCACACGTCTGAACTCTTTCCCTACACGACGCTCTTCCGATCTNNNAAAAGTTGGGTAGATGGAGTC
AGCACACGTCTGAACTCTTTCCCTACACGACGCTCTTCCGATCTNNNAAAAGTTGGGTAGATGGAGTC
GCACACGTCTGAACTCTTTCCCTACACGACGCTCTTCCGATCTNNNAGCCTCTCCAGTAGCTGAGATA
GCACACGTCTGAACTCTTTCCCTACACGACGCTCTTCCGATCTNNNAGCCTCTCCAGTAGCTGAGATA
AGCACACGTCTGAACTCTTTCCCTACACGACGCTCTTCCGATCTNNNGCCTCCTTCTCTCTGATGTTCA
AGCACACGTCTGAACTCTTTCCCTACACGACGCTCTTCCGATCTNNNGCCTCCTTCTCTCTGATGTTCA
AGCACACGTCTGAACTCTTTCCCTACACGACGCTCTTCCGATCTNNNCACTACTTCTGGCTAACGTCTC
AGCACACGTCTGAACTCTTTCCCTACACGACGCTCTTCCGATCTNNNCACTACTTCTGGCTAACGTCTC
AGCACACGTCTGAACTCTTTCCCTACACGACGCTCTTCCGATCTNNNAAATTAGTTGCCCAAGGTTGC
AGCACACGTCTGAACTCTTTCCCTACACGACGCTCTTCCGATCTNNNAAATTAGTTGCCCAAGGTTGC
AGCACACGTCTGAACTCTTTCCCTACACGACGCTCTTCCGATCTNNNCTACTGCCCTTACCTTCTGGTTT
AGCACACGTCTGAACTCTTTCCCTACACGACGCTCTTCCGATCTNNNCTACTGCCCTTACCTTCTGGTTT
AGAGCACACGTCTGAACTCTTTCCCTACACGACGCTCTTCCGATCTNNNCTGAGATTGAGCCATTGC
AGAGCACACGTCTGAACTCTTTCCCTACACGACGCTCTTCCGATCTNNNCTGAGATTGAGCCATTGC
GGAAGAGCACACGTCTGAACTCTTTCCCTACACGACGCTCTTCCGATCTNNNATCATGAGTTCTGAGA
GGAAGAGCACACGTCTGAACTCTTTCCCTACACGACGCTCTTCCGATCTNNNATCATGAGTTCTGAGA
AGCACACGTCTGAACTCTTTCCCTACACGACGCTCTTCCGATCTNNNTATAAACAGAATGGGCGGATC
AGCACACGTCTGAACTCTTTCCCTACACGACGCTCTTCCGATCTNNNTATAAACAGAATGGGCGGATC
AGCACACGTCTGAACTCTTTCCCTACACGACGCTCTTCCGATCTNNNCTCTGGTTATTATCCCTGCCCT
AGCACACGTCTGAACTCTTTCCCTACACGACGCTCTTCCGATCTNNNCTCTGGTTATTATCCCTGCCCT
AGCACACGTCTGAACTCTTTCCCTACACGACGCTCTTCCGATCTNNNAAGGCTAATGGATTCCAAGAA
AGCACACGTCTGAACTCTTTCCCTACACGACGCTCTTCCGATCTNNNAAGGCTAATGGATTCCAAGAA

AGAGCACACGTCTGAACTCTTTCCCTACACGACGCTCTTCCGATCTNNNTGCCACAACCTATAATCTTAC

AGAGCACACGTCTGAACTCTTTCCCTACACGACGCTCTTCCGATCTNNNTGCCACAACCTATAATCTTAC

GAGCACACGTCTGAACTCTTTCCCTACACGACGCTCTTCCGATCTNNNGGTGAGACCTGGTCTATAGCA

GAGCACACGTCTGAACTCTTTCCCTACACGACGCTCTTCCGATCTNNNGGTGAGACCTGGTCTATAGCA

AGAGCACACGTCTGAACTCTTTCCCTACACGACGCTCTTCCGATCTNNNGTCAGGGTGGTGGTTGTTA

AGAGCACACGTCTGAACTCTTTCCCTACACGACGCTCTTCCGATCTNNNGTCAGGGTGGTGGTTGTTA

GAGCACACGTCTGAACTCTTTCCCTACACGACGCTCTTCCGATCTNNNCTGTAGTCCCAGCTACTCAGGA

GAGCACACGTCTGAACTCTTTCCCTACACGACGCTCTTCCGATCTNNNCTGTAGTCCCAGCTACTCAGGA

AGCACACGTCTGAACTCTTTCCCTACACGACGCTCTTCCGATCTNNNGCCTCTGTATTCACTTGCATGTT

AGCACACGTCTGAACTCTTTCCCTACACGACGCTCTTCCGATCTNNNGCCTCTGTATTCACTTGCATGTT

AGCACACGTCTGAACTCTTTCCCTACACGACGCTCTTCCGATCTNNNTAGGCTGTGTAGGTAAAATTTA

AGCACACGTCTGAACTCTTTCCCTACACGACGCTCTTCCGATCTNNNTAGGCTGTGTAGGTAAAATTTA

GCACACGTCTGAACTCTTTCCCTACACGACGCTCTTCCGATCTNNNAGGCAGGAGAATCACTTGAACTT

GCACACGTCTGAACTCTTTCCCTACACGACGCTCTTCCGATCTNNNAGGCAGGAGAATCACTTGAACTT

AGCACACGTCTGAACTCTTTCCCTACACGACGCTCTTCCGATCTNNNTCCCAGAGAAACAGAACCAATGT

AGCACACGTCTGAACTCTTTCCCTACACGACGCTCTTCCGATCTNNNTCCCAGAGAAACAGAACCAATGT

AGCACACGTCTGAACTCTTTCCCTACACGACGCTCTTCCGATCTNNNTCCTGACCTCAGATGAGATGAC

AGCACACGTCTGAACTCTTTCCCTACACGACGCTCTTCCGATCTNNNTCCTGACCTCAGATGAGATGAC

AGCACACGTCTGAACTCTTTCCCTACACGACGCTCTTCCGATCTNNNATTACAGAAGGGTGAGGTGAT

AGCACACGTCTGAACTCTTTCCCTACACGACGCTCTTCCGATCTNNNATTACAGAAGGGTGAGGTGAT

AGCACACGTCTGAACTCTTTCCCTACACGACGCTCTTCCGATCTNNNGTAATCCCAGCTACTCAGGAGT

AGCACACGTCTGAACTCTTTCCCTACACGACGCTCTTCCGATCTNNNGTAATCCCAGCTACTCAGGAGT

GCACACGTCTGAACTCTTTCCCTACACGACGCTCTTCCGATCTNNNAGCTCAGGAATTGTGAGTGTTCA

GCACACGTCTGAACTCTTTCCCTACACGACGCTCTTCCGATCTNNNAGCTCAGGAATTGTGAGTGTTCA

CACACGTCTGAACTCTTTCCCTACACGACGCTCTTCCGATCTNNNTTTGGAGAAGCAAGATGTGTGTG

CACACGTCTGAACTCTTTCCCTACACGACGCTCTTCCGATCTNNNTTTGGAGAAGCAAGATGTGTGTG

AGCACACGTCTGAACTCTTTCCCTACACGACGCTCTTCCGATCTNNNCCTATAGTCCCAGCTACTCGGG

AGCACACGTCTGAACTCTTTCCCTACACGACGCTCTTCCGATCTNNNCCTATAGTCCCAGCTACTCGGG

GAGCACACGTCTGAACTCTTTCCCTACACGACGCTCTTCCGATCTNNNAATACAAAAATTAGCCGGGCG

GAGCACACGTCTGAACTCTTTCCCTACACGACGCTCTTCCGATCTNNNAATACAAAAATTAGCCGGGCG

GAGCACACGTCTGAACTCTTTCCCTACACGACGCTCTTCCGATCTNNNAAATCCCAACTATGTGGAAG

GAGCACACGTCTGAACTCTTTCCCTACACGACGCTCTTCCGATCTNNNAAATCCCAACTATGTGGAAG

AGAGCACACGTCTGAACTCTTTCCCTACACGACGCTCTTCCGATCTNNNGTTTCAAGCGATTCTTCTGCC

AGAGCACACGTCTGAACTCTTTCCCTACACGACGCTCTTCCGATCTNNNGTTTCAAGCGATTCTTCTGCC

AGCACACGTCTGAACTCTTTCCCTACACGACGCTCTTCCGATCTNNNATATCGCATGGAAGACTGTTGG

AGCACACGTCTGAACTCTTTCCCTACACGACGCTCTTCCGATCTNNNATATCGCATGGAAGACTGTTGG

AGCACACGTCTGAACTCTTTCCCTACACGACGCTCTTCCGATCTNNNCTCAGCATATTTTTTCATCTCTTC

AGCACACGTCTGAACTCTTTCCCTACACGACGCTCTTCCGATCTNNNCTCAGCATATTTTTTCATCTCTTC

GAGCACACGTCTGAACTCTTTCCCTACACGACGCTCTTCCGATCTNNNATGCTTGTAATCCCAGCTCCTC

GAGCACACGTCTGAACTCTTTCCCTACACGACGCTCTTCCGATCTNNNATGCTTGTAATCCCAGCTCCTC

AGCACACGTCTGAACTCTTTCCCTACACGACGCTCTTCCGATCTNNNAAATGATGGTGGGATAACTGG

AGCACACGTCTGAACTCTTTCCCTACACGACGCTCTTCCGATCTNNNAAATGATGGTGGGATAACTGG

AGCACACGTCTGAACTCTTTCCCTACACGACGCTCTTCCGATCTNNNTGGAGGAAGACAAGAAAACAC

AGCACACGTCTGAACTCTTTCCCTACACGACGCTCTTCCGATCTNNNTGGAGGAAGACAAGAAAACA
GAGCACACGTCTGAACTCTTTCCCTACACGACGCTCTTCCGATCTNNNCCACCACACCCAGCTAATTTTT
GAGCACACGTCTGAACTCTTTCCCTACACGACGCTCTTCCGATCTNNNCCACCACACCCAGCTAATTTTT
AGCACACGTCTGAACTCTTTCCCTACACGACGCTCTTCCGATCTNNNTCAGCAATTGGTAGAAACAGT
AGCACACGTCTGAACTCTTTCCCTACACGACGCTCTTCCGATCTNNNTCAGCAATTGGTAGAAACAGT
AGCACACGTCTGAACTCTTTCCCTACACGACGCTCTTCCGATCTNNNCTGCTCTCTCTAGGAGTCTCAG
AGCACACGTCTGAACTCTTTCCCTACACGACGCTCTTCCGATCTNNNCTGCTCTCTCTAGGAGTCTCAG
AGCACACGTCTGAACTCTTTCCCTACACGACGCTCTTCCGATCTNNNCATGCCTGTAATCCCAGCTACT
AGCACACGTCTGAACTCTTTCCCTACACGACGCTCTTCCGATCTNNNCATGCCTGTAATCCCAGCTACT
GAGCACACGTCTGAACTCTTTCCCTACACGACGCTCTTCCGATCTNNNGCTAGAATCAGGTGTTAGGT
GAGCACACGTCTGAACTCTTTCCCTACACGACGCTCTTCCGATCTNNNGCTAGAATCAGGTGTTAGGT
GAGCACACGTCTGAACTCTTTCCCTACACGACGCTCTTCCGATCTNNNAAACATCCTATTTCATGGGGG
GAGCACACGTCTGAACTCTTTCCCTACACGACGCTCTTCCGATCTNNNAAACATCCTATTTCATGGGGG
AGCACACGTCTGAACTCTTTCCCTACACGACGCTCTTCCGATCTNNNCTCCCAGGTTCATGCCATTCTC
AGCACACGTCTGAACTCTTTCCCTACACGACGCTCTTCCGATCTNNNCTCCCAGGTTCATGCCATTCTC
AGCACACGTCTGAACTCTTTCCCTACACGACGCTCTTCCGATCTNNNTGAGGATTTTCACAACACAACC
AGCACACGTCTGAACTCTTTCCCTACACGACGCTCTTCCGATCTNNNTGAGGATTTTCACAACACAACC
AGCACACGTCTGAACTCTTTCCCTACACGACGCTCTTCCGATCTNNNACCTATGTAGTCCCAGCTACTCA
AGCACACGTCTGAACTCTTTCCCTACACGACGCTCTTCCGATCTNNNACCTATGTAGTCCCAGCTACTCA
AGCACACGTCTGAACTCTTTCCCTACACGACGCTCTTCCGATCTNNNCTGAAGCAGGAGAATCACTGG
AGCACACGTCTGAACTCTTTCCCTACACGACGCTCTTCCGATCTNNNCTGAAGCAGGAGAATCACTGG
AGCACACGTCTGAACTCTTTCCCTACACGACGCTCTTCCGATCTNNNGACCATCGACAAAGCAAAAAG
AGCACACGTCTGAACTCTTTCCCTACACGACGCTCTTCCGATCTNNNGACCATCGACAAAGCAAAAAG
AGCACACGTCTGAACTCTTTCCCTACACGACGCTCTTCCGATCTNNNTGCATAGAATAGGGAATGGGA
AGCACACGTCTGAACTCTTTCCCTACACGACGCTCTTCCGATCTNNNTGCATAGAATAGGGAATGGGA
AGCACACGTCTGAACTCTTTCCCTACACGACGCTCTTCCGATCTNNNTTCCCTAATTGAAGAGAGCCAC
AGCACACGTCTGAACTCTTTCCCTACACGACGCTCTTCCGATCTNNNTTCCCTAATTGAAGAGAGCCAC
AGCACACGTCTGAACTCTTTCCCTACACGACGCTCTTCCGATCTNNNAATTGCTGGGATTACAGGCGT
AGCACACGTCTGAACTCTTTCCCTACACGACGCTCTTCCGATCTNNNAATTGCTGGGATTACAGGCGT
AGAGCACACGTCTGAACTCTTTCCCTACACGACGCTCTTCCGATCTNNNCTGTGCTGGAGGATGTTGA
AGAGCACACGTCTGAACTCTTTCCCTACACGACGCTCTTCCGATCTNNNCTGTGCTGGAGGATGTTGA
ACACGTCTGAACTCTTTCCCTACACGACGCTCTTCCGATCTNNNCCACATTTCATGCACCAGGTATGCA
ACACGTCTGAACTCTTTCCCTACACGACGCTCTTCCGATCTNNNCCACATTTCATGCACCAGGTATGCA
AGCACACGTCTGAACTCTTTCCCTACACGACGCTCTTCCGATCTNNNGCAACTGTGAGAACAAAAGG
AGCACACGTCTGAACTCTTTCCCTACACGACGCTCTTCCGATCTNNNGCAACTGTGAGAACAAAAGG
AGCACACGTCTGAACTCTTTCCCTACACGACGCTCTTCCGATCTNNNAAAAGAAGGAGGCCCGTATG
AGCACACGTCTGAACTCTTTCCCTACACGACGCTCTTCCGATCTNNNAAAAGAAGGAGGCCCGTATG
AGCACACGTCTGAACTCTTTCCCTACACGACGCTCTTCCGATCTNNNGTGGTTCTATCCATGCATTGTG
AGCACACGTCTGAACTCTTTCCCTACACGACGCTCTTCCGATCTNNNGTGGTTCTATCCATGCATTGTG
AGCACACGTCTGAACTCTTTCCCTACACGACGCTCTTCCGATCTNNNGGTTGGTGTCACTTTGGTGTA
AGCACACGTCTGAACTCTTTCCCTACACGACGCTCTTCCGATCTNNNGGTTGGTGTCACTTTGGTGTA
GAGCACACGTCTGAACTCTTTCCCTACACGACGCTCTTCCGATCTNNNGTACTGATTTTCTTTCTTTTGG
AGCACACGTCTGAACTCTTTCCCTACACGACGCTCTTCCGATCTNNNGTACTGATTTTCTTTCTTTTGG

GAGCACACGTCTGAACTCTTTCCCTACACGACGCTCTTCCGATCTNNNATGCCTGTAATCCCAGCTATT
GAGCACACGTCTGAACTCTTTCCCTACACGACGCTCTTCCGATCTNNNATGCCTGTAATCCCAGCTATT
GAGCACACGTCTGAACTCTTTCCCTACACGACGCTCTTCCGATCTNNNCAAGAGCGAATCTCCGTCTCAT
GAGCACACGTCTGAACTCTTTCCCTACACGACGCTCTTCCGATCTNNNCAAGAGCGAATCTCCGTCTCAT
AGCACACGTCTGAACTCTTTCCCTACACGACGCTCTTCCGATCTNNNAACATTGTTTGCACACTTTTGCC
AGCACACGTCTGAACTCTTTCCCTACACGACGCTCTTCCGATCTNNNAACATTGTTTGCACACTTTTGCC
AGAGCACACGTCTGAACTCTTTCCCTACACGACGCTCTTCCGATCTNNNCTGAGGCAGAGAATTGCTT
AGAGCACACGTCTGAACTCTTTCCCTACACGACGCTCTTCCGATCTNNNCTGAGGCAGAGAATTGCTT
GAGCACACGTCTGAACTCTTTCCCTACACGACGCTCTTCCGATCTNNNAGACTGCCCAGATGCTATTGA
GAGCACACGTCTGAACTCTTTCCCTACACGACGCTCTTCCGATCTNNNAGACTGCCCAGATGCTATTGA
AGCACACGTCTGAACTCTTTCCCTACACGACGCTCTTCCGATCTNNNAAGTGAAGAGACAACCCACAGA
AGCACACGTCTGAACTCTTTCCCTACACGACGCTCTTCCGATCTNNNAAGTGAAGAGACAACCCACAGA
AGAGCACACGTCTGAACTCTTTCCCTACACGACGCTCTTCCGATCTNNNGTGTGCATCTCTCCTTGTATG
AGAGCACACGTCTGAACTCTTTCCCTACACGACGCTCTTCCGATCTNNNGTGTGCATCTCTCCTTGTATG
AGCACACGTCTGAACTCTTTCCCTACACGACGCTCTTCCGATCTNNNCACAGTGAAGGAAGATGATCC
AGCACACGTCTGAACTCTTTCCCTACACGACGCTCTTCCGATCTNNNCACAGTGAAGGAAGATGATCC
GAGCACACGTCTGAACTCTTTCCCTACACGACGCTCTTCCGATCTNNNGCATATAATCCCAGCTACTCG
GAGCACACGTCTGAACTCTTTCCCTACACGACGCTCTTCCGATCTNNNGCATATAATCCCAGCTACTCG
GAGCACACGTCTGAACTCTTTCCCTACACGACGCTCTTCCGATCTNNNACTTGCCCTACATGTGCTAACT
GAGCACACGTCTGAACTCTTTCCCTACACGACGCTCTTCCGATCTNNNACTTGCCCTACATGTGCTAACT
AGCACACGTCTGAACTCTTTCCCTACACGACGCTCTTCCGATCTNNNGTTGTCCAAACCTGTTACCTCCA
AGCACACGTCTGAACTCTTTCCCTACACGACGCTCTTCCGATCTNNNGTTGTCCAAACCTGTTACCTCCA
AGCACACGTCTGAACTCTTTCCCTACACGACGCTCTTCCGATCTNNNCAACAAGGAGAGAGATGCTGT
AGCACACGTCTGAACTCTTTCCCTACACGACGCTCTTCCGATCTNNNCAACAAGGAGAGAGATGCTGT
GAGCACACGTCTGAACTCTTTCCCTACACGACGCTCTTCCGATCTNNNTGGGAGAGTCAAAGGAAGAT
GAGCACACGTCTGAACTCTTTCCCTACACGACGCTCTTCCGATCTNNNTGGGAGAGTCAAAGGAAGAT
GAGCACACGTCTGAACTCTTTCCCTACACGACGCTCTTCCGATCTNNNTGAGGCAGGAGAATGACTTGA
GAGCACACGTCTGAACTCTTTCCCTACACGACGCTCTTCCGATCTNNNTGAGGCAGGAGAATGACTTGA
GAGCACACGTCTGAACTCTTTCCCTACACGACGCTCTTCCGATCTNNNGAAGCTCTGGGATATGATGTG
GAGCACACGTCTGAACTCTTTCCCTACACGACGCTCTTCCGATCTNNNGAAGCTCTGGGATATGATGTG
GCACACGTCTGAACTCTTTCCCTACACGACGCTCTTCCGATCTNNNTTTATATCCCTGTAGAACTGAAG
GCACACGTCTGAACTCTTTCCCTACACGACGCTCTTCCGATCTNNNTTTATATCCCTGTAGAACTGAAG
AGCACACGTCTGAACTCTTTCCCTACACGACGCTCTTCCGATCTNNNAGTCAGGGTTTTCCAGAGAAA
AGCACACGTCTGAACTCTTTCCCTACACGACGCTCTTCCGATCTNNNAGTCAGGGTTTTCCAGAGAAA
AGCACACGTCTGAACTCTTTCCCTACACGACGCTCTTCCGATCTNNNTGAGAATTACTTGAACCCGGGA
AGCACACGTCTGAACTCTTTCCCTACACGACGCTCTTCCGATCTNNNTGAGAATTACTTGAACCCGGGA
AGCACACGTCTGAACTCTTTCCCTACACGACGCTCTTCCGATCTNNNGCACAGATCAATGCACAGCCATA
AGCACACGTCTGAACTCTTTCCCTACACGACGCTCTTCCGATCTNNNGCACAGATCAATGCACAGCCATA
GAGCACACGTCTGAACTCTTTCCCTACACGACGCTCTTCCGATCTNNNTTGTTAGCCTGGACTTCACTGT
GAGCACACGTCTGAACTCTTTCCCTACACGACGCTCTTCCGATCTNNNTTGTTAGCCTGGACTTCACTGT
GAGCACACGTCTGAACTCTTTCCCTACACGACGCTCTTCCGATCTNNNAAAGATCAATCTGAATGTGTT
GAGCACACGTCTGAACTCTTTCCCTACACGACGCTCTTCCGATCTNNNAAAGATCAATCTGAATGTGTT
AGAGCACACGTCTGAACTCTTTCCCTACACGACGCTCTTCCGATCTNNNGGCTGAGACAGAAGAATTG

GAGCACACGTCTGAACTCTTTCCCTACACGACGCTCTTCCGATCTNNNGGCTGAGACAGAAGAATTG
GCACACGTCTGAACTCTTTCCCTACACGACGCTCTTCCGATCTNNNGTTATTTAGATCACCTTTTGGCC
GCACACGTCTGAACTCTTTCCCTACACGACGCTCTTCCGATCTNNNGTTATTTAGATCACCTTTTGGCC
GCACACGTCTGAACTCTTTCCCTACACGACGCTCTTCCGATCTNNNAGGCATGTGGTCAATCTTAGAC
GCACACGTCTGAACTCTTTCCCTACACGACGCTCTTCCGATCTNNNAGGCATGTGGTCAATCTTAGAC
AGAGCACACGTCTGAACTCTTTCCCTACACGACGCTCTTCCGATCTNNNCTGAGGCAGGAGAATTGC
AGAGCACACGTCTGAACTCTTTCCCTACACGACGCTCTTCCGATCTNNNCTGAGGCAGGAGAATTGC
AGCACACGTCTGAACTCTTTCCCTACACGACGCTCTTCCGATCTNNNCCACATAACACCTTCTGTCTCC
AGCACACGTCTGAACTCTTTCCCTACACGACGCTCTTCCGATCTNNNCCACATAACACCTTCTGTCTCC
CACACGTCTGAACTCTTTCCCTACACGACGCTCTTCCGATCTNNNGAGCCATTGTTCTTTCCTCTTCGC
CACACGTCTGAACTCTTTCCCTACACGACGCTCTTCCGATCTNNNGAGCCATTGTTCTTTCCTCTTCGC
AGCACACGTCTGAACTCTTTCCCTACACGACGCTCTTCCGATCTNNNGCACCCTTCTCACCAGAAAATT
AGCACACGTCTGAACTCTTTCCCTACACGACGCTCTTCCGATCTNNNGCACCCTTCTCACCAGAAAATT
AGCACACGTCTGAACTCTTTCCCTACACGACGCTCTTCCGATCTNNNCTTGCAGTGAGCTGAGATCG
AGCACACGTCTGAACTCTTTCCCTACACGACGCTCTTCCGATCTNNNCTTGCAGTGAGCTGAGATCG
AGCACACGTCTGAACTCTTTCCCTACACGACGCTCTTCCGATCTNNNGTGAAACCCCGTCTCTACTGA
AGCACACGTCTGAACTCTTTCCCTACACGACGCTCTTCCGATCTNNNGTGAAACCCCGTCTCTACTGA
AGCACACGTCTGAACTCTTTCCCTACACGACGCTCTTCCGATCTNNNGCTCCCTGACATTCTTTGGAAA
AGCACACGTCTGAACTCTTTCCCTACACGACGCTCTTCCGATCTNNNGCTCCCTGACATTCTTTGGAAA
AGCACACGTCTGAACTCTTTCCCTACACGACGCTCTTCCGATCTNNNTGCTAATGGTAGATTTGGCTG
AGCACACGTCTGAACTCTTTCCCTACACGACGCTCTTCCGATCTNNNTGCTAATGGTAGATTTGGCTG
AGCACACGTCTGAACTCTTTCCCTACACGACGCTCTTCCGATCTNNNATGCCTTTAGTCCCAGCTACTT
AGCACACGTCTGAACTCTTTCCCTACACGACGCTCTTCCGATCTNNNATGCCTTTAGTCCCAGCTACTT
GCACACGTCTGAACTCTTTCCCTACACGACGCTCTTCCGATCTNNNGGGAACAGTGTTGAGCTGTAT
GCACACGTCTGAACTCTTTCCCTACACGACGCTCTTCCGATCTNNNGGGAACAGTGTTGAGCTGTAT
GAGCACACGTCTGAACTCTTTCCCTACACGACGCTCTTCCGATCTNNNAAAACCACTTGTAGCCAGAC
GAGCACACGTCTGAACTCTTTCCCTACACGACGCTCTTCCGATCTNNNAAAACCACTTGTAGCCAGAC
AGCACACGTCTGAACTCTTTCCCTACACGACGCTCTTCCGATCTNNNGTGGTAGTTTTGGCGCACTTG
AGCACACGTCTGAACTCTTTCCCTACACGACGCTCTTCCGATCTNNNGTGGTAGTTTTGGCGCACTTG
GCACACGTCTGAACTCTTTCCCTACACGACGCTCTTCCGATCTNNNGTTGCAGTGAGCCAAGATTGTG
GCACACGTCTGAACTCTTTCCCTACACGACGCTCTTCCGATCTNNNGTTGCAGTGAGCCAAGATTGTG
AGCACACGTCTGAACTCTTTCCCTACACGACGCTCTTCCGATCTNNNGTTCCAGAGCTTCTTCCACCTT
AGCACACGTCTGAACTCTTTCCCTACACGACGCTCTTCCGATCTNNNGTTCCAGAGCTTCTTCCACCTT
AGCACACGTCTGAACTCTTTCCCTACACGACGCTCTTCCGATCTNNNGGGAGTCTGAGATTTCCTGT
AGCACACGTCTGAACTCTTTCCCTACACGACGCTCTTCCGATCTNNNGGGAGTCTGAGATTTCCTGT
GCACACGTCTGAACTCTTTCCCTACACGACGCTCTTCCGATCTNNNGTTAACATTCATTCTCCTTTGAC
GCACACGTCTGAACTCTTTCCCTACACGACGCTCTTCCGATCTNNNGTTAACATTCATTCTCCTTTGAC
AGCACACGTCTGAACTCTTTCCCTACACGACGCTCTTCCGATCTNNNCCTGTAGTCTCAGCTACTCAGG
AGCACACGTCTGAACTCTTTCCCTACACGACGCTCTTCCGATCTNNNCCTGTAGTCTCAGCTACTCAGG
AGCACACGTCTGAACTCTTTCCCTACACGACGCTCTTCCGATCTNNNCCATGAGAACAGATAGAGTC
AGCACACGTCTGAACTCTTTCCCTACACGACGCTCTTCCGATCTNNNCCATGAGAACAGATAGAGTC
AGCACACGTCTGAACTCTTTCCCTACACGACGCTCTTCCGATCTNNNATATAAGTTCCTTGGGGTGG
AGCACACGTCTGAACTCTTTCCCTACACGACGCTCTTCCGATCTNNNATATAAGTTCCTTGGGGTGG

AGCACACGTCTGAACTCTTTCCCTACACGACGCTCTTCCGATCTNNNGAGAATGGTGTGAACCTGGGA
AGCACACGTCTGAACTCTTTCCCTACACGACGCTCTTCCGATCTNNNGAGAATGGTGTGAACCTGGGA
CACACGTCTGAACTCTTTCCCTACACGACGCTCTTCCGATCTNNNCTCACTTGATCAACTGCGAGAACCA
CACACGTCTGAACTCTTTCCCTACACGACGCTCTTCCGATCTNNNCTCACTTGATCAACTGCGAGAACCA
AGCACACGTCTGAACTCTTTCCCTACACGACGCTCTTCCGATCTNNNTATCTCCTAATGCTACCCCTCCCC
AGCACACGTCTGAACTCTTTCCCTACACGACGCTCTTCCGATCTNNNTATCTCCTAATGCTACCCCTCCCC
GCACACGTCTGAACTCTTTCCCTACACGACGCTCTTCCGATCTNNNGGGAAAGAGCAATGGAAGCAAT
GCACACGTCTGAACTCTTTCCCTACACGACGCTCTTCCGATCTNNNGGGAAAGAGCAATGGAAGCAAT
AGCACACGTCTGAACTCTTTCCCTACACGACGCTCTTCCGATCTNNNATATCAAGCTCCTGTTCTCTGCC
AGCACACGTCTGAACTCTTTCCCTACACGACGCTCTTCCGATCTNNNATATCAAGCTCCTGTTCTCTGCC
GCACACGTCTGAACTCTTTCCCTACACGACGCTCTTCCGATCTNNNTGCATTTCTTATGTGGCTCTTGG
GCACACGTCTGAACTCTTTCCCTACACGACGCTCTTCCGATCTNNNTGCATTTCTTATGTGGCTCTTGG
AGAGCACACGTCTGAACTCTTTCCCTACACGACGCTCTTCCGATCTNNNTTTGAGGCTTTGTTGGAGG
AGAGCACACGTCTGAACTCTTTCCCTACACGACGCTCTTCCGATCTNNNTTTGAGGCTTTGTTGGAGG
AGCACACGTCTGAACTCTTTCCCTACACGACGCTCTTCCGATCTNNNTTACACTGAGAGCAGCCAGAAT
AGCACACGTCTGAACTCTTTCCCTACACGACGCTCTTCCGATCTNNNTTACACTGAGAGCAGCCAGAAT
CACACGTCTGAACTCTTTCCCTACACGACGCTCTTCCGATCTNNNGTTTCTGTGTCTCCGGGTAAGTCCA
CACACGTCTGAACTCTTTCCCTACACGACGCTCTTCCGATCTNNNGTTTCTGTGTCTCCGGGTAAGTCCA
AGCACACGTCTGAACTCTTTCCCTACACGACGCTCTTCCGATCTNNNCACCTGTAGTCCCAGCTACTTG
AGCACACGTCTGAACTCTTTCCCTACACGACGCTCTTCCGATCTNNNCACCTGTAGTCCCAGCTACTTG
AGCACACGTCTGAACTCTTTCCCTACACGACGCTCTTCCGATCTNNNCAAGGGCAGAAAGGCTAATGT
AGCACACGTCTGAACTCTTTCCCTACACGACGCTCTTCCGATCTNNNCAAGGGCAGAAAGGCTAATGT
AGCACACGTCTGAACTCTTTCCCTACACGACGCTCTTCCGATCTNNNTGCTGTTTTTTATCCCCAGAGC
AGCACACGTCTGAACTCTTTCCCTACACGACGCTCTTCCGATCTNNNTGCTGTTTTTTATCCCCAGAGC
AGCACACGTCTGAACTCTTTCCCTACACGACGCTCTTCCGATCTNNNGATCCTGGTGGGGTGATTCCTA
AGCACACGTCTGAACTCTTTCCCTACACGACGCTCTTCCGATCTNNNGATCCTGGTGGGGTGATTCCTA
AGCACACGTCTGAACTCTTTCCCTACACGACGCTCTTCCGATCTNNNGGACATGGTAGCATGTGCCTAT
AGCACACGTCTGAACTCTTTCCCTACACGACGCTCTTCCGATCTNNNGGACATGGTAGCATGTGCCTAT
AGCACACGTCTGAACTCTTTCCCTACACGACGCTCTTCCGATCTNNNGTGTGTGTGTGATTTAGAGCA
AGCACACGTCTGAACTCTTTCCCTACACGACGCTCTTCCGATCTNNNGTGTGTGTGTGATTTAGAGCA
AGCACACGTCTGAACTCTTTCCCTACACGACGCTCTTCCGATCTNNNAGAGATACACACACACACACAC
AGCACACGTCTGAACTCTTTCCCTACACGACGCTCTTCCGATCTNNNAGAGATACACACACACACACAC
AGAGCACACGTCTGAACTCTTTCCCTACACGACGCTCTTCCGATCTNNNACCTTTTGGACTTACATTGG
AGAGCACACGTCTGAACTCTTTCCCTACACGACGCTCTTCCGATCTNNNACCTTTTGGACTTACATTGG
GAGCACACGTCTGAACTCTTTCCCTACACGACGCTCTTCCGATCTNNNCTCCTTTGACTCCATGTCTCACA
GAGCACACGTCTGAACTCTTTCCCTACACGACGCTCTTCCGATCTNNNCTCCTTTGACTCCATGTCTCACA
AGCACACGTCTGAACTCTTTCCCTACACGACGCTCTTCCGATCTNNNCATGCCTATGGTCCCATCTACAC
AGCACACGTCTGAACTCTTTCCCTACACGACGCTCTTCCGATCTNNNCATGCCTATGGTCCCATCTACAC
AGCACACGTCTGAACTCTTTCCCTACACGACGCTCTTCCGATCTNNNCCTTTTTCTCACTTTCCTCCCCTC
AGCACACGTCTGAACTCTTTCCCTACACGACGCTCTTCCGATCTNNNCCTTTTTCTCACTTTCCTCCCCTC
GAGCACACGTCTGAACTCTTTCCCTACACGACGCTCTTCCGATCTNNNTGGCTGTGCTAATTTACACTCC
GAGCACACGTCTGAACTCTTTCCCTACACGACGCTCTTCCGATCTNNNTGGCTGTGCTAATTTACACTCC
AGCACACGTCTGAACTCTTTCCCTACACGACGCTCTTCCGATCTNNNCTTGGTGAGTGAAAGGCTGATT

GCACACGTCTGAACTCTTTCCCTACACGACGCTCTTCCGATCTNNNCTTGGTGAGTGAAAGGCTGATT
AGCACACGTCTGAACTCTTTCCCTACACGACGCTCTTCCGATCTNNNCAAAGAAAAGTCCTCTGTGCA
AGCACACGTCTGAACTCTTTCCCTACACGACGCTCTTCCGATCTNNNCAAAGAAAAGTCCTCTGTGCA
AGCACACGTCTGAACTCTTTCCCTACACGACGCTCTTCCGATCTNNNTGATTTCTTCTGGATTGTCCCA
AGCACACGTCTGAACTCTTTCCCTACACGACGCTCTTCCGATCTNNNTGATTTCTTCTGGATTGTCCCA
AGCACACGTCTGAACTCTTTCCCTACACGACGCTCTTCCGATCTNNNTGGTTTCGATTTGTGACCCTAG
AGCACACGTCTGAACTCTTTCCCTACACGACGCTCTTCCGATCTNNNTGGTTTCGATTTGTGACCCTAG
AGCACACGTCTGAACTCTTTCCCTACACGACGCTCTTCCGATCTNNNTACTCACACAGTTCTGGAGTAC
AGCACACGTCTGAACTCTTTCCCTACACGACGCTCTTCCGATCTNNNTACTCACACAGTTCTGGAGTAC
AGCACACGTCTGAACTCTTTCCCTACACGACGCTCTTCCGATCTNNNAGAGAAAAGGACGATGTAGC
AGCACACGTCTGAACTCTTTCCCTACACGACGCTCTTCCGATCTNNNAGAGAAAAGGACGATGTAGC
AGCACACGTCTGAACTCTTTCCCTACACGACGCTCTTCCGATCTNNNGCTTTCTTGTTTGGTGGATGGT
AGCACACGTCTGAACTCTTTCCCTACACGACGCTCTTCCGATCTNNNGCTTTCTTGTTTGGTGGATGGT
GAGCACACGTCTGAACTCTTTCCCTACACGACGCTCTTCCGATCTNNNGAAGCAGGAGAATGGCATGA
GAGCACACGTCTGAACTCTTTCCCTACACGACGCTCTTCCGATCTNNNGAAGCAGGAGAATGGCATGA
GAGCACACGTCTGAACTCTTTCCCTACACGACGCTCTTCCGATCTNNNGTAGCTGGTACTACAAGGGC
GAGCACACGTCTGAACTCTTTCCCTACACGACGCTCTTCCGATCTNNNGTAGCTGGTACTACAAGGGC
AGCACACGTCTGAACTCTTTCCCTACACGACGCTCTTCCGATCTNNNACAAATAGTGAAAGGTGCCAAC
AGCACACGTCTGAACTCTTTCCCTACACGACGCTCTTCCGATCTNNNACAAATAGTGAAAGGTGCCAAC
AGCACACGTCTGAACTCTTTCCCTACACGACGCTCTTCCGATCTNNNTGTCTGTGCATATGTTCATGTG
AGCACACGTCTGAACTCTTTCCCTACACGACGCTCTTCCGATCTNNNTGTCTGTGCATATGTTCATGTG
GCACACGTCTGAACTCTTTCCCTACACGACGCTCTTCCGATCTNNNGTGAGTGTAGTGTCTGACTCTGG
GCACACGTCTGAACTCTTTCCCTACACGACGCTCTTCCGATCTNNNGTGAGTGTAGTGTCTGACTCTGG
AGCACACGTCTGAACTCTTTCCCTACACGACGCTCTTCCGATCTNNNCCATGATGGGAAGTTGACGTT
AGCACACGTCTGAACTCTTTCCCTACACGACGCTCTTCCGATCTNNNCCATGATGGGAAGTTGACGTT
GCACACGTCTGAACTCTTTCCCTACACGACGCTCTTCCGATCTNNNGGCTGGAATACTACGAAAATGC
GCACACGTCTGAACTCTTTCCCTACACGACGCTCTTCCGATCTNNNGGCTGGAATACTACGAAAATGC
AGCACACGTCTGAACTCTTTCCCTACACGACGCTCTTCCGATCTNNNTCGAAGTTCAACCGAGTGACAA
AGCACACGTCTGAACTCTTTCCCTACACGACGCTCTTCCGATCTNNNTCGAAGTTCAACCGAGTGACAA
GAGCACACGTCTGAACTCTTTCCCTACACGACGCTCTTCCGATCTNNNAAATGCCCATCAATGGTGAAC
GAGCACACGTCTGAACTCTTTCCCTACACGACGCTCTTCCGATCTNNNAAATGCCCATCAATGGTGAAC
GAGCACACGTCTGAACTCTTTCCCTACACGACGCTCTTCCGATCTNNNCCAAAGAGGCATGCAACCTAA
GAGCACACGTCTGAACTCTTTCCCTACACGACGCTCTTCCGATCTNNNCCAAAGAGGCATGCAACCTAA
AGCACACGTCTGAACTCTTTCCCTACACGACGCTCTTCCGATCTNNNTCACAGATGAGGAAACTAACG
AGCACACGTCTGAACTCTTTCCCTACACGACGCTCTTCCGATCTNNNTCACAGATGAGGAAACTAACG
AAGAGCACACGTCTGAACTCTTTCCCTACACGACGCTCTTCCGATCTNNNTATGATTACATCGGAACGG
AAGAGCACACGTCTGAACTCTTTCCCTACACGACGCTCTTCCGATCTNNNTATGATTACATCGGAACGG
AGCACACGTCTGAACTCTTTCCCTACACGACGCTCTTCCGATCTNNNCCACGCTGACTTGATCATATTCA
AGCACACGTCTGAACTCTTTCCCTACACGACGCTCTTCCGATCTNNNCCACGCTGACTTGATCATATTCA
AGCACACGTCTGAACTCTTTCCCTACACGACGCTCTTCCGATCTNNNGTAATCACAGCACTTTGGGATC
AGCACACGTCTGAACTCTTTCCCTACACGACGCTCTTCCGATCTNNNGTAATCACAGCACTTTGGGATC
GAGCACACGTCTGAACTCTTTCCCTACACGACGCTCTTCCGATCTNNNCAGTGAGCCATAGTCGTATCA
GAGCACACGTCTGAACTCTTTCCCTACACGACGCTCTTCCGATCTNNNCAGTGAGCCATAGTCGTATCA

GAGCACACGTCTGAACTCTTTCCCTACACGACGCTCTTCCGATCTNNNTCAATGACAAATTTGATCTG
GAGCACACGTCTGAACTCTTTCCCTACACGACGCTCTTCCGATCTNNNTCAATGACAAATTTGATCTG
CACACGTCTGAACTCTTTCCCTACACGACGCTCTTCCGATCTNNNTACTCATTTGGCTCCGCATTTCAC
CACACGTCTGAACTCTTTCCCTACACGACGCTCTTCCGATCTNNNTACTCATTTGGCTCCGCATTTCAC
AGCACACGTCTGAACTCTTTCCCTACACGACGCTCTTCCGATCTNNNCTGTAGTCCCAGCTACTCAGG
AGCACACGTCTGAACTCTTTCCCTACACGACGCTCTTCCGATCTNNNCTGTAGTCCCAGCTACTCAGG
AAGAGCACACGTCTGAACTCTTTCCCTACACGACGCTCTTCCGATCTNNNCTGAGGCAGGAGAATCAC
AAGAGCACACGTCTGAACTCTTTCCCTACACGACGCTCTTCCGATCTNNNCTGAGGCAGGAGAATCAC
CACACGTCTGAACTCTTTCCCTACACGACGCTCTTCCGATCTNNNAATGAGCTGTTGAGTACACCTCCC
CACACGTCTGAACTCTTTCCCTACACGACGCTCTTCCGATCTNNNAATGAGCTGTTGAGTACACCTCCC
AGCACACGTCTGAACTCTTTCCCTACACGACGCTCTTCCGATCTNNNTCTACCCAGGGAAGAAGAAGT
AGCACACGTCTGAACTCTTTCCCTACACGACGCTCTTCCGATCTNNNTCTACCCAGGGAAGAAGAAGT
CACACGTCTGAACTCTTTCCCTACACGACGCTCTTCCGATCTNNNCTCAGCGTGGAAAATATGGAACC
CACACGTCTGAACTCTTTCCCTACACGACGCTCTTCCGATCTNNNCTCAGCGTGGAAAATATGGAACC
GCACACGTCTGAACTCTTTCCCTACACGACGCTCTTCCGATCTNNNATTCAATTACCTCCCACTGGGTC
GCACACGTCTGAACTCTTTCCCTACACGACGCTCTTCCGATCTNNNATTCAATTACCTCCCACTGGGTC
AGCACACGTCTGAACTCTTTCCCTACACGACGCTCTTCCGATCTNNNACCATGAACCTGACTTTCCTTG
AGCACACGTCTGAACTCTTTCCCTACACGACGCTCTTCCGATCTNNNACCATGAACCTGACTTTCCTTG
AGCACACGTCTGAACTCTTTCCCTACACGACGCTCTTCCGATCTNNNATTCTCACATCCCAAGTAGTCC
AGCACACGTCTGAACTCTTTCCCTACACGACGCTCTTCCGATCTNNNATTCTCACATCCCAAGTAGTCC
GCACACGTCTGAACTCTTTCCCTACACGACGCTCTTCCGATCTNNNCTCCTGGACTCAAGTGATCTAC
GCACACGTCTGAACTCTTTCCCTACACGACGCTCTTCCGATCTNNNCTCCTGGACTCAAGTGATCTAC
AGCACACGTCTGAACTCTTTCCCTACACGACGCTCTTCCGATCTNNNGAGGCTGGTCCAAAGCTAGTA
AGCACACGTCTGAACTCTTTCCCTACACGACGCTCTTCCGATCTNNNGAGGCTGGTCCAAAGCTAGTA
GCACACGTCTGAACTCTTTCCCTACACGACGCTCTTCCGATCTNNNTTTGACAATGTTCTTAGTAACTG
GCACACGTCTGAACTCTTTCCCTACACGACGCTCTTCCGATCTNNNTTTGACAATGTTCTTAGTAACTG
GCACACGTCTGAACTCTTTCCCTACACGACGCTCTTCCGATCTNNNTGGGAGACTGAGGCAGAAAAT
GCACACGTCTGAACTCTTTCCCTACACGACGCTCTTCCGATCTNNNTGGGAGACTGAGGCAGAAAAT
GCACACGTCTGAACTCTTTCCCTACACGACGCTCTTCCGATCTNNNGATTGCTTGAGCTCAGGAGTTT
GCACACGTCTGAACTCTTTCCCTACACGACGCTCTTCCGATCTNNNGATTGCTTGAGCTCAGGAGTTT
GAGCACACGTCTGAACTCTTTCCCTACACGACGCTCTTCCGATCTNNNAGTGAGTGGAGATCGTGCCA
GAGCACACGTCTGAACTCTTTCCCTACACGACGCTCTTCCGATCTNNNAGTGAGTGGAGATCGTGCCA
GAGCACACGTCTGAACTCTTTCCCTACACGACGCTCTTCCGATCTNNNGGAAGACGACCTAGGCAATA
GAGCACACGTCTGAACTCTTTCCCTACACGACGCTCTTCCGATCTNNNGGAAGACGACCTAGGCAATA
GAGCACACGTCTGAACTCTTTCCCTACACGACGCTCTTCCGATCTNNNGGAAGGCCCTGTCTCAAAAA
GAGCACACGTCTGAACTCTTTCCCTACACGACGCTCTTCCGATCTNNNGGAAGGCCCTGTCTCAAAAA
GCACACGTCTGAACTCTTTCCCTACACGACGCTCTTCCGATCTNNNCTGCAAGTTGATAAGGAGCATG
GCACACGTCTGAACTCTTTCCCTACACGACGCTCTTCCGATCTNNNCTGCAAGTTGATAAGGAGCATG
GAGCACACGTCTGAACTCTTTCCCTACACGACGCTCTTCCGATCTNNNCAGGATCTAGTGGTCCATAT
GAGCACACGTCTGAACTCTTTCCCTACACGACGCTCTTCCGATCTNNNCAGGATCTAGTGGTCCATAT
AGCACACGTCTGAACTCTTTCCCTACACGACGCTCTTCCGATCTNNNCTTTGCCTGGGTATAGAGTACA
AGCACACGTCTGAACTCTTTCCCTACACGACGCTCTTCCGATCTNNNCTTTGCCTGGGTATAGAGTACA
AGCACACGTCTGAACTCTTTCCCTACACGACGCTCTTCCGATCTNNNTGAGACTGTTTTTCCAGCCCA

AGCACACGTCTGAACTCTTTCCCTACACGACGCTCTTCCGATCTNNNTGAGACTGTTTTTCCAGCCCA

AGCACACGTCTGAACTCTTTCCCTACACGACGCTCTTCCGATCTNNNATGTCTAGCCCTCAGTTTCCATC

AGCACACGTCTGAACTCTTTCCCTACACGACGCTCTTCCGATCTNNNATGTCTAGCCCTCAGTTTCCATC

AGCACACGTCTGAACTCTTTCCCTACACGACGCTCTTCCGATCTNNNGTACTTTTGTGATTCTGATGTA

AGCACACGTCTGAACTCTTTCCCTACACGACGCTCTTCCGATCTNNNGTACTTTTGTGATTCTGATGTA

AGCACACGTCTGAACTCTTTCCCTACACGACGCTCTTCCGATCTNNNCAGAGGTTACAGTGAGCAGA

AGCACACGTCTGAACTCTTTCCCTACACGACGCTCTTCCGATCTNNNCAGAGGTTACAGTGAGCAGA

GAGCACACGTCTGAACTCTTTCCCTACACGACGCTCTTCCGATCTNNNTGGGCAACAGAGTGAGACTT

GAGCACACGTCTGAACTCTTTCCCTACACGACGCTCTTCCGATCTNNNTGGGCAACAGAGTGAGACTT

AGCACACGTCTGAACTCTTTCCCTACACGACGCTCTTCCGATCTNNNGCACAAAGTAGACACTGAGCA

AGCACACGTCTGAACTCTTTCCCTACACGACGCTCTTCCGATCTNNNGCACAAAGTAGACACTGAGCA

AGCACACGTCTGAACTCTTTCCCTACACGACGCTCTTCCGATCTNNNTTCCCCAGCAGTTCATTATCCAA

AGCACACGTCTGAACTCTTTCCCTACACGACGCTCTTCCGATCTNNNTTCCCCAGCAGTTCATTATCCAA

GAGCACACGTCTGAACTCTTTCCCTACACGACGCTCTTCCGATCTNNNGAAGTCAGGAGATCGAGAC

GAGCACACGTCTGAACTCTTTCCCTACACGACGCTCTTCCGATCTNNNGAAGTCAGGAGATCGAGAC

AGCACACGTCTGAACTCTTTCCCTACACGACGCTCTTCCGATCTNNNGACTGGAGTGTCTCATCTCTG

AGCACACGTCTGAACTCTTTCCCTACACGACGCTCTTCCGATCTNNNGACTGGAGTGTCTCATCTCTG

GAGCACACGTCTGAACTCTTTCCCTACACGACGCTCTTCCGATCTNNNGTGATGAACGCAACAGCTTA

GAGCACACGTCTGAACTCTTTCCCTACACGACGCTCTTCCGATCTNNNGTGATGAACGCAACAGCTTA

AGCACACGTCTGAACTCTTTCCCTACACGACGCTCTTCCGATCTNNNTGCTTTTTTCCAGTTGCTCACT

AGCACACGTCTGAACTCTTTCCCTACACGACGCTCTTCCGATCTNNNTGCTTTTTTCCAGTTGCTCACT

GAGCACACGTCTGAACTCTTTCCCTACACGACGCTCTTCCGATCTNNNATGGATTTGGGCACTGGTAG

GAGCACACGTCTGAACTCTTTCCCTACACGACGCTCTTCCGATCTNNNATGGATTTGGGCACTGGTAG

AGCACACGTCTGAACTCTTTCCCTACACGACGCTCTTCCGATCTNNNGAACTGAGATTGCATACCATC

AGCACACGTCTGAACTCTTTCCCTACACGACGCTCTTCCGATCTNNNGAACTGAGATTGCATACCATC

AGCACACGTCTGAACTCTTTCCCTACACGACGCTCTTCCGATCTNNNCTACCCTGTCAAGAGGAAAGA

AGCACACGTCTGAACTCTTTCCCTACACGACGCTCTTCCGATCTNNNCTACCCTGTCAAGAGGAAAGA

GCACACGTCTGAACTCTTTCCCTACACGACGCTCTTCCGATCTNNNTGGAGAGACATGTAGGCTACA

GCACACGTCTGAACTCTTTCCCTACACGACGCTCTTCCGATCTNNNTGGAGAGACATGTAGGCTACA

GCACACGTCTGAACTCTTTCCCTACACGACGCTCTTCCGATCTNNNTGTCCCCAGTATCTGTTTTGTGC

GCACACGTCTGAACTCTTTCCCTACACGACGCTCTTCCGATCTNNNTGTCCCCAGTATCTGTTTTGTGC

GAGCACACGTCTGAACTCTTTCCCTACACGACGCTCTTCCGATCTNNNCCTGGGAGGTTGGTTCTGTAT

GAGCACACGTCTGAACTCTTTCCCTACACGACGCTCTTCCGATCTNNNCCTGGGAGGTTGGTTCTGTAT

AGCACACGTCTGAACTCTTTCCCTACACGACGCTCTTCCGATCTNNNTCACGATCAAGTCCAAACTCCTT

AGCACACGTCTGAACTCTTTCCCTACACGACGCTCTTCCGATCTNNNTCACGATCAAGTCCAAACTCCTT

GAGCACACGTCTGAACTCTTTCCCTACACGACGCTCTTCCGATCTNNNCCACCTATTCTCTCTGAAGCCT

GAGCACACGTCTGAACTCTTTCCCTACACGACGCTCTTCCGATCTNNNCCACCTATTCTCTCTGAAGCCT

GAGCACACGTCTGAACTCTTTCCCTACACGACGCTCTTCCGATCTNNNACAATCCGTCTATTGAGGACA

GAGCACACGTCTGAACTCTTTCCCTACACGACGCTCTTCCGATCTNNNACAATCCGTCTATTGAGGACA

GAGCACACGTCTGAACTCTTTCCCTACACGACGCTCTTCCGATCTNNNTCACCATTGATTCAACAATAC

GAGCACACGTCTGAACTCTTTCCCTACACGACGCTCTTCCGATCTNNNTCACCATTGATTCAACAATAC

AGCACACGTCTGAACTCTTTCCCTACACGACGCTCTTCCGATCTNNNCCCCATCGCATCCTTTCCTATAG

AGCACACGTCTGAACTCTTTCCCTACACGACGCTCTTCCGATCTNNNCCCCATCGCATCCTTTCCTATAG

GCACACGTCTGAACTCTTTCCCTACACGACGCTCTTCCGATCTNNNACATTGGTCTGAATCTCTTGGCT

GCACACGTCTGAACTCTTTCCCTACACGACGCTCTTCCGATCTNNNACATTGGTCTGAATCTCTTGGCT

GAGCACACGTCTGAACTCTTTCCCTACACGACGCTCTTCCGATCTNNNCCTCAGAGAGCTGACACAGAA

GAGCACACGTCTGAACTCTTTCCCTACACGACGCTCTTCCGATCTNNNCCTCAGAGAGCTGACACAGAA

GAGCACACGTCTGAACTCTTTCCCTACACGACGCTCTTCCGATCTNNNGGTTGGGAGATTGAGAACAT

GAGCACACGTCTGAACTCTTTCCCTACACGACGCTCTTCCGATCTNNNGGTTGGGAGATTGAGAACAT

AGCACACGTCTGAACTCTTTCCCTACACGACGCTCTTCCGATCTNNNGTATCATAGCTGGGACTACAG

AGCACACGTCTGAACTCTTTCCCTACACGACGCTCTTCCGATCTNNNGTATCATAGCTGGGACTACAG

GAAGAGCACACGTCTGAACTCTTTCCCTACACGACGCTCTTCCGATCTNNNGCCATCCCCAGACTTCCCC

GAAGAGCACACGTCTGAACTCTTTCCCTACACGACGCTCTTCCGATCTNNNGCCATCCCCAGACTTCCCC

CACACGTCTGAACTCTTTCCCTACACGACGCTCTTCCGATCTNNNGCTAGTATTCGATACTTCAGTAGGC

CACACGTCTGAACTCTTTCCCTACACGACGCTCTTCCGATCTNNNGCTAGTATTCGATACTTCAGTAGGC

GAGCACACGTCTGAACTCTTTCCCTACACGACGCTCTTCCGATCTNNNCATAGTCCCAGCTACTCAGAA

GAGCACACGTCTGAACTCTTTCCCTACACGACGCTCTTCCGATCTNNNCATAGTCCCAGCTACTCAGAA

GAGCACACGTCTGAACTCTTTCCCTACACGACGCTCTTCCGATCTNNNTGTCATTCTAGCAGCCCTTTCA

GAGCACACGTCTGAACTCTTTCCCTACACGACGCTCTTCCGATCTNNNTGTCATTCTAGCAGCCCTTTCA

AGCACACGTCTGAACTCTTTCCCTACACGACGCTCTTCCGATCTNNNGGACAGTGCTGAAGAGAAGT

AGCACACGTCTGAACTCTTTCCCTACACGACGCTCTTCCGATCTNNNGGACAGTGCTGAAGAGAAGT

GAGCACACGTCTGAACTCTTTCCCTACACGACGCTCTTCCGATCTNNNCTCTTGAGTTGGTATTGTA

GAGCACACGTCTGAACTCTTTCCCTACACGACGCTCTTCCGATCTNNNCTCTTGAGTTGGTATTGTA

GCACACGTCTGAACTCTTTCCCTACACGACGCTCTTCCGATCTNNNAGAAGTGCTTGGATGAAACTTG

GCACACGTCTGAACTCTTTCCCTACACGACGCTCTTCCGATCTNNNAGAAGTGCTTGGATGAAACTTG

GCACACGTCTGAACTCTTTCCCTACACGACGCTCTTCCGATCTNNNGTTTTTCTTGTGTGTGTGTGTGT

GCACACGTCTGAACTCTTTCCCTACACGACGCTCTTCCGATCTNNNGTTTTTCTTGTGTGTGTGTGTGT

GAGCACACGTCTGAACTCTTTCCCTACACGACGCTCTTCCGATCTNNNGAGGTTGAGAGTTAGAGAC

GAGCACACGTCTGAACTCTTTCCCTACACGACGCTCTTCCGATCTNNNGAGGTTGAGAGTTAGAGAC

AGCACACGTCTGAACTCTTTCCCTACACGACGCTCTTCCGATCTNNNGTTGATCACTGCCTGTTCCAAA

AGCACACGTCTGAACTCTTTCCCTACACGACGCTCTTCCGATCTNNNGTTGATCACTGCCTGTTCCAAA

GAGCACACGTCTGAACTCTTTCCCTACACGACGCTCTTCCGATCTNNNCAAACTCCCAAGCTTGAATCA

GAGCACACGTCTGAACTCTTTCCCTACACGACGCTCTTCCGATCTNNNCAAACTCCCAAGCTTGAATCA

AGCACACGTCTGAACTCTTTCCCTACACGACGCTCTTCCGATCTNNNTGCTCGATGGAGACTAAAATG

AGCACACGTCTGAACTCTTTCCCTACACGACGCTCTTCCGATCTNNNTGCTCGATGGAGACTAAAATG

AGCACACGTCTGAACTCTTTCCCTACACGACGCTCTTCCGATCTNNNACTTGAGGTCAGAAGTTCGAG

AGCACACGTCTGAACTCTTTCCCTACACGACGCTCTTCCGATCTNNNACTTGAGGTCAGAAGTTCGAG

AGCACACGTCTGAACTCTTTCCCTACACGACGCTCTTCCGATCTNNNTTAGTTGGCTCCCATGTCTCTC

AGCACACGTCTGAACTCTTTCCCTACACGACGCTCTTCCGATCTNNNTTAGTTGGCTCCCATGTCTCTC

AGCACACGTCTGAACTCTTTCCCTACACGACGCTCTTCCGATCTNNNGGTTGATTTGAAGGCCTTGTCT

AGCACACGTCTGAACTCTTTCCCTACACGACGCTCTTCCGATCTNNNGGTTGATTTGAAGGCCTTGTCT

AGCACACGTCTGAACTCTTTCCCTACACGACGCTCTTCCGATCTNNNTTGTGTGACACGGAAATGCAAT

AGCACACGTCTGAACTCTTTCCCTACACGACGCTCTTCCGATCTNNNTTGTGTGACACGGAAATGCAAT

GAGCACACGTCTGAACTCTTTCCCTACACGACGCTCTTCCGATCTNNNTGTGATGCTTCTAGCCTTGTT

GAGCACACGTCTGAACTCTTTCCCTACACGACGCTCTTCCGATCTNNNTGTGATGCTTCTAGCCTTGTT

AGCACACGTCTGAACTCTTTCCCTACACGACGCTCTTCCGATCTNNNGGGCACAGGAACTAGAAAAT

AGCACACGTCTGAACTCTTTCCCTACACGACGCTCTTCCGATCTNNNGGGCACAGGAACTAGAAAAAT

GAGCACACGTCTGAACTCTTTCCCTACACGACGCTCTTCCGATCTNNNCTGTGTGGTTAGACGGCTTCC

GAGCACACGTCTGAACTCTTTCCCTACACGACGCTCTTCCGATCTNNNCTGTGTGGTTAGACGGCTTCC

AGCACACGTCTGAACTCTTTCCCTACACGACGCTCTTCCGATCTNNNTATATGCAGCTTTCCAGGGACA

AGCACACGTCTGAACTCTTTCCCTACACGACGCTCTTCCGATCTNNNTATATGCAGCTTTCCAGGGACA

AGCACACGTCTGAACTCTTTCCCTACACGACGCTCTTCCGATCTNNNCTGAGGCAGGAGAATGACATG

AGCACACGTCTGAACTCTTTCCCTACACGACGCTCTTCCGATCTNNNCTGAGGCAGGAGAATGACATG

GAGCACACGTCTGAACTCTTTCCCTACACGACGCTCTTCCGATCTNNNTCACTTCTATGTGAGAGAGG

GAGCACACGTCTGAACTCTTTCCCTACACGACGCTCTTCCGATCTNNNTCACTTCTATGTGAGAGAGG

AGCACACGTCTGAACTCTTTCCCTACACGACGCTCTTCCGATCTNNNCCACAGGAAGAAAAGGAGAGA

AGCACACGTCTGAACTCTTTCCCTACACGACGCTCTTCCGATCTNNNCCACAGGAAGAAAAGGAGAGA

GCACACGTCTGAACTCTTTCCCTACACGACGCTCTTCCGATCTNNNATCATAGCTCATTGCAGCCTTGAC

GCACACGTCTGAACTCTTTCCCTACACGACGCTCTTCCGATCTNNNATCATAGCTCATTGCAGCCTTGAC

AGCACACGTCTGAACTCTTTCCCTACACGACGCTCTTCCGATCTNNNGCAGTGTTGTGCTGTTACTCAA

AGCACACGTCTGAACTCTTTCCCTACACGACGCTCTTCCGATCTNNNGCAGTGTTGTGCTGTTACTCAA

AGCACACGTCTGAACTCTTTCCCTACACGACGCTCTTCCGATCTNNNTGTTGCTAATTTTGGAACCTGT

AGCACACGTCTGAACTCTTTCCCTACACGACGCTCTTCCGATCTNNNTGTTGCTAATTTTGGAACCTGT

ACACGTCTGAACTCTTTCCCTACACGACGCTCTTCCGATCTNNNTGAATCATTGCAATAACCACGTTCAT

ACACGTCTGAACTCTTTCCCTACACGACGCTCTTCCGATCTNNNTGAATCATTGCAATAACCACGTTCAT

GAGCACACGTCTGAACTCTTTCCCTACACGACGCTCTTCCGATCTNNNGAGGAGTGGTCTCCTCACTCT

GAGCACACGTCTGAACTCTTTCCCTACACGACGCTCTTCCGATCTNNNGAGGAGTGGTCTCCTCACTCT

GAGCACACGTCTGAACTCTTTCCCTACACGACGCTCTTCCGATCTNNNGCTTGAATCAAGGAAGGTGCT

GAGCACACGTCTGAACTCTTTCCCTACACGACGCTCTTCCGATCTNNNGCTTGAATCAAGGAAGGTGCT

GCACACGTCTGAACTCTTTCCCTACACGACGCTCTTCCGATCTNNNTATGCCTGTAATCCCAGCACTTTC

GCACACGTCTGAACTCTTTCCCTACACGACGCTCTTCCGATCTNNNTATGCCTGTAATCCCAGCACTTTC

GAGCACACGTCTGAACTCTTTCCCTACACGACGCTCTTCCGATCTNNNAAAAAGACCCTGGAGTCCTGC

GAGCACACGTCTGAACTCTTTCCCTACACGACGCTCTTCCGATCTNNNAAAAAGACCCTGGAGTCCTGC

GAGCACACGTCTGAACTCTTTCCCTACACGACGCTCTTCCGATCTNNNACAAAGGAGAGACACCTAAC

GAGCACACGTCTGAACTCTTTCCCTACACGACGCTCTTCCGATCTNNNACAAAGGAGAGACACCTAAC

AGCACACGTCTGAACTCTTTCCCTACACGACGCTCTTCCGATCTNNNTGTCTTGGGTTTCTAAGTCTTCT

AGCACACGTCTGAACTCTTTCCCTACACGACGCTCTTCCGATCTNNNTGTCTTGGGTTTCTAAGTCTTCT

AGCACACGTCTGAACTCTTTCCCTACACGACGCTCTTCCGATCTNNNGTAATGAAGTTCTTCTAGTTCA

AGCACACGTCTGAACTCTTTCCCTACACGACGCTCTTCCGATCTNNNGTAATGAAGTTCTTCTAGTTCA

GAGCACACGTCTGAACTCTTTCCCTACACGACGCTCTTCCGATCTNNNGTGTTTGTATCCCCAGCTACTT

GAGCACACGTCTGAACTCTTTCCCTACACGACGCTCTTCCGATCTNNNGTGTTTGTATCCCCAGCTACTT

GCACACGTCTGAACTCTTTCCCTACACGACGCTCTTCCGATCTNNNCACCACACCCAGCTAATTTTTGTC

GCACACGTCTGAACTCTTTCCCTACACGACGCTCTTCCGATCTNNNCACCACACCCAGCTAATTTTTGTC

GAGCACACGTCTGAACTCTTTCCCTACACGACGCTCTTCCGATCTNNNCTGTAGTCTCAGCTACTGGGC

GAGCACACGTCTGAACTCTTTCCCTACACGACGCTCTTCCGATCTNNNCTGTAGTCTCAGCTACTGGGC

AGCACACGTCTGAACTCTTTCCCTACACGACGCTCTTCCGATCTNNNAAATGAATTTCAGTGCCACCCTC

AGCACACGTCTGAACTCTTTCCCTACACGACGCTCTTCCGATCTNNNAAATGAATTTCAGTGCCACCCTC

GAGCACACGTCTGAACTCTTTCCCTACACGACGCTCTTCCGATCTNNNCAGCTGGTTGTATATCCAATAT

GAGCACACGTCTGAACTCTTTCCCTACACGACGCTCTTCCGATCTNNNCAGCTGGTTGTATATCCAATAT

AGCACACGTCTGAACTCTTTCCCTACACGACGCTCTTCCGATCTNNNTTACTGCTTGTGAGTCACCCTA
AGCACACGTCTGAACTCTTTCCCTACACGACGCTCTTCCGATCTNNNTTACTGCTTGTGAGTCACCCTA
AGCACACGTCTGAACTCTTTCCCTACACGACGCTCTTCCGATCTNNNTTCCTTCCTTCCTTCCTTCCTTCC
AGCACACGTCTGAACTCTTTCCCTACACGACGCTCTTCCGATCTNNNTTCCTTCCTTCCTTCCTTCCTTCC
GAGCACACGTCTGAACTCTTTCCCTACACGACGCTCTTCCGATCTNNNTGTTCTTGAGCAAATCCAAAC
GAGCACACGTCTGAACTCTTTCCCTACACGACGCTCTTCCGATCTNNNTGTTCTTGAGCAAATCCAAAC
GAGCACACGTCTGAACTCTTTCCCTACACGACGCTCTTCCGATCTNNNCTGAGGCAGGAGAATCACTT
GAGCACACGTCTGAACTCTTTCCCTACACGACGCTCTTCCGATCTNNNCTGAGGCAGGAGAATCACTT
AGCACACGTCTGAACTCTTTCCCTACACGACGCTCTTCCGATCTNNNAAAAAAGCCCAGCCTTAGCAAA
AGCACACGTCTGAACTCTTTCCCTACACGACGCTCTTCCGATCTNNNAAAAAAGCCCAGCCTTAGCAAA
GAGCACACGTCTGAACTCTTTCCCTACACGACGCTCTTCCGATCTNNNCTTGTCTCCCATCTTACCACAC
GAGCACACGTCTGAACTCTTTCCCTACACGACGCTCTTCCGATCTNNNCTTGTCTCCCATCTTACCACAC
AGCACACGTCTGAACTCTTTCCCTACACGACGCTCTTCCGATCTNNNCCAGCATCTCCACCATCTACAAA
AGCACACGTCTGAACTCTTTCCCTACACGACGCTCTTCCGATCTNNNCCAGCATCTCCACCATCTACAAA
GAGCACACGTCTGAACTCTTTCCCTACACGACGCTCTTCCGATCTNNNGATGGAATCCATGATAGGGG
GAGCACACGTCTGAACTCTTTCCCTACACGACGCTCTTCCGATCTNNNGATGGAATCCATGATAGGGG
AGCACACGTCTGAACTCTTTCCCTACACGACGCTCTTCCGATCTNNNCAGGGAACAAGAAAGTAAGC
AGCACACGTCTGAACTCTTTCCCTACACGACGCTCTTCCGATCTNNNCAGGGAACAAGAAAGTAAGC
GAGCACACGTCTGAACTCTTTCCCTACACGACGCTCTTCCGATCTNNNCAGCCTGAGTGATAGAGTGA
GAGCACACGTCTGAACTCTTTCCCTACACGACGCTCTTCCGATCTNNNCAGCCTGAGTGATAGAGTGA
AGCACACGTCTGAACTCTTTCCCTACACGACGCTCTTCCGATCTNNNACAGGAAGCACAGTTCTAGCAT
AGCACACGTCTGAACTCTTTCCCTACACGACGCTCTTCCGATCTNNNACAGGAAGCACAGTTCTAGCAT
GAGCACACGTCTGAACTCTTTCCCTACACGACGCTCTTCCGATCTNNNTACTTAGAACTGGGTCCAGGT
GAGCACACGTCTGAACTCTTTCCCTACACGACGCTCTTCCGATCTNNNTACTTAGAACTGGGTCCAGGT
AGCACACGTCTGAACTCTTTCCCTACACGACGCTCTTCCGATCTNNNAACCCACGTCTTGTTTTCCTTTCC
AGCACACGTCTGAACTCTTTCCCTACACGACGCTCTTCCGATCTNNNAACCCACGTCTTGTTTTCCTTTCC
GAGCACACGTCTGAACTCTTTCCCTACACGACGCTCTTCCGATCTNNNTGAGGCAGGAGAATCTCTTG
GAGCACACGTCTGAACTCTTTCCCTACACGACGCTCTTCCGATCTNNNTGAGGCAGGAGAATCTCTTG
AGCACACGTCTGAACTCTTTCCCTACACGACGCTCTTCCGATCTNNNAACAAAATTAGCTGGGTGTGG
AGCACACGTCTGAACTCTTTCCCTACACGACGCTCTTCCGATCTNNNAACAAAATTAGCTGGGTGTGG
GAGCACACGTCTGAACTCTTTCCCTACACGACGCTCTTCCGATCTNNNTTTTTGAGACAGGGTCTTGC
GAGCACACGTCTGAACTCTTTCCCTACACGACGCTCTTCCGATCTNNNTTTTTGAGACAGGGTCTTGC
GAGCACACGTCTGAACTCTTTCCCTACACGACGCTCTTCCGATCTNNNTAAGGTGGGAGGATCAATT
GAGCACACGTCTGAACTCTTTCCCTACACGACGCTCTTCCGATCTNNNTAAGGTGGGAGGATCAATT
GAGCACACGTCTGAACTCTTTCCCTACACGACGCTCTTCCGATCTNNNTGGCATGTATTCAACAACAA
GAGCACACGTCTGAACTCTTTCCCTACACGACGCTCTTCCGATCTNNNTGGCATGTATTCAACAACAA
GCACACGTCTGAACTCTTTCCCTACACGACGCTCTTCCGATCTNNNGCTCCACACATACTTCATGAGGA
GCACACGTCTGAACTCTTTCCCTACACGACGCTCTTCCGATCTNNNGCTCCACACATACTTCATGAGGA
GCACACGTCTGAACTCTTTCCCTACACGACGCTCTTCCGATCTNNNCCATGTCAATTCCTCTCACTGGG
GCACACGTCTGAACTCTTTCCCTACACGACGCTCTTCCGATCTNNNCCATGTCAATTCCTCTCACTGGG
AGCACACGTCTGAACTCTTTCCCTACACGACGCTCTTCCGATCTNNNGCAGCTACTCCCACATCTACAT
AGCACACGTCTGAACTCTTTCCCTACACGACGCTCTTCCGATCTNNNGCAGCTACTCCCACATCTACAT
GAGCACACGTCTGAACTCTTTCCCTACACGACGCTCTTCCGATCTNNNAGACAGAGGCAATTTGGTTG

AGCACACGTCTGAACTCTTTCCCTACACGACGCTCTTCCGATCTNNNAGACAGAGGCAATTTGGTTG
AGCACACGTCTGAACTCTTTCCCTACACGACGCTCTTCCGATCTNNNCCTGTTGTCCCAGCTACTTGGC
AGCACACGTCTGAACTCTTTCCCTACACGACGCTCTTCCGATCTNNNCCTGTTGTCCCAGCTACTTGGC
AGCACACGTCTGAACTCTTTCCCTACACGACGCTCTTCCGATCTNNNCTCTAGCCTGGGCAATGAGAG
AGCACACGTCTGAACTCTTTCCCTACACGACGCTCTTCCGATCTNNNCTCTAGCCTGGGCAATGAGAG
GAGCACACGTCTGAACTCTTTCCCTACACGACGCTCTTCCGATCTNNNAATACAAAATTAGCTGGGCG
GAGCACACGTCTGAACTCTTTCCCTACACGACGCTCTTCCGATCTNNNAATACAAAATTAGCTGGGCG
GCACACGTCTGAACTCTTTCCCTACACGACGCTCTTCCGATCTNNNTGAAATCTTTAACTGCATTTTCT
GCACACGTCTGAACTCTTTCCCTACACGACGCTCTTCCGATCTNNNTGAAATCTTTAACTGCATTTTCT
CACACGTCTGAACTCTTTCCCTACACGACGCTCTTCCGATCTNNNTGCCAGAGAGTTTTCAATACACAC
CACACGTCTGAACTCTTTCCCTACACGACGCTCTTCCGATCTNNNTGCCAGAGAGTTTTCAATACACAC
GAGCACACGTCTGAACTCTTTCCCTACACGACGCTCTTCCGATCTNNNTTAGCATAATGCCCATGGGT
GAGCACACGTCTGAACTCTTTCCCTACACGACGCTCTTCCGATCTNNNTTAGCATAATGCCCATGGGT
AGCACACGTCTGAACTCTTTCCCTACACGACGCTCTTCCGATCTNNNGCAGTGTTTTGTAGTTCTCCTT
AGCACACGTCTGAACTCTTTCCCTACACGACGCTCTTCCGATCTNNNGCAGTGTTTTGTAGTTCTCCTT
GAGCACACGTCTGAACTCTTTCCCTACACGACGCTCTTCCGATCTNNNATACACACACATCCTCCCAAGT
GAGCACACGTCTGAACTCTTTCCCTACACGACGCTCTTCCGATCTNNNATACACACACATCCTCCCAAGT
GCACACGTCTGAACTCTTTCCCTACACGACGCTCTTCCGATCTNNNCACCTTTTTATTGGCCTGTTTGCC
GCACACGTCTGAACTCTTTCCCTACACGACGCTCTTCCGATCTNNNCACCTTTTTATTGGCCTGTTTGCC
GCACACGTCTGAACTCTTTCCCTACACGACGCTCTTCCGATCTNNNGGATGGTCTTGATTTCCTGACC
GCACACGTCTGAACTCTTTCCCTACACGACGCTCTTCCGATCTNNNGGATGGTCTTGATTTCCTGACC
GAGCACACGTCTGAACTCTTTCCCTACACGACGCTCTTCCGATCTNNNCTCACCAGTTTCCCAATGAGG
GAGCACACGTCTGAACTCTTTCCCTACACGACGCTCTTCCGATCTNNNCTCACCAGTTTCCCAATGAGG
GAGCACACGTCTGAACTCTTTCCCTACACGACGCTCTTCCGATCTNNNGAGGCTGAGGCAGGAGAAT
GAGCACACGTCTGAACTCTTTCCCTACACGACGCTCTTCCGATCTNNNGAGGCTGAGGCAGGAGAAT
AGAGCACACGTCTGAACTCTTTCCCTACACGACGCTCTTCCGATCTNNNATGGGGAAGAAAAGGAGA
AGAGCACACGTCTGAACTCTTTCCCTACACGACGCTCTTCCGATCTNNNATGGGGAAGAAAAGGAGA
GAGCACACGTCTGAACTCTTTCCCTACACGACGCTCTTCCGATCTNNNAAGGGGAAGAAGGGTGAA
GAGCACACGTCTGAACTCTTTCCCTACACGACGCTCTTCCGATCTNNNAAGGGGAAGAAGGGTGAA
GAGCACACGTCTGAACTCTTTCCCTACACGACGCTCTTCCGATCTNNNAGACAAAGTCTCACTCTGTTG
GAGCACACGTCTGAACTCTTTCCCTACACGACGCTCTTCCGATCTNNNAGACAAAGTCTCACTCTGTTG
GCACACGTCTGAACTCTTTCCCTACACGACGCTCTTCCGATCTNNNTTCCCTTTCCTTCATGCCTACTCC
GCACACGTCTGAACTCTTTCCCTACACGACGCTCTTCCGATCTNNNTTCCCTTTCCTTCATGCCTACTCC
GCACACGTCTGAACTCTTTCCCTACACGACGCTCTTCCGATCTNNNGCGCCTGGCCATAAATAAACTAT
GCACACGTCTGAACTCTTTCCCTACACGACGCTCTTCCGATCTNNNGCGCCTGGCCATAAATAAACTAT
AGCACACGTCTGAACTCTTTCCCTACACGACGCTCTTCCGATCTNNNGGTTGCAGTAAGCTGATATCA
AGCACACGTCTGAACTCTTTCCCTACACGACGCTCTTCCGATCTNNNGGTTGCAGTAAGCTGATATCA
AGCACACGTCTGAACTCTTTCCCTACACGACGCTCTTCCGATCTNNNCATCCTCTTTTGTGCTCTTCAG
AGCACACGTCTGAACTCTTTCCCTACACGACGCTCTTCCGATCTNNNCATCCTCTTTTGTGCTCTTCAG
GAGCACACGTCTGAACTCTTTCCCTACACGACGCTCTTCCGATCTNNNGGTTGGGACATCCTTACACA
GAGCACACGTCTGAACTCTTTCCCTACACGACGCTCTTCCGATCTNNNGGTTGGGACATCCTTACACA
GAGCACACGTCTGAACTCTTTCCCTACACGACGCTCTTCCGATCTNNNGAGGCTCTGAAGTTTTCCCTG
GAGCACACGTCTGAACTCTTTCCCTACACGACGCTCTTCCGATCTNNNGAGGCTCTGAAGTTTTCCCTG

AGCACACGTCTGAACTCTTTCCCTACACGACGCTCTTCCGATCTNNNTTGTGTTATGGCTTTACCCCAC
AGCACACGTCTGAACTCTTTCCCTACACGACGCTCTTCCGATCTNNNTTGTGTTATGGCTTTACCCCAC
AGCACACGTCTGAACTCTTTCCCTACACGACGCTCTTCCGATCTNNNAGTGCTAGGATTACAGGCATG
AGCACACGTCTGAACTCTTTCCCTACACGACGCTCTTCCGATCTNNNAGTGCTAGGATTACAGGCATG
AGAGCACACGTCTGAACTCTTTCCCTACACGACGCTCTTCCGATCTNNNTAACTTGGGGAGTGGGATC
AGAGCACACGTCTGAACTCTTTCCCTACACGACGCTCTTCCGATCTNNNTAACTTGGGGAGTGGGATC
GAGCACACGTCTGAACTCTTTCCCTACACGACGCTCTTCCGATCTNNNTGTGTGCATCTAGCTTTGTAA
GAGCACACGTCTGAACTCTTTCCCTACACGACGCTCTTCCGATCTNNNTGTGTGCATCTAGCTTTGTAA
GAGCACACGTCTGAACTCTTTCCCTACACGACGCTCTTCCGATCTNNNGAGTTTGTATTAAAAGGTACT
GAGCACACGTCTGAACTCTTTCCCTACACGACGCTCTTCCGATCTNNNGAGTTTGTATTAAAAGGTACT
GAGCACACGTCTGAACTCTTTCCCTACACGACGCTCTTCCGATCTNNNTCTGCCTATGAAAAAGATGGG
GAGCACACGTCTGAACTCTTTCCCTACACGACGCTCTTCCGATCTNNNTCTGCCTATGAAAAAGATGGG
GCACACGTCTGAACTCTTTCCCTACACGACGCTCTTCCGATCTNNNACAGGGAGGTTTCTGCTATGTT
GCACACGTCTGAACTCTTTCCCTACACGACGCTCTTCCGATCTNNNACAGGGAGGTTTCTGCTATGTT
GCACACGTCTGAACTCTTTCCCTACACGACGCTCTTCCGATCTNNNCCTTGGTCTCTTCGGTCTCCTCAT
GCACACGTCTGAACTCTTTCCCTACACGACGCTCTTCCGATCTNNNCCTTGGTCTCTTCGGTCTCCTCAT
GAGCACACGTCTGAACTCTTTCCCTACACGACGCTCTTCCGATCTNNNCTACTGCACTCCAGCCTATTTG
GAGCACACGTCTGAACTCTTTCCCTACACGACGCTCTTCCGATCTNNNCTACTGCACTCCAGCCTATTTG
AGAGCACACGTCTGAACTCTTTCCCTACACGACGCTCTTCCGATCTNNNACACAGAACTGACTCTACGT
AGAGCACACGTCTGAACTCTTTCCCTACACGACGCTCTTCCGATCTNNNACACAGAACTGACTCTACGT
AGAGCACACGTCTGAACTCTTTCCCTACACGACGCTCTTCCGATCTNNNCATGCCTCTAATCCCAGCTAC
AGAGCACACGTCTGAACTCTTTCCCTACACGACGCTCTTCCGATCTNNNCATGCCTCTAATCCCAGCTAC
GAGCACACGTCTGAACTCTTTCCCTACACGACGCTCTTCCGATCTNNNGGTACCTCAGATGGAAATGC
GAGCACACGTCTGAACTCTTTCCCTACACGACGCTCTTCCGATCTNNNGGTACCTCAGATGGAAATGC
GCACACGTCTGAACTCTTTCCCTACACGACGCTCTTCCGATCTNNNCTGTAGTACCAGCTACTTGGGA
GCACACGTCTGAACTCTTTCCCTACACGACGCTCTTCCGATCTNNNCTGTAGTACCAGCTACTTGGGA
GCACACGTCTGAACTCTTTCCCTACACGACGCTCTTCCGATCTNNNAGTCCAACTCTTATGCAGGGAA
GCACACGTCTGAACTCTTTCCCTACACGACGCTCTTCCGATCTNNNAGTCCAACTCTTATGCAGGGAA
GCACACGTCTGAACTCTTTCCCTACACGACGCTCTTCCGATCTNNNTTGTGAGGAGATTTCTGAGTGC
GCACACGTCTGAACTCTTTCCCTACACGACGCTCTTCCGATCTNNNTTGTGAGGAGATTTCTGAGTGC
GAGCACACGTCTGAACTCTTTCCCTACACGACGCTCTTCCGATCTNNNGGCACTGGGGATAGAGATGT
GAGCACACGTCTGAACTCTTTCCCTACACGACGCTCTTCCGATCTNNNGGCACTGGGGATAGAGATGT
GAGCACACGTCTGAACTCTTTCCCTACACGACGCTCTTCCGATCTNNNTGAATACCTCTGCCCCACTTCA
GAGCACACGTCTGAACTCTTTCCCTACACGACGCTCTTCCGATCTNNNTGAATACCTCTGCCCCACTTCA
GCACACGTCTGAACTCTTTCCCTACACGACGCTCTTCCGATCTNNNAATGCCCTTGATCCCAACTACAT
GCACACGTCTGAACTCTTTCCCTACACGACGCTCTTCCGATCTNNNAATGCCCTTGATCCCAACTACAT
AGCACACGTCTGAACTCTTTCCCTACACGACGCTCTTCCGATCTNNNGATCGAGACCATCCTGGCTAA
AGCACACGTCTGAACTCTTTCCCTACACGACGCTCTTCCGATCTNNNGATCGAGACCATCCTGGCTAA
AGCACACGTCTGAACTCTTTCCCTACACGACGCTCTTCCGATCTNNNCTCTTAGCTGCCTACCCACATAG
AGCACACGTCTGAACTCTTTCCCTACACGACGCTCTTCCGATCTNNNCTCTTAGCTGCCTACCCACATAG
GCACACGTCTGAACTCTTTCCCTACACGACGCTCTTCCGATCTNNNTTTAGTACAGACGGGGTTTCAC
GCACACGTCTGAACTCTTTCCCTACACGACGCTCTTCCGATCTNNNTTTAGTACAGACGGGGTTTCAC
GAGCACACGTCTGAACTCTTTCCCTACACGACGCTCTTCCGATCTNNNGTTCCCAGCGTAGTCCATTTTT

GAGCACACGTCTGAACTCTTTCCCTACACGACGCTCTTCCGATCTNNNGTTCCCAGCGTAGTCCATTTTT
AGAGCACACGTCTGAACTCTTTCCCTACACGACGCTCTTCCGATCTNNNCTGAGGCAGGAGAATCACT
AGAGCACACGTCTGAACTCTTTCCCTACACGACGCTCTTCCGATCTNNNCTGAGGCAGGAGAATCACT
AGCACACGTCTGAACTCTTTCCCTACACGACGCTCTTCCGATCTNNNGTGTGAAATGCTTCTGGCTTTCT
AGCACACGTCTGAACTCTTTCCCTACACGACGCTCTTCCGATCTNNNGTGTGAAATGCTTCTGGCTTTCT
AGAGCACACGTCTGAACTCTTTCCCTACACGACGCTCTTCCGATCTNNNACAAGGAAGGAGGAAGGC
AGAGCACACGTCTGAACTCTTTCCCTACACGACGCTCTTCCGATCTNNNACAAGGAAGGAGGAAGGC
AGAGCACACGTCTGAACTCTTTCCCTACACGACGCTCTTCCGATCTNNNCCTCTGCTAGCATTTTATTGG
AGAGCACACGTCTGAACTCTTTCCCTACACGACGCTCTTCCGATCTNNNCCTCTGCTAGCATTTTATTGG
AAGAGCACACGTCTGAACTCTTTCCCTACACGACGCTCTTCCGATCTNNNTAAGAGAAGGAAGAGGCA
AAGAGCACACGTCTGAACTCTTTCCCTACACGACGCTCTTCCGATCTNNNTAAGAGAAGGAAGAGGCA
GAGCACACGTCTGAACTCTTTCCCTACACGACGCTCTTCCGATCTNNNTGTGTCATTATACTCCAGCCTG
GAGCACACGTCTGAACTCTTTCCCTACACGACGCTCTTCCGATCTNNNTGTGTCATTATACTCCAGCCTG
AGCACACGTCTGAACTCTTTCCCTACACGACGCTCTTCCGATCTNNNGCCTGCAGTTCCAATTGGTAAAT
AGCACACGTCTGAACTCTTTCCCTACACGACGCTCTTCCGATCTNNNGCCTGCAGTTCCAATTGGTAAAT
GAGCACACGTCTGAACTCTTTCCCTACACGACGCTCTTCCGATCTNNNGGCTGTTACCTTGGAGTGGTT
GAGCACACGTCTGAACTCTTTCCCTACACGACGCTCTTCCGATCTNNNGGCTGTTACCTTGGAGTGGTT
GAGCACACGTCTGAACTCTTTCCCTACACGACGCTCTTCCGATCTNNNAAGCCGGCCAATTCAATAACA
GAGCACACGTCTGAACTCTTTCCCTACACGACGCTCTTCCGATCTNNNAAGCCGGCCAATTCAATAACA
AGCACACGTCTGAACTCTTTCCCTACACGACGCTCTTCCGATCTNNNGAGGAGCAGCTTCTAGACTCTT
AGCACACGTCTGAACTCTTTCCCTACACGACGCTCTTCCGATCTNNNGAGGAGCAGCTTCTAGACTCTT
GAGCACACGTCTGAACTCTTTCCCTACACGACGCTCTTCCGATCTNNNTCGCAAAAGGAGAGAGTGAA
GAGCACACGTCTGAACTCTTTCCCTACACGACGCTCTTCCGATCTNNNTCGCAAAAGGAGAGAGTGAA
AGCACACGTCTGAACTCTTTCCCTACACGACGCTCTTCCGATCTNNNTCCATGAACATTGGATTTATCTA
AGCACACGTCTGAACTCTTTCCCTACACGACGCTCTTCCGATCTNNNTCCATGAACATTGGATTTATCTA
GAGCACACGTCTGAACTCTTTCCCTACACGACGCTCTTCCGATCTNNNAATTTCTTCCACCAGGCAACCT
GAGCACACGTCTGAACTCTTTCCCTACACGACGCTCTTCCGATCTNNNAATTTCTTCCACCAGGCAACCT
AGCACACGTCTGAACTCTTTCCCTACACGACGCTCTTCCGATCTNNNCCCAGGTGGCCTAATTAGTCAAA
AGCACACGTCTGAACTCTTTCCCTACACGACGCTCTTCCGATCTNNNCCCAGGTGGCCTAATTAGTCAAA
GAGCACACGTCTGAACTCTTTCCCTACACGACGCTCTTCCGATCTNNNCTTTGACAAGACACAGGCACTG
GAGCACACGTCTGAACTCTTTCCCTACACGACGCTCTTCCGATCTNNNCTTTGACAAGACACAGGCACTG
GAGCACACGTCTGAACTCTTTCCCTACACGACGCTCTTCCGATCTNNNGTGGTCCTGAGGGGAGAAGA
GAGCACACGTCTGAACTCTTTCCCTACACGACGCTCTTCCGATCTNNNGTGGTCCTGAGGGGAGAAGA
AGAGCACACGTCTGAACTCTTTCCCTACACGACGCTCTTCCGATCTNNNTGATGCCCAGTAGGTAAGGA
AGAGCACACGTCTGAACTCTTTCCCTACACGACGCTCTTCCGATCTNNNTGATGCCCAGTAGGTAAGGA
GAGCACACGTCTGAACTCTTTCCCTACACGACGCTCTTCCGATCTNNNTTTGGCAGAACTGTTTTCTTGG
GAGCACACGTCTGAACTCTTTCCCTACACGACGCTCTTCCGATCTNNNTTTGGCAGAACTGTTTTCTTGG
GAGCACACGTCTGAACTCTTTCCCTACACGACGCTCTTCCGATCTNNNCTTCTGGAAAATGGTGTGGG
GAGCACACGTCTGAACTCTTTCCCTACACGACGCTCTTCCGATCTNNNCTTCTGGAAAATGGTGTGGG
AGCACACGTCTGAACTCTTTCCCTACACGACGCTCTTCCGATCTNNNAGAGGCTCTTGTTTTCTTCCCTT
AGCACACGTCTGAACTCTTTCCCTACACGACGCTCTTCCGATCTNNNAGAGGCTCTTGTTTTCTTCCCTT
GAGCACACGTCTGAACTCTTTCCCTACACGACGCTCTTCCGATCTNNNCATTGGTTGGACAGTGGTAG
GAGCACACGTCTGAACTCTTTCCCTACACGACGCTCTTCCGATCTNNNCATTGGTTGGACAGTGGTAG

GAGCACACGTCTGAACTCTTTCCCTACACGACGCTCTTCCGATCTNNNGCAATATACATCTGTATGCA
GAGCACACGTCTGAACTCTTTCCCTACACGACGCTCTTCCGATCTNNNGCAATATACATCTGTATGCA
AGCACACGTCTGAACTCTTTCCCTACACGACGCTCTTCCGATCTNNNTGTAATCCCAGCACTTTGAGA
AGCACACGTCTGAACTCTTTCCCTACACGACGCTCTTCCGATCTNNNTGTAATCCCAGCACTTTGAGA
AGCACACGTCTGAACTCTTTCCCTACACGACGCTCTTCCGATCTNNNCCGAAAGAGGCAATCATCACA
AGCACACGTCTGAACTCTTTCCCTACACGACGCTCTTCCGATCTNNNCCGAAAGAGGCAATCATCACA
AGCACACGTCTGAACTCTTTCCCTACACGACGCTCTTCCGATCTNNNACCATGTGACCAGCTACAGAA
AGCACACGTCTGAACTCTTTCCCTACACGACGCTCTTCCGATCTNNNACCATGTGACCAGCTACAGAA
GAGCACACGTCTGAACTCTTTCCCTACACGACGCTCTTCCGATCTNNNAAATTAGCTATGTGTGGTGA
GAGCACACGTCTGAACTCTTTCCCTACACGACGCTCTTCCGATCTNNNAAATTAGCTATGTGTGGTGA
GCACACGTCTGAACTCTTTCCCTACACGACGCTCTTCCGATCTNNNAGCTGAGATTACAGGTTCACAC
GCACACGTCTGAACTCTTTCCCTACACGACGCTCTTCCGATCTNNNAGCTGAGATTACAGGTTCACAC
GCACACGTCTGAACTCTTTCCCTACACGACGCTCTTCCGATCTNNNTGAAAGAGTTGAGAGGTGAAA
GCACACGTCTGAACTCTTTCCCTACACGACGCTCTTCCGATCTNNNTGAAAGAGTTGAGAGGTGAAA
AGCACACGTCTGAACTCTTTCCCTACACGACGCTCTTCCGATCTNNNTGAAATAGAGACGTCTGGGTT
AGCACACGTCTGAACTCTTTCCCTACACGACGCTCTTCCGATCTNNNTGAAATAGAGACGTCTGGGTT
GCACACGTCTGAACTCTTTCCCTACACGACGCTCTTCCGATCTNNNCCTGGCTAATTTTGTGTGTGTGG
GCACACGTCTGAACTCTTTCCCTACACGACGCTCTTCCGATCTNNNCCTGGCTAATTTTGTGTGTGTGG
AGCACACGTCTGAACTCTTTCCCTACACGACGCTCTTCCGATCTNNNCGAGATGCTACAGAATGCAG
AGCACACGTCTGAACTCTTTCCCTACACGACGCTCTTCCGATCTNNNCGAGATGCTACAGAATGCAG
GCACACGTCTGAACTCTTTCCCTACACGACGCTCTTCCGATCTNNNGTGGTCAACAGCATCTTCAAAG
GCACACGTCTGAACTCTTTCCCTACACGACGCTCTTCCGATCTNNNGTGGTCAACAGCATCTTCAAAG
GCACACGTCTGAACTCTTTCCCTACACGACGCTCTTCCGATCTNNNACAGCATGAAGAACAAAACCA
GCACACGTCTGAACTCTTTCCCTACACGACGCTCTTCCGATCTNNNACAGCATGAAGAACAAAACCA
GAGCACACGTCTGAACTCTTTCCCTACACGACGCTCTTCCGATCTNNNTATTCCAGTCTAGCCGCTCTT
GAGCACACGTCTGAACTCTTTCCCTACACGACGCTCTTCCGATCTNNNTATTCCAGTCTAGCCGCTCTT
AGCACACGTCTGAACTCTTTCCCTACACGACGCTCTTCCGATCTNNNACACTGGTTATTTGGTGCATG
AGCACACGTCTGAACTCTTTCCCTACACGACGCTCTTCCGATCTNNNACACTGGTTATTTGGTGCATG
GCACACGTCTGAACTCTTTCCCTACACGACGCTCTTCCGATCTNNNTAAATGTGGAGCTAGCTCTGA
GCACACGTCTGAACTCTTTCCCTACACGACGCTCTTCCGATCTNNNTAAATGTGGAGCTAGCTCTGA
AGCACACGTCTGAACTCTTTCCCTACACGACGCTCTTCCGATCTNNNGGTGCAGTTATGCCTCAGATT
AGCACACGTCTGAACTCTTTCCCTACACGACGCTCTTCCGATCTNNNGGTGCAGTTATGCCTCAGATT
AGCACACGTCTGAACTCTTTCCCTACACGACGCTCTTCCGATCTNNNCAGATACTCGGGAAGCTGAG
AGCACACGTCTGAACTCTTTCCCTACACGACGCTCTTCCGATCTNNNCAGATACTCGGGAAGCTGAG
AGCACACGTCTGAACTCTTTCCCTACACGACGCTCTTCCGATCTNNNAGGGACTTGTGCTTTTCTGAG
AGCACACGTCTGAACTCTTTCCCTACACGACGCTCTTCCGATCTNNNAGGGACTTGTGCTTTTCTGAG
GCACACGTCTGAACTCTTTCCCTACACGACGCTCTTCCGATCTNNNAGTAATACAATGTTCTTGTCAT
GCACACGTCTGAACTCTTTCCCTACACGACGCTCTTCCGATCTNNNAGTAATACAATGTTCTTGTCAT
AGCACACGTCTGAACTCTTTCCCTACACGACGCTCTTCCGATCTNNNGATCACAAGGTCAGGAGATC
AGCACACGTCTGAACTCTTTCCCTACACGACGCTCTTCCGATCTNNNGATCACAAGGTCAGGAGATC
GCACACGTCTGAACTCTTTCCCTACACGACGCTCTTCCGATCTNNNTTTTATTGTTACGAAGGACTTTT
GCACACGTCTGAACTCTTTCCCTACACGACGCTCTTCCGATCTNNNTTTTATTGTTACGAAGGACTTTT
GAGCACACGTCTGAACTCTTTCCCTACACGACGCTCTTCCGATCTNNNAATGCAGATGGTTTAGGGTT

GAGCACACGTCTGAACTCTTTCCCTACACGACGCTCTTCCGATCTNNNAATGCAGATGGTTTAGGGTT
AGCACACGTCTGAACTCTTTCCCTACACGACGCTCTTCCGATCTNNNCAGTTGACCCAGGAATTACAG
AGCACACGTCTGAACTCTTTCCCTACACGACGCTCTTCCGATCTNNNCAGTTGACCCAGGAATTACAG
GAGCACACGTCTGAACTCTTTCCCTACACGACGCTCTTCCGATCTNNNGAGATGGAGCCAAGAAGG
GAGCACACGTCTGAACTCTTTCCCTACACGACGCTCTTCCGATCTNNNGAGATGGAGCCAAGAAGG
AGCACACGTCTGAACTCTTTCCCTACACGACGCTCTTCCGATCTNNNCCCTCCCCCATTTAGAAAG
AGCACACGTCTGAACTCTTTCCCTACACGACGCTCTTCCGATCTNNNCCCTCCCCCATTTAGAAAG
AGAGCACACGTCTGAACTCTTTCCCTACACGACGCTCTTCCGATCTNNNCGCCTATAGTCCCAGCTACT
AGAGCACACGTCTGAACTCTTTCCCTACACGACGCTCTTCCGATCTNNNCGCCTATAGTCCCAGCTACT
AGCACACGTCTGAACTCTTTCCCTACACGACGCTCTTCCGATCTNNNTCAATAACTGTGGCTGCATTTG
AGCACACGTCTGAACTCTTTCCCTACACGACGCTCTTCCGATCTNNNTCAATAACTGTGGCTGCATTTG
GCACACGTCTGAACTCTTTCCCTACACGACGCTCTTCCGATCTNNNCCCACAAAAAGTCATCAGGGTT
GCACACGTCTGAACTCTTTCCCTACACGACGCTCTTCCGATCTNNNCCCACAAAAAGTCATCAGGGTT
AGCACACGTCTGAACTCTTTCCCTACACGACGCTCTTCCGATCTNNNTTTGCATTCCTTTGTGTCTCCCA
AGCACACGTCTGAACTCTTTCCCTACACGACGCTCTTCCGATCTNNNTTTGCATTCCTTTGTGTCTCCCA
GCACACGTCTGAACTCTTTCCCTACACGACGCTCTTCCGATCTNNNCTCATGGTGGTTGTGATTTGCAT
GCACACGTCTGAACTCTTTCCCTACACGACGCTCTTCCGATCTNNNCTCATGGTGGTTGTGATTTGCAT
GAGCACACGTCTGAACTCTTTCCCTACACGACGCTCTTCCGATCTNNNTGGTCAGCTATTTTGCAGAA
GAGCACACGTCTGAACTCTTTCCCTACACGACGCTCTTCCGATCTNNNTGGTCAGCTATTTTGCAGAA
CACACGTCTGAACTCTTTCCCTACACGACGCTCTTCCGATCTNNNCAAGCAATTCTCTCTCCCACCTCCA
CACACGTCTGAACTCTTTCCCTACACGACGCTCTTCCGATCTNNNCAAGCAATTCTCTCTCCCACCTCCA
GCACACGTCTGAACTCTTTCCCTACACGACGCTCTTCCGATCTNNNGGGATCCAAGACTGATTCACCAT
GCACACGTCTGAACTCTTTCCCTACACGACGCTCTTCCGATCTNNNGGGATCCAAGACTGATTCACCAT
AGCACACGTCTGAACTCTTTCCCTACACGACGCTCTTCCGATCTNNNCGACAGAGTGAGAATCCGTCT
AGCACACGTCTGAACTCTTTCCCTACACGACGCTCTTCCGATCTNNNCGACAGAGTGAGAATCCGTCT
CACACGTCTGAACTCTTTCCCTACACGACGCTCTTCCGATCTNNNCTTTGGTGGCTTATCTGATGCAT
CACACGTCTGAACTCTTTCCCTACACGACGCTCTTCCGATCTNNNCTTTGGTGGCTTATCTGATGCAT
CACACGTCTGAACTCTTTCCCTACACGACGCTCTTCCGATCTNNNTTCAGGTCCACTTTCTCCACAAACA
CACACGTCTGAACTCTTTCCCTACACGACGCTCTTCCGATCTNNNTTCAGGTCCACTTTCTCCACAAACA
GAGCACACGTCTGAACTCTTTCCCTACACGACGCTCTTCCGATCTNNNTTTAAAGATTGGTTTCTGTTT
GAGCACACGTCTGAACTCTTTCCCTACACGACGCTCTTCCGATCTNNNTTTAAAGATTGGTTTCTGTTT
AGCACACGTCTGAACTCTTTCCCTACACGACGCTCTTCCGATCTNNNTGGGAACTAGTGACATTTTGC
AGCACACGTCTGAACTCTTTCCCTACACGACGCTCTTCCGATCTNNNTGGGAACTAGTGACATTTTGC
GCACACGTCTGAACTCTTTCCCTACACGACGCTCTTCCGATCTNNNACAATCTATTCTGTGAATCATTT
GCACACGTCTGAACTCTTTCCCTACACGACGCTCTTCCGATCTNNNACAATCTATTCTGTGAATCATTT
AGCACACGTCTGAACTCTTTCCCTACACGACGCTCTTCCGATCTNNNGTGCATCTCCAGACTCACAGA
AGCACACGTCTGAACTCTTTCCCTACACGACGCTCTTCCGATCTNNNGTGCATCTCCAGACTCACAGA
AGCACACGTCTGAACTCTTTCCCTACACGACGCTCTTCCGATCTNNNTGTTACTACAGCACAAAAGCA
AGCACACGTCTGAACTCTTTCCCTACACGACGCTCTTCCGATCTNNNTGTTACTACAGCACAAAAGCA
AGCACACGTCTGAACTCTTTCCCTACACGACGCTCTTCCGATCTNNNGTGGAGTGCTGTTTGTGGT
AGCACACGTCTGAACTCTTTCCCTACACGACGCTCTTCCGATCTNNNGTGGAGTGCTGTTTGTGGT
GAGCACACGTCTGAACTCTTTCCCTACACGACGCTCTTCCGATCTNNNGTTACAATGAGCCAAGATCC
GAGCACACGTCTGAACTCTTTCCCTACACGACGCTCTTCCGATCTNNNGTTACAATGAGCCAAGATCC

GCACACGTCTGAACTCTTTCCCTACACGACGCTCTTCCGATCTNNNGAGAGACAGAGCCACACACTGC
GCACACGTCTGAACTCTTTCCCTACACGACGCTCTTCCGATCTNNNGAGAGACAGAGCCACACACTGC
AGCACACGTCTGAACTCTTTCCCTACACGACGCTCTTCCGATCTNNNACAAGAGAAAATGTAAGCCTC
AGCACACGTCTGAACTCTTTCCCTACACGACGCTCTTCCGATCTNNNACAAGAGAAAATGTAAGCCTC
GAGCACACGTCTGAACTCTTTCCCTACACGACGCTCTTCCGATCTNNNCTGCTCACTATAACCTCCACCT
GAGCACACGTCTGAACTCTTTCCCTACACGACGCTCTTCCGATCTNNNCTGCTCACTATAACCTCCACCT
AGCACACGTCTGAACTCTTTCCCTACACGACGCTCTTCCGATCTNNNACTCCTCTAATGACCTCTTCCCA
AGCACACGTCTGAACTCTTTCCCTACACGACGCTCTTCCGATCTNNNACTCCTCTAATGACCTCTTCCCA
CACACGTCTGAACTCTTTCCCTACACGACGCTCTTCCGATCTNNNAGCATTGAAATAAAAGCACAGCA
CACACGTCTGAACTCTTTCCCTACACGACGCTCTTCCGATCTNNNAGCATTGAAATAAAAGCACAGCA
AGCACACGTCTGAACTCTTTCCCTACACGACGCTCTTCCGATCTNNNGTCACATATCCCACTGCTTCAG
AGCACACGTCTGAACTCTTTCCCTACACGACGCTCTTCCGATCTNNNGTCACATATCCCACTGCTTCAG
AGCACACGTCTGAACTCTTTCCCTACACGACGCTCTTCCGATCTNNNAAACACAAAAATTAGCTGGGC
AGCACACGTCTGAACTCTTTCCCTACACGACGCTCTTCCGATCTNNNAAACACAAAAATTAGCTGGGC
GAGCACACGTCTGAACTCTTTCCCTACACGACGCTCTTCCGATCTNNNCTGCACCCACTAACTCATCAT
GAGCACACGTCTGAACTCTTTCCCTACACGACGCTCTTCCGATCTNNNCTGCACCCACTAACTCATCAT
AGCACACGTCTGAACTCTTTCCCTACACGACGCTCTTCCGATCTNNNGTTGCAGTGACCCAAGATCATG
AGCACACGTCTGAACTCTTTCCCTACACGACGCTCTTCCGATCTNNNGTTGCAGTGACCCAAGATCATG
GAGCACACGTCTGAACTCTTTCCCTACACGACGCTCTTCCGATCTNNNTTACGGTTGGCCCATTTCTTTT
GAGCACACGTCTGAACTCTTTCCCTACACGACGCTCTTCCGATCTNNNTTACGGTTGGCCCATTTCTTTT
GAGCACACGTCTGAACTCTTTCCCTACACGACGCTCTTCCGATCTNNNAACATGGCAAAACCCCATCTCT
GAGCACACGTCTGAACTCTTTCCCTACACGACGCTCTTCCGATCTNNNAACATGGCAAAACCCCATCTCT
GAGCACACGTCTGAACTCTTTCCCTACACGACGCTCTTCCGATCTNNNTGATGATTAGTGGTGTTGAG
GAGCACACGTCTGAACTCTTTCCCTACACGACGCTCTTCCGATCTNNNTGATGATTAGTGGTGTTGAG
GAGCACACGTCTGAACTCTTTCCCTACACGACGCTCTTCCGATCTNNNTAAGGAAAGGAGAGCCGTTT
GAGCACACGTCTGAACTCTTTCCCTACACGACGCTCTTCCGATCTNNNTAAGGAAAGGAGAGCCGTTT
GCACACGTCTGAACTCTTTCCCTACACGACGCTCTTCCGATCTNNNCACCCAATCATAGTGGCGGAGCA
GCACACGTCTGAACTCTTTCCCTACACGACGCTCTTCCGATCTNNNCACCCAATCATAGTGGCGGAGCA
GAGCACACGTCTGAACTCTTTCCCTACACGACGCTCTTCCGATCTNNNCTGCATGAGCTGTGCTATTTC
GAGCACACGTCTGAACTCTTTCCCTACACGACGCTCTTCCGATCTNNNCTGCATGAGCTGTGCTATTTC
AGCACACGTCTGAACTCTTTCCCTACACGACGCTCTTCCGATCTNNNTAGCCTTTATTGCCTACTCCACC
AGCACACGTCTGAACTCTTTCCCTACACGACGCTCTTCCGATCTNNNTAGCCTTTATTGCCTACTCCACC
AGCACACGTCTGAACTCTTTCCCTACACGACGCTCTTCCGATCTNNNCTGATGCCTGTAATCCTAGCAC
AGCACACGTCTGAACTCTTTCCCTACACGACGCTCTTCCGATCTNNNCTGATGCCTGTAATCCTAGCAC
AGCACACGTCTGAACTCTTTCCCTACACGACGCTCTTCCGATCTNNNTCTTCACCTTTGACATCTGTGT
AGCACACGTCTGAACTCTTTCCCTACACGACGCTCTTCCGATCTNNNTCTTCACCTTTGACATCTGTGT
GAGCACACGTCTGAACTCTTTCCCTACACGACGCTCTTCCGATCTNNNTACTTGGGTTTCTCAGGTGAT
GAGCACACGTCTGAACTCTTTCCCTACACGACGCTCTTCCGATCTNNNTACTTGGGTTTCTCAGGTGAT
AAGAGCACACGTCTGAACTCTTTCCCTACACGACGCTCTTCCGATCTNNNGCCTATAATCCTAGCTACT
AAGAGCACACGTCTGAACTCTTTCCCTACACGACGCTCTTCCGATCTNNNGCCTATAATCCTAGCTACT
GCACACGTCTGAACTCTTTCCCTACACGACGCTCTTCCGATCTNNNTCTCACTGCTGACTACTTGCTTTC
GCACACGTCTGAACTCTTTCCCTACACGACGCTCTTCCGATCTNNNTCTCACTGCTGACTACTTGCTTTC
CACACGTCTGAACTCTTTCCCTACACGACGCTCTTCCGATCTNNNGCTGGAGAATTGCTTGAACCTGCA

CACACGTCTGAACTCTTTCCCTACACGACGCTCTTCCGATCTNNNGCTGGAGAATTGCTTGAACCTGCA
AGCACACGTCTGAACTCTTTCCCTACACGACGCTCTTCCGATCTNNNTCAGGAGAATCACTTGAACCCA
AGCACACGTCTGAACTCTTTCCCTACACGACGCTCTTCCGATCTNNNTCAGGAGAATCACTTGAACCCA
GAGCACACGTCTGAACTCTTTCCCTACACGACGCTCTTCCGATCTNNNATCTTGCCATTGTGAGATTGT
GAGCACACGTCTGAACTCTTTCCCTACACGACGCTCTTCCGATCTNNNATCTTGCCATTGTGAGATTGT
GCACACGTCTGAACTCTTTCCCTACACGACGCTCTTCCGATCTNNNCAATGGCTTCGCTCAGACCCCATC
GCACACGTCTGAACTCTTTCCCTACACGACGCTCTTCCGATCTNNNCAATGGCTTCGCTCAGACCCCATC
AGCACACGTCTGAACTCTTTCCCTACACGACGCTCTTCCGATCTNNNGGCAGAATGGGAGAAACGTTTA
AGCACACGTCTGAACTCTTTCCCTACACGACGCTCTTCCGATCTNNNGGCAGAATGGGAGAAACGTTTA
AGCACACGTCTGAACTCTTTCCCTACACGACGCTCTTCCGATCTNNNTGAGATTACAGATGTGAGCCAC
AGCACACGTCTGAACTCTTTCCCTACACGACGCTCTTCCGATCTNNNTGAGATTACAGATGTGAGCCAC
AGAGCACACGTCTGAACTCTTTCCCTACACGACGCTCTTCCGATCTNNNGGCATAGCATCACCAAGTTG
AGAGCACACGTCTGAACTCTTTCCCTACACGACGCTCTTCCGATCTNNNGGCATAGCATCACCAAGTTG
AGAGCACACGTCTGAACTCTTTCCCTACACGACGCTCTTCCGATCTNNNCACCTGTAATCCCAGCTACTT
AGAGCACACGTCTGAACTCTTTCCCTACACGACGCTCTTCCGATCTNNNCACCTGTAATCCCAGCTACTT
AGCACACGTCTGAACTCTTTCCCTACACGACGCTCTTCCGATCTNNNGAACCCAGGAGACAGAGCTTG
AGCACACGTCTGAACTCTTTCCCTACACGACGCTCTTCCGATCTNNNGAACCCAGGAGACAGAGCTTG
CACACGTCTGAACTCTTTCCCTACACGACGCTCTTCCGATCTNNNGGCTCAAATTCTGCCTCATTGTTCA
CACACGTCTGAACTCTTTCCCTACACGACGCTCTTCCGATCTNNNGGCTCAAATTCTGCCTCATTGTTCA
GAGCACACGTCTGAACTCTTTCCCTACACGACGCTCTTCCGATCTNNNGATGACTCACAGCCCATCCCAT
GAGCACACGTCTGAACTCTTTCCCTACACGACGCTCTTCCGATCTNNNGATGACTCACAGCCCATCCCAT
AGCACACGTCTGAACTCTTTCCCTACACGACGCTCTTCCGATCTNNNGTTGTACAGAACCAATGGTACT
AGCACACGTCTGAACTCTTTCCCTACACGACGCTCTTCCGATCTNNNGTTGTACAGAACCAATGGTACT
AGCACACGTCTGAACTCTTTCCCTACACGACGCTCTTCCGATCTNNNATTTGCTTCACAGGATGGTTGT
AGCACACGTCTGAACTCTTTCCCTACACGACGCTCTTCCGATCTNNNATTTGCTTCACAGGATGGTTGT
AGCACACGTCTGAACTCTTTCCCTACACGACGCTCTTCCGATCTNNNAAACACTCCCTCCCTTTTCCTTTCA
AGCACACGTCTGAACTCTTTCCCTACACGACGCTCTTCCGATCTNNNAAACACTCCCTCCCTTTTCCTTTCA
AAGAGCACACGTCTGAACTCTTTCCCTACACGACGCTCTTCCGATCTNNNGTACCAAAAAGCAGAGGG
AAGAGCACACGTCTGAACTCTTTCCCTACACGACGCTCTTCCGATCTNNNGTACCAAAAAGCAGAGGG
AGAGCACACGTCTGAACTCTTTCCCTACACGACGCTCTTCCGATCTNNNGTGAACGCATATATTGGGCT
AGAGCACACGTCTGAACTCTTTCCCTACACGACGCTCTTCCGATCTNNNGTGAACGCATATATTGGGCT
AGAGCACACGTCTGAACTCTTTCCCTACACGACGCTCTTCCGATCTNNNATGATCTATTGCAGTCAGTG
AGAGCACACGTCTGAACTCTTTCCCTACACGACGCTCTTCCGATCTNNNATGATCTATTGCAGTCAGTG
AGCACACGTCTGAACTCTTTCCCTACACGACGCTCTTCCGATCTNNNACCGTTGATACGGTTACAGATA
AGCACACGTCTGAACTCTTTCCCTACACGACGCTCTTCCGATCTNNNACCGTTGATACGGTTACAGATA
AGCACACGTCTGAACTCTTTCCCTACACGACGCTCTTCCGATCTNNNTGAGTTTGAACCTCTGGTCTGA
AGCACACGTCTGAACTCTTTCCCTACACGACGCTCTTCCGATCTNNNTGAGTTTGAACCTCTGGTCTGA
GAGCACACGTCTGAACTCTTTCCCTACACGACGCTCTTCCGATCTNNNAAAGGCAGATAGTGTTAGTG
GAGCACACGTCTGAACTCTTTCCCTACACGACGCTCTTCCGATCTNNNAAAGGCAGATAGTGTTAGTG
AGAGCACACGTCTGAACTCTTTCCCTACACGACGCTCTTCCGATCTNNNATGTTCACGATATGCATGGC
AGAGCACACGTCTGAACTCTTTCCCTACACGACGCTCTTCCGATCTNNNATGTTCACGATATGCATGGC
AGCACACGTCTGAACTCTTTCCCTACACGACGCTCTTCCGATCTNNNCACTCATAGGATTTCCTTGGGC
AGCACACGTCTGAACTCTTTCCCTACACGACGCTCTTCCGATCTNNNCACTCATAGGATTTCCTTGGGC

GAGCACACGTCTGAACTCTTTCCCTACACGACGCTCTTCCGATCTNNNCTCAGTAAAACAGCACAACCGA
GAGCACACGTCTGAACTCTTTCCCTACACGACGCTCTTCCGATCTNNNCTCAGTAAAACAGCACAACCGA
AGCACACGTCTGAACTCTTTCCCTACACGACGCTCTTCCGATCTNNNACTTTGAGAGACAACATCTGCC
AGCACACGTCTGAACTCTTTCCCTACACGACGCTCTTCCGATCTNNNACTTTGAGAGACAACATCTGCC
AGAGCACACGTCTGAACTCTTTCCCTACACGACGCTCTTCCGATCTNNNAGTGAATGGTTCTGTCTCACT
AGAGCACACGTCTGAACTCTTTCCCTACACGACGCTCTTCCGATCTNNNAGTGAATGGTTCTGTCTCACT
CACACGTCTGAACTCTTTCCCTACACGACGCTCTTCCGATCTNNNAGTGCTGAGATTACAGGTGTGAG
CACACGTCTGAACTCTTTCCCTACACGACGCTCTTCCGATCTNNNAGTGCTGAGATTACAGGTGTGAG
AGCACACGTCTGAACTCTTTCCCTACACGACGCTCTTCCGATCTNNNGTTTCACTGTGTTAGCCAGGATC
AGCACACGTCTGAACTCTTTCCCTACACGACGCTCTTCCGATCTNNNGTTTCACTGTGTTAGCCAGGATC
GCACACGTCTGAACTCTTTCCCTACACGACGCTCTTCCGATCTNNNAGAAGCAGAAGGAGACCCAGAG
GCACACGTCTGAACTCTTTCCCTACACGACGCTCTTCCGATCTNNNAGAAGCAGAAGGAGACCCAGAG
GAGCACACGTCTGAACTCTTTCCCTACACGACGCTCTTCCGATCTNNNGTTGGGGTCCTGAGTATGTAT
GAGCACACGTCTGAACTCTTTCCCTACACGACGCTCTTCCGATCTNNNGTTGGGGTCCTGAGTATGTAT
CACACGTCTGAACTCTTTCCCTACACGACGCTCTTCCGATCTNNNGGGTTCCCCTAGACTCCCACCATCG
CACACGTCTGAACTCTTTCCCTACACGACGCTCTTCCGATCTNNNGGGTTCCCCTAGACTCCCACCATCG
GCACACGTCTGAACTCTTTCCCTACACGACGCTCTTCCGATCTNNNCTCAAGCAAAACCGTGAAGTAGC
GCACACGTCTGAACTCTTTCCCTACACGACGCTCTTCCGATCTNNNCTCAAGCAAAACCGTGAAGTAGC
GAGCACACGTCTGAACTCTTTCCCTACACGACGCTCTTCCGATCTNNNGTACAGCAAAGTCCTTGTCCC
GAGCACACGTCTGAACTCTTTCCCTACACGACGCTCTTCCGATCTNNNGTACAGCAAAGTCCTTGTCCC
AGAGCACACGTCTGAACTCTTTCCCTACACGACGCTCTTCCGATCTNNNCAATAGCTAAGCTCCTCACTG
AGAGCACACGTCTGAACTCTTTCCCTACACGACGCTCTTCCGATCTNNNCAATAGCTAAGCTCCTCACTG
GAGCACACGTCTGAACTCTTTCCCTACACGACGCTCTTCCGATCTNNNCTCCCTGGTTCATGCCATTCTCC
GAGCACACGTCTGAACTCTTTCCCTACACGACGCTCTTCCGATCTNNNCTCCCTGGTTCATGCCATTCTCC
AGAGCACACGTCTGAACTCTTTCCCTACACGACGCTCTTCCGATCTNNNCCGGGTTGACTTTCACAGCAA
AGAGCACACGTCTGAACTCTTTCCCTACACGACGCTCTTCCGATCTNNNCCGGGTTGACTTTCACAGCAA
AGCACACGTCTGAACTCTTTCCCTACACGACGCTCTTCCGATCTNNNTGAATTCAGCTTTATGTGGCAG
AGCACACGTCTGAACTCTTTCCCTACACGACGCTCTTCCGATCTNNNTGAATTCAGCTTTATGTGGCAG
AGCACACGTCTGAACTCTTTCCCTACACGACGCTCTTCCGATCTNNNGAACAACAGCCTTTAGCCCTAG
AGCACACGTCTGAACTCTTTCCCTACACGACGCTCTTCCGATCTNNNGAACAACAGCCTTTAGCCCTAG
GAGCACACGTCTGAACTCTTTCCCTACACGACGCTCTTCCGATCTNNNCAAGTGTTGTAGTTGGTCTTG
GAGCACACGTCTGAACTCTTTCCCTACACGACGCTCTTCCGATCTNNNCAAGTGTTGTAGTTGGTCTTG
AGCACACGTCTGAACTCTTTCCCTACACGACGCTCTTCCGATCTNNNATTGCTTTGTTTTGGATCGTGG
AGCACACGTCTGAACTCTTTCCCTACACGACGCTCTTCCGATCTNNNATTGCTTTGTTTTGGATCGTGG
GAGCACACGTCTGAACTCTTTCCCTACACGACGCTCTTCCGATCTNNNGACTACAGGTGTCCTTCTGTG
GAGCACACGTCTGAACTCTTTCCCTACACGACGCTCTTCCGATCTNNNGACTACAGGTGTCCTTCTGTG
AGCACACGTCTGAACTCTTTCCCTACACGACGCTCTTCCGATCTNNNGCACAGTGGCTCATACCTCTAAT
AGCACACGTCTGAACTCTTTCCCTACACGACGCTCTTCCGATCTNNNGCACAGTGGCTCATACCTCTAAT
AGAGCACACGTCTGAACTCTTTCCCTACACGACGCTCTTCCGATCTNNNCAGGAAGGATGTACTAAGAC
AGAGCACACGTCTGAACTCTTTCCCTACACGACGCTCTTCCGATCTNNNCAGGAAGGATGTACTAAGAC
CACACGTCTGAACTCTTTCCCTACACGACGCTCTTCCGATCTNNNATCTCCCAAAGGATATGCTCCATCG
CACACGTCTGAACTCTTTCCCTACACGACGCTCTTCCGATCTNNNATCTCCCAAAGGATATGCTCCATCG
GCACACGTCTGAACTCTTTCCCTACACGACGCTCTTCCGATCTNNNCTCATTGCCTTATGCTGTTGCTTC

GCACACGTCTGAACTCTTTCCCTACACGACGCTCTTCCGATCTNNNCTCATTGCCTTATGCTGTTGCTT
GCACACGTCTGAACTCTTTCCCTACACGACGCTCTTCCGATCTNNNAAATATACACAGGGCAGAGATA
GCACACGTCTGAACTCTTTCCCTACACGACGCTCTTCCGATCTNNNAAATATACACAGGGCAGAGATA
AGCACACGTCTGAACTCTTTCCCTACACGACGCTCTTCCGATCTNNNCATCAGTTTGCATCATGGAGCA
AGCACACGTCTGAACTCTTTCCCTACACGACGCTCTTCCGATCTNNNCATCAGTTTGCATCATGGAGCA
AGCACACGTCTGAACTCTTTCCCTACACGACGCTCTTCCGATCTNNNCAGGACACTCTTACACAGGAG
AGCACACGTCTGAACTCTTTCCCTACACGACGCTCTTCCGATCTNNNCAGGACACTCTTACACAGGAG
AGCACACGTCTGAACTCTTTCCCTACACGACGCTCTTCCGATCTNNNAGATCGTCCCAGTGATTAGAA
AGCACACGTCTGAACTCTTTCCCTACACGACGCTCTTCCGATCTNNNAGATCGTCCCAGTGATTAGAA
AGCACACGTCTGAACTCTTTCCCTACACGACGCTCTTCCGATCTNNNATCTGAGGCAGGAGAATCACTT
AGCACACGTCTGAACTCTTTCCCTACACGACGCTCTTCCGATCTNNNATCTGAGGCAGGAGAATCACTT
AGCACACGTCTGAACTCTTTCCCTACACGACGCTCTTCCGATCTNNNAAAAAATTAGCTGGGTGTGG
AGCACACGTCTGAACTCTTTCCCTACACGACGCTCTTCCGATCTNNNAAAAAATTAGCTGGGTGTGG
GCACACGTCTGAACTCTTTCCCTACACGACGCTCTTCCGATCTNNNACCTGGACATTTTGAAAATGGC
GCACACGTCTGAACTCTTTCCCTACACGACGCTCTTCCGATCTNNNACCTGGACATTTTGAAAATGGC
AGCACACGTCTGAACTCTTTCCCTACACGACGCTCTTCCGATCTNNNTTTGGACAGCATCAAGGAGCT
AGCACACGTCTGAACTCTTTCCCTACACGACGCTCTTCCGATCTNNNTTTGGACAGCATCAAGGAGCT
AGCACACGTCTGAACTCTTTCCCTACACGACGCTCTTCCGATCTNNNCAGAGTCCAGACCTAACACATC
AGCACACGTCTGAACTCTTTCCCTACACGACGCTCTTCCGATCTNNNCAGAGTCCAGACCTAACACATC
GAGCACACGTCTGAACTCTTTCCCTACACGACGCTCTTCCGATCTNNNACACACTCTTCTTTTCCCACCA
GAGCACACGTCTGAACTCTTTCCCTACACGACGCTCTTCCGATCTNNNACACACTCTTCTTTTCCCACCA
AGCACACGTCTGAACTCTTTCCCTACACGACGCTCTTCCGATCTNNNGGCAACAAGAGTGTAACAGT
AGCACACGTCTGAACTCTTTCCCTACACGACGCTCTTCCGATCTNNNGGCAACAAGAGTGTAACAGT
AGCACACGTCTGAACTCTTTCCCTACACGACGCTCTTCCGATCTNNNTGTGCTGACATGATCTTCTCAC
AGCACACGTCTGAACTCTTTCCCTACACGACGCTCTTCCGATCTNNNTGTGCTGACATGATCTTCTCAC
GCACACGTCTGAACTCTTTCCCTACACGACGCTCTTCCGATCTNNNTGATCACTGTGCAATGGTATCT
GCACACGTCTGAACTCTTTCCCTACACGACGCTCTTCCGATCTNNNTGATCACTGTGCAATGGTATCT
GCACACGTCTGAACTCTTTCCCTACACGACGCTCTTCCGATCTNNNCACAGAAAGCCAAAGCAAGAT
GCACACGTCTGAACTCTTTCCCTACACGACGCTCTTCCGATCTNNNCACAGAAAGCCAAAGCAAGAT
AGCACACGTCTGAACTCTTTCCCTACACGACGCTCTTCCGATCTNNNCTGTAATCCAAGCTACTGGGG
AGCACACGTCTGAACTCTTTCCCTACACGACGCTCTTCCGATCTNNNCTGTAATCCAAGCTACTGGGG
GAGCACACGTCTGAACTCTTTCCCTACACGACGCTCTTCCGATCTNNNATCTATGTGCAGGCCATCAA
GAGCACACGTCTGAACTCTTTCCCTACACGACGCTCTTCCGATCTNNNATCTATGTGCAGGCCATCAA
ACACGTCTGAACTCTTTCCCTACACGACGCTCTTCCGATCTNNNATGTTATAGTCCTTTGCATCCCATCG
ACACGTCTGAACTCTTTCCCTACACGACGCTCTTCCGATCTNNNATGTTATAGTCCTTTGCATCCCATCG
GAGCACACGTCTGAACTCTTTCCCTACACGACGCTCTTCCGATCTNNNAGACTGGATGACAAAGTGAG
GAGCACACGTCTGAACTCTTTCCCTACACGACGCTCTTCCGATCTNNNAGACTGGATGACAAAGTGAG
ACACGTCTGAACTCTTTCCCTACACGACGCTCTTCCGATCTNNNCCCAAGTGACTTAGGGAAGAGAG
ACACGTCTGAACTCTTTCCCTACACGACGCTCTTCCGATCTNNNCCCAAGTGACTTAGGGAAGAGAG
AGCACACGTCTGAACTCTTTCCCTACACGACGCTCTTCCGATCTNNNCTGGATAAGTCTGGGCTCATGT
AGCACACGTCTGAACTCTTTCCCTACACGACGCTCTTCCGATCTNNNCTGGATAAGTCTGGGCTCATGT
AGCACACGTCTGAACTCTTTCCCTACACGACGCTCTTCCGATCTNNNTCCCTCATGTTCTGTCAGTTAG
AGCACACGTCTGAACTCTTTCCCTACACGACGCTCTTCCGATCTNNNTCCCTCATGTTCTGTCAGTTAG

AGAGCACACGTCTGAACTCTTTCCCTACACGACGCTCTTCCGATCTNNNAGACTGCAGTGAGCTATGAT

AGAGCACACGTCTGAACTCTTTCCCTACACGACGCTCTTCCGATCTNNNAGACTGCAGTGAGCTATGAT

GAGCACACGTCTGAACTCTTTCCCTACACGACGCTCTTCCGATCTNNNTTCCACTTGGACAGACAGAAG

GAGCACACGTCTGAACTCTTTCCCTACACGACGCTCTTCCGATCTNNNTTCCACTTGGACAGACAGAAG

AGCACACGTCTGAACTCTTTCCCTACACGACGCTCTTCCGATCTNNNAAATTGTGGGTATAGACCTGGC

AGCACACGTCTGAACTCTTTCCCTACACGACGCTCTTCCGATCTNNNAAATTGTGGGTATAGACCTGGC

AGCACACGTCTGAACTCTTTCCCTACACGACGCTCTTCCGATCTNNNCCACCATGCCCAGCTAATTTATT

AGCACACGTCTGAACTCTTTCCCTACACGACGCTCTTCCGATCTNNNCCACCATGCCCAGCTAATTTATT

AAGAGCACACGTCTGAACTCTTTCCCTACACGACGCTCTTCCGATCTNNNAGGAGAATGGCTTGAACC

AAGAGCACACGTCTGAACTCTTTCCCTACACGACGCTCTTCCGATCTNNNAGGAGAATGGCTTGAACC

AGCACACGTCTGAACTCTTTCCCTACACGACGCTCTTCCGATCTNNNATGCACTAATGGCTGTCTCAGAA

AGCACACGTCTGAACTCTTTCCCTACACGACGCTCTTCCGATCTNNNATGCACTAATGGCTGTCTCAGAA

GAGCACACGTCTGAACTCTTTCCCTACACGACGCTCTTCCGATCTNNNTTGCAAGGATCAAGGAGGAAA

GAGCACACGTCTGAACTCTTTCCCTACACGACGCTCTTCCGATCTNNNTTGCAAGGATCAAGGAGGAAA

GAGCACACGTCTGAACTCTTTCCCTACACGACGCTCTTCCGATCTNNNCTGGCCTCTTTACAGTTTTGCA

GAGCACACGTCTGAACTCTTTCCCTACACGACGCTCTTCCGATCTNNNCTGGCCTCTTTACAGTTTTGCA

GCACACGTCTGAACTCTTTCCCTACACGACGCTCTTCCGATCTNNNACACAGCATCATCTAAACCACCAC

GCACACGTCTGAACTCTTTCCCTACACGACGCTCTTCCGATCTNNNACACAGCATCATCTAAACCACCAC

GAGCACACGTCTGAACTCTTTCCCTACACGACGCTCTTCCGATCTNNNTCAATCAATCCTCCCACCTTAGC

GAGCACACGTCTGAACTCTTTCCCTACACGACGCTCTTCCGATCTNNNTCAATCAATCCTCCCACCTTAGC

GAGCACACGTCTGAACTCTTTCCCTACACGACGCTCTTCCGATCTNNNCTTGCAGTGAGCCGAGATCGC

GAGCACACGTCTGAACTCTTTCCCTACACGACGCTCTTCCGATCTNNNCTTGCAGTGAGCCGAGATCGC

AGCACACGTCTGAACTCTTTCCCTACACGACGCTCTTCCGATCTNNNTTCTTTCTAGCCGAGGGAAATC

AGCACACGTCTGAACTCTTTCCCTACACGACGCTCTTCCGATCTNNNTTCTTTCTAGCCGAGGGAAATC

AGAGCACACGTCTGAACTCTTTCCCTACACGACGCTCTTCCGATCTNNNTGAGGAGTTACCCTGCAAAA

AGAGCACACGTCTGAACTCTTTCCCTACACGACGCTCTTCCGATCTNNNTGAGGAGTTACCCTGCAAAA

AGAGCACACGTCTGAACTCTTTCCCTACACGACGCTCTTCCGATCTNNNCAGGAGAACGGTGTGAACCT

AGAGCACACGTCTGAACTCTTTCCCTACACGACGCTCTTCCGATCTNNNCAGGAGAACGGTGTGAACCT

AGCACACGTCTGAACTCTTTCCCTACACGACGCTCTTCCGATCTNNNTTAATTTCTGAAAAGCACCATTA

AGCACACGTCTGAACTCTTTCCCTACACGACGCTCTTCCGATCTNNNTTAATTTCTGAAAAGCACCATTA

GAGCACACGTCTGAACTCTTTCCCTACACGACGCTCTTCCGATCTNNNACAGTTCTGGAGAGCAAAGAG

GAGCACACGTCTGAACTCTTTCCCTACACGACGCTCTTCCGATCTNNNACAGTTCTGGAGAGCAAAGAG

AGCACACGTCTGAACTCTTTCCCTACACGACGCTCTTCCGATCTNNNCCAAAAACCCAAAAGCAAATGC

AGCACACGTCTGAACTCTTTCCCTACACGACGCTCTTCCGATCTNNNCCAAAAACCCAAAAGCAAATGC

GAGCACACGTCTGAACTCTTTCCCTACACGACGCTCTTCCGATCTNNNGTATCACCTCTTCACATCCCGA

GAGCACACGTCTGAACTCTTTCCCTACACGACGCTCTTCCGATCTNNNGTATCACCTCTTCACATCCCGA

GAGCACACGTCTGAACTCTTTCCCTACACGACGCTCTTCCGATCTNNNGAGGCAGAAGAATCGCTTGAA

GAGCACACGTCTGAACTCTTTCCCTACACGACGCTCTTCCGATCTNNNGAGGCAGAAGAATCGCTTGAA

AGCACACGTCTGAACTCTTTCCCTACACGACGCTCTTCCGATCTNNNCATTAGGGAAGTCCTCCGCTAC

AGCACACGTCTGAACTCTTTCCCTACACGACGCTCTTCCGATCTNNNCATTAGGGAAGTCCTCCGCTAC

AGAGCACACGTCTGAACTCTTTCCCTACACGACGCTCTTCCGATCTNNNAGACGAGCAGAAAGAGGAA

AGAGCACACGTCTGAACTCTTTCCCTACACGACGCTCTTCCGATCTNNNAGACGAGCAGAAAGAGGAA

GAGCACACGTCTGAACTCTTTCCCTACACGACGCTCTTCCGATCTNNNTGAAACCCCGTCTCCACTAAAA

AGCACACGTCTGAACTCTTTCCCTACACGACGCTCTTCCGATCTNNNTGAAACCCCGTCTCCACTAAAA
GCACACGTCTGAACTCTTTCCCTACACGACGCTCTTCCGATCTNNNCTTCATGAATGTGTGCATCTCCC
GCACACGTCTGAACTCTTTCCCTACACGACGCTCTTCCGATCTNNNCTTCATGAATGTGTGCATCTCCC
GAGCACACGTCTGAACTCTTTCCCTACACGACGCTCTTCCGATCTNNNTCACCTTGGAAGGTAGAGAG
GAGCACACGTCTGAACTCTTTCCCTACACGACGCTCTTCCGATCTNNNTCACCTTGGAAGGTAGAGAG
AGCACACGTCTGAACTCTTTCCCTACACGACGCTCTTCCGATCTNNNGGGGTGACAGAACAAGACTC
AGCACACGTCTGAACTCTTTCCCTACACGACGCTCTTCCGATCTNNNGGGGTGACAGAACAAGACTC
AGCACACGTCTGAACTCTTTCCCTACACGACGCTCTTCCGATCTNNNACACAGCTGGAGATCTGAGA
AGCACACGTCTGAACTCTTTCCCTACACGACGCTCTTCCGATCTNNNACACAGCTGGAGATCTGAGA
AGCACACGTCTGAACTCTTTCCCTACACGACGCTCTTCCGATCTNNNTCCATGGTGGCTTGGATTTTT
AGCACACGTCTGAACTCTTTCCCTACACGACGCTCTTCCGATCTNNNTCCATGGTGGCTTGGATTTTT
GAGCACACGTCTGAACTCTTTCCCTACACGACGCTCTTCCGATCTNNNGAGGCAGAAGAATCGCTTG
GAGCACACGTCTGAACTCTTTCCCTACACGACGCTCTTCCGATCTNNNGAGGCAGAAGAATCGCTTG
AGCACACGTCTGAACTCTTTCCCTACACGACGCTCTTCCGATCTNNNCAGGATAATACACATCACAGT
AGCACACGTCTGAACTCTTTCCCTACACGACGCTCTTCCGATCTNNNCAGGATAATACACATCACAGT
AGCACACGTCTGAACTCTTTCCCTACACGACGCTCTTCCGATCTNNNAGGCCCTCACAGTAATCAAGG
AGCACACGTCTGAACTCTTTCCCTACACGACGCTCTTCCGATCTNNNAGGCCCTCACAGTAATCAAGG
AGCACACGTCTGAACTCTTTCCCTACACGACGCTCTTCCGATCTNNNTGCTGTAATCAATAACCTTGTG
AGCACACGTCTGAACTCTTTCCCTACACGACGCTCTTCCGATCTNNNTGCTGTAATCAATAACCTTGTG
GCACACGTCTGAACTCTTTCCCTACACGACGCTCTTCCGATCTNNNACTCCCTCACAAAACATCTACCCC
GCACACGTCTGAACTCTTTCCCTACACGACGCTCTTCCGATCTNNNACTCCCTCACAAAACATCTACCCC
AGCACACGTCTGAACTCTTTCCCTACACGACGCTCTTCCGATCTNNNTCTAGTCCCAGCTACTCAGAAG
AGCACACGTCTGAACTCTTTCCCTACACGACGCTCTTCCGATCTNNNTCTAGTCCCAGCTACTCAGAAG
GCACACGTCTGAACTCTTTCCCTACACGACGCTCTTCCGATCTNNNTGGACAAAGTTTGGGGGTTTTTT
GCACACGTCTGAACTCTTTCCCTACACGACGCTCTTCCGATCTNNNTGGACAAAGTTTGGGGGTTTTTT
GAGCACACGTCTGAACTCTTTCCCTACACGACGCTCTTCCGATCTNNNTTGAATTAGGCCCAGAGAC
GAGCACACGTCTGAACTCTTTCCCTACACGACGCTCTTCCGATCTNNNTTGAATTAGGCCCAGAGAC
GCACACGTCTGAACTCTTTCCCTACACGACGCTCTTCCGATCTNNNGAGGCAGGAGAATGGTGTGAA
GCACACGTCTGAACTCTTTCCCTACACGACGCTCTTCCGATCTNNNGAGGCAGGAGAATGGTGTGAA
GAGCACACGTCTGAACTCTTTCCCTACACGACGCTCTTCCGATCTNNNACCTTTCTCTTGCTGTCCTTGA
GAGCACACGTCTGAACTCTTTCCCTACACGACGCTCTTCCGATCTNNNACCTTTCTCTTGCTGTCCTTGA
CACACGTCTGAACTCTTTCCCTACACGACGCTCTTCCGATCTNNNCGAGATTACACCACTTCACTCCAC
CACACGTCTGAACTCTTTCCCTACACGACGCTCTTCCGATCTNNNCGAGATTACACCACTTCACTCCAC
AGCACACGTCTGAACTCTTTCCCTACACGACGCTCTTCCGATCTNNNAAGCCAGAATCTTTGTTCAGC
AGCACACGTCTGAACTCTTTCCCTACACGACGCTCTTCCGATCTNNNAAGCCAGAATCTTTGTTCAGC
CACACGTCTGAACTCTTTCCCTACACGACGCTCTTCCGATCTNNNTGATGTGAACAGCATATTCGCATC
CACACGTCTGAACTCTTTCCCTACACGACGCTCTTCCGATCTNNNTGATGTGAACAGCATATTCGCATC
CACACGTCTGAACTCTTTCCCTACACGACGCTCTTCCGATCTNNNGACTACAGGTCTACACTACCACGC
CACACGTCTGAACTCTTTCCCTACACGACGCTCTTCCGATCTNNNGACTACAGGTCTACACTACCACGC
AGCACACGTCTGAACTCTTTCCCTACACGACGCTCTTCCGATCTNNNCTCAAGTGTAGAGCTGCCTAA
AGCACACGTCTGAACTCTTTCCCTACACGACGCTCTTCCGATCTNNNCTCAAGTGTAGAGCTGCCTAA
ACACGTCTGAACTCTTTCCCTACACGACGCTCTTCCGATCTNNNTTACCAGTCCCCTCAGCGATCATCG
ACACGTCTGAACTCTTTCCCTACACGACGCTCTTCCGATCTNNNTTACCAGTCCCCTCAGCGATCATCG

GAGCACACGTCTGAACTCTTTCCCTACACGACGCTCTTCCGATCTNNNCTTGCAGTGAGCCGAGATTGT
GAGCACACGTCTGAACTCTTTCCCTACACGACGCTCTTCCGATCTNNNCTTGCAGTGAGCCGAGATTGT
AGAGCACACGTCTGAACTCTTTCCCTACACGACGCTCTTCCGATCTNNNCCAGTCTTTCAACAGCAGAC
AGAGCACACGTCTGAACTCTTTCCCTACACGACGCTCTTCCGATCTNNNCCAGTCTTTCAACAGCAGAC
AGCACACGTCTGAACTCTTTCCCTACACGACGCTCTTCCGATCTNNNGCAGGAGAAATCACTTGAACCT
AGCACACGTCTGAACTCTTTCCCTACACGACGCTCTTCCGATCTNNNGCAGGAGAAATCACTTGAACCT
GCACACGTCTGAACTCTTTCCCTACACGACGCTCTTCCGATCTNNNAGAACCAGTTCTTTGTCTCATTGA
GCACACGTCTGAACTCTTTCCCTACACGACGCTCTTCCGATCTNNNAGAACCAGTTCTTTGTCTCATTGA
AGCACACGTCTGAACTCTTTCCCTACACGACGCTCTTCCGATCTNNNCTGGCTGGCTTTCTTTCTTTCTCC
AGCACACGTCTGAACTCTTTCCCTACACGACGCTCTTCCGATCTNNNCTGGCTGGCTTTCTTTCTTTCTCC
AGCACACGTCTGAACTCTTTCCCTACACGACGCTCTTCCGATCTNNNCATTAGAAAACATCAGCACTCCA
AGCACACGTCTGAACTCTTTCCCTACACGACGCTCTTCCGATCTNNNCATTAGAAAACATCAGCACTCCA
AGCACACGTCTGAACTCTTTCCCTACACGACGCTCTTCCGATCTNNNACCGCAACTCTTAGGTGAATTCT
AGCACACGTCTGAACTCTTTCCCTACACGACGCTCTTCCGATCTNNNACCGCAACTCTTAGGTGAATTCT
AGCACACGTCTGAACTCTTTCCCTACACGACGCTCTTCCGATCTNNNGTGCTGGCAGTGAGAATTTCAA
AGCACACGTCTGAACTCTTTCCCTACACGACGCTCTTCCGATCTNNNGTGCTGGCAGTGAGAATTTCAA
GCACACGTCTGAACTCTTTCCCTACACGACGCTCTTCCGATCTNNNCTGTAACTCTTCCCAGTGCTCATC
GCACACGTCTGAACTCTTTCCCTACACGACGCTCTTCCGATCTNNNCTGTAACTCTTCCCAGTGCTCATC
AGCACACGTCTGAACTCTTTCCCTACACGACGCTCTTCCGATCTNNNGTCTCCGTTGCGACAGAATTTC
AGCACACGTCTGAACTCTTTCCCTACACGACGCTCTTCCGATCTNNNGTCTCCGTTGCGACAGAATTTC
AGCACACGTCTGAACTCTTTCCCTACACGACGCTCTTCCGATCTNNNCCTCCAGGACATGATTTCTGTGA
AGCACACGTCTGAACTCTTTCCCTACACGACGCTCTTCCGATCTNNNCCTCCAGGACATGATTTCTGTGA
AGAGCACACGTCTGAACTCTTTCCCTACACGACGCTCTTCCGATCTNNNAATTAGTGCTGAATGAAAGA
AGAGCACACGTCTGAACTCTTTCCCTACACGACGCTCTTCCGATCTNNNAATTAGTGCTGAATGAAAGA
AGCACACGTCTGAACTCTTTCCCTACACGACGCTCTTCCGATCTNNNTGAGCTGAGATCATGCCATTGT
AGCACACGTCTGAACTCTTTCCCTACACGACGCTCTTCCGATCTNNNTGAGCTGAGATCATGCCATTGT
GAGCACACGTCTGAACTCTTTCCCTACACGACGCTCTTCCGATCTNNNGAAAGGAGAAAGAACCAAAC
GAGCACACGTCTGAACTCTTTCCCTACACGACGCTCTTCCGATCTNNNGAAAGGAGAAAGAACCAAAC
AGCACACGTCTGAACTCTTTCCCTACACGACGCTCTTCCGATCTNNNCCATGTCTGACTTGTTTCACTTG
AGCACACGTCTGAACTCTTTCCCTACACGACGCTCTTCCGATCTNNNCCATGTCTGACTTGTTTCACTTG
AGCACACGTCTGAACTCTTTCCCTACACGACGCTCTTCCGATCTNNNTGTCCTTTCTTAGAGGAACCAGG
AGCACACGTCTGAACTCTTTCCCTACACGACGCTCTTCCGATCTNNNTGTCCTTTCTTAGAGGAACCAGG
ACACGTCTGAACTCTTTCCCTACACGACGCTCTTCCGATCTNNNTCCCAACCATTCCTTCATTCCTCCATCG
ACACGTCTGAACTCTTTCCCTACACGACGCTCTTCCGATCTNNNTCCCAACCATTCCTTCATTCCTCCATCG
GCACACGTCTGAACTCTTTCCCTACACGACGCTCTTCCGATCTNNNGTTGTAGAGGATGGGGTTTACAT
GCACACGTCTGAACTCTTTCCCTACACGACGCTCTTCCGATCTNNNGTTGTAGAGGATGGGGTTTACAT
GAGCACACGTCTGAACTCTTTCCCTACACGACGCTCTTCCGATCTNNNTGTGTGTATGTATATCCCCAGC
GAGCACACGTCTGAACTCTTTCCCTACACGACGCTCTTCCGATCTNNNTGTGTGTATGTATATCCCCAGC
AGCACACGTCTGAACTCTTTCCCTACACGACGCTCTTCCGATCTNNNCTGAGGCAGGAGAATCACTTG
AGCACACGTCTGAACTCTTTCCCTACACGACGCTCTTCCGATCTNNNCTGAGGCAGGAGAATCACTTG
GCACACGTCTGAACTCTTTCCCTACACGACGCTCTTCCGATCTNNNTGAGGCATGATTAGCAGTTTCCA
GCACACGTCTGAACTCTTTCCCTACACGACGCTCTTCCGATCTNNNTGAGGCATGATTAGCAGTTTCCA
AGCACACGTCTGAACTCTTTCCCTACACGACGCTCTTCCGATCTNNNAGAAATGCAGCTGATTTTTGTCT

AGCACACGTCTGAACTCTTTCCCTACACGACGCTCTTCCGATCTNNNAGAAATGCAGCTGATTTTTGTCT
AGAGCACACGTCTGAACTCTTTCCCTACACGACGCTCTTCCGATCTNNNAGAAGGCTCCTTTTCATCCAG
AGAGCACACGTCTGAACTCTTTCCCTACACGACGCTCTTCCGATCTNNNAGAAGGCTCCTTTTCATCCAG
GAGCACACGTCTGAACTCTTTCCCTACACGACGCTCTTCCGATCTNNNAGGTAGACTGGATTGCGAATC
GAGCACACGTCTGAACTCTTTCCCTACACGACGCTCTTCCGATCTNNNAGGTAGACTGGATTGCGAATC
CACACGTCTGAACTCTTTCCCTACACGACGCTCTTCCGATCTNNNAAAATAACATTGTCTTTGAGCCCAT
CACACGTCTGAACTCTTTCCCTACACGACGCTCTTCCGATCTNNNAAAATAACATTGTCTTTGAGCCCAT
GCACACGTCTGAACTCTTTCCCTACACGACGCTCTTCCGATCTNNNGGTCTTGACACCATAGATTATGCA
GCACACGTCTGAACTCTTTCCCTACACGACGCTCTTCCGATCTNNNGGTCTTGACACCATAGATTATGCA
AGCACACGTCTGAACTCTTTCCCTACACGACGCTCTTCCGATCTNNNTGGGTAATTGGGATATCCATTG
AGCACACGTCTGAACTCTTTCCCTACACGACGCTCTTCCGATCTNNNTGGGTAATTGGGATATCCATTG
AGCACACGTCTGAACTCTTTCCCTACACGACGCTCTTCCGATCTNNNTTTCCCCTTCCTGTTCTTGTGATC
AGCACACGTCTGAACTCTTTCCCTACACGACGCTCTTCCGATCTNNNTTTCCCCTTCCTGTTCTTGTGATC
AGCACACGTCTGAACTCTTTCCCTACACGACGCTCTTCCGATCTNNNGGCATAGATCCAAGTGTAGGCT
AGCACACGTCTGAACTCTTTCCCTACACGACGCTCTTCCGATCTNNNGGCATAGATCCAAGTGTAGGCT
AGCACACGTCTGAACTCTTTCCCTACACGACGCTCTTCCGATCTNNNCCATGTGAGGATTGCATACTTG
AGCACACGTCTGAACTCTTTCCCTACACGACGCTCTTCCGATCTNNNCCATGTGAGGATTGCATACTTG
AGCACACGTCTGAACTCTTTCCCTACACGACGCTCTTCCGATCTNNNACTCCTATGTTTGCTTCTCAGGA
AGCACACGTCTGAACTCTTTCCCTACACGACGCTCTTCCGATCTNNNACTCCTATGTTTGCTTCTCAGGA
AGCACACGTCTGAACTCTTTCCCTACACGACGCTCTTCCGATCTNNNGTGTTTGAGGCAAGTCAGTAAC
AGCACACGTCTGAACTCTTTCCCTACACGACGCTCTTCCGATCTNNNGTGTTTGAGGCAAGTCAGTAAC
AGCACACGTCTGAACTCTTTCCCTACACGACGCTCTTCCGATCTNNNCCAAGTAGCTGGTACTGACAGT
AGCACACGTCTGAACTCTTTCCCTACACGACGCTCTTCCGATCTNNNCCAAGTAGCTGGTACTGACAGT
GAGCACACGTCTGAACTCTTTCCCTACACGACGCTCTTCCGATCTNNNGCCTGACAAGATACAAAACCTC
GAGCACACGTCTGAACTCTTTCCCTACACGACGCTCTTCCGATCTNNNGCCTGACAAGATACAAAACCTC
GAGCACACGTCTGAACTCTTTCCCTACACGACGCTCTTCCGATCTNNNAAACTAGAAGGCCTCAGAAGA
GAGCACACGTCTGAACTCTTTCCCTACACGACGCTCTTCCGATCTNNNAAACTAGAAGGCCTCAGAAGA
AGCACACGTCTGAACTCTTTCCCTACACGACGCTCTTCCGATCTNNNATTCTGTAGGTTGCCTGTTCACT
AGCACACGTCTGAACTCTTTCCCTACACGACGCTCTTCCGATCTNNNATTCTGTAGGTTGCCTGTTCACT
AGCACACGTCTGAACTCTTTCCCTACACGACGCTCTTCCGATCTNNNTGTGTGAATGTGTGAGCCAATTC
AGCACACGTCTGAACTCTTTCCCTACACGACGCTCTTCCGATCTNNNTGTGTGAATGTGTGAGCCAATTC
AGCACACGTCTGAACTCTTTCCCTACACGACGCTCTTCCGATCTNNNATGCCTGTGGGTAATGCAGATA
AGCACACGTCTGAACTCTTTCCCTACACGACGCTCTTCCGATCTNNNATGCCTGTGGGTAATGCAGATA
AGCACACGTCTGAACTCTTTCCCTACACGACGCTCTTCCGATCTNNNGTGTCCTCTTCCATTTCTTGCATC
AGCACACGTCTGAACTCTTTCCCTACACGACGCTCTTCCGATCTNNNGTGTCCTCTTCCATTTCTTGCATC
AGCACACGTCTGAACTCTTTCCCTACACGACGCTCTTCCGATCTNNNGGATCACTTGAGTCCAGGAAGT
AGCACACGTCTGAACTCTTTCCCTACACGACGCTCTTCCGATCTNNNGGATCACTTGAGTCCAGGAAGT
AGCACACGTCTGAACTCTTTCCCTACACGACGCTCTTCCGATCTNNNCTGTAATCCCAGCTACTCAGGA
AGCACACGTCTGAACTCTTTCCCTACACGACGCTCTTCCGATCTNNNCTGTAATCCCAGCTACTCAGGA
GCACACGTCTGAACTCTTTCCCTACACGACGCTCTTCCGATCTNNNTTCATAATGGTGCTTTCAAACCATC
GCACACGTCTGAACTCTTTCCCTACACGACGCTCTTCCGATCTNNNTTCATAATGGTGCTTTCAAACCATC
AGAGCACACGTCTGAACTCTTTCCCTACACGACGCTCTTCCGATCTNNNACAGAAGGGACATCACTCCA
AGAGCACACGTCTGAACTCTTTCCCTACACGACGCTCTTCCGATCTNNNACAGAAGGGACATCACTCCA

GAGCACACGTCTGAACTCTTTCCCTACACGACGCTCTTCCGATCTNNNGTCAAGAGATCGAGACCATC
GAGCACACGTCTGAACTCTTTCCCTACACGACGCTCTTCCGATCTNNNGTCAAGAGATCGAGACCATC
AGCACACGTCTGAACTCTTTCCCTACACGACGCTCTTCCGATCTNNNGTAAGTTGTCACATGTCACGG
AGCACACGTCTGAACTCTTTCCCTACACGACGCTCTTCCGATCTNNNGTAAGTTGTCACATGTCACGG
GCACACGTCTGAACTCTTTCCCTACACGACGCTCTTCCGATCTNNNTTTCTCTCCTCCTCCTTCTCCTTCA
GCACACGTCTGAACTCTTTCCCTACACGACGCTCTTCCGATCTNNNTTTCTCTCCTCCTCCTTCTCCTTCA
CACACGTCTGAACTCTTTCCCTACACGACGCTCTTCCGATCTNNNTTGCCTTCTATCAGTAACCCCAC
CACACGTCTGAACTCTTTCCCTACACGACGCTCTTCCGATCTNNNTTGCCTTCTATCAGTAACCCCAC
AGCACACGTCTGAACTCTTTCCCTACACGACGCTCTTCCGATCTNNNTTGCACACGCAATTGCAAAAA
AGCACACGTCTGAACTCTTTCCCTACACGACGCTCTTCCGATCTNNNTTGCACACGCAATTGCAAAAA
GCACACGTCTGAACTCTTTCCCTACACGACGCTCTTCCGATCTNNNAAGATACTTGTGTGGTGTGGT
GCACACGTCTGAACTCTTTCCCTACACGACGCTCTTCCGATCTNNNAAGATACTTGTGTGGTGTGGT
AGCACACGTCTGAACTCTTTCCCTACACGACGCTCTTCCGATCTNNNATGATCCTTGGTTGGGCTTGT
AGCACACGTCTGAACTCTTTCCCTACACGACGCTCTTCCGATCTNNNATGATCCTTGGTTGGGCTTGT
GCACACGTCTGAACTCTTTCCCTACACGACGCTCTTCCGATCTNNNAAAGCCTCTGGGAATGTATGTG
GCACACGTCTGAACTCTTTCCCTACACGACGCTCTTCCGATCTNNNAAAGCCTCTGGGAATGTATGTG
GCACACGTCTGAACTCTTTCCCTACACGACGCTCTTCCGATCTNNNAAAAAATTAGCTGGGCGTAGTG
GCACACGTCTGAACTCTTTCCCTACACGACGCTCTTCCGATCTNNNAAAAAATTAGCTGGGCGTAGTG
ACACGTCTGAACTCTTTCCCTACACGACGCTCTTCCGATCTNNNACAGGGGAAACTACCAAACACTTC
ACACGTCTGAACTCTTTCCCTACACGACGCTCTTCCGATCTNNNACAGGGGAAACTACCAAACACTTC
AGCACACGTCTGAACTCTTTCCCTACACGACGCTCTTCCGATCTNNNAAGTGCTGGGATTACAGGCAT
AGCACACGTCTGAACTCTTTCCCTACACGACGCTCTTCCGATCTNNNAAGTGCTGGGATTACAGGCAT
AGCACACGTCTGAACTCTTTCCCTACACGACGCTCTTCCGATCTNNNGATAAGCAGTAGCAAGCAAA
AGCACACGTCTGAACTCTTTCCCTACACGACGCTCTTCCGATCTNNNGATAAGCAGTAGCAAGCAAA
AGCACACGTCTGAACTCTTTCCCTACACGACGCTCTTCCGATCTNNNATGCTGTTCTCATGGTAGTGA
AGCACACGTCTGAACTCTTTCCCTACACGACGCTCTTCCGATCTNNNATGCTGTTCTCATGGTAGTGA
AGCACACGTCTGAACTCTTTCCCTACACGACGCTCTTCCGATCTNNNGCTTCAATTTGTTCTCCCACAG
AGCACACGTCTGAACTCTTTCCCTACACGACGCTCTTCCGATCTNNNGCTTCAATTTGTTCTCCCACAG
AGCACACGTCTGAACTCTTTCCCTACACGACGCTCTTCCGATCTNNNACACTGTAGGCTGTATCTAGG
AGCACACGTCTGAACTCTTTCCCTACACGACGCTCTTCCGATCTNNNACACTGTAGGCTGTATCTAGG
GAGCACACGTCTGAACTCTTTCCCTACACGACGCTCTTCCGATCTNNNATAAATGGGCAAAACCTGG
GAGCACACGTCTGAACTCTTTCCCTACACGACGCTCTTCCGATCTNNNATAAATGGGCAAAACCTGG
AGCACACGTCTGAACTCTTTCCCTACACGACGCTCTTCCGATCTNNNTATTTGTGACAAGGGTAGGT
AGCACACGTCTGAACTCTTTCCCTACACGACGCTCTTCCGATCTNNNTATTTGTGACAAGGGTAGGT
AGCACACGTCTGAACTCTTTCCCTACACGACGCTCTTCCGATCTNNNGTAATCCTAGCTACTTGGGAG
AGCACACGTCTGAACTCTTTCCCTACACGACGCTCTTCCGATCTNNNGTAATCCTAGCTACTTGGGAG
AGCACACGTCTGAACTCTTTCCCTACACGACGCTCTTCCGATCTNNNCTCTAGATCAATCACTGCCTGC
AGCACACGTCTGAACTCTTTCCCTACACGACGCTCTTCCGATCTNNNCTCTAGATCAATCACTGCCTGC
GAGCACACGTCTGAACTCTTTCCCTACACGACGCTCTTCCGATCTNNNGGCTTGGAACCTAGTTTACC
GAGCACACGTCTGAACTCTTTCCCTACACGACGCTCTTCCGATCTNNNGGCTTGGAACCTAGTTTACC
AGCACACGTCTGAACTCTTTCCCTACACGACGCTCTTCCGATCTNNNAGAGGTTGCAGTAAGCCAGA
AGCACACGTCTGAACTCTTTCCCTACACGACGCTCTTCCGATCTNNNAGAGGTTGCAGTAAGCCAGA
GCACACGTCTGAACTCTTTCCCTACACGACGCTCTTCCGATCTNNNAGAGGCAGGAGAATCACTTGA

GCACACGTCTGAACTCTTTCCCTACACGACGCTCTTCCGATCTNNNAGAGGCAGGAGAATCACTTGAA
AGCACACGTCTGAACTCTTTCCCTACACGACGCTCTTCCGATCTNNNGACTTCTGGCTCACCACTAATCA
AGCACACGTCTGAACTCTTTCCCTACACGACGCTCTTCCGATCTNNNGACTTCTGGCTCACCACTAATCA
GAGCACACGTCTGAACTCTTTCCCTACACGACGCTCTTCCGATCTNNNCTGTGCCTTCTTCATAGGAGTC
GAGCACACGTCTGAACTCTTTCCCTACACGACGCTCTTCCGATCTNNNCTGTGCCTTCTTCATAGGAGTC
AGCACACGTCTGAACTCTTTCCCTACACGACGCTCTTCCGATCTNNNAGCAACAGAACAAGATCCCATC
AGCACACGTCTGAACTCTTTCCCTACACGACGCTCTTCCGATCTNNNAGCAACAGAACAAGATCCCATC
GAGCACACGTCTGAACTCTTTCCCTACACGACGCTCTTCCGATCTNNNCTTGAGGTCAGGAGTTCGAG
GAGCACACGTCTGAACTCTTTCCCTACACGACGCTCTTCCGATCTNNNCTTGAGGTCAGGAGTTCGAG
AGCACACGTCTGAACTCTTTCCCTACACGACGCTCTTCCGATCTNNNGCCTTCTTCTGTCCATGGTTTTC
AGCACACGTCTGAACTCTTTCCCTACACGACGCTCTTCCGATCTNNNGCCTTCTTCTGTCCATGGTTTTC
GCACACGTCTGAACTCTTTCCCTACACGACGCTCTTCCGATCTNNNCCACATAGCGAGATCTCTGTCTTC
GCACACGTCTGAACTCTTTCCCTACACGACGCTCTTCCGATCTNNNCCACATAGCGAGATCTCTGTCTTC
GAGCACACGTCTGAACTCTTTCCCTACACGACGCTCTTCCGATCTNNNGGGTTCTGATAGAGGTGGCT
GAGCACACGTCTGAACTCTTTCCCTACACGACGCTCTTCCGATCTNNNGGGTTCTGATAGAGGTGGCT
AGCACACGTCTGAACTCTTTCCCTACACGACGCTCTTCCGATCTNNNCTTGCAGTGAGCCAAGATTGC
AGCACACGTCTGAACTCTTTCCCTACACGACGCTCTTCCGATCTNNNCTTGCAGTGAGCCAAGATTGC
AAGAGCACACGTCTGAACTCTTTCCCTACACGACGCTCTTCCGATCTNNNAGAGGACTGACTATCGGA
AAGAGCACACGTCTGAACTCTTTCCCTACACGACGCTCTTCCGATCTNNNAGAGGACTGACTATCGGA
GAGCACACGTCTGAACTCTTTCCCTACACGACGCTCTTCCGATCTNNNAGCTGAGAACAAAGACAAGC
GAGCACACGTCTGAACTCTTTCCCTACACGACGCTCTTCCGATCTNNNAGCTGAGAACAAAGACAAGC
AGCACACGTCTGAACTCTTTCCCTACACGACGCTCTTCCGATCTNNNACGCCCGGCTATTTTTTTGTA
AGCACACGTCTGAACTCTTTCCCTACACGACGCTCTTCCGATCTNNNACGCCCGGCTATTTTTTTGTA
GAGCACACGTCTGAACTCTTTCCCTACACGACGCTCTTCCGATCTNNNGACCTCTCTCCCCATCAATCTTC
GAGCACACGTCTGAACTCTTTCCCTACACGACGCTCTTCCGATCTNNNGACCTCTCTCCCCATCAATCTTC
GAGCACACGTCTGAACTCTTTCCCTACACGACGCTCTTCCGATCTNNNTCAGAGAGCAGAGGAGAAGA
GAGCACACGTCTGAACTCTTTCCCTACACGACGCTCTTCCGATCTNNNTCAGAGAGCAGAGGAGAAGA
AGCACACGTCTGAACTCTTTCCCTACACGACGCTCTTCCGATCTNNNAATCACCTCTCAAAGACCCCATC
AGCACACGTCTGAACTCTTTCCCTACACGACGCTCTTCCGATCTNNNAATCACCTCTCAAAGACCCCATC
AGCACACGTCTGAACTCTTTCCCTACACGACGCTCTTCCGATCTNNNTTTCATGTGTCTTTTGGCTGCAT
AGCACACGTCTGAACTCTTTCCCTACACGACGCTCTTCCGATCTNNNTTTCATGTGTCTTTTGGCTGCAT
AGAGCACACGTCTGAACTCTTTCCCTACACGACGCTCTTCCGATCTNNNGTATGCCTGTAATCCCAGCTA
AGAGCACACGTCTGAACTCTTTCCCTACACGACGCTCTTCCGATCTNNNGTATGCCTGTAATCCCAGCTA
GAGCACACGTCTGAACTCTTTCCCTACACGACGCTCTTCCGATCTNNNCCAATTCCAGGGCCTTTTGAAA
GAGCACACGTCTGAACTCTTTCCCTACACGACGCTCTTCCGATCTNNNCCAATTCCAGGGCCTTTTGAAA
GAGCACACGTCTGAACTCTTTCCCTACACGACGCTCTTCCGATCTNNNTGAGCAAAGGAATGAATGAC
GAGCACACGTCTGAACTCTTTCCCTACACGACGCTCTTCCGATCTNNNTGAGCAAAGGAATGAATGAC
AGCACACGTCTGAACTCTTTCCCTACACGACGCTCTTCCGATCTNNNAACTGGATAAACCACATGGATC
AGCACACGTCTGAACTCTTTCCCTACACGACGCTCTTCCGATCTNNNAACTGGATAAACCACATGGATC
AGCACACGTCTGAACTCTTTCCCTACACGACGCTCTTCCGATCTNNNCTGAGGCAGGAGAATCACTTGA
AGCACACGTCTGAACTCTTTCCCTACACGACGCTCTTCCGATCTNNNCTGAGGCAGGAGAATCACTTGA
AGCACACGTCTGAACTCTTTCCCTACACGACGCTCTTCCGATCTNNNGAACACGGTATGGAGGTTCCTA
AGCACACGTCTGAACTCTTTCCCTACACGACGCTCTTCCGATCTNNNGAACACGGTATGGAGGTTCCTA

AGCACACGTCTGAACTCTTTCCCTACACGACGCTCTTCCGATCTNNNATTGAAGCACGCAGAAATCCAA
AGCACACGTCTGAACTCTTTCCCTACACGACGCTCTTCCGATCTNNNATTGAAGCACGCAGAAATCCAA
AGCACACGTCTGAACTCTTTCCCTACACGACGCTCTTCCGATCTNNNCACCTGTAATCCCAGCTACTCA
AGCACACGTCTGAACTCTTTCCCTACACGACGCTCTTCCGATCTNNNCACCTGTAATCCCAGCTACTCA
AGCACACGTCTGAACTCTTTCCCTACACGACGCTCTTCCGATCTNNNTCCATTTGCATTTTCTCCTTGG
AGCACACGTCTGAACTCTTTCCCTACACGACGCTCTTCCGATCTNNNTCCATTTGCATTTTCTCCTTGG
AGCACACGTCTGAACTCTTTCCCTACACGACGCTCTTCCGATCTNNNTTAGTCCTGCCCTGGTTTTTC
AGCACACGTCTGAACTCTTTCCCTACACGACGCTCTTCCGATCTNNNTTAGTCCTGCCCTGGTTTTTC
GCACACGTCTGAACTCTTTCCCTACACGACGCTCTTCCGATCTNNNAAACCCAGGCTTTTGGAGAAA
GCACACGTCTGAACTCTTTCCCTACACGACGCTCTTCCGATCTNNNAAACCCAGGCTTTTGGAGAAA
AGCACACGTCTGAACTCTTTCCCTACACGACGCTCTTCCGATCTNNNATCAGCAACTTCGCAGACTTAG
AGCACACGTCTGAACTCTTTCCCTACACGACGCTCTTCCGATCTNNNATCAGCAACTTCGCAGACTTAG
GCACACGTCTGAACTCTTTCCCTACACGACGCTCTTCCGATCTNNNCCCTTTACTTGGACCACCTTTAC
GCACACGTCTGAACTCTTTCCCTACACGACGCTCTTCCGATCTNNNCCCTTTACTTGGACCACCTTTAC
AGCACACGTCTGAACTCTTTCCCTACACGACGCTCTTCCGATCTNNNTTGTGAAGGGGCAAAATTTAC
AGCACACGTCTGAACTCTTTCCCTACACGACGCTCTTCCGATCTNNNTTGTGAAGGGGCAAAATTTAC
AGCACACGTCTGAACTCTTTCCCTACACGACGCTCTTCCGATCTNNNTGGTTAAGAAAACTGTTCCAA
AGCACACGTCTGAACTCTTTCCCTACACGACGCTCTTCCGATCTNNNTGGTTAAGAAAACTGTTCCAA
GAGCACACGTCTGAACTCTTTCCCTACACGACGCTCTTCCGATCTNNNAGGTGACACATTCTGGTTTCC
GAGCACACGTCTGAACTCTTTCCCTACACGACGCTCTTCCGATCTNNNAGGTGACACATTCTGGTTTCC
CACACGTCTGAACTCTTTCCCTACACGACGCTCTTCCGATCTNNNCTACATGATGTACATTTGGTCTAC
CACACGTCTGAACTCTTTCCCTACACGACGCTCTTCCGATCTNNNCTACATGATGTACATTTGGTCTAC
AGCACACGTCTGAACTCTTTCCCTACACGACGCTCTTCCGATCTNNNAACAAACCTGACAAGCCATAA
AGCACACGTCTGAACTCTTTCCCTACACGACGCTCTTCCGATCTNNNAACAAACCTGACAAGCCATAA
CACACGTCTGAACTCTTTCCCTACACGACGCTCTTCCGATCTNNNGTGAGATGATGTGTGTGTGTGT
CACACGTCTGAACTCTTTCCCTACACGACGCTCTTCCGATCTNNNGTGAGATGATGTGTGTGTGTGT
AGCACACGTCTGAACTCTTTCCCTACACGACGCTCTTCCGATCTNNNGGAGACTGAGGCAGAAGAAT
AGCACACGTCTGAACTCTTTCCCTACACGACGCTCTTCCGATCTNNNGGAGACTGAGGCAGAAGAAT
AGCACACGTCTGAACTCTTTCCCTACACGACGCTCTTCCGATCTNNNTCTGAGAATGTAGCATCGTTC
AGCACACGTCTGAACTCTTTCCCTACACGACGCTCTTCCGATCTNNNTCTGAGAATGTAGCATCGTTC
GCACACGTCTGAACTCTTTCCCTACACGACGCTCTTCCGATCTNNNTCTCTGCAGATAAAGGCACTGT
GCACACGTCTGAACTCTTTCCCTACACGACGCTCTTCCGATCTNNNTCTCTGCAGATAAAGGCACTGT
AGCACACGTCTGAACTCTTTCCCTACACGACGCTCTTCCGATCTNNNCTGACCTCAGTTGATCCACCTG
AGCACACGTCTGAACTCTTTCCCTACACGACGCTCTTCCGATCTNNNCTGACCTCAGTTGATCCACCTG
GAGCACACGTCTGAACTCTTTCCCTACACGACGCTCTTCCGATCTNNNAGATAATTTCCCAGGCTGCTT
GAGCACACGTCTGAACTCTTTCCCTACACGACGCTCTTCCGATCTNNNAGATAATTTCCCAGGCTGCTT
AGCACACGTCTGAACTCTTTCCCTACACGACGCTCTTCCGATCTNNNGATTGAGACCATCCTGGCTAA
AGCACACGTCTGAACTCTTTCCCTACACGACGCTCTTCCGATCTNNNGATTGAGACCATCCTGGCTAA
AGCACACGTCTGAACTCTTTCCCTACACGACGCTCTTCCGATCTNNNTAATCCTAGCACTTTTGGAGGC
AGCACACGTCTGAACTCTTTCCCTACACGACGCTCTTCCGATCTNNNTAATCCTAGCACTTTTGGAGGC
AGCACACGTCTGAACTCTTTCCCTACACGACGCTCTTCCGATCTNNNTTTCCTGCTGAAATGACCTAAG
AGCACACGTCTGAACTCTTTCCCTACACGACGCTCTTCCGATCTNNNTTTCCTGCTGAAATGACCTAAG
GCACACGTCTGAACTCTTTCCCTACACGACGCTCTTCCGATCTNNNTACAAAAAATTAGCCGGGCATG

GCACACGTCTGAACTCTTTCCCTACACGACGCTCTTCCGATCTNNNTACAAAAAATTAGCCGGGCATG
AGCACACGTCTGAACTCTTTCCCTACACGACGCTCTTCCGATCTNNNTTCCAGAACTGTGTCAGAGAA
AGCACACGTCTGAACTCTTTCCCTACACGACGCTCTTCCGATCTNNNTTCCAGAACTGTGTCAGAGAA
GCACACGTCTGAACTCTTTCCCTACACGACGCTCTTCCGATCTNNNAGACCCCTGTTTTTGGTGTTTTG
GCACACGTCTGAACTCTTTCCCTACACGACGCTCTTCCGATCTNNNAGACCCCTGTTTTTGGTGTTTTG
AGCACACGTCTGAACTCTTTCCCTACACGACGCTCTTCCGATCTNNNTAGTCTTTTTTCCCTCACCACC
AGCACACGTCTGAACTCTTTCCCTACACGACGCTCTTCCGATCTNNNTAGTCTTTTTTCCCTCACCACC
GCACACGTCTGAACTCTTTCCCTACACGACGCTCTTCCGATCTNNNACATGGTGCATTATTCAAAACAA
GCACACGTCTGAACTCTTTCCCTACACGACGCTCTTCCGATCTNNNACATGGTGCATTATTCAAAACAA
AGCACACGTCTGAACTCTTTCCCTACACGACGCTCTTCCGATCTNNNTGAGGTTAGAGGGAGGAGA
AGCACACGTCTGAACTCTTTCCCTACACGACGCTCTTCCGATCTNNNTGAGGTTAGAGGGAGGAGA
ACACGTCTGAACTCTTTCCCTACACGACGCTCTTCCGATCTNNNCCTCCCCAGGCCATAATAGTTTTCAT
ACACGTCTGAACTCTTTCCCTACACGACGCTCTTCCGATCTNNNCCTCCCCAGGCCATAATAGTTTTCAT
AGCACACGTCTGAACTCTTTCCCTACACGACGCTCTTCCGATCTNNNTGAGGTGGTAAGTAGATGGG
AGCACACGTCTGAACTCTTTCCCTACACGACGCTCTTCCGATCTNNNTGAGGTGGTAAGTAGATGGG
AGCACACGTCTGAACTCTTTCCCTACACGACGCTCTTCCGATCTNNNTTGAGAAGGGCAAAAGCATG
AGCACACGTCTGAACTCTTTCCCTACACGACGCTCTTCCGATCTNNNTTGAGAAGGGCAAAAGCATG
AGCACACGTCTGAACTCTTTCCCTACACGACGCTCTTCCGATCTNNNTTCCTGATGCCAACTTTGACCT
AGCACACGTCTGAACTCTTTCCCTACACGACGCTCTTCCGATCTNNNTTCCTGATGCCAACTTTGACCT
GCACACGTCTGAACTCTTTCCCTACACGACGCTCTTCCGATCTNNNCCTGTGCAATGGAAGACAAACT
GCACACGTCTGAACTCTTTCCCTACACGACGCTCTTCCGATCTNNNCCTGTGCAATGGAAGACAAACT
GCACACGTCTGAACTCTTTCCCTACACGACGCTCTTCCGATCTNNNACGCTTATGATTGCATCACATGG
GCACACGTCTGAACTCTTTCCCTACACGACGCTCTTCCGATCTNNNACGCTTATGATTGCATCACATGG
GAGCACACGTCTGAACTCTTTCCCTACACGACGCTCTTCCGATCTNNNGAGGCAGGAGAATGGTGTG
GAGCACACGTCTGAACTCTTTCCCTACACGACGCTCTTCCGATCTNNNGAGGCAGGAGAATGGTGTG
AGCACACGTCTGAACTCTTTCCCTACACGACGCTCTTCCGATCTNNNATTGTGTGTGTTTGTGTGTGT
AGCACACGTCTGAACTCTTTCCCTACACGACGCTCTTCCGATCTNNNATTGTGTGTGTTTGTGTGTGT
AGCACACGTCTGAACTCTTTCCCTACACGACGCTCTTCCGATCTNNNTTCAGCTATAGTTGAGGCAGT
AGCACACGTCTGAACTCTTTCCCTACACGACGCTCTTCCGATCTNNNTTCAGCTATAGTTGAGGCAGT
GCACACGTCTGAACTCTTTCCCTACACGACGCTCTTCCGATCTNNNTTTTGGGGCAATTGTGAGGAA
GCACACGTCTGAACTCTTTCCCTACACGACGCTCTTCCGATCTNNNTTTTGGGGCAATTGTGAGGAA
AGCACACGTCTGAACTCTTTCCCTACACGACGCTCTTCCGATCTNNNTTTTGAGACAAAGTCACGCTC
AGCACACGTCTGAACTCTTTCCCTACACGACGCTCTTCCGATCTNNNTTTTGAGACAAAGTCACGCTC
GCACACGTCTGAACTCTTTCCCTACACGACGCTCTTCCGATCTNNNAATTTTGATTCTGTTACCTTGTT
GCACACGTCTGAACTCTTTCCCTACACGACGCTCTTCCGATCTNNNAATTTTGATTCTGTTACCTTGTT
GCACACGTCTGAACTCTTTCCCTACACGACGCTCTTCCGATCTNNNTTACCAACCTCATTTCTTTACTTC
GCACACGTCTGAACTCTTTCCCTACACGACGCTCTTCCGATCTNNNTTACCAACCTCATTTCTTTACTTC
AGCACACGTCTGAACTCTTTCCCTACACGACGCTCTTCCGATCTNNNAAAATAATTAGCCGGGCATGG
AGCACACGTCTGAACTCTTTCCCTACACGACGCTCTTCCGATCTNNNAAAATAATTAGCCGGGCATGG
AGCACACGTCTGAACTCTTTCCCTACACGACGCTCTTCCGATCTNNNGAGGCATAGCACCCTGAAGA
AGCACACGTCTGAACTCTTTCCCTACACGACGCTCTTCCGATCTNNNGAGGCATAGCACCCTGAAGA
AGCACACGTCTGAACTCTTTCCCTACACGACGCTCTTCCGATCTNNNGGGGATACAGAGCCAAACCA
AGCACACGTCTGAACTCTTTCCCTACACGACGCTCTTCCGATCTNNNGGGGATACAGAGCCAAACCA

GCACACGTCTGAACTCTTTCCCTACACGACGCTCTTCCGATCTNNNTTAGGATGGGAGGATGGTGGA
GCACACGTCTGAACTCTTTCCCTACACGACGCTCTTCCGATCTNNNTTAGGATGGGAGGATGGTGGA
AGCACACGTCTGAACTCTTTCCCTACACGACGCTCTTCCGATCTNNNGTGTGTATCTGTGTGTGCATG
AGCACACGTCTGAACTCTTTCCCTACACGACGCTCTTCCGATCTNNNGTGTGTATCTGTGTGTGCATG
AGCACACGTCTGAACTCTTTCCCTACACGACGCTCTTCCGATCTNNNCTTGCAGTGAGTGGAGATTG
AGCACACGTCTGAACTCTTTCCCTACACGACGCTCTTCCGATCTNNNCTTGCAGTGAGTGGAGATTG
GCACACGTCTGAACTCTTTCCCTACACGACGCTCTTCCGATCTNNNATCGCATCATGCTTTTCCACATTC
GCACACGTCTGAACTCTTTCCCTACACGACGCTCTTCCGATCTNNNATCGCATCATGCTTTTCCACATTC
GCACACGTCTGAACTCTTTCCCTACACGACGCTCTTCCGATCTNNNTCACAAGCTTCATGGTTCCAGTA
GCACACGTCTGAACTCTTTCCCTACACGACGCTCTTCCGATCTNNNTCACAAGCTTCATGGTTCCAGTA
GAGCACACGTCTGAACTCTTTCCCTACACGACGCTCTTCCGATCTNNNGCCTATAGTTCCAGCTACTCG
GAGCACACGTCTGAACTCTTTCCCTACACGACGCTCTTCCGATCTNNNGCCTATAGTTCCAGCTACTCG
AGCACACGTCTGAACTCTTTCCCTACACGACGCTCTTCCGATCTNNNCTAAGAATGCCTAACCTCCTGG
AGCACACGTCTGAACTCTTTCCCTACACGACGCTCTTCCGATCTNNNCTAAGAATGCCTAACCTCCTGG
GAGCACACGTCTGAACTCTTTCCCTACACGACGCTCTTCCGATCTNNNTATTGTACACAACTGACGCTC
GAGCACACGTCTGAACTCTTTCCCTACACGACGCTCTTCCGATCTNNNTATTGTACACAACTGACGCTC
AGCACACGTCTGAACTCTTTCCCTACACGACGCTCTTCCGATCTNNNTCCCGCAGTCTTATTTATACCCG
AGCACACGTCTGAACTCTTTCCCTACACGACGCTCTTCCGATCTNNNTCCCGCAGTCTTATTTATACCCG
AGCACACGTCTGAACTCTTTCCCTACACGACGCTCTTCCGATCTNNNTATTGGCACAGATGAGCTCACA
AGCACACGTCTGAACTCTTTCCCTACACGACGCTCTTCCGATCTNNNTATTGGCACAGATGAGCTCACA
GAGCACACGTCTGAACTCTTTCCCTACACGACGCTCTTCCGATCTNNNATATCAAGACCTTGCCACTCC
GAGCACACGTCTGAACTCTTTCCCTACACGACGCTCTTCCGATCTNNNATATCAAGACCTTGCCACTCC
GAGCACACGTCTGAACTCTTTCCCTACACGACGCTCTTCCGATCTNNNAAGCAAGGAAGAACTCAAGC
GAGCACACGTCTGAACTCTTTCCCTACACGACGCTCTTCCGATCTNNNAAGCAAGGAAGAACTCAAGC
AGCACACGTCTGAACTCTTTCCCTACACGACGCTCTTCCGATCTNNNGGAGGTGCTTCTTATCTGTCA
AGCACACGTCTGAACTCTTTCCCTACACGACGCTCTTCCGATCTNNNGGAGGTGCTTCTTATCTGTCA
GCACACGTCTGAACTCTTTCCCTACACGACGCTCTTCCGATCTNNNAAATCACTGCACAACTGAGGAAG
GCACACGTCTGAACTCTTTCCCTACACGACGCTCTTCCGATCTNNNAAATCACTGCACAACTGAGGAAG
GAGCACACGTCTGAACTCTTTCCCTACACGACGCTCTTCCGATCTNNNCCAATCTGGACAACAAGAGCA
GAGCACACGTCTGAACTCTTTCCCTACACGACGCTCTTCCGATCTNNNCCAATCTGGACAACAAGAGCA
AGCACACGTCTGAACTCTTTCCCTACACGACGCTCTTCCGATCTNNNAATGCAAAGTTACAGGTTGAG
AGCACACGTCTGAACTCTTTCCCTACACGACGCTCTTCCGATCTNNNAATGCAAAGTTACAGGTTGAG
AGCACACGTCTGAACTCTTTCCCTACACGACGCTCTTCCGATCTNNNTAAGGTTAGGAGTTCGAGACC
AGCACACGTCTGAACTCTTTCCCTACACGACGCTCTTCCGATCTNNNTAAGGTTAGGAGTTCGAGACC
AGCACACGTCTGAACTCTTTCCCTACACGACGCTCTTCCGATCTNNNTTTCCTCCTGTTACTGGTCATGG
AGCACACGTCTGAACTCTTTCCCTACACGACGCTCTTCCGATCTNNNTTTCCTCCTGTTACTGGTCATGG
AGCACACGTCTGAACTCTTTCCCTACACGACGCTCTTCCGATCTNNNTGAGCTGAGATCATACCATTG
AGCACACGTCTGAACTCTTTCCCTACACGACGCTCTTCCGATCTNNNTGAGCTGAGATCATACCATTG
AGCACACGTCTGAACTCTTTCCCTACACGACGCTCTTCCGATCTNNNGGTGATCACTGTGGACACTAA
AGCACACGTCTGAACTCTTTCCCTACACGACGCTCTTCCGATCTNNNGGTGATCACTGTGGACACTAA
AGCACACGTCTGAACTCTTTCCCTACACGACGCTCTTCCGATCTNNNTTAGTTTACTTGCCTGGCATCC
AGCACACGTCTGAACTCTTTCCCTACACGACGCTCTTCCGATCTNNNTTAGTTTACTTGCCTGGCATCC
AGCACACGTCTGAACTCTTTCCCTACACGACGCTCTTCCGATCTNNNCTAGGGGATGTAGAGGTGAA

AGCACACGTCTGAACTCTTTCCCTACACGACGCTCTTCCGATCTNNNCTAGGGGATGTAGAGGTGAAT
GAGCACACGTCTGAACTCTTTCCCTACACGACGCTCTTCCGATCTNNNAACTACTCGGGAAGTTGAGGT
GAGCACACGTCTGAACTCTTTCCCTACACGACGCTCTTCCGATCTNNNAACTACTCGGGAAGTTGAGGT
AGAGCACACGTCTGAACTCTTTCCCTACACGACGCTCTTCCGATCTNNNCAGGGCAACATAGTGAGAAC
AGAGCACACGTCTGAACTCTTTCCCTACACGACGCTCTTCCGATCTNNNCAGGGCAACATAGTGAGAAC
GAGCACACGTCTGAACTCTTTCCCTACACGACGCTCTTCCGATCTNNNGGCATGAGAATCTTTTGAGCCT
GAGCACACGTCTGAACTCTTTCCCTACACGACGCTCTTCCGATCTNNNGGCATGAGAATCTTTTGAGCCT
GCACACGTCTGAACTCTTTCCCTACACGACGCTCTTCCGATCTNNNTAAGAAACTGCCAAATTGCCTTCC
GCACACGTCTGAACTCTTTCCCTACACGACGCTCTTCCGATCTNNNTAAGAAACTGCCAAATTGCCTTCC
GAGCACACGTCTGAACTCTTTCCCTACACGACGCTCTTCCGATCTNNNTCACTGGTTTCATTTTGATTAA
GAGCACACGTCTGAACTCTTTCCCTACACGACGCTCTTCCGATCTNNNTCACTGGTTTCATTTTGATTAA
AGCACACGTCTGAACTCTTTCCCTACACGACGCTCTTCCGATCTNNNCAAGAAAGAATTCAGGACGGCT
AGCACACGTCTGAACTCTTTCCCTACACGACGCTCTTCCGATCTNNNCAAGAAAGAATTCAGGACGGCT
CACACGTCTGAACTCTTTCCCTACACGACGCTCTTCCGATCTNNNATGAGTTCTTAAATTCTCAGTCTCA
CACACGTCTGAACTCTTTCCCTACACGACGCTCTTCCGATCTNNNATGAGTTCTTAAATTCTCAGTCTCA
AGCACACGTCTGAACTCTTTCCCTACACGACGCTCTTCCGATCTNNNACGAAGTCCTTGCCTCTGTAATT
AGCACACGTCTGAACTCTTTCCCTACACGACGCTCTTCCGATCTNNNACGAAGTCCTTGCCTCTGTAATT
AGCACACGTCTGAACTCTTTCCCTACACGACGCTCTTCCGATCTNNNGATTACATGAATTTACCCATACA
AGCACACGTCTGAACTCTTTCCCTACACGACGCTCTTCCGATCTNNNGATTACATGAATTTACCCATACA
AGCACACGTCTGAACTCTTTCCCTACACGACGCTCTTCCGATCTNNNATATACATCAAACACCTACTTCTC
AGCACACGTCTGAACTCTTTCCCTACACGACGCTCTTCCGATCTNNNATATACATCAAACACCTACTTCTC
GAGCACACGTCTGAACTCTTTCCCTACACGACGCTCTTCCGATCTNNNCATTATCCTGTCTATAGATCCAC
GAGCACACGTCTGAACTCTTTCCCTACACGACGCTCTTCCGATCTNNNCATTATCCTGTCTATAGATCCAC
AGCACACGTCTGAACTCTTTCCCTACACGACGCTCTTCCGATCTNNNAAGATTGTGCCACTACACTCCAC
AGCACACGTCTGAACTCTTTCCCTACACGACGCTCTTCCGATCTNNNAAGATTGTGCCACTACACTCCAC
AGCACACGTCTGAACTCTTTCCCTACACGACGCTCTTCCGATCTNNNCCTCCTCCTGTTGCTTGTTAGAT
AGCACACGTCTGAACTCTTTCCCTACACGACGCTCTTCCGATCTNNNCCTCCTCCTGTTGCTTGTTAGAT
AGCACACGTCTGAACTCTTTCCCTACACGACGCTCTTCCGATCTNNNAACTTTCACAAAGGAGGGGAGA
AGCACACGTCTGAACTCTTTCCCTACACGACGCTCTTCCGATCTNNNAACTTTCACAAAGGAGGGGAGA
AGCACACGTCTGAACTCTTTCCCTACACGACGCTCTTCCGATCTNNNTAGGAGATTCCCTCAAGTGCCT
AGCACACGTCTGAACTCTTTCCCTACACGACGCTCTTCCGATCTNNNTAGGAGATTCCCTCAAGTGCCT
GAGCACACGTCTGAACTCTTTCCCTACACGACGCTCTTCCGATCTNNNTACCAAAGTGCTGGGATTACA
GAGCACACGTCTGAACTCTTTCCCTACACGACGCTCTTCCGATCTNNNTACCAAAGTGCTGGGATTACA
GAGCACACGTCTGAACTCTTTCCCTACACGACGCTCTTCCGATCTNNNATAGAATCTGTCTCCCCTCCCT
GAGCACACGTCTGAACTCTTTCCCTACACGACGCTCTTCCGATCTNNNATAGAATCTGTCTCCCCTCCCT
AGCACACGTCTGAACTCTTTCCCTACACGACGCTCTTCCGATCTNNNCTGGGACTATAGGCATCTGTCAC
AGCACACGTCTGAACTCTTTCCCTACACGACGCTCTTCCGATCTNNNCTGGGACTATAGGCATCTGTCAC
AGCACACGTCTGAACTCTTTCCCTACACGACGCTCTTCCGATCTNNNGCCATTATCAGTTGTCTTCCTTTC
AGCACACGTCTGAACTCTTTCCCTACACGACGCTCTTCCGATCTNNNGCCATTATCAGTTGTCTTCCTTTC
GAGCACACGTCTGAACTCTTTCCCTACACGACGCTCTTCCGATCTNNNAGATGCACATCACTCACACAG
GAGCACACGTCTGAACTCTTTCCCTACACGACGCTCTTCCGATCTNNNAGATGCACATCACTCACACAG
GCACACGTCTGAACTCTTTCCCTACACGACGCTCTTCCGATCTNNNCCAGTGCCCTATAACTCCATCTGC
GCACACGTCTGAACTCTTTCCCTACACGACGCTCTTCCGATCTNNNCCAGTGCCCTATAACTCCATCTGC

GCACACGTCTGAACTCTTTCCCTACACGACGCTCTTCCGATCTNNNTTTAGTTCAATGGATAGATTCCT

GCACACGTCTGAACTCTTTCCCTACACGACGCTCTTCCGATCTNNNTTTAGTTCAATGGATAGATTCCT

ACACGTCTGAACTCTTTCCCTACACGACGCTCTTCCGATCTNNNCACTCGGGGTAGCCGATTAGCATC

ACACGTCTGAACTCTTTCCCTACACGACGCTCTTCCGATCTNNNCACTCGGGGTAGCCGATTAGCATC

AGCACACGTCTGAACTCTTTCCCTACACGACGCTCTTCCGATCTNNNGCCTATAATCCCAGCTACTCAG

AGCACACGTCTGAACTCTTTCCCTACACGACGCTCTTCCGATCTNNNGCCTATAATCCCAGCTACTCAG

AGCACACGTCTGAACTCTTTCCCTACACGACGCTCTTCCGATCTNNNAAGATTGGGTGTTCATGCACA

AGCACACGTCTGAACTCTTTCCCTACACGACGCTCTTCCGATCTNNNAAGATTGGGTGTTCATGCACA

AGCACACGTCTGAACTCTTTCCCTACACGACGCTCTTCCGATCTNNNGCAGATAATCCTCTTCCTTCGG

AGCACACGTCTGAACTCTTTCCCTACACGACGCTCTTCCGATCTNNNGCAGATAATCCTCTTCCTTCGG

AGCACACGTCTGAACTCTTTCCCTACACGACGCTCTTCCGATCTNNNGTCATCTTCTCTCTCCTGTTGCA

AGCACACGTCTGAACTCTTTCCCTACACGACGCTCTTCCGATCTNNNGTCATCTTCTCTCTCCTGTTGCA

GAGCACACGTCTGAACTCTTTCCCTACACGACGCTCTTCCGATCTNNNTAATTTGTTACCTGTCTGGGT

GAGCACACGTCTGAACTCTTTCCCTACACGACGCTCTTCCGATCTNNNTAATTTGTTACCTGTCTGGGT

GCACACGTCTGAACTCTTTCCCTACACGACGCTCTTCCGATCTNNNAGCTACTGTTGATCCCCTCTAGT

GCACACGTCTGAACTCTTTCCCTACACGACGCTCTTCCGATCTNNNAGCTACTGTTGATCCCCTCTAGT

GAGCACACGTCTGAACTCTTTCCCTACACGACGCTCTTCCGATCTNNNGCACCAACATTTACTCACTCAC

GAGCACACGTCTGAACTCTTTCCCTACACGACGCTCTTCCGATCTNNNGCACCAACATTTACTCACTCAC

GAGCACACGTCTGAACTCTTTCCCTACACGACGCTCTTCCGATCTNNNAATGGTGAGTGAGTACCCCA

GAGCACACGTCTGAACTCTTTCCCTACACGACGCTCTTCCGATCTNNNAATGGTGAGTGAGTACCCCA

AGCACACGTCTGAACTCTTTCCCTACACGACGCTCTTCCGATCTNNNGAAAGTACACTTGGAAAAGGG

AGCACACGTCTGAACTCTTTCCCTACACGACGCTCTTCCGATCTNNNGAAAGTACACTTGGAAAAGGG

AGCACACGTCTGAACTCTTTCCCTACACGACGCTCTTCCGATCTNNNCTGCTGGAACCATAGTCTCCAA

AGCACACGTCTGAACTCTTTCCCTACACGACGCTCTTCCGATCTNNNCTGCTGGAACCATAGTCTCCAA

GCACACGTCTGAACTCTTTCCCTACACGACGCTCTTCCGATCTNNNGGTTTGGGTGACTGTGACAATT

GCACACGTCTGAACTCTTTCCCTACACGACGCTCTTCCGATCTNNNGGTTTGGGTGACTGTGACAATT

AGCACACGTCTGAACTCTTTCCCTACACGACGCTCTTCCGATCTNNNAAACCCCGACGCTACAAAAAA

AGCACACGTCTGAACTCTTTCCCTACACGACGCTCTTCCGATCTNNNAAACCCCGACGCTACAAAAAA

GAGCACACGTCTGAACTCTTTCCCTACACGACGCTCTTCCGATCTNNNTGCACCTATAGTCCCAGCTAC

GAGCACACGTCTGAACTCTTTCCCTACACGACGCTCTTCCGATCTNNNTGCACCTATAGTCCCAGCTAC

AGAGCACACGTCTGAACTCTTTCCCTACACGACGCTCTTCCGATCTNNNTGAAGGTACATCCTCAGAA

AGAGCACACGTCTGAACTCTTTCCCTACACGACGCTCTTCCGATCTNNNTGAAGGTACATCCTCAGAA

GCACACGTCTGAACTCTTTCCCTACACGACGCTCTTCCGATCTNNNCTGAAGCAGGAGAATGGCTTG

GCACACGTCTGAACTCTTTCCCTACACGACGCTCTTCCGATCTNNNCTGAAGCAGGAGAATGGCTTG

AGCACACGTCTGAACTCTTTCCCTACACGACGCTCTTCCGATCTNNNCTCCACACACCCCAACATACATA

AGCACACGTCTGAACTCTTTCCCTACACGACGCTCTTCCGATCTNNNCTCCACACACCCCAACATACATA

AGCACACGTCTGAACTCTTTCCCTACACGACGCTCTTCCGATCTNNNCCTCAGCTTCCCAAGTATCTGGC

AGCACACGTCTGAACTCTTTCCCTACACGACGCTCTTCCGATCTNNNCCTCAGCTTCCCAAGTATCTGGC

GAGCACACGTCTGAACTCTTTCCCTACACGACGCTCTTCCGATCTNNNCCTCAGTCTCCTGAATAGTTG

GAGCACACGTCTGAACTCTTTCCCTACACGACGCTCTTCCGATCTNNNCCTCAGTCTCCTGAATAGTTG

AGCACACGTCTGAACTCTTTCCCTACACGACGCTCTTCCGATCTNNNATGAATTTCTGTTTCCTGCAGC

AGCACACGTCTGAACTCTTTCCCTACACGACGCTCTTCCGATCTNNNATGAATTTCTGTTTCCTGCAGC

GAGCACACGTCTGAACTCTTTCCCTACACGACGCTCTTCCGATCTNNNTTCAAGCAATTCTCCTGCCTCA

GAGCACACGTCTGAACTCTTTCCCTACACGACGCTCTTCCGATCTNNNTTCAAGCAATTCTCCTGCCTCA
GAGCACACGTCTGAACTCTTTCCCTACACGACGCTCTTCCGATCTNNNAGTGAACTACGATTTTGCCACT
GAGCACACGTCTGAACTCTTTCCCTACACGACGCTCTTCCGATCTNNNAGTGAACTACGATTTTGCCACT
AGAGCACACGTCTGAACTCTTTCCCTACACGACGCTCTTCCGATCTNNNGAGGATTAGGACGAGGACA
AGAGCACACGTCTGAACTCTTTCCCTACACGACGCTCTTCCGATCTNNNGAGGATTAGGACGAGGACA
AAGAGCACACGTCTGAACTCTTTCCCTACACGACGCTCTTCCGATCTNNNAAGATACATATGGCCAATA
AAGAGCACACGTCTGAACTCTTTCCCTACACGACGCTCTTCCGATCTNNNAAGATACATATGGCCAATA
AGCACACGTCTGAACTCTTTCCCTACACGACGCTCTTCCGATCTNNNTATCCACCAGTGCCCAATGTTTA
AGCACACGTCTGAACTCTTTCCCTACACGACGCTCTTCCGATCTNNNTATCCACCAGTGCCCAATGTTTA
GCACACGTCTGAACTCTTTCCCTACACGACGCTCTTCCGATCTNNNGCTAGGTAGGAGAATCGCTTGAA
GCACACGTCTGAACTCTTTCCCTACACGACGCTCTTCCGATCTNNNGCTAGGTAGGAGAATCGCTTGAA
AGCACACGTCTGAACTCTTTCCCTACACGACGCTCTTCCGATCTNNNTTCAGGTGATTCTGCCTCACTCC
AGCACACGTCTGAACTCTTTCCCTACACGACGCTCTTCCGATCTNNNTTCAGGTGATTCTGCCTCACTCC
GAGCACACGTCTGAACTCTTTCCCTACACGACGCTCTTCCGATCTNNNCCCACCACTATCTCTCCAACAA
GAGCACACGTCTGAACTCTTTCCCTACACGACGCTCTTCCGATCTNNNCCCACCACTATCTCTCCAACAA
AGCACACGTCTGAACTCTTTCCCTACACGACGCTCTTCCGATCTNNNACTTCTTCCTTTCCTATCTGCATT
AGCACACGTCTGAACTCTTTCCCTACACGACGCTCTTCCGATCTNNNACTTCTTCCTTTCCTATCTGCATT
GAGCACACGTCTGAACTCTTTCCCTACACGACGCTCTTCCGATCTNNNTAATCCCAGCTACTCAGGAGGA
GAGCACACGTCTGAACTCTTTCCCTACACGACGCTCTTCCGATCTNNNTAATCCCAGCTACTCAGGAGGA
GCACACGTCTGAACTCTTTCCCTACACGACGCTCTTCCGATCTNNNCTCCCATCCTCAAGATTCCTTCACA
GCACACGTCTGAACTCTTTCCCTACACGACGCTCTTCCGATCTNNNCTCCCATCCTCAAGATTCCTTCACA
AGCACACGTCTGAACTCTTTCCCTACACGACGCTCTTCCGATCTNNNTTCAAAACATCGCATTGGCAGAT
AGCACACGTCTGAACTCTTTCCCTACACGACGCTCTTCCGATCTNNNTTCAAAACATCGCATTGGCAGAT
AGCACACGTCTGAACTCTTTCCCTACACGACGCTCTTCCGATCTNNNCAGATGTGCACTGCTTTAGGAT
AGCACACGTCTGAACTCTTTCCCTACACGACGCTCTTCCGATCTNNNCAGATGTGCACTGCTTTAGGAT
GAGCACACGTCTGAACTCTTTCCCTACACGACGCTCTTCCGATCTNNNATCTTCAGCTCTTGGAACTCAG
GAGCACACGTCTGAACTCTTTCCCTACACGACGCTCTTCCGATCTNNNATCTTCAGCTCTTGGAACTCAG
AGAGCACACGTCTGAACTCTTTCCCTACACGACGCTCTTCCGATCTNNNGTAACAGAGCTAGATGCTGT
AGAGCACACGTCTGAACTCTTTCCCTACACGACGCTCTTCCGATCTNNNGTAACAGAGCTAGATGCTGT
GAGCACACGTCTGAACTCTTTCCCTACACGACGCTCTTCCGATCTNNNGTACTTCCTCAGCTTCTGTCTG
GAGCACACGTCTGAACTCTTTCCCTACACGACGCTCTTCCGATCTNNNGTACTTCCTCAGCTTCTGTCTG
GCACACGTCTGAACTCTTTCCCTACACGACGCTCTTCCGATCTNNNGATCCTGCAGAAGAGGAGTTCTG
GCACACGTCTGAACTCTTTCCCTACACGACGCTCTTCCGATCTNNNGATCCTGCAGAAGAGGAGTTCTG
AGCACACGTCTGAACTCTTTCCCTACACGACGCTCTTCCGATCTNNNTGAAGCTGGATCTCTATCTCACT
AGCACACGTCTGAACTCTTTCCCTACACGACGCTCTTCCGATCTNNNTGAAGCTGGATCTCTATCTCACT
GCACACGTCTGAACTCTTTCCCTACACGACGCTCTTCCGATCTNNNCAGACAGACACGCAGAACTCCCAT
GCACACGTCTGAACTCTTTCCCTACACGACGCTCTTCCGATCTNNNCAGACAGACACGCAGAACTCCCAT
AGCACACGTCTGAACTCTTTCCCTACACGACGCTCTTCCGATCTNNNAATTTCACGATGGCAGTTTCACC
AGCACACGTCTGAACTCTTTCCCTACACGACGCTCTTCCGATCTNNNAATTTCACGATGGCAGTTTCACC
GCACACGTCTGAACTCTTTCCCTACACGACGCTCTTCCGATCTNNNCAGCGATGGCCTCTTCTGTAAGC
GCACACGTCTGAACTCTTTCCCTACACGACGCTCTTCCGATCTNNNCAGCGATGGCCTCTTCTGTAAGC
GCACACGTCTGAACTCTTTCCCTACACGACGCTCTTCCGATCTNNNGGGTTCCTCTACACTCATCTGACC
GCACACGTCTGAACTCTTTCCCTACACGACGCTCTTCCGATCTNNNGGGTTCCTCTACACTCATCTGACC

AGCACACGTCTGAACTCTTTCCCTACACGACGCTCTTCCGATCTNNNCTCAAACTCCTGACCTCGTGAT
AGCACACGTCTGAACTCTTTCCCTACACGACGCTCTTCCGATCTNNNCTCAAACTCCTGACCTCGTGAT
AGCACACGTCTGAACTCTTTCCCTACACGACGCTCTTCCGATCTNNNAGTGCTGAGACTCGAATTCTGA
AGCACACGTCTGAACTCTTTCCCTACACGACGCTCTTCCGATCTNNNAGTGCTGAGACTCGAATTCTGA
AAGAGCACACGTCTGAACTCTTTCCCTACACGACGCTCTTCCGATCTNNNATCAGCAGCCATGTGTAG
AAGAGCACACGTCTGAACTCTTTCCCTACACGACGCTCTTCCGATCTNNNATCAGCAGCCATGTGTAG
AGCACACGTCTGAACTCTTTCCCTACACGACGCTCTTCCGATCTNNNAGACTTGTGATAATGGCCTCCA
AGCACACGTCTGAACTCTTTCCCTACACGACGCTCTTCCGATCTNNNAGACTTGTGATAATGGCCTCCA
AGCACACGTCTGAACTCTTTCCCTACACGACGCTCTTCCGATCTNNNCAACCTCTTTGGGGTCCTCAAT
AGCACACGTCTGAACTCTTTCCCTACACGACGCTCTTCCGATCTNNNCAACCTCTTTGGGGTCCTCAAT
AGCACACGTCTGAACTCTTTCCCTACACGACGCTCTTCCGATCTNNNATACAGGATGCACCCTTTTGTG
AGCACACGTCTGAACTCTTTCCCTACACGACGCTCTTCCGATCTNNNATACAGGATGCACCCTTTTGTG
CACGTCTGAACTCTTTCCCTACACGACGCTCTTCCGATCTNNNAGATGTGTCTCTTGAATATGGCACCA
CACGTCTGAACTCTTTCCCTACACGACGCTCTTCCGATCTNNNAGATGTGTCTCTTGAATATGGCACCA
AGCACACGTCTGAACTCTTTCCCTACACGACGCTCTTCCGATCTNNNTGGTCAGATTTCCTAGCCTCTG
AGCACACGTCTGAACTCTTTCCCTACACGACGCTCTTCCGATCTNNNTGGTCAGATTTCCTAGCCTCTG
AGCACACGTCTGAACTCTTTCCCTACACGACGCTCTTCCGATCTNNNGCCTATTATACACCTGGGTCTG
AGCACACGTCTGAACTCTTTCCCTACACGACGCTCTTCCGATCTNNNGCCTATTATACACCTGGGTCTG
GCACACGTCTGAACTCTTTCCCTACACGACGCTCTTCCGATCTNNNCTCTCCCCCCTCTCTCCCCATCG
GCACACGTCTGAACTCTTTCCCTACACGACGCTCTTCCGATCTNNNCTCTCCCCCCTCTCTCCCCATCG
GAGCACACGTCTGAACTCTTTCCCTACACGACGCTCTTCCGATCTNNNGGAGCTTGCAGTGAGACTAG
GAGCACACGTCTGAACTCTTTCCCTACACGACGCTCTTCCGATCTNNNGGAGCTTGCAGTGAGACTAG
AAGAGCACACGTCTGAACTCTTTCCCTACACGACGCTCTTCCGATCTNNNCTGATGACCTTTGGAGGA
AAGAGCACACGTCTGAACTCTTTCCCTACACGACGCTCTTCCGATCTNNNCTGATGACCTTTGGAGGA
GCACACGTCTGAACTCTTTCCCTACACGACGCTCTTCCGATCTNNNTGGTATCAGTCAGTCAATAAGC
GCACACGTCTGAACTCTTTCCCTACACGACGCTCTTCCGATCTNNNTGGTATCAGTCAGTCAATAAGC
AGCACACGTCTGAACTCTTTCCCTACACGACGCTCTTCCGATCTNNNTTGCTGTGCAAGATCAGTCATT
AGCACACGTCTGAACTCTTTCCCTACACGACGCTCTTCCGATCTNNNTTGCTGTGCAAGATCAGTCATT
GCACACGTCTGAACTCTTTCCCTACACGACGCTCTTCCGATCTNNNCATGCCTGTAATCCCAGCTACTC
GCACACGTCTGAACTCTTTCCCTACACGACGCTCTTCCGATCTNNNCATGCCTGTAATCCCAGCTACTC
CACACGTCTGAACTCTTTCCCTACACGACGCTCTTCCGATCTNNNAGCGCCATGAGCAACAAATGCAT
CACACGTCTGAACTCTTTCCCTACACGACGCTCTTCCGATCTNNNAGCGCCATGAGCAACAAATGCAT
AGCACACGTCTGAACTCTTTCCCTACACGACGCTCTTCCGATCTNNNGTGTGAAGGAGAAGAAAGTG
AGCACACGTCTGAACTCTTTCCCTACACGACGCTCTTCCGATCTNNNGTGTGAAGGAGAAGAAAGTG
GCACACGTCTGAACTCTTTCCCTACACGACGCTCTTCCGATCTNNNCAATTCTTCCTTTAGCCCTGCAG
GCACACGTCTGAACTCTTTCCCTACACGACGCTCTTCCGATCTNNNCAATTCTTCCTTTAGCCCTGCAG
GAGCACACGTCTGAACTCTTTCCCTACACGACGCTCTTCCGATCTNNNACAACACTGCACTCTAGATCT
GAGCACACGTCTGAACTCTTTCCCTACACGACGCTCTTCCGATCTNNNACAACACTGCACTCTAGATCT
AGCACACGTCTGAACTCTTTCCCTACACGACGCTCTTCCGATCTNNNGCAGAAGAATCACTTGAATCG
AGCACACGTCTGAACTCTTTCCCTACACGACGCTCTTCCGATCTNNNGCAGAAGAATCACTTGAATCG
GAGCACACGTCTGAACTCTTTCCCTACACGACGCTCTTCCGATCTNNNCTCCTTTCCAGATGAGTGAA
GAGCACACGTCTGAACTCTTTCCCTACACGACGCTCTTCCGATCTNNNCTCCTTTCCAGATGAGTGAA
GAGCACACGTCTGAACTCTTTCCCTACACGACGCTCTTCCGATCTNNNAAGTAGCTGGGATTACATGC

GAGCACACGTCTGAACTCTTTCCCTACACGACGCTCTTCCGATCTNNNAAGTAGCTGGGATTACATGC
GAGCACACGTCTGAACTCTTTCCCTACACGACGCTCTTCCGATCTNNNCATCTCATGGGCAGAGACAC
GAGCACACGTCTGAACTCTTTCCCTACACGACGCTCTTCCGATCTNNNCATCTCATGGGCAGAGACAC
AGCACACGTCTGAACTCTTTCCCTACACGACGCTCTTCCGATCTNNNACTTGATGTCCAGAGTTCGAG
AGCACACGTCTGAACTCTTTCCCTACACGACGCTCTTCCGATCTNNNACTTGATGTCCAGAGTTCGAG
GCACACGTCTGAACTCTTTCCCTACACGACGCTCTTCCGATCTNNNCTGAGGCAGGAGAATCACTTGA
GCACACGTCTGAACTCTTTCCCTACACGACGCTCTTCCGATCTNNNCTGAGGCAGGAGAATCACTTGA
AGCACACGTCTGAACTCTTTCCCTACACGACGCTCTTCCGATCTNNNCTGTTGAATGGTTAACTGTAAA
AGCACACGTCTGAACTCTTTCCCTACACGACGCTCTTCCGATCTNNNCTGTTGAATGGTTAACTGTAAA
ACACGTCTGAACTCTTTCCCTACACGACGCTCTTCCGATCTNNNTGTCTCTCTCTCTCTCTCTCTCATC
ACACGTCTGAACTCTTTCCCTACACGACGCTCTTCCGATCTNNNTGTCTCTCTCTCTCTCTCTCTCATC
GAGCACACGTCTGAACTCTTTCCCTACACGACGCTCTTCCGATCTNNNGGGCCAAATGAGGTATTTCC
GAGCACACGTCTGAACTCTTTCCCTACACGACGCTCTTCCGATCTNNNGGGCCAAATGAGGTATTTCC
GAGCACACGTCTGAACTCTTTCCCTACACGACGCTCTTCCGATCTNNNTCAGACTCCTGAGTAGCTGA
GAGCACACGTCTGAACTCTTTCCCTACACGACGCTCTTCCGATCTNNNTCAGACTCCTGAGTAGCTGA
GAGCACACGTCTGAACTCTTTCCCTACACGACGCTCTTCCGATCTNNNTAGAGTGGAGTCTAGCATTG
GAGCACACGTCTGAACTCTTTCCCTACACGACGCTCTTCCGATCTNNNTAGAGTGGAGTCTAGCATTG
AGCACACGTCTGAACTCTTTCCCTACACGACGCTCTTCCGATCTNNNTGACAGAGTGAGACTCCCTCT
AGCACACGTCTGAACTCTTTCCCTACACGACGCTCTTCCGATCTNNNTGACAGAGTGAGACTCCCTCT
GAGCACACGTCTGAACTCTTTCCCTACACGACGCTCTTCCGATCTNNNATGGACTGGTTTTTCTTTCAG
GAGCACACGTCTGAACTCTTTCCCTACACGACGCTCTTCCGATCTNNNATGGACTGGTTTTTCTTTCAG
AGCACACGTCTGAACTCTTTCCCTACACGACGCTCTTCCGATCTNNNGACCCTGCATTACTGTACCACT
AGCACACGTCTGAACTCTTTCCCTACACGACGCTCTTCCGATCTNNNGACCCTGCATTACTGTACCACT
AGCACACGTCTGAACTCTTTCCCTACACGACGCTCTTCCGATCTNNNTACGATCTTTGGACAGCTACC
AGCACACGTCTGAACTCTTTCCCTACACGACGCTCTTCCGATCTNNNTACGATCTTTGGACAGCTACC
GAGCACACGTCTGAACTCTTTCCCTACACGACGCTCTTCCGATCTNNNCCTCTGTTTCTTGGTGGGTTA
GAGCACACGTCTGAACTCTTTCCCTACACGACGCTCTTCCGATCTNNNCCTCTGTTTCTTGGTGGGTTA
AGCACACGTCTGAACTCTTTCCCTACACGACGCTCTTCCGATCTNNNCTCATCTTCTAGCTCCTTCTTCAT
AGCACACGTCTGAACTCTTTCCCTACACGACGCTCTTCCGATCTNNNCTCATCTTCTAGCTCCTTCTTCAT
GCACACGTCTGAACTCTTTCCCTACACGACGCTCTTCCGATCTNNNGGAATCTGCCCTGTGTCGTCCAT
GCACACGTCTGAACTCTTTCCCTACACGACGCTCTTCCGATCTNNNGGAATCTGCCCTGTGTCGTCCAT
AGCACACGTCTGAACTCTTTCCCTACACGACGCTCTTCCGATCTNNNACCACCTTCATGTTCATCACAG
AGCACACGTCTGAACTCTTTCCCTACACGACGCTCTTCCGATCTNNNACCACCTTCATGTTCATCACAG
AGCACACGTCTGAACTCTTTCCCTACACGACGCTCTTCCGATCTNNNCCTGTGAAGAAAATTTGGAGC
AGCACACGTCTGAACTCTTTCCCTACACGACGCTCTTCCGATCTNNNCCTGTGAAGAAAATTTGGAGC
AGCACACGTCTGAACTCTTTCCCTACACGACGCTCTTCCGATCTNNNGCCTGGCCTAACTTGCTTTATT
AGCACACGTCTGAACTCTTTCCCTACACGACGCTCTTCCGATCTNNNGCCTGGCCTAACTTGCTTTATT
AGCACACGTCTGAACTCTTTCCCTACACGACGCTCTTCCGATCTNNNGAATTGCTTGAACCAGATGGA
AGCACACGTCTGAACTCTTTCCCTACACGACGCTCTTCCGATCTNNNGAATTGCTTGAACCAGATGGA
AGCACACGTCTGAACTCTTTCCCTACACGACGCTCTTCCGATCTNNNTCACTGGGCTGTGCTTACTTTT
AGCACACGTCTGAACTCTTTCCCTACACGACGCTCTTCCGATCTNNNTCACTGGGCTGTGCTTACTTTT
AGCACACGTCTGAACTCTTTCCCTACACGACGCTCTTCCGATCTNNNGGGAAGAGACTTCTGGAACA
AGCACACGTCTGAACTCTTTCCCTACACGACGCTCTTCCGATCTNNNGGGAAGAGACTTCTGGAACA

AGCACACGTCTGAACTCTTTCCCTACACGACGCTCTTCCGATCTNNNCAAGTTGCTGCAGAAGACGTT

AGCACACGTCTGAACTCTTTCCCTACACGACGCTCTTCCGATCTNNNCAAGTTGCTGCAGAAGACGTT

GCACACGTCTGAACTCTTTCCCTACACGACGCTCTTCCGATCTNNNCATGGTGTGGTTCCTTCCTCTTA

GCACACGTCTGAACTCTTTCCCTACACGACGCTCTTCCGATCTNNNCATGGTGTGGTTCCTTCCTCTTA

GAGCACACGTCTGAACTCTTTCCCTACACGACGCTCTTCCGATCTNNNGTGGTTGAAGTGAAAACGA

GAGCACACGTCTGAACTCTTTCCCTACACGACGCTCTTCCGATCTNNNGTGGTTGAAGTGAAAACGA

GCACACGTCTGAACTCTTTCCCTACACGACGCTCTTCCGATCTNNNAACTGCAGAGCTTGAGACTTCT

GCACACGTCTGAACTCTTTCCCTACACGACGCTCTTCCGATCTNNNAACTGCAGAGCTTGAGACTTCT

AGCACACGTCTGAACTCTTTCCCTACACGACGCTCTTCCGATCTNNNAAAATGCTGGGATTACAGGC

AGCACACGTCTGAACTCTTTCCCTACACGACGCTCTTCCGATCTNNNAAAATGCTGGGATTACAGGC

AGCACACGTCTGAACTCTTTCCCTACACGACGCTCTTCCGATCTNNNAGATATGGCCAAACACTACCCA

AGCACACGTCTGAACTCTTTCCCTACACGACGCTCTTCCGATCTNNNAGATATGGCCAAACACTACCCA

AGCACACGTCTGAACTCTTTCCCTACACGACGCTCTTCCGATCTNNNAGCCAGAGAGATGTAACTTCA

AGCACACGTCTGAACTCTTTCCCTACACGACGCTCTTCCGATCTNNNAGCCAGAGAGATGTAACTTCA

AAGAGCACACGTCTGAACTCTTTCCCTACACGACGCTCTTCCGATCTNNNCCCAGAAACCTCTGATTAT

AAGAGCACACGTCTGAACTCTTTCCCTACACGACGCTCTTCCGATCTNNNCCCAGAAACCTCTGATTAT

GAGCACACGTCTGAACTCTTTCCCTACACGACGCTCTTCCGATCTNNNGTGGATCACAAGGTCAGGA

GAGCACACGTCTGAACTCTTTCCCTACACGACGCTCTTCCGATCTNNNGTGGATCACAAGGTCAGGA

GAGCACACGTCTGAACTCTTTCCCTACACGACGCTCTTCCGATCTNNNTGAGAAGTGCTCAATGCTCT

GAGCACACGTCTGAACTCTTTCCCTACACGACGCTCTTCCGATCTNNNTGAGAAGTGCTCAATGCTCT

GCACACGTCTGAACTCTTTCCCTACACGACGCTCTTCCGATCTNNNTGAGCTCTGAGTCAAATCAAGC

GCACACGTCTGAACTCTTTCCCTACACGACGCTCTTCCGATCTNNNTGAGCTCTGAGTCAAATCAAGC

AGCACACGTCTGAACTCTTTCCCTACACGACGCTCTTCCGATCTNNNGTGTGTTTGTAAGAGTTCCTG

AGCACACGTCTGAACTCTTTCCCTACACGACGCTCTTCCGATCTNNNGTGTGTTTGTAAGAGTTCCTG

GCACACGTCTGAACTCTTTCCCTACACGACGCTCTTCCGATCTNNNAATTCCATATCTTGCGAGGCTG

GCACACGTCTGAACTCTTTCCCTACACGACGCTCTTCCGATCTNNNAATTCCATATCTTGCGAGGCTG

AGAGCACACGTCTGAACTCTTTCCCTACACGACGCTCTTCCGATCTNNNTCATTCTGTCAGTGGGAAA

AGAGCACACGTCTGAACTCTTTCCCTACACGACGCTCTTCCGATCTNNNTCATTCTGTCAGTGGGAAA

GCACACGTCTGAACTCTTTCCCTACACGACGCTCTTCCGATCTNNNACAGCACTGATTACATGGGTAC

GCACACGTCTGAACTCTTTCCCTACACGACGCTCTTCCGATCTNNNACAGCACTGATTACATGGGTAC

CACACGTCTGAACTCTTTCCCTACACGACGCTCTTCCGATCTNNNGAGGTTGTGGTGAGCTGAGATC

CACACGTCTGAACTCTTTCCCTACACGACGCTCTTCCGATCTNNNGAGGTTGTGGTGAGCTGAGATC

CACACGTCTGAACTCTTTCCCTACACGACGCTCTTCCGATCTNNNTGACTCAGATGACCACGTTTGAT

CACACGTCTGAACTCTTTCCCTACACGACGCTCTTCCGATCTNNNTGACTCAGATGACCACGTTTGAT

AGCACACGTCTGAACTCTTTCCCTACACGACGCTCTTCCGATCTNNNCAGTCTGGACTTTTGCTCCAAA

AGCACACGTCTGAACTCTTTCCCTACACGACGCTCTTCCGATCTNNNCAGTCTGGACTTTTGCTCCAAA

GAGCACACGTCTGAACTCTTTCCCTACACGACGCTCTTCCGATCTNNNGCTCACACCTGTAAAACCAAC

GAGCACACGTCTGAACTCTTTCCCTACACGACGCTCTTCCGATCTNNNGCTCACACCTGTAAAACCAAC

AGCACACGTCTGAACTCTTTCCCTACACGACGCTCTTCCGATCTNNNAGAATGCTAGCTCAGGTGTTA

AGCACACGTCTGAACTCTTTCCCTACACGACGCTCTTCCGATCTNNNAGAATGCTAGCTCAGGTGTTA

AGCACACGTCTGAACTCTTTCCCTACACGACGCTCTTCCGATCTNNNATAGTCCTAGCTATGTGGGG

AGCACACGTCTGAACTCTTTCCCTACACGACGCTCTTCCGATCTNNNATAGTCCTAGCTATGTGGGG

GAGCACACGTCTGAACTCTTTCCCTACACGACGCTCTTCCGATCTNNNTGGTTTCACTGTGACTTAGA

GAGCACACGTCTGAACTCTTTCCCTACACGACGCTCTTCCGATCTNNNTGGTTTCACTGTGACTTAGA
AGCACACGTCTGAACTCTTTCCCTACACGACGCTCTTCCGATCTNNNACATACTAAGACAGGAGCACCT
AGCACACGTCTGAACTCTTTCCCTACACGACGCTCTTCCGATCTNNNACATACTAAGACAGGAGCACCT
GCACACGTCTGAACTCTTTCCCTACACGACGCTCTTCCGATCTNNNCCTTCCACTCTCTCAACCTTTGTC
GCACACGTCTGAACTCTTTCCCTACACGACGCTCTTCCGATCTNNNCCTTCCACTCTCTCAACCTTTGTC
AGCACACGTCTGAACTCTTTCCCTACACGACGCTCTTCCGATCTNNNTACTCTGCACTACTCCTGTGA
AGCACACGTCTGAACTCTTTCCCTACACGACGCTCTTCCGATCTNNNTACTCTGCACTACTCCTGTGA
AGCACACGTCTGAACTCTTTCCCTACACGACGCTCTTCCGATCTNNNCTGTAAGCCCAGCTACTCAAGA
AGCACACGTCTGAACTCTTTCCCTACACGACGCTCTTCCGATCTNNNCTGTAAGCCCAGCTACTCAAGA
AGCACACGTCTGAACTCTTTCCCTACACGACGCTCTTCCGATCTNNNATCCAATTCCTGCCTCTCTGTCT
AGCACACGTCTGAACTCTTTCCCTACACGACGCTCTTCCGATCTNNNATCCAATTCCTGCCTCTCTGTCT
GCACACGTCTGAACTCTTTCCCTACACGACGCTCTTCCGATCTNNNTCTGTAATCCCAGCTACTCAGG
GCACACGTCTGAACTCTTTCCCTACACGACGCTCTTCCGATCTNNNTCTGTAATCCCAGCTACTCAGG
AGCACACGTCTGAACTCTTTCCCTACACGACGCTCTTCCGATCTNNNCTACCTTCTCCCAGAGCTCACTT
AGCACACGTCTGAACTCTTTCCCTACACGACGCTCTTCCGATCTNNNCTACCTTCTCCCAGAGCTCACTT
GCACACGTCTGAACTCTTTCCCTACACGACGCTCTTCCGATCTNNNAGTAACTGGGACTATAAGCGCG
GCACACGTCTGAACTCTTTCCCTACACGACGCTCTTCCGATCTNNNAGTAACTGGGACTATAAGCGCG
AGCACACGTCTGAACTCTTTCCCTACACGACGCTCTTCCGATCTNNNTGTGTTAGCCAGGATGTTCTT
AGCACACGTCTGAACTCTTTCCCTACACGACGCTCTTCCGATCTNNNTGTGTTAGCCAGGATGTTCTT
AGCACACGTCTGAACTCTTTCCCTACACGACGCTCTTCCGATCTNNNACCTTAAGATCAACCACCATGG
AGCACACGTCTGAACTCTTTCCCTACACGACGCTCTTCCGATCTNNNACCTTAAGATCAACCACCATGG
AGAGCACACGTCTGAACTCTTTCCCTACACGACGCTCTTCCGATCTNNNAGCCCTCAACAAAATTACGG
AGAGCACACGTCTGAACTCTTTCCCTACACGACGCTCTTCCGATCTNNNAGCCCTCAACAAAATTACGG
CACACGTCTGAACTCTTTCCCTACACGACGCTCTTCCGATCTNNNGCCAGTGTATTAGTCTGCTCTCAC
CACACGTCTGAACTCTTTCCCTACACGACGCTCTTCCGATCTNNNGCCAGTGTATTAGTCTGCTCTCAC
AGCACACGTCTGAACTCTTTCCCTACACGACGCTCTTCCGATCTNNNTGGAATGTGCCAAATGTTTCTG
AGCACACGTCTGAACTCTTTCCCTACACGACGCTCTTCCGATCTNNNTGGAATGTGCCAAATGTTTCTG
GCACACGTCTGAACTCTTTCCCTACACGACGCTCTTCCGATCTNNNGAAAGTGGGGGCCAATATTCA
GCACACGTCTGAACTCTTTCCCTACACGACGCTCTTCCGATCTNNNGAAAGTGGGGGCCAATATTCA
AGCACACGTCTGAACTCTTTCCCTACACGACGCTCTTCCGATCTNNNCTAAGTGGCTTCTGAAGGGG
AGCACACGTCTGAACTCTTTCCCTACACGACGCTCTTCCGATCTNNNCTAAGTGGCTTCTGAAGGGG
GCACACGTCTGAACTCTTTCCCTACACGACGCTCTTCCGATCTNNNGGGTGTTCTAAGTACACTGAG
GCACACGTCTGAACTCTTTCCCTACACGACGCTCTTCCGATCTNNNGGGTGTTCTAAGTACACTGAG
CACACGTCTGAACTCTTTCCCTACACGACGCTCTTCCGATCTNNNTAAGGATCGCTTGAACCTGAGAA
CACACGTCTGAACTCTTTCCCTACACGACGCTCTTCCGATCTNNNTAAGGATCGCTTGAACCTGAGAA
AGAGCACACGTCTGAACTCTTTCCCTACACGACGCTCTTCCGATCTNNNGACAGAGCGAGACTCCGTC
AGAGCACACGTCTGAACTCTTTCCCTACACGACGCTCTTCCGATCTNNNGACAGAGCGAGACTCCGTC
AGCACACGTCTGAACTCTTTCCCTACACGACGCTCTTCCGATCTNNNTTGCTGACAATCTTTGGTGTT
AGCACACGTCTGAACTCTTTCCCTACACGACGCTCTTCCGATCTNNNTTGCTGACAATCTTTGGTGTT
GAGCACACGTCTGAACTCTTTCCCTACACGACGCTCTTCCGATCTNNNTGTATTTGTTTGGGGCAGTT
GAGCACACGTCTGAACTCTTTCCCTACACGACGCTCTTCCGATCTNNNTGTATTTGTTTGGGGCAGTT
AGCACACGTCTGAACTCTTTCCCTACACGACGCTCTTCCGATCTNNNCCTGGGTTCAAGCAATTCTTCA
AGCACACGTCTGAACTCTTTCCCTACACGACGCTCTTCCGATCTNNNCCTGGGTTCAAGCAATTCTTCA

GAGCACACGTCTGAACTCTTTCCCTACACGACGCTCTTCCGATCTNNNAGACAGGAGAATTGCTTGAA
GAGCACACGTCTGAACTCTTTCCCTACACGACGCTCTTCCGATCTNNNAGACAGGAGAATTGCTTGAA
GCACACGTCTGAACTCTTTCCCTACACGACGCTCTTCCGATCTNNNGCTATCCAAAAGCTGTCACAACA
GCACACGTCTGAACTCTTTCCCTACACGACGCTCTTCCGATCTNNNGCTATCCAAAAGCTGTCACAACA
AGCACACGTCTGAACTCTTTCCCTACACGACGCTCTTCCGATCTNNNTTCAAAACGTTATCCAGCTGTG
AGCACACGTCTGAACTCTTTCCCTACACGACGCTCTTCCGATCTNNNTTCAAAACGTTATCCAGCTGTG
GAGCACACGTCTGAACTCTTTCCCTACACGACGCTCTTCCGATCTNNNAAGTATCTTCTTCCCAAGGTG
GAGCACACGTCTGAACTCTTTCCCTACACGACGCTCTTCCGATCTNNNAAGTATCTTCTTCCCAAGGTG
AGCACACGTCTGAACTCTTTCCCTACACGACGCTCTTCCGATCTNNNGGTCTAATACAGATGCAGCAG
AGCACACGTCTGAACTCTTTCCCTACACGACGCTCTTCCGATCTNNNGGTCTAATACAGATGCAGCAG
GAGCACACGTCTGAACTCTTTCCCTACACGACGCTCTTCCGATCTNNNACCAAACCCGGCTAAATTTTTG
GAGCACACGTCTGAACTCTTTCCCTACACGACGCTCTTCCGATCTNNNACCAAACCCGGCTAAATTTTTG
GAGCACACGTCTGAACTCTTTCCCTACACGACGCTCTTCCGATCTNNNCTGTAATCCCAGCTACTCAGG
GAGCACACGTCTGAACTCTTTCCCTACACGACGCTCTTCCGATCTNNNCTGTAATCCCAGCTACTCAGG
GAGCACACGTCTGAACTCTTTCCCTACACGACGCTCTTCCGATCTNNNTTGTGTGATTCACTTCATTGT
GAGCACACGTCTGAACTCTTTCCCTACACGACGCTCTTCCGATCTNNNTTGTGTGATTCACTTCATTGT
AGCACACGTCTGAACTCTTTCCCTACACGACGCTCTTCCGATCTNNNCACCTGTAGTCCCAGCTACTTG
AGCACACGTCTGAACTCTTTCCCTACACGACGCTCTTCCGATCTNNNCACCTGTAGTCCCAGCTACTTG
GCACACGTCTGAACTCTTTCCCTACACGACGCTCTTCCGATCTNNNATCAGTAGTGGGATTGCTGGAT
GCACACGTCTGAACTCTTTCCCTACACGACGCTCTTCCGATCTNNNATCAGTAGTGGGATTGCTGGAT
GCACACGTCTGAACTCTTTCCCTACACGACGCTCTTCCGATCTNNNTCACAAGGACTGAAATACTGGG
GCACACGTCTGAACTCTTTCCCTACACGACGCTCTTCCGATCTNNNTCACAAGGACTGAAATACTGGG
GCACACGTCTGAACTCTTTCCCTACACGACGCTCTTCCGATCTNNNTCCTGTTTCTAGGACCCATGTTG
GCACACGTCTGAACTCTTTCCCTACACGACGCTCTTCCGATCTNNNTCCTGTTTCTAGGACCCATGTTG
AGCACACGTCTGAACTCTTTCCCTACACGACGCTCTTCCGATCTNNNATTCAAGTGGATGCCATTTGCT
AGCACACGTCTGAACTCTTTCCCTACACGACGCTCTTCCGATCTNNNATTCAAGTGGATGCCATTTGCT
GAGCACACGTCTGAACTCTTTCCCTACACGACGCTCTTCCGATCTNNNGAAATGCAGGCTTGAGTTCT
GAGCACACGTCTGAACTCTTTCCCTACACGACGCTCTTCCGATCTNNNGAAATGCAGGCTTGAGTTCT
GAGCACACGTCTGAACTCTTTCCCTACACGACGCTCTTCCGATCTNNNAGCCTAGGGAGAAATCTTTG
GAGCACACGTCTGAACTCTTTCCCTACACGACGCTCTTCCGATCTNNNAGCCTAGGGAGAAATCTTTG
AGCACACGTCTGAACTCTTTCCCTACACGACGCTCTTCCGATCTNNNTTGGCTGCCAGATCTAAAGAG
AGCACACGTCTGAACTCTTTCCCTACACGACGCTCTTCCGATCTNNNTTGGCTGCCAGATCTAAAGAG
AGCACACGTCTGAACTCTTTCCCTACACGACGCTCTTCCGATCTNNNTCATCAAGACATACCCCTCCTCT
AGCACACGTCTGAACTCTTTCCCTACACGACGCTCTTCCGATCTNNNTCATCAAGACATACCCCTCCTCT
GAGCACACGTCTGAACTCTTTCCCTACACGACGCTCTTCCGATCTNNNCTATCTCATCCTCCTGCCTTGA
GAGCACACGTCTGAACTCTTTCCCTACACGACGCTCTTCCGATCTNNNCTATCTCATCCTCCTGCCTTGA
GAGCACACGTCTGAACTCTTTCCCTACACGACGCTCTTCCGATCTNNNCTCTTTTGAACTTAGAGGCGT
GAGCACACGTCTGAACTCTTTCCCTACACGACGCTCTTCCGATCTNNNCTCTTTTGAACTTAGAGGCGT
GCACACGTCTGAACTCTTTCCCTACACGACGCTCTTCCGATCTNNNCAGGACAATCGCTTGAACACAG
GCACACGTCTGAACTCTTTCCCTACACGACGCTCTTCCGATCTNNNCAGGACAATCGCTTGAACACAG
GAGCACACGTCTGAACTCTTTCCCTACACGACGCTCTTCCGATCTNNNAATGGTGTGATCTCAGCTCACT
GAGCACACGTCTGAACTCTTTCCCTACACGACGCTCTTCCGATCTNNNAATGGTGTGATCTCAGCTCACT
GAGCACACGTCTGAACTCTTTCCCTACACGACGCTCTTCCGATCTNNNCGCTTAAGGATGGAAGAGG

GAGCACACGTCTGAACTCTTTCCCTACACGACGCTCTTCCGATCTNNNCGCTTAAGGATGGAAGAGG
GCACACGTCTGAACTCTTTCCCTACACGACGCTCTTCCGATCTNNNTCTGTTCCTGAAATGAACGACC
GCACACGTCTGAACTCTTTCCCTACACGACGCTCTTCCGATCTNNNTCTGTTCCTGAAATGAACGACC
AGCACACGTCTGAACTCTTTCCCTACACGACGCTCTTCCGATCTNNNCTGTGACAGTTTCTTGTGTGG
AGCACACGTCTGAACTCTTTCCCTACACGACGCTCTTCCGATCTNNNCTGTGACAGTTTCTTGTGTGG
CACACGTCTGAACTCTTTCCCTACACGACGCTCTTCCGATCTNNNACTTATGGGCACCATGTCAACTC
CACACGTCTGAACTCTTTCCCTACACGACGCTCTTCCGATCTNNNACTTATGGGCACCATGTCAACTC
AGCACACGTCTGAACTCTTTCCCTACACGACGCTCTTCCGATCTNNNAAATAGACTCGGATGTGGCTG
AGCACACGTCTGAACTCTTTCCCTACACGACGCTCTTCCGATCTNNNAAATAGACTCGGATGTGGCTG
AGCACACGTCTGAACTCTTTCCCTACACGACGCTCTTCCGATCTNNNTGAATCCGAGGGCTGATAGTA
AGCACACGTCTGAACTCTTTCCCTACACGACGCTCTTCCGATCTNNNTGAATCCGAGGGCTGATAGTA
GCACACGTCTGAACTCTTTCCCTACACGACGCTCTTCCGATCTNNNATCCAGAATTTCTTCCCCCAAATC
GCACACGTCTGAACTCTTTCCCTACACGACGCTCTTCCGATCTNNNATCCAGAATTTCTTCCCCCAAATC
GCACACGTCTGAACTCTTTCCCTACACGACGCTCTTCCGATCTNNNTAGATATCTTGCTGATGTGGCC
GCACACGTCTGAACTCTTTCCCTACACGACGCTCTTCCGATCTNNNTAGATATCTTGCTGATGTGGCC
GCACACGTCTGAACTCTTTCCCTACACGACGCTCTTCCGATCTNNNGCTGAGATAGGAGAATCGCTT
GCACACGTCTGAACTCTTTCCCTACACGACGCTCTTCCGATCTNNNGCTGAGATAGGAGAATCGCTT
AGCACACGTCTGAACTCTTTCCCTACACGACGCTCTTCCGATCTNNNGACCCACTCTGCACAGTATCAG
AGCACACGTCTGAACTCTTTCCCTACACGACGCTCTTCCGATCTNNNGACCCACTCTGCACAGTATCAG
GAGCACACGTCTGAACTCTTTCCCTACACGACGCTCTTCCGATCTNNNACAAAAAATTAGCCGTGTGT
GAGCACACGTCTGAACTCTTTCCCTACACGACGCTCTTCCGATCTNNNACAAAAAATTAGCCGTGTGT
GCACACGTCTGAACTCTTTCCCTACACGACGCTCTTCCGATCTNNNCCTGGCCCTCTTTCAGTTACTAA
GCACACGTCTGAACTCTTTCCCTACACGACGCTCTTCCGATCTNNNCCTGGCCCTCTTTCAGTTACTAA
AGCACACGTCTGAACTCTTTCCCTACACGACGCTCTTCCGATCTNNNGGCACCTGTAATCCTAGCTACT
AGCACACGTCTGAACTCTTTCCCTACACGACGCTCTTCCGATCTNNNGGCACCTGTAATCCTAGCTACT
AGCACACGTCTGAACTCTTTCCCTACACGACGCTCTTCCGATCTNNNCACACATCACCACACCCTACTA
AGCACACGTCTGAACTCTTTCCCTACACGACGCTCTTCCGATCTNNNCACACATCACCACACCCTACTA
AGCACACGTCTGAACTCTTTCCCTACACGACGCTCTTCCGATCTNNNGGAGAACTCTGAACATGGAC
AGCACACGTCTGAACTCTTTCCCTACACGACGCTCTTCCGATCTNNNGGAGAACTCTGAACATGGAC
AGCACACGTCTGAACTCTTTCCCTACACGACGCTCTTCCGATCTNNNTCATTCATTCATTCCCCCACAT
AGCACACGTCTGAACTCTTTCCCTACACGACGCTCTTCCGATCTNNNTCATTCATTCATTCCCCCACAT
AGCACACGTCTGAACTCTTTCCCTACACGACGCTCTTCCGATCTNNNAAGAGAATTTCTTGAACCCGG
AGCACACGTCTGAACTCTTTCCCTACACGACGCTCTTCCGATCTNNNAAGAGAATTTCTTGAACCCGG
AGCACACGTCTGAACTCTTTCCCTACACGACGCTCTTCCGATCTNNNGTGCTCTAGTCACACAGAGTC
AGCACACGTCTGAACTCTTTCCCTACACGACGCTCTTCCGATCTNNNGTGCTCTAGTCACACAGAGTC
AGCACACGTCTGAACTCTTTCCCTACACGACGCTCTTCCGATCTNNNGCCCCACTCCCTCTACTCTATAT
AGCACACGTCTGAACTCTTTCCCTACACGACGCTCTTCCGATCTNNNGCCCCACTCCCTCTACTCTATAT
GAGCACACGTCTGAACTCTTTCCCTACACGACGCTCTTCCGATCTNNNTAGTCCCAGCTATACTCGGGA
GAGCACACGTCTGAACTCTTTCCCTACACGACGCTCTTCCGATCTNNNTAGTCCCAGCTATACTCGGGA
AGCACACGTCTGAACTCTTTCCCTACACGACGCTCTTCCGATCTNNNGGATAAACTGGTTGCATAATA
AGCACACGTCTGAACTCTTTCCCTACACGACGCTCTTCCGATCTNNNGGATAAACTGGTTGCATAATA
AGCACACGTCTGAACTCTTTCCCTACACGACGCTCTTCCGATCTNNNGCCAGTTCCTTGTTTTGTCCTA
AGCACACGTCTGAACTCTTTCCCTACACGACGCTCTTCCGATCTNNNGCCAGTTCCTTGTTTTGTCCTA

GAGCACACGTCTGAACTCTTTCCCTACACGACGCTCTTCCGATCTNNNTCACACCTGTAATTCCAGCACT
GAGCACACGTCTGAACTCTTTCCCTACACGACGCTCTTCCGATCTNNNTCACACCTGTAATTCCAGCACT
GCACACGTCTGAACTCTTTCCCTACACGACGCTCTTCCGATCTNNNTAGAATGAGGAAGAACCACTCAT
GCACACGTCTGAACTCTTTCCCTACACGACGCTCTTCCGATCTNNNTAGAATGAGGAAGAACCACTCAT
GAGCACACGTCTGAACTCTTTCCCTACACGACGCTCTTCCGATCTNNNTTGTCCATCTCAGTTGGAGG
GAGCACACGTCTGAACTCTTTCCCTACACGACGCTCTTCCGATCTNNNTTGTCCATCTCAGTTGGAGG
GAGCACACGTCTGAACTCTTTCCCTACACGACGCTCTTCCGATCTNNNGAGGATCACTTTAGCGTAGG
GAGCACACGTCTGAACTCTTTCCCTACACGACGCTCTTCCGATCTNNNGAGGATCACTTTAGCGTAGG
GAGCACACGTCTGAACTCTTTCCCTACACGACGCTCTTCCGATCTNNNCACCTACTCAGGAGGTTGAG
GAGCACACGTCTGAACTCTTTCCCTACACGACGCTCTTCCGATCTNNNCACCTACTCAGGAGGTTGAG
GAGCACACGTCTGAACTCTTTCCCTACACGACGCTCTTCCGATCTNNNGCCCATAAGAGACTCTGTTT
GAGCACACGTCTGAACTCTTTCCCTACACGACGCTCTTCCGATCTNNNGCCCATAAGAGACTCTGTTT
GCACACGTCTGAACTCTTTCCCTACACGACGCTCTTCCGATCTNNNAGTGGCATGATATAGGTGTGCT
GCACACGTCTGAACTCTTTCCCTACACGACGCTCTTCCGATCTNNNAGTGGCATGATATAGGTGTGCT
AGCACACGTCTGAACTCTTTCCCTACACGACGCTCTTCCGATCTNNNACCTATAGTCCCAGCTACTCGG
AGCACACGTCTGAACTCTTTCCCTACACGACGCTCTTCCGATCTNNNACCTATAGTCCCAGCTACTCGG
AGCACACGTCTGAACTCTTTCCCTACACGACGCTCTTCCGATCTNNNAGAGCTTGCAGTTAGTTGAGA
AGCACACGTCTGAACTCTTTCCCTACACGACGCTCTTCCGATCTNNNAGAGCTTGCAGTTAGTTGAGA
GCACACGTCTGAACTCTTTCCCTACACGACGCTCTTCCGATCTNNNTGTGAGACCTACATTGCATCCA
GCACACGTCTGAACTCTTTCCCTACACGACGCTCTTCCGATCTNNNTGTGAGACCTACATTGCATCCA
AGCACACGTCTGAACTCTTTCCCTACACGACGCTCTTCCGATCTNNNAAAACAGGATGGTGCTGGTG
AGCACACGTCTGAACTCTTTCCCTACACGACGCTCTTCCGATCTNNNAAAACAGGATGGTGCTGGTG
AGCACACGTCTGAACTCTTTCCCTACACGACGCTCTTCCGATCTNNNCTGGGCAATAAGAGCGAAACT
AGCACACGTCTGAACTCTTTCCCTACACGACGCTCTTCCGATCTNNNCTGGGCAATAAGAGCGAAACT
AGCACACGTCTGAACTCTTTCCCTACACGACGCTCTTCCGATCTNNNTAAAAGCGGGAAGTTCTTTTG
AGCACACGTCTGAACTCTTTCCCTACACGACGCTCTTCCGATCTNNNTAAAAGCGGGAAGTTCTTTTG
AGCACACGTCTGAACTCTTTCCCTACACGACGCTCTTCCGATCTNNNACAGACACAGAAGACAATAGG
AGCACACGTCTGAACTCTTTCCCTACACGACGCTCTTCCGATCTNNNACAGACACAGAAGACAATAGG
GAGCACACGTCTGAACTCTTTCCCTACACGACGCTCTTCCGATCTNNNAACTGGTTGGTTGTCTCTGAG
GAGCACACGTCTGAACTCTTTCCCTACACGACGCTCTTCCGATCTNNNAACTGGTTGGTTGTCTCTGAG
GCACACGTCTGAACTCTTTCCCTACACGACGCTCTTCCGATCTNNNCTGAGGTGGGAAGATCATCTG
GCACACGTCTGAACTCTTTCCCTACACGACGCTCTTCCGATCTNNNCTGAGGTGGGAAGATCATCTG
AGCACACGTCTGAACTCTTTCCCTACACGACGCTCTTCCGATCTNNNACTCTATTCTGTTGCACTGGTCT
AGCACACGTCTGAACTCTTTCCCTACACGACGCTCTTCCGATCTNNNACTCTATTCTGTTGCACTGGTCT
AGCACACGTCTGAACTCTTTCCCTACACGACGCTCTTCCGATCTNNNCCTCCCCCTCTCTCTTTCTTTTTC
AGCACACGTCTGAACTCTTTCCCTACACGACGCTCTTCCGATCTNNNCCTCCCCCTCTCTCTTTCTTTTTC
AGCACACGTCTGAACTCTTTCCCTACACGACGCTCTTCCGATCTNNNACAGGGCTGGAATAACTAAAG
AGCACACGTCTGAACTCTTTCCCTACACGACGCTCTTCCGATCTNNNACAGGGCTGGAATAACTAAAG
GAGCACACGTCTGAACTCTTTCCCTACACGACGCTCTTCCGATCTNNNAGGGTGGGTAGAGAAAAAC
GAGCACACGTCTGAACTCTTTCCCTACACGACGCTCTTCCGATCTNNNAGGGTGGGTAGAGAAAAAC
GCACACGTCTGAACTCTTTCCCTACACGACGCTCTTCCGATCTNNNTTCTTGAAGTTGGGCTCCTAGT
GCACACGTCTGAACTCTTTCCCTACACGACGCTCTTCCGATCTNNNTTCTTGAAGTTGGGCTCCTAGT
AGCACACGTCTGAACTCTTTCCCTACACGACGCTCTTCCGATCTNNNCAGTTTTTATGATGGTGCAGG

AGCACACGTCTGAACTCTTTCCCTACACGACGCTCTTCCGATCTNNNCAGTTTTTATGATGGTGCAGG
AGCACACGTCTGAACTCTTTCCCTACACGACGCTCTTCCGATCTNNNGTGGGAGAAGGCCATTTCTTTT
AGCACACGTCTGAACTCTTTCCCTACACGACGCTCTTCCGATCTNNNGTGGGAGAAGGCCATTTCTTTT
GCACACGTCTGAACTCTTTCCCTACACGACGCTCTTCCGATCTNNNTACTGCTGTTCCCTTTGCCTTATC
GCACACGTCTGAACTCTTTCCCTACACGACGCTCTTCCGATCTNNNTACTGCTGTTCCCTTTGCCTTATC
CACACGTCTGAACTCTTTCCCTACACGACGCTCTTCCGATCTNNNGCCGATGTTTTGTCTTTCTTTCCC
CACACGTCTGAACTCTTTCCCTACACGACGCTCTTCCGATCTNNNGCCGATGTTTTGTCTTTCTTTCCC
AGCACACGTCTGAACTCTTTCCCTACACGACGCTCTTCCGATCTNNNAACTGTTGATACTCATTCAATT
AGCACACGTCTGAACTCTTTCCCTACACGACGCTCTTCCGATCTNNNAACTGTTGATACTCATTCAATT
GCACACGTCTGAACTCTTTCCCTACACGACGCTCTTCCGATCTNNNATCACTCCCTGGGCTTTTTTTCTC
GCACACGTCTGAACTCTTTCCCTACACGACGCTCTTCCGATCTNNNATCACTCCCTGGGCTTTTTTTCTC
AGCACACGTCTGAACTCTTTCCCTACACGACGCTCTTCCGATCTNNNGTTGACTACATTCTCCTGCCTC
AGCACACGTCTGAACTCTTTCCCTACACGACGCTCTTCCGATCTNNNGTTGACTACATTCTCCTGCCTC
AGCACACGTCTGAACTCTTTCCCTACACGACGCTCTTCCGATCTNNNCCTTGGCTCCATACTCTAAACCA
AGCACACGTCTGAACTCTTTCCCTACACGACGCTCTTCCGATCTNNNCCTTGGCTCCATACTCTAAACCA
AGCACACGTCTGAACTCTTTCCCTACACGACGCTCTTCCGATCTNNNAAAGACAGGCAGGGGAAAA
AGCACACGTCTGAACTCTTTCCCTACACGACGCTCTTCCGATCTNNNAAAGACAGGCAGGGGAAAA
AGCACACGTCTGAACTCTTTCCCTACACGACGCTCTTCCGATCTNNNTCAGCAATTGGTAGAAACAGT
AGCACACGTCTGAACTCTTTCCCTACACGACGCTCTTCCGATCTNNNTCAGCAATTGGTAGAAACAGT
AGCACACGTCTGAACTCTTTCCCTACACGACGCTCTTCCGATCTNNNTGTAATCCTAGCTACTCGGGA
AGCACACGTCTGAACTCTTTCCCTACACGACGCTCTTCCGATCTNNNTGTAATCCTAGCTACTCGGGA
AGAGCACACGTCTGAACTCTTTCCCTACACGACGCTCTTCCGATCTNNNGCCCTAGAGGAAGTTGGAA
AGAGCACACGTCTGAACTCTTTCCCTACACGACGCTCTTCCGATCTNNNGCCCTAGAGGAAGTTGGAA
AGCACACGTCTGAACTCTTTCCCTACACGACGCTCTTCCGATCTNNNCAGCCTTCCCTCTCACTACTTT
AGCACACGTCTGAACTCTTTCCCTACACGACGCTCTTCCGATCTNNNCAGCCTTCCCTCTCACTACTTT
CACACGTCTGAACTCTTTCCCTACACGACGCTCTTCCGATCTNNNCCTCTCAGAGCCAAAAGTGGAAA
CACACGTCTGAACTCTTTCCCTACACGACGCTCTTCCGATCTNNNCCTCTCAGAGCCAAAAGTGGAAA
GAGCACACGTCTGAACTCTTTCCCTACACGACGCTCTTCCGATCTNNNCCCTGTGGGTTCTTGAAGAG
GAGCACACGTCTGAACTCTTTCCCTACACGACGCTCTTCCGATCTNNNCCCTGTGGGTTCTTGAAGAG
GAGCACACGTCTGAACTCTTTCCCTACACGACGCTCTTCCGATCTNNNGCTAGAGTACAGTGGCATGA
GAGCACACGTCTGAACTCTTTCCCTACACGACGCTCTTCCGATCTNNNGCTAGAGTACAGTGGCATGA
AGCACACGTCTGAACTCTTTCCCTACACGACGCTCTTCCGATCTNNNGCTCTCATTTACATGGTTCCTG
AGCACACGTCTGAACTCTTTCCCTACACGACGCTCTTCCGATCTNNNGCTCTCATTTACATGGTTCCTG
AGCACACGTCTGAACTCTTTCCCTACACGACGCTCTTCCGATCTNNNGAGGTTGCAGTGAGCTAAGA
AGCACACGTCTGAACTCTTTCCCTACACGACGCTCTTCCGATCTNNNGAGGTTGCAGTGAGCTAAGA
AGCACACGTCTGAACTCTTTCCCTACACGACGCTCTTCCGATCTNNNTCAGTAGGAATGGTACCAGC
AGCACACGTCTGAACTCTTTCCCTACACGACGCTCTTCCGATCTNNNTCAGTAGGAATGGTACCAGC
AGCACACGTCTGAACTCTTTCCCTACACGACGCTCTTCCGATCTNNNGTGGTCTTTCTGGTGATGGAT
AGCACACGTCTGAACTCTTTCCCTACACGACGCTCTTCCGATCTNNNGTGGTCTTTCTGGTGATGGAT
GAGCACACGTCTGAACTCTTTCCCTACACGACGCTCTTCCGATCTNNNAAACCTTGGAATTTGAGACC
GAGCACACGTCTGAACTCTTTCCCTACACGACGCTCTTCCGATCTNNNAAACCTTGGAATTTGAGACC
AGCACACGTCTGAACTCTTTCCCTACACGACGCTCTTCCGATCTNNNAGGCAAAAGAATCGCTTGAAC
AGCACACGTCTGAACTCTTTCCCTACACGACGCTCTTCCGATCTNNNAGGCAAAAGAATCGCTTGAAC

CACACGTCTGAACTCTTTCCCTACACGACGCTCTTCCGATCTNNNGGGGAAGTGTGGGCATAAAATT
CACACGTCTGAACTCTTTCCCTACACGACGCTCTTCCGATCTNNNGGGGAAGTGTGGGCATAAAATT
GAGCACACGTCTGAACTCTTTCCCTACACGACGCTCTTCCGATCTNNNCATCTCCTTTCTTCATCACCCA
GAGCACACGTCTGAACTCTTTCCCTACACGACGCTCTTCCGATCTNNNCATCTCCTTTCTTCATCACCCA
GAGCACACGTCTGAACTCTTTCCCTACACGACGCTCTTCCGATCTNNNTGGTTGAATAAGACAAGCC
GAGCACACGTCTGAACTCTTTCCCTACACGACGCTCTTCCGATCTNNNTGGTTGAATAAGACAAGCC
AGCACACGTCTGAACTCTTTCCCTACACGACGCTCTTCCGATCTNNNGCTTATTTGCTGTCTGCTTG
AGCACACGTCTGAACTCTTTCCCTACACGACGCTCTTCCGATCTNNNGCTTATTTGCTGTCTGCTTG
AGCACACGTCTGAACTCTTTCCCTACACGACGCTCTTCCGATCTNNNTGATAGTTAAATCTGGGCCAG
AGCACACGTCTGAACTCTTTCCCTACACGACGCTCTTCCGATCTNNNTGATAGTTAAATCTGGGCCAG
GCACACGTCTGAACTCTTTCCCTACACGACGCTCTTCCGATCTNNNTCTCCAAGGTACATCTCAACTG
GCACACGTCTGAACTCTTTCCCTACACGACGCTCTTCCGATCTNNNTCTCCAAGGTACATCTCAACTG
AGCACACGTCTGAACTCTTTCCCTACACGACGCTCTTCCGATCTNNNCACCTGTAATCCCAGCTACTTG
AGCACACGTCTGAACTCTTTCCCTACACGACGCTCTTCCGATCTNNNCACCTGTAATCCCAGCTACTTG
GAGCACACGTCTGAACTCTTTCCCTACACGACGCTCTTCCGATCTNNNGAGAGCAGGAAAGGAGAAG
GAGCACACGTCTGAACTCTTTCCCTACACGACGCTCTTCCGATCTNNNGAGAGCAGGAAAGGAGAAG
GAGCACACGTCTGAACTCTTTCCCTACACGACGCTCTTCCGATCTNNNCTGAGGCAGGAGAATCACTT
GAGCACACGTCTGAACTCTTTCCCTACACGACGCTCTTCCGATCTNNNCTGAGGCAGGAGAATCACTT
AGCACACGTCTGAACTCTTTCCCTACACGACGCTCTTCCGATCTNNNGAATGATGACAATACCACTG
AGCACACGTCTGAACTCTTTCCCTACACGACGCTCTTCCGATCTNNNGAATGATGACAATACCACTG
CACACGTCTGAACTCTTTCCCTACACGACGCTCTTCCGATCTNNNCAGGGACCTTACCTTATACACATC
CACACGTCTGAACTCTTTCCCTACACGACGCTCTTCCGATCTNNNCAGGGACCTTACCTTATACACATC
GAGCACACGTCTGAACTCTTTCCCTACACGACGCTCTTCCGATCTNNNTGCCTATAATCGCAGCTACTC
GAGCACACGTCTGAACTCTTTCCCTACACGACGCTCTTCCGATCTNNNTGCCTATAATCGCAGCTACTC
AGCACACGTCTGAACTCTTTCCCTACACGACGCTCTTCCGATCTNNNTCTAACCTCATGCAAAATAAAA
AGCACACGTCTGAACTCTTTCCCTACACGACGCTCTTCCGATCTNNNTCTAACCTCATGCAAAATAAAA
AGCACACGTCTGAACTCTTTCCCTACACGACGCTCTTCCGATCTNNNAAAGAGGGAATGAATGCCAG
AGCACACGTCTGAACTCTTTCCCTACACGACGCTCTTCCGATCTNNNAAAGAGGGAATGAATGCCAG
AGCACACGTCTGAACTCTTTCCCTACACGACGCTCTTCCGATCTNNNTGCCTGTAGTCCCAGCTACTCC
AGCACACGTCTGAACTCTTTCCCTACACGACGCTCTTCCGATCTNNNTGCCTGTAGTCCCAGCTACTCC
AAGAGCACACGTCTGAACTCTTTCCCTACACGACGCTCTTCCGATCTNNNGAGGCTGAGGCAGGAG
AAGAGCACACGTCTGAACTCTTTCCCTACACGACGCTCTTCCGATCTNNNGAGGCTGAGGCAGGAG
AGCACACGTCTGAACTCTTTCCCTACACGACGCTCTTCCGATCTNNNAGCACTCCATTCCTTTCCTACA
AGCACACGTCTGAACTCTTTCCCTACACGACGCTCTTCCGATCTNNNAGCACTCCATTCCTTTCCTACA
AGCACACGTCTGAACTCTTTCCCTACACGACGCTCTTCCGATCTNNNATTTCTGGTAGGCGAGAGAT
AGCACACGTCTGAACTCTTTCCCTACACGACGCTCTTCCGATCTNNNATTTCTGGTAGGCGAGAGAT
AGCACACGTCTGAACTCTTTCCCTACACGACGCTCTTCCGATCTNNNTATAAGTAAAAGTGTGGGCAT
AGCACACGTCTGAACTCTTTCCCTACACGACGCTCTTCCGATCTNNNTATAAGTAAAAGTGTGGGCAT
CACACGTCTGAACTCTTTCCCTACACGACGCTCTTCCGATCTNNNTAGTCCCAGCTACTTGAGGATGA
CACACGTCTGAACTCTTTCCCTACACGACGCTCTTCCGATCTNNNTAGTCCCAGCTACTTGAGGATGA
AGCACACGTCTGAACTCTTTCCCTACACGACGCTCTTCCGATCTNNNTGTTTCTCAGATTCCCAGACTC
AGCACACGTCTGAACTCTTTCCCTACACGACGCTCTTCCGATCTNNNTGTTTCTCAGATTCCCAGACTC
AGCACACGTCTGAACTCTTTCCCTACACGACGCTCTTCCGATCTNNNTGGCATACCCCTCTGAAATGA

AGCACACGTCTGAACTCTTTCCCTACACGACGCTCTTCCGATCTNNNTGGCATACCCCTCTGAAATGAA
GAGCACACGTCTGAACTCTTTCCCTACACGACGCTCTTCCGATCTNNNGAGAATGGTGTGAATCCAGG
GAGCACACGTCTGAACTCTTTCCCTACACGACGCTCTTCCGATCTNNNGAGAATGGTGTGAATCCAGG
GAGCACACGTCTGAACTCTTTCCCTACACGACGCTCTTCCGATCTNNNCATTCTCACAGGCTCTTTTTGC
GAGCACACGTCTGAACTCTTTCCCTACACGACGCTCTTCCGATCTNNNCATTCTCACAGGCTCTTTTTGC
GAGCACACGTCTGAACTCTTTCCCTACACGACGCTCTTCCGATCTNNNGCCTGTAGTCTCATCTACTTC
GAGCACACGTCTGAACTCTTTCCCTACACGACGCTCTTCCGATCTNNNGCCTGTAGTCTCATCTACTTC
AGCACACGTCTGAACTCTTTCCCTACACGACGCTCTTCCGATCTNNNAGACCAAAACCACATGCAAAA
AGCACACGTCTGAACTCTTTCCCTACACGACGCTCTTCCGATCTNNNAGACCAAAACCACATGCAAAA
AGCACACGTCTGAACTCTTTCCCTACACGACGCTCTTCCGATCTNNNTCTGAACATCCAAGGGAATGAC
AGCACACGTCTGAACTCTTTCCCTACACGACGCTCTTCCGATCTNNNTCTGAACATCCAAGGGAATGAC
AGCACACGTCTGAACTCTTTCCCTACACGACGCTCTTCCGATCTNNNTGACAAGGCCATGATACCATCA
AGCACACGTCTGAACTCTTTCCCTACACGACGCTCTTCCGATCTNNNTGACAAGGCCATGATACCATCA
AGCACACGTCTGAACTCTTTCCCTACACGACGCTCTTCCGATCTNNNCGACTTTTCCTGCTGAAATGAC
AGCACACGTCTGAACTCTTTCCCTACACGACGCTCTTCCGATCTNNNCGACTTTTCCTGCTGAAATGAC
AGCACACGTCTGAACTCTTTCCCTACACGACGCTCTTCCGATCTNNNCAAAAATTAGCCGGGTATGGT
AGCACACGTCTGAACTCTTTCCCTACACGACGCTCTTCCGATCTNNNCAAAAATTAGCCGGGTATGGT
AGCACACGTCTGAACTCTTTCCCTACACGACGCTCTTCCGATCTNNNGAAGGTTTGGGGTTCAAGAC
AGCACACGTCTGAACTCTTTCCCTACACGACGCTCTTCCGATCTNNNGAAGGTTTGGGGTTCAAGAC
AGCACACGTCTGAACTCTTTCCCTACACGACGCTCTTCCGATCTNNNGGTGCCTGTAATCCTACCTACT
AGCACACGTCTGAACTCTTTCCCTACACGACGCTCTTCCGATCTNNNGGTGCCTGTAATCCTACCTACT
GAGCACACGTCTGAACTCTTTCCCTACACGACGCTCTTCCGATCTNNNGTAATCCTAGCTACTTGGGA
GAGCACACGTCTGAACTCTTTCCCTACACGACGCTCTTCCGATCTNNNGTAATCCTAGCTACTTGGGA
ACACGTCTGAACTCTTTCCCTACACGACGCTCTTCCGATCTNNNATAACCTTCCTAGTACCTGAGCATC
ACACGTCTGAACTCTTTCCCTACACGACGCTCTTCCGATCTNNNATAACCTTCCTAGTACCTGAGCATC
AGCACACGTCTGAACTCTTTCCCTACACGACGCTCTTCCGATCTNNNCCTGTTGGCTTTGTTTACACTG
AGCACACGTCTGAACTCTTTCCCTACACGACGCTCTTCCGATCTNNNCCTGTTGGCTTTGTTTACACTG
AGCACACGTCTGAACTCTTTCCCTACACGACGCTCTTCCGATCTNNNTCTGGATGAGTCTTTGGTTCT
AGCACACGTCTGAACTCTTTCCCTACACGACGCTCTTCCGATCTNNNTCTGGATGAGTCTTTGGTTCT
CACACGTCTGAACTCTTTCCCTACACGACGCTCTTCCGATCTNNNACAGGCCAAAGAATGTCTGTACT
CACACGTCTGAACTCTTTCCCTACACGACGCTCTTCCGATCTNNNACAGGCCAAAGAATGTCTGTACT
AGCACACGTCTGAACTCTTTCCCTACACGACGCTCTTCCGATCTNNNGGCCTGAATGATGGTGGTGA
AGCACACGTCTGAACTCTTTCCCTACACGACGCTCTTCCGATCTNNNGGCCTGAATGATGGTGGTGA
AGCACACGTCTGAACTCTTTCCCTACACGACGCTCTTCCGATCTNNNTGTCCATGTGTTTACAATGCTG
AGCACACGTCTGAACTCTTTCCCTACACGACGCTCTTCCGATCTNNNTGTCCATGTGTTTACAATGCTG
GAGCACACGTCTGAACTCTTTCCCTACACGACGCTCTTCCGATCTNNNACCTCCACTAAAAGGAAGAC
GAGCACACGTCTGAACTCTTTCCCTACACGACGCTCTTCCGATCTNNNACCTCCACTAAAAGGAAGAC
AGCACACGTCTGAACTCTTTCCCTACACGACGCTCTTCCGATCTNNNGCTGAGATAGGAGAATCGCTT
AGCACACGTCTGAACTCTTTCCCTACACGACGCTCTTCCGATCTNNNGCTGAGATAGGAGAATCGCTT
GAGCACACGTCTGAACTCTTTCCCTACACGACGCTCTTCCGATCTNNNTTGGAATGGGAATGCCTACT
GAGCACACGTCTGAACTCTTTCCCTACACGACGCTCTTCCGATCTNNNTTGGAATGGGAATGCCTACT
CACACGTCTGAACTCTTTCCCTACACGACGCTCTTCCGATCTNNNTTCTCTTTGCTGATCATTGTTCAT
CACACGTCTGAACTCTTTCCCTACACGACGCTCTTCCGATCTNNNTTCTCTTTGCTGATCATTGTTCAT

AGCACACGTCTGAACTCTTTCCCTACACGACGCTCTTCCGATCTNNNATCATGTGCTCTAGGTTGTGT
AGCACACGTCTGAACTCTTTCCCTACACGACGCTCTTCCGATCTNNNATCATGTGCTCTAGGTTGTGT
GAGCACACGTCTGAACTCTTTCCCTACACGACGCTCTTCCGATCTNNNAGCTGAGTCCTTTCTCCACTAG
GAGCACACGTCTGAACTCTTTCCCTACACGACGCTCTTCCGATCTNNNAGCTGAGTCCTTTCTCCACTAG
GAGCACACGTCTGAACTCTTTCCCTACACGACGCTCTTCCGATCTNNNTGGACACAGGATGGCAACAA
GAGCACACGTCTGAACTCTTTCCCTACACGACGCTCTTCCGATCTNNNTGGACACAGGATGGCAACAA
CACACGTCTGAACTCTTTCCCTACACGACGCTCTTCCGATCTNNNAGAGGAATTCTTAGACAGATAAC
CACACGTCTGAACTCTTTCCCTACACGACGCTCTTCCGATCTNNNAGAGGAATTCTTAGACAGATAAC
GAGCACACGTCTGAACTCTTTCCCTACACGACGCTCTTCCGATCTNNNTGCCTGTAATCCCAGTGAAAG
GAGCACACGTCTGAACTCTTTCCCTACACGACGCTCTTCCGATCTNNNTGCCTGTAATCCCAGTGAAAG
GAGCACACGTCTGAACTCTTTCCCTACACGACGCTCTTCCGATCTNNNGGAAATGCTGAGAATTATGG
GAGCACACGTCTGAACTCTTTCCCTACACGACGCTCTTCCGATCTNNNGGAAATGCTGAGAATTATGG
GCACACGTCTGAACTCTTTCCCTACACGACGCTCTTCCGATCTNNNAGCACAGTTGAGGAAGAAGTA
GCACACGTCTGAACTCTTTCCCTACACGACGCTCTTCCGATCTNNNAGCACAGTTGAGGAAGAAGTA
GAGCACACGTCTGAACTCTTTCCCTACACGACGCTCTTCCGATCTNNNCTTGTGAAATTACCTGGCCCT
GAGCACACGTCTGAACTCTTTCCCTACACGACGCTCTTCCGATCTNNNCTTGTGAAATTACCTGGCCCT
AGCACACGTCTGAACTCTTTCCCTACACGACGCTCTTCCGATCTNNNCTGAGGCAGGAGAATCACTTG
AGCACACGTCTGAACTCTTTCCCTACACGACGCTCTTCCGATCTNNNCTGAGGCAGGAGAATCACTTG
GCACACGTCTGAACTCTTTCCCTACACGACGCTCTTCCGATCTNNNACTTGAATAAGGGGCTTGGCAT
GCACACGTCTGAACTCTTTCCCTACACGACGCTCTTCCGATCTNNNACTTGAATAAGGGGCTTGGCAT
AGCACACGTCTGAACTCTTTCCCTACACGACGCTCTTCCGATCTNNNGCTGGGGCTACTGTATGTACA
AGCACACGTCTGAACTCTTTCCCTACACGACGCTCTTCCGATCTNNNGCTGGGGCTACTGTATGTACA
GCACACGTCTGAACTCTTTCCCTACACGACGCTCTTCCGATCTNNNGGTGAACCTTAGTACCTAAAAG
GCACACGTCTGAACTCTTTCCCTACACGACGCTCTTCCGATCTNNNGGTGAACCTTAGTACCTAAAAG
AGCACACGTCTGAACTCTTTCCCTACACGACGCTCTTCCGATCTNNNAGGGGTAGTCCAGTGCTAAAT
AGCACACGTCTGAACTCTTTCCCTACACGACGCTCTTCCGATCTNNNAGGGGTAGTCCAGTGCTAAAT
AGCACACGTCTGAACTCTTTCCCTACACGACGCTCTTCCGATCTNNNCACACACACAAACACACACACT
AGCACACGTCTGAACTCTTTCCCTACACGACGCTCTTCCGATCTNNNCACACACACAAACACACACACT
GAGCACACGTCTGAACTCTTTCCCTACACGACGCTCTTCCGATCTNNNCGATGAGGTGTCCGTTTGGA
GAGCACACGTCTGAACTCTTTCCCTACACGACGCTCTTCCGATCTNNNCGATGAGGTGTCCGTTTGGA
AGCACACGTCTGAACTCTTTCCCTACACGACGCTCTTCCGATCTNNNTCCATCAGGAAAATACCACATT
AGCACACGTCTGAACTCTTTCCCTACACGACGCTCTTCCGATCTNNNTCCATCAGGAAAATACCACATT
AGCACACGTCTGAACTCTTTCCCTACACGACGCTCTTCCGATCTNNNTTACATGGATTATCCCTGAGGC
AGCACACGTCTGAACTCTTTCCCTACACGACGCTCTTCCGATCTNNNTTACATGGATTATCCCTGAGGC
GAGCACACGTCTGAACTCTTTCCCTACACGACGCTCTTCCGATCTNNNCAACTGTAGTCCCAGCTACTT
GAGCACACGTCTGAACTCTTTCCCTACACGACGCTCTTCCGATCTNNNCAACTGTAGTCCCAGCTACTT
AGCACACGTCTGAACTCTTTCCCTACACGACGCTCTTCCGATCTNNNAGTGCTGGGATTACAGGTGTA
AGCACACGTCTGAACTCTTTCCCTACACGACGCTCTTCCGATCTNNNAGTGCTGGGATTACAGGTGTA
GCACACGTCTGAACTCTTTCCCTACACGACGCTCTTCCGATCTNNNAACTCGTAAGTCGTTTCTCAGG
GCACACGTCTGAACTCTTTCCCTACACGACGCTCTTCCGATCTNNNAACTCGTAAGTCGTTTCTCAGG
AGCACACGTCTGAACTCTTTCCCTACACGACGCTCTTCCGATCTNNNGCCATAGCTCCTTATGCACTGA
AGCACACGTCTGAACTCTTTCCCTACACGACGCTCTTCCGATCTNNNGCCATAGCTCCTTATGCACTGA
CACGTCTGAACTCTTTCCCTACACGACGCTCTTCCGATCTNNNTCTTTACCCTCTAAGCTGAACATCGGA

ACGTCTGAACTCTTTCCCTACACGACGCTCTTCCGATCTNNNTCTTTACCCTCTAAGCTGAACATCGGA
AGCACACGTCTGAACTCTTTCCCTACACGACGCTCTTCCGATCTNNNTCCCCTGACCTATCAACTCAGA
AGCACACGTCTGAACTCTTTCCCTACACGACGCTCTTCCGATCTNNNTCCCCTGACCTATCAACTCAGA
AGCACACGTCTGAACTCTTTCCCTACACGACGCTCTTCCGATCTNNNATTCATTTTCCCCACAACCCTCC
AGCACACGTCTGAACTCTTTCCCTACACGACGCTCTTCCGATCTNNNATTCATTTTCCCCACAACCCTCC
GAGCACACGTCTGAACTCTTTCCCTACACGACGCTCTTCCGATCTNNNTATCTTGGAAGGTTGAGACG
GAGCACACGTCTGAACTCTTTCCCTACACGACGCTCTTCCGATCTNNNTATCTTGGAAGGTTGAGACG
CACGTCTGAACTCTTTCCCTACACGACGCTCTTCCGATCTNNNACAGGTACAATTTAGCGCGTTTCCAT
CACGTCTGAACTCTTTCCCTACACGACGCTCTTCCGATCTNNNACAGGTACAATTTAGCGCGTTTCCAT
AGCACACGTCTGAACTCTTTCCCTACACGACGCTCTTCCGATCTNNNCAGCAGTGGATTTGGTTTTCTC
AGCACACGTCTGAACTCTTTCCCTACACGACGCTCTTCCGATCTNNNCAGCAGTGGATTTGGTTTTCTC
AGCACACGTCTGAACTCTTTCCCTACACGACGCTCTTCCGATCTNNNAAAAAAAATTAGCCGGGCGTG
AGCACACGTCTGAACTCTTTCCCTACACGACGCTCTTCCGATCTNNNAAAAAAAATTAGCCGGGCGTG
AGCACACGTCTGAACTCTTTCCCTACACGACGCTCTTCCGATCTNNNAGCTAGGACTAAAAGCCCTGT
AGCACACGTCTGAACTCTTTCCCTACACGACGCTCTTCCGATCTNNNAGCTAGGACTAAAAGCCCTGT
AGCACACGTCTGAACTCTTTCCCTACACGACGCTCTTCCGATCTNNNGTGTATCCTTTTATTCTGGGC
AGCACACGTCTGAACTCTTTCCCTACACGACGCTCTTCCGATCTNNNGTGTATCCTTTTATTCTGGGC
AGCACACGTCTGAACTCTTTCCCTACACGACGCTCTTCCGATCTNNNCTGAGGCAGGAGAATCGTTTG
AGCACACGTCTGAACTCTTTCCCTACACGACGCTCTTCCGATCTNNNCTGAGGCAGGAGAATCGTTTG
GCACACGTCTGAACTCTTTCCCTACACGACGCTCTTCCGATCTNNNTTCTGTGATTTGAGAGTAAGAG
GCACACGTCTGAACTCTTTCCCTACACGACGCTCTTCCGATCTNNNTTCTGTGATTTGAGAGTAAGAG
GAGCACACGTCTGAACTCTTTCCCTACACGACGCTCTTCCGATCTNNNGAAACCAGGTCAGTTGGGA
GAGCACACGTCTGAACTCTTTCCCTACACGACGCTCTTCCGATCTNNNGAAACCAGGTCAGTTGGGA
AGCACACGTCTGAACTCTTTCCCTACACGACGCTCTTCCGATCTNNNTTGGGAAGGGGGTGTGTATAT
AGCACACGTCTGAACTCTTTCCCTACACGACGCTCTTCCGATCTNNNTTGGGAAGGGGGTGTGTATAT
AGCACACGTCTGAACTCTTTCCCTACACGACGCTCTTCCGATCTNNNTTTACCATGCACATTTGTGGC
AGCACACGTCTGAACTCTTTCCCTACACGACGCTCTTCCGATCTNNNTTTACCATGCACATTTGTGGC
GAGCACACGTCTGAACTCTTTCCCTACACGACGCTCTTCCGATCTNNNCCACTCTTCCCTGTCTCCAAAT
GAGCACACGTCTGAACTCTTTCCCTACACGACGCTCTTCCGATCTNNNCCACTCTTCCCTGTCTCCAAAT
AGCACACGTCTGAACTCTTTCCCTACACGACGCTCTTCCGATCTNNNCTGACTCTTGAACCAAACACGA
AGCACACGTCTGAACTCTTTCCCTACACGACGCTCTTCCGATCTNNNCTGACTCTTGAACCAAACACGA
AGCACACGTCTGAACTCTTTCCCTACACGACGCTCTTCCGATCTNNNCAAGCGGAGAACAGAAAGAA
AGCACACGTCTGAACTCTTTCCCTACACGACGCTCTTCCGATCTNNNCAAGCGGAGAACAGAAAGAA
AGCACACGTCTGAACTCTTTCCCTACACGACGCTCTTCCGATCTNNNTCGGTTTCGACAGTCTTCTGA
AGCACACGTCTGAACTCTTTCCCTACACGACGCTCTTCCGATCTNNNTCGGTTTCGACAGTCTTCTGA
AGCACACGTCTGAACTCTTTCCCTACACGACGCTCTTCCGATCTNNNGGTTAGTCTCAAACTCCTGGGT
AGCACACGTCTGAACTCTTTCCCTACACGACGCTCTTCCGATCTNNNGGTTAGTCTCAAACTCCTGGGT
AGCACACGTCTGAACTCTTTCCCTACACGACGCTCTTCCGATCTNNNCTCACCGATGGGTCCTGAAAA
AGCACACGTCTGAACTCTTTCCCTACACGACGCTCTTCCGATCTNNNCTCACCGATGGGTCCTGAAAA
GAGCACACGTCTGAACTCTTTCCCTACACGACGCTCTTCCGATCTNNNCTGTAGTCCCAGCTACTCAG
GAGCACACGTCTGAACTCTTTCCCTACACGACGCTCTTCCGATCTNNNCTGTAGTCCCAGCTACTCAG
AGCACACGTCTGAACTCTTTCCCTACACGACGCTCTTCCGATCTNNNAAGCTAGCTCTGTTGTACACCA
AGCACACGTCTGAACTCTTTCCCTACACGACGCTCTTCCGATCTNNNAAGCTAGCTCTGTTGTACACCA

GCACACGTCTGAACTCTTTCCCTACACGACGCTCTTCCGATCTNNNCACCGGGAAGTAGAACATTGGT

GCACACGTCTGAACTCTTTCCCTACACGACGCTCTTCCGATCTNNNCACCGGGAAGTAGAACATTGGT

AGCACACGTCTGAACTCTTTCCCTACACGACGCTCTTCCGATCTNNNGTTGCAGTAAGCCAAGGTTGT

AGCACACGTCTGAACTCTTTCCCTACACGACGCTCTTCCGATCTNNNGTTGCAGTAAGCCAAGGTTGT

AGCACACGTCTGAACTCTTTCCCTACACGACGCTCTTCCGATCTNNNCTCCTTTGACTTGTTGGTTACG

AGCACACGTCTGAACTCTTTCCCTACACGACGCTCTTCCGATCTNNNCTCCTTTGACTTGTTGGTTACG

AGCACACGTCTGAACTCTTTCCCTACACGACGCTCTTCCGATCTNNNTGAATTAGAGTTGGAAGGAAC

AGCACACGTCTGAACTCTTTCCCTACACGACGCTCTTCCGATCTNNNTGAATTAGAGTTGGAAGGAAC

AGCACACGTCTGAACTCTTTCCCTACACGACGCTCTTCCGATCTNNNACTTTGTGTGACAAGAGGAA

AGCACACGTCTGAACTCTTTCCCTACACGACGCTCTTCCGATCTNNNACTTTGTGTGACAAGAGGAA

AGCACACGTCTGAACTCTTTCCCTACACGACGCTCTTCCGATCTNNNATGAGAACAGCATGGGAAAGA

AGCACACGTCTGAACTCTTTCCCTACACGACGCTCTTCCGATCTNNNATGAGAACAGCATGGGAAAGA

AGCACACGTCTGAACTCTTTCCCTACACGACGCTCTTCCGATCTNNNTGCCTGGTATGAACTCAGTAAG

AGCACACGTCTGAACTCTTTCCCTACACGACGCTCTTCCGATCTNNNTGCCTGGTATGAACTCAGTAAG

AGCACACGTCTGAACTCTTTCCCTACACGACGCTCTTCCGATCTNNNTCTCCAGTGGTGGCATTAAT

AGCACACGTCTGAACTCTTTCCCTACACGACGCTCTTCCGATCTNNNTCTCCAGTGGTGGCATTAAT

GAGCACACGTCTGAACTCTTTCCCTACACGACGCTCTTCCGATCTNNNGCCTGTAGTCCCAGCTACTTGC

GAGCACACGTCTGAACTCTTTCCCTACACGACGCTCTTCCGATCTNNNGCCTGTAGTCCCAGCTACTTGC

AGCACACGTCTGAACTCTTTCCCTACACGACGCTCTTCCGATCTNNNCGAATGATGAGAGATTTCCCAC

AGCACACGTCTGAACTCTTTCCCTACACGACGCTCTTCCGATCTNNNCGAATGATGAGAGATTTCCCAC

AGCACACGTCTGAACTCTTTCCCTACACGACGCTCTTCCGATCTNNNCATAATCCCAGCTACTCAGGAGC

AGCACACGTCTGAACTCTTTCCCTACACGACGCTCTTCCGATCTNNNCATAATCCCAGCTACTCAGGAGC

AGAGCACACGTCTGAACTCTTTCCCTACACGACGCTCTTCCGATCTNNNCATGAGAATTGCTTGAACCT

AGAGCACACGTCTGAACTCTTTCCCTACACGACGCTCTTCCGATCTNNNCATGAGAATTGCTTGAACCT

AGAGCACACGTCTGAACTCTTTCCCTACACGACGCTCTTCCGATCTNNNTGTTGACTGTGTGTTGTGTA

AGAGCACACGTCTGAACTCTTTCCCTACACGACGCTCTTCCGATCTNNNTGTTGACTGTGTGTTGTGTA

GCACACGTCTGAACTCTTTCCCTACACGACGCTCTTCCGATCTNNNGATACCCTATAGGCGGTTTCCAG

GCACACGTCTGAACTCTTTCCCTACACGACGCTCTTCCGATCTNNNGATACCCTATAGGCGGTTTCCAG

AGCACACGTCTGAACTCTTTCCCTACACGACGCTCTTCCGATCTNNNGAGCTTGCATGAGCCAAGATCC

AGCACACGTCTGAACTCTTTCCCTACACGACGCTCTTCCGATCTNNNGAGCTTGCATGAGCCAAGATCC

AGCACACGTCTGAACTCTTTCCCTACACGACGCTCTTCCGATCTNNNCAAAGAAAGATCAGGGACAGC

AGCACACGTCTGAACTCTTTCCCTACACGACGCTCTTCCGATCTNNNCAAAGAAAGATCAGGGACAGC

GCACACGTCTGAACTCTTTCCCTACACGACGCTCTTCCGATCTNNNAGAACTACCATTCAACCCAGCAAC

GCACACGTCTGAACTCTTTCCCTACACGACGCTCTTCCGATCTNNNAGAACTACCATTCAACCCAGCAAC

AGCACACGTCTGAACTCTTTCCCTACACGACGCTCTTCCGATCTNNNCCACGCCCAGCTAATTTTTGTAT

AGCACACGTCTGAACTCTTTCCCTACACGACGCTCTTCCGATCTNNNCCACGCCCAGCTAATTTTTGTAT

AGCACACGTCTGAACTCTTTCCCTACACGACGCTCTTCCGATCTNNNAGTAGGTAGGATTACAGGCAC

AGCACACGTCTGAACTCTTTCCCTACACGACGCTCTTCCGATCTNNNAGTAGGTAGGATTACAGGCAC

CACACGTCTGAACTCTTTCCCTACACGACGCTCTTCCGATCTNNNTACGATTTCCTGTGTCCTATGCGCA

CACACGTCTGAACTCTTTCCCTACACGACGCTCTTCCGATCTNNNTACGATTTCCTGTGTCCTATGCGCA

AGCACACGTCTGAACTCTTTCCCTACACGACGCTCTTCCGATCTNNNAGGAAGGAGAATTGCTTGAAC

AGCACACGTCTGAACTCTTTCCCTACACGACGCTCTTCCGATCTNNNAGGAAGGAGAATTGCTTGAAC

CACACGTCTGAACTCTTTCCCTACACGACGCTCTTCCGATCTNNNCTGAGGTGGAGCTGGAATACTCCA

CACACGTCTGAACTCTTTCCCTACACGACGCTCTTCCGATCTNNNCTGAGGTGGAGCTGGAATACTCCA
GCACACGTCTGAACTCTTTCCCTACACGACGCTCTTCCGATCTNNNTCCTGCCACAAGTTTATGGATGA
GCACACGTCTGAACTCTTTCCCTACACGACGCTCTTCCGATCTNNNTCCTGCCACAAGTTTATGGATGA
GCACACGTCTGAACTCTTTCCCTACACGACGCTCTTCCGATCTNNNGGAGCATGATCCCGTATAAACCA
GCACACGTCTGAACTCTTTCCCTACACGACGCTCTTCCGATCTNNNGGAGCATGATCCCGTATAAACCA
GAGCACACGTCTGAACTCTTTCCCTACACGACGCTCTTCCGATCTNNNAGAGTTAGTGTGAGCATGTG
GAGCACACGTCTGAACTCTTTCCCTACACGACGCTCTTCCGATCTNNNAGAGTTAGTGTGAGCATGTG
GAGCACACGTCTGAACTCTTTCCCTACACGACGCTCTTCCGATCTNNNTCCAAGTAGCCCAAATGACGT
GAGCACACGTCTGAACTCTTTCCCTACACGACGCTCTTCCGATCTNNNTCCAAGTAGCCCAAATGACGT
GAGCACACGTCTGAACTCTTTCCCTACACGACGCTCTTCCGATCTNNNGAGGTTGCAGTGAGCTGAG
GAGCACACGTCTGAACTCTTTCCCTACACGACGCTCTTCCGATCTNNNGAGGTTGCAGTGAGCTGAG
GAGCACACGTCTGAACTCTTTCCCTACACGACGCTCTTCCGATCTNNNTCCCAAAGTGCTAGGATTAC
GAGCACACGTCTGAACTCTTTCCCTACACGACGCTCTTCCGATCTNNNTCCCAAAGTGCTAGGATTAC
AGCACACGTCTGAACTCTTTCCCTACACGACGCTCTTCCGATCTNNNGTTGTTCCCCTCTATGTGTCCAT
AGCACACGTCTGAACTCTTTCCCTACACGACGCTCTTCCGATCTNNNGTTGTTCCCCTCTATGTGTCCAT
GCACACGTCTGAACTCTTTCCCTACACGACGCTCTTCCGATCTNNNACATCAGGTTTTCTTGACGTTCT
GCACACGTCTGAACTCTTTCCCTACACGACGCTCTTCCGATCTNNNACATCAGGTTTTCTTGACGTTCT
AGAGCACACGTCTGAACTCTTTCCCTACACGACGCTCTTCCGATCTNNNTGTAGTACCAGCTACTCAG
AGAGCACACGTCTGAACTCTTTCCCTACACGACGCTCTTCCGATCTNNNTGTAGTACCAGCTACTCAG
GCACACGTCTGAACTCTTTCCCTACACGACGCTCTTCCGATCTNNNCTCCCGAGTAGCTAGGACTACA
GCACACGTCTGAACTCTTTCCCTACACGACGCTCTTCCGATCTNNNCTCCCGAGTAGCTAGGACTACA
GAGCACACGTCTGAACTCTTTCCCTACACGACGCTCTTCCGATCTNNNGTGGTCAAGTTTTCCTCCATG
GAGCACACGTCTGAACTCTTTCCCTACACGACGCTCTTCCGATCTNNNGTGGTCAAGTTTTCCTCCATG
GCACACGTCTGAACTCTTTCCCTACACGACGCTCTTCCGATCTNNNTGGAAAAGATCAAAAGGAAAC
GCACACGTCTGAACTCTTTCCCTACACGACGCTCTTCCGATCTNNNTGGAAAAGATCAAAAGGAAAC
GAGCACACGTCTGAACTCTTTCCCTACACGACGCTCTTCCGATCTNNNTAGTTGCCGATTTGAGTGGA
GAGCACACGTCTGAACTCTTTCCCTACACGACGCTCTTCCGATCTNNNTAGTTGCCGATTTGAGTGGA
AGCACACGTCTGAACTCTTTCCCTACACGACGCTCTTCCGATCTNNNTGTGTATCTAATTTTTGGGGGG
AGCACACGTCTGAACTCTTTCCCTACACGACGCTCTTCCGATCTNNNTGTGTATCTAATTTTTGGGGGG
GAGCACACGTCTGAACTCTTTCCCTACACGACGCTCTTCCGATCTNNNAATCACTTGAACCTAGGAGG
GAGCACACGTCTGAACTCTTTCCCTACACGACGCTCTTCCGATCTNNNAATCACTTGAACCTAGGAGG
CACACGTCTGAACTCTTTCCCTACACGACGCTCTTCCGATCTNNNCCAGGCTGATCTCAAACTCCTAGC
CACACGTCTGAACTCTTTCCCTACACGACGCTCTTCCGATCTNNNCCAGGCTGATCTCAAACTCCTAGC
GCACACGTCTGAACTCTTTCCCTACACGACGCTCTTCCGATCTNNNAAGGAAAAAAAGTGGGCTAGCA
GCACACGTCTGAACTCTTTCCCTACACGACGCTCTTCCGATCTNNNAAGGAAAAAAAGTGGGCTAGCA
AGAGCACACGTCTGAACTCTTTCCCTACACGACGCTCTTCCGATCTNNNCCATGTTCTCACCTGAAGGG
AGAGCACACGTCTGAACTCTTTCCCTACACGACGCTCTTCCGATCTNNNCCATGTTCTCACCTGAAGGG
GCACACGTCTGAACTCTTTCCCTACACGACGCTCTTCCGATCTNNNGTTATGGGTTTGAGGTCCTGTCT
GCACACGTCTGAACTCTTTCCCTACACGACGCTCTTCCGATCTNNNGTTATGGGTTTGAGGTCCTGTCT
GAGCACACGTCTGAACTCTTTCCCTACACGACGCTCTTCCGATCTNNNAGCCCAGGGACATAATTGAAT
GAGCACACGTCTGAACTCTTTCCCTACACGACGCTCTTCCGATCTNNNAGCCCAGGGACATAATTGAAT
AGCACACGTCTGAACTCTTTCCCTACACGACGCTCTTCCGATCTNNNGGAAACCATGCTTTTCCCATTG
AGCACACGTCTGAACTCTTTCCCTACACGACGCTCTTCCGATCTNNNGGAAACCATGCTTTTCCCATTG

GAGCACACGTCTGAACTCTTTCCCTACACGACGCTCTTCCGATCTNNNGAGAAAGGTTGTTAGGGAAC
GAGCACACGTCTGAACTCTTTCCCTACACGACGCTCTTCCGATCTNNNGAGAAAGGTTGTTAGGGAAC
AGCACACGTCTGAACTCTTTCCCTACACGACGCTCTTCCGATCTNNNCTAGGGAAAAGCTGGTGTCACT
AGCACACGTCTGAACTCTTTCCCTACACGACGCTCTTCCGATCTNNNCTAGGGAAAAGCTGGTGTCACT
AGCACACGTCTGAACTCTTTCCCTACACGACGCTCTTCCGATCTNNNTTGCACCTGAACACTAAAGGAC
AGCACACGTCTGAACTCTTTCCCTACACGACGCTCTTCCGATCTNNNTTGCACCTGAACACTAAAGGAC
AGCACACGTCTGAACTCTTTCCCTACACGACGCTCTTCCGATCTNNNCCCACTCACCACTTCAAGGATTA
AGCACACGTCTGAACTCTTTCCCTACACGACGCTCTTCCGATCTNNNCCCACTCACCACTTCAAGGATTA
GAGCACACGTCTGAACTCTTTCCCTACACGACGCTCTTCCGATCTNNNCTCCATTCTCACTCGATACTCCC
GAGCACACGTCTGAACTCTTTCCCTACACGACGCTCTTCCGATCTNNNCTCCATTCTCACTCGATACTCCC
AGCACACGTCTGAACTCTTTCCCTACACGACGCTCTTCCGATCTNNNCTGTTGCCTTCTGCCAAATACTTC
AGCACACGTCTGAACTCTTTCCCTACACGACGCTCTTCCGATCTNNNCTGTTGCCTTCTGCCAAATACTTC
AGCACACGTCTGAACTCTTTCCCTACACGACGCTCTTCCGATCTNNNTGTTATAGCAGCCCAAACAGAC
AGCACACGTCTGAACTCTTTCCCTACACGACGCTCTTCCGATCTNNNTGTTATAGCAGCCCAAACAGAC
AGAGCACACGTCTGAACTCTTTCCCTACACGACGCTCTTCCGATCTNNNCCTCTATGGGTCTGACTTGG
AGAGCACACGTCTGAACTCTTTCCCTACACGACGCTCTTCCGATCTNNNCCTCTATGGGTCTGACTTGG
AGCACACGTCTGAACTCTTTCCCTACACGACGCTCTTCCGATCTNNNCTGAGGCAGGAGAATCACTTGA
AGCACACGTCTGAACTCTTTCCCTACACGACGCTCTTCCGATCTNNNCTGAGGCAGGAGAATCACTTGA
AGCACACGTCTGAACTCTTTCCCTACACGACGCTCTTCCGATCTNNNAAGGACAAGATTTCCAGGAAC
AGCACACGTCTGAACTCTTTCCCTACACGACGCTCTTCCGATCTNNNAAGGACAAGATTTCCAGGAAC
GAGCACACGTCTGAACTCTTTCCCTACACGACGCTCTTCCGATCTNNNGACTCTTTTCAGCACATAAGG
GAGCACACGTCTGAACTCTTTCCCTACACGACGCTCTTCCGATCTNNNGACTCTTTTCAGCACATAAGG
GCACACGTCTGAACTCTTTCCCTACACGACGCTCTTCCGATCTNNNAGGATGGCTACTGATCCTAGCAC
GCACACGTCTGAACTCTTTCCCTACACGACGCTCTTCCGATCTNNNAGGATGGCTACTGATCCTAGCAC
GAGCACACGTCTGAACTCTTTCCCTACACGACGCTCTTCCGATCTNNNGAGGCTGAGGCAGGAGAATC
GAGCACACGTCTGAACTCTTTCCCTACACGACGCTCTTCCGATCTNNNGAGGCTGAGGCAGGAGAATC
GAGCACACGTCTGAACTCTTTCCCTACACGACGCTCTTCCGATCTNNNTGTGTAGAGGTGTGAGTGTTC
GAGCACACGTCTGAACTCTTTCCCTACACGACGCTCTTCCGATCTNNNTGTGTAGAGGTGTGAGTGTTC
AGCACACGTCTGAACTCTTTCCCTACACGACGCTCTTCCGATCTNNNGAGGCTGAGGCAGGAGAATG
AGCACACGTCTGAACTCTTTCCCTACACGACGCTCTTCCGATCTNNNGAGGCTGAGGCAGGAGAATG
GCACACGTCTGAACTCTTTCCCTACACGACGCTCTTCCGATCTNNNTCCAACGGCAATATGATTGGAAC
GCACACGTCTGAACTCTTTCCCTACACGACGCTCTTCCGATCTNNNTCCAACGGCAATATGATTGGAAC
GAGCACACGTCTGAACTCTTTCCCTACACGACGCTCTTCCGATCTNNNTCGTCATCGTGGAAGACATAG
GAGCACACGTCTGAACTCTTTCCCTACACGACGCTCTTCCGATCTNNNTCGTCATCGTGGAAGACATAG
GAGCACACGTCTGAACTCTTTCCCTACACGACGCTCTTCCGATCTNNNAAAAGCATGCGGTCAGAGTTT
GAGCACACGTCTGAACTCTTTCCCTACACGACGCTCTTCCGATCTNNNAAAAGCATGCGGTCAGAGTTT
GAGCACACGTCTGAACTCTTTCCCTACACGACGCTCTTCCGATCTNNNAAATACAAAAATTAGCCGGGC
GAGCACACGTCTGAACTCTTTCCCTACACGACGCTCTTCCGATCTNNNAAATACAAAAATTAGCCGGGC
GAGCACACGTCTGAACTCTTTCCCTACACGACGCTCTTCCGATCTNNNGAGAATCGCTTGAATCCAGGA
GAGCACACGTCTGAACTCTTTCCCTACACGACGCTCTTCCGATCTNNNGAGAATCGCTTGAATCCAGGA
AGCACACGTCTGAACTCTTTCCCTACACGACGCTCTTCCGATCTNNNCCAAGTGCAGGACATACTAAAT
AGCACACGTCTGAACTCTTTCCCTACACGACGCTCTTCCGATCTNNNCCAAGTGCAGGACATACTAAAT
AGAGCACACGTCTGAACTCTTTCCCTACACGACGCTCTTCCGATCTNNNGTCTCTTCTGGTGTGTAGGG

GAGCACACGTCTGAACTCTTTCCCTACACGACGCTCTTCCGATCTNNNGTCTCTTCTGGTGTGTAGGG

AGCACACGTCTGAACTCTTTCCCTACACGACGCTCTTCCGATCTNNNACTTCAGTCCTCTTCCATATGG

AGCACACGTCTGAACTCTTTCCCTACACGACGCTCTTCCGATCTNNNACTTCAGTCCTCTTCCATATGG

GCACACGTCTGAACTCTTTCCCTACACGACGCTCTTCCGATCTNNNTCCTCAACCTTGGCAAAATAAG

GCACACGTCTGAACTCTTTCCCTACACGACGCTCTTCCGATCTNNNTCCTCAACCTTGGCAAAATAAG

AGCACACGTCTGAACTCTTTCCCTACACGACGCTCTTCCGATCTNNNGTGTCCTGTTGTTGCTGCTATT

AGCACACGTCTGAACTCTTTCCCTACACGACGCTCTTCCGATCTNNNGTGTCCTGTTGTTGCTGCTATT

AGCACACGTCTGAACTCTTTCCCTACACGACGCTCTTCCGATCTNNNATCCATGTGTCTGTGCTTCTT

AGCACACGTCTGAACTCTTTCCCTACACGACGCTCTTCCGATCTNNNATCCATGTGTCTGTGCTTCTT

GCACACGTCTGAACTCTTTCCCTACACGACGCTCTTCCGATCTNNNATTCCAGAAGTCCGGCCTAATT

GCACACGTCTGAACTCTTTCCCTACACGACGCTCTTCCGATCTNNNATTCCAGAAGTCCGGCCTAATT

GCACACGTCTGAACTCTTTCCCTACACGACGCTCTTCCGATCTNNNTTGGGGCCCAAAATAACATTCT

GCACACGTCTGAACTCTTTCCCTACACGACGCTCTTCCGATCTNNNTTGGGGCCCAAAATAACATTCT

CACACGTCTGAACTCTTTCCCTACACGACGCTCTTCCGATCTNNNCTGAAGGAGTTATCTCTCTTTAA

CACACGTCTGAACTCTTTCCCTACACGACGCTCTTCCGATCTNNNCTGAAGGAGTTATCTCTCTTTAA

AGCACACGTCTGAACTCTTTCCCTACACGACGCTCTTCCGATCTNNNTAGTCCCAGCTACTCAAAAGG

AGCACACGTCTGAACTCTTTCCCTACACGACGCTCTTCCGATCTNNNTAGTCCCAGCTACTCAAAAGG

AGCACACGTCTGAACTCTTTCCCTACACGACGCTCTTCCGATCTNNNCATCATCAAGTGTGTCACTCAG

AGCACACGTCTGAACTCTTTCCCTACACGACGCTCTTCCGATCTNNNCATCATCAAGTGTGTCACTCAG

GAGCACACGTCTGAACTCTTTCCCTACACGACGCTCTTCCGATCTNNNTGAATTCTTCCCGGTTCTCT

GAGCACACGTCTGAACTCTTTCCCTACACGACGCTCTTCCGATCTNNNTGAATTCTTCCCGGTTCTCT

GAGCACACGTCTGAACTCTTTCCCTACACGACGCTCTTCCGATCTNNNGAGGTCCCAGCTACTCAGGA

GAGCACACGTCTGAACTCTTTCCCTACACGACGCTCTTCCGATCTNNNGAGGTCCCAGCTACTCAGGA

AGCACACGTCTGAACTCTTTCCCTACACGACGCTCTTCCGATCTNNNCTGCACTCCTCAGCCTTTGGCAT

AGCACACGTCTGAACTCTTTCCCTACACGACGCTCTTCCGATCTNNNCTGCACTCCTCAGCCTTTGGCAT

AGCACACGTCTGAACTCTTTCCCTACACGACGCTCTTCCGATCTNNNGTGTCCAGTAAAGTGCCATGA

AGCACACGTCTGAACTCTTTCCCTACACGACGCTCTTCCGATCTNNNGTGTCCAGTAAAGTGCCATGA

GCACACGTCTGAACTCTTTCCCTACACGACGCTCTTCCGATCTNNNCCTCCTGAGTAGCTAGTACACCA

GCACACGTCTGAACTCTTTCCCTACACGACGCTCTTCCGATCTNNNCCTCCTGAGTAGCTAGTACACCA

GAGCACACGTCTGAACTCTTTCCCTACACGACGCTCTTCCGATCTNNNGCCCCCTCTTCCTCTAGAAAT

GAGCACACGTCTGAACTCTTTCCCTACACGACGCTCTTCCGATCTNNNGCCCCCTCTTCCTCTAGAAAT

AGCACACGTCTGAACTCTTTCCCTACACGACGCTCTTCCGATCTNNNGAGAATGGCATGAACCCCAGA

AGCACACGTCTGAACTCTTTCCCTACACGACGCTCTTCCGATCTNNNGAGAATGGCATGAACCCCAGA

AGCACACGTCTGAACTCTTTCCCTACACGACGCTCTTCCGATCTNNNACCTGGCAGTGTTTCTGTTGA

AGCACACGTCTGAACTCTTTCCCTACACGACGCTCTTCCGATCTNNNACCTGGCAGTGTTTCTGTTGA

AGCACACGTCTGAACTCTTTCCCTACACGACGCTCTTCCGATCTNNNTTCCACCTCAACCGACTGAATA

AGCACACGTCTGAACTCTTTCCCTACACGACGCTCTTCCGATCTNNNTTCCACCTCAACCGACTGAATA

AGCACACGTCTGAACTCTTTCCCTACACGACGCTCTTCCGATCTNNNACTTCCTCCTCCCATTCAGACTA

AGCACACGTCTGAACTCTTTCCCTACACGACGCTCTTCCGATCTNNNACTTCCTCCTCCCATTCAGACTA

GCACACGTCTGAACTCTTTCCCTACACGACGCTCTTCCGATCTNNNCTGAGGCAGGAGAATAGCTTG

GCACACGTCTGAACTCTTTCCCTACACGACGCTCTTCCGATCTNNNCTGAGGCAGGAGAATAGCTTG

AGCACACGTCTGAACTCTTTCCCTACACGACGCTCTTCCGATCTNNNGGATTCCATATTGTTCATAACA

AGCACACGTCTGAACTCTTTCCCTACACGACGCTCTTCCGATCTNNNGGATTCCATATTGTTCATAACA

AGCACACGTCTGAACTCTTTCCCTACACGACGCTCTTCCGATCTNNNTGCTTTCCACATTCAGAATTGGT
AGCACACGTCTGAACTCTTTCCCTACACGACGCTCTTCCGATCTNNNTGCTTTCCACATTCAGAATTGGT
AGCACACGTCTGAACTCTTTCCCTACACGACGCTCTTCCGATCTNNNACCAAGGCTAAAGTAGAAAGG
AGCACACGTCTGAACTCTTTCCCTACACGACGCTCTTCCGATCTNNNACCAAGGCTAAAGTAGAAAGG
AGCACACGTCTGAACTCTTTCCCTACACGACGCTCTTCCGATCTNNNCCAGCCATCATCTTGTCCATTTT
AGCACACGTCTGAACTCTTTCCCTACACGACGCTCTTCCGATCTNNNCCAGCCATCATCTTGTCCATTTT
GAGCACACGTCTGAACTCTTTCCCTACACGACGCTCTTCCGATCTNNNCACACACAATTGCTACACCTC
GAGCACACGTCTGAACTCTTTCCCTACACGACGCTCTTCCGATCTNNNCACACACAATTGCTACACCTC
AGCACACGTCTGAACTCTTTCCCTACACGACGCTCTTCCGATCTNNNTGTGTTTATCCTGTGGTCCAGT
AGCACACGTCTGAACTCTTTCCCTACACGACGCTCTTCCGATCTNNNTGTGTTTATCCTGTGGTCCAGT
AGCACACGTCTGAACTCTTTCCCTACACGACGCTCTTCCGATCTNNNGAGTTCAGGAGATCGAGACCA
AGCACACGTCTGAACTCTTTCCCTACACGACGCTCTTCCGATCTNNNGAGTTCAGGAGATCGAGACCA
AGCACACGTCTGAACTCTTTCCCTACACGACGCTCTTCCGATCTNNNTGTAGATCACAACACCCAAGTC
AGCACACGTCTGAACTCTTTCCCTACACGACGCTCTTCCGATCTNNNTGTAGATCACAACACCCAAGTC
AGCACACGTCTGAACTCTTTCCCTACACGACGCTCTTCCGATCTNNNCTGTTTTAGGAGACTGGTGGA
AGCACACGTCTGAACTCTTTCCCTACACGACGCTCTTCCGATCTNNNCTGTTTTAGGAGACTGGTGGA
AGCACACGTCTGAACTCTTTCCCTACACGACGCTCTTCCGATCTNNNGAGCTACCTCATCAGCATCCAT
AGCACACGTCTGAACTCTTTCCCTACACGACGCTCTTCCGATCTNNNGAGCTACCTCATCAGCATCCAT
AGCACACGTCTGAACTCTTTCCCTACACGACGCTCTTCCGATCTNNNCTGTAATCCCAGCTACTCAGGA
AGCACACGTCTGAACTCTTTCCCTACACGACGCTCTTCCGATCTNNNCTGTAATCCCAGCTACTCAGGA
AGCACACGTCTGAACTCTTTCCCTACACGACGCTCTTCCGATCTNNNGTAGCGTTTAGAGGAGTCTTC
AGCACACGTCTGAACTCTTTCCCTACACGACGCTCTTCCGATCTNNNGTAGCGTTTAGAGGAGTCTTC
AGAGCACACGTCTGAACTCTTTCCCTACACGACGCTCTTCCGATCTNNNTTTCATTTTCTAGCTCCTCCC
AGAGCACACGTCTGAACTCTTTCCCTACACGACGCTCTTCCGATCTNNNTTTCATTTTCTAGCTCCTCCC
AGCACACGTCTGAACTCTTTCCCTACACGACGCTCTTCCGATCTNNNGTCGGGATCTAAATTCCTCCTC
AGCACACGTCTGAACTCTTTCCCTACACGACGCTCTTCCGATCTNNNGTCGGGATCTAAATTCCTCCTC
GAGCACACGTCTGAACTCTTTCCCTACACGACGCTCTTCCGATCTNNNCATTTGTGTACAAGTGTGTGT
GAGCACACGTCTGAACTCTTTCCCTACACGACGCTCTTCCGATCTNNNCATTTGTGTACAAGTGTGTGT
AGCACACGTCTGAACTCTTTCCCTACACGACGCTCTTCCGATCTNNNACCATTACATCACTGCAGCCAA
AGCACACGTCTGAACTCTTTCCCTACACGACGCTCTTCCGATCTNNNACCATTACATCACTGCAGCCAA
AGCACACGTCTGAACTCTTTCCCTACACGACGCTCTTCCGATCTNNNGAGGCTGAAGTGAGCTATGAT
AGCACACGTCTGAACTCTTTCCCTACACGACGCTCTTCCGATCTNNNGAGGCTGAAGTGAGCTATGAT
AGCACACGTCTGAACTCTTTCCCTACACGACGCTCTTCCGATCTNNNATCCTGTTCTCTCTCAGCCAAA
AGCACACGTCTGAACTCTTTCCCTACACGACGCTCTTCCGATCTNNNATCCTGTTCTCTCTCAGCCAAA
GAGCACACGTCTGAACTCTTTCCCTACACGACGCTCTTCCGATCTNNNCTTCTTTCCCAAGAGCCTCTCT
GAGCACACGTCTGAACTCTTTCCCTACACGACGCTCTTCCGATCTNNNCTTCTTTCCCAAGAGCCTCTCT
AGCACACGTCTGAACTCTTTCCCTACACGACGCTCTTCCGATCTNNNACCTGTAGTCCCAGCTACTCGC
AGCACACGTCTGAACTCTTTCCCTACACGACGCTCTTCCGATCTNNNACCTGTAGTCCCAGCTACTCGC
GAGCACACGTCTGAACTCTTTCCCTACACGACGCTCTTCCGATCTNNNGCAACAGAGTGAGACTCCAT
GAGCACACGTCTGAACTCTTTCCCTACACGACGCTCTTCCGATCTNNNGCAACAGAGTGAGACTCCAT
AGCACACGTCTGAACTCTTTCCCTACACGACGCTCTTCCGATCTNNNTAATCTCAGCTACTCAGGAGCC
AGCACACGTCTGAACTCTTTCCCTACACGACGCTCTTCCGATCTNNNTAATCTCAGCTACTCAGGAGCC
AGCACACGTCTGAACTCTTTCCCTACACGACGCTCTTCCGATCTNNNCAAGATCTCAGCTCACTGCAAG

AGCACACGTCTGAACTCTTTCCCTACACGACGCTCTTCCGATCTNNNCAAGATCTCAGCTCACTGCAAG
AGCACACGTCTGAACTCTTTCCCTACACGACGCTCTTCCGATCTNNNATTAGTACAAGTGTGAGCAGG
AGCACACGTCTGAACTCTTTCCCTACACGACGCTCTTCCGATCTNNNATTAGTACAAGTGTGAGCAGG
GAGCACACGTCTGAACTCTTTCCCTACACGACGCTCTTCCGATCTNNNGCTTGGTAAAAAGCCACATCA
GAGCACACGTCTGAACTCTTTCCCTACACGACGCTCTTCCGATCTNNNGCTTGGTAAAAAGCCACATCA
GAGCACACGTCTGAACTCTTTCCCTACACGACGCTCTTCCGATCTNNNGGAGAAAAGCAGGAGAGGA
GAGCACACGTCTGAACTCTTTCCCTACACGACGCTCTTCCGATCTNNNGGAGAAAAGCAGGAGAGGA
GAGCACACGTCTGAACTCTTTCCCTACACGACGCTCTTCCGATCTNNNGCCATGTACTAATCAATGGTG
GAGCACACGTCTGAACTCTTTCCCTACACGACGCTCTTCCGATCTNNNGCCATGTACTAATCAATGGTG
GAGCACACGTCTGAACTCTTTCCCTACACGACGCTCTTCCGATCTNNNCTCCACTCTTGCTGTCCAGTG
GAGCACACGTCTGAACTCTTTCCCTACACGACGCTCTTCCGATCTNNNCTCCACTCTTGCTGTCCAGTG
AGCACACGTCTGAACTCTTTCCCTACACGACGCTCTTCCGATCTNNNACACACACACACACATACACGTA
AGCACACGTCTGAACTCTTTCCCTACACGACGCTCTTCCGATCTNNNACACACACACACACATACACGTA
AGCACACGTCTGAACTCTTTCCCTACACGACGCTCTTCCGATCTNNNCACCTGATACATGTTCCCAGAC
AGCACACGTCTGAACTCTTTCCCTACACGACGCTCTTCCGATCTNNNCACCTGATACATGTTCCCAGAC
GAGCACACGTCTGAACTCTTTCCCTACACGACGCTCTTCCGATCTNNNTGAAACCATTCCAGTGTAAAA
GAGCACACGTCTGAACTCTTTCCCTACACGACGCTCTTCCGATCTNNNTGAAACCATTCCAGTGTAAAA
AGCACACGTCTGAACTCTTTCCCTACACGACGCTCTTCCGATCTNNNGAGAATGGCATGATCCCAGGA
AGCACACGTCTGAACTCTTTCCCTACACGACGCTCTTCCGATCTNNNGAGAATGGCATGATCCCAGGA
GAGCACACGTCTGAACTCTTTCCCTACACGACGCTCTTCCGATCTNNNAGAATCACTTGAATCCAGGA
GAGCACACGTCTGAACTCTTTCCCTACACGACGCTCTTCCGATCTNNNAGAATCACTTGAATCCAGGA
GCACACGTCTGAACTCTTTCCCTACACGACGCTCTTCCGATCTNNNACACACACACACACATATATACG
GCACACGTCTGAACTCTTTCCCTACACGACGCTCTTCCGATCTNNNACACACACACACACATATATACG
GCACACGTCTGAACTCTTTCCCTACACGACGCTCTTCCGATCTNNNCATGTGCTGCTCTGATCATTCTG
GCACACGTCTGAACTCTTTCCCTACACGACGCTCTTCCGATCTNNNCATGTGCTGCTCTGATCATTCTG
GAGCACACGTCTGAACTCTTTCCCTACACGACGCTCTTCCGATCTNNNCCTGGGAGATGGAGATTACA
GAGCACACGTCTGAACTCTTTCCCTACACGACGCTCTTCCGATCTNNNCCTGGGAGATGGAGATTACA
AGCACACGTCTGAACTCTTTCCCTACACGACGCTCTTCCGATCTNNNCTGTAGTCCCAGCTACTTGAGA
AGCACACGTCTGAACTCTTTCCCTACACGACGCTCTTCCGATCTNNNCTGTAGTCCCAGCTACTTGAGA
GCACACGTCTGAACTCTTTCCCTACACGACGCTCTTCCGATCTNNNGATCTCCTGACCTCATGATCCAC
GCACACGTCTGAACTCTTTCCCTACACGACGCTCTTCCGATCTNNNGATCTCCTGACCTCATGATCCAC
GAGCACACGTCTGAACTCTTTCCCTACACGACGCTCTTCCGATCTNNNCTGCCCACCTTTGTTACTCTT
GAGCACACGTCTGAACTCTTTCCCTACACGACGCTCTTCCGATCTNNNCTGCCCACCTTTGTTACTCTT
GAGCACACGTCTGAACTCTTTCCCTACACGACGCTCTTCCGATCTNNNACTGAGTCAATTCAATCAGCA
GAGCACACGTCTGAACTCTTTCCCTACACGACGCTCTTCCGATCTNNNACTGAGTCAATTCAATCAGCA
AGCACACGTCTGAACTCTTTCCCTACACGACGCTCTTCCGATCTNNNATTTCAGTCTTCAGGTCCAGGA
AGCACACGTCTGAACTCTTTCCCTACACGACGCTCTTCCGATCTNNNATTTCAGTCTTCAGGTCCAGGA
GAGCACACGTCTGAACTCTTTCCCTACACGACGCTCTTCCGATCTNNNGAGGCTGAGGCAGGAGAAT
GAGCACACGTCTGAACTCTTTCCCTACACGACGCTCTTCCGATCTNNNGAGGCTGAGGCAGGAGAAT
AGCACACGTCTGAACTCTTTCCCTACACGACGCTCTTCCGATCTNNNGTGTTTCTGATGGGAGAAAAG
AGCACACGTCTGAACTCTTTCCCTACACGACGCTCTTCCGATCTNNNGTGTTTCTGATGGGAGAAAAG
GCACACGTCTGAACTCTTTCCCTACACGACGCTCTTCCGATCTNNNAATAGTGTACCTCCCCATCAGTG
GCACACGTCTGAACTCTTTCCCTACACGACGCTCTTCCGATCTNNNAATAGTGTACCTCCCCATCAGTG

GAGCACACGTCTGAACTCTTTCCCTACACGACGCTCTTCCGATCTNNNAATTACGTATCACTGGGCTT
GAGCACACGTCTGAACTCTTTCCCTACACGACGCTCTTCCGATCTNNNAATTACGTATCACTGGGCTT
GAGCACACGTCTGAACTCTTTCCCTACACGACGCTCTTCCGATCTNNNATGTAGCAAAATCGCTTGAA
GAGCACACGTCTGAACTCTTTCCCTACACGACGCTCTTCCGATCTNNNATGTAGCAAAATCGCTTGAA
AGCACACGTCTGAACTCTTTCCCTACACGACGCTCTTCCGATCTNNNATTGCATCCTTTTGCTCCCATCT
AGCACACGTCTGAACTCTTTCCCTACACGACGCTCTTCCGATCTNNNATTGCATCCTTTTGCTCCCATCT
GAGCACACGTCTGAACTCTTTCCCTACACGACGCTCTTCCGATCTNNNTGGGGTTCCAAGTTTTCTCTG
GAGCACACGTCTGAACTCTTTCCCTACACGACGCTCTTCCGATCTNNNTGGGGTTCCAAGTTTTCTCTG
GAGCACACGTCTGAACTCTTTCCCTACACGACGCTCTTCCGATCTNNNTTCCAGAGGAGTGAATGGTT
GAGCACACGTCTGAACTCTTTCCCTACACGACGCTCTTCCGATCTNNNTTCCAGAGGAGTGAATGGTT
GCACACGTCTGAACTCTTTCCCTACACGACGCTCTTCCGATCTNNNGCCGGCACTTTGTTCTGTATATC
GCACACGTCTGAACTCTTTCCCTACACGACGCTCTTCCGATCTNNNGCCGGCACTTTGTTCTGTATATC
GAGCACACGTCTGAACTCTTTCCCTACACGACGCTCTTCCGATCTNNNCACCTGTAATTGCAGCTACTT
GAGCACACGTCTGAACTCTTTCCCTACACGACGCTCTTCCGATCTNNNCACCTGTAATTGCAGCTACTT
AGCACACGTCTGAACTCTTTCCCTACACGACGCTCTTCCGATCTNNNCACGTCATTTTCTGTGGTTTCCA
AGCACACGTCTGAACTCTTTCCCTACACGACGCTCTTCCGATCTNNNCACGTCATTTTCTGTGGTTTCCA
GAGCACACGTCTGAACTCTTTCCCTACACGACGCTCTTCCGATCTNNNAGGAGTACTATCATGGCTCCT
GAGCACACGTCTGAACTCTTTCCCTACACGACGCTCTTCCGATCTNNNAGGAGTACTATCATGGCTCCT
AGCACACGTCTGAACTCTTTCCCTACACGACGCTCTTCCGATCTNNNATAGTCAAAGGGGAAATGAG
AGCACACGTCTGAACTCTTTCCCTACACGACGCTCTTCCGATCTNNNATAGTCAAAGGGGAAATGAG
AGCACACGTCTGAACTCTTTCCCTACACGACGCTCTTCCGATCTNNNCTTGCCTTCTGCTAGCTTTTGA
AGCACACGTCTGAACTCTTTCCCTACACGACGCTCTTCCGATCTNNNCTTGCCTTCTGCTAGCTTTTGA
AGCACACGTCTGAACTCTTTCCCTACACGACGCTCTTCCGATCTNNNGAATATAAGTTATGGGCTAG
AGCACACGTCTGAACTCTTTCCCTACACGACGCTCTTCCGATCTNNNGAATATAAGTTATGGGCTAG
ACACGTCTGAACTCTTTCCCTACACGACGCTCTTCCGATCTNNNAACTGTAGCCCTCTGTAGTACATCG
ACACGTCTGAACTCTTTCCCTACACGACGCTCTTCCGATCTNNNAACTGTAGCCCTCTGTAGTACATCG
AGCACACGTCTGAACTCTTTCCCTACACGACGCTCTTCCGATCTNNNTAGGAGGCAAGGATCGGGAA
AGCACACGTCTGAACTCTTTCCCTACACGACGCTCTTCCGATCTNNNTAGGAGGCAAGGATCGGGAA
GCACACGTCTGAACTCTTTCCCTACACGACGCTCTTCCGATCTNNNCTCAGATGAGGTGGCGCTTATCA
GCACACGTCTGAACTCTTTCCCTACACGACGCTCTTCCGATCTNNNCTCAGATGAGGTGGCGCTTATCA
AGCACACGTCTGAACTCTTTCCCTACACGACGCTCTTCCGATCTNNNCCTGGGAAGTCAAGGCTATAG
AGCACACGTCTGAACTCTTTCCCTACACGACGCTCTTCCGATCTNNNCCTGGGAAGTCAAGGCTATAG
GAGCACACGTCTGAACTCTTTCCCTACACGACGCTCTTCCGATCTNNNGAGATCATGCCACTTCACTCC
GAGCACACGTCTGAACTCTTTCCCTACACGACGCTCTTCCGATCTNNNGAGATCATGCCACTTCACTCC
AGCACACGTCTGAACTCTTTCCCTACACGACGCTCTTCCGATCTNNNTGCTGAAGATGGTTGAACTGG
AGCACACGTCTGAACTCTTTCCCTACACGACGCTCTTCCGATCTNNNTGCTGAAGATGGTTGAACTGG
AGCACACGTCTGAACTCTTTCCCTACACGACGCTCTTCCGATCTNNNCAAGGAAGCTCATGTTTTCCAG
AGCACACGTCTGAACTCTTTCCCTACACGACGCTCTTCCGATCTNNNCAAGGAAGCTCATGTTTTCCAG
ACACGTCTGAACTCTTTCCCTACACGACGCTCTTCCGATCTNNNATCCCATCCAGATTGAAAATGCATCG
ACACGTCTGAACTCTTTCCCTACACGACGCTCTTCCGATCTNNNATCCCATCCAGATTGAAAATGCATCG
GCACACGTCTGAACTCTTTCCCTACACGACGCTCTTCCGATCTNNNACTCCCAGTCTGTGATTTGTCTTC
GCACACGTCTGAACTCTTTCCCTACACGACGCTCTTCCGATCTNNNACTCCCAGTCTGTGATTTGTCTTC
CACACGTCTGAACTCTTTCCCTACACGACGCTCTTCCGATCTNNNTTTAGTAGAGACGGGGTTTCACCC

CACACGTCTGAACTCTTTCCCTACACGACGCTCTTCCGATCTNNNTTTAGTAGAGACGGGGTTTCACCC
AGCACACGTCTGAACTCTTTCCCTACACGACGCTCTTCCGATCTNNNAAGGTTACATTTGCCTGAGTGG
AGCACACGTCTGAACTCTTTCCCTACACGACGCTCTTCCGATCTNNNAAGGTTACATTTGCCTGAGTGG
GAGCACACGTCTGAACTCTTTCCCTACACGACGCTCTTCCGATCTNNNAGAAATAGTTGAATGACAGG
GAGCACACGTCTGAACTCTTTCCCTACACGACGCTCTTCCGATCTNNNAGAAATAGTTGAATGACAGG
AGCACACGTCTGAACTCTTTCCCTACACGACGCTCTTCCGATCTNNNTAAAGGAAGGAAGGAAGAGG
AGCACACGTCTGAACTCTTTCCCTACACGACGCTCTTCCGATCTNNNTAAAGGAAGGAAGGAAGAGG
GCACACGTCTGAACTCTTTCCCTACACGACGCTCTTCCGATCTNNNCATCTCAAATCCCACGCAATCTCC
GCACACGTCTGAACTCTTTCCCTACACGACGCTCTTCCGATCTNNNCATCTCAAATCCCACGCAATCTCC
AGCACACGTCTGAACTCTTTCCCTACACGACGCTCTTCCGATCTNNNATTCAAGTGATTCTCCTGCCTC
AGCACACGTCTGAACTCTTTCCCTACACGACGCTCTTCCGATCTNNNATTCAAGTGATTCTCCTGCCTC
GAGCACACGTCTGAACTCTTTCCCTACACGACGCTCTTCCGATCTNNNCGGTTTCTAAGGGACTAGGT
GAGCACACGTCTGAACTCTTTCCCTACACGACGCTCTTCCGATCTNNNCGGTTTCTAAGGGACTAGGT
GAGCACACGTCTGAACTCTTTCCCTACACGACGCTCTTCCGATCTNNNCTGATATGACCCTTCGTTTTC
GAGCACACGTCTGAACTCTTTCCCTACACGACGCTCTTCCGATCTNNNCTGATATGACCCTTCGTTTTC
AGCACACGTCTGAACTCTTTCCCTACACGACGCTCTTCCGATCTNNNGAGCTCCAGTGTTAGGTGCAT
AGCACACGTCTGAACTCTTTCCCTACACGACGCTCTTCCGATCTNNNGAGCTCCAGTGTTAGGTGCAT
AGCACACGTCTGAACTCTTTCCCTACACGACGCTCTTCCGATCTNNNATGATTTCTCGCTACAAACCCC
AGCACACGTCTGAACTCTTTCCCTACACGACGCTCTTCCGATCTNNNATGATTTCTCGCTACAAACCCC
AGCACACGTCTGAACTCTTTCCCTACACGACGCTCTTCCGATCTNNNGCTATGGAGCCCTTTTATGCAT
AGCACACGTCTGAACTCTTTCCCTACACGACGCTCTTCCGATCTNNNGCTATGGAGCCCTTTTATGCAT
AGCACACGTCTGAACTCTTTCCCTACACGACGCTCTTCCGATCTNNNTTGGGTGGGTTTAGGAATCTC
AGCACACGTCTGAACTCTTTCCCTACACGACGCTCTTCCGATCTNNNTTGGGTGGGTTTAGGAATCTC
AGCACACGTCTGAACTCTTTCCCTACACGACGCTCTTCCGATCTNNNCAGGAGCTGTGATGAAGACC
AGCACACGTCTGAACTCTTTCCCTACACGACGCTCTTCCGATCTNNNCAGGAGCTGTGATGAAGACC
AGCACACGTCTGAACTCTTTCCCTACACGACGCTCTTCCGATCTNNNCTCTGTCTCTCTCTCTCCCTCTC
AGCACACGTCTGAACTCTTTCCCTACACGACGCTCTTCCGATCTNNNCTCTGTCTCTCTCTCTCCCTCTC
GAGCACACGTCTGAACTCTTTCCCTACACGACGCTCTTCCGATCTNNNTCCTTTCCCTGTCAGTCTCTCT
GAGCACACGTCTGAACTCTTTCCCTACACGACGCTCTTCCGATCTNNNTCCTTTCCCTGTCAGTCTCTCT
GCACACGTCTGAACTCTTTCCCTACACGACGCTCTTCCGATCTNNNCCTTTTCATCCCACTTCCCTACAC
GCACACGTCTGAACTCTTTCCCTACACGACGCTCTTCCGATCTNNNCCTTTTCATCCCACTTCCCTACAC
AGCACACGTCTGAACTCTTTCCCTACACGACGCTCTTCCGATCTNNNGAAAACTCTACGGCAGTCTCTC
AGCACACGTCTGAACTCTTTCCCTACACGACGCTCTTCCGATCTNNNGAAAACTCTACGGCAGTCTCTC
AGCACACGTCTGAACTCTTTCCCTACACGACGCTCTTCCGATCTNNNGGCCTGAATGATGGTGGTGAG
AGCACACGTCTGAACTCTTTCCCTACACGACGCTCTTCCGATCTNNNGGCCTGAATGATGGTGGTGAG
GCACACGTCTGAACTCTTTCCCTACACGACGCTCTTCCGATCTNNNGTTTTCCATTCCAGAGGTTTGCA
GCACACGTCTGAACTCTTTCCCTACACGACGCTCTTCCGATCTNNNGTTTTCCATTCCAGAGGTTTGCA
GAGCACACGTCTGAACTCTTTCCCTACACGACGCTCTTCCGATCTNNNAAGGCCTCCTGCATGTATAA
GAGCACACGTCTGAACTCTTTCCCTACACGACGCTCTTCCGATCTNNNAAGGCCTCCTGCATGTATAA
AGCACACGTCTGAACTCTTTCCCTACACGACGCTCTTCCGATCTNNNCACCAGAGGCCACAACAAATAT
AGCACACGTCTGAACTCTTTCCCTACACGACGCTCTTCCGATCTNNNCACCAGAGGCCACAACAAATAT
GAGCACACGTCTGAACTCTTTCCCTACACGACGCTCTTCCGATCTNNNCATGCAGGATTATGACTGGC
GAGCACACGTCTGAACTCTTTCCCTACACGACGCTCTTCCGATCTNNNCATGCAGGATTATGACTGGC

GAGCACACGTCTGAACTCTTTCCCTACACGACGCTCTTCCGATCTNNNTTCCTACTTTTGTTGGTTCCC

GAGCACACGTCTGAACTCTTTCCCTACACGACGCTCTTCCGATCTNNNTTCCTACTTTTGTTGGTTCCC

AGCACACGTCTGAACTCTTTCCCTACACGACGCTCTTCCGATCTNNNAAGATCATGCCACTGTACTCCA

AGCACACGTCTGAACTCTTTCCCTACACGACGCTCTTCCGATCTNNNAAGATCATGCCACTGTACTCCA

AGAGCACACGTCTGAACTCTTTCCCTACACGACGCTCTTCCGATCTNNNCTGTAGTCAGTCCCAGCTAC

AGAGCACACGTCTGAACTCTTTCCCTACACGACGCTCTTCCGATCTNNNCTGTAGTCAGTCCCAGCTAC

AGCACACGTCTGAACTCTTTCCCTACACGACGCTCTTCCGATCTNNNAGCAAGCACATTTGGGATTAC

AGCACACGTCTGAACTCTTTCCCTACACGACGCTCTTCCGATCTNNNAGCAAGCACATTTGGGATTAC

GCACACGTCTGAACTCTTTCCCTACACGACGCTCTTCCGATCTNNNCACCTGTAATCCCAGCTACTTGG

GCACACGTCTGAACTCTTTCCCTACACGACGCTCTTCCGATCTNNNCACCTGTAATCCCAGCTACTTGG

AGCACACGTCTGAACTCTTTCCCTACACGACGCTCTTCCGATCTNNNTAATCTGATAGGGACAGGGCT

AGCACACGTCTGAACTCTTTCCCTACACGACGCTCTTCCGATCTNNNTAATCTGATAGGGACAGGGCT

AGCACACGTCTGAACTCTTTCCCTACACGACGCTCTTCCGATCTNNNTTCACACCATTCTCCTGTCTCA

AGCACACGTCTGAACTCTTTCCCTACACGACGCTCTTCCGATCTNNNTTCACACCATTCTCCTGTCTCA

GAGCACACGTCTGAACTCTTTCCCTACACGACGCTCTTCCGATCTNNNCTGAGGCCTAGGTGGTAGA

GAGCACACGTCTGAACTCTTTCCCTACACGACGCTCTTCCGATCTNNNCTGAGGCCTAGGTGGTAGA

AGCACACGTCTGAACTCTTTCCCTACACGACGCTCTTCCGATCTNNNTGATGGAGCACTCACTTTAGGA

AGCACACGTCTGAACTCTTTCCCTACACGACGCTCTTCCGATCTNNNTGATGGAGCACTCACTTTAGGA

AGCACACGTCTGAACTCTTTCCCTACACGACGCTCTTCCGATCTNNNTCTGAGATTTTTTCCCATGTGC

AGCACACGTCTGAACTCTTTCCCTACACGACGCTCTTCCGATCTNNNTCTGAGATTTTTTCCCATGTGC

AGCACACGTCTGAACTCTTTCCCTACACGACGCTCTTCCGATCTNNNGGATCTGGTTTCCATTGCCTCT

AGCACACGTCTGAACTCTTTCCCTACACGACGCTCTTCCGATCTNNNGGATCTGGTTTCCATTGCCTCT

AGCACACGTCTGAACTCTTTCCCTACACGACGCTCTTCCGATCTNNNAGGTTTCAGTAAGCCAAGATC

AGCACACGTCTGAACTCTTTCCCTACACGACGCTCTTCCGATCTNNNAGGTTTCAGTAAGCCAAGATC

GCACACGTCTGAACTCTTTCCCTACACGACGCTCTTCCGATCTNNNGTCAAGTCTCCAGAGAGCAACA

GCACACGTCTGAACTCTTTCCCTACACGACGCTCTTCCGATCTNNNGTCAAGTCTCCAGAGAGCAACA

GCACACGTCTGAACTCTTTCCCTACACGACGCTCTTCCGATCTNNNTTCTTTCCTGAGTTCAGTCAGCA

GCACACGTCTGAACTCTTTCCCTACACGACGCTCTTCCGATCTNNNTTCTTTCCTGAGTTCAGTCAGCA

GCACACGTCTGAACTCTTTCCCTACACGACGCTCTTCCGATCTNNNGAGGTTGCAGTGAGCTGAGAT

GCACACGTCTGAACTCTTTCCCTACACGACGCTCTTCCGATCTNNNGAGGTTGCAGTGAGCTGAGAT

GAGCACACGTCTGAACTCTTTCCCTACACGACGCTCTTCCGATCTNNNATCAATCCAAGTGTCCGTCAT

GAGCACACGTCTGAACTCTTTCCCTACACGACGCTCTTCCGATCTNNNATCAATCCAAGTGTCCGTCAT

GCACACGTCTGAACTCTTTCCCTACACGACGCTCTTCCGATCTNNNGCCAGAAAAAAAACAGGGGAAC

GCACACGTCTGAACTCTTTCCCTACACGACGCTCTTCCGATCTNNNGCCAGAAAAAAAACAGGGGAAC

AGCACACGTCTGAACTCTTTCCCTACACGACGCTCTTCCGATCTNNNATAGTGAGTTCAGAAAGCCCA

AGCACACGTCTGAACTCTTTCCCTACACGACGCTCTTCCGATCTNNNATAGTGAGTTCAGAAAGCCCA

AGCACACGTCTGAACTCTTTCCCTACACGACGCTCTTCCGATCTNNNCACAGTGGCTCACAACTGTAA

AGCACACGTCTGAACTCTTTCCCTACACGACGCTCTTCCGATCTNNNCACAGTGGCTCACAACTGTAA

GCACACGTCTGAACTCTTTCCCTACACGACGCTCTTCCGATCTNNNCTGAATCCTGGTCTACTGCTAC

GCACACGTCTGAACTCTTTCCCTACACGACGCTCTTCCGATCTNNNCTGAATCCTGGTCTACTGCTAC

GCACACGTCTGAACTCTTTCCCTACACGACGCTCTTCCGATCTNNNAAAATTAGCCAGGCGTGGTGA

GCACACGTCTGAACTCTTTCCCTACACGACGCTCTTCCGATCTNNNAAAATTAGCCAGGCGTGGTGA

GCACACGTCTGAACTCTTTCCCTACACGACGCTCTTCCGATCTNNNTAGTCCCAGCTACTCCGGAAGC

AGCACACGTCTGAACTCTTTCCCTACACGACGCTCTTCCGATCTNNNTAGTCCCAGCTACTCCGGAAGCA
GCACACGTCTGAACTCTTTCCCTACACGACGCTCTTCCGATCTNNNGGACCTACCAAGCAGTCCTAATT
GCACACGTCTGAACTCTTTCCCTACACGACGCTCTTCCGATCTNNNGGACCTACCAAGCAGTCCTAATT
GAGCACACGTCTGAACTCTTTCCCTACACGACGCTCTTCCGATCTNNNCAGATCACGTGAGATCAGGA
GAGCACACGTCTGAACTCTTTCCCTACACGACGCTCTTCCGATCTNNNCAGATCACGTGAGATCAGGA
GCACACGTCTGAACTCTTTCCCTACACGACGCTCTTCCGATCTNNNTCAGCTGCCAGCAAATATAAAGC
GCACACGTCTGAACTCTTTCCCTACACGACGCTCTTCCGATCTNNNTCAGCTGCCAGCAAATATAAAGC
AGCACACGTCTGAACTCTTTCCCTACACGACGCTCTTCCGATCTNNNTCACACATCTTTCTCTCCTCTGCC
AGCACACGTCTGAACTCTTTCCCTACACGACGCTCTTCCGATCTNNNTCACACATCTTTCTCTCCTCTGCC
GAGCACACGTCTGAACTCTTTCCCTACACGACGCTCTTCCGATCTNNNTGCCTGTAGTCCCAGCTACTC
GAGCACACGTCTGAACTCTTTCCCTACACGACGCTCTTCCGATCTNNNTGCCTGTAGTCCCAGCTACTC
AGCACACGTCTGAACTCTTTCCCTACACGACGCTCTTCCGATCTNNNTATATGCAGCTTTCCAGGGACA
AGCACACGTCTGAACTCTTTCCCTACACGACGCTCTTCCGATCTNNNTATATGCAGCTTTCCAGGGACA
AGCACACGTCTGAACTCTTTCCCTACACGACGCTCTTCCGATCTNNNATCTTGGCTTTTTGCAGGATGA
AGCACACGTCTGAACTCTTTCCCTACACGACGCTCTTCCGATCTNNNATCTTGGCTTTTTGCAGGATGA
AGCACACGTCTGAACTCTTTCCCTACACGACGCTCTTCCGATCTNNNTTCATATTCACGGCTGATCCAG
AGCACACGTCTGAACTCTTTCCCTACACGACGCTCTTCCGATCTNNNTTCATATTCACGGCTGATCCAG
AGCACACGTCTGAACTCTTTCCCTACACGACGCTCTTCCGATCTNNNTTGTTTCATCCTTGCAGAAGAG
AGCACACGTCTGAACTCTTTCCCTACACGACGCTCTTCCGATCTNNNTTGTTTCATCCTTGCAGAAGAG
GCACACGTCTGAACTCTTTCCCTACACGACGCTCTTCCGATCTNNNTTTGCCTCTGTCTCTTCCTCTCCA
GCACACGTCTGAACTCTTTCCCTACACGACGCTCTTCCGATCTNNNTTTGCCTCTGTCTCTTCCTCTCCA
GCACACGTCTGAACTCTTTCCCTACACGACGCTCTTCCGATCTNNNGACTATTGTCAACCAACACACGCC
GCACACGTCTGAACTCTTTCCCTACACGACGCTCTTCCGATCTNNNGACTATTGTCAACCAACACACGCC
AGCACACGTCTGAACTCTTTCCCTACACGACGCTCTTCCGATCTNNNAGGTACTGGGGATTAGGACTTC
AGCACACGTCTGAACTCTTTCCCTACACGACGCTCTTCCGATCTNNNAGGTACTGGGGATTAGGACTTC
AGCACACGTCTGAACTCTTTCCCTACACGACGCTCTTCCGATCTNNNTATGAGGAAGAGCTGACCGACA
AGCACACGTCTGAACTCTTTCCCTACACGACGCTCTTCCGATCTNNNTATGAGGAAGAGCTGACCGACA
AGCACACGTCTGAACTCTTTCCCTACACGACGCTCTTCCGATCTNNNTTAGCCTTGTGAGTAGTTGGGA
AGCACACGTCTGAACTCTTTCCCTACACGACGCTCTTCCGATCTNNNTTAGCCTTGTGAGTAGTTGGGA
AGCACACGTCTGAACTCTTTCCCTACACGACGCTCTTCCGATCTNNNATGTTGGTCAGTCTGGTCTTGA
AGCACACGTCTGAACTCTTTCCCTACACGACGCTCTTCCGATCTNNNATGTTGGTCAGTCTGGTCTTGA
AGAGCACACGTCTGAACTCTTTCCCTACACGACGCTCTTCCGATCTNNNCTGAGGCAGGAGAATCACTT
AGAGCACACGTCTGAACTCTTTCCCTACACGACGCTCTTCCGATCTNNNCTGAGGCAGGAGAATCACTT
GAGCACACGTCTGAACTCTTTCCCTACACGACGCTCTTCCGATCTNNNACTACTTGCTGTTGTTACACTT
GAGCACACGTCTGAACTCTTTCCCTACACGACGCTCTTCCGATCTNNNACTACTTGCTGTTGTTACACTT
AGCACACGTCTGAACTCTTTCCCTACACGACGCTCTTCCGATCTNNNTCTCTTGATGTTTGCAGTTTTCA
AGCACACGTCTGAACTCTTTCCCTACACGACGCTCTTCCGATCTNNNTCTCTTGATGTTTGCAGTTTTCA
GAGCACACGTCTGAACTCTTTCCCTACACGACGCTCTTCCGATCTNNNAAGGAGAGGAAAGAGTTTAG
GAGCACACGTCTGAACTCTTTCCCTACACGACGCTCTTCCGATCTNNNAAGGAGAGGAAAGAGTTTAG
GAGCACACGTCTGAACTCTTTCCCTACACGACGCTCTTCCGATCTNNNAAGATTTGAGAAGGGCCATTC
GAGCACACGTCTGAACTCTTTCCCTACACGACGCTCTTCCGATCTNNNAAGATTTGAGAAGGGCCATTC
AGCACACGTCTGAACTCTTTCCCTACACGACGCTCTTCCGATCTNNNCTCCTGAGTAGCTGGGACTACA
AGCACACGTCTGAACTCTTTCCCTACACGACGCTCTTCCGATCTNNNCTCCTGAGTAGCTGGGACTACA

GAGCACACGTCTGAACTCTTTCCCTACACGACGCTCTTCCGATCTNNNTCTGCTTCCTGAGTTATCTGCA
GAGCACACGTCTGAACTCTTTCCCTACACGACGCTCTTCCGATCTNNNTCTGCTTCCTGAGTTATCTGCA
GAGCACACGTCTGAACTCTTTCCCTACACGACGCTCTTCCGATCTNNNATATAGGAGGGAGGGAAGA
GAGCACACGTCTGAACTCTTTCCCTACACGACGCTCTTCCGATCTNNNATATAGGAGGGAGGGAAGA
AGCACACGTCTGAACTCTTTCCCTACACGACGCTCTTCCGATCTNNNCTCTTCCCTGACCTTGGCTTCCA
AGCACACGTCTGAACTCTTTCCCTACACGACGCTCTTCCGATCTNNNCTCTTCCCTGACCTTGGCTTCCA
AGAGCACACGTCTGAACTCTTTCCCTACACGACGCTCTTCCGATCTNNNTTACATTTTGAAAGACTGGTT
AGAGCACACGTCTGAACTCTTTCCCTACACGACGCTCTTCCGATCTNNNTTACATTTTGAAAGACTGGTT
GAGCACACGTCTGAACTCTTTCCCTACACGACGCTCTTCCGATCTNNNAGGCCACTCACATCACACAAT
GAGCACACGTCTGAACTCTTTCCCTACACGACGCTCTTCCGATCTNNNAGGCCACTCACATCACACAAT
GAGCACACGTCTGAACTCTTTCCCTACACGACGCTCTTCCGATCTNNNGACGAAACTAAGGTGTGGTTT
GAGCACACGTCTGAACTCTTTCCCTACACGACGCTCTTCCGATCTNNNGACGAAACTAAGGTGTGGTTT
GAGCACACGTCTGAACTCTTTCCCTACACGACGCTCTTCCGATCTNNNTGTCTTTTCTTGTCATTAATAGC
GAGCACACGTCTGAACTCTTTCCCTACACGACGCTCTTCCGATCTNNNTGTCTTTTCTTGTCATTAATAGC
GAGCACACGTCTGAACTCTTTCCCTACACGACGCTCTTCCGATCTNNNGGAAACCTAGCAAGACTCTGTC
GAGCACACGTCTGAACTCTTTCCCTACACGACGCTCTTCCGATCTNNNGGAAACCTAGCAAGACTCTGTC
AGCACACGTCTGAACTCTTTCCCTACACGACGCTCTTCCGATCTNNNCAGCTCTGAGATAAGACTGCTG
AGCACACGTCTGAACTCTTTCCCTACACGACGCTCTTCCGATCTNNNCAGCTCTGAGATAAGACTGCTG
GCACACGTCTGAACTCTTTCCCTACACGACGCTCTTCCGATCTNNNCTCATTCCACCTCAGCTCGAGCAT
GCACACGTCTGAACTCTTTCCCTACACGACGCTCTTCCGATCTNNNCTCATTCCACCTCAGCTCGAGCAT
AGCACACGTCTGAACTCTTTCCCTACACGACGCTCTTCCGATCTNNNAAATCCGAGGTCTGAAACCTGA
AGCACACGTCTGAACTCTTTCCCTACACGACGCTCTTCCGATCTNNNAAATCCGAGGTCTGAAACCTGA
AGAGCACACGTCTGAACTCTTTCCCTACACGACGCTCTTCCGATCTNNNTCACTAAAGGAAGACAGGA
AGAGCACACGTCTGAACTCTTTCCCTACACGACGCTCTTCCGATCTNNNTCACTAAAGGAAGACAGGA
GAGCACACGTCTGAACTCTTTCCCTACACGACGCTCTTCCGATCTNNNAGTTCCCCAACAAGTACTACCA
GAGCACACGTCTGAACTCTTTCCCTACACGACGCTCTTCCGATCTNNNAGTTCCCCAACAAGTACTACCA
GCACACGTCTGAACTCTTTCCCTACACGACGCTCTTCCGATCTNNNCATCACCATGCACAGCTAACATAC
GCACACGTCTGAACTCTTTCCCTACACGACGCTCTTCCGATCTNNNCATCACCATGCACAGCTAACATAC
GCACACGTCTGAACTCTTTCCCTACACGACGCTCTTCCGATCTNNNGGCTAGCTTATTTTGCTGAGCATC
GCACACGTCTGAACTCTTTCCCTACACGACGCTCTTCCGATCTNNNGGCTAGCTTATTTTGCTGAGCATC
GAGCACACGTCTGAACTCTTTCCCTACACGACGCTCTTCCGATCTNNNGTGAAACCCCTTCTCTACCAAA
GAGCACACGTCTGAACTCTTTCCCTACACGACGCTCTTCCGATCTNNNGTGAAACCCCTTCTCTACCAAA
GAGCACACGTCTGAACTCTTTCCCTACACGACGCTCTTCCGATCTNNNGGACCCAAAATCAATGCACAAC
GAGCACACGTCTGAACTCTTTCCCTACACGACGCTCTTCCGATCTNNNGGACCCAAAATCAATGCACAAC
GAGCACACGTCTGAACTCTTTCCCTACACGACGCTCTTCCGATCTNNNTAATCCCAGCTACTCAGGAGA
GAGCACACGTCTGAACTCTTTCCCTACACGACGCTCTTCCGATCTNNNTAATCCCAGCTACTCAGGAGA
AGCACACGTCTGAACTCTTTCCCTACACGACGCTCTTCCGATCTNNNCTTCCTTCCTTCCTTCTTCCCTCCA
AGCACACGTCTGAACTCTTTCCCTACACGACGCTCTTCCGATCTNNNCTTCCTTCCTTCCTTCTTCCCTCCA
GAGCACACGTCTGAACTCTTTCCCTACACGACGCTCTTCCGATCTNNNCCTTCTCCGTCTCCTTCTTCTTC
GAGCACACGTCTGAACTCTTTCCCTACACGACGCTCTTCCGATCTNNNCCTTCTCCGTCTCCTTCTTCTTC
AGCACACGTCTGAACTCTTTCCCTACACGACGCTCTTCCGATCTNNNCAAATGGCTTAGCTGACTGCCCA
AGCACACGTCTGAACTCTTTCCCTACACGACGCTCTTCCGATCTNNNCAAATGGCTTAGCTGACTGCCCA
GAGCACACGTCTGAACTCTTTCCCTACACGACGCTCTTCCGATCTNNNCATACATGCACCCATTCATTCCC

GAGCACACGTCTGAACTCTTTCCCTACACGACGCTCTTCCGATCTNNNCATACATGCACCCATTCATTCCC
AGCACACGTCTGAACTCTTTCCCTACACGACGCTCTTCCGATCTNNNCTGAGGCAGAGAATTGCTTGA
AGCACACGTCTGAACTCTTTCCCTACACGACGCTCTTCCGATCTNNNCTGAGGCAGAGAATTGCTTGA
GCACACGTCTGAACTCTTTCCCTACACGACGCTCTTCCGATCTNNNTTTTCTTTTTTTCAGACAGAGTCA
GCACACGTCTGAACTCTTTCCCTACACGACGCTCTTCCGATCTNNNTTTTCTTTTTTTCAGACAGAGTCA
GAGCACACGTCTGAACTCTTTCCCTACACGACGCTCTTCCGATCTNNNTACTCATCATCCTTGCCAGCAT
GAGCACACGTCTGAACTCTTTCCCTACACGACGCTCTTCCGATCTNNNTACTCATCATCCTTGCCAGCAT
AGCACACGTCTGAACTCTTTCCCTACACGACGCTCTTCCGATCTNNNTCTGAGTCTCAGCCTTACTCTGA
AGCACACGTCTGAACTCTTTCCCTACACGACGCTCTTCCGATCTNNNTCTGAGTCTCAGCCTTACTCTGA
GAGCACACGTCTGAACTCTTTCCCTACACGACGCTCTTCCGATCTNNNAGGCCAATTATTTCTCCCCTGA
GAGCACACGTCTGAACTCTTTCCCTACACGACGCTCTTCCGATCTNNNAGGCCAATTATTTCTCCCCTGA
AGAGCACACGTCTGAACTCTTTCCCTACACGACGCTCTTCCGATCTNNNCCCTCTTTGAAGTGAAGACAC
AGAGCACACGTCTGAACTCTTTCCCTACACGACGCTCTTCCGATCTNNNCCCTCTTTGAAGTGAAGACAC
AGCACACGTCTGAACTCTTTCCCTACACGACGCTCTTCCGATCTNNNCAAGCTGGAGTTCAGTGGTATC
AGCACACGTCTGAACTCTTTCCCTACACGACGCTCTTCCGATCTNNNCAAGCTGGAGTTCAGTGGTATC
GCACACGTCTGAACTCTTTCCCTACACGACGCTCTTCCGATCTNNNTTCTAGGAAGTGGGTTGAGAAG
GCACACGTCTGAACTCTTTCCCTACACGACGCTCTTCCGATCTNNNTTCTAGGAAGTGGGTTGAGAAG
GAGCACACGTCTGAACTCTTTCCCTACACGACGCTCTTCCGATCTNNNTTAACATTTGATGGTTTGAAA
GAGCACACGTCTGAACTCTTTCCCTACACGACGCTCTTCCGATCTNNNTTAACATTTGATGGTTTGAAA
AGCACACGTCTGAACTCTTTCCCTACACGACGCTCTTCCGATCTNNNTAAACTCCTTGCCTCTCACATCCC
AGCACACGTCTGAACTCTTTCCCTACACGACGCTCTTCCGATCTNNNTAAACTCCTTGCCTCTCACATCCC
ACACGTCTGAACTCTTTCCCTACACGACGCTCTTCCGATCTNNNAAAGAGACAGAAGCCTTCCCAGCAT
ACACGTCTGAACTCTTTCCCTACACGACGCTCTTCCGATCTNNNAAAGAGACAGAAGCCTTCCCAGCAT
AAGAGCACACGTCTGAACTCTTTCCCTACACGACGCTCTTCCGATCTNNNCTTGCAGTGAGTCGAGTTT
AAGAGCACACGTCTGAACTCTTTCCCTACACGACGCTCTTCCGATCTNNNCTTGCAGTGAGTCGAGTTT
GCACACGTCTGAACTCTTTCCCTACACGACGCTCTTCCGATCTNNNAGCGTAATCTAACGGAGGAGTGA
GCACACGTCTGAACTCTTTCCCTACACGACGCTCTTCCGATCTNNNAGCGTAATCTAACGGAGGAGTGA
GAGCACACGTCTGAACTCTTTCCCTACACGACGCTCTTCCGATCTNNNGTTGCAGTGAGCTGAGATCA
GAGCACACGTCTGAACTCTTTCCCTACACGACGCTCTTCCGATCTNNNGTTGCAGTGAGCTGAGATCA
GCACACGTCTGAACTCTTTCCCTACACGACGCTCTTCCGATCTNNNCGCAGCATGGTGGAGGTGCTCAT
GCACACGTCTGAACTCTTTCCCTACACGACGCTCTTCCGATCTNNNCGCAGCATGGTGGAGGTGCTCAT
AGCACACGTCTGAACTCTTTCCCTACACGACGCTCTTCCGATCTNNNCACCTGTAATCCCAGCTACTTGG
AGCACACGTCTGAACTCTTTCCCTACACGACGCTCTTCCGATCTNNNCACCTGTAATCCCAGCTACTTGG
AGCACACGTCTGAACTCTTTCCCTACACGACGCTCTTCCGATCTNNNTGCGGCTAAGATGTCAAAGTAC
AGCACACGTCTGAACTCTTTCCCTACACGACGCTCTTCCGATCTNNNTGCGGCTAAGATGTCAAAGTAC
AGCACACGTCTGAACTCTTTCCCTACACGACGCTCTTCCGATCTNNNAAGGTTTTGGTGAGCCATGAT
AGCACACGTCTGAACTCTTTCCCTACACGACGCTCTTCCGATCTNNNAAGGTTTTGGTGAGCCATGAT
GCACACGTCTGAACTCTTTCCCTACACGACGCTCTTCCGATCTNNNTCTAAATACACACACACGCATGCC
GCACACGTCTGAACTCTTTCCCTACACGACGCTCTTCCGATCTNNNTCTAAATACACACACACGCATGCC
GAGCACACGTCTGAACTCTTTCCCTACACGACGCTCTTCCGATCTNNNCACAATCTCGGCTCATTGCAAG
GAGCACACGTCTGAACTCTTTCCCTACACGACGCTCTTCCGATCTNNNCACAATCTCGGCTCATTGCAAG
AGAGCACACGTCTGAACTCTTTCCCTACACGACGCTCTTCCGATCTNNNGCCTGTAATCCCAGCTACTCT
AGAGCACACGTCTGAACTCTTTCCCTACACGACGCTCTTCCGATCTNNNGCCTGTAATCCCAGCTACTCT

AGCACACGTCTGAACTCTTTCCCTACACGACGCTCTTCCGATCTNNNAAGACTGTCCTTTCAGTGCTCAT

AGCACACGTCTGAACTCTTTCCCTACACGACGCTCTTCCGATCTNNNAAGACTGTCCTTTCAGTGCTCAT

AGCACACGTCTGAACTCTTTCCCTACACGACGCTCTTCCGATCTNNNCACCCTATGGCTGAAGATGTCT

AGCACACGTCTGAACTCTTTCCCTACACGACGCTCTTCCGATCTNNNCACCCTATGGCTGAAGATGTCT

GAGCACACGTCTGAACTCTTTCCCTACACGACGCTCTTCCGATCTNNNCATACCCACCACTACCAGACAC

GAGCACACGTCTGAACTCTTTCCCTACACGACGCTCTTCCGATCTNNNCATACCCACCACTACCAGACAC

AGAGCACACGTCTGAACTCTTTCCCTACACGACGCTCTTCCGATCTNNNGAGAATTCCTTGAACCTGGG

AGAGCACACGTCTGAACTCTTTCCCTACACGACGCTCTTCCGATCTNNNGAGAATTCCTTGAACCTGGG

GAGCACACGTCTGAACTCTTTCCCTACACGACGCTCTTCCGATCTNNNTACGTATACACACACACACACA

GAGCACACGTCTGAACTCTTTCCCTACACGACGCTCTTCCGATCTNNNTACGTATACACACACACACACA

AGCACACGTCTGAACTCTTTCCCTACACGACGCTCTTCCGATCTNNNTTGAGGCAGGAGAATCACTTG

AGCACACGTCTGAACTCTTTCCCTACACGACGCTCTTCCGATCTNNNTTGAGGCAGGAGAATCACTTG

GCACACGTCTGAACTCTTTCCCTACACGACGCTCTTCCGATCTNNNTCCAGATCCTCACCAATCTCTCCCA

GCACACGTCTGAACTCTTTCCCTACACGACGCTCTTCCGATCTNNNTCCAGATCCTCACCAATCTCTCCCA

AGAGCACACGTCTGAACTCTTTCCCTACACGACGCTCTTCCGATCTNNNAGTACAAGTGGAGGCCATC

AGAGCACACGTCTGAACTCTTTCCCTACACGACGCTCTTCCGATCTNNNAGTACAAGTGGAGGCCATC

AGAGCACACGTCTGAACTCTTTCCCTACACGACGCTCTTCCGATCTNNNGAGGCTGAATCCCATAAGCA

AGAGCACACGTCTGAACTCTTTCCCTACACGACGCTCTTCCGATCTNNNGAGGCTGAATCCCATAAGCA

AGAGCACACGTCTGAACTCTTTCCCTACACGACGCTCTTCCGATCTNNNATGAACATGAGGTCATCCCA

AGAGCACACGTCTGAACTCTTTCCCTACACGACGCTCTTCCGATCTNNNATGAACATGAGGTCATCCCA

GCACACGTCTGAACTCTTTCCCTACACGACGCTCTTCCGATCTNNNTAGAGTAGGAATCAGCTTGCCAC

GCACACGTCTGAACTCTTTCCCTACACGACGCTCTTCCGATCTNNNTAGAGTAGGAATCAGCTTGCCAC

GAGCACACGTCTGAACTCTTTCCCTACACGACGCTCTTCCGATCTNNNCTGAGGCAGGAGAATAGCTTC

GAGCACACGTCTGAACTCTTTCCCTACACGACGCTCTTCCGATCTNNNCTGAGGCAGGAGAATAGCTTC

GAGCACACGTCTGAACTCTTTCCCTACACGACGCTCTTCCGATCTNNNTCCCTCAAGACCTACGTGAATC

GAGCACACGTCTGAACTCTTTCCCTACACGACGCTCTTCCGATCTNNNTCCCTCAAGACCTACGTGAATC

GAGCACACGTCTGAACTCTTTCCCTACACGACGCTCTTCCGATCTNNNGTGTGTGTGTGTGTGTGTGTA

GAGCACACGTCTGAACTCTTTCCCTACACGACGCTCTTCCGATCTNNNGTGTGTGTGTGTGTGTGTGTA

GAGCACACGTCTGAACTCTTTCCCTACACGACGCTCTTCCGATCTNNNCACCTGTAATCCCAGCTACTCAC

GAGCACACGTCTGAACTCTTTCCCTACACGACGCTCTTCCGATCTNNNCACCTGTAATCCCAGCTACTCAC

AGCACACGTCTGAACTCTTTCCCTACACGACGCTCTTCCGATCTNNNGTAATCCTAGCTACTTGGGAGC

AGCACACGTCTGAACTCTTTCCCTACACGACGCTCTTCCGATCTNNNGTAATCCTAGCTACTTGGGAGC

AGCACACGTCTGAACTCTTTCCCTACACGACGCTCTTCCGATCTNNNATCAGCTCCTCTTTGTACCTGTC

AGCACACGTCTGAACTCTTTCCCTACACGACGCTCTTCCGATCTNNNATCAGCTCCTCTTTGTACCTGTC

GAGCACACGTCTGAACTCTTTCCCTACACGACGCTCTTCCGATCTNNNGCAATTAGAGAAGAGAGAGG

GAGCACACGTCTGAACTCTTTCCCTACACGACGCTCTTCCGATCTNNNGCAATTAGAGAAGAGAGAGG

AGCACACGTCTGAACTCTTTCCCTACACGACGCTCTTCCGATCTNNNAGGAAGTCATGTGCATGTCATC

AGCACACGTCTGAACTCTTTCCCTACACGACGCTCTTCCGATCTNNNAGGAAGTCATGTGCATGTCATC

AGAGCACACGTCTGAACTCTTTCCCTACACGACGCTCTTCCGATCTNNNACACCTGTAATCCCAGCTACT

AGAGCACACGTCTGAACTCTTTCCCTACACGACGCTCTTCCGATCTNNNACACCTGTAATCCCAGCTACT

GAGCACACGTCTGAACTCTTTCCCTACACGACGCTCTTCCGATCTNNNCTTTCAGGGTGACGTTGAGGC

GAGCACACGTCTGAACTCTTTCCCTACACGACGCTCTTCCGATCTNNNCTTTCAGGGTGACGTTGAGGC

AGCACACGTCTGAACTCTTTCCCTACACGACGCTCTTCCGATCTNNNACACCTCAGATATATTTCTTCATC

AGCACACGTCTGAACTCTTTCCCTACACGACGCTCTTCCGATCTNNNACACCTCAGATATATTTCTTCAT
GAGCACACGTCTGAACTCTTTCCCTACACGACGCTCTTCCGATCTNNNGCTTATATGCACATTGGGAG
GAGCACACGTCTGAACTCTTTCCCTACACGACGCTCTTCCGATCTNNNGCTTATATGCACATTGGGAG
GAGCACACGTCTGAACTCTTTCCCTACACGACGCTCTTCCGATCTNNNCTCTGCCTCAGAATTAGGTGG
GAGCACACGTCTGAACTCTTTCCCTACACGACGCTCTTCCGATCTNNNCTCTGCCTCAGAATTAGGTGG
AGAGCACACGTCTGAACTCTTTCCCTACACGACGCTCTTCCGATCTNNNGAGGCTGAGGCAGGAGAA
AGAGCACACGTCTGAACTCTTTCCCTACACGACGCTCTTCCGATCTNNNGAGGCTGAGGCAGGAGAA
GAGCACACGTCTGAACTCTTTCCCTACACGACGCTCTTCCGATCTNNNGTGGTCTGGAAATGAGGATG
GAGCACACGTCTGAACTCTTTCCCTACACGACGCTCTTCCGATCTNNNGTGGTCTGGAAATGAGGATG
AGCACACGTCTGAACTCTTTCCCTACACGACGCTCTTCCGATCTNNNTCAGGTATGCTTCCATTTAATAG
AGCACACGTCTGAACTCTTTCCCTACACGACGCTCTTCCGATCTNNNTCAGGTATGCTTCCATTTAATAG
AGCACACGTCTGAACTCTTTCCCTACACGACGCTCTTCCGATCTNNNTGTCTCTTTACTTGCCTGTGCTC
AGCACACGTCTGAACTCTTTCCCTACACGACGCTCTTCCGATCTNNNTGTCTCTTTACTTGCCTGTGCTC
GCACACGTCTGAACTCTTTCCCTACACGACGCTCTTCCGATCTNNNTGTTTGTTGGCTGCATAAATGAC
GCACACGTCTGAACTCTTTCCCTACACGACGCTCTTCCGATCTNNNTGTTTGTTGGCTGCATAAATGAC
GAGCACACGTCTGAACTCTTTCCCTACACGACGCTCTTCCGATCTNNNTAACCCCAGATACTCAGGATG
GAGCACACGTCTGAACTCTTTCCCTACACGACGCTCTTCCGATCTNNNTAACCCCAGATACTCAGGATG
AGCACACGTCTGAACTCTTTCCCTACACGACGCTCTTCCGATCTNNNTCTTACCCAGCACAGTCATAGT
AGCACACGTCTGAACTCTTTCCCTACACGACGCTCTTCCGATCTNNNTCTTACCCAGCACAGTCATAGT
GAGCACACGTCTGAACTCTTTCCCTACACGACGCTCTTCCGATCTNNNGAGATCACGCCACTGCACTTT
GAGCACACGTCTGAACTCTTTCCCTACACGACGCTCTTCCGATCTNNNGAGATCACGCCACTGCACTTT
GAGCACACGTCTGAACTCTTTCCCTACACGACGCTCTTCCGATCTNNNGGACGTACTCTCCTTTTATGG
GAGCACACGTCTGAACTCTTTCCCTACACGACGCTCTTCCGATCTNNNGGACGTACTCTCCTTTTATGG
ACACGTCTGAACTCTTTCCCTACACGACGCTCTTCCGATCTNNNTTCTCAATGAGCTGTTGGGTACACA
ACACGTCTGAACTCTTTCCCTACACGACGCTCTTCCGATCTNNNTTCTCAATGAGCTGTTGGGTACACA
AGCACACGTCTGAACTCTTTCCCTACACGACGCTCTTCCGATCTNNNACACACAAGTCACAAATTAGCC
AGCACACGTCTGAACTCTTTCCCTACACGACGCTCTTCCGATCTNNNACACACAAGTCACAAATTAGCC
GAAGAGCACACGTCTGAACTCTTTCCCTACACGACGCTCTTCCGATCTNNNCAGCTATAATACCCCTCCC
GAAGAGCACACGTCTGAACTCTTTCCCTACACGACGCTCTTCCGATCTNNNCAGCTATAATACCCCTCCC
AGCACACGTCTGAACTCTTTCCCTACACGACGCTCTTCCGATCTNNNGTTCACCTTGCTCAGTAAACCA
AGCACACGTCTGAACTCTTTCCCTACACGACGCTCTTCCGATCTNNNGTTCACCTTGCTCAGTAAACCA
GCACACGTCTGAACTCTTTCCCTACACGACGCTCTTCCGATCTNNNGATGGTGCAACAGTTTCAGTAC
GCACACGTCTGAACTCTTTCCCTACACGACGCTCTTCCGATCTNNNGATGGTGCAACAGTTTCAGTAC
GAGCACACGTCTGAACTCTTTCCCTACACGACGCTCTTCCGATCTNNNACCCCAATTTCTAGACCTGCAT
GAGCACACGTCTGAACTCTTTCCCTACACGACGCTCTTCCGATCTNNNACCCCAATTTCTAGACCTGCAT
AGAGCACACGTCTGAACTCTTTCCCTACACGACGCTCTTCCGATCTNNNAAGAGATCTCTGGTTGGTT
AGAGCACACGTCTGAACTCTTTCCCTACACGACGCTCTTCCGATCTNNNAAGAGATCTCTGGTTGGTT
GAGCACACGTCTGAACTCTTTCCCTACACGACGCTCTTCCGATCTNNNCAGGAAGGAAGGAAGGAAG
GAGCACACGTCTGAACTCTTTCCCTACACGACGCTCTTCCGATCTNNNCAGGAAGGAAGGAAGGAAG
AGAGCACACGTCTGAACTCTTTCCCTACACGACGCTCTTCCGATCTNNNTCCCCAATTATATGCTTATAT
AGAGCACACGTCTGAACTCTTTCCCTACACGACGCTCTTCCGATCTNNNTCCCCAATTATATGCTTATAT
AGCACACGTCTGAACTCTTTCCCTACACGACGCTCTTCCGATCTNNNGTCGTCACATTACAGAAGGGGT
AGCACACGTCTGAACTCTTTCCCTACACGACGCTCTTCCGATCTNNNGTCGTCACATTACAGAAGGGGT

GAGCACACGTCTGAACTCTTTCCCTACACGACGCTCTTCCGATCTNNNAAAAATTAGCCAGGCTTGGT
GAGCACACGTCTGAACTCTTTCCCTACACGACGCTCTTCCGATCTNNNAAAAATTAGCCAGGCTTGGT
AGCACACGTCTGAACTCTTTCCCTACACGACGCTCTTCCGATCTNNNGAGATTGCTCCATTACGCTCCA
AGCACACGTCTGAACTCTTTCCCTACACGACGCTCTTCCGATCTNNNGAGATTGCTCCATTACGCTCCA
GCACACGTCTGAACTCTTTCCCTACACGACGCTCTTCCGATCTNNNTGAGAATCACAATCACTGCACAT
GCACACGTCTGAACTCTTTCCCTACACGACGCTCTTCCGATCTNNNTGAGAATCACAATCACTGCACAT
ACACGTCTGAACTCTTTCCCTACACGACGCTCTTCCGATCTNNNTATTCCATGGTGTATGTGTGCCACA
ACACGTCTGAACTCTTTCCCTACACGACGCTCTTCCGATCTNNNTATTCCATGGTGTATGTGTGCCACA
GAGCACACGTCTGAACTCTTTCCCTACACGACGCTCTTCCGATCTNNNACCTGAAAACTCTTGGTGGT
GAGCACACGTCTGAACTCTTTCCCTACACGACGCTCTTCCGATCTNNNACCTGAAAACTCTTGGTGGT
GCACACGTCTGAACTCTTTCCCTACACGACGCTCTTCCGATCTNNNTTGGGATCTCAACTCATCTGACC
GCACACGTCTGAACTCTTTCCCTACACGACGCTCTTCCGATCTNNNTTGGGATCTCAACTCATCTGACC
GAGCACACGTCTGAACTCTTTCCCTACACGACGCTCTTCCGATCTNNNCTGTAATCCCAGCTACTCAGG
GAGCACACGTCTGAACTCTTTCCCTACACGACGCTCTTCCGATCTNNNCTGTAATCCCAGCTACTCAGG
GAGCACACGTCTGAACTCTTTCCCTACACGACGCTCTTCCGATCTNNNATGCTGCCACCATCTTAGAGA
GAGCACACGTCTGAACTCTTTCCCTACACGACGCTCTTCCGATCTNNNATGCTGCCACCATCTTAGAGA
GCACACGTCTGAACTCTTTCCCTACACGACGCTCTTCCGATCTNNNTGGAGGGAAGAATCTCAGAGCT
GCACACGTCTGAACTCTTTCCCTACACGACGCTCTTCCGATCTNNNTGGAGGGAAGAATCTCAGAGCT
AGCACACGTCTGAACTCTTTCCCTACACGACGCTCTTCCGATCTNNNAGCCAATATGACTTATTTACAA
AGCACACGTCTGAACTCTTTCCCTACACGACGCTCTTCCGATCTNNNAGCCAATATGACTTATTTACAA
GAGCACACGTCTGAACTCTTTCCCTACACGACGCTCTTCCGATCTNNNCTGTAGTCCCAGCTACTTGAG
GAGCACACGTCTGAACTCTTTCCCTACACGACGCTCTTCCGATCTNNNCTGTAGTCCCAGCTACTTGAG
GAGCACACGTCTGAACTCTTTCCCTACACGACGCTCTTCCGATCTNNNCCTGATTACCTGCCCCTACAAT
GAGCACACGTCTGAACTCTTTCCCTACACGACGCTCTTCCGATCTNNNCCTGATTACCTGCCCCTACAAT
AGCACACGTCTGAACTCTTTCCCTACACGACGCTCTTCCGATCTNNNAAGTGGATTAGACCAGCCTTCT
AGCACACGTCTGAACTCTTTCCCTACACGACGCTCTTCCGATCTNNNAAGTGGATTAGACCAGCCTTCT
GAGCACACGTCTGAACTCTTTCCCTACACGACGCTCTTCCGATCTNNNAAAAAAAAATTAGCTGGGCGT
GAGCACACGTCTGAACTCTTTCCCTACACGACGCTCTTCCGATCTNNNAAAAAAAAATTAGCTGGGCGT
AGCACACGTCTGAACTCTTTCCCTACACGACGCTCTTCCGATCTNNNTTTACCCTCTATTTTCACTTCCCA
AGCACACGTCTGAACTCTTTCCCTACACGACGCTCTTCCGATCTNNNTTTACCCTCTATTTTCACTTCCCA
AGCACACGTCTGAACTCTTTCCCTACACGACGCTCTTCCGATCTNNNTTACATTCCAGTCCAGTGAGGT
AGCACACGTCTGAACTCTTTCCCTACACGACGCTCTTCCGATCTNNNTTACATTCCAGTCCAGTGAGGT
AGCACACGTCTGAACTCTTTCCCTACACGACGCTCTTCCGATCTNNNCTGTAGTCCCAGCTACTTGGAA
AGCACACGTCTGAACTCTTTCCCTACACGACGCTCTTCCGATCTNNNCTGTAGTCCCAGCTACTTGGAA
AGCACACGTCTGAACTCTTTCCCTACACGACGCTCTTCCGATCTNNNTTCCTCTTTCAGTGGATGGCTA
AGCACACGTCTGAACTCTTTCCCTACACGACGCTCTTCCGATCTNNNTTCCTCTTTCAGTGGATGGCTA
GAGCACACGTCTGAACTCTTTCCCTACACGACGCTCTTCCGATCTNNNGAGTCTTTGCTTTTAACAAAAT
GAGCACACGTCTGAACTCTTTCCCTACACGACGCTCTTCCGATCTNNNGAGTCTTTGCTTTTAACAAAAT
GAGCACACGTCTGAACTCTTTCCCTACACGACGCTCTTCCGATCTNNNGATCACAAGGTCTGGAGAT
GAGCACACGTCTGAACTCTTTCCCTACACGACGCTCTTCCGATCTNNNGATCACAAGGTCTGGAGAT
AGCACACGTCTGAACTCTTTCCCTACACGACGCTCTTCCGATCTNNNGCTCATGCTTATAATCCCAGCA
AGCACACGTCTGAACTCTTTCCCTACACGACGCTCTTCCGATCTNNNGCTCATGCTTATAATCCCAGCA
GAGCACACGTCTGAACTCTTTCCCTACACGACGCTCTTCCGATCTNNNCTACTCTGGAGAAACAGCGT

GAGCACACGTCTGAACTCTTTCCCTACACGACGCTCTTCCGATCTNNNCTACTCTGGAGAAACAGCGT
GAGCACACGTCTGAACTCTTTCCCTACACGACGCTCTTCCGATCTNNNAAAAAAAAATAGCCGGGTG
GAGCACACGTCTGAACTCTTTCCCTACACGACGCTCTTCCGATCTNNNAAAAAAAAATAGCCGGGTG
GCACACGTCTGAACTCTTTCCCTACACGACGCTCTTCCGATCTNNNGGTTGAGGTCATAGTTGAGGA
GCACACGTCTGAACTCTTTCCCTACACGACGCTCTTCCGATCTNNNGGTTGAGGTCATAGTTGAGGA
AGCACACGTCTGAACTCTTTCCCTACACGACGCTCTTCCGATCTNNNAAGCTCTCATCTTGACGTCTTG
AGCACACGTCTGAACTCTTTCCCTACACGACGCTCTTCCGATCTNNNAAGCTCTCATCTTGACGTCTTG
AGAGCACACGTCTGAACTCTTTCCCTACACGACGCTCTTCCGATCTNNNAGGAAGCAAACTCTTCTCA
AGAGCACACGTCTGAACTCTTTCCCTACACGACGCTCTTCCGATCTNNNAGGAAGCAAACTCTTCTCA
AGCACACGTCTGAACTCTTTCCCTACACGACGCTCTTCCGATCTNNNTGTAGTACCAGCTGCTCAGAA
AGCACACGTCTGAACTCTTTCCCTACACGACGCTCTTCCGATCTNNNTGTAGTACCAGCTGCTCAGAA
GCACACGTCTGAACTCTTTCCCTACACGACGCTCTTCCGATCTNNNTGTATCCATGTCGTAGCATGTG
GCACACGTCTGAACTCTTTCCCTACACGACGCTCTTCCGATCTNNNTGTATCCATGTCGTAGCATGTG
AGCACACGTCTGAACTCTTTCCCTACACGACGCTCTTCCGATCTNNNTGTTCTTTGGGTCCCAGAAAG
AGCACACGTCTGAACTCTTTCCCTACACGACGCTCTTCCGATCTNNNTGTTCTTTGGGTCCCAGAAAG
AGCACACGTCTGAACTCTTTCCCTACACGACGCTCTTCCGATCTNNNTACACAATTCTCCTTGCCACTCT
AGCACACGTCTGAACTCTTTCCCTACACGACGCTCTTCCGATCTNNNTACACAATTCTCCTTGCCACTCT
CACACGTCTGAACTCTTTCCCTACACGACGCTCTTCCGATCTNNNCTGAGGTGGGAGTATCAGCTAA
CACACGTCTGAACTCTTTCCCTACACGACGCTCTTCCGATCTNNNCTGAGGTGGGAGTATCAGCTAA
AGCACACGTCTGAACTCTTTCCCTACACGACGCTCTTCCGATCTNNNGTGCTTGTGTCATTCCACCAAT
AGCACACGTCTGAACTCTTTCCCTACACGACGCTCTTCCGATCTNNNGTGCTTGTGTCATTCCACCAAT
AGCACACGTCTGAACTCTTTCCCTACACGACGCTCTTCCGATCTNNNAGTAGCACCTACCCTTTTTGAG
AGCACACGTCTGAACTCTTTCCCTACACGACGCTCTTCCGATCTNNNAGTAGCACCTACCCTTTTTGAG
CACACGTCTGAACTCTTTCCCTACACGACGCTCTTCCGATCTNNNCTGGCGGTTGTAGATGTCATACT
CACACGTCTGAACTCTTTCCCTACACGACGCTCTTCCGATCTNNNCTGGCGGTTGTAGATGTCATACT
GCACACGTCTGAACTCTTTCCCTACACGACGCTCTTCCGATCTNNNGAAGTTGCTGGGAACATGTGT
GCACACGTCTGAACTCTTTCCCTACACGACGCTCTTCCGATCTNNNGAAGTTGCTGGGAACATGTGT
GCACACGTCTGAACTCTTTCCCTACACGACGCTCTTCCGATCTNNNGTACAGTCTACAAGAGCCTGCT
GCACACGTCTGAACTCTTTCCCTACACGACGCTCTTCCGATCTNNNGTACAGTCTACAAGAGCCTGCT
GCACACGTCTGAACTCTTTCCCTACACGACGCTCTTCCGATCTNNNCTCCACCATAACTCCCACAGTAG
GCACACGTCTGAACTCTTTCCCTACACGACGCTCTTCCGATCTNNNCTCCACCATAACTCCCACAGTAG
AGCACACGTCTGAACTCTTTCCCTACACGACGCTCTTCCGATCTNNNTAGTGAGCTGAAATCATGCCA
AGCACACGTCTGAACTCTTTCCCTACACGACGCTCTTCCGATCTNNNTAGTGAGCTGAAATCATGCCA
AGCACACGTCTGAACTCTTTCCCTACACGACGCTCTTCCGATCTNNNGAGAATCTCTTGAGCCTGGG
AGCACACGTCTGAACTCTTTCCCTACACGACGCTCTTCCGATCTNNNGAGAATCTCTTGAGCCTGGG
GAGCACACGTCTGAACTCTTTCCCTACACGACGCTCTTCCGATCTNNNGAGGACAAGATGTGAGCTCC
GAGCACACGTCTGAACTCTTTCCCTACACGACGCTCTTCCGATCTNNNGAGGACAAGATGTGAGCTCC
CACACGTCTGAACTCTTTCCCTACACGACGCTCTTCCGATCTNNNTTCTCAATGAGCTGTTGGGTACAC
CACACGTCTGAACTCTTTCCCTACACGACGCTCTTCCGATCTNNNTTCTCAATGAGCTGTTGGGTACAC
GCACACGTCTGAACTCTTTCCCTACACGACGCTCTTCCGATCTNNNCTGTAATCCCAGCTACTCAGGA
GCACACGTCTGAACTCTTTCCCTACACGACGCTCTTCCGATCTNNNCTGTAATCCCAGCTACTCAGGA
GCACACGTCTGAACTCTTTCCCTACACGACGCTCTTCCGATCTNNNCAACTGTGACAGCCAAAACTCT
GCACACGTCTGAACTCTTTCCCTACACGACGCTCTTCCGATCTNNNCAACTGTGACAGCCAAAACTCT

AGCACACGTCTGAACTCTTTCCCTACACGACGCTCTTCCGATCTNNNCTGTAATCCCAGCTACTCAGGA
AGCACACGTCTGAACTCTTTCCCTACACGACGCTCTTCCGATCTNNNCTGTAATCCCAGCTACTCAGGA
AGCACACGTCTGAACTCTTTCCCTACACGACGCTCTTCCGATCTNNNAGTAAGCCAAGATCATGCCATT
AGCACACGTCTGAACTCTTTCCCTACACGACGCTCTTCCGATCTNNNAGTAAGCCAAGATCATGCCATT
GCACACGTCTGAACTCTTTCCCTACACGACGCTCTTCCGATCTNNNTATGTGCCAGTAGTCCCAGCTAC
GCACACGTCTGAACTCTTTCCCTACACGACGCTCTTCCGATCTNNNTATGTGCCAGTAGTCCCAGCTAC
GAGCACACGTCTGAACTCTTTCCCTACACGACGCTCTTCCGATCTNNNAGTAAGCTGAGATCACACCAC
GAGCACACGTCTGAACTCTTTCCCTACACGACGCTCTTCCGATCTNNNAGTAAGCTGAGATCACACCAC
GAGCACACGTCTGAACTCTTTCCCTACACGACGCTCTTCCGATCTNNNTGAGGAGAGTGGATTGCTTG
GAGCACACGTCTGAACTCTTTCCCTACACGACGCTCTTCCGATCTNNNTGAGGAGAGTGGATTGCTTG
GAGCACACGTCTGAACTCTTTCCCTACACGACGCTCTTCCGATCTNNNAGTGAACCAAGATCATGCCAT
GAGCACACGTCTGAACTCTTTCCCTACACGACGCTCTTCCGATCTNNNAGTGAACCAAGATCATGCCAT
AGCACACGTCTGAACTCTTTCCCTACACGACGCTCTTCCGATCTNNNTTGGGGGTTAGCACTTCAACA
AGCACACGTCTGAACTCTTTCCCTACACGACGCTCTTCCGATCTNNNTTGGGGGTTAGCACTTCAACA
GCACACGTCTGAACTCTTTCCCTACACGACGCTCTTCCGATCTNNNTGTGTCTCTGTCTCTGTCTCTC
GCACACGTCTGAACTCTTTCCCTACACGACGCTCTTCCGATCTNNNTGTGTCTCTGTCTCTGTCTCTC
GCACACGTCTGAACTCTTTCCCTACACGACGCTCTTCCGATCTNNNCTGAGGCAGGAGAATTGCTTG
GCACACGTCTGAACTCTTTCCCTACACGACGCTCTTCCGATCTNNNCTGAGGCAGGAGAATTGCTTG
GAGCACACGTCTGAACTCTTTCCCTACACGACGCTCTTCCGATCTNNNTCTGGAGTGGGCCTAAGATTT
GAGCACACGTCTGAACTCTTTCCCTACACGACGCTCTTCCGATCTNNNTCTGGAGTGGGCCTAAGATTT
AGCACACGTCTGAACTCTTTCCCTACACGACGCTCTTCCGATCTNNNTCTGGAAGTGGGTCAAAAGGA
AGCACACGTCTGAACTCTTTCCCTACACGACGCTCTTCCGATCTNNNTCTGGAAGTGGGTCAAAAGGA
GAGCACACGTCTGAACTCTTTCCCTACACGACGCTCTTCCGATCTNNNTAACTCTGCACCAAAGAGGGA
GAGCACACGTCTGAACTCTTTCCCTACACGACGCTCTTCCGATCTNNNTAACTCTGCACCAAAGAGGGA
GAGCACACGTCTGAACTCTTTCCCTACACGACGCTCTTCCGATCTNNNTGTTGCAATGTATTCTGATATA
GAGCACACGTCTGAACTCTTTCCCTACACGACGCTCTTCCGATCTNNNTGTTGCAATGTATTCTGATATA
GCACACGTCTGAACTCTTTCCCTACACGACGCTCTTCCGATCTNNNATAATCCAATCACTTCTTTCTCTC
GCACACGTCTGAACTCTTTCCCTACACGACGCTCTTCCGATCTNNNATAATCCAATCACTTCTTTCTCTC
GAGCACACGTCTGAACTCTTTCCCTACACGACGCTCTTCCGATCTNNNTACAAAAAATTAGCCGGGCAT
GAGCACACGTCTGAACTCTTTCCCTACACGACGCTCTTCCGATCTNNNTACAAAAAATTAGCCGGGCAT
AGAGCACACGTCTGAACTCTTTCCCTACACGACGCTCTTCCGATCTNNNGATTATTTGCTCGATACTGA
AGAGCACACGTCTGAACTCTTTCCCTACACGACGCTCTTCCGATCTNNNGATTATTTGCTCGATACTGA
ACACGTCTGAACTCTTTCCCTACACGACGCTCTTCCGATCTNNNAAGGGGAGTCTCAGTCTGGTGCAT
ACACGTCTGAACTCTTTCCCTACACGACGCTCTTCCGATCTNNNAAGGGGAGTCTCAGTCTGGTGCAT
AGCACACGTCTGAACTCTTTCCCTACACGACGCTCTTCCGATCTNNNCAATTCATACACCCACCTTTCCG
AGCACACGTCTGAACTCTTTCCCTACACGACGCTCTTCCGATCTNNNCAATTCATACACCCACCTTTCCG
GCACACGTCTGAACTCTTTCCCTACACGACGCTCTTCCGATCTNNNAGAAAACATGGGTGAATTCCTCT
GCACACGTCTGAACTCTTTCCCTACACGACGCTCTTCCGATCTNNNAGAAAACATGGGTGAATTCCTCT
GAGCACACGTCTGAACTCTTTCCCTACACGACGCTCTTCCGATCTNNNTTTTACATACCTCATCCTCATTT
GAGCACACGTCTGAACTCTTTCCCTACACGACGCTCTTCCGATCTNNNTTTTACATACCTCATCCTCATTT
GCACACGTCTGAACTCTTTCCCTACACGACGCTCTTCCGATCTNNNATCATACTGCTAAGAATTCAAAC
GCACACGTCTGAACTCTTTCCCTACACGACGCTCTTCCGATCTNNNATCATACTGCTAAGAATTCAAAC
GAGCACACGTCTGAACTCTTTCCCTACACGACGCTCTTCCGATCTNNNTTGTTTGCTCGTTTGCATCTAT

GAGCACACGTCTGAACTCTTTCCCTACACGACGCTCTTCCGATCTNNNTTGTTTGCTCGTTTGCATCTAT
AGCACACGTCTGAACTCTTTCCCTACACGACGCTCTTCCGATCTNNNCTACTGGGCTCAAGAGATTCTC
AGCACACGTCTGAACTCTTTCCCTACACGACGCTCTTCCGATCTNNNCTACTGGGCTCAAGAGATTCTC
AGCACACGTCTGAACTCTTTCCCTACACGACGCTCTTCCGATCTNNNAATAGGCAGCTTTTCCCATACC
AGCACACGTCTGAACTCTTTCCCTACACGACGCTCTTCCGATCTNNNAATAGGCAGCTTTTCCCATACC
GAGCACACGTCTGAACTCTTTCCCTACACGACGCTCTTCCGATCTNNNGAGATCACACCACTACACTCC
GAGCACACGTCTGAACTCTTTCCCTACACGACGCTCTTCCGATCTNNNGAGATCACACCACTACACTCC
GAGCACACGTCTGAACTCTTTCCCTACACGACGCTCTTCCGATCTNNNCTCCCAAGCAGCTGAAATTAC
GAGCACACGTCTGAACTCTTTCCCTACACGACGCTCTTCCGATCTNNNCTCCCAAGCAGCTGAAATTAC
GAGCACACGTCTGAACTCTTTCCCTACACGACGCTCTTCCGATCTNNNAGGAGAATGGCGTGAACCT
GAGCACACGTCTGAACTCTTTCCCTACACGACGCTCTTCCGATCTNNNAGGAGAATGGCGTGAACCT
AGCACACGTCTGAACTCTTTCCCTACACGACGCTCTTCCGATCTNNNAGCACCCTTGTCTTCTCATGTTA
AGCACACGTCTGAACTCTTTCCCTACACGACGCTCTTCCGATCTNNNAGCACCCTTGTCTTCTCATGTTA
AGCACACGTCTGAACTCTTTCCCTACACGACGCTCTTCCGATCTNNNATCCTAGCCAACATGGTGAAAC
AGCACACGTCTGAACTCTTTCCCTACACGACGCTCTTCCGATCTNNNATCCTAGCCAACATGGTGAAAC
AGAGCACACGTCTGAACTCTTTCCCTACACGACGCTCTTCCGATCTNNNCTGTTATGCCACTTAGCGTA
AGAGCACACGTCTGAACTCTTTCCCTACACGACGCTCTTCCGATCTNNNCTGTTATGCCACTTAGCGTA
GAAGAGCACACGTCTGAACTCTTTCCCTACACGACGCTCTTCCGATCTNNNGAAAATGCTGGCAAATC
GAAGAGCACACGTCTGAACTCTTTCCCTACACGACGCTCTTCCGATCTNNNGAAAATGCTGGCAAATC
AGCACACGTCTGAACTCTTTCCCTACACGACGCTCTTCCGATCTNNNGTAGCTGGGATTACAGGCACA
AGCACACGTCTGAACTCTTTCCCTACACGACGCTCTTCCGATCTNNNGTAGCTGGGATTACAGGCACA
GAGCACACGTCTGAACTCTTTCCCTACACGACGCTCTTCCGATCTNNNTTGCAACAACTGAAGAACTTG
GAGCACACGTCTGAACTCTTTCCCTACACGACGCTCTTCCGATCTNNNTTGCAACAACTGAAGAACTTG
AGCACACGTCTGAACTCTTTCCCTACACGACGCTCTTCCGATCTNNNCATTTTGGAATCAATATGTCTC
AGCACACGTCTGAACTCTTTCCCTACACGACGCTCTTCCGATCTNNNCATTTTGGAATCAATATGTCTC
AGCACACGTCTGAACTCTTTCCCTACACGACGCTCTTCCGATCTNNNAATCTGAATGAAGTGAGGGAG
AGCACACGTCTGAACTCTTTCCCTACACGACGCTCTTCCGATCTNNNAATCTGAATGAAGTGAGGGAG
GAGCACACGTCTGAACTCTTTCCCTACACGACGCTCTTCCGATCTNNNTTTGTGGGTCTCCAGGATCTTT
GAGCACACGTCTGAACTCTTTCCCTACACGACGCTCTTCCGATCTNNNTTTGTGGGTCTCCAGGATCTTT
AGAGCACACGTCTGAACTCTTTCCCTACACGACGCTCTTCCGATCTNNNATCTTAGCACTTTGGGAGCC
AGAGCACACGTCTGAACTCTTTCCCTACACGACGCTCTTCCGATCTNNNATCTTAGCACTTTGGGAGCC
AGCACACGTCTGAACTCTTTCCCTACACGACGCTCTTCCGATCTNNNGCCTGAAACTCACACAATCAAC
AGCACACGTCTGAACTCTTTCCCTACACGACGCTCTTCCGATCTNNNGCCTGAAACTCACACAATCAAC
GAGCACACGTCTGAACTCTTTCCCTACACGACGCTCTTCCGATCTNNNTGAGGCAGGAGAATCACTTG
GAGCACACGTCTGAACTCTTTCCCTACACGACGCTCTTCCGATCTNNNTGAGGCAGGAGAATCACTTG
AGCACACGTCTGAACTCTTTCCCTACACGACGCTCTTCCGATCTNNNATGAGTTCTGATGTGCTGTTTG
AGCACACGTCTGAACTCTTTCCCTACACGACGCTCTTCCGATCTNNNATGAGTTCTGATGTGCTGTTTG
AGAGCACACGTCTGAACTCTTTCCCTACACGACGCTCTTCCGATCTNNNGGGGTATTCCTGGAGACTCA
AGAGCACACGTCTGAACTCTTTCCCTACACGACGCTCTTCCGATCTNNNGGGGTATTCCTGGAGACTCA
AGCACACGTCTGAACTCTTTCCCTACACGACGCTCTTCCGATCTNNNTCAGCCTTCTGAGTAGCTAGGA
AGCACACGTCTGAACTCTTTCCCTACACGACGCTCTTCCGATCTNNNTCAGCCTTCTGAGTAGCTAGGA
GAGCACACGTCTGAACTCTTTCCCTACACGACGCTCTTCCGATCTNNNGGAGATTGCAGTGAGTGAG
GAGCACACGTCTGAACTCTTTCCCTACACGACGCTCTTCCGATCTNNNGGAGATTGCAGTGAGTGAG

AGAGCACACGTCTGAACTCTTTCCCTACACGACGCTCTTCCGATCTNNNACTCCAGCCTAGTCTACAGAG

AGAGCACACGTCTGAACTCTTTCCCTACACGACGCTCTTCCGATCTNNNACTCCAGCCTAGTCTACAGAG

AGCACACGTCTGAACTCTTTCCCTACACGACGCTCTTCCGATCTNNNGAGATCGCACCACTGTACTCTAG

AGCACACGTCTGAACTCTTTCCCTACACGACGCTCTTCCGATCTNNNGAGATCGCACCACTGTACTCTAG

AGCACACGTCTGAACTCTTTCCCTACACGACGCTCTTCCGATCTNNNTGTTTGCAAGCTTTGACAACAGA

AGCACACGTCTGAACTCTTTCCCTACACGACGCTCTTCCGATCTNNNTGTTTGCAAGCTTTGACAACAGA

AGCACACGTCTGAACTCTTTCCCTACACGACGCTCTTCCGATCTNNNACTGGAATTATGAGCATGAGCC

AGCACACGTCTGAACTCTTTCCCTACACGACGCTCTTCCGATCTNNNACTGGAATTATGAGCATGAGCC

GCACACGTCTGAACTCTTTCCCTACACGACGCTCTTCCGATCTNNNTCCCCAAGAGTTCAAGGTTGTAG

GCACACGTCTGAACTCTTTCCCTACACGACGCTCTTCCGATCTNNNTCCCCAAGAGTTCAAGGTTGTAG

GAGCACACGTCTGAACTCTTTCCCTACACGACGCTCTTCCGATCTNNNTCCTTATGAAAGAGGACCTCC

GAGCACACGTCTGAACTCTTTCCCTACACGACGCTCTTCCGATCTNNNTCCTTATGAAAGAGGACCTCC

AGCACACGTCTGAACTCTTTCCCTACACGACGCTCTTCCGATCTNNNAATGCAGAAAATTAGCCGGGCC

AGCACACGTCTGAACTCTTTCCCTACACGACGCTCTTCCGATCTNNNAATGCAGAAAATTAGCCGGGCC

AGCACACGTCTGAACTCTTTCCCTACACGACGCTCTTCCGATCTNNNGAGAGATGCCACCACTGTAGAT

AGCACACGTCTGAACTCTTTCCCTACACGACGCTCTTCCGATCTNNNGAGAGATGCCACCACTGTAGAT

AGCACACGTCTGAACTCTTTCCCTACACGACGCTCTTCCGATCTNNNTTTTCATGTCACCCCATCTCCTCA

AGCACACGTCTGAACTCTTTCCCTACACGACGCTCTTCCGATCTNNNTTTTCATGTCACCCCATCTCCTCA

GCACACGTCTGAACTCTTTCCCTACACGACGCTCTTCCGATCTNNNGCTGCACTACAGAGTACAGTGGC

GCACACGTCTGAACTCTTTCCCTACACGACGCTCTTCCGATCTNNNGCTGCACTACAGAGTACAGTGGC

AGCACACGTCTGAACTCTTTCCCTACACGACGCTCTTCCGATCTNNNCTGAGGTAGGAGGATCACTTG

AGCACACGTCTGAACTCTTTCCCTACACGACGCTCTTCCGATCTNNNCTGAGGTAGGAGGATCACTTG

AGCACACGTCTGAACTCTTTCCCTACACGACGCTCTTCCGATCTNNNGCTTCTAGGGAATGCAAAGCAA

AGCACACGTCTGAACTCTTTCCCTACACGACGCTCTTCCGATCTNNNGCTTCTAGGGAATGCAAAGCAA

AGCACACGTCTGAACTCTTTCCCTACACGACGCTCTTCCGATCTNNNGTGGTCCTACCCAAAGAATCACT

AGCACACGTCTGAACTCTTTCCCTACACGACGCTCTTCCGATCTNNNGTGGTCCTACCCAAAGAATCACT

AGCACACGTCTGAACTCTTTCCCTACACGACGCTCTTCCGATCTNNNATTTTGTATTTTTAGTAGAGAC

AGCACACGTCTGAACTCTTTCCCTACACGACGCTCTTCCGATCTNNNATTTTGTATTTTTAGTAGAGAC

AGCACACGTCTGAACTCTTTCCCTACACGACGCTCTTCCGATCTNNNTCTACGTGAAGGTGAATGGTGG

AGCACACGTCTGAACTCTTTCCCTACACGACGCTCTTCCGATCTNNNTCTACGTGAAGGTGAATGGTGG

AGCACACGTCTGAACTCTTTCCCTACACGACGCTCTTCCGATCTNNNTTGTCTCTACAGGTCAGTGAAGC

AGCACACGTCTGAACTCTTTCCCTACACGACGCTCTTCCGATCTNNNTTGTCTCTACAGGTCAGTGAAGC

GAGCACACGTCTGAACTCTTTCCCTACACGACGCTCTTCCGATCTNNNCCAACATGGTGAAACTCCATCT

GAGCACACGTCTGAACTCTTTCCCTACACGACGCTCTTCCGATCTNNNCCAACATGGTGAAACTCCATCT

AGCACACGTCTGAACTCTTTCCCTACACGACGCTCTTCCGATCTNNNGAAAAGAGAGTAAAACGCCGG

AGCACACGTCTGAACTCTTTCCCTACACGACGCTCTTCCGATCTNNNGAAAAGAGAGTAAAACGCCGG

GCACACGTCTGAACTCTTTCCCTACACGACGCTCTTCCGATCTNNNAGCCATGGGAGACACTGATGTA

GCACACGTCTGAACTCTTTCCCTACACGACGCTCTTCCGATCTNNNAGCCATGGGAGACACTGATGTA

AGCACACGTCTGAACTCTTTCCCTACACGACGCTCTTCCGATCTNNNTTCAAGCAATTCTCCTGCCTCAG

AGCACACGTCTGAACTCTTTCCCTACACGACGCTCTTCCGATCTNNNTTCAAGCAATTCTCCTGCCTCAG

GAGCACACGTCTGAACTCTTTCCCTACACGACGCTCTTCCGATCTNNNGACAACTCACTAGATAGGCCCA

GAGCACACGTCTGAACTCTTTCCCTACACGACGCTCTTCCGATCTNNNGACAACTCACTAGATAGGCCCA

AGCACACGTCTGAACTCTTTCCCTACACGACGCTCTTCCGATCTNNNGAGGCAGGAGAATCGCTTGAA

AGCACACGTCTGAACTCTTTCCCTACACGACGCTCTTCCGATCTNNNGAGGCAGGAGAATCGCTTGAA
AGCACACGTCTGAACTCTTTCCCTACACGACGCTCTTCCGATCTNNNCTACAGCACAACAAAGACAACG
AGCACACGTCTGAACTCTTTCCCTACACGACGCTCTTCCGATCTNNNCTACAGCACAACAAAGACAACG
GAGCACACGTCTGAACTCTTTCCCTACACGACGCTCTTCCGATCTNNNTTTTGGATTGTTCACTGTTG
GAGCACACGTCTGAACTCTTTCCCTACACGACGCTCTTCCGATCTNNNTTTTGGATTGTTCACTGTTG
GAGCACACGTCTGAACTCTTTCCCTACACGACGCTCTTCCGATCTNNNGCCTCACTAGTAGCTGGGATT
GAGCACACGTCTGAACTCTTTCCCTACACGACGCTCTTCCGATCTNNNGCCTCACTAGTAGCTGGGATT
GAGCACACGTCTGAACTCTTTCCCTACACGACGCTCTTCCGATCTNNNAGGCACAATAATTGCTTGAAC
GAGCACACGTCTGAACTCTTTCCCTACACGACGCTCTTCCGATCTNNNAGGCACAATAATTGCTTGAAC
ACACGTCTGAACTCTTTCCCTACACGACGCTCTTCCGATCTNNNCTCTTTGTTTCTCATCAACTGCATCGG
ACACGTCTGAACTCTTTCCCTACACGACGCTCTTCCGATCTNNNCTCTTTGTTTCTCATCAACTGCATCGG
AGCACACGTCTGAACTCTTTCCCTACACGACGCTCTTCCGATCTNNNCCATGGAATGTCATTTTGGAGG
AGCACACGTCTGAACTCTTTCCCTACACGACGCTCTTCCGATCTNNNCCATGGAATGTCATTTTGGAGG
AGAGCACACGTCTGAACTCTTTCCCTACACGACGCTCTTCCGATCTNNNTGTAATCCTAGCACTTTGGG
AGAGCACACGTCTGAACTCTTTCCCTACACGACGCTCTTCCGATCTNNNTGTAATCCTAGCACTTTGGG
GAGCACACGTCTGAACTCTTTCCCTACACGACGCTCTTCCGATCTNNNCACAGTCTTCAAAACCCACAGA
GAGCACACGTCTGAACTCTTTCCCTACACGACGCTCTTCCGATCTNNNCACAGTCTTCAAAACCCACAGA
GCACACGTCTGAACTCTTTCCCTACACGACGCTCTTCCGATCTNNNGAGAATTGACCCATGAGTTGCAT
GCACACGTCTGAACTCTTTCCCTACACGACGCTCTTCCGATCTNNNGAGAATTGACCCATGAGTTGCAT
AGAGCACACGTCTGAACTCTTTCCCTACACGACGCTCTTCCGATCTNNNACCATCCTAGGCTTGTCAAAC
AGAGCACACGTCTGAACTCTTTCCCTACACGACGCTCTTCCGATCTNNNACCATCCTAGGCTTGTCAAAC
AGCACACGTCTGAACTCTTTCCCTACACGACGCTCTTCCGATCTNNNCCGTCTCCTTTCTCCTTTCCTTTCA
AGCACACGTCTGAACTCTTTCCCTACACGACGCTCTTCCGATCTNNNCCGTCTCCTTTCTCCTTTCCTTTCA
GAGCACACGTCTGAACTCTTTCCCTACACGACGCTCTTCCGATCTNNNACACCTGTAATCCCAGCACTTT
GAGCACACGTCTGAACTCTTTCCCTACACGACGCTCTTCCGATCTNNNACACCTGTAATCCCAGCACTTT
GAGCACACGTCTGAACTCTTTCCCTACACGACGCTCTTCCGATCTNNNAATCTCGACTCACTTCAAGCTCC
GAGCACACGTCTGAACTCTTTCCCTACACGACGCTCTTCCGATCTNNNAATCTCGACTCACTTCAAGCTCC
AGCACACGTCTGAACTCTTTCCCTACACGACGCTCTTCCGATCTNNNCACCTGTAATCTCAGCTACTCGG
AGCACACGTCTGAACTCTTTCCCTACACGACGCTCTTCCGATCTNNNCACCTGTAATCTCAGCTACTCGG
AGCACACGTCTGAACTCTTTCCCTACACGACGCTCTTCCGATCTNNNATTACTTCCTATGCATATGCGCC
AGCACACGTCTGAACTCTTTCCCTACACGACGCTCTTCCGATCTNNNATTACTTCCTATGCATATGCGCC
AGCACACGTCTGAACTCTTTCCCTACACGACGCTCTTCCGATCTNNNAGGAATGGAGGTCAGTAGGA
AGCACACGTCTGAACTCTTTCCCTACACGACGCTCTTCCGATCTNNNAGGAATGGAGGTCAGTAGGA
GAGCACACGTCTGAACTCTTTCCCTACACGACGCTCTTCCGATCTNNNAATGAACACACAATGCCAAGA
GAGCACACGTCTGAACTCTTTCCCTACACGACGCTCTTCCGATCTNNNAATGAACACACAATGCCAAGA
GAGCACACGTCTGAACTCTTTCCCTACACGACGCTCTTCCGATCTNNNATGAGGTTTCACCATATTGGC
GAGCACACGTCTGAACTCTTTCCCTACACGACGCTCTTCCGATCTNNNATGAGGTTTCACCATATTGGC
AAGAGCACACGTCTGAACTCTTTCCCTACACGACGCTCTTCCGATCTNNNATACATTATGGTTGGGTTG
AAGAGCACACGTCTGAACTCTTTCCCTACACGACGCTCTTCCGATCTNNNATACATTATGGTTGGGTTG
AGCACACGTCTGAACTCTTTCCCTACACGACGCTCTTCCGATCTNNNTCATGCCTGTAATCCCAACACTT
AGCACACGTCTGAACTCTTTCCCTACACGACGCTCTTCCGATCTNNNTCATGCCTGTAATCCCAACACTT
CACACGTCTGAACTCTTTCCCTACACGACGCTCTTCCGATCTNNNGTTCAAACGACTCTCTGCTTCAGCA
CACACGTCTGAACTCTTTCCCTACACGACGCTCTTCCGATCTNNNGTTCAAACGACTCTCTGCTTCAGCA

GCACACGTCTGAACTCTTTCCCTACACGACGCTCTTCCGATCTNNNAGGGTTTATGATCCAGGCTGAA
GCACACGTCTGAACTCTTTCCCTACACGACGCTCTTCCGATCTNNNAGGGTTTATGATCCAGGCTGAA
AGCACACGTCTGAACTCTTTCCCTACACGACGCTCTTCCGATCTNNNCTCTAAGAAGCGGTTCAGTCTG
AGCACACGTCTGAACTCTTTCCCTACACGACGCTCTTCCGATCTNNNCTCTAAGAAGCGGTTCAGTCTG
AGCACACGTCTGAACTCTTTCCCTACACGACGCTCTTCCGATCTNNNCAGCCTGGCTAACATAGTAAA
AGCACACGTCTGAACTCTTTCCCTACACGACGCTCTTCCGATCTNNNCAGCCTGGCTAACATAGTAAA
GCACACGTCTGAACTCTTTCCCTACACGACGCTCTTCCGATCTNNNTAATCACTTGAACCCAGGAAGC
GCACACGTCTGAACTCTTTCCCTACACGACGCTCTTCCGATCTNNNTAATCACTTGAACCCAGGAAGC
GAGCACACGTCTGAACTCTTTCCCTACACGACGCTCTTCCGATCTNNNGCTATTGAGATGGCATTGTC
GAGCACACGTCTGAACTCTTTCCCTACACGACGCTCTTCCGATCTNNNGCTATTGAGATGGCATTGTC
AGCACACGTCTGAACTCTTTCCCTACACGACGCTCTTCCGATCTNNNGTCTGTTCATGTACTTTGCCCA
AGCACACGTCTGAACTCTTTCCCTACACGACGCTCTTCCGATCTNNNGTCTGTTCATGTACTTTGCCCA
GAGCACACGTCTGAACTCTTTCCCTACACGACGCTCTTCCGATCTNNNTTCAAGCAAGGGGGAAAAA
GAGCACACGTCTGAACTCTTTCCCTACACGACGCTCTTCCGATCTNNNTTCAAGCAAGGGGGAAAAA
GAGCACACGTCTGAACTCTTTCCCTACACGACGCTCTTCCGATCTNNNTACGTAGATTGGGTCAAGAT
GAGCACACGTCTGAACTCTTTCCCTACACGACGCTCTTCCGATCTNNNTACGTAGATTGGGTCAAGAT
GCACACGTCTGAACTCTTTCCCTACACGACGCTCTTCCGATCTNNNGTTTCTTCTGTGGGCAGTTACA
GCACACGTCTGAACTCTTTCCCTACACGACGCTCTTCCGATCTNNNGTTTCTTCTGTGGGCAGTTACA
GCACACGTCTGAACTCTTTCCCTACACGACGCTCTTCCGATCTNNNGAAGGAGGATCGCTTGAGACT
GCACACGTCTGAACTCTTTCCCTACACGACGCTCTTCCGATCTNNNGAAGGAGGATCGCTTGAGACT
GCACACGTCTGAACTCTTTCCCTACACGACGCTCTTCCGATCTNNNGCACGCTTCAACAATAACACCTT
GCACACGTCTGAACTCTTTCCCTACACGACGCTCTTCCGATCTNNNGCACGCTTCAACAATAACACCTT
CACACGTCTGAACTCTTTCCCTACACGACGCTCTTCCGATCTNNNGATGGGTCAGGTGTTACTCTCTG
CACACGTCTGAACTCTTTCCCTACACGACGCTCTTCCGATCTNNNGATGGGTCAGGTGTTACTCTCTG
GAGCACACGTCTGAACTCTTTCCCTACACGACGCTCTTCCGATCTNNNTCAGATGTCGCCAAGTTTCAA
GAGCACACGTCTGAACTCTTTCCCTACACGACGCTCTTCCGATCTNNNTCAGATGTCGCCAAGTTTCAA
GAGCACACGTCTGAACTCTTTCCCTACACGACGCTCTTCCGATCTNNNGGCTTTTTGTCCTTGCGATAG
GAGCACACGTCTGAACTCTTTCCCTACACGACGCTCTTCCGATCTNNNGGCTTTTTGTCCTTGCGATAG
AGCACACGTCTGAACTCTTTCCCTACACGACGCTCTTCCGATCTNNNAGAGCTACAAGGGGATGAGT
AGCACACGTCTGAACTCTTTCCCTACACGACGCTCTTCCGATCTNNNAGAGCTACAAGGGGATGAGT
GCACACGTCTGAACTCTTTCCCTACACGACGCTCTTCCGATCTNNNGCCATTTCATATATGGAGGTGC
GCACACGTCTGAACTCTTTCCCTACACGACGCTCTTCCGATCTNNNGCCATTTCATATATGGAGGTGC
GCACACGTCTGAACTCTTTCCCTACACGACGCTCTTCCGATCTNNNAGGCCACACTAACTTTGAAAGC
GCACACGTCTGAACTCTTTCCCTACACGACGCTCTTCCGATCTNNNAGGCCACACTAACTTTGAAAGC
GCACACGTCTGAACTCTTTCCCTACACGACGCTCTTCCGATCTNNNGCTGGTGAGATCCTTTTGCTTTT
GCACACGTCTGAACTCTTTCCCTACACGACGCTCTTCCGATCTNNNGCTGGTGAGATCCTTTTGCTTTT
AGCACACGTCTGAACTCTTTCCCTACACGACGCTCTTCCGATCTNNNGCTTTGCTATGGAAGAGAGAG
AGCACACGTCTGAACTCTTTCCCTACACGACGCTCTTCCGATCTNNNGCTTTGCTATGGAAGAGAGAG
AGCACACGTCTGAACTCTTTCCCTACACGACGCTCTTCCGATCTNNNGATGCCTGTAATCCCAGCTACT
AGCACACGTCTGAACTCTTTCCCTACACGACGCTCTTCCGATCTNNNGATGCCTGTAATCCCAGCTACT
CACACGTCTGAACTCTTTCCCTACACGACGCTCTTCCGATCTNNNGCAGGTGCTAGTAAACCTGATCC
CACACGTCTGAACTCTTTCCCTACACGACGCTCTTCCGATCTNNNGCAGGTGCTAGTAAACCTGATCC
AGCACACGTCTGAACTCTTTCCCTACACGACGCTCTTCCGATCTNNNTCCCAAACCAAACATGTCTTTC

GAGCACACGTCTGAACTCTTTCCCTACACGACGCTCTTCCGATCTNNNTCCCAAACCAAACATGTCTTTC

GAGCACACGTCTGAACTCTTTCCCTACACGACGCTCTTCCGATCTNNNCAACAGAAGAAAGGTTGGTG

GAGCACACGTCTGAACTCTTTCCCTACACGACGCTCTTCCGATCTNNNCAACAGAAGAAAGGTTGGTG

AGAGCACACGTCTGAACTCTTTCCCTACACGACGCTCTTCCGATCTNNNGAATGAAAAGCCAGTTGGA

AGAGCACACGTCTGAACTCTTTCCCTACACGACGCTCTTCCGATCTNNNGAATGAAAAGCCAGTTGGA

CACACGTCTGAACTCTTTCCCTACACGACGCTCTTCCGATCTNNNGGCCTCTGCTGGATTTTGAGATTCA

CACACGTCTGAACTCTTTCCCTACACGACGCTCTTCCGATCTNNNGGCCTCTGCTGGATTTTGAGATTCA

GAGCACACGTCTGAACTCTTTCCCTACACGACGCTCTTCCGATCTNNNTGGAGTATAGTGGCGTGATC

GAGCACACGTCTGAACTCTTTCCCTACACGACGCTCTTCCGATCTNNNTGGAGTATAGTGGCGTGATC

GAGCACACGTCTGAACTCTTTCCCTACACGACGCTCTTCCGATCTNNNATGGGGACGAGAAGGGATTT

GAGCACACGTCTGAACTCTTTCCCTACACGACGCTCTTCCGATCTNNNATGGGGACGAGAAGGGATTT

GAGCACACGTCTGAACTCTTTCCCTACACGACGCTCTTCCGATCTNNNGAGTCTCCAGCCTCATACACCT

GAGCACACGTCTGAACTCTTTCCCTACACGACGCTCTTCCGATCTNNNGAGTCTCCAGCCTCATACACCT

AGAGCACACGTCTGAACTCTTTCCCTACACGACGCTCTTCCGATCTNNNGAGCAAGACTCCGTCTCAAAA

AGAGCACACGTCTGAACTCTTTCCCTACACGACGCTCTTCCGATCTNNNGAGCAAGACTCCGTCTCAAAA

AGAGCACACGTCTGAACTCTTTCCCTACACGACGCTCTTCCGATCTNNNCTGAGGCAGGAGAATCACTT

AGAGCACACGTCTGAACTCTTTCCCTACACGACGCTCTTCCGATCTNNNCTGAGGCAGGAGAATCACTT

AGCACACGTCTGAACTCTTTCCCTACACGACGCTCTTCCGATCTNNNGTAATCCCAGCATTTTGGTAGG

AGCACACGTCTGAACTCTTTCCCTACACGACGCTCTTCCGATCTNNNGTAATCCCAGCATTTTGGTAGG

GAGCACACGTCTGAACTCTTTCCCTACACGACGCTCTTCCGATCTNNNGATTCTTCAATAGACTGGAAG

GAGCACACGTCTGAACTCTTTCCCTACACGACGCTCTTCCGATCTNNNGATTCTTCAATAGACTGGAAG

AGCACACGTCTGAACTCTTTCCCTACACGACGCTCTTCCGATCTNNNGAATTCACGACGTCACACTTGT

AGCACACGTCTGAACTCTTTCCCTACACGACGCTCTTCCGATCTNNNGAATTCACGACGTCACACTTGT

GAGCACACGTCTGAACTCTTTCCCTACACGACGCTCTTCCGATCTNNNGCCTCCAGTATATGTTAGCAGC

GAGCACACGTCTGAACTCTTTCCCTACACGACGCTCTTCCGATCTNNNGCCTCCAGTATATGTTAGCAGC

AGCACACGTCTGAACTCTTTCCCTACACGACGCTCTTCCGATCTNNNCACCTCTAATCCCAGCTACTCAG

AGCACACGTCTGAACTCTTTCCCTACACGACGCTCTTCCGATCTNNNCACCTCTAATCCCAGCTACTCAG

GCACACGTCTGAACTCTTTCCCTACACGACGCTCTTCCGATCTNNNCTGTAATCCCAGCTACTCAGGAG

GCACACGTCTGAACTCTTTCCCTACACGACGCTCTTCCGATCTNNNCTGTAATCCCAGCTACTCAGGAG

AGCACACGTCTGAACTCTTTCCCTACACGACGCTCTTCCGATCTNNNCCTGGAGAACTTGTTCAAAGCA

AGCACACGTCTGAACTCTTTCCCTACACGACGCTCTTCCGATCTNNNCCTGGAGAACTTGTTCAAAGCA

GAGCACACGTCTGAACTCTTTCCCTACACGACGCTCTTCCGATCTNNNTCAAGAGCCCCTAGTTTGAAAG

GAGCACACGTCTGAACTCTTTCCCTACACGACGCTCTTCCGATCTNNNTCAAGAGCCCCTAGTTTGAAAG

GAGCACACGTCTGAACTCTTTCCCTACACGACGCTCTTCCGATCTNNNCTGAGGTAGGAGGATTGCTT

GAGCACACGTCTGAACTCTTTCCCTACACGACGCTCTTCCGATCTNNNCTGAGGTAGGAGGATTGCTT

AGCACACGTCTGAACTCTTTCCCTACACGACGCTCTTCCGATCTNNNGTCCACATTTTAAAAACTAAGCT

AGCACACGTCTGAACTCTTTCCCTACACGACGCTCTTCCGATCTNNNGTCCACATTTTAAAAACTAAGCT

ACACGTCTGAACTCTTTCCCTACACGACGCTCTTCCGATCTNNNTCTCTCAAGTAGCTGGGACTACACAT

ACACGTCTGAACTCTTTCCCTACACGACGCTCTTCCGATCTNNNTCTCTCAAGTAGCTGGGACTACACAT

AGCACACGTCTGAACTCTTTCCCTACACGACGCTCTTCCGATCTNNNGGAGACAAGAATGAAACGCCA

AGCACACGTCTGAACTCTTTCCCTACACGACGCTCTTCCGATCTNNNGGAGACAAGAATGAAACGCCA

AGCACACGTCTGAACTCTTTCCCTACACGACGCTCTTCCGATCTNNNTATGTCTGCAATCCCAGCTACTC

AGCACACGTCTGAACTCTTTCCCTACACGACGCTCTTCCGATCTNNNTATGTCTGCAATCCCAGCTACTC

GAGCACACGTCTGAACTCTTTCCCTACACGACGCTCTTCCGATCTNNNCAGGAGAATTGCTTGAACAC
GAGCACACGTCTGAACTCTTTCCCTACACGACGCTCTTCCGATCTNNNCAGGAGAATTGCTTGAACAC
CACACGTCTGAACTCTTTCCCTACACGACGCTCTTCCGATCTNNNCTGATAGTCCTTGCCTCTAACCCC
CACACGTCTGAACTCTTTCCCTACACGACGCTCTTCCGATCTNNNCTGATAGTCCTTGCCTCTAACCCC
AGCACACGTCTGAACTCTTTCCCTACACGACGCTCTTCCGATCTNNNCTTCTCCTTTCCCTGGACTAACC
AGCACACGTCTGAACTCTTTCCCTACACGACGCTCTTCCGATCTNNNCTTCTCCTTTCCCTGGACTAACC
GCACACGTCTGAACTCTTTCCCTACACGACGCTCTTCCGATCTNNNCCACTTTCTTCCACCAAAAGGACC
GCACACGTCTGAACTCTTTCCCTACACGACGCTCTTCCGATCTNNNCCACTTTCTTCCACCAAAAGGACC
GCACACGTCTGAACTCTTTCCCTACACGACGCTCTTCCGATCTNNNTCCGAACTCTAACTGGTGTTCTG
GCACACGTCTGAACTCTTTCCCTACACGACGCTCTTCCGATCTNNNTCCGAACTCTAACTGGTGTTCTG
GCACACGTCTGAACTCTTTCCCTACACGACGCTCTTCCGATCTNNNTATGACGTTGTCAATATACAGTA
GCACACGTCTGAACTCTTTCCCTACACGACGCTCTTCCGATCTNNNTATGACGTTGTCAATATACAGTA
GAGCACACGTCTGAACTCTTTCCCTACACGACGCTCTTCCGATCTNNNTGTCCCAAAAATGTATCATGC
GAGCACACGTCTGAACTCTTTCCCTACACGACGCTCTTCCGATCTNNNTGTCCCAAAAATGTATCATGC
AGAGCACACGTCTGAACTCTTTCCCTACACGACGCTCTTCCGATCTNNNAATCACTTGAACCCAGGAG
AGAGCACACGTCTGAACTCTTTCCCTACACGACGCTCTTCCGATCTNNNAATCACTTGAACCCAGGAG
GCACACGTCTGAACTCTTTCCCTACACGACGCTCTTCCGATCTNNNCGGTGTGAAGTTTCCGCTTGCAT
GCACACGTCTGAACTCTTTCCCTACACGACGCTCTTCCGATCTNNNCGGTGTGAAGTTTCCGCTTGCAT
AGCACACGTCTGAACTCTTTCCCTACACGACGCTCTTCCGATCTNNNTACAGACATGAGGAAATGGTG
AGCACACGTCTGAACTCTTTCCCTACACGACGCTCTTCCGATCTNNNTACAGACATGAGGAAATGGTG
ACACGTCTGAACTCTTTCCCTACACGACGCTCTTCCGATCTNNNCCAAATGGGTACATGACAGACCTC
ACACGTCTGAACTCTTTCCCTACACGACGCTCTTCCGATCTNNNCCAAATGGGTACATGACAGACCTC
AGCACACGTCTGAACTCTTTCCCTACACGACGCTCTTCCGATCTNNNGGCTTTTACCTCTAGGGCTTCT
AGCACACGTCTGAACTCTTTCCCTACACGACGCTCTTCCGATCTNNNGGCTTTTACCTCTAGGGCTTCT
GCACACGTCTGAACTCTTTCCCTACACGACGCTCTTCCGATCTNNNAATGAGCTTGTTCTTGGGAAA
GCACACGTCTGAACTCTTTCCCTACACGACGCTCTTCCGATCTNNNAATGAGCTTGTTCTTGGGAAA
GAGCACACGTCTGAACTCTTTCCCTACACGACGCTCTTCCGATCTNNNCAGGCATAAGACTCCAACCTC
GAGCACACGTCTGAACTCTTTCCCTACACGACGCTCTTCCGATCTNNNCAGGCATAAGACTCCAACCTC
GCACACGTCTGAACTCTTTCCCTACACGACGCTCTTCCGATCTNNNGATTGTGTCCAGGTCCTTGTTCA
GCACACGTCTGAACTCTTTCCCTACACGACGCTCTTCCGATCTNNNGATTGTGTCCAGGTCCTTGTTCA
GAGCACACGTCTGAACTCTTTCCCTACACGACGCTCTTCCGATCTNNNCCCCCCCTTTTTTCTAGAACTG
GAGCACACGTCTGAACTCTTTCCCTACACGACGCTCTTCCGATCTNNNCCCCCCCTTTTTTCTAGAACTG
GCACACGTCTGAACTCTTTCCCTACACGACGCTCTTCCGATCTNNNTGTGTCAGAGAGAGAGCAAGGA
GCACACGTCTGAACTCTTTCCCTACACGACGCTCTTCCGATCTNNNTGTGTCAGAGAGAGAGCAAGGA
AGCACACGTCTGAACTCTTTCCCTACACGACGCTCTTCCGATCTNNNTCTCTCTTTCCTTCCTTTCTTGCT
AGCACACGTCTGAACTCTTTCCCTACACGACGCTCTTCCGATCTNNNTCTCTCTTTCCTTCCTTTCTTGCT
AGCACACGTCTGAACTCTTTCCCTACACGACGCTCTTCCGATCTNNNCTATGGCAAGACTCAGTTTGCA
AGCACACGTCTGAACTCTTTCCCTACACGACGCTCTTCCGATCTNNNCTATGGCAAGACTCAGTTTGCA
GAGCACACGTCTGAACTCTTTCCCTACACGACGCTCTTCCGATCTNNNTCGCTTCTGCAAATGAGGAA
GAGCACACGTCTGAACTCTTTCCCTACACGACGCTCTTCCGATCTNNNTCGCTTCTGCAAATGAGGAA
AGCACACGTCTGAACTCTTTCCCTACACGACGCTCTTCCGATCTNNNAAGCCACCTGATGCACAGATA
AGCACACGTCTGAACTCTTTCCCTACACGACGCTCTTCCGATCTNNNAAGCCACCTGATGCACAGATA
GCACACGTCTGAACTCTTTCCCTACACGACGCTCTTCCGATCTNNNAGATTGCAGTGAGCTGAGATC

AGCACACGTCTGAACTCTTTCCCTACACGACGCTCTTCCGATCTNNNAGATTGCAGTGAGCTGAGATCA
AGCACACGTCTGAACTCTTTCCCTACACGACGCTCTTCCGATCTNNNTCTCCATGGAGTTTTCATGCAGA
AGCACACGTCTGAACTCTTTCCCTACACGACGCTCTTCCGATCTNNNTCTCCATGGAGTTTTCATGCAGA
GAGCACACGTCTGAACTCTTTCCCTACACGACGCTCTTCCGATCTNNNCTGCAAAGGACATGAACTCAT
GAGCACACGTCTGAACTCTTTCCCTACACGACGCTCTTCCGATCTNNNCTGCAAAGGACATGAACTCAT
GCACACGTCTGAACTCTTTCCCTACACGACGCTCTTCCGATCTNNNACTATATGCCTAGTTGTGTGCAT
GCACACGTCTGAACTCTTTCCCTACACGACGCTCTTCCGATCTNNNACTATATGCCTAGTTGTGTGCAT
AGCACACGTCTGAACTCTTTCCCTACACGACGCTCTTCCGATCTNNNATTTCCAATAAACCAGCACCAG
AGCACACGTCTGAACTCTTTCCCTACACGACGCTCTTCCGATCTNNNATTTCCAATAAACCAGCACCAG
GCACACGTCTGAACTCTTTCCCTACACGACGCTCTTCCGATCTNNNCCCACATAAATGCACCCAACTGA
GCACACGTCTGAACTCTTTCCCTACACGACGCTCTTCCGATCTNNNCCCACATAAATGCACCCAACTGA
GAGCACACGTCTGAACTCTTTCCCTACACGACGCTCTTCCGATCTNNNCTTTGCTATACTGCAGGCCATT
GAGCACACGTCTGAACTCTTTCCCTACACGACGCTCTTCCGATCTNNNCTTTGCTATACTGCAGGCCATT
GAGCACACGTCTGAACTCTTTCCCTACACGACGCTCTTCCGATCTNNNGTTTCACTTCCTCTCCTCTGTG
GAGCACACGTCTGAACTCTTTCCCTACACGACGCTCTTCCGATCTNNNGTTTCACTTCCTCTCCTCTGTG
GAGCACACGTCTGAACTCTTTCCCTACACGACGCTCTTCCGATCTNNNGGGCCTGACTGTTAGAAGGA
GAGCACACGTCTGAACTCTTTCCCTACACGACGCTCTTCCGATCTNNNGGGCCTGACTGTTAGAAGGA
GAGCACACGTCTGAACTCTTTCCCTACACGACGCTCTTCCGATCTNNNGGTGACTGTGGTTTTAGAAA
GAGCACACGTCTGAACTCTTTCCCTACACGACGCTCTTCCGATCTNNNGGTGACTGTGGTTTTAGAAA
AGCACACGTCTGAACTCTTTCCCTACACGACGCTCTTCCGATCTNNNTAGACAGGATCTTCTCAGGAGC
AGCACACGTCTGAACTCTTTCCCTACACGACGCTCTTCCGATCTNNNTAGACAGGATCTTCTCAGGAGC
AGCACACGTCTGAACTCTTTCCCTACACGACGCTCTTCCGATCTNNNTCTCTCTCTGTGTCACTCTCTCTC
AGCACACGTCTGAACTCTTTCCCTACACGACGCTCTTCCGATCTNNNTCTCTCTCTGTGTCACTCTCTCTC
AGCACACGTCTGAACTCTTTCCCTACACGACGCTCTTCCGATCTNNNACTAAAACTCAGCGATACTGCCA
AGCACACGTCTGAACTCTTTCCCTACACGACGCTCTTCCGATCTNNNACTAAAACTCAGCGATACTGCCA
GCACACGTCTGAACTCTTTCCCTACACGACGCTCTTCCGATCTNNNGTTTAAATTCTCCACAGATTGTAC
GCACACGTCTGAACTCTTTCCCTACACGACGCTCTTCCGATCTNNNGTTTAAATTCTCCACAGATTGTAC
AGAGCACACGTCTGAACTCTTTCCCTACACGACGCTCTTCCGATCTNNNCTTTTTGCTTAGGATTGTCG
AGAGCACACGTCTGAACTCTTTCCCTACACGACGCTCTTCCGATCTNNNCTTTTTGCTTAGGATTGTCG
AGAGCACACGTCTGAACTCTTTCCCTACACGACGCTCTTCCGATCTNNNGAAAAAATTAGCCGGGCAT
AGAGCACACGTCTGAACTCTTTCCCTACACGACGCTCTTCCGATCTNNNGAAAAAATTAGCCGGGCAT
AGAGCACACGTCTGAACTCTTTCCCTACACGACGCTCTTCCGATCTNNNCTTGGGTAGCAGCTCATATT
AGAGCACACGTCTGAACTCTTTCCCTACACGACGCTCTTCCGATCTNNNCTTGGGTAGCAGCTCATATT
GAGCACACGTCTGAACTCTTTCCCTACACGACGCTCTTCCGATCTNNNCTCAGGAAGACAGAAGGAGA
GAGCACACGTCTGAACTCTTTCCCTACACGACGCTCTTCCGATCTNNNCTCAGGAAGACAGAAGGAGA
GAGCACACGTCTGAACTCTTTCCCTACACGACGCTCTTCCGATCTNNNACAAACAAGTGAGGAGAGGG
GAGCACACGTCTGAACTCTTTCCCTACACGACGCTCTTCCGATCTNNNACAAACAAGTGAGGAGAGGG
GCACACGTCTGAACTCTTTCCCTACACGACGCTCTTCCGATCTNNNGGGAAGCTTACTGTGTCCAAGA
GCACACGTCTGAACTCTTTCCCTACACGACGCTCTTCCGATCTNNNGGGAAGCTTACTGTGTCCAAGA
GAGCACACGTCTGAACTCTTTCCCTACACGACGCTCTTCCGATCTNNNCTTCACTGTGGTCCTGCCTCCA
GAGCACACGTCTGAACTCTTTCCCTACACGACGCTCTTCCGATCTNNNCTTCACTGTGGTCCTGCCTCCA
AGAGCACACGTCTGAACTCTTTCCCTACACGACGCTCTTCCGATCTNNNAGAATCACTTGAACCTGGGG
AGAGCACACGTCTGAACTCTTTCCCTACACGACGCTCTTCCGATCTNNNAGAATCACTTGAACCTGGGG

GAGCACACGTCTGAACTCTTTCCCTACACGACGCTCTTCCGATCTNNNATACTGTGGCTCCTATTTGG

GAGCACACGTCTGAACTCTTTCCCTACACGACGCTCTTCCGATCTNNNATACTGTGGCTCCTATTTGG

AGCACACGTCTGAACTCTTTCCCTACACGACGCTCTTCCGATCTNNNTCAATCTCAGCTTACTGCAACC

AGCACACGTCTGAACTCTTTCCCTACACGACGCTCTTCCGATCTNNNTCAATCTCAGCTTACTGCAACC

GAGCACACGTCTGAACTCTTTCCCTACACGACGCTCTTCCGATCTNNNTGCCCATTTTCTAATTGGACT

GAGCACACGTCTGAACTCTTTCCCTACACGACGCTCTTCCGATCTNNNTGCCCATTTTCTAATTGGACT

GAGCACACGTCTGAACTCTTTCCCTACACGACGCTCTTCCGATCTNNNGAATGGGAGTTGCAGTCGAG

GAGCACACGTCTGAACTCTTTCCCTACACGACGCTCTTCCGATCTNNNGAATGGGAGTTGCAGTCGAG

GCACACGTCTGAACTCTTTCCCTACACGACGCTCTTCCGATCTNNNCCATCCCTGACTCATGTGAACAT

GCACACGTCTGAACTCTTTCCCTACACGACGCTCTTCCGATCTNNNCCATCCCTGACTCATGTGAACAT

AGAGCACACGTCTGAACTCTTTCCCTACACGACGCTCTTCCGATCTNNNTGTTTGCTCATTTGCTCCTTT

AGAGCACACGTCTGAACTCTTTCCCTACACGACGCTCTTCCGATCTNNNTGTTTGCTCATTTGCTCCTTT

GAGCACACGTCTGAACTCTTTCCCTACACGACGCTCTTCCGATCTNNNGTTGCAGTGAGCCAAGATTG

GAGCACACGTCTGAACTCTTTCCCTACACGACGCTCTTCCGATCTNNNGTTGCAGTGAGCCAAGATTG

GAGCACACGTCTGAACTCTTTCCCTACACGACGCTCTTCCGATCTNNNAAAGAACCTCTTCACACACAC

GAGCACACGTCTGAACTCTTTCCCTACACGACGCTCTTCCGATCTNNNAAAGAACCTCTTCACACACAC

GAGCACACGTCTGAACTCTTTCCCTACACGACGCTCTTCCGATCTNNNCTTGCAGTGAACCAAGATCTC

GAGCACACGTCTGAACTCTTTCCCTACACGACGCTCTTCCGATCTNNNCTTGCAGTGAACCAAGATCTC

AGAGCACACGTCTGAACTCTTTCCCTACACGACGCTCTTCCGATCTNNNCTCTAAAGACACAGGAGCTT

AGAGCACACGTCTGAACTCTTTCCCTACACGACGCTCTTCCGATCTNNNCTCTAAAGACACAGGAGCTT

AGCACACGTCTGAACTCTTTCCCTACACGACGCTCTTCCGATCTNNNCCACAGCCTTCCTCATTTTTGTT

AGCACACGTCTGAACTCTTTCCCTACACGACGCTCTTCCGATCTNNNCCACAGCCTTCCTCATTTTTGTT

GCACACGTCTGAACTCTTTCCCTACACGACGCTCTTCCGATCTNNNAAAGGAAAAGCGAATCTCCGGA

GCACACGTCTGAACTCTTTCCCTACACGACGCTCTTCCGATCTNNNAAAGGAAAAGCGAATCTCCGGA

AGCACACGTCTGAACTCTTTCCCTACACGACGCTCTTCCGATCTNNNCGCTTGGATGTTCATTGGGAAT

AGCACACGTCTGAACTCTTTCCCTACACGACGCTCTTCCGATCTNNNCGCTTGGATGTTCATTGGGAAT

GAGCACACGTCTGAACTCTTTCCCTACACGACGCTCTTCCGATCTNNNTCCCAAGTTCCGTACATTCCTT

GAGCACACGTCTGAACTCTTTCCCTACACGACGCTCTTCCGATCTNNNTCCCAAGTTCCGTACATTCCTT

AGAGCACACGTCTGAACTCTTTCCCTACACGACGCTCTTCCGATCTNNNGGAACAGAAAACCAAACACC

AGAGCACACGTCTGAACTCTTTCCCTACACGACGCTCTTCCGATCTNNNGGAACAGAAAACCAAACACC

AGAGCACACGTCTGAACTCTTTCCCTACACGACGCTCTTCCGATCTNNNTGAGGCAAGAGAATTGCTT

AGAGCACACGTCTGAACTCTTTCCCTACACGACGCTCTTCCGATCTNNNTGAGGCAAGAGAATTGCTT

GAGCACACGTCTGAACTCTTTCCCTACACGACGCTCTTCCGATCTNNNTGAGCCTTGTGGACGATTAT

GAGCACACGTCTGAACTCTTTCCCTACACGACGCTCTTCCGATCTNNNTGAGCCTTGTGGACGATTAT

CACACGTCTGAACTCTTTCCCTACACGACGCTCTTCCGATCTNNNCACAATGTGCCGCTTGTGTGCATC

CACACGTCTGAACTCTTTCCCTACACGACGCTCTTCCGATCTNNNCACAATGTGCCGCTTGTGTGCATC

GAGCACACGTCTGAACTCTTTCCCTACACGACGCTCTTCCGATCTNNNACATTTTCATGCTCACAAGGA

GAGCACACGTCTGAACTCTTTCCCTACACGACGCTCTTCCGATCTNNNACATTTTCATGCTCACAAGGA

AGCACACGTCTGAACTCTTTCCCTACACGACGCTCTTCCGATCTNNNCACCTCCCAGCTAAAGTGATTC

AGCACACGTCTGAACTCTTTCCCTACACGACGCTCTTCCGATCTNNNCACCTCCCAGCTAAAGTGATTC

GAGCACACGTCTGAACTCTTTCCCTACACGACGCTCTTCCGATCTNNNCAGGCATTCCATTCTGTGTTC

GAGCACACGTCTGAACTCTTTCCCTACACGACGCTCTTCCGATCTNNNCAGGCATTCCATTCTGTGTTC

AGCACACGTCTGAACTCTTTCCCTACACGACGCTCTTCCGATCTNNNTGTAATTCCAGCTACTTGGGA

AGCACACGTCTGAACTCTTTCCCTACACGACGCTCTTCCGATCTNNNTGTAATTCCAGCTACTTGGGA
GAGCACACGTCTGAACTCTTTCCCTACACGACGCTCTTCCGATCTNNNATGTGTGTGTGTGTGTGTTT
GAGCACACGTCTGAACTCTTTCCCTACACGACGCTCTTCCGATCTNNNATGTGTGTGTGTGTGTGTTT
GCACACGTCTGAACTCTTTCCCTACACGACGCTCTTCCGATCTNNNACAAAAATTAGCCAGGTGTGGT
GCACACGTCTGAACTCTTTCCCTACACGACGCTCTTCCGATCTNNNACAAAAATTAGCCAGGTGTGGT
AGCACACGTCTGAACTCTTTCCCTACACGACGCTCTTCCGATCTNNNGGGAACTTGGACCTTGTTTCAA
AGCACACGTCTGAACTCTTTCCCTACACGACGCTCTTCCGATCTNNNGGGAACTTGGACCTTGTTTCAA
AGCACACGTCTGAACTCTTTCCCTACACGACGCTCTTCCGATCTNNNTTCATCTTGCAGAAACCCTCTCA
AGCACACGTCTGAACTCTTTCCCTACACGACGCTCTTCCGATCTNNNTTCATCTTGCAGAAACCCTCTCA
AGCACACGTCTGAACTCTTTCCCTACACGACGCTCTTCCGATCTNNNCTCTGAACACGACTTGGGCTTT
AGCACACGTCTGAACTCTTTCCCTACACGACGCTCTTCCGATCTNNNCTCTGAACACGACTTGGGCTTT
GCACACGTCTGAACTCTTTCCCTACACGACGCTCTTCCGATCTNNNCCTCAGAAGTTGTTTGCTTCCTG
GCACACGTCTGAACTCTTTCCCTACACGACGCTCTTCCGATCTNNNCCTCAGAAGTTGTTTGCTTCCTG
AGCACACGTCTGAACTCTTTCCCTACACGACGCTCTTCCGATCTNNNAACTGATTCTGAAATCCAGCA
AGCACACGTCTGAACTCTTTCCCTACACGACGCTCTTCCGATCTNNNAACTGATTCTGAAATCCAGCA
GCACACGTCTGAACTCTTTCCCTACACGACGCTCTTCCGATCTNNNGAGCCCTCTGGAAGACTTGTTTA
GCACACGTCTGAACTCTTTCCCTACACGACGCTCTTCCGATCTNNNGAGCCCTCTGGAAGACTTGTTTA
AGCACACGTCTGAACTCTTTCCCTACACGACGCTCTTCCGATCTNNNGGTCCAGGGATGGTGTTCTGC
AGCACACGTCTGAACTCTTTCCCTACACGACGCTCTTCCGATCTNNNGGTCCAGGGATGGTGTTCTGC
AGCACACGTCTGAACTCTTTCCCTACACGACGCTCTTCCGATCTNNNAAAACTAAATCAACTGAAACA
AGCACACGTCTGAACTCTTTCCCTACACGACGCTCTTCCGATCTNNNAAAACTAAATCAACTGAAACA
AGCACACGTCTGAACTCTTTCCCTACACGACGCTCTTCCGATCTNNNGTGGATCACGAGGTCAGGAT
AGCACACGTCTGAACTCTTTCCCTACACGACGCTCTTCCGATCTNNNGTGGATCACGAGGTCAGGAT
AGCACACGTCTGAACTCTTTCCCTACACGACGCTCTTCCGATCTNNNGCTTTTTCGATTACGGAGTCTG
AGCACACGTCTGAACTCTTTCCCTACACGACGCTCTTCCGATCTNNNGCTTTTTCGATTACGGAGTCTG
GAGCACACGTCTGAACTCTTTCCCTACACGACGCTCTTCCGATCTNNNATTGAGCCGAGATCAACCCA
GAGCACACGTCTGAACTCTTTCCCTACACGACGCTCTTCCGATCTNNNATTGAGCCGAGATCAACCCA
AGCACACGTCTGAACTCTTTCCCTACACGACGCTCTTCCGATCTNNNCATTCTATTCAGTCCCCCCTCTG
AGCACACGTCTGAACTCTTTCCCTACACGACGCTCTTCCGATCTNNNCATTCTATTCAGTCCCCCCTCTG
AGCACACGTCTGAACTCTTTCCCTACACGACGCTCTTCCGATCTNNNCTACTGGGAGAAATGTGTGTC
AGCACACGTCTGAACTCTTTCCCTACACGACGCTCTTCCGATCTNNNCTACTGGGAGAAATGTGTGTC
AGCACACGTCTGAACTCTTTCCCTACACGACGCTCTTCCGATCTNNNCAGAGGTGCATATACACTCCCT
AGCACACGTCTGAACTCTTTCCCTACACGACGCTCTTCCGATCTNNNCAGAGGTGCATATACACTCCCT
AGCACACGTCTGAACTCTTTCCCTACACGACGCTCTTCCGATCTNNNTCTTCCTTTTGCTTACTGCACC
AGCACACGTCTGAACTCTTTCCCTACACGACGCTCTTCCGATCTNNNTCTTCCTTTTGCTTACTGCACC
AGCACACGTCTGAACTCTTTCCCTACACGACGCTCTTCCGATCTNNNTGGAACACTCTGCCATATTGTC
AGCACACGTCTGAACTCTTTCCCTACACGACGCTCTTCCGATCTNNNTGGAACACTCTGCCATATTGTC
AGCACACGTCTGAACTCTTTCCCTACACGACGCTCTTCCGATCTNNNCCTTGTCGATGGCTTTGGTAA
AGCACACGTCTGAACTCTTTCCCTACACGACGCTCTTCCGATCTNNNCCTTGTCGATGGCTTTGGTAA
GAGCACACGTCTGAACTCTTTCCCTACACGACGCTCTTCCGATCTNNNTACAAAAAATTAGCCGGGCAT
GAGCACACGTCTGAACTCTTTCCCTACACGACGCTCTTCCGATCTNNNTACAAAAAATTAGCCGGGCAT
AGCACACGTCTGAACTCTTTCCCTACACGACGCTCTTCCGATCTNNNGTGGCTATCTATGAATGGTGG
AGCACACGTCTGAACTCTTTCCCTACACGACGCTCTTCCGATCTNNNGTGGCTATCTATGAATGGTGG

GCACACGTCTGAACTCTTTCCCTACACGACGCTCTTCCGATCTNNNTCACTGAACAGACCAAGGATGA
GCACACGTCTGAACTCTTTCCCTACACGACGCTCTTCCGATCTNNNTCACTGAACAGACCAAGGATGA
AGCACACGTCTGAACTCTTTCCCTACACGACGCTCTTCCGATCTNNNAGTGCTGGGATTATAGGCATG
AGCACACGTCTGAACTCTTTCCCTACACGACGCTCTTCCGATCTNNNAGTGCTGGGATTATAGGCATG
AGCACACGTCTGAACTCTTTCCCTACACGACGCTCTTCCGATCTNNNAGCTTTGAATGCATCAAGGTCT
AGCACACGTCTGAACTCTTTCCCTACACGACGCTCTTCCGATCTNNNAGCTTTGAATGCATCAAGGTCT
AGCACACGTCTGAACTCTTTCCCTACACGACGCTCTTCCGATCTNNNACTGATCTTGCCCACATTTCTC
AGCACACGTCTGAACTCTTTCCCTACACGACGCTCTTCCGATCTNNNACTGATCTTGCCCACATTTCTC
GAGCACACGTCTGAACTCTTTCCCTACACGACGCTCTTCCGATCTNNNCTGCAATCTCAGCTACTCAGG
GAGCACACGTCTGAACTCTTTCCCTACACGACGCTCTTCCGATCTNNNCTGCAATCTCAGCTACTCAGG
AGCACACGTCTGAACTCTTTCCCTACACGACGCTCTTCCGATCTNNNCATTCTCTGCTCCTCTCGATGA
AGCACACGTCTGAACTCTTTCCCTACACGACGCTCTTCCGATCTNNNCATTCTCTGCTCCTCTCGATGA
GAGCACACGTCTGAACTCTTTCCCTACACGACGCTCTTCCGATCTNNNGACCCAAATAGAGATGAGGT
GAGCACACGTCTGAACTCTTTCCCTACACGACGCTCTTCCGATCTNNNGACCCAAATAGAGATGAGGT
AGCACACGTCTGAACTCTTTCCCTACACGACGCTCTTCCGATCTNNNGAGAACAACAAACCACTGCTCA
AGCACACGTCTGAACTCTTTCCCTACACGACGCTCTTCCGATCTNNNGAGAACAACAAACCACTGCTCA
AGCACACGTCTGAACTCTTTCCCTACACGACGCTCTTCCGATCTNNNAACTACTACAGGTTGAGCATCC
AGCACACGTCTGAACTCTTTCCCTACACGACGCTCTTCCGATCTNNNAACTACTACAGGTTGAGCATCC
AGCACACGTCTGAACTCTTTCCCTACACGACGCTCTTCCGATCTNNNCTCCTACAGAGTGCTAACAGCC
AGCACACGTCTGAACTCTTTCCCTACACGACGCTCTTCCGATCTNNNCTCCTACAGAGTGCTAACAGCC
AGCACACGTCTGAACTCTTTCCCTACACGACGCTCTTCCGATCTNNNTGTCTCACATCTAGGTCATGCT
AGCACACGTCTGAACTCTTTCCCTACACGACGCTCTTCCGATCTNNNTGTCTCACATCTAGGTCATGCT
AGAGCACACGTCTGAACTCTTTCCCTACACGACGCTCTTCCGATCTNNNGCAGAGGTTGAAATGAGC
AGAGCACACGTCTGAACTCTTTCCCTACACGACGCTCTTCCGATCTNNNGCAGAGGTTGAAATGAGC
AGAGCACACGTCTGAACTCTTTCCCTACACGACGCTCTTCCGATCTNNNATTTGTACCTAGCAGGTGT
AGAGCACACGTCTGAACTCTTTCCCTACACGACGCTCTTCCGATCTNNNATTTGTACCTAGCAGGTGT
GAGCACACGTCTGAACTCTTTCCCTACACGACGCTCTTCCGATCTNNNAGACCAGGATGATGATGACG
GAGCACACGTCTGAACTCTTTCCCTACACGACGCTCTTCCGATCTNNNAGACCAGGATGATGATGACG
GCACACGTCTGAACTCTTTCCCTACACGACGCTCTTCCGATCTNNNTGGTTGGATTCTAATGAGGGCT
GCACACGTCTGAACTCTTTCCCTACACGACGCTCTTCCGATCTNNNTGGTTGGATTCTAATGAGGGCT
AGCACACGTCTGAACTCTTTCCCTACACGACGCTCTTCCGATCTNNNGCTAGTCTTACAGACCCATTGG
AGCACACGTCTGAACTCTTTCCCTACACGACGCTCTTCCGATCTNNNGCTAGTCTTACAGACCCATTGG
AGCACACGTCTGAACTCTTTCCCTACACGACGCTCTTCCGATCTNNNGTTTCACTATATTGGCCAGGCT
AGCACACGTCTGAACTCTTTCCCTACACGACGCTCTTCCGATCTNNNGTTTCACTATATTGGCCAGGCT
GAGCACACGTCTGAACTCTTTCCCTACACGACGCTCTTCCGATCTNNNCATGCCTGTAATCCCAGCTAC
GAGCACACGTCTGAACTCTTTCCCTACACGACGCTCTTCCGATCTNNNCATGCCTGTAATCCCAGCTAC
GAGCACACGTCTGAACTCTTTCCCTACACGACGCTCTTCCGATCTNNNCTGAGGCAGAGGAATCACTT
GAGCACACGTCTGAACTCTTTCCCTACACGACGCTCTTCCGATCTNNNCTGAGGCAGAGGAATCACTT
GAGCACACGTCTGAACTCTTTCCCTACACGACGCTCTTCCGATCTNNNGCAGGCATCTCCGCTATAAAA
GAGCACACGTCTGAACTCTTTCCCTACACGACGCTCTTCCGATCTNNNGCAGGCATCTCCGCTATAAAA
AGAGCACACGTCTGAACTCTTTCCCTACACGACGCTCTTCCGATCTNNNGCCTGTAGTCCCAGCTATGA
AGAGCACACGTCTGAACTCTTTCCCTACACGACGCTCTTCCGATCTNNNGCCTGTAGTCCCAGCTATGA
AGCACACGTCTGAACTCTTTCCCTACACGACGCTCTTCCGATCTNNNTTCACACCATTCTTCTGCCTCAG

GAGCACACGTCTGAACTCTTTCCCTACACGACGCTCTTCCGATCTNNNTTCACACCATTCTTCTGCCTCAG
GAGCACACGTCTGAACTCTTTCCCTACACGACGCTCTTCCGATCTNNNCTACGGTCACCAAGTGCTCTG
GAGCACACGTCTGAACTCTTTCCCTACACGACGCTCTTCCGATCTNNNCTACGGTCACCAAGTGCTCTG
AGCACACGTCTGAACTCTTTCCCTACACGACGCTCTTCCGATCTNNNTGATTTCAGTTAGGGCCTACAC
AGCACACGTCTGAACTCTTTCCCTACACGACGCTCTTCCGATCTNNNTGATTTCAGTTAGGGCCTACAC
AGCACACGTCTGAACTCTTTCCCTACACGACGCTCTTCCGATCTNNNTTCCTAAAAAGTCCACACCTGGC
AGCACACGTCTGAACTCTTTCCCTACACGACGCTCTTCCGATCTNNNTTCCTAAAAAGTCCACACCTGGC
AGCACACGTCTGAACTCTTTCCCTACACGACGCTCTTCCGATCTNNNGCAACTCCCTTTCCTTACCTAGCC
AGCACACGTCTGAACTCTTTCCCTACACGACGCTCTTCCGATCTNNNGCAACTCCCTTTCCTTACCTAGCC
GAGCACACGTCTGAACTCTTTCCCTACACGACGCTCTTCCGATCTNNNACACCTGGCTGAAACTCACTA
GAGCACACGTCTGAACTCTTTCCCTACACGACGCTCTTCCGATCTNNNACACCTGGCTGAAACTCACTA
AGCACACGTCTGAACTCTTTCCCTACACGACGCTCTTCCGATCTNNNGTAAGTGCACATTCTAAGAACT
AGCACACGTCTGAACTCTTTCCCTACACGACGCTCTTCCGATCTNNNGTAAGTGCACATTCTAAGAACT
GAGCACACGTCTGAACTCTTTCCCTACACGACGCTCTTCCGATCTNNNAGCTCTCATTTCCTTTTAGCG
GAGCACACGTCTGAACTCTTTCCCTACACGACGCTCTTCCGATCTNNNAGCTCTCATTTCCTTTTAGCG
GCACACGTCTGAACTCTTTCCCTACACGACGCTCTTCCGATCTNNNAGCATTTACTGGTGCTCATGTCAC
GCACACGTCTGAACTCTTTCCCTACACGACGCTCTTCCGATCTNNNAGCATTTACTGGTGCTCATGTCAC
GAGCACACGTCTGAACTCTTTCCCTACACGACGCTCTTCCGATCTNNNCAGTGACCTGGTTGATAGCAA
GAGCACACGTCTGAACTCTTTCCCTACACGACGCTCTTCCGATCTNNNCAGTGACCTGGTTGATAGCAA
AGCACACGTCTGAACTCTTTCCCTACACGACGCTCTTCCGATCTNNNCTTTGTGGATGTTGCAGGGGA
AGCACACGTCTGAACTCTTTCCCTACACGACGCTCTTCCGATCTNNNCTTTGTGGATGTTGCAGGGGA
GAGCACACGTCTGAACTCTTTCCCTACACGACGCTCTTCCGATCTNNNGAGATGTCAGGAGACTTTTAA
GAGCACACGTCTGAACTCTTTCCCTACACGACGCTCTTCCGATCTNNNGAGATGTCAGGAGACTTTTAA
GAGCACACGTCTGAACTCTTTCCCTACACGACGCTCTTCCGATCTNNNATGGAGAAAATTAGACCGGCC
GAGCACACGTCTGAACTCTTTCCCTACACGACGCTCTTCCGATCTNNNATGGAGAAAATTAGACCGGCC
GAGCACACGTCTGAACTCTTTCCCTACACGACGCTCTTCCGATCTNNNACAGGCCCAGAAGTTCATAAC
GAGCACACGTCTGAACTCTTTCCCTACACGACGCTCTTCCGATCTNNNACAGGCCCAGAAGTTCATAAC
AGAGCACACGTCTGAACTCTTTCCCTACACGACGCTCTTCCGATCTNNNGAAAGGTATCCCTACAGAGC
AGAGCACACGTCTGAACTCTTTCCCTACACGACGCTCTTCCGATCTNNNGAAAGGTATCCCTACAGAGC
AGCACACGTCTGAACTCTTTCCCTACACGACGCTCTTCCGATCTNNNATTTGGAGGCTCTAATGTTGGG
AGCACACGTCTGAACTCTTTCCCTACACGACGCTCTTCCGATCTNNNATTTGGAGGCTCTAATGTTGGG
ACACGTCTGAACTCTTTCCCTACACGACGCTCTTCCGATCTNNNTTTCCTTTTGGTAATGGCAGACTCAT
ACACGTCTGAACTCTTTCCCTACACGACGCTCTTCCGATCTNNNTTTCCTTTTGGTAATGGCAGACTCAT
GAGCACACGTCTGAACTCTTTCCCTACACGACGCTCTTCCGATCTNNNAAAGTGACACCCTGTCTCTCTC
GAGCACACGTCTGAACTCTTTCCCTACACGACGCTCTTCCGATCTNNNAAAGTGACACCCTGTCTCTCTC
GAGCACACGTCTGAACTCTTTCCCTACACGACGCTCTTCCGATCTNNNTGTGATGTTGAAAATGGCATC
GAGCACACGTCTGAACTCTTTCCCTACACGACGCTCTTCCGATCTNNNTGTGATGTTGAAAATGGCATC
AGCACACGTCTGAACTCTTTCCCTACACGACGCTCTTCCGATCTNNNATAGGCCCATTATACCAAGCAGC
AGCACACGTCTGAACTCTTTCCCTACACGACGCTCTTCCGATCTNNNATAGGCCCATTATACCAAGCAGC
AGAGCACACGTCTGAACTCTTTCCCTACACGACGCTCTTCCGATCTNNNTAGAGTTTGAGTGTGTGTGC
AGAGCACACGTCTGAACTCTTTCCCTACACGACGCTCTTCCGATCTNNNTAGAGTTTGAGTGTGTGTGC
GAGCACACGTCTGAACTCTTTCCCTACACGACGCTCTTCCGATCTNNNAATGTCTTGCTGCTAGGATTGC
GAGCACACGTCTGAACTCTTTCCCTACACGACGCTCTTCCGATCTNNNAATGTCTTGCTGCTAGGATTGC

GCACACGTCTGAACTCTTTCCCTACACGACGCTCTTCCGATCTNNNGTTTAGGCAATCCTCTCACCTCA
GCACACGTCTGAACTCTTTCCCTACACGACGCTCTTCCGATCTNNNGTTTAGGCAATCCTCTCACCTCA
GAGCACACGTCTGAACTCTTTCCCTACACGACGCTCTTCCGATCTNNNAGGCTGCCAAAAGTAGGTTT
GAGCACACGTCTGAACTCTTTCCCTACACGACGCTCTTCCGATCTNNNAGGCTGCCAAAAGTAGGTTT
AGCACACGTCTGAACTCTTTCCCTACACGACGCTCTTCCGATCTNNNAAGAACCAGGGGCACATTACT
AGCACACGTCTGAACTCTTTCCCTACACGACGCTCTTCCGATCTNNNAAGAACCAGGGGCACATTACT
GCACACGTCTGAACTCTTTCCCTACACGACGCTCTTCCGATCTNNNCCACGAAAACCCTAGTGAATCAC
GCACACGTCTGAACTCTTTCCCTACACGACGCTCTTCCGATCTNNNCCACGAAAACCCTAGTGAATCAC
AGCACACGTCTGAACTCTTTCCCTACACGACGCTCTTCCGATCTNNNTCACACATACTACTCCCTGTCCT
AGCACACGTCTGAACTCTTTCCCTACACGACGCTCTTCCGATCTNNNTCACACATACTACTCCCTGTCCT
AGCACACGTCTGAACTCTTTCCCTACACGACGCTCTTCCGATCTNNNCCTTACTTGAAGCCAGAGGTCA
AGCACACGTCTGAACTCTTTCCCTACACGACGCTCTTCCGATCTNNNCCTTACTTGAAGCCAGAGGTCA
GAGCACACGTCTGAACTCTTTCCCTACACGACGCTCTTCCGATCTNNNCTTGCAACCAAGTCTTCCTAT
GAGCACACGTCTGAACTCTTTCCCTACACGACGCTCTTCCGATCTNNNCTTGCAACCAAGTCTTCCTAT
CACGTCTGAACTCTTTCCCTACACGACGCTCTTCCGATCTNNNACCATGGCAGCTACTGTAAATCACAT
CACGTCTGAACTCTTTCCCTACACGACGCTCTTCCGATCTNNNACCATGGCAGCTACTGTAAATCACAT
GAGCACACGTCTGAACTCTTTCCCTACACGACGCTCTTCCGATCTNNNACAACTATGACTCCACCTGGA
GAGCACACGTCTGAACTCTTTCCCTACACGACGCTCTTCCGATCTNNNACAACTATGACTCCACCTGGA
AGCACACGTCTGAACTCTTTCCCTACACGACGCTCTTCCGATCTNNNCCCTGGTAACCACCTTTCTACTC
AGCACACGTCTGAACTCTTTCCCTACACGACGCTCTTCCGATCTNNNCCCTGGTAACCACCTTTCTACTC
AGCACACGTCTGAACTCTTTCCCTACACGACGCTCTTCCGATCTNNNTGACCTCTGAGCTTCAATTCGA
AGCACACGTCTGAACTCTTTCCCTACACGACGCTCTTCCGATCTNNNTGACCTCTGAGCTTCAATTCGA
AGCACACGTCTGAACTCTTTCCCTACACGACGCTCTTCCGATCTNNNCTATGGTCGTTATTTTCTCATT
AGCACACGTCTGAACTCTTTCCCTACACGACGCTCTTCCGATCTNNNCTATGGTCGTTATTTTCTCATT
AGCACACGTCTGAACTCTTTCCCTACACGACGCTCTTCCGATCTNNNCAAGGATCCCAGTTGAGATTCC
AGCACACGTCTGAACTCTTTCCCTACACGACGCTCTTCCGATCTNNNCAAGGATCCCAGTTGAGATTCC
AGCACACGTCTGAACTCTTTCCCTACACGACGCTCTTCCGATCTNNNAGCTGTCCCTTTGTGATTCATCT
AGCACACGTCTGAACTCTTTCCCTACACGACGCTCTTCCGATCTNNNAGCTGTCCCTTTGTGATTCATCT
AGCACACGTCTGAACTCTTTCCCTACACGACGCTCTTCCGATCTNNNGGTTTACCCTGGGTGTCTTCTT
AGCACACGTCTGAACTCTTTCCCTACACGACGCTCTTCCGATCTNNNGGTTTACCCTGGGTGTCTTCTT
GCACACGTCTGAACTCTTTCCCTACACGACGCTCTTCCGATCTNNNGGCAGCAAGTGAGGAAGTAAG
GCACACGTCTGAACTCTTTCCCTACACGACGCTCTTCCGATCTNNNGGCAGCAAGTGAGGAAGTAAG
GCACACGTCTGAACTCTTTCCCTACACGACGCTCTTCCGATCTNNNCTTCTCCCTTAGAACCGCGTGCAT
GCACACGTCTGAACTCTTTCCCTACACGACGCTCTTCCGATCTNNNCTTCTCCCTTAGAACCGCGTGCAT
GCACACGTCTGAACTCTTTCCCTACACGACGCTCTTCCGATCTNNNCGCTTTCCCTTTCTGAGATCTGG
GCACACGTCTGAACTCTTTCCCTACACGACGCTCTTCCGATCTNNNCGCTTTCCCTTTCTGAGATCTGG
AGCACACGTCTGAACTCTTTCCCTACACGACGCTCTTCCGATCTNNNGGATCATTCCCATATAGAAGA
AGCACACGTCTGAACTCTTTCCCTACACGACGCTCTTCCGATCTNNNGGATCATTCCCATATAGAAGA
GCACACGTCTGAACTCTTTCCCTACACGACGCTCTTCCGATCTNNNTTGCAGTGAGTCGAGATCTTACC
GCACACGTCTGAACTCTTTCCCTACACGACGCTCTTCCGATCTNNNTTGCAGTGAGTCGAGATCTTACC
AGCACACGTCTGAACTCTTTCCCTACACGACGCTCTTCCGATCTNNNAATTAGACAACAACTCCTGGGC
AGCACACGTCTGAACTCTTTCCCTACACGACGCTCTTCCGATCTNNNAATTAGACAACAACTCCTGGGC
AGCACACGTCTGAACTCTTTCCCTACACGACGCTCTTCCGATCTNNNATGGCCTGGGATTTGTTTTT

AGCACACGTCTGAACTCTTTCCCTACACGACGCTCTTCCGATCTNNNATGGCCTGGGATTTGTTTTTC
CACACGTCTGAACTCTTTCCCTACACGACGCTCTTCCGATCTNNNATACTTTTCACACTCTTCCTCCATCG
CACACGTCTGAACTCTTTCCCTACACGACGCTCTTCCGATCTNNNATACTTTTCACACTCTTCCTCCATCG
AGCACACGTCTGAACTCTTTCCCTACACGACGCTCTTCCGATCTNNNATTCTAACTAGGGTGAGTTGGC
AGCACACGTCTGAACTCTTTCCCTACACGACGCTCTTCCGATCTNNNATTCTAACTAGGGTGAGTTGGC
GCACACGTCTGAACTCTTTCCCTACACGACGCTCTTCCGATCTNNNCTCCTGAGTAGCTGGGACTACAC
GCACACGTCTGAACTCTTTCCCTACACGACGCTCTTCCGATCTNNNCTCCTGAGTAGCTGGGACTACAC
GAGCACACGTCTGAACTCTTTCCCTACACGACGCTCTTCCGATCTNNNTAGTTGTTGGGCATTTGGGTT
GAGCACACGTCTGAACTCTTTCCCTACACGACGCTCTTCCGATCTNNNTAGTTGTTGGGCATTTGGGTT
GCACACGTCTGAACTCTTTCCCTACACGACGCTCTTCCGATCTNNNTCAGCCATTGAAACTATTGATCT
GCACACGTCTGAACTCTTTCCCTACACGACGCTCTTCCGATCTNNNTCAGCCATTGAAACTATTGATCT
AGCACACGTCTGAACTCTTTCCCTACACGACGCTCTTCCGATCTNNNGCCTCCCAAGTAGCTAGGATTA
AGCACACGTCTGAACTCTTTCCCTACACGACGCTCTTCCGATCTNNNGCCTCCCAAGTAGCTAGGATTA
AGAGCACACGTCTGAACTCTTTCCCTACACGACGCTCTTCCGATCTNNNCTGAGCATGACCCTGAAAAG
AGAGCACACGTCTGAACTCTTTCCCTACACGACGCTCTTCCGATCTNNNCTGAGCATGACCCTGAAAAG
AGCACACGTCTGAACTCTTTCCCTACACGACGCTCTTCCGATCTNNNCCATAGATTGAGGGCTCAGTA
AGCACACGTCTGAACTCTTTCCCTACACGACGCTCTTCCGATCTNNNCCATAGATTGAGGGCTCAGTA
AGCACACGTCTGAACTCTTTCCCTACACGACGCTCTTCCGATCTNNNCCCATTTGAACCAGTTCAGGAC
AGCACACGTCTGAACTCTTTCCCTACACGACGCTCTTCCGATCTNNNCCCATTTGAACCAGTTCAGGAC
GAGCACACGTCTGAACTCTTTCCCTACACGACGCTCTTCCGATCTNNNTGGTTGGCCAAGAACTGATAA
GAGCACACGTCTGAACTCTTTCCCTACACGACGCTCTTCCGATCTNNNTGGTTGGCCAAGAACTGATAA
AGAGCACACGTCTGAACTCTTTCCCTACACGACGCTCTTCCGATCTNNNCCAAAGGCTTCAACAAGAAC
AGAGCACACGTCTGAACTCTTTCCCTACACGACGCTCTTCCGATCTNNNCCAAAGGCTTCAACAAGAAC
AGAGCACACGTCTGAACTCTTTCCCTACACGACGCTCTTCCGATCTNNNGAGGTTACGGTGAGTGGA
AGAGCACACGTCTGAACTCTTTCCCTACACGACGCTCTTCCGATCTNNNGAGGTTACGGTGAGTGGA
AGCACACGTCTGAACTCTTTCCCTACACGACGCTCTTCCGATCTNNNATTCTAGGCTGTGTTCTAGAG
AGCACACGTCTGAACTCTTTCCCTACACGACGCTCTTCCGATCTNNNATTCTAGGCTGTGTTCTAGAG
AGCACACGTCTGAACTCTTTCCCTACACGACGCTCTTCCGATCTNNNCAGGAAAATGGCATGAACCCAC
AGCACACGTCTGAACTCTTTCCCTACACGACGCTCTTCCGATCTNNNCAGGAAAATGGCATGAACCCAC
AGCACACGTCTGAACTCTTTCCCTACACGACGCTCTTCCGATCTNNNGGATCTCTTCCTGCTGGATCTT
AGCACACGTCTGAACTCTTTCCCTACACGACGCTCTTCCGATCTNNNGGATCTCTTCCTGCTGGATCTT
ACACGTCTGAACTCTTTCCCTACACGACGCTCTTCCGATCTNNNATTGTGACGTCTCTGGCAGCCATCG
ACACGTCTGAACTCTTTCCCTACACGACGCTCTTCCGATCTNNNATTGTGACGTCTCTGGCAGCCATCG
AGCACACGTCTGAACTCTTTCCCTACACGACGCTCTTCCGATCTNNNTGCTGTTCCCACTTGTACTTCTT
AGCACACGTCTGAACTCTTTCCCTACACGACGCTCTTCCGATCTNNNTGCTGTTCCCACTTGTACTTCTT
AGCACACGTCTGAACTCTTTCCCTACACGACGCTCTTCCGATCTNNNCTGAGAAGGGAGAATGGTGTG
AGCACACGTCTGAACTCTTTCCCTACACGACGCTCTTCCGATCTNNNCTGAGAAGGGAGAATGGTGTG
AGCACACGTCTGAACTCTTTCCCTACACGACGCTCTTCCGATCTNNNGTGGAACTCTCTCGCTCTTTCTC
AGCACACGTCTGAACTCTTTCCCTACACGACGCTCTTCCGATCTNNNGTGGAACTCTCTCGCTCTTTCTC
GAGCACACGTCTGAACTCTTTCCCTACACGACGCTCTTCCGATCTNNNCAATTGCAAAAACGTGGAAC
GAGCACACGTCTGAACTCTTTCCCTACACGACGCTCTTCCGATCTNNNCAATTGCAAAAACGTGGAAC
AGCACACGTCTGAACTCTTTCCCTACACGACGCTCTTCCGATCTNNNACAGGGGTCTCACTATGTTTGA
AGCACACGTCTGAACTCTTTCCCTACACGACGCTCTTCCGATCTNNNACAGGGGTCTCACTATGTTTGA

GAGCACACGTCTGAACTCTTTCCCTACACGACGCTCTTCCGATCTNNNGAGAATGGCATGAACCCAGG

GAGCACACGTCTGAACTCTTTCCCTACACGACGCTCTTCCGATCTNNNGAGAATGGCATGAACCCAGG

AGCACACGTCTGAACTCTTTCCCTACACGACGCTCTTCCGATCTNNNTGTAATCCAGCTACTTGGAAG

AGCACACGTCTGAACTCTTTCCCTACACGACGCTCTTCCGATCTNNNTGTAATCCAGCTACTTGGAAG

GAGCACACGTCTGAACTCTTTCCCTACACGACGCTCTTCCGATCTNNNGCAAGGTCATTCTGGAGATCT

GAGCACACGTCTGAACTCTTTCCCTACACGACGCTCTTCCGATCTNNNGCAAGGTCATTCTGGAGATCT

AGCACACGTCTGAACTCTTTCCCTACACGACGCTCTTCCGATCTNNNAACAAAGCAATGAGGTTGGCA

AGCACACGTCTGAACTCTTTCCCTACACGACGCTCTTCCGATCTNNNAACAAAGCAATGAGGTTGGCA

GAGCACACGTCTGAACTCTTTCCCTACACGACGCTCTTCCGATCTNNNGCAGTGTTTTGTGTGTGCTTT

GAGCACACGTCTGAACTCTTTCCCTACACGACGCTCTTCCGATCTNNNGCAGTGTTTTGTGTGTGCTTT

AGCACACGTCTGAACTCTTTCCCTACACGACGCTCTTCCGATCTNNNATGAACACACTCTTTAATTTCAT

AGCACACGTCTGAACTCTTTCCCTACACGACGCTCTTCCGATCTNNNATGAACACACTCTTTAATTTCAT

GCACACGTCTGAACTCTTTCCCTACACGACGCTCTTCCGATCTNNNCCATGCCCGGCTAATTTTTTGTA

GCACACGTCTGAACTCTTTCCCTACACGACGCTCTTCCGATCTNNNCCATGCCCGGCTAATTTTTTGTA

GAGCACACGTCTGAACTCTTTCCCTACACGACGCTCTTCCGATCTNNNCCTGTAGTCCCAGCTACTTGG

GAGCACACGTCTGAACTCTTTCCCTACACGACGCTCTTCCGATCTNNNCCTGTAGTCCCAGCTACTTGG

AGCACACGTCTGAACTCTTTCCCTACACGACGCTCTTCCGATCTNNNTATTTACTGGCCAAGCTGTTG

AGCACACGTCTGAACTCTTTCCCTACACGACGCTCTTCCGATCTNNNTATTTACTGGCCAAGCTGTTG

AGAGCACACGTCTGAACTCTTTCCCTACACGACGCTCTTCCGATCTNNNTCTATACCAGGGTTCATTCC

AGAGCACACGTCTGAACTCTTTCCCTACACGACGCTCTTCCGATCTNNNTCTATACCAGGGTTCATTCC

GAGCACACGTCTGAACTCTTTCCCTACACGACGCTCTTCCGATCTNNNCGAGATGATGCCATTGTACTC

GAGCACACGTCTGAACTCTTTCCCTACACGACGCTCTTCCGATCTNNNCGAGATGATGCCATTGTACTC

AGCACACGTCTGAACTCTTTCCCTACACGACGCTCTTCCGATCTNNNCTCCTGAGTAGCTGGGATTACA

AGCACACGTCTGAACTCTTTCCCTACACGACGCTCTTCCGATCTNNNCTCCTGAGTAGCTGGGATTACA

AGCACACGTCTGAACTCTTTCCCTACACGACGCTCTTCCGATCTNNNGAAGAGATTGATCATGTAATT

AGCACACGTCTGAACTCTTTCCCTACACGACGCTCTTCCGATCTNNNGAAGAGATTGATCATGTAATT

GCACACGTCTGAACTCTTTCCCTACACGACGCTCTTCCGATCTNNNCAACATGAGCGAAACTCTGTCT

GCACACGTCTGAACTCTTTCCCTACACGACGCTCTTCCGATCTNNNCAACATGAGCGAAACTCTGTCT

AGCACACGTCTGAACTCTTTCCCTACACGACGCTCTTCCGATCTNNNCTGAAAGGCGGTTCTATTTGCA

AGCACACGTCTGAACTCTTTCCCTACACGACGCTCTTCCGATCTNNNCTGAAAGGCGGTTCTATTTGCA

CACACGTCTGAACTCTTTCCCTACACGACGCTCTTCCGATCTNNNGCCCTATGGAATTGTAACTTGCCC

CACACGTCTGAACTCTTTCCCTACACGACGCTCTTCCGATCTNNNGCCCTATGGAATTGTAACTTGCCC

CACACGTCTGAACTCTTTCCCTACACGACGCTCTTCCGATCTNNNGCATGCCTGTAATCCTAGCTACTC

CACACGTCTGAACTCTTTCCCTACACGACGCTCTTCCGATCTNNNGCATGCCTGTAATCCTAGCTACTC

AGCACACGTCTGAACTCTTTCCCTACACGACGCTCTTCCGATCTNNNTCTATCTTATCTGCTATCCCATA

AGCACACGTCTGAACTCTTTCCCTACACGACGCTCTTCCGATCTNNNTCTATCTTATCTGCTATCCCATA

AGCACACGTCTGAACTCTTTCCCTACACGACGCTCTTCCGATCTNNNAGTGTGAGGAACCCAGTGATT

AGCACACGTCTGAACTCTTTCCCTACACGACGCTCTTCCGATCTNNNAGTGTGAGGAACCCAGTGATT

GAGCACACGTCTGAACTCTTTCCCTACACGACGCTCTTCCGATCTNNNGTTCTCCTGGTTTTCTTTCTG

GAGCACACGTCTGAACTCTTTCCCTACACGACGCTCTTCCGATCTNNNGTTCTCCTGGTTTTCTTTCTG

GAGCACACGTCTGAACTCTTTCCCTACACGACGCTCTTCCGATCTNNNAGCTTCAATTGTGAATGCAG

GAGCACACGTCTGAACTCTTTCCCTACACGACGCTCTTCCGATCTNNNAGCTTCAATTGTGAATGCAG

GAGCACACGTCTGAACTCTTTCCCTACACGACGCTCTTCCGATCTNNNGCTGAGACAAGGGAGTGTT

GAGCACACGTCTGAACTCTTTCCCTACACGACGCTCTTCCGATCTNNNGCTGAGACAAGGGAGTGTT
AGCACACGTCTGAACTCTTTCCCTACACGACGCTCTTCCGATCTNNNATGGTACTTCCCTCCAAACTGG
AGCACACGTCTGAACTCTTTCCCTACACGACGCTCTTCCGATCTNNNATGGTACTTCCCTCCAAACTGG
AGAGCACACGTCTGAACTCTTTCCCTACACGACGCTCTTCCGATCTNNNAGAATTGTGTGTATCAGCT
AGAGCACACGTCTGAACTCTTTCCCTACACGACGCTCTTCCGATCTNNNAGAATTGTGTGTATCAGCT
AGCACACGTCTGAACTCTTTCCCTACACGACGCTCTTCCGATCTNNNTTGCTGAATCCAAACCCATTTG
AGCACACGTCTGAACTCTTTCCCTACACGACGCTCTTCCGATCTNNNTTGCTGAATCCAAACCCATTTG
AGCACACGTCTGAACTCTTTCCCTACACGACGCTCTTCCGATCTNNNGGACATGTAAATCATTTGTGA
AGCACACGTCTGAACTCTTTCCCTACACGACGCTCTTCCGATCTNNNGGACATGTAAATCATTTGTGA
GCACACGTCTGAACTCTTTCCCTACACGACGCTCTTCCGATCTNNNGTGGTAGCTCTGCTGATGAACCC
GCACACGTCTGAACTCTTTCCCTACACGACGCTCTTCCGATCTNNNGTGGTAGCTCTGCTGATGAACCC
GCACACGTCTGAACTCTTTCCCTACACGACGCTCTTCCGATCTNNNAGTCCTCCATAAATTATGTAACC
GCACACGTCTGAACTCTTTCCCTACACGACGCTCTTCCGATCTNNNAGTCCTCCATAAATTATGTAACC
AGCACACGTCTGAACTCTTTCCCTACACGACGCTCTTCCGATCTNNNTGTCAGTGGATGTACCTTTCTG
AGCACACGTCTGAACTCTTTCCCTACACGACGCTCTTCCGATCTNNNTGTCAGTGGATGTACCTTTCTG
AGCACACGTCTGAACTCTTTCCCTACACGACGCTCTTCCGATCTNNNCTGTAGTCCCAGCTACTCAGG
AGCACACGTCTGAACTCTTTCCCTACACGACGCTCTTCCGATCTNNNCTGTAGTCCCAGCTACTCAGG
CACACGTCTGAACTCTTTCCCTACACGACGCTCTTCCGATCTNNNTGTTGCCATCTCTAACCTACCAGC
CACACGTCTGAACTCTTTCCCTACACGACGCTCTTCCGATCTNNNTGTTGCCATCTCTAACCTACCAGC
GCACACGTCTGAACTCTTTCCCTACACGACGCTCTTCCGATCTNNNATGGTCTTGATCTCCTGAACTCG
GCACACGTCTGAACTCTTTCCCTACACGACGCTCTTCCGATCTNNNATGGTCTTGATCTCCTGAACTCG
AGCACACGTCTGAACTCTTTCCCTACACGACGCTCTTCCGATCTNNNACATGACCATCAGATTGAAGCC
AGCACACGTCTGAACTCTTTCCCTACACGACGCTCTTCCGATCTNNNACATGACCATCAGATTGAAGCC
AGCACACGTCTGAACTCTTTCCCTACACGACGCTCTTCCGATCTNNNAGGGTTGAAGAGGATGTAGT
AGCACACGTCTGAACTCTTTCCCTACACGACGCTCTTCCGATCTNNNAGGGTTGAAGAGGATGTAGT
CACACGTCTGAACTCTTTCCCTACACGACGCTCTTCCGATCTNNNGTTTACCTGTCTGCCTCCTCATTCAT
CACACGTCTGAACTCTTTCCCTACACGACGCTCTTCCGATCTNNNGTTTACCTGTCTGCCTCCTCATTCAT
AGCACACGTCTGAACTCTTTCCCTACACGACGCTCTTCCGATCTNNNGTGGTGAATTCCCTAAGCTTT
AGCACACGTCTGAACTCTTTCCCTACACGACGCTCTTCCGATCTNNNGTGGTGAATTCCCTAAGCTTT
GAGCACACGTCTGAACTCTTTCCCTACACGACGCTCTTCCGATCTNNNCCCATGCCTGTAATTTCAGCT
GAGCACACGTCTGAACTCTTTCCCTACACGACGCTCTTCCGATCTNNNCCCATGCCTGTAATTTCAGCT
AGCACACGTCTGAACTCTTTCCCTACACGACGCTCTTCCGATCTNNNAGAGGAAGGCAGACTGACTA
AGCACACGTCTGAACTCTTTCCCTACACGACGCTCTTCCGATCTNNNAGAGGAAGGCAGACTGACTA
GAGCACACGTCTGAACTCTTTCCCTACACGACGCTCTTCCGATCTNNNCCGTTCCCTTCCTTTCTCTTCTC
GAGCACACGTCTGAACTCTTTCCCTACACGACGCTCTTCCGATCTNNNCCGTTCCCTTCCTTTCTCTTCTC
GCACACGTCTGAACTCTTTCCCTACACGACGCTCTTCCGATCTNNNGAGTGGAAGTTGAAGTTTGTG
GCACACGTCTGAACTCTTTCCCTACACGACGCTCTTCCGATCTNNNGAGTGGAAGTTGAAGTTTGTG
GCACACGTCTGAACTCTTTCCCTACACGACGCTCTTCCGATCTNNNAACCTGTTCTTCCATGAGGCATT
GCACACGTCTGAACTCTTTCCCTACACGACGCTCTTCCGATCTNNNAACCTGTTCTTCCATGAGGCATT
GAGCACACGTCTGAACTCTTTCCCTACACGACGCTCTTCCGATCTNNNTTATAGCTCAGTGCAGCCTTG
GAGCACACGTCTGAACTCTTTCCCTACACGACGCTCTTCCGATCTNNNTTATAGCTCAGTGCAGCCTTG
GAGCACACGTCTGAACTCTTTCCCTACACGACGCTCTTCCGATCTNNNAGTTCATGCTTTATCCCTGGC
GAGCACACGTCTGAACTCTTTCCCTACACGACGCTCTTCCGATCTNNNAGTTCATGCTTTATCCCTGGC

AGCACACGTCTGAACTCTTTCCCTACACGACGCTCTTCCGATCTNNNCTCAACCCCATTTACAACAGCTG
AGCACACGTCTGAACTCTTTCCCTACACGACGCTCTTCCGATCTNNNCTCAACCCCATTTACAACAGCTG
GAGCACACGTCTGAACTCTTTCCCTACACGACGCTCTTCCGATCTNNNATATGCCAACAACAGTCAAG
GAGCACACGTCTGAACTCTTTCCCTACACGACGCTCTTCCGATCTNNNATATGCCAACAACAGTCAAG
AGCACACGTCTGAACTCTTTCCCTACACGACGCTCTTCCGATCTNNNCTGTGAAAAATCACCTGGTGA
AGCACACGTCTGAACTCTTTCCCTACACGACGCTCTTCCGATCTNNNCTGTGAAAAATCACCTGGTGA
AGCACACGTCTGAACTCTTTCCCTACACGACGCTCTTCCGATCTNNNGCGCTTATAATCCCAGCTACTC
AGCACACGTCTGAACTCTTTCCCTACACGACGCTCTTCCGATCTNNNGCGCTTATAATCCCAGCTACTC
AGCACACGTCTGAACTCTTTCCCTACACGACGCTCTTCCGATCTNNNGATGGAGGACATTGAGTGTG
AGCACACGTCTGAACTCTTTCCCTACACGACGCTCTTCCGATCTNNNGATGGAGGACATTGAGTGTG
CACACGTCTGAACTCTTTCCCTACACGACGCTCTTCCGATCTNNNCCATAATCCCATGACAAACACCAC
CACACGTCTGAACTCTTTCCCTACACGACGCTCTTCCGATCTNNNCCATAATCCCATGACAAACACCAC
GCACACGTCTGAACTCTTTCCCTACACGACGCTCTTCCGATCTNNNATAGGAGGTTTGGAGATCAGGC
GCACACGTCTGAACTCTTTCCCTACACGACGCTCTTCCGATCTNNNATAGGAGGTTTGGAGATCAGGC
GAGCACACGTCTGAACTCTTTCCCTACACGACGCTCTTCCGATCTNNNAAGTCCCTTTTGGGGAAGAA
GAGCACACGTCTGAACTCTTTCCCTACACGACGCTCTTCCGATCTNNNAAGTCCCTTTTGGGGAAGAA
AGCACACGTCTGAACTCTTTCCCTACACGACGCTCTTCCGATCTNNNTCAACCCATTACCCTTTAGCTCC
AGCACACGTCTGAACTCTTTCCCTACACGACGCTCTTCCGATCTNNNTCAACCCATTACCCTTTAGCTCC
AGAGCACACGTCTGAACTCTTTCCCTACACGACGCTCTTCCGATCTNNNACAAGCTCAAACTGCACATT
AGAGCACACGTCTGAACTCTTTCCCTACACGACGCTCTTCCGATCTNNNACAAGCTCAAACTGCACATT
AGCACACGTCTGAACTCTTTCCCTACACGACGCTCTTCCGATCTNNNTTCTGAAACACACACACACACA
AGCACACGTCTGAACTCTTTCCCTACACGACGCTCTTCCGATCTNNNTTCTGAAACACACACACACACA
GAGCACACGTCTGAACTCTTTCCCTACACGACGCTCTTCCGATCTNNNATTTACCAAAGGTGTGGGGT
GAGCACACGTCTGAACTCTTTCCCTACACGACGCTCTTCCGATCTNNNATTTACCAAAGGTGTGGGGT
AAGAGCACACGTCTGAACTCTTTCCCTACACGACGCTCTTCCGATCTNNNGGATGCCCTTTTGTATTTG
AAGAGCACACGTCTGAACTCTTTCCCTACACGACGCTCTTCCGATCTNNNGGATGCCCTTTTGTATTTG
GAGCACACGTCTGAACTCTTTCCCTACACGACGCTCTTCCGATCTNNNTGGGCACTTGTTATAGAACCA
GAGCACACGTCTGAACTCTTTCCCTACACGACGCTCTTCCGATCTNNNTGGGCACTTGTTATAGAACCA
GAGCACACGTCTGAACTCTTTCCCTACACGACGCTCTTCCGATCTNNNCTTACGCCTGTAATCCTAGCA
GAGCACACGTCTGAACTCTTTCCCTACACGACGCTCTTCCGATCTNNNCTTACGCCTGTAATCCTAGCA
AGAGCACACGTCTGAACTCTTTCCCTACACGACGCTCTTCCGATCTNNNTCTTCACAAAACACTGAGCT
AGAGCACACGTCTGAACTCTTTCCCTACACGACGCTCTTCCGATCTNNNTCTTCACAAAACACTGAGCT
GAGCACACGTCTGAACTCTTTCCCTACACGACGCTCTTCCGATCTNNNGAGATCTCGCCACTGCACTCC
GAGCACACGTCTGAACTCTTTCCCTACACGACGCTCTTCCGATCTNNNGAGATCTCGCCACTGCACTCC
GCACACGTCTGAACTCTTTCCCTACACGACGCTCTTCCGATCTNNNTAATCTGGCAGGTAGGTGAGG
GCACACGTCTGAACTCTTTCCCTACACGACGCTCTTCCGATCTNNNTAATCTGGCAGGTAGGTGAGG
AGAGCACACGTCTGAACTCTTTCCCTACACGACGCTCTTCCGATCTNNNTACATTTCACAGTCCTTGCC
AGAGCACACGTCTGAACTCTTTCCCTACACGACGCTCTTCCGATCTNNNTACATTTCACAGTCCTTGCC
AGCACACGTCTGAACTCTTTCCCTACACGACGCTCTTCCGATCTNNNGGCTACAGAGGGAGACTACAT
AGCACACGTCTGAACTCTTTCCCTACACGACGCTCTTCCGATCTNNNGGCTACAGAGGGAGACTACAT
GAGCACACGTCTGAACTCTTTCCCTACACGACGCTCTTCCGATCTNNNTGGGGTAACAAATGGTGAAC
GAGCACACGTCTGAACTCTTTCCCTACACGACGCTCTTCCGATCTNNNTGGGGTAACAAATGGTGAAC
GCACACGTCTGAACTCTTTCCCTACACGACGCTCTTCCGATCTNNNAGTTGGTTGTCCTGTTATCCTCC

AGCACACGTCTGAACTCTTTCCCTACACGACGCTCTTCCGATCTNNNAGTTGGTTGTCCTGTTATCCTCC
AGCACACGTCTGAACTCTTTCCCTACACGACGCTCTTCCGATCTNNNAGATGGTACTGACTCTTACCTG
AGCACACGTCTGAACTCTTTCCCTACACGACGCTCTTCCGATCTNNNAGATGGTACTGACTCTTACCTG
GAGCACACGTCTGAACTCTTTCCCTACACGACGCTCTTCCGATCTNNNATATGGATCAAGTGCAGGAG
GAGCACACGTCTGAACTCTTTCCCTACACGACGCTCTTCCGATCTNNNATATGGATCAAGTGCAGGAG
GAGCACACGTCTGAACTCTTTCCCTACACGACGCTCTTCCGATCTNNNAACACATGGCCTAAACCAGAG
GAGCACACGTCTGAACTCTTTCCCTACACGACGCTCTTCCGATCTNNNAACACATGGCCTAAACCAGAG
GAGCACACGTCTGAACTCTTTCCCTACACGACGCTCTTCCGATCTNNNACCCATAGCACTGGGAAAAAC
GAGCACACGTCTGAACTCTTTCCCTACACGACGCTCTTCCGATCTNNNACCCATAGCACTGGGAAAAAC
AGCACACGTCTGAACTCTTTCCCTACACGACGCTCTTCCGATCTNNNCAGTAGTTTCCATTTGTTGTCAC
AGCACACGTCTGAACTCTTTCCCTACACGACGCTCTTCCGATCTNNNCAGTAGTTTCCATTTGTTGTCAC
AGCACACGTCTGAACTCTTTCCCTACACGACGCTCTTCCGATCTNNNTGGCAAGACATAAACTTGTAG
AGCACACGTCTGAACTCTTTCCCTACACGACGCTCTTCCGATCTNNNTGGCAAGACATAAACTTGTAG
AGCACACGTCTGAACTCTTTCCCTACACGACGCTCTTCCGATCTNNNAGCTGAAAGTCCAAGACTACTG
AGCACACGTCTGAACTCTTTCCCTACACGACGCTCTTCCGATCTNNNAGCTGAAAGTCCAAGACTACTG
AGCACACGTCTGAACTCTTTCCCTACACGACGCTCTTCCGATCTNNNGTGAACCACTCTAGTACGACAG
AGCACACGTCTGAACTCTTTCCCTACACGACGCTCTTCCGATCTNNNGTGAACCACTCTAGTACGACAG
AGCACACGTCTGAACTCTTTCCCTACACGACGCTCTTCCGATCTNNNAAATCATCCCTACTTGGTCCCAG
AGCACACGTCTGAACTCTTTCCCTACACGACGCTCTTCCGATCTNNNAAATCATCCCTACTTGGTCCCAG
AGCACACGTCTGAACTCTTTCCCTACACGACGCTCTTCCGATCTNNNGAGATTGTGCCACTCTACTCCA
AGCACACGTCTGAACTCTTTCCCTACACGACGCTCTTCCGATCTNNNGAGATTGTGCCACTCTACTCCA
AGCACACGTCTGAACTCTTTCCCTACACGACGCTCTTCCGATCTNNNTGTACAAGATGCAGCAAAGATG
AGCACACGTCTGAACTCTTTCCCTACACGACGCTCTTCCGATCTNNNTGTACAAGATGCAGCAAAGATG
AGCACACGTCTGAACTCTTTCCCTACACGACGCTCTTCCGATCTNNNCAAGATCCTGTGGTCTGAAGT
AGCACACGTCTGAACTCTTTCCCTACACGACGCTCTTCCGATCTNNNCAAGATCCTGTGGTCTGAAGT
GCACACGTCTGAACTCTTTCCCTACACGACGCTCTTCCGATCTNNNTATTGTTCTACAAAGGCAGAAAC
GCACACGTCTGAACTCTTTCCCTACACGACGCTCTTCCGATCTNNNTATTGTTCTACAAAGGCAGAAAC
AGCACACGTCTGAACTCTTTCCCTACACGACGCTCTTCCGATCTNNNTTGTGTCAGCAGGAGAATCAG
AGCACACGTCTGAACTCTTTCCCTACACGACGCTCTTCCGATCTNNNTTGTGTCAGCAGGAGAATCAG
GCACACGTCTGAACTCTTTCCCTACACGACGCTCTTCCGATCTNNNTGAAACTGAAAAGTGCTGTGTGA
GCACACGTCTGAACTCTTTCCCTACACGACGCTCTTCCGATCTNNNTGAAACTGAAAAGTGCTGTGTGA
GAGCACACGTCTGAACTCTTTCCCTACACGACGCTCTTCCGATCTNNNCTGTAATCCCAGCTACTCGAG
GAGCACACGTCTGAACTCTTTCCCTACACGACGCTCTTCCGATCTNNNCTGTAATCCCAGCTACTCGAG
GAGCACACGTCTGAACTCTTTCCCTACACGACGCTCTTCCGATCTNNNTACTAGGCAATGGGGATAGG
GAGCACACGTCTGAACTCTTTCCCTACACGACGCTCTTCCGATCTNNNTACTAGGCAATGGGGATAGG
GCACACGTCTGAACTCTTTCCCTACACGACGCTCTTCCGATCTNNNGGCAAGGACTTCATGTCTGAAA
GCACACGTCTGAACTCTTTCCCTACACGACGCTCTTCCGATCTNNNGGCAAGGACTTCATGTCTGAAA
AGCACACGTCTGAACTCTTTCCCTACACGACGCTCTTCCGATCTNNNCTCCTCTTCCACTCACATTCACAC
AGCACACGTCTGAACTCTTTCCCTACACGACGCTCTTCCGATCTNNNCTCCTCTTCCACTCACATTCACAC
GAGCACACGTCTGAACTCTTTCCCTACACGACGCTCTTCCGATCTNNNAGTACTTCTGCCCCAGAGTTA
GAGCACACGTCTGAACTCTTTCCCTACACGACGCTCTTCCGATCTNNNAGTACTTCTGCCCCAGAGTTA
GAGCACACGTCTGAACTCTTTCCCTACACGACGCTCTTCCGATCTNNNCAACAACAGCGAAACTCCATC
GAGCACACGTCTGAACTCTTTCCCTACACGACGCTCTTCCGATCTNNNCAACAACAGCGAAACTCCATC

AGCACACGTCTGAACTCTTTCCCTACACGACGCTCTTCCGATCTNNNTGGGATTTGACTTAGGTGAT
AGCACACGTCTGAACTCTTTCCCTACACGACGCTCTTCCGATCTNNNTGGGATTTGACTTAGGTGAT
GCACACGTCTGAACTCTTTCCCTACACGACGCTCTTCCGATCTNNNACAAACATCATGGTGGGCTTTT
GCACACGTCTGAACTCTTTCCCTACACGACGCTCTTCCGATCTNNNACAAACATCATGGTGGGCTTTT
AGCACACGTCTGAACTCTTTCCCTACACGACGCTCTTCCGATCTNNNAATGCAGGAGAATTGCTTGAA
AGCACACGTCTGAACTCTTTCCCTACACGACGCTCTTCCGATCTNNNAATGCAGGAGAATTGCTTGAA
AGCACACGTCTGAACTCTTTCCCTACACGACGCTCTTCCGATCTNNNGAGACAGGAGAATTGCTTGA
AGCACACGTCTGAACTCTTTCCCTACACGACGCTCTTCCGATCTNNNGAGACAGGAGAATTGCTTGA
AGCACACGTCTGAACTCTTTCCCTACACGACGCTCTTCCGATCTNNNTGGTTTCAAACTCCTGACCTCA
AGCACACGTCTGAACTCTTTCCCTACACGACGCTCTTCCGATCTNNNTGGTTTCAAACTCCTGACCTCA
AGCACACGTCTGAACTCTTTCCCTACACGACGCTCTTCCGATCTNNNGTGTAGACTTGGGAGGATGA
AGCACACGTCTGAACTCTTTCCCTACACGACGCTCTTCCGATCTNNNGTGTAGACTTGGGAGGATGA
AGCACACGTCTGAACTCTTTCCCTACACGACGCTCTTCCGATCTNNNTGTGGGATAATGTGTGGAAA
AGCACACGTCTGAACTCTTTCCCTACACGACGCTCTTCCGATCTNNNTGTGGGATAATGTGTGGAAA
AGCACACGTCTGAACTCTTTCCCTACACGACGCTCTTCCGATCTNNNTGCACATCCAGACTGACTTAC
AGCACACGTCTGAACTCTTTCCCTACACGACGCTCTTCCGATCTNNNTGCACATCCAGACTGACTTAC
AGCACACGTCTGAACTCTTTCCCTACACGACGCTCTTCCGATCTNNNTGGCCATGAATCAATGAATGC
AGCACACGTCTGAACTCTTTCCCTACACGACGCTCTTCCGATCTNNNTGGCCATGAATCAATGAATGC
GCACACGTCTGAACTCTTTCCCTACACGACGCTCTTCCGATCTNNNCCTTATTGGACTTCCCTCTTGGTC
GCACACGTCTGAACTCTTTCCCTACACGACGCTCTTCCGATCTNNNCCTTATTGGACTTCCCTCTTGGTC
GAGCACACGTCTGAACTCTTTCCCTACACGACGCTCTTCCGATCTNNNTCCCTTTTCCAGAGTCTCCAG
GAGCACACGTCTGAACTCTTTCCCTACACGACGCTCTTCCGATCTNNNTCCCTTTTCCAGAGTCTCCAG
AGCACACGTCTGAACTCTTTCCCTACACGACGCTCTTCCGATCTNNNCCTCAAGCAGACCTAATAGGC
AGCACACGTCTGAACTCTTTCCCTACACGACGCTCTTCCGATCTNNNCCTCAAGCAGACCTAATAGGC
GCACACGTCTGAACTCTTTCCCTACACGACGCTCTTCCGATCTNNNCTGGAGAGACAGTGCGATTTA
GCACACGTCTGAACTCTTTCCCTACACGACGCTCTTCCGATCTNNNCTGGAGAGACAGTGCGATTTA
CACACGTCTGAACTCTTTCCCTACACGACGCTCTTCCGATCTNNNTCCACTGTCCCCAAAATTTCTCTCA
CACACGTCTGAACTCTTTCCCTACACGACGCTCTTCCGATCTNNNTCCACTGTCCCCAAAATTTCTCTCA
GCACACGTCTGAACTCTTTCCCTACACGACGCTCTTCCGATCTNNNTGACAGAGCAAGATTCCGTCTCC
GCACACGTCTGAACTCTTTCCCTACACGACGCTCTTCCGATCTNNNTGACAGAGCAAGATTCCGTCTCC
AGCACACGTCTGAACTCTTTCCCTACACGACGCTCTTCCGATCTNNNACAAAAATTAGCCAGGTGTG
AGCACACGTCTGAACTCTTTCCCTACACGACGCTCTTCCGATCTNNNACAAAAATTAGCCAGGTGTG
GCACACGTCTGAACTCTTTCCCTACACGACGCTCTTCCGATCTNNNTCATTGATGAAGGTGGCTACA
GCACACGTCTGAACTCTTTCCCTACACGACGCTCTTCCGATCTNNNTCATTGATGAAGGTGGCTACA
GAGCACACGTCTGAACTCTTTCCCTACACGACGCTCTTCCGATCTNNNGGATCACTTGAGGTCAGGA
GAGCACACGTCTGAACTCTTTCCCTACACGACGCTCTTCCGATCTNNNGGATCACTTGAGGTCAGGA
AGCACACGTCTGAACTCTTTCCCTACACGACGCTCTTCCGATCTNNNGAGCTTGCAGTGAGTTGAGA
AGCACACGTCTGAACTCTTTCCCTACACGACGCTCTTCCGATCTNNNGAGCTTGCAGTGAGTTGAGA
AGCACACGTCTGAACTCTTTCCCTACACGACGCTCTTCCGATCTNNNTCCATGTTCCTCTACCAGTAAG
AGCACACGTCTGAACTCTTTCCCTACACGACGCTCTTCCGATCTNNNTCCATGTTCCTCTACCAGTAAG
GCACACGTCTGAACTCTTTCCCTACACGACGCTCTTCCGATCTNNNTGCAAACCCTAACTGATGTGGA
GCACACGTCTGAACTCTTTCCCTACACGACGCTCTTCCGATCTNNNTGCAAACCCTAACTGATGTGGA
GAGCACACGTCTGAACTCTTTCCCTACACGACGCTCTTCCGATCTNNNTGAACAGATATCACACCACTG

GAGCACACGTCTGAACTCTTTCCCTACACGACGCTCTTCCGATCTNNNTGAACAGATATCACACCACTG
GAGCACACGTCTGAACTCTTTCCCTACACGACGCTCTTCCGATCTNNNGCACGACATATGGGTATGCTC
GAGCACACGTCTGAACTCTTTCCCTACACGACGCTCTTCCGATCTNNNGCACGACATATGGGTATGCTC
AGCACACGTCTGAACTCTTTCCCTACACGACGCTCTTCCGATCTNNNACTTTGAAGCTAGGAACCTCAG
AGCACACGTCTGAACTCTTTCCCTACACGACGCTCTTCCGATCTNNNACTTTGAAGCTAGGAACCTCAG
AGCACACGTCTGAACTCTTTCCCTACACGACGCTCTTCCGATCTNNNACAATAAAAATGCCACAAGTAG
AGCACACGTCTGAACTCTTTCCCTACACGACGCTCTTCCGATCTNNNACAATAAAAATGCCACAAGTAG
GAGCACACGTCTGAACTCTTTCCCTACACGACGCTCTTCCGATCTNNNCATTTGTGATTTCCAACCTCGC
GAGCACACGTCTGAACTCTTTCCCTACACGACGCTCTTCCGATCTNNNCATTTGTGATTTCCAACCTCGC
GCACACGTCTGAACTCTTTCCCTACACGACGCTCTTCCGATCTNNNTTAGGATGGGAGGATGGTGGA
GCACACGTCTGAACTCTTTCCCTACACGACGCTCTTCCGATCTNNNTTAGGATGGGAGGATGGTGGA
AGCACACGTCTGAACTCTTTCCCTACACGACGCTCTTCCGATCTNNNCTGAGATGGGAGGATTGCTAG
AGCACACGTCTGAACTCTTTCCCTACACGACGCTCTTCCGATCTNNNCTGAGATGGGAGGATTGCTAG
GAGCACACGTCTGAACTCTTTCCCTACACGACGCTCTTCCGATCTNNNAGATTGTGCCACTGTACTCCA
GAGCACACGTCTGAACTCTTTCCCTACACGACGCTCTTCCGATCTNNNAGATTGTGCCACTGTACTCCA
GCACACGTCTGAACTCTTTCCCTACACGACGCTCTTCCGATCTNNNCTCACCTCTGCCTCTGTCTCAT
GCACACGTCTGAACTCTTTCCCTACACGACGCTCTTCCGATCTNNNCTCACCTCTGCCTCTGTCTCAT
GAGCACACGTCTGAACTCTTTCCCTACACGACGCTCTTCCGATCTNNNCTTGAACCTGGGAGGCAGAG
GAGCACACGTCTGAACTCTTTCCCTACACGACGCTCTTCCGATCTNNNCTTGAACCTGGGAGGCAGAG
GCACACGTCTGAACTCTTTCCCTACACGACGCTCTTCCGATCTNNNACTTTCCCACTTCACAGCACTATC
GCACACGTCTGAACTCTTTCCCTACACGACGCTCTTCCGATCTNNNACTTTCCCACTTCACAGCACTATC
GAGCACACGTCTGAACTCTTTCCCTACACGACGCTCTTCCGATCTNNNTGACTATGGAGTGAAGATGA
GAGCACACGTCTGAACTCTTTCCCTACACGACGCTCTTCCGATCTNNNTGACTATGGAGTGAAGATGA
AGCACACGTCTGAACTCTTTCCCTACACGACGCTCTTCCGATCTNNNTGTTGGACAGAGCACCTCTAGA
AGCACACGTCTGAACTCTTTCCCTACACGACGCTCTTCCGATCTNNNTGTTGGACAGAGCACCTCTAGA
AGCACACGTCTGAACTCTTTCCCTACACGACGCTCTTCCGATCTNNNCTAGGAAATGAAGCGAGGCTG
AGCACACGTCTGAACTCTTTCCCTACACGACGCTCTTCCGATCTNNNCTAGGAAATGAAGCGAGGCTG
AGCACACGTCTGAACTCTTTCCCTACACGACGCTCTTCCGATCTNNNGCCTGGCCCATAGCTTTCTTTT
AGCACACGTCTGAACTCTTTCCCTACACGACGCTCTTCCGATCTNNNGCCTGGCCCATAGCTTTCTTTT
AGCACACGTCTGAACTCTTTCCCTACACGACGCTCTTCCGATCTNNNCTGAGGCAGGAGAATCACTTG
AGCACACGTCTGAACTCTTTCCCTACACGACGCTCTTCCGATCTNNNCTGAGGCAGGAGAATCACTTG
GAGCACACGTCTGAACTCTTTCCCTACACGACGCTCTTCCGATCTNNNATCTATGGTTTCAGGCATCCA
GAGCACACGTCTGAACTCTTTCCCTACACGACGCTCTTCCGATCTNNNATCTATGGTTTCAGGCATCCA
CACACGTCTGAACTCTTTCCCTACACGACGCTCTTCCGATCTNNNGGTACAGAGCAAACAATGTGGG
CACACGTCTGAACTCTTTCCCTACACGACGCTCTTCCGATCTNNNGGTACAGAGCAAACAATGTGGG
GCACACGTCTGAACTCTTTCCCTACACGACGCTCTTCCGATCTNNNTTAGCCTGAAATTGCTTGAACCC
GCACACGTCTGAACTCTTTCCCTACACGACGCTCTTCCGATCTNNNTTAGCCTGAAATTGCTTGAACCC
GAGCACACGTCTGAACTCTTTCCCTACACGACGCTCTTCCGATCTNNNATTGGACAATTCTCCAGGAG
GAGCACACGTCTGAACTCTTTCCCTACACGACGCTCTTCCGATCTNNNATTGGACAATTCTCCAGGAG
AGCACACGTCTGAACTCTTTCCCTACACGACGCTCTTCCGATCTNNNTGTTTCAATTTCTTGATCTGAAG
AGCACACGTCTGAACTCTTTCCCTACACGACGCTCTTCCGATCTNNNTGTTTCAATTTCTTGATCTGAAG
GAGCACACGTCTGAACTCTTTCCCTACACGACGCTCTTCCGATCTNNNTTCTGACCTCAAGTAATCCACC
GAGCACACGTCTGAACTCTTTCCCTACACGACGCTCTTCCGATCTNNNTTCTGACCTCAAGTAATCCACC

GCACACGTCTGAACTCTTTCCCTACACGACGCTCTTCCGATCTNNNAGTGCTGGGATTATAGGCTTGA
GCACACGTCTGAACTCTTTCCCTACACGACGCTCTTCCGATCTNNNAGTGCTGGGATTATAGGCTTGA
GAGCACACGTCTGAACTCTTTCCCTACACGACGCTCTTCCGATCTNNNTCTTTGTCAGTCCTATGCAGC
GAGCACACGTCTGAACTCTTTCCCTACACGACGCTCTTCCGATCTNNNTCTTTGTCAGTCCTATGCAGC
GAGCACACGTCTGAACTCTTTCCCTACACGACGCTCTTCCGATCTNNNTTTGCAGAGCTATCCACACT
GAGCACACGTCTGAACTCTTTCCCTACACGACGCTCTTCCGATCTNNNTTTGCAGAGCTATCCACACT
GCACACGTCTGAACTCTTTCCCTACACGACGCTCTTCCGATCTNNNGATCACGAGGTCAGGAATTCGA
GCACACGTCTGAACTCTTTCCCTACACGACGCTCTTCCGATCTNNNGATCACGAGGTCAGGAATTCGA
GAGCACACGTCTGAACTCTTTCCCTACACGACGCTCTTCCGATCTNNNCCAAATAGCTAGTACCACAG
GAGCACACGTCTGAACTCTTTCCCTACACGACGCTCTTCCGATCTNNNCCAAATAGCTAGTACCACAG
GAGCACACGTCTGAACTCTTTCCCTACACGACGCTCTTCCGATCTNNNACACACACACATCATATGTACA
GAGCACACGTCTGAACTCTTTCCCTACACGACGCTCTTCCGATCTNNNACACACACACATCATATGTACA
AGCACACGTCTGAACTCTTTCCCTACACGACGCTCTTCCGATCTNNNCATTCTCCTCATGCCAAAGAAG
AGCACACGTCTGAACTCTTTCCCTACACGACGCTCTTCCGATCTNNNCATTCTCCTCATGCCAAAGAAG
AGCACACGTCTGAACTCTTTCCCTACACGACGCTCTTCCGATCTNNNTGGGTTTCCCTGAATTGAATTG
AGCACACGTCTGAACTCTTTCCCTACACGACGCTCTTCCGATCTNNNTGGGTTTCCCTGAATTGAATTG
GAGCACACGTCTGAACTCTTTCCCTACACGACGCTCTTCCGATCTNNNATCTTTGTGTGTGTGTGTGT
GAGCACACGTCTGAACTCTTTCCCTACACGACGCTCTTCCGATCTNNNATCTTTGTGTGTGTGTGTGT
GCACACGTCTGAACTCTTTCCCTACACGACGCTCTTCCGATCTNNNACCTGTAGTAGTCCCAGCTACTT
GCACACGTCTGAACTCTTTCCCTACACGACGCTCTTCCGATCTNNNACCTGTAGTAGTCCCAGCTACTT
GAGCACACGTCTGAACTCTTTCCCTACACGACGCTCTTCCGATCTNNNTCTGAATCTGCCAACACATCTG
GAGCACACGTCTGAACTCTTTCCCTACACGACGCTCTTCCGATCTNNNTCTGAATCTGCCAACACATCTG
GAGCACACGTCTGAACTCTTTCCCTACACGACGCTCTTCCGATCTNNNACAGAGGGGGTACAATGAA
GAGCACACGTCTGAACTCTTTCCCTACACGACGCTCTTCCGATCTNNNACAGAGGGGGTACAATGAA
GCACACGTCTGAACTCTTTCCCTACACGACGCTCTTCCGATCTNNNGGTTCGTTTTACGTTCCGTGAAG
GCACACGTCTGAACTCTTTCCCTACACGACGCTCTTCCGATCTNNNGGTTCGTTTTACGTTCCGTGAAG
GCACACGTCTGAACTCTTTCCCTACACGACGCTCTTCCGATCTNNNCTTTAACCAGCTATAAACAAGAG
GCACACGTCTGAACTCTTTCCCTACACGACGCTCTTCCGATCTNNNCTTTAACCAGCTATAAACAAGAG
AGCACACGTCTGAACTCTTTCCCTACACGACGCTCTTCCGATCTNNNCCTCTTCTGAAATAGCCTGGTG
AGCACACGTCTGAACTCTTTCCCTACACGACGCTCTTCCGATCTNNNCCTCTTCTGAAATAGCCTGGTG
AGCACACGTCTGAACTCTTTCCCTACACGACGCTCTTCCGATCTNNNCCCACTGATGTAGTCCATGTAG
AGCACACGTCTGAACTCTTTCCCTACACGACGCTCTTCCGATCTNNNCCCACTGATGTAGTCCATGTAG
AGAGCACACGTCTGAACTCTTTCCCTACACGACGCTCTTCCGATCTNNNAGAGGTGTATTTCCCCCTCC
AGAGCACACGTCTGAACTCTTTCCCTACACGACGCTCTTCCGATCTNNNAGAGGTGTATTTCCCCCTCC
AGAGCACACGTCTGAACTCTTTCCCTACACGACGCTCTTCCGATCTNNNGAAGTCTGTCTCCAACTCTGT
AGAGCACACGTCTGAACTCTTTCCCTACACGACGCTCTTCCGATCTNNNGAAGTCTGTCTCCAACTCTGT
AGCACACGTCTGAACTCTTTCCCTACACGACGCTCTTCCGATCTNNNAAGAATGCATCTCAGCCTGTGT
AGCACACGTCTGAACTCTTTCCCTACACGACGCTCTTCCGATCTNNNAAGAATGCATCTCAGCCTGTGT
GAGCACACGTCTGAACTCTTTCCCTACACGACGCTCTTCCGATCTNNNGTACAGGAACAGACACATAG
GAGCACACGTCTGAACTCTTTCCCTACACGACGCTCTTCCGATCTNNNGTACAGGAACAGACACATAG
AGCACACGTCTGAACTCTTTCCCTACACGACGCTCTTCCGATCTNNNTAATCTCAGCACTTTGGGAGGT
AGCACACGTCTGAACTCTTTCCCTACACGACGCTCTTCCGATCTNNNTAATCTCAGCACTTTGGGAGGT
AGCACACGTCTGAACTCTTTCCCTACACGACGCTCTTCCGATCTNNNACTCACCTGGTTGCAGTCTGCAT

AGCACACGTCTGAACTCTTTCCCTACACGACGCTCTTCCGATCTNNNACTCACCTGGTTGCAGTCTGCAT
AGCACACGTCTGAACTCTTTCCCTACACGACGCTCTTCCGATCTNNNCAGGAGAGAAGGCTCAAGAATC
AGCACACGTCTGAACTCTTTCCCTACACGACGCTCTTCCGATCTNNNCAGGAGAGAAGGCTCAAGAATC
GAGCACACGTCTGAACTCTTTCCCTACACGACGCTCTTCCGATCTNNNCTGGGTTCAAGCAATTCTCGTC
GAGCACACGTCTGAACTCTTTCCCTACACGACGCTCTTCCGATCTNNNCTGGGTTCAAGCAATTCTCGTC
GAGCACACGTCTGAACTCTTTCCCTACACGACGCTCTTCCGATCTNNNCAAAGTGCACAGGATGATTCCA
GAGCACACGTCTGAACTCTTTCCCTACACGACGCTCTTCCGATCTNNNCAAAGTGCACAGGATGATTCCA
AGCACACGTCTGAACTCTTTCCCTACACGACGCTCTTCCGATCTNNNTTAGGAGGTATACAGCCACATC
AGCACACGTCTGAACTCTTTCCCTACACGACGCTCTTCCGATCTNNNTTAGGAGGTATACAGCCACATC
AGCACACGTCTGAACTCTTTCCCTACACGACGCTCTTCCGATCTNNNTTCGGGTGGAAGTGAGTTCTTA
AGCACACGTCTGAACTCTTTCCCTACACGACGCTCTTCCGATCTNNNTTCGGGTGGAAGTGAGTTCTTA
AGAGCACACGTCTGAACTCTTTCCCTACACGACGCTCTTCCGATCTNNNCACCACGCCCAGCTAATTTTA
AGAGCACACGTCTGAACTCTTTCCCTACACGACGCTCTTCCGATCTNNNCACCACGCCCAGCTAATTTTA
AGCACACGTCTGAACTCTTTCCCTACACGACGCTCTTCCGATCTNNNCCACCCATCTAAAATCATCCCCAC
AGCACACGTCTGAACTCTTTCCCTACACGACGCTCTTCCGATCTNNNCCACCCATCTAAAATCATCCCCAC
AGCACACGTCTGAACTCTTTCCCTACACGACGCTCTTCCGATCTNNNTGTGTGGGTGATCCTAGTTTGT
AGCACACGTCTGAACTCTTTCCCTACACGACGCTCTTCCGATCTNNNTGTGTGGGTGATCCTAGTTTGT
GAGCACACGTCTGAACTCTTTCCCTACACGACGCTCTTCCGATCTNNNAGTAGCTAGGAGTACAGGCTC
GAGCACACGTCTGAACTCTTTCCCTACACGACGCTCTTCCGATCTNNNAGTAGCTAGGAGTACAGGCTC
GAGCACACGTCTGAACTCTTTCCCTACACGACGCTCTTCCGATCTNNNAACGTACAAGGGCTCCAATGT
GAGCACACGTCTGAACTCTTTCCCTACACGACGCTCTTCCGATCTNNNAACGTACAAGGGCTCCAATGT
GCACACGTCTGAACTCTTTCCCTACACGACGCTCTTCCGATCTNNNCAAGTCAGCTTGCATCTTTGTGAC
GCACACGTCTGAACTCTTTCCCTACACGACGCTCTTCCGATCTNNNCAAGTCAGCTTGCATCTTTGTGAC
GAGCACACGTCTGAACTCTTTCCCTACACGACGCTCTTCCGATCTNNNTAAAAAATTAGCCGGGTGTGG
GAGCACACGTCTGAACTCTTTCCCTACACGACGCTCTTCCGATCTNNNTAAAAAATTAGCCGGGTGTGG
GAGCACACGTCTGAACTCTTTCCCTACACGACGCTCTTCCGATCTNNNTTAGGACTTTACTGTTGGGCT
GAGCACACGTCTGAACTCTTTCCCTACACGACGCTCTTCCGATCTNNNTTAGGACTTTACTGTTGGGCT
GAGCACACGTCTGAACTCTTTCCCTACACGACGCTCTTCCGATCTNNNTCTCCTGTGAACTGAATTCGA
GAGCACACGTCTGAACTCTTTCCCTACACGACGCTCTTCCGATCTNNNTCTCCTGTGAACTGAATTCGA
AGCACACGTCTGAACTCTTTCCCTACACGACGCTCTTCCGATCTNNNTGTGGCTGTCTACTTGTGTCTAC
AGCACACGTCTGAACTCTTTCCCTACACGACGCTCTTCCGATCTNNNTGTGGCTGTCTACTTGTGTCTAC
GAGCACACGTCTGAACTCTTTCCCTACACGACGCTCTTCCGATCTNNNTACAAAAAATTAGCCTGGCGT
GAGCACACGTCTGAACTCTTTCCCTACACGACGCTCTTCCGATCTNNNTACAAAAAATTAGCCTGGCGT
GAGCACACGTCTGAACTCTTTCCCTACACGACGCTCTTCCGATCTNNNTGGGAGAACAATACAACCAAG
GAGCACACGTCTGAACTCTTTCCCTACACGACGCTCTTCCGATCTNNNTGGGAGAACAATACAACCAAG
AGCACACGTCTGAACTCTTTCCCTACACGACGCTCTTCCGATCTNNNCCCATACCAGGTGGCATTATTCA
AGCACACGTCTGAACTCTTTCCCTACACGACGCTCTTCCGATCTNNNCCCATACCAGGTGGCATTATTCA
ACACGTCTGAACTCTTTCCCTACACGACGCTCTTCCGATCTNNNACAGTTCAGTGATCAAACTCAGGAC
ACACGTCTGAACTCTTTCCCTACACGACGCTCTTCCGATCTNNNACAGTTCAGTGATCAAACTCAGGAC
AGCACACGTCTGAACTCTTTCCCTACACGACGCTCTTCCGATCTNNNTAGTGGTGCATGCCTATAATCCC
AGCACACGTCTGAACTCTTTCCCTACACGACGCTCTTCCGATCTNNNTAGTGGTGCATGCCTATAATCCC
AGCACACGTCTGAACTCTTTCCCTACACGACGCTCTTCCGATCTNNNCTTGCAGTGAGCTGAGATTGCC
AGCACACGTCTGAACTCTTTCCCTACACGACGCTCTTCCGATCTNNNCTTGCAGTGAGCTGAGATTGCC

AGCACACGTCTGAACTCTTTCCCTACACGACGCTCTTCCGATCTNNNCGCCTGGCATTTTTTGGTGT

AGCACACGTCTGAACTCTTTCCCTACACGACGCTCTTCCGATCTNNNCGCCTGGCATTTTTTGGTGT

AGCACACGTCTGAACTCTTTCCCTACACGACGCTCTTCCGATCTNNNAAACTCCTGGCCTCAAGTGATT

AGCACACGTCTGAACTCTTTCCCTACACGACGCTCTTCCGATCTNNNAAACTCCTGGCCTCAAGTGATT

GAGCACACGTCTGAACTCTTTCCCTACACGACGCTCTTCCGATCTNNNCCTTTCTCTGTCCCAATCCTCT

GAGCACACGTCTGAACTCTTTCCCTACACGACGCTCTTCCGATCTNNNCCTTTCTCTGTCCCAATCCTCT

AGCACACGTCTGAACTCTTTCCCTACACGACGCTCTTCCGATCTNNNCTGTTGTCCCTGGGAACTGTG

AGCACACGTCTGAACTCTTTCCCTACACGACGCTCTTCCGATCTNNNCTGTTGTCCCTGGGAACTGTG

GCACACGTCTGAACTCTTTCCCTACACGACGCTCTTCCGATCTNNNGAGGCTCCAGAATGATTTACCC

GCACACGTCTGAACTCTTTCCCTACACGACGCTCTTCCGATCTNNNGAGGCTCCAGAATGATTTACCC

GCACACGTCTGAACTCTTTCCCTACACGACGCTCTTCCGATCTNNNGAGGATCACTTGAGCTCAGGTC

GCACACGTCTGAACTCTTTCCCTACACGACGCTCTTCCGATCTNNNGAGGATCACTTGAGCTCAGGTC

AGCACACGTCTGAACTCTTTCCCTACACGACGCTCTTCCGATCTNNNTCATCCCTAATCCAAACAAGCC

AGCACACGTCTGAACTCTTTCCCTACACGACGCTCTTCCGATCTNNNTCATCCCTAATCCAAACAAGCC

AGCACACGTCTGAACTCTTTCCCTACACGACGCTCTTCCGATCTNNNCTCACATCTGTAATCCCAGTGC

AGCACACGTCTGAACTCTTTCCCTACACGACGCTCTTCCGATCTNNNCTCACATCTGTAATCCCAGTGC

GAGCACACGTCTGAACTCTTTCCCTACACGACGCTCTTCCGATCTNNNCCACTCTGGGTAAGGGAAG

GAGCACACGTCTGAACTCTTTCCCTACACGACGCTCTTCCGATCTNNNCCACTCTGGGTAAGGGAAG

AGCACACGTCTGAACTCTTTCCCTACACGACGCTCTTCCGATCTNNNTTTCAAAACACAACCATGCCCTT

AGCACACGTCTGAACTCTTTCCCTACACGACGCTCTTCCGATCTNNNTTTCAAAACACAACCATGCCCTT

GAGCACACGTCTGAACTCTTTCCCTACACGACGCTCTTCCGATCTNNNTAAGGGAGAAAAGCAAGTG

GAGCACACGTCTGAACTCTTTCCCTACACGACGCTCTTCCGATCTNNNTAAGGGAGAAAAGCAAGTG

GCACACGTCTGAACTCTTTCCCTACACGACGCTCTTCCGATCTNNNAAATCCATGCTTCAGAAGGCCT

GCACACGTCTGAACTCTTTCCCTACACGACGCTCTTCCGATCTNNNAAATCCATGCTTCAGAAGGCCT

AGCACACGTCTGAACTCTTTCCCTACACGACGCTCTTCCGATCTNNNGAGGCAGAAGAATCGCTTGA

AGCACACGTCTGAACTCTTTCCCTACACGACGCTCTTCCGATCTNNNGAGGCAGAAGAATCGCTTGA

GAGCACACGTCTGAACTCTTTCCCTACACGACGCTCTTCCGATCTNNNCCTTTGACCCATCCCATGTTT

GAGCACACGTCTGAACTCTTTCCCTACACGACGCTCTTCCGATCTNNNCCTTTGACCCATCCCATGTTT

GAGCACACGTCTGAACTCTTTCCCTACACGACGCTCTTCCGATCTNNNAATGGGATCAGGGTGTGGT

GAGCACACGTCTGAACTCTTTCCCTACACGACGCTCTTCCGATCTNNNAATGGGATCAGGGTGTGGT

GCACACGTCTGAACTCTTTCCCTACACGACGCTCTTCCGATCTNNNGCCATGGTTTTCCGGAGTTTTAG

GCACACGTCTGAACTCTTTCCCTACACGACGCTCTTCCGATCTNNNGCCATGGTTTTCCGGAGTTTTAG

AGCACACGTCTGAACTCTTTCCCTACACGACGCTCTTCCGATCTNNNTAGGCATTTGGGTTGGTTCCAT

AGCACACGTCTGAACTCTTTCCCTACACGACGCTCTTCCGATCTNNNTAGGCATTTGGGTTGGTTCCAT

GCACACGTCTGAACTCTTTCCCTACACGACGCTCTTCCGATCTNNNTTCTTTTCCTTCCTTCCTTCCCCCA

GCACACGTCTGAACTCTTTCCCTACACGACGCTCTTCCGATCTNNNTTCTTTTCCTTCCTTCCTTCCCCCA

GAGCACACGTCTGAACTCTTTCCCTACACGACGCTCTTCCGATCTNNNGGATCATCTGAGGTCAGGA

GAGCACACGTCTGAACTCTTTCCCTACACGACGCTCTTCCGATCTNNNGGATCATCTGAGGTCAGGA

GCACACGTCTGAACTCTTTCCCTACACGACGCTCTTCCGATCTNNNGGAAAGCCAGTGTGACACAAA

GCACACGTCTGAACTCTTTCCCTACACGACGCTCTTCCGATCTNNNGGAAAGCCAGTGTGACACAAA

AGCACACGTCTGAACTCTTTCCCTACACGACGCTCTTCCGATCTNNNAAATACAAAAATTAGCCGGG

AGCACACGTCTGAACTCTTTCCCTACACGACGCTCTTCCGATCTNNNAAATACAAAAATTAGCCGGG

GCACACGTCTGAACTCTTTCCCTACACGACGCTCTTCCGATCTNNNCCTCATAAAGAATGCCAGCCTG

GCACACGTCTGAACTCTTTCCCTACACGACGCTCTTCCGATCTNNNCCTCATAAAGAATGCCAGCCTGT
AGCACACGTCTGAACTCTTTCCCTACACGACGCTCTTCCGATCTNNNCACACTCAAAGTTCCTGTCCTC
AGCACACGTCTGAACTCTTTCCCTACACGACGCTCTTCCGATCTNNNCACACTCAAAGTTCCTGTCCTC
GCACACGTCTGAACTCTTTCCCTACACGACGCTCTTCCGATCTNNNTTACCTCCCTGACCTGACCTTAT
GCACACGTCTGAACTCTTTCCCTACACGACGCTCTTCCGATCTNNNTTACCTCCCTGACCTGACCTTAT
AGCACACGTCTGAACTCTTTCCCTACACGACGCTCTTCCGATCTNNNCACAGGACCTTCTTCCTCTAAC
AGCACACGTCTGAACTCTTTCCCTACACGACGCTCTTCCGATCTNNNCACAGGACCTTCTTCCTCTAAC
AGCACACGTCTGAACTCTTTCCCTACACGACGCTCTTCCGATCTNNNGAGAATCGCTTGAACCCAGG
AGCACACGTCTGAACTCTTTCCCTACACGACGCTCTTCCGATCTNNNGAGAATCGCTTGAACCCAGG
GAGCACACGTCTGAACTCTTTCCCTACACGACGCTCTTCCGATCTNNNCCTGTAACTTCCTCTTCTTCCC
GAGCACACGTCTGAACTCTTTCCCTACACGACGCTCTTCCGATCTNNNCCTGTAACTTCCTCTTCTTCCC
AGAGCACACGTCTGAACTCTTTCCCTACACGACGCTCTTCCGATCTNNNCCTATAATCTCCCTTTGCCAC
AGAGCACACGTCTGAACTCTTTCCCTACACGACGCTCTTCCGATCTNNNCCTATAATCTCCCTTTGCCAC
CACACGTCTGAACTCTTTCCCTACACGACGCTCTTCCGATCTNNNTTCAGACATGCCAGTGAGAATGT
CACACGTCTGAACTCTTTCCCTACACGACGCTCTTCCGATCTNNNTTCAGACATGCCAGTGAGAATGT
GAGCACACGTCTGAACTCTTTCCCTACACGACGCTCTTCCGATCTNNNAACTTCTGGTGTCAGTCTCC
GAGCACACGTCTGAACTCTTTCCCTACACGACGCTCTTCCGATCTNNNAACTTCTGGTGTCAGTCTCC
GAGCACACGTCTGAACTCTTTCCCTACACGACGCTCTTCCGATCTNNNCAATGGCTCATGACTGTAAT
GAGCACACGTCTGAACTCTTTCCCTACACGACGCTCTTCCGATCTNNNCAATGGCTCATGACTGTAAT
GAGCACACGTCTGAACTCTTTCCCTACACGACGCTCTTCCGATCTNNNGTTTTGGATGTCGGGGTACA
GAGCACACGTCTGAACTCTTTCCCTACACGACGCTCTTCCGATCTNNNGTTTTGGATGTCGGGGTACA
AGCACACGTCTGAACTCTTTCCCTACACGACGCTCTTCCGATCTNNNATGGAGCAGGGAGTAAAGCTT
AGCACACGTCTGAACTCTTTCCCTACACGACGCTCTTCCGATCTNNNATGGAGCAGGGAGTAAAGCTT
GAGCACACGTCTGAACTCTTTCCCTACACGACGCTCTTCCGATCTNNNGAGGTTGCAGTGAGCTGAG
GAGCACACGTCTGAACTCTTTCCCTACACGACGCTCTTCCGATCTNNNGAGGTTGCAGTGAGCTGAG
CACACGTCTGAACTCTTTCCCTACACGACGCTCTTCCGATCTNNNGAAATGTGACTTTGGACAAGGGG
CACACGTCTGAACTCTTTCCCTACACGACGCTCTTCCGATCTNNNGAAATGTGACTTTGGACAAGGGG
GAGCACACGTCTGAACTCTTTCCCTACACGACGCTCTTCCGATCTNNNAACTGCTTGGTGTTTTCAGTC
GAGCACACGTCTGAACTCTTTCCCTACACGACGCTCTTCCGATCTNNNAACTGCTTGGTGTTTTCAGTC
AGCACACGTCTGAACTCTTTCCCTACACGACGCTCTTCCGATCTNNNCTTTAGATGAGCAACCATGGT
AGCACACGTCTGAACTCTTTCCCTACACGACGCTCTTCCGATCTNNNCTTTAGATGAGCAACCATGGT
GAGCACACGTCTGAACTCTTTCCCTACACGACGCTCTTCCGATCTNNNACAGAGTTTTGTTCTTGTTGC
GAGCACACGTCTGAACTCTTTCCCTACACGACGCTCTTCCGATCTNNNACAGAGTTTTGTTCTTGTTGC
GAGCACACGTCTGAACTCTTTCCCTACACGACGCTCTTCCGATCTNNNAAACTTTTTGATGTGCTGCTG
GAGCACACGTCTGAACTCTTTCCCTACACGACGCTCTTCCGATCTNNNAAACTTTTTGATGTGCTGCTG
GAGCACACGTCTGAACTCTTTCCCTACACGACGCTCTTCCGATCTNNNCTGTAATCCCAGCTACTCAGG
GAGCACACGTCTGAACTCTTTCCCTACACGACGCTCTTCCGATCTNNNCTGTAATCCCAGCTACTCAGG
AGCACACGTCTGAACTCTTTCCCTACACGACGCTCTTCCGATCTNNNGCACCATTTTACATTCCCACCAA
AGCACACGTCTGAACTCTTTCCCTACACGACGCTCTTCCGATCTNNNGCACCATTTTACATTCCCACCAA
AGCACACGTCTGAACTCTTTCCCTACACGACGCTCTTCCGATCTNNNGAGTAGCTGGGTTTACAGACA
AGCACACGTCTGAACTCTTTCCCTACACGACGCTCTTCCGATCTNNNGAGTAGCTGGGTTTACAGACA
AGCACACGTCTGAACTCTTTCCCTACACGACGCTCTTCCGATCTNNNCTCTCTCTGGTGTGACAATTGG
AGCACACGTCTGAACTCTTTCCCTACACGACGCTCTTCCGATCTNNNCTCTCTCTGGTGTGACAATTGG

GAGCACACGTCTGAACTCTTTCCCTACACGACGCTCTTCCGATCTNNNCTTGCAGTGAGCCAAGATCAT
GAGCACACGTCTGAACTCTTTCCCTACACGACGCTCTTCCGATCTNNNCTTGCAGTGAGCCAAGATCAT
AGCACACGTCTGAACTCTTTCCCTACACGACGCTCTTCCGATCTNNNTGTCTTTGTCTCTTCATGGCCTT
AGCACACGTCTGAACTCTTTCCCTACACGACGCTCTTCCGATCTNNNTGTCTTTGTCTCTTCATGGCCTT
AGCACACGTCTGAACTCTTTCCCTACACGACGCTCTTCCGATCTNNNGGATTCGGCAGTTTTTATTAGC
AGCACACGTCTGAACTCTTTCCCTACACGACGCTCTTCCGATCTNNNGGATTCGGCAGTTTTTATTAGC
AGCACACGTCTGAACTCTTTCCCTACACGACGCTCTTCCGATCTNNNTGACTCCCTTTTCTGCAAATTCA
AGCACACGTCTGAACTCTTTCCCTACACGACGCTCTTCCGATCTNNNTGACTCCCTTTTCTGCAAATTCA
GAGCACACGTCTGAACTCTTTCCCTACACGACGCTCTTCCGATCTNNNATGTCAACATCCCTTTCCCACTT
GAGCACACGTCTGAACTCTTTCCCTACACGACGCTCTTCCGATCTNNNATGTCAACATCCCTTTCCCACTT
GAGCACACGTCTGAACTCTTTCCCTACACGACGCTCTTCCGATCTNNNAATCCTAGCACTTTGGGAGA
GAGCACACGTCTGAACTCTTTCCCTACACGACGCTCTTCCGATCTNNNAATCCTAGCACTTTGGGAGA
AGCACACGTCTGAACTCTTTCCCTACACGACGCTCTTCCGATCTNNNCACTATGCTGGTTTCAGGTCTG
AGCACACGTCTGAACTCTTTCCCTACACGACGCTCTTCCGATCTNNNCACTATGCTGGTTTCAGGTCTG
AGCACACGTCTGAACTCTTTCCCTACACGACGCTCTTCCGATCTNNNTCCATAAACTGACTTCACAGGCA
AGCACACGTCTGAACTCTTTCCCTACACGACGCTCTTCCGATCTNNNTCCATAAACTGACTTCACAGGCA
GCACACGTCTGAACTCTTTCCCTACACGACGCTCTTCCGATCTNNNACAGGGTATTCCATAATAACTTTC
GCACACGTCTGAACTCTTTCCCTACACGACGCTCTTCCGATCTNNNACAGGGTATTCCATAATAACTTTC
AGCACACGTCTGAACTCTTTCCCTACACGACGCTCTTCCGATCTNNNAAGAAAGGGAACAGAGGAAG
AGCACACGTCTGAACTCTTTCCCTACACGACGCTCTTCCGATCTNNNAAGAAAGGGAACAGAGGAAG
AGCACACGTCTGAACTCTTTCCCTACACGACGCTCTTCCGATCTNNNCTTCTTTCTCCACCTCCCTGTTTCA
AGCACACGTCTGAACTCTTTCCCTACACGACGCTCTTCCGATCTNNNCTTCTTTCTCCACCTCCCTGTTTCA
GAGCACACGTCTGAACTCTTTCCCTACACGACGCTCTTCCGATCTNNNCATTCAGTAATGGTTCTGGGG
GAGCACACGTCTGAACTCTTTCCCTACACGACGCTCTTCCGATCTNNNCATTCAGTAATGGTTCTGGGG
GAGCACACGTCTGAACTCTTTCCCTACACGACGCTCTTCCGATCTNNNCAGGTGACAGAGTGAGATTCC
GAGCACACGTCTGAACTCTTTCCCTACACGACGCTCTTCCGATCTNNNCAGGTGACAGAGTGAGATTCC
AGCACACGTCTGAACTCTTTCCCTACACGACGCTCTTCCGATCTNNNAAGAGGTCCTTCACATCTGTTG
AGCACACGTCTGAACTCTTTCCCTACACGACGCTCTTCCGATCTNNNAAGAGGTCCTTCACATCTGTTG
GAGCACACGTCTGAACTCTTTCCCTACACGACGCTCTTCCGATCTNNNGTGCTTCCATCATGACACCTTT
GAGCACACGTCTGAACTCTTTCCCTACACGACGCTCTTCCGATCTNNNGTGCTTCCATCATGACACCTTT
AGCACACGTCTGAACTCTTTCCCTACACGACGCTCTTCCGATCTNNNCAAAGGGGACTCAATCAGCAA
AGCACACGTCTGAACTCTTTCCCTACACGACGCTCTTCCGATCTNNNCAAAGGGGACTCAATCAGCAA
GAGCACACGTCTGAACTCTTTCCCTACACGACGCTCTTCCGATCTNNNAGTAGCTGAGATTACAAGTG
GAGCACACGTCTGAACTCTTTCCCTACACGACGCTCTTCCGATCTNNNAGTAGCTGAGATTACAAGTG
AGCACACGTCTGAACTCTTTCCCTACACGACGCTCTTCCGATCTNNNTCTGTCTGAGACTTAGGGGACA
AGCACACGTCTGAACTCTTTCCCTACACGACGCTCTTCCGATCTNNNTCTGTCTGAGACTTAGGGGACA
AGAGCACACGTCTGAACTCTTTCCCTACACGACGCTCTTCCGATCTNNNAAAAAATTAGCTGGGCACAG
AGAGCACACGTCTGAACTCTTTCCCTACACGACGCTCTTCCGATCTNNNAAAAAATTAGCTGGGCACAG
GAGCACACGTCTGAACTCTTTCCCTACACGACGCTCTTCCGATCTNNNCCTTTCCCTTGTGTTAGTGCCT
GAGCACACGTCTGAACTCTTTCCCTACACGACGCTCTTCCGATCTNNNCCTTTCCCTTGTGTTAGTGCCT
AGCACACGTCTGAACTCTTTCCCTACACGACGCTCTTCCGATCTNNNTTTTATAGACTGGCTCACTGGG
AGCACACGTCTGAACTCTTTCCCTACACGACGCTCTTCCGATCTNNNTTTTATAGACTGGCTCACTGGG
AGCACACGTCTGAACTCTTTCCCTACACGACGCTCTTCCGATCTNNNCTCTAGGGGATGCTGGGAATC

AGCACACGTCTGAACTCTTTCCCTACACGACGCTCTTCCGATCTNNNCTCTAGGGGATGCTGGGAATC
GAGCACACGTCTGAACTCTTTCCCTACACGACGCTCTTCCGATCTNNNGGGGATTGATGAAGTTGTTC
GAGCACACGTCTGAACTCTTTCCCTACACGACGCTCTTCCGATCTNNNGGGGATTGATGAAGTTGTTC
AGCACACGTCTGAACTCTTTCCCTACACGACGCTCTTCCGATCTNNNAGGGTTGTCTGCTTGTGATATC
AGCACACGTCTGAACTCTTTCCCTACACGACGCTCTTCCGATCTNNNAGGGTTGTCTGCTTGTGATATC
AGAGCACACGTCTGAACTCTTTCCCTACACGACGCTCTTCCGATCTNNNTATAATTCTTCCTTGTCCTTTC
AGAGCACACGTCTGAACTCTTTCCCTACACGACGCTCTTCCGATCTNNNTATAATTCTTCCTTGTCCTTTC
AGAGCACACGTCTGAACTCTTTCCCTACACGACGCTCTTCCGATCTNNNTGTTTTGGGACTACCATGGAT
AGAGCACACGTCTGAACTCTTTCCCTACACGACGCTCTTCCGATCTNNNTGTTTTGGGACTACCATGGAT
GAGCACACGTCTGAACTCTTTCCCTACACGACGCTCTTCCGATCTNNNCACACCAGACTAGGTAACAGA
GAGCACACGTCTGAACTCTTTCCCTACACGACGCTCTTCCGATCTNNNCACACCAGACTAGGTAACAGA
GAGCACACGTCTGAACTCTTTCCCTACACGACGCTCTTCCGATCTNNNCCAGGAGGTTGAAGCTGTAA
GAGCACACGTCTGAACTCTTTCCCTACACGACGCTCTTCCGATCTNNNCCAGGAGGTTGAAGCTGTAA
AGCACACGTCTGAACTCTTTCCCTACACGACGCTCTTCCGATCTNNNATACAAAAATTAGCCGGGTGTG
AGCACACGTCTGAACTCTTTCCCTACACGACGCTCTTCCGATCTNNNATACAAAAATTAGCCGGGTGTG
GAGCACACGTCTGAACTCTTTCCCTACACGACGCTCTTCCGATCTNNNGCGCCTAGCCATATGTTTCTT
GAGCACACGTCTGAACTCTTTCCCTACACGACGCTCTTCCGATCTNNNGCGCCTAGCCATATGTTTCTT
AGAGCACACGTCTGAACTCTTTCCCTACACGACGCTCTTCCGATCTNNNCAAAGTGTTGGGATTACAGC
AGAGCACACGTCTGAACTCTTTCCCTACACGACGCTCTTCCGATCTNNNCAAAGTGTTGGGATTACAGC
GAGCACACGTCTGAACTCTTTCCCTACACGACGCTCTTCCGATCTNNNACCACGGAAAGAAAGAAAGG
GAGCACACGTCTGAACTCTTTCCCTACACGACGCTCTTCCGATCTNNNACCACGGAAAGAAAGAAAGG
GAGCACACGTCTGAACTCTTTCCCTACACGACGCTCTTCCGATCTNNNTCCATTTACTGAATTTATGGAG
GAGCACACGTCTGAACTCTTTCCCTACACGACGCTCTTCCGATCTNNNTCCATTTACTGAATTTATGGAG
AGCACACGTCTGAACTCTTTCCCTACACGACGCTCTTCCGATCTNNNGACAGAGCAAGACTCCGTCTCC
AGCACACGTCTGAACTCTTTCCCTACACGACGCTCTTCCGATCTNNNGACAGAGCAAGACTCCGTCTCC
GAGCACACGTCTGAACTCTTTCCCTACACGACGCTCTTCCGATCTNNNCTCAGGGGTTGGTGGAATCC
GAGCACACGTCTGAACTCTTTCCCTACACGACGCTCTTCCGATCTNNNCTCAGGGGTTGGTGGAATCC
AGCACACGTCTGAACTCTTTCCCTACACGACGCTCTTCCGATCTNNNTGTAGTCCCAGCTACTTAGGAG
AGCACACGTCTGAACTCTTTCCCTACACGACGCTCTTCCGATCTNNNTGTAGTCCCAGCTACTTAGGAG
GAGCACACGTCTGAACTCTTTCCCTACACGACGCTCTTCCGATCTNNNATCTCTTGTTCATTCACCGCCTC
GAGCACACGTCTGAACTCTTTCCCTACACGACGCTCTTCCGATCTNNNATCTCTTGTTCATTCACCGCCTC
AGCACACGTCTGAACTCTTTCCCTACACGACGCTCTTCCGATCTNNNACCTGTAGTCCTAGCTACTCCAG
AGCACACGTCTGAACTCTTTCCCTACACGACGCTCTTCCGATCTNNNACCTGTAGTCCTAGCTACTCCAG
GAGCACACGTCTGAACTCTTTCCCTACACGACGCTCTTCCGATCTNNNATGCCTGTAATCCCAGCTACTT
GAGCACACGTCTGAACTCTTTCCCTACACGACGCTCTTCCGATCTNNNATGCCTGTAATCCCAGCTACTT
AGCACACGTCTGAACTCTTTCCCTACACGACGCTCTTCCGATCTNNNACTTCATTGCAACTCCCTGATCTC
AGCACACGTCTGAACTCTTTCCCTACACGACGCTCTTCCGATCTNNNACTTCATTGCAACTCCCTGATCTC
GCACACGTCTGAACTCTTTCCCTACACGACGCTCTTCCGATCTNNNTAACTCAGTGTTTACCAGTGGCC
GCACACGTCTGAACTCTTTCCCTACACGACGCTCTTCCGATCTNNNTAACTCAGTGTTTACCAGTGGCC
GCACACGTCTGAACTCTTTCCCTACACGACGCTCTTCCGATCTNNNCTTTAACCAGCTATAAACAAGAG
GCACACGTCTGAACTCTTTCCCTACACGACGCTCTTCCGATCTNNNCTTTAACCAGCTATAAACAAGAG
GAGCACACGTCTGAACTCTTTCCCTACACGACGCTCTTCCGATCTNNNGCTTGAATTTATGCACATTCTT
GAGCACACGTCTGAACTCTTTCCCTACACGACGCTCTTCCGATCTNNNGCTTGAATTTATGCACATTCTT

GCACACGTCTGAACTCTTTCCCTACACGACGCTCTTCCGATCTNNNGGCTGTCTTAGTCCATTTTCTGC
GCACACGTCTGAACTCTTTCCCTACACGACGCTCTTCCGATCTNNNGGCTGTCTTAGTCCATTTTCTGC
AGCACACGTCTGAACTCTTTCCCTACACGACGCTCTTCCGATCTNNNAGTAAACCACCTTTCACTGAGG
AGCACACGTCTGAACTCTTTCCCTACACGACGCTCTTCCGATCTNNNAGTAAACCACCTTTCACTGAGG
GCACACGTCTGAACTCTTTCCCTACACGACGCTCTTCCGATCTNNNAGACAAGCTGACTCTGCCTACTT
GCACACGTCTGAACTCTTTCCCTACACGACGCTCTTCCGATCTNNNAGACAAGCTGACTCTGCCTACTT
AGCACACGTCTGAACTCTTTCCCTACACGACGCTCTTCCGATCTNNNGAGCTCCAGCAAGAAATCAAA
AGCACACGTCTGAACTCTTTCCCTACACGACGCTCTTCCGATCTNNNGAGCTCCAGCAAGAAATCAAA
AGCACACGTCTGAACTCTTTCCCTACACGACGCTCTTCCGATCTNNNATGGCTGCATAGTATTCCATGG
AGCACACGTCTGAACTCTTTCCCTACACGACGCTCTTCCGATCTNNNATGGCTGCATAGTATTCCATGG
GCACACGTCTGAACTCTTTCCCTACACGACGCTCTTCCGATCTNNNTAATGGAAGTCACTGCTGCTTCT
GCACACGTCTGAACTCTTTCCCTACACGACGCTCTTCCGATCTNNNTAATGGAAGTCACTGCTGCTTCT
AGCACACGTCTGAACTCTTTCCCTACACGACGCTCTTCCGATCTNNNGCAGGAGAATTGCTTGAACCTG
AGCACACGTCTGAACTCTTTCCCTACACGACGCTCTTCCGATCTNNNGCAGGAGAATTGCTTGAACCTG
GAGCACACGTCTGAACTCTTTCCCTACACGACGCTCTTCCGATCTNNNCGTGAATTTGGAAGTGGAA
GAGCACACGTCTGAACTCTTTCCCTACACGACGCTCTTCCGATCTNNNCGTGAATTTGGAAGTGGAA
AGAGCACACGTCTGAACTCTTTCCCTACACGACGCTCTTCCGATCTNNNCTGAGGTGGGAGAATCAC
AGAGCACACGTCTGAACTCTTTCCCTACACGACGCTCTTCCGATCTNNNCTGAGGTGGGAGAATCAC
AGCACACGTCTGAACTCTTTCCCTACACGACGCTCTTCCGATCTNNNTAGTTTTACACAGAGGTTGGC
AGCACACGTCTGAACTCTTTCCCTACACGACGCTCTTCCGATCTNNNTAGTTTTACACAGAGGTTGGC
AGCACACGTCTGAACTCTTTCCCTACACGACGCTCTTCCGATCTNNNGAGGCAGACCAGACCAACTGC
AGCACACGTCTGAACTCTTTCCCTACACGACGCTCTTCCGATCTNNNGAGGCAGACCAGACCAACTGC
AGCACACGTCTGAACTCTTTCCCTACACGACGCTCTTCCGATCTNNNATTGAGTACTTTCTGGAGGCT
AGCACACGTCTGAACTCTTTCCCTACACGACGCTCTTCCGATCTNNNATTGAGTACTTTCTGGAGGCT
AGCACACGTCTGAACTCTTTCCCTACACGACGCTCTTCCGATCTNNNATCTCTTGACCTTATGATCCGC
AGCACACGTCTGAACTCTTTCCCTACACGACGCTCTTCCGATCTNNNATCTCTTGACCTTATGATCCGC
GAGCACACGTCTGAACTCTTTCCCTACACGACGCTCTTCCGATCTNNNGAGTCACATTTCCAGACCTCA
GAGCACACGTCTGAACTCTTTCCCTACACGACGCTCTTCCGATCTNNNGAGTCACATTTCCAGACCTCA
GAGCACACGTCTGAACTCTTTCCCTACACGACGCTCTTCCGATCTNNNATCTTGGCTCACTACAACCTCT
GAGCACACGTCTGAACTCTTTCCCTACACGACGCTCTTCCGATCTNNNATCTTGGCTCACTACAACCTCT
GAGCACACGTCTGAACTCTTTCCCTACACGACGCTCTTCCGATCTNNNTTTTTAGCAGGAGGATGTGA
GAGCACACGTCTGAACTCTTTCCCTACACGACGCTCTTCCGATCTNNNTTTTTAGCAGGAGGATGTGA
AGCACACGTCTGAACTCTTTCCCTACACGACGCTCTTCCGATCTNNNGGGCTGGACCTAATAGCTTTCT
AGCACACGTCTGAACTCTTTCCCTACACGACGCTCTTCCGATCTNNNGGGCTGGACCTAATAGCTTTCT
AGCACACGTCTGAACTCTTTCCCTACACGACGCTCTTCCGATCTNNNCCGTTTTGTGGTTACAGGAGG
AGCACACGTCTGAACTCTTTCCCTACACGACGCTCTTCCGATCTNNNCCGTTTTGTGGTTACAGGAGG
GCACACGTCTGAACTCTTTCCCTACACGACGCTCTTCCGATCTNNNTGACAGGATATTAGAACAGCAG
GCACACGTCTGAACTCTTTCCCTACACGACGCTCTTCCGATCTNNNTGACAGGATATTAGAACAGCAG
GAGCACACGTCTGAACTCTTTCCCTACACGACGCTCTTCCGATCTNNNGCGACAAAGCAAGATTGTGT
GAGCACACGTCTGAACTCTTTCCCTACACGACGCTCTTCCGATCTNNNGCGACAAAGCAAGATTGTGT
AGCACACGTCTGAACTCTTTCCCTACACGACGCTCTTCCGATCTNNNCCAGTTTCATGCTTTTATTTCAT
AGCACACGTCTGAACTCTTTCCCTACACGACGCTCTTCCGATCTNNNCCAGTTTCATGCTTTTATTTCAT
AGCACACGTCTGAACTCTTTCCCTACACGACGCTCTTCCGATCTNNNGCCTGTAGTCCCAGCTACTTGC

AGCACACGTCTGAACTCTTTCCCTACACGACGCTCTTCCGATCTNNNGCCTGTAGTCCCAGCTACTTGC
AGCACACGTCTGAACTCTTTCCCTACACGACGCTCTTCCGATCTNNNAAGAAAGCTCTGAGTTATGGG
AGCACACGTCTGAACTCTTTCCCTACACGACGCTCTTCCGATCTNNNAAGAAAGCTCTGAGTTATGGG
AGCACACGTCTGAACTCTTTCCCTACACGACGCTCTTCCGATCTNNNTTCTGAGTTGCTGGGATTACA
AGCACACGTCTGAACTCTTTCCCTACACGACGCTCTTCCGATCTNNNTTCTGAGTTGCTGGGATTACA
AGCACACGTCTGAACTCTTTCCCTACACGACGCTCTTCCGATCTNNNGTTGGTATATCCGTGTGTGCT
AGCACACGTCTGAACTCTTTCCCTACACGACGCTCTTCCGATCTNNNGTTGGTATATCCGTGTGTGCT
GCACACGTCTGAACTCTTTCCCTACACGACGCTCTTCCGATCTNNNGACTTTGGGGATTTGGGGAAA
GCACACGTCTGAACTCTTTCCCTACACGACGCTCTTCCGATCTNNNGACTTTGGGGATTTGGGGAAA
AGCACACGTCTGAACTCTTTCCCTACACGACGCTCTTCCGATCTNNNCTGAGGCAGGAGAATTGCTT
AGCACACGTCTGAACTCTTTCCCTACACGACGCTCTTCCGATCTNNNCTGAGGCAGGAGAATTGCTT
AGCACACGTCTGAACTCTTTCCCTACACGACGCTCTTCCGATCTNNNTTTTTAACTTACGGTTGATGAA
AGCACACGTCTGAACTCTTTCCCTACACGACGCTCTTCCGATCTNNNTTTTTAACTTACGGTTGATGAA
AGCACACGTCTGAACTCTTTCCCTACACGACGCTCTTCCGATCTNNNGAGCGAGGATGTAGCTTGTC
AGCACACGTCTGAACTCTTTCCCTACACGACGCTCTTCCGATCTNNNGAGCGAGGATGTAGCTTGTC
GCACACGTCTGAACTCTTTCCCTACACGACGCTCTTCCGATCTNNNTCCATCAATCCCATGTAAGCCAA
GCACACGTCTGAACTCTTTCCCTACACGACGCTCTTCCGATCTNNNTCCATCAATCCCATGTAAGCCAA
GAGCACACGTCTGAACTCTTTCCCTACACGACGCTCTTCCGATCTNNNGAGTGCTGTCTTGGAGGGT
GAGCACACGTCTGAACTCTTTCCCTACACGACGCTCTTCCGATCTNNNGAGTGCTGTCTTGGAGGGT
GCACACGTCTGAACTCTTTCCCTACACGACGCTCTTCCGATCTNNNGAGCTTGCAGTGAGCCAAGAT
GCACACGTCTGAACTCTTTCCCTACACGACGCTCTTCCGATCTNNNGAGCTTGCAGTGAGCCAAGAT
GAGCACACGTCTGAACTCTTTCCCTACACGACGCTCTTCCGATCTNNNACTTGGAATGCTGAAGTAGG
GAGCACACGTCTGAACTCTTTCCCTACACGACGCTCTTCCGATCTNNNACTTGGAATGCTGAAGTAGG
AGCACACGTCTGAACTCTTTCCCTACACGACGCTCTTCCGATCTNNNAAAATGTGTGTGTGTGTGTG
AGCACACGTCTGAACTCTTTCCCTACACGACGCTCTTCCGATCTNNNAAAATGTGTGTGTGTGTGTG
GCACACGTCTGAACTCTTTCCCTACACGACGCTCTTCCGATCTNNNTCCTTAGTGTCTGGCTTAAATCA
GCACACGTCTGAACTCTTTCCCTACACGACGCTCTTCCGATCTNNNTCCTTAGTGTCTGGCTTAAATCA
AGCACACGTCTGAACTCTTTCCCTACACGACGCTCTTCCGATCTNNNACATCATGGAGAAACGGGTA
AGCACACGTCTGAACTCTTTCCCTACACGACGCTCTTCCGATCTNNNACATCATGGAGAAACGGGTA
AGCACACGTCTGAACTCTTTCCCTACACGACGCTCTTCCGATCTNNNGTGCACACACCTCCATATACAG
AGCACACGTCTGAACTCTTTCCCTACACGACGCTCTTCCGATCTNNNGTGCACACACCTCCATATACAG
AGAGCACACGTCTGAACTCTTTCCCTACACGACGCTCTTCCGATCTNNNACAAGGAAAAGGAACATG
AGAGCACACGTCTGAACTCTTTCCCTACACGACGCTCTTCCGATCTNNNACAAGGAAAAGGAACATG
ACACGTCTGAACTCTTTCCCTACACGACGCTCTTCCGATCTNNNTACCACAGTTTCTTCTGTAAATCAT
ACACGTCTGAACTCTTTCCCTACACGACGCTCTTCCGATCTNNNTACCACAGTTTCTTCTGTAAATCAT
AGCACACGTCTGAACTCTTTCCCTACACGACGCTCTTCCGATCTNNNTAGGAATGTCATGTGCTCTCG
AGCACACGTCTGAACTCTTTCCCTACACGACGCTCTTCCGATCTNNNTAGGAATGTCATGTGCTCTCG
AGCACACGTCTGAACTCTTTCCCTACACGACGCTCTTCCGATCTNNNTAATTCCAGCTATTCAGGAGG
AGCACACGTCTGAACTCTTTCCCTACACGACGCTCTTCCGATCTNNNTAATTCCAGCTATTCAGGAGG
GCACACGTCTGAACTCTTTCCCTACACGACGCTCTTCCGATCTNNNGAGGTTGAAGTGGGAGAATCA
GCACACGTCTGAACTCTTTCCCTACACGACGCTCTTCCGATCTNNNGAGGTTGAAGTGGGAGAATCA
AGCACACGTCTGAACTCTTTCCCTACACGACGCTCTTCCGATCTNNNGTATTTCTGCAGTCCCAGCTAC
AGCACACGTCTGAACTCTTTCCCTACACGACGCTCTTCCGATCTNNNGTATTTCTGCAGTCCCAGCTAC

GAGCACACGTCTGAACTCTTTCCCTACACGACGCTCTTCCGATCTNNNACGTTCTCACATGATGAAGAG

GAGCACACGTCTGAACTCTTTCCCTACACGACGCTCTTCCGATCTNNNACGTTCTCACATGATGAAGAG

GAGCACACGTCTGAACTCTTTCCCTACACGACGCTCTTCCGATCTNNNACTTGCTCACTGGGGACAAAT

GAGCACACGTCTGAACTCTTTCCCTACACGACGCTCTTCCGATCTNNNACTTGCTCACTGGGGACAAAT

AGCACACGTCTGAACTCTTTCCCTACACGACGCTCTTCCGATCTNNNAAGCTCCCTCTGAAAAGTTTCAT

AGCACACGTCTGAACTCTTTCCCTACACGACGCTCTTCCGATCTNNNAAGCTCCCTCTGAAAAGTTTCAT

AGCACACGTCTGAACTCTTTCCCTACACGACGCTCTTCCGATCTNNNGTACAGGTTGGTTGGAGGTTC

AGCACACGTCTGAACTCTTTCCCTACACGACGCTCTTCCGATCTNNNGTACAGGTTGGTTGGAGGTTC

AGCACACGTCTGAACTCTTTCCCTACACGACGCTCTTCCGATCTNNNGCTCCGTAGAGAAAGGATGTA

AGCACACGTCTGAACTCTTTCCCTACACGACGCTCTTCCGATCTNNNGCTCCGTAGAGAAAGGATGTA

GAGCACACGTCTGAACTCTTTCCCTACACGACGCTCTTCCGATCTNNNTCTGGAGACTGAGACAGGA

GAGCACACGTCTGAACTCTTTCCCTACACGACGCTCTTCCGATCTNNNTCTGGAGACTGAGACAGGA

AGCACACGTCTGAACTCTTTCCCTACACGACGCTCTTCCGATCTNNNAGGAGATTTGCTTGAACCTGA

AGCACACGTCTGAACTCTTTCCCTACACGACGCTCTTCCGATCTNNNAGGAGATTTGCTTGAACCTGA

AGCACACGTCTGAACTCTTTCCCTACACGACGCTCTTCCGATCTNNNCAAGCAGGTGAGGGAGAAAA

AGCACACGTCTGAACTCTTTCCCTACACGACGCTCTTCCGATCTNNNCAAGCAGGTGAGGGAGAAAA

AGCACACGTCTGAACTCTTTCCCTACACGACGCTCTTCCGATCTNNNGAGAATCACTCGAACCCAGGA

AGCACACGTCTGAACTCTTTCCCTACACGACGCTCTTCCGATCTNNNGAGAATCACTCGAACCCAGGA

CACACGTCTGAACTCTTTCCCTACACGACGCTCTTCCGATCTNNNGCGTCATGTCTTTATTACGGTGG

CACACGTCTGAACTCTTTCCCTACACGACGCTCTTCCGATCTNNNGCGTCATGTCTTTATTACGGTGG

AGCACACGTCTGAACTCTTTCCCTACACGACGCTCTTCCGATCTNNNGGATCTCAGCTCACTACAACCTC

AGCACACGTCTGAACTCTTTCCCTACACGACGCTCTTCCGATCTNNNGGATCTCAGCTCACTACAACCTC

GCACACGTCTGAACTCTTTCCCTACACGACGCTCTTCCGATCTNNNGGCTTCTTCACTCAGCATAGTCT

GCACACGTCTGAACTCTTTCCCTACACGACGCTCTTCCGATCTNNNGGCTTCTTCACTCAGCATAGTCT

GAGCACACGTCTGAACTCTTTCCCTACACGACGCTCTTCCGATCTNNNCAAGACCATGACTGCAGTGTT

GAGCACACGTCTGAACTCTTTCCCTACACGACGCTCTTCCGATCTNNNCAAGACCATGACTGCAGTGTT

GAGCACACGTCTGAACTCTTTCCCTACACGACGCTCTTCCGATCTNNNTGCTTAGGCCACTCTAGGACT

GAGCACACGTCTGAACTCTTTCCCTACACGACGCTCTTCCGATCTNNNTGCTTAGGCCACTCTAGGACT

GAGCACACGTCTGAACTCTTTCCCTACACGACGCTCTTCCGATCTNNNCTGCTTTACCACTTCAACACA

GAGCACACGTCTGAACTCTTTCCCTACACGACGCTCTTCCGATCTNNNCTGCTTTACCACTTCAACACA

AGCACACGTCTGAACTCTTTCCCTACACGACGCTCTTCCGATCTNNNTTAGCCTAGGTGACAGAGTGA

AGCACACGTCTGAACTCTTTCCCTACACGACGCTCTTCCGATCTNNNTTAGCCTAGGTGACAGAGTGA

GCACACGTCTGAACTCTTTCCCTACACGACGCTCTTCCGATCTNNNGAGATGCTAGTCAGGCTGTACA

GCACACGTCTGAACTCTTTCCCTACACGACGCTCTTCCGATCTNNNGAGATGCTAGTCAGGCTGTACA

GAGCACACGTCTGAACTCTTTCCCTACACGACGCTCTTCCGATCTNNNCTGGTGGGTTTTATCTTTTTT

GAGCACACGTCTGAACTCTTTCCCTACACGACGCTCTTCCGATCTNNNCTGGTGGGTTTTATCTTTTTT

GCACACGTCTGAACTCTTTCCCTACACGACGCTCTTCCGATCTNNNCTTGCTGTCAAGTTGTAGCAGTC

GCACACGTCTGAACTCTTTCCCTACACGACGCTCTTCCGATCTNNNCTTGCTGTCAAGTTGTAGCAGTC

GCACACGTCTGAACTCTTTCCCTACACGACGCTCTTCCGATCTNNNTGAGAAAGGTACGCATGAAGT

GCACACGTCTGAACTCTTTCCCTACACGACGCTCTTCCGATCTNNNTGAGAAAGGTACGCATGAAGT

AGCACACGTCTGAACTCTTTCCCTACACGACGCTCTTCCGATCTNNNAGAATCACTTGAACTCAGGAG

AGCACACGTCTGAACTCTTTCCCTACACGACGCTCTTCCGATCTNNNAGAATCACTTGAACTCAGGAG

AGCACACGTCTGAACTCTTTCCCTACACGACGCTCTTCCGATCTNNNACAAAAATTAGCCAGGCTTGG

AGCACACGTCTGAACTCTTTCCCTACACGACGCTCTTCCGATCTNNNACAAAAATTAGCCAGGCTTGG
GAGCACACGTCTGAACTCTTTCCCTACACGACGCTCTTCCGATCTNNNGGCACTCTAGCAAGTCCATAT
GAGCACACGTCTGAACTCTTTCCCTACACGACGCTCTTCCGATCTNNNGGCACTCTAGCAAGTCCATAT
GAGCACACGTCTGAACTCTTTCCCTACACGACGCTCTTCCGATCTNNNACCTGAATGAGCAAGAAGAT
GAGCACACGTCTGAACTCTTTCCCTACACGACGCTCTTCCGATCTNNNACCTGAATGAGCAAGAAGAT
GAGCACACGTCTGAACTCTTTCCCTACACGACGCTCTTCCGATCTNNNTCTCAAAGTGGCATCTTGGTG
GAGCACACGTCTGAACTCTTTCCCTACACGACGCTCTTCCGATCTNNNTCTCAAAGTGGCATCTTGGTG
GAGCACACGTCTGAACTCTTTCCCTACACGACGCTCTTCCGATCTNNNGCAAAATCTGGCTCCTAGTG
GAGCACACGTCTGAACTCTTTCCCTACACGACGCTCTTCCGATCTNNNGCAAAATCTGGCTCCTAGTG
GCACACGTCTGAACTCTTTCCCTACACGACGCTCTTCCGATCTNNNCATCCTGGCTAACATGGTGAAAC
GCACACGTCTGAACTCTTTCCCTACACGACGCTCTTCCGATCTNNNCATCCTGGCTAACATGGTGAAAC
ACACGTCTGAACTCTTTCCCTACACGACGCTCTTCCGATCTNNNAAGTTCAAGTGATTCTCCTGCCTCA
ACACGTCTGAACTCTTTCCCTACACGACGCTCTTCCGATCTNNNAAGTTCAAGTGATTCTCCTGCCTCA
AGCACACGTCTGAACTCTTTCCCTACACGACGCTCTTCCGATCTNNNCTGCAGTGAGCCAAAATCATAC
AGCACACGTCTGAACTCTTTCCCTACACGACGCTCTTCCGATCTNNNCTGCAGTGAGCCAAAATCATAC
AGCACACGTCTGAACTCTTTCCCTACACGACGCTCTTCCGATCTNNNCAAGATCAAGATCACGTCACTG
AGCACACGTCTGAACTCTTTCCCTACACGACGCTCTTCCGATCTNNNCAAGATCAAGATCACGTCACTG
GAGCACACGTCTGAACTCTTTCCCTACACGACGCTCTTCCGATCTNNNGCAAACTCTTCAGGCTCCTTT
GAGCACACGTCTGAACTCTTTCCCTACACGACGCTCTTCCGATCTNNNGCAAACTCTTCAGGCTCCTTT
AGCACACGTCTGAACTCTTTCCCTACACGACGCTCTTCCGATCTNNNTGCAGAGATTGGGCTCTCAAT
AGCACACGTCTGAACTCTTTCCCTACACGACGCTCTTCCGATCTNNNTGCAGAGATTGGGCTCTCAAT
AGCACACGTCTGAACTCTTTCCCTACACGACGCTCTTCCGATCTNNNAATCACCTGGGGATCTTGTTAC
AGCACACGTCTGAACTCTTTCCCTACACGACGCTCTTCCGATCTNNNAATCACCTGGGGATCTTGTTAC
AGCACACGTCTGAACTCTTTCCCTACACGACGCTCTTCCGATCTNNNAAGATTAGATCTGGTGCACTG
AGCACACGTCTGAACTCTTTCCCTACACGACGCTCTTCCGATCTNNNAAGATTAGATCTGGTGCACTG
GAGCACACGTCTGAACTCTTTCCCTACACGACGCTCTTCCGATCTNNNGGAGGACAAAGTGAGACTCC
GAGCACACGTCTGAACTCTTTCCCTACACGACGCTCTTCCGATCTNNNGGAGGACAAAGTGAGACTCC
GAGCACACGTCTGAACTCTTTCCCTACACGACGCTCTTCCGATCTNNNAAATGAACATCCAAGGGCAA
GAGCACACGTCTGAACTCTTTCCCTACACGACGCTCTTCCGATCTNNNAAATGAACATCCAAGGGCAA
AGCACACGTCTGAACTCTTTCCCTACACGACGCTCTTCCGATCTNNNTGTCTGCATTCAACTCACAGAG
AGCACACGTCTGAACTCTTTCCCTACACGACGCTCTTCCGATCTNNNTGTCTGCATTCAACTCACAGAG
GAGCACACGTCTGAACTCTTTCCCTACACGACGCTCTTCCGATCTNNNTATACACTTGCTACTTCCAGCC
GAGCACACGTCTGAACTCTTTCCCTACACGACGCTCTTCCGATCTNNNTATACACTTGCTACTTCCAGCC
GAGCACACGTCTGAACTCTTTCCCTACACGACGCTCTTCCGATCTNNNGAGGCTGAGGCAGGAGAAT
GAGCACACGTCTGAACTCTTTCCCTACACGACGCTCTTCCGATCTNNNGAGGCTGAGGCAGGAGAAT
GCACACGTCTGAACTCTTTCCCTACACGACGCTCTTCCGATCTNNNGAAGAGTCTGTTAGCTCCAGGA
GCACACGTCTGAACTCTTTCCCTACACGACGCTCTTCCGATCTNNNGAAGAGTCTGTTAGCTCCAGGA
AGAGCACACGTCTGAACTCTTTCCCTACACGACGCTCTTCCGATCTNNNGAGGCAGAAGAATCGCTTG
AGAGCACACGTCTGAACTCTTTCCCTACACGACGCTCTTCCGATCTNNNGAGGCAGAAGAATCGCTTG
GAGCACACGTCTGAACTCTTTCCCTACACGACGCTCTTCCGATCTNNNATTTATTGAGGGCTGTGTGTG
GAGCACACGTCTGAACTCTTTCCCTACACGACGCTCTTCCGATCTNNNATTTATTGAGGGCTGTGTGTG
AGCACACGTCTGAACTCTTTCCCTACACGACGCTCTTCCGATCTNNNTGAGGTAGGAGAATCGCTTGA
AGCACACGTCTGAACTCTTTCCCTACACGACGCTCTTCCGATCTNNNTGAGGTAGGAGAATCGCTTGA

AGCACACGTCTGAACTCTTTCCCTACACGACGCTCTTCCGATCTNNNGCAGGAGAATTGCTTGAACCT
AGCACACGTCTGAACTCTTTCCCTACACGACGCTCTTCCGATCTNNNGCAGGAGAATTGCTTGAACCT
AGCACACGTCTGAACTCTTTCCCTACACGACGCTCTTCCGATCTNNNTATTGAAAGAGGATGTGGGA
AGCACACGTCTGAACTCTTTCCCTACACGACGCTCTTCCGATCTNNNTATTGAAAGAGGATGTGGGA
AGCACACGTCTGAACTCTTTCCCTACACGACGCTCTTCCGATCTNNNTTAGAGAAGTTTAAGAGGCA
AGCACACGTCTGAACTCTTTCCCTACACGACGCTCTTCCGATCTNNNTTAGAGAAGTTTAAGAGGCA
GCACACGTCTGAACTCTTTCCCTACACGACGCTCTTCCGATCTNNNTCCTGACCACATTCATCTCAAGG
GCACACGTCTGAACTCTTTCCCTACACGACGCTCTTCCGATCTNNNTCCTGACCACATTCATCTCAAGG
AGCACACGTCTGAACTCTTTCCCTACACGACGCTCTTCCGATCTNNNAGAGCAAGACCCTGTCTCAAA
AGCACACGTCTGAACTCTTTCCCTACACGACGCTCTTCCGATCTNNNAGAGCAAGACCCTGTCTCAAA
GCACACGTCTGAACTCTTTCCCTACACGACGCTCTTCCGATCTNNNTACTGGCATTATCAGCATCATCA
GCACACGTCTGAACTCTTTCCCTACACGACGCTCTTCCGATCTNNNTACTGGCATTATCAGCATCATCA
GCACACGTCTGAACTCTTTCCCTACACGACGCTCTTCCGATCTNNNCTTTCCCCATTACATTGGCCTTCC
GCACACGTCTGAACTCTTTCCCTACACGACGCTCTTCCGATCTNNNCTTTCCCCATTACATTGGCCTTCC
AGCACACGTCTGAACTCTTTCCCTACACGACGCTCTTCCGATCTNNNTTTAGATGGGATTCCGCTTGG
AGCACACGTCTGAACTCTTTCCCTACACGACGCTCTTCCGATCTNNNTTTAGATGGGATTCCGCTTGG
GCACACGTCTGAACTCTTTCCCTACACGACGCTCTTCCGATCTNNNTTCAAGCAATTCTCCTGCCTCAGC
GCACACGTCTGAACTCTTTCCCTACACGACGCTCTTCCGATCTNNNTTCAAGCAATTCTCCTGCCTCAGC
AGCACACGTCTGAACTCTTTCCCTACACGACGCTCTTCCGATCTNNNACCACAGCTGCATTATTTGCTT
AGCACACGTCTGAACTCTTTCCCTACACGACGCTCTTCCGATCTNNNACCACAGCTGCATTATTTGCTT
AGCACACGTCTGAACTCTTTCCCTACACGACGCTCTTCCGATCTNNNTACAAAAAATTAGCTGGGCGT
AGCACACGTCTGAACTCTTTCCCTACACGACGCTCTTCCGATCTNNNTACAAAAAATTAGCTGGGCGT
GCACACGTCTGAACTCTTTCCCTACACGACGCTCTTCCGATCTNNNGGGTACATCAACTCTCCAACACT
GCACACGTCTGAACTCTTTCCCTACACGACGCTCTTCCGATCTNNNGGGTACATCAACTCTCCAACACT
AGCACACGTCTGAACTCTTTCCCTACACGACGCTCTTCCGATCTNNNAAATTTTCAAGACCAAGGCCA
AGCACACGTCTGAACTCTTTCCCTACACGACGCTCTTCCGATCTNNNAAATTTTCAAGACCAAGGCCA
AGCACACGTCTGAACTCTTTCCCTACACGACGCTCTTCCGATCTNNNATGAAGGCCAATGTGAGAGA
AGCACACGTCTGAACTCTTTCCCTACACGACGCTCTTCCGATCTNNNATGAAGGCCAATGTGAGAGA
GCACACGTCTGAACTCTTTCCCTACACGACGCTCTTCCGATCTNNNGCCGTCTTATGAATGAAAGGA
GCACACGTCTGAACTCTTTCCCTACACGACGCTCTTCCGATCTNNNGCCGTCTTATGAATGAAAGGA
AGCACACGTCTGAACTCTTTCCCTACACGACGCTCTTCCGATCTNNNATATGGATGCAGTTTCCCCCTT
AGCACACGTCTGAACTCTTTCCCTACACGACGCTCTTCCGATCTNNNATATGGATGCAGTTTCCCCCTT
GCACACGTCTGAACTCTTTCCCTACACGACGCTCTTCCGATCTNNNGAGGCTGAGGCAGGAGAATG
GCACACGTCTGAACTCTTTCCCTACACGACGCTCTTCCGATCTNNNGAGGCTGAGGCAGGAGAATG
AGAGCACACGTCTGAACTCTTTCCCTACACGACGCTCTTCCGATCTNNNCCACCATGTCTTGGTTCCTA
AGAGCACACGTCTGAACTCTTTCCCTACACGACGCTCTTCCGATCTNNNCCACCATGTCTTGGTTCCTA
GAGCACACGTCTGAACTCTTTCCCTACACGACGCTCTTCCGATCTNNNTTCTCATGAATTGGTCCTCGT
GAGCACACGTCTGAACTCTTTCCCTACACGACGCTCTTCCGATCTNNNTTCTCATGAATTGGTCCTCGT
AGCACACGTCTGAACTCTTTCCCTACACGACGCTCTTCCGATCTNNNGGCAGGGACTCTGTTCTGTAT
AGCACACGTCTGAACTCTTTCCCTACACGACGCTCTTCCGATCTNNNGGCAGGGACTCTGTTCTGTAT
AGCACACGTCTGAACTCTTTCCCTACACGACGCTCTTCCGATCTNNNATCCTGGCTAACATGGTGAAA
AGCACACGTCTGAACTCTTTCCCTACACGACGCTCTTCCGATCTNNNATCCTGGCTAACATGGTGAAA
GCACACGTCTGAACTCTTTCCCTACACGACGCTCTTCCGATCTNNNCTTACATAACCCCCATCTCCCAC

GCACACGTCTGAACTCTTTCCCTACACGACGCTCTTCCGATCTNNNCTTACATAACCCCATCTCCCAC
GAGCACACGTCTGAACTCTTTCCCTACACGACGCTCTTCCGATCTNNNGTGGGGAGGGAATAGAAA
GAGCACACGTCTGAACTCTTTCCCTACACGACGCTCTTCCGATCTNNNGTGGGGAGGGAATAGAAA
AGCACACGTCTGAACTCTTTCCCTACACGACGCTCTTCCGATCTNNNTTACCTTGATGTGGATGACTG
AGCACACGTCTGAACTCTTTCCCTACACGACGCTCTTCCGATCTNNNTTACCTTGATGTGGATGACTG
CACACGTCTGAACTCTTTCCCTACACGACGCTCTTCCGATCTNNNCCCACAAAGGCAGAGAGATCTCC
CACACGTCTGAACTCTTTCCCTACACGACGCTCTTCCGATCTNNNCCCACAAAGGCAGAGAGATCTCC
AGCACACGTCTGAACTCTTTCCCTACACGACGCTCTTCCGATCTNNNGTAGCCTGCATATCCAAAGCA
AGCACACGTCTGAACTCTTTCCCTACACGACGCTCTTCCGATCTNNNGTAGCCTGCATATCCAAAGCA
GAGCACACGTCTGAACTCTTTCCCTACACGACGCTCTTCCGATCTNNNGTCTTACCATATTGCCCATGC
GAGCACACGTCTGAACTCTTTCCCTACACGACGCTCTTCCGATCTNNNGTCTTACCATATTGCCCATGC
GCACACGTCTGAACTCTTTCCCTACACGACGCTCTTCCGATCTNNNAGGATCACATGTTCGGTGTTCA
GCACACGTCTGAACTCTTTCCCTACACGACGCTCTTCCGATCTNNNAGGATCACATGTTCGGTGTTCA
GAGCACACGTCTGAACTCTTTCCCTACACGACGCTCTTCCGATCTNNNCTCTGTGGTGAGGATGTGTT
GAGCACACGTCTGAACTCTTTCCCTACACGACGCTCTTCCGATCTNNNCTCTGTGGTGAGGATGTGTT
AGCACACGTCTGAACTCTTTCCCTACACGACGCTCTTCCGATCTNNNGGTGTATCTGTATGTCCGTGT
AGCACACGTCTGAACTCTTTCCCTACACGACGCTCTTCCGATCTNNNGGTGTATCTGTATGTCCGTGT
AGCACACGTCTGAACTCTTTCCCTACACGACGCTCTTCCGATCTNNNGAGGCTGAGGCAGGAGAAT
AGCACACGTCTGAACTCTTTCCCTACACGACGCTCTTCCGATCTNNNGAGGCTGAGGCAGGAGAAT
AGCACACGTCTGAACTCTTTCCCTACACGACGCTCTTCCGATCTNNNGCACATGCTACCATAGATGGC
AGCACACGTCTGAACTCTTTCCCTACACGACGCTCTTCCGATCTNNNGCACATGCTACCATAGATGGC
GCACACGTCTGAACTCTTTCCCTACACGACGCTCTTCCGATCTNNNAGTTTCTTCTTGATTTTTTGGAC
GCACACGTCTGAACTCTTTCCCTACACGACGCTCTTCCGATCTNNNAGTTTCTTCTTGATTTTTTGGAC
GCACACGTCTGAACTCTTTCCCTACACGACGCTCTTCCGATCTNNNCTAATTCTCTGGCAAGCACATG
GCACACGTCTGAACTCTTTCCCTACACGACGCTCTTCCGATCTNNNCTAATTCTCTGGCAAGCACATG
AGCACACGTCTGAACTCTTTCCCTACACGACGCTCTTCCGATCTNNNTGAGGAGGGAATCCAAACTC
AGCACACGTCTGAACTCTTTCCCTACACGACGCTCTTCCGATCTNNNTGAGGAGGGAATCCAAACTC
AGAGCACACGTCTGAACTCTTTCCCTACACGACGCTCTTCCGATCTNNNCAACACACAGTGGTTGAG
AGAGCACACGTCTGAACTCTTTCCCTACACGACGCTCTTCCGATCTNNNCAACACACAGTGGTTGAG
AGCACACGTCTGAACTCTTTCCCTACACGACGCTCTTCCGATCTNNNAGAGGAGATAGTGAGGGTCA
AGCACACGTCTGAACTCTTTCCCTACACGACGCTCTTCCGATCTNNNAGAGGAGATAGTGAGGGTCA
GCACACGTCTGAACTCTTTCCCTACACGACGCTCTTCCGATCTNNNGCAGTCAAAGCAAAGGCAAAG
GCACACGTCTGAACTCTTTCCCTACACGACGCTCTTCCGATCTNNNGCAGTCAAAGCAAAGGCAAAG
GAGCACACGTCTGAACTCTTTCCCTACACGACGCTCTTCCGATCTNNNAGCAGAGAAAAGCTGATCTG
GAGCACACGTCTGAACTCTTTCCCTACACGACGCTCTTCCGATCTNNNAGCAGAGAAAAGCTGATCTG
AGCACACGTCTGAACTCTTTCCCTACACGACGCTCTTCCGATCTNNNCTGAGGTGGGAGGATCAATT
AGCACACGTCTGAACTCTTTCCCTACACGACGCTCTTCCGATCTNNNCTGAGGTGGGAGGATCAATT
GAGCACACGTCTGAACTCTTTCCCTACACGACGCTCTTCCGATCTNNNGACACCTGTATTCCCAGCTAC
GAGCACACGTCTGAACTCTTTCCCTACACGACGCTCTTCCGATCTNNNGACACCTGTATTCCCAGCTAC
AGCACACGTCTGAACTCTTTCCCTACACGACGCTCTTCCGATCTNNNGATGCAGGAGAATCGCTTGAA
AGCACACGTCTGAACTCTTTCCCTACACGACGCTCTTCCGATCTNNNGATGCAGGAGAATCGCTTGAA
AGCACACGTCTGAACTCTTTCCCTACACGACGCTCTTCCGATCTNNNTGTTGTTGTTGTTGCTGACAG
AGCACACGTCTGAACTCTTTCCCTACACGACGCTCTTCCGATCTNNNTGTTGTTGTTGTTGCTGACAG

GCACACGTCTGAACTCTTTCCCTACACGACGCTCTTCCGATCTNNNGATCTCCTGACCTCATGATCCACC
GCACACGTCTGAACTCTTTCCCTACACGACGCTCTTCCGATCTNNNGATCTCCTGACCTCATGATCCACC
AGCACACGTCTGAACTCTTTCCCTACACGACGCTCTTCCGATCTNNNTTAGAGTTTCCTAGCTCTTGGCC
AGCACACGTCTGAACTCTTTCCCTACACGACGCTCTTCCGATCTNNNTTAGAGTTTCCTAGCTCTTGGCC
AGCACACGTCTGAACTCTTTCCCTACACGACGCTCTTCCGATCTNNNTACATCTGGCTGTCTGAAAGTGT
AGCACACGTCTGAACTCTTTCCCTACACGACGCTCTTCCGATCTNNNTACATCTGGCTGTCTGAAAGTGT
GCACACGTCTGAACTCTTTCCCTACACGACGCTCTTCCGATCTNNNAGTCCTTCTTTAGTATGCATTGCT
GCACACGTCTGAACTCTTTCCCTACACGACGCTCTTCCGATCTNNNAGTCCTTCTTTAGTATGCATTGCT
AGCACACGTCTGAACTCTTTCCCTACACGACGCTCTTCCGATCTNNNTGGTCTCTGGTTCTCTCTCATGA
AGCACACGTCTGAACTCTTTCCCTACACGACGCTCTTCCGATCTNNNTGGTCTCTGGTTCTCTCTCATGA
GAGCACACGTCTGAACTCTTTCCCTACACGACGCTCTTCCGATCTNNNCTTGAACTCCTGAACTTGCGAT
GAGCACACGTCTGAACTCTTTCCCTACACGACGCTCTTCCGATCTNNNCTTGAACTCCTGAACTTGCGAT
GAGCACACGTCTGAACTCTTTCCCTACACGACGCTCTTCCGATCTNNNGCATTTCTGATTGGTTGCCTTC
GAGCACACGTCTGAACTCTTTCCCTACACGACGCTCTTCCGATCTNNNGCATTTCTGATTGGTTGCCTTC
AGCACACGTCTGAACTCTTTCCCTACACGACGCTCTTCCGATCTNNNTAGAAAAGTTAGGAAGACAGA
AGCACACGTCTGAACTCTTTCCCTACACGACGCTCTTCCGATCTNNNTAGAAAAGTTAGGAAGACAGA
AGCACACGTCTGAACTCTTTCCCTACACGACGCTCTTCCGATCTNNNAATTCAAGCCATGGAAATTAAG
AGCACACGTCTGAACTCTTTCCCTACACGACGCTCTTCCGATCTNNNAATTCAAGCCATGGAAATTAAG
GAGCACACGTCTGAACTCTTTCCCTACACGACGCTCTTCCGATCTNNNCCTCCCAGGTTCAAGCAATTCT
GAGCACACGTCTGAACTCTTTCCCTACACGACGCTCTTCCGATCTNNNCCTCCCAGGTTCAAGCAATTCT
AGAGCACACGTCTGAACTCTTTCCCTACACGACGCTCTTCCGATCTNNNGAAGCGAGAGGAATGTGAC
AGAGCACACGTCTGAACTCTTTCCCTACACGACGCTCTTCCGATCTNNNGAAGCGAGAGGAATGTGAC
AGCACACGTCTGAACTCTTTCCCTACACGACGCTCTTCCGATCTNNNACATACTTAGAACAGTGCCTGG
AGCACACGTCTGAACTCTTTCCCTACACGACGCTCTTCCGATCTNNNACATACTTAGAACAGTGCCTGG
GAGCACACGTCTGAACTCTTTCCCTACACGACGCTCTTCCGATCTNNNGAAAAGCAGAGCCAACGTTGA
GAGCACACGTCTGAACTCTTTCCCTACACGACGCTCTTCCGATCTNNNGAAAAGCAGAGCCAACGTTGA
GAGCACACGTCTGAACTCTTTCCCTACACGACGCTCTTCCGATCTNNNCTGAGGCAGGAGAATCACTT
GAGCACACGTCTGAACTCTTTCCCTACACGACGCTCTTCCGATCTNNNCTGAGGCAGGAGAATCACTT
AGCACACGTCTGAACTCTTTCCCTACACGACGCTCTTCCGATCTNNNCAACCGCTCATTCCTCGCAGCAT
AGCACACGTCTGAACTCTTTCCCTACACGACGCTCTTCCGATCTNNNCAACCGCTCATTCCTCGCAGCAT
AGCACACGTCTGAACTCTTTCCCTACACGACGCTCTTCCGATCTNNNCTTGAAATTCTCACTCCCAGCAC
AGCACACGTCTGAACTCTTTCCCTACACGACGCTCTTCCGATCTNNNCTTGAAATTCTCACTCCCAGCAC
AGCACACGTCTGAACTCTTTCCCTACACGACGCTCTTCCGATCTNNNACCCCACAATAATGGAGAAAGA
AGCACACGTCTGAACTCTTTCCCTACACGACGCTCTTCCGATCTNNNACCCCACAATAATGGAGAAAGA
CACACGTCTGAACTCTTTCCCTACACGACGCTCTTCCGATCTNNNGGTACAACTGGACAAGCAGAGAG
CACACGTCTGAACTCTTTCCCTACACGACGCTCTTCCGATCTNNNGGTACAACTGGACAAGCAGAGAG
AGCACACGTCTGAACTCTTTCCCTACACGACGCTCTTCCGATCTNNNACACCTCAGATATATTTCTTCATC
AGCACACGTCTGAACTCTTTCCCTACACGACGCTCTTCCGATCTNNNACACCTCAGATATATTTCTTCATC
AGCACACGTCTGAACTCTTTCCCTACACGACGCTCTTCCGATCTNNNTCACAAGGACTCTCTCAGGATC
AGCACACGTCTGAACTCTTTCCCTACACGACGCTCTTCCGATCTNNNTCACAAGGACTCTCTCAGGATC
AGCACACGTCTGAACTCTTTCCCTACACGACGCTCTTCCGATCTNNNGGCAGCCTTTGTTCTGAACTATC
AGCACACGTCTGAACTCTTTCCCTACACGACGCTCTTCCGATCTNNNGGCAGCCTTTGTTCTGAACTATC
AGCACACGTCTGAACTCTTTCCCTACACGACGCTCTTCCGATCTNNNAGATTGCAGTGAGCCGAGATT

AGCACACGTCTGAACTCTTTCCCTACACGACGCTCTTCCGATCTNNNAGATTGCAGTGAGCCGAGATT
GCACACGTCTGAACTCTTTCCCTACACGACGCTCTTCCGATCTNNNTGATTCCTGCCTTCTTCAGAGTC
GCACACGTCTGAACTCTTTCCCTACACGACGCTCTTCCGATCTNNNTGATTCCTGCCTTCTTCAGAGTC
AGCACACGTCTGAACTCTTTCCCTACACGACGCTCTTCCGATCTNNNTAGAAACAATACGTATTTGCTT
AGCACACGTCTGAACTCTTTCCCTACACGACGCTCTTCCGATCTNNNTAGAAACAATACGTATTTGCTT
AGCACACGTCTGAACTCTTTCCCTACACGACGCTCTTCCGATCTNNNCATCCTGGCTAACAAGGTGAA
AGCACACGTCTGAACTCTTTCCCTACACGACGCTCTTCCGATCTNNNCATCCTGGCTAACAAGGTGAA
GAGCACACGTCTGAACTCTTTCCCTACACGACGCTCTTCCGATCTNNNAAGTGGCAGAGCTCTGAATA
GAGCACACGTCTGAACTCTTTCCCTACACGACGCTCTTCCGATCTNNNAAGTGGCAGAGCTCTGAATA
AGCACACGTCTGAACTCTTTCCCTACACGACGCTCTTCCGATCTNNNGCATCGCCCTGATCATTCTATTC
AGCACACGTCTGAACTCTTTCCCTACACGACGCTCTTCCGATCTNNNGCATCGCCCTGATCATTCTATTC
AGCACACGTCTGAACTCTTTCCCTACACGACGCTCTTCCGATCTNNNCACCCTTTGCATTTCCATCAGTT
AGCACACGTCTGAACTCTTTCCCTACACGACGCTCTTCCGATCTNNNCACCCTTTGCATTTCCATCAGTT
AGCACACGTCTGAACTCTTTCCCTACACGACGCTCTTCCGATCTNNNACATGCAAAGTGCTTAGGGTA
AGCACACGTCTGAACTCTTTCCCTACACGACGCTCTTCCGATCTNNNACATGCAAAGTGCTTAGGGTA
GCACACGTCTGAACTCTTTCCCTACACGACGCTCTTCCGATCTNNNTACAGCTGCCCCTATTTACTGACC
GCACACGTCTGAACTCTTTCCCTACACGACGCTCTTCCGATCTNNNTACAGCTGCCCCTATTTACTGACC
GAGCACACGTCTGAACTCTTTCCCTACACGACGCTCTTCCGATCTNNNAGTGAACCGAGATTGCATCA
GAGCACACGTCTGAACTCTTTCCCTACACGACGCTCTTCCGATCTNNNAGTGAACCGAGATTGCATCA
AGCACACGTCTGAACTCTTTCCCTACACGACGCTCTTCCGATCTNNNCGTGGACAAAATCACTTGAAGC
AGCACACGTCTGAACTCTTTCCCTACACGACGCTCTTCCGATCTNNNCGTGGACAAAATCACTTGAAGC
AGCACACGTCTGAACTCTTTCCCTACACGACGCTCTTCCGATCTNNNCTTGCAGTGAGCTGAGACTGT
AGCACACGTCTGAACTCTTTCCCTACACGACGCTCTTCCGATCTNNNCTTGCAGTGAGCTGAGACTGT
GAGCACACGTCTGAACTCTTTCCCTACACGACGCTCTTCCGATCTNNNCTCATGCCTATAATCCCAGCA
GAGCACACGTCTGAACTCTTTCCCTACACGACGCTCTTCCGATCTNNNCTCATGCCTATAATCCCAGCA
AGCACACGTCTGAACTCTTTCCCTACACGACGCTCTTCCGATCTNNNATGCTTCCTGCTATCCAACTGA
AGCACACGTCTGAACTCTTTCCCTACACGACGCTCTTCCGATCTNNNATGCTTCCTGCTATCCAACTGA
AGCACACGTCTGAACTCTTTCCCTACACGACGCTCTTCCGATCTNNNGAAGTATCCATGGTCAGGGGT
AGCACACGTCTGAACTCTTTCCCTACACGACGCTCTTCCGATCTNNNGAAGTATCCATGGTCAGGGGT
GCACACGTCTGAACTCTTTCCCTACACGACGCTCTTCCGATCTNNNTCAGAGTATAACCAAGCTGCTG
GCACACGTCTGAACTCTTTCCCTACACGACGCTCTTCCGATCTNNNTCAGAGTATAACCAAGCTGCTG
GAGCACACGTCTGAACTCTTTCCCTACACGACGCTCTTCCGATCTNNNTTTGGGCAAGTCACAATCTCT
GAGCACACGTCTGAACTCTTTCCCTACACGACGCTCTTCCGATCTNNNTTTGGGCAAGTCACAATCTCT
CACGTCTGAACTCTTTCCCTACACGACGCTCTTCCGATCTNNNCCGGTGATTTGTTCAACTCTAGGCAT
CACGTCTGAACTCTTTCCCTACACGACGCTCTTCCGATCTNNNCCGGTGATTTGTTCAACTCTAGGCAT
GCACACGTCTGAACTCTTTCCCTACACGACGCTCTTCCGATCTNNNTTTGTGAGATGACCTCGTGTGA
GCACACGTCTGAACTCTTTCCCTACACGACGCTCTTCCGATCTNNNTTTGTGAGATGACCTCGTGTGA
AGAGCACACGTCTGAACTCTTTCCCTACACGACGCTCTTCCGATCTNNNCTCGAAACCACTTGGAGAC
AGAGCACACGTCTGAACTCTTTCCCTACACGACGCTCTTCCGATCTNNNCTCGAAACCACTTGGAGAC
AGCACACGTCTGAACTCTTTCCCTACACGACGCTCTTCCGATCTNNNGTTCATAAGATGTTTTGGCCC
AGCACACGTCTGAACTCTTTCCCTACACGACGCTCTTCCGATCTNNNGTTCATAAGATGTTTTGGCCC
GAGCACACGTCTGAACTCTTTCCCTACACGACGCTCTTCCGATCTNNNCTTGCAGTCAGCCGAGATCAT
GAGCACACGTCTGAACTCTTTCCCTACACGACGCTCTTCCGATCTNNNCTTGCAGTCAGCCGAGATCAT

GAGCACACGTCTGAACTCTTTCCCTACACGACGCTCTTCCGATCTNNNCCAGGTTCAAGCGATCCTCTTC

GAGCACACGTCTGAACTCTTTCCCTACACGACGCTCTTCCGATCTNNNCCAGGTTCAAGCGATCCTCTTC

GAGCACACGTCTGAACTCTTTCCCTACACGACGCTCTTCCGATCTNNNCTCCATGTGAGCTCCTCCAT

GAGCACACGTCTGAACTCTTTCCCTACACGACGCTCTTCCGATCTNNNCTCCATGTGAGCTCCTCCAT

AGAGCACACGTCTGAACTCTTTCCCTACACGACGCTCTTCCGATCTNNNGAGGGTGGAATTCAAAGCC

AGAGCACACGTCTGAACTCTTTCCCTACACGACGCTCTTCCGATCTNNNGAGGGTGGAATTCAAAGCC

GAGCACACGTCTGAACTCTTTCCCTACACGACGCTCTTCCGATCTNNNCGTGCTTGTAATCCCAGCTACT

GAGCACACGTCTGAACTCTTTCCCTACACGACGCTCTTCCGATCTNNNCGTGCTTGTAATCCCAGCTACT

AGCACACGTCTGAACTCTTTCCCTACACGACGCTCTTCCGATCTNNNAGATTTCCACCAAGAAAACCAG

AGCACACGTCTGAACTCTTTCCCTACACGACGCTCTTCCGATCTNNNAGATTTCCACCAAGAAAACCAG

AGCACACGTCTGAACTCTTTCCCTACACGACGCTCTTCCGATCTNNNGACAACTACCACAGACTGGGAA

AGCACACGTCTGAACTCTTTCCCTACACGACGCTCTTCCGATCTNNNGACAACTACCACAGACTGGGAA

AGCACACGTCTGAACTCTTTCCCTACACGACGCTCTTCCGATCTNNNCAGAGTAGCTGGGACTACAGAT

AGCACACGTCTGAACTCTTTCCCTACACGACGCTCTTCCGATCTNNNCAGAGTAGCTGGGACTACAGAT

GAGCACACGTCTGAACTCTTTCCCTACACGACGCTCTTCCGATCTNNNAAATCCCAACTATGTGGAAGC

GAGCACACGTCTGAACTCTTTCCCTACACGACGCTCTTCCGATCTNNNAAATCCCAACTATGTGGAAGC

AGCACACGTCTGAACTCTTTCCCTACACGACGCTCTTCCGATCTNNNGACTCACTTGCTCCAAAACAGTC

AGCACACGTCTGAACTCTTTCCCTACACGACGCTCTTCCGATCTNNNGACTCACTTGCTCCAAAACAGTC

AGAGCACACGTCTGAACTCTTTCCCTACACGACGCTCTTCCGATCTNNNGAGGTTGCAGTGAGCTGAGA

AGAGCACACGTCTGAACTCTTTCCCTACACGACGCTCTTCCGATCTNNNGAGGTTGCAGTGAGCTGAGA

GAGCACACGTCTGAACTCTTTCCCTACACGACGCTCTTCCGATCTNNNTCATTCCGTCTCTATGAACAGG

GAGCACACGTCTGAACTCTTTCCCTACACGACGCTCTTCCGATCTNNNTCATTCCGTCTCTATGAACAGG

AGCACACGTCTGAACTCTTTCCCTACACGACGCTCTTCCGATCTNNNTTTAGACGGCCTTGCCTCATAA

AGCACACGTCTGAACTCTTTCCCTACACGACGCTCTTCCGATCTNNNTTTAGACGGCCTTGCCTCATAA

ACACGTCTGAACTCTTTCCCTACACGACGCTCTTCCGATCTNNNCCACGGCTTTTAATTCCCTACCGCATC

ACACGTCTGAACTCTTTCCCTACACGACGCTCTTCCGATCTNNNCCACGGCTTTTAATTCCCTACCGCATC

AGCACACGTCTGAACTCTTTCCCTACACGACGCTCTTCCGATCTNNNACAAAAATTAGCTGGGTGTGG

AGCACACGTCTGAACTCTTTCCCTACACGACGCTCTTCCGATCTNNNACAAAAATTAGCTGGGTGTGG

AGAGCACACGTCTGAACTCTTTCCCTACACGACGCTCTTCCGATCTNNNCCTAAGTGGCCCATATTGGA

AGAGCACACGTCTGAACTCTTTCCCTACACGACGCTCTTCCGATCTNNNCCTAAGTGGCCCATATTGGA

GCACACGTCTGAACTCTTTCCCTACACGACGCTCTTCCGATCTNNNGCCCCTAGGATTGTCTTTCAGAA

GCACACGTCTGAACTCTTTCCCTACACGACGCTCTTCCGATCTNNNGCCCCTAGGATTGTCTTTCAGAA

AGCACACGTCTGAACTCTTTCCCTACACGACGCTCTTCCGATCTNNNAAGAAAGTCATGCACCCAAACT

AGCACACGTCTGAACTCTTTCCCTACACGACGCTCTTCCGATCTNNNAAGAAAGTCATGCACCCAAACT

AGCACACGTCTGAACTCTTTCCCTACACGACGCTCTTCCGATCTNNNGGTACCTTGCTTTTACAATCTGT

AGCACACGTCTGAACTCTTTCCCTACACGACGCTCTTCCGATCTNNNGGTACCTTGCTTTTACAATCTGT

AGAGCACACGTCTGAACTCTTTCCCTACACGACGCTCTTCCGATCTNNNGCCTGTAGTCTGAGCTACTAA

AGAGCACACGTCTGAACTCTTTCCCTACACGACGCTCTTCCGATCTNNNGCCTGTAGTCTGAGCTACTAA

GAGCACACGTCTGAACTCTTTCCCTACACGACGCTCTTCCGATCTNNNGGATACGCTGACAATTTGAAC

GAGCACACGTCTGAACTCTTTCCCTACACGACGCTCTTCCGATCTNNNGGATACGCTGACAATTTGAAC

AGCACACGTCTGAACTCTTTCCCTACACGACGCTCTTCCGATCTNNNTTGCCATTGTGTCTTGATGTCAC

AGCACACGTCTGAACTCTTTCCCTACACGACGCTCTTCCGATCTNNNTTGCCATTGTGTCTTGATGTCAC

GCACACGTCTGAACTCTTTCCCTACACGACGCTCTTCCGATCTNNNAGCTAGTTATTTGGGGTCACTTTC

GCACACGTCTGAACTCTTTCCCTACACGACGCTCTTCCGATCTNNNAGCTAGTTATTTGGGGTCACTTTC
AGCACACGTCTGAACTCTTTCCCTACACGACGCTCTTCCGATCTNNNAGTTGCCAAAAATGGTTACCTG
AGCACACGTCTGAACTCTTTCCCTACACGACGCTCTTCCGATCTNNNAGTTGCCAAAAATGGTTACCTG
AGCACACGTCTGAACTCTTTCCCTACACGACGCTCTTCCGATCTNNNTTGCCTTCTCTTTCAGATCTCAA
AGCACACGTCTGAACTCTTTCCCTACACGACGCTCTTCCGATCTNNNTTGCCTTCTCTTTCAGATCTCAA
AGAGCACACGTCTGAACTCTTTCCCTACACGACGCTCTTCCGATCTNNNGCAGTTCATGTCCACCATATT
AGAGCACACGTCTGAACTCTTTCCCTACACGACGCTCTTCCGATCTNNNGCAGTTCATGTCCACCATATT
AGCACACGTCTGAACTCTTTCCCTACACGACGCTCTTCCGATCTNNNCATAGCCCGTGTCTATCACATCA
AGCACACGTCTGAACTCTTTCCCTACACGACGCTCTTCCGATCTNNNCATAGCCCGTGTCTATCACATCA
GAGCACACGTCTGAACTCTTTCCCTACACGACGCTCTTCCGATCTNNNTGATTCCACTTTATTAAAAAGC
GAGCACACGTCTGAACTCTTTCCCTACACGACGCTCTTCCGATCTNNNTGATTCCACTTTATTAAAAAGC
AAGAGCACACGTCTGAACTCTTTCCCTACACGACGCTCTTCCGATCTNNNTGAGCTGTGAACTTGTGTG
AAGAGCACACGTCTGAACTCTTTCCCTACACGACGCTCTTCCGATCTNNNTGAGCTGTGAACTTGTGTG
GAGCACACGTCTGAACTCTTTCCCTACACGACGCTCTTCCGATCTNNNGGGGGTTATCCTGCATCTCTTT
GAGCACACGTCTGAACTCTTTCCCTACACGACGCTCTTCCGATCTNNNGGGGGTTATCCTGCATCTCTTT
AGCACACGTCTGAACTCTTTCCCTACACGACGCTCTTCCGATCTNNNAGGTACAGAAATACTCGCACTG
AGCACACGTCTGAACTCTTTCCCTACACGACGCTCTTCCGATCTNNNAGGTACAGAAATACTCGCACTG
GAGCACACGTCTGAACTCTTTCCCTACACGACGCTCTTCCGATCTNNNAAGTTCAACATCAAGATGCCA
GAGCACACGTCTGAACTCTTTCCCTACACGACGCTCTTCCGATCTNNNAAGTTCAACATCAAGATGCCA
AGCACACGTCTGAACTCTTTCCCTACACGACGCTCTTCCGATCTNNNGGATTCTGATTCCCACTCCAAGC
AGCACACGTCTGAACTCTTTCCCTACACGACGCTCTTCCGATCTNNNGGATTCTGATTCCCACTCCAAGC
AGCACACGTCTGAACTCTTTCCCTACACGACGCTCTTCCGATCTNNNAGCCATCTAGCAAAAAGCCAGA
AGCACACGTCTGAACTCTTTCCCTACACGACGCTCTTCCGATCTNNNAGCCATCTAGCAAAAAGCCAGA
AGAGCACACGTCTGAACTCTTTCCCTACACGACGCTCTTCCGATCTNNNTGCCTCTAGTCCCAGCTACTC
AGAGCACACGTCTGAACTCTTTCCCTACACGACGCTCTTCCGATCTNNNTGCCTCTAGTCCCAGCTACTC
GAGCACACGTCTGAACTCTTTCCCTACACGACGCTCTTCCGATCTNNNGATTTGATGCTTTACCCTCTGC
GAGCACACGTCTGAACTCTTTCCCTACACGACGCTCTTCCGATCTNNNGATTTGATGCTTTACCCTCTGC
AGAGCACACGTCTGAACTCTTTCCCTACACGACGCTCTTCCGATCTNNNCTTTGTAGTTGCCTTCGTGGA
AGAGCACACGTCTGAACTCTTTCCCTACACGACGCTCTTCCGATCTNNNCTTTGTAGTTGCCTTCGTGGA
GAGCACACGTCTGAACTCTTTCCCTACACGACGCTCTTCCGATCTNNNAGAGGCAGACAAATACCAGAC
GAGCACACGTCTGAACTCTTTCCCTACACGACGCTCTTCCGATCTNNNAGAGGCAGACAAATACCAGAC
AGCACACGTCTGAACTCTTTCCCTACACGACGCTCTTCCGATCTNNNCCCTCTAGCAGCCAATCAAAAAC
AGCACACGTCTGAACTCTTTCCCTACACGACGCTCTTCCGATCTNNNCCCTCTAGCAGCCAATCAAAAAC
AGCACACGTCTGAACTCTTTCCCTACACGACGCTCTTCCGATCTNNNGTAATCCCAGCTACTTAGGAGG
AGCACACGTCTGAACTCTTTCCCTACACGACGCTCTTCCGATCTNNNGTAATCCCAGCTACTTAGGAGG
AGCACACGTCTGAACTCTTTCCCTACACGACGCTCTTCCGATCTNNNGCTAAGCTGCAAACTACATAGCC
AGCACACGTCTGAACTCTTTCCCTACACGACGCTCTTCCGATCTNNNGCTAAGCTGCAAACTACATAGCC
GCACACGTCTGAACTCTTTCCCTACACGACGCTCTTCCGATCTNNNTGATGGATGCACAGCTCTATGAA
GCACACGTCTGAACTCTTTCCCTACACGACGCTCTTCCGATCTNNNTGATGGATGCACAGCTCTATGAA
AGCACACGTCTGAACTCTTTCCCTACACGACGCTCTTCCGATCTNNNCTAAAATGCCTGTGTGTGTGTC
AGCACACGTCTGAACTCTTTCCCTACACGACGCTCTTCCGATCTNNNCTAAAATGCCTGTGTGTGTGTC
GAGCACACGTCTGAACTCTTTCCCTACACGACGCTCTTCCGATCTNNNACTGGGTGTTGCAGTATCCTA
GAGCACACGTCTGAACTCTTTCCCTACACGACGCTCTTCCGATCTNNNACTGGGTGTTGCAGTATCCTA

GAGCACACGTCTGAACTCTTTCCCTACACGACGCTCTTCCGATCTNNNTAGTAGATAGCACCCCAACAC
GAGCACACGTCTGAACTCTTTCCCTACACGACGCTCTTCCGATCTNNNTAGTAGATAGCACCCCAACAC
GAGCACACGTCTGAACTCTTTCCCTACACGACGCTCTTCCGATCTNNNGCGCCCAGCTATTATTTCTTTC
GAGCACACGTCTGAACTCTTTCCCTACACGACGCTCTTCCGATCTNNNGCGCCCAGCTATTATTTCTTTC
GAGCACACGTCTGAACTCTTTCCCTACACGACGCTCTTCCGATCTNNNTCATTCATTCATTCCCCCCACAT
GAGCACACGTCTGAACTCTTTCCCTACACGACGCTCTTCCGATCTNNNTCATTCATTCATTCCCCCCACAT
GAGCACACGTCTGAACTCTTTCCCTACACGACGCTCTTCCGATCTNNNACCCCTCACCAAGCTATGCTACA
GAGCACACGTCTGAACTCTTTCCCTACACGACGCTCTTCCGATCTNNNACCCCTCACCAAGCTATGCTACA
AGCACACGTCTGAACTCTTTCCCTACACGACGCTCTTCCGATCTNNNCCTGAGACTGGGTAATTTTGGG
AGCACACGTCTGAACTCTTTCCCTACACGACGCTCTTCCGATCTNNNCCTGAGACTGGGTAATTTTGGG
AGAGCACACGTCTGAACTCTTTCCCTACACGACGCTCTTCCGATCTNNNCAGTTGGCTCAGATGGGAG
AGAGCACACGTCTGAACTCTTTCCCTACACGACGCTCTTCCGATCTNNNCAGTTGGCTCAGATGGGAG
GAGCACACGTCTGAACTCTTTCCCTACACGACGCTCTTCCGATCTNNNCAAGAACGGCATTCCCCTTTTT
GAGCACACGTCTGAACTCTTTCCCTACACGACGCTCTTCCGATCTNNNCAAGAACGGCATTCCCCTTTTT
AGCACACGTCTGAACTCTTTCCCTACACGACGCTCTTCCGATCTNNNTCAGTTGGATTGTTACATACCTG
AGCACACGTCTGAACTCTTTCCCTACACGACGCTCTTCCGATCTNNNTCAGTTGGATTGTTACATACCTG
AGCACACGTCTGAACTCTTTCCCTACACGACGCTCTTCCGATCTNNNGATCATTAGATCTGGGGTCTTGA
AGCACACGTCTGAACTCTTTCCCTACACGACGCTCTTCCGATCTNNNGATCATTAGATCTGGGGTCTTGA
AGCACACGTCTGAACTCTTTCCCTACACGACGCTCTTCCGATCTNNNGGATAACCCCCAGTTGGTTGTA
AGCACACGTCTGAACTCTTTCCCTACACGACGCTCTTCCGATCTNNNGGATAACCCCCAGTTGGTTGTA
GAGCACACGTCTGAACTCTTTCCCTACACGACGCTCTTCCGATCTNNNCTCTAACCTGACTCAGCGACTA
GAGCACACGTCTGAACTCTTTCCCTACACGACGCTCTTCCGATCTNNNCTCTAACCTGACTCAGCGACTA
GAGCACACGTCTGAACTCTTTCCCTACACGACGCTCTTCCGATCTNNNGGTGACAGAACGAGACTTCAT
GAGCACACGTCTGAACTCTTTCCCTACACGACGCTCTTCCGATCTNNNGGTGACAGAACGAGACTTCAT
AGCACACGTCTGAACTCTTTCCCTACACGACGCTCTTCCGATCTNNNGTCATCCTAGCTACTCAGAAGG
AGCACACGTCTGAACTCTTTCCCTACACGACGCTCTTCCGATCTNNNGTCATCCTAGCTACTCAGAAGG
GAGCACACGTCTGAACTCTTTCCCTACACGACGCTCTTCCGATCTNNNCAGGAGAATTGCTTGAACCAA
GAGCACACGTCTGAACTCTTTCCCTACACGACGCTCTTCCGATCTNNNCAGGAGAATTGCTTGAACCAA
GAGCACACGTCTGAACTCTTTCCCTACACGACGCTCTTCCGATCTNNNAGAAGTCTGTACCCTTTGTTCC
GAGCACACGTCTGAACTCTTTCCCTACACGACGCTCTTCCGATCTNNNAGAAGTCTGTACCCTTTGTTCC
CACACGTCTGAACTCTTTCCCTACACGACGCTCTTCCGATCTNNNTGCAACTGGAATGGTATCTCTCTCA
CACACGTCTGAACTCTTTCCCTACACGACGCTCTTCCGATCTNNNTGCAACTGGAATGGTATCTCTCTCA
AGCACACGTCTGAACTCTTTCCCTACACGACGCTCTTCCGATCTNNNCTGCCCTCACCTTCCTTTCTTCCA
AGCACACGTCTGAACTCTTTCCCTACACGACGCTCTTCCGATCTNNNCTGCCCTCACCTTCCTTTCTTCCA
AGCACACGTCTGAACTCTTTCCCTACACGACGCTCTTCCGATCTNNNAATGCACAGAAGAGAAACCTG
AGCACACGTCTGAACTCTTTCCCTACACGACGCTCTTCCGATCTNNNAATGCACAGAAGAGAAACCTG
GAGCACACGTCTGAACTCTTTCCCTACACGACGCTCTTCCGATCTNNNGGAGCAGGGAAGAAATTTTG
GAGCACACGTCTGAACTCTTTCCCTACACGACGCTCTTCCGATCTNNNGGAGCAGGGAAGAAATTTTG
GCACACGTCTGAACTCTTTCCCTACACGACGCTCTTCCGATCTNNNTGACAATAACAAGCCCTGGTTCTC
GCACACGTCTGAACTCTTTCCCTACACGACGCTCTTCCGATCTNNNTGACAATAACAAGCCCTGGTTCTC
AGCACACGTCTGAACTCTTTCCCTACACGACGCTCTTCCGATCTNNNATTGGCACACATTTTTGAGGGA
AGCACACGTCTGAACTCTTTCCCTACACGACGCTCTTCCGATCTNNNATTGGCACACATTTTTGAGGGA
GAGCACACGTCTGAACTCTTTCCCTACACGACGCTCTTCCGATCTNNNCTGTAGTCCCAGCTACTCAGGA

GAGCACACGTCTGAACTCTTTCCCTACACGACGCTCTTCCGATCTNNNCTGTAGTCCCAGCTACTCAGG
AGCACACGTCTGAACTCTTTCCCTACACGACGCTCTTCCGATCTNNNAGTGACCAGAGCAAGTATGAA
AGCACACGTCTGAACTCTTTCCCTACACGACGCTCTTCCGATCTNNNAGTGACCAGAGCAAGTATGAA
GCACACGTCTGAACTCTTTCCCTACACGACGCTCTTCCGATCTNNNGTTTTCTGCACGACCTGTACTTC
GCACACGTCTGAACTCTTTCCCTACACGACGCTCTTCCGATCTNNNGTTTTCTGCACGACCTGTACTTC
AGCACACGTCTGAACTCTTTCCCTACACGACGCTCTTCCGATCTNNNAGGAGCTAAGACTACAGGAGT
AGCACACGTCTGAACTCTTTCCCTACACGACGCTCTTCCGATCTNNNAGGAGCTAAGACTACAGGAGT
AGCACACGTCTGAACTCTTTCCCTACACGACGCTCTTCCGATCTNNNAGTATACAGTGCAACAAGGGG
AGCACACGTCTGAACTCTTTCCCTACACGACGCTCTTCCGATCTNNNAGTATACAGTGCAACAAGGGG
GCACACGTCTGAACTCTTTCCCTACACGACGCTCTTCCGATCTNNNGCTCATGCCTGTAATCTCAACAC
GCACACGTCTGAACTCTTTCCCTACACGACGCTCTTCCGATCTNNNGCTCATGCCTGTAATCTCAACAC
AGCACACGTCTGAACTCTTTCCCTACACGACGCTCTTCCGATCTNNNTTCAGAGGGCACATGAAAGTTC
AGCACACGTCTGAACTCTTTCCCTACACGACGCTCTTCCGATCTNNNTTCAGAGGGCACATGAAAGTTC
GAGCACACGTCTGAACTCTTTCCCTACACGACGCTCTTCCGATCTNNNGATGCCTGTAATCCCAGCTAC
GAGCACACGTCTGAACTCTTTCCCTACACGACGCTCTTCCGATCTNNNGATGCCTGTAATCCCAGCTAC
AGCACACGTCTGAACTCTTTCCCTACACGACGCTCTTCCGATCTNNNCTTTTCCCATGGTCTTCACAACC
AGCACACGTCTGAACTCTTTCCCTACACGACGCTCTTCCGATCTNNNCTTTTCCCATGGTCTTCACAACC
GAGCACACGTCTGAACTCTTTCCCTACACGACGCTCTTCCGATCTNNNCTGTAGTCCCAGCTACTCAGG
GAGCACACGTCTGAACTCTTTCCCTACACGACGCTCTTCCGATCTNNNCTGTAGTCCCAGCTACTCAGG
GCACACGTCTGAACTCTTTCCCTACACGACGCTCTTCCGATCTNNNAGCCTGGGCAACATAAGTTAGA
GCACACGTCTGAACTCTTTCCCTACACGACGCTCTTCCGATCTNNNAGCCTGGGCAACATAAGTTAGA
AGCACACGTCTGAACTCTTTCCCTACACGACGCTCTTCCGATCTNNNAAAGGAAAGGGCTTCTTGCTG
AGCACACGTCTGAACTCTTTCCCTACACGACGCTCTTCCGATCTNNNAAAGGAAAGGGCTTCTTGCTG
AGCACACGTCTGAACTCTTTCCCTACACGACGCTCTTCCGATCTNNNGAGGCTGAGGCAGGAGAATG
AGCACACGTCTGAACTCTTTCCCTACACGACGCTCTTCCGATCTNNNGAGGCTGAGGCAGGAGAATG
GAGCACACGTCTGAACTCTTTCCCTACACGACGCTCTTCCGATCTNNNTGGGTAACAAAGCGAGATCTT
GAGCACACGTCTGAACTCTTTCCCTACACGACGCTCTTCCGATCTNNNTGGGTAACAAAGCGAGATCTT
GAGCACACGTCTGAACTCTTTCCCTACACGACGCTCTTCCGATCTNNNGTTCACACACACACATACACAC
GAGCACACGTCTGAACTCTTTCCCTACACGACGCTCTTCCGATCTNNNGTTCACACACACACATACACAC
GCACACGTCTGAACTCTTTCCCTACACGACGCTCTTCCGATCTNNNGCCTGTAGTCCCAGCTACTTGCA
GCACACGTCTGAACTCTTTCCCTACACGACGCTCTTCCGATCTNNNGCCTGTAGTCCCAGCTACTTGCA
AGCACACGTCTGAACTCTTTCCCTACACGACGCTCTTCCGATCTNNNCCGTGGAGCTTTCTATTCTGCT
AGCACACGTCTGAACTCTTTCCCTACACGACGCTCTTCCGATCTNNNCCGTGGAGCTTTCTATTCTGCT
AGCACACGTCTGAACTCTTTCCCTACACGACGCTCTTCCGATCTNNNTTCCACTCTATCCCCCTTCTTCTC
AGCACACGTCTGAACTCTTTCCCTACACGACGCTCTTCCGATCTNNNTTCCACTCTATCCCCCTTCTTCTC
GAGCACACGTCTGAACTCTTTCCCTACACGACGCTCTTCCGATCTNNNAATTGGATGACCCTTGTGCTTT
GAGCACACGTCTGAACTCTTTCCCTACACGACGCTCTTCCGATCTNNNAATTGGATGACCCTTGTGCTTT
AGAGCACACGTCTGAACTCTTTCCCTACACGACGCTCTTCCGATCTNNNCTATGTGCTGATGCTGGGTA
AGAGCACACGTCTGAACTCTTTCCCTACACGACGCTCTTCCGATCTNNNCTATGTGCTGATGCTGGGTA
GAGCACACGTCTGAACTCTTTCCCTACACGACGCTCTTCCGATCTNNNCCTATAGTCCCAGCTACTCGG
GAGCACACGTCTGAACTCTTTCCCTACACGACGCTCTTCCGATCTNNNCCTATAGTCCCAGCTACTCGG
AGCACACGTCTGAACTCTTTCCCTACACGACGCTCTTCCGATCTNNNGAGAGAGAGAGTGTGCAAGAA
AGCACACGTCTGAACTCTTTCCCTACACGACGCTCTTCCGATCTNNNGAGAGAGAGAGTGTGCAAGAA

AGCACACGTCTGAACTCTTTCCCTACACGACGCTCTTCCGATCTNNNACTCAATGGTTTCTGTGAAGG
AGCACACGTCTGAACTCTTTCCCTACACGACGCTCTTCCGATCTNNNACTCAATGGTTTCTGTGAAGG
GAGCACACGTCTGAACTCTTTCCCTACACGACGCTCTTCCGATCTNNNCCCATGCCTGGCTAATTTTGT
GAGCACACGTCTGAACTCTTTCCCTACACGACGCTCTTCCGATCTNNNCCCATGCCTGGCTAATTTTGT
GCACACGTCTGAACTCTTTCCCTACACGACGCTCTTCCGATCTNNNCTGAAACAGGGTTAGGGCTTTT
GCACACGTCTGAACTCTTTCCCTACACGACGCTCTTCCGATCTNNNCTGAAACAGGGTTAGGGCTTTT
GAGCACACGTCTGAACTCTTTCCCTACACGACGCTCTTCCGATCTNNNCTGAGGTAGGAGAATCGCTT
GAGCACACGTCTGAACTCTTTCCCTACACGACGCTCTTCCGATCTNNNCTGAGGTAGGAGAATCGCTT
GCACACGTCTGAACTCTTTCCCTACACGACGCTCTTCCGATCTNNNCAGGAGAATTGCTTGAACTTGG
GCACACGTCTGAACTCTTTCCCTACACGACGCTCTTCCGATCTNNNCAGGAGAATTGCTTGAACTTGG
AGCACACGTCTGAACTCTTTCCCTACACGACGCTCTTCCGATCTNNNATGATGTAGCTGTCTTCTGCAC
AGCACACGTCTGAACTCTTTCCCTACACGACGCTCTTCCGATCTNNNATGATGTAGCTGTCTTCTGCAC
GAGCACACGTCTGAACTCTTTCCCTACACGACGCTCTTCCGATCTNNNTGTAATCTCAGCACTTTGGG
GAGCACACGTCTGAACTCTTTCCCTACACGACGCTCTTCCGATCTNNNTGTAATCTCAGCACTTTGGG
AGCACACGTCTGAACTCTTTCCCTACACGACGCTCTTCCGATCTNNNAGCATGAAAAGCTACCAGACTC
AGCACACGTCTGAACTCTTTCCCTACACGACGCTCTTCCGATCTNNNAGCATGAAAAGCTACCAGACTC
AGCACACGTCTGAACTCTTTCCCTACACGACGCTCTTCCGATCTNNNTCTGGTACCTCAGTTGGGAAA
AGCACACGTCTGAACTCTTTCCCTACACGACGCTCTTCCGATCTNNNTCTGGTACCTCAGTTGGGAAA
GAGCACACGTCTGAACTCTTTCCCTACACGACGCTCTTCCGATCTNNNCTGGTGCCTTACTCTGACATTC
GAGCACACGTCTGAACTCTTTCCCTACACGACGCTCTTCCGATCTNNNCTGGTGCCTTACTCTGACATTC
AGCACACGTCTGAACTCTTTCCCTACACGACGCTCTTCCGATCTNNNCCTGGCTTTTTTACTGTCGTG
AGCACACGTCTGAACTCTTTCCCTACACGACGCTCTTCCGATCTNNNCCTGGCTTTTTTACTGTCGTG
GAGCACACGTCTGAACTCTTTCCCTACACGACGCTCTTCCGATCTNNNCTGTAATTCCAGCTACTCGGG
GAGCACACGTCTGAACTCTTTCCCTACACGACGCTCTTCCGATCTNNNCTGTAATTCCAGCTACTCGGG
GCACACGTCTGAACTCTTTCCCTACACGACGCTCTTCCGATCTNNNGTCTTGACTTGCCAGCCATATTG
GCACACGTCTGAACTCTTTCCCTACACGACGCTCTTCCGATCTNNNGTCTTGACTTGCCAGCCATATTG
GAGCACACGTCTGAACTCTTTCCCTACACGACGCTCTTCCGATCTNNNACAAGATGGGAGAGTGCTAG
GAGCACACGTCTGAACTCTTTCCCTACACGACGCTCTTCCGATCTNNNACAAGATGGGAGAGTGCTAG
GCACACGTCTGAACTCTTTCCCTACACGACGCTCTTCCGATCTNNNAGACTTGAAGTATGCTGCCAAG
GCACACGTCTGAACTCTTTCCCTACACGACGCTCTTCCGATCTNNNAGACTTGAAGTATGCTGCCAAG
GCACACGTCTGAACTCTTTCCCTACACGACGCTCTTCCGATCTNNNGGTCCATTCTGCAATCTGTGATG
GCACACGTCTGAACTCTTTCCCTACACGACGCTCTTCCGATCTNNNGGTCCATTCTGCAATCTGTGATG
AGCACACGTCTGAACTCTTTCCCTACACGACGCTCTTCCGATCTNNNTTTTTTTCAAAGACCCTCCTGGC
AGCACACGTCTGAACTCTTTCCCTACACGACGCTCTTCCGATCTNNNTTTTTTTCAAAGACCCTCCTGGC
AGCACACGTCTGAACTCTTTCCCTACACGACGCTCTTCCGATCTNNNAAATCTGTGTGTGTGTGTGTG
AGCACACGTCTGAACTCTTTCCCTACACGACGCTCTTCCGATCTNNNAAATCTGTGTGTGTGTGTGTG
AGCACACGTCTGAACTCTTTCCCTACACGACGCTCTTCCGATCTNNNAAAAGCAACAAGGTTTCACAG
AGCACACGTCTGAACTCTTTCCCTACACGACGCTCTTCCGATCTNNNAAAAGCAACAAGGTTTCACAG
GAGCACACGTCTGAACTCTTTCCCTACACGACGCTCTTCCGATCTNNNGGGGCTCAGCAAAGTTTTCTT
GAGCACACGTCTGAACTCTTTCCCTACACGACGCTCTTCCGATCTNNNGGGGCTCAGCAAAGTTTTCTT
AGCACACGTCTGAACTCTTTCCCTACACGACGCTCTTCCGATCTNNNAATCCCAGAACTTTGAGAGGC
AGCACACGTCTGAACTCTTTCCCTACACGACGCTCTTCCGATCTNNNAATCCCAGAACTTTGAGAGGC
CACGTCTGAACTCTTTCCCTACACGACGCTCTTCCGATCTNNNTTTCACACACTCCAGTACCATCCATC

CACGTCTGAACTCTTTCCCTACACGACGCTCTTCCGATCTNNNTTTCACACACTCCAGTACCATCCCATC
ACACGTCTGAACTCTTTCCCTACACGACGCTCTTCCGATCTNNNCCAAGACTCATTGGGTCTCCAGCAT
ACACGTCTGAACTCTTTCCCTACACGACGCTCTTCCGATCTNNNCCAAGACTCATTGGGTCTCCAGCAT
GCACACGTCTGAACTCTTTCCCTACACGACGCTCTTCCGATCTNNNAACCTCACTGTCTTCTCTGTTGG
GCACACGTCTGAACTCTTTCCCTACACGACGCTCTTCCGATCTNNNAACCTCACTGTCTTCTCTGTTGG
AGCACACGTCTGAACTCTTTCCCTACACGACGCTCTTCCGATCTNNNGAGACAGAAGCCAAGAGATT
AGCACACGTCTGAACTCTTTCCCTACACGACGCTCTTCCGATCTNNNGAGACAGAAGCCAAGAGATT
AGCACACGTCTGAACTCTTTCCCTACACGACGCTCTTCCGATCTNNNCCATTGCCAAACTTCGGACTTT
AGCACACGTCTGAACTCTTTCCCTACACGACGCTCTTCCGATCTNNNCCATTGCCAAACTTCGGACTTT
ACACGTCTGAACTCTTTCCCTACACGACGCTCTTCCGATCTNNNAGAATGAGATCCTGTCATTGGCAC
ACACGTCTGAACTCTTTCCCTACACGACGCTCTTCCGATCTNNNAGAATGAGATCCTGTCATTGGCAC
AGCACACGTCTGAACTCTTTCCCTACACGACGCTCTTCCGATCTNNNCCTTTGCCTCCTTCTAGCTGATT
AGCACACGTCTGAACTCTTTCCCTACACGACGCTCTTCCGATCTNNNCCTTTGCCTCCTTCTAGCTGATT
GCACACGTCTGAACTCTTTCCCTACACGACGCTCTTCCGATCTNNNAAGACCATCTTGGCTAACATGG
GCACACGTCTGAACTCTTTCCCTACACGACGCTCTTCCGATCTNNNAAGACCATCTTGGCTAACATGG
AGCACACGTCTGAACTCTTTCCCTACACGACGCTCTTCCGATCTNNNAGACAAACCATATCCCTGCCA
AGCACACGTCTGAACTCTTTCCCTACACGACGCTCTTCCGATCTNNNAGACAAACCATATCCCTGCCA
AGCACACGTCTGAACTCTTTCCCTACACGACGCTCTTCCGATCTNNNTCCTAAACTGCTGGGATTACA
AGCACACGTCTGAACTCTTTCCCTACACGACGCTCTTCCGATCTNNNTCCTAAACTGCTGGGATTACA
AGCACACGTCTGAACTCTTTCCCTACACGACGCTCTTCCGATCTNNNGCCAGTTGAACATTATAGACC
AGCACACGTCTGAACTCTTTCCCTACACGACGCTCTTCCGATCTNNNGCCAGTTGAACATTATAGACC
AGCACACGTCTGAACTCTTTCCCTACACGACGCTCTTCCGATCTNNNGTATTTTTGTACGGCTGCACA
AGCACACGTCTGAACTCTTTCCCTACACGACGCTCTTCCGATCTNNNGTATTTTTGTACGGCTGCACA
AGCACACGTCTGAACTCTTTCCCTACACGACGCTCTTCCGATCTNNNTTCAGGAAGGTATAGAGAGCC
AGCACACGTCTGAACTCTTTCCCTACACGACGCTCTTCCGATCTNNNTTCAGGAAGGTATAGAGAGCC
GCACACGTCTGAACTCTTTCCCTACACGACGCTCTTCCGATCTNNNCATGGTGGTTCTTGACTGTCATC
GCACACGTCTGAACTCTTTCCCTACACGACGCTCTTCCGATCTNNNCATGGTGGTTCTTGACTGTCATC
AGCACACGTCTGAACTCTTTCCCTACACGACGCTCTTCCGATCTNNNATTAGGAGGAGACGCTGTTTC
AGCACACGTCTGAACTCTTTCCCTACACGACGCTCTTCCGATCTNNNATTAGGAGGAGACGCTGTTTC
AGAGCACACGTCTGAACTCTTTCCCTACACGACGCTCTTCCGATCTNNNCAAGTCTGTAGCATGATCCT
AGAGCACACGTCTGAACTCTTTCCCTACACGACGCTCTTCCGATCTNNNCAAGTCTGTAGCATGATCCT
AGCACACGTCTGAACTCTTTCCCTACACGACGCTCTTCCGATCTNNNTAGGAAGAAGCAAAAGCGGA
AGCACACGTCTGAACTCTTTCCCTACACGACGCTCTTCCGATCTNNNTAGGAAGAAGCAAAAGCGGA
GAGCACACGTCTGAACTCTTTCCCTACACGACGCTCTTCCGATCTNNNGACCAGCTCCTACTCATTGTG
GAGCACACGTCTGAACTCTTTCCCTACACGACGCTCTTCCGATCTNNNGACCAGCTCCTACTCATTGTG
CACACGTCTGAACTCTTTCCCTACACGACGCTCTTCCGATCTNNNTTCACCATTCTCCTTCTCAGCCA
CACACGTCTGAACTCTTTCCCTACACGACGCTCTTCCGATCTNNNTTCACCATTCTCCTTCTCAGCCA
AGCACACGTCTGAACTCTTTCCCTACACGACGCTCTTCCGATCTNNNGGCACCTGTAATCCCAGATAC
AGCACACGTCTGAACTCTTTCCCTACACGACGCTCTTCCGATCTNNNGGCACCTGTAATCCCAGATAC
GCACACGTCTGAACTCTTTCCCTACACGACGCTCTTCCGATCTNNNCTCCAGGTCCTTATCTGTGTCAC
GCACACGTCTGAACTCTTTCCCTACACGACGCTCTTCCGATCTNNNCTCCAGGTCCTTATCTGTGTCAC
AGCACACGTCTGAACTCTTTCCCTACACGACGCTCTTCCGATCTNNNCCATGCATACAAAAGCCAGCA
AGCACACGTCTGAACTCTTTCCCTACACGACGCTCTTCCGATCTNNNCCATGCATACAAAAGCCAGCA

GAGCACACGTCTGAACTCTTTCCCTACACGACGCTCTTCCGATCTNNNAAAAAATTAGCCTGGCATGG
GAGCACACGTCTGAACTCTTTCCCTACACGACGCTCTTCCGATCTNNNAAAAAATTAGCCTGGCATGG
GAGCACACGTCTGAACTCTTTCCCTACACGACGCTCTTCCGATCTNNNCTCCCTTAGACTGTAATAGCCC
GAGCACACGTCTGAACTCTTTCCCTACACGACGCTCTTCCGATCTNNNCTCCCTTAGACTGTAATAGCCC
AGCACACGTCTGAACTCTTTCCCTACACGACGCTCTTCCGATCTNNNACAAGATGCTGTTTTTGGCAGA
AGCACACGTCTGAACTCTTTCCCTACACGACGCTCTTCCGATCTNNNACAAGATGCTGTTTTTGGCAGA
GAGCACACGTCTGAACTCTTTCCCTACACGACGCTCTTCCGATCTNNNGTATACTCCCTACCCCAGCAA
GAGCACACGTCTGAACTCTTTCCCTACACGACGCTCTTCCGATCTNNNGTATACTCCCTACCCCAGCAA
AGCACACGTCTGAACTCTTTCCCTACACGACGCTCTTCCGATCTNNNCAATCCCCTCCCTGCTTAGAAA
AGCACACGTCTGAACTCTTTCCCTACACGACGCTCTTCCGATCTNNNCAATCCCCTCCCTGCTTAGAAA
GCACACGTCTGAACTCTTTCCCTACACGACGCTCTTCCGATCTNNNTTTTGCCTCTGTCTCTTCCTCTCCA
GCACACGTCTGAACTCTTTCCCTACACGACGCTCTTCCGATCTNNNTTTTGCCTCTGTCTCTTCCTCTCCA
AGCACACGTCTGAACTCTTTCCCTACACGACGCTCTTCCGATCTNNNAAGGAATTGTGTGTGTGTGT
AGCACACGTCTGAACTCTTTCCCTACACGACGCTCTTCCGATCTNNNAAGGAATTGTGTGTGTGTGT
CACACGTCTGAACTCTTTCCCTACACGACGCTCTTCCGATCTNNNTTTCTGTGTCTTTTCCCTCTGCTCAT
CACACGTCTGAACTCTTTCCCTACACGACGCTCTTCCGATCTNNNTTTCTGTGTCTTTTCCCTCTGCTCAT
AGCACACGTCTGAACTCTTTCCCTACACGACGCTCTTCCGATCTNNNCTCAACCCAGAAAACCAGTCACA
AGCACACGTCTGAACTCTTTCCCTACACGACGCTCTTCCGATCTNNNCTCAACCCAGAAAACCAGTCACA
GAGCACACGTCTGAACTCTTTCCCTACACGACGCTCTTCCGATCTNNNGAAAGAGAATTCAGCCAGAG
GAGCACACGTCTGAACTCTTTCCCTACACGACGCTCTTCCGATCTNNNGAAAGAGAATTCAGCCAGAG
GCACACGTCTGAACTCTTTCCCTACACGACGCTCTTCCGATCTNNNCAACCCAACTCTCACTCATACCAC
GCACACGTCTGAACTCTTTCCCTACACGACGCTCTTCCGATCTNNNCAACCCAACTCTCACTCATACCAC
GCACACGTCTGAACTCTTTCCCTACACGACGCTCTTCCGATCTNNNCCTACCAAGTAGCTGGGACTTAC
GCACACGTCTGAACTCTTTCCCTACACGACGCTCTTCCGATCTNNNCCTACCAAGTAGCTGGGACTTAC
AGCACACGTCTGAACTCTTTCCCTACACGACGCTCTTCCGATCTNNNCAGTGAGCTGTGATTGTGTCC
AGCACACGTCTGAACTCTTTCCCTACACGACGCTCTTCCGATCTNNNCAGTGAGCTGTGATTGTGTCC
AGCACACGTCTGAACTCTTTCCCTACACGACGCTCTTCCGATCTNNNTCACTTGAATCCTGGAGGTCA
AGCACACGTCTGAACTCTTTCCCTACACGACGCTCTTCCGATCTNNNTCACTTGAATCCTGGAGGTCA
GAGCACACGTCTGAACTCTTTCCCTACACGACGCTCTTCCGATCTNNNAATGTGCCGAAAATCCTAACA
GAGCACACGTCTGAACTCTTTCCCTACACGACGCTCTTCCGATCTNNNAATGTGCCGAAAATCCTAACA
GAGCACACGTCTGAACTCTTTCCCTACACGACGCTCTTCCGATCTNNNGCAGAGGTCCCTGAGGAATT
GAGCACACGTCTGAACTCTTTCCCTACACGACGCTCTTCCGATCTNNNGCAGAGGTCCCTGAGGAATT
GAGCACACGTCTGAACTCTTTCCCTACACGACGCTCTTCCGATCTNNNGCTAACCATCCAAGTTCCTGG
GAGCACACGTCTGAACTCTTTCCCTACACGACGCTCTTCCGATCTNNNGCTAACCATCCAAGTTCCTGG
AGCACACGTCTGAACTCTTTCCCTACACGACGCTCTTCCGATCTNNNGTAAGGTCACAGGTCAACAGG
AGCACACGTCTGAACTCTTTCCCTACACGACGCTCTTCCGATCTNNNGTAAGGTCACAGGTCAACAGG
GCACACGTCTGAACTCTTTCCCTACACGACGCTCTTCCGATCTNNNCACCCGGTTGCAATCTTACATTTC
GCACACGTCTGAACTCTTTCCCTACACGACGCTCTTCCGATCTNNNCACCCGGTTGCAATCTTACATTTC
GCACACGTCTGAACTCTTTCCCTACACGACGCTCTTCCGATCTNNNGCAGCAGCGTAACTTTTCTTTTT
GCACACGTCTGAACTCTTTCCCTACACGACGCTCTTCCGATCTNNNGCAGCAGCGTAACTTTTCTTTTT
AGCACACGTCTGAACTCTTTCCCTACACGACGCTCTTCCGATCTNNNCTCAAGCAATCTTCTTGCCTCAG
AGCACACGTCTGAACTCTTTCCCTACACGACGCTCTTCCGATCTNNNCTCAAGCAATCTTCTTGCCTCAG
GAGCACACGTCTGAACTCTTTCCCTACACGACGCTCTTCCGATCTNNNATTCCTCTTTTCAGACCGGAT

GAGCACACGTCTGAACTCTTTCCCTACACGACGCTCTTCCGATCTNNNATTCCTCTTTTCAGACCGGATC
GAGCACACGTCTGAACTCTTTCCCTACACGACGCTCTTCCGATCTNNNTATTTCTCCTCCGCGATCTCTG
GAGCACACGTCTGAACTCTTTCCCTACACGACGCTCTTCCGATCTNNNTATTTCTCCTCCGCGATCTCTG
GCACACGTCTGAACTCTTTCCCTACACGACGCTCTTCCGATCTNNNCTAGAATGAAGCTGGAGAATGCA
GCACACGTCTGAACTCTTTCCCTACACGACGCTCTTCCGATCTNNNCTAGAATGAAGCTGGAGAATGCA
GAGCACACGTCTGAACTCTTTCCCTACACGACGCTCTTCCGATCTNNNTTTGATCTGGGTGGTGGTTA
GAGCACACGTCTGAACTCTTTCCCTACACGACGCTCTTCCGATCTNNNTTTGATCTGGGTGGTGGTTA
GAGCACACGTCTGAACTCTTTCCCTACACGACGCTCTTCCGATCTNNNAATGTGAGGTGTGTGTGAGA
GAGCACACGTCTGAACTCTTTCCCTACACGACGCTCTTCCGATCTNNNAATGTGAGGTGTGTGTGAGA
GAGCACACGTCTGAACTCTTTCCCTACACGACGCTCTTCCGATCTNNNTGAGGTAGGAGAATGGTGTG
GAGCACACGTCTGAACTCTTTCCCTACACGACGCTCTTCCGATCTNNNTGAGGTAGGAGAATGGTGTG
AGCACACGTCTGAACTCTTTCCCTACACGACGCTCTTCCGATCTNNNCTCCTGAATGACACTAGTGCCAT
AGCACACGTCTGAACTCTTTCCCTACACGACGCTCTTCCGATCTNNNCTCCTGAATGACACTAGTGCCAT
AGCACACGTCTGAACTCTTTCCCTACACGACGCTCTTCCGATCTNNNCGATCTCCTGACCTCATGATCTG
AGCACACGTCTGAACTCTTTCCCTACACGACGCTCTTCCGATCTNNNCGATCTCCTGACCTCATGATCTG
AGCACACGTCTGAACTCTTTCCCTACACGACGCTCTTCCGATCTNNNACCTACCACATCTGTCAACGTAC
AGCACACGTCTGAACTCTTTCCCTACACGACGCTCTTCCGATCTNNNACCTACCACATCTGTCAACGTAC
GAGCACACGTCTGAACTCTTTCCCTACACGACGCTCTTCCGATCTNNNCCCAAAGTGCTAGGATTACAG
GAGCACACGTCTGAACTCTTTCCCTACACGACGCTCTTCCGATCTNNNCCCAAAGTGCTAGGATTACAG
AGCACACGTCTGAACTCTTTCCCTACACGACGCTCTTCCGATCTNNNTGTTCATCAGGGATTTTGGCCTA
AGCACACGTCTGAACTCTTTCCCTACACGACGCTCTTCCGATCTNNNTGTTCATCAGGGATTTTGGCCTA
AGCACACGTCTGAACTCTTTCCCTACACGACGCTCTTCCGATCTNNNGAGAACAGTTTGGAGGTTCCTG
AGCACACGTCTGAACTCTTTCCCTACACGACGCTCTTCCGATCTNNNGAGAACAGTTTGGAGGTTCCTG
AGCACACGTCTGAACTCTTTCCCTACACGACGCTCTTCCGATCTNNNGAGAATCACTTGAACCAGGGA
AGCACACGTCTGAACTCTTTCCCTACACGACGCTCTTCCGATCTNNNGAGAATCACTTGAACCAGGGA
GAGCACACGTCTGAACTCTTTCCCTACACGACGCTCTTCCGATCTNNNCTCCTCACCTTGTGATCTTCCAC
GAGCACACGTCTGAACTCTTTCCCTACACGACGCTCTTCCGATCTNNNCTCCTCACCTTGTGATCTTCCAC
AGCACACGTCTGAACTCTTTCCCTACACGACGCTCTTCCGATCTNNNCTGAATCAGCGTCTAGGCTACT
AGCACACGTCTGAACTCTTTCCCTACACGACGCTCTTCCGATCTNNNCTGAATCAGCGTCTAGGCTACT
GAGCACACGTCTGAACTCTTTCCCTACACGACGCTCTTCCGATCTNNNGATCACGAGGTCAGGAGATTC
GAGCACACGTCTGAACTCTTTCCCTACACGACGCTCTTCCGATCTNNNGATCACGAGGTCAGGAGATTC
GCACACGTCTGAACTCTTTCCCTACACGACGCTCTTCCGATCTNNNGTGAACCTTAGTACCTAAAAGGCA
GCACACGTCTGAACTCTTTCCCTACACGACGCTCTTCCGATCTNNNGTGAACCTTAGTACCTAAAAGGCA
AGCACACGTCTGAACTCTTTCCCTACACGACGCTCTTCCGATCTNNNACAGTACTGTTGTAGCAGCAGAT
AGCACACGTCTGAACTCTTTCCCTACACGACGCTCTTCCGATCTNNNACAGTACTGTTGTAGCAGCAGAT
GAGCACACGTCTGAACTCTTTCCCTACACGACGCTCTTCCGATCTNNNGTCATCGCCAGCATCATCAAAC
GAGCACACGTCTGAACTCTTTCCCTACACGACGCTCTTCCGATCTNNNGTCATCGCCAGCATCATCAAAC
GAGCACACGTCTGAACTCTTTCCCTACACGACGCTCTTCCGATCTNNNTATCACCACCCTACAGCAAACT
GAGCACACGTCTGAACTCTTTCCCTACACGACGCTCTTCCGATCTNNNTATCACCACCCTACAGCAAACT
AGCACACGTCTGAACTCTTTCCCTACACGACGCTCTTCCGATCTNNNCACCTGTAGTCCCAGCTACTTGC
AGCACACGTCTGAACTCTTTCCCTACACGACGCTCTTCCGATCTNNNCACCTGTAGTCCCAGCTACTTGC
AGAGCACACGTCTGAACTCTTTCCCTACACGACGCTCTTCCGATCTNNNGTTCTGCTCTGGGGTACTCTC
AGAGCACACGTCTGAACTCTTTCCCTACACGACGCTCTTCCGATCTNNNGTTCTGCTCTGGGGTACTCTC

AGCACACGTCTGAACTCTTTCCCTACACGACGCTCTTCCGATCTNNNACTGTCAGTCTCCTCGCCTCCA
AGCACACGTCTGAACTCTTTCCCTACACGACGCTCTTCCGATCTNNNACTGTCAGTCTCCTCGCCTCCA
AGCACACGTCTGAACTCTTTCCCTACACGACGCTCTTCCGATCTNNNTGTAGGAAGATGGAATTGCA
AGCACACGTCTGAACTCTTTCCCTACACGACGCTCTTCCGATCTNNNTGTAGGAAGATGGAATTGCA
AGCACACGTCTGAACTCTTTCCCTACACGACGCTCTTCCGATCTNNNCATAGACTCAGGTACCAGGCTT
AGCACACGTCTGAACTCTTTCCCTACACGACGCTCTTCCGATCTNNNCATAGACTCAGGTACCAGGCTT
GCACACGTCTGAACTCTTTCCCTACACGACGCTCTTCCGATCTNNNCCATGCCCGGCTAATTTTTGTAT
GCACACGTCTGAACTCTTTCCCTACACGACGCTCTTCCGATCTNNNCCATGCCCGGCTAATTTTTGTAT
AGCACACGTCTGAACTCTTTCCCTACACGACGCTCTTCCGATCTNNNCAGACAGCAGGGTAGAGAAA
AGCACACGTCTGAACTCTTTCCCTACACGACGCTCTTCCGATCTNNNCAGACAGCAGGGTAGAGAAA
AGAGCACACGTCTGAACTCTTTCCCTACACGACGCTCTTCCGATCTNNNGGTTTCCTTGAACGTATTCT
AGAGCACACGTCTGAACTCTTTCCCTACACGACGCTCTTCCGATCTNNNGGTTTCCTTGAACGTATTCT
GAGCACACGTCTGAACTCTTTCCCTACACGACGCTCTTCCGATCTNNNCTGCAGCAGGAAAATCACTT
GAGCACACGTCTGAACTCTTTCCCTACACGACGCTCTTCCGATCTNNNCTGCAGCAGGAAAATCACTT
CACACGTCTGAACTCTTTCCCTACACGACGCTCTTCCGATCTNNNCGATCTCCTAACCTTGTGATCCG
CACACGTCTGAACTCTTTCCCTACACGACGCTCTTCCGATCTNNNCGATCTCCTAACCTTGTGATCCG
GAGCACACGTCTGAACTCTTTCCCTACACGACGCTCTTCCGATCTNNNTTAGACTTCTGGGCAGCTAT
GAGCACACGTCTGAACTCTTTCCCTACACGACGCTCTTCCGATCTNNNTTAGACTTCTGGGCAGCTAT
GAGCACACGTCTGAACTCTTTCCCTACACGACGCTCTTCCGATCTNNNCCTGGGCAATAAGAGCAAAA
GAGCACACGTCTGAACTCTTTCCCTACACGACGCTCTTCCGATCTNNNCCTGGGCAATAAGAGCAAAA
AGCACACGTCTGAACTCTTTCCCTACACGACGCTCTTCCGATCTNNNCGCCCGGCTTTTCTTTTTTTTT
AGCACACGTCTGAACTCTTTCCCTACACGACGCTCTTCCGATCTNNNCGCCCGGCTTTTCTTTTTTTTT
AGCACACGTCTGAACTCTTTCCCTACACGACGCTCTTCCGATCTNNNGTCAGAGGTTGTAGTGAGCC
AGCACACGTCTGAACTCTTTCCCTACACGACGCTCTTCCGATCTNNNGTCAGAGGTTGTAGTGAGCC
GAGCACACGTCTGAACTCTTTCCCTACACGACGCTCTTCCGATCTNNNAAAAAATTAGCCAGGTGTG
GAGCACACGTCTGAACTCTTTCCCTACACGACGCTCTTCCGATCTNNNAAAAAATTAGCCAGGTGTG
AGCACACGTCTGAACTCTTTCCCTACACGACGCTCTTCCGATCTNNNTTGAGGTTTGGAAATCTTGCA
AGCACACGTCTGAACTCTTTCCCTACACGACGCTCTTCCGATCTNNNTTGAGGTTTGGAAATCTTGCA
AGCACACGTCTGAACTCTTTCCCTACACGACGCTCTTCCGATCTNNNCCGCTCATGATACTTGGTGTA
AGCACACGTCTGAACTCTTTCCCTACACGACGCTCTTCCGATCTNNNCCGCTCATGATACTTGGTGTA
CACACGTCTGAACTCTTTCCCTACACGACGCTCTTCCGATCTNNNTACGATTTCCTGTGTCCTATGCGC
CACACGTCTGAACTCTTTCCCTACACGACGCTCTTCCGATCTNNNTACGATTTCCTGTGTCCTATGCGC
GAGCACACGTCTGAACTCTTTCCCTACACGACGCTCTTCCGATCTNNNTTCAAGTGATTCTCCTGCCTC
GAGCACACGTCTGAACTCTTTCCCTACACGACGCTCTTCCGATCTNNNTTCAAGTGATTCTCCTGCCTC
AGCACACGTCTGAACTCTTTCCCTACACGACGCTCTTCCGATCTNNNCTCATCTCTTCTCTTTTCCTCCAT
AGCACACGTCTGAACTCTTTCCCTACACGACGCTCTTCCGATCTNNNCTCATCTCTTCTCTTTTCCTCCAT
AGCACACGTCTGAACTCTTTCCCTACACGACGCTCTTCCGATCTNNNGAAGATTGCTTGAACCTGGGA
AGCACACGTCTGAACTCTTTCCCTACACGACGCTCTTCCGATCTNNNGAAGATTGCTTGAACCTGGGA
GCACACGTCTGAACTCTTTCCCTACACGACGCTCTTCCGATCTNNNTCAGCAACTGGAATTCCTCAAGT
GCACACGTCTGAACTCTTTCCCTACACGACGCTCTTCCGATCTNNNTCAGCAACTGGAATTCCTCAAGT
AGCACACGTCTGAACTCTTTCCCTACACGACGCTCTTCCGATCTNNNACACTCTCCTTTTCTGAAGCCTG
AGCACACGTCTGAACTCTTTCCCTACACGACGCTCTTCCGATCTNNNACACTCTCCTTTTCTGAAGCCTG
AGCACACGTCTGAACTCTTTCCCTACACGACGCTCTTCCGATCTNNNCTCACTGTCTCCCTACCGAAAA

AGCACACGTCTGAACTCTTTCCCTACACGACGCTCTTCCGATCTNNNCTCACTGTCTCCCTACCGAAAAC
GCACACGTCTGAACTCTTTCCCTACACGACGCTCTTCCGATCTNNNTGGTTGAATCCTCATCCTGGATG
GCACACGTCTGAACTCTTTCCCTACACGACGCTCTTCCGATCTNNNTGGTTGAATCCTCATCCTGGATG
GAGCACACGTCTGAACTCTTTCCCTACACGACGCTCTTCCGATCTNNNTACAAAAAATTAGCTGGGCGT
GAGCACACGTCTGAACTCTTTCCCTACACGACGCTCTTCCGATCTNNNTACAAAAAATTAGCTGGGCGT
GAGCACACGTCTGAACTCTTTCCCTACACGACGCTCTTCCGATCTNNNCAGAATCTGTTTTCCGAGGCTT
GAGCACACGTCTGAACTCTTTCCCTACACGACGCTCTTCCGATCTNNNCAGAATCTGTTTTCCGAGGCTT
AGCACACGTCTGAACTCTTTCCCTACACGACGCTCTTCCGATCTNNNTTCTGTCTCTATGAACAGGGCAG
AGCACACGTCTGAACTCTTTCCCTACACGACGCTCTTCCGATCTNNNTTCTGTCTCTATGAACAGGGCAG
GAGCACACGTCTGAACTCTTTCCCTACACGACGCTCTTCCGATCTNNNCTGAGGCACGAGAATCACTTC
GAGCACACGTCTGAACTCTTTCCCTACACGACGCTCTTCCGATCTNNNCTGAGGCACGAGAATCACTTC
GAGCACACGTCTGAACTCTTTCCCTACACGACGCTCTTCCGATCTNNNTCTGTTTCCCCGCTCTTTTCTTAC
GAGCACACGTCTGAACTCTTTCCCTACACGACGCTCTTCCGATCTNNNTCTGTTTCCCCGCTCTTTTCTTAC
AGCACACGTCTGAACTCTTTCCCTACACGACGCTCTTCCGATCTNNNTAAAAAACATCCCCGTTTGGCAA
AGCACACGTCTGAACTCTTTCCCTACACGACGCTCTTCCGATCTNNNTAAAAAACATCCCCGTTTGGCAA
GAGCACACGTCTGAACTCTTTCCCTACACGACGCTCTTCCGATCTNNNCTGTAATCCCAGCTACTCAGG
GAGCACACGTCTGAACTCTTTCCCTACACGACGCTCTTCCGATCTNNNCTGTAATCCCAGCTACTCAGG
AGCACACGTCTGAACTCTTTCCCTACACGACGCTCTTCCGATCTNNNTTACACACACACACACACACACC
AGCACACGTCTGAACTCTTTCCCTACACGACGCTCTTCCGATCTNNNTTACACACACACACACACACACC
AGCACACGTCTGAACTCTTTCCCTACACGACGCTCTTCCGATCTNNNACAGATTATTTCATCACCCAGGT
AGCACACGTCTGAACTCTTTCCCTACACGACGCTCTTCCGATCTNNNACAGATTATTTCATCACCCAGGT
GAGCACACGTCTGAACTCTTTCCCTACACGACGCTCTTCCGATCTNNNAAAAATTAGCTGGATGTGGT
GAGCACACGTCTGAACTCTTTCCCTACACGACGCTCTTCCGATCTNNNAAAAATTAGCTGGATGTGGT
GAGCACACGTCTGAACTCTTTCCCTACACGACGCTCTTCCGATCTNNNGTAGTTCCAGGTATTCAGGAC
GAGCACACGTCTGAACTCTTTCCCTACACGACGCTCTTCCGATCTNNNGTAGTTCCAGGTATTCAGGAC
GAGCACACGTCTGAACTCTTTCCCTACACGACGCTCTTCCGATCTNNNTGGGGAAAGACAGTTTGTTCC
GAGCACACGTCTGAACTCTTTCCCTACACGACGCTCTTCCGATCTNNNTGGGGAAAGACAGTTTGTTCC
AAGAGCACACGTCTGAACTCTTTCCCTACACGACGCTCTTCCGATCTNNNTTCCCCTAATCTCACTAACCA
AAGAGCACACGTCTGAACTCTTTCCCTACACGACGCTCTTCCGATCTNNNTTCCCCTAATCTCACTAACCA
GAGCACACGTCTGAACTCTTTCCCTACACGACGCTCTTCCGATCTNNNGCACCTGTAATCCCAGCTATTT
GAGCACACGTCTGAACTCTTTCCCTACACGACGCTCTTCCGATCTNNNGCACCTGTAATCCCAGCTATTT
GAGCACACGTCTGAACTCTTTCCCTACACGACGCTCTTCCGATCTNNNTACATATTCATGGGATACATCA
GAGCACACGTCTGAACTCTTTCCCTACACGACGCTCTTCCGATCTNNNTACATATTCATGGGATACATCA
GAGCACACGTCTGAACTCTTTCCCTACACGACGCTCTTCCGATCTNNNTGGTACATGACTGTAATCCCAG
GAGCACACGTCTGAACTCTTTCCCTACACGACGCTCTTCCGATCTNNNTGGTACATGACTGTAATCCCAG
AGCACACGTCTGAACTCTTTCCCTACACGACGCTCTTCCGATCTNNNGCTGTGGCAGACATGTTTTACA
AGCACACGTCTGAACTCTTTCCCTACACGACGCTCTTCCGATCTNNNGCTGTGGCAGACATGTTTTACA
GCACACGTCTGAACTCTTTCCCTACACGACGCTCTTCCGATCTNNNAAAAAATTAGCTGGGCATGGTG
GCACACGTCTGAACTCTTTCCCTACACGACGCTCTTCCGATCTNNNAAAAAATTAGCTGGGCATGGTG
AGCACACGTCTGAACTCTTTCCCTACACGACGCTCTTCCGATCTNNNGCTGTGGTTTTTTGCTTACATGC
AGCACACGTCTGAACTCTTTCCCTACACGACGCTCTTCCGATCTNNNGCTGTGGTTTTTTGCTTACATGC
GAGCACACGTCTGAACTCTTTCCCTACACGACGCTCTTCCGATCTNNNACCAACATGATGAAACCCCTTC
GAGCACACGTCTGAACTCTTTCCCTACACGACGCTCTTCCGATCTNNNACCAACATGATGAAACCCCTTC

GAGCACACGTCTGAACTCTTTCCCTACACGACGCTCTTCCGATCTNNNAAATTGGGAGTAAGGAGATG

GAGCACACGTCTGAACTCTTTCCCTACACGACGCTCTTCCGATCTNNNAAATTGGGAGTAAGGAGATG

GAGCACACGTCTGAACTCTTTCCCTACACGACGCTCTTCCGATCTNNNACATACTGACAGAGATGTCCA

GAGCACACGTCTGAACTCTTTCCCTACACGACGCTCTTCCGATCTNNNACATACTGACAGAGATGTCCA

GAGCACACGTCTGAACTCTTTCCCTACACGACGCTCTTCCGATCTNNNCACTGGCCTACAGAGCATTTC

GAGCACACGTCTGAACTCTTTCCCTACACGACGCTCTTCCGATCTNNNCACTGGCCTACAGAGCATTTC

GAGCACACGTCTGAACTCTTTCCCTACACGACGCTCTTCCGATCTNNNCAGGAGAATTGCTTGAACCTT

GAGCACACGTCTGAACTCTTTCCCTACACGACGCTCTTCCGATCTNNNCAGGAGAATTGCTTGAACCTT

GCACACGTCTGAACTCTTTCCCTACACGACGCTCTTCCGATCTNNNGTGAGTGGAGATTATGCCATTGC

GCACACGTCTGAACTCTTTCCCTACACGACGCTCTTCCGATCTNNNGTGAGTGGAGATTATGCCATTGC

AGCACACGTCTGAACTCTTTCCCTACACGACGCTCTTCCGATCTNNNATCCTGCATTCAGAGTCCATGTT

AGCACACGTCTGAACTCTTTCCCTACACGACGCTCTTCCGATCTNNNATCCTGCATTCAGAGTCCATGTT

GAGCACACGTCTGAACTCTTTCCCTACACGACGCTCTTCCGATCTNNNGAGAGTTACTACAAGCTGGGC

GAGCACACGTCTGAACTCTTTCCCTACACGACGCTCTTCCGATCTNNNGAGAGTTACTACAAGCTGGGC

AGCACACGTCTGAACTCTTTCCCTACACGACGCTCTTCCGATCTNNNATGCCTGTAATCCCAACTACTCG

AGCACACGTCTGAACTCTTTCCCTACACGACGCTCTTCCGATCTNNNATGCCTGTAATCCCAACTACTCG

GAGCACACGTCTGAACTCTTTCCCTACACGACGCTCTTCCGATCTNNNCGCCTTTTCCAAAGACAAAGCT

GAGCACACGTCTGAACTCTTTCCCTACACGACGCTCTTCCGATCTNNNCGCCTTTTCCAAAGACAAAGCT

AGCACACGTCTGAACTCTTTCCCTACACGACGCTCTTCCGATCTNNNTTGTCCCCTTTTGTATCTGGGAG

AGCACACGTCTGAACTCTTTCCCTACACGACGCTCTTCCGATCTNNNTTGTCCCCTTTTGTATCTGGGAG

AGCACACGTCTGAACTCTTTCCCTACACGACGCTCTTCCGATCTNNNTAGGCAAATCACTTGAACCTGG

AGCACACGTCTGAACTCTTTCCCTACACGACGCTCTTCCGATCTNNNTAGGCAAATCACTTGAACCTGG

AGCACACGTCTGAACTCTTTCCCTACACGACGCTCTTCCGATCTNNNAGGGGAATCACAGTGGCATAT

AGCACACGTCTGAACTCTTTCCCTACACGACGCTCTTCCGATCTNNNAGGGGAATCACAGTGGCATAT

GAGCACACGTCTGAACTCTTTCCCTACACGACGCTCTTCCGATCTNNNCCCTGTTTTTCCAATGCTATGG

GAGCACACGTCTGAACTCTTTCCCTACACGACGCTCTTCCGATCTNNNCCCTGTTTTTCCAATGCTATGG

AGCACACGTCTGAACTCTTTCCCTACACGACGCTCTTCCGATCTNNNCTGTAGTCCCAGCTACTCAGGA

AGCACACGTCTGAACTCTTTCCCTACACGACGCTCTTCCGATCTNNNCTGTAGTCCCAGCTACTCAGGA

GAGCACACGTCTGAACTCTTTCCCTACACGACGCTCTTCCGATCTNNNCTGAGTGAGGAGAGAAACAC

GAGCACACGTCTGAACTCTTTCCCTACACGACGCTCTTCCGATCTNNNCTGAGTGAGGAGAGAAACAC

GAGCACACGTCTGAACTCTTTCCCTACACGACGCTCTTCCGATCTNNNCTTGCAGTGAGCTGAGATCCT

GAGCACACGTCTGAACTCTTTCCCTACACGACGCTCTTCCGATCTNNNCTTGCAGTGAGCTGAGATCCT

AGCACACGTCTGAACTCTTTCCCTACACGACGCTCTTCCGATCTNNNGACCCTTCATTCCAGGACATTCT

AGCACACGTCTGAACTCTTTCCCTACACGACGCTCTTCCGATCTNNNGACCCTTCATTCCAGGACATTCT

GAGCACACGTCTGAACTCTTTCCCTACACGACGCTCTTCCGATCTNNNAGGCAGGAGAATTTCTTGAAC

GAGCACACGTCTGAACTCTTTCCCTACACGACGCTCTTCCGATCTNNNAGGCAGGAGAATTTCTTGAAC

AGCACACGTCTGAACTCTTTCCCTACACGACGCTCTTCCGATCTNNNGAAGACTCTTTGTTGGTTCTGC

AGCACACGTCTGAACTCTTTCCCTACACGACGCTCTTCCGATCTNNNGAAGACTCTTTGTTGGTTCTGC

AGCACACGTCTGAACTCTTTCCCTACACGACGCTCTTCCGATCTNNNGATGTGCAAGGATGTCTGTGAA

AGCACACGTCTGAACTCTTTCCCTACACGACGCTCTTCCGATCTNNNGATGTGCAAGGATGTCTGTGAA

AGCACACGTCTGAACTCTTTCCCTACACGACGCTCTTCCGATCTNNNGGGTCTGCTACAATGAGATAGC

AGCACACGTCTGAACTCTTTCCCTACACGACGCTCTTCCGATCTNNNGGGTCTGCTACAATGAGATAGC

GAGCACACGTCTGAACTCTTTCCCTACACGACGCTCTTCCGATCTNNNCTCTGGGAAGCTTTCTGACACT

GAGCACACGTCTGAACTCTTTCCCTACACGACGCTCTTCCGATCTNNNCTCTGGGAAGCTTTCTGACACT

AGCACACGTCTGAACTCTTTCCCTACACGACGCTCTTCCGATCTNNNCAAAGAAGCATGTGGAATTCCC

AGCACACGTCTGAACTCTTTCCCTACACGACGCTCTTCCGATCTNNNCAAAGAAGCATGTGGAATTCCC

AGCACACGTCTGAACTCTTTCCCTACACGACGCTCTTCCGATCTNNNAGGGAGGAGTATCTCGTTCTCA

AGCACACGTCTGAACTCTTTCCCTACACGACGCTCTTCCGATCTNNNAGGGAGGAGTATCTCGTTCTCA

GAGCACACGTCTGAACTCTTTCCCTACACGACGCTCTTCCGATCTNNNCAACAAAAGCGAGACTGTCTC

GAGCACACGTCTGAACTCTTTCCCTACACGACGCTCTTCCGATCTNNNCAACAAAAGCGAGACTGTCTC

GCACACGTCTGAACTCTTTCCCTACACGACGCTCTTCCGATCTNNNTGACACCATGCTTCTCAGACTTCC

GCACACGTCTGAACTCTTTCCCTACACGACGCTCTTCCGATCTNNNTGACACCATGCTTCTCAGACTTCC

AGCACACGTCTGAACTCTTTCCCTACACGACGCTCTTCCGATCTNNNAACATATTTCTGGAGTGTGGGC

AGCACACGTCTGAACTCTTTCCCTACACGACGCTCTTCCGATCTNNNAACATATTTCTGGAGTGTGGGC

AGCACACGTCTGAACTCTTTCCCTACACGACGCTCTTCCGATCTNNNAATGGCTCTGAAATGTCACTTGC

AGCACACGTCTGAACTCTTTCCCTACACGACGCTCTTCCGATCTNNNAATGGCTCTGAAATGTCACTTGC

GAGCACACGTCTGAACTCTTTCCCTACACGACGCTCTTCCGATCTNNNACGCAAATTCCACTAGGTACTC

GAGCACACGTCTGAACTCTTTCCCTACACGACGCTCTTCCGATCTNNNACGCAAATTCCACTAGGTACTC

AAGAGCACACGTCTGAACTCTTTCCCTACACGACGCTCTTCCGATCTNNNACAAGTTTTTTGTCCTGGT

AAGAGCACACGTCTGAACTCTTTCCCTACACGACGCTCTTCCGATCTNNNACAAGTTTTTTGTCCTGGT

GAGCACACGTCTGAACTCTTTCCCTACACGACGCTCTTCCGATCTNNNGACTATACCAGTCCACGGATA

GAGCACACGTCTGAACTCTTTCCCTACACGACGCTCTTCCGATCTNNNGACTATACCAGTCCACGGATA

AGAGCACACGTCTGAACTCTTTCCCTACACGACGCTCTTCCGATCTNNNCATGGATGAAGCTGGAAAC

AGAGCACACGTCTGAACTCTTTCCCTACACGACGCTCTTCCGATCTNNNCATGGATGAAGCTGGAAAC

GAGCACACGTCTGAACTCTTTCCCTACACGACGCTCTTCCGATCTNNNTCAATGTAGGTCCTGGAAGCA

GAGCACACGTCTGAACTCTTTCCCTACACGACGCTCTTCCGATCTNNNTCAATGTAGGTCCTGGAAGCA

AGCACACGTCTGAACTCTTTCCCTACACGACGCTCTTCCGATCTNNNACCCTGTCTCCACTAAAAACACA

AGCACACGTCTGAACTCTTTCCCTACACGACGCTCTTCCGATCTNNNACCCTGTCTCCACTAAAAACACA

AGCACACGTCTGAACTCTTTCCCTACACGACGCTCTTCCGATCTNNNAGGAGGTTTGGGGTTTTTTGA

AGCACACGTCTGAACTCTTTCCCTACACGACGCTCTTCCGATCTNNNAGGAGGTTTGGGGTTTTTTGA

CACACGTCTGAACTCTTTCCCTACACGACGCTCTTCCGATCTNNNTAGAACTTACATAAGGATTGCCCAT

CACACGTCTGAACTCTTTCCCTACACGACGCTCTTCCGATCTNNNTAGAACTTACATAAGGATTGCCCAT

AGCACACGTCTGAACTCTTTCCCTACACGACGCTCTTCCGATCTNNNACACACCTTACTGCTAATCATGG

AGCACACGTCTGAACTCTTTCCCTACACGACGCTCTTCCGATCTNNNACACACCTTACTGCTAATCATGG

ACACGTCTGAACTCTTTCCCTACACGACGCTCTTCCGATCTNNNAAATGCTAGTTGCTTGTTTGCATCGG

ACACGTCTGAACTCTTTCCCTACACGACGCTCTTCCGATCTNNNAAATGCTAGTTGCTTGTTTGCATCGG

AGCACACGTCTGAACTCTTTCCCTACACGACGCTCTTCCGATCTNNNGAGGCTGAGGCAGGAGAATGC

AGCACACGTCTGAACTCTTTCCCTACACGACGCTCTTCCGATCTNNNGAGGCTGAGGCAGGAGAATGC

AGCACACGTCTGAACTCTTTCCCTACACGACGCTCTTCCGATCTNNNGTAGAGACAGGATTTGACCGTC

AGCACACGTCTGAACTCTTTCCCTACACGACGCTCTTCCGATCTNNNGTAGAGACAGGATTTGACCGTC

AGCACACGTCTGAACTCTTTCCCTACACGACGCTCTTCCGATCTNNNTCCCTGGAATGAAACCAACTTGA

AGCACACGTCTGAACTCTTTCCCTACACGACGCTCTTCCGATCTNNNTCCCTGGAATGAAACCAACTTGA

AGCACACGTCTGAACTCTTTCCCTACACGACGCTCTTCCGATCTNNNAGCCTTCTGAGTAGCTGAGACT

AGCACACGTCTGAACTCTTTCCCTACACGACGCTCTTCCGATCTNNNAGCCTTCTGAGTAGCTGAGACT

AGCACACGTCTGAACTCTTTCCCTACACGACGCTCTTCCGATCTNNNGTGAGCAGAGATCATGCCACTC

AGCACACGTCTGAACTCTTTCCCTACACGACGCTCTTCCGATCTNNNGTGAGCAGAGATCATGCCACTC

GAGCACACGTCTGAACTCTTTCCCTACACGACGCTCTTCCGATCTNNNTTGGAAAGCTAAGGCAGGA
GAGCACACGTCTGAACTCTTTCCCTACACGACGCTCTTCCGATCTNNNTTGGAAAGCTAAGGCAGGA
AGCACACGTCTGAACTCTTTCCCTACACGACGCTCTTCCGATCTNNNTGGGCTTTCAGCACAACAGAT
AGCACACGTCTGAACTCTTTCCCTACACGACGCTCTTCCGATCTNNNTGGGCTTTCAGCACAACAGAT
GCACACGTCTGAACTCTTTCCCTACACGACGCTCTTCCGATCTNNNCATCTTCTACAGTGGCTCCCAAT
GCACACGTCTGAACTCTTTCCCTACACGACGCTCTTCCGATCTNNNCATCTTCTACAGTGGCTCCCAAT
GAGCACACGTCTGAACTCTTTCCCTACACGACGCTCTTCCGATCTNNNCACCTGTAGTCCCAGTTACTT
GAGCACACGTCTGAACTCTTTCCCTACACGACGCTCTTCCGATCTNNNCACCTGTAGTCCCAGTTACTT
AGCACACGTCTGAACTCTTTCCCTACACGACGCTCTTCCGATCTNNNGTGAGTCTTTATCATGCCACTG
AGCACACGTCTGAACTCTTTCCCTACACGACGCTCTTCCGATCTNNNGTGAGTCTTTATCATGCCACTG
GCACACGTCTGAACTCTTTCCCTACACGACGCTCTTCCGATCTNNNTAGGATTACAGGCATGAACCAC
GCACACGTCTGAACTCTTTCCCTACACGACGCTCTTCCGATCTNNNTAGGATTACAGGCATGAACCAC
GCACACGTCTGAACTCTTTCCCTACACGACGCTCTTCCGATCTNNNAAATGAACTCACAAGGTTCCAG
GCACACGTCTGAACTCTTTCCCTACACGACGCTCTTCCGATCTNNNAAATGAACTCACAAGGTTCCAG
CACACGTCTGAACTCTTTCCCTACACGACGCTCTTCCGATCTNNNAGTGACACGATCTTGGTTCACTG
CACACGTCTGAACTCTTTCCCTACACGACGCTCTTCCGATCTNNNAGTGACACGATCTTGGTTCACTG
AGCACACGTCTGAACTCTTTCCCTACACGACGCTCTTCCGATCTNNNCCCCAGGTGAGACAGTAAACT
AGCACACGTCTGAACTCTTTCCCTACACGACGCTCTTCCGATCTNNNCCCCAGGTGAGACAGTAAACT
AGCACACGTCTGAACTCTTTCCCTACACGACGCTCTTCCGATCTNNNAAATACAGGTCTGCACCAGTA
AGCACACGTCTGAACTCTTTCCCTACACGACGCTCTTCCGATCTNNNAAATACAGGTCTGCACCAGTA
GCACACGTCTGAACTCTTTCCCTACACGACGCTCTTCCGATCTNNNTGCAGTAGGCTTTACCATCTAG
GCACACGTCTGAACTCTTTCCCTACACGACGCTCTTCCGATCTNNNTGCAGTAGGCTTTACCATCTAG
GAGCACACGTCTGAACTCTTTCCCTACACGACGCTCTTCCGATCTNNNAAGTCAAAGCCCTGAATAGA
GAGCACACGTCTGAACTCTTTCCCTACACGACGCTCTTCCGATCTNNNAAGTCAAAGCCCTGAATAGA
GCACACGTCTGAACTCTTTCCCTACACGACGCTCTTCCGATCTNNNACTGGCTTGCAGACTCAAAATTT
GCACACGTCTGAACTCTTTCCCTACACGACGCTCTTCCGATCTNNNACTGGCTTGCAGACTCAAAATTT
GCACACGTCTGAACTCTTTCCCTACACGACGCTCTTCCGATCTNNNTTTTGTGTTTGGTGTGATGCTA
GCACACGTCTGAACTCTTTCCCTACACGACGCTCTTCCGATCTNNNTTTTGTGTTTGGTGTGATGCTA
AGCACACGTCTGAACTCTTTCCCTACACGACGCTCTTCCGATCTNNNTGTGTTGCAACTAAGAGAGA
AGCACACGTCTGAACTCTTTCCCTACACGACGCTCTTCCGATCTNNNTGTGTTGCAACTAAGAGAGA
GCACACGTCTGAACTCTTTCCCTACACGACGCTCTTCCGATCTNNNCTGGGTTCACGCCATTCTTCTCA
GCACACGTCTGAACTCTTTCCCTACACGACGCTCTTCCGATCTNNNCTGGGTTCACGCCATTCTTCTCA
GAGCACACGTCTGAACTCTTTCCCTACACGACGCTCTTCCGATCTNNNAATGTGGAAACAGCAAAGTG
GAGCACACGTCTGAACTCTTTCCCTACACGACGCTCTTCCGATCTNNNAATGTGGAAACAGCAAAGTG
AGCACACGTCTGAACTCTTTCCCTACACGACGCTCTTCCGATCTNNNGTGATCACACCACTACACTCCA
AGCACACGTCTGAACTCTTTCCCTACACGACGCTCTTCCGATCTNNNGTGATCACACCACTACACTCCA
GCACACGTCTGAACTCTTTCCCTACACGACGCTCTTCCGATCTNNNGGCCCCAGTGTCTATTGTTCTAT
GCACACGTCTGAACTCTTTCCCTACACGACGCTCTTCCGATCTNNNGGCCCCAGTGTCTATTGTTCTAT
AGCACACGTCTGAACTCTTTCCCTACACGACGCTCTTCCGATCTNNNCACCAGGGAATCTGAATAACC
AGCACACGTCTGAACTCTTTCCCTACACGACGCTCTTCCGATCTNNNCACCAGGGAATCTGAATAACC
GCACACGTCTGAACTCTTTCCCTACACGACGCTCTTCCGATCTNNNGAAACTTCAATCACCAGCAGTCC
GCACACGTCTGAACTCTTTCCCTACACGACGCTCTTCCGATCTNNNGAAACTTCAATCACCAGCAGTCC
AGCACACGTCTGAACTCTTTCCCTACACGACGCTCTTCCGATCTNNNACAACAGAAGCAGAGGAACA

AGCACACGTCTGAACTCTTTCCCTACACGACGCTCTTCCGATCTNNNACAACAGAAGCAGAGGAACA
GAGCACACGTCTGAACTCTTTCCCTACACGACGCTCTTCCGATCTNNNATGAAAGCAGCAAAGGTCCA
GAGCACACGTCTGAACTCTTTCCCTACACGACGCTCTTCCGATCTNNNATGAAAGCAGCAAAGGTCCA
GCACACGTCTGAACTCTTTCCCTACACGACGCTCTTCCGATCTNNNCACCTGTAGTCCCAGCTTAGTTG
GCACACGTCTGAACTCTTTCCCTACACGACGCTCTTCCGATCTNNNCACCTGTAGTCCCAGCTTAGTTG
GCACACGTCTGAACTCTTTCCCTACACGACGCTCTTCCGATCTNNNGCCTGGCCCAAAAGCTATTATTA
GCACACGTCTGAACTCTTTCCCTACACGACGCTCTTCCGATCTNNNGCCTGGCCCAAAAGCTATTATTA
GCACACGTCTGAACTCTTTCCCTACACGACGCTCTTCCGATCTNNNGCATCTTTGTTCCCTTTGCCTAG
GCACACGTCTGAACTCTTTCCCTACACGACGCTCTTCCGATCTNNNGCATCTTTGTTCCCTTTGCCTAG
GCACACGTCTGAACTCTTTCCCTACACGACGCTCTTCCGATCTNNNTCTAACAAGCTCTCTGGTGTCTC
GCACACGTCTGAACTCTTTCCCTACACGACGCTCTTCCGATCTNNNTCTAACAAGCTCTCTGGTGTCTC
GCACACGTCTGAACTCTTTCCCTACACGACGCTCTTCCGATCTNNNTTTGTGAAGAGAGGAGTCAGT
GCACACGTCTGAACTCTTTCCCTACACGACGCTCTTCCGATCTNNNTTTGTGAAGAGAGGAGTCAGT
AGCACACGTCTGAACTCTTTCCCTACACGACGCTCTTCCGATCTNNNGGAGATTGCAGTGAGCTGAG
AGCACACGTCTGAACTCTTTCCCTACACGACGCTCTTCCGATCTNNNGGAGATTGCAGTGAGCTGAG
GAGCACACGTCTGAACTCTTTCCCTACACGACGCTCTTCCGATCTNNNTAGACCGGAAAAGAGAAGC
GAGCACACGTCTGAACTCTTTCCCTACACGACGCTCTTCCGATCTNNNTAGACCGGAAAAGAGAAGC
AGCACACGTCTGAACTCTTTCCCTACACGACGCTCTTCCGATCTNNNTTTATTTGCAACCACGGAAGA
AGCACACGTCTGAACTCTTTCCCTACACGACGCTCTTCCGATCTNNNTTTATTTGCAACCACGGAAGA
AGCACACGTCTGAACTCTTTCCCTACACGACGCTCTTCCGATCTNNNAGGTCACAAAATGTTCCCCTA
AGCACACGTCTGAACTCTTTCCCTACACGACGCTCTTCCGATCTNNNAGGTCACAAAATGTTCCCCTA
AGCACACGTCTGAACTCTTTCCCTACACGACGCTCTTCCGATCTNNNCTGAGGCAGGAGAATTGCTTG
AGCACACGTCTGAACTCTTTCCCTACACGACGCTCTTCCGATCTNNNCTGAGGCAGGAGAATTGCTTG
GCACACGTCTGAACTCTTTCCCTACACGACGCTCTTCCGATCTNNNTGAGAGACAGAGTGAGAACCT
GCACACGTCTGAACTCTTTCCCTACACGACGCTCTTCCGATCTNNNTGAGAGACAGAGTGAGAACCT
GCACACGTCTGAACTCTTTCCCTACACGACGCTCTTCCGATCTNNNCCCCCTGTCTACCCCTTTAGATAC
GCACACGTCTGAACTCTTTCCCTACACGACGCTCTTCCGATCTNNNCCCCCTGTCTACCCCTTTAGATAC
AGCACACGTCTGAACTCTTTCCCTACACGACGCTCTTCCGATCTNNNGCACACCACACTCTCAATACCA
AGCACACGTCTGAACTCTTTCCCTACACGACGCTCTTCCGATCTNNNGCACACCACACTCTCAATACCA
AGCACACGTCTGAACTCTTTCCCTACACGACGCTCTTCCGATCTNNNGGTGGGACAAGCATAGGATA
AGCACACGTCTGAACTCTTTCCCTACACGACGCTCTTCCGATCTNNNGGTGGGACAAGCATAGGATA
AGCACACGTCTGAACTCTTTCCCTACACGACGCTCTTCCGATCTNNNCTCTGTACGCGAAGGTTTTGTT
AGCACACGTCTGAACTCTTTCCCTACACGACGCTCTTCCGATCTNNNCTCTGTACGCGAAGGTTTTGTT
AGCACACGTCTGAACTCTTTCCCTACACGACGCTCTTCCGATCTNNNCTGTAATCCCAGCTACTCAGG
AGCACACGTCTGAACTCTTTCCCTACACGACGCTCTTCCGATCTNNNCTGTAATCCCAGCTACTCAGG
AGCACACGTCTGAACTCTTTCCCTACACGACGCTCTTCCGATCTNNNAGAGAGAGAGAGAGAGAGG
AGCACACGTCTGAACTCTTTCCCTACACGACGCTCTTCCGATCTNNNAGAGAGAGAGAGAGAGAGG
GAGCACACGTCTGAACTCTTTCCCTACACGACGCTCTTCCGATCTNNNACAATGCCTAGTGCTGGTAG
GAGCACACGTCTGAACTCTTTCCCTACACGACGCTCTTCCGATCTNNNACAATGCCTAGTGCTGGTAG
GCACACGTCTGAACTCTTTCCCTACACGACGCTCTTCCGATCTNNNTGTTTCTTTATTGGGCAGGCAA
GCACACGTCTGAACTCTTTCCCTACACGACGCTCTTCCGATCTNNNTGTTTCTTTATTGGGCAGGCAA
GAGCACACGTCTGAACTCTTTCCCTACACGACGCTCTTCCGATCTNNNGTTCTCATTTCCAGGGTTGCA
GAGCACACGTCTGAACTCTTTCCCTACACGACGCTCTTCCGATCTNNNGTTCTCATTTCCAGGGTTGCA

AGCACACGTCTGAACTCTTTCCCTACACGACGCTCTTCCGATCTNNNTGAACACAAGTCTTCCACTCCT
AGCACACGTCTGAACTCTTTCCCTACACGACGCTCTTCCGATCTNNNTGAACACAAGTCTTCCACTCCT
AGCACACGTCTGAACTCTTTCCCTACACGACGCTCTTCCGATCTNNNTGGACACAAAGAGAGGAACAA
AGCACACGTCTGAACTCTTTCCCTACACGACGCTCTTCCGATCTNNNTGGACACAAAGAGAGGAACAA
AGCACACGTCTGAACTCTTTCCCTACACGACGCTCTTCCGATCTNNNCGTCTAACCTTCACAGTCTTGG
AGCACACGTCTGAACTCTTTCCCTACACGACGCTCTTCCGATCTNNNCGTCTAACCTTCACAGTCTTGG
AGCACACGTCTGAACTCTTTCCCTACACGACGCTCTTCCGATCTNNNATCCTTCACACCTTCAGCTCAAT
AGCACACGTCTGAACTCTTTCCCTACACGACGCTCTTCCGATCTNNNATCCTTCACACCTTCAGCTCAAT
AGCACACGTCTGAACTCTTTCCCTACACGACGCTCTTCCGATCTNNNTCTCTCTCTCTCTCTCTCCCTC
AGCACACGTCTGAACTCTTTCCCTACACGACGCTCTTCCGATCTNNNTCTCTCTCTCTCTCTCTCCCTC
AGCACACGTCTGAACTCTTTCCCTACACGACGCTCTTCCGATCTNNNTACTAAATCAAACTTCTCGACTT
AGCACACGTCTGAACTCTTTCCCTACACGACGCTCTTCCGATCTNNNTACTAAATCAAACTTCTCGACTT
GCACACGTCTGAACTCTTTCCCTACACGACGCTCTTCCGATCTNNNAATTTCCTCCCAGTCCTTGTCAG
GCACACGTCTGAACTCTTTCCCTACACGACGCTCTTCCGATCTNNNAATTTCCTCCCAGTCCTTGTCAG
GAGCACACGTCTGAACTCTTTCCCTACACGACGCTCTTCCGATCTNNNTCACACCTGTAATGCCAGCTA
GAGCACACGTCTGAACTCTTTCCCTACACGACGCTCTTCCGATCTNNNTCACACCTGTAATGCCAGCTA
AGCACACGTCTGAACTCTTTCCCTACACGACGCTCTTCCGATCTNNNCCAGTGTTTTAGCCCAGAACCT
AGCACACGTCTGAACTCTTTCCCTACACGACGCTCTTCCGATCTNNNCCAGTGTTTTAGCCCAGAACCT
AGCACACGTCTGAACTCTTTCCCTACACGACGCTCTTCCGATCTNNNGGAATTTGAGACCAGCCTGAG
AGCACACGTCTGAACTCTTTCCCTACACGACGCTCTTCCGATCTNNNGGAATTTGAGACCAGCCTGAG
AGCACACGTCTGAACTCTTTCCCTACACGACGCTCTTCCGATCTNNNGGAGAATCTGTTGAACCCAGG
AGCACACGTCTGAACTCTTTCCCTACACGACGCTCTTCCGATCTNNNGGAGAATCTGTTGAACCCAGG
AGCACACGTCTGAACTCTTTCCCTACACGACGCTCTTCCGATCTNNNAACTTCACCACAGGTCCTTGAA
AGCACACGTCTGAACTCTTTCCCTACACGACGCTCTTCCGATCTNNNAACTTCACCACAGGTCCTTGAA
AGCACACGTCTGAACTCTTTCCCTACACGACGCTCTTCCGATCTNNNACAAGAAGCAAAATCAACAGC
AGCACACGTCTGAACTCTTTCCCTACACGACGCTCTTCCGATCTNNNACAAGAAGCAAAATCAACAGC
AGCACACGTCTGAACTCTTTCCCTACACGACGCTCTTCCGATCTNNNGGAGTTTGAGGCTGTAGTGA
AGCACACGTCTGAACTCTTTCCCTACACGACGCTCTTCCGATCTNNNGGAGTTTGAGGCTGTAGTGA
AGCACACGTCTGAACTCTTTCCCTACACGACGCTCTTCCGATCTNNNGAGAGTAGCTTGGCAGGAGA
AGCACACGTCTGAACTCTTTCCCTACACGACGCTCTTCCGATCTNNNGAGAGTAGCTTGGCAGGAGA
GCACACGTCTGAACTCTTTCCCTACACGACGCTCTTCCGATCTNNNATCTACACCTTTCCTTCCTGCAGC
GCACACGTCTGAACTCTTTCCCTACACGACGCTCTTCCGATCTNNNATCTACACCTTTCCTTCCTGCAGC
AGCACACGTCTGAACTCTTTCCCTACACGACGCTCTTCCGATCTNNNCATTTTAATTCAGTTCCGTTTAC
AGCACACGTCTGAACTCTTTCCCTACACGACGCTCTTCCGATCTNNNCATTTTAATTCAGTTCCGTTTAC
AGCACACGTCTGAACTCTTTCCCTACACGACGCTCTTCCGATCTNNNTGAATGGGTATTTGTGCCTATG
AGCACACGTCTGAACTCTTTCCCTACACGACGCTCTTCCGATCTNNNTGAATGGGTATTTGTGCCTATG
AGCACACGTCTGAACTCTTTCCCTACACGACGCTCTTCCGATCTNNNTCCCTCTGTCTCTCCTTCTTTCT
AGCACACGTCTGAACTCTTTCCCTACACGACGCTCTTCCGATCTNNNTCCCTCTGTCTCTCCTTCTTTCT
AGCACACGTCTGAACTCTTTCCCTACACGACGCTCTTCCGATCTNNNAGCATAAGCTCCTTTGCCTTTG
AGCACACGTCTGAACTCTTTCCCTACACGACGCTCTTCCGATCTNNNAGCATAAGCTCCTTTGCCTTTG
GCACACGTCTGAACTCTTTCCCTACACGACGCTCTTCCGATCTNNNTCACAGATAAGGAAGTTGAGC
GCACACGTCTGAACTCTTTCCCTACACGACGCTCTTCCGATCTNNNTCACAGATAAGGAAGTTGAGC
AGCACACGTCTGAACTCTTTCCCTACACGACGCTCTTCCGATCTNNNTACTGAGAAAAAAAAGGGGG

AGCACACGTCTGAACTCTTTCCCTACACGACGCTCTTCCGATCTNNNTACTGAGAAAAAAAAGGGGGC
AAGAGCACACGTCTGAACTCTTTCCCTACACGACGCTCTTCCGATCTNNNTGAGCTGAGATTGAGCCA
AAGAGCACACGTCTGAACTCTTTCCCTACACGACGCTCTTCCGATCTNNNTGAGCTGAGATTGAGCCA
ACACGTCTGAACTCTTTCCCTACACGACGCTCTTCCGATCTNNNTGATGCCTTTGTCTGATTTTGGTCA
ACACGTCTGAACTCTTTCCCTACACGACGCTCTTCCGATCTNNNTGATGCCTTTGTCTGATTTTGGTCA
AGCACACGTCTGAACTCTTTCCCTACACGACGCTCTTCCGATCTNNNTCCTTTCTACTGCATCCTTCCCC
AGCACACGTCTGAACTCTTTCCCTACACGACGCTCTTCCGATCTNNNTCCTTTCTACTGCATCCTTCCCC
AGCACACGTCTGAACTCTTTCCCTACACGACGCTCTTCCGATCTNNNGGTCTCTCAAAGTGCTGGGAT
AGCACACGTCTGAACTCTTTCCCTACACGACGCTCTTCCGATCTNNNGGTCTCTCAAAGTGCTGGGAT
GAGCACACGTCTGAACTCTTTCCCTACACGACGCTCTTCCGATCTNNNAAGCCACTCACTGAGAATTGG
GAGCACACGTCTGAACTCTTTCCCTACACGACGCTCTTCCGATCTNNNAAGCCACTCACTGAGAATTGG
GCACACGTCTGAACTCTTTCCCTACACGACGCTCTTCCGATCTNNNGTTATCTCTGCTACTCAGGAGGC
GCACACGTCTGAACTCTTTCCCTACACGACGCTCTTCCGATCTNNNGTTATCTCTGCTACTCAGGAGGC
AGAGCACACGTCTGAACTCTTTCCCTACACGACGCTCTTCCGATCTNNNTAGGACAGCATTTCCTTAGA
AGAGCACACGTCTGAACTCTTTCCCTACACGACGCTCTTCCGATCTNNNTAGGACAGCATTTCCTTAGA
GAGCACACGTCTGAACTCTTTCCCTACACGACGCTCTTCCGATCTNNNACCCACATTACCTGGAGAGTC
GAGCACACGTCTGAACTCTTTCCCTACACGACGCTCTTCCGATCTNNNACCCACATTACCTGGAGAGTC
AGCACACGTCTGAACTCTTTCCCTACACGACGCTCTTCCGATCTNNNCTGGAGCATTTCCCCAATGTTTT
AGCACACGTCTGAACTCTTTCCCTACACGACGCTCTTCCGATCTNNNCTGGAGCATTTCCCCAATGTTTT
AGCACACGTCTGAACTCTTTCCCTACACGACGCTCTTCCGATCTNNNCAAGTCAGGAGGAATGGGATC
AGCACACGTCTGAACTCTTTCCCTACACGACGCTCTTCCGATCTNNNCAAGTCAGGAGGAATGGGATC
AGCACACGTCTGAACTCTTTCCCTACACGACGCTCTTCCGATCTNNNTGGCTACATTGTATTCCATGGT
AGCACACGTCTGAACTCTTTCCCTACACGACGCTCTTCCGATCTNNNTGGCTACATTGTATTCCATGGT
GCACACGTCTGAACTCTTTCCCTACACGACGCTCTTCCGATCTNNNATATCTCCCTTTTTTTCTCTCTTCAT
GCACACGTCTGAACTCTTTCCCTACACGACGCTCTTCCGATCTNNNATATCTCCCTTTTTTTCTCTCTTCAT
GAGCACACGTCTGAACTCTTTCCCTACACGACGCTCTTCCGATCTNNNCATTTAGTGTTCACCCTTGGCT
GAGCACACGTCTGAACTCTTTCCCTACACGACGCTCTTCCGATCTNNNCATTTAGTGTTCACCCTTGGCT
GCACACGTCTGAACTCTTTCCCTACACGACGCTCTTCCGATCTNNNCCGTGTGAAGATTCATGAACCTC
GCACACGTCTGAACTCTTTCCCTACACGACGCTCTTCCGATCTNNNCCGTGTGAAGATTCATGAACCTC
GAGCACACGTCTGAACTCTTTCCCTACACGACGCTCTTCCGATCTNNNCACAGTGAGACCCCCATTTTTT
GAGCACACGTCTGAACTCTTTCCCTACACGACGCTCTTCCGATCTNNNCACAGTGAGACCCCCATTTTTT
GAGCACACGTCTGAACTCTTTCCCTACACGACGCTCTTCCGATCTNNNCTGAGGCAGGAGAATGAACCT
GAGCACACGTCTGAACTCTTTCCCTACACGACGCTCTTCCGATCTNNNCTGAGGCAGGAGAATGAACCT
AGCACACGTCTGAACTCTTTCCCTACACGACGCTCTTCCGATCTNNNTGTTGGCCTTTGGTATCTAGGT
AGCACACGTCTGAACTCTTTCCCTACACGACGCTCTTCCGATCTNNNTGTTGGCCTTTGGTATCTAGGT
GAGCACACGTCTGAACTCTTTCCCTACACGACGCTCTTCCGATCTNNNTCATGGACTTTACCTTGGGCAA
GAGCACACGTCTGAACTCTTTCCCTACACGACGCTCTTCCGATCTNNNTCATGGACTTTACCTTGGGCAA
AGCACACGTCTGAACTCTTTCCCTACACGACGCTCTTCCGATCTNNNGTAGGCAGGAAGTAGCTGTGT
AGCACACGTCTGAACTCTTTCCCTACACGACGCTCTTCCGATCTNNNGTAGGCAGGAAGTAGCTGTGT
GAGCACACGTCTGAACTCTTTCCCTACACGACGCTCTTCCGATCTNNNCATGCTGAAACCCGATCTCTA
GAGCACACGTCTGAACTCTTTCCCTACACGACGCTCTTCCGATCTNNNCATGCTGAAACCCGATCTCTA
AGCACACGTCTGAACTCTTTCCCTACACGACGCTCTTCCGATCTNNNCTCTAACCTGCACATGGTTCTCT
AGCACACGTCTGAACTCTTTCCCTACACGACGCTCTTCCGATCTNNNCTCTAACCTGCACATGGTTCTCT

AGCACACGTCTGAACTCTTTCCCTACACGACGCTCTTCCGATCTNNNCTGAGGCAGGAGAATTGCTA

AGCACACGTCTGAACTCTTTCCCTACACGACGCTCTTCCGATCTNNNCTGAGGCAGGAGAATTGCTA

GCACACGTCTGAACTCTTTCCCTACACGACGCTCTTCCGATCTNNNCTGAGTGTCCTCCAGATCAGTTC

GCACACGTCTGAACTCTTTCCCTACACGACGCTCTTCCGATCTNNNCTGAGTGTCCTCCAGATCAGTTC

GAAGAGCACACGTCTGAACTCTTTCCCTACACGACGCTCTTCCGATCTNNNCCCAGTCTTATTCAGG

GAAGAGCACACGTCTGAACTCTTTCCCTACACGACGCTCTTCCGATCTNNNCCCAGTCTTATTCAGG

AGCACACGTCTGAACTCTTTCCCTACACGACGCTCTTCCGATCTNNNGAGCTGGCAATGAGTCCTATC

AGCACACGTCTGAACTCTTTCCCTACACGACGCTCTTCCGATCTNNNGAGCTGGCAATGAGTCCTATC

AGCACACGTCTGAACTCTTTCCCTACACGACGCTCTTCCGATCTNNNATCAAGTGGGTTTCATCCTTG

AGCACACGTCTGAACTCTTTCCCTACACGACGCTCTTCCGATCTNNNATCAAGTGGGTTTCATCCTTG

AGCACACGTCTGAACTCTTTCCCTACACGACGCTCTTCCGATCTNNNAGCCTGGGAGGATGGACTATT

AGCACACGTCTGAACTCTTTCCCTACACGACGCTCTTCCGATCTNNNAGCCTGGGAGGATGGACTATT

AGCACACGTCTGAACTCTTTCCCTACACGACGCTCTTCCGATCTNNNGATGCCGAAACCTAAACTTTG

AGCACACGTCTGAACTCTTTCCCTACACGACGCTCTTCCGATCTNNNGATGCCGAAACCTAAACTTTG

ACACGTCTGAACTCTTTCCCTACACGACGCTCTTCCGATCTNNNAGAGAAACAGCCTGAGTCTTTCCC

ACACGTCTGAACTCTTTCCCTACACGACGCTCTTCCGATCTNNNAGAGAAACAGCCTGAGTCTTTCCC

GAGCACACGTCTGAACTCTTTCCCTACACGACGCTCTTCCGATCTNNNAGAGCTGCCTGTGATTTGAT

GAGCACACGTCTGAACTCTTTCCCTACACGACGCTCTTCCGATCTNNNAGAGCTGCCTGTGATTTGAT

AGCACACGTCTGAACTCTTTCCCTACACGACGCTCTTCCGATCTNNNAAGAGTGAACATGGTCCCTTCT

AGCACACGTCTGAACTCTTTCCCTACACGACGCTCTTCCGATCTNNNAAGAGTGAACATGGTCCCTTCT

AGCACACGTCTGAACTCTTTCCCTACACGACGCTCTTCCGATCTNNNAATCATATTGGCTGCAGCAAA

AGCACACGTCTGAACTCTTTCCCTACACGACGCTCTTCCGATCTNNNAATCATATTGGCTGCAGCAAA

GAGCACACGTCTGAACTCTTTCCCTACACGACGCTCTTCCGATCTNNNGAGGTTGCAGTGAGCTGAG

GAGCACACGTCTGAACTCTTTCCCTACACGACGCTCTTCCGATCTNNNGAGGTTGCAGTGAGCTGAG

AGAGCACACGTCTGAACTCTTTCCCTACACGACGCTCTTCCGATCTNNNCATGCGATGAAGAAAGCAG

AGAGCACACGTCTGAACTCTTTCCCTACACGACGCTCTTCCGATCTNNNCATGCGATGAAGAAAGCAG

GAGCACACGTCTGAACTCTTTCCCTACACGACGCTCTTCCGATCTNNNTGGGAAGGTCTTCGGTCTAA

GAGCACACGTCTGAACTCTTTCCCTACACGACGCTCTTCCGATCTNNNTGGGAAGGTCTTCGGTCTAA

AGCACACGTCTGAACTCTTTCCCTACACGACGCTCTTCCGATCTNNNGGTTGCAGTGAGTGGAGATC

AGCACACGTCTGAACTCTTTCCCTACACGACGCTCTTCCGATCTNNNGGTTGCAGTGAGTGGAGATC

AGCACACGTCTGAACTCTTTCCCTACACGACGCTCTTCCGATCTNNNCAGGGCAAATAGGCTCCTAGG

AGCACACGTCTGAACTCTTTCCCTACACGACGCTCTTCCGATCTNNNCAGGGCAAATAGGCTCCTAGG

GAGCACACGTCTGAACTCTTTCCCTACACGACGCTCTTCCGATCTNNNGTAGCCATCCTAACAGGTAT

GAGCACACGTCTGAACTCTTTCCCTACACGACGCTCTTCCGATCTNNNGTAGCCATCCTAACAGGTAT

GCACACGTCTGAACTCTTTCCCTACACGACGCTCTTCCGATCTNNNCATGGTGGTTCTTGACTGTCAT

GCACACGTCTGAACTCTTTCCCTACACGACGCTCTTCCGATCTNNNCATGGTGGTTCTTGACTGTCAT

GCACACGTCTGAACTCTTTCCCTACACGACGCTCTTCCGATCTNNNGTTTCTTTTGAGTGGCCTTCCTC

GCACACGTCTGAACTCTTTCCCTACACGACGCTCTTCCGATCTNNNGTTTCTTTTGAGTGGCCTTCCTC

GAGCACACGTCTGAACTCTTTCCCTACACGACGCTCTTCCGATCTNNNCAGAGCGAGATTCTGTCTAA

GAGCACACGTCTGAACTCTTTCCCTACACGACGCTCTTCCGATCTNNNCAGAGCGAGATTCTGTCTAA

GCACACGTCTGAACTCTTTCCCTACACGACGCTCTTCCGATCTNNNTCCTGTGTCCATGTGTTCTCATTC

GCACACGTCTGAACTCTTTCCCTACACGACGCTCTTCCGATCTNNNTCCTGTGTCCATGTGTTCTCATTC

AGCACACGTCTGAACTCTTTCCCTACACGACGCTCTTCCGATCTNNNCTTCAGGACTAACTCCATCTGG

AGCACACGTCTGAACTCTTTCCCTACACGACGCTCTTCCGATCTNNNCTTCAGGACTAACTCCATCTGG

GCACACGTCTGAACTCTTTCCCTACACGACGCTCTTCCGATCTNNNTCAACCACAATGCAGGTCTTGAG

GCACACGTCTGAACTCTTTCCCTACACGACGCTCTTCCGATCTNNNTCAACCACAATGCAGGTCTTGAG

GCACACGTCTGAACTCTTTCCCTACACGACGCTCTTCCGATCTNNNGCCCTTGAAAGATCACACCAAA

GCACACGTCTGAACTCTTTCCCTACACGACGCTCTTCCGATCTNNNGCCCTTGAAAGATCACACCAAA

AGCACACGTCTGAACTCTTTCCCTACACGACGCTCTTCCGATCTNNNTCTCCCATTGCATAGGTTGTCTT

AGCACACGTCTGAACTCTTTCCCTACACGACGCTCTTCCGATCTNNNTCTCCCATTGCATAGGTTGTCTT

AGCACACGTCTGAACTCTTTCCCTACACGACGCTCTTCCGATCTNNNAGAATCTCTTGAATCTGGGAG

AGCACACGTCTGAACTCTTTCCCTACACGACGCTCTTCCGATCTNNNAGAATCTCTTGAATCTGGGAG

AGCACACGTCTGAACTCTTTCCCTACACGACGCTCTTCCGATCTNNNAGGCTAATCAGGGGAAAACA

AGCACACGTCTGAACTCTTTCCCTACACGACGCTCTTCCGATCTNNNAGGCTAATCAGGGGAAAACA

CACACGTCTGAACTCTTTCCCTACACGACGCTCTTCCGATCTNNNCTACGCCTTCTAAACGTGGTTCCC

CACACGTCTGAACTCTTTCCCTACACGACGCTCTTCCGATCTNNNCTACGCCTTCTAAACGTGGTTCCC

GCACACGTCTGAACTCTTTCCCTACACGACGCTCTTCCGATCTNNNTCCTTCTTTCCTTCTTTTCTTTTTCT

GCACACGTCTGAACTCTTTCCCTACACGACGCTCTTCCGATCTNNNTCCTTCTTTCCTTCTTTTCTTTTTCT

GAGCACACGTCTGAACTCTTTCCCTACACGACGCTCTTCCGATCTNNNATGTTGCAGTGAGCCGAGAT

GAGCACACGTCTGAACTCTTTCCCTACACGACGCTCTTCCGATCTNNNATGTTGCAGTGAGCCGAGAT

GAGCACACGTCTGAACTCTTTCCCTACACGACGCTCTTCCGATCTNNNCATCTGGTCCTGGGCTTTTTT

GAGCACACGTCTGAACTCTTTCCCTACACGACGCTCTTCCGATCTNNNCATCTGGTCCTGGGCTTTTTT

AGAGCACACGTCTGAACTCTTTCCCTACACGACGCTCTTCCGATCTNNNGAGGCTGAGGCAGGAGAA

AGAGCACACGTCTGAACTCTTTCCCTACACGACGCTCTTCCGATCTNNNGAGGCTGAGGCAGGAGAA

GCACACGTCTGAACTCTTTCCCTACACGACGCTCTTCCGATCTNNNCAATAGACCGTCTCTTTGTGGGA

GCACACGTCTGAACTCTTTCCCTACACGACGCTCTTCCGATCTNNNCAATAGACCGTCTCTTTGTGGGA

AGCACACGTCTGAACTCTTTCCCTACACGACGCTCTTCCGATCTNNNGCTGGGTCAAATGGTATTCAC

AGCACACGTCTGAACTCTTTCCCTACACGACGCTCTTCCGATCTNNNGCTGGGTCAAATGGTATTCAC

GCACACGTCTGAACTCTTTCCCTACACGACGCTCTTCCGATCTNNNGGAGAATCGCTTGAACCTGGAA

GCACACGTCTGAACTCTTTCCCTACACGACGCTCTTCCGATCTNNNGGAGAATCGCTTGAACCTGGAA

GCACACGTCTGAACTCTTTCCCTACACGACGCTCTTCCGATCTNNNTGGGATTGCCTTTTGATTTCTCT

GCACACGTCTGAACTCTTTCCCTACACGACGCTCTTCCGATCTNNNTGGGATTGCCTTTTGATTTCTCT

AGCACACGTCTGAACTCTTTCCCTACACGACGCTCTTCCGATCTNNNGAGAATCGCTTGAACCCAGG

AGCACACGTCTGAACTCTTTCCCTACACGACGCTCTTCCGATCTNNNGAGAATCGCTTGAACCCAGG

GAGCACACGTCTGAACTCTTTCCCTACACGACGCTCTTCCGATCTNNNGCACAGATACACAGACAGAC

GAGCACACGTCTGAACTCTTTCCCTACACGACGCTCTTCCGATCTNNNGCACAGATACACAGACAGAC

GAGCACACGTCTGAACTCTTTCCCTACACGACGCTCTTCCGATCTNNNTTTCATCAACAAACGAAGGA

GAGCACACGTCTGAACTCTTTCCCTACACGACGCTCTTCCGATCTNNNTTTCATCAACAAACGAAGGA

GAGCACACGTCTGAACTCTTTCCCTACACGACGCTCTTCCGATCTNNNATACACCTGTAGTCCCAGCTA

GAGCACACGTCTGAACTCTTTCCCTACACGACGCTCTTCCGATCTNNNATACACCTGTAGTCCCAGCTA

AGCACACGTCTGAACTCTTTCCCTACACGACGCTCTTCCGATCTNNNATCTAACAGATTTCCTAAAATG

AGCACACGTCTGAACTCTTTCCCTACACGACGCTCTTCCGATCTNNNATCTAACAGATTTCCTAAAATG

AGAGCACACGTCTGAACTCTTTCCCTACACGACGCTCTTCCGATCTNNNAGAGGTTGCAGTGAGTCAA

AGAGCACACGTCTGAACTCTTTCCCTACACGACGCTCTTCCGATCTNNNAGAGGTTGCAGTGAGTCAA

AGCACACGTCTGAACTCTTTCCCTACACGACGCTCTTCCGATCTNNNAGGAGGCTGAGACAGAAGAAT

AGCACACGTCTGAACTCTTTCCCTACACGACGCTCTTCCGATCTNNNAGGAGGCTGAGACAGAAGAAT

GCACACGTCTGAACTCTTTCCCTACACGACGCTCTTCCGATCTNNNCATGCAACTCCTGAATCTGGTTG

GCACACGTCTGAACTCTTTCCCTACACGACGCTCTTCCGATCTNNNCATGCAACTCCTGAATCTGGTTG

GAGCACACGTCTGAACTCTTTCCCTACACGACGCTCTTCCGATCTNNNGCCTCCTGAGTAACTGGGATT

GAGCACACGTCTGAACTCTTTCCCTACACGACGCTCTTCCGATCTNNNGCCTCCTGAGTAACTGGGATT

AGCACACGTCTGAACTCTTTCCCTACACGACGCTCTTCCGATCTNNNCAGAACTGGAAGCAGTCTTCT

AGCACACGTCTGAACTCTTTCCCTACACGACGCTCTTCCGATCTNNNCAGAACTGGAAGCAGTCTTCT

AGCACACGTCTGAACTCTTTCCCTACACGACGCTCTTCCGATCTNNNTCTTTTCCCGATCTATCATCCA

AGCACACGTCTGAACTCTTTCCCTACACGACGCTCTTCCGATCTNNNTCTTTTCCCGATCTATCATCCA

AGCACACGTCTGAACTCTTTCCCTACACGACGCTCTTCCGATCTNNNGCATATTGCATGATGCTGAGGT

AGCACACGTCTGAACTCTTTCCCTACACGACGCTCTTCCGATCTNNNGCATATTGCATGATGCTGAGGT

GAGCACACGTCTGAACTCTTTCCCTACACGACGCTCTTCCGATCTNNNCTTTGCATCTTGGAAGGGA

GAGCACACGTCTGAACTCTTTCCCTACACGACGCTCTTCCGATCTNNNCTTTGCATCTTGGAAGGGA

AGCACACGTCTGAACTCTTTCCCTACACGACGCTCTTCCGATCTNNNGCTCTCTTGTTGCTTCTCCTTTG

AGCACACGTCTGAACTCTTTCCCTACACGACGCTCTTCCGATCTNNNGCTCTCTTGTTGCTTCTCCTTTG

AGCACACGTCTGAACTCTTTCCCTACACGACGCTCTTCCGATCTNNNATGTATTTTGTGTGTGTGTGG

AGCACACGTCTGAACTCTTTCCCTACACGACGCTCTTCCGATCTNNNATGTATTTTGTGTGTGTGTGG

GAGCACACGTCTGAACTCTTTCCCTACACGACGCTCTTCCGATCTNNNGGGCCTCCTGGAACTGTAAAA

GAGCACACGTCTGAACTCTTTCCCTACACGACGCTCTTCCGATCTNNNGGGCCTCCTGGAACTGTAAAA

GAGCACACGTCTGAACTCTTTCCCTACACGACGCTCTTCCGATCTNNNTGAGGCTACAATGAGCTGTG

GAGCACACGTCTGAACTCTTTCCCTACACGACGCTCTTCCGATCTNNNTGAGGCTACAATGAGCTGTG

GAGCACACGTCTGAACTCTTTCCCTACACGACGCTCTTCCGATCTNNNGCATGCCTATAATCCCAGCTA

GAGCACACGTCTGAACTCTTTCCCTACACGACGCTCTTCCGATCTNNNGCATGCCTATAATCCCAGCTA

AAGAGCACACGTCTGAACTCTTTCCCTACACGACGCTCTTCCGATCTNNNGTGGAGGATCGCTTGAA

AAGAGCACACGTCTGAACTCTTTCCCTACACGACGCTCTTCCGATCTNNNGTGGAGGATCGCTTGAA

AGCACACGTCTGAACTCTTTCCCTACACGACGCTCTTCCGATCTNNNATAAATAAATAGGCCGTGCGC

AGCACACGTCTGAACTCTTTCCCTACACGACGCTCTTCCGATCTNNNATAAATAAATAGGCCGTGCGC

AGCACACGTCTGAACTCTTTCCCTACACGACGCTCTTCCGATCTNNNTACAGCTCAGGAATCTGTAGGG

AGCACACGTCTGAACTCTTTCCCTACACGACGCTCTTCCGATCTNNNTACAGCTCAGGAATCTGTAGGG

AGCACACGTCTGAACTCTTTCCCTACACGACGCTCTTCCGATCTNNNGTGGTGTACTCCCAGTTACTCA

AGCACACGTCTGAACTCTTTCCCTACACGACGCTCTTCCGATCTNNNGTGGTGTACTCCCAGTTACTCA

GAGCACACGTCTGAACTCTTTCCCTACACGACGCTCTTCCGATCTNNNTGAGGCAGGAGAATGGTAT

GAGCACACGTCTGAACTCTTTCCCTACACGACGCTCTTCCGATCTNNNTGAGGCAGGAGAATGGTAT

AGCACACGTCTGAACTCTTTCCCTACACGACGCTCTTCCGATCTNNNACTTGTGATCCCAGCTATTCAG

AGCACACGTCTGAACTCTTTCCCTACACGACGCTCTTCCGATCTNNNACTTGTGATCCCAGCTATTCAG

AGAGCACACGTCTGAACTCTTTCCCTACACGACGCTCTTCCGATCTNNNAAGGAAAAGCAAAGGACTG

AGAGCACACGTCTGAACTCTTTCCCTACACGACGCTCTTCCGATCTNNNAAGGAAAAGCAAAGGACTG

AGCACACGTCTGAACTCTTTCCCTACACGACGCTCTTCCGATCTNNNCTAAATCTGCTTACCCTGCCTCA

AGCACACGTCTGAACTCTTTCCCTACACGACGCTCTTCCGATCTNNNCTAAATCTGCTTACCCTGCCTCA

GAGCACACGTCTGAACTCTTTCCCTACACGACGCTCTTCCGATCTNNNGAGGAATGGTTTGCATGGAT

GAGCACACGTCTGAACTCTTTCCCTACACGACGCTCTTCCGATCTNNNGAGGAATGGTTTGCATGGAT

AGCACACGTCTGAACTCTTTCCCTACACGACGCTCTTCCGATCTNNNCCTCCTGTTCCTCTTCTTCCTTTC

AGCACACGTCTGAACTCTTTCCCTACACGACGCTCTTCCGATCTNNNCCTCCTGTTCCTCTTCTTCCTTTC

GAGCACACGTCTGAACTCTTTCCCTACACGACGCTCTTCCGATCTNNNAAGAATTCTCTGGAAGGCCC

GAGCACACGTCTGAACTCTTTCCCTACACGACGCTCTTCCGATCTNNNAAGAATTCTCTGGAAGGCCCA
AGCACACGTCTGAACTCTTTCCCTACACGACGCTCTTCCGATCTNNNCTGCTCTCTGTCAATCTCTCTGGC
AGCACACGTCTGAACTCTTTCCCTACACGACGCTCTTCCGATCTNNNCTGCTCTCTGTCAATCTCTCTGGC
AGCACACGTCTGAACTCTTTCCCTACACGACGCTCTTCCGATCTNNNGGCACCTGTAATCCTAGCTACTC
AGCACACGTCTGAACTCTTTCCCTACACGACGCTCTTCCGATCTNNNGGCACCTGTAATCCTAGCTACTC
AGAGCACACGTCTGAACTCTTTCCCTACACGACGCTCTTCCGATCTNNNTTCCCATGTTCTGGGTTGACA
AGAGCACACGTCTGAACTCTTTCCCTACACGACGCTCTTCCGATCTNNNTTCCCATGTTCTGGGTTGACA
GAGCACACGTCTGAACTCTTTCCCTACACGACGCTCTTCCGATCTNNNGTACTACAGCCTGGGCGAGTA
GAGCACACGTCTGAACTCTTTCCCTACACGACGCTCTTCCGATCTNNNGTACTACAGCCTGGGCGAGTA
AGCACACGTCTGAACTCTTTCCCTACACGACGCTCTTCCGATCTNNNTTCTATGTTGCTCAGGTTGCTCT
AGCACACGTCTGAACTCTTTCCCTACACGACGCTCTTCCGATCTNNNTTCTATGTTGCTCAGGTTGCTCT
ACACGTCTGAACTCTTTCCCTACACGACGCTCTTCCGATCTNNNTTACTGTCTTCCAAGATGGCCTCCAT
ACACGTCTGAACTCTTTCCCTACACGACGCTCTTCCGATCTNNNTTACTGTCTTCCAAGATGGCCTCCAT
GAGCACACGTCTGAACTCTTTCCCTACACGACGCTCTTCCGATCTNNNTGCAAATTTGGAGGGAGGAA
GAGCACACGTCTGAACTCTTTCCCTACACGACGCTCTTCCGATCTNNNTGCAAATTTGGAGGGAGGAA
GAGCACACGTCTGAACTCTTTCCCTACACGACGCTCTTCCGATCTNNNGGATCACTTGAGCCTAGGAG
GAGCACACGTCTGAACTCTTTCCCTACACGACGCTCTTCCGATCTNNNGGATCACTTGAGCCTAGGAG
GAGCACACGTCTGAACTCTTTCCCTACACGACGCTCTTCCGATCTNNNGGGTGACAGAGTGAGATTCCA
GAGCACACGTCTGAACTCTTTCCCTACACGACGCTCTTCCGATCTNNNGGGTGACAGAGTGAGATTCCA
GAGCACACGTCTGAACTCTTTCCCTACACGACGCTCTTCCGATCTNNNGTCTTGAATTCCTGACCTCGTG
GAGCACACGTCTGAACTCTTTCCCTACACGACGCTCTTCCGATCTNNNGTCTTGAATTCCTGACCTCGTG
CACACGTCTGAACTCTTTCCCTACACGACGCTCTTCCGATCTNNNAGACACTTCTCTATCCTTGTGCTCAT
CACACGTCTGAACTCTTTCCCTACACGACGCTCTTCCGATCTNNNAGACACTTCTCTATCCTTGTGCTCAT
AGCACACGTCTGAACTCTTTCCCTACACGACGCTCTTCCGATCTNNNGACCTGGCTTGTGTAGTAACTCA
AGCACACGTCTGAACTCTTTCCCTACACGACGCTCTTCCGATCTNNNGACCTGGCTTGTGTAGTAACTCA
GAGCACACGTCTGAACTCTTTCCCTACACGACGCTCTTCCGATCTNNNTGTTGACCTGAACAGAAGTGC
GAGCACACGTCTGAACTCTTTCCCTACACGACGCTCTTCCGATCTNNNTGTTGACCTGAACAGAAGTGC
AGCACACGTCTGAACTCTTTCCCTACACGACGCTCTTCCGATCTNNNTGAGACAGGAGAATCACTCGA
AGCACACGTCTGAACTCTTTCCCTACACGACGCTCTTCCGATCTNNNTGAGACAGGAGAATCACTCGA
AAGAGCACACGTCTGAACTCTTTCCCTACACGACGCTCTTCCGATCTNNNTGTTTCTGTTTCCAGCCAGT
AAGAGCACACGTCTGAACTCTTTCCCTACACGACGCTCTTCCGATCTNNNTGTTTCTGTTTCCAGCCAGT
AGCACACGTCTGAACTCTTTCCCTACACGACGCTCTTCCGATCTNNNCCTGGACTCCTGATTTTCTCTCAC
AGCACACGTCTGAACTCTTTCCCTACACGACGCTCTTCCGATCTNNNCCTGGACTCCTGATTTTCTCTCAC
AGCACACGTCTGAACTCTTTCCCTACACGACGCTCTTCCGATCTNNNACCTTGAGTGCACTTTTTTCCTAC
AGCACACGTCTGAACTCTTTCCCTACACGACGCTCTTCCGATCTNNNACCTTGAGTGCACTTTTTTCCTAC
AGAGCACACGTCTGAACTCTTTCCCTACACGACGCTCTTCCGATCTNNNTGGCTGAGTAGTATTCCATG
AGAGCACACGTCTGAACTCTTTCCCTACACGACGCTCTTCCGATCTNNNTGGCTGAGTAGTATTCCATG
GAGCACACGTCTGAACTCTTTCCCTACACGACGCTCTTCCGATCTNNNGTGCTCACTAGAGAGAACTGC
GAGCACACGTCTGAACTCTTTCCCTACACGACGCTCTTCCGATCTNNNGTGCTCACTAGAGAGAACTGC
AGCACACGTCTGAACTCTTTCCCTACACGACGCTCTTCCGATCTNNNGGAAAGGAAAAAGGAAAGACT
AGCACACGTCTGAACTCTTTCCCTACACGACGCTCTTCCGATCTNNNGGAAAGGAAAAAGGAAAGACT
GCACACGTCTGAACTCTTTCCCTACACGACGCTCTTCCGATCTNNNCAGTTGTTCAAAGAGTAGCTGCC
GCACACGTCTGAACTCTTTCCCTACACGACGCTCTTCCGATCTNNNCAGTTGTTCAAAGAGTAGCTGCC

GCACACGTCTGAACTCTTTCCCTACACGACGCTCTTCCGATCTNNNGGTTTGTTCTTCATGTAGTGGC
GCACACGTCTGAACTCTTTCCCTACACGACGCTCTTCCGATCTNNNGGTTTGTTCTTCATGTAGTGGC
AGCACACGTCTGAACTCTTTCCCTACACGACGCTCTTCCGATCTNNNGAATCCACCAAGGCAATTTAC
AGCACACGTCTGAACTCTTTCCCTACACGACGCTCTTCCGATCTNNNGAATCCACCAAGGCAATTTAC
AGCACACGTCTGAACTCTTTCCCTACACGACGCTCTTCCGATCTNNNAAGTCCAGACTGAATAATACTA
AGCACACGTCTGAACTCTTTCCCTACACGACGCTCTTCCGATCTNNNAAGTCCAGACTGAATAATACTA
AGCACACGTCTGAACTCTTTCCCTACACGACGCTCTTCCGATCTNNNTGTCCTCCAGAAAGAACACCAA
AGCACACGTCTGAACTCTTTCCCTACACGACGCTCTTCCGATCTNNNTGTCCTCCAGAAAGAACACCAA
GAGCACACGTCTGAACTCTTTCCCTACACGACGCTCTTCCGATCTNNNTGGTGCATACAGAGAGAGA
GAGCACACGTCTGAACTCTTTCCCTACACGACGCTCTTCCGATCTNNNTGGTGCATACAGAGAGAGA
AGCACACGTCTGAACTCTTTCCCTACACGACGCTCTTCCGATCTNNNCACACACCTACAGTACCAGCTAC
AGCACACGTCTGAACTCTTTCCCTACACGACGCTCTTCCGATCTNNNCACACACCTACAGTACCAGCTAC
AAGAGCACACGTCTGAACTCTTTCCCTACACGACGCTCTTCCGATCTNNNCCTGAGGTATATCTGTGCT
AAGAGCACACGTCTGAACTCTTTCCCTACACGACGCTCTTCCGATCTNNNCCTGAGGTATATCTGTGCT
AGCACACGTCTGAACTCTTTCCCTACACGACGCTCTTCCGATCTNNNCTTGCAGTGAGCCGAGATTGTC
AGCACACGTCTGAACTCTTTCCCTACACGACGCTCTTCCGATCTNNNCTTGCAGTGAGCCGAGATTGTC
GAGCACACGTCTGAACTCTTTCCCTACACGACGCTCTTCCGATCTNNNCAGTCTCCATAATTGTGAATC
GAGCACACGTCTGAACTCTTTCCCTACACGACGCTCTTCCGATCTNNNCAGTCTCCATAATTGTGAATC
AGCACACGTCTGAACTCTTTCCCTACACGACGCTCTTCCGATCTNNNGAGGCATGAGAATCGCTTGAA
AGCACACGTCTGAACTCTTTCCCTACACGACGCTCTTCCGATCTNNNGAGGCATGAGAATCGCTTGAA
GAGCACACGTCTGAACTCTTTCCCTACACGACGCTCTTCCGATCTNNNGAGAACTGCTTGACCCAGGA
GAGCACACGTCTGAACTCTTTCCCTACACGACGCTCTTCCGATCTNNNGAGAACTGCTTGACCCAGGA
AGAGCACACGTCTGAACTCTTTCCCTACACGACGCTCTTCCGATCTNNNAATAGCCACATTTCTGGGT
AGAGCACACGTCTGAACTCTTTCCCTACACGACGCTCTTCCGATCTNNNAATAGCCACATTTCTGGGT
AGCACACGTCTGAACTCTTTCCCTACACGACGCTCTTCCGATCTNNNCCCTCATAGAGGACACATCCA
AGCACACGTCTGAACTCTTTCCCTACACGACGCTCTTCCGATCTNNNCCCTCATAGAGGACACATCCA
AGCACACGTCTGAACTCTTTCCCTACACGACGCTCTTCCGATCTNNNAATGTCAATCCCAACTGGTCTG
AGCACACGTCTGAACTCTTTCCCTACACGACGCTCTTCCGATCTNNNAATGTCAATCCCAACTGGTCTG
GAGCACACGTCTGAACTCTTTCCCTACACGACGCTCTTCCGATCTNNNCATAAGCACCCTATGAAACTG
GAGCACACGTCTGAACTCTTTCCCTACACGACGCTCTTCCGATCTNNNCATAAGCACCCTATGAAACTG
GCACACGTCTGAACTCTTTCCCTACACGACGCTCTTCCGATCTNNNTCTCTGGGGCAGAAAGATCTTTC
GCACACGTCTGAACTCTTTCCCTACACGACGCTCTTCCGATCTNNNTCTCTGGGGCAGAAAGATCTTTC
CACACGTCTGAACTCTTTCCCTACACGACGCTCTTCCGATCTNNNTTTGGTTTTATTTTTGTCCTACTCAT
CACACGTCTGAACTCTTTCCCTACACGACGCTCTTCCGATCTNNNTTTGGTTTTATTTTTGTCCTACTCAT
AGCACACGTCTGAACTCTTTCCCTACACGACGCTCTTCCGATCTNNNCACGTCACAGTTATGCAGCTAT
AGCACACGTCTGAACTCTTTCCCTACACGACGCTCTTCCGATCTNNNCACGTCACAGTTATGCAGCTAT
AGCACACGTCTGAACTCTTTCCCTACACGACGCTCTTCCGATCTNNNGTGATGCCCTGTTGCAGATATT
AGCACACGTCTGAACTCTTTCCCTACACGACGCTCTTCCGATCTNNNGTGATGCCCTGTTGCAGATATT
AGCACACGTCTGAACTCTTTCCCTACACGACGCTCTTCCGATCTNNNGGAGGCAAGGATACAATGGTG
AGCACACGTCTGAACTCTTTCCCTACACGACGCTCTTCCGATCTNNNGGAGGCAAGGATACAATGGTG
AGCACACGTCTGAACTCTTTCCCTACACGACGCTCTTCCGATCTNNNCCTGAGTATTGGGTGTGTTTGC
AGCACACGTCTGAACTCTTTCCCTACACGACGCTCTTCCGATCTNNNCCTGAGTATTGGGTGTGTTTGC
GCACACGTCTGAACTCTTTCCCTACACGACGCTCTTCCGATCTNNNTCACTGAGACCACTGACAAAACA

GCACACGTCTGAACTCTTTCCCTACACGACGCTCTTCCGATCTNNNTCACTGAGACCACTGACAAAACA

AGCACACGTCTGAACTCTTTCCCTACACGACGCTCTTCCGATCTNNNAGGCTGAATTTCTCAAACACAG

AGCACACGTCTGAACTCTTTCCCTACACGACGCTCTTCCGATCTNNNAGGCTGAATTTCTCAAACACAG

AGCACACGTCTGAACTCTTTCCCTACACGACGCTCTTCCGATCTNNNCCACTTCTCTTAGGAGCCTGTTT

AGCACACGTCTGAACTCTTTCCCTACACGACGCTCTTCCGATCTNNNCCACTTCTCTTAGGAGCCTGTTT

AGCACACGTCTGAACTCTTTCCCTACACGACGCTCTTCCGATCTNNNACACATACATGCTGATGGTTTC

AGCACACGTCTGAACTCTTTCCCTACACGACGCTCTTCCGATCTNNNACACATACATGCTGATGGTTTC

GAGCACACGTCTGAACTCTTTCCCTACACGACGCTCTTCCGATCTNNNTGCATCTCTCTCTCTCTCTCTCT

GAGCACACGTCTGAACTCTTTCCCTACACGACGCTCTTCCGATCTNNNTGCATCTCTCTCTCTCTCTCTCT

GAGCACACGTCTGAACTCTTTCCCTACACGACGCTCTTCCGATCTNNNGGAAAGGTTTCATATTTCCTT

GAGCACACGTCTGAACTCTTTCCCTACACGACGCTCTTCCGATCTNNNGGAAAGGTTTCATATTTCCTT

AGCACACGTCTGAACTCTTTCCCTACACGACGCTCTTCCGATCTNNNTACAAAAAATTAGCTGGGCGT

AGCACACGTCTGAACTCTTTCCCTACACGACGCTCTTCCGATCTNNNTACAAAAAATTAGCTGGGCGT

GAGCACACGTCTGAACTCTTTCCCTACACGACGCTCTTCCGATCTNNNTAGTCCCAGCTACTCAGGAGG

GAGCACACGTCTGAACTCTTTCCCTACACGACGCTCTTCCGATCTNNNTAGTCCCAGCTACTCAGGAGG

GCACACGTCTGAACTCTTTCCCTACACGACGCTCTTCCGATCTNNNATGACTTCCTGCTCAAGTTCCTG

GCACACGTCTGAACTCTTTCCCTACACGACGCTCTTCCGATCTNNNATGACTTCCTGCTCAAGTTCCTG

GAGCACACGTCTGAACTCTTTCCCTACACGACGCTCTTCCGATCTNNNACCGCGCCCGACCTATATATA

GAGCACACGTCTGAACTCTTTCCCTACACGACGCTCTTCCGATCTNNNACCGCGCCCGACCTATATATA

AGCACACGTCTGAACTCTTTCCCTACACGACGCTCTTCCGATCTNNNTTAATTTCTGAAAAGCACCATT

AGCACACGTCTGAACTCTTTCCCTACACGACGCTCTTCCGATCTNNNTTAATTTCTGAAAAGCACCATT

AGCACACGTCTGAACTCTTTCCCTACACGACGCTCTTCCGATCTNNNACCCTGCCTTGACTGAAAATAC

AGCACACGTCTGAACTCTTTCCCTACACGACGCTCTTCCGATCTNNNACCCTGCCTTGACTGAAAATAC

AGCACACGTCTGAACTCTTTCCCTACACGACGCTCTTCCGATCTNNNCTCCAAGGATGAGCATTCCTTC

AGCACACGTCTGAACTCTTTCCCTACACGACGCTCTTCCGATCTNNNCTCCAAGGATGAGCATTCCTTC

GCACACGTCTGAACTCTTTCCCTACACGACGCTCTTCCGATCTNNNGAGAATCACTTGAACCTGGGAG

GCACACGTCTGAACTCTTTCCCTACACGACGCTCTTCCGATCTNNNGAGAATCACTTGAACCTGGGAG

GAGCACACGTCTGAACTCTTTCCCTACACGACGCTCTTCCGATCTNNNTCAGAAAGGAACATTGCATGT

GAGCACACGTCTGAACTCTTTCCCTACACGACGCTCTTCCGATCTNNNTCAGAAAGGAACATTGCATGT

GCACACGTCTGAACTCTTTCCCTACACGACGCTCTTCCGATCTNNNCGTTAGTCCAGGAGTTCGAGAT

GCACACGTCTGAACTCTTTCCCTACACGACGCTCTTCCGATCTNNNCGTTAGTCCAGGAGTTCGAGAT

AGCACACGTCTGAACTCTTTCCCTACACGACGCTCTTCCGATCTNNNTTTTGGAAAGGGAGCATCGGT

AGCACACGTCTGAACTCTTTCCCTACACGACGCTCTTCCGATCTNNNTTTTGGAAAGGGAGCATCGGT

GCACACGTCTGAACTCTTTCCCTACACGACGCTCTTCCGATCTNNNAAAAAAACTAGCAGGGCATGGT

GCACACGTCTGAACTCTTTCCCTACACGACGCTCTTCCGATCTNNNAAAAAAACTAGCAGGGCATGGT

GAGCACACGTCTGAACTCTTTCCCTACACGACGCTCTTCCGATCTNNNAGCATTACTTAGTGCTGGGTC

GAGCACACGTCTGAACTCTTTCCCTACACGACGCTCTTCCGATCTNNNAGCATTACTTAGTGCTGGGTC

GAGCACACGTCTGAACTCTTTCCCTACACGACGCTCTTCCGATCTNNNCCTTCAGGAAGACTCACTTGCT

GAGCACACGTCTGAACTCTTTCCCTACACGACGCTCTTCCGATCTNNNCCTTCAGGAAGACTCACTTGCT

AGAGCACACGTCTGAACTCTTTCCCTACACGACGCTCTTCCGATCTNNNCCATCTAAGCACCTTGGGAT

AGAGCACACGTCTGAACTCTTTCCCTACACGACGCTCTTCCGATCTNNNCCATCTAAGCACCTTGGGAT

GCACACGTCTGAACTCTTTCCCTACACGACGCTCTTCCGATCTNNNCCTGATCACTCCACTCTTTTCCACA

GCACACGTCTGAACTCTTTCCCTACACGACGCTCTTCCGATCTNNNCCTGATCACTCCACTCTTTTCCACA

GAGCACACGTCTGAACTCTTTCCCTACACGACGCTCTTCCGATCTNNNAAGAATTGCTTGAACCGAGA
GAGCACACGTCTGAACTCTTTCCCTACACGACGCTCTTCCGATCTNNNAAGAATTGCTTGAACCGAGA
GAGCACACGTCTGAACTCTTTCCCTACACGACGCTCTTCCGATCTNNNAGGAAATGTGAACCCCCAAT
GAGCACACGTCTGAACTCTTTCCCTACACGACGCTCTTCCGATCTNNNAGGAAATGTGAACCCCCAAT
CACACGTCTGAACTCTTTCCCTACACGACGCTCTTCCGATCTNNNCTGAGGCTGATCCCTTCCTAGTCA
CACACGTCTGAACTCTTTCCCTACACGACGCTCTTCCGATCTNNNCTGAGGCTGATCCCTTCCTAGTCA
AGCACACGTCTGAACTCTTTCCCTACACGACGCTCTTCCGATCTNNNGTGCTTGGGAATAGGGAAGA
AGCACACGTCTGAACTCTTTCCCTACACGACGCTCTTCCGATCTNNNGTGCTTGGGAATAGGGAAGA
AGCACACGTCTGAACTCTTTCCCTACACGACGCTCTTCCGATCTNNNTCTCTGTTCCCTTCTACTGTCCT
AGCACACGTCTGAACTCTTTCCCTACACGACGCTCTTCCGATCTNNNTCTCTGTTCCCTTCTACTGTCCT
AGCACACGTCTGAACTCTTTCCCTACACGACGCTCTTCCGATCTNNNACGCCCAGGTTCTTTAGATGAA
AGCACACGTCTGAACTCTTTCCCTACACGACGCTCTTCCGATCTNNNACGCCCAGGTTCTTTAGATGAA
ACACGTCTGAACTCTTTCCCTACACGACGCTCTTCCGATCTNNNAAATCTCGAGACCCCAGATCTCCCA
ACACGTCTGAACTCTTTCCCTACACGACGCTCTTCCGATCTNNNAAATCTCGAGACCCCAGATCTCCCA
GCACACGTCTGAACTCTTTCCCTACACGACGCTCTTCCGATCTNNNGCCTCCCAAATAGCTAGGACTAC
GCACACGTCTGAACTCTTTCCCTACACGACGCTCTTCCGATCTNNNGCCTCCCAAATAGCTAGGACTAC
GAGCACACGTCTGAACTCTTTCCCTACACGACGCTCTTCCGATCTNNNGCTCATACCTGTAATCCCAG
GAGCACACGTCTGAACTCTTTCCCTACACGACGCTCTTCCGATCTNNNGCTCATACCTGTAATCCCAG
AGCACACGTCTGAACTCTTTCCCTACACGACGCTCTTCCGATCTNNNCACACAAAGCTTCTCAGTGACT
AGCACACGTCTGAACTCTTTCCCTACACGACGCTCTTCCGATCTNNNCACACAAAGCTTCTCAGTGACT
GCACACGTCTGAACTCTTTCCCTACACGACGCTCTTCCGATCTNNNATTGCCCAAAAAGAAACTCCAT
GCACACGTCTGAACTCTTTCCCTACACGACGCTCTTCCGATCTNNNATTGCCCAAAAAGAAACTCCAT
AGCACACGTCTGAACTCTTTCCCTACACGACGCTCTTCCGATCTNNNCTCACAGGGGAGGAAATGCA
AGCACACGTCTGAACTCTTTCCCTACACGACGCTCTTCCGATCTNNNCTCACAGGGGAGGAAATGCA
GAGCACACGTCTGAACTCTTTCCCTACACGACGCTCTTCCGATCTNNNTATCTGTTCAACTCCCTGCCTT
GAGCACACGTCTGAACTCTTTCCCTACACGACGCTCTTCCGATCTNNNTATCTGTTCAACTCCCTGCCTT
AGCACACGTCTGAACTCTTTCCCTACACGACGCTCTTCCGATCTNNNTCAACAACAGTAGTAGGAGACT
AGCACACGTCTGAACTCTTTCCCTACACGACGCTCTTCCGATCTNNNTCAACAACAGTAGTAGGAGACT
AGCACACGTCTGAACTCTTTCCCTACACGACGCTCTTCCGATCTNNNGGCAGAGTCACAAACTTTAGC
AGCACACGTCTGAACTCTTTCCCTACACGACGCTCTTCCGATCTNNNGGCAGAGTCACAAACTTTAGC
AGCACACGTCTGAACTCTTTCCCTACACGACGCTCTTCCGATCTNNNAGGCATCACATAGACACACAC
AGCACACGTCTGAACTCTTTCCCTACACGACGCTCTTCCGATCTNNNAGGCATCACATAGACACACAC
AGCACACGTCTGAACTCTTTCCCTACACGACGCTCTTCCGATCTNNNGGGTAGCTACTCTGTCTGGTT
AGCACACGTCTGAACTCTTTCCCTACACGACGCTCTTCCGATCTNNNGGGTAGCTACTCTGTCTGGTT
GAGCACACGTCTGAACTCTTTCCCTACACGACGCTCTTCCGATCTNNNGCGAAACCCCCTCTCTACTAAA
GAGCACACGTCTGAACTCTTTCCCTACACGACGCTCTTCCGATCTNNNGCGAAACCCCCTCTCTACTAAA
AGCACACGTCTGAACTCTTTCCCTACACGACGCTCTTCCGATCTNNNCCATGACTTCTCCTTCACTGACA
AGCACACGTCTGAACTCTTTCCCTACACGACGCTCTTCCGATCTNNNCCATGACTTCTCCTTCACTGACA
AGCACACGTCTGAACTCTTTCCCTACACGACGCTCTTCCGATCTNNNCTCATCTCTTCTCTTTTCCTCCAT
AGCACACGTCTGAACTCTTTCCCTACACGACGCTCTTCCGATCTNNNCTCATCTCTTCTCTTTTCCTCCAT
AGCACACGTCTGAACTCTTTCCCTACACGACGCTCTTCCGATCTNNNTCAGGTTTCACAGTGACATCG
AGCACACGTCTGAACTCTTTCCCTACACGACGCTCTTCCGATCTNNNTCAGGTTTCACAGTGACATCG
GCACACGTCTGAACTCTTTCCCTACACGACGCTCTTCCGATCTNNNTCTCAGCTCAGCCTATTTTCCTGC

GCACACGTCTGAACTCTTTCCCTACACGACGCTCTTCCGATCTNNNTCTCAGCTCAGCCTATTTTCCTGC
AGCACACGTCTGAACTCTTTCCCTACACGACGCTCTTCCGATCTNNNCAGAAGTGGAGAAGGATGGT
AGCACACGTCTGAACTCTTTCCCTACACGACGCTCTTCCGATCTNNNCAGAAGTGGAGAAGGATGGT
GCACACGTCTGAACTCTTTCCCTACACGACGCTCTTCCGATCTNNNCACCCATAATCCCACCTACTCGCA
GCACACGTCTGAACTCTTTCCCTACACGACGCTCTTCCGATCTNNNCACCCATAATCCCACCTACTCGCA
GAGCACACGTCTGAACTCTTTCCCTACACGACGCTCTTCCGATCTNNNTGTGAGGTTTTGTTCCCTCCAT
GAGCACACGTCTGAACTCTTTCCCTACACGACGCTCTTCCGATCTNNNTGTGAGGTTTTGTTCCCTCCAT
AAGAGCACACGTCTGAACTCTTTCCCTACACGACGCTCTTCCGATCTNNNCGTCTCAACAGAAGAAAG
AAGAGCACACGTCTGAACTCTTTCCCTACACGACGCTCTTCCGATCTNNNCGTCTCAACAGAAGAAAG
GAGCACACGTCTGAACTCTTTCCCTACACGACGCTCTTCCGATCTNNNGAGCTTGCAGTGAGCCAAGAT
GAGCACACGTCTGAACTCTTTCCCTACACGACGCTCTTCCGATCTNNNGAGCTTGCAGTGAGCCAAGAT
AGCACACGTCTGAACTCTTTCCCTACACGACGCTCTTCCGATCTNNNCGGATCGCGGAGTTGAGAATA
AGCACACGTCTGAACTCTTTCCCTACACGACGCTCTTCCGATCTNNNCGGATCGCGGAGTTGAGAATA
AGCACACGTCTGAACTCTTTCCCTACACGACGCTCTTCCGATCTNNNCATGCACCTGTAAACCCACCTA
AGCACACGTCTGAACTCTTTCCCTACACGACGCTCTTCCGATCTNNNCATGCACCTGTAAACCCACCTA
GCACACGTCTGAACTCTTTCCCTACACGACGCTCTTCCGATCTNNNAAGATCACACAACTGCACTCCAC
GCACACGTCTGAACTCTTTCCCTACACGACGCTCTTCCGATCTNNNAAGATCACACAACTGCACTCCAC
AGCACACGTCTGAACTCTTTCCCTACACGACGCTCTTCCGATCTNNNCTCTTCTAAATCAGCCTTTTTCC
AGCACACGTCTGAACTCTTTCCCTACACGACGCTCTTCCGATCTNNNCTCTTCTAAATCAGCCTTTTTCC
GAGCACACGTCTGAACTCTTTCCCTACACGACGCTCTTCCGATCTNNNCAGAGTGCTGGGATTACAGG
GAGCACACGTCTGAACTCTTTCCCTACACGACGCTCTTCCGATCTNNNCAGAGTGCTGGGATTACAGG
AGCACACGTCTGAACTCTTTCCCTACACGACGCTCTTCCGATCTNNNGATCAAGACCATCCTGGCTAAC
AGCACACGTCTGAACTCTTTCCCTACACGACGCTCTTCCGATCTNNNGATCAAGACCATCCTGGCTAAC
AGCACACGTCTGAACTCTTTCCCTACACGACGCTCTTCCGATCTNNNTCTCTTGATGCTTTTGCCAACTT
AGCACACGTCTGAACTCTTTCCCTACACGACGCTCTTCCGATCTNNNTCTCTTGATGCTTTTGCCAACTT
AGCACACGTCTGAACTCTTTCCCTACACGACGCTCTTCCGATCTNNNGAAAATGATTCTGGGGTCTAT
AGCACACGTCTGAACTCTTTCCCTACACGACGCTCTTCCGATCTNNNGAAAATGATTCTGGGGTCTAT
GCACACGTCTGAACTCTTTCCCTACACGACGCTCTTCCGATCTNNNTCTTGCTCCTCAATGTGTGTGTG
GCACACGTCTGAACTCTTTCCCTACACGACGCTCTTCCGATCTNNNTCTTGCTCCTCAATGTGTGTGTG
AGCACACGTCTGAACTCTTTCCCTACACGACGCTCTTCCGATCTNNNACCACACCCGGCTAATTTTTTT
AGCACACGTCTGAACTCTTTCCCTACACGACGCTCTTCCGATCTNNNACCACACCCGGCTAATTTTTTT
GCACACGTCTGAACTCTTTCCCTACACGACGCTCTTCCGATCTNNNAGTTCCAGCGAATACACTCAACT
GCACACGTCTGAACTCTTTCCCTACACGACGCTCTTCCGATCTNNNAGTTCCAGCGAATACACTCAACT
GAGCACACGTCTGAACTCTTTCCCTACACGACGCTCTTCCGATCTNNNGGAAATTGGTCAGTGAAAGA
GAGCACACGTCTGAACTCTTTCCCTACACGACGCTCTTCCGATCTNNNGGAAATTGGTCAGTGAAAGA
AGCACACGTCTGAACTCTTTCCCTACACGACGCTCTTCCGATCTNNNTAGCTGGAACTACAGGCATAT
AGCACACGTCTGAACTCTTTCCCTACACGACGCTCTTCCGATCTNNNTAGCTGGAACTACAGGCATAT
GAGCACACGTCTGAACTCTTTCCCTACACGACGCTCTTCCGATCTNNNGTCTCAAGGTCTGTTTAGGG
GAGCACACGTCTGAACTCTTTCCCTACACGACGCTCTTCCGATCTNNNGTCTCAAGGTCTGTTTAGGG
AGCACACGTCTGAACTCTTTCCCTACACGACGCTCTTCCGATCTNNNCAGTCTGTTTTCTGTGTGTGTG
AGCACACGTCTGAACTCTTTCCCTACACGACGCTCTTCCGATCTNNNCAGTCTGTTTTCTGTGTGTGTG
GAGCACACGTCTGAACTCTTTCCCTACACGACGCTCTTCCGATCTNNNCAATTCTAGCCAGGCACATTG
GAGCACACGTCTGAACTCTTTCCCTACACGACGCTCTTCCGATCTNNNCAATTCTAGCCAGGCACATTG

GAGCACACGTCTGAACTCTTTCCCTACACGACGCTCTTCCGATCTNNNCACATTAGGAACAGAAGGGA
GAGCACACGTCTGAACTCTTTCCCTACACGACGCTCTTCCGATCTNNNCACATTAGGAACAGAAGGGA
AGCACACGTCTGAACTCTTTCCCTACACGACGCTCTTCCGATCTNNNTCAGCAGCATGAAAATGGACAA
AGCACACGTCTGAACTCTTTCCCTACACGACGCTCTTCCGATCTNNNTCAGCAGCATGAAAATGGACAA
AGCACACGTCTGAACTCTTTCCCTACACGACGCTCTTCCGATCTNNNAAAAAAATTAGCCAGGCGTGG
AGCACACGTCTGAACTCTTTCCCTACACGACGCTCTTCCGATCTNNNAAAAAAATTAGCCAGGCGTGG
AGCACACGTCTGAACTCTTTCCCTACACGACGCTCTTCCGATCTNNNTCTTCTGAAAGATTTTTACAAA
AGCACACGTCTGAACTCTTTCCCTACACGACGCTCTTCCGATCTNNNTCTTCTGAAAGATTTTTACAAA
AGCACACGTCTGAACTCTTTCCCTACACGACGCTCTTCCGATCTNNNGTGGGAATCACATGATTTTGG
AGCACACGTCTGAACTCTTTCCCTACACGACGCTCTTCCGATCTNNNGTGGGAATCACATGATTTTGG
AGCACACGTCTGAACTCTTTCCCTACACGACGCTCTTCCGATCTNNNCTGCGTCTTCCTATGTAACTTGC
AGCACACGTCTGAACTCTTTCCCTACACGACGCTCTTCCGATCTNNNCTGCGTCTTCCTATGTAACTTGC
GAGCACACGTCTGAACTCTTTCCCTACACGACGCTCTTCCGATCTNNNCTCCTGAGTAGCTGGGATTAC
GAGCACACGTCTGAACTCTTTCCCTACACGACGCTCTTCCGATCTNNNCTCCTGAGTAGCTGGGATTAC
GAGCACACGTCTGAACTCTTTCCCTACACGACGCTCTTCCGATCTNNNAGAAATATGAACTGCACAGCC
GAGCACACGTCTGAACTCTTTCCCTACACGACGCTCTTCCGATCTNNNAGAAATATGAACTGCACAGCC
GAGCACACGTCTGAACTCTTTCCCTACACGACGCTCTTCCGATCTNNNGAGTTTTTGCTCTTGTTGCC
GAGCACACGTCTGAACTCTTTCCCTACACGACGCTCTTCCGATCTNNNGAGTTTTTGCTCTTGTTGCC
GAGCACACGTCTGAACTCTTTCCCTACACGACGCTCTTCCGATCTNNNTTGGGGCATGGAAATAAGGG
GAGCACACGTCTGAACTCTTTCCCTACACGACGCTCTTCCGATCTNNNTTGGGGCATGGAAATAAGGG
GAGCACACGTCTGAACTCTTTCCCTACACGACGCTCTTCCGATCTNNNGAGTTTGCAGTGAGCCGAGA
GAGCACACGTCTGAACTCTTTCCCTACACGACGCTCTTCCGATCTNNNGAGTTTGCAGTGAGCCGAGA
GAGCACACGTCTGAACTCTTTCCCTACACGACGCTCTTCCGATCTNNNAATACAAAAAATTAGCCGGG
GAGCACACGTCTGAACTCTTTCCCTACACGACGCTCTTCCGATCTNNNAATACAAAAAATTAGCCGGG
GAGCACACGTCTGAACTCTTTCCCTACACGACGCTCTTCCGATCTNNNGTGTCCTACCCCTTCGACACCA
GAGCACACGTCTGAACTCTTTCCCTACACGACGCTCTTCCGATCTNNNGTGTCCTACCCCTTCGACACCA
AGCACACGTCTGAACTCTTTCCCTACACGACGCTCTTCCGATCTNNNAGCTAGTATTACAGGCACACAC
AGCACACGTCTGAACTCTTTCCCTACACGACGCTCTTCCGATCTNNNAGCTAGTATTACAGGCACACAC
AGCACACGTCTGAACTCTTTCCCTACACGACGCTCTTCCGATCTNNNCACCATAAAGAGCATGGGTGT
AGCACACGTCTGAACTCTTTCCCTACACGACGCTCTTCCGATCTNNNCACCATAAAGAGCATGGGTGT
GAGCACACGTCTGAACTCTTTCCCTACACGACGCTCTTCCGATCTNNNTAGGTGACAGACTGAGACTGA
GAGCACACGTCTGAACTCTTTCCCTACACGACGCTCTTCCGATCTNNNTAGGTGACAGACTGAGACTGA
AGCACACGTCTGAACTCTTTCCCTACACGACGCTCTTCCGATCTNNNTGTAAACATATAACACCCCACA
AGCACACGTCTGAACTCTTTCCCTACACGACGCTCTTCCGATCTNNNTGTAAACATATAACACCCCACA
GAGCACACGTCTGAACTCTTTCCCTACACGACGCTCTTCCGATCTNNNCTTGAGGTCGGGAGTTCTAG
GAGCACACGTCTGAACTCTTTCCCTACACGACGCTCTTCCGATCTNNNCTTGAGGTCGGGAGTTCTAG
AGCACACGTCTGAACTCTTTCCCTACACGACGCTCTTCCGATCTNNNCTGTAATCCCAGCTACTCAGGAG
AGCACACGTCTGAACTCTTTCCCTACACGACGCTCTTCCGATCTNNNCTGTAATCCCAGCTACTCAGGAG
AGCACACGTCTGAACTCTTTCCCTACACGACGCTCTTCCGATCTNNNAGAGAAAAAAAAAGAATCTAG
AGCACACGTCTGAACTCTTTCCCTACACGACGCTCTTCCGATCTNNNAGAGAAAAAAAAAGAATCTAG
AGCACACGTCTGAACTCTTTCCCTACACGACGCTCTTCCGATCTNNNATCATGAGGTCAGGAGATCGA
AGCACACGTCTGAACTCTTTCCCTACACGACGCTCTTCCGATCTNNNATCATGAGGTCAGGAGATCGA
CACACGTCTGAACTCTTTCCCTACACGACGCTCTTCCGATCTNNNTTGCTTCAACCTAGTAGGCAGAGC

CACACGTCTGAACTCTTTCCCTACACGACGCTCTTCCGATCTNNNTTGCTTCAACCTAGTAGGCAGAGC

AGCACACGTCTGAACTCTTTCCCTACACGACGCTCTTCCGATCTNNNAAAAAAATTAGCCAGGCGTGG

AGCACACGTCTGAACTCTTTCCCTACACGACGCTCTTCCGATCTNNNAAAAAAATTAGCCAGGCGTGG

GCACACGTCTGAACTCTTTCCCTACACGACGCTCTTCCGATCTNNNGTGTGAGGAGTCATTAGCAGTG

GCACACGTCTGAACTCTTTCCCTACACGACGCTCTTCCGATCTNNNGTGTGAGGAGTCATTAGCAGTG

AGCACACGTCTGAACTCTTTCCCTACACGACGCTCTTCCGATCTNNNAGGAGAATTGTTTGAACCTGGC

AGCACACGTCTGAACTCTTTCCCTACACGACGCTCTTCCGATCTNNNAGGAGAATTGTTTGAACCTGGC

AGAGCACACGTCTGAACTCTTTCCCTACACGACGCTCTTCCGATCTNNNAGAATAGCTTGAACTCAGGA

AGAGCACACGTCTGAACTCTTTCCCTACACGACGCTCTTCCGATCTNNNAGAATAGCTTGAACTCAGGA

AGAGCACACGTCTGAACTCTTTCCCTACACGACGCTCTTCCGATCTNNNCTGAGGCAGGAGAATTGCTT

AGAGCACACGTCTGAACTCTTTCCCTACACGACGCTCTTCCGATCTNNNCTGAGGCAGGAGAATTGCTT

AGCACACGTCTGAACTCTTTCCCTACACGACGCTCTTCCGATCTNNNGAAAACCAACGGATCCCTGAGA

AGCACACGTCTGAACTCTTTCCCTACACGACGCTCTTCCGATCTNNNGAAAACCAACGGATCCCTGAGA

AGCACACGTCTGAACTCTTTCCCTACACGACGCTCTTCCGATCTNNNTTTCCCTGTTTCTGACCACAACCC

AGCACACGTCTGAACTCTTTCCCTACACGACGCTCTTCCGATCTNNNTTTCCCTGTTTCTGACCACAACCC

AGCACACGTCTGAACTCTTTCCCTACACGACGCTCTTCCGATCTNNNTACAGTTAGCGATGATCATGCCA

AGCACACGTCTGAACTCTTTCCCTACACGACGCTCTTCCGATCTNNNTACAGTTAGCGATGATCATGCCA

AGCACACGTCTGAACTCTTTCCCTACACGACGCTCTTCCGATCTNNNGTTATTTGGGTCAAGGGCTGTT

AGCACACGTCTGAACTCTTTCCCTACACGACGCTCTTCCGATCTNNNGTTATTTGGGTCAAGGGCTGTT

GCACACGTCTGAACTCTTTCCCTACACGACGCTCTTCCGATCTNNNGATCAGGCTTATTTTATTGAACTC

GCACACGTCTGAACTCTTTCCCTACACGACGCTCTTCCGATCTNNNGATCAGGCTTATTTTATTGAACTC

AGCACACGTCTGAACTCTTTCCCTACACGACGCTCTTCCGATCTNNNCCCCAGTGTATATTCTTGGCACTC

AGCACACGTCTGAACTCTTTCCCTACACGACGCTCTTCCGATCTNNNCCCCAGTGTATATTCTTGGCACTC

GAGCACACGTCTGAACTCTTTCCCTACACGACGCTCTTCCGATCTNNNAGACGAGGTCAAAGGAATCCA

GAGCACACGTCTGAACTCTTTCCCTACACGACGCTCTTCCGATCTNNNAGACGAGGTCAAAGGAATCCA

AGAGCACACGTCTGAACTCTTTCCCTACACGACGCTCTTCCGATCTNNNGCAACAAGAGTGAGACTCCA

AGAGCACACGTCTGAACTCTTTCCCTACACGACGCTCTTCCGATCTNNNGCAACAAGAGTGAGACTCCA

AGCACACGTCTGAACTCTTTCCCTACACGACGCTCTTCCGATCTNNNTCTCAATCCTTTACTTTGCCACAC

AGCACACGTCTGAACTCTTTCCCTACACGACGCTCTTCCGATCTNNNTCTCAATCCTTTACTTTGCCACAC

AGCACACGTCTGAACTCTTTCCCTACACGACGCTCTTCCGATCTNNNCTGTAATCCCAGCTACTCAGGAC

AGCACACGTCTGAACTCTTTCCCTACACGACGCTCTTCCGATCTNNNCTGTAATCCCAGCTACTCAGGAC

GAGCACACGTCTGAACTCTTTCCCTACACGACGCTCTTCCGATCTNNNCCAGTCTTGCTGAGAGAGAGA

GAGCACACGTCTGAACTCTTTCCCTACACGACGCTCTTCCGATCTNNNCCAGTCTTGCTGAGAGAGAGA

GAGCACACGTCTGAACTCTTTCCCTACACGACGCTCTTCCGATCTNNNTACATGTTGTGTGTGGCTGTA

GAGCACACGTCTGAACTCTTTCCCTACACGACGCTCTTCCGATCTNNNTACATGTTGTGTGTGGCTGTA

GAGCACACGTCTGAACTCTTTCCCTACACGACGCTCTTCCGATCTNNNATAAAAGGTCAGCCAGTTTCCC

GAGCACACGTCTGAACTCTTTCCCTACACGACGCTCTTCCGATCTNNNATAAAAGGTCAGCCAGTTTCCC

GAGCACACGTCTGAACTCTTTCCCTACACGACGCTCTTCCGATCTNNNCTGGGCAACAAAAGCAAAATT

GAGCACACGTCTGAACTCTTTCCCTACACGACGCTCTTCCGATCTNNNCTGGGCAACAAAAGCAAAATT

CACACGTCTGAACTCTTTCCCTACACGACGCTCTTCCGATCTNNNTCTCTCTCTCTCTCTCCCTCTCATC

CACACGTCTGAACTCTTTCCCTACACGACGCTCTTCCGATCTNNNTCTCTCTCTCTCTCTCCCTCTCATC

AGCACACGTCTGAACTCTTTCCCTACACGACGCTCTTCCGATCTNNNTGCAGTTATAACTCATATCAGA

AGCACACGTCTGAACTCTTTCCCTACACGACGCTCTTCCGATCTNNNTGCAGTTATAACTCATATCAGA

AGCACACGTCTGAACTCTTTCCCTACACGACGCTCTTCCGATCTNNNGCTGTGTTCAAGTGGGATGTG

AGCACACGTCTGAACTCTTTCCCTACACGACGCTCTTCCGATCTNNNGCTGTGTTCAAGTGGGATGTG

AGCACACGTCTGAACTCTTTCCCTACACGACGCTCTTCCGATCTNNNCTGAGGCAGGAGAATAGCTTG

AGCACACGTCTGAACTCTTTCCCTACACGACGCTCTTCCGATCTNNNCTGAGGCAGGAGAATAGCTTG

GAGCACACGTCTGAACTCTTTCCCTACACGACGCTCTTCCGATCTNNNCCATCTAGGTTGCTGCAAATA

GAGCACACGTCTGAACTCTTTCCCTACACGACGCTCTTCCGATCTNNNCCATCTAGGTTGCTGCAAATA

CACACGTCTGAACTCTTTCCCTACACGACGCTCTTCCGATCTNNNGAGAGAAAGTCCGGGCAGAAGA

CACACGTCTGAACTCTTTCCCTACACGACGCTCTTCCGATCTNNNGAGAGAAAGTCCGGGCAGAAGA

GCACACGTCTGAACTCTTTCCCTACACGACGCTCTTCCGATCTNNNCAGAATCTGTCTCTGTCACCCAA

GCACACGTCTGAACTCTTTCCCTACACGACGCTCTTCCGATCTNNNCAGAATCTGTCTCTGTCACCCAA

AGCACACGTCTGAACTCTTTCCCTACACGACGCTCTTCCGATCTNNNGATGTGCCTAGTTTTTGCAGA

AGCACACGTCTGAACTCTTTCCCTACACGACGCTCTTCCGATCTNNNGATGTGCCTAGTTTTTGCAGA

GCACACGTCTGAACTCTTTCCCTACACGACGCTCTTCCGATCTNNNCTGCCTCCTCTTCAGCTTGATTTC

GCACACGTCTGAACTCTTTCCCTACACGACGCTCTTCCGATCTNNNCTGCCTCCTCTTCAGCTTGATTTC

AGCACACGTCTGAACTCTTTCCCTACACGACGCTCTTCCGATCTNNNTCAGGAGAAGTTGGGCAAAGT

AGCACACGTCTGAACTCTTTCCCTACACGACGCTCTTCCGATCTNNNTCAGGAGAAGTTGGGCAAAGT

AGCACACGTCTGAACTCTTTCCCTACACGACGCTCTTCCGATCTNNNGCTTCTCAATTTCATGGCCATC

AGCACACGTCTGAACTCTTTCCCTACACGACGCTCTTCCGATCTNNNGCTTCTCAATTTCATGGCCATC

GCACACGTCTGAACTCTTTCCCTACACGACGCTCTTCCGATCTNNNCTGTAATCCCAGCTACTCAGGAG

GCACACGTCTGAACTCTTTCCCTACACGACGCTCTTCCGATCTNNNCTGTAATCCCAGCTACTCAGGAG

GCACACGTCTGAACTCTTTCCCTACACGACGCTCTTCCGATCTNNNGGGGCTTCTTGCAAAAACATCA

GCACACGTCTGAACTCTTTCCCTACACGACGCTCTTCCGATCTNNNGGGGCTTCTTGCAAAAACATCA

GCACACGTCTGAACTCTTTCCCTACACGACGCTCTTCCGATCTNNNGCCATGTAAAAGCAGAAGAGC

GCACACGTCTGAACTCTTTCCCTACACGACGCTCTTCCGATCTNNNGCCATGTAAAAGCAGAAGAGC

AGCACACGTCTGAACTCTTTCCCTACACGACGCTCTTCCGATCTNNNCAGAAGAATTGCTTGAACCTG

AGCACACGTCTGAACTCTTTCCCTACACGACGCTCTTCCGATCTNNNCAGAAGAATTGCTTGAACCTG

CACACGTCTGAACTCTTTCCCTACACGACGCTCTTCCGATCTNNNTTCTCAATGAGCTGTTGGGTACAC

CACACGTCTGAACTCTTTCCCTACACGACGCTCTTCCGATCTNNNTTCTCAATGAGCTGTTGGGTACAC

GAGCACACGTCTGAACTCTTTCCCTACACGACGCTCTTCCGATCTNNNACAGAGTGAAACCCTGTCTC

GAGCACACGTCTGAACTCTTTCCCTACACGACGCTCTTCCGATCTNNNACAGAGTGAAACCCTGTCTC

GCACACGTCTGAACTCTTTCCCTACACGACGCTCTTCCGATCTNNNTTCACCTGTTAGGGAATTCAGA

GCACACGTCTGAACTCTTTCCCTACACGACGCTCTTCCGATCTNNNTTCACCTGTTAGGGAATTCAGA

AGCACACGTCTGAACTCTTTCCCTACACGACGCTCTTCCGATCTNNNCATGTCCAACCATCTGAATTCCG

AGCACACGTCTGAACTCTTTCCCTACACGACGCTCTTCCGATCTNNNCATGTCCAACCATCTGAATTCCG

AGCACACGTCTGAACTCTTTCCCTACACGACGCTCTTCCGATCTNNNCGATCTCCAATTACTGCAAGCT

AGCACACGTCTGAACTCTTTCCCTACACGACGCTCTTCCGATCTNNNCGATCTCCAATTACTGCAAGCT

GCACACGTCTGAACTCTTTCCCTACACGACGCTCTTCCGATCTNNNGAAGGACACTGAGTGGAGATCA

GCACACGTCTGAACTCTTTCCCTACACGACGCTCTTCCGATCTNNNGAAGGACACTGAGTGGAGATCA

AGCACACGTCTGAACTCTTTCCCTACACGACGCTCTTCCGATCTNNNGCACAAATGCCCATTGATCAAT

AGCACACGTCTGAACTCTTTCCCTACACGACGCTCTTCCGATCTNNNGCACAAATGCCCATTGATCAAT

GCACACGTCTGAACTCTTTCCCTACACGACGCTCTTCCGATCTNNNGCTGAAAAGGTGTTCCAATCTCA

GCACACGTCTGAACTCTTTCCCTACACGACGCTCTTCCGATCTNNNGCTGAAAAGGTGTTCCAATCTCA

AGCACACGTCTGAACTCTTTCCCTACACGACGCTCTTCCGATCTNNNGAACTGAGATTGCATACCATC

AGCACACGTCTGAACTCTTTCCCTACACGACGCTCTTCCGATCTNNNGAACTGAGATTGCATACCATC
AGCACACGTCTGAACTCTTTCCCTACACGACGCTCTTCCGATCTNNNCACCTGTAATCCCAGCTACTCA
AGCACACGTCTGAACTCTTTCCCTACACGACGCTCTTCCGATCTNNNCACCTGTAATCCCAGCTACTCA
GAGCACACGTCTGAACTCTTTCCCTACACGACGCTCTTCCGATCTNNNCATTTTACATGGCAGCAGGA
GAGCACACGTCTGAACTCTTTCCCTACACGACGCTCTTCCGATCTNNNCATTTTACATGGCAGCAGGA
AGCACACGTCTGAACTCTTTCCCTACACGACGCTCTTCCGATCTNNNGTTTGCCTTTACTTCCCTGTTCC
AGCACACGTCTGAACTCTTTCCCTACACGACGCTCTTCCGATCTNNNGTTTGCCTTTACTTCCCTGTTCC
AGCACACGTCTGAACTCTTTCCCTACACGACGCTCTTCCGATCTNNNACCATTATCTCTTCCAGTTCCTG
AGCACACGTCTGAACTCTTTCCCTACACGACGCTCTTCCGATCTNNNACCATTATCTCTTCCAGTTCCTG
AGAGCACACGTCTGAACTCTTTCCCTACACGACGCTCTTCCGATCTNNNCTGAGCACAGCGGTTGATA
AGAGCACACGTCTGAACTCTTTCCCTACACGACGCTCTTCCGATCTNNNCTGAGCACAGCGGTTGATA
AGCACACGTCTGAACTCTTTCCCTACACGACGCTCTTCCGATCTNNNCCCTAACCTGCTCTCTGTTCTTCC
AGCACACGTCTGAACTCTTTCCCTACACGACGCTCTTCCGATCTNNNCCCTAACCTGCTCTCTGTTCTTCC
GCACACGTCTGAACTCTTTCCCTACACGACGCTCTTCCGATCTNNNTATTCTTCAGGGTTATTGAATCA
GCACACGTCTGAACTCTTTCCCTACACGACGCTCTTCCGATCTNNNTATTCTTCAGGGTTATTGAATCA
GAGCACACGTCTGAACTCTTTCCCTACACGACGCTCTTCCGATCTNNNGAGCTTACAGTGAGCCGAGA
GAGCACACGTCTGAACTCTTTCCCTACACGACGCTCTTCCGATCTNNNGAGCTTACAGTGAGCCGAGA
AAGAGCACACGTCTGAACTCTTTCCCTACACGACGCTCTTCCGATCTNNNTGTACCACACGTCTGTGA
AAGAGCACACGTCTGAACTCTTTCCCTACACGACGCTCTTCCGATCTNNNTGTACCACACGTCTGTGA
AGCACACGTCTGAACTCTTTCCCTACACGACGCTCTTCCGATCTNNNACATCTGTAGTCCCAGCTACTC
AGCACACGTCTGAACTCTTTCCCTACACGACGCTCTTCCGATCTNNNACATCTGTAGTCCCAGCTACTC
AGCACACGTCTGAACTCTTTCCCTACACGACGCTCTTCCGATCTNNNTGAAACAGGATCTCACTTGGG
AGCACACGTCTGAACTCTTTCCCTACACGACGCTCTTCCGATCTNNNTGAAACAGGATCTCACTTGGG
GAGCACACGTCTGAACTCTTTCCCTACACGACGCTCTTCCGATCTNNNACCTCAGAACATGTAATTCCT
GAGCACACGTCTGAACTCTTTCCCTACACGACGCTCTTCCGATCTNNNACCTCAGAACATGTAATTCCT
AGCACACGTCTGAACTCTTTCCCTACACGACGCTCTTCCGATCTNNNCTGTAGTCCCAGCTACACGGA
AGCACACGTCTGAACTCTTTCCCTACACGACGCTCTTCCGATCTNNNCTGTAGTCCCAGCTACACGGA
AGCACACGTCTGAACTCTTTCCCTACACGACGCTCTTCCGATCTNNNCTGGGCTAAGTTCAAGGTGTG
AGCACACGTCTGAACTCTTTCCCTACACGACGCTCTTCCGATCTNNNCTGGGCTAAGTTCAAGGTGTG
AGCACACGTCTGAACTCTTTCCCTACACGACGCTCTTCCGATCTNNNAGACACACATATCATATGCACA
AGCACACGTCTGAACTCTTTCCCTACACGACGCTCTTCCGATCTNNNAGACACACATATCATATGCACA
GAGCACACGTCTGAACTCTTTCCCTACACGACGCTCTTCCGATCTNNNCAAGATCACACTACCACACTC
GAGCACACGTCTGAACTCTTTCCCTACACGACGCTCTTCCGATCTNNNCAAGATCACACTACCACACTC
GAGCACACGTCTGAACTCTTTCCCTACACGACGCTCTTCCGATCTNNNGGTTTCCCTCTCTGTCTTTGA
GAGCACACGTCTGAACTCTTTCCCTACACGACGCTCTTCCGATCTNNNGGTTTCCCTCTCTGTCTTTGA
GCACACGTCTGAACTCTTTCCCTACACGACGCTCTTCCGATCTNNNTCTGTTCATGTAAGCAGGAAGG
GCACACGTCTGAACTCTTTCCCTACACGACGCTCTTCCGATCTNNNTCTGTTCATGTAAGCAGGAAGG
AGCACACGTCTGAACTCTTTCCCTACACGACGCTCTTCCGATCTNNNCTAGCAATGGAGTGATGAGA
AGCACACGTCTGAACTCTTTCCCTACACGACGCTCTTCCGATCTNNNCTAGCAATGGAGTGATGAGA
GCACACGTCTGAACTCTTTCCCTACACGACGCTCTTCCGATCTNNNCCTCACTACCCATCGGAATTAGG
GCACACGTCTGAACTCTTTCCCTACACGACGCTCTTCCGATCTNNNCCTCACTACCCATCGGAATTAGG
AGCACACGTCTGAACTCTTTCCCTACACGACGCTCTTCCGATCTNNNAATGTGGATGAACTGAGACCC
AGCACACGTCTGAACTCTTTCCCTACACGACGCTCTTCCGATCTNNNAATGTGGATGAACTGAGACCC

GCACACGTCTGAACTCTTTCCCTACACGACGCTCTTCCGATCTNNNAGACTATTTGGCCAATGGGGAA
GCACACGTCTGAACTCTTTCCCTACACGACGCTCTTCCGATCTNNNAGACTATTTGGCCAATGGGGAA
AGCACACGTCTGAACTCTTTCCCTACACGACGCTCTTCCGATCTNNNGAGGCAGGAGAATGGTGTG
AGCACACGTCTGAACTCTTTCCCTACACGACGCTCTTCCGATCTNNNGAGGCAGGAGAATGGTGTG
AGCACACGTCTGAACTCTTTCCCTACACGACGCTCTTCCGATCTNNNGAGATCATGCCACTACTCTCCA
AGCACACGTCTGAACTCTTTCCCTACACGACGCTCTTCCGATCTNNNGAGATCATGCCACTACTCTCCA
AGCACACGTCTGAACTCTTTCCCTACACGACGCTCTTCCGATCTNNNCAGATGGTTATGTTCCTCCCT
AGCACACGTCTGAACTCTTTCCCTACACGACGCTCTTCCGATCTNNNCAGATGGTTATGTTCCTCCCT
AGCACACGTCTGAACTCTTTCCCTACACGACGCTCTTCCGATCTNNNTGGCATCTGGTGATTTTTCGA
AGCACACGTCTGAACTCTTTCCCTACACGACGCTCTTCCGATCTNNNTGGCATCTGGTGATTTTTCGA
CACACGTCTGAACTCTTTCCCTACACGACGCTCTTCCGATCTNNNGAAAGTTAAGGTATCAATGAGA
CACACGTCTGAACTCTTTCCCTACACGACGCTCTTCCGATCTNNNGAAAGTTAAGGTATCAATGAGA
AGCACACGTCTGAACTCTTTCCCTACACGACGCTCTTCCGATCTNNNGTATTCCAGTCGATGAGGAGT
AGCACACGTCTGAACTCTTTCCCTACACGACGCTCTTCCGATCTNNNGTATTCCAGTCGATGAGGAGT
GAGCACACGTCTGAACTCTTTCCCTACACGACGCTCTTCCGATCTNNNCCAGCTACCTGAGAGAAGT
GAGCACACGTCTGAACTCTTTCCCTACACGACGCTCTTCCGATCTNNNCCAGCTACCTGAGAGAAGT
GAGCACACGTCTGAACTCTTTCCCTACACGACGCTCTTCCGATCTNNNTAATCATGGTGGAAGGTGA
GAGCACACGTCTGAACTCTTTCCCTACACGACGCTCTTCCGATCTNNNTAATCATGGTGGAAGGTGA
AGCACACGTCTGAACTCTTTCCCTACACGACGCTCTTCCGATCTNNNCTGGACTAAAATCAAGGGGAC
AGCACACGTCTGAACTCTTTCCCTACACGACGCTCTTCCGATCTNNNCTGGACTAAAATCAAGGGGAC
GCACACGTCTGAACTCTTTCCCTACACGACGCTCTTCCGATCTNNNCTCCCAGGTTCAAGCGATTCTCC
GCACACGTCTGAACTCTTTCCCTACACGACGCTCTTCCGATCTNNNCTCCCAGGTTCAAGCGATTCTCC
GAGCACACGTCTGAACTCTTTCCCTACACGACGCTCTTCCGATCTNNNATTGCACTCTAGCCTGAATGA
GAGCACACGTCTGAACTCTTTCCCTACACGACGCTCTTCCGATCTNNNATTGCACTCTAGCCTGAATGA
GAGCACACGTCTGAACTCTTTCCCTACACGACGCTCTTCCGATCTNNNAGTCTAGGAAGATTTGCCG
GAGCACACGTCTGAACTCTTTCCCTACACGACGCTCTTCCGATCTNNNAGTCTAGGAAGATTTGCCG
AGCACACGTCTGAACTCTTTCCCTACACGACGCTCTTCCGATCTNNNAAGTCCAACATGCTGGGAATA
AGCACACGTCTGAACTCTTTCCCTACACGACGCTCTTCCGATCTNNNAAGTCCAACATGCTGGGAATA
AGAGCACACGTCTGAACTCTTTCCCTACACGACGCTCTTCCGATCTNNNGGGGGTGTCTGTGAATAC
AGAGCACACGTCTGAACTCTTTCCCTACACGACGCTCTTCCGATCTNNNGGGGGTGTCTGTGAATAC
GCACACGTCTGAACTCTTTCCCTACACGACGCTCTTCCGATCTNNNGACAAACAGGAACCAGAACAG
GCACACGTCTGAACTCTTTCCCTACACGACGCTCTTCCGATCTNNNGACAAACAGGAACCAGAACAG
AGCACACGTCTGAACTCTTTCCCTACACGACGCTCTTCCGATCTNNNTGAGTCATGCATTCTCCAAGAC
AGCACACGTCTGAACTCTTTCCCTACACGACGCTCTTCCGATCTNNNTGAGTCATGCATTCTCCAAGAC
GAGCACACGTCTGAACTCTTTCCCTACACGACGCTCTTCCGATCTNNNGGTAACCATGTGAAGGGCTC
GAGCACACGTCTGAACTCTTTCCCTACACGACGCTCTTCCGATCTNNNGGTAACCATGTGAAGGGCTC
AGAGCACACGTCTGAACTCTTTCCCTACACGACGCTCTTCCGATCTNNNCATGCCTGTAATCCCAGCTA
AGAGCACACGTCTGAACTCTTTCCCTACACGACGCTCTTCCGATCTNNNCATGCCTGTAATCCCAGCTA
GCACACGTCTGAACTCTTTCCCTACACGACGCTCTTCCGATCTNNNGGATGGTCTCTATCTCCTGACCT
GCACACGTCTGAACTCTTTCCCTACACGACGCTCTTCCGATCTNNNGGATGGTCTCTATCTCCTGACCT
AGCACACGTCTGAACTCTTTCCCTACACGACGCTCTTCCGATCTNNNTCCTTCATCTCTCAAAAGCTGC
AGCACACGTCTGAACTCTTTCCCTACACGACGCTCTTCCGATCTNNNTCCTTCATCTCTCAAAAGCTGC
GAGCACACGTCTGAACTCTTTCCCTACACGACGCTCTTCCGATCTNNNTCAAGCAGTCATGTGAGGTA

GAGCACACGTCTGAACTCTTTCCCTACACGACGCTCTTCCGATCTNNNTCAAGCAGTCATGTGAGGTA
AGCACACGTCTGAACTCTTTCCCTACACGACGCTCTTCCGATCTNNNTTTCCTGTCCATTCGTATCAGCA
AGCACACGTCTGAACTCTTTCCCTACACGACGCTCTTCCGATCTNNNTTTCCTGTCCATTCGTATCAGCA
AGCACACGTCTGAACTCTTTCCCTACACGACGCTCTTCCGATCTNNNACCACTGCCAGATTCAAAAAAG
AGCACACGTCTGAACTCTTTCCCTACACGACGCTCTTCCGATCTNNNACCACTGCCAGATTCAAAAAAG
AGCACACGTCTGAACTCTTTCCCTACACGACGCTCTTCCGATCTNNNTCTACCACCCAAAGGAAACCAA
AGCACACGTCTGAACTCTTTCCCTACACGACGCTCTTCCGATCTNNNTCTACCACCCAAAGGAAACCAA
AGCACACGTCTGAACTCTTTCCCTACACGACGCTCTTCCGATCTNNNTCAGAAAAATAGAAGCCCAGA
AGCACACGTCTGAACTCTTTCCCTACACGACGCTCTTCCGATCTNNNTCAGAAAAATAGAAGCCCAGA
AGCACACGTCTGAACTCTTTCCCTACACGACGCTCTTCCGATCTNNNGCTTGCAGTGAGTGGAGATCA
AGCACACGTCTGAACTCTTTCCCTACACGACGCTCTTCCGATCTNNNGCTTGCAGTGAGTGGAGATCA
GCACACGTCTGAACTCTTTCCCTACACGACGCTCTTCCGATCTNNNCTAGGAGCCCGGTCTCTGAGCAT
GCACACGTCTGAACTCTTTCCCTACACGACGCTCTTCCGATCTNNNCTAGGAGCCCGGTCTCTGAGCAT
GAGCACACGTCTGAACTCTTTCCCTACACGACGCTCTTCCGATCTNNNCCCTGCATGTCTGTATTGTTAC
GAGCACACGTCTGAACTCTTTCCCTACACGACGCTCTTCCGATCTNNNCCCTGCATGTCTGTATTGTTAC
GAGCACACGTCTGAACTCTTTCCCTACACGACGCTCTTCCGATCTNNNTCATTCATTCATTCCCCCACAT
GAGCACACGTCTGAACTCTTTCCCTACACGACGCTCTTCCGATCTNNNTCATTCATTCATTCCCCCACAT
GAGCACACGTCTGAACTCTTTCCCTACACGACGCTCTTCCGATCTNNNCGAGGGGATGAGAGAGAGA
GAGCACACGTCTGAACTCTTTCCCTACACGACGCTCTTCCGATCTNNNCGAGGGGATGAGAGAGAGA
GAGCACACGTCTGAACTCTTTCCCTACACGACGCTCTTCCGATCTNNNCTTTAATGTCTGCGAGGGTCC
GAGCACACGTCTGAACTCTTTCCCTACACGACGCTCTTCCGATCTNNNCTTTAATGTCTGCGAGGGTCC
AGCACACGTCTGAACTCTTTCCCTACACGACGCTCTTCCGATCTNNNTGATTTGGCCCCACTCATTCATA
AGCACACGTCTGAACTCTTTCCCTACACGACGCTCTTCCGATCTNNNTGATTTGGCCCCACTCATTCATA
AGCACACGTCTGAACTCTTTCCCTACACGACGCTCTTCCGATCTNNNCACATGAGGTCAGGAGTTCAT
AGCACACGTCTGAACTCTTTCCCTACACGACGCTCTTCCGATCTNNNCACATGAGGTCAGGAGTTCAT
AGCACACGTCTGAACTCTTTCCCTACACGACGCTCTTCCGATCTNNNTCACTTCTGCGACTCTTTGGAG
AGCACACGTCTGAACTCTTTCCCTACACGACGCTCTTCCGATCTNNNTCACTTCTGCGACTCTTTGGAG
GAGCACACGTCTGAACTCTTTCCCTACACGACGCTCTTCCGATCTNNNTCTGTAGAGGAGAGGAGAAG
GAGCACACGTCTGAACTCTTTCCCTACACGACGCTCTTCCGATCTNNNTCTGTAGAGGAGAGGAGAAG
GAGCACACGTCTGAACTCTTTCCCTACACGACGCTCTTCCGATCTNNNTCAAACAATTCTCTTGCCTCA
GAGCACACGTCTGAACTCTTTCCCTACACGACGCTCTTCCGATCTNNNTCAAACAATTCTCTTGCCTCA
AGAGCACACGTCTGAACTCTTTCCCTACACGACGCTCTTCCGATCTNNNGGATTCCGGCTATGCTAATG
AGAGCACACGTCTGAACTCTTTCCCTACACGACGCTCTTCCGATCTNNNGGATTCCGGCTATGCTAATG
GAGCACACGTCTGAACTCTTTCCCTACACGACGCTCTTCCGATCTNNNTGACCATGTATTGAGCATGAC
GAGCACACGTCTGAACTCTTTCCCTACACGACGCTCTTCCGATCTNNNTGACCATGTATTGAGCATGAC
GAGCACACGTCTGAACTCTTTCCCTACACGACGCTCTTCCGATCTNNNTGTTTGCCACGTGTATGTTTT
GAGCACACGTCTGAACTCTTTCCCTACACGACGCTCTTCCGATCTNNNTGTTTGCCACGTGTATGTTTT
GAGCACACGTCTGAACTCTTTCCCTACACGACGCTCTTCCGATCTNNNAGGAAGTCCGACAAAGTCCT
GAGCACACGTCTGAACTCTTTCCCTACACGACGCTCTTCCGATCTNNNAGGAAGTCCGACAAAGTCCT
AGCACACGTCTGAACTCTTTCCCTACACGACGCTCTTCCGATCTNNNATGGAAACTAAGCCCCAGAGA
AGCACACGTCTGAACTCTTTCCCTACACGACGCTCTTCCGATCTNNNATGGAAACTAAGCCCCAGAGA
AGCACACGTCTGAACTCTTTCCCTACACGACGCTCTTCCGATCTNNNGTTGAAGAAAAGAAAATTGTT
AGCACACGTCTGAACTCTTTCCCTACACGACGCTCTTCCGATCTNNNGTTGAAGAAAAGAAAATTGTT

AGCACACGTCTGAACTCTTTCCCTACACGACGCTCTTCCGATCTNNNCCTACTTCAAACTCAGAGTCCCC

AGCACACGTCTGAACTCTTTCCCTACACGACGCTCTTCCGATCTNNNCCTACTTCAAACTCAGAGTCCCC

GAGCACACGTCTGAACTCTTTCCCTACACGACGCTCTTCCGATCTNNNTAGGAGGCAAAGGTTACAGT

GAGCACACGTCTGAACTCTTTCCCTACACGACGCTCTTCCGATCTNNNTAGGAGGCAAAGGTTACAGT

GAGCACACGTCTGAACTCTTTCCCTACACGACGCTCTTCCGATCTNNNACAGGAGAATAGTTTGAGCC

GAGCACACGTCTGAACTCTTTCCCTACACGACGCTCTTCCGATCTNNNACAGGAGAATAGTTTGAGCC

AGCACACGTCTGAACTCTTTCCCTACACGACGCTCTTCCGATCTNNNGCCTAAGAGTTTGAGACCAGTG

AGCACACGTCTGAACTCTTTCCCTACACGACGCTCTTCCGATCTNNNGCCTAAGAGTTTGAGACCAGTG

AGCACACGTCTGAACTCTTTCCCTACACGACGCTCTTCCGATCTNNNCCAGTGGGATATTATCTAACTTT

AGCACACGTCTGAACTCTTTCCCTACACGACGCTCTTCCGATCTNNNCCAGTGGGATATTATCTAACTTT

AGAGCACACGTCTGAACTCTTTCCCTACACGACGCTCTTCCGATCTNNNGCAGTGGTTGTGAATGTGTC

AGAGCACACGTCTGAACTCTTTCCCTACACGACGCTCTTCCGATCTNNNGCAGTGGTTGTGAATGTGTC

AGAGCACACGTCTGAACTCTTTCCCTACACGACGCTCTTCCGATCTNNNTCTCTCCCAGGTGATTGAC

AGAGCACACGTCTGAACTCTTTCCCTACACGACGCTCTTCCGATCTNNNTCTCTCCCAGGTGATTGAC

GAGCACACGTCTGAACTCTTTCCCTACACGACGCTCTTCCGATCTNNNGTCTCCCGAGTTCTAAGCGAT

GAGCACACGTCTGAACTCTTTCCCTACACGACGCTCTTCCGATCTNNNGTCTCCCGAGTTCTAAGCGAT

GAGCACACGTCTGAACTCTTTCCCTACACGACGCTCTTCCGATCTNNNGCACCTGTAGTCCCAGATACT

GAGCACACGTCTGAACTCTTTCCCTACACGACGCTCTTCCGATCTNNNGCACCTGTAGTCCCAGATACT

AGCACACGTCTGAACTCTTTCCCTACACGACGCTCTTCCGATCTNNNCTAATGTTGGCAAAAATTGAAT

AGCACACGTCTGAACTCTTTCCCTACACGACGCTCTTCCGATCTNNNCTAATGTTGGCAAAAATTGAAT

GAGCACACGTCTGAACTCTTTCCCTACACGACGCTCTTCCGATCTNNNACCTCTGTGTATGCCTGTCTGA

GAGCACACGTCTGAACTCTTTCCCTACACGACGCTCTTCCGATCTNNNACCTCTGTGTATGCCTGTCTGA

GCACACGTCTGAACTCTTTCCCTACACGACGCTCTTCCGATCTNNNCCACAAATGGCAGGATTTCACTC

GCACACGTCTGAACTCTTTCCCTACACGACGCTCTTCCGATCTNNNCCACAAATGGCAGGATTTCACTC

AGCACACGTCTGAACTCTTTCCCTACACGACGCTCTTCCGATCTNNNAGGAGAATGGCGTGAACTGGC

AGCACACGTCTGAACTCTTTCCCTACACGACGCTCTTCCGATCTNNNAGGAGAATGGCGTGAACTGGC

GAGCACACGTCTGAACTCTTTCCCTACACGACGCTCTTCCGATCTNNNGAGACCAGCTCCTCCCCCTACA

GAGCACACGTCTGAACTCTTTCCCTACACGACGCTCTTCCGATCTNNNGAGACCAGCTCCTCCCCCTACA

AGCACACGTCTGAACTCTTTCCCTACACGACGCTCTTCCGATCTNNNGGTGCTCAACTCCTCAGTGGCA

AGCACACGTCTGAACTCTTTCCCTACACGACGCTCTTCCGATCTNNNGGTGCTCAACTCCTCAGTGGCA

AGCACACGTCTGAACTCTTTCCCTACACGACGCTCTTCCGATCTNNNGTGAATAGTGCCTGTGATGAAC

AGCACACGTCTGAACTCTTTCCCTACACGACGCTCTTCCGATCTNNNGTGAATAGTGCCTGTGATGAAC

AGAGCACACGTCTGAACTCTTTCCCTACACGACGCTCTTCCGATCTNNNTTCAGTTCAAGTGGTGTGTC

AGAGCACACGTCTGAACTCTTTCCCTACACGACGCTCTTCCGATCTNNNTTCAGTTCAAGTGGTGTGTC

AGCACACGTCTGAACTCTTTCCCTACACGACGCTCTTCCGATCTNNNTGGGCAACATGGCGAAATAAA

AGCACACGTCTGAACTCTTTCCCTACACGACGCTCTTCCGATCTNNNTGGGCAACATGGCGAAATAAA

AGCACACGTCTGAACTCTTTCCCTACACGACGCTCTTCCGATCTNNNAGGTTCAAATGATTCTCCTGCCT

AGCACACGTCTGAACTCTTTCCCTACACGACGCTCTTCCGATCTNNNAGGTTCAAATGATTCTCCTGCCT

AGCACACGTCTGAACTCTTTCCCTACACGACGCTCTTCCGATCTNNNACCATGTTTTTGCACACTCTTCAC

AGCACACGTCTGAACTCTTTCCCTACACGACGCTCTTCCGATCTNNNACCATGTTTTTGCACACTCTTCAC

GCACACGTCTGAACTCTTTCCCTACACGACGCTCTTCCGATCTNNNTGTGGATCAAAACACGTACCTCA

GCACACGTCTGAACTCTTTCCCTACACGACGCTCTTCCGATCTNNNTGTGGATCAAAACACGTACCTCA

GAGCACACGTCTGAACTCTTTCCCTACACGACGCTCTTCCGATCTNNNCAGCTGGGGTTCTTTATTGTG

GAGCACACGTCTGAACTCTTTCCCTACACGACGCTCTTCCGATCTNNNCAGCTGGGGTTCTTTATTGTG
GCACACGTCTGAACTCTTTCCCTACACGACGCTCTTCCGATCTNNNAAGACACAAGACAGACACAGAG
GCACACGTCTGAACTCTTTCCCTACACGACGCTCTTCCGATCTNNNAAGACACAAGACAGACACAGAG
GAGCACACGTCTGAACTCTTTCCCTACACGACGCTCTTCCGATCTNNNCCCACATTCTTTCCAGTCTCTC
GAGCACACGTCTGAACTCTTTCCCTACACGACGCTCTTCCGATCTNNNCCCACATTCTTTCCAGTCTCTC
AGCACACGTCTGAACTCTTTCCCTACACGACGCTCTTCCGATCTNNNTAGGTTACTTCAGCAGGGTGT
AGCACACGTCTGAACTCTTTCCCTACACGACGCTCTTCCGATCTNNNTAGGTTACTTCAGCAGGGTGT
AGCACACGTCTGAACTCTTTCCCTACACGACGCTCTTCCGATCTNNNCTGGCTGTCTATGAACTAGGG
AGCACACGTCTGAACTCTTTCCCTACACGACGCTCTTCCGATCTNNNCTGGCTGTCTATGAACTAGGG
AGCACACGTCTGAACTCTTTCCCTACACGACGCTCTTCCGATCTNNNTAAATGGAGTAACACACCCTGG
AGCACACGTCTGAACTCTTTCCCTACACGACGCTCTTCCGATCTNNNTAAATGGAGTAACACACCCTGG
GCACACGTCTGAACTCTTTCCCTACACGACGCTCTTCCGATCTNNNTATTTGTCTTCGCCGCTTGTTG
GCACACGTCTGAACTCTTTCCCTACACGACGCTCTTCCGATCTNNNTATTTGTCTTCGCCGCTTGTTG
GAGCACACGTCTGAACTCTTTCCCTACACGACGCTCTTCCGATCTNNNCAAATCTCAAAGACTCAGCCC
GAGCACACGTCTGAACTCTTTCCCTACACGACGCTCTTCCGATCTNNNCAAATCTCAAAGACTCAGCCC
GAGCACACGTCTGAACTCTTTCCCTACACGACGCTCTTCCGATCTNNNTAAAATGGAAAGTAGGAGCA
GAGCACACGTCTGAACTCTTTCCCTACACGACGCTCTTCCGATCTNNNTAAAATGGAAAGTAGGAGCA
AGCACACGTCTGAACTCTTTCCCTACACGACGCTCTTCCGATCTNNNCTGGGTGAAAGACTGAGACTT
AGCACACGTCTGAACTCTTTCCCTACACGACGCTCTTCCGATCTNNNCTGGGTGAAAGACTGAGACTT
GAGCACACGTCTGAACTCTTTCCCTACACGACGCTCTTCCGATCTNNNAAAAATTAGCTGGTTGTGGT
GAGCACACGTCTGAACTCTTTCCCTACACGACGCTCTTCCGATCTNNNAAAAATTAGCTGGTTGTGGT
GCACACGTCTGAACTCTTTCCCTACACGACGCTCTTCCGATCTNNNCATCTGCAGACAAATGTGAGGGA
GCACACGTCTGAACTCTTTCCCTACACGACGCTCTTCCGATCTNNNCATCTGCAGACAAATGTGAGGGA
AGCACACGTCTGAACTCTTTCCCTACACGACGCTCTTCCGATCTNNNATTGGGAGCTCTTTCAGTTGAC
AGCACACGTCTGAACTCTTTCCCTACACGACGCTCTTCCGATCTNNNATTGGGAGCTCTTTCAGTTGAC
AGCACACGTCTGAACTCTTTCCCTACACGACGCTCTTCCGATCTNNNTGTGTACAGACTCTCCAGGAAA
AGCACACGTCTGAACTCTTTCCCTACACGACGCTCTTCCGATCTNNNTGTGTACAGACTCTCCAGGAAA
AGCACACGTCTGAACTCTTTCCCTACACGACGCTCTTCCGATCTNNNAAATTTGCCCAGGAACCCAAAA
AGCACACGTCTGAACTCTTTCCCTACACGACGCTCTTCCGATCTNNNAAATTTGCCCAGGAACCCAAAA
GCACACGTCTGAACTCTTTCCCTACACGACGCTCTTCCGATCTNNNCAGTTACTTGAGAGATGGAGGCA
GCACACGTCTGAACTCTTTCCCTACACGACGCTCTTCCGATCTNNNCAGTTACTTGAGAGATGGAGGCA
GCACACGTCTGAACTCTTTCCCTACACGACGCTCTTCCGATCTNNNGTTGTAGTGAGCTGAGATTGTG
GCACACGTCTGAACTCTTTCCCTACACGACGCTCTTCCGATCTNNNGTTGTAGTGAGCTGAGATTGTG
AGCACACGTCTGAACTCTTTCCCTACACGACGCTCTTCCGATCTNNNGTGCCTATCACACAGTAACCAC
AGCACACGTCTGAACTCTTTCCCTACACGACGCTCTTCCGATCTNNNGTGCCTATCACACAGTAACCAC
GCACACGTCTGAACTCTTTCCCTACACGACGCTCTTCCGATCTNNNGGTGCTTCTGTCTCTGCCTGCAT
GCACACGTCTGAACTCTTTCCCTACACGACGCTCTTCCGATCTNNNGGTGCTTCTGTCTCTGCCTGCAT
AGCACACGTCTGAACTCTTTCCCTACACGACGCTCTTCCGATCTNNNAATCGCTCTCTCTCTGTCTCTCT
AGCACACGTCTGAACTCTTTCCCTACACGACGCTCTTCCGATCTNNNAATCGCTCTCTCTCTGTCTCTCT
AGAGCACACGTCTGAACTCTTTCCCTACACGACGCTCTTCCGATCTNNNGCACTTACTGTCTTCAACCCC
AGAGCACACGTCTGAACTCTTTCCCTACACGACGCTCTTCCGATCTNNNGCACTTACTGTCTTCAACCCC
GAGCACACGTCTGAACTCTTTCCCTACACGACGCTCTTCCGATCTNNNCTGAGGCAGGAGAATCACTTC
GAGCACACGTCTGAACTCTTTCCCTACACGACGCTCTTCCGATCTNNNCTGAGGCAGGAGAATCACTTC

GAGCACACGTCTGAACTCTTTCCCTACACGACGCTCTTCCGATCTNNNGGGCAACAGAGTGAGAATGT
GAGCACACGTCTGAACTCTTTCCCTACACGACGCTCTTCCGATCTNNNGGGCAACAGAGTGAGAATGT
GAGCACACGTCTGAACTCTTTCCCTACACGACGCTCTTCCGATCTNNNGACTGGAGGTGGTGCTATCA
GAGCACACGTCTGAACTCTTTCCCTACACGACGCTCTTCCGATCTNNNGACTGGAGGTGGTGCTATCA
AGCACACGTCTGAACTCTTTCCCTACACGACGCTCTTCCGATCTNNNGATTGCAGTGAGCCAAGATCAT
AGCACACGTCTGAACTCTTTCCCTACACGACGCTCTTCCGATCTNNNGATTGCAGTGAGCCAAGATCAT
GCACACGTCTGAACTCTTTCCCTACACGACGCTCTTCCGATCTNNNACTACACCCTAAACTGCCCTCTAC
GCACACGTCTGAACTCTTTCCCTACACGACGCTCTTCCGATCTNNNACTACACCCTAAACTGCCCTCTAC
GAGCACACGTCTGAACTCTTTCCCTACACGACGCTCTTCCGATCTNNNTTGAGGTTCAGCCACATAACA
GAGCACACGTCTGAACTCTTTCCCTACACGACGCTCTTCCGATCTNNNTTGAGGTTCAGCCACATAACA
AGCACACGTCTGAACTCTTTCCCTACACGACGCTCTTCCGATCTNNNTGAGGCAGGAGAATCACTTGT
AGCACACGTCTGAACTCTTTCCCTACACGACGCTCTTCCGATCTNNNTGAGGCAGGAGAATCACTTGT
GAGCACACGTCTGAACTCTTTCCCTACACGACGCTCTTCCGATCTNNNTACGACTTTCTCCCTAGGCAA
GAGCACACGTCTGAACTCTTTCCCTACACGACGCTCTTCCGATCTNNNTACGACTTTCTCCCTAGGCAA
AGCACACGTCTGAACTCTTTCCCTACACGACGCTCTTCCGATCTNNNTCCTAAATCCAGCCCTGTAATAC
AGCACACGTCTGAACTCTTTCCCTACACGACGCTCTTCCGATCTNNNTCCTAAATCCAGCCCTGTAATAC
AGCACACGTCTGAACTCTTTCCCTACACGACGCTCTTCCGATCTNNNGCAACAAGAGCAAAATTCCGT
AGCACACGTCTGAACTCTTTCCCTACACGACGCTCTTCCGATCTNNNGCAACAAGAGCAAAATTCCGT
GCACACGTCTGAACTCTTTCCCTACACGACGCTCTTCCGATCTNNNTACCCAACTCCTGTACCCTCATTC
GCACACGTCTGAACTCTTTCCCTACACGACGCTCTTCCGATCTNNNTACCCAACTCCTGTACCCTCATTC
AGCACACGTCTGAACTCTTTCCCTACACGACGCTCTTCCGATCTNNNATTAGCCTTGCTAGGGGTCTTT
AGCACACGTCTGAACTCTTTCCCTACACGACGCTCTTCCGATCTNNNATTAGCCTTGCTAGGGGTCTTT
AGAGCACACGTCTGAACTCTTTCCCTACACGACGCTCTTCCGATCTNNNAACATCAGGGAAACACTAC
AGAGCACACGTCTGAACTCTTTCCCTACACGACGCTCTTCCGATCTNNNAACATCAGGGAAACACTAC
AGCACACGTCTGAACTCTTTCCCTACACGACGCTCTTCCGATCTNNNGATTTTCCCATTAGATGCCCAG
AGCACACGTCTGAACTCTTTCCCTACACGACGCTCTTCCGATCTNNNGATTTTCCCATTAGATGCCCAG
AGCACACGTCTGAACTCTTTCCCTACACGACGCTCTTCCGATCTNNNAAGTATAAGAAAGCTCTCCATT
AGCACACGTCTGAACTCTTTCCCTACACGACGCTCTTCCGATCTNNNAAGTATAAGAAAGCTCTCCATT
GAGCACACGTCTGAACTCTTTCCCTACACGACGCTCTTCCGATCTNNNGCCTGTAGTCCCAGCTACTTG
GAGCACACGTCTGAACTCTTTCCCTACACGACGCTCTTCCGATCTNNNGCCTGTAGTCCCAGCTACTTG
GCACACGTCTGAACTCTTTCCCTACACGACGCTCTTCCGATCTNNNTCTTGGTTCCTCCTTCTAGGGAA
GCACACGTCTGAACTCTTTCCCTACACGACGCTCTTCCGATCTNNNTCTTGGTTCCTCCTTCTAGGGAA
AGCACACGTCTGAACTCTTTCCCTACACGACGCTCTTCCGATCTNNNGGTCTTTTTGTGTCCCATCCTT
AGCACACGTCTGAACTCTTTCCCTACACGACGCTCTTCCGATCTNNNGGTCTTTTTGTGTCCCATCCTT
GCACACGTCTGAACTCTTTCCCTACACGACGCTCTTCCGATCTNNNTTAGAGTCAGAAGTTGGGGAA
GCACACGTCTGAACTCTTTCCCTACACGACGCTCTTCCGATCTNNNTTAGAGTCAGAAGTTGGGGAA
GAGCACACGTCTGAACTCTTTCCCTACACGACGCTCTTCCGATCTNNNGGCCAACCATGCTGTTCTTATT
GAGCACACGTCTGAACTCTTTCCCTACACGACGCTCTTCCGATCTNNNGGCCAACCATGCTGTTCTTATT
AGAGCACACGTCTGAACTCTTTCCCTACACGACGCTCTTCCGATCTNNNCAGTCTGAAGGGCACATCTT
AGAGCACACGTCTGAACTCTTTCCCTACACGACGCTCTTCCGATCTNNNCAGTCTGAAGGGCACATCTT
GAGCACACGTCTGAACTCTTTCCCTACACGACGCTCTTCCGATCTNNNTGTAGTCCTCTCTACTTGGGA
GAGCACACGTCTGAACTCTTTCCCTACACGACGCTCTTCCGATCTNNNTGTAGTCCTCTCTACTTGGGA
CACACGTCTGAACTCTTTCCCTACACGACGCTCTTCCGATCTNNNGAGTGACGACTGCAACGAAGGCA

CACACGTCTGAACTCTTTCCCTACACGACGCTCTTCCGATCTNNNGAGTGACGACTGCAACGAAGGCAT
AGCACACGTCTGAACTCTTTCCCTACACGACGCTCTTCCGATCTNNNGCTCCAGTTCAGTTAGCTAGAG
AGCACACGTCTGAACTCTTTCCCTACACGACGCTCTTCCGATCTNNNGCTCCAGTTCAGTTAGCTAGAG
AGCACACGTCTGAACTCTTTCCCTACACGACGCTCTTCCGATCTNNNAAAGTGCTGGGATTACAGGTG
AGCACACGTCTGAACTCTTTCCCTACACGACGCTCTTCCGATCTNNNAAAGTGCTGGGATTACAGGTG
AGAGCACACGTCTGAACTCTTTCCCTACACGACGCTCTTCCGATCTNNNGACTCCTTGTTGTGCAGGACT
AGAGCACACGTCTGAACTCTTTCCCTACACGACGCTCTTCCGATCTNNNGACTCCTTGTTGTGCAGGACT
AGCACACGTCTGAACTCTTTCCCTACACGACGCTCTTCCGATCTNNNTTGTAACCCCTCACCAGTTTTGA
AGCACACGTCTGAACTCTTTCCCTACACGACGCTCTTCCGATCTNNNTTGTAACCCCTCACCAGTTTTGA
AGCACACGTCTGAACTCTTTCCCTACACGACGCTCTTCCGATCTNNNTGCCTAAAGTACCATCTTTGCCTC
AGCACACGTCTGAACTCTTTCCCTACACGACGCTCTTCCGATCTNNNTGCCTAAAGTACCATCTTTGCCTC
CACACGTCTGAACTCTTTCCCTACACGACGCTCTTCCGATCTNNNCTTTGCTTCAGTAGGATCTGCACCA
CACACGTCTGAACTCTTTCCCTACACGACGCTCTTCCGATCTNNNCTTTGCTTCAGTAGGATCTGCACCA
ACACGTCTGAACTCTTTCCCTACACGACGCTCTTCCGATCTNNNCTCCCCGTTTCTGTCCCTTCCATCGGAC
ACACGTCTGAACTCTTTCCCTACACGACGCTCTTCCGATCTNNNCTCCCCGTTTCTGTCCCTTCCATCGGAC
AGCACACGTCTGAACTCTTTCCCTACACGACGCTCTTCCGATCTNNNATGCACGTGGTAGACACTCAGTA
AGCACACGTCTGAACTCTTTCCCTACACGACGCTCTTCCGATCTNNNATGCACGTGGTAGACACTCAGTA
GAGCACACGTCTGAACTCTTTCCCTACACGACGCTCTTCCGATCTNNNACATATACCTATCCTGTGACGC
GAGCACACGTCTGAACTCTTTCCCTACACGACGCTCTTCCGATCTNNNACATATACCTATCCTGTGACGC
GAGCACACGTCTGAACTCTTTCCCTACACGACGCTCTTCCGATCTNNNGTTGTGGCATGTGCTTATAGT
GAGCACACGTCTGAACTCTTTCCCTACACGACGCTCTTCCGATCTNNNGTTGTGGCATGTGCTTATAGT
AGCACACGTCTGAACTCTTTCCCTACACGACGCTCTTCCGATCTNNNCAGTCCAGTTTTTCAGCCTTCAG
AGCACACGTCTGAACTCTTTCCCTACACGACGCTCTTCCGATCTNNNCAGTCCAGTTTTTCAGCCTTCAG
GAGCACACGTCTGAACTCTTTCCCTACACGACGCTCTTCCGATCTNNNCGTTGTCACAAAAGGCAAGAT
GAGCACACGTCTGAACTCTTTCCCTACACGACGCTCTTCCGATCTNNNCGTTGTCACAAAAGGCAAGAT
AGAGCACACGTCTGAACTCTTTCCCTACACGACGCTCTTCCGATCTNNNCATGCCTGTAATCCCAGCTAC
AGAGCACACGTCTGAACTCTTTCCCTACACGACGCTCTTCCGATCTNNNCATGCCTGTAATCCCAGCTAC
GAGCACACGTCTGAACTCTTTCCCTACACGACGCTCTTCCGATCTNNNTAGGAGTTTGAGGCTGCATGA
GAGCACACGTCTGAACTCTTTCCCTACACGACGCTCTTCCGATCTNNNTAGGAGTTTGAGGCTGCATGA
GAGCACACGTCTGAACTCTTTCCCTACACGACGCTCTTCCGATCTNNNCCTCTCCTATACTCTGCTCCATC
GAGCACACGTCTGAACTCTTTCCCTACACGACGCTCTTCCGATCTNNNCCTCTCCTATACTCTGCTCCATC
AGCACACGTCTGAACTCTTTCCCTACACGACGCTCTTCCGATCTNNNGACAAGACCAAAACACTAGGGC
AGCACACGTCTGAACTCTTTCCCTACACGACGCTCTTCCGATCTNNNGACAAGACCAAAACACTAGGGC
AGCACACGTCTGAACTCTTTCCCTACACGACGCTCTTCCGATCTNNNGTTTCCATGCACACTGTGAAGT
AGCACACGTCTGAACTCTTTCCCTACACGACGCTCTTCCGATCTNNNGTTTCCATGCACACTGTGAAGT
GAGCACACGTCTGAACTCTTTCCCTACACGACGCTCTTCCGATCTNNNTCTCCAGAAGGTGAGGAAGGA
GAGCACACGTCTGAACTCTTTCCCTACACGACGCTCTTCCGATCTNNNTCTCCAGAAGGTGAGGAAGGA
GAGCACACGTCTGAACTCTTTCCCTACACGACGCTCTTCCGATCTNNNGTGTATCTCACGCATATGCACA
GAGCACACGTCTGAACTCTTTCCCTACACGACGCTCTTCCGATCTNNNGTGTATCTCACGCATATGCACA
AGCACACGTCTGAACTCTTTCCCTACACGACGCTCTTCCGATCTNNNGCCTTCTGTTCACATGACAATCC
AGCACACGTCTGAACTCTTTCCCTACACGACGCTCTTCCGATCTNNNGCCTTCTGTTCACATGACAATCC
GCACACGTCTGAACTCTTTCCCTACACGACGCTCTTCCGATCTNNNGTAAAGGTGCCTCTTAAAGAACA
GCACACGTCTGAACTCTTTCCCTACACGACGCTCTTCCGATCTNNNGTAAAGGTGCCTCTTAAAGAACA

AGCACACGTCTGAACTCTTTCCCTACACGACGCTCTTCCGATCTNNNAGAGCTTGTATCATTCCTTCTG

AGCACACGTCTGAACTCTTTCCCTACACGACGCTCTTCCGATCTNNNAGAGCTTGTATCATTCCTTCTG

GAGCACACGTCTGAACTCTTTCCCTACACGACGCTCTTCCGATCTNNNGATAATGTGAGGGCCCTTTG

GAGCACACGTCTGAACTCTTTCCCTACACGACGCTCTTCCGATCTNNNGATAATGTGAGGGCCCTTTG

GCACACGTCTGAACTCTTTCCCTACACGACGCTCTTCCGATCTNNNACCAATGAGGGGATGAATGTG

GCACACGTCTGAACTCTTTCCCTACACGACGCTCTTCCGATCTNNNACCAATGAGGGGATGAATGTG

GAGCACACGTCTGAACTCTTTCCCTACACGACGCTCTTCCGATCTNNNGGATTTGCAGTGAGTCAAGA

GAGCACACGTCTGAACTCTTTCCCTACACGACGCTCTTCCGATCTNNNGGATTTGCAGTGAGTCAAGA

AGCACACGTCTGAACTCTTTCCCTACACGACGCTCTTCCGATCTNNNGGTCTTTGATGATGGTGACGA

AGCACACGTCTGAACTCTTTCCCTACACGACGCTCTTCCGATCTNNNGGTCTTTGATGATGGTGACGA

GAGCACACGTCTGAACTCTTTCCCTACACGACGCTCTTCCGATCTNNNAGTCTGTCTTGCTCTCTATAT

GAGCACACGTCTGAACTCTTTCCCTACACGACGCTCTTCCGATCTNNNAGTCTGTCTTGCTCTCTATAT

AGCACACGTCTGAACTCTTTCCCTACACGACGCTCTTCCGATCTNNNGAGCAAGACTCCATCTCAAAA

AGCACACGTCTGAACTCTTTCCCTACACGACGCTCTTCCGATCTNNNGAGCAAGACTCCATCTCAAAA

AGAGCACACGTCTGAACTCTTTCCCTACACGACGCTCTTCCGATCTNNNTGAGCCCAGAAGTTCAAGA

AGAGCACACGTCTGAACTCTTTCCCTACACGACGCTCTTCCGATCTNNNTGAGCCCAGAAGTTCAAGA

GCACACGTCTGAACTCTTTCCCTACACGACGCTCTTCCGATCTNNNTATGATTTGACTCTTGCCTGCCT

GCACACGTCTGAACTCTTTCCCTACACGACGCTCTTCCGATCTNNNTATGATTTGACTCTTGCCTGCCT

GAGCACACGTCTGAACTCTTTCCCTACACGACGCTCTTCCGATCTNNNTGATGCCTTTCTTAGCTCCCA

GAGCACACGTCTGAACTCTTTCCCTACACGACGCTCTTCCGATCTNNNTGATGCCTTTCTTAGCTCCCA

GAGCACACGTCTGAACTCTTTCCCTACACGACGCTCTTCCGATCTNNNCGGCCCTCACTCTCCTATAGAC

GAGCACACGTCTGAACTCTTTCCCTACACGACGCTCTTCCGATCTNNNCGGCCCTCACTCTCCTATAGAC

GAGCACACGTCTGAACTCTTTCCCTACACGACGCTCTTCCGATCTNNNTGTAATCCCAGCACTTTGAGA

GAGCACACGTCTGAACTCTTTCCCTACACGACGCTCTTCCGATCTNNNTGTAATCCCAGCACTTTGAGA

AGCACACGTCTGAACTCTTTCCCTACACGACGCTCTTCCGATCTNNNTGCTCACCATAGTGTCATTTCC

AGCACACGTCTGAACTCTTTCCCTACACGACGCTCTTCCGATCTNNNTGCTCACCATAGTGTCATTTCC

GCACACGTCTGAACTCTTTCCCTACACGACGCTCTTCCGATCTNNNCTTCGAACTTCTACTTTAACAAA

GCACACGTCTGAACTCTTTCCCTACACGACGCTCTTCCGATCTNNNCTTCGAACTTCTACTTTAACAAA

AGCACACGTCTGAACTCTTTCCCTACACGACGCTCTTCCGATCTNNNGAGGCATGAGAATCGCTTGAA

AGCACACGTCTGAACTCTTTCCCTACACGACGCTCTTCCGATCTNNNGAGGCATGAGAATCGCTTGAA

GCACACGTCTGAACTCTTTCCCTACACGACGCTCTTCCGATCTNNNGAACGGATTCCTTCTCCTCTACCC

GCACACGTCTGAACTCTTTCCCTACACGACGCTCTTCCGATCTNNNGAACGGATTCCTTCTCCTCTACCC

GAGCACACGTCTGAACTCTTTCCCTACACGACGCTCTTCCGATCTNNNCTTCCTTCCTTCCTTCCTTCCTC

GAGCACACGTCTGAACTCTTTCCCTACACGACGCTCTTCCGATCTNNNCTTCCTTCCTTCCTTCCTTCCTC

GCACACGTCTGAACTCTTTCCCTACACGACGCTCTTCCGATCTNNNGAGGTTTTACTGAGCCCAGATC

GCACACGTCTGAACTCTTTCCCTACACGACGCTCTTCCGATCTNNNGAGGTTTTACTGAGCCCAGATC

GAGCACACGTCTGAACTCTTTCCCTACACGACGCTCTTCCGATCTNNNATCAAGGACATCAAGGAGG

GAGCACACGTCTGAACTCTTTCCCTACACGACGCTCTTCCGATCTNNNATCAAGGACATCAAGGAGG

AGAGCACACGTCTGAACTCTTTCCCTACACGACGCTCTTCCGATCTNNNTAACACAGGGACAGGAAAC

AGAGCACACGTCTGAACTCTTTCCCTACACGACGCTCTTCCGATCTNNNTAACACAGGGACAGGAAAC

AGCACACGTCTGAACTCTTTCCCTACACGACGCTCTTCCGATCTNNNACTTCTAATACCACAGGGCACA

AGCACACGTCTGAACTCTTTCCCTACACGACGCTCTTCCGATCTNNNACTTCTAATACCACAGGGCACA

GAGCACACGTCTGAACTCTTTCCCTACACGACGCTCTTCCGATCTNNNTCCATTCTCTCCAGCCTATCTC

GAGCACACGTCTGAACTCTTTCCCTACACGACGCTCTTCCGATCTNNNTCCATTCTCTCCAGCCTATCTC
GCACACGTCTGAACTCTTTCCCTACACGACGCTCTTCCGATCTNNNACTAGCCAGCACACTTAGGTAT
GCACACGTCTGAACTCTTTCCCTACACGACGCTCTTCCGATCTNNNACTAGCCAGCACACTTAGGTAT
AGCACACGTCTGAACTCTTTCCCTACACGACGCTCTTCCGATCTNNNCGACAGAGTGAGACTCCGTCT
AGCACACGTCTGAACTCTTTCCCTACACGACGCTCTTCCGATCTNNNCGACAGAGTGAGACTCCGTCT
AGCACACGTCTGAACTCTTTCCCTACACGACGCTCTTCCGATCTNNNAAGAAATCTCAGAGACTCAGC
AGCACACGTCTGAACTCTTTCCCTACACGACGCTCTTCCGATCTNNNAAGAAATCTCAGAGACTCAGC
GCACACGTCTGAACTCTTTCCCTACACGACGCTCTTCCGATCTNNNCTGTAGTCCCAGCTACTTGGAA
GCACACGTCTGAACTCTTTCCCTACACGACGCTCTTCCGATCTNNNCTGTAGTCCCAGCTACTTGGAA
GAGCACACGTCTGAACTCTTTCCCTACACGACGCTCTTCCGATCTNNNATTAGTCAGGGTTCCTTAGA
GAGCACACGTCTGAACTCTTTCCCTACACGACGCTCTTCCGATCTNNNATTAGTCAGGGTTCCTTAGA
GCACACGTCTGAACTCTTTCCCTACACGACGCTCTTCCGATCTNNNTCTCAGCACCTTAGAACAAACCA
GCACACGTCTGAACTCTTTCCCTACACGACGCTCTTCCGATCTNNNTCTCAGCACCTTAGAACAAACCA
ACACGTCTGAACTCTTTCCCTACACGACGCTCTTCCGATCTNNNCACCAGTTTCTCTGCTTTCCACCAT
ACACGTCTGAACTCTTTCCCTACACGACGCTCTTCCGATCTNNNCACCAGTTTCTCTGCTTTCCACCAT
GCACACGTCTGAACTCTTTCCCTACACGACGCTCTTCCGATCTNNNCAGGCATTTTGCTAGGCACTTT
GCACACGTCTGAACTCTTTCCCTACACGACGCTCTTCCGATCTNNNCAGGCATTTTGCTAGGCACTTT
CACACGTCTGAACTCTTTCCCTACACGACGCTCTTCCGATCTNNNCCTGAGCAACAGAGTGATACTCC
CACACGTCTGAACTCTTTCCCTACACGACGCTCTTCCGATCTNNNCCTGAGCAACAGAGTGATACTCC
AGCACACGTCTGAACTCTTTCCCTACACGACGCTCTTCCGATCTNNNGTGAAAGAAATTCTAAGGGCC
AGCACACGTCTGAACTCTTTCCCTACACGACGCTCTTCCGATCTNNNGTGAAAGAAATTCTAAGGGCC
GAGCACACGTCTGAACTCTTTCCCTACACGACGCTCTTCCGATCTNNNGTTCCATGGTCTTGGCTGG
GAGCACACGTCTGAACTCTTTCCCTACACGACGCTCTTCCGATCTNNNGTTCCATGGTCTTGGCTGG
AGAGCACACGTCTGAACTCTTTCCCTACACGACGCTCTTCCGATCTNNNGAGATTGCAGTGAGCCTA
AGAGCACACGTCTGAACTCTTTCCCTACACGACGCTCTTCCGATCTNNNGAGATTGCAGTGAGCCTA
GCACACGTCTGAACTCTTTCCCTACACGACGCTCTTCCGATCTNNNCTGATGGGGAAGGGTGTGTAG
GCACACGTCTGAACTCTTTCCCTACACGACGCTCTTCCGATCTNNNCTGATGGGGAAGGGTGTGTAG
GAGCACACGTCTGAACTCTTTCCCTACACGACGCTCTTCCGATCTNNNTATGACTTGATGGGGTCTAG
GAGCACACGTCTGAACTCTTTCCCTACACGACGCTCTTCCGATCTNNNTATGACTTGATGGGGTCTAG
GAGCACACGTCTGAACTCTTTCCCTACACGACGCTCTTCCGATCTNNNTCTCCTTTCCCATCCTGCACA
GAGCACACGTCTGAACTCTTTCCCTACACGACGCTCTTCCGATCTNNNTCTCCTTTCCCATCCTGCACA
AGAGCACACGTCTGAACTCTTTCCCTACACGACGCTCTTCCGATCTNNNAAAAAAAATTAGCCGGGC
AGAGCACACGTCTGAACTCTTTCCCTACACGACGCTCTTCCGATCTNNNAAAAAAAATTAGCCGGGC
GAGCACACGTCTGAACTCTTTCCCTACACGACGCTCTTCCGATCTNNNTGGGAAGTGCCATCATTTTC
GAGCACACGTCTGAACTCTTTCCCTACACGACGCTCTTCCGATCTNNNTGGGAAGTGCCATCATTTTC
GAGCACACGTCTGAACTCTTTCCCTACACGACGCTCTTCCGATCTNNNCCAGGCAGGATTAGCTACAA
GAGCACACGTCTGAACTCTTTCCCTACACGACGCTCTTCCGATCTNNNCCAGGCAGGATTAGCTACAA
AGCACACGTCTGAACTCTTTCCCTACACGACGCTCTTCCGATCTNNNCTGGGCAACAAGAGCAAAATT
AGCACACGTCTGAACTCTTTCCCTACACGACGCTCTTCCGATCTNNNCTGGGCAACAAGAGCAAAATT
GAGCACACGTCTGAACTCTTTCCCTACACGACGCTCTTCCGATCTNNNTCTGATGTTACTTGGCAACCA
GAGCACACGTCTGAACTCTTTCCCTACACGACGCTCTTCCGATCTNNNTCTGATGTTACTTGGCAACCA
GAGCACACGTCTGAACTCTTTCCCTACACGACGCTCTTCCGATCTNNNGCATGGTAGAAGGGAGAA
GAGCACACGTCTGAACTCTTTCCCTACACGACGCTCTTCCGATCTNNNGCATGGTAGAAGGGAGAA

GAGCACACGTCTGAACTCTTTCCCTACACGACGCTCTTCCGATCTNNNTGTAGTACCAGCTACTCAGGA

GAGCACACGTCTGAACTCTTTCCCTACACGACGCTCTTCCGATCTNNNTGTAGTACCAGCTACTCAGGA

GCACACGTCTGAACTCTTTCCCTACACGACGCTCTTCCGATCTNNNATGAAAAAGACATCTCAGGCCCTC

GCACACGTCTGAACTCTTTCCCTACACGACGCTCTTCCGATCTNNNATGAAAAAGACATCTCAGGCCCTC

GAGCACACGTCTGAACTCTTTCCCTACACGACGCTCTTCCGATCTNNNCATTGCATTGATATGATGGAT

GAGCACACGTCTGAACTCTTTCCCTACACGACGCTCTTCCGATCTNNNCATTGCATTGATATGATGGAT

AGCACACGTCTGAACTCTTTCCCTACACGACGCTCTTCCGATCTNNNCAGCCAACCACAAGTGAGAATTT

AGCACACGTCTGAACTCTTTCCCTACACGACGCTCTTCCGATCTNNNCAGCCAACCACAAGTGAGAATTT

AGAGCACACGTCTGAACTCTTTCCCTACACGACGCTCTTCCGATCTNNNACAAAGCCACAGGACTAGTT

AGAGCACACGTCTGAACTCTTTCCCTACACGACGCTCTTCCGATCTNNNACAAAGCCACAGGACTAGTT

GAGCACACGTCTGAACTCTTTCCCTACACGACGCTCTTCCGATCTNNNTTTTTTTTTTTGCAGGGGAGG

GAGCACACGTCTGAACTCTTTCCCTACACGACGCTCTTCCGATCTNNNTTTTTTTTTTTGCAGGGGAGG

AGAGCACACGTCTGAACTCTTTCCCTACACGACGCTCTTCCGATCTNNNTGGTATCCCGGTCATTTTTGC

AGAGCACACGTCTGAACTCTTTCCCTACACGACGCTCTTCCGATCTNNNTGGTATCCCGGTCATTTTTGC

GAGCACACGTCTGAACTCTTTCCCTACACGACGCTCTTCCGATCTNNNCCGTCTTTTCTGTGACTCATCCT

GAGCACACGTCTGAACTCTTTCCCTACACGACGCTCTTCCGATCTNNNCCGTCTTTTCTGTGACTCATCCT

GCACACGTCTGAACTCTTTCCCTACACGACGCTCTTCCGATCTNNNTGCCTGTAATCTCAGCTACTCAGC

GCACACGTCTGAACTCTTTCCCTACACGACGCTCTTCCGATCTNNNTGCCTGTAATCTCAGCTACTCAGC

AGCACACGTCTGAACTCTTTCCCTACACGACGCTCTTCCGATCTNNNCTCTTGGCTCCCCATTCATCATCA

AGCACACGTCTGAACTCTTTCCCTACACGACGCTCTTCCGATCTNNNCTCTTGGCTCCCCATTCATCATCA

AGCACACGTCTGAACTCTTTCCCTACACGACGCTCTTCCGATCTNNNAAACTTACAGTCATAGCAGAAG

AGCACACGTCTGAACTCTTTCCCTACACGACGCTCTTCCGATCTNNNAAACTTACAGTCATAGCAGAAG

AGCACACGTCTGAACTCTTTCCCTACACGACGCTCTTCCGATCTNNNCTCCTGAGTAGCTGGGACTACA

AGCACACGTCTGAACTCTTTCCCTACACGACGCTCTTCCGATCTNNNCTCCTGAGTAGCTGGGACTACA

GAGCACACGTCTGAACTCTTTCCCTACACGACGCTCTTCCGATCTNNNAGGCACAGACTCTTGTTCTCT

GAGCACACGTCTGAACTCTTTCCCTACACGACGCTCTTCCGATCTNNNAGGCACAGACTCTTGTTCTCT

AGCACACGTCTGAACTCTTTCCCTACACGACGCTCTTCCGATCTNNNTAATCACAGCTACTTGGGAAGG

AGCACACGTCTGAACTCTTTCCCTACACGACGCTCTTCCGATCTNNNTAATCACAGCTACTTGGGAAGG

AGAGCACACGTCTGAACTCTTTCCCTACACGACGCTCTTCCGATCTNNNGCCTGTAATCCCAGCTACTTG

AGAGCACACGTCTGAACTCTTTCCCTACACGACGCTCTTCCGATCTNNNGCCTGTAATCCCAGCTACTTG

AGCACACGTCTGAACTCTTTCCCTACACGACGCTCTTCCGATCTNNNTAGTACACACACACACACACACA

AGCACACGTCTGAACTCTTTCCCTACACGACGCTCTTCCGATCTNNNTAGTACACACACACACACACACA

GAGCACACGTCTGAACTCTTTCCCTACACGACGCTCTTCCGATCTNNNAAAAGTAGTCAGGCATGGTG

GAGCACACGTCTGAACTCTTTCCCTACACGACGCTCTTCCGATCTNNNAAAAGTAGTCAGGCATGGTG

AGCACACGTCTGAACTCTTTCCCTACACGACGCTCTTCCGATCTNNNTGTTCCACTGTTTGTTTTGGAGC

AGCACACGTCTGAACTCTTTCCCTACACGACGCTCTTCCGATCTNNNTGTTCCACTGTTTGTTTTGGAGC

AGCACACGTCTGAACTCTTTCCCTACACGACGCTCTTCCGATCTNNNTGGGAAGCTCCACCAAGAGATT

AGCACACGTCTGAACTCTTTCCCTACACGACGCTCTTCCGATCTNNNTGGGAAGCTCCACCAAGAGATT

AGCACACGTCTGAACTCTTTCCCTACACGACGCTCTTCCGATCTNNNTTAGGCAGGAGAATCGCTTGA

AGCACACGTCTGAACTCTTTCCCTACACGACGCTCTTCCGATCTNNNTTAGGCAGGAGAATCGCTTGA

AGCACACGTCTGAACTCTTTCCCTACACGACGCTCTTCCGATCTNNNCCGTCGGTGGCTTTTTTACAAAA

AGCACACGTCTGAACTCTTTCCCTACACGACGCTCTTCCGATCTNNNCCGTCGGTGGCTTTTTTACAAAA

GAGCACACGTCTGAACTCTTTCCCTACACGACGCTCTTCCGATCTNNNGCTACTCAAACCTTCCTCACAGA

AGCACACGTCTGAACTCTTTCCCTACACGACGCTCTTCCGATCTNNNGCTACTCAAACCTTCCTCACAG
AGCACACGTCTGAACTCTTTCCCTACACGACGCTCTTCCGATCTNNNTGCAAGAAACTGAATTCTGCCA
AGCACACGTCTGAACTCTTTCCCTACACGACGCTCTTCCGATCTNNNTGCAAGAAACTGAATTCTGCCA
AGCACACGTCTGAACTCTTTCCCTACACGACGCTCTTCCGATCTNNNAGACAATTAGTGGAAGGAGC
AGCACACGTCTGAACTCTTTCCCTACACGACGCTCTTCCGATCTNNNAGACAATTAGTGGAAGGAGC
GCACACGTCTGAACTCTTTCCCTACACGACGCTCTTCCGATCTNNNGCAGAAGAATCACTTGAACCCA
GCACACGTCTGAACTCTTTCCCTACACGACGCTCTTCCGATCTNNNGCAGAAGAATCACTTGAACCCA
AGCACACGTCTGAACTCTTTCCCTACACGACGCTCTTCCGATCTNNNAAAGAAAGAGAGAGAAGCGA
AGCACACGTCTGAACTCTTTCCCTACACGACGCTCTTCCGATCTNNNAAAGAAAGAGAGAGAAGCGA
AGCACACGTCTGAACTCTTTCCCTACACGACGCTCTTCCGATCTNNNTGAGCTATAAATGGGCCCCTTT
AGCACACGTCTGAACTCTTTCCCTACACGACGCTCTTCCGATCTNNNTGAGCTATAAATGGGCCCCTTT
GCACACGTCTGAACTCTTTCCCTACACGACGCTCTTCCGATCTNNNAAAGGCCACCACAAAGTCTATC
GCACACGTCTGAACTCTTTCCCTACACGACGCTCTTCCGATCTNNNAAAGGCCACCACAAAGTCTATC
GAGCACACGTCTGAACTCTTTCCCTACACGACGCTCTTCCGATCTNNNAAAGCCCATATTCAAGCAG
GAGCACACGTCTGAACTCTTTCCCTACACGACGCTCTTCCGATCTNNNAAAGCCCATATTCAAGCAG
CACACGTCTGAACTCTTTCCCTACACGACGCTCTTCCGATCTNNNTGTAGTCCCAGTTACTTGTGGTT
CACACGTCTGAACTCTTTCCCTACACGACGCTCTTCCGATCTNNNTGTAGTCCCAGTTACTTGTGGTT
GCACACGTCTGAACTCTTTCCCTACACGACGCTCTTCCGATCTNNNGATGGTCTTGATCTCCTGACCTG
GCACACGTCTGAACTCTTTCCCTACACGACGCTCTTCCGATCTNNNGATGGTCTTGATCTCCTGACCTG
AGCACACGTCTGAACTCTTTCCCTACACGACGCTCTTCCGATCTNNNACCTTGGCAGAGTGAATTCGT
AGCACACGTCTGAACTCTTTCCCTACACGACGCTCTTCCGATCTNNNACCTTGGCAGAGTGAATTCGT
AGCACACGTCTGAACTCTTTCCCTACACGACGCTCTTCCGATCTNNNTGCACCCAGTAACTCGTCATTT
AGCACACGTCTGAACTCTTTCCCTACACGACGCTCTTCCGATCTNNNTGCACCCAGTAACTCGTCATTT
GAGCACACGTCTGAACTCTTTCCCTACACGACGCTCTTCCGATCTNNNGTACAAGCTACTTGAGAGGC
GAGCACACGTCTGAACTCTTTCCCTACACGACGCTCTTCCGATCTNNNGTACAAGCTACTTGAGAGGC
GAGCACACGTCTGAACTCTTTCCCTACACGACGCTCTTCCGATCTNNNAATTATTTGGGGGTAAAGGG
GAGCACACGTCTGAACTCTTTCCCTACACGACGCTCTTCCGATCTNNNAATTATTTGGGGGTAAAGGG
GAGCACACGTCTGAACTCTTTCCCTACACGACGCTCTTCCGATCTNNNTCTCCAATTCTTCCTCTCCCTCT
GAGCACACGTCTGAACTCTTTCCCTACACGACGCTCTTCCGATCTNNNTCTCCAATTCTTCCTCTCCCTCT
CACGTCTGAACTCTTTCCCTACACGACGCTCTTCCGATCTNNNCTTTCCTTAGCCTCCTGAGTAGCCAT
CACGTCTGAACTCTTTCCCTACACGACGCTCTTCCGATCTNNNCTTTCCTTAGCCTCCTGAGTAGCCAT
CACACGTCTGAACTCTTTCCCTACACGACGCTCTTCCGATCTNNNACCTCTTCTTCCTCCTCGTTTTCCAT
CACACGTCTGAACTCTTTCCCTACACGACGCTCTTCCGATCTNNNACCTCTTCTTCCTCCTCGTTTTCCAT
AGCACACGTCTGAACTCTTTCCCTACACGACGCTCTTCCGATCTNNNTGTGTGAATTTCTCCTGTAGG
AGCACACGTCTGAACTCTTTCCCTACACGACGCTCTTCCGATCTNNNTGTGTGAATTTCTCCTGTAGG
GAGCACACGTCTGAACTCTTTCCCTACACGACGCTCTTCCGATCTNNNTAGATGTGGCTCGCCAATTA
GAGCACACGTCTGAACTCTTTCCCTACACGACGCTCTTCCGATCTNNNTAGATGTGGCTCGCCAATTA
GCACACGTCTGAACTCTTTCCCTACACGACGCTCTTCCGATCTNNNTTGTGCACCTTTCATTCTTGGTC
GCACACGTCTGAACTCTTTCCCTACACGACGCTCTTCCGATCTNNNTTGTGCACCTTTCATTCTTGGTC
AGCACACGTCTGAACTCTTTCCCTACACGACGCTCTTCCGATCTNNNTGTTCATCTGGGAGTTCATCTG
AGCACACGTCTGAACTCTTTCCCTACACGACGCTCTTCCGATCTNNNTGTTCATCTGGGAGTTCATCTG
GAGCACACGTCTGAACTCTTTCCCTACACGACGCTCTTCCGATCTNNNAGCCCACTGAGAATATACCCT
GAGCACACGTCTGAACTCTTTCCCTACACGACGCTCTTCCGATCTNNNAGCCCACTGAGAATATACCCT

AGCACACGTCTGAACTCTTTCCCTACACGACGCTCTTCCGATCTNNNCCACTCCTAGATTACTCGAACCG
AGCACACGTCTGAACTCTTTCCCTACACGACGCTCTTCCGATCTNNNCCACTCCTAGATTACTCGAACCG
GCACACGTCTGAACTCTTTCCCTACACGACGCTCTTCCGATCTNNNTTTTGCTTTCAGAAATCAACCCAC
GCACACGTCTGAACTCTTTCCCTACACGACGCTCTTCCGATCTNNNTTTTGCTTTCAGAAATCAACCCAC
AGCACACGTCTGAACTCTTTCCCTACACGACGCTCTTCCGATCTNNNCAATGCCCGGCTAATTTTCTGT
AGCACACGTCTGAACTCTTTCCCTACACGACGCTCTTCCGATCTNNNCAATGCCCGGCTAATTTTCTGT
AGCACACGTCTGAACTCTTTCCCTACACGACGCTCTTCCGATCTNNNTCCTTCCCCCACATTTCATGAAA
AGCACACGTCTGAACTCTTTCCCTACACGACGCTCTTCCGATCTNNNTCCTTCCCCCACATTTCATGAAA
GAGCACACGTCTGAACTCTTTCCCTACACGACGCTCTTCCGATCTNNNCTGAGGCAGGAGAATCACTTG
GAGCACACGTCTGAACTCTTTCCCTACACGACGCTCTTCCGATCTNNNCTGAGGCAGGAGAATCACTTG
AGCACACGTCTGAACTCTTTCCCTACACGACGCTCTTCCGATCTNNNCCTGCCTTTAGTCTTGGCTTAGA
AGCACACGTCTGAACTCTTTCCCTACACGACGCTCTTCCGATCTNNNCCTGCCTTTAGTCTTGGCTTAGA
GAGCACACGTCTGAACTCTTTCCCTACACGACGCTCTTCCGATCTNNNACAGCAACATCCCACTTCTAGT
GAGCACACGTCTGAACTCTTTCCCTACACGACGCTCTTCCGATCTNNNACAGCAACATCCCACTTCTAGT
AGCACACGTCTGAACTCTTTCCCTACACGACGCTCTTCCGATCTNNNGCCTGTAGTCCCAGCTACTTGC
AGCACACGTCTGAACTCTTTCCCTACACGACGCTCTTCCGATCTNNNGCCTGTAGTCCCAGCTACTTGC
AGCACACGTCTGAACTCTTTCCCTACACGACGCTCTTCCGATCTNNNGATTGTAGCTACTCAGGAGGC
AGCACACGTCTGAACTCTTTCCCTACACGACGCTCTTCCGATCTNNNGATTGTAGCTACTCAGGAGGC
GCACACGTCTGAACTCTTTCCCTACACGACGCTCTTCCGATCTNNNTCACCATTGCGCATGAAGTAATC
GCACACGTCTGAACTCTTTCCCTACACGACGCTCTTCCGATCTNNNTCACCATTGCGCATGAAGTAATC
AGAGCACACGTCTGAACTCTTTCCCTACACGACGCTCTTCCGATCTNNNGCTTAGGGATTCTTAGTTGG
AGAGCACACGTCTGAACTCTTTCCCTACACGACGCTCTTCCGATCTNNNGCTTAGGGATTCTTAGTTGG
GAGCACACGTCTGAACTCTTTCCCTACACGACGCTCTTCCGATCTNNNGTAATCCTACTTGGGAGGCTG
GAGCACACGTCTGAACTCTTTCCCTACACGACGCTCTTCCGATCTNNNGTAATCCTACTTGGGAGGCTG
GAGCACACGTCTGAACTCTTTCCCTACACGACGCTCTTCCGATCTNNNAAAGTCAGGTTCTGCCAGTGA
GAGCACACGTCTGAACTCTTTCCCTACACGACGCTCTTCCGATCTNNNAAAGTCAGGTTCTGCCAGTGA
AGCACACGTCTGAACTCTTTCCCTACACGACGCTCTTCCGATCTNNNCTAATGTTGGCAAAAATTGAAT
AGCACACGTCTGAACTCTTTCCCTACACGACGCTCTTCCGATCTNNNCTAATGTTGGCAAAAATTGAAT
AGCACACGTCTGAACTCTTTCCCTACACGACGCTCTTCCGATCTNNNCTGGGCAACAAGAGCAAAACTC
AGCACACGTCTGAACTCTTTCCCTACACGACGCTCTTCCGATCTNNNCTGGGCAACAAGAGCAAAACTC
AGCACACGTCTGAACTCTTTCCCTACACGACGCTCTTCCGATCTNNNCTTCTTCGCACTAGACAATGAG
AGCACACGTCTGAACTCTTTCCCTACACGACGCTCTTCCGATCTNNNCTTCTTCGCACTAGACAATGAG
GAGCACACGTCTGAACTCTTTCCCTACACGACGCTCTTCCGATCTNNNACCAACATGATGAAACCCTGT
GAGCACACGTCTGAACTCTTTCCCTACACGACGCTCTTCCGATCTNNNACCAACATGATGAAACCCTGT
GAGCACACGTCTGAACTCTTTCCCTACACGACGCTCTTCCGATCTNNNTTCAGGGATTCCACTTCTTCCT
GAGCACACGTCTGAACTCTTTCCCTACACGACGCTCTTCCGATCTNNNTTCAGGGATTCCACTTCTTCCT
GAGCACACGTCTGAACTCTTTCCCTACACGACGCTCTTCCGATCTNNNTTGACTTTTTAATGACATGAG
GAGCACACGTCTGAACTCTTTCCCTACACGACGCTCTTCCGATCTNNNTTGACTTTTTAATGACATGAG
GAGCACACGTCTGAACTCTTTCCCTACACGACGCTCTTCCGATCTNNNCAGAAGGTTTTGCTATATGTA
GAGCACACGTCTGAACTCTTTCCCTACACGACGCTCTTCCGATCTNNNCAGAAGGTTTTGCTATATGTA
AGCACACGTCTGAACTCTTTCCCTACACGACGCTCTTCCGATCTNNNCACTTGAGTCCAGGAGTTCAAG
AGCACACGTCTGAACTCTTTCCCTACACGACGCTCTTCCGATCTNNNCACTTGAGTCCAGGAGTTCAAG
AGCACACGTCTGAACTCTTTCCCTACACGACGCTCTTCCGATCTNNNCTCCTGGGTTCAAGAGATTCTC

AGCACACGTCTGAACTCTTTCCCTACACGACGCTCTTCCGATCTNNNCTCCTGGGTTCAAGAGATTCTC
GAGCACACGTCTGAACTCTTTCCCTACACGACGCTCTTCCGATCTNNNCTGCCCAAAACAGGAAAAGTC
GAGCACACGTCTGAACTCTTTCCCTACACGACGCTCTTCCGATCTNNNCTGCCCAAAACAGGAAAAGTC
AGCACACGTCTGAACTCTTTCCCTACACGACGCTCTTCCGATCTNNNCACTTCTGAGGGGATAGTGTCA
AGCACACGTCTGAACTCTTTCCCTACACGACGCTCTTCCGATCTNNNCACTTCTGAGGGGATAGTGTCA
AGCACACGTCTGAACTCTTTCCCTACACGACGCTCTTCCGATCTNNNTTCCATTGTATGTCTATATCACA
AGCACACGTCTGAACTCTTTCCCTACACGACGCTCTTCCGATCTNNNTTCCATTGTATGTCTATATCACA
GAGCACACGTCTGAACTCTTTCCCTACACGACGCTCTTCCGATCTNNNACTGTGTTATCTGTCTTGAGAA
GAGCACACGTCTGAACTCTTTCCCTACACGACGCTCTTCCGATCTNNNACTGTGTTATCTGTCTTGAGAA
GAGCACACGTCTGAACTCTTTCCCTACACGACGCTCTTCCGATCTNNNGTCAATTGCAACATGGAGCA
GAGCACACGTCTGAACTCTTTCCCTACACGACGCTCTTCCGATCTNNNGTCAATTGCAACATGGAGCA
AGCACACGTCTGAACTCTTTCCCTACACGACGCTCTTCCGATCTNNNACTCAATCCCCTTTACAACAGCT
AGCACACGTCTGAACTCTTTCCCTACACGACGCTCTTCCGATCTNNNACTCAATCCCCTTTACAACAGCT
GAGCACACGTCTGAACTCTTTCCCTACACGACGCTCTTCCGATCTNNNCAGTGCCACTGGTCTATAATC
GAGCACACGTCTGAACTCTTTCCCTACACGACGCTCTTCCGATCTNNNCAGTGCCACTGGTCTATAATC
GAGCACACGTCTGAACTCTTTCCCTACACGACGCTCTTCCGATCTNNNTGAGGTAGGAGAATGGCATG
GAGCACACGTCTGAACTCTTTCCCTACACGACGCTCTTCCGATCTNNNTGAGGTAGGAGAATGGCATG
AGCACACGTCTGAACTCTTTCCCTACACGACGCTCTTCCGATCTNNNCTGGGTCCTTTCCTTCAAGAGAG
AGCACACGTCTGAACTCTTTCCCTACACGACGCTCTTCCGATCTNNNCTGGGTCCTTTCCTTCAAGAGAG
GAGCACACGTCTGAACTCTTTCCCTACACGACGCTCTTCCGATCTNNNACTCACCTGACAACCTTTGACA
GAGCACACGTCTGAACTCTTTCCCTACACGACGCTCTTCCGATCTNNNACTCACCTGACAACCTTTGACA
GAGCACACGTCTGAACTCTTTCCCTACACGACGCTCTTCCGATCTNNNGGGTTGCATTGAGTCACAAT
GAGCACACGTCTGAACTCTTTCCCTACACGACGCTCTTCCGATCTNNNGGGTTGCATTGAGTCACAAT
AGCACACGTCTGAACTCTTTCCCTACACGACGCTCTTCCGATCTNNNCACGTTTGACCCCAGGTAATAG
AGCACACGTCTGAACTCTTTCCCTACACGACGCTCTTCCGATCTNNNCACGTTTGACCCCAGGTAATAG
GAGCACACGTCTGAACTCTTTCCCTACACGACGCTCTTCCGATCTNNNGCAACAAGAACAAAACTCCGT
GAGCACACGTCTGAACTCTTTCCCTACACGACGCTCTTCCGATCTNNNGCAACAAGAACAAAACTCCGT
AGAGCACACGTCTGAACTCTTTCCCTACACGACGCTCTTCCGATCTNNNTCTCTTGCCTAATTGCTATGG
AGAGCACACGTCTGAACTCTTTCCCTACACGACGCTCTTCCGATCTNNNTCTCTTGCCTAATTGCTATGG
GAGCACACGTCTGAACTCTTTCCCTACACGACGCTCTTCCGATCTNNNATGGCTAAAATCACACACACA
GAGCACACGTCTGAACTCTTTCCCTACACGACGCTCTTCCGATCTNNNATGGCTAAAATCACACACACA
AGAGCACACGTCTGAACTCTTTCCCTACACGACGCTCTTCCGATCTNNNGAAACGCAAAGTCTCAGGT
AGAGCACACGTCTGAACTCTTTCCCTACACGACGCTCTTCCGATCTNNNGAAACGCAAAGTCTCAGGT
AGCACACGTCTGAACTCTTTCCCTACACGACGCTCTTCCGATCTNNNGAGGAGTAGAAACAGTCCAGC
AGCACACGTCTGAACTCTTTCCCTACACGACGCTCTTCCGATCTNNNGAGGAGTAGAAACAGTCCAGC
GAGCACACGTCTGAACTCTTTCCCTACACGACGCTCTTCCGATCTNNNGAAGCAGGAGAATCGCTTGA
GAGCACACGTCTGAACTCTTTCCCTACACGACGCTCTTCCGATCTNNNGAAGCAGGAGAATCGCTTGA
AGCACACGTCTGAACTCTTTCCCTACACGACGCTCTTCCGATCTNNNCCACTTGGGTTACAGGATCATT
AGCACACGTCTGAACTCTTTCCCTACACGACGCTCTTCCGATCTNNNCCACTTGGGTTACAGGATCATT
GAGCACACGTCTGAACTCTTTCCCTACACGACGCTCTTCCGATCTNNNGATTTGCTCTAGATGGTGGGA
GAGCACACGTCTGAACTCTTTCCCTACACGACGCTCTTCCGATCTNNNGATTTGCTCTAGATGGTGGGA
AGCACACGTCTGAACTCTTTCCCTACACGACGCTCTTCCGATCTNNNCTCTCCCCATCTCTCTCTCTCTTCA
AGCACACGTCTGAACTCTTTCCCTACACGACGCTCTTCCGATCTNNNCTCTCCCCATCTCTCTCTCTCTTCA

AGCACACGTCTGAACTCTTTCCCTACACGACGCTCTTCCGATCTNNNTCCCCTTTGTCCTTTTTATTTGTG
AGCACACGTCTGAACTCTTTCCCTACACGACGCTCTTCCGATCTNNNTCCCCTTTGTCCTTTTTATTTGTG
AGCACACGTCTGAACTCTTTCCCTACACGACGCTCTTCCGATCTNNNTCCCAAGCAAAAAACACAGTTG
AGCACACGTCTGAACTCTTTCCCTACACGACGCTCTTCCGATCTNNNTCCCAAGCAAAAAACACAGTTG
GAGCACACGTCTGAACTCTTTCCCTACACGACGCTCTTCCGATCTNNNGAGGAAACTGGCCTGGTATC
GAGCACACGTCTGAACTCTTTCCCTACACGACGCTCTTCCGATCTNNNGAGGAAACTGGCCTGGTATC
AGCACACGTCTGAACTCTTTCCCTACACGACGCTCTTCCGATCTNNNGAGTAGCTGGTACTACAGGCA
AGCACACGTCTGAACTCTTTCCCTACACGACGCTCTTCCGATCTNNNGAGTAGCTGGTACTACAGGCA
AGCACACGTCTGAACTCTTTCCCTACACGACGCTCTTCCGATCTNNNCGCATCAGTTTCCTGTAATTGG
AGCACACGTCTGAACTCTTTCCCTACACGACGCTCTTCCGATCTNNNCGCATCAGTTTCCTGTAATTGG
AGCACACGTCTGAACTCTTTCCCTACACGACGCTCTTCCGATCTNNNATTTTGGGTAGAGACAGAGCC
AGCACACGTCTGAACTCTTTCCCTACACGACGCTCTTCCGATCTNNNATTTTGGGTAGAGACAGAGCC
GCACACGTCTGAACTCTTTCCCTACACGACGCTCTTCCGATCTNNNGGCCCAAAAGTACACGTGAATA
GCACACGTCTGAACTCTTTCCCTACACGACGCTCTTCCGATCTNNNGGCCCAAAAGTACACGTGAATA
AGCACACGTCTGAACTCTTTCCCTACACGACGCTCTTCCGATCTNNNCCCACAAACCTGTCATGAGAAA
AGCACACGTCTGAACTCTTTCCCTACACGACGCTCTTCCGATCTNNNCCCACAAACCTGTCATGAGAAA
AGCACACGTCTGAACTCTTTCCCTACACGACGCTCTTCCGATCTNNNCCTGACCAAAATCAATCTCGCAA
AGCACACGTCTGAACTCTTTCCCTACACGACGCTCTTCCGATCTNNNCCTGACCAAAATCAATCTCGCAA
AAGAGCACACGTCTGAACTCTTTCCCTACACGACGCTCTTCCGATCTNNNAGGGAGAGAAAGAAAGA
AAGAGCACACGTCTGAACTCTTTCCCTACACGACGCTCTTCCGATCTNNNAGGGAGAGAAAGAAAGA
GAGCACACGTCTGAACTCTTTCCCTACACGACGCTCTTCCGATCTNNNAGAATCGCTTGAACCTGGGA
GAGCACACGTCTGAACTCTTTCCCTACACGACGCTCTTCCGATCTNNNAGAATCGCTTGAACCTGGGA
AGCACACGTCTGAACTCTTTCCCTACACGACGCTCTTCCGATCTNNNCTTTCTTTCTCCTTCCCTCCCTCC
AGCACACGTCTGAACTCTTTCCCTACACGACGCTCTTCCGATCTNNNCTTTCTTTCTCCTTCCCTCCCTCC
CACGTCTGAACTCTTTCCCTACACGACGCTCTTCCGATCTNNNGATCAAAGGGGAATACTGGGGAAC
CACGTCTGAACTCTTTCCCTACACGACGCTCTTCCGATCTNNNGATCAAAGGGGAATACTGGGGAAC
GAGCACACGTCTGAACTCTTTCCCTACACGACGCTCTTCCGATCTNNNTACACCACTTTCTCAGCAGGAT
GAGCACACGTCTGAACTCTTTCCCTACACGACGCTCTTCCGATCTNNNTACACCACTTTCTCAGCAGGAT
GAGCACACGTCTGAACTCTTTCCCTACACGACGCTCTTCCGATCTNNNTTCCGGGCTTAGAAAGTTAG
GAGCACACGTCTGAACTCTTTCCCTACACGACGCTCTTCCGATCTNNNTTCCGGGCTTAGAAAGTTAG
GAGCACACGTCTGAACTCTTTCCCTACACGACGCTCTTCCGATCTNNNAAGCTCACTATTGTCCTCTGCA
GAGCACACGTCTGAACTCTTTCCCTACACGACGCTCTTCCGATCTNNNAAGCTCACTATTGTCCTCTGCA
AGCACACGTCTGAACTCTTTCCCTACACGACGCTCTTCCGATCTNNNACTTGACCAGCTCAACATTAGGT
AGCACACGTCTGAACTCTTTCCCTACACGACGCTCTTCCGATCTNNNACTTGACCAGCTCAACATTAGGT
AGAGCACACGTCTGAACTCTTTCCCTACACGACGCTCTTCCGATCTNNNAGAGAAATCTGTCTAAGCTC
AGAGCACACGTCTGAACTCTTTCCCTACACGACGCTCTTCCGATCTNNNAGAGAAATCTGTCTAAGCTC
GAGCACACGTCTGAACTCTTTCCCTACACGACGCTCTTCCGATCTNNNATTTGGTAGAGAGAAAGAGG
GAGCACACGTCTGAACTCTTTCCCTACACGACGCTCTTCCGATCTNNNATTTGGTAGAGAGAAAGAGG
AGCACACGTCTGAACTCTTTCCCTACACGACGCTCTTCCGATCTNNNCTGCTCTCCATTTTCTACCCAGAC
AGCACACGTCTGAACTCTTTCCCTACACGACGCTCTTCCGATCTNNNCTGCTCTCCATTTTCTACCCAGAC
AGCACACGTCTGAACTCTTTCCCTACACGACGCTCTTCCGATCTNNNGATCAACTCTGAAACATACATA
AGCACACGTCTGAACTCTTTCCCTACACGACGCTCTTCCGATCTNNNGATCAACTCTGAAACATACATA
GAGCACACGTCTGAACTCTTTCCCTACACGACGCTCTTCCGATCTNNNAATGCCCTTGTTTATTGCCTGT

AGCACACGTCTGAACTCTTTCCCTACACGACGCTCTTCCGATCTNNNAATGCCCTTGTTTATTGCCTGT
CACACGTCTGAACTCTTTCCCTACACGACGCTCTTCCGATCTNNNATTTGGTACCAGACACTGAGCTTC
CACACGTCTGAACTCTTTCCCTACACGACGCTCTTCCGATCTNNNATTTGGTACCAGACACTGAGCTTC
AGAGCACACGTCTGAACTCTTTCCCTACACGACGCTCTTCCGATCTNNNTCTGCTTTATACTCTCCTAA
AGAGCACACGTCTGAACTCTTTCCCTACACGACGCTCTTCCGATCTNNNTCTGCTTTATACTCTCCTAA
AGCACACGTCTGAACTCTTTCCCTACACGACGCTCTTCCGATCTNNNATGTACAACCTCTTCCCAAACCC
AGCACACGTCTGAACTCTTTCCCTACACGACGCTCTTCCGATCTNNNATGTACAACCTCTTCCCAAACCC
AGCACACGTCTGAACTCTTTCCCTACACGACGCTCTTCCGATCTNNNGTACCTCTTCCTCTGTGTGCTGC
AGCACACGTCTGAACTCTTTCCCTACACGACGCTCTTCCGATCTNNNGTACCTCTTCCTCTGTGTGCTGC
GCACACGTCTGAACTCTTTCCCTACACGACGCTCTTCCGATCTNNNCGTCTGTACTAGGTACCCAAAG
GCACACGTCTGAACTCTTTCCCTACACGACGCTCTTCCGATCTNNNCGTCTGTACTAGGTACCCAAAG
AGCACACGTCTGAACTCTTTCCCTACACGACGCTCTTCCGATCTNNNTTCAAGCAATTTTCCTGCCTCAG
AGCACACGTCTGAACTCTTTCCCTACACGACGCTCTTCCGATCTNNNTTCAAGCAATTTTCCTGCCTCAG
GAGCACACGTCTGAACTCTTTCCCTACACGACGCTCTTCCGATCTNNNGCTTGTAATCCTAGCTACTCG
GAGCACACGTCTGAACTCTTTCCCTACACGACGCTCTTCCGATCTNNNGCTTGTAATCCTAGCTACTCG
GAGCACACGTCTGAACTCTTTCCCTACACGACGCTCTTCCGATCTNNNGCAGTGAATTCCCATTGCACT
GAGCACACGTCTGAACTCTTTCCCTACACGACGCTCTTCCGATCTNNNGCAGTGAATTCCCATTGCACT
GAGCACACGTCTGAACTCTTTCCCTACACGACGCTCTTCCGATCTNNNTTCAAGTGATTCTCCTGCCTC
GAGCACACGTCTGAACTCTTTCCCTACACGACGCTCTTCCGATCTNNNTTCAAGTGATTCTCCTGCCTC
GAGCACACGTCTGAACTCTTTCCCTACACGACGCTCTTCCGATCTNNNTTTTGGAGATTGAGGCAAG
GAGCACACGTCTGAACTCTTTCCCTACACGACGCTCTTCCGATCTNNNTTTTGGAGATTGAGGCAAG
AGCACACGTCTGAACTCTTTCCCTACACGACGCTCTTCCGATCTNNNAGAATTGCTTGAACCTGGGAGT
AGCACACGTCTGAACTCTTTCCCTACACGACGCTCTTCCGATCTNNNAGAATTGCTTGAACCTGGGAGT
AGAGCACACGTCTGAACTCTTTCCCTACACGACGCTCTTCCGATCTNNNGAGTAGGATTGGTAGAAG
AGAGCACACGTCTGAACTCTTTCCCTACACGACGCTCTTCCGATCTNNNGAGTAGGATTGGTAGAAG
GAGCACACGTCTGAACTCTTTCCCTACACGACGCTCTTCCGATCTNNNATAGTTACTCTTCCCCATCCC
GAGCACACGTCTGAACTCTTTCCCTACACGACGCTCTTCCGATCTNNNATAGTTACTCTTCCCCATCCC
GCACACGTCTGAACTCTTTCCCTACACGACGCTCTTCCGATCTNNNACCATTTTATGCCCTCAGTAGCA
GCACACGTCTGAACTCTTTCCCTACACGACGCTCTTCCGATCTNNNACCATTTTATGCCCTCAGTAGCA
GAGCACACGTCTGAACTCTTTCCCTACACGACGCTCTTCCGATCTNNNCAGGGGTGTGTATGGTTTTT
GAGCACACGTCTGAACTCTTTCCCTACACGACGCTCTTCCGATCTNNNCAGGGGTGTGTATGGTTTTT
GAGCACACGTCTGAACTCTTTCCCTACACGACGCTCTTCCGATCTNNNCTGTAGTCCCAGGTACTCAGG
GAGCACACGTCTGAACTCTTTCCCTACACGACGCTCTTCCGATCTNNNCTGTAGTCCCAGGTACTCAGG
AGCACACGTCTGAACTCTTTCCCTACACGACGCTCTTCCGATCTNNNAGTAGTTGGGACTAAGGTGC
AGCACACGTCTGAACTCTTTCCCTACACGACGCTCTTCCGATCTNNNAGTAGTTGGGACTAAGGTGC
AGAGCACACGTCTGAACTCTTTCCCTACACGACGCTCTTCCGATCTNNNGATCACTTGATGCCAGGAG
AGAGCACACGTCTGAACTCTTTCCCTACACGACGCTCTTCCGATCTNNNGATCACTTGATGCCAGGAG
GAGCACACGTCTGAACTCTTTCCCTACACGACGCTCTTCCGATCTNNNCATGCCTGTAATCCCAGCTAC
GAGCACACGTCTGAACTCTTTCCCTACACGACGCTCTTCCGATCTNNNCATGCCTGTAATCCCAGCTAC
AGCACACGTCTGAACTCTTTCCCTACACGACGCTCTTCCGATCTNNNATAAGCTAATTTTGGCTGGGT
AGCACACGTCTGAACTCTTTCCCTACACGACGCTCTTCCGATCTNNNATAAGCTAATTTTGGCTGGGT
AGCACACGTCTGAACTCTTTCCCTACACGACGCTCTTCCGATCTNNNGAGGCAGGAGAATCGCTTGAA
AGCACACGTCTGAACTCTTTCCCTACACGACGCTCTTCCGATCTNNNGAGGCAGGAGAATCGCTTGAA

AGCACACGTCTGAACTCTTTCCCTACACGACGCTCTTCCGATCTNNNGTTGGGAAGGGTTTCAATAGA
AGCACACGTCTGAACTCTTTCCCTACACGACGCTCTTCCGATCTNNNGTTGGGAAGGGTTTCAATAGA
GAGCACACGTCTGAACTCTTTCCCTACACGACGCTCTTCCGATCTNNNGAGGTTGCAGTGAGCTGAGA
GAGCACACGTCTGAACTCTTTCCCTACACGACGCTCTTCCGATCTNNNGAGGTTGCAGTGAGCTGAGA
CACACGTCTGAACTCTTTCCCTACACGACGCTCTTCCGATCTNNNTCCCTCTGGATAGACTTCCAAGTCA
CACACGTCTGAACTCTTTCCCTACACGACGCTCTTCCGATCTNNNTCCCTCTGGATAGACTTCCAAGTCA
AGCACACGTCTGAACTCTTTCCCTACACGACGCTCTTCCGATCTNNNAGAGCATGGGGAAGTCTTTTTC
AGCACACGTCTGAACTCTTTCCCTACACGACGCTCTTCCGATCTNNNAGAGCATGGGGAAGTCTTTTTC
AGAGCACACGTCTGAACTCTTTCCCTACACGACGCTCTTCCGATCTNNNGTGCTCATATCCTCAGCTACT
AGAGCACACGTCTGAACTCTTTCCCTACACGACGCTCTTCCGATCTNNNGTGCTCATATCCTCAGCTACT
ACACGTCTGAACTCTTTCCCTACACGACGCTCTTCCGATCTNNNGAAGGGTCTTTCTCTGTCTCCTGCAT
ACACGTCTGAACTCTTTCCCTACACGACGCTCTTCCGATCTNNNGAAGGGTCTTTCTCTGTCTCCTGCAT
GCACACGTCTGAACTCTTTCCCTACACGACGCTCTTCCGATCTNNNGCAGACTCACCATGCATTCTACAC
GCACACGTCTGAACTCTTTCCCTACACGACGCTCTTCCGATCTNNNGCAGACTCACCATGCATTCTACAC
GAGCACACGTCTGAACTCTTTCCCTACACGACGCTCTTCCGATCTNNNCACCACCATGCCTGACTAATTTT
GAGCACACGTCTGAACTCTTTCCCTACACGACGCTCTTCCGATCTNNNCACCACCATGCCTGACTAATTTT
AGCACACGTCTGAACTCTTTCCCTACACGACGCTCTTCCGATCTNNNCTGTAATCCCAGCTACTCAGGA
AGCACACGTCTGAACTCTTTCCCTACACGACGCTCTTCCGATCTNNNCTGTAATCCCAGCTACTCAGGA
AGCACACGTCTGAACTCTTTCCCTACACGACGCTCTTCCGATCTNNNGCAACAGAGTGAGACTCCATCT
AGCACACGTCTGAACTCTTTCCCTACACGACGCTCTTCCGATCTNNNGCAACAGAGTGAGACTCCATCT
GAGCACACGTCTGAACTCTTTCCCTACACGACGCTCTTCCGATCTNNNAGTACACGGACACATCCTCAA
GAGCACACGTCTGAACTCTTTCCCTACACGACGCTCTTCCGATCTNNNAGTACACGGACACATCCTCAA
GAGCACACGTCTGAACTCTTTCCCTACACGACGCTCTTCCGATCTNNNTAATTTCTAAGGCAAGAATCTT
GAGCACACGTCTGAACTCTTTCCCTACACGACGCTCTTCCGATCTNNNTAATTTCTAAGGCAAGAATCTT
AGAGCACACGTCTGAACTCTTTCCCTACACGACGCTCTTCCGATCTNNNCCCAAGATTGCCTTCATTCCA
AGAGCACACGTCTGAACTCTTTCCCTACACGACGCTCTTCCGATCTNNNCCCAAGATTGCCTTCATTCCA
AGCACACGTCTGAACTCTTTCCCTACACGACGCTCTTCCGATCTNNNCATGAGGGTATAAGTCTGGGCT
AGCACACGTCTGAACTCTTTCCCTACACGACGCTCTTCCGATCTNNNCATGAGGGTATAAGTCTGGGCT
GAGCACACGTCTGAACTCTTTCCCTACACGACGCTCTTCCGATCTNNNAGAAGAATGGTGTGAACCCA
GAGCACACGTCTGAACTCTTTCCCTACACGACGCTCTTCCGATCTNNNAGAAGAATGGTGTGAACCCA
GAGCACACGTCTGAACTCTTTCCCTACACGACGCTCTTCCGATCTNNNTAACACTGCCTTGCTTGCTACTA
GAGCACACGTCTGAACTCTTTCCCTACACGACGCTCTTCCGATCTNNNTAACACTGCCTTGCTTGCTACTA
GAGCACACGTCTGAACTCTTTCCCTACACGACGCTCTTCCGATCTNNNGGCCATCAAGGTGGTACTTCT
GAGCACACGTCTGAACTCTTTCCCTACACGACGCTCTTCCGATCTNNNGGCCATCAAGGTGGTACTTCT
GAGCACACGTCTGAACTCTTTCCCTACACGACGCTCTTCCGATCTNNNATGTCCCTCTGTCTCTCTCTCTC
GAGCACACGTCTGAACTCTTTCCCTACACGACGCTCTTCCGATCTNNNATGTCCCTCTGTCTCTCTCTCTC
GAGCACACGTCTGAACTCTTTCCCTACACGACGCTCTTCCGATCTNNNGCCAAATCTGAGGTGGCTATG
GAGCACACGTCTGAACTCTTTCCCTACACGACGCTCTTCCGATCTNNNGCCAAATCTGAGGTGGCTATG
GAGCACACGTCTGAACTCTTTCCCTACACGACGCTCTTCCGATCTNNNATCTCACTCTGTTGCACAGGTT
GAGCACACGTCTGAACTCTTTCCCTACACGACGCTCTTCCGATCTNNNATCTCACTCTGTTGCACAGGTT
AGCACACGTCTGAACTCTTTCCCTACACGACGCTCTTCCGATCTNNNGGCTTCAGAGAGAACAGGTTG
AGCACACGTCTGAACTCTTTCCCTACACGACGCTCTTCCGATCTNNNGGCTTCAGAGAGAACAGGTTG
GCACACGTCTGAACTCTTTCCCTACACGACGCTCTTCCGATCTNNNTATACTGACGTCTCCAACATCATC

GCACACGTCTGAACTCTTTCCCTACACGACGCTCTTCCGATCTNNNTATACTGACGTCTCCAACATCATC

AGCACACGTCTGAACTCTTTCCCTACACGACGCTCTTCCGATCTNNNAGGATTACAACTCTCCTGCTTTG

AGCACACGTCTGAACTCTTTCCCTACACGACGCTCTTCCGATCTNNNAGGATTACAACTCTCCTGCTTTG

GAGCACACGTCTGAACTCTTTCCCTACACGACGCTCTTCCGATCTNNNGTTTTCCTTGTGGAGGTCATT

GAGCACACGTCTGAACTCTTTCCCTACACGACGCTCTTCCGATCTNNNGTTTTCCTTGTGGAGGTCATT

AGCACACGTCTGAACTCTTTCCCTACACGACGCTCTTCCGATCTNNNATCACTATACTCTAGCCTGGGCA

AGCACACGTCTGAACTCTTTCCCTACACGACGCTCTTCCGATCTNNNATCACTATACTCTAGCCTGGGCA

GAGCACACGTCTGAACTCTTTCCCTACACGACGCTCTTCCGATCTNNNAGCCAAGATCACACCACTGTAA

GAGCACACGTCTGAACTCTTTCCCTACACGACGCTCTTCCGATCTNNNAGCCAAGATCACACCACTGTAA

AAGAGCACACGTCTGAACTCTTTCCCTACACGACGCTCTTCCGATCTNNNTGGCCATTGACTAGGTCAA

AAGAGCACACGTCTGAACTCTTTCCCTACACGACGCTCTTCCGATCTNNNTGGCCATTGACTAGGTCAA

GAGCACACGTCTGAACTCTTTCCCTACACGACGCTCTTCCGATCTNNNGAGATTGCACCATTGTACTCCA

GAGCACACGTCTGAACTCTTTCCCTACACGACGCTCTTCCGATCTNNNGAGATTGCACCATTGTACTCCA

AGCACACGTCTGAACTCTTTCCCTACACGACGCTCTTCCGATCTNNNTCATAGGCAATGGCATTTTGTAA

AGCACACGTCTGAACTCTTTCCCTACACGACGCTCTTCCGATCTNNNTCATAGGCAATGGCATTTTGTAA

AGCACACGTCTGAACTCTTTCCCTACACGACGCTCTTCCGATCTNNNCACTGTCAGCTTCTCTGGTTTTG

AGCACACGTCTGAACTCTTTCCCTACACGACGCTCTTCCGATCTNNNCACTGTCAGCTTCTCTGGTTTTG

GCACACGTCTGAACTCTTTCCCTACACGACGCTCTTCCGATCTNNNTTCAAGCAATTTTCCTGCCTCAGCA

GCACACGTCTGAACTCTTTCCCTACACGACGCTCTTCCGATCTNNNTTCAAGCAATTTTCCTGCCTCAGCA

GAGCACACGTCTGAACTCTTTCCCTACACGACGCTCTTCCGATCTNNNCTTCATGAGGAATGGAGGTA

GAGCACACGTCTGAACTCTTTCCCTACACGACGCTCTTCCGATCTNNNCTTCATGAGGAATGGAGGTA

AGCACACGTCTGAACTCTTTCCCTACACGACGCTCTTCCGATCTNNNTTTGCCCAGGTTGACATTGATTC

AGCACACGTCTGAACTCTTTCCCTACACGACGCTCTTCCGATCTNNNTTTGCCCAGGTTGACATTGATTC

GCACACGTCTGAACTCTTTCCCTACACGACGCTCTTCCGATCTNNNCACCACACCCAGCTAATTTTTGTC

GCACACGTCTGAACTCTTTCCCTACACGACGCTCTTCCGATCTNNNCACCACACCCAGCTAATTTTTGTC

GAGCACACGTCTGAACTCTTTCCCTACACGACGCTCTTCCGATCTNNNCCTCCTGGGTTCAAGTGATTCT

GAGCACACGTCTGAACTCTTTCCCTACACGACGCTCTTCCGATCTNNNCCTCCTGGGTTCAAGTGATTCT

GCACACGTCTGAACTCTTTCCCTACACGACGCTCTTCCGATCTNNNAGAAAGGCCTGACTTTGGAATCA

GCACACGTCTGAACTCTTTCCCTACACGACGCTCTTCCGATCTNNNAGAAAGGCCTGACTTTGGAATCA

AGCACACGTCTGAACTCTTTCCCTACACGACGCTCTTCCGATCTNNNTTGCAGACTCCCTGAAAAGTTCT

AGCACACGTCTGAACTCTTTCCCTACACGACGCTCTTCCGATCTNNNTTGCAGACTCCCTGAAAAGTTCT

GCACACGTCTGAACTCTTTCCCTACACGACGCTCTTCCGATCTNNNGATGAGGTAGGGAGAGGGAGG

GCACACGTCTGAACTCTTTCCCTACACGACGCTCTTCCGATCTNNNGATGAGGTAGGGAGAGGGAGG

AGCACACGTCTGAACTCTTTCCCTACACGACGCTCTTCCGATCTNNNAGAATCACTCGAACCCAGAAGT

AGCACACGTCTGAACTCTTTCCCTACACGACGCTCTTCCGATCTNNNAGAATCACTCGAACCCAGAAGT

GCACACGTCTGAACTCTTTCCCTACACGACGCTCTTCCGATCTNNNAAAGTGACTTGTCCAAGGTCACA

GCACACGTCTGAACTCTTTCCCTACACGACGCTCTTCCGATCTNNNAAAGTGACTTGTCCAAGGTCACA

AGCACACGTCTGAACTCTTTCCCTACACGACGCTCTTCCGATCTNNNGCAGATTGGGAACAGAGGAGT

AGCACACGTCTGAACTCTTTCCCTACACGACGCTCTTCCGATCTNNNGCAGATTGGGAACAGAGGAGT

AGCACACGTCTGAACTCTTTCCCTACACGACGCTCTTCCGATCTNNNGTCTCCTTTCTGACACCCTGAAA

AGCACACGTCTGAACTCTTTCCCTACACGACGCTCTTCCGATCTNNNGTCTCCTTTCTGACACCCTGAAA

GCACACGTCTGAACTCTTTCCCTACACGACGCTCTTCCGATCTNNNTAGATCAAGGGTTGCAAGACGAT

GCACACGTCTGAACTCTTTCCCTACACGACGCTCTTCCGATCTNNNTAGATCAAGGGTTGCAAGACGAT

AGCACACGTCTGAACTCTTTCCCTACACGACGCTCTTCCGATCTNNNAAATTCAGAGGCTACTGTCTG
AGCACACGTCTGAACTCTTTCCCTACACGACGCTCTTCCGATCTNNNAAATTCAGAGGCTACTGTCTG
AGCACACGTCTGAACTCTTTCCCTACACGACGCTCTTCCGATCTNNNTTCAAGAAGATCACACCCAGG
AGCACACGTCTGAACTCTTTCCCTACACGACGCTCTTCCGATCTNNNTTCAAGAAGATCACACCCAGG
AGCACACGTCTGAACTCTTTCCCTACACGACGCTCTTCCGATCTNNNACCTCAGTACACTCTCTCTCTCT
AGCACACGTCTGAACTCTTTCCCTACACGACGCTCTTCCGATCTNNNACCTCAGTACACTCTCTCTCTCT
AGCACACGTCTGAACTCTTTCCCTACACGACGCTCTTCCGATCTNNNCGACAGAGTGAGACTCCATCT
AGCACACGTCTGAACTCTTTCCCTACACGACGCTCTTCCGATCTNNNCGACAGAGTGAGACTCCATCT
AGCACACGTCTGAACTCTTTCCCTACACGACGCTCTTCCGATCTNNNAACACTGTTTTGCTAGGGGGT
AGCACACGTCTGAACTCTTTCCCTACACGACGCTCTTCCGATCTNNNAACACTGTTTTGCTAGGGGGT
AGCACACGTCTGAACTCTTTCCCTACACGACGCTCTTCCGATCTNNNCAGGAGAATTGCTTGAACAC
AGCACACGTCTGAACTCTTTCCCTACACGACGCTCTTCCGATCTNNNCAGGAGAATTGCTTGAACAC
GCACACGTCTGAACTCTTTCCCTACACGACGCTCTTCCGATCTNNNCTTCAGCTCAAGAATCAAGCACC
GCACACGTCTGAACTCTTTCCCTACACGACGCTCTTCCGATCTNNNCTTCAGCTCAAGAATCAAGCACC
GCACACGTCTGAACTCTTTCCCTACACGACGCTCTTCCGATCTNNNAGTAACTTGTTTCTTCTCTCTGTC
GCACACGTCTGAACTCTTTCCCTACACGACGCTCTTCCGATCTNNNAGTAACTTGTTTCTTCTCTCTGTC
AGCACACGTCTGAACTCTTTCCCTACACGACGCTCTTCCGATCTNNNCTGAGGCAGGAGAATCACTT
AGCACACGTCTGAACTCTTTCCCTACACGACGCTCTTCCGATCTNNNCTGAGGCAGGAGAATCACTT
AGCACACGTCTGAACTCTTTCCCTACACGACGCTCTTCCGATCTNNNATCATAACCTGAAACCCCTGGA
AGCACACGTCTGAACTCTTTCCCTACACGACGCTCTTCCGATCTNNNATCATAACCTGAAACCCCTGGA
GCACACGTCTGAACTCTTTCCCTACACGACGCTCTTCCGATCTNNNGTTCCAGGCTGCTAATCACTTTG
GCACACGTCTGAACTCTTTCCCTACACGACGCTCTTCCGATCTNNNGTTCCAGGCTGCTAATCACTTTG
GAGCACACGTCTGAACTCTTTCCCTACACGACGCTCTTCCGATCTNNNCTGTTTGTCTAGAGGAGCTG
GAGCACACGTCTGAACTCTTTCCCTACACGACGCTCTTCCGATCTNNNCTGTTTGTCTAGAGGAGCTG
GAGCACACGTCTGAACTCTTTCCCTACACGACGCTCTTCCGATCTNNNAGCCTAATGGTCTGAGTGGA
GAGCACACGTCTGAACTCTTTCCCTACACGACGCTCTTCCGATCTNNNAGCCTAATGGTCTGAGTGGA
AGAGCACACGTCTGAACTCTTTCCCTACACGACGCTCTTCCGATCTNNNACGGAAGTGGCTTTGAAA
AGAGCACACGTCTGAACTCTTTCCCTACACGACGCTCTTCCGATCTNNNACGGAAGTGGCTTTGAAA
GCACACGTCTGAACTCTTTCCCTACACGACGCTCTTCCGATCTNNNCTTTGCATACTTCCAGGTCCTCT
GCACACGTCTGAACTCTTTCCCTACACGACGCTCTTCCGATCTNNNCTTTGCATACTTCCAGGTCCTCT
GAGCACACGTCTGAACTCTTTCCCTACACGACGCTCTTCCGATCTNNNGGTCTCACAGACGATACATG
GAGCACACGTCTGAACTCTTTCCCTACACGACGCTCTTCCGATCTNNNGGTCTCACAGACGATACATG
GCACACGTCTGAACTCTTTCCCTACACGACGCTCTTCCGATCTNNNACTCCTGGACTCAAGTGATTGT
GCACACGTCTGAACTCTTTCCCTACACGACGCTCTTCCGATCTNNNACTCCTGGACTCAAGTGATTGT
GAGCACACGTCTGAACTCTTTCCCTACACGACGCTCTTCCGATCTNNNAATATCTGGGGTGCTGCCAT
GAGCACACGTCTGAACTCTTTCCCTACACGACGCTCTTCCGATCTNNNAATATCTGGGGTGCTGCCAT
AGCACACGTCTGAACTCTTTCCCTACACGACGCTCTTCCGATCTNNNACTTAGGCTACTGGTCTAGCTG
AGCACACGTCTGAACTCTTTCCCTACACGACGCTCTTCCGATCTNNNACTTAGGCTACTGGTCTAGCTG
AGCACACGTCTGAACTCTTTCCCTACACGACGCTCTTCCGATCTNNNTCTGTAGTCTCAGCTACTTGGG
AGCACACGTCTGAACTCTTTCCCTACACGACGCTCTTCCGATCTNNNTCTGTAGTCTCAGCTACTTGGG
GAGCACACGTCTGAACTCTTTCCCTACACGACGCTCTTCCGATCTNNNGAGAGAACAGGCAACCTAC
GAGCACACGTCTGAACTCTTTCCCTACACGACGCTCTTCCGATCTNNNGAGAGAACAGGCAACCTAC
AGCACACGTCTGAACTCTTTCCCTACACGACGCTCTTCCGATCTNNNCAGGAAGGGATCTACGTAGT

AGCACACGTCTGAACTCTTTCCCTACACGACGCTCTTCCGATCTNNNCAGGAAGGGATCTACGTAGT
AGCACACGTCTGAACTCTTTCCCTACACGACGCTCTTCCGATCTNNNATCTTTGCCACCACTTCTCTGAT
AGCACACGTCTGAACTCTTTCCCTACACGACGCTCTTCCGATCTNNNATCTTTGCCACCACTTCTCTGAT
AGCACACGTCTGAACTCTTTCCCTACACGACGCTCTTCCGATCTNNNGAGAATCGCTTGAACCCAGG
AGCACACGTCTGAACTCTTTCCCTACACGACGCTCTTCCGATCTNNNGAGAATCGCTTGAACCCAGG
GCACACGTCTGAACTCTTTCCCTACACGACGCTCTTCCGATCTNNNATGATAAGTTTCCTGAGGCCTCC
GCACACGTCTGAACTCTTTCCCTACACGACGCTCTTCCGATCTNNNATGATAAGTTTCCTGAGGCCTCC
AGCACACGTCTGAACTCTTTCCCTACACGACGCTCTTCCGATCTNNNATATTACATTTCAGAGTTCACC
AGCACACGTCTGAACTCTTTCCCTACACGACGCTCTTCCGATCTNNNATATTACATTTCAGAGTTCACC
GAGCACACGTCTGAACTCTTTCCCTACACGACGCTCTTCCGATCTNNNCTTGCAGTGAGCCAAGATCA
GAGCACACGTCTGAACTCTTTCCCTACACGACGCTCTTCCGATCTNNNCTTGCAGTGAGCCAAGATCA
GAGCACACGTCTGAACTCTTTCCCTACACGACGCTCTTCCGATCTNNNCATTTACTGCAGCTGGGAA
GAGCACACGTCTGAACTCTTTCCCTACACGACGCTCTTCCGATCTNNNCATTTACTGCAGCTGGGAA
GAGCACACGTCTGAACTCTTTCCCTACACGACGCTCTTCCGATCTNNNTAAAAGGTAAGGGGGGAA
GAGCACACGTCTGAACTCTTTCCCTACACGACGCTCTTCCGATCTNNNTAAAAGGTAAGGGGGGAA
AGAGCACACGTCTGAACTCTTTCCCTACACGACGCTCTTCCGATCTNNNTAGGGAGGCTGAGGTACG
AGAGCACACGTCTGAACTCTTTCCCTACACGACGCTCTTCCGATCTNNNTAGGGAGGCTGAGGTACG
AGCACACGTCTGAACTCTTTCCCTACACGACGCTCTTCCGATCTNNNGACAGACATCGGGAAAACAA
AGCACACGTCTGAACTCTTTCCCTACACGACGCTCTTCCGATCTNNNGACAGACATCGGGAAAACAA
AGCACACGTCTGAACTCTTTCCCTACACGACGCTCTTCCGATCTNNNAAAAAGGATTTGCTTAGGTG
AGCACACGTCTGAACTCTTTCCCTACACGACGCTCTTCCGATCTNNNAAAAAGGATTTGCTTAGGTG
GCACACGTCTGAACTCTTTCCCTACACGACGCTCTTCCGATCTNNNATTGTGGAACCTAAGATTGGCC
GCACACGTCTGAACTCTTTCCCTACACGACGCTCTTCCGATCTNNNATTGTGGAACCTAAGATTGGCC
GCACACGTCTGAACTCTTTCCCTACACGACGCTCTTCCGATCTNNNTCACTTGGTTGTTGGCAGAATT
GCACACGTCTGAACTCTTTCCCTACACGACGCTCTTCCGATCTNNNTCACTTGGTTGTTGGCAGAATT
GAGCACACGTCTGAACTCTTTCCCTACACGACGCTCTTCCGATCTNNNTGAGAGAGTATGAGGTTGT
GAGCACACGTCTGAACTCTTTCCCTACACGACGCTCTTCCGATCTNNNTGAGAGAGTATGAGGTTGT
GAGCACACGTCTGAACTCTTTCCCTACACGACGCTCTTCCGATCTNNNGGGGTCAGAACAGTAGGAA
GAGCACACGTCTGAACTCTTTCCCTACACGACGCTCTTCCGATCTNNNGGGGTCAGAACAGTAGGAA
CACACGTCTGAACTCTTTCCCTACACGACGCTCTTCCGATCTNNNTGAATGATGTTCACCATTTTTATC
CACACGTCTGAACTCTTTCCCTACACGACGCTCTTCCGATCTNNNTGAATGATGTTCACCATTTTTATC
GAGCACACGTCTGAACTCTTTCCCTACACGACGCTCTTCCGATCTNNNCATGTATCTGCAGGGTGTC
GAGCACACGTCTGAACTCTTTCCCTACACGACGCTCTTCCGATCTNNNCATGTATCTGCAGGGTGTC
GCACACGTCTGAACTCTTTCCCTACACGACGCTCTTCCGATCTNNNGGAGAGTCAATTTGTATGCAG
GCACACGTCTGAACTCTTTCCCTACACGACGCTCTTCCGATCTNNNGGAGAGTCAATTTGTATGCAG
AGCACACGTCTGAACTCTTTCCCTACACGACGCTCTTCCGATCTNNNAAAGTGCTGGGATTACAGGC
AGCACACGTCTGAACTCTTTCCCTACACGACGCTCTTCCGATCTNNNAAAGTGCTGGGATTACAGGC
AGCACACGTCTGAACTCTTTCCCTACACGACGCTCTTCCGATCTNNNGTGTGGAGCAGGAAAAACTG
AGCACACGTCTGAACTCTTTCCCTACACGACGCTCTTCCGATCTNNNGTGTGGAGCAGGAAAAACTG
AGCACACGTCTGAACTCTTTCCCTACACGACGCTCTTCCGATCTNNNCAATGGACCCTCATGTGGTAG
AGCACACGTCTGAACTCTTTCCCTACACGACGCTCTTCCGATCTNNNCAATGGACCCTCATGTGGTAG
AGCACACGTCTGAACTCTTTCCCTACACGACGCTCTTCCGATCTNNNACCAAAATCCTGCTGTGGTTT
AGCACACGTCTGAACTCTTTCCCTACACGACGCTCTTCCGATCTNNNACCAAAATCCTGCTGTGGTTT

GAGCACACGTCTGAACTCTTTCCCTACACGACGCTCTTCCGATCTNNNGTCCTCACACTTCCTTCCCAGC
GAGCACACGTCTGAACTCTTTCCCTACACGACGCTCTTCCGATCTNNNGTCCTCACACTTCCTTCCCAGC
GAGCACACGTCTGAACTCTTTCCCTACACGACGCTCTTCCGATCTNNNGTAGCACTGGAAGCAGGAA
GAGCACACGTCTGAACTCTTTCCCTACACGACGCTCTTCCGATCTNNNGTAGCACTGGAAGCAGGAA
AGCACACGTCTGAACTCTTTCCCTACACGACGCTCTTCCGATCTNNNCTGTGAGCCATGCTGAGGTAT
AGCACACGTCTGAACTCTTTCCCTACACGACGCTCTTCCGATCTNNNCTGTGAGCCATGCTGAGGTAT
GCACACGTCTGAACTCTTTCCCTACACGACGCTCTTCCGATCTNNNAGGCTAGGACATAAATCTGCAT
GCACACGTCTGAACTCTTTCCCTACACGACGCTCTTCCGATCTNNNAGGCTAGGACATAAATCTGCAT
AGCACACGTCTGAACTCTTTCCCTACACGACGCTCTTCCGATCTNNNATGAATCTTTCTATAATTGCAG
AGCACACGTCTGAACTCTTTCCCTACACGACGCTCTTCCGATCTNNNATGAATCTTTCTATAATTGCAG
GCACACGTCTGAACTCTTTCCCTACACGACGCTCTTCCGATCTNNNCCGGATTCAAACGATTCTCCTCT
GCACACGTCTGAACTCTTTCCCTACACGACGCTCTTCCGATCTNNNCCGGATTCAAACGATTCTCCTCT
GCACACGTCTGAACTCTTTCCCTACACGACGCTCTTCCGATCTNNNAATCCAGCCTTTACCACTCCAATC
GCACACGTCTGAACTCTTTCCCTACACGACGCTCTTCCGATCTNNNAATCCAGCCTTTACCACTCCAATC
AGCACACGTCTGAACTCTTTCCCTACACGACGCTCTTCCGATCTNNNCAATCTGCACATCTGACAAAG
AGCACACGTCTGAACTCTTTCCCTACACGACGCTCTTCCGATCTNNNCAATCTGCACATCTGACAAAG
CACACGTCTGAACTCTTTCCCTACACGACGCTCTTCCGATCTNNNCAGGCTTCCCATCAGAACTTACCC
CACACGTCTGAACTCTTTCCCTACACGACGCTCTTCCGATCTNNNCAGGCTTCCCATCAGAACTTACCC
GCACACGTCTGAACTCTTTCCCTACACGACGCTCTTCCGATCTNNNATATTCATAGGATTTGTTTCCCT
GCACACGTCTGAACTCTTTCCCTACACGACGCTCTTCCGATCTNNNATATTCATAGGATTTGTTTCCCT
GAGCACACGTCTGAACTCTTTCCCTACACGACGCTCTTCCGATCTNNNTAGGGAGAGAGAAGGCTAT
GAGCACACGTCTGAACTCTTTCCCTACACGACGCTCTTCCGATCTNNNTAGGGAGAGAGAAGGCTAT
AGCACACGTCTGAACTCTTTCCCTACACGACGCTCTTCCGATCTNNNCTTGCAGTGAGCCGAGATTGC
AGCACACGTCTGAACTCTTTCCCTACACGACGCTCTTCCGATCTNNNCTTGCAGTGAGCCGAGATTGC
AGCACACGTCTGAACTCTTTCCCTACACGACGCTCTTCCGATCTNNNTACCCAGCCAATGAGGACACAT
AGCACACGTCTGAACTCTTTCCCTACACGACGCTCTTCCGATCTNNNTACCCAGCCAATGAGGACACAT
AGCACACGTCTGAACTCTTTCCCTACACGACGCTCTTCCGATCTNNNCATGATGGTGCATGCCTGTAAT
AGCACACGTCTGAACTCTTTCCCTACACGACGCTCTTCCGATCTNNNCATGATGGTGCATGCCTGTAAT
GAGCACACGTCTGAACTCTTTCCCTACACGACGCTCTTCCGATCTNNNCTGAGGGAGGAGAATCGTTT
GAGCACACGTCTGAACTCTTTCCCTACACGACGCTCTTCCGATCTNNNCTGAGGGAGGAGAATCGTTT
GCACACGTCTGAACTCTTTCCCTACACGACGCTCTTCCGATCTNNNACAGGGTGCTCAGATATTTGGT
GCACACGTCTGAACTCTTTCCCTACACGACGCTCTTCCGATCTNNNACAGGGTGCTCAGATATTTGGT
GCACACGTCTGAACTCTTTCCCTACACGACGCTCTTCCGATCTNNNCACTCAAAAGGTCCCTGTGTTTG
GCACACGTCTGAACTCTTTCCCTACACGACGCTCTTCCGATCTNNNCACTCAAAAGGTCCCTGTGTTTG
CACACGTCTGAACTCTTTCCCTACACGACGCTCTTCCGATCTNNNTTTTCTTAATAGCTGAGTTCATGC
CACACGTCTGAACTCTTTCCCTACACGACGCTCTTCCGATCTNNNTTTTCTTAATAGCTGAGTTCATGC
AGCACACGTCTGAACTCTTTCCCTACACGACGCTCTTCCGATCTNNNTTACATTCTTGCTTCAAGCTG
AGCACACGTCTGAACTCTTTCCCTACACGACGCTCTTCCGATCTNNNTTACATTCTTGCTTCAAGCTG
GCACACGTCTGAACTCTTTCCCTACACGACGCTCTTCCGATCTNNNGACATTTTTCCCCCATGGTGTTT
GCACACGTCTGAACTCTTTCCCTACACGACGCTCTTCCGATCTNNNGACATTTTTCCCCCATGGTGTTT
GAGCACACGTCTGAACTCTTTCCCTACACGACGCTCTTCCGATCTNNNGAGGCAGGAGAATGGTGT
GAGCACACGTCTGAACTCTTTCCCTACACGACGCTCTTCCGATCTNNNGAGGCAGGAGAATGGTGT
GAGCACACGTCTGAACTCTTTCCCTACACGACGCTCTTCCGATCTNNNGACTTCATCCAAGACGTGCA

GAGCACACGTCTGAACTCTTTCCCTACACGACGCTCTTCCGATCTNNNGACTTCATCCAAGACGTGCA
AGCACACGTCTGAACTCTTTCCCTACACGACGCTCTTCCGATCTNNNTACTCTGACTTCTTGGTCTCCCT
AGCACACGTCTGAACTCTTTCCCTACACGACGCTCTTCCGATCTNNNTACTCTGACTTCTTGGTCTCCCT
AGCACACGTCTGAACTCTTTCCCTACACGACGCTCTTCCGATCTNNNGCCTGTAGTCCCAGCTACTTGC
AGCACACGTCTGAACTCTTTCCCTACACGACGCTCTTCCGATCTNNNGCCTGTAGTCCCAGCTACTTGC
AGCACACGTCTGAACTCTTTCCCTACACGACGCTCTTCCGATCTNNNCTCCAATAAGAGCATCCATGC
AGCACACGTCTGAACTCTTTCCCTACACGACGCTCTTCCGATCTNNNCTCCAATAAGAGCATCCATGC
GCACACGTCTGAACTCTTTCCCTACACGACGCTCTTCCGATCTNNNCTAGAGTGCAGTGGTGTGATCT
GCACACGTCTGAACTCTTTCCCTACACGACGCTCTTCCGATCTNNNCTAGAGTGCAGTGGTGTGATCT
AGCACACGTCTGAACTCTTTCCCTACACGACGCTCTTCCGATCTNNNTAGGAGAATTGCTTGAACCCC
AGCACACGTCTGAACTCTTTCCCTACACGACGCTCTTCCGATCTNNNTAGGAGAATTGCTTGAACCCC
GAGCACACGTCTGAACTCTTTCCCTACACGACGCTCTTCCGATCTNNNTGGAAATTTTTAAAAACTGTG
GAGCACACGTCTGAACTCTTTCCCTACACGACGCTCTTCCGATCTNNNTGGAAATTTTTAAAAACTGTG
AGCACACGTCTGAACTCTTTCCCTACACGACGCTCTTCCGATCTNNNGCATGTATAACACAAAGTAGG
AGCACACGTCTGAACTCTTTCCCTACACGACGCTCTTCCGATCTNNNGCATGTATAACACAAAGTAGG
GAGCACACGTCTGAACTCTTTCCCTACACGACGCTCTTCCGATCTNNNCTGAGGCAGGAGAATCACTT
GAGCACACGTCTGAACTCTTTCCCTACACGACGCTCTTCCGATCTNNNCTGAGGCAGGAGAATCACTT
GCACACGTCTGAACTCTTTCCCTACACGACGCTCTTCCGATCTNNNTTCTCAGTTCTCTCTCCCTGCTAC
GCACACGTCTGAACTCTTTCCCTACACGACGCTCTTCCGATCTNNNTTCTCAGTTCTCTCTCCCTGCTAC
AGCACACGTCTGAACTCTTTCCCTACACGACGCTCTTCCGATCTNNNCAATGTGGGAAACGAGATCAC
AGCACACGTCTGAACTCTTTCCCTACACGACGCTCTTCCGATCTNNNCAATGTGGGAAACGAGATCAC
GAGCACACGTCTGAACTCTTTCCCTACACGACGCTCTTCCGATCTNNNTGAATGTTCTTTGAGCCTTGC
GAGCACACGTCTGAACTCTTTCCCTACACGACGCTCTTCCGATCTNNNTGAATGTTCTTTGAGCCTTGC
AGCACACGTCTGAACTCTTTCCCTACACGACGCTCTTCCGATCTNNNCTGAAGTGAGAGGATCATTG
AGCACACGTCTGAACTCTTTCCCTACACGACGCTCTTCCGATCTNNNCTGAAGTGAGAGGATCATTG
AGCACACGTCTGAACTCTTTCCCTACACGACGCTCTTCCGATCTNNNATAAAGCAGGTGGGAAGGAG
AGCACACGTCTGAACTCTTTCCCTACACGACGCTCTTCCGATCTNNNATAAAGCAGGTGGGAAGGAG
GCACACGTCTGAACTCTTTCCCTACACGACGCTCTTCCGATCTNNNAGGAAGGAGAATCACTTGAAC
GCACACGTCTGAACTCTTTCCCTACACGACGCTCTTCCGATCTNNNAGGAAGGAGAATCACTTGAAC
AGCACACGTCTGAACTCTTTCCCTACACGACGCTCTTCCGATCTNNNATCAGAAATGACAGCCTGGTG
AGCACACGTCTGAACTCTTTCCCTACACGACGCTCTTCCGATCTNNNATCAGAAATGACAGCCTGGTG
CACACGTCTGAACTCTTTCCCTACACGACGCTCTTCCGATCTNNNTTTCAAATAAAACTTGTACTGGG
CACACGTCTGAACTCTTTCCCTACACGACGCTCTTCCGATCTNNNTTTCAAATAAAACTTGTACTGGG
CACGTCTGAACTCTTTCCCTACACGACGCTCTTCCGATCTNNNAGCCAGGCTCTTTCTTTTTCTTTCATC
CACGTCTGAACTCTTTCCCTACACGACGCTCTTCCGATCTNNNAGCCAGGCTCTTTCTTTTTCTTTCATC
GAGCACACGTCTGAACTCTTTCCCTACACGACGCTCTTCCGATCTNNNCCTGGAAAGATGAGTGGG
GAGCACACGTCTGAACTCTTTCCCTACACGACGCTCTTCCGATCTNNNCCTGGAAAGATGAGTGGG
AGAGCACACGTCTGAACTCTTTCCCTACACGACGCTCTTCCGATCTNNNGATCTTTGGGTCTCGGAGG
AGAGCACACGTCTGAACTCTTTCCCTACACGACGCTCTTCCGATCTNNNGATCTTTGGGTCTCGGAGG
ACACGTCTGAACTCTTTCCCTACACGACGCTCTTCCGATCTNNNAAGGCATATGGAGATGTAGTTAC
ACACGTCTGAACTCTTTCCCTACACGACGCTCTTCCGATCTNNNAAGGCATATGGAGATGTAGTTAC
CACACGTCTGAACTCTTTCCCTACACGACGCTCTTCCGATCTNNNGTTGCAGTGAGCTGAGATGATG
CACACGTCTGAACTCTTTCCCTACACGACGCTCTTCCGATCTNNNGTTGCAGTGAGCTGAGATGATG

GAGCACACGTCTGAACTCTTTCCCTACACGACGCTCTTCCGATCTNNNTCCACTAGTTGATTGATGGG
GAGCACACGTCTGAACTCTTTCCCTACACGACGCTCTTCCGATCTNNNTCCACTAGTTGATTGATGGG
GCACACGTCTGAACTCTTTCCCTACACGACGCTCTTCCGATCTNNNTTTACAACCTCCTGGATCCACAG
GCACACGTCTGAACTCTTTCCCTACACGACGCTCTTCCGATCTNNNTTTACAACCTCCTGGATCCACAG
AGCACACGTCTGAACTCTTTCCCTACACGACGCTCTTCCGATCTNNNTTGCTGCAAATTCTAGAAGCCT
AGCACACGTCTGAACTCTTTCCCTACACGACGCTCTTCCGATCTNNNTTGCTGCAAATTCTAGAAGCCT
AGCACACGTCTGAACTCTTTCCCTACACGACGCTCTTCCGATCTNNNTGGAAGTTTCAGTCAGCCAAGA
AGCACACGTCTGAACTCTTTCCCTACACGACGCTCTTCCGATCTNNNTGGAAGTTTCAGTCAGCCAAGA
AGCACACGTCTGAACTCTTTCCCTACACGACGCTCTTCCGATCTNNNGAGTTACAAATAACAGTGGAA
AGCACACGTCTGAACTCTTTCCCTACACGACGCTCTTCCGATCTNNNGAGTTACAAATAACAGTGGAA
CACACGTCTGAACTCTTTCCCTACACGACGCTCTTCCGATCTNNNCTGCAGGCTCTTTTCCACAATGCAT
CACACGTCTGAACTCTTTCCCTACACGACGCTCTTCCGATCTNNNCTGCAGGCTCTTTTCCACAATGCAT
GAGCACACGTCTGAACTCTTTCCCTACACGACGCTCTTCCGATCTNNNGAGACTACCTAACATCCAACC
GAGCACACGTCTGAACTCTTTCCCTACACGACGCTCTTCCGATCTNNNGAGACTACCTAACATCCAACC
AGCACACGTCTGAACTCTTTCCCTACACGACGCTCTTCCGATCTNNNAGAGACGGGGTTTCACTATGT
AGCACACGTCTGAACTCTTTCCCTACACGACGCTCTTCCGATCTNNNAGAGACGGGGTTTCACTATGT
AGCACACGTCTGAACTCTTTCCCTACACGACGCTCTTCCGATCTNNNGAAAGGAGGAGCAAGGAAAG
AGCACACGTCTGAACTCTTTCCCTACACGACGCTCTTCCGATCTNNNGAAAGGAGGAGCAAGGAAAG
AGCACACGTCTGAACTCTTTCCCTACACGACGCTCTTCCGATCTNNNAGTCCCTACTGCCTATCTCTACC
AGCACACGTCTGAACTCTTTCCCTACACGACGCTCTTCCGATCTNNNAGTCCCTACTGCCTATCTCTACC
AGCACACGTCTGAACTCTTTCCCTACACGACGCTCTTCCGATCTNNNCCTTGTCTGTCTGTCTGTCTCTC
AGCACACGTCTGAACTCTTTCCCTACACGACGCTCTTCCGATCTNNNCCTTGTCTGTCTGTCTGTCTCTC
AGCACACGTCTGAACTCTTTCCCTACACGACGCTCTTCCGATCTNNNCTGGAGGCTAAGACAGGAGAA
AGCACACGTCTGAACTCTTTCCCTACACGACGCTCTTCCGATCTNNNCTGGAGGCTAAGACAGGAGAA
AGCACACGTCTGAACTCTTTCCCTACACGACGCTCTTCCGATCTNNNCTCGTAGCTTGAGAAAAGGGA
AGCACACGTCTGAACTCTTTCCCTACACGACGCTCTTCCGATCTNNNCTCGTAGCTTGAGAAAAGGGA
AGAGCACACGTCTGAACTCTTTCCCTACACGACGCTCTTCCGATCTNNNTAGAGTTTGAGTGTGTGTG
AGAGCACACGTCTGAACTCTTTCCCTACACGACGCTCTTCCGATCTNNNTAGAGTTTGAGTGTGTGTG
GCACACGTCTGAACTCTTTCCCTACACGACGCTCTTCCGATCTNNNGGTGCTCTGCTCTTCTGTATCTTC
GCACACGTCTGAACTCTTTCCCTACACGACGCTCTTCCGATCTNNNGGTGCTCTGCTCTTCTGTATCTTC
GAGCACACGTCTGAACTCTTTCCCTACACGACGCTCTTCCGATCTNNNACCGTTGATACGGTTACAGAT
GAGCACACGTCTGAACTCTTTCCCTACACGACGCTCTTCCGATCTNNNACCGTTGATACGGTTACAGAT
AGCACACGTCTGAACTCTTTCCCTACACGACGCTCTTCCGATCTNNNTGGAGTACAGTGCCATGATTTC
AGCACACGTCTGAACTCTTTCCCTACACGACGCTCTTCCGATCTNNNTGGAGTACAGTGCCATGATTTC
AGCACACGTCTGAACTCTTTCCCTACACGACGCTCTTCCGATCTNNNGTTTTTTCATGTTCTTTGCCCAC
AGCACACGTCTGAACTCTTTCCCTACACGACGCTCTTCCGATCTNNNGTTTTTTCATGTTCTTTGCCCAC
GCACACGTCTGAACTCTTTCCCTACACGACGCTCTTCCGATCTNNNGTATTTTGGTGTTTGTAACCATA
GCACACGTCTGAACTCTTTCCCTACACGACGCTCTTCCGATCTNNNGTATTTTGGTGTTTGTAACCATA
GAGCACACGTCTGAACTCTTTCCCTACACGACGCTCTTCCGATCTNNNTGTAATTCCAGCTACTTGGGA
GAGCACACGTCTGAACTCTTTCCCTACACGACGCTCTTCCGATCTNNNTGTAATTCCAGCTACTTGGGA
GCACACGTCTGAACTCTTTCCCTACACGACGCTCTTCCGATCTNNNTGAGAAGAAGGCTTGCATCTCAT
GCACACGTCTGAACTCTTTCCCTACACGACGCTCTTCCGATCTNNNTGAGAAGAAGGCTTGCATCTCAT
GCACACGTCTGAACTCTTTCCCTACACGACGCTCTTCCGATCTNNNTTGGCTAGACTGGTCTCAAACTC

GCACACGTCTGAACTCTTTCCCTACACGACGCTCTTCCGATCTNNNTTGGCTAGACTGGTCTCAAACTC
GAGCACACGTCTGAACTCTTTCCCTACACGACGCTCTTCCGATCTNNNTAGCAGCCTGAAGGAACTAA
GAGCACACGTCTGAACTCTTTCCCTACACGACGCTCTTCCGATCTNNNTAGCAGCCTGAAGGAACTAA
AGCACACGTCTGAACTCTTTCCCTACACGACGCTCTTCCGATCTNNNGGTGTGATGTTCTTTCTTTGTT
AGCACACGTCTGAACTCTTTCCCTACACGACGCTCTTCCGATCTNNNGGTGTGATGTTCTTTCTTTGTT
AGCACACGTCTGAACTCTTTCCCTACACGACGCTCTTCCGATCTNNNTACAACAAAGCAGTTCCAATGG
AGCACACGTCTGAACTCTTTCCCTACACGACGCTCTTCCGATCTNNNTACAACAAAGCAGTTCCAATGG
AGCACACGTCTGAACTCTTTCCCTACACGACGCTCTTCCGATCTNNNACTGGAAAGGCAGTCTCCCTAT
AGCACACGTCTGAACTCTTTCCCTACACGACGCTCTTCCGATCTNNNACTGGAAAGGCAGTCTCCCTAT
GAGCACACGTCTGAACTCTTTCCCTACACGACGCTCTTCCGATCTNNNCCCTCACCATACTGATGCTTTC
GAGCACACGTCTGAACTCTTTCCCTACACGACGCTCTTCCGATCTNNNCCCTCACCATACTGATGCTTTC
AGCACACGTCTGAACTCTTTCCCTACACGACGCTCTTCCGATCTNNNAAAAAGGAAAAAGCTCCCTCT
AGCACACGTCTGAACTCTTTCCCTACACGACGCTCTTCCGATCTNNNAAAAAGGAAAAAGCTCCCTCT
GCACACGTCTGAACTCTTTCCCTACACGACGCTCTTCCGATCTNNNCATGGATGCTTGGACAACATGA
GCACACGTCTGAACTCTTTCCCTACACGACGCTCTTCCGATCTNNNCATGGATGCTTGGACAACATGA
AGCACACGTCTGAACTCTTTCCCTACACGACGCTCTTCCGATCTNNNGGTTACAGTGATGTGCCCATTT
AGCACACGTCTGAACTCTTTCCCTACACGACGCTCTTCCGATCTNNNGGTTACAGTGATGTGCCCATTT
AGCACACGTCTGAACTCTTTCCCTACACGACGCTCTTCCGATCTNNNCTGATGTTTTCACAGTGAATTT
AGCACACGTCTGAACTCTTTCCCTACACGACGCTCTTCCGATCTNNNCTGATGTTTTCACAGTGAATTT
AGCACACGTCTGAACTCTTTCCCTACACGACGCTCTTCCGATCTNNNCTAAGAGGAGGAGCAGGGGG
AGCACACGTCTGAACTCTTTCCCTACACGACGCTCTTCCGATCTNNNCTAAGAGGAGGAGCAGGGGG
AGAGCACACGTCTGAACTCTTTCCCTACACGACGCTCTTCCGATCTNNNCCCACCTCAGCTTCTTGAGTA
AGAGCACACGTCTGAACTCTTTCCCTACACGACGCTCTTCCGATCTNNNCCCACCTCAGCTTCTTGAGTA
GAGCACACGTCTGAACTCTTTCCCTACACGACGCTCTTCCGATCTNNNGCGGCCTCTTGTTGAATTTTCT
GAGCACACGTCTGAACTCTTTCCCTACACGACGCTCTTCCGATCTNNNGCGGCCTCTTGTTGAATTTTCT
AGCACACGTCTGAACTCTTTCCCTACACGACGCTCTTCCGATCTNNNCCCAAATCAAACACTCTAGCCC
AGCACACGTCTGAACTCTTTCCCTACACGACGCTCTTCCGATCTNNNCCCAAATCAAACACTCTAGCCC
GAGCACACGTCTGAACTCTTTCCCTACACGACGCTCTTCCGATCTNNNTGGTCACAAGTTTCTGAGGA
GAGCACACGTCTGAACTCTTTCCCTACACGACGCTCTTCCGATCTNNNTGGTCACAAGTTTCTGAGGA
AGCACACGTCTGAACTCTTTCCCTACACGACGCTCTTCCGATCTNNNTACACACACTACATTGTAGGTG
AGCACACGTCTGAACTCTTTCCCTACACGACGCTCTTCCGATCTNNNTACACACACTACATTGTAGGTG
GAGCACACGTCTGAACTCTTTCCCTACACGACGCTCTTCCGATCTNNNGAGGTTGCAGTGAGCTGAGA
GAGCACACGTCTGAACTCTTTCCCTACACGACGCTCTTCCGATCTNNNGAGGTTGCAGTGAGCTGAGA
AGCACACGTCTGAACTCTTTCCCTACACGACGCTCTTCCGATCTNNNCAGACCTGTTGAATCAGCATCT
AGCACACGTCTGAACTCTTTCCCTACACGACGCTCTTCCGATCTNNNCAGACCTGTTGAATCAGCATCT
GAGCACACGTCTGAACTCTTTCCCTACACGACGCTCTTCCGATCTNNNAAAAAATTAGCCAGGTGTGG
GAGCACACGTCTGAACTCTTTCCCTACACGACGCTCTTCCGATCTNNNAAAAAATTAGCCAGGTGTGG
AGCACACGTCTGAACTCTTTCCCTACACGACGCTCTTCCGATCTNNNTCTGAAGCCAGAAAACGTATG
AGCACACGTCTGAACTCTTTCCCTACACGACGCTCTTCCGATCTNNNTCTGAAGCCAGAAAACGTATG
GAGCACACGTCTGAACTCTTTCCCTACACGACGCTCTTCCGATCTNNNGCAGAGACACTTGTTTTCTTC
GAGCACACGTCTGAACTCTTTCCCTACACGACGCTCTTCCGATCTNNNGCAGAGACACTTGTTTTCTTC
GAGCACACGTCTGAACTCTTTCCCTACACGACGCTCTTCCGATCTNNNCAGAAACAGGAAGCCAGTAA
GAGCACACGTCTGAACTCTTTCCCTACACGACGCTCTTCCGATCTNNNCAGAAACAGGAAGCCAGTAA

GCACACGTCTGAACTCTTTCCCTACACGACGCTCTTCCGATCTNNNGCCAATATTGTGTGAGTGTGTG
GCACACGTCTGAACTCTTTCCCTACACGACGCTCTTCCGATCTNNNGCCAATATTGTGTGAGTGTGTG
AGCACACGTCTGAACTCTTTCCCTACACGACGCTCTTCCGATCTNNNTGATTGGTCAACACTCCACTACC
AGCACACGTCTGAACTCTTTCCCTACACGACGCTCTTCCGATCTNNNTGATTGGTCAACACTCCACTACC
GCACACGTCTGAACTCTTTCCCTACACGACGCTCTTCCGATCTNNNACGTGCTACACATTTCCATTTGGC
GCACACGTCTGAACTCTTTCCCTACACGACGCTCTTCCGATCTNNNACGTGCTACACATTTCCATTTGGC
GCACACGTCTGAACTCTTTCCCTACACGACGCTCTTCCGATCTNNNGTTCACATGACAGCAATGTTCCA
GCACACGTCTGAACTCTTTCCCTACACGACGCTCTTCCGATCTNNNGTTCACATGACAGCAATGTTCCA
AGAGCACACGTCTGAACTCTTTCCCTACACGACGCTCTTCCGATCTNNNTAGCCTGGGTGAAACTCCAT
AGAGCACACGTCTGAACTCTTTCCCTACACGACGCTCTTCCGATCTNNNTAGCCTGGGTGAAACTCCAT
AGCACACGTCTGAACTCTTTCCCTACACGACGCTCTTCCGATCTNNNGCTGAGATCATTCCATTGCACTC
AGCACACGTCTGAACTCTTTCCCTACACGACGCTCTTCCGATCTNNNGCTGAGATCATTCCATTGCACTC
GCACACGTCTGAACTCTTTCCCTACACGACGCTCTTCCGATCTNNNGAGAGAGAGAGAGCAAGCTGA
AGCACACGTCTGAACTCTTTCCCTACACGACGCTCTTCCGATCTNNNGAGAGAGAGAGAGCAAGCTGA
AAGAGCACACGTCTGAACTCTTTCCCTACACGACGCTCTTCCGATCTNNNGTTACACGGAACCCAGTAC
AAGAGCACACGTCTGAACTCTTTCCCTACACGACGCTCTTCCGATCTNNNGTTACACGGAACCCAGTAC
AGCACACGTCTGAACTCTTTCCCTACACGACGCTCTTCCGATCTNNNTAGTATACTCCTGTCCAGGCTCC
AGCACACGTCTGAACTCTTTCCCTACACGACGCTCTTCCGATCTNNNTAGTATACTCCTGTCCAGGCTCC
GAGCACACGTCTGAACTCTTTCCCTACACGACGCTCTTCCGATCTNNNGAAAAGCTCAAATGAGAGGC
GAGCACACGTCTGAACTCTTTCCCTACACGACGCTCTTCCGATCTNNNGAAAAGCTCAAATGAGAGGC
GAGCACACGTCTGAACTCTTTCCCTACACGACGCTCTTCCGATCTNNNGAGATCACACCACTGCACTCC
GAGCACACGTCTGAACTCTTTCCCTACACGACGCTCTTCCGATCTNNNGAGATCACACCACTGCACTCC
AGCACACGTCTGAACTCTTTCCCTACACGACGCTCTTCCGATCTNNNTGCATACCTGTAATCCCAGCTA
AGCACACGTCTGAACTCTTTCCCTACACGACGCTCTTCCGATCTNNNTGCATACCTGTAATCCCAGCTA
GAGCACACGTCTGAACTCTTTCCCTACACGACGCTCTTCCGATCTNNNTGAGCCTGAATAGCCAAAGGA
GAGCACACGTCTGAACTCTTTCCCTACACGACGCTCTTCCGATCTNNNTGAGCCTGAATAGCCAAAGGA
GCACACGTCTGAACTCTTTCCCTACACGACGCTCTTCCGATCTNNNCAGAACTGCCCTTGCTTTTTAGCCA
GCACACGTCTGAACTCTTTCCCTACACGACGCTCTTCCGATCTNNNCAGAACTGCCCTTGCTTTTTAGCCA
GCACACGTCTGAACTCTTTCCCTACACGACGCTCTTCCGATCTNNNGACAAGAGGAGAACGGGAGACT
GCACACGTCTGAACTCTTTCCCTACACGACGCTCTTCCGATCTNNNGACAAGAGGAGAACGGGAGACT
GCACACGTCTGAACTCTTTCCCTACACGACGCTCTTCCGATCTNNNTATCATGTGTGTGTCACTTGGCTC
GCACACGTCTGAACTCTTTCCCTACACGACGCTCTTCCGATCTNNNTATCATGTGTGTGTCACTTGGCTC
AGCACACGTCTGAACTCTTTCCCTACACGACGCTCTTCCGATCTNNNAAACAGGCAGTGGTCATTTCAA
AGCACACGTCTGAACTCTTTCCCTACACGACGCTCTTCCGATCTNNNAAACAGGCAGTGGTCATTTCAA
AGCACACGTCTGAACTCTTTCCCTACACGACGCTCTTCCGATCTNNNAATCCAAAGCGAACTCTAGACC
AGCACACGTCTGAACTCTTTCCCTACACGACGCTCTTCCGATCTNNNAATCCAAAGCGAACTCTAGACC
AGCACACGTCTGAACTCTTTCCCTACACGACGCTCTTCCGATCTNNNCTCCCAGGTTCAAGTGATCCTCC
AGCACACGTCTGAACTCTTTCCCTACACGACGCTCTTCCGATCTNNNCTCCCAGGTTCAAGTGATCCTCC
GCACACGTCTGAACTCTTTCCCTACACGACGCTCTTCCGATCTNNNGCCACTAGCATATCACTTAGCACCA
GCACACGTCTGAACTCTTTCCCTACACGACGCTCTTCCGATCTNNNGCCACTAGCATATCACTTAGCACCA
GCACACGTCTGAACTCTTTCCCTACACGACGCTCTTCCGATCTNNNAGAAAATTGCTTGAACCGGGAT
GCACACGTCTGAACTCTTTCCCTACACGACGCTCTTCCGATCTNNNAGAAAATTGCTTGAACCGGGAT
GCACACGTCTGAACTCTTTCCCTACACGACGCTCTTCCGATCTNNNTTCTAATGCTGGTACCGTCTGACC

GCACACGTCTGAACTCTTTCCCTACACGACGCTCTTCCGATCTNNNTTCTAATGCTGGTACCGTCTGACC

AGCACACGTCTGAACTCTTTCCCTACACGACGCTCTTCCGATCTNNNGCCTGAATGCAAAGCTCTTGTTT

AGCACACGTCTGAACTCTTTCCCTACACGACGCTCTTCCGATCTNNNGCCTGAATGCAAAGCTCTTGTTT

GAGCACACGTCTGAACTCTTTCCCTACACGACGCTCTTCCGATCTNNNTGGATCTGTAGTCCCACCTACT

GAGCACACGTCTGAACTCTTTCCCTACACGACGCTCTTCCGATCTNNNTGGATCTGTAGTCCCACCTACT

AGCACACGTCTGAACTCTTTCCCTACACGACGCTCTTCCGATCTNNNCCATCCATCCAAAGGTTGCATTT

AGCACACGTCTGAACTCTTTCCCTACACGACGCTCTTCCGATCTNNNCCATCCATCCAAAGGTTGCATTT

AGAGCACACGTCTGAACTCTTTCCCTACACGACGCTCTTCCGATCTNNNTGGCCACATCTTTCTACTTCC

AGAGCACACGTCTGAACTCTTTCCCTACACGACGCTCTTCCGATCTNNNTGGCCACATCTTTCTACTTCC

AGCACACGTCTGAACTCTTTCCCTACACGACGCTCTTCCGATCTNNNGACTCCTGAGTAGCTGGGACTA

AGCACACGTCTGAACTCTTTCCCTACACGACGCTCTTCCGATCTNNNGACTCCTGAGTAGCTGGGACTA

AGCACACGTCTGAACTCTTTCCCTACACGACGCTCTTCCGATCTNNNCTACTTACCCACCTGGAGTCAAC

AGCACACGTCTGAACTCTTTCCCTACACGACGCTCTTCCGATCTNNNCTACTTACCCACCTGGAGTCAAC

AGCACACGTCTGAACTCTTTCCCTACACGACGCTCTTCCGATCTNNNGAGTAGCTGGGACTACAGATGC

AGCACACGTCTGAACTCTTTCCCTACACGACGCTCTTCCGATCTNNNGAGTAGCTGGGACTACAGATGC

GAGCACACGTCTGAACTCTTTCCCTACACGACGCTCTTCCGATCTNNNTGAGAGAGAGAGAGCAAGAC

GAGCACACGTCTGAACTCTTTCCCTACACGACGCTCTTCCGATCTNNNTGAGAGAGAGAGAGCAAGAC

AGCACACGTCTGAACTCTTTCCCTACACGACGCTCTTCCGATCTNNNGAGAAAAAAACCACCCCAACACA

AGCACACGTCTGAACTCTTTCCCTACACGACGCTCTTCCGATCTNNNGAGAAAAAAACCACCCCAACACA

AAGAGCACACGTCTGAACTCTTTCCCTACACGACGCTCTTCCGATCTNNNGAAGGAAGGAAGGAAGG

AAGAGCACACGTCTGAACTCTTTCCCTACACGACGCTCTTCCGATCTNNNGAAGGAAGGAAGGAAGG

GAGCACACGTCTGAACTCTTTCCCTACACGACGCTCTTCCGATCTNNNTAGGAAGCTGGATAAGGGGA

GAGCACACGTCTGAACTCTTTCCCTACACGACGCTCTTCCGATCTNNNTAGGAAGCTGGATAAGGGGA

AGCACACGTCTGAACTCTTTCCCTACACGACGCTCTTCCGATCTNNNGATCTAGAAGCAGCGAAGGACT

AGCACACGTCTGAACTCTTTCCCTACACGACGCTCTTCCGATCTNNNGATCTAGAAGCAGCGAAGGACT

AGAGCACACGTCTGAACTCTTTCCCTACACGACGCTCTTCCGATCTNNNGAGGCAAGAGAATCGCTTG

AGAGCACACGTCTGAACTCTTTCCCTACACGACGCTCTTCCGATCTNNNGAGGCAAGAGAATCGCTTG

GAGCACACGTCTGAACTCTTTCCCTACACGACGCTCTTCCGATCTNNNGGCCCAGTGCAAAATAAAAAT

GAGCACACGTCTGAACTCTTTCCCTACACGACGCTCTTCCGATCTNNNGGCCCAGTGCAAAATAAAAAT

GAGCACACGTCTGAACTCTTTCCCTACACGACGCTCTTCCGATCTNNNATTTTGAAGGCTGTGGAATGC

GAGCACACGTCTGAACTCTTTCCCTACACGACGCTCTTCCGATCTNNNATTTTGAAGGCTGTGGAATGC

AGCACACGTCTGAACTCTTTCCCTACACGACGCTCTTCCGATCTNNNCATGACACAGTGAGACTCTGTC

AGCACACGTCTGAACTCTTTCCCTACACGACGCTCTTCCGATCTNNNCATGACACAGTGAGACTCTGTC

AGCACACGTCTGAACTCTTTCCCTACACGACGCTCTTCCGATCTNNNTGAGGCAGGAGAATACCTTGAA

AGCACACGTCTGAACTCTTTCCCTACACGACGCTCTTCCGATCTNNNTGAGGCAGGAGAATACCTTGAA

AGCACACGTCTGAACTCTTTCCCTACACGACGCTCTTCCGATCTNNNAGAATCTTGCTCTGTCACTCAGG

AGCACACGTCTGAACTCTTTCCCTACACGACGCTCTTCCGATCTNNNAGAATCTTGCTCTGTCACTCAGG

GAGCACACGTCTGAACTCTTTCCCTACACGACGCTCTTCCGATCTNNNGAGAATAGGTTGAACGCAGG

GAGCACACGTCTGAACTCTTTCCCTACACGACGCTCTTCCGATCTNNNGAGAATAGGTTGAACGCAGG

AGCACACGTCTGAACTCTTTCCCTACACGACGCTCTTCCGATCTNNNCTGTAGTCCTAGCTACTCGGGA

AGCACACGTCTGAACTCTTTCCCTACACGACGCTCTTCCGATCTNNNCTGTAGTCCTAGCTACTCGGGA

AGCACACGTCTGAACTCTTTCCCTACACGACGCTCTTCCGATCTNNNCCAGCCTGGATGATGGAGTAAA

AGCACACGTCTGAACTCTTTCCCTACACGACGCTCTTCCGATCTNNNCCAGCCTGGATGATGGAGTAAA

AGCACACGTCTGAACTCTTTCCCTACACGACGCTCTTCCGATCTNNNATATTACATTTCAGAGTTCACCT
AGCACACGTCTGAACTCTTTCCCTACACGACGCTCTTCCGATCTNNNATATTACATTTCAGAGTTCACCT
GAGCACACGTCTGAACTCTTTCCCTACACGACGCTCTTCCGATCTNNNCTCCCCAGGGTAAAGATGAAG
GAGCACACGTCTGAACTCTTTCCCTACACGACGCTCTTCCGATCTNNNCTCCCCAGGGTAAAGATGAAG
GAGCACACGTCTGAACTCTTTCCCTACACGACGCTCTTCCGATCTNNNCTGTAGTGCCAGCTACTCAGGA
GAGCACACGTCTGAACTCTTTCCCTACACGACGCTCTTCCGATCTNNNCTGTAGTGCCAGCTACTCAGGA
GAGCACACGTCTGAACTCTTTCCCTACACGACGCTCTTCCGATCTNNNTAGGTGCAGTACAGTCAAACT
GAGCACACGTCTGAACTCTTTCCCTACACGACGCTCTTCCGATCTNNNTAGGTGCAGTACAGTCAAACT
GAGCACACGTCTGAACTCTTTCCCTACACGACGCTCTTCCGATCTNNNGTATGCCCAAGTTTATGCAGC
GAGCACACGTCTGAACTCTTTCCCTACACGACGCTCTTCCGATCTNNNGTATGCCCAAGTTTATGCAGC
GCACACGTCTGAACTCTTTCCCTACACGACGCTCTTCCGATCTNNNTTGCCTTTGCCAGAATGTCATATC
GCACACGTCTGAACTCTTTCCCTACACGACGCTCTTCCGATCTNNNTTGCCTTTGCCAGAATGTCATATC
GAGCACACGTCTGAACTCTTTCCCTACACGACGCTCTTCCGATCTNNNGTGAGCTGAGATCCGAGATCA
GAGCACACGTCTGAACTCTTTCCCTACACGACGCTCTTCCGATCTNNNGTGAGCTGAGATCCGAGATCA
GCACACGTCTGAACTCTTTCCCTACACGACGCTCTTCCGATCTNNNGGGTGAACCACTCTCTTTCTTTCCA
GCACACGTCTGAACTCTTTCCCTACACGACGCTCTTCCGATCTNNNGGGTGAACCACTCTCTTTCTTTCCA
GCACACGTCTGAACTCTTTCCCTACACGACGCTCTTCCGATCTNNNTAGTGCTGCTATGGACATCTGTG
GCACACGTCTGAACTCTTTCCCTACACGACGCTCTTCCGATCTNNNTAGTGCTGCTATGGACATCTGTG
GAGCACACGTCTGAACTCTTTCCCTACACGACGCTCTTCCGATCTNNNATGGAATCAACCTAACTGCCCA
GAGCACACGTCTGAACTCTTTCCCTACACGACGCTCTTCCGATCTNNNATGGAATCAACCTAACTGCCCA
GCACACGTCTGAACTCTTTCCCTACACGACGCTCTTCCGATCTNNNCTTACCTGAGAGAGAGGTTCCATC
GCACACGTCTGAACTCTTTCCCTACACGACGCTCTTCCGATCTNNNCTTACCTGAGAGAGAGGTTCCATC
AGCACACGTCTGAACTCTTTCCCTACACGACGCTCTTCCGATCTNNNTAAGAGAATGCTGGGATGCAG
AGCACACGTCTGAACTCTTTCCCTACACGACGCTCTTCCGATCTNNNTAAGAGAATGCTGGGATGCAG
GAGCACACGTCTGAACTCTTTCCCTACACGACGCTCTTCCGATCTNNNAGGCTGAAGCAGGTAGATAG
GAGCACACGTCTGAACTCTTTCCCTACACGACGCTCTTCCGATCTNNNAGGCTGAAGCAGGTAGATAG
AGAGCACACGTCTGAACTCTTTCCCTACACGACGCTCTTCCGATCTNNNGAGCTTGCAGTTAGCCGAGA
AGAGCACACGTCTGAACTCTTTCCCTACACGACGCTCTTCCGATCTNNNGAGCTTGCAGTTAGCCGAGA
GAGCACACGTCTGAACTCTTTCCCTACACGACGCTCTTCCGATCTNNNAATGGCCCGATCTAGATTCAC
GAGCACACGTCTGAACTCTTTCCCTACACGACGCTCTTCCGATCTNNNAATGGCCCGATCTAGATTCAC
AGCACACGTCTGAACTCTTTCCCTACACGACGCTCTTCCGATCTNNNGAAATTGCAGTGAGCCGAGAT
AGCACACGTCTGAACTCTTTCCCTACACGACGCTCTTCCGATCTNNNGAAATTGCAGTGAGCCGAGAT
AGAGCACACGTCTGAACTCTTTCCCTACACGACGCTCTTCCGATCTNNNTACAGAAATTAGCTGGGCAT
AGAGCACACGTCTGAACTCTTTCCCTACACGACGCTCTTCCGATCTNNNTACAGAAATTAGCTGGGCAT
AGAGCACACGTCTGAACTCTTTCCCTACACGACGCTCTTCCGATCTNNNATGCCTGTAGTCCCAGTTACT
AGAGCACACGTCTGAACTCTTTCCCTACACGACGCTCTTCCGATCTNNNATGCCTGTAGTCCCAGTTACT
GAGCACACGTCTGAACTCTTTCCCTACACGACGCTCTTCCGATCTNNNTTTGCTGCTCAAACCTCCAAA
GAGCACACGTCTGAACTCTTTCCCTACACGACGCTCTTCCGATCTNNNTTTGCTGCTCAAACCTCCAAA
GAGCACACGTCTGAACTCTTTCCCTACACGACGCTCTTCCGATCTNNNGAGATCGCGCTACTGTACTCC
GAGCACACGTCTGAACTCTTTCCCTACACGACGCTCTTCCGATCTNNNGAGATCGCGCTACTGTACTCC
GAGCACACGTCTGAACTCTTTCCCTACACGACGCTCTTCCGATCTNNNCACAAACAGATTGCATATTATC
GAGCACACGTCTGAACTCTTTCCCTACACGACGCTCTTCCGATCTNNNCACAAACAGATTGCATATTATC
GAGCACACGTCTGAACTCTTTCCCTACACGACGCTCTTCCGATCTNNNATGAGCAGATAGCCTGAACTC

GAGCACACGTCTGAACTCTTTCCCTACACGACGCTCTTCCGATCTNNNATGAGCAGATAGCCTGAACTC
GCACACGTCTGAACTCTTTCCCTACACGACGCTCTTCCGATCTNNNCGATCCTGACTAAGACGGTGAAA
GCACACGTCTGAACTCTTTCCCTACACGACGCTCTTCCGATCTNNNCGATCCTGACTAAGACGGTGAAA
AGCACACGTCTGAACTCTTTCCCTACACGACGCTCTTCCGATCTNNNCTCTTTCTATTCTCTGAGTGATA
AGCACACGTCTGAACTCTTTCCCTACACGACGCTCTTCCGATCTNNNCTCTTTCTATTCTCTGAGTGATA
AGCACACGTCTGAACTCTTTCCCTACACGACGCTCTTCCGATCTNNNACCCTTATCTCTTTGCCTCCTTTCA
AGCACACGTCTGAACTCTTTCCCTACACGACGCTCTTCCGATCTNNNACCCTTATCTCTTTGCCTCCTTTCA
AGCACACGTCTGAACTCTTTCCCTACACGACGCTCTTCCGATCTNNNGGCAACAGAGAGAGATCTTGT
AGCACACGTCTGAACTCTTTCCCTACACGACGCTCTTCCGATCTNNNGGCAACAGAGAGAGATCTTGT
AGCACACGTCTGAACTCTTTCCCTACACGACGCTCTTCCGATCTNNNCCTTTCCTAGTGCTTTGGGTTT
AGCACACGTCTGAACTCTTTCCCTACACGACGCTCTTCCGATCTNNNCCTTTCCTAGTGCTTTGGGTTT
AGAGCACACGTCTGAACTCTTTCCCTACACGACGCTCTTCCGATCTNNNTGATTGAGTAAGACCCTGCC
AGAGCACACGTCTGAACTCTTTCCCTACACGACGCTCTTCCGATCTNNNTGATTGAGTAAGACCCTGCC
GCACACGTCTGAACTCTTTCCCTACACGACGCTCTTCCGATCTNNNCACCTGTAATCCCAGCTACTCAG
GCACACGTCTGAACTCTTTCCCTACACGACGCTCTTCCGATCTNNNCACCTGTAATCCCAGCTACTCAG
AGCACACGTCTGAACTCTTTCCCTACACGACGCTCTTCCGATCTNNNCAGGAGGCTGAGGTATGAGA
AGCACACGTCTGAACTCTTTCCCTACACGACGCTCTTCCGATCTNNNCAGGAGGCTGAGGTATGAGA
AGCACACGTCTGAACTCTTTCCCTACACGACGCTCTTCCGATCTNNNGAGGCCCAGACATTATAGAGCA
AGCACACGTCTGAACTCTTTCCCTACACGACGCTCTTCCGATCTNNNGAGGCCCAGACATTATAGAGCA
AGCACACGTCTGAACTCTTTCCCTACACGACGCTCTTCCGATCTNNNGCTATAGTCAGTCTGAGCACACA
AGCACACGTCTGAACTCTTTCCCTACACGACGCTCTTCCGATCTNNNGCTATAGTCAGTCTGAGCACACA
AGCACACGTCTGAACTCTTTCCCTACACGACGCTCTTCCGATCTNNNTTTTAGCAGCAAAAGCCAGGT
AGCACACGTCTGAACTCTTTCCCTACACGACGCTCTTCCGATCTNNNTTTTAGCAGCAAAAGCCAGGT
AGAGCACACGTCTGAACTCTTTCCCTACACGACGCTCTTCCGATCTNNNGACTGGAGTGAAACCCTGTC
AGAGCACACGTCTGAACTCTTTCCCTACACGACGCTCTTCCGATCTNNNGACTGGAGTGAAACCCTGTC
AGCACACGTCTGAACTCTTTCCCTACACGACGCTCTTCCGATCTNNNTGAAATTTAGGTGGAGGTTCC
AGCACACGTCTGAACTCTTTCCCTACACGACGCTCTTCCGATCTNNNTGAAATTTAGGTGGAGGTTCC
AGCACACGTCTGAACTCTTTCCCTACACGACGCTCTTCCGATCTNNNGGGAGGTGAGCAATTACAAAT
AGCACACGTCTGAACTCTTTCCCTACACGACGCTCTTCCGATCTNNNGGGAGGTGAGCAATTACAAAT
AGCACACGTCTGAACTCTTTCCCTACACGACGCTCTTCCGATCTNNNACAACAGACCACATACAGTCTCA
AGCACACGTCTGAACTCTTTCCCTACACGACGCTCTTCCGATCTNNNACAACAGACCACATACAGTCTCA
AGCACACGTCTGAACTCTTTCCCTACACGACGCTCTTCCGATCTNNNACTCACCGTGCAATCTTGATGTA
AGCACACGTCTGAACTCTTTCCCTACACGACGCTCTTCCGATCTNNNACTCACCGTGCAATCTTGATGTA
AGCACACGTCTGAACTCTTTCCCTACACGACGCTCTTCCGATCTNNNACAGTGCCTGACAAGTTGCTAT
AGCACACGTCTGAACTCTTTCCCTACACGACGCTCTTCCGATCTNNNACAGTGCCTGACAAGTTGCTAT
GAGCACACGTCTGAACTCTTTCCCTACACGACGCTCTTCCGATCTNNNATCAGCAATGACTTTGACCAA
GAGCACACGTCTGAACTCTTTCCCTACACGACGCTCTTCCGATCTNNNATCAGCAATGACTTTGACCAA
GAGCACACGTCTGAACTCTTTCCCTACACGACGCTCTTCCGATCTNNNTGCAGATAACCTTGGATGAGA
GAGCACACGTCTGAACTCTTTCCCTACACGACGCTCTTCCGATCTNNNTGCAGATAACCTTGGATGAGA
AGAGCACACGTCTGAACTCTTTCCCTACACGACGCTCTTCCGATCTNNNATAATGCTACACAAAGGAG
AGAGCACACGTCTGAACTCTTTCCCTACACGACGCTCTTCCGATCTNNNATAATGCTACACAAAGGAG
AGCACACGTCTGAACTCTTTCCCTACACGACGCTCTTCCGATCTNNNCAGCTGTGGTTATTTGGCTCCTA
AGCACACGTCTGAACTCTTTCCCTACACGACGCTCTTCCGATCTNNNCAGCTGTGGTTATTTGGCTCCTA

AGCACACGTCTGAACTCTTTCCCTACACGACGCTCTTCCGATCTNNNGCATATGGCTGCTATCTACCCTT
AGCACACGTCTGAACTCTTTCCCTACACGACGCTCTTCCGATCTNNNGCATATGGCTGCTATCTACCCTT
AGCACACGTCTGAACTCTTTCCCTACACGACGCTCTTCCGATCTNNNTTCAGACTCCCAATCACACCAT
AGCACACGTCTGAACTCTTTCCCTACACGACGCTCTTCCGATCTNNNTTCAGACTCCCAATCACACCAT
AGCACACGTCTGAACTCTTTCCCTACACGACGCTCTTCCGATCTNNNTTTCAGGGAGGATTGCTGATG
AGCACACGTCTGAACTCTTTCCCTACACGACGCTCTTCCGATCTNNNTTTCAGGGAGGATTGCTGATG
AGAGCACACGTCTGAACTCTTTCCCTACACGACGCTCTTCCGATCTNNNCCCGCCAAAAGAGAGCATT
AGAGCACACGTCTGAACTCTTTCCCTACACGACGCTCTTCCGATCTNNNCCCGCCAAAAGAGAGCATT
GCACACGTCTGAACTCTTTCCCTACACGACGCTCTTCCGATCTNNNTACAGCTATACTCCCACCTCAGCC
GCACACGTCTGAACTCTTTCCCTACACGACGCTCTTCCGATCTNNNTACAGCTATACTCCCACCTCAGCC
CACACGTCTGAACTCTTTCCCTACACGACGCTCTTCCGATCTNNNGTGTGGTCTTTTGTCCCTCATCAC
CACACGTCTGAACTCTTTCCCTACACGACGCTCTTCCGATCTNNNGTGTGGTCTTTTGTCCCTCATCAC
AGCACACGTCTGAACTCTTTCCCTACACGACGCTCTTCCGATCTNNNCAAATCACTGGGTTCTGCCAA
AGCACACGTCTGAACTCTTTCCCTACACGACGCTCTTCCGATCTNNNCAAATCACTGGGTTCTGCCAA
AGCACACGTCTGAACTCTTTCCCTACACGACGCTCTTCCGATCTNNNTGCAGAAAAATGATGGCAGT
AGCACACGTCTGAACTCTTTCCCTACACGACGCTCTTCCGATCTNNNTGCAGAAAAATGATGGCAGT
AGCACACGTCTGAACTCTTTCCCTACACGACGCTCTTCCGATCTNNNTATAAAAGGGTGTGGGAGAG
AGCACACGTCTGAACTCTTTCCCTACACGACGCTCTTCCGATCTNNNTATAAAAGGGTGTGGGAGAG
AGCACACGTCTGAACTCTTTCCCTACACGACGCTCTTCCGATCTNNNACAGGAGAGAGAGAGAGAG
AGCACACGTCTGAACTCTTTCCCTACACGACGCTCTTCCGATCTNNNACAGGAGAGAGAGAGAGAG
AGCACACGTCTGAACTCTTTCCCTACACGACGCTCTTCCGATCTNNNCCAGGAAAGAAGCCAGTTCAT
AGCACACGTCTGAACTCTTTCCCTACACGACGCTCTTCCGATCTNNNCCAGGAAAGAAGCCAGTTCAT
GCACACGTCTGAACTCTTTCCCTACACGACGCTCTTCCGATCTNNNCTTAGGTCCTGGGAATCCTGACC
GCACACGTCTGAACTCTTTCCCTACACGACGCTCTTCCGATCTNNNCTTAGGTCCTGGGAATCCTGACC
AGCACACGTCTGAACTCTTTCCCTACACGACGCTCTTCCGATCTNNNTGAAGTGGCATGATCATAGCT
AGCACACGTCTGAACTCTTTCCCTACACGACGCTCTTCCGATCTNNNTGAAGTGGCATGATCATAGCT
AGCACACGTCTGAACTCTTTCCCTACACGACGCTCTTCCGATCTNNNTTTGCTGAAGATATTTGCCAC
AGCACACGTCTGAACTCTTTCCCTACACGACGCTCTTCCGATCTNNNTTTGCTGAAGATATTTGCCAC
AGCACACGTCTGAACTCTTTCCCTACACGACGCTCTTCCGATCTNNNTCTACACATCACATCTGCCCTTCC
AGCACACGTCTGAACTCTTTCCCTACACGACGCTCTTCCGATCTNNNTCTACACATCACATCTGCCCTTCC
AGCACACGTCTGAACTCTTTCCCTACACGACGCTCTTCCGATCTNNNCAACATAGTCTCCTGTCACTGC
AGCACACGTCTGAACTCTTTCCCTACACGACGCTCTTCCGATCTNNNCAACATAGTCTCCTGTCACTGC
AGCACACGTCTGAACTCTTTCCCTACACGACGCTCTTCCGATCTNNNCACTCACTCTAGGCAAACAGGA
AGCACACGTCTGAACTCTTTCCCTACACGACGCTCTTCCGATCTNNNCACTCACTCTAGGCAAACAGGA
AGCACACGTCTGAACTCTTTCCCTACACGACGCTCTTCCGATCTNNNGGAGACAGTGAGACTCCGTCT
AGCACACGTCTGAACTCTTTCCCTACACGACGCTCTTCCGATCTNNNGGAGACAGTGAGACTCCGTCT
AGCACACGTCTGAACTCTTTCCCTACACGACGCTCTTCCGATCTNNNACCTGCATCTCAAGCATTCAGT
AGCACACGTCTGAACTCTTTCCCTACACGACGCTCTTCCGATCTNNNACCTGCATCTCAAGCATTCAGT
AGCACACGTCTGAACTCTTTCCCTACACGACGCTCTTCCGATCTNNNTTGTAACCCTAGCCTACACTACC
AGCACACGTCTGAACTCTTTCCCTACACGACGCTCTTCCGATCTNNNTTGTAACCCTAGCCTACACTACC
AGCACACGTCTGAACTCTTTCCCTACACGACGCTCTTCCGATCTNNNAGACAGGAGAATTGCTTGAA
AGCACACGTCTGAACTCTTTCCCTACACGACGCTCTTCCGATCTNNNAGACAGGAGAATTGCTTGAA
AGCACACGTCTGAACTCTTTCCCTACACGACGCTCTTCCGATCTNNNCTGTGGTTAGAAGCCTGGGGC

AGCACACGTCTGAACTCTTTCCCTACACGACGCTCTTCCGATCTNNNCTGTGGTTAGAAGCCTGGGGC
AGCACACGTCTGAACTCTTTCCCTACACGACGCTCTTCCGATCTNNNGGAGTCTGCAGTAACCTGAGA
AGCACACGTCTGAACTCTTTCCCTACACGACGCTCTTCCGATCTNNNGGAGTCTGCAGTAACCTGAGA
AGCACACGTCTGAACTCTTTCCCTACACGACGCTCTTCCGATCTNNNCTTTCACCTAGGCTGGAGTGTA
AGCACACGTCTGAACTCTTTCCCTACACGACGCTCTTCCGATCTNNNCTTTCACCTAGGCTGGAGTGTA
AGCACACGTCTGAACTCTTTCCCTACACGACGCTCTTCCGATCTNNNTATCCAGGAGTGTGTTTGACAG
AGCACACGTCTGAACTCTTTCCCTACACGACGCTCTTCCGATCTNNNTATCCAGGAGTGTGTTTGACAG
AGCACACGTCTGAACTCTTTCCCTACACGACGCTCTTCCGATCTNNNGAGGCTGAGGCAAGAGAATTA
AGCACACGTCTGAACTCTTTCCCTACACGACGCTCTTCCGATCTNNNGAGGCTGAGGCAAGAGAATTA
AGCACACGTCTGAACTCTTTCCCTACACGACGCTCTTCCGATCTNNNGGCCCAGTTCATACACCTAGAT
AGCACACGTCTGAACTCTTTCCCTACACGACGCTCTTCCGATCTNNNGGCCCAGTTCATACACCTAGAT
CACACGTCTGAACTCTTTCCCTACACGACGCTCTTCCGATCTNNNTTCCTCTTTATCTCAGATATGTGCA
CACACGTCTGAACTCTTTCCCTACACGACGCTCTTCCGATCTNNNTTCCTCTTTATCTCAGATATGTGCA
GAGCACACGTCTGAACTCTTTCCCTACACGACGCTCTTCCGATCTNNNTAGGAACAAAAGTGGGAGG
GAGCACACGTCTGAACTCTTTCCCTACACGACGCTCTTCCGATCTNNNTAGGAACAAAAGTGGGAGG
GAGCACACGTCTGAACTCTTTCCCTACACGACGCTCTTCCGATCTNNNTGTAGTCCCAGCCATTTTAGA
GAGCACACGTCTGAACTCTTTCCCTACACGACGCTCTTCCGATCTNNNTGTAGTCCCAGCCATTTTAGA
GAGCACACGTCTGAACTCTTTCCCTACACGACGCTCTTCCGATCTNNNTCCGAAAGTACTGGGATTAT
GAGCACACGTCTGAACTCTTTCCCTACACGACGCTCTTCCGATCTNNNTCCGAAAGTACTGGGATTAT
AGCACACGTCTGAACTCTTTCCCTACACGACGCTCTTCCGATCTNNNCACACTCACACACACAACACATT
AGCACACGTCTGAACTCTTTCCCTACACGACGCTCTTCCGATCTNNNCACACTCACACACACAACACATT
AGCACACGTCTGAACTCTTTCCCTACACGACGCTCTTCCGATCTNNNGTTTCTCTTTTTCACGCTCCTCC
AGCACACGTCTGAACTCTTTCCCTACACGACGCTCTTCCGATCTNNNGTTTCTCTTTTTCACGCTCCTCC
AGAGCACACGTCTGAACTCTTTCCCTACACGACGCTCTTCCGATCTNNNTCTGGCCAAACTTTTGTATG
AGAGCACACGTCTGAACTCTTTCCCTACACGACGCTCTTCCGATCTNNNTCTGGCCAAACTTTTGTATG
AGCACACGTCTGAACTCTTTCCCTACACGACGCTCTTCCGATCTNNNAGTGAATCACTACCTCTCCTCGA
AGCACACGTCTGAACTCTTTCCCTACACGACGCTCTTCCGATCTNNNAGTGAATCACTACCTCTCCTCGA
AGCACACGTCTGAACTCTTTCCCTACACGACGCTCTTCCGATCTNNNATTAGAAAGGAAATGAATGGT
AGCACACGTCTGAACTCTTTCCCTACACGACGCTCTTCCGATCTNNNATTAGAAAGGAAATGAATGGT
GAGCACACGTCTGAACTCTTTCCCTACACGACGCTCTTCCGATCTNNNCTGCCTTCCATACTTCCTTTGG
GAGCACACGTCTGAACTCTTTCCCTACACGACGCTCTTCCGATCTNNNCTGCCTTCCATACTTCCTTTGG
AGAGCACACGTCTGAACTCTTTCCCTACACGACGCTCTTCCGATCTNNNAGAACAGAAAATGAAATGC
AGAGCACACGTCTGAACTCTTTCCCTACACGACGCTCTTCCGATCTNNNAGAACAGAAAATGAAATGC
AGCACACGTCTGAACTCTTTCCCTACACGACGCTCTTCCGATCTNNNCTATGCTTGCCTGACTCTGGAA
AGCACACGTCTGAACTCTTTCCCTACACGACGCTCTTCCGATCTNNNCTATGCTTGCCTGACTCTGGAA
GAGCACACGTCTGAACTCTTTCCCTACACGACGCTCTTCCGATCTNNNTCTTTGTAGGTGTCAGTGCC
GAGCACACGTCTGAACTCTTTCCCTACACGACGCTCTTCCGATCTNNNTCTTTGTAGGTGTCAGTGCC
GCACACGTCTGAACTCTTTCCCTACACGACGCTCTTCCGATCTNNNAAGTAGGTGACCTTGCAGTTCTC
GCACACGTCTGAACTCTTTCCCTACACGACGCTCTTCCGATCTNNNAAGTAGGTGACCTTGCAGTTCTC
AGAGCACACGTCTGAACTCTTTCCCTACACGACGCTCTTCCGATCTNNNGCAGGAGCCCAGAGGGATG
AGAGCACACGTCTGAACTCTTTCCCTACACGACGCTCTTCCGATCTNNNGCAGGAGCCCAGAGGGATG
GCACACGTCTGAACTCTTTCCCTACACGACGCTCTTCCGATCTNNNGGCACCAATTCACCATTCTACAG
GCACACGTCTGAACTCTTTCCCTACACGACGCTCTTCCGATCTNNNGGCACCAATTCACCATTCTACAG

AGCACACGTCTGAACTCTTTCCCTACACGACGCTCTTCCGATCTNNNACCTCAAGTATCTGACACATGG
AGCACACGTCTGAACTCTTTCCCTACACGACGCTCTTCCGATCTNNNACCTCAAGTATCTGACACATGG
AGCACACGTCTGAACTCTTTCCCTACACGACGCTCTTCCGATCTNNNATCCCTCACTTCTGCTTCCACTT
AGCACACGTCTGAACTCTTTCCCTACACGACGCTCTTCCGATCTNNNATCCCTCACTTCTGCTTCCACTT
GAGCACACGTCTGAACTCTTTCCCTACACGACGCTCTTCCGATCTNNNCCCAGTAGAAAGTTGGTGGA
GAGCACACGTCTGAACTCTTTCCCTACACGACGCTCTTCCGATCTNNNCCCAGTAGAAAGTTGGTGGA
AGCACACGTCTGAACTCTTTCCCTACACGACGCTCTTCCGATCTNNNACCTGAACCTGAACTCAATACTT
AGCACACGTCTGAACTCTTTCCCTACACGACGCTCTTCCGATCTNNNACCTGAACCTGAACTCAATACTT
GCACACGTCTGAACTCTTTCCCTACACGACGCTCTTCCGATCTNNNACACTGAGATGCTTCTTGTGCTA
GCACACGTCTGAACTCTTTCCCTACACGACGCTCTTCCGATCTNNNACACTGAGATGCTTCTTGTGCTA
AGCACACGTCTGAACTCTTTCCCTACACGACGCTCTTCCGATCTNNNGGAAACCAGTGCTCTAGAGAT
AGCACACGTCTGAACTCTTTCCCTACACGACGCTCTTCCGATCTNNNGGAAACCAGTGCTCTAGAGAT
AGCACACGTCTGAACTCTTTCCCTACACGACGCTCTTCCGATCTNNNGTGAGCTGAGATCACCACT
AGCACACGTCTGAACTCTTTCCCTACACGACGCTCTTCCGATCTNNNGTGAGCTGAGATCACCACT
AGCACACGTCTGAACTCTTTCCCTACACGACGCTCTTCCGATCTNNNTGCAATACATCATGGTGCACAA
AGCACACGTCTGAACTCTTTCCCTACACGACGCTCTTCCGATCTNNNTGCAATACATCATGGTGCACAA
GAGCACACGTCTGAACTCTTTCCCTACACGACGCTCTTCCGATCTNNNGCTGGGGACTTTCCAAAACTA
GAGCACACGTCTGAACTCTTTCCCTACACGACGCTCTTCCGATCTNNNGCTGGGGACTTTCCAAAACTA
AGCACACGTCTGAACTCTTTCCCTACACGACGCTCTTCCGATCTNNNCTTATGAACTCTCCAGATTGGCA
AGCACACGTCTGAACTCTTTCCCTACACGACGCTCTTCCGATCTNNNCTTATGAACTCTCCAGATTGGCA
AGCACACGTCTGAACTCTTTCCCTACACGACGCTCTTCCGATCTNNNTCCTTCCTTTCTTTCTTTCTCCCCA
AGCACACGTCTGAACTCTTTCCCTACACGACGCTCTTCCGATCTNNNTCCTTCCTTTCTTTCTTTCTCCCCA
GAGCACACGTCTGAACTCTTTCCCTACACGACGCTCTTCCGATCTNNNCTCCCAGTGGTAGCCTAGTCT
GAGCACACGTCTGAACTCTTTCCCTACACGACGCTCTTCCGATCTNNNCTCCCAGTGGTAGCCTAGTCT
GCACACGTCTGAACTCTTTCCCTACACGACGCTCTTCCGATCTNNNCTGGGTTCAAGAAATTCTCCT
GCACACGTCTGAACTCTTTCCCTACACGACGCTCTTCCGATCTNNNCTGGGTTCAAGAAATTCTCCT
CACACGTCTGAACTCTTTCCCTACACGACGCTCTTCCGATCTNNNCACAACGACCACATCCTCATAGACA
CACACGTCTGAACTCTTTCCCTACACGACGCTCTTCCGATCTNNNCACAACGACCACATCCTCATAGACA
AGCACACGTCTGAACTCTTTCCCTACACGACGCTCTTCCGATCTNNNACTGATCCCACAGACATACGAA
AGCACACGTCTGAACTCTTTCCCTACACGACGCTCTTCCGATCTNNNACTGATCCCACAGACATACGAA
AGCACACGTCTGAACTCTTTCCCTACACGACGCTCTTCCGATCTNNNGTGTGCTGTCTCTTGTAAAGCA
AGCACACGTCTGAACTCTTTCCCTACACGACGCTCTTCCGATCTNNNGTGTGCTGTCTCTTGTAAAGCA
GAGCACACGTCTGAACTCTTTCCCTACACGACGCTCTTCCGATCTNNNTGTTGTTGTTGTTGCTGACAG
GAGCACACGTCTGAACTCTTTCCCTACACGACGCTCTTCCGATCTNNNTGTTGTTGTTGTTGCTGACAG
AGCACACGTCTGAACTCTTTCCCTACACGACGCTCTTCCGATCTNNNGCACTTTCGACATCTTCTACACG
AGCACACGTCTGAACTCTTTCCCTACACGACGCTCTTCCGATCTNNNGCACTTTCGACATCTTCTACACG
AGCACACGTCTGAACTCTTTCCCTACACGACGCTCTTCCGATCTNNNGCCTCCACAAACTAAACCTTCA
AGCACACGTCTGAACTCTTTCCCTACACGACGCTCTTCCGATCTNNNGCCTCCACAAACTAAACCTTCA
GAGCACACGTCTGAACTCTTTCCCTACACGACGCTCTTCCGATCTNNNTTCTCTGCAAGGAATTCTCAG
GAGCACACGTCTGAACTCTTTCCCTACACGACGCTCTTCCGATCTNNNTTCTCTGCAAGGAATTCTCAG
GCACACGTCTGAACTCTTTCCCTACACGACGCTCTTCCGATCTNNNGTCCTTCCTTCCTTCCCTTCTTTCA
GCACACGTCTGAACTCTTTCCCTACACGACGCTCTTCCGATCTNNNGTCCTTCCTTCCTTCCCTTCTTTCA
AGCACACGTCTGAACTCTTTCCCTACACGACGCTCTTCCGATCTNNNTCCACTTGTTCCAGTCAGTCATT

AGCACACGTCTGAACTCTTTCCCTACACGACGCTCTTCCGATCTNNNTCCACTTGTTCCAGTCAGTCATT
AGCACACGTCTGAACTCTTTCCCTACACGACGCTCTTCCGATCTNNNTTCATGCCATTCTCCTGCCTCCA
AGCACACGTCTGAACTCTTTCCCTACACGACGCTCTTCCGATCTNNNTTCATGCCATTCTCCTGCCTCCA
AGCACACGTCTGAACTCTTTCCCTACACGACGCTCTTCCGATCTNNNTGAGAGCAATGCAGAGTGAT
AGCACACGTCTGAACTCTTTCCCTACACGACGCTCTTCCGATCTNNNTGAGAGCAATGCAGAGTGAT
GCACACGTCTGAACTCTTTCCCTACACGACGCTCTTCCGATCTNNNTGTGGAAAATGCTACCATGTTG
GCACACGTCTGAACTCTTTCCCTACACGACGCTCTTCCGATCTNNNTGTGGAAAATGCTACCATGTTG
GAGCACACGTCTGAACTCTTTCCCTACACGACGCTCTTCCGATCTNNNATCATTAGGATGGCAGAGTC
GAGCACACGTCTGAACTCTTTCCCTACACGACGCTCTTCCGATCTNNNATCATTAGGATGGCAGAGTC
AGCACACGTCTGAACTCTTTCCCTACACGACGCTCTTCCGATCTNNNAAAAGAACAACCTGTATGTGG
AGCACACGTCTGAACTCTTTCCCTACACGACGCTCTTCCGATCTNNNAAAAGAACAACCTGTATGTGG
GAGCACACGTCTGAACTCTTTCCCTACACGACGCTCTTCCGATCTNNNACACACACAGAGAGAGAGAG
GAGCACACGTCTGAACTCTTTCCCTACACGACGCTCTTCCGATCTNNNACACACACAGAGAGAGAGAG
AGCACACGTCTGAACTCTTTCCCTACACGACGCTCTTCCGATCTNNNCCAAGTAGCTGGGATTACAG
AGCACACGTCTGAACTCTTTCCCTACACGACGCTCTTCCGATCTNNNCCAAGTAGCTGGGATTACAG
AGAGCACACGTCTGAACTCTTTCCCTACACGACGCTCTTCCGATCTNNNTAAGAGAGAAGGGGGTCG
AGAGCACACGTCTGAACTCTTTCCCTACACGACGCTCTTCCGATCTNNNTAAGAGAGAAGGGGGTCG
GCACACGTCTGAACTCTTTCCCTACACGACGCTCTTCCGATCTNNNGTCTGGGCAACAAGAGTGAAAC
GCACACGTCTGAACTCTTTCCCTACACGACGCTCTTCCGATCTNNNGTCTGGGCAACAAGAGTGAAAC
GCACACGTCTGAACTCTTTCCCTACACGACGCTCTTCCGATCTNNNCCACCACACCAGGCTAATTTTTCC
GCACACGTCTGAACTCTTTCCCTACACGACGCTCTTCCGATCTNNNCCACCACACCAGGCTAATTTTTCC
GAGCACACGTCTGAACTCTTTCCCTACACGACGCTCTTCCGATCTNNNCTGAGGCAGGAGAATCACTT
GAGCACACGTCTGAACTCTTTCCCTACACGACGCTCTTCCGATCTNNNCTGAGGCAGGAGAATCACTT
AGCACACGTCTGAACTCTTTCCCTACACGACGCTCTTCCGATCTNNNAGCTGAAAGTTCACCAGAGT
AGCACACGTCTGAACTCTTTCCCTACACGACGCTCTTCCGATCTNNNAGCTGAAAGTTCACCAGAGT
AGCACACGTCTGAACTCTTTCCCTACACGACGCTCTTCCGATCTNNNACACATTGCACTTCTTTTCAGG
AGCACACGTCTGAACTCTTTCCCTACACGACGCTCTTCCGATCTNNNACACATTGCACTTCTTTTCAGG
AGCACACGTCTGAACTCTTTCCCTACACGACGCTCTTCCGATCTNNNTTGGATACCTATACAGATCCA
AGCACACGTCTGAACTCTTTCCCTACACGACGCTCTTCCGATCTNNNTTGGATACCTATACAGATCCA
GAGCACACGTCTGAACTCTTTCCCTACACGACGCTCTTCCGATCTNNNGAGAATCTCTTGAGCCCAGT
GAGCACACGTCTGAACTCTTTCCCTACACGACGCTCTTCCGATCTNNNGAGAATCTCTTGAGCCCAGT
AGCACACGTCTGAACTCTTTCCCTACACGACGCTCTTCCGATCTNNNGGAACCAGTGTACCTCTCAACA
AGCACACGTCTGAACTCTTTCCCTACACGACGCTCTTCCGATCTNNNGGAACCAGTGTACCTCTCAACA
GAGCACACGTCTGAACTCTTTCCCTACACGACGCTCTTCCGATCTNNNCCACAGAACTGAAGTCTACCA
GAGCACACGTCTGAACTCTTTCCCTACACGACGCTCTTCCGATCTNNNCCACAGAACTGAAGTCTACCA
GCACACGTCTGAACTCTTTCCCTACACGACGCTCTTCCGATCTNNNCTCTACTTGATGGAGTTCTTAAT
GCACACGTCTGAACTCTTTCCCTACACGACGCTCTTCCGATCTNNNCTCTACTTGATGGAGTTCTTAAT
AGCACACGTCTGAACTCTTTCCCTACACGACGCTCTTCCGATCTNNNGTTGCAGTGAGCTGAGATCAT
AGCACACGTCTGAACTCTTTCCCTACACGACGCTCTTCCGATCTNNNGTTGCAGTGAGCTGAGATCAT
AGCACACGTCTGAACTCTTTCCCTACACGACGCTCTTCCGATCTNNNATCAAGTTCCTCAAAATGTGCC
AGCACACGTCTGAACTCTTTCCCTACACGACGCTCTTCCGATCTNNNATCAAGTTCCTCAAAATGTGCC
AGCACACGTCTGAACTCTTTCCCTACACGACGCTCTTCCGATCTNNNCATACCTGTGATCCCAGCTACT
AGCACACGTCTGAACTCTTTCCCTACACGACGCTCTTCCGATCTNNNCATACCTGTGATCCCAGCTACT

GAGCACACGTCTGAACTCTTTCCCTACACGACGCTCTTCCGATCTNNNGGAACAGAAAACCAAACACC
GAGCACACGTCTGAACTCTTTCCCTACACGACGCTCTTCCGATCTNNNGGAACAGAAAACCAAACACC
AGAGCACACGTCTGAACTCTTTCCCTACACGACGCTCTTCCGATCTNNNTGTGTGTGTGTGTGTGG
AGAGCACACGTCTGAACTCTTTCCCTACACGACGCTCTTCCGATCTNNNTGTGTGTGTGTGTGTGG
AGCACACGTCTGAACTCTTTCCCTACACGACGCTCTTCCGATCTNNNACAGTAATGGATGCCCACTCTT
AGCACACGTCTGAACTCTTTCCCTACACGACGCTCTTCCGATCTNNNACAGTAATGGATGCCCACTCTT
AGCACACGTCTGAACTCTTTCCCTACACGACGCTCTTCCGATCTNNNTCAAGACACTGAAGGGAAATG
AGCACACGTCTGAACTCTTTCCCTACACGACGCTCTTCCGATCTNNNTCAAGACACTGAAGGGAAATG
AGCACACGTCTGAACTCTTTCCCTACACGACGCTCTTCCGATCTNNNAGATATCCACCACCGTTTCCAT
AGCACACGTCTGAACTCTTTCCCTACACGACGCTCTTCCGATCTNNNAGATATCCACCACCGTTTCCAT
GCACACGTCTGAACTCTTTCCCTACACGACGCTCTTCCGATCTNNNGACAGAGCAAGACTCCGTCTCC
GCACACGTCTGAACTCTTTCCCTACACGACGCTCTTCCGATCTNNNGACAGAGCAAGACTCCGTCTCC
AGCACACGTCTGAACTCTTTCCCTACACGACGCTCTTCCGATCTNNNACAGGACAGCACCAAATTCAAT
AGCACACGTCTGAACTCTTTCCCTACACGACGCTCTTCCGATCTNNNACAGGACAGCACCAAATTCAAT
GAGCACACGTCTGAACTCTTTCCCTACACGACGCTCTTCCGATCTNNNGAGGTTGTAGTGAGCTGAGA
GAGCACACGTCTGAACTCTTTCCCTACACGACGCTCTTCCGATCTNNNGAGGTTGTAGTGAGCTGAGA
GCACACGTCTGAACTCTTTCCCTACACGACGCTCTTCCGATCTNNNGGATGCTTCCAGGAGAAATGGA
GCACACGTCTGAACTCTTTCCCTACACGACGCTCTTCCGATCTNNNGGATGCTTCCAGGAGAAATGGA
AGCACACGTCTGAACTCTTTCCCTACACGACGCTCTTCCGATCTNNNCTTCCAGTCTGGTGGGTAAGTA
AGCACACGTCTGAACTCTTTCCCTACACGACGCTCTTCCGATCTNNNCTTCCAGTCTGGTGGGTAAGTA
GCACACGTCTGAACTCTTTCCCTACACGACGCTCTTCCGATCTNNNGAATGGCGTGAACCCAGGAGC
GCACACGTCTGAACTCTTTCCCTACACGACGCTCTTCCGATCTNNNGAATGGCGTGAACCCAGGAGC
AGCACACGTCTGAACTCTTTCCCTACACGACGCTCTTCCGATCTNNNTATTTTCCCAGACTGGAGTGGC
AGCACACGTCTGAACTCTTTCCCTACACGACGCTCTTCCGATCTNNNTATTTTCCCAGACTGGAGTGGC
AGCACACGTCTGAACTCTTTCCCTACACGACGCTCTTCCGATCTNNNTTCACACCATTCTCCTGCCTCCA
AGCACACGTCTGAACTCTTTCCCTACACGACGCTCTTCCGATCTNNNTTCACACCATTCTCCTGCCTCCA
GCACACGTCTGAACTCTTTCCCTACACGACGCTCTTCCGATCTNNNCCCTGGAGCAAAGACTACAACAG
GCACACGTCTGAACTCTTTCCCTACACGACGCTCTTCCGATCTNNNCCCTGGAGCAAAGACTACAACAG
AGCACACGTCTGAACTCTTTCCCTACACGACGCTCTTCCGATCTNNNGAATGTAAGGGCATCCATCCAC
AGCACACGTCTGAACTCTTTCCCTACACGACGCTCTTCCGATCTNNNGAATGTAAGGGCATCCATCCAC
GAGCACACGTCTGAACTCTTTCCCTACACGACGCTCTTCCGATCTNNNAACTTAAAGACAGGCACGAA
GAGCACACGTCTGAACTCTTTCCCTACACGACGCTCTTCCGATCTNNNAACTTAAAGACAGGCACGAA
GAGCACACGTCTGAACTCTTTCCCTACACGACGCTCTTCCGATCTNNNGAGGCAGGATAATTGCTTGA
GAGCACACGTCTGAACTCTTTCCCTACACGACGCTCTTCCGATCTNNNGAGGCAGGATAATTGCTTGA
GCACACGTCTGAACTCTTTCCCTACACGACGCTCTTCCGATCTNNNAAAAAAAAAAAGCGGCAAAGCAC
GCACACGTCTGAACTCTTTCCCTACACGACGCTCTTCCGATCTNNNAAAAAAAAAAAGCGGCAAAGCAC
GAGCACACGTCTGAACTCTTTCCCTACACGACGCTCTTCCGATCTNNNCCTCTATTCCACCTTCACCATCC
GAGCACACGTCTGAACTCTTTCCCTACACGACGCTCTTCCGATCTNNNCCTCTATTCCACCTTCACCATCC
GCACACGTCTGAACTCTTTCCCTACACGACGCTCTTCCGATCTNNNGGGTTCCTCAGAGAAGGAGAA
GCACACGTCTGAACTCTTTCCCTACACGACGCTCTTCCGATCTNNNGGGTTCCTCAGAGAAGGAGAA
AGCACACGTCTGAACTCTTTCCCTACACGACGCTCTTCCGATCTNNNTGAAGATGGCCAAATAGTGAG
AGCACACGTCTGAACTCTTTCCCTACACGACGCTCTTCCGATCTNNNTGAAGATGGCCAAATAGTGAG
AGCACACGTCTGAACTCTTTCCCTACACGACGCTCTTCCGATCTNNNTTCAGAAGGAATGGTACCAGC

AGCACACGTCTGAACTCTTTCCCTACACGACGCTCTTCCGATCTNNNTTCAGAAGGAATGGTACCAG
GCACACGTCTGAACTCTTTCCCTACACGACGCTCTTCCGATCTNNNCGCCTATAGTCCCAGCTACTTG
GCACACGTCTGAACTCTTTCCCTACACGACGCTCTTCCGATCTNNNCGCCTATAGTCCCAGCTACTTG
GCACACGTCTGAACTCTTTCCCTACACGACGCTCTTCCGATCTNNNAGGCACAGAGACACTTCTTTGT
GCACACGTCTGAACTCTTTCCCTACACGACGCTCTTCCGATCTNNNAGGCACAGAGACACTTCTTTGT
GCACACGTCTGAACTCTTTCCCTACACGACGCTCTTCCGATCTNNNCAGGAGGGGCACTGTAGCTGC
GCACACGTCTGAACTCTTTCCCTACACGACGCTCTTCCGATCTNNNCAGGAGGGGCACTGTAGCTGC
AGCACACGTCTGAACTCTTTCCCTACACGACGCTCTTCCGATCTNNNGAGGTTGCAGTGAGCTGAGA
AGCACACGTCTGAACTCTTTCCCTACACGACGCTCTTCCGATCTNNNGAGGTTGCAGTGAGCTGAGA
AGAGCACACGTCTGAACTCTTTCCCTACACGACGCTCTTCCGATCTNNNGCTAAGGCAGGAGAATCA
AGAGCACACGTCTGAACTCTTTCCCTACACGACGCTCTTCCGATCTNNNGCTAAGGCAGGAGAATCA
AGCACACGTCTGAACTCTTTCCCTACACGACGCTCTTCCGATCTNNNTGAGGCATCGAAACAGGAAA
AGCACACGTCTGAACTCTTTCCCTACACGACGCTCTTCCGATCTNNNTGAGGCATCGAAACAGGAAA
GCACACGTCTGAACTCTTTCCCTACACGACGCTCTTCCGATCTNNNGGTCAACATAGCGAAACCCCAT
GCACACGTCTGAACTCTTTCCCTACACGACGCTCTTCCGATCTNNNGGTCAACATAGCGAAACCCCAT
GCACACGTCTGAACTCTTTCCCTACACGACGCTCTTCCGATCTNNNCTGACATTATCTGCTAAGAGAG
GCACACGTCTGAACTCTTTCCCTACACGACGCTCTTCCGATCTNNNCTGACATTATCTGCTAAGAGAG
AGCACACGTCTGAACTCTTTCCCTACACGACGCTCTTCCGATCTNNNAGGAGAGAGAGAGAGGAGA
AGCACACGTCTGAACTCTTTCCCTACACGACGCTCTTCCGATCTNNNAGGAGAGAGAGAGAGGAGA
AGCACACGTCTGAACTCTTTCCCTACACGACGCTCTTCCGATCTNNNGACTGCCCTTATTTGCCTGTTT
AGCACACGTCTGAACTCTTTCCCTACACGACGCTCTTCCGATCTNNNGACTGCCCTTATTTGCCTGTTT
AGCACACGTCTGAACTCTTTCCCTACACGACGCTCTTCCGATCTNNNGTAGTACCAGCTACTCAGGAA
AGCACACGTCTGAACTCTTTCCCTACACGACGCTCTTCCGATCTNNNGTAGTACCAGCTACTCAGGAA
GCACACGTCTGAACTCTTTCCCTACACGACGCTCTTCCGATCTNNNAAGCCCAGTTGTACTCTCCAGAT
GCACACGTCTGAACTCTTTCCCTACACGACGCTCTTCCGATCTNNNAAGCCCAGTTGTACTCTCCAGAT
GCACACGTCTGAACTCTTTCCCTACACGACGCTCTTCCGATCTNNNTGAGGGAAACAGGAACATACCA
GCACACGTCTGAACTCTTTCCCTACACGACGCTCTTCCGATCTNNNTGAGGGAAACAGGAACATACCA
GCACACGTCTGAACTCTTTCCCTACACGACGCTCTTCCGATCTNNNTATGGGTAAGGAAACTGAGGC
GCACACGTCTGAACTCTTTCCCTACACGACGCTCTTCCGATCTNNNTATGGGTAAGGAAACTGAGGC
AAGAGCACACGTCTGAACTCTTTCCCTACACGACGCTCTTCCGATCTNNNGCAATGGATGTCACAACT
AAGAGCACACGTCTGAACTCTTTCCCTACACGACGCTCTTCCGATCTNNNGCAATGGATGTCACAACT
AGCACACGTCTGAACTCTTTCCCTACACGACGCTCTTCCGATCTNNNTTCTTTCTCTCTCTCTCCCTCCCC
AGCACACGTCTGAACTCTTTCCCTACACGACGCTCTTCCGATCTNNNTTCTTTCTCTCTCTCTCCCTCCCC
GCACACGTCTGAACTCTTTCCCTACACGACGCTCTTCCGATCTNNNACTCCTGACCTCAAGAGATTTGG
GCACACGTCTGAACTCTTTCCCTACACGACGCTCTTCCGATCTNNNACTCCTGACCTCAAGAGATTTGG
GCACACGTCTGAACTCTTTCCCTACACGACGCTCTTCCGATCTNNNGGGGAAAACAGTGGGAAATGA
GCACACGTCTGAACTCTTTCCCTACACGACGCTCTTCCGATCTNNNGGGGAAAACAGTGGGAAATGA
AGCACACGTCTGAACTCTTTCCCTACACGACGCTCTTCCGATCTNNNACAAGCAGCAAGATTGAAAT
AGCACACGTCTGAACTCTTTCCCTACACGACGCTCTTCCGATCTNNNACAAGCAGCAAGATTGAAAT
CACACGTCTGAACTCTTTCCCTACACGACGCTCTTCCGATCTNNNTGGTACATCTTCTTTGGTGGAGT
CACACGTCTGAACTCTTTCCCTACACGACGCTCTTCCGATCTNNNTGGTACATCTTCTTTGGTGGAGT
GCACACGTCTGAACTCTTTCCCTACACGACGCTCTTCCGATCTNNNCGTGGCCAACAAGCATATTGTA
GCACACGTCTGAACTCTTTCCCTACACGACGCTCTTCCGATCTNNNCGTGGCCAACAAGCATATTGTA

GAGCACACGTCTGAACTCTTTCCCTACACGACGCTCTTCCGATCTNNNCTGCTGGGTTCTCTTATGACAC
GAGCACACGTCTGAACTCTTTCCCTACACGACGCTCTTCCGATCTNNNCTGCTGGGTTCTCTTATGACAC
AGCACACGTCTGAACTCTTTCCCTACACGACGCTCTTCCGATCTNNNAGGTGTTGGTATCAATCCACGA
AGCACACGTCTGAACTCTTTCCCTACACGACGCTCTTCCGATCTNNNAGGTGTTGGTATCAATCCACGA
AGCACACGTCTGAACTCTTTCCCTACACGACGCTCTTCCGATCTNNNTGGGATGTTTTGTGTCATCTGA
AGCACACGTCTGAACTCTTTCCCTACACGACGCTCTTCCGATCTNNNTGGGATGTTTTGTGTCATCTGA
AGCACACGTCTGAACTCTTTCCCTACACGACGCTCTTCCGATCTNNNGTGTTACTCCATCCCCAGCTTTA
AGCACACGTCTGAACTCTTTCCCTACACGACGCTCTTCCGATCTNNNGTGTTACTCCATCCCCAGCTTTA
AGCACACGTCTGAACTCTTTCCCTACACGACGCTCTTCCGATCTNNNGGCCACTTTGCTCTCATTTACCT
AGCACACGTCTGAACTCTTTCCCTACACGACGCTCTTCCGATCTNNNGGCCACTTTGCTCTCATTTACCT
AGAGCACACGTCTGAACTCTTTCCCTACACGACGCTCTTCCGATCTNNNGAGGCTGAGGCACAAGAAT
AGAGCACACGTCTGAACTCTTTCCCTACACGACGCTCTTCCGATCTNNNGAGGCTGAGGCACAAGAAT
AGCACACGTCTGAACTCTTTCCCTACACGACGCTCTTCCGATCTNNNCACCATGCCAGGCTAAATTTTGT
AGCACACGTCTGAACTCTTTCCCTACACGACGCTCTTCCGATCTNNNCACCATGCCAGGCTAAATTTTGT
GAGCACACGTCTGAACTCTTTCCCTACACGACGCTCTTCCGATCTNNNGAGTTGGGGTGTCTACATTGT
GAGCACACGTCTGAACTCTTTCCCTACACGACGCTCTTCCGATCTNNNGAGTTGGGGTGTCTACATTGT
GCACACGTCTGAACTCTTTCCCTACACGACGCTCTTCCGATCTNNNCGCATCTGGGAAGTGAGGAGCA
GCACACGTCTGAACTCTTTCCCTACACGACGCTCTTCCGATCTNNNCGCATCTGGGAAGTGAGGAGCA
GAGCACACGTCTGAACTCTTTCCCTACACGACGCTCTTCCGATCTNNNGAGAGTGAGCCATCTCCAATCA
GAGCACACGTCTGAACTCTTTCCCTACACGACGCTCTTCCGATCTNNNGAGAGTGAGCCATCTCCAATCA
GAGCACACGTCTGAACTCTTTCCCTACACGACGCTCTTCCGATCTNNNGGGATAAGTATGGAGAGGCA
GAGCACACGTCTGAACTCTTTCCCTACACGACGCTCTTCCGATCTNNNGGGATAAGTATGGAGAGGCA
AGAGCACACGTCTGAACTCTTTCCCTACACGACGCTCTTCCGATCTNNNGAGGCTGAGGCAGGAGAAT
AGAGCACACGTCTGAACTCTTTCCCTACACGACGCTCTTCCGATCTNNNGAGGCTGAGGCAGGAGAAT
GCACACGTCTGAACTCTTTCCCTACACGACGCTCTTCCGATCTNNNCATACACCTGTGGTCCCAGATAC
GCACACGTCTGAACTCTTTCCCTACACGACGCTCTTCCGATCTNNNCATACACCTGTGGTCCCAGATAC
AGAGCACACGTCTGAACTCTTTCCCTACACGACGCTCTTCCGATCTNNNCCAGAAGAAGCAGTTTGCAC
AGAGCACACGTCTGAACTCTTTCCCTACACGACGCTCTTCCGATCTNNNCCAGAAGAAGCAGTTTGCAC
AGCACACGTCTGAACTCTTTCCCTACACGACGCTCTTCCGATCTNNNGCACCTGGCCCTTTTTTTTTCTT
AGCACACGTCTGAACTCTTTCCCTACACGACGCTCTTCCGATCTNNNGCACCTGGCCCTTTTTTTTTCTT
GAGCACACGTCTGAACTCTTTCCCTACACGACGCTCTTCCGATCTNNNTTCAAGCAACTCTCTAGCCTCA
GAGCACACGTCTGAACTCTTTCCCTACACGACGCTCTTCCGATCTNNNTTCAAGCAACTCTCTAGCCTCA
CACACGTCTGAACTCTTTCCCTACACGACGCTCTTCCGATCTNNNCACAACGACCACATCCTCATAGACA
CACACGTCTGAACTCTTTCCCTACACGACGCTCTTCCGATCTNNNCACAACGACCACATCCTCATAGACA
AGCACACGTCTGAACTCTTTCCCTACACGACGCTCTTCCGATCTNNNAGTAAGCCATGTTCACTCCAGT
AGCACACGTCTGAACTCTTTCCCTACACGACGCTCTTCCGATCTNNNAGTAAGCCATGTTCACTCCAGT
GAGCACACGTCTGAACTCTTTCCCTACACGACGCTCTTCCGATCTNNNAACACAGACTGCACACATTTCT
GAGCACACGTCTGAACTCTTTCCCTACACGACGCTCTTCCGATCTNNNAACACAGACTGCACACATTTCT
GAGCACACGTCTGAACTCTTTCCCTACACGACGCTCTTCCGATCTNNNGAAGGTTGCAGTGAGTGGAA
GAGCACACGTCTGAACTCTTTCCCTACACGACGCTCTTCCGATCTNNNGAAGGTTGCAGTGAGTGGAA
AGCACACGTCTGAACTCTTTCCCTACACGACGCTCTTCCGATCTNNNTGCTTTCTCTGGTTCTCTGAGTC
AGCACACGTCTGAACTCTTTCCCTACACGACGCTCTTCCGATCTNNNTGCTTTCTCTGGTTCTCTGAGTC
AGCACACGTCTGAACTCTTTCCCTACACGACGCTCTTCCGATCTNNNGACAAGGCCCTCTATGTTCAGA

GCACACGTCTGAACTCTTTCCCTACACGACGCTCTTCCGATCTNNNGACAAGGCCCTCTATGTTCAGA
GCACACGTCTGAACTCTTTCCCTACACGACGCTCTTCCGATCTNNNGGCACCTGTAATCCCAGTTACTT
GCACACGTCTGAACTCTTTCCCTACACGACGCTCTTCCGATCTNNNGGCACCTGTAATCCCAGTTACTT
AGCACACGTCTGAACTCTTTCCCTACACGACGCTCTTCCGATCTNNNCATGCCTGTAATCCCAGCTAC
AGCACACGTCTGAACTCTTTCCCTACACGACGCTCTTCCGATCTNNNCATGCCTGTAATCCCAGCTAC
GCACACGTCTGAACTCTTTCCCTACACGACGCTCTTCCGATCTNNNGATGCTTCACCCGGAAATACAC
GCACACGTCTGAACTCTTTCCCTACACGACGCTCTTCCGATCTNNNGATGCTTCACCCGGAAATACAC
GAGCACACGTCTGAACTCTTTCCCTACACGACGCTCTTCCGATCTNNNGATGGAGGACATTGAGTGT
GAGCACACGTCTGAACTCTTTCCCTACACGACGCTCTTCCGATCTNNNGATGGAGGACATTGAGTGT
GCACACGTCTGAACTCTTTCCCTACACGACGCTCTTCCGATCTNNNTCTGATGGTGTTGACTTCTCTG
GCACACGTCTGAACTCTTTCCCTACACGACGCTCTTCCGATCTNNNTCTGATGGTGTTGACTTCTCTG
AGCACACGTCTGAACTCTTTCCCTACACGACGCTCTTCCGATCTNNNGGGTGTCGTTTCCCTATGTAT
AGCACACGTCTGAACTCTTTCCCTACACGACGCTCTTCCGATCTNNNGGGTGTCGTTTCCCTATGTAT
GCACACGTCTGAACTCTTTCCCTACACGACGCTCTTCCGATCTNNNAGATAGACACACACACACACA
GCACACGTCTGAACTCTTTCCCTACACGACGCTCTTCCGATCTNNNAGATAGACACACACACACACA
AGCACACGTCTGAACTCTTTCCCTACACGACGCTCTTCCGATCTNNNCCACGCCCAGCTAATTTTTCTTA
AGCACACGTCTGAACTCTTTCCCTACACGACGCTCTTCCGATCTNNNCCACGCCCAGCTAATTTTTCTTA
GAGCACACGTCTGAACTCTTTCCCTACACGACGCTCTTCCGATCTNNNGCTCACCTTGGAAAGTGACA
GAGCACACGTCTGAACTCTTTCCCTACACGACGCTCTTCCGATCTNNNGCTCACCTTGGAAAGTGACA
AGCACACGTCTGAACTCTTTCCCTACACGACGCTCTTCCGATCTNNNAAGAGGAACCACCAACTGAA
AGCACACGTCTGAACTCTTTCCCTACACGACGCTCTTCCGATCTNNNAAGAGGAACCACCAACTGAA
GAGCACACGTCTGAACTCTTTCCCTACACGACGCTCTTCCGATCTNNNCTTGCAGTGAGCTGAGATC
GAGCACACGTCTGAACTCTTTCCCTACACGACGCTCTTCCGATCTNNNCTTGCAGTGAGCTGAGATC
GCACACGTCTGAACTCTTTCCCTACACGACGCTCTTCCGATCTNNNGCTCACAGCTCTTACCATAAACA
GCACACGTCTGAACTCTTTCCCTACACGACGCTCTTCCGATCTNNNGCTCACAGCTCTTACCATAAACA
AGCACACGTCTGAACTCTTTCCCTACACGACGCTCTTCCGATCTNNNCTGAGGCAGGAGAATCGATTG
AGCACACGTCTGAACTCTTTCCCTACACGACGCTCTTCCGATCTNNNCTGAGGCAGGAGAATCGATTG
ACACGTCTGAACTCTTTCCCTACACGACGCTCTTCCGATCTNNNCTCCCCTCATCCCACTAAGACATCG
ACACGTCTGAACTCTTTCCCTACACGACGCTCTTCCGATCTNNNCTCCCCTCATCCCACTAAGACATCG
GCACACGTCTGAACTCTTTCCCTACACGACGCTCTTCCGATCTNNNTACTAGGATTGCAGGCATGAGA
GCACACGTCTGAACTCTTTCCCTACACGACGCTCTTCCGATCTNNNTACTAGGATTGCAGGCATGAGA
GCACACGTCTGAACTCTTTCCCTACACGACGCTCTTCCGATCTNNNGGGAACTCTTGTGTCAGTCTA
GCACACGTCTGAACTCTTTCCCTACACGACGCTCTTCCGATCTNNNGGGAACTCTTGTGTCAGTCTA
AGCACACGTCTGAACTCTTTCCCTACACGACGCTCTTCCGATCTNNNGACTGAAGTGATCCTCCTGTC
AGCACACGTCTGAACTCTTTCCCTACACGACGCTCTTCCGATCTNNNGACTGAAGTGATCCTCCTGTC
CACGTCTGAACTCTTTCCCTACACGACGCTCTTCCGATCTNNNATTTTCCTGTCTTCATGAATGACATC
CACGTCTGAACTCTTTCCCTACACGACGCTCTTCCGATCTNNNATTTTCCTGTCTTCATGAATGACATC
GCACACGTCTGAACTCTTTCCCTACACGACGCTCTTCCGATCTNNNTCCTCTTCCTGCTTCTTTCTCTTC
GCACACGTCTGAACTCTTTCCCTACACGACGCTCTTCCGATCTNNNTCCTCTTCCTGCTTCTTTCTCTTC
AGCACACGTCTGAACTCTTTCCCTACACGACGCTCTTCCGATCTNNNCTGTAGTCCCAGCTACTCAGG
AGCACACGTCTGAACTCTTTCCCTACACGACGCTCTTCCGATCTNNNCTGTAGTCCCAGCTACTCAGG
AGCACACGTCTGAACTCTTTCCCTACACGACGCTCTTCCGATCTNNNGAGAATTGCTTGAACCCAGG
AGCACACGTCTGAACTCTTTCCCTACACGACGCTCTTCCGATCTNNNGAGAATTGCTTGAACCCAGG

AGAGCACACGTCTGAACTCTTTCCCTACACGACGCTCTTCCGATCTNNNTTGCTGGAGACAAATTGCGA
AGAGCACACGTCTGAACTCTTTCCCTACACGACGCTCTTCCGATCTNNNTTGCTGGAGACAAATTGCGA
GAGCACACGTCTGAACTCTTTCCCTACACGACGCTCTTCCGATCTNNNTACAGACACACAAATTCCCCAC
GAGCACACGTCTGAACTCTTTCCCTACACGACGCTCTTCCGATCTNNNTACAGACACACAAATTCCCCAC
AGCACACGTCTGAACTCTTTCCCTACACGACGCTCTTCCGATCTNNNAAGTCTGTAGTCCCAGCTACTCA
AGCACACGTCTGAACTCTTTCCCTACACGACGCTCTTCCGATCTNNNAAGTCTGTAGTCCCAGCTACTCA
AGCACACGTCTGAACTCTTTCCCTACACGACGCTCTTCCGATCTNNNGCATAGTCTAGTGCAGGACTGA
AGCACACGTCTGAACTCTTTCCCTACACGACGCTCTTCCGATCTNNNGCATAGTCTAGTGCAGGACTGA
AGCACACGTCTGAACTCTTTCCCTACACGACGCTCTTCCGATCTNNNTGTGTGAGGATAATGGCTTTG
AGCACACGTCTGAACTCTTTCCCTACACGACGCTCTTCCGATCTNNNTGTGTGAGGATAATGGCTTTG
AGCACACGTCTGAACTCTTTCCCTACACGACGCTCTTCCGATCTNNNATATGGACCAAAAAATGTGCCC
AGCACACGTCTGAACTCTTTCCCTACACGACGCTCTTCCGATCTNNNATATGGACCAAAAAATGTGCCC
AGAGCACACGTCTGAACTCTTTCCCTACACGACGCTCTTCCGATCTNNNCTGCCGGAGAAAAGGAACCT
AGAGCACACGTCTGAACTCTTTCCCTACACGACGCTCTTCCGATCTNNNCTGCCGGAGAAAAGGAACCT
GCACACGTCTGAACTCTTTCCCTACACGACGCTCTTCCGATCTNNNCCCTAAGCAGAAGGAAAGGGAAT
GCACACGTCTGAACTCTTTCCCTACACGACGCTCTTCCGATCTNNNCCCTAAGCAGAAGGAAAGGGAAT
GAGCACACGTCTGAACTCTTTCCCTACACGACGCTCTTCCGATCTNNNAAATAGGCCCCTTTGTGAGTT
GAGCACACGTCTGAACTCTTTCCCTACACGACGCTCTTCCGATCTNNNAAATAGGCCCCTTTGTGAGTT
AGCACACGTCTGAACTCTTTCCCTACACGACGCTCTTCCGATCTNNNATCTCTCCTGTGGATTCCTACCAC
AGCACACGTCTGAACTCTTTCCCTACACGACGCTCTTCCGATCTNNNATCTCTCCTGTGGATTCCTACCAC
GAGCACACGTCTGAACTCTTTCCCTACACGACGCTCTTCCGATCTNNNTAGTGTCTCAGAGGAGTGTGT
GAGCACACGTCTGAACTCTTTCCCTACACGACGCTCTTCCGATCTNNNTAGTGTCTCAGAGGAGTGTGT
AGCACACGTCTGAACTCTTTCCCTACACGACGCTCTTCCGATCTNNNCTTGCAGTGAGCCGAGATCCCA
AGCACACGTCTGAACTCTTTCCCTACACGACGCTCTTCCGATCTNNNCTTGCAGTGAGCCGAGATCCCA
AGCACACGTCTGAACTCTTTCCCTACACGACGCTCTTCCGATCTNNNGACAGCACACAGATTTATCAGG
AGCACACGTCTGAACTCTTTCCCTACACGACGCTCTTCCGATCTNNNGACAGCACACAGATTTATCAGG
GAGCACACGTCTGAACTCTTTCCCTACACGACGCTCTTCCGATCTNNNCTGAGGCAAGAGAATCCCTTG
GAGCACACGTCTGAACTCTTTCCCTACACGACGCTCTTCCGATCTNNNCTGAGGCAAGAGAATCCCTTG
GCACACGTCTGAACTCTTTCCCTACACGACGCTCTTCCGATCTNNNCATGAGAAGCTGGGTAGGCATAT
GCACACGTCTGAACTCTTTCCCTACACGACGCTCTTCCGATCTNNNCATGAGAAGCTGGGTAGGCATAT
GAGCACACGTCTGAACTCTTTCCCTACACGACGCTCTTCCGATCTNNNGAGAATCACTTGAACCTGGGA
GAGCACACGTCTGAACTCTTTCCCTACACGACGCTCTTCCGATCTNNNGAGAATCACTTGAACCTGGGA
AGCACACGTCTGAACTCTTTCCCTACACGACGCTCTTCCGATCTNNNAGCACTCAGTGGTAGAGACTTC
AGCACACGTCTGAACTCTTTCCCTACACGACGCTCTTCCGATCTNNNAGCACTCAGTGGTAGAGACTTC
GAGCACACGTCTGAACTCTTTCCCTACACGACGCTCTTCCGATCTNNNTCTGTCTTGCCCAATCTCATGT
GAGCACACGTCTGAACTCTTTCCCTACACGACGCTCTTCCGATCTNNNTCTGTCTTGCCCAATCTCATGT
AGAGCACACGTCTGAACTCTTTCCCTACACGACGCTCTTCCGATCTNNNCTGAGGCAGGAAAATTGCTT
AGAGCACACGTCTGAACTCTTTCCCTACACGACGCTCTTCCGATCTNNNCTGAGGCAGGAAAATTGCTT
GAGCACACGTCTGAACTCTTTCCCTACACGACGCTCTTCCGATCTNNNACACTCCTCAAGTCTCATCCTCT
GAGCACACGTCTGAACTCTTTCCCTACACGACGCTCTTCCGATCTNNNACACTCCTCAAGTCTCATCCTCT
GAGCACACGTCTGAACTCTTTCCCTACACGACGCTCTTCCGATCTNNNATAGAATTCAGAATCAACCAA
GAGCACACGTCTGAACTCTTTCCCTACACGACGCTCTTCCGATCTNNNATAGAATTCAGAATCAACCAA
AGAGCACACGTCTGAACTCTTTCCCTACACGACGCTCTTCCGATCTNNNTAATACCCACAAGATGAGGC

AGAGCACACGTCTGAACTCTTTCCCTACACGACGCTCTTCCGATCTNNNTAATACCCACAAGATGAGG
GCACACGTCTGAACTCTTTCCCTACACGACGCTCTTCCGATCTNNNATTTAGATCATCTTAAAGGCTGT
GCACACGTCTGAACTCTTTCCCTACACGACGCTCTTCCGATCTNNNATTTAGATCATCTTAAAGGCTGT
AGCACACGTCTGAACTCTTTCCCTACACGACGCTCTTCCGATCTNNNCTGAGGCAGGAGAATTGCTT
AGCACACGTCTGAACTCTTTCCCTACACGACGCTCTTCCGATCTNNNCTGAGGCAGGAGAATTGCTT
AGCACACGTCTGAACTCTTTCCCTACACGACGCTCTTCCGATCTNNNGGAAAAAGAGGGAGGTTCTG
AGCACACGTCTGAACTCTTTCCCTACACGACGCTCTTCCGATCTNNNGGAAAAAGAGGGAGGTTCTG
GCACACGTCTGAACTCTTTCCCTACACGACGCTCTTCCGATCTNNNAAAAACCAAGCTGTGAGTGGAT
GCACACGTCTGAACTCTTTCCCTACACGACGCTCTTCCGATCTNNNAAAAACCAAGCTGTGAGTGGAT
AGCACACGTCTGAACTCTTTCCCTACACGACGCTCTTCCGATCTNNNCACGCTGCTGATAAAGACATA
AGCACACGTCTGAACTCTTTCCCTACACGACGCTCTTCCGATCTNNNCACGCTGCTGATAAAGACATA
GAGCACACGTCTGAACTCTTTCCCTACACGACGCTCTTCCGATCTNNNTGAGATCAGGAGTTCGAGAC
GAGCACACGTCTGAACTCTTTCCCTACACGACGCTCTTCCGATCTNNNTGAGATCAGGAGTTCGAGAC
GCACACGTCTGAACTCTTTCCCTACACGACGCTCTTCCGATCTNNNCCGTCGGTGGCTTTTTACAAA
GCACACGTCTGAACTCTTTCCCTACACGACGCTCTTCCGATCTNNNCCGTCGGTGGCTTTTTACAAA
AGCACACGTCTGAACTCTTTCCCTACACGACGCTCTTCCGATCTNNNGTTTGAGCTCAGCAGTTCAAG
AGCACACGTCTGAACTCTTTCCCTACACGACGCTCTTCCGATCTNNNGTTTGAGCTCAGCAGTTCAAG
GCACACGTCTGAACTCTTTCCCTACACGACGCTCTTCCGATCTNNNTGAGAAAGAGGAAAATGGCTG
GCACACGTCTGAACTCTTTCCCTACACGACGCTCTTCCGATCTNNNTGAGAAAGAGGAAAATGGCTG
AGAGCACACGTCTGAACTCTTTCCCTACACGACGCTCTTCCGATCTNNNTCTTTCACCATTCTGGATGC
AGAGCACACGTCTGAACTCTTTCCCTACACGACGCTCTTCCGATCTNNNTCTTTCACCATTCTGGATGC
AGCACACGTCTGAACTCTTTCCCTACACGACGCTCTTCCGATCTNNNTGTTGAACCTTTTGAGGAACT
AGCACACGTCTGAACTCTTTCCCTACACGACGCTCTTCCGATCTNNNTGTTGAACCTTTTGAGGAACT
AGCACACGTCTGAACTCTTTCCCTACACGACGCTCTTCCGATCTNNNTGACACTATGAACCCTGGATG
AGCACACGTCTGAACTCTTTCCCTACACGACGCTCTTCCGATCTNNNTGACACTATGAACCCTGGATG
AGCACACGTCTGAACTCTTTCCCTACACGACGCTCTTCCGATCTNNNGAGGTTGCAGTAAGCCGAGA
AGCACACGTCTGAACTCTTTCCCTACACGACGCTCTTCCGATCTNNNGAGGTTGCAGTAAGCCGAGA
GCACACGTCTGAACTCTTTCCCTACACGACGCTCTTCCGATCTNNNCTGAGGCAGGAGAATCACTTGA
GCACACGTCTGAACTCTTTCCCTACACGACGCTCTTCCGATCTNNNCTGAGGCAGGAGAATCACTTGA
AGCACACGTCTGAACTCTTTCCCTACACGACGCTCTTCCGATCTNNNTGAGGCAGGAGAATCACTTGA
AGCACACGTCTGAACTCTTTCCCTACACGACGCTCTTCCGATCTNNNTGAGGCAGGAGAATCACTTGA
AGCACACGTCTGAACTCTTTCCCTACACGACGCTCTTCCGATCTNNNAATTGGAGATTGTGAGAGGG
AGCACACGTCTGAACTCTTTCCCTACACGACGCTCTTCCGATCTNNNAATTGGAGATTGTGAGAGGG
GAGCACACGTCTGAACTCTTTCCCTACACGACGCTCTTCCGATCTNNNTAAGGTGGGAGGATCAATT
GAGCACACGTCTGAACTCTTTCCCTACACGACGCTCTTCCGATCTNNNTAAGGTGGGAGGATCAATT
GCACACGTCTGAACTCTTTCCCTACACGACGCTCTTCCGATCTNNNAATCATGGTGGAAAGCAAAGT
GCACACGTCTGAACTCTTTCCCTACACGACGCTCTTCCGATCTNNNAATCATGGTGGAAAGCAAAGT
AGCACACGTCTGAACTCTTTCCCTACACGACGCTCTTCCGATCTNNNTCTGCATTCGCTGAGATTACCT
AGCACACGTCTGAACTCTTTCCCTACACGACGCTCTTCCGATCTNNNTCTGCATTCGCTGAGATTACCT
GCACACGTCTGAACTCTTTCCCTACACGACGCTCTTCCGATCTNNNTCCCAAAGTGCTAGGATTACAG
GCACACGTCTGAACTCTTTCCCTACACGACGCTCTTCCGATCTNNNTCCCAAAGTGCTAGGATTACAG
GCACACGTCTGAACTCTTTCCCTACACGACGCTCTTCCGATCTNNNAGGAAATCAGAGAGAGTGGG
GCACACGTCTGAACTCTTTCCCTACACGACGCTCTTCCGATCTNNNAGGAAATCAGAGAGAGTGGG

AGCACACGTCTGAACTCTTTCCCTACACGACGCTCTTCCGATCTNNNCCACTGAACACACTCAGATCTG
AGCACACGTCTGAACTCTTTCCCTACACGACGCTCTTCCGATCTNNNCCACTGAACACACTCAGATCTG
GAGCACACGTCTGAACTCTTTCCCTACACGACGCTCTTCCGATCTNNNGTGTGTGTGTGTGTGTAATG
GAGCACACGTCTGAACTCTTTCCCTACACGACGCTCTTCCGATCTNNNGTGTGTGTGTGTGTGTAATG
CACACGTCTGAACTCTTTCCCTACACGACGCTCTTCCGATCTNNNAGTGTGCCATGGAGGAGAACTC
CACACGTCTGAACTCTTTCCCTACACGACGCTCTTCCGATCTNNNAGTGTGCCATGGAGGAGAACTC
GCACACGTCTGAACTCTTTCCCTACACGACGCTCTTCCGATCTNNNTACCTCTTCCCCAAGATCTTCCTC
GCACACGTCTGAACTCTTTCCCTACACGACGCTCTTCCGATCTNNNTACCTCTTCCCCAAGATCTTCCTC
GCACACGTCTGAACTCTTTCCCTACACGACGCTCTTCCGATCTNNNGTGGCATTTTGTAACTGGCTTCT
GCACACGTCTGAACTCTTTCCCTACACGACGCTCTTCCGATCTNNNGTGGCATTTTGTAACTGGCTTCT
GAGCACACGTCTGAACTCTTTCCCTACACGACGCTCTTCCGATCTNNNAGATTGCTGTGAGTCAAGA
GAGCACACGTCTGAACTCTTTCCCTACACGACGCTCTTCCGATCTNNNAGATTGCTGTGAGTCAAGA
CACACGTCTGAACTCTTTCCCTACACGACGCTCTTCCGATCTNNNTGCCAGATGTTTCACTTTCATCAC
CACACGTCTGAACTCTTTCCCTACACGACGCTCTTCCGATCTNNNTGCCAGATGTTTCACTTTCATCAC
AGCACACGTCTGAACTCTTTCCCTACACGACGCTCTTCCGATCTNNNCTGTCTACTGTTTTAGTGCCTGC
AGCACACGTCTGAACTCTTTCCCTACACGACGCTCTTCCGATCTNNNCTGTCTACTGTTTTAGTGCCTGC
AGCACACGTCTGAACTCTTTCCCTACACGACGCTCTTCCGATCTNNNAGGAGACTCTAACAGAAACGC
AGCACACGTCTGAACTCTTTCCCTACACGACGCTCTTCCGATCTNNNAGGAGACTCTAACAGAAACGC
AGCACACGTCTGAACTCTTTCCCTACACGACGCTCTTCCGATCTNNNCTACAGCATGGGGAAGTCTCT
AGCACACGTCTGAACTCTTTCCCTACACGACGCTCTTCCGATCTNNNCTACAGCATGGGGAAGTCTCT
AGCACACGTCTGAACTCTTTCCCTACACGACGCTCTTCCGATCTNNNCTTAGATCATAGCCATGAAAGA
AGCACACGTCTGAACTCTTTCCCTACACGACGCTCTTCCGATCTNNNCTTAGATCATAGCCATGAAAGA
AGCACACGTCTGAACTCTTTCCCTACACGACGCTCTTCCGATCTNNNGCAACAAGAGTGAAAATCTGC
AGCACACGTCTGAACTCTTTCCCTACACGACGCTCTTCCGATCTNNNGCAACAAGAGTGAAAATCTGC
CACACGTCTGAACTCTTTCCCTACACGACGCTCTTCCGATCTNNNCCCATATGCTATCCCTTATATTTCA
CACACGTCTGAACTCTTTCCCTACACGACGCTCTTCCGATCTNNNCCCATATGCTATCCCTTATATTTCA
CACACGTCTGAACTCTTTCCCTACACGACGCTCTTCCGATCTNNNTCTTCCGAGAGATCCTGCCCCATC
CACACGTCTGAACTCTTTCCCTACACGACGCTCTTCCGATCTNNNTCTTCCGAGAGATCCTGCCCCATC
AGCACACGTCTGAACTCTTTCCCTACACGACGCTCTTCCGATCTNNNCCTGGTGCATGCTACCTACATA
AGCACACGTCTGAACTCTTTCCCTACACGACGCTCTTCCGATCTNNNCCTGGTGCATGCTACCTACATA
AGCACACGTCTGAACTCTTTCCCTACACGACGCTCTTCCGATCTNNNACATACTCCTGCTGTGTAACCAG
AGCACACGTCTGAACTCTTTCCCTACACGACGCTCTTCCGATCTNNNACATACTCCTGCTGTGTAACCAG
AGCACACGTCTGAACTCTTTCCCTACACGACGCTCTTCCGATCTNNNAAGATGGCACCACTGCACTCC
AGCACACGTCTGAACTCTTTCCCTACACGACGCTCTTCCGATCTNNNAAGATGGCACCACTGCACTCC
AGAGCACACGTCTGAACTCTTTCCCTACACGACGCTCTTCCGATCTNNNATTTCAGATGAAAGACCAG
AGAGCACACGTCTGAACTCTTTCCCTACACGACGCTCTTCCGATCTNNNATTTCAGATGAAAGACCAG
GAGCACACGTCTGAACTCTTTCCCTACACGACGCTCTTCCGATCTNNNAAGCAACAGTGGTGTTTTCC
GAGCACACGTCTGAACTCTTTCCCTACACGACGCTCTTCCGATCTNNNAAGCAACAGTGGTGTTTTCC
AGCACACGTCTGAACTCTTTCCCTACACGACGCTCTTCCGATCTNNNTCTCTCTCTCTCACTCTCTCAC
AGCACACGTCTGAACTCTTTCCCTACACGACGCTCTTCCGATCTNNNTCTCTCTCTCTCACTCTCTCAC
AGCACACGTCTGAACTCTTTCCCTACACGACGCTCTTCCGATCTNNNGGAGTTTGCAGTGAGCTGAGA
AGCACACGTCTGAACTCTTTCCCTACACGACGCTCTTCCGATCTNNNGGAGTTTGCAGTGAGCTGAGA
GAGCACACGTCTGAACTCTTTCCCTACACGACGCTCTTCCGATCTNNNCATGAGAATTGCTTCAACCTG

GAGCACACGTCTGAACTCTTTCCCTACACGACGCTCTTCCGATCTNNNCATGAGAATTGCTTCAACCTG
AAGAGCACACGTCTGAACTCTTTCCCTACACGACGCTCTTCCGATCTNNNCTGAGACAGGAGAATCGC
AAGAGCACACGTCTGAACTCTTTCCCTACACGACGCTCTTCCGATCTNNNCTGAGACAGGAGAATCGC
GAGCACACGTCTGAACTCTTTCCCTACACGACGCTCTTCCGATCTNNNAAAAAATTAGCAGGGTGTG
GAGCACACGTCTGAACTCTTTCCCTACACGACGCTCTTCCGATCTNNNAAAAAATTAGCAGGGTGTG
AGCACACGTCTGAACTCTTTCCCTACACGACGCTCTTCCGATCTNNNGAGGCAGGAGAATTGCTTGAA
AGCACACGTCTGAACTCTTTCCCTACACGACGCTCTTCCGATCTNNNGAGGCAGGAGAATTGCTTGAA
GAGCACACGTCTGAACTCTTTCCCTACACGACGCTCTTCCGATCTNNNCACTCTGCCTTTTGTGTTTCCT
GAGCACACGTCTGAACTCTTTCCCTACACGACGCTCTTCCGATCTNNNCACTCTGCCTTTTGTGTTTCCT
AGCACACGTCTGAACTCTTTCCCTACACGACGCTCTTCCGATCTNNNAATCCCAGTTACTCAGGAAGCT
AGCACACGTCTGAACTCTTTCCCTACACGACGCTCTTCCGATCTNNNAATCCCAGTTACTCAGGAAGCT
GAGCACACGTCTGAACTCTTTCCCTACACGACGCTCTTCCGATCTNNNGCTGCCAGGATCCTATGTCAG
GAGCACACGTCTGAACTCTTTCCCTACACGACGCTCTTCCGATCTNNNGCTGCCAGGATCCTATGTCAG
GCACACGTCTGAACTCTTTCCCTACACGACGCTCTTCCGATCTNNNACTACATAGCCTCCTCGCTTTTCC
GCACACGTCTGAACTCTTTCCCTACACGACGCTCTTCCGATCTNNNACTACATAGCCTCCTCGCTTTTCC
AGCACACGTCTGAACTCTTTCCCTACACGACGCTCTTCCGATCTNNNCTGGGTTCCCAGTTTCCTTAGAG
AGCACACGTCTGAACTCTTTCCCTACACGACGCTCTTCCGATCTNNNCTGGGTTCCCAGTTTCCTTAGAG
CACACGTCTGAACTCTTTCCCTACACGACGCTCTTCCGATCTNNNGAGGTTTTCTGTCTGTGTTGAGCC
CACACGTCTGAACTCTTTCCCTACACGACGCTCTTCCGATCTNNNGAGGTTTTCTGTCTGTGTTGAGCC
GAGCACACGTCTGAACTCTTTCCCTACACGACGCTCTTCCGATCTNNNATTTCTCCCTTTCCCTTACACCC
GAGCACACGTCTGAACTCTTTCCCTACACGACGCTCTTCCGATCTNNNATTTCTCCCTTTCCCTTACACCC
AGCACACGTCTGAACTCTTTCCCTACACGACGCTCTTCCGATCTNNNATTTTTTATGCAGCTCCCTGGTC
AGCACACGTCTGAACTCTTTCCCTACACGACGCTCTTCCGATCTNNNATTTTTTATGCAGCTCCCTGGTC
GAGCACACGTCTGAACTCTTTCCCTACACGACGCTCTTCCGATCTNNNGAAATCATCCTAACAGCAGCT
GAGCACACGTCTGAACTCTTTCCCTACACGACGCTCTTCCGATCTNNNGAAATCATCCTAACAGCAGCT
AGCACACGTCTGAACTCTTTCCCTACACGACGCTCTTCCGATCTNNNTGTGTGTGTGTGAGAGAG
AGCACACGTCTGAACTCTTTCCCTACACGACGCTCTTCCGATCTNNNTGTGTGTGTGTGAGAGAG
GAGCACACGTCTGAACTCTTTCCCTACACGACGCTCTTCCGATCTNNNCCATAAGCTCCTCTTTTCCCCAT
GAGCACACGTCTGAACTCTTTCCCTACACGACGCTCTTCCGATCTNNNCCATAAGCTCCTCTTTTCCCCAT
GAGCACACGTCTGAACTCTTTCCCTACACGACGCTCTTCCGATCTNNNCACAGGAGAACCACTAAGGT
GAGCACACGTCTGAACTCTTTCCCTACACGACGCTCTTCCGATCTNNNCACAGGAGAACCACTAAGGT
AGCACACGTCTGAACTCTTTCCCTACACGACGCTCTTCCGATCTNNNCATGGTGTGGTTCCTTCCTCTTA
AGCACACGTCTGAACTCTTTCCCTACACGACGCTCTTCCGATCTNNNCATGGTGTGGTTCCTTCCTCTTA
GAGCACACGTCTGAACTCTTTCCCTACACGACGCTCTTCCGATCTNNNAAAATGCAAGCCAGAAGACA
GAGCACACGTCTGAACTCTTTCCCTACACGACGCTCTTCCGATCTNNNAAAATGCAAGCCAGAAGACA
GAAGAGCACACGTCTGAACTCTTTCCCTACACGACGCTCTTCCGATCTNNNATCCAAGTAAAAACTGA
GAAGAGCACACGTCTGAACTCTTTCCCTACACGACGCTCTTCCGATCTNNNATCCAAGTAAAAACTGA
AGCACACGTCTGAACTCTTTCCCTACACGACGCTCTTCCGATCTNNNGGGTGCTACTGCTGAGATTTT
AGCACACGTCTGAACTCTTTCCCTACACGACGCTCTTCCGATCTNNNGGGTGCTACTGCTGAGATTTT
GAGCACACGTCTGAACTCTTTCCCTACACGACGCTCTTCCGATCTNNNTCCTGAGTAGCTGAGACTAG
GAGCACACGTCTGAACTCTTTCCCTACACGACGCTCTTCCGATCTNNNTCCTGAGTAGCTGAGACTAG
GCACACGTCTGAACTCTTTCCCTACACGACGCTCTTCCGATCTNNNTCACCATTCACTCAAGAGAGAAA
GCACACGTCTGAACTCTTTCCCTACACGACGCTCTTCCGATCTNNNTCACCATTCACTCAAGAGAGAAA

GCACACGTCTGAACTCTTTCCCTACACGACGCTCTTCCGATCTNNNCTCACATCGAGGAGCAAGACGCA
GCACACGTCTGAACTCTTTCCCTACACGACGCTCTTCCGATCTNNNCTCACATCGAGGAGCAAGACGCA
GCACACGTCTGAACTCTTTCCCTACACGACGCTCTTCCGATCTNNNGAGGCAGGAGAATTGCTTGAAT
GCACACGTCTGAACTCTTTCCCTACACGACGCTCTTCCGATCTNNNGAGGCAGGAGAATTGCTTGAAT
GAGCACACGTCTGAACTCTTTCCCTACACGACGCTCTTCCGATCTNNNGGCAAAACCCTGTCCCTACTAA
GAGCACACGTCTGAACTCTTTCCCTACACGACGCTCTTCCGATCTNNNGGCAAAACCCTGTCCCTACTAA
GCACACGTCTGAACTCTTTCCCTACACGACGCTCTTCCGATCTNNNGAGCACTGTGGGGCATTCAGCAT
GCACACGTCTGAACTCTTTCCCTACACGACGCTCTTCCGATCTNNNGAGCACTGTGGGGCATTCAGCAT
CACACGTCTGAACTCTTTCCCTACACGACGCTCTTCCGATCTNNNAATACAAAAAATTAGCCGGGCGCCA
CACACGTCTGAACTCTTTCCCTACACGACGCTCTTCCGATCTNNNAATACAAAAAATTAGCCGGGCGCCA
AGCACACGTCTGAACTCTTTCCCTACACGACGCTCTTCCGATCTNNNACAACAGACCACATACAGTCTCA
AGCACACGTCTGAACTCTTTCCCTACACGACGCTCTTCCGATCTNNNACAACAGACCACATACAGTCTCA
GAGCACACGTCTGAACTCTTTCCCTACACGACGCTCTTCCGATCTNNNCTTGGCTCTAGACTGTTACTCC
GAGCACACGTCTGAACTCTTTCCCTACACGACGCTCTTCCGATCTNNNCTTGGCTCTAGACTGTTACTCC
GAGCACACGTCTGAACTCTTTCCCTACACGACGCTCTTCCGATCTNNNTGAAAGAGCAGGCATCCCTAA
GAGCACACGTCTGAACTCTTTCCCTACACGACGCTCTTCCGATCTNNNTGAAAGAGCAGGCATCCCTAA
AGCACACGTCTGAACTCTTTCCCTACACGACGCTCTTCCGATCTNNNACTTCCATCCAGCCTTGAAGAAT
AGCACACGTCTGAACTCTTTCCCTACACGACGCTCTTCCGATCTNNNACTTCCATCCAGCCTTGAAGAAT
GCACACGTCTGAACTCTTTCCCTACACGACGCTCTTCCGATCTNNNGGACTTTGGTTCCCTAAGAGGACC
GCACACGTCTGAACTCTTTCCCTACACGACGCTCTTCCGATCTNNNGGACTTTGGTTCCCTAAGAGGACC
AGCACACGTCTGAACTCTTTCCCTACACGACGCTCTTCCGATCTNNNAAGCGCATCTAGGGTACAGATT
AGCACACGTCTGAACTCTTTCCCTACACGACGCTCTTCCGATCTNNNAAGCGCATCTAGGGTACAGATT
AGCACACGTCTGAACTCTTTCCCTACACGACGCTCTTCCGATCTNNNAGGATTGGAGCAGACAAAGTTC
AGCACACGTCTGAACTCTTTCCCTACACGACGCTCTTCCGATCTNNNAGGATTGGAGCAGACAAAGTTC
AGAGCACACGTCTGAACTCTTTCCCTACACGACGCTCTTCCGATCTNNNTTCCTTCCCCACCTCTTTTTCCC
AGAGCACACGTCTGAACTCTTTCCCTACACGACGCTCTTCCGATCTNNNTTCCTTCCCCACCTCTTTTTCCC
GAGCACACGTCTGAACTCTTTCCCTACACGACGCTCTTCCGATCTNNNTGTTTAGCCTACCTGAATTCGC
GAGCACACGTCTGAACTCTTTCCCTACACGACGCTCTTCCGATCTNNNTGTTTAGCCTACCTGAATTCGC
GCACACGTCTGAACTCTTTCCCTACACGACGCTCTTCCGATCTNNNGTGAGAGAGGGCCATATAGAGC
GCACACGTCTGAACTCTTTCCCTACACGACGCTCTTCCGATCTNNNGTGAGAGAGGGCCATATAGAGC
AGCACACGTCTGAACTCTTTCCCTACACGACGCTCTTCCGATCTNNNAAAAAATAAGGCAGAATCACAT
AGCACACGTCTGAACTCTTTCCCTACACGACGCTCTTCCGATCTNNNAAAAAATAAGGCAGAATCACAT
AGAGCACACGTCTGAACTCTTTCCCTACACGACGCTCTTCCGATCTNNNGGTCTATGGTAATGGAGGA
AGAGCACACGTCTGAACTCTTTCCCTACACGACGCTCTTCCGATCTNNNGGTCTATGGTAATGGAGGA
AGCACACGTCTGAACTCTTTCCCTACACGACGCTCTTCCGATCTNNNGATGTTGCAGTGAGCCGAGAT
AGCACACGTCTGAACTCTTTCCCTACACGACGCTCTTCCGATCTNNNGATGTTGCAGTGAGCCGAGAT
GAGCACACGTCTGAACTCTTTCCCTACACGACGCTCTTCCGATCTNNNTGATGGCTTCCTTCCATTCTGTC
GAGCACACGTCTGAACTCTTTCCCTACACGACGCTCTTCCGATCTNNNTGATGGCTTCCTTCCATTCTGTC
AGCACACGTCTGAACTCTTTCCCTACACGACGCTCTTCCGATCTNNNATGTTTCCTCTGTACCCAGCTTT
AGCACACGTCTGAACTCTTTCCCTACACGACGCTCTTCCGATCTNNNATGTTTCCTCTGTACCCAGCTTT
AGCACACGTCTGAACTCTTTCCCTACACGACGCTCTTCCGATCTNNNTCTGCAGTTAGTCCATTTCAGCT
AGCACACGTCTGAACTCTTTCCCTACACGACGCTCTTCCGATCTNNNTCTGCAGTTAGTCCATTTCAGCT
AGCACACGTCTGAACTCTTTCCCTACACGACGCTCTTCCGATCTNNNAGAGCTGAGATAAACGCCCTTT

AGCACACGTCTGAACTCTTTCCCTACACGACGCTCTTCCGATCTNNNAGAGCTGAGATAAACGCCCTTT
CACACGTCTGAACTCTTTCCCTACACGACGCTCTTCCGATCTNNNTCTCCAAAGGCTGTTTTCAAAGCC
CACACGTCTGAACTCTTTCCCTACACGACGCTCTTCCGATCTNNNTCTCCAAAGGCTGTTTTCAAAGCC
AGCACACGTCTGAACTCTTTCCCTACACGACGCTCTTCCGATCTNNNAGGCAGGAGAATTACTTGAAC
AGCACACGTCTGAACTCTTTCCCTACACGACGCTCTTCCGATCTNNNAGGCAGGAGAATTACTTGAAC
AGCACACGTCTGAACTCTTTCCCTACACGACGCTCTTCCGATCTNNNACTAGACAGGCCAGTCTCTAG
AGCACACGTCTGAACTCTTTCCCTACACGACGCTCTTCCGATCTNNNACTAGACAGGCCAGTCTCTAG
AGCACACGTCTGAACTCTTTCCCTACACGACGCTCTTCCGATCTNNNGCTGTTGAGTTGTCATTGCTTCT
AGCACACGTCTGAACTCTTTCCCTACACGACGCTCTTCCGATCTNNNGCTGTTGAGTTGTCATTGCTTCT
GAGCACACGTCTGAACTCTTTCCCTACACGACGCTCTTCCGATCTNNNAGTTGTTTTGCATCGGGTCA
GAGCACACGTCTGAACTCTTTCCCTACACGACGCTCTTCCGATCTNNNAGTTGTTTTGCATCGGGTCA
AGCACACGTCTGAACTCTTTCCCTACACGACGCTCTTCCGATCTNNNCTTGAAACTGAAGGCAGAGGT
AGCACACGTCTGAACTCTTTCCCTACACGACGCTCTTCCGATCTNNNCTTGAAACTGAAGGCAGAGGT
GAGCACACGTCTGAACTCTTTCCCTACACGACGCTCTTCCGATCTNNNGGCAGGAGAATTGAGCCAA
GAGCACACGTCTGAACTCTTTCCCTACACGACGCTCTTCCGATCTNNNGGCAGGAGAATTGAGCCAA
AGAGCACACGTCTGAACTCTTTCCCTACACGACGCTCTTCCGATCTNNNTTGAGCCCAGGAATTCAAGG
AGAGCACACGTCTGAACTCTTTCCCTACACGACGCTCTTCCGATCTNNNTTGAGCCCAGGAATTCAAGG
AGAGCACACGTCTGAACTCTTTCCCTACACGACGCTCTTCCGATCTNNNAAAGTGCTGGGATTACAGG
AGAGCACACGTCTGAACTCTTTCCCTACACGACGCTCTTCCGATCTNNNAAAGTGCTGGGATTACAGG
AGCACACGTCTGAACTCTTTCCCTACACGACGCTCTTCCGATCTNNNGTAATCCTAGCTACTCGGAAG
AGCACACGTCTGAACTCTTTCCCTACACGACGCTCTTCCGATCTNNNGTAATCCTAGCTACTCGGAAG
GAGCACACGTCTGAACTCTTTCCCTACACGACGCTCTTCCGATCTNNNAAGTGCTGAGATTACAGGCA
GAGCACACGTCTGAACTCTTTCCCTACACGACGCTCTTCCGATCTNNNAAGTGCTGAGATTACAGGCA
AGCACACGTCTGAACTCTTTCCCTACACGACGCTCTTCCGATCTNNNGTAATAGGCTGGCCTCCAAGAT
AGCACACGTCTGAACTCTTTCCCTACACGACGCTCTTCCGATCTNNNGTAATAGGCTGGCCTCCAAGAT
GAGCACACGTCTGAACTCTTTCCCTACACGACGCTCTTCCGATCTNNNTCTTCCCAGACATCACTTTCCT
GAGCACACGTCTGAACTCTTTCCCTACACGACGCTCTTCCGATCTNNNTCTTCCCAGACATCACTTTCCT
AGCACACGTCTGAACTCTTTCCCTACACGACGCTCTTCCGATCTNNNGCTTCAGGCTTTCTCTGTGTAAG
AGCACACGTCTGAACTCTTTCCCTACACGACGCTCTTCCGATCTNNNGCTTCAGGCTTTCTCTGTGTAAG
GCACACGTCTGAACTCTTTCCCTACACGACGCTCTTCCGATCTNNNCCATGCCTGGCAAATTTTTGTAA
GCACACGTCTGAACTCTTTCCCTACACGACGCTCTTCCGATCTNNNCCATGCCTGGCAAATTTTTGTAA
AGCACACGTCTGAACTCTTTCCCTACACGACGCTCTTCCGATCTNNNCCTTTCTCAGCACCCTCATCAG
AGCACACGTCTGAACTCTTTCCCTACACGACGCTCTTCCGATCTNNNCCTTTCTCAGCACCCTCATCAG
AGCACACGTCTGAACTCTTTCCCTACACGACGCTCTTCCGATCTNNNAATTGCTGGGCCATATGGTAG
AGCACACGTCTGAACTCTTTCCCTACACGACGCTCTTCCGATCTNNNAATTGCTGGGCCATATGGTAG
AGCACACGTCTGAACTCTTTCCCTACACGACGCTCTTCCGATCTNNNGGGAAACATTTCATAGTGGTT
AGCACACGTCTGAACTCTTTCCCTACACGACGCTCTTCCGATCTNNNGGGAAACATTTCATAGTGGTT
GAGCACACGTCTGAACTCTTTCCCTACACGACGCTCTTCCGATCTNNNCTCATGTTTTGGAATGGTTCG
GAGCACACGTCTGAACTCTTTCCCTACACGACGCTCTTCCGATCTNNNCTCATGTTTTGGAATGGTTCG
AGAGCACACGTCTGAACTCTTTCCCTACACGACGCTCTTCCGATCTNNNAGCAAGAGGAGGAGTTTCA
AGAGCACACGTCTGAACTCTTTCCCTACACGACGCTCTTCCGATCTNNNAGCAAGAGGAGGAGTTTCA
GCACACGTCTGAACTCTTTCCCTACACGACGCTCTTCCGATCTNNNTCCTCACATCCAGAACAGTAAGC
GCACACGTCTGAACTCTTTCCCTACACGACGCTCTTCCGATCTNNNTCCTCACATCCAGAACAGTAAGC

AGCACACGTCTGAACTCTTTCCCTACACGACGCTCTTCCGATCTNNNCCAAAAGAGCAAGACATTCCAG
AGCACACGTCTGAACTCTTTCCCTACACGACGCTCTTCCGATCTNNNCCAAAAGAGCAAGACATTCCAG
GCACACGTCTGAACTCTTTCCCTACACGACGCTCTTCCGATCTNNNACTGCCTTTAGAGCTGGTCATTT
GCACACGTCTGAACTCTTTCCCTACACGACGCTCTTCCGATCTNNNACTGCCTTTAGAGCTGGTCATTT
GCACACGTCTGAACTCTTTCCCTACACGACGCTCTTCCGATCTNNNTCAGATGTGCTAGTTGCTCAAG
GCACACGTCTGAACTCTTTCCCTACACGACGCTCTTCCGATCTNNNTCAGATGTGCTAGTTGCTCAAG
AGCACACGTCTGAACTCTTTCCCTACACGACGCTCTTCCGATCTNNNTCTCATGGAAAGTCAGCTGTTG
AGCACACGTCTGAACTCTTTCCCTACACGACGCTCTTCCGATCTNNNTCTCATGGAAAGTCAGCTGTTG
AGCACACGTCTGAACTCTTTCCCTACACGACGCTCTTCCGATCTNNNTCATCAGCTCCAACTTTACCTG
AGCACACGTCTGAACTCTTTCCCTACACGACGCTCTTCCGATCTNNNTCATCAGCTCCAACTTTACCTG
AGCACACGTCTGAACTCTTTCCCTACACGACGCTCTTCCGATCTNNNTGCAGCTTTCTTCTTCCCAATTG
AGCACACGTCTGAACTCTTTCCCTACACGACGCTCTTCCGATCTNNNTGCAGCTTTCTTCTTCCCAATTG
GAGCACACGTCTGAACTCTTTCCCTACACGACGCTCTTCCGATCTNNNCAGGGCCCTTGGAAAAATGT
GAGCACACGTCTGAACTCTTTCCCTACACGACGCTCTTCCGATCTNNNCAGGGCCCTTGGAAAAATGT
GCACACGTCTGAACTCTTTCCCTACACGACGCTCTTCCGATCTNNNTAAAGCTGGCTCTAATCGGTGT
GCACACGTCTGAACTCTTTCCCTACACGACGCTCTTCCGATCTNNNTAAAGCTGGCTCTAATCGGTGT
GCACACGTCTGAACTCTTTCCCTACACGACGCTCTTCCGATCTNNNCTTCTCCCTCTCTAGAGCCTGTAC
GCACACGTCTGAACTCTTTCCCTACACGACGCTCTTCCGATCTNNNCTTCTCCCTCTCTAGAGCCTGTAC
AGCACACGTCTGAACTCTTTCCCTACACGACGCTCTTCCGATCTNNNCTGTAGTCCCAGCTACTCAGGA
AGCACACGTCTGAACTCTTTCCCTACACGACGCTCTTCCGATCTNNNCTGTAGTCCCAGCTACTCAGGA
AGCACACGTCTGAACTCTTTCCCTACACGACGCTCTTCCGATCTNNNCATCAGCAAACTGAACAAACCC
AGCACACGTCTGAACTCTTTCCCTACACGACGCTCTTCCGATCTNNNCATCAGCAAACTGAACAAACCC
AGCACACGTCTGAACTCTTTCCCTACACGACGCTCTTCCGATCTNNNTATTTGGAAAATATTGCATGT
AGCACACGTCTGAACTCTTTCCCTACACGACGCTCTTCCGATCTNNNTATTTGGAAAATATTGCATGT
AGCACACGTCTGAACTCTTTCCCTACACGACGCTCTTCCGATCTNNNTGTAGAGAGAGACAGGGTCT
AGCACACGTCTGAACTCTTTCCCTACACGACGCTCTTCCGATCTNNNTGTAGAGAGAGACAGGGTCT
GCACACGTCTGAACTCTTTCCCTACACGACGCTCTTCCGATCTNNNCAGACTGGCATTCTCAAAACTGG
GCACACGTCTGAACTCTTTCCCTACACGACGCTCTTCCGATCTNNNCAGACTGGCATTCTCAAAACTGG
GCACACGTCTGAACTCTTTCCCTACACGACGCTCTTCCGATCTNNNTTCACCATTCTCCTGCCTCCAT
GCACACGTCTGAACTCTTTCCCTACACGACGCTCTTCCGATCTNNNTTCACCATTCTCCTGCCTCCAT
AGCACACGTCTGAACTCTTTCCCTACACGACGCTCTTCCGATCTNNNCTGGGGATGGTCCTAAGAGG
AGCACACGTCTGAACTCTTTCCCTACACGACGCTCTTCCGATCTNNNCTGGGGATGGTCCTAAGAGG
AAGAGCACACGTCTGAACTCTTTCCCTACACGACGCTCTTCCGATCTNNNTCAAACACCCATCACTTTCC
AAGAGCACACGTCTGAACTCTTTCCCTACACGACGCTCTTCCGATCTNNNTCAAACACCCATCACTTTCC
GAGCACACGTCTGAACTCTTTCCCTACACGACGCTCTTCCGATCTNNNGCTGATGTGGGAAAGAAGT
GAGCACACGTCTGAACTCTTTCCCTACACGACGCTCTTCCGATCTNNNGCTGATGTGGGAAAGAAGT
AGCACACGTCTGAACTCTTTCCCTACACGACGCTCTTCCGATCTNNNATTCTTCACCAATGTCAGGAGC
AGCACACGTCTGAACTCTTTCCCTACACGACGCTCTTCCGATCTNNNATTCTTCACCAATGTCAGGAGC
GAGCACACGTCTGAACTCTTTCCCTACACGACGCTCTTCCGATCTNNNGGTCAGATGCTATTTATTCA
GAGCACACGTCTGAACTCTTTCCCTACACGACGCTCTTCCGATCTNNNGGTCAGATGCTATTTATTCA
CACACGTCTGAACTCTTTCCCTACACGACGCTCTTCCGATCTNNNAGGAACTATGACAGTGTAAAATA
CACACGTCTGAACTCTTTCCCTACACGACGCTCTTCCGATCTNNNAGGAACTATGACAGTGTAAAATA
AGCACACGTCTGAACTCTTTCCCTACACGACGCTCTTCCGATCTNNNCAAACCATATCAGTGGCACGT

AGCACACGTCTGAACTCTTTCCCTACACGACGCTCTTCCGATCTNNNCAAACCATATCAGTGGCACGT
AGCACACGTCTGAACTCTTTCCCTACACGACGCTCTTCCGATCTNNNTCTTGTCTAAACTGGGGGAG
AGCACACGTCTGAACTCTTTCCCTACACGACGCTCTTCCGATCTNNNTCTTGTCTAAACTGGGGGAG
GAGCACACGTCTGAACTCTTTCCCTACACGACGCTCTTCCGATCTNNNGGGTTACCAAGCAAAGACTG
GAGCACACGTCTGAACTCTTTCCCTACACGACGCTCTTCCGATCTNNNGGGTTACCAAGCAAAGACTG
GCACACGTCTGAACTCTTTCCCTACACGACGCTCTTCCGATCTNNNGGAAGGAAATGAGGAGATGGA
GCACACGTCTGAACTCTTTCCCTACACGACGCTCTTCCGATCTNNNGGAAGGAAATGAGGAGATGGA
AGCACACGTCTGAACTCTTTCCCTACACGACGCTCTTCCGATCTNNNGCACATGCAACCAATCATTAG
AGCACACGTCTGAACTCTTTCCCTACACGACGCTCTTCCGATCTNNNGCACATGCAACCAATCATTAG
GAGCACACGTCTGAACTCTTTCCCTACACGACGCTCTTCCGATCTNNNACAGACACCCAGACTTCTATT
GAGCACACGTCTGAACTCTTTCCCTACACGACGCTCTTCCGATCTNNNACAGACACCCAGACTTCTATT
AGCACACGTCTGAACTCTTTCCCTACACGACGCTCTTCCGATCTNNNGCCCTGCAAATACAAGGAGTT
AGCACACGTCTGAACTCTTTCCCTACACGACGCTCTTCCGATCTNNNGCCCTGCAAATACAAGGAGTT
AGCACACGTCTGAACTCTTTCCCTACACGACGCTCTTCCGATCTNNNATGGTTACCAGATGCTGAGAA
AGCACACGTCTGAACTCTTTCCCTACACGACGCTCTTCCGATCTNNNATGGTTACCAGATGCTGAGAA
AGCACACGTCTGAACTCTTTCCCTACACGACGCTCTTCCGATCTNNNCTGCTTTCCTCTGGGGGCTCCAT
AGCACACGTCTGAACTCTTTCCCTACACGACGCTCTTCCGATCTNNNCTGCTTTCCTCTGGGGGCTCCAT
AGCACACGTCTGAACTCTTTCCCTACACGACGCTCTTCCGATCTNNNCACTGTGCCTGGTGACTTTATT
AGCACACGTCTGAACTCTTTCCCTACACGACGCTCTTCCGATCTNNNCACTGTGCCTGGTGACTTTATT
AGCACACGTCTGAACTCTTTCCCTACACGACGCTCTTCCGATCTNNNAGGTACACACATACACACAC
AGCACACGTCTGAACTCTTTCCCTACACGACGCTCTTCCGATCTNNNAGGTACACACATACACACAC
AGAGCACACGTCTGAACTCTTTCCCTACACGACGCTCTTCCGATCTNNNACACTGAATCTTGTGGTTCC
AGAGCACACGTCTGAACTCTTTCCCTACACGACGCTCTTCCGATCTNNNACACTGAATCTTGTGGTTCC
AGCACACGTCTGAACTCTTTCCCTACACGACGCTCTTCCGATCTNNNTATTACATATCGGCCTGCAGT
AGCACACGTCTGAACTCTTTCCCTACACGACGCTCTTCCGATCTNNNTATTACATATCGGCCTGCAGT
GAGCACACGTCTGAACTCTTTCCCTACACGACGCTCTTCCGATCTNNNGCTACTTGCAGAATGAAGCC
GAGCACACGTCTGAACTCTTTCCCTACACGACGCTCTTCCGATCTNNNGCTACTTGCAGAATGAAGCC
GCACACGTCTGAACTCTTTCCCTACACGACGCTCTTCCGATCTNNNATCTCCATTTATTGCTTCTGTTAC
GCACACGTCTGAACTCTTTCCCTACACGACGCTCTTCCGATCTNNNATCTCCATTTATTGCTTCTGTTAC
AAGAGCACACGTCTGAACTCTTTCCCTACACGACGCTCTTCCGATCTNNNCTAGGGTCCCTTCCTTCCAG
AAGAGCACACGTCTGAACTCTTTCCCTACACGACGCTCTTCCGATCTNNNCTAGGGTCCCTTCCTTCCAG
AGCACACGTCTGAACTCTTTCCCTACACGACGCTCTTCCGATCTNNNCTGACATCCAAGGCCCTTTTTTT
AGCACACGTCTGAACTCTTTCCCTACACGACGCTCTTCCGATCTNNNCTGACATCCAAGGCCCTTTTTTT
GAGCACACGTCTGAACTCTTTCCCTACACGACGCTCTTCCGATCTNNNAGTAGGAGCTAGACTATGGG
GAGCACACGTCTGAACTCTTTCCCTACACGACGCTCTTCCGATCTNNNAGTAGGAGCTAGACTATGGG
AGCACACGTCTGAACTCTTTCCCTACACGACGCTCTTCCGATCTNNNACAGCTCCTGGATTTGTTGATCT
AGCACACGTCTGAACTCTTTCCCTACACGACGCTCTTCCGATCTNNNACAGCTCCTGGATTTGTTGATCT
CACACGTCTGAACTCTTTCCCTACACGACGCTCTTCCGATCTNNNGGCACAAGCTTGAACACTAACAT
CACACGTCTGAACTCTTTCCCTACACGACGCTCTTCCGATCTNNNGGCACAAGCTTGAACACTAACAT
AGCACACGTCTGAACTCTTTCCCTACACGACGCTCTTCCGATCTNNNTCACTTTCCTTCTGGATGTTCC
AGCACACGTCTGAACTCTTTCCCTACACGACGCTCTTCCGATCTNNNTCACTTTCCTTCTGGATGTTCC
AGCACACGTCTGAACTCTTTCCCTACACGACGCTCTTCCGATCTNNNTCACTGAGGTGACAGAAACTGT
AGCACACGTCTGAACTCTTTCCCTACACGACGCTCTTCCGATCTNNNTCACTGAGGTGACAGAAACTGT

AGCACACGTCTGAACTCTTTCCCTACACGACGCTCTTCCGATCTNNNTAGCTAGTGATGTTTTTTTGTT
AGCACACGTCTGAACTCTTTCCCTACACGACGCTCTTCCGATCTNNNTAGCTAGTGATGTTTTTTTGTT
AGCACACGTCTGAACTCTTTCCCTACACGACGCTCTTCCGATCTNNNAAATTGGAGAAAGAGCATGCC
AGCACACGTCTGAACTCTTTCCCTACACGACGCTCTTCCGATCTNNNAAATTGGAGAAAGAGCATGCC
CACACGTCTGAACTCTTTCCCTACACGACGCTCTTCCGATCTNNNTGACCCAGGAAAACAAGAGAACA
CACACGTCTGAACTCTTTCCCTACACGACGCTCTTCCGATCTNNNTGACCCAGGAAAACAAGAGAACA
AGCACACGTCTGAACTCTTTCCCTACACGACGCTCTTCCGATCTNNNAGAACTGAGGTTTGCTTTTGC
AGCACACGTCTGAACTCTTTCCCTACACGACGCTCTTCCGATCTNNNAGAACTGAGGTTTGCTTTTGC
AGCACACGTCTGAACTCTTTCCCTACACGACGCTCTTCCGATCTNNNTTACTTTACCCACACCACCACTC
AGCACACGTCTGAACTCTTTCCCTACACGACGCTCTTCCGATCTNNNTTACTTTACCCACACCACCACTC
AGCACACGTCTGAACTCTTTCCCTACACGACGCTCTTCCGATCTNNNATAAATTGCTCTCAGGTCCTGG
AGCACACGTCTGAACTCTTTCCCTACACGACGCTCTTCCGATCTNNNATAAATTGCTCTCAGGTCCTGG
AGCACACGTCTGAACTCTTTCCCTACACGACGCTCTTCCGATCTNNNTCCAGGAGATTCAGAAGAGGC
AGCACACGTCTGAACTCTTTCCCTACACGACGCTCTTCCGATCTNNNTCCAGGAGATTCAGAAGAGGC
AGCACACGTCTGAACTCTTTCCCTACACGACGCTCTTCCGATCTNNNAGATTTCTTGAAGATGACACA
AGCACACGTCTGAACTCTTTCCCTACACGACGCTCTTCCGATCTNNNAGATTTCTTGAAGATGACACA
GCACACGTCTGAACTCTTTCCCTACACGACGCTCTTCCGATCTNNNCTCATGATCTTAGACACCCCCAG
GCACACGTCTGAACTCTTTCCCTACACGACGCTCTTCCGATCTNNNCTCATGATCTTAGACACCCCCAG
AGCACACGTCTGAACTCTTTCCCTACACGACGCTCTTCCGATCTNNNGGGAATTTGGGTAGGGAAG
AGCACACGTCTGAACTCTTTCCCTACACGACGCTCTTCCGATCTNNNGGGAATTTGGGTAGGGAAG
GAGCACACGTCTGAACTCTTTCCCTACACGACGCTCTTCCGATCTNNNTACCTGTGGCCATATCAAAGA
GAGCACACGTCTGAACTCTTTCCCTACACGACGCTCTTCCGATCTNNNTACCTGTGGCCATATCAAAGA
AGCACACGTCTGAACTCTTTCCCTACACGACGCTCTTCCGATCTNNNAATAGTCTCCAATTCCATCCGG
AGCACACGTCTGAACTCTTTCCCTACACGACGCTCTTCCGATCTNNNAATAGTCTCCAATTCCATCCGG
AGCACACGTCTGAACTCTTTCCCTACACGACGCTCTTCCGATCTNNNCACTTGTAGTCCCAGCTACTCGC
AGCACACGTCTGAACTCTTTCCCTACACGACGCTCTTCCGATCTNNNCACTTGTAGTCCCAGCTACTCGC
AGCACACGTCTGAACTCTTTCCCTACACGACGCTCTTCCGATCTNNNGGCCACGTGTATGTCTTTTGAA
AGCACACGTCTGAACTCTTTCCCTACACGACGCTCTTCCGATCTNNNGGCCACGTGTATGTCTTTTGAA
AGCACACGTCTGAACTCTTTCCCTACACGACGCTCTTCCGATCTNNNGGCTTTCTTGGAATCTTGATCCA
AGCACACGTCTGAACTCTTTCCCTACACGACGCTCTTCCGATCTNNNGGCTTTCTTGGAATCTTGATCCA
AGCACACGTCTGAACTCTTTCCCTACACGACGCTCTTCCGATCTNNNGTTCACACCATTCTCCTACCTCAC
AGCACACGTCTGAACTCTTTCCCTACACGACGCTCTTCCGATCTNNNGTTCACACCATTCTCCTACCTCAC
GCACACGTCTGAACTCTTTCCCTACACGACGCTCTTCCGATCTNNNCTGCCCCAAATGAAGACAAATGT
GCACACGTCTGAACTCTTTCCCTACACGACGCTCTTCCGATCTNNNCTGCCCCAAATGAAGACAAATGT
AGCACACGTCTGAACTCTTTCCCTACACGACGCTCTTCCGATCTNNNGGCCATAGTAAAGAGTCTCCCA
AGCACACGTCTGAACTCTTTCCCTACACGACGCTCTTCCGATCTNNNGGCCATAGTAAAGAGTCTCCCA
AGCACACGTCTGAACTCTTTCCCTACACGACGCTCTTCCGATCTNNNTCGATCTCCTGGGATCAAGTGA
AGCACACGTCTGAACTCTTTCCCTACACGACGCTCTTCCGATCTNNNTCGATCTCCTGGGATCAAGTGA
GCACACGTCTGAACTCTTTCCCTACACGACGCTCTTCCGATCTNNNCATCTTGGCTCACTGCAACTTCCA
GCACACGTCTGAACTCTTTCCCTACACGACGCTCTTCCGATCTNNNCATCTTGGCTCACTGCAACTTCCA
AGAGCACACGTCTGAACTCTTTCCCTACACGACGCTCTTCCGATCTNNNTATACAAACAAAGCTTCAACA
AGAGCACACGTCTGAACTCTTTCCCTACACGACGCTCTTCCGATCTNNNTATACAAACAAAGCTTCAACA
GAGCACACGTCTGAACTCTTTCCCTACACGACGCTCTTCCGATCTNNNATGCTGGCACATGTTTGTAAT

AGCACACGTCTGAACTCTTTCCCTACACGACGCTCTTCCGATCTNNNATGCTGGCACATGTTTGTAAT
AGCACACGTCTGAACTCTTTCCCTACACGACGCTCTTCCGATCTNNNCTGTACCCTTGCTACCTGATTG
AGCACACGTCTGAACTCTTTCCCTACACGACGCTCTTCCGATCTNNNCTGTACCCTTGCTACCTGATTG
AGCACACGTCTGAACTCTTTCCCTACACGACGCTCTTCCGATCTNNNCTGTCTTCTGCATGGCTGTTAT
AGCACACGTCTGAACTCTTTCCCTACACGACGCTCTTCCGATCTNNNCTGTCTTCTGCATGGCTGTTAT
AGCACACGTCTGAACTCTTTCCCTACACGACGCTCTTCCGATCTNNNGACCTGCCAGCCCACAATACAT
AGCACACGTCTGAACTCTTTCCCTACACGACGCTCTTCCGATCTNNNGACCTGCCAGCCCACAATACAT
GCACACGTCTGAACTCTTTCCCTACACGACGCTCTTCCGATCTNNNGCTCTCTCTCTTTGCCTCTTTCTC
GCACACGTCTGAACTCTTTCCCTACACGACGCTCTTCCGATCTNNNGCTCTCTCTCTTTGCCTCTTTCTC
GAGCACACGTCTGAACTCTTTCCCTACACGACGCTCTTCCGATCTNNNCATCACTCCATAATTGCTGCC
GAGCACACGTCTGAACTCTTTCCCTACACGACGCTCTTCCGATCTNNNCATCACTCCATAATTGCTGCC
AGCACACGTCTGAACTCTTTCCCTACACGACGCTCTTCCGATCTNNNTTCTGACATCTCTTCTTTTCTTT
AGCACACGTCTGAACTCTTTCCCTACACGACGCTCTTCCGATCTNNNTTCTGACATCTCTTCTTTTCTTT
GAGCACACGTCTGAACTCTTTCCCTACACGACGCTCTTCCGATCTNNNGTGGCACATACCTGTAGTCCT
GAGCACACGTCTGAACTCTTTCCCTACACGACGCTCTTCCGATCTNNNGTGGCACATACCTGTAGTCCT
GCACACGTCTGAACTCTTTCCCTACACGACGCTCTTCCGATCTNNNTTGAGATCTGAGAACAGGCAG
GCACACGTCTGAACTCTTTCCCTACACGACGCTCTTCCGATCTNNNTTGAGATCTGAGAACAGGCAG
AGCACACGTCTGAACTCTTTCCCTACACGACGCTCTTCCGATCTNNNACACATATTCCCATTGCAATGCC
AGCACACGTCTGAACTCTTTCCCTACACGACGCTCTTCCGATCTNNNACACATATTCCCATTGCAATGCC
AGCACACGTCTGAACTCTTTCCCTACACGACGCTCTTCCGATCTNNNAGGCTTCCACTCAAAAGTAGGT
AGCACACGTCTGAACTCTTTCCCTACACGACGCTCTTCCGATCTNNNAGGCTTCCACTCAAAAGTAGGT
GAGCACACGTCTGAACTCTTTCCCTACACGACGCTCTTCCGATCTNNNGGGAAGCTGAGACCAGAGA
GAGCACACGTCTGAACTCTTTCCCTACACGACGCTCTTCCGATCTNNNGGGAAGCTGAGACCAGAGA
AGAGCACACGTCTGAACTCTTTCCCTACACGACGCTCTTCCGATCTNNNTGAGATGAATGAGTTCTGG
AGAGCACACGTCTGAACTCTTTCCCTACACGACGCTCTTCCGATCTNNNTGAGATGAATGAGTTCTGG
GAGCACACGTCTGAACTCTTTCCCTACACGACGCTCTTCCGATCTNNNTAAGGACAGAGAGATGGAG
GAGCACACGTCTGAACTCTTTCCCTACACGACGCTCTTCCGATCTNNNTAAGGACAGAGAGATGGAG
GAGCACACGTCTGAACTCTTTCCCTACACGACGCTCTTCCGATCTNNNGTAAATAGTCTCAGCTCCTCG
GAGCACACGTCTGAACTCTTTCCCTACACGACGCTCTTCCGATCTNNNGTAAATAGTCTCAGCTCCTCG
GAGCACACGTCTGAACTCTTTCCCTACACGACGCTCTTCCGATCTNNNCTGAGGCAGGAGAATCACTT
GAGCACACGTCTGAACTCTTTCCCTACACGACGCTCTTCCGATCTNNNCTGAGGCAGGAGAATCACTT
GAGCACACGTCTGAACTCTTTCCCTACACGACGCTCTTCCGATCTNNNTGTGTTCATAGCTGAGTGCA
GAGCACACGTCTGAACTCTTTCCCTACACGACGCTCTTCCGATCTNNNTGTGTTCATAGCTGAGTGCA
GAGCACACGTCTGAACTCTTTCCCTACACGACGCTCTTCCGATCTNNNTTTGAGATCAGTCTGGGCAA
GAGCACACGTCTGAACTCTTTCCCTACACGACGCTCTTCCGATCTNNNTTTGAGATCAGTCTGGGCAA
AGCACACGTCTGAACTCTTTCCCTACACGACGCTCTTCCGATCTNNNTTTCTTTTCTCAGTCTCTCGCCC
AGCACACGTCTGAACTCTTTCCCTACACGACGCTCTTCCGATCTNNNTTTCTTTTCTCAGTCTCTCGCCC
AGCACACGTCTGAACTCTTTCCCTACACGACGCTCTTCCGATCTNNNGGTGTGGATGTCCTTTCTGTTT
AGCACACGTCTGAACTCTTTCCCTACACGACGCTCTTCCGATCTNNNGGTGTGGATGTCCTTTCTGTTT
AGCACACGTCTGAACTCTTTCCCTACACGACGCTCTTCCGATCTNNNCTGGAAGGTAATGATGGTCAG
AGCACACGTCTGAACTCTTTCCCTACACGACGCTCTTCCGATCTNNNCTGGAAGGTAATGATGGTCAG
GAGCACACGTCTGAACTCTTTCCCTACACGACGCTCTTCCGATCTNNNCACTCAACACTAAGTGGGTGA
GAGCACACGTCTGAACTCTTTCCCTACACGACGCTCTTCCGATCTNNNCACTCAACACTAAGTGGGTGA

CACACGTCTGAACTCTTTCCCTACACGACGCTCTTCCGATCTNNNGGGTCGATCTCAAACAGGTGCAT
CACACGTCTGAACTCTTTCCCTACACGACGCTCTTCCGATCTNNNGGGTCGATCTCAAACAGGTGCAT
AGCACACGTCTGAACTCTTTCCCTACACGACGCTCTTCCGATCTNNNCACACACACACACACACATACT
AGCACACGTCTGAACTCTTTCCCTACACGACGCTCTTCCGATCTNNNCACACACACACACACACATACT
GAAGAGCACACGTCTGAACTCTTTCCCTACACGACGCTCTTCCGATCTNNNTAAGCATCCAAAGCACA
GAAGAGCACACGTCTGAACTCTTTCCCTACACGACGCTCTTCCGATCTNNNTAAGCATCCAAAGCACA
AGAGCACACGTCTGAACTCTTTCCCTACACGACGCTCTTCCGATCTNNNAGTTTTGGGGAGTGTTGTT
AGAGCACACGTCTGAACTCTTTCCCTACACGACGCTCTTCCGATCTNNNAGTTTTGGGGAGTGTTGTT
GCACACGTCTGAACTCTTTCCCTACACGACGCTCTTCCGATCTNNNCACAGCATTTGTTCCAGAGTCAG
GCACACGTCTGAACTCTTTCCCTACACGACGCTCTTCCGATCTNNNCACAGCATTTGTTCCAGAGTCAG
AGAGCACACGTCTGAACTCTTTCCCTACACGACGCTCTTCCGATCTNNNTGTCTTTCCAGCTCCATAGA
AGAGCACACGTCTGAACTCTTTCCCTACACGACGCTCTTCCGATCTNNNTGTCTTTCCAGCTCCATAGA
GCACACGTCTGAACTCTTTCCCTACACGACGCTCTTCCGATCTNNNCTAAGTATCTCCACCAGGATGC
GCACACGTCTGAACTCTTTCCCTACACGACGCTCTTCCGATCTNNNCTAAGTATCTCCACCAGGATGC
AGCACACGTCTGAACTCTTTCCCTACACGACGCTCTTCCGATCTNNNTTGGATGGAAGGTGAGAGAG
AGCACACGTCTGAACTCTTTCCCTACACGACGCTCTTCCGATCTNNNTTGGATGGAAGGTGAGAGAG
GCACACGTCTGAACTCTTTCCCTACACGACGCTCTTCCGATCTNNNCCTGCCAGGTAACCTATAATCCC
GCACACGTCTGAACTCTTTCCCTACACGACGCTCTTCCGATCTNNNCCTGCCAGGTAACCTATAATCCC
AGCACACGTCTGAACTCTTTCCCTACACGACGCTCTTCCGATCTNNNACAAGTCAAGGGTTACTGTGT
AGCACACGTCTGAACTCTTTCCCTACACGACGCTCTTCCGATCTNNNACAAGTCAAGGGTTACTGTGT
GCACACGTCTGAACTCTTTCCCTACACGACGCTCTTCCGATCTNNNTCCAGGCTCATGGATTTCCTGCA
GCACACGTCTGAACTCTTTCCCTACACGACGCTCTTCCGATCTNNNTCCAGGCTCATGGATTTCCTGCA
AGCACACGTCTGAACTCTTTCCCTACACGACGCTCTTCCGATCTNNNTCCTGAAGAACACGGAATCTG
AGCACACGTCTGAACTCTTTCCCTACACGACGCTCTTCCGATCTNNNTCCTGAAGAACACGGAATCTG
GCACACGTCTGAACTCTTTCCCTACACGACGCTCTTCCGATCTNNNTATTCTGGGATGTTGGAAGCAC
GCACACGTCTGAACTCTTTCCCTACACGACGCTCTTCCGATCTNNNTATTCTGGGATGTTGGAAGCAC
AGCACACGTCTGAACTCTTTCCCTACACGACGCTCTTCCGATCTNNNTGCCTATAAGTCCCAGCTACTC
AGCACACGTCTGAACTCTTTCCCTACACGACGCTCTTCCGATCTNNNTGCCTATAAGTCCCAGCTACTC
AGCACACGTCTGAACTCTTTCCCTACACGACGCTCTTCCGATCTNNNCTGAGGCAGGAGAATCACTTG
AGCACACGTCTGAACTCTTTCCCTACACGACGCTCTTCCGATCTNNNCTGAGGCAGGAGAATCACTTG
AGCACACGTCTGAACTCTTTCCCTACACGACGCTCTTCCGATCTNNNTGTAGATGAAACAAGCCCCAT
AGCACACGTCTGAACTCTTTCCCTACACGACGCTCTTCCGATCTNNNTGTAGATGAAACAAGCCCCAT
GAGCACACGTCTGAACTCTTTCCCTACACGACGCTCTTCCGATCTNNNGCACAGAAAGGCAAATACCA
GAGCACACGTCTGAACTCTTTCCCTACACGACGCTCTTCCGATCTNNNGCACAGAAAGGCAAATACCA
AGCACACGTCTGAACTCTTTCCCTACACGACGCTCTTCCGATCTNNNCTCTGGGCAAGTTACTCATCTG
AGCACACGTCTGAACTCTTTCCCTACACGACGCTCTTCCGATCTNNNCTCTGGGCAAGTTACTCATCTG
AGCACACGTCTGAACTCTTTCCCTACACGACGCTCTTCCGATCTNNNTGGGTGACAGAGTGAGATAGT
AGCACACGTCTGAACTCTTTCCCTACACGACGCTCTTCCGATCTNNNTGGGTGACAGAGTGAGATAGT
AGCACACGTCTGAACTCTTTCCCTACACGACGCTCTTCCGATCTNNNGATTACAGGTGTGAGCTACTG
AGCACACGTCTGAACTCTTTCCCTACACGACGCTCTTCCGATCTNNNGATTACAGGTGTGAGCTACTG
GAGCACACGTCTGAACTCTTTCCCTACACGACGCTCTTCCGATCTNNNCACAGGAGAATTGCTTGAAC
GAGCACACGTCTGAACTCTTTCCCTACACGACGCTCTTCCGATCTNNNCACAGGAGAATTGCTTGAAC
CACACGTCTGAACTCTTTCCCTACACGACGCTCTTCCGATCTNNNTGGAAACACAATCTGGATAATATC

CACACGTCTGAACTCTTTCCCTACACGACGCTCTTCCGATCTNNNTGGAAACACAATCTGGATAATATC
AGCACACGTCTGAACTCTTTCCCTACACGACGCTCTTCCGATCTNNNGATGCCTGTAGTCTCAGCTACT
AGCACACGTCTGAACTCTTTCCCTACACGACGCTCTTCCGATCTNNNGATGCCTGTAGTCTCAGCTACT
AGCACACGTCTGAACTCTTTCCCTACACGACGCTCTTCCGATCTNNNGCCTAAGCTACTGCAAAGGAAA
AGCACACGTCTGAACTCTTTCCCTACACGACGCTCTTCCGATCTNNNGCCTAAGCTACTGCAAAGGAAA
GAGCACACGTCTGAACTCTTTCCCTACACGACGCTCTTCCGATCTNNNATACAGAGAGACCCTTGTGCA
GAGCACACGTCTGAACTCTTTCCCTACACGACGCTCTTCCGATCTNNNATACAGAGAGACCCTTGTGCA
GAGCACACGTCTGAACTCTTTCCCTACACGACGCTCTTCCGATCTNNNTCAAATGGCCATCTAGCAGAC
GAGCACACGTCTGAACTCTTTCCCTACACGACGCTCTTCCGATCTNNNTCAAATGGCCATCTAGCAGAC
CACACGTCTGAACTCTTTCCCTACACGACGCTCTTCCGATCTNNNGAGCGACCCTTGATCTTTTTTGTCA
CACACGTCTGAACTCTTTCCCTACACGACGCTCTTCCGATCTNNNGAGCGACCCTTGATCTTTTTTGTCA
GAGCACACGTCTGAACTCTTTCCCTACACGACGCTCTTCCGATCTNNNAGAATAGCATGAACCTGAGAC
GAGCACACGTCTGAACTCTTTCCCTACACGACGCTCTTCCGATCTNNNAGAATAGCATGAACCTGAGAC
AGAGCACACGTCTGAACTCTTTCCCTACACGACGCTCTTCCGATCTNNNCCAAATCTTGACACTTGACCA
AGAGCACACGTCTGAACTCTTTCCCTACACGACGCTCTTCCGATCTNNNCCAAATCTTGACACTTGACCA
GAGCACACGTCTGAACTCTTTCCCTACACGACGCTCTTCCGATCTNNNTTCGAGACCATTCTGACCAACA
GAGCACACGTCTGAACTCTTTCCCTACACGACGCTCTTCCGATCTNNNTTCGAGACCATTCTGACCAACA
CACACGTCTGAACTCTTTCCCTACACGACGCTCTTCCGATCTNNNAAGATAAACAATAATTGGGTTCCCA
CACACGTCTGAACTCTTTCCCTACACGACGCTCTTCCGATCTNNNAAGATAAACAATAATTGGGTTCCCA
AGAGCACACGTCTGAACTCTTTCCCTACACGACGCTCTTCCGATCTNNNTTTTTCAAGAAGCCATCTCCTG
AGAGCACACGTCTGAACTCTTTCCCTACACGACGCTCTTCCGATCTNNNTTTTTCAAGAAGCCATCTCCTG
GAGCACACGTCTGAACTCTTTCCCTACACGACGCTCTTCCGATCTNNNATTTGACTGACAGTAGGAGG
GAGCACACGTCTGAACTCTTTCCCTACACGACGCTCTTCCGATCTNNNATTTGACTGACAGTAGGAGG
ACACGTCTGAACTCTTTCCCTACACGACGCTCTTCCGATCTNNNCAAAAATTAGCCGGACATGGTGTCA
ACACGTCTGAACTCTTTCCCTACACGACGCTCTTCCGATCTNNNCAAAAATTAGCCGGACATGGTGTCA
GAGCACACGTCTGAACTCTTTCCCTACACGACGCTCTTCCGATCTNNNAAGCAATGGTGAGAAGTGCTA
GAGCACACGTCTGAACTCTTTCCCTACACGACGCTCTTCCGATCTNNNAAGCAATGGTGAGAAGTGCTA
CACACGTCTGAACTCTTTCCCTACACGACGCTCTTCCGATCTNNNGTTTTTCCTCCCACCCACTTTTGCAT
CACACGTCTGAACTCTTTCCCTACACGACGCTCTTCCGATCTNNNGTTTTTCCTCCCACCCACTTTTGCAT
AGCACACGTCTGAACTCTTTCCCTACACGACGCTCTTCCGATCTNNNTGAATGCTATTCCTCCAATGGCT
AGCACACGTCTGAACTCTTTCCCTACACGACGCTCTTCCGATCTNNNTGAATGCTATTCCTCCAATGGCT
GAGCACACGTCTGAACTCTTTCCCTACACGACGCTCTTCCGATCTNNNTGTGAATTCTGATGTACCCAG
GAGCACACGTCTGAACTCTTTCCCTACACGACGCTCTTCCGATCTNNNTGTGAATTCTGATGTACCCAG
GAGCACACGTCTGAACTCTTTCCCTACACGACGCTCTTCCGATCTNNNCTTGGGTTTATTTGGATGGGA
GAGCACACGTCTGAACTCTTTCCCTACACGACGCTCTTCCGATCTNNNCTTGGGTTTATTTGGATGGGA
AGCACACGTCTGAACTCTTTCCCTACACGACGCTCTTCCGATCTNNNCTCTTCTCAGACAACGTTTGCTGG
AGCACACGTCTGAACTCTTTCCCTACACGACGCTCTTCCGATCTNNNCTCTTCTCAGACAACGTTTGCTGG
AGCACACGTCTGAACTCTTTCCCTACACGACGCTCTTCCGATCTNNNTCCTCTCCTCCTTTCCTCTTCTCCA
AGCACACGTCTGAACTCTTTCCCTACACGACGCTCTTCCGATCTNNNTCCTCTCCTCCTTTCCTCTTCTCCA
GCACACGTCTGAACTCTTTCCCTACACGACGCTCTTCCGATCTNNNGGGAATCATTGTCTATGGCAGCT
GCACACGTCTGAACTCTTTCCCTACACGACGCTCTTCCGATCTNNNGGGAATCATTGTCTATGGCAGCT
AGCACACGTCTGAACTCTTTCCCTACACGACGCTCTTCCGATCTNNNGCTGAGGAAGGAGAATGGTTT
AGCACACGTCTGAACTCTTTCCCTACACGACGCTCTTCCGATCTNNNGCTGAGGAAGGAGAATGGTTT

AGCACACGTCTGAACTCTTTCCCTACACGACGCTCTTCCGATCTNNNTCCCTCCCATGATATGTGGAGAT

AGCACACGTCTGAACTCTTTCCCTACACGACGCTCTTCCGATCTNNNTCCCTCCCATGATATGTGGAGAT

CACACGTCTGAACTCTTTCCCTACACGACGCTCTTCCGATCTNNNGGCCATGCCTATGTACAGGTTTAC

CACACGTCTGAACTCTTTCCCTACACGACGCTCTTCCGATCTNNNGGCCATGCCTATGTACAGGTTTAC

AGCACACGTCTGAACTCTTTCCCTACACGACGCTCTTCCGATCTNNNCACCATTTGTTTCCACTTCTGCAC

AGCACACGTCTGAACTCTTTCCCTACACGACGCTCTTCCGATCTNNNCACCATTTGTTTCCACTTCTGCAC

ACACGTCTGAACTCTTTCCCTACACGACGCTCTTCCGATCTNNNGTGGCTTTCACTGTAAATTTGCATCG

ACACGTCTGAACTCTTTCCCTACACGACGCTCTTCCGATCTNNNGTGGCTTTCACTGTAAATTTGCATCG

AGCACACGTCTGAACTCTTTCCCTACACGACGCTCTTCCGATCTNNNTGAACATTACTTGGACTCCAGCA

AGCACACGTCTGAACTCTTTCCCTACACGACGCTCTTCCGATCTNNNTGAACATTACTTGGACTCCAGCA

CACACGTCTGAACTCTTTCCCTACACGACGCTCTTCCGATCTNNNCACAACGACCACATCCTCATAGACA

CACACGTCTGAACTCTTTCCCTACACGACGCTCTTCCGATCTNNNCACAACGACCACATCCTCATAGACA

AGCACACGTCTGAACTCTTTCCCTACACGACGCTCTTCCGATCTNNNGAAGCTGACCATAGAGGAGTT

AGCACACGTCTGAACTCTTTCCCTACACGACGCTCTTCCGATCTNNNGAAGCTGACCATAGAGGAGTT

GAGCACACGTCTGAACTCTTTCCCTACACGACGCTCTTCCGATCTNNNTAAGCTTTTTGATGTGCTGCTG

GAGCACACGTCTGAACTCTTTCCCTACACGACGCTCTTCCGATCTNNNTAAGCTTTTTGATGTGCTGCTG

AGAGCACACGTCTGAACTCTTTCCCTACACGACGCTCTTCCGATCTNNNGAGAGGCAGAGAAGGAGA

AGAGCACACGTCTGAACTCTTTCCCTACACGACGCTCTTCCGATCTNNNGAGAGGCAGAGAAGGAGA

AGCACACGTCTGAACTCTTTCCCTACACGACGCTCTTCCGATCTNNNACTCACTTTCCACAAAGGTACCA

AGCACACGTCTGAACTCTTTCCCTACACGACGCTCTTCCGATCTNNNACTCACTTTCCACAAAGGTACCA

GCACACGTCTGAACTCTTTCCCTACACGACGCTCTTCCGATCTNNNTGCATAGGCTTTGTATGGTAGGC

GCACACGTCTGAACTCTTTCCCTACACGACGCTCTTCCGATCTNNNTGCATAGGCTTTGTATGGTAGGC

GCACACGTCTGAACTCTTTCCCTACACGACGCTCTTCCGATCTNNNCATTCTGCTTGTTCTGTTCCACACA

GCACACGTCTGAACTCTTTCCCTACACGACGCTCTTCCGATCTNNNCATTCTGCTTGTTCTGTTCCACACA

AGAGCACACGTCTGAACTCTTTCCCTACACGACGCTCTTCCGATCTNNNATAGTTGCAGGTGCCTGTAG

AGAGCACACGTCTGAACTCTTTCCCTACACGACGCTCTTCCGATCTNNNATAGTTGCAGGTGCCTGTAG

GAGCACACGTCTGAACTCTTTCCCTACACGACGCTCTTCCGATCTNNNCACCGATTTCCACAATGGTTGA

GAGCACACGTCTGAACTCTTTCCCTACACGACGCTCTTCCGATCTNNNCACCGATTTCCACAATGGTTGA

GCACACGTCTGAACTCTTTCCCTACACGACGCTCTTCCGATCTNNNGGCAGTAAGCTCCATTCTGAGTT

GCACACGTCTGAACTCTTTCCCTACACGACGCTCTTCCGATCTNNNGGCAGTAAGCTCCATTCTGAGTT

GAGCACACGTCTGAACTCTTTCCCTACACGACGCTCTTCCGATCTNNNCGGGTGAATCAATTGAGTTCA

GAGCACACGTCTGAACTCTTTCCCTACACGACGCTCTTCCGATCTNNNCGGGTGAATCAATTGAGTTCA

GAGCACACGTCTGAACTCTTTCCCTACACGACGCTCTTCCGATCTNNNGCCATTCATCTCGGGACTTTCC

GAGCACACGTCTGAACTCTTTCCCTACACGACGCTCTTCCGATCTNNNGCCATTCATCTCGGGACTTTCC

GAGCACACGTCTGAACTCTTTCCCTACACGACGCTCTTCCGATCTNNNGAACTTCCCAGGTGTGAAATG

GAGCACACGTCTGAACTCTTTCCCTACACGACGCTCTTCCGATCTNNNGAACTTCCCAGGTGTGAAATG

AGAGCACACGTCTGAACTCTTTCCCTACACGACGCTCTTCCGATCTNNNGCTTTTGGTGTCTTCCATCTG

AGAGCACACGTCTGAACTCTTTCCCTACACGACGCTCTTCCGATCTNNNGCTTTTGGTGTCTTCCATCTG

AGCACACGTCTGAACTCTTTCCCTACACGACGCTCTTCCGATCTNNNGCATTGAATGTGCCTTTCCTTTC

AGCACACGTCTGAACTCTTTCCCTACACGACGCTCTTCCGATCTNNNGCATTGAATGTGCCTTTCCTTTC

AGAGCACACGTCTGAACTCTTTCCCTACACGACGCTCTTCCGATCTNNNACAAAATTTAGCTGGGTGTG

AGAGCACACGTCTGAACTCTTTCCCTACACGACGCTCTTCCGATCTNNNACAAAATTTAGCTGGGTGTG

AGCACACGTCTGAACTCTTTCCCTACACGACGCTCTTCCGATCTNNNAGCCTTCCAAGAAGCTGAAACT

AGCACACGTCTGAACTCTTTCCCTACACGACGCTCTTCCGATCTNNNAGCCTTCCAAGAAGCTGAAACT
GCACACGTCTGAACTCTTTCCCTACACGACGCTCTTCCGATCTNNNTGGGAGAGAGGTGTGTGATTTT
GCACACGTCTGAACTCTTTCCCTACACGACGCTCTTCCGATCTNNNTGGGAGAGAGGTGTGTGATTTT
AGCACACGTCTGAACTCTTTCCCTACACGACGCTCTTCCGATCTNNNTCTCAGTTTTGTTTGTCTGGGA
AGCACACGTCTGAACTCTTTCCCTACACGACGCTCTTCCGATCTNNNTCTCAGTTTTGTTTGTCTGGGA
GCACACGTCTGAACTCTTTCCCTACACGACGCTCTTCCGATCTNNNAGCTAGGATGGTCTTGTACTCCT
GCACACGTCTGAACTCTTTCCCTACACGACGCTCTTCCGATCTNNNAGCTAGGATGGTCTTGTACTCCT
AGCACACGTCTGAACTCTTTCCCTACACGACGCTCTTCCGATCTNNNCACACACACACATAGCACAGTTT
AGCACACGTCTGAACTCTTTCCCTACACGACGCTCTTCCGATCTNNNCACACACACACATAGCACAGTTT
AGCACACGTCTGAACTCTTTCCCTACACGACGCTCTTCCGATCTNNNCTGAGGCAGGAAAATTGCTTGA
AGCACACGTCTGAACTCTTTCCCTACACGACGCTCTTCCGATCTNNNCTGAGGCAGGAAAATTGCTTGA
GAGCACACGTCTGAACTCTTTCCCTACACGACGCTCTTCCGATCTNNNCACCACTCCTTGTATCTGAGGT
GAGCACACGTCTGAACTCTTTCCCTACACGACGCTCTTCCGATCTNNNCACCACTCCTTGTATCTGAGGT
AGCACACGTCTGAACTCTTTCCCTACACGACGCTCTTCCGATCTNNNCTCCTGGGTTCAAGCCATTCTCCA
AGCACACGTCTGAACTCTTTCCCTACACGACGCTCTTCCGATCTNNNCTCCTGGGTTCAAGCCATTCTCCA
AGCACACGTCTGAACTCTTTCCCTACACGACGCTCTTCCGATCTNNNTAACCAACCCAGCTGTTTCTGTA
AGCACACGTCTGAACTCTTTCCCTACACGACGCTCTTCCGATCTNNNTAACCAACCCAGCTGTTTCTGTA
AGCACACGTCTGAACTCTTTCCCTACACGACGCTCTTCCGATCTNNNGCTGTCATGAACAATGCTGCTAT
AGCACACGTCTGAACTCTTTCCCTACACGACGCTCTTCCGATCTNNNGCTGTCATGAACAATGCTGCTAT
AGCACACGTCTGAACTCTTTCCCTACACGACGCTCTTCCGATCTNNNGCAACAGAGCAAAAACTCCATC
AGCACACGTCTGAACTCTTTCCCTACACGACGCTCTTCCGATCTNNNGCAACAGAGCAAAAACTCCATC
GCACACGTCTGAACTCTTTCCCTACACGACGCTCTTCCGATCTNNNGGGAAGAACAATCTACCCAATGCC
GCACACGTCTGAACTCTTTCCCTACACGACGCTCTTCCGATCTNNNGGGAAGAACAATCTACCCAATGCC
AGAGCACACGTCTGAACTCTTTCCCTACACGACGCTCTTCCGATCTNNNTGGAAGAAAAGGCTAAGGC
AGAGCACACGTCTGAACTCTTTCCCTACACGACGCTCTTCCGATCTNNNTGGAAGAAAAGGCTAAGGC
GAGCACACGTCTGAACTCTTTCCCTACACGACGCTCTTCCGATCTNNNCCCCAAAGTCCATTGTGTCATT
GAGCACACGTCTGAACTCTTTCCCTACACGACGCTCTTCCGATCTNNNCCCCAAAGTCCATTGTGTCATT
GAGCACACGTCTGAACTCTTTCCCTACACGACGCTCTTCCGATCTNNNCCAGCAGGGATGTCAGACTTTA
GAGCACACGTCTGAACTCTTTCCCTACACGACGCTCTTCCGATCTNNNCCAGCAGGGATGTCAGACTTTA
AGCACACGTCTGAACTCTTTCCCTACACGACGCTCTTCCGATCTNNNGAACTCCTGACCTCGTGATCTG
AGCACACGTCTGAACTCTTTCCCTACACGACGCTCTTCCGATCTNNNGAACTCCTGACCTCGTGATCTG
AGCACACGTCTGAACTCTTTCCCTACACGACGCTCTTCCGATCTNNNCATATGGCAATTTGGGGGACTC
AGCACACGTCTGAACTCTTTCCCTACACGACGCTCTTCCGATCTNNNCATATGGCAATTTGGGGGACTC
AGAGCACACGTCTGAACTCTTTCCCTACACGACGCTCTTCCGATCTNNNGCTTGGAAGATCAGCAAGCT
AGAGCACACGTCTGAACTCTTTCCCTACACGACGCTCTTCCGATCTNNNGCTTGGAAGATCAGCAAGCT
AGCACACGTCTGAACTCTTTCCCTACACGACGCTCTTCCGATCTNNNGAATAGGAATGGCCAAGTGGA
AGCACACGTCTGAACTCTTTCCCTACACGACGCTCTTCCGATCTNNNGAATAGGAATGGCCAAGTGGA
GAGCACACGTCTGAACTCTTTCCCTACACGACGCTCTTCCGATCTNNNGAAATCCCGGGAGTTGGAAA
GAGCACACGTCTGAACTCTTTCCCTACACGACGCTCTTCCGATCTNNNGAAATCCCGGGAGTTGGAAA
AGCACACGTCTGAACTCTTTCCCTACACGACGCTCTTCCGATCTNNNTGCATACCTGTAATCCCAGCTAC
AGCACACGTCTGAACTCTTTCCCTACACGACGCTCTTCCGATCTNNNTGCATACCTGTAATCCCAGCTAC
AGCACACGTCTGAACTCTTTCCCTACACGACGCTCTTCCGATCTNNNTAAGAGTGTAGTATGAATGATC
AGCACACGTCTGAACTCTTTCCCTACACGACGCTCTTCCGATCTNNNTAAGAGTGTAGTATGAATGATC

AGCACACGTCTGAACTCTTTCCCTACACGACGCTCTTCCGATCTNNNCTGGGCAAGAGACTGAGACT
AGCACACGTCTGAACTCTTTCCCTACACGACGCTCTTCCGATCTNNNCTGGGCAAGAGACTGAGACT
CACACGTCTGAACTCTTTCCCTACACGACGCTCTTCCGATCTNNNTTGTATTTCTAGCAGAGACGCATC
CACACGTCTGAACTCTTTCCCTACACGACGCTCTTCCGATCTNNNTTGTATTTCTAGCAGAGACGCATC
GAGCACACGTCTGAACTCTTTCCCTACACGACGCTCTTCCGATCTNNNGAGATCCCACCACTGCACTCC
GAGCACACGTCTGAACTCTTTCCCTACACGACGCTCTTCCGATCTNNNGAGATCCCACCACTGCACTCC
GAGCACACGTCTGAACTCTTTCCCTACACGACGCTCTTCCGATCTNNNCTGAGGCAGGAGAATCACT
GAGCACACGTCTGAACTCTTTCCCTACACGACGCTCTTCCGATCTNNNCTGAGGCAGGAGAATCACT
AGCACACGTCTGAACTCTTTCCCTACACGACGCTCTTCCGATCTNNNCCATGCAAATGGAAACCAAAA
AGCACACGTCTGAACTCTTTCCCTACACGACGCTCTTCCGATCTNNNCCATGCAAATGGAAACCAAAA
GCACACGTCTGAACTCTTTCCCTACACGACGCTCTTCCGATCTNNNCCATCATGACATTGAGACTCTG
GCACACGTCTGAACTCTTTCCCTACACGACGCTCTTCCGATCTNNNCCATCATGACATTGAGACTCTG
GCACACGTCTGAACTCTTTCCCTACACGACGCTCTTCCGATCTNNNGAAGTACATGGACAGGCAGACA
GCACACGTCTGAACTCTTTCCCTACACGACGCTCTTCCGATCTNNNGAAGTACATGGACAGGCAGACA
GCACACGTCTGAACTCTTTCCCTACACGACGCTCTTCCGATCTNNNTACAAAAAATTAGCCGGGCGTG
GCACACGTCTGAACTCTTTCCCTACACGACGCTCTTCCGATCTNNNTACAAAAAATTAGCCGGGCGTG
GCACACGTCTGAACTCTTTCCCTACACGACGCTCTTCCGATCTNNNCAGATTCATCTAACATTGCAATA
GCACACGTCTGAACTCTTTCCCTACACGACGCTCTTCCGATCTNNNCAGATTCATCTAACATTGCAATA
GCACACGTCTGAACTCTTTCCCTACACGACGCTCTTCCGATCTNNNGAGGCAGAAGAATCGCTTGAA
GCACACGTCTGAACTCTTTCCCTACACGACGCTCTTCCGATCTNNNGAGGCAGAAGAATCGCTTGAA
AGCACACGTCTGAACTCTTTCCCTACACGACGCTCTTCCGATCTNNNAATGTCCCAGTTTCTTTCCCACT
AGCACACGTCTGAACTCTTTCCCTACACGACGCTCTTCCGATCTNNNAATGTCCCAGTTTCTTTCCCACT
AGCACACGTCTGAACTCTTTCCCTACACGACGCTCTTCCGATCTNNNGGGACTTTCAGGTTGGATGAG
AGCACACGTCTGAACTCTTTCCCTACACGACGCTCTTCCGATCTNNNGGGACTTTCAGGTTGGATGAG
GAGCACACGTCTGAACTCTTTCCCTACACGACGCTCTTCCGATCTNNNAAAAAAAATTAGCCGGGTGT
GAGCACACGTCTGAACTCTTTCCCTACACGACGCTCTTCCGATCTNNNAAAAAAAATTAGCCGGGTGT
AGCACACGTCTGAACTCTTTCCCTACACGACGCTCTTCCGATCTNNNCATATTCACTGCCATGCTTGGT
AGCACACGTCTGAACTCTTTCCCTACACGACGCTCTTCCGATCTNNNCATATTCACTGCCATGCTTGGT
ACACGTCTGAACTCTTTCCCTACACGACGCTCTTCCGATCTNNNATCAACTCTTCTCTGGTTCTCGGCA
ACACGTCTGAACTCTTTCCCTACACGACGCTCTTCCGATCTNNNATCAACTCTTCTCTGGTTCTCGGCA
ACACGTCTGAACTCTTTCCCTACACGACGCTCTTCCGATCTNNNGATGTAGTTACTGTGGAGTCTCAT
ACACGTCTGAACTCTTTCCCTACACGACGCTCTTCCGATCTNNNGATGTAGTTACTGTGGAGTCTCAT
AGCACACGTCTGAACTCTTTCCCTACACGACGCTCTTCCGATCTNNNCATGATGGCGAGTGCTTGTAA
AGCACACGTCTGAACTCTTTCCCTACACGACGCTCTTCCGATCTNNNCATGATGGCGAGTGCTTGTAA
GAGCACACGTCTGAACTCTTTCCCTACACGACGCTCTTCCGATCTNNNACATGTTGCCGAACAGACATG
GAGCACACGTCTGAACTCTTTCCCTACACGACGCTCTTCCGATCTNNNACATGTTGCCGAACAGACATG
GCACACGTCTGAACTCTTTCCCTACACGACGCTCTTCCGATCTNNNATCAGGTATCCCGGCATCTTTTG
GCACACGTCTGAACTCTTTCCCTACACGACGCTCTTCCGATCTNNNATCAGGTATCCCGGCATCTTTTG
GAGCACACGTCTGAACTCTTTCCCTACACGACGCTCTTCCGATCTNNNTTCCACCATTCCAGATGTTTTG
GAGCACACGTCTGAACTCTTTCCCTACACGACGCTCTTCCGATCTNNNTTCCACCATTCCAGATGTTTTG
GCACACGTCTGAACTCTTTCCCTACACGACGCTCTTCCGATCTNNNAATAACCGAGAGAGAGGAGG
GCACACGTCTGAACTCTTTCCCTACACGACGCTCTTCCGATCTNNNAATAACCGAGAGAGAGGAGG
GAGCACACGTCTGAACTCTTTCCCTACACGACGCTCTTCCGATCTNNNTCATCTCCCACAATGTAACCTC

GAGCACACGTCTGAACTCTTTCCCTACACGACGCTCTTCCGATCTNNNTCATCTCCCACAATGTAACCTC
CACACGTCTGAACTCTTTCCCTACACGACGCTCTTCCGATCTNNNGGAGGAGGGTAAGGGTTGAAAA
CACACGTCTGAACTCTTTCCCTACACGACGCTCTTCCGATCTNNNGGAGGAGGGTAAGGGTTGAAAA
AGCACACGTCTGAACTCTTTCCCTACACGACGCTCTTCCGATCTNNNTTTTGAGTCAGGGTCTCACTCT
AGCACACGTCTGAACTCTTTCCCTACACGACGCTCTTCCGATCTNNNTTTTGAGTCAGGGTCTCACTCT
AGCACACGTCTGAACTCTTTCCCTACACGACGCTCTTCCGATCTNNNTGGCAACAAAACAAAACTCTC
AGCACACGTCTGAACTCTTTCCCTACACGACGCTCTTCCGATCTNNNTGGCAACAAAACAAAACTCTC
AGCACACGTCTGAACTCTTTCCCTACACGACGCTCTTCCGATCTNNNAGGGAAGCATTTTTCTCATTGG
AGCACACGTCTGAACTCTTTCCCTACACGACGCTCTTCCGATCTNNNAGGGAAGCATTTTTCTCATTGG
AGCACACGTCTGAACTCTTTCCCTACACGACGCTCTTCCGATCTNNNCAGGATTCTTACAGAAAACAAG
AGCACACGTCTGAACTCTTTCCCTACACGACGCTCTTCCGATCTNNNCAGGATTCTTACAGAAAACAAG
AGCACACGTCTGAACTCTTTCCCTACACGACGCTCTTCCGATCTNNNCAGGATTCTTACAGAAAACAAG
AGCACACGTCTGAACTCTTTCCCTACACGACGCTCTTCCGATCTNNNCAGGATTCTTACAGAAAACAAG
GAGCACACGTCTGAACTCTTTCCCTACACGACGCTCTTCCGATCTNNNTGGGAGTGGGAGGGGAAAA
GAGCACACGTCTGAACTCTTTCCCTACACGACGCTCTTCCGATCTNNNTGGGAGTGGGAGGGGAAAA
AGCACACGTCTGAACTCTTTCCCTACACGACGCTCTTCCGATCTNNNCCTATTCCGTAGCATTTTGCCTG
AGCACACGTCTGAACTCTTTCCCTACACGACGCTCTTCCGATCTNNNCCTATTCCGTAGCATTTTGCCTG
GCACACGTCTGAACTCTTTCCCTACACGACGCTCTTCCGATCTNNNCTCCTGGGTTCAAGAGATTCTCC
GCACACGTCTGAACTCTTTCCCTACACGACGCTCTTCCGATCTNNNCTCCTGGGTTCAAGAGATTCTCC
GAGCACACGTCTGAACTCTTTCCCTACACGACGCTCTTCCGATCTNNNTGGAGTGCAGTGGTGAATCA
GAGCACACGTCTGAACTCTTTCCCTACACGACGCTCTTCCGATCTNNNTGGAGTGCAGTGGTGAATCA
GCACACGTCTGAACTCTTTCCCTACACGACGCTCTTCCGATCTNNNGGTAAGGAAAGCAAAACCCTGT
GCACACGTCTGAACTCTTTCCCTACACGACGCTCTTCCGATCTNNNGGTAAGGAAAGCAAAACCCTGT
AGCACACGTCTGAACTCTTTCCCTACACGACGCTCTTCCGATCTNNNTCTCTCTCTCTCTCTCCCTCTC
AGCACACGTCTGAACTCTTTCCCTACACGACGCTCTTCCGATCTNNNTCTCTCTCTCTCTCTCCCTCTC
GAGCACACGTCTGAACTCTTTCCCTACACGACGCTCTTCCGATCTNNNCCTTGCATATCAGACAAACCA
GAGCACACGTCTGAACTCTTTCCCTACACGACGCTCTTCCGATCTNNNCCTTGCATATCAGACAAACCA
GCACACGTCTGAACTCTTTCCCTACACGACGCTCTTCCGATCTNNNTTGATGGAGATGGGTTGCTGAA
GCACACGTCTGAACTCTTTCCCTACACGACGCTCTTCCGATCTNNNTTGATGGAGATGGGTTGCTGAA
AGCACACGTCTGAACTCTTTCCCTACACGACGCTCTTCCGATCTNNNCATGCATGCTGAGAAAGAGAG
AGCACACGTCTGAACTCTTTCCCTACACGACGCTCTTCCGATCTNNNCATGCATGCTGAGAAAGAGAG
GAGCACACGTCTGAACTCTTTCCCTACACGACGCTCTTCCGATCTNNNATGTCTTGTGTTCTCCATCCA
GAGCACACGTCTGAACTCTTTCCCTACACGACGCTCTTCCGATCTNNNATGTCTTGTGTTCTCCATCCA
GCACACGTCTGAACTCTTTCCCTACACGACGCTCTTCCGATCTNNNAAGTAGCTGAGATTACAGGTGC
GCACACGTCTGAACTCTTTCCCTACACGACGCTCTTCCGATCTNNNAAGTAGCTGAGATTACAGGTGC
AGCACACGTCTGAACTCTTTCCCTACACGACGCTCTTCCGATCTNNNGCAATCCTAGCACTTTGGGAG
AGCACACGTCTGAACTCTTTCCCTACACGACGCTCTTCCGATCTNNNGCAATCCTAGCACTTTGGGAG
GAGCACACGTCTGAACTCTTTCCCTACACGACGCTCTTCCGATCTNNNTGAACTGTGATTGAGCCACTG
GAGCACACGTCTGAACTCTTTCCCTACACGACGCTCTTCCGATCTNNNTGAACTGTGATTGAGCCACTG
GAGCACACGTCTGAACTCTTTCCCTACACGACGCTCTTCCGATCTNNNGGCTTTCAAACCGGAAAAGA
GAGCACACGTCTGAACTCTTTCCCTACACGACGCTCTTCCGATCTNNNGGCTTTCAAACCGGAAAAGA
AGCACACGTCTGAACTCTTTCCCTACACGACGCTCTTCCGATCTNNNGCATAGCTCACTTATTATGCGG
AGCACACGTCTGAACTCTTTCCCTACACGACGCTCTTCCGATCTNNNGCATAGCTCACTTATTATGCGG

AAGAGCACACGTCTGAACTCTTTCCCTACACGACGCTCTTCCGATCTNNNTGGGTGACAGAGTGAGAC

AAGAGCACACGTCTGAACTCTTTCCCTACACGACGCTCTTCCGATCTNNNTGGGTGACAGAGTGAGAC

AGAGCACACGTCTGAACTCTTTCCCTACACGACGCTCTTCCGATCTNNNTGGAGACCATGAAAGTAAT

AGAGCACACGTCTGAACTCTTTCCCTACACGACGCTCTTCCGATCTNNNTGGAGACCATGAAAGTAAT

GAGCACACGTCTGAACTCTTTCCCTACACGACGCTCTTCCGATCTNNNAGTTGCAGTGAGCTGAGATTC

GAGCACACGTCTGAACTCTTTCCCTACACGACGCTCTTCCGATCTNNNAGTTGCAGTGAGCTGAGATTC

AGCACACGTCTGAACTCTTTCCCTACACGACGCTCTTCCGATCTNNNTTGCTAAAGACATCTGTCCCTCC

AGCACACGTCTGAACTCTTTCCCTACACGACGCTCTTCCGATCTNNNTTGCTAAAGACATCTGTCCCTCC

GAGCACACGTCTGAACTCTTTCCCTACACGACGCTCTTCCGATCTNNNTCAAGGACGCCAGGAAAATCA

GAGCACACGTCTGAACTCTTTCCCTACACGACGCTCTTCCGATCTNNNTCAAGGACGCCAGGAAAATCA

AGCACACGTCTGAACTCTTTCCCTACACGACGCTCTTCCGATCTNNNTGAGATAAAGCCACTGCACTTCA

AGCACACGTCTGAACTCTTTCCCTACACGACGCTCTTCCGATCTNNNTGAGATAAAGCCACTGCACTTCA

AGCACACGTCTGAACTCTTTCCCTACACGACGCTCTTCCGATCTNNNATACAAAAATTAGCCGGGCATG

AGCACACGTCTGAACTCTTTCCCTACACGACGCTCTTCCGATCTNNNATACAAAAATTAGCCGGGCATG

AGAGCACACGTCTGAACTCTTTCCCTACACGACGCTCTTCCGATCTNNNGCATCTTGACCATCCCTTTAC

AGAGCACACGTCTGAACTCTTTCCCTACACGACGCTCTTCCGATCTNNNGCATCTTGACCATCCCTTTAC

AGCACACGTCTGAACTCTTTCCCTACACGACGCTCTTCCGATCTNNNCATCCTGGGCAGAAGACATAGTA

AGCACACGTCTGAACTCTTTCCCTACACGACGCTCTTCCGATCTNNNCATCCTGGGCAGAAGACATAGTA

AGAGCACACGTCTGAACTCTTTCCCTACACGACGCTCTTCCGATCTNNNTCCAGTAAGTTCAGAGAACAC

AGAGCACACGTCTGAACTCTTTCCCTACACGACGCTCTTCCGATCTNNNTCCAGTAAGTTCAGAGAACAC

GAGCACACGTCTGAACTCTTTCCCTACACGACGCTCTTCCGATCTNNNAGAGAGAGAGCAAGAGATAG

GAGCACACGTCTGAACTCTTTCCCTACACGACGCTCTTCCGATCTNNNAGAGAGAGAGCAAGAGATAG

AGCACACGTCTGAACTCTTTCCCTACACGACGCTCTTCCGATCTNNNCTGTGCCTGTCCAGTTTCTTTTT

AGCACACGTCTGAACTCTTTCCCTACACGACGCTCTTCCGATCTNNNCTGTGCCTGTCCAGTTTCTTTTT

AGCACACGTCTGAACTCTTTCCCTACACGACGCTCTTCCGATCTNNNAACCAACACCCATTCATGAGGTA

AGCACACGTCTGAACTCTTTCCCTACACGACGCTCTTCCGATCTNNNAACCAACACCCATTCATGAGGTA

AGAGCACACGTCTGAACTCTTTCCCTACACGACGCTCTTCCGATCTNNNGTAATCCCAGCTATACAGGAC

AGAGCACACGTCTGAACTCTTTCCCTACACGACGCTCTTCCGATCTNNNGTAATCCCAGCTATACAGGAC

GAGCACACGTCTGAACTCTTTCCCTACACGACGCTCTTCCGATCTNNNCCATGGTGGAAGTAGATGTGT

GAGCACACGTCTGAACTCTTTCCCTACACGACGCTCTTCCGATCTNNNCCATGGTGGAAGTAGATGTGT

GAGCACACGTCTGAACTCTTTCCCTACACGACGCTCTTCCGATCTNNNCATTTGTAGTCCCAGCTACTCG

GAGCACACGTCTGAACTCTTTCCCTACACGACGCTCTTCCGATCTNNNCATTTGTAGTCCCAGCTACTCG

AGCACACGTCTGAACTCTTTCCCTACACGACGCTCTTCCGATCTNNNGCTGGTTGCCCATTTTTTTATG

AGCACACGTCTGAACTCTTTCCCTACACGACGCTCTTCCGATCTNNNGCTGGTTGCCCATTTTTTTATG

AGCACACGTCTGAACTCTTTCCCTACACGACGCTCTTCCGATCTNNNCAATCTCCTGAAGACAGATGCA

AGCACACGTCTGAACTCTTTCCCTACACGACGCTCTTCCGATCTNNNCAATCTCCTGAAGACAGATGCA

AGCACACGTCTGAACTCTTTCCCTACACGACGCTCTTCCGATCTNNNTAGAAACAATACGTATTTGCTTC

AGCACACGTCTGAACTCTTTCCCTACACGACGCTCTTCCGATCTNNNTAGAAACAATACGTATTTGCTTC

AGCACACGTCTGAACTCTTTCCCTACACGACGCTCTTCCGATCTNNNACACAAAAAAACAGTGAGCTGC

AGCACACGTCTGAACTCTTTCCCTACACGACGCTCTTCCGATCTNNNACACAAAAAAACAGTGAGCTGC

AGCACACGTCTGAACTCTTTCCCTACACGACGCTCTTCCGATCTNNNTGGTGAATCCAGGCAGTTAGA

AGCACACGTCTGAACTCTTTCCCTACACGACGCTCTTCCGATCTNNNTGGTGAATCCAGGCAGTTAGA

AGCACACGTCTGAACTCTTTCCCTACACGACGCTCTTCCGATCTNNNCTTACCAAGTAGATGCCCCAAG

AGCACACGTCTGAACTCTTTCCCTACACGACGCTCTTCCGATCTNNNCTTACCAAGTAGATGCCCCAAG
AGAGCACACGTCTGAACTCTTTCCCTACACGACGCTCTTCCGATCTNNNGAGATTCGCTTGAACTCAG
AGAGCACACGTCTGAACTCTTTCCCTACACGACGCTCTTCCGATCTNNNGAGATTCGCTTGAACTCAG
AGCACACGTCTGAACTCTTTCCCTACACGACGCTCTTCCGATCTNNNTTGATTTAGGAACCAGGCTGT
AGCACACGTCTGAACTCTTTCCCTACACGACGCTCTTCCGATCTNNNTTGATTTAGGAACCAGGCTGT
ACACGTCTGAACTCTTTCCCTACACGACGCTCTTCCGATCTNNNATATATGTGCCATCTTTCTACATCAT
ACACGTCTGAACTCTTTCCCTACACGACGCTCTTCCGATCTNNNATATATGTGCCATCTTTCTACATCAT
AGCACACGTCTGAACTCTTTCCCTACACGACGCTCTTCCGATCTNNNCACCTGTATTCCCAGCTACTCA
AGCACACGTCTGAACTCTTTCCCTACACGACGCTCTTCCGATCTNNNCACCTGTATTCCCAGCTACTCA
AGCACACGTCTGAACTCTTTCCCTACACGACGCTCTTCCGATCTNNNCAGAAGGTTTTGCTATATGTA
AGCACACGTCTGAACTCTTTCCCTACACGACGCTCTTCCGATCTNNNCAGAAGGTTTTGCTATATGTA
AGCACACGTCTGAACTCTTTCCCTACACGACGCTCTTCCGATCTNNNTTTGCTATTGCCTGGTTGACTT
AGCACACGTCTGAACTCTTTCCCTACACGACGCTCTTCCGATCTNNNTTTGCTATTGCCTGGTTGACTT
GCACACGTCTGAACTCTTTCCCTACACGACGCTCTTCCGATCTNNNCCCGGCCTAAACTCTAGTTTTGA
GCACACGTCTGAACTCTTTCCCTACACGACGCTCTTCCGATCTNNNCCCGGCCTAAACTCTAGTTTTGA
GAGCACACGTCTGAACTCTTTCCCTACACGACGCTCTTCCGATCTNNNTCTTTAGCAAAGAGACAGCC
GAGCACACGTCTGAACTCTTTCCCTACACGACGCTCTTCCGATCTNNNTCTTTAGCAAAGAGACAGCC
AGCACACGTCTGAACTCTTTCCCTACACGACGCTCTTCCGATCTNNNGCAACAGAGCAAGAAACTGTC
AGCACACGTCTGAACTCTTTCCCTACACGACGCTCTTCCGATCTNNNGCAACAGAGCAAGAAACTGTC
AGCACACGTCTGAACTCTTTCCCTACACGACGCTCTTCCGATCTNNNGGAAACCTAGCAAGACTCTGT
AGCACACGTCTGAACTCTTTCCCTACACGACGCTCTTCCGATCTNNNGGAAACCTAGCAAGACTCTGT
AGCACACGTCTGAACTCTTTCCCTACACGACGCTCTTCCGATCTNNNCTTCCTGAAGCTCAGGTCAATG
AGCACACGTCTGAACTCTTTCCCTACACGACGCTCTTCCGATCTNNNCTTCCTGAAGCTCAGGTCAATG
GAGCACACGTCTGAACTCTTTCCCTACACGACGCTCTTCCGATCTNNNCAACATAAGGAGACCCCATC
GAGCACACGTCTGAACTCTTTCCCTACACGACGCTCTTCCGATCTNNNCAACATAAGGAGACCCCATC
AGCACACGTCTGAACTCTTTCCCTACACGACGCTCTTCCGATCTNNNAAGACACATACCGGCTGTAAC
AGCACACGTCTGAACTCTTTCCCTACACGACGCTCTTCCGATCTNNNAAGACACATACCGGCTGTAAC
AGCACACGTCTGAACTCTTTCCCTACACGACGCTCTTCCGATCTNNNCTGTAGTTCCAGCTACTCAGG
AGCACACGTCTGAACTCTTTCCCTACACGACGCTCTTCCGATCTNNNCTGTAGTTCCAGCTACTCAGG
AGAGCACACGTCTGAACTCTTTCCCTACACGACGCTCTTCCGATCTNNNAACAGACCCACAGCCATTAT
AGAGCACACGTCTGAACTCTTTCCCTACACGACGCTCTTCCGATCTNNNAACAGACCCACAGCCATTAT
AGCACACGTCTGAACTCTTTCCCTACACGACGCTCTTCCGATCTNNNCTGAACTACAGGGTGGATGAC
AGCACACGTCTGAACTCTTTCCCTACACGACGCTCTTCCGATCTNNNCTGAACTACAGGGTGGATGAC
AGCACACGTCTGAACTCTTTCCCTACACGACGCTCTTCCGATCTNNNTTTCTTCTCTTTTCCACCCCACCC
AGCACACGTCTGAACTCTTTCCCTACACGACGCTCTTCCGATCTNNNTTTCTTCTCTTTTCCACCCCACCC
GCACACGTCTGAACTCTTTCCCTACACGACGCTCTTCCGATCTNNNGAAAAGGAAGGGGAGGAAGA
GCACACGTCTGAACTCTTTCCCTACACGACGCTCTTCCGATCTNNNGAAAAGGAAGGGGAGGAAGA
AGCACACGTCTGAACTCTTTCCCTACACGACGCTCTTCCGATCTNNNGTAGCTGGGACTACAGGCATA
AGCACACGTCTGAACTCTTTCCCTACACGACGCTCTTCCGATCTNNNGTAGCTGGGACTACAGGCATA
AGCACACGTCTGAACTCTTTCCCTACACGACGCTCTTCCGATCTNNNCATGAGAATTGCTTGAACCAGG
AGCACACGTCTGAACTCTTTCCCTACACGACGCTCTTCCGATCTNNNCATGAGAATTGCTTGAACCAGG
AGCACACGTCTGAACTCTTTCCCTACACGACGCTCTTCCGATCTNNNGGTGAGATTATAGGCGTGAG
AGCACACGTCTGAACTCTTTCCCTACACGACGCTCTTCCGATCTNNNGGTGAGATTATAGGCGTGAG

AGCACACGTCTGAACTCTTTCCCTACACGACGCTCTTCCGATCTNNNCAGGGTTTTACCATGTTAGCC

AGCACACGTCTGAACTCTTTCCCTACACGACGCTCTTCCGATCTNNNCAGGGTTTTACCATGTTAGCC

ACACGTCTGAACTCTTTCCCTACACGACGCTCTTCCGATCTNNNATAGTGGGGCCCTTGTCACAATAC

ACACGTCTGAACTCTTTCCCTACACGACGCTCTTCCGATCTNNNATAGTGGGGCCCTTGTCACAATAC

CACACGTCTGAACTCTTTCCCTACACGACGCTCTTCCGATCTNNNCGCTTGGCCTGCAATGTACTCCAT

CACACGTCTGAACTCTTTCCCTACACGACGCTCTTCCGATCTNNNCGCTTGGCCTGCAATGTACTCCAT

AGCACACGTCTGAACTCTTTCCCTACACGACGCTCTTCCGATCTNNNTTGCAGTGAGCTGAGATCGTA

AGCACACGTCTGAACTCTTTCCCTACACGACGCTCTTCCGATCTNNNTTGCAGTGAGCTGAGATCGTA

GCACACGTCTGAACTCTTTCCCTACACGACGCTCTTCCGATCTNNNATACCTGCAAGATCTTCCTTCAT

GCACACGTCTGAACTCTTTCCCTACACGACGCTCTTCCGATCTNNNATACCTGCAAGATCTTCCTTCAT

GAGCACACGTCTGAACTCTTTCCCTACACGACGCTCTTCCGATCTNNNCATCACTTGAGGACAGGAGT

GAGCACACGTCTGAACTCTTTCCCTACACGACGCTCTTCCGATCTNNNCATCACTTGAGGACAGGAGT

GCACACGTCTGAACTCTTTCCCTACACGACGCTCTTCCGATCTNNNACAGTGGGGCAAGTTACAAAAA

GCACACGTCTGAACTCTTTCCCTACACGACGCTCTTCCGATCTNNNACAGTGGGGCAAGTTACAAAAA

AGCACACGTCTGAACTCTTTCCCTACACGACGCTCTTCCGATCTNNNTCTTGAAACAGTTTGTGCCAC

AGCACACGTCTGAACTCTTTCCCTACACGACGCTCTTCCGATCTNNNTCTTGAAACAGTTTGTGCCAC

AGCACACGTCTGAACTCTTTCCCTACACGACGCTCTTCCGATCTNNNGCAATGAGTGAGACTCCGTCT

AGCACACGTCTGAACTCTTTCCCTACACGACGCTCTTCCGATCTNNNGCAATGAGTGAGACTCCGTCT

AGCACACGTCTGAACTCTTTCCCTACACGACGCTCTTCCGATCTNNNACCTGTAGTCCCAGCTACTCGC

AGCACACGTCTGAACTCTTTCCCTACACGACGCTCTTCCGATCTNNNACCTGTAGTCCCAGCTACTCGC

GCACACGTCTGAACTCTTTCCCTACACGACGCTCTTCCGATCTNNNGATTCAAGCATTTCTCCTGCCTCC

GCACACGTCTGAACTCTTTCCCTACACGACGCTCTTCCGATCTNNNGATTCAAGCATTTCTCCTGCCTCC

AGCACACGTCTGAACTCTTTCCCTACACGACGCTCTTCCGATCTNNNCAGAAAGTTGAGGTTGCAGT

AGCACACGTCTGAACTCTTTCCCTACACGACGCTCTTCCGATCTNNNCAGAAAGTTGAGGTTGCAGT

CACACGTCTGAACTCTTTCCCTACACGACGCTCTTCCGATCTNNNATTCTTTCTTATCACTCAAATCGCA

CACACGTCTGAACTCTTTCCCTACACGACGCTCTTCCGATCTNNNATTCTTTCTTATCACTCAAATCGCA

GAGCACACGTCTGAACTCTTTCCCTACACGACGCTCTTCCGATCTNNNTAGAGCCTTGTAGGTGGAA

GAGCACACGTCTGAACTCTTTCCCTACACGACGCTCTTCCGATCTNNNTAGAGCCTTGTAGGTGGAA

GCACACGTCTGAACTCTTTCCCTACACGACGCTCTTCCGATCTNNNCACGCTTGTAATTCCAGCAGTTT

GCACACGTCTGAACTCTTTCCCTACACGACGCTCTTCCGATCTNNNCACGCTTGTAATTCCAGCAGTTT

AGCACACGTCTGAACTCTTTCCCTACACGACGCTCTTCCGATCTNNNACAAGGTATAAGGTGCCTGG

AGCACACGTCTGAACTCTTTCCCTACACGACGCTCTTCCGATCTNNNACAAGGTATAAGGTGCCTGG

AGCACACGTCTGAACTCTTTCCCTACACGACGCTCTTCCGATCTNNNGCCTGTAGTCCCAGCTACTTGC

AGCACACGTCTGAACTCTTTCCCTACACGACGCTCTTCCGATCTNNNGCCTGTAGTCCCAGCTACTTGC

ACACGTCTGAACTCTTTCCCTACACGACGCTCTTCCGATCTNNNCAACTCTCCTTCCACCTCCTTCCATC

ACACGTCTGAACTCTTTCCCTACACGACGCTCTTCCGATCTNNNCAACTCTCCTTCCACCTCCTTCCATC

GCACACGTCTGAACTCTTTCCCTACACGACGCTCTTCCGATCTNNNGAGGTTGAGGCAGTAGAATCA

GCACACGTCTGAACTCTTTCCCTACACGACGCTCTTCCGATCTNNNGAGGTTGAGGCAGTAGAATCA

AGCACACGTCTGAACTCTTTCCCTACACGACGCTCTTCCGATCTNNNAAGATTGCTTGAGCTCAGGA

AGCACACGTCTGAACTCTTTCCCTACACGACGCTCTTCCGATCTNNNAAGATTGCTTGAGCTCAGGA

AGCACACGTCTGAACTCTTTCCCTACACGACGCTCTTCCGATCTNNNCTATCTTTCTTCCCCACCCCCTC

AGCACACGTCTGAACTCTTTCCCTACACGACGCTCTTCCGATCTNNNCTATCTTTCTTCCCCACCCCCTC

AGCACACGTCTGAACTCTTTCCCTACACGACGCTCTTCCGATCTNNNGAAGAATGGTGTGAACCCAG

GAGCACACGTCTGAACTCTTTCCCTACACGACGCTCTTCCGATCTNNNGAAGAATGGTGTGAACCCAG
GCACACGTCTGAACTCTTTCCCTACACGACGCTCTTCCGATCTNNNCCATGCCCGGCTAATTTTTGTATC
GCACACGTCTGAACTCTTTCCCTACACGACGCTCTTCCGATCTNNNCCATGCCCGGCTAATTTTTGTATC
AGCACACGTCTGAACTCTTTCCCTACACGACGCTCTTCCGATCTNNNCCCAGGTTCAAGCAATTCTCTTG
AGCACACGTCTGAACTCTTTCCCTACACGACGCTCTTCCGATCTNNNCCCAGGTTCAAGCAATTCTCTTG
GAGCACACGTCTGAACTCTTTCCCTACACGACGCTCTTCCGATCTNNNTCAGACTCCAAAGAGAAAACC
GAGCACACGTCTGAACTCTTTCCCTACACGACGCTCTTCCGATCTNNNTCAGACTCCAAAGAGAAAACC
AGCACACGTCTGAACTCTTTCCCTACACGACGCTCTTCCGATCTNNNTTTGGAGGACAAAGGGAGCAC
AGCACACGTCTGAACTCTTTCCCTACACGACGCTCTTCCGATCTNNNTTTGGAGGACAAAGGGAGCAC
GAGCACACGTCTGAACTCTTTCCCTACACGACGCTCTTCCGATCTNNNCTGAGGCAGGAGAATCACTT
GAGCACACGTCTGAACTCTTTCCCTACACGACGCTCTTCCGATCTNNNCTGAGGCAGGAGAATCACTT
GCACACGTCTGAACTCTTTCCCTACACGACGCTCTTCCGATCTNNNGCATCACATTCCTCAAGAAGCATC
GCACACGTCTGAACTCTTTCCCTACACGACGCTCTTCCGATCTNNNGCATCACATTCCTCAAGAAGCATC
AGCACACGTCTGAACTCTTTCCCTACACGACGCTCTTCCGATCTNNNTTGGATGAGTGGGCACATAAAC
AGCACACGTCTGAACTCTTTCCCTACACGACGCTCTTCCGATCTNNNTTGGATGAGTGGGCACATAAAC
AGAGCACACGTCTGAACTCTTTCCCTACACGACGCTCTTCCGATCTNNNGTGAATGAGAGCTTCCCAAA
AGAGCACACGTCTGAACTCTTTCCCTACACGACGCTCTTCCGATCTNNNGTGAATGAGAGCTTCCCAAA
AGCACACGTCTGAACTCTTTCCCTACACGACGCTCTTCCGATCTNNNGGGTGATGAAGTGAGATTTTG
AGCACACGTCTGAACTCTTTCCCTACACGACGCTCTTCCGATCTNNNGGGTGATGAAGTGAGATTTTG
AGCACACGTCTGAACTCTTTCCCTACACGACGCTCTTCCGATCTNNNGCTCCCAGCCAAAATACTCATTT
AGCACACGTCTGAACTCTTTCCCTACACGACGCTCTTCCGATCTNNNGCTCCCAGCCAAAATACTCATTT
GAGCACACGTCTGAACTCTTTCCCTACACGACGCTCTTCCGATCTNNNAAGCAATTCAAAGCCAGTGAA
GAGCACACGTCTGAACTCTTTCCCTACACGACGCTCTTCCGATCTNNNAAGCAATTCAAAGCCAGTGAA
CACACGTCTGAACTCTTTCCCTACACGACGCTCTTCCGATCTNNNGGCCACATATTCAGTTTGGTTGCCA
CACACGTCTGAACTCTTTCCCTACACGACGCTCTTCCGATCTNNNGGCCACATATTCAGTTTGGTTGCCA
GAGCACACGTCTGAACTCTTTCCCTACACGACGCTCTTCCGATCTNNNTGGTTTTCTGTACTGCCAGAG
GAGCACACGTCTGAACTCTTTCCCTACACGACGCTCTTCCGATCTNNNTGGTTTTCTGTACTGCCAGAG
AGCACACGTCTGAACTCTTTCCCTACACGACGCTCTTCCGATCTNNNGGATTCCTGGTTTGTTCTTGCAC
AGCACACGTCTGAACTCTTTCCCTACACGACGCTCTTCCGATCTNNNGGATTCCTGGTTTGTTCTTGCAC
AGCACACGTCTGAACTCTTTCCCTACACGACGCTCTTCCGATCTNNNTGAGATAAAGCCACTGCACTTCA
AGCACACGTCTGAACTCTTTCCCTACACGACGCTCTTCCGATCTNNNTGAGATAAAGCCACTGCACTTCA
GAGCACACGTCTGAACTCTTTCCCTACACGACGCTCTTCCGATCTNNNGAGATCACACCACTGCACTCCC
GAGCACACGTCTGAACTCTTTCCCTACACGACGCTCTTCCGATCTNNNGAGATCACACCACTGCACTCCC
AGCACACGTCTGAACTCTTTCCCTACACGACGCTCTTCCGATCTNNNCAGTGTTTTGTGGTTCTCCTTGT
AGCACACGTCTGAACTCTTTCCCTACACGACGCTCTTCCGATCTNNNCAGTGTTTTGTGGTTCTCCTTGT
GAGCACACGTCTGAACTCTTTCCCTACACGACGCTCTTCCGATCTNNNAACAGTAAAAGCACTTGCCCA
GAGCACACGTCTGAACTCTTTCCCTACACGACGCTCTTCCGATCTNNNAACAGTAAAAGCACTTGCCCA
GAGCACACGTCTGAACTCTTTCCCTACACGACGCTCTTCCGATCTNNNCTTCTCAGGCTCAAGGTAACAC
GAGCACACGTCTGAACTCTTTCCCTACACGACGCTCTTCCGATCTNNNCTTCTCAGGCTCAAGGTAACAC
AGCACACGTCTGAACTCTTTCCCTACACGACGCTCTTCCGATCTNNNTCTCAAACTGCTGGGATTACAG
AGCACACGTCTGAACTCTTTCCCTACACGACGCTCTTCCGATCTNNNTCTCAAACTGCTGGGATTACAG
ACACGTCTGAACTCTTTCCCTACACGACGCTCTTCCGATCTNNNTAGTTCAATTGCGGGTGGAACTACAT
ACACGTCTGAACTCTTTCCCTACACGACGCTCTTCCGATCTNNNTAGTTCAATTGCGGGTGGAACTACAT

GCACACGTCTGAACTCTTTCCCTACACGACGCTCTTCCGATCTNNNACACACAAAGAGAAGGATACAG
GCACACGTCTGAACTCTTTCCCTACACGACGCTCTTCCGATCTNNNACACACAAAGAGAAGGATACAG
GCACACGTCTGAACTCTTTCCCTACACGACGCTCTTCCGATCTNNNCCTGACTGAGGACTGATGGTAT
GCACACGTCTGAACTCTTTCCCTACACGACGCTCTTCCGATCTNNNCCTGACTGAGGACTGATGGTAT
GCACACGTCTGAACTCTTTCCCTACACGACGCTCTTCCGATCTNNNTATGTCTACCCTCTCCCGAGAAA
GCACACGTCTGAACTCTTTCCCTACACGACGCTCTTCCGATCTNNNTATGTCTACCCTCTCCCGAGAAA
GCACACGTCTGAACTCTTTCCCTACACGACGCTCTTCCGATCTNNNCTCTCAAGTAGCTGGGACTACA
GCACACGTCTGAACTCTTTCCCTACACGACGCTCTTCCGATCTNNNCTCTCAAGTAGCTGGGACTACA
AGAGCACACGTCTGAACTCTTTCCCTACACGACGCTCTTCCGATCTNNNCTGAGGCAGGAAAATTGCT
AGAGCACACGTCTGAACTCTTTCCCTACACGACGCTCTTCCGATCTNNNCTGAGGCAGGAAAATTGCT
AGCACACGTCTGAACTCTTTCCCTACACGACGCTCTTCCGATCTNNNAATCTGACGGATCGAAGGCAT
AGCACACGTCTGAACTCTTTCCCTACACGACGCTCTTCCGATCTNNNAATCTGACGGATCGAAGGCAT
GCACACGTCTGAACTCTTTCCCTACACGACGCTCTTCCGATCTNNNTGGGCAGTCAGTAAAGAAAACC
GCACACGTCTGAACTCTTTCCCTACACGACGCTCTTCCGATCTNNNTGGGCAGTCAGTAAAGAAAACC
AGCACACGTCTGAACTCTTTCCCTACACGACGCTCTTCCGATCTNNNCGACAGAGTGAGACTCCGTCT
AGCACACGTCTGAACTCTTTCCCTACACGACGCTCTTCCGATCTNNNCGACAGAGTGAGACTCCGTCT
GCACACGTCTGAACTCTTTCCCTACACGACGCTCTTCCGATCTNNNCTGAGATGGGAGGATCACTTGA
GCACACGTCTGAACTCTTTCCCTACACGACGCTCTTCCGATCTNNNCTGAGATGGGAGGATCACTTGA
AGCACACGTCTGAACTCTTTCCCTACACGACGCTCTTCCGATCTNNNGGATGGTCCTAAGAGGAGG
AGCACACGTCTGAACTCTTTCCCTACACGACGCTCTTCCGATCTNNNGGATGGTCCTAAGAGGAGG
GAGCACACGTCTGAACTCTTTCCCTACACGACGCTCTTCCGATCTNNNGTGTATATTGAACCAGCCTTG
GAGCACACGTCTGAACTCTTTCCCTACACGACGCTCTTCCGATCTNNNGTGTATATTGAACCAGCCTTG
AGCACACGTCTGAACTCTTTCCCTACACGACGCTCTTCCGATCTNNNCGGTGACTCCATCTTAGTTGG
AGCACACGTCTGAACTCTTTCCCTACACGACGCTCTTCCGATCTNNNCGGTGACTCCATCTTAGTTGG
AGCACACGTCTGAACTCTTTCCCTACACGACGCTCTTCCGATCTNNNGAACTCCCGACTTCAGTTGATC
AGCACACGTCTGAACTCTTTCCCTACACGACGCTCTTCCGATCTNNNGAACTCCCGACTTCAGTTGATC
GAGCACACGTCTGAACTCTTTCCCTACACGACGCTCTTCCGATCTNNNTAAGGCAGCCAGTTACCACA
GAGCACACGTCTGAACTCTTTCCCTACACGACGCTCTTCCGATCTNNNTAAGGCAGCCAGTTACCACA
AGCACACGTCTGAACTCTTTCCCTACACGACGCTCTTCCGATCTNNNCCAAAAAAAGCCATGGACCAG
AGCACACGTCTGAACTCTTTCCCTACACGACGCTCTTCCGATCTNNNCCAAAAAAAGCCATGGACCAG
AGCACACGTCTGAACTCTTTCCCTACACGACGCTCTTCCGATCTNNNTCCTTCTGCTTCTGCTTCTTCTT
AGCACACGTCTGAACTCTTTCCCTACACGACGCTCTTCCGATCTNNNTCCTTCTGCTTCTGCTTCTTCTT
AGAGCACACGTCTGAACTCTTTCCCTACACGACGCTCTTCCGATCTNNNTCATGCAAACTAGATAACTA
AGAGCACACGTCTGAACTCTTTCCCTACACGACGCTCTTCCGATCTNNNTCATGCAAACTAGATAACTA
AGAGCACACGTCTGAACTCTTTCCCTACACGACGCTCTTCCGATCTNNNTTCAAGATCAGGAGAGGGA
AGAGCACACGTCTGAACTCTTTCCCTACACGACGCTCTTCCGATCTNNNTTCAAGATCAGGAGAGGGA
AGCACACGTCTGAACTCTTTCCCTACACGACGCTCTTCCGATCTNNNTCTGCTTGCATCTCAGTCTACAT
AGCACACGTCTGAACTCTTTCCCTACACGACGCTCTTCCGATCTNNNTCTGCTTGCATCTCAGTCTACAT
AGCACACGTCTGAACTCTTTCCCTACACGACGCTCTTCCGATCTNNNACTAAAATAACCACTGTCAACA
AGCACACGTCTGAACTCTTTCCCTACACGACGCTCTTCCGATCTNNNACTAAAATAACCACTGTCAACA
GCACACGTCTGAACTCTTTCCCTACACGACGCTCTTCCGATCTNNNTCTCCCTATTGCTTGCTCTTTCTCA
GCACACGTCTGAACTCTTTCCCTACACGACGCTCTTCCGATCTNNNTCTCCCTATTGCTTGCTCTTTCTCA
AGAGCACACGTCTGAACTCTTTCCCTACACGACGCTCTTCCGATCTNNNAATCAGGTAGAACCGGAG

AAGAGCACACGTCTGAACTCTTTCCCTACACGACGCTCTTCCGATCTNNNAATCAGGTAGAACCGGAGA
AGAGCACACGTCTGAACTCTTTCCCTACACGACGCTCTTCCGATCTNNNGTAATCTCAGCTACTCAGGAG
AGAGCACACGTCTGAACTCTTTCCCTACACGACGCTCTTCCGATCTNNNGTAATCTCAGCTACTCAGGAG
GAGCACACGTCTGAACTCTTTCCCTACACGACGCTCTTCCGATCTNNNAGTCAAATTACCTGGCTGAGC
GAGCACACGTCTGAACTCTTTCCCTACACGACGCTCTTCCGATCTNNNAGTCAAATTACCTGGCTGAGC
AGCACACGTCTGAACTCTTTCCCTACACGACGCTCTTCCGATCTNNNAAAGTAGTTCTGGCTGTCTGGA
AGCACACGTCTGAACTCTTTCCCTACACGACGCTCTTCCGATCTNNNAAAGTAGTTCTGGCTGTCTGGA
AGCACACGTCTGAACTCTTTCCCTACACGACGCTCTTCCGATCTNNNGCCTGTAGTCCCAGCTACTTGCA
AGCACACGTCTGAACTCTTTCCCTACACGACGCTCTTCCGATCTNNNGCCTGTAGTCCCAGCTACTTGCA
GAGCACACGTCTGAACTCTTTCCCTACACGACGCTCTTCCGATCTNNNTGCTAGTAGACCTGCATTACA
GAGCACACGTCTGAACTCTTTCCCTACACGACGCTCTTCCGATCTNNNTGCTAGTAGACCTGCATTACA
AGCACACGTCTGAACTCTTTCCCTACACGACGCTCTTCCGATCTNNNTGCCTGTAATCCCAGTACTTTGG
AGCACACGTCTGAACTCTTTCCCTACACGACGCTCTTCCGATCTNNNTGCCTGTAATCCCAGTACTTTGG
AGAGCACACGTCTGAACTCTTTCCCTACACGACGCTCTTCCGATCTNNNCCCTATGTCTTGGCCATAAGA
AGAGCACACGTCTGAACTCTTTCCCTACACGACGCTCTTCCGATCTNNNCCCTATGTCTTGGCCATAAGA
AAGAGCACACGTCTGAACTCTTTCCCTACACGACGCTCTTCCGATCTNNNAAAATTAGCCAGGTGTGGT
AAGAGCACACGTCTGAACTCTTTCCCTACACGACGCTCTTCCGATCTNNNAAAATTAGCCAGGTGTGGT
GCACACGTCTGAACTCTTTCCCTACACGACGCTCTTCCGATCTNNNGAGTGACGACTGCACCGAAGCAT
GCACACGTCTGAACTCTTTCCCTACACGACGCTCTTCCGATCTNNNGAGTGACGACTGCACCGAAGCAT
AGAGCACACGTCTGAACTCTTTCCCTACACGACGCTCTTCCGATCTNNNGATGTTTTCAAGGTCACCTGC
AGAGCACACGTCTGAACTCTTTCCCTACACGACGCTCTTCCGATCTNNNGATGTTTTCAAGGTCACCTGC
GAGCACACGTCTGAACTCTTTCCCTACACGACGCTCTTCCGATCTNNNCTGTCACCATGCCATGAATTCA
GAGCACACGTCTGAACTCTTTCCCTACACGACGCTCTTCCGATCTNNNCTGTCACCATGCCATGAATTCA
AGCACACGTCTGAACTCTTTCCCTACACGACGCTCTTCCGATCTNNNGTTCGTAGCTCAAGATCCCGAA
AGCACACGTCTGAACTCTTTCCCTACACGACGCTCTTCCGATCTNNNGTTCGTAGCTCAAGATCCCGAA
GAGCACACGTCTGAACTCTTTCCCTACACGACGCTCTTCCGATCTNNNAGTATCTAAAGGGGACACTG
GAGCACACGTCTGAACTCTTTCCCTACACGACGCTCTTCCGATCTNNNAGTATCTAAAGGGGACACTG
AGCACACGTCTGAACTCTTTCCCTACACGACGCTCTTCCGATCTNNNCAGACTCTTACCAGCATATATAC
AGCACACGTCTGAACTCTTTCCCTACACGACGCTCTTCCGATCTNNNCAGACTCTTACCAGCATATATAC
AGCACACGTCTGAACTCTTTCCCTACACGACGCTCTTCCGATCTNNNCTGTGTTATCTCCATGCCAGACTC
AGCACACGTCTGAACTCTTTCCCTACACGACGCTCTTCCGATCTNNNCTGTGTTATCTCCATGCCAGACTC
AGCACACGTCTGAACTCTTTCCCTACACGACGCTCTTCCGATCTNNNTGTGCAACTTCTTTTCATTGTGA
AGCACACGTCTGAACTCTTTCCCTACACGACGCTCTTCCGATCTNNNTGTGCAACTTCTTTTCATTGTGA
AGAGCACACGTCTGAACTCTTTCCCTACACGACGCTCTTCCGATCTNNNATCTTGCAGTGAGCCAAGAT
AGAGCACACGTCTGAACTCTTTCCCTACACGACGCTCTTCCGATCTNNNATCTTGCAGTGAGCCAAGAT
GAGCACACGTCTGAACTCTTTCCCTACACGACGCTCTTCCGATCTNNNGGGGCTTCTTGCAAAAACATCA
GAGCACACGTCTGAACTCTTTCCCTACACGACGCTCTTCCGATCTNNNGGGGCTTCTTGCAAAAACATCA
AGCACACGTCTGAACTCTTTCCCTACACGACGCTCTTCCGATCTNNNTAAGGAAGTGTGTGTGTGTGTC
AGCACACGTCTGAACTCTTTCCCTACACGACGCTCTTCCGATCTNNNTAAGGAAGTGTGTGTGTGTGTC
GAGCACACGTCTGAACTCTTTCCCTACACGACGCTCTTCCGATCTNNNGACTGGCTTATTTCACATCACA
GAGCACACGTCTGAACTCTTTCCCTACACGACGCTCTTCCGATCTNNNGACTGGCTTATTTCACATCACA
GAGCACACGTCTGAACTCTTTCCCTACACGACGCTCTTCCGATCTNNNAGTAGTCCCCCCTCTGAGTAAA
GAGCACACGTCTGAACTCTTTCCCTACACGACGCTCTTCCGATCTNNNAGTAGTCCCCCCTCTGAGTAAA

AGCACACGTCTGAACTCTTTCCCTACACGACGCTCTTCCGATCTNNNTTCTGCTACCTCCTTGCAAATCT

AGCACACGTCTGAACTCTTTCCCTACACGACGCTCTTCCGATCTNNNTTCTGCTACCTCCTTGCAAATCT

CACACGTCTGAACTCTTTCCCTACACGACGCTCTTCCGATCTNNNCACATCTGACCCAAAATGCCATTC

CACACGTCTGAACTCTTTCCCTACACGACGCTCTTCCGATCTNNNCACATCTGACCCAAAATGCCATTC

GCACACGTCTGAACTCTTTCCCTACACGACGCTCTTCCGATCTNNNCTCTAGAGAGGAAGTGACATG

GCACACGTCTGAACTCTTTCCCTACACGACGCTCTTCCGATCTNNNCTCTAGAGAGGAAGTGACATG

GCACACGTCTGAACTCTTTCCCTACACGACGCTCTTCCGATCTNNNTGTGTGAATGTGTGAGCCAATT

GCACACGTCTGAACTCTTTCCCTACACGACGCTCTTCCGATCTNNNTGTGTGAATGTGTGAGCCAATT

AGCACACGTCTGAACTCTTTCCCTACACGACGCTCTTCCGATCTNNNCAGGAAAATCGCTTGAACCTG

AGCACACGTCTGAACTCTTTCCCTACACGACGCTCTTCCGATCTNNNCAGGAAAATCGCTTGAACCTG

AGCACACGTCTGAACTCTTTCCCTACACGACGCTCTTCCGATCTNNNTTCTTCTAAGCAGATCACAGC

AGCACACGTCTGAACTCTTTCCCTACACGACGCTCTTCCGATCTNNNTTCTTCTAAGCAGATCACAGC

AGCACACGTCTGAACTCTTTCCCTACACGACGCTCTTCCGATCTNNNGATCTGGTTTCTCTTCTGCCAC

AGCACACGTCTGAACTCTTTCCCTACACGACGCTCTTCCGATCTNNNGATCTGGTTTCTCTTCTGCCAC

GCACACGTCTGAACTCTTTCCCTACACGACGCTCTTCCGATCTNNNTCAGAACAACAGGAAAAAGAC

GCACACGTCTGAACTCTTTCCCTACACGACGCTCTTCCGATCTNNNTCAGAACAACAGGAAAAAGAC

GAGCACACGTCTGAACTCTTTCCCTACACGACGCTCTTCCGATCTNNNTGTGAGGGACGAATGGAA

GAGCACACGTCTGAACTCTTTCCCTACACGACGCTCTTCCGATCTNNNTGTGAGGGACGAATGGAA

GAGCACACGTCTGAACTCTTTCCCTACACGACGCTCTTCCGATCTNNNTTTGCAAGTCTCTTCAAGTCT

GAGCACACGTCTGAACTCTTTCCCTACACGACGCTCTTCCGATCTNNNTTTGCAAGTCTCTTCAAGTCT

AGCACACGTCTGAACTCTTTCCCTACACGACGCTCTTCCGATCTNNNCCTTCTGGCAAAACTGATGTG

AGCACACGTCTGAACTCTTTCCCTACACGACGCTCTTCCGATCTNNNCCTTCTGGCAAAACTGATGTG

AGCACACGTCTGAACTCTTTCCCTACACGACGCTCTTCCGATCTNNNCAGCACCTACCCAGATCGAAGC

AGCACACGTCTGAACTCTTTCCCTACACGACGCTCTTCCGATCTNNNCAGCACCTACCCAGATCGAAGC

AGCACACGTCTGAACTCTTTCCCTACACGACGCTCTTCCGATCTNNNGAATCCAACACCTTCCCTGAGA

AGCACACGTCTGAACTCTTTCCCTACACGACGCTCTTCCGATCTNNNGAATCCAACACCTTCCCTGAGA

AGCACACGTCTGAACTCTTTCCCTACACGACGCTCTTCCGATCTNNNCAGTGGTCTTCAGAAACTCAC

AGCACACGTCTGAACTCTTTCCCTACACGACGCTCTTCCGATCTNNNCAGTGGTCTTCAGAAACTCAC

AGCACACGTCTGAACTCTTTCCCTACACGACGCTCTTCCGATCTNNNGGGGTCTACTTGAGGGTAAA

AGCACACGTCTGAACTCTTTCCCTACACGACGCTCTTCCGATCTNNNGGGGTCTACTTGAGGGTAAA

GAGCACACGTCTGAACTCTTTCCCTACACGACGCTCTTCCGATCTNNNCACACCTGTAGTCCCAGTTAC

GAGCACACGTCTGAACTCTTTCCCTACACGACGCTCTTCCGATCTNNNCACACCTGTAGTCCCAGTTAC

GCACACGTCTGAACTCTTTCCCTACACGACGCTCTTCCGATCTNNNCTAGGGAGATGTACCTGACCTG

GCACACGTCTGAACTCTTTCCCTACACGACGCTCTTCCGATCTNNNCTAGGGAGATGTACCTGACCTG

GAGCACACGTCTGAACTCTTTCCCTACACGACGCTCTTCCGATCTNNNATTATTTGGGACATATTGAT

GAGCACACGTCTGAACTCTTTCCCTACACGACGCTCTTCCGATCTNNNATTATTTGGGACATATTGAT

AGCACACGTCTGAACTCTTTCCCTACACGACGCTCTTCCGATCTNNNTGGAAAATCCTATGGTGCCA

AGCACACGTCTGAACTCTTTCCCTACACGACGCTCTTCCGATCTNNNTGGAAAATCCTATGGTGCCA

AGCACACGTCTGAACTCTTTCCCTACACGACGCTCTTCCGATCTNNNTGTTTCTCGAGACCCTGACAAT

AGCACACGTCTGAACTCTTTCCCTACACGACGCTCTTCCGATCTNNNTGTTTCTCGAGACCCTGACAAT

GCACACGTCTGAACTCTTTCCCTACACGACGCTCTTCCGATCTNNNATTAGGAGGAGACGCTGTTTC

GCACACGTCTGAACTCTTTCCCTACACGACGCTCTTCCGATCTNNNATTAGGAGGAGACGCTGTTTC

GCACACGTCTGAACTCTTTCCCTACACGACGCTCTTCCGATCTNNNTCTGCACATGCTTGCTTTCTTTT

AGCACACGTCTGAACTCTTTCCCTACACGACGCTCTTCCGATCTNNNTCTGCACATGCTTGCTTTCTTTC
AGCACACGTCTGAACTCTTTCCCTACACGACGCTCTTCCGATCTNNNCACATCCCAACTTGAATAAAAG
AGCACACGTCTGAACTCTTTCCCTACACGACGCTCTTCCGATCTNNNCACATCCCAACTTGAATAAAAG
AGCACACGTCTGAACTCTTTCCCTACACGACGCTCTTCCGATCTNNNCCTGTCTGAGAAGTGAGGAGCC
AGCACACGTCTGAACTCTTTCCCTACACGACGCTCTTCCGATCTNNNCCTGTCTGAGAAGTGAGGAGCC
AGCACACGTCTGAACTCTTTCCCTACACGACGCTCTTCCGATCTNNNTGGGCCAGAATTTAGTTAGGGA
AGCACACGTCTGAACTCTTTCCCTACACGACGCTCTTCCGATCTNNNTGGGCCAGAATTTAGTTAGGGA
AGCACACGTCTGAACTCTTTCCCTACACGACGCTCTTCCGATCTNNNGTGGGGGCCATAGTCTTTTTTT
AGCACACGTCTGAACTCTTTCCCTACACGACGCTCTTCCGATCTNNNGTGGGGGCCATAGTCTTTTTTT
GAGCACACGTCTGAACTCTTTCCCTACACGACGCTCTTCCGATCTNNNCAGATTGTATGAGGGTGTGG
GAGCACACGTCTGAACTCTTTCCCTACACGACGCTCTTCCGATCTNNNCAGATTGTATGAGGGTGTGG
AGAGCACACGTCTGAACTCTTTCCCTACACGACGCTCTTCCGATCTNNNAAAGTAAGCCTGAGTGTCTC
AGAGCACACGTCTGAACTCTTTCCCTACACGACGCTCTTCCGATCTNNNAAAGTAAGCCTGAGTGTCTC
AGCACACGTCTGAACTCTTTCCCTACACGACGCTCTTCCGATCTNNNCCATGGAGGGGTAGTTTCAAA
AGCACACGTCTGAACTCTTTCCCTACACGACGCTCTTCCGATCTNNNCCATGGAGGGGTAGTTTCAAA
GAGCACACGTCTGAACTCTTTCCCTACACGACGCTCTTCCGATCTNNNAGCCTTCTGCCTCTTCCTTCATA
GAGCACACGTCTGAACTCTTTCCCTACACGACGCTCTTCCGATCTNNNAGCCTTCTGCCTCTTCCTTCATA
CACACGTCTGAACTCTTTCCCTACACGACGCTCTTCCGATCTNNNCACACCTGATTGTTGGAAAGGTCCA
CACACGTCTGAACTCTTTCCCTACACGACGCTCTTCCGATCTNNNCACACCTGATTGTTGGAAAGGTCCA
AGCACACGTCTGAACTCTTTCCCTACACGACGCTCTTCCGATCTNNNAAGATGTGTAGCAGCTGTGGT
AGCACACGTCTGAACTCTTTCCCTACACGACGCTCTTCCGATCTNNNAAGATGTGTAGCAGCTGTGGT
AGCACACGTCTGAACTCTTTCCCTACACGACGCTCTTCCGATCTNNNGTGAGCTGAGATCATGCCACTC
AGCACACGTCTGAACTCTTTCCCTACACGACGCTCTTCCGATCTNNNGTGAGCTGAGATCATGCCACTC
AGCACACGTCTGAACTCTTTCCCTACACGACGCTCTTCCGATCTNNNATCTGTGTCTGAAAAGTGGCTC
AGCACACGTCTGAACTCTTTCCCTACACGACGCTCTTCCGATCTNNNATCTGTGTCTGAAAAGTGGCTC
GCACACGTCTGAACTCTTTCCCTACACGACGCTCTTCCGATCTNNNCCAACATGCCTGGCTAATTTTTGC
GCACACGTCTGAACTCTTTCCCTACACGACGCTCTTCCGATCTNNNCCAACATGCCTGGCTAATTTTTGC
GAGCACACGTCTGAACTCTTTCCCTACACGACGCTCTTCCGATCTNNNAAAAATTAGTCAGGCATGGTC
GAGCACACGTCTGAACTCTTTCCCTACACGACGCTCTTCCGATCTNNNAAAAATTAGTCAGGCATGGTC
GAGCACACGTCTGAACTCTTTCCCTACACGACGCTCTTCCGATCTNNNGAGCAAGATGGGCAGATGTA
GAGCACACGTCTGAACTCTTTCCCTACACGACGCTCTTCCGATCTNNNGAGCAAGATGGGCAGATGTA
GAGCACACGTCTGAACTCTTTCCCTACACGACGCTCTTCCGATCTNNNTAGTGAGCTGAGATTGTGCC
GAGCACACGTCTGAACTCTTTCCCTACACGACGCTCTTCCGATCTNNNTAGTGAGCTGAGATTGTGCC
GAGCACACGTCTGAACTCTTTCCCTACACGACGCTCTTCCGATCTNNNGGAGAAATCACTTGAATCCGC
GAGCACACGTCTGAACTCTTTCCCTACACGACGCTCTTCCGATCTNNNGGAGAAATCACTTGAATCCGC
AGCACACGTCTGAACTCTTTCCCTACACGACGCTCTTCCGATCTNNNAAAGTGCTGGGATTACAGGTG
AGCACACGTCTGAACTCTTTCCCTACACGACGCTCTTCCGATCTNNNAAAGTGCTGGGATTACAGGTG
GAGCACACGTCTGAACTCTTTCCCTACACGACGCTCTTCCGATCTNNNCTTGCAGTGAGCCGAGATCAT
GAGCACACGTCTGAACTCTTTCCCTACACGACGCTCTTCCGATCTNNNCTTGCAGTGAGCCGAGATCAT
AGCACACGTCTGAACTCTTTCCCTACACGACGCTCTTCCGATCTNNNTTTCTGTCTGCCTCAGGAGATTC
AGCACACGTCTGAACTCTTTCCCTACACGACGCTCTTCCGATCTNNNTTTCTGTCTGCCTCAGGAGATTC
CACACGTCTGAACTCTTTCCCTACACGACGCTCTTCCGATCTNNNGAGACCACATGAATACACACCATCC
CACACGTCTGAACTCTTTCCCTACACGACGCTCTTCCGATCTNNNGAGACCACATGAATACACACCATCC

CACACGTCTGAACTCTTTCCCTACACGACGCTCTTCCGATCTNNNCCTCTCCTTCTCCCTTTTTCCTCCATC

CACACGTCTGAACTCTTTCCCTACACGACGCTCTTCCGATCTNNNCCTCTCCTTCTCCCTTTTTCCTCCATC

GAGCACACGTCTGAACTCTTTCCCTACACGACGCTCTTCCGATCTNNNTGGGGAAATGGCAGATGCT

GAGCACACGTCTGAACTCTTTCCCTACACGACGCTCTTCCGATCTNNNTGGGGAAATGGCAGATGCT

GAGCACACGTCTGAACTCTTTCCCTACACGACGCTCTTCCGATCTNNNAGTGCAGCCAAATTCATCAA

GAGCACACGTCTGAACTCTTTCCCTACACGACGCTCTTCCGATCTNNNAGTGCAGCCAAATTCATCAA

AGCACACGTCTGAACTCTTTCCCTACACGACGCTCTTCCGATCTNNNGCATGGTTTTTGAGGAGTTGG

AGCACACGTCTGAACTCTTTCCCTACACGACGCTCTTCCGATCTNNNGCATGGTTTTTGAGGAGTTGG

AGCACACGTCTGAACTCTTTCCCTACACGACGCTCTTCCGATCTNNNAGGGATTTGCACATAAAGACC

AGCACACGTCTGAACTCTTTCCCTACACGACGCTCTTCCGATCTNNNAGGGATTTGCACATAAAGACC

GCACACGTCTGAACTCTTTCCCTACACGACGCTCTTCCGATCTNNNGTTACTGCAGAATTGACGTAGC

GCACACGTCTGAACTCTTTCCCTACACGACGCTCTTCCGATCTNNNGTTACTGCAGAATTGACGTAGC

GAGCACACGTCTGAACTCTTTCCCTACACGACGCTCTTCCGATCTNNNAGTGAAGTGTGAGTGAGTG

GAGCACACGTCTGAACTCTTTCCCTACACGACGCTCTTCCGATCTNNNAGTGAAGTGTGAGTGAGTG

GCACACGTCTGAACTCTTTCCCTACACGACGCTCTTCCGATCTNNNCTCCTGGGTTCATGCCATTCTCC

GCACACGTCTGAACTCTTTCCCTACACGACGCTCTTCCGATCTNNNCTCCTGGGTTCATGCCATTCTCC

AGCACACGTCTGAACTCTTTCCCTACACGACGCTCTTCCGATCTNNNACTTCCTAGGTGGATTCCTAGC

AGCACACGTCTGAACTCTTTCCCTACACGACGCTCTTCCGATCTNNNACTTCCTAGGTGGATTCCTAGC

GCACACGTCTGAACTCTTTCCCTACACGACGCTCTTCCGATCTNNNGAGGTTGCAGTGAGCTGAGAT

GCACACGTCTGAACTCTTTCCCTACACGACGCTCTTCCGATCTNNNGAGGTTGCAGTGAGCTGAGAT

CACACGTCTGAACTCTTTCCCTACACGACGCTCTTCCGATCTNNNACTGTGCAGCAAAGAAGACTGAA

CACACGTCTGAACTCTTTCCCTACACGACGCTCTTCCGATCTNNNACTGTGCAGCAAAGAAGACTGAA

GAGCACACGTCTGAACTCTTTCCCTACACGACGCTCTTCCGATCTNNNAGAGGGACAGGAGGAGAG

GAGCACACGTCTGAACTCTTTCCCTACACGACGCTCTTCCGATCTNNNAGAGGGACAGGAGGAGAG

AGCACACGTCTGAACTCTTTCCCTACACGACGCTCTTCCGATCTNNNTAAGTAAGGTACATCAGAGGC

AGCACACGTCTGAACTCTTTCCCTACACGACGCTCTTCCGATCTNNNTAAGTAAGGTACATCAGAGGC

GCACACGTCTGAACTCTTTCCCTACACGACGCTCTTCCGATCTNNNCTACTGGGAAGTGAGGAGCCCC

GCACACGTCTGAACTCTTTCCCTACACGACGCTCTTCCGATCTNNNCTACTGGGAAGTGAGGAGCCCC

GAGCACACGTCTGAACTCTTTCCCTACACGACGCTCTTCCGATCTNNNCCCCAGCCAAAATGACACATA

GAGCACACGTCTGAACTCTTTCCCTACACGACGCTCTTCCGATCTNNNCCCCAGCCAAAATGACACATA

GAGCACACGTCTGAACTCTTTCCCTACACGACGCTCTTCCGATCTNNNTCTCAGGTTCCACCAAAGGTT

GAGCACACGTCTGAACTCTTTCCCTACACGACGCTCTTCCGATCTNNNTCTCAGGTTCCACCAAAGGTT

GCACACGTCTGAACTCTTTCCCTACACGACGCTCTTCCGATCTNNNAAGCCATTATCTGCAAAAATAT

GCACACGTCTGAACTCTTTCCCTACACGACGCTCTTCCGATCTNNNAAGCCATTATCTGCAAAAATAT

AGCACACGTCTGAACTCTTTCCCTACACGACGCTCTTCCGATCTNNNAGAGCAGAAGACTGACTGTCA

AGCACACGTCTGAACTCTTTCCCTACACGACGCTCTTCCGATCTNNNAGAGCAGAAGACTGACTGTCA

GAGCACACGTCTGAACTCTTTCCCTACACGACGCTCTTCCGATCTNNNCTGTAACGTGGTTTTGGTTTG

GAGCACACGTCTGAACTCTTTCCCTACACGACGCTCTTCCGATCTNNNCTGTAACGTGGTTTTGGTTTG

GCACACGTCTGAACTCTTTCCCTACACGACGCTCTTCCGATCTNNNTGACAACATCAAGAAGGTGGT

GCACACGTCTGAACTCTTTCCCTACACGACGCTCTTCCGATCTNNNTGACAACATCAAGAAGGTGGT

AGCACACGTCTGAACTCTTTCCCTACACGACGCTCTTCCGATCTNNNTGTAGTCTCAGCTACTAGGGA

AGCACACGTCTGAACTCTTTCCCTACACGACGCTCTTCCGATCTNNNTGTAGTCTCAGCTACTAGGGA

AGCACACGTCTGAACTCTTTCCCTACACGACGCTCTTCCGATCTNNNGCATGCTTTCTTGTTCCTCCATT

AGCACACGTCTGAACTCTTTCCCTACACGACGCTCTTCCGATCTNNNGCATGCTTTCTTGTTCCTCCATT

AGCACACGTCTGAACTCTTTCCCTACACGACGCTCTTCCGATCTNNNTTGAGGCAAATCCCCGAAAGTA

AGCACACGTCTGAACTCTTTCCCTACACGACGCTCTTCCGATCTNNNTTGAGGCAAATCCCCGAAAGTA

GAGCACACGTCTGAACTCTTTCCCTACACGACGCTCTTCCGATCTNNNGGTTGGTACCCCAAACCACTAT

GAGCACACGTCTGAACTCTTTCCCTACACGACGCTCTTCCGATCTNNNGGTTGGTACCCCAAACCACTAT

GAAGAGCACACGTCTGAACTCTTTCCCTACACGACGCTCTTCCGATCTNNNGAAGTTGAGGGGGGAAT

GAAGAGCACACGTCTGAACTCTTTCCCTACACGACGCTCTTCCGATCTNNNGAAGTTGAGGGGGGAAT

AGCACACGTCTGAACTCTTTCCCTACACGACGCTCTTCCGATCTNNNCTGTAGTCCCAGCTACTCAGGAC

AGCACACGTCTGAACTCTTTCCCTACACGACGCTCTTCCGATCTNNNCTGTAGTCCCAGCTACTCAGGAC

GAGCACACGTCTGAACTCTTTCCCTACACGACGCTCTTCCGATCTNNNTCTGAGATCTGAGCCTGAAGA

GAGCACACGTCTGAACTCTTTCCCTACACGACGCTCTTCCGATCTNNNTCTGAGATCTGAGCCTGAAGA

AGAGCACACGTCTGAACTCTTTCCCTACACGACGCTCTTCCGATCTNNNCTGAGGTAGGAAGATGGCTT

AGAGCACACGTCTGAACTCTTTCCCTACACGACGCTCTTCCGATCTNNNCTGAGGTAGGAAGATGGCTT

GAGCACACGTCTGAACTCTTTCCCTACACGACGCTCTTCCGATCTNNNTCAGGAACTCTCTTTGCTGAG

GAGCACACGTCTGAACTCTTTCCCTACACGACGCTCTTCCGATCTNNNTCAGGAACTCTCTTTGCTGAG

AGCACACGTCTGAACTCTTTCCCTACACGACGCTCTTCCGATCTNNNCTGAAGTCCCAGCTACCAGGAG

AGCACACGTCTGAACTCTTTCCCTACACGACGCTCTTCCGATCTNNNCTGAAGTCCCAGCTACCAGGAG

GAGCACACGTCTGAACTCTTTCCCTACACGACGCTCTTCCGATCTNNNCTGTTGCCTTCTGTCTGCAATTA

GAGCACACGTCTGAACTCTTTCCCTACACGACGCTCTTCCGATCTNNNCTGTTGCCTTCTGTCTGCAATTA

GAGCACACGTCTGAACTCTTTCCCTACACGACGCTCTTCCGATCTNNNACTCTCACTTTGTCATCCATGCT

GAGCACACGTCTGAACTCTTTCCCTACACGACGCTCTTCCGATCTNNNACTCTCACTTTGTCATCCATGCT

GAGCACACGTCTGAACTCTTTCCCTACACGACGCTCTTCCGATCTNNNAGACTGCTTTCGAACTGTAGCT

GAGCACACGTCTGAACTCTTTCCCTACACGACGCTCTTCCGATCTNNNAGACTGCTTTCGAACTGTAGCT

AGCACACGTCTGAACTCTTTCCCTACACGACGCTCTTCCGATCTNNNTCTTGAGAGTTCATGCAGGTGG

AGCACACGTCTGAACTCTTTCCCTACACGACGCTCTTCCGATCTNNNTCTTGAGAGTTCATGCAGGTGG

GAGCACACGTCTGAACTCTTTCCCTACACGACGCTCTTCCGATCTNNNGGCAACAAGAATGAAACTCCT

GAGCACACGTCTGAACTCTTTCCCTACACGACGCTCTTCCGATCTNNNGGCAACAAGAATGAAACTCCT

GCACACGTCTGAACTCTTTCCCTACACGACGCTCTTCCGATCTNNNAATGCTAAGCACTGTGGACCATA

GCACACGTCTGAACTCTTTCCCTACACGACGCTCTTCCGATCTNNNAATGCTAAGCACTGTGGACCATA

AGAGCACACGTCTGAACTCTTTCCCTACACGACGCTCTTCCGATCTNNNGTCAGTAGATCGAGACCATC

AGAGCACACGTCTGAACTCTTTCCCTACACGACGCTCTTCCGATCTNNNGTCAGTAGATCGAGACCATC

GAGCACACGTCTGAACTCTTTCCCTACACGACGCTCTTCCGATCTNNNAAGAAATGTTTTTAGTTATGG

GAGCACACGTCTGAACTCTTTCCCTACACGACGCTCTTCCGATCTNNNAAGAAATGTTTTTAGTTATGG

GAGCACACGTCTGAACTCTTTCCCTACACGACGCTCTTCCGATCTNNNCAGAGGGTCCTACACCCTGAC

GAGCACACGTCTGAACTCTTTCCCTACACGACGCTCTTCCGATCTNNNCAGAGGGTCCTACACCCTGAC

AGCACACGTCTGAACTCTTTCCCTACACGACGCTCTTCCGATCTNNNCTCTGGGCTCTAACATGCTAC

AGCACACGTCTGAACTCTTTCCCTACACGACGCTCTTCCGATCTNNNCTCTGGGCTCTAACATGCTAC

GCACACGTCTGAACTCTTTCCCTACACGACGCTCTTCCGATCTNNNAGGGAACACTAATTTTCTTTCCTG

GCACACGTCTGAACTCTTTCCCTACACGACGCTCTTCCGATCTNNNAGGGAACACTAATTTTCTTTCCTG

AGCACACGTCTGAACTCTTTCCCTACACGACGCTCTTCCGATCTNNNACAACAAGCTCTTGAGACTTGA

AGCACACGTCTGAACTCTTTCCCTACACGACGCTCTTCCGATCTNNNACAACAAGCTCTTGAGACTTGA

CACACGTCTGAACTCTTTCCCTACACGACGCTCTTCCGATCTNNNGAATTTCCAGTGTATTGTTTGGCAT

CACACGTCTGAACTCTTTCCCTACACGACGCTCTTCCGATCTNNNGAATTTCCAGTGTATTGTTTGGCAT

GCACACGTCTGAACTCTTTCCCTACACGACGCTCTTCCGATCTNNNGTACTCTCTCCGCAGTGAACTGC
GCACACGTCTGAACTCTTTCCCTACACGACGCTCTTCCGATCTNNNGTACTCTCTCCGCAGTGAACTGC
AGCACACGTCTGAACTCTTTCCCTACACGACGCTCTTCCGATCTNNNCGAACCTCTTCCACCTCCATCC
AGCACACGTCTGAACTCTTTCCCTACACGACGCTCTTCCGATCTNNNCGAACCTCTTCCACCTCCATCC
GCACACGTCTGAACTCTTTCCCTACACGACGCTCTTCCGATCTNNNTTCTTTTCCTCTTCCTCGTCGCC
GCACACGTCTGAACTCTTTCCCTACACGACGCTCTTCCGATCTNNNTTCTTTTCCTCTTCCTCGTCGCC
GCACACGTCTGAACTCTTTCCCTACACGACGCTCTTCCGATCTNNNACTTTTGGTTTTGGTTCACATGG
GCACACGTCTGAACTCTTTCCCTACACGACGCTCTTCCGATCTNNNACTTTTGGTTTTGGTTCACATGG
GCACACGTCTGAACTCTTTCCCTACACGACGCTCTTCCGATCTNNNTGAAGCAGAAGAATCGCTTGAA
GCACACGTCTGAACTCTTTCCCTACACGACGCTCTTCCGATCTNNNTGAAGCAGAAGAATCGCTTGAA
GAGCACACGTCTGAACTCTTTCCCTACACGACGCTCTTCCGATCTNNNCTTCCCAGCATGTGTGAGGT
GAGCACACGTCTGAACTCTTTCCCTACACGACGCTCTTCCGATCTNNNCTTCCCAGCATGTGTGAGGT
AGCACACGTCTGAACTCTTTCCCTACACGACGCTCTTCCGATCTNNNTCTACATGCGCAGTCTCTACTTC
AGCACACGTCTGAACTCTTTCCCTACACGACGCTCTTCCGATCTNNNTCTACATGCGCAGTCTCTACTTC
AGCACACGTCTGAACTCTTTCCCTACACGACGCTCTTCCGATCTNNNCTAGGAACATGGCTCACCTCTT
AGCACACGTCTGAACTCTTTCCCTACACGACGCTCTTCCGATCTNNNCTAGGAACATGGCTCACCTCTT
GAGCACACGTCTGAACTCTTTCCCTACACGACGCTCTTCCGATCTNNNTACAGCAGGACTTGGACAAG
GAGCACACGTCTGAACTCTTTCCCTACACGACGCTCTTCCGATCTNNNTACAGCAGGACTTGGACAAG
GCACACGTCTGAACTCTTTCCCTACACGACGCTCTTCCGATCTNNNACATCATTTTAGCCCTTACTCCAT
GCACACGTCTGAACTCTTTCCCTACACGACGCTCTTCCGATCTNNNACATCATTTTAGCCCTTACTCCAT
AGCACACGTCTGAACTCTTTCCCTACACGACGCTCTTCCGATCTNNNGGTAATGGTTTAGGGTGGTGG
AGCACACGTCTGAACTCTTTCCCTACACGACGCTCTTCCGATCTNNNGGTAATGGTTTAGGGTGGTGG
GAGCACACGTCTGAACTCTTTCCCTACACGACGCTCTTCCGATCTNNNTGCTATTTTTCCCAGGTGAAG
GAGCACACGTCTGAACTCTTTCCCTACACGACGCTCTTCCGATCTNNNTGCTATTTTTCCCAGGTGAAG
GCACACGTCTGAACTCTTTCCCTACACGACGCTCTTCCGATCTNNNTAATAATCCTGGGAGTTTGGCC
GCACACGTCTGAACTCTTTCCCTACACGACGCTCTTCCGATCTNNNTAATAATCCTGGGAGTTTGGCC
CACACGTCTGAACTCTTTCCCTACACGACGCTCTTCCGATCTNNNTGTCTCCCTTCCTCTCTTCTGTCCA
CACACGTCTGAACTCTTTCCCTACACGACGCTCTTCCGATCTNNNTGTCTCCCTTCCTCTCTTCTGTCCA
AGCACACGTCTGAACTCTTTCCCTACACGACGCTCTTCCGATCTNNNACCTTGAAATTGGCCATGATG
AGCACACGTCTGAACTCTTTCCCTACACGACGCTCTTCCGATCTNNNACCTTGAAATTGGCCATGATG
AGCACACGTCTGAACTCTTTCCCTACACGACGCTCTTCCGATCTNNNTGCCTGGTACACGTTTGTCAT
AGCACACGTCTGAACTCTTTCCCTACACGACGCTCTTCCGATCTNNNTGCCTGGTACACGTTTGTCAT
CACACGTCTGAACTCTTTCCCTACACGACGCTCTTCCGATCTNNNGACTTCATAGGGTCATTGTGGGT
CACACGTCTGAACTCTTTCCCTACACGACGCTCTTCCGATCTNNNGACTTCATAGGGTCATTGTGGGT
AGCACACGTCTGAACTCTTTCCCTACACGACGCTCTTCCGATCTNNNAATTCGTCCAGCTATATGGCA
AGCACACGTCTGAACTCTTTCCCTACACGACGCTCTTCCGATCTNNNAATTCGTCCAGCTATATGGCA
GCACACGTCTGAACTCTTTCCCTACACGACGCTCTTCCGATCTNNNTGGTGTATCAGACAGGGAAACA
GCACACGTCTGAACTCTTTCCCTACACGACGCTCTTCCGATCTNNNTGGTGTATCAGACAGGGAAACA
AGCACACGTCTGAACTCTTTCCCTACACGACGCTCTTCCGATCTNNNTCTACACTTTTCCCCTGGTCTTCC
AGCACACGTCTGAACTCTTTCCCTACACGACGCTCTTCCGATCTNNNTCTACACTTTTCCCCTGGTCTTCC
AGCACACGTCTGAACTCTTTCCCTACACGACGCTCTTCCGATCTNNNCACCTGTAGTCCCAGCTACTTG
AGCACACGTCTGAACTCTTTCCCTACACGACGCTCTTCCGATCTNNNCACCTGTAGTCCCAGCTACTTG
GCACACGTCTGAACTCTTTCCCTACACGACGCTCTTCCGATCTNNNGCTGAGCCCTAACCTCAAGTAAG

GCACACGTCTGAACTCTTTCCCTACACGACGCTCTTCCGATCTNNNGCTGAGCCCTAACCTCAAGTAAG
AGCACACGTCTGAACTCTTTCCCTACACGACGCTCTTCCGATCTNNNTGTGTTGTCAGTGGTCTTCAGA
AGCACACGTCTGAACTCTTTCCCTACACGACGCTCTTCCGATCTNNNTGTGTTGTCAGTGGTCTTCAGA
GCACACGTCTGAACTCTTTCCCTACACGACGCTCTTCCGATCTNNNGTGAAAATAAAGAAGATTGTCC
GCACACGTCTGAACTCTTTCCCTACACGACGCTCTTCCGATCTNNNGTGAAAATAAAGAAGATTGTCC
AGCACACGTCTGAACTCTTTCCCTACACGACGCTCTTCCGATCTNNNTGCAAGTAACACAAGTGGCAAT
AGCACACGTCTGAACTCTTTCCCTACACGACGCTCTTCCGATCTNNNTGCAAGTAACACAAGTGGCAAT
AGCACACGTCTGAACTCTTTCCCTACACGACGCTCTTCCGATCTNNNCTGGAATGCAATGGTGTGATC
AGCACACGTCTGAACTCTTTCCCTACACGACGCTCTTCCGATCTNNNCTGGAATGCAATGGTGTGATC
AGCACACGTCTGAACTCTTTCCCTACACGACGCTCTTCCGATCTNNNGATCCTTGAGGTCAGAGACAC
AGCACACGTCTGAACTCTTTCCCTACACGACGCTCTTCCGATCTNNNGATCCTTGAGGTCAGAGACAC
AGCACACGTCTGAACTCTTTCCCTACACGACGCTCTTCCGATCTNNNTACCTCACAGAAAAGTGCTCAC
AGCACACGTCTGAACTCTTTCCCTACACGACGCTCTTCCGATCTNNNTACCTCACAGAAAAGTGCTCAC
AGCACACGTCTGAACTCTTTCCCTACACGACGCTCTTCCGATCTNNNGACTGATGTTTGATCACAATGC
AGCACACGTCTGAACTCTTTCCCTACACGACGCTCTTCCGATCTNNNGACTGATGTTTGATCACAATGC
GCACACGTCTGAACTCTTTCCCTACACGACGCTCTTCCGATCTNNNGATGCAGGAGAATCACTTGAAC
GCACACGTCTGAACTCTTTCCCTACACGACGCTCTTCCGATCTNNNGATGCAGGAGAATCACTTGAAC
AGAGCACACGTCTGAACTCTTTCCCTACACGACGCTCTTCCGATCTNNNTAGGCAACACAGTGAGAAT
AGAGCACACGTCTGAACTCTTTCCCTACACGACGCTCTTCCGATCTNNNTAGGCAACACAGTGAGAAT
GCACACGTCTGAACTCTTTCCCTACACGACGCTCTTCCGATCTNNNGTATGTAGCTTTCCTTTTTCTTAC
GCACACGTCTGAACTCTTTCCCTACACGACGCTCTTCCGATCTNNNGTATGTAGCTTTCCTTTTTCTTAC
GAGCACACGTCTGAACTCTTTCCCTACACGACGCTCTTCCGATCTNNNGTACTGGTACAAAAACAGGT
GAGCACACGTCTGAACTCTTTCCCTACACGACGCTCTTCCGATCTNNNGTACTGGTACAAAAACAGGT
GAGCACACGTCTGAACTCTTTCCCTACACGACGCTCTTCCGATCTNNNTTGAACTCAGGAGATGGAGG
GAGCACACGTCTGAACTCTTTCCCTACACGACGCTCTTCCGATCTNNNTTGAACTCAGGAGATGGAGG
GAGCACACGTCTGAACTCTTTCCCTACACGACGCTCTTCCGATCTNNNTGCTGGCTCAGGAGAAAAAA
GAGCACACGTCTGAACTCTTTCCCTACACGACGCTCTTCCGATCTNNNTGCTGGCTCAGGAGAAAAAA
GCACACGTCTGAACTCTTTCCCTACACGACGCTCTTCCGATCTNNNGGAGCCCCTTGAGATTCTATCTG
GCACACGTCTGAACTCTTTCCCTACACGACGCTCTTCCGATCTNNNGGAGCCCCTTGAGATTCTATCTG
AGCACACGTCTGAACTCTTTCCCTACACGACGCTCTTCCGATCTNNNCCAGCTGGCTATTTGCAAAACTA
AGCACACGTCTGAACTCTTTCCCTACACGACGCTCTTCCGATCTNNNCCAGCTGGCTATTTGCAAAACTA
GAGCACACGTCTGAACTCTTTCCCTACACGACGCTCTTCCGATCTNNNTTGCTCTCTGTGACATGAACG
GAGCACACGTCTGAACTCTTTCCCTACACGACGCTCTTCCGATCTNNNTTGCTCTCTGTGACATGAACG
AGCACACGTCTGAACTCTTTCCCTACACGACGCTCTTCCGATCTNNNGAGAGAGTGAGGACCTACTGG
AGCACACGTCTGAACTCTTTCCCTACACGACGCTCTTCCGATCTNNNGAGAGAGTGAGGACCTACTGG
AGCACACGTCTGAACTCTTTCCCTACACGACGCTCTTCCGATCTNNNGTATCTAGCAGTCACCACCCTTT
AGCACACGTCTGAACTCTTTCCCTACACGACGCTCTTCCGATCTNNNGTATCTAGCAGTCACCACCCTTT
GCACACGTCTGAACTCTTTCCCTACACGACGCTCTTCCGATCTNNNCACGCTCTGTGTATTCACTTCAAC
GCACACGTCTGAACTCTTTCCCTACACGACGCTCTTCCGATCTNNNCACGCTCTGTGTATTCACTTCAAC
AGAGCACACGTCTGAACTCTTTCCCTACACGACGCTCTTCCGATCTNNNTGGCTTTTTAGAGATCAGGC
AGAGCACACGTCTGAACTCTTTCCCTACACGACGCTCTTCCGATCTNNNTGGCTTTTTAGAGATCAGGC
GAGCACACGTCTGAACTCTTTCCCTACACGACGCTCTTCCGATCTNNNTGTGCATTTTCACATGTATGA
GAGCACACGTCTGAACTCTTTCCCTACACGACGCTCTTCCGATCTNNNTGTGCATTTTCACATGTATGA

AGCACACGTCTGAACTCTTTCCCTACACGACGCTCTTCCGATCTNNNGAGTGTGTGTTGAGTTTGGTC
AGCACACGTCTGAACTCTTTCCCTACACGACGCTCTTCCGATCTNNNGAGTGTGTGTTGAGTTTGGTC
AGCACACGTCTGAACTCTTTCCCTACACGACGCTCTTCCGATCTNNNCAGAACCTTGTGGAGATAGCC
AGCACACGTCTGAACTCTTTCCCTACACGACGCTCTTCCGATCTNNNCAGAACCTTGTGGAGATAGCC
GCACACGTCTGAACTCTTTCCCTACACGACGCTCTTCCGATCTNNNTCTCTTGGAAATGAGGGAAAG
GCACACGTCTGAACTCTTTCCCTACACGACGCTCTTCCGATCTNNNTCTCTTGGAAATGAGGGAAAG
AGCACACGTCTGAACTCTTTCCCTACACGACGCTCTTCCGATCTNNNCTGTATTCCCAGCTACTCGGA
AGCACACGTCTGAACTCTTTCCCTACACGACGCTCTTCCGATCTNNNCTGTATTCCCAGCTACTCGGA
GAGCACACGTCTGAACTCTTTCCCTACACGACGCTCTTCCGATCTNNNCAGCCCTGCAACTCCTGTAGC
GAGCACACGTCTGAACTCTTTCCCTACACGACGCTCTTCCGATCTNNNCAGCCCTGCAACTCCTGTAGC
GAGCACACGTCTGAACTCTTTCCCTACACGACGCTCTTCCGATCTNNNTGAGGCAGGAGAATCACTT
GAGCACACGTCTGAACTCTTTCCCTACACGACGCTCTTCCGATCTNNNTGAGGCAGGAGAATCACTT
AGCACACGTCTGAACTCTTTCCCTACACGACGCTCTTCCGATCTNNNTTTGAACTCCTGGTTTCAAGC
AGCACACGTCTGAACTCTTTCCCTACACGACGCTCTTCCGATCTNNNTTTGAACTCCTGGTTTCAAGC
GCACACGTCTGAACTCTTTCCCTACACGACGCTCTTCCGATCTNNNTGTGATCTTGGCTCACTGCAAG
GCACACGTCTGAACTCTTTCCCTACACGACGCTCTTCCGATCTNNNTGTGATCTTGGCTCACTGCAAG
GAGCACACGTCTGAACTCTTTCCCTACACGACGCTCTTCCGATCTNNNCCCATCTCTAAGGATGTGTG
GAGCACACGTCTGAACTCTTTCCCTACACGACGCTCTTCCGATCTNNNCCCATCTCTAAGGATGTGTG
GAGCACACGTCTGAACTCTTTCCCTACACGACGCTCTTCCGATCTNNNCTTGGAGCGAGACACTCTTG
GAGCACACGTCTGAACTCTTTCCCTACACGACGCTCTTCCGATCTNNNCTTGGAGCGAGACACTCTTG
AGCACACGTCTGAACTCTTTCCCTACACGACGCTCTTCCGATCTNNNAAACAAATGTCAAAGACAGCCC
AGCACACGTCTGAACTCTTTCCCTACACGACGCTCTTCCGATCTNNNAAACAAATGTCAAAGACAGCCC
AGCACACGTCTGAACTCTTTCCCTACACGACGCTCTTCCGATCTNNNGTAATCCCAGCTACTCAAGAG
AGCACACGTCTGAACTCTTTCCCTACACGACGCTCTTCCGATCTNNNGTAATCCCAGCTACTCAAGAG
AGCACACGTCTGAACTCTTTCCCTACACGACGCTCTTCCGATCTNNNCCTCTCTCCTTGGTCCAATTCA
AGCACACGTCTGAACTCTTTCCCTACACGACGCTCTTCCGATCTNNNCCTCTCTCCTTGGTCCAATTCA
AGCACACGTCTGAACTCTTTCCCTACACGACGCTCTTCCGATCTNNNGGGAATGATTTGGGGAACAAG
AGCACACGTCTGAACTCTTTCCCTACACGACGCTCTTCCGATCTNNNGGGAATGATTTGGGGAACAAG
AGCACACGTCTGAACTCTTTCCCTACACGACGCTCTTCCGATCTNNNCTCTTCCCCCAGTGGCAAATCCA
AGCACACGTCTGAACTCTTTCCCTACACGACGCTCTTCCGATCTNNNCTCTTCCCCCAGTGGCAAATCCA
AGCACACGTCTGAACTCTTTCCCTACACGACGCTCTTCCGATCTNNNCATCCTGGCTAACATGGTGAAA
AGCACACGTCTGAACTCTTTCCCTACACGACGCTCTTCCGATCTNNNCATCCTGGCTAACATGGTGAAA
GCACACGTCTGAACTCTTTCCCTACACGACGCTCTTCCGATCTNNNTACCCTAGACTGTGACCACCATA
GCACACGTCTGAACTCTTTCCCTACACGACGCTCTTCCGATCTNNNTACCCTAGACTGTGACCACCATA
CACACGTCTGAACTCTTTCCCTACACGACGCTCTTCCGATCTNNNTAAGAAACATCTGACCAGCCGGG
CACACGTCTGAACTCTTTCCCTACACGACGCTCTTCCGATCTNNNTAAGAAACATCTGACCAGCCGGG
AGCACACGTCTGAACTCTTTCCCTACACGACGCTCTTCCGATCTNNNCTGTGTCATGGATTCCAGAGTG
AGCACACGTCTGAACTCTTTCCCTACACGACGCTCTTCCGATCTNNNCTGTGTCATGGATTCCAGAGTG
CACGTCTGAACTCTTTCCCTACACGACGCTCTTCCGATCTNNNGTTCTGGGCATCCTCTTCCTCCATCGG
CACGTCTGAACTCTTTCCCTACACGACGCTCTTCCGATCTNNNGTTCTGGGCATCCTCTTCCTCCATCGG
AGCACACGTCTGAACTCTTTCCCTACACGACGCTCTTCCGATCTNNNGCCAATAGACACACAAAAGAT
AGCACACGTCTGAACTCTTTCCCTACACGACGCTCTTCCGATCTNNNGCCAATAGACACACAAAAGAT
AGCACACGTCTGAACTCTTTCCCTACACGACGCTCTTCCGATCTNNNGAAACGAGTGTTGGACAAAG

GAGCACACGTCTGAACTCTTTCCCTACACGACGCTCTTCCGATCTNNNGAAACGAGTGTTGGACAAAG
GCACACGTCTGAACTCTTTCCCTACACGACGCTCTTCCGATCTNNNTCCTTCTCTTGCTCCAAAAAGGT
GCACACGTCTGAACTCTTTCCCTACACGACGCTCTTCCGATCTNNNTCCTTCTCTTGCTCCAAAAAGGT
AGCACACGTCTGAACTCTTTCCCTACACGACGCTCTTCCGATCTNNNGGAAGGTTGAAGTGGGTGGA
AGCACACGTCTGAACTCTTTCCCTACACGACGCTCTTCCGATCTNNNGGAAGGTTGAAGTGGGTGGA
GAGCACACGTCTGAACTCTTTCCCTACACGACGCTCTTCCGATCTNNNCTGAGACGGGAGAATCACTT
GAGCACACGTCTGAACTCTTTCCCTACACGACGCTCTTCCGATCTNNNCTGAGACGGGAGAATCACTT
AGCACACGTCTGAACTCTTTCCCTACACGACGCTCTTCCGATCTNNNGGGAGGGATAGCATTGGGAG
AGCACACGTCTGAACTCTTTCCCTACACGACGCTCTTCCGATCTNNNGGGAGGGATAGCATTGGGAG
GCACACGTCTGAACTCTTTCCCTACACGACGCTCTTCCGATCTNNNTCACATGGCCCTCTTTCCTAAATC
GCACACGTCTGAACTCTTTCCCTACACGACGCTCTTCCGATCTNNNTCACATGGCCCTCTTTCCTAAATC
AGCACACGTCTGAACTCTTTCCCTACACGACGCTCTTCCGATCTNNNTGGGGTAATTTGTTATGCAGT
AGCACACGTCTGAACTCTTTCCCTACACGACGCTCTTCCGATCTNNNTGGGGTAATTTGTTATGCAGT
GAGCACACGTCTGAACTCTTTCCCTACACGACGCTCTTCCGATCTNNNAGAGGGAGAGGGAGACAAT
GAGCACACGTCTGAACTCTTTCCCTACACGACGCTCTTCCGATCTNNNAGAGGGAGAGGGAGACAAT
AAGAGCACACGTCTGAACTCTTTCCCTACACGACGCTCTTCCGATCTNNNTTTGAGGTTGTAGTCAGT
AAGAGCACACGTCTGAACTCTTTCCCTACACGACGCTCTTCCGATCTNNNTTTGAGGTTGTAGTCAGT
GAGCACACGTCTGAACTCTTTCCCTACACGACGCTCTTCCGATCTNNNACTTCTTCAGGACTTTTTGGG
GAGCACACGTCTGAACTCTTTCCCTACACGACGCTCTTCCGATCTNNNACTTCTTCAGGACTTTTTGGG
GAGCACACGTCTGAACTCTTTCCCTACACGACGCTCTTCCGATCTNNNTGCAGTGATTCATTGGCTGTC
GAGCACACGTCTGAACTCTTTCCCTACACGACGCTCTTCCGATCTNNNTGCAGTGATTCATTGGCTGTC
GAGCACACGTCTGAACTCTTTCCCTACACGACGCTCTTCCGATCTNNNTGAGGCAGGAGAATCACTTG
GAGCACACGTCTGAACTCTTTCCCTACACGACGCTCTTCCGATCTNNNTGAGGCAGGAGAATCACTTG
CACACGTCTGAACTCTTTCCCTACACGACGCTCTTCCGATCTNNNAGTTCTAACATGTATGTATGCAAA
CACACGTCTGAACTCTTTCCCTACACGACGCTCTTCCGATCTNNNAGTTCTAACATGTATGTATGCAAA
GAGCACACGTCTGAACTCTTTCCCTACACGACGCTCTTCCGATCTNNNATGATCACACCACTGCACTCTA
GAGCACACGTCTGAACTCTTTCCCTACACGACGCTCTTCCGATCTNNNATGATCACACCACTGCACTCTA
GAGCACACGTCTGAACTCTTTCCCTACACGACGCTCTTCCGATCTNNNTTATTACACAACTTGCTGGGG
GAGCACACGTCTGAACTCTTTCCCTACACGACGCTCTTCCGATCTNNNTTATTACACAACTTGCTGGGG
GAGCACACGTCTGAACTCTTTCCCTACACGACGCTCTTCCGATCTNNNGGACACCCTTCTTCTTCTTCTC
GAGCACACGTCTGAACTCTTTCCCTACACGACGCTCTTCCGATCTNNNGGACACCCTTCTTCTTCTTCTC
AGCACACGTCTGAACTCTTTCCCTACACGACGCTCTTCCGATCTNNNTCTCAGACTTAGACTTCTTCCTG
AGCACACGTCTGAACTCTTTCCCTACACGACGCTCTTCCGATCTNNNTCTCAGACTTAGACTTCTTCCTG
GAGCACACGTCTGAACTCTTTCCCTACACGACGCTCTTCCGATCTNNNTTTGCTGATCCTCTTCCCTAAC
GAGCACACGTCTGAACTCTTTCCCTACACGACGCTCTTCCGATCTNNNTTTGCTGATCCTCTTCCCTAAC
CACGTCTGAACTCTTTCCCTACACGACGCTCTTCCGATCTNNNACAAGGACCCCCACCTTCTCCATCGGA
CACGTCTGAACTCTTTCCCTACACGACGCTCTTCCGATCTNNNACAAGGACCCCCACCTTCTCCATCGGA
AGCACACGTCTGAACTCTTTCCCTACACGACGCTCTTCCGATCTNNNTTTTCTTTTAGTTGGTCGAATT
AGCACACGTCTGAACTCTTTCCCTACACGACGCTCTTCCGATCTNNNTTTTCTTTTAGTTGGTCGAATT
AGCACACGTCTGAACTCTTTCCCTACACGACGCTCTTCCGATCTNNNTCCCAAAGTGCTAGGATTACAG
AGCACACGTCTGAACTCTTTCCCTACACGACGCTCTTCCGATCTNNNTCCCAAAGTGCTAGGATTACAG
AGCACACGTCTGAACTCTTTCCCTACACGACGCTCTTCCGATCTNNNCTCCTTCTGGTGTGGGTCTTGC
AGCACACGTCTGAACTCTTTCCCTACACGACGCTCTTCCGATCTNNNCTCCTTCTGGTGTGGGTCTTGC

AGCACACGTCTGAACTCTTTCCCTACACGACGCTCTTCCGATCTNNNAGTCCGTTATCAGTTGGCAAT
AGCACACGTCTGAACTCTTTCCCTACACGACGCTCTTCCGATCTNNNAGTCCGTTATCAGTTGGCAAT
GCACACGTCTGAACTCTTTCCCTACACGACGCTCTTCCGATCTNNNAGGATAATCGCTTGAAACTGGG
GCACACGTCTGAACTCTTTCCCTACACGACGCTCTTCCGATCTNNNAGGATAATCGCTTGAAACTGGG
AGCACACGTCTGAACTCTTTCCCTACACGACGCTCTTCCGATCTNNNGAAAGGCCATGGAATCTGCAT
AGCACACGTCTGAACTCTTTCCCTACACGACGCTCTTCCGATCTNNNGAAAGGCCATGGAATCTGCAT
GAGCACACGTCTGAACTCTTTCCCTACACGACGCTCTTCCGATCTNNNAAGCAAAGCAGGACCACAAA
GAGCACACGTCTGAACTCTTTCCCTACACGACGCTCTTCCGATCTNNNAAGCAAAGCAGGACCACAAA
GAGCACACGTCTGAACTCTTTCCCTACACGACGCTCTTCCGATCTNNNAAAAGTCTGCCCACTCAACAC
GAGCACACGTCTGAACTCTTTCCCTACACGACGCTCTTCCGATCTNNNAAAAGTCTGCCCACTCAACAC
GCACACGTCTGAACTCTTTCCCTACACGACGCTCTTCCGATCTNNNAAGAACAGACCATGAGCAGAGG
GCACACGTCTGAACTCTTTCCCTACACGACGCTCTTCCGATCTNNNAAGAACAGACCATGAGCAGAGG
AGAGCACACGTCTGAACTCTTTCCCTACACGACGCTCTTCCGATCTNNNAAACACTATTCACGGGTACT
AGAGCACACGTCTGAACTCTTTCCCTACACGACGCTCTTCCGATCTNNNAAACACTATTCACGGGTACT
GCACACGTCTGAACTCTTTCCCTACACGACGCTCTTCCGATCTNNNCTATATTTTCCAAAATCCCTACCC
GCACACGTCTGAACTCTTTCCCTACACGACGCTCTTCCGATCTNNNCTATATTTTCCAAAATCCCTACCC
AGCACACGTCTGAACTCTTTCCCTACACGACGCTCTTCCGATCTNNNGGAGACTGCAGTAAGCCATG
AGCACACGTCTGAACTCTTTCCCTACACGACGCTCTTCCGATCTNNNGGAGACTGCAGTAAGCCATG
AGAGCACACGTCTGAACTCTTTCCCTACACGACGCTCTTCCGATCTNNNGTAATCCCAGCTACTTAGG
AGAGCACACGTCTGAACTCTTTCCCTACACGACGCTCTTCCGATCTNNNGTAATCCCAGCTACTTAGG
AGCACACGTCTGAACTCTTTCCCTACACGACGCTCTTCCGATCTNNNGGTATCAAAATGATGCTGGCC
AGCACACGTCTGAACTCTTTCCCTACACGACGCTCTTCCGATCTNNNGGTATCAAAATGATGCTGGCC
GAGCACACGTCTGAACTCTTTCCCTACACGACGCTCTTCCGATCTNNNGAAGGTAGGGGTCTCTTTTG
GAGCACACGTCTGAACTCTTTCCCTACACGACGCTCTTCCGATCTNNNGAAGGTAGGGGTCTCTTTTG
AGCACACGTCTGAACTCTTTCCCTACACGACGCTCTTCCGATCTNNNCAGTAGCTCCCCATTTACACACC
AGCACACGTCTGAACTCTTTCCCTACACGACGCTCTTCCGATCTNNNCAGTAGCTCCCCATTTACACACC
AGCACACGTCTGAACTCTTTCCCTACACGACGCTCTTCCGATCTNNNCTGTAGTCCCAGCTACTCAGGA
AGCACACGTCTGAACTCTTTCCCTACACGACGCTCTTCCGATCTNNNCTGTAGTCCCAGCTACTCAGGA
GCACACGTCTGAACTCTTTCCCTACACGACGCTCTTCCGATCTNNNAATCTCCTGATCTCATGATCCGC
GCACACGTCTGAACTCTTTCCCTACACGACGCTCTTCCGATCTNNNAATCTCCTGATCTCATGATCCGC
CACACGTCTGAACTCTTTCCCTACACGACGCTCTTCCGATCTNNNAAATTCTCGATTGTCTTCTCTAGC
CACACGTCTGAACTCTTTCCCTACACGACGCTCTTCCGATCTNNNAAATTCTCGATTGTCTTCTCTAGC
GCACACGTCTGAACTCTTTCCCTACACGACGCTCTTCCGATCTNNNGACATCTAGGCAGGTTCTGGTG
GCACACGTCTGAACTCTTTCCCTACACGACGCTCTTCCGATCTNNNGACATCTAGGCAGGTTCTGGTG
GAGCACACGTCTGAACTCTTTCCCTACACGACGCTCTTCCGATCTNNNCCTGGGTGACATAGAACTGG
GAGCACACGTCTGAACTCTTTCCCTACACGACGCTCTTCCGATCTNNNCCTGGGTGACATAGAACTGG
GCACACGTCTGAACTCTTTCCCTACACGACGCTCTTCCGATCTNNNGGTATGTATCCCCTCAGGCATTT
GCACACGTCTGAACTCTTTCCCTACACGACGCTCTTCCGATCTNNNGGTATGTATCCCCTCAGGCATTT
AGCACACGTCTGAACTCTTTCCCTACACGACGCTCTTCCGATCTNNNGTTGCAATGAGCTGAGACTGT
AGCACACGTCTGAACTCTTTCCCTACACGACGCTCTTCCGATCTNNNGTTGCAATGAGCTGAGACTGT
AGCACACGTCTGAACTCTTTCCCTACACGACGCTCTTCCGATCTNNNACAAAAATTAGCTGGGTGTGG
AGCACACGTCTGAACTCTTTCCCTACACGACGCTCTTCCGATCTNNNACAAAAATTAGCTGGGTGTGG
GCACACGTCTGAACTCTTTCCCTACACGACGCTCTTCCGATCTNNNGAGGTTGCAGTGAGCTGAGAT

AGCACACGTCTGAACTCTTTCCCTACACGACGCTCTTCCGATCTNNNGAGGTTGCAGTGAGCTGAGATC
AGAGCACACGTCTGAACTCTTTCCCTACACGACGCTCTTCCGATCTNNNTGTGAGGCCTGGTTCCTATTT
AGAGCACACGTCTGAACTCTTTCCCTACACGACGCTCTTCCGATCTNNNTGTGAGGCCTGGTTCCTATTT
AGCACACGTCTGAACTCTTTCCCTACACGACGCTCTTCCGATCTNNNAAGGTCCATGGCCAAATCTACA
AGCACACGTCTGAACTCTTTCCCTACACGACGCTCTTCCGATCTNNNAAGGTCCATGGCCAAATCTACA
GAGCACACGTCTGAACTCTTTCCCTACACGACGCTCTTCCGATCTNNNCATGAACAACCAAAAGAGAAG
GAGCACACGTCTGAACTCTTTCCCTACACGACGCTCTTCCGATCTNNNCATGAACAACCAAAAGAGAAG
AGCACACGTCTGAACTCTTTCCCTACACGACGCTCTTCCGATCTNNNTGTAATCTCAGCACTTTGGGAG
AGCACACGTCTGAACTCTTTCCCTACACGACGCTCTTCCGATCTNNNTGTAATCTCAGCACTTTGGGAG
GCACACGTCTGAACTCTTTCCCTACACGACGCTCTTCCGATCTNNNTTTGTTCCCTCATCCTCTCCTGCA
GCACACGTCTGAACTCTTTCCCTACACGACGCTCTTCCGATCTNNNTTTGTTCCCTCATCCTCTCCTGCA
AGCACACGTCTGAACTCTTTCCCTACACGACGCTCTTCCGATCTNNNTTGACAACCCTGACTACTGGAAC
AGCACACGTCTGAACTCTTTCCCTACACGACGCTCTTCCGATCTNNNTTGACAACCCTGACTACTGGAAC
AGCACACGTCTGAACTCTTTCCCTACACGACGCTCTTCCGATCTNNNACTCTATCTCCACTCACACTTGCC
AGCACACGTCTGAACTCTTTCCCTACACGACGCTCTTCCGATCTNNNACTCTATCTCCACTCACACTTGCC
CACACGTCTGAACTCTTTCCCTACACGACGCTCTTCCGATCTNNNGACTGAAAAGGCCAGAGCAAATG
CACACGTCTGAACTCTTTCCCTACACGACGCTCTTCCGATCTNNNGACTGAAAAGGCCAGAGCAAATG
GAGCACACGTCTGAACTCTTTCCCTACACGACGCTCTTCCGATCTNNNCACTTGAGTACGAAGATCGTC
GAGCACACGTCTGAACTCTTTCCCTACACGACGCTCTTCCGATCTNNNCACTTGAGTACGAAGATCGTC
GAGCACACGTCTGAACTCTTTCCCTACACGACGCTCTTCCGATCTNNNGCAAGAGAATCACTTGAACCC
GAGCACACGTCTGAACTCTTTCCCTACACGACGCTCTTCCGATCTNNNGCAAGAGAATCACTTGAACCC
AGCACACGTCTGAACTCTTTCCCTACACGACGCTCTTCCGATCTNNNAGGTTCGATAATCCTCAGCATC
AGCACACGTCTGAACTCTTTCCCTACACGACGCTCTTCCGATCTNNNAGGTTCGATAATCCTCAGCATC
AGCACACGTCTGAACTCTTTCCCTACACGACGCTCTTCCGATCTNNNCTGGGTTCAAGCAATTCTACTG
AGCACACGTCTGAACTCTTTCCCTACACGACGCTCTTCCGATCTNNNCTGGGTTCAAGCAATTCTACTG
GAGCACACGTCTGAACTCTTTCCCTACACGACGCTCTTCCGATCTNNNTCAGGTACTACGCTTCAGAGAC
GAGCACACGTCTGAACTCTTTCCCTACACGACGCTCTTCCGATCTNNNTCAGGTACTACGCTTCAGAGAC
AGCACACGTCTGAACTCTTTCCCTACACGACGCTCTTCCGATCTNNNGTAATCCCAGCTACTCAGTAGGC
AGCACACGTCTGAACTCTTTCCCTACACGACGCTCTTCCGATCTNNNGTAATCCCAGCTACTCAGTAGGC
AGCACACGTCTGAACTCTTTCCCTACACGACGCTCTTCCGATCTNNNATATGTGTGTGTGTGTGTGTGT
AGCACACGTCTGAACTCTTTCCCTACACGACGCTCTTCCGATCTNNNATATGTGTGTGTGTGTGTGTGT
GAGCACACGTCTGAACTCTTTCCCTACACGACGCTCTTCCGATCTNNNTGGCTAAACTTTTCTAGGCAT
GAGCACACGTCTGAACTCTTTCCCTACACGACGCTCTTCCGATCTNNNTGGCTAAACTTTTCTAGGCAT
GAGCACACGTCTGAACTCTTTCCCTACACGACGCTCTTCCGATCTNNNTCTGAAGTCAGGTAGCGTGAT
GAGCACACGTCTGAACTCTTTCCCTACACGACGCTCTTCCGATCTNNNTCTGAAGTCAGGTAGCGTGAT
AGAGCACACGTCTGAACTCTTTCCCTACACGACGCTCTTCCGATCTNNNTCCAATTATTCCCATTGCAGA
AGAGCACACGTCTGAACTCTTTCCCTACACGACGCTCTTCCGATCTNNNTCCAATTATTCCCATTGCAGA
AGCACACGTCTGAACTCTTTCCCTACACGACGCTCTTCCGATCTNNNCATCTCTTCCCTGTTCTGTTCGAC
AGCACACGTCTGAACTCTTTCCCTACACGACGCTCTTCCGATCTNNNCATCTCTTCCCTGTTCTGTTCGAC
AGAGCACACGTCTGAACTCTTTCCCTACACGACGCTCTTCCGATCTNNNTGGATTATAGGAGGGCAAA
AGAGCACACGTCTGAACTCTTTCCCTACACGACGCTCTTCCGATCTNNNTGGATTATAGGAGGGCAAA
GAGCACACGTCTGAACTCTTTCCCTACACGACGCTCTTCCGATCTNNNACTAAACACCCCCATCCAAAAG
GAGCACACGTCTGAACTCTTTCCCTACACGACGCTCTTCCGATCTNNNACTAAACACCCCCATCCAAAAG

AGCACACGTCTGAACTCTTTCCCTACACGACGCTCTTCCGATCTNNNGATGCTGCAAAAGGCAAGAT
AGCACACGTCTGAACTCTTTCCCTACACGACGCTCTTCCGATCTNNNGATGCTGCAAAAGGCAAGAT
GAGCACACGTCTGAACTCTTTCCCTACACGACGCTCTTCCGATCTNNNGAGGCTGAGGCAGGAGAAT
GAGCACACGTCTGAACTCTTTCCCTACACGACGCTCTTCCGATCTNNNGAGGCTGAGGCAGGAGAAT
AGCACACGTCTGAACTCTTTCCCTACACGACGCTCTTCCGATCTNNNAGTCAAGAGGGGTTGTCACT
AGCACACGTCTGAACTCTTTCCCTACACGACGCTCTTCCGATCTNNNAGTCAAGAGGGGTTGTCACT
AGCACACGTCTGAACTCTTTCCCTACACGACGCTCTTCCGATCTNNNGAACAATAACAAAAAGGGGG
AGCACACGTCTGAACTCTTTCCCTACACGACGCTCTTCCGATCTNNNGAACAATAACAAAAAGGGGG
AGCACACGTCTGAACTCTTTCCCTACACGACGCTCTTCCGATCTNNNCGGTGGAAATTGTAAACAAGG
AGCACACGTCTGAACTCTTTCCCTACACGACGCTCTTCCGATCTNNNCGGTGGAAATTGTAAACAAGG
GAGCACACGTCTGAACTCTTTCCCTACACGACGCTCTTCCGATCTNNNTGTTGTCTTTTCTGCCAGTTCT
GAGCACACGTCTGAACTCTTTCCCTACACGACGCTCTTCCGATCTNNNTGTTGTCTTTTCTGCCAGTTCT
GAGCACACGTCTGAACTCTTTCCCTACACGACGCTCTTCCGATCTNNNCTCTAGAGAGACACCCTTGTT
GAGCACACGTCTGAACTCTTTCCCTACACGACGCTCTTCCGATCTNNNCTCTAGAGAGACACCCTTGTT
AAGAGCACACGTCTGAACTCTTTCCCTACACGACGCTCTTCCGATCTNNNCTTGCAGTGAGCCAAGAT
AAGAGCACACGTCTGAACTCTTTCCCTACACGACGCTCTTCCGATCTNNNCTTGCAGTGAGCCAAGAT
AGCACACGTCTGAACTCTTTCCCTACACGACGCTCTTCCGATCTNNNATATAACATGTAACGCCCTCCC
AGCACACGTCTGAACTCTTTCCCTACACGACGCTCTTCCGATCTNNNATATAACATGTAACGCCCTCCC
AGCACACGTCTGAACTCTTTCCCTACACGACGCTCTTCCGATCTNNNATGCCTATATTCCCAGCACTGA
AGCACACGTCTGAACTCTTTCCCTACACGACGCTCTTCCGATCTNNNATGCCTATATTCCCAGCACTGA
CACGTCTGAACTCTTTCCCTACACGACGCTCTTCCGATCTNNNTTGGGTAGCTTGCACCTAGAATTCAT
CACGTCTGAACTCTTTCCCTACACGACGCTCTTCCGATCTNNNTTGGGTAGCTTGCACCTAGAATTCAT
AGCACACGTCTGAACTCTTTCCCTACACGACGCTCTTCCGATCTNNNATGAGACAGCAGCCCATATTT
AGCACACGTCTGAACTCTTTCCCTACACGACGCTCTTCCGATCTNNNATGAGACAGCAGCCCATATTT
GAGCACACGTCTGAACTCTTTCCCTACACGACGCTCTTCCGATCTNNNAGAATAACTTGAACCTGGGA
GAGCACACGTCTGAACTCTTTCCCTACACGACGCTCTTCCGATCTNNNAGAATAACTTGAACCTGGGA
AGAGCACACGTCTGAACTCTTTCCCTACACGACGCTCTTCCGATCTNNNTGTGTGGAATCAGAAAGAA
AGAGCACACGTCTGAACTCTTTCCCTACACGACGCTCTTCCGATCTNNNTGTGTGGAATCAGAAAGAA
AGCACACGTCTGAACTCTTTCCCTACACGACGCTCTTCCGATCTNNNGATGTTCTCGATCTCCTGACCTC
AGCACACGTCTGAACTCTTTCCCTACACGACGCTCTTCCGATCTNNNGATGTTCTCGATCTCCTGACCTC
GAGCACACGTCTGAACTCTTTCCCTACACGACGCTCTTCCGATCTNNNCCCAGCTAACTAAGTGCACA
GAGCACACGTCTGAACTCTTTCCCTACACGACGCTCTTCCGATCTNNNCCCAGCTAACTAAGTGCACA
AGCACACGTCTGAACTCTTTCCCTACACGACGCTCTTCCGATCTNNNCACAGTTATAGTGGTGGTGGT
AGCACACGTCTGAACTCTTTCCCTACACGACGCTCTTCCGATCTNNNCACAGTTATAGTGGTGGTGGT
AGCACACGTCTGAACTCTTTCCCTACACGACGCTCTTCCGATCTNNNCTGAAAACAAGGTATTGGCTG
AGCACACGTCTGAACTCTTTCCCTACACGACGCTCTTCCGATCTNNNCTGAAAACAAGGTATTGGCTG
GCACACGTCTGAACTCTTTCCCTACACGACGCTCTTCCGATCTNNNAACACTCAGGGCATGAATTTGT
GCACACGTCTGAACTCTTTCCCTACACGACGCTCTTCCGATCTNNNAACACTCAGGGCATGAATTTGT
AGCACACGTCTGAACTCTTTCCCTACACGACGCTCTTCCGATCTNNNACCTGGGTAATATAGCAAGA
AGCACACGTCTGAACTCTTTCCCTACACGACGCTCTTCCGATCTNNNACCTGGGTAATATAGCAAGA
AGCACACGTCTGAACTCTTTCCCTACACGACGCTCTTCCGATCTNNNTTTTTGGAGAGGGCATCTTCTT
AGCACACGTCTGAACTCTTTCCCTACACGACGCTCTTCCGATCTNNNTTTTTGGAGAGGGCATCTTCTT
GCACACGTCTGAACTCTTTCCCTACACGACGCTCTTCCGATCTNNNATATTTCCCCGCATCTTTCCTGAC

GCACACGTCTGAACTCTTTCCCTACACGACGCTCTTCCGATCTNNNATATTTCCCCGCATCTTTCCTGAC
GCACACGTCTGAACTCTTTCCCTACACGACGCTCTTCCGATCTNNNCAATTAACCTAAGCTTGATGATG
GCACACGTCTGAACTCTTTCCCTACACGACGCTCTTCCGATCTNNNCAATTAACCTAAGCTTGATGATG
AGCACACGTCTGAACTCTTTCCCTACACGACGCTCTTCCGATCTNNNTAACCGCTGGACTTTGTTGAA
AGCACACGTCTGAACTCTTTCCCTACACGACGCTCTTCCGATCTNNNTAACCGCTGGACTTTGTTGAA
GCACACGTCTGAACTCTTTCCCTACACGACGCTCTTCCGATCTNNNGAGAATGGCATGAACCCAGAA
GCACACGTCTGAACTCTTTCCCTACACGACGCTCTTCCGATCTNNNGAGAATGGCATGAACCCAGAA
AGCACACGTCTGAACTCTTTCCCTACACGACGCTCTTCCGATCTNNNACACTTCTGGTCTCAAAGCTTG
AGCACACGTCTGAACTCTTTCCCTACACGACGCTCTTCCGATCTNNNACACTTCTGGTCTCAAAGCTTG
AGAGCACACGTCTGAACTCTTTCCCTACACGACGCTCTTCCGATCTNNNGCAAGACCTTGTGCTAATTG
AGAGCACACGTCTGAACTCTTTCCCTACACGACGCTCTTCCGATCTNNNGCAAGACCTTGTGCTAATTG
AGCACACGTCTGAACTCTTTCCCTACACGACGCTCTTCCGATCTNNNTCTGTCGAGTTATACCACTGCA
AGCACACGTCTGAACTCTTTCCCTACACGACGCTCTTCCGATCTNNNTCTGTCGAGTTATACCACTGCA
AGAGCACACGTCTGAACTCTTTCCCTACACGACGCTCTTCCGATCTNNNGGAGGTGGAACTTGAAATA
AGAGCACACGTCTGAACTCTTTCCCTACACGACGCTCTTCCGATCTNNNGGAGGTGGAACTTGAAATA
AGCACACGTCTGAACTCTTTCCCTACACGACGCTCTTCCGATCTNNNTGTAGTCCCAGCTACTTAGGAG
AGCACACGTCTGAACTCTTTCCCTACACGACGCTCTTCCGATCTNNNTGTAGTCCCAGCTACTTAGGAG
AGCACACGTCTGAACTCTTTCCCTACACGACGCTCTTCCGATCTNNNATAGAAAAAATTAGCCGGGCG
AGCACACGTCTGAACTCTTTCCCTACACGACGCTCTTCCGATCTNNNATAGAAAAAATTAGCCGGGCG
AGCACACGTCTGAACTCTTTCCCTACACGACGCTCTTCCGATCTNNNCGCCCCTTCACATAAGGGATAT
AGCACACGTCTGAACTCTTTCCCTACACGACGCTCTTCCGATCTNNNCGCCCCTTCACATAAGGGATAT
GAGCACACGTCTGAACTCTTTCCCTACACGACGCTCTTCCGATCTNNNTCAAGTGATTTTCTTGCGTCG
GAGCACACGTCTGAACTCTTTCCCTACACGACGCTCTTCCGATCTNNNTCAAGTGATTTTCTTGCGTCG
AGCACACGTCTGAACTCTTTCCCTACACGACGCTCTTCCGATCTNNNCTGTAGTCCCAGCTACTCAGG
AGCACACGTCTGAACTCTTTCCCTACACGACGCTCTTCCGATCTNNNCTGTAGTCCCAGCTACTCAGG
GCACACGTCTGAACTCTTTCCCTACACGACGCTCTTCCGATCTNNNGCAGCTCTTCATCAAGTTCAGTT
GCACACGTCTGAACTCTTTCCCTACACGACGCTCTTCCGATCTNNNGCAGCTCTTCATCAAGTTCAGTT
AGCACACGTCTGAACTCTTTCCCTACACGACGCTCTTCCGATCTNNNTGTTGTATTTGCTAGCTAAGG
AGCACACGTCTGAACTCTTTCCCTACACGACGCTCTTCCGATCTNNNTGTTGTATTTGCTAGCTAAGG
AGCACACGTCTGAACTCTTTCCCTACACGACGCTCTTCCGATCTNNNAACTGGAGGCCATTATCCTAA
AGCACACGTCTGAACTCTTTCCCTACACGACGCTCTTCCGATCTNNNAACTGGAGGCCATTATCCTAA
AGAGCACACGTCTGAACTCTTTCCCTACACGACGCTCTTCCGATCTNNNTGCAGTGAACCAAAATTGTG
AGAGCACACGTCTGAACTCTTTCCCTACACGACGCTCTTCCGATCTNNNTGCAGTGAACCAAAATTGTG
GCACACGTCTGAACTCTTTCCCTACACGACGCTCTTCCGATCTNNNAAACTCATGGAGCGTATTCCCAC
GCACACGTCTGAACTCTTTCCCTACACGACGCTCTTCCGATCTNNNAAACTCATGGAGCGTATTCCCAC
AGCACACGTCTGAACTCTTTCCCTACACGACGCTCTTCCGATCTNNNCGACGTTTCATGGAGTTGTCA
AGCACACGTCTGAACTCTTTCCCTACACGACGCTCTTCCGATCTNNNCGACGTTTCATGGAGTTGTCA
AGCACACGTCTGAACTCTTTCCCTACACGACGCTCTTCCGATCTNNNCCAGATGCAGTCACCTATCAA
AGCACACGTCTGAACTCTTTCCCTACACGACGCTCTTCCGATCTNNNCCAGATGCAGTCACCTATCAA
GAGCACACGTCTGAACTCTTTCCCTACACGACGCTCTTCCGATCTNNNGTTCCAGTGAGCCAATATCA
GAGCACACGTCTGAACTCTTTCCCTACACGACGCTCTTCCGATCTNNNGTTCCAGTGAGCCAATATCA
AGCACACGTCTGAACTCTTTCCCTACACGACGCTCTTCCGATCTNNNTTAGCTGCGTAAGGCCTCTTAT
AGCACACGTCTGAACTCTTTCCCTACACGACGCTCTTCCGATCTNNNTTAGCTGCGTAAGGCCTCTTAT

GCACACGTCTGAACTCTTTCCCTACACGACGCTCTTCCGATCTNNNCTCTAACCCTCATCCCATTCCTGC
GCACACGTCTGAACTCTTTCCCTACACGACGCTCTTCCGATCTNNNCTCTAACCCTCATCCCATTCCTGC
AGCACACGTCTGAACTCTTTCCCTACACGACGCTCTTCCGATCTNNNGCTGTCACTGATCTGTTTCACA
AGCACACGTCTGAACTCTTTCCCTACACGACGCTCTTCCGATCTNNNGCTGTCACTGATCTGTTTCACA
AGAGCACACGTCTGAACTCTTTCCCTACACGACGCTCTTCCGATCTNNNCTAGCCAGACAGAGAGAAA
AGAGCACACGTCTGAACTCTTTCCCTACACGACGCTCTTCCGATCTNNNCTAGCCAGACAGAGAGAAA
AGCACACGTCTGAACTCTTTCCCTACACGACGCTCTTCCGATCTNNNTCCAGGAGATTCAGAAGAGGC
AGCACACGTCTGAACTCTTTCCCTACACGACGCTCTTCCGATCTNNNTCCAGGAGATTCAGAAGAGGC
AAGAGCACACGTCTGAACTCTTTCCCTACACGACGCTCTTCCGATCTNNNGAAAAATGCTCAAGTGGT
AAGAGCACACGTCTGAACTCTTTCCCTACACGACGCTCTTCCGATCTNNNGAAAAATGCTCAAGTGGT
AGCACACGTCTGAACTCTTTCCCTACACGACGCTCTTCCGATCTNNNGTTTCTCTATGTTGTTGCCTCCA
AGCACACGTCTGAACTCTTTCCCTACACGACGCTCTTCCGATCTNNNGTTTCTCTATGTTGTTGCCTCCA
AGCACACGTCTGAACTCTTTCCCTACACGACGCTCTTCCGATCTNNNGTTCTGCATCGGGACAGGACC
AGCACACGTCTGAACTCTTTCCCTACACGACGCTCTTCCGATCTNNNGTTCTGCATCGGGACAGGACC
AGCACACGTCTGAACTCTTTCCCTACACGACGCTCTTCCGATCTNNNGCAGATGTCCTTGAAGAGAGTG
AGCACACGTCTGAACTCTTTCCCTACACGACGCTCTTCCGATCTNNNGCAGATGTCCTTGAAGAGAGTG
AGCACACGTCTGAACTCTTTCCCTACACGACGCTCTTCCGATCTNNNAGGTGGTGTTTGGTTACATGA
AGCACACGTCTGAACTCTTTCCCTACACGACGCTCTTCCGATCTNNNAGGTGGTGTTTGGTTACATGA
AGCACACGTCTGAACTCTTTCCCTACACGACGCTCTTCCGATCTNNNCATTCTCCTCCTACCAAGCCAGC
AGCACACGTCTGAACTCTTTCCCTACACGACGCTCTTCCGATCTNNNCATTCTCCTCCTACCAAGCCAGC
GCACACGTCTGAACTCTTTCCCTACACGACGCTCTTCCGATCTNNNCCTGGCCCCATTTTTCTTCTTTTCA
GCACACGTCTGAACTCTTTCCCTACACGACGCTCTTCCGATCTNNNCCTGGCCCCATTTTTCTTCTTTTCA
AGCACACGTCTGAACTCTTTCCCTACACGACGCTCTTCCGATCTNNNTCTTTCCCATGCTGTTCTCATGA
AGCACACGTCTGAACTCTTTCCCTACACGACGCTCTTCCGATCTNNNTCTTTCCCATGCTGTTCTCATGA
GAGCACACGTCTGAACTCTTTCCCTACACGACGCTCTTCCGATCTNNNTAACCCACTGAGCTCACAGAA
GAGCACACGTCTGAACTCTTTCCCTACACGACGCTCTTCCGATCTNNNTAACCCACTGAGCTCACAGAA
AGAGCACACGTCTGAACTCTTTCCCTACACGACGCTCTTCCGATCTNNNCTGAGGCAGGAGAATCACT
AGAGCACACGTCTGAACTCTTTCCCTACACGACGCTCTTCCGATCTNNNCTGAGGCAGGAGAATCACT
AGCACACGTCTGAACTCTTTCCCTACACGACGCTCTTCCGATCTNNNTATGAAAGACTGGCTTGTTTGC
AGCACACGTCTGAACTCTTTCCCTACACGACGCTCTTCCGATCTNNNTATGAAAGACTGGCTTGTTTGC
GAGCACACGTCTGAACTCTTTCCCTACACGACGCTCTTCCGATCTNNNAAACACACAACATTCCCTGCAA
GAGCACACGTCTGAACTCTTTCCCTACACGACGCTCTTCCGATCTNNNAAACACACAACATTCCCTGCAA
GAGCACACGTCTGAACTCTTTCCCTACACGACGCTCTTCCGATCTNNNAACATTCTTGACATTGCTGGT
GAGCACACGTCTGAACTCTTTCCCTACACGACGCTCTTCCGATCTNNNAACATTCTTGACATTGCTGGT
GAGCACACGTCTGAACTCTTTCCCTACACGACGCTCTTCCGATCTNNNTGGCCATTTTCTTTCTGGGTA
GAGCACACGTCTGAACTCTTTCCCTACACGACGCTCTTCCGATCTNNNTGGCCATTTTCTTTCTGGGTA
AGCACACGTCTGAACTCTTTCCCTACACGACGCTCTTCCGATCTNNNCACAGTGCCTACAATTACAACA
AGCACACGTCTGAACTCTTTCCCTACACGACGCTCTTCCGATCTNNNCACAGTGCCTACAATTACAACA
GAGCACACGTCTGAACTCTTTCCCTACACGACGCTCTTCCGATCTNNNCTTTGCTTACCACCGTCTCTCT
GAGCACACGTCTGAACTCTTTCCCTACACGACGCTCTTCCGATCTNNNCTTTGCTTACCACCGTCTCTCT
CACACGTCTGAACTCTTTCCCTACACGACGCTCTTCCGATCTNNNTAGTGTGGGAGGGACTTAGAGA
CACACGTCTGAACTCTTTCCCTACACGACGCTCTTCCGATCTNNNTAGTGTGGGAGGGACTTAGAGA
GAGCACACGTCTGAACTCTTTCCCTACACGACGCTCTTCCGATCTNNNCAGGAGGTCAAGGCTACAGT

GAGCACACGTCTGAACTCTTTCCCTACACGACGCTCTTCCGATCTNNNCAGGAGGTCAAGGCTACAGT
AGAGCACACGTCTGAACTCTTTCCCTACACGACGCTCTTCCGATCTNNNTGAGGCAGGAGAATTGCTT
AGAGCACACGTCTGAACTCTTTCCCTACACGACGCTCTTCCGATCTNNNTGAGGCAGGAGAATTGCTT
GAGCACACGTCTGAACTCTTTCCCTACACGACGCTCTTCCGATCTNNNTACCTCCCTCCTACTAGGTTGT
GAGCACACGTCTGAACTCTTTCCCTACACGACGCTCTTCCGATCTNNNTACCTCCCTCCTACTAGGTTGT
AGCACACGTCTGAACTCTTTCCCTACACGACGCTCTTCCGATCTNNNTTGTATGTGGTGGAGCACAACT
AGCACACGTCTGAACTCTTTCCCTACACGACGCTCTTCCGATCTNNNTTGTATGTGGTGGAGCACAACT
GAGCACACGTCTGAACTCTTTCCCTACACGACGCTCTTCCGATCTNNNCAGAGGTTACAGTGAGCCAA
GAGCACACGTCTGAACTCTTTCCCTACACGACGCTCTTCCGATCTNNNCAGAGGTTACAGTGAGCCAA
AGCACACGTCTGAACTCTTTCCCTACACGACGCTCTTCCGATCTNNNCTCACACACACACAAATAGCCAC
AGCACACGTCTGAACTCTTTCCCTACACGACGCTCTTCCGATCTNNNCTCACACACACACAAATAGCCAC
GAGCACACGTCTGAACTCTTTCCCTACACGACGCTCTTCCGATCTNNNATGCCAGTTTTGGCTTACTTC
GAGCACACGTCTGAACTCTTTCCCTACACGACGCTCTTCCGATCTNNNATGCCAGTTTTGGCTTACTTC
AAGAGCACACGTCTGAACTCTTTCCCTACACGACGCTCTTCCGATCTNNNTCATTACACCAGTTCGTCCC
AAGAGCACACGTCTGAACTCTTTCCCTACACGACGCTCTTCCGATCTNNNTCATTACACCAGTTCGTCCC
AGCACACGTCTGAACTCTTTCCCTACACGACGCTCTTCCGATCTNNNCCACCTTTCAAAATCACAGTCCC
AGCACACGTCTGAACTCTTTCCCTACACGACGCTCTTCCGATCTNNNCCACCTTTCAAAATCACAGTCCC
GAGCACACGTCTGAACTCTTTCCCTACACGACGCTCTTCCGATCTNNNATCTATTTTCTCAACTGCCATA
GAGCACACGTCTGAACTCTTTCCCTACACGACGCTCTTCCGATCTNNNATCTATTTTCTCAACTGCCATA
GCACACGTCTGAACTCTTTCCCTACACGACGCTCTTCCGATCTNNNGTGAGAAGGACACAAATTTCGG
GCACACGTCTGAACTCTTTCCCTACACGACGCTCTTCCGATCTNNNGTGAGAAGGACACAAATTTCGG
ACACGTCTGAACTCTTTCCCTACACGACGCTCTTCCGATCTNNNAAATGCGTCTCCCTGTGAATCTACAT
ACACGTCTGAACTCTTTCCCTACACGACGCTCTTCCGATCTNNNAAATGCGTCTCCCTGTGAATCTACAT
GAGCACACGTCTGAACTCTTTCCCTACACGACGCTCTTCCGATCTNNNTCAATGACCTCTGGAACTATG
GAGCACACGTCTGAACTCTTTCCCTACACGACGCTCTTCCGATCTNNNTCAATGACCTCTGGAACTATG
GAGCACACGTCTGAACTCTTTCCCTACACGACGCTCTTCCGATCTNNNAACCTTTTTCCCAGGTAAGGAC
GAGCACACGTCTGAACTCTTTCCCTACACGACGCTCTTCCGATCTNNNAACCTTTTTCCCAGGTAAGGAC
GAGCACACGTCTGAACTCTTTCCCTACACGACGCTCTTCCGATCTNNNATATGTGTGTGTGTGTGTGTG
GAGCACACGTCTGAACTCTTTCCCTACACGACGCTCTTCCGATCTNNNATATGTGTGTGTGTGTGTGTG
AGCACACGTCTGAACTCTTTCCCTACACGACGCTCTTCCGATCTNNNCAGGAATGATGTAACTTGGTG
AGCACACGTCTGAACTCTTTCCCTACACGACGCTCTTCCGATCTNNNCAGGAATGATGTAACTTGGTG
GAGCACACGTCTGAACTCTTTCCCTACACGACGCTCTTCCGATCTNNNCCTGGCCTGGGATTGTTATTT
GAGCACACGTCTGAACTCTTTCCCTACACGACGCTCTTCCGATCTNNNCCTGGCCTGGGATTGTTATTT
AGCACACGTCTGAACTCTTTCCCTACACGACGCTCTTCCGATCTNNNGTCCTGTTGCAAAAGGAAATG
AGCACACGTCTGAACTCTTTCCCTACACGACGCTCTTCCGATCTNNNGTCCTGTTGCAAAAGGAAATG
AAGAGCACACGTCTGAACTCTTTCCCTACACGACGCTCTTCCGATCTNNNTAGGCCTGTAATCCCAGCT
AAGAGCACACGTCTGAACTCTTTCCCTACACGACGCTCTTCCGATCTNNNTAGGCCTGTAATCCCAGCT
AGCACACGTCTGAACTCTTTCCCTACACGACGCTCTTCCGATCTNNNCAAGGTGGGAGGATCACTTGA
AGCACACGTCTGAACTCTTTCCCTACACGACGCTCTTCCGATCTNNNCAAGGTGGGAGGATCACTTGA
GAGCACACGTCTGAACTCTTTCCCTACACGACGCTCTTCCGATCTNNNGCATACACCTGTAATCCCAGC
GAGCACACGTCTGAACTCTTTCCCTACACGACGCTCTTCCGATCTNNNGCATACACCTGTAATCCCAGC
GAGCACACGTCTGAACTCTTTCCCTACACGACGCTCTTCCGATCTNNNAGACGAGTGACTTCCCTGAAT
GAGCACACGTCTGAACTCTTTCCCTACACGACGCTCTTCCGATCTNNNAGACGAGTGACTTCCCTGAAT

AGCACACGTCTGAACTCTTTCCCTACACGACGCTCTTCCGATCTNNNCACACACAATTGCTACTACTGCC

AGCACACGTCTGAACTCTTTCCCTACACGACGCTCTTCCGATCTNNNCACACACAATTGCTACTACTGCC

AGCACACGTCTGAACTCTTTCCCTACACGACGCTCTTCCGATCTNNNATGTTCCAGGCCTTGTCTTGTCC

AGCACACGTCTGAACTCTTTCCCTACACGACGCTCTTCCGATCTNNNATGTTCCAGGCCTTGTCTTGTCC

GAGCACACGTCTGAACTCTTTCCCTACACGACGCTCTTCCGATCTNNNAATGCCTGTAATCCCAGCTACT

GAGCACACGTCTGAACTCTTTCCCTACACGACGCTCTTCCGATCTNNNAATGCCTGTAATCCCAGCTACT

AGCACACGTCTGAACTCTTTCCCTACACGACGCTCTTCCGATCTNNNTGCCCTGATATCAAAACCAGACA

AGCACACGTCTGAACTCTTTCCCTACACGACGCTCTTCCGATCTNNNTGCCCTGATATCAAAACCAGACA

CACGTCTGAACTCTTTCCCTACACGACGCTCTTCCGATCTNNNTCTAAGAGGTGATTAGGTCAACATCG

CACGTCTGAACTCTTTCCCTACACGACGCTCTTCCGATCTNNNTCTAAGAGGTGATTAGGTCAACATCG

AGCACACGTCTGAACTCTTTCCCTACACGACGCTCTTCCGATCTNNNCATGCCACTATAGTCCAGCCTA

AGCACACGTCTGAACTCTTTCCCTACACGACGCTCTTCCGATCTNNNCATGCCACTATAGTCCAGCCTA

GCACACGTCTGAACTCTTTCCCTACACGACGCTCTTCCGATCTNNNATGGATGGATTCAATCGGCAGA

GCACACGTCTGAACTCTTTCCCTACACGACGCTCTTCCGATCTNNNATGGATGGATTCAATCGGCAGA

AGCACACGTCTGAACTCTTTCCCTACACGACGCTCTTCCGATCTNNNACTCTTGTGTCTGCAGTTTCTCA

AGCACACGTCTGAACTCTTTCCCTACACGACGCTCTTCCGATCTNNNACTCTTGTGTCTGCAGTTTCTCA

GCACACGTCTGAACTCTTTCCCTACACGACGCTCTTCCGATCTNNNTGCAGCACTACTCTTTTCCTCTACA

GCACACGTCTGAACTCTTTCCCTACACGACGCTCTTCCGATCTNNNTGCAGCACTACTCTTTTCCTCTACA

AGCACACGTCTGAACTCTTTCCCTACACGACGCTCTTCCGATCTNNNGCCTTTGGCTGATTTCCAATCTTC

AGCACACGTCTGAACTCTTTCCCTACACGACGCTCTTCCGATCTNNNGCCTTTGGCTGATTTCCAATCTTC

GAGCACACGTCTGAACTCTTTCCCTACACGACGCTCTTCCGATCTNNNGATTATGCCACTACACTCCAGC

GAGCACACGTCTGAACTCTTTCCCTACACGACGCTCTTCCGATCTNNNGATTATGCCACTACACTCCAGC

GAGCACACGTCTGAACTCTTTCCCTACACGACGCTCTTCCGATCTNNNAATGACACAAAGAACACAGAC

GAGCACACGTCTGAACTCTTTCCCTACACGACGCTCTTCCGATCTNNNAATGACACAAAGAACACAGAC

AGCACACGTCTGAACTCTTTCCCTACACGACGCTCTTCCGATCTNNNGAGTCTTCACTTTTTTCCGAGG

AGCACACGTCTGAACTCTTTCCCTACACGACGCTCTTCCGATCTNNNGAGTCTTCACTTTTTTCCGAGG

GAGCACACGTCTGAACTCTTTCCCTACACGACGCTCTTCCGATCTNNNGCCAGAGCTTGAGCCTAGTAA

GAGCACACGTCTGAACTCTTTCCCTACACGACGCTCTTCCGATCTNNNGCCAGAGCTTGAGCCTAGTAA

AGAGCACACGTCTGAACTCTTTCCCTACACGACGCTCTTCCGATCTNNNAAGAACATTGCAACACATCTC

AGAGCACACGTCTGAACTCTTTCCCTACACGACGCTCTTCCGATCTNNNAAGAACATTGCAACACATCTC

AGCACACGTCTGAACTCTTTCCCTACACGACGCTCTTCCGATCTNNNCTGAGGCAGGAGAATCACTTGA

AGCACACGTCTGAACTCTTTCCCTACACGACGCTCTTCCGATCTNNNCTGAGGCAGGAGAATCACTTGA

AGCACACGTCTGAACTCTTTCCCTACACGACGCTCTTCCGATCTNNNGTTCTCACCCATATATTGATTTT

AGCACACGTCTGAACTCTTTCCCTACACGACGCTCTTCCGATCTNNNGTTCTCACCCATATATTGATTTT

GAGCACACGTCTGAACTCTTTCCCTACACGACGCTCTTCCGATCTNNNATTCTTATAGGGGGGTGTGTC

GAGCACACGTCTGAACTCTTTCCCTACACGACGCTCTTCCGATCTNNNATTCTTATAGGGGGGTGTGTC

GAGCACACGTCTGAACTCTTTCCCTACACGACGCTCTTCCGATCTNNNCTGTAATCCCAGCTACTCAGG

GAGCACACGTCTGAACTCTTTCCCTACACGACGCTCTTCCGATCTNNNCTGTAATCCCAGCTACTCAGG

AGAGCACACGTCTGAACTCTTTCCCTACACGACGCTCTTCCGATCTNNNACACATACACACACACACAG

AGAGCACACGTCTGAACTCTTTCCCTACACGACGCTCTTCCGATCTNNNACACATACACACACACACAG

AGCACACGTCTGAACTCTTTCCCTACACGACGCTCTTCCGATCTNNNCTGAGTAACTGGGAATACAGG

AGCACACGTCTGAACTCTTTCCCTACACGACGCTCTTCCGATCTNNNCTGAGTAACTGGGAATACAGG

AGCACACGTCTGAACTCTTTCCCTACACGACGCTCTTCCGATCTNNNCATGCCTGTAATCCCAGCTACTC

AGCACACGTCTGAACTCTTTCCCTACACGACGCTCTTCCGATCTNNNCATGCCTGTAATCCCAGCTACTC
AGCACACGTCTGAACTCTTTCCCTACACGACGCTCTTCCGATCTNNNGACGACAGGACCAAGAATTAG
AGCACACGTCTGAACTCTTTCCCTACACGACGCTCTTCCGATCTNNNGACGACAGGACCAAGAATTAG
GAGCACACGTCTGAACTCTTTCCCTACACGACGCTCTTCCGATCTNNNGAGATGCATGCATACACATCC
GAGCACACGTCTGAACTCTTTCCCTACACGACGCTCTTCCGATCTNNNGAGATGCATGCATACACATCC
AGCACACGTCTGAACTCTTTCCCTACACGACGCTCTTCCGATCTNNNTACAAATCAACTCCTGGGATCC
AGCACACGTCTGAACTCTTTCCCTACACGACGCTCTTCCGATCTNNNTACAAATCAACTCCTGGGATCC
AGCACACGTCTGAACTCTTTCCCTACACGACGCTCTTCCGATCTNNNCAATGAGAACACATGGACACA
AGCACACGTCTGAACTCTTTCCCTACACGACGCTCTTCCGATCTNNNCAATGAGAACACATGGACACA
GCACACGTCTGAACTCTTTCCCTACACGACGCTCTTCCGATCTNNNGTAATCCTAGCTACTCGGAAGGC
GCACACGTCTGAACTCTTTCCCTACACGACGCTCTTCCGATCTNNNGTAATCCTAGCTACTCGGAAGGC
AGCACACGTCTGAACTCTTTCCCTACACGACGCTCTTCCGATCTNNNAATCACTAGCGTCATCCTCACAG
AGCACACGTCTGAACTCTTTCCCTACACGACGCTCTTCCGATCTNNNAATCACTAGCGTCATCCTCACAG
AGCACACGTCTGAACTCTTTCCCTACACGACGCTCTTCCGATCTNNNTCTTTTGATCTTTGTTGGGCGA
AGCACACGTCTGAACTCTTTCCCTACACGACGCTCTTCCGATCTNNNTCTTTTGATCTTTGTTGGGCGA
AGCACACGTCTGAACTCTTTCCCTACACGACGCTCTTCCGATCTNNNGCAAGAGAATCACTTGAACCCA
AGCACACGTCTGAACTCTTTCCCTACACGACGCTCTTCCGATCTNNNGCAAGAGAATCACTTGAACCCA
AGCACACGTCTGAACTCTTTCCCTACACGACGCTCTTCCGATCTNNNAGCCACAGCACCTGACATATAT
AGCACACGTCTGAACTCTTTCCCTACACGACGCTCTTCCGATCTNNNAGCCACAGCACCTGACATATAT
AGCACACGTCTGAACTCTTTCCCTACACGACGCTCTTCCGATCTNNNACTGCTTTTGAAAAGTTGCACC
AGCACACGTCTGAACTCTTTCCCTACACGACGCTCTTCCGATCTNNNACTGCTTTTGAAAAGTTGCACC
AGCACACGTCTGAACTCTTTCCCTACACGACGCTCTTCCGATCTNNNCTGTGATCACCACTGCACTTCAG
AGCACACGTCTGAACTCTTTCCCTACACGACGCTCTTCCGATCTNNNCTGTGATCACCACTGCACTTCAG
GAGCACACGTCTGAACTCTTTCCCTACACGACGCTCTTCCGATCTNNNAAAAGAACCCACATAGCCAAA
GAGCACACGTCTGAACTCTTTCCCTACACGACGCTCTTCCGATCTNNNAAAAGAACCCACATAGCCAAA
GCACACGTCTGAACTCTTTCCCTACACGACGCTCTTCCGATCTNNNGTGTCCCCTTGGAATCTTCTTCTC
GCACACGTCTGAACTCTTTCCCTACACGACGCTCTTCCGATCTNNNGTGTCCCCTTGGAATCTTCTTCTC
GAGCACACGTCTGAACTCTTTCCCTACACGACGCTCTTCCGATCTNNNCTACAGACTTGTCCCTCTGATG
GAGCACACGTCTGAACTCTTTCCCTACACGACGCTCTTCCGATCTNNNCTACAGACTTGTCCCTCTGATG
AGCACACGTCTGAACTCTTTCCCTACACGACGCTCTTCCGATCTNNNGTTTCTTACTTACACTTCCGGCGC
AGCACACGTCTGAACTCTTTCCCTACACGACGCTCTTCCGATCTNNNGTTTCTTACTTACACTTCCGGCGC
AGCACACGTCTGAACTCTTTCCCTACACGACGCTCTTCCGATCTNNNCTCCTGGTCACATGAGATCTACC
AGCACACGTCTGAACTCTTTCCCTACACGACGCTCTTCCGATCTNNNCTCCTGGTCACATGAGATCTACC
AGCACACGTCTGAACTCTTTCCCTACACGACGCTCTTCCGATCTNNNGTGGCTCACTCCTATAATCCCAG
AGCACACGTCTGAACTCTTTCCCTACACGACGCTCTTCCGATCTNNNGTGGCTCACTCCTATAATCCCAG
CACACGTCTGAACTCTTTCCCTACACGACGCTCTTCCGATCTNNNGAATATGGCAAGCATTTTGGCATC
CACACGTCTGAACTCTTTCCCTACACGACGCTCTTCCGATCTNNNGAATATGGCAAGCATTTTGGCATC
AGAGCACACGTCTGAACTCTTTCCCTACACGACGCTCTTCCGATCTNNNGAAGTGAAACCAAGAAACAG
AGAGCACACGTCTGAACTCTTTCCCTACACGACGCTCTTCCGATCTNNNGAAGTGAAACCAAGAAACAG
AGCACACGTCTGAACTCTTTCCCTACACGACGCTCTTCCGATCTNNNACACCTCCCTAGTCTCAAAAACTC
AGCACACGTCTGAACTCTTTCCCTACACGACGCTCTTCCGATCTNNNACACCTCCCTAGTCTCAAAAACTC
GAGCACACGTCTGAACTCTTTCCCTACACGACGCTCTTCCGATCTNNNAATCACTTGAACTCAGGAGGT
GAGCACACGTCTGAACTCTTTCCCTACACGACGCTCTTCCGATCTNNNAATCACTTGAACTCAGGAGGT

GCACACGTCTGAACTCTTTCCCTACACGACGCTCTTCCGATCTNNNGAAGTTGTGAGCACTTGGTAAC
GCACACGTCTGAACTCTTTCCCTACACGACGCTCTTCCGATCTNNNGAAGTTGTGAGCACTTGGTAAC
AGAGCACACGTCTGAACTCTTTCCCTACACGACGCTCTTCCGATCTNNNGTGATGTGTGGGAGATGAT
AGAGCACACGTCTGAACTCTTTCCCTACACGACGCTCTTCCGATCTNNNGTGATGTGTGGGAGATGAT
GAGCACACGTCTGAACTCTTTCCCTACACGACGCTCTTCCGATCTNNNTCCTAATGCTATCCCTCCCCTA
GAGCACACGTCTGAACTCTTTCCCTACACGACGCTCTTCCGATCTNNNTCCTAATGCTATCCCTCCCCTA
AGAGCACACGTCTGAACTCTTTCCCTACACGACGCTCTTCCGATCTNNNCATGGAAAAATTGTGTTCCG
AGAGCACACGTCTGAACTCTTTCCCTACACGACGCTCTTCCGATCTNNNCATGGAAAAATTGTGTTCCG
AGCACACGTCTGAACTCTTTCCCTACACGACGCTCTTCCGATCTNNNGGGAAAGCGGGAAAAACAAA
AGCACACGTCTGAACTCTTTCCCTACACGACGCTCTTCCGATCTNNNGGGAAAGCGGGAAAAACAAA
GAGCACACGTCTGAACTCTTTCCCTACACGACGCTCTTCCGATCTNNNCTGAGATTGCCCTACTGTACT
GAGCACACGTCTGAACTCTTTCCCTACACGACGCTCTTCCGATCTNNNCTGAGATTGCCCTACTGTACT
GAGCACACGTCTGAACTCTTTCCCTACACGACGCTCTTCCGATCTNNNAGATTATAAATCAGAATAGCG
GAGCACACGTCTGAACTCTTTCCCTACACGACGCTCTTCCGATCTNNNAGATTATAAATCAGAATAGCG
AGCACACGTCTGAACTCTTTCCCTACACGACGCTCTTCCGATCTNNNTGTCCCCTTTTCTTATTCTCTGGA
AGCACACGTCTGAACTCTTTCCCTACACGACGCTCTTCCGATCTNNNTGTCCCCTTTTCTTATTCTCTGGA
GCACACGTCTGAACTCTTTCCCTACACGACGCTCTTCCGATCTNNNCCCACCATACAAAATTACAGGGG
GCACACGTCTGAACTCTTTCCCTACACGACGCTCTTCCGATCTNNNCCCACCATACAAAATTACAGGGG
AGCACACGTCTGAACTCTTTCCCTACACGACGCTCTTCCGATCTNNNTCACTTAGCTTTTCCCTAAAATA
AGCACACGTCTGAACTCTTTCCCTACACGACGCTCTTCCGATCTNNNTCACTTAGCTTTTCCCTAAAATA
GAGCACACGTCTGAACTCTTTCCCTACACGACGCTCTTCCGATCTNNNGAGGCTGAGGCAGGAGAATG
GAGCACACGTCTGAACTCTTTCCCTACACGACGCTCTTCCGATCTNNNGAGGCTGAGGCAGGAGAATG
GAGCACACGTCTGAACTCTTTCCCTACACGACGCTCTTCCGATCTNNNGACCGGTCTGTCCATTTCTACT
GAGCACACGTCTGAACTCTTTCCCTACACGACGCTCTTCCGATCTNNNGACCGGTCTGTCCATTTCTACT
GAGCACACGTCTGAACTCTTTCCCTACACGACGCTCTTCCGATCTNNNTCTAACCTCAGTACCATTGCCA
GAGCACACGTCTGAACTCTTTCCCTACACGACGCTCTTCCGATCTNNNTCTAACCTCAGTACCATTGCCA
AGAGCACACGTCTGAACTCTTTCCCTACACGACGCTCTTCCGATCTNNNGCACTACAGACTGGACAATG
AGAGCACACGTCTGAACTCTTTCCCTACACGACGCTCTTCCGATCTNNNGCACTACAGACTGGACAATG
GAGCACACGTCTGAACTCTTTCCCTACACGACGCTCTTCCGATCTNNNTGATGTCCTCGTAAGCAAACT
GAGCACACGTCTGAACTCTTTCCCTACACGACGCTCTTCCGATCTNNNTGATGTCCTCGTAAGCAAACT
GAGCACACGTCTGAACTCTTTCCCTACACGACGCTCTTCCGATCTNNNCTGTGCCTTGTGAAACTTACGT
GAGCACACGTCTGAACTCTTTCCCTACACGACGCTCTTCCGATCTNNNCTGTGCCTTGTGAAACTTACGT
GAGCACACGTCTGAACTCTTTCCCTACACGACGCTCTTCCGATCTNNNAATTTTCCCCCATTTCCATGCT
GAGCACACGTCTGAACTCTTTCCCTACACGACGCTCTTCCGATCTNNNAATTTTCCCCCATTTCCATGCT
GAGCACACGTCTGAACTCTTTCCCTACACGACGCTCTTCCGATCTNNNTATTTTCTTTTTCAGTTCTTTC
GAGCACACGTCTGAACTCTTTCCCTACACGACGCTCTTCCGATCTNNNTATTTTCTTTTTCAGTTCTTTC
AGCACACGTCTGAACTCTTTCCCTACACGACGCTCTTCCGATCTNNNCCAAACCAAGATGGAAAGGAC
AGCACACGTCTGAACTCTTTCCCTACACGACGCTCTTCCGATCTNNNCCAAACCAAGATGGAAAGGAC
GAAGAGCACACGTCTGAACTCTTTCCCTACACGACGCTCTTCCGATCTNNNGTGATTTGAGGTGTCAC
GAAGAGCACACGTCTGAACTCTTTCCCTACACGACGCTCTTCCGATCTNNNGTGATTTGAGGTGTCAC
AGAGCACACGTCTGAACTCTTTCCCTACACGACGCTCTTCCGATCTNNNGATGTCAGGAGATCGAGAC
AGAGCACACGTCTGAACTCTTTCCCTACACGACGCTCTTCCGATCTNNNGATGTCAGGAGATCGAGAC
GAGCACACGTCTGAACTCTTTCCCTACACGACGCTCTTCCGATCTNNNTGAACACCCACTCCATGAAGA

GAGCACACGTCTGAACTCTTTCCCTACACGACGCTCTTCCGATCTNNNTGAACACCCACTCCATGAAGA
AGCACACGTCTGAACTCTTTCCCTACACGACGCTCTTCCGATCTNNNCTTTCTCCCAGACAGATGGAGT
AGCACACGTCTGAACTCTTTCCCTACACGACGCTCTTCCGATCTNNNCTTTCTCCCAGACAGATGGAGT
GAGCACACGTCTGAACTCTTTCCCTACACGACGCTCTTCCGATCTNNNGAGATGGAGTCTCGCTCTGT
GAGCACACGTCTGAACTCTTTCCCTACACGACGCTCTTCCGATCTNNNGAGATGGAGTCTCGCTCTGT
GAGCACACGTCTGAACTCTTTCCCTACACGACGCTCTTCCGATCTNNNAAAAGGATCCTGCATCTTGAG
GAGCACACGTCTGAACTCTTTCCCTACACGACGCTCTTCCGATCTNNNAAAAGGATCCTGCATCTTGAG
GAGCACACGTCTGAACTCTTTCCCTACACGACGCTCTTCCGATCTNNNTGACTGAATCTCTGACCTCTCC
GAGCACACGTCTGAACTCTTTCCCTACACGACGCTCTTCCGATCTNNNTGACTGAATCTCTGACCTCTCC
GCACACGTCTGAACTCTTTCCCTACACGACGCTCTTCCGATCTNNNGGAGATCGAGACCATGGTGAAA
GCACACGTCTGAACTCTTTCCCTACACGACGCTCTTCCGATCTNNNGGAGATCGAGACCATGGTGAAA
GAGCACACGTCTGAACTCTTTCCCTACACGACGCTCTTCCGATCTNNNTGGTTAGCTGCTTCCTCTCTT
GAGCACACGTCTGAACTCTTTCCCTACACGACGCTCTTCCGATCTNNNTGGTTAGCTGCTTCCTCTCTT
AGAGCACACGTCTGAACTCTTTCCCTACACGACGCTCTTCCGATCTNNNCAGGAAAAGTTTGACAACCC
AGAGCACACGTCTGAACTCTTTCCCTACACGACGCTCTTCCGATCTNNNCAGGAAAAGTTTGACAACCC
AGCACACGTCTGAACTCTTTCCCTACACGACGCTCTTCCGATCTNNNCATTGAAAATGAAGGAAAACA
AGCACACGTCTGAACTCTTTCCCTACACGACGCTCTTCCGATCTNNNCATTGAAAATGAAGGAAAACA
AGCACACGTCTGAACTCTTTCCCTACACGACGCTCTTCCGATCTNNNTACTCTGAAGGCTGAGGTAGG
AGCACACGTCTGAACTCTTTCCCTACACGACGCTCTTCCGATCTNNNTACTCTGAAGGCTGAGGTAGG
AGCACACGTCTGAACTCTTTCCCTACACGACGCTCTTCCGATCTNNNGAGTCTGAGGAACTGTAGCCT
AGCACACGTCTGAACTCTTTCCCTACACGACGCTCTTCCGATCTNNNGAGTCTGAGGAACTGTAGCCT
AGAGCACACGTCTGAACTCTTTCCCTACACGACGCTCTTCCGATCTNNNTGTAGTCCCAGCTACTCAGG
AGAGCACACGTCTGAACTCTTTCCCTACACGACGCTCTTCCGATCTNNNTGTAGTCCCAGCTACTCAGG
AGAGCACACGTCTGAACTCTTTCCCTACACGACGCTCTTCCGATCTNNNCTGAGGCAGGAGAATTGCT
AGAGCACACGTCTGAACTCTTTCCCTACACGACGCTCTTCCGATCTNNNCTGAGGCAGGAGAATTGCT
GCACACGTCTGAACTCTTTCCCTACACGACGCTCTTCCGATCTNNNTTTCAAATAAAACTTGTACTGGG
GCACACGTCTGAACTCTTTCCCTACACGACGCTCTTCCGATCTNNNTTTCAAATAAAACTTGTACTGGG
AGCACACGTCTGAACTCTTTCCCTACACGACGCTCTTCCGATCTNNNATGCCTCTAGTACCAGCTACTC
AGCACACGTCTGAACTCTTTCCCTACACGACGCTCTTCCGATCTNNNATGCCTCTAGTACCAGCTACTC
GAGCACACGTCTGAACTCTTTCCCTACACGACGCTCTTCCGATCTNNNGAGATGAGATACGCGAGTCT
GAGCACACGTCTGAACTCTTTCCCTACACGACGCTCTTCCGATCTNNNGAGATGAGATACGCGAGTCT
GCACACGTCTGAACTCTTTCCCTACACGACGCTCTTCCGATCTNNNCTTTCAGCCTGTGTGAGTGGGCA
GCACACGTCTGAACTCTTTCCCTACACGACGCTCTTCCGATCTNNNCTTTCAGCCTGTGTGAGTGGGCA
GCACACGTCTGAACTCTTTCCCTACACGACGCTCTTCCGATCTNNNCACCTGTAGTCCCAGCTACTTGC
GCACACGTCTGAACTCTTTCCCTACACGACGCTCTTCCGATCTNNNCACCTGTAGTCCCAGCTACTTGC
AGCACACGTCTGAACTCTTTCCCTACACGACGCTCTTCCGATCTNNNGGACTTTAGACACACAGAACA
AGCACACGTCTGAACTCTTTCCCTACACGACGCTCTTCCGATCTNNNGGACTTTAGACACACAGAACA
GCACACGTCTGAACTCTTTCCCTACACGACGCTCTTCCGATCTNNNATCACCCAACCTGTGCTTCTCTAC
GCACACGTCTGAACTCTTTCCCTACACGACGCTCTTCCGATCTNNNATCACCCAACCTGTGCTTCTCTAC
GCACACGTCTGAACTCTTTCCCTACACGACGCTCTTCCGATCTNNNCCTGACTGGTAGAAAACTCCAG
GCACACGTCTGAACTCTTTCCCTACACGACGCTCTTCCGATCTNNNCCTGACTGGTAGAAAACTCCAG
CACGTCTGAACTCTTTCCCTACACGACGCTCTTCCGATCTNNNGCAGGACTAGTTGATTATCCCCACAT
CACGTCTGAACTCTTTCCCTACACGACGCTCTTCCGATCTNNNGCAGGACTAGTTGATTATCCCCACAT

AGCACACGTCTGAACTCTTTCCCTACACGACGCTCTTCCGATCTNNNGCCTCCTAAGTAGCTGGGATT
AGCACACGTCTGAACTCTTTCCCTACACGACGCTCTTCCGATCTNNNGCCTCCTAAGTAGCTGGGATT
GCACACGTCTGAACTCTTTCCCTACACGACGCTCTTCCGATCTNNNGGAATTCCTTTGTGCAGGCTTTC
GCACACGTCTGAACTCTTTCCCTACACGACGCTCTTCCGATCTNNNGGAATTCCTTTGTGCAGGCTTTC
GCACACGTCTGAACTCTTTCCCTACACGACGCTCTTCCGATCTNNNCACAACAGCTCTGCTTTTAGCTT
GCACACGTCTGAACTCTTTCCCTACACGACGCTCTTCCGATCTNNNCACAACAGCTCTGCTTTTAGCTT
GCACACGTCTGAACTCTTTCCCTACACGACGCTCTTCCGATCTNNNAAGACAGGGGTTAGCAAACTAG
GCACACGTCTGAACTCTTTCCCTACACGACGCTCTTCCGATCTNNNAAGACAGGGGTTAGCAAACTAG
AGCACACGTCTGAACTCTTTCCCTACACGACGCTCTTCCGATCTNNNCTCTTGGCTTGAAGGGAATG
AGCACACGTCTGAACTCTTTCCCTACACGACGCTCTTCCGATCTNNNCTCTTGGCTTGAAGGGAATG
GCACACGTCTGAACTCTTTCCCTACACGACGCTCTTCCGATCTNNNAGAAGAATCGTTTGAACCTGG
GCACACGTCTGAACTCTTTCCCTACACGACGCTCTTCCGATCTNNNAGAAGAATCGTTTGAACCTGG
GCACACGTCTGAACTCTTTCCCTACACGACGCTCTTCCGATCTNNNCTGTTACACTAGCCCCCATTTCT
GCACACGTCTGAACTCTTTCCCTACACGACGCTCTTCCGATCTNNNCTGTTACACTAGCCCCCATTTCT
GCACACGTCTGAACTCTTTCCCTACACGACGCTCTTCCGATCTNNNATGTGGATACTGACAGGACGA
GCACACGTCTGAACTCTTTCCCTACACGACGCTCTTCCGATCTNNNATGTGGATACTGACAGGACGA
AGCACACGTCTGAACTCTTTCCCTACACGACGCTCTTCCGATCTNNNTGGATCCTGCAGTTGTTACCAT
AGCACACGTCTGAACTCTTTCCCTACACGACGCTCTTCCGATCTNNNTGGATCCTGCAGTTGTTACCAT
AGCACACGTCTGAACTCTTTCCCTACACGACGCTCTTCCGATCTNNNTGGCCTCTGTGGGATAGAAA
AGCACACGTCTGAACTCTTTCCCTACACGACGCTCTTCCGATCTNNNTGGCCTCTGTGGGATAGAAA
GAGCACACGTCTGAACTCTTTCCCTACACGACGCTCTTCCGATCTNNNAGAATTACTTGAACCCAGGA
GAGCACACGTCTGAACTCTTTCCCTACACGACGCTCTTCCGATCTNNNAGAATTACTTGAACCCAGGA
GCACACGTCTGAACTCTTTCCCTACACGACGCTCTTCCGATCTNNNAACACCTGGAAACTTGGAAATG
GCACACGTCTGAACTCTTTCCCTACACGACGCTCTTCCGATCTNNNAACACCTGGAAACTTGGAAATG
GAGCACACGTCTGAACTCTTTCCCTACACGACGCTCTTCCGATCTNNNGAAGTCTGTCTCCAACTCTGT
GAGCACACGTCTGAACTCTTTCCCTACACGACGCTCTTCCGATCTNNNGAAGTCTGTCTCCAACTCTGT
GCACACGTCTGAACTCTTTCCCTACACGACGCTCTTCCGATCTNNNTCTCTCTCTCTCTCTCTCCCTCA
GCACACGTCTGAACTCTTTCCCTACACGACGCTCTTCCGATCTNNNTCTCTCTCTCTCTCTCTCCCTCA
GCACACGTCTGAACTCTTTCCCTACACGACGCTCTTCCGATCTNNNCCTGGCCAAAAGCAGATAGTTT
GCACACGTCTGAACTCTTTCCCTACACGACGCTCTTCCGATCTNNNCCTGGCCAAAAGCAGATAGTTT
GCACACGTCTGAACTCTTTCCCTACACGACGCTCTTCCGATCTNNNATAAACCATGACCTCAGCCGACC
GCACACGTCTGAACTCTTTCCCTACACGACGCTCTTCCGATCTNNNATAAACCATGACCTCAGCCGACC
AGCACACGTCTGAACTCTTTCCCTACACGACGCTCTTCCGATCTNNNCTAGGTGACACAGCAAGACTC
AGCACACGTCTGAACTCTTTCCCTACACGACGCTCTTCCGATCTNNNCTAGGTGACACAGCAAGACTC
AGCACACGTCTGAACTCTTTCCCTACACGACGCTCTTCCGATCTNNNAGTAAGGCAGCAAACACAAA
AGCACACGTCTGAACTCTTTCCCTACACGACGCTCTTCCGATCTNNNAGTAAGGCAGCAAACACAAA
AGAGCACACGTCTGAACTCTTTCCCTACACGACGCTCTTCCGATCTNNNCTGAGGCAGAGAATTGCTT
AGAGCACACGTCTGAACTCTTTCCCTACACGACGCTCTTCCGATCTNNNCTGAGGCAGAGAATTGCTT
AGCACACGTCTGAACTCTTTCCCTACACGACGCTCTTCCGATCTNNNGCAATAGTTTTGTTCAGCCCAG
AGCACACGTCTGAACTCTTTCCCTACACGACGCTCTTCCGATCTNNNGCAATAGTTTTGTTCAGCCCAG
AGCACACGTCTGAACTCTTTCCCTACACGACGCTCTTCCGATCTNNNCAGATTCATGAGATCGTACAT
AGCACACGTCTGAACTCTTTCCCTACACGACGCTCTTCCGATCTNNNCAGATTCATGAGATCGTACAT
AGCACACGTCTGAACTCTTTCCCTACACGACGCTCTTCCGATCTNNNAAAAAAAAGTAGCCAGGCGAT

AGCACACGTCTGAACTCTTTCCCTACACGACGCTCTTCCGATCTNNNAAAAAAAAGTAGCCAGGCGAT
GCACACGTCTGAACTCTTTCCCTACACGACGCTCTTCCGATCTNNNCTGGAAGGCCTTTTGTCACTAGG
GCACACGTCTGAACTCTTTCCCTACACGACGCTCTTCCGATCTNNNCTGGAAGGCCTTTTGTCACTAGG
AGAGCACACGTCTGAACTCTTTCCCTACACGACGCTCTTCCGATCTNNNAAAATTAGCCAGACGTGGTG
AGAGCACACGTCTGAACTCTTTCCCTACACGACGCTCTTCCGATCTNNNAAAATTAGCCAGACGTGGTG
AGCACACGTCTGAACTCTTTCCCTACACGACGCTCTTCCGATCTNNNAGACAAGGGGGAAATGTATGC
AGCACACGTCTGAACTCTTTCCCTACACGACGCTCTTCCGATCTNNNAGACAAGGGGGAAATGTATGC
AGCACACGTCTGAACTCTTTCCCTACACGACGCTCTTCCGATCTNNNTTGTTTCTTTAACTAGCATGTG
AGCACACGTCTGAACTCTTTCCCTACACGACGCTCTTCCGATCTNNNTTGTTTCTTTAACTAGCATGTG
AGCACACGTCTGAACTCTTTCCCTACACGACGCTCTTCCGATCTNNNGCTGGACCAAACCGATGTATTT
AGCACACGTCTGAACTCTTTCCCTACACGACGCTCTTCCGATCTNNNGCTGGACCAAACCGATGTATTT
GAGCACACGTCTGAACTCTTTCCCTACACGACGCTCTTCCGATCTNNNCCATCCTGGGAAACAAGAGTG
GAGCACACGTCTGAACTCTTTCCCTACACGACGCTCTTCCGATCTNNNCCATCCTGGGAAACAAGAGTG
GCACACGTCTGAACTCTTTCCCTACACGACGCTCTTCCGATCTNNNCTCTCTAGCTATCCCCCTCTCCA
GCACACGTCTGAACTCTTTCCCTACACGACGCTCTTCCGATCTNNNCTCTCTAGCTATCCCCCTCTCCA
GAGCACACGTCTGAACTCTTTCCCTACACGACGCTCTTCCGATCTNNNCCAACACATCCTCCATTTCCTCA
GAGCACACGTCTGAACTCTTTCCCTACACGACGCTCTTCCGATCTNNNCCAACACATCCTCCATTTCCTCA
AGCACACGTCTGAACTCTTTCCCTACACGACGCTCTTCCGATCTNNNGAGAATACCTTGAACCTGGGAG
AGCACACGTCTGAACTCTTTCCCTACACGACGCTCTTCCGATCTNNNGAGAATACCTTGAACCTGGGAG
GCACACGTCTGAACTCTTTCCCTACACGACGCTCTTCCGATCTNNNTGGCCTTCTCCTCTATGTCTCTTCA
GCACACGTCTGAACTCTTTCCCTACACGACGCTCTTCCGATCTNNNTGGCCTTCTCCTCTATGTCTCTTCA
GAGCACACGTCTGAACTCTTTCCCTACACGACGCTCTTCCGATCTNNNAGAGAATTGCTTGAACCCAGC
GAGCACACGTCTGAACTCTTTCCCTACACGACGCTCTTCCGATCTNNNAGAGAATTGCTTGAACCCAGC
GAGCACACGTCTGAACTCTTTCCCTACACGACGCTCTTCCGATCTNNNTGGTGGGAAGGGAAAGAAA
GAGCACACGTCTGAACTCTTTCCCTACACGACGCTCTTCCGATCTNNNTGGTGGGAAGGGAAAGAAA
GAGCACACGTCTGAACTCTTTCCCTACACGACGCTCTTCCGATCTNNNCTGAGGCAGGAGAATCACTTG
GAGCACACGTCTGAACTCTTTCCCTACACGACGCTCTTCCGATCTNNNCTGAGGCAGGAGAATCACTTG
AGCACACGTCTGAACTCTTTCCCTACACGACGCTCTTCCGATCTNNNCTGGGAGAAAGAGCGAGACTC
AGCACACGTCTGAACTCTTTCCCTACACGACGCTCTTCCGATCTNNNCTGGGAGAAAGAGCGAGACTC
AGCACACGTCTGAACTCTTTCCCTACACGACGCTCTTCCGATCTNNNCCAAATCCAGTGGGCTAAATGT
AGCACACGTCTGAACTCTTTCCCTACACGACGCTCTTCCGATCTNNNCCAAATCCAGTGGGCTAAATGT
AGCACACGTCTGAACTCTTTCCCTACACGACGCTCTTCCGATCTNNNTATCCATGGAAGTCAGACCCTCT
AGCACACGTCTGAACTCTTTCCCTACACGACGCTCTTCCGATCTNNNTATCCATGGAAGTCAGACCCTCT
AGCACACGTCTGAACTCTTTCCCTACACGACGCTCTTCCGATCTNNNTGTGACCAGGTTTCATCCAATCA
AGCACACGTCTGAACTCTTTCCCTACACGACGCTCTTCCGATCTNNNTGTGACCAGGTTTCATCCAATCA
AGAGCACACGTCTGAACTCTTTCCCTACACGACGCTCTTCCGATCTNNNGAGGCTGAGGCAGGAGAAT
AGAGCACACGTCTGAACTCTTTCCCTACACGACGCTCTTCCGATCTNNNGAGGCTGAGGCAGGAGAAT
GCACACGTCTGAACTCTTTCCCTACACGACGCTCTTCCGATCTNNNGTGCCTGTAGTCTCAGCTACTCCA
GCACACGTCTGAACTCTTTCCCTACACGACGCTCTTCCGATCTNNNGTGCCTGTAGTCTCAGCTACTCCA
GAGCACACGTCTGAACTCTTTCCCTACACGACGCTCTTCCGATCTNNNCCCTGTGGTCAAGCTACAAAAA
GAGCACACGTCTGAACTCTTTCCCTACACGACGCTCTTCCGATCTNNNCCCTGTGGTCAAGCTACAAAAA
GCACACGTCTGAACTCTTTCCCTACACGACGCTCTTCCGATCTNNNCCTTCAAACCTCCACTCCAAGTTCA
GCACACGTCTGAACTCTTTCCCTACACGACGCTCTTCCGATCTNNNCCTTCAAACCTCCACTCCAAGTTCA

GCACACGTCTGAACTCTTTCCCTACACGACGCTCTTCCGATCTNNNTTTTTCTGTGCATGTGTGTGTGT
GCACACGTCTGAACTCTTTCCCTACACGACGCTCTTCCGATCTNNNTTTTTCTGTGCATGTGTGTGTGT
GCACACGTCTGAACTCTTTCCCTACACGACGCTCTTCCGATCTNNNTGTGGATGCTTGCTGTTTTTGTT
GCACACGTCTGAACTCTTTCCCTACACGACGCTCTTCCGATCTNNNTGTGGATGCTTGCTGTTTTTGTT
GCACACGTCTGAACTCTTTCCCTACACGACGCTCTTCCGATCTNNNACATAATGAGGCTTTCTGGGG
GCACACGTCTGAACTCTTTCCCTACACGACGCTCTTCCGATCTNNNACATAATGAGGCTTTCTGGGG
AAGAGCACACGTCTGAACTCTTTCCCTACACGACGCTCTTCCGATCTNNNACTGCCTTTTTAAAATTC
AAGAGCACACGTCTGAACTCTTTCCCTACACGACGCTCTTCCGATCTNNNACTGCCTTTTTAAAATTC
AGCACACGTCTGAACTCTTTCCCTACACGACGCTCTTCCGATCTNNNAACATGGCAAAACCCCATCTCT
AGCACACGTCTGAACTCTTTCCCTACACGACGCTCTTCCGATCTNNNAACATGGCAAAACCCCATCTCT
AGCACACGTCTGAACTCTTTCCCTACACGACGCTCTTCCGATCTNNNTTACCGTTCCACACAGTCAGA
AGCACACGTCTGAACTCTTTCCCTACACGACGCTCTTCCGATCTNNNTTACCGTTCCACACAGTCAGA
AGCACACGTCTGAACTCTTTCCCTACACGACGCTCTTCCGATCTNNNTTCAAGTGATTCTCCTGCCTCA
AGCACACGTCTGAACTCTTTCCCTACACGACGCTCTTCCGATCTNNNTTCAAGTGATTCTCCTGCCTCA
GCACACGTCTGAACTCTTTCCCTACACGACGCTCTTCCGATCTNNNGAACTCCCAACCTCAGGTGATC
GCACACGTCTGAACTCTTTCCCTACACGACGCTCTTCCGATCTNNNGAACTCCCAACCTCAGGTGATC
GCACACGTCTGAACTCTTTCCCTACACGACGCTCTTCCGATCTNNNCTTGCAGTGAGTCGAGATTGTG
GCACACGTCTGAACTCTTTCCCTACACGACGCTCTTCCGATCTNNNCTTGCAGTGAGTCGAGATTGTG
AGCACACGTCTGAACTCTTTCCCTACACGACGCTCTTCCGATCTNNNCTGTGATTGGCTGGAGTTTTG
AGCACACGTCTGAACTCTTTCCCTACACGACGCTCTTCCGATCTNNNCTGTGATTGGCTGGAGTTTTG
AGCACACGTCTGAACTCTTTCCCTACACGACGCTCTTCCGATCTNNNGGCAATGATGAAAACTACTCG
AGCACACGTCTGAACTCTTTCCCTACACGACGCTCTTCCGATCTNNNGGCAATGATGAAAACTACTCG
GCACACGTCTGAACTCTTTCCCTACACGACGCTCTTCCGATCTNNNTGGTATTCCTGTACAGCAACAC
GCACACGTCTGAACTCTTTCCCTACACGACGCTCTTCCGATCTNNNTGGTATTCCTGTACAGCAACAC
AGCACACGTCTGAACTCTTTCCCTACACGACGCTCTTCCGATCTNNNAAAATCGCAAGGAGTTTGGA
AGCACACGTCTGAACTCTTTCCCTACACGACGCTCTTCCGATCTNNNAAAATCGCAAGGAGTTTGGA
GCACACGTCTGAACTCTTTCCCTACACGACGCTCTTCCGATCTNNNAGGTGAATAAAACAGCCCTCAC
GCACACGTCTGAACTCTTTCCCTACACGACGCTCTTCCGATCTNNNAGGTGAATAAAACAGCCCTCAC
GCACACGTCTGAACTCTTTCCCTACACGACGCTCTTCCGATCTNNNCCAAAGTGCTGGAATTATAGG
GCACACGTCTGAACTCTTTCCCTACACGACGCTCTTCCGATCTNNNCCAAAGTGCTGGAATTATAGG
GCACACGTCTGAACTCTTTCCCTACACGACGCTCTTCCGATCTNNNGGAAGGAGACAGGAAAGGTG
GCACACGTCTGAACTCTTTCCCTACACGACGCTCTTCCGATCTNNNGGAAGGAGACAGGAAAGGTG
AGCACACGTCTGAACTCTTTCCCTACACGACGCTCTTCCGATCTNNNTCCCAGACTGATGAGAAAGCC
AGCACACGTCTGAACTCTTTCCCTACACGACGCTCTTCCGATCTNNNTCCCAGACTGATGAGAAAGCC
GAGCACACGTCTGAACTCTTTCCCTACACGACGCTCTTCCGATCTNNNCTATGCAGGGTCCCTTGCTAT
GAGCACACGTCTGAACTCTTTCCCTACACGACGCTCTTCCGATCTNNNCTATGCAGGGTCCCTTGCTAT
AGCACACGTCTGAACTCTTTCCCTACACGACGCTCTTCCGATCTNNNCATGCCTGTAGTACCAGCTACT
AGCACACGTCTGAACTCTTTCCCTACACGACGCTCTTCCGATCTNNNCATGCCTGTAGTACCAGCTACT
GCACACGTCTGAACTCTTTCCCTACACGACGCTCTTCCGATCTNNNGGCTATTTGGGGCAATGAAAA
GCACACGTCTGAACTCTTTCCCTACACGACGCTCTTCCGATCTNNNGGCTATTTGGGGCAATGAAAA
AGCACACGTCTGAACTCTTTCCCTACACGACGCTCTTCCGATCTNNNTAGTTTTCCGAACACACACAC
AGCACACGTCTGAACTCTTTCCCTACACGACGCTCTTCCGATCTNNNTAGTTTTCCGAACACACACAC
AGCACACGTCTGAACTCTTTCCCTACACGACGCTCTTCCGATCTNNNTTCTCACTCCTGGACTACAGC

AGCACACGTCTGAACTCTTTCCCTACACGACGCTCTTCCGATCTNNNTTCTCACTCCTGGACTACAGC
GCACACGTCTGAACTCTTTCCCTACACGACGCTCTTCCGATCTNNNGCATTGGGCTTCAGATTAGCA
GCACACGTCTGAACTCTTTCCCTACACGACGCTCTTCCGATCTNNNGCATTGGGCTTCAGATTAGCA
GAGCACACGTCTGAACTCTTTCCCTACACGACGCTCTTCCGATCTNNNCAGCCTCCAGAACAGCAAAA
GAGCACACGTCTGAACTCTTTCCCTACACGACGCTCTTCCGATCTNNNCAGCCTCCAGAACAGCAAAA
GCACACGTCTGAACTCTTTCCCTACACGACGCTCTTCCGATCTNNNGGGAATGTAGCAGTGAAGTA
GCACACGTCTGAACTCTTTCCCTACACGACGCTCTTCCGATCTNNNGGGAATGTAGCAGTGAAGTA
GAGCACACGTCTGAACTCTTTCCCTACACGACGCTCTTCCGATCTNNNAGCTTGTAGTGAGCCTATAT
GAGCACACGTCTGAACTCTTTCCCTACACGACGCTCTTCCGATCTNNNAGCTTGTAGTGAGCCTATAT
GCACACGTCTGAACTCTTTCCCTACACGACGCTCTTCCGATCTNNNTCCTCTCCTTCTGGGATTCTCAT
GCACACGTCTGAACTCTTTCCCTACACGACGCTCTTCCGATCTNNNTCCTCTCCTTCTGGGATTCTCAT
AGCACACGTCTGAACTCTTTCCCTACACGACGCTCTTCCGATCTNNNCATATCCTCAGAGCCTTCGGA
AGCACACGTCTGAACTCTTTCCCTACACGACGCTCTTCCGATCTNNNCATATCCTCAGAGCCTTCGGA
GAGCACACGTCTGAACTCTTTCCCTACACGACGCTCTTCCGATCTNNNCAGAGCCGAGATCACGTTCA
GAGCACACGTCTGAACTCTTTCCCTACACGACGCTCTTCCGATCTNNNCAGAGCCGAGATCACGTTCA
GAGCACACGTCTGAACTCTTTCCCTACACGACGCTCTTCCGATCTNNNGTGTTCAGATTGAGCCATTG
GAGCACACGTCTGAACTCTTTCCCTACACGACGCTCTTCCGATCTNNNGTGTTCAGATTGAGCCATTG
GCACACGTCTGAACTCTTTCCCTACACGACGCTCTTCCGATCTNNNGCCTTTGTATTTCTGTGATGTCA
GCACACGTCTGAACTCTTTCCCTACACGACGCTCTTCCGATCTNNNGCCTTTGTATTTCTGTGATGTCA
GCACACGTCTGAACTCTTTCCCTACACGACGCTCTTCCGATCTNNNTGCAATGTTCAAAAAACCTTAG
GCACACGTCTGAACTCTTTCCCTACACGACGCTCTTCCGATCTNNNTGCAATGTTCAAAAAACCTTAG
AGCACACGTCTGAACTCTTTCCCTACACGACGCTCTTCCGATCTNNNTGAGGCAGGAGAATCTCTTGA
AGCACACGTCTGAACTCTTTCCCTACACGACGCTCTTCCGATCTNNNTGAGGCAGGAGAATCTCTTGA
GCACACGTCTGAACTCTTTCCCTACACGACGCTCTTCCGATCTNNNCAGCAAGACCCCATCTCTATACA
GCACACGTCTGAACTCTTTCCCTACACGACGCTCTTCCGATCTNNNCAGCAAGACCCCATCTCTATACA
AGCACACGTCTGAACTCTTTCCCTACACGACGCTCTTCCGATCTNNNACACCCAGCCTGTTTTTTTGTT
AGCACACGTCTGAACTCTTTCCCTACACGACGCTCTTCCGATCTNNNACACCCAGCCTGTTTTTTTGTT
AGAGCACACGTCTGAACTCTTTCCCTACACGACGCTCTTCCGATCTNNNGATCAGTGATGTACACGCA
AGAGCACACGTCTGAACTCTTTCCCTACACGACGCTCTTCCGATCTNNNGATCAGTGATGTACACGCA
AGCACACGTCTGAACTCTTTCCCTACACGACGCTCTTCCGATCTNNNTAGTTACTGGTAGCCGTGTCT
AGCACACGTCTGAACTCTTTCCCTACACGACGCTCTTCCGATCTNNNTAGTTACTGGTAGCCGTGTCT
AGAGCACACGTCTGAACTCTTTCCCTACACGACGCTCTTCCGATCTNNNAGATCTGGCAGGGTTAGT
AGAGCACACGTCTGAACTCTTTCCCTACACGACGCTCTTCCGATCTNNNAGATCTGGCAGGGTTAGT
GCACACGTCTGAACTCTTTCCCTACACGACGCTCTTCCGATCTNNNAAACAAATGTCAAAGACAGCCC
GCACACGTCTGAACTCTTTCCCTACACGACGCTCTTCCGATCTNNNAAACAAATGTCAAAGACAGCCC
GCACACGTCTGAACTCTTTCCCTACACGACGCTCTTCCGATCTNNNACATGTGGTGTCTGAAAGGAC
GCACACGTCTGAACTCTTTCCCTACACGACGCTCTTCCGATCTNNNACATGTGGTGTCTGAAAGGAC
GCACACGTCTGAACTCTTTCCCTACACGACGCTCTTCCGATCTNNNATGGAGTTTCACTCTTTCACCCA
GCACACGTCTGAACTCTTTCCCTACACGACGCTCTTCCGATCTNNNATGGAGTTTCACTCTTTCACCCA
GAGCACACGTCTGAACTCTTTCCCTACACGACGCTCTTCCGATCTNNNCAGTGAGATATACGCTGGTG
GAGCACACGTCTGAACTCTTTCCCTACACGACGCTCTTCCGATCTNNNCAGTGAGATATACGCTGGTG
AGCACACGTCTGAACTCTTTCCCTACACGACGCTCTTCCGATCTNNNTATCTGAGGTTTTGACCTTGG
AGCACACGTCTGAACTCTTTCCCTACACGACGCTCTTCCGATCTNNNTATCTGAGGTTTTGACCTTGG

AGCACACGTCTGAACTCTTTCCCTACACGACGCTCTTCCGATCTNNNGCATCCTTGTAATCCCAGCTAC
AGCACACGTCTGAACTCTTTCCCTACACGACGCTCTTCCGATCTNNNGCATCCTTGTAATCCCAGCTAC
GAGCACACGTCTGAACTCTTTCCCTACACGACGCTCTTCCGATCTNNNCCTCCTGGAGACACACTTTG
GAGCACACGTCTGAACTCTTTCCCTACACGACGCTCTTCCGATCTNNNCCTCCTGGAGACACACTTTG
AGCACACGTCTGAACTCTTTCCCTACACGACGCTCTTCCGATCTNNNGAAGGAGGTGGAAGAGAAG
AGCACACGTCTGAACTCTTTCCCTACACGACGCTCTTCCGATCTNNNGAAGGAGGTGGAAGAGAAG
AGCACACGTCTGAACTCTTTCCCTACACGACGCTCTTCCGATCTNNNCTCTGTGTCACTTCCTCACTGT
AGCACACGTCTGAACTCTTTCCCTACACGACGCTCTTCCGATCTNNNCTCTGTGTCACTTCCTCACTGT
AGAGCACACGTCTGAACTCTTTCCCTACACGACGCTCTTCCGATCTNNNCTAGGCTGCAGTGAGCTGA
AGAGCACACGTCTGAACTCTTTCCCTACACGACGCTCTTCCGATCTNNNCTAGGCTGCAGTGAGCTGA
AGCACACGTCTGAACTCTTTCCCTACACGACGCTCTTCCGATCTNNNTGGACAGTACCACAGATCGAT
AGCACACGTCTGAACTCTTTCCCTACACGACGCTCTTCCGATCTNNNTGGACAGTACCACAGATCGAT
AGCACACGTCTGAACTCTTTCCCTACACGACGCTCTTCCGATCTNNNACACACAAGTAGTAAGGGCCA
AGCACACGTCTGAACTCTTTCCCTACACGACGCTCTTCCGATCTNNNACACACAAGTAGTAAGGGCCA
AGCACACGTCTGAACTCTTTCCCTACACGACGCTCTTCCGATCTNNNAGTGAGGAGTGAGACTCTAC
AGCACACGTCTGAACTCTTTCCCTACACGACGCTCTTCCGATCTNNNAGTGAGGAGTGAGACTCTAC
GAGCACACGTCTGAACTCTTTCCCTACACGACGCTCTTCCGATCTNNNAATAACATTCAGAGAGGCC
GAGCACACGTCTGAACTCTTTCCCTACACGACGCTCTTCCGATCTNNNAATAACATTCAGAGAGGCC
GAGCACACGTCTGAACTCTTTCCCTACACGACGCTCTTCCGATCTNNNTATCCTGCCACACCATTCATGA
GAGCACACGTCTGAACTCTTTCCCTACACGACGCTCTTCCGATCTNNNTATCCTGCCACACCATTCATGA
AGCACACGTCTGAACTCTTTCCCTACACGACGCTCTTCCGATCTNNNCTCCCAGGTTCAAGCAATTCTC
AGCACACGTCTGAACTCTTTCCCTACACGACGCTCTTCCGATCTNNNCTCCCAGGTTCAAGCAATTCTC
CACACGTCTGAACTCTTTCCCTACACGACGCTCTTCCGATCTNNNGATCAAGACCATCCTGGCTAACAC
CACACGTCTGAACTCTTTCCCTACACGACGCTCTTCCGATCTNNNGATCAAGACCATCCTGGCTAACAC
AGAGCACACGTCTGAACTCTTTCCCTACACGACGCTCTTCCGATCTNNNAGCAGGGAAAAGAGGAAA
AGAGCACACGTCTGAACTCTTTCCCTACACGACGCTCTTCCGATCTNNNAGCAGGGAAAAGAGGAAA
GAGCACACGTCTGAACTCTTTCCCTACACGACGCTCTTCCGATCTNNNAGGAGGGAAATTTGTCCAC
GAGCACACGTCTGAACTCTTTCCCTACACGACGCTCTTCCGATCTNNNAGGAGGGAAATTTGTCCAC
GAGCACACGTCTGAACTCTTTCCCTACACGACGCTCTTCCGATCTNNNCCACAGAATAGGCCAGTTAG
GAGCACACGTCTGAACTCTTTCCCTACACGACGCTCTTCCGATCTNNNCCACAGAATAGGCCAGTTAG
AGCACACGTCTGAACTCTTTCCCTACACGACGCTCTTCCGATCTNNNAAAAAATTAGCCAGGTGTGG
AGCACACGTCTGAACTCTTTCCCTACACGACGCTCTTCCGATCTNNNAAAAAATTAGCCAGGTGTGG
AGCACACGTCTGAACTCTTTCCCTACACGACGCTCTTCCGATCTNNNCGGATCACTTGAGATCAGGAG
AGCACACGTCTGAACTCTTTCCCTACACGACGCTCTTCCGATCTNNNCGGATCACTTGAGATCAGGAG
AGCACACGTCTGAACTCTTTCCCTACACGACGCTCTTCCGATCTNNNTACTAGAGAGACTGAGGCAG
AGCACACGTCTGAACTCTTTCCCTACACGACGCTCTTCCGATCTNNNTACTAGAGAGACTGAGGCAG
GCACACGTCTGAACTCTTTCCCTACACGACGCTCTTCCGATCTNNNGGTTGCATCCTTTTTACCCTCACC
GCACACGTCTGAACTCTTTCCCTACACGACGCTCTTCCGATCTNNNGGTTGCATCCTTTTTACCCTCACC
GCACACGTCTGAACTCTTTCCCTACACGACGCTCTTCCGATCTNNNAAAGTGCTGGGATTACAGGCA
GCACACGTCTGAACTCTTTCCCTACACGACGCTCTTCCGATCTNNNAAAGTGCTGGGATTACAGGCA
AGCACACGTCTGAACTCTTTCCCTACACGACGCTCTTCCGATCTNNNACTTTGGACCATGTACCTCCAT
AGCACACGTCTGAACTCTTTCCCTACACGACGCTCTTCCGATCTNNNACTTTGGACCATGTACCTCCAT
GCACACGTCTGAACTCTTTCCCTACACGACGCTCTTCCGATCTNNNGGTGAGAACAGCCATATTCATG

AGCACACGTCTGAACTCTTTCCCTACACGACGCTCTTCCGATCTNNNGGTGAGAACAGCCATATTCATG
AGCACACGTCTGAACTCTTTCCCTACACGACGCTCTTCCGATCTNNNAGTCCCATCTATGTCAGAGGT
AGCACACGTCTGAACTCTTTCCCTACACGACGCTCTTCCGATCTNNNAGTCCCATCTATGTCAGAGGT
AGCACACGTCTGAACTCTTTCCCTACACGACGCTCTTCCGATCTNNNTGAGAGTGTCTCTAGGATGTG
AGCACACGTCTGAACTCTTTCCCTACACGACGCTCTTCCGATCTNNNTGAGAGTGTCTCTAGGATGTG
AGCACACGTCTGAACTCTTTCCCTACACGACGCTCTTCCGATCTNNNAAAGTGCTGGGATTACAGGT
AGCACACGTCTGAACTCTTTCCCTACACGACGCTCTTCCGATCTNNNAAAGTGCTGGGATTACAGGT
GCACACGTCTGAACTCTTTCCCTACACGACGCTCTTCCGATCTNNNTTGTTTCTCCCACAATTCTCCAC
GCACACGTCTGAACTCTTTCCCTACACGACGCTCTTCCGATCTNNNTTGTTTCTCCCACAATTCTCCAC
GAGCACACGTCTGAACTCTTTCCCTACACGACGCTCTTCCGATCTNNNATTGTCACCAGTTCCATGGCT
GAGCACACGTCTGAACTCTTTCCCTACACGACGCTCTTCCGATCTNNNATTGTCACCAGTTCCATGGCT
AGCACACGTCTGAACTCTTTCCCTACACGACGCTCTTCCGATCTNNNGTAATCCCAGCTACTAGTGAG
AGCACACGTCTGAACTCTTTCCCTACACGACGCTCTTCCGATCTNNNGTAATCCCAGCTACTAGTGAG
AGCACACGTCTGAACTCTTTCCCTACACGACGCTCTTCCGATCTNNNCGAAACCATCCATCTTCCTCCTC
AGCACACGTCTGAACTCTTTCCCTACACGACGCTCTTCCGATCTNNNCGAAACCATCCATCTTCCTCCTC
ACACGTCTGAACTCTTTCCCTACACGACGCTCTTCCGATCTNNNCCCTCTTGCAGACGGTACTTGCATC
ACACGTCTGAACTCTTTCCCTACACGACGCTCTTCCGATCTNNNCCCTCTTGCAGACGGTACTTGCATC
AGCACACGTCTGAACTCTTTCCCTACACGACGCTCTTCCGATCTNNNCTGTGAGGAAATGGCCCCAGC
AGCACACGTCTGAACTCTTTCCCTACACGACGCTCTTCCGATCTNNNCTGTGAGGAAATGGCCCCAGC
GCACACGTCTGAACTCTTTCCCTACACGACGCTCTTCCGATCTNNNGGTGATTTCTCTCTTGGCAGGT
GCACACGTCTGAACTCTTTCCCTACACGACGCTCTTCCGATCTNNNGGTGATTTCTCTCTTGGCAGGT
AGAGCACACGTCTGAACTCTTTCCCTACACGACGCTCTTCCGATCTNNNGAGGTTGCAGTGAGCTGA
AGAGCACACGTCTGAACTCTTTCCCTACACGACGCTCTTCCGATCTNNNGAGGTTGCAGTGAGCTGA
AGCACACGTCTGAACTCTTTCCCTACACGACGCTCTTCCGATCTNNNTACCTGACCAGTGAAGTGTGT
AGCACACGTCTGAACTCTTTCCCTACACGACGCTCTTCCGATCTNNNTACCTGACCAGTGAAGTGTGT
AGCACACGTCTGAACTCTTTCCCTACACGACGCTCTTCCGATCTNNNCTCTTCCACTTCACCAGCTGGCA
AGCACACGTCTGAACTCTTTCCCTACACGACGCTCTTCCGATCTNNNCTCTTCCACTTCACCAGCTGGCA
GCACACGTCTGAACTCTTTCCCTACACGACGCTCTTCCGATCTNNNCAACTGCAAATGGCTTTCGACAT
GCACACGTCTGAACTCTTTCCCTACACGACGCTCTTCCGATCTNNNCAACTGCAAATGGCTTTCGACAT
AGCACACGTCTGAACTCTTTCCCTACACGACGCTCTTCCGATCTNNNCAAACAAGCAAACAAAAGCC
AGCACACGTCTGAACTCTTTCCCTACACGACGCTCTTCCGATCTNNNCAAACAAGCAAACAAAAGCC
AGCACACGTCTGAACTCTTTCCCTACACGACGCTCTTCCGATCTNNNGCTGCACACACTAGCTACTTCT
AGCACACGTCTGAACTCTTTCCCTACACGACGCTCTTCCGATCTNNNGCTGCACACACTAGCTACTTCT
AGCACACGTCTGAACTCTTTCCCTACACGACGCTCTTCCGATCTNNNATGAGGAATGAATGAGCGGA
AGCACACGTCTGAACTCTTTCCCTACACGACGCTCTTCCGATCTNNNATGAGGAATGAATGAGCGGA
AGCACACGTCTGAACTCTTTCCCTACACGACGCTCTTCCGATCTNNNCCTTTCCTGTACTTTCTGGGTG
AGCACACGTCTGAACTCTTTCCCTACACGACGCTCTTCCGATCTNNNCCTTTCCTGTACTTTCTGGGTG
AGCACACGTCTGAACTCTTTCCCTACACGACGCTCTTCCGATCTNNNCATCTGTAGTCCCAGCTACTC
AGCACACGTCTGAACTCTTTCCCTACACGACGCTCTTCCGATCTNNNCATCTGTAGTCCCAGCTACTC
GCACACGTCTGAACTCTTTCCCTACACGACGCTCTTCCGATCTNNNACCTGGTTTCTGTGCTTCAGTAT
GCACACGTCTGAACTCTTTCCCTACACGACGCTCTTCCGATCTNNNACCTGGTTTCTGTGCTTCAGTAT
GCACACGTCTGAACTCTTTCCCTACACGACGCTCTTCCGATCTNNNGGAGTTTGCAGTGAGCTGAGAT
GCACACGTCTGAACTCTTTCCCTACACGACGCTCTTCCGATCTNNNGGAGTTTGCAGTGAGCTGAGAT

AGAGCACACGTCTGAACTCTTTCCCTACACGACGCTCTTCCGATCTNNNCTGTAATCCCAGCTACTCAG
AGAGCACACGTCTGAACTCTTTCCCTACACGACGCTCTTCCGATCTNNNCTGTAATCCCAGCTACTCAG
GAGCACACGTCTGAACTCTTTCCCTACACGACGCTCTTCCGATCTNNNATTGTCCCCTTCTTTGTGACCAT
GAGCACACGTCTGAACTCTTTCCCTACACGACGCTCTTCCGATCTNNNATTGTCCCCTTCTTTGTGACCAT
AGCACACGTCTGAACTCTTTCCCTACACGACGCTCTTCCGATCTNNNCACCTGTAGTCCCAGCTACTTGC
AGCACACGTCTGAACTCTTTCCCTACACGACGCTCTTCCGATCTNNNCACCTGTAGTCCCAGCTACTTGC
GCACACGTCTGAACTCTTTCCCTACACGACGCTCTTCCGATCTNNNCCAGTTTACTCCCCCAGCAATACA
GCACACGTCTGAACTCTTTCCCTACACGACGCTCTTCCGATCTNNNCCAGTTTACTCCCCCAGCAATACA
AGAGCACACGTCTGAACTCTTTCCCTACACGACGCTCTTCCGATCTNNNCATGCCTGTAATCCCAGCTA
AGAGCACACGTCTGAACTCTTTCCCTACACGACGCTCTTCCGATCTNNNCATGCCTGTAATCCCAGCTA
AGAGCACACGTCTGAACTCTTTCCCTACACGACGCTCTTCCGATCTNNNAGCAACAGCGTGATCTAAGG
AGAGCACACGTCTGAACTCTTTCCCTACACGACGCTCTTCCGATCTNNNAGCAACAGCGTGATCTAAGG
AGCACACGTCTGAACTCTTTCCCTACACGACGCTCTTCCGATCTNNNGCCTAAATAGCCAGAGCAATTG
AGCACACGTCTGAACTCTTTCCCTACACGACGCTCTTCCGATCTNNNGCCTAAATAGCCAGAGCAATTG
AGCACACGTCTGAACTCTTTCCCTACACGACGCTCTTCCGATCTNNNCAGGGAGTCTGGAAGGAGGTA
AGCACACGTCTGAACTCTTTCCCTACACGACGCTCTTCCGATCTNNNCAGGGAGTCTGGAAGGAGGTA
GAGCACACGTCTGAACTCTTTCCCTACACGACGCTCTTCCGATCTNNNGCAGGAGAATTGCTTGAACT
GAGCACACGTCTGAACTCTTTCCCTACACGACGCTCTTCCGATCTNNNGCAGGAGAATTGCTTGAACT
GAGCACACGTCTGAACTCTTTCCCTACACGACGCTCTTCCGATCTNNNGGTGACATAGTGAGACCCCAT
GAGCACACGTCTGAACTCTTTCCCTACACGACGCTCTTCCGATCTNNNGGTGACATAGTGAGACCCCAT
AGCACACGTCTGAACTCTTTCCCTACACGACGCTCTTCCGATCTNNNTAGTTATGGATTATGTGAGAAG
AGCACACGTCTGAACTCTTTCCCTACACGACGCTCTTCCGATCTNNNTAGTTATGGATTATGTGAGAAG
GAGCACACGTCTGAACTCTTTCCCTACACGACGCTCTTCCGATCTNNNGAGAATCTCTTGAACCCAGGA
GAGCACACGTCTGAACTCTTTCCCTACACGACGCTCTTCCGATCTNNNGAGAATCTCTTGAACCCAGGA
CACACGTCTGAACTCTTTCCCTACACGACGCTCTTCCGATCTNNNAAGTAAACAGAGAAAGGGATCAT
CACACGTCTGAACTCTTTCCCTACACGACGCTCTTCCGATCTNNNAAGTAAACAGAGAAAGGGATCAT
GCACACGTCTGAACTCTTTCCCTACACGACGCTCTTCCGATCTNNNTAAGAACTCTCAGCAAACGGGAA
GCACACGTCTGAACTCTTTCCCTACACGACGCTCTTCCGATCTNNNTAAGAACTCTCAGCAAACGGGAA
AGAGCACACGTCTGAACTCTTTCCCTACACGACGCTCTTCCGATCTNNNTGAGGCAGGAGAAATCACT
AGAGCACACGTCTGAACTCTTTCCCTACACGACGCTCTTCCGATCTNNNTGAGGCAGGAGAAATCACT
AGCACACGTCTGAACTCTTTCCCTACACGACGCTCTTCCGATCTNNNCAGAGGTTGCAGTGAGCTAAG
AGCACACGTCTGAACTCTTTCCCTACACGACGCTCTTCCGATCTNNNCAGAGGTTGCAGTGAGCTAAG
CACACGTCTGAACTCTTTCCCTACACGACGCTCTTCCGATCTNNNCTAGAGCTTTTCCTTTTAAGAGACA
CACACGTCTGAACTCTTTCCCTACACGACGCTCTTCCGATCTNNNCTAGAGCTTTTCCTTTTAAGAGACA
GAGCACACGTCTGAACTCTTTCCCTACACGACGCTCTTCCGATCTNNNGAGGCATGAGAATTGCTTGCT
GAGCACACGTCTGAACTCTTTCCCTACACGACGCTCTTCCGATCTNNNGAGGCATGAGAATTGCTTGCT
GAGCACACGTCTGAACTCTTTCCCTACACGACGCTCTTCCGATCTNNNTTCAAGCGATTCTCCTGCCTTA
GAGCACACGTCTGAACTCTTTCCCTACACGACGCTCTTCCGATCTNNNTTCAAGCGATTCTCCTGCCTTA
AGCACACGTCTGAACTCTTTCCCTACACGACGCTCTTCCGATCTNNNTTGAACATGGGGAGTTTAGGA
AGCACACGTCTGAACTCTTTCCCTACACGACGCTCTTCCGATCTNNNTTGAACATGGGGAGTTTAGGA
AGAGCACACGTCTGAACTCTTTCCCTACACGACGCTCTTCCGATCTNNNGTGAACAAGTCAGACAAAG
AGAGCACACGTCTGAACTCTTTCCCTACACGACGCTCTTCCGATCTNNNGTGAACAAGTCAGACAAAG
AGCACACGTCTGAACTCTTTCCCTACACGACGCTCTTCCGATCTNNNTGGATTTTCAATGCTATGCCAG

AGCACACGTCTGAACTCTTTCCCTACACGACGCTCTTCCGATCTNNNTGGATTTTCAATGCTATGCCAG
AGCACACGTCTGAACTCTTTCCCTACACGACGCTCTTCCGATCTNNNGAGAATCGCTTGAACCAAGGA
AGCACACGTCTGAACTCTTTCCCTACACGACGCTCTTCCGATCTNNNGAGAATCGCTTGAACCAAGGA
GCACACGTCTGAACTCTTTCCCTACACGACGCTCTTCCGATCTNNNCCAGGAGATTGAGAGTGGCTTC
GCACACGTCTGAACTCTTTCCCTACACGACGCTCTTCCGATCTNNNCCAGGAGATTGAGAGTGGCTTC
GAGCACACGTCTGAACTCTTTCCCTACACGACGCTCTTCCGATCTNNNCAGGAGAATTGCTTGAAACC
GAGCACACGTCTGAACTCTTTCCCTACACGACGCTCTTCCGATCTNNNCAGGAGAATTGCTTGAAACC
GAGCACACGTCTGAACTCTTTCCCTACACGACGCTCTTCCGATCTNNNATATAAAAGACTACTGGCCG
GAGCACACGTCTGAACTCTTTCCCTACACGACGCTCTTCCGATCTNNNATATAAAAGACTACTGGCCG
ACACGTCTGAACTCTTTCCCTACACGACGCTCTTCCGATCTNNNAGCCAAGTTGGTAATCCCTTTCTCAT
ACACGTCTGAACTCTTTCCCTACACGACGCTCTTCCGATCTNNNAGCCAAGTTGGTAATCCCTTTCTCAT
CACACGTCTGAACTCTTTCCCTACACGACGCTCTTCCGATCTNNNTGTGCATGTGTTGGGGGGAGCAT
CACACGTCTGAACTCTTTCCCTACACGACGCTCTTCCGATCTNNNTGTGCATGTGTTGGGGGGAGCAT
AGCACACGTCTGAACTCTTTCCCTACACGACGCTCTTCCGATCTNNNGGAATAGTGATTGGGAAGTGG
AGCACACGTCTGAACTCTTTCCCTACACGACGCTCTTCCGATCTNNNGGAATAGTGATTGGGAAGTGG
GCACACGTCTGAACTCTTTCCCTACACGACGCTCTTCCGATCTNNNCACACACACACACACACACATAT
GCACACGTCTGAACTCTTTCCCTACACGACGCTCTTCCGATCTNNNCACACACACACACACACACATAT
AGCACACGTCTGAACTCTTTCCCTACACGACGCTCTTCCGATCTNNNAAGGAATTGTGTGTGTGTGTG
AGCACACGTCTGAACTCTTTCCCTACACGACGCTCTTCCGATCTNNNAAGGAATTGTGTGTGTGTGTG
AGCACACGTCTGAACTCTTTCCCTACACGACGCTCTTCCGATCTNNNCACCTGGTCACAGTATGCAATT
AGCACACGTCTGAACTCTTTCCCTACACGACGCTCTTCCGATCTNNNCACCTGGTCACAGTATGCAATT
AGCACACGTCTGAACTCTTTCCCTACACGACGCTCTTCCGATCTNNNCTTAGGGTGGAGGAAGGGGA
AGCACACGTCTGAACTCTTTCCCTACACGACGCTCTTCCGATCTNNNCTTAGGGTGGAGGAAGGGGA
GAGCACACGTCTGAACTCTTTCCCTACACGACGCTCTTCCGATCTNNNGATTCCACTAATGACTGACAT
GAGCACACGTCTGAACTCTTTCCCTACACGACGCTCTTCCGATCTNNNGATTCCACTAATGACTGACAT
AGCACACGTCTGAACTCTTTCCCTACACGACGCTCTTCCGATCTNNNATCAAACAAGCCTGTGAATGT
AGCACACGTCTGAACTCTTTCCCTACACGACGCTCTTCCGATCTNNNATCAAACAAGCCTGTGAATGT
GAGCACACGTCTGAACTCTTTCCCTACACGACGCTCTTCCGATCTNNNAGAATCACTTGAGAACACGGG
AGCACACGTCTGAACTCTTTCCCTACACGACGCTCTTCCGATCTNNNAGAATCACTTGAGAACACGGG
AGCACACGTCTGAACTCTTTCCCTACACGACGCTCTTCCGATCTNNNGTGCAGTTCCAAACAAAATCC
AGCACACGTCTGAACTCTTTCCCTACACGACGCTCTTCCGATCTNNNGTGCAGTTCCAAACAAAATCC
AGCACACGTCTGAACTCTTTCCCTACACGACGCTCTTCCGATCTNNNAATCAGAAGACACTCCTCAACC
AGCACACGTCTGAACTCTTTCCCTACACGACGCTCTTCCGATCTNNNAATCAGAAGACACTCCTCAACC
GAGCACACGTCTGAACTCTTTCCCTACACGACGCTCTTCCGATCTNNNCTCGAACTCCTGACCTTGTGA
GAGCACACGTCTGAACTCTTTCCCTACACGACGCTCTTCCGATCTNNNCTCGAACTCCTGACCTTGTGA
GAGCACACGTCTGAACTCTTTCCCTACACGACGCTCTTCCGATCTNNNAAGTGCTCTTGACTGACGACT
AGCACACGTCTGAACTCTTTCCCTACACGACGCTCTTCCGATCTNNNAAGTGCTCTTGACTGACGACT
AGCACACGTCTGAACTCTTTCCCTACACGACGCTCTTCCGATCTNNNATTAGGGAGCTGAAAGACTG
AGCACACGTCTGAACTCTTTCCCTACACGACGCTCTTCCGATCTNNNATTAGGGAGCTGAAAGACTG
GCACACGTCTGAACTCTTTCCCTACACGACGCTCTTCCGATCTNNNTGGGCATAAGCTATCTGTTTGTT
CACACGTCTGAACTCTTTCCCTACACGACGCTCTTCCGATCTNNNTGGGCATAAGCTATCTGTTTGTT
GAGCACACGTCTGAACTCTTTCCCTACACGACGCTCTTCCGATCTNNNGGTAGGTTCTGATGGCATCT
GAGCACACGTCTGAACTCTTTCCCTACACGACGCTCTTCCGATCTNNNGGTAGGTTCTGATGGCATCT

AGCACACGTCTGAACTCTTTCCCTACACGACGCTCTTCCGATCTNNNCATGGTGGTAGGCACCTGTA
AGCACACGTCTGAACTCTTTCCCTACACGACGCTCTTCCGATCTNNNCATGGTGGTAGGCACCTGTA
GCACACGTCTGAACTCTTTCCCTACACGACGCTCTTCCGATCTNNNATTCCATCCAGGTTGCTACGAAT
GCACACGTCTGAACTCTTTCCCTACACGACGCTCTTCCGATCTNNNATTCCATCCAGGTTGCTACGAAT
AGCACACGTCTGAACTCTTTCCCTACACGACGCTCTTCCGATCTNNNAATTCAGTTTCTTCACGTCCCTG
AGCACACGTCTGAACTCTTTCCCTACACGACGCTCTTCCGATCTNNNAATTCAGTTTCTTCACGTCCCTG
GAGCACACGTCTGAACTCTTTCCCTACACGACGCTCTTCCGATCTNNNAAAAAATTAGCCAGGTGTGG
GAGCACACGTCTGAACTCTTTCCCTACACGACGCTCTTCCGATCTNNNAAAAAATTAGCCAGGTGTGG
AGCACACGTCTGAACTCTTTCCCTACACGACGCTCTTCCGATCTNNNGCCTTTCAGGTTCAAGCAATTC
AGCACACGTCTGAACTCTTTCCCTACACGACGCTCTTCCGATCTNNNGCCTTTCAGGTTCAAGCAATTC
AAGAGCACACGTCTGAACTCTTTCCCTACACGACGCTCTTCCGATCTNNNCTGGAGAATGGCGTGAA
AAGAGCACACGTCTGAACTCTTTCCCTACACGACGCTCTTCCGATCTNNNCTGGAGAATGGCGTGAA
AGCACACGTCTGAACTCTTTCCCTACACGACGCTCTTCCGATCTNNNCCTGAGGATAAGCAGGGAGT
AGCACACGTCTGAACTCTTTCCCTACACGACGCTCTTCCGATCTNNNCCTGAGGATAAGCAGGGAGT
GAGCACACGTCTGAACTCTTTCCCTACACGACGCTCTTCCGATCTNNNAGCTTTATTCGCCCTAACCAA
GAGCACACGTCTGAACTCTTTCCCTACACGACGCTCTTCCGATCTNNNAGCTTTATTCGCCCTAACCAA
AGCACACGTCTGAACTCTTTCCCTACACGACGCTCTTCCGATCTNNNCTGGCTTTGTGCTAGGGGAGA
AGCACACGTCTGAACTCTTTCCCTACACGACGCTCTTCCGATCTNNNCTGGCTTTGTGCTAGGGGAGA
GCACACGTCTGAACTCTTTCCCTACACGACGCTCTTCCGATCTNNNCAGCCACTCAAATTCTCACGTTT
GCACACGTCTGAACTCTTTCCCTACACGACGCTCTTCCGATCTNNNCAGCCACTCAAATTCTCACGTTT
AGCACACGTCTGAACTCTTTCCCTACACGACGCTCTTCCGATCTNNNGCCAACATGTTGAAATCCTGTC
AGCACACGTCTGAACTCTTTCCCTACACGACGCTCTTCCGATCTNNNGCCAACATGTTGAAATCCTGTC
GCACACGTCTGAACTCTTTCCCTACACGACGCTCTTCCGATCTNNNAAGCAAGATGAGTGGGGTGAT
GCACACGTCTGAACTCTTTCCCTACACGACGCTCTTCCGATCTNNNAAGCAAGATGAGTGGGGTGAT
GCACACGTCTGAACTCTTTCCCTACACGACGCTCTTCCGATCTNNNTCTGGCAAAACCCATCTTACTCA
GCACACGTCTGAACTCTTTCCCTACACGACGCTCTTCCGATCTNNNTCTGGCAAAACCCATCTTACTCA
AGCACACGTCTGAACTCTTTCCCTACACGACGCTCTTCCGATCTNNNCTGAGGCAGGAGAATCACTTG
AGCACACGTCTGAACTCTTTCCCTACACGACGCTCTTCCGATCTNNNCTGAGGCAGGAGAATCACTTG
AGCACACGTCTGAACTCTTTCCCTACACGACGCTCTTCCGATCTNNNGGTCAAAAGGTGAAGCAACTC
AGCACACGTCTGAACTCTTTCCCTACACGACGCTCTTCCGATCTNNNGGTCAAAAGGTGAAGCAACTC
GAGCACACGTCTGAACTCTTTCCCTACACGACGCTCTTCCGATCTNNNTACTCTCAGGTTCTCTGCTGTC
GAGCACACGTCTGAACTCTTTCCCTACACGACGCTCTTCCGATCTNNNTACTCTCAGGTTCTCTGCTGTC
AGCACACGTCTGAACTCTTTCCCTACACGACGCTCTTCCGATCTNNNACATGGTGAAACTCCGTCTCTA
AGCACACGTCTGAACTCTTTCCCTACACGACGCTCTTCCGATCTNNNACATGGTGAAACTCCGTCTCTA
GAGCACACGTCTGAACTCTTTCCCTACACGACGCTCTTCCGATCTNNNATGTAGTCCCACCTACTCTGG
GAGCACACGTCTGAACTCTTTCCCTACACGACGCTCTTCCGATCTNNNATGTAGTCCCACCTACTCTGG
GAGCACACGTCTGAACTCTTTCCCTACACGACGCTCTTCCGATCTNNNGCTCAGAAGACTCCTGTCAGG
GAGCACACGTCTGAACTCTTTCCCTACACGACGCTCTTCCGATCTNNNGCTCAGAAGACTCCTGTCAGG
AGCACACGTCTGAACTCTTTCCCTACACGACGCTCTTCCGATCTNNNTGACATTCTTTCCCGCTGAACTA
AGCACACGTCTGAACTCTTTCCCTACACGACGCTCTTCCGATCTNNNTGACATTCTTTCCCGCTGAACTA
AGCACACGTCTGAACTCTTTCCCTACACGACGCTCTTCCGATCTNNNCTTTGCTGGGGGTTGACAGGC
AGCACACGTCTGAACTCTTTCCCTACACGACGCTCTTCCGATCTNNNCTTTGCTGGGGGTTGACAGGC
AGCACACGTCTGAACTCTTTCCCTACACGACGCTCTTCCGATCTNNNCCTGGTGCATGCTACCTACATA

AGCACACGTCTGAACTCTTTCCCTACACGACGCTCTTCCGATCTNNNCCTGGTGCATGCTACCTACATA

AGCACACGTCTGAACTCTTTCCCTACACGACGCTCTTCCGATCTNNNCAGTGATCATGCCAATCGATGA

AGCACACGTCTGAACTCTTTCCCTACACGACGCTCTTCCGATCTNNNCAGTGATCATGCCAATCGATGA

GAGCACACGTCTGAACTCTTTCCCTACACGACGCTCTTCCGATCTNNNGTGGAATCTCAGAAACCCTGA

GAGCACACGTCTGAACTCTTTCCCTACACGACGCTCTTCCGATCTNNNGTGGAATCTCAGAAACCCTGA

CACACGTCTGAACTCTTTCCCTACACGACGCTCTTCCGATCTNNNACCCCACTAGGCAGTGTTCTGCATC

CACACGTCTGAACTCTTTCCCTACACGACGCTCTTCCGATCTNNNACCCCACTAGGCAGTGTTCTGCATC

GAGCACACGTCTGAACTCTTTCCCTACACGACGCTCTTCCGATCTNNNTACAGAAGACGGGAGAAGA

GAGCACACGTCTGAACTCTTTCCCTACACGACGCTCTTCCGATCTNNNTACAGAAGACGGGAGAAGA

AGAGCACACGTCTGAACTCTTTCCCTACACGACGCTCTTCCGATCTNNNGGGATTGTGTCTCTTGCAGA

AGAGCACACGTCTGAACTCTTTCCCTACACGACGCTCTTCCGATCTNNNGGGATTGTGTCTCTTGCAGA

GAGCACACGTCTGAACTCTTTCCCTACACGACGCTCTTCCGATCTNNNAAAAATTAGTCGGATGTGGTG

GAGCACACGTCTGAACTCTTTCCCTACACGACGCTCTTCCGATCTNNNAAAAATTAGTCGGATGTGGTG

AGCACACGTCTGAACTCTTTCCCTACACGACGCTCTTCCGATCTNNNAGTTGGATCTAGGTCACCTTTCC

AGCACACGTCTGAACTCTTTCCCTACACGACGCTCTTCCGATCTNNNAGTTGGATCTAGGTCACCTTTCC

GCACACGTCTGAACTCTTTCCCTACACGACGCTCTTCCGATCTNNNCCTCAACCCTGCCAGAAAAGTAAC

GCACACGTCTGAACTCTTTCCCTACACGACGCTCTTCCGATCTNNNCCTCAACCCTGCCAGAAAAGTAAC

AGCACACGTCTGAACTCTTTCCCTACACGACGCTCTTCCGATCTNNNATGTAGCTTGCACCATTCACTGT

AGCACACGTCTGAACTCTTTCCCTACACGACGCTCTTCCGATCTNNNATGTAGCTTGCACCATTCACTGT

GCACACGTCTGAACTCTTTCCCTACACGACGCTCTTCCGATCTNNNCACAGACCCTCAACATTGTGACAG

GCACACGTCTGAACTCTTTCCCTACACGACGCTCTTCCGATCTNNNCACAGACCCTCAACATTGTGACAG

CACACGTCTGAACTCTTTCCCTACACGACGCTCTTCCGATCTNNNTGCAGGAAGTAGGAGGGTTTATG

CACACGTCTGAACTCTTTCCCTACACGACGCTCTTCCGATCTNNNTGCAGGAAGTAGGAGGGTTTATG

AAGAGCACACGTCTGAACTCTTTCCCTACACGACGCTCTTCCGATCTNNNTTTCTGCGCAGGTTTTCAT

AAGAGCACACGTCTGAACTCTTTCCCTACACGACGCTCTTCCGATCTNNNTTTCTGCGCAGGTTTTCAT

CACACGTCTGAACTCTTTCCCTACACGACGCTCTTCCGATCTNNNCACATCAAGCATGACAGATGCATC

CACACGTCTGAACTCTTTCCCTACACGACGCTCTTCCGATCTNNNCACATCAAGCATGACAGATGCATC

AGCACACGTCTGAACTCTTTCCCTACACGACGCTCTTCCGATCTNNNAGCACTCCCTTATCTGCTTTGTG

AGCACACGTCTGAACTCTTTCCCTACACGACGCTCTTCCGATCTNNNAGCACTCCCTTATCTGCTTTGTG

GAGCACACGTCTGAACTCTTTCCCTACACGACGCTCTTCCGATCTNNNTACTAAGGCTGTTCCTCTTCTC

GAGCACACGTCTGAACTCTTTCCCTACACGACGCTCTTCCGATCTNNNTACTAAGGCTGTTCCTCTTCTC

CACACGTCTGAACTCTTTCCCTACACGACGCTCTTCCGATCTNNNCCATGCCTTGCATCCATTTATCCAT

CACACGTCTGAACTCTTTCCCTACACGACGCTCTTCCGATCTNNNCCATGCCTTGCATCCATTTATCCAT

AGAGCACACGTCTGAACTCTTTCCCTACACGACGCTCTTCCGATCTNNNCTTTTTCCCCTTCCTGGCTGA

AGAGCACACGTCTGAACTCTTTCCCTACACGACGCTCTTCCGATCTNNNCTTTTTCCCCTTCCTGGCTGA

AGCACACGTCTGAACTCTTTCCCTACACGACGCTCTTCCGATCTNNNTGGTTTTCGGCATTTTCAACCTTG

AGCACACGTCTGAACTCTTTCCCTACACGACGCTCTTCCGATCTNNNTGGTTTTCGGCATTTTCAACCTTG

AGAGCACACGTCTGAACTCTTTCCCTACACGACGCTCTTCCGATCTNNNACCGTAGTATACCCCCGGTCC

AGAGCACACGTCTGAACTCTTTCCCTACACGACGCTCTTCCGATCTNNNACCGTAGTATACCCCCGGTCC

AGAGCACACGTCTGAACTCTTTCCCTACACGACGCTCTTCCGATCTNNNAGCATTCTCCAGAGTTGTCC

AGAGCACACGTCTGAACTCTTTCCCTACACGACGCTCTTCCGATCTNNNAGCATTCTCCAGAGTTGTCC

GAGCACACGTCTGAACTCTTTCCCTACACGACGCTCTTCCGATCTNNNACATTCATATGTACGGCTCGG

GAGCACACGTCTGAACTCTTTCCCTACACGACGCTCTTCCGATCTNNNACATTCATATGTACGGCTCGG

GAGCACACGTCTGAACTCTTTCCCTACACGACGCTCTTCCGATCTNNNAGATGGCCATTATTGGGATCC

GAGCACACGTCTGAACTCTTTCCCTACACGACGCTCTTCCGATCTNNNAGATGGCCATTATTGGGATCC

AGAGCACACGTCTGAACTCTTTCCCTACACGACGCTCTTCCGATCTNNNAAGCTTCAATCTACTGCTGTT

AGAGCACACGTCTGAACTCTTTCCCTACACGACGCTCTTCCGATCTNNNAAGCTTCAATCTACTGCTGTT

AGCACACGTCTGAACTCTTTCCCTACACGACGCTCTTCCGATCTNNNTGAGGCAGGAGAATCACTTGA

AGCACACGTCTGAACTCTTTCCCTACACGACGCTCTTCCGATCTNNNTGAGGCAGGAGAATCACTTGA

CACACGTCTGAACTCTTTCCCTACACGACGCTCTTCCGATCTNNNACCATGATTTCCCAAATGGCATCCA

CACACGTCTGAACTCTTTCCCTACACGACGCTCTTCCGATCTNNNACCATGATTTCCCAAATGGCATCCA

AGCACACGTCTGAACTCTTTCCCTACACGACGCTCTTCCGATCTNNNCTCCAATCCTGACACTTCCATCAC

AGCACACGTCTGAACTCTTTCCCTACACGACGCTCTTCCGATCTNNNCTCCAATCCTGACACTTCCATCAC

GAGCACACGTCTGAACTCTTTCCCTACACGACGCTCTTCCGATCTNNNAGGCATCCAGAAAACTCTCTCT

GAGCACACGTCTGAACTCTTTCCCTACACGACGCTCTTCCGATCTNNNAGGCATCCAGAAAACTCTCTCT

AGAGCACACGTCTGAACTCTTTCCCTACACGACGCTCTTCCGATCTNNNTGGGTGACAGAGCAAGACT

AGAGCACACGTCTGAACTCTTTCCCTACACGACGCTCTTCCGATCTNNNTGGGTGACAGAGCAAGACT

AGCACACGTCTGAACTCTTTCCCTACACGACGCTCTTCCGATCTNNNTGTCTCTCTCACTCTCTCTCTCCCA

AGCACACGTCTGAACTCTTTCCCTACACGACGCTCTTCCGATCTNNNTGTCTCTCTCACTCTCTCTCTCCCA

AGAGCACACGTCTGAACTCTTTCCCTACACGACGCTCTTCCGATCTNNNCAATGGTACCTCCCGATCTAA

AGAGCACACGTCTGAACTCTTTCCCTACACGACGCTCTTCCGATCTNNNCAATGGTACCTCCCGATCTAA

GAGCACACGTCTGAACTCTTTCCCTACACGACGCTCTTCCGATCTNNNAAAAAAAATTAGCTGGGCGTG

GAGCACACGTCTGAACTCTTTCCCTACACGACGCTCTTCCGATCTNNNAAAAAAAATTAGCTGGGCGTG

AGAGCACACGTCTGAACTCTTTCCCTACACGACGCTCTTCCGATCTNNNGAGACAGACCAAGACTCCAT

AGAGCACACGTCTGAACTCTTTCCCTACACGACGCTCTTCCGATCTNNNGAGACAGACCAAGACTCCAT

AGCACACGTCTGAACTCTTTCCCTACACGACGCTCTTCCGATCTNNNCTTGCAAATCCTCATCCTAGGGT

AGCACACGTCTGAACTCTTTCCCTACACGACGCTCTTCCGATCTNNNCTTGCAAATCCTCATCCTAGGGT

GAGCACACGTCTGAACTCTTTCCCTACACGACGCTCTTCCGATCTNNNCAGGCAGACTCTTGTTCTCTTC

GAGCACACGTCTGAACTCTTTCCCTACACGACGCTCTTCCGATCTNNNCAGGCAGACTCTTGTTCTCTTC

AGCACACGTCTGAACTCTTTCCCTACACGACGCTCTTCCGATCTNNNTGTGCCTCCTTCCTGAATTCTAG

AGCACACGTCTGAACTCTTTCCCTACACGACGCTCTTCCGATCTNNNTGTGCCTCCTTCCTGAATTCTAG

AGCACACGTCTGAACTCTTTCCCTACACGACGCTCTTCCGATCTNNNGAGATCAAATCATGCCTCAGTG

AGCACACGTCTGAACTCTTTCCCTACACGACGCTCTTCCGATCTNNNGAGATCAAATCATGCCTCAGTG

AGCACACGTCTGAACTCTTTCCCTACACGACGCTCTTCCGATCTNNNGAGGTTGCAGTGAGCCTAGAC

AGCACACGTCTGAACTCTTTCCCTACACGACGCTCTTCCGATCTNNNGAGGTTGCAGTGAGCCTAGAC

GCACACGTCTGAACTCTTTCCCTACACGACGCTCTTCCGATCTNNNCCTGGACATGTCTTTTGGCAAATC

GCACACGTCTGAACTCTTTCCCTACACGACGCTCTTCCGATCTNNNCCTGGACATGTCTTTTGGCAAATC

GAGCACACGTCTGAACTCTTTCCCTACACGACGCTCTTCCGATCTNNNGCAAAGGGGAAAAAAAGCAC

GAGCACACGTCTGAACTCTTTCCCTACACGACGCTCTTCCGATCTNNNGCAAAGGGGAAAAAAAGCAC

CACACGTCTGAACTCTTTCCCTACACGACGCTCTTCCGATCTNNNCCTCCACCCAGAATTCTAACACTCAT

CACACGTCTGAACTCTTTCCCTACACGACGCTCTTCCGATCTNNNCCTCCACCCAGAATTCTAACACTCAT

AGCACACGTCTGAACTCTTTCCCTACACGACGCTCTTCCGATCTNNNTTTGATCACAATGGGAAAATTA

AGCACACGTCTGAACTCTTTCCCTACACGACGCTCTTCCGATCTNNNTTTGATCACAATGGGAAAATTA

AGCACACGTCTGAACTCTTTCCCTACACGACGCTCTTCCGATCTNNNCACCTGTAATCCCAGCTACTTGG

AGCACACGTCTGAACTCTTTCCCTACACGACGCTCTTCCGATCTNNNCACCTGTAATCCCAGCTACTTGG

AGCACACGTCTGAACTCTTTCCCTACACGACGCTCTTCCGATCTNNNGGCTCGCAACTAATTTCCTGTTT

AGCACACGTCTGAACTCTTTCCCTACACGACGCTCTTCCGATCTNNNGGCTCGCAACTAATTTCCTGTTT
AGCACACGTCTGAACTCTTTCCCTACACGACGCTCTTCCGATCTNNNGAAGCAAGCACCTTCTTCACAA
AGCACACGTCTGAACTCTTTCCCTACACGACGCTCTTCCGATCTNNNGAAGCAAGCACCTTCTTCACAA
AGCACACGTCTGAACTCTTTCCCTACACGACGCTCTTCCGATCTNNNCACCTCGTAAGCTTTGCCTTAA
AGCACACGTCTGAACTCTTTCCCTACACGACGCTCTTCCGATCTNNNCACCTCGTAAGCTTTGCCTTAA
GCACACGTCTGAACTCTTTCCCTACACGACGCTCTTCCGATCTNNNGCTGATCCAGAAAGAGGTAAGCA
GCACACGTCTGAACTCTTTCCCTACACGACGCTCTTCCGATCTNNNGCTGATCCAGAAAGAGGTAAGCA
AGAGCACACGTCTGAACTCTTTCCCTACACGACGCTCTTCCGATCTNNNGTAATCCCAGCTACTCAGAA
AGAGCACACGTCTGAACTCTTTCCCTACACGACGCTCTTCCGATCTNNNGTAATCCCAGCTACTCAGAA
GCACACGTCTGAACTCTTTCCCTACACGACGCTCTTCCGATCTNNNTTGCTGATAATCTCATGGGAGCG
GCACACGTCTGAACTCTTTCCCTACACGACGCTCTTCCGATCTNNNTTGCTGATAATCTCATGGGAGCG
GAGCACACGTCTGAACTCTTTCCCTACACGACGCTCTTCCGATCTNNNAAAAAAAAATTTAGCCGGGC
GAGCACACGTCTGAACTCTTTCCCTACACGACGCTCTTCCGATCTNNNAAAAAAAAATTTAGCCGGGC
GAGCACACGTCTGAACTCTTTCCCTACACGACGCTCTTCCGATCTNNNAGATTAGAATGCTGCCTTGGA
GAGCACACGTCTGAACTCTTTCCCTACACGACGCTCTTCCGATCTNNNAGATTAGAATGCTGCCTTGGA
GAGCACACGTCTGAACTCTTTCCCTACACGACGCTCTTCCGATCTNNNCTGAGGCAGGAGAATCACTTG
GAGCACACGTCTGAACTCTTTCCCTACACGACGCTCTTCCGATCTNNNCTGAGGCAGGAGAATCACTTG
CACGTCTGAACTCTTTCCCTACACGACGCTCTTCCGATCTNNNCATCCCACTCCTTCTGACATTCACATCC
CACGTCTGAACTCTTTCCCTACACGACGCTCTTCCGATCTNNNCATCCCACTCCTTCTGACATTCACATCC
AGCACACGTCTGAACTCTTTCCCTACACGACGCTCTTCCGATCTNNNGCTCCAAGTGTCTTCAGCAAAA
AGCACACGTCTGAACTCTTTCCCTACACGACGCTCTTCCGATCTNNNGCTCCAAGTGTCTTCAGCAAAA
AGCACACGTCTGAACTCTTTCCCTACACGACGCTCTTCCGATCTNNNTACTGGAGACACATCAAAGCAC
AGCACACGTCTGAACTCTTTCCCTACACGACGCTCTTCCGATCTNNNTACTGGAGACACATCAAAGCAC
AGCACACGTCTGAACTCTTTCCCTACACGACGCTCTTCCGATCTNNNCCTCCCTCTCTCTCTCTCTTTCTCA
AGCACACGTCTGAACTCTTTCCCTACACGACGCTCTTCCGATCTNNNCCTCCCTCTCTCTCTCTCTTTCTCA
GCACACGTCTGAACTCTTTCCCTACACGACGCTCTTCCGATCTNNNCTTTGGAGAATAAGGATGGAGC
GCACACGTCTGAACTCTTTCCCTACACGACGCTCTTCCGATCTNNNCTTTGGAGAATAAGGATGGAGC
GAGCACACGTCTGAACTCTTTCCCTACACGACGCTCTTCCGATCTNNNGTGAAAACAAGAGATGAGGG
GAGCACACGTCTGAACTCTTTCCCTACACGACGCTCTTCCGATCTNNNGTGAAAACAAGAGATGAGGG
AGCACACGTCTGAACTCTTTCCCTACACGACGCTCTTCCGATCTNNNTGTTCTGCAGGGAACATATGTG
AGCACACGTCTGAACTCTTTCCCTACACGACGCTCTTCCGATCTNNNTGTTCTGCAGGGAACATATGTG
GAGCACACGTCTGAACTCTTTCCCTACACGACGCTCTTCCGATCTNNNAAAAAATTAGCAGGGCATGG
GAGCACACGTCTGAACTCTTTCCCTACACGACGCTCTTCCGATCTNNNAAAAAATTAGCAGGGCATGG
CACACGTCTGAACTCTTTCCCTACACGACGCTCTTCCGATCTNNNGGGTGGAAGTAAACAGAAAAAGC
CACACGTCTGAACTCTTTCCCTACACGACGCTCTTCCGATCTNNNGGGTGGAAGTAAACAGAAAAAGC
GAGCACACGTCTGAACTCTTTCCCTACACGACGCTCTTCCGATCTNNNTGCGAGGCTATTACTGTCAGT
GAGCACACGTCTGAACTCTTTCCCTACACGACGCTCTTCCGATCTNNNTGCGAGGCTATTACTGTCAGT
AGCACACGTCTGAACTCTTTCCCTACACGACGCTCTTCCGATCTNNNGAAGCTCCGATATTCCCCAGAA
AGCACACGTCTGAACTCTTTCCCTACACGACGCTCTTCCGATCTNNNGAAGCTCCGATATTCCCCAGAA
GAGCACACGTCTGAACTCTTTCCCTACACGACGCTCTTCCGATCTNNNTAGGACCCCATTCTTGACAAAG
GAGCACACGTCTGAACTCTTTCCCTACACGACGCTCTTCCGATCTNNNTAGGACCCCATTCTTGACAAAG
GAGCACACGTCTGAACTCTTTCCCTACACGACGCTCTTCCGATCTNNNTTGGAACTGATTTGGAGAGTG
GAGCACACGTCTGAACTCTTTCCCTACACGACGCTCTTCCGATCTNNNTTGGAACTGATTTGGAGAGTG

AGCACACGTCTGAACTCTTTCCCTACACGACGCTCTTCCGATCTNNNTTAAAGCTATACGACTAAGTCT
AGCACACGTCTGAACTCTTTCCCTACACGACGCTCTTCCGATCTNNNTTAAAGCTATACGACTAAGTCT
GAGCACACGTCTGAACTCTTTCCCTACACGACGCTCTTCCGATCTNNNCTCTTCTCGCTCCGTCTATCTCT
GAGCACACGTCTGAACTCTTTCCCTACACGACGCTCTTCCGATCTNNNCTCTTCTCGCTCCGTCTATCTCT
AGAGCACACGTCTGAACTCTTTCCCTACACGACGCTCTTCCGATCTNNNTTACTGGGCCTCCATCTTGCC
AGAGCACACGTCTGAACTCTTTCCCTACACGACGCTCTTCCGATCTNNNTTACTGGGCCTCCATCTTGCC
GAGCACACGTCTGAACTCTTTCCCTACACGACGCTCTTCCGATCTNNNTGTTTGATACAGGCATGCAA
AGCACACGTCTGAACTCTTTCCCTACACGACGCTCTTCCGATCTNNNTGTTTGATACAGGCATGCAA
AGCACACGTCTGAACTCTTTCCCTACACGACGCTCTTCCGATCTNNNATTTCGGGAATCTTGAATGTAT
AGCACACGTCTGAACTCTTTCCCTACACGACGCTCTTCCGATCTNNNATTTCGGGAATCTTGAATGTAT
GCACACGTCTGAACTCTTTCCCTACACGACGCTCTTCCGATCTNNNGTGTGAGGAGTCATTAGCAGTG
GCACACGTCTGAACTCTTTCCCTACACGACGCTCTTCCGATCTNNNGTGTGAGGAGTCATTAGCAGTG
GAGCACACGTCTGAACTCTTTCCCTACACGACGCTCTTCCGATCTNNNTAAGGCATCTAGAAACCAGC
GAGCACACGTCTGAACTCTTTCCCTACACGACGCTCTTCCGATCTNNNTAAGGCATCTAGAAACCAGC
GAGCACACGTCTGAACTCTTTCCCTACACGACGCTCTTCCGATCTNNNCAGTGAGCTAAGATCCTGCC
GAGCACACGTCTGAACTCTTTCCCTACACGACGCTCTTCCGATCTNNNCAGTGAGCTAAGATCCTGCC
AGCACACGTCTGAACTCTTTCCCTACACGACGCTCTTCCGATCTNNNGGAACTGTTTGAGCCTAGGAG
AGCACACGTCTGAACTCTTTCCCTACACGACGCTCTTCCGATCTNNNGGAACTGTTTGAGCCTAGGAG
AGAGCACACGTCTGAACTCTTTCCCTACACGACGCTCTTCCGATCTNNNCACAGTGGGTCTTCCTCACC
AGAGCACACGTCTGAACTCTTTCCCTACACGACGCTCTTCCGATCTNNNCACAGTGGGTCTTCCTCACC
GAGCACACGTCTGAACTCTTTCCCTACACGACGCTCTTCCGATCTNNNAATAGTCCAAGAAAGGCTGA
GAGCACACGTCTGAACTCTTTCCCTACACGACGCTCTTCCGATCTNNNAATAGTCCAAGAAAGGCTGA
GCACACGTCTGAACTCTTTCCCTACACGACGCTCTTCCGATCTNNNCTATTCATACCTTGGAACCCCCC
GCACACGTCTGAACTCTTTCCCTACACGACGCTCTTCCGATCTNNNCTATTCATACCTTGGAACCCCCC
AGCACACGTCTGAACTCTTTCCCTACACGACGCTCTTCCGATCTNNNGTTACTTGGGAGGCTGAGAC
AGCACACGTCTGAACTCTTTCCCTACACGACGCTCTTCCGATCTNNNGTTACTTGGGAGGCTGAGAC
AGCACACGTCTGAACTCTTTCCCTACACGACGCTCTTCCGATCTNNNTGAGGCAGGAGAATAGTGTG
AGCACACGTCTGAACTCTTTCCCTACACGACGCTCTTCCGATCTNNNTGAGGCAGGAGAATAGTGTG
AGCACACGTCTGAACTCTTTCCCTACACGACGCTCTTCCGATCTNNNATCTGGTCCGTTCTTCACACTCC
AGCACACGTCTGAACTCTTTCCCTACACGACGCTCTTCCGATCTNNNATCTGGTCCGTTCTTCACACTCC
GAGCACACGTCTGAACTCTTTCCCTACACGACGCTCTTCCGATCTNNNGAGGATCACTTGAGCTCGAG
GAGCACACGTCTGAACTCTTTCCCTACACGACGCTCTTCCGATCTNNNGAGGATCACTTGAGCTCGAG
GAGCACACGTCTGAACTCTTTCCCTACACGACGCTCTTCCGATCTNNNAAATAGAAGTAGGCCAAGGT
GAGCACACGTCTGAACTCTTTCCCTACACGACGCTCTTCCGATCTNNNAAATAGAAGTAGGCCAAGGT
GCACACGTCTGAACTCTTTCCCTACACGACGCTCTTCCGATCTNNNCAGGAAAATCGCTTGAACCAGG
GCACACGTCTGAACTCTTTCCCTACACGACGCTCTTCCGATCTNNNCAGGAAAATCGCTTGAACCAGG
AGCACACGTCTGAACTCTTTCCCTACACGACGCTCTTCCGATCTNNNTGGATGTGATGGAGTGCGAA
AGCACACGTCTGAACTCTTTCCCTACACGACGCTCTTCCGATCTNNNTGGATGTGATGGAGTGCGAA
AGCACACGTCTGAACTCTTTCCCTACACGACGCTCTTCCGATCTNNNACTTATGAGTGAGAACATGCG
AGCACACGTCTGAACTCTTTCCCTACACGACGCTCTTCCGATCTNNNACTTATGAGTGAGAACATGCG
GAGCACACGTCTGAACTCTTTCCCTACACGACGCTCTTCCGATCTNNNGTCTTCCTTGGTCTTTTGCTGT
GAGCACACGTCTGAACTCTTTCCCTACACGACGCTCTTCCGATCTNNNGTCTTCCTTGGTCTTTTGCTGT
GAGCACACGTCTGAACTCTTTCCCTACACGACGCTCTTCCGATCTNNNCTCACTCTCATTGTCCAGTGG

GAGCACACGTCTGAACTCTTTCCCTACACGACGCTCTTCCGATCTNNNCTCACTCTCATTGTCCAGTGGA
AGCACACGTCTGAACTCTTTCCCTACACGACGCTCTTCCGATCTNNNCTCCACGCATTCCTTACCTATGAC
AGCACACGTCTGAACTCTTTCCCTACACGACGCTCTTCCGATCTNNNCTCCACGCATTCCTTACCTATGAC
AGAGCACACGTCTGAACTCTTTCCCTACACGACGCTCTTCCGATCTNNNGTAATCTCAGCTACTTGGGAC
AGAGCACACGTCTGAACTCTTTCCCTACACGACGCTCTTCCGATCTNNNGTAATCTCAGCTACTTGGGAC
AGCACACGTCTGAACTCTTTCCCTACACGACGCTCTTCCGATCTNNNATGGTTGAATGAATGGTTAAA
AGCACACGTCTGAACTCTTTCCCTACACGACGCTCTTCCGATCTNNNATGGTTGAATGAATGGTTAAA
AGAGCACACGTCTGAACTCTTTCCCTACACGACGCTCTTCCGATCTNNNCAAAAATTACCCAGGTGTGG
AGAGCACACGTCTGAACTCTTTCCCTACACGACGCTCTTCCGATCTNNNCAAAAATTACCCAGGTGTGG
AGCACACGTCTGAACTCTTTCCCTACACGACGCTCTTCCGATCTNNNCTTCTTCTTCCTGGTGCACTGAC
AGCACACGTCTGAACTCTTTCCCTACACGACGCTCTTCCGATCTNNNCTTCTTCTTCCTGGTGCACTGAC
AGCACACGTCTGAACTCTTTCCCTACACGACGCTCTTCCGATCTNNNAGCAGATGGGAAGTCATGAGA
AGCACACGTCTGAACTCTTTCCCTACACGACGCTCTTCCGATCTNNNAGCAGATGGGAAGTCATGAGA
CACACGTCTGAACTCTTTCCCTACACGACGCTCTTCCGATCTNNNCACAACGACCACATCCTCATAGACA
CACACGTCTGAACTCTTTCCCTACACGACGCTCTTCCGATCTNNNCACAACGACCACATCCTCATAGACA
GAGCACACGTCTGAACTCTTTCCCTACACGACGCTCTTCCGATCTNNNCAGGAGAATCACTTGAACCCAC
GAGCACACGTCTGAACTCTTTCCCTACACGACGCTCTTCCGATCTNNNCAGGAGAATCACTTGAACCCAC
AGCACACGTCTGAACTCTTTCCCTACACGACGCTCTTCCGATCTNNNCAAAAAGCTAGACTCCTGGGAG
AGCACACGTCTGAACTCTTTCCCTACACGACGCTCTTCCGATCTNNNCAAAAAGCTAGACTCCTGGGAG
GAGCACACGTCTGAACTCTTTCCCTACACGACGCTCTTCCGATCTNNNAGTGAGCTGAGACCACTGACT
GAGCACACGTCTGAACTCTTTCCCTACACGACGCTCTTCCGATCTNNNAGTGAGCTGAGACCACTGACT
AGCACACGTCTGAACTCTTTCCCTACACGACGCTCTTCCGATCTNNNAATCTCGATTCACTGCAACCTCT
AGCACACGTCTGAACTCTTTCCCTACACGACGCTCTTCCGATCTNNNAATCTCGATTCACTGCAACCTCT
GAGCACACGTCTGAACTCTTTCCCTACACGACGCTCTTCCGATCTNNNAGTGTCTTGAACTTGATGGAC
GAGCACACGTCTGAACTCTTTCCCTACACGACGCTCTTCCGATCTNNNAGTGTCTTGAACTTGATGGAC
GCACACGTCTGAACTCTTTCCCTACACGACGCTCTTCCGATCTNNNACTTCTGTACTCTAGTCCAGCCTC
GCACACGTCTGAACTCTTTCCCTACACGACGCTCTTCCGATCTNNNACTTCTGTACTCTAGTCCAGCCTC
GAGCACACGTCTGAACTCTTTCCCTACACGACGCTCTTCCGATCTNNNCTGCCTTCTCATTTGACAGACC
GAGCACACGTCTGAACTCTTTCCCTACACGACGCTCTTCCGATCTNNNCTGCCTTCTCATTTGACAGACC
GAGCACACGTCTGAACTCTTTCCCTACACGACGCTCTTCCGATCTNNNCTTGACGTAGGAAACCCAGGT
GAGCACACGTCTGAACTCTTTCCCTACACGACGCTCTTCCGATCTNNNCTTGACGTAGGAAACCCAGGT
AGCACACGTCTGAACTCTTTCCCTACACGACGCTCTTCCGATCTNNNACACACACACTATGGAATACTAC
AGCACACGTCTGAACTCTTTCCCTACACGACGCTCTTCCGATCTNNNACACACACACTATGGAATACTAC
AGCACACGTCTGAACTCTTTCCCTACACGACGCTCTTCCGATCTNNNGCTGCTGGTCATCTTCACTTTAC
AGCACACGTCTGAACTCTTTCCCTACACGACGCTCTTCCGATCTNNNGCTGCTGGTCATCTTCACTTTAC
GAGCACACGTCTGAACTCTTTCCCTACACGACGCTCTTCCGATCTNNNAGAGAGATCCTATGCACCCCAT
GAGCACACGTCTGAACTCTTTCCCTACACGACGCTCTTCCGATCTNNNAGAGAGATCCTATGCACCCCAT
GAGCACACGTCTGAACTCTTTCCCTACACGACGCTCTTCCGATCTNNNCATTCCCACTTTGCATTCTGCTC
GAGCACACGTCTGAACTCTTTCCCTACACGACGCTCTTCCGATCTNNNCATTCCCACTTTGCATTCTGCTC
AGCACACGTCTGAACTCTTTCCCTACACGACGCTCTTCCGATCTNNNTATGTCCTGCTTGGGAAAGTTG
AGCACACGTCTGAACTCTTTCCCTACACGACGCTCTTCCGATCTNNNTATGTCCTGCTTGGGAAAGTTG
GCACACGTCTGAACTCTTTCCCTACACGACGCTCTTCCGATCTNNNATTATACTTTAATGACCTCCCCAT
GCACACGTCTGAACTCTTTCCCTACACGACGCTCTTCCGATCTNNNATTATACTTTAATGACCTCCCCAT

AGCACACGTCTGAACTCTTTCCCTACACGACGCTCTTCCGATCTNNNCATTTGATCCTGCAGTTCCAGT
AGCACACGTCTGAACTCTTTCCCTACACGACGCTCTTCCGATCTNNNCATTTGATCCTGCAGTTCCAGT
GAGCACACGTCTGAACTCTTTCCCTACACGACGCTCTTCCGATCTNNNTGCCAATGATGCCAAGAGTTC
GAGCACACGTCTGAACTCTTTCCCTACACGACGCTCTTCCGATCTNNNTGCCAATGATGCCAAGAGTTC
AGAGCACACGTCTGAACTCTTTCCCTACACGACGCTCTTCCGATCTNNNAGGCAAGAGAATCACTTGA
AGAGCACACGTCTGAACTCTTTCCCTACACGACGCTCTTCCGATCTNNNAGGCAAGAGAATCACTTGA
GCACACGTCTGAACTCTTTCCCTACACGACGCTCTTCCGATCTNNNATCTTGGGCCTGTGTTATCTCCTC
GCACACGTCTGAACTCTTTCCCTACACGACGCTCTTCCGATCTNNNATCTTGGGCCTGTGTTATCTCCTC
AGCACACGTCTGAACTCTTTCCCTACACGACGCTCTTCCGATCTNNNAATGTGCACCAAAGACAGACTT
AGCACACGTCTGAACTCTTTCCCTACACGACGCTCTTCCGATCTNNNAATGTGCACCAAAGACAGACTT
GAGCACACGTCTGAACTCTTTCCCTACACGACGCTCTTCCGATCTNNNAGAATCACTTGAATCCAGGAG
GAGCACACGTCTGAACTCTTTCCCTACACGACGCTCTTCCGATCTNNNAGAATCACTTGAATCCAGGAG
GCACACGTCTGAACTCTTTCCCTACACGACGCTCTTCCGATCTNNNTATTTCTCTTTCTGTGCCTGGCTC
GCACACGTCTGAACTCTTTCCCTACACGACGCTCTTCCGATCTNNNTATTTCTCTTTCTGTGCCTGGCTC
AGCACACGTCTGAACTCTTTCCCTACACGACGCTCTTCCGATCTNNNATGGATCACAAGGTCAGGAGT
AGCACACGTCTGAACTCTTTCCCTACACGACGCTCTTCCGATCTNNNATGGATCACAAGGTCAGGAGT
AGCACACGTCTGAACTCTTTCCCTACACGACGCTCTTCCGATCTNNNCTGAGGAAGGAGATTGGCTTG
AGCACACGTCTGAACTCTTTCCCTACACGACGCTCTTCCGATCTNNNCTGAGGAAGGAGATTGGCTTG
GAGCACACGTCTGAACTCTTTCCCTACACGACGCTCTTCCGATCTNNNGCCTACACAGTAAGAGCAAG
GAGCACACGTCTGAACTCTTTCCCTACACGACGCTCTTCCGATCTNNNGCCTACACAGTAAGAGCAAG
GAGCACACGTCTGAACTCTTTCCCTACACGACGCTCTTCCGATCTNNNAGTTTTGCATGATAAAAGTGG
GAGCACACGTCTGAACTCTTTCCCTACACGACGCTCTTCCGATCTNNNAGTTTTGCATGATAAAAGTGG
AGCACACGTCTGAACTCTTTCCCTACACGACGCTCTTCCGATCTNNNTCAGCTGCAAATGAGACTTCTC
AGCACACGTCTGAACTCTTTCCCTACACGACGCTCTTCCGATCTNNNTCAGCTGCAAATGAGACTTCTC
AGCACACGTCTGAACTCTTTCCCTACACGACGCTCTTCCGATCTNNNACTGGATCATGATGGCAGTTTC
AGCACACGTCTGAACTCTTTCCCTACACGACGCTCTTCCGATCTNNNACTGGATCATGATGGCAGTTTC
GAGCACACGTCTGAACTCTTTCCCTACACGACGCTCTTCCGATCTNNNCCTCACCTCTCTGACTTCTTTCC
GAGCACACGTCTGAACTCTTTCCCTACACGACGCTCTTCCGATCTNNNCCTCACCTCTCTGACTTCTTTCC
AGCACACGTCTGAACTCTTTCCCTACACGACGCTCTTCCGATCTNNNACTCCAAAAGTGCTGGGATTA
AGCACACGTCTGAACTCTTTCCCTACACGACGCTCTTCCGATCTNNNACTCCAAAAGTGCTGGGATTA
GAGCACACGTCTGAACTCTTTCCCTACACGACGCTCTTCCGATCTNNNACCCCACAGATTTGCCTAAAT
GAGCACACGTCTGAACTCTTTCCCTACACGACGCTCTTCCGATCTNNNACCCCACAGATTTGCCTAAAT
AGCACACGTCTGAACTCTTTCCCTACACGACGCTCTTCCGATCTNNNGCCATGAGAATTTGGGCTACT
AGCACACGTCTGAACTCTTTCCCTACACGACGCTCTTCCGATCTNNNGCCATGAGAATTTGGGCTACT
GAGCACACGTCTGAACTCTTTCCCTACACGACGCTCTTCCGATCTNNNGGTCAGGTGGCCTAATACTG
GAGCACACGTCTGAACTCTTTCCCTACACGACGCTCTTCCGATCTNNNGGTCAGGTGGCCTAATACTG
GAGCACACGTCTGAACTCTTTCCCTACACGACGCTCTTCCGATCTNNNTGCCCATGAGTCAGAATCTAC
GAGCACACGTCTGAACTCTTTCCCTACACGACGCTCTTCCGATCTNNNTGCCCATGAGTCAGAATCTAC
GAGCACACGTCTGAACTCTTTCCCTACACGACGCTCTTCCGATCTNNNTGCATGTTTCAAAGAACTCCA
GAGCACACGTCTGAACTCTTTCCCTACACGACGCTCTTCCGATCTNNNTGCATGTTTCAAAGAACTCCA
GAGCACACGTCTGAACTCTTTCCCTACACGACGCTCTTCCGATCTNNNTTGTTGTTGTCATGGATGTG
GAGCACACGTCTGAACTCTTTCCCTACACGACGCTCTTCCGATCTNNNTTGTTGTTGTCATGGATGTG
AGCACACGTCTGAACTCTTTCCCTACACGACGCTCTTCCGATCTNNNGGCCCCAGTAGCTTCTTTGTAA

AGCACACGTCTGAACTCTTTCCCTACACGACGCTCTTCCGATCTNNNGGCCCCAGTAGCTTCTTTGTAAA
GAGCACACGTCTGAACTCTTTCCCTACACGACGCTCTTCCGATCTNNNCCCAGGTTGAAGAGTTTTGG
GAGCACACGTCTGAACTCTTTCCCTACACGACGCTCTTCCGATCTNNNCCCAGGTTGAAGAGTTTTGG
ACACGTCTGAACTCTTTCCCTACACGACGCTCTTCCGATCTNNNGCAAGATTCTGAGGGGAAGTGTAC
ACACGTCTGAACTCTTTCCCTACACGACGCTCTTCCGATCTNNNGCAAGATTCTGAGGGGAAGTGTAC
AGCACACGTCTGAACTCTTTCCCTACACGACGCTCTTCCGATCTNNNGTGACAGAGCCAGATTCCATC
AGCACACGTCTGAACTCTTTCCCTACACGACGCTCTTCCGATCTNNNGTGACAGAGCCAGATTCCATC
AGCACACGTCTGAACTCTTTCCCTACACGACGCTCTTCCGATCTNNNTTTTTCTTCAAAGAGCGTGAGC
AGCACACGTCTGAACTCTTTCCCTACACGACGCTCTTCCGATCTNNNTTTTTCTTCAAAGAGCGTGAGC
AGCACACGTCTGAACTCTTTCCCTACACGACGCTCTTCCGATCTNNNGAAAGTTGCAGTGAGCTGAGA
AGCACACGTCTGAACTCTTTCCCTACACGACGCTCTTCCGATCTNNNGAAAGTTGCAGTGAGCTGAGA
GAGCACACGTCTGAACTCTTTCCCTACACGACGCTCTTCCGATCTNNNAGAACAAGAACATCACCTGAC
GAGCACACGTCTGAACTCTTTCCCTACACGACGCTCTTCCGATCTNNNAGAACAAGAACATCACCTGAC
GCACACGTCTGAACTCTTTCCCTACACGACGCTCTTCCGATCTNNNGGAAATCTCTGGAGTGGGAAAC
GCACACGTCTGAACTCTTTCCCTACACGACGCTCTTCCGATCTNNNGGAAATCTCTGGAGTGGGAAAC
AGCACACGTCTGAACTCTTTCCCTACACGACGCTCTTCCGATCTNNNTGCCTGGCCAGCTTTTTATTATC
AGCACACGTCTGAACTCTTTCCCTACACGACGCTCTTCCGATCTNNNTGCCTGGCCAGCTTTTTATTATC
GAGCACACGTCTGAACTCTTTCCCTACACGACGCTCTTCCGATCTNNNTGGTGAAGATGGCGGTAAA
GAGCACACGTCTGAACTCTTTCCCTACACGACGCTCTTCCGATCTNNNTGGTGAAGATGGCGGTAAA
GAGCACACGTCTGAACTCTTTCCCTACACGACGCTCTTCCGATCTNNNTGGTGTTTTAGTGTAGGCGAC
GAGCACACGTCTGAACTCTTTCCCTACACGACGCTCTTCCGATCTNNNTGGTGTTTTAGTGTAGGCGAC
AGCACACGTCTGAACTCTTTCCCTACACGACGCTCTTCCGATCTNNNATGCCTGTAATCCAAGCACTTTG
AGCACACGTCTGAACTCTTTCCCTACACGACGCTCTTCCGATCTNNNATGCCTGTAATCCAAGCACTTTG
GAGCACACGTCTGAACTCTTTCCCTACACGACGCTCTTCCGATCTNNNGTAATCCCAGCTACTCAAGAG
GAGCACACGTCTGAACTCTTTCCCTACACGACGCTCTTCCGATCTNNNGTAATCCCAGCTACTCAAGAG
AGCACACGTCTGAACTCTTTCCCTACACGACGCTCTTCCGATCTNNNTAGTCTTTTCCCATTCCTCACCG
AGCACACGTCTGAACTCTTTCCCTACACGACGCTCTTCCGATCTNNNTAGTCTTTTCCCATTCCTCACCG
GAGCACACGTCTGAACTCTTTCCCTACACGACGCTCTTCCGATCTNNNGCTCCGATCGTGTACAAATGA
GAGCACACGTCTGAACTCTTTCCCTACACGACGCTCTTCCGATCTNNNGCTCCGATCGTGTACAAATGA
GCACACGTCTGAACTCTTTCCCTACACGACGCTCTTCCGATCTNNNATCCACCCCAATGCACATACTAAC
GCACACGTCTGAACTCTTTCCCTACACGACGCTCTTCCGATCTNNNATCCACCCCAATGCACATACTAAC
AGCACACGTCTGAACTCTTTCCCTACACGACGCTCTTCCGATCTNNNGAGGCAGGAAAATGGTGTGAA
AGCACACGTCTGAACTCTTTCCCTACACGACGCTCTTCCGATCTNNNGAGGCAGGAAAATGGTGTGAA
GAGCACACGTCTGAACTCTTTCCCTACACGACGCTCTTCCGATCTNNNCTTTGCATTCTTTGGGTCTGG
GAGCACACGTCTGAACTCTTTCCCTACACGACGCTCTTCCGATCTNNNCTTTGCATTCTTTGGGTCTGG
AGAGCACACGTCTGAACTCTTTCCCTACACGACGCTCTTCCGATCTNNNTGTCACTCAATGAACCTCCTC
AGAGCACACGTCTGAACTCTTTCCCTACACGACGCTCTTCCGATCTNNNTGTCACTCAATGAACCTCCTC
AGCACACGTCTGAACTCTTTCCCTACACGACGCTCTTCCGATCTNNNCTGTAATCCCAGCTACTCAGGAG
AGCACACGTCTGAACTCTTTCCCTACACGACGCTCTTCCGATCTNNNCTGTAATCCCAGCTACTCAGGAG
AGAGCACACGTCTGAACTCTTTCCCTACACGACGCTCTTCCGATCTNNNATTAATGCACAGCAAATCAC
AGAGCACACGTCTGAACTCTTTCCCTACACGACGCTCTTCCGATCTNNNATTAATGCACAGCAAATCAC
AGCACACGTCTGAACTCTTTCCCTACACGACGCTCTTCCGATCTNNNTCTCCTAATGCTATCCCTCTCCC
AGCACACGTCTGAACTCTTTCCCTACACGACGCTCTTCCGATCTNNNTCTCCTAATGCTATCCCTCTCCC

AGAGCACACGTCTGAACTCTTTCCCTACACGACGCTCTTCCGATCTNNNAGGAATTCAAGACCAGCCTG
AGAGCACACGTCTGAACTCTTTCCCTACACGACGCTCTTCCGATCTNNNAGGAATTCAAGACCAGCCTG
AAGAGCACACGTCTGAACTCTTTCCCTACACGACGCTCTTCCGATCTNNNAAACGTTCAAACCATCGCAC
AAGAGCACACGTCTGAACTCTTTCCCTACACGACGCTCTTCCGATCTNNNAAACGTTCAAACCATCGCAC
GAGCACACGTCTGAACTCTTTCCCTACACGACGCTCTTCCGATCTNNNAGCCCTTACACTTTGGTCTTTG
GAGCACACGTCTGAACTCTTTCCCTACACGACGCTCTTCCGATCTNNNAGCCCTTACACTTTGGTCTTTG
AGCACACGTCTGAACTCTTTCCCTACACGACGCTCTTCCGATCTNNNGGATACTTGACACAGTTTTGGT
AGCACACGTCTGAACTCTTTCCCTACACGACGCTCTTCCGATCTNNNGGATACTTGACACAGTTTTGGT
GAGCACACGTCTGAACTCTTTCCCTACACGACGCTCTTCCGATCTNNNGATAGCAAGATTGGCCTCCCTA
GAGCACACGTCTGAACTCTTTCCCTACACGACGCTCTTCCGATCTNNNGATAGCAAGATTGGCCTCCCTA
AGCACACGTCTGAACTCTTTCCCTACACGACGCTCTTCCGATCTNNNCTATAGTCCCAGCTACTTGGCA
AGCACACGTCTGAACTCTTTCCCTACACGACGCTCTTCCGATCTNNNCTATAGTCCCAGCTACTTGGCA
AGCACACGTCTGAACTCTTTCCCTACACGACGCTCTTCCGATCTNNNGTGTGTGTGTGTGTGTTTTGA
AGCACACGTCTGAACTCTTTCCCTACACGACGCTCTTCCGATCTNNNGTGTGTGTGTGTGTGTTTTGA
AGAGCACACGTCTGAACTCTTTCCCTACACGACGCTCTTCCGATCTNNNTTGCCAGGTGTAACTGCAAG
AGAGCACACGTCTGAACTCTTTCCCTACACGACGCTCTTCCGATCTNNNTTGCCAGGTGTAACTGCAAG
AGCACACGTCTGAACTCTTTCCCTACACGACGCTCTTCCGATCTNNNTAGTAGTACTACCAGTATAGGA
AGCACACGTCTGAACTCTTTCCCTACACGACGCTCTTCCGATCTNNNTAGTAGTACTACCAGTATAGGA
AGCACACGTCTGAACTCTTTCCCTACACGACGCTCTTCCGATCTNNNTGTCTGTGTGTTAGGTGGTCAA
AGCACACGTCTGAACTCTTTCCCTACACGACGCTCTTCCGATCTNNNTGTCTGTGTGTTAGGTGGTCAA
AGCACACGTCTGAACTCTTTCCCTACACGACGCTCTTCCGATCTNNNAACTTTCAGAGTCTGAGGTGGT
AGCACACGTCTGAACTCTTTCCCTACACGACGCTCTTCCGATCTNNNAACTTTCAGAGTCTGAGGTGGT
AGCACACGTCTGAACTCTTTCCCTACACGACGCTCTTCCGATCTNNNAGAATGGTTCCAAGAAATGTGC
AGCACACGTCTGAACTCTTTCCCTACACGACGCTCTTCCGATCTNNNAGAATGGTTCCAAGAAATGTGC
AGAGCACACGTCTGAACTCTTTCCCTACACGACGCTCTTCCGATCTNNNGCCATTGCACTTCCTTCTACA
AGAGCACACGTCTGAACTCTTTCCCTACACGACGCTCTTCCGATCTNNNGCCATTGCACTTCCTTCTACA
GAGCACACGTCTGAACTCTTTCCCTACACGACGCTCTTCCGATCTNNNACTGGTTGGGTTGGAGATGA
GAGCACACGTCTGAACTCTTTCCCTACACGACGCTCTTCCGATCTNNNACTGGTTGGGTTGGAGATGA
AGCACACGTCTGAACTCTTTCCCTACACGACGCTCTTCCGATCTNNNGGCAAAATGGTGAAATCCCATC
AGCACACGTCTGAACTCTTTCCCTACACGACGCTCTTCCGATCTNNNGGCAAAATGGTGAAATCCCATC
ACACGTCTGAACTCTTTCCCTACACGACGCTCTTCCGATCTNNNCGTATAATTTTTTGGGAAGTGAACAT
ACACGTCTGAACTCTTTCCCTACACGACGCTCTTCCGATCTNNNCGTATAATTTTTTGGGAAGTGAACAT
CACACGTCTGAACTCTTTCCCTACACGACGCTCTTCCGATCTNNNCACAAGACATTTTTCTACAGTATCA
CACACGTCTGAACTCTTTCCCTACACGACGCTCTTCCGATCTNNNCACAAGACATTTTTCTACAGTATCA
AGCACACGTCTGAACTCTTTCCCTACACGACGCTCTTCCGATCTNNNTTACTGCCCCTTTCCATTTCAGTC
AGCACACGTCTGAACTCTTTCCCTACACGACGCTCTTCCGATCTNNNTTACTGCCCCTTTCCATTTCAGTC
GAGCACACGTCTGAACTCTTTCCCTACACGACGCTCTTCCGATCTNNNTTTTACACCCCCATGAAGCTACA
GAGCACACGTCTGAACTCTTTCCCTACACGACGCTCTTCCGATCTNNNTTTTACACCCCCATGAAGCTACA
AGAGCACACGTCTGAACTCTTTCCCTACACGACGCTCTTCCGATCTNNNATCCGACTGTCGAAATGAGC
AGAGCACACGTCTGAACTCTTTCCCTACACGACGCTCTTCCGATCTNNNATCCGACTGTCGAAATGAGC
AGCACACGTCTGAACTCTTTCCCTACACGACGCTCTTCCGATCTNNNCGAGAATACTTTATTGCTATGTC
AGCACACGTCTGAACTCTTTCCCTACACGACGCTCTTCCGATCTNNNCGAGAATACTTTATTGCTATGTC
AGCACACGTCTGAACTCTTTCCCTACACGACGCTCTTCCGATCTNNNAGATGCATTCTTAGACGCCAAG

AGCACACGTCTGAACTCTTTCCCTACACGACGCTCTTCCGATCTNNNAGATGCATTCTTAGACGCCAAG
AGCACACGTCTGAACTCTTTCCCTACACGACGCTCTTCCGATCTNNNCCAGGAAGAAGTTGAATCCCT
AGCACACGTCTGAACTCTTTCCCTACACGACGCTCTTCCGATCTNNNCCAGGAAGAAGTTGAATCCCT
AGCACACGTCTGAACTCTTTCCCTACACGACGCTCTTCCGATCTNNNTATTGTTCTACCTTTCAGATGAA
AGCACACGTCTGAACTCTTTCCCTACACGACGCTCTTCCGATCTNNNTATTGTTCTACCTTTCAGATGAA
AGCACACGTCTGAACTCTTTCCCTACACGACGCTCTTCCGATCTNNNGGCTTCTGCTCACAGTAGGATA
AGCACACGTCTGAACTCTTTCCCTACACGACGCTCTTCCGATCTNNNGGCTTCTGCTCACAGTAGGATA
GCACACGTCTGAACTCTTTCCCTACACGACGCTCTTCCGATCTNNNTAGATCAAGGGTTGCAAGACGAT
GCACACGTCTGAACTCTTTCCCTACACGACGCTCTTCCGATCTNNNTAGATCAAGGGTTGCAAGACGAT
AGAGCACACGTCTGAACTCTTTCCCTACACGACGCTCTTCCGATCTNNNTGGATCAAGGCACCTTTTCA
AGAGCACACGTCTGAACTCTTTCCCTACACGACGCTCTTCCGATCTNNNTGGATCAAGGCACCTTTTCA
GAGCACACGTCTGAACTCTTTCCCTACACGACGCTCTTCCGATCTNNNCTGAAATCCAGGGACCACTAG
GAGCACACGTCTGAACTCTTTCCCTACACGACGCTCTTCCGATCTNNNCTGAAATCCAGGGACCACTAG
AGAGCACACGTCTGAACTCTTTCCCTACACGACGCTCTTCCGATCTNNNGAATTCAGATCAAACGCCCTG
AGAGCACACGTCTGAACTCTTTCCCTACACGACGCTCTTCCGATCTNNNGAATTCAGATCAAACGCCCTG
GAGCACACGTCTGAACTCTTTCCCTACACGACGCTCTTCCGATCTNNNTAATGTGTGTGTGTGTGTGT
GAGCACACGTCTGAACTCTTTCCCTACACGACGCTCTTCCGATCTNNNTAATGTGTGTGTGTGTGTGT
AGAGCACACGTCTGAACTCTTTCCCTACACGACGCTCTTCCGATCTNNNCCATGTATTTGCTGTGTTTG
AGAGCACACGTCTGAACTCTTTCCCTACACGACGCTCTTCCGATCTNNNCCATGTATTTGCTGTGTTTG
AGCACACGTCTGAACTCTTTCCCTACACGACGCTCTTCCGATCTNNNGATTTTGGGTGTTCGGTTTAG
AGCACACGTCTGAACTCTTTCCCTACACGACGCTCTTCCGATCTNNNGATTTTGGGTGTTCGGTTTAG
AGCACACGTCTGAACTCTTTCCCTACACGACGCTCTTCCGATCTNNNGGTTGCAGTGAGCCAAGATTG
AGCACACGTCTGAACTCTTTCCCTACACGACGCTCTTCCGATCTNNNGGTTGCAGTGAGCCAAGATTG
GAGCACACGTCTGAACTCTTTCCCTACACGACGCTCTTCCGATCTNNNAAAGCTCAAGAAAAATCCTGG
GAGCACACGTCTGAACTCTTTCCCTACACGACGCTCTTCCGATCTNNNAAAGCTCAAGAAAAATCCTGG
AGCACACGTCTGAACTCTTTCCCTACACGACGCTCTTCCGATCTNNNAAAATCAGCTGAGTGCCTGTAG
AGCACACGTCTGAACTCTTTCCCTACACGACGCTCTTCCGATCTNNNAAAATCAGCTGAGTGCCTGTAG
AGAGCACACGTCTGAACTCTTTCCCTACACGACGCTCTTCCGATCTNNNAAGCAGGAGAATCACTTGAA
AGAGCACACGTCTGAACTCTTTCCCTACACGACGCTCTTCCGATCTNNNAAGCAGGAGAATCACTTGAA
GAGCACACGTCTGAACTCTTTCCCTACACGACGCTCTTCCGATCTNNNAATCCTTACCTTGGGTTCTGCT
GAGCACACGTCTGAACTCTTTCCCTACACGACGCTCTTCCGATCTNNNAATCCTTACCTTGGGTTCTGCT
AGCACACGTCTGAACTCTTTCCCTACACGACGCTCTTCCGATCTNNNTTTTAGTAGAGGCGGGGTTT
AGCACACGTCTGAACTCTTTCCCTACACGACGCTCTTCCGATCTNNNTTTTAGTAGAGGCGGGGTTT
AGCACACGTCTGAACTCTTTCCCTACACGACGCTCTTCCGATCTNNNGGCCAGAGAAGTTCAGTATGG
AGCACACGTCTGAACTCTTTCCCTACACGACGCTCTTCCGATCTNNNGGCCAGAGAAGTTCAGTATGG
GAGCACACGTCTGAACTCTTTCCCTACACGACGCTCTTCCGATCTNNNCAGGAGACAGAGAAAGACCCT
GAGCACACGTCTGAACTCTTTCCCTACACGACGCTCTTCCGATCTNNNCAGGAGACAGAGAAAGACCCT
GAGCACACGTCTGAACTCTTTCCCTACACGACGCTCTTCCGATCTNNNTGGATTCAGTGGAGTCATCCT
GAGCACACGTCTGAACTCTTTCCCTACACGACGCTCTTCCGATCTNNNTGGATTCAGTGGAGTCATCCT
AGCACACGTCTGAACTCTTTCCCTACACGACGCTCTTCCGATCTNNNACCTGTTATGATTGACCCAGGAG
AGCACACGTCTGAACTCTTTCCCTACACGACGCTCTTCCGATCTNNNACCTGTTATGATTGACCCAGGAG
GAGCACACGTCTGAACTCTTTCCCTACACGACGCTCTTCCGATCTNNNTGAGACTCTGATTGCAGTGGA
GAGCACACGTCTGAACTCTTTCCCTACACGACGCTCTTCCGATCTNNNTGAGACTCTGATTGCAGTGGA

AGCACACGTCTGAACTCTTTCCCTACACGACGCTCTTCCGATCTNNNGAAAAACCCCAAAGCTTCAGG
AGCACACGTCTGAACTCTTTCCCTACACGACGCTCTTCCGATCTNNNGAAAAACCCCAAAGCTTCAGG
ACACGTCTGAACTCTTTCCCTACACGACGCTCTTCCGATCTNNNTGACAACTTCTCTCCCCTTCAGACAT
ACACGTCTGAACTCTTTCCCTACACGACGCTCTTCCGATCTNNNTGACAACTTCTCTCCCCTTCAGACAT
AGCACACGTCTGAACTCTTTCCCTACACGACGCTCTTCCGATCTNNNTTCATGCCATTCTCCTGCCTCCA
AGCACACGTCTGAACTCTTTCCCTACACGACGCTCTTCCGATCTNNNTTCATGCCATTCTCCTGCCTCCA
GAGCACACGTCTGAACTCTTTCCCTACACGACGCTCTTCCGATCTNNNGAGAGGGACTTGGAGCTGAT
GAGCACACGTCTGAACTCTTTCCCTACACGACGCTCTTCCGATCTNNNGAGAGGGACTTGGAGCTGAT
AGCACACGTCTGAACTCTTTCCCTACACGACGCTCTTCCGATCTNNNCTGAGGTAGGAGGATTGCTT
AGCACACGTCTGAACTCTTTCCCTACACGACGCTCTTCCGATCTNNNCTGAGGTAGGAGGATTGCTT
GCACACGTCTGAACTCTTTCCCTACACGACGCTCTTCCGATCTNNNTCAAGTCCTCTCTTCCAGCTACTC
GCACACGTCTGAACTCTTTCCCTACACGACGCTCTTCCGATCTNNNTCAAGTCCTCTCTTCCAGCTACTC
AGCACACGTCTGAACTCTTTCCCTACACGACGCTCTTCCGATCTNNNACGATCCATTGGTTTCCAGACAT
AGCACACGTCTGAACTCTTTCCCTACACGACGCTCTTCCGATCTNNNACGATCCATTGGTTTCCAGACAT
AGCACACGTCTGAACTCTTTCCCTACACGACGCTCTTCCGATCTNNNCCTCTTTGGTTTGGTATATTCC
AGCACACGTCTGAACTCTTTCCCTACACGACGCTCTTCCGATCTNNNCCTCTTTGGTTTGGTATATTCC
GCACACGTCTGAACTCTTTCCCTACACGACGCTCTTCCGATCTNNNTCCTGAAATCTTATGCAAATGGG
GCACACGTCTGAACTCTTTCCCTACACGACGCTCTTCCGATCTNNNTCCTGAAATCTTATGCAAATGGG
GAGCACACGTCTGAACTCTTTCCCTACACGACGCTCTTCCGATCTNNNCAGGATGCTGAGGTAGGAG
GAGCACACGTCTGAACTCTTTCCCTACACGACGCTCTTCCGATCTNNNCAGGATGCTGAGGTAGGAG
AGCACACGTCTGAACTCTTTCCCTACACGACGCTCTTCCGATCTNNNGATGGTCTCAATCTCCTGACCT
AGCACACGTCTGAACTCTTTCCCTACACGACGCTCTTCCGATCTNNNGATGGTCTCAATCTCCTGACCT
AGCACACGTCTGAACTCTTTCCCTACACGACGCTCTTCCGATCTNNNCTGCCTACATGTCCCAAGAGG
AGCACACGTCTGAACTCTTTCCCTACACGACGCTCTTCCGATCTNNNCTGCCTACATGTCCCAAGAGG
AAGAGCACACGTCTGAACTCTTTCCCTACACGACGCTCTTCCGATCTNNNTCAGCAGCTGAGGAAGA
AAGAGCACACGTCTGAACTCTTTCCCTACACGACGCTCTTCCGATCTNNNTCAGCAGCTGAGGAAGA
GCACACGTCTGAACTCTTTCCCTACACGACGCTCTTCCGATCTNNNTGCCTTCTCCTGTCCTTCTCAATC
GCACACGTCTGAACTCTTTCCCTACACGACGCTCTTCCGATCTNNNTGCCTTCTCCTGTCCTTCTCAATC
AGCACACGTCTGAACTCTTTCCCTACACGACGCTCTTCCGATCTNNNGCTTGCAGTGAGCTGAGATAG
AGCACACGTCTGAACTCTTTCCCTACACGACGCTCTTCCGATCTNNNGCTTGCAGTGAGCTGAGATAG
AGCACACGTCTGAACTCTTTCCCTACACGACGCTCTTCCGATCTNNNCTGAGGCAGGAGAATAGCTT
AGCACACGTCTGAACTCTTTCCCTACACGACGCTCTTCCGATCTNNNCTGAGGCAGGAGAATAGCTT
GAGCACACGTCTGAACTCTTTCCCTACACGACGCTCTTCCGATCTNNNTTGCCTGGCAAGAACAACTAT
GAGCACACGTCTGAACTCTTTCCCTACACGACGCTCTTCCGATCTNNNTTGCCTGGCAAGAACAACTAT
GAGCACACGTCTGAACTCTTTCCCTACACGACGCTCTTCCGATCTNNNCCTGGGCAATAGAGTAAGAC
GAGCACACGTCTGAACTCTTTCCCTACACGACGCTCTTCCGATCTNNNCCTGGGCAATAGAGTAAGAC
AGCACACGTCTGAACTCTTTCCCTACACGACGCTCTTCCGATCTNNNAGAAGAATGGTGTGAACCCAG
AGCACACGTCTGAACTCTTTCCCTACACGACGCTCTTCCGATCTNNNAGAAGAATGGTGTGAACCCAG
AGCACACGTCTGAACTCTTTCCCTACACGACGCTCTTCCGATCTNNNACTTCTCCTTTGCTGTTCTTGTG
AGCACACGTCTGAACTCTTTCCCTACACGACGCTCTTCCGATCTNNNACTTCTCCTTTGCTGTTCTTGTG
AGCACACGTCTGAACTCTTTCCCTACACGACGCTCTTCCGATCTNNNGTTTTTCCCCACGTCAGTGCCAT
AGCACACGTCTGAACTCTTTCCCTACACGACGCTCTTCCGATCTNNNGTTTTTCCCCACGTCAGTGCCAT
AGCACACGTCTGAACTCTTTCCCTACACGACGCTCTTCCGATCTNNNGAGTGTGAGTCTAGGGAGTG

AGCACACGTCTGAACTCTTTCCCTACACGACGCTCTTCCGATCTNNNGAGTGTGAGTCTAGGGAGTGA
GAGCACACGTCTGAACTCTTTCCCTACACGACGCTCTTCCGATCTNNNGAGTTTGAGACCAGACTAGG
GAGCACACGTCTGAACTCTTTCCCTACACGACGCTCTTCCGATCTNNNGAGTTTGAGACCAGACTAGG
AGCACACGTCTGAACTCTTTCCCTACACGACGCTCTTCCGATCTNNNTGTAGTCCCAGCTACTTAGGAG
AGCACACGTCTGAACTCTTTCCCTACACGACGCTCTTCCGATCTNNNTGTAGTCCCAGCTACTTAGGAG
AGCACACGTCTGAACTCTTTCCCTACACGACGCTCTTCCGATCTNNNGAGAGGAGAGGTGCTACACTT
AGCACACGTCTGAACTCTTTCCCTACACGACGCTCTTCCGATCTNNNGAGAGGAGAGGTGCTACACTT
GAGCACACGTCTGAACTCTTTCCCTACACGACGCTCTTCCGATCTNNNAATGTAAAGGAGCAGTCTCA
GAGCACACGTCTGAACTCTTTCCCTACACGACGCTCTTCCGATCTNNNAATGTAAAGGAGCAGTCTCA
GAGCACACGTCTGAACTCTTTCCCTACACGACGCTCTTCCGATCTNNNGGTGAAGATGTTGATGAATA
GAGCACACGTCTGAACTCTTTCCCTACACGACGCTCTTCCGATCTNNNGGTGAAGATGTTGATGAATA
AGAGCACACGTCTGAACTCTTTCCCTACACGACGCTCTTCCGATCTNNNAGCCCCAGATTTTATATCAGA
AGAGCACACGTCTGAACTCTTTCCCTACACGACGCTCTTCCGATCTNNNAGCCCCAGATTTTATATCAGA
GAGCACACGTCTGAACTCTTTCCCTACACGACGCTCTTCCGATCTNNNGGACAGGCTACAAAATAACAG
GAGCACACGTCTGAACTCTTTCCCTACACGACGCTCTTCCGATCTNNNGGACAGGCTACAAAATAACAG
AGCACACGTCTGAACTCTTTCCCTACACGACGCTCTTCCGATCTNNNCTCCTGAGTAGCTGGGATTACA
AGCACACGTCTGAACTCTTTCCCTACACGACGCTCTTCCGATCTNNNCTCCTGAGTAGCTGGGATTACA
AGAGCACACGTCTGAACTCTTTCCCTACACGACGCTCTTCCGATCTNNNACGAGAATCACTTGAACCCA
AGAGCACACGTCTGAACTCTTTCCCTACACGACGCTCTTCCGATCTNNNACGAGAATCACTTGAACCCA
AGCACACGTCTGAACTCTTTCCCTACACGACGCTCTTCCGATCTNNNCAGCACTTTAAGTAAAGAATCT
AGCACACGTCTGAACTCTTTCCCTACACGACGCTCTTCCGATCTNNNCAGCACTTTAAGTAAAGAATCT
GCACACGTCTGAACTCTTTCCCTACACGACGCTCTTCCGATCTNNNACAAGGAGAGTACAGCTCAGTTG
GCACACGTCTGAACTCTTTCCCTACACGACGCTCTTCCGATCTNNNACAAGGAGAGTACAGCTCAGTTG
GAGCACACGTCTGAACTCTTTCCCTACACGACGCTCTTCCGATCTNNNCCAAGTAGCTGGGACTACAG
GAGCACACGTCTGAACTCTTTCCCTACACGACGCTCTTCCGATCTNNNCCAAGTAGCTGGGACTACAG
AGCACACGTCTGAACTCTTTCCCTACACGACGCTCTTCCGATCTNNNGCTTTCTTTTATGTGCCCCCTTT
AGCACACGTCTGAACTCTTTCCCTACACGACGCTCTTCCGATCTNNNGCTTTCTTTTATGTGCCCCCTTT
AGCACACGTCTGAACTCTTTCCCTACACGACGCTCTTCCGATCTNNNAGAATCACTCGAACCCAGAAGT
AGCACACGTCTGAACTCTTTCCCTACACGACGCTCTTCCGATCTNNNAGAATCACTCGAACCCAGAAGT
CACACGTCTGAACTCTTTCCCTACACGACGCTCTTCCGATCTNNNAAACTCATGCGTGGTCCTGTCCAT
CACACGTCTGAACTCTTTCCCTACACGACGCTCTTCCGATCTNNNAAACTCATGCGTGGTCCTGTCCAT
GAGCACACGTCTGAACTCTTTCCCTACACGACGCTCTTCCGATCTNNNAAAGCCACATACCTACAGCCTA
GAGCACACGTCTGAACTCTTTCCCTACACGACGCTCTTCCGATCTNNNAAAGCCACATACCTACAGCCTA
AGCACACGTCTGAACTCTTTCCCTACACGACGCTCTTCCGATCTNNNCTGAGGCAGGAGAATCACTTGA
AGCACACGTCTGAACTCTTTCCCTACACGACGCTCTTCCGATCTNNNCTGAGGCAGGAGAATCACTTGA
GAGCACACGTCTGAACTCTTTCCCTACACGACGCTCTTCCGATCTNNNCCTTACCTTCACTCTCCTCTCCT
GAGCACACGTCTGAACTCTTTCCCTACACGACGCTCTTCCGATCTNNNCCTTACCTTCACTCTCCTCTCCT
AGCACACGTCTGAACTCTTTCCCTACACGACGCTCTTCCGATCTNNNGAGGCATGAGAATCGCTTGAT
AGCACACGTCTGAACTCTTTCCCTACACGACGCTCTTCCGATCTNNNGAGGCATGAGAATCGCTTGAT
GAGCACACGTCTGAACTCTTTCCCTACACGACGCTCTTCCGATCTNNNAACCCTGTCTCTACCAAAAAT
GAGCACACGTCTGAACTCTTTCCCTACACGACGCTCTTCCGATCTNNNAACCCTGTCTCTACCAAAAAT
AGCACACGTCTGAACTCTTTCCCTACACGACGCTCTTCCGATCTNNNCTCAGCTTCAAGTGGCTCAGAA
AGCACACGTCTGAACTCTTTCCCTACACGACGCTCTTCCGATCTNNNCTCAGCTTCAAGTGGCTCAGAA

AGAGCACACGTCTGAACTCTTTCCCTACACGACGCTCTTCCGATCTNNNTTCCAGCAGGTGAAGGAAA

AGAGCACACGTCTGAACTCTTTCCCTACACGACGCTCTTCCGATCTNNNTTCCAGCAGGTGAAGGAAA

AGAGCACACGTCTGAACTCTTTCCCTACACGACGCTCTTCCGATCTNNNAAGGATGCAGCTGGAGAAC

AGAGCACACGTCTGAACTCTTTCCCTACACGACGCTCTTCCGATCTNNNAAGGATGCAGCTGGAGAAC

AGCACACGTCTGAACTCTTTCCCTACACGACGCTCTTCCGATCTNNNCACAAACAGCGCCTCAAAGAATA

AGCACACGTCTGAACTCTTTCCCTACACGACGCTCTTCCGATCTNNNCACAAACAGCGCCTCAAAGAATA

GAGCACACGTCTGAACTCTTTCCCTACACGACGCTCTTCCGATCTNNNGAGAATCTCTTGAACCCAGGA

GAGCACACGTCTGAACTCTTTCCCTACACGACGCTCTTCCGATCTNNNGAGAATCTCTTGAACCCAGGA

AGCACACGTCTGAACTCTTTCCCTACACGACGCTCTTCCGATCTNNNTCTGAGAGCTGCAGTTAGAGG

AGCACACGTCTGAACTCTTTCCCTACACGACGCTCTTCCGATCTNNNTCTGAGAGCTGCAGTTAGAGG

GAGCACACGTCTGAACTCTTTCCCTACACGACGCTCTTCCGATCTNNNAGAGGTTGCAGTAAGCTGAG

GAGCACACGTCTGAACTCTTTCCCTACACGACGCTCTTCCGATCTNNNAGAGGTTGCAGTAAGCTGAG

AGAGCACACGTCTGAACTCTTTCCCTACACGACGCTCTTCCGATCTNNNAAAACACCACAAATCAGCTG

AGAGCACACGTCTGAACTCTTTCCCTACACGACGCTCTTCCGATCTNNNAAAACACCACAAATCAGCTG

GAGCACACGTCTGAACTCTTTCCCTACACGACGCTCTTCCGATCTNNNGCTGCCTGATACAGTAGGATG

GAGCACACGTCTGAACTCTTTCCCTACACGACGCTCTTCCGATCTNNNGCTGCCTGATACAGTAGGATG

AGCACACGTCTGAACTCTTTCCCTACACGACGCTCTTCCGATCTNNNTGAGGTTTGTGAATCTCCTGACA

AGCACACGTCTGAACTCTTTCCCTACACGACGCTCTTCCGATCTNNNTGAGGTTTGTGAATCTCCTGACA

AGCACACGTCTGAACTCTTTCCCTACACGACGCTCTTCCGATCTNNNCCACAGAAAGCACAGGAAGAAA

AGCACACGTCTGAACTCTTTCCCTACACGACGCTCTTCCGATCTNNNCCACAGAAAGCACAGGAAGAAA

AGAGCACACGTCTGAACTCTTTCCCTACACGACGCTCTTCCGATCTNNNCCTGAATGGTGGTCACTCTAC

AGAGCACACGTCTGAACTCTTTCCCTACACGACGCTCTTCCGATCTNNNCCTGAATGGTGGTCACTCTAC

GAGCACACGTCTGAACTCTTTCCCTACACGACGCTCTTCCGATCTNNNCCTTCCTCCCTTCCTTCTTCATGC

GAGCACACGTCTGAACTCTTTCCCTACACGACGCTCTTCCGATCTNNNCCTTCCTCCCTTCCTTCTTCATGC

GCACACGTCTGAACTCTTTCCCTACACGACGCTCTTCCGATCTNNNAACTGTGGATGCCCTTGTTACAG

GCACACGTCTGAACTCTTTCCCTACACGACGCTCTTCCGATCTNNNAACTGTGGATGCCCTTGTTACAG

AGCACACGTCTGAACTCTTTCCCTACACGACGCTCTTCCGATCTNNNGCAGGAGAATCACTTGAACCTT

AGCACACGTCTGAACTCTTTCCCTACACGACGCTCTTCCGATCTNNNGCAGGAGAATCACTTGAACCTT

AGCACACGTCTGAACTCTTTCCCTACACGACGCTCTTCCGATCTNNNACAAAAAAATTAGCTGGGCAGC

AGCACACGTCTGAACTCTTTCCCTACACGACGCTCTTCCGATCTNNNACAAAAAAATTAGCTGGGCAGC

AGCACACGTCTGAACTCTTTCCCTACACGACGCTCTTCCGATCTNNNACTCTTTTTTGGTTTCCATTGGCA

AGCACACGTCTGAACTCTTTCCCTACACGACGCTCTTCCGATCTNNNACTCTTTTTTGGTTTCCATTGGCA

GAGCACACGTCTGAACTCTTTCCCTACACGACGCTCTTCCGATCTNNNCCCCTAAAATTGGTGTGGCAA

GAGCACACGTCTGAACTCTTTCCCTACACGACGCTCTTCCGATCTNNNCCCCTAAAATTGGTGTGGCAA

AGAGCACACGTCTGAACTCTTTCCCTACACGACGCTCTTCCGATCTNNNGAGGCTGAGGCAGGAGAAT

AGAGCACACGTCTGAACTCTTTCCCTACACGACGCTCTTCCGATCTNNNGAGGCTGAGGCAGGAGAAT

GCACACGTCTGAACTCTTTCCCTACACGACGCTCTTCCGATCTNNNGAAATATCTGTCTTATCCTCTGTCA

GCACACGTCTGAACTCTTTCCCTACACGACGCTCTTCCGATCTNNNGAAATATCTGTCTTATCCTCTGTCA

AGCACACGTCTGAACTCTTTCCCTACACGACGCTCTTCCGATCTNNNCACATTGTGCACATGTACCCTAA

AGCACACGTCTGAACTCTTTCCCTACACGACGCTCTTCCGATCTNNNCACATTGTGCACATGTACCCTAA

AGCACACGTCTGAACTCTTTCCCTACACGACGCTCTTCCGATCTNNNCAGGACAATCGCTTGAACACAGC

AGCACACGTCTGAACTCTTTCCCTACACGACGCTCTTCCGATCTNNNCAGGACAATCGCTTGAACACAGC

GAGCACACGTCTGAACTCTTTCCCTACACGACGCTCTTCCGATCTNNNCTCAGCTCAGACCAGAGTTTC

GAGCACACGTCTGAACTCTTTCCCTACACGACGCTCTTCCGATCTNNNCTCAGCTCAGACCAGAGTTTC

AGCACACGTCTGAACTCTTTCCCTACACGACGCTCTTCCGATCTNNNTCCTTTACACTGATGCAGAGTTT

AGCACACGTCTGAACTCTTTCCCTACACGACGCTCTTCCGATCTNNNTCCTTTACACTGATGCAGAGTTT

AGCACACGTCTGAACTCTTTCCCTACACGACGCTCTTCCGATCTNNNGTTTTTGCGTGTAGAGGTGTT

AGCACACGTCTGAACTCTTTCCCTACACGACGCTCTTCCGATCTNNNGTTTTTGCGTGTAGAGGTGTT

AGCACACGTCTGAACTCTTTCCCTACACGACGCTCTTCCGATCTNNNATGCCTGTAGTCCTAGCTACTC

AGCACACGTCTGAACTCTTTCCCTACACGACGCTCTTCCGATCTNNNATGCCTGTAGTCCTAGCTACTC

GAGCACACGTCTGAACTCTTTCCCTACACGACGCTCTTCCGATCTNNNTCTACATTTGGAAGGGGAGA

GAGCACACGTCTGAACTCTTTCCCTACACGACGCTCTTCCGATCTNNNTCTACATTTGGAAGGGGAGA

AGCACACGTCTGAACTCTTTCCCTACACGACGCTCTTCCGATCTNNNTTCAAGTGATTCTCCTGCCTCAG

AGCACACGTCTGAACTCTTTCCCTACACGACGCTCTTCCGATCTNNNTTCAAGTGATTCTCCTGCCTCAG

GCACACGTCTGAACTCTTTCCCTACACGACGCTCTTCCGATCTNNNGTGATCAGGTTCCTCCTCTCCAC

GCACACGTCTGAACTCTTTCCCTACACGACGCTCTTCCGATCTNNNGTGATCAGGTTCCTCCTCTCCAC

GAGCACACGTCTGAACTCTTTCCCTACACGACGCTCTTCCGATCTNNNTGGGAGCTCTAAGAACAAGA

GAGCACACGTCTGAACTCTTTCCCTACACGACGCTCTTCCGATCTNNNTGGGAGCTCTAAGAACAAGA

AGCACACGTCTGAACTCTTTCCCTACACGACGCTCTTCCGATCTNNNGGAATGTCAGAAACCTTCATG

AGCACACGTCTGAACTCTTTCCCTACACGACGCTCTTCCGATCTNNNGGAATGTCAGAAACCTTCATG

GAGCACACGTCTGAACTCTTTCCCTACACGACGCTCTTCCGATCTNNNGATCATGAGGTCAGGAGAT

GAGCACACGTCTGAACTCTTTCCCTACACGACGCTCTTCCGATCTNNNGATCATGAGGTCAGGAGAT

AGCACACGTCTGAACTCTTTCCCTACACGACGCTCTTCCGATCTNNNGGGATGAATAGGAGTTCACCT

AGCACACGTCTGAACTCTTTCCCTACACGACGCTCTTCCGATCTNNNGGGATGAATAGGAGTTCACCT

GCACACGTCTGAACTCTTTCCCTACACGACGCTCTTCCGATCTNNNAGTAAACCCAAGGAAACCCCAAT

GCACACGTCTGAACTCTTTCCCTACACGACGCTCTTCCGATCTNNNAGTAAACCCAAGGAAACCCCAAT

AGCACACGTCTGAACTCTTTCCCTACACGACGCTCTTCCGATCTNNNATGTATACATGCACCAGGTTGG

AGCACACGTCTGAACTCTTTCCCTACACGACGCTCTTCCGATCTNNNATGTATACATGCACCAGGTTGG

GAGCACACGTCTGAACTCTTTCCCTACACGACGCTCTTCCGATCTNNNGCAGGAGATGACTCACATGG

GAGCACACGTCTGAACTCTTTCCCTACACGACGCTCTTCCGATCTNNNGCAGGAGATGACTCACATGG

AGCACACGTCTGAACTCTTTCCCTACACGACGCTCTTCCGATCTNNNGGTTCTCAACCCTGGCTATACA

AGCACACGTCTGAACTCTTTCCCTACACGACGCTCTTCCGATCTNNNGGTTCTCAACCCTGGCTATACA

AGCACACGTCTGAACTCTTTCCCTACACGACGCTCTTCCGATCTNNNCACTGCATCCGACCTTTCTTTTC

AGCACACGTCTGAACTCTTTCCCTACACGACGCTCTTCCGATCTNNNCACTGCATCCGACCTTTCTTTTC

AGCACACGTCTGAACTCTTTCCCTACACGACGCTCTTCCGATCTNNNCTGTAATCCCAGCTACTCAGGA

AGCACACGTCTGAACTCTTTCCCTACACGACGCTCTTCCGATCTNNNCTGTAATCCCAGCTACTCAGGA

AGCACACGTCTGAACTCTTTCCCTACACGACGCTCTTCCGATCTNNNGAACCGAGATCAAGTCATTGC

AGCACACGTCTGAACTCTTTCCCTACACGACGCTCTTCCGATCTNNNGAACCGAGATCAAGTCATTGC

AGCACACGTCTGAACTCTTTCCCTACACGACGCTCTTCCGATCTNNNAGTTGTCTAAACTGGCCCTTCA

AGCACACGTCTGAACTCTTTCCCTACACGACGCTCTTCCGATCTNNNAGTTGTCTAAACTGGCCCTTCA

AGCACACGTCTGAACTCTTTCCCTACACGACGCTCTTCCGATCTNNNTGTAGAAGTGGGTTTGGTCTC

AGCACACGTCTGAACTCTTTCCCTACACGACGCTCTTCCGATCTNNNTGTAGAAGTGGGTTTGGTCTC

GAGCACACGTCTGAACTCTTTCCCTACACGACGCTCTTCCGATCTNNNGGAGAAGCAAGAAAGAAGC

GAGCACACGTCTGAACTCTTTCCCTACACGACGCTCTTCCGATCTNNNGGAGAAGCAAGAAAGAAGC

AGCACACGTCTGAACTCTTTCCCTACACGACGCTCTTCCGATCTNNNTGTCCTGATTTCTGGCATCTTCT

AGCACACGTCTGAACTCTTTCCCTACACGACGCTCTTCCGATCTNNNTGTCCTGATTTCTGGCATCTTCT

AGCACACGTCTGAACTCTTTCCCTACACGACGCTCTTCCGATCTNNNCACCTGTAATCCCAGCTACTCGC
AGCACACGTCTGAACTCTTTCCCTACACGACGCTCTTCCGATCTNNNCACCTGTAATCCCAGCTACTCGC
GCACACGTCTGAACTCTTTCCCTACACGACGCTCTTCCGATCTNNNAGCCAGCTAACAACCTCACTAAAC
GCACACGTCTGAACTCTTTCCCTACACGACGCTCTTCCGATCTNNNAGCCAGCTAACAACCTCACTAAAC
GAGCACACGTCTGAACTCTTTCCCTACACGACGCTCTTCCGATCTNNNTTACAGAAATCTAGGCCTCCAA
GAGCACACGTCTGAACTCTTTCCCTACACGACGCTCTTCCGATCTNNNTTACAGAAATCTAGGCCTCCAA
AGCACACGTCTGAACTCTTTCCCTACACGACGCTCTTCCGATCTNNNGGCTCATTCCAGATCTCTCCTTT
AGCACACGTCTGAACTCTTTCCCTACACGACGCTCTTCCGATCTNNNGGCTCATTCCAGATCTCTCCTTT
AGCACACGTCTGAACTCTTTCCCTACACGACGCTCTTCCGATCTNNNGGACATCTGAATTGGAAAGGAA
AGCACACGTCTGAACTCTTTCCCTACACGACGCTCTTCCGATCTNNNGGACATCTGAATTGGAAAGGAA
GAGCACACGTCTGAACTCTTTCCCTACACGACGCTCTTCCGATCTNNNAGTGAGCAAATCCTCTCTATGC
GAGCACACGTCTGAACTCTTTCCCTACACGACGCTCTTCCGATCTNNNAGTGAGCAAATCCTCTCTATGC
AGCACACGTCTGAACTCTTTCCCTACACGACGCTCTTCCGATCTNNNTTGGAGAAGGGAGAAAAAGG
AGCACACGTCTGAACTCTTTCCCTACACGACGCTCTTCCGATCTNNNTTGGAGAAGGGAGAAAAAGG
AGCACACGTCTGAACTCTTTCCCTACACGACGCTCTTCCGATCTNNNTTTGTTTCCTGTGCTTTTGTGGT
AGCACACGTCTGAACTCTTTCCCTACACGACGCTCTTCCGATCTNNNTTTGTTTCCTGTGCTTTTGTGGT
GAGCACACGTCTGAACTCTTTCCCTACACGACGCTCTTCCGATCTNNNGTTGCAGTGAACCGAGATCAC
GAGCACACGTCTGAACTCTTTCCCTACACGACGCTCTTCCGATCTNNNGTTGCAGTGAACCGAGATCAC
AGCACACGTCTGAACTCTTTCCCTACACGACGCTCTTCCGATCTNNNTCAGACAACACGTTCTGACCTAC
AGCACACGTCTGAACTCTTTCCCTACACGACGCTCTTCCGATCTNNNTCAGACAACACGTTCTGACCTAC
GAGCACACGTCTGAACTCTTTCCCTACACGACGCTCTTCCGATCTNNNGCTCATGCTTGTAATCCCAGA
GAGCACACGTCTGAACTCTTTCCCTACACGACGCTCTTCCGATCTNNNGCTCATGCTTGTAATCCCAGA
GAGCACACGTCTGAACTCTTTCCCTACACGACGCTCTTCCGATCTNNNCACCTCCTTCCTCTGCTTAGCCA
GAGCACACGTCTGAACTCTTTCCCTACACGACGCTCTTCCGATCTNNNCACCTCCTTCCTCTGCTTAGCCA
AGCACACGTCTGAACTCTTTCCCTACACGACGCTCTTCCGATCTNNNGCCACACCCAGCTTTTTTTTTTC
AGCACACGTCTGAACTCTTTCCCTACACGACGCTCTTCCGATCTNNNGCCACACCCAGCTTTTTTTTTTC
AGCACACGTCTGAACTCTTTCCCTACACGACGCTCTTCCGATCTNNNGGTGACAGAGCAAGACTTCATC
AGCACACGTCTGAACTCTTTCCCTACACGACGCTCTTCCGATCTNNNGGTGACAGAGCAAGACTTCATC
AGCACACGTCTGAACTCTTTCCCTACACGACGCTCTTCCGATCTNNNTACAAAAAATTAGCCGAGCGTG
AGCACACGTCTGAACTCTTTCCCTACACGACGCTCTTCCGATCTNNNTACAAAAAATTAGCCGAGCGTG
GCACACGTCTGAACTCTTTCCCTACACGACGCTCTTCCGATCTNNNCATGAGAATTGCTTGAACCTGGC
GCACACGTCTGAACTCTTTCCCTACACGACGCTCTTCCGATCTNNNCATGAGAATTGCTTGAACCTGGC
GCACACGTCTGAACTCTTTCCCTACACGACGCTCTTCCGATCTNNNGGAGAAATGGGAGACAGGTTAG
GCACACGTCTGAACTCTTTCCCTACACGACGCTCTTCCGATCTNNNGGAGAAATGGGAGACAGGTTAG
AGAGCACACGTCTGAACTCTTTCCCTACACGACGCTCTTCCGATCTNNNAAGCTACCTGCACTCATATGT
AGAGCACACGTCTGAACTCTTTCCCTACACGACGCTCTTCCGATCTNNNAAGCTACCTGCACTCATATGT
GAGCACACGTCTGAACTCTTTCCCTACACGACGCTCTTCCGATCTNNNAGTAGTCCCCCCTCTGAGTAAA
GAGCACACGTCTGAACTCTTTCCCTACACGACGCTCTTCCGATCTNNNAGTAGTCCCCCCTCTGAGTAAA
GAGCACACGTCTGAACTCTTTCCCTACACGACGCTCTTCCGATCTNNNCTGGGTTGCTGTTGAAGTTTT
GAGCACACGTCTGAACTCTTTCCCTACACGACGCTCTTCCGATCTNNNCTGGGTTGCTGTTGAAGTTTT
AGCACACGTCTGAACTCTTTCCCTACACGACGCTCTTCCGATCTNNNATTATAAAGGGCTTGGGGCTGT
AGCACACGTCTGAACTCTTTCCCTACACGACGCTCTTCCGATCTNNNATTATAAAGGGCTTGGGGCTGT
AGAGCACACGTCTGAACTCTTTCCCTACACGACGCTCTTCCGATCTNNNGAATCCCCAGCTCTCCTGCTA

AGAGCACACGTCTGAACTCTTTCCCTACACGACGCTCTTCCGATCTNNNGAATCCCCAGCTCTCCTGCTA

ACACGTCTGAACTCTTTCCCTACACGACGCTCTTCCGATCTNNNCAGATTTAGGGTGGATCTGCCTTCA

ACACGTCTGAACTCTTTCCCTACACGACGCTCTTCCGATCTNNNCAGATTTAGGGTGGATCTGCCTTCA

AGCACACGTCTGAACTCTTTCCCTACACGACGCTCTTCCGATCTNNNCTACCTGACTACTGGACATGGT

AGCACACGTCTGAACTCTTTCCCTACACGACGCTCTTCCGATCTNNNCTACCTGACTACTGGACATGGT

AGCACACGTCTGAACTCTTTCCCTACACGACGCTCTTCCGATCTNNNCTTTGATGATCTTCAGTATCAG

AGCACACGTCTGAACTCTTTCCCTACACGACGCTCTTCCGATCTNNNCTTTGATGATCTTCAGTATCAG

GAGCACACGTCTGAACTCTTTCCCTACACGACGCTCTTCCGATCTNNNACGATTGGTCAGTTTACCCAA

GAGCACACGTCTGAACTCTTTCCCTACACGACGCTCTTCCGATCTNNNACGATTGGTCAGTTTACCCAA

GAGCACACGTCTGAACTCTTTCCCTACACGACGCTCTTCCGATCTNNNGCTTTTAGAAAAAGGCCTTG

GAGCACACGTCTGAACTCTTTCCCTACACGACGCTCTTCCGATCTNNNGCTTTTAGAAAAAGGCCTTG

AGCACACGTCTGAACTCTTTCCCTACACGACGCTCTTCCGATCTNNNATGAAACTCAGTGGCAGATCC

AGCACACGTCTGAACTCTTTCCCTACACGACGCTCTTCCGATCTNNNATGAAACTCAGTGGCAGATCC

AGCACACGTCTGAACTCTTTCCCTACACGACGCTCTTCCGATCTNNNGTCAAGACCCATCAGTGTACT

AGCACACGTCTGAACTCTTTCCCTACACGACGCTCTTCCGATCTNNNGTCAAGACCCATCAGTGTACT

GAGCACACGTCTGAACTCTTTCCCTACACGACGCTCTTCCGATCTNNNTCACCTGATCCACAAATACTG

GAGCACACGTCTGAACTCTTTCCCTACACGACGCTCTTCCGATCTNNNTCACCTGATCCACAAATACTG

GAGCACACGTCTGAACTCTTTCCCTACACGACGCTCTTCCGATCTNNNTGGTGACTTGAGAGGTTTGA

GAGCACACGTCTGAACTCTTTCCCTACACGACGCTCTTCCGATCTNNNTGGTGACTTGAGAGGTTTGA

GCACACGTCTGAACTCTTTCCCTACACGACGCTCTTCCGATCTNNNTAACTCGTTATCTCCCATCGTCCC

GCACACGTCTGAACTCTTTCCCTACACGACGCTCTTCCGATCTNNNTAACTCGTTATCTCCCATCGTCCC

AGCACACGTCTGAACTCTTTCCCTACACGACGCTCTTCCGATCTNNNAAGAACTTTCAGGGTTTTGTGC

AGCACACGTCTGAACTCTTTCCCTACACGACGCTCTTCCGATCTNNNAAGAACTTTCAGGGTTTTGTGC

GAGCACACGTCTGAACTCTTTCCCTACACGACGCTCTTCCGATCTNNNCACCTGAAGTCTCAGCTACTC

GAGCACACGTCTGAACTCTTTCCCTACACGACGCTCTTCCGATCTNNNCACCTGAAGTCTCAGCTACTC

GAGCACACGTCTGAACTCTTTCCCTACACGACGCTCTTCCGATCTNNNTGGCCTACACAAATGTACCCA

GAGCACACGTCTGAACTCTTTCCCTACACGACGCTCTTCCGATCTNNNTGGCCTACACAAATGTACCCA

GAGCACACGTCTGAACTCTTTCCCTACACGACGCTCTTCCGATCTNNNGTAGAAGCTGCAGTGAGTT

GAGCACACGTCTGAACTCTTTCCCTACACGACGCTCTTCCGATCTNNNGTAGAAGCTGCAGTGAGTT

GAGCACACGTCTGAACTCTTTCCCTACACGACGCTCTTCCGATCTNNNGATCATGCCATTACACTTCAGC

GAGCACACGTCTGAACTCTTTCCCTACACGACGCTCTTCCGATCTNNNGATCATGCCATTACACTTCAGC

AGCACACGTCTGAACTCTTTCCCTACACGACGCTCTTCCGATCTNNNCTGAGGGCTGTAAGTAGGAGT

AGCACACGTCTGAACTCTTTCCCTACACGACGCTCTTCCGATCTNNNCTGAGGGCTGTAAGTAGGAGT

GAGCACACGTCTGAACTCTTTCCCTACACGACGCTCTTCCGATCTNNNCACCTGTAATCCCAGCTACTCA

GAGCACACGTCTGAACTCTTTCCCTACACGACGCTCTTCCGATCTNNNCACCTGTAATCCCAGCTACTCA

GAGCACACGTCTGAACTCTTTCCCTACACGACGCTCTTCCGATCTNNNCTGAGGCAGGAGAATCACTT

GAGCACACGTCTGAACTCTTTCCCTACACGACGCTCTTCCGATCTNNNCTGAGGCAGGAGAATCACTT

GAGCACACGTCTGAACTCTTTCCCTACACGACGCTCTTCCGATCTNNNGAGCTTGCAGTGAGCCAAGAT

GAGCACACGTCTGAACTCTTTCCCTACACGACGCTCTTCCGATCTNNNGAGCTTGCAGTGAGCCAAGAT

GAGCACACGTCTGAACTCTTTCCCTACACGACGCTCTTCCGATCTNNNGTGACAAAGCAAGACTCCAT

GAGCACACGTCTGAACTCTTTCCCTACACGACGCTCTTCCGATCTNNNGTGACAAAGCAAGACTCCAT

AGCACACGTCTGAACTCTTTCCCTACACGACGCTCTTCCGATCTNNNCTACTCAGAAGCTGATGTGGGA

AGCACACGTCTGAACTCTTTCCCTACACGACGCTCTTCCGATCTNNNCTACTCAGAAGCTGATGTGGGA

AGCACACGTCTGAACTCTTTCCCTACACGACGCTCTTCCGATCTNNNCCTTCTCCCCATAAAAATGCTCT
AGCACACGTCTGAACTCTTTCCCTACACGACGCTCTTCCGATCTNNNCCTTCTCCCCATAAAAATGCTCT
AGCACACGTCTGAACTCTTTCCCTACACGACGCTCTTCCGATCTNNNCAGGAGCAATATGTGTTTGTGG
AGCACACGTCTGAACTCTTTCCCTACACGACGCTCTTCCGATCTNNNCAGGAGCAATATGTGTTTGTGG
GAGCACACGTCTGAACTCTTTCCCTACACGACGCTCTTCCGATCTNNNAAAATGCAAAAAATTAGCCGG
GAGCACACGTCTGAACTCTTTCCCTACACGACGCTCTTCCGATCTNNNAAAATGCAAAAAATTAGCCGG
GAGCACACGTCTGAACTCTTTCCCTACACGACGCTCTTCCGATCTNNNCTCAGGAGTTTGAGACCAGCC
GAGCACACGTCTGAACTCTTTCCCTACACGACGCTCTTCCGATCTNNNCTCAGGAGTTTGAGACCAGCC
GAGCACACGTCTGAACTCTTTCCCTACACGACGCTCTTCCGATCTNNNCTGAAATAAGGTGAGGATGCC
GAGCACACGTCTGAACTCTTTCCCTACACGACGCTCTTCCGATCTNNNCTGAAATAAGGTGAGGATGCC
GAGCACACGTCTGAACTCTTTCCCTACACGACGCTCTTCCGATCTNNNGTATTGCCAGCTATTTTGGAG
GAGCACACGTCTGAACTCTTTCCCTACACGACGCTCTTCCGATCTNNNGTATTGCCAGCTATTTTGGAG
GAGCACACGTCTGAACTCTTTCCCTACACGACGCTCTTCCGATCTNNNGCAGTCTTGTAATCACATGAG
GAGCACACGTCTGAACTCTTTCCCTACACGACGCTCTTCCGATCTNNNGCAGTCTTGTAATCACATGAG
AGCACACGTCTGAACTCTTTCCCTACACGACGCTCTTCCGATCTNNNTCAGAGACTTTGAGCATGGAAG
AGCACACGTCTGAACTCTTTCCCTACACGACGCTCTTCCGATCTNNNTCAGAGACTTTGAGCATGGAAG
AGCACACGTCTGAACTCTTTCCCTACACGACGCTCTTCCGATCTNNNTATGGCCATTCTTGCAGGAGTA
AGCACACGTCTGAACTCTTTCCCTACACGACGCTCTTCCGATCTNNNTATGGCCATTCTTGCAGGAGTA
GAGCACACGTCTGAACTCTTTCCCTACACGACGCTCTTCCGATCTNNNATTCACCCCAGAGTTAGTTGCA
GAGCACACGTCTGAACTCTTTCCCTACACGACGCTCTTCCGATCTNNNATTCACCCCAGAGTTAGTTGCA
CACACGTCTGAACTCTTTCCCTACACGACGCTCTTCCGATCTNNNCCATTTGCTTCAGTAGGATCTGCCA
CACACGTCTGAACTCTTTCCCTACACGACGCTCTTCCGATCTNNNCCATTTGCTTCAGTAGGATCTGCCA
AGCACACGTCTGAACTCTTTCCCTACACGACGCTCTTCCGATCTNNNACACATTGATTTAGAGCTGGGG
AGCACACGTCTGAACTCTTTCCCTACACGACGCTCTTCCGATCTNNNACACATTGATTTAGAGCTGGGG
GAGCACACGTCTGAACTCTTTCCCTACACGACGCTCTTCCGATCTNNNACTTTCATTCTGTATGCTTTGGT
GAGCACACGTCTGAACTCTTTCCCTACACGACGCTCTTCCGATCTNNNACTTTCATTCTGTATGCTTTGGT
CACACGTCTGAACTCTTTCCCTACACGACGCTCTTCCGATCTNNNGCAGGTTTGTAGAGTGATATATAC
CACACGTCTGAACTCTTTCCCTACACGACGCTCTTCCGATCTNNNGCAGGTTTGTAGAGTGATATATAC
GAGCACACGTCTGAACTCTTTCCCTACACGACGCTCTTCCGATCTNNNAGGAGAGTTCTGCTTTTAGCT
GAGCACACGTCTGAACTCTTTCCCTACACGACGCTCTTCCGATCTNNNAGGAGAGTTCTGCTTTTAGCT
GAGCACACGTCTGAACTCTTTCCCTACACGACGCTCTTCCGATCTNNNCCCAGGCATTCAGCTGGATAT
GAGCACACGTCTGAACTCTTTCCCTACACGACGCTCTTCCGATCTNNNCCCAGGCATTCAGCTGGATAT
AGCACACGTCTGAACTCTTTCCCTACACGACGCTCTTCCGATCTNNNTTCATTCCTGTAATCCCAGTGGC
AGCACACGTCTGAACTCTTTCCCTACACGACGCTCTTCCGATCTNNNTTCATTCCTGTAATCCCAGTGGC
AGCACACGTCTGAACTCTTTCCCTACACGACGCTCTTCCGATCTNNNTTGATGTGCTCCTATAGAAGGC
AGCACACGTCTGAACTCTTTCCCTACACGACGCTCTTCCGATCTNNNTTGATGTGCTCCTATAGAAGGC
GAGCACACGTCTGAACTCTTTCCCTACACGACGCTCTTCCGATCTNNNGTGTACCCAGTAGTCATTCAG
GAGCACACGTCTGAACTCTTTCCCTACACGACGCTCTTCCGATCTNNNGTGTACCCAGTAGTCATTCAG
GAGCACACGTCTGAACTCTTTCCCTACACGACGCTCTTCCGATCTNNNGGATAGTCAGTCTTCCATATTT
GAGCACACGTCTGAACTCTTTCCCTACACGACGCTCTTCCGATCTNNNGGATAGTCAGTCTTCCATATTT
AGCACACGTCTGAACTCTTTCCCTACACGACGCTCTTCCGATCTNNNTTCAAAACCAGCCTGGGTAAGAT
AGCACACGTCTGAACTCTTTCCCTACACGACGCTCTTCCGATCTNNNTTCAAAACCAGCCTGGGTAAGAT
GAGCACACGTCTGAACTCTTTCCCTACACGACGCTCTTCCGATCTNNNGACAAAATGAGACTCCACCTCC

AGCACACGTCTGAACTCTTTCCCTACACGACGCTCTTCCGATCTNNNGACAAAATGAGACTCCACCTC
GAGCACACGTCTGAACTCTTTCCCTACACGACGCTCTTCCGATCTNNNGCAAGACCCTGGCTCTACTAA
GAGCACACGTCTGAACTCTTTCCCTACACGACGCTCTTCCGATCTNNNGCAAGACCCTGGCTCTACTAA
AGCACACGTCTGAACTCTTTCCCTACACGACGCTCTTCCGATCTNNNGAAGGGGAGAAGGGAAAAG
AGCACACGTCTGAACTCTTTCCCTACACGACGCTCTTCCGATCTNNNGAAGGGGAGAAGGGAAAAG
GCACACGTCTGAACTCTTTCCCTACACGACGCTCTTCCGATCTNNNAATACAAAAAATTAGCCGGGCG
GCACACGTCTGAACTCTTTCCCTACACGACGCTCTTCCGATCTNNNAATACAAAAAATTAGCCGGGCG
GAGCACACGTCTGAACTCTTTCCCTACACGACGCTCTTCCGATCTNNNCCTACCTGAACTGCCCTTAC
GAGCACACGTCTGAACTCTTTCCCTACACGACGCTCTTCCGATCTNNNCCTACCTGAACTGCCCTTAC
AGCACACGTCTGAACTCTTTCCCTACACGACGCTCTTCCGATCTNNNCTCTGCACCTTATTCAGAGCT
AGCACACGTCTGAACTCTTTCCCTACACGACGCTCTTCCGATCTNNNCTCTGCACCTTATTCAGAGCT
GCACACGTCTGAACTCTTTCCCTACACGACGCTCTTCCGATCTNNNCACTTGTAGTCCCAGCTACTTG
GCACACGTCTGAACTCTTTCCCTACACGACGCTCTTCCGATCTNNNCACTTGTAGTCCCAGCTACTTG
AGCACACGTCTGAACTCTTTCCCTACACGACGCTCTTCCGATCTNNNTAGCAAAGTAGGCAGCTCTTG
AGCACACGTCTGAACTCTTTCCCTACACGACGCTCTTCCGATCTNNNTAGCAAAGTAGGCAGCTCTTG
AGCACACGTCTGAACTCTTTCCCTACACGACGCTCTTCCGATCTNNNGCTTTTGATAAGCTCCTGACAC
AGCACACGTCTGAACTCTTTCCCTACACGACGCTCTTCCGATCTNNNGCTTTTGATAAGCTCCTGACAC
GAGCACACGTCTGAACTCTTTCCCTACACGACGCTCTTCCGATCTNNNATGGATGGGGATGAGGAG
GAGCACACGTCTGAACTCTTTCCCTACACGACGCTCTTCCGATCTNNNATGGATGGGGATGAGGAG
AGCACACGTCTGAACTCTTTCCCTACACGACGCTCTTCCGATCTNNNTCATGGGAGAAAGGAAACTG
AGCACACGTCTGAACTCTTTCCCTACACGACGCTCTTCCGATCTNNNTCATGGGAGAAAGGAAACTG
AGCACACGTCTGAACTCTTTCCCTACACGACGCTCTTCCGATCTNNNGCATGGACATTATCAAGCCTT
AGCACACGTCTGAACTCTTTCCCTACACGACGCTCTTCCGATCTNNNGCATGGACATTATCAAGCCTT
GAGCACACGTCTGAACTCTTTCCCTACACGACGCTCTTCCGATCTNNNGAGCTTGCAGTGAGACGAG
GAGCACACGTCTGAACTCTTTCCCTACACGACGCTCTTCCGATCTNNNGAGCTTGCAGTGAGACGAG
AGCACACGTCTGAACTCTTTCCCTACACGACGCTCTTCCGATCTNNNTGTAATTTCCTCCTGAGTAGCG
AGCACACGTCTGAACTCTTTCCCTACACGACGCTCTTCCGATCTNNNTGTAATTTCCTCCTGAGTAGCG
GAGCACACGTCTGAACTCTTTCCCTACACGACGCTCTTCCGATCTNNNGACAGAGCTCTCAAATCGAG
GAGCACACGTCTGAACTCTTTCCCTACACGACGCTCTTCCGATCTNNNGACAGAGCTCTCAAATCGAG
AGCACACGTCTGAACTCTTTCCCTACACGACGCTCTTCCGATCTNNNGTTTCAAGCGATTCTCGTGTCT
AGCACACGTCTGAACTCTTTCCCTACACGACGCTCTTCCGATCTNNNGTTTCAAGCGATTCTCGTGTCT
AGCACACGTCTGAACTCTTTCCCTACACGACGCTCTTCCGATCTNNNCCTATAGTCCCAGCTACTCAGG
AGCACACGTCTGAACTCTTTCCCTACACGACGCTCTTCCGATCTNNNCCTATAGTCCCAGCTACTCAGG
AGCACACGTCTGAACTCTTTCCCTACACGACGCTCTTCCGATCTNNNGGTGTTTGATTACATGAATAA
AGCACACGTCTGAACTCTTTCCCTACACGACGCTCTTCCGATCTNNNGGTGTTTGATTACATGAATAA
GAGCACACGTCTGAACTCTTTCCCTACACGACGCTCTTCCGATCTNNNCACATATATGCATGCACACAC
GAGCACACGTCTGAACTCTTTCCCTACACGACGCTCTTCCGATCTNNNCACATATATGCATGCACACAC
AAGAGCACACGTCTGAACTCTTTCCCTACACGACGCTCTTCCGATCTNNNTGTCCCATTCACTGCTGG
AAGAGCACACGTCTGAACTCTTTCCCTACACGACGCTCTTCCGATCTNNNTGTCCCATTCACTGCTGG
CACACGTCTGAACTCTTTCCCTACACGACGCTCTTCCGATCTNNNGCTGAGAGTAAGGACCAGACAAA
CACACGTCTGAACTCTTTCCCTACACGACGCTCTTCCGATCTNNNGCTGAGAGTAAGGACCAGACAAA
GCACACGTCTGAACTCTTTCCCTACACGACGCTCTTCCGATCTNNNGCAGACAGAAATCACACCTCAA
GCACACGTCTGAACTCTTTCCCTACACGACGCTCTTCCGATCTNNNGCAGACAGAAATCACACCTCAA

GCACACGTCTGAACTCTTTCCCTACACGACGCTCTTCCGATCTNNNTTCTCGGTACCCACTCTTAGAGA
GCACACGTCTGAACTCTTTCCCTACACGACGCTCTTCCGATCTNNNTTCTCGGTACCCACTCTTAGAGA
AGCACACGTCTGAACTCTTTCCCTACACGACGCTCTTCCGATCTNNNAATCACAAAAACTGCACTCCAC
AGCACACGTCTGAACTCTTTCCCTACACGACGCTCTTCCGATCTNNNAATCACAAAAACTGCACTCCAC
GAGCACACGTCTGAACTCTTTCCCTACACGACGCTCTTCCGATCTNNNTGGAGAAGACAATGTGTGT
GAGCACACGTCTGAACTCTTTCCCTACACGACGCTCTTCCGATCTNNNTGGAGAAGACAATGTGTGT
AGAGCACACGTCTGAACTCTTTCCCTACACGACGCTCTTCCGATCTNNNAGCCAAATAGTTCAGACAG
AGAGCACACGTCTGAACTCTTTCCCTACACGACGCTCTTCCGATCTNNNAGCCAAATAGTTCAGACAG
AGCACACGTCTGAACTCTTTCCCTACACGACGCTCTTCCGATCTNNNTGTGCTGGCATCATAGTTCTCA
AGCACACGTCTGAACTCTTTCCCTACACGACGCTCTTCCGATCTNNNTGTGCTGGCATCATAGTTCTCA
AGCACACGTCTGAACTCTTTCCCTACACGACGCTCTTCCGATCTNNNGCAACAGAGGAATGGTGAAC
AGCACACGTCTGAACTCTTTCCCTACACGACGCTCTTCCGATCTNNNGCAACAGAGGAATGGTGAAC
GAGCACACGTCTGAACTCTTTCCCTACACGACGCTCTTCCGATCTNNNTGTTCATCAGGGATATTGAC
GAGCACACGTCTGAACTCTTTCCCTACACGACGCTCTTCCGATCTNNNTGTTCATCAGGGATATTGAC
GAGCACACGTCTGAACTCTTTCCCTACACGACGCTCTTCCGATCTNNNGTGCCTCAGGCTGCTATCTT
GAGCACACGTCTGAACTCTTTCCCTACACGACGCTCTTCCGATCTNNNGTGCCTCAGGCTGCTATCTT
GCACACGTCTGAACTCTTTCCCTACACGACGCTCTTCCGATCTNNNAATTTAGCTACTCGGGAGGCTG
GCACACGTCTGAACTCTTTCCCTACACGACGCTCTTCCGATCTNNNAATTTAGCTACTCGGGAGGCTG
GCACACGTCTGAACTCTTTCCCTACACGACGCTCTTCCGATCTNNNACTCTGTCCATTGCTCTCCATTTC
GCACACGTCTGAACTCTTTCCCTACACGACGCTCTTCCGATCTNNNACTCTGTCCATTGCTCTCCATTTC
AGCACACGTCTGAACTCTTTCCCTACACGACGCTCTTCCGATCTNNNGGGAGTCAGGAACTTTGGGAT
AGCACACGTCTGAACTCTTTCCCTACACGACGCTCTTCCGATCTNNNGGGAGTCAGGAACTTTGGGAT
GAGCACACGTCTGAACTCTTTCCCTACACGACGCTCTTCCGATCTNNNGCACTTTAGTTATGGAAGGC
GAGCACACGTCTGAACTCTTTCCCTACACGACGCTCTTCCGATCTNNNGCACTTTAGTTATGGAAGGC
AGCACACGTCTGAACTCTTTCCCTACACGACGCTCTTCCGATCTNNNGGAGACTGAGGCATGAGAAT
AGCACACGTCTGAACTCTTTCCCTACACGACGCTCTTCCGATCTNNNGGAGACTGAGGCATGAGAAT
AGCACACGTCTGAACTCTTTCCCTACACGACGCTCTTCCGATCTNNNAGAGGCAGGAGAATCACTTG
AGCACACGTCTGAACTCTTTCCCTACACGACGCTCTTCCGATCTNNNAGAGGCAGGAGAATCACTTG
AGCACACGTCTGAACTCTTTCCCTACACGACGCTCTTCCGATCTNNNAGTGGAATTGCTGGATCATAT
AGCACACGTCTGAACTCTTTCCCTACACGACGCTCTTCCGATCTNNNAGTGGAATTGCTGGATCATAT
AGCACACGTCTGAACTCTTTCCCTACACGACGCTCTTCCGATCTNNNAAACACAGTAGCTTCTCTTCCA
AGCACACGTCTGAACTCTTTCCCTACACGACGCTCTTCCGATCTNNNAAACACAGTAGCTTCTCTTCCA
GCACACGTCTGAACTCTTTCCCTACACGACGCTCTTCCGATCTNNNCTTCATCCCCACGGTGCTTCCATC
GCACACGTCTGAACTCTTTCCCTACACGACGCTCTTCCGATCTNNNCTTCATCCCCACGGTGCTTCCATC
AGCACACGTCTGAACTCTTTCCCTACACGACGCTCTTCCGATCTNNNGGCAACATAGCAAAACCTCAT
AGCACACGTCTGAACTCTTTCCCTACACGACGCTCTTCCGATCTNNNGGCAACATAGCAAAACCTCAT
GCACACGTCTGAACTCTTTCCCTACACGACGCTCTTCCGATCTNNNTGTGGCTTTGTGGACTAAGTTT
GCACACGTCTGAACTCTTTCCCTACACGACGCTCTTCCGATCTNNNTGTGGCTTTGTGGACTAAGTTT
GAGCACACGTCTGAACTCTTTCCCTACACGACGCTCTTCCGATCTNNNCCTTCAGCCAACAAATTACTC
GAGCACACGTCTGAACTCTTTCCCTACACGACGCTCTTCCGATCTNNNCCTTCAGCCAACAAATTACTC
AGAGCACACGTCTGAACTCTTTCCCTACACGACGCTCTTCCGATCTNNNCGACAGAGTGAAAGTCCAT
AGAGCACACGTCTGAACTCTTTCCCTACACGACGCTCTTCCGATCTNNNCGACAGAGTGAAAGTCCAT
AGCACACGTCTGAACTCTTTCCCTACACGACGCTCTTCCGATCTNNNTGGATTTGTTTGCTGCTTCTG

GAGCACACGTCTGAACTCTTTCCCTACACGACGCTCTTCCGATCTNNNTGGATTTGTTTGCTGCTTCTGT
GAGCACACGTCTGAACTCTTTCCCTACACGACGCTCTTCCGATCTNNNACCCCAAAGTGATACATCATTC
GAGCACACGTCTGAACTCTTTCCCTACACGACGCTCTTCCGATCTNNNACCCCAAAGTGATACATCATTC
AGCACACGTCTGAACTCTTTCCCTACACGACGCTCTTCCGATCTNNNTGAAGCCATCTGGACATACAGT
AGCACACGTCTGAACTCTTTCCCTACACGACGCTCTTCCGATCTNNNTGAAGCCATCTGGACATACAGT
GAGCACACGTCTGAACTCTTTCCCTACACGACGCTCTTCCGATCTNNNCAGCCTGGATCCTGAAGATACT
GAGCACACGTCTGAACTCTTTCCCTACACGACGCTCTTCCGATCTNNNCAGCCTGGATCCTGAAGATACT
GAGCACACGTCTGAACTCTTTCCCTACACGACGCTCTTCCGATCTNNNACATTTGAGTAAGTGCTTTGC
GAGCACACGTCTGAACTCTTTCCCTACACGACGCTCTTCCGATCTNNNACATTTGAGTAAGTGCTTTGC
AGCACACGTCTGAACTCTTTCCCTACACGACGCTCTTCCGATCTNNNGCCTCTGGAGTACCTAGTCTTTC
AGCACACGTCTGAACTCTTTCCCTACACGACGCTCTTCCGATCTNNNGCCTCTGGAGTACCTAGTCTTTC
AGCACACGTCTGAACTCTTTCCCTACACGACGCTCTTCCGATCTNNNTAGAAAAGTTAGGAAGACAGAA
AGCACACGTCTGAACTCTTTCCCTACACGACGCTCTTCCGATCTNNNTAGAAAAGTTAGGAAGACAGAA
AGCACACGTCTGAACTCTTTCCCTACACGACGCTCTTCCGATCTNNNCCTGGTACATCCCCTACATCAGTC
AGCACACGTCTGAACTCTTTCCCTACACGACGCTCTTCCGATCTNNNCCTGGTACATCCCCTACATCAGTC
AGCACACGTCTGAACTCTTTCCCTACACGACGCTCTTCCGATCTNNNGCCTCTTGAGTAGCTGGAACTA
AGCACACGTCTGAACTCTTTCCCTACACGACGCTCTTCCGATCTNNNGCCTCTTGAGTAGCTGGAACTA
AGAGCACACGTCTGAACTCTTTCCCTACACGACGCTCTTCCGATCTNNNACCCAACTCCTATGCAAACTG
AGAGCACACGTCTGAACTCTTTCCCTACACGACGCTCTTCCGATCTNNNACCCAACTCCTATGCAAACTG
GAGCACACGTCTGAACTCTTTCCCTACACGACGCTCTTCCGATCTNNNGAGGCTGAGGCAGGAGAATC
GAGCACACGTCTGAACTCTTTCCCTACACGACGCTCTTCCGATCTNNNGAGGCTGAGGCAGGAGAATC
GAGCACACGTCTGAACTCTTTCCCTACACGACGCTCTTCCGATCTNNNTGAGGTAGGAGAATCGCTTG
GAGCACACGTCTGAACTCTTTCCCTACACGACGCTCTTCCGATCTNNNTGAGGTAGGAGAATCGCTTG
AGCACACGTCTGAACTCTTTCCCTACACGACGCTCTTCCGATCTNNNTACGAGCTACTTAGTGCAATCC
AGCACACGTCTGAACTCTTTCCCTACACGACGCTCTTCCGATCTNNNTACGAGCTACTTAGTGCAATCC
AGCACACGTCTGAACTCTTTCCCTACACGACGCTCTTCCGATCTNNNTGCTATTGCACACTTAGTATGCA
AGCACACGTCTGAACTCTTTCCCTACACGACGCTCTTCCGATCTNNNTGCTATTGCACACTTAGTATGCA
GCACACGTCTGAACTCTTTCCCTACACGACGCTCTTCCGATCTNNNGGCAATATTTTAACCATACTCAACA
GCACACGTCTGAACTCTTTCCCTACACGACGCTCTTCCGATCTNNNGGCAATATTTTAACCATACTCAACA
GCACACGTCTGAACTCTTTCCCTACACGACGCTCTTCCGATCTNNNTAAACACACACACACACACACACC
GCACACGTCTGAACTCTTTCCCTACACGACGCTCTTCCGATCTNNNTAAACACACACACACACACACACC
GAGCACACGTCTGAACTCTTTCCCTACACGACGCTCTTCCGATCTNNNATCTGCAGAAGAGGGGTGTG
GAGCACACGTCTGAACTCTTTCCCTACACGACGCTCTTCCGATCTNNNATCTGCAGAAGAGGGGTGTG
AGCACACGTCTGAACTCTTTCCCTACACGACGCTCTTCCGATCTNNNGCCTGTAGTCCCAGCTACTTGCA
AGCACACGTCTGAACTCTTTCCCTACACGACGCTCTTCCGATCTNNNGCCTGTAGTCCCAGCTACTTGCA
AGCACACGTCTGAACTCTTTCCCTACACGACGCTCTTCCGATCTNNNCCGACCCGGTCTTTCTATTTTCTC
AGCACACGTCTGAACTCTTTCCCTACACGACGCTCTTCCGATCTNNNCCGACCCGGTCTTTCTATTTTCTC
AGCACACGTCTGAACTCTTTCCCTACACGACGCTCTTCCGATCTNNNGAGCAAGACTCCGTCACAAAAA
AGCACACGTCTGAACTCTTTCCCTACACGACGCTCTTCCGATCTNNNGAGCAAGACTCCGTCACAAAAA
AGCACACGTCTGAACTCTTTCCCTACACGACGCTCTTCCGATCTNNNAAAGAGAGAGAGAGAGAGAGGG
AGCACACGTCTGAACTCTTTCCCTACACGACGCTCTTCCGATCTNNNAAAGAGAGAGAGAGAGAGAGGG
GAGCACACGTCTGAACTCTTTCCCTACACGACGCTCTTCCGATCTNNNAAATGAGATGAAACCGTGGAA
GAGCACACGTCTGAACTCTTTCCCTACACGACGCTCTTCCGATCTNNNAAATGAGATGAAACCGTGGAA

GAGCACACGTCTGAACTCTTTCCCTACACGACGCTCTTCCGATCTNNNCTGTAATCCCAGCTACTCAGG
GAGCACACGTCTGAACTCTTTCCCTACACGACGCTCTTCCGATCTNNNCTGTAATCCCAGCTACTCAGG
AGCACACGTCTGAACTCTTTCCCTACACGACGCTCTTCCGATCTNNNTTGTTTGCTTGTTGACCAGTTG
AGCACACGTCTGAACTCTTTCCCTACACGACGCTCTTCCGATCTNNNTTGTTTGCTTGTTGACCAGTTG
AGCACACGTCTGAACTCTTTCCCTACACGACGCTCTTCCGATCTNNNCTGTTCTATCTGGGGAGGGGA
AGCACACGTCTGAACTCTTTCCCTACACGACGCTCTTCCGATCTNNNCTGTTCTATCTGGGGAGGGGA
ACACGTCTGAACTCTTTCCCTACACGACGCTCTTCCGATCTNNNGTCCTCGTCAATATGGCGGATCCAT
ACACGTCTGAACTCTTTCCCTACACGACGCTCTTCCGATCTNNNGTCCTCGTCAATATGGCGGATCCAT
GAGCACACGTCTGAACTCTTTCCCTACACGACGCTCTTCCGATCTNNNTCTGATGGAGAGAACTTGGT
GAGCACACGTCTGAACTCTTTCCCTACACGACGCTCTTCCGATCTNNNTCTGATGGAGAGAACTTGGT
GCACACGTCTGAACTCTTTCCCTACACGACGCTCTTCCGATCTNNNAGGACAATGAACTCTTTAGAAT
GCACACGTCTGAACTCTTTCCCTACACGACGCTCTTCCGATCTNNNAGGACAATGAACTCTTTAGAAT
GAGCACACGTCTGAACTCTTTCCCTACACGACGCTCTTCCGATCTNNNGAGAATCGCTTGAACCCAGGA
GAGCACACGTCTGAACTCTTTCCCTACACGACGCTCTTCCGATCTNNNGAGAATCGCTTGAACCCAGGA
GCACACGTCTGAACTCTTTCCCTACACGACGCTCTTCCGATCTNNNTCAGACTCCCAAGTAGCTAGGAT
GCACACGTCTGAACTCTTTCCCTACACGACGCTCTTCCGATCTNNNTCAGACTCCCAAGTAGCTAGGAT
AGCACACGTCTGAACTCTTTCCCTACACGACGCTCTTCCGATCTNNNAGGCGGCATATGATAGTGACAA
AGCACACGTCTGAACTCTTTCCCTACACGACGCTCTTCCGATCTNNNAGGCGGCATATGATAGTGACAA
AGCACACGTCTGAACTCTTTCCCTACACGACGCTCTTCCGATCTNNNTGGCTGACTAGTACTCCATTGT
AGCACACGTCTGAACTCTTTCCCTACACGACGCTCTTCCGATCTNNNTGGCTGACTAGTACTCCATTGT
GAGCACACGTCTGAACTCTTTCCCTACACGACGCTCTTCCGATCTNNNTTGTAAGGCCACTGAGGTTCT
GAGCACACGTCTGAACTCTTTCCCTACACGACGCTCTTCCGATCTNNNTTGTAAGGCCACTGAGGTTCT
GAGCACACGTCTGAACTCTTTCCCTACACGACGCTCTTCCGATCTNNNTTTTTTTTGAGACAGAGTCCC
GAGCACACGTCTGAACTCTTTCCCTACACGACGCTCTTCCGATCTNNNTTTTTTTTGAGACAGAGTCCC
GAGCACACGTCTGAACTCTTTCCCTACACGACGCTCTTCCGATCTNNNGAGTATCATAAGGGACCCTG
GAGCACACGTCTGAACTCTTTCCCTACACGACGCTCTTCCGATCTNNNGAGTATCATAAGGGACCCTG
GAGCACACGTCTGAACTCTTTCCCTACACGACGCTCTTCCGATCTNNNTATAGCCAAGGAGCTGGAAA
GAGCACACGTCTGAACTCTTTCCCTACACGACGCTCTTCCGATCTNNNTATAGCCAAGGAGCTGGAAA
AGCACACGTCTGAACTCTTTCCCTACACGACGCTCTTCCGATCTNNNCCGACCTCTTGAAGCTTTTCTTT
AGCACACGTCTGAACTCTTTCCCTACACGACGCTCTTCCGATCTNNNCCGACCTCTTGAAGCTTTTCTTT
GCACACGTCTGAACTCTTTCCCTACACGACGCTCTTCCGATCTNNNGACTTCCGTGACCCTCTTATGGCA
GCACACGTCTGAACTCTTTCCCTACACGACGCTCTTCCGATCTNNNGACTTCCGTGACCCTCTTATGGCA
GAGCACACGTCTGAACTCTTTCCCTACACGACGCTCTTCCGATCTNNNTGGAAACAGAGAGAGACTCCA
GAGCACACGTCTGAACTCTTTCCCTACACGACGCTCTTCCGATCTNNNTGGAAACAGAGAGAGACTCCA
AGCACACGTCTGAACTCTTTCCCTACACGACGCTCTTCCGATCTNNNCATTGAGTTGGTGTGTGTGTG
AGCACACGTCTGAACTCTTTCCCTACACGACGCTCTTCCGATCTNNNCATTGAGTTGGTGTGTGTGTG
AGCACACGTCTGAACTCTTTCCCTACACGACGCTCTTCCGATCTNNNGATTCCCATTCTCACCACTCCTTC
AGCACACGTCTGAACTCTTTCCCTACACGACGCTCTTCCGATCTNNNGATTCCCATTCTCACCACTCCTTC
AGCACACGTCTGAACTCTTTCCCTACACGACGCTCTTCCGATCTNNNGTGGCCAGTTTCAGCTTGTATT
AGCACACGTCTGAACTCTTTCCCTACACGACGCTCTTCCGATCTNNNGTGGCCAGTTTCAGCTTGTATT
GAGCACACGTCTGAACTCTTTCCCTACACGACGCTCTTCCGATCTNNNGGAAAGGGAAACGTTGTACA
GAGCACACGTCTGAACTCTTTCCCTACACGACGCTCTTCCGATCTNNNGGAAAGGGAAACGTTGTACA
GCACACGTCTGAACTCTTTCCCTACACGACGCTCTTCCGATCTNNNAGCATTTTACTTGTGGCTTTCCC

AGCACACGTCTGAACTCTTTCCCTACACGACGCTCTTCCGATCTNNNAGCATTTTACTTGTGGCTTTCCCC

GAGCACACGTCTGAACTCTTTCCCTACACGACGCTCTTCCGATCTNNNGGCAAGAGAGAACCTTTTCAT

GAGCACACGTCTGAACTCTTTCCCTACACGACGCTCTTCCGATCTNNNGGCAAGAGAGAACCTTTTCAT

AGCACACGTCTGAACTCTTTCCCTACACGACGCTCTTCCGATCTNNNACACACACACACACACACAAATTC

AGCACACGTCTGAACTCTTTCCCTACACGACGCTCTTCCGATCTNNNACACACACACACACACACAAATTC

AGCACACGTCTGAACTCTTTCCCTACACGACGCTCTTCCGATCTNNNAGGGCCTTCTTTATCCATTCCAG

AGCACACGTCTGAACTCTTTCCCTACACGACGCTCTTCCGATCTNNNAGGGCCTTCTTTATCCATTCCAG

GAGCACACGTCTGAACTCTTTCCCTACACGACGCTCTTCCGATCTNNNAGGTAGGAGAATTGCTTGAA

GAGCACACGTCTGAACTCTTTCCCTACACGACGCTCTTCCGATCTNNNAGGTAGGAGAATTGCTTGAA

AGAGCACACGTCTGAACTCTTTCCCTACACGACGCTCTTCCGATCTNNNGCAGGTCCTAACAGAAGAAA

AGAGCACACGTCTGAACTCTTTCCCTACACGACGCTCTTCCGATCTNNNGCAGGTCCTAACAGAAGAAA

GAGCACACGTCTGAACTCTTTCCCTACACGACGCTCTTCCGATCTNNNTCACAAGGTCAGGAGATCAA

GAGCACACGTCTGAACTCTTTCCCTACACGACGCTCTTCCGATCTNNNTCACAAGGTCAGGAGATCAA

GAGCACACGTCTGAACTCTTTCCCTACACGACGCTCTTCCGATCTNNNGAGCAAACTGAAAACAGTGG

GAGCACACGTCTGAACTCTTTCCCTACACGACGCTCTTCCGATCTNNNGAGCAAACTGAAAACAGTGG

GAGCACACGTCTGAACTCTTTCCCTACACGACGCTCTTCCGATCTNNNCGACTTTTCCTGCTGAAATGAC

GAGCACACGTCTGAACTCTTTCCCTACACGACGCTCTTCCGATCTNNNCGACTTTTCCTGCTGAAATGAC

GCACACGTCTGAACTCTTTCCCTACACGACGCTCTTCCGATCTNNNGGGTCATCCTGGAAGATACCATC

GCACACGTCTGAACTCTTTCCCTACACGACGCTCTTCCGATCTNNNGGGTCATCCTGGAAGATACCATC

GAGCACACGTCTGAACTCTTTCCCTACACGACGCTCTTCCGATCTNNNACACCAAGCATCTGGAAATCTC

GAGCACACGTCTGAACTCTTTCCCTACACGACGCTCTTCCGATCTNNNACACCAAGCATCTGGAAATCTC

AGCACACGTCTGAACTCTTTCCCTACACGACGCTCTTCCGATCTNNNCTTGCAGTGAGCCGAGATCGCA

AGCACACGTCTGAACTCTTTCCCTACACGACGCTCTTCCGATCTNNNCTTGCAGTGAGCCGAGATCGCA

AGCACACGTCTGAACTCTTTCCCTACACGACGCTCTTCCGATCTNNNACCCTTTAGTACGGAATTGGGG

AGCACACGTCTGAACTCTTTCCCTACACGACGCTCTTCCGATCTNNNACCCTTTAGTACGGAATTGGGG

AGCACACGTCTGAACTCTTTCCCTACACGACGCTCTTCCGATCTNNNAGTGGATGGGTATATATGGTGT

AGCACACGTCTGAACTCTTTCCCTACACGACGCTCTTCCGATCTNNNAGTGGATGGGTATATATGGTGT

GAGCACACGTCTGAACTCTTTCCCTACACGACGCTCTTCCGATCTNNNAGGCAAGAGCAAACAAATTCG

GAGCACACGTCTGAACTCTTTCCCTACACGACGCTCTTCCGATCTNNNAGGCAAGAGCAAACAAATTCG

AGCACACGTCTGAACTCTTTCCCTACACGACGCTCTTCCGATCTNNNAAATGAGAACTTTACCCACAATC

AGCACACGTCTGAACTCTTTCCCTACACGACGCTCTTCCGATCTNNNAAATGAGAACTTTACCCACAATC

AGCACACGTCTGAACTCTTTCCCTACACGACGCTCTTCCGATCTNNNCTGAGGCAGGAGAATAGCTTG

AGCACACGTCTGAACTCTTTCCCTACACGACGCTCTTCCGATCTNNNCTGAGGCAGGAGAATAGCTTG

GAGCACACGTCTGAACTCTTTCCCTACACGACGCTCTTCCGATCTNNNCTGTCTGTGTCTCCTCTTCTGT

GAGCACACGTCTGAACTCTTTCCCTACACGACGCTCTTCCGATCTNNNCTGTCTGTGTCTCCTCTTCTGT

AGCACACGTCTGAACTCTTTCCCTACACGACGCTCTTCCGATCTNNNGATCTTTGCGATGCTTTGGGTT

AGCACACGTCTGAACTCTTTCCCTACACGACGCTCTTCCGATCTNNNGATCTTTGCGATGCTTTGGGTT

GAGCACACGTCTGAACTCTTTCCCTACACGACGCTCTTCCGATCTNNNATTTCTTACCATCTGGAGGCTG

GAGCACACGTCTGAACTCTTTCCCTACACGACGCTCTTCCGATCTNNNATTTCTTACCATCTGGAGGCTG

GAGCACACGTCTGAACTCTTTCCCTACACGACGCTCTTCCGATCTNNNCTGTAATCCCAGCTACTCAGG

GAGCACACGTCTGAACTCTTTCCCTACACGACGCTCTTCCGATCTNNNCTGTAATCCCAGCTACTCAGG

AGAGCACACGTCTGAACTCTTTCCCTACACGACGCTCTTCCGATCTNNNCACTCACAGTTTGGGTAGAG

AGAGCACACGTCTGAACTCTTTCCCTACACGACGCTCTTCCGATCTNNNCACTCACAGTTTGGGTAGAG

AGAGCACACGTCTGAACTCTTTCCCTACACGACGCTCTTCCGATCTNNNTCAGTCCTGCCTAGAAATGT

AGAGCACACGTCTGAACTCTTTCCCTACACGACGCTCTTCCGATCTNNNTCAGTCCTGCCTAGAAATGT

GAGCACACGTCTGAACTCTTTCCCTACACGACGCTCTTCCGATCTNNNCATCTGACCTGTTGTGCTCACT

GAGCACACGTCTGAACTCTTTCCCTACACGACGCTCTTCCGATCTNNNCATCTGACCTGTTGTGCTCACT

GCACACGTCTGAACTCTTTCCCTACACGACGCTCTTCCGATCTNNNAGTGCTTTTTCCTTCTGAAGCATC

GCACACGTCTGAACTCTTTCCCTACACGACGCTCTTCCGATCTNNNAGTGCTTTTTCCTTCTGAAGCATC

AGAGCACACGTCTGAACTCTTTCCCTACACGACGCTCTTCCGATCTNNNTTGGAGTGTGCTCTCTCTCTC

AGAGCACACGTCTGAACTCTTTCCCTACACGACGCTCTTCCGATCTNNNTTGGAGTGTGCTCTCTCTCTC

AGAGCACACGTCTGAACTCTTTCCCTACACGACGCTCTTCCGATCTNNNGATAGGAATGAGATTGCGG

AGAGCACACGTCTGAACTCTTTCCCTACACGACGCTCTTCCGATCTNNNGATAGGAATGAGATTGCGG

GAGCACACGTCTGAACTCTTTCCCTACACGACGCTCTTCCGATCTNNNTCATTTGGGTGTACAGTAAAG

GAGCACACGTCTGAACTCTTTCCCTACACGACGCTCTTCCGATCTNNNTCATTTGGGTGTACAGTAAAG

AGCACACGTCTGAACTCTTTCCCTACACGACGCTCTTCCGATCTNNNGTGAAACCCCACCTCTACAAAAA

AGCACACGTCTGAACTCTTTCCCTACACGACGCTCTTCCGATCTNNNGTGAAACCCCACCTCTACAAAAA

GCACACGTCTGAACTCTTTCCCTACACGACGCTCTTCCGATCTNNNTCTCCTGAATAGCTGGGACTACA

GCACACGTCTGAACTCTTTCCCTACACGACGCTCTTCCGATCTNNNTCTCCTGAATAGCTGGGACTACA

AGCACACGTCTGAACTCTTTCCCTACACGACGCTCTTCCGATCTNNNCTGCAACTGGGTACATTTTGGA

AGCACACGTCTGAACTCTTTCCCTACACGACGCTCTTCCGATCTNNNCTGCAACTGGGTACATTTTGGA

AGCACACGTCTGAACTCTTTCCCTACACGACGCTCTTCCGATCTNNNTTCCTACTGAATGCTCCAGGAA

AGCACACGTCTGAACTCTTTCCCTACACGACGCTCTTCCGATCTNNNTTCCTACTGAATGCTCCAGGAA

GCACACGTCTGAACTCTTTCCCTACACGACGCTCTTCCGATCTNNNGACGGTGTCTCTTCAGCTTTCTA

GCACACGTCTGAACTCTTTCCCTACACGACGCTCTTCCGATCTNNNGACGGTGTCTCTTCAGCTTTCTA

GAGCACACGTCTGAACTCTTTCCCTACACGACGCTCTTCCGATCTNNNTGCAAATGGCCAACAAGTCT

GAGCACACGTCTGAACTCTTTCCCTACACGACGCTCTTCCGATCTNNNTGCAAATGGCCAACAAGTCT

GAGCACACGTCTGAACTCTTTCCCTACACGACGCTCTTCCGATCTNNNCCTGTTCAACCTCGTACACTGT

GAGCACACGTCTGAACTCTTTCCCTACACGACGCTCTTCCGATCTNNNCCTGTTCAACCTCGTACACTGT

GAGCACACGTCTGAACTCTTTCCCTACACGACGCTCTTCCGATCTNNNAGTGAGCCAAGATAACACCAC

GAGCACACGTCTGAACTCTTTCCCTACACGACGCTCTTCCGATCTNNNAGTGAGCCAAGATAACACCAC

AGAGCACACGTCTGAACTCTTTCCCTACACGACGCTCTTCCGATCTNNNGTCAGGGTGCTTGACCTAA

AGAGCACACGTCTGAACTCTTTCCCTACACGACGCTCTTCCGATCTNNNGTCAGGGTGCTTGACCTAA

AGCACACGTCTGAACTCTTTCCCTACACGACGCTCTTCCGATCTNNNATCTAGTATAACCTGGTGGGGG

AGCACACGTCTGAACTCTTTCCCTACACGACGCTCTTCCGATCTNNNATCTAGTATAACCTGGTGGGGG

GCACACGTCTGAACTCTTTCCCTACACGACGCTCTTCCGATCTNNNGCTGATGTGGACAAAGTGTTTC

GCACACGTCTGAACTCTTTCCCTACACGACGCTCTTCCGATCTNNNGCTGATGTGGACAAAGTGTTTC

GAGCACACGTCTGAACTCTTTCCCTACACGACGCTCTTCCGATCTNNNGGTTGCAGTGAGCCAAGATT

GAGCACACGTCTGAACTCTTTCCCTACACGACGCTCTTCCGATCTNNNGGTTGCAGTGAGCCAAGATT

GCACACGTCTGAACTCTTTCCCTACACGACGCTCTTCCGATCTNNNGGATCAAACCAGGCAATTCATGT

GCACACGTCTGAACTCTTTCCCTACACGACGCTCTTCCGATCTNNNGGATCAAACCAGGCAATTCATGT

AGCACACGTCTGAACTCTTTCCCTACACGACGCTCTTCCGATCTNNNACTATAGTTGTGTGCCACCATG

AGCACACGTCTGAACTCTTTCCCTACACGACGCTCTTCCGATCTNNNACTATAGTTGTGTGCCACCATG

GAGCACACGTCTGAACTCTTTCCCTACACGACGCTCTTCCGATCTNNNGAGGCTGAGGCAGGAGAAT

GAGCACACGTCTGAACTCTTTCCCTACACGACGCTCTTCCGATCTNNNGAGGCTGAGGCAGGAGAAT

AGCACACGTCTGAACTCTTTCCCTACACGACGCTCTTCCGATCTNNNCAGACATTGTTACCCACGTGAA

AGCACACGTCTGAACTCTTTCCCTACACGACGCTCTTCCGATCTNNNCAGACATTGTTACCCACGTGA
CACACGTCTGAACTCTTTCCCTACACGACGCTCTTCCGATCTNNNTGTGTGTGTGTGTGTATCTCA
CACACGTCTGAACTCTTTCCCTACACGACGCTCTTCCGATCTNNNTGTGTGTGTGTGTGTATCTCA
CACACGTCTGAACTCTTTCCCTACACGACGCTCTTCCGATCTNNNTCCACTGTTTCTTATCGCAAGGA
CACACGTCTGAACTCTTTCCCTACACGACGCTCTTCCGATCTNNNTCCACTGTTTCTTATCGCAAGGA
GCACACGTCTGAACTCTTTCCCTACACGACGCTCTTCCGATCTNNNTGGAGGCATAGACAAGAAAGC
GCACACGTCTGAACTCTTTCCCTACACGACGCTCTTCCGATCTNNNTGGAGGCATAGACAAGAAAGC
GCACACGTCTGAACTCTTTCCCTACACGACGCTCTTCCGATCTNNNAGTAAGCACATAGTGTCACGAG
GCACACGTCTGAACTCTTTCCCTACACGACGCTCTTCCGATCTNNNAGTAAGCACATAGTGTCACGAG
AGCACACGTCTGAACTCTTTCCCTACACGACGCTCTTCCGATCTNNNGCTTTGCATCTAACAGTCAGC
AGCACACGTCTGAACTCTTTCCCTACACGACGCTCTTCCGATCTNNNGCTTTGCATCTAACAGTCAGC
CACACGTCTGAACTCTTTCCCTACACGACGCTCTTCCGATCTNNNAGAAAGAAAAGTCAAAGTGTAA
CACACGTCTGAACTCTTTCCCTACACGACGCTCTTCCGATCTNNNAGAAAGAAAAGTCAAAGTGTAA
GAGCACACGTCTGAACTCTTTCCCTACACGACGCTCTTCCGATCTNNNTACCCCCACCTTCTTTTATTG
GAGCACACGTCTGAACTCTTTCCCTACACGACGCTCTTCCGATCTNNNTACCCCCACCTTCTTTTATTG
AGCACACGTCTGAACTCTTTCCCTACACGACGCTCTTCCGATCTNNNAGGCCTGTGAGCATATGGTAA
AGCACACGTCTGAACTCTTTCCCTACACGACGCTCTTCCGATCTNNNAGGCCTGTGAGCATATGGTAA
AGCACACGTCTGAACTCTTTCCCTACACGACGCTCTTCCGATCTNNNAGACCAAGGTTCTGACTCCTA
AGCACACGTCTGAACTCTTTCCCTACACGACGCTCTTCCGATCTNNNAGACCAAGGTTCTGACTCCTA
GCACACGTCTGAACTCTTTCCCTACACGACGCTCTTCCGATCTNNNCACCACACCCAGCTAATTTTTGT
GCACACGTCTGAACTCTTTCCCTACACGACGCTCTTCCGATCTNNNCACCACACCCAGCTAATTTTTGT
GCACACGTCTGAACTCTTTCCCTACACGACGCTCTTCCGATCTNNNGTGGAACCCCGTCTCTACTAAA
GCACACGTCTGAACTCTTTCCCTACACGACGCTCTTCCGATCTNNNGTGGAACCCCGTCTCTACTAAA
AGCACACGTCTGAACTCTTTCCCTACACGACGCTCTTCCGATCTNNNCACTTGTAATCCCAGCACTTTG
AGCACACGTCTGAACTCTTTCCCTACACGACGCTCTTCCGATCTNNNCACTTGTAATCCCAGCACTTTG
AGCACACGTCTGAACTCTTTCCCTACACGACGCTCTTCCGATCTNNNGTGAGCCAAGATCACACCACT
AGCACACGTCTGAACTCTTTCCCTACACGACGCTCTTCCGATCTNNNGTGAGCCAAGATCACACCACT
CACACGTCTGAACTCTTTCCCTACACGACGCTCTTCCGATCTNNNCCTGAACACAAGGACTCCTGTAA
CACACGTCTGAACTCTTTCCCTACACGACGCTCTTCCGATCTNNNCCTGAACACAAGGACTCCTGTAA
GCACACGTCTGAACTCTTTCCCTACACGACGCTCTTCCGATCTNNNGGTACAATCATGGCTCATTGCA
GCACACGTCTGAACTCTTTCCCTACACGACGCTCTTCCGATCTNNNGGTACAATCATGGCTCATTGCA
GCACACGTCTGAACTCTTTCCCTACACGACGCTCTTCCGATCTNNNGCCAAGATCATACCATTGCACT
GCACACGTCTGAACTCTTTCCCTACACGACGCTCTTCCGATCTNNNGCCAAGATCATACCATTGCACT
GAGCACACGTCTGAACTCTTTCCCTACACGACGCTCTTCCGATCTNNNGGCCTTGGAAGTGTTAGAT
GAGCACACGTCTGAACTCTTTCCCTACACGACGCTCTTCCGATCTNNNGGCCTTGGAAGTGTTAGAT
ACACGTCTGAACTCTTTCCCTACACGACGCTCTTCCGATCTNNNGAAAAGAACTCAGTACCTGAATCAT
ACACGTCTGAACTCTTTCCCTACACGACGCTCTTCCGATCTNNNGAAAAGAACTCAGTACCTGAATCAT
AGAGCACACGTCTGAACTCTTTCCCTACACGACGCTCTTCCGATCTNNNGACGCCGTGAAAAGAAAG
AGAGCACACGTCTGAACTCTTTCCCTACACGACGCTCTTCCGATCTNNNGACGCCGTGAAAAGAAAG
AGCACACGTCTGAACTCTTTCCCTACACGACGCTCTTCCGATCTNNNTTTTGCAAACCATCAATATCTA
AGCACACGTCTGAACTCTTTCCCTACACGACGCTCTTCCGATCTNNNTTTTGCAAACCATCAATATCTA
AGCACACGTCTGAACTCTTTCCCTACACGACGCTCTTCCGATCTNNNAAACAAACAACCCCCCCAAAAA
AGCACACGTCTGAACTCTTTCCCTACACGACGCTCTTCCGATCTNNNAAACAAACAACCCCCCCAAAAA

AGCACACGTCTGAACTCTTTCCCTACACGACGCTCTTCCGATCTNNNTATTCTTTCAGTGTGGTCCAGG
AGCACACGTCTGAACTCTTTCCCTACACGACGCTCTTCCGATCTNNNTATTCTTTCAGTGTGGTCCAGG
GCACACGTCTGAACTCTTTCCCTACACGACGCTCTTCCGATCTNNNTTCCTTCCTTCCTTCCTTCCTTCC
GCACACGTCTGAACTCTTTCCCTACACGACGCTCTTCCGATCTNNNTTCCTTCCTTCCTTCCTTCCTTCC
GCACACGTCTGAACTCTTTCCCTACACGACGCTCTTCCGATCTNNNTACTCCAGACTGGGCAATATGA
GCACACGTCTGAACTCTTTCCCTACACGACGCTCTTCCGATCTNNNTACTCCAGACTGGGCAATATGA
AGCACACGTCTGAACTCTTTCCCTACACGACGCTCTTCCGATCTNNNCATCCTGGCTAACACAGTGAA
AGCACACGTCTGAACTCTTTCCCTACACGACGCTCTTCCGATCTNNNCATCCTGGCTAACACAGTGAA
GCACACGTCTGAACTCTTTCCCTACACGACGCTCTTCCGATCTNNNTATGTATTAGAGAGGGGGCCAC
GCACACGTCTGAACTCTTTCCCTACACGACGCTCTTCCGATCTNNNTATGTATTAGAGAGGGGGCCAC
AGCACACGTCTGAACTCTTTCCCTACACGACGCTCTTCCGATCTNNNTCCTGAACTAGACACTGGTAG
AGCACACGTCTGAACTCTTTCCCTACACGACGCTCTTCCGATCTNNNTCCTGAACTAGACACTGGTAG
AGCACACGTCTGAACTCTTTCCCTACACGACGCTCTTCCGATCTNNNTGAGGCAGGAGAATCTCTTGA
AGCACACGTCTGAACTCTTTCCCTACACGACGCTCTTCCGATCTNNNTGAGGCAGGAGAATCTCTTGA
AGCACACGTCTGAACTCTTTCCCTACACGACGCTCTTCCGATCTNNNGCCAAGATCACACCATTTCACT
AGCACACGTCTGAACTCTTTCCCTACACGACGCTCTTCCGATCTNNNGCCAAGATCACACCATTTCACT
AGCACACGTCTGAACTCTTTCCCTACACGACGCTCTTCCGATCTNNNAGGAGTTGCACCTGATGTTAG
AGCACACGTCTGAACTCTTTCCCTACACGACGCTCTTCCGATCTNNNAGGAGTTGCACCTGATGTTAG
AGCACACGTCTGAACTCTTTCCCTACACGACGCTCTTCCGATCTNNNAACCCTGAACAGAAGCACTAC
AGCACACGTCTGAACTCTTTCCCTACACGACGCTCTTCCGATCTNNNAACCCTGAACAGAAGCACTAC
AGCACACGTCTGAACTCTTTCCCTACACGACGCTCTTCCGATCTNNNGAGAATCGCTTGAACCCAGG
AGCACACGTCTGAACTCTTTCCCTACACGACGCTCTTCCGATCTNNNGAGAATCGCTTGAACCCAGG
AGCACACGTCTGAACTCTTTCCCTACACGACGCTCTTCCGATCTNNNGTCCCAGCTACTTGAAAGTCT
AGCACACGTCTGAACTCTTTCCCTACACGACGCTCTTCCGATCTNNNGTCCCAGCTACTTGAAAGTCT
GAGCACACGTCTGAACTCTTTCCCTACACGACGCTCTTCCGATCTNNNAAAACAGTGAAATGCTGTCC
GAGCACACGTCTGAACTCTTTCCCTACACGACGCTCTTCCGATCTNNNAAAACAGTGAAATGCTGTCC
GCACACGTCTGAACTCTTTCCCTACACGACGCTCTTCCGATCTNNNTCCAGCCTGATACTGAAGTAGG
GCACACGTCTGAACTCTTTCCCTACACGACGCTCTTCCGATCTNNNTCCAGCCTGATACTGAAGTAGG
AGCACACGTCTGAACTCTTTCCCTACACGACGCTCTTCCGATCTNNNTGAAAATGAGGCCAGCTAAAA
AGCACACGTCTGAACTCTTTCCCTACACGACGCTCTTCCGATCTNNNTGAAAATGAGGCCAGCTAAAA
AGCACACGTCTGAACTCTTTCCCTACACGACGCTCTTCCGATCTNNNACAGTTGGTATTTTTAATTCTC
AGCACACGTCTGAACTCTTTCCCTACACGACGCTCTTCCGATCTNNNACAGTTGGTATTTTTAATTCTC
GAGCACACGTCTGAACTCTTTCCCTACACGACGCTCTTCCGATCTNNNCCAACATGGTGAAATCCCAT
GAGCACACGTCTGAACTCTTTCCCTACACGACGCTCTTCCGATCTNNNCCAACATGGTGAAATCCCAT
AGCACACGTCTGAACTCTTTCCCTACACGACGCTCTTCCGATCTNNNGGGGAAGGAGAAAAGGA
AGCACACGTCTGAACTCTTTCCCTACACGACGCTCTTCCGATCTNNNGGGGAAGGAGAAAAGGA
GAGCACACGTCTGAACTCTTTCCCTACACGACGCTCTTCCGATCTNNNGTACATACGTATGTGCGCAC
GAGCACACGTCTGAACTCTTTCCCTACACGACGCTCTTCCGATCTNNNGTACATACGTATGTGCGCAC
GAGCACACGTCTGAACTCTTTCCCTACACGACGCTCTTCCGATCTNNNCAGTGGAAGAAGAGGAGG
GAGCACACGTCTGAACTCTTTCCCTACACGACGCTCTTCCGATCTNNNCAGTGGAAGAAGAGGAGG
CACACGTCTGAACTCTTTCCCTACACGACGCTCTTCCGATCTNNNGAAACAAAGAAAGGCATGCGTG
CACACGTCTGAACTCTTTCCCTACACGACGCTCTTCCGATCTNNNGAAACAAAGAAAGGCATGCGTG
AGCACACGTCTGAACTCTTTCCCTACACGACGCTCTTCCGATCTNNNGCATCACACACTTACACACTCA

AGCACACGTCTGAACTCTTTCCCTACACGACGCTCTTCCGATCTNNNGCATCACACACTTACACACTCA
AGCACACGTCTGAACTCTTTCCCTACACGACGCTCTTCCGATCTNNNACAGATTGTTTACAGTGGGAG
AGCACACGTCTGAACTCTTTCCCTACACGACGCTCTTCCGATCTNNNACAGATTGTTTACAGTGGGAG
AGCACACGTCTGAACTCTTTCCCTACACGACGCTCTTCCGATCTNNNCCAGGGGAGTGAATAGTTCTG
AGCACACGTCTGAACTCTTTCCCTACACGACGCTCTTCCGATCTNNNCCAGGGGAGTGAATAGTTCTG
AGCACACGTCTGAACTCTTTCCCTACACGACGCTCTTCCGATCTNNNCTCACAAACCTTCCCTCTCTTCCC
AGCACACGTCTGAACTCTTTCCCTACACGACGCTCTTCCGATCTNNNCTCACAAACCTTCCCTCTCTTCCC
GAGCACACGTCTGAACTCTTTCCCTACACGACGCTCTTCCGATCTNNNTGTCAGCTGCTGTTGACTTT
GAGCACACGTCTGAACTCTTTCCCTACACGACGCTCTTCCGATCTNNNTGTCAGCTGCTGTTGACTTT
GAGCACACGTCTGAACTCTTTCCCTACACGACGCTCTTCCGATCTNNNAGAATCACTTGAACCCGAGA
GAGCACACGTCTGAACTCTTTCCCTACACGACGCTCTTCCGATCTNNNAGAATCACTTGAACCCGAGA
AGCACACGTCTGAACTCTTTCCCTACACGACGCTCTTCCGATCTNNNGAGTGTCGACATCATGCTCAA
AGCACACGTCTGAACTCTTTCCCTACACGACGCTCTTCCGATCTNNNGAGTGTCGACATCATGCTCAA
GCACACGTCTGAACTCTTTCCCTACACGACGCTCTTCCGATCTNNNTAGAGATCACAATCTGGGCACT
GCACACGTCTGAACTCTTTCCCTACACGACGCTCTTCCGATCTNNNTAGAGATCACAATCTGGGCACT
AGCACACGTCTGAACTCTTTCCCTACACGACGCTCTTCCGATCTNNNTATCCACTCGTTGATTGTTGG
AGCACACGTCTGAACTCTTTCCCTACACGACGCTCTTCCGATCTNNNTATCCACTCGTTGATTGTTGG
AGCACACGTCTGAACTCTTTCCCTACACGACGCTCTTCCGATCTNNNGCTACAGTGAGTTGTGTTCAC
AGCACACGTCTGAACTCTTTCCCTACACGACGCTCTTCCGATCTNNNGCTACAGTGAGTTGTGTTCAC
AGCACACGTCTGAACTCTTTCCCTACACGACGCTCTTCCGATCTNNNTGCTGAGATTACAGGCTTGAAC
AGCACACGTCTGAACTCTTTCCCTACACGACGCTCTTCCGATCTNNNTGCTGAGATTACAGGCTTGAAC
AGCACACGTCTGAACTCTTTCCCTACACGACGCTCTTCCGATCTNNNGGGAAGGCATGATTGTGTTTT
AGCACACGTCTGAACTCTTTCCCTACACGACGCTCTTCCGATCTNNNGGGAAGGCATGATTGTGTTTT
AGCACACGTCTGAACTCTTTCCCTACACGACGCTCTTCCGATCTNNNGCACATGCCTATAATCCCAGCT
AGCACACGTCTGAACTCTTTCCCTACACGACGCTCTTCCGATCTNNNGCACATGCCTATAATCCCAGCT
AGCACACGTCTGAACTCTTTCCCTACACGACGCTCTTCCGATCTNNNTGAGGGAAAGGAGTCTAAGA
AGCACACGTCTGAACTCTTTCCCTACACGACGCTCTTCCGATCTNNNTGAGGGAAAGGAGTCTAAGA
AGCACACGTCTGAACTCTTTCCCTACACGACGCTCTTCCGATCTNNNTGGATTACCAGGGAGAGACT
AGCACACGTCTGAACTCTTTCCCTACACGACGCTCTTCCGATCTNNNTGGATTACCAGGGAGAGACT
GAGCACACGTCTGAACTCTTTCCCTACACGACGCTCTTCCGATCTNNNCGGGTGGGAGATAACGTGT
GAGCACACGTCTGAACTCTTTCCCTACACGACGCTCTTCCGATCTNNNCGGGTGGGAGATAACGTGT
GAGCACACGTCTGAACTCTTTCCCTACACGACGCTCTTCCGATCTNNNCTTTTCTCTACAGCCTTGCCA
GAGCACACGTCTGAACTCTTTCCCTACACGACGCTCTTCCGATCTNNNCTTTTCTCTACAGCCTTGCCA
GAGCACACGTCTGAACTCTTTCCCTACACGACGCTCTTCCGATCTNNNGCCATCATTTGAGTTGGAAG
GAGCACACGTCTGAACTCTTTCCCTACACGACGCTCTTCCGATCTNNNGCCATCATTTGAGTTGGAAG
GCACACGTCTGAACTCTTTCCCTACACGACGCTCTTCCGATCTNNNATGCCTGTAATCCCAACTACTCGC
GCACACGTCTGAACTCTTTCCCTACACGACGCTCTTCCGATCTNNNATGCCTGTAATCCCAACTACTCGC
AGCACACGTCTGAACTCTTTCCCTACACGACGCTCTTCCGATCTNNNAAAGCCCAAGCATATCACCATT
AGCACACGTCTGAACTCTTTCCCTACACGACGCTCTTCCGATCTNNNAAAGCCCAAGCATATCACCATT
AGCACACGTCTGAACTCTTTCCCTACACGACGCTCTTCCGATCTNNNCCCTGTAAAGAAACACTGTACC
AGCACACGTCTGAACTCTTTCCCTACACGACGCTCTTCCGATCTNNNCCCTGTAAAGAAACACTGTACC
AGCACACGTCTGAACTCTTTCCCTACACGACGCTCTTCCGATCTNNNGCTTCCGGAATTTGACTAGATG
AGCACACGTCTGAACTCTTTCCCTACACGACGCTCTTCCGATCTNNNGCTTCCGGAATTTGACTAGATG

AGCACACGTCTGAACTCTTTCCCTACACGACGCTCTTCCGATCTNNNTTTTGCATGGCAGACTCTGAT
AGCACACGTCTGAACTCTTTCCCTACACGACGCTCTTCCGATCTNNNTTTTGCATGGCAGACTCTGAT
GAGCACACGTCTGAACTCTTTCCCTACACGACGCTCTTCCGATCTNNNTTTATGTGACTGTTGCCAGGA
GAGCACACGTCTGAACTCTTTCCCTACACGACGCTCTTCCGATCTNNNTTTATGTGACTGTTGCCAGGA
GAGCACACGTCTGAACTCTTTCCCTACACGACGCTCTTCCGATCTNNNCTTACTGTGCTGGGATGGCTG
GAGCACACGTCTGAACTCTTTCCCTACACGACGCTCTTCCGATCTNNNCTTACTGTGCTGGGATGGCTG
AGCACACGTCTGAACTCTTTCCCTACACGACGCTCTTCCGATCTNNNTGAGGCAAGAGAATTGCTTGA
AGCACACGTCTGAACTCTTTCCCTACACGACGCTCTTCCGATCTNNNTGAGGCAAGAGAATTGCTTGA
GAGCACACGTCTGAACTCTTTCCCTACACGACGCTCTTCCGATCTNNNAGGCCTAGCAAATAAGTGTTC
GAGCACACGTCTGAACTCTTTCCCTACACGACGCTCTTCCGATCTNNNAGGCCTAGCAAATAAGTGTTC
AGCACACGTCTGAACTCTTTCCCTACACGACGCTCTTCCGATCTNNNTTTTACCTAAGCCAGTGGACGA
AGCACACGTCTGAACTCTTTCCCTACACGACGCTCTTCCGATCTNNNTTTTACCTAAGCCAGTGGACGA
AGAGCACACGTCTGAACTCTTTCCCTACACGACGCTCTTCCGATCTNNNAAGGTTATGGTGAGCTATG
AGAGCACACGTCTGAACTCTTTCCCTACACGACGCTCTTCCGATCTNNNAAGGTTATGGTGAGCTATG
CACGTCTGAACTCTTTCCCTACACGACGCTCTTCCGATCTNNNAAGTGTTGGGATTACAGGTGTGACA
CACGTCTGAACTCTTTCCCTACACGACGCTCTTCCGATCTNNNAAGTGTTGGGATTACAGGTGTGACA
GAGCACACGTCTGAACTCTTTCCCTACACGACGCTCTTCCGATCTNNNCTGAGACAGGAGAATGGCATC
GAGCACACGTCTGAACTCTTTCCCTACACGACGCTCTTCCGATCTNNNCTGAGACAGGAGAATGGCATC
AGCACACGTCTGAACTCTTTCCCTACACGACGCTCTTCCGATCTNNNAAGTAAAGGCTGCAGTGAGCTA
AGCACACGTCTGAACTCTTTCCCTACACGACGCTCTTCCGATCTNNNAAGTAAAGGCTGCAGTGAGCTA
AGCACACGTCTGAACTCTTTCCCTACACGACGCTCTTCCGATCTNNNAAGAGCCATCTGTGTGCTGTAT
AGCACACGTCTGAACTCTTTCCCTACACGACGCTCTTCCGATCTNNNAAGAGCCATCTGTGTGCTGTAT
GAGCACACGTCTGAACTCTTTCCCTACACGACGCTCTTCCGATCTNNNAGAATCACTTGAACCCAGTAG
GAGCACACGTCTGAACTCTTTCCCTACACGACGCTCTTCCGATCTNNNAGAATCACTTGAACCCAGTAG
GAGCACACGTCTGAACTCTTTCCCTACACGACGCTCTTCCGATCTNNNAGAGGCTTGGCAACTAGAATG
GAGCACACGTCTGAACTCTTTCCCTACACGACGCTCTTCCGATCTNNNAGAGGCTTGGCAACTAGAATG
GCACACGTCTGAACTCTTTCCCTACACGACGCTCTTCCGATCTNNNCCCATATGCTATCCCTTATATTTCA
GCACACGTCTGAACTCTTTCCCTACACGACGCTCTTCCGATCTNNNCCCATATGCTATCCCTTATATTTCA
GAGCACACGTCTGAACTCTTTCCCTACACGACGCTCTTCCGATCTNNNGCCAATGGCGTAACTATTCTG
GAGCACACGTCTGAACTCTTTCCCTACACGACGCTCTTCCGATCTNNNGCCAATGGCGTAACTATTCTG
AGCACACGTCTGAACTCTTTCCCTACACGACGCTCTTCCGATCTNNNCAGGTAGCTGGGACTACATACA
AGCACACGTCTGAACTCTTTCCCTACACGACGCTCTTCCGATCTNNNCAGGTAGCTGGGACTACATACA
AGCACACGTCTGAACTCTTTCCCTACACGACGCTCTTCCGATCTNNNTAGAATGAAGAGGAGCACACC
AGCACACGTCTGAACTCTTTCCCTACACGACGCTCTTCCGATCTNNNTAGAATGAAGAGGAGCACACC
AGCACACGTCTGAACTCTTTCCCTACACGACGCTCTTCCGATCTNNNGTTTCAGTGAGCCGAGATTGTC
AGCACACGTCTGAACTCTTTCCCTACACGACGCTCTTCCGATCTNNNGTTTCAGTGAGCCGAGATTGTC
AGCACACGTCTGAACTCTTTCCCTACACGACGCTCTTCCGATCTNNNGTACACCGCGGCTTTTGCTGCAT
AGCACACGTCTGAACTCTTTCCCTACACGACGCTCTTCCGATCTNNNGTACACCGCGGCTTTTGCTGCAT
GAGCACACGTCTGAACTCTTTCCCTACACGACGCTCTTCCGATCTNNNCTGTAGTCCCAGCTACTCAGAA
GAGCACACGTCTGAACTCTTTCCCTACACGACGCTCTTCCGATCTNNNCTGTAGTCCCAGCTACTCAGAA
GAGCACACGTCTGAACTCTTTCCCTACACGACGCTCTTCCGATCTNNNTACACATTTTTGTGGGGACAG
GAGCACACGTCTGAACTCTTTCCCTACACGACGCTCTTCCGATCTNNNTACACATTTTTGTGGGGACAG
AGCACACGTCTGAACTCTTTCCCTACACGACGCTCTTCCGATCTNNNGGAATCACTTGAACCTGGGAGG

AGCACACGTCTGAACTCTTTCCCTACACGACGCTCTTCCGATCTNNNGGAATCACTTGAACCTGGGAGG
GCACACGTCTGAACTCTTTCCCTACACGACGCTCTTCCGATCTNNNGAGAATGAGCTGTCACCATACAT
GCACACGTCTGAACTCTTTCCCTACACGACGCTCTTCCGATCTNNNGAGAATGAGCTGTCACCATACAT
GAGCACACGTCTGAACTCTTTCCCTACACGACGCTCTTCCGATCTNNNCGCTTTGCAATTTTATTAGGTC
GAGCACACGTCTGAACTCTTTCCCTACACGACGCTCTTCCGATCTNNNCGCTTTGCAATTTTATTAGGTC
ACACGTCTGAACTCTTTCCCTACACGACGCTCTTCCGATCTNNNTAGAAAATCCAGGCCCATAACATCGG
ACACGTCTGAACTCTTTCCCTACACGACGCTCTTCCGATCTNNNTAGAAAATCCAGGCCCATAACATCGG
GAGCACACGTCTGAACTCTTTCCCTACACGACGCTCTTCCGATCTNNNGATCGAGACCATCCTGGCTAA
GAGCACACGTCTGAACTCTTTCCCTACACGACGCTCTTCCGATCTNNNGATCGAGACCATCCTGGCTAA
AGCACACGTCTGAACTCTTTCCCTACACGACGCTCTTCCGATCTNNNGTGTTCATGCCATTCTCCTGCCA
AGCACACGTCTGAACTCTTTCCCTACACGACGCTCTTCCGATCTNNNGTGTTCATGCCATTCTCCTGCCA
AGCACACGTCTGAACTCTTTCCCTACACGACGCTCTTCCGATCTNNNGAGATCATGCCACTACACTCCA
AGCACACGTCTGAACTCTTTCCCTACACGACGCTCTTCCGATCTNNNGAGATCATGCCACTACACTCCA
GAGCACACGTCTGAACTCTTTCCCTACACGACGCTCTTCCGATCTNNNGTGGTGCATTCCTGTAATTCCA
GAGCACACGTCTGAACTCTTTCCCTACACGACGCTCTTCCGATCTNNNGTGGTGCATTCCTGTAATTCCA
AGCACACGTCTGAACTCTTTCCCTACACGACGCTCTTCCGATCTNNNCTGGGCAACAAGAGCAAAACT
AGCACACGTCTGAACTCTTTCCCTACACGACGCTCTTCCGATCTNNNCTGGGCAACAAGAGCAAAACT
AGCACACGTCTGAACTCTTTCCCTACACGACGCTCTTCCGATCTNNNGCACCCGGCTGATAACTTTTTTT
AGCACACGTCTGAACTCTTTCCCTACACGACGCTCTTCCGATCTNNNGCACCCGGCTGATAACTTTTTTT
AGCACACGTCTGAACTCTTTCCCTACACGACGCTCTTCCGATCTNNNCCCATTATGAATACAAAACCAAA
AGCACACGTCTGAACTCTTTCCCTACACGACGCTCTTCCGATCTNNNCCCATTATGAATACAAAACCAAA
GAGCACACGTCTGAACTCTTTCCCTACACGACGCTCTTCCGATCTNNNGAAAGAGTTGTCAGTTTGGG
GAGCACACGTCTGAACTCTTTCCCTACACGACGCTCTTCCGATCTNNNGAAAGAGTTGTCAGTTTGGG
AGCACACGTCTGAACTCTTTCCCTACACGACGCTCTTCCGATCTNNNCTAGAGTGCAGTGGTGTGATCT
AGCACACGTCTGAACTCTTTCCCTACACGACGCTCTTCCGATCTNNNCTAGAGTGCAGTGGTGTGATCT
AGCACACGTCTGAACTCTTTCCCTACACGACGCTCTTCCGATCTNNNAAAATGCTCCTCTCTCCTCCCTTCA
AGCACACGTCTGAACTCTTTCCCTACACGACGCTCTTCCGATCTNNNAAAATGCTCCTCTCTCCTCCCTTCA
ACACGTCTGAACTCTTTCCCTACACGACGCTCTTCCGATCTNNNCTACGATCAGTCCTACCAATACATCG
ACACGTCTGAACTCTTTCCCTACACGACGCTCTTCCGATCTNNNCTACGATCAGTCCTACCAATACATCG
GCACACGTCTGAACTCTTTCCCTACACGACGCTCTTCCGATCTNNNGAGATCGTGTCACTGCACTCTAG
GCACACGTCTGAACTCTTTCCCTACACGACGCTCTTCCGATCTNNNGAGATCGTGTCACTGCACTCTAG
AGCACACGTCTGAACTCTTTCCCTACACGACGCTCTTCCGATCTNNNCCCTCCTCCACAAAGCAAAAGAT
AGCACACGTCTGAACTCTTTCCCTACACGACGCTCTTCCGATCTNNNCCCTCCTCCACAAAGCAAAAGAT
GAGCACACGTCTGAACTCTTTCCCTACACGACGCTCTTCCGATCTNNNGGATGGTCTTGAACTCCTGAC
GAGCACACGTCTGAACTCTTTCCCTACACGACGCTCTTCCGATCTNNNGGATGGTCTTGAACTCCTGAC
GAGCACACGTCTGAACTCTTTCCCTACACGACGCTCTTCCGATCTNNNAAAGTTGTCATGGTCCAAGAG
GAGCACACGTCTGAACTCTTTCCCTACACGACGCTCTTCCGATCTNNNAAAGTTGTCATGGTCCAAGAG
AGCACACGTCTGAACTCTTTCCCTACACGACGCTCTTCCGATCTNNNTAAGGGCATCATTTCAGCAAGA
AGCACACGTCTGAACTCTTTCCCTACACGACGCTCTTCCGATCTNNNTAAGGGCATCATTTCAGCAAGA
GAGCACACGTCTGAACTCTTTCCCTACACGACGCTCTTCCGATCTNNNGCAATTAGAGAAGAGAGAGG
GAGCACACGTCTGAACTCTTTCCCTACACGACGCTCTTCCGATCTNNNGCAATTAGAGAAGAGAGAGG
AGCACACGTCTGAACTCTTTCCCTACACGACGCTCTTCCGATCTNNNGGAACTGTGAGTCCACTAAAC
AGCACACGTCTGAACTCTTTCCCTACACGACGCTCTTCCGATCTNNNGGAACTGTGAGTCCACTAAAC

GAAGAGCACACGTCTGAACTCTTTCCCTACACGACGCTCTTCCGATCTNNNTTTGGATAATTGACTAGC

GAAGAGCACACGTCTGAACTCTTTCCCTACACGACGCTCTTCCGATCTNNNTTTGGATAATTGACTAGC

GAGCACACGTCTGAACTCTTTCCCTACACGACGCTCTTCCGATCTNNNTGTGCCTGGCTAATTTTTGTGT

GAGCACACGTCTGAACTCTTTCCCTACACGACGCTCTTCCGATCTNNNTGTGCCTGGCTAATTTTTGTGT

GAGCACACGTCTGAACTCTTTCCCTACACGACGCTCTTCCGATCTNNNGAGGCAGGAGAATAGTGGGA

GAGCACACGTCTGAACTCTTTCCCTACACGACGCTCTTCCGATCTNNNGAGGCAGGAGAATAGTGGGA

AGCACACGTCTGAACTCTTTCCCTACACGACGCTCTTCCGATCTNNNCTTGAACTCCTGACCTCCTGATC

AGCACACGTCTGAACTCTTTCCCTACACGACGCTCTTCCGATCTNNNCTTGAACTCCTGACCTCCTGATC

AGCACACGTCTGAACTCTTTCCCTACACGACGCTCTTCCGATCTNNNTTGTGTCATGACAATTTGAGCCA

AGCACACGTCTGAACTCTTTCCCTACACGACGCTCTTCCGATCTNNNTTGTGTCATGACAATTTGAGCCA

GAGCACACGTCTGAACTCTTTCCCTACACGACGCTCTTCCGATCTNNNGAGATTGCAGTGAGCCGAGA

GAGCACACGTCTGAACTCTTTCCCTACACGACGCTCTTCCGATCTNNNGAGATTGCAGTGAGCCGAGA

GAGCACACGTCTGAACTCTTTCCCTACACGACGCTCTTCCGATCTNNNCAATCTTCGCTCACTGCAAGCCA

GAGCACACGTCTGAACTCTTTCCCTACACGACGCTCTTCCGATCTNNNCAATCTTCGCTCACTGCAAGCCA

CACACGTCTGAACTCTTTCCCTACACGACGCTCTTCCGATCTNNNCTGATTGAGAAGACTCTGTACCATC

CACACGTCTGAACTCTTTCCCTACACGACGCTCTTCCGATCTNNNCTGATTGAGAAGACTCTGTACCATC

AGCACACGTCTGAACTCTTTCCCTACACGACGCTCTTCCGATCTNNNCACAATGTGCACATGTACCCTAA

AGCACACGTCTGAACTCTTTCCCTACACGACGCTCTTCCGATCTNNNCACAATGTGCACATGTACCCTAA

GCACACGTCTGAACTCTTTCCCTACACGACGCTCTTCCGATCTNNNCACAGTATGACTCCAGTGACTCCCA

GCACACGTCTGAACTCTTTCCCTACACGACGCTCTTCCGATCTNNNCACAGTATGACTCCAGTGACTCCCA

AGAGCACACGTCTGAACTCTTTCCCTACACGACGCTCTTCCGATCTNNNGACTGAATCCTGGCACTTTG

AGAGCACACGTCTGAACTCTTTCCCTACACGACGCTCTTCCGATCTNNNGACTGAATCCTGGCACTTTG

AGCACACGTCTGAACTCTTTCCCTACACGACGCTCTTCCGATCTNNNTAGTTTGGGTCCTGTTGGTTTG

AGCACACGTCTGAACTCTTTCCCTACACGACGCTCTTCCGATCTNNNTAGTTTGGGTCCTGTTGGTTTG

AGCACACGTCTGAACTCTTTCCCTACACGACGCTCTTCCGATCTNNNTAGCTAATGTGTGCCTGTAGTCC

AGCACACGTCTGAACTCTTTCCCTACACGACGCTCTTCCGATCTNNNTAGCTAATGTGTGCCTGTAGTCC

AGCACACGTCTGAACTCTTTCCCTACACGACGCTCTTCCGATCTNNNGAGCTTGCAGTGAGCCAAGATC

AGCACACGTCTGAACTCTTTCCCTACACGACGCTCTTCCGATCTNNNGAGCTTGCAGTGAGCCAAGATC

GAGCACACGTCTGAACTCTTTCCCTACACGACGCTCTTCCGATCTNNNCAACAGAGCGAGACCCTGTCCA

GAGCACACGTCTGAACTCTTTCCCTACACGACGCTCTTCCGATCTNNNCAACAGAGCGAGACCCTGTCCA

AGCACACGTCTGAACTCTTTCCCTACACGACGCTCTTCCGATCTNNNATTTCCAGTCTCGCCATTATCCAG

AGCACACGTCTGAACTCTTTCCCTACACGACGCTCTTCCGATCTNNNATTTCCAGTCTCGCCATTATCCAG

CACACGTCTGAACTCTTTCCCTACACGACGCTCTTCCGATCTNNNATAATGAGTACAAGTAAGATGGTC

CACACGTCTGAACTCTTTCCCTACACGACGCTCTTCCGATCTNNNATAATGAGTACAAGTAAGATGGTC

AGAGCACACGTCTGAACTCTTTCCCTACACGACGCTCTTCCGATCTNNNTATGACCTGGCCAGTGCTAA

AGAGCACACGTCTGAACTCTTTCCCTACACGACGCTCTTCCGATCTNNNTATGACCTGGCCAGTGCTAA

AGCACACGTCTGAACTCTTTCCCTACACGACGCTCTTCCGATCTNNNCAGGTTGTCGTGAATGCCATTA

AGCACACGTCTGAACTCTTTCCCTACACGACGCTCTTCCGATCTNNNCAGGTTGTCGTGAATGCCATTA

GCACACGTCTGAACTCTTTCCCTACACGACGCTCTTCCGATCTNNNTCCTGTTCCCCCATTATCTTTCCAT

GCACACGTCTGAACTCTTTCCCTACACGACGCTCTTCCGATCTNNNTCCTGTTCCCCCATTATCTTTCCAT

AGCACACGTCTGAACTCTTTCCCTACACGACGCTCTTCCGATCTNNNGACAGTTCTACAAAGGAGGCTG

AGCACACGTCTGAACTCTTTCCCTACACGACGCTCTTCCGATCTNNNGACAGTTCTACAAAGGAGGCTG

AGCACACGTCTGAACTCTTTCCCTACACGACGCTCTTCCGATCTNNNATGCTCCAGAACACGTTACACTC

AGCACACGTCTGAACTCTTTCCCTACACGACGCTCTTCCGATCTNNNATGCTCCAGAACACGTTACACTC
GAGCACACGTCTGAACTCTTTCCCTACACGACGCTCTTCCGATCTNNNAGGCAGTAGAATCACTTGAAC
GAGCACACGTCTGAACTCTTTCCCTACACGACGCTCTTCCGATCTNNNAGGCAGTAGAATCACTTGAAC
AGCACACGTCTGAACTCTTTCCCTACACGACGCTCTTCCGATCTNNNCAACATTTGGGTGTGCTACTTCA
AGCACACGTCTGAACTCTTTCCCTACACGACGCTCTTCCGATCTNNNCAACATTTGGGTGTGCTACTTCA
AGAGCACACGTCTGAACTCTTTCCCTACACGACGCTCTTCCGATCTNNNCTACCCCTGTTTTTCACTTGGC
AGAGCACACGTCTGAACTCTTTCCCTACACGACGCTCTTCCGATCTNNNCTACCCCTGTTTTTCACTTGGC
AGCACACGTCTGAACTCTTTCCCTACACGACGCTCTTCCGATCTNNNCTGCAGTGAACTATGATTGTGC
AGCACACGTCTGAACTCTTTCCCTACACGACGCTCTTCCGATCTNNNCTGCAGTGAACTATGATTGTGC
AGCACACGTCTGAACTCTTTCCCTACACGACGCTCTTCCGATCTNNNAAAAAGTGCTAGCCCATGGTTA
AGCACACGTCTGAACTCTTTCCCTACACGACGCTCTTCCGATCTNNNAAAAAGTGCTAGCCCATGGTTA
AGCACACGTCTGAACTCTTTCCCTACACGACGCTCTTCCGATCTNNNGAGGTTGCAGTGAGCTGAGAT
AGCACACGTCTGAACTCTTTCCCTACACGACGCTCTTCCGATCTNNNGAGGTTGCAGTGAGCTGAGAT
CACACGTCTGAACTCTTTCCCTACACGACGCTCTTCCGATCTNNNAAGGGACGGGGAAGACTTTGTTA
CACACGTCTGAACTCTTTCCCTACACGACGCTCTTCCGATCTNNNAAGGGACGGGGAAGACTTTGTTA
AGCACACGTCTGAACTCTTTCCCTACACGACGCTCTTCCGATCTNNNCGGATTCCAGTCTGTCTCTCTCTC
AGCACACGTCTGAACTCTTTCCCTACACGACGCTCTTCCGATCTNNNCGGATTCCAGTCTGTCTCTCTCTC
GAGCACACGTCTGAACTCTTTCCCTACACGACGCTCTTCCGATCTNNNGAGAATCGTGTGAACCCAGGA
GAGCACACGTCTGAACTCTTTCCCTACACGACGCTCTTCCGATCTNNNGAGAATCGTGTGAACCCAGGA
GAGCACACGTCTGAACTCTTTCCCTACACGACGCTCTTCCGATCTNNNAGAGGTTTCAGTGACCCAAGA
GAGCACACGTCTGAACTCTTTCCCTACACGACGCTCTTCCGATCTNNNAGAGGTTTCAGTGACCCAAGA
AAGAGCACACGTCTGAACTCTTTCCCTACACGACGCTCTTCCGATCTNNNCAGCTAGCATACAAGGAG
AAGAGCACACGTCTGAACTCTTTCCCTACACGACGCTCTTCCGATCTNNNCAGCTAGCATACAAGGAG
AGCACACGTCTGAACTCTTTCCCTACACGACGCTCTTCCGATCTNNNGCCTCATAAGTAGCTGGGACTA
AGCACACGTCTGAACTCTTTCCCTACACGACGCTCTTCCGATCTNNNGCCTCATAAGTAGCTGGGACTA
GCACACGTCTGAACTCTTTCCCTACACGACGCTCTTCCGATCTNNNTCATGCATAAAGTGATCCACTCTC
GCACACGTCTGAACTCTTTCCCTACACGACGCTCTTCCGATCTNNNTCATGCATAAAGTGATCCACTCTC
AGCACACGTCTGAACTCTTTCCCTACACGACGCTCTTCCGATCTNNNCGGTAGAGAAATGAGACTCTG
AGCACACGTCTGAACTCTTTCCCTACACGACGCTCTTCCGATCTNNNCGGTAGAGAAATGAGACTCTG
AGCACACGTCTGAACTCTTTCCCTACACGACGCTCTTCCGATCTNNNAATGTTGGGATTACAGGCATGA
AGCACACGTCTGAACTCTTTCCCTACACGACGCTCTTCCGATCTNNNAATGTTGGGATTACAGGCATGA
GAGCACACGTCTGAACTCTTTCCCTACACGACGCTCTTCCGATCTNNNCAGCCACAAAGGGAAGAAAAC
GAGCACACGTCTGAACTCTTTCCCTACACGACGCTCTTCCGATCTNNNCAGCCACAAAGGGAAGAAAAC
GAGCACACGTCTGAACTCTTTCCCTACACGACGCTCTTCCGATCTNNNCAGATCGCACCACTGGACTCCA
GAGCACACGTCTGAACTCTTTCCCTACACGACGCTCTTCCGATCTNNNCAGATCGCACCACTGGACTCCA
AGCACACGTCTGAACTCTTTCCCTACACGACGCTCTTCCGATCTNNNAAAAACAGAAAGAAGCCCTCTG
AGCACACGTCTGAACTCTTTCCCTACACGACGCTCTTCCGATCTNNNAAAAACAGAAAGAAGCCCTCTG
GAGCACACGTCTGAACTCTTTCCCTACACGACGCTCTTCCGATCTNNNCCTGGCTGCCTTTCTTTTTTTCT
GAGCACACGTCTGAACTCTTTCCCTACACGACGCTCTTCCGATCTNNNCCTGGCTGCCTTTCTTTTTTTCT
AGCACACGTCTGAACTCTTTCCCTACACGACGCTCTTCCGATCTNNNGCACACACACACTTTCTAATCTC
AGCACACGTCTGAACTCTTTCCCTACACGACGCTCTTCCGATCTNNNGCACACACACACTTTCTAATCTC
GAGCACACGTCTGAACTCTTTCCCTACACGACGCTCTTCCGATCTNNNTTCTACTGAATGTCTGCATGG
GAGCACACGTCTGAACTCTTTCCCTACACGACGCTCTTCCGATCTNNNTTCTACTGAATGTCTGCATGG

GAGCACACGTCTGAACTCTTTCCCTACACGACGCTCTTCCGATCTNNNCTAACCCTTCATGCCGTTCATT
GAGCACACGTCTGAACTCTTTCCCTACACGACGCTCTTCCGATCTNNNCTAACCCTTCATGCCGTTCATT
AGCACACGTCTGAACTCTTTCCCTACACGACGCTCTTCCGATCTNNNAACTTATGGCAGAAGGAGAAG
AGCACACGTCTGAACTCTTTCCCTACACGACGCTCTTCCGATCTNNNAACTTATGGCAGAAGGAGAAG
AGCACACGTCTGAACTCTTTCCCTACACGACGCTCTTCCGATCTNNNCCTTGATTTGCACCCCTTGATTT
AGCACACGTCTGAACTCTTTCCCTACACGACGCTCTTCCGATCTNNNCCTTGATTTGCACCCCTTGATTT
AGCACACGTCTGAACTCTTTCCCTACACGACGCTCTTCCGATCTNNNTCCTGTTTCTAGGACCCATGTTG
AGCACACGTCTGAACTCTTTCCCTACACGACGCTCTTCCGATCTNNNTCCTGTTTCTAGGACCCATGTTG
GCACACGTCTGAACTCTTTCCCTACACGACGCTCTTCCGATCTNNNCAAGCCTATTTCCTTTCCCTCCTC
GCACACGTCTGAACTCTTTCCCTACACGACGCTCTTCCGATCTNNNCAAGCCTATTTCCTTTCCCTCCTC
AGCACACGTCTGAACTCTTTCCCTACACGACGCTCTTCCGATCTNNNTGGAGATAAAATGGGGAAAG
AGCACACGTCTGAACTCTTTCCCTACACGACGCTCTTCCGATCTNNNTGGAGATAAAATGGGGAAAG
GAGCACACGTCTGAACTCTTTCCCTACACGACGCTCTTCCGATCTNNNGCTAGGGTTGCTCATTGCTA
GAGCACACGTCTGAACTCTTTCCCTACACGACGCTCTTCCGATCTNNNGCTAGGGTTGCTCATTGCTA
CACACGTCTGAACTCTTTCCCTACACGACGCTCTTCCGATCTNNNATGCTCTTATGCTGGTAAAAGCAT
CACACGTCTGAACTCTTTCCCTACACGACGCTCTTCCGATCTNNNATGCTCTTATGCTGGTAAAAGCAT
AGCACACGTCTGAACTCTTTCCCTACACGACGCTCTTCCGATCTNNNGCAGCAGGCAAGATTTACTGA
AGCACACGTCTGAACTCTTTCCCTACACGACGCTCTTCCGATCTNNNGCAGCAGGCAAGATTTACTGA
GAGCACACGTCTGAACTCTTTCCCTACACGACGCTCTTCCGATCTNNNGTACTTCCTCAGCTTCTGTCTG
GAGCACACGTCTGAACTCTTTCCCTACACGACGCTCTTCCGATCTNNNGTACTTCCTCAGCTTCTGTCTG
AGCACACGTCTGAACTCTTTCCCTACACGACGCTCTTCCGATCTNNNATGATGACCCAAAGCAATCTCC
AGCACACGTCTGAACTCTTTCCCTACACGACGCTCTTCCGATCTNNNATGATGACCCAAAGCAATCTCC
GCACACGTCTGAACTCTTTCCCTACACGACGCTCTTCCGATCTNNNTAGATCTGGGTTCCCTGGAGTTG
GCACACGTCTGAACTCTTTCCCTACACGACGCTCTTCCGATCTNNNTAGATCTGGGTTCCCTGGAGTTG
GAGCACACGTCTGAACTCTTTCCCTACACGACGCTCTTCCGATCTNNNTGATCTATACGTGGACCAGCA
GAGCACACGTCTGAACTCTTTCCCTACACGACGCTCTTCCGATCTNNNTGATCTATACGTGGACCAGCA
GAGCACACGTCTGAACTCTTTCCCTACACGACGCTCTTCCGATCTNNNTATACACACACACACACACACA
GAGCACACGTCTGAACTCTTTCCCTACACGACGCTCTTCCGATCTNNNTATACACACACACACACACACA
GAGCACACGTCTGAACTCTTTCCCTACACGACGCTCTTCCGATCTNNNCACCTGTGTATTAGCTCTGGG
GAGCACACGTCTGAACTCTTTCCCTACACGACGCTCTTCCGATCTNNNCACCTGTGTATTAGCTCTGGG
GAGCACACGTCTGAACTCTTTCCCTACACGACGCTCTTCCGATCTNNNGACGTCTGTAATCCCAGCTAC
GAGCACACGTCTGAACTCTTTCCCTACACGACGCTCTTCCGATCTNNNGACGTCTGTAATCCCAGCTAC
GAGCACACGTCTGAACTCTTTCCCTACACGACGCTCTTCCGATCTNNNGCCCCTGCCCAGTTCTATAAAT
GAGCACACGTCTGAACTCTTTCCCTACACGACGCTCTTCCGATCTNNNGCCCCTGCCCAGTTCTATAAAT
AGCACACGTCTGAACTCTTTCCCTACACGACGCTCTTCCGATCTNNNAGAGTTCTGTGCAGTTTTACCC
AGCACACGTCTGAACTCTTTCCCTACACGACGCTCTTCCGATCTNNNAGAGTTCTGTGCAGTTTTACCC
GCACACGTCTGAACTCTTTCCCTACACGACGCTCTTCCGATCTNNNAATCAATGGAGAGGGAAGGATC
GCACACGTCTGAACTCTTTCCCTACACGACGCTCTTCCGATCTNNNAATCAATGGAGAGGGAAGGATC
AGAGCACACGTCTGAACTCTTTCCCTACACGACGCTCTTCCGATCTNNNCTGAGGCAGGAGAATTGCT
AGAGCACACGTCTGAACTCTTTCCCTACACGACGCTCTTCCGATCTNNNCTGAGGCAGGAGAATTGCT
AGCACACGTCTGAACTCTTTCCCTACACGACGCTCTTCCGATCTNNNGCATATTGTCCAGTCTCATTACA
AGCACACGTCTGAACTCTTTCCCTACACGACGCTCTTCCGATCTNNNGCATATTGTCCAGTCTCATTACA
AGAGCACACGTCTGAACTCTTTCCCTACACGACGCTCTTCCGATCTNNNATAGCAGGAACACAAAAGG

AGAGCACACGTCTGAACTCTTTCCCTACACGACGCTCTTCCGATCTNNNATAGCAGGAACACAAAAGG

GCACACGTCTGAACTCTTTCCCTACACGACGCTCTTCCGATCTNNNCCATGCCTGAGATGCAAGAATT

GCACACGTCTGAACTCTTTCCCTACACGACGCTCTTCCGATCTNNNCCATGCCTGAGATGCAAGAATT

AGCACACGTCTGAACTCTTTCCCTACACGACGCTCTTCCGATCTNNNAGAATCAACATAAGGCCCACA

AGCACACGTCTGAACTCTTTCCCTACACGACGCTCTTCCGATCTNNNAGAATCAACATAAGGCCCACA

CACACGTCTGAACTCTTTCCCTACACGACGCTCTTCCGATCTNNNGCTCCAGTAGTGCCCTTTATCTGC

CACACGTCTGAACTCTTTCCCTACACGACGCTCTTCCGATCTNNNGCTCCAGTAGTGCCCTTTATCTGC

GAGCACACGTCTGAACTCTTTCCCTACACGACGCTCTTCCGATCTNNNGCAGGAGAATCGCTTGAACA

GAGCACACGTCTGAACTCTTTCCCTACACGACGCTCTTCCGATCTNNNGCAGGAGAATCGCTTGAACA

AGCACACGTCTGAACTCTTTCCCTACACGACGCTCTTCCGATCTNNNACAAAAATTAGCTGGGCATGG

AGCACACGTCTGAACTCTTTCCCTACACGACGCTCTTCCGATCTNNNACAAAAATTAGCTGGGCATGG

AGCACACGTCTGAACTCTTTCCCTACACGACGCTCTTCCGATCTNNNCTGGATTCTGTGGTCCCTCTAT

AGCACACGTCTGAACTCTTTCCCTACACGACGCTCTTCCGATCTNNNCTGGATTCTGTGGTCCCTCTAT

GAGCACACGTCTGAACTCTTTCCCTACACGACGCTCTTCCGATCTNNNCCATCAGGAAAGGCAAATGA

GAGCACACGTCTGAACTCTTTCCCTACACGACGCTCTTCCGATCTNNNCCATCAGGAAAGGCAAATGA

AGCACACGTCTGAACTCTTTCCCTACACGACGCTCTTCCGATCTNNNAAATCCACTTTCCTTCTGCCTGA

AGCACACGTCTGAACTCTTTCCCTACACGACGCTCTTCCGATCTNNNAAATCCACTTTCCTTCTGCCTGA

GAGCACACGTCTGAACTCTTTCCCTACACGACGCTCTTCCGATCTNNNGCATGGCTAGATTTGAAATG

GAGCACACGTCTGAACTCTTTCCCTACACGACGCTCTTCCGATCTNNNGCATGGCTAGATTTGAAATG

GAGCACACGTCTGAACTCTTTCCCTACACGACGCTCTTCCGATCTNNNATCTCTCTATGTTGCCTAGGC

GAGCACACGTCTGAACTCTTTCCCTACACGACGCTCTTCCGATCTNNNATCTCTCTATGTTGCCTAGGC

GAGCACACGTCTGAACTCTTTCCCTACACGACGCTCTTCCGATCTNNNTCAAGAATTAGGGACGCTGC

GAGCACACGTCTGAACTCTTTCCCTACACGACGCTCTTCCGATCTNNNTCAAGAATTAGGGACGCTGC

GAAGAGCACACGTCTGAACTCTTTCCCTACACGACGCTCTTCCGATCTNNNGGACTGTCGCATCATGG

GAAGAGCACACGTCTGAACTCTTTCCCTACACGACGCTCTTCCGATCTNNNGGACTGTCGCATCATGG

AGCACACGTCTGAACTCTTTCCCTACACGACGCTCTTCCGATCTNNNCCCAAATGATGAAGTGATGCA

AGCACACGTCTGAACTCTTTCCCTACACGACGCTCTTCCGATCTNNNCCCAAATGATGAAGTGATGCA

GAGCACACGTCTGAACTCTTTCCCTACACGACGCTCTTCCGATCTNNNGAGAATTGCTTGAACCTGGG

GAGCACACGTCTGAACTCTTTCCCTACACGACGCTCTTCCGATCTNNNGAGAATTGCTTGAACCTGGG

AGCACACGTCTGAACTCTTTCCCTACACGACGCTCTTCCGATCTNNNGAGTACAGAGGTGGATATGCT

AGCACACGTCTGAACTCTTTCCCTACACGACGCTCTTCCGATCTNNNGAGTACAGAGGTGGATATGCT

GCACACGTCTGAACTCTTTCCCTACACGACGCTCTTCCGATCTNNNATCCACTGTCCCCAAACTTTTTG

GCACACGTCTGAACTCTTTCCCTACACGACGCTCTTCCGATCTNNNATCCACTGTCCCCAAACTTTTTG

AGCACACGTCTGAACTCTTTCCCTACACGACGCTCTTCCGATCTNNNTAAGCTTCCACCCACTAAGGTT

AGCACACGTCTGAACTCTTTCCCTACACGACGCTCTTCCGATCTNNNTAAGCTTCCACCCACTAAGGTT

GAGCACACGTCTGAACTCTTTCCCTACACGACGCTCTTCCGATCTNNNCACCTGGCCTCGTTAGTCTTTT

GAGCACACGTCTGAACTCTTTCCCTACACGACGCTCTTCCGATCTNNNCACCTGGCCTCGTTAGTCTTTT

AGCACACGTCTGAACTCTTTCCCTACACGACGCTCTTCCGATCTNNNGGCAACTTCCTGATGCTTCTAT

AGCACACGTCTGAACTCTTTCCCTACACGACGCTCTTCCGATCTNNNGGCAACTTCCTGATGCTTCTAT

GAGCACACGTCTGAACTCTTTCCCTACACGACGCTCTTCCGATCTNNNGAGGCAAGAGAATCGCTTGA

GAGCACACGTCTGAACTCTTTCCCTACACGACGCTCTTCCGATCTNNNGAGGCAAGAGAATCGCTTGA

AGCACACGTCTGAACTCTTTCCCTACACGACGCTCTTCCGATCTNNNTACTGATCTAGGATGCCTTGTG

AGCACACGTCTGAACTCTTTCCCTACACGACGCTCTTCCGATCTNNNTACTGATCTAGGATGCCTTGTG

AGCACACGTCTGAACTCTTTCCCTACACGACGCTCTTCCGATCTNNNTTGTCACCTCATTTGGAACCTCA

AGCACACGTCTGAACTCTTTCCCTACACGACGCTCTTCCGATCTNNNTTGTCACCTCATTTGGAACCTCA

AGCACACGTCTGAACTCTTTCCCTACACGACGCTCTTCCGATCTNNNTGGATGTCCATTCAGAGGAAG

AGCACACGTCTGAACTCTTTCCCTACACGACGCTCTTCCGATCTNNNTGGATGTCCATTCAGAGGAAG

AGAGCACACGTCTGAACTCTTTCCCTACACGACGCTCTTCCGATCTNNNATGTACCTGTACTCCCAGCTA

AGAGCACACGTCTGAACTCTTTCCCTACACGACGCTCTTCCGATCTNNNATGTACCTGTACTCCCAGCTA

AGCACACGTCTGAACTCTTTCCCTACACGACGCTCTTCCGATCTNNNCTTGTTTTGGGCCATTCAGCTAA

AGCACACGTCTGAACTCTTTCCCTACACGACGCTCTTCCGATCTNNNCTTGTTTTGGGCCATTCAGCTAA

AGCACACGTCTGAACTCTTTCCCTACACGACGCTCTTCCGATCTNNNAAATATCTCAAAAAGTCAGTGA

AGCACACGTCTGAACTCTTTCCCTACACGACGCTCTTCCGATCTNNNAAATATCTCAAAAAGTCAGTGA

GAGCACACGTCTGAACTCTTTCCCTACACGACGCTCTTCCGATCTNNNCCTACGCCAATCCCAAGGAGTC

GAGCACACGTCTGAACTCTTTCCCTACACGACGCTCTTCCGATCTNNNCCTACGCCAATCCCAAGGAGTC

GAGCACACGTCTGAACTCTTTCCCTACACGACGCTCTTCCGATCTNNNGTGAAACCCCGTCTCTACTAAA

GAGCACACGTCTGAACTCTTTCCCTACACGACGCTCTTCCGATCTNNNGTGAAACCCCGTCTCTACTAAA

GCACACGTCTGAACTCTTTCCCTACACGACGCTCTTCCGATCTNNNACTGGCTCTTTCTCATAACGACTC

GCACACGTCTGAACTCTTTCCCTACACGACGCTCTTCCGATCTNNNACTGGCTCTTTCTCATAACGACTC

AGCACACGTCTGAACTCTTTCCCTACACGACGCTCTTCCGATCTNNNGTGACAAAGTGAGACCCAGTCT

AGCACACGTCTGAACTCTTTCCCTACACGACGCTCTTCCGATCTNNNGTGACAAAGTGAGACCCAGTCT

GAGCACACGTCTGAACTCTTTCCCTACACGACGCTCTTCCGATCTNNNTGTATCAGTCAAGGTTCTCCAC

GAGCACACGTCTGAACTCTTTCCCTACACGACGCTCTTCCGATCTNNNTGTATCAGTCAAGGTTCTCCAC

AGCACACGTCTGAACTCTTTCCCTACACGACGCTCTTCCGATCTNNNACATGGCAAAAGCCCATTCATAC

AGCACACGTCTGAACTCTTTCCCTACACGACGCTCTTCCGATCTNNNACATGGCAAAAGCCCATTCATAC

AGCACACGTCTGAACTCTTTCCCTACACGACGCTCTTCCGATCTNNNAAGTACATGGGACTACAGGCA

AGCACACGTCTGAACTCTTTCCCTACACGACGCTCTTCCGATCTNNNAAGTACATGGGACTACAGGCA

GAGCACACGTCTGAACTCTTTCCCTACACGACGCTCTTCCGATCTNNNGCTTGTAGTCCCAGCTACTCTG

GAGCACACGTCTGAACTCTTTCCCTACACGACGCTCTTCCGATCTNNNGCTTGTAGTCCCAGCTACTCTG

AGCACACGTCTGAACTCTTTCCCTACACGACGCTCTTCCGATCTNNNGAGACAGGAGAATGGCGTGAA

AGCACACGTCTGAACTCTTTCCCTACACGACGCTCTTCCGATCTNNNGAGACAGGAGAATGGCGTGAA

GAGCACACGTCTGAACTCTTTCCCTACACGACGCTCTTCCGATCTNNNAACAGAGGAACCAAACTGCCTT

GAGCACACGTCTGAACTCTTTCCCTACACGACGCTCTTCCGATCTNNNAACAGAGGAACCAAACTGCCTT

GAGCACACGTCTGAACTCTTTCCCTACACGACGCTCTTCCGATCTNNNGGGCTACAGAGTGAGATGTTC

GAGCACACGTCTGAACTCTTTCCCTACACGACGCTCTTCCGATCTNNNGGGCTACAGAGTGAGATGTTC

AGCACACGTCTGAACTCTTTCCCTACACGACGCTCTTCCGATCTNNNAGGGCAATATAAAGGGAAAAT

AGCACACGTCTGAACTCTTTCCCTACACGACGCTCTTCCGATCTNNNAGGGCAATATAAAGGGAAAAT

GAGCACACGTCTGAACTCTTTCCCTACACGACGCTCTTCCGATCTNNNGATCTCTGATGATCCGCTCTCC

GAGCACACGTCTGAACTCTTTCCCTACACGACGCTCTTCCGATCTNNNGATCTCTGATGATCCGCTCTCC

AGCACACGTCTGAACTCTTTCCCTACACGACGCTCTTCCGATCTNNNCACAGTGTCTGAAACATGCTGA

AGCACACGTCTGAACTCTTTCCCTACACGACGCTCTTCCGATCTNNNCACAGTGTCTGAAACATGCTGA

AGAGCACACGTCTGAACTCTTTCCCTACACGACGCTCTTCCGATCTNNNAGTTCCCTGGTCCCATTTTCA

AGAGCACACGTCTGAACTCTTTCCCTACACGACGCTCTTCCGATCTNNNAGTTCCCTGGTCCCATTTTCA

GAGCACACGTCTGAACTCTTTCCCTACACGACGCTCTTCCGATCTNNNTCTGCATCTCTTGAGATAATTC

GAGCACACGTCTGAACTCTTTCCCTACACGACGCTCTTCCGATCTNNNTCTGCATCTCTTGAGATAATTC

GAGCACACGTCTGAACTCTTTCCCTACACGACGCTCTTCCGATCTNNNCAAGATCGCACTACTGTACTCC

GAGCACACGTCTGAACTCTTTCCCTACACGACGCTCTTCCGATCTNNNCAAGATCGCACTACTGTACTC

AGCACACGTCTGAACTCTTTCCCTACACGACGCTCTTCCGATCTNNNTGGCCAGTTCTCTAGTAAGGA

AGCACACGTCTGAACTCTTTCCCTACACGACGCTCTTCCGATCTNNNTGGCCAGTTCTCTAGTAAGGA

GAGCACACGTCTGAACTCTTTCCCTACACGACGCTCTTCCGATCTNNNACCTCGTGAAACTGGAGAA

GAGCACACGTCTGAACTCTTTCCCTACACGACGCTCTTCCGATCTNNNACCTCGTGAAACTGGAGAA

AGCACACGTCTGAACTCTTTCCCTACACGACGCTCTTCCGATCTNNNGACAAGTTCCATAGCTTGAGG

AGCACACGTCTGAACTCTTTCCCTACACGACGCTCTTCCGATCTNNNGACAAGTTCCATAGCTTGAGG

AGCACACGTCTGAACTCTTTCCCTACACGACGCTCTTCCGATCTNNNTTAGACCTGAAATCGTGCCAC

AGCACACGTCTGAACTCTTTCCCTACACGACGCTCTTCCGATCTNNNTTAGACCTGAAATCGTGCCAC

AGCACACGTCTGAACTCTTTCCCTACACGACGCTCTTCCGATCTNNNCAGCAGATCCTCATCCCAGAGC

AGCACACGTCTGAACTCTTTCCCTACACGACGCTCTTCCGATCTNNNCAGCAGATCCTCATCCCAGAGC

AGCACACGTCTGAACTCTTTCCCTACACGACGCTCTTCCGATCTNNNTATCATCAATGGCTGCTAAAA

AGCACACGTCTGAACTCTTTCCCTACACGACGCTCTTCCGATCTNNNTATCATCAATGGCTGCTAAAA

GAGCACACGTCTGAACTCTTTCCCTACACGACGCTCTTCCGATCTNNNCTGAGGCAGGAGAATCACTT

GAGCACACGTCTGAACTCTTTCCCTACACGACGCTCTTCCGATCTNNNCTGAGGCAGGAGAATCACTT

AGCACACGTCTGAACTCTTTCCCTACACGACGCTCTTCCGATCTNNNTCAGATGGCTATGTGTGTGTG

AGCACACGTCTGAACTCTTTCCCTACACGACGCTCTTCCGATCTNNNTCAGATGGCTATGTGTGTGTG

GAGCACACGTCTGAACTCTTTCCCTACACGACGCTCTTCCGATCTNNNCTGTGCATGCGTGTTTTTCAT

GAGCACACGTCTGAACTCTTTCCCTACACGACGCTCTTCCGATCTNNNCTGTGCATGCGTGTTTTTCAT

GCACACGTCTGAACTCTTTCCCTACACGACGCTCTTCCGATCTNNNGCACCTGTAATCCCAGCTACTAG

GCACACGTCTGAACTCTTTCCCTACACGACGCTCTTCCGATCTNNNGCACCTGTAATCCCAGCTACTAG

GCACACGTCTGAACTCTTTCCCTACACGACGCTCTTCCGATCTNNNTGGAGTACTCAGTTTAATCCAG

GCACACGTCTGAACTCTTTCCCTACACGACGCTCTTCCGATCTNNNTGGAGTACTCAGTTTAATCCAG

AGCACACGTCTGAACTCTTTCCCTACACGACGCTCTTCCGATCTNNNCCAGGGAGAAAATCAGGTTCT

AGCACACGTCTGAACTCTTTCCCTACACGACGCTCTTCCGATCTNNNCCAGGGAGAAAATCAGGTTCT

AGCACACGTCTGAACTCTTTCCCTACACGACGCTCTTCCGATCTNNNTACCACCTTTCACCATGGATAC

AGCACACGTCTGAACTCTTTCCCTACACGACGCTCTTCCGATCTNNNTACCACCTTTCACCATGGATAC

AGAGCACACGTCTGAACTCTTTCCCTACACGACGCTCTTCCGATCTNNNCAATCTCGGCTCACTACAAC

AGAGCACACGTCTGAACTCTTTCCCTACACGACGCTCTTCCGATCTNNNCAATCTCGGCTCACTACAAC

GCACACGTCTGAACTCTTTCCCTACACGACGCTCTTCCGATCTNNNCATGCATGCATACAATTCCCCTCC

GCACACGTCTGAACTCTTTCCCTACACGACGCTCTTCCGATCTNNNCATGCATGCATACAATTCCCCTCC

GCACACGTCTGAACTCTTTCCCTACACGACGCTCTTCCGATCTNNNAAACATCTTCCCCTTCTCCCTTCC

GCACACGTCTGAACTCTTTCCCTACACGACGCTCTTCCGATCTNNNAAACATCTTCCCCTTCTCCCTTCC

AGCACACGTCTGAACTCTTTCCCTACACGACGCTCTTCCGATCTNNNATTAGGAGCTCTGTGTCAGAA

AGCACACGTCTGAACTCTTTCCCTACACGACGCTCTTCCGATCTNNNATTAGGAGCTCTGTGTCAGAA

GAGCACACGTCTGAACTCTTTCCCTACACGACGCTCTTCCGATCTNNNCTGCACATGTATCCCTGAACC

GAGCACACGTCTGAACTCTTTCCCTACACGACGCTCTTCCGATCTNNNCTGCACATGTATCCCTGAACC

AGCACACGTCTGAACTCTTTCCCTACACGACGCTCTTCCGATCTNNNCTCAAGTTATTTGCCCACCTTG

AGCACACGTCTGAACTCTTTCCCTACACGACGCTCTTCCGATCTNNNCTCAAGTTATTTGCCCACCTTG

AGCACACGTCTGAACTCTTTCCCTACACGACGCTCTTCCGATCTNNNTTGGGCTGTCTAGGTAACAGG

AGCACACGTCTGAACTCTTTCCCTACACGACGCTCTTCCGATCTNNNTTGGGCTGTCTAGGTAACAGG

GAGCACACGTCTGAACTCTTTCCCTACACGACGCTCTTCCGATCTNNNGAGAATGGTGTGAACTCAGG

GAGCACACGTCTGAACTCTTTCCCTACACGACGCTCTTCCGATCTNNNGAGAATGGTGTGAACTCAGG

GAGCACACGTCTGAACTCTTTCCCTACACGACGCTCTTCCGATCTNNNTAAAATCCAAAAACCAGCGGT
GAGCACACGTCTGAACTCTTTCCCTACACGACGCTCTTCCGATCTNNNTAAAATCCAAAAACCAGCGGT
GAGCACACGTCTGAACTCTTTCCCTACACGACGCTCTTCCGATCTNNNGAGGTTTCCAGACTGATTTGC
GAGCACACGTCTGAACTCTTTCCCTACACGACGCTCTTCCGATCTNNNGAGGTTTCCAGACTGATTTGC
AGCACACGTCTGAACTCTTTCCCTACACGACGCTCTTCCGATCTNNNGCCAGATTGCACTACTGTACTT
AGCACACGTCTGAACTCTTTCCCTACACGACGCTCTTCCGATCTNNNGCCAGATTGCACTACTGTACTT
GAGCACACGTCTGAACTCTTTCCCTACACGACGCTCTTCCGATCTNNNAGTCTCCCGAGTAGCATGAGT
GAGCACACGTCTGAACTCTTTCCCTACACGACGCTCTTCCGATCTNNNAGTCTCCCGAGTAGCATGAGT
GAGCACACGTCTGAACTCTTTCCCTACACGACGCTCTTCCGATCTNNNAGTCTCAAACTCCTGAGCTGA
GAGCACACGTCTGAACTCTTTCCCTACACGACGCTCTTCCGATCTNNNAGTCTCAAACTCCTGAGCTGA
GAGCACACGTCTGAACTCTTTCCCTACACGACGCTCTTCCGATCTNNNTTTCCATCTGCTGTCCATTGGT
GAGCACACGTCTGAACTCTTTCCCTACACGACGCTCTTCCGATCTNNNTTTCCATCTGCTGTCCATTGGT
GAGCACACGTCTGAACTCTTTCCCTACACGACGCTCTTCCGATCTNNNTGCAAATGGACAAATGGGAAC
GAGCACACGTCTGAACTCTTTCCCTACACGACGCTCTTCCGATCTNNNTGCAAATGGACAAATGGGAAC
GAGCACACGTCTGAACTCTTTCCCTACACGACGCTCTTCCGATCTNNNGAATCGGATGAGGGGAGTAC
GAGCACACGTCTGAACTCTTTCCCTACACGACGCTCTTCCGATCTNNNGAATCGGATGAGGGGAGTAC
AGCACACGTCTGAACTCTTTCCCTACACGACGCTCTTCCGATCTNNNTTCACACCATTCTCCTGCCTCCAT
AGCACACGTCTGAACTCTTTCCCTACACGACGCTCTTCCGATCTNNNTTCACACCATTCTCCTGCCTCCAT
GAGCACACGTCTGAACTCTTTCCCTACACGACGCTCTTCCGATCTNNNCAAGGGAGTTGCCAATCTAGC
GAGCACACGTCTGAACTCTTTCCCTACACGACGCTCTTCCGATCTNNNCAAGGGAGTTGCCAATCTAGC
GAGCACACGTCTGAACTCTTTCCCTACACGACGCTCTTCCGATCTNNNGCTCTGTCATTGCCCTCATACT
GAGCACACGTCTGAACTCTTTCCCTACACGACGCTCTTCCGATCTNNNGCTCTGTCATTGCCCTCATACT
GAGCACACGTCTGAACTCTTTCCCTACACGACGCTCTTCCGATCTNNNTAGACTGGTTCATGGTCACAG
GAGCACACGTCTGAACTCTTTCCCTACACGACGCTCTTCCGATCTNNNTAGACTGGTTCATGGTCACAG
GAGCACACGTCTGAACTCTTTCCCTACACGACGCTCTTCCGATCTNNNGAGCTGAGATCATACCACTCC
GAGCACACGTCTGAACTCTTTCCCTACACGACGCTCTTCCGATCTNNNGAGCTGAGATCATACCACTCC
AAGAGCACACGTCTGAACTCTTTCCCTACACGACGCTCTTCCGATCTNNNTTGAGCAGTATTGCGCTCT
AAGAGCACACGTCTGAACTCTTTCCCTACACGACGCTCTTCCGATCTNNNTTGAGCAGTATTGCGCTCT
AGCACACGTCTGAACTCTTTCCCTACACGACGCTCTTCCGATCTNNNCCACACTGTTTTCCATGATGGT
AGCACACGTCTGAACTCTTTCCCTACACGACGCTCTTCCGATCTNNNCCACACTGTTTTCCATGATGGT
AGCACACGTCTGAACTCTTTCCCTACACGACGCTCTTCCGATCTNNNCCAAAGTGTTGAGATTACAGG
AGCACACGTCTGAACTCTTTCCCTACACGACGCTCTTCCGATCTNNNCCAAAGTGTTGAGATTACAGG
GCACACGTCTGAACTCTTTCCCTACACGACGCTCTTCCGATCTNNNTGAAAATTAAGTGCTATGGTAAG
GCACACGTCTGAACTCTTTCCCTACACGACGCTCTTCCGATCTNNNTGAAAATTAAGTGCTATGGTAAG
GAGCACACGTCTGAACTCTTTCCCTACACGACGCTCTTCCGATCTNNNTCCATCTATAGTGCAACATGT
GAGCACACGTCTGAACTCTTTCCCTACACGACGCTCTTCCGATCTNNNTCCATCTATAGTGCAACATGT
AGCACACGTCTGAACTCTTTCCCTACACGACGCTCTTCCGATCTNNNGGAGAGGACTGCTTGAGAAGA
AGCACACGTCTGAACTCTTTCCCTACACGACGCTCTTCCGATCTNNNGGAGAGGACTGCTTGAGAAGA
GCACACGTCTGAACTCTTTCCCTACACGACGCTCTTCCGATCTNNNTGCCTTGAAAGAAAATGCACACA
GCACACGTCTGAACTCTTTCCCTACACGACGCTCTTCCGATCTNNNTGCCTTGAAAGAAAATGCACACA
AGCACACGTCTGAACTCTTTCCCTACACGACGCTCTTCCGATCTNNNGCAAAACTCCAGTAAATCGCTCA
AGCACACGTCTGAACTCTTTCCCTACACGACGCTCTTCCGATCTNNNGCAAAACTCCAGTAAATCGCTCA
ACACGTCTGAACTCTTTCCCTACACGACGCTCTTCCGATCTNNNCAGAGTTGTTCAACCATCACCACCATC

CACGTCTGAACTCTTTCCCTACACGACGCTCTTCCGATCTNNNCAGAGTTGTTCAACCATCACCACCAT
AGCACACGTCTGAACTCTTTCCCTACACGACGCTCTTCCGATCTNNNTTTTCCTCCCCAACACCTACTTCC
AGCACACGTCTGAACTCTTTCCCTACACGACGCTCTTCCGATCTNNNTTTTCCTCCCCAACACCTACTTCC
AGCACACGTCTGAACTCTTTCCCTACACGACGCTCTTCCGATCTNNNATAAGCCAGACCAATGTGACAG
AGCACACGTCTGAACTCTTTCCCTACACGACGCTCTTCCGATCTNNNATAAGCCAGACCAATGTGACAG
GCACACGTCTGAACTCTTTCCCTACACGACGCTCTTCCGATCTNNNAACATAATGAGACCCCCATCTCT
GCACACGTCTGAACTCTTTCCCTACACGACGCTCTTCCGATCTNNNAACATAATGAGACCCCCATCTCT
GCACACGTCTGAACTCTTTCCCTACACGACGCTCTTCCGATCTNNNTGGGTGATGTTTACTCTTCTCCT
GCACACGTCTGAACTCTTTCCCTACACGACGCTCTTCCGATCTNNNTGGGTGATGTTTACTCTTCTCCT
AGCACACGTCTGAACTCTTTCCCTACACGACGCTCTTCCGATCTNNNGACAGGAACAATCACACAACTG
AGCACACGTCTGAACTCTTTCCCTACACGACGCTCTTCCGATCTNNNGACAGGAACAATCACACAACTG
AGCACACGTCTGAACTCTTTCCCTACACGACGCTCTTCCGATCTNNNAAAATGGTAGAAGGAGAGAG
AGCACACGTCTGAACTCTTTCCCTACACGACGCTCTTCCGATCTNNNAAAATGGTAGAAGGAGAGAG
AGCACACGTCTGAACTCTTTCCCTACACGACGCTCTTCCGATCTNNNGCCTATGTCTGCTTTCATATGG
AGCACACGTCTGAACTCTTTCCCTACACGACGCTCTTCCGATCTNNNGCCTATGTCTGCTTTCATATGG
GAGCACACGTCTGAACTCTTTCCCTACACGACGCTCTTCCGATCTNNNAATGGTGTGTTTTCATCTGCA
GAGCACACGTCTGAACTCTTTCCCTACACGACGCTCTTCCGATCTNNNAATGGTGTGTTTTCATCTGCA
GAGCACACGTCTGAACTCTTTCCCTACACGACGCTCTTCCGATCTNNNTCAGTAGGAATGGTACCAGC
GAGCACACGTCTGAACTCTTTCCCTACACGACGCTCTTCCGATCTNNNTCAGTAGGAATGGTACCAGC
GAGCACACGTCTGAACTCTTTCCCTACACGACGCTCTTCCGATCTNNNGGCACCTCTAATCTCAGCTAC
GAGCACACGTCTGAACTCTTTCCCTACACGACGCTCTTCCGATCTNNNGGCACCTCTAATCTCAGCTAC
AGCACACGTCTGAACTCTTTCCCTACACGACGCTCTTCCGATCTNNNCTGGTAATCCTTTGGGACCAGT
AGCACACGTCTGAACTCTTTCCCTACACGACGCTCTTCCGATCTNNNCTGGTAATCCTTTGGGACCAGT
GAGCACACGTCTGAACTCTTTCCCTACACGACGCTCTTCCGATCTNNNATTACACCCAGCCTCCAATTCA
GAGCACACGTCTGAACTCTTTCCCTACACGACGCTCTTCCGATCTNNNATTACACCCAGCCTCCAATTCA
GCACACGTCTGAACTCTTTCCCTACACGACGCTCTTCCGATCTNNNTTCTTTCCTTCCTTCCTTCCCTCCA
GCACACGTCTGAACTCTTTCCCTACACGACGCTCTTCCGATCTNNNTTCTTTCCTTCCTTCCTTCCCTCCA
AGCACACGTCTGAACTCTTTCCCTACACGACGCTCTTCCGATCTNNNTATTGAACTCCTGGGCTCAAGT
AGCACACGTCTGAACTCTTTCCCTACACGACGCTCTTCCGATCTNNNTATTGAACTCCTGGGCTCAAGT
GAGCACACGTCTGAACTCTTTCCCTACACGACGCTCTTCCGATCTNNNATTTGCAGAGCTACTCAAGG
GAGCACACGTCTGAACTCTTTCCCTACACGACGCTCTTCCGATCTNNNATTTGCAGAGCTACTCAAGG
GCACACGTCTGAACTCTTTCCCTACACGACGCTCTTCCGATCTNNNCCAACAAGCCTTCCTCCAAAATTC
GCACACGTCTGAACTCTTTCCCTACACGACGCTCTTCCGATCTNNNCCAACAAGCCTTCCTCCAAAATTC
GCACACGTCTGAACTCTTTCCCTACACGACGCTCTTCCGATCTNNNTCCAGGGAATTTGTTTGTTCGT
GCACACGTCTGAACTCTTTCCCTACACGACGCTCTTCCGATCTNNNTCCAGGGAATTTGTTTGTTCGT
GAGCACACGTCTGAACTCTTTCCCTACACGACGCTCTTCCGATCTNNNTGCAAATAGGCATTCTGAGC
GAGCACACGTCTGAACTCTTTCCCTACACGACGCTCTTCCGATCTNNNTGCAAATAGGCATTCTGAGC
GCACACGTCTGAACTCTTTCCCTACACGACGCTCTTCCGATCTNNNCCAACAAGCAGTCATGACATGA
GCACACGTCTGAACTCTTTCCCTACACGACGCTCTTCCGATCTNNNCCAACAAGCAGTCATGACATGA
GAAGAGCACACGTCTGAACTCTTTCCCTACACGACGCTCTTCCGATCTNNNTAACGCCTGGATTTATCT
GAAGAGCACACGTCTGAACTCTTTCCCTACACGACGCTCTTCCGATCTNNNTAACGCCTGGATTTATCT
GAGCACACGTCTGAACTCTTTCCCTACACGACGCTCTTCCGATCTNNNATAGGGACTGTTCAAAAGCT
GAGCACACGTCTGAACTCTTTCCCTACACGACGCTCTTCCGATCTNNNATAGGGACTGTTCAAAAGCT

AGCACACGTCTGAACTCTTTCCCTACACGACGCTCTTCCGATCTNNNCCCACAAGCAGAACTACTTAG
AGCACACGTCTGAACTCTTTCCCTACACGACGCTCTTCCGATCTNNNCCCACAAGCAGAACTACTTAG
GAGCACACGTCTGAACTCTTTCCCTACACGACGCTCTTCCGATCTNNNGCAGTTGGAGAGCATGAAAC
GAGCACACGTCTGAACTCTTTCCCTACACGACGCTCTTCCGATCTNNNGCAGTTGGAGAGCATGAAAC
AGCACACGTCTGAACTCTTTCCCTACACGACGCTCTTCCGATCTNNNATAAAAAGTTGCCGCATATTC
AGCACACGTCTGAACTCTTTCCCTACACGACGCTCTTCCGATCTNNNATAAAAAGTTGCCGCATATTC
AGCACACGTCTGAACTCTTTCCCTACACGACGCTCTTCCGATCTNNNCCCTGAGGAAATGTTTGCACT
AGCACACGTCTGAACTCTTTCCCTACACGACGCTCTTCCGATCTNNNCCCTGAGGAAATGTTTGCACT
GCACACGTCTGAACTCTTTCCCTACACGACGCTCTTCCGATCTNNNCGCATCAGGCTTTCTTATGTCTG
GCACACGTCTGAACTCTTTCCCTACACGACGCTCTTCCGATCTNNNCGCATCAGGCTTTCTTATGTCTG
AGCACACGTCTGAACTCTTTCCCTACACGACGCTCTTCCGATCTNNNGCCCTAGTTGACATATGCTTCG
AGCACACGTCTGAACTCTTTCCCTACACGACGCTCTTCCGATCTNNNGCCCTAGTTGACATATGCTTCG
ACACGTCTGAACTCTTTCCCTACACGACGCTCTTCCGATCTNNNTATAACACATCTACTGAAAAAGCC
ACACGTCTGAACTCTTTCCCTACACGACGCTCTTCCGATCTNNNTATAACACATCTACTGAAAAAGCC
AGCACACGTCTGAACTCTTTCCCTACACGACGCTCTTCCGATCTNNNGAGCCTGTGCCCATGAGTATA
AGCACACGTCTGAACTCTTTCCCTACACGACGCTCTTCCGATCTNNNGAGCCTGTGCCCATGAGTATA
AGCACACGTCTGAACTCTTTCCCTACACGACGCTCTTCCGATCTNNNGTCACAGTACTGATAGGCCTA
AGCACACGTCTGAACTCTTTCCCTACACGACGCTCTTCCGATCTNNNGTCACAGTACTGATAGGCCTA
AGCACACGTCTGAACTCTTTCCCTACACGACGCTCTTCCGATCTNNNCTGTAGTCTCAGCCACTAGGG
AGCACACGTCTGAACTCTTTCCCTACACGACGCTCTTCCGATCTNNNCTGTAGTCTCAGCCACTAGGG
CACACGTCTGAACTCTTTCCCTACACGACGCTCTTCCGATCTNNNAACAGGGAGAAGAAAGACAGCA
CACACGTCTGAACTCTTTCCCTACACGACGCTCTTCCGATCTNNNAACAGGGAGAAGAAAGACAGCA
GAGCACACGTCTGAACTCTTTCCCTACACGACGCTCTTCCGATCTNNNCTCACTGCAACCTCCACTTCCC
GAGCACACGTCTGAACTCTTTCCCTACACGACGCTCTTCCGATCTNNNCTCACTGCAACCTCCACTTCCC
GAGCACACGTCTGAACTCTTTCCCTACACGACGCTCTTCCGATCTNNNTGCCTAAAAAAAAACCTGGAT
GAGCACACGTCTGAACTCTTTCCCTACACGACGCTCTTCCGATCTNNNTGCCTAAAAAAAAACCTGGAT
AGCACACGTCTGAACTCTTTCCCTACACGACGCTCTTCCGATCTNNNCTGTAATCCCAGCTACTCAGG
AGCACACGTCTGAACTCTTTCCCTACACGACGCTCTTCCGATCTNNNCTGTAATCCCAGCTACTCAGG
AGCACACGTCTGAACTCTTTCCCTACACGACGCTCTTCCGATCTNNNTGAGACTCTGATTGCAGTGG
AGCACACGTCTGAACTCTTTCCCTACACGACGCTCTTCCGATCTNNNTGAGACTCTGATTGCAGTGG
AGCACACGTCTGAACTCTTTCCCTACACGACGCTCTTCCGATCTNNNAGCTTTTGTTGGCTATAACAG
AGCACACGTCTGAACTCTTTCCCTACACGACGCTCTTCCGATCTNNNAGCTTTTGTTGGCTATAACAG
AGCACACGTCTGAACTCTTTCCCTACACGACGCTCTTCCGATCTNNNAGTGTTGGTCTCCCAAGAGGT
AGCACACGTCTGAACTCTTTCCCTACACGACGCTCTTCCGATCTNNNAGTGTTGGTCTCCCAAGAGGT
AGCACACGTCTGAACTCTTTCCCTACACGACGCTCTTCCGATCTNNNACTGGCAGAGAATGGTACTT
AGCACACGTCTGAACTCTTTCCCTACACGACGCTCTTCCGATCTNNNACTGGCAGAGAATGGTACTT
GCACACGTCTGAACTCTTTCCCTACACGACGCTCTTCCGATCTNNNTGAGTAACATGTGCTCACCATC
GCACACGTCTGAACTCTTTCCCTACACGACGCTCTTCCGATCTNNNTGAGTAACATGTGCTCACCATC
AGCACACGTCTGAACTCTTTCCCTACACGACGCTCTTCCGATCTNNNTCCGAAAAATGTAGGCAGTGA
AGCACACGTCTGAACTCTTTCCCTACACGACGCTCTTCCGATCTNNNTCCGAAAAATGTAGGCAGTGA
AGCACACGTCTGAACTCTTTCCCTACACGACGCTCTTCCGATCTNNNCAAGGTCAACACAATGCATGA
AGCACACGTCTGAACTCTTTCCCTACACGACGCTCTTCCGATCTNNNCAAGGTCAACACAATGCATGA
GCACACGTCTGAACTCTTTCCCTACACGACGCTCTTCCGATCTNNNTGCCCTCTTTTTATTCACCACA

AGCACACGTCTGAACTCTTTCCCTACACGACGCTCTTCCGATCTNNNTGCCCTCTTTTTATTCACCACCAC

CACACGTCTGAACTCTTTCCCTACACGACGCTCTTCCGATCTNNNGTAATCGTAGCTACTCAGGAGGCC

CACACGTCTGAACTCTTTCCCTACACGACGCTCTTCCGATCTNNNGTAATCGTAGCTACTCAGGAGGCC

GAGCACACGTCTGAACTCTTTCCCTACACGACGCTCTTCCGATCTNNNGTTGCAGTGAGCTGAGATTG

GAGCACACGTCTGAACTCTTTCCCTACACGACGCTCTTCCGATCTNNNGTTGCAGTGAGCTGAGATTG

GAGCACACGTCTGAACTCTTTCCCTACACGACGCTCTTCCGATCTNNNAAGGTGTTTTTGTGTGTGTGT

GAGCACACGTCTGAACTCTTTCCCTACACGACGCTCTTCCGATCTNNNAAGGTGTTTTTGTGTGTGTGT

AGCACACGTCTGAACTCTTTCCCTACACGACGCTCTTCCGATCTNNNGGAGTCTGAGGCACAAAAATCA

AGCACACGTCTGAACTCTTTCCCTACACGACGCTCTTCCGATCTNNNGGAGTCTGAGGCACAAAAATCA

GAGCACACGTCTGAACTCTTTCCCTACACGACGCTCTTCCGATCTNNNGAGAGAGAGAGAGAGAGAGA

GAGCACACGTCTGAACTCTTTCCCTACACGACGCTCTTCCGATCTNNNGAGAGAGAGAGAGAGAGAGA

AGCACACGTCTGAACTCTTTCCCTACACGACGCTCTTCCGATCTNNNGAGGCTGAGGCAGGAGAATGC

AGCACACGTCTGAACTCTTTCCCTACACGACGCTCTTCCGATCTNNNGAGGCTGAGGCAGGAGAATGC

GAGCACACGTCTGAACTCTTTCCCTACACGACGCTCTTCCGATCTNNNATCTTCCATTTGTGGCCTGAGA

GAGCACACGTCTGAACTCTTTCCCTACACGACGCTCTTCCGATCTNNNATCTTCCATTTGTGGCCTGAGA

GAGCACACGTCTGAACTCTTTCCCTACACGACGCTCTTCCGATCTNNNATGTTCATTGCAGCCGTATTCA

GAGCACACGTCTGAACTCTTTCCCTACACGACGCTCTTCCGATCTNNNATGTTCATTGCAGCCGTATTCA

AGAGCACACGTCTGAACTCTTTCCCTACACGACGCTCTTCCGATCTNNNGTTCCATGTGGCATTCGACCT

AGAGCACACGTCTGAACTCTTTCCCTACACGACGCTCTTCCGATCTNNNGTTCCATGTGGCATTCGACCT

AGCACACGTCTGAACTCTTTCCCTACACGACGCTCTTCCGATCTNNNTCAGGGTTATCTAGGGATGGCA

AGCACACGTCTGAACTCTTTCCCTACACGACGCTCTTCCGATCTNNNTCAGGGTTATCTAGGGATGGCA

AGAGCACACGTCTGAACTCTTTCCCTACACGACGCTCTTCCGATCTNNNAGAGTCAGCTGTCTCCTCAA

AGAGCACACGTCTGAACTCTTTCCCTACACGACGCTCTTCCGATCTNNNAGAGTCAGCTGTCTCCTCAA

AGCACACGTCTGAACTCTTTCCCTACACGACGCTCTTCCGATCTNNNCTATCCAGGTCACGTGGTTAGA

AGCACACGTCTGAACTCTTTCCCTACACGACGCTCTTCCGATCTNNNCTATCCAGGTCACGTGGTTAGA

AAGAGCACACGTCTGAACTCTTTCCCTACACGACGCTCTTCCGATCTNNNCAGTAGTTCAGCAGGCAGT

AAGAGCACACGTCTGAACTCTTTCCCTACACGACGCTCTTCCGATCTNNNCAGTAGTTCAGCAGGCAGT

AGCACACGTCTGAACTCTTTCCCTACACGACGCTCTTCCGATCTNNNCCTTTTTTTGCTCAGCATTGTGA

AGCACACGTCTGAACTCTTTCCCTACACGACGCTCTTCCGATCTNNNCCTTTTTTTGCTCAGCATTGTGA

GAGCACACGTCTGAACTCTTTCCCTACACGACGCTCTTCCGATCTNNNAAGGTCTTGCTCTATTGCCCAG

GAGCACACGTCTGAACTCTTTCCCTACACGACGCTCTTCCGATCTNNNAAGGTCTTGCTCTATTGCCCAG

GAGCACACGTCTGAACTCTTTCCCTACACGACGCTCTTCCGATCTNNNACATGGCTGTTCATACTGAGG

GAGCACACGTCTGAACTCTTTCCCTACACGACGCTCTTCCGATCTNNNACATGGCTGTTCATACTGAGG

GAGCACACGTCTGAACTCTTTCCCTACACGACGCTCTTCCGATCTNNNGATGGCTTATGATCTTGGTCCC

GAGCACACGTCTGAACTCTTTCCCTACACGACGCTCTTCCGATCTNNNGATGGCTTATGATCTTGGTCCC

CACACGTCTGAACTCTTTCCCTACACGACGCTCTTCCGATCTNNNCTATCAGAATGGCAAAGACGCGCA

CACACGTCTGAACTCTTTCCCTACACGACGCTCTTCCGATCTNNNCTATCAGAATGGCAAAGACGCGCA

GCACACGTCTGAACTCTTTCCCTACACGACGCTCTTCCGATCTNNNGGTGACAGAGAGAGATTCCGTC

GCACACGTCTGAACTCTTTCCCTACACGACGCTCTTCCGATCTNNNGGTGACAGAGAGAGATTCCGTC

CACACGTCTGAACTCTTTCCCTACACGACGCTCTTCCGATCTNNNTCCGAAGGAAAGGCTTATTCACGC

CACACGTCTGAACTCTTTCCCTACACGACGCTCTTCCGATCTNNNTCCGAAGGAAAGGCTTATTCACGC

GAGCACACGTCTGAACTCTTTCCCTACACGACGCTCTTCCGATCTNNNACTGGGGAAGTATGCCTGTTT

GAGCACACGTCTGAACTCTTTCCCTACACGACGCTCTTCCGATCTNNNACTGGGGAAGTATGCCTGTTT

AGCACACGTCTGAACTCTTTCCCTACACGACGCTCTTCCGATCTNNNTGTAGTCCCAGCTACTTAGGA
AGCACACGTCTGAACTCTTTCCCTACACGACGCTCTTCCGATCTNNNTGTAGTCCCAGCTACTTAGGA
GAGCACACGTCTGAACTCTTTCCCTACACGACGCTCTTCCGATCTNNNGGTTGTAGGGAGCCAAGATT
GAGCACACGTCTGAACTCTTTCCCTACACGACGCTCTTCCGATCTNNNGGTTGTAGGGAGCCAAGATT
GAGCACACGTCTGAACTCTTTCCCTACACGACGCTCTTCCGATCTNNNACCCAGACTGGTTTTGTGGT
GAGCACACGTCTGAACTCTTTCCCTACACGACGCTCTTCCGATCTNNNACCCAGACTGGTTTTGTGGT
AGCACACGTCTGAACTCTTTCCCTACACGACGCTCTTCCGATCTNNNAATTCACAGATCAAGAGCCCCT
AGCACACGTCTGAACTCTTTCCCTACACGACGCTCTTCCGATCTNNNAATTCACAGATCAAGAGCCCCT
GAGCACACGTCTGAACTCTTTCCCTACACGACGCTCTTCCGATCTNNNAGGAGGCTGAGACAAAAGA
GAGCACACGTCTGAACTCTTTCCCTACACGACGCTCTTCCGATCTNNNAGGAGGCTGAGACAAAAGA
GAGCACACGTCTGAACTCTTTCCCTACACGACGCTCTTCCGATCTNNNTCTCTGCATTTTCAGGAACGA
GAGCACACGTCTGAACTCTTTCCCTACACGACGCTCTTCCGATCTNNNTCTCTGCATTTTCAGGAACGA
GAGCACACGTCTGAACTCTTTCCCTACACGACGCTCTTCCGATCTNNNAGAATGGTGTGAACCCAGA
GAGCACACGTCTGAACTCTTTCCCTACACGACGCTCTTCCGATCTNNNAGAATGGTGTGAACCCAGA
CACACGTCTGAACTCTTTCCCTACACGACGCTCTTCCGATCTNNNGAAACGTCTAGATGCTCAACCATC
CACACGTCTGAACTCTTTCCCTACACGACGCTCTTCCGATCTNNNGAAACGTCTAGATGCTCAACCATC
GAGCACACGTCTGAACTCTTTCCCTACACGACGCTCTTCCGATCTNNNCTGCTGTTTTACTAAGAACGA
GAGCACACGTCTGAACTCTTTCCCTACACGACGCTCTTCCGATCTNNNCTGCTGTTTTACTAAGAACGA
AGCACACGTCTGAACTCTTTCCCTACACGACGCTCTTCCGATCTNNNTCAAGCAGAAGGATGTTCCAA
AGCACACGTCTGAACTCTTTCCCTACACGACGCTCTTCCGATCTNNNTCAAGCAGAAGGATGTTCCAA
AGCACACGTCTGAACTCTTTCCCTACACGACGCTCTTCCGATCTNNNCCCAAATGATGAAGTGATGCA
AGCACACGTCTGAACTCTTTCCCTACACGACGCTCTTCCGATCTNNNCCCAAATGATGAAGTGATGCA
AGCACACGTCTGAACTCTTTCCCTACACGACGCTCTTCCGATCTNNNGCACATGCCTGTAATCCTAGCT
AGCACACGTCTGAACTCTTTCCCTACACGACGCTCTTCCGATCTNNNGCACATGCCTGTAATCCTAGCT
CACACGTCTGAACTCTTTCCCTACACGACGCTCTTCCGATCTNNNGATTAAATGGACTTTACCTTCAGC
CACACGTCTGAACTCTTTCCCTACACGACGCTCTTCCGATCTNNNGATTAAATGGACTTTACCTTCAGC
AGCACACGTCTGAACTCTTTCCCTACACGACGCTCTTCCGATCTNNNTCCTCCATCTCACACAGGAAGT
AGCACACGTCTGAACTCTTTCCCTACACGACGCTCTTCCGATCTNNNTCCTCCATCTCACACAGGAAGT
AGCACACGTCTGAACTCTTTCCCTACACGACGCTCTTCCGATCTNNNTTCAAACTCCTGGGGCTCATG
AGCACACGTCTGAACTCTTTCCCTACACGACGCTCTTCCGATCTNNNTTCAAACTCCTGGGGCTCATG
GCACACGTCTGAACTCTTTCCCTACACGACGCTCTTCCGATCTNNNGCTGGTAAATTGAAGAGCTGG
GCACACGTCTGAACTCTTTCCCTACACGACGCTCTTCCGATCTNNNGCTGGTAAATTGAAGAGCTGG
GCACACGTCTGAACTCTTTCCCTACACGACGCTCTTCCGATCTNNNAATGTGGATCAACTATGTGTGT
GCACACGTCTGAACTCTTTCCCTACACGACGCTCTTCCGATCTNNNAATGTGGATCAACTATGTGTGT
AGCACACGTCTGAACTCTTTCCCTACACGACGCTCTTCCGATCTNNNTCTAGCTTCATCCATGTACCTGC
AGCACACGTCTGAACTCTTTCCCTACACGACGCTCTTCCGATCTNNNTCTAGCTTCATCCATGTACCTGC
GAGCACACGTCTGAACTCTTTCCCTACACGACGCTCTTCCGATCTNNNCTGAAATCCCAGCTACTCAGG
GAGCACACGTCTGAACTCTTTCCCTACACGACGCTCTTCCGATCTNNNCTGAAATCCCAGCTACTCAGG
GCACACGTCTGAACTCTTTCCCTACACGACGCTCTTCCGATCTNNNAGGGTTCAAGACTGATTGGTG
GCACACGTCTGAACTCTTTCCCTACACGACGCTCTTCCGATCTNNNAGGGTTCAAGACTGATTGGTG
GCACACGTCTGAACTCTTTCCCTACACGACGCTCTTCCGATCTNNNAGAGCTTGCAGTCAGATGAGAT
GCACACGTCTGAACTCTTTCCCTACACGACGCTCTTCCGATCTNNNAGAGCTTGCAGTCAGATGAGAT
GAGCACACGTCTGAACTCTTTCCCTACACGACGCTCTTCCGATCTNNNACTAGAGAACCAAGAGCAAG

GAGCACACGTCTGAACTCTTTCCCTACACGACGCTCTTCCGATCTNNNACTAGAGAACCAAGAGCAAG
AGCACACGTCTGAACTCTTTCCCTACACGACGCTCTTCCGATCTNNNTATACAGTTCAGGTACTTTGT
AGCACACGTCTGAACTCTTTCCCTACACGACGCTCTTCCGATCTNNNTATACAGTTCAGGTACTTTGT
GCACACGTCTGAACTCTTTCCCTACACGACGCTCTTCCGATCTNNNCCCAAAATCCAACGTGTCATTC
GCACACGTCTGAACTCTTTCCCTACACGACGCTCTTCCGATCTNNNCCCAAAATCCAACGTGTCATTC
AGCACACGTCTGAACTCTTTCCCTACACGACGCTCTTCCGATCTNNNGGGTGACAGACTGAGACACT
AGCACACGTCTGAACTCTTTCCCTACACGACGCTCTTCCGATCTNNNGGGTGACAGACTGAGACACT
GCACACGTCTGAACTCTTTCCCTACACGACGCTCTTCCGATCTNNNGTGGCTGGTTTGTCTTTGATTC
GCACACGTCTGAACTCTTTCCCTACACGACGCTCTTCCGATCTNNNGTGGCTGGTTTGTCTTTGATTC
GCACACGTCTGAACTCTTTCCCTACACGACGCTCTTCCGATCTNNNCTAGTGAGGTTAATGATCCTTC
GCACACGTCTGAACTCTTTCCCTACACGACGCTCTTCCGATCTNNNCTAGTGAGGTTAATGATCCTTC
AGCACACGTCTGAACTCTTTCCCTACACGACGCTCTTCCGATCTNNNCTGTAGTCCCAGCTACTCAGG
AGCACACGTCTGAACTCTTTCCCTACACGACGCTCTTCCGATCTNNNCTGTAGTCCCAGCTACTCAGG
AGCACACGTCTGAACTCTTTCCCTACACGACGCTCTTCCGATCTNNNACACGGAATAAGAACTTCCTG
AGCACACGTCTGAACTCTTTCCCTACACGACGCTCTTCCGATCTNNNACACGGAATAAGAACTTCCTG
GCACACGTCTGAACTCTTTCCCTACACGACGCTCTTCCGATCTNNNCTCCTAGGGATTTTGTGGTTTGC
GCACACGTCTGAACTCTTTCCCTACACGACGCTCTTCCGATCTNNNCTCCTAGGGATTTTGTGGTTTGC
AGCACACGTCTGAACTCTTTCCCTACACGACGCTCTTCCGATCTNNNCTCTCTTTCTTTTCCCTGCAAG
AGCACACGTCTGAACTCTTTCCCTACACGACGCTCTTCCGATCTNNNCTCTCTTTCTTTTCCCTGCAAG
AGCACACGTCTGAACTCTTTCCCTACACGACGCTCTTCCGATCTNNNCAAGAGAATCGCTTGAACCTT
AGCACACGTCTGAACTCTTTCCCTACACGACGCTCTTCCGATCTNNNCAAGAGAATCGCTTGAACCTT
GAGCACACGTCTGAACTCTTTCCCTACACGACGCTCTTCCGATCTNNNCTGAGGCAGGAGATTCAGT
GAGCACACGTCTGAACTCTTTCCCTACACGACGCTCTTCCGATCTNNNCTGAGGCAGGAGATTCAGT
AGCACACGTCTGAACTCTTTCCCTACACGACGCTCTTCCGATCTNNNACAGGAACCCTAATTCGACCA
AGCACACGTCTGAACTCTTTCCCTACACGACGCTCTTCCGATCTNNNACAGGAACCCTAATTCGACCA
AGCACACGTCTGAACTCTTTCCCTACACGACGCTCTTCCGATCTNNNAGAACAGTCATTGTAGGCATC
AGCACACGTCTGAACTCTTTCCCTACACGACGCTCTTCCGATCTNNNAGAACAGTCATTGTAGGCATC
GAGCACACGTCTGAACTCTTTCCCTACACGACGCTCTTCCGATCTNNNGAGCTTGCAGTGAGGTGAG
GAGCACACGTCTGAACTCTTTCCCTACACGACGCTCTTCCGATCTNNNGAGCTTGCAGTGAGGTGAG
AGCACACGTCTGAACTCTTTCCCTACACGACGCTCTTCCGATCTNNNGCATTTTCCCAGCTTCCACTTT
AGCACACGTCTGAACTCTTTCCCTACACGACGCTCTTCCGATCTNNNGCATTTTCCCAGCTTCCACTTT
AGCACACGTCTGAACTCTTTCCCTACACGACGCTCTTCCGATCTNNNCTGCAATCTAGGAAAGGCCTC
AGCACACGTCTGAACTCTTTCCCTACACGACGCTCTTCCGATCTNNNCTGCAATCTAGGAAAGGCCTC
GCACACGTCTGAACTCTTTCCCTACACGACGCTCTTCCGATCTNNNTGTTCTTGCCTCTCTAGTTCCTTC
GCACACGTCTGAACTCTTTCCCTACACGACGCTCTTCCGATCTNNNTGTTCTTGCCTCTCTAGTTCCTTC
AGCACACGTCTGAACTCTTTCCCTACACGACGCTCTTCCGATCTNNNACCACGCCTGGCTAATTTTTGT
AGCACACGTCTGAACTCTTTCCCTACACGACGCTCTTCCGATCTNNNACCACGCCTGGCTAATTTTTGT
AGAGCACACGTCTGAACTCTTTCCCTACACGACGCTCTTCCGATCTNNNGCAGGAGAATTGCTTGAA
AGAGCACACGTCTGAACTCTTTCCCTACACGACGCTCTTCCGATCTNNNGCAGGAGAATTGCTTGAA
GCACACGTCTGAACTCTTTCCCTACACGACGCTCTTCCGATCTNNNTGTTCTCGTGACATTGAGTGAG
GCACACGTCTGAACTCTTTCCCTACACGACGCTCTTCCGATCTNNNTGTTCTCGTGACATTGAGTGAG
AGCACACGTCTGAACTCTTTCCCTACACGACGCTCTTCCGATCTNNNCCTCTATTTGAATGGGCCTTTG
AGCACACGTCTGAACTCTTTCCCTACACGACGCTCTTCCGATCTNNNCCTCTATTTGAATGGGCCTTTG

GCACACGTCTGAACTCTTTCCCTACACGACGCTCTTCCGATCTNNNCATGAGAATTGCTTGAACTCGG

GCACACGTCTGAACTCTTTCCCTACACGACGCTCTTCCGATCTNNNCATGAGAATTGCTTGAACTCGG

AGAGCACACGTCTGAACTCTTTCCCTACACGACGCTCTTCCGATCTNNNCTTAGGTGGGAGGATTGC

AGAGCACACGTCTGAACTCTTTCCCTACACGACGCTCTTCCGATCTNNNCTTAGGTGGGAGGATTGC

CACACGTCTGAACTCTTTCCCTACACGACGCTCTTCCGATCTNNNGCTAAAAGCATGGACACTGGAG

CACACGTCTGAACTCTTTCCCTACACGACGCTCTTCCGATCTNNNGCTAAAAGCATGGACACTGGAG

ACACGTCTGAACTCTTTCCCTACACGACGCTCTTCCGATCTNNNTAGCTATCCTGGCTGAAAGAACCC

ACACGTCTGAACTCTTTCCCTACACGACGCTCTTCCGATCTNNNTAGCTATCCTGGCTGAAAGAACCC

AGCACACGTCTGAACTCTTTCCCTACACGACGCTCTTCCGATCTNNNCTCAAGTGCTTACATATGCTG

AGCACACGTCTGAACTCTTTCCCTACACGACGCTCTTCCGATCTNNNCTCAAGTGCTTACATATGCTG

GAGCACACGTCTGAACTCTTTCCCTACACGACGCTCTTCCGATCTNNNATGCCTGTAATCTCAGCTACT

GAGCACACGTCTGAACTCTTTCCCTACACGACGCTCTTCCGATCTNNNATGCCTGTAATCTCAGCTACT

AGCACACGTCTGAACTCTTTCCCTACACGACGCTCTTCCGATCTNNNCTAGGCATCCAGAGTCTTGTGT

AGCACACGTCTGAACTCTTTCCCTACACGACGCTCTTCCGATCTNNNCTAGGCATCCAGAGTCTTGTGT

GCACACGTCTGAACTCTTTCCCTACACGACGCTCTTCCGATCTNNNTATTTGGAGTGTTGCCTCAACT

GCACACGTCTGAACTCTTTCCCTACACGACGCTCTTCCGATCTNNNTATTTGGAGTGTTGCCTCAACT

AGCACACGTCTGAACTCTTTCCCTACACGACGCTCTTCCGATCTNNNTTTGAGGCTAGAAGTTCGAG

AGCACACGTCTGAACTCTTTCCCTACACGACGCTCTTCCGATCTNNNTTTGAGGCTAGAAGTTCGAG

GAGCACACGTCTGAACTCTTTCCCTACACGACGCTCTTCCGATCTNNNCCAGATGGCAATGTTGTCTCT

GAGCACACGTCTGAACTCTTTCCCTACACGACGCTCTTCCGATCTNNNCCAGATGGCAATGTTGTCTCT

GCACACGTCTGAACTCTTTCCCTACACGACGCTCTTCCGATCTNNNTTCCCTTTCTGCTCTAACAGGACC

GCACACGTCTGAACTCTTTCCCTACACGACGCTCTTCCGATCTNNNTTCCCTTTCTGCTCTAACAGGACC

AGCACACGTCTGAACTCTTTCCCTACACGACGCTCTTCCGATCTNNNATCAAGATCTGACCTTTGGACC

AGCACACGTCTGAACTCTTTCCCTACACGACGCTCTTCCGATCTNNNATCAAGATCTGACCTTTGGACC

AGCACACGTCTGAACTCTTTCCCTACACGACGCTCTTCCGATCTNNNTGGCAAAGGAAAAAAGTTGC

AGCACACGTCTGAACTCTTTCCCTACACGACGCTCTTCCGATCTNNNTGGCAAAGGAAAAAAGTTGC

AGCACACGTCTGAACTCTTTCCCTACACGACGCTCTTCCGATCTNNNTGTGAACACTCAGTGAGGTAA

AGCACACGTCTGAACTCTTTCCCTACACGACGCTCTTCCGATCTNNNTGTGAACACTCAGTGAGGTAA

AGCACACGTCTGAACTCTTTCCCTACACGACGCTCTTCCGATCTNNNAATCAGACAACTCTTGCCAAAG

AGCACACGTCTGAACTCTTTCCCTACACGACGCTCTTCCGATCTNNNAATCAGACAACTCTTGCCAAAG

AGCACACGTCTGAACTCTTTCCCTACACGACGCTCTTCCGATCTNNNGCAGGAAGAAAACTTTGCTA

AGCACACGTCTGAACTCTTTCCCTACACGACGCTCTTCCGATCTNNNGCAGGAAGAAAACTTTGCTA

AGCACACGTCTGAACTCTTTCCCTACACGACGCTCTTCCGATCTNNNGAAGCAAAATGGGTCTCCCAT

AGCACACGTCTGAACTCTTTCCCTACACGACGCTCTTCCGATCTNNNGAAGCAAAATGGGTCTCCCAT

AGAGCACACGTCTGAACTCTTTCCCTACACGACGCTCTTCCGATCTNNNAGCCAATGGATAGCTGTAG

AGAGCACACGTCTGAACTCTTTCCCTACACGACGCTCTTCCGATCTNNNAGCCAATGGATAGCTGTAG

AGCACACGTCTGAACTCTTTCCCTACACGACGCTCTTCCGATCTNNNCAGGAGAATTGCTTGAACAC

AGCACACGTCTGAACTCTTTCCCTACACGACGCTCTTCCGATCTNNNCAGGAGAATTGCTTGAACAC

CACACGTCTGAACTCTTTCCCTACACGACGCTCTTCCGATCTNNNCCTCTCCTGTTCTTACTAAGCCCC

CACACGTCTGAACTCTTTCCCTACACGACGCTCTTCCGATCTNNNCCTCTCCTGTTCTTACTAAGCCCC

GCACACGTCTGAACTCTTTCCCTACACGACGCTCTTCCGATCTNNNGTTCAAGGTTGCAGTGAGTTAC

GCACACGTCTGAACTCTTTCCCTACACGACGCTCTTCCGATCTNNNGTTCAAGGTTGCAGTGAGTTAC

ACACGTCTGAACTCTTTCCCTACACGACGCTCTTCCGATCTNNNACCATAGCACCATAGCAGTTTGACA

ACACGTCTGAACTCTTTCCCTACACGACGCTCTTCCGATCTNNNACCATAGCACCATAGCAGTTTGACA
GCACACGTCTGAACTCTTTCCCTACACGACGCTCTTCCGATCTNNNTCCTCAAGCCATGTCCTCAATTCC
GCACACGTCTGAACTCTTTCCCTACACGACGCTCTTCCGATCTNNNTCCTCAAGCCATGTCCTCAATTCC
AGCACACGTCTGAACTCTTTCCCTACACGACGCTCTTCCGATCTNNNGCTCAATCTTCCTTTGCCATCT
AGCACACGTCTGAACTCTTTCCCTACACGACGCTCTTCCGATCTNNNGCTCAATCTTCCTTTGCCATCT
GCACACGTCTGAACTCTTTCCCTACACGACGCTCTTCCGATCTNNNCACTGGCATCCCTACTTCTCCCAT
GCACACGTCTGAACTCTTTCCCTACACGACGCTCTTCCGATCTNNNCACTGGCATCCCTACTTCTCCCAT
GAGCACACGTCTGAACTCTTTCCCTACACGACGCTCTTCCGATCTNNNAATCTCATGGGAGCACTGTT
GAGCACACGTCTGAACTCTTTCCCTACACGACGCTCTTCCGATCTNNNAATCTCATGGGAGCACTGTT
AGCACACGTCTGAACTCTTTCCCTACACGACGCTCTTCCGATCTNNNCCCTGTACCAAGATCATTCCGA
AGCACACGTCTGAACTCTTTCCCTACACGACGCTCTTCCGATCTNNNCCCTGTACCAAGATCATTCCGA
AGCACACGTCTGAACTCTTTCCCTACACGACGCTCTTCCGATCTNNNAACATATGCATGCCAGATGGT
AGCACACGTCTGAACTCTTTCCCTACACGACGCTCTTCCGATCTNNNAACATATGCATGCCAGATGGT
GCACACGTCTGAACTCTTTCCCTACACGACGCTCTTCCGATCTNNNACACATATGTACACACGGTGGA
GCACACGTCTGAACTCTTTCCCTACACGACGCTCTTCCGATCTNNNACACATATGTACACACGGTGGA
AGCACACGTCTGAACTCTTTCCCTACACGACGCTCTTCCGATCTNNNCTACCTCTTCACCTTCCTCAACC
AGCACACGTCTGAACTCTTTCCCTACACGACGCTCTTCCGATCTNNNCTACCTCTTCACCTTCCTCAACC
GAGCACACGTCTGAACTCTTTCCCTACACGACGCTCTTCCGATCTNNNATGCCTGTAATCCCAGCTAC
GAGCACACGTCTGAACTCTTTCCCTACACGACGCTCTTCCGATCTNNNATGCCTGTAATCCCAGCTAC
AGCACACGTCTGAACTCTTTCCCTACACGACGCTCTTCCGATCTNNNTGAAAAGACAGCCATCTGCAA
AGCACACGTCTGAACTCTTTCCCTACACGACGCTCTTCCGATCTNNNTGAAAAGACAGCCATCTGCAA
AGCACACGTCTGAACTCTTTCCCTACACGACGCTCTTCCGATCTNNNGTAATCCGGCCCAGGTTTTTT
AGCACACGTCTGAACTCTTTCCCTACACGACGCTCTTCCGATCTNNNGTAATCCGGCCCAGGTTTTTT
AGCACACGTCTGAACTCTTTCCCTACACGACGCTCTTCCGATCTNNNCAGAAGAATTGCTTGAACCTG
AGCACACGTCTGAACTCTTTCCCTACACGACGCTCTTCCGATCTNNNCAGAAGAATTGCTTGAACCTG
GCACACGTCTGAACTCTTTCCCTACACGACGCTCTTCCGATCTNNNGCTCCCTATATCTCCTCTCCCCTC
GCACACGTCTGAACTCTTTCCCTACACGACGCTCTTCCGATCTNNNGCTCCCTATATCTCCTCTCCCCTC
AGCACACGTCTGAACTCTTTCCCTACACGACGCTCTTCCGATCTNNNGAGAATCGCTTGAACCAGGAA
AGCACACGTCTGAACTCTTTCCCTACACGACGCTCTTCCGATCTNNNGAGAATCGCTTGAACCAGGAA
AGCACACGTCTGAACTCTTTCCCTACACGACGCTCTTCCGATCTNNNCTCGAACTCCTGACCTTGTGAT
AGCACACGTCTGAACTCTTTCCCTACACGACGCTCTTCCGATCTNNNCTCGAACTCCTGACCTTGTGAT
AGCACACGTCTGAACTCTTTCCCTACACGACGCTCTTCCGATCTNNNTCCCCATCTCCATAGTATTCCC
AGCACACGTCTGAACTCTTTCCCTACACGACGCTCTTCCGATCTNNNTCCCCATCTCCATAGTATTCCC
AGCACACGTCTGAACTCTTTCCCTACACGACGCTCTTCCGATCTNNNGGATGGATGAAGCAAGTTGT
AGCACACGTCTGAACTCTTTCCCTACACGACGCTCTTCCGATCTNNNGGATGGATGAAGCAAGTTGT
AGCACACGTCTGAACTCTTTCCCTACACGACGCTCTTCCGATCTNNNGTACATGTACATCCTCAGCCCA
AGCACACGTCTGAACTCTTTCCCTACACGACGCTCTTCCGATCTNNNGTACATGTACATCCTCAGCCCA
AGCACACGTCTGAACTCTTTCCCTACACGACGCTCTTCCGATCTNNNCAAGATCACGTGACTGCACTT
AGCACACGTCTGAACTCTTTCCCTACACGACGCTCTTCCGATCTNNNCAAGATCACGTGACTGCACTT
GCACACGTCTGAACTCTTTCCCTACACGACGCTCTTCCGATCTNNNCGTCTGTAATCCCAGCTACTCAA
GCACACGTCTGAACTCTTTCCCTACACGACGCTCTTCCGATCTNNNCGTCTGTAATCCCAGCTACTCAA
GAGCACACGTCTGAACTCTTTCCCTACACGACGCTCTTCCGATCTNNNCCTGGCCCTTCTTTTCTTTTCT
GAGCACACGTCTGAACTCTTTCCCTACACGACGCTCTTCCGATCTNNNCCTGGCCCTTCTTTTCTTTTCT

GAGCACACGTCTGAACTCTTTCCCTACACGACGCTCTTCCGATCTNNNTACATGCCTGTAATCCCAGCT

GAGCACACGTCTGAACTCTTTCCCTACACGACGCTCTTCCGATCTNNNTACATGCCTGTAATCCCAGCT

AGCACACGTCTGAACTCTTTCCCTACACGACGCTCTTCCGATCTNNNTCCTTTGCCCAGTTTTTAGTAG

AGCACACGTCTGAACTCTTTCCCTACACGACGCTCTTCCGATCTNNNTCCTTTGCCCAGTTTTTAGTAG

GAGCACACGTCTGAACTCTTTCCCTACACGACGCTCTTCCGATCTNNNCTCCTGTAGTCCCAGCTACTT

GAGCACACGTCTGAACTCTTTCCCTACACGACGCTCTTCCGATCTNNNCTCCTGTAGTCCCAGCTACTT

AGCACACGTCTGAACTCTTTCCCTACACGACGCTCTTCCGATCTNNNAATGCCAGAATAATCGCTGCA

AGCACACGTCTGAACTCTTTCCCTACACGACGCTCTTCCGATCTNNNAATGCCAGAATAATCGCTGCA

AGCACACGTCTGAACTCTTTCCCTACACGACGCTCTTCCGATCTNNNCGGTTAGGAACTGAGAGCAA

AGCACACGTCTGAACTCTTTCCCTACACGACGCTCTTCCGATCTNNNCGGTTAGGAACTGAGAGCAA

GCACACGTCTGAACTCTTTCCCTACACGACGCTCTTCCGATCTNNNTGCAGAAAAGGAAGTCTCTGT

GCACACGTCTGAACTCTTTCCCTACACGACGCTCTTCCGATCTNNNTGCAGAAAAGGAAGTCTCTGT

AGAGCACACGTCTGAACTCTTTCCCTACACGACGCTCTTCCGATCTNNNGAGCAGGAGGATTGAGG

AGAGCACACGTCTGAACTCTTTCCCTACACGACGCTCTTCCGATCTNNNGAGCAGGAGGATTGAGG

AGCACACGTCTGAACTCTTTCCCTACACGACGCTCTTCCGATCTNNNCTTAATGTGAGAGAGACAAA

AGCACACGTCTGAACTCTTTCCCTACACGACGCTCTTCCGATCTNNNCTTAATGTGAGAGAGACAAA

GCACACGTCTGAACTCTTTCCCTACACGACGCTCTTCCGATCTNNNCAGAGACACACACACATGAACA

GCACACGTCTGAACTCTTTCCCTACACGACGCTCTTCCGATCTNNNCAGAGACACACACACATGAACA

AGCACACGTCTGAACTCTTTCCCTACACGACGCTCTTCCGATCTNNNGAGAATCGCTTGAATCCGAGA

AGCACACGTCTGAACTCTTTCCCTACACGACGCTCTTCCGATCTNNNGAGAATCGCTTGAATCCGAGA

ACACGTCTGAACTCTTTCCCTACACGACGCTCTTCCGATCTNNNATTCAGGATTTCAATGACTTTTACA

ACACGTCTGAACTCTTTCCCTACACGACGCTCTTCCGATCTNNNATTCAGGATTTCAATGACTTTTACA

AGCACACGTCTGAACTCTTTCCCTACACGACGCTCTTCCGATCTNNNTGCTTGTGTGTTTGTGTTCAT

AGCACACGTCTGAACTCTTTCCCTACACGACGCTCTTCCGATCTNNNTGCTTGTGTGTTTGTGTTCAT

AGCACACGTCTGAACTCTTTCCCTACACGACGCTCTTCCGATCTNNNGAGGCAGGATAATTGCTTGA

AGCACACGTCTGAACTCTTTCCCTACACGACGCTCTTCCGATCTNNNGAGGCAGGATAATTGCTTGA

AGCACACGTCTGAACTCTTTCCCTACACGACGCTCTTCCGATCTNNNCTCCCCACTCTAGGAGTCTGTA

AGCACACGTCTGAACTCTTTCCCTACACGACGCTCTTCCGATCTNNNCTCCCCACTCTAGGAGTCTGTA

GAGCACACGTCTGAACTCTTTCCCTACACGACGCTCTTCCGATCTNNNTGTAGTGACACCAAGAGGCT

GAGCACACGTCTGAACTCTTTCCCTACACGACGCTCTTCCGATCTNNNTGTAGTGACACCAAGAGGCT

GAGCACACGTCTGAACTCTTTCCCTACACGACGCTCTTCCGATCTNNNTAGCACATCCTCGGTACTAGA

GAGCACACGTCTGAACTCTTTCCCTACACGACGCTCTTCCGATCTNNNTAGCACATCCTCGGTACTAGA

AGCACACGTCTGAACTCTTTCCCTACACGACGCTCTTCCGATCTNNNCTTGCAGTAAGCTGAGATTGC

AGCACACGTCTGAACTCTTTCCCTACACGACGCTCTTCCGATCTNNNCTTGCAGTAAGCTGAGATTGC

AGCACACGTCTGAACTCTTTCCCTACACGACGCTCTTCCGATCTNNNAGAACTTGCAGTGAGTCGAGA

AGCACACGTCTGAACTCTTTCCCTACACGACGCTCTTCCGATCTNNNAGAACTTGCAGTGAGTCGAGA

AGCACACGTCTGAACTCTTTCCCTACACGACGCTCTTCCGATCTNNNCTGAGACAGGAGAATCGCTTG

AGCACACGTCTGAACTCTTTCCCTACACGACGCTCTTCCGATCTNNNCTGAGACAGGAGAATCGCTTG

GAGCACACGTCTGAACTCTTTCCCTACACGACGCTCTTCCGATCTNNNGAGACATACAGTTGCCCAGA

GAGCACACGTCTGAACTCTTTCCCTACACGACGCTCTTCCGATCTNNNGAGACATACAGTTGCCCAGA

GCACACGTCTGAACTCTTTCCCTACACGACGCTCTTCCGATCTNNNGCTGCCATGTGAAAATGAGGAA

GCACACGTCTGAACTCTTTCCCTACACGACGCTCTTCCGATCTNNNGCTGCCATGTGAAAATGAGGAA

AGCACACGTCTGAACTCTTTCCCTACACGACGCTCTTCCGATCTNNNGCTTCATTGCCATTGACACCAT

GAGCACACGTCTGAACTCTTTCCCTACACGACGCTCTTCCGATCTNNNGCTTCATTGCCATTGACACCAT

AGCACACGTCTGAACTCTTTCCCTACACGACGCTCTTCCGATCTNNNCTTGCAGTGAGCTGAGATTGCC

AGCACACGTCTGAACTCTTTCCCTACACGACGCTCTTCCGATCTNNNCTTGCAGTGAGCTGAGATTGCC

GAGCACACGTCTGAACTCTTTCCCTACACGACGCTCTTCCGATCTNNNGGTCTTGAACTCCTGACCTCAG

GAGCACACGTCTGAACTCTTTCCCTACACGACGCTCTTCCGATCTNNNGGTCTTGAACTCCTGACCTCAG

AGCACACGTCTGAACTCTTTCCCTACACGACGCTCTTCCGATCTNNNACTGTTGGACAATTGCAAAAGG

AGCACACGTCTGAACTCTTTCCCTACACGACGCTCTTCCGATCTNNNACTGTTGGACAATTGCAAAAGG

GCACACGTCTGAACTCTTTCCCTACACGACGCTCTTCCGATCTNNNATATTTCCCAGTGATGGAGAGCT

GCACACGTCTGAACTCTTTCCCTACACGACGCTCTTCCGATCTNNNATATTTCCCAGTGATGGAGAGCT

AGAGCACACGTCTGAACTCTTTCCCTACACGACGCTCTTCCGATCTNNNGCAGAATCCTGGCCTCTATCT

AGAGCACACGTCTGAACTCTTTCCCTACACGACGCTCTTCCGATCTNNNGCAGAATCCTGGCCTCTATCT

GCACACGTCTGAACTCTTTCCCTACACGACGCTCTTCCGATCTNNNGTAATCTCAGCTACTCAGGAGGC

GCACACGTCTGAACTCTTTCCCTACACGACGCTCTTCCGATCTNNNGTAATCTCAGCTACTCAGGAGGC

GAGCACACGTCTGAACTCTTTCCCTACACGACGCTCTTCCGATCTNNNTCGACCAATATGAGATCCAGC

GAGCACACGTCTGAACTCTTTCCCTACACGACGCTCTTCCGATCTNNNTCGACCAATATGAGATCCAGC

AGAGCACACGTCTGAACTCTTTCCCTACACGACGCTCTTCCGATCTNNNATTGCTTGAGACCAGGAGTT

AGAGCACACGTCTGAACTCTTTCCCTACACGACGCTCTTCCGATCTNNNATTGCTTGAGACCAGGAGTT

AGCACACGTCTGAACTCTTTCCCTACACGACGCTCTTCCGATCTNNNGCTTCTAAAACACCATTTAAAAT

AGCACACGTCTGAACTCTTTCCCTACACGACGCTCTTCCGATCTNNNGCTTCTAAAACACCATTTAAAAT

GCACACGTCTGAACTCTTTCCCTACACGACGCTCTTCCGATCTNNNGGGGATCTGTACCATTTTAATATC

GCACACGTCTGAACTCTTTCCCTACACGACGCTCTTCCGATCTNNNGGGGATCTGTACCATTTTAATATC

AGCACACGTCTGAACTCTTTCCCTACACGACGCTCTTCCGATCTNNNGTGTTACTGTGTTAGCTTCTGG

AGCACACGTCTGAACTCTTTCCCTACACGACGCTCTTCCGATCTNNNGTGTTACTGTGTTAGCTTCTGG

GAGCACACGTCTGAACTCTTTCCCTACACGACGCTCTTCCGATCTNNNTCTAGGGAGCAATGGGAAAC

GAGCACACGTCTGAACTCTTTCCCTACACGACGCTCTTCCGATCTNNNTCTAGGGAGCAATGGGAAAC

AGCACACGTCTGAACTCTTTCCCTACACGACGCTCTTCCGATCTNNNCACAATTATAGCTCACTGCAGCC

AGCACACGTCTGAACTCTTTCCCTACACGACGCTCTTCCGATCTNNNCACAATTATAGCTCACTGCAGCC

AGAGCACACGTCTGAACTCTTTCCCTACACGACGCTCTTCCGATCTNNNGAGGCTGAGGCAGGAGAAT

AGAGCACACGTCTGAACTCTTTCCCTACACGACGCTCTTCCGATCTNNNGAGGCTGAGGCAGGAGAAT

GAGCACACGTCTGAACTCTTTCCCTACACGACGCTCTTCCGATCTNNNCCAGAAGTTTACCAGAGCATC

GAGCACACGTCTGAACTCTTTCCCTACACGACGCTCTTCCGATCTNNNCCAGAAGTTTACCAGAGCATC

AGCACACGTCTGAACTCTTTCCCTACACGACGCTCTTCCGATCTNNNAGTGATGGGATTACAGGTGTG

AGCACACGTCTGAACTCTTTCCCTACACGACGCTCTTCCGATCTNNNAGTGATGGGATTACAGGTGTG

AGCACACGTCTGAACTCTTTCCCTACACGACGCTCTTCCGATCTNNNCGAGTCCCAACCGTATTACAGAA

AGCACACGTCTGAACTCTTTCCCTACACGACGCTCTTCCGATCTNNNCGAGTCCCAACCGTATTACAGAA

GCACACGTCTGAACTCTTTCCCTACACGACGCTCTTCCGATCTNNNGTGTACTGAAACAGTTGGTGCAA

GCACACGTCTGAACTCTTTCCCTACACGACGCTCTTCCGATCTNNNGTGTACTGAAACAGTTGGTGCAA

GAGCACACGTCTGAACTCTTTCCCTACACGACGCTCTTCCGATCTNNNCTGTAGTCCCAGCTACTCAGGA

GAGCACACGTCTGAACTCTTTCCCTACACGACGCTCTTCCGATCTNNNCTGTAGTCCCAGCTACTCAGGA

CACACGTCTGAACTCTTTCCCTACACGACGCTCTTCCGATCTNNNATATCAACGTAGGGTATTACACCA

CACACGTCTGAACTCTTTCCCTACACGACGCTCTTCCGATCTNNNATATCAACGTAGGGTATTACACCA

GAGCACACGTCTGAACTCTTTCCCTACACGACGCTCTTCCGATCTNNNTGAGATTGTGCCATTGTACTCC

GAGCACACGTCTGAACTCTTTCCCTACACGACGCTCTTCCGATCTNNNTGAGATTGTGCCATTGTACTCC

AGCACACGTCTGAACTCTTTCCCTACACGACGCTCTTCCGATCTNNNGGAAAAAATTACTGTCTTCTAC
AGCACACGTCTGAACTCTTTCCCTACACGACGCTCTTCCGATCTNNNGGAAAAAATTACTGTCTTCTAC
GAGCACACGTCTGAACTCTTTCCCTACACGACGCTCTTCCGATCTNNNAGGCAATCAAAAGTGTGTGCA
GAGCACACGTCTGAACTCTTTCCCTACACGACGCTCTTCCGATCTNNNAGGCAATCAAAAGTGTGTGCA
GAGCACACGTCTGAACTCTTTCCCTACACGACGCTCTTCCGATCTNNNAGGGAGAATTACCTGAACCCA
GAGCACACGTCTGAACTCTTTCCCTACACGACGCTCTTCCGATCTNNNAGGGAGAATTACCTGAACCCA
GAGCACACGTCTGAACTCTTTCCCTACACGACGCTCTTCCGATCTNNNTTTTTTTTTTGCAGGGGAGG
GAGCACACGTCTGAACTCTTTCCCTACACGACGCTCTTCCGATCTNNNTTTTTTTTTTGCAGGGGAGG
GAGCACACGTCTGAACTCTTTCCCTACACGACGCTCTTCCGATCTNNNCAGATATTGGGGAGTATGGG
GAGCACACGTCTGAACTCTTTCCCTACACGACGCTCTTCCGATCTNNNCAGATATTGGGGAGTATGGG
AGAGCACACGTCTGAACTCTTTCCCTACACGACGCTCTTCCGATCTNNNTGTAGTCCCATCTACTCAGGA
AGAGCACACGTCTGAACTCTTTCCCTACACGACGCTCTTCCGATCTNNNTGTAGTCCCATCTACTCAGGA
AGAGCACACGTCTGAACTCTTTCCCTACACGACGCTCTTCCGATCTNNNGGATCTCTGCTTCACTTAGCT
AGAGCACACGTCTGAACTCTTTCCCTACACGACGCTCTTCCGATCTNNNGGATCTCTGCTTCACTTAGCT
AGCACACGTCTGAACTCTTTCCCTACACGACGCTCTTCCGATCTNNNTAATCCCAGCTACTCAGGAGGTT
AGCACACGTCTGAACTCTTTCCCTACACGACGCTCTTCCGATCTNNNTAATCCCAGCTACTCAGGAGGTT
AGCACACGTCTGAACTCTTTCCCTACACGACGCTCTTCCGATCTNNNGCATACGATCCAGAACAACGTTT
AGCACACGTCTGAACTCTTTCCCTACACGACGCTCTTCCGATCTNNNGCATACGATCCAGAACAACGTTT
AGCACACGTCTGAACTCTTTCCCTACACGACGCTCTTCCGATCTNNNAAACAGCATTCCTAGACCATGGT
AGCACACGTCTGAACTCTTTCCCTACACGACGCTCTTCCGATCTNNNAAACAGCATTCCTAGACCATGGT
AGCACACGTCTGAACTCTTTCCCTACACGACGCTCTTCCGATCTNNNAGTAGAGACAAGGCTTCACCAT
AGCACACGTCTGAACTCTTTCCCTACACGACGCTCTTCCGATCTNNNAGTAGAGACAAGGCTTCACCAT
AGCACACGTCTGAACTCTTTCCCTACACGACGCTCTTCCGATCTNNNCCAGGTTCCTCCTACTTCTAGGG
AGCACACGTCTGAACTCTTTCCCTACACGACGCTCTTCCGATCTNNNCCAGGTTCCTCCTACTTCTAGGG
AGAGCACACGTCTGAACTCTTTCCCTACACGACGCTCTTCCGATCTNNNTTGTGAAGGTGGAACAGTG
AGAGCACACGTCTGAACTCTTTCCCTACACGACGCTCTTCCGATCTNNNTTGTGAAGGTGGAACAGTG
AGCACACGTCTGAACTCTTTCCCTACACGACGCTCTTCCGATCTNNNTCATACGCACAATACTCAGGGA
AGCACACGTCTGAACTCTTTCCCTACACGACGCTCTTCCGATCTNNNTCATACGCACAATACTCAGGGA
GAGCACACGTCTGAACTCTTTCCCTACACGACGCTCTTCCGATCTNNNTTGCTGCAAATTCTAGAAGCCT
GAGCACACGTCTGAACTCTTTCCCTACACGACGCTCTTCCGATCTNNNTTGCTGCAAATTCTAGAAGCCT
AGCACACGTCTGAACTCTTTCCCTACACGACGCTCTTCCGATCTNNNACTTCTTGAAATATCTACTCTGT
AGCACACGTCTGAACTCTTTCCCTACACGACGCTCTTCCGATCTNNNACTTCTTGAAATATCTACTCTGT
AGCACACGTCTGAACTCTTTCCCTACACGACGCTCTTCCGATCTNNNCTATTTCTGGGGCTACGCTAGAG
AGCACACGTCTGAACTCTTTCCCTACACGACGCTCTTCCGATCTNNNCTATTTCTGGGGCTACGCTAGAG
GAGCACACGTCTGAACTCTTTCCCTACACGACGCTCTTCCGATCTNNNAAGCAGGAGAATCACTTGAAC
GAGCACACGTCTGAACTCTTTCCCTACACGACGCTCTTCCGATCTNNNAAGCAGGAGAATCACTTGAAC
AGCACACGTCTGAACTCTTTCCCTACACGACGCTCTTCCGATCTNNNCTGAGGCAGGAGAATAGCTTG
AGCACACGTCTGAACTCTTTCCCTACACGACGCTCTTCCGATCTNNNCTGAGGCAGGAGAATAGCTTG
GAGCACACGTCTGAACTCTTTCCCTACACGACGCTCTTCCGATCTNNNACAAAAATTAGCTGGGCATGC
GAGCACACGTCTGAACTCTTTCCCTACACGACGCTCTTCCGATCTNNNACAAAAATTAGCTGGGCATGC
GAGCACACGTCTGAACTCTTTCCCTACACGACGCTCTTCCGATCTNNNTGTTTCTGAAGCAGAGCATGT
GAGCACACGTCTGAACTCTTTCCCTACACGACGCTCTTCCGATCTNNNTGTTTCTGAAGCAGAGCATGT
AGCACACGTCTGAACTCTTTCCCTACACGACGCTCTTCCGATCTNNNTGAGAGCTAAAGTGTTCTGGG

AGCACACGTCTGAACTCTTTCCCTACACGACGCTCTTCCGATCTNNNTGAGAGCTAAAGTGTTCTGGG
AGCACACGTCTGAACTCTTTCCCTACACGACGCTCTTCCGATCTNNNCTACTGGGGTAGATCTGTTTC
AGCACACGTCTGAACTCTTTCCCTACACGACGCTCTTCCGATCTNNNCTACTGGGGTAGATCTGTTTC
AGCACACGTCTGAACTCTTTCCCTACACGACGCTCTTCCGATCTNNNGAAAGAAAGAGAGAGGGAG
AGCACACGTCTGAACTCTTTCCCTACACGACGCTCTTCCGATCTNNNGAAAGAAAGAGAGAGGGAG
GCACACGTCTGAACTCTTTCCCTACACGACGCTCTTCCGATCTNNNTTTACCTGCTCAAAGAACCCGTC
GCACACGTCTGAACTCTTTCCCTACACGACGCTCTTCCGATCTNNNTTTACCTGCTCAAAGAACCCGTC
GAGCACACGTCTGAACTCTTTCCCTACACGACGCTCTTCCGATCTNNNCAGGAGGCTGAGACATGAGA
GAGCACACGTCTGAACTCTTTCCCTACACGACGCTCTTCCGATCTNNNCAGGAGGCTGAGACATGAGA
AGCACACGTCTGAACTCTTTCCCTACACGACGCTCTTCCGATCTNNNTTCACACCTTCAGACTCAGACT
AGCACACGTCTGAACTCTTTCCCTACACGACGCTCTTCCGATCTNNNTTCACACCTTCAGACTCAGACT
AGCACACGTCTGAACTCTTTCCCTACACGACGCTCTTCCGATCTNNNGTCACCCGACATGATCAATTTC
AGCACACGTCTGAACTCTTTCCCTACACGACGCTCTTCCGATCTNNNGTCACCCGACATGATCAATTTC
GAGCACACGTCTGAACTCTTTCCCTACACGACGCTCTTCCGATCTNNNCCCGTTTCAAGCAACCAACTA
GAGCACACGTCTGAACTCTTTCCCTACACGACGCTCTTCCGATCTNNNCCCGTTTCAAGCAACCAACTA
GCACACGTCTGAACTCTTTCCCTACACGACGCTCTTCCGATCTNNNAGGATGCTTTCATGTCTAGTGCA
GCACACGTCTGAACTCTTTCCCTACACGACGCTCTTCCGATCTNNNAGGATGCTTTCATGTCTAGTGCA
AGCACACGTCTGAACTCTTTCCCTACACGACGCTCTTCCGATCTNNNAGTGAGCCATGATCATGCTAC
AGCACACGTCTGAACTCTTTCCCTACACGACGCTCTTCCGATCTNNNAGTGAGCCATGATCATGCTAC
AGCACACGTCTGAACTCTTTCCCTACACGACGCTCTTCCGATCTNNNTCCCCAAGGATTAGTTTTGAG
AGCACACGTCTGAACTCTTTCCCTACACGACGCTCTTCCGATCTNNNTCCCCAAGGATTAGTTTTGAG
GAGCACACGTCTGAACTCTTTCCCTACACGACGCTCTTCCGATCTNNNCACCTGTAATCCCAGCTACTC
GAGCACACGTCTGAACTCTTTCCCTACACGACGCTCTTCCGATCTNNNCACCTGTAATCCCAGCTACTC
AGCACACGTCTGAACTCTTTCCCTACACGACGCTCTTCCGATCTNNNTGAAAAAGCTCTTGAAGACGC
AGCACACGTCTGAACTCTTTCCCTACACGACGCTCTTCCGATCTNNNTGAAAAAGCTCTTGAAGACGC
AGCACACGTCTGAACTCTTTCCCTACACGACGCTCTTCCGATCTNNNATTCTCTCCACTCCTTTCCCGAT
AGCACACGTCTGAACTCTTTCCCTACACGACGCTCTTCCGATCTNNNATTCTCTCCACTCCTTTCCCGAT
AGCACACGTCTGAACTCTTTCCCTACACGACGCTCTTCCGATCTNNNAGTAGCTGGAATTACTGACA
AGCACACGTCTGAACTCTTTCCCTACACGACGCTCTTCCGATCTNNNAGTAGCTGGAATTACTGACA
AGCACACGTCTGAACTCTTTCCCTACACGACGCTCTTCCGATCTNNNCTCTTTGTCTTCTGGGTCCTTG
AGCACACGTCTGAACTCTTTCCCTACACGACGCTCTTCCGATCTNNNCTCTTTGTCTTCTGGGTCCTTG
AGCACACGTCTGAACTCTTTCCCTACACGACGCTCTTCCGATCTNNNGGAAAGAAGGGAATGGAAGA
AGCACACGTCTGAACTCTTTCCCTACACGACGCTCTTCCGATCTNNNGGAAAGAAGGGAATGGAAGA
GAGCACACGTCTGAACTCTTTCCCTACACGACGCTCTTCCGATCTNNNCTGAGGGAGGAGATTTGCT
GAGCACACGTCTGAACTCTTTCCCTACACGACGCTCTTCCGATCTNNNCTGAGGGAGGAGATTTGCT
AGCACACGTCTGAACTCTTTCCCTACACGACGCTCTTCCGATCTNNNATAGTGTAATCTCATGGGCTC
AGCACACGTCTGAACTCTTTCCCTACACGACGCTCTTCCGATCTNNNATAGTGTAATCTCATGGGCTC
GAGCACACGTCTGAACTCTTTCCCTACACGACGCTCTTCCGATCTNNNACTCCAAATTGAGGTTTCTTC
GAGCACACGTCTGAACTCTTTCCCTACACGACGCTCTTCCGATCTNNNACTCCAAATTGAGGTTTCTTC
GAGCACACGTCTGAACTCTTTCCCTACACGACGCTCTTCCGATCTNNNGATAGAGCAAGACGGCGTCT
GAGCACACGTCTGAACTCTTTCCCTACACGACGCTCTTCCGATCTNNNGATAGAGCAAGACGGCGTCT
AGCACACGTCTGAACTCTTTCCCTACACGACGCTCTTCCGATCTNNNTACAGTGGCATTGTCATAGCTC
AGCACACGTCTGAACTCTTTCCCTACACGACGCTCTTCCGATCTNNNTACAGTGGCATTGTCATAGCTC

GAGCACACGTCTGAACTCTTTCCCTACACGACGCTCTTCCGATCTNNNCAAATCTCAAAGACTCAGCCC
GAGCACACGTCTGAACTCTTTCCCTACACGACGCTCTTCCGATCTNNNCAAATCTCAAAGACTCAGCCC
GAGCACACGTCTGAACTCTTTCCCTACACGACGCTCTTCCGATCTNNNCCTCTTCTCTCTCTCAGGGCCCA
GAGCACACGTCTGAACTCTTTCCCTACACGACGCTCTTCCGATCTNNNCCTCTTCTCTCTCTCAGGGCCCA
AGCACACGTCTGAACTCTTTCCCTACACGACGCTCTTCCGATCTNNNAAGGCAACAGACATCTTCCTAG
AGCACACGTCTGAACTCTTTCCCTACACGACGCTCTTCCGATCTNNNAAGGCAACAGACATCTTCCTAG
ACACGTCTGAACTCTTTCCCTACACGACGCTCTTCCGATCTNNNACGCGGATATCCTTCCAAAGAATCAT
ACACGTCTGAACTCTTTCCCTACACGACGCTCTTCCGATCTNNNACGCGGATATCCTTCCAAAGAATCAT
AGCACACGTCTGAACTCTTTCCCTACACGACGCTCTTCCGATCTNNNGTTATCTACTTGGCCTTTGCTGG
AGCACACGTCTGAACTCTTTCCCTACACGACGCTCTTCCGATCTNNNGTTATCTACTTGGCCTTTGCTGG
AGCACACGTCTGAACTCTTTCCCTACACGACGCTCTTCCGATCTNNNGAGGACATCCACATACCTGACT
AGCACACGTCTGAACTCTTTCCCTACACGACGCTCTTCCGATCTNNNGAGGACATCCACATACCTGACT
GAGCACACGTCTGAACTCTTTCCCTACACGACGCTCTTCCGATCTNNNAGGCAATCAGAGGTATGTTTC
GAGCACACGTCTGAACTCTTTCCCTACACGACGCTCTTCCGATCTNNNAGGCAATCAGAGGTATGTTTC
AAGAGCACACGTCTGAACTCTTTCCCTACACGACGCTCTTCCGATCTNNNCCCAAGGCTTCCCCATATTT
AAGAGCACACGTCTGAACTCTTTCCCTACACGACGCTCTTCCGATCTNNNCCCAAGGCTTCCCCATATTT
AGCACACGTCTGAACTCTTTCCCTACACGACGCTCTTCCGATCTNNNTCTCTGCTATACCTGCTTTCGTCC
AGCACACGTCTGAACTCTTTCCCTACACGACGCTCTTCCGATCTNNNTCTCTGCTATACCTGCTTTCGTCC
GAGCACACGTCTGAACTCTTTCCCTACACGACGCTCTTCCGATCTNNNTCTATTCCTTGCTACCCACAAG
GAGCACACGTCTGAACTCTTTCCCTACACGACGCTCTTCCGATCTNNNTCTATTCCTTGCTACCCACAAG
AGCACACGTCTGAACTCTTTCCCTACACGACGCTCTTCCGATCTNNNAGCTGGAATGATGATGCTGAGT
AGCACACGTCTGAACTCTTTCCCTACACGACGCTCTTCCGATCTNNNAGCTGGAATGATGATGCTGAGT
GAGCACACGTCTGAACTCTTTCCCTACACGACGCTCTTCCGATCTNNNGAGGTTGCAATGAGTCGAGA
GAGCACACGTCTGAACTCTTTCCCTACACGACGCTCTTCCGATCTNNNGAGGTTGCAATGAGTCGAGA
AGCACACGTCTGAACTCTTTCCCTACACGACGCTCTTCCGATCTNNNGATGAAATTTTAGAGAAAAGG
AGCACACGTCTGAACTCTTTCCCTACACGACGCTCTTCCGATCTNNNGATGAAATTTTAGAGAAAAGG
GAGCACACGTCTGAACTCTTTCCCTACACGACGCTCTTCCGATCTNNNCCTGAGCTACCTTCAGAAGTGT
GAGCACACGTCTGAACTCTTTCCCTACACGACGCTCTTCCGATCTNNNCCTGAGCTACCTTCAGAAGTGT
AGCACACGTCTGAACTCTTTCCCTACACGACGCTCTTCCGATCTNNNGTAGTGAGTTGTGAGTCCTCCA
AGCACACGTCTGAACTCTTTCCCTACACGACGCTCTTCCGATCTNNNGTAGTGAGTTGTGAGTCCTCCA
GAGCACACGTCTGAACTCTTTCCCTACACGACGCTCTTCCGATCTNNNCTGTAATCCCAGCTACTCAGGA
GAGCACACGTCTGAACTCTTTCCCTACACGACGCTCTTCCGATCTNNNCTGTAATCCCAGCTACTCAGGA
AGCACACGTCTGAACTCTTTCCCTACACGACGCTCTTCCGATCTNNNGGGAGTATTGTGAGGGTTGGA
AGCACACGTCTGAACTCTTTCCCTACACGACGCTCTTCCGATCTNNNGGGAGTATTGTGAGGGTTGGA
AGAGCACACGTCTGAACTCTTTCCCTACACGACGCTCTTCCGATCTNNNCCTGTAGTCCCAGCTACTTGG
AGAGCACACGTCTGAACTCTTTCCCTACACGACGCTCTTCCGATCTNNNCCTGTAGTCCCAGCTACTTGG
AGCACACGTCTGAACTCTTTCCCTACACGACGCTCTTCCGATCTNNNCAGCTCCATTTCTTCTAAAATAG
AGCACACGTCTGAACTCTTTCCCTACACGACGCTCTTCCGATCTNNNCAGCTCCATTTCTTCTAAAATAG
GAGCACACGTCTGAACTCTTTCCCTACACGACGCTCTTCCGATCTNNNCATGCCTGTAATCCCAGCTACT
GAGCACACGTCTGAACTCTTTCCCTACACGACGCTCTTCCGATCTNNNCATGCCTGTAATCCCAGCTACT
AGCACACGTCTGAACTCTTTCCCTACACGACGCTCTTCCGATCTNNNACTTACATGGTAAGGCCCAGAA
AGCACACGTCTGAACTCTTTCCCTACACGACGCTCTTCCGATCTNNNACTTACATGGTAAGGCCCAGAA
AGCACACGTCTGAACTCTTTCCCTACACGACGCTCTTCCGATCTNNNCCAAGTTGAGGTGGTGGGTTT

AGCACACGTCTGAACTCTTTCCCTACACGACGCTCTTCCGATCTNNNCCAAGTTGAGGTGGTGGGTTT
AGCACACGTCTGAACTCTTTCCCTACACGACGCTCTTCCGATCTNNNGGAATACCAAGGCAGAGAAA
AGCACACGTCTGAACTCTTTCCCTACACGACGCTCTTCCGATCTNNNGGAATACCAAGGCAGAGAAA
AGCACACGTCTGAACTCTTTCCCTACACGACGCTCTTCCGATCTNNNTGTGATGGACAGACAGAGTCT
AGCACACGTCTGAACTCTTTCCCTACACGACGCTCTTCCGATCTNNNTGTGATGGACAGACAGAGTCT
GAGCACACGTCTGAACTCTTTCCCTACACGACGCTCTTCCGATCTNNNGAGGCTAAGTCAGGAGAATT
GAGCACACGTCTGAACTCTTTCCCTACACGACGCTCTTCCGATCTNNNGAGGCTAAGTCAGGAGAATT
GAGCACACGTCTGAACTCTTTCCCTACACGACGCTCTTCCGATCTNNNGAGAATTGCTTGAACTCGAAG
GAGCACACGTCTGAACTCTTTCCCTACACGACGCTCTTCCGATCTNNNGAGAATTGCTTGAACTCGAAG
GAGCACACGTCTGAACTCTTTCCCTACACGACGCTCTTCCGATCTNNNGGGGTAAAGCTCCTTGGAAT
GAGCACACGTCTGAACTCTTTCCCTACACGACGCTCTTCCGATCTNNNGGGGTAAAGCTCCTTGGAAT
AGCACACGTCTGAACTCTTTCCCTACACGACGCTCTTCCGATCTNNNACATTCAAACAAGGCAGGTCAA
AGCACACGTCTGAACTCTTTCCCTACACGACGCTCTTCCGATCTNNNACATTCAAACAAGGCAGGTCAA
GCACACGTCTGAACTCTTTCCCTACACGACGCTCTTCCGATCTNNNAACCTCAAATTCTGGCTCCTCTAC
GCACACGTCTGAACTCTTTCCCTACACGACGCTCTTCCGATCTNNNAACCTCAAATTCTGGCTCCTCTAC
AGCACACGTCTGAACTCTTTCCCTACACGACGCTCTTCCGATCTNNNATAACAAAAGTGGAAACCTGG
AGCACACGTCTGAACTCTTTCCCTACACGACGCTCTTCCGATCTNNNATAACAAAAGTGGAAACCTGG
GAGCACACGTCTGAACTCTTTCCCTACACGACGCTCTTCCGATCTNNNGGTGATTGTATTCTGTGTACT
GAGCACACGTCTGAACTCTTTCCCTACACGACGCTCTTCCGATCTNNNGGTGATTGTATTCTGTGTACT
AGCACACGTCTGAACTCTTTCCCTACACGACGCTCTTCCGATCTNNNCTGTGGTCCTAGCTACTCAGAA
AGCACACGTCTGAACTCTTTCCCTACACGACGCTCTTCCGATCTNNNCTGTGGTCCTAGCTACTCAGAA
AGCACACGTCTGAACTCTTTCCCTACACGACGCTCTTCCGATCTNNNCCTGATGCCCTATTGCAGATCTT
AGCACACGTCTGAACTCTTTCCCTACACGACGCTCTTCCGATCTNNNCCTGATGCCCTATTGCAGATCTT
AGCACACGTCTGAACTCTTTCCCTACACGACGCTCTTCCGATCTNNNGCGTTTCAGTGGTTTAGTTAG
AGCACACGTCTGAACTCTTTCCCTACACGACGCTCTTCCGATCTNNNGCGTTTCAGTGGTTTAGTTAG
GAGCACACGTCTGAACTCTTTCCCTACACGACGCTCTTCCGATCTNNNCCACCATACCCGGCTAATTTTT
GAGCACACGTCTGAACTCTTTCCCTACACGACGCTCTTCCGATCTNNNCCACCATACCCGGCTAATTTTT
CACACGTCTGAACTCTTTCCCTACACGACGCTCTTCCGATCTNNNCAGAGAACAGCCCCTCTGCGCATC
CACACGTCTGAACTCTTTCCCTACACGACGCTCTTCCGATCTNNNCAGAGAACAGCCCCTCTGCGCATC
AGCACACGTCTGAACTCTTTCCCTACACGACGCTCTTCCGATCTNNNAATTTCTCTAACCTGGGAGGCA
AGCACACGTCTGAACTCTTTCCCTACACGACGCTCTTCCGATCTNNNAATTTCTCTAACCTGGGAGGCA
GAGCACACGTCTGAACTCTTTCCCTACACGACGCTCTTCCGATCTNNNCTGAGGCACGAGAATCACTTG
GAGCACACGTCTGAACTCTTTCCCTACACGACGCTCTTCCGATCTNNNCTGAGGCACGAGAATCACTTG
AGCACACGTCTGAACTCTTTCCCTACACGACGCTCTTCCGATCTNNNATACAAAAATTAGCCGGGTGTG
AGCACACGTCTGAACTCTTTCCCTACACGACGCTCTTCCGATCTNNNATACAAAAATTAGCCGGGTGTG
AGAGCACACGTCTGAACTCTTTCCCTACACGACGCTCTTCCGATCTNNNGCCTGTAGTCCCAGCTACTCT
AGAGCACACGTCTGAACTCTTTCCCTACACGACGCTCTTCCGATCTNNNGCCTGTAGTCCCAGCTACTCT
AGCACACGTCTGAACTCTTTCCCTACACGACGCTCTTCCGATCTNNNATGGAACGCATTTTGGGTTGTT
AGCACACGTCTGAACTCTTTCCCTACACGACGCTCTTCCGATCTNNNATGGAACGCATTTTGGGTTGTT
AGCACACGTCTGAACTCTTTCCCTACACGACGCTCTTCCGATCTNNNTACTGAAGAGGCTGAGGTAG
AGCACACGTCTGAACTCTTTCCCTACACGACGCTCTTCCGATCTNNNTACTGAAGAGGCTGAGGTAG
AGCACACGTCTGAACTCTTTCCCTACACGACGCTCTTCCGATCTNNNTCAACCAGAATACCTCCACATG
AGCACACGTCTGAACTCTTTCCCTACACGACGCTCTTCCGATCTNNNTCAACCAGAATACCTCCACATG

GAGCACACGTCTGAACTCTTTCCCTACACGACGCTCTTCCGATCTNNNAAAAAATTAGCCAGGCGTAGT
GAGCACACGTCTGAACTCTTTCCCTACACGACGCTCTTCCGATCTNNNAAAAAATTAGCCAGGCGTAGT
AGAGCACACGTCTGAACTCTTTCCCTACACGACGCTCTTCCGATCTNNNTCTCCCTCATTTTCTCTTGCTC
AGAGCACACGTCTGAACTCTTTCCCTACACGACGCTCTTCCGATCTNNNTCTCCCTCATTTTCTCTTGCTC
GCACACGTCTGAACTCTTTCCCTACACGACGCTCTTCCGATCTNNNAGAATTGCTTGAACCCAGAAAG
GCACACGTCTGAACTCTTTCCCTACACGACGCTCTTCCGATCTNNNAGAATTGCTTGAACCCAGAAAG
GAGCACACGTCTGAACTCTTTCCCTACACGACGCTCTTCCGATCTNNNGAGACAGAGCAAACAAACGA
GAGCACACGTCTGAACTCTTTCCCTACACGACGCTCTTCCGATCTNNNGAGACAGAGCAAACAAACGA
AGCACACGTCTGAACTCTTTCCCTACACGACGCTCTTCCGATCTNNNGTGGTGCAATCCTAGCTCAGTA
AGCACACGTCTGAACTCTTTCCCTACACGACGCTCTTCCGATCTNNNGTGGTGCAATCCTAGCTCAGTA
GAGCACACGTCTGAACTCTTTCCCTACACGACGCTCTTCCGATCTNNNGGAGAATGGCGTAAACCTGGA
GAGCACACGTCTGAACTCTTTCCCTACACGACGCTCTTCCGATCTNNNGGAGAATGGCGTAAACCTGGA
AGCACACGTCTGAACTCTTTCCCTACACGACGCTCTTCCGATCTNNNCCTGAGTAGCTGGGATTATAGG
AGCACACGTCTGAACTCTTTCCCTACACGACGCTCTTCCGATCTNNNCCTGAGTAGCTGGGATTATAGG
GAGCACACGTCTGAACTCTTTCCCTACACGACGCTCTTCCGATCTNNNCAGATCACAAGGTCAGGAGTT
GAGCACACGTCTGAACTCTTTCCCTACACGACGCTCTTCCGATCTNNNCAGATCACAAGGTCAGGAGTT
AGCACACGTCTGAACTCTTTCCCTACACGACGCTCTTCCGATCTNNNACCTAGGGGAGTACTTCTCAGT
AGCACACGTCTGAACTCTTTCCCTACACGACGCTCTTCCGATCTNNNACCTAGGGGAGTACTTCTCAGT
GAGCACACGTCTGAACTCTTTCCCTACACGACGCTCTTCCGATCTNNNGCCTGGCTACCTGAATCATCTT
GAGCACACGTCTGAACTCTTTCCCTACACGACGCTCTTCCGATCTNNNGCCTGGCTACCTGAATCATCTT
AGCACACGTCTGAACTCTTTCCCTACACGACGCTCTTCCGATCTNNNACTCCTCATCTCAAGTGATCTG
AGCACACGTCTGAACTCTTTCCCTACACGACGCTCTTCCGATCTNNNACTCCTCATCTCAAGTGATCTG
GAGCACACGTCTGAACTCTTTCCCTACACGACGCTCTTCCGATCTNNNGAAGAAGAAAGAGGGAAGG
GAGCACACGTCTGAACTCTTTCCCTACACGACGCTCTTCCGATCTNNNGAAGAAGAAAGAGGGAAGG
GAGCACACGTCTGAACTCTTTCCCTACACGACGCTCTTCCGATCTNNNGTAATCTCAGCTACTCAGGAG
GAGCACACGTCTGAACTCTTTCCCTACACGACGCTCTTCCGATCTNNNGTAATCTCAGCTACTCAGGAG
GAGCACACGTCTGAACTCTTTCCCTACACGACGCTCTTCCGATCTNNNTGCTTGTAATCCCAGCTACTCA
GAGCACACGTCTGAACTCTTTCCCTACACGACGCTCTTCCGATCTNNNTGCTTGTAATCCCAGCTACTCA
GCACACGTCTGAACTCTTTCCCTACACGACGCTCTTCCGATCTNNNAGACATGCAGTTCCAGCTACCCAT
GCACACGTCTGAACTCTTTCCCTACACGACGCTCTTCCGATCTNNNAGACATGCAGTTCCAGCTACCCAT
GAGCACACGTCTGAACTCTTTCCCTACACGACGCTCTTCCGATCTNNNCTGAGGCAGGAGAATAGCTTG
GAGCACACGTCTGAACTCTTTCCCTACACGACGCTCTTCCGATCTNNNCTGAGGCAGGAGAATAGCTTG
GAGCACACGTCTGAACTCTTTCCCTACACGACGCTCTTCCGATCTNNNCTACAGCCTGTCTTTTCTACCGT
GAGCACACGTCTGAACTCTTTCCCTACACGACGCTCTTCCGATCTNNNCTACAGCCTGTCTTTTCTACCGT
GAGCACACGTCTGAACTCTTTCCCTACACGACGCTCTTCCGATCTNNNAGTTGGGGACTGTCTGTAAAC
GAGCACACGTCTGAACTCTTTCCCTACACGACGCTCTTCCGATCTNNNAGTTGGGGACTGTCTGTAAAC
AGCACACGTCTGAACTCTTTCCCTACACGACGCTCTTCCGATCTNNNGTAACTCCACGTAGAAGCCTCAT
AGCACACGTCTGAACTCTTTCCCTACACGACGCTCTTCCGATCTNNNGTAACTCCACGTAGAAGCCTCAT
AGCACACGTCTGAACTCTTTCCCTACACGACGCTCTTCCGATCTNNNACCCTTGAGTTTTGGGACTTTGA
AGCACACGTCTGAACTCTTTCCCTACACGACGCTCTTCCGATCTNNNACCCTTGAGTTTTGGGACTTTGA
AGAGCACACGTCTGAACTCTTTCCCTACACGACGCTCTTCCGATCTNNNGCCTTGCACTTGCTATTCTCTT
AGAGCACACGTCTGAACTCTTTCCCTACACGACGCTCTTCCGATCTNNNGCCTTGCACTTGCTATTCTCTT
AGCACACGTCTGAACTCTTTCCCTACACGACGCTCTTCCGATCTNNNATTTCTGCAGCTGCCTTGAATTT

GCACACGTCTGAACTCTTTCCCTACACGACGCTCTTCCGATCTNNNATTTCTGCAGCTGCCTTGAATTT
AGAGCACACGTCTGAACTCTTTCCCTACACGACGCTCTTCCGATCTNNNTGAGGCATTGGAGATGGTG
AGAGCACACGTCTGAACTCTTTCCCTACACGACGCTCTTCCGATCTNNNTGAGGCATTGGAGATGGTG
AGCACACGTCTGAACTCTTTCCCTACACGACGCTCTTCCGATCTNNNCTCTACCTCATCCCAGAAGGTT
AGCACACGTCTGAACTCTTTCCCTACACGACGCTCTTCCGATCTNNNCTCTACCTCATCCCAGAAGGTT
AGCACACGTCTGAACTCTTTCCCTACACGACGCTCTTCCGATCTNNNTCTCTAGAGTCAGAATCCTGC
AGCACACGTCTGAACTCTTTCCCTACACGACGCTCTTCCGATCTNNNTCTCTAGAGTCAGAATCCTGC
AGCACACGTCTGAACTCTTTCCCTACACGACGCTCTTCCGATCTNNNCCCGAAAAGTAATGAGCTCTCC
AGCACACGTCTGAACTCTTTCCCTACACGACGCTCTTCCGATCTNNNCCCGAAAAGTAATGAGCTCTCC
AGAGCACACGTCTGAACTCTTTCCCTACACGACGCTCTTCCGATCTNNNACAACTGACATACCATCTCTG
AGAGCACACGTCTGAACTCTTTCCCTACACGACGCTCTTCCGATCTNNNACAACTGACATACCATCTCTG
GAGCACACGTCTGAACTCTTTCCCTACACGACGCTCTTCCGATCTNNNAACAAGAAGGTACAGAACAC
GAGCACACGTCTGAACTCTTTCCCTACACGACGCTCTTCCGATCTNNNAACAAGAAGGTACAGAACAC
AGCACACGTCTGAACTCTTTCCCTACACGACGCTCTTCCGATCTNNNGGATTTGGGCTCTCATATCAC
AGCACACGTCTGAACTCTTTCCCTACACGACGCTCTTCCGATCTNNNGGATTTGGGCTCTCATATCAC
AGCACACGTCTGAACTCTTTCCCTACACGACGCTCTTCCGATCTNNNGTCCTGGACAAGACCTTTCACT
AGCACACGTCTGAACTCTTTCCCTACACGACGCTCTTCCGATCTNNNGTCCTGGACAAGACCTTTCACT
CACACGTCTGAACTCTTTCCCTACACGACGCTCTTCCGATCTNNNCCAGGCTGATCTCAAACTCCTAGC
CACACGTCTGAACTCTTTCCCTACACGACGCTCTTCCGATCTNNNCCAGGCTGATCTCAAACTCCTAGC
GAGCACACGTCTGAACTCTTTCCCTACACGACGCTCTTCCGATCTNNNACCATATTTCTTCAGCAGCGTT
GAGCACACGTCTGAACTCTTTCCCTACACGACGCTCTTCCGATCTNNNACCATATTTCTTCAGCAGCGTT
AGCACACGTCTGAACTCTTTCCCTACACGACGCTCTTCCGATCTNNNATGTTTTTGCATAGGCTTGTCG
AGCACACGTCTGAACTCTTTCCCTACACGACGCTCTTCCGATCTNNNATGTTTTTGCATAGGCTTGTCG
GCACACGTCTGAACTCTTTCCCTACACGACGCTCTTCCGATCTNNNCTCACTTTTTGTCCTGTCTGCAG
GCACACGTCTGAACTCTTTCCCTACACGACGCTCTTCCGATCTNNNCTCACTTTTTGTCCTGTCTGCAG
GCACACGTCTGAACTCTTTCCCTACACGACGCTCTTCCGATCTNNNATGCTGGCAACTTCTCTCCTTTCC
GCACACGTCTGAACTCTTTCCCTACACGACGCTCTTCCGATCTNNNATGCTGGCAACTTCTCTCCTTTCC
GCACACGTCTGAACTCTTTCCCTACACGACGCTCTTCCGATCTNNNCCATCCGACTCAACCCTATAACCC
GCACACGTCTGAACTCTTTCCCTACACGACGCTCTTCCGATCTNNNCCATCCGACTCAACCCTATAACCC
GCACACGTCTGAACTCTTTCCCTACACGACGCTCTTCCGATCTNNNTTGTGAGGGTAGAAGTTTAGG
GCACACGTCTGAACTCTTTCCCTACACGACGCTCTTCCGATCTNNNTTGTGAGGGTAGAAGTTTAGG
ACACGTCTGAACTCTTTCCCTACACGACGCTCTTCCGATCTNNNGTCCTGGTTGTGGATAGTGACTGC
ACACGTCTGAACTCTTTCCCTACACGACGCTCTTCCGATCTNNNGTCCTGGTTGTGGATAGTGACTGC
AGCACACGTCTGAACTCTTTCCCTACACGACGCTCTTCCGATCTNNNCTGTAGTCCCAGCTATTCAGGA
AGCACACGTCTGAACTCTTTCCCTACACGACGCTCTTCCGATCTNNNCTGTAGTCCCAGCTATTCAGGA
ACACGTCTGAACTCTTTCCCTACACGACGCTCTTCCGATCTNNNATGTGTCTCCCGTCTGCTTTCCATCG
ACACGTCTGAACTCTTTCCCTACACGACGCTCTTCCGATCTNNNATGTGTCTCCCGTCTGCTTTCCATCG
AGCACACGTCTGAACTCTTTCCCTACACGACGCTCTTCCGATCTNNNAAATAAAGAGAGAGGACCTGC
AGCACACGTCTGAACTCTTTCCCTACACGACGCTCTTCCGATCTNNNAAATAAAGAGAGAGGACCTGC
GAGCACACGTCTGAACTCTTTCCCTACACGACGCTCTTCCGATCTNNNCTGGGCAAAAGAGCAAGACT
GAGCACACGTCTGAACTCTTTCCCTACACGACGCTCTTCCGATCTNNNCTGGGCAAAAGAGCAAGACT
AGCACACGTCTGAACTCTTTCCCTACACGACGCTCTTCCGATCTNNNTACGTACTGGGTCCAATACTTCG
AGCACACGTCTGAACTCTTTCCCTACACGACGCTCTTCCGATCTNNNTACGTACTGGGTCCAATACTTCG

AGCACACGTCTGAACTCTTTCCCTACACGACGCTCTTCCGATCTNNNTTTAGCACCCAGCTGTCCTGAA

AGCACACGTCTGAACTCTTTCCCTACACGACGCTCTTCCGATCTNNNTTTAGCACCCAGCTGTCCTGAA

GAGCACACGTCTGAACTCTTTCCCTACACGACGCTCTTCCGATCTNNNTCTGCCTCTCAGATTCAAGCAA

GAGCACACGTCTGAACTCTTTCCCTACACGACGCTCTTCCGATCTNNNTCTGCCTCTCAGATTCAAGCAA

GAGCACACGTCTGAACTCTTTCCCTACACGACGCTCTTCCGATCTNNNCCTGGGTTATGGAGTGAGA

GAGCACACGTCTGAACTCTTTCCCTACACGACGCTCTTCCGATCTNNNCCTGGGTTATGGAGTGAGA

GCACACGTCTGAACTCTTTCCCTACACGACGCTCTTCCGATCTNNNAAAGAAGTGGAGAGGTGCTTCA

GCACACGTCTGAACTCTTTCCCTACACGACGCTCTTCCGATCTNNNAAAGAAGTGGAGAGGTGCTTCA

GAGCACACGTCTGAACTCTTTCCCTACACGACGCTCTTCCGATCTNNNCTGTAATCCCAGCTACTCGAGA

GAGCACACGTCTGAACTCTTTCCCTACACGACGCTCTTCCGATCTNNNCTGTAATCCCAGCTACTCGAGA

AGCACACGTCTGAACTCTTTCCCTACACGACGCTCTTCCGATCTNNNCACAATGTGCACATGTACCCTA

AGCACACGTCTGAACTCTTTCCCTACACGACGCTCTTCCGATCTNNNCACAATGTGCACATGTACCCTA

AGCACACGTCTGAACTCTTTCCCTACACGACGCTCTTCCGATCTNNNTGTTCCTATGCTGAAAGCACCTC

AGCACACGTCTGAACTCTTTCCCTACACGACGCTCTTCCGATCTNNNTGTTCCTATGCTGAAAGCACCTC

AGCACACGTCTGAACTCTTTCCCTACACGACGCTCTTCCGATCTNNNGGCCTTGGCTTCTGTATCTACTT

AGCACACGTCTGAACTCTTTCCCTACACGACGCTCTTCCGATCTNNNGGCCTTGGCTTCTGTATCTACTT

GAGCACACGTCTGAACTCTTTCCCTACACGACGCTCTTCCGATCTNNNCTCCAATAAGAGCATCCATGCA

GAGCACACGTCTGAACTCTTTCCCTACACGACGCTCTTCCGATCTNNNCTCCAATAAGAGCATCCATGCA

AGCACACGTCTGAACTCTTTCCCTACACGACGCTCTTCCGATCTNNNCTAGCCAAAAACCAGGAAAAGC

AGCACACGTCTGAACTCTTTCCCTACACGACGCTCTTCCGATCTNNNCTAGCCAAAAACCAGGAAAAGC

GCACACGTCTGAACTCTTTCCCTACACGACGCTCTTCCGATCTNNNATACTGTGACCACCCTACCCTAACA

GCACACGTCTGAACTCTTTCCCTACACGACGCTCTTCCGATCTNNNATACTGTGACCACCCTACCCTAACA

AGCACACGTCTGAACTCTTTCCCTACACGACGCTCTTCCGATCTNNNAGCAGAGCCTGATGATGTAAA

AGCACACGTCTGAACTCTTTCCCTACACGACGCTCTTCCGATCTNNNAGCAGAGCCTGATGATGTAAA

GAGCACACGTCTGAACTCTTTCCCTACACGACGCTCTTCCGATCTNNNTAGAAACATGCCAGTGTTGGC

GAGCACACGTCTGAACTCTTTCCCTACACGACGCTCTTCCGATCTNNNTAGAAACATGCCAGTGTTGGC

GCACACGTCTGAACTCTTTCCCTACACGACGCTCTTCCGATCTNNNGAGGCTGAGATGGGAAACTAGA

GCACACGTCTGAACTCTTTCCCTACACGACGCTCTTCCGATCTNNNGAGGCTGAGATGGGAAACTAGA

GAGCACACGTCTGAACTCTTTCCCTACACGACGCTCTTCCGATCTNNNCCGCCTTGAGAGTGTATGGA

GAGCACACGTCTGAACTCTTTCCCTACACGACGCTCTTCCGATCTNNNCCGCCTTGAGAGTGTATGGA

AGCACACGTCTGAACTCTTTCCCTACACGACGCTCTTCCGATCTNNNCTGTAATCCCAGCTACTCAGGA

AGCACACGTCTGAACTCTTTCCCTACACGACGCTCTTCCGATCTNNNCTGTAATCCCAGCTACTCAGGA

ACACGTCTGAACTCTTTCCCTACACGACGCTCTTCCGATCTNNNCATGTTGGAAGTGCTTGGATTGACA

ACACGTCTGAACTCTTTCCCTACACGACGCTCTTCCGATCTNNNCATGTTGGAAGTGCTTGGATTGACA

AGCACACGTCTGAACTCTTTCCCTACACGACGCTCTTCCGATCTNNNATCCAGTTGTCACTCTGCATCTT

AGCACACGTCTGAACTCTTTCCCTACACGACGCTCTTCCGATCTNNNATCCAGTTGTCACTCTGCATCTT

GCACACGTCTGAACTCTTTCCCTACACGACGCTCTTCCGATCTNNNTATGCTGGGATTACAGGCATGAG

GCACACGTCTGAACTCTTTCCCTACACGACGCTCTTCCGATCTNNNTATGCTGGGATTACAGGCATGAG

AGCACACGTCTGAACTCTTTCCCTACACGACGCTCTTCCGATCTNNNAAATCTTCAACTGGAGGGATG

AGCACACGTCTGAACTCTTTCCCTACACGACGCTCTTCCGATCTNNNAAATCTTCAACTGGAGGGATG

AGCACACGTCTGAACTCTTTCCCTACACGACGCTCTTCCGATCTNNNGCCTGTAGTCCCAGCTACTTGCA

AGCACACGTCTGAACTCTTTCCCTACACGACGCTCTTCCGATCTNNNGCCTGTAGTCCCAGCTACTTGCA

AGAGCACACGTCTGAACTCTTTCCCTACACGACGCTCTTCCGATCTNNNGCAAAAACATGGATGAACC

AGAGCACACGTCTGAACTCTTTCCCTACACGACGCTCTTCCGATCTNNNGCAAAAACATGGATGAACC
GAGCACACGTCTGAACTCTTTCCCTACACGACGCTCTTCCGATCTNNNCCACGCCTGGCTAATTTTTTTC
GAGCACACGTCTGAACTCTTTCCCTACACGACGCTCTTCCGATCTNNNCCACGCCTGGCTAATTTTTTTC
AGCACACGTCTGAACTCTTTCCCTACACGACGCTCTTCCGATCTNNNGTCAGTGGAGTGTGCTTGAAA
AGCACACGTCTGAACTCTTTCCCTACACGACGCTCTTCCGATCTNNNGTCAGTGGAGTGTGCTTGAAA
AGCACACGTCTGAACTCTTTCCCTACACGACGCTCTTCCGATCTNNNGATCCGAATGGCAAAGACGAGT
AGCACACGTCTGAACTCTTTCCCTACACGACGCTCTTCCGATCTNNNGATCCGAATGGCAAAGACGAGT
GAGCACACGTCTGAACTCTTTCCCTACACGACGCTCTTCCGATCTNNNAGGTGATGTGGAAACATGGA
GAGCACACGTCTGAACTCTTTCCCTACACGACGCTCTTCCGATCTNNNAGGTGATGTGGAAACATGGA
AGCACACGTCTGAACTCTTTCCCTACACGACGCTCTTCCGATCTNNNCTTTCTCCAGTGCCGCTCTCCATC
AGCACACGTCTGAACTCTTTCCCTACACGACGCTCTTCCGATCTNNNCTTTCTCCAGTGCCGCTCTCCATC
AGCACACGTCTGAACTCTTTCCCTACACGACGCTCTTCCGATCTNNNTACCAAATGTAAAAATAACTGG
AGCACACGTCTGAACTCTTTCCCTACACGACGCTCTTCCGATCTNNNTACCAAATGTAAAAATAACTGG
GAGCACACGTCTGAACTCTTTCCCTACACGACGCTCTTCCGATCTNNNGCAATGAACATGGAGGTGCA
GAGCACACGTCTGAACTCTTTCCCTACACGACGCTCTTCCGATCTNNNGCAATGAACATGGAGGTGCA
GAGCACACGTCTGAACTCTTTCCCTACACGACGCTCTTCCGATCTNNNAAATAGAGCAGGGCTGTTTGC
GAGCACACGTCTGAACTCTTTCCCTACACGACGCTCTTCCGATCTNNNAAATAGAGCAGGGCTGTTTGC
AGCACACGTCTGAACTCTTTCCCTACACGACGCTCTTCCGATCTNNNATACAAAATTACCCAGGCGTGC
AGCACACGTCTGAACTCTTTCCCTACACGACGCTCTTCCGATCTNNNATACAAAATTACCCAGGCGTGC
GAGCACACGTCTGAACTCTTTCCCTACACGACGCTCTTCCGATCTNNNGCTTTCAGTCCTGTGAGCTTTT
GAGCACACGTCTGAACTCTTTCCCTACACGACGCTCTTCCGATCTNNNGCTTTCAGTCCTGTGAGCTTTT
AGCACACGTCTGAACTCTTTCCCTACACGACGCTCTTCCGATCTNNNAGATTTGTACAGGCATGGAGA
AGCACACGTCTGAACTCTTTCCCTACACGACGCTCTTCCGATCTNNNAGATTTGTACAGGCATGGAGA
AGCACACGTCTGAACTCTTTCCCTACACGACGCTCTTCCGATCTNNNCTTGTCTGAGTGGAAGGCTGTA
AGCACACGTCTGAACTCTTTCCCTACACGACGCTCTTCCGATCTNNNCTTGTCTGAGTGGAAGGCTGTA
GAGCACACGTCTGAACTCTTTCCCTACACGACGCTCTTCCGATCTNNNACTTCCCATGTCCAGACTAGGT
GAGCACACGTCTGAACTCTTTCCCTACACGACGCTCTTCCGATCTNNNACTTCCCATGTCCAGACTAGGT
GAGCACACGTCTGAACTCTTTCCCTACACGACGCTCTTCCGATCTNNNGTCGTATCAGCCCTCTTAGAGC
GAGCACACGTCTGAACTCTTTCCCTACACGACGCTCTTCCGATCTNNNGTCGTATCAGCCCTCTTAGAGC
GAGCACACGTCTGAACTCTTTCCCTACACGACGCTCTTCCGATCTNNNGAGAGCACGTGTTGGAGTGG
GAGCACACGTCTGAACTCTTTCCCTACACGACGCTCTTCCGATCTNNNGAGAGCACGTGTTGGAGTGG
GAGCACACGTCTGAACTCTTTCCCTACACGACGCTCTTCCGATCTNNNACTTAAGGCTGACTTGAGCGA
GAGCACACGTCTGAACTCTTTCCCTACACGACGCTCTTCCGATCTNNNACTTAAGGCTGACTTGAGCGA
GCACACGTCTGAACTCTTTCCCTACACGACGCTCTTCCGATCTNNNACCAACTTTGTGTCTTTTTCTTCTC
GCACACGTCTGAACTCTTTCCCTACACGACGCTCTTCCGATCTNNNACCAACTTTGTGTCTTTTTCTTCTC
AGCACACGTCTGAACTCTTTCCCTACACGACGCTCTTCCGATCTNNNGGAACTTCCAAATAATGTGTTTC
AGCACACGTCTGAACTCTTTCCCTACACGACGCTCTTCCGATCTNNNGGAACTTCCAAATAATGTGTTTC
AGCACACGTCTGAACTCTTTCCCTACACGACGCTCTTCCGATCTNNNCTCCCTCAAGCGTACCAAAAAAA
AGCACACGTCTGAACTCTTTCCCTACACGACGCTCTTCCGATCTNNNCTCCCTCAAGCGTACCAAAAAAA
GAGCACACGTCTGAACTCTTTCCCTACACGACGCTCTTCCGATCTNNNATGCCTGTAATCCCAGCACTTT
GAGCACACGTCTGAACTCTTTCCCTACACGACGCTCTTCCGATCTNNNATGCCTGTAATCCCAGCACTTT
AGCACACGTCTGAACTCTTTCCCTACACGACGCTCTTCCGATCTNNNATGAAAAGGCAACCCACAGAAT
AGCACACGTCTGAACTCTTTCCCTACACGACGCTCTTCCGATCTNNNATGAAAAGGCAACCCACAGAAT

AGCACACGTCTGAACTCTTTCCCTACACGACGCTCTTCCGATCTNNNTCTCTCCTCTCTCTCTCTCCTCTC
AGCACACGTCTGAACTCTTTCCCTACACGACGCTCTTCCGATCTNNNTCTCTCCTCTCTCTCTCTCCTCTC
GCACACGTCTGAACTCTTTCCCTACACGACGCTCTTCCGATCTNNNGTAATCTCAGCTACTTGGGAGG
GCACACGTCTGAACTCTTTCCCTACACGACGCTCTTCCGATCTNNNGTAATCTCAGCTACTTGGGAGG
GCACACGTCTGAACTCTTTCCCTACACGACGCTCTTCCGATCTNNNGTTGAGATTTGGGTTAGCAGC
GCACACGTCTGAACTCTTTCCCTACACGACGCTCTTCCGATCTNNNGTTGAGATTTGGGTTAGCAGC
CACACGTCTGAACTCTTTCCCTACACGACGCTCTTCCGATCTNNNCTAACTGGTTTCTCTCTCCCCACCA
CACACGTCTGAACTCTTTCCCTACACGACGCTCTTCCGATCTNNNCTAACTGGTTTCTCTCTCCCCACCA
GCACACGTCTGAACTCTTTCCCTACACGACGCTCTTCCGATCTNNNTGTGGTGGACTGAAGGATACA
GCACACGTCTGAACTCTTTCCCTACACGACGCTCTTCCGATCTNNNTGTGGTGGACTGAAGGATACA
GCACACGTCTGAACTCTTTCCCTACACGACGCTCTTCCGATCTNNNGCTGGGATTATAGTCATGAGCC
GCACACGTCTGAACTCTTTCCCTACACGACGCTCTTCCGATCTNNNGCTGGGATTATAGTCATGAGCC
AGAGCACACGTCTGAACTCTTTCCCTACACGACGCTCTTCCGATCTNNNCCCAGCTACTTAGGAGGTT
AGAGCACACGTCTGAACTCTTTCCCTACACGACGCTCTTCCGATCTNNNCCCAGCTACTTAGGAGGTT
GCACACGTCTGAACTCTTTCCCTACACGACGCTCTTCCGATCTNNNCTGTAATTCCAGCTACTCGGGA
GCACACGTCTGAACTCTTTCCCTACACGACGCTCTTCCGATCTNNNCTGTAATTCCAGCTACTCGGGA
ACACGTCTGAACTCTTTCCCTACACGACGCTCTTCCGATCTNNNCTATCTGGTCCTCCCCTCACCCATCG
ACACGTCTGAACTCTTTCCCTACACGACGCTCTTCCGATCTNNNCTATCTGGTCCTCCCCTCACCCATCG
AGAGCACACGTCTGAACTCTTTCCCTACACGACGCTCTTCCGATCTNNNTCTCTGTTCCTCTTACCC
AGAGCACACGTCTGAACTCTTTCCCTACACGACGCTCTTCCGATCTNNNTCTCTGTTCCTCTTACCC
AGCACACGTCTGAACTCTTTCCCTACACGACGCTCTTCCGATCTNNNATGCCTCTAATCCCAGCTACTTG
AGCACACGTCTGAACTCTTTCCCTACACGACGCTCTTCCGATCTNNNATGCCTCTAATCCCAGCTACTTG
GCACACGTCTGAACTCTTTCCCTACACGACGCTCTTCCGATCTNNNCTGATCAGTACAGAAGAGTGG
GCACACGTCTGAACTCTTTCCCTACACGACGCTCTTCCGATCTNNNCTGATCAGTACAGAAGAGTGG
GAGCACACGTCTGAACTCTTTCCCTACACGACGCTCTTCCGATCTNNNATGAGGATTCGTGATGGTGC
GAGCACACGTCTGAACTCTTTCCCTACACGACGCTCTTCCGATCTNNNATGAGGATTCGTGATGGTGC
GCACACGTCTGAACTCTTTCCCTACACGACGCTCTTCCGATCTNNNTAATGCTGCTTCTTGAAGGCTTG
GCACACGTCTGAACTCTTTCCCTACACGACGCTCTTCCGATCTNNNTAATGCTGCTTCTTGAAGGCTTG
AGCACACGTCTGAACTCTTTCCCTACACGACGCTCTTCCGATCTNNNCCACGCCTGGCTAATTTTTGTA
AGCACACGTCTGAACTCTTTCCCTACACGACGCTCTTCCGATCTNNNCCACGCCTGGCTAATTTTTGTA
GAGCACACGTCTGAACTCTTTCCCTACACGACGCTCTTCCGATCTNNNTGTCTGAGTGTAGGCAGAAC
GAGCACACGTCTGAACTCTTTCCCTACACGACGCTCTTCCGATCTNNNTGTCTGAGTGTAGGCAGAAC
GCACACGTCTGAACTCTTTCCCTACACGACGCTCTTCCGATCTNNNGTTCAAGAGATTCTCCTGCCTCA
GCACACGTCTGAACTCTTTCCCTACACGACGCTCTTCCGATCTNNNGTTCAAGAGATTCTCCTGCCTCA
GCACACGTCTGAACTCTTTCCCTACACGACGCTCTTCCGATCTNNNTGACATAGGGCCTGTGAGAAA
GCACACGTCTGAACTCTTTCCCTACACGACGCTCTTCCGATCTNNNTGACATAGGGCCTGTGAGAAA
GAGCACACGTCTGAACTCTTTCCCTACACGACGCTCTTCCGATCTNNNGAGGCTGAGGCAGGAGAAT
GAGCACACGTCTGAACTCTTTCCCTACACGACGCTCTTCCGATCTNNNGAGGCTGAGGCAGGAGAAT
AGCACACGTCTGAACTCTTTCCCTACACGACGCTCTTCCGATCTNNNGAGGCAAGAGAATCGCTTGA
AGCACACGTCTGAACTCTTTCCCTACACGACGCTCTTCCGATCTNNNGAGGCAAGAGAATCGCTTGA
GAGCACACGTCTGAACTCTTTCCCTACACGACGCTCTTCCGATCTNNNAGAGGTTGCAGTGAACAGA
GAGCACACGTCTGAACTCTTTCCCTACACGACGCTCTTCCGATCTNNNAGAGGTTGCAGTGAACAGA
GCACACGTCTGAACTCTTTCCCTACACGACGCTCTTCCGATCTNNNTCTGGTACACTTTTCTCACATGC

AGCACACGTCTGAACTCTTTCCCTACACGACGCTCTTCCGATCTNNNTCTGGTACACTTTTCTCACATGCC
GAGCACACGTCTGAACTCTTTCCCTACACGACGCTCTTCCGATCTNNNGGCACCTATAATCCCAGCTAC
GAGCACACGTCTGAACTCTTTCCCTACACGACGCTCTTCCGATCTNNNGGCACCTATAATCCCAGCTAC
AGCACACGTCTGAACTCTTTCCCTACACGACGCTCTTCCGATCTNNNGGAGGCTTCCTTTGTGGTTAGA
AGCACACGTCTGAACTCTTTCCCTACACGACGCTCTTCCGATCTNNNGGAGGCTTCCTTTGTGGTTAGA
AGCACACGTCTGAACTCTTTCCCTACACGACGCTCTTCCGATCTNNNTTTAAAATATCAAGAGATCGTC
AGCACACGTCTGAACTCTTTCCCTACACGACGCTCTTCCGATCTNNNTTTAAAATATCAAGAGATCGTC
AGCACACGTCTGAACTCTTTCCCTACACGACGCTCTTCCGATCTNNNTCGGTAAGAGAAAGAGGTAGC
AGCACACGTCTGAACTCTTTCCCTACACGACGCTCTTCCGATCTNNNTCGGTAAGAGAAAGAGGTAGC
GAGCACACGTCTGAACTCTTTCCCTACACGACGCTCTTCCGATCTNNNTACCTGAGCACTCCTTCATTTG
GAGCACACGTCTGAACTCTTTCCCTACACGACGCTCTTCCGATCTNNNTACCTGAGCACTCCTTCATTTG
GAGCACACGTCTGAACTCTTTCCCTACACGACGCTCTTCCGATCTNNNGGACACATACACAGTCCCAAG
GAGCACACGTCTGAACTCTTTCCCTACACGACGCTCTTCCGATCTNNNGGACACATACACAGTCCCAAG
AGCACACGTCTGAACTCTTTCCCTACACGACGCTCTTCCGATCTNNNTCCTCTTCCTAATAAGCCTGGGA
AGCACACGTCTGAACTCTTTCCCTACACGACGCTCTTCCGATCTNNNTCCTCTTCCTAATAAGCCTGGGA
AGCACACGTCTGAACTCTTTCCCTACACGACGCTCTTCCGATCTNNNAAGGGTTTACTGCTGCCATTTT
AGCACACGTCTGAACTCTTTCCCTACACGACGCTCTTCCGATCTNNNAAGGGTTTACTGCTGCCATTTT
AGCACACGTCTGAACTCTTTCCCTACACGACGCTCTTCCGATCTNNNCTTACTGACTTGGAGTCCCTCA
AGCACACGTCTGAACTCTTTCCCTACACGACGCTCTTCCGATCTNNNCTTACTGACTTGGAGTCCCTCA
AGCACACGTCTGAACTCTTTCCCTACACGACGCTCTTCCGATCTNNNAGTTTATTTGCATGGAGGTGTT
AGCACACGTCTGAACTCTTTCCCTACACGACGCTCTTCCGATCTNNNAGTTTATTTGCATGGAGGTGTT
AGAGCACACGTCTGAACTCTTTCCCTACACGACGCTCTTCCGATCTNNNTGGGGGTGGTGAAGAAGGA
AGAGCACACGTCTGAACTCTTTCCCTACACGACGCTCTTCCGATCTNNNTGGGGGTGGTGAAGAAGGA
GCACACGTCTGAACTCTTTCCCTACACGACGCTCTTCCGATCTNNNCCCCAAACATGGCAATGTGATTA
GCACACGTCTGAACTCTTTCCCTACACGACGCTCTTCCGATCTNNNCCCCAAACATGGCAATGTGATTA
GAGCACACGTCTGAACTCTTTCCCTACACGACGCTCTTCCGATCTNNNGGATCTCGTACATGACTTCTG
GAGCACACGTCTGAACTCTTTCCCTACACGACGCTCTTCCGATCTNNNGGATCTCGTACATGACTTCTG
AGCACACGTCTGAACTCTTTCCCTACACGACGCTCTTCCGATCTNNNCATTTTTGTACCTGTGGCTGTCC
AGCACACGTCTGAACTCTTTCCCTACACGACGCTCTTCCGATCTNNNCATTTTTGTACCTGTGGCTGTCC
AGCACACGTCTGAACTCTTTCCCTACACGACGCTCTTCCGATCTNNNAGTGAGCTGAAATCATGCCATT
AGCACACGTCTGAACTCTTTCCCTACACGACGCTCTTCCGATCTNNNAGTGAGCTGAAATCATGCCATT
GAGCACACGTCTGAACTCTTTCCCTACACGACGCTCTTCCGATCTNNNATGTGTGTGTGCTACCTAGAC
GAGCACACGTCTGAACTCTTTCCCTACACGACGCTCTTCCGATCTNNNATGTGTGTGTGCTACCTAGAC
AGCACACGTCTGAACTCTTTCCCTACACGACGCTCTTCCGATCTNNNTTAGGATGGGAGGATGGTGGA
AGCACACGTCTGAACTCTTTCCCTACACGACGCTCTTCCGATCTNNNTTAGGATGGGAGGATGGTGGA
AGCACACGTCTGAACTCTTTCCCTACACGACGCTCTTCCGATCTNNNCCCGAACTAATCCTCTCTGTCACC
AGCACACGTCTGAACTCTTTCCCTACACGACGCTCTTCCGATCTNNNCCCGAACTAATCCTCTCTGTCACC
AGCACACGTCTGAACTCTTTCCCTACACGACGCTCTTCCGATCTNNNGTGGAAAGGGCTGAGAGAAG
AGCACACGTCTGAACTCTTTCCCTACACGACGCTCTTCCGATCTNNNGTGGAAAGGGCTGAGAGAAG
GCACACGTCTGAACTCTTTCCCTACACGACGCTCTTCCGATCTNNNGAGAAGGCTGTGAGTGCTGTCCA
GCACACGTCTGAACTCTTTCCCTACACGACGCTCTTCCGATCTNNNGAGAAGGCTGTGAGTGCTGTCCA
AGAGCACACGTCTGAACTCTTTCCCTACACGACGCTCTTCCGATCTNNNATTTGTCCCTGTGTGACATAG
AGAGCACACGTCTGAACTCTTTCCCTACACGACGCTCTTCCGATCTNNNATTTGTCCCTGTGTGACATAG

AGCACACGTCTGAACTCTTTCCCTACACGACGCTCTTCCGATCTNNNGCCACCTTGATTTTGAGAGTGT
AGCACACGTCTGAACTCTTTCCCTACACGACGCTCTTCCGATCTNNNGCCACCTTGATTTTGAGAGTGT
GAGCACACGTCTGAACTCTTTCCCTACACGACGCTCTTCCGATCTNNNAAAAAATTAGCCAGACGTGGT
GAGCACACGTCTGAACTCTTTCCCTACACGACGCTCTTCCGATCTNNNAAAAAATTAGCCAGACGTGGT
AGCACACGTCTGAACTCTTTCCCTACACGACGCTCTTCCGATCTNNNGCTGGGATTATAGGCATGAGA
AGCACACGTCTGAACTCTTTCCCTACACGACGCTCTTCCGATCTNNNGCTGGGATTATAGGCATGAGA
GAGCACACGTCTGAACTCTTTCCCTACACGACGCTCTTCCGATCTNNNGCATTTTGAAGTCCCAACCAAG
GAGCACACGTCTGAACTCTTTCCCTACACGACGCTCTTCCGATCTNNNGCATTTTGAAGTCCCAACCAAG
GAGCACACGTCTGAACTCTTTCCCTACACGACGCTCTTCCGATCTNNNAGATTGCAGTGAGCTGAGAT
GAGCACACGTCTGAACTCTTTCCCTACACGACGCTCTTCCGATCTNNNAGATTGCAGTGAGCTGAGAT
GAGCACACGTCTGAACTCTTTCCCTACACGACGCTCTTCCGATCTNNNATTGCACAGTAGGGTGGCTAT
GAGCACACGTCTGAACTCTTTCCCTACACGACGCTCTTCCGATCTNNNATTGCACAGTAGGGTGGCTAT
AGCACACGTCTGAACTCTTTCCCTACACGACGCTCTTCCGATCTNNNAACATGTCAAAACCCCGTCTCTA
AGCACACGTCTGAACTCTTTCCCTACACGACGCTCTTCCGATCTNNNAACATGTCAAAACCCCGTCTCTA
AGCACACGTCTGAACTCTTTCCCTACACGACGCTCTTCCGATCTNNNCTGTAATCCCAGCTACTCAGGA
AGCACACGTCTGAACTCTTTCCCTACACGACGCTCTTCCGATCTNNNCTGTAATCCCAGCTACTCAGGA
AGCACACGTCTGAACTCTTTCCCTACACGACGCTCTTCCGATCTNNNGTGGTTGTTATTGCCACTCTGTG
AGCACACGTCTGAACTCTTTCCCTACACGACGCTCTTCCGATCTNNNGTGGTTGTTATTGCCACTCTGTG
AGAGCACACGTCTGAACTCTTTCCCTACACGACGCTCTTCCGATCTNNNCCTTGAGAGAGGTTTCTGGA
AGAGCACACGTCTGAACTCTTTCCCTACACGACGCTCTTCCGATCTNNNCCTTGAGAGAGGTTTCTGGA
GAGCACACGTCTGAACTCTTTCCCTACACGACGCTCTTCCGATCTNNNGGTGTCTGTAATCCCAGCTACT
GAGCACACGTCTGAACTCTTTCCCTACACGACGCTCTTCCGATCTNNNGGTGTCTGTAATCCCAGCTACT
GAGCACACGTCTGAACTCTTTCCCTACACGACGCTCTTCCGATCTNNNACTCAACCCCTTTCACAATAGCT
GAGCACACGTCTGAACTCTTTCCCTACACGACGCTCTTCCGATCTNNNACTCAACCCCTTTCACAATAGCT
GAGCACACGTCTGAACTCTTTCCCTACACGACGCTCTTCCGATCTNNNAAAATGTCTTCTATACCCCAGA
GAGCACACGTCTGAACTCTTTCCCTACACGACGCTCTTCCGATCTNNNAAAATGTCTTCTATACCCCAGA
AGCACACGTCTGAACTCTTTCCCTACACGACGCTCTTCCGATCTNNNGCTTTTCACAAGTAAACAGGCCC
AGCACACGTCTGAACTCTTTCCCTACACGACGCTCTTCCGATCTNNNGCTTTTCACAAGTAAACAGGCCC
GAGCACACGTCTGAACTCTTTCCCTACACGACGCTCTTCCGATCTNNNTGAGGCAAGAGAATTGCTTGA
GAGCACACGTCTGAACTCTTTCCCTACACGACGCTCTTCCGATCTNNNTGAGGCAAGAGAATTGCTTGA
AGCACACGTCTGAACTCTTTCCCTACACGACGCTCTTCCGATCTNNNTGTATGATCGTTTTCCCCAGGTT
AGCACACGTCTGAACTCTTTCCCTACACGACGCTCTTCCGATCTNNNTGTATGATCGTTTTCCCCAGGTT
AGCACACGTCTGAACTCTTTCCCTACACGACGCTCTTCCGATCTNNNATCGCTTGAGTTCAGGTGATCA
AGCACACGTCTGAACTCTTTCCCTACACGACGCTCTTCCGATCTNNNATCGCTTGAGTTCAGGTGATCA
AGAGCACACGTCTGAACTCTTTCCCTACACGACGCTCTTCCGATCTNNNGAGAATCACTTGAACCTGGG
AGAGCACACGTCTGAACTCTTTCCCTACACGACGCTCTTCCGATCTNNNGAGAATCACTTGAACCTGGG
AGCACACGTCTGAACTCTTTCCCTACACGACGCTCTTCCGATCTNNNGCCAGGTATGATTCCAGCTACTT
AGCACACGTCTGAACTCTTTCCCTACACGACGCTCTTCCGATCTNNNGCCAGGTATGATTCCAGCTACTT
AGAGCACACGTCTGAACTCTTTCCCTACACGACGCTCTTCCGATCTNNNGAAGTTTCATGGTGGAAGG
AGAGCACACGTCTGAACTCTTTCCCTACACGACGCTCTTCCGATCTNNNGAAGTTTCATGGTGGAAGG
AGAGCACACGTCTGAACTCTTTCCCTACACGACGCTCTTCCGATCTNNNCTGACACAGGAGAATCGCTTG
AGAGCACACGTCTGAACTCTTTCCCTACACGACGCTCTTCCGATCTNNNCTGACACAGGAGAATCGCTTG
AGCACACGTCTGAACTCTTTCCCTACACGACGCTCTTCCGATCTNNNTTGCTGTCCAAACAAGAGGAAA

GCACACGTCTGAACTCTTTCCCTACACGACGCTCTTCCGATCTNNNTTGCTGTCCAAACAAGAGGAAA
AGCACACGTCTGAACTCTTTCCCTACACGACGCTCTTCCGATCTNNNATAACTCAGCTCCTGGCATATG
AGCACACGTCTGAACTCTTTCCCTACACGACGCTCTTCCGATCTNNNATAACTCAGCTCCTGGCATATG
CACACGTCTGAACTCTTTCCCTACACGACGCTCTTCCGATCTNNNTTCAAGTCCCTGCTTTCCATTCTCA
CACACGTCTGAACTCTTTCCCTACACGACGCTCTTCCGATCTNNNTTCAAGTCCCTGCTTTCCATTCTCA
GAGCACACGTCTGAACTCTTTCCCTACACGACGCTCTTCCGATCTNNNCACTCAAACCACAAGATGCA
GAGCACACGTCTGAACTCTTTCCCTACACGACGCTCTTCCGATCTNNNCACTCAAACCACAAGATGCA
GCACACGTCTGAACTCTTTCCCTACACGACGCTCTTCCGATCTNNNTCTCACTGTAGAGGTCAATTGG
GCACACGTCTGAACTCTTTCCCTACACGACGCTCTTCCGATCTNNNTCTCACTGTAGAGGTCAATTGG
AGCACACGTCTGAACTCTTTCCCTACACGACGCTCTTCCGATCTNNNTACACACACACTCACACACAAA
AGCACACGTCTGAACTCTTTCCCTACACGACGCTCTTCCGATCTNNNTACACACACACTCACACACAAA
AGCACACGTCTGAACTCTTTCCCTACACGACGCTCTTCCGATCTNNNCTAGCATCCAAATCAGCTGCT
AGCACACGTCTGAACTCTTTCCCTACACGACGCTCTTCCGATCTNNNCTAGCATCCAAATCAGCTGCT
GAGCACACGTCTGAACTCTTTCCCTACACGACGCTCTTCCGATCTNNNGCACTGATAGCTTTAGGCCA
GAGCACACGTCTGAACTCTTTCCCTACACGACGCTCTTCCGATCTNNNGCACTGATAGCTTTAGGCCA
AGCACACGTCTGAACTCTTTCCCTACACGACGCTCTTCCGATCTNNNACTCCCATCTCATCTCCTAGGA
AGCACACGTCTGAACTCTTTCCCTACACGACGCTCTTCCGATCTNNNACTCCCATCTCATCTCCTAGGA
AGCACACGTCTGAACTCTTTCCCTACACGACGCTCTTCCGATCTNNNACTGACCTCATGATTCACTCAC
AGCACACGTCTGAACTCTTTCCCTACACGACGCTCTTCCGATCTNNNACTGACCTCATGATTCACTCAC
AGAGCACACGTCTGAACTCTTTCCCTACACGACGCTCTTCCGATCTNNNTCAGGCTTCCTAAGATGCA
AGAGCACACGTCTGAACTCTTTCCCTACACGACGCTCTTCCGATCTNNNTCAGGCTTCCTAAGATGCA
GAGCACACGTCTGAACTCTTTCCCTACACGACGCTCTTCCGATCTNNNCACACATCTAGCTCAATGCC
GAGCACACGTCTGAACTCTTTCCCTACACGACGCTCTTCCGATCTNNNCACACATCTAGCTCAATGCC
AGCACACGTCTGAACTCTTTCCCTACACGACGCTCTTCCGATCTNNNGCATTCCACTCCACATCTGAAT
AGCACACGTCTGAACTCTTTCCCTACACGACGCTCTTCCGATCTNNNGCATTCCACTCCACATCTGAAT
AGCACACGTCTGAACTCTTTCCCTACACGACGCTCTTCCGATCTNNNGAAACTAGGTGTGTATTGTTT
AGCACACGTCTGAACTCTTTCCCTACACGACGCTCTTCCGATCTNNNGAAACTAGGTGTGTATTGTTT
GCACACGTCTGAACTCTTTCCCTACACGACGCTCTTCCGATCTNNNTCTCAATTGCAGGTGACACAAA
GCACACGTCTGAACTCTTTCCCTACACGACGCTCTTCCGATCTNNNTCTCAATTGCAGGTGACACAAA
AGCACACGTCTGAACTCTTTCCCTACACGACGCTCTTCCGATCTNNNAGGAGAATGGCGTGAACCTG
AGCACACGTCTGAACTCTTTCCCTACACGACGCTCTTCCGATCTNNNAGGAGAATGGCGTGAACCTG
CACACGTCTGAACTCTTTCCCTACACGACGCTCTTCCGATCTNNNTTTAGACTTTGTACCTGTTTTACCA
CACACGTCTGAACTCTTTCCCTACACGACGCTCTTCCGATCTNNNTTTAGACTTTGTACCTGTTTTACCA
AGCACACGTCTGAACTCTTTCCCTACACGACGCTCTTCCGATCTNNNTTTGTTTTTGCCTGGAAGAGTG
AGCACACGTCTGAACTCTTTCCCTACACGACGCTCTTCCGATCTNNNTTTGTTTTTGCCTGGAAGAGTG
AGAGCACACGTCTGAACTCTTTCCCTACACGACGCTCTTCCGATCTNNNTGGAGGGTGGTTGCCTAT
AGAGCACACGTCTGAACTCTTTCCCTACACGACGCTCTTCCGATCTNNNTGGAGGGTGGTTGCCTAT
GCACACGTCTGAACTCTTTCCCTACACGACGCTCTTCCGATCTNNNAATCTGTCCTCTGTTGGGGTTTT
GCACACGTCTGAACTCTTTCCCTACACGACGCTCTTCCGATCTNNNAATCTGTCCTCTGTTGGGGTTTT
AGCACACGTCTGAACTCTTTCCCTACACGACGCTCTTCCGATCTNNNGGTCACATTTACAGAAAGCAA
AGCACACGTCTGAACTCTTTCCCTACACGACGCTCTTCCGATCTNNNGGTCACATTTACAGAAAGCAA
AGCACACGTCTGAACTCTTTCCCTACACGACGCTCTTCCGATCTNNNGTGGCACATGTCTGTAATTCCA
AGCACACGTCTGAACTCTTTCCCTACACGACGCTCTTCCGATCTNNNGTGGCACATGTCTGTAATTCCA

AGCACACGTCTGAACTCTTTCCCTACACGACGCTCTTCCGATCTNNNCAGTCTGGGAGTGAGATGAA
AGCACACGTCTGAACTCTTTCCCTACACGACGCTCTTCCGATCTNNNCAGTCTGGGAGTGAGATGAA
AGCACACGTCTGAACTCTTTCCCTACACGACGCTCTTCCGATCTNNNAAATTAGCTGAGCAAAGTGGC
AGCACACGTCTGAACTCTTTCCCTACACGACGCTCTTCCGATCTNNNAAATTAGCTGAGCAAAGTGGC
AGCACACGTCTGAACTCTTTCCCTACACGACGCTCTTCCGATCTNNNTACAAGCTCTTCTTTGTGCCTCT
AGCACACGTCTGAACTCTTTCCCTACACGACGCTCTTCCGATCTNNNTACAAGCTCTTCTTTGTGCCTCT
AGCACACGTCTGAACTCTTTCCCTACACGACGCTCTTCCGATCTNNNTGCCTGTAATTCCAGCTACTCT
AGCACACGTCTGAACTCTTTCCCTACACGACGCTCTTCCGATCTNNNTGCCTGTAATTCCAGCTACTCT
GCACACGTCTGAACTCTTTCCCTACACGACGCTCTTCCGATCTNNNGAAATTTCATGTTTCTGCGGGG
GCACACGTCTGAACTCTTTCCCTACACGACGCTCTTCCGATCTNNNGAAATTTCATGTTTCTGCGGGG
GCACACGTCTGAACTCTTTCCCTACACGACGCTCTTCCGATCTNNNATTGAAAAGCTGGAGGAAGGA
GCACACGTCTGAACTCTTTCCCTACACGACGCTCTTCCGATCTNNNATTGAAAAGCTGGAGGAAGGA
GAGCACACGTCTGAACTCTTTCCCTACACGACGCTCTTCCGATCTNNNAGTACATCATGTTGGGGTTC
GAGCACACGTCTGAACTCTTTCCCTACACGACGCTCTTCCGATCTNNNAGTACATCATGTTGGGGTTC
AGCACACGTCTGAACTCTTTCCCTACACGACGCTCTTCCGATCTNNNGCTAAGGAGCTGGCATAGTAC
AGCACACGTCTGAACTCTTTCCCTACACGACGCTCTTCCGATCTNNNGCTAAGGAGCTGGCATAGTAC
GAGCACACGTCTGAACTCTTTCCCTACACGACGCTCTTCCGATCTNNNCCTGTAATCCCAGCTACTCGA
GAGCACACGTCTGAACTCTTTCCCTACACGACGCTCTTCCGATCTNNNCCTGTAATCCCAGCTACTCGA
GCACACGTCTGAACTCTTTCCCTACACGACGCTCTTCCGATCTNNNGGGGCTTGTAATCTCAGCTACT
GCACACGTCTGAACTCTTTCCCTACACGACGCTCTTCCGATCTNNNGGGGCTTGTAATCTCAGCTACT
AGCACACGTCTGAACTCTTTCCCTACACGACGCTCTTCCGATCTNNNTGTGCTGCTGATGAGATCATA
AGCACACGTCTGAACTCTTTCCCTACACGACGCTCTTCCGATCTNNNTGTGCTGCTGATGAGATCATA
AGCACACGTCTGAACTCTTTCCCTACACGACGCTCTTCCGATCTNNNACGAATTATGTCATGCACAGTC
AGCACACGTCTGAACTCTTTCCCTACACGACGCTCTTCCGATCTNNNACGAATTATGTCATGCACAGTC
GCACACGTCTGAACTCTTTCCCTACACGACGCTCTTCCGATCTNNNCTGTTGAACAACCATGTCTGGA
GCACACGTCTGAACTCTTTCCCTACACGACGCTCTTCCGATCTNNNCTGTTGAACAACCATGTCTGGA
AGCACACGTCTGAACTCTTTCCCTACACGACGCTCTTCCGATCTNNNATTCAGCCGCTTGTCTAATGGT
AGCACACGTCTGAACTCTTTCCCTACACGACGCTCTTCCGATCTNNNATTCAGCCGCTTGTCTAATGGT
GAGCACACGTCTGAACTCTTTCCCTACACGACGCTCTTCCGATCTNNNCTGAGATCACATGACTGCACT
GAGCACACGTCTGAACTCTTTCCCTACACGACGCTCTTCCGATCTNNNCTGAGATCACATGACTGCACT
AGCACACGTCTGAACTCTTTCCCTACACGACGCTCTTCCGATCTNNNCTCAGGTGATCCTCTCATCTCAG
AGCACACGTCTGAACTCTTTCCCTACACGACGCTCTTCCGATCTNNNCTCAGGTGATCCTCTCATCTCAG
AGCACACGTCTGAACTCTTTCCCTACACGACGCTCTTCCGATCTNNNGTTGGGAACTGTGGACAAAA
AGCACACGTCTGAACTCTTTCCCTACACGACGCTCTTCCGATCTNNNGTTGGGAACTGTGGACAAAA
GAGCACACGTCTGAACTCTTTCCCTACACGACGCTCTTCCGATCTNNNTGCTTTGAGGCAGACTGAGT
GAGCACACGTCTGAACTCTTTCCCTACACGACGCTCTTCCGATCTNNNTGCTTTGAGGCAGACTGAGT
GCACACGTCTGAACTCTTTCCCTACACGACGCTCTTCCGATCTNNNATTCTTTTGTGCGTCCGATTGTT
GCACACGTCTGAACTCTTTCCCTACACGACGCTCTTCCGATCTNNNATTCTTTTGTGCGTCCGATTGTT
AGCACACGTCTGAACTCTTTCCCTACACGACGCTCTTCCGATCTNNNACTGATGGGAAGGGTGATTCT
AGCACACGTCTGAACTCTTTCCCTACACGACGCTCTTCCGATCTNNNACTGATGGGAAGGGTGATTCT
GCACACGTCTGAACTCTTTCCCTACACGACGCTCTTCCGATCTNNNTCAAGATAATGGCATCAGAGTA
GCACACGTCTGAACTCTTTCCCTACACGACGCTCTTCCGATCTNNNTCAAGATAATGGCATCAGAGTA
GCACACGTCTGAACTCTTTCCCTACACGACGCTCTTCCGATCTNNNTTGGACAAGGGACAGTAAAGC

GCACACGTCTGAACTCTTTCCCTACACGACGCTCTTCCGATCTNNNTTGGACAAGGGACAGTAAAGC
AGCACACGTCTGAACTCTTTCCCTACACGACGCTCTTCCGATCTNNNGCGAAACCCCGTCTCTACTAAAA
AGCACACGTCTGAACTCTTTCCCTACACGACGCTCTTCCGATCTNNNGCGAAACCCCGTCTCTACTAAAA
AGCACACGTCTGAACTCTTTCCCTACACGACGCTCTTCCGATCTNNNGCTTCTCCAGAGAAACAGAAC
AGCACACGTCTGAACTCTTTCCCTACACGACGCTCTTCCGATCTNNNGCTTCTCCAGAGAAACAGAAC
GAGCACACGTCTGAACTCTTTCCCTACACGACGCTCTTCCGATCTNNNTTCAGTTCCCTTTCTCTGTCAG
GAGCACACGTCTGAACTCTTTCCCTACACGACGCTCTTCCGATCTNNNTTCAGTTCCCTTTCTCTGTCAG
GCACACGTCTGAACTCTTTCCCTACACGACGCTCTTCCGATCTNNNGAGAATGGTGTGACCCCAGAA
GCACACGTCTGAACTCTTTCCCTACACGACGCTCTTCCGATCTNNNGAGAATGGTGTGACCCCAGAA
AGCACACGTCTGAACTCTTTCCCTACACGACGCTCTTCCGATCTNNNTCTCATTTTTGGCATGGTTTCC
AGCACACGTCTGAACTCTTTCCCTACACGACGCTCTTCCGATCTNNNTCTCATTTTTGGCATGGTTTCC
AGCACACGTCTGAACTCTTTCCCTACACGACGCTCTTCCGATCTNNNAATCTTGACTCACTGCAACCTC
AGCACACGTCTGAACTCTTTCCCTACACGACGCTCTTCCGATCTNNNAATCTTGACTCACTGCAACCTC
AGCACACGTCTGAACTCTTTCCCTACACGACGCTCTTCCGATCTNNNTGGGAATGTGTTATGATGCCT
AGCACACGTCTGAACTCTTTCCCTACACGACGCTCTTCCGATCTNNNTGGGAATGTGTTATGATGCCT
AGCACACGTCTGAACTCTTTCCCTACACGACGCTCTTCCGATCTNNNTGTGTGGATGTGGGAGACTAA
AGCACACGTCTGAACTCTTTCCCTACACGACGCTCTTCCGATCTNNNTGTGTGGATGTGGGAGACTAA
GCACACGTCTGAACTCTTTCCCTACACGACGCTCTTCCGATCTNNNGCCAAAGCAAGACTAGGCAAA
GCACACGTCTGAACTCTTTCCCTACACGACGCTCTTCCGATCTNNNGCCAAAGCAAGACTAGGCAAA
GAGCACACGTCTGAACTCTTTCCCTACACGACGCTCTTCCGATCTNNNGTGTGTCTGTAGTCCCAGCTA
GAGCACACGTCTGAACTCTTTCCCTACACGACGCTCTTCCGATCTNNNGTGTGTCTGTAGTCCCAGCTA
AGCACACGTCTGAACTCTTTCCCTACACGACGCTCTTCCGATCTNNNCTTCTCATGAGGCCATTGAGA
AGCACACGTCTGAACTCTTTCCCTACACGACGCTCTTCCGATCTNNNCTTCTCATGAGGCCATTGAGA
AGCACACGTCTGAACTCTTTCCCTACACGACGCTCTTCCGATCTNNNGGTTGCAGTGAGACGAGATCT
AGCACACGTCTGAACTCTTTCCCTACACGACGCTCTTCCGATCTNNNGGTTGCAGTGAGACGAGATCT
GAGCACACGTCTGAACTCTTTCCCTACACGACGCTCTTCCGATCTNNNGAAGTTTGCAGTGAGCTGAG
GAGCACACGTCTGAACTCTTTCCCTACACGACGCTCTTCCGATCTNNNGAAGTTTGCAGTGAGCTGAG
GCACACGTCTGAACTCTTTCCCTACACGACGCTCTTCCGATCTNNNTTAGGCACTGTGAGCTACATAT
GCACACGTCTGAACTCTTTCCCTACACGACGCTCTTCCGATCTNNNTTAGGCACTGTGAGCTACATAT
CACACGTCTGAACTCTTTCCCTACACGACGCTCTTCCGATCTNNNATTTGCTCATCGTGGCGGAGCAT
CACACGTCTGAACTCTTTCCCTACACGACGCTCTTCCGATCTNNNATTTGCTCATCGTGGCGGAGCAT
AGCACACGTCTGAACTCTTTCCCTACACGACGCTCTTCCGATCTNNNGTTTTAGCGTCACTGTGTCAA
AGCACACGTCTGAACTCTTTCCCTACACGACGCTCTTCCGATCTNNNGTTTTAGCGTCACTGTGTCAA
AGCACACGTCTGAACTCTTTCCCTACACGACGCTCTTCCGATCTNNNAGTCTTGCTCTGTTGCCCAGCAT
AGCACACGTCTGAACTCTTTCCCTACACGACGCTCTTCCGATCTNNNAGTCTTGCTCTGTTGCCCAGCAT
AGCACACGTCTGAACTCTTTCCCTACACGACGCTCTTCCGATCTNNNGCTGGGATTACAGGCGTGAG
AGCACACGTCTGAACTCTTTCCCTACACGACGCTCTTCCGATCTNNNGCTGGGATTACAGGCGTGAG
GAGCACACGTCTGAACTCTTTCCCTACACGACGCTCTTCCGATCTNNNAGCAACATAGCACATGGACA
GAGCACACGTCTGAACTCTTTCCCTACACGACGCTCTTCCGATCTNNNAGCAACATAGCACATGGACA
GAGCACACGTCTGAACTCTTTCCCTACACGACGCTCTTCCGATCTNNNAAAAAATTAGCCAGGCGTGG
GAGCACACGTCTGAACTCTTTCCCTACACGACGCTCTTCCGATCTNNNAAAAAATTAGCCAGGCGTGG
GAGCACACGTCTGAACTCTTTCCCTACACGACGCTCTTCCGATCTNNNGCTCACAGGTTCTCACTTTTG
GAGCACACGTCTGAACTCTTTCCCTACACGACGCTCTTCCGATCTNNNGCTCACAGGTTCTCACTTTTG

AGCACACGTCTGAACTCTTTCCCTACACGACGCTCTTCCGATCTNNNTGTGGATGTATATCTGGGTTCT
AGCACACGTCTGAACTCTTTCCCTACACGACGCTCTTCCGATCTNNNTGTGGATGTATATCTGGGTTCT
AGCACACGTCTGAACTCTTTCCCTACACGACGCTCTTCCGATCTNNNTAAGAGCCTGTCTCATTTCTGG
AGCACACGTCTGAACTCTTTCCCTACACGACGCTCTTCCGATCTNNNTAAGAGCCTGTCTCATTTCTGG
GCACACGTCTGAACTCTTTCCCTACACGACGCTCTTCCGATCTNNNGAGCTAATTCCACTCCTGACAGC
GCACACGTCTGAACTCTTTCCCTACACGACGCTCTTCCGATCTNNNGAGCTAATTCCACTCCTGACAGC
AGCACACGTCTGAACTCTTTCCCTACACGACGCTCTTCCGATCTNNNGAGGCAGGAGAATGGTGTGA
AGCACACGTCTGAACTCTTTCCCTACACGACGCTCTTCCGATCTNNNGAGGCAGGAGAATGGTGTGA
AGCACACGTCTGAACTCTTTCCCTACACGACGCTCTTCCGATCTNNNTGCCTTGGTGATCTTGGATAAT
AGCACACGTCTGAACTCTTTCCCTACACGACGCTCTTCCGATCTNNNTGCCTTGGTGATCTTGGATAAT
GCACACGTCTGAACTCTTTCCCTACACGACGCTCTTCCGATCTNNNTTCTGTGATTTGAGAGTAAGAG
GCACACGTCTGAACTCTTTCCCTACACGACGCTCTTCCGATCTNNNTTCTGTGATTTGAGAGTAAGAG
GCACACGTCTGAACTCTTTCCCTACACGACGCTCTTCCGATCTNNNTGCAAAGTTCAGAAGTTGAAG
GCACACGTCTGAACTCTTTCCCTACACGACGCTCTTCCGATCTNNNTGCAAAGTTCAGAAGTTGAAG
GCACACGTCTGAACTCTTTCCCTACACGACGCTCTTCCGATCTNNNGGAACAGGCTTTTTCGTGGAAT
GCACACGTCTGAACTCTTTCCCTACACGACGCTCTTCCGATCTNNNGGAACAGGCTTTTTCGTGGAAT
GAGCACACGTCTGAACTCTTTCCCTACACGACGCTCTTCCGATCTNNNCAGGTGCCTATAATCCCAGCT
GAGCACACGTCTGAACTCTTTCCCTACACGACGCTCTTCCGATCTNNNCAGGTGCCTATAATCCCAGCT
GAGCACACGTCTGAACTCTTTCCCTACACGACGCTCTTCCGATCTNNNACCAAAGGAAAACCTTAGGC
GAGCACACGTCTGAACTCTTTCCCTACACGACGCTCTTCCGATCTNNNACCAAAGGAAAACCTTAGGC
AGCACACGTCTGAACTCTTTCCCTACACGACGCTCTTCCGATCTNNNTTGCCATCTGTAAAACCTCTTCG
AGCACACGTCTGAACTCTTTCCCTACACGACGCTCTTCCGATCTNNNTTGCCATCTGTAAAACCTCTTCG
GAGCACACGTCTGAACTCTTTCCCTACACGACGCTCTTCCGATCTNNNGAAACCTTGCCATTCCTGTCA
GAGCACACGTCTGAACTCTTTCCCTACACGACGCTCTTCCGATCTNNNGAAACCTTGCCATTCCTGTCA
GCACACGTCTGAACTCTTTCCCTACACGACGCTCTTCCGATCTNNNGAGAATCACTTGAACTCGGGAG
GCACACGTCTGAACTCTTTCCCTACACGACGCTCTTCCGATCTNNNGAGAATCACTTGAACTCGGGAG
AGCACACGTCTGAACTCTTTCCCTACACGACGCTCTTCCGATCTNNNAGAATTGCTTGAACCCAGGAG
AGCACACGTCTGAACTCTTTCCCTACACGACGCTCTTCCGATCTNNNAGAATTGCTTGAACCCAGGAG
AGCACACGTCTGAACTCTTTCCCTACACGACGCTCTTCCGATCTNNNTGCCTTTGCATCCTCATAGCTTA
AGCACACGTCTGAACTCTTTCCCTACACGACGCTCTTCCGATCTNNNTGCCTTTGCATCCTCATAGCTTA
AGCACACGTCTGAACTCTTTCCCTACACGACGCTCTTCCGATCTNNNCTTGCAGTGAGCCGAGATCGC
AGCACACGTCTGAACTCTTTCCCTACACGACGCTCTTCCGATCTNNNCTTGCAGTGAGCCGAGATCGC
GAGCACACGTCTGAACTCTTTCCCTACACGACGCTCTTCCGATCTNNNGTCAGAAGTTTGAGACCAGC
GAGCACACGTCTGAACTCTTTCCCTACACGACGCTCTTCCGATCTNNNGTCAGAAGTTTGAGACCAGC
CACACGTCTGAACTCTTTCCCTACACGACGCTCTTCCGATCTNNNGGCCCTCTTCAAGTTTTCTTTCTCAT
CACACGTCTGAACTCTTTCCCTACACGACGCTCTTCCGATCTNNNGGCCCTCTTCAAGTTTTCTTTCTCAT
AGCACACGTCTGAACTCTTTCCCTACACGACGCTCTTCCGATCTNNNCCTCAGCTTCTAAAGGACACAG
AGCACACGTCTGAACTCTTTCCCTACACGACGCTCTTCCGATCTNNNCCTCAGCTTCTAAAGGACACAG
GAGCACACGTCTGAACTCTTTCCCTACACGACGCTCTTCCGATCTNNNCATCCTGGCTAACATGGTGAA
GAGCACACGTCTGAACTCTTTCCCTACACGACGCTCTTCCGATCTNNNCATCCTGGCTAACATGGTGAA
GCACACGTCTGAACTCTTTCCCTACACGACGCTCTTCCGATCTNNNTAAGAGTGTCAGCATTATCATTC
GCACACGTCTGAACTCTTTCCCTACACGACGCTCTTCCGATCTNNNTAAGAGTGTCAGCATTATCATTC
GAGCACACGTCTGAACTCTTTCCCTACACGACGCTCTTCCGATCTNNNCACTGAATGTGGCCTGAAAG

GAGCACACGTCTGAACTCTTTCCCTACACGACGCTCTTCCGATCTNNNCACTGAATGTGGCCTGAAAG
AGCACACGTCTGAACTCTTTCCCTACACGACGCTCTTCCGATCTNNNAAATTCTTATGTCCCAGGGCCA
AGCACACGTCTGAACTCTTTCCCTACACGACGCTCTTCCGATCTNNNAAATTCTTATGTCCCAGGGCCA
GCACACGTCTGAACTCTTTCCCTACACGACGCTCTTCCGATCTNNNGGCTGATATGACAACACTTTGC
GCACACGTCTGAACTCTTTCCCTACACGACGCTCTTCCGATCTNNNGGCTGATATGACAACACTTTGC
AGAGCACACGTCTGAACTCTTTCCCTACACGACGCTCTTCCGATCTNNNTGAGGTAGGAGAAACGCT
AGAGCACACGTCTGAACTCTTTCCCTACACGACGCTCTTCCGATCTNNNTGAGGTAGGAGAAACGCT
GCACACGTCTGAACTCTTTCCCTACACGACGCTCTTCCGATCTNNNCACCACACCCAGCTATTTTTCTTC
GCACACGTCTGAACTCTTTCCCTACACGACGCTCTTCCGATCTNNNCACCACACCCAGCTATTTTTCTTC
AGCACACGTCTGAACTCTTTCCCTACACGACGCTCTTCCGATCTNNNGAGGCAGGAGAATCGCTTGA
AGCACACGTCTGAACTCTTTCCCTACACGACGCTCTTCCGATCTNNNGAGGCAGGAGAATCGCTTGA
GAGCACACGTCTGAACTCTTTCCCTACACGACGCTCTTCCGATCTNNNGATGGAGGACATTGAGTGT
GAGCACACGTCTGAACTCTTTCCCTACACGACGCTCTTCCGATCTNNNGATGGAGGACATTGAGTGT
AGCACACGTCTGAACTCTTTCCCTACACGACGCTCTTCCGATCTNNNTCTCCAAAGAAACTGTCTTGCCT
AGCACACGTCTGAACTCTTTCCCTACACGACGCTCTTCCGATCTNNNTCTCCAAAGAAACTGTCTTGCCT
GAGCACACGTCTGAACTCTTTCCCTACACGACGCTCTTCCGATCTNNNCAGCCTGCTCTCAGATGTTG
GAGCACACGTCTGAACTCTTTCCCTACACGACGCTCTTCCGATCTNNNCAGCCTGCTCTCAGATGTTG
GAGCACACGTCTGAACTCTTTCCCTACACGACGCTCTTCCGATCTNNNTCAAGTGTGCCCTCATATGTC
GAGCACACGTCTGAACTCTTTCCCTACACGACGCTCTTCCGATCTNNNTCAAGTGTGCCCTCATATGTC
AGCACACGTCTGAACTCTTTCCCTACACGACGCTCTTCCGATCTNNNGCAATGGCATATTCTCAGGTGA
AGCACACGTCTGAACTCTTTCCCTACACGACGCTCTTCCGATCTNNNGCAATGGCATATTCTCAGGTGA
GCACACGTCTGAACTCTTTCCCTACACGACGCTCTTCCGATCTNNNCCACAGTTTTTGGACATTCTGGG
GCACACGTCTGAACTCTTTCCCTACACGACGCTCTTCCGATCTNNNCCACAGTTTTTGGACATTCTGGG
AGCACACGTCTGAACTCTTTCCCTACACGACGCTCTTCCGATCTNNNTCAGTAGCAAACCTAAAACCA
AGCACACGTCTGAACTCTTTCCCTACACGACGCTCTTCCGATCTNNNTCAGTAGCAAACCTAAAACCA
GAGCACACGTCTGAACTCTTTCCCTACACGACGCTCTTCCGATCTNNNGAGAGAACACACAAAAGAAC
GAGCACACGTCTGAACTCTTTCCCTACACGACGCTCTTCCGATCTNNNGAGAGAACACACAAAAGAAC
GAGCACACGTCTGAACTCTTTCCCTACACGACGCTCTTCCGATCTNNNCTCCCAAGTAGCTGGAACTAC
GAGCACACGTCTGAACTCTTTCCCTACACGACGCTCTTCCGATCTNNNCTCCCAAGTAGCTGGAACTAC
GAGCACACGTCTGAACTCTTTCCCTACACGACGCTCTTCCGATCTNNNTAAGATGGCTGTTTGCTTCAC
GAGCACACGTCTGAACTCTTTCCCTACACGACGCTCTTCCGATCTNNNTAAGATGGCTGTTTGCTTCAC
AGCACACGTCTGAACTCTTTCCCTACACGACGCTCTTCCGATCTNNNAGAGGTTGCAGTAAGCTGAGA
AGCACACGTCTGAACTCTTTCCCTACACGACGCTCTTCCGATCTNNNAGAGGTTGCAGTAAGCTGAGA
AGCACACGTCTGAACTCTTTCCCTACACGACGCTCTTCCGATCTNNNGAAGTGAGGACAAAAGGCAT
AGCACACGTCTGAACTCTTTCCCTACACGACGCTCTTCCGATCTNNNGAAGTGAGGACAAAAGGCAT
GCACACGTCTGAACTCTTTCCCTACACGACGCTCTTCCGATCTNNNTCCGAGGTGTCCCTGAGTATGC
GCACACGTCTGAACTCTTTCCCTACACGACGCTCTTCCGATCTNNNTCCGAGGTGTCCCTGAGTATGC
CACACGTCTGAACTCTTTCCCTACACGACGCTCTTCCGATCTNNNAAGAGCCCCTTTGAGCAGGTATCA
CACACGTCTGAACTCTTTCCCTACACGACGCTCTTCCGATCTNNNAAGAGCCCCTTTGAGCAGGTATCA
GAGCACACGTCTGAACTCTTTCCCTACACGACGCTCTTCCGATCTNNNTGGTTTTCCATTCCCGAGTTAC
GAGCACACGTCTGAACTCTTTCCCTACACGACGCTCTTCCGATCTNNNTGGTTTTCCATTCCCGAGTTAC
GAGCACACGTCTGAACTCTTTCCCTACACGACGCTCTTCCGATCTNNNATCAGGCTGAGCAACATAGA
GAGCACACGTCTGAACTCTTTCCCTACACGACGCTCTTCCGATCTNNNATCAGGCTGAGCAACATAGA

GAGCACACGTCTGAACTCTTTCCCTACACGACGCTCTTCCGATCTNNNGTAATTACCCTTCAGCTCTTAG

GAGCACACGTCTGAACTCTTTCCCTACACGACGCTCTTCCGATCTNNNGTAATTACCCTTCAGCTCTTAG

GAGCACACGTCTGAACTCTTTCCCTACACGACGCTCTTCCGATCTNNNAAAAACTCCAAAGAAGAGCCA

GAGCACACGTCTGAACTCTTTCCCTACACGACGCTCTTCCGATCTNNNAAAAACTCCAAAGAAGAGCCA

GAGCACACGTCTGAACTCTTTCCCTACACGACGCTCTTCCGATCTNNNGGACAGGAGTTCTTGGGATT

GAGCACACGTCTGAACTCTTTCCCTACACGACGCTCTTCCGATCTNNNGGACAGGAGTTCTTGGGATT

AGCACACGTCTGAACTCTTTCCCTACACGACGCTCTTCCGATCTNNNACCTATGCATCCATTTTTCTTTA

AGCACACGTCTGAACTCTTTCCCTACACGACGCTCTTCCGATCTNNNACCTATGCATCCATTTTTCTTTA

GAGCACACGTCTGAACTCTTTCCCTACACGACGCTCTTCCGATCTNNNATACATGAGAGTGCAGGTAC

GAGCACACGTCTGAACTCTTTCCCTACACGACGCTCTTCCGATCTNNNATACATGAGAGTGCAGGTAC

AGCACACGTCTGAACTCTTTCCCTACACGACGCTCTTCCGATCTNNNACTGTGCTTTACAACCTCTCCAA

AGCACACGTCTGAACTCTTTCCCTACACGACGCTCTTCCGATCTNNNACTGTGCTTTACAACCTCTCCAA

AGCACACGTCTGAACTCTTTCCCTACACGACGCTCTTCCGATCTNNNGACCCTTGCATTCAACCTCTTTCC

AGCACACGTCTGAACTCTTTCCCTACACGACGCTCTTCCGATCTNNNGACCCTTGCATTCAACCTCTTTCC

AGCACACGTCTGAACTCTTTCCCTACACGACGCTCTTCCGATCTNNNTGTGGCACATATACACCATGGA

AGCACACGTCTGAACTCTTTCCCTACACGACGCTCTTCCGATCTNNNTGTGGCACATATACACCATGGA

GAGCACACGTCTGAACTCTTTCCCTACACGACGCTCTTCCGATCTNNNATCCAATTACCTCCCACAAGA

GAGCACACGTCTGAACTCTTTCCCTACACGACGCTCTTCCGATCTNNNATCCAATTACCTCCCACAAGA

AGAGCACACGTCTGAACTCTTTCCCTACACGACGCTCTTCCGATCTNNNGGTTTGTGGGAAAGGGAGA

AGAGCACACGTCTGAACTCTTTCCCTACACGACGCTCTTCCGATCTNNNGGTTTGTGGGAAAGGGAGA

AGCACACGTCTGAACTCTTTCCCTACACGACGCTCTTCCGATCTNNNCAGAGAGGTTACAGTGAGCAG

AGCACACGTCTGAACTCTTTCCCTACACGACGCTCTTCCGATCTNNNCAGAGAGGTTACAGTGAGCAG

GCACACGTCTGAACTCTTTCCCTACACGACGCTCTTCCGATCTNNNTTAGGTCTCAGCTCAAGTGTCAC

GCACACGTCTGAACTCTTTCCCTACACGACGCTCTTCCGATCTNNNTTAGGTCTCAGCTCAAGTGTCAC

AGCACACGTCTGAACTCTTTCCCTACACGACGCTCTTCCGATCTNNNTGCTCCTGGATAGCATAGCATT

AGCACACGTCTGAACTCTTTCCCTACACGACGCTCTTCCGATCTNNNTGCTCCTGGATAGCATAGCATT

AGAGCACACGTCTGAACTCTTTCCCTACACGACGCTCTTCCGATCTNNNACAAAAATTAGCCGGATGT

AGAGCACACGTCTGAACTCTTTCCCTACACGACGCTCTTCCGATCTNNNACAAAAATTAGCCGGATGT

AGAGCACACGTCTGAACTCTTTCCCTACACGACGCTCTTCCGATCTNNNTTCCCATTCTCCCATCCCTATC

AGAGCACACGTCTGAACTCTTTCCCTACACGACGCTCTTCCGATCTNNNTTCCCATTCTCCCATCCCTATC

GAGCACACGTCTGAACTCTTTCCCTACACGACGCTCTTCCGATCTNNNGGGTAACAGCTTCACACCCTA

GAGCACACGTCTGAACTCTTTCCCTACACGACGCTCTTCCGATCTNNNGGGTAACAGCTTCACACCCTA

AGCACACGTCTGAACTCTTTCCCTACACGACGCTCTTCCGATCTNNNCTGAGGTGGGAGGATCACTTG

AGCACACGTCTGAACTCTTTCCCTACACGACGCTCTTCCGATCTNNNCTGAGGTGGGAGGATCACTTG

AGCACACGTCTGAACTCTTTCCCTACACGACGCTCTTCCGATCTNNNTGCCCAGCTAATGTTTGTGTTT

AGCACACGTCTGAACTCTTTCCCTACACGACGCTCTTCCGATCTNNNTGCCCAGCTAATGTTTGTGTTT

GAGCACACGTCTGAACTCTTTCCCTACACGACGCTCTTCCGATCTNNNCACACCTGTAGTCTCAGCTACT

GAGCACACGTCTGAACTCTTTCCCTACACGACGCTCTTCCGATCTNNNCACACCTGTAGTCTCAGCTACT

GAGCACACGTCTGAACTCTTTCCCTACACGACGCTCTTCCGATCTNNNTGGTGACAGATCAAGACTTCC

GAGCACACGTCTGAACTCTTTCCCTACACGACGCTCTTCCGATCTNNNTGGTGACAGATCAAGACTTCC

AGCACACGTCTGAACTCTTTCCCTACACGACGCTCTTCCGATCTNNNGGTACAGAGCCAACCTGTAGAA

AGCACACGTCTGAACTCTTTCCCTACACGACGCTCTTCCGATCTNNNGGTACAGAGCCAACCTGTAGAA

GAGCACACGTCTGAACTCTTTCCCTACACGACGCTCTTCCGATCTNNNCCTTAGCCTCATCACCTTCATCT

GAGCACACGTCTGAACTCTTTCCCTACACGACGCTCTTCCGATCTNNNCCTTAGCCTCATCACCTTCATCT
AGCACACGTCTGAACTCTTTCCCTACACGACGCTCTTCCGATCTNNNTGCCACCAAATAATCCAGGAGTT
AGCACACGTCTGAACTCTTTCCCTACACGACGCTCTTCCGATCTNNNTGCCACCAAATAATCCAGGAGTT
AGAGCACACGTCTGAACTCTTTCCCTACACGACGCTCTTCCGATCTNNNCTATTCCACAAAGCCACAACTC
AGAGCACACGTCTGAACTCTTTCCCTACACGACGCTCTTCCGATCTNNNCTATTCCACAAAGCCACAACTC
CACACGTCTGAACTCTTTCCCTACACGACGCTCTTCCGATCTNNNTGTCTTTTGTAGCAGAATTCAACTCA
CACACGTCTGAACTCTTTCCCTACACGACGCTCTTCCGATCTNNNTGTCTTTTGTAGCAGAATTCAACTCA
GAGCACACGTCTGAACTCTTTCCCTACACGACGCTCTTCCGATCTNNNTGATCATAGCTCACTTCAGCCT
GAGCACACGTCTGAACTCTTTCCCTACACGACGCTCTTCCGATCTNNNTGATCATAGCTCACTTCAGCCT
AGAGCACACGTCTGAACTCTTTCCCTACACGACGCTCTTCCGATCTNNNACAGAATGAGGCCTTGTCTG
AGAGCACACGTCTGAACTCTTTCCCTACACGACGCTCTTCCGATCTNNNACAGAATGAGGCCTTGTCTG
AGAGCACACGTCTGAACTCTTTCCCTACACGACGCTCTTCCGATCTNNNCTCATGAGTCCTTTTCCCTGA
AGAGCACACGTCTGAACTCTTTCCCTACACGACGCTCTTCCGATCTNNNCTCATGAGTCCTTTTCCCTGA
AGCACACGTCTGAACTCTTTCCCTACACGACGCTCTTCCGATCTNNNATGTGTGTGTGTGTCTCTCTCTC
AGCACACGTCTGAACTCTTTCCCTACACGACGCTCTTCCGATCTNNNATGTGTGTGTGTGTCTCTCTCTC
GAGCACACGTCTGAACTCTTTCCCTACACGACGCTCTTCCGATCTNNNGTGTGTGTGTGAGAGAGAGA
GAGCACACGTCTGAACTCTTTCCCTACACGACGCTCTTCCGATCTNNNGTGTGTGTGTGAGAGAGAGA
GAGCACACGTCTGAACTCTTTCCCTACACGACGCTCTTCCGATCTNNNACGTGAGATCAGGAGTTCAAG
GAGCACACGTCTGAACTCTTTCCCTACACGACGCTCTTCCGATCTNNNACGTGAGATCAGGAGTTCAAG
GAGCACACGTCTGAACTCTTTCCCTACACGACGCTCTTCCGATCTNNNGTTTCTGAACTACAGGCTGCAA
GAGCACACGTCTGAACTCTTTCCCTACACGACGCTCTTCCGATCTNNNGTTTCTGAACTACAGGCTGCAA
GAGCACACGTCTGAACTCTTTCCCTACACGACGCTCTTCCGATCTNNNCAAGGCAGAAGGATCATTTGA
GAGCACACGTCTGAACTCTTTCCCTACACGACGCTCTTCCGATCTNNNCAAGGCAGAAGGATCATTTGA
AGCACACGTCTGAACTCTTTCCCTACACGACGCTCTTCCGATCTNNNTCCTCTTCATGCTGACATCTG
AGCACACGTCTGAACTCTTTCCCTACACGACGCTCTTCCGATCTNNNTCCTCTTCATGCTGACATCTG
AGCACACGTCTGAACTCTTTCCCTACACGACGCTCTTCCGATCTNNNACTGGGATGTTTTCCCCTCTGTA
AGCACACGTCTGAACTCTTTCCCTACACGACGCTCTTCCGATCTNNNACTGGGATGTTTTCCCCTCTGTA
GAGCACACGTCTGAACTCTTTCCCTACACGACGCTCTTCCGATCTNNNTGTCACTTCTGAAGCTCCGTTTA
GAGCACACGTCTGAACTCTTTCCCTACACGACGCTCTTCCGATCTNNNTGTCACTTCTGAAGCTCCGTTTA
GAGCACACGTCTGAACTCTTTCCCTACACGACGCTCTTCCGATCTNNNTCATGAGGCTGAAGCAAGAG
GAGCACACGTCTGAACTCTTTCCCTACACGACGCTCTTCCGATCTNNNTCATGAGGCTGAAGCAAGAG
GAGCACACGTCTGAACTCTTTCCCTACACGACGCTCTTCCGATCTNNNCTCAGAGTTAGAGGTCTGTGC
GAGCACACGTCTGAACTCTTTCCCTACACGACGCTCTTCCGATCTNNNCTCAGAGTTAGAGGTCTGTGC
AGCACACGTCTGAACTCTTTCCCTACACGACGCTCTTCCGATCTNNNTCTTCTTCAGTTCCTCTGGCTTCC
AGCACACGTCTGAACTCTTTCCCTACACGACGCTCTTCCGATCTNNNTCTTCTTCAGTTCCTCTGGCTTCC
AGCACACGTCTGAACTCTTTCCCTACACGACGCTCTTCCGATCTNNNGTTGCAAAAGATGATGGTGGT
AGCACACGTCTGAACTCTTTCCCTACACGACGCTCTTCCGATCTNNNGTTGCAAAAGATGATGGTGGT
AGCACACGTCTGAACTCTTTCCCTACACGACGCTCTTCCGATCTNNNGCATGGAGGATAAGGCACTCA
AGCACACGTCTGAACTCTTTCCCTACACGACGCTCTTCCGATCTNNNGCATGGAGGATAAGGCACTCA
AGAGCACACGTCTGAACTCTTTCCCTACACGACGCTCTTCCGATCTNNNAGACGAGAGAGAAGTACCG
AGAGCACACGTCTGAACTCTTTCCCTACACGACGCTCTTCCGATCTNNNAGACGAGAGAGAAGTACCG
AGCACACGTCTGAACTCTTTCCCTACACGACGCTCTTCCGATCTNNNCTGAGTCAGGAGAATCGCTTG
AGCACACGTCTGAACTCTTTCCCTACACGACGCTCTTCCGATCTNNNCTGAGTCAGGAGAATCGCTTG

GAGCACACGTCTGAACTCTTTCCCTACACGACGCTCTTCCGATCTNNNAGACACGGGAATGTATTTACC
GAGCACACGTCTGAACTCTTTCCCTACACGACGCTCTTCCGATCTNNNAGACACGGGAATGTATTTACC
AGCACACGTCTGAACTCTTTCCCTACACGACGCTCTTCCGATCTNNNAGTGCTGAGACTCGAATTCTGA
AGCACACGTCTGAACTCTTTCCCTACACGACGCTCTTCCGATCTNNNAGTGCTGAGACTCGAATTCTGA
GAGCACACGTCTGAACTCTTTCCCTACACGACGCTCTTCCGATCTNNNGATCCGTGCTTTGAACTCCTTC
GAGCACACGTCTGAACTCTTTCCCTACACGACGCTCTTCCGATCTNNNGATCCGTGCTTTGAACTCCTTC
GCACACGTCTGAACTCTTTCCCTACACGACGCTCTTCCGATCTNNNGCTTTTTCTTGCATCACATTGAG
GCACACGTCTGAACTCTTTCCCTACACGACGCTCTTCCGATCTNNNGCTTTTTCTTGCATCACATTGAG
AGCACACGTCTGAACTCTTTCCCTACACGACGCTCTTCCGATCTNNNGATCCACAGTGCACGAAGAAAT
AGCACACGTCTGAACTCTTTCCCTACACGACGCTCTTCCGATCTNNNGATCCACAGTGCACGAAGAAAT
AGCACACGTCTGAACTCTTTCCCTACACGACGCTCTTCCGATCTNNNGCATCTCCTCTGCACCTCTATTA
AGCACACGTCTGAACTCTTTCCCTACACGACGCTCTTCCGATCTNNNGCATCTCCTCTGCACCTCTATTA
AGCACACGTCTGAACTCTTTCCCTACACGACGCTCTTCCGATCTNNNTGACAGATCTCAGTGTGTGTTG
AGCACACGTCTGAACTCTTTCCCTACACGACGCTCTTCCGATCTNNNTGACAGATCTCAGTGTGTGTTG
AGCACACGTCTGAACTCTTTCCCTACACGACGCTCTTCCGATCTNNNGAGAACTCGCTCATTACTGCAA
AGCACACGTCTGAACTCTTTCCCTACACGACGCTCTTCCGATCTNNNGAGAACTCGCTCATTACTGCAA
AGCACACGTCTGAACTCTTTCCCTACACGACGCTCTTCCGATCTNNNAGTGTAGAAGTCTCTCTCCACC
AGCACACGTCTGAACTCTTTCCCTACACGACGCTCTTCCGATCTNNNAGTGTAGAAGTCTCTCTCCACC
GAGCACACGTCTGAACTCTTTCCCTACACGACGCTCTTCCGATCTNNNGTATACACCTAGCAGAAGGG
GAGCACACGTCTGAACTCTTTCCCTACACGACGCTCTTCCGATCTNNNGTATACACCTAGCAGAAGGG
GAGCACACGTCTGAACTCTTTCCCTACACGACGCTCTTCCGATCTNNNGATACATGCCAGAGAACTGG
GAGCACACGTCTGAACTCTTTCCCTACACGACGCTCTTCCGATCTNNNGATACATGCCAGAGAACTGG
GAGCACACGTCTGAACTCTTTCCCTACACGACGCTCTTCCGATCTNNNTTGGCCACAAACAACTAGTGG
GAGCACACGTCTGAACTCTTTCCCTACACGACGCTCTTCCGATCTNNNTTGGCCACAAACAACTAGTGG
GCACACGTCTGAACTCTTTCCCTACACGACGCTCTTCCGATCTNNNAGATATTCCTCATATGTTACCAT
GCACACGTCTGAACTCTTTCCCTACACGACGCTCTTCCGATCTNNNAGATATTCCTCATATGTTACCAT
GCACACGTCTGAACTCTTTCCCTACACGACGCTCTTCCGATCTNNNTGGATCCCAAAGATTTTCTCCTG
GCACACGTCTGAACTCTTTCCCTACACGACGCTCTTCCGATCTNNNTGGATCCCAAAGATTTTCTCCTG
GAGCACACGTCTGAACTCTTTCCCTACACGACGCTCTTCCGATCTNNNCTGAGGCAGAGAATTGCTTG
GAGCACACGTCTGAACTCTTTCCCTACACGACGCTCTTCCGATCTNNNCTGAGGCAGAGAATTGCTTG
AGCACACGTCTGAACTCTTTCCCTACACGACGCTCTTCCGATCTNNNGATTGCAGTGAGCCAGGATCA
AGCACACGTCTGAACTCTTTCCCTACACGACGCTCTTCCGATCTNNNGATTGCAGTGAGCCAGGATCA
GAGCACACGTCTGAACTCTTTCCCTACACGACGCTCTTCCGATCTNNNCTCTCAGCTACTCGGGACTTT
GAGCACACGTCTGAACTCTTTCCCTACACGACGCTCTTCCGATCTNNNCTCTCAGCTACTCGGGACTTT
GAGCACACGTCTGAACTCTTTCCCTACACGACGCTCTTCCGATCTNNNCACATTTTCAGGCCAGTCATG
GAGCACACGTCTGAACTCTTTCCCTACACGACGCTCTTCCGATCTNNNCACATTTTCAGGCCAGTCATG
AGCACACGTCTGAACTCTTTCCCTACACGACGCTCTTCCGATCTNNNTGCACAAATCTTCTGTTTTCCCC
AGCACACGTCTGAACTCTTTCCCTACACGACGCTCTTCCGATCTNNNTGCACAAATCTTCTGTTTTCCCC
AGCACACGTCTGAACTCTTTCCCTACACGACGCTCTTCCGATCTNNNTTGGGGGAACAGAAGGAATA
AGCACACGTCTGAACTCTTTCCCTACACGACGCTCTTCCGATCTNNNTTGGGGGAACAGAAGGAATA
GAGCACACGTCTGAACTCTTTCCCTACACGACGCTCTTCCGATCTNNNAGGGAGGCAAATGTCACTAA
GAGCACACGTCTGAACTCTTTCCCTACACGACGCTCTTCCGATCTNNNAGGGAGGCAAATGTCACTAA
AGCACACGTCTGAACTCTTTCCCTACACGACGCTCTTCCGATCTNNNTTCTACTGGAGCCTGACACTAG

AGCACACGTCTGAACTCTTTCCCTACACGACGCTCTTCCGATCTNNNTTCTACTGGAGCCTGACACTAG
AGCACACGTCTGAACTCTTTCCCTACACGACGCTCTTCCGATCTNNNGGCTTGCTCTCTGTGTCTCTTAG
AGCACACGTCTGAACTCTTTCCCTACACGACGCTCTTCCGATCTNNNGGCTTGCTCTCTGTGTCTCTTAG
AGCACACGTCTGAACTCTTTCCCTACACGACGCTCTTCCGATCTNNNCCTCTCTGGTCTGGCCTCGCCATC
AGCACACGTCTGAACTCTTTCCCTACACGACGCTCTTCCGATCTNNNCCTCTCTGGTCTGGCCTCGCCATC
GAGCACACGTCTGAACTCTTTCCCTACACGACGCTCTTCCGATCTNNNAGAATCACTTGAACCTGGGAG
GAGCACACGTCTGAACTCTTTCCCTACACGACGCTCTTCCGATCTNNNAGAATCACTTGAACCTGGGAG
CACACGTCTGAACTCTTTCCCTACACGACGCTCTTCCGATCTNNNCTCAAGTAGCTGGGAGTACAGC
CACACGTCTGAACTCTTTCCCTACACGACGCTCTTCCGATCTNNNCTCAAGTAGCTGGGAGTACAGC
GAGCACACGTCTGAACTCTTTCCCTACACGACGCTCTTCCGATCTNNNGCCTCAAGGTATTATCCCCAGA
GAGCACACGTCTGAACTCTTTCCCTACACGACGCTCTTCCGATCTNNNGCCTCAAGGTATTATCCCCAGA
AGCACACGTCTGAACTCTTTCCCTACACGACGCTCTTCCGATCTNNNGTGTAGACTTGGGAGGATGAC
AGCACACGTCTGAACTCTTTCCCTACACGACGCTCTTCCGATCTNNNGTGTAGACTTGGGAGGATGAC
AGAGCACACGTCTGAACTCTTTCCCTACACGACGCTCTTCCGATCTNNNAACAAAGCTCAAGAATACCT
AGAGCACACGTCTGAACTCTTTCCCTACACGACGCTCTTCCGATCTNNNAACAAAGCTCAAGAATACCT
GAGCACACGTCTGAACTCTTTCCCTACACGACGCTCTTCCGATCTNNNTCTTTCCTGCTCTCTTCTCCTCC
GAGCACACGTCTGAACTCTTTCCCTACACGACGCTCTTCCGATCTNNNTCTTTCCTGCTCTCTTCTCCTCC
AGCACACGTCTGAACTCTTTCCCTACACGACGCTCTTCCGATCTNNNGAGAATGGTGTGAACCTGGGA
AGCACACGTCTGAACTCTTTCCCTACACGACGCTCTTCCGATCTNNNGAGAATGGTGTGAACCTGGGA
GCACACGTCTGAACTCTTTCCCTACACGACGCTCTTCCGATCTNNNTTCAGGGACTCTTCAGGTAAGGA
GCACACGTCTGAACTCTTTCCCTACACGACGCTCTTCCGATCTNNNTTCAGGGACTCTTCAGGTAAGGA
GAGCACACGTCTGAACTCTTTCCCTACACGACGCTCTTCCGATCTNNNCTGTAGTCCCAGCTACTCAGGA
GAGCACACGTCTGAACTCTTTCCCTACACGACGCTCTTCCGATCTNNNCTGTAGTCCCAGCTACTCAGGA
GAGCACACGTCTGAACTCTTTCCCTACACGACGCTCTTCCGATCTNNNATATCTGGCAATCACTGGAGCA
GAGCACACGTCTGAACTCTTTCCCTACACGACGCTCTTCCGATCTNNNATATCTGGCAATCACTGGAGCA
GCACACGTCTGAACTCTTTCCCTACACGACGCTCTTCCGATCTNNNGAGAATCACTTGAACCCAGGAGG
GCACACGTCTGAACTCTTTCCCTACACGACGCTCTTCCGATCTNNNGAGAATCACTTGAACCCAGGAGG
GCACACGTCTGAACTCTTTCCCTACACGACGCTCTTCCGATCTNNNACATTTGCAGGGAATATCTGGGT
GCACACGTCTGAACTCTTTCCCTACACGACGCTCTTCCGATCTNNNACATTTGCAGGGAATATCTGGGT
GAGCACACGTCTGAACTCTTTCCCTACACGACGCTCTTCCGATCTNNNCATTGCAGTGAGCTCAGATCAA
GAGCACACGTCTGAACTCTTTCCCTACACGACGCTCTTCCGATCTNNNCATTGCAGTGAGCTCAGATCAA
GAGCACACGTCTGAACTCTTTCCCTACACGACGCTCTTCCGATCTNNNCAGCCTCCGAAGTAGCTAGAA
GAGCACACGTCTGAACTCTTTCCCTACACGACGCTCTTCCGATCTNNNCAGCCTCCGAAGTAGCTAGAA
GAGCACACGTCTGAACTCTTTCCCTACACGACGCTCTTCCGATCTNNNTCTAGGATTTTTCCAAGTTGG
GAGCACACGTCTGAACTCTTTCCCTACACGACGCTCTTCCGATCTNNNTCTAGGATTTTTCCAAGTTGG
AGAGCACACGTCTGAACTCTTTCCCTACACGACGCTCTTCCGATCTNNNACACCTGTAGTCCTTGCTACT
AGAGCACACGTCTGAACTCTTTCCCTACACGACGCTCTTCCGATCTNNNACACCTGTAGTCCTTGCTACT
AGCACACGTCTGAACTCTTTCCCTACACGACGCTCTTCCGATCTNNNGGAGTGATGCAATTGCTGTCTT
AGCACACGTCTGAACTCTTTCCCTACACGACGCTCTTCCGATCTNNNGGAGTGATGCAATTGCTGTCTT
GAGCACACGTCTGAACTCTTTCCCTACACGACGCTCTTCCGATCTNNNCCGTTCCTCCCAACTCTGAATA
GAGCACACGTCTGAACTCTTTCCCTACACGACGCTCTTCCGATCTNNNCCGTTCCTCCCAACTCTGAATA
GAGCACACGTCTGAACTCTTTCCCTACACGACGCTCTTCCGATCTNNNCAAAGAGCAGACCTACCTTCCA
GAGCACACGTCTGAACTCTTTCCCTACACGACGCTCTTCCGATCTNNNCAAAGAGCAGACCTACCTTCCA

AGCACACGTCTGAACTCTTTCCCTACACGACGCTCTTCCGATCTNNNCCTATCTCTGGTATCCAAACGCA
AGCACACGTCTGAACTCTTTCCCTACACGACGCTCTTCCGATCTNNNCCTATCTCTGGTATCCAAACGCA
GAGCACACGTCTGAACTCTTTCCCTACACGACGCTCTTCCGATCTNNNCATATGATCCCACAGCTATGCC
GAGCACACGTCTGAACTCTTTCCCTACACGACGCTCTTCCGATCTNNNCATATGATCCCACAGCTATGCC
AGCACACGTCTGAACTCTTTCCCTACACGACGCTCTTCCGATCTNNNCAACTGAGATTGGGGAGAGAG
AGCACACGTCTGAACTCTTTCCCTACACGACGCTCTTCCGATCTNNNCAACTGAGATTGGGGAGAGAG
GAGCACACGTCTGAACTCTTTCCCTACACGACGCTCTTCCGATCTNNNGGACTACACTTCCCAGTAATG
GAGCACACGTCTGAACTCTTTCCCTACACGACGCTCTTCCGATCTNNNGGACTACACTTCCCAGTAATG
AGCACACGTCTGAACTCTTTCCCTACACGACGCTCTTCCGATCTNNNAGCTTCTGATTTAGCAGGTGTG
AGCACACGTCTGAACTCTTTCCCTACACGACGCTCTTCCGATCTNNNAGCTTCTGATTTAGCAGGTGTG
GAGCACACGTCTGAACTCTTTCCCTACACGACGCTCTTCCGATCTNNNTAGGATAAGCTCACACCACCA
GAGCACACGTCTGAACTCTTTCCCTACACGACGCTCTTCCGATCTNNNTAGGATAAGCTCACACCACCA
GAGCACACGTCTGAACTCTTTCCCTACACGACGCTCTTCCGATCTNNNAAAAGAAGAAATTCTTTCCAGT
GAGCACACGTCTGAACTCTTTCCCTACACGACGCTCTTCCGATCTNNNAAAAGAAGAAATTCTTTCCAGT
AGCACACGTCTGAACTCTTTCCCTACACGACGCTCTTCCGATCTNNNCCCACGATCCTTCTCGCCAACATC
AGCACACGTCTGAACTCTTTCCCTACACGACGCTCTTCCGATCTNNNCCCACGATCCTTCTCGCCAACATC
ACACGTCTGAACTCTTTCCCTACACGACGCTCTTCCGATCTNNNGAGTCTCACTCTGTCATCCAGGCATC
ACACGTCTGAACTCTTTCCCTACACGACGCTCTTCCGATCTNNNGAGTCTCACTCTGTCATCCAGGCATC
AGCACACGTCTGAACTCTTTCCCTACACGACGCTCTTCCGATCTNNNTGTAGTTTCAGCTACTTGGGAG
AGCACACGTCTGAACTCTTTCCCTACACGACGCTCTTCCGATCTNNNTGTAGTTTCAGCTACTTGGGAG
GAGCACACGTCTGAACTCTTTCCCTACACGACGCTCTTCCGATCTNNNCAAGTCAGTCTGGTACCAGTA
GAGCACACGTCTGAACTCTTTCCCTACACGACGCTCTTCCGATCTNNNCAAGTCAGTCTGGTACCAGTA
ACACGTCTGAACTCTTTCCCTACACGACGCTCTTCCGATCTNNNCTCTTTGTTTCTCATCAACTGCATCGG
ACACGTCTGAACTCTTTCCCTACACGACGCTCTTCCGATCTNNNCTCTTTGTTTCTCATCAACTGCATCGG
GCACACGTCTGAACTCTTTCCCTACACGACGCTCTTCCGATCTNNNATCCAAATAGCTGGGATCACAGC
GCACACGTCTGAACTCTTTCCCTACACGACGCTCTTCCGATCTNNNATCCAAATAGCTGGGATCACAGC
GAGCACACGTCTGAACTCTTTCCCTACACGACGCTCTTCCGATCTNNNAGGCCAGGAGAGAGGAAATC
GAGCACACGTCTGAACTCTTTCCCTACACGACGCTCTTCCGATCTNNNAGGCCAGGAGAGAGGAAATC
GAGCACACGTCTGAACTCTTTCCCTACACGACGCTCTTCCGATCTNNNAACTGTTTTCCAGAGTGGCTG
GAGCACACGTCTGAACTCTTTCCCTACACGACGCTCTTCCGATCTNNNAACTGTTTTCCAGAGTGGCTG
AGCACACGTCTGAACTCTTTCCCTACACGACGCTCTTCCGATCTNNNAGAGAGCTTTTTTCCCTTGAGGT
AGCACACGTCTGAACTCTTTCCCTACACGACGCTCTTCCGATCTNNNAGAGAGCTTTTTTCCCTTGAGGT
AGAGCACACGTCTGAACTCTTTCCCTACACGACGCTCTTCCGATCTNNNGGACCCCATAGCCTGAAACAT
AGAGCACACGTCTGAACTCTTTCCCTACACGACGCTCTTCCGATCTNNNGGACCCCATAGCCTGAAACAT
GAGCACACGTCTGAACTCTTTCCCTACACGACGCTCTTCCGATCTNNNAACTGTAGTCCCAGCTGCTTGC
GAGCACACGTCTGAACTCTTTCCCTACACGACGCTCTTCCGATCTNNNAACTGTAGTCCCAGCTGCTTGC
AGCACACGTCTGAACTCTTTCCCTACACGACGCTCTTCCGATCTNNNTGTAGTCACAGCTACTAGGGAC
AGCACACGTCTGAACTCTTTCCCTACACGACGCTCTTCCGATCTNNNTGTAGTCACAGCTACTAGGGAC
AGCACACGTCTGAACTCTTTCCCTACACGACGCTCTTCCGATCTNNNTAGGAGCAGTGGGTGTATAGC
AGCACACGTCTGAACTCTTTCCCTACACGACGCTCTTCCGATCTNNNTAGGAGCAGTGGGTGTATAGC
AGCACACGTCTGAACTCTTTCCCTACACGACGCTCTTCCGATCTNNNACTACATAGCCTCCTCGCTTTTCC
AGCACACGTCTGAACTCTTTCCCTACACGACGCTCTTCCGATCTNNNACTACATAGCCTCCTCGCTTTTCC
AGCACACGTCTGAACTCTTTCCCTACACGACGCTCTTCCGATCTNNNTTTACCAGATTGCTAATGCCCCT

AGCACACGTCTGAACTCTTTCCCTACACGACGCTCTTCCGATCTNNNTTTACCAGATTGCTAATGCCCCT

GAGCACACGTCTGAACTCTTTCCCTACACGACGCTCTTCCGATCTNNNGGGTGAGACCGTAATGAGAA

GAGCACACGTCTGAACTCTTTCCCTACACGACGCTCTTCCGATCTNNNGGGTGAGACCGTAATGAGAA

GAGCACACGTCTGAACTCTTTCCCTACACGACGCTCTTCCGATCTNNNATTGTCAAGAGAGCAGACACC

GAGCACACGTCTGAACTCTTTCCCTACACGACGCTCTTCCGATCTNNNATTGTCAAGAGAGCAGACACC

CACGTCTGAACTCTTTCCCTACACGACGCTCTTCCGATCTNNNTTTCTCTCTCCCTCCCTCTTTCTCATCGG

CACGTCTGAACTCTTTCCCTACACGACGCTCTTCCGATCTNNNTTTCTCTCTCCCTCCCTCTTTCTCATCGG

AGCACACGTCTGAACTCTTTCCCTACACGACGCTCTTCCGATCTNNNGCAAATCAGCCTCCAGAGATCTA

AGCACACGTCTGAACTCTTTCCCTACACGACGCTCTTCCGATCTNNNGCAAATCAGCCTCCAGAGATCTA

GAGCACACGTCTGAACTCTTTCCCTACACGACGCTCTTCCGATCTNNNGGTACCGAATATTGTACTAGCC

GAGCACACGTCTGAACTCTTTCCCTACACGACGCTCTTCCGATCTNNNGGTACCGAATATTGTACTAGCC

GAGCACACGTCTGAACTCTTTCCCTACACGACGCTCTTCCGATCTNNNGTGTCATCACAGTAGCAATGA

GAGCACACGTCTGAACTCTTTCCCTACACGACGCTCTTCCGATCTNNNGTGTCATCACAGTAGCAATGA

AGAGCACACGTCTGAACTCTTTCCCTACACGACGCTCTTCCGATCTNNNTGCAGACTTCTGACTCTGAAC

AGAGCACACGTCTGAACTCTTTCCCTACACGACGCTCTTCCGATCTNNNTGCAGACTTCTGACTCTGAAC

AGCACACGTCTGAACTCTTTCCCTACACGACGCTCTTCCGATCTNNNGGATGACCTGAAGCTAAGAAC

AGCACACGTCTGAACTCTTTCCCTACACGACGCTCTTCCGATCTNNNGGATGACCTGAAGCTAAGAAC

AGCACACGTCTGAACTCTTTCCCTACACGACGCTCTTCCGATCTNNNTCTGGGTTCTCTATTCTGTTCCA

AGCACACGTCTGAACTCTTTCCCTACACGACGCTCTTCCGATCTNNNTCTGGGTTCTCTATTCTGTTCCA

AGCACACGTCTGAACTCTTTCCCTACACGACGCTCTTCCGATCTNNNTTGCTATGTCTTGTGGCCTTACTC

AGCACACGTCTGAACTCTTTCCCTACACGACGCTCTTCCGATCTNNNTTGCTATGTCTTGTGGCCTTACTC

AGCACACGTCTGAACTCTTTCCCTACACGACGCTCTTCCGATCTNNNGAGACATCTAGGCAGGTTCTGG

AGCACACGTCTGAACTCTTTCCCTACACGACGCTCTTCCGATCTNNNGAGACATCTAGGCAGGTTCTGG

GCACACGTCTGAACTCTTTCCCTACACGACGCTCTTCCGATCTNNNTCTTGCCTAAAACCTCCACAGTTCA

GCACACGTCTGAACTCTTTCCCTACACGACGCTCTTCCGATCTNNNTCTTGCCTAAAACCTCCACAGTTCA

GAGCACACGTCTGAACTCTTTCCCTACACGACGCTCTTCCGATCTNNNAATTCAGCTAATGGCCAGGTA

GAGCACACGTCTGAACTCTTTCCCTACACGACGCTCTTCCGATCTNNNAATTCAGCTAATGGCCAGGTA

AGCACACGTCTGAACTCTTTCCCTACACGACGCTCTTCCGATCTNNNAATTTTGATTCTGTTACCTTGTTC

AGCACACGTCTGAACTCTTTCCCTACACGACGCTCTTCCGATCTNNNAATTTTGATTCTGTTACCTTGTTC

AGAGCACACGTCTGAACTCTTTCCCTACACGACGCTCTTCCGATCTNNNGGAGGTTTTGACATTCTGAG

AGAGCACACGTCTGAACTCTTTCCCTACACGACGCTCTTCCGATCTNNNGGAGGTTTTGACATTCTGAG

AGCACACGTCTGAACTCTTTCCCTACACGACGCTCTTCCGATCTNNNAGAAAAACGCTTCCAAATCCAAT

AGCACACGTCTGAACTCTTTCCCTACACGACGCTCTTCCGATCTNNNAGAAAAACGCTTCCAAATCCAAT

GCACACGTCTGAACTCTTTCCCTACACGACGCTCTTCCGATCTNNNTGCAGTCTCTCACTTCAGATCTAC

GCACACGTCTGAACTCTTTCCCTACACGACGCTCTTCCGATCTNNNTGCAGTCTCTCACTTCAGATCTAC

AGAGCACACGTCTGAACTCTTTCCCTACACGACGCTCTTCCGATCTNNNGACTGAGGCAACAAGACTG

AGAGCACACGTCTGAACTCTTTCCCTACACGACGCTCTTCCGATCTNNNGACTGAGGCAACAAGACTG

AGCACACGTCTGAACTCTTTCCCTACACGACGCTCTTCCGATCTNNNAGCAGCACTATTCACAATAGCC

AGCACACGTCTGAACTCTTTCCCTACACGACGCTCTTCCGATCTNNNAGCAGCACTATTCACAATAGCC

AGCACACGTCTGAACTCTTTCCCTACACGACGCTCTTCCGATCTNNNCTGGAGATTACAGTGGCTGAG

AGCACACGTCTGAACTCTTTCCCTACACGACGCTCTTCCGATCTNNNCTGGAGATTACAGTGGCTGAG

AGCACACGTCTGAACTCTTTCCCTACACGACGCTCTTCCGATCTNNNTGGAGGGAGAAAGTTGGTTTA

AGCACACGTCTGAACTCTTTCCCTACACGACGCTCTTCCGATCTNNNTGGAGGGAGAAAGTTGGTTTA

AGCACACGTCTGAACTCTTTCCCTACACGACGCTCTTCCGATCTNNNAACATCAGAAAATACCATTTCAT
AGCACACGTCTGAACTCTTTCCCTACACGACGCTCTTCCGATCTNNNAACATCAGAAAATACCATTTCAT
GCACACGTCTGAACTCTTTCCCTACACGACGCTCTTCCGATCTNNNCTCATGCCTGTAATCCCAGCTATC
GCACACGTCTGAACTCTTTCCCTACACGACGCTCTTCCGATCTNNNCTCATGCCTGTAATCCCAGCTATC
AGCACACGTCTGAACTCTTTCCCTACACGACGCTCTTCCGATCTNNNGCAACTAGAAGCAAAGGCAGTA
AGCACACGTCTGAACTCTTTCCCTACACGACGCTCTTCCGATCTNNNGCAACTAGAAGCAAAGGCAGTA
AGCACACGTCTGAACTCTTTCCCTACACGACGCTCTTCCGATCTNNNGTGCTGGTATTACAAGTGTGAC
AGCACACGTCTGAACTCTTTCCCTACACGACGCTCTTCCGATCTNNNGTGCTGGTATTACAAGTGTGAC
AGCACACGTCTGAACTCTTTCCCTACACGACGCTCTTCCGATCTNNNACAGACCAATAACAAGGAGAG
AGCACACGTCTGAACTCTTTCCCTACACGACGCTCTTCCGATCTNNNACAGACCAATAACAAGGAGAG
AGCACACGTCTGAACTCTTTCCCTACACGACGCTCTTCCGATCTNNNGTCTCAGAATGGAAAGGAGCAC
AGCACACGTCTGAACTCTTTCCCTACACGACGCTCTTCCGATCTNNNGTCTCAGAATGGAAAGGAGCAC
GAGCACACGTCTGAACTCTTTCCCTACACGACGCTCTTCCGATCTNNNTGGTGCAATGCTTCATCTGTT
GAGCACACGTCTGAACTCTTTCCCTACACGACGCTCTTCCGATCTNNNTGGTGCAATGCTTCATCTGTT
AGCACACGTCTGAACTCTTTCCCTACACGACGCTCTTCCGATCTNNNAATTGCTGGGATTATAGGTGCC
AGCACACGTCTGAACTCTTTCCCTACACGACGCTCTTCCGATCTNNNAATTGCTGGGATTATAGGTGCC
AGAGCACACGTCTGAACTCTTTCCCTACACGACGCTCTTCCGATCTNNNAGTTTCAATGAGACCCAAGC
AGAGCACACGTCTGAACTCTTTCCCTACACGACGCTCTTCCGATCTNNNAGTTTCAATGAGACCCAAGC
GCACACGTCTGAACTCTTTCCCTACACGACGCTCTTCCGATCTNNNCCAGGACCTAGTGGAATCATGTT
GCACACGTCTGAACTCTTTCCCTACACGACGCTCTTCCGATCTNNNCCAGGACCTAGTGGAATCATGTT
AGCACACGTCTGAACTCTTTCCCTACACGACGCTCTTCCGATCTNNNTGCGATAGTTGCTATGCTCAGA
AGCACACGTCTGAACTCTTTCCCTACACGACGCTCTTCCGATCTNNNTGCGATAGTTGCTATGCTCAGA
AGCACACGTCTGAACTCTTTCCCTACACGACGCTCTTCCGATCTNNNTTGTTAGAAAATGTTGGCCAGG
AGCACACGTCTGAACTCTTTCCCTACACGACGCTCTTCCGATCTNNNTTGTTAGAAAATGTTGGCCAGG
AGCACACGTCTGAACTCTTTCCCTACACGACGCTCTTCCGATCTNNNTGGAGAAAGAGAAGCCATAGC
AGCACACGTCTGAACTCTTTCCCTACACGACGCTCTTCCGATCTNNNTGGAGAAAGAGAAGCCATAGC
CACGTCTGAACTCTTTCCCTACACGACGCTCTTCCGATCTNNNTAGTTTGCTCTTTTTCCCAGGGTCATC
CACGTCTGAACTCTTTCCCTACACGACGCTCTTCCGATCTNNNTAGTTTGCTCTTTTTCCCAGGGTCATC
AGCACACGTCTGAACTCTTTCCCTACACGACGCTCTTCCGATCTNNNTATGGTGTTGACGGACCTGTA
AGCACACGTCTGAACTCTTTCCCTACACGACGCTCTTCCGATCTNNNTATGGTGTTGACGGACCTGTA
GCACACGTCTGAACTCTTTCCCTACACGACGCTCTTCCGATCTNNNTGGTTGTGTGTGAATGATGCAAT
GCACACGTCTGAACTCTTTCCCTACACGACGCTCTTCCGATCTNNNTGGTTGTGTGTGAATGATGCAAT
GAGCACACGTCTGAACTCTTTCCCTACACGACGCTCTTCCGATCTNNNTGAGAATAGCTTCAACCCAGC
GAGCACACGTCTGAACTCTTTCCCTACACGACGCTCTTCCGATCTNNNTGAGAATAGCTTCAACCCAGC
GCACACGTCTGAACTCTTTCCCTACACGACGCTCTTCCGATCTNNNGTGGACTTTAGAAAATGTTAAGT
GCACACGTCTGAACTCTTTCCCTACACGACGCTCTTCCGATCTNNNGTGGACTTTAGAAAATGTTAAGT
GAGCACACGTCTGAACTCTTTCCCTACACGACGCTCTTCCGATCTNNNTTCTCAAACAAATTTGGAGCG
GAGCACACGTCTGAACTCTTTCCCTACACGACGCTCTTCCGATCTNNNTTCTCAAACAAATTTGGAGCG
CACACGTCTGAACTCTTTCCCTACACGACGCTCTTCCGATCTNNNCTTCTTTATGATGTCGCATTATGCAT
CACACGTCTGAACTCTTTCCCTACACGACGCTCTTCCGATCTNNNCTTCTTTATGATGTCGCATTATGCAT
GAGCACACGTCTGAACTCTTTCCCTACACGACGCTCTTCCGATCTNNNTACCCTTTTGTCTGTCTTCCACC
GAGCACACGTCTGAACTCTTTCCCTACACGACGCTCTTCCGATCTNNNTACCCTTTTGTCTGTCTTCCACC
CACGTCTGAACTCTTTCCCTACACGACGCTCTTCCGATCTNNNAGTTATGTTAGGTATACCCGACATCG

CACGTCTGAACTCTTTCCCTACACGACGCTCTTCCGATCTNNNAGTTATGTTAGGTATACCCGACATCG
AGCACACGTCTGAACTCTTTCCCTACACGACGCTCTTCCGATCTNNNACAGCAAGTATTTTAAGTTTTG
AGCACACGTCTGAACTCTTTCCCTACACGACGCTCTTCCGATCTNNNACAGCAAGTATTTTAAGTTTTG
AGCACACGTCTGAACTCTTTCCCTACACGACGCTCTTCCGATCTNNNCTGCAGGACCAAAGAATGAGT
AGCACACGTCTGAACTCTTTCCCTACACGACGCTCTTCCGATCTNNNCTGCAGGACCAAAGAATGAGT
AGCACACGTCTGAACTCTTTCCCTACACGACGCTCTTCCGATCTNNNATGTTGCAGTGAGCCAAGATC
AGCACACGTCTGAACTCTTTCCCTACACGACGCTCTTCCGATCTNNNATGTTGCAGTGAGCCAAGATC
AAGAGCACACGTCTGAACTCTTTCCCTACACGACGCTCTTCCGATCTNNNAGACTCTGTCTGAGGGTG
AAGAGCACACGTCTGAACTCTTTCCCTACACGACGCTCTTCCGATCTNNNAGACTCTGTCTGAGGGTG
GAGCACACGTCTGAACTCTTTCCCTACACGACGCTCTTCCGATCTNNNTGCTTTACTTGTTGGACTGCA
GAGCACACGTCTGAACTCTTTCCCTACACGACGCTCTTCCGATCTNNNTGCTTTACTTGTTGGACTGCA
AGAGCACACGTCTGAACTCTTTCCCTACACGACGCTCTTCCGATCTNNNCGTCTCAACAGAAGAAAGG
AGAGCACACGTCTGAACTCTTTCCCTACACGACGCTCTTCCGATCTNNNCGTCTCAACAGAAGAAAGG
AGCACACGTCTGAACTCTTTCCCTACACGACGCTCTTCCGATCTNNNCTTTTGGGTTCAAGCAGTTCT
AGCACACGTCTGAACTCTTTCCCTACACGACGCTCTTCCGATCTNNNCTTTTGGGTTCAAGCAGTTCT
ACACGTCTGAACTCTTTCCCTACACGACGCTCTTCCGATCTNNNTACTTACTTCTCCCCCTCCTCTGCATC
ACACGTCTGAACTCTTTCCCTACACGACGCTCTTCCGATCTNNNTACTTACTTCTCCCCCTCCTCTGCATC
AGCACACGTCTGAACTCTTTCCCTACACGACGCTCTTCCGATCTNNNACCTGTAGTCCCAGCTACTCGC
AGCACACGTCTGAACTCTTTCCCTACACGACGCTCTTCCGATCTNNNACCTGTAGTCCCAGCTACTCGC
AGCACACGTCTGAACTCTTTCCCTACACGACGCTCTTCCGATCTNNNGTAGAGAAGGGGTTTCACCAT
AGCACACGTCTGAACTCTTTCCCTACACGACGCTCTTCCGATCTNNNGTAGAGAAGGGGTTTCACCAT
AGCACACGTCTGAACTCTTTCCCTACACGACGCTCTTCCGATCTNNNTTGAATCCGGGAAGTAGAGG
AGCACACGTCTGAACTCTTTCCCTACACGACGCTCTTCCGATCTNNNTTGAATCCGGGAAGTAGAGG
AGCACACGTCTGAACTCTTTCCCTACACGACGCTCTTCCGATCTNNNCCCATTACCATTATGTGATGCC
AGCACACGTCTGAACTCTTTCCCTACACGACGCTCTTCCGATCTNNNCCCATTACCATTATGTGATGCC
GCACACGTCTGAACTCTTTCCCTACACGACGCTCTTCCGATCTNNNGTAGGCTGAGGCAGGAGAATG
GCACACGTCTGAACTCTTTCCCTACACGACGCTCTTCCGATCTNNNGTAGGCTGAGGCAGGAGAATG
AGCACACGTCTGAACTCTTTCCCTACACGACGCTCTTCCGATCTNNNCTCTTTCTCAGTGAAGGCAATG
AGCACACGTCTGAACTCTTTCCCTACACGACGCTCTTCCGATCTNNNCTCTTTCTCAGTGAAGGCAATG
AGCACACGTCTGAACTCTTTCCCTACACGACGCTCTTCCGATCTNNNGGGGATGAGCAAGGAGATAA
AGCACACGTCTGAACTCTTTCCCTACACGACGCTCTTCCGATCTNNNGGGGATGAGCAAGGAGATAA
GCACACGTCTGAACTCTTTCCCTACACGACGCTCTTCCGATCTNNNTCCTCCGGGGGAAAAAAAAAAA
GCACACGTCTGAACTCTTTCCCTACACGACGCTCTTCCGATCTNNNTCCTCCGGGGGAAAAAAAAAAA
AGCACACGTCTGAACTCTTTCCCTACACGACGCTCTTCCGATCTNNNATGAACTTGAGCACCTCCATG
AGCACACGTCTGAACTCTTTCCCTACACGACGCTCTTCCGATCTNNNATGAACTTGAGCACCTCCATG
AGCACACGTCTGAACTCTTTCCCTACACGACGCTCTTCCGATCTNNNGTGCCTGTAATCCCAGCTACTT
AGCACACGTCTGAACTCTTTCCCTACACGACGCTCTTCCGATCTNNNGTGCCTGTAATCCCAGCTACTT
AGCACACGTCTGAACTCTTTCCCTACACGACGCTCTTCCGATCTNNNAGGTACTGTCAGAATTTGCAG
AGCACACGTCTGAACTCTTTCCCTACACGACGCTCTTCCGATCTNNNAGGTACTGTCAGAATTTGCAG
AGCACACGTCTGAACTCTTTCCCTACACGACGCTCTTCCGATCTNNNCCCAAAAAGGTTCTGACTGCAA
AGCACACGTCTGAACTCTTTCCCTACACGACGCTCTTCCGATCTNNNCCCAAAAAGGTTCTGACTGCAA
AGAGCACACGTCTGAACTCTTTCCCTACACGACGCTCTTCCGATCTNNNCTGAGGCAGGAGAATCACT
AGAGCACACGTCTGAACTCTTTCCCTACACGACGCTCTTCCGATCTNNNCTGAGGCAGGAGAATCACT

GCACACGTCTGAACTCTTTCCCTACACGACGCTCTTCCGATCTNNNAATCCCCATTGTTCTTATTCCGC
GCACACGTCTGAACTCTTTCCCTACACGACGCTCTTCCGATCTNNNAATCCCCATTGTTCTTATTCCGC
AGCACACGTCTGAACTCTTTCCCTACACGACGCTCTTCCGATCTNNNCTTGCAGTGAACCGAGATTGT
AGCACACGTCTGAACTCTTTCCCTACACGACGCTCTTCCGATCTNNNCTTGCAGTGAACCGAGATTGT
GAGCACACGTCTGAACTCTTTCCCTACACGACGCTCTTCCGATCTNNNGTGACTGAGCAAGACTCCAT
GAGCACACGTCTGAACTCTTTCCCTACACGACGCTCTTCCGATCTNNNGTGACTGAGCAAGACTCCAT
GCACACGTCTGAACTCTTTCCCTACACGACGCTCTTCCGATCTNNNCCAGGTTGCTGCAAATGTCATT
GCACACGTCTGAACTCTTTCCCTACACGACGCTCTTCCGATCTNNNCCAGGTTGCTGCAAATGTCATT
AGCACACGTCTGAACTCTTTCCCTACACGACGCTCTTCCGATCTNNNCTGAGGTAGGAAGATCGCTTG
AGCACACGTCTGAACTCTTTCCCTACACGACGCTCTTCCGATCTNNNCTGAGGTAGGAAGATCGCTTG
GCACACGTCTGAACTCTTTCCCTACACGACGCTCTTCCGATCTNNNTGAGGCACGAGAATCACTGAA
GCACACGTCTGAACTCTTTCCCTACACGACGCTCTTCCGATCTNNNTGAGGCACGAGAATCACTGAA
GAGCACACGTCTGAACTCTTTCCCTACACGACGCTCTTCCGATCTNNNAAAAATTAGCTGGTTGTGGT
GAGCACACGTCTGAACTCTTTCCCTACACGACGCTCTTCCGATCTNNNAAAAATTAGCTGGTTGTGGT
AGCACACGTCTGAACTCTTTCCCTACACGACGCTCTTCCGATCTNNNCGTTCTGGATGACATTAGGAG
AGCACACGTCTGAACTCTTTCCCTACACGACGCTCTTCCGATCTNNNCGTTCTGGATGACATTAGGAG
GAGCACACGTCTGAACTCTTTCCCTACACGACGCTCTTCCGATCTNNNACATTCAGGAGTCACTGACG
GAGCACACGTCTGAACTCTTTCCCTACACGACGCTCTTCCGATCTNNNACATTCAGGAGTCACTGACG
GCACACGTCTGAACTCTTTCCCTACACGACGCTCTTCCGATCTNNNCAAGTTCAAGCAATTCTCCTGCC
GCACACGTCTGAACTCTTTCCCTACACGACGCTCTTCCGATCTNNNCAAGTTCAAGCAATTCTCCTGCC
AGCACACGTCTGAACTCTTTCCCTACACGACGCTCTTCCGATCTNNNCTGATCCCAGTGAAGCCTTGAA
AGCACACGTCTGAACTCTTTCCCTACACGACGCTCTTCCGATCTNNNCTGATCCCAGTGAAGCCTTGAA
AGCACACGTCTGAACTCTTTCCCTACACGACGCTCTTCCGATCTNNNTGCATTTTATCCCACTAGTGCCA
AGCACACGTCTGAACTCTTTCCCTACACGACGCTCTTCCGATCTNNNTGCATTTTATCCCACTAGTGCCA
GCACACGTCTGAACTCTTTCCCTACACGACGCTCTTCCGATCTNNNGGAAGTATCCCCAGCTAGATA
GCACACGTCTGAACTCTTTCCCTACACGACGCTCTTCCGATCTNNNGGAAGTATCCCCAGCTAGATA
GCACACGTCTGAACTCTTTCCCTACACGACGCTCTTCCGATCTNNNGTTACTTGCTCTGCCTGAAATCC
GCACACGTCTGAACTCTTTCCCTACACGACGCTCTTCCGATCTNNNGTTACTTGCTCTGCCTGAAATCC
GCACACGTCTGAACTCTTTCCCTACACGACGCTCTTCCGATCTNNNCTGAGGCAGGAGAATCACTTGA
GCACACGTCTGAACTCTTTCCCTACACGACGCTCTTCCGATCTNNNCTGAGGCAGGAGAATCACTTGA
AGCACACGTCTGAACTCTTTCCCTACACGACGCTCTTCCGATCTNNNCGGTAAGCTCCGAGTAGAGT
AGCACACGTCTGAACTCTTTCCCTACACGACGCTCTTCCGATCTNNNCGGTAAGCTCCGAGTAGAGT
GCACACGTCTGAACTCTTTCCCTACACGACGCTCTTCCGATCTNNNGGGAGCCTGTGAAAATTTATG
GCACACGTCTGAACTCTTTCCCTACACGACGCTCTTCCGATCTNNNGGGAGCCTGTGAAAATTTATG
GCACACGTCTGAACTCTTTCCCTACACGACGCTCTTCCGATCTNNNTCTGTGTCTCTGTCTCTCTCCATC
GCACACGTCTGAACTCTTTCCCTACACGACGCTCTTCCGATCTNNNTCTGTGTCTCTGTCTCTCTCCATC
GCACACGTCTGAACTCTTTCCCTACACGACGCTCTTCCGATCTNNNGTGAAGGGGAGAAAGAAAGC
GCACACGTCTGAACTCTTTCCCTACACGACGCTCTTCCGATCTNNNGTGAAGGGGAGAAAGAAAGC
AGCACACGTCTGAACTCTTTCCCTACACGACGCTCTTCCGATCTNNNATCAGGCATGACATCCTAAGT
AGCACACGTCTGAACTCTTTCCCTACACGACGCTCTTCCGATCTNNNATCAGGCATGACATCCTAAGT
GCACACGTCTGAACTCTTTCCCTACACGACGCTCTTCCGATCTNNNAAAGATGCTCCCAAGTGTAGTC
GCACACGTCTGAACTCTTTCCCTACACGACGCTCTTCCGATCTNNNAAAGATGCTCCCAAGTGTAGTC
AGCACACGTCTGAACTCTTTCCCTACACGACGCTCTTCCGATCTNNNTCCATCTGCATTATGTGACCT

GAGCACACGTCTGAACTCTTTCCCTACACGACGCTCTTCCGATCTNNNTCCATCTGCATTATGTGACCTC
AGCACACGTCTGAACTCTTTCCCTACACGACGCTCTTCCGATCTNNNTAGGGACAAACTCAGACACAC
AGCACACGTCTGAACTCTTTCCCTACACGACGCTCTTCCGATCTNNNTAGGGACAAACTCAGACACAC
AGCACACGTCTGAACTCTTTCCCTACACGACGCTCTTCCGATCTNNNTGCAAGTAGTCACTCAGATCCA
AGCACACGTCTGAACTCTTTCCCTACACGACGCTCTTCCGATCTNNNTGCAAGTAGTCACTCAGATCCA
AGCACACGTCTGAACTCTTTCCCTACACGACGCTCTTCCGATCTNNNAGGCAGGAGAATCATTTGAA
AGCACACGTCTGAACTCTTTCCCTACACGACGCTCTTCCGATCTNNNAGGCAGGAGAATCATTTGAA
AGCACACGTCTGAACTCTTTCCCTACACGACGCTCTTCCGATCTNNNTGTTTAGGATACTGCACTGGG
AGCACACGTCTGAACTCTTTCCCTACACGACGCTCTTCCGATCTNNNTGTTTAGGATACTGCACTGGG
AGCACACGTCTGAACTCTTTCCCTACACGACGCTCTTCCGATCTNNNGAGCAAGCCCAGAACTTGAAA
AGCACACGTCTGAACTCTTTCCCTACACGACGCTCTTCCGATCTNNNGAGCAAGCCCAGAACTTGAAA
GAGCACACGTCTGAACTCTTTCCCTACACGACGCTCTTCCGATCTNNNGTAACTGAGCACTTCACAACC
GAGCACACGTCTGAACTCTTTCCCTACACGACGCTCTTCCGATCTNNNGTAACTGAGCACTTCACAACC
AGCACACGTCTGAACTCTTTCCCTACACGACGCTCTTCCGATCTNNNCTGTGGGATGATCTCAGCTCAT
AGCACACGTCTGAACTCTTTCCCTACACGACGCTCTTCCGATCTNNNCTGTGGGATGATCTCAGCTCAT
AGCACACGTCTGAACTCTTTCCCTACACGACGCTCTTCCGATCTNNNTGAGAAGTGTGGAGGAAACTT
AGCACACGTCTGAACTCTTTCCCTACACGACGCTCTTCCGATCTNNNTGAGAAGTGTGGAGGAAACTT
AGCACACGTCTGAACTCTTTCCCTACACGACGCTCTTCCGATCTNNNTGTGATCTTTGATGTGTCTATT
AGCACACGTCTGAACTCTTTCCCTACACGACGCTCTTCCGATCTNNNTGTGATCTTTGATGTGTCTATT
AGCACACGTCTGAACTCTTTCCCTACACGACGCTCTTCCGATCTNNNCGTACACTGAAACAGAACTGC
AGCACACGTCTGAACTCTTTCCCTACACGACGCTCTTCCGATCTNNNCGTACACTGAAACAGAACTGC
AGCACACGTCTGAACTCTTTCCCTACACGACGCTCTTCCGATCTNNNAAGAATCACTTGAACCTGGGAG
AGCACACGTCTGAACTCTTTCCCTACACGACGCTCTTCCGATCTNNNAAGAATCACTTGAACCTGGGAG
AGCACACGTCTGAACTCTTTCCCTACACGACGCTCTTCCGATCTNNNGAATCTGTCCTCTCCCCTTCTTG
AGCACACGTCTGAACTCTTTCCCTACACGACGCTCTTCCGATCTNNNGAATCTGTCCTCTCCCCTTCTTG
AGCACACGTCTGAACTCTTTCCCTACACGACGCTCTTCCGATCTNNNATGCTCAGTCTAGGCCAAATGT
AGCACACGTCTGAACTCTTTCCCTACACGACGCTCTTCCGATCTNNNATGCTCAGTCTAGGCCAAATGT
AGCACACGTCTGAACTCTTTCCCTACACGACGCTCTTCCGATCTNNNCGACAAGAACGAAACTCTGTCT
AGCACACGTCTGAACTCTTTCCCTACACGACGCTCTTCCGATCTNNNCGACAAGAACGAAACTCTGTCT
GCACACGTCTGAACTCTTTCCCTACACGACGCTCTTCCGATCTNNNATTAATCCTTCACAAAGTCTTTGC
GCACACGTCTGAACTCTTTCCCTACACGACGCTCTTCCGATCTNNNATTAATCCTTCACAAAGTCTTTGC
AGCACACGTCTGAACTCTTTCCCTACACGACGCTCTTCCGATCTNNNGAGGCTGAGGCAGGAGAATG
AGCACACGTCTGAACTCTTTCCCTACACGACGCTCTTCCGATCTNNNGAGGCTGAGGCAGGAGAATG
AGCACACGTCTGAACTCTTTCCCTACACGACGCTCTTCCGATCTNNNGCGGGAGTTGAACAATGAGAA
AGCACACGTCTGAACTCTTTCCCTACACGACGCTCTTCCGATCTNNNGCGGGAGTTGAACAATGAGAA
GCACACGTCTGAACTCTTTCCCTACACGACGCTCTTCCGATCTNNNACCAATTGACCATAGATGCATAG
GCACACGTCTGAACTCTTTCCCTACACGACGCTCTTCCGATCTNNNACCAATTGACCATAGATGCATAG
GAGCACACGTCTGAACTCTTTCCCTACACGACGCTCTTCCGATCTNNNGAGGCAAGAGAATCGCTTGA
GAGCACACGTCTGAACTCTTTCCCTACACGACGCTCTTCCGATCTNNNGAGGCAAGAGAATCGCTTGA
GAGCACACGTCTGAACTCTTTCCCTACACGACGCTCTTCCGATCTNNNAAAGTCCAAATCTCTTGCAAT
GAGCACACGTCTGAACTCTTTCCCTACACGACGCTCTTCCGATCTNNNAAAGTCCAAATCTCTTGCAAT
GCACACGTCTGAACTCTTTCCCTACACGACGCTCTTCCGATCTNNNCCTGCCATCATGTTCAGCTCATA
GCACACGTCTGAACTCTTTCCCTACACGACGCTCTTCCGATCTNNNCCTGCCATCATGTTCAGCTCATA

AGCACACGTCTGAACTCTTTCCCTACACGACGCTCTTCCGATCTNNNTGGAGGCTACAATTCAAGAGG

AGCACACGTCTGAACTCTTTCCCTACACGACGCTCTTCCGATCTNNNTGGAGGCTACAATTCAAGAGG

AGCACACGTCTGAACTCTTTCCCTACACGACGCTCTTCCGATCTNNNCCCTACAATTGCAGACTTACCCT

AGCACACGTCTGAACTCTTTCCCTACACGACGCTCTTCCGATCTNNNCCCTACAATTGCAGACTTACCCT

AGAGCACACGTCTGAACTCTTTCCCTACACGACGCTCTTCCGATCTNNNAAAAATTAGCTGGGTTGGTC

AGAGCACACGTCTGAACTCTTTCCCTACACGACGCTCTTCCGATCTNNNAAAAATTAGCTGGGTTGGTC

GAGCACACGTCTGAACTCTTTCCCTACACGACGCTCTTCCGATCTNNNTGCTGAAGAGAGAGAGGAT

GAGCACACGTCTGAACTCTTTCCCTACACGACGCTCTTCCGATCTNNNTGCTGAAGAGAGAGAGGAT

AGCACACGTCTGAACTCTTTCCCTACACGACGCTCTTCCGATCTNNNAGAAATGCTCGGAGTCAGACAC

AGCACACGTCTGAACTCTTTCCCTACACGACGCTCTTCCGATCTNNNAGAAATGCTCGGAGTCAGACAC

GAGCACACGTCTGAACTCTTTCCCTACACGACGCTCTTCCGATCTNNNCATTATTACCCCCTTCCCCCTTT

GAGCACACGTCTGAACTCTTTCCCTACACGACGCTCTTCCGATCTNNNCATTATTACCCCCTTCCCCCTTT

GCACACGTCTGAACTCTTTCCCTACACGACGCTCTTCCGATCTNNNTGAAATCTTTAACTGCATTTTCTC

GCACACGTCTGAACTCTTTCCCTACACGACGCTCTTCCGATCTNNNTGAAATCTTTAACTGCATTTTCTC

AGCACACGTCTGAACTCTTTCCCTACACGACGCTCTTCCGATCTNNNCGCAACTAACATCTCAGGCTTTT

AGCACACGTCTGAACTCTTTCCCTACACGACGCTCTTCCGATCTNNNCGCAACTAACATCTCAGGCTTTT

GAGCACACGTCTGAACTCTTTCCCTACACGACGCTCTTCCGATCTNNNCTGAGGCAGGAGAATCACTT

GAGCACACGTCTGAACTCTTTCCCTACACGACGCTCTTCCGATCTNNNCTGAGGCAGGAGAATCACTT

GAGCACACGTCTGAACTCTTTCCCTACACGACGCTCTTCCGATCTNNNTCTCTCTTCCCTTTCCCCTTCTG

GAGCACACGTCTGAACTCTTTCCCTACACGACGCTCTTCCGATCTNNNTCTCTCTTCCCTTTCCCCTTCTG

AGCACACGTCTGAACTCTTTCCCTACACGACGCTCTTCCGATCTNNNACCTGTAGTCCCAGCTACTAGG

AGCACACGTCTGAACTCTTTCCCTACACGACGCTCTTCCGATCTNNNACCTGTAGTCCCAGCTACTAGG

GAGCACACGTCTGAACTCTTTCCCTACACGACGCTCTTCCGATCTNNNTTTTCGTCTCTAAGGATGGGGT

GAGCACACGTCTGAACTCTTTCCCTACACGACGCTCTTCCGATCTNNNTTTTCGTCTCTAAGGATGGGGT

GAGCACACGTCTGAACTCTTTCCCTACACGACGCTCTTCCGATCTNNNGCTCTTTTTTGGTTTCCATTGG

GAGCACACGTCTGAACTCTTTCCCTACACGACGCTCTTCCGATCTNNNGCTCTTTTTTGGTTTCCATTGG

AGCACACGTCTGAACTCTTTCCCTACACGACGCTCTTCCGATCTNNNGACAGAGTAACACTCCGTCTCG

AGCACACGTCTGAACTCTTTCCCTACACGACGCTCTTCCGATCTNNNGACAGAGTAACACTCCGTCTCG

GAGCACACGTCTGAACTCTTTCCCTACACGACGCTCTTCCGATCTNNNAAGGTTATGCTGACGTAGAAC

GAGCACACGTCTGAACTCTTTCCCTACACGACGCTCTTCCGATCTNNNAAGGTTATGCTGACGTAGAAC

AGCACACGTCTGAACTCTTTCCCTACACGACGCTCTTCCGATCTNNNGCGCTGTCTTTGCTCTTTTTTTT

AGCACACGTCTGAACTCTTTCCCTACACGACGCTCTTCCGATCTNNNGCGCTGTCTTTGCTCTTTTTTTT

GCACACGTCTGAACTCTTTCCCTACACGACGCTCTTCCGATCTNNNCAAAATCAAGAAGCTGGAAGCAT

GCACACGTCTGAACTCTTTCCCTACACGACGCTCTTCCGATCTNNNCAAAATCAAGAAGCTGGAAGCAT

GCACACGTCTGAACTCTTTCCCTACACGACGCTCTTCCGATCTNNNCCTTCATCTCCATACCCTCGACACA

GCACACGTCTGAACTCTTTCCCTACACGACGCTCTTCCGATCTNNNCCTTCATCTCCATACCCTCGACACA

AGCACACGTCTGAACTCTTTCCCTACACGACGCTCTTCCGATCTNNNGTATTAACCTTATGTGTGACAT

AGCACACGTCTGAACTCTTTCCCTACACGACGCTCTTCCGATCTNNNGTATTAACCTTATGTGTGACAT

GCACACGTCTGAACTCTTTCCCTACACGACGCTCTTCCGATCTNNNGTGCTCTGTTAAATCTTTCCCTTG

GCACACGTCTGAACTCTTTCCCTACACGACGCTCTTCCGATCTNNNGTGCTCTGTTAAATCTTTCCCTTG

AGCACACGTCTGAACTCTTTCCCTACACGACGCTCTTCCGATCTNNNTGCAGCTAAACTTTATTGGGCA

AGCACACGTCTGAACTCTTTCCCTACACGACGCTCTTCCGATCTNNNTGCAGCTAAACTTTATTGGGCA

AGAGCACACGTCTGAACTCTTTCCCTACACGACGCTCTTCCGATCTNNNAGCCTGTAATCCCAGTACTTT

AGAGCACACGTCTGAACTCTTTCCCTACACGACGCTCTTCCGATCTNNNAGCCTGTAATCCCAGTACTTT
AGCACACGTCTGAACTCTTTCCCTACACGACGCTCTTCCGATCTNNNGTTGAGTGATAGTGATATGGG
AGCACACGTCTGAACTCTTTCCCTACACGACGCTCTTCCGATCTNNNGTTGAGTGATAGTGATATGGG
GAGCACACGTCTGAACTCTTTCCCTACACGACGCTCTTCCGATCTNNNCTGCCTTCTAGTTGCCGGATAT
GAGCACACGTCTGAACTCTTTCCCTACACGACGCTCTTCCGATCTNNNCTGCCTTCTAGTTGCCGGATAT
AGCACACGTCTGAACTCTTTCCCTACACGACGCTCTTCCGATCTNNNCTAATTCGGAAGGTGGCACATT
AGCACACGTCTGAACTCTTTCCCTACACGACGCTCTTCCGATCTNNNCTAATTCGGAAGGTGGCACATT
AGAGCACACGTCTGAACTCTTTCCCTACACGACGCTCTTCCGATCTNNNGCGACAGAGTGAGACTCCAT
AGAGCACACGTCTGAACTCTTTCCCTACACGACGCTCTTCCGATCTNNNGCGACAGAGTGAGACTCCAT
ACACGTCTGAACTCTTTCCCTACACGACGCTCTTCCGATCTNNNCAGAGTTGTTCAACCATCACCACCAT
ACACGTCTGAACTCTTTCCCTACACGACGCTCTTCCGATCTNNNCAGAGTTGTTCAACCATCACCACCAT
AGCACACGTCTGAACTCTTTCCCTACACGACGCTCTTCCGATCTNNNAAACGTAATGCTTTTTCCCCCAG
AGCACACGTCTGAACTCTTTCCCTACACGACGCTCTTCCGATCTNNNAAACGTAATGCTTTTTCCCCCAG
AGCACACGTCTGAACTCTTTCCCTACACGACGCTCTTCCGATCTNNNGGTGACAAAGTGAGACTCTGT
AGCACACGTCTGAACTCTTTCCCTACACGACGCTCTTCCGATCTNNNGGTGACAAAGTGAGACTCTGT
GCACACGTCTGAACTCTTTCCCTACACGACGCTCTTCCGATCTNNNCTGGGAGACAAAGCGAGAATCT
GCACACGTCTGAACTCTTTCCCTACACGACGCTCTTCCGATCTNNNCTGGGAGACAAAGCGAGAATCT
CACACGTCTGAACTCTTTCCCTACACGACGCTCTTCCGATCTNNNAAAATGTGGAAGTTACGTGGCTG
CACACGTCTGAACTCTTTCCCTACACGACGCTCTTCCGATCTNNNAAAATGTGGAAGTTACGTGGCTG
AGAGCACACGTCTGAACTCTTTCCCTACACGACGCTCTTCCGATCTNNNGTACCTGTAATCCCAGCTAC
AGAGCACACGTCTGAACTCTTTCCCTACACGACGCTCTTCCGATCTNNNGTACCTGTAATCCCAGCTAC
AGCACACGTCTGAACTCTTTCCCTACACGACGCTCTTCCGATCTNNNATGCTTACGTGACCATCTTCTTG
AGCACACGTCTGAACTCTTTCCCTACACGACGCTCTTCCGATCTNNNATGCTTACGTGACCATCTTCTTG
AGCACACGTCTGAACTCTTTCCCTACACGACGCTCTTCCGATCTNNNTGCAACTGACTTTCTAATGTTGA
AGCACACGTCTGAACTCTTTCCCTACACGACGCTCTTCCGATCTNNNTGCAACTGACTTTCTAATGTTGA
GAGCACACGTCTGAACTCTTTCCCTACACGACGCTCTTCCGATCTNNNATGCCAGCAGTGTCAATAGAG
GAGCACACGTCTGAACTCTTTCCCTACACGACGCTCTTCCGATCTNNNATGCCAGCAGTGTCAATAGAG
GCACACGTCTGAACTCTTTCCCTACACGACGCTCTTCCGATCTNNNGTGGAGAAGAGAATGAGTGCG
GCACACGTCTGAACTCTTTCCCTACACGACGCTCTTCCGATCTNNNGTGGAGAAGAGAATGAGTGCG
AGCACACGTCTGAACTCTTTCCCTACACGACGCTCTTCCGATCTNNNTCGTCCATAGTCACACTACCCAA
AGCACACGTCTGAACTCTTTCCCTACACGACGCTCTTCCGATCTNNNTCGTCCATAGTCACACTACCCAA
GCACACGTCTGAACTCTTTCCCTACACGACGCTCTTCCGATCTNNNTTGATCTTCTCCTGCTTGGCTTCCA
GCACACGTCTGAACTCTTTCCCTACACGACGCTCTTCCGATCTNNNTTGATCTTCTCCTGCTTGGCTTCCA
GAGCACACGTCTGAACTCTTTCCCTACACGACGCTCTTCCGATCTNNNTGAAACCCCGTCTCCACTAAAA
GAGCACACGTCTGAACTCTTTCCCTACACGACGCTCTTCCGATCTNNNTGAAACCCCGTCTCCACTAAAA
GAGCACACGTCTGAACTCTTTCCCTACACGACGCTCTTCCGATCTNNNGGACTCTACACACACACTTTCC
GAGCACACGTCTGAACTCTTTCCCTACACGACGCTCTTCCGATCTNNNGGACTCTACACACACACTTTCC
AGAGCACACGTCTGAACTCTTTCCCTACACGACGCTCTTCCGATCTNNNGGTAATCTAGAGGTGGCTC
AGAGCACACGTCTGAACTCTTTCCCTACACGACGCTCTTCCGATCTNNNGGTAATCTAGAGGTGGCTC
AGCACACGTCTGAACTCTTTCCCTACACGACGCTCTTCCGATCTNNNTACATTACCTGTTGCTCCAGTCC
AGCACACGTCTGAACTCTTTCCCTACACGACGCTCTTCCGATCTNNNTACATTACCTGTTGCTCCAGTCC
GAGCACACGTCTGAACTCTTTCCCTACACGACGCTCTTCCGATCTNNNTGAGGAGGGAGAATTGCTTG
GAGCACACGTCTGAACTCTTTCCCTACACGACGCTCTTCCGATCTNNNTGAGGAGGGAGAATTGCTTG

AGCACACGTCTGAACTCTTTCCCTACACGACGCTCTTCCGATCTNNNATGGGAAAGAAAGGCAATTT
AGCACACGTCTGAACTCTTTCCCTACACGACGCTCTTCCGATCTNNNATGGGAAAGAAAGGCAATTT
AGCACACGTCTGAACTCTTTCCCTACACGACGCTCTTCCGATCTNNNACAGCTACAGCAACAGGAACT
AGCACACGTCTGAACTCTTTCCCTACACGACGCTCTTCCGATCTNNNACAGCTACAGCAACAGGAACT
GAGCACACGTCTGAACTCTTTCCCTACACGACGCTCTTCCGATCTNNNCTAAGGCAGGAGAATCGCTT
GAGCACACGTCTGAACTCTTTCCCTACACGACGCTCTTCCGATCTNNNCTAAGGCAGGAGAATCGCTT
GAGCACACGTCTGAACTCTTTCCCTACACGACGCTCTTCCGATCTNNNGAAGTTGCAGTGAGCTGAGA
GAGCACACGTCTGAACTCTTTCCCTACACGACGCTCTTCCGATCTNNNGAAGTTGCAGTGAGCTGAGA
GAGCACACGTCTGAACTCTTTCCCTACACGACGCTCTTCCGATCTNNNCTGAGGCAGGAGAATCACTT
GAGCACACGTCTGAACTCTTTCCCTACACGACGCTCTTCCGATCTNNNCTGAGGCAGGAGAATCACTT
AGCACACGTCTGAACTCTTTCCCTACACGACGCTCTTCCGATCTNNNCTGAGGCAGGAGAATTGCTTG
AGCACACGTCTGAACTCTTTCCCTACACGACGCTCTTCCGATCTNNNCTGAGGCAGGAGAATTGCTTG
AGCACACGTCTGAACTCTTTCCCTACACGACGCTCTTCCGATCTNNNTCGTAGAGAAGAAGGAGGAG
AGCACACGTCTGAACTCTTTCCCTACACGACGCTCTTCCGATCTNNNTCGTAGAGAAGAAGGAGGAG
AGCACACGTCTGAACTCTTTCCCTACACGACGCTCTTCCGATCTNNNAGTGGCATATTCTCAGCTCTCT
AGCACACGTCTGAACTCTTTCCCTACACGACGCTCTTCCGATCTNNNAGTGGCATATTCTCAGCTCTCT
GAGCACACGTCTGAACTCTTTCCCTACACGACGCTCTTCCGATCTNNNCTGAGGCAAGAGAATCCCTT
GAGCACACGTCTGAACTCTTTCCCTACACGACGCTCTTCCGATCTNNNCTGAGGCAAGAGAATCCCTT
GAGCACACGTCTGAACTCTTTCCCTACACGACGCTCTTCCGATCTNNNAGCGGACAAAGTGTAATGTC
GAGCACACGTCTGAACTCTTTCCCTACACGACGCTCTTCCGATCTNNNAGCGGACAAAGTGTAATGTC
AGCACACGTCTGAACTCTTTCCCTACACGACGCTCTTCCGATCTNNNCTGAGGCAGGAGAATTGCTTG
AGCACACGTCTGAACTCTTTCCCTACACGACGCTCTTCCGATCTNNNCTGAGGCAGGAGAATTGCTTG
CACACGTCTGAACTCTTTCCCTACACGACGCTCTTCCGATCTNNNTAGCTGGGATTACAGGCACATGC
CACACGTCTGAACTCTTTCCCTACACGACGCTCTTCCGATCTNNNTAGCTGGGATTACAGGCACATGC
AGCACACGTCTGAACTCTTTCCCTACACGACGCTCTTCCGATCTNNNATTCCTGACCTCAGATGATCTG
AGCACACGTCTGAACTCTTTCCCTACACGACGCTCTTCCGATCTNNNATTCCTGACCTCAGATGATCTG
AAGAGCACACGTCTGAACTCTTTCCCTACACGACGCTCTTCCGATCTNNNCCCGGCTCCAAGCATCTTT
AAGAGCACACGTCTGAACTCTTTCCCTACACGACGCTCTTCCGATCTNNNCCCGGCTCCAAGCATCTTT
CACGTCTGAACTCTTTCCCTACACGACGCTCTTCCGATCTNNNAAATGCTAGTTGCTTGTTTGCATCGG
CACGTCTGAACTCTTTCCCTACACGACGCTCTTCCGATCTNNNAAATGCTAGTTGCTTGTTTGCATCGG
GAGCACACGTCTGAACTCTTTCCCTACACGACGCTCTTCCGATCTNNNTTATCTTTCCTGTGGCAAGG
GAGCACACGTCTGAACTCTTTCCCTACACGACGCTCTTCCGATCTNNNTTATCTTTCCTGTGGCAAGG
AGCACACGTCTGAACTCTTTCCCTACACGACGCTCTTCCGATCTNNNTAAAGGGAGTAGGGGAGACT
AGCACACGTCTGAACTCTTTCCCTACACGACGCTCTTCCGATCTNNNTAAAGGGAGTAGGGGAGACT
GCACACGTCTGAACTCTTTCCCTACACGACGCTCTTCCGATCTNNNAATTGGTATCCTCGCCAACTCTT
GCACACGTCTGAACTCTTTCCCTACACGACGCTCTTCCGATCTNNNAATTGGTATCCTCGCCAACTCTT
CACACGTCTGAACTCTTTCCCTACACGACGCTCTTCCGATCTNNNTCTCCCGAGTAGCTGAGATTACAC
CACACGTCTGAACTCTTTCCCTACACGACGCTCTTCCGATCTNNNTCTCCCGAGTAGCTGAGATTACAC
AGCACACGTCTGAACTCTTTCCCTACACGACGCTCTTCCGATCTNNNGCGACAGAGTGAGACTCCATC
AGCACACGTCTGAACTCTTTCCCTACACGACGCTCTTCCGATCTNNNGCGACAGAGTGAGACTCCATC
CACGTCTGAACTCTTTCCCTACACGACGCTCTTCCGATCTNNNCAAAGTGTTGGGATTACAGGCGCAT
CACGTCTGAACTCTTTCCCTACACGACGCTCTTCCGATCTNNNCAAAGTGTTGGGATTACAGGCGCAT
GCACACGTCTGAACTCTTTCCCTACACGACGCTCTTCCGATCTNNNTGGGATTTCCACACGGTATATCC

GCACACGTCTGAACTCTTTCCCTACACGACGCTCTTCCGATCTNNNTGGGATTTCCACACGGTATATCC
GAGCACACGTCTGAACTCTTTCCCTACACGACGCTCTTCCGATCTNNNTCACCATCAATATGTCCTGCA
GAGCACACGTCTGAACTCTTTCCCTACACGACGCTCTTCCGATCTNNNTCACCATCAATATGTCCTGCA
AGCACACGTCTGAACTCTTTCCCTACACGACGCTCTTCCGATCTNNNATTTGTGCCTTTTTCACCACCTT
AGCACACGTCTGAACTCTTTCCCTACACGACGCTCTTCCGATCTNNNATTTGTGCCTTTTTCACCACCTT
AGCACACGTCTGAACTCTTTCCCTACACGACGCTCTTCCGATCTNNNAGACAGGAGAATTGCTTGAAC
AGCACACGTCTGAACTCTTTCCCTACACGACGCTCTTCCGATCTNNNAGACAGGAGAATTGCTTGAAC
AGCACACGTCTGAACTCTTTCCCTACACGACGCTCTTCCGATCTNNNACAGTTCCTCAGCCTTTCTTTG
AGCACACGTCTGAACTCTTTCCCTACACGACGCTCTTCCGATCTNNNACAGTTCCTCAGCCTTTCTTTG
GCACACGTCTGAACTCTTTCCCTACACGACGCTCTTCCGATCTNNNAATCACTCCACCTCAGTAATGGC
GCACACGTCTGAACTCTTTCCCTACACGACGCTCTTCCGATCTNNNAATCACTCCACCTCAGTAATGGC
ACACGTCTGAACTCTTTCCCTACACGACGCTCTTCCGATCTNNNTCCCATGAAAATTCCTTGAAAATAA
ACACGTCTGAACTCTTTCCCTACACGACGCTCTTCCGATCTNNNTCCCATGAAAATTCCTTGAAAATAA
GAGCACACGTCTGAACTCTTTCCCTACACGACGCTCTTCCGATCTNNNAAATGGGTCAGCACCTCCAA
GAGCACACGTCTGAACTCTTTCCCTACACGACGCTCTTCCGATCTNNNAAATGGGTCAGCACCTCCAA
AGCACACGTCTGAACTCTTTCCCTACACGACGCTCTTCCGATCTNNNCTTTGTGTGTCTGTGTGTGT
AGCACACGTCTGAACTCTTTCCCTACACGACGCTCTTCCGATCTNNNCTTTGTGTGTCTGTGTGTGT
AGCACACGTCTGAACTCTTTCCCTACACGACGCTCTTCCGATCTNNNGGAATCACTTGAACCTGGGA
AGCACACGTCTGAACTCTTTCCCTACACGACGCTCTTCCGATCTNNNGGAATCACTTGAACCTGGGA
GCACACGTCTGAACTCTTTCCCTACACGACGCTCTTCCGATCTNNNCTGGAATCTTTACAGGAGAACC
GCACACGTCTGAACTCTTTCCCTACACGACGCTCTTCCGATCTNNNCTGGAATCTTTACAGGAGAACC
AGCACACGTCTGAACTCTTTCCCTACACGACGCTCTTCCGATCTNNNTATCTAACAGCTGCAATGAGC
AGCACACGTCTGAACTCTTTCCCTACACGACGCTCTTCCGATCTNNNTATCTAACAGCTGCAATGAGC
CACACGTCTGAACTCTTTCCCTACACGACGCTCTTCCGATCTNNNCAGGCATAGTCAAACAATAGATCA
CACACGTCTGAACTCTTTCCCTACACGACGCTCTTCCGATCTNNNCAGGCATAGTCAAACAATAGATCA
GCACACGTCTGAACTCTTTCCCTACACGACGCTCTTCCGATCTNNNCACCACACCTGGCTAATTTTCTT
GCACACGTCTGAACTCTTTCCCTACACGACGCTCTTCCGATCTNNNCACCACACCTGGCTAATTTTCTT
GCACACGTCTGAACTCTTTCCCTACACGACGCTCTTCCGATCTNNNTCTCTTTAGACAAGGGCCCTTTC
GCACACGTCTGAACTCTTTCCCTACACGACGCTCTTCCGATCTNNNTCTCTTTAGACAAGGGCCCTTTC
AGCACACGTCTGAACTCTTTCCCTACACGACGCTCTTCCGATCTNNNTAAATCGCCCCTTCAGAAGTC
AGCACACGTCTGAACTCTTTCCCTACACGACGCTCTTCCGATCTNNNTAAATCGCCCCTTCAGAAGTC
GAGCACACGTCTGAACTCTTTCCCTACACGACGCTCTTCCGATCTNNNAGAAAGGAAGCATCACATA
GAGCACACGTCTGAACTCTTTCCCTACACGACGCTCTTCCGATCTNNNAGAAAGGAAGCATCACATA
AGCACACGTCTGAACTCTTTCCCTACACGACGCTCTTCCGATCTNNNATCACCTTCAACACGGTTTTGA
AGCACACGTCTGAACTCTTTCCCTACACGACGCTCTTCCGATCTNNNATCACCTTCAACACGGTTTTGA
AGCACACGTCTGAACTCTTTCCCTACACGACGCTCTTCCGATCTNNNAAACTCCTGGGCTCAAGTGAT
AGCACACGTCTGAACTCTTTCCCTACACGACGCTCTTCCGATCTNNNAAACTCCTGGGCTCAAGTGAT
CACACGTCTGAACTCTTTCCCTACACGACGCTCTTCCGATCTNNNTTACCCTTACTCCAACACTGCAACA
CACACGTCTGAACTCTTTCCCTACACGACGCTCTTCCGATCTNNNTTACCCTTACTCCAACACTGCAACA
GCACACGTCTGAACTCTTTCCCTACACGACGCTCTTCCGATCTNNNAGATCCAGTCTTCCACCACTCATC
GCACACGTCTGAACTCTTTCCCTACACGACGCTCTTCCGATCTNNNAGATCCAGTCTTCCACCACTCATC
AGCACACGTCTGAACTCTTTCCCTACACGACGCTCTTCCGATCTNNNGCTACACCTGCTCTTTTTTGGT
AGCACACGTCTGAACTCTTTCCCTACACGACGCTCTTCCGATCTNNNGCTACACCTGCTCTTTTTTGGT

AGCACACGTCTGAACTCTTTCCCTACACGACGCTCTTCCGATCTNNNTTCAGGGACTATTCCAGAAGC
AGCACACGTCTGAACTCTTTCCCTACACGACGCTCTTCCGATCTNNNTTCAGGGACTATTCCAGAAGC
CACACGTCTGAACTCTTTCCCTACACGACGCTCTTCCGATCTNNNCTTCACACGGTCATCCCCAAAGCA
CACACGTCTGAACTCTTTCCCTACACGACGCTCTTCCGATCTNNNCTTCACACGGTCATCCCCAAAGCA
GCACACGTCTGAACTCTTTCCCTACACGACGCTCTTCCGATCTNNNTATACAACGACTCTTGAGCCCA
GCACACGTCTGAACTCTTTCCCTACACGACGCTCTTCCGATCTNNNTATACAACGACTCTTGAGCCCA
AGCACACGTCTGAACTCTTTCCCTACACGACGCTCTTCCGATCTNNNTGTCTGAAGGAAAGAGAACCC
AGCACACGTCTGAACTCTTTCCCTACACGACGCTCTTCCGATCTNNNTGTCTGAAGGAAAGAGAACCC
GCACACGTCTGAACTCTTTCCCTACACGACGCTCTTCCGATCTNNNCAACTAACCTCACTTTCTAGAAA
GCACACGTCTGAACTCTTTCCCTACACGACGCTCTTCCGATCTNNNCAACTAACCTCACTTTCTAGAAA
GCACACGTCTGAACTCTTTCCCTACACGACGCTCTTCCGATCTNNNTATGCAGTGAACCCAGTTGAGT
GCACACGTCTGAACTCTTTCCCTACACGACGCTCTTCCGATCTNNNTATGCAGTGAACCCAGTTGAGT
AGCACACGTCTGAACTCTTTCCCTACACGACGCTCTTCCGATCTNNNAGTGCCACATTTTTATGCAG
AGCACACGTCTGAACTCTTTCCCTACACGACGCTCTTCCGATCTNNNAGTGCCACATTTTTATGCAG
AGCACACGTCTGAACTCTTTCCCTACACGACGCTCTTCCGATCTNNNTAAGTGCTCCAGAAACCTACCC
AGCACACGTCTGAACTCTTTCCCTACACGACGCTCTTCCGATCTNNNTAAGTGCTCCAGAAACCTACCC
GCACACGTCTGAACTCTTTCCCTACACGACGCTCTTCCGATCTNNNATCATAGATGTTTTCACCTCTAC
GCACACGTCTGAACTCTTTCCCTACACGACGCTCTTCCGATCTNNNATCATAGATGTTTTCACCTCTAC
AGCACACGTCTGAACTCTTTCCCTACACGACGCTCTTCCGATCTNNNGTTTTCCAGGGTAGCCACACT
AGCACACGTCTGAACTCTTTCCCTACACGACGCTCTTCCGATCTNNNGTTTTCCAGGGTAGCCACACT
AGCACACGTCTGAACTCTTTCCCTACACGACGCTCTTCCGATCTNNNAATATGAAACCAGCCCAAATG
AGCACACGTCTGAACTCTTTCCCTACACGACGCTCTTCCGATCTNNNAATATGAAACCAGCCCAAATG
AGCACACGTCTGAACTCTTTCCCTACACGACGCTCTTCCGATCTNNNAAGCACAAATACTCCCCAAATG
AGCACACGTCTGAACTCTTTCCCTACACGACGCTCTTCCGATCTNNNAAGCACAAATACTCCCCAAATG
AGCACACGTCTGAACTCTTTCCCTACACGACGCTCTTCCGATCTNNNCACTCAGCTGATTTACCTAAG
AGCACACGTCTGAACTCTTTCCCTACACGACGCTCTTCCGATCTNNNCACTCAGCTGATTTACCTAAG
AGCACACGTCTGAACTCTTTCCCTACACGACGCTCTTCCGATCTNNNATTCAAGGACAGAGCTTCTCC
AGCACACGTCTGAACTCTTTCCCTACACGACGCTCTTCCGATCTNNNATTCAAGGACAGAGCTTCTCC
GCACACGTCTGAACTCTTTCCCTACACGACGCTCTTCCGATCTNNNTTTGCAGAAGTGAATCTGGGGT
GCACACGTCTGAACTCTTTCCCTACACGACGCTCTTCCGATCTNNNTTTGCAGAAGTGAATCTGGGGT
GCACACGTCTGAACTCTTTCCCTACACGACGCTCTTCCGATCTNNNGAACAGAATTACAGGAAGTTT
GCACACGTCTGAACTCTTTCCCTACACGACGCTCTTCCGATCTNNNGAACAGAATTACAGGAAGTTT
GCACACGTCTGAACTCTTTCCCTACACGACGCTCTTCCGATCTNNNCAGGAGAATCACTTGAACCAGG
GCACACGTCTGAACTCTTTCCCTACACGACGCTCTTCCGATCTNNNCAGGAGAATCACTTGAACCAGG
AGCACACGTCTGAACTCTTTCCCTACACGACGCTCTTCCGATCTNNNCTCCGAAAGTGCTGGGATTAC
AGCACACGTCTGAACTCTTTCCCTACACGACGCTCTTCCGATCTNNNCTCCGAAAGTGCTGGGATTAC
GAGCACACGTCTGAACTCTTTCCCTACACGACGCTCTTCCGATCTNNNCTTGAACCTGGGAGATGGA
GAGCACACGTCTGAACTCTTTCCCTACACGACGCTCTTCCGATCTNNNCTTGAACCTGGGAGATGGA
AGCACACGTCTGAACTCTTTCCCTACACGACGCTCTTCCGATCTNNNTTTAGGCCATCTGTCTGTTTCC
AGCACACGTCTGAACTCTTTCCCTACACGACGCTCTTCCGATCTNNNTTTAGGCCATCTGTCTGTTTCC
GAGCACACGTCTGAACTCTTTCCCTACACGACGCTCTTCCGATCTNNNCACCTGTAGTCCCAGCTACTT
GAGCACACGTCTGAACTCTTTCCCTACACGACGCTCTTCCGATCTNNNCACCTGTAGTCCCAGCTACTT
GCACACGTCTGAACTCTTTCCCTACACGACGCTCTTCCGATCTNNNAGAAGACGTTCATCAGCATCCT

GCACACGTCTGAACTCTTTCCCTACACGACGCTCTTCCGATCTNNNAGAAGACGTTCATCAGCATCCTT

AGCACACGTCTGAACTCTTTCCCTACACGACGCTCTTCCGATCTNNNTATAATGTGGCCAGCTAGGTG

AGCACACGTCTGAACTCTTTCCCTACACGACGCTCTTCCGATCTNNNTATAATGTGGCCAGCTAGGTG

AGCACACGTCTGAACTCTTTCCCTACACGACGCTCTTCCGATCTNNNTGCTCTGTTCTGTTCTCGTCTT

AGCACACGTCTGAACTCTTTCCCTACACGACGCTCTTCCGATCTNNNTGCTCTGTTCTGTTCTCGTCTT

GAGCACACGTCTGAACTCTTTCCCTACACGACGCTCTTCCGATCTNNNGGCTGAGGTAGGAGAATCA

GAGCACACGTCTGAACTCTTTCCCTACACGACGCTCTTCCGATCTNNNGGCTGAGGTAGGAGAATCA

AGCACACGTCTGAACTCTTTCCCTACACGACGCTCTTCCGATCTNNNTTTGTCCTAATGCTCTCCCTCT

AGCACACGTCTGAACTCTTTCCCTACACGACGCTCTTCCGATCTNNNTTTGTCCTAATGCTCTCCCTCT

AGCACACGTCTGAACTCTTTCCCTACACGACGCTCTTCCGATCTNNNATATTTCTTGCATTGGGCTCAC

AGCACACGTCTGAACTCTTTCCCTACACGACGCTCTTCCGATCTNNNATATTTCTTGCATTGGGCTCAC

GCACACGTCTGAACTCTTTCCCTACACGACGCTCTTCCGATCTNNNCAGCACATGGGACTTTTTCCAGA

GCACACGTCTGAACTCTTTCCCTACACGACGCTCTTCCGATCTNNNCAGCACATGGGACTTTTTCCAGA

GCACACGTCTGAACTCTTTCCCTACACGACGCTCTTCCGATCTNNNTAACCTCCAGTTCCATCCAGTTG

GCACACGTCTGAACTCTTTCCCTACACGACGCTCTTCCGATCTNNNTAACCTCCAGTTCCATCCAGTTG

AGCACACGTCTGAACTCTTTCCCTACACGACGCTCTTCCGATCTNNNTTATCAGACAGTTGCCCCAGAT

AGCACACGTCTGAACTCTTTCCCTACACGACGCTCTTCCGATCTNNNTTATCAGACAGTTGCCCCAGAT

GCACACGTCTGAACTCTTTCCCTACACGACGCTCTTCCGATCTNNNCTGCAGAGGGAGTACAGCAAAC

GCACACGTCTGAACTCTTTCCCTACACGACGCTCTTCCGATCTNNNCTGCAGAGGGAGTACAGCAAAC

AGCACACGTCTGAACTCTTTCCCTACACGACGCTCTTCCGATCTNNNCATTGGCTGGACAGTGGTAG

AGCACACGTCTGAACTCTTTCCCTACACGACGCTCTTCCGATCTNNNCATTGGCTGGACAGTGGTAG

GCACACGTCTGAACTCTTTCCCTACACGACGCTCTTCCGATCTNNNATGGCTTTGCATGTGTGTATGT

GCACACGTCTGAACTCTTTCCCTACACGACGCTCTTCCGATCTNNNATGGCTTTGCATGTGTGTATGT

GCACACGTCTGAACTCTTTCCCTACACGACGCTCTTCCGATCTNNNCCCATGAGAAGAACTCAGTAG

GCACACGTCTGAACTCTTTCCCTACACGACGCTCTTCCGATCTNNNCCCATGAGAAGAACTCAGTAG

GCACACGTCTGAACTCTTTCCCTACACGACGCTCTTCCGATCTNNNTTTTCTTTGTCCTCCTCTCCCAC

GCACACGTCTGAACTCTTTCCCTACACGACGCTCTTCCGATCTNNNTTTTCTTTGTCCTCCTCTCCCAC

GAGCACACGTCTGAACTCTTTCCCTACACGACGCTCTTCCGATCTNNNCGACTAGCTGAGATTACGG

GAGCACACGTCTGAACTCTTTCCCTACACGACGCTCTTCCGATCTNNNCGACTAGCTGAGATTACGG

AGCACACGTCTGAACTCTTTCCCTACACGACGCTCTTCCGATCTNNNCAAAATGGCAAAACCCCATCTC

AGCACACGTCTGAACTCTTTCCCTACACGACGCTCTTCCGATCTNNNCAAAATGGCAAAACCCCATCTC

AGCACACGTCTGAACTCTTTCCCTACACGACGCTCTTCCGATCTNNNACCTTCATGTCTATCTCGCTCT

AGCACACGTCTGAACTCTTTCCCTACACGACGCTCTTCCGATCTNNNACCTTCATGTCTATCTCGCTCT

GCACACGTCTGAACTCTTTCCCTACACGACGCTCTTCCGATCTNNNTCCATAACCTAAGTATTTAGATG

GCACACGTCTGAACTCTTTCCCTACACGACGCTCTTCCGATCTNNNTCCATAACCTAAGTATTTAGATG

AGCACACGTCTGAACTCTTTCCCTACACGACGCTCTTCCGATCTNNNGACTGATGACAGAGTGAGAC

AGCACACGTCTGAACTCTTTCCCTACACGACGCTCTTCCGATCTNNNGACTGATGACAGAGTGAGAC

GCACACGTCTGAACTCTTTCCCTACACGACGCTCTTCCGATCTNNNTCTTCACATCTTATTGCTCTTGCT

GCACACGTCTGAACTCTTTCCCTACACGACGCTCTTCCGATCTNNNTCTTCACATCTTATTGCTCTTGCT

GCACACGTCTGAACTCTTTCCCTACACGACGCTCTTCCGATCTNNNAGATTGACTGGCTTTGGTTTTT

GCACACGTCTGAACTCTTTCCCTACACGACGCTCTTCCGATCTNNNAGATTGACTGGCTTTGGTTTTT

AGCACACGTCTGAACTCTTTCCCTACACGACGCTCTTCCGATCTNNNTCTTACTTTGGGGCATCTTGTG

AGCACACGTCTGAACTCTTTCCCTACACGACGCTCTTCCGATCTNNNTCTTACTTTGGGGCATCTTGTG

GAGCACACGTCTGAACTCTTTCCCTACACGACGCTCTTCCGATCTNNNTAGCTGTGCTTTCAGTTGAG
GAGCACACGTCTGAACTCTTTCCCTACACGACGCTCTTCCGATCTNNNTAGCTGTGCTTTCAGTTGAG
AGCACACGTCTGAACTCTTTCCCTACACGACGCTCTTCCGATCTNNNCTCTCCTGTTCTGGATAATGC
AGCACACGTCTGAACTCTTTCCCTACACGACGCTCTTCCGATCTNNNCTCTCCTGTTCTGGATAATGC
GCACACGTCTGAACTCTTTCCCTACACGACGCTCTTCCGATCTNNNAACAGTGCAATTACCTCTTTGG
GCACACGTCTGAACTCTTTCCCTACACGACGCTCTTCCGATCTNNNAACAGTGCAATTACCTCTTTGG
GCACACGTCTGAACTCTTTCCCTACACGACGCTCTTCCGATCTNNNACTAGTCTGATAGCTATTGGTG
GCACACGTCTGAACTCTTTCCCTACACGACGCTCTTCCGATCTNNNACTAGTCTGATAGCTATTGGTG
AGCACACGTCTGAACTCTTTCCCTACACGACGCTCTTCCGATCTNNNCCATGCTGAAGGTGTTCATGA
AGCACACGTCTGAACTCTTTCCCTACACGACGCTCTTCCGATCTNNNCCATGCTGAAGGTGTTCATGA
GCACACGTCTGAACTCTTTCCCTACACGACGCTCTTCCGATCTNNNGCCACCAACCTCATTCTGTTTTG
GCACACGTCTGAACTCTTTCCCTACACGACGCTCTTCCGATCTNNNGCCACCAACCTCATTCTGTTTTG
GAGCACACGTCTGAACTCTTTCCCTACACGACGCTCTTCCGATCTNNNCTGAGTAGGTGGGATTACAG
GAGCACACGTCTGAACTCTTTCCCTACACGACGCTCTTCCGATCTNNNCTGAGTAGGTGGGATTACAG
GCACACGTCTGAACTCTTTCCCTACACGACGCTCTTCCGATCTNNNAGAAAATGCCTACATGTGATGA
GCACACGTCTGAACTCTTTCCCTACACGACGCTCTTCCGATCTNNNAGAAAATGCCTACATGTGATGA
CACACGTCTGAACTCTTTCCCTACACGACGCTCTTCCGATCTNNNGAAAGTTAAGGTATCAATGAGA
CACACGTCTGAACTCTTTCCCTACACGACGCTCTTCCGATCTNNNGAAAGTTAAGGTATCAATGAGA
GCACACGTCTGAACTCTTTCCCTACACGACGCTCTTCCGATCTNNNGGTTGCATAACAGGAAGCCAA
GCACACGTCTGAACTCTTTCCCTACACGACGCTCTTCCGATCTNNNGGTTGCATAACAGGAAGCCAA
GAGCACACGTCTGAACTCTTTCCCTACACGACGCTCTTCCGATCTNNNCAGAAAGGTAGGAGGCGAC
GAGCACACGTCTGAACTCTTTCCCTACACGACGCTCTTCCGATCTNNNCAGAAAGGTAGGAGGCGAC
AGCACACGTCTGAACTCTTTCCCTACACGACGCTCTTCCGATCTNNNTTTTGAATAGCTGGATCTGGC
AGCACACGTCTGAACTCTTTCCCTACACGACGCTCTTCCGATCTNNNTTTTGAATAGCTGGATCTGGC
GCACACGTCTGAACTCTTTCCCTACACGACGCTCTTCCGATCTNNNCACAGCTTGGACCCCCTCTCCAT
GCACACGTCTGAACTCTTTCCCTACACGACGCTCTTCCGATCTNNNCACAGCTTGGACCCCCTCTCCAT
GCACACGTCTGAACTCTTTCCCTACACGACGCTCTTCCGATCTNNNGAGGATTGCTTGAGCCTAGGA
GCACACGTCTGAACTCTTTCCCTACACGACGCTCTTCCGATCTNNNGAGGATTGCTTGAGCCTAGGA
GCACACGTCTGAACTCTTTCCCTACACGACGCTCTTCCGATCTNNNAAAAGTAGGCACTTGTCCATTG
GCACACGTCTGAACTCTTTCCCTACACGACGCTCTTCCGATCTNNNAAAAGTAGGCACTTGTCCATTG
AGCACACGTCTGAACTCTTTCCCTACACGACGCTCTTCCGATCTNNNCAGAAGAATTGCTTGAACCCA
AGCACACGTCTGAACTCTTTCCCTACACGACGCTCTTCCGATCTNNNCAGAAGAATTGCTTGAACCCA
AGCACACGTCTGAACTCTTTCCCTACACGACGCTCTTCCGATCTNNNCGGTTTGCTAATCATGGTTTTG
AGCACACGTCTGAACTCTTTCCCTACACGACGCTCTTCCGATCTNNNCGGTTTGCTAATCATGGTTTTG
GAGCACACGTCTGAACTCTTTCCCTACACGACGCTCTTCCGATCTNNNGGAGGCTAAGACAGGAGAA
GAGCACACGTCTGAACTCTTTCCCTACACGACGCTCTTCCGATCTNNNGGAGGCTAAGACAGGAGAA
AGCACACGTCTGAACTCTTTCCCTACACGACGCTCTTCCGATCTNNNGTCTGGCTTAGTGTGATTCAT
AGCACACGTCTGAACTCTTTCCCTACACGACGCTCTTCCGATCTNNNGTCTGGCTTAGTGTGATTCAT
AGCACACGTCTGAACTCTTTCCCTACACGACGCTCTTCCGATCTNNNAGCTGAGGAATGAGAAGCTT
AGCACACGTCTGAACTCTTTCCCTACACGACGCTCTTCCGATCTNNNAGCTGAGGAATGAGAAGCTT
GCACACGTCTGAACTCTTTCCCTACACGACGCTCTTCCGATCTNNNTGGATAACTCGCCTTCATGGAA
GCACACGTCTGAACTCTTTCCCTACACGACGCTCTTCCGATCTNNNTGGATAACTCGCCTTCATGGAA
GCACACGTCTGAACTCTTTCCCTACACGACGCTCTTCCGATCTNNNGTGGCTCATTCCTGGAAGATGT

GCACACGTCTGAACTCTTTCCCTACACGACGCTCTTCCGATCTNNNGTGGCTCATTCCTGGAAGATGT

GAGCACACGTCTGAACTCTTTCCCTACACGACGCTCTTCCGATCTNNNAAAATTAGCCAGACGTGGT

GAGCACACGTCTGAACTCTTTCCCTACACGACGCTCTTCCGATCTNNNAAAATTAGCCAGACGTGGT

AGCACACGTCTGAACTCTTTCCCTACACGACGCTCTTCCGATCTNNNCCAAACCATATGAGCCCCTTAT

AGCACACGTCTGAACTCTTTCCCTACACGACGCTCTTCCGATCTNNNCCAAACCATATGAGCCCCTTAT

AGCACACGTCTGAACTCTTTCCCTACACGACGCTCTTCCGATCTNNNTCCAGGAACTCCAACATGATT

AGCACACGTCTGAACTCTTTCCCTACACGACGCTCTTCCGATCTNNNTCCAGGAACTCCAACATGATT

AGCACACGTCTGAACTCTTTCCCTACACGACGCTCTTCCGATCTNNNGAGAATCGCTTGAACTTGGGA

AGCACACGTCTGAACTCTTTCCCTACACGACGCTCTTCCGATCTNNNGAGAATCGCTTGAACTTGGGA

AGCACACGTCTGAACTCTTTCCCTACACGACGCTCTTCCGATCTNNNCAGTGACCTATGACTGGACTA

AGCACACGTCTGAACTCTTTCCCTACACGACGCTCTTCCGATCTNNNCAGTGACCTATGACTGGACTA

AGCACACGTCTGAACTCTTTCCCTACACGACGCTCTTCCGATCTNNNGGTATTGTGTGCCTGTAGTCT

AGCACACGTCTGAACTCTTTCCCTACACGACGCTCTTCCGATCTNNNGGTATTGTGTGCCTGTAGTCT

AGCACACGTCTGAACTCTTTCCCTACACGACGCTCTTCCGATCTNNNTCTCAGTCTTACACACACAAGCA

AGCACACGTCTGAACTCTTTCCCTACACGACGCTCTTCCGATCTNNNTCTCAGTCTTACACACACAAGCA

AGCACACGTCTGAACTCTTTCCCTACACGACGCTCTTCCGATCTNNNGAAAGGATGGAGAAGTTGTTG

AGCACACGTCTGAACTCTTTCCCTACACGACGCTCTTCCGATCTNNNGAAAGGATGGAGAAGTTGTTG

GCACACGTCTGAACTCTTTCCCTACACGACGCTCTTCCGATCTNNNTGTAATTTATTGGCAGATCTCTA

GCACACGTCTGAACTCTTTCCCTACACGACGCTCTTCCGATCTNNNTGTAATTTATTGGCAGATCTCTA

AGCACACGTCTGAACTCTTTCCCTACACGACGCTCTTCCGATCTNNNTATCCTAGCACAAACACCCACA

AGCACACGTCTGAACTCTTTCCCTACACGACGCTCTTCCGATCTNNNTATCCTAGCACAAACACCCACA

GAGCACACGTCTGAACTCTTTCCCTACACGACGCTCTTCCGATCTNNNCAGTCCTGTAATCCCAGCTAC

GAGCACACGTCTGAACTCTTTCCCTACACGACGCTCTTCCGATCTNNNCAGTCCTGTAATCCCAGCTAC

CACACGTCTGAACTCTTTCCCTACACGACGCTCTTCCGATCTNNNACAGGGTATTCCATAATAACTTT

CACACGTCTGAACTCTTTCCCTACACGACGCTCTTCCGATCTNNNACAGGGTATTCCATAATAACTTT

GCACACGTCTGAACTCTTTCCCTACACGACGCTCTTCCGATCTNNNTGGCCCCATAATCATCTGATTCT

GCACACGTCTGAACTCTTTCCCTACACGACGCTCTTCCGATCTNNNTGGCCCCATAATCATCTGATTCT

GAGCACACGTCTGAACTCTTTCCCTACACGACGCTCTTCCGATCTNNNAGTGAGCCAAGATTGCATCA

GAGCACACGTCTGAACTCTTTCCCTACACGACGCTCTTCCGATCTNNNAGTGAGCCAAGATTGCATCA

GAGCACACGTCTGAACTCTTTCCCTACACGACGCTCTTCCGATCTNNNAAAGCCAAGAAAAGCAGCAC

GAGCACACGTCTGAACTCTTTCCCTACACGACGCTCTTCCGATCTNNNAAAGCCAAGAAAAGCAGCAC

AGCACACGTCTGAACTCTTTCCCTACACGACGCTCTTCCGATCTNNNATGCCATCTAGTACCATAAGCG

AGCACACGTCTGAACTCTTTCCCTACACGACGCTCTTCCGATCTNNNATGCCATCTAGTACCATAAGCG

AGCACACGTCTGAACTCTTTCCCTACACGACGCTCTTCCGATCTNNNAGAGCTGAAAGACAGAAAGAT

AGCACACGTCTGAACTCTTTCCCTACACGACGCTCTTCCGATCTNNNAGAGCTGAAAGACAGAAAGAT

AGCACACGTCTGAACTCTTTCCCTACACGACGCTCTTCCGATCTNNNCCATTTTATTACCTCCCCCTCCCC

AGCACACGTCTGAACTCTTTCCCTACACGACGCTCTTCCGATCTNNNCCATTTTATTACCTCCCCCTCCCC

GAGCACACGTCTGAACTCTTTCCCTACACGACGCTCTTCCGATCTNNNATGATCTCTGGTTGTGTGTTT

GAGCACACGTCTGAACTCTTTCCCTACACGACGCTCTTCCGATCTNNNATGATCTCTGGTTGTGTGTTT

AGCACACGTCTGAACTCTTTCCCTACACGACGCTCTTCCGATCTNNNAGCTACTTGGAAGGTTGAGG

AGCACACGTCTGAACTCTTTCCCTACACGACGCTCTTCCGATCTNNNAGCTACTTGGAAGGTTGAGG

CACACGTCTGAACTCTTTCCCTACACGACGCTCTTCCGATCTNNNATGATCTCTGCAAAGACCCCTTTCA

CACACGTCTGAACTCTTTCCCTACACGACGCTCTTCCGATCTNNNATGATCTCTGCAAAGACCCCTTTCA

AGCACACGTCTGAACTCTTTCCCTACACGACGCTCTTCCGATCTNNNGTGCATTGAAAGGAGAACACG
AGCACACGTCTGAACTCTTTCCCTACACGACGCTCTTCCGATCTNNNGTGCATTGAAAGGAGAACACG
AGCACACGTCTGAACTCTTTCCCTACACGACGCTCTTCCGATCTNNNCATTTCTTTGGTGTTGGCTCCA
AGCACACGTCTGAACTCTTTCCCTACACGACGCTCTTCCGATCTNNNCATTTCTTTGGTGTTGGCTCCA
AGCACACGTCTGAACTCTTTCCCTACACGACGCTCTTCCGATCTNNNAAGAACTTGCTGAAGAGCTTT
AGCACACGTCTGAACTCTTTCCCTACACGACGCTCTTCCGATCTNNNAAGAACTTGCTGAAGAGCTTT
AGCACACGTCTGAACTCTTTCCCTACACGACGCTCTTCCGATCTNNNTACTCATGTGAGAAAGGAAGG
AGCACACGTCTGAACTCTTTCCCTACACGACGCTCTTCCGATCTNNNTACTCATGTGAGAAAGGAAGG
GAGCACACGTCTGAACTCTTTCCCTACACGACGCTCTTCCGATCTNNNTGCTTTGTTTTGTTTTGGTTT
GAGCACACGTCTGAACTCTTTCCCTACACGACGCTCTTCCGATCTNNNTGCTTTGTTTTGTTTTGGTTT
CACACGTCTGAACTCTTTCCCTACACGACGCTCTTCCGATCTNNNCTGCTGGTGATGCTGCTCAGCAT
CACACGTCTGAACTCTTTCCCTACACGACGCTCTTCCGATCTNNNCTGCTGGTGATGCTGCTCAGCAT
AGAGCACACGTCTGAACTCTTTCCCTACACGACGCTCTTCCGATCTNNNCTTGCAGTGAGCCGAGATT
AGAGCACACGTCTGAACTCTTTCCCTACACGACGCTCTTCCGATCTNNNCTTGCAGTGAGCCGAGATT
GCACACGTCTGAACTCTTTCCCTACACGACGCTCTTCCGATCTNNNACTTTCTGCTCATCTCACTTTCTC
GCACACGTCTGAACTCTTTCCCTACACGACGCTCTTCCGATCTNNNACTTTCTGCTCATCTCACTTTCTC
GCACACGTCTGAACTCTTTCCCTACACGACGCTCTTCCGATCTNNNTTTGCAGAAGTGAATCTGGGGT
GCACACGTCTGAACTCTTTCCCTACACGACGCTCTTCCGATCTNNNTTTGCAGAAGTGAATCTGGGGT
GCACACGTCTGAACTCTTTCCCTACACGACGCTCTTCCGATCTNNNATGGGGTGATGTGACATATTG
GCACACGTCTGAACTCTTTCCCTACACGACGCTCTTCCGATCTNNNATGGGGTGATGTGACATATTG
GCACACGTCTGAACTCTTTCCCTACACGACGCTCTTCCGATCTNNNCTTAGTGCTGCCCTCATGATAGT
GCACACGTCTGAACTCTTTCCCTACACGACGCTCTTCCGATCTNNNCTTAGTGCTGCCCTCATGATAGT
GAGCACACGTCTGAACTCTTTCCCTACACGACGCTCTTCCGATCTNNNACAGCTCTGGATTCTGTCTCA
GAGCACACGTCTGAACTCTTTCCCTACACGACGCTCTTCCGATCTNNNACAGCTCTGGATTCTGTCTCA
GAGCACACGTCTGAACTCTTTCCCTACACGACGCTCTTCCGATCTNNNAGGCTGGAGTACAATGTCA
GAGCACACGTCTGAACTCTTTCCCTACACGACGCTCTTCCGATCTNNNAGGCTGGAGTACAATGTCA
GAGCACACGTCTGAACTCTTTCCCTACACGACGCTCTTCCGATCTNNNGTCAGGAGATCCAGACCATC
GAGCACACGTCTGAACTCTTTCCCTACACGACGCTCTTCCGATCTNNNGTCAGGAGATCCAGACCATC
AGCACACGTCTGAACTCTTTCCCTACACGACGCTCTTCCGATCTNNNGGGGCACTGAGAACTTGATAG
AGCACACGTCTGAACTCTTTCCCTACACGACGCTCTTCCGATCTNNNGGGGCACTGAGAACTTGATAG
GAGCACACGTCTGAACTCTTTCCCTACACGACGCTCTTCCGATCTNNNCACTTGAGGTCAGGAGTTTG
GAGCACACGTCTGAACTCTTTCCCTACACGACGCTCTTCCGATCTNNNCACTTGAGGTCAGGAGTTTG
GAGCACACGTCTGAACTCTTTCCCTACACGACGCTCTTCCGATCTNNNGCTGCTTTGTCATAGATGCT
GAGCACACGTCTGAACTCTTTCCCTACACGACGCTCTTCCGATCTNNNGCTGCTTTGTCATAGATGCT
AGCACACGTCTGAACTCTTTCCCTACACGACGCTCTTCCGATCTNNNATATATACACACACACACACAC
AGCACACGTCTGAACTCTTTCCCTACACGACGCTCTTCCGATCTNNNATATATACACACACACACACAC
GCACACGTCTGAACTCTTTCCCTACACGACGCTCTTCCGATCTNNNCTGCTGTCTGTCAGAGAGGATC
GCACACGTCTGAACTCTTTCCCTACACGACGCTCTTCCGATCTNNNCTGCTGTCTGTCAGAGAGGATC
GAGCACACGTCTGAACTCTTTCCCTACACGACGCTCTTCCGATCTNNNGAATTGCTTGAACCCAGGAA
GAGCACACGTCTGAACTCTTTCCCTACACGACGCTCTTCCGATCTNNNGAATTGCTTGAACCCAGGAA
GCACACGTCTGAACTCTTTCCCTACACGACGCTCTTCCGATCTNNNCACTGTTTTGGACTAAGGCAAG
GCACACGTCTGAACTCTTTCCCTACACGACGCTCTTCCGATCTNNNCACTGTTTTGGACTAAGGCAAG
AGCACACGTCTGAACTCTTTCCCTACACGACGCTCTTCCGATCTNNNTTTACCTGTTTGGGCATCTCGA

AGCACACGTCTGAACTCTTTCCCTACACGACGCTCTTCCGATCTNNNTTTACCTGTTTGGGCATCTCGA
GAGCACACGTCTGAACTCTTTCCCTACACGACGCTCTTCCGATCTNNNCACAGCCTTCTTCCTTTTCATG
GAGCACACGTCTGAACTCTTTCCCTACACGACGCTCTTCCGATCTNNNCACAGCCTTCTTCCTTTTCATG
GAGCACACGTCTGAACTCTTTCCCTACACGACGCTCTTCCGATCTNNNGTGTCCAACCAATGCAAAGTT
GAGCACACGTCTGAACTCTTTCCCTACACGACGCTCTTCCGATCTNNNGTGTCCAACCAATGCAAAGTT
AGCACACGTCTGAACTCTTTCCCTACACGACGCTCTTCCGATCTNNNTCCAAGGGGTGATGCAATGAA
AGCACACGTCTGAACTCTTTCCCTACACGACGCTCTTCCGATCTNNNTCCAAGGGGTGATGCAATGAA
GCACACGTCTGAACTCTTTCCCTACACGACGCTCTTCCGATCTNNNAGAACCAGTATGTTCACCTATGC
GCACACGTCTGAACTCTTTCCCTACACGACGCTCTTCCGATCTNNNAGAACCAGTATGTTCACCTATGC
GAGCACACGTCTGAACTCTTTCCCTACACGACGCTCTTCCGATCTNNNTCCTGTAACAGCAGACACACTA
GAGCACACGTCTGAACTCTTTCCCTACACGACGCTCTTCCGATCTNNNTCCTGTAACAGCAGACACACTA
GAGCACACGTCTGAACTCTTTCCCTACACGACGCTCTTCCGATCTNNNGAGGCTGAGGCAGGAGAATG
GAGCACACGTCTGAACTCTTTCCCTACACGACGCTCTTCCGATCTNNNGAGGCTGAGGCAGGAGAATG
AGCACACGTCTGAACTCTTTCCCTACACGACGCTCTTCCGATCTNNNCAGGCACCTATAGTCCCAGATTC
AGCACACGTCTGAACTCTTTCCCTACACGACGCTCTTCCGATCTNNNCAGGCACCTATAGTCCCAGATTC
AGCACACGTCTGAACTCTTTCCCTACACGACGCTCTTCCGATCTNNNAGAACTGGTAAACAGCACTATC
AGCACACGTCTGAACTCTTTCCCTACACGACGCTCTTCCGATCTNNNAGAACTGGTAAACAGCACTATC
AGCACACGTCTGAACTCTTTCCCTACACGACGCTCTTCCGATCTNNNGTCTCCATCAATGCCTCATACCT
AGCACACGTCTGAACTCTTTCCCTACACGACGCTCTTCCGATCTNNNGTCTCCATCAATGCCTCATACCT
GAGCACACGTCTGAACTCTTTCCCTACACGACGCTCTTCCGATCTNNNGGAGACCATCCTGGCTAATAT
GAGCACACGTCTGAACTCTTTCCCTACACGACGCTCTTCCGATCTNNNGGAGACCATCCTGGCTAATAT
AGCACACGTCTGAACTCTTTCCCTACACGACGCTCTTCCGATCTNNNCACACACACACACATCAAAG
AGCACACGTCTGAACTCTTTCCCTACACGACGCTCTTCCGATCTNNNCACACACACACACATCAAAG
AGCACACGTCTGAACTCTTTCCCTACACGACGCTCTTCCGATCTNNNCAGTCCCTGTCCAATGTACTTCA
AGCACACGTCTGAACTCTTTCCCTACACGACGCTCTTCCGATCTNNNCAGTCCCTGTCCAATGTACTTCA
AGCACACGTCTGAACTCTTTCCCTACACGACGCTCTTCCGATCTNNNCTCAAGTAGCTGGGACTATAGG
AGCACACGTCTGAACTCTTTCCCTACACGACGCTCTTCCGATCTNNNCTCAAGTAGCTGGGACTATAGG
AGCACACGTCTGAACTCTTTCCCTACACGACGCTCTTCCGATCTNNNCACACAGACAAGGTCTAGGTC
AGCACACGTCTGAACTCTTTCCCTACACGACGCTCTTCCGATCTNNNCACACAGACAAGGTCTAGGTC
GAGCACACGTCTGAACTCTTTCCCTACACGACGCTCTTCCGATCTNNNCAGTATGTTCTGCTCCATCCAC
GAGCACACGTCTGAACTCTTTCCCTACACGACGCTCTTCCGATCTNNNCAGTATGTTCTGCTCCATCCAC
AGCACACGTCTGAACTCTTTCCCTACACGACGCTCTTCCGATCTNNNGCATATGGATGATACTCTAAGA
AGCACACGTCTGAACTCTTTCCCTACACGACGCTCTTCCGATCTNNNGCATATGGATGATACTCTAAGA
GCACACGTCTGAACTCTTTCCCTACACGACGCTCTTCCGATCTNNNGGCTTCATCTTCAATATCATCCCT
GCACACGTCTGAACTCTTTCCCTACACGACGCTCTTCCGATCTNNNGGCTTCATCTTCAATATCATCCCT
AGCACACGTCTGAACTCTTTCCCTACACGACGCTCTTCCGATCTNNNGCATCCAGCCTTGAAATGTTTT
AGCACACGTCTGAACTCTTTCCCTACACGACGCTCTTCCGATCTNNNGCATCCAGCCTTGAAATGTTTT
AGCACACGTCTGAACTCTTTCCCTACACGACGCTCTTCCGATCTNNNAGTGAGCTGAGATTGTACCACT
AGCACACGTCTGAACTCTTTCCCTACACGACGCTCTTCCGATCTNNNAGTGAGCTGAGATTGTACCACT
AGAGCACACGTCTGAACTCTTTCCCTACACGACGCTCTTCCGATCTNNNGATCCCCATTACTCTGCATGC
AGAGCACACGTCTGAACTCTTTCCCTACACGACGCTCTTCCGATCTNNNGATCCCCATTACTCTGCATGC
GAGCACACGTCTGAACTCTTTCCCTACACGACGCTCTTCCGATCTNNNGAAACCACTTCTGTGTCATCTC
GAGCACACGTCTGAACTCTTTCCCTACACGACGCTCTTCCGATCTNNNGAAACCACTTCTGTGTCATCTC

AGCACACGTCTGAACTCTTTCCCTACACGACGCTCTTCCGATCTNNNCTGTCCCTCTAGAAAACCCTGAC
AGCACACGTCTGAACTCTTTCCCTACACGACGCTCTTCCGATCTNNNCTGTCCCTCTAGAAAACCCTGAC
AGCACACGTCTGAACTCTTTCCCTACACGACGCTCTTCCGATCTNNNAAAAACAGAAAGAAGCCCTCTG
AGCACACGTCTGAACTCTTTCCCTACACGACGCTCTTCCGATCTNNNAAAAACAGAAAGAAGCCCTCTG
CACACGTCTGAACTCTTTCCCTACACGACGCTCTTCCGATCTNNNGTTTTGGGCTGAAAGATGGCGCA
CACACGTCTGAACTCTTTCCCTACACGACGCTCTTCCGATCTNNNGTTTTGGGCTGAAAGATGGCGCA
AGCACACGTCTGAACTCTTTCCCTACACGACGCTCTTCCGATCTNNNTGAGGTTACAGTGAGCCAAGA
AGCACACGTCTGAACTCTTTCCCTACACGACGCTCTTCCGATCTNNNTGAGGTTACAGTGAGCCAAGA
GAGCACACGTCTGAACTCTTTCCCTACACGACGCTCTTCCGATCTNNNAAAACAAATTCCTAACCAGT
GAGCACACGTCTGAACTCTTTCCCTACACGACGCTCTTCCGATCTNNNAAAACAAATTCCTAACCAGT
AGCACACGTCTGAACTCTTTCCCTACACGACGCTCTTCCGATCTNNNTCCTCCTTTTCTGTTAGCAGCTT
AGCACACGTCTGAACTCTTTCCCTACACGACGCTCTTCCGATCTNNNTCCTCCTTTTCTGTTAGCAGCTT
AGCACACGTCTGAACTCTTTCCCTACACGACGCTCTTCCGATCTNNNAAAACAGAAACAACAACCAGG
AGCACACGTCTGAACTCTTTCCCTACACGACGCTCTTCCGATCTNNNAAAACAGAAACAACAACCAGG
AGCACACGTCTGAACTCTTTCCCTACACGACGCTCTTCCGATCTNNNTGGTCTAGAAACCTGCGATAA
AGCACACGTCTGAACTCTTTCCCTACACGACGCTCTTCCGATCTNNNTGGTCTAGAAACCTGCGATAA
AGCACACGTCTGAACTCTTTCCCTACACGACGCTCTTCCGATCTNNNCTGGATCCCTGTAAAAAACGA
AGCACACGTCTGAACTCTTTCCCTACACGACGCTCTTCCGATCTNNNCTGGATCCCTGTAAAAAACGA
GAGCACACGTCTGAACTCTTTCCCTACACGACGCTCTTCCGATCTNNNCCTCCACATTGATGGCCTTTA
GAGCACACGTCTGAACTCTTTCCCTACACGACGCTCTTCCGATCTNNNCCTCCACATTGATGGCCTTTA
GAGCACACGTCTGAACTCTTTCCCTACACGACGCTCTTCCGATCTNNNCCTGGAATGTGAAGGTTGC
GAGCACACGTCTGAACTCTTTCCCTACACGACGCTCTTCCGATCTNNNCCTGGAATGTGAAGGTTGC
AGAGCACACGTCTGAACTCTTTCCCTACACGACGCTCTTCCGATCTNNNATCGTCCTTCATAACTGGG
AGAGCACACGTCTGAACTCTTTCCCTACACGACGCTCTTCCGATCTNNNATCGTCCTTCATAACTGGG
AGCACACGTCTGAACTCTTTCCCTACACGACGCTCTTCCGATCTNNNAGGGACTTGGGGTGATCATTT
AGCACACGTCTGAACTCTTTCCCTACACGACGCTCTTCCGATCTNNNAGGGACTTGGGGTGATCATTT
AGCACACGTCTGAACTCTTTCCCTACACGACGCTCTTCCGATCTNNNAAGGTTTACAAAATAATTTTTC
AGCACACGTCTGAACTCTTTCCCTACACGACGCTCTTCCGATCTNNNAAGGTTTACAAAATAATTTTTC
GCACACGTCTGAACTCTTTCCCTACACGACGCTCTTCCGATCTNNNCTCCCTCATGCTACTGTATCAGCC
GCACACGTCTGAACTCTTTCCCTACACGACGCTCTTCCGATCTNNNCTCCCTCATGCTACTGTATCAGCC
AGCACACGTCTGAACTCTTTCCCTACACGACGCTCTTCCGATCTNNNGGGGTGAAGAGATGGATATG
AGCACACGTCTGAACTCTTTCCCTACACGACGCTCTTCCGATCTNNNGGGGTGAAGAGATGGATATG
AGCACACGTCTGAACTCTTTCCCTACACGACGCTCTTCCGATCTNNNAAGAGACACCATGAAAGCAA
AGCACACGTCTGAACTCTTTCCCTACACGACGCTCTTCCGATCTNNNAAGAGACACCATGAAAGCAA
AGCACACGTCTGAACTCTTTCCCTACACGACGCTCTTCCGATCTNNNTTACTTGTCCTTAGACCCAAGCC
AGCACACGTCTGAACTCTTTCCCTACACGACGCTCTTCCGATCTNNNTTACTTGTCCTTAGACCCAAGCC
ACACGTCTGAACTCTTTCCCTACACGACGCTCTTCCGATCTNNNTGGTCTTGAACTCCTGACTTCAGCA
ACACGTCTGAACTCTTTCCCTACACGACGCTCTTCCGATCTNNNTGGTCTTGAACTCCTGACTTCAGCA
AGCACACGTCTGAACTCTTTCCCTACACGACGCTCTTCCGATCTNNNTGGCAGGCCAAAAAAAGAAAG
AGCACACGTCTGAACTCTTTCCCTACACGACGCTCTTCCGATCTNNNTGGCAGGCCAAAAAAAGAAAG
GCACACGTCTGAACTCTTTCCCTACACGACGCTCTTCCGATCTNNNACATCTCCATTCTTCTCTTTTAAC
GCACACGTCTGAACTCTTTCCCTACACGACGCTCTTCCGATCTNNNACATCTCCATTCTTCTCTTTTAAC
AGAGCACACGTCTGAACTCTTTCCCTACACGACGCTCTTCCGATCTNNNGGAACTAATCAGCAGGATG

GAGCACACGTCTGAACTCTTTCCCTACACGACGCTCTTCCGATCTNNNGGAACTAATCAGCAGGATG
AGCACACGTCTGAACTCTTTCCCTACACGACGCTCTTCCGATCTNNNTTTCATCAGAATCTGGAGGCAC
AGCACACGTCTGAACTCTTTCCCTACACGACGCTCTTCCGATCTNNNTTTCATCAGAATCTGGAGGCAC
AGCACACGTCTGAACTCTTTCCCTACACGACGCTCTTCCGATCTNNNGCTGAGGTAGGAGAATGGAA
AGCACACGTCTGAACTCTTTCCCTACACGACGCTCTTCCGATCTNNNGCTGAGGTAGGAGAATGGAA
AGCACACGTCTGAACTCTTTCCCTACACGACGCTCTTCCGATCTNNNTTTTATGAGCCTTTACAAATT
AGCACACGTCTGAACTCTTTCCCTACACGACGCTCTTCCGATCTNNNTTTTATGAGCCTTTACAAATT
AGCACACGTCTGAACTCTTTCCCTACACGACGCTCTTCCGATCTNNNTGGGGTCTCTGATGTTAGTAC
AGCACACGTCTGAACTCTTTCCCTACACGACGCTCTTCCGATCTNNNTGGGGTCTCTGATGTTAGTAC
GCACACGTCTGAACTCTTTCCCTACACGACGCTCTTCCGATCTNNNTGTCACAGCCTAACATTACCCTCC
GCACACGTCTGAACTCTTTCCCTACACGACGCTCTTCCGATCTNNNTGTCACAGCCTAACATTACCCTCC
GCACACGTCTGAACTCTTTCCCTACACGACGCTCTTCCGATCTNNNTGCTTGTATCAGCAGTCCTCAAT
GCACACGTCTGAACTCTTTCCCTACACGACGCTCTTCCGATCTNNNTGCTTGTATCAGCAGTCCTCAAT
AGCACACGTCTGAACTCTTTCCCTACACGACGCTCTTCCGATCTNNNAGAGATGGTTCATGCTCTCAA
AGCACACGTCTGAACTCTTTCCCTACACGACGCTCTTCCGATCTNNNAGAGATGGTTCATGCTCTCAA
AGCACACGTCTGAACTCTTTCCCTACACGACGCTCTTCCGATCTNNNGACTATCTAAGGCAAGCATGC
AGCACACGTCTGAACTCTTTCCCTACACGACGCTCTTCCGATCTNNNGACTATCTAAGGCAAGCATGC
GAGCACACGTCTGAACTCTTTCCCTACACGACGCTCTTCCGATCTNNNCCCATGAAACTAGGGACATT
GAGCACACGTCTGAACTCTTTCCCTACACGACGCTCTTCCGATCTNNNCCCATGAAACTAGGGACATT
GCACACGTCTGAACTCTTTCCCTACACGACGCTCTTCCGATCTNNNGAGGCAGCTAGAGATAAGAAG
GCACACGTCTGAACTCTTTCCCTACACGACGCTCTTCCGATCTNNNGAGGCAGCTAGAGATAAGAAG
AGCACACGTCTGAACTCTTTCCCTACACGACGCTCTTCCGATCTNNNCCATATGCCGGGCTAATTTT
AGCACACGTCTGAACTCTTTCCCTACACGACGCTCTTCCGATCTNNNCCATATGCCGGGCTAATTTT
GCACACGTCTGAACTCTTTCCCTACACGACGCTCTTCCGATCTNNNCCACCAGATGCCAGTAGTAGTT
GCACACGTCTGAACTCTTTCCCTACACGACGCTCTTCCGATCTNNNCCACCAGATGCCAGTAGTAGTT
CACGTCTGAACTCTTTCCCTACACGACGCTCTTCCGATCTNNNTGATTTGGTGTTGATCGCACATTCAT
CACGTCTGAACTCTTTCCCTACACGACGCTCTTCCGATCTNNNTGATTTGGTGTTGATCGCACATTCAT
AGCACACGTCTGAACTCTTTCCCTACACGACGCTCTTCCGATCTNNNAGGCTCCTCATCACTAATCATT
AGCACACGTCTGAACTCTTTCCCTACACGACGCTCTTCCGATCTNNNAGGCTCCTCATCACTAATCATT
AGAGCACACGTCTGAACTCTTTCCCTACACGACGCTCTTCCGATCTNNNCAGGAGGCAGAGCTTACAG
AGAGCACACGTCTGAACTCTTTCCCTACACGACGCTCTTCCGATCTNNNCAGGAGGCAGAGCTTACAG
CACACGTCTGAACTCTTTCCCTACACGACGCTCTTCCGATCTNNNTTCTCTTTGGAATGGGGAGAAGC
CACACGTCTGAACTCTTTCCCTACACGACGCTCTTCCGATCTNNNTTCTCTTTGGAATGGGGAGAAGC
ACACGTCTGAACTCTTTCCCTACACGACGCTCTTCCGATCTNNNCTCCACCCTAGATCACTTTGCATCG
ACACGTCTGAACTCTTTCCCTACACGACGCTCTTCCGATCTNNNCTCCACCCTAGATCACTTTGCATCG
CACACGTCTGAACTCTTTCCCTACACGACGCTCTTCCGATCTNNNCCTCTGTGACGTTTGTGTTTTGTC
CACACGTCTGAACTCTTTCCCTACACGACGCTCTTCCGATCTNNNCCTCTGTGACGTTTGTGTTTTGTC
GAGCACACGTCTGAACTCTTTCCCTACACGACGCTCTTCCGATCTNNNTCCTCTAAGGACAGACTTGG
GAGCACACGTCTGAACTCTTTCCCTACACGACGCTCTTCCGATCTNNNTCCTCTAAGGACAGACTTGG
AGCACACGTCTGAACTCTTTCCCTACACGACGCTCTTCCGATCTNNNCAATCTGAGAACTGTCTGGACC
AGCACACGTCTGAACTCTTTCCCTACACGACGCTCTTCCGATCTNNNCAATCTGAGAACTGTCTGGACC
GCACACGTCTGAACTCTTTCCCTACACGACGCTCTTCCGATCTNNNATGCCGGTAATCCTAGCTACTT
GCACACGTCTGAACTCTTTCCCTACACGACGCTCTTCCGATCTNNNATGCCGGTAATCCTAGCTACTT

GCACACGTCTGAACTCTTTCCCTACACGACGCTCTTCCGATCTNNNGTCAGAAGGGGTTCTAAAGCCA
GCACACGTCTGAACTCTTTCCCTACACGACGCTCTTCCGATCTNNNGTCAGAAGGGGTTCTAAAGCCA
GAGCACACGTCTGAACTCTTTCCCTACACGACGCTCTTCCGATCTNNNTGGAAAAACCTAAAGACTCCA
GAGCACACGTCTGAACTCTTTCCCTACACGACGCTCTTCCGATCTNNNTGGAAAAACCTAAAGACTCCA
GAGCACACGTCTGAACTCTTTCCCTACACGACGCTCTTCCGATCTNNNTGAGGTCAGGAGTTTGAGAC
GAGCACACGTCTGAACTCTTTCCCTACACGACGCTCTTCCGATCTNNNTGAGGTCAGGAGTTTGAGAC
GCACACGTCTGAACTCTTTCCCTACACGACGCTCTTCCGATCTNNNGCTCTCTCTCTCCTCTCTCTCCA
GCACACGTCTGAACTCTTTCCCTACACGACGCTCTTCCGATCTNNNGCTCTCTCTCTCCTCTCTCTCCA
AGCACACGTCTGAACTCTTTCCCTACACGACGCTCTTCCGATCTNNNCTGTAATCCCAGCTACTCGGAA
AGCACACGTCTGAACTCTTTCCCTACACGACGCTCTTCCGATCTNNNCTGTAATCCCAGCTACTCGGAA
GCACACGTCTGAACTCTTTCCCTACACGACGCTCTTCCGATCTNNNTATCCTTTCTCTCCCACCACTCTCA
GCACACGTCTGAACTCTTTCCCTACACGACGCTCTTCCGATCTNNNTATCCTTTCTCTCCCACCACTCTCA
AGCACACGTCTGAACTCTTTCCCTACACGACGCTCTTCCGATCTNNNCAACTAGCATTGCATCAGGGTT
AGCACACGTCTGAACTCTTTCCCTACACGACGCTCTTCCGATCTNNNCAACTAGCATTGCATCAGGGTT
GAGCACACGTCTGAACTCTTTCCCTACACGACGCTCTTCCGATCTNNNTGGAAGAGCACAAAAATGAG
GAGCACACGTCTGAACTCTTTCCCTACACGACGCTCTTCCGATCTNNNTGGAAGAGCACAAAAATGAG
AGCACACGTCTGAACTCTTTCCCTACACGACGCTCTTCCGATCTNNNGGCCTCTGTTAGATTCACTCCA
AGCACACGTCTGAACTCTTTCCCTACACGACGCTCTTCCGATCTNNNGGCCTCTGTTAGATTCACTCCA
GAGCACACGTCTGAACTCTTTCCCTACACGACGCTCTTCCGATCTNNNCAAGCCCCTAAAATCCTGGAA
GAGCACACGTCTGAACTCTTTCCCTACACGACGCTCTTCCGATCTNNNCAAGCCCCTAAAATCCTGGAA
AGCACACGTCTGAACTCTTTCCCTACACGACGCTCTTCCGATCTNNNAGCAAACTCATTGAATCATGGG
AGCACACGTCTGAACTCTTTCCCTACACGACGCTCTTCCGATCTNNNAGCAAACTCATTGAATCATGGG
AGCACACGTCTGAACTCTTTCCCTACACGACGCTCTTCCGATCTNNNGCTGTGTTGTGCTTCATCTGAT
AGCACACGTCTGAACTCTTTCCCTACACGACGCTCTTCCGATCTNNNGCTGTGTTGTGCTTCATCTGAT
GCACACGTCTGAACTCTTTCCCTACACGACGCTCTTCCGATCTNNNGAGAATTGACCCATGAGTTGCAT
GCACACGTCTGAACTCTTTCCCTACACGACGCTCTTCCGATCTNNNGAGAATTGACCCATGAGTTGCAT
AGCACACGTCTGAACTCTTTCCCTACACGACGCTCTTCCGATCTNNNCATTCTATGACCTAGTGGGCCTG
AGCACACGTCTGAACTCTTTCCCTACACGACGCTCTTCCGATCTNNNCATTCTATGACCTAGTGGGCCTG
AGCACACGTCTGAACTCTTTCCCTACACGACGCTCTTCCGATCTNNNGAGGCAAGCATTTCTGACACTT
AGCACACGTCTGAACTCTTTCCCTACACGACGCTCTTCCGATCTNNNGAGGCAAGCATTTCTGACACTT
AAGAGCACACGTCTGAACTCTTTCCCTACACGACGCTCTTCCGATCTNNNGTAGGTGGGAGACAGAAA
AAGAGCACACGTCTGAACTCTTTCCCTACACGACGCTCTTCCGATCTNNNGTAGGTGGGAGACAGAAA
AGCACACGTCTGAACTCTTTCCCTACACGACGCTCTTCCGATCTNNNGGTAATGGGGATACTGCTGTGA
AGCACACGTCTGAACTCTTTCCCTACACGACGCTCTTCCGATCTNNNGGTAATGGGGATACTGCTGTGA
GAGCACACGTCTGAACTCTTTCCCTACACGACGCTCTTCCGATCTNNNGAGCAAGACTCCGTCTCAAAA
GAGCACACGTCTGAACTCTTTCCCTACACGACGCTCTTCCGATCTNNNGAGCAAGACTCCGTCTCAAAA
AGCACACGTCTGAACTCTTTCCCTACACGACGCTCTTCCGATCTNNNTCTCAGATACTGGAGAGGCTGA
AGCACACGTCTGAACTCTTTCCCTACACGACGCTCTTCCGATCTNNNTCTCAGATACTGGAGAGGCTGA
AGCACACGTCTGAACTCTTTCCCTACACGACGCTCTTCCGATCTNNNTGATATAAAGGCAAATCTTCTTG
AGCACACGTCTGAACTCTTTCCCTACACGACGCTCTTCCGATCTNNNTGATATAAAGGCAAATCTTCTTG
GAGCACACGTCTGAACTCTTTCCCTACACGACGCTCTTCCGATCTNNNTCACCTGGGTATAGCCTCAAGT
GAGCACACGTCTGAACTCTTTCCCTACACGACGCTCTTCCGATCTNNNTCACCTGGGTATAGCCTCAAGT
GAGCACACGTCTGAACTCTTTCCCTACACGACGCTCTTCCGATCTNNNTTCAGAACGTGAACATGGACA

GAGCACACGTCTGAACTCTTTCCCTACACGACGCTCTTCCGATCTNNNTTCAGAACGTGAACATGGACA
AGAGCACACGTCTGAACTCTTTCCCTACACGACGCTCTTCCGATCTNNNCCCAAGTGCTTACAAGGATCA
AGAGCACACGTCTGAACTCTTTCCCTACACGACGCTCTTCCGATCTNNNCCCAAGTGCTTACAAGGATCA
GAAGAGCACACGTCTGAACTCTTTCCCTACACGACGCTCTTCCGATCTNNNGAATGGCGTGAACCCAGA
GAAGAGCACACGTCTGAACTCTTTCCCTACACGACGCTCTTCCGATCTNNNGAATGGCGTGAACCCAGA
GAGCACACGTCTGAACTCTTTCCCTACACGACGCTCTTCCGATCTNNNCCTGTAATCCCAGCTCCTTGGC
GAGCACACGTCTGAACTCTTTCCCTACACGACGCTCTTCCGATCTNNNCCTGTAATCCCAGCTCCTTGGC
GAGCACACGTCTGAACTCTTTCCCTACACGACGCTCTTCCGATCTNNNTCACTATTGGTGGAATTCAGG
GAGCACACGTCTGAACTCTTTCCCTACACGACGCTCTTCCGATCTNNNTCACTATTGGTGGAATTCAGG
GAGCACACGTCTGAACTCTTTCCCTACACGACGCTCTTCCGATCTNNNTGTATCTTCTTGCAAGAAAAC
GAGCACACGTCTGAACTCTTTCCCTACACGACGCTCTTCCGATCTNNNTGTATCTTCTTGCAAGAAAAC
AGCACACGTCTGAACTCTTTCCCTACACGACGCTCTTCCGATCTNNNGAAAGTCACTACTCCTGCTTGCT
AGCACACGTCTGAACTCTTTCCCTACACGACGCTCTTCCGATCTNNNGAAAGTCACTACTCCTGCTTGCT
GAGCACACGTCTGAACTCTTTCCCTACACGACGCTCTTCCGATCTNNNCTCTGTGCGTGTGTGTGTGCC
GAGCACACGTCTGAACTCTTTCCCTACACGACGCTCTTCCGATCTNNNCTCTGTGCGTGTGTGTGTGCC
GAGCACACGTCTGAACTCTTTCCCTACACGACGCTCTTCCGATCTNNNTCGTGATCCTTAGTACCCTGAG
GAGCACACGTCTGAACTCTTTCCCTACACGACGCTCTTCCGATCTNNNTCGTGATCCTTAGTACCCTGAG
GCACACGTCTGAACTCTTTCCCTACACGACGCTCTTCCGATCTNNNTGAAGCAGTAGGATCACTTGAG
GCACACGTCTGAACTCTTTCCCTACACGACGCTCTTCCGATCTNNNTGAAGCAGTAGGATCACTTGAG
GAGCACACGTCTGAACTCTTTCCCTACACGACGCTCTTCCGATCTNNNGTGCCTGTAGTCCCATCTACTC
GAGCACACGTCTGAACTCTTTCCCTACACGACGCTCTTCCGATCTNNNGTGCCTGTAGTCCCATCTACTC
GAGCACACGTCTGAACTCTTTCCCTACACGACGCTCTTCCGATCTNNNTTACCAGCCAATGACAAACACC
GAGCACACGTCTGAACTCTTTCCCTACACGACGCTCTTCCGATCTNNNTTACCAGCCAATGACAAACACC
GAGCACACGTCTGAACTCTTTCCCTACACGACGCTCTTCCGATCTNNNCACTTGAGGTCAGGAGTTCAA
GAGCACACGTCTGAACTCTTTCCCTACACGACGCTCTTCCGATCTNNNCACTTGAGGTCAGGAGTTCAA
GAGCACACGTCTGAACTCTTTCCCTACACGACGCTCTTCCGATCTNNNTATTTAGGCCCTGCTTTACCTG
GAGCACACGTCTGAACTCTTTCCCTACACGACGCTCTTCCGATCTNNNTATTTAGGCCCTGCTTTACCTG
GAGCACACGTCTGAACTCTTTCCCTACACGACGCTCTTCCGATCTNNNAGTTTTGCATGATAAAAGTGG
GAGCACACGTCTGAACTCTTTCCCTACACGACGCTCTTCCGATCTNNNAGTTTTGCATGATAAAAGTGG
GAGCACACGTCTGAACTCTTTCCCTACACGACGCTCTTCCGATCTNNNTCACTTTAGCTACCCCCAAGTG
GAGCACACGTCTGAACTCTTTCCCTACACGACGCTCTTCCGATCTNNNTCACTTTAGCTACCCCCAAGTG
GAGCACACGTCTGAACTCTTTCCCTACACGACGCTCTTCCGATCTNNNAAGTTACGAGTCTCAGAAATT
GAGCACACGTCTGAACTCTTTCCCTACACGACGCTCTTCCGATCTNNNAAGTTACGAGTCTCAGAAATT
CACACGTCTGAACTCTTTCCCTACACGACGCTCTTCCGATCTNNNGTCTTGAACTTCTGACCTCGTGACA
CACACGTCTGAACTCTTTCCCTACACGACGCTCTTCCGATCTNNNGTCTTGAACTTCTGACCTCGTGACA
GCACACGTCTGAACTCTTTCCCTACACGACGCTCTTCCGATCTNNNGAGACTGAGGTGAGAGTTTTGCC
GCACACGTCTGAACTCTTTCCCTACACGACGCTCTTCCGATCTNNNGAGACTGAGGTGAGAGTTTTGCC
GCACACGTCTGAACTCTTTCCCTACACGACGCTCTTCCGATCTNNNCTTTTGCATGTTTTGTAGAGATCA
GCACACGTCTGAACTCTTTCCCTACACGACGCTCTTCCGATCTNNNCTTTTGCATGTTTTGTAGAGATCA
GAGCACACGTCTGAACTCTTTCCCTACACGACGCTCTTCCGATCTNNNAAGAAGCACCTCAGTCAGTAG
GAGCACACGTCTGAACTCTTTCCCTACACGACGCTCTTCCGATCTNNNAAGAAGCACCTCAGTCAGTAG
GCACACGTCTGAACTCTTTCCCTACACGACGCTCTTCCGATCTNNNCTGGGCAACAAGAGCAAAACTCC
GCACACGTCTGAACTCTTTCCCTACACGACGCTCTTCCGATCTNNNCTGGGCAACAAGAGCAAAACTCC

AGCACACGTCTGAACTCTTTCCCTACACGACGCTCTTCCGATCTNNNTGCTTTTAACTGAGCCCAAGGA
AGCACACGTCTGAACTCTTTCCCTACACGACGCTCTTCCGATCTNNNTGCTTTTAACTGAGCCCAAGGA
AGCACACGTCTGAACTCTTTCCCTACACGACGCTCTTCCGATCTNNNTTCAACCCTTGTCTCCATCTCTC
AGCACACGTCTGAACTCTTTCCCTACACGACGCTCTTCCGATCTNNNTTCAACCCTTGTCTCCATCTCTC
AGAGCACACGTCTGAACTCTTTCCCTACACGACGCTCTTCCGATCTNNNCAGGGTGGGGTTATGAATG
AGAGCACACGTCTGAACTCTTTCCCTACACGACGCTCTTCCGATCTNNNCAGGGTGGGGTTATGAATG
AGCACACGTCTGAACTCTTTCCCTACACGACGCTCTTCCGATCTNNNAGCAAGCTATCAAAAGAAATTT
AGCACACGTCTGAACTCTTTCCCTACACGACGCTCTTCCGATCTNNNAGCAAGCTATCAAAAGAAATTT
AGCACACGTCTGAACTCTTTCCCTACACGACGCTCTTCCGATCTNNNGGACTTTTTGAGATCAGTGCGT
AGCACACGTCTGAACTCTTTCCCTACACGACGCTCTTCCGATCTNNNGGACTTTTTGAGATCAGTGCGT
AGCACACGTCTGAACTCTTTCCCTACACGACGCTCTTCCGATCTNNNCATTTCAACAACCACAGCAGACT
AGCACACGTCTGAACTCTTTCCCTACACGACGCTCTTCCGATCTNNNCATTTCAACAACCACAGCAGACT
GAGCACACGTCTGAACTCTTTCCCTACACGACGCTCTTCCGATCTNNNAAAAGAATGGTGGGCGGGA
GAGCACACGTCTGAACTCTTTCCCTACACGACGCTCTTCCGATCTNNNAAAAGAATGGTGGGCGGGA
AGAGCACACGTCTGAACTCTTTCCCTACACGACGCTCTTCCGATCTNNNACAAGGAAGGAGGAAGGC
AGAGCACACGTCTGAACTCTTTCCCTACACGACGCTCTTCCGATCTNNNACAAGGAAGGAGGAAGGC
GAGCACACGTCTGAACTCTTTCCCTACACGACGCTCTTCCGATCTNNNCTCCTAGTAGTGGCTCCAGAA
GAGCACACGTCTGAACTCTTTCCCTACACGACGCTCTTCCGATCTNNNCTCCTAGTAGTGGCTCCAGAA
AGCACACGTCTGAACTCTTTCCCTACACGACGCTCTTCCGATCTNNNACCCTGTCTCCAGAAAAACTGTT
AGCACACGTCTGAACTCTTTCCCTACACGACGCTCTTCCGATCTNNNACCCTGTCTCCAGAAAAACTGTT
GAGCACACGTCTGAACTCTTTCCCTACACGACGCTCTTCCGATCTNNNTAGAGCTGGTGGTTTTGCTT
GAGCACACGTCTGAACTCTTTCCCTACACGACGCTCTTCCGATCTNNNTAGAGCTGGTGGTTTTGCTT
GAGCACACGTCTGAACTCTTTCCCTACACGACGCTCTTCCGATCTNNNGCCAACAGGAAATGAACTCTT
GAGCACACGTCTGAACTCTTTCCCTACACGACGCTCTTCCGATCTNNNGCCAACAGGAAATGAACTCTT
GAGCACACGTCTGAACTCTTTCCCTACACGACGCTCTTCCGATCTNNNTCAGGTGTTACAATACTGTCA
GAGCACACGTCTGAACTCTTTCCCTACACGACGCTCTTCCGATCTNNNTCAGGTGTTACAATACTGTCA
AGCACACGTCTGAACTCTTTCCCTACACGACGCTCTTCCGATCTNNNAGTGAGCTGAGATTATACCGCT
AGCACACGTCTGAACTCTTTCCCTACACGACGCTCTTCCGATCTNNNAGTGAGCTGAGATTATACCGCT
AGCACACGTCTGAACTCTTTCCCTACACGACGCTCTTCCGATCTNNNAGAAAACCTAGACAACCTTGGG
AGCACACGTCTGAACTCTTTCCCTACACGACGCTCTTCCGATCTNNNAGAAAACCTAGACAACCTTGGG
AGCACACGTCTGAACTCTTTCCCTACACGACGCTCTTCCGATCTNNNCCTAGTTTTTTAGTTAAACAGCA
AGCACACGTCTGAACTCTTTCCCTACACGACGCTCTTCCGATCTNNNCCTAGTTTTTTAGTTAAACAGCA
AGCACACGTCTGAACTCTTTCCCTACACGACGCTCTTCCGATCTNNNAAATACAAAAATCAGCCAGGCG
AGCACACGTCTGAACTCTTTCCCTACACGACGCTCTTCCGATCTNNNAAATACAAAAATCAGCCAGGCG
GAGCACACGTCTGAACTCTTTCCCTACACGACGCTCTTCCGATCTNNNTACATGCCCTCGTATGTTTCCT
GAGCACACGTCTGAACTCTTTCCCTACACGACGCTCTTCCGATCTNNNTACATGCCCTCGTATGTTTCCT
GAGCACACGTCTGAACTCTTTCCCTACACGACGCTCTTCCGATCTNNNATACCCAAAGAAGCAAAGAG
GAGCACACGTCTGAACTCTTTCCCTACACGACGCTCTTCCGATCTNNNATACCCAAAGAAGCAAAGAG
AGCACACGTCTGAACTCTTTCCCTACACGACGCTCTTCCGATCTNNNTCCTGATGTTCTGAGCAAGAGT
AGCACACGTCTGAACTCTTTCCCTACACGACGCTCTTCCGATCTNNNTCCTGATGTTCTGAGCAAGAGT
AGCACACGTCTGAACTCTTTCCCTACACGACGCTCTTCCGATCTNNNCACACACGTACACACACTTACA
AGCACACGTCTGAACTCTTTCCCTACACGACGCTCTTCCGATCTNNNCACACACGTACACACACTTACA
AGCACACGTCTGAACTCTTTCCCTACACGACGCTCTTCCGATCTNNNCAGAGAGCAGAGATCACTTGG

AGCACACGTCTGAACTCTTTCCCTACACGACGCTCTTCCGATCTNNNCAGAGAGCAGAGATCACTTGG
AGCACACGTCTGAACTCTTTCCCTACACGACGCTCTTCCGATCTNNNCACCTGGCCCCTGCAATATATA
AGCACACGTCTGAACTCTTTCCCTACACGACGCTCTTCCGATCTNNNCACCTGGCCCCTGCAATATATA
GAGCACACGTCTGAACTCTTTCCCTACACGACGCTCTTCCGATCTNNNAGCAGTGTACAGGAATTCCAG
GAGCACACGTCTGAACTCTTTCCCTACACGACGCTCTTCCGATCTNNNAGCAGTGTACAGGAATTCCAG
CACACGTCTGAACTCTTTCCCTACACGACGCTCTTCCGATCTNNNCCCCTCCAAACTTCTAGAAGCTTCA
CACACGTCTGAACTCTTTCCCTACACGACGCTCTTCCGATCTNNNCCCCTCCAAACTTCTAGAAGCTTCA
AGCACACGTCTGAACTCTTTCCCTACACGACGCTCTTCCGATCTNNNTCCATTCATCTTTCCTCAGGTGG
AGCACACGTCTGAACTCTTTCCCTACACGACGCTCTTCCGATCTNNNTCCATTCATCTTTCCTCAGGTGG
GCACACGTCTGAACTCTTTCCCTACACGACGCTCTTCCGATCTNNNAAAAACTAAGCTTGTGTAAGAAT
GCACACGTCTGAACTCTTTCCCTACACGACGCTCTTCCGATCTNNNAAAAACTAAGCTTGTGTAAGAAT
AGCACACGTCTGAACTCTTTCCCTACACGACGCTCTTCCGATCTNNNTTCCTATTTGGCCATCTTGGGAA
AGCACACGTCTGAACTCTTTCCCTACACGACGCTCTTCCGATCTNNNTTCCTATTTGGCCATCTTGGGAA
GCACACGTCTGAACTCTTTCCCTACACGACGCTCTTCCGATCTNNNTAACAGAAGCAAAAGTGGGCATC
GCACACGTCTGAACTCTTTCCCTACACGACGCTCTTCCGATCTNNNTAACAGAAGCAAAAGTGGGCATC
GCACACGTCTGAACTCTTTCCCTACACGACGCTCTTCCGATCTNNNTGTTCATCTCCTCGTATCATTAGC
GCACACGTCTGAACTCTTTCCCTACACGACGCTCTTCCGATCTNNNTGTTCATCTCCTCGTATCATTAGC
GAGCACACGTCTGAACTCTTTCCCTACACGACGCTCTTCCGATCTNNNCAGAGGCAGGAGAATCACTTC
GAGCACACGTCTGAACTCTTTCCCTACACGACGCTCTTCCGATCTNNNCAGAGGCAGGAGAATCACTTC
AGCACACGTCTGAACTCTTTCCCTACACGACGCTCTTCCGATCTNNNTGGAATCTTCTGTGTTGTCTTT
AGCACACGTCTGAACTCTTTCCCTACACGACGCTCTTCCGATCTNNNTGGAATCTTCTGTGTTGTCTTT
AGCACACGTCTGAACTCTTTCCCTACACGACGCTCTTCCGATCTNNNGAGGTTGCTATGAGCCGAGAT
AGCACACGTCTGAACTCTTTCCCTACACGACGCTCTTCCGATCTNNNGAGGTTGCTATGAGCCGAGAT
AGCACACGTCTGAACTCTTTCCCTACACGACGCTCTTCCGATCTNNNATTGCTGGGGAACTAGTGTGAT
AGCACACGTCTGAACTCTTTCCCTACACGACGCTCTTCCGATCTNNNATTGCTGGGGAACTAGTGTGAT
GAGCACACGTCTGAACTCTTTCCCTACACGACGCTCTTCCGATCTNNNGCAGATGTCATCCTGATCAGT
GAGCACACGTCTGAACTCTTTCCCTACACGACGCTCTTCCGATCTNNNGCAGATGTCATCCTGATCAGT
AGCACACGTCTGAACTCTTTCCCTACACGACGCTCTTCCGATCTNNNCCCAGAGCAAGCAAAATCAAGA
AGCACACGTCTGAACTCTTTCCCTACACGACGCTCTTCCGATCTNNNCCCAGAGCAAGCAAAATCAAGA
GAGCACACGTCTGAACTCTTTCCCTACACGACGCTCTTCCGATCTNNNGAGCTGAGATCACGCCACTGC
GAGCACACGTCTGAACTCTTTCCCTACACGACGCTCTTCCGATCTNNNGAGCTGAGATCACGCCACTGC
CACACGTCTGAACTCTTTCCCTACACGACGCTCTTCCGATCTNNNCTTGGTAATGATCCTACAGATCATC
CACACGTCTGAACTCTTTCCCTACACGACGCTCTTCCGATCTNNNCTTGGTAATGATCCTACAGATCATC
GAGCACACGTCTGAACTCTTTCCCTACACGACGCTCTTCCGATCTNNNCACGTGGGTGTCTGTAAAGAA
GAGCACACGTCTGAACTCTTTCCCTACACGACGCTCTTCCGATCTNNNCACGTGGGTGTCTGTAAAGAA
AGAGCACACGTCTGAACTCTTTCCCTACACGACGCTCTTCCGATCTNNNGCAGACTTGTTTGCTTTCAGT
AGAGCACACGTCTGAACTCTTTCCCTACACGACGCTCTTCCGATCTNNNGCAGACTTGTTTGCTTTCAGT
AGCACACGTCTGAACTCTTTCCCTACACGACGCTCTTCCGATCTNNNTAATTACAGCAACCAGTTTTAT
AGCACACGTCTGAACTCTTTCCCTACACGACGCTCTTCCGATCTNNNTAATTACAGCAACCAGTTTTAT
GAGCACACGTCTGAACTCTTTCCCTACACGACGCTCTTCCGATCTNNNGTGAGGGAACTGGGAATCAA
GAGCACACGTCTGAACTCTTTCCCTACACGACGCTCTTCCGATCTNNNGTGAGGGAACTGGGAATCAA
GCACACGTCTGAACTCTTTCCCTACACGACGCTCTTCCGATCTNNNTCAAATTTCCTGCACACAGTTCCCA
GCACACGTCTGAACTCTTTCCCTACACGACGCTCTTCCGATCTNNNTCAAATTTCCTGCACACAGTTCCCA

GAGCACACGTCTGAACTCTTTCCCTACACGACGCTCTTCCGATCTNNNAGCATACATAGTCATGGACA
GAGCACACGTCTGAACTCTTTCCCTACACGACGCTCTTCCGATCTNNNAGCATACATAGTCATGGACA
CACACGTCTGAACTCTTTCCCTACACGACGCTCTTCCGATCTNNNTCCAGGAGCTAGCTTTTTTGACAC
CACACGTCTGAACTCTTTCCCTACACGACGCTCTTCCGATCTNNNTCCAGGAGCTAGCTTTTTTGACAC
AGAGCACACGTCTGAACTCTTTCCCTACACGACGCTCTTCCGATCTNNNGTAACATCTAAGCTACGGCC
AGAGCACACGTCTGAACTCTTTCCCTACACGACGCTCTTCCGATCTNNNGTAACATCTAAGCTACGGCC
GAGCACACGTCTGAACTCTTTCCCTACACGACGCTCTTCCGATCTNNNTGCACACCTATAATCCCAGCT
GAGCACACGTCTGAACTCTTTCCCTACACGACGCTCTTCCGATCTNNNTGCACACCTATAATCCCAGCT
AGCACACGTCTGAACTCTTTCCCTACACGACGCTCTTCCGATCTNNNGGTGACTTGTCCATGTTCAACA
AGCACACGTCTGAACTCTTTCCCTACACGACGCTCTTCCGATCTNNNGGTGACTTGTCCATGTTCAACA
AGCACACGTCTGAACTCTTTCCCTACACGACGCTCTTCCGATCTNNNTCCATTCTTTAGGTTGTCTCTTC
AGCACACGTCTGAACTCTTTCCCTACACGACGCTCTTCCGATCTNNNTCCATTCTTTAGGTTGTCTCTTC
AGCACACGTCTGAACTCTTTCCCTACACGACGCTCTTCCGATCTNNNTTTATCCACGAATGTAGCCTGG
AGCACACGTCTGAACTCTTTCCCTACACGACGCTCTTCCGATCTNNNTTTATCCACGAATGTAGCCTGG
ACACGTCTGAACTCTTTCCCTACACGACGCTCTTCCGATCTNNNTAAGAGAAGGTCATCCTCTCCCACA
ACACGTCTGAACTCTTTCCCTACACGACGCTCTTCCGATCTNNNTAAGAGAAGGTCATCCTCTCCCACA
AGCACACGTCTGAACTCTTTCCCTACACGACGCTCTTCCGATCTNNNTTCTGTTAGAGCAGTGTGGAAG
AGCACACGTCTGAACTCTTTCCCTACACGACGCTCTTCCGATCTNNNTTCTGTTAGAGCAGTGTGGAAG
AGCACACGTCTGAACTCTTTCCCTACACGACGCTCTTCCGATCTNNNCACATATGTCCTCCCTCCATAGC
AGCACACGTCTGAACTCTTTCCCTACACGACGCTCTTCCGATCTNNNCACATATGTCCTCCCTCCATAGC
AGCACACGTCTGAACTCTTTCCCTACACGACGCTCTTCCGATCTNNNACTGATGCTACTGGAGAATTG
AGCACACGTCTGAACTCTTTCCCTACACGACGCTCTTCCGATCTNNNACTGATGCTACTGGAGAATTG
ACACGTCTGAACTCTTTCCCTACACGACGCTCTTCCGATCTNNNTTCACCCCTCAAAAATATTGTAACAT
ACACGTCTGAACTCTTTCCCTACACGACGCTCTTCCGATCTNNNTTCACCCCTCAAAAATATTGTAACAT
GCACACGTCTGAACTCTTTCCCTACACGACGCTCTTCCGATCTNNNTTCGGCGAAATGATTTTCCACAC
GCACACGTCTGAACTCTTTCCCTACACGACGCTCTTCCGATCTNNNTTCGGCGAAATGATTTTCCACAC
AGCACACGTCTGAACTCTTTCCCTACACGACGCTCTTCCGATCTNNNACCTGTAGTCCCAGCTACTCGC
AGCACACGTCTGAACTCTTTCCCTACACGACGCTCTTCCGATCTNNNACCTGTAGTCCCAGCTACTCGC
AGCACACGTCTGAACTCTTTCCCTACACGACGCTCTTCCGATCTNNNAGAGGTTGCAGTAAGCTGAG
AGCACACGTCTGAACTCTTTCCCTACACGACGCTCTTCCGATCTNNNAGAGGTTGCAGTAAGCTGAG
AGCACACGTCTGAACTCTTTCCCTACACGACGCTCTTCCGATCTNNNCTGGAGTGCAGTGGTACAATC
AGCACACGTCTGAACTCTTTCCCTACACGACGCTCTTCCGATCTNNNCTGGAGTGCAGTGGTACAATC
ACACGTCTGAACTCTTTCCCTACACGACGCTCTTCCGATCTNNNGGAATTCAACAGTGAGGAGTGCAC
ACACGTCTGAACTCTTTCCCTACACGACGCTCTTCCGATCTNNNGGAATTCAACAGTGAGGAGTGCAC
GAGCACACGTCTGAACTCTTTCCCTACACGACGCTCTTCCGATCTNNNAGAAGTTGCAGTGAGCTGAG
GAGCACACGTCTGAACTCTTTCCCTACACGACGCTCTTCCGATCTNNNAGAAGTTGCAGTGAGCTGAG
AGCACACGTCTGAACTCTTTCCCTACACGACGCTCTTCCGATCTNNNTTGTTTTCTTCTTTCCCATTTAAC
AGCACACGTCTGAACTCTTTCCCTACACGACGCTCTTCCGATCTNNNTTGTTTTCTTCTTTCCCATTTAAC
AGCACACGTCTGAACTCTTTCCCTACACGACGCTCTTCCGATCTNNNCCAAAAGAGTAAAATCTCAATG
AGCACACGTCTGAACTCTTTCCCTACACGACGCTCTTCCGATCTNNNCCAAAAGAGTAAAATCTCAATG
AGCACACGTCTGAACTCTTTCCCTACACGACGCTCTTCCGATCTNNNGCTCCTTCTTATTTGGGTAGGC
AGCACACGTCTGAACTCTTTCCCTACACGACGCTCTTCCGATCTNNNGCTCCTTCTTATTTGGGTAGGC
AGCACACGTCTGAACTCTTTCCCTACACGACGCTCTTCCGATCTNNNTGTAAATACAGGGAAAGGCAG

AGCACACGTCTGAACTCTTTCCCTACACGACGCTCTTCCGATCTNNNTGTAAATACAGGGAAAGGCAG

CACACGTCTGAACTCTTTCCCTACACGACGCTCTTCCGATCTNNNCTTCCAAGTCCCACCTCAGCCATCGG

CACACGTCTGAACTCTTTCCCTACACGACGCTCTTCCGATCTNNNCTTCCAAGTCCCACCTCAGCCATCGG

AGCACACGTCTGAACTCTTTCCCTACACGACGCTCTTCCGATCTNNNGAAAGAAGGAAGGAAGGAGG

AGCACACGTCTGAACTCTTTCCCTACACGACGCTCTTCCGATCTNNNGAAAGAAGGAAGGAAGGAGG

AGCACACGTCTGAACTCTTTCCCTACACGACGCTCTTCCGATCTNNNCCTTAGTGTCTGGCTTAAATTCA

AGCACACGTCTGAACTCTTTCCCTACACGACGCTCTTCCGATCTNNNCCTTAGTGTCTGGCTTAAATTCA

AGCACACGTCTGAACTCTTTCCCTACACGACGCTCTTCCGATCTNNNTATTCGTGTGTGTGTACATGG

AGCACACGTCTGAACTCTTTCCCTACACGACGCTCTTCCGATCTNNNTATTCGTGTGTGTGTACATGG

GAGCACACGTCTGAACTCTTTCCCTACACGACGCTCTTCCGATCTNNNAGAAGAATGGCGAGAACCTG

GAGCACACGTCTGAACTCTTTCCCTACACGACGCTCTTCCGATCTNNNAGAAGAATGGCGAGAACCTG

GAGCACACGTCTGAACTCTTTCCCTACACGACGCTCTTCCGATCTNNNGCTATTTTGGCCTCTTGGGAA

GAGCACACGTCTGAACTCTTTCCCTACACGACGCTCTTCCGATCTNNNGCTATTTTGGCCTCTTGGGAA

GAGCACACGTCTGAACTCTTTCCCTACACGACGCTCTTCCGATCTNNNGGAGGATTTCTTGAGCCTAG

GAGCACACGTCTGAACTCTTTCCCTACACGACGCTCTTCCGATCTNNNGGAGGATTTCTTGAGCCTAG

AGCACACGTCTGAACTCTTTCCCTACACGACGCTCTTCCGATCTNNNCAAAGACACAAGTCCAGCTGAA

AGCACACGTCTGAACTCTTTCCCTACACGACGCTCTTCCGATCTNNNCAAAGACACAAGTCCAGCTGAA

GAGCACACGTCTGAACTCTTTCCCTACACGACGCTCTTCCGATCTNNNCAAGATCGTGCCACTACACTCT

GAGCACACGTCTGAACTCTTTCCCTACACGACGCTCTTCCGATCTNNNCAAGATCGTGCCACTACACTCT

AGCACACGTCTGAACTCTTTCCCTACACGACGCTCTTCCGATCTNNNCGCGCACTTTATTTTCCAGTCTA

AGCACACGTCTGAACTCTTTCCCTACACGACGCTCTTCCGATCTNNNCGCGCACTTTATTTTCCAGTCTA

AGAGCACACGTCTGAACTCTTTCCCTACACGACGCTCTTCCGATCTNNNCAAGGAGGCAGGAGATGGT

AGAGCACACGTCTGAACTCTTTCCCTACACGACGCTCTTCCGATCTNNNCAAGGAGGCAGGAGATGGT

AGCACACGTCTGAACTCTTTCCCTACACGACGCTCTTCCGATCTNNNTGCAGTGAGCCAAGATTATGCT

AGCACACGTCTGAACTCTTTCCCTACACGACGCTCTTCCGATCTNNNTGCAGTGAGCCAAGATTATGCT

AGCACACGTCTGAACTCTTTCCCTACACGACGCTCTTCCGATCTNNNTCTGTAATCCCAGCTACTCAGG

AGCACACGTCTGAACTCTTTCCCTACACGACGCTCTTCCGATCTNNNTCTGTAATCCCAGCTACTCAGG

AGCACACGTCTGAACTCTTTCCCTACACGACGCTCTTCCGATCTNNNTTTCTGCAACAAAGAGTGGTTC

AGCACACGTCTGAACTCTTTCCCTACACGACGCTCTTCCGATCTNNNTTTCTGCAACAAAGAGTGGTTC

GAGCACACGTCTGAACTCTTTCCCTACACGACGCTCTTCCGATCTNNNCTCCGGTAGGACAGGAGGG

GAGCACACGTCTGAACTCTTTCCCTACACGACGCTCTTCCGATCTNNNCTCCGGTAGGACAGGAGGG

GAGCACACGTCTGAACTCTTTCCCTACACGACGCTCTTCCGATCTNNNTATTCTGTTCTTATCCATGAG

GAGCACACGTCTGAACTCTTTCCCTACACGACGCTCTTCCGATCTNNNTATTCTGTTCTTATCCATGAG

GAGCACACGTCTGAACTCTTTCCCTACACGACGCTCTTCCGATCTNNNCCTTAAAAGGGAGTAACAATT

GAGCACACGTCTGAACTCTTTCCCTACACGACGCTCTTCCGATCTNNNCCTTAAAAGGGAGTAACAATT

AGCACACGTCTGAACTCTTTCCCTACACGACGCTCTTCCGATCTNNNTTTATTTCTTCTTTCACAACCCC

AGCACACGTCTGAACTCTTTCCCTACACGACGCTCTTCCGATCTNNNTTTATTTCTTCTTTCACAACCCC

GAGCACACGTCTGAACTCTTTCCCTACACGACGCTCTTCCGATCTNNNGGTGTGAGCAAGACCCCCAG

GAGCACACGTCTGAACTCTTTCCCTACACGACGCTCTTCCGATCTNNNGGTGTGAGCAAGACCCCCAG

GCACACGTCTGAACTCTTTCCCTACACGACGCTCTTCCGATCTNNNTCTTCTTTCCTCCCTCTTTCTGCCA

GCACACGTCTGAACTCTTTCCCTACACGACGCTCTTCCGATCTNNNTCTTCTTTCCTCCCTCTTTCTGCCA

AGCACACGTCTGAACTCTTTCCCTACACGACGCTCTTCCGATCTNNNTGAAATCAACATGTCAACAGGG

AGCACACGTCTGAACTCTTTCCCTACACGACGCTCTTCCGATCTNNNTGAAATCAACATGTCAACAGGG

AGAGCACACGTCTGAACTCTTTCCCTACACGACGCTCTTCCGATCTNNNACCTTCCTGACTTAGGCCAAT

AGAGCACACGTCTGAACTCTTTCCCTACACGACGCTCTTCCGATCTNNNACCTTCCTGACTTAGGCCAAT

GAGCACACGTCTGAACTCTTTCCCTACACGACGCTCTTCCGATCTNNNCTTAGGAGTTTGATGCTGCAG

GAGCACACGTCTGAACTCTTTCCCTACACGACGCTCTTCCGATCTNNNCTTAGGAGTTTGATGCTGCAG

AGAGCACACGTCTGAACTCTTTCCCTACACGACGCTCTTCCGATCTNNNGTTGCAGTGAGCTGAGATTC

AGAGCACACGTCTGAACTCTTTCCCTACACGACGCTCTTCCGATCTNNNGTTGCAGTGAGCTGAGATTC

GAGCACACGTCTGAACTCTTTCCCTACACGACGCTCTTCCGATCTNNNAGTGAATGGGTTTGTACAATA

GAGCACACGTCTGAACTCTTTCCCTACACGACGCTCTTCCGATCTNNNAGTGAATGGGTTTGTACAATA

AGCACACGTCTGAACTCTTTCCCTACACGACGCTCTTCCGATCTNNNTTTGGTGTGCATTCTTTGTGTG

AGCACACGTCTGAACTCTTTCCCTACACGACGCTCTTCCGATCTNNNTTTGGTGTGCATTCTTTGTGTG

AGAGCACACGTCTGAACTCTTTCCCTACACGACGCTCTTCCGATCTNNNTGCACATGAACTACCATACCA

AGAGCACACGTCTGAACTCTTTCCCTACACGACGCTCTTCCGATCTNNNTGCACATGAACTACCATACCA

AGCACACGTCTGAACTCTTTCCCTACACGACGCTCTTCCGATCTNNNCCATACCACTGACCCAAACTGTA

AGCACACGTCTGAACTCTTTCCCTACACGACGCTCTTCCGATCTNNNCCATACCACTGACCCAAACTGTA

GAGCACACGTCTGAACTCTTTCCCTACACGACGCTCTTCCGATCTNNNAGGTTATGAAGACTAACATG

GAGCACACGTCTGAACTCTTTCCCTACACGACGCTCTTCCGATCTNNNAGGTTATGAAGACTAACATG

AGCACACGTCTGAACTCTTTCCCTACACGACGCTCTTCCGATCTNNNACCTACTCATTTATCACCTTTGCA

AGCACACGTCTGAACTCTTTCCCTACACGACGCTCTTCCGATCTNNNACCTACTCATTTATCACCTTTGCA

CACACGTCTGAACTCTTTCCCTACACGACGCTCTTCCGATCTNNNTTCTTCCTTCTGGTGAGTTCCTGCAT

CACACGTCTGAACTCTTTCCCTACACGACGCTCTTCCGATCTNNNTTCTTCCTTCTGGTGAGTTCCTGCAT

GAGCACACGTCTGAACTCTTTCCCTACACGACGCTCTTCCGATCTNNNCCCACCACTACACCTAGCTAATC

GAGCACACGTCTGAACTCTTTCCCTACACGACGCTCTTCCGATCTNNNCCCACCACTACACCTAGCTAATC

AGAGCACACGTCTGAACTCTTTCCCTACACGACGCTCTTCCGATCTNNNGTTGCAGTGAGCTGAGATTC

AGAGCACACGTCTGAACTCTTTCCCTACACGACGCTCTTCCGATCTNNNGTTGCAGTGAGCTGAGATTC

GAGCACACGTCTGAACTCTTTCCCTACACGACGCTCTTCCGATCTNNNCTAGGAGCTCTTTGGGAGTCA

GAGCACACGTCTGAACTCTTTCCCTACACGACGCTCTTCCGATCTNNNCTAGGAGCTCTTTGGGAGTCA

AGCACACGTCTGAACTCTTTCCCTACACGACGCTCTTCCGATCTNNNGACACAGGTGGCATTTTGCTTA

AGCACACGTCTGAACTCTTTCCCTACACGACGCTCTTCCGATCTNNNGACACAGGTGGCATTTTGCTTA

AGCACACGTCTGAACTCTTTCCCTACACGACGCTCTTCCGATCTNNNCAGACATCTCCCCTTTACACCAA

AGCACACGTCTGAACTCTTTCCCTACACGACGCTCTTCCGATCTNNNCAGACATCTCCCCTTTACACCAA

GCACACGTCTGAACTCTTTCCCTACACGACGCTCTTCCGATCTNNNCATGCTCTTACCTGGGTGATGTCC

GCACACGTCTGAACTCTTTCCCTACACGACGCTCTTCCGATCTNNNCATGCTCTTACCTGGGTGATGTCC

AGCACACGTCTGAACTCTTTCCCTACACGACGCTCTTCCGATCTNNNAAGCACATCAGACTAAAACACG

AGCACACGTCTGAACTCTTTCCCTACACGACGCTCTTCCGATCTNNNAAGCACATCAGACTAAAACACG

AGAGCACACGTCTGAACTCTTTCCCTACACGACGCTCTTCCGATCTNNNTGAGACTCCACCTCAAAACAA

AGAGCACACGTCTGAACTCTTTCCCTACACGACGCTCTTCCGATCTNNNTGAGACTCCACCTCAAAACAA

GAGCACACGTCTGAACTCTTTCCCTACACGACGCTCTTCCGATCTNNNACTTTCTGGGAAATGCAACCTC

GAGCACACGTCTGAACTCTTTCCCTACACGACGCTCTTCCGATCTNNNACTTTCTGGGAAATGCAACCTC

GAGCACACGTCTGAACTCTTTCCCTACACGACGCTCTTCCGATCTNNNAAGAGATGCGGTTTCACCATCT

GAGCACACGTCTGAACTCTTTCCCTACACGACGCTCTTCCGATCTNNNAAGAGATGCGGTTTCACCATCT

GAGCACACGTCTGAACTCTTTCCCTACACGACGCTCTTCCGATCTNNNGGAGCTCAGGGACCGTGTAG

GAGCACACGTCTGAACTCTTTCCCTACACGACGCTCTTCCGATCTNNNGGAGCTCAGGGACCGTGTAG

AGCACACGTCTGAACTCTTTCCCTACACGACGCTCTTCCGATCTNNNACAGAGGTTTGTTCCCAAAGGA

AGCACACGTCTGAACTCTTTCCCTACACGACGCTCTTCCGATCTNNNACAGAGGTTTGTTCCCAAAGGA
GCACACGTCTGAACTCTTTCCCTACACGACGCTCTTCCGATCTNNNCGTAAGGAGAGGGATTTGGAGA
GCACACGTCTGAACTCTTTCCCTACACGACGCTCTTCCGATCTNNNCGTAAGGAGAGGGATTTGGAGA
GCACACGTCTGAACTCTTTCCCTACACGACGCTCTTCCGATCTNNNGGAAGGAAGAGAGGCAAAACAA
GCACACGTCTGAACTCTTTCCCTACACGACGCTCTTCCGATCTNNNGGAAGGAAGAGAGGCAAAACAA
AGCACACGTCTGAACTCTTTCCCTACACGACGCTCTTCCGATCTNNNGCCACAGTGGAAGAAGAATTG
AGCACACGTCTGAACTCTTTCCCTACACGACGCTCTTCCGATCTNNNGCCACAGTGGAAGAAGAATTG
AGCACACGTCTGAACTCTTTCCCTACACGACGCTCTTCCGATCTNNNGAGGATCTTTGAAGTGGGAGA
AGCACACGTCTGAACTCTTTCCCTACACGACGCTCTTCCGATCTNNNGAGGATCTTTGAAGTGGGAGA
GAGCACACGTCTGAACTCTTTCCCTACACGACGCTCTTCCGATCTNNNATAGGTGCAGGGATTGTGGA
GAGCACACGTCTGAACTCTTTCCCTACACGACGCTCTTCCGATCTNNNATAGGTGCAGGGATTGTGGA
AGCACACGTCTGAACTCTTTCCCTACACGACGCTCTTCCGATCTNNNCACCTGTAATCCCAGCTACTCAGC
AGCACACGTCTGAACTCTTTCCCTACACGACGCTCTTCCGATCTNNNCACCTGTAATCCCAGCTACTCAGC
AGCACACGTCTGAACTCTTTCCCTACACGACGCTCTTCCGATCTNNNAAACTCAACTCTGTAATAAGAAA
AGCACACGTCTGAACTCTTTCCCTACACGACGCTCTTCCGATCTNNNAAACTCAACTCTGTAATAAGAAA
AGCACACGTCTGAACTCTTTCCCTACACGACGCTCTTCCGATCTNNNGTTTCCTGTTTGTTTTGCCTTGCC
AGCACACGTCTGAACTCTTTCCCTACACGACGCTCTTCCGATCTNNNGTTTCCTGTTTGTTTTGCCTTGCC
GAGCACACGTCTGAACTCTTTCCCTACACGACGCTCTTCCGATCTNNNCCCGACAGGTGAGGGGTAAC
GAGCACACGTCTGAACTCTTTCCCTACACGACGCTCTTCCGATCTNNNCCCGACAGGTGAGGGGTAAC
GAGCACACGTCTGAACTCTTTCCCTACACGACGCTCTTCCGATCTNNNATCCTAGCTACTCAAGAGGCT
GAGCACACGTCTGAACTCTTTCCCTACACGACGCTCTTCCGATCTNNNATCCTAGCTACTCAAGAGGCT
GCACACGTCTGAACTCTTTCCCTACACGACGCTCTTCCGATCTNNNACTGTATAGAAGGTGGTCTCCATC
GCACACGTCTGAACTCTTTCCCTACACGACGCTCTTCCGATCTNNNACTGTATAGAAGGTGGTCTCCATC
AGCACACGTCTGAACTCTTTCCCTACACGACGCTCTTCCGATCTNNNATCCAATAGTTCCTCAACCCTCG
AGCACACGTCTGAACTCTTTCCCTACACGACGCTCTTCCGATCTNNNATCCAATAGTTCCTCAACCCTCG
GCACACGTCTGAACTCTTTCCCTACACGACGCTCTTCCGATCTNNNGAAAACAATGTGTGGCCTCAAGT
GCACACGTCTGAACTCTTTCCCTACACGACGCTCTTCCGATCTNNNGAAAACAATGTGTGGCCTCAAGT
GCACACGTCTGAACTCTTTCCCTACACGACGCTCTTCCGATCTNNNTTCTGTTGCTCCTTCTGTCTCTCCAT
GCACACGTCTGAACTCTTTCCCTACACGACGCTCTTCCGATCTNNNTTCTGTTGCTCCTTCTGTCTCTCCAT
AGAGCACACGTCTGAACTCTTTCCCTACACGACGCTCTTCCGATCTNNNCCTTCAGCAATGGTGTATGCA
AGAGCACACGTCTGAACTCTTTCCCTACACGACGCTCTTCCGATCTNNNCCTTCAGCAATGGTGTATGCA
AGCACACGTCTGAACTCTTTCCCTACACGACGCTCTTCCGATCTNNNATAATTTACCCCCAGAAGCTCC
AGCACACGTCTGAACTCTTTCCCTACACGACGCTCTTCCGATCTNNNATAATTTACCCCCAGAAGCTCC
GAGCACACGTCTGAACTCTTTCCCTACACGACGCTCTTCCGATCTNNNTCTCCTTCTCCCCTAGGCAATTC
GAGCACACGTCTGAACTCTTTCCCTACACGACGCTCTTCCGATCTNNNTCTCCTTCTCCCCTAGGCAATTC
GAGCACACGTCTGAACTCTTTCCCTACACGACGCTCTTCCGATCTNNNTGGGGATGATGGAAGTGTTG
GAGCACACGTCTGAACTCTTTCCCTACACGACGCTCTTCCGATCTNNNTGGGGATGATGGAAGTGTTG
AGCACACGTCTGAACTCTTTCCCTACACGACGCTCTTCCGATCTNNNTCAAACAATTCTCCACAACTGGC
AGCACACGTCTGAACTCTTTCCCTACACGACGCTCTTCCGATCTNNNTCAAACAATTCTCCACAACTGGC
AGCACACGTCTGAACTCTTTCCCTACACGACGCTCTTCCGATCTNNNACAGAGAGAGACCCTGTATCCA
AGCACACGTCTGAACTCTTTCCCTACACGACGCTCTTCCGATCTNNNACAGAGAGAGACCCTGTATCCA
AGAGCACACGTCTGAACTCTTTCCCTACACGACGCTCTTCCGATCTNNNTGGTGTACATCTGTAATCCCA
AGAGCACACGTCTGAACTCTTTCCCTACACGACGCTCTTCCGATCTNNNTGGTGTACATCTGTAATCCCA

GAGCACACGTCTGAACTCTTTCCCTACACGACGCTCTTCCGATCTNNNGAGGTTGCAGTAAGCCAAGA
GAGCACACGTCTGAACTCTTTCCCTACACGACGCTCTTCCGATCTNNNGAGGTTGCAGTAAGCCAAGA
GAGCACACGTCTGAACTCTTTCCCTACACGACGCTCTTCCGATCTNNNAGGCTTTCATCTTTTCCCACTC
GAGCACACGTCTGAACTCTTTCCCTACACGACGCTCTTCCGATCTNNNAGGCTTTCATCTTTTCCCACTC
ACACGTCTGAACTCTTTCCCTACACGACGCTCTTCCGATCTNNNTTCTCAATGAGCTGCTGGGTACCATC
ACACGTCTGAACTCTTTCCCTACACGACGCTCTTCCGATCTNNNTTCTCAATGAGCTGCTGGGTACCATC
AGCACACGTCTGAACTCTTTCCCTACACGACGCTCTTCCGATCTNNNGAGATTGCAGTGAGCCAAGAT
AGCACACGTCTGAACTCTTTCCCTACACGACGCTCTTCCGATCTNNNGAGATTGCAGTGAGCCAAGAT
GAGCACACGTCTGAACTCTTTCCCTACACGACGCTCTTCCGATCTNNNGAGAATCCCTTGAACCCAGGA
GAGCACACGTCTGAACTCTTTCCCTACACGACGCTCTTCCGATCTNNNGAGAATCCCTTGAACCCAGGA
GAGCACACGTCTGAACTCTTTCCCTACACGACGCTCTTCCGATCTNNNGAGCTTGTAGTGAGCCGAGAT
GAGCACACGTCTGAACTCTTTCCCTACACGACGCTCTTCCGATCTNNNGAGCTTGTAGTGAGCCGAGAT
GAGCACACGTCTGAACTCTTTCCCTACACGACGCTCTTCCGATCTNNNCCACCCAGTACTTCCCTTCTAAC
GAGCACACGTCTGAACTCTTTCCCTACACGACGCTCTTCCGATCTNNNCCACCCAGTACTTCCCTTCTAAC
AGAGCACACGTCTGAACTCTTTCCCTACACGACGCTCTTCCGATCTNNNTATCCAGGAGCCCACCAAAA
AGAGCACACGTCTGAACTCTTTCCCTACACGACGCTCTTCCGATCTNNNTATCCAGGAGCCCACCAAAA
AGCACACGTCTGAACTCTTTCCCTACACGACGCTCTTCCGATCTNNNCTGTAGTCCCAGCTACTCAGGA
AGCACACGTCTGAACTCTTTCCCTACACGACGCTCTTCCGATCTNNNCTGTAGTCCCAGCTACTCAGGA
GAGCACACGTCTGAACTCTTTCCCTACACGACGCTCTTCCGATCTNNNACTCCCTGGCTACTACTGATGA
GAGCACACGTCTGAACTCTTTCCCTACACGACGCTCTTCCGATCTNNNACTCCCTGGCTACTACTGATGA
GAGCACACGTCTGAACTCTTTCCCTACACGACGCTCTTCCGATCTNNNACCGCATGTTCTCACTCATAAC
GAGCACACGTCTGAACTCTTTCCCTACACGACGCTCTTCCGATCTNNNACCGCATGTTCTCACTCATAAC
GAGCACACGTCTGAACTCTTTCCCTACACGACGCTCTTCCGATCTNNNAAAAGAGGAGCAAGCATGAA
GAGCACACGTCTGAACTCTTTCCCTACACGACGCTCTTCCGATCTNNNAAAAGAGGAGCAAGCATGAA
GCACACGTCTGAACTCTTTCCCTACACGACGCTCTTCCGATCTNNNCAAAGACAGAACTTAACCTCTCAT
GCACACGTCTGAACTCTTTCCCTACACGACGCTCTTCCGATCTNNNCAAAGACAGAACTTAACCTCTCAT
GCACACGTCTGAACTCTTTCCCTACACGACGCTCTTCCGATCTNNNGTTTCTCACTTTAGCCAATTCCATC
GCACACGTCTGAACTCTTTCCCTACACGACGCTCTTCCGATCTNNNGTTTCTCACTTTAGCCAATTCCATC
GCACACGTCTGAACTCTTTCCCTACACGACGCTCTTCCGATCTNNNCTGGGCAACAAGAGCAAAACTCC
GCACACGTCTGAACTCTTTCCCTACACGACGCTCTTCCGATCTNNNCTGGGCAACAAGAGCAAAACTCC
GAGCACACGTCTGAACTCTTTCCCTACACGACGCTCTTCCGATCTNNNTGAGTGTTTCCCATCTCTCTTG
GAGCACACGTCTGAACTCTTTCCCTACACGACGCTCTTCCGATCTNNNTGAGTGTTTCCCATCTCTCTTG
AGCACACGTCTGAACTCTTTCCCTACACGACGCTCTTCCGATCTNNNTGCTGAGCACATTTTCATGTACC
AGCACACGTCTGAACTCTTTCCCTACACGACGCTCTTCCGATCTNNNTGCTGAGCACATTTTCATGTACC
AGCACACGTCTGAACTCTTTCCCTACACGACGCTCTTCCGATCTNNNACCATCTCTGACAGTTTCCCTTCC
AGCACACGTCTGAACTCTTTCCCTACACGACGCTCTTCCGATCTNNNACCATCTCTGACAGTTTCCCTTCC
GAGCACACGTCTGAACTCTTTCCCTACACGACGCTCTTCCGATCTNNNATCCCTCATTCATCTACCACGTC
GAGCACACGTCTGAACTCTTTCCCTACACGACGCTCTTCCGATCTNNNATCCCTCATTCATCTACCACGTC
GAGCACACGTCTGAACTCTTTCCCTACACGACGCTCTTCCGATCTNNNTAGCTTGTCTTCTCCTTCTGTG
GAGCACACGTCTGAACTCTTTCCCTACACGACGCTCTTCCGATCTNNNTAGCTTGTCTTCTCCTTCTGTG
GAGCACACGTCTGAACTCTTTCCCTACACGACGCTCTTCCGATCTNNNGAGGCCTGGTTGTCTTTAGTA
GAGCACACGTCTGAACTCTTTCCCTACACGACGCTCTTCCGATCTNNNGAGGCCTGGTTGTCTTTAGTA
AGCACACGTCTGAACTCTTTCCCTACACGACGCTCTTCCGATCTNNNATGCCTGTAATCCCAGCTACTTC

AGCACACGTCTGAACTCTTTCCCTACACGACGCTCTTCCGATCTNNNATGCCTGTAATCCCAGCTACTTG
GAGCACACGTCTGAACTCTTTCCCTACACGACGCTCTTCCGATCTNNNACAAAATTTGGTTGTGGTGC
GAGCACACGTCTGAACTCTTTCCCTACACGACGCTCTTCCGATCTNNNACAAAATTTGGTTGTGGTGC
GAGCACACGTCTGAACTCTTTCCCTACACGACGCTCTTCCGATCTNNNCATTGCATTACAGCCTGGTTG
GAGCACACGTCTGAACTCTTTCCCTACACGACGCTCTTCCGATCTNNNCATTGCATTACAGCCTGGTTG
GAGCACACGTCTGAACTCTTTCCCTACACGACGCTCTTCCGATCTNNNATTTATTCCAGTGAGTGCCCA
GAGCACACGTCTGAACTCTTTCCCTACACGACGCTCTTCCGATCTNNNATTTATTCCAGTGAGTGCCCA
GCACACGTCTGAACTCTTTCCCTACACGACGCTCTTCCGATCTNNNCTCTTGCACTGAATTGTTCCTGG
GCACACGTCTGAACTCTTTCCCTACACGACGCTCTTCCGATCTNNNCTCTTGCACTGAATTGTTCCTGG
AGCACACGTCTGAACTCTTTCCCTACACGACGCTCTTCCGATCTNNNTGAGAACTTTTGGGGGTCTGT
AGCACACGTCTGAACTCTTTCCCTACACGACGCTCTTCCGATCTNNNTGAGAACTTTTGGGGGTCTGT
AGCACACGTCTGAACTCTTTCCCTACACGACGCTCTTCCGATCTNNNTGAGCTGAGATCACACTATTG
AGCACACGTCTGAACTCTTTCCCTACACGACGCTCTTCCGATCTNNNTGAGCTGAGATCACACTATTG
GAGCACACGTCTGAACTCTTTCCCTACACGACGCTCTTCCGATCTNNNATTTATGAGGCTGTGCAATG
GAGCACACGTCTGAACTCTTTCCCTACACGACGCTCTTCCGATCTNNNATTTATGAGGCTGTGCAATG
GAGCACACGTCTGAACTCTTTCCCTACACGACGCTCTTCCGATCTNNNGCAATATGGCAAAACCCTGTC
GAGCACACGTCTGAACTCTTTCCCTACACGACGCTCTTCCGATCTNNNGCAATATGGCAAAACCCTGTC
GCACACGTCTGAACTCTTTCCCTACACGACGCTCTTCCGATCTNNNCCTGTAGTCCCAGCTACTTGGCAT
GCACACGTCTGAACTCTTTCCCTACACGACGCTCTTCCGATCTNNNCCTGTAGTCCCAGCTACTTGGCAT
AGCACACGTCTGAACTCTTTCCCTACACGACGCTCTTCCGATCTNNNCTCACCTGTAATCTCAGCACT
AGCACACGTCTGAACTCTTTCCCTACACGACGCTCTTCCGATCTNNNCTCACCTGTAATCTCAGCACT
GAGCACACGTCTGAACTCTTTCCCTACACGACGCTCTTCCGATCTNNNCACACACACACACACACACATA
GAGCACACGTCTGAACTCTTTCCCTACACGACGCTCTTCCGATCTNNNCACACACACACACACACACATA
GCACACGTCTGAACTCTTTCCCTACACGACGCTCTTCCGATCTNNNCTTGGCAAAAGAGCAAGACCTTG
GCACACGTCTGAACTCTTTCCCTACACGACGCTCTTCCGATCTNNNCTTGGCAAAAGAGCAAGACCTTG
GAGCACACGTCTGAACTCTTTCCCTACACGACGCTCTTCCGATCTNNNCAGGACTACACAGGCTACACT
GAGCACACGTCTGAACTCTTTCCCTACACGACGCTCTTCCGATCTNNNCAGGACTACACAGGCTACACT
GAGCACACGTCTGAACTCTTTCCCTACACGACGCTCTTCCGATCTNNNATAGGCCCCAAAGCAGTAGTA
GAGCACACGTCTGAACTCTTTCCCTACACGACGCTCTTCCGATCTNNNATAGGCCCCAAAGCAGTAGTA
GCACACGTCTGAACTCTTTCCCTACACGACGCTCTTCCGATCTNNNGCAAACCTGCCTCCCATTTTATTCA
GCACACGTCTGAACTCTTTCCCTACACGACGCTCTTCCGATCTNNNGCAAACCTGCCTCCCATTTTATTCA
AGCACACGTCTGAACTCTTTCCCTACACGACGCTCTTCCGATCTNNNACAGTGAAACCCCCTCTCTACTA
AGCACACGTCTGAACTCTTTCCCTACACGACGCTCTTCCGATCTNNNACAGTGAAACCCCCTCTCTACTA
AGCACACGTCTGAACTCTTTCCCTACACGACGCTCTTCCGATCTNNNATCTTTATACCTATGCTGGAAA
AGCACACGTCTGAACTCTTTCCCTACACGACGCTCTTCCGATCTNNNATCTTTATACCTATGCTGGAAA
AGCACACGTCTGAACTCTTTCCCTACACGACGCTCTTCCGATCTNNNATTAATGTGGATGATGAGATTA
AGCACACGTCTGAACTCTTTCCCTACACGACGCTCTTCCGATCTNNNATTAATGTGGATGATGAGATTA
AGAGCACACGTCTGAACTCTTTCCCTACACGACGCTCTTCCGATCTNNNGAAGAAGAAGCAGGCACA
AGAGCACACGTCTGAACTCTTTCCCTACACGACGCTCTTCCGATCTNNNGAAGAAGAAGCAGGCACA
GAGCACACGTCTGAACTCTTTCCCTACACGACGCTCTTCCGATCTNNNTTGGAGTAGGCTGAGATTATC
GAGCACACGTCTGAACTCTTTCCCTACACGACGCTCTTCCGATCTNNNTTGGAGTAGGCTGAGATTATC
AGCACACGTCTGAACTCTTTCCCTACACGACGCTCTTCCGATCTNNNCGCCTGTAATTTCAGCTACTTC
AGCACACGTCTGAACTCTTTCCCTACACGACGCTCTTCCGATCTNNNCGCCTGTAATTTCAGCTACTTC

AGAGCACACGTCTGAACTCTTTCCCTACACGACGCTCTTCCGATCTNNNGAGGTGGTAGGGAGTAAAT
AGAGCACACGTCTGAACTCTTTCCCTACACGACGCTCTTCCGATCTNNNGAGGTGGTAGGGAGTAAAT
CACACGTCTGAACTCTTTCCCTACACGACGCTCTTCCGATCTNNNCTGTTCTTGGTTTCACAGATCATCG
CACACGTCTGAACTCTTTCCCTACACGACGCTCTTCCGATCTNNNCTGTTCTTGGTTTCACAGATCATCG
AGCACACGTCTGAACTCTTTCCCTACACGACGCTCTTCCGATCTNNNATATACCCAACTACTCTGCATAT
AGCACACGTCTGAACTCTTTCCCTACACGACGCTCTTCCGATCTNNNATATACCCAACTACTCTGCATAT
AGCACACGTCTGAACTCTTTCCCTACACGACGCTCTTCCGATCTNNNTTGGAGAAAGGTCTATCTGCTG
AGCACACGTCTGAACTCTTTCCCTACACGACGCTCTTCCGATCTNNNTTGGAGAAAGGTCTATCTGCTG
AGCACACGTCTGAACTCTTTCCCTACACGACGCTCTTCCGATCTNNNCTTGCAGTGAGCCGAGATAGCC
AGCACACGTCTGAACTCTTTCCCTACACGACGCTCTTCCGATCTNNNCTTGCAGTGAGCCGAGATAGCC
GCACACGTCTGAACTCTTTCCCTACACGACGCTCTTCCGATCTNNNAGGATGTAGGGTAACAGGGAAG
GCACACGTCTGAACTCTTTCCCTACACGACGCTCTTCCGATCTNNNAGGATGTAGGGTAACAGGGAAG
AGAGCACACGTCTGAACTCTTTCCCTACACGACGCTCTTCCGATCTNNNCTGTAGTTCCAGCTACTCGGG
AGAGCACACGTCTGAACTCTTTCCCTACACGACGCTCTTCCGATCTNNNCTGTAGTTCCAGCTACTCGGG
AGCACACGTCTGAACTCTTTCCCTACACGACGCTCTTCCGATCTNNNTGTTTCAATTTCTTGATCTGAAG
AGCACACGTCTGAACTCTTTCCCTACACGACGCTCTTCCGATCTNNNTGTTTCAATTTCTTGATCTGAAG
AGCACACGTCTGAACTCTTTCCCTACACGACGCTCTTCCGATCTNNNATGTAAAATGTTTGCTTAGCAT
AGCACACGTCTGAACTCTTTCCCTACACGACGCTCTTCCGATCTNNNATGTAAAATGTTTGCTTAGCAT
AGCACACGTCTGAACTCTTTCCCTACACGACGCTCTTCCGATCTNNNCAGAACAGAGACTGACATCTCC
AGCACACGTCTGAACTCTTTCCCTACACGACGCTCTTCCGATCTNNNCAGAACAGAGACTGACATCTCC
AGAGCACACGTCTGAACTCTTTCCCTACACGACGCTCTTCCGATCTNNNCTGTAGTCCCAGCTACTCAGG
AGAGCACACGTCTGAACTCTTTCCCTACACGACGCTCTTCCGATCTNNNCTGTAGTCCCAGCTACTCAGG
GAGCACACGTCTGAACTCTTTCCCTACACGACGCTCTTCCGATCTNNNAGGAGAATGGCATGAACCCGC
GAGCACACGTCTGAACTCTTTCCCTACACGACGCTCTTCCGATCTNNNAGGAGAATGGCATGAACCCGC
GAGCACACGTCTGAACTCTTTCCCTACACGACGCTCTTCCGATCTNNNCATATGCTAGTTGGAGGTGA
GAGCACACGTCTGAACTCTTTCCCTACACGACGCTCTTCCGATCTNNNCATATGCTAGTTGGAGGTGA
GAGCACACGTCTGAACTCTTTCCCTACACGACGCTCTTCCGATCTNNNGAGGCTGAGGCAGGAGAATG
GAGCACACGTCTGAACTCTTTCCCTACACGACGCTCTTCCGATCTNNNGAGGCTGAGGCAGGAGAATG
ACACGTCTGAACTCTTTCCCTACACGACGCTCTTCCGATCTNNNCACCTCTCCGCTCAAGGAAGCATCGC
ACACGTCTGAACTCTTTCCCTACACGACGCTCTTCCGATCTNNNCACCTCTCCGCTCAAGGAAGCATCGC
GAGCACACGTCTGAACTCTTTCCCTACACGACGCTCTTCCGATCTNNNGTGTTTCACTATAGGCAGCTG
GAGCACACGTCTGAACTCTTTCCCTACACGACGCTCTTCCGATCTNNNGTGTTTCACTATAGGCAGCTG
GCACACGTCTGAACTCTTTCCCTACACGACGCTCTTCCGATCTNNNGTATTACCAGCTACTTGGGAGGC
GCACACGTCTGAACTCTTTCCCTACACGACGCTCTTCCGATCTNNNGTATTACCAGCTACTTGGGAGGC
AGCACACGTCTGAACTCTTTCCCTACACGACGCTCTTCCGATCTNNNAGTGAGCCAGAAGTCAGACATT
AGCACACGTCTGAACTCTTTCCCTACACGACGCTCTTCCGATCTNNNAGTGAGCCAGAAGTCAGACATT
AGAGCACACGTCTGAACTCTTTCCCTACACGACGCTCTTCCGATCTNNNTAAATGGTTCTGTCTAGCTGC
AGAGCACACGTCTGAACTCTTTCCCTACACGACGCTCTTCCGATCTNNNTAAATGGTTCTGTCTAGCTGC
AGAGCACACGTCTGAACTCTTTCCCTACACGACGCTCTTCCGATCTNNNGTGTGTGTATGGTGTGTAT
AGAGCACACGTCTGAACTCTTTCCCTACACGACGCTCTTCCGATCTNNNGTGTGTGTATGGTGTGTAT
AGCACACGTCTGAACTCTTTCCCTACACGACGCTCTTCCGATCTNNNAGAATCACTTGAACCTGGGAGA
AGCACACGTCTGAACTCTTTCCCTACACGACGCTCTTCCGATCTNNNAGAATCACTTGAACCTGGGAGA
GCACACGTCTGAACTCTTTCCCTACACGACGCTCTTCCGATCTNNNAATACAAAAAATTAGCCGGGCGT

GCACACGTCTGAACTCTTTCCCTACACGACGCTCTTCCGATCTNNNAATACAAAAAATTAGCCGGGCG
AGCACACGTCTGAACTCTTTCCCTACACGACGCTCTTCCGATCTNNNTGTTAGGATCTACACATCTGCC
AGCACACGTCTGAACTCTTTCCCTACACGACGCTCTTCCGATCTNNNTGTTAGGATCTACACATCTGCC
GCACACGTCTGAACTCTTTCCCTACACGACGCTCTTCCGATCTNNNAGTTTGGAGGTACAAAGGGTTG
GCACACGTCTGAACTCTTTCCCTACACGACGCTCTTCCGATCTNNNAGTTTGGAGGTACAAAGGGTTG
AGCACACGTCTGAACTCTTTCCCTACACGACGCTCTTCCGATCTNNNAAAGGATCAAGGGAATTGGGG
AGCACACGTCTGAACTCTTTCCCTACACGACGCTCTTCCGATCTNNNAAAGGATCAAGGGAATTGGGG
AGCACACGTCTGAACTCTTTCCCTACACGACGCTCTTCCGATCTNNNTACTTCTGCCATGGAGAGAATG
AGCACACGTCTGAACTCTTTCCCTACACGACGCTCTTCCGATCTNNNTACTTCTGCCATGGAGAGAATG
AGCACACGTCTGAACTCTTTCCCTACACGACGCTCTTCCGATCTNNNACACAAAAAATTAGCCACGTGT
AGCACACGTCTGAACTCTTTCCCTACACGACGCTCTTCCGATCTNNNACACAAAAAATTAGCCACGTGT
GAGCACACGTCTGAACTCTTTCCCTACACGACGCTCTTCCGATCTNNNTCAAGTTGAGAGCCAAATCAG
GAGCACACGTCTGAACTCTTTCCCTACACGACGCTCTTCCGATCTNNNTCAAGTTGAGAGCCAAATCAG
GAGCACACGTCTGAACTCTTTCCCTACACGACGCTCTTCCGATCTNNNACCAACAACCAGCAATCATTG
GAGCACACGTCTGAACTCTTTCCCTACACGACGCTCTTCCGATCTNNNACCAACAACCAGCAATCATTG
GAGCACACGTCTGAACTCTTTCCCTACACGACGCTCTTCCGATCTNNNGGAAGCCTGAAGGAAAGAG
GAGCACACGTCTGAACTCTTTCCCTACACGACGCTCTTCCGATCTNNNGGAAGCCTGAAGGAAAGAG
AGCACACGTCTGAACTCTTTCCCTACACGACGCTCTTCCGATCTNNNGGAGGTAGAGCTTGCAGTGAG
AGCACACGTCTGAACTCTTTCCCTACACGACGCTCTTCCGATCTNNNGGAGGTAGAGCTTGCAGTGAG
AGAGCACACGTCTGAACTCTTTCCCTACACGACGCTCTTCCGATCTNNNACCCAGTTGCAGTACACATAT
AGAGCACACGTCTGAACTCTTTCCCTACACGACGCTCTTCCGATCTNNNACCCAGTTGCAGTACACATAT
GAGCACACGTCTGAACTCTTTCCCTACACGACGCTCTTCCGATCTNNNTGAGTTGAATCAGCTTTCCAG
GAGCACACGTCTGAACTCTTTCCCTACACGACGCTCTTCCGATCTNNNTGAGTTGAATCAGCTTTCCAG
GAGCACACGTCTGAACTCTTTCCCTACACGACGCTCTTCCGATCTNNNCTAGGTAGTGCTGGGCTTATG
GAGCACACGTCTGAACTCTTTCCCTACACGACGCTCTTCCGATCTNNNCTAGGTAGTGCTGGGCTTATG
AGAGCACACGTCTGAACTCTTTCCCTACACGACGCTCTTCCGATCTNNNTGAGGCAGGAGAATCACTT
AGAGCACACGTCTGAACTCTTTCCCTACACGACGCTCTTCCGATCTNNNTGAGGCAGGAGAATCACTT
AGAGCACACGTCTGAACTCTTTCCCTACACGACGCTCTTCCGATCTNNNTTATTGCACTGGAGAGATGG
AGAGCACACGTCTGAACTCTTTCCCTACACGACGCTCTTCCGATCTNNNTTATTGCACTGGAGAGATGG
AGCACACGTCTGAACTCTTTCCCTACACGACGCTCTTCCGATCTNNNGGTCAGTCTAAGCCTCCTGTTTT
AGCACACGTCTGAACTCTTTCCCTACACGACGCTCTTCCGATCTNNNGGTCAGTCTAAGCCTCCTGTTTT
AGCACACGTCTGAACTCTTTCCCTACACGACGCTCTTCCGATCTNNNAGATGGCTGTGAAATAGGAGT
AGCACACGTCTGAACTCTTTCCCTACACGACGCTCTTCCGATCTNNNAGATGGCTGTGAAATAGGAGT
GAGCACACGTCTGAACTCTTTCCCTACACGACGCTCTTCCGATCTNNNCTGTGATTTTGCCACTGTACTC
GAGCACACGTCTGAACTCTTTCCCTACACGACGCTCTTCCGATCTNNNCTGTGATTTTGCCACTGTACTC
GCACACGTCTGAACTCTTTCCCTACACGACGCTCTTCCGATCTNNNGTTTGCAGATCACATGTACACCC
GCACACGTCTGAACTCTTTCCCTACACGACGCTCTTCCGATCTNNNGTTTGCAGATCACATGTACACCC
AGCACACGTCTGAACTCTTTCCCTACACGACGCTCTTCCGATCTNNNCAATCATGGTGGAAGGTGAAA
AGCACACGTCTGAACTCTTTCCCTACACGACGCTCTTCCGATCTNNNCAATCATGGTGGAAGGTGAAA
GAGCACACGTCTGAACTCTTTCCCTACACGACGCTCTTCCGATCTNNNACGTTTGAGAACCTCACATGG
GAGCACACGTCTGAACTCTTTCCCTACACGACGCTCTTCCGATCTNNNACGTTTGAGAACCTCACATGG
GAGCACACGTCTGAACTCTTTCCCTACACGACGCTCTTCCGATCTNNNCCTGGTCTCTCTCTCTCTCTC
GAGCACACGTCTGAACTCTTTCCCTACACGACGCTCTTCCGATCTNNNCCTGGTCTCTCTCTCTCTCTC

AGCACACGTCTGAACTCTTTCCCTACACGACGCTCTTCCGATCTNNNTGGACTGGCAAAGCAGAATA

AGCACACGTCTGAACTCTTTCCCTACACGACGCTCTTCCGATCTNNNTGGACTGGCAAAGCAGAATA

GCACACGTCTGAACTCTTTCCCTACACGACGCTCTTCCGATCTNNNCTTCTGGCTCTTTCTGCTTCAAC

GCACACGTCTGAACTCTTTCCCTACACGACGCTCTTCCGATCTNNNCTTCTGGCTCTTTCTGCTTCAAC

GCACACGTCTGAACTCTTTCCCTACACGACGCTCTTCCGATCTNNNTCTACTTGCTGCTCCAAGAACAA

GCACACGTCTGAACTCTTTCCCTACACGACGCTCTTCCGATCTNNNTCTACTTGCTGCTCCAAGAACAA

AGCACACGTCTGAACTCTTTCCCTACACGACGCTCTTCCGATCTNNNAGGCAGGAGAATTACTTGAA

AGCACACGTCTGAACTCTTTCCCTACACGACGCTCTTCCGATCTNNNAGGCAGGAGAATTACTTGAA

AGCACACGTCTGAACTCTTTCCCTACACGACGCTCTTCCGATCTNNNCCACTCTGTTGACTAGGATGG

AGCACACGTCTGAACTCTTTCCCTACACGACGCTCTTCCGATCTNNNCCACTCTGTTGACTAGGATGG

AGCACACGTCTGAACTCTTTCCCTACACGACGCTCTTCCGATCTNNNTGGTGCTGACCTCATGAAGTA

AGCACACGTCTGAACTCTTTCCCTACACGACGCTCTTCCGATCTNNNTGGTGCTGACCTCATGAAGTA

GCACACGTCTGAACTCTTTCCCTACACGACGCTCTTCCGATCTNNNCCAGCCATTGTTCAACAAACCTA

GCACACGTCTGAACTCTTTCCCTACACGACGCTCTTCCGATCTNNNCCAGCCATTGTTCAACAAACCTA

GAGCACACGTCTGAACTCTTTCCCTACACGACGCTCTTCCGATCTNNNTAGGAATCGACTGTGTCTTC

GAGCACACGTCTGAACTCTTTCCCTACACGACGCTCTTCCGATCTNNNTAGGAATCGACTGTGTCTTC

GAGCACACGTCTGAACTCTTTCCCTACACGACGCTCTTCCGATCTNNNTCCTTGGAGTCTCACCTTGG

GAGCACACGTCTGAACTCTTTCCCTACACGACGCTCTTCCGATCTNNNTCCTTGGAGTCTCACCTTGG

GAGCACACGTCTGAACTCTTTCCCTACACGACGCTCTTCCGATCTNNNCTCATTTCATCACTGTTTGGC

GAGCACACGTCTGAACTCTTTCCCTACACGACGCTCTTCCGATCTNNNCTCATTTCATCACTGTTTGGC

AGCACACGTCTGAACTCTTTCCCTACACGACGCTCTTCCGATCTNNNCAAAGGAGACACTCATGGCA

AGCACACGTCTGAACTCTTTCCCTACACGACGCTCTTCCGATCTNNNCAAAGGAGACACTCATGGCA

CACGTCTGAACTCTTTCCCTACACGACGCTCTTCCGATCTNNNAAAAATGTTCTCTAGTGCCAAACTCA

CACGTCTGAACTCTTTCCCTACACGACGCTCTTCCGATCTNNNAAAAATGTTCTCTAGTGCCAAACTCA

AGCACACGTCTGAACTCTTTCCCTACACGACGCTCTTCCGATCTNNNAAGCGACTCTATGCAAAGAGA

AGCACACGTCTGAACTCTTTCCCTACACGACGCTCTTCCGATCTNNNAAGCGACTCTATGCAAAGAGA

AGCACACGTCTGAACTCTTTCCCTACACGACGCTCTTCCGATCTNNNTGGATTACAGGTGTGAGTCACT

AGCACACGTCTGAACTCTTTCCCTACACGACGCTCTTCCGATCTNNNTGGATTACAGGTGTGAGTCACT

AGCACACGTCTGAACTCTTTCCCTACACGACGCTCTTCCGATCTNNNATCCTTCACTTCCTCACCCCTAC

AGCACACGTCTGAACTCTTTCCCTACACGACGCTCTTCCGATCTNNNATCCTTCACTTCCTCACCCCTAC

GAGCACACGTCTGAACTCTTTCCCTACACGACGCTCTTCCGATCTNNNGTAATCCCAGATACTCGGAA

GAGCACACGTCTGAACTCTTTCCCTACACGACGCTCTTCCGATCTNNNGTAATCCCAGATACTCGGAA

GCACACGTCTGAACTCTTTCCCTACACGACGCTCTTCCGATCTNNNCCACCTCCTCCACAAAACATCTA

GCACACGTCTGAACTCTTTCCCTACACGACGCTCTTCCGATCTNNNCCACCTCCTCCACAAAACATCTA

GAGCACACGTCTGAACTCTTTCCCTACACGACGCTCTTCCGATCTNNNTGTCCCAAATGGTCTGATTTT

GAGCACACGTCTGAACTCTTTCCCTACACGACGCTCTTCCGATCTNNNTGTCCCAAATGGTCTGATTTT

AGCACACGTCTGAACTCTTTCCCTACACGACGCTCTTCCGATCTNNNCTCTTTCTGGGACTCAGGTTC

AGCACACGTCTGAACTCTTTCCCTACACGACGCTCTTCCGATCTNNNCTCTTTCTGGGACTCAGGTTC

GCACACGTCTGAACTCTTTCCCTACACGACGCTCTTCCGATCTNNNAATGAATGTGGGGCTGAAAAG

GCACACGTCTGAACTCTTTCCCTACACGACGCTCTTCCGATCTNNNAATGAATGTGGGGCTGAAAAG

GAGCACACGTCTGAACTCTTTCCCTACACGACGCTCTTCCGATCTNNNCTGAGGCAGGAGAATCACTT

GAGCACACGTCTGAACTCTTTCCCTACACGACGCTCTTCCGATCTNNNCTGAGGCAGGAGAATCACTT

AGCACACGTCTGAACTCTTTCCCTACACGACGCTCTTCCGATCTNNNTTTCTGTCAGTCCACTTCACCA

AGCACACGTCTGAACTCTTTCCCTACACGACGCTCTTCCGATCTNNNTTTCTGTCAGTCCACTTCACCA
AGCACACGTCTGAACTCTTTCCCTACACGACGCTCTTCCGATCTNNNTTTCTGTGGTGCCAGTGGTAAT
AGCACACGTCTGAACTCTTTCCCTACACGACGCTCTTCCGATCTNNNTTTCTGTGGTGCCAGTGGTAAT
AGCACACGTCTGAACTCTTTCCCTACACGACGCTCTTCCGATCTNNNCCTTTGCAGCTTGGCATCATAT
AGCACACGTCTGAACTCTTTCCCTACACGACGCTCTTCCGATCTNNNCCTTTGCAGCTTGGCATCATAT
GAGCACACGTCTGAACTCTTTCCCTACACGACGCTCTTCCGATCTNNNCTGCAGTGAGCCAAGATCGT
GAGCACACGTCTGAACTCTTTCCCTACACGACGCTCTTCCGATCTNNNCTGCAGTGAGCCAAGATCGT
AGCACACGTCTGAACTCTTTCCCTACACGACGCTCTTCCGATCTNNNAGCTGATGACTATGTGCCTAT
AGCACACGTCTGAACTCTTTCCCTACACGACGCTCTTCCGATCTNNNAGCTGATGACTATGTGCCTAT
AGCACACGTCTGAACTCTTTCCCTACACGACGCTCTTCCGATCTNNNCAAGGATGCAGCAACTAAGCA
AGCACACGTCTGAACTCTTTCCCTACACGACGCTCTTCCGATCTNNNCAAGGATGCAGCAACTAAGCA
AGCACACGTCTGAACTCTTTCCCTACACGACGCTCTTCCGATCTNNNCGGCCATACTGCTTTCAGGAAT
AGCACACGTCTGAACTCTTTCCCTACACGACGCTCTTCCGATCTNNNCGGCCATACTGCTTTCAGGAAT
AGCACACGTCTGAACTCTTTCCCTACACGACGCTCTTCCGATCTNNNAAGAGCTGTAAGAGGAACTG
AGCACACGTCTGAACTCTTTCCCTACACGACGCTCTTCCGATCTNNNAAGAGCTGTAAGAGGAACTG
AGCACACGTCTGAACTCTTTCCCTACACGACGCTCTTCCGATCTNNNCCCACGTCTTATTGTAGAGGCT
AGCACACGTCTGAACTCTTTCCCTACACGACGCTCTTCCGATCTNNNCCCACGTCTTATTGTAGAGGCT
AGCACACGTCTGAACTCTTTCCCTACACGACGCTCTTCCGATCTNNNCCACAACCTGGAATTCCCTATC
AGCACACGTCTGAACTCTTTCCCTACACGACGCTCTTCCGATCTNNNCCACAACCTGGAATTCCCTATC
GAGCACACGTCTGAACTCTTTCCCTACACGACGCTCTTCCGATCTNNNGTATTTCTGCAGTCCCAGCTA
GAGCACACGTCTGAACTCTTTCCCTACACGACGCTCTTCCGATCTNNNGTATTTCTGCAGTCCCAGCTA
AGCACACGTCTGAACTCTTTCCCTACACGACGCTCTTCCGATCTNNNATACACCTGTAGTCCCAGCTAC
AGCACACGTCTGAACTCTTTCCCTACACGACGCTCTTCCGATCTNNNATACACCTGTAGTCCCAGCTAC
GAGCACACGTCTGAACTCTTTCCCTACACGACGCTCTTCCGATCTNNNCTGGGCCTGAACAAATAGGT
GAGCACACGTCTGAACTCTTTCCCTACACGACGCTCTTCCGATCTNNNCTGGGCCTGAACAAATAGGT
GAGCACACGTCTGAACTCTTTCCCTACACGACGCTCTTCCGATCTNNNTCTCTCCATTCCTCCCTATCC
GAGCACACGTCTGAACTCTTTCCCTACACGACGCTCTTCCGATCTNNNTCTCTCCATTCCTCCCTATCC
GCACACGTCTGAACTCTTTCCCTACACGACGCTCTTCCGATCTNNNGAATCGGGTTCCTGGGTCTCCAT
GCACACGTCTGAACTCTTTCCCTACACGACGCTCTTCCGATCTNNNGAATCGGGTTCCTGGGTCTCCAT
AGCACACGTCTGAACTCTTTCCCTACACGACGCTCTTCCGATCTNNNGGAAGGAGGAGTTTGCAGTG
AGCACACGTCTGAACTCTTTCCCTACACGACGCTCTTCCGATCTNNNGGAAGGAGGAGTTTGCAGTG
AGCACACGTCTGAACTCTTTCCCTACACGACGCTCTTCCGATCTNNNCTCCACATCAGAACCAAAAGCTG
AGCACACGTCTGAACTCTTTCCCTACACGACGCTCTTCCGATCTNNNCTCCACATCAGAACCAAAAGCTG
GCACACGTCTGAACTCTTTCCCTACACGACGCTCTTCCGATCTNNNTTAGGATGGGAGGATGGTGGA
GCACACGTCTGAACTCTTTCCCTACACGACGCTCTTCCGATCTNNNTTAGGATGGGAGGATGGTGGA
GCACACGTCTGAACTCTTTCCCTACACGACGCTCTTCCGATCTNNNGATGGTTACACACACACACACA
GCACACGTCTGAACTCTTTCCCTACACGACGCTCTTCCGATCTNNNGATGGTTACACACACACACACA
GAGCACACGTCTGAACTCTTTCCCTACACGACGCTCTTCCGATCTNNNCGAGTAGCTGAGATTACAGG
GAGCACACGTCTGAACTCTTTCCCTACACGACGCTCTTCCGATCTNNNCGAGTAGCTGAGATTACAGG
AGCACACGTCTGAACTCTTTCCCTACACGACGCTCTTCCGATCTNNNCATTTGGGCAGAATATGTGCT
AGCACACGTCTGAACTCTTTCCCTACACGACGCTCTTCCGATCTNNNCATTTGGGCAGAATATGTGCT
AGCACACGTCTGAACTCTTTCCCTACACGACGCTCTTCCGATCTNNNTGCCATCAGTAACACAAACCCA
AGCACACGTCTGAACTCTTTCCCTACACGACGCTCTTCCGATCTNNNTGCCATCAGTAACACAAACCCA

GCACACGTCTGAACTCTTTCCCTACACGACGCTCTTCCGATCTNNNAAGAAAATTCACTTCCCAGGCA
AGCACACGTCTGAACTCTTTCCCTACACGACGCTCTTCCGATCTNNNAAGAAAATTCACTTCCCAGGCA
GAGCACACGTCTGAACTCTTTCCCTACACGACGCTCTTCCGATCTNNNCCTCCAAAATGCTGTGAAACT
GAGCACACGTCTGAACTCTTTCCCTACACGACGCTCTTCCGATCTNNNCCTCCAAAATGCTGTGAAACT
AGCACACGTCTGAACTCTTTCCCTACACGACGCTCTTCCGATCTNNNCTGACATCCTAGCTCCTGACTG
AGCACACGTCTGAACTCTTTCCCTACACGACGCTCTTCCGATCTNNNCTGACATCCTAGCTCCTGACTG
AGCACACGTCTGAACTCTTTCCCTACACGACGCTCTTCCGATCTNNNTGGTGTCAGTTGTAGTATCTCC
AGCACACGTCTGAACTCTTTCCCTACACGACGCTCTTCCGATCTNNNTGGTGTCAGTTGTAGTATCTCC
GCACACGTCTGAACTCTTTCCCTACACGACGCTCTTCCGATCTNNNATTCCCAAATGAAGCTTCAATGT
GCACACGTCTGAACTCTTTCCCTACACGACGCTCTTCCGATCTNNNATTCCCAAATGAAGCTTCAATGT
AGCACACGTCTGAACTCTTTCCCTACACGACGCTCTTCCGATCTNNNGCCTGATCTTTCCTGCAATCAA
AGCACACGTCTGAACTCTTTCCCTACACGACGCTCTTCCGATCTNNNGCCTGATCTTTCCTGCAATCAA
GAGCACACGTCTGAACTCTTTCCCTACACGACGCTCTTCCGATCTNNNCTGGAGGATTGTAGCTAAC
GAGCACACGTCTGAACTCTTTCCCTACACGACGCTCTTCCGATCTNNNCTGGAGGATTGTAGCTAAC
GAGCACACGTCTGAACTCTTTCCCTACACGACGCTCTTCCGATCTNNNACTTCACTAAATGCTCCGTGT
GAGCACACGTCTGAACTCTTTCCCTACACGACGCTCTTCCGATCTNNNACTTCACTAAATGCTCCGTGT
AGCACACGTCTGAACTCTTTCCCTACACGACGCTCTTCCGATCTNNNTGGACAGTGGCATAGAAGTTG
AGCACACGTCTGAACTCTTTCCCTACACGACGCTCTTCCGATCTNNNTGGACAGTGGCATAGAAGTTG
GAGCACACGTCTGAACTCTTTCCCTACACGACGCTCTTCCGATCTNNNGTCAAGACCCTTCGGTATGCT
GAGCACACGTCTGAACTCTTTCCCTACACGACGCTCTTCCGATCTNNNGTCAAGACCCTTCGGTATGCT
GCACACGTCTGAACTCTTTCCCTACACGACGCTCTTCCGATCTNNNCAGGGACCTTACCTTATACACATC
GCACACGTCTGAACTCTTTCCCTACACGACGCTCTTCCGATCTNNNCAGGGACCTTACCTTATACACATC
GAGCACACGTCTGAACTCTTTCCCTACACGACGCTCTTCCGATCTNNNTCCTGGAGAAATAAAACCTG
GAGCACACGTCTGAACTCTTTCCCTACACGACGCTCTTCCGATCTNNNTCCTGGAGAAATAAAACCTG
GCACACGTCTGAACTCTTTCCCTACACGACGCTCTTCCGATCTNNNATGTAGGGCAATTTGGCAGTTT
GCACACGTCTGAACTCTTTCCCTACACGACGCTCTTCCGATCTNNNATGTAGGGCAATTTGGCAGTTT
GAGCACACGTCTGAACTCTTTCCCTACACGACGCTCTTCCGATCTNNNGCACAATCACAGTTCAACCTCA
GAGCACACGTCTGAACTCTTTCCCTACACGACGCTCTTCCGATCTNNNGCACAATCACAGTTCAACCTCA
AGCACACGTCTGAACTCTTTCCCTACACGACGCTCTTCCGATCTNNNTGGTCATTAGGTCTCAAATACG
AGCACACGTCTGAACTCTTTCCCTACACGACGCTCTTCCGATCTNNNTGGTCATTAGGTCTCAAATACG
GAGCACACGTCTGAACTCTTTCCCTACACGACGCTCTTCCGATCTNNNTATCCCCCAACTCAGTTTTGA
GAGCACACGTCTGAACTCTTTCCCTACACGACGCTCTTCCGATCTNNNTATCCCCCAACTCAGTTTTGA
GAGCACACGTCTGAACTCTTTCCCTACACGACGCTCTTCCGATCTNNNGTTCTATCCTGTTCCATCAGC
GAGCACACGTCTGAACTCTTTCCCTACACGACGCTCTTCCGATCTNNNGTTCTATCCTGTTCCATCAGC
AGCACACGTCTGAACTCTTTCCCTACACGACGCTCTTCCGATCTNNNTCATCTTAGTCCATGCTCAGCAG
AGCACACGTCTGAACTCTTTCCCTACACGACGCTCTTCCGATCTNNNTCATCTTAGTCCATGCTCAGCAG
AGCACACGTCTGAACTCTTTCCCTACACGACGCTCTTCCGATCTNNNCATGCCCGGCTAGTTTTTGTATT
AGCACACGTCTGAACTCTTTCCCTACACGACGCTCTTCCGATCTNNNCATGCCCGGCTAGTTTTTGTATT
AGCACACGTCTGAACTCTTTCCCTACACGACGCTCTTCCGATCTNNNGAGAGAGGAGAATCGCTTGA
AGCACACGTCTGAACTCTTTCCCTACACGACGCTCTTCCGATCTNNNGAGAGAGGAGAATCGCTTGA
GAGCACACGTCTGAACTCTTTCCCTACACGACGCTCTTCCGATCTNNNGCACTGTTCGAAACACTATTA
GAGCACACGTCTGAACTCTTTCCCTACACGACGCTCTTCCGATCTNNNGCACTGTTCGAAACACTATTA
CACACGTCTGAACTCTTTCCCTACACGACGCTCTTCCGATCTNNNCTAGAAATGTGGTTCTCGTGCATC

CACACGTCTGAACTCTTTCCCTACACGACGCTCTTCCGATCTNNNCTAGAAATGTGGTTCTCGTGCATC

AGCACACGTCTGAACTCTTTCCCTACACGACGCTCTTCCGATCTNNNAACAACAAACTACTGATGAGG

AGCACACGTCTGAACTCTTTCCCTACACGACGCTCTTCCGATCTNNNAACAACAAACTACTGATGAGG

AGAGCACACGTCTGAACTCTTTCCCTACACGACGCTCTTCCGATCTNNNAGACTCTGCGGAAGAAAGG

AGAGCACACGTCTGAACTCTTTCCCTACACGACGCTCTTCCGATCTNNNAGACTCTGCGGAAGAAAGG

AGCACACGTCTGAACTCTTTCCCTACACGACGCTCTTCCGATCTNNNAGGCAATACAGTAGGGTGTG

AGCACACGTCTGAACTCTTTCCCTACACGACGCTCTTCCGATCTNNNAGGCAATACAGTAGGGTGTG

AGCACACGTCTGAACTCTTTCCCTACACGACGCTCTTCCGATCTNNNTGGAGGAAGGTGATTTCATGT

AGCACACGTCTGAACTCTTTCCCTACACGACGCTCTTCCGATCTNNNTGGAGGAAGGTGATTTCATGT

GAGCACACGTCTGAACTCTTTCCCTACACGACGCTCTTCCGATCTNNNTGTAGTCCCAGCTACTCAAAA

GAGCACACGTCTGAACTCTTTCCCTACACGACGCTCTTCCGATCTNNNTGTAGTCCCAGCTACTCAAAA

GAGCACACGTCTGAACTCTTTCCCTACACGACGCTCTTCCGATCTNNNCAACAGAGCAAAAACTCCGT

GAGCACACGTCTGAACTCTTTCCCTACACGACGCTCTTCCGATCTNNNCAACAGAGCAAAAACTCCGT

GCACACGTCTGAACTCTTTCCCTACACGACGCTCTTCCGATCTNNNTTGACTACTCCAGGTACCTCAGT

GCACACGTCTGAACTCTTTCCCTACACGACGCTCTTCCGATCTNNNTTGACTACTCCAGGTACCTCAGT

AGCACACGTCTGAACTCTTTCCCTACACGACGCTCTTCCGATCTNNNAGGAGACTGAGATGGGAGAA

AGCACACGTCTGAACTCTTTCCCTACACGACGCTCTTCCGATCTNNNAGGAGACTGAGATGGGAGAA

GCACACGTCTGAACTCTTTCCCTACACGACGCTCTTCCGATCTNNNACATGTGACTCCCCAGAATACAC

GCACACGTCTGAACTCTTTCCCTACACGACGCTCTTCCGATCTNNNACATGTGACTCCCCAGAATACAC

AGCACACGTCTGAACTCTTTCCCTACACGACGCTCTTCCGATCTNNNCTGGGTTTCTAGTAAGTAGCC

AGCACACGTCTGAACTCTTTCCCTACACGACGCTCTTCCGATCTNNNCTGGGTTTCTAGTAAGTAGCC

GAGCACACGTCTGAACTCTTTCCCTACACGACGCTCTTCCGATCTNNNCCTGAGCAATACAGTGAGAC

GAGCACACGTCTGAACTCTTTCCCTACACGACGCTCTTCCGATCTNNNCCTGAGCAATACAGTGAGAC

AGCACACGTCTGAACTCTTTCCCTACACGACGCTCTTCCGATCTNNNTGGAACTACACCATCAACTCTCC

AGCACACGTCTGAACTCTTTCCCTACACGACGCTCTTCCGATCTNNNTGGAACTACACCATCAACTCTCC

GCACACGTCTGAACTCTTTCCCTACACGACGCTCTTCCGATCTNNNGCTTTAGGATCTGGAACAACCT

GCACACGTCTGAACTCTTTCCCTACACGACGCTCTTCCGATCTNNNGCTTTAGGATCTGGAACAACCT

GCACACGTCTGAACTCTTTCCCTACACGACGCTCTTCCGATCTNNNGCTTTCGTTTTTGATCCCAGGA

GCACACGTCTGAACTCTTTCCCTACACGACGCTCTTCCGATCTNNNGCTTTCGTTTTTGATCCCAGGA

AGCACACGTCTGAACTCTTTCCCTACACGACGCTCTTCCGATCTNNNATCTGGACCTCTACTGTGAATG

AGCACACGTCTGAACTCTTTCCCTACACGACGCTCTTCCGATCTNNNATCTGGACCTCTACTGTGAATG

AAGAGCACACGTCTGAACTCTTTCCCTACACGACGCTCTTCCGATCTNNNCGAAGTCCTGAGTAGGGT

AAGAGCACACGTCTGAACTCTTTCCCTACACGACGCTCTTCCGATCTNNNCGAAGTCCTGAGTAGGGT

AGCACACGTCTGAACTCTTTCCCTACACGACGCTCTTCCGATCTNNNCTGTAGTCCCAGCTACTCAGG

AGCACACGTCTGAACTCTTTCCCTACACGACGCTCTTCCGATCTNNNCTGTAGTCCCAGCTACTCAGG

GCACACGTCTGAACTCTTTCCCTACACGACGCTCTTCCGATCTNNNGAAATTGTGGGGGAAAGAGCT

GCACACGTCTGAACTCTTTCCCTACACGACGCTCTTCCGATCTNNNGAAATTGTGGGGGAAAGAGCT

AGCACACGTCTGAACTCTTTCCCTACACGACGCTCTTCCGATCTNNNGACATAAGCAAGGCAGATGT

AGCACACGTCTGAACTCTTTCCCTACACGACGCTCTTCCGATCTNNNGACATAAGCAAGGCAGATGT

AAGAGCACACGTCTGAACTCTTTCCCTACACGACGCTCTTCCGATCTNNNCGTCTCTGGCCTATGTGA

AAGAGCACACGTCTGAACTCTTTCCCTACACGACGCTCTTCCGATCTNNNCGTCTCTGGCCTATGTGA

AGCACACGTCTGAACTCTTTCCCTACACGACGCTCTTCCGATCTNNNCCTTGTCGATGGCTTTGGTAA

AGCACACGTCTGAACTCTTTCCCTACACGACGCTCTTCCGATCTNNNCCTTGTCGATGGCTTTGGTAA

GCACACGTCTGAACTCTTTCCCTACACGACGCTCTTCCGATCTNNNTCCTTCATTCTCACCTTTCCCTGCA
GCACACGTCTGAACTCTTTCCCTACACGACGCTCTTCCGATCTNNNTCCTTCATTCTCACCTTTCCCTGCA
GAGCACACGTCTGAACTCTTTCCCTACACGACGCTCTTCCGATCTNNNGAGACTGAGGTGGGAGAATT
GAGCACACGTCTGAACTCTTTCCCTACACGACGCTCTTCCGATCTNNNGAGACTGAGGTGGGAGAATT
GCACACGTCTGAACTCTTTCCCTACACGACGCTCTTCCGATCTNNNTCTTTCCCCTCTCCCTACCTTAGCAT
GCACACGTCTGAACTCTTTCCCTACACGACGCTCTTCCGATCTNNNTCTTTCCCCTCTCCCTACCTTAGCAT
GAGCACACGTCTGAACTCTTTCCCTACACGACGCTCTTCCGATCTNNNCTGAGGCAGAGAATTGCTTG
GAGCACACGTCTGAACTCTTTCCCTACACGACGCTCTTCCGATCTNNNCTGAGGCAGAGAATTGCTTG
GAGCACACGTCTGAACTCTTTCCCTACACGACGCTCTTCCGATCTNNNGTAGAGACGGGGTTTACCAT
GAGCACACGTCTGAACTCTTTCCCTACACGACGCTCTTCCGATCTNNNGTAGAGACGGGGTTTACCAT
AGCACACGTCTGAACTCTTTCCCTACACGACGCTCTTCCGATCTNNNTGGTATAATGCCTCCAGCTTTGA
AGCACACGTCTGAACTCTTTCCCTACACGACGCTCTTCCGATCTNNNTGGTATAATGCCTCCAGCTTTGA
GAGCACACGTCTGAACTCTTTCCCTACACGACGCTCTTCCGATCTNNNGCCTGTCTGGTTGTCATAGTT
GAGCACACGTCTGAACTCTTTCCCTACACGACGCTCTTCCGATCTNNNGCCTGTCTGGTTGTCATAGTT
GAGCACACGTCTGAACTCTTTCCCTACACGACGCTCTTCCGATCTNNNGGTTGCAGTGATCCGATATTG
GAGCACACGTCTGAACTCTTTCCCTACACGACGCTCTTCCGATCTNNNGGTTGCAGTGATCCGATATTG
AGCACACGTCTGAACTCTTTCCCTACACGACGCTCTTCCGATCTNNNAAGAATCACTTGAACCCAGGAG
AGCACACGTCTGAACTCTTTCCCTACACGACGCTCTTCCGATCTNNNAAGAATCACTTGAACCCAGGAG
AGCACACGTCTGAACTCTTTCCCTACACGACGCTCTTCCGATCTNNNGGCCTTTGAGTCTACCTTTCTGA
AGCACACGTCTGAACTCTTTCCCTACACGACGCTCTTCCGATCTNNNGGCCTTTGAGTCTACCTTTCTGA
GAGCACACGTCTGAACTCTTTCCCTACACGACGCTCTTCCGATCTNNNTCTCAATGAGCATTTTCTTCTG
GAGCACACGTCTGAACTCTTTCCCTACACGACGCTCTTCCGATCTNNNTCTCAATGAGCATTTTCTTCTG
AGCACACGTCTGAACTCTTTCCCTACACGACGCTCTTCCGATCTNNNATCTGTGTCTGCTTTTGTGCTTC
AGCACACGTCTGAACTCTTTCCCTACACGACGCTCTTCCGATCTNNNATCTGTGTCTGCTTTTGTGCTTC
GAGCACACGTCTGAACTCTTTCCCTACACGACGCTCTTCCGATCTNNNGAGGTTGTAGTGAGCCGAGA
GAGCACACGTCTGAACTCTTTCCCTACACGACGCTCTTCCGATCTNNNGAGGTTGTAGTGAGCCGAGA
GAGCACACGTCTGAACTCTTTCCCTACACGACGCTCTTCCGATCTNNNGGTGTGCTGGTAAAACATGA
GAGCACACGTCTGAACTCTTTCCCTACACGACGCTCTTCCGATCTNNNGGTGTGCTGGTAAAACATGA
GAGCACACGTCTGAACTCTTTCCCTACACGACGCTCTTCCGATCTNNNACACAAGATGGAGTCAAAGA
GAGCACACGTCTGAACTCTTTCCCTACACGACGCTCTTCCGATCTNNNACACAAGATGGAGTCAAAGA
AGCACACGTCTGAACTCTTTCCCTACACGACGCTCTTCCGATCTNNNCTGTCATCTCTCCTCAACCAACA
AGCACACGTCTGAACTCTTTCCCTACACGACGCTCTTCCGATCTNNNCTGTCATCTCTCCTCAACCAACA
AGCACACGTCTGAACTCTTTCCCTACACGACGCTCTTCCGATCTNNNCCAGACTAGATGCAAATGTCAG
AGCACACGTCTGAACTCTTTCCCTACACGACGCTCTTCCGATCTNNNCCAGACTAGATGCAAATGTCAG
GAGCACACGTCTGAACTCTTTCCCTACACGACGCTCTTCCGATCTNNNTGTTGCTGCCTGATTATTCCTC
GAGCACACGTCTGAACTCTTTCCCTACACGACGCTCTTCCGATCTNNNTGTTGCTGCCTGATTATTCCTC
AGCACACGTCTGAACTCTTTCCCTACACGACGCTCTTCCGATCTNNNCAAGGATTAGAGGAGGCATG
AGCACACGTCTGAACTCTTTCCCTACACGACGCTCTTCCGATCTNNNCAAGGATTAGAGGAGGCATG
AGCACACGTCTGAACTCTTTCCCTACACGACGCTCTTCCGATCTNNNAGCCTATCCACAGAGGTTCAAT
AGCACACGTCTGAACTCTTTCCCTACACGACGCTCTTCCGATCTNNNAGCCTATCCACAGAGGTTCAAT
GAGCACACGTCTGAACTCTTTCCCTACACGACGCTCTTCCGATCTNNNCCTGGCAATTTTTCTTGGGGA
GAGCACACGTCTGAACTCTTTCCCTACACGACGCTCTTCCGATCTNNNCCTGGCAATTTTTCTTGGGGA
AGCACACGTCTGAACTCTTTCCCTACACGACGCTCTTCCGATCTNNNAGGGAGTGGGGATGGTCAATA

AGCACACGTCTGAACTCTTTCCCTACACGACGCTCTTCCGATCTNNNAGGGAGTGGGGATGGTCAATA
GAGCACACGTCTGAACTCTTTCCCTACACGACGCTCTTCCGATCTNNNGTGAGGCCCTGTCTCAAAAAA
GAGCACACGTCTGAACTCTTTCCCTACACGACGCTCTTCCGATCTNNNGTGAGGCCCTGTCTCAAAAAA
GAGCACACGTCTGAACTCTTTCCCTACACGACGCTCTTCCGATCTNNNTACAAAAAATTAGCTGGGCGT
GAGCACACGTCTGAACTCTTTCCCTACACGACGCTCTTCCGATCTNNNTACAAAAAATTAGCTGGGCGT
AGCACACGTCTGAACTCTTTCCCTACACGACGCTCTTCCGATCTNNNCTGAGGCAGGAGAATTGCTTG
AGCACACGTCTGAACTCTTTCCCTACACGACGCTCTTCCGATCTNNNCTGAGGCAGGAGAATTGCTTG
AGCACACGTCTGAACTCTTTCCCTACACGACGCTCTTCCGATCTNNNAGAATTGCTTGAACCCAGGAGA
AGCACACGTCTGAACTCTTTCCCTACACGACGCTCTTCCGATCTNNNAGAATTGCTTGAACCCAGGAGA
GAGCACACGTCTGAACTCTTTCCCTACACGACGCTCTTCCGATCTNNNAGAATTGCTTGAACTCAGGAG
GAGCACACGTCTGAACTCTTTCCCTACACGACGCTCTTCCGATCTNNNAGAATTGCTTGAACTCAGGAG
AGCACACGTCTGAACTCTTTCCCTACACGACGCTCTTCCGATCTNNNAGGCAGAAGAATCACTTGAAC
AGCACACGTCTGAACTCTTTCCCTACACGACGCTCTTCCGATCTNNNAGGCAGAAGAATCACTTGAAC
AGCACACGTCTGAACTCTTTCCCTACACGACGCTCTTCCGATCTNNNCCAGGGATGAAGCCTACTTGAT
AGCACACGTCTGAACTCTTTCCCTACACGACGCTCTTCCGATCTNNNCCAGGGATGAAGCCTACTTGAT
GAGCACACGTCTGAACTCTTTCCCTACACGACGCTCTTCCGATCTNNNGAGGCAGGAGAATGGTGTG
GAGCACACGTCTGAACTCTTTCCCTACACGACGCTCTTCCGATCTNNNGAGGCAGGAGAATGGTGTG
GAGCACACGTCTGAACTCTTTCCCTACACGACGCTCTTCCGATCTNNNCTCCTTTCCTTCACCTTCTCCTC
GAGCACACGTCTGAACTCTTTCCCTACACGACGCTCTTCCGATCTNNNCTCCTTTCCTTCACCTTCTCCTC
AGAGCACACGTCTGAACTCTTTCCCTACACGACGCTCTTCCGATCTNNNCTGAGATCACACCACTGCAGT
AGAGCACACGTCTGAACTCTTTCCCTACACGACGCTCTTCCGATCTNNNCTGAGATCACACCACTGCAGT
GAGCACACGTCTGAACTCTTTCCCTACACGACGCTCTTCCGATCTNNNCAGATCATGAGGTCAGGAGAT
GAGCACACGTCTGAACTCTTTCCCTACACGACGCTCTTCCGATCTNNNCAGATCATGAGGTCAGGAGAT
AGCACACGTCTGAACTCTTTCCCTACACGACGCTCTTCCGATCTNNNTAGTAAGCACAATGGGCTGAG
AGCACACGTCTGAACTCTTTCCCTACACGACGCTCTTCCGATCTNNNTAGTAAGCACAATGGGCTGAG
AGAGCACACGTCTGAACTCTTTCCCTACACGACGCTCTTCCGATCTNNNACCGATGTGATGTGTGTGGT
AGAGCACACGTCTGAACTCTTTCCCTACACGACGCTCTTCCGATCTNNNACCGATGTGATGTGTGTGGT
GAGCACACGTCTGAACTCTTTCCCTACACGACGCTCTTCCGATCTNNNAAGCATTTTATTCTTCCCGACG
GAGCACACGTCTGAACTCTTTCCCTACACGACGCTCTTCCGATCTNNNAAGCATTTTATTCTTCCCGACG
GAGCACACGTCTGAACTCTTTCCCTACACGACGCTCTTCCGATCTNNNAGCAAGGTTGTGGTTTTTGAA
GAGCACACGTCTGAACTCTTTCCCTACACGACGCTCTTCCGATCTNNNAGCAAGGTTGTGGTTTTTGAA
GAGCACACGTCTGAACTCTTTCCCTACACGACGCTCTTCCGATCTNNNAAACAAAGGAAATCAGGCCAC
GAGCACACGTCTGAACTCTTTCCCTACACGACGCTCTTCCGATCTNNNAAACAAAGGAAATCAGGCCAC
GAGCACACGTCTGAACTCTTTCCCTACACGACGCTCTTCCGATCTNNNTTTGAAGAGTCTGCCCTGTTTC
GAGCACACGTCTGAACTCTTTCCCTACACGACGCTCTTCCGATCTNNNTTTGAAGAGTCTGCCCTGTTTC
AGCACACGTCTGAACTCTTTCCCTACACGACGCTCTTCCGATCTNNNCACAGACACCTGCAATTGCTAAA
AGCACACGTCTGAACTCTTTCCCTACACGACGCTCTTCCGATCTNNNCACAGACACCTGCAATTGCTAAA
GAGCACACGTCTGAACTCTTTCCCTACACGACGCTCTTCCGATCTNNNCCAGGGTTCAAGCAATTCTTCT
GAGCACACGTCTGAACTCTTTCCCTACACGACGCTCTTCCGATCTNNNCCAGGGTTCAAGCAATTCTTCT
GAGCACACGTCTGAACTCTTTCCCTACACGACGCTCTTCCGATCTNNNCCATTCCCTCCAGTCTCCTAAAC
GAGCACACGTCTGAACTCTTTCCCTACACGACGCTCTTCCGATCTNNNCCATTCCCTCCAGTCTCCTAAAC
AGCACACGTCTGAACTCTTTCCCTACACGACGCTCTTCCGATCTNNNACAAAAATTAGCTGGGTGTGGT
AGCACACGTCTGAACTCTTTCCCTACACGACGCTCTTCCGATCTNNNACAAAAATTAGCTGGGTGTGGT

AAGAGCACACGTCTGAACTCTTTCCCTACACGACGCTCTTCCGATCTNNNGGAGACAGAGGTTGCAG
AAGAGCACACGTCTGAACTCTTTCCCTACACGACGCTCTTCCGATCTNNNGGAGACAGAGGTTGCAG
GAGCACACGTCTGAACTCTTTCCCTACACGACGCTCTTCCGATCTNNNTTTGGTGCTGAGATCTTACCT
GAGCACACGTCTGAACTCTTTCCCTACACGACGCTCTTCCGATCTNNNTTTGGTGCTGAGATCTTACCT
ACACGTCTGAACTCTTTCCCTACACGACGCTCTTCCGATCTNNNTTGGGCGTTTATCATGATCATGGC
ACACGTCTGAACTCTTTCCCTACACGACGCTCTTCCGATCTNNNTTGGGCGTTTATCATGATCATGGC
ACACGTCTGAACTCTTTCCCTACACGACGCTCTTCCGATCTNNNTTATTTCTGGTCAGAGATATTGTCA
ACACGTCTGAACTCTTTCCCTACACGACGCTCTTCCGATCTNNNTTATTTCTGGTCAGAGATATTGTCA
GAGCACACGTCTGAACTCTTTCCCTACACGACGCTCTTCCGATCTNNNGAGGCTGAGGCAGGAGAAT
GAGCACACGTCTGAACTCTTTCCCTACACGACGCTCTTCCGATCTNNNGAGGCTGAGGCAGGAGAAT
AGCACACGTCTGAACTCTTTCCCTACACGACGCTCTTCCGATCTNNNTTCGAACCATAGCAAGAAGGC
AGCACACGTCTGAACTCTTTCCCTACACGACGCTCTTCCGATCTNNNTTCGAACCATAGCAAGAAGGC
GCACACGTCTGAACTCTTTCCCTACACGACGCTCTTCCGATCTNNNTGACATGATACCAAGCAACAGG
GCACACGTCTGAACTCTTTCCCTACACGACGCTCTTCCGATCTNNNTGACATGATACCAAGCAACAGG
AAGAGCACACGTCTGAACTCTTTCCCTACACGACGCTCTTCCGATCTNNNAAGCAATGCTAAGAGCGA
AAGAGCACACGTCTGAACTCTTTCCCTACACGACGCTCTTCCGATCTNNNAAGCAATGCTAAGAGCGA
AGCACACGTCTGAACTCTTTCCCTACACGACGCTCTTCCGATCTNNNATCATGCCTGGCCAAGATATCT
AGCACACGTCTGAACTCTTTCCCTACACGACGCTCTTCCGATCTNNNATCATGCCTGGCCAAGATATCT
GAGCACACGTCTGAACTCTTTCCCTACACGACGCTCTTCCGATCTNNNGACTGTAGTCCCAGCTACTCC
GAGCACACGTCTGAACTCTTTCCCTACACGACGCTCTTCCGATCTNNNGACTGTAGTCCCAGCTACTCC
AGCACACGTCTGAACTCTTTCCCTACACGACGCTCTTCCGATCTNNNTTCAAGTGATTCTCCTGCCTCAG
AGCACACGTCTGAACTCTTTCCCTACACGACGCTCTTCCGATCTNNNTTCAAGTGATTCTCCTGCCTCAG
ACACGTCTGAACTCTTTCCCTACACGACGCTCTTCCGATCTNNNCCTCCTCTCCTCCTCTCAGGCATCGG
ACACGTCTGAACTCTTTCCCTACACGACGCTCTTCCGATCTNNNCCTCCTCTCCTCCTCTCAGGCATCGG
GCACACGTCTGAACTCTTTCCCTACACGACGCTCTTCCGATCTNNNCAATCACTACGCTTTGTTGGGT
GCACACGTCTGAACTCTTTCCCTACACGACGCTCTTCCGATCTNNNCAATCACTACGCTTTGTTGGGT
AGCACACGTCTGAACTCTTTCCCTACACGACGCTCTTCCGATCTNNNGTTGGAACAGACTCACACACAT
AGCACACGTCTGAACTCTTTCCCTACACGACGCTCTTCCGATCTNNNGTTGGAACAGACTCACACACAT
AGCACACGTCTGAACTCTTTCCCTACACGACGCTCTTCCGATCTNNNTTTCTTTTCCCAGTGCAAATGC
AGCACACGTCTGAACTCTTTCCCTACACGACGCTCTTCCGATCTNNNTTTCTTTTCCCAGTGCAAATGC
AGCACACGTCTGAACTCTTTCCCTACACGACGCTCTTCCGATCTNNNACCTAGAGTGTGACAAACCAA
AGCACACGTCTGAACTCTTTCCCTACACGACGCTCTTCCGATCTNNNACCTAGAGTGTGACAAACCAA
AGCACACGTCTGAACTCTTTCCCTACACGACGCTCTTCCGATCTNNNTGCTGATGGACTTATTGCTTCC
AGCACACGTCTGAACTCTTTCCCTACACGACGCTCTTCCGATCTNNNTGCTGATGGACTTATTGCTTCC
AGCACACGTCTGAACTCTTTCCCTACACGACGCTCTTCCGATCTNNNGCTAAAGCTGAAAAGCTGACT
AGCACACGTCTGAACTCTTTCCCTACACGACGCTCTTCCGATCTNNNGCTAAAGCTGAAAAGCTGACT
GAGCACACGTCTGAACTCTTTCCCTACACGACGCTCTTCCGATCTNNNGGGTCTCCAGCTTTCAGGTGC
GAGCACACGTCTGAACTCTTTCCCTACACGACGCTCTTCCGATCTNNNGGGTCTCCAGCTTTCAGGTGC
ACACGTCTGAACTCTTTCCCTACACGACGCTCTTCCGATCTNNNGTAAAACGTCATTAGCACCAAGCCA
ACACGTCTGAACTCTTTCCCTACACGACGCTCTTCCGATCTNNNGTAAAACGTCATTAGCACCAAGCCA
GAGCACACGTCTGAACTCTTTCCCTACACGACGCTCTTCCGATCTNNNCAATTCTGAGTTGGGTCATT
GAGCACACGTCTGAACTCTTTCCCTACACGACGCTCTTCCGATCTNNNCAATTCTGAGTTGGGTCATT
AGAGCACACGTCTGAACTCTTTCCCTACACGACGCTCTTCCGATCTNNNAGCAAAAGAAGAAGCAGT

AGAGCACACGTCTGAACTCTTTCCCTACACGACGCTCTTCCGATCTNNNAGCAAAAGAAGAAGCAGT
GAGCACACGTCTGAACTCTTTCCCTACACGACGCTCTTCCGATCTNNNTTGGCCTCCTCTTTCTCACAAA
GAGCACACGTCTGAACTCTTTCCCTACACGACGCTCTTCCGATCTNNNTTGGCCTCCTCTTTCTCACAAA
AGCACACGTCTGAACTCTTTCCCTACACGACGCTCTTCCGATCTNNNTAGAGAGTTCTGTGCCCCAATC
AGCACACGTCTGAACTCTTTCCCTACACGACGCTCTTCCGATCTNNNTAGAGAGTTCTGTGCCCCAATC
GAGCACACGTCTGAACTCTTTCCCTACACGACGCTCTTCCGATCTNNNCACGCTCAACAATTATCCACC
GAGCACACGTCTGAACTCTTTCCCTACACGACGCTCTTCCGATCTNNNCACGCTCAACAATTATCCACC
AGCACACGTCTGAACTCTTTCCCTACACGACGCTCTTCCGATCTNNNCCCTAGTGCAGTGTAGATGAT
AGCACACGTCTGAACTCTTTCCCTACACGACGCTCTTCCGATCTNNNCCCTAGTGCAGTGTAGATGAT
GAGCACACGTCTGAACTCTTTCCCTACACGACGCTCTTCCGATCTNNNGAGACAGAAGGATCTCTTGA
GAGCACACGTCTGAACTCTTTCCCTACACGACGCTCTTCCGATCTNNNGAGACAGAAGGATCTCTTGA
AGCACACGTCTGAACTCTTTCCCTACACGACGCTCTTCCGATCTNNNCTCATGCCTGTAATCCTAGCAC
AGCACACGTCTGAACTCTTTCCCTACACGACGCTCTTCCGATCTNNNCTCATGCCTGTAATCCTAGCAC
CACACGTCTGAACTCTTTCCCTACACGACGCTCTTCCGATCTNNNGGATTACTGGCTGGAATGGTAGT
CACACGTCTGAACTCTTTCCCTACACGACGCTCTTCCGATCTNNNGGATTACTGGCTGGAATGGTAGT
AGCACACGTCTGAACTCTTTCCCTACACGACGCTCTTCCGATCTNNNATCAGACAGCCATCAGAATCAC
AGCACACGTCTGAACTCTTTCCCTACACGACGCTCTTCCGATCTNNNATCAGACAGCCATCAGAATCAC
GAGCACACGTCTGAACTCTTTCCCTACACGACGCTCTTCCGATCTNNNCACAATCTTGGCTCACTGCAA
GAGCACACGTCTGAACTCTTTCCCTACACGACGCTCTTCCGATCTNNNCACAATCTTGGCTCACTGCAA
GCACACGTCTGAACTCTTTCCCTACACGACGCTCTTCCGATCTNNNGTTGCAGTGAGCTGAGATCATG
GCACACGTCTGAACTCTTTCCCTACACGACGCTCTTCCGATCTNNNGTTGCAGTGAGCTGAGATCATG
AGCACACGTCTGAACTCTTTCCCTACACGACGCTCTTCCGATCTNNNCCTGAGGACAAACTCATGAGGT
AGCACACGTCTGAACTCTTTCCCTACACGACGCTCTTCCGATCTNNNCCTGAGGACAAACTCATGAGGT
GCACACGTCTGAACTCTTTCCCTACACGACGCTCTTCCGATCTNNNACCTGCAAACTGTTCTCCATAGTC
GCACACGTCTGAACTCTTTCCCTACACGACGCTCTTCCGATCTNNNACCTGCAAACTGTTCTCCATAGTC
AGCACACGTCTGAACTCTTTCCCTACACGACGCTCTTCCGATCTNNNATCCTGGATGTCCAATGTTTCCA
AGCACACGTCTGAACTCTTTCCCTACACGACGCTCTTCCGATCTNNNATCCTGGATGTCCAATGTTTCCA
AGCACACGTCTGAACTCTTTCCCTACACGACGCTCTTCCGATCTNNNTGCCACGAGGATTTTGTGAAAT
AGCACACGTCTGAACTCTTTCCCTACACGACGCTCTTCCGATCTNNNTGCCACGAGGATTTTGTGAAAT
AGCACACGTCTGAACTCTTTCCCTACACGACGCTCTTCCGATCTNNNGGTTCCAAGTCACTGCAGAAG
AGCACACGTCTGAACTCTTTCCCTACACGACGCTCTTCCGATCTNNNGGTTCCAAGTCACTGCAGAAG
AGCACACGTCTGAACTCTTTCCCTACACGACGCTCTTCCGATCTNNNTATGTGTCCTCAACCTCTGCAAA
AGCACACGTCTGAACTCTTTCCCTACACGACGCTCTTCCGATCTNNNTATGTGTCCTCAACCTCTGCAAA
GAGCACACGTCTGAACTCTTTCCCTACACGACGCTCTTCCGATCTNNNTTCTTCAGTGCTGCATGTCAA
GAGCACACGTCTGAACTCTTTCCCTACACGACGCTCTTCCGATCTNNNTTCTTCAGTGCTGCATGTCAA
AGCACACGTCTGAACTCTTTCCCTACACGACGCTCTTCCGATCTNNNGGGGTAACAGCTTTACACCCTA
AGCACACGTCTGAACTCTTTCCCTACACGACGCTCTTCCGATCTNNNGGGGTAACAGCTTTACACCCTA
GAGCACACGTCTGAACTCTTTCCCTACACGACGCTCTTCCGATCTNNNACATACTGGGGGTTGAAATGT
GAGCACACGTCTGAACTCTTTCCCTACACGACGCTCTTCCGATCTNNNACATACTGGGGGTTGAAATGT
AGCACACGTCTGAACTCTTTCCCTACACGACGCTCTTCCGATCTNNNGGAGGACACATACCATCTCTGT
AGCACACGTCTGAACTCTTTCCCTACACGACGCTCTTCCGATCTNNNGGAGGACACATACCATCTCTGT
CACACGTCTGAACTCTTTCCCTACACGACGCTCTTCCGATCTNNNGATGGTGGAGGTGGTGTACAGC
CACACGTCTGAACTCTTTCCCTACACGACGCTCTTCCGATCTNNNGATGGTGGAGGTGGTGTACAGC

AGCACACGTCTGAACTCTTTCCCTACACGACGCTCTTCCGATCTNNNGTAGCTGGGATTAGAGGCATGT
AGCACACGTCTGAACTCTTTCCCTACACGACGCTCTTCCGATCTNNNGTAGCTGGGATTAGAGGCATGT
AGCACACGTCTGAACTCTTTCCCTACACGACGCTCTTCCGATCTNNNCACATCTCAGCAACAGTGTTACA
AGCACACGTCTGAACTCTTTCCCTACACGACGCTCTTCCGATCTNNNCACATCTCAGCAACAGTGTTACA
AGCACACGTCTGAACTCTTTCCCTACACGACGCTCTTCCGATCTNNNATTATAGCCACCCGGGTTTTCTT
AGCACACGTCTGAACTCTTTCCCTACACGACGCTCTTCCGATCTNNNATTATAGCCACCCGGGTTTTCTT
GAGCACACGTCTGAACTCTTTCCCTACACGACGCTCTTCCGATCTNNNAGCTTTGAGGGTGTAAAGGCA
GAGCACACGTCTGAACTCTTTCCCTACACGACGCTCTTCCGATCTNNNAGCTTTGAGGGTGTAAAGGCA
GAGCACACGTCTGAACTCTTTCCCTACACGACGCTCTTCCGATCTNNNATGCCTGTAATCCCAGCTACTT
GAGCACACGTCTGAACTCTTTCCCTACACGACGCTCTTCCGATCTNNNATGCCTGTAATCCCAGCTACTT
AGAGCACACGTCTGAACTCTTTCCCTACACGACGCTCTTCCGATCTNNNGAATATTCAGGTTTCGCTCTG
AGAGCACACGTCTGAACTCTTTCCCTACACGACGCTCTTCCGATCTNNNGAATATTCAGGTTTCGCTCTG
AGCACACGTCTGAACTCTTTCCCTACACGACGCTCTTCCGATCTNNNATTTTTCAGCACACTGCCATCAT
AGCACACGTCTGAACTCTTTCCCTACACGACGCTCTTCCGATCTNNNATTTTTCAGCACACTGCCATCAT
AGCACACGTCTGAACTCTTTCCCTACACGACGCTCTTCCGATCTNNNGAATACTCACCATTCCCCACACT
AGCACACGTCTGAACTCTTTCCCTACACGACGCTCTTCCGATCTNNNGAATACTCACCATTCCCCACACT
AGAGCACACGTCTGAACTCTTTCCCTACACGACGCTCTTCCGATCTNNNCTTTGAGATGGGCAGTAAG
AGAGCACACGTCTGAACTCTTTCCCTACACGACGCTCTTCCGATCTNNNCTTTGAGATGGGCAGTAAG
GCACACGTCTGAACTCTTTCCCTACACGACGCTCTTCCGATCTNNNTGTACCTCCATTACAGTATGAGA
GCACACGTCTGAACTCTTTCCCTACACGACGCTCTTCCGATCTNNNTGTACCTCCATTACAGTATGAGA
GAGCACACGTCTGAACTCTTTCCCTACACGACGCTCTTCCGATCTNNNGATCGAGACCATCCTGGCTAA
GAGCACACGTCTGAACTCTTTCCCTACACGACGCTCTTCCGATCTNNNGATCGAGACCATCCTGGCTAA
GAGCACACGTCTGAACTCTTTCCCTACACGACGCTCTTCCGATCTNNNCTGAGGCAGTAGAACCACTTG
GAGCACACGTCTGAACTCTTTCCCTACACGACGCTCTTCCGATCTNNNCTGAGGCAGTAGAACCACTTG
GAGCACACGTCTGAACTCTTTCCCTACACGACGCTCTTCCGATCTNNNTGAGGTGAACTGAAGAGCAT
GAGCACACGTCTGAACTCTTTCCCTACACGACGCTCTTCCGATCTNNNTGAGGTGAACTGAAGAGCAT
GAGCACACGTCTGAACTCTTTCCCTACACGACGCTCTTCCGATCTNNNCACCTGTAATTCCAGCAGTTTG
GAGCACACGTCTGAACTCTTTCCCTACACGACGCTCTTCCGATCTNNNCACCTGTAATTCCAGCAGTTTG
AGCACACGTCTGAACTCTTTCCCTACACGACGCTCTTCCGATCTNNNCCTTTCCCTTCCTTTCCTTTCCTCA
AGCACACGTCTGAACTCTTTCCCTACACGACGCTCTTCCGATCTNNNCCTTTCCCTTCCTTTCCTTTCCTCA
GAGCACACGTCTGAACTCTTTCCCTACACGACGCTCTTCCGATCTNNNGGATAGATCTGATGCTCCATC
GAGCACACGTCTGAACTCTTTCCCTACACGACGCTCTTCCGATCTNNNGGATAGATCTGATGCTCCATC
CACACGTCTGAACTCTTTCCCTACACGACGCTCTTCCGATCTNNNTAAAAAAACATACAAAACGTACTGC
CACACGTCTGAACTCTTTCCCTACACGACGCTCTTCCGATCTNNNTAAAAAAACATACAAAACGTACTGC
GAGCACACGTCTGAACTCTTTCCCTACACGACGCTCTTCCGATCTNNNCAGAGACTACAGTGAGCCAA
GAGCACACGTCTGAACTCTTTCCCTACACGACGCTCTTCCGATCTNNNCAGAGACTACAGTGAGCCAA
CACACGTCTGAACTCTTTCCCTACACGACGCTCTTCCGATCTNNNGAGTCTGTGGGAAGGGACAGCATC
CACACGTCTGAACTCTTTCCCTACACGACGCTCTTCCGATCTNNNGAGTCTGTGGGAAGGGACAGCATC
CACACGTCTGAACTCTTTCCCTACACGACGCTCTTCCGATCTNNNGCCATCATCATCACTCTCTTCCTCATC
CACACGTCTGAACTCTTTCCCTACACGACGCTCTTCCGATCTNNNGCCATCATCATCACTCTCTTCCTCATC
GAGCACACGTCTGAACTCTTTCCCTACACGACGCTCTTCCGATCTNNNTTTACCTGGAACTCCTGATTCA
GAGCACACGTCTGAACTCTTTCCCTACACGACGCTCTTCCGATCTNNNTTTACCTGGAACTCCTGATTCA
AGCACACGTCTGAACTCTTTCCCTACACGACGCTCTTCCGATCTNNNCTGTAATCCCAGCTACTCAGGA

AGCACACGTCTGAACTCTTTCCCTACACGACGCTCTTCCGATCTNNNCTGTAATCCCAGCTACTCAGGA
AGCACACGTCTGAACTCTTTCCCTACACGACGCTCTTCCGATCTNNNATTCCCAGCTATTCAGGTGTCTG
AGCACACGTCTGAACTCTTTCCCTACACGACGCTCTTCCGATCTNNNATTCCCAGCTATTCAGGTGTCTG
GAGCACACGTCTGAACTCTTTCCCTACACGACGCTCTTCCGATCTNNNGATAGAGAATAGGCTGGGCA
GAGCACACGTCTGAACTCTTTCCCTACACGACGCTCTTCCGATCTNNNGATAGAGAATAGGCTGGGCA
AGCACACGTCTGAACTCTTTCCCTACACGACGCTCTTCCGATCTNNNCTTCTATCTCTATTGGGCTTTTA
AGCACACGTCTGAACTCTTTCCCTACACGACGCTCTTCCGATCTNNNCTTCTATCTCTATTGGGCTTTTA
AGCACACGTCTGAACTCTTTCCCTACACGACGCTCTTCCGATCTNNNGAGGGATAATTTCACACAGCA
AGCACACGTCTGAACTCTTTCCCTACACGACGCTCTTCCGATCTNNNGAGGGATAATTTCACACAGCA
CACGTCTGAACTCTTTCCCTACACGACGCTCTTCCGATCTNNNTAAACCGTGTGCTTCCCAATTTCCATC
CACGTCTGAACTCTTTCCCTACACGACGCTCTTCCGATCTNNNTAAACCGTGTGCTTCCCAATTTCCATC
AGCACACGTCTGAACTCTTTCCCTACACGACGCTCTTCCGATCTNNNGAAAGGACAAAGCTCAGGTTTC
AGCACACGTCTGAACTCTTTCCCTACACGACGCTCTTCCGATCTNNNGAAAGGACAAAGCTCAGGTTTC
AGCACACGTCTGAACTCTTTCCCTACACGACGCTCTTCCGATCTNNNCCTCCCCAATGTTGACATCTTA
AGCACACGTCTGAACTCTTTCCCTACACGACGCTCTTCCGATCTNNNCCTCCCCAATGTTGACATCTTA
GCACACGTCTGAACTCTTTCCCTACACGACGCTCTTCCGATCTNNNGTAGGGACTGGAAGCAGCAGC
GCACACGTCTGAACTCTTTCCCTACACGACGCTCTTCCGATCTNNNGTAGGGACTGGAAGCAGCAGC
GAGCACACGTCTGAACTCTTTCCCTACACGACGCTCTTCCGATCTNNNGATCACGTCACTGCACTCCAAC
GAGCACACGTCTGAACTCTTTCCCTACACGACGCTCTTCCGATCTNNNGATCACGTCACTGCACTCCAAC
GAGCACACGTCTGAACTCTTTCCCTACACGACGCTCTTCCGATCTNNNATGATTACTTAGATGCTATCG
GAGCACACGTCTGAACTCTTTCCCTACACGACGCTCTTCCGATCTNNNATGATTACTTAGATGCTATCG
AGCACACGTCTGAACTCTTTCCCTACACGACGCTCTTCCGATCTNNNGCCATCACACCAGGCTAATTTT
AGCACACGTCTGAACTCTTTCCCTACACGACGCTCTTCCGATCTNNNGCCATCACACCAGGCTAATTTT
AGCACACGTCTGAACTCTTTCCCTACACGACGCTCTTCCGATCTNNNATTGAAGCAGGGTCAGGGTTT
AGCACACGTCTGAACTCTTTCCCTACACGACGCTCTTCCGATCTNNNATTGAAGCAGGGTCAGGGTTT
GAGCACACGTCTGAACTCTTTCCCTACACGACGCTCTTCCGATCTNNNGGGAGGGATAGCATTGGGAC
GAGCACACGTCTGAACTCTTTCCCTACACGACGCTCTTCCGATCTNNNGGGAGGGATAGCATTGGGAC
AGCACACGTCTGAACTCTTTCCCTACACGACGCTCTTCCGATCTNNNTACTCGGCTGGATTCAGTTTG
AGCACACGTCTGAACTCTTTCCCTACACGACGCTCTTCCGATCTNNNTACTCGGCTGGATTCAGTTTG
GCACACGTCTGAACTCTTTCCCTACACGACGCTCTTCCGATCTNNNATTTGTAAGGCGTGACTCCTGA
GCACACGTCTGAACTCTTTCCCTACACGACGCTCTTCCGATCTNNNATTTGTAAGGCGTGACTCCTGA
GAGCACACGTCTGAACTCTTTCCCTACACGACGCTCTTCCGATCTNNNAAGCAAGGAAATCTGCCTTTA
GAGCACACGTCTGAACTCTTTCCCTACACGACGCTCTTCCGATCTNNNAAGCAAGGAAATCTGCCTTTA
AGAGCACACGTCTGAACTCTTTCCCTACACGACGCTCTTCCGATCTNNNTGAGGCAGGAGAATCTCTT
AGAGCACACGTCTGAACTCTTTCCCTACACGACGCTCTTCCGATCTNNNTGAGGCAGGAGAATCTCTT
GAGCACACGTCTGAACTCTTTCCCTACACGACGCTCTTCCGATCTNNNCCTGCCTTGTCTACATCCTGA
GAGCACACGTCTGAACTCTTTCCCTACACGACGCTCTTCCGATCTNNNCCTGCCTTGTCTACATCCTGA
AGCACACGTCTGAACTCTTTCCCTACACGACGCTCTTCCGATCTNNNGGGACTATTTTTGCCCCAAACTT
AGCACACGTCTGAACTCTTTCCCTACACGACGCTCTTCCGATCTNNNGGGACTATTTTTGCCCCAAACTT
AGAGCACACGTCTGAACTCTTTCCCTACACGACGCTCTTCCGATCTNNNGGAGATGTGTGTCTGTGTG
AGAGCACACGTCTGAACTCTTTCCCTACACGACGCTCTTCCGATCTNNNGGAGATGTGTGTCTGTGTG
AGCACACGTCTGAACTCTTTCCCTACACGACGCTCTTCCGATCTNNNCATCTGGGTTGTTTCCACCTTT
AGCACACGTCTGAACTCTTTCCCTACACGACGCTCTTCCGATCTNNNCATCTGGGTTGTTTCCACCTTT

AGCACACGTCTGAACTCTTTCCCTACACGACGCTCTTCCGATCTNNNGCTGTGGTCTTCTGCTCCTAAA
AGCACACGTCTGAACTCTTTCCCTACACGACGCTCTTCCGATCTNNNGCTGTGGTCTTCTGCTCCTAAA
GCACACGTCTGAACTCTTTCCCTACACGACGCTCTTCCGATCTNNNTCCCAACAAGGACATAGAGCTTC
GCACACGTCTGAACTCTTTCCCTACACGACGCTCTTCCGATCTNNNTCCCAACAAGGACATAGAGCTTC
AGCACACGTCTGAACTCTTTCCCTACACGACGCTCTTCCGATCTNNNAAAATGGAGATGGTGTGTGG
AGCACACGTCTGAACTCTTTCCCTACACGACGCTCTTCCGATCTNNNAAAATGGAGATGGTGTGTGG
GAGCACACGTCTGAACTCTTTCCCTACACGACGCTCTTCCGATCTNNNAAGGTGATGGCCTCCATTTC
GAGCACACGTCTGAACTCTTTCCCTACACGACGCTCTTCCGATCTNNNAAGGTGATGGCCTCCATTTC
GAGCACACGTCTGAACTCTTTCCCTACACGACGCTCTTCCGATCTNNNTCGTGACTTTGTGGTTTTTGT
GAGCACACGTCTGAACTCTTTCCCTACACGACGCTCTTCCGATCTNNNTCGTGACTTTGTGGTTTTTGT
GAGCACACGTCTGAACTCTTTCCCTACACGACGCTCTTCCGATCTNNNTGGGTTAGTGGGCAAAGTG
GAGCACACGTCTGAACTCTTTCCCTACACGACGCTCTTCCGATCTNNNTGGGTTAGTGGGCAAAGTG
GAGCACACGTCTGAACTCTTTCCCTACACGACGCTCTTCCGATCTNNNCAGGATTATGCTGGCCTCATA
GAGCACACGTCTGAACTCTTTCCCTACACGACGCTCTTCCGATCTNNNCAGGATTATGCTGGCCTCATA
GAGCACACGTCTGAACTCTTTCCCTACACGACGCTCTTCCGATCTNNNGGACTTGTCAGGCTCCATAA
GAGCACACGTCTGAACTCTTTCCCTACACGACGCTCTTCCGATCTNNNGGACTTGTCAGGCTCCATAA
GAGCACACGTCTGAACTCTTTCCCTACACGACGCTCTTCCGATCTNNNTGTCTGTATTGTGGCAGTGG
GAGCACACGTCTGAACTCTTTCCCTACACGACGCTCTTCCGATCTNNNTGTCTGTATTGTGGCAGTGG
GAGCACACGTCTGAACTCTTTCCCTACACGACGCTCTTCCGATCTNNNAGTACCCCTATCATCTACTAG
GAGCACACGTCTGAACTCTTTCCCTACACGACGCTCTTCCGATCTNNNAGTACCCCTATCATCTACTAG
GAGCACACGTCTGAACTCTTTCCCTACACGACGCTCTTCCGATCTNNNGGAGTGGGAGAGAAGAGAG
GAGCACACGTCTGAACTCTTTCCCTACACGACGCTCTTCCGATCTNNNGGAGTGGGAGAGAAGAGAG
GAGCACACGTCTGAACTCTTTCCCTACACGACGCTCTTCCGATCTNNNCATCTGTAGTCCCAGCTACTC
GAGCACACGTCTGAACTCTTTCCCTACACGACGCTCTTCCGATCTNNNCATCTGTAGTCCCAGCTACTC
GAGCACACGTCTGAACTCTTTCCCTACACGACGCTCTTCCGATCTNNNGCCACACATACTGACACCTAA
GAGCACACGTCTGAACTCTTTCCCTACACGACGCTCTTCCGATCTNNNGCCACACATACTGACACCTAA
AGCACACGTCTGAACTCTTTCCCTACACGACGCTCTTCCGATCTNNNGGCAGGAGAATCACTTGAACC
AGCACACGTCTGAACTCTTTCCCTACACGACGCTCTTCCGATCTNNNGGCAGGAGAATCACTTGAACC
GAGCACACGTCTGAACTCTTTCCCTACACGACGCTCTTCCGATCTNNNAGTTGGGAACTAAATGGGAG
GAGCACACGTCTGAACTCTTTCCCTACACGACGCTCTTCCGATCTNNNAGTTGGGAACTAAATGGGAG
GAGCACACGTCTGAACTCTTTCCCTACACGACGCTCTTCCGATCTNNNGGTGTCCAAGGGAGAGAATA
GAGCACACGTCTGAACTCTTTCCCTACACGACGCTCTTCCGATCTNNNGGTGTCCAAGGGAGAGAATA
GAGCACACGTCTGAACTCTTTCCCTACACGACGCTCTTCCGATCTNNNCTAGGCACAGTTTCCTTCTCTG
GAGCACACGTCTGAACTCTTTCCCTACACGACGCTCTTCCGATCTNNNCTAGGCACAGTTTCCTTCTCTG
AGCACACGTCTGAACTCTTTCCCTACACGACGCTCTTCCGATCTNNNAGACAATACTCGCAGGATCAA
AGCACACGTCTGAACTCTTTCCCTACACGACGCTCTTCCGATCTNNNAGACAATACTCGCAGGATCAA
GAGCACACGTCTGAACTCTTTCCCTACACGACGCTCTTCCGATCTNNNCCTCAGCCACAAAACAAAGAC
GAGCACACGTCTGAACTCTTTCCCTACACGACGCTCTTCCGATCTNNNCCTCAGCCACAAAACAAAGAC
GCACACGTCTGAACTCTTTCCCTACACGACGCTCTTCCGATCTNNNACCCCTACTGTTTTAGTTTGGGC
GCACACGTCTGAACTCTTTCCCTACACGACGCTCTTCCGATCTNNNACCCCTACTGTTTTAGTTTGGGC
GAGCACACGTCTGAACTCTTTCCCTACACGACGCTCTTCCGATCTNNNAATGTCAGGTATTGAGGAGG
GAGCACACGTCTGAACTCTTTCCCTACACGACGCTCTTCCGATCTNNNAATGTCAGGTATTGAGGAGG
GAGCACACGTCTGAACTCTTTCCCTACACGACGCTCTTCCGATCTNNNATTGTGAAAATGACCAGACTG

GAGCACACGTCTGAACTCTTTCCCTACACGACGCTCTTCCGATCTNNNATTGTGAAAATGACCAGACTG
AGCACACGTCTGAACTCTTTCCCTACACGACGCTCTTCCGATCTNNNATGTGCTTGTAATCCCAGCTAC
AGCACACGTCTGAACTCTTTCCCTACACGACGCTCTTCCGATCTNNNATGTGCTTGTAATCCCAGCTAC
GCACACGTCTGAACTCTTTCCCTACACGACGCTCTTCCGATCTNNNCAGCAGCAATATTGGAAGGCTTC
GCACACGTCTGAACTCTTTCCCTACACGACGCTCTTCCGATCTNNNCAGCAGCAATATTGGAAGGCTTC
GAGCACACGTCTGAACTCTTTCCCTACACGACGCTCTTCCGATCTNNNAAGCATGAAGGAGATTTCCA
GAGCACACGTCTGAACTCTTTCCCTACACGACGCTCTTCCGATCTNNNAAGCATGAAGGAGATTTCCA
AGCACACGTCTGAACTCTTTCCCTACACGACGCTCTTCCGATCTNNNGAGACTGAGGTGGGAGAATCA
AGCACACGTCTGAACTCTTTCCCTACACGACGCTCTTCCGATCTNNNGAGACTGAGGTGGGAGAATCA
GAGCACACGTCTGAACTCTTTCCCTACACGACGCTCTTCCGATCTNNNGCTGATTGCATGTGTGATACA
GAGCACACGTCTGAACTCTTTCCCTACACGACGCTCTTCCGATCTNNNGCTGATTGCATGTGTGATACA
GAGCACACGTCTGAACTCTTTCCCTACACGACGCTCTTCCGATCTNNNGGCACATGGCTGTATATGGCT
GAGCACACGTCTGAACTCTTTCCCTACACGACGCTCTTCCGATCTNNNGGCACATGGCTGTATATGGCT
GAGCACACGTCTGAACTCTTTCCCTACACGACGCTCTTCCGATCTNNNACCTTATTTGGCACCTGCTAAG
GAGCACACGTCTGAACTCTTTCCCTACACGACGCTCTTCCGATCTNNNACCTTATTTGGCACCTGCTAAG
CACACGTCTGAACTCTTTCCCTACACGACGCTCTTCCGATCTNNNTGGCTTTTCTCAAGATGACAGAAC
CACACGTCTGAACTCTTTCCCTACACGACGCTCTTCCGATCTNNNTGGCTTTTCTCAAGATGACAGAAC
AGAGCACACGTCTGAACTCTTTCCCTACACGACGCTCTTCCGATCTNNNTCTGGTTAGACAGAATCCAGA
AGAGCACACGTCTGAACTCTTTCCCTACACGACGCTCTTCCGATCTNNNTCTGGTTAGACAGAATCCAGA
AGCACACGTCTGAACTCTTTCCCTACACGACGCTCTTCCGATCTNNNCTGAGGCAGGAGAATCACTTCA
AGCACACGTCTGAACTCTTTCCCTACACGACGCTCTTCCGATCTNNNCTGAGGCAGGAGAATCACTTCA
AGAGCACACGTCTGAACTCTTTCCCTACACGACGCTCTTCCGATCTNNNCTTTGATCCACAAATCGGCCT
AGAGCACACGTCTGAACTCTTTCCCTACACGACGCTCTTCCGATCTNNNCTTTGATCCACAAATCGGCCT
AGCACACGTCTGAACTCTTTCCCTACACGACGCTCTTCCGATCTNNNGAGACACAGTTCAGTCCATAGCA
AGCACACGTCTGAACTCTTTCCCTACACGACGCTCTTCCGATCTNNNGAGACACAGTTCAGTCCATAGCA
AGCACACGTCTGAACTCTTTCCCTACACGACGCTCTTCCGATCTNNNGTCTGATCCAGCACAGTCATAG
AGCACACGTCTGAACTCTTTCCCTACACGACGCTCTTCCGATCTNNNGTCTGATCCAGCACAGTCATAG
AGAGCACACGTCTGAACTCTTTCCCTACACGACGCTCTTCCGATCTNNNAAGGAGCAGCAAGTCACATC
AGAGCACACGTCTGAACTCTTTCCCTACACGACGCTCTTCCGATCTNNNAAGGAGCAGCAAGTCACATC
AGCACACGTCTGAACTCTTTCCCTACACGACGCTCTTCCGATCTNNNTTTCTTGTGGGGTTTAGGGTTC
AGCACACGTCTGAACTCTTTCCCTACACGACGCTCTTCCGATCTNNNTTTCTTGTGGGGTTTAGGGTTC
AGCACACGTCTGAACTCTTTCCCTACACGACGCTCTTCCGATCTNNNTGCCAAAGAATTCAGCCACAATT
AGCACACGTCTGAACTCTTTCCCTACACGACGCTCTTCCGATCTNNNTGCCAAAGAATTCAGCCACAATT
GAGCACACGTCTGAACTCTTTCCCTACACGACGCTCTTCCGATCTNNNGGAGGTTGCAGTGAGCTAAGA
GAGCACACGTCTGAACTCTTTCCCTACACGACGCTCTTCCGATCTNNNGGAGGTTGCAGTGAGCTAAGA
AGCACACGTCTGAACTCTTTCCCTACACGACGCTCTTCCGATCTNNNCCACCATTCCTTTCTTTGCATCTC
AGCACACGTCTGAACTCTTTCCCTACACGACGCTCTTCCGATCTNNNCCACCATTCCTTTCTTTGCATCTC
AAGAGCACACGTCTGAACTCTTTCCCTACACGACGCTCTTCCGATCTNNNCAAGACCAGGTCAGCCCTTA
AAGAGCACACGTCTGAACTCTTTCCCTACACGACGCTCTTCCGATCTNNNCAAGACCAGGTCAGCCCTTA
AGAGCACACGTCTGAACTCTTTCCCTACACGACGCTCTTCCGATCTNNNCTTGCAGTGAGCCGAGATCA
AGAGCACACGTCTGAACTCTTTCCCTACACGACGCTCTTCCGATCTNNNCTTGCAGTGAGCCGAGATCA
GCACACGTCTGAACTCTTTCCCTACACGACGCTCTTCCGATCTNNNCATGGTAGTGTGCACCTGTAGTC
GCACACGTCTGAACTCTTTCCCTACACGACGCTCTTCCGATCTNNNCATGGTAGTGTGCACCTGTAGTC

GCACACGTCTGAACTCTTTCCCTACACGACGCTCTTCCGATCTNNNTATCCTGGGCTGTCTCACTCTAACA
GCACACGTCTGAACTCTTTCCCTACACGACGCTCTTCCGATCTNNNTATCCTGGGCTGTCTCACTCTAACA
GAGCACACGTCTGAACTCTTTCCCTACACGACGCTCTTCCGATCTNNNTCAGGCATTTGGGAAAAGAC
GAGCACACGTCTGAACTCTTTCCCTACACGACGCTCTTCCGATCTNNNTCAGGCATTTGGGAAAAGAC
AGAGCACACGTCTGAACTCTTTCCCTACACGACGCTCTTCCGATCTNNNGTCTGCTTCTGAGAGAGTCC
AGAGCACACGTCTGAACTCTTTCCCTACACGACGCTCTTCCGATCTNNNGTCTGCTTCTGAGAGAGTCC
AGCACACGTCTGAACTCTTTCCCTACACGACGCTCTTCCGATCTNNNATTCTCCACACGTCTTTTTGACTC
AGCACACGTCTGAACTCTTTCCCTACACGACGCTCTTCCGATCTNNNATTCTCCACACGTCTTTTTGACTC
AGCACACGTCTGAACTCTTTCCCTACACGACGCTCTTCCGATCTNNNTCAATATTTGGTTCCAGGAGGC
AGCACACGTCTGAACTCTTTCCCTACACGACGCTCTTCCGATCTNNNTCAATATTTGGTTCCAGGAGGC
AGCACACGTCTGAACTCTTTCCCTACACGACGCTCTTCCGATCTNNNAGAGGCTATGTCTGCTTTGTTCA
AGCACACGTCTGAACTCTTTCCCTACACGACGCTCTTCCGATCTNNNAGAGGCTATGTCTGCTTTGTTCA
AGCACACGTCTGAACTCTTTCCCTACACGACGCTCTTCCGATCTNNNAAAGCCTTCAGAGTCACAGGAA
AGCACACGTCTGAACTCTTTCCCTACACGACGCTCTTCCGATCTNNNAAAGCCTTCAGAGTCACAGGAA
GAGCACACGTCTGAACTCTTTCCCTACACGACGCTCTTCCGATCTNNNCTTCAGTCTGCTTTTGAGTGGA
GAGCACACGTCTGAACTCTTTCCCTACACGACGCTCTTCCGATCTNNNCTTCAGTCTGCTTTTGAGTGGA
AGCACACGTCTGAACTCTTTCCCTACACGACGCTCTTCCGATCTNNNCGAGATCACACCATTACACTCCA
AGCACACGTCTGAACTCTTTCCCTACACGACGCTCTTCCGATCTNNNCGAGATCACACCATTACACTCCA
GCACACGTCTGAACTCTTTCCCTACACGACGCTCTTCCGATCTNNNCTGAGGCAGGAGAATCACTTGA
GCACACGTCTGAACTCTTTCCCTACACGACGCTCTTCCGATCTNNNCTGAGGCAGGAGAATCACTTGA
GAGCACACGTCTGAACTCTTTCCCTACACGACGCTCTTCCGATCTNNNGAGAATGGTGTGAACCTAGG
GAGCACACGTCTGAACTCTTTCCCTACACGACGCTCTTCCGATCTNNNGAGAATGGTGTGAACCTAGG
AGCACACGTCTGAACTCTTTCCCTACACGACGCTCTTCCGATCTNNNAAAGTGCTGGGATTACAGGCAT
AGCACACGTCTGAACTCTTTCCCTACACGACGCTCTTCCGATCTNNNAAAGTGCTGGGATTACAGGCAT
GAGCACACGTCTGAACTCTTTCCCTACACGACGCTCTTCCGATCTNNNACCTGTAATCCCAACTGCTAAC
GAGCACACGTCTGAACTCTTTCCCTACACGACGCTCTTCCGATCTNNNACCTGTAATCCCAACTGCTAAC
AGAGCACACGTCTGAACTCTTTCCCTACACGACGCTCTTCCGATCTNNNATACAGGGAGGAAAAGGG
AGAGCACACGTCTGAACTCTTTCCCTACACGACGCTCTTCCGATCTNNNATACAGGGAGGAAAAGGG
GAGCACACGTCTGAACTCTTTCCCTACACGACGCTCTTCCGATCTNNNACAACCACACAAGCTTCAATCA
GAGCACACGTCTGAACTCTTTCCCTACACGACGCTCTTCCGATCTNNNACAACCACACAAGCTTCAATCA
GAGCACACGTCTGAACTCTTTCCCTACACGACGCTCTTCCGATCTNNNCTGTTTTTGGCATGCTTGGGA
GAGCACACGTCTGAACTCTTTCCCTACACGACGCTCTTCCGATCTNNNCTGTTTTTGGCATGCTTGGGA
GAGCACACGTCTGAACTCTTTCCCTACACGACGCTCTTCCGATCTNNNATCACGAGGTCAGGAGATCGA
GAGCACACGTCTGAACTCTTTCCCTACACGACGCTCTTCCGATCTNNNATCACGAGGTCAGGAGATCGA
GAGCACACGTCTGAACTCTTTCCCTACACGACGCTCTTCCGATCTNNNGCAGAAGAATCACTTGAACCC
GAGCACACGTCTGAACTCTTTCCCTACACGACGCTCTTCCGATCTNNNGCAGAAGAATCACTTGAACCC
GCACACGTCTGAACTCTTTCCCTACACGACGCTCTTCCGATCTNNNCTTGGATGGTTGCCATTTTTCCTC
GCACACGTCTGAACTCTTTCCCTACACGACGCTCTTCCGATCTNNNCTTGGATGGTTGCCATTTTTCCTC
GAGCACACGTCTGAACTCTTTCCCTACACGACGCTCTTCCGATCTNNNTCCTCTACCTCATAGTGCCTGAT
GAGCACACGTCTGAACTCTTTCCCTACACGACGCTCTTCCGATCTNNNTCCTCTACCTCATAGTGCCTGAT
AGCACACGTCTGAACTCTTTCCCTACACGACGCTCTTCCGATCTNNNTAGTACACACACACACACACACA
AGCACACGTCTGAACTCTTTCCCTACACGACGCTCTTCCGATCTNNNTAGTACACACACACACACACACA
GAGCACACGTCTGAACTCTTTCCCTACACGACGCTCTTCCGATCTNNNCCCTTTTTTGCCATTCACCGGCA

AGCACACGTCTGAACTCTTTCCCTACACGACGCTCTTCCGATCTNNNCCCTTTTTTGCCATTCACCGGC
GCACACGTCTGAACTCTTTCCCTACACGACGCTCTTCCGATCTNNNCCCATACCAGGTGGCATTATTC
GCACACGTCTGAACTCTTTCCCTACACGACGCTCTTCCGATCTNNNCCCATACCAGGTGGCATTATTC
GAGCACACGTCTGAACTCTTTCCCTACACGACGCTCTTCCGATCTNNNGCTGTTCTTGGTTGTTGAAG
GAGCACACGTCTGAACTCTTTCCCTACACGACGCTCTTCCGATCTNNNGCTGTTCTTGGTTGTTGAAG
ACACGTCTGAACTCTTTCCCTACACGACGCTCTTCCGATCTNNNTGCAAATCACAAACATGGCTTGTCA
ACACGTCTGAACTCTTTCCCTACACGACGCTCTTCCGATCTNNNTGCAAATCACAAACATGGCTTGTCA
AGCACACGTCTGAACTCTTTCCCTACACGACGCTCTTCCGATCTNNNAGCTATTTGGGAGACAAGACA
AGCACACGTCTGAACTCTTTCCCTACACGACGCTCTTCCGATCTNNNAGCTATTTGGGAGACAAGACA
AGAGCACACGTCTGAACTCTTTCCCTACACGACGCTCTTCCGATCTNNNCCAAGGTGTGACTGTATTT
AGAGCACACGTCTGAACTCTTTCCCTACACGACGCTCTTCCGATCTNNNCCAAGGTGTGACTGTATTT
AGCACACGTCTGAACTCTTTCCCTACACGACGCTCTTCCGATCTNNNGGATTACAGACGTGAGCCACT
AGCACACGTCTGAACTCTTTCCCTACACGACGCTCTTCCGATCTNNNGGATTACAGACGTGAGCCACT
GCACACGTCTGAACTCTTTCCCTACACGACGCTCTTCCGATCTNNNCGTAGTAACTTTTATTCGTGAA
GCACACGTCTGAACTCTTTCCCTACACGACGCTCTTCCGATCTNNNCGTAGTAACTTTTATTCGTGAA
AGCACACGTCTGAACTCTTTCCCTACACGACGCTCTTCCGATCTNNNTCCTTGTTTTTCTGCAACTTGC
AGCACACGTCTGAACTCTTTCCCTACACGACGCTCTTCCGATCTNNNTCCTTGTTTTTCTGCAACTTGC
GCACACGTCTGAACTCTTTCCCTACACGACGCTCTTCCGATCTNNNGGATACCTGTAGTCCCAGCTACT
GCACACGTCTGAACTCTTTCCCTACACGACGCTCTTCCGATCTNNNGGATACCTGTAGTCCCAGCTACT
AGCACACGTCTGAACTCTTTCCCTACACGACGCTCTTCCGATCTNNNTTTCTCTCCCTCTGCCTCTTTCC
AGCACACGTCTGAACTCTTTCCCTACACGACGCTCTTCCGATCTNNNTTTCTCTCCCTCTGCCTCTTTCC
AGCACACGTCTGAACTCTTTCCCTACACGACGCTCTTCCGATCTNNNAAAGCTGACTGTAAACTCCCCT
AGCACACGTCTGAACTCTTTCCCTACACGACGCTCTTCCGATCTNNNAAAGCTGACTGTAAACTCCCCT
AGCACACGTCTGAACTCTTTCCCTACACGACGCTCTTCCGATCTNNNAACCTAAGTGCCCATCAACTG
AGCACACGTCTGAACTCTTTCCCTACACGACGCTCTTCCGATCTNNNAACCTAAGTGCCCATCAACTG
AGCACACGTCTGAACTCTTTCCCTACACGACGCTCTTCCGATCTNNNGAAATCGCCTAACAACGCATT
AGCACACGTCTGAACTCTTTCCCTACACGACGCTCTTCCGATCTNNNGAAATCGCCTAACAACGCATT
AGCACACGTCTGAACTCTTTCCCTACACGACGCTCTTCCGATCTNNNAGATTTGTACAGGCATGGAGA
AGCACACGTCTGAACTCTTTCCCTACACGACGCTCTTCCGATCTNNNAGATTTGTACAGGCATGGAGA
AGCACACGTCTGAACTCTTTCCCTACACGACGCTCTTCCGATCTNNNACAGAAACCAGGATTTCAGG
AGCACACGTCTGAACTCTTTCCCTACACGACGCTCTTCCGATCTNNNACAGAAACCAGGATTTCAGG
GCACACGTCTGAACTCTTTCCCTACACGACGCTCTTCCGATCTNNNTGAGGCAGGAGAATAGCTTGA
GCACACGTCTGAACTCTTTCCCTACACGACGCTCTTCCGATCTNNNTGAGGCAGGAGAATAGCTTGA
AGCACACGTCTGAACTCTTTCCCTACACGACGCTCTTCCGATCTNNNTGTCCTTTCCTTCTCACTGCTT
AGCACACGTCTGAACTCTTTCCCTACACGACGCTCTTCCGATCTNNNTGTCCTTTCCTTCTCACTGCTT
AGAGCACACGTCTGAACTCTTTCCCTACACGACGCTCTTCCGATCTNNNCTCCATTCTGGTCCAACTC
AGAGCACACGTCTGAACTCTTTCCCTACACGACGCTCTTCCGATCTNNNCTCCATTCTGGTCCAACTC
AGCACACGTCTGAACTCTTTCCCTACACGACGCTCTTCCGATCTNNNAATATGCCTACGTGCTGAGAG
AGCACACGTCTGAACTCTTTCCCTACACGACGCTCTTCCGATCTNNNAATATGCCTACGTGCTGAGAG
AGCACACGTCTGAACTCTTTCCCTACACGACGCTCTTCCGATCTNNNCTGGGCCTGCATATACTCATC
AGCACACGTCTGAACTCTTTCCCTACACGACGCTCTTCCGATCTNNNCTGGGCCTGCATATACTCATC
GCACACGTCTGAACTCTTTCCCTACACGACGCTCTTCCGATCTNNNCACCGTGTCCAGCTAATTTTTG
GCACACGTCTGAACTCTTTCCCTACACGACGCTCTTCCGATCTNNNCACCGTGTCCAGCTAATTTTTG

AGCACACGTCTGAACTCTTTCCCTACACGACGCTCTTCCGATCTNNNCAAGATGGAGGGAAGTCGTA
AGCACACGTCTGAACTCTTTCCCTACACGACGCTCTTCCGATCTNNNCAAGATGGAGGGAAGTCGTA
GAGCACACGTCTGAACTCTTTCCCTACACGACGCTCTTCCGATCTNNNTTCCAGAAAGCTTCCTCTAAC
GAGCACACGTCTGAACTCTTTCCCTACACGACGCTCTTCCGATCTNNNTTCCAGAAAGCTTCCTCTAAC
AGCACACGTCTGAACTCTTTCCCTACACGACGCTCTTCCGATCTNNNTTTCACTGTGTTAGCCAGGATG
AGCACACGTCTGAACTCTTTCCCTACACGACGCTCTTCCGATCTNNNTTTCACTGTGTTAGCCAGGATG
GAGCACACGTCTGAACTCTTTCCCTACACGACGCTCTTCCGATCTNNNTTGAGACAGGAGAATCGCTT
GAGCACACGTCTGAACTCTTTCCCTACACGACGCTCTTCCGATCTNNNTTGAGACAGGAGAATCGCTT
GAGCACACGTCTGAACTCTTTCCCTACACGACGCTCTTCCGATCTNNNTGGTCTAAATGGAGCCAAAGA
GAGCACACGTCTGAACTCTTTCCCTACACGACGCTCTTCCGATCTNNNTGGTCTAAATGGAGCCAAAGA
GAGCACACGTCTGAACTCTTTCCCTACACGACGCTCTTCCGATCTNNNCTCCATCCAGTTTTGTTGCCTT
GAGCACACGTCTGAACTCTTTCCCTACACGACGCTCTTCCGATCTNNNCTCCATCCAGTTTTGTTGCCTT
AGCACACGTCTGAACTCTTTCCCTACACGACGCTCTTCCGATCTNNNTATTCTTTGACTGTGGTGCTG
AGCACACGTCTGAACTCTTTCCCTACACGACGCTCTTCCGATCTNNNTATTCTTTGACTGTGGTGCTG
GAGCACACGTCTGAACTCTTTCCCTACACGACGCTCTTCCGATCTNNNCAGGATTTGCACGTGTGAACTA
GAGCACACGTCTGAACTCTTTCCCTACACGACGCTCTTCCGATCTNNNCAGGATTTGCACGTGTGAACTA
AGCACACGTCTGAACTCTTTCCCTACACGACGCTCTTCCGATCTNNNCTCCTGTGTAGCTGGGATTACA
AGCACACGTCTGAACTCTTTCCCTACACGACGCTCTTCCGATCTNNNCTCCTGTGTAGCTGGGATTACA
GAGCACACGTCTGAACTCTTTCCCTACACGACGCTCTTCCGATCTNNNGATCACACTACTACACTCCAGC
GAGCACACGTCTGAACTCTTTCCCTACACGACGCTCTTCCGATCTNNNGATCACACTACTACACTCCAGC
AGCACACGTCTGAACTCTTTCCCTACACGACGCTCTTCCGATCTNNNGAGAATGGCATGAACCTGGGA
AGCACACGTCTGAACTCTTTCCCTACACGACGCTCTTCCGATCTNNNGAGAATGGCATGAACCTGGGA
GCACACGTCTGAACTCTTTCCCTACACGACGCTCTTCCGATCTNNNGAGACAGGGTTTCAACATGTTG
GCACACGTCTGAACTCTTTCCCTACACGACGCTCTTCCGATCTNNNGAGACAGGGTTTCAACATGTTG
GAGCACACGTCTGAACTCTTTCCCTACACGACGCTCTTCCGATCTNNNACGCCTGTAAATCCAACACTTT
GAGCACACGTCTGAACTCTTTCCCTACACGACGCTCTTCCGATCTNNNACGCCTGTAAATCCAACACTTT
AGCACACGTCTGAACTCTTTCCCTACACGACGCTCTTCCGATCTNNNCTCCCTTTCCTTCCTTCCTTTCTCA
AGCACACGTCTGAACTCTTTCCCTACACGACGCTCTTCCGATCTNNNCTCCCTTTCCTTCCTTCCTTTCTCA
AGCACACGTCTGAACTCTTTCCCTACACGACGCTCTTCCGATCTNNNGGAGCTTACAGTGAGTGGAGA
AGCACACGTCTGAACTCTTTCCCTACACGACGCTCTTCCGATCTNNNGGAGCTTACAGTGAGTGGAGA
GCACACGTCTGAACTCTTTCCCTACACGACGCTCTTCCGATCTNNNGGTAACCTTTGATTGGATGCTGG
GCACACGTCTGAACTCTTTCCCTACACGACGCTCTTCCGATCTNNNGGTAACCTTTGATTGGATGCTGG
GCACACGTCTGAACTCTTTCCCTACACGACGCTCTTCCGATCTNNNTAGAATTTCTGCCGAGAGATCC
GCACACGTCTGAACTCTTTCCCTACACGACGCTCTTCCGATCTNNNTAGAATTTCTGCCGAGAGATCC
GAGCACACGTCTGAACTCTTTCCCTACACGACGCTCTTCCGATCTNNNAGAGAGACCAGGGAGAAAA
GAGCACACGTCTGAACTCTTTCCCTACACGACGCTCTTCCGATCTNNNAGAGAGACCAGGGAGAAAA
AGCACACGTCTGAACTCTTTCCCTACACGACGCTCTTCCGATCTNNNAGCTGAATAAAAGAGGGAGGT
AGCACACGTCTGAACTCTTTCCCTACACGACGCTCTTCCGATCTNNNAGCTGAATAAAAGAGGGAGGT
AGCACACGTCTGAACTCTTTCCCTACACGACGCTCTTCCGATCTNNNTTTTTTTGTTGGGATTTCAGATA
AGCACACGTCTGAACTCTTTCCCTACACGACGCTCTTCCGATCTNNNTTTTTTTGTTGGGATTTCAGATA
GCACACGTCTGAACTCTTTCCCTACACGACGCTCTTCCGATCTNNNGGTGTGGTGAAATCAAACAGCTT
GCACACGTCTGAACTCTTTCCCTACACGACGCTCTTCCGATCTNNNGGTGTGGTGAAATCAAACAGCTT
AGCACACGTCTGAACTCTTTCCCTACACGACGCTCTTCCGATCTNNNTGAATTACTGTAACAGGAGTG

AGCACACGTCTGAACTCTTTCCCTACACGACGCTCTTCCGATCTNNNTGAATTACTGTAACAGGAGTGC

GAGCACACGTCTGAACTCTTTCCCTACACGACGCTCTTCCGATCTNNNAGACTGGCTGTAAGGAATGCA

GAGCACACGTCTGAACTCTTTCCCTACACGACGCTCTTCCGATCTNNNAGACTGGCTGTAAGGAATGCA

AGCACACGTCTGAACTCTTTCCCTACACGACGCTCTTCCGATCTNNNGAGCTTGCAGTGAGCCAAGATC

AGCACACGTCTGAACTCTTTCCCTACACGACGCTCTTCCGATCTNNNGAGCTTGCAGTGAGCCAAGATC

AGCACACGTCTGAACTCTTTCCCTACACGACGCTCTTCCGATCTNNNGGGAGAATTGCTTGAATCTGG

AGCACACGTCTGAACTCTTTCCCTACACGACGCTCTTCCGATCTNNNGGGAGAATTGCTTGAATCTGG

AGCACACGTCTGAACTCTTTCCCTACACGACGCTCTTCCGATCTNNNAGTTCTAATAAGCTTGTGACTG

AGCACACGTCTGAACTCTTTCCCTACACGACGCTCTTCCGATCTNNNAGTTCTAATAAGCTTGTGACTG

GAGCACACGTCTGAACTCTTTCCCTACACGACGCTCTTCCGATCTNNNGTATGTGTGTGTGTGTGTGTC

GAGCACACGTCTGAACTCTTTCCCTACACGACGCTCTTCCGATCTNNNGTATGTGTGTGTGTGTGTGTC

AGAGCACACGTCTGAACTCTTTCCCTACACGACGCTCTTCCGATCTNNNAGCAGAAATCCCTGATAAAC

AGAGCACACGTCTGAACTCTTTCCCTACACGACGCTCTTCCGATCTNNNAGCAGAAATCCCTGATAAAC

GAGCACACGTCTGAACTCTTTCCCTACACGACGCTCTTCCGATCTNNNAAATCTCAGTGTGTGTCTGTCG

GAGCACACGTCTGAACTCTTTCCCTACACGACGCTCTTCCGATCTNNNAAATCTCAGTGTGTGTCTGTCG

AGCACACGTCTGAACTCTTTCCCTACACGACGCTCTTCCGATCTNNNTATTAGAGGCTGGGTACTCTCC

AGCACACGTCTGAACTCTTTCCCTACACGACGCTCTTCCGATCTNNNTATTAGAGGCTGGGTACTCTCC

GAGCACACGTCTGAACTCTTTCCCTACACGACGCTCTTCCGATCTNNNGAGCAAGACTCCATCTCAAAA

GAGCACACGTCTGAACTCTTTCCCTACACGACGCTCTTCCGATCTNNNGAGCAAGACTCCATCTCAAAA

AGCACACGTCTGAACTCTTTCCCTACACGACGCTCTTCCGATCTNNNAAAAATTGGCCTGGCATCATGA

AGCACACGTCTGAACTCTTTCCCTACACGACGCTCTTCCGATCTNNNAAAAATTGGCCTGGCATCATGA

AGAGCACACGTCTGAACTCTTTCCCTACACGACGCTCTTCCGATCTNNNGAGAATCTCGCTGTCCTCTGA

AGAGCACACGTCTGAACTCTTTCCCTACACGACGCTCTTCCGATCTNNNGAGAATCTCGCTGTCCTCTGA

AGCACACGTCTGAACTCTTTCCCTACACGACGCTCTTCCGATCTNNNCTGGGTGACACAATGAGTCTCT

AGCACACGTCTGAACTCTTTCCCTACACGACGCTCTTCCGATCTNNNCTGGGTGACACAATGAGTCTCT

AGCACACGTCTGAACTCTTTCCCTACACGACGCTCTTCCGATCTNNNACCTACACTTGTCACCTTTGTGT

AGCACACGTCTGAACTCTTTCCCTACACGACGCTCTTCCGATCTNNNACCTACACTTGTCACCTTTGTGT

AGCACACGTCTGAACTCTTTCCCTACACGACGCTCTTCCGATCTNNNGTGAGCCGAGATCATGTCACTA

AGCACACGTCTGAACTCTTTCCCTACACGACGCTCTTCCGATCTNNNGTGAGCCGAGATCATGTCACTA

AGCACACGTCTGAACTCTTTCCCTACACGACGCTCTTCCGATCTNNNGAAATCATCTCCACTGAATGCC

AGCACACGTCTGAACTCTTTCCCTACACGACGCTCTTCCGATCTNNNGAAATCATCTCCACTGAATGCC

AGCACACGTCTGAACTCTTTCCCTACACGACGCTCTTCCGATCTNNNCTAAGGATAATGGCCTCCACCTC

AGCACACGTCTGAACTCTTTCCCTACACGACGCTCTTCCGATCTNNNCTAAGGATAATGGCCTCCACCTC

GAGCACACGTCTGAACTCTTTCCCTACACGACGCTCTTCCGATCTNNNAAAAATTTAGCCAGGCGTGGT

GAGCACACGTCTGAACTCTTTCCCTACACGACGCTCTTCCGATCTNNNAAAAATTTAGCCAGGCGTGGT

GAGCACACGTCTGAACTCTTTCCCTACACGACGCTCTTCCGATCTNNNTCTGCATTTCCAACTGAGGTAC

GAGCACACGTCTGAACTCTTTCCCTACACGACGCTCTTCCGATCTNNNTCTGCATTTCCAACTGAGGTAC

GAGCACACGTCTGAACTCTTTCCCTACACGACGCTCTTCCGATCTNNNCATGAAAGCCATCAACCCTGA

GAGCACACGTCTGAACTCTTTCCCTACACGACGCTCTTCCGATCTNNNCATGAAAGCCATCAACCCTGA

AGAGCACACGTCTGAACTCTTTCCCTACACGACGCTCTTCCGATCTNNNTCAGTTGAATTGGAGAGAAC

AGAGCACACGTCTGAACTCTTTCCCTACACGACGCTCTTCCGATCTNNNTCAGTTGAATTGGAGAGAAC

AGCACACGTCTGAACTCTTTCCCTACACGACGCTCTTCCGATCTNNNTCACAGACACGGGTGTTCTTAC

AGCACACGTCTGAACTCTTTCCCTACACGACGCTCTTCCGATCTNNNTCACAGACACGGGTGTTCTTAC

GAGCACACGTCTGAACTCTTTCCCTACACGACGCTCTTCCGATCTNNNATATGCTGCCAGCCTCAGAT

GAGCACACGTCTGAACTCTTTCCCTACACGACGCTCTTCCGATCTNNNATATGCTGCCAGCCTCAGAT

AGCACACGTCTGAACTCTTTCCCTACACGACGCTCTTCCGATCTNNNCAGGACCCTTTTGAGAGTGAA

AGCACACGTCTGAACTCTTTCCCTACACGACGCTCTTCCGATCTNNNCAGGACCCTTTTGAGAGTGAA

AGCACACGTCTGAACTCTTTCCCTACACGACGCTCTTCCGATCTNNNTTGTTGGGAAATGGTAGTGA

AGCACACGTCTGAACTCTTTCCCTACACGACGCTCTTCCGATCTNNNTTGTTGGGAAATGGTAGTGA

AGCACACGTCTGAACTCTTTCCCTACACGACGCTCTTCCGATCTNNNCTTAGGCAGGAGAATCGCTTG

AGCACACGTCTGAACTCTTTCCCTACACGACGCTCTTCCGATCTNNNCTTAGGCAGGAGAATCGCTTG

GAGCACACGTCTGAACTCTTTCCCTACACGACGCTCTTCCGATCTNNNTAGGTGCTGTTATACCGCAG

GAGCACACGTCTGAACTCTTTCCCTACACGACGCTCTTCCGATCTNNNTAGGTGCTGTTATACCGCAG

GAGCACACGTCTGAACTCTTTCCCTACACGACGCTCTTCCGATCTNNNGTGGTACAGTCACAGCTCAC

GAGCACACGTCTGAACTCTTTCCCTACACGACGCTCTTCCGATCTNNNGTGGTACAGTCACAGCTCAC

AGCACACGTCTGAACTCTTTCCCTACACGACGCTCTTCCGATCTNNNCCCAAATGATGAAGTGATGCA

AGCACACGTCTGAACTCTTTCCCTACACGACGCTCTTCCGATCTNNNCCCAAATGATGAAGTGATGCA

AGCACACGTCTGAACTCTTTCCCTACACGACGCTCTTCCGATCTNNNAAAAAATTAGCCGGGCATAGT

AGCACACGTCTGAACTCTTTCCCTACACGACGCTCTTCCGATCTNNNAAAAAATTAGCCGGGCATAGT

CACACGTCTGAACTCTTTCCCTACACGACGCTCTTCCGATCTNNNAATTCAGATTCGCTTCAGGAGGTC

CACACGTCTGAACTCTTTCCCTACACGACGCTCTTCCGATCTNNNAATTCAGATTCGCTTCAGGAGGTC

GCACACGTCTGAACTCTTTCCCTACACGACGCTCTTCCGATCTNNNTCTTTTTGGTTGTGTCTCTGCC

GCACACGTCTGAACTCTTTCCCTACACGACGCTCTTCCGATCTNNNTCTTTTTGGTTGTGTCTCTGCC

GAGCACACGTCTGAACTCTTTCCCTACACGACGCTCTTCCGATCTNNNGAGATCACACCATTGCACTCC

GAGCACACGTCTGAACTCTTTCCCTACACGACGCTCTTCCGATCTNNNGAGATCACACCATTGCACTCC

GAGCACACGTCTGAACTCTTTCCCTACACGACGCTCTTCCGATCTNNNTCTCCTGCTATACACACACAC

GAGCACACGTCTGAACTCTTTCCCTACACGACGCTCTTCCGATCTNNNTCTCCTGCTATACACACACAC

AGCACACGTCTGAACTCTTTCCCTACACGACGCTCTTCCGATCTNNNCTGTAATCCCAGCAACACTTTG

AGCACACGTCTGAACTCTTTCCCTACACGACGCTCTTCCGATCTNNNCTGTAATCCCAGCAACACTTTG

GCACACGTCTGAACTCTTTCCCTACACGACGCTCTTCCGATCTNNNAATAGCAGAATTCCTAGTCAAAA

GCACACGTCTGAACTCTTTCCCTACACGACGCTCTTCCGATCTNNNAATAGCAGAATTCCTAGTCAAAA

AGCACACGTCTGAACTCTTTCCCTACACGACGCTCTTCCGATCTNNNCTGTAATCCCAGCTACTCAGGA

AGCACACGTCTGAACTCTTTCCCTACACGACGCTCTTCCGATCTNNNCTGTAATCCCAGCTACTCAGGA

GAGCACACGTCTGAACTCTTTCCCTACACGACGCTCTTCCGATCTNNNTAATAACAGGCCAGCAAACAT

GAGCACACGTCTGAACTCTTTCCCTACACGACGCTCTTCCGATCTNNNTAATAACAGGCCAGCAAACAT

GAGCACACGTCTGAACTCTTTCCCTACACGACGCTCTTCCGATCTNNNGAGATCGTGCCACTACACTCT

GAGCACACGTCTGAACTCTTTCCCTACACGACGCTCTTCCGATCTNNNGAGATCGTGCCACTACACTCT

GCACACGTCTGAACTCTTTCCCTACACGACGCTCTTCCGATCTNNNGCCACAGTTGAAAAACCCTGTTC

GCACACGTCTGAACTCTTTCCCTACACGACGCTCTTCCGATCTNNNGCCACAGTTGAAAAACCCTGTTC

AGCACACGTCTGAACTCTTTCCCTACACGACGCTCTTCCGATCTNNNTGTAGTCCTAGCTACTTGGGAG

AGCACACGTCTGAACTCTTTCCCTACACGACGCTCTTCCGATCTNNNTGTAGTCCTAGCTACTTGGGAG

AGCACACGTCTGAACTCTTTCCCTACACGACGCTCTTCCGATCTNNNGAGGGAGGTAGGGGGGTCA

AGCACACGTCTGAACTCTTTCCCTACACGACGCTCTTCCGATCTNNNGAGGGAGGTAGGGGGGTCA

AGCACACGTCTGAACTCTTTCCCTACACGACGCTCTTCCGATCTNNNATGTAGTTGCCAAATATCCTTA

AGCACACGTCTGAACTCTTTCCCTACACGACGCTCTTCCGATCTNNNATGTAGTTGCCAAATATCCTTA

AGCACACGTCTGAACTCTTTCCCTACACGACGCTCTTCCGATCTNNNCTGGGAAACAAGAGTGAAAA

AGCACACGTCTGAACTCTTTCCCTACACGACGCTCTTCCGATCTNNNCTGGGAAACAAGAGTGAAAA
GCACACGTCTGAACTCTTTCCCTACACGACGCTCTTCCGATCTNNNTCACCTTGTGATCATGTGTCAGT
GCACACGTCTGAACTCTTTCCCTACACGACGCTCTTCCGATCTNNNTCACCTTGTGATCATGTGTCAGT
GAGCACACGTCTGAACTCTTTCCCTACACGACGCTCTTCCGATCTNNNGCACATGTAGTCCCAGATACT
GAGCACACGTCTGAACTCTTTCCCTACACGACGCTCTTCCGATCTNNNGCACATGTAGTCCCAGATACT
GCACACGTCTGAACTCTTTCCCTACACGACGCTCTTCCGATCTNNNTCCATCACAGTCTTGACAAGAGG
GCACACGTCTGAACTCTTTCCCTACACGACGCTCTTCCGATCTNNNTCCATCACAGTCTTGACAAGAGG
AGCACACGTCTGAACTCTTTCCCTACACGACGCTCTTCCGATCTNNNGTTCACAGTCTGAGTTCCATTG
AGCACACGTCTGAACTCTTTCCCTACACGACGCTCTTCCGATCTNNNGTTCACAGTCTGAGTTCCATTG
GCACACGTCTGAACTCTTTCCCTACACGACGCTCTTCCGATCTNNNCTTCTCCTGTTCTTGGGTGGTAA
GCACACGTCTGAACTCTTTCCCTACACGACGCTCTTCCGATCTNNNCTTCTCCTGTTCTTGGGTGGTAA
AGCACACGTCTGAACTCTTTCCCTACACGACGCTCTTCCGATCTNNNCTGCTCAGAGCGCAGTACTTAC
AGCACACGTCTGAACTCTTTCCCTACACGACGCTCTTCCGATCTNNNCTGCTCAGAGCGCAGTACTTAC
AGAGCACACGTCTGAACTCTTTCCCTACACGACGCTCTTCCGATCTNNNCCTAGCAAAAGTGGAAGGA
AGAGCACACGTCTGAACTCTTTCCCTACACGACGCTCTTCCGATCTNNNCCTAGCAAAAGTGGAAGGA
AGCACACGTCTGAACTCTTTCCCTACACGACGCTCTTCCGATCTNNNGACGGGGTTTTCCTTGTACAG
AGCACACGTCTGAACTCTTTCCCTACACGACGCTCTTCCGATCTNNNGACGGGGTTTTCCTTGTACAG
CACGTCTGAACTCTTTCCCTACACGACGCTCTTCCGATCTNNNACTCACCCTTACCTTGTCCTTTCCATCG
CACGTCTGAACTCTTTCCCTACACGACGCTCTTCCGATCTNNNACTCACCCTTACCTTGTCCTTTCCATCG
GAGCACACGTCTGAACTCTTTCCCTACACGACGCTCTTCCGATCTNNNTGGACACAGCTGAGGAAAA
GAGCACACGTCTGAACTCTTTCCCTACACGACGCTCTTCCGATCTNNNTGGACACAGCTGAGGAAAA
AGAGCACACGTCTGAACTCTTTCCCTACACGACGCTCTTCCGATCTNNNTCAGACACCCAATATGGATC
AGAGCACACGTCTGAACTCTTTCCCTACACGACGCTCTTCCGATCTNNNTCAGACACCCAATATGGATC
AGCACACGTCTGAACTCTTTCCCTACACGACGCTCTTCCGATCTNNNATCACCTGGAGACTTGGACAA
AGCACACGTCTGAACTCTTTCCCTACACGACGCTCTTCCGATCTNNNATCACCTGGAGACTTGGACAA
AGCACACGTCTGAACTCTTTCCCTACACGACGCTCTTCCGATCTNNNTGCCTGTAATCCCAGCACTTTC
AGCACACGTCTGAACTCTTTCCCTACACGACGCTCTTCCGATCTNNNTGCCTGTAATCCCAGCACTTTC
AGCACACGTCTGAACTCTTTCCCTACACGACGCTCTTCCGATCTNNNTTGTCATTCAGTCTTGGGAGG
AGCACACGTCTGAACTCTTTCCCTACACGACGCTCTTCCGATCTNNNTTGTCATTCAGTCTTGGGAGG
AGCACACGTCTGAACTCTTTCCCTACACGACGCTCTTCCGATCTNNNGCTGTTTTAACTTTATTGTGAT
AGCACACGTCTGAACTCTTTCCCTACACGACGCTCTTCCGATCTNNNGCTGTTTTAACTTTATTGTGAT
GAGCACACGTCTGAACTCTTTCCCTACACGACGCTCTTCCGATCTNNNTCCTCCTTCTTTTCTCCCTAGGT
GAGCACACGTCTGAACTCTTTCCCTACACGACGCTCTTCCGATCTNNNTCCTCCTTCTTTTCTCCCTAGGT
AGAGCACACGTCTGAACTCTTTCCCTACACGACGCTCTTCCGATCTNNNCTGAGGCAGGAGAATCACT
AGAGCACACGTCTGAACTCTTTCCCTACACGACGCTCTTCCGATCTNNNCTGAGGCAGGAGAATCACT
AGCACACGTCTGAACTCTTTCCCTACACGACGCTCTTCCGATCTNNNCTATGGCAAGACTCAGTTTGC
AGCACACGTCTGAACTCTTTCCCTACACGACGCTCTTCCGATCTNNNCTATGGCAAGACTCAGTTTGC
AGCACACGTCTGAACTCTTTCCCTACACGACGCTCTTCCGATCTNNNTAGCCTCTGGTTTCTCCTTTTCC
AGCACACGTCTGAACTCTTTCCCTACACGACGCTCTTCCGATCTNNNTAGCCTCTGGTTTCTCCTTTTCC
GAGCACACGTCTGAACTCTTTCCCTACACGACGCTCTTCCGATCTNNNGTGATGGTGGTAGGAAGAT
GAGCACACGTCTGAACTCTTTCCCTACACGACGCTCTTCCGATCTNNNGTGATGGTGGTAGGAAGAT
AGAGCACACGTCTGAACTCTTTCCCTACACGACGCTCTTCCGATCTNNNGAAGACTCCAGGAGCGGGA
AGAGCACACGTCTGAACTCTTTCCCTACACGACGCTCTTCCGATCTNNNGAAGACTCCAGGAGCGGGA

AGCACACGTCTGAACTCTTTCCCTACACGACGCTCTTCCGATCTNNNGCTCCTGTATTGGTTGGTTCAA

AGCACACGTCTGAACTCTTTCCCTACACGACGCTCTTCCGATCTNNNGCTCCTGTATTGGTTGGTTCAA

CACACGTCTGAACTCTTTCCCTACACGACGCTCTTCCGATCTNNNCAGAAACACTTTTCGACATAGCAT

CACACGTCTGAACTCTTTCCCTACACGACGCTCTTCCGATCTNNNCAGAAACACTTTTCGACATAGCAT

CACACGTCTGAACTCTTTCCCTACACGACGCTCTTCCGATCTNNNTCCTTTCAGTGGGCAGAAATAGTC

CACACGTCTGAACTCTTTCCCTACACGACGCTCTTCCGATCTNNNTCCTTTCAGTGGGCAGAAATAGTC

AGCACACGTCTGAACTCTTTCCCTACACGACGCTCTTCCGATCTNNNTCAGATATGGTGGCATCAGAGT

AGCACACGTCTGAACTCTTTCCCTACACGACGCTCTTCCGATCTNNNTCAGATATGGTGGCATCAGAGT

GAGCACACGTCTGAACTCTTTCCCTACACGACGCTCTTCCGATCTNNNTAATTTCAGCACTTTGGGAGG

GAGCACACGTCTGAACTCTTTCCCTACACGACGCTCTTCCGATCTNNNTAATTTCAGCACTTTGGGAGG

GCACACGTCTGAACTCTTTCCCTACACGACGCTCTTCCGATCTNNNGCCTAGAAGAAAATGTACTTAGT

GCACACGTCTGAACTCTTTCCCTACACGACGCTCTTCCGATCTNNNGCCTAGAAGAAAATGTACTTAGT

AGCACACGTCTGAACTCTTTCCCTACACGACGCTCTTCCGATCTNNNCTCACAGAACACCAGTGTCCTAA

AGCACACGTCTGAACTCTTTCCCTACACGACGCTCTTCCGATCTNNNCTCACAGAACACCAGTGTCCTAA

GAGCACACGTCTGAACTCTTTCCCTACACGACGCTCTTCCGATCTNNNGCATTGTGCAAGTGTATAGTG

GAGCACACGTCTGAACTCTTTCCCTACACGACGCTCTTCCGATCTNNNGCATTGTGCAAGTGTATAGTG

GAGCACACGTCTGAACTCTTTCCCTACACGACGCTCTTCCGATCTNNNCTGTAATCCCAGCTACTCAGG

GAGCACACGTCTGAACTCTTTCCCTACACGACGCTCTTCCGATCTNNNCTGTAATCCCAGCTACTCAGG

GCACACGTCTGAACTCTTTCCCTACACGACGCTCTTCCGATCTNNNATAATGAGGTTTTTAACTTGTGC

GCACACGTCTGAACTCTTTCCCTACACGACGCTCTTCCGATCTNNNATAATGAGGTTTTTAACTTGTGC

GAGCACACGTCTGAACTCTTTCCCTACACGACGCTCTTCCGATCTNNNGAGATCGTACCACTGCACTCT

GAGCACACGTCTGAACTCTTTCCCTACACGACGCTCTTCCGATCTNNNGAGATCGTACCACTGCACTCT

AGCACACGTCTGAACTCTTTCCCTACACGACGCTCTTCCGATCTNNNAAAAGGAGAGAGAGCGGCAA

AGCACACGTCTGAACTCTTTCCCTACACGACGCTCTTCCGATCTNNNAAAAGGAGAGAGAGCGGCAA

GAGCACACGTCTGAACTCTTTCCCTACACGACGCTCTTCCGATCTNNNCTCTTACCTCCAGGCCATTTC

GAGCACACGTCTGAACTCTTTCCCTACACGACGCTCTTCCGATCTNNNCTCTTACCTCCAGGCCATTTC

AGCACACGTCTGAACTCTTTCCCTACACGACGCTCTTCCGATCTNNNTGCGCTTTTTCTGGTCTACATTC

AGCACACGTCTGAACTCTTTCCCTACACGACGCTCTTCCGATCTNNNTGCGCTTTTTCTGGTCTACATTC

GCACACGTCTGAACTCTTTCCCTACACGACGCTCTTCCGATCTNNNGCACATGTAGTCCCAGCTACTAG

GCACACGTCTGAACTCTTTCCCTACACGACGCTCTTCCGATCTNNNGCACATGTAGTCCCAGCTACTAG

GCACACGTCTGAACTCTTTCCCTACACGACGCTCTTCCGATCTNNNTTTCCACAGTATCTTGCCTTCCAC

GCACACGTCTGAACTCTTTCCCTACACGACGCTCTTCCGATCTNNNTTTCCACAGTATCTTGCCTTCCAC

AGCACACGTCTGAACTCTTTCCCTACACGACGCTCTTCCGATCTNNNTGGATAAAGGTTAGGTGTGGT

AGCACACGTCTGAACTCTTTCCCTACACGACGCTCTTCCGATCTNNNTGGATAAAGGTTAGGTGTGGT

AGCACACGTCTGAACTCTTTCCCTACACGACGCTCTTCCGATCTNNNATGCCTGTAATCCCAGCTACTTG

AGCACACGTCTGAACTCTTTCCCTACACGACGCTCTTCCGATCTNNNATGCCTGTAATCCCAGCTACTTG

AGAGCACACGTCTGAACTCTTTCCCTACACGACGCTCTTCCGATCTNNNGAGGCAGGAGAATCATTTG

AGAGCACACGTCTGAACTCTTTCCCTACACGACGCTCTTCCGATCTNNNGAGGCAGGAGAATCATTTG

AGAGCACACGTCTGAACTCTTTCCCTACACGACGCTCTTCCGATCTNNNGAGATGCGGAAATGGGGAT

AGAGCACACGTCTGAACTCTTTCCCTACACGACGCTCTTCCGATCTNNNGAGATGCGGAAATGGGGAT

GAGCACACGTCTGAACTCTTTCCCTACACGACGCTCTTCCGATCTNNNTCTCGGCCTTAGCATGTAGTT

GAGCACACGTCTGAACTCTTTCCCTACACGACGCTCTTCCGATCTNNNTCTCGGCCTTAGCATGTAGTT

GAGCACACGTCTGAACTCTTTCCCTACACGACGCTCTTCCGATCTNNNTGAGGCAGCAAAATCACTTGA

AGCACACGTCTGAACTCTTTCCCTACACGACGCTCTTCCGATCTNNNTGAGGCAGCAAAATCACTTG
AGCACACGTCTGAACTCTTTCCCTACACGACGCTCTTCCGATCTNNNTAAGGCAGGAGAATCGCTTG
AGCACACGTCTGAACTCTTTCCCTACACGACGCTCTTCCGATCTNNNTAAGGCAGGAGAATCGCTTG
AGCACACGTCTGAACTCTTTCCCTACACGACGCTCTTCCGATCTNNNGCCATTTCCACCTCTCCATTTTC
AGCACACGTCTGAACTCTTTCCCTACACGACGCTCTTCCGATCTNNNGCCATTTCCACCTCTCCATTTTC
AGCACACGTCTGAACTCTTTCCCTACACGACGCTCTTCCGATCTNNNAAAAAATTAGCTGGGTGTGGT
AGCACACGTCTGAACTCTTTCCCTACACGACGCTCTTCCGATCTNNNAAAAAATTAGCTGGGTGTGGT
GCACACGTCTGAACTCTTTCCCTACACGACGCTCTTCCGATCTNNNAACACCTACATGTGCCAAACAGT
GCACACGTCTGAACTCTTTCCCTACACGACGCTCTTCCGATCTNNNAACACCTACATGTGCCAAACAGT
AGAGCACACGTCTGAACTCTTTCCCTACACGACGCTCTTCCGATCTNNNGCTCGCCCTCTTTCTATTCT
AGAGCACACGTCTGAACTCTTTCCCTACACGACGCTCTTCCGATCTNNNGCTCGCCCTCTTTCTATTCT
GCACACGTCTGAACTCTTTCCCTACACGACGCTCTTCCGATCTNNNTTAGCTGCTAGTGGCTGAAAGA
GCACACGTCTGAACTCTTTCCCTACACGACGCTCTTCCGATCTNNNTTAGCTGCTAGTGGCTGAAAGA
AGCACACGTCTGAACTCTTTCCCTACACGACGCTCTTCCGATCTNNNATTGCAACCCTAGGCTGATCT
AGCACACGTCTGAACTCTTTCCCTACACGACGCTCTTCCGATCTNNNATTGCAACCCTAGGCTGATCT
GCACACGTCTGAACTCTTTCCCTACACGACGCTCTTCCGATCTNNNATTTGGACACTTTTGCTGGGGT
GCACACGTCTGAACTCTTTCCCTACACGACGCTCTTCCGATCTNNNATTTGGACACTTTTGCTGGGGT
AGCACACGTCTGAACTCTTTCCCTACACGACGCTCTTCCGATCTNNNGTCAGTCGAAAACTCCCTTACC
AGCACACGTCTGAACTCTTTCCCTACACGACGCTCTTCCGATCTNNNGTCAGTCGAAAACTCCCTTACC
CACACGTCTGAACTCTTTCCCTACACGACGCTCTTCCGATCTNNNGTTCCAGCCACCTCCATTATGCCA
CACACGTCTGAACTCTTTCCCTACACGACGCTCTTCCGATCTNNNGTTCCAGCCACCTCCATTATGCCA
AGCACACGTCTGAACTCTTTCCCTACACGACGCTCTTCCGATCTNNNTGTGTGTGTGTGTGTGTGAA
AGCACACGTCTGAACTCTTTCCCTACACGACGCTCTTCCGATCTNNNTGTGTGTGTGTGTGTGTGAA
AGCACACGTCTGAACTCTTTCCCTACACGACGCTCTTCCGATCTNNNACCTGTAGTCCCAGCTACTCGC
AGCACACGTCTGAACTCTTTCCCTACACGACGCTCTTCCGATCTNNNACCTGTAGTCCCAGCTACTCGC
GCACACGTCTGAACTCTTTCCCTACACGACGCTCTTCCGATCTNNNGCAGCTCTCTTCTTCCTTCTCTC
GCACACGTCTGAACTCTTTCCCTACACGACGCTCTTCCGATCTNNNGCAGCTCTCTTCTTCCTTCTCTC
AGCACACGTCTGAACTCTTTCCCTACACGACGCTCTTCCGATCTNNNCTAGGCAACAAAGAGAGACT
AGCACACGTCTGAACTCTTTCCCTACACGACGCTCTTCCGATCTNNNCTAGGCAACAAAGAGAGACT
AGCACACGTCTGAACTCTTTCCCTACACGACGCTCTTCCGATCTNNNATGTGTGTCCTAGAGATCCTG
AGCACACGTCTGAACTCTTTCCCTACACGACGCTCTTCCGATCTNNNATGTGTGTCCTAGAGATCCTG
GAGCACACGTCTGAACTCTTTCCCTACACGACGCTCTTCCGATCTNNNCAAGTGTCTTTGTGTGTGTGT
GAGCACACGTCTGAACTCTTTCCCTACACGACGCTCTTCCGATCTNNNCAAGTGTCTTTGTGTGTGTGT
AGAGCACACGTCTGAACTCTTTCCCTACACGACGCTCTTCCGATCTNNNCTTGCAGTGAGCTGAGATT
AGAGCACACGTCTGAACTCTTTCCCTACACGACGCTCTTCCGATCTNNNCTTGCAGTGAGCTGAGATT
GAGCACACGTCTGAACTCTTTCCCTACACGACGCTCTTCCGATCTNNNCCTCTCAAAGTGCTGGGATT
GAGCACACGTCTGAACTCTTTCCCTACACGACGCTCTTCCGATCTNNNCCTCTCAAAGTGCTGGGATT
AGCACACGTCTGAACTCTTTCCCTACACGACGCTCTTCCGATCTNNNAGTCCCCAAATTAGTGCCATA
AGCACACGTCTGAACTCTTTCCCTACACGACGCTCTTCCGATCTNNNAGTCCCCAAATTAGTGCCATA
GAGCACACGTCTGAACTCTTTCCCTACACGACGCTCTTCCGATCTNNNCTGAGGCTGGAGAATCACTT
GAGCACACGTCTGAACTCTTTCCCTACACGACGCTCTTCCGATCTNNNCTGAGGCTGGAGAATCACTT
GAGCACACGTCTGAACTCTTTCCCTACACGACGCTCTTCCGATCTNNNGAGTGAACAGTTCTGTCTTG
GAGCACACGTCTGAACTCTTTCCCTACACGACGCTCTTCCGATCTNNNGAGTGAACAGTTCTGTCTTG

AGCACACGTCTGAACTCTTTCCCTACACGACGCTCTTCCGATCTNNNGCAGACACCTGTAATCCCAGTT
AGCACACGTCTGAACTCTTTCCCTACACGACGCTCTTCCGATCTNNNGCAGACACCTGTAATCCCAGTT
AGCACACGTCTGAACTCTTTCCCTACACGACGCTCTTCCGATCTNNNAGGGCTTTGTTCTTTGAACACT
AGCACACGTCTGAACTCTTTCCCTACACGACGCTCTTCCGATCTNNNAGGGCTTTGTTCTTTGAACACT
GAGCACACGTCTGAACTCTTTCCCTACACGACGCTCTTCCGATCTNNNCGCCTGTAATCCCATCTACTG
GAGCACACGTCTGAACTCTTTCCCTACACGACGCTCTTCCGATCTNNNCGCCTGTAATCCCATCTACTG
GAGCACACGTCTGAACTCTTTCCCTACACGACGCTCTTCCGATCTNNNGCTTGTGAACTCTCAGCATT
GAGCACACGTCTGAACTCTTTCCCTACACGACGCTCTTCCGATCTNNNGCTTGTGAACTCTCAGCATT
AGCACACGTCTGAACTCTTTCCCTACACGACGCTCTTCCGATCTNNNTGAAGCTACTCATGAATGTGG
AGCACACGTCTGAACTCTTTCCCTACACGACGCTCTTCCGATCTNNNTGAAGCTACTCATGAATGTGG
AGAGCACACGTCTGAACTCTTTCCCTACACGACGCTCTTCCGATCTNNNACAACAGAGAAAATCACCT
AGAGCACACGTCTGAACTCTTTCCCTACACGACGCTCTTCCGATCTNNNACAACAGAGAAAATCACCT
CACACGTCTGAACTCTTTCCCTACACGACGCTCTTCCGATCTNNNTCATTGCATGCTACCAAGAAACCC
CACACGTCTGAACTCTTTCCCTACACGACGCTCTTCCGATCTNNNTCATTGCATGCTACCAAGAAACCC
GCACACGTCTGAACTCTTTCCCTACACGACGCTCTTCCGATCTNNNCAGAGGTTTCCCATCGCTCCCATC
GCACACGTCTGAACTCTTTCCCTACACGACGCTCTTCCGATCTNNNCAGAGGTTTCCCATCGCTCCCATC
GAGCACACGTCTGAACTCTTTCCCTACACGACGCTCTTCCGATCTNNNGCATGGATTGCAGGGAAACT
GAGCACACGTCTGAACTCTTTCCCTACACGACGCTCTTCCGATCTNNNGCATGGATTGCAGGGAAACT
AGAGCACACGTCTGAACTCTTTCCCTACACGACGCTCTTCCGATCTNNNCAGGGAGTGAGGAAAGGA
AGAGCACACGTCTGAACTCTTTCCCTACACGACGCTCTTCCGATCTNNNCAGGGAGTGAGGAAAGGA
GCACACGTCTGAACTCTTTCCCTACACGACGCTCTTCCGATCTNNNCTGAGGCTTTGGATATGAGAGC
GCACACGTCTGAACTCTTTCCCTACACGACGCTCTTCCGATCTNNNCTGAGGCTTTGGATATGAGAGC
GAGCACACGTCTGAACTCTTTCCCTACACGACGCTCTTCCGATCTNNNCCTGTTTTGAACACATGGCTG
GAGCACACGTCTGAACTCTTTCCCTACACGACGCTCTTCCGATCTNNNCCTGTTTTGAACACATGGCTG
GCACACGTCTGAACTCTTTCCCTACACGACGCTCTTCCGATCTNNNGTTCTCATCACTTAGCTCCCACTC
GCACACGTCTGAACTCTTTCCCTACACGACGCTCTTCCGATCTNNNGTTCTCATCACTTAGCTCCCACTC
GCACACGTCTGAACTCTTTCCCTACACGACGCTCTTCCGATCTNNNCTGTAATCCCAGCTACTCAGGA
GCACACGTCTGAACTCTTTCCCTACACGACGCTCTTCCGATCTNNNCTGTAATCCCAGCTACTCAGGA
GCACACGTCTGAACTCTTTCCCTACACGACGCTCTTCCGATCTNNNCTGAAACAGGGTTAGGGCTTTT
GCACACGTCTGAACTCTTTCCCTACACGACGCTCTTCCGATCTNNNCTGAAACAGGGTTAGGGCTTTT
AGCACACGTCTGAACTCTTTCCCTACACGACGCTCTTCCGATCTNNNACTTCCAGTGCTTCCTGTTGTA
AGCACACGTCTGAACTCTTTCCCTACACGACGCTCTTCCGATCTNNNACTTCCAGTGCTTCCTGTTGTA
AGCACACGTCTGAACTCTTTCCCTACACGACGCTCTTCCGATCTNNNTCAATACTGCATTAGCTCTGG
AGCACACGTCTGAACTCTTTCCCTACACGACGCTCTTCCGATCTNNNTCAATACTGCATTAGCTCTGG
AGCACACGTCTGAACTCTTTCCCTACACGACGCTCTTCCGATCTNNNCACACCAGACATGAGTGAGA
AGCACACGTCTGAACTCTTTCCCTACACGACGCTCTTCCGATCTNNNCACACCAGACATGAGTGAGA
GAGCACACGTCTGAACTCTTTCCCTACACGACGCTCTTCCGATCTNNNAAAAAATTAGCTGGACGTG
GAGCACACGTCTGAACTCTTTCCCTACACGACGCTCTTCCGATCTNNNAAAAAATTAGCTGGACGTG
GCACACGTCTGAACTCTTTCCCTACACGACGCTCTTCCGATCTNNNATACAAAAAATTAGCCGGGCGT
GCACACGTCTGAACTCTTTCCCTACACGACGCTCTTCCGATCTNNNATACAAAAAATTAGCCGGGCGT
GAGCACACGTCTGAACTCTTTCCCTACACGACGCTCTTCCGATCTNNNTCCATTTAGTTGGTTCTGGG
GAGCACACGTCTGAACTCTTTCCCTACACGACGCTCTTCCGATCTNNNTCCATTTAGTTGGTTCTGGG
GCACACGTCTGAACTCTTTCCCTACACGACGCTCTTCCGATCTNNNTCCTCATCCTCTACTCTCCATGTC

AGCACACGTCTGAACTCTTTCCCTACACGACGCTCTTCCGATCTNNNTCCTCATCCTCTACTCTCCATGTC
AGCACACGTCTGAACTCTTTCCCTACACGACGCTCTTCCGATCTNNNAGACATACACTGACAGAAGAG
AGCACACGTCTGAACTCTTTCCCTACACGACGCTCTTCCGATCTNNNAGACATACACTGACAGAAGAG
AGAGCACACGTCTGAACTCTTTCCCTACACGACGCTCTTCCGATCTNNNACCGTGACAATAAAGCCATA
AGAGCACACGTCTGAACTCTTTCCCTACACGACGCTCTTCCGATCTNNNACCGTGACAATAAAGCCATA
GAGCACACGTCTGAACTCTTTCCCTACACGACGCTCTTCCGATCTNNNCAGGCTCATGATTCACAACAG
GAGCACACGTCTGAACTCTTTCCCTACACGACGCTCTTCCGATCTNNNCAGGCTCATGATTCACAACAG
AGCACACGTCTGAACTCTTTCCCTACACGACGCTCTTCCGATCTNNNGGGGTAGAGAGGGAGAAAAG
AGCACACGTCTGAACTCTTTCCCTACACGACGCTCTTCCGATCTNNNGGGGTAGAGAGGGAGAAAAG
GAGCACACGTCTGAACTCTTTCCCTACACGACGCTCTTCCGATCTNNNTGATGAAACCCCGTCGTTACT
GAGCACACGTCTGAACTCTTTCCCTACACGACGCTCTTCCGATCTNNNTGATGAAACCCCGTCGTTACT
AGCACACGTCTGAACTCTTTCCCTACACGACGCTCTTCCGATCTNNNGTCACCAGGAACCAGGAAGTAT
AGCACACGTCTGAACTCTTTCCCTACACGACGCTCTTCCGATCTNNNGTCACCAGGAACCAGGAAGTAT
AGCACACGTCTGAACTCTTTCCCTACACGACGCTCTTCCGATCTNNNCGCGTCTATAATCCCAGCTACT
AGCACACGTCTGAACTCTTTCCCTACACGACGCTCTTCCGATCTNNNCGCGTCTATAATCCCAGCTACT
AGAGCACACGTCTGAACTCTTTCCCTACACGACGCTCTTCCGATCTNNNGAGAGGACAGAGCAGGAG
AGAGCACACGTCTGAACTCTTTCCCTACACGACGCTCTTCCGATCTNNNGAGAGGACAGAGCAGGAG
GCACACGTCTGAACTCTTTCCCTACACGACGCTCTTCCGATCTNNNTGCATAAAGAAAGAGGCTTTGC
GCACACGTCTGAACTCTTTCCCTACACGACGCTCTTCCGATCTNNNTGCATAAAGAAAGAGGCTTTGC
GAGCACACGTCTGAACTCTTTCCCTACACGACGCTCTTCCGATCTNNNTTCTATACCACAACAGCGACCA
GAGCACACGTCTGAACTCTTTCCCTACACGACGCTCTTCCGATCTNNNTTCTATACCACAACAGCGACCA
AGCACACGTCTGAACTCTTTCCCTACACGACGCTCTTCCGATCTNNNCTCAAACTCCTGACCTCAAGTGA
AGCACACGTCTGAACTCTTTCCCTACACGACGCTCTTCCGATCTNNNCTCAAACTCCTGACCTCAAGTGA
AGCACACGTCTGAACTCTTTCCCTACACGACGCTCTTCCGATCTNNNTTGTAGTGATTGGAGCTGTGT
AGCACACGTCTGAACTCTTTCCCTACACGACGCTCTTCCGATCTNNNTTGTAGTGATTGGAGCTGTGT
GCACACGTCTGAACTCTTTCCCTACACGACGCTCTTCCGATCTNNNAGGATCTTGTTCTCTGAAGTCCA
GCACACGTCTGAACTCTTTCCCTACACGACGCTCTTCCGATCTNNNAGGATCTTGTTCTCTGAAGTCCA
AGCACACGTCTGAACTCTTTCCCTACACGACGCTCTTCCGATCTNNNACTCAATGTCATTCAGGGCTGG
AGCACACGTCTGAACTCTTTCCCTACACGACGCTCTTCCGATCTNNNACTCAATGTCATTCAGGGCTGG
GCACACGTCTGAACTCTTTCCCTACACGACGCTCTTCCGATCTNNNTAGGAGTTTTTGGCCCTCTTACC
GCACACGTCTGAACTCTTTCCCTACACGACGCTCTTCCGATCTNNNTAGGAGTTTTTGGCCCTCTTACC
AGCACACGTCTGAACTCTTTCCCTACACGACGCTCTTCCGATCTNNNAATACAAAAAATTAGCCGGGC
AGCACACGTCTGAACTCTTTCCCTACACGACGCTCTTCCGATCTNNNAATACAAAAAATTAGCCGGGC
GCACACGTCTGAACTCTTTCCCTACACGACGCTCTTCCGATCTNNNTGTACATGGATGGAAAGTGGTG
GCACACGTCTGAACTCTTTCCCTACACGACGCTCTTCCGATCTNNNTGTACATGGATGGAAAGTGGTG
AGCACACGTCTGAACTCTTTCCCTACACGACGCTCTTCCGATCTNNNGCCGATGAAACATCTCAGTCCT
AGCACACGTCTGAACTCTTTCCCTACACGACGCTCTTCCGATCTNNNGCCGATGAAACATCTCAGTCCT
AGCACACGTCTGAACTCTTTCCCTACACGACGCTCTTCCGATCTNNNATAATTTACCCCCCAGAAGCTC
AGCACACGTCTGAACTCTTTCCCTACACGACGCTCTTCCGATCTNNNATAATTTACCCCCCAGAAGCTC
GAGCACACGTCTGAACTCTTTCCCTACACGACGCTCTTCCGATCTNNNATGGATGTTTTCTGGAGAGG
GAGCACACGTCTGAACTCTTTCCCTACACGACGCTCTTCCGATCTNNNATGGATGTTTTCTGGAGAGG
AGCACACGTCTGAACTCTTTCCCTACACGACGCTCTTCCGATCTNNNTTTGGAACACAAAGGTGGTTT
AGCACACGTCTGAACTCTTTCCCTACACGACGCTCTTCCGATCTNNNTTTGGAACACAAAGGTGGTTT

AGCACACGTCTGAACTCTTTCCCTACACGACGCTCTTCCGATCTNNNTCAACAGAAATGGCAGGTACT
AGCACACGTCTGAACTCTTTCCCTACACGACGCTCTTCCGATCTNNNTCAACAGAAATGGCAGGTACT
AGAGCACACGTCTGAACTCTTTCCCTACACGACGCTCTTCCGATCTNNNGAAAATTGGTGCCCAGATA
AGAGCACACGTCTGAACTCTTTCCCTACACGACGCTCTTCCGATCTNNNGAAAATTGGTGCCCAGATA
AGCACACGTCTGAACTCTTTCCCTACACGACGCTCTTCCGATCTNNNTGCTAAAAATACAATAGCCTG
AGCACACGTCTGAACTCTTTCCCTACACGACGCTCTTCCGATCTNNNTGCTAAAAATACAATAGCCTG
AGCACACGTCTGAACTCTTTCCCTACACGACGCTCTTCCGATCTNNNACATCCTTTCCATGTCAAGGGA
AGCACACGTCTGAACTCTTTCCCTACACGACGCTCTTCCGATCTNNNACATCCTTTCCATGTCAAGGGA
GCACACGTCTGAACTCTTTCCCTACACGACGCTCTTCCGATCTNNNTTCAAAAATTGGCTCCCATACCG
GCACACGTCTGAACTCTTTCCCTACACGACGCTCTTCCGATCTNNNTTCAAAAATTGGCTCCCATACCG
AGCACACGTCTGAACTCTTTCCCTACACGACGCTCTTCCGATCTNNNATGCCAAAAATGCAAAGTCCT
AGCACACGTCTGAACTCTTTCCCTACACGACGCTCTTCCGATCTNNNATGCCAAAAATGCAAAGTCCT
GAGCACACGTCTGAACTCTTTCCCTACACGACGCTCTTCCGATCTNNNTGACGAAACAAGGGCCTTTA
GAGCACACGTCTGAACTCTTTCCCTACACGACGCTCTTCCGATCTNNNTGACGAAACAAGGGCCTTTA
AGCACACGTCTGAACTCTTTCCCTACACGACGCTCTTCCGATCTNNNGCGGTGAAATGCTGTCTCTACT
AGCACACGTCTGAACTCTTTCCCTACACGACGCTCTTCCGATCTNNNGCGGTGAAATGCTGTCTCTACT
GCACACGTCTGAACTCTTTCCCTACACGACGCTCTTCCGATCTNNNTTAGAGAGCAGGTTGGTCAACA
GCACACGTCTGAACTCTTTCCCTACACGACGCTCTTCCGATCTNNNTTAGAGAGCAGGTTGGTCAACA
CACACGTCTGAACTCTTTCCCTACACGACGCTCTTCCGATCTNNNTCACAAAACATCCTGAAAATTCCAT
CACACGTCTGAACTCTTTCCCTACACGACGCTCTTCCGATCTNNNTCACAAAACATCCTGAAAATTCCAT
GCACACGTCTGAACTCTTTCCCTACACGACGCTCTTCCGATCTNNNAAAGTGCTGGGATTACAGGCAT
GCACACGTCTGAACTCTTTCCCTACACGACGCTCTTCCGATCTNNNAAAGTGCTGGGATTACAGGCAT
AGCACACGTCTGAACTCTTTCCCTACACGACGCTCTTCCGATCTNNNAGTCAATTGGCTTCATTTCTGG
AGCACACGTCTGAACTCTTTCCCTACACGACGCTCTTCCGATCTNNNAGTCAATTGGCTTCATTTCTGG
GAGCACACGTCTGAACTCTTTCCCTACACGACGCTCTTCCGATCTNNNCCTGGCCTGGGATTGTTATTT
GAGCACACGTCTGAACTCTTTCCCTACACGACGCTCTTCCGATCTNNNCCTGGCCTGGGATTGTTATTT
GCACACGTCTGAACTCTTTCCCTACACGACGCTCTTCCGATCTNNNTGCAAAACTGCCTAATTCCCAAG
GCACACGTCTGAACTCTTTCCCTACACGACGCTCTTCCGATCTNNNTGCAAAACTGCCTAATTCCCAAG
GAGCACACGTCTGAACTCTTTCCCTACACGACGCTCTTCCGATCTNNNCACTGCATCTATCCTGGGTGA
GAGCACACGTCTGAACTCTTTCCCTACACGACGCTCTTCCGATCTNNNCACTGCATCTATCCTGGGTGA
GCACACGTCTGAACTCTTTCCCTACACGACGCTCTTCCGATCTNNNTGAGAAATGCTATGGAAAGGG
GCACACGTCTGAACTCTTTCCCTACACGACGCTCTTCCGATCTNNNTGAGAAATGCTATGGAAAGGG
AGCACACGTCTGAACTCTTTCCCTACACGACGCTCTTCCGATCTNNNCATGATGCTAGATTGCACCTGT
AGCACACGTCTGAACTCTTTCCCTACACGACGCTCTTCCGATCTNNNCATGATGCTAGATTGCACCTGT
CACACGTCTGAACTCTTTCCCTACACGACGCTCTTCCGATCTNNNGAGACTGGACACCTGAGTCTTG
CACACGTCTGAACTCTTTCCCTACACGACGCTCTTCCGATCTNNNGAGACTGGACACCTGAGTCTTG
AGCACACGTCTGAACTCTTTCCCTACACGACGCTCTTCCGATCTNNNCTGTAATCCCAGCTACTCAGG
AGCACACGTCTGAACTCTTTCCCTACACGACGCTCTTCCGATCTNNNCTGTAATCCCAGCTACTCAGG
GAGCACACGTCTGAACTCTTTCCCTACACGACGCTCTTCCGATCTNNNCTCCCCATGACATGTAGATAG
GAGCACACGTCTGAACTCTTTCCCTACACGACGCTCTTCCGATCTNNNCTCCCCATGACATGTAGATAG
AGCACACGTCTGAACTCTTTCCCTACACGACGCTCTTCCGATCTNNNCAGGAGAACTGCTACAACCTA
AGCACACGTCTGAACTCTTTCCCTACACGACGCTCTTCCGATCTNNNCAGGAGAACTGCTACAACCTA
GAGCACACGTCTGAACTCTTTCCCTACACGACGCTCTTCCGATCTNNNGAGAATTGCTTGAACCCAGG

GAGCACACGTCTGAACTCTTTCCCTACACGACGCTCTTCCGATCTNNNGAGAATTGCTTGAACCCAGG

AGCACACGTCTGAACTCTTTCCCTACACGACGCTCTTCCGATCTNNNGGTAAAAGGGCAAGACTCTGT

AGCACACGTCTGAACTCTTTCCCTACACGACGCTCTTCCGATCTNNNGGTAAAAGGGCAAGACTCTGT

ACACGTCTGAACTCTTTCCCTACACGACGCTCTTCCGATCTNNNTCTCAATGAGCTGTTGGGTACACCA

ACACGTCTGAACTCTTTCCCTACACGACGCTCTTCCGATCTNNNTCTCAATGAGCTGTTGGGTACACCA

GAGCACACGTCTGAACTCTTTCCCTACACGACGCTCTTCCGATCTNNNAGTGAGCTGAGATTACACCA

GAGCACACGTCTGAACTCTTTCCCTACACGACGCTCTTCCGATCTNNNAGTGAGCTGAGATTACACCA

AGCACACGTCTGAACTCTTTCCCTACACGACGCTCTTCCGATCTNNNTTAGTACTGGATCTAACTCATTG

AGCACACGTCTGAACTCTTTCCCTACACGACGCTCTTCCGATCTNNNTTAGTACTGGATCTAACTCATTG

GCACACGTCTGAACTCTTTCCCTACACGACGCTCTTCCGATCTNNNAACTAGCCAAGTCTCTTTCTCCCC

GCACACGTCTGAACTCTTTCCCTACACGACGCTCTTCCGATCTNNNAACTAGCCAAGTCTCTTTCTCCCC

GCACACGTCTGAACTCTTTCCCTACACGACGCTCTTCCGATCTNNNCCTCGCCTGGCTAATTTTTTTGTC

GCACACGTCTGAACTCTTTCCCTACACGACGCTCTTCCGATCTNNNCCTCGCCTGGCTAATTTTTTTGTC

GAGCACACGTCTGAACTCTTTCCCTACACGACGCTCTTCCGATCTNNNCAAGTATCTGGGACTACAGG

GAGCACACGTCTGAACTCTTTCCCTACACGACGCTCTTCCGATCTNNNCAAGTATCTGGGACTACAGG

GAGCACACGTCTGAACTCTTTCCCTACACGACGCTCTTCCGATCTNNNTGGCATAGAGTTATTTCTCCC

GAGCACACGTCTGAACTCTTTCCCTACACGACGCTCTTCCGATCTNNNTGGCATAGAGTTATTTCTCCC

AGAGCACACGTCTGAACTCTTTCCCTACACGACGCTCTTCCGATCTNNNCACACACACACATACACACA

AGAGCACACGTCTGAACTCTTTCCCTACACGACGCTCTTCCGATCTNNNCACACACACACATACACACA

AGCACACGTCTGAACTCTTTCCCTACACGACGCTCTTCCGATCTNNNGAGAGACTTGAGCATCCCAGA

AGCACACGTCTGAACTCTTTCCCTACACGACGCTCTTCCGATCTNNNGAGAGACTTGAGCATCCCAGA

GCACACGTCTGAACTCTTTCCCTACACGACGCTCTTCCGATCTNNNCGATGGGAGTTCTGGAAGCTAT

GCACACGTCTGAACTCTTTCCCTACACGACGCTCTTCCGATCTNNNCGATGGGAGTTCTGGAAGCTAT

AGCACACGTCTGAACTCTTTCCCTACACGACGCTCTTCCGATCTNNNAGAAGTGTTGGGAGACTGGAA

AGCACACGTCTGAACTCTTTCCCTACACGACGCTCTTCCGATCTNNNAGAAGTGTTGGGAGACTGGAA

GCACACGTCTGAACTCTTTCCCTACACGACGCTCTTCCGATCTNNNGATAATTTGGATCTGTGGCTGCC

GCACACGTCTGAACTCTTTCCCTACACGACGCTCTTCCGATCTNNNGATAATTTGGATCTGTGGCTGCC

AGCACACGTCTGAACTCTTTCCCTACACGACGCTCTTCCGATCTNNNCTGCATCCATGAACTCGTCATTT

AGCACACGTCTGAACTCTTTCCCTACACGACGCTCTTCCGATCTNNNCTGCATCCATGAACTCGTCATTT

GCACACGTCTGAACTCTTTCCCTACACGACGCTCTTCCGATCTNNNGCTTTCCTGGTTCTCCAACTTTCC

GCACACGTCTGAACTCTTTCCCTACACGACGCTCTTCCGATCTNNNGCTTTCCTGGTTCTCCAACTTTCC

GAGCACACGTCTGAACTCTTTCCCTACACGACGCTCTTCCGATCTNNNAAGCAGAAAGGGTGGTCTAA

GAGCACACGTCTGAACTCTTTCCCTACACGACGCTCTTCCGATCTNNNAAGCAGAAAGGGTGGTCTAA

AGCACACGTCTGAACTCTTTCCCTACACGACGCTCTTCCGATCTNNNGAGACAGGAGAATTGCTCAAA

AGCACACGTCTGAACTCTTTCCCTACACGACGCTCTTCCGATCTNNNGAGACAGGAGAATTGCTCAAA

AGCACACGTCTGAACTCTTTCCCTACACGACGCTCTTCCGATCTNNNTCAACAGAGTCTTACATTGGCC

AGCACACGTCTGAACTCTTTCCCTACACGACGCTCTTCCGATCTNNNTCAACAGAGTCTTACATTGGCC

AGCACACGTCTGAACTCTTTCCCTACACGACGCTCTTCCGATCTNNNCGAGATCTCTGAAATCCGCACTA

AGCACACGTCTGAACTCTTTCCCTACACGACGCTCTTCCGATCTNNNCGAGATCTCTGAAATCCGCACTA

GAGCACACGTCTGAACTCTTTCCCTACACGACGCTCTTCCGATCTNNNCAAGAGAATTGCTTGAACCCA

GAGCACACGTCTGAACTCTTTCCCTACACGACGCTCTTCCGATCTNNNCAAGAGAATTGCTTGAACCCA

GAGCACACGTCTGAACTCTTTCCCTACACGACGCTCTTCCGATCTNNNTTGTTGCAAGAAACAAGAGG

GAGCACACGTCTGAACTCTTTCCCTACACGACGCTCTTCCGATCTNNNTTGTTGCAAGAAACAAGAGG

AGCACACGTCTGAACTCTTTCCCTACACGACGCTCTTCCGATCTNNNCGTTTCATGGTTCCGTTTTCTCA

AGCACACGTCTGAACTCTTTCCCTACACGACGCTCTTCCGATCTNNNCGTTTCATGGTTCCGTTTTCTCA

AGCACACGTCTGAACTCTTTCCCTACACGACGCTCTTCCGATCTNNNGATCTTCCTGGCACCATTTGATG

AGCACACGTCTGAACTCTTTCCCTACACGACGCTCTTCCGATCTNNNGATCTTCCTGGCACCATTTGATG

CACACGTCTGAACTCTTTCCCTACACGACGCTCTTCCGATCTNNNCTGGGAAGTGAGGAGCGTCTCCAT

CACACGTCTGAACTCTTTCCCTACACGACGCTCTTCCGATCTNNNCTGGGAAGTGAGGAGCGTCTCCAT

GCACACGTCTGAACTCTTTCCCTACACGACGCTCTTCCGATCTNNNCCTTACTCTCAACCTCCTGGAACCA

GCACACGTCTGAACTCTTTCCCTACACGACGCTCTTCCGATCTNNNCCTTACTCTCAACCTCCTGGAACCA

AGAGCACACGTCTGAACTCTTTCCCTACACGACGCTCTTCCGATCTNNNGAAGGAGATTGAGACACAC

AGAGCACACGTCTGAACTCTTTCCCTACACGACGCTCTTCCGATCTNNNGAAGGAGATTGAGACACAC

AGCACACGTCTGAACTCTTTCCCTACACGACGCTCTTCCGATCTNNNATATTACATTTCAGAGTTCACCT

AGCACACGTCTGAACTCTTTCCCTACACGACGCTCTTCCGATCTNNNATATTACATTTCAGAGTTCACCT

AGCACACGTCTGAACTCTTTCCCTACACGACGCTCTTCCGATCTNNNGCAGTAGCAAGACCTTGTCTCA

AGCACACGTCTGAACTCTTTCCCTACACGACGCTCTTCCGATCTNNNGCAGTAGCAAGACCTTGTCTCA

AGCACACGTCTGAACTCTTTCCCTACACGACGCTCTTCCGATCTNNNAGCGATTCTTCTACTTCAGCCTC

AGCACACGTCTGAACTCTTTCCCTACACGACGCTCTTCCGATCTNNNAGCGATTCTTCTACTTCAGCCTC

AGCACACGTCTGAACTCTTTCCCTACACGACGCTCTTCCGATCTNNNCTAATCCCAGTTACTCAGGAGG

AGCACACGTCTGAACTCTTTCCCTACACGACGCTCTTCCGATCTNNNCTAATCCCAGTTACTCAGGAGG

GCACACGTCTGAACTCTTTCCCTACACGACGCTCTTCCGATCTNNNATTCCAGCCTGAGCAATTGAGAG

GCACACGTCTGAACTCTTTCCCTACACGACGCTCTTCCGATCTNNNATTCCAGCCTGAGCAATTGAGAG

AGCACACGTCTGAACTCTTTCCCTACACGACGCTCTTCCGATCTNNNCAAAATGCAAACCAATGTGGCA

AGCACACGTCTGAACTCTTTCCCTACACGACGCTCTTCCGATCTNNNCAAAATGCAAACCAATGTGGCA

GAGCACACGTCTGAACTCTTTCCCTACACGACGCTCTTCCGATCTNNNCAAACCCTGAGTGTGGGTGTT

GAGCACACGTCTGAACTCTTTCCCTACACGACGCTCTTCCGATCTNNNCAAACCCTGAGTGTGGGTGTT

AGCACACGTCTGAACTCTTTCCCTACACGACGCTCTTCCGATCTNNNGAAGATGGGTTGGAAGGAAGA

AGCACACGTCTGAACTCTTTCCCTACACGACGCTCTTCCGATCTNNNGAAGATGGGTTGGAAGGAAGA

GAGCACACGTCTGAACTCTTTCCCTACACGACGCTCTTCCGATCTNNNGGCAACAGTAAGACTCCATG

GAGCACACGTCTGAACTCTTTCCCTACACGACGCTCTTCCGATCTNNNGGCAACAGTAAGACTCCATG

AGCACACGTCTGAACTCTTTCCCTACACGACGCTCTTCCGATCTNNNGCTTCTGGCTTGGATGATGTAA

AGCACACGTCTGAACTCTTTCCCTACACGACGCTCTTCCGATCTNNNGCTTCTGGCTTGGATGATGTAA

AGCACACGTCTGAACTCTTTCCCTACACGACGCTCTTCCGATCTNNNTAAGAACACATGGACACAGGG

AGCACACGTCTGAACTCTTTCCCTACACGACGCTCTTCCGATCTNNNTAAGAACACATGGACACAGGG

AGCACACGTCTGAACTCTTTCCCTACACGACGCTCTTCCGATCTNNNTCCTTCCTTCTTTCCTTCCTTCCCA

AGCACACGTCTGAACTCTTTCCCTACACGACGCTCTTCCGATCTNNNTCCTTCCTTCTTTCCTTCCTTCCCA

AGCACACGTCTGAACTCTTTCCCTACACGACGCTCTTCCGATCTNNNCTGTAATCCCAGCTACTCGGAA

AGCACACGTCTGAACTCTTTCCCTACACGACGCTCTTCCGATCTNNNCTGTAATCCCAGCTACTCGGAA

GAGCACACGTCTGAACTCTTTCCCTACACGACGCTCTTCCGATCTNNNAGTTTGCCTGAATGAGAACAA

GAGCACACGTCTGAACTCTTTCCCTACACGACGCTCTTCCGATCTNNNAGTTTGCCTGAATGAGAACAA

AGCACACGTCTGAACTCTTTCCCTACACGACGCTCTTCCGATCTNNNGAGTCAGAGCAAGGCTGTATC

AGCACACGTCTGAACTCTTTCCCTACACGACGCTCTTCCGATCTNNNGAGTCAGAGCAAGGCTGTATC

GCACACGTCTGAACTCTTTCCCTACACGACGCTCTTCCGATCTNNNAGGTACATCTGTGTTGTCACCAA

GCACACGTCTGAACTCTTTCCCTACACGACGCTCTTCCGATCTNNNAGGTACATCTGTGTTGTCACCAA

AGCACACGTCTGAACTCTTTCCCTACACGACGCTCTTCCGATCTNNNAGACATATGCAAACCTCCCAAA

AGCACACGTCTGAACTCTTTCCCTACACGACGCTCTTCCGATCTNNNAGACATATGCAAACCTCCCAAA
AGCACACGTCTGAACTCTTTCCCTACACGACGCTCTTCCGATCTNNNCAACAAGAGTGAGACTCCACC
AGCACACGTCTGAACTCTTTCCCTACACGACGCTCTTCCGATCTNNNCAACAAGAGTGAGACTCCACC
AGCACACGTCTGAACTCTTTCCCTACACGACGCTCTTCCGATCTNNNGAAAGGTGGGAGGAATGGG
AGCACACGTCTGAACTCTTTCCCTACACGACGCTCTTCCGATCTNNNGAAAGGTGGGAGGAATGGG
GCACACGTCTGAACTCTTTCCCTACACGACGCTCTTCCGATCTNNNATAAAGCATGAGCAAATCCCAG
GCACACGTCTGAACTCTTTCCCTACACGACGCTCTTCCGATCTNNNATAAAGCATGAGCAAATCCCAG
GCACACGTCTGAACTCTTTCCCTACACGACGCTCTTCCGATCTNNNCTGTGACAGGTTGTAGAACCCA
GCACACGTCTGAACTCTTTCCCTACACGACGCTCTTCCGATCTNNNCTGTGACAGGTTGTAGAACCCA
AGCACACGTCTGAACTCTTTCCCTACACGACGCTCTTCCGATCTNNNCAGAGTGTAAGCAAGCAGAC
AGCACACGTCTGAACTCTTTCCCTACACGACGCTCTTCCGATCTNNNCAGAGTGTAAGCAAGCAGAC
AGCACACGTCTGAACTCTTTCCCTACACGACGCTCTTCCGATCTNNNATCACTTGAGCCTACTTGGAG
AGCACACGTCTGAACTCTTTCCCTACACGACGCTCTTCCGATCTNNNATCACTTGAGCCTACTTGGAG
AGCACACGTCTGAACTCTTTCCCTACACGACGCTCTTCCGATCTNNNGAAAATGGCATAAACCCGGGA
AGCACACGTCTGAACTCTTTCCCTACACGACGCTCTTCCGATCTNNNGAAAATGGCATAAACCCGGGA
AGAGCACACGTCTGAACTCTTTCCCTACACGACGCTCTTCCGATCTNNNGGCAACAAGAATGTCTACA
AGAGCACACGTCTGAACTCTTTCCCTACACGACGCTCTTCCGATCTNNNGGCAACAAGAATGTCTACA
AGCACACGTCTGAACTCTTTCCCTACACGACGCTCTTCCGATCTNNNCTGTGCTGGCTTCTGTCTATTC
AGCACACGTCTGAACTCTTTCCCTACACGACGCTCTTCCGATCTNNNCTGTGCTGGCTTCTGTCTATTC
CACACGTCTGAACTCTTTCCCTACACGACGCTCTTCCGATCTNNNCTACCCCCTTCTCCCCTCCGCATCG
CACACGTCTGAACTCTTTCCCTACACGACGCTCTTCCGATCTNNNCTACCCCCTTCTCCCCTCCGCATCG
AGCACACGTCTGAACTCTTTCCCTACACGACGCTCTTCCGATCTNNNTGGAGTGCAGTGGTATGATA
AGCACACGTCTGAACTCTTTCCCTACACGACGCTCTTCCGATCTNNNTGGAGTGCAGTGGTATGATA
AGCACACGTCTGAACTCTTTCCCTACACGACGCTCTTCCGATCTNNNCTCAAAACTTTGGGAGGCTGA
AGCACACGTCTGAACTCTTTCCCTACACGACGCTCTTCCGATCTNNNCTCAAAACTTTGGGAGGCTGA
AGCACACGTCTGAACTCTTTCCCTACACGACGCTCTTCCGATCTNNNTAAAAACACGTCTTTGTGCAC
AGCACACGTCTGAACTCTTTCCCTACACGACGCTCTTCCGATCTNNNTAAAAACACGTCTTTGTGCAC
GCACACGTCTGAACTCTTTCCCTACACGACGCTCTTCCGATCTNNNTCTCTCTCTCTCTCTCTTTCTTC
GCACACGTCTGAACTCTTTCCCTACACGACGCTCTTCCGATCTNNNTCTCTCTCTCTCTCTCTTTCTTC
ACACGTCTGAACTCTTTCCCTACACGACGCTCTTCCGATCTNNNAGTGGTGAAGTCAGGGCTTTTATC
ACACGTCTGAACTCTTTCCCTACACGACGCTCTTCCGATCTNNNAGTGGTGAAGTCAGGGCTTTTATC
AGCACACGTCTGAACTCTTTCCCTACACGACGCTCTTCCGATCTNNNGAGATGAGATACGCGAGTCT
AGCACACGTCTGAACTCTTTCCCTACACGACGCTCTTCCGATCTNNNGAGATGAGATACGCGAGTCT
GAGCACACGTCTGAACTCTTTCCCTACACGACGCTCTTCCGATCTNNNAATCCTAGCTACTTTGGAGG
GAGCACACGTCTGAACTCTTTCCCTACACGACGCTCTTCCGATCTNNNAATCCTAGCTACTTTGGAGG
CACACGTCTGAACTCTTTCCCTACACGACGCTCTTCCGATCTNNNCCACGAAAACCCTAGTGAATCAC
CACACGTCTGAACTCTTTCCCTACACGACGCTCTTCCGATCTNNNCCACGAAAACCCTAGTGAATCAC
AGAGCACACGTCTGAACTCTTTCCCTACACGACGCTCTTCCGATCTNNNCTCCAGTAGTGCCCTTTATC
AGAGCACACGTCTGAACTCTTTCCCTACACGACGCTCTTCCGATCTNNNCTCCAGTAGTGCCCTTTATC
AGCACACGTCTGAACTCTTTCCCTACACGACGCTCTTCCGATCTNNNGAGTATCACTCTGTTGCCCAG
AGCACACGTCTGAACTCTTTCCCTACACGACGCTCTTCCGATCTNNNGAGTATCACTCTGTTGCCCAG
AGCACACGTCTGAACTCTTTCCCTACACGACGCTCTTCCGATCTNNNGGGGTGAGATCGAAGGATGA
AGCACACGTCTGAACTCTTTCCCTACACGACGCTCTTCCGATCTNNNGGGGTGAGATCGAAGGATGA

AGCACACGTCTGAACTCTTTCCCTACACGACGCTCTTCCGATCTNNNGTCTGTGGCACCTCCTCTATTTT
AGCACACGTCTGAACTCTTTCCCTACACGACGCTCTTCCGATCTNNNGTCTGTGGCACCTCCTCTATTTT
AGCACACGTCTGAACTCTTTCCCTACACGACGCTCTTCCGATCTNNNATCACACCTGTAATCCCAGCTA
AGCACACGTCTGAACTCTTTCCCTACACGACGCTCTTCCGATCTNNNATCACACCTGTAATCCCAGCTA
GCACACGTCTGAACTCTTTCCCTACACGACGCTCTTCCGATCTNNNGACTGCCTCGTATTACATGCAAC
GCACACGTCTGAACTCTTTCCCTACACGACGCTCTTCCGATCTNNNGACTGCCTCGTATTACATGCAAC
GCACACGTCTGAACTCTTTCCCTACACGACGCTCTTCCGATCTNNNGTTTCACTGTGTTAGCCAGGATG
GCACACGTCTGAACTCTTTCCCTACACGACGCTCTTCCGATCTNNNGTTTCACTGTGTTAGCCAGGATG
CACACGTCTGAACTCTTTCCCTACACGACGCTCTTCCGATCTNNNACTTTCAGTCCTATGCTCCTTGACAT
CACACGTCTGAACTCTTTCCCTACACGACGCTCTTCCGATCTNNNACTTTCAGTCCTATGCTCCTTGACAT
AGCACACGTCTGAACTCTTTCCCTACACGACGCTCTTCCGATCTNNNTACAAAAAATTAGCTGGGCGTG
AGCACACGTCTGAACTCTTTCCCTACACGACGCTCTTCCGATCTNNNTACAAAAAATTAGCTGGGCGTG
GCACACGTCTGAACTCTTTCCCTACACGACGCTCTTCCGATCTNNNGATCCATGCGCAGAAAACTACA
GCACACGTCTGAACTCTTTCCCTACACGACGCTCTTCCGATCTNNNGATCCATGCGCAGAAAACTACA
AGCACACGTCTGAACTCTTTCCCTACACGACGCTCTTCCGATCTNNNTGAGAATCACTTGAACCCAGGA
AGCACACGTCTGAACTCTTTCCCTACACGACGCTCTTCCGATCTNNNTGAGAATCACTTGAACCCAGGA
GCACACGTCTGAACTCTTTCCCTACACGACGCTCTTCCGATCTNNNACAGGGTATTCCATAATAACTTTC
GCACACGTCTGAACTCTTTCCCTACACGACGCTCTTCCGATCTNNNACAGGGTATTCCATAATAACTTTC
GAGCACACGTCTGAACTCTTTCCCTACACGACGCTCTTCCGATCTNNNATCTGCTGGTATCTTGATCTCG
GAGCACACGTCTGAACTCTTTCCCTACACGACGCTCTTCCGATCTNNNATCTGCTGGTATCTTGATCTCG
ACACGTCTGAACTCTTTCCCTACACGACGCTCTTCCGATCTNNNCCACTGCATACTGTTTATTCTCACAC
ACACGTCTGAACTCTTTCCCTACACGACGCTCTTCCGATCTNNNCCACTGCATACTGTTTATTCTCACAC
AGCACACGTCTGAACTCTTTCCCTACACGACGCTCTTCCGATCTNNNAACTCACGTGTGCCAGAACTAT
AGCACACGTCTGAACTCTTTCCCTACACGACGCTCTTCCGATCTNNNAACTCACGTGTGCCAGAACTAT
AGCACACGTCTGAACTCTTTCCCTACACGACGCTCTTCCGATCTNNNGGGTCTCCTGTAACATGAATGC
AGCACACGTCTGAACTCTTTCCCTACACGACGCTCTTCCGATCTNNNGGGTCTCCTGTAACATGAATGC
AGAGCACACGTCTGAACTCTTTCCCTACACGACGCTCTTCCGATCTNNNGAGGTTGAGGTTGGAGAA
AGAGCACACGTCTGAACTCTTTCCCTACACGACGCTCTTCCGATCTNNNGAGGTTGAGGTTGGAGAA
GAGCACACGTCTGAACTCTTTCCCTACACGACGCTCTTCCGATCTNNNTAAGGGAGATGCATCAACAA
GAGCACACGTCTGAACTCTTTCCCTACACGACGCTCTTCCGATCTNNNTAAGGGAGATGCATCAACAA
CACGTCTGAACTCTTTCCCTACACGACGCTCTTCCGATCTNNNCTGGCCCTACTTGTCCATTTTTCCATC
CACGTCTGAACTCTTTCCCTACACGACGCTCTTCCGATCTNNNCTGGCCCTACTTGTCCATTTTTCCATC
GCACACGTCTGAACTCTTTCCCTACACGACGCTCTTCCGATCTNNNCTAGCTTTGTATTGCCAGTGAG
GCACACGTCTGAACTCTTTCCCTACACGACGCTCTTCCGATCTNNNCTAGCTTTGTATTGCCAGTGAG
GAGCACACGTCTGAACTCTTTCCCTACACGACGCTCTTCCGATCTNNNGAGGCTGAGGCAGGAGAAT
GAGCACACGTCTGAACTCTTTCCCTACACGACGCTCTTCCGATCTNNNGAGGCTGAGGCAGGAGAAT
GAGCACACGTCTGAACTCTTTCCCTACACGACGCTCTTCCGATCTNNNTTTGGTGAATCTGCAACTGG
GAGCACACGTCTGAACTCTTTCCCTACACGACGCTCTTCCGATCTNNNTTTGGTGAATCTGCAACTGG
GCACACGTCTGAACTCTTTCCCTACACGACGCTCTTCCGATCTNNNGGAATCTGCCCTGTGTCGTCCAT
GCACACGTCTGAACTCTTTCCCTACACGACGCTCTTCCGATCTNNNGGAATCTGCCCTGTGTCGTCCAT
AGCACACGTCTGAACTCTTTCCCTACACGACGCTCTTCCGATCTNNNAAAAAAAAAAAAAGGTCGGG
AGCACACGTCTGAACTCTTTCCCTACACGACGCTCTTCCGATCTNNNAAAAAAAAAAAAAGGTCGGG
AGCACACGTCTGAACTCTTTCCCTACACGACGCTCTTCCGATCTNNNAGAAAGATCAAGGTGTGAGT

AGCACACGTCTGAACTCTTTCCCTACACGACGCTCTTCCGATCTNNNAGAAAGATCAAGGTGTGAGT
AGCACACGTCTGAACTCTTTCCCTACACGACGCTCTTCCGATCTNNNCATTTAGCACAAGTCAAGGCT
AGCACACGTCTGAACTCTTTCCCTACACGACGCTCTTCCGATCTNNNCATTTAGCACAAGTCAAGGCT
AGCACACGTCTGAACTCTTTCCCTACACGACGCTCTTCCGATCTNNNATTGTTGCTCAAAGTGGTGG
AGCACACGTCTGAACTCTTTCCCTACACGACGCTCTTCCGATCTNNNATTGTTGCTCAAAGTGGTGG
AGCACACGTCTGAACTCTTTCCCTACACGACGCTCTTCCGATCTNNNTGGGAAGCTAATAACCTACAG
AGCACACGTCTGAACTCTTTCCCTACACGACGCTCTTCCGATCTNNNTGGGAAGCTAATAACCTACAG
AGCACACGTCTGAACTCTTTCCCTACACGACGCTCTTCCGATCTNNNGCATTTGCTTGTCTGGGAAAG
AGCACACGTCTGAACTCTTTCCCTACACGACGCTCTTCCGATCTNNNGCATTTGCTTGTCTGGGAAAG
AGCACACGTCTGAACTCTTTCCCTACACGACGCTCTTCCGATCTNNNGTGTTTCCACATGTCATGCAG
AGCACACGTCTGAACTCTTTCCCTACACGACGCTCTTCCGATCTNNNGTGTTTCCACATGTCATGCAG
GCACACGTCTGAACTCTTTCCCTACACGACGCTCTTCCGATCTNNNCTTGTCTTAGGTTTGGGTTCTG
GCACACGTCTGAACTCTTTCCCTACACGACGCTCTTCCGATCTNNNCTTGTCTTAGGTTTGGGTTCTG
AGCACACGTCTGAACTCTTTCCCTACACGACGCTCTTCCGATCTNNNCTACGCCTTCTAAACGTGGTT
AGCACACGTCTGAACTCTTTCCCTACACGACGCTCTTCCGATCTNNNCTACGCCTTCTAAACGTGGTT
GCACACGTCTGAACTCTTTCCCTACACGACGCTCTTCCGATCTNNNTAGCCTGGGCAAAAAGAGTGA
GCACACGTCTGAACTCTTTCCCTACACGACGCTCTTCCGATCTNNNTAGCCTGGGCAAAAAGAGTGA
GAGCACACGTCTGAACTCTTTCCCTACACGACGCTCTTCCGATCTNNNGTTCCTGGCTGTCCTTTCAC
GAGCACACGTCTGAACTCTTTCCCTACACGACGCTCTTCCGATCTNNNGTTCCTGGCTGTCCTTTCAC
GCACACGTCTGAACTCTTTCCCTACACGACGCTCTTCCGATCTNNNCCTTCAAAATAAGAGGGTACTT
GCACACGTCTGAACTCTTTCCCTACACGACGCTCTTCCGATCTNNNCCTTCAAAATAAGAGGGTACTT
AGCACACGTCTGAACTCTTTCCCTACACGACGCTCTTCCGATCTNNNAGAGGCAAGCAAACTTCAACC
AGCACACGTCTGAACTCTTTCCCTACACGACGCTCTTCCGATCTNNNAGAGGCAAGCAAACTTCAACC
GCACACGTCTGAACTCTTTCCCTACACGACGCTCTTCCGATCTNNNTCAGGTAGGCTCAGTGTCTCC
GCACACGTCTGAACTCTTTCCCTACACGACGCTCTTCCGATCTNNNTCAGGTAGGCTCAGTGTCTCC
AGAGCACACGTCTGAACTCTTTCCCTACACGACGCTCTTCCGATCTNNNGTGAGGCAGGAGAATAGCT
AGAGCACACGTCTGAACTCTTTCCCTACACGACGCTCTTCCGATCTNNNGTGAGGCAGGAGAATAGCT
AGCACACGTCTGAACTCTTTCCCTACACGACGCTCTTCCGATCTNNNTTCCAAATGTTGCCTCTGTTGA
AGCACACGTCTGAACTCTTTCCCTACACGACGCTCTTCCGATCTNNNTTCCAAATGTTGCCTCTGTTGA
AGCACACGTCTGAACTCTTTCCCTACACGACGCTCTTCCGATCTNNNGTGTTGGAAAGTTGGGCACAA
AGCACACGTCTGAACTCTTTCCCTACACGACGCTCTTCCGATCTNNNGTGTTGGAAAGTTGGGCACAA
ACACGTCTGAACTCTTTCCCTACACGACGCTCTTCCGATCTNNNGAAGAGAATTTGAGAGAAAAGAG
ACACGTCTGAACTCTTTCCCTACACGACGCTCTTCCGATCTNNNGAAGAGAATTTGAGAGAAAAGAG
AGCACACGTCTGAACTCTTTCCCTACACGACGCTCTTCCGATCTNNNCTGGTCATTTGAGGCCAAGA
AGCACACGTCTGAACTCTTTCCCTACACGACGCTCTTCCGATCTNNNCTGGTCATTTGAGGCCAAGA
AGCACACGTCTGAACTCTTTCCCTACACGACGCTCTTCCGATCTNNNACAATTAGCTAAAAGCCAGCAG
AGCACACGTCTGAACTCTTTCCCTACACGACGCTCTTCCGATCTNNNACAATTAGCTAAAAGCCAGCAG
AGCACACGTCTGAACTCTTTCCCTACACGACGCTCTTCCGATCTNNNGCCTGTTATCCCAGCTACATG
AGCACACGTCTGAACTCTTTCCCTACACGACGCTCTTCCGATCTNNNGCCTGTTATCCCAGCTACATG
AGAGCACACGTCTGAACTCTTTCCCTACACGACGCTCTTCCGATCTNNNTTAGGGGTCACTACTCAAC
AGAGCACACGTCTGAACTCTTTCCCTACACGACGCTCTTCCGATCTNNNTTAGGGGTCACTACTCAAC
AGCACACGTCTGAACTCTTTCCCTACACGACGCTCTTCCGATCTNNNCGCCAGACATTGAGCCATAATT
AGCACACGTCTGAACTCTTTCCCTACACGACGCTCTTCCGATCTNNNCGCCAGACATTGAGCCATAATT

CACACGTCTGAACTCTTTCCCTACACGACGCTCTTCCGATCTNNNTTCACCTCACGATCTAGCCTTCTCA
CACACGTCTGAACTCTTTCCCTACACGACGCTCTTCCGATCTNNNTTCACCTCACGATCTAGCCTTCTCA
GCACACGTCTGAACTCTTTCCCTACACGACGCTCTTCCGATCTNNNGAAGAACTGCTTGAACCTGGGA
GCACACGTCTGAACTCTTTCCCTACACGACGCTCTTCCGATCTNNNGAAGAACTGCTTGAACCTGGGA
AGCACACGTCTGAACTCTTTCCCTACACGACGCTCTTCCGATCTNNNAGCATGCATGAAGATGAAAAC
AGCACACGTCTGAACTCTTTCCCTACACGACGCTCTTCCGATCTNNNAGCATGCATGAAGATGAAAAC
AGCACACGTCTGAACTCTTTCCCTACACGACGCTCTTCCGATCTNNNCTCTCCCTCCCTTTTTCCTCTTCC
AGCACACGTCTGAACTCTTTCCCTACACGACGCTCTTCCGATCTNNNCTCTCCCTCCCTTTTTCCTCTTCC
GCACACGTCTGAACTCTTTCCCTACACGACGCTCTTCCGATCTNNNGGCACCAACACCTCCTTCTACCA
GCACACGTCTGAACTCTTTCCCTACACGACGCTCTTCCGATCTNNNGGCACCAACACCTCCTTCTACCA
GAGCACACGTCTGAACTCTTTCCCTACACGACGCTCTTCCGATCTNNNCCAGCCTCGATTCTCTACACT
GAGCACACGTCTGAACTCTTTCCCTACACGACGCTCTTCCGATCTNNNCCAGCCTCGATTCTCTACACT
GCACACGTCTGAACTCTTTCCCTACACGACGCTCTTCCGATCTNNNGGACACCTTGTCTGCTCTCTATT
GCACACGTCTGAACTCTTTCCCTACACGACGCTCTTCCGATCTNNNGGACACCTTGTCTGCTCTCTATT
AGCACACGTCTGAACTCTTTCCCTACACGACGCTCTTCCGATCTNNNCCCCAGCTACCCTTCTAACATA
AGCACACGTCTGAACTCTTTCCCTACACGACGCTCTTCCGATCTNNNCCCCAGCTACCCTTCTAACATA
AGCACACGTCTGAACTCTTTCCCTACACGACGCTCTTCCGATCTNNNTATAGTCCCAGCTACTCAGGA
AGCACACGTCTGAACTCTTTCCCTACACGACGCTCTTCCGATCTNNNTATAGTCCCAGCTACTCAGGA
GAGCACACGTCTGAACTCTTTCCCTACACGACGCTCTTCCGATCTNNNTCCAAGATTGAATCGGGAAG
GAGCACACGTCTGAACTCTTTCCCTACACGACGCTCTTCCGATCTNNNTCCAAGATTGAATCGGGAAG
AGCACACGTCTGAACTCTTTCCCTACACGACGCTCTTCCGATCTNNNCCCATCGTTGTCATTGTGAGTT
AGCACACGTCTGAACTCTTTCCCTACACGACGCTCTTCCGATCTNNNCCCATCGTTGTCATTGTGAGTT
GCACACGTCTGAACTCTTTCCCTACACGACGCTCTTCCGATCTNNNTAAAGTGCTGGGATTACAGGC
GCACACGTCTGAACTCTTTCCCTACACGACGCTCTTCCGATCTNNNTAAAGTGCTGGGATTACAGGC
CACGTCTGAACTCTTTCCCTACACGACGCTCTTCCGATCTNNNTAAGGTGTGTCCAAACTCCCATCCAT
CACGTCTGAACTCTTTCCCTACACGACGCTCTTCCGATCTNNNTAAGGTGTGTCCAAACTCCCATCCAT
GAGCACACGTCTGAACTCTTTCCCTACACGACGCTCTTCCGATCTNNNCCTTTTTTTCATACTCATGCTA
GAGCACACGTCTGAACTCTTTCCCTACACGACGCTCTTCCGATCTNNNCCTTTTTTTCATACTCATGCTA
AGCACACGTCTGAACTCTTTCCCTACACGACGCTCTTCCGATCTNNNCTCTAAATTATCCAGACTGAAG
AGCACACGTCTGAACTCTTTCCCTACACGACGCTCTTCCGATCTNNNCTCTAAATTATCCAGACTGAAG
AGCACACGTCTGAACTCTTTCCCTACACGACGCTCTTCCGATCTNNNCCAAGCTACAATCCACAGATG
AGCACACGTCTGAACTCTTTCCCTACACGACGCTCTTCCGATCTNNNCCAAGCTACAATCCACAGATG
AGCACACGTCTGAACTCTTTCCCTACACGACGCTCTTCCGATCTNNNTTCCTCCACTATACACTCCCACT
AGCACACGTCTGAACTCTTTCCCTACACGACGCTCTTCCGATCTNNNTTCCTCCACTATACACTCCCACT
GAGCACACGTCTGAACTCTTTCCCTACACGACGCTCTTCCGATCTNNNAAAAAGGTCTATCCCCAGCAA
GAGCACACGTCTGAACTCTTTCCCTACACGACGCTCTTCCGATCTNNNAAAAAGGTCTATCCCCAGCAA
ACACGTCTGAACTCTTTCCCTACACGACGCTCTTCCGATCTNNNAGAAACCTTCATTTGTGTCAGCTCAT
ACACGTCTGAACTCTTTCCCTACACGACGCTCTTCCGATCTNNNAGAAACCTTCATTTGTGTCAGCTCAT
AGCACACGTCTGAACTCTTTCCCTACACGACGCTCTTCCGATCTNNNTGCCTGTAATCCCAGCTATTCA
AGCACACGTCTGAACTCTTTCCCTACACGACGCTCTTCCGATCTNNNTGCCTGTAATCCCAGCTATTCA
GCACACGTCTGAACTCTTTCCCTACACGACGCTCTTCCGATCTNNNCTGGGATCATGGTATCCAGGCC
GCACACGTCTGAACTCTTTCCCTACACGACGCTCTTCCGATCTNNNCTGGGATCATGGTATCCAGGCC
GCACACGTCTGAACTCTTTCCCTACACGACGCTCTTCCGATCTNNNAGTATACATGGGATGACGAACG

GCACACGTCTGAACTCTTTCCCTACACGACGCTCTTCCGATCTNNNAGTATACATGGGATGACGAACG

GCACACGTCTGAACTCTTTCCCTACACGACGCTCTTCCGATCTNNNGACTATCACCAGGCTCTGCAATA

GCACACGTCTGAACTCTTTCCCTACACGACGCTCTTCCGATCTNNNGACTATCACCAGGCTCTGCAATA

AGCACACGTCTGAACTCTTTCCCTACACGACGCTCTTCCGATCTNNNTGGGACAAATCATGCACCTGTA

AGCACACGTCTGAACTCTTTCCCTACACGACGCTCTTCCGATCTNNNTGGGACAAATCATGCACCTGTA

GAGCACACGTCTGAACTCTTTCCCTACACGACGCTCTTCCGATCTNNNGTAAGCTGTGATTGTGCCACT

GAGCACACGTCTGAACTCTTTCCCTACACGACGCTCTTCCGATCTNNNGTAAGCTGTGATTGTGCCACT

GCACACGTCTGAACTCTTTCCCTACACGACGCTCTTCCGATCTNNNCTTCTTACCCTCTCCTGCAGAAGC

GCACACGTCTGAACTCTTTCCCTACACGACGCTCTTCCGATCTNNNCTTCTTACCCTCTCCTGCAGAAGC

AGCACACGTCTGAACTCTTTCCCTACACGACGCTCTTCCGATCTNNNATTCTGTCAGCCTCTTGAGTAGC

AGCACACGTCTGAACTCTTTCCCTACACGACGCTCTTCCGATCTNNNATTCTGTCAGCCTCTTGAGTAGC

GAGCACACGTCTGAACTCTTTCCCTACACGACGCTCTTCCGATCTNNNAGGTGGGAGGATTGGTTTATT

GAGCACACGTCTGAACTCTTTCCCTACACGACGCTCTTCCGATCTNNNAGGTGGGAGGATTGGTTTATT

GAGCACACGTCTGAACTCTTTCCCTACACGACGCTCTTCCGATCTNNNTCAACTGCCCAAAACAAGTTAC

GAGCACACGTCTGAACTCTTTCCCTACACGACGCTCTTCCGATCTNNNTCAACTGCCCAAAACAAGTTAC

AGCACACGTCTGAACTCTTTCCCTACACGACGCTCTTCCGATCTNNNTCCCTTTCCCTTGCTGATTTTGG

AGCACACGTCTGAACTCTTTCCCTACACGACGCTCTTCCGATCTNNNTCCCTTTCCCTTGCTGATTTTGG

GCACACGTCTGAACTCTTTCCCTACACGACGCTCTTCCGATCTNNNCTCCTCTGACCCACTCACAGCATC

GCACACGTCTGAACTCTTTCCCTACACGACGCTCTTCCGATCTNNNCTCCTCTGACCCACTCACAGCATC

AGCACACGTCTGAACTCTTTCCCTACACGACGCTCTTCCGATCTNNNAGGGTTAGAGAAAGGGAGGA

AGCACACGTCTGAACTCTTTCCCTACACGACGCTCTTCCGATCTNNNAGGGTTAGAGAAAGGGAGGA

AGCACACGTCTGAACTCTTTCCCTACACGACGCTCTTCCGATCTNNNAGAGGAATCATGCACACAGCTT

AGCACACGTCTGAACTCTTTCCCTACACGACGCTCTTCCGATCTNNNAGAGGAATCATGCACACAGCTT

GAGCACACGTCTGAACTCTTTCCCTACACGACGCTCTTCCGATCTNNNTAGCTCACTGAAGCCTCAATCT

GAGCACACGTCTGAACTCTTTCCCTACACGACGCTCTTCCGATCTNNNTAGCTCACTGAAGCCTCAATCT

GCACACGTCTGAACTCTTTCCCTACACGACGCTCTTCCGATCTNNNGGAAGGATGATACAGTGTGAGC

GCACACGTCTGAACTCTTTCCCTACACGACGCTCTTCCGATCTNNNGGAAGGATGATACAGTGTGAGC

AGCACACGTCTGAACTCTTTCCCTACACGACGCTCTTCCGATCTNNNCAAATGGACCAGTCAGCAGGAT

AGCACACGTCTGAACTCTTTCCCTACACGACGCTCTTCCGATCTNNNCAAATGGACCAGTCAGCAGGAT

GCACACGTCTGAACTCTTTCCCTACACGACGCTCTTCCGATCTNNNCTTAGGCTGGAGTTTACAATAAC

GCACACGTCTGAACTCTTTCCCTACACGACGCTCTTCCGATCTNNNCTTAGGCTGGAGTTTACAATAAC

AGCACACGTCTGAACTCTTTCCCTACACGACGCTCTTCCGATCTNNNCTGTCAGTTTGCCATTGCCATTT

AGCACACGTCTGAACTCTTTCCCTACACGACGCTCTTCCGATCTNNNCTGTCAGTTTGCCATTGCCATTT

GAGCACACGTCTGAACTCTTTCCCTACACGACGCTCTTCCGATCTNNNTGTAGTCCCAGCTACTAGAGA

GAGCACACGTCTGAACTCTTTCCCTACACGACGCTCTTCCGATCTNNNTGTAGTCCCAGCTACTAGAGA

AGCACACGTCTGAACTCTTTCCCTACACGACGCTCTTCCGATCTNNNGAGCGAGTTCATCCGTGTTATT

AGCACACGTCTGAACTCTTTCCCTACACGACGCTCTTCCGATCTNNNGAGCGAGTTCATCCGTGTTATT

AGCACACGTCTGAACTCTTTCCCTACACGACGCTCTTCCGATCTNNNGGTCCCAGAAATCTCTTGGCAT

AGCACACGTCTGAACTCTTTCCCTACACGACGCTCTTCCGATCTNNNGGTCCCAGAAATCTCTTGGCAT

GAGCACACGTCTGAACTCTTTCCCTACACGACGCTCTTCCGATCTNNNGGATGGGAGAGAAGGCTGA

GAGCACACGTCTGAACTCTTTCCCTACACGACGCTCTTCCGATCTNNNGGATGGGAGAGAAGGCTGA

AGAGCACACGTCTGAACTCTTTCCCTACACGACGCTCTTCCGATCTNNNTTCCAGGCTTATTAGGCTTC

AGAGCACACGTCTGAACTCTTTCCCTACACGACGCTCTTCCGATCTNNNTTCCAGGCTTATTAGGCTTC

AGCACACGTCTGAACTCTTTCCCTACACGACGCTCTTCCGATCTNNNGCAGCATTACTAGGCTTTAGGC
AGCACACGTCTGAACTCTTTCCCTACACGACGCTCTTCCGATCTNNNGCAGCATTACTAGGCTTTAGGC
GAGCACACGTCTGAACTCTTTCCCTACACGACGCTCTTCCGATCTNNNGAATGCCTTGGAGTTCTTTTG
GAGCACACGTCTGAACTCTTTCCCTACACGACGCTCTTCCGATCTNNNGAATGCCTTGGAGTTCTTTTG
GAGCACACGTCTGAACTCTTTCCCTACACGACGCTCTTCCGATCTNNNACCTGTAGTCCCAGCTACTCG
GAGCACACGTCTGAACTCTTTCCCTACACGACGCTCTTCCGATCTNNNACCTGTAGTCCCAGCTACTCG
GAGCACACGTCTGAACTCTTTCCCTACACGACGCTCTTCCGATCTNNNAGGCAGGAAAATTGCTTGAA
GAGCACACGTCTGAACTCTTTCCCTACACGACGCTCTTCCGATCTNNNAGGCAGGAAAATTGCTTGAA
GAGCACACGTCTGAACTCTTTCCCTACACGACGCTCTTCCGATCTNNNGTGTGTGTGTGTGTGTAT
GAGCACACGTCTGAACTCTTTCCCTACACGACGCTCTTCCGATCTNNNGTGTGTGTGTGTGTGTAT
ACACGTCTGAACTCTTTCCCTACACGACGCTCTTCCGATCTNNNCTATGAAATACAGTGACAGTGTCAT
ACACGTCTGAACTCTTTCCCTACACGACGCTCTTCCGATCTNNNCTATGAAATACAGTGACAGTGTCAT
GAGCACACGTCTGAACTCTTTCCCTACACGACGCTCTTCCGATCTNNNTTCCCCAATTTCACTCCATTTG
GAGCACACGTCTGAACTCTTTCCCTACACGACGCTCTTCCGATCTNNNTTCCCCAATTTCACTCCATTTG
GCACACGTCTGAACTCTTTCCCTACACGACGCTCTTCCGATCTNNNGAGGCAGGAGAATTGCTTGAAT
GCACACGTCTGAACTCTTTCCCTACACGACGCTCTTCCGATCTNNNGAGGCAGGAGAATTGCTTGAAT
GAGCACACGTCTGAACTCTTTCCCTACACGACGCTCTTCCGATCTNNNATCTTGGACGTAGATAAGTG
GAGCACACGTCTGAACTCTTTCCCTACACGACGCTCTTCCGATCTNNNATCTTGGACGTAGATAAGTG
AGCACACGTCTGAACTCTTTCCCTACACGACGCTCTTCCGATCTNNNGTAGGGAAGACAACACAACAG
AGCACACGTCTGAACTCTTTCCCTACACGACGCTCTTCCGATCTNNNGTAGGGAAGACAACACAACAG
AGCACACGTCTGAACTCTTTCCCTACACGACGCTCTTCCGATCTNNNAATTCAATCCTGCAGCTTTCCAC
AGCACACGTCTGAACTCTTTCCCTACACGACGCTCTTCCGATCTNNNAATTCAATCCTGCAGCTTTCCAC
AGCACACGTCTGAACTCTTTCCCTACACGACGCTCTTCCGATCTNNNATTCTTTTGCCTTGAAAGACGG
AGCACACGTCTGAACTCTTTCCCTACACGACGCTCTTCCGATCTNNNATTCTTTTGCCTTGAAAGACGG
GCACACGTCTGAACTCTTTCCCTACACGACGCTCTTCCGATCTNNNGCCTCTCCAACAATTTCAGTCAGC
GCACACGTCTGAACTCTTTCCCTACACGACGCTCTTCCGATCTNNNGCCTCTCCAACAATTTCAGTCAGC
AGCACACGTCTGAACTCTTTCCCTACACGACGCTCTTCCGATCTNNNGGAAGAGGAGCTTGCAGTGAG
AGCACACGTCTGAACTCTTTCCCTACACGACGCTCTTCCGATCTNNNGGAAGAGGAGCTTGCAGTGAG
GAGCACACGTCTGAACTCTTTCCCTACACGACGCTCTTCCGATCTNNNCACAATGTGCACATGTACCCT
GAGCACACGTCTGAACTCTTTCCCTACACGACGCTCTTCCGATCTNNNCACAATGTGCACATGTACCCT
AGCACACGTCTGAACTCTTTCCCTACACGACGCTCTTCCGATCTNNNTTGTTTATCTGTGCAGCCATGT
AGCACACGTCTGAACTCTTTCCCTACACGACGCTCTTCCGATCTNNNTTGTTTATCTGTGCAGCCATGT
AGCACACGTCTGAACTCTTTCCCTACACGACGCTCTTCCGATCTNNNCTTTTCAGCAAGGCCTTTTCTG
AGCACACGTCTGAACTCTTTCCCTACACGACGCTCTTCCGATCTNNNCTTTTCAGCAAGGCCTTTTCTG
GAGCACACGTCTGAACTCTTTCCCTACACGACGCTCTTCCGATCTNNNCTTCTTTGTGTGTGTGTGTGT
GAGCACACGTCTGAACTCTTTCCCTACACGACGCTCTTCCGATCTNNNCTTCTTTGTGTGTGTGTGTGT
AGCACACGTCTGAACTCTTTCCCTACACGACGCTCTTCCGATCTNNNTTCAGGAAGGTGGTACTTGG
AGCACACGTCTGAACTCTTTCCCTACACGACGCTCTTCCGATCTNNNTTCAGGAAGGTGGTACTTGG
AGCACACGTCTGAACTCTTTCCCTACACGACGCTCTTCCGATCTNNNCTATATGCAGCCGGATTCTCTG
AGCACACGTCTGAACTCTTTCCCTACACGACGCTCTTCCGATCTNNNCTATATGCAGCCGGATTCTCTG
AGCACACGTCTGAACTCTTTCCCTACACGACGCTCTTCCGATCTNNNTTGAACATCTTCAGGTGTCCCA
AGCACACGTCTGAACTCTTTCCCTACACGACGCTCTTCCGATCTNNNTTGAACATCTTCAGGTGTCCCA
AGCACACGTCTGAACTCTTTCCCTACACGACGCTCTTCCGATCTNNNGTAATCTCAGCTACTCAGGAG

AGCACACGTCTGAACTCTTTCCCTACACGACGCTCTTCCGATCTNNNGTAATCTCAGCTACTCAGGAG
GAGCACACGTCTGAACTCTTTCCCTACACGACGCTCTTCCGATCTNNNAATCACTTGAATCCTAGGGG
GAGCACACGTCTGAACTCTTTCCCTACACGACGCTCTTCCGATCTNNNAATCACTTGAATCCTAGGGG
AGCACACGTCTGAACTCTTTCCCTACACGACGCTCTTCCGATCTNNNTGTAATGCCAGCTACTCTACA
AGCACACGTCTGAACTCTTTCCCTACACGACGCTCTTCCGATCTNNNTGTAATGCCAGCTACTCTACA
GAGCACACGTCTGAACTCTTTCCCTACACGACGCTCTTCCGATCTNNNACAACTGCATGAACATAGTCC
GAGCACACGTCTGAACTCTTTCCCTACACGACGCTCTTCCGATCTNNNACAACTGCATGAACATAGTCC
GCACACGTCTGAACTCTTTCCCTACACGACGCTCTTCCGATCTNNNCTGAGGCAGGAGAATCACTTGA
GCACACGTCTGAACTCTTTCCCTACACGACGCTCTTCCGATCTNNNCTGAGGCAGGAGAATCACTTGA
AGCACACGTCTGAACTCTTTCCCTACACGACGCTCTTCCGATCTNNNCATATGCCTGTAGTCCTAGCCA
AGCACACGTCTGAACTCTTTCCCTACACGACGCTCTTCCGATCTNNNCATATGCCTGTAGTCCTAGCCA
AGCACACGTCTGAACTCTTTCCCTACACGACGCTCTTCCGATCTNNNATATTACATTTCAGAGTTCACCT
AGCACACGTCTGAACTCTTTCCCTACACGACGCTCTTCCGATCTNNNATATTACATTTCAGAGTTCACCT
GAGCACACGTCTGAACTCTTTCCCTACACGACGCTCTTCCGATCTNNNTTTAGTAGAGACGGGGTTT
GAGCACACGTCTGAACTCTTTCCCTACACGACGCTCTTCCGATCTNNNTTTAGTAGAGACGGGGTTT
GCACACGTCTGAACTCTTTCCCTACACGACGCTCTTCCGATCTNNNGACACACTGATTCCTACACCACT
GCACACGTCTGAACTCTTTCCCTACACGACGCTCTTCCGATCTNNNGACACACTGATTCCTACACCACT
AGCACACGTCTGAACTCTTTCCCTACACGACGCTCTTCCGATCTNNNAGACATGGAATCAACCTAGGT
AGCACACGTCTGAACTCTTTCCCTACACGACGCTCTTCCGATCTNNNAGACATGGAATCAACCTAGGT
CACACGTCTGAACTCTTTCCCTACACGACGCTCTTCCGATCTNNNGCCTATAGTTTTCTGCGTGTGTG
CACACGTCTGAACTCTTTCCCTACACGACGCTCTTCCGATCTNNNGCCTATAGTTTTCTGCGTGTGTG
CACACGTCTGAACTCTTTCCCTACACGACGCTCTTCCGATCTNNNCTCTCTCTCTGTCTCCCTGTCCAT
CACACGTCTGAACTCTTTCCCTACACGACGCTCTTCCGATCTNNNCTCTCTCTCTGTCTCCCTGTCCAT
AGCACACGTCTGAACTCTTTCCCTACACGACGCTCTTCCGATCTNNNCAAGGAGTTTGAGAACAGCTT
AGCACACGTCTGAACTCTTTCCCTACACGACGCTCTTCCGATCTNNNCAAGGAGTTTGAGAACAGCTT
GCACACGTCTGAACTCTTTCCCTACACGACGCTCTTCCGATCTNNNCCAAAATCCAAAGTGCTGGGAA
GCACACGTCTGAACTCTTTCCCTACACGACGCTCTTCCGATCTNNNCCAAAATCCAAAGTGCTGGGAA
GAGCACACGTCTGAACTCTTTCCCTACACGACGCTCTTCCGATCTNNNCACAGTCCATTTTCTGAGCC
GAGCACACGTCTGAACTCTTTCCCTACACGACGCTCTTCCGATCTNNNCACAGTCCATTTTCTGAGCC
AGCACACGTCTGAACTCTTTCCCTACACGACGCTCTTCCGATCTNNNGAGGATCATTTGAGTCCAGGA
AGCACACGTCTGAACTCTTTCCCTACACGACGCTCTTCCGATCTNNNGAGGATCATTTGAGTCCAGGA
AGCACACGTCTGAACTCTTTCCCTACACGACGCTCTTCCGATCTNNNCGAGGTCTGGAATTAGACACC
AGCACACGTCTGAACTCTTTCCCTACACGACGCTCTTCCGATCTNNNCGAGGTCTGGAATTAGACACC
GAGCACACGTCTGAACTCTTTCCCTACACGACGCTCTTCCGATCTNNNTTCACCATGCCGGTGATAAAC
GAGCACACGTCTGAACTCTTTCCCTACACGACGCTCTTCCGATCTNNNTTCACCATGCCGGTGATAAAC
GCACACGTCTGAACTCTTTCCCTACACGACGCTCTTCCGATCTNNNACGCCACCATACCTGTCTAATTT
GCACACGTCTGAACTCTTTCCCTACACGACGCTCTTCCGATCTNNNACGCCACCATACCTGTCTAATTT
AGAGCACACGTCTGAACTCTTTCCCTACACGACGCTCTTCCGATCTNNNGAGGCTGAGGCAGGAGAA
AGAGCACACGTCTGAACTCTTTCCCTACACGACGCTCTTCCGATCTNNNGAGGCTGAGGCAGGAGAA
GCACACGTCTGAACTCTTTCCCTACACGACGCTCTTCCGATCTNNNAGATAGCGAAGTCCAAGGTCA
GCACACGTCTGAACTCTTTCCCTACACGACGCTCTTCCGATCTNNNAGATAGCGAAGTCCAAGGTCA
GCACACGTCTGAACTCTTTCCCTACACGACGCTCTTCCGATCTNNNACCAACCCTAGATCCCTCTACAA
GCACACGTCTGAACTCTTTCCCTACACGACGCTCTTCCGATCTNNNACCAACCCTAGATCCCTCTACAA

CACACGTCTGAACTCTTTCCCTACACGACGCTCTTCCGATCTNNNCATGGCCTGCTTCCTCACAGCATC
CACACGTCTGAACTCTTTCCCTACACGACGCTCTTCCGATCTNNNCATGGCCTGCTTCCTCACAGCATC
GAGCACACGTCTGAACTCTTTCCCTACACGACGCTCTTCCGATCTNNNTGGTTTCAAGGAGGAAGTAT
GAGCACACGTCTGAACTCTTTCCCTACACGACGCTCTTCCGATCTNNNTGGTTTCAAGGAGGAAGTAT
AGAGCACACGTCTGAACTCTTTCCCTACACGACGCTCTTCCGATCTNNNTCAGGTTGATTTTGTGCCT
AGAGCACACGTCTGAACTCTTTCCCTACACGACGCTCTTCCGATCTNNNTCAGGTTGATTTTGTGCCT
GAGCACACGTCTGAACTCTTTCCCTACACGACGCTCTTCCGATCTNNNTCCCTACACCACGAAATATGC
GAGCACACGTCTGAACTCTTTCCCTACACGACGCTCTTCCGATCTNNNTCCCTACACCACGAAATATGC
GCACACGTCTGAACTCTTTCCCTACACGACGCTCTTCCGATCTNNNAATCCTGGGCTCCTTTGATCTTT
GCACACGTCTGAACTCTTTCCCTACACGACGCTCTTCCGATCTNNNAATCCTGGGCTCCTTTGATCTTT
GAGCACACGTCTGAACTCTTTCCCTACACGACGCTCTTCCGATCTNNNGAGGCTGAGGCAGGAGAA
GAGCACACGTCTGAACTCTTTCCCTACACGACGCTCTTCCGATCTNNNGAGGCTGAGGCAGGAGAA
GAGCACACGTCTGAACTCTTTCCCTACACGACGCTCTTCCGATCTNNNTGAGATACCACTTCACATCCA
GAGCACACGTCTGAACTCTTTCCCTACACGACGCTCTTCCGATCTNNNTGAGATACCACTTCACATCCA
AGCACACGTCTGAACTCTTTCCCTACACGACGCTCTTCCGATCTNNNCTGTAATCCCAGCTACTCAGG
AGCACACGTCTGAACTCTTTCCCTACACGACGCTCTTCCGATCTNNNCTGTAATCCCAGCTACTCAGG
GAGCACACGTCTGAACTCTTTCCCTACACGACGCTCTTCCGATCTNNNAGCTCCAATAACTTTTGGTG
GAGCACACGTCTGAACTCTTTCCCTACACGACGCTCTTCCGATCTNNNAGCTCCAATAACTTTTGGTG
AGCACACGTCTGAACTCTTTCCCTACACGACGCTCTTCCGATCTNNNATGCAATGTTGAGACCTGGA
AGCACACGTCTGAACTCTTTCCCTACACGACGCTCTTCCGATCTNNNATGCAATGTTGAGACCTGGA
AGCACACGTCTGAACTCTTTCCCTACACGACGCTCTTCCGATCTNNNTCGGCAAGTACCCCAAGGAGC
AGCACACGTCTGAACTCTTTCCCTACACGACGCTCTTCCGATCTNNNTCGGCAAGTACCCCAAGGAGC
GCACACGTCTGAACTCTTTCCCTACACGACGCTCTTCCGATCTNNNCGTGATCACACCACTGCATTGC
GCACACGTCTGAACTCTTTCCCTACACGACGCTCTTCCGATCTNNNCGTGATCACACCACTGCATTGC
CACACGTCTGAACTCTTTCCCTACACGACGCTCTTCCGATCTNNNCCAGACACCCTAACTCATCACTCCA
CACACGTCTGAACTCTTTCCCTACACGACGCTCTTCCGATCTNNNCCAGACACCCTAACTCATCACTCCA
AGCACACGTCTGAACTCTTTCCCTACACGACGCTCTTCCGATCTNNNAGGCATTGTATTGAAGGATG
AGCACACGTCTGAACTCTTTCCCTACACGACGCTCTTCCGATCTNNNAGGCATTGTATTGAAGGATG
AGCACACGTCTGAACTCTTTCCCTACACGACGCTCTTCCGATCTNNNGGAGAATCACTTGAACCTGG
AGCACACGTCTGAACTCTTTCCCTACACGACGCTCTTCCGATCTNNNGGAGAATCACTTGAACCTGG
GCACACGTCTGAACTCTTTCCCTACACGACGCTCTTCCGATCTNNNAGGCATGAAGAAAAAAAAAAA
GCACACGTCTGAACTCTTTCCCTACACGACGCTCTTCCGATCTNNNAGGCATGAAGAAAAAAAAAAA
GCACACGTCTGAACTCTTTCCCTACACGACGCTCTTCCGATCTNNNCCAGAAAGTCAAAAGCAGTGG
GCACACGTCTGAACTCTTTCCCTACACGACGCTCTTCCGATCTNNNCCAGAAAGTCAAAAGCAGTGG
AGCACACGTCTGAACTCTTTCCCTACACGACGCTCTTCCGATCTNNNTTCTACCAAGTGGCTGAGTAC
AGCACACGTCTGAACTCTTTCCCTACACGACGCTCTTCCGATCTNNNTTCTACCAAGTGGCTGAGTAC
AGCACACGTCTGAACTCTTTCCCTACACGACGCTCTTCCGATCTNNNTTCTAGTGGGCAGCAGAGAA
AGCACACGTCTGAACTCTTTCCCTACACGACGCTCTTCCGATCTNNNTTCTAGTGGGCAGCAGAGAA
GAGCACACGTCTGAACTCTTTCCCTACACGACGCTCTTCCGATCTNNNCCCTCTCATCCCAGAGCTCAC
GAGCACACGTCTGAACTCTTTCCCTACACGACGCTCTTCCGATCTNNNCCCTCTCATCCCAGAGCTCAC
AGAGCACACGTCTGAACTCTTTCCCTACACGACGCTCTTCCGATCTNNNCTGGAGTGTTAGGTTTTGG
AGAGCACACGTCTGAACTCTTTCCCTACACGACGCTCTTCCGATCTNNNCTGGAGTGTTAGGTTTTGG
AGCACACGTCTGAACTCTTTCCCTACACGACGCTCTTCCGATCTNNNCTCCATCTCCAAGGTCAAGTGA

AGCACACGTCTGAACTCTTTCCCTACACGACGCTCTTCCGATCTNNNCTCCATCTCCAAGGTCAAGTGA

AGCACACGTCTGAACTCTTTCCCTACACGACGCTCTTCCGATCTNNNCTGTCAGTCAACAGTGAGCTT

AGCACACGTCTGAACTCTTTCCCTACACGACGCTCTTCCGATCTNNNCTGTCAGTCAACAGTGAGCTT

GCACACGTCTGAACTCTTTCCCTACACGACGCTCTTCCGATCTNNNTAATGCAAAGACAGCACCAAACC

GCACACGTCTGAACTCTTTCCCTACACGACGCTCTTCCGATCTNNNTAATGCAAAGACAGCACCAAACC

AGCACACGTCTGAACTCTTTCCCTACACGACGCTCTTCCGATCTNNNGCACCAGCTGAGGTTACAATTT

AGCACACGTCTGAACTCTTTCCCTACACGACGCTCTTCCGATCTNNNGCACCAGCTGAGGTTACAATTT

AGCACACGTCTGAACTCTTTCCCTACACGACGCTCTTCCGATCTNNNAGGATCTCTTGAGCTCAGGGA

AGCACACGTCTGAACTCTTTCCCTACACGACGCTCTTCCGATCTNNNAGGATCTCTTGAGCTCAGGGA

AGCACACGTCTGAACTCTTTCCCTACACGACGCTCTTCCGATCTNNNCCATGACAGGATGTAGTGACC

AGCACACGTCTGAACTCTTTCCCTACACGACGCTCTTCCGATCTNNNCCATGACAGGATGTAGTGACC

GAGCACACGTCTGAACTCTTTCCCTACACGACGCTCTTCCGATCTNNNTTAATCAAGCCATTCATCGCT

GAGCACACGTCTGAACTCTTTCCCTACACGACGCTCTTCCGATCTNNNTTAATCAAGCCATTCATCGCT

AGCACACGTCTGAACTCTTTCCCTACACGACGCTCTTCCGATCTNNNCTCTTCCCCTAATTTCCCCAAAC

AGCACACGTCTGAACTCTTTCCCTACACGACGCTCTTCCGATCTNNNCTCTTCCCCTAATTTCCCCAAAC

AGCACACGTCTGAACTCTTTCCCTACACGACGCTCTTCCGATCTNNNTTCTAGATGGGGGCAGGAAA

AGCACACGTCTGAACTCTTTCCCTACACGACGCTCTTCCGATCTNNNTTCTAGATGGGGGCAGGAAA

GCACACGTCTGAACTCTTTCCCTACACGACGCTCTTCCGATCTNNNTAACTATATTCGGTCCTTCATAC

GCACACGTCTGAACTCTTTCCCTACACGACGCTCTTCCGATCTNNNTAACTATATTCGGTCCTTCATAC

AGCACACGTCTGAACTCTTTCCCTACACGACGCTCTTCCGATCTNNNCACACACACACTCTCTCTCTT

AGCACACGTCTGAACTCTTTCCCTACACGACGCTCTTCCGATCTNNNCACACACACACTCTCTCTCTT

GCACACGTCTGAACTCTTTCCCTACACGACGCTCTTCCGATCTNNNGAAGGAGCAAAATGAAGCCTTG

GCACACGTCTGAACTCTTTCCCTACACGACGCTCTTCCGATCTNNNGAAGGAGCAAAATGAAGCCTTG

GAGCACACGTCTGAACTCTTTCCCTACACGACGCTCTTCCGATCTNNNGGTTGTAGTGAGCCAAGTTT

GAGCACACGTCTGAACTCTTTCCCTACACGACGCTCTTCCGATCTNNNGGTTGTAGTGAGCCAAGTTT

AGCACACGTCTGAACTCTTTCCCTACACGACGCTCTTCCGATCTNNNTGAGGCAGGAGAATCTCTTGA

AGCACACGTCTGAACTCTTTCCCTACACGACGCTCTTCCGATCTNNNTGAGGCAGGAGAATCTCTTGA

AGCACACGTCTGAACTCTTTCCCTACACGACGCTCTTCCGATCTNNNAGAAGTTGCAGTGAGCTGAG

AGCACACGTCTGAACTCTTTCCCTACACGACGCTCTTCCGATCTNNNAGAAGTTGCAGTGAGCTGAG

AGCACACGTCTGAACTCTTTCCCTACACGACGCTCTTCCGATCTNNNATCGTAGGGTGTCACTTTTCCA

AGCACACGTCTGAACTCTTTCCCTACACGACGCTCTTCCGATCTNNNATCGTAGGGTGTCACTTTTCCA

GAGCACACGTCTGAACTCTTTCCCTACACGACGCTCTTCCGATCTNNNACAAAAAATAGCCAGGTGTG

GAGCACACGTCTGAACTCTTTCCCTACACGACGCTCTTCCGATCTNNNACAAAAAATAGCCAGGTGTG

AGCACACGTCTGAACTCTTTCCCTACACGACGCTCTTCCGATCTNNNCCGACTCAACTCCACAGAAATCT

AGCACACGTCTGAACTCTTTCCCTACACGACGCTCTTCCGATCTNNNCCGACTCAACTCCACAGAAATCT

AGAGCACACGTCTGAACTCTTTCCCTACACGACGCTCTTCCGATCTNNNCGCCGCATATTCTAGAAAAC

AGAGCACACGTCTGAACTCTTTCCCTACACGACGCTCTTCCGATCTNNNCGCCGCATATTCTAGAAAAC

AGCACACGTCTGAACTCTTTCCCTACACGACGCTCTTCCGATCTNNNACCAGCACTAAATCATCCCAAG

AGCACACGTCTGAACTCTTTCCCTACACGACGCTCTTCCGATCTNNNACCAGCACTAAATCATCCCAAG

AGCACACGTCTGAACTCTTTCCCTACACGACGCTCTTCCGATCTNNNAGCAATGGACAGGAGAAGGA

AGCACACGTCTGAACTCTTTCCCTACACGACGCTCTTCCGATCTNNNAGCAATGGACAGGAGAAGGA

AGCACACGTCTGAACTCTTTCCCTACACGACGCTCTTCCGATCTNNNCTCATATGACCACTCCCATCTGG

AGCACACGTCTGAACTCTTTCCCTACACGACGCTCTTCCGATCTNNNCTCATATGACCACTCCCATCTGG

GAGCACACGTCTGAACTCTTTCCCTACACGACGCTCTTCCGATCTNNNTGGAGATTGAACAAAAGGT
GAGCACACGTCTGAACTCTTTCCCTACACGACGCTCTTCCGATCTNNNTGGAGATTGAACAAAAGGT
GCACACGTCTGAACTCTTTCCCTACACGACGCTCTTCCGATCTNNNTCCTCCTCCTTCTTCTCCTTCTTCA
GCACACGTCTGAACTCTTTCCCTACACGACGCTCTTCCGATCTNNNTCCTCCTCCTTCTTCTCCTTCTTCA
GAGCACACGTCTGAACTCTTTCCCTACACGACGCTCTTCCGATCTNNNATGTGTGTGTGTATGTGTG
GAGCACACGTCTGAACTCTTTCCCTACACGACGCTCTTCCGATCTNNNATGTGTGTGTGTATGTGTG
GAGCACACGTCTGAACTCTTTCCCTACACGACGCTCTTCCGATCTNNNCACCTGTAATCCCAGCTACTC
GAGCACACGTCTGAACTCTTTCCCTACACGACGCTCTTCCGATCTNNNCACCTGTAATCCCAGCTACTC
AGCACACGTCTGAACTCTTTCCCTACACGACGCTCTTCCGATCTNNNCCAGGCATGACAATCAAAACT
AGCACACGTCTGAACTCTTTCCCTACACGACGCTCTTCCGATCTNNNCCAGGCATGACAATCAAAACT
ACACGTCTGAACTCTTTCCCTACACGACGCTCTTCCGATCTNNNGTGCTACTGGCCCCTGATATTTACA
ACACGTCTGAACTCTTTCCCTACACGACGCTCTTCCGATCTNNNGTGCTACTGGCCCCTGATATTTACA
AGCACACGTCTGAACTCTTTCCCTACACGACGCTCTTCCGATCTNNNAAGGACAGTGAATGGGGATA
AGCACACGTCTGAACTCTTTCCCTACACGACGCTCTTCCGATCTNNNAAGGACAGTGAATGGGGATA
GAGCACACGTCTGAACTCTTTCCCTACACGACGCTCTTCCGATCTNNNACCCATTTCATAATCAGTTTC
GAGCACACGTCTGAACTCTTTCCCTACACGACGCTCTTCCGATCTNNNACCCATTTCATAATCAGTTTC
GCACACGTCTGAACTCTTTCCCTACACGACGCTCTTCCGATCTNNNGTGGAGGTTACAGTGAGTTGA
GCACACGTCTGAACTCTTTCCCTACACGACGCTCTTCCGATCTNNNGTGGAGGTTACAGTGAGTTGA
GCACACGTCTGAACTCTTTCCCTACACGACGCTCTTCCGATCTNNNTTTAGTAGAGGCGAGGTTTCAC
GCACACGTCTGAACTCTTTCCCTACACGACGCTCTTCCGATCTNNNTTTAGTAGAGGCGAGGTTTCAC
CACACGTCTGAACTCTTTCCCTACACGACGCTCTTCCGATCTNNNATGTTCAGTAGCTTTCCTCTAAGT
CACACGTCTGAACTCTTTCCCTACACGACGCTCTTCCGATCTNNNATGTTCAGTAGCTTTCCTCTAAGT
GCACACGTCTGAACTCTTTCCCTACACGACGCTCTTCCGATCTNNNGTAGGATTCTGAGCCATCTGAC
GCACACGTCTGAACTCTTTCCCTACACGACGCTCTTCCGATCTNNNGTAGGATTCTGAGCCATCTGAC
GCACACGTCTGAACTCTTTCCCTACACGACGCTCTTCCGATCTNNNAGTCTCATTTCATCCTGCTCTGG
GCACACGTCTGAACTCTTTCCCTACACGACGCTCTTCCGATCTNNNAGTCTCATTTCATCCTGCTCTGG
CACACGTCTGAACTCTTTCCCTACACGACGCTCTTCCGATCTNNNATCAAAGTTCTGAGACTTAATCTC
CACACGTCTGAACTCTTTCCCTACACGACGCTCTTCCGATCTNNNATCAAAGTTCTGAGACTTAATCTC
GAGCACACGTCTGAACTCTTTCCCTACACGACGCTCTTCCGATCTNNNGAGCTTGCAGTGAGTCGAG
GAGCACACGTCTGAACTCTTTCCCTACACGACGCTCTTCCGATCTNNNGAGCTTGCAGTGAGTCGAG
AGCACACGTCTGAACTCTTTCCCTACACGACGCTCTTCCGATCTNNNTCCTGGAAGCTTTTTGTAGCA
AGCACACGTCTGAACTCTTTCCCTACACGACGCTCTTCCGATCTNNNTCCTGGAAGCTTTTTGTAGCA
AGCACACGTCTGAACTCTTTCCCTACACGACGCTCTTCCGATCTNNNACTCTGTCTTCTGGACTATCCC
AGCACACGTCTGAACTCTTTCCCTACACGACGCTCTTCCGATCTNNNACTCTGTCTTCTGGACTATCCC
GCACACGTCTGAACTCTTTCCCTACACGACGCTCTTCCGATCTNNNCTAAGGTGTCAACAGGGCTATG
GCACACGTCTGAACTCTTTCCCTACACGACGCTCTTCCGATCTNNNCTAAGGTGTCAACAGGGCTATG
AGCACACGTCTGAACTCTTTCCCTACACGACGCTCTTCCGATCTNNNGAGATCTCACCATTGCACTCCA
AGCACACGTCTGAACTCTTTCCCTACACGACGCTCTTCCGATCTNNNGAGATCTCACCATTGCACTCCA
AGCACACGTCTGAACTCTTTCCCTACACGACGCTCTTCCGATCTNNNCAGAAGCAGGAGGATCACTT
AGCACACGTCTGAACTCTTTCCCTACACGACGCTCTTCCGATCTNNNCAGAAGCAGGAGGATCACTT
AGCACACGTCTGAACTCTTTCCCTACACGACGCTCTTCCGATCTNNNAGACTGTGTGTGTCTGTGCAT
AGCACACGTCTGAACTCTTTCCCTACACGACGCTCTTCCGATCTNNNAGACTGTGTGTGTCTGTGCAT
GAGCACACGTCTGAACTCTTTCCCTACACGACGCTCTTCCGATCTNNNGATGAGGACCTCACGGAAA

GAGCACACGTCTGAACTCTTTCCCTACACGACGCTCTTCCGATCTNNNGATGAGGACCTCACGGAAAT
GCACACGTCTGAACTCTTTCCCTACACGACGCTCTTCCGATCTNNNCTTTGACCTCAGGTTTCTAACGC
GCACACGTCTGAACTCTTTCCCTACACGACGCTCTTCCGATCTNNNCTTTGACCTCAGGTTTCTAACGC
GCACACGTCTGAACTCTTTCCCTACACGACGCTCTTCCGATCTNNNAAGGGTTTTTACCTTGAAGCGAT
AGCACACGTCTGAACTCTTTCCCTACACGACGCTCTTCCGATCTNNNAAGGGTTTTTACCTTGAAGCGAT
AGCACACGTCTGAACTCTTTCCCTACACGACGCTCTTCCGATCTNNNGTGTGTGTGTGTGTGTGTGTA
AGCACACGTCTGAACTCTTTCCCTACACGACGCTCTTCCGATCTNNNGTGTGTGTGTGTGTGTGTGTA
GAGCACACGTCTGAACTCTTTCCCTACACGACGCTCTTCCGATCTNNNACCATGTCATCCTTCCCTTTCT
GAGCACACGTCTGAACTCTTTCCCTACACGACGCTCTTCCGATCTNNNACCATGTCATCCTTCCCTTTCT
AGCACACGTCTGAACTCTTTCCCTACACGACGCTCTTCCGATCTNNNAATCAGAGGAAAGCATACAGG
AGCACACGTCTGAACTCTTTCCCTACACGACGCTCTTCCGATCTNNNAATCAGAGGAAAGCATACAGG
GCACACGTCTGAACTCTTTCCCTACACGACGCTCTTCCGATCTNNNTTAGGATGGGAGGATGGTGGA
GCACACGTCTGAACTCTTTCCCTACACGACGCTCTTCCGATCTNNNTTAGGATGGGAGGATGGTGGA
AGCACACGTCTGAACTCTTTCCCTACACGACGCTCTTCCGATCTNNNCTCCCTGCTTCTGTCTCCTACAC
AGCACACGTCTGAACTCTTTCCCTACACGACGCTCTTCCGATCTNNNCTCCCTGCTTCTGTCTCCTACAC
GAGCACACGTCTGAACTCTTTCCCTACACGACGCTCTTCCGATCTNNNTTTGAGTGTCGAACTTGTGG
GAGCACACGTCTGAACTCTTTCCCTACACGACGCTCTTCCGATCTNNNTTTGAGTGTCGAACTTGTGG
GAGCACACGTCTGAACTCTTTCCCTACACGACGCTCTTCCGATCTNNNACATTTTGTAGCCATTTGACA
GAGCACACGTCTGAACTCTTTCCCTACACGACGCTCTTCCGATCTNNNACATTTTGTAGCCATTTGACA
GCACACGTCTGAACTCTTTCCCTACACGACGCTCTTCCGATCTNNNCCAAATCTCCATCCCTCAGAGAG
GCACACGTCTGAACTCTTTCCCTACACGACGCTCTTCCGATCTNNNCCAAATCTCCATCCCTCAGAGAG
GAGCACACGTCTGAACTCTTTCCCTACACGACGCTCTTCCGATCTNNNACTGGGAGTATGCAGTAAG
GAGCACACGTCTGAACTCTTTCCCTACACGACGCTCTTCCGATCTNNNACTGGGAGTATGCAGTAAG
GCACACGTCTGAACTCTTTCCCTACACGACGCTCTTCCGATCTNNNTGATCATTTCCCCAAGTTTTAGC
GCACACGTCTGAACTCTTTCCCTACACGACGCTCTTCCGATCTNNNTGATCATTTCCCCAAGTTTTAGC
GAGCACACGTCTGAACTCTTTCCCTACACGACGCTCTTCCGATCTNNNACTCGTTTACTCTGGCCTTCA
GAGCACACGTCTGAACTCTTTCCCTACACGACGCTCTTCCGATCTNNNACTCGTTTACTCTGGCCTTCA
AGCACACGTCTGAACTCTTTCCCTACACGACGCTCTTCCGATCTNNNCTCCTGGATTCACGCCATTCTC
AGCACACGTCTGAACTCTTTCCCTACACGACGCTCTTCCGATCTNNNCTCCTGGATTCACGCCATTCTC
GAGCACACGTCTGAACTCTTTCCCTACACGACGCTCTTCCGATCTNNNGAGAATGACGTGAACCCAGG
GAGCACACGTCTGAACTCTTTCCCTACACGACGCTCTTCCGATCTNNNGAGAATGACGTGAACCCAGG
GAGCACACGTCTGAACTCTTTCCCTACACGACGCTCTTCCGATCTNNNCAGCTGTTTTGTCTTCTCCCA
GAGCACACGTCTGAACTCTTTCCCTACACGACGCTCTTCCGATCTNNNCAGCTGTTTTGTCTTCTCCCA
AGCACACGTCTGAACTCTTTCCCTACACGACGCTCTTCCGATCTNNNTTTCCCCCAGTCTCCATTACTTG
AGCACACGTCTGAACTCTTTCCCTACACGACGCTCTTCCGATCTNNNTTTCCCCCAGTCTCCATTACTTG
GCACACGTCTGAACTCTTTCCCTACACGACGCTCTTCCGATCTNNNAAAGCCCAGGGAGACTATTTAG
GCACACGTCTGAACTCTTTCCCTACACGACGCTCTTCCGATCTNNNAAAGCCCAGGGAGACTATTTAG
AGCACACGTCTGAACTCTTTCCCTACACGACGCTCTTCCGATCTNNNTCCCCTTCAAACATATACATTA
AGCACACGTCTGAACTCTTTCCCTACACGACGCTCTTCCGATCTNNNTCCCCTTCAAACATATACATTA
AGCACACGTCTGAACTCTTTCCCTACACGACGCTCTTCCGATCTNNNACAAAAATTAGCCAGGTGTGG
AGCACACGTCTGAACTCTTTCCCTACACGACGCTCTTCCGATCTNNNACAAAAATTAGCCAGGTGTGG
AGCACACGTCTGAACTCTTTCCCTACACGACGCTCTTCCGATCTNNNACAATGGAAAATGGGCAAATG
AGCACACGTCTGAACTCTTTCCCTACACGACGCTCTTCCGATCTNNNACAATGGAAAATGGGCAAATG

AGCACACGTCTGAACTCTTTCCCTACACGACGCTCTTCCGATCTNNNTTCCCCGATGCTATTGTTCCTAG
AGCACACGTCTGAACTCTTTCCCTACACGACGCTCTTCCGATCTNNNTTCCCCGATGCTATTGTTCCTAG
GCACACGTCTGAACTCTTTCCCTACACGACGCTCTTCCGATCTNNNACCATCTCCTTGTTTCTGTCACTCA
GCACACGTCTGAACTCTTTCCCTACACGACGCTCTTCCGATCTNNNACCATCTCCTTGTTTCTGTCACTCA
GAGCACACGTCTGAACTCTTTCCCTACACGACGCTCTTCCGATCTNNNATCTTCCCCTCATTTTGGTGAG
GAGCACACGTCTGAACTCTTTCCCTACACGACGCTCTTCCGATCTNNNATCTTCCCCTCATTTTGGTGAG
AGCACACGTCTGAACTCTTTCCCTACACGACGCTCTTCCGATCTNNNTTCAGAAGACTGCTCCTTCAGA
AGCACACGTCTGAACTCTTTCCCTACACGACGCTCTTCCGATCTNNNTTCAGAAGACTGCTCCTTCAGA
GAGCACACGTCTGAACTCTTTCCCTACACGACGCTCTTCCGATCTNNNGGCCTATAATTCCAGCCACTT
GAGCACACGTCTGAACTCTTTCCCTACACGACGCTCTTCCGATCTNNNGGCCTATAATTCCAGCCACTT
GAGCACACGTCTGAACTCTTTCCCTACACGACGCTCTTCCGATCTNNNCGCTTTGCAATTTTATTAGGTC
GAGCACACGTCTGAACTCTTTCCCTACACGACGCTCTTCCGATCTNNNCGCTTTGCAATTTTATTAGGTC
AGCACACGTCTGAACTCTTTCCCTACACGACGCTCTTCCGATCTNNNCTGCCACCTTCCAGGATCATTTT
AGCACACGTCTGAACTCTTTCCCTACACGACGCTCTTCCGATCTNNNCTGCCACCTTCCAGGATCATTTT
AGCACACGTCTGAACTCTTTCCCTACACGACGCTCTTCCGATCTNNNGCTTTGCCATAAATTGGGATCG
AGCACACGTCTGAACTCTTTCCCTACACGACGCTCTTCCGATCTNNNGCTTTGCCATAAATTGGGATCG
GAGCACACGTCTGAACTCTTTCCCTACACGACGCTCTTCCGATCTNNNGGGGTTATTGTGAGGACTGA
GAGCACACGTCTGAACTCTTTCCCTACACGACGCTCTTCCGATCTNNNGGGGTTATTGTGAGGACTGA
GCACACGTCTGAACTCTTTCCCTACACGACGCTCTTCCGATCTNNNTCACTCTACCTTTATGCCCCATCCA
GCACACGTCTGAACTCTTTCCCTACACGACGCTCTTCCGATCTNNNTCACTCTACCTTTATGCCCCATCCA
AGCACACGTCTGAACTCTTTCCCTACACGACGCTCTTCCGATCTNNNCTACCAAGTAGCCAGGACTACA
AGCACACGTCTGAACTCTTTCCCTACACGACGCTCTTCCGATCTNNNCTACCAAGTAGCCAGGACTACA
AGCACACGTCTGAACTCTTTCCCTACACGACGCTCTTCCGATCTNNNTTTTCTCACAGTTCTGGAGGCT
AGCACACGTCTGAACTCTTTCCCTACACGACGCTCTTCCGATCTNNNTTTTCTCACAGTTCTGGAGGCT
AGAGCACACGTCTGAACTCTTTCCCTACACGACGCTCTTCCGATCTNNNTATTTTGAGACAGAGTCCC
AGAGCACACGTCTGAACTCTTTCCCTACACGACGCTCTTCCGATCTNNNTATTTTGAGACAGAGTCCC
GAGCACACGTCTGAACTCTTTCCCTACACGACGCTCTTCCGATCTNNNTCCACAGCTATATGTAAACCG
GAGCACACGTCTGAACTCTTTCCCTACACGACGCTCTTCCGATCTNNNTCCACAGCTATATGTAAACCG
AGCACACGTCTGAACTCTTTCCCTACACGACGCTCTTCCGATCTNNNCAAAAGACACCTGCACACCTAT
AGCACACGTCTGAACTCTTTCCCTACACGACGCTCTTCCGATCTNNNCAAAAGACACCTGCACACCTAT
AGCACACGTCTGAACTCTTTCCCTACACGACGCTCTTCCGATCTNNNTCAGTTGGTGTCTCAGTAGGTT
AGCACACGTCTGAACTCTTTCCCTACACGACGCTCTTCCGATCTNNNTCAGTTGGTGTCTCAGTAGGTT
AGCACACGTCTGAACTCTTTCCCTACACGACGCTCTTCCGATCTNNNAGGACTGCTAGTTTTTCTCCCTG
AGCACACGTCTGAACTCTTTCCCTACACGACGCTCTTCCGATCTNNNAGGACTGCTAGTTTTTCTCCCTG
GAGCACACGTCTGAACTCTTTCCCTACACGACGCTCTTCCGATCTNNNCCATGATGAGGGATTCCATGA
GAGCACACGTCTGAACTCTTTCCCTACACGACGCTCTTCCGATCTNNNCCATGATGAGGGATTCCATGA
GAGCACACGTCTGAACTCTTTCCCTACACGACGCTCTTCCGATCTNNNGGGGGGTACTTACTCTCTAA
GAGCACACGTCTGAACTCTTTCCCTACACGACGCTCTTCCGATCTNNNGGGGGGTACTTACTCTCTAA
AGCACACGTCTGAACTCTTTCCCTACACGACGCTCTTCCGATCTNNNCTGTTTGTGACCTGAAAACCCT
AGCACACGTCTGAACTCTTTCCCTACACGACGCTCTTCCGATCTNNNCTGTTTGTGACCTGAAAACCCT
GCACACGTCTGAACTCTTTCCCTACACGACGCTCTTCCGATCTNNNATGGTAGCTTTTGTGGCTTGATG
GCACACGTCTGAACTCTTTCCCTACACGACGCTCTTCCGATCTNNNATGGTAGCTTTTGTGGCTTGATG
AGCACACGTCTGAACTCTTTCCCTACACGACGCTCTTCCGATCTNNNTTTGAAATGGAGTCTCACTTTC

GCACACGTCTGAACTCTTTCCCTACACGACGCTCTTCCGATCTNNNTTTGAAATGGAGTCTCACTTTC
GCACACGTCTGAACTCTTTCCCTACACGACGCTCTTCCGATCTNNNAAGACGGATCTTCTGTCCTCTGG
GCACACGTCTGAACTCTTTCCCTACACGACGCTCTTCCGATCTNNNAAGACGGATCTTCTGTCCTCTGG
CACACGTCTGAACTCTTTCCCTACACGACGCTCTTCCGATCTNNNCCCTGAACCACGGATTTGACTATC
CACACGTCTGAACTCTTTCCCTACACGACGCTCTTCCGATCTNNNCCCTGAACCACGGATTTGACTATC
GAGCACACGTCTGAACTCTTTCCCTACACGACGCTCTTCCGATCTNNNAAAAAATTAGCTGGGTGTGG
GAGCACACGTCTGAACTCTTTCCCTACACGACGCTCTTCCGATCTNNNAAAAAATTAGCTGGGTGTGG
AGCACACGTCTGAACTCTTTCCCTACACGACGCTCTTCCGATCTNNNCACAATCTCAGCTCACTGCAAC
AGCACACGTCTGAACTCTTTCCCTACACGACGCTCTTCCGATCTNNNCACAATCTCAGCTCACTGCAAC
AGAGCACACGTCTGAACTCTTTCCCTACACGACGCTCTTCCGATCTNNNGGAAAAGATCAGAAGTGAC
AGAGCACACGTCTGAACTCTTTCCCTACACGACGCTCTTCCGATCTNNNGGAAAAGATCAGAAGTGAC
AGCACACGTCTGAACTCTTTCCCTACACGACGCTCTTCCGATCTNNNCTGGCCTTGAACTTTTCTGGTT
AGCACACGTCTGAACTCTTTCCCTACACGACGCTCTTCCGATCTNNNCTGGCCTTGAACTTTTCTGGTT
GCACACGTCTGAACTCTTTCCCTACACGACGCTCTTCCGATCTNNNAATGAATGACTGTGTGTGTCTG
GCACACGTCTGAACTCTTTCCCTACACGACGCTCTTCCGATCTNNNAATGAATGACTGTGTGTGTCTG
ACACGTCTGAACTCTTTCCCTACACGACGCTCTTCCGATCTNNNTTGAACATCAGAGAGGCAATCATC
ACACGTCTGAACTCTTTCCCTACACGACGCTCTTCCGATCTNNNTTGAACATCAGAGAGGCAATCATC
GAGCACACGTCTGAACTCTTTCCCTACACGACGCTCTTCCGATCTNNNGGAGATTTTCCTATTTCAGAG
GAGCACACGTCTGAACTCTTTCCCTACACGACGCTCTTCCGATCTNNNGGAGATTTTCCTATTTCAGAG
GAGCACACGTCTGAACTCTTTCCCTACACGACGCTCTTCCGATCTNNNGGAGATTGTAGTGAGCCGAC
GAGCACACGTCTGAACTCTTTCCCTACACGACGCTCTTCCGATCTNNNGGAGATTGTAGTGAGCCGAC
GCACACGTCTGAACTCTTTCCCTACACGACGCTCTTCCGATCTNNNGCTCGGAGAAGTAGAGCGCACA
GCACACGTCTGAACTCTTTCCCTACACGACGCTCTTCCGATCTNNNGCTCGGAGAAGTAGAGCGCACA
GAGCACACGTCTGAACTCTTTCCCTACACGACGCTCTTCCGATCTNNNCTTGAGTCCTGGAGTTTGAG
GAGCACACGTCTGAACTCTTTCCCTACACGACGCTCTTCCGATCTNNNCTTGAGTCCTGGAGTTTGAG
AGCACACGTCTGAACTCTTTCCCTACACGACGCTCTTCCGATCTNNNCCCATGACTCAAACATCACCAAC
AGCACACGTCTGAACTCTTTCCCTACACGACGCTCTTCCGATCTNNNCCCATGACTCAAACATCACCAAC
GCACACGTCTGAACTCTTTCCCTACACGACGCTCTTCCGATCTNNNTGAGCACAGTGAAGTTTAGCTC
GCACACGTCTGAACTCTTTCCCTACACGACGCTCTTCCGATCTNNNTGAGCACAGTGAAGTTTAGCTC
CACACGTCTGAACTCTTTCCCTACACGACGCTCTTCCGATCTNNNTGAGAGAGAAATGCATCCATCATC
CACACGTCTGAACTCTTTCCCTACACGACGCTCTTCCGATCTNNNTGAGAGAGAAATGCATCCATCATC
AGAGCACACGTCTGAACTCTTTCCCTACACGACGCTCTTCCGATCTNNNGAACAGCTTTGGTTCTGCTT
AGAGCACACGTCTGAACTCTTTCCCTACACGACGCTCTTCCGATCTNNNGAACAGCTTTGGTTCTGCTT
GAGCACACGTCTGAACTCTTTCCCTACACGACGCTCTTCCGATCTNNNGGGTAGCACCTTTCCCTTTA
GAGCACACGTCTGAACTCTTTCCCTACACGACGCTCTTCCGATCTNNNGGGTAGCACCTTTCCCTTTA
GAGCACACGTCTGAACTCTTTCCCTACACGACGCTCTTCCGATCTNNNAGCTTTACCTATAACACAGGG
GAGCACACGTCTGAACTCTTTCCCTACACGACGCTCTTCCGATCTNNNAGCTTTACCTATAACACAGGG
AGCACACGTCTGAACTCTTTCCCTACACGACGCTCTTCCGATCTNNNATATAGCGAGAACCCTGCCTC
AGCACACGTCTGAACTCTTTCCCTACACGACGCTCTTCCGATCTNNNATATAGCGAGAACCCTGCCTC
GCACACGTCTGAACTCTTTCCCTACACGACGCTCTTCCGATCTNNNTTTCAACATGAGGTTTGAGGGG
GCACACGTCTGAACTCTTTCCCTACACGACGCTCTTCCGATCTNNNTTTCAACATGAGGTTTGAGGGG
AGCACACGTCTGAACTCTTTCCCTACACGACGCTCTTCCGATCTNNNGGGAGACTCCTTTCCAACTCAT
AGCACACGTCTGAACTCTTTCCCTACACGACGCTCTTCCGATCTNNNGGGAGACTCCTTTCCAACTCAT

ACACGTCTGAACTCTTTCCCTACACGACGCTCTTCCGATCTNNNTGAACAGGTTAAAGACGAAACATC
ACACGTCTGAACTCTTTCCCTACACGACGCTCTTCCGATCTNNNTGAACAGGTTAAAGACGAAACATC
AGCACACGTCTGAACTCTTTCCCTACACGACGCTCTTCCGATCTNNNGAGGCTGAGGCAGGAGAATG
AGCACACGTCTGAACTCTTTCCCTACACGACGCTCTTCCGATCTNNNGAGGCTGAGGCAGGAGAATG
AGCACACGTCTGAACTCTTTCCCTACACGACGCTCTTCCGATCTNNNTCGTTACAATTGCTTTTGGGG
AGCACACGTCTGAACTCTTTCCCTACACGACGCTCTTCCGATCTNNNTCGTTACAATTGCTTTTGGGG
AGCACACGTCTGAACTCTTTCCCTACACGACGCTCTTCCGATCTNNNCTCCTCGGCTCAAGGAAAGTAC
AGCACACGTCTGAACTCTTTCCCTACACGACGCTCTTCCGATCTNNNCTCCTCGGCTCAAGGAAAGTAC
GCACACGTCTGAACTCTTTCCCTACACGACGCTCTTCCGATCTNNNATACATACACAAGCACATGCACG
GCACACGTCTGAACTCTTTCCCTACACGACGCTCTTCCGATCTNNNATACATACACAAGCACATGCACG
GCACACGTCTGAACTCTTTCCCTACACGACGCTCTTCCGATCTNNNATACTCTTTCTGACTGATGACTA
GCACACGTCTGAACTCTTTCCCTACACGACGCTCTTCCGATCTNNNATACTCTTTCTGACTGATGACTA
AGCACACGTCTGAACTCTTTCCCTACACGACGCTCTTCCGATCTNNNAGAGAGAGAGAGAGAGGGAG
AGCACACGTCTGAACTCTTTCCCTACACGACGCTCTTCCGATCTNNNAGAGAGAGAGAGAGAGGGAG
AGCACACGTCTGAACTCTTTCCCTACACGACGCTCTTCCGATCTNNNTCTTTCTCGTCATTGTGGGCTT
AGCACACGTCTGAACTCTTTCCCTACACGACGCTCTTCCGATCTNNNTCTTTCTCGTCATTGTGGGCTT
GCACACGTCTGAACTCTTTCCCTACACGACGCTCTTCCGATCTNNNGTTGTACGATCTTGGCTAACTGC
GCACACGTCTGAACTCTTTCCCTACACGACGCTCTTCCGATCTNNNGTTGTACGATCTTGGCTAACTGC
AGCACACGTCTGAACTCTTTCCCTACACGACGCTCTTCCGATCTNNNGTTGCAGTGAGCTGAGATCAT
AGCACACGTCTGAACTCTTTCCCTACACGACGCTCTTCCGATCTNNNGTTGCAGTGAGCTGAGATCAT
GAGCACACGTCTGAACTCTTTCCCTACACGACGCTCTTCCGATCTNNNGAGACATGAGAATCGCTTGA
GAGCACACGTCTGAACTCTTTCCCTACACGACGCTCTTCCGATCTNNNGAGACATGAGAATCGCTTGA
AGCACACGTCTGAACTCTTTCCCTACACGACGCTCTTCCGATCTNNNGAATGCAGAACAGTCTATGCT
AGCACACGTCTGAACTCTTTCCCTACACGACGCTCTTCCGATCTNNNGAATGCAGAACAGTCTATGCT
CACACGTCTGAACTCTTTCCCTACACGACGCTCTTCCGATCTNNNGTCTGTGAAGGCTTAGGGTACT
CACACGTCTGAACTCTTTCCCTACACGACGCTCTTCCGATCTNNNGTCTGTGAAGGCTTAGGGTACT
GAGCACACGTCTGAACTCTTTCCCTACACGACGCTCTTCCGATCTNNNTTTATGTGAGCAGGGAGGA
GAGCACACGTCTGAACTCTTTCCCTACACGACGCTCTTCCGATCTNNNTTTATGTGAGCAGGGAGGA
GCACACGTCTGAACTCTTTCCCTACACGACGCTCTTCCGATCTNNNCTTCTCTGGAAGCAGTGAAGCT
GCACACGTCTGAACTCTTTCCCTACACGACGCTCTTCCGATCTNNNCTTCTCTGGAAGCAGTGAAGCT
GCACACGTCTGAACTCTTTCCCTACACGACGCTCTTCCGATCTNNNTCTAAGTCCCTCTCGAAGTCTCA
GCACACGTCTGAACTCTTTCCCTACACGACGCTCTTCCGATCTNNNTCTAAGTCCCTCTCGAAGTCTCA
AGCACACGTCTGAACTCTTTCCCTACACGACGCTCTTCCGATCTNNNCATGCCTGTAATCCCACCTACT
AGCACACGTCTGAACTCTTTCCCTACACGACGCTCTTCCGATCTNNNCATGCCTGTAATCCCACCTACT
ACACGTCTGAACTCTTTCCCTACACGACGCTCTTCCGATCTNNNACAAAAGCATCAGAATTATCTTTCA
ACACGTCTGAACTCTTTCCCTACACGACGCTCTTCCGATCTNNNACAAAAGCATCAGAATTATCTTTCA
AGCACACGTCTGAACTCTTTCCCTACACGACGCTCTTCCGATCTNNNTGACAGGATTTGGTACCAGGA
AGCACACGTCTGAACTCTTTCCCTACACGACGCTCTTCCGATCTNNNTGACAGGATTTGGTACCAGGA
AGCACACGTCTGAACTCTTTCCCTACACGACGCTCTTCCGATCTNNNGGTTTCTCCTGAGGCCTCTTT
AGCACACGTCTGAACTCTTTCCCTACACGACGCTCTTCCGATCTNNNGGTTTCTCCTGAGGCCTCTTT
AGCACACGTCTGAACTCTTTCCCTACACGACGCTCTTCCGATCTNNNGCATCCTGAATAGCTGGGACT
AGCACACGTCTGAACTCTTTCCCTACACGACGCTCTTCCGATCTNNNGCATCCTGAATAGCTGGGACT
GAGCACACGTCTGAACTCTTTCCCTACACGACGCTCTTCCGATCTNNNATGCTGTGTTTTATTGTGTG

AGAGCACACGTCTGAACTCTTTCCCTACACGACGCTCTTCCGATCTNNNATGCTGTGTTTTATTGTGTGT
AGCACACGTCTGAACTCTTTCCCTACACGACGCTCTTCCGATCTNNNAACCAGAAGCCAGAAAGCAAAA
AGCACACGTCTGAACTCTTTCCCTACACGACGCTCTTCCGATCTNNNAACCAGAAGCCAGAAAGCAAAA
AGCACACGTCTGAACTCTTTCCCTACACGACGCTCTTCCGATCTNNNAAAAGGACTAGGAAACACCAAC
AGCACACGTCTGAACTCTTTCCCTACACGACGCTCTTCCGATCTNNNAAAAGGACTAGGAAACACCAAC
AGCACACGTCTGAACTCTTTCCCTACACGACGCTCTTCCGATCTNNNCTCCAACGCAGACTGGTTATTT
AGCACACGTCTGAACTCTTTCCCTACACGACGCTCTTCCGATCTNNNCTCCAACGCAGACTGGTTATTT
AGCACACGTCTGAACTCTTTCCCTACACGACGCTCTTCCGATCTNNNGCAGCCAAACATAAGCTCATGA
AGCACACGTCTGAACTCTTTCCCTACACGACGCTCTTCCGATCTNNNGCAGCCAAACATAAGCTCATGA
AGCACACGTCTGAACTCTTTCCCTACACGACGCTCTTCCGATCTNNNTTCTCTGTTGGTAGTGTGGGAT
AGCACACGTCTGAACTCTTTCCCTACACGACGCTCTTCCGATCTNNNTTCTCTGTTGGTAGTGTGGGAT
AGCACACGTCTGAACTCTTTCCCTACACGACGCTCTTCCGATCTNNNCCGCCCTGAAAAATCTGTTTCTT
AGCACACGTCTGAACTCTTTCCCTACACGACGCTCTTCCGATCTNNNCCGCCCTGAAAAATCTGTTTCTT
GAGCACACGTCTGAACTCTTTCCCTACACGACGCTCTTCCGATCTNNNATTACCACTGACCCCACAGAAA
GAGCACACGTCTGAACTCTTTCCCTACACGACGCTCTTCCGATCTNNNATTACCACTGACCCCACAGAAA
AGAGCACACGTCTGAACTCTTTCCCTACACGACGCTCTTCCGATCTNNNCGAAGGTCACCAACATCAAAC
AGAGCACACGTCTGAACTCTTTCCCTACACGACGCTCTTCCGATCTNNNCGAAGGTCACCAACATCAAAC
AGAGCACACGTCTGAACTCTTTCCCTACACGACGCTCTTCCGATCTNNNAGCAAGGAAGGTCATTCAGC
AGAGCACACGTCTGAACTCTTTCCCTACACGACGCTCTTCCGATCTNNNAGCAAGGAAGGTCATTCAGC
AGCACACGTCTGAACTCTTTCCCTACACGACGCTCTTCCGATCTNNNTTTGGTGGCATAAAGGAGCTTT
AGCACACGTCTGAACTCTTTCCCTACACGACGCTCTTCCGATCTNNNTTTGGTGGCATAAAGGAGCTTT
GCACACGTCTGAACTCTTTCCCTACACGACGCTCTTCCGATCTNNNGGAATTTCTCCTTCTTCCCCACTCA
GCACACGTCTGAACTCTTTCCCTACACGACGCTCTTCCGATCTNNNGGAATTTCTCCTTCTTCCCCACTCA
GGAAGAGCACACGTCTGAACTCTTTCCCTACACGACGCTCTTCCGATCTNNNTCAAAATCAAGTTGTGT
GGAAGAGCACACGTCTGAACTCTTTCCCTACACGACGCTCTTCCGATCTNNNTCAAAATCAAGTTGTGT
AGCACACGTCTGAACTCTTTCCCTACACGACGCTCTTCCGATCTNNNGAAGAAAAAGGAGAGAGGGAC
AGCACACGTCTGAACTCTTTCCCTACACGACGCTCTTCCGATCTNNNGAAGAAAAAGGAGAGAGGGAC
AGCACACGTCTGAACTCTTTCCCTACACGACGCTCTTCCGATCTNNNAACCAGCCTGCTCTTCATTCATAC
AGCACACGTCTGAACTCTTTCCCTACACGACGCTCTTCCGATCTNNNAACCAGCCTGCTCTTCATTCATAC
GCACACGTCTGAACTCTTTCCCTACACGACGCTCTTCCGATCTNNNCGCATCAGGCTTTCTTATGTCTGC
GCACACGTCTGAACTCTTTCCCTACACGACGCTCTTCCGATCTNNNCGCATCAGGCTTTCTTATGTCTGC
GAGCACACGTCTGAACTCTTTCCCTACACGACGCTCTTCCGATCTNNNAGTCTGAGATCAAACTGCAAC
GAGCACACGTCTGAACTCTTTCCCTACACGACGCTCTTCCGATCTNNNAGTCTGAGATCAAACTGCAAC
GAGCACACGTCTGAACTCTTTCCCTACACGACGCTCTTCCGATCTNNNGGCCTTTTTTTTCTATTATGAGA
GAGCACACGTCTGAACTCTTTCCCTACACGACGCTCTTCCGATCTNNNGGCCTTTTTTTTCTATTATGAGA
AGCACACGTCTGAACTCTTTCCCTACACGACGCTCTTCCGATCTNNNCTTGGTAGAGATGGCAGTAC
AGCACACGTCTGAACTCTTTCCCTACACGACGCTCTTCCGATCTNNNCTTGGTAGAGATGGCAGTAC
GCACACGTCTGAACTCTTTCCCTACACGACGCTCTTCCGATCTNNNTCATCTGAAGAGCCAGCATACAA
GCACACGTCTGAACTCTTTCCCTACACGACGCTCTTCCGATCTNNNTCATCTGAAGAGCCAGCATACAA
GAGCACACGTCTGAACTCTTTCCCTACACGACGCTCTTCCGATCTNNNTTGCATATGTTCAACCAGCCTT
GAGCACACGTCTGAACTCTTTCCCTACACGACGCTCTTCCGATCTNNNTTGCATATGTTCAACCAGCCTT
AGCACACGTCTGAACTCTTTCCCTACACGACGCTCTTCCGATCTNNNCTCCTGAGTAGCTGGGACTACA
AGCACACGTCTGAACTCTTTCCCTACACGACGCTCTTCCGATCTNNNCTCCTGAGTAGCTGGGACTACA

GAGCACACGTCTGAACTCTTTCCCTACACGACGCTCTTCCGATCTNNNTGGGACTGTGTTTATGTGAG

GAGCACACGTCTGAACTCTTTCCCTACACGACGCTCTTCCGATCTNNNTGGGACTGTGTTTATGTGAG

GAGCACACGTCTGAACTCTTTCCCTACACGACGCTCTTCCGATCTNNNACCTTGCTCCGAAGATTTGTT

GAGCACACGTCTGAACTCTTTCCCTACACGACGCTCTTCCGATCTNNNACCTTGCTCCGAAGATTTGTT

AGCACACGTCTGAACTCTTTCCCTACACGACGCTCTTCCGATCTNNNGGATTAGTGGCAGTTACAAGT

AGCACACGTCTGAACTCTTTCCCTACACGACGCTCTTCCGATCTNNNGGATTAGTGGCAGTTACAAGT

GAGCACACGTCTGAACTCTTTCCCTACACGACGCTCTTCCGATCTNNNTCATCAAATACCCGGCAGTCT

GAGCACACGTCTGAACTCTTTCCCTACACGACGCTCTTCCGATCTNNNTCATCAAATACCCGGCAGTCT

GAGCACACGTCTGAACTCTTTCCCTACACGACGCTCTTCCGATCTNNNCATAGAGAGGGGAAGATGC

GAGCACACGTCTGAACTCTTTCCCTACACGACGCTCTTCCGATCTNNNCATAGAGAGGGGAAGATGC

GCACACGTCTGAACTCTTTCCCTACACGACGCTCTTCCGATCTNNNCTCATGGCGAAACACACATACAC

GCACACGTCTGAACTCTTTCCCTACACGACGCTCTTCCGATCTNNNCTCATGGCGAAACACACATACAC

AGCACACGTCTGAACTCTTTCCCTACACGACGCTCTTCCGATCTNNNTGCACGCCGAAATTTCTGTTTA

AGCACACGTCTGAACTCTTTCCCTACACGACGCTCTTCCGATCTNNNTGCACGCCGAAATTTCTGTTTA

AGCACACGTCTGAACTCTTTCCCTACACGACGCTCTTCCGATCTNNNGTACTTGGGTTTCAGCTCCAGG

AGCACACGTCTGAACTCTTTCCCTACACGACGCTCTTCCGATCTNNNGTACTTGGGTTTCAGCTCCAGG

AGCACACGTCTGAACTCTTTCCCTACACGACGCTCTTCCGATCTNNNTAAGGCAGGAGAATCGCTTGA

AGCACACGTCTGAACTCTTTCCCTACACGACGCTCTTCCGATCTNNNTAAGGCAGGAGAATCGCTTGA

GAGCACACGTCTGAACTCTTTCCCTACACGACGCTCTTCCGATCTNNNTTGAATCCGGGAAGTAGAGG

GAGCACACGTCTGAACTCTTTCCCTACACGACGCTCTTCCGATCTNNNTTGAATCCGGGAAGTAGAGG

AGCACACGTCTGAACTCTTTCCCTACACGACGCTCTTCCGATCTNNNTGAGGAAAAGTAAAAGCGCT

AGCACACGTCTGAACTCTTTCCCTACACGACGCTCTTCCGATCTNNNTGAGGAAAAGTAAAAGCGCT

GAGCACACGTCTGAACTCTTTCCCTACACGACGCTCTTCCGATCTNNNTCCGTTCTCTTCTACAGCCAAC

GAGCACACGTCTGAACTCTTTCCCTACACGACGCTCTTCCGATCTNNNTCCGTTCTCTTCTACAGCCAAC

AGCACACGTCTGAACTCTTTCCCTACACGACGCTCTTCCGATCTNNNTGAGCCATGTTTCCAATTTTCCT

AGCACACGTCTGAACTCTTTCCCTACACGACGCTCTTCCGATCTNNNTGAGCCATGTTTCCAATTTTCCT

AGCACACGTCTGAACTCTTTCCCTACACGACGCTCTTCCGATCTNNNTAATCCTAGATACTCGGGAGGC

AGCACACGTCTGAACTCTTTCCCTACACGACGCTCTTCCGATCTNNNTAATCCTAGATACTCGGGAGGC

GAGCACACGTCTGAACTCTTTCCCTACACGACGCTCTTCCGATCTNNNTCACTTGAGTCCCAGAGTTCA

GAGCACACGTCTGAACTCTTTCCCTACACGACGCTCTTCCGATCTNNNTCACTTGAGTCCCAGAGTTCA

GAGCACACGTCTGAACTCTTTCCCTACACGACGCTCTTCCGATCTNNNACGGAGCAAGACTCTTTCTC

GAGCACACGTCTGAACTCTTTCCCTACACGACGCTCTTCCGATCTNNNACGGAGCAAGACTCTTTCTC

AGCACACGTCTGAACTCTTTCCCTACACGACGCTCTTCCGATCTNNNAGCGAAACTCCGTCTCTCTCTAT

AGCACACGTCTGAACTCTTTCCCTACACGACGCTCTTCCGATCTNNNAGCGAAACTCCGTCTCTCTCTAT

AGCACACGTCTGAACTCTTTCCCTACACGACGCTCTTCCGATCTNNNAGTGGCTATCATTATGTTTAAG

AGCACACGTCTGAACTCTTTCCCTACACGACGCTCTTCCGATCTNNNAGTGGCTATCATTATGTTTAAG

AGCACACGTCTGAACTCTTTCCCTACACGACGCTCTTCCGATCTNNNTGGCCCCTCAAAACCTCATTATT

AGCACACGTCTGAACTCTTTCCCTACACGACGCTCTTCCGATCTNNNTGGCCCCTCAAAACCTCATTATT

AAGAGCACACGTCTGAACTCTTTCCCTACACGACGCTCTTCCGATCTNNNCATCTTGGAAGTAGTACCC

AAGAGCACACGTCTGAACTCTTTCCCTACACGACGCTCTTCCGATCTNNNCATCTTGGAAGTAGTACCC

GAGCACACGTCTGAACTCTTTCCCTACACGACGCTCTTCCGATCTNNNAGCATTGCCAGTATGACAAG

GAGCACACGTCTGAACTCTTTCCCTACACGACGCTCTTCCGATCTNNNAGCATTGCCAGTATGACAAG

GAGCACACGTCTGAACTCTTTCCCTACACGACGCTCTTCCGATCTNNNTTCCTTTCTTTCGTCTCTCCCTC

AGCACACGTCTGAACTCTTTCCCTACACGACGCTCTTCCGATCTNNNTTCCTTTCTTTCGTCTCTCCCTC

GAGCACACGTCTGAACTCTTTCCCTACACGACGCTCTTCCGATCTNNNCTGTAGTCCCAGCTACTCAG

GAGCACACGTCTGAACTCTTTCCCTACACGACGCTCTTCCGATCTNNNCTGTAGTCCCAGCTACTCAG

AGCACACGTCTGAACTCTTTCCCTACACGACGCTCTTCCGATCTNNNAAAAAAAATTAGCCAGGCGTG

AGCACACGTCTGAACTCTTTCCCTACACGACGCTCTTCCGATCTNNNAAAAAAAATTAGCCAGGCGTG

AGCACACGTCTGAACTCTTTCCCTACACGACGCTCTTCCGATCTNNNAGAGAGAGAGAGAGAGAGA

AGCACACGTCTGAACTCTTTCCCTACACGACGCTCTTCCGATCTNNNAGAGAGAGAGAGAGAGAGA

GCACACGTCTGAACTCTTTCCCTACACGACGCTCTTCCGATCTNNNGAACCTTTGCCTTTGAGGTCAT

GCACACGTCTGAACTCTTTCCCTACACGACGCTCTTCCGATCTNNNGAACCTTTGCCTTTGAGGTCAT

GCACACGTCTGAACTCTTTCCCTACACGACGCTCTTCCGATCTNNNCTTGGGACAGAGTCTGGAATCA

GCACACGTCTGAACTCTTTCCCTACACGACGCTCTTCCGATCTNNNCTTGGGACAGAGTCTGGAATCA

AGCACACGTCTGAACTCTTTCCCTACACGACGCTCTTCCGATCTNNNGTAATCCCAGCTACTAGGAAG

AGCACACGTCTGAACTCTTTCCCTACACGACGCTCTTCCGATCTNNNGTAATCCCAGCTACTAGGAAG

GCACACGTCTGAACTCTTTCCCTACACGACGCTCTTCCGATCTNNNATTTTGTGCCTATTCCAGTTGCC

GCACACGTCTGAACTCTTTCCCTACACGACGCTCTTCCGATCTNNNATTTTGTGCCTATTCCAGTTGCC

GCACACGTCTGAACTCTTTCCCTACACGACGCTCTTCCGATCTNNNCCTACAGAACAGTGAGCAACAA

GCACACGTCTGAACTCTTTCCCTACACGACGCTCTTCCGATCTNNNCCTACAGAACAGTGAGCAACAA

AGCACACGTCTGAACTCTTTCCCTACACGACGCTCTTCCGATCTNNNATGCTTTCTATGCTGGTTGGTT

AGCACACGTCTGAACTCTTTCCCTACACGACGCTCTTCCGATCTNNNATGCTTTCTATGCTGGTTGGTT

GCACACGTCTGAACTCTTTCCCTACACGACGCTCTTCCGATCTNNNAATATAGTGAGACCGAGACCCC

GCACACGTCTGAACTCTTTCCCTACACGACGCTCTTCCGATCTNNNAATATAGTGAGACCGAGACCCC

AGCACACGTCTGAACTCTTTCCCTACACGACGCTCTTCCGATCTNNNTAGGAAAATTGTGAGACCGG

AGCACACGTCTGAACTCTTTCCCTACACGACGCTCTTCCGATCTNNNTAGGAAAATTGTGAGACCGG

GCACACGTCTGAACTCTTTCCCTACACGACGCTCTTCCGATCTNNNAGTGAGCCAAGATAGAACCAC

GCACACGTCTGAACTCTTTCCCTACACGACGCTCTTCCGATCTNNNAGTGAGCCAAGATAGAACCAC

GCACACGTCTGAACTCTTTCCCTACACGACGCTCTTCCGATCTNNNAGTGTTTTGCTTCTCTAGGTACC

GCACACGTCTGAACTCTTTCCCTACACGACGCTCTTCCGATCTNNNAGTGTTTTGCTTCTCTAGGTACC

AGCACACGTCTGAACTCTTTCCCTACACGACGCTCTTCCGATCTNNNGTAAATAGTCTCAGCTCCTCG

AGCACACGTCTGAACTCTTTCCCTACACGACGCTCTTCCGATCTNNNGTAAATAGTCTCAGCTCCTCG

GCACACGTCTGAACTCTTTCCCTACACGACGCTCTTCCGATCTNNNACAGTGACACAACACACACATC

GCACACGTCTGAACTCTTTCCCTACACGACGCTCTTCCGATCTNNNACAGTGACACAACACACACATC

AGCACACGTCTGAACTCTTTCCCTACACGACGCTCTTCCGATCTNNNATGTCCTCTGGCATCTCATGGA

AGCACACGTCTGAACTCTTTCCCTACACGACGCTCTTCCGATCTNNNATGTCCTCTGGCATCTCATGGA

GCACACGTCTGAACTCTTTCCCTACACGACGCTCTTCCGATCTNNNCTCAGAAGCACTACAAGAAGCGT

GCACACGTCTGAACTCTTTCCCTACACGACGCTCTTCCGATCTNNNCTCAGAAGCACTACAAGAAGCGT

AGCACACGTCTGAACTCTTTCCCTACACGACGCTCTTCCGATCTNNNCACAAGTAGCTGGGACTACA

AGCACACGTCTGAACTCTTTCCCTACACGACGCTCTTCCGATCTNNNCACAAGTAGCTGGGACTACA

GAGCACACGTCTGAACTCTTTCCCTACACGACGCTCTTCCGATCTNNNGAGAGATGAAAGAAGCTTC

GAGCACACGTCTGAACTCTTTCCCTACACGACGCTCTTCCGATCTNNNGAGAGATGAAAGAAGCTTC

AGCACACGTCTGAACTCTTTCCCTACACGACGCTCTTCCGATCTNNNACTGCATGTTCTCACTCATCAG

AGCACACGTCTGAACTCTTTCCCTACACGACGCTCTTCCGATCTNNNACTGCATGTTCTCACTCATCAG

GCACACGTCTGAACTCTTTCCCTACACGACGCTCTTCCGATCTNNNTCAACATGTCAGTGATGCTCTC

GCACACGTCTGAACTCTTTCCCTACACGACGCTCTTCCGATCTNNNTCAACATGTCAGTGATGCTCTC

AGCACACGTCTGAACTCTTTCCCTACACGACGCTCTTCCGATCTNNNTTCTCATAGAGGCTCCTGACA
AGCACACGTCTGAACTCTTTCCCTACACGACGCTCTTCCGATCTNNNTTCTCATAGAGGCTCCTGACA
GAGCACACGTCTGAACTCTTTCCCTACACGACGCTCTTCCGATCTNNNTTCAGCTGTGAATCCATCAGG
GAGCACACGTCTGAACTCTTTCCCTACACGACGCTCTTCCGATCTNNNTTCAGCTGTGAATCCATCAGG
AGCACACGTCTGAACTCTTTCCCTACACGACGCTCTTCCGATCTNNNAACTATTTGAGGGCTGAGTT
AGCACACGTCTGAACTCTTTCCCTACACGACGCTCTTCCGATCTNNNAACTATTTGAGGGCTGAGTT
CACACGTCTGAACTCTTTCCCTACACGACGCTCTTCCGATCTNNNCTCTCTATCTTGCCTCTTCATCG
CACACGTCTGAACTCTTTCCCTACACGACGCTCTTCCGATCTNNNCTCTCTATCTTGCCTCTTCATCG
GCACACGTCTGAACTCTTTCCCTACACGACGCTCTTCCGATCTNNNGGCTGGTATACTGTCTCCATAG
GCACACGTCTGAACTCTTTCCCTACACGACGCTCTTCCGATCTNNNGGCTGGTATACTGTCTCCATAG
AGCACACGTCTGAACTCTTTCCCTACACGACGCTCTTCCGATCTNNNGTTGTGGTGAGCTGAGATTGT
AGCACACGTCTGAACTCTTTCCCTACACGACGCTCTTCCGATCTNNNGTTGTGGTGAGCTGAGATTGT
AGCACACGTCTGAACTCTTTCCCTACACGACGCTCTTCCGATCTNNNAAAGCATAAAGGCACAAGAC
AGCACACGTCTGAACTCTTTCCCTACACGACGCTCTTCCGATCTNNNAAAGCATAAAGGCACAAGAC
AGCACACGTCTGAACTCTTTCCCTACACGACGCTCTTCCGATCTNNNAAAAATTAGCTGGGCATAGTG
AGCACACGTCTGAACTCTTTCCCTACACGACGCTCTTCCGATCTNNNAAAAATTAGCTGGGCATAGTG
GCACACGTCTGAACTCTTTCCCTACACGACGCTCTTCCGATCTNNNAACAGCAGCACAGAAACAAAAC
GCACACGTCTGAACTCTTTCCCTACACGACGCTCTTCCGATCTNNNAACAGCAGCACAGAAACAAAAC
GCACACGTCTGAACTCTTTCCCTACACGACGCTCTTCCGATCTNNNAGGATTCTCTGAGTTCTTGCCA
GCACACGTCTGAACTCTTTCCCTACACGACGCTCTTCCGATCTNNNAGGATTCTCTGAGTTCTTGCCA
GCACACGTCTGAACTCTTTCCCTACACGACGCTCTTCCGATCTNNNCCCTTCATCCCACCCTACTAATGC
GCACACGTCTGAACTCTTTCCCTACACGACGCTCTTCCGATCTNNNCCCTTCATCCCACCCTACTAATGC
GCACACGTCTGAACTCTTTCCCTACACGACGCTCTTCCGATCTNNNGCACAAAATAGAACCCACTGCT
GCACACGTCTGAACTCTTTCCCTACACGACGCTCTTCCGATCTNNNGCACAAAATAGAACCCACTGCT
GAGCACACGTCTGAACTCTTTCCCTACACGACGCTCTTCCGATCTNNNTCATCTTCCAACTTCCAGTCCT
GAGCACACGTCTGAACTCTTTCCCTACACGACGCTCTTCCGATCTNNNTCATCTTCCAACTTCCAGTCCT
GCACACGTCTGAACTCTTTCCCTACACGACGCTCTTCCGATCTNNNCCTACCACAAACACCTCTCTCTCC
GCACACGTCTGAACTCTTTCCCTACACGACGCTCTTCCGATCTNNNCCTACCACAAACACCTCTCTCTCC
AGCACACGTCTGAACTCTTTCCCTACACGACGCTCTTCCGATCTNNNCCTCCAGCTAGTTGAACATCAC
AGCACACGTCTGAACTCTTTCCCTACACGACGCTCTTCCGATCTNNNCCTCCAGCTAGTTGAACATCAC
GCACACGTCTGAACTCTTTCCCTACACGACGCTCTTCCGATCTNNNGAACCCAGAAATAAAGCAGCA
GCACACGTCTGAACTCTTTCCCTACACGACGCTCTTCCGATCTNNNGAACCCAGAAATAAAGCAGCA
AGCACACGTCTGAACTCTTTCCCTACACGACGCTCTTCCGATCTNNNGCAATTTCTTTTCTAATTAGTG
AGCACACGTCTGAACTCTTTCCCTACACGACGCTCTTCCGATCTNNNGCAATTTCTTTTCTAATTAGTG
GAGCACACGTCTGAACTCTTTCCCTACACGACGCTCTTCCGATCTNNNCAAAATACTCACCTACCCACC
GAGCACACGTCTGAACTCTTTCCCTACACGACGCTCTTCCGATCTNNNCAAAATACTCACCTACCCACC
GAGCACACGTCTGAACTCTTTCCCTACACGACGCTCTTCCGATCTNNNCTGGGTGACAAGAGCAAAA
GAGCACACGTCTGAACTCTTTCCCTACACGACGCTCTTCCGATCTNNNCTGGGTGACAAGAGCAAAA
CACACGTCTGAACTCTTTCCCTACACGACGCTCTTCCGATCTNNNGGACCACTGTAATCCTGAAGACA
CACACGTCTGAACTCTTTCCCTACACGACGCTCTTCCGATCTNNNGGACCACTGTAATCCTGAAGACA
GCACACGTCTGAACTCTTTCCCTACACGACGCTCTTCCGATCTNNNTGTGCTTGTAGTCACAGCTACT
GCACACGTCTGAACTCTTTCCCTACACGACGCTCTTCCGATCTNNNTGTGCTTGTAGTCACAGCTACT
GAGCACACGTCTGAACTCTTTCCCTACACGACGCTCTTCCGATCTNNNGTAATCCTAGCTACTTGGGA

GAGCACACGTCTGAACTCTTTCCCTACACGACGCTCTTCCGATCTNNNGTAATCCTAGCTACTTGGGA
AGCACACGTCTGAACTCTTTCCCTACACGACGCTCTTCCGATCTNNNGTGAAAGCCTCTTAGGTTGCA
AGCACACGTCTGAACTCTTTCCCTACACGACGCTCTTCCGATCTNNNGTGAAAGCCTCTTAGGTTGCA
AGCACACGTCTGAACTCTTTCCCTACACGACGCTCTTCCGATCTNNNCAACAGCAAGATCCTGTCTCA
AGCACACGTCTGAACTCTTTCCCTACACGACGCTCTTCCGATCTNNNCAACAGCAAGATCCTGTCTCA
GCACACGTCTGAACTCTTTCCCTACACGACGCTCTTCCGATCTNNNGAGAGCTGAACTAAACACAGG
GCACACGTCTGAACTCTTTCCCTACACGACGCTCTTCCGATCTNNNGAGAGCTGAACTAAACACAGG
GAGCACACGTCTGAACTCTTTCCCTACACGACGCTCTTCCGATCTNNNAAAAAATTAGCCAGGTGTGG
GAGCACACGTCTGAACTCTTTCCCTACACGACGCTCTTCCGATCTNNNAAAAAATTAGCCAGGTGTGG
GAGCACACGTCTGAACTCTTTCCCTACACGACGCTCTTCCGATCTNNNGAAAACCTGTTGTAACCCTCA
GAGCACACGTCTGAACTCTTTCCCTACACGACGCTCTTCCGATCTNNNGAAAACCTGTTGTAACCCTCA
AGCACACGTCTGAACTCTTTCCCTACACGACGCTCTTCCGATCTNNNCATTCAGAGAGTCCACTGTGTG
AGCACACGTCTGAACTCTTTCCCTACACGACGCTCTTCCGATCTNNNCATTCAGAGAGTCCACTGTGTG
AGCACACGTCTGAACTCTTTCCCTACACGACGCTCTTCCGATCTNNNACCAGAATTCACATGGTGGAA
AGCACACGTCTGAACTCTTTCCCTACACGACGCTCTTCCGATCTNNNACCAGAATTCACATGGTGGAA
AGCACACGTCTGAACTCTTTCCCTACACGACGCTCTTCCGATCTNNNCACACATACCAACAGACAGGC
AGCACACGTCTGAACTCTTTCCCTACACGACGCTCTTCCGATCTNNNCACACATACCAACAGACAGGC
AGCACACGTCTGAACTCTTTCCCTACACGACGCTCTTCCGATCTNNNTTGAACCCAGGAGTCATAGG
AGCACACGTCTGAACTCTTTCCCTACACGACGCTCTTCCGATCTNNNTTGAACCCAGGAGTCATAGG
AGCACACGTCTGAACTCTTTCCCTACACGACGCTCTTCCGATCTNNNTGACCTGGCCTTTTTTCTTTTC
AGCACACGTCTGAACTCTTTCCCTACACGACGCTCTTCCGATCTNNNTGACCTGGCCTTTTTTCTTTTC
AGCACACGTCTGAACTCTTTCCCTACACGACGCTCTTCCGATCTNNNGCAGGCTTCCTCCAATTTTCTTT
AGCACACGTCTGAACTCTTTCCCTACACGACGCTCTTCCGATCTNNNGCAGGCTTCCTCCAATTTTCTTT
AGAGCACACGTCTGAACTCTTTCCCTACACGACGCTCTTCCGATCTNNNCAGCTGAGTGAAAGAGTG
AGAGCACACGTCTGAACTCTTTCCCTACACGACGCTCTTCCGATCTNNNCAGCTGAGTGAAAGAGTG
GCACACGTCTGAACTCTTTCCCTACACGACGCTCTTCCGATCTNNNCCTGTTTTCCTCGGTGGGAACA
GCACACGTCTGAACTCTTTCCCTACACGACGCTCTTCCGATCTNNNCCTGTTTTCCTCGGTGGGAACA
GAGCACACGTCTGAACTCTTTCCCTACACGACGCTCTTCCGATCTNNNATTGTTCATTTTCTTGGGCGT
GAGCACACGTCTGAACTCTTTCCCTACACGACGCTCTTCCGATCTNNNATTGTTCATTTTCTTGGGCGT
GCACACGTCTGAACTCTTTCCCTACACGACGCTCTTCCGATCTNNNGCAGAGCTGATACTGGCTTGA
GCACACGTCTGAACTCTTTCCCTACACGACGCTCTTCCGATCTNNNGCAGAGCTGATACTGGCTTGA
AGCACACGTCTGAACTCTTTCCCTACACGACGCTCTTCCGATCTNNNTACAGTCCTCATTTCTGCAGCT
AGCACACGTCTGAACTCTTTCCCTACACGACGCTCTTCCGATCTNNNTACAGTCCTCATTTCTGCAGCT
GCACACGTCTGAACTCTTTCCCTACACGACGCTCTTCCGATCTNNNTGTCCCCAAAATCATCCTCACTTC
GCACACGTCTGAACTCTTTCCCTACACGACGCTCTTCCGATCTNNNTGTCCCCAAAATCATCCTCACTTC
AGCACACGTCTGAACTCTTTCCCTACACGACGCTCTTCCGATCTNNNCCAGAAACAGATGGATTCACA
AGCACACGTCTGAACTCTTTCCCTACACGACGCTCTTCCGATCTNNNCCAGAAACAGATGGATTCACA
GCACACGTCTGAACTCTTTCCCTACACGACGCTCTTCCGATCTNNNGGAGGCAAGGATACAATGGTG
GCACACGTCTGAACTCTTTCCCTACACGACGCTCTTCCGATCTNNNGGAGGCAAGGATACAATGGTG
GAGCACACGTCTGAACTCTTTCCCTACACGACGCTCTTCCGATCTNNNGACCAAGGAAACCCAGAGAC
GAGCACACGTCTGAACTCTTTCCCTACACGACGCTCTTCCGATCTNNNGACCAAGGAAACCCAGAGAC
AGCACACGTCTGAACTCTTTCCCTACACGACGCTCTTCCGATCTNNNGAGAGAAGAGACAGCAGGG
AGCACACGTCTGAACTCTTTCCCTACACGACGCTCTTCCGATCTNNNGAGAGAAGAGACAGCAGGG

CACACGTCTGAACTCTTTCCCTACACGACGCTCTTCCGATCTNNNAGACAGGAGAATTGCTTGAACCAC
CACACGTCTGAACTCTTTCCCTACACGACGCTCTTCCGATCTNNNAGACAGGAGAATTGCTTGAACCAC
GAGCACACGTCTGAACTCTTTCCCTACACGACGCTCTTCCGATCTNNNTGGTTTCGATCCATTACTGG
GAGCACACGTCTGAACTCTTTCCCTACACGACGCTCTTCCGATCTNNNTGGTTTCGATCCATTACTGG
AAGAGCACACGTCTGAACTCTTTCCCTACACGACGCTCTTCCGATCTNNNGAGGTAGAGGTTGCAGT
AAGAGCACACGTCTGAACTCTTTCCCTACACGACGCTCTTCCGATCTNNNGAGGTAGAGGTTGCAGT
CACACGTCTGAACTCTTTCCCTACACGACGCTCTTCCGATCTNNNCCCTCATTAACTTAATGGAATTGC
CACACGTCTGAACTCTTTCCCTACACGACGCTCTTCCGATCTNNNCCCTCATTAACTTAATGGAATTGC
AGCACACGTCTGAACTCTTTCCCTACACGACGCTCTTCCGATCTNNNACAAAAATTAGCTGGGCATGG
AGCACACGTCTGAACTCTTTCCCTACACGACGCTCTTCCGATCTNNNACAAAAATTAGCTGGGCATGG
GAGCACACGTCTGAACTCTTTCCCTACACGACGCTCTTCCGATCTNNNACATTTCTATCTGCTGACAGC
GAGCACACGTCTGAACTCTTTCCCTACACGACGCTCTTCCGATCTNNNACATTTCTATCTGCTGACAGC
GAGCACACGTCTGAACTCTTTCCCTACACGACGCTCTTCCGATCTNNNGGTGAGGGGGATAGGGAAA
GAGCACACGTCTGAACTCTTTCCCTACACGACGCTCTTCCGATCTNNNGGTGAGGGGGATAGGGAAA
AAGAGCACACGTCTGAACTCTTTCCCTACACGACGCTCTTCCGATCTNNNACACAGTGAGGTCTGTAT
AAGAGCACACGTCTGAACTCTTTCCCTACACGACGCTCTTCCGATCTNNNACACAGTGAGGTCTGTAT
GAAGAGCACACGTCTGAACTCTTTCCCTACACGACGCTCTTCCGATCTNNNTCACAGATGAACCACCC
GAAGAGCACACGTCTGAACTCTTTCCCTACACGACGCTCTTCCGATCTNNNTCACAGATGAACCACCC
CACACGTCTGAACTCTTTCCCTACACGACGCTCTTCCGATCTNNNTTGTGTCCTTTCAGAGAACCACAC
CACACGTCTGAACTCTTTCCCTACACGACGCTCTTCCGATCTNNNTTGTGTCCTTTCAGAGAACCACAC
GCACACGTCTGAACTCTTTCCCTACACGACGCTCTTCCGATCTNNNAGAGGCATTCCATAATCAGTGCT
GCACACGTCTGAACTCTTTCCCTACACGACGCTCTTCCGATCTNNNAGAGGCATTCCATAATCAGTGCT
GAGCACACGTCTGAACTCTTTCCCTACACGACGCTCTTCCGATCTNNNCGAGACCAGCCTGGGAAATA
GAGCACACGTCTGAACTCTTTCCCTACACGACGCTCTTCCGATCTNNNCGAGACCAGCCTGGGAAATA
AGCACACGTCTGAACTCTTTCCCTACACGACGCTCTTCCGATCTNNNTGGAGAGCTTCACTGATTCTCTG
AGCACACGTCTGAACTCTTTCCCTACACGACGCTCTTCCGATCTNNNTGGAGAGCTTCACTGATTCTCTG
GAGCACACGTCTGAACTCTTTCCCTACACGACGCTCTTCCGATCTNNNTTTTCTTGTGGGCTCCTACTG
GAGCACACGTCTGAACTCTTTCCCTACACGACGCTCTTCCGATCTNNNTTTTCTTGTGGGCTCCTACTG
AGCACACGTCTGAACTCTTTCCCTACACGACGCTCTTCCGATCTNNNGCTGACTGTGAAGGGAGTCAT
AGCACACGTCTGAACTCTTTCCCTACACGACGCTCTTCCGATCTNNNGCTGACTGTGAAGGGAGTCAT
AGCACACGTCTGAACTCTTTCCCTACACGACGCTCTTCCGATCTNNNTTTATACACACACACACGCACAC
AGCACACGTCTGAACTCTTTCCCTACACGACGCTCTTCCGATCTNNNTTTATACACACACACACGCACAC
GCACACGTCTGAACTCTTTCCCTACACGACGCTCTTCCGATCTNNNTGAAGTAAGAAGATGTGGGCTG
GCACACGTCTGAACTCTTTCCCTACACGACGCTCTTCCGATCTNNNTGAAGTAAGAAGATGTGGGCTG
GAGCACACGTCTGAACTCTTTCCCTACACGACGCTCTTCCGATCTNNNTACTATGCTGTCCTCTTCTGCA
GAGCACACGTCTGAACTCTTTCCCTACACGACGCTCTTCCGATCTNNNTACTATGCTGTCCTCTTCTGCA
AGCACACGTCTGAACTCTTTCCCTACACGACGCTCTTCCGATCTNNNACCCTAGGAAACCAAAGAACG
AGCACACGTCTGAACTCTTTCCCTACACGACGCTCTTCCGATCTNNNACCCTAGGAAACCAAAGAACG
GAGCACACGTCTGAACTCTTTCCCTACACGACGCTCTTCCGATCTNNNTTGCTCAGCATAGCTAATAGG
GAGCACACGTCTGAACTCTTTCCCTACACGACGCTCTTCCGATCTNNNTTGCTCAGCATAGCTAATAGG
AGCACACGTCTGAACTCTTTCCCTACACGACGCTCTTCCGATCTNNNAGACTTTCAAGCTTGCCAACTTC
AGCACACGTCTGAACTCTTTCCCTACACGACGCTCTTCCGATCTNNNAGACTTTCAAGCTTGCCAACTTC
AGCACACGTCTGAACTCTTTCCCTACACGACGCTCTTCCGATCTNNNGTCATTGTTGCTTTAGGGGCA

AGCACACGTCTGAACTCTTTCCCTACACGACGCTCTTCCGATCTNNNGTCATTGTTGCTTTAGGGGCA
AGCACACGTCTGAACTCTTTCCCTACACGACGCTCTTCCGATCTNNNCTGGCAAAATAGTGAAACC
AGCACACGTCTGAACTCTTTCCCTACACGACGCTCTTCCGATCTNNNCTGGCAAAATAGTGAAACC
AGCACACGTCTGAACTCTTTCCCTACACGACGCTCTTCCGATCTNNNTTGGAGAAATCCAGATGTCC
AGCACACGTCTGAACTCTTTCCCTACACGACGCTCTTCCGATCTNNNTTGGAGAAATCCAGATGTCC
AGCACACGTCTGAACTCTTTCCCTACACGACGCTCTTCCGATCTNNNCAGTCTGTATGTCATTCCTGCA
AGCACACGTCTGAACTCTTTCCCTACACGACGCTCTTCCGATCTNNNCAGTCTGTATGTCATTCCTGCA
AGCACACGTCTGAACTCTTTCCCTACACGACGCTCTTCCGATCTNNNATGTGTAATGATCAAGCCAGG
AGCACACGTCTGAACTCTTTCCCTACACGACGCTCTTCCGATCTNNNATGTGTAATGATCAAGCCAGG
GAGCACACGTCTGAACTCTTTCCCTACACGACGCTCTTCCGATCTNNNGCCAGATCTTCATTCCCTATG
GAGCACACGTCTGAACTCTTTCCCTACACGACGCTCTTCCGATCTNNNGCCAGATCTTCATTCCCTATG
AGCACACGTCTGAACTCTTTCCCTACACGACGCTCTTCCGATCTNNNTTGAACTCTTGACCTCAGGTG
AGCACACGTCTGAACTCTTTCCCTACACGACGCTCTTCCGATCTNNNTTGAACTCTTGACCTCAGGTG
AGCACACGTCTGAACTCTTTCCCTACACGACGCTCTTCCGATCTNNNCTGATGATCTGTCACTGTCTC
AGCACACGTCTGAACTCTTTCCCTACACGACGCTCTTCCGATCTNNNCTGATGATCTGTCACTGTCTC
GAGCACACGTCTGAACTCTTTCCCTACACGACGCTCTTCCGATCTNNNCTGTGCAAATGCGTGTGTGT
GAGCACACGTCTGAACTCTTTCCCTACACGACGCTCTTCCGATCTNNNCTGTGCAAATGCGTGTGTGT
AGCACACGTCTGAACTCTTTCCCTACACGACGCTCTTCCGATCTNNNAGTACATACACACAGACGCAA
AGCACACGTCTGAACTCTTTCCCTACACGACGCTCTTCCGATCTNNNAGTACATACACACAGACGCAA
AGCACACGTCTGAACTCTTTCCCTACACGACGCTCTTCCGATCTNNNTCAGAGAATGTTCCTTCAGCTC
AGCACACGTCTGAACTCTTTCCCTACACGACGCTCTTCCGATCTNNNTCAGAGAATGTTCCTTCAGCTC
AGCACACGTCTGAACTCTTTCCCTACACGACGCTCTTCCGATCTNNNTGGAGTTGTACACACTTCCAA
AGCACACGTCTGAACTCTTTCCCTACACGACGCTCTTCCGATCTNNNTGGAGTTGTACACACTTCCAA
GCACACGTCTGAACTCTTTCCCTACACGACGCTCTTCCGATCTNNNCTGTAATCCCAGCTACTCAGGA
GCACACGTCTGAACTCTTTCCCTACACGACGCTCTTCCGATCTNNNCTGTAATCCCAGCTACTCAGGA
GCACACGTCTGAACTCTTTCCCTACACGACGCTCTTCCGATCTNNNAGCTAGAGGCACTAGGTTCTGT
GCACACGTCTGAACTCTTTCCCTACACGACGCTCTTCCGATCTNNNAGCTAGAGGCACTAGGTTCTGT
GCACACGTCTGAACTCTTTCCCTACACGACGCTCTTCCGATCTNNNAGCTTGCCATTTCTTTTCAGATG
GCACACGTCTGAACTCTTTCCCTACACGACGCTCTTCCGATCTNNNAGCTTGCCATTTCTTTTCAGATG
GCACACGTCTGAACTCTTTCCCTACACGACGCTCTTCCGATCTNNNGGGCCTTGATGAGCTAGTAGCC
GCACACGTCTGAACTCTTTCCCTACACGACGCTCTTCCGATCTNNNGGGCCTTGATGAGCTAGTAGCC
GAGCACACGTCTGAACTCTTTCCCTACACGACGCTCTTCCGATCTNNNAAAGAATGTTTGTATACAAA
GAGCACACGTCTGAACTCTTTCCCTACACGACGCTCTTCCGATCTNNNAAAGAATGTTTGTATACAAA
AGAGCACACGTCTGAACTCTTTCCCTACACGACGCTCTTCCGATCTNNNTACGTTGAACCACTACTCA
AGAGCACACGTCTGAACTCTTTCCCTACACGACGCTCTTCCGATCTNNNTACGTTGAACCACTACTCA
GAGCACACGTCTGAACTCTTTCCCTACACGACGCTCTTCCGATCTNNNCTACAAGAGGAGAAGGACA
GAGCACACGTCTGAACTCTTTCCCTACACGACGCTCTTCCGATCTNNNCTACAAGAGGAGAAGGACA
AGCACACGTCTGAACTCTTTCCCTACACGACGCTCTTCCGATCTNNNCTTCCACCCAACCAATACAACAC
AGCACACGTCTGAACTCTTTCCCTACACGACGCTCTTCCGATCTNNNCTTCCACCCAACCAATACAACAC
GAGCACACGTCTGAACTCTTTCCCTACACGACGCTCTTCCGATCTNNNGTTGAGGTAGGAGGATCGC
GAGCACACGTCTGAACTCTTTCCCTACACGACGCTCTTCCGATCTNNNGTTGAGGTAGGAGGATCGC
AGAGCACACGTCTGAACTCTTTCCCTACACGACGCTCTTCCGATCTNNNCTACAAGCTACTCTGAAGG
AGAGCACACGTCTGAACTCTTTCCCTACACGACGCTCTTCCGATCTNNNCTACAAGCTACTCTGAAGG

ACACGTCTGAACTCTTTCCCTACACGACGCTCTTCCGATCTNNNGAATTTCGATGCTGGAGACACGCAT

ACACGTCTGAACTCTTTCCCTACACGACGCTCTTCCGATCTNNNGAATTTCGATGCTGGAGACACGCAT

GCACACGTCTGAACTCTTTCCCTACACGACGCTCTTCCGATCTNNNTGGTCTCAAATTCCTGACCTCAG

GCACACGTCTGAACTCTTTCCCTACACGACGCTCTTCCGATCTNNNTGGTCTCAAATTCCTGACCTCAG

GAGCACACGTCTGAACTCTTTCCCTACACGACGCTCTTCCGATCTNNNAGCTTACCAAGATTGAACCAT

GAGCACACGTCTGAACTCTTTCCCTACACGACGCTCTTCCGATCTNNNAGCTTACCAAGATTGAACCAT

GAGCACACGTCTGAACTCTTTCCCTACACGACGCTCTTCCGATCTNNNTGGGCTTTCCAAGAATACCA

GAGCACACGTCTGAACTCTTTCCCTACACGACGCTCTTCCGATCTNNNTGGGCTTTCCAAGAATACCA

AGCACACGTCTGAACTCTTTCCCTACACGACGCTCTTCCGATCTNNNGCTTGAGACCAGGAGTTTGAA

AGCACACGTCTGAACTCTTTCCCTACACGACGCTCTTCCGATCTNNNGCTTGAGACCAGGAGTTTGAA

AGCACACGTCTGAACTCTTTCCCTACACGACGCTCTTCCGATCTNNNGTATTCCCAGCTACTAAGGAG

AGCACACGTCTGAACTCTTTCCCTACACGACGCTCTTCCGATCTNNNGTATTCCCAGCTACTAAGGAG

AGCACACGTCTGAACTCTTTCCCTACACGACGCTCTTCCGATCTNNNCTGCAACCTGCCTGACCAATAT

AGCACACGTCTGAACTCTTTCCCTACACGACGCTCTTCCGATCTNNNCTGCAACCTGCCTGACCAATAT

GAGCACACGTCTGAACTCTTTCCCTACACGACGCTCTTCCGATCTNNNCTTGCAGTGAGCCGAGATCA

GAGCACACGTCTGAACTCTTTCCCTACACGACGCTCTTCCGATCTNNNCTTGCAGTGAGCCGAGATCA

GAGCACACGTCTGAACTCTTTCCCTACACGACGCTCTTCCGATCTNNNTGCAGAGATGCAGGTATTGT

GAGCACACGTCTGAACTCTTTCCCTACACGACGCTCTTCCGATCTNNNTGCAGAGATGCAGGTATTGT

AGCACACGTCTGAACTCTTTCCCTACACGACGCTCTTCCGATCTNNNAAAAAAAATCAGCTGGGTGT

AGCACACGTCTGAACTCTTTCCCTACACGACGCTCTTCCGATCTNNNAAAAAAAATCAGCTGGGTGT

GCACACGTCTGAACTCTTTCCCTACACGACGCTCTTCCGATCTNNNCAATCAATCTCCTGCCTCAGCTCC

GCACACGTCTGAACTCTTTCCCTACACGACGCTCTTCCGATCTNNNCAATCAATCTCCTGCCTCAGCTCC

AGCACACGTCTGAACTCTTTCCCTACACGACGCTCTTCCGATCTNNNCTGTAGTCCCAGCTACTCAGAA

AGCACACGTCTGAACTCTTTCCCTACACGACGCTCTTCCGATCTNNNCTGTAGTCCCAGCTACTCAGAA

AGCACACGTCTGAACTCTTTCCCTACACGACGCTCTTCCGATCTNNNGTGAGCCTTCTTTGCTTCTTGAG

AGCACACGTCTGAACTCTTTCCCTACACGACGCTCTTCCGATCTNNNGTGAGCCTTCTTTGCTTCTTGAG

AGCACACGTCTGAACTCTTTCCCTACACGACGCTCTTCCGATCTNNNAGGCAAGAGAATGGTGTGAA

AGCACACGTCTGAACTCTTTCCCTACACGACGCTCTTCCGATCTNNNAGGCAAGAGAATGGTGTGAA

AGCACACGTCTGAACTCTTTCCCTACACGACGCTCTTCCGATCTNNNTCCTTCCTTCTGCATGTAGATG

AGCACACGTCTGAACTCTTTCCCTACACGACGCTCTTCCGATCTNNNTCCTTCCTTCTGCATGTAGATG

GCACACGTCTGAACTCTTTCCCTACACGACGCTCTTCCGATCTNNNTAGGCAGGAGAATCACTTGAAC

GCACACGTCTGAACTCTTTCCCTACACGACGCTCTTCCGATCTNNNTAGGCAGGAGAATCACTTGAAC

GCACACGTCTGAACTCTTTCCCTACACGACGCTCTTCCGATCTNNNACCTTGAAGAAAATCAGTTTCCA

GCACACGTCTGAACTCTTTCCCTACACGACGCTCTTCCGATCTNNNACCTTGAAGAAAATCAGTTTCCA

CACACGTCTGAACTCTTTCCCTACACGACGCTCTTCCGATCTNNNCTACCAGGCCGTTACACTTAGTTC

CACACGTCTGAACTCTTTCCCTACACGACGCTCTTCCGATCTNNNCTACCAGGCCGTTACACTTAGTTC

AGCACACGTCTGAACTCTTTCCCTACACGACGCTCTTCCGATCTNNNGAGATTGCAGTGAGCCGAGAT

AGCACACGTCTGAACTCTTTCCCTACACGACGCTCTTCCGATCTNNNGAGATTGCAGTGAGCCGAGAT

AGCACACGTCTGAACTCTTTCCCTACACGACGCTCTTCCGATCTNNNCTGAACCCATGAGGCAAATGT

AGCACACGTCTGAACTCTTTCCCTACACGACGCTCTTCCGATCTNNNCTGAACCCATGAGGCAAATGT

AGCACACGTCTGAACTCTTTCCCTACACGACGCTCTTCCGATCTNNNCTTCCACAAATTCTGCCCACAAT

AGCACACGTCTGAACTCTTTCCCTACACGACGCTCTTCCGATCTNNNCTTCCACAAATTCTGCCCACAAT

GAGCACACGTCTGAACTCTTTCCCTACACGACGCTCTTCCGATCTNNNAAAGAGGCAGAGAGACAAA

GAGCACACGTCTGAACTCTTTCCCTACACGACGCTCTTCCGATCTNNNAAAGAGGCAGAGAGACAAA
GCACACGTCTGAACTCTTTCCCTACACGACGCTCTTCCGATCTNNNGGAGTTGTACTCTGCTCAATCC
GCACACGTCTGAACTCTTTCCCTACACGACGCTCTTCCGATCTNNNGGAGTTGTACTCTGCTCAATCC
AGCACACGTCTGAACTCTTTCCCTACACGACGCTCTTCCGATCTNNNCACTCCCAACTTAGACAATCCC
AGCACACGTCTGAACTCTTTCCCTACACGACGCTCTTCCGATCTNNNCACTCCCAACTTAGACAATCCC
GAGCACACGTCTGAACTCTTTCCCTACACGACGCTCTTCCGATCTNNNACCTGTAATTCCAGCTACTCA
GAGCACACGTCTGAACTCTTTCCCTACACGACGCTCTTCCGATCTNNNACCTGTAATTCCAGCTACTCA
AGAGCACACGTCTGAACTCTTTCCCTACACGACGCTCTTCCGATCTNNNTTTATACAGGGGACGCCCA
AGAGCACACGTCTGAACTCTTTCCCTACACGACGCTCTTCCGATCTNNNTTTATACAGGGGACGCCCA
AGCACACGTCTGAACTCTTTCCCTACACGACGCTCTTCCGATCTNNNATAGAATTCAGAATCAACCAA
AGCACACGTCTGAACTCTTTCCCTACACGACGCTCTTCCGATCTNNNATAGAATTCAGAATCAACCAA
CACACGTCTGAACTCTTTCCCTACACGACGCTCTTCCGATCTNNNTCTGTTCAAGTCCTTGTTTGTGAC
CACACGTCTGAACTCTTTCCCTACACGACGCTCTTCCGATCTNNNTCTGTTCAAGTCCTTGTTTGTGAC
ACACGTCTGAACTCTTTCCCTACACGACGCTCTTCCGATCTNNNCTTGGTAATGATCCTACAGATCAT
ACACGTCTGAACTCTTTCCCTACACGACGCTCTTCCGATCTNNNCTTGGTAATGATCCTACAGATCAT
GAGCACACGTCTGAACTCTTTCCCTACACGACGCTCTTCCGATCTNNNAGTCAGGTTCCAAGTCCTTG
GAGCACACGTCTGAACTCTTTCCCTACACGACGCTCTTCCGATCTNNNAGTCAGGTTCCAAGTCCTTG
GAGCACACGTCTGAACTCTTTCCCTACACGACGCTCTTCCGATCTNNNCACTCTCCCAGTCTGTCAAGT
GAGCACACGTCTGAACTCTTTCCCTACACGACGCTCTTCCGATCTNNNCACTCTCCCAGTCTGTCAAGT
GCACACGTCTGAACTCTTTCCCTACACGACGCTCTTCCGATCTNNNTAGACACTGGGGCCAACTTGAT
GCACACGTCTGAACTCTTTCCCTACACGACGCTCTTCCGATCTNNNTAGACACTGGGGCCAACTTGAT
GCACACGTCTGAACTCTTTCCCTACACGACGCTCTTCCGATCTNNNACTGTAATTGATCGTGTGTGTG
GCACACGTCTGAACTCTTTCCCTACACGACGCTCTTCCGATCTNNNACTGTAATTGATCGTGTGTGTG
CACACGTCTGAACTCTTTCCCTACACGACGCTCTTCCGATCTNNNGGCTGAGATAGGAGAATCACCT
CACACGTCTGAACTCTTTCCCTACACGACGCTCTTCCGATCTNNNGGCTGAGATAGGAGAATCACCT
AGCACACGTCTGAACTCTTTCCCTACACGACGCTCTTCCGATCTNNNCTGATCTCAAGTGATCCTCCCA
AGCACACGTCTGAACTCTTTCCCTACACGACGCTCTTCCGATCTNNNCTGATCTCAAGTGATCCTCCCA
AGCACACGTCTGAACTCTTTCCCTACACGACGCTCTTCCGATCTNNNCCAGCCAGATAACCCTGACAAT
AGCACACGTCTGAACTCTTTCCCTACACGACGCTCTTCCGATCTNNNCCAGCCAGATAACCCTGACAAT
AGCACACGTCTGAACTCTTTCCCTACACGACGCTCTTCCGATCTNNNGAATCCTCGGGTCAGAAGAG
AGCACACGTCTGAACTCTTTCCCTACACGACGCTCTTCCGATCTNNNGAATCCTCGGGTCAGAAGAG
GAGCACACGTCTGAACTCTTTCCCTACACGACGCTCTTCCGATCTNNNAGTGAGGCTGCAGTTTACGA
GAGCACACGTCTGAACTCTTTCCCTACACGACGCTCTTCCGATCTNNNAGTGAGGCTGCAGTTTACGA
GCACACGTCTGAACTCTTTCCCTACACGACGCTCTTCCGATCTNNNTTCTGAATGGGCTGCTCATATCA
GCACACGTCTGAACTCTTTCCCTACACGACGCTCTTCCGATCTNNNTTCTGAATGGGCTGCTCATATCA
GCACACGTCTGAACTCTTTCCCTACACGACGCTCTTCCGATCTNNNTGGGTGACAGAGTGAGACTCCC
GCACACGTCTGAACTCTTTCCCTACACGACGCTCTTCCGATCTNNNTGGGTGACAGAGTGAGACTCCC
AGCACACGTCTGAACTCTTTCCCTACACGACGCTCTTCCGATCTNNNAACAGTATTGCACATTCAGGC
AGCACACGTCTGAACTCTTTCCCTACACGACGCTCTTCCGATCTNNNAACAGTATTGCACATTCAGGC
GCACACGTCTGAACTCTTTCCCTACACGACGCTCTTCCGATCTNNNCACAGTCCCCATTCCTGCTATAC
GCACACGTCTGAACTCTTTCCCTACACGACGCTCTTCCGATCTNNNCACAGTCCCCATTCCTGCTATAC
GCACACGTCTGAACTCTTTCCCTACACGACGCTCTTCCGATCTNNNATCCAGCACTTCACCAGATTCTT
GCACACGTCTGAACTCTTTCCCTACACGACGCTCTTCCGATCTNNNATCCAGCACTTCACCAGATTCTT

AGCACACGTCTGAACTCTTTCCCTACACGACGCTCTTCCGATCTNNNGAGGTTACAGTGAGCTGAGA
AGCACACGTCTGAACTCTTTCCCTACACGACGCTCTTCCGATCTNNNGAGGTTACAGTGAGCTGAGA
GAGCACACGTCTGAACTCTTTCCCTACACGACGCTCTTCCGATCTNNNGGAGGTGGAACTTGAAATA
GAGCACACGTCTGAACTCTTTCCCTACACGACGCTCTTCCGATCTNNNGGAGGTGGAACTTGAAATA
GCACACGTCTGAACTCTTTCCCTACACGACGCTCTTCCGATCTNNNCAACTACCACCACCTCCAGTCCAT
GCACACGTCTGAACTCTTTCCCTACACGACGCTCTTCCGATCTNNNCAACTACCACCACCTCCAGTCCAT
AGCACACGTCTGAACTCTTTCCCTACACGACGCTCTTCCGATCTNNNCAAGTGGCTTATGGGATAGG
AGCACACGTCTGAACTCTTTCCCTACACGACGCTCTTCCGATCTNNNCAAGTGGCTTATGGGATAGG
GAGCACACGTCTGAACTCTTTCCCTACACGACGCTCTTCCGATCTNNNGCCCAGGAATTGTTAGAGG
GAGCACACGTCTGAACTCTTTCCCTACACGACGCTCTTCCGATCTNNNGCCCAGGAATTGTTAGAGG
AGCACACGTCTGAACTCTTTCCCTACACGACGCTCTTCCGATCTNNNTGGGAGACAGAGAGAGGCTC
AGCACACGTCTGAACTCTTTCCCTACACGACGCTCTTCCGATCTNNNTGGGAGACAGAGAGAGGCTC
GAGCACACGTCTGAACTCTTTCCCTACACGACGCTCTTCCGATCTNNNAGAGAGAGAGAGAGAGAG
GAGCACACGTCTGAACTCTTTCCCTACACGACGCTCTTCCGATCTNNNAGAGAGAGAGAGAGAGAG
GAGCACACGTCTGAACTCTTTCCCTACACGACGCTCTTCCGATCTNNNTGTTCATTGCAACACTGTTC
GAGCACACGTCTGAACTCTTTCCCTACACGACGCTCTTCCGATCTNNNTGTTCATTGCAACACTGTTC
AGCACACGTCTGAACTCTTTCCCTACACGACGCTCTTCCGATCTNNNGGGAAAGAACTGGGGTGATG
AGCACACGTCTGAACTCTTTCCCTACACGACGCTCTTCCGATCTNNNGGGAAAGAACTGGGGTGATG
GCACACGTCTGAACTCTTTCCCTACACGACGCTCTTCCGATCTNNNCCCTGATGGTGTCTATGTCACAT
GCACACGTCTGAACTCTTTCCCTACACGACGCTCTTCCGATCTNNNCCCTGATGGTGTCTATGTCACAT
GAGCACACGTCTGAACTCTTTCCCTACACGACGCTCTTCCGATCTNNNTGAGGCAAGAGAATTGCTTG
GAGCACACGTCTGAACTCTTTCCCTACACGACGCTCTTCCGATCTNNNTGAGGCAAGAGAATTGCTTG
AGCACACGTCTGAACTCTTTCCCTACACGACGCTCTTCCGATCTNNNAAAAGGTAGCTTCCCCTTGTT
AGCACACGTCTGAACTCTTTCCCTACACGACGCTCTTCCGATCTNNNAAAAGGTAGCTTCCCCTTGTT
AGCACACGTCTGAACTCTTTCCCTACACGACGCTCTTCCGATCTNNNCTTGAACTCCTGACCTCAAGTG
AGCACACGTCTGAACTCTTTCCCTACACGACGCTCTTCCGATCTNNNCTTGAACTCCTGACCTCAAGTG
AGCACACGTCTGAACTCTTTCCCTACACGACGCTCTTCCGATCTNNNGACACAAATGGCTCAGATTCTG
AGCACACGTCTGAACTCTTTCCCTACACGACGCTCTTCCGATCTNNNGACACAAATGGCTCAGATTCTG
AGCACACGTCTGAACTCTTTCCCTACACGACGCTCTTCCGATCTNNNCATGCCTGTAATCCCAGCTACT
AGCACACGTCTGAACTCTTTCCCTACACGACGCTCTTCCGATCTNNNCATGCCTGTAATCCCAGCTACT
AGCACACGTCTGAACTCTTTCCCTACACGACGCTCTTCCGATCTNNNGGACCATTTAGATAACAGCCC
AGCACACGTCTGAACTCTTTCCCTACACGACGCTCTTCCGATCTNNNGGACCATTTAGATAACAGCCC
GAGCACACGTCTGAACTCTTTCCCTACACGACGCTCTTCCGATCTNNNGGTTGATGCACAAGAATCG
GAGCACACGTCTGAACTCTTTCCCTACACGACGCTCTTCCGATCTNNNGGTTGATGCACAAGAATCG
AGCACACGTCTGAACTCTTTCCCTACACGACGCTCTTCCGATCTNNNTACCGTGGTGTGTTGTTTTGT
AGCACACGTCTGAACTCTTTCCCTACACGACGCTCTTCCGATCTNNNTACCGTGGTGTGTTGTTTTGT
AGCACACGTCTGAACTCTTTCCCTACACGACGCTCTTCCGATCTNNNATCAGAGACTAGGATTGCAAC
AGCACACGTCTGAACTCTTTCCCTACACGACGCTCTTCCGATCTNNNATCAGAGACTAGGATTGCAAC
AGCACACGTCTGAACTCTTTCCCTACACGACGCTCTTCCGATCTNNNTCAATGCTACCTTTGATGTGGA
AGCACACGTCTGAACTCTTTCCCTACACGACGCTCTTCCGATCTNNNTCAATGCTACCTTTGATGTGGA
AGCACACGTCTGAACTCTTTCCCTACACGACGCTCTTCCGATCTNNNGGATAATGAGTCTGCTAGAGG
AGCACACGTCTGAACTCTTTCCCTACACGACGCTCTTCCGATCTNNNGGATAATGAGTCTGCTAGAGG
AGCACACGTCTGAACTCTTTCCCTACACGACGCTCTTCCGATCTNNNTGTTTTTCCCCTAAGATCAGGC

GCACACGTCTGAACTCTTTCCCTACACGACGCTCTTCCGATCTNNNTGTTTTTCCCCTAAGATCAGGC
CACACGTCTGAACTCTTTCCCTACACGACGCTCTTCCGATCTNNNCAGGCCCTAATCCAGGATGATCT
CACACGTCTGAACTCTTTCCCTACACGACGCTCTTCCGATCTNNNCAGGCCCTAATCCAGGATGATCT
GCACACGTCTGAACTCTTTCCCTACACGACGCTCTTCCGATCTNNNAAATCCTCCGTACAGACAAAGC
GCACACGTCTGAACTCTTTCCCTACACGACGCTCTTCCGATCTNNNAAATCCTCCGTACAGACAAAGC
GCACACGTCTGAACTCTTTCCCTACACGACGCTCTTCCGATCTNNNGGAGGGTCCAGTTTCAATCTTC
GCACACGTCTGAACTCTTTCCCTACACGACGCTCTTCCGATCTNNNGGAGGGTCCAGTTTCAATCTTC
AGCACACGTCTGAACTCTTTCCCTACACGACGCTCTTCCGATCTNNNGCACTTAGCAAAACTTCCTCCA
AGCACACGTCTGAACTCTTTCCCTACACGACGCTCTTCCGATCTNNNGCACTTAGCAAAACTTCCTCCA
AGCACACGTCTGAACTCTTTCCCTACACGACGCTCTTCCGATCTNNNGAGAATGGTGTGAACCCAGG
AGCACACGTCTGAACTCTTTCCCTACACGACGCTCTTCCGATCTNNNGAGAATGGTGTGAACCCAGG
AGCACACGTCTGAACTCTTTCCCTACACGACGCTCTTCCGATCTNNNTGTAATCACAGCACTTTGGGA
AGCACACGTCTGAACTCTTTCCCTACACGACGCTCTTCCGATCTNNNTGTAATCACAGCACTTTGGGA
AGCACACGTCTGAACTCTTTCCCTACACGACGCTCTTCCGATCTNNNAAATTAGCCGAGCCTAGTGGT
AGCACACGTCTGAACTCTTTCCCTACACGACGCTCTTCCGATCTNNNAAATTAGCCGAGCCTAGTGGT
AGCACACGTCTGAACTCTTTCCCTACACGACGCTCTTCCGATCTNNNTGATAGCTGATAAGCCATGTC
AGCACACGTCTGAACTCTTTCCCTACACGACGCTCTTCCGATCTNNNTGATAGCTGATAAGCCATGTC
AGCACACGTCTGAACTCTTTCCCTACACGACGCTCTTCCGATCTNNNTGCAATCCTACAATTCTGTGTG
AGCACACGTCTGAACTCTTTCCCTACACGACGCTCTTCCGATCTNNNTGCAATCCTACAATTCTGTGTG
CACACGTCTGAACTCTTTCCCTACACGACGCTCTTCCGATCTNNNGTTGGGGATCATAGTGATGGTGT
CACACGTCTGAACTCTTTCCCTACACGACGCTCTTCCGATCTNNNGTTGGGGATCATAGTGATGGTGT
AGCACACGTCTGAACTCTTTCCCTACACGACGCTCTTCCGATCTNNNGTTTGCAGTGAGCCAAGATCA
AGCACACGTCTGAACTCTTTCCCTACACGACGCTCTTCCGATCTNNNGTTTGCAGTGAGCCAAGATCA
GCACACGTCTGAACTCTTTCCCTACACGACGCTCTTCCGATCTNNNCTGTAATCCCAGCTACTCCAGAG
GCACACGTCTGAACTCTTTCCCTACACGACGCTCTTCCGATCTNNNCTGTAATCCCAGCTACTCCAGAG
AGCACACGTCTGAACTCTTTCCCTACACGACGCTCTTCCGATCTNNNGAGGTAAGAGGATGGATTG
AGCACACGTCTGAACTCTTTCCCTACACGACGCTCTTCCGATCTNNNGAGGTAAGAGGATGGATTG
CACACGTCTGAACTCTTTCCCTACACGACGCTCTTCCGATCTNNNATCACCATTTTTACCCTCCCCTCCAT
CACACGTCTGAACTCTTTCCCTACACGACGCTCTTCCGATCTNNNATCACCATTTTTACCCTCCCCTCCAT
ACACGTCTGAACTCTTTCCCTACACGACGCTCTTCCGATCTNNNTGTTTTTTCTTCTAAAGGCTTTTCAT
ACACGTCTGAACTCTTTCCCTACACGACGCTCTTCCGATCTNNNTGTTTTTTCTTCTAAAGGCTTTTCAT
AGCACACGTCTGAACTCTTTCCCTACACGACGCTCTTCCGATCTNNNGCCACCATGCCCAACTAATTTT
AGCACACGTCTGAACTCTTTCCCTACACGACGCTCTTCCGATCTNNNGCCACCATGCCCAACTAATTTT
AGCACACGTCTGAACTCTTTCCCTACACGACGCTCTTCCGATCTNNNCATCAGAAAGACACCTGCACAT
AGCACACGTCTGAACTCTTTCCCTACACGACGCTCTTCCGATCTNNNCATCAGAAAGACACCTGCACAT
AGCACACGTCTGAACTCTTTCCCTACACGACGCTCTTCCGATCTNNNGAGGCAGGAGAATGGTGTGA
AGCACACGTCTGAACTCTTTCCCTACACGACGCTCTTCCGATCTNNNGAGGCAGGAGAATGGTGTGA
GCACACGTCTGAACTCTTTCCCTACACGACGCTCTTCCGATCTNNNGAAAGACCCAATGTGGTTGTCT
GCACACGTCTGAACTCTTTCCCTACACGACGCTCTTCCGATCTNNNGAAAGACCCAATGTGGTTGTCT
GAGCACACGTCTGAACTCTTTCCCTACACGACGCTCTTCCGATCTNNNCATCTGCACACACCATCATCA
GAGCACACGTCTGAACTCTTTCCCTACACGACGCTCTTCCGATCTNNNCATCTGCACACACCATCATCA
GCACACGTCTGAACTCTTTCCCTACACGACGCTCTTCCGATCTNNNACATGACGAAACCCCATCTCTA
GCACACGTCTGAACTCTTTCCCTACACGACGCTCTTCCGATCTNNNACATGACGAAACCCCATCTCTA

AGCACACGTCTGAACTCTTTCCCTACACGACGCTCTTCCGATCTNNNCATGCCTGTAATCCCAGCTACTC

AGCACACGTCTGAACTCTTTCCCTACACGACGCTCTTCCGATCTNNNCATGCCTGTAATCCCAGCTACTC

GAGCACACGTCTGAACTCTTTCCCTACACGACGCTCTTCCGATCTNNNTCATTTCATGGATGCCAGTCTG

GAGCACACGTCTGAACTCTTTCCCTACACGACGCTCTTCCGATCTNNNTCATTTCATGGATGCCAGTCTG

GAGCACACGTCTGAACTCTTTCCCTACACGACGCTCTTCCGATCTNNNGACCTCAAGACCAAATGGAAA

GAGCACACGTCTGAACTCTTTCCCTACACGACGCTCTTCCGATCTNNNGACCTCAAGACCAAATGGAAA

AGCACACGTCTGAACTCTTTCCCTACACGACGCTCTTCCGATCTNNNAAAGACTTCTTGCCCTCTATGCAC

AGCACACGTCTGAACTCTTTCCCTACACGACGCTCTTCCGATCTNNNAAAGACTTCTTGCCCTCTATGCAC

GAGCACACGTCTGAACTCTTTCCCTACACGACGCTCTTCCGATCTNNNCTGGCCAACTTTCCCATTTTTGA

GAGCACACGTCTGAACTCTTTCCCTACACGACGCTCTTCCGATCTNNNCTGGCCAACTTTCCCATTTTTGA

GAGCACACGTCTGAACTCTTTCCCTACACGACGCTCTTCCGATCTNNNAGAGACATCACATGTGCTTCCT

GAGCACACGTCTGAACTCTTTCCCTACACGACGCTCTTCCGATCTNNNAGAGACATCACATGTGCTTCCT

GAGCACACGTCTGAACTCTTTCCCTACACGACGCTCTTCCGATCTNNNGAAGGCAGGAACCAGGATTC

GAGCACACGTCTGAACTCTTTCCCTACACGACGCTCTTCCGATCTNNNGAAGGCAGGAACCAGGATTC

AAGAGCACACGTCTGAACTCTTTCCCTACACGACGCTCTTCCGATCTNNNTGAATGGCCTTATTGGATC

AAGAGCACACGTCTGAACTCTTTCCCTACACGACGCTCTTCCGATCTNNNTGAATGGCCTTATTGGATC

GAGCACACGTCTGAACTCTTTCCCTACACGACGCTCTTCCGATCTNNNGGAGTGTCATGTAAGTGGAC

GAGCACACGTCTGAACTCTTTCCCTACACGACGCTCTTCCGATCTNNNGGAGTGTCATGTAAGTGGAC

GAGCACACGTCTGAACTCTTTCCCTACACGACGCTCTTCCGATCTNNNCCAAATCTCCATCCCTCAGAGA

GAGCACACGTCTGAACTCTTTCCCTACACGACGCTCTTCCGATCTNNNCCAAATCTCCATCCCTCAGAGA

AGAGCACACGTCTGAACTCTTTCCCTACACGACGCTCTTCCGATCTNNNATCTTCCTTTGCTGTTGCTTG

AGAGCACACGTCTGAACTCTTTCCCTACACGACGCTCTTCCGATCTNNNATCTTCCTTTGCTGTTGCTTG

AGCACACGTCTGAACTCTTTCCCTACACGACGCTCTTCCGATCTNNNATCCAGTGATCCTCTCTCCTCAA

AGCACACGTCTGAACTCTTTCCCTACACGACGCTCTTCCGATCTNNNATCCAGTGATCCTCTCTCCTCAA

GAGCACACGTCTGAACTCTTTCCCTACACGACGCTCTTCCGATCTNNNTTCTAACCTTACCACCACTGCTC

GAGCACACGTCTGAACTCTTTCCCTACACGACGCTCTTCCGATCTNNNTTCTAACCTTACCACCACTGCTC

AGCACACGTCTGAACTCTTTCCCTACACGACGCTCTTCCGATCTNNNCTATGTAAGGTCAGTGGCAAC

AGCACACGTCTGAACTCTTTCCCTACACGACGCTCTTCCGATCTNNNCTATGTAAGGTCAGTGGCAAC

GAGCACACGTCTGAACTCTTTCCCTACACGACGCTCTTCCGATCTNNNGTGAGACCCTGCCTCTAAAGA

GAGCACACGTCTGAACTCTTTCCCTACACGACGCTCTTCCGATCTNNNGTGAGACCCTGCCTCTAAAGA

GCACACGTCTGAACTCTTTCCCTACACGACGCTCTTCCGATCTNNNATTGTGTCATTTGCTGTGCTGAA

GCACACGTCTGAACTCTTTCCCTACACGACGCTCTTCCGATCTNNNATTGTGTCATTTGCTGTGCTGAA

AGCACACGTCTGAACTCTTTCCCTACACGACGCTCTTCCGATCTNNNAAGACCCATCAGTGTGCTGTAT

AGCACACGTCTGAACTCTTTCCCTACACGACGCTCTTCCGATCTNNNAAGACCCATCAGTGTGCTGTAT

AGCACACGTCTGAACTCTTTCCCTACACGACGCTCTTCCGATCTNNNTTGTCACCCTGAAAGAGGAACT

AGCACACGTCTGAACTCTTTCCCTACACGACGCTCTTCCGATCTNNNTTGTCACCCTGAAAGAGGAACT

GAGCACACGTCTGAACTCTTTCCCTACACGACGCTCTTCCGATCTNNNAGAAAGAGAGAGAGAGAGG

GAGCACACGTCTGAACTCTTTCCCTACACGACGCTCTTCCGATCTNNNAGAAAGAGAGAGAGAGAGG

AGCACACGTCTGAACTCTTTCCCTACACGACGCTCTTCCGATCTNNNGTGCTTTTACCTAGTAACTGCTCA

AGCACACGTCTGAACTCTTTCCCTACACGACGCTCTTCCGATCTNNNGTGCTTTTACCTAGTAACTGCTCA

GAGCACACGTCTGAACTCTTTCCCTACACGACGCTCTTCCGATCTNNNCTGTAGTCCCAGCTACTCAGG

GAGCACACGTCTGAACTCTTTCCCTACACGACGCTCTTCCGATCTNNNCTGTAGTCCCAGCTACTCAGG

AGCACACGTCTGAACTCTTTCCCTACACGACGCTCTTCCGATCTNNNCCTCCTTCCTCTTGTGAATATAAC

AGCACACGTCTGAACTCTTTCCCTACACGACGCTCTTCCGATCTNNNCCTCCTTCCTCTTGTGAATATAA
AGCACACGTCTGAACTCTTTCCCTACACGACGCTCTTCCGATCTNNNGTTGCAGTGAGCCGAGATCAC
AGCACACGTCTGAACTCTTTCCCTACACGACGCTCTTCCGATCTNNNGTTGCAGTGAGCCGAGATCAC
GCACACGTCTGAACTCTTTCCCTACACGACGCTCTTCCGATCTNNNTATCTTTGAGGCCTGCAAAACTC
GCACACGTCTGAACTCTTTCCCTACACGACGCTCTTCCGATCTNNNTATCTTTGAGGCCTGCAAAACTC
AGCACACGTCTGAACTCTTTCCCTACACGACGCTCTTCCGATCTNNNTCCTTGACACACTCCTGGTTAT
AGCACACGTCTGAACTCTTTCCCTACACGACGCTCTTCCGATCTNNNTCCTTGACACACTCCTGGTTAT
GCACACGTCTGAACTCTTTCCCTACACGACGCTCTTCCGATCTNNNCACCATCCATCTTCATTCCAGGG
GCACACGTCTGAACTCTTTCCCTACACGACGCTCTTCCGATCTNNNCACCATCCATCTTCATTCCAGGG
GAGCACACGTCTGAACTCTTTCCCTACACGACGCTCTTCCGATCTNNNACACCTGTATCCTAGCACTTT
GAGCACACGTCTGAACTCTTTCCCTACACGACGCTCTTCCGATCTNNNACACCTGTATCCTAGCACTTT
GCACACGTCTGAACTCTTTCCCTACACGACGCTCTTCCGATCTNNNCTCGGACAGCAACCCAGATTACC
GCACACGTCTGAACTCTTTCCCTACACGACGCTCTTCCGATCTNNNCTCGGACAGCAACCCAGATTACC
AGCACACGTCTGAACTCTTTCCCTACACGACGCTCTTCCGATCTNNNGAATGGTTTTGGTGTCCTTGT
AGCACACGTCTGAACTCTTTCCCTACACGACGCTCTTCCGATCTNNNGAATGGTTTTGGTGTCCTTGT
AGCACACGTCTGAACTCTTTCCCTACACGACGCTCTTCCGATCTNNNCAAAAAAAGGAGTACTATCAT
AGCACACGTCTGAACTCTTTCCCTACACGACGCTCTTCCGATCTNNNCAAAAAAAGGAGTACTATCAT
AGCACACGTCTGAACTCTTTCCCTACACGACGCTCTTCCGATCTNNNAGACTTTCCCAAAAAGCGGATT
AGCACACGTCTGAACTCTTTCCCTACACGACGCTCTTCCGATCTNNNAGACTTTCCCAAAAAGCGGATT
AGCACACGTCTGAACTCTTTCCCTACACGACGCTCTTCCGATCTNNNGGTCTGCATGTGTTCAGCTTTA
AGCACACGTCTGAACTCTTTCCCTACACGACGCTCTTCCGATCTNNNGGTCTGCATGTGTTCAGCTTTA
AGCACACGTCTGAACTCTTTCCCTACACGACGCTCTTCCGATCTNNNCTGAGACAGGAGAATCGCTTG
AGCACACGTCTGAACTCTTTCCCTACACGACGCTCTTCCGATCTNNNCTGAGACAGGAGAATCGCTTG
ACACGTCTGAACTCTTTCCCTACACGACGCTCTTCCGATCTNNNAATGGTGCTGAAATGAAGATCATC
ACACGTCTGAACTCTTTCCCTACACGACGCTCTTCCGATCTNNNAATGGTGCTGAAATGAAGATCATC
AGCACACGTCTGAACTCTTTCCCTACACGACGCTCTTCCGATCTNNNCCAGACACACTATCCAGATCCA
AGCACACGTCTGAACTCTTTCCCTACACGACGCTCTTCCGATCTNNNCCAGACACACTATCCAGATCCA
AGCACACGTCTGAACTCTTTCCCTACACGACGCTCTTCCGATCTNNNTAATACACATGAACCATTTTTG
AGCACACGTCTGAACTCTTTCCCTACACGACGCTCTTCCGATCTNNNTAATACACATGAACCATTTTTG
AGCACACGTCTGAACTCTTTCCCTACACGACGCTCTTCCGATCTNNNTCCCATTCTCTCTCTCTCTCTCT
AGCACACGTCTGAACTCTTTCCCTACACGACGCTCTTCCGATCTNNNTCCCATTCTCTCTCTCTCTCTCT
AGCACACGTCTGAACTCTTTCCCTACACGACGCTCTTCCGATCTNNNGAGAATGGCATGAACCTGGG
AGCACACGTCTGAACTCTTTCCCTACACGACGCTCTTCCGATCTNNNGAGAATGGCATGAACCTGGG
AGAGCACACGTCTGAACTCTTTCCCTACACGACGCTCTTCCGATCTNNNAGCATCAAGCAAGTCCAAAA
AGAGCACACGTCTGAACTCTTTCCCTACACGACGCTCTTCCGATCTNNNAGCATCAAGCAAGTCCAAAA
AGCACACGTCTGAACTCTTTCCCTACACGACGCTCTTCCGATCTNNNCCTACACATCCAAGACTTTCCCA
AGCACACGTCTGAACTCTTTCCCTACACGACGCTCTTCCGATCTNNNCCTACACATCCAAGACTTTCCCA
GAGCACACGTCTGAACTCTTTCCCTACACGACGCTCTTCCGATCTNNNCTCTTCCCTGGACTTGAGTGA
GAGCACACGTCTGAACTCTTTCCCTACACGACGCTCTTCCGATCTNNNCTCTTCCCTGGACTTGAGTGA
AGCACACGTCTGAACTCTTTCCCTACACGACGCTCTTCCGATCTNNNAGATCATGTCACTACACTCCAG
AGCACACGTCTGAACTCTTTCCCTACACGACGCTCTTCCGATCTNNNAGATCATGTCACTACACTCCAG
GAGCACACGTCTGAACTCTTTCCCTACACGACGCTCTTCCGATCTNNNTGTAATCTCAGCACTTTGGGA
GAGCACACGTCTGAACTCTTTCCCTACACGACGCTCTTCCGATCTNNNTGTAATCTCAGCACTTTGGGA

GAGCACACGTCTGAACTCTTTCCCTACACGACGCTCTTCCGATCTNNNTAGCCAGGTGTCTCAAATTAG
GAGCACACGTCTGAACTCTTTCCCTACACGACGCTCTTCCGATCTNNNTAGCCAGGTGTCTCAAATTAG
AGAGCACACGTCTGAACTCTTTCCCTACACGACGCTCTTCCGATCTNNNTAGGCAGACACTAAGCTCCC
AGAGCACACGTCTGAACTCTTTCCCTACACGACGCTCTTCCGATCTNNNTAGGCAGACACTAAGCTCCC
AAGAGCACACGTCTGAACTCTTTCCCTACACGACGCTCTTCCGATCTNNNAAAATTAGTTGGGCATGG
AAGAGCACACGTCTGAACTCTTTCCCTACACGACGCTCTTCCGATCTNNNAAAATTAGTTGGGCATGG
AGCACACGTCTGAACTCTTTCCCTACACGACGCTCTTCCGATCTNNNAACCAGTACATGAGGCAATCCA
AGCACACGTCTGAACTCTTTCCCTACACGACGCTCTTCCGATCTNNNAACCAGTACATGAGGCAATCCA
AGAGCACACGTCTGAACTCTTTCCCTACACGACGCTCTTCCGATCTNNNGATGGAGGATTCTGGTGGA
AGAGCACACGTCTGAACTCTTTCCCTACACGACGCTCTTCCGATCTNNNGATGGAGGATTCTGGTGGA
GAGCACACGTCTGAACTCTTTCCCTACACGACGCTCTTCCGATCTNNNCTGCTGTTAGTTAGCTGGCTA
GAGCACACGTCTGAACTCTTTCCCTACACGACGCTCTTCCGATCTNNNCTGCTGTTAGTTAGCTGGCTA
AGCACACGTCTGAACTCTTTCCCTACACGACGCTCTTCCGATCTNNNCTTTCTCCACAACCTTGCTAGCC
AGCACACGTCTGAACTCTTTCCCTACACGACGCTCTTCCGATCTNNNCTTTCTCCACAACCTTGCTAGCC
AGAGCACACGTCTGAACTCTTTCCCTACACGACGCTCTTCCGATCTNNNGTTGTTAGAGCATAAGCTG
AGAGCACACGTCTGAACTCTTTCCCTACACGACGCTCTTCCGATCTNNNGTTGTTAGAGCATAAGCTG
AGCACACGTCTGAACTCTTTCCCTACACGACGCTCTTCCGATCTNNNTGTGTGTGTGTGTGTTTCAA
AGCACACGTCTGAACTCTTTCCCTACACGACGCTCTTCCGATCTNNNTGTGTGTGTGTGTGTTTCAA
GAGCACACGTCTGAACTCTTTCCCTACACGACGCTCTTCCGATCTNNNGTCCAACCTCTCATCACTACCTG
GAGCACACGTCTGAACTCTTTCCCTACACGACGCTCTTCCGATCTNNNGTCCAACCTCTCATCACTACCTG
GAGCACACGTCTGAACTCTTTCCCTACACGACGCTCTTCCGATCTNNNTCACAGCAGATTTGGAGAGAC
GAGCACACGTCTGAACTCTTTCCCTACACGACGCTCTTCCGATCTNNNTCACAGCAGATTTGGAGAGAC
ACACGTCTGAACTCTTTCCCTACACGACGCTCTTCCGATCTNNNTCCCTGCCCTATCACTACTTATCATCG
ACACGTCTGAACTCTTTCCCTACACGACGCTCTTCCGATCTNNNTCCCTGCCCTATCACTACTTATCATCG
AGAGCACACGTCTGAACTCTTTCCCTACACGACGCTCTTCCGATCTNNNAGCAATGGGGAGCATTTTTT
AGAGCACACGTCTGAACTCTTTCCCTACACGACGCTCTTCCGATCTNNNAGCAATGGGGAGCATTTTTT
AGCACACGTCTGAACTCTTTCCCTACACGACGCTCTTCCGATCTNNNGTGGAAAGGGCTGAGAGAAG
AGCACACGTCTGAACTCTTTCCCTACACGACGCTCTTCCGATCTNNNGTGGAAAGGGCTGAGAGAAG
GAGCACACGTCTGAACTCTTTCCCTACACGACGCTCTTCCGATCTNNNTCCTTCCCAGGATCTACCCAAT
GAGCACACGTCTGAACTCTTTCCCTACACGACGCTCTTCCGATCTNNNTCCTTCCCAGGATCTACCCAAT
AGCACACGTCTGAACTCTTTCCCTACACGACGCTCTTCCGATCTNNNTGTGTGGTAGGTGGAATTCTGA
AGCACACGTCTGAACTCTTTCCCTACACGACGCTCTTCCGATCTNNNTGTGTGGTAGGTGGAATTCTGA
AGCACACGTCTGAACTCTTTCCCTACACGACGCTCTTCCGATCTNNNCCTGACATTTGTGGCAATCAACT
AGCACACGTCTGAACTCTTTCCCTACACGACGCTCTTCCGATCTNNNCCTGACATTTGTGGCAATCAACT
AGCACACGTCTGAACTCTTTCCCTACACGACGCTCTTCCGATCTNNNTAAAGAGTCAAGACCCATCCGT
AGCACACGTCTGAACTCTTTCCCTACACGACGCTCTTCCGATCTNNNTAAAGAGTCAAGACCCATCCGT
AGCACACGTCTGAACTCTTTCCCTACACGACGCTCTTCCGATCTNNNCCTGGCCCCTATAAAGACTCAAG
AGCACACGTCTGAACTCTTTCCCTACACGACGCTCTTCCGATCTNNNCCTGGCCCCTATAAAGACTCAAG
CACACGTCTGAACTCTTTCCCTACACGACGCTCTTCCGATCTNNNGAACACTGTGGCACAAGCTCAGCA
CACACGTCTGAACTCTTTCCCTACACGACGCTCTTCCGATCTNNNGAACACTGTGGCACAAGCTCAGCA
AGCACACGTCTGAACTCTTTCCCTACACGACGCTCTTCCGATCTNNNCTCCTGAAGTGCGGTTATCCTTA
AGCACACGTCTGAACTCTTTCCCTACACGACGCTCTTCCGATCTNNNCTCCTGAAGTGCGGTTATCCTTA
GAGCACACGTCTGAACTCTTTCCCTACACGACGCTCTTCCGATCTNNNGACCTACCCCATCCGAATCTAA

GAGCACACGTCTGAACTCTTTCCCTACACGACGCTCTTCCGATCTNNNGACCTACCCCATCCGAATCTAA

GAGCACACGTCTGAACTCTTTCCCTACACGACGCTCTTCCGATCTNNNATAGACCAAAGGGAAAGCGA

GAGCACACGTCTGAACTCTTTCCCTACACGACGCTCTTCCGATCTNNNATAGACCAAAGGGAAAGCGA

GAGCACACGTCTGAACTCTTTCCCTACACGACGCTCTTCCGATCTNNNATCATGAGGTCAGGAGATCG

GAGCACACGTCTGAACTCTTTCCCTACACGACGCTCTTCCGATCTNNNATCATGAGGTCAGGAGATCG

AGCACACGTCTGAACTCTTTCCCTACACGACGCTCTTCCGATCTNNNCAGATGCCTTTTGTGCTGTCTA

AGCACACGTCTGAACTCTTTCCCTACACGACGCTCTTCCGATCTNNNCAGATGCCTTTTGTGCTGTCTA

AGCACACGTCTGAACTCTTTCCCTACACGACGCTCTTCCGATCTNNNTGCATAGCATGACTGAGTTGTC

AGCACACGTCTGAACTCTTTCCCTACACGACGCTCTTCCGATCTNNNTGCATAGCATGACTGAGTTGTC

GCACACGTCTGAACTCTTTCCCTACACGACGCTCTTCCGATCTNNNCAACTTATCAAGCATGTTTGGCA

GCACACGTCTGAACTCTTTCCCTACACGACGCTCTTCCGATCTNNNCAACTTATCAAGCATGTTTGGCA

GCACACGTCTGAACTCTTTCCCTACACGACGCTCTTCCGATCTNNNCTGGGCAACAAGAGAAAAACTC

GCACACGTCTGAACTCTTTCCCTACACGACGCTCTTCCGATCTNNNCTGGGCAACAAGAGAAAAACTC

AGCACACGTCTGAACTCTTTCCCTACACGACGCTCTTCCGATCTNNNTGTGCCAAGAAATAAAAGCTG

AGCACACGTCTGAACTCTTTCCCTACACGACGCTCTTCCGATCTNNNTGTGCCAAGAAATAAAAGCTG

AGCACACGTCTGAACTCTTTCCCTACACGACGCTCTTCCGATCTNNNGGTCAAACCCAAACAGAAAAC

AGCACACGTCTGAACTCTTTCCCTACACGACGCTCTTCCGATCTNNNGGTCAAACCCAAACAGAAAAC

AGAGCACACGTCTGAACTCTTTCCCTACACGACGCTCTTCCGATCTNNNGGTAGAATATCAGCCTGGA

AGAGCACACGTCTGAACTCTTTCCCTACACGACGCTCTTCCGATCTNNNGGTAGAATATCAGCCTGGA

GAGCACACGTCTGAACTCTTTCCCTACACGACGCTCTTCCGATCTNNNGACACGTAGGCTCAGATTCCT

GAGCACACGTCTGAACTCTTTCCCTACACGACGCTCTTCCGATCTNNNGACACGTAGGCTCAGATTCCT

AGCACACGTCTGAACTCTTTCCCTACACGACGCTCTTCCGATCTNNNGTGTGTGTCTGTCTCTCCATCAC

AGCACACGTCTGAACTCTTTCCCTACACGACGCTCTTCCGATCTNNNGTGTGTGTCTGTCTCTCCATCAC

GAGCACACGTCTGAACTCTTTCCCTACACGACGCTCTTCCGATCTNNNATGCAATCTCAGCTTACTGCAA

GAGCACACGTCTGAACTCTTTCCCTACACGACGCTCTTCCGATCTNNNATGCAATCTCAGCTTACTGCAA

CACACGTCTGAACTCTTTCCCTACACGACGCTCTTCCGATCTNNNCTGGTGCAGACGCTCATAGTGCATC

CACACGTCTGAACTCTTTCCCTACACGACGCTCTTCCGATCTNNNCTGGTGCAGACGCTCATAGTGCATC

AGCACACGTCTGAACTCTTTCCCTACACGACGCTCTTCCGATCTNNNTGTTAAAAATCATGCCACGGTG

AGCACACGTCTGAACTCTTTCCCTACACGACGCTCTTCCGATCTNNNTGTTAAAAATCATGCCACGGTG

GAGCACACGTCTGAACTCTTTCCCTACACGACGCTCTTCCGATCTNNNCTATATCCTTGAGCCAGGAGA

GAGCACACGTCTGAACTCTTTCCCTACACGACGCTCTTCCGATCTNNNCTATATCCTTGAGCCAGGAGA

AGCACACGTCTGAACTCTTTCCCTACACGACGCTCTTCCGATCTNNNTGTAGCAAACGTGTATCTCCCTC

AGCACACGTCTGAACTCTTTCCCTACACGACGCTCTTCCGATCTNNNTGTAGCAAACGTGTATCTCCCTC

AGAGCACACGTCTGAACTCTTTCCCTACACGACGCTCTTCCGATCTNNNGAAGATAAACTAGGGGGAT

AGAGCACACGTCTGAACTCTTTCCCTACACGACGCTCTTCCGATCTNNNGAAGATAAACTAGGGGGAT

AGAGCACACGTCTGAACTCTTTCCCTACACGACGCTCTTCCGATCTNNNGAGGCTGAGGCAGGAGAA

AGAGCACACGTCTGAACTCTTTCCCTACACGACGCTCTTCCGATCTNNNGAGGCTGAGGCAGGAGAA

GAGCACACGTCTGAACTCTTTCCCTACACGACGCTCTTCCGATCTNNNGGAGGGAATTGAGGAGAA

GAGCACACGTCTGAACTCTTTCCCTACACGACGCTCTTCCGATCTNNNGGAGGGAATTGAGGAGAA

GAGCACACGTCTGAACTCTTTCCCTACACGACGCTCTTCCGATCTNNNACTTGAATCTGGGAAGCAGA

GAGCACACGTCTGAACTCTTTCCCTACACGACGCTCTTCCGATCTNNNACTTGAATCTGGGAAGCAGA

GAGCACACGTCTGAACTCTTTCCCTACACGACGCTCTTCCGATCTNNNATAATGACGGCTGTGATCTGC

GAGCACACGTCTGAACTCTTTCCCTACACGACGCTCTTCCGATCTNNNATAATGACGGCTGTGATCTGC

GCACACGTCTGAACTCTTTCCCTACACGACGCTCTTCCGATCTNNNGGAGTCAATCCTCTCAAACCCTG
GCACACGTCTGAACTCTTTCCCTACACGACGCTCTTCCGATCTNNNGGAGTCAATCCTCTCAAACCCTG
AGCACACGTCTGAACTCTTTCCCTACACGACGCTCTTCCGATCTNNNGAGAATTGCTTGAACCGAGG
AGCACACGTCTGAACTCTTTCCCTACACGACGCTCTTCCGATCTNNNGAGAATTGCTTGAACCGAGG
GCACACGTCTGAACTCTTTCCCTACACGACGCTCTTCCGATCTNNNTAGGATTGCTCACTGTTGGGAT
GCACACGTCTGAACTCTTTCCCTACACGACGCTCTTCCGATCTNNNTAGGATTGCTCACTGTTGGGAT
GAGCACACGTCTGAACTCTTTCCCTACACGACGCTCTTCCGATCTNNNTGAGGCACGAGAATTGTTT
GAGCACACGTCTGAACTCTTTCCCTACACGACGCTCTTCCGATCTNNNTGAGGCACGAGAATTGTTT
GAGCACACGTCTGAACTCTTTCCCTACACGACGCTCTTCCGATCTNNNATAATAACCGTCCCCAACA
GAGCACACGTCTGAACTCTTTCCCTACACGACGCTCTTCCGATCTNNNATAATAACCGTCCCCAACA
AGCACACGTCTGAACTCTTTCCCTACACGACGCTCTTCCGATCTNNNCTAAGAGGGGAGATGACATG
AGCACACGTCTGAACTCTTTCCCTACACGACGCTCTTCCGATCTNNNCTAAGAGGGGAGATGACATG
AGCACACGTCTGAACTCTTTCCCTACACGACGCTCTTCCGATCTNNNATGTGTGTATATGTGTGTGT
AGCACACGTCTGAACTCTTTCCCTACACGACGCTCTTCCGATCTNNNATGTGTGTATATGTGTGTGT
GCACACGTCTGAACTCTTTCCCTACACGACGCTCTTCCGATCTNNNCTGTTTGCAGTAGAAAGAGAC
GCACACGTCTGAACTCTTTCCCTACACGACGCTCTTCCGATCTNNNCTGTTTGCAGTAGAAAGAGAC
CACACGTCTGAACTCTTTCCCTACACGACGCTCTTCCGATCTNNNTAGAACCCCCTTGAGAAAACTGG
CACACGTCTGAACTCTTTCCCTACACGACGCTCTTCCGATCTNNNTAGAACCCCCTTGAGAAAACTGG
AGCACACGTCTGAACTCTTTCCCTACACGACGCTCTTCCGATCTNNNCTTCACATTCTGGGCTTGTTTG
AGCACACGTCTGAACTCTTTCCCTACACGACGCTCTTCCGATCTNNNCTTCACATTCTGGGCTTGTTTG
GAGCACACGTCTGAACTCTTTCCCTACACGACGCTCTTCCGATCTNNNCTCTGCTTGATGCTGCCTCTA
GAGCACACGTCTGAACTCTTTCCCTACACGACGCTCTTCCGATCTNNNCTCTGCTTGATGCTGCCTCTA
AGCACACGTCTGAACTCTTTCCCTACACGACGCTCTTCCGATCTNNNAAAGTGATCCTCCCTGTCTTAG
AGCACACGTCTGAACTCTTTCCCTACACGACGCTCTTCCGATCTNNNAAAGTGATCCTCCCTGTCTTAG
AGCACACGTCTGAACTCTTTCCCTACACGACGCTCTTCCGATCTNNNGAAAGGCAAGAGGACTGTCG
AGCACACGTCTGAACTCTTTCCCTACACGACGCTCTTCCGATCTNNNGAAAGGCAAGAGGACTGTCG
AGCACACGTCTGAACTCTTTCCCTACACGACGCTCTTCCGATCTNNNTATCTAACAGAGGTCCTTGGG
AGCACACGTCTGAACTCTTTCCCTACACGACGCTCTTCCGATCTNNNTATCTAACAGAGGTCCTTGGG
AGCACACGTCTGAACTCTTTCCCTACACGACGCTCTTCCGATCTNNNCAAAAGCCAAGGGTATGGGA
AGCACACGTCTGAACTCTTTCCCTACACGACGCTCTTCCGATCTNNNCAAAAGCCAAGGGTATGGGA
GCACACGTCTGAACTCTTTCCCTACACGACGCTCTTCCGATCTNNNACAGAGAGTTCTCATGTTTCCC
GCACACGTCTGAACTCTTTCCCTACACGACGCTCTTCCGATCTNNNACAGAGAGTTCTCATGTTTCCC
AGCACACGTCTGAACTCTTTCCCTACACGACGCTCTTCCGATCTNNNCTGAGGCAGGAGAATGACAT
AGCACACGTCTGAACTCTTTCCCTACACGACGCTCTTCCGATCTNNNCTGAGGCAGGAGAATGACAT
GCACACGTCTGAACTCTTTCCCTACACGACGCTCTTCCGATCTNNNCCCTAATCCAATATGGCTGGTG
GCACACGTCTGAACTCTTTCCCTACACGACGCTCTTCCGATCTNNNCCCTAATCCAATATGGCTGGTG
ACACGTCTGAACTCTTTCCCTACACGACGCTCTTCCGATCTNNNCTCTCAGCTGGTCTCATCTTCTCCAT
ACACGTCTGAACTCTTTCCCTACACGACGCTCTTCCGATCTNNNCTCTCAGCTGGTCTCATCTTCTCCAT
AGCACACGTCTGAACTCTTTCCCTACACGACGCTCTTCCGATCTNNNCTGTAGTCCCAGCTACTCGGG
AGCACACGTCTGAACTCTTTCCCTACACGACGCTCTTCCGATCTNNNCTGTAGTCCCAGCTACTCGGG
AGCACACGTCTGAACTCTTTCCCTACACGACGCTCTTCCGATCTNNNTGGGCAATAAAGTGACACTCT
AGCACACGTCTGAACTCTTTCCCTACACGACGCTCTTCCGATCTNNNTGGGCAATAAAGTGACACTCT
AGCACACGTCTGAACTCTTTCCCTACACGACGCTCTTCCGATCTNNNCAAAATCACACCACTGCACTCC

AGCACACGTCTGAACTCTTTCCCTACACGACGCTCTTCCGATCTNNNCAAAATCACACCACTGCACTCC
GCACACGTCTGAACTCTTTCCCTACACGACGCTCTTCCGATCTNNNAATTTGTGTGTGTGTGTGTGTG
GCACACGTCTGAACTCTTTCCCTACACGACGCTCTTCCGATCTNNNAATTTGTGTGTGTGTGTGTGTG
CACACGTCTGAACTCTTTCCCTACACGACGCTCTTCCGATCTNNNCTGCTATCTGTGGGTTTGGTTTG
CACACGTCTGAACTCTTTCCCTACACGACGCTCTTCCGATCTNNNCTGCTATCTGTGGGTTTGGTTTG
AGCACACGTCTGAACTCTTTCCCTACACGACGCTCTTCCGATCTNNNCTGAGGATAAGGATGACTTAA
AGCACACGTCTGAACTCTTTCCCTACACGACGCTCTTCCGATCTNNNCTGAGGATAAGGATGACTTAA
AGCACACGTCTGAACTCTTTCCCTACACGACGCTCTTCCGATCTNNNGCTTTTGGGTTCTTGGTCATG
AGCACACGTCTGAACTCTTTCCCTACACGACGCTCTTCCGATCTNNNGCTTTTGGGTTCTTGGTCATG
AGCACACGTCTGAACTCTTTCCCTACACGACGCTCTTCCGATCTNNNCTCTTCTTTTGACTCTTTTTCTT
AGCACACGTCTGAACTCTTTCCCTACACGACGCTCTTCCGATCTNNNCTCTTCTTTTGACTCTTTTTCTT
AGCACACGTCTGAACTCTTTCCCTACACGACGCTCTTCCGATCTNNNGCAAGAGAGTGCATTTCTGTT
AGCACACGTCTGAACTCTTTCCCTACACGACGCTCTTCCGATCTNNNGCAAGAGAGTGCATTTCTGTT
GAGCACACGTCTGAACTCTTTCCCTACACGACGCTCTTCCGATCTNNNATAGGTGGCTGTTCAAGTGT
GAGCACACGTCTGAACTCTTTCCCTACACGACGCTCTTCCGATCTNNNATAGGTGGCTGTTCAAGTGT
GAGCACACGTCTGAACTCTTTCCCTACACGACGCTCTTCCGATCTNNNCACCAAGTGGAAAGATACCT
GAGCACACGTCTGAACTCTTTCCCTACACGACGCTCTTCCGATCTNNNCACCAAGTGGAAAGATACCT
AGCACACGTCTGAACTCTTTCCCTACACGACGCTCTTCCGATCTNNNGAGCTTGCAGTGAACCAAGAT
AGCACACGTCTGAACTCTTTCCCTACACGACGCTCTTCCGATCTNNNGAGCTTGCAGTGAACCAAGAT
AGCACACGTCTGAACTCTTTCCCTACACGACGCTCTTCCGATCTNNNCTCAACCCCTGTAGTGGATTG
AGCACACGTCTGAACTCTTTCCCTACACGACGCTCTTCCGATCTNNNCTCAACCCCTGTAGTGGATTG
AGCACACGTCTGAACTCTTTCCCTACACGACGCTCTTCCGATCTNNNAAACTTTCCCCATCATTCACAGT
AGCACACGTCTGAACTCTTTCCCTACACGACGCTCTTCCGATCTNNNAAACTTTCCCCATCATTCACAGT
AGCACACGTCTGAACTCTTTCCCTACACGACGCTCTTCCGATCTNNNGGAGTTTTCAATAAGCTCTTTT
AGCACACGTCTGAACTCTTTCCCTACACGACGCTCTTCCGATCTNNNGGAGTTTTCAATAAGCTCTTTT
AGCACACGTCTGAACTCTTTCCCTACACGACGCTCTTCCGATCTNNNCACTAGATGTGGGTTCTGTGG
AGCACACGTCTGAACTCTTTCCCTACACGACGCTCTTCCGATCTNNNCACTAGATGTGGGTTCTGTGG
GAGCACACGTCTGAACTCTTTCCCTACACGACGCTCTTCCGATCTNNNCATCTTCAGTACAGGTGCAG
GAGCACACGTCTGAACTCTTTCCCTACACGACGCTCTTCCGATCTNNNCATCTTCAGTACAGGTGCAG
AGCACACGTCTGAACTCTTTCCCTACACGACGCTCTTCCGATCTNNNTCGAACTCCTGGGTTTGAATG
AGCACACGTCTGAACTCTTTCCCTACACGACGCTCTTCCGATCTNNNTCGAACTCCTGGGTTTGAATG
AGCACACGTCTGAACTCTTTCCCTACACGACGCTCTTCCGATCTNNNGGACCTCCAACAATCACAAAT
AGCACACGTCTGAACTCTTTCCCTACACGACGCTCTTCCGATCTNNNGGACCTCCAACAATCACAAAT
AGCACACGTCTGAACTCTTTCCCTACACGACGCTCTTCCGATCTNNNCTGTAGTCCCAGCTACTCAGG
AGCACACGTCTGAACTCTTTCCCTACACGACGCTCTTCCGATCTNNNCTGTAGTCCCAGCTACTCAGG
GCACACGTCTGAACTCTTTCCCTACACGACGCTCTTCCGATCTNNNCGACATGCAGAATTTCTTGAGCT
GCACACGTCTGAACTCTTTCCCTACACGACGCTCTTCCGATCTNNNCGACATGCAGAATTTCTTGAGCT
AGCACACGTCTGAACTCTTTCCCTACACGACGCTCTTCCGATCTNNNGCCTTGCAGGTTTACAGTTTTC
AGCACACGTCTGAACTCTTTCCCTACACGACGCTCTTCCGATCTNNNGCCTTGCAGGTTTACAGTTTTC
AGCACACGTCTGAACTCTTTCCCTACACGACGCTCTTCCGATCTNNNCAAAGTGCTGGGATTACAGG
AGCACACGTCTGAACTCTTTCCCTACACGACGCTCTTCCGATCTNNNCAAAGTGCTGGGATTACAGG
AGCACACGTCTGAACTCTTTCCCTACACGACGCTCTTCCGATCTNNNATAGTTTGAAAAAGCCCCTCAG
AGCACACGTCTGAACTCTTTCCCTACACGACGCTCTTCCGATCTNNNATAGTTTGAAAAAGCCCCTCAG

AGAGCACACGTCTGAACTCTTTCCCTACACGACGCTCTTCCGATCTNNNGATTGCGCCACTGCACTCTA
AGAGCACACGTCTGAACTCTTTCCCTACACGACGCTCTTCCGATCTNNNGATTGCGCCACTGCACTCTA
GAGCACACGTCTGAACTCTTTCCCTACACGACGCTCTTCCGATCTNNNAATGATTGGCTGTAGTCCTGT
GAGCACACGTCTGAACTCTTTCCCTACACGACGCTCTTCCGATCTNNNAATGATTGGCTGTAGTCCTGT
GAGCACACGTCTGAACTCTTTCCCTACACGACGCTCTTCCGATCTNNNGAGATAGTGTGTGTGTGCAT
GAGCACACGTCTGAACTCTTTCCCTACACGACGCTCTTCCGATCTNNNGAGATAGTGTGTGTGTGCAT
AGAGCACACGTCTGAACTCTTTCCCTACACGACGCTCTTCCGATCTNNNTAAAGTGGCTATTTGCACA
AGAGCACACGTCTGAACTCTTTCCCTACACGACGCTCTTCCGATCTNNNTAAAGTGGCTATTTGCACA
GAGCACACGTCTGAACTCTTTCCCTACACGACGCTCTTCCGATCTNNNCAGCAGTTGCAGTCATTCAAT
GAGCACACGTCTGAACTCTTTCCCTACACGACGCTCTTCCGATCTNNNCAGCAGTTGCAGTCATTCAAT
AGCACACGTCTGAACTCTTTCCCTACACGACGCTCTTCCGATCTNNNAAAGGTTGCAGTGAGCCAAGAT
AGCACACGTCTGAACTCTTTCCCTACACGACGCTCTTCCGATCTNNNAAAGGTTGCAGTGAGCCAAGAT
GAGCACACGTCTGAACTCTTTCCCTACACGACGCTCTTCCGATCTNNNCTCGCTTTGTACCGTATGGAG
GAGCACACGTCTGAACTCTTTCCCTACACGACGCTCTTCCGATCTNNNCTCGCTTTGTACCGTATGGAG
GAGCACACGTCTGAACTCTTTCCCTACACGACGCTCTTCCGATCTNNNGGACCATTTTAGGGCTAGACA
GAGCACACGTCTGAACTCTTTCCCTACACGACGCTCTTCCGATCTNNNGGACCATTTTAGGGCTAGACA
GAGCACACGTCTGAACTCTTTCCCTACACGACGCTCTTCCGATCTNNNAAGGGAGTTTTATACACAGG
GAGCACACGTCTGAACTCTTTCCCTACACGACGCTCTTCCGATCTNNNAAGGGAGTTTTATACACAGG
GAGCACACGTCTGAACTCTTTCCCTACACGACGCTCTTCCGATCTNNNCCAAGATCAGGCCACTATACT
GAGCACACGTCTGAACTCTTTCCCTACACGACGCTCTTCCGATCTNNNCCAAGATCAGGCCACTATACT
AGCACACGTCTGAACTCTTTCCCTACACGACGCTCTTCCGATCTNNNAGGCCCTGTTTACTTGGTGATT
AGCACACGTCTGAACTCTTTCCCTACACGACGCTCTTCCGATCTNNNAGGCCCTGTTTACTTGGTGATT
GAGCACACGTCTGAACTCTTTCCCTACACGACGCTCTTCCGATCTNNNCCACCCGGGTCAACTATTTTTA
GAGCACACGTCTGAACTCTTTCCCTACACGACGCTCTTCCGATCTNNNCCACCCGGGTCAACTATTTTTA
GAGCACACGTCTGAACTCTTTCCCTACACGACGCTCTTCCGATCTNNNAAAAAATTAGCCAGGTGTGG
GAGCACACGTCTGAACTCTTTCCCTACACGACGCTCTTCCGATCTNNNAAAAAATTAGCCAGGTGTGG
AGCACACGTCTGAACTCTTTCCCTACACGACGCTCTTCCGATCTNNNACGTGTATTCAACACCATAGTC
AGCACACGTCTGAACTCTTTCCCTACACGACGCTCTTCCGATCTNNNACGTGTATTCAACACCATAGTC
CACACGTCTGAACTCTTTCCCTACACGACGCTCTTCCGATCTNNNCCTCAGTTATTTGATGGGAGGCT
CACACGTCTGAACTCTTTCCCTACACGACGCTCTTCCGATCTNNNCCTCAGTTATTTGATGGGAGGCT
GCACACGTCTGAACTCTTTCCCTACACGACGCTCTTCCGATCTNNNCACAAGACATTTTTCTACAGTATC
GCACACGTCTGAACTCTTTCCCTACACGACGCTCTTCCGATCTNNNCACAAGACATTTTTCTACAGTATC
GAGCACACGTCTGAACTCTTTCCCTACACGACGCTCTTCCGATCTNNNATTACAAGCATGAGCCATGTG
GAGCACACGTCTGAACTCTTTCCCTACACGACGCTCTTCCGATCTNNNATTACAAGCATGAGCCATGTG
AGAGCACACGTCTGAACTCTTTCCCTACACGACGCTCTTCCGATCTNNNGTCTGTGTGATTTTAATTCA
AGAGCACACGTCTGAACTCTTTCCCTACACGACGCTCTTCCGATCTNNNGTCTGTGTGATTTTAATTCA
GCACACGTCTGAACTCTTTCCCTACACGACGCTCTTCCGATCTNNNTTTCTGTCCTGCTAGCTTTGGATC
GCACACGTCTGAACTCTTTCCCTACACGACGCTCTTCCGATCTNNNTTTCTGTCCTGCTAGCTTTGGATC
AGAGCACACGTCTGAACTCTTTCCCTACACGACGCTCTTCCGATCTNNNGAAGAATGGTTAGTGGTAG
AGAGCACACGTCTGAACTCTTTCCCTACACGACGCTCTTCCGATCTNNNGAAGAATGGTTAGTGGTAG
AGCACACGTCTGAACTCTTTCCCTACACGACGCTCTTCCGATCTNNNATGCCTGTAATACCAGCTACTCG
AGCACACGTCTGAACTCTTTCCCTACACGACGCTCTTCCGATCTNNNATGCCTGTAATACCAGCTACTCG
AGAGCACACGTCTGAACTCTTTCCCTACACGACGCTCTTCCGATCTNNNGAGGCAGGAGAATCGCTTG

GAGCACACGTCTGAACTCTTTCCCTACACGACGCTCTTCCGATCTNNNGAGGCAGGAGAATCGCTTG

AGCACACGTCTGAACTCTTTCCCTACACGACGCTCTTCCGATCTNNNTGTCTATGAGAAGGTCACCCTC

AGCACACGTCTGAACTCTTTCCCTACACGACGCTCTTCCGATCTNNNTGTCTATGAGAAGGTCACCCTC

AGCACACGTCTGAACTCTTTCCCTACACGACGCTCTTCCGATCTNNNAAATGGGAGGATCACTTGAGT

AGCACACGTCTGAACTCTTTCCCTACACGACGCTCTTCCGATCTNNNAAATGGGAGGATCACTTGAGT

AGCACACGTCTGAACTCTTTCCCTACACGACGCTCTTCCGATCTNNNACCAGCTTTTCCCCCTTTTCATAC

AGCACACGTCTGAACTCTTTCCCTACACGACGCTCTTCCGATCTNNNACCAGCTTTTCCCCCTTTTCATAC

GAGCACACGTCTGAACTCTTTCCCTACACGACGCTCTTCCGATCTNNNACTGATGGGAAATGCACAAG

GAGCACACGTCTGAACTCTTTCCCTACACGACGCTCTTCCGATCTNNNACTGATGGGAAATGCACAAG

GAGCACACGTCTGAACTCTTTCCCTACACGACGCTCTTCCGATCTNNNTGACTTGCAGAAGGCTTCCTA

GAGCACACGTCTGAACTCTTTCCCTACACGACGCTCTTCCGATCTNNNTGACTTGCAGAAGGCTTCCTA

AGCACACGTCTGAACTCTTTCCCTACACGACGCTCTTCCGATCTNNNAAATTCATCTTAAATCAGGAA

AGCACACGTCTGAACTCTTTCCCTACACGACGCTCTTCCGATCTNNNAAATTCATCTTAAATCAGGAA

GCACACGTCTGAACTCTTTCCCTACACGACGCTCTTCCGATCTNNNCATCAATTCTCTTGGGGGTCCTG

GCACACGTCTGAACTCTTTCCCTACACGACGCTCTTCCGATCTNNNCATCAATTCTCTTGGGGGTCCTG

AGCACACGTCTGAACTCTTTCCCTACACGACGCTCTTCCGATCTNNNGGTAGGAACCGTAAAATGTTT

AGCACACGTCTGAACTCTTTCCCTACACGACGCTCTTCCGATCTNNNGGTAGGAACCGTAAAATGTTT

GAGCACACGTCTGAACTCTTTCCCTACACGACGCTCTTCCGATCTNNNTCCTCTCTTCTTTCCTTTCTTTCT

GAGCACACGTCTGAACTCTTTCCCTACACGACGCTCTTCCGATCTNNNTCCTCTCTTCTTTCCTTTCTTTCT

GAGCACACGTCTGAACTCTTTCCCTACACGACGCTCTTCCGATCTNNNCTGAGTCAGGAGAATCGCTTG

GAGCACACGTCTGAACTCTTTCCCTACACGACGCTCTTCCGATCTNNNCTGAGTCAGGAGAATCGCTTG

GAGCACACGTCTGAACTCTTTCCCTACACGACGCTCTTCCGATCTNNNTGAGCTGAGATCGCACCATTG

GAGCACACGTCTGAACTCTTTCCCTACACGACGCTCTTCCGATCTNNNTGAGCTGAGATCGCACCATTG

AGCACACGTCTGAACTCTTTCCCTACACGACGCTCTTCCGATCTNNNGTCCACTGTTACAGCTCAGAG

AGCACACGTCTGAACTCTTTCCCTACACGACGCTCTTCCGATCTNNNGTCCACTGTTACAGCTCAGAG

AGCACACGTCTGAACTCTTTCCCTACACGACGCTCTTCCGATCTNNNACCTGTAGTCCCAGCTACTAGG

AGCACACGTCTGAACTCTTTCCCTACACGACGCTCTTCCGATCTNNNACCTGTAGTCCCAGCTACTAGG

AGCACACGTCTGAACTCTTTCCCTACACGACGCTCTTCCGATCTNNNTCCTCTTCCTAATAAGCCTGGGA

AGCACACGTCTGAACTCTTTCCCTACACGACGCTCTTCCGATCTNNNTCCTCTTCCTAATAAGCCTGGGA

GAGCACACGTCTGAACTCTTTCCCTACACGACGCTCTTCCGATCTNNNGTTCTCCAAGTTCTTCCCTTTG

GAGCACACGTCTGAACTCTTTCCCTACACGACGCTCTTCCGATCTNNNGTTCTCCAAGTTCTTCCCTTTG

CACGTCTGAACTCTTTCCCTACACGACGCTCTTCCGATCTNNNCCCTTTATTCAAGCCCACCTTTCCATC

CACGTCTGAACTCTTTCCCTACACGACGCTCTTCCGATCTNNNCCCTTTATTCAAGCCCACCTTTCCATC

AGCACACGTCTGAACTCTTTCCCTACACGACGCTCTTCCGATCTNNNTTCAAGTGATTCTCCTGCCTCA

AGCACACGTCTGAACTCTTTCCCTACACGACGCTCTTCCGATCTNNNTTCAAGTGATTCTCCTGCCTCA

AGAGCACACGTCTGAACTCTTTCCCTACACGACGCTCTTCCGATCTNNNCCTGCAAACATCAGATTAGG

AGAGCACACGTCTGAACTCTTTCCCTACACGACGCTCTTCCGATCTNNNCCTGCAAACATCAGATTAGG

GAGCACACGTCTGAACTCTTTCCCTACACGACGCTCTTCCGATCTNNNTGCCTGTTTGCTCTGATGATA

GAGCACACGTCTGAACTCTTTCCCTACACGACGCTCTTCCGATCTNNNTGCCTGTTTGCTCTGATGATA

AGCACACGTCTGAACTCTTTCCCTACACGACGCTCTTCCGATCTNNNCTGAGCGACCTTGTTGAGATTT

AGCACACGTCTGAACTCTTTCCCTACACGACGCTCTTCCGATCTNNNCTGAGCGACCTTGTTGAGATTT

AGCACACGTCTGAACTCTTTCCCTACACGACGCTCTTCCGATCTNNNGCACTGGGTTCAGAATTTCCTC

AGCACACGTCTGAACTCTTTCCCTACACGACGCTCTTCCGATCTNNNGCACTGGGTTCAGAATTTCCTC

GCACACGTCTGAACTCTTTCCCTACACGACGCTCTTCCGATCTNNNGTGAGTAGTGCTGCAGTTCTTT
GCACACGTCTGAACTCTTTCCCTACACGACGCTCTTCCGATCTNNNGTGAGTAGTGCTGCAGTTCTTT
AGCACACGTCTGAACTCTTTCCCTACACGACGCTCTTCCGATCTNNNTCTGAAACTAAGGAGGCAATG
AGCACACGTCTGAACTCTTTCCCTACACGACGCTCTTCCGATCTNNNTCTGAAACTAAGGAGGCAATG
GAGCACACGTCTGAACTCTTTCCCTACACGACGCTCTTCCGATCTNNNGTTTCCTACCCTTTGAGTCTG
GAGCACACGTCTGAACTCTTTCCCTACACGACGCTCTTCCGATCTNNNGTTTCCTACCCTTTGAGTCTG
AGAGCACACGTCTGAACTCTTTCCCTACACGACGCTCTTCCGATCTNNNCTGTAATCCCAGCTACTCAG
AGAGCACACGTCTGAACTCTTTCCCTACACGACGCTCTTCCGATCTNNNCTGTAATCCCAGCTACTCAG
CACACGTCTGAACTCTTTCCCTACACGACGCTCTTCCGATCTNNNCTGGAGACGAAGAAGCTAGAAG
CACACGTCTGAACTCTTTCCCTACACGACGCTCTTCCGATCTNNNCTGGAGACGAAGAAGCTAGAAG
GAGCACACGTCTGAACTCTTTCCCTACACGACGCTCTTCCGATCTNNNAGGCTGACTAACATGGTGAA
GAGCACACGTCTGAACTCTTTCCCTACACGACGCTCTTCCGATCTNNNAGGCTGACTAACATGGTGAA
AGCACACGTCTGAACTCTTTCCCTACACGACGCTCTTCCGATCTNNNTGCGGTTTTTGTCACTACTTTC
AGCACACGTCTGAACTCTTTCCCTACACGACGCTCTTCCGATCTNNNTGCGGTTTTTGTCACTACTTTC
AGCACACGTCTGAACTCTTTCCCTACACGACGCTCTTCCGATCTNNNTCTAGTACTGGACGATTTGTG
AGCACACGTCTGAACTCTTTCCCTACACGACGCTCTTCCGATCTNNNTCTAGTACTGGACGATTTGTG
GAGCACACGTCTGAACTCTTTCCCTACACGACGCTCTTCCGATCTNNNGCACATGCCTATAATCCCAGC
GAGCACACGTCTGAACTCTTTCCCTACACGACGCTCTTCCGATCTNNNGCACATGCCTATAATCCCAGC
GCACACGTCTGAACTCTTTCCCTACACGACGCTCTTCCGATCTNNNCTTGCCTGGCTCATAATATGCAC
GCACACGTCTGAACTCTTTCCCTACACGACGCTCTTCCGATCTNNNCTTGCCTGGCTCATAATATGCAC
CACACGTCTGAACTCTTTCCCTACACGACGCTCTTCCGATCTNNNGGAGTCTTGTTCTATAGCCCAGGC
CACACGTCTGAACTCTTTCCCTACACGACGCTCTTCCGATCTNNNGGAGTCTTGTTCTATAGCCCAGGC
AGCACACGTCTGAACTCTTTCCCTACACGACGCTCTTCCGATCTNNNCCCAAATGATGAAGTGATGCA
AGCACACGTCTGAACTCTTTCCCTACACGACGCTCTTCCGATCTNNNCCCAAATGATGAAGTGATGCA
GCACACGTCTGAACTCTTTCCCTACACGACGCTCTTCCGATCTNNNTTCCATGGTGTATATGTGCCACA
GCACACGTCTGAACTCTTTCCCTACACGACGCTCTTCCGATCTNNNTTCCATGGTGTATATGTGCCACA
GCACACGTCTGAACTCTTTCCCTACACGACGCTCTTCCGATCTNNNCACCACACCCAGCTAATTTTTGTC
GCACACGTCTGAACTCTTTCCCTACACGACGCTCTTCCGATCTNNNCACCACACCCAGCTAATTTTTGTC
AGCACACGTCTGAACTCTTTCCCTACACGACGCTCTTCCGATCTNNNGGTGAGACTCCGCCTCTAAAAA
AGCACACGTCTGAACTCTTTCCCTACACGACGCTCTTCCGATCTNNNGGTGAGACTCCGCCTCTAAAAA
GCACACGTCTGAACTCTTTCCCTACACGACGCTCTTCCGATCTNNNGAGGCAGGAGAATCGCTTGAA
GCACACGTCTGAACTCTTTCCCTACACGACGCTCTTCCGATCTNNNGAGGCAGGAGAATCGCTTGAA
AGCACACGTCTGAACTCTTTCCCTACACGACGCTCTTCCGATCTNNNGAAAATCGCTTGAATCCAGGA
AGCACACGTCTGAACTCTTTCCCTACACGACGCTCTTCCGATCTNNNGAAAATCGCTTGAATCCAGGA
GAGCACACGTCTGAACTCTTTCCCTACACGACGCTCTTCCGATCTNNNACATTGACCTCTCTCTTTCCCC
GAGCACACGTCTGAACTCTTTCCCTACACGACGCTCTTCCGATCTNNNACATTGACCTCTCTCTTTCCCC
GAGCACACGTCTGAACTCTTTCCCTACACGACGCTCTTCCGATCTNNNGATTGCATGAGCTCAGGAGT
GAGCACACGTCTGAACTCTTTCCCTACACGACGCTCTTCCGATCTNNNGATTGCATGAGCTCAGGAGT
GCACACGTCTGAACTCTTTCCCTACACGACGCTCTTCCGATCTNNNATTTGTTCTTTTTTGCACCCTGCC
GCACACGTCTGAACTCTTTCCCTACACGACGCTCTTCCGATCTNNNATTTGTTCTTTTTTGCACCCTGCC
GAGCACACGTCTGAACTCTTTCCCTACACGACGCTCTTCCGATCTNNNGCAAGAGAATGGCATGAAC
GAGCACACGTCTGAACTCTTTCCCTACACGACGCTCTTCCGATCTNNNGCAAGAGAATGGCATGAAC
GCACACGTCTGAACTCTTTCCCTACACGACGCTCTTCCGATCTNNNCTGAACAAACAGGCTTTGCTACA

AGCACACGTCTGAACTCTTTCCCTACACGACGCTCTTCCGATCTNNNCTGAACAAACAGGCTTTGCTACA
GAGCACACGTCTGAACTCTTTCCCTACACGACGCTCTTCCGATCTNNNGCTTCCCTTTCTACCATGACTG
GAGCACACGTCTGAACTCTTTCCCTACACGACGCTCTTCCGATCTNNNGCTTCCCTTTCTACCATGACTG
GAGCACACGTCTGAACTCTTTCCCTACACGACGCTCTTCCGATCTNNNGTAGAGAGCATGGCGACTAA
GAGCACACGTCTGAACTCTTTCCCTACACGACGCTCTTCCGATCTNNNGTAGAGAGCATGGCGACTAA
GAGCACACGTCTGAACTCTTTCCCTACACGACGCTCTTCCGATCTNNNAGAAAAATTGCTTGAACCGGG
GAGCACACGTCTGAACTCTTTCCCTACACGACGCTCTTCCGATCTNNNAGAAAAATTGCTTGAACCGGG
AGAGCACACGTCTGAACTCTTTCCCTACACGACGCTCTTCCGATCTNNNGCACTATGCTGAGACACATG
AGAGCACACGTCTGAACTCTTTCCCTACACGACGCTCTTCCGATCTNNNGCACTATGCTGAGACACATG
GAGCACACGTCTGAACTCTTTCCCTACACGACGCTCTTCCGATCTNNNGAGATTGCAGTGAGCCAAGAT
GAGCACACGTCTGAACTCTTTCCCTACACGACGCTCTTCCGATCTNNNGAGATTGCAGTGAGCCAAGAT
AGCACACGTCTGAACTCTTTCCCTACACGACGCTCTTCCGATCTNNNAAAACTTACTTACTAGGCACTAT
AGCACACGTCTGAACTCTTTCCCTACACGACGCTCTTCCGATCTNNNAAAACTTACTTACTAGGCACTAT
AGCACACGTCTGAACTCTTTCCCTACACGACGCTCTTCCGATCTNNNTATCTCCCGATGCTATCTCTCCC
AGCACACGTCTGAACTCTTTCCCTACACGACGCTCTTCCGATCTNNNTATCTCCCGATGCTATCTCTCCC
GAGCACACGTCTGAACTCTTTCCCTACACGACGCTCTTCCGATCTNNNAAAGGATAGTCTGGTCTACGG
GAGCACACGTCTGAACTCTTTCCCTACACGACGCTCTTCCGATCTNNNAAAGGATAGTCTGGTCTACGG
AGCACACGTCTGAACTCTTTCCCTACACGACGCTCTTCCGATCTNNNTCTAAAGTGCAGTGGCGTAATC
AGCACACGTCTGAACTCTTTCCCTACACGACGCTCTTCCGATCTNNNTCTAAAGTGCAGTGGCGTAATC
AGCACACGTCTGAACTCTTTCCCTACACGACGCTCTTCCGATCTNNNGGAAATCCCAGGATTATCCTCC
AGCACACGTCTGAACTCTTTCCCTACACGACGCTCTTCCGATCTNNNGGAAATCCCAGGATTATCCTCC
AGAGCACACGTCTGAACTCTTTCCCTACACGACGCTCTTCCGATCTNNNCTGTAATCTCAGCTACTCGGG
AGAGCACACGTCTGAACTCTTTCCCTACACGACGCTCTTCCGATCTNNNCTGTAATCTCAGCTACTCGGG
GAGCACACGTCTGAACTCTTTCCCTACACGACGCTCTTCCGATCTNNNACCTCAGTATCATGAGCAGTT
GAGCACACGTCTGAACTCTTTCCCTACACGACGCTCTTCCGATCTNNNACCTCAGTATCATGAGCAGTT
ACACGTCTGAACTCTTTCCCTACACGACGCTCTTCCGATCTNNNTCAACATCCACCCAGTTACTCAGCATC
ACACGTCTGAACTCTTTCCCTACACGACGCTCTTCCGATCTNNNTCAACATCCACCCAGTTACTCAGCATC
GAGCACACGTCTGAACTCTTTCCCTACACGACGCTCTTCCGATCTNNNTAATCTCAGCACTTTAGGAGG
GAGCACACGTCTGAACTCTTTCCCTACACGACGCTCTTCCGATCTNNNTAATCTCAGCACTTTAGGAGG
AGAGCACACGTCTGAACTCTTTCCCTACACGACGCTCTTCCGATCTNNNAGGTCATGCCAATAGACACA
AGAGCACACGTCTGAACTCTTTCCCTACACGACGCTCTTCCGATCTNNNAGGTCATGCCAATAGACACA
AGCACACGTCTGAACTCTTTCCCTACACGACGCTCTTCCGATCTNNNAGAGAGGTAGGGTCTCAGCTTA
AGCACACGTCTGAACTCTTTCCCTACACGACGCTCTTCCGATCTNNNAGAGAGGTAGGGTCTCAGCTTA
AGAGCACACGTCTGAACTCTTTCCCTACACGACGCTCTTCCGATCTNNNCAGTCCCAATGAACAAACAT
AGAGCACACGTCTGAACTCTTTCCCTACACGACGCTCTTCCGATCTNNNCAGTCCCAATGAACAAACAT
AGAGCACACGTCTGAACTCTTTCCCTACACGACGCTCTTCCGATCTNNNGACATCTACAGAACTCTCCAC
AGAGCACACGTCTGAACTCTTTCCCTACACGACGCTCTTCCGATCTNNNGACATCTACAGAACTCTCCAC
AGCACACGTCTGAACTCTTTCCCTACACGACGCTCTTCCGATCTNNNGAGTGACAGAGTGAGACTCCA
AGCACACGTCTGAACTCTTTCCCTACACGACGCTCTTCCGATCTNNNGAGTGACAGAGTGAGACTCCA
AGCACACGTCTGAACTCTTTCCCTACACGACGCTCTTCCGATCTNNNCTCAAGAGTCCTACCCTCTGGTA
AGCACACGTCTGAACTCTTTCCCTACACGACGCTCTTCCGATCTNNNCTCAAGAGTCCTACCCTCTGGTA
AGCACACGTCTGAACTCTTTCCCTACACGACGCTCTTCCGATCTNNNGAGACTTTTTCAAGGCAGGCAA
AGCACACGTCTGAACTCTTTCCCTACACGACGCTCTTCCGATCTNNNGAGACTTTTTCAAGGCAGGCAA

AGAGCACACGTCTGAACTCTTTCCCTACACGACGCTCTTCCGATCTNNNACCCCATCTCTACTAAACACAA

AGAGCACACGTCTGAACTCTTTCCCTACACGACGCTCTTCCGATCTNNNACCCCATCTCTACTAAACACAA

AGAGCACACGTCTGAACTCTTTCCCTACACGACGCTCTTCCGATCTNNNCTGCACAGAACTGAACTTCAA

AGAGCACACGTCTGAACTCTTTCCCTACACGACGCTCTTCCGATCTNNNCTGCACAGAACTGAACTTCAA

AGCACACGTCTGAACTCTTTCCCTACACGACGCTCTTCCGATCTNNNGATTGCAGTGAGCTGAGATTG

AGCACACGTCTGAACTCTTTCCCTACACGACGCTCTTCCGATCTNNNGATTGCAGTGAGCTGAGATTG

GCACACGTCTGAACTCTTTCCCTACACGACGCTCTTCCGATCTNNNTGAGTTTCTTTGTGCCAGCTTTCC

GCACACGTCTGAACTCTTTCCCTACACGACGCTCTTCCGATCTNNNTGAGTTTCTTTGTGCCAGCTTTCC

GAGCACACGTCTGAACTCTTTCCCTACACGACGCTCTTCCGATCTNNNGAGATCATGCCACTTCACTCCA

GAGCACACGTCTGAACTCTTTCCCTACACGACGCTCTTCCGATCTNNNGAGATCATGCCACTTCACTCCA

AGCACACGTCTGAACTCTTTCCCTACACGACGCTCTTCCGATCTNNNCTCCTGAGTAGCCGAGATTACAC

AGCACACGTCTGAACTCTTTCCCTACACGACGCTCTTCCGATCTNNNCTCCTGAGTAGCCGAGATTACAC

AGCACACGTCTGAACTCTTTCCCTACACGACGCTCTTCCGATCTNNNCTCTCCTTCATAAGCGCATCCTCC

AGCACACGTCTGAACTCTTTCCCTACACGACGCTCTTCCGATCTNNNCTCTCCTTCATAAGCGCATCCTCC

AGCACACGTCTGAACTCTTTCCCTACACGACGCTCTTCCGATCTNNNACACAACTTCCCAAAATTGAACC

AGCACACGTCTGAACTCTTTCCCTACACGACGCTCTTCCGATCTNNNACACAACTTCCCAAAATTGAACC

GAGCACACGTCTGAACTCTTTCCCTACACGACGCTCTTCCGATCTNNNGATGTGAGGTCCCCGTTCTTT

GAGCACACGTCTGAACTCTTTCCCTACACGACGCTCTTCCGATCTNNNGATGTGAGGTCCCCGTTCTTT

ACACGTCTGAACTCTTTCCCTACACGACGCTCTTCCGATCTNNNCACGCCTGGCTATTATTATTATCATCC

ACACGTCTGAACTCTTTCCCTACACGACGCTCTTCCGATCTNNNCACGCCTGGCTATTATTATTATCATCC

AGCACACGTCTGAACTCTTTCCCTACACGACGCTCTTCCGATCTNNNCAAGTAATTCTTGTGCCTCAGCC

AGCACACGTCTGAACTCTTTCCCTACACGACGCTCTTCCGATCTNNNCAAGTAATTCTTGTGCCTCAGCC

GCACACGTCTGAACTCTTTCCCTACACGACGCTCTTCCGATCTNNNCAGGAGAGAGCACAGATCAGAT

GCACACGTCTGAACTCTTTCCCTACACGACGCTCTTCCGATCTNNNCAGGAGAGAGCACAGATCAGAT

GAGCACACGTCTGAACTCTTTCCCTACACGACGCTCTTCCGATCTNNNGTCCCCGGTTCCTGTATTTTCTA

GAGCACACGTCTGAACTCTTTCCCTACACGACGCTCTTCCGATCTNNNGTCCCCGGTTCCTGTATTTTCTA

GAGCACACGTCTGAACTCTTTCCCTACACGACGCTCTTCCGATCTNNNACGCCTGGCTAATTTTCACATT

GAGCACACGTCTGAACTCTTTCCCTACACGACGCTCTTCCGATCTNNNACGCCTGGCTAATTTTCACATT

AGAGCACACGTCTGAACTCTTTCCCTACACGACGCTCTTCCGATCTNNNCATCCCTGTAATCCCAGCTACT

AGAGCACACGTCTGAACTCTTTCCCTACACGACGCTCTTCCGATCTNNNCATCCCTGTAATCCCAGCTACT

GAGCACACGTCTGAACTCTTTCCCTACACGACGCTCTTCCGATCTNNNTACAAGGGACTGTTATGCTGA

GAGCACACGTCTGAACTCTTTCCCTACACGACGCTCTTCCGATCTNNNTACAAGGGACTGTTATGCTGA

AGCACACGTCTGAACTCTTTCCCTACACGACGCTCTTCCGATCTNNNACAGGGATGTTTATAGCAGCAC

AGCACACGTCTGAACTCTTTCCCTACACGACGCTCTTCCGATCTNNNACAGGGATGTTTATAGCAGCAC

AGCACACGTCTGAACTCTTTCCCTACACGACGCTCTTCCGATCTNNNTGTCATTTTTTCCTTTATAGTCT

AGCACACGTCTGAACTCTTTCCCTACACGACGCTCTTCCGATCTNNNTGTCATTTTTTCCTTTATAGTCT

GAGCACACGTCTGAACTCTTTCCCTACACGACGCTCTTCCGATCTNNNCATCTGTAGTCCCAGCTACTCTC

GAGCACACGTCTGAACTCTTTCCCTACACGACGCTCTTCCGATCTNNNCATCTGTAGTCCCAGCTACTCTC

AGCACACGTCTGAACTCTTTCCCTACACGACGCTCTTCCGATCTNNNATGATCTCAACTCACTGCAACCT

AGCACACGTCTGAACTCTTTCCCTACACGACGCTCTTCCGATCTNNNATGATCTCAACTCACTGCAACCT

AGCACACGTCTGAACTCTTTCCCTACACGACGCTCTTCCGATCTNNNAAATCTTTCTTTTAAAATGCTGC

AGCACACGTCTGAACTCTTTCCCTACACGACGCTCTTCCGATCTNNNAAATCTTTCTTTTAAAATGCTGC

GCACACGTCTGAACTCTTTCCCTACACGACGCTCTTCCGATCTNNNTTCCAAAGCGGTTTCTTTTCTCCCA

GCACACGTCTGAACTCTTTCCCTACACGACGCTCTTCCGATCTNNNTTCCAAAGCGGTTTCTTTTCTCCCA
GAGCACACGTCTGAACTCTTTCCCTACACGACGCTCTTCCGATCTNNNTCACTCTCACAAGCAAAACAAC
GAGCACACGTCTGAACTCTTTCCCTACACGACGCTCTTCCGATCTNNNTCACTCTCACAAGCAAAACAAC
GAAGAGCACACGTCTGAACTCTTTCCCTACACGACGCTCTTCCGATCTNNNGCAGGAGAATTGCTTGAA
GAAGAGCACACGTCTGAACTCTTTCCCTACACGACGCTCTTCCGATCTNNNGCAGGAGAATTGCTTGAA
GAGCACACGTCTGAACTCTTTCCCTACACGACGCTCTTCCGATCTNNNTGATCACCTCTACTCCCTCCTAC
GAGCACACGTCTGAACTCTTTCCCTACACGACGCTCTTCCGATCTNNNTGATCACCTCTACTCCCTCCTAC
AGCACACGTCTGAACTCTTTCCCTACACGACGCTCTTCCGATCTNNNAACCAGGACTACAAATTGGGGT
AGCACACGTCTGAACTCTTTCCCTACACGACGCTCTTCCGATCTNNNAACCAGGACTACAAATTGGGGT
GCACACGTCTGAACTCTTTCCCTACACGACGCTCTTCCGATCTNNNGAGTAGCTGGGACTATAGGCAA
GCACACGTCTGAACTCTTTCCCTACACGACGCTCTTCCGATCTNNNGAGTAGCTGGGACTATAGGCAA
AGCACACGTCTGAACTCTTTCCCTACACGACGCTCTTCCGATCTNNNGGCTGTACCATTTTGCATTTCCA
AGCACACGTCTGAACTCTTTCCCTACACGACGCTCTTCCGATCTNNNGGCTGTACCATTTTGCATTTCCA
GAGCACACGTCTGAACTCTTTCCCTACACGACGCTCTTCCGATCTNNNTTGAGGAGGCATCAGAGTCTT
GAGCACACGTCTGAACTCTTTCCCTACACGACGCTCTTCCGATCTNNNTTGAGGAGGCATCAGAGTCTT
AGCACACGTCTGAACTCTTTCCCTACACGACGCTCTTCCGATCTNNNTATATGGAACCACAAAAGGCCC
AGCACACGTCTGAACTCTTTCCCTACACGACGCTCTTCCGATCTNNNTATATGGAACCACAAAAGGCCC
GCACACGTCTGAACTCTTTCCCTACACGACGCTCTTCCGATCTNNNGATTTACTGAATACATCCATGACC
GCACACGTCTGAACTCTTTCCCTACACGACGCTCTTCCGATCTNNNGATTTACTGAATACATCCATGACC
GAGCACACGTCTGAACTCTTTCCCTACACGACGCTCTTCCGATCTNNNGAGACGTGAGAATCGCTTGAA
GAGCACACGTCTGAACTCTTTCCCTACACGACGCTCTTCCGATCTNNNGAGACGTGAGAATCGCTTGAA
AGCACACGTCTGAACTCTTTCCCTACACGACGCTCTTCCGATCTNNNCCTTTGGATGTATACCCAGGAG
AGCACACGTCTGAACTCTTTCCCTACACGACGCTCTTCCGATCTNNNCCTTTGGATGTATACCCAGGAG
GAGCACACGTCTGAACTCTTTCCCTACACGACGCTCTTCCGATCTNNNCTCTTCCCTCCCACCGCTACCAT
GAGCACACGTCTGAACTCTTTCCCTACACGACGCTCTTCCGATCTNNNCTCTTCCCTCCCACCGCTACCAT
AGCACACGTCTGAACTCTTTCCCTACACGACGCTCTTCCGATCTNNNAAAGACAGTACATGCACAAGGC
AGCACACGTCTGAACTCTTTCCCTACACGACGCTCTTCCGATCTNNNAAAGACAGTACATGCACAAGGC
AGCACACGTCTGAACTCTTTCCCTACACGACGCTCTTCCGATCTNNNTGTTCCCAAAGTTCTAGCTCCTCC
AGCACACGTCTGAACTCTTTCCCTACACGACGCTCTTCCGATCTNNNTGTTCCCAAAGTTCTAGCTCCTCC
AGCACACGTCTGAACTCTTTCCCTACACGACGCTCTTCCGATCTNNNAGTGGCTGAGAGTATGGATTTC
AGCACACGTCTGAACTCTTTCCCTACACGACGCTCTTCCGATCTNNNAGTGGCTGAGAGTATGGATTTC
GAGCACACGTCTGAACTCTTTCCCTACACGACGCTCTTCCGATCTNNNAGGAAGGAGCCATAGCCTAG
GAGCACACGTCTGAACTCTTTCCCTACACGACGCTCTTCCGATCTNNNAGGAAGGAGCCATAGCCTAG
GAGCACACGTCTGAACTCTTTCCCTACACGACGCTCTTCCGATCTNNNTGAGGCAGGAGAATCACTTGA
GAGCACACGTCTGAACTCTTTCCCTACACGACGCTCTTCCGATCTNNNTGAGGCAGGAGAATCACTTGA
GAGCACACGTCTGAACTCTTTCCCTACACGACGCTCTTCCGATCTNNNAGTTGTTCTGTGGTCTTCCTTT
GAGCACACGTCTGAACTCTTTCCCTACACGACGCTCTTCCGATCTNNNAGTTGTTCTGTGGTCTTCCTTT
GCACACGTCTGAACTCTTTCCCTACACGACGCTCTTCCGATCTNNNCTACTGGGAAGTGAGGAGCCCCA
GCACACGTCTGAACTCTTTCCCTACACGACGCTCTTCCGATCTNNNCTACTGGGAAGTGAGGAGCCCCA
AGCACACGTCTGAACTCTTTCCCTACACGACGCTCTTCCGATCTNNNCACTGCACCTGGTCTTTTTTCTTC
AGCACACGTCTGAACTCTTTCCCTACACGACGCTCTTCCGATCTNNNCACTGCACCTGGTCTTTTTTCTTC
AGCACACGTCTGAACTCTTTCCCTACACGACGCTCTTCCGATCTNNNGGGTTGGCAATGGCTTTCTTTTT
AGCACACGTCTGAACTCTTTCCCTACACGACGCTCTTCCGATCTNNNGGGTTGGCAATGGCTTTCTTTTT

AGCACACGTCTGAACTCTTTCCCTACACGACGCTCTTCCGATCTNNNCTCAGGCAAAAAGGGGGAAA
AGCACACGTCTGAACTCTTTCCCTACACGACGCTCTTCCGATCTNNNCTCAGGCAAAAAGGGGGAAA
AGCACACGTCTGAACTCTTTCCCTACACGACGCTCTTCCGATCTNNNAGCACCCTACCCAACCAAAATT
AGCACACGTCTGAACTCTTTCCCTACACGACGCTCTTCCGATCTNNNAGCACCCTACCCAACCAAAATT
GCACACGTCTGAACTCTTTCCCTACACGACGCTCTTCCGATCTNNNTAGTAGCAGACTGATTGTGTG
GCACACGTCTGAACTCTTTCCCTACACGACGCTCTTCCGATCTNNNTAGTAGCAGACTGATTGTGTG
GAGCACACGTCTGAACTCTTTCCCTACACGACGCTCTTCCGATCTNNNGAATCGCTTGAACCCAGGA
GAGCACACGTCTGAACTCTTTCCCTACACGACGCTCTTCCGATCTNNNGAATCGCTTGAACCCAGGA
AGCACACGTCTGAACTCTTTCCCTACACGACGCTCTTCCGATCTNNNGAAGACAGAAGAACACAGGG
AGCACACGTCTGAACTCTTTCCCTACACGACGCTCTTCCGATCTNNNGAAGACAGAAGAACACAGGG
AGAGCACACGTCTGAACTCTTTCCCTACACGACGCTCTTCCGATCTNNNAGAAGACTTGTCCGGAGG
AGAGCACACGTCTGAACTCTTTCCCTACACGACGCTCTTCCGATCTNNNAGAAGACTTGTCCGGAGG
GCACACGTCTGAACTCTTTCCCTACACGACGCTCTTCCGATCTNNNCTGAGGCAGGAGAATGATGTG
GCACACGTCTGAACTCTTTCCCTACACGACGCTCTTCCGATCTNNNCTGAGGCAGGAGAATGATGTG
GCACACGTCTGAACTCTTTCCCTACACGACGCTCTTCCGATCTNNNAGCATCTCCTTGATATGTGTATG
GCACACGTCTGAACTCTTTCCCTACACGACGCTCTTCCGATCTNNNAGCATCTCCTTGATATGTGTATG
GCACACGTCTGAACTCTTTCCCTACACGACGCTCTTCCGATCTNNNGGCCCTTTCGTTTACAATGTCTC
GCACACGTCTGAACTCTTTCCCTACACGACGCTCTTCCGATCTNNNGGCCCTTTCGTTTACAATGTCTC
GCACACGTCTGAACTCTTTCCCTACACGACGCTCTTCCGATCTNNNACTGCTAGTGTCTCTCTCTGGAT
GCACACGTCTGAACTCTTTCCCTACACGACGCTCTTCCGATCTNNNACTGCTAGTGTCTCTCTCTGGAT
AGAGCACACGTCTGAACTCTTTCCCTACACGACGCTCTTCCGATCTNNNCACAGGATCCTACTCTTTG
AGAGCACACGTCTGAACTCTTTCCCTACACGACGCTCTTCCGATCTNNNCACAGGATCCTACTCTTTG
GAGCACACGTCTGAACTCTTTCCCTACACGACGCTCTTCCGATCTNNNTCTTTACCCATCTGTCCTCCTG
GAGCACACGTCTGAACTCTTTCCCTACACGACGCTCTTCCGATCTNNNTCTTTACCCATCTGTCCTCCTG
AGCACACGTCTGAACTCTTTCCCTACACGACGCTCTTCCGATCTNNNTGCATGACATACTGCAATTGA
AGCACACGTCTGAACTCTTTCCCTACACGACGCTCTTCCGATCTNNNTGCATGACATACTGCAATTGA
AGCACACGTCTGAACTCTTTCCCTACACGACGCTCTTCCGATCTNNNTTTACATCACAGGACTCCATGC
AGCACACGTCTGAACTCTTTCCCTACACGACGCTCTTCCGATCTNNNTTTACATCACAGGACTCCATGC
GAGCACACGTCTGAACTCTTTCCCTACACGACGCTCTTCCGATCTNNNCTGTAGACTGCTCTCTACCAA
GAGCACACGTCTGAACTCTTTCCCTACACGACGCTCTTCCGATCTNNNCTGTAGACTGCTCTCTACCAA
GCACACGTCTGAACTCTTTCCCTACACGACGCTCTTCCGATCTNNNTCCTTCTTTCTCTGTCTTTCTCCT
GCACACGTCTGAACTCTTTCCCTACACGACGCTCTTCCGATCTNNNTCCTTCTTTCTCTGTCTTTCTCCT
CACACGTCTGAACTCTTTCCCTACACGACGCTCTTCCGATCTNNNCTCTCTGTCTCTCCCTCATACATCG
CACACGTCTGAACTCTTTCCCTACACGACGCTCTTCCGATCTNNNCTCTCTGTCTCTCCCTCATACATCG
CACACGTCTGAACTCTTTCCCTACACGACGCTCTTCCGATCTNNNTATTGCCATTCACACTCCACCAGC
CACACGTCTGAACTCTTTCCCTACACGACGCTCTTCCGATCTNNNTATTGCCATTCACACTCCACCAGC
GCACACGTCTGAACTCTTTCCCTACACGACGCTCTTCCGATCTNNNAGGCAGAAGAATTGCTTGAAC
GCACACGTCTGAACTCTTTCCCTACACGACGCTCTTCCGATCTNNNAGGCAGAAGAATTGCTTGAAC
AGCACACGTCTGAACTCTTTCCCTACACGACGCTCTTCCGATCTNNNATCTGAAACCAAACACCTCTCC
AGCACACGTCTGAACTCTTTCCCTACACGACGCTCTTCCGATCTNNNATCTGAAACCAAACACCTCTCC
CACACGTCTGAACTCTTTCCCTACACGACGCTCTTCCGATCTNNNCTCATATCAAAAAGGCCCTGCATC
CACACGTCTGAACTCTTTCCCTACACGACGCTCTTCCGATCTNNNCTCATATCAAAAAGGCCCTGCATC
GAGCACACGTCTGAACTCTTTCCCTACACGACGCTCTTCCGATCTNNNACAAGCACAATAGAAGCAGG

GAGCACACGTCTGAACTCTTTCCCTACACGACGCTCTTCCGATCTNNNACAAGCACAATAGAAGCAGG
GCACACGTCTGAACTCTTTCCCTACACGACGCTCTTCCGATCTNNNAGGGACTTGGCATCCATGAATT
GCACACGTCTGAACTCTTTCCCTACACGACGCTCTTCCGATCTNNNAGGGACTTGGCATCCATGAATT
AGAGCACACGTCTGAACTCTTTCCCTACACGACGCTCTTCCGATCTNNNGTGTGGCAGGACAGTGTTA
AGAGCACACGTCTGAACTCTTTCCCTACACGACGCTCTTCCGATCTNNNGTGTGGCAGGACAGTGTTA
GAGCACACGTCTGAACTCTTTCCCTACACGACGCTCTTCCGATCTNNNGCCAGTGATTGACTTGGAATG
GAGCACACGTCTGAACTCTTTCCCTACACGACGCTCTTCCGATCTNNNGCCAGTGATTGACTTGGAATG
AGCACACGTCTGAACTCTTTCCCTACACGACGCTCTTCCGATCTNNNAAAATTAGCTGGGTGTGGTAG
AGCACACGTCTGAACTCTTTCCCTACACGACGCTCTTCCGATCTNNNAAAATTAGCTGGGTGTGGTAG
GAGCACACGTCTGAACTCTTTCCCTACACGACGCTCTTCCGATCTNNNGTGGATGAAAAGCAGTTGG
GAGCACACGTCTGAACTCTTTCCCTACACGACGCTCTTCCGATCTNNNGTGGATGAAAAGCAGTTGG
AGCACACGTCTGAACTCTTTCCCTACACGACGCTCTTCCGATCTNNNGGATCAAACTCCCCTTCTTGTC
AGCACACGTCTGAACTCTTTCCCTACACGACGCTCTTCCGATCTNNNGGATCAAACTCCCCTTCTTGTC
AGCACACGTCTGAACTCTTTCCCTACACGACGCTCTTCCGATCTNNNCTGAGTTCCTTAGAGGCAGCA
AGCACACGTCTGAACTCTTTCCCTACACGACGCTCTTCCGATCTNNNCTGAGTTCCTTAGAGGCAGCA
AGCACACGTCTGAACTCTTTCCCTACACGACGCTCTTCCGATCTNNNGCGATGCTCATTTCTCTTCAAC
AGCACACGTCTGAACTCTTTCCCTACACGACGCTCTTCCGATCTNNNGCGATGCTCATTTCTCTTCAAC
AGCACACGTCTGAACTCTTTCCCTACACGACGCTCTTCCGATCTNNNTGGCCAAATCCAAAGGAAAGT
AGCACACGTCTGAACTCTTTCCCTACACGACGCTCTTCCGATCTNNNTGGCCAAATCCAAAGGAAAGT
GAGCACACGTCTGAACTCTTTCCCTACACGACGCTCTTCCGATCTNNNAGCATCAATAGGTGACAGA
GAGCACACGTCTGAACTCTTTCCCTACACGACGCTCTTCCGATCTNNNAGCATCAATAGGTGACAGA
AGCACACGTCTGAACTCTTTCCCTACACGACGCTCTTCCGATCTNNNCATCAACTCAGAAGGGAGCAG
AGCACACGTCTGAACTCTTTCCCTACACGACGCTCTTCCGATCTNNNCATCAACTCAGAAGGGAGCAG
CACACGTCTGAACTCTTTCCCTACACGACGCTCTTCCGATCTNNNAGCATGTGAGGTGAGAGATATGT
CACACGTCTGAACTCTTTCCCTACACGACGCTCTTCCGATCTNNNAGCATGTGAGGTGAGAGATATGT
AGCACACGTCTGAACTCTTTCCCTACACGACGCTCTTCCGATCTNNNCTCCTGGATTCAAGCCATCTTC
AGCACACGTCTGAACTCTTTCCCTACACGACGCTCTTCCGATCTNNNCTCCTGGATTCAAGCCATCTTC
GAGCACACGTCTGAACTCTTTCCCTACACGACGCTCTTCCGATCTNNNAGCATGGGATGTGTTTCCAT
GAGCACACGTCTGAACTCTTTCCCTACACGACGCTCTTCCGATCTNNNAGCATGGGATGTGTTTCCAT
GAGCACACGTCTGAACTCTTTCCCTACACGACGCTCTTCCGATCTNNNAGCATGAAAGCTACTGTTCC
GAGCACACGTCTGAACTCTTTCCCTACACGACGCTCTTCCGATCTNNNAGCATGAAAGCTACTGTTCC
GAGCACACGTCTGAACTCTTTCCCTACACGACGCTCTTCCGATCTNNNGGATCACTTGAGGTCAGGAG
GAGCACACGTCTGAACTCTTTCCCTACACGACGCTCTTCCGATCTNNNGGATCACTTGAGGTCAGGAG
AGCACACGTCTGAACTCTTTCCCTACACGACGCTCTTCCGATCTNNNGCTTTAGGTACAAGGCCAAAG
AGCACACGTCTGAACTCTTTCCCTACACGACGCTCTTCCGATCTNNNGCTTTAGGTACAAGGCCAAAG
GCACACGTCTGAACTCTTTCCCTACACGACGCTCTTCCGATCTNNNCAGAGGAAGAGAAGAGCCAAA
GCACACGTCTGAACTCTTTCCCTACACGACGCTCTTCCGATCTNNNCAGAGGAAGAGAAGAGCCAAA
CACGTCTGAACTCTTTCCCTACACGACGCTCTTCCGATCTNNNTCAAATCTCCTGACCTCGTGATCCATC
CACGTCTGAACTCTTTCCCTACACGACGCTCTTCCGATCTNNNTCAAATCTCCTGACCTCGTGATCCATC
AGCACACGTCTGAACTCTTTCCCTACACGACGCTCTTCCGATCTNNNGCCTGTTTCTGGTGCTTTTCTA
AGCACACGTCTGAACTCTTTCCCTACACGACGCTCTTCCGATCTNNNGCCTGTTTCTGGTGCTTTTCTA
AGCACACGTCTGAACTCTTTCCCTACACGACGCTCTTCCGATCTNNNAAAAATTCATTGGTGCTCGCTT
AGCACACGTCTGAACTCTTTCCCTACACGACGCTCTTCCGATCTNNNAAAAATTCATTGGTGCTCGCTT

AGCACACGTCTGAACTCTTTCCCTACACGACGCTCTTCCGATCTNNNAATCAATCTGTCTTTCTCCAGGG
AGCACACGTCTGAACTCTTTCCCTACACGACGCTCTTCCGATCTNNNAATCAATCTGTCTTTCTCCAGGG
GAGCACACGTCTGAACTCTTTCCCTACACGACGCTCTTCCGATCTNNNAATCAAGGGAATGATCAACAC
GAGCACACGTCTGAACTCTTTCCCTACACGACGCTCTTCCGATCTNNNAATCAAGGGAATGATCAACAC
AGCACACGTCTGAACTCTTTCCCTACACGACGCTCTTCCGATCTNNNGCACGATGCAGCTTTTTTCAAA
AGCACACGTCTGAACTCTTTCCCTACACGACGCTCTTCCGATCTNNNGCACGATGCAGCTTTTTTCAAA
GAGCACACGTCTGAACTCTTTCCCTACACGACGCTCTTCCGATCTNNNCTGTAGTCCCAGCTACTTCAG
GAGCACACGTCTGAACTCTTTCCCTACACGACGCTCTTCCGATCTNNNCTGTAGTCCCAGCTACTTCAG
GAGCACACGTCTGAACTCTTTCCCTACACGACGCTCTTCCGATCTNNNCTGTTATCCCAGCTACTCGGAA
GAGCACACGTCTGAACTCTTTCCCTACACGACGCTCTTCCGATCTNNNCTGTTATCCCAGCTACTCGGAA
GAGCACACGTCTGAACTCTTTCCCTACACGACGCTCTTCCGATCTNNNATTACTTGAACCCGGGAAATGG
GAGCACACGTCTGAACTCTTTCCCTACACGACGCTCTTCCGATCTNNNATTACTTGAACCCGGGAAATGG
GAGCACACGTCTGAACTCTTTCCCTACACGACGCTCTTCCGATCTNNNCAAAATCGCTTGAACTGAGGA
GAGCACACGTCTGAACTCTTTCCCTACACGACGCTCTTCCGATCTNNNCAAAATCGCTTGAACTGAGGA
AGCACACGTCTGAACTCTTTCCCTACACGACGCTCTTCCGATCTNNNAGTCAATTTTTGGTTGCCAGGA
AGCACACGTCTGAACTCTTTCCCTACACGACGCTCTTCCGATCTNNNAGTCAATTTTTGGTTGCCAGGA
AGCACACGTCTGAACTCTTTCCCTACACGACGCTCTTCCGATCTNNNATGTTTTAGCAGCACTGGGGAT
AGCACACGTCTGAACTCTTTCCCTACACGACGCTCTTCCGATCTNNNATGTTTTAGCAGCACTGGGGAT
AGCACACGTCTGAACTCTTTCCCTACACGACGCTCTTCCGATCTNNNCTGGAATGCAATGGTGTGATCT
AGCACACGTCTGAACTCTTTCCCTACACGACGCTCTTCCGATCTNNNCTGGAATGCAATGGTGTGATCT
AGAGCACACGTCTGAACTCTTTCCCTACACGACGCTCTTCCGATCTNNNTTCACAAATGGCCATTTCTCA
AGAGCACACGTCTGAACTCTTTCCCTACACGACGCTCTTCCGATCTNNNTTCACAAATGGCCATTTCTCA
AGCACACGTCTGAACTCTTTCCCTACACGACGCTCTTCCGATCTNNNGAAAATTTGAGAATGGCCTGCC
AGCACACGTCTGAACTCTTTCCCTACACGACGCTCTTCCGATCTNNNGAAAATTTGAGAATGGCCTGCC
AGCACACGTCTGAACTCTTTCCCTACACGACGCTCTTCCGATCTNNNGTAGCGTTTAGAGGAGTCTTCC
AGCACACGTCTGAACTCTTTCCCTACACGACGCTCTTCCGATCTNNNGTAGCGTTTAGAGGAGTCTTCC
AGAGCACACGTCTGAACTCTTTCCCTACACGACGCTCTTCCGATCTNNNGCTTCTTCTGCAGGATGGAA
AGAGCACACGTCTGAACTCTTTCCCTACACGACGCTCTTCCGATCTNNNGCTTCTTCTGCAGGATGGAA
GAGCACACGTCTGAACTCTTTCCCTACACGACGCTCTTCCGATCTNNNTGGTTTTCCATTCTTGAGTTACT
GAGCACACGTCTGAACTCTTTCCCTACACGACGCTCTTCCGATCTNNNTGGTTTTCCATTCTTGAGTTACT
CACACGTCTGAACTCTTTCCCTACACGACGCTCTTCCGATCTNNNACCATGGGTTGTTTTTGTTTTGCCA
CACACGTCTGAACTCTTTCCCTACACGACGCTCTTCCGATCTNNNACCATGGGTTGTTTTTGTTTTGCCA
GAGCACACGTCTGAACTCTTTCCCTACACGACGCTCTTCCGATCTNNNTCCTGACTCCTGACTTGATGTT
GAGCACACGTCTGAACTCTTTCCCTACACGACGCTCTTCCGATCTNNNTCCTGACTCCTGACTTGATGTT
AGCACACGTCTGAACTCTTTCCCTACACGACGCTCTTCCGATCTNNNGGGCTTGCTTATTCCATGGAAT
AGCACACGTCTGAACTCTTTCCCTACACGACGCTCTTCCGATCTNNNGGGCTTGCTTATTCCATGGAAT
GGAAGAGCACACGTCTGAACTCTTTCCCTACACGACGCTCTTCCGATCTNNNCGTACCTGGCCTAAGAC
GGAAGAGCACACGTCTGAACTCTTTCCCTACACGACGCTCTTCCGATCTNNNCGTACCTGGCCTAAGAC
AGCACACGTCTGAACTCTTTCCCTACACGACGCTCTTCCGATCTNNNGGAGACTCCTTTCAATTGACCAT
AGCACACGTCTGAACTCTTTCCCTACACGACGCTCTTCCGATCTNNNGGAGACTCCTTTCAATTGACCAT
AGCACACGTCTGAACTCTTTCCCTACACGACGCTCTTCCGATCTNNNAGTCTGGAACACTTCTATTGCCT
AGCACACGTCTGAACTCTTTCCCTACACGACGCTCTTCCGATCTNNNAGTCTGGAACACTTCTATTGCCT
GAGCACACGTCTGAACTCTTTCCCTACACGACGCTCTTCCGATCTNNNGCAGTATTTTGAGAGCTGAAC

GAGCACACGTCTGAACTCTTTCCCTACACGACGCTCTTCCGATCTNNNGCAGTATTTGAGAGCTGAAC
GCACACGTCTGAACTCTTTCCCTACACGACGCTCTTCCGATCTNNNCTAATGCAGGCTGAAATGCTAGC
GCACACGTCTGAACTCTTTCCCTACACGACGCTCTTCCGATCTNNNCTAATGCAGGCTGAAATGCTAGC
GCACACGTCTGAACTCTTTCCCTACACGACGCTCTTCCGATCTNNNGAAAAGCACCACTTTCTTGGTGG
GCACACGTCTGAACTCTTTCCCTACACGACGCTCTTCCGATCTNNNGAAAAGCACCACTTTCTTGGTGG
GCACACGTCTGAACTCTTTCCCTACACGACGCTCTTCCGATCTNNNGGTGGTGCTGGTATAAACAAAC
GCACACGTCTGAACTCTTTCCCTACACGACGCTCTTCCGATCTNNNGGTGGTGCTGGTATAAACAAAC
AGCACACGTCTGAACTCTTTCCCTACACGACGCTCTTCCGATCTNNNACAACAAGAGATTTGGAGGGAC
AGCACACGTCTGAACTCTTTCCCTACACGACGCTCTTCCGATCTNNNACAACAAGAGATTTGGAGGGAC
GCACACGTCTGAACTCTTTCCCTACACGACGCTCTTCCGATCTNNNTGACCACATTGCTAATCACAGGA
GCACACGTCTGAACTCTTTCCCTACACGACGCTCTTCCGATCTNNNTGACCACATTGCTAATCACAGGA
AGCACACGTCTGAACTCTTTCCCTACACGACGCTCTTCCGATCTNNNCTTCCTATCCCCACATATGCACA
AGCACACGTCTGAACTCTTTCCCTACACGACGCTCTTCCGATCTNNNCTTCCTATCCCCACATATGCACA
AGCACACGTCTGAACTCTTTCCCTACACGACGCTCTTCCGATCTNNNCATTCTCTATCCCCATTCCACCAC
AGCACACGTCTGAACTCTTTCCCTACACGACGCTCTTCCGATCTNNNCATTCTCTATCCCCATTCCACCAC
AGCACACGTCTGAACTCTTTCCCTACACGACGCTCTTCCGATCTNNNCACCTGTAATCCCAGCTACTTGG
AGCACACGTCTGAACTCTTTCCCTACACGACGCTCTTCCGATCTNNNCACCTGTAATCCCAGCTACTTGG
AGCACACGTCTGAACTCTTTCCCTACACGACGCTCTTCCGATCTNNNGCCATAGACTCAGCTCTGTTCAA
AGCACACGTCTGAACTCTTTCCCTACACGACGCTCTTCCGATCTNNNGCCATAGACTCAGCTCTGTTCAA
GAGCACACGTCTGAACTCTTTCCCTACACGACGCTCTTCCGATCTNNNAGAGGTTGCAGTGAGCTAAG
GAGCACACGTCTGAACTCTTTCCCTACACGACGCTCTTCCGATCTNNNAGAGGTTGCAGTGAGCTAAG
AGCACACGTCTGAACTCTTTCCCTACACGACGCTCTTCCGATCTNNNCTGTAGTCCCAGCTACTCCTGAC
AGCACACGTCTGAACTCTTTCCCTACACGACGCTCTTCCGATCTNNNCTGTAGTCCCAGCTACTCCTGAC
GAGCACACGTCTGAACTCTTTCCCTACACGACGCTCTTCCGATCTNNNCTTTTGGACCAAAGGCCTACA
GAGCACACGTCTGAACTCTTTCCCTACACGACGCTCTTCCGATCTNNNCTTTTGGACCAAAGGCCTACA
GAGCACACGTCTGAACTCTTTCCCTACACGACGCTCTTCCGATCTNNNCTGTAGTCCCAGCTATTCAGGA
GAGCACACGTCTGAACTCTTTCCCTACACGACGCTCTTCCGATCTNNNCTGTAGTCCCAGCTATTCAGGA
CACACGTCTGAACTCTTTCCCTACACGACGCTCTTCCGATCTNNNATCTGTCAATTTAAGCTGGTTCATC
CACACGTCTGAACTCTTTCCCTACACGACGCTCTTCCGATCTNNNATCTGTCAATTTAAGCTGGTTCATC
ACACGTCTGAACTCTTTCCCTACACGACGCTCTTCCGATCTNNNACAGTCAATCTTTGTGTGTGCACCAT
ACACGTCTGAACTCTTTCCCTACACGACGCTCTTCCGATCTNNNACAGTCAATCTTTGTGTGTGCACCAT
AGCACACGTCTGAACTCTTTCCCTACACGACGCTCTTCCGATCTNNNTGGCAACCACAATTCCACTTTTT
AGCACACGTCTGAACTCTTTCCCTACACGACGCTCTTCCGATCTNNNTGGCAACCACAATTCCACTTTTT
GAGCACACGTCTGAACTCTTTCCCTACACGACGCTCTTCCGATCTNNNTGTAGCTTGCAGTGAGTTGAC
GAGCACACGTCTGAACTCTTTCCCTACACGACGCTCTTCCGATCTNNNTGTAGCTTGCAGTGAGTTGAC
GAGCACACGTCTGAACTCTTTCCCTACACGACGCTCTTCCGATCTNNNATTGCAATGAGTCGAGATCAT
GAGCACACGTCTGAACTCTTTCCCTACACGACGCTCTTCCGATCTNNNATTGCAATGAGTCGAGATCAT
AGCACACGTCTGAACTCTTTCCCTACACGACGCTCTTCCGATCTNNNACCAAATTGAAGCCAGTTGTCTC
AGCACACGTCTGAACTCTTTCCCTACACGACGCTCTTCCGATCTNNNACCAAATTGAAGCCAGTTGTCTC
AGCACACGTCTGAACTCTTTCCCTACACGACGCTCTTCCGATCTNNNGGACCCCCTCTAATCTGTTCCATC
AGCACACGTCTGAACTCTTTCCCTACACGACGCTCTTCCGATCTNNNGGACCCCCTCTAATCTGTTCCATC
AGCACACGTCTGAACTCTTTCCCTACACGACGCTCTTCCGATCTNNNTCTTCACTCCCACTAGAAGCAAC
AGCACACGTCTGAACTCTTTCCCTACACGACGCTCTTCCGATCTNNNTCTTCACTCCCACTAGAAGCAAC

GAGCACACGTCTGAACTCTTTCCCTACACGACGCTCTTCCGATCTNNNATGAGTTGGAAAAGATGTGC
GAGCACACGTCTGAACTCTTTCCCTACACGACGCTCTTCCGATCTNNNATGAGTTGGAAAAGATGTGC
GAGCACACGTCTGAACTCTTTCCCTACACGACGCTCTTCCGATCTNNNGGTCAGACATGGACTATTGG
GAGCACACGTCTGAACTCTTTCCCTACACGACGCTCTTCCGATCTNNNGGTCAGACATGGACTATTGG
AGCACACGTCTGAACTCTTTCCCTACACGACGCTCTTCCGATCTNNNAAAAAATAATTACCCGGGCGT
AGCACACGTCTGAACTCTTTCCCTACACGACGCTCTTCCGATCTNNNAAAAAATAATTACCCGGGCGT
AGCACACGTCTGAACTCTTTCCCTACACGACGCTCTTCCGATCTNNNGAGAATTGCTTGAACTTGGGA
AGCACACGTCTGAACTCTTTCCCTACACGACGCTCTTCCGATCTNNNGAGAATTGCTTGAACTTGGGA
GCACACGTCTGAACTCTTTCCCTACACGACGCTCTTCCGATCTNNNCAAACTATACACACGCACACACA
GCACACGTCTGAACTCTTTCCCTACACGACGCTCTTCCGATCTNNNCAAACTATACACACGCACACACA
GAGCACACGTCTGAACTCTTTCCCTACACGACGCTCTTCCGATCTNNNCAGATTCCAGAGGAGAGAG
GAGCACACGTCTGAACTCTTTCCCTACACGACGCTCTTCCGATCTNNNCAGATTCCAGAGGAGAGAG
GCACACGTCTGAACTCTTTCCCTACACGACGCTCTTCCGATCTNNNGCTTCCTCCATCCTTCTTTCTCTCA
GCACACGTCTGAACTCTTTCCCTACACGACGCTCTTCCGATCTNNNGCTTCCTCCATCCTTCTTTCTCTCA
GCACACGTCTGAACTCTTTCCCTACACGACGCTCTTCCGATCTNNNAGAGACTCAGACACGTACCTTC
GCACACGTCTGAACTCTTTCCCTACACGACGCTCTTCCGATCTNNNAGAGACTCAGACACGTACCTTC
GCACACGTCTGAACTCTTTCCCTACACGACGCTCTTCCGATCTNNNCCCAACACCCTTCCTTTCCTTATC
GCACACGTCTGAACTCTTTCCCTACACGACGCTCTTCCGATCTNNNCCCAACACCCTTCCTTTCCTTATC
GCACACGTCTGAACTCTTTCCCTACACGACGCTCTTCCGATCTNNNCCAGCTTCCTCACTGTCAAACTTC
GCACACGTCTGAACTCTTTCCCTACACGACGCTCTTCCGATCTNNNCCAGCTTCCTCACTGTCAAACTTC
GAGCACACGTCTGAACTCTTTCCCTACACGACGCTCTTCCGATCTNNNTGTACACCTGTTTATGCATGT
GAGCACACGTCTGAACTCTTTCCCTACACGACGCTCTTCCGATCTNNNTGTACACCTGTTTATGCATGT
GCACACGTCTGAACTCTTTCCCTACACGACGCTCTTCCGATCTNNNCTTCCTTATTGGCCTCTGAGCTG
GCACACGTCTGAACTCTTTCCCTACACGACGCTCTTCCGATCTNNNCTTCCTTATTGGCCTCTGAGCTG
GAGCACACGTCTGAACTCTTTCCCTACACGACGCTCTTCCGATCTNNNATATATACACACACACACACA
GAGCACACGTCTGAACTCTTTCCCTACACGACGCTCTTCCGATCTNNNATATATACACACACACACACA
AGCACACGTCTGAACTCTTTCCCTACACGACGCTCTTCCGATCTNNNGGTGGATGGATTATGAGGTC
AGCACACGTCTGAACTCTTTCCCTACACGACGCTCTTCCGATCTNNNGGTGGATGGATTATGAGGTC
GAGCACACGTCTGAACTCTTTCCCTACACGACGCTCTTCCGATCTNNNCCCGTTCTGTGTTCAAACTTG
GAGCACACGTCTGAACTCTTTCCCTACACGACGCTCTTCCGATCTNNNCCCGTTCTGTGTTCAAACTTG
AAGAGCACACGTCTGAACTCTTTCCCTACACGACGCTCTTCCGATCTNNNAGGCCATAAAACCTGCCC
AAGAGCACACGTCTGAACTCTTTCCCTACACGACGCTCTTCCGATCTNNNAGGCCATAAAACCTGCCC
GCACACGTCTGAACTCTTTCCCTACACGACGCTCTTCCGATCTNNNCCATCCCATGGTGGAAAGCATA
GCACACGTCTGAACTCTTTCCCTACACGACGCTCTTCCGATCTNNNCCATCCCATGGTGGAAAGCATA
AGCACACGTCTGAACTCTTTCCCTACACGACGCTCTTCCGATCTNNNTAGCTGGGATTACAGGCATAT
AGCACACGTCTGAACTCTTTCCCTACACGACGCTCTTCCGATCTNNNTAGCTGGGATTACAGGCATAT
AGCACACGTCTGAACTCTTTCCCTACACGACGCTCTTCCGATCTNNNATGCGCGCATGTATACAGATA
AGCACACGTCTGAACTCTTTCCCTACACGACGCTCTTCCGATCTNNNATGCGCGCATGTATACAGATA
AGCACACGTCTGAACTCTTTCCCTACACGACGCTCTTCCGATCTNNNGGGCCTAACTCTAGCTCTTTGA
AGCACACGTCTGAACTCTTTCCCTACACGACGCTCTTCCGATCTNNNGGGCCTAACTCTAGCTCTTTGA
AGCACACGTCTGAACTCTTTCCCTACACGACGCTCTTCCGATCTNNNGGGACATGGAGCTTGCAGTAA
AGCACACGTCTGAACTCTTTCCCTACACGACGCTCTTCCGATCTNNNGGGACATGGAGCTTGCAGTAA
AGCACACGTCTGAACTCTTTCCCTACACGACGCTCTTCCGATCTNNNAAGTATACTCAGTCAAGCAGC

AGCACACGTCTGAACTCTTTCCCTACACGACGCTCTTCCGATCTNNNAAGTATACTCAGTCAAGCAGCC
GAGCACACGTCTGAACTCTTTCCCTACACGACGCTCTTCCGATCTNNNCCACATGAGGCAAGGAGTTCA
GAGCACACGTCTGAACTCTTTCCCTACACGACGCTCTTCCGATCTNNNCCACATGAGGCAAGGAGTTCA
AGCACACGTCTGAACTCTTTCCCTACACGACGCTCTTCCGATCTNNNCCTGGAAGAAAGACAGATGGT
AGCACACGTCTGAACTCTTTCCCTACACGACGCTCTTCCGATCTNNNCCTGGAAGAAAGACAGATGGT
AGAGCACACGTCTGAACTCTTTCCCTACACGACGCTCTTCCGATCTNNNCAGATGGACAAGGATGAAG
AGAGCACACGTCTGAACTCTTTCCCTACACGACGCTCTTCCGATCTNNNCAGATGGACAAGGATGAAG
AGCACACGTCTGAACTCTTTCCCTACACGACGCTCTTCCGATCTNNNACTTCAGTACAAGGAAGTTCTC
AGCACACGTCTGAACTCTTTCCCTACACGACGCTCTTCCGATCTNNNACTTCAGTACAAGGAAGTTCTC
AGCACACGTCTGAACTCTTTCCCTACACGACGCTCTTCCGATCTNNNTTGGGATTACAGGTATGAGCCA
AGCACACGTCTGAACTCTTTCCCTACACGACGCTCTTCCGATCTNNNTTGGGATTACAGGTATGAGCCA
AGCACACGTCTGAACTCTTTCCCTACACGACGCTCTTCCGATCTNNNACCATGTTTCTCACTGCCATTTG
AGCACACGTCTGAACTCTTTCCCTACACGACGCTCTTCCGATCTNNNACCATGTTTCTCACTGCCATTTG
GCACACGTCTGAACTCTTTCCCTACACGACGCTCTTCCGATCTNNNGAATCTCACTCTGTCACCAAGCTCA
GCACACGTCTGAACTCTTTCCCTACACGACGCTCTTCCGATCTNNNGAATCTCACTCTGTCACCAAGCTCA
GAGCACACGTCTGAACTCTTTCCCTACACGACGCTCTTCCGATCTNNNGACCTTAGGAATTCCCACATTC
GAGCACACGTCTGAACTCTTTCCCTACACGACGCTCTTCCGATCTNNNGACCTTAGGAATTCCCACATTC
AGCACACGTCTGAACTCTTTCCCTACACGACGCTCTTCCGATCTNNNTCCAGGGATAAACAGTCTCCTGA
AGCACACGTCTGAACTCTTTCCCTACACGACGCTCTTCCGATCTNNNTCCAGGGATAAACAGTCTCCTGA
GAGCACACGTCTGAACTCTTTCCCTACACGACGCTCTTCCGATCTNNNTCACTGTGAACACGTAGAAAA
GAGCACACGTCTGAACTCTTTCCCTACACGACGCTCTTCCGATCTNNNTCACTGTGAACACGTAGAAAA
AGCACACGTCTGAACTCTTTCCCTACACGACGCTCTTCCGATCTNNNGCTAAGCTGCAAACTACATAGCC
AGCACACGTCTGAACTCTTTCCCTACACGACGCTCTTCCGATCTNNNGCTAAGCTGCAAACTACATAGCC
AGCACACGTCTGAACTCTTTCCCTACACGACGCTCTTCCGATCTNNNTATGAAAATGGCCTCCAACTCCA
AGCACACGTCTGAACTCTTTCCCTACACGACGCTCTTCCGATCTNNNTATGAAAATGGCCTCCAACTCCA
GAGCACACGTCTGAACTCTTTCCCTACACGACGCTCTTCCGATCTNNNTTGACCTTCTGACTCCATCACT
GAGCACACGTCTGAACTCTTTCCCTACACGACGCTCTTCCGATCTNNNTTGACCTTCTGACTCCATCACT
ACACGTCTGAACTCTTTCCCTACACGACGCTCTTCCGATCTNNNTTCACTCACTACAAGGCTTCCTCCATG
ACACGTCTGAACTCTTTCCCTACACGACGCTCTTCCGATCTNNNTTCACTCACTACAAGGCTTCCTCCATG
AGAGCACACGTCTGAACTCTTTCCCTACACGACGCTCTTCCGATCTNNNGATAGCTCTAGAGACACCGA
AGAGCACACGTCTGAACTCTTTCCCTACACGACGCTCTTCCGATCTNNNGATAGCTCTAGAGACACCGA
AGCACACGTCTGAACTCTTTCCCTACACGACGCTCTTCCGATCTNNNATATTCTTGCACATAAATTTTGC
AGCACACGTCTGAACTCTTTCCCTACACGACGCTCTTCCGATCTNNNATATTCTTGCACATAAATTTTGC
GCACACGTCTGAACTCTTTCCCTACACGACGCTCTTCCGATCTNNNCAAAATTGCTTGGTCTCAGGGAG
GCACACGTCTGAACTCTTTCCCTACACGACGCTCTTCCGATCTNNNCAAAATTGCTTGGTCTCAGGGAG
AGCACACGTCTGAACTCTTTCCCTACACGACGCTCTTCCGATCTNNNGTAGAGACGGGGTTTCATCAT
AGCACACGTCTGAACTCTTTCCCTACACGACGCTCTTCCGATCTNNNGTAGAGACGGGGTTTCATCAT
GAGCACACGTCTGAACTCTTTCCCTACACGACGCTCTTCCGATCTNNNGAGATTTCCGTGAGTGTGTCC
GAGCACACGTCTGAACTCTTTCCCTACACGACGCTCTTCCGATCTNNNGAGATTTCCGTGAGTGTGTCC
AGCACACGTCTGAACTCTTTCCCTACACGACGCTCTTCCGATCTNNNATTTTCCTGGTATTCAGAGGGG
AGCACACGTCTGAACTCTTTCCCTACACGACGCTCTTCCGATCTNNNATTTTCCTGGTATTCAGAGGGG
GCACACGTCTGAACTCTTTCCCTACACGACGCTCTTCCGATCTNNNGAGCTTGCAGTGAGCCAAGATAC
GCACACGTCTGAACTCTTTCCCTACACGACGCTCTTCCGATCTNNNGAGCTTGCAGTGAGCCAAGATAC

AGCACACGTCTGAACTCTTTCCCTACACGACGCTCTTCCGATCTNNNGCCTTTCTTTCCTACCACTCTGA

AGCACACGTCTGAACTCTTTCCCTACACGACGCTCTTCCGATCTNNNGCCTTTCTTTCCTACCACTCTGA

CACACGTCTGAACTCTTTCCCTACACGACGCTCTTCCGATCTNNNCACCTGGCTCGTGGCTTTATCCAT

CACACGTCTGAACTCTTTCCCTACACGACGCTCTTCCGATCTNNNCACCTGGCTCGTGGCTTTATCCAT

AGCACACGTCTGAACTCTTTCCCTACACGACGCTCTTCCGATCTNNNACACCCTTGCTGATCTTAATAT

AGCACACGTCTGAACTCTTTCCCTACACGACGCTCTTCCGATCTNNNACACCCTTGCTGATCTTAATAT

AGCACACGTCTGAACTCTTTCCCTACACGACGCTCTTCCGATCTNNNATCTGTGTGGAGGCAGCAAT

AGCACACGTCTGAACTCTTTCCCTACACGACGCTCTTCCGATCTNNNATCTGTGTGGAGGCAGCAAT

AGCACACGTCTGAACTCTTTCCCTACACGACGCTCTTCCGATCTNNNAGGAGTGAAAATTTCAGAGG

AGCACACGTCTGAACTCTTTCCCTACACGACGCTCTTCCGATCTNNNAGGAGTGAAAATTTCAGAGG

AGCACACGTCTGAACTCTTTCCCTACACGACGCTCTTCCGATCTNNNCTGTATTCTGCTAACAGGAGG

AGCACACGTCTGAACTCTTTCCCTACACGACGCTCTTCCGATCTNNNCTGTATTCTGCTAACAGGAGG

AGCACACGTCTGAACTCTTTCCCTACACGACGCTCTTCCGATCTNNNGCATTGACCCAGCCAATACAAA

AGCACACGTCTGAACTCTTTCCCTACACGACGCTCTTCCGATCTNNNGCATTGACCCAGCCAATACAAA

AGCACACGTCTGAACTCTTTCCCTACACGACGCTCTTCCGATCTNNNGCCTAGGCCTTTAAATAACTCA

AGCACACGTCTGAACTCTTTCCCTACACGACGCTCTTCCGATCTNNNGCCTAGGCCTTTAAATAACTCA

GAGCACACGTCTGAACTCTTTCCCTACACGACGCTCTTCCGATCTNNNGCAGTGGCAGCTTGGATTTA

GAGCACACGTCTGAACTCTTTCCCTACACGACGCTCTTCCGATCTNNNGCAGTGGCAGCTTGGATTTA

AGCACACGTCTGAACTCTTTCCCTACACGACGCTCTTCCGATCTNNNTTCTCCAGAATGTTGCTGAGT

AGCACACGTCTGAACTCTTTCCCTACACGACGCTCTTCCGATCTNNNTTCTCCAGAATGTTGCTGAGT

AGCACACGTCTGAACTCTTTCCCTACACGACGCTCTTCCGATCTNNNTCCTAATGCTATCCCTCCCCTAG

AGCACACGTCTGAACTCTTTCCCTACACGACGCTCTTCCGATCTNNNTCCTAATGCTATCCCTCCCCTAG

CACACGTCTGAACTCTTTCCCTACACGACGCTCTTCCGATCTNNNCCGCTTTGTCTCTTCACTTCTTGCA

CACACGTCTGAACTCTTTCCCTACACGACGCTCTTCCGATCTNNNCCGCTTTGTCTCTTCACTTCTTGCA

AGCACACGTCTGAACTCTTTCCCTACACGACGCTCTTCCGATCTNNNCTTGCCTAGATTAGCTGCTGC

AGCACACGTCTGAACTCTTTCCCTACACGACGCTCTTCCGATCTNNNCTTGCCTAGATTAGCTGCTGC

AGCACACGTCTGAACTCTTTCCCTACACGACGCTCTTCCGATCTNNNTGTGAAAGGCAGCTTTGCAA

AGCACACGTCTGAACTCTTTCCCTACACGACGCTCTTCCGATCTNNNTGTGAAAGGCAGCTTTGCAA

CACACGTCTGAACTCTTTCCCTACACGACGCTCTTCCGATCTNNNGATTTGCCCTGTGACTTCACTTCCA

CACACGTCTGAACTCTTTCCCTACACGACGCTCTTCCGATCTNNNGATTTGCCCTGTGACTTCACTTCCA

AGCACACGTCTGAACTCTTTCCCTACACGACGCTCTTCCGATCTNNNACAGAGTCAGTGAACCAATC

AGCACACGTCTGAACTCTTTCCCTACACGACGCTCTTCCGATCTNNNACAGAGTCAGTGAACCAATC

GCACACGTCTGAACTCTTTCCCTACACGACGCTCTTCCGATCTNNNCCCATTCAGGAAGATCTATCCAC

GCACACGTCTGAACTCTTTCCCTACACGACGCTCTTCCGATCTNNNCCCATTCAGGAAGATCTATCCAC

AGCACACGTCTGAACTCTTTCCCTACACGACGCTCTTCCGATCTNNNAGACCACGGAGAAGTGACTTT

AGCACACGTCTGAACTCTTTCCCTACACGACGCTCTTCCGATCTNNNAGACCACGGAGAAGTGACTTT

AGCACACGTCTGAACTCTTTCCCTACACGACGCTCTTCCGATCTNNNGGAAAACAGAGAGGGAACAA

AGCACACGTCTGAACTCTTTCCCTACACGACGCTCTTCCGATCTNNNGGAAAACAGAGAGGGAACAA

GAGCACACGTCTGAACTCTTTCCCTACACGACGCTCTTCCGATCTNNNCCAAACACCGCATATTCTCAC

GAGCACACGTCTGAACTCTTTCCCTACACGACGCTCTTCCGATCTNNNCCAAACACCGCATATTCTCAC

AGCACACGTCTGAACTCTTTCCCTACACGACGCTCTTCCGATCTNNNGAATGGCATAAACCCGGGAG

AGCACACGTCTGAACTCTTTCCCTACACGACGCTCTTCCGATCTNNNGAATGGCATAAACCCGGGAG

GAGCACACGTCTGAACTCTTTCCCTACACGACGCTCTTCCGATCTNNNGAGAAGCTGGGATTACAAG

GAGCACACGTCTGAACTCTTTCCCTACACGACGCTCTTCCGATCTNNNGAGAAGCTGGGATTACAAGC
AGCACACGTCTGAACTCTTTCCCTACACGACGCTCTTCCGATCTNNNTGTTTCTGGTCACAGCCTATTG
AGCACACGTCTGAACTCTTTCCCTACACGACGCTCTTCCGATCTNNNTGTTTCTGGTCACAGCCTATTG
GAGCACACGTCTGAACTCTTTCCCTACACGACGCTCTTCCGATCTNNNGTCGTCTACCTCACTTCACAGG
GAGCACACGTCTGAACTCTTTCCCTACACGACGCTCTTCCGATCTNNNGTCGTCTACCTCACTTCACAGG
GCACACGTCTGAACTCTTTCCCTACACGACGCTCTTCCGATCTNNNTGACAACCGACTTGAGTTTGAGA
GCACACGTCTGAACTCTTTCCCTACACGACGCTCTTCCGATCTNNNTGACAACCGACTTGAGTTTGAGA
AGCACACGTCTGAACTCTTTCCCTACACGACGCTCTTCCGATCTNNNGGAGATCCTGTAGTGCTGTAAG
AGCACACGTCTGAACTCTTTCCCTACACGACGCTCTTCCGATCTNNNGGAGATCCTGTAGTGCTGTAAG
AGCACACGTCTGAACTCTTTCCCTACACGACGCTCTTCCGATCTNNNGGTTGCAGTGAGCCAAGATTGC
AGCACACGTCTGAACTCTTTCCCTACACGACGCTCTTCCGATCTNNNGGTTGCAGTGAGCCAAGATTGC
GCACACGTCTGAACTCTTTCCCTACACGACGCTCTTCCGATCTNNNCAGCAAAGGAAAGTGGGGAAC
GCACACGTCTGAACTCTTTCCCTACACGACGCTCTTCCGATCTNNNCAGCAAAGGAAAGTGGGGAAC
GAGCACACGTCTGAACTCTTTCCCTACACGACGCTCTTCCGATCTNNNTAACAACATAAAGGGGGCAT
GAGCACACGTCTGAACTCTTTCCCTACACGACGCTCTTCCGATCTNNNTAACAACATAAAGGGGGCAT
AGAGCACACGTCTGAACTCTTTCCCTACACGACGCTCTTCCGATCTNNNAGAATGTGGGGCATTTCAAC
AGAGCACACGTCTGAACTCTTTCCCTACACGACGCTCTTCCGATCTNNNAGAATGTGGGGCATTTCAAC
CACACGTCTGAACTCTTTCCCTACACGACGCTCTTCCGATCTNNNTTACACACATATTCATCATTTCTGGT
CACACGTCTGAACTCTTTCCCTACACGACGCTCTTCCGATCTNNNTTACACACATATTCATCATTTCTGGT
GAGCACACGTCTGAACTCTTTCCCTACACGACGCTCTTCCGATCTNNNAAAGTACTGGGATTACAGGC
GAGCACACGTCTGAACTCTTTCCCTACACGACGCTCTTCCGATCTNNNAAAGTACTGGGATTACAGGC
AGCACACGTCTGAACTCTTTCCCTACACGACGCTCTTCCGATCTNNNGTGCGGGCTTAGACTTTTTGTA
AGCACACGTCTGAACTCTTTCCCTACACGACGCTCTTCCGATCTNNNGTGCGGGCTTAGACTTTTTGTA
AGCACACGTCTGAACTCTTTCCCTACACGACGCTCTTCCGATCTNNNCCTAACTCTCATTCCTCACCCACC
AGCACACGTCTGAACTCTTTCCCTACACGACGCTCTTCCGATCTNNNCCTAACTCTCATTCCTCACCCACC
GCACACGTCTGAACTCTTTCCCTACACGACGCTCTTCCGATCTNNNACGTCTGTCTTTGTAGGCATTCAC
GCACACGTCTGAACTCTTTCCCTACACGACGCTCTTCCGATCTNNNACGTCTGTCTTTGTAGGCATTCAC
AGCACACGTCTGAACTCTTTCCCTACACGACGCTCTTCCGATCTNNNCTTTTCAGTACATCCACTTAAG
AGCACACGTCTGAACTCTTTCCCTACACGACGCTCTTCCGATCTNNNCTTTTCAGTACATCCACTTAAG
GCACACGTCTGAACTCTTTCCCTACACGACGCTCTTCCGATCTNNNCAGAATTCACCCTTCATACACCAT
GCACACGTCTGAACTCTTTCCCTACACGACGCTCTTCCGATCTNNNCAGAATTCACCCTTCATACACCAT
AGCACACGTCTGAACTCTTTCCCTACACGACGCTCTTCCGATCTNNNGTGTCTCTCCATTGTCCCCATAA
AGCACACGTCTGAACTCTTTCCCTACACGACGCTCTTCCGATCTNNNGTGTCTCTCCATTGTCCCCATAA
AGCACACGTCTGAACTCTTTCCCTACACGACGCTCTTCCGATCTNNNTGGGGTTTAGGAGAGGGAAT
AGCACACGTCTGAACTCTTTCCCTACACGACGCTCTTCCGATCTNNNTGGGGTTTAGGAGAGGGAAT
GCACACGTCTGAACTCTTTCCCTACACGACGCTCTTCCGATCTNNNATGAGGACAGTGAAGCAACA
GCACACGTCTGAACTCTTTCCCTACACGACGCTCTTCCGATCTNNNATGAGGACAGTGAAGCAACA
GAGCACACGTCTGAACTCTTTCCCTACACGACGCTCTTCCGATCTNNNAGCGACTCCTGCTATATTTTG
GAGCACACGTCTGAACTCTTTCCCTACACGACGCTCTTCCGATCTNNNAGCGACTCCTGCTATATTTTG
AGCACACGTCTGAACTCTTTCCCTACACGACGCTCTTCCGATCTNNNAAGTCCTCTTCTCTTGGTTTGCA
AGCACACGTCTGAACTCTTTCCCTACACGACGCTCTTCCGATCTNNNAAGTCCTCTTCTCTTGGTTTGCA
AGCACACGTCTGAACTCTTTCCCTACACGACGCTCTTCCGATCTNNNGGGAATTGGTGTTTCATGGGA
AGCACACGTCTGAACTCTTTCCCTACACGACGCTCTTCCGATCTNNNGGGAATTGGTGTTTCATGGGA

AGCACACGTCTGAACTCTTTCCCTACACGACGCTCTTCCGATCTNNNAAAAAATTAGCTGGGCATGGTC
AGCACACGTCTGAACTCTTTCCCTACACGACGCTCTTCCGATCTNNNAAAAAATTAGCTGGGCATGGTC
AGCACACGTCTGAACTCTTTCCCTACACGACGCTCTTCCGATCTNNNAGTCCAATTTCTTCAGAGGCTGA
AGCACACGTCTGAACTCTTTCCCTACACGACGCTCTTCCGATCTNNNAGTCCAATTTCTTCAGAGGCTGA
AGCACACGTCTGAACTCTTTCCCTACACGACGCTCTTCCGATCTNNNAGCTTTTGCATGTACTACTGGG
AGCACACGTCTGAACTCTTTCCCTACACGACGCTCTTCCGATCTNNNAGCTTTTGCATGTACTACTGGG
GAGCACACGTCTGAACTCTTTCCCTACACGACGCTCTTCCGATCTNNNGAAACAGTTGGGTGGCTAACT
GAGCACACGTCTGAACTCTTTCCCTACACGACGCTCTTCCGATCTNNNGAAACAGTTGGGTGGCTAACT
AGCACACGTCTGAACTCTTTCCCTACACGACGCTCTTCCGATCTNNNCAGTTTGCTGTTTGTCTCCAAGT
AGCACACGTCTGAACTCTTTCCCTACACGACGCTCTTCCGATCTNNNCAGTTTGCTGTTTGTCTCCAAGT
GAGCACACGTCTGAACTCTTTCCCTACACGACGCTCTTCCGATCTNNNCGTATCTGTAACCCCAGCTACT
GAGCACACGTCTGAACTCTTTCCCTACACGACGCTCTTCCGATCTNNNCGTATCTGTAACCCCAGCTACT
AGCACACGTCTGAACTCTTTCCCTACACGACGCTCTTCCGATCTNNNTGGCACGGTACTAGGTGTTTTA
AGCACACGTCTGAACTCTTTCCCTACACGACGCTCTTCCGATCTNNNTGGCACGGTACTAGGTGTTTTA
GAGCACACGTCTGAACTCTTTCCCTACACGACGCTCTTCCGATCTNNNGAGGAGACGAGCAGGATCAT
GAGCACACGTCTGAACTCTTTCCCTACACGACGCTCTTCCGATCTNNNGAGGAGACGAGCAGGATCAT
AGCACACGTCTGAACTCTTTCCCTACACGACGCTCTTCCGATCTNNNAATCACAGTAGCTTCCCCTTTGT
AGCACACGTCTGAACTCTTTCCCTACACGACGCTCTTCCGATCTNNNAATCACAGTAGCTTCCCCTTTGT
AGCACACGTCTGAACTCTTTCCCTACACGACGCTCTTCCGATCTNNNCCCTTCCAAAATGCCACATTACAC
AGCACACGTCTGAACTCTTTCCCTACACGACGCTCTTCCGATCTNNNCCCTTCCAAAATGCCACATTACAC
AGCACACGTCTGAACTCTTTCCCTACACGACGCTCTTCCGATCTNNNAAAAAATTAGCAGGCGTAGTG
AGCACACGTCTGAACTCTTTCCCTACACGACGCTCTTCCGATCTNNNAAAAAATTAGCAGGCGTAGTG
AGCACACGTCTGAACTCTTTCCCTACACGACGCTCTTCCGATCTNNNAGTCACATCTTACAAGGTGGCA
AGCACACGTCTGAACTCTTTCCCTACACGACGCTCTTCCGATCTNNNAGTCACATCTTACAAGGTGGCA
AGCACACGTCTGAACTCTTTCCCTACACGACGCTCTTCCGATCTNNNCAGCCTTTTTTTCGCTGGTTTTCC
AGCACACGTCTGAACTCTTTCCCTACACGACGCTCTTCCGATCTNNNCAGCCTTTTTTTCGCTGGTTTTCC
AAGAGCACACGTCTGAACTCTTTCCCTACACGACGCTCTTCCGATCTNNNATTGCTCAAGCTCAGGAGT
AAGAGCACACGTCTGAACTCTTTCCCTACACGACGCTCTTCCGATCTNNNATTGCTCAAGCTCAGGAGT
GAGCACACGTCTGAACTCTTTCCCTACACGACGCTCTTCCGATCTNNNCATCTTACATGGATGGCAGCA
GAGCACACGTCTGAACTCTTTCCCTACACGACGCTCTTCCGATCTNNNCATCTTACATGGATGGCAGCA
AGCACACGTCTGAACTCTTTCCCTACACGACGCTCTTCCGATCTNNNGGAAACCTCTAACCTGGAAGCT
AGCACACGTCTGAACTCTTTCCCTACACGACGCTCTTCCGATCTNNNGGAAACCTCTAACCTGGAAGCT
GAGCACACGTCTGAACTCTTTCCCTACACGACGCTCTTCCGATCTNNNGAGGATTGCTTGATCTCAGGA
GAGCACACGTCTGAACTCTTTCCCTACACGACGCTCTTCCGATCTNNNGAGGATTGCTTGATCTCAGGA
GAAGAGCACACGTCTGAACTCTTTCCCTACACGACGCTCTTCCGATCTNNNAAAAAATTAGCCAGGCGT
GAAGAGCACACGTCTGAACTCTTTCCCTACACGACGCTCTTCCGATCTNNNAAAAAATTAGCCAGGCGT
AGCACACGTCTGAACTCTTTCCCTACACGACGCTCTTCCGATCTNNNCACCTGTAATCCCAGCTACTCAG
AGCACACGTCTGAACTCTTTCCCTACACGACGCTCTTCCGATCTNNNCACCTGTAATCCCAGCTACTCAG
GAGCACACGTCTGAACTCTTTCCCTACACGACGCTCTTCCGATCTNNNGAGAATTCTTCTGCCAAGCAC
GAGCACACGTCTGAACTCTTTCCCTACACGACGCTCTTCCGATCTNNNGAGAATTCTTCTGCCAAGCAC
AGCACACGTCTGAACTCTTTCCCTACACGACGCTCTTCCGATCTNNNTTGAACCTCAAGAACTGCTCTGA
AGCACACGTCTGAACTCTTTCCCTACACGACGCTCTTCCGATCTNNNTTGAACCTCAAGAACTGCTCTGA
GAGCACACGTCTGAACTCTTTCCCTACACGACGCTCTTCCGATCTNNNTGCAGTGATATTTACCCGACA

GAGCACACGTCTGAACTCTTTCCCTACACGACGCTCTTCCGATCTNNNTGCAGTGATATTTACCCGACA
AGCACACGTCTGAACTCTTTCCCTACACGACGCTCTTCCGATCTNNNCTGCAGGTAGTTGAGTTGGAA
AGCACACGTCTGAACTCTTTCCCTACACGACGCTCTTCCGATCTNNNCTGCAGGTAGTTGAGTTGGAA
AGCACACGTCTGAACTCTTTCCCTACACGACGCTCTTCCGATCTNNNAGTGACTTCAGAGCTGTGTAAG
AGCACACGTCTGAACTCTTTCCCTACACGACGCTCTTCCGATCTNNNAGTGACTTCAGAGCTGTGTAAG
GAGCACACGTCTGAACTCTTTCCCTACACGACGCTCTTCCGATCTNNNGATCAGTGGGACTCCGTGTAT
GAGCACACGTCTGAACTCTTTCCCTACACGACGCTCTTCCGATCTNNNGATCAGTGGGACTCCGTGTAT
AGCACACGTCTGAACTCTTTCCCTACACGACGCTCTTCCGATCTNNNCAAGCTGATATGCAGTTGCTAG
AGCACACGTCTGAACTCTTTCCCTACACGACGCTCTTCCGATCTNNNCAAGCTGATATGCAGTTGCTAG
AGCACACGTCTGAACTCTTTCCCTACACGACGCTCTTCCGATCTNNNTGTACCTCCCTAACTGGTCTTCT
AGCACACGTCTGAACTCTTTCCCTACACGACGCTCTTCCGATCTNNNTGTACCTCCCTAACTGGTCTTCT
GCACACGTCTGAACTCTTTCCCTACACGACGCTCTTCCGATCTNNNACACATACGCGTACACACTTCTCC
GCACACGTCTGAACTCTTTCCCTACACGACGCTCTTCCGATCTNNNACACATACGCGTACACACTTCTCC
AGCACACGTCTGAACTCTTTCCCTACACGACGCTCTTCCGATCTNNNATTTTGAGACAGGGTCTTGCTC
AGCACACGTCTGAACTCTTTCCCTACACGACGCTCTTCCGATCTNNNATTTTGAGACAGGGTCTTGCTC
GCACACGTCTGAACTCTTTCCCTACACGACGCTCTTCCGATCTNNNGCTCAGAAAGTTTTGATATGTTAC
GCACACGTCTGAACTCTTTCCCTACACGACGCTCTTCCGATCTNNNGCTCAGAAAGTTTTGATATGTTAC
AGAGCACACGTCTGAACTCTTTCCCTACACGACGCTCTTCCGATCTNNNTAATGAGGCATCAGTGGAG
AGAGCACACGTCTGAACTCTTTCCCTACACGACGCTCTTCCGATCTNNNTAATGAGGCATCAGTGGAG
AGCACACGTCTGAACTCTTTCCCTACACGACGCTCTTCCGATCTNNNCCGGTTAGCTCAGTTGGTAAGA
AGCACACGTCTGAACTCTTTCCCTACACGACGCTCTTCCGATCTNNNCCGGTTAGCTCAGTTGGTAAGA
AAGAGCACACGTCTGAACTCTTTCCCTACACGACGCTCTTCCGATCTNNNCAGTGCCTGTAATCCCAGC
AAGAGCACACGTCTGAACTCTTTCCCTACACGACGCTCTTCCGATCTNNNCAGTGCCTGTAATCCCAGC
GAGCACACGTCTGAACTCTTTCCCTACACGACGCTCTTCCGATCTNNNTGTTGTCATGCCATTTTGTTTC
GAGCACACGTCTGAACTCTTTCCCTACACGACGCTCTTCCGATCTNNNTGTTGTCATGCCATTTTGTTTC
AGAGCACACGTCTGAACTCTTTCCCTACACGACGCTCTTCCGATCTNNNGCCTGTAATCCCAGCTACTCC
AGAGCACACGTCTGAACTCTTTCCCTACACGACGCTCTTCCGATCTNNNGCCTGTAATCCCAGCTACTCC
GAGCACACGTCTGAACTCTTTCCCTACACGACGCTCTTCCGATCTNNNGACCTACAATTCTGAGGCACC
GAGCACACGTCTGAACTCTTTCCCTACACGACGCTCTTCCGATCTNNNGACCTACAATTCTGAGGCACC
GAGCACACGTCTGAACTCTTTCCCTACACGACGCTCTTCCGATCTNNNAGAATTGCTTGAATCTGGGAG
GAGCACACGTCTGAACTCTTTCCCTACACGACGCTCTTCCGATCTNNNAGAATTGCTTGAATCTGGGAG
GAGCACACGTCTGAACTCTTTCCCTACACGACGCTCTTCCGATCTNNNTCCAATAGCCTTGAACAAGCTC
GAGCACACGTCTGAACTCTTTCCCTACACGACGCTCTTCCGATCTNNNTCCAATAGCCTTGAACAAGCTC
GAGCACACGTCTGAACTCTTTCCCTACACGACGCTCTTCCGATCTNNNTTACATCTTTTCAGGTCCCGAG
GAGCACACGTCTGAACTCTTTCCCTACACGACGCTCTTCCGATCTNNNTTACATCTTTTCAGGTCCCGAG
AGCACACGTCTGAACTCTTTCCCTACACGACGCTCTTCCGATCTNNNACTAGCAGCTTCACTCTTTCACA
AGCACACGTCTGAACTCTTTCCCTACACGACGCTCTTCCGATCTNNNACTAGCAGCTTCACTCTTTCACA
AGCACACGTCTGAACTCTTTCCCTACACGACGCTCTTCCGATCTNNNGGGAGAATTGCTTGAATCTGG
AGCACACGTCTGAACTCTTTCCCTACACGACGCTCTTCCGATCTNNNGGGAGAATTGCTTGAATCTGG
GAGCACACGTCTGAACTCTTTCCCTACACGACGCTCTTCCGATCTNNNCTCAAGGCATTGTCACCATCTT
GAGCACACGTCTGAACTCTTTCCCTACACGACGCTCTTCCGATCTNNNCTCAAGGCATTGTCACCATCTT
AGAGCACACGTCTGAACTCTTTCCCTACACGACGCTCTTCCGATCTNNNTTTTACAAAACAGGATGCTG
AGAGCACACGTCTGAACTCTTTCCCTACACGACGCTCTTCCGATCTNNNTTTTACAAAACAGGATGCTG

GAGCACACGTCTGAACTCTTTCCCTACACGACGCTCTTCCGATCTNNNGTAGGACAAAGGAGGAGTA
GAGCACACGTCTGAACTCTTTCCCTACACGACGCTCTTCCGATCTNNNGTAGGACAAAGGAGGAGTA
AGAGCACACGTCTGAACTCTTTCCCTACACGACGCTCTTCCGATCTNNNCACAGCCTTGAGTTTGATTC
AGAGCACACGTCTGAACTCTTTCCCTACACGACGCTCTTCCGATCTNNNCACAGCCTTGAGTTTGATTC
AGAGCACACGTCTGAACTCTTTCCCTACACGACGCTCTTCCGATCTNNNGAAAGAGAATCGCTTGAAC
AGAGCACACGTCTGAACTCTTTCCCTACACGACGCTCTTCCGATCTNNNGAAAGAGAATCGCTTGAAC
AGCACACGTCTGAACTCTTTCCCTACACGACGCTCTTCCGATCTNNNCTGGGTGATAGAGAGAGACC
AGCACACGTCTGAACTCTTTCCCTACACGACGCTCTTCCGATCTNNNCTGGGTGATAGAGAGAGACC
AGCACACGTCTGAACTCTTTCCCTACACGACGCTCTTCCGATCTNNNCTGTAATCCCAGCTACTCCAGAG
AGCACACGTCTGAACTCTTTCCCTACACGACGCTCTTCCGATCTNNNCTGTAATCCCAGCTACTCCAGAG
GCACACGTCTGAACTCTTTCCCTACACGACGCTCTTCCGATCTNNNAGTGCTAGGATTACAGGCATGA
GCACACGTCTGAACTCTTTCCCTACACGACGCTCTTCCGATCTNNNAGTGCTAGGATTACAGGCATGA
AGCACACGTCTGAACTCTTTCCCTACACGACGCTCTTCCGATCTNNNACAACAGTTGGTTCAGAGGTG
AGCACACGTCTGAACTCTTTCCCTACACGACGCTCTTCCGATCTNNNACAACAGTTGGTTCAGAGGTG
GAGCACACGTCTGAACTCTTTCCCTACACGACGCTCTTCCGATCTNNNTCTGTAGTCCCAGCTACTCAA
GAGCACACGTCTGAACTCTTTCCCTACACGACGCTCTTCCGATCTNNNTCTGTAGTCCCAGCTACTCAA
GAGCACACGTCTGAACTCTTTCCCTACACGACGCTCTTCCGATCTNNNTTGAGACAGAGTCTTGCACTG
GAGCACACGTCTGAACTCTTTCCCTACACGACGCTCTTCCGATCTNNNTTGAGACAGAGTCTTGCACTG
GCACACGTCTGAACTCTTTCCCTACACGACGCTCTTCCGATCTNNNTTTGACCTCTGCAAAATGGGTTG
GCACACGTCTGAACTCTTTCCCTACACGACGCTCTTCCGATCTNNNTTTGACCTCTGCAAAATGGGTTG
GCACACGTCTGAACTCTTTCCCTACACGACGCTCTTCCGATCTNNNAGGTCATTCAAAGTTTCACAGCG
GCACACGTCTGAACTCTTTCCCTACACGACGCTCTTCCGATCTNNNAGGTCATTCAAAGTTTCACAGCG
GCACACGTCTGAACTCTTTCCCTACACGACGCTCTTCCGATCTNNNCTAATGCAGGCTGAAATGCTAG
GCACACGTCTGAACTCTTTCCCTACACGACGCTCTTCCGATCTNNNCTAATGCAGGCTGAAATGCTAG
AGCACACGTCTGAACTCTTTCCCTACACGACGCTCTTCCGATCTNNNACCTCCATCATTCCATATCTGGG
AGCACACGTCTGAACTCTTTCCCTACACGACGCTCTTCCGATCTNNNACCTCCATCATTCCATATCTGGG
AGCACACGTCTGAACTCTTTCCCTACACGACGCTCTTCCGATCTNNNGATAGAGTCAAGACCCATCGG
AGCACACGTCTGAACTCTTTCCCTACACGACGCTCTTCCGATCTNNNGATAGAGTCAAGACCCATCGG
AGCACACGTCTGAACTCTTTCCCTACACGACGCTCTTCCGATCTNNNTGTGGGCAGCAGAAATGTAAA
AGCACACGTCTGAACTCTTTCCCTACACGACGCTCTTCCGATCTNNNTGTGGGCAGCAGAAATGTAAA
AGAGCACACGTCTGAACTCTTTCCCTACACGACGCTCTTCCGATCTNNNGATCAAGACCATCCTGGCTA
AGAGCACACGTCTGAACTCTTTCCCTACACGACGCTCTTCCGATCTNNNGATCAAGACCATCCTGGCTA
ACACGTCTGAACTCTTTCCCTACACGACGCTCTTCCGATCTNNNGCCCCTTTGTAATTCCCTTCCTACAT
ACACGTCTGAACTCTTTCCCTACACGACGCTCTTCCGATCTNNNGCCCCTTTGTAATTCCCTTCCTACAT
AGCACACGTCTGAACTCTTTCCCTACACGACGCTCTTCCGATCTNNNGAAACAGCTACAGACCTTCTCC
AGCACACGTCTGAACTCTTTCCCTACACGACGCTCTTCCGATCTNNNGAAACAGCTACAGACCTTCTCC
GAGCACACGTCTGAACTCTTTCCCTACACGACGCTCTTCCGATCTNNNAAAAAATTAGCCGGGCATGG
GAGCACACGTCTGAACTCTTTCCCTACACGACGCTCTTCCGATCTNNNAAAAAATTAGCCGGGCATGG
AGCACACGTCTGAACTCTTTCCCTACACGACGCTCTTCCGATCTNNNCTGAACACGCTCATGTAACCAT
AGCACACGTCTGAACTCTTTCCCTACACGACGCTCTTCCGATCTNNNCTGAACACGCTCATGTAACCAT
GAGCACACGTCTGAACTCTTTCCCTACACGACGCTCTTCCGATCTNNNAGGTACATTTTGACCCCATGG
GAGCACACGTCTGAACTCTTTCCCTACACGACGCTCTTCCGATCTNNNAGGTACATTTTGACCCCATGG
GAGCACACGTCTGAACTCTTTCCCTACACGACGCTCTTCCGATCTNNNGGGAGGTTGCAATGAACTGA

GAGCACACGTCTGAACTCTTTCCCTACACGACGCTCTTCCGATCTNNNGGGAGGTTGCAATGAACTGA
GAGCACACGTCTGAACTCTTTCCCTACACGACGCTCTTCCGATCTNNNGACTTTCCACTTGTTCTCCTCCC
GAGCACACGTCTGAACTCTTTCCCTACACGACGCTCTTCCGATCTNNNGACTTTCCACTTGTTCTCCTCCC
GCACACGTCTGAACTCTTTCCCTACACGACGCTCTTCCGATCTNNNACACGTACTTCTTCCAACAGTGAC
GCACACGTCTGAACTCTTTCCCTACACGACGCTCTTCCGATCTNNNACACGTACTTCTTCCAACAGTGAC
AGCACACGTCTGAACTCTTTCCCTACACGACGCTCTTCCGATCTNNNTTGGCATAGCACTCTGACAGTT
AGCACACGTCTGAACTCTTTCCCTACACGACGCTCTTCCGATCTNNNTTGGCATAGCACTCTGACAGTT
GCACACGTCTGAACTCTTTCCCTACACGACGCTCTTCCGATCTNNNATGGGTACAAAGCATAGTGCTCC
GCACACGTCTGAACTCTTTCCCTACACGACGCTCTTCCGATCTNNNATGGGTACAAAGCATAGTGCTCC
AGCACACGTCTGAACTCTTTCCCTACACGACGCTCTTCCGATCTNNNTTCACAGCTGATGGGAGCATAA
AGCACACGTCTGAACTCTTTCCCTACACGACGCTCTTCCGATCTNNNTTCACAGCTGATGGGAGCATAA
GAGCACACGTCTGAACTCTTTCCCTACACGACGCTCTTCCGATCTNNNCCTCGGAAGATACGGCGTAG
GAGCACACGTCTGAACTCTTTCCCTACACGACGCTCTTCCGATCTNNNCCTCGGAAGATACGGCGTAG
GAGCACACGTCTGAACTCTTTCCCTACACGACGCTCTTCCGATCTNNNCTGTAGTCCCAGCTACTCAGG
GAGCACACGTCTGAACTCTTTCCCTACACGACGCTCTTCCGATCTNNNCTGTAGTCCCAGCTACTCAGG
GAGCACACGTCTGAACTCTTTCCCTACACGACGCTCTTCCGATCTNNNGTAATCCCAGCTACTCAGGAA
GAGCACACGTCTGAACTCTTTCCCTACACGACGCTCTTCCGATCTNNNGTAATCCCAGCTACTCAGGAA
GAGCACACGTCTGAACTCTTTCCCTACACGACGCTCTTCCGATCTNNNAAAGTTAGGTGGAGACATGG
GAGCACACGTCTGAACTCTTTCCCTACACGACGCTCTTCCGATCTNNNAAAGTTAGGTGGAGACATGG
GCACACGTCTGAACTCTTTCCCTACACGACGCTCTTCCGATCTNNNATTGCTGCAATAAACATGGGAGT
GCACACGTCTGAACTCTTTCCCTACACGACGCTCTTCCGATCTNNNATTGCTGCAATAAACATGGGAGT
GAGCACACGTCTGAACTCTTTCCCTACACGACGCTCTTCCGATCTNNNCTCGTTTTCCAATCATTTTATTC
GAGCACACGTCTGAACTCTTTCCCTACACGACGCTCTTCCGATCTNNNCTCGTTTTCCAATCATTTTATTC
GAGCACACGTCTGAACTCTTTCCCTACACGACGCTCTTCCGATCTNNNTCCGTCCTAGTTGGGTTTTGAT
GAGCACACGTCTGAACTCTTTCCCTACACGACGCTCTTCCGATCTNNNTCCGTCCTAGTTGGGTTTTGAT
GAGCACACGTCTGAACTCTTTCCCTACACGACGCTCTTCCGATCTNNNAGTAAAAGATTTGGGATGTT
GAGCACACGTCTGAACTCTTTCCCTACACGACGCTCTTCCGATCTNNNAGTAAAAGATTTGGGATGTT
GAGCACACGTCTGAACTCTTTCCCTACACGACGCTCTTCCGATCTNNNTCAACCTAAGTGTTCATCAGTC
GAGCACACGTCTGAACTCTTTCCCTACACGACGCTCTTCCGATCTNNNTCAACCTAAGTGTTCATCAGTC
GAGCACACGTCTGAACTCTTTCCCTACACGACGCTCTTCCGATCTNNNATGTTTGGGGTTTCTGTTATC
GAGCACACGTCTGAACTCTTTCCCTACACGACGCTCTTCCGATCTNNNATGTTTGGGGTTTCTGTTATC
AGAGCACACGTCTGAACTCTTTCCCTACACGACGCTCTTCCGATCTNNNGGCTCTAGGCAAACCCATAAT
AGAGCACACGTCTGAACTCTTTCCCTACACGACGCTCTTCCGATCTNNNGGCTCTAGGCAAACCCATAAT
AGCACACGTCTGAACTCTTTCCCTACACGACGCTCTTCCGATCTNNNCTGGAGGCACAGCATCTATCAT
AGCACACGTCTGAACTCTTTCCCTACACGACGCTCTTCCGATCTNNNCTGGAGGCACAGCATCTATCAT
AGAGCACACGTCTGAACTCTTTCCCTACACGACGCTCTTCCGATCTNNNATCCAACTCCGTGTGTGGTTT
AGAGCACACGTCTGAACTCTTTCCCTACACGACGCTCTTCCGATCTNNNATCCAACTCCGTGTGTGGTTT
CACACGTCTGAACTCTTTCCCTACACGACGCTCTTCCGATCTNNNCAGTCTGGGAAGTGAGGAGCCAT
CACACGTCTGAACTCTTTCCCTACACGACGCTCTTCCGATCTNNNCAGTCTGGGAAGTGAGGAGCCAT
GCACACGTCTGAACTCTTTCCCTACACGACGCTCTTCCGATCTNNNCACTGTGACCTCTTGAGCTCAACA
GCACACGTCTGAACTCTTTCCCTACACGACGCTCTTCCGATCTNNNCACTGTGACCTCTTGAGCTCAACA
GAGCACACGTCTGAACTCTTTCCCTACACGACGCTCTTCCGATCTNNNAAGGCAAAAACAACTCTGGAG
GAGCACACGTCTGAACTCTTTCCCTACACGACGCTCTTCCGATCTNNNAAGGCAAAAACAACTCTGGAG

AGCACACGTCTGAACTCTTTCCCTACACGACGCTCTTCCGATCTNNNGCGAAACCCCGTCTCTACTAAAA
AGCACACGTCTGAACTCTTTCCCTACACGACGCTCTTCCGATCTNNNGCGAAACCCCGTCTCTACTAAAA
CACACGTCTGAACTCTTTCCCTACACGACGCTCTTCCGATCTNNNAGCTGCTGATCACTCATTTCAGGCA
CACACGTCTGAACTCTTTCCCTACACGACGCTCTTCCGATCTNNNAGCTGCTGATCACTCATTTCAGGCA
AGAGCACACGTCTGAACTCTTTCCCTACACGACGCTCTTCCGATCTNNNAAAACAAGTGATGACGGTG
AGAGCACACGTCTGAACTCTTTCCCTACACGACGCTCTTCCGATCTNNNAAAACAAGTGATGACGGTG
GAGCACACGTCTGAACTCTTTCCCTACACGACGCTCTTCCGATCTNNNATGAACCATGGCAGGAGAATC
GAGCACACGTCTGAACTCTTTCCCTACACGACGCTCTTCCGATCTNNNATGAACCATGGCAGGAGAATC
GAGCACACGTCTGAACTCTTTCCCTACACGACGCTCTTCCGATCTNNNTCACCCACCTGACTAGTCATAG
GAGCACACGTCTGAACTCTTTCCCTACACGACGCTCTTCCGATCTNNNTCACCCACCTGACTAGTCATAG
GAGCACACGTCTGAACTCTTTCCCTACACGACGCTCTTCCGATCTNNNCCTGGTCTGTTGGAACTCCATT
GAGCACACGTCTGAACTCTTTCCCTACACGACGCTCTTCCGATCTNNNCCTGGTCTGTTGGAACTCCATT
GCACACGTCTGAACTCTTTCCCTACACGACGCTCTTCCGATCTNNNAGATCTTACCACTGCACTCTAGCC
GCACACGTCTGAACTCTTTCCCTACACGACGCTCTTCCGATCTNNNAGATCTTACCACTGCACTCTAGCC
GAGCACACGTCTGAACTCTTTCCCTACACGACGCTCTTCCGATCTNNNCATGCCTGTAATCCCAGCTACT
GAGCACACGTCTGAACTCTTTCCCTACACGACGCTCTTCCGATCTNNNCATGCCTGTAATCCCAGCTACT
GAGCACACGTCTGAACTCTTTCCCTACACGACGCTCTTCCGATCTNNNTTTGTTCTCGCAACCTGAAACA
GAGCACACGTCTGAACTCTTTCCCTACACGACGCTCTTCCGATCTNNNTTTGTTCTCGCAACCTGAAACA
AGCACACGTCTGAACTCTTTCCCTACACGACGCTCTTCCGATCTNNNTCTCCTGGGTTCAAGCAATTCTC
AGCACACGTCTGAACTCTTTCCCTACACGACGCTCTTCCGATCTNNNTCTCCTGGGTTCAAGCAATTCTC
AGCACACGTCTGAACTCTTTCCCTACACGACGCTCTTCCGATCTNNNTCTACTAATTAGAGGGGGTGG
AGCACACGTCTGAACTCTTTCCCTACACGACGCTCTTCCGATCTNNNTCTACTAATTAGAGGGGGTGG
GAGCACACGTCTGAACTCTTTCCCTACACGACGCTCTTCCGATCTNNNAGACCCTCCTGAAGTCACTCAT
GAGCACACGTCTGAACTCTTTCCCTACACGACGCTCTTCCGATCTNNNAGACCCTCCTGAAGTCACTCAT
GAGCACACGTCTGAACTCTTTCCCTACACGACGCTCTTCCGATCTNNNAGTTCGAGATCAGTCTGACCAA
GAGCACACGTCTGAACTCTTTCCCTACACGACGCTCTTCCGATCTNNNAGTTCGAGATCAGTCTGACCAA
AGCACACGTCTGAACTCTTTCCCTACACGACGCTCTTCCGATCTNNNATTAACCTGATTCACTGCTCCAT
AGCACACGTCTGAACTCTTTCCCTACACGACGCTCTTCCGATCTNNNATTAACCTGATTCACTGCTCCAT
AGCACACGTCTGAACTCTTTCCCTACACGACGCTCTTCCGATCTNNNATAGGGAGTCCTTTCTGCATTGC
AGCACACGTCTGAACTCTTTCCCTACACGACGCTCTTCCGATCTNNNATAGGGAGTCCTTTCTGCATTGC
AGAGCACACGTCTGAACTCTTTCCCTACACGACGCTCTTCCGATCTNNNCTGTAGTCCCAGCTACTCAGC
AGAGCACACGTCTGAACTCTTTCCCTACACGACGCTCTTCCGATCTNNNCTGTAGTCCCAGCTACTCAGC
GAGCACACGTCTGAACTCTTTCCCTACACGACGCTCTTCCGATCTNNNACACAAGTAGGAAGCAGCCA
GAGCACACGTCTGAACTCTTTCCCTACACGACGCTCTTCCGATCTNNNACACAAGTAGGAAGCAGCCA
GAGCACACGTCTGAACTCTTTCCCTACACGACGCTCTTCCGATCTNNNTTGAGGCAGGAGAATTGCTTG
GAGCACACGTCTGAACTCTTTCCCTACACGACGCTCTTCCGATCTNNNTTGAGGCAGGAGAATTGCTTG
AGCACACGTCTGAACTCTTTCCCTACACGACGCTCTTCCGATCTNNNCTTCTCAGGACCTAGCACAGTAC
AGCACACGTCTGAACTCTTTCCCTACACGACGCTCTTCCGATCTNNNCTTCTCAGGACCTAGCACAGTAC
AGCACACGTCTGAACTCTTTCCCTACACGACGCTCTTCCGATCTNNNCCATCAAGGGATATGTGTAGG
AGCACACGTCTGAACTCTTTCCCTACACGACGCTCTTCCGATCTNNNCCATCAAGGGATATGTGTAGG
AGCACACGTCTGAACTCTTTCCCTACACGACGCTCTTCCGATCTNNNCTCCCATGTTCATTGCAGCATAA
AGCACACGTCTGAACTCTTTCCCTACACGACGCTCTTCCGATCTNNNCTCCCATGTTCATTGCAGCATAA
AGCACACGTCTGAACTCTTTCCCTACACGACGCTCTTCCGATCTNNNACTTAGTGCTTCTCTTGTCCCAGC

GCACACGTCTGAACTCTTTCCCTACACGACGCTCTTCCGATCTNNNACTTAGTGCTTCTCTTGTCCCAG

AGCACACGTCTGAACTCTTTCCCTACACGACGCTCTTCCGATCTNNNAGAGAGAGACTTGTTCTGTTG

AGCACACGTCTGAACTCTTTCCCTACACGACGCTCTTCCGATCTNNNAGAGAGAGACTTGTTCTGTTG

GCACACGTCTGAACTCTTTCCCTACACGACGCTCTTCCGATCTNNNCCTCAGTGGTCGGAAACTCCCAT

GCACACGTCTGAACTCTTTCCCTACACGACGCTCTTCCGATCTNNNCCTCAGTGGTCGGAAACTCCCAT

GAGCACACGTCTGAACTCTTTCCCTACACGACGCTCTTCCGATCTNNNGTTAGGAGTTCAAAACCAGC

GAGCACACGTCTGAACTCTTTCCCTACACGACGCTCTTCCGATCTNNNGTTAGGAGTTCAAAACCAGC

GAGCACACGTCTGAACTCTTTCCCTACACGACGCTCTTCCGATCTNNNGAGGCTGAGGCAGGAGAAT

GAGCACACGTCTGAACTCTTTCCCTACACGACGCTCTTCCGATCTNNNGAGGCTGAGGCAGGAGAAT

AGCACACGTCTGAACTCTTTCCCTACACGACGCTCTTCCGATCTNNNTCCAAGTTCTAACTAGGCCCAA

AGCACACGTCTGAACTCTTTCCCTACACGACGCTCTTCCGATCTNNNTCCAAGTTCTAACTAGGCCCAA

AGCACACGTCTGAACTCTTTCCCTACACGACGCTCTTCCGATCTNNNCAAAGGGTGACTCTCCTTTCTG

AGCACACGTCTGAACTCTTTCCCTACACGACGCTCTTCCGATCTNNNCAAAGGGTGACTCTCCTTTCTG

AGCACACGTCTGAACTCTTTCCCTACACGACGCTCTTCCGATCTNNNGTGGGGATTATGGGAGCTACA

AGCACACGTCTGAACTCTTTCCCTACACGACGCTCTTCCGATCTNNNGTGGGGATTATGGGAGCTACA

AGCACACGTCTGAACTCTTTCCCTACACGACGCTCTTCCGATCTNNNCAGCTTGGGAGGCTCAGTAAA

AGCACACGTCTGAACTCTTTCCCTACACGACGCTCTTCCGATCTNNNCAGCTTGGGAGGCTCAGTAAA

AGCACACGTCTGAACTCTTTCCCTACACGACGCTCTTCCGATCTNNNGATCATGCCATTGTACTTCAG

AGCACACGTCTGAACTCTTTCCCTACACGACGCTCTTCCGATCTNNNGATCATGCCATTGTACTTCAG

AGCACACGTCTGAACTCTTTCCCTACACGACGCTCTTCCGATCTNNNCTGATCATTTGGCTCTTTTTCA

AGCACACGTCTGAACTCTTTCCCTACACGACGCTCTTCCGATCTNNNCTGATCATTTGGCTCTTTTTCA

AGCACACGTCTGAACTCTTTCCCTACACGACGCTCTTCCGATCTNNNGTTGAGGCTGTAGCTGTAGTG

AGCACACGTCTGAACTCTTTCCCTACACGACGCTCTTCCGATCTNNNGTTGAGGCTGTAGCTGTAGTG

AGCACACGTCTGAACTCTTTCCCTACACGACGCTCTTCCGATCTNNNTGATGCACACCTGTAATTCCAG

AGCACACGTCTGAACTCTTTCCCTACACGACGCTCTTCCGATCTNNNTGATGCACACCTGTAATTCCAG

AGAGCACACGTCTGAACTCTTTCCCTACACGACGCTCTTCCGATCTNNNTCCCAGCAAAATAATCAGCT

AGAGCACACGTCTGAACTCTTTCCCTACACGACGCTCTTCCGATCTNNNTCCCAGCAAAATAATCAGCT

AGAGCACACGTCTGAACTCTTTCCCTACACGACGCTCTTCCGATCTNNNGCTAACACGGTGAAACTCCA

AGAGCACACGTCTGAACTCTTTCCCTACACGACGCTCTTCCGATCTNNNGCTAACACGGTGAAACTCCA

GCACACGTCTGAACTCTTTCCCTACACGACGCTCTTCCGATCTNNNCCACGCCTGGCTAATTTTTTGTA

GCACACGTCTGAACTCTTTCCCTACACGACGCTCTTCCGATCTNNNCCACGCCTGGCTAATTTTTTGTA

AGCACACGTCTGAACTCTTTCCCTACACGACGCTCTTCCGATCTNNNATGCCTGTAATCCCAGCTACTT

AGCACACGTCTGAACTCTTTCCCTACACGACGCTCTTCCGATCTNNNATGCCTGTAATCCCAGCTACTT

AGAGCACACGTCTGAACTCTTTCCCTACACGACGCTCTTCCGATCTNNNCTGAGGCAGGAGAATCACT

AGAGCACACGTCTGAACTCTTTCCCTACACGACGCTCTTCCGATCTNNNCTGAGGCAGGAGAATCACT

GAGCACACGTCTGAACTCTTTCCCTACACGACGCTCTTCCGATCTNNNGCCTGTAGTCCCAGCTACTTG

GAGCACACGTCTGAACTCTTTCCCTACACGACGCTCTTCCGATCTNNNGCCTGTAGTCCCAGCTACTTG

AGCACACGTCTGAACTCTTTCCCTACACGACGCTCTTCCGATCTNNNTCCCCACCAATAGTAAAATGCT

AGCACACGTCTGAACTCTTTCCCTACACGACGCTCTTCCGATCTNNNTCCCCACCAATAGTAAAATGCT

AGCACACGTCTGAACTCTTTCCCTACACGACGCTCTTCCGATCTNNNGTTCCAAGTAAGCCCAAAACAC

AGCACACGTCTGAACTCTTTCCCTACACGACGCTCTTCCGATCTNNNGTTCCAAGTAAGCCCAAAACAC

AGAGCACACGTCTGAACTCTTTCCCTACACGACGCTCTTCCGATCTNNNGCGAGACTCTGCCACAAAAA

AGAGCACACGTCTGAACTCTTTCCCTACACGACGCTCTTCCGATCTNNNGCGAGACTCTGCCACAAAAA

GAGCACACGTCTGAACTCTTTCCCTACACGACGCTCTTCCGATCTNNNATGGAAAAGGCAGTCACCCA
GAGCACACGTCTGAACTCTTTCCCTACACGACGCTCTTCCGATCTNNNATGGAAAAGGCAGTCACCCA
GAGCACACGTCTGAACTCTTTCCCTACACGACGCTCTTCCGATCTNNNCACACACATACACACACACACA
GAGCACACGTCTGAACTCTTTCCCTACACGACGCTCTTCCGATCTNNNCACACACATACACACACACACA
GAGCACACGTCTGAACTCTTTCCCTACACGACGCTCTTCCGATCTNNNGCCAATATCACATCACTGCACT
GAGCACACGTCTGAACTCTTTCCCTACACGACGCTCTTCCGATCTNNNGCCAATATCACATCACTGCACT
GCACACGTCTGAACTCTTTCCCTACACGACGCTCTTCCGATCTNNNCTGCTACTGGTAAACCCATAGCTC
GCACACGTCTGAACTCTTTCCCTACACGACGCTCTTCCGATCTNNNCTGCTACTGGTAAACCCATAGCTC
GAGCACACGTCTGAACTCTTTCCCTACACGACGCTCTTCCGATCTNNNAAAATTAGCTGGGCATGCTAC
GAGCACACGTCTGAACTCTTTCCCTACACGACGCTCTTCCGATCTNNNAAAATTAGCTGGGCATGCTAC
AGAGCACACGTCTGAACTCTTTCCCTACACGACGCTCTTCCGATCTNNNAGACAGGAGAATCACTTGAA
AGAGCACACGTCTGAACTCTTTCCCTACACGACGCTCTTCCGATCTNNNAGACAGGAGAATCACTTGAA
GCACACGTCTGAACTCTTTCCCTACACGACGCTCTTCCGATCTNNNCTCCCTTTCATTCACCTCCATCACA
GCACACGTCTGAACTCTTTCCCTACACGACGCTCTTCCGATCTNNNCTCCCTTTCATTCACCTCCATCACA
AGCACACGTCTGAACTCTTTCCCTACACGACGCTCTTCCGATCTNNNAACAACCCAAGTGTCCATCAATG
AGCACACGTCTGAACTCTTTCCCTACACGACGCTCTTCCGATCTNNNAACAACCCAAGTGTCCATCAATG
GAGCACACGTCTGAACTCTTTCCCTACACGACGCTCTTCCGATCTNNNTAGAGATGCCCTTGCTTGACAT
GAGCACACGTCTGAACTCTTTCCCTACACGACGCTCTTCCGATCTNNNTAGAGATGCCCTTGCTTGACAT
AGCACACGTCTGAACTCTTTCCCTACACGACGCTCTTCCGATCTNNNACTGCCTTCTCAGAGAAAAGAG
AGCACACGTCTGAACTCTTTCCCTACACGACGCTCTTCCGATCTNNNACTGCCTTCTCAGAGAAAAGAG
AGCACACGTCTGAACTCTTTCCCTACACGACGCTCTTCCGATCTNNNGAGTCCGAGACAAGCCTGTAA
AGCACACGTCTGAACTCTTTCCCTACACGACGCTCTTCCGATCTNNNGAGTCCGAGACAAGCCTGTAA
GAGCACACGTCTGAACTCTTTCCCTACACGACGCTCTTCCGATCTNNNGAGGAGGCTAAGGAAGGACA
GAGCACACGTCTGAACTCTTTCCCTACACGACGCTCTTCCGATCTNNNGAGGAGGCTAAGGAAGGACA
AGCACACGTCTGAACTCTTTCCCTACACGACGCTCTTCCGATCTNNNGCTGGGAGTAGATGGGATTAC
AGCACACGTCTGAACTCTTTCCCTACACGACGCTCTTCCGATCTNNNGCTGGGAGTAGATGGGATTAC
GCACACGTCTGAACTCTTTCCCTACACGACGCTCTTCCGATCTNNNTCCAAAGCCCAGAGATAAAGCAT
GCACACGTCTGAACTCTTTCCCTACACGACGCTCTTCCGATCTNNNTCCAAAGCCCAGAGATAAAGCAT
GAGCACACGTCTGAACTCTTTCCCTACACGACGCTCTTCCGATCTNNNGCACATGTAGTCCCAGATACT
GAGCACACGTCTGAACTCTTTCCCTACACGACGCTCTTCCGATCTNNNGCACATGTAGTCCCAGATACT
AGCACACGTCTGAACTCTTTCCCTACACGACGCTCTTCCGATCTNNNCGCCAGACCTCACTACTTTTTTG
AGCACACGTCTGAACTCTTTCCCTACACGACGCTCTTCCGATCTNNNCGCCAGACCTCACTACTTTTTTG
GAGCACACGTCTGAACTCTTTCCCTACACGACGCTCTTCCGATCTNNNATTTGTCAAGGAAGAAGGTCC
GAGCACACGTCTGAACTCTTTCCCTACACGACGCTCTTCCGATCTNNNATTTGTCAAGGAAGAAGGTCC
GCACACGTCTGAACTCTTTCCCTACACGACGCTCTTCCGATCTNNNCACGCCTGTAATCCCTTCTACTCC
GCACACGTCTGAACTCTTTCCCTACACGACGCTCTTCCGATCTNNNCACGCCTGTAATCCCTTCTACTCC
GAGCACACGTCTGAACTCTTTCCCTACACGACGCTCTTCCGATCTNNNCATATGATGGTTTGCGCCTGT
GAGCACACGTCTGAACTCTTTCCCTACACGACGCTCTTCCGATCTNNNCATATGATGGTTTGCGCCTGT
AGAGCACACGTCTGAACTCTTTCCCTACACGACGCTCTTCCGATCTNNNTAGCTAGGATTACAGGTGTG
AGAGCACACGTCTGAACTCTTTCCCTACACGACGCTCTTCCGATCTNNNTAGCTAGGATTACAGGTGTG
AGCACACGTCTGAACTCTTTCCCTACACGACGCTCTTCCGATCTNNNCCACGCCTGGCTAATTTTTTGTA
AGCACACGTCTGAACTCTTTCCCTACACGACGCTCTTCCGATCTNNNCCACGCCTGGCTAATTTTTTGTA
GCACACGTCTGAACTCTTTCCCTACACGACGCTCTTCCGATCTNNNTTTCCTTCCTTGGGTCCTCTCTCCA

GCACACGTCTGAACTCTTTCCCTACACGACGCTCTTCCGATCTNNNTTTCCTTCCTTGGGTCCTCTCTCC

AGCACACGTCTGAACTCTTTCCCTACACGACGCTCTTCCGATCTNNNAATGTTTGGGCTTCCTCAGAG

AGCACACGTCTGAACTCTTTCCCTACACGACGCTCTTCCGATCTNNNAATGTTTGGGCTTCCTCAGAG

AGCACACGTCTGAACTCTTTCCCTACACGACGCTCTTCCGATCTNNNAATGAAGGAGGAGAGTTCAT

AGCACACGTCTGAACTCTTTCCCTACACGACGCTCTTCCGATCTNNNAATGAAGGAGGAGAGTTCAT

GCACACGTCTGAACTCTTTCCCTACACGACGCTCTTCCGATCTNNNGTTTCACTGTGTTAGCCAGGATG

GCACACGTCTGAACTCTTTCCCTACACGACGCTCTTCCGATCTNNNGTTTCACTGTGTTAGCCAGGATG

AGCACACGTCTGAACTCTTTCCCTACACGACGCTCTTCCGATCTNNNCTACTCCCACCATTCCCATCTTCC

AGCACACGTCTGAACTCTTTCCCTACACGACGCTCTTCCGATCTNNNCTACTCCCACCATTCCCATCTTCC

GCACACGTCTGAACTCTTTCCCTACACGACGCTCTTCCGATCTNNNTGAGCCTTTGTAAGAGGGCAATT

GCACACGTCTGAACTCTTTCCCTACACGACGCTCTTCCGATCTNNNTGAGCCTTTGTAAGAGGGCAATT

GAGCACACGTCTGAACTCTTTCCCTACACGACGCTCTTCCGATCTNNNGTAATCCCAGCTATACAGGA

GAGCACACGTCTGAACTCTTTCCCTACACGACGCTCTTCCGATCTNNNGTAATCCCAGCTATACAGGA

AGCACACGTCTGAACTCTTTCCCTACACGACGCTCTTCCGATCTNNNAAAAGTTACTCCCATGGCTCCTC

AGCACACGTCTGAACTCTTTCCCTACACGACGCTCTTCCGATCTNNNAAAAGTTACTCCCATGGCTCCTC

GAGCACACGTCTGAACTCTTTCCCTACACGACGCTCTTCCGATCTNNNCTCCCAAGTTGCTGAGACTAC

GAGCACACGTCTGAACTCTTTCCCTACACGACGCTCTTCCGATCTNNNCTCCCAAGTTGCTGAGACTAC

AGCACACGTCTGAACTCTTTCCCTACACGACGCTCTTCCGATCTNNNCTCCCTTCAGTGTCTGCTAGAA

AGCACACGTCTGAACTCTTTCCCTACACGACGCTCTTCCGATCTNNNCTCCCTTCAGTGTCTGCTAGAA

GAGCACACGTCTGAACTCTTTCCCTACACGACGCTCTTCCGATCTNNNTCCAAAGGAGAGAAGTTTGC

GAGCACACGTCTGAACTCTTTCCCTACACGACGCTCTTCCGATCTNNNTCCAAAGGAGAGAAGTTTGC

AGCACACGTCTGAACTCTTTCCCTACACGACGCTCTTCCGATCTNNNCCTTCCTGAAAGTAGGCGAGG

AGCACACGTCTGAACTCTTTCCCTACACGACGCTCTTCCGATCTNNNCCTTCCTGAAAGTAGGCGAGG

GAGCACACGTCTGAACTCTTTCCCTACACGACGCTCTTCCGATCTNNNCATTTTGTCACCCCAACCAGTA

GAGCACACGTCTGAACTCTTTCCCTACACGACGCTCTTCCGATCTNNNCATTTTGTCACCCCAACCAGTA

AGAGCACACGTCTGAACTCTTTCCCTACACGACGCTCTTCCGATCTNNNCTGAGGCAGGAGAATTGCT

AGAGCACACGTCTGAACTCTTTCCCTACACGACGCTCTTCCGATCTNNNCTGAGGCAGGAGAATTGCT

AGCACACGTCTGAACTCTTTCCCTACACGACGCTCTTCCGATCTNNNGAGAATCACTTGAACCCAGGA

AGCACACGTCTGAACTCTTTCCCTACACGACGCTCTTCCGATCTNNNGAGAATCACTTGAACCCAGGA

AGCACACGTCTGAACTCTTTCCCTACACGACGCTCTTCCGATCTNNNTCCCTCCCATGATATGTGGAGA

AGCACACGTCTGAACTCTTTCCCTACACGACGCTCTTCCGATCTNNNTCCCTCCCATGATATGTGGAGA

GAGCACACGTCTGAACTCTTTCCCTACACGACGCTCTTCCGATCTNNNGGTTGCAGTAAGCTGAGATT

GAGCACACGTCTGAACTCTTTCCCTACACGACGCTCTTCCGATCTNNNGGTTGCAGTAAGCTGAGATT

AGCACACGTCTGAACTCTTTCCCTACACGACGCTCTTCCGATCTNNNGGCAATGATAGCAGTGACTAT

AGCACACGTCTGAACTCTTTCCCTACACGACGCTCTTCCGATCTNNNGGCAATGATAGCAGTGACTAT

GAGCACACGTCTGAACTCTTTCCCTACACGACGCTCTTCCGATCTNNNTGCTTGGAGAGAGAGAGAG

GAGCACACGTCTGAACTCTTTCCCTACACGACGCTCTTCCGATCTNNNTGCTTGGAGAGAGAGAGAG

AGCACACGTCTGAACTCTTTCCCTACACGACGCTCTTCCGATCTNNNTAGCAAGACCTCGTATCTACC

AGCACACGTCTGAACTCTTTCCCTACACGACGCTCTTCCGATCTNNNTAGCAAGACCTCGTATCTACC

AGCACACGTCTGAACTCTTTCCCTACACGACGCTCTTCCGATCTNNNCAAAATAATTAGCTGGGCGTG

AGCACACGTCTGAACTCTTTCCCTACACGACGCTCTTCCGATCTNNNCAAAATAATTAGCTGGGCGTG

AGCACACGTCTGAACTCTTTCCCTACACGACGCTCTTCCGATCTNNNCAGCATATTTGGGATGTGACA

AGCACACGTCTGAACTCTTTCCCTACACGACGCTCTTCCGATCTNNNCAGCATATTTGGGATGTGACA

AGCACACGTCTGAACTCTTTCCCTACACGACGCTCTTCCGATCTNNNAGGTTGAGGCAGGAGAATCATT
AGCACACGTCTGAACTCTTTCCCTACACGACGCTCTTCCGATCTNNNAGGTTGAGGCAGGAGAATCATT
GAGCACACGTCTGAACTCTTTCCCTACACGACGCTCTTCCGATCTNNNTGGAAGACTCTCACGGCTTTT
GAGCACACGTCTGAACTCTTTCCCTACACGACGCTCTTCCGATCTNNNTGGAAGACTCTCACGGCTTTT
GCACACGTCTGAACTCTTTCCCTACACGACGCTCTTCCGATCTNNNCTCTCCTTCATAAGCGCATCCTCCA
GCACACGTCTGAACTCTTTCCCTACACGACGCTCTTCCGATCTNNNCTCTCCTTCATAAGCGCATCCTCCA
AGCACACGTCTGAACTCTTTCCCTACACGACGCTCTTCCGATCTNNNGAGAGAGAGAGAGTGTGTGTG
AGCACACGTCTGAACTCTTTCCCTACACGACGCTCTTCCGATCTNNNGAGAGAGAGAGAGTGTGTGTG
AGCACACGTCTGAACTCTTTCCCTACACGACGCTCTTCCGATCTNNNTCTTTATAGGGAATGGGCTGAC
AGCACACGTCTGAACTCTTTCCCTACACGACGCTCTTCCGATCTNNNTCTTTATAGGGAATGGGCTGAC
AGCACACGTCTGAACTCTTTCCCTACACGACGCTCTTCCGATCTNNNGTCAACGGTACTTGGATCTGTCT
AGCACACGTCTGAACTCTTTCCCTACACGACGCTCTTCCGATCTNNNGTCAACGGTACTTGGATCTGTCT
AGCACACGTCTGAACTCTTTCCCTACACGACGCTCTTCCGATCTNNNCTTGTGAGGGGGATTACAAGGA
AGCACACGTCTGAACTCTTTCCCTACACGACGCTCTTCCGATCTNNNCTTGTGAGGGGGATTACAAGGA
GAGCACACGTCTGAACTCTTTCCCTACACGACGCTCTTCCGATCTNNNTGAATTCAGTCCCCAAACTAGC
GAGCACACGTCTGAACTCTTTCCCTACACGACGCTCTTCCGATCTNNNTGAATTCAGTCCCCAAACTAGC
AGAGCACACGTCTGAACTCTTTCCCTACACGACGCTCTTCCGATCTNNNTATAGTCCCAGCTACTCAGGA
AGAGCACACGTCTGAACTCTTTCCCTACACGACGCTCTTCCGATCTNNNTATAGTCCCAGCTACTCAGGA
GAGCACACGTCTGAACTCTTTCCCTACACGACGCTCTTCCGATCTNNNCAGTCCTCACCATACGCTAATC
GAGCACACGTCTGAACTCTTTCCCTACACGACGCTCTTCCGATCTNNNCAGTCCTCACCATACGCTAATC
GAGCACACGTCTGAACTCTTTCCCTACACGACGCTCTTCCGATCTNNNAGAATCTTTTGAGCATGGGAC
GAGCACACGTCTGAACTCTTTCCCTACACGACGCTCTTCCGATCTNNNAGAATCTTTTGAGCATGGGAC
GAGCACACGTCTGAACTCTTTCCCTACACGACGCTCTTCCGATCTNNNTGGAAAGTGGGAAGAACAGA
GAGCACACGTCTGAACTCTTTCCCTACACGACGCTCTTCCGATCTNNNTGGAAAGTGGGAAGAACAGA
AGCACACGTCTGAACTCTTTCCCTACACGACGCTCTTCCGATCTNNNCACCACACCTGGCTAATTTTTGT
AGCACACGTCTGAACTCTTTCCCTACACGACGCTCTTCCGATCTNNNCACCACACCTGGCTAATTTTTGT
GAGCACACGTCTGAACTCTTTCCCTACACGACGCTCTTCCGATCTNNNGAGAGAGAGGTTGCAGTGAA
GAGCACACGTCTGAACTCTTTCCCTACACGACGCTCTTCCGATCTNNNGAGAGAGAGGTTGCAGTGAA
AGCACACGTCTGAACTCTTTCCCTACACGACGCTCTTCCGATCTNNNGACCTCATTTTGCCTGAATCACC
AGCACACGTCTGAACTCTTTCCCTACACGACGCTCTTCCGATCTNNNGACCTCATTTTGCCTGAATCACC
GAAGAGCACACGTCTGAACTCTTTCCCTACACGACGCTCTTCCGATCTNNNGAGGTTGTAGTGAGCCG
GAAGAGCACACGTCTGAACTCTTTCCCTACACGACGCTCTTCCGATCTNNNGAGGTTGTAGTGAGCCG
GAGCACACGTCTGAACTCTTTCCCTACACGACGCTCTTCCGATCTNNNGCTTCTCAAGAGTAGCTGGGA
GAGCACACGTCTGAACTCTTTCCCTACACGACGCTCTTCCGATCTNNNGCTTCTCAAGAGTAGCTGGGA
GAGCACACGTCTGAACTCTTTCCCTACACGACGCTCTTCCGATCTNNNATGCCTATAACCCCAGCACTTT
GAGCACACGTCTGAACTCTTTCCCTACACGACGCTCTTCCGATCTNNNATGCCTATAACCCCAGCACTTT
GCACACGTCTGAACTCTTTCCCTACACGACGCTCTTCCGATCTNNNGCTTATGTGACCTCTGAGGAGAC
GCACACGTCTGAACTCTTTCCCTACACGACGCTCTTCCGATCTNNNGCTTATGTGACCTCTGAGGAGAC
GAGCACACGTCTGAACTCTTTCCCTACACGACGCTCTTCCGATCTNNNGCCTGTAATCCCATCTACTCAC
GAGCACACGTCTGAACTCTTTCCCTACACGACGCTCTTCCGATCTNNNGCCTGTAATCCCATCTACTCAC
AGCACACGTCTGAACTCTTTCCCTACACGACGCTCTTCCGATCTNNNTGCTAGCTCTCTATTTTGCCTGT
AGCACACGTCTGAACTCTTTCCCTACACGACGCTCTTCCGATCTNNNTGCTAGCTCTCTATTTTGCCTGT
GCACACGTCTGAACTCTTTCCCTACACGACGCTCTTCCGATCTNNNTTGAATCCGTCTCATTGAAGGGT

GCACACGTCTGAACTCTTTCCCTACACGACGCTCTTCCGATCTNNNTTGAATCCGTCTCATTGAAGGGT
AGCACACGTCTGAACTCTTTCCCTACACGACGCTCTTCCGATCTNNNCTCAGCAAATGGAAGTAGTGC
AGCACACGTCTGAACTCTTTCCCTACACGACGCTCTTCCGATCTNNNCTCAGCAAATGGAAGTAGTGC
GAGCACACGTCTGAACTCTTTCCCTACACGACGCTCTTCCGATCTNNNTCTCTACCAATATTAAGCCTT
GAGCACACGTCTGAACTCTTTCCCTACACGACGCTCTTCCGATCTNNNTCTCTACCAATATTAAGCCTT
CACACGTCTGAACTCTTTCCCTACACGACGCTCTTCCGATCTNNNTCACTGATGTCTCCCTTCTCTCTCAT
CACACGTCTGAACTCTTTCCCTACACGACGCTCTTCCGATCTNNNTCACTGATGTCTCCCTTCTCTCTCAT
AGCACACGTCTGAACTCTTTCCCTACACGACGCTCTTCCGATCTNNNGCAGAAACATGCCTCCAGAAT
AGCACACGTCTGAACTCTTTCCCTACACGACGCTCTTCCGATCTNNNGCAGAAACATGCCTCCAGAAT
AGCACACGTCTGAACTCTTTCCCTACACGACGCTCTTCCGATCTNNNCACCTGTAATCCCAGCTACTTG
AGCACACGTCTGAACTCTTTCCCTACACGACGCTCTTCCGATCTNNNCACCTGTAATCCCAGCTACTTG
AGCACACGTCTGAACTCTTTCCCTACACGACGCTCTTCCGATCTNNNCTGTAGTCCCAGCTACTTGTG
AGCACACGTCTGAACTCTTTCCCTACACGACGCTCTTCCGATCTNNNCTGTAGTCCCAGCTACTTGTG
AGAGCACACGTCTGAACTCTTTCCCTACACGACGCTCTTCCGATCTNNNTGATGTTCATCAGGGGCTA
AGAGCACACGTCTGAACTCTTTCCCTACACGACGCTCTTCCGATCTNNNTGATGTTCATCAGGGGCTA
AGCACACGTCTGAACTCTTTCCCTACACGACGCTCTTCCGATCTNNNAGGTTTCAGTGAGCCAAGATC
AGCACACGTCTGAACTCTTTCCCTACACGACGCTCTTCCGATCTNNNAGGTTTCAGTGAGCCAAGATC
GCACACGTCTGAACTCTTTCCCTACACGACGCTCTTCCGATCTNNNATCACACCTCACTGATGAAACC
GCACACGTCTGAACTCTTTCCCTACACGACGCTCTTCCGATCTNNNATCACACCTCACTGATGAAACC
AGCACACGTCTGAACTCTTTCCCTACACGACGCTCTTCCGATCTNNNAAAGAGAAAGATGTCACTGG
AGCACACGTCTGAACTCTTTCCCTACACGACGCTCTTCCGATCTNNNAAAGAGAAAGATGTCACTGG
AGCACACGTCTGAACTCTTTCCCTACACGACGCTCTTCCGATCTNNNGCCACTACTGTTTTCCAAAGCA
AGCACACGTCTGAACTCTTTCCCTACACGACGCTCTTCCGATCTNNNGCCACTACTGTTTTCCAAAGCA
AGCACACGTCTGAACTCTTTCCCTACACGACGCTCTTCCGATCTNNNGTTGTAGTCAGAAGCCATGCA
AGCACACGTCTGAACTCTTTCCCTACACGACGCTCTTCCGATCTNNNGTTGTAGTCAGAAGCCATGCA
GCACACGTCTGAACTCTTTCCCTACACGACGCTCTTCCGATCTNNNTTTGCTTTTCTATCCCGCCCTAC
GCACACGTCTGAACTCTTTCCCTACACGACGCTCTTCCGATCTNNNTTTGCTTTTCTATCCCGCCCTAC
GAGCACACGTCTGAACTCTTTCCCTACACGACGCTCTTCCGATCTNNNGGCTGCAGTGAGCTATAATT
GAGCACACGTCTGAACTCTTTCCCTACACGACGCTCTTCCGATCTNNNGGCTGCAGTGAGCTATAATT
AGCACACGTCTGAACTCTTTCCCTACACGACGCTCTTCCGATCTNNNTCTTAGCCTGACTTCTAGCTTG
AGCACACGTCTGAACTCTTTCCCTACACGACGCTCTTCCGATCTNNNTCTTAGCCTGACTTCTAGCTTG
AGCACACGTCTGAACTCTTTCCCTACACGACGCTCTTCCGATCTNNNCCTTCAAATACCATCACCTTGA
AGCACACGTCTGAACTCTTTCCCTACACGACGCTCTTCCGATCTNNNCCTTCAAATACCATCACCTTGA
GAGCACACGTCTGAACTCTTTCCCTACACGACGCTCTTCCGATCTNNNCTCACTCCTGTAATCCCAACA
GAGCACACGTCTGAACTCTTTCCCTACACGACGCTCTTCCGATCTNNNCTCACTCCTGTAATCCCAACA
AGCACACGTCTGAACTCTTTCCCTACACGACGCTCTTCCGATCTNNNCACCACAGGGCTTTTCTTTCTTT
AGCACACGTCTGAACTCTTTCCCTACACGACGCTCTTCCGATCTNNNCACCACAGGGCTTTTCTTTCTTT
AGAGCACACGTCTGAACTCTTTCCCTACACGACGCTCTTCCGATCTNNNGCCACCATTTTCTTCTCTCTC
AGAGCACACGTCTGAACTCTTTCCCTACACGACGCTCTTCCGATCTNNNGCCACCATTTTCTTCTCTCTC
GCACACGTCTGAACTCTTTCCCTACACGACGCTCTTCCGATCTNNNTACCAGTGAATTTTTGCAATAAT
GCACACGTCTGAACTCTTTCCCTACACGACGCTCTTCCGATCTNNNTACCAGTGAATTTTTGCAATAAT
GAGCACACGTCTGAACTCTTTCCCTACACGACGCTCTTCCGATCTNNNCTGAGGCAGGAGAATCACTT
GAGCACACGTCTGAACTCTTTCCCTACACGACGCTCTTCCGATCTNNNCTGAGGCAGGAGAATCACTT

AGCACACGTCTGAACTCTTTCCCTACACGACGCTCTTCCGATCTNNNATCCTCCCACCTAAGACTTCTG
AGCACACGTCTGAACTCTTTCCCTACACGACGCTCTTCCGATCTNNNATCCTCCCACCTAAGACTTCTG
AGCACACGTCTGAACTCTTTCCCTACACGACGCTCTTCCGATCTNNNATATAAGGTACTGTCCTGGGC
AGCACACGTCTGAACTCTTTCCCTACACGACGCTCTTCCGATCTNNNATATAAGGTACTGTCCTGGGC
GCACACGTCTGAACTCTTTCCCTACACGACGCTCTTCCGATCTNNNAGTCCAACCAACATATTTTAAAT
GCACACGTCTGAACTCTTTCCCTACACGACGCTCTTCCGATCTNNNAGTCCAACCAACATATTTTAAAT
AGCACACGTCTGAACTCTTTCCCTACACGACGCTCTTCCGATCTNNNGAGGCTGAGGCAGGAGAAT
AGCACACGTCTGAACTCTTTCCCTACACGACGCTCTTCCGATCTNNNGAGGCTGAGGCAGGAGAAT
AGCACACGTCTGAACTCTTTCCCTACACGACGCTCTTCCGATCTNNNCTAAGAAAAGTGCCTAACTGC
AGCACACGTCTGAACTCTTTCCCTACACGACGCTCTTCCGATCTNNNCTAAGAAAAGTGCCTAACTGC
GAGCACACGTCTGAACTCTTTCCCTACACGACGCTCTTCCGATCTNNNTGTTTTCCCAAGAATGTCACC
GAGCACACGTCTGAACTCTTTCCCTACACGACGCTCTTCCGATCTNNNTGTTTTCCCAAGAATGTCACC
GAGCACACGTCTGAACTCTTTCCCTACACGACGCTCTTCCGATCTNNNTTTCATTTCTGTGTGGATTGT
GAGCACACGTCTGAACTCTTTCCCTACACGACGCTCTTCCGATCTNNNTTTCATTTCTGTGTGGATTGT
AGCACACGTCTGAACTCTTTCCCTACACGACGCTCTTCCGATCTNNNAAAAAATTAGCCAGGTGTGG
AGCACACGTCTGAACTCTTTCCCTACACGACGCTCTTCCGATCTNNNAAAAAATTAGCCAGGTGTGG
GAGCACACGTCTGAACTCTTTCCCTACACGACGCTCTTCCGATCTNNNACAGTCTCGGTACAGTACAC
GAGCACACGTCTGAACTCTTTCCCTACACGACGCTCTTCCGATCTNNNACAGTCTCGGTACAGTACAC
GAGCACACGTCTGAACTCTTTCCCTACACGACGCTCTTCCGATCTNNNCGAAGCTGAGTCTTACCCCTT
GAGCACACGTCTGAACTCTTTCCCTACACGACGCTCTTCCGATCTNNNCGAAGCTGAGTCTTACCCCTT
CACACGTCTGAACTCTTTCCCTACACGACGCTCTTCCGATCTNNNTCAACATACAAAAATCAGTAGCAT
CACACGTCTGAACTCTTTCCCTACACGACGCTCTTCCGATCTNNNTCAACATACAAAAATCAGTAGCAT
GAGCACACGTCTGAACTCTTTCCCTACACGACGCTCTTCCGATCTNNNTCAAAGAAGCTCAGAGAACA
GAGCACACGTCTGAACTCTTTCCCTACACGACGCTCTTCCGATCTNNNTCAAAGAAGCTCAGAGAACA
GAGCACACGTCTGAACTCTTTCCCTACACGACGCTCTTCCGATCTNNNGAAGCTGATTAGAGGCTCTT
GAGCACACGTCTGAACTCTTTCCCTACACGACGCTCTTCCGATCTNNNGAAGCTGATTAGAGGCTCTT
GAGCACACGTCTGAACTCTTTCCCTACACGACGCTCTTCCGATCTNNNCGGTAAGCTGAGATCATGTC
GAGCACACGTCTGAACTCTTTCCCTACACGACGCTCTTCCGATCTNNNCGGTAAGCTGAGATCATGTC
AGCACACGTCTGAACTCTTTCCCTACACGACGCTCTTCCGATCTNNNATTATGTCCAACACTCCACCTCC
AGCACACGTCTGAACTCTTTCCCTACACGACGCTCTTCCGATCTNNNATTATGTCCAACACTCCACCTCC
GCACACGTCTGAACTCTTTCCCTACACGACGCTCTTCCGATCTNNNCCCGGCCAAGACTTTTTTCTGCA
GCACACGTCTGAACTCTTTCCCTACACGACGCTCTTCCGATCTNNNCCCGGCCAAGACTTTTTTCTGCA
GCACACGTCTGAACTCTTTCCCTACACGACGCTCTTCCGATCTNNNAAGCAACAAACAGTATAATGG
GCACACGTCTGAACTCTTTCCCTACACGACGCTCTTCCGATCTNNNAAGCAACAAACAGTATAATGG
AGCACACGTCTGAACTCTTTCCCTACACGACGCTCTTCCGATCTNNNTCCCTTCCTTCCTTCCTTCTTTCC
AGCACACGTCTGAACTCTTTCCCTACACGACGCTCTTCCGATCTNNNTCCCTTCCTTCCTTCCTTCTTTCC
GAGCACACGTCTGAACTCTTTCCCTACACGACGCTCTTCCGATCTNNNTGAGGTTGGAGGTCACTAA
GAGCACACGTCTGAACTCTTTCCCTACACGACGCTCTTCCGATCTNNNTGAGGTTGGAGGTCACTAA
GCACACGTCTGAACTCTTTCCCTACACGACGCTCTTCCGATCTNNNAGTGCTGGGATTACAGACATG
GCACACGTCTGAACTCTTTCCCTACACGACGCTCTTCCGATCTNNNAGTGCTGGGATTACAGACATG
GAGCACACGTCTGAACTCTTTCCCTACACGACGCTCTTCCGATCTNNNTGACTTGCTATTCCTTGCCTTC
GAGCACACGTCTGAACTCTTTCCCTACACGACGCTCTTCCGATCTNNNTGACTTGCTATTCCTTGCCTTC
GAGCACACGTCTGAACTCTTTCCCTACACGACGCTCTTCCGATCTNNNACCAAGTTGTAGAAGGACTG

GAGCACACGTCTGAACTCTTTCCCTACACGACGCTCTTCCGATCTNNNACCAAGTTGTAGAAGGACTG
GAGCACACGTCTGAACTCTTTCCCTACACGACGCTCTTCCGATCTNNNGAGGTTAGGATGTGTGTGTG
GAGCACACGTCTGAACTCTTTCCCTACACGACGCTCTTCCGATCTNNNGAGGTTAGGATGTGTGTGTG
AGCACACGTCTGAACTCTTTCCCTACACGACGCTCTTCCGATCTNNNGAGATCACACCATTGCACTCCA
AGCACACGTCTGAACTCTTTCCCTACACGACGCTCTTCCGATCTNNNGAGATCACACCATTGCACTCCA
GCACACGTCTGAACTCTTTCCCTACACGACGCTCTTCCGATCTNNNCTTGCAGTGAGCCGAGATTGTC
GCACACGTCTGAACTCTTTCCCTACACGACGCTCTTCCGATCTNNNCTTGCAGTGAGCCGAGATTGTC
GCACACGTCTGAACTCTTTCCCTACACGACGCTCTTCCGATCTNNNCCAGAAATCAGCTCATTTTCAGG
GCACACGTCTGAACTCTTTCCCTACACGACGCTCTTCCGATCTNNNCCAGAAATCAGCTCATTTTCAGG
GAGCACACGTCTGAACTCTTTCCCTACACGACGCTCTTCCGATCTNNNATGAACTCTGGCAAGTAGAG
GAGCACACGTCTGAACTCTTTCCCTACACGACGCTCTTCCGATCTNNNATGAACTCTGGCAAGTAGAG
AGCACACGTCTGAACTCTTTCCCTACACGACGCTCTTCCGATCTNNNCTTGTAATTCCAGCTACTCGGG
AGCACACGTCTGAACTCTTTCCCTACACGACGCTCTTCCGATCTNNNCTTGTAATTCCAGCTACTCGGG
GCACACGTCTGAACTCTTTCCCTACACGACGCTCTTCCGATCTNNNGAGAGAGAGGAAGAGAGCTTG
GCACACGTCTGAACTCTTTCCCTACACGACGCTCTTCCGATCTNNNGAGAGAGAGGAAGAGAGCTTG
AGCACACGTCTGAACTCTTTCCCTACACGACGCTCTTCCGATCTNNNGTCAAAGATGTGCGGCCTTAT
AGCACACGTCTGAACTCTTTCCCTACACGACGCTCTTCCGATCTNNNGTCAAAGATGTGCGGCCTTAT
GCACACGTCTGAACTCTTTCCCTACACGACGCTCTTCCGATCTNNNCTGAGGTGGGAAGATCACTTGA
GCACACGTCTGAACTCTTTCCCTACACGACGCTCTTCCGATCTNNNCTGAGGTGGGAAGATCACTTGA
AGCACACGTCTGAACTCTTTCCCTACACGACGCTCTTCCGATCTNNNATCCAGACCTCCTCCCACTACCA
AGCACACGTCTGAACTCTTTCCCTACACGACGCTCTTCCGATCTNNNATCCAGACCTCCTCCCACTACCA
GAGCACACGTCTGAACTCTTTCCCTACACGACGCTCTTCCGATCTNNNCTCATAGGGTTAGACCACATC
GAGCACACGTCTGAACTCTTTCCCTACACGACGCTCTTCCGATCTNNNCTCATAGGGTTAGACCACATC
AGCACACGTCTGAACTCTTTCCCTACACGACGCTCTTCCGATCTNNNAAGTCATCTCAGCTTTGAGGAG
AGCACACGTCTGAACTCTTTCCCTACACGACGCTCTTCCGATCTNNNAAGTCATCTCAGCTTTGAGGAG
AGCACACGTCTGAACTCTTTCCCTACACGACGCTCTTCCGATCTNNNGAAGTGTGTTTCCTGTAGGCA
AGCACACGTCTGAACTCTTTCCCTACACGACGCTCTTCCGATCTNNNGAAGTGTGTTTCCTGTAGGCA
GCACACGTCTGAACTCTTTCCCTACACGACGCTCTTCCGATCTNNNTAGAAAGAGGACACCCTTACCC
GCACACGTCTGAACTCTTTCCCTACACGACGCTCTTCCGATCTNNNTAGAAAGAGGACACCCTTACCC
GCACACGTCTGAACTCTTTCCCTACACGACGCTCTTCCGATCTNNNTAAGTAGAGGGGGAGAAAGGG
GCACACGTCTGAACTCTTTCCCTACACGACGCTCTTCCGATCTNNNTAAGTAGAGGGGGAGAAAGGG
CACACGTCTGAACTCTTTCCCTACACGACGCTCTTCCGATCTNNNCATCCTGGCTAACACAGTGAAACC
CACACGTCTGAACTCTTTCCCTACACGACGCTCTTCCGATCTNNNCATCCTGGCTAACACAGTGAAACC
AGCACACGTCTGAACTCTTTCCCTACACGACGCTCTTCCGATCTNNNAGGGTCACAGCTGGGAAGAG
AGCACACGTCTGAACTCTTTCCCTACACGACGCTCTTCCGATCTNNNAGGGTCACAGCTGGGAAGAG
GCACACGTCTGAACTCTTTCCCTACACGACGCTCTTCCGATCTNNNTAGATTCACCTAGCCTCCACAAC
GCACACGTCTGAACTCTTTCCCTACACGACGCTCTTCCGATCTNNNTAGATTCACCTAGCCTCCACAAC
CACACGTCTGAACTCTTTCCCTACACGACGCTCTTCCGATCTNNNTTCAGTCAAGTTACTGCACCTCGC
CACACGTCTGAACTCTTTCCCTACACGACGCTCTTCCGATCTNNNTTCAGTCAAGTTACTGCACCTCGC
GCACACGTCTGAACTCTTTCCCTACACGACGCTCTTCCGATCTNNNGGCTTTTGTTACATTCCAGCCTTC
GCACACGTCTGAACTCTTTCCCTACACGACGCTCTTCCGATCTNNNGGCTTTTGTTACATTCCAGCCTTC
AGCACACGTCTGAACTCTTTCCCTACACGACGCTCTTCCGATCTNNNCTCAGTGGTCACTTCTCTGTCTT
AGCACACGTCTGAACTCTTTCCCTACACGACGCTCTTCCGATCTNNNCTCAGTGGTCACTTCTCTGTCTT

AGCACACGTCTGAACTCTTTCCCTACACGACGCTCTTCCGATCTNNNACTTCCGGCCTCTTTCTCTCTTT
AGCACACGTCTGAACTCTTTCCCTACACGACGCTCTTCCGATCTNNNACTTCCGGCCTCTTTCTCTCTTT
GAGCACACGTCTGAACTCTTTCCCTACACGACGCTCTTCCGATCTNNNGAATGCGACCTCTTCTGGGA
GAGCACACGTCTGAACTCTTTCCCTACACGACGCTCTTCCGATCTNNNGAATGCGACCTCTTCTGGGA
AGCACACGTCTGAACTCTTTCCCTACACGACGCTCTTCCGATCTNNNCCAACTGGTCTATGCCTCTAGA
AGCACACGTCTGAACTCTTTCCCTACACGACGCTCTTCCGATCTNNNCCAACTGGTCTATGCCTCTAGA
AGCACACGTCTGAACTCTTTCCCTACACGACGCTCTTCCGATCTNNNTCATGTAAACAAGAAGGGAGC
AGCACACGTCTGAACTCTTTCCCTACACGACGCTCTTCCGATCTNNNTCATGTAAACAAGAAGGGAGC
GAGCACACGTCTGAACTCTTTCCCTACACGACGCTCTTCCGATCTNNNGCAAAAAACAGACCTCAACCA
GAGCACACGTCTGAACTCTTTCCCTACACGACGCTCTTCCGATCTNNNGCAAAAAACAGACCTCAACCA
AGCACACGTCTGAACTCTTTCCCTACACGACGCTCTTCCGATCTNNNGATAATTGCTTGAACCCAGAGG
AGCACACGTCTGAACTCTTTCCCTACACGACGCTCTTCCGATCTNNNGATAATTGCTTGAACCCAGAGG
GAGCACACGTCTGAACTCTTTCCCTACACGACGCTCTTCCGATCTNNNGCCTGTAGTCCCAGCTATTCTG
GAGCACACGTCTGAACTCTTTCCCTACACGACGCTCTTCCGATCTNNNGCCTGTAGTCCCAGCTATTCTG
GAGCACACGTCTGAACTCTTTCCCTACACGACGCTCTTCCGATCTNNNGGGACCCCAAAATCACTAAG
GAGCACACGTCTGAACTCTTTCCCTACACGACGCTCTTCCGATCTNNNGGGACCCCAAAATCACTAAG
GAGCACACGTCTGAACTCTTTCCCTACACGACGCTCTTCCGATCTNNNTTTATCCTCTCTCCTTGTTGCA
GAGCACACGTCTGAACTCTTTCCCTACACGACGCTCTTCCGATCTNNNTTTATCCTCTCTCCTTGTTGCA
AGCACACGTCTGAACTCTTTCCCTACACGACGCTCTTCCGATCTNNNAGTTATACATATGCCCCTTGGG
AGCACACGTCTGAACTCTTTCCCTACACGACGCTCTTCCGATCTNNNAGTTATACATATGCCCCTTGGG
GAGCACACGTCTGAACTCTTTCCCTACACGACGCTCTTCCGATCTNNNGGTGTCTGTAATCCCAGCTAC
GAGCACACGTCTGAACTCTTTCCCTACACGACGCTCTTCCGATCTNNNGGTGTCTGTAATCCCAGCTAC
GAGCACACGTCTGAACTCTTTCCCTACACGACGCTCTTCCGATCTNNNTCATGTGCCGGTATGGATCT
GAGCACACGTCTGAACTCTTTCCCTACACGACGCTCTTCCGATCTNNNTCATGTGCCGGTATGGATCT
GAGCACACGTCTGAACTCTTTCCCTACACGACGCTCTTCCGATCTNNNTCCAAGGCAGTCGTAAACAC
GAGCACACGTCTGAACTCTTTCCCTACACGACGCTCTTCCGATCTNNNTCCAAGGCAGTCGTAAACAC
AGCACACGTCTGAACTCTTTCCCTACACGACGCTCTTCCGATCTNNNTCTGGCAACCCTATTCTTGATCC
AGCACACGTCTGAACTCTTTCCCTACACGACGCTCTTCCGATCTNNNTCTGGCAACCCTATTCTTGATCC
GCACACGTCTGAACTCTTTCCCTACACGACGCTCTTCCGATCTNNNCTGGGTTCATGCCATTTTCCTGC
GCACACGTCTGAACTCTTTCCCTACACGACGCTCTTCCGATCTNNNCTGGGTTCATGCCATTTTCCTGC
AGCACACGTCTGAACTCTTTCCCTACACGACGCTCTTCCGATCTNNNCACACACACATGCACACACATAT
AGCACACGTCTGAACTCTTTCCCTACACGACGCTCTTCCGATCTNNNCACACACACATGCACACACATAT
GCACACGTCTGAACTCTTTCCCTACACGACGCTCTTCCGATCTNNNATCTTCCCCTCATTTTGGTGAGT
GCACACGTCTGAACTCTTTCCCTACACGACGCTCTTCCGATCTNNNATCTTCCCCTCATTTTGGTGAGT
AGCACACGTCTGAACTCTTTCCCTACACGACGCTCTTCCGATCTNNNACTCTGTCCTTTTGTCCCTAGAA
AGCACACGTCTGAACTCTTTCCCTACACGACGCTCTTCCGATCTNNNACTCTGTCCTTTTGTCCCTAGAA
GCACACGTCTGAACTCTTTCCCTACACGACGCTCTTCCGATCTNNNGCATGTAGGACTTACCCGGTAA
GCACACGTCTGAACTCTTTCCCTACACGACGCTCTTCCGATCTNNNGCATGTAGGACTTACCCGGTAA
GAGCACACGTCTGAACTCTTTCCCTACACGACGCTCTTCCGATCTNNNAGAATTGTCTTGGGCCACACA
GAGCACACGTCTGAACTCTTTCCCTACACGACGCTCTTCCGATCTNNNAGAATTGTCTTGGGCCACACA
GAGCACACGTCTGAACTCTTTCCCTACACGACGCTCTTCCGATCTNNNCCTCTCAGGTTCACGCTATTCT
GAGCACACGTCTGAACTCTTTCCCTACACGACGCTCTTCCGATCTNNNCCTCTCAGGTTCACGCTATTCT
GAGCACACGTCTGAACTCTTTCCCTACACGACGCTCTTCCGATCTNNNTCACACCACTACATTCCAGAG

GAGCACACGTCTGAACTCTTTCCCTACACGACGCTCTTCCGATCTNNNTCACACCACTACATTCCAGAG
AGCACACGTCTGAACTCTTTCCCTACACGACGCTCTTCCGATCTNNNTACTGATAGATAGATAGGGTC
AGCACACGTCTGAACTCTTTCCCTACACGACGCTCTTCCGATCTNNNTACTGATAGATAGATAGGGTC
AGCACACGTCTGAACTCTTTCCCTACACGACGCTCTTCCGATCTNNNCCTGGCCAGAAAGGGAAAAA
AGCACACGTCTGAACTCTTTCCCTACACGACGCTCTTCCGATCTNNNCCTGGCCAGAAAGGGAAAAA
GCACACGTCTGAACTCTTTCCCTACACGACGCTCTTCCGATCTNNNAAGGAGGGCAGACTAACTAGA
GCACACGTCTGAACTCTTTCCCTACACGACGCTCTTCCGATCTNNNAAGGAGGGCAGACTAACTAGA
GCACACGTCTGAACTCTTTCCCTACACGACGCTCTTCCGATCTNNNAGTCATCAGAGAGGAATACTG
GCACACGTCTGAACTCTTTCCCTACACGACGCTCTTCCGATCTNNNAGTCATCAGAGAGGAATACTG
GCACACGTCTGAACTCTTTCCCTACACGACGCTCTTCCGATCTNNNTCAAAGATTCTCAAGGCATCCCA
GCACACGTCTGAACTCTTTCCCTACACGACGCTCTTCCGATCTNNNTCAAAGATTCTCAAGGCATCCCA
CACACGTCTGAACTCTTTCCCTACACGACGCTCTTCCGATCTNNNTCCAAGGAAATCTGCTCCAAAGAC
CACACGTCTGAACTCTTTCCCTACACGACGCTCTTCCGATCTNNNTCCAAGGAAATCTGCTCCAAAGAC
CACACGTCTGAACTCTTTCCCTACACGACGCTCTTCCGATCTNNNTTAATTAGCTTAGAATCCCTGAA
CACACGTCTGAACTCTTTCCCTACACGACGCTCTTCCGATCTNNNTTAATTAGCTTAGAATCCCTGAA
GCACACGTCTGAACTCTTTCCCTACACGACGCTCTTCCGATCTNNNCATTTGCCACTATTTTTGGGGGT
GCACACGTCTGAACTCTTTCCCTACACGACGCTCTTCCGATCTNNNCATTTGCCACTATTTTTGGGGGT
GAGCACACGTCTGAACTCTTTCCCTACACGACGCTCTTCCGATCTNNNGAGGCAGGAGAATGGTGT
GAGCACACGTCTGAACTCTTTCCCTACACGACGCTCTTCCGATCTNNNGAGGCAGGAGAATGGTGT
GCACACGTCTGAACTCTTTCCCTACACGACGCTCTTCCGATCTNNNAGCACTATGCCAGAAAGACTCTC
GCACACGTCTGAACTCTTTCCCTACACGACGCTCTTCCGATCTNNNAGCACTATGCCAGAAAGACTCTC
GCACACGTCTGAACTCTTTCCCTACACGACGCTCTTCCGATCTNNNCGAACAAAGTTGGAAGGGGA
GCACACGTCTGAACTCTTTCCCTACACGACGCTCTTCCGATCTNNNCGAACAAAGTTGGAAGGGGA
GCACACGTCTGAACTCTTTCCCTACACGACGCTCTTCCGATCTNNNCTTGATCTTGGCTCACTGCAATC
GCACACGTCTGAACTCTTTCCCTACACGACGCTCTTCCGATCTNNNCTTGATCTTGGCTCACTGCAATC
GAGCACACGTCTGAACTCTTTCCCTACACGACGCTCTTCCGATCTNNNCTCCTCTTTGGTGACTTGCTT
GAGCACACGTCTGAACTCTTTCCCTACACGACGCTCTTCCGATCTNNNCTCCTCTTTGGTGACTTGCTT
AGCACACGTCTGAACTCTTTCCCTACACGACGCTCTTCCGATCTNNNAGTCTCAGTGGACTCATTTGA
AGCACACGTCTGAACTCTTTCCCTACACGACGCTCTTCCGATCTNNNAGTCTCAGTGGACTCATTTGA
AGCACACGTCTGAACTCTTTCCCTACACGACGCTCTTCCGATCTNNNGTGAAACCCCGTCTCTACTAAA
AGCACACGTCTGAACTCTTTCCCTACACGACGCTCTTCCGATCTNNNGTGAAACCCCGTCTCTACTAAA
AGCACACGTCTGAACTCTTTCCCTACACGACGCTCTTCCGATCTNNNTGGAGGAAGCAGAGATAACC
AGCACACGTCTGAACTCTTTCCCTACACGACGCTCTTCCGATCTNNNTGGAGGAAGCAGAGATAACC
AGCACACGTCTGAACTCTTTCCCTACACGACGCTCTTCCGATCTNNNTATTGCAGTGCCTTCTCTCTCTC
AGCACACGTCTGAACTCTTTCCCTACACGACGCTCTTCCGATCTNNNTATTGCAGTGCCTTCTCTCTCTC
AGCACACGTCTGAACTCTTTCCCTACACGACGCTCTTCCGATCTNNNGTTGTTGGAAAACATGAGGG
AGCACACGTCTGAACTCTTTCCCTACACGACGCTCTTCCGATCTNNNGTTGTTGGAAAACATGAGGG
GAGCACACGTCTGAACTCTTTCCCTACACGACGCTCTTCCGATCTNNNTTTTTCCAGTTGGGCTTGAA
GAGCACACGTCTGAACTCTTTCCCTACACGACGCTCTTCCGATCTNNNTTTTTCCAGTTGGGCTTGAA
AGCACACGTCTGAACTCTTTCCCTACACGACGCTCTTCCGATCTNNNACAGGAATTTCAGGACAGTAT
AGCACACGTCTGAACTCTTTCCCTACACGACGCTCTTCCGATCTNNNACAGGAATTTCAGGACAGTAT
AGCACACGTCTGAACTCTTTCCCTACACGACGCTCTTCCGATCTNNNAAAGACACAGACTGGCAAATT
AGCACACGTCTGAACTCTTTCCCTACACGACGCTCTTCCGATCTNNNAAAGACACAGACTGGCAAATT

GAGCACACGTCTGAACTCTTTCCCTACACGACGCTCTTCCGATCTNNNATGCAATGTCCTTCTTAGTGC
GAGCACACGTCTGAACTCTTTCCCTACACGACGCTCTTCCGATCTNNNATGCAATGTCCTTCTTAGTGC
AGCACACGTCTGAACTCTTTCCCTACACGACGCTCTTCCGATCTNNNCCCTTTCATTCCAGGAGGATG
AGCACACGTCTGAACTCTTTCCCTACACGACGCTCTTCCGATCTNNNCCCTTTCATTCCAGGAGGATG
GCACACGTCTGAACTCTTTCCCTACACGACGCTCTTCCGATCTNNNTCACTCTCTCTTGCTCTCTCTCTCA
GCACACGTCTGAACTCTTTCCCTACACGACGCTCTTCCGATCTNNNTCACTCTCTCTTGCTCTCTCTCTCA
AGAGCACACGTCTGAACTCTTTCCCTACACGACGCTCTTCCGATCTNNNAGGGGTAAATCTCACTTGG
AGAGCACACGTCTGAACTCTTTCCCTACACGACGCTCTTCCGATCTNNNAGGGGTAAATCTCACTTGG
AGCACACGTCTGAACTCTTTCCCTACACGACGCTCTTCCGATCTNNNGGTCTTGAACTCCTGACCTCA
AGCACACGTCTGAACTCTTTCCCTACACGACGCTCTTCCGATCTNNNGGTCTTGAACTCCTGACCTCA
AGCACACGTCTGAACTCTTTCCCTACACGACGCTCTTCCGATCTNNNAGGTCCATGTCACAGAATTGCT
AGCACACGTCTGAACTCTTTCCCTACACGACGCTCTTCCGATCTNNNAGGTCCATGTCACAGAATTGCT
AGCACACGTCTGAACTCTTTCCCTACACGACGCTCTTCCGATCTNNNATCACTTGAAATCGGAAGGTG
AGCACACGTCTGAACTCTTTCCCTACACGACGCTCTTCCGATCTNNNATCACTTGAAATCGGAAGGTG
AGCACACGTCTGAACTCTTTCCCTACACGACGCTCTTCCGATCTNNNCTCCTGGGTTTACGCCATTCTC
AGCACACGTCTGAACTCTTTCCCTACACGACGCTCTTCCGATCTNNNCTCCTGGGTTTACGCCATTCTC
AGCACACGTCTGAACTCTTTCCCTACACGACGCTCTTCCGATCTNNNACTTGGGAGGTTGAGACAGGA
AGCACACGTCTGAACTCTTTCCCTACACGACGCTCTTCCGATCTNNNACTTGGGAGGTTGAGACAGGA
CACACGTCTGAACTCTTTCCCTACACGACGCTCTTCCGATCTNNNTATGGTTTGAATCATTTTTGTGTC
CACACGTCTGAACTCTTTCCCTACACGACGCTCTTCCGATCTNNNTATGGTTTGAATCATTTTTGTGTC
AGCACACGTCTGAACTCTTTCCCTACACGACGCTCTTCCGATCTNNNGGAGTACAGTGGCACGATCTC
AGCACACGTCTGAACTCTTTCCCTACACGACGCTCTTCCGATCTNNNGGAGTACAGTGGCACGATCTC
AGCACACGTCTGAACTCTTTCCCTACACGACGCTCTTCCGATCTNNNAAGTACAGGTGGCATGAGATT
AGCACACGTCTGAACTCTTTCCCTACACGACGCTCTTCCGATCTNNNAAGTACAGGTGGCATGAGATT
AGCACACGTCTGAACTCTTTCCCTACACGACGCTCTTCCGATCTNNNATGGAGAAACCCCGTCTCTACT
AGCACACGTCTGAACTCTTTCCCTACACGACGCTCTTCCGATCTNNNATGGAGAAACCCCGTCTCTACT
AGCACACGTCTGAACTCTTTCCCTACACGACGCTCTTCCGATCTNNNGAGTTTGAGACCATCCTGGCT
AGCACACGTCTGAACTCTTTCCCTACACGACGCTCTTCCGATCTNNNGAGTTTGAGACCATCCTGGCT
AGCACACGTCTGAACTCTTTCCCTACACGACGCTCTTCCGATCTNNNTCTAATGCTGCCAAAGTTCTTG
AGCACACGTCTGAACTCTTTCCCTACACGACGCTCTTCCGATCTNNNTCTAATGCTGCCAAAGTTCTTG
GCACACGTCTGAACTCTTTCCCTACACGACGCTCTTCCGATCTNNNGATCACGAGGTCAGGAGTTCAA
GCACACGTCTGAACTCTTTCCCTACACGACGCTCTTCCGATCTNNNGATCACGAGGTCAGGAGTTCAA
GAGCACACGTCTGAACTCTTTCCCTACACGACGCTCTTCCGATCTNNNGAATCATCGCTTGAACCTGG
GAGCACACGTCTGAACTCTTTCCCTACACGACGCTCTTCCGATCTNNNGAATCATCGCTTGAACCTGG
AGCACACGTCTGAACTCTTTCCCTACACGACGCTCTTCCGATCTNNNATACCTAGGGTAAGGCTGGAT
AGCACACGTCTGAACTCTTTCCCTACACGACGCTCTTCCGATCTNNNATACCTAGGGTAAGGCTGGAT
AGCACACGTCTGAACTCTTTCCCTACACGACGCTCTTCCGATCTNNNCTCAAGCAGTCCTCCTACTTCAG
AGCACACGTCTGAACTCTTTCCCTACACGACGCTCTTCCGATCTNNNCTCAAGCAGTCCTCCTACTTCAG
AGCACACGTCTGAACTCTTTCCCTACACGACGCTCTTCCGATCTNNNACAACACAGTTGCTTTATCCAC
AGCACACGTCTGAACTCTTTCCCTACACGACGCTCTTCCGATCTNNNACAACACAGTTGCTTTATCCAC
CACACGTCTGAACTCTTTCCCTACACGACGCTCTTCCGATCTNNNACCGGTGTTGCAAATAACGTTTCC
CACACGTCTGAACTCTTTCCCTACACGACGCTCTTCCGATCTNNNACCGGTGTTGCAAATAACGTTTCC
GAGCACACGTCTGAACTCTTTCCCTACACGACGCTCTTCCGATCTNNNCTGTCACAGCCTTCAAATTCA

GAGCACACGTCTGAACTCTTTCCCTACACGACGCTCTTCCGATCTNNNCTGTCACAGCCTTCAAATTCAC
GAGCACACGTCTGAACTCTTTCCCTACACGACGCTCTTCCGATCTNNNACGAGACCCCATCACTACAAAA
GAGCACACGTCTGAACTCTTTCCCTACACGACGCTCTTCCGATCTNNNACGAGACCCCATCACTACAAAA
GAGCACACGTCTGAACTCTTTCCCTACACGACGCTCTTCCGATCTNNNCATACACAGACACACACACACA
GAGCACACGTCTGAACTCTTTCCCTACACGACGCTCTTCCGATCTNNNCATACACAGACACACACACACA
GAGCACACGTCTGAACTCTTTCCCTACACGACGCTCTTCCGATCTNNNTTCTCTTCTTTTGCTCTGTGGG
GAGCACACGTCTGAACTCTTTCCCTACACGACGCTCTTCCGATCTNNNTTCTCTTCTTTTGCTCTGTGGG
GAGCACACGTCTGAACTCTTTCCCTACACGACGCTCTTCCGATCTNNNCTCCTCTGTCACCTTCTCTTAGC
GAGCACACGTCTGAACTCTTTCCCTACACGACGCTCTTCCGATCTNNNCTCCTCTGTCACCTTCTCTTAGC
AGCACACGTCTGAACTCTTTCCCTACACGACGCTCTTCCGATCTNNNTGGACAATAGGCATTATTCTAG
AGCACACGTCTGAACTCTTTCCCTACACGACGCTCTTCCGATCTNNNTGGACAATAGGCATTATTCTAG
AGCACACGTCTGAACTCTTTCCCTACACGACGCTCTTCCGATCTNNNTGAGGCATGAGAATTGCTTGAA
AGCACACGTCTGAACTCTTTCCCTACACGACGCTCTTCCGATCTNNNTGAGGCATGAGAATTGCTTGAA
GAGCACACGTCTGAACTCTTTCCCTACACGACGCTCTTCCGATCTNNNTCAACTATTGACCTGAATGCCC
GAGCACACGTCTGAACTCTTTCCCTACACGACGCTCTTCCGATCTNNNTCAACTATTGACCTGAATGCCC
GAGCACACGTCTGAACTCTTTCCCTACACGACGCTCTTCCGATCTNNNTTACCTGGCCTTTTCCTTTTCGT
GAGCACACGTCTGAACTCTTTCCCTACACGACGCTCTTCCGATCTNNNTTACCTGGCCTTTTCCTTTTCGT
AGCACACGTCTGAACTCTTTCCCTACACGACGCTCTTCCGATCTNNNCGCTACCAGCCTGTCTCTTTATC
AGCACACGTCTGAACTCTTTCCCTACACGACGCTCTTCCGATCTNNNCGCTACCAGCCTGTCTCTTTATC
AGCACACGTCTGAACTCTTTCCCTACACGACGCTCTTCCGATCTNNNGTGAATCCTTGAAACCAAGAGG
AGCACACGTCTGAACTCTTTCCCTACACGACGCTCTTCCGATCTNNNGTGAATCCTTGAAACCAAGAGG
AGCACACGTCTGAACTCTTTCCCTACACGACGCTCTTCCGATCTNNNTACTGACTCCAATGGCTGACTTC
AGCACACGTCTGAACTCTTTCCCTACACGACGCTCTTCCGATCTNNNTACTGACTCCAATGGCTGACTTC
AGCACACGTCTGAACTCTTTCCCTACACGACGCTCTTCCGATCTNNNGCCTTCTTCTGTCCATGGTTTTC
AGCACACGTCTGAACTCTTTCCCTACACGACGCTCTTCCGATCTNNNGCCTTCTTCTGTCCATGGTTTTC
AGCACACGTCTGAACTCTTTCCCTACACGACGCTCTTCCGATCTNNNGATTACGCCACTGCACTCCACAT
AGCACACGTCTGAACTCTTTCCCTACACGACGCTCTTCCGATCTNNNGATTACGCCACTGCACTCCACAT
GAGCACACGTCTGAACTCTTTCCCTACACGACGCTCTTCCGATCTNNNGTAGTCCCAGCTACTTGGACAC
GAGCACACGTCTGAACTCTTTCCCTACACGACGCTCTTCCGATCTNNNGTAGTCCCAGCTACTTGGACAC
GAGCACACGTCTGAACTCTTTCCCTACACGACGCTCTTCCGATCTNNNTTCTGAGAAGTCTCTGGTTTGG
GAGCACACGTCTGAACTCTTTCCCTACACGACGCTCTTCCGATCTNNNTTCTGAGAAGTCTCTGGTTTGG
AAGAGCACACGTCTGAACTCTTTCCCTACACGACGCTCTTCCGATCTNNNTTTATGTTGGAGCGACAGA
AAGAGCACACGTCTGAACTCTTTCCCTACACGACGCTCTTCCGATCTNNNTTTATGTTGGAGCGACAGA
GAGCACACGTCTGAACTCTTTCCCTACACGACGCTCTTCCGATCTNNNCAGCCCTATGCATTTGTCAAAC
GAGCACACGTCTGAACTCTTTCCCTACACGACGCTCTTCCGATCTNNNCAGCCCTATGCATTTGTCAAAC
GAGCACACGTCTGAACTCTTTCCCTACACGACGCTCTTCCGATCTNNNTAACTACCTGTTGCCTTCTTCCC
GAGCACACGTCTGAACTCTTTCCCTACACGACGCTCTTCCGATCTNNNTAACTACCTGTTGCCTTCTTCCC
GAGCACACGTCTGAACTCTTTCCCTACACGACGCTCTTCCGATCTNNNAAGCTCCCCTAGAGACAAGAT
GAGCACACGTCTGAACTCTTTCCCTACACGACGCTCTTCCGATCTNNNAAGCTCCCCTAGAGACAAGAT
AGCACACGTCTGAACTCTTTCCCTACACGACGCTCTTCCGATCTNNNGAGGTTGCAGTGAGCTGAGAT
AGCACACGTCTGAACTCTTTCCCTACACGACGCTCTTCCGATCTNNNGAGGTTGCAGTGAGCTGAGAT
GAGCACACGTCTGAACTCTTTCCCTACACGACGCTCTTCCGATCTNNNAGGAGTCTACTCTTGGGGACT
GAGCACACGTCTGAACTCTTTCCCTACACGACGCTCTTCCGATCTNNNAGGAGTCTACTCTTGGGGACT

AGCACACGTCTGAACTCTTTCCCTACACGACGCTCTTCCGATCTNNNTTCCCTAAACTTACCGGCCACTT
AGCACACGTCTGAACTCTTTCCCTACACGACGCTCTTCCGATCTNNNTTCCCTAAACTTACCGGCCACTT
AGCACACGTCTGAACTCTTTCCCTACACGACGCTCTTCCGATCTNNNCCCAAATGATGAAGTGATGCA
AGCACACGTCTGAACTCTTTCCCTACACGACGCTCTTCCGATCTNNNCCCAAATGATGAAGTGATGCA
GAGCACACGTCTGAACTCTTTCCCTACACGACGCTCTTCCGATCTNNNGAGATCGCACCACTGTACTTG
GAGCACACGTCTGAACTCTTTCCCTACACGACGCTCTTCCGATCTNNNGAGATCGCACCACTGTACTTG
GAGCACACGTCTGAACTCTTTCCCTACACGACGCTCTTCCGATCTNNNGTAAGCAGTGGAGATGACAG
GAGCACACGTCTGAACTCTTTCCCTACACGACGCTCTTCCGATCTNNNGTAAGCAGTGGAGATGACAG
AGAGCACACGTCTGAACTCTTTCCCTACACGACGCTCTTCCGATCTNNNAGGCTATGACAAATGAGCA
AGAGCACACGTCTGAACTCTTTCCCTACACGACGCTCTTCCGATCTNNNAGGCTATGACAAATGAGCA
GCACACGTCTGAACTCTTTCCCTACACGACGCTCTTCCGATCTNNNAAAGCACAGTTTTAGCCACAGTC
GCACACGTCTGAACTCTTTCCCTACACGACGCTCTTCCGATCTNNNAAAGCACAGTTTTAGCCACAGTC
GCACACGTCTGAACTCTTTCCCTACACGACGCTCTTCCGATCTNNNAAAGGAGAAAAAGCTGTAGGGG
GCACACGTCTGAACTCTTTCCCTACACGACGCTCTTCCGATCTNNNAAAGGAGAAAAAGCTGTAGGGG
GAGCACACGTCTGAACTCTTTCCCTACACGACGCTCTTCCGATCTNNNGGTTGCAGTGAGCCAAGATT
GAGCACACGTCTGAACTCTTTCCCTACACGACGCTCTTCCGATCTNNNGGTTGCAGTGAGCCAAGATT
GAGCACACGTCTGAACTCTTTCCCTACACGACGCTCTTCCGATCTNNNTAATCCCAGCTACTCTACTCGG
GAGCACACGTCTGAACTCTTTCCCTACACGACGCTCTTCCGATCTNNNTAATCCCAGCTACTCTACTCGG
AGCACACGTCTGAACTCTTTCCCTACACGACGCTCTTCCGATCTNNNAGGAACACAGACATGGAGAAA
AGCACACGTCTGAACTCTTTCCCTACACGACGCTCTTCCGATCTNNNAGGAACACAGACATGGAGAAA
GCACACGTCTGAACTCTTTCCCTACACGACGCTCTTCCGATCTNNNCGTCCTCAACCTTAGCAAAACACC
GCACACGTCTGAACTCTTTCCCTACACGACGCTCTTCCGATCTNNNCGTCCTCAACCTTAGCAAAACACC
GCACACGTCTGAACTCTTTCCCTACACGACGCTCTTCCGATCTNNNGACGTGCAGGTTTGTAACATAG
GCACACGTCTGAACTCTTTCCCTACACGACGCTCTTCCGATCTNNNGACGTGCAGGTTTGTAACATAG
GAGCACACGTCTGAACTCTTTCCCTACACGACGCTCTTCCGATCTNNNACAGGACCACAGGCTTCTCTA
GAGCACACGTCTGAACTCTTTCCCTACACGACGCTCTTCCGATCTNNNACAGGACCACAGGCTTCTCTA
AGCACACGTCTGAACTCTTTCCCTACACGACGCTCTTCCGATCTNNNAGAGTTGAGCAATGCCTTGTTT
AGCACACGTCTGAACTCTTTCCCTACACGACGCTCTTCCGATCTNNNAGAGTTGAGCAATGCCTTGTTT
GAGCACACGTCTGAACTCTTTCCCTACACGACGCTCTTCCGATCTNNNGAAGTCAGGAGTTCGAGACC
GAGCACACGTCTGAACTCTTTCCCTACACGACGCTCTTCCGATCTNNNGAAGTCAGGAGTTCGAGACC
GAGCACACGTCTGAACTCTTTCCCTACACGACGCTCTTCCGATCTNNNCACCTTCGAAGACCTCAGTGA
GAGCACACGTCTGAACTCTTTCCCTACACGACGCTCTTCCGATCTNNNCACCTTCGAAGACCTCAGTGA
AGCACACGTCTGAACTCTTTCCCTACACGACGCTCTTCCGATCTNNNTATCCACCTTTCTGCCTACCTTG
AGCACACGTCTGAACTCTTTCCCTACACGACGCTCTTCCGATCTNNNTATCCACCTTTCTGCCTACCTTG
AGCACACGTCTGAACTCTTTCCCTACACGACGCTCTTCCGATCTNNNCTCCTCTGTGCCATCCCTGGCAT
AGCACACGTCTGAACTCTTTCCCTACACGACGCTCTTCCGATCTNNNCTCCTCTGTGCCATCCCTGGCAT
GCACACGTCTGAACTCTTTCCCTACACGACGCTCTTCCGATCTNNNATGTGTGGTTGTGTGCATATGT
GCACACGTCTGAACTCTTTCCCTACACGACGCTCTTCCGATCTNNNATGTGTGGTTGTGTGCATATGT
AGCACACGTCTGAACTCTTTCCCTACACGACGCTCTTCCGATCTNNNGTCATTACGCTAGGAGTCCCA
AGCACACGTCTGAACTCTTTCCCTACACGACGCTCTTCCGATCTNNNGTCATTACGCTAGGAGTCCCA
GAGCACACGTCTGAACTCTTTCCCTACACGACGCTCTTCCGATCTNNNTCCAGTATCTTCTTGGACAGT
GAGCACACGTCTGAACTCTTTCCCTACACGACGCTCTTCCGATCTNNNTCCAGTATCTTCTTGGACAGT
GAGCACACGTCTGAACTCTTTCCCTACACGACGCTCTTCCGATCTNNNTGTGGGGTTGTTTTTCGTGT

AGCACACGTCTGAACTCTTTCCCTACACGACGCTCTTCCGATCTNNNTGTGGGGTTGTTTTTCGTGT
AGCACACGTCTGAACTCTTTCCCTACACGACGCTCTTCCGATCTNNNAATTCAGACATTAGGCCATCG
AGCACACGTCTGAACTCTTTCCCTACACGACGCTCTTCCGATCTNNNAATTCAGACATTAGGCCATCG
AGCACACGTCTGAACTCTTTCCCTACACGACGCTCTTCCGATCTNNNGAGCTGAGATTGCGCCATTGC
AGCACACGTCTGAACTCTTTCCCTACACGACGCTCTTCCGATCTNNNGAGCTGAGATTGCGCCATTGC
AGCACACGTCTGAACTCTTTCCCTACACGACGCTCTTCCGATCTNNNCTGAGGCAGGAGAATCACTTG
AGCACACGTCTGAACTCTTTCCCTACACGACGCTCTTCCGATCTNNNCTGAGGCAGGAGAATCACTTG
AGCACACGTCTGAACTCTTTCCCTACACGACGCTCTTCCGATCTNNNTCTCTGGGTGTCTTTCTCCGAA
AGCACACGTCTGAACTCTTTCCCTACACGACGCTCTTCCGATCTNNNTCTCTGGGTGTCTTTCTCCGAA
AGCACACGTCTGAACTCTTTCCCTACACGACGCTCTTCCGATCTNNNGCAGGGTAGATTTCTTTCCCC
AGCACACGTCTGAACTCTTTCCCTACACGACGCTCTTCCGATCTNNNGCAGGGTAGATTTCTTTCCCC
AGCACACGTCTGAACTCTTTCCCTACACGACGCTCTTCCGATCTNNNTCCTGATTCACCATCAATCCCTG
AGCACACGTCTGAACTCTTTCCCTACACGACGCTCTTCCGATCTNNNTCCTGATTCACCATCAATCCCTG
AGCACACGTCTGAACTCTTTCCCTACACGACGCTCTTCCGATCTNNNATTTTTCACCACCCTCATAGGC
AGCACACGTCTGAACTCTTTCCCTACACGACGCTCTTCCGATCTNNNATTTTTCACCACCCTCATAGGC
AGCACACGTCTGAACTCTTTCCCTACACGACGCTCTTCCGATCTNNNTGAAATAAGGACCCCAAACGC
AGCACACGTCTGAACTCTTTCCCTACACGACGCTCTTCCGATCTNNNTGAAATAAGGACCCCAAACGC
AGCACACGTCTGAACTCTTTCCCTACACGACGCTCTTCCGATCTNNNGAGGAGCTGGGTTCAGAGG
AGCACACGTCTGAACTCTTTCCCTACACGACGCTCTTCCGATCTNNNGAGGAGCTGGGTTCAGAGG
AGCACACGTCTGAACTCTTTCCCTACACGACGCTCTTCCGATCTNNNAGGTGTATGACCATCACCCAA
AGCACACGTCTGAACTCTTTCCCTACACGACGCTCTTCCGATCTNNNAGGTGTATGACCATCACCCAA
AGCACACGTCTGAACTCTTTCCCTACACGACGCTCTTCCGATCTNNNGTGGGAATGAAAAGCAGCAC
AGCACACGTCTGAACTCTTTCCCTACACGACGCTCTTCCGATCTNNNGTGGGAATGAAAAGCAGCAC
AGCACACGTCTGAACTCTTTCCCTACACGACGCTCTTCCGATCTNNNGAAGCTAATGCCGTCCATTCTT
AGCACACGTCTGAACTCTTTCCCTACACGACGCTCTTCCGATCTNNNGAAGCTAATGCCGTCCATTCTT
AGCACACGTCTGAACTCTTTCCCTACACGACGCTCTTCCGATCTNNNGAAGGGTCTTATCCTCTCTGC
AGCACACGTCTGAACTCTTTCCCTACACGACGCTCTTCCGATCTNNNGAAGGGTCTTATCCTCTCTGC
GAGCACACGTCTGAACTCTTTCCCTACACGACGCTCTTCCGATCTNNNGAGGAAGGAAATGATTCGG
GAGCACACGTCTGAACTCTTTCCCTACACGACGCTCTTCCGATCTNNNGAGGAAGGAAATGATTCGG
GAGCACACGTCTGAACTCTTTCCCTACACGACGCTCTTCCGATCTNNNCTGTTTGAGCCCAGGAATTC
GAGCACACGTCTGAACTCTTTCCCTACACGACGCTCTTCCGATCTNNNCTGTTTGAGCCCAGGAATTC
GAGCACACGTCTGAACTCTTTCCCTACACGACGCTCTTCCGATCTNNNGCAGATCACTTGAGGCTAAG
GAGCACACGTCTGAACTCTTTCCCTACACGACGCTCTTCCGATCTNNNGCAGATCACTTGAGGCTAAG
GCACACGTCTGAACTCTTTCCCTACACGACGCTCTTCCGATCTNNNCCAAGTAGCTGAGATTACAGGC
GCACACGTCTGAACTCTTTCCCTACACGACGCTCTTCCGATCTNNNCCAAGTAGCTGAGATTACAGGC
GAGCACACGTCTGAACTCTTTCCCTACACGACGCTCTTCCGATCTNNNGATTGCATGAGCCTGAGAGA
GAGCACACGTCTGAACTCTTTCCCTACACGACGCTCTTCCGATCTNNNGATTGCATGAGCCTGAGAGA
GCACACGTCTGAACTCTTTCCCTACACGACGCTCTTCCGATCTNNNAGCCTCAGAATATCACCCAAAGG
GCACACGTCTGAACTCTTTCCCTACACGACGCTCTTCCGATCTNNNAGCCTCAGAATATCACCCAAAGG
AGCACACGTCTGAACTCTTTCCCTACACGACGCTCTTCCGATCTNNNGAGAATCTGAAGGTTTGGGG
AGCACACGTCTGAACTCTTTCCCTACACGACGCTCTTCCGATCTNNNGAGAATCTGAAGGTTTGGGG
AGCACACGTCTGAACTCTTTCCCTACACGACGCTCTTCCGATCTNNNGTGTGTGTGTGTGTGTGT
AGCACACGTCTGAACTCTTTCCCTACACGACGCTCTTCCGATCTNNNGTGTGTGTGTGTGTGTGT

AGAGCACACGTCTGAACTCTTTCCCTACACGACGCTCTTCCGATCTNNNCACTACCCAACGACCACTCT
AGAGCACACGTCTGAACTCTTTCCCTACACGACGCTCTTCCGATCTNNNCACTACCCAACGACCACTCT
GCACACGTCTGAACTCTTTCCCTACACGACGCTCTTCCGATCTNNNGTAGGGCCACAGGTCGAAATA
GCACACGTCTGAACTCTTTCCCTACACGACGCTCTTCCGATCTNNNGTAGGGCCACAGGTCGAAATA
AGCACACGTCTGAACTCTTTCCCTACACGACGCTCTTCCGATCTNNNGCTGAGAGAGGACATCACTT
AGCACACGTCTGAACTCTTTCCCTACACGACGCTCTTCCGATCTNNNGCTGAGAGAGGACATCACTT
AGCACACGTCTGAACTCTTTCCCTACACGACGCTCTTCCGATCTNNNCACCTGTAATCCCAGCTACTCA
AGCACACGTCTGAACTCTTTCCCTACACGACGCTCTTCCGATCTNNNCACCTGTAATCCCAGCTACTCA
AGCACACGTCTGAACTCTTTCCCTACACGACGCTCTTCCGATCTNNNATTGATTGGTTGTGAGGGTG
AGCACACGTCTGAACTCTTTCCCTACACGACGCTCTTCCGATCTNNNATTGATTGGTTGTGAGGGTG
AGCACACGTCTGAACTCTTTCCCTACACGACGCTCTTCCGATCTNNNACACTTCCACAGTTATCATCAG
AGCACACGTCTGAACTCTTTCCCTACACGACGCTCTTCCGATCTNNNACACTTCCACAGTTATCATCAG
GCACACGTCTGAACTCTTTCCCTACACGACGCTCTTCCGATCTNNNACTGATAAGTGATGTGCCAAGG
GCACACGTCTGAACTCTTTCCCTACACGACGCTCTTCCGATCTNNNACTGATAAGTGATGTGCCAAGG
AGAGCACACGTCTGAACTCTTTCCCTACACGACGCTCTTCCGATCTNNNTGAGGCAGAAGAATTGCTT
AGAGCACACGTCTGAACTCTTTCCCTACACGACGCTCTTCCGATCTNNNTGAGGCAGAAGAATTGCTT
AGCACACGTCTGAACTCTTTCCCTACACGACGCTCTTCCGATCTNNNAGACAATTACTTGAACCCGGG
AGCACACGTCTGAACTCTTTCCCTACACGACGCTCTTCCGATCTNNNAGACAATTACTTGAACCCGGG
AGCACACGTCTGAACTCTTTCCCTACACGACGCTCTTCCGATCTNNNCAAAGTGCTGGGATTACAGG
AGCACACGTCTGAACTCTTTCCCTACACGACGCTCTTCCGATCTNNNCAAAGTGCTGGGATTACAGG
GCACACGTCTGAACTCTTTCCCTACACGACGCTCTTCCGATCTNNNTCTTTTCCATTGGTCTATGTGCC
GCACACGTCTGAACTCTTTCCCTACACGACGCTCTTCCGATCTNNNTCTTTTCCATTGGTCTATGTGCC
AGCACACGTCTGAACTCTTTCCCTACACGACGCTCTTCCGATCTNNNCTGCAGAACTGACAGCTAAAC
AGCACACGTCTGAACTCTTTCCCTACACGACGCTCTTCCGATCTNNNCTGCAGAACTGACAGCTAAAC
AGCACACGTCTGAACTCTTTCCCTACACGACGCTCTTCCGATCTNNNTTGTAGTGAGCCAAAATCATG
AGCACACGTCTGAACTCTTTCCCTACACGACGCTCTTCCGATCTNNNTTGTAGTGAGCCAAAATCATG
GAGCACACGTCTGAACTCTTTCCCTACACGACGCTCTTCCGATCTNNNAAAATCTGTGTTTGTCGGGG
GAGCACACGTCTGAACTCTTTCCCTACACGACGCTCTTCCGATCTNNNAAAATCTGTGTTTGTCGGGG
AGCACACGTCTGAACTCTTTCCCTACACGACGCTCTTCCGATCTNNNGATCACGAGGTCAGGAGATT
AGCACACGTCTGAACTCTTTCCCTACACGACGCTCTTCCGATCTNNNGATCACGAGGTCAGGAGATT
GCACACGTCTGAACTCTTTCCCTACACGACGCTCTTCCGATCTNNNAGGAGAAAATCTGAGAATAAG
GCACACGTCTGAACTCTTTCCCTACACGACGCTCTTCCGATCTNNNAGGAGAAAATCTGAGAATAAG
AGCACACGTCTGAACTCTTTCCCTACACGACGCTCTTCCGATCTNNNGCCTGTAGTCCCAGCTACTTGC
AGCACACGTCTGAACTCTTTCCCTACACGACGCTCTTCCGATCTNNNGCCTGTAGTCCCAGCTACTTGC
GAGCACACGTCTGAACTCTTTCCCTACACGACGCTCTTCCGATCTNNNGAGATACATGTGGGAGTCA
GAGCACACGTCTGAACTCTTTCCCTACACGACGCTCTTCCGATCTNNNGAGATACATGTGGGAGTCA
AGCACACGTCTGAACTCTTTCCCTACACGACGCTCTTCCGATCTNNNTTTTAACTGGCAGTCTTCCAAG
AGCACACGTCTGAACTCTTTCCCTACACGACGCTCTTCCGATCTNNNTTTTAACTGGCAGTCTTCCAAG
GCACACGTCTGAACTCTTTCCCTACACGACGCTCTTCCGATCTNNNTGTTGGGACTTTTTAGGCTGTT
GCACACGTCTGAACTCTTTCCCTACACGACGCTCTTCCGATCTNNNTGTTGGGACTTTTTAGGCTGTT
AGCACACGTCTGAACTCTTTCCCTACACGACGCTCTTCCGATCTNNNTATTATCTGTGTCACTGCAGCC
AGCACACGTCTGAACTCTTTCCCTACACGACGCTCTTCCGATCTNNNTATTATCTGTGTCACTGCAGCC
AGCACACGTCTGAACTCTTTCCCTACACGACGCTCTTCCGATCTNNNCAAAGGCATTGAACACTTGTT

AGCACACGTCTGAACTCTTTCCCTACACGACGCTCTTCCGATCTNNNCAAAGGCATTGAACACTTGTTG
GCACACGTCTGAACTCTTTCCCTACACGACGCTCTTCCGATCTNNNCCCAGGTTCAAGCAATTCTCATGC
GCACACGTCTGAACTCTTTCCCTACACGACGCTCTTCCGATCTNNNCCCAGGTTCAAGCAATTCTCATGC
AGCACACGTCTGAACTCTTTCCCTACACGACGCTCTTCCGATCTNNNGCACCTGTAATCCCAGCTACTA
AGCACACGTCTGAACTCTTTCCCTACACGACGCTCTTCCGATCTNNNGCACCTGTAATCCCAGCTACTA
GAGCACACGTCTGAACTCTTTCCCTACACGACGCTCTTCCGATCTNNNGACTTTACCGCCACCTTCCTG
GAGCACACGTCTGAACTCTTTCCCTACACGACGCTCTTCCGATCTNNNGACTTTACCGCCACCTTCCTG
GAGCACACGTCTGAACTCTTTCCCTACACGACGCTCTTCCGATCTNNNTTGAGTTCAAGACTAGCCTGG
GAGCACACGTCTGAACTCTTTCCCTACACGACGCTCTTCCGATCTNNNTTGAGTTCAAGACTAGCCTGG
AGCACACGTCTGAACTCTTTCCCTACACGACGCTCTTCCGATCTNNNTGATCCAGACATTCCCTCTTTGG
AGCACACGTCTGAACTCTTTCCCTACACGACGCTCTTCCGATCTNNNTGATCCAGACATTCCCTCTTTGG
AGCACACGTCTGAACTCTTTCCCTACACGACGCTCTTCCGATCTNNNATAGTCCCAGCTACTTAGGAG
AGCACACGTCTGAACTCTTTCCCTACACGACGCTCTTCCGATCTNNNATAGTCCCAGCTACTTAGGAG
AGCACACGTCTGAACTCTTTCCCTACACGACGCTCTTCCGATCTNNNTGAGGCAGGAGAATCTCTTGA
AGCACACGTCTGAACTCTTTCCCTACACGACGCTCTTCCGATCTNNNTGAGGCAGGAGAATCTCTTGA
GCACACGTCTGAACTCTTTCCCTACACGACGCTCTTCCGATCTNNNACAAGCTCCAAGTCCACTTTCTGC
GCACACGTCTGAACTCTTTCCCTACACGACGCTCTTCCGATCTNNNACAAGCTCCAAGTCCACTTTCTGC
AGCACACGTCTGAACTCTTTCCCTACACGACGCTCTTCCGATCTNNNAACTAAAAAAATCAAGAAACTA
AGCACACGTCTGAACTCTTTCCCTACACGACGCTCTTCCGATCTNNNAACTAAAAAAATCAAGAAACTA
GAGCACACGTCTGAACTCTTTCCCTACACGACGCTCTTCCGATCTNNNCCAGAGATCAGTTTCGTGGA
GAGCACACGTCTGAACTCTTTCCCTACACGACGCTCTTCCGATCTNNNCCAGAGATCAGTTTCGTGGA
AGCACACGTCTGAACTCTTTCCCTACACGACGCTCTTCCGATCTNNNCTCATGCCTGTAATCCCAGAAC
AGCACACGTCTGAACTCTTTCCCTACACGACGCTCTTCCGATCTNNNCTCATGCCTGTAATCCCAGAAC
AGCACACGTCTGAACTCTTTCCCTACACGACGCTCTTCCGATCTNNNCCCATGAGTATAGGTGGCGTA
AGCACACGTCTGAACTCTTTCCCTACACGACGCTCTTCCGATCTNNNCCCATGAGTATAGGTGGCGTA
AGAGCACACGTCTGAACTCTTTCCCTACACGACGCTCTTCCGATCTNNNAAAAAAATTAGCTGGGCGT
AGAGCACACGTCTGAACTCTTTCCCTACACGACGCTCTTCCGATCTNNNAAAAAAATTAGCTGGGCGT
AGCACACGTCTGAACTCTTTCCCTACACGACGCTCTTCCGATCTNNNATGGCTTAGGGTGTAGGGTA
AGCACACGTCTGAACTCTTTCCCTACACGACGCTCTTCCGATCTNNNATGGCTTAGGGTGTAGGGTA
GAGCACACGTCTGAACTCTTTCCCTACACGACGCTCTTCCGATCTNNNTGAGGTAAAGTGGCATGATT
GAGCACACGTCTGAACTCTTTCCCTACACGACGCTCTTCCGATCTNNNTGAGGTAAAGTGGCATGATT
AGCACACGTCTGAACTCTTTCCCTACACGACGCTCTTCCGATCTNNNCTAAGAGCTGGGATTACAGGC
AGCACACGTCTGAACTCTTTCCCTACACGACGCTCTTCCGATCTNNNCTAAGAGCTGGGATTACAGGC
GCACACGTCTGAACTCTTTCCCTACACGACGCTCTTCCGATCTNNNGAGAATCGCTTGAACCCAGGAG
GCACACGTCTGAACTCTTTCCCTACACGACGCTCTTCCGATCTNNNGAGAATCGCTTGAACCCAGGAG
ACACGTCTGAACTCTTTCCCTACACGACGCTCTTCCGATCTNNNCTTGGAAGAGGATCCCTGTGATGC
ACACGTCTGAACTCTTTCCCTACACGACGCTCTTCCGATCTNNNCTTGGAAGAGGATCCCTGTGATGC
CACGTCTGAACTCTTTCCCTACACGACGCTCTTCCGATCTNNNAGCTAGGACTACAGGACTACAGGCA
CACGTCTGAACTCTTTCCCTACACGACGCTCTTCCGATCTNNNAGCTAGGACTACAGGACTACAGGCA
AGAGCACACGTCTGAACTCTTTCCCTACACGACGCTCTTCCGATCTNNNACTCTGGGATTTGAGTTGG
AGAGCACACGTCTGAACTCTTTCCCTACACGACGCTCTTCCGATCTNNNACTCTGGGATTTGAGTTGG
AGCACACGTCTGAACTCTTTCCCTACACGACGCTCTTCCGATCTNNNTGGAACACTGTCCTTATCACAC
AGCACACGTCTGAACTCTTTCCCTACACGACGCTCTTCCGATCTNNNTGGAACACTGTCCTTATCACAC

AGCACACGTCTGAACTCTTTCCCTACACGACGCTCTTCCGATCTNNNCTCTCAGCTACTCGGGACTTTA
AGCACACGTCTGAACTCTTTCCCTACACGACGCTCTTCCGATCTNNNCTCTCAGCTACTCGGGACTTTA
GAGCACACGTCTGAACTCTTTCCCTACACGACGCTCTTCCGATCTNNNTATTGTTACACACACACACAC
GAGCACACGTCTGAACTCTTTCCCTACACGACGCTCTTCCGATCTNNNTATTGTTACACACACACACAC
ACACGTCTGAACTCTTTCCCTACACGACGCTCTTCCGATCTNNNACTCTAGGGGTAAAGGGTGAGAG
ACACGTCTGAACTCTTTCCCTACACGACGCTCTTCCGATCTNNNACTCTAGGGGTAAAGGGTGAGAG
AGCACACGTCTGAACTCTTTCCCTACACGACGCTCTTCCGATCTNNNCTGAGGCAGGAGAATCACTTG
AGCACACGTCTGAACTCTTTCCCTACACGACGCTCTTCCGATCTNNNCTGAGGCAGGAGAATCACTTG
AGCACACGTCTGAACTCTTTCCCTACACGACGCTCTTCCGATCTNNNAACAAATAAACTCCGCGGAAC
AGCACACGTCTGAACTCTTTCCCTACACGACGCTCTTCCGATCTNNNAACAAATAAACTCCGCGGAAC
AGCACACGTCTGAACTCTTTCCCTACACGACGCTCTTCCGATCTNNNCATGTAACTCCAGGCAGCAAT
AGCACACGTCTGAACTCTTTCCCTACACGACGCTCTTCCGATCTNNNCATGTAACTCCAGGCAGCAAT
GCACACGTCTGAACTCTTTCCCTACACGACGCTCTTCCGATCTNNNGATGTATGCACGTTGGAGAGTT
GCACACGTCTGAACTCTTTCCCTACACGACGCTCTTCCGATCTNNNGATGTATGCACGTTGGAGAGTT
GCACACGTCTGAACTCTTTCCCTACACGACGCTCTTCCGATCTNNNTGCATCTGAGTCTCTCTCTCTCC
GCACACGTCTGAACTCTTTCCCTACACGACGCTCTTCCGATCTNNNTGCATCTGAGTCTCTCTCTCTCC
GCACACGTCTGAACTCTTTCCCTACACGACGCTCTTCCGATCTNNNGGGTCAAACATCCAAACTGTAG
GCACACGTCTGAACTCTTTCCCTACACGACGCTCTTCCGATCTNNNGGGTCAAACATCCAAACTGTAG
AGCACACGTCTGAACTCTTTCCCTACACGACGCTCTTCCGATCTNNNCTGCTCTCTCATAACCACACCTT
AGCACACGTCTGAACTCTTTCCCTACACGACGCTCTTCCGATCTNNNCTGCTCTCTCATAACCACACCTT
CACACGTCTGAACTCTTTCCCTACACGACGCTCTTCCGATCTNNNTGGTGCAGGTCACTGTCTATTTGC
CACACGTCTGAACTCTTTCCCTACACGACGCTCTTCCGATCTNNNTGGTGCAGGTCACTGTCTATTTGC
AGCACACGTCTGAACTCTTTCCCTACACGACGCTCTTCCGATCTNNNAACATCACACCACTGCACTCTT
AGCACACGTCTGAACTCTTTCCCTACACGACGCTCTTCCGATCTNNNAACATCACACCACTGCACTCTT
GCACACGTCTGAACTCTTTCCCTACACGACGCTCTTCCGATCTNNNATAATGCCAGCTACTTAGGAGG
GCACACGTCTGAACTCTTTCCCTACACGACGCTCTTCCGATCTNNNATAATGCCAGCTACTTAGGAGG
CACACGTCTGAACTCTTTCCCTACACGACGCTCTTCCGATCTNNNAAGCTGCTGTATTTTAGTGAGCAT
CACACGTCTGAACTCTTTCCCTACACGACGCTCTTCCGATCTNNNAAGCTGCTGTATTTTAGTGAGCAT
AGCACACGTCTGAACTCTTTCCCTACACGACGCTCTTCCGATCTNNNAACTCAGTGAAGATGCCTTGTC
AGCACACGTCTGAACTCTTTCCCTACACGACGCTCTTCCGATCTNNNAACTCAGTGAAGATGCCTTGTC
AGCACACGTCTGAACTCTTTCCCTACACGACGCTCTTCCGATCTNNNCTTGCAGTGACCCATGAGAAT
AGCACACGTCTGAACTCTTTCCCTACACGACGCTCTTCCGATCTNNNCTTGCAGTGACCCATGAGAAT
GAGCACACGTCTGAACTCTTTCCCTACACGACGCTCTTCCGATCTNNNCGCTGCATGTTCTCACTCATA
GAGCACACGTCTGAACTCTTTCCCTACACGACGCTCTTCCGATCTNNNCGCTGCATGTTCTCACTCATA
GCACACGTCTGAACTCTTTCCCTACACGACGCTCTTCCGATCTNNNCTCTTCCTTCCTTCCTTCCTTCCCA
GCACACGTCTGAACTCTTTCCCTACACGACGCTCTTCCGATCTNNNCTCTTCCTTCCTTCCTTCCTTCCCA
AGCACACGTCTGAACTCTTTCCCTACACGACGCTCTTCCGATCTNNNTATAGCAGTGTGTGGCCAAGA
AGCACACGTCTGAACTCTTTCCCTACACGACGCTCTTCCGATCTNNNTATAGCAGTGTGTGGCCAAGA
GCACACGTCTGAACTCTTTCCCTACACGACGCTCTTCCGATCTNNNGAACTTTTTCTCTGAGGGGGCT
GCACACGTCTGAACTCTTTCCCTACACGACGCTCTTCCGATCTNNNGAACTTTTTCTCTGAGGGGGCT
GCACACGTCTGAACTCTTTCCCTACACGACGCTCTTCCGATCTNNNTCCTACAATTGAGGGTTGTGTGA
GCACACGTCTGAACTCTTTCCCTACACGACGCTCTTCCGATCTNNNTCCTACAATTGAGGGTTGTGTGA
GCACACGTCTGAACTCTTTCCCTACACGACGCTCTTCCGATCTNNNACCTCTGAAAATGAAAGGCTGTC

GCACACGTCTGAACTCTTTCCCTACACGACGCTCTTCCGATCTNNNACCTCTGAAAATGAAAGGCTGT
GAGCACACGTCTGAACTCTTTCCCTACACGACGCTCTTCCGATCTNNNCTCAGCTGTGTTGACTACAGT
GAGCACACGTCTGAACTCTTTCCCTACACGACGCTCTTCCGATCTNNNCTCAGCTGTGTTGACTACAGT
AGAGCACACGTCTGAACTCTTTCCCTACACGACGCTCTTCCGATCTNNNGGATAAACCGATTTAATGGG
AGAGCACACGTCTGAACTCTTTCCCTACACGACGCTCTTCCGATCTNNNGGATAAACCGATTTAATGGG
AGCACACGTCTGAACTCTTTCCCTACACGACGCTCTTCCGATCTNNNAGAGCTAACTGCCTCCAATGAA
AGCACACGTCTGAACTCTTTCCCTACACGACGCTCTTCCGATCTNNNAGAGCTAACTGCCTCCAATGAA
AGAGCACACGTCTGAACTCTTTCCCTACACGACGCTCTTCCGATCTNNNACAAAGGAGACGCAAGAAT
AGAGCACACGTCTGAACTCTTTCCCTACACGACGCTCTTCCGATCTNNNACAAAGGAGACGCAAGAAT
GAGCACACGTCTGAACTCTTTCCCTACACGACGCTCTTCCGATCTNNNAGGAAAAAGGGCTAGATGAG
GAGCACACGTCTGAACTCTTTCCCTACACGACGCTCTTCCGATCTNNNAGGAAAAAGGGCTAGATGAG
GAGCACACGTCTGAACTCTTTCCCTACACGACGCTCTTCCGATCTNNNCACGCCATCTAAGTTTGTGAT
GAGCACACGTCTGAACTCTTTCCCTACACGACGCTCTTCCGATCTNNNCACGCCATCTAAGTTTGTGAT
AGCACACGTCTGAACTCTTTCCCTACACGACGCTCTTCCGATCTNNNAAGTTGTTCATTGGCCCCTCTAT
AGCACACGTCTGAACTCTTTCCCTACACGACGCTCTTCCGATCTNNNAAGTTGTTCATTGGCCCCTCTAT
AGAGCACACGTCTGAACTCTTTCCCTACACGACGCTCTTCCGATCTNNNGGGATTTGGCAAATGACTGT
AGAGCACACGTCTGAACTCTTTCCCTACACGACGCTCTTCCGATCTNNNGGGATTTGGCAAATGACTGT
AGCACACGTCTGAACTCTTTCCCTACACGACGCTCTTCCGATCTNNNCCAAGATTACACCATTGCACTCC
AGCACACGTCTGAACTCTTTCCCTACACGACGCTCTTCCGATCTNNNCCAAGATTACACCATTGCACTCC
GAGCACACGTCTGAACTCTTTCCCTACACGACGCTCTTCCGATCTNNNCACCTGTAATCCCAGCTACTCA
GAGCACACGTCTGAACTCTTTCCCTACACGACGCTCTTCCGATCTNNNCACCTGTAATCCCAGCTACTCA
GAGCACACGTCTGAACTCTTTCCCTACACGACGCTCTTCCGATCTNNNTAACCCAGTTCCATCCTCTACCT
GAGCACACGTCTGAACTCTTTCCCTACACGACGCTCTTCCGATCTNNNTAACCCAGTTCCATCCTCTACCT
AGCACACGTCTGAACTCTTTCCCTACACGACGCTCTTCCGATCTNNNTACACCGAACTATTAGTTGGCC
AGCACACGTCTGAACTCTTTCCCTACACGACGCTCTTCCGATCTNNNTACACCGAACTATTAGTTGGCC
AGCACACGTCTGAACTCTTTCCCTACACGACGCTCTTCCGATCTNNNCCTCATTCTCCTCAGCCATAGAG
AGCACACGTCTGAACTCTTTCCCTACACGACGCTCTTCCGATCTNNNCCTCATTCTCCTCAGCCATAGAG
AGCACACGTCTGAACTCTTTCCCTACACGACGCTCTTCCGATCTNNNCAAAAGTTATCTGGGCATGGT
AGCACACGTCTGAACTCTTTCCCTACACGACGCTCTTCCGATCTNNNCAAAAGTTATCTGGGCATGGT
GAGCACACGTCTGAACTCTTTCCCTACACGACGCTCTTCCGATCTNNNCTCTACAAGCGTCAGGATTCT
GAGCACACGTCTGAACTCTTTCCCTACACGACGCTCTTCCGATCTNNNCTCTACAAGCGTCAGGATTCT
GAGCACACGTCTGAACTCTTTCCCTACACGACGCTCTTCCGATCTNNNCCCTAACTTGCATGACCAATTC
GAGCACACGTCTGAACTCTTTCCCTACACGACGCTCTTCCGATCTNNNCCCTAACTTGCATGACCAATTC
AAGAGCACACGTCTGAACTCTTTCCCTACACGACGCTCTTCCGATCTNNNTGTGTGTGTGTGTAGTGT
AAGAGCACACGTCTGAACTCTTTCCCTACACGACGCTCTTCCGATCTNNNTGTGTGTGTGTGTAGTGT
AGCACACGTCTGAACTCTTTCCCTACACGACGCTCTTCCGATCTNNNAGAGAGAGTGTTTGTGTGTGT
AGCACACGTCTGAACTCTTTCCCTACACGACGCTCTTCCGATCTNNNAGAGAGAGTGTTTGTGTGTGT
GAGCACACGTCTGAACTCTTTCCCTACACGACGCTCTTCCGATCTNNNCAAACAGTTGGAGGAAAGAG
GAGCACACGTCTGAACTCTTTCCCTACACGACGCTCTTCCGATCTNNNCAAACAGTTGGAGGAAAGAG
GCACACGTCTGAACTCTTTCCCTACACGACGCTCTTCCGATCTNNNTGGTGACTAACTGTTCAGACTGG
GCACACGTCTGAACTCTTTCCCTACACGACGCTCTTCCGATCTNNNTGGTGACTAACTGTTCAGACTGG
CACACGTCTGAACTCTTTCCCTACACGACGCTCTTCCGATCTNNNATGCAGTCTCATAAACACTCCATCG
CACACGTCTGAACTCTTTCCCTACACGACGCTCTTCCGATCTNNNATGCAGTCTCATAAACACTCCATCG

AGCACACGTCTGAACTCTTTCCCTACACGACGCTCTTCCGATCTNNNGGCCATGCCTATGTACAGGTTTA
AGCACACGTCTGAACTCTTTCCCTACACGACGCTCTTCCGATCTNNNGGCCATGCCTATGTACAGGTTTA
AGCACACGTCTGAACTCTTTCCCTACACGACGCTCTTCCGATCTNNNTTCTGCCAGCTTGTTACATAGGT
AGCACACGTCTGAACTCTTTCCCTACACGACGCTCTTCCGATCTNNNTTCTGCCAGCTTGTTACATAGGT
AGCACACGTCTGAACTCTTTCCCTACACGACGCTCTTCCGATCTNNNAATTGGGAGAGTGGCTTGAGG
AGCACACGTCTGAACTCTTTCCCTACACGACGCTCTTCCGATCTNNNAATTGGGAGAGTGGCTTGAGG
AGCACACGTCTGAACTCTTTCCCTACACGACGCTCTTCCGATCTNNNCCTGAGCCATAGCACCTTTTCAT
AGCACACGTCTGAACTCTTTCCCTACACGACGCTCTTCCGATCTNNNCCTGAGCCATAGCACCTTTTCAT
AGCACACGTCTGAACTCTTTCCCTACACGACGCTCTTCCGATCTNNNTGGGGATTTCTCACCTCTTTCTG
AGCACACGTCTGAACTCTTTCCCTACACGACGCTCTTCCGATCTNNNTGGGGATTTCTCACCTCTTTCTG
GCACACGTCTGAACTCTTTCCCTACACGACGCTCTTCCGATCTNNNCTTAAAGAATGAGAACTTCGACCA
GCACACGTCTGAACTCTTTCCCTACACGACGCTCTTCCGATCTNNNCTTAAAGAATGAGAACTTCGACCA
AGCACACGTCTGAACTCTTTCCCTACACGACGCTCTTCCGATCTNNNTAGTCCCAACTACTCAGGAGGTT
AGCACACGTCTGAACTCTTTCCCTACACGACGCTCTTCCGATCTNNNTAGTCCCAACTACTCAGGAGGTT
AGCACACGTCTGAACTCTTTCCCTACACGACGCTCTTCCGATCTNNNCTGGCCTGTTTCAGATCCTACAT
AGCACACGTCTGAACTCTTTCCCTACACGACGCTCTTCCGATCTNNNCTGGCCTGTTTCAGATCCTACAT
GAGCACACGTCTGAACTCTTTCCCTACACGACGCTCTTCCGATCTNNNCCTTGAATCCCAGGAATAAGC
GAGCACACGTCTGAACTCTTTCCCTACACGACGCTCTTCCGATCTNNNCCTTGAATCCCAGGAATAAGC
AGCACACGTCTGAACTCTTTCCCTACACGACGCTCTTCCGATCTNNNCAACAAGGAGAGAGATGCTGT
AGCACACGTCTGAACTCTTTCCCTACACGACGCTCTTCCGATCTNNNCAACAAGGAGAGAGATGCTGT
AGCACACGTCTGAACTCTTTCCCTACACGACGCTCTTCCGATCTNNNCCCTGTTAGTCATGTGAAGGTG
AGCACACGTCTGAACTCTTTCCCTACACGACGCTCTTCCGATCTNNNCCCTGTTAGTCATGTGAAGGTG
AGCACACGTCTGAACTCTTTCCCTACACGACGCTCTTCCGATCTNNNCATGCCAACATCCAAATCACCTC
AGCACACGTCTGAACTCTTTCCCTACACGACGCTCTTCCGATCTNNNCATGCCAACATCCAAATCACCTC
AGAGCACACGTCTGAACTCTTTCCCTACACGACGCTCTTCCGATCTNNNGAAGCGAGAAAGGTCTGA
AGAGCACACGTCTGAACTCTTTCCCTACACGACGCTCTTCCGATCTNNNGAAGCGAGAAAGGTCTGA
AGAGCACACGTCTGAACTCTTTCCCTACACGACGCTCTTCCGATCTNNNTGAGCTGATATCATGCCACT
AGAGCACACGTCTGAACTCTTTCCCTACACGACGCTCTTCCGATCTNNNTGAGCTGATATCATGCCACT
AGCACACGTCTGAACTCTTTCCCTACACGACGCTCTTCCGATCTNNNGGCTCATCAAAGTACATGTTG
AGCACACGTCTGAACTCTTTCCCTACACGACGCTCTTCCGATCTNNNGGCTCATCAAAGTACATGTTG
GCACACGTCTGAACTCTTTCCCTACACGACGCTCTTCCGATCTNNNTAGACCCTCCATAACCAGTCCATC
GCACACGTCTGAACTCTTTCCCTACACGACGCTCTTCCGATCTNNNTAGACCCTCCATAACCAGTCCATC
GAGCACACGTCTGAACTCTTTCCCTACACGACGCTCTTCCGATCTNNNATTTCTCACCTCCTTCAAGCCTT
GAGCACACGTCTGAACTCTTTCCCTACACGACGCTCTTCCGATCTNNNATTTCTCACCTCCTTCAAGCCTT
AGCACACGTCTGAACTCTTTCCCTACACGACGCTCTTCCGATCTNNNTCTCTCTCTCTCTGTGTGTGT
AGCACACGTCTGAACTCTTTCCCTACACGACGCTCTTCCGATCTNNNTCTCTCTCTCTCTGTGTGTGT
AGCACACGTCTGAACTCTTTCCCTACACGACGCTCTTCCGATCTNNNCCAATTTGGTGGGGCAAGTAA
AGCACACGTCTGAACTCTTTCCCTACACGACGCTCTTCCGATCTNNNCCAATTTGGTGGGGCAAGTAA
AGCACACGTCTGAACTCTTTCCCTACACGACGCTCTTCCGATCTNNNTCACACTTTTCCTTTGAGACTCT
AGCACACGTCTGAACTCTTTCCCTACACGACGCTCTTCCGATCTNNNTCACACTTTTCCTTTGAGACTCT
AGAGCACACGTCTGAACTCTTTCCCTACACGACGCTCTTCCGATCTNNNCGCCTATAGTCCCAGCTATTC
AGAGCACACGTCTGAACTCTTTCCCTACACGACGCTCTTCCGATCTNNNCGCCTATAGTCCCAGCTATTC
AGAGCACACGTCTGAACTCTTTCCCTACACGACGCTCTTCCGATCTNNNGAGATCACATCACTGCACTCC

GAGCACACGTCTGAACTCTTTCCCTACACGACGCTCTTCCGATCTNNNGAGATCACATCACTGCACTCC

ACACGTCTGAACTCTTTCCCTACACGACGCTCTTCCGATCTNNNACACCAAGAATTGTGTCAGAGGAC

ACACGTCTGAACTCTTTCCCTACACGACGCTCTTCCGATCTNNNACACCAAGAATTGTGTCAGAGGAC

AAGAGCACACGTCTGAACTCTTTCCCTACACGACGCTCTTCCGATCTNNNAATGTGACTGTTGTTTCA

AAGAGCACACGTCTGAACTCTTTCCCTACACGACGCTCTTCCGATCTNNNAATGTGACTGTTGTTTCA

AGCACACGTCTGAACTCTTTCCCTACACGACGCTCTTCCGATCTNNNATTTCCAAGTTTGGCCACTGAG

AGCACACGTCTGAACTCTTTCCCTACACGACGCTCTTCCGATCTNNNATTTCCAAGTTTGGCCACTGAG

AGCACACGTCTGAACTCTTTCCCTACACGACGCTCTTCCGATCTNNNGAGACAGGAAAATGGCATGAA

AGCACACGTCTGAACTCTTTCCCTACACGACGCTCTTCCGATCTNNNGAGACAGGAAAATGGCATGAA

AGCACACGTCTGAACTCTTTCCCTACACGACGCTCTTCCGATCTNNNCCTGGTGTTTTCATCATTCTGG

AGCACACGTCTGAACTCTTTCCCTACACGACGCTCTTCCGATCTNNNCCTGGTGTTTTCATCATTCTGG

AGCACACGTCTGAACTCTTTCCCTACACGACGCTCTTCCGATCTNNNATCATCTCCACTAAAAGGCGGA

AGCACACGTCTGAACTCTTTCCCTACACGACGCTCTTCCGATCTNNNATCATCTCCACTAAAAGGCGGA

AGCACACGTCTGAACTCTTTCCCTACACGACGCTCTTCCGATCTNNNTGAAGAACACTTTGGAGGTTC

AGCACACGTCTGAACTCTTTCCCTACACGACGCTCTTCCGATCTNNNTGAAGAACACTTTGGAGGTTC

AGCACACGTCTGAACTCTTTCCCTACACGACGCTCTTCCGATCTNNNGCACGATGTAGCTCTTTCTTG

AGCACACGTCTGAACTCTTTCCCTACACGACGCTCTTCCGATCTNNNGCACGATGTAGCTCTTTCTTG

GCACACGTCTGAACTCTTTCCCTACACGACGCTCTTCCGATCTNNNGCCTCCTGAGTAGTTGGGACTA

GCACACGTCTGAACTCTTTCCCTACACGACGCTCTTCCGATCTNNNGCCTCCTGAGTAGTTGGGACTA

GCACACGTCTGAACTCTTTCCCTACACGACGCTCTTCCGATCTNNNTCAGGGACTCCTCCTAACTCATT

GCACACGTCTGAACTCTTTCCCTACACGACGCTCTTCCGATCTNNNTCAGGGACTCCTCCTAACTCATT

GCACACGTCTGAACTCTTTCCCTACACGACGCTCTTCCGATCTNNNGGATCACCTGAGGTCAAGAGTT

GCACACGTCTGAACTCTTTCCCTACACGACGCTCTTCCGATCTNNNGGATCACCTGAGGTCAAGAGTT

GAGCACACGTCTGAACTCTTTCCCTACACGACGCTCTTCCGATCTNNNGAAAATCCTTGAAAGTCAACA

GAGCACACGTCTGAACTCTTTCCCTACACGACGCTCTTCCGATCTNNNGAAAATCCTTGAAAGTCAACA

GAGCACACGTCTGAACTCTTTCCCTACACGACGCTCTTCCGATCTNNNAATACATACACTGTTCTTTGT

GAGCACACGTCTGAACTCTTTCCCTACACGACGCTCTTCCGATCTNNNAATACATACACTGTTCTTTGT

GCACACGTCTGAACTCTTTCCCTACACGACGCTCTTCCGATCTNNNACCTGGGAAAAGCTTCTGTTAGT

GCACACGTCTGAACTCTTTCCCTACACGACGCTCTTCCGATCTNNNACCTGGGAAAAGCTTCTGTTAGT

GCACACGTCTGAACTCTTTCCCTACACGACGCTCTTCCGATCTNNNCAGCCTGGATGATAGAGTGAGA

GCACACGTCTGAACTCTTTCCCTACACGACGCTCTTCCGATCTNNNCAGCCTGGATGATAGAGTGAGA

GAGCACACGTCTGAACTCTTTCCCTACACGACGCTCTTCCGATCTNNNTACACATATGCACAGACACAT

GAGCACACGTCTGAACTCTTTCCCTACACGACGCTCTTCCGATCTNNNTACACATATGCACAGACACAT

GAGCACACGTCTGAACTCTTTCCCTACACGACGCTCTTCCGATCTNNNCCTTATCACTACTTCCAGAAG

GAGCACACGTCTGAACTCTTTCCCTACACGACGCTCTTCCGATCTNNNCCTTATCACTACTTCCAGAAG

AGAGCACACGTCTGAACTCTTTCCCTACACGACGCTCTTCCGATCTNNNTGAAATAGGTGTGACAGGA

AGAGCACACGTCTGAACTCTTTCCCTACACGACGCTCTTCCGATCTNNNTGAAATAGGTGTGACAGGA

GAGCACACGTCTGAACTCTTTCCCTACACGACGCTCTTCCGATCTNNNAGAAAAGCACTACACAAAACG

GAGCACACGTCTGAACTCTTTCCCTACACGACGCTCTTCCGATCTNNNAGAAAAGCACTACACAAAACG

AGCACACGTCTGAACTCTTTCCCTACACGACGCTCTTCCGATCTNNNATTCATAAAAGGCCATCCCCAC

AGCACACGTCTGAACTCTTTCCCTACACGACGCTCTTCCGATCTNNNATTCATAAAAGGCCATCCCCAC

AGCACACGTCTGAACTCTTTCCCTACACGACGCTCTTCCGATCTNNNTGACCTGCTGCTAAAGTGTGTA

AGCACACGTCTGAACTCTTTCCCTACACGACGCTCTTCCGATCTNNNTGACCTGCTGCTAAAGTGTGTA

GAGCACACGTCTGAACTCTTTCCCTACACGACGCTCTTCCGATCTNNNAAATACAAAAATTGCCTGGGC
GAGCACACGTCTGAACTCTTTCCCTACACGACGCTCTTCCGATCTNNNAAATACAAAAATTGCCTGGGC
AGAGCACACGTCTGAACTCTTTCCCTACACGACGCTCTTCCGATCTNNNAACATGCCAATGAAACCTCA
AGAGCACACGTCTGAACTCTTTCCCTACACGACGCTCTTCCGATCTNNNAACATGCCAATGAAACCTCA
GAGCACACGTCTGAACTCTTTCCCTACACGACGCTCTTCCGATCTNNNCCATCATACACACACACACACA
GAGCACACGTCTGAACTCTTTCCCTACACGACGCTCTTCCGATCTNNNCCATCATACACACACACACACA
AGCACACGTCTGAACTCTTTCCCTACACGACGCTCTTCCGATCTNNNATTTGCAAGGGACAAACAACC
AGCACACGTCTGAACTCTTTCCCTACACGACGCTCTTCCGATCTNNNATTTGCAAGGGACAAACAACC
AGAGCACACGTCTGAACTCTTTCCCTACACGACGCTCTTCCGATCTNNNCCATCTACCTGCCTACCTACC
AGAGCACACGTCTGAACTCTTTCCCTACACGACGCTCTTCCGATCTNNNCCATCTACCTGCCTACCTACC
AGAGCACACGTCTGAACTCTTTCCCTACACGACGCTCTTCCGATCTNNNAAACTCCTGAGACTGGCTTT
AGAGCACACGTCTGAACTCTTTCCCTACACGACGCTCTTCCGATCTNNNAAACTCCTGAGACTGGCTTT
AGCACACGTCTGAACTCTTTCCCTACACGACGCTCTTCCGATCTNNNGAGATCACACCACTGCACTCTAG
AGCACACGTCTGAACTCTTTCCCTACACGACGCTCTTCCGATCTNNNGAGATCACACCACTGCACTCTAG
GAGCACACGTCTGAACTCTTTCCCTACACGACGCTCTTCCGATCTNNNTCAGTTACTGTGTGTGGCCTA
GAGCACACGTCTGAACTCTTTCCCTACACGACGCTCTTCCGATCTNNNTCAGTTACTGTGTGTGGCCTA
GAGCACACGTCTGAACTCTTTCCCTACACGACGCTCTTCCGATCTNNNGTGGAAGCCTCCAAAAGCAG
GAGCACACGTCTGAACTCTTTCCCTACACGACGCTCTTCCGATCTNNNGTGGAAGCCTCCAAAAGCAG
GAGCACACGTCTGAACTCTTTCCCTACACGACGCTCTTCCGATCTNNNCTTGCAGTGAGCCCAGATTGT
GAGCACACGTCTGAACTCTTTCCCTACACGACGCTCTTCCGATCTNNNCTTGCAGTGAGCCCAGATTGT
GAGCACACGTCTGAACTCTTTCCCTACACGACGCTCTTCCGATCTNNNGAGTGCCCTACCATGCTACAG
GAGCACACGTCTGAACTCTTTCCCTACACGACGCTCTTCCGATCTNNNGAGTGCCCTACCATGCTACAG
GCACACGTCTGAACTCTTTCCCTACACGACGCTCTTCCGATCTNNNTCAATGAAGACCCAGTGAGTGA
GCACACGTCTGAACTCTTTCCCTACACGACGCTCTTCCGATCTNNNTCAATGAAGACCCAGTGAGTGA
GAGCACACGTCTGAACTCTTTCCCTACACGACGCTCTTCCGATCTNNNGACAGAGAAAGACCCCATCTC
GAGCACACGTCTGAACTCTTTCCCTACACGACGCTCTTCCGATCTNNNGACAGAGAAAGACCCCATCTC
AGAGCACACGTCTGAACTCTTTCCCTACACGACGCTCTTCCGATCTNNNGGCTCTAGGCAAACCCATAA
AGAGCACACGTCTGAACTCTTTCCCTACACGACGCTCTTCCGATCTNNNGGCTCTAGGCAAACCCATAA
AGAGCACACGTCTGAACTCTTTCCCTACACGACGCTCTTCCGATCTNNNCCTCTCTCTTGGGTAGACTG
AGAGCACACGTCTGAACTCTTTCCCTACACGACGCTCTTCCGATCTNNNCCTCTCTCTTGGGTAGACTG
AGAGCACACGTCTGAACTCTTTCCCTACACGACGCTCTTCCGATCTNNNTGGTCCTCCCCTTTTGTCTTTT
AGAGCACACGTCTGAACTCTTTCCCTACACGACGCTCTTCCGATCTNNNTGGTCCTCCCCTTTTGTCTTTT
AGCACACGTCTGAACTCTTTCCCTACACGACGCTCTTCCGATCTNNNCTGCTTCACTTGTGAGGTCTTTG
AGCACACGTCTGAACTCTTTCCCTACACGACGCTCTTCCGATCTNNNCTGCTTCACTTGTGAGGTCTTTG
CACACGTCTGAACTCTTTCCCTACACGACGCTCTTCCGATCTNNNCAGATGCTCGCGAACCAGAGCATC
CACACGTCTGAACTCTTTCCCTACACGACGCTCTTCCGATCTNNNCAGATGCTCGCGAACCAGAGCATC
AGCACACGTCTGAACTCTTTCCCTACACGACGCTCTTCCGATCTNNNCCCTTCTCAATCAAGGTCCAGAA
AGCACACGTCTGAACTCTTTCCCTACACGACGCTCTTCCGATCTNNNCCCTTCTCAATCAAGGTCCAGAA
AGCACACGTCTGAACTCTTTCCCTACACGACGCTCTTCCGATCTNNNGTCATTTCTTTTTCCCTGGCACA
AGCACACGTCTGAACTCTTTCCCTACACGACGCTCTTCCGATCTNNNGTCATTTCTTTTTCCCTGGCACA
GAGCACACGTCTGAACTCTTTCCCTACACGACGCTCTTCCGATCTNNNTGCCACCTACTTAACCTCTTTG
GAGCACACGTCTGAACTCTTTCCCTACACGACGCTCTTCCGATCTNNNTGCCACCTACTTAACCTCTTTG
CACACGTCTGAACTCTTTCCCTACACGACGCTCTTCCGATCTNNNGAAGTATTGAATAAGCCATTTCTC

CACACGTCTGAACTCTTTCCCTACACGACGCTCTTCCGATCTNNNGAAGTATTGAATAAGCCATTTCTC
GAGCACACGTCTGAACTCTTTCCCTACACGACGCTCTTCCGATCTNNNCTGAGACAGGAGAATCGCTT
GAGCACACGTCTGAACTCTTTCCCTACACGACGCTCTTCCGATCTNNNCTGAGACAGGAGAATCGCTT
AGCACACGTCTGAACTCTTTCCCTACACGACGCTCTTCCGATCTNNNGAAAAGCTTCAACCAGGCTTAG
AGCACACGTCTGAACTCTTTCCCTACACGACGCTCTTCCGATCTNNNGAAAAGCTTCAACCAGGCTTAG
AGCACACGTCTGAACTCTTTCCCTACACGACGCTCTTCCGATCTNNNCAAGAAGCATTTTTAAAGCATTT
AGCACACGTCTGAACTCTTTCCCTACACGACGCTCTTCCGATCTNNNCAAGAAGCATTTTTAAAGCATTT
AGCACACGTCTGAACTCTTTCCCTACACGACGCTCTTCCGATCTNNNCACTGAAACTTTAGGGGCACAA
AGCACACGTCTGAACTCTTTCCCTACACGACGCTCTTCCGATCTNNNCACTGAAACTTTAGGGGCACAA
AAGAGCACACGTCTGAACTCTTTCCCTACACGACGCTCTTCCGATCTNNNGGTTACCGTGACCTATGAT
AAGAGCACACGTCTGAACTCTTTCCCTACACGACGCTCTTCCGATCTNNNGGTTACCGTGACCTATGAT
AGCACACGTCTGAACTCTTTCCCTACACGACGCTCTTCCGATCTNNNTTAGGATGGGAGGATGGTGGA
AGCACACGTCTGAACTCTTTCCCTACACGACGCTCTTCCGATCTNNNTTAGGATGGGAGGATGGTGGA
GAGCACACGTCTGAACTCTTTCCCTACACGACGCTCTTCCGATCTNNNCAAGCTTACAGTAGTCCAACCC
GAGCACACGTCTGAACTCTTTCCCTACACGACGCTCTTCCGATCTNNNCAAGCTTACAGTAGTCCAACCC
AGCACACGTCTGAACTCTTTCCCTACACGACGCTCTTCCGATCTNNNATTAGAAAGGAAATGAATGGT
AGCACACGTCTGAACTCTTTCCCTACACGACGCTCTTCCGATCTNNNATTAGAAAGGAAATGAATGGT
AGCACACGTCTGAACTCTTTCCCTACACGACGCTCTTCCGATCTNNNCACCCCATTCTCACCAACACATAC
AGCACACGTCTGAACTCTTTCCCTACACGACGCTCTTCCGATCTNNNCACCCCATTCTCACCAACACATAC
GCACACGTCTGAACTCTTTCCCTACACGACGCTCTTCCGATCTNNNAAACTGCTGGGATTACAGGCACA
GCACACGTCTGAACTCTTTCCCTACACGACGCTCTTCCGATCTNNNAAACTGCTGGGATTACAGGCACA
GAGCACACGTCTGAACTCTTTCCCTACACGACGCTCTTCCGATCTNNNGTGAGCTTCAGCAGGACCAAT
GAGCACACGTCTGAACTCTTTCCCTACACGACGCTCTTCCGATCTNNNGTGAGCTTCAGCAGGACCAAT
AGCACACGTCTGAACTCTTTCCCTACACGACGCTCTTCCGATCTNNNTGGTTGGGAGAGATTTTTTCCTT
AGCACACGTCTGAACTCTTTCCCTACACGACGCTCTTCCGATCTNNNTGGTTGGGAGAGATTTTTTCCTT
AGAGCACACGTCTGAACTCTTTCCCTACACGACGCTCTTCCGATCTNNNTGGTCTACTCTGGATACCTTC
AGAGCACACGTCTGAACTCTTTCCCTACACGACGCTCTTCCGATCTNNNTGGTCTACTCTGGATACCTTC
GAGCACACGTCTGAACTCTTTCCCTACACGACGCTCTTCCGATCTNNNCTGATCAATGAGGAGTCTGAC
GAGCACACGTCTGAACTCTTTCCCTACACGACGCTCTTCCGATCTNNNCTGATCAATGAGGAGTCTGAC
AGAGCACACGTCTGAACTCTTTCCCTACACGACGCTCTTCCGATCTNNNAACATCCTGTCTTGCTTGGCA
AGAGCACACGTCTGAACTCTTTCCCTACACGACGCTCTTCCGATCTNNNAACATCCTGTCTTGCTTGGCA
GAGCACACGTCTGAACTCTTTCCCTACACGACGCTCTTCCGATCTNNNCCTTCCATACATCCCTGTGGAA
GAGCACACGTCTGAACTCTTTCCCTACACGACGCTCTTCCGATCTNNNCCTTCCATACATCCCTGTGGAA
GAGCACACGTCTGAACTCTTTCCCTACACGACGCTCTTCCGATCTNNNACTTCGCACACACACACACATA
GAGCACACGTCTGAACTCTTTCCCTACACGACGCTCTTCCGATCTNNNACTTCGCACACACACACACATA
AGCACACGTCTGAACTCTTTCCCTACACGACGCTCTTCCGATCTNNNAATTGGGTTCCCGTGTGTATTTC
AGCACACGTCTGAACTCTTTCCCTACACGACGCTCTTCCGATCTNNNAATTGGGTTCCCGTGTGTATTTC
GAGCACACGTCTGAACTCTTTCCCTACACGACGCTCTTCCGATCTNNNCCACTGAAAACCCTGAGAACCA
GAGCACACGTCTGAACTCTTTCCCTACACGACGCTCTTCCGATCTNNNCCACTGAAAACCCTGAGAACCA
AAGAGCACACGTCTGAACTCTTTCCCTACACGACGCTCTTCCGATCTNNNCATTCCCTATGCTGCCTAATC
AAGAGCACACGTCTGAACTCTTTCCCTACACGACGCTCTTCCGATCTNNNCATTCCCTATGCTGCCTAATC
AGCACACGTCTGAACTCTTTCCCTACACGACGCTCTTCCGATCTNNNCTACCACTTTCTTTATGCCCCCTCA
AGCACACGTCTGAACTCTTTCCCTACACGACGCTCTTCCGATCTNNNCTACCACTTTCTTTATGCCCCCTCA

AGCACACGTCTGAACTCTTTCCCTACACGACGCTCTTCCGATCTNNNCTGAGATGGGAGGATCATTGA

AGCACACGTCTGAACTCTTTCCCTACACGACGCTCTTCCGATCTNNNCTGAGATGGGAGGATCATTGA

AGAGCACACGTCTGAACTCTTTCCCTACACGACGCTCTTCCGATCTNNNATGAGATGCTATGATGGAA

AGAGCACACGTCTGAACTCTTTCCCTACACGACGCTCTTCCGATCTNNNATGAGATGCTATGATGGAA

AGCACACGTCTGAACTCTTTCCCTACACGACGCTCTTCCGATCTNNNCTTCCAGGTTCATGCCATTCTCC

AGCACACGTCTGAACTCTTTCCCTACACGACGCTCTTCCGATCTNNNCTTCCAGGTTCATGCCATTCTCC

AGCACACGTCTGAACTCTTTCCCTACACGACGCTCTTCCGATCTNNNCCACAATGAGACTTCACCTTGTG

AGCACACGTCTGAACTCTTTCCCTACACGACGCTCTTCCGATCTNNNCCACAATGAGACTTCACCTTGTG

GAGCACACGTCTGAACTCTTTCCCTACACGACGCTCTTCCGATCTNNNAGACAACTCACAGAATGGGGA

GAGCACACGTCTGAACTCTTTCCCTACACGACGCTCTTCCGATCTNNNAGACAACTCACAGAATGGGGA

AGAGCACACGTCTGAACTCTTTCCCTACACGACGCTCTTCCGATCTNNNAGGCCCAAGAGGAACATTTA

AGAGCACACGTCTGAACTCTTTCCCTACACGACGCTCTTCCGATCTNNNAGGCCCAAGAGGAACATTTA

GAGCACACGTCTGAACTCTTTCCCTACACGACGCTCTTCCGATCTNNNTGAACTCCTGACCTCAAGTGAT

GAGCACACGTCTGAACTCTTTCCCTACACGACGCTCTTCCGATCTNNNTGAACTCCTGACCTCAAGTGAT

GAGCACACGTCTGAACTCTTTCCCTACACGACGCTCTTCCGATCTNNNTGTGGGGAATTCAGGAAAGA

GAGCACACGTCTGAACTCTTTCCCTACACGACGCTCTTCCGATCTNNNTGTGGGGAATTCAGGAAAGA

AGCACACGTCTGAACTCTTTCCCTACACGACGCTCTTCCGATCTNNNCTGAGGCAGGAGAATCACTTG

AGCACACGTCTGAACTCTTTCCCTACACGACGCTCTTCCGATCTNNNCTGAGGCAGGAGAATCACTTG

AGCACACGTCTGAACTCTTTCCCTACACGACGCTCTTCCGATCTNNNGACAAAGTGAGACCCTGACTCA

AGCACACGTCTGAACTCTTTCCCTACACGACGCTCTTCCGATCTNNNGACAAAGTGAGACCCTGACTCA

GAGCACACGTCTGAACTCTTTCCCTACACGACGCTCTTCCGATCTNNNTACACACACAGACAGACAGAC

GAGCACACGTCTGAACTCTTTCCCTACACGACGCTCTTCCGATCTNNNTACACACACAGACAGACAGAC

GAGCACACGTCTGAACTCTTTCCCTACACGACGCTCTTCCGATCTNNNACACAGACACACACTATCACT

GAGCACACGTCTGAACTCTTTCCCTACACGACGCTCTTCCGATCTNNNACACAGACACACACTATCACT

CACACGTCTGAACTCTTTCCCTACACGACGCTCTTCCGATCTNNNAGTACAGTCAGCATTATTTTACACT

CACACGTCTGAACTCTTTCCCTACACGACGCTCTTCCGATCTNNNAGTACAGTCAGCATTATTTTACACT

GAGCACACGTCTGAACTCTTTCCCTACACGACGCTCTTCCGATCTNNNAGAGGTTGCAGTGAGCTATGA

GAGCACACGTCTGAACTCTTTCCCTACACGACGCTCTTCCGATCTNNNAGAGGTTGCAGTGAGCTATGA

AGCACACGTCTGAACTCTTTCCCTACACGACGCTCTTCCGATCTNNNGGAGTTTGCAGTGAGCTGAGA

AGCACACGTCTGAACTCTTTCCCTACACGACGCTCTTCCGATCTNNNGGAGTTTGCAGTGAGCTGAGA

AGCACACGTCTGAACTCTTTCCCTACACGACGCTCTTCCGATCTNNNATTCACATCATCTTTGTAGGTAT

AGCACACGTCTGAACTCTTTCCCTACACGACGCTCTTCCGATCTNNNATTCACATCATCTTTGTAGGTAT

GAGCACACGTCTGAACTCTTTCCCTACACGACGCTCTTCCGATCTNNNCTGAGGCAGGAGAATTGCTT

GAGCACACGTCTGAACTCTTTCCCTACACGACGCTCTTCCGATCTNNNCTGAGGCAGGAGAATTGCTT

AGCACACGTCTGAACTCTTTCCCTACACGACGCTCTTCCGATCTNNNCCACTGATATTTGTAGCTGGGC

AGCACACGTCTGAACTCTTTCCCTACACGACGCTCTTCCGATCTNNNCCACTGATATTTGTAGCTGGGC

GAGCACACGTCTGAACTCTTTCCCTACACGACGCTCTTCCGATCTNNNCAGAATGGCCCACAAAAGAAA

GAGCACACGTCTGAACTCTTTCCCTACACGACGCTCTTCCGATCTNNNCAGAATGGCCCACAAAAGAAA

AGCACACGTCTGAACTCTTTCCCTACACGACGCTCTTCCGATCTNNNGAGAATGACATGAACCTGGGA

AGCACACGTCTGAACTCTTTCCCTACACGACGCTCTTCCGATCTNNNGAGAATGACATGAACCTGGGA

AGCACACGTCTGAACTCTTTCCCTACACGACGCTCTTCCGATCTNNNCTCCTGAGTTCCAGTGATTCTCC

AGCACACGTCTGAACTCTTTCCCTACACGACGCTCTTCCGATCTNNNCTCCTGAGTTCCAGTGATTCTCC

AGCACACGTCTGAACTCTTTCCCTACACGACGCTCTTCCGATCTNNNCTTCCTTCATCCCTCTTTCCCTCCA

AGCACACGTCTGAACTCTTTCCCTACACGACGCTCTTCCGATCTNNNCTTCCTTCATCCCTCTTTCCCTCC

AGCACACGTCTGAACTCTTTCCCTACACGACGCTCTTCCGATCTNNNGCACTTTGTTTTCACACCAATG

AGCACACGTCTGAACTCTTTCCCTACACGACGCTCTTCCGATCTNNNGCACTTTGTTTTCACACCAATG

GAGCACACGTCTGAACTCTTTCCCTACACGACGCTCTTCCGATCTNNNGAGCAGGACTCTGTCTCAAA

GAGCACACGTCTGAACTCTTTCCCTACACGACGCTCTTCCGATCTNNNGAGCAGGACTCTGTCTCAAA

AGCACACGTCTGAACTCTTTCCCTACACGACGCTCTTCCGATCTNNNAAGGCTGCTGTGAGTTATGAT

AGCACACGTCTGAACTCTTTCCCTACACGACGCTCTTCCGATCTNNNAAGGCTGCTGTGAGTTATGAT

AGCACACGTCTGAACTCTTTCCCTACACGACGCTCTTCCGATCTNNNTCACTTCCTCTCCTATGGTAGC

AGCACACGTCTGAACTCTTTCCCTACACGACGCTCTTCCGATCTNNNTCACTTCCTCTCCTATGGTAGC

GAGCACACGTCTGAACTCTTTCCCTACACGACGCTCTTCCGATCTNNNGCTTTTCATAGGGAAAGGCA

GAGCACACGTCTGAACTCTTTCCCTACACGACGCTCTTCCGATCTNNNGCTTTTCATAGGGAAAGGCA

AGCACACGTCTGAACTCTTTCCCTACACGACGCTCTTCCGATCTNNNGATAATCACTTGAGGCCAGG

AGCACACGTCTGAACTCTTTCCCTACACGACGCTCTTCCGATCTNNNGATAATCACTTGAGGCCAGG

AGCACACGTCTGAACTCTTTCCCTACACGACGCTCTTCCGATCTNNNCATACATTTCTGTGATGGTCT

AGCACACGTCTGAACTCTTTCCCTACACGACGCTCTTCCGATCTNNNCATACATTTCTGTGATGGTCT

CACACGTCTGAACTCTTTCCCTACACGACGCTCTTCCGATCTNNNGCTCTTATGCTGGTAAAAGACATC

CACACGTCTGAACTCTTTCCCTACACGACGCTCTTCCGATCTNNNGCTCTTATGCTGGTAAAAGACATC

AGCACACGTCTGAACTCTTTCCCTACACGACGCTCTTCCGATCTNNNAAAAATCCTTGCCCAGACCAA

AGCACACGTCTGAACTCTTTCCCTACACGACGCTCTTCCGATCTNNNAAAAATCCTTGCCCAGACCAA

AGCACACGTCTGAACTCTTTCCCTACACGACGCTCTTCCGATCTNNNGATGGTCTCAATCTCCTGACCT

AGCACACGTCTGAACTCTTTCCCTACACGACGCTCTTCCGATCTNNNGATGGTCTCAATCTCCTGACCT

ACACGTCTGAACTCTTTCCCTACACGACGCTCTTCCGATCTNNNCTTCAGAATGGTGTGGAACATGGC

ACACGTCTGAACTCTTTCCCTACACGACGCTCTTCCGATCTNNNCTTCAGAATGGTGTGGAACATGGC

AGCACACGTCTGAACTCTTTCCCTACACGACGCTCTTCCGATCTNNNCAATGAGTCACACACACATTC

AGCACACGTCTGAACTCTTTCCCTACACGACGCTCTTCCGATCTNNNCAATGAGTCACACACACATTC

GCACACGTCTGAACTCTTTCCCTACACGACGCTCTTCCGATCTNNNGAGAGCTTGTGCAGGGAAACT

GCACACGTCTGAACTCTTTCCCTACACGACGCTCTTCCGATCTNNNGAGAGCTTGTGCAGGGAAACT

AGCACACGTCTGAACTCTTTCCCTACACGACGCTCTTCCGATCTNNNTAGTCTCAACTACTCGGGAGG

AGCACACGTCTGAACTCTTTCCCTACACGACGCTCTTCCGATCTNNNTAGTCTCAACTACTCGGGAGG

GAGCACACGTCTGAACTCTTTCCCTACACGACGCTCTTCCGATCTNNNTGTAGTCCCAGCTACTCAGGA

GAGCACACGTCTGAACTCTTTCCCTACACGACGCTCTTCCGATCTNNNTGTAGTCCCAGCTACTCAGGA

GCACACGTCTGAACTCTTTCCCTACACGACGCTCTTCCGATCTNNNCAAGGCTCCATCTCAAACAAAC

GCACACGTCTGAACTCTTTCCCTACACGACGCTCTTCCGATCTNNNCAAGGCTCCATCTCAAACAAAC

GAGCACACGTCTGAACTCTTTCCCTACACGACGCTCTTCCGATCTNNNGCATGAGAATGGCTTGAAC

GAGCACACGTCTGAACTCTTTCCCTACACGACGCTCTTCCGATCTNNNGCATGAGAATGGCTTGAAC

AGAGCACACGTCTGAACTCTTTCCCTACACGACGCTCTTCCGATCTNNNGGAGAAGGAGCTAGCGTT

AGAGCACACGTCTGAACTCTTTCCCTACACGACGCTCTTCCGATCTNNNGGAGAAGGAGCTAGCGTT

GCACACGTCTGAACTCTTTCCCTACACGACGCTCTTCCGATCTNNNTCGTCCCTGGTAGATTACTCCTTC

GCACACGTCTGAACTCTTTCCCTACACGACGCTCTTCCGATCTNNNTCGTCCCTGGTAGATTACTCCTTC

GAGCACACGTCTGAACTCTTTCCCTACACGACGCTCTTCCGATCTNNNAAATTGTGACCTCTGCATTGT

GAGCACACGTCTGAACTCTTTCCCTACACGACGCTCTTCCGATCTNNNAAATTGTGACCTCTGCATTGT

AGCACACGTCTGAACTCTTTCCCTACACGACGCTCTTCCGATCTNNNGTAATCCCAGCTACTCAGGCAG

AGCACACGTCTGAACTCTTTCCCTACACGACGCTCTTCCGATCTNNNGTAATCCCAGCTACTCAGGCAG

GCACACGTCTGAACTCTTTCCCTACACGACGCTCTTCCGATCTNNNGCTGGAAAGGAAGGTGAAAG
GCACACGTCTGAACTCTTTCCCTACACGACGCTCTTCCGATCTNNNGCTGGAAAGGAAGGTGAAAG
AGCACACGTCTGAACTCTTTCCCTACACGACGCTCTTCCGATCTNNNAGTTTTTCTGGTGTCTTTGGGG
AGCACACGTCTGAACTCTTTCCCTACACGACGCTCTTCCGATCTNNNAGTTTTTCTGGTGTCTTTGGGG
AGCACACGTCTGAACTCTTTCCCTACACGACGCTCTTCCGATCTNNNACATTGTGATGACCAAAGCAAG
AGCACACGTCTGAACTCTTTCCCTACACGACGCTCTTCCGATCTNNNACATTGTGATGACCAAAGCAAG
GCACACGTCTGAACTCTTTCCCTACACGACGCTCTTCCGATCTNNNGATTTGAACCAGGTCAAGCTGA
GCACACGTCTGAACTCTTTCCCTACACGACGCTCTTCCGATCTNNNGATTTGAACCAGGTCAAGCTGA
AGCACACGTCTGAACTCTTTCCCTACACGACGCTCTTCCGATCTNNNTCATTTTTCCTGAACCCCATG
AGCACACGTCTGAACTCTTTCCCTACACGACGCTCTTCCGATCTNNNTCATTTTTCCTGAACCCCATG
GAGCACACGTCTGAACTCTTTCCCTACACGACGCTCTTCCGATCTNNNCATGGTAGAGTTTGAGGTC
GAGCACACGTCTGAACTCTTTCCCTACACGACGCTCTTCCGATCTNNNCATGGTAGAGTTTGAGGTC
AGCACACGTCTGAACTCTTTCCCTACACGACGCTCTTCCGATCTNNNGGAGAAAAGGGAAGGAACC
AGCACACGTCTGAACTCTTTCCCTACACGACGCTCTTCCGATCTNNNGGAGAAAAGGGAAGGAACC
GCACACGTCTGAACTCTTTCCCTACACGACGCTCTTCCGATCTNNNAAAAACTAAGCTTGTGTAAGAA
GCACACGTCTGAACTCTTTCCCTACACGACGCTCTTCCGATCTNNNAAAAACTAAGCTTGTGTAAGAA
GCACACGTCTGAACTCTTTCCCTACACGACGCTCTTCCGATCTNNNCGCATCAGGCTTTCTTATGTCTG
GCACACGTCTGAACTCTTTCCCTACACGACGCTCTTCCGATCTNNNCGCATCAGGCTTTCTTATGTCTG
AGCACACGTCTGAACTCTTTCCCTACACGACGCTCTTCCGATCTNNNCGAGGGGCAGATGTGTATCA
AGCACACGTCTGAACTCTTTCCCTACACGACGCTCTTCCGATCTNNNCGAGGGGCAGATGTGTATCA
GCACACGTCTGAACTCTTTCCCTACACGACGCTCTTCCGATCTNNNATGTAATTTAGGGCATGGGAGG
GCACACGTCTGAACTCTTTCCCTACACGACGCTCTTCCGATCTNNNATGTAATTTAGGGCATGGGAGG
GCACACGTCTGAACTCTTTCCCTACACGACGCTCTTCCGATCTNNNCATGGGAGGGTAAGGTAGCTT
GCACACGTCTGAACTCTTTCCCTACACGACGCTCTTCCGATCTNNNCATGGGAGGGTAAGGTAGCTT
AGCACACGTCTGAACTCTTTCCCTACACGACGCTCTTCCGATCTNNNTCAGTGGTAATCTCTCCTTTGC
AGCACACGTCTGAACTCTTTCCCTACACGACGCTCTTCCGATCTNNNTCAGTGGTAATCTCTCCTTTGC
AGCACACGTCTGAACTCTTTCCCTACACGACGCTCTTCCGATCTNNNCCTGACTTTCTCCTGCCTTACTT
AGCACACGTCTGAACTCTTTCCCTACACGACGCTCTTCCGATCTNNNCCTGACTTTCTCCTGCCTTACTT
GCACACGTCTGAACTCTTTCCCTACACGACGCTCTTCCGATCTNNNGAGGGGAATGTTGTGGAGGA
GCACACGTCTGAACTCTTTCCCTACACGACGCTCTTCCGATCTNNNGAGGGGAATGTTGTGGAGGA
GAGCACACGTCTGAACTCTTTCCCTACACGACGCTCTTCCGATCTNNNACTCATCTGTAGTCCCAGCTA
GAGCACACGTCTGAACTCTTTCCCTACACGACGCTCTTCCGATCTNNNACTCATCTGTAGTCCCAGCTA
CACACGTCTGAACTCTTTCCCTACACGACGCTCTTCCGATCTNNNCCCTTGAGTGTTCTCCTGAGATTCA
CACACGTCTGAACTCTTTCCCTACACGACGCTCTTCCGATCTNNNCCCTTGAGTGTTCTCCTGAGATTCA
AGCACACGTCTGAACTCTTTCCCTACACGACGCTCTTCCGATCTNNNATGGTGTGTAGATGTGTCTCT
AGCACACGTCTGAACTCTTTCCCTACACGACGCTCTTCCGATCTNNNATGGTGTGTAGATGTGTCTCT
AGCACACGTCTGAACTCTTTCCCTACACGACGCTCTTCCGATCTNNNTTGAAGAAAACACTGTGACCAG
AGCACACGTCTGAACTCTTTCCCTACACGACGCTCTTCCGATCTNNNTTGAAGAAAACACTGTGACCAG
AGCACACGTCTGAACTCTTTCCCTACACGACGCTCTTCCGATCTNNNTTGCTGTTTCCCAACACAAATG
AGCACACGTCTGAACTCTTTCCCTACACGACGCTCTTCCGATCTNNNTTGCTGTTTCCCAACACAAATG
GAGCACACGTCTGAACTCTTTCCCTACACGACGCTCTTCCGATCTNNNAGTGTTTGCATCAGTACCTCA
GAGCACACGTCTGAACTCTTTCCCTACACGACGCTCTTCCGATCTNNNAGTGTTTGCATCAGTACCTCA
GCACACGTCTGAACTCTTTCCCTACACGACGCTCTTCCGATCTNNNACCTGGATGACTGTTTCACACTT

GCACACGTCTGAACTCTTTCCCTACACGACGCTCTTCCGATCTNNNACCTGGATGACTGTTTCACACTT
GCACACGTCTGAACTCTTTCCCTACACGACGCTCTTCCGATCTNNNACACTGTCCTTTCTAAAGTGGC
GCACACGTCTGAACTCTTTCCCTACACGACGCTCTTCCGATCTNNNACACTGTCCTTTCTAAAGTGGC
GAGCACACGTCTGAACTCTTTCCCTACACGACGCTCTTCCGATCTNNNTGGACTTTTCTATGAGTTCTT
GAGCACACGTCTGAACTCTTTCCCTACACGACGCTCTTCCGATCTNNNTGGACTTTTCTATGAGTTCTT
AGCACACGTCTGAACTCTTTCCCTACACGACGCTCTTCCGATCTNNNGTGAAACCCCGTCTCTACTAAA
AGCACACGTCTGAACTCTTTCCCTACACGACGCTCTTCCGATCTNNNGTGAAACCCCGTCTCTACTAAA
GAGCACACGTCTGAACTCTTTCCCTACACGACGCTCTTCCGATCTNNNAAAGTACTGGGATTACAGGC
GAGCACACGTCTGAACTCTTTCCCTACACGACGCTCTTCCGATCTNNNAAAGTACTGGGATTACAGGC
AGCACACGTCTGAACTCTTTCCCTACACGACGCTCTTCCGATCTNNNAAACTCCTGACCTCAAGTGATC
AGCACACGTCTGAACTCTTTCCCTACACGACGCTCTTCCGATCTNNNAAACTCCTGACCTCAAGTGATC
AGCACACGTCTGAACTCTTTCCCTACACGACGCTCTTCCGATCTNNNCGGAGCTTGACACCTAGTAAA
AGCACACGTCTGAACTCTTTCCCTACACGACGCTCTTCCGATCTNNNCGGAGCTTGACACCTAGTAAA
AGCACACGTCTGAACTCTTTCCCTACACGACGCTCTTCCGATCTNNNATGAGCAGTTGTAGTCCTTGT
AGCACACGTCTGAACTCTTTCCCTACACGACGCTCTTCCGATCTNNNATGAGCAGTTGTAGTCCTTGT
CACGTCTGAACTCTTTCCCTACACGACGCTCTTCCGATCTNNNCCAAGCTCATTATGTCTTCAGCCCATC
CACGTCTGAACTCTTTCCCTACACGACGCTCTTCCGATCTNNNCCAAGCTCATTATGTCTTCAGCCCATC
GCACACGTCTGAACTCTTTCCCTACACGACGCTCTTCCGATCTNNNGTATTTTTAGTAGAGACAGGGT
GCACACGTCTGAACTCTTTCCCTACACGACGCTCTTCCGATCTNNNGTATTTTTAGTAGAGACAGGGT
AGCACACGTCTGAACTCTTTCCCTACACGACGCTCTTCCGATCTNNNCTGAGGCAGGAGAATTGCTT
AGCACACGTCTGAACTCTTTCCCTACACGACGCTCTTCCGATCTNNNCTGAGGCAGGAGAATTGCTT
GAGCACACGTCTGAACTCTTTCCCTACACGACGCTCTTCCGATCTNNNGTGTCACTTGTCCCCATAGA
GAGCACACGTCTGAACTCTTTCCCTACACGACGCTCTTCCGATCTNNNGTGTCACTTGTCCCCATAGA
AGCACACGTCTGAACTCTTTCCCTACACGACGCTCTTCCGATCTNNNTGCTCTAAACAAGGCCAAATGA
AGCACACGTCTGAACTCTTTCCCTACACGACGCTCTTCCGATCTNNNTGCTCTAAACAAGGCCAAATGA
GAGCACACGTCTGAACTCTTTCCCTACACGACGCTCTTCCGATCTNNNAAGGAGGTATACATCTTGCA
GAGCACACGTCTGAACTCTTTCCCTACACGACGCTCTTCCGATCTNNNAAGGAGGTATACATCTTGCA
AGCACACGTCTGAACTCTTTCCCTACACGACGCTCTTCCGATCTNNNTGAGGCAGGAGAATCTCTTGA
AGCACACGTCTGAACTCTTTCCCTACACGACGCTCTTCCGATCTNNNTGAGGCAGGAGAATCTCTTGA
AGCACACGTCTGAACTCTTTCCCTACACGACGCTCTTCCGATCTNNNTGAAGAAGCAGAAACTGTCC
AGCACACGTCTGAACTCTTTCCCTACACGACGCTCTTCCGATCTNNNTGAAGAAGCAGAAACTGTCC
AGCACACGTCTGAACTCTTTCCCTACACGACGCTCTTCCGATCTNNNAGGTTCAGAATGTACTCCTCA
AGCACACGTCTGAACTCTTTCCCTACACGACGCTCTTCCGATCTNNNAGGTTCAGAATGTACTCCTCA
AGCACACGTCTGAACTCTTTCCCTACACGACGCTCTTCCGATCTNNNGAGGAGTACAGAATGCAGGA
AGCACACGTCTGAACTCTTTCCCTACACGACGCTCTTCCGATCTNNNGAGGAGTACAGAATGCAGGA
CACGTCTGAACTCTTTCCCTACACGACGCTCTTCCGATCTNNNGGCAGACTAGAAAACCCTGACTTCA
CACGTCTGAACTCTTTCCCTACACGACGCTCTTCCGATCTNNNGGCAGACTAGAAAACCCTGACTTCA
GAGCACACGTCTGAACTCTTTCCCTACACGACGCTCTTCCGATCTNNNAGGCTCAAAATACAGGGAT
GAGCACACGTCTGAACTCTTTCCCTACACGACGCTCTTCCGATCTNNNAGGCTCAAAATACAGGGAT
AGCACACGTCTGAACTCTTTCCCTACACGACGCTCTTCCGATCTNNNAAAGCCACCACAATTACCAAAC
AGCACACGTCTGAACTCTTTCCCTACACGACGCTCTTCCGATCTNNNAAAGCCACCACAATTACCAAAC
AGAGCACACGTCTGAACTCTTTCCCTACACGACGCTCTTCCGATCTNNNAAAAAATTAGCTGGGCATG
AGAGCACACGTCTGAACTCTTTCCCTACACGACGCTCTTCCGATCTNNNAAAAAATTAGCTGGGCATG

AGCACACGTCTGAACTCTTTCCCTACACGACGCTCTTCCGATCTNNNATGCTTGACCTCCAAAGCATA

AGCACACGTCTGAACTCTTTCCCTACACGACGCTCTTCCGATCTNNNATGCTTGACCTCCAAAGCATA

GCACACGTCTGAACTCTTTCCCTACACGACGCTCTTCCGATCTNNNCACTCAGAGAAATTCAAGACTG

GCACACGTCTGAACTCTTTCCCTACACGACGCTCTTCCGATCTNNNCACTCAGAGAAATTCAAGACTG

AGCACACGTCTGAACTCTTTCCCTACACGACGCTCTTCCGATCTNNNACCTGCAGTCCCAGCTACTAGC

AGCACACGTCTGAACTCTTTCCCTACACGACGCTCTTCCGATCTNNNACCTGCAGTCCCAGCTACTAGC

GCACACGTCTGAACTCTTTCCCTACACGACGCTCTTCCGATCTNNNGTAGGAGGATCACTTGAACCTG

GCACACGTCTGAACTCTTTCCCTACACGACGCTCTTCCGATCTNNNGTAGGAGGATCACTTGAACCTG

GCACACGTCTGAACTCTTTCCCTACACGACGCTCTTCCGATCTNNNCCCACTCGCCCTAAATGATTAGT

GCACACGTCTGAACTCTTTCCCTACACGACGCTCTTCCGATCTNNNCCCACTCGCCCTAAATGATTAGT

CACACGTCTGAACTCTTTCCCTACACGACGCTCTTCCGATCTNNNGATCCTCTCCTACAGCCCTGATAC

CACACGTCTGAACTCTTTCCCTACACGACGCTCTTCCGATCTNNNGATCCTCTCCTACAGCCCTGATAC

GCACACGTCTGAACTCTTTCCCTACACGACGCTCTTCCGATCTNNNAACACTAAGGCAGGAAACAAG

GCACACGTCTGAACTCTTTCCCTACACGACGCTCTTCCGATCTNNNAACACTAAGGCAGGAAACAAG

AGCACACGTCTGAACTCTTTCCCTACACGACGCTCTTCCGATCTNNNCTGAGGCAGGAGAATCGTTT

AGCACACGTCTGAACTCTTTCCCTACACGACGCTCTTCCGATCTNNNCTGAGGCAGGAGAATCGTTT

AGCACACGTCTGAACTCTTTCCCTACACGACGCTCTTCCGATCTNNNCTCCTGCTCTCTACCATCCTAG

AGCACACGTCTGAACTCTTTCCCTACACGACGCTCTTCCGATCTNNNCTCCTGCTCTCTACCATCCTAG

AGCACACGTCTGAACTCTTTCCCTACACGACGCTCTTCCGATCTNNNTCTTGAGGGTTTTGAGTGAG

AGCACACGTCTGAACTCTTTCCCTACACGACGCTCTTCCGATCTNNNTCTTGAGGGTTTTGAGTGAG

AGCACACGTCTGAACTCTTTCCCTACACGACGCTCTTCCGATCTNNNAGGAGAATGGCATGAACCCGC

AGCACACGTCTGAACTCTTTCCCTACACGACGCTCTTCCGATCTNNNAGGAGAATGGCATGAACCCGC

CACACGTCTGAACTCTTTCCCTACACGACGCTCTTCCGATCTNNNAATGTTGTCCACTGCTACCCTAGC

CACACGTCTGAACTCTTTCCCTACACGACGCTCTTCCGATCTNNNAATGTTGTCCACTGCTACCCTAGC

GAGCACACGTCTGAACTCTTTCCCTACACGACGCTCTTCCGATCTNNNATGGTGACAAGCACTACACA

GAGCACACGTCTGAACTCTTTCCCTACACGACGCTCTTCCGATCTNNNATGGTGACAAGCACTACACA

GAGCACACGTCTGAACTCTTTCCCTACACGACGCTCTTCCGATCTNNNGAGACAGACCAAGACTCCAT

GAGCACACGTCTGAACTCTTTCCCTACACGACGCTCTTCCGATCTNNNGAGACAGACCAAGACTCCAT

AGCACACGTCTGAACTCTTTCCCTACACGACGCTCTTCCGATCTNNNAGATGGATGGTGAAGCAATG

AGCACACGTCTGAACTCTTTCCCTACACGACGCTCTTCCGATCTNNNAGATGGATGGTGAAGCAATG

AGCACACGTCTGAACTCTTTCCCTACACGACGCTCTTCCGATCTNNNGGGCAGTAGTTGTTCTCTGTA

AGCACACGTCTGAACTCTTTCCCTACACGACGCTCTTCCGATCTNNNGGGCAGTAGTTGTTCTCTGTA

AGCACACGTCTGAACTCTTTCCCTACACGACGCTCTTCCGATCTNNNCTGAGGTCAGGAGTTCGAGAC

AGCACACGTCTGAACTCTTTCCCTACACGACGCTCTTCCGATCTNNNCTGAGGTCAGGAGTTCGAGAC

GAGCACACGTCTGAACTCTTTCCCTACACGACGCTCTTCCGATCTNNNCACCTGTAATCCCAGCTACTCT

GAGCACACGTCTGAACTCTTTCCCTACACGACGCTCTTCCGATCTNNNCACCTGTAATCCCAGCTACTCT

AGCACACGTCTGAACTCTTTCCCTACACGACGCTCTTCCGATCTNNNTCCAAGACATCGAAAAGGAGA

AGCACACGTCTGAACTCTTTCCCTACACGACGCTCTTCCGATCTNNNTCCAAGACATCGAAAAGGAGA

GAGCACACGTCTGAACTCTTTCCCTACACGACGCTCTTCCGATCTNNNAGGAGAATTGCTTGAACCTG

GAGCACACGTCTGAACTCTTTCCCTACACGACGCTCTTCCGATCTNNNAGGAGAATTGCTTGAACCTG

GAGCACACGTCTGAACTCTTTCCCTACACGACGCTCTTCCGATCTNNNTGCTTTGTGTTTTTGGTTTAC

GAGCACACGTCTGAACTCTTTCCCTACACGACGCTCTTCCGATCTNNNTGCTTTGTGTTTTTGGTTTAC

GAGCACACGTCTGAACTCTTTCCCTACACGACGCTCTTCCGATCTNNNATTTCACTGTCACACAGCAG

AGAGCACACGTCTGAACTCTTTCCCTACACGACGCTCTTCCGATCTNNNATTTCACTGTCACACAGCAGC

GAGCACACGTCTGAACTCTTTCCCTACACGACGCTCTTCCGATCTNNNTTACTGTTTGAGGAACTGCCA

GAGCACACGTCTGAACTCTTTCCCTACACGACGCTCTTCCGATCTNNNTTACTGTTTGAGGAACTGCCA

AGCACACGTCTGAACTCTTTCCCTACACGACGCTCTTCCGATCTNNNTCATTTTTGTGGGCAGATGTTC

AGCACACGTCTGAACTCTTTCCCTACACGACGCTCTTCCGATCTNNNTCATTTTTGTGGGCAGATGTTC

GAGCACACGTCTGAACTCTTTCCCTACACGACGCTCTTCCGATCTNNNTGAGGCAGGAGAATCACTTGA

GAGCACACGTCTGAACTCTTTCCCTACACGACGCTCTTCCGATCTNNNTGAGGCAGGAGAATCACTTGA

AGCACACGTCTGAACTCTTTCCCTACACGACGCTCTTCCGATCTNNNTTAAAGCAAATACATTGTTCTG

AGCACACGTCTGAACTCTTTCCCTACACGACGCTCTTCCGATCTNNNTTAAAGCAAATACATTGTTCTG

GCACACGTCTGAACTCTTTCCCTACACGACGCTCTTCCGATCTNNNTTCCTGATGACTACCCTCTCCATCA

GCACACGTCTGAACTCTTTCCCTACACGACGCTCTTCCGATCTNNNTTCCTGATGACTACCCTCTCCATCA

AGAGCACACGTCTGAACTCTTTCCCTACACGACGCTCTTCCGATCTNNNGTGGAGAACGAGATCGCAG

AGAGCACACGTCTGAACTCTTTCCCTACACGACGCTCTTCCGATCTNNNGTGGAGAACGAGATCGCAG

GAGCACACGTCTGAACTCTTTCCCTACACGACGCTCTTCCGATCTNNNCCCGCCTTTATGGACCAAACTA

GAGCACACGTCTGAACTCTTTCCCTACACGACGCTCTTCCGATCTNNNCCCGCCTTTATGGACCAAACTA

CACACGTCTGAACTCTTTCCCTACACGACGCTCTTCCGATCTNNNTTTTGTGGGTTGATCTTGTCCCCCAT

CACACGTCTGAACTCTTTCCCTACACGACGCTCTTCCGATCTNNNTTTTGTGGGTTGATCTTGTCCCCCAT

CACACGTCTGAACTCTTTCCCTACACGACGCTCTTCCGATCTNNNGAAGGAGAAAACTGCCTTGGTTGC

CACACGTCTGAACTCTTTCCCTACACGACGCTCTTCCGATCTNNNGAAGGAGAAAACTGCCTTGGTTGC

AGCACACGTCTGAACTCTTTCCCTACACGACGCTCTTCCGATCTNNNGCTGAGACAGGAGAATCACTTG

AGCACACGTCTGAACTCTTTCCCTACACGACGCTCTTCCGATCTNNNGCTGAGACAGGAGAATCACTTG

AGAGCACACGTCTGAACTCTTTCCCTACACGACGCTCTTCCGATCTNNNGAGGCTCTGACCACTCTTGGA

AGAGCACACGTCTGAACTCTTTCCCTACACGACGCTCTTCCGATCTNNNGAGGCTCTGACCACTCTTGGA

AGAGCACACGTCTGAACTCTTTCCCTACACGACGCTCTTCCGATCTNNNCTAGGGTAACTGCTCTTCAAC

AGAGCACACGTCTGAACTCTTTCCCTACACGACGCTCTTCCGATCTNNNCTAGGGTAACTGCTCTTCAAC

GAGCACACGTCTGAACTCTTTCCCTACACGACGCTCTTCCGATCTNNNTAATTCAACTCCACTGTGGCCA

GAGCACACGTCTGAACTCTTTCCCTACACGACGCTCTTCCGATCTNNNTAATTCAACTCCACTGTGGCCA

GAGCACACGTCTGAACTCTTTCCCTACACGACGCTCTTCCGATCTNNNGAGAATCACTTGAACCTGCGAA

GAGCACACGTCTGAACTCTTTCCCTACACGACGCTCTTCCGATCTNNNGAGAATCACTTGAACCTGCGAA

AGCACACGTCTGAACTCTTTCCCTACACGACGCTCTTCCGATCTNNNAGGGAAATCACTTGAACATGGC

AGCACACGTCTGAACTCTTTCCCTACACGACGCTCTTCCGATCTNNNAGGGAAATCACTTGAACATGGC

GAGCACACGTCTGAACTCTTTCCCTACACGACGCTCTTCCGATCTNNNAGTAAAGCACAGTGTGAGTA

GAGCACACGTCTGAACTCTTTCCCTACACGACGCTCTTCCGATCTNNNAGTAAAGCACAGTGTGAGTA

GAGCACACGTCTGAACTCTTTCCCTACACGACGCTCTTCCGATCTNNNGGGATATGTAGACAAGGACA

GAGCACACGTCTGAACTCTTTCCCTACACGACGCTCTTCCGATCTNNNGGGATATGTAGACAAGGACA

AGCACACGTCTGAACTCTTTCCCTACACGACGCTCTTCCGATCTNNNATGCCTGTAGTCCCAGCTAATTG

AGCACACGTCTGAACTCTTTCCCTACACGACGCTCTTCCGATCTNNNATGCCTGTAGTCCCAGCTAATTG

AGCACACGTCTGAACTCTTTCCCTACACGACGCTCTTCCGATCTNNNAGTGCTTAGCTTGTTCAGTGAC

AGCACACGTCTGAACTCTTTCCCTACACGACGCTCTTCCGATCTNNNAGTGCTTAGCTTGTTCAGTGAC

CACACGTCTGAACTCTTTCCCTACACGACGCTCTTCCGATCTNNNTATACGTTTATACCTTCCGCGCTCAT

CACACGTCTGAACTCTTTCCCTACACGACGCTCTTCCGATCTNNNTATACGTTTATACCTTCCGCGCTCAT

AGCACACGTCTGAACTCTTTCCCTACACGACGCTCTTCCGATCTNNNTCCTGAGTAGCTGGGACTATAC

AGCACACGTCTGAACTCTTTCCCTACACGACGCTCTTCCGATCTNNNTCCTGAGTAGCTGGGACTATAC

AGCACACGTCTGAACTCTTTCCCTACACGACGCTCTTCCGATCTNNNTCATCATCTTGATCTTCAAATTC
AGCACACGTCTGAACTCTTTCCCTACACGACGCTCTTCCGATCTNNNTCATCATCTTGATCTTCAAATTC
AGCACACGTCTGAACTCTTTCCCTACACGACGCTCTTCCGATCTNNNCCGCGTTGGAGTAATTGACAAA
AGCACACGTCTGAACTCTTTCCCTACACGACGCTCTTCCGATCTNNNCCGCGTTGGAGTAATTGACAAA
AGCACACGTCTGAACTCTTTCCCTACACGACGCTCTTCCGATCTNNNTAATCACAGCTACTTGGGAAGG
AGCACACGTCTGAACTCTTTCCCTACACGACGCTCTTCCGATCTNNNTAATCACAGCTACTTGGGAAGG
AGCACACGTCTGAACTCTTTCCCTACACGACGCTCTTCCGATCTNNNACAATTTGGGGATAGAAGCCAC
AGCACACGTCTGAACTCTTTCCCTACACGACGCTCTTCCGATCTNNNACAATTTGGGGATAGAAGCCAC
CACACGTCTGAACTCTTTCCCTACACGACGCTCTTCCGATCTNNNCTGAAGAAATAAAGTTCTTTTTTTT
CACACGTCTGAACTCTTTCCCTACACGACGCTCTTCCGATCTNNNCTGAAGAAATAAAGTTCTTTTTTTT
AGCACACGTCTGAACTCTTTCCCTACACGACGCTCTTCCGATCTNNNCTCTTGGGAATACTCCCCTGGTA
AGCACACGTCTGAACTCTTTCCCTACACGACGCTCTTCCGATCTNNNCTCTTGGGAATACTCCCCTGGTA
GAGCACACGTCTGAACTCTTTCCCTACACGACGCTCTTCCGATCTNNNGAGATCATGCCACTGTACTCC
GAGCACACGTCTGAACTCTTTCCCTACACGACGCTCTTCCGATCTNNNGAGATCATGCCACTGTACTCC
GAGCACACGTCTGAACTCTTTCCCTACACGACGCTCTTCCGATCTNNNGCATCTGGGCTAAGAAATGA
GAGCACACGTCTGAACTCTTTCCCTACACGACGCTCTTCCGATCTNNNGCATCTGGGCTAAGAAATGA
AGCACACGTCTGAACTCTTTCCCTACACGACGCTCTTCCGATCTNNNATCAAGTGGGTTTCATCCTTGC
AGCACACGTCTGAACTCTTTCCCTACACGACGCTCTTCCGATCTNNNATCAAGTGGGTTTCATCCTTGC
AGCACACGTCTGAACTCTTTCCCTACACGACGCTCTTCCGATCTNNNAACCGACCTAACAGGATTCTAG
AGCACACGTCTGAACTCTTTCCCTACACGACGCTCTTCCGATCTNNNAACCGACCTAACAGGATTCTAG
GCACACGTCTGAACTCTTTCCCTACACGACGCTCTTCCGATCTNNNCCTGTAATCCCAGCTTCTCAAGTC
GCACACGTCTGAACTCTTTCCCTACACGACGCTCTTCCGATCTNNNCCTGTAATCCCAGCTTCTCAAGTC
GCACACGTCTGAACTCTTTCCCTACACGACGCTCTTCCGATCTNNNAAATGCCCTCCAGGTTTATCCAT
GCACACGTCTGAACTCTTTCCCTACACGACGCTCTTCCGATCTNNNAAATGCCCTCCAGGTTTATCCAT
GCACACGTCTGAACTCTTTCCCTACACGACGCTCTTCCGATCTNNNGCAGATTCCTTGGCCTTTCATTT
GCACACGTCTGAACTCTTTCCCTACACGACGCTCTTCCGATCTNNNGCAGATTCCTTGGCCTTTCATTT
GAGCACACGTCTGAACTCTTTCCCTACACGACGCTCTTCCGATCTNNNTCAATGGGCTCCAAAATGTCT
GAGCACACGTCTGAACTCTTTCCCTACACGACGCTCTTCCGATCTNNNTCAATGGGCTCCAAAATGTCT
GAGCACACGTCTGAACTCTTTCCCTACACGACGCTCTTCCGATCTNNNGCTAAGATAGTAGTGGTGGC
GAGCACACGTCTGAACTCTTTCCCTACACGACGCTCTTCCGATCTNNNGCTAAGATAGTAGTGGTGGC
AGCACACGTCTGAACTCTTTCCCTACACGACGCTCTTCCGATCTNNNGGTAATAGAGAGGTTGAGGC
AGCACACGTCTGAACTCTTTCCCTACACGACGCTCTTCCGATCTNNNGGTAATAGAGAGGTTGAGGC
AGCACACGTCTGAACTCTTTCCCTACACGACGCTCTTCCGATCTNNNCGAAGTAGAGCATCCTGGAGT
AGCACACGTCTGAACTCTTTCCCTACACGACGCTCTTCCGATCTNNNCGAAGTAGAGCATCCTGGAGT
AGCACACGTCTGAACTCTTTCCCTACACGACGCTCTTCCGATCTNNNGTCTCAAGCTCCTGACCAAGTA
AGCACACGTCTGAACTCTTTCCCTACACGACGCTCTTCCGATCTNNNGTCTCAAGCTCCTGACCAAGTA
CACACGTCTGAACTCTTTCCCTACACGACGCTCTTCCGATCTNNNTGTCCCAGATTTGGTCATTGGAAC
CACACGTCTGAACTCTTTCCCTACACGACGCTCTTCCGATCTNNNTGTCCCAGATTTGGTCATTGGAAC
GCACACGTCTGAACTCTTTCCCTACACGACGCTCTTCCGATCTNNNCAGCAACAACAGCAATTTGGTTC
GCACACGTCTGAACTCTTTCCCTACACGACGCTCTTCCGATCTNNNCAGCAACAACAGCAATTTGGTTC
AGCACACGTCTGAACTCTTTCCCTACACGACGCTCTTCCGATCTNNNAGTGATGGGATTACAGGTGTC
AGCACACGTCTGAACTCTTTCCCTACACGACGCTCTTCCGATCTNNNAGTGATGGGATTACAGGTGTC
GAGCACACGTCTGAACTCTTTCCCTACACGACGCTCTTCCGATCTNNNTTGTTTGTTTTTGAGACGGGG

GAGCACACGTCTGAACTCTTTCCCTACACGACGCTCTTCCGATCTNNNTTGTTTGTTTTTGAGACGGG
GAGCACACGTCTGAACTCTTTCCCTACACGACGCTCTTCCGATCTNNNTTAGTCCATGGCCATACTTTGC
GAGCACACGTCTGAACTCTTTCCCTACACGACGCTCTTCCGATCTNNNTTAGTCCATGGCCATACTTTGC
AGCACACGTCTGAACTCTTTCCCTACACGACGCTCTTCCGATCTNNNGAGCTTGCAGTGAACTGAGAT
AGCACACGTCTGAACTCTTTCCCTACACGACGCTCTTCCGATCTNNNGAGCTTGCAGTGAACTGAGAT
AGCACACGTCTGAACTCTTTCCCTACACGACGCTCTTCCGATCTNNNGGTCAAGGAGGAATTGAGTGA
AGCACACGTCTGAACTCTTTCCCTACACGACGCTCTTCCGATCTNNNGGTCAAGGAGGAATTGAGTGA
AGCACACGTCTGAACTCTTTCCCTACACGACGCTCTTCCGATCTNNNATCTTTGGATACCAGAACCCCA
AGCACACGTCTGAACTCTTTCCCTACACGACGCTCTTCCGATCTNNNATCTTTGGATACCAGAACCCCA
AGCACACGTCTGAACTCTTTCCCTACACGACGCTCTTCCGATCTNNNGCCTGTAGTCCCAGCTACTTGC
AGCACACGTCTGAACTCTTTCCCTACACGACGCTCTTCCGATCTNNNGCCTGTAGTCCCAGCTACTTGC
GAGCACACGTCTGAACTCTTTCCCTACACGACGCTCTTCCGATCTNNNCAAGTAGCTGGGACTATAGG
GAGCACACGTCTGAACTCTTTCCCTACACGACGCTCTTCCGATCTNNNCAAGTAGCTGGGACTATAGG
GAGCACACGTCTGAACTCTTTCCCTACACGACGCTCTTCCGATCTNNNTGTAGTCCCAGCTACTTGAAA
GAGCACACGTCTGAACTCTTTCCCTACACGACGCTCTTCCGATCTNNNTGTAGTCCCAGCTACTTGAAA
GAGCACACGTCTGAACTCTTTCCCTACACGACGCTCTTCCGATCTNNNATCTATTTTCTCAACTGCCATA
GAGCACACGTCTGAACTCTTTCCCTACACGACGCTCTTCCGATCTNNNATCTATTTTCTCAACTGCCATA
GAGCACACGTCTGAACTCTTTCCCTACACGACGCTCTTCCGATCTNNNTTTGAGTTAAGTGGAGCAAT
GAGCACACGTCTGAACTCTTTCCCTACACGACGCTCTTCCGATCTNNNTTTGAGTTAAGTGGAGCAAT
AGCACACGTCTGAACTCTTTCCCTACACGACGCTCTTCCGATCTNNNCATTCGCAGCACCAGAATAACA
AGCACACGTCTGAACTCTTTCCCTACACGACGCTCTTCCGATCTNNNCATTCGCAGCACCAGAATAACA
GAGCACACGTCTGAACTCTTTCCCTACACGACGCTCTTCCGATCTNNNGCAGGAGAATTGCTTGAACT
GAGCACACGTCTGAACTCTTTCCCTACACGACGCTCTTCCGATCTNNNGCAGGAGAATTGCTTGAACT
AGCACACGTCTGAACTCTTTCCCTACACGACGCTCTTCCGATCTNNNGTGTGTGTGTGTGTGTGTGTA
AGCACACGTCTGAACTCTTTCCCTACACGACGCTCTTCCGATCTNNNGTGTGTGTGTGTGTGTGTGTA
CACACGTCTGAACTCTTTCCCTACACGACGCTCTTCCGATCTNNNCTGCGCCTCTGTCCCTACTACATCG
CACACGTCTGAACTCTTTCCCTACACGACGCTCTTCCGATCTNNNCTGCGCCTCTGTCCCTACTACATCG
GAGCACACGTCTGAACTCTTTCCCTACACGACGCTCTTCCGATCTNNNTAGATTGAATGGGATGTTGG
GAGCACACGTCTGAACTCTTTCCCTACACGACGCTCTTCCGATCTNNNTAGATTGAATGGGATGTTGG
GAGCACACGTCTGAACTCTTTCCCTACACGACGCTCTTCCGATCTNNNAAGCACATCTTACCATGGAGG
GAGCACACGTCTGAACTCTTTCCCTACACGACGCTCTTCCGATCTNNNAAGCACATCTTACCATGGAGG
AGCACACGTCTGAACTCTTTCCCTACACGACGCTCTTCCGATCTNNNGGTAGGACTACTAGTGGTTTG
AGCACACGTCTGAACTCTTTCCCTACACGACGCTCTTCCGATCTNNNGGTAGGACTACTAGTGGTTTG
GCACACGTCTGAACTCTTTCCCTACACGACGCTCTTCCGATCTNNNTTTCCATCCAGCCATCAACAGTA
GCACACGTCTGAACTCTTTCCCTACACGACGCTCTTCCGATCTNNNTTTCCATCCAGCCATCAACAGTA
AGCACACGTCTGAACTCTTTCCCTACACGACGCTCTTCCGATCTNNNTTCAAATTAGGCACAGATCCA
AGCACACGTCTGAACTCTTTCCCTACACGACGCTCTTCCGATCTNNNTTCAAATTAGGCACAGATCCA
AGAGCACACGTCTGAACTCTTTCCCTACACGACGCTCTTCCGATCTNNNTAAAGGGAGATTGGCAGAC
AGAGCACACGTCTGAACTCTTTCCCTACACGACGCTCTTCCGATCTNNNTAAAGGGAGATTGGCAGAC
GAGCACACGTCTGAACTCTTTCCCTACACGACGCTCTTCCGATCTNNNAAAGCCACTCTCATAATGAG
GAGCACACGTCTGAACTCTTTCCCTACACGACGCTCTTCCGATCTNNNAAAGCCACTCTCATAATGAG
GAGCACACGTCTGAACTCTTTCCCTACACGACGCTCTTCCGATCTNNNGGTTGATCCACTTTGTTGAGT
GAGCACACGTCTGAACTCTTTCCCTACACGACGCTCTTCCGATCTNNNGGTTGATCCACTTTGTTGAGT

AGCACACGTCTGAACTCTTTCCCTACACGACGCTCTTCCGATCTNNNGTGAAGCAGTGTGAAAACTCGA

AGCACACGTCTGAACTCTTTCCCTACACGACGCTCTTCCGATCTNNNGTGAAGCAGTGTGAAAACTCGA

GAGCACACGTCTGAACTCTTTCCCTACACGACGCTCTTCCGATCTNNNTTCATTCCATTCTCCTGCCTCAG

GAGCACACGTCTGAACTCTTTCCCTACACGACGCTCTTCCGATCTNNNTTCATTCCATTCTCCTGCCTCAG

AGCACACGTCTGAACTCTTTCCCTACACGACGCTCTTCCGATCTNNNCTGTAGTCCCAGCTACTCAGGA

AGCACACGTCTGAACTCTTTCCCTACACGACGCTCTTCCGATCTNNNCTGTAGTCCCAGCTACTCAGGA

AGCACACGTCTGAACTCTTTCCCTACACGACGCTCTTCCGATCTNNNCTGTAATCCCAGCTACTCAGGA

AGCACACGTCTGAACTCTTTCCCTACACGACGCTCTTCCGATCTNNNCTGTAATCCCAGCTACTCAGGA

AGAGCACACGTCTGAACTCTTTCCCTACACGACGCTCTTCCGATCTNNNCAACAGATGTTTCCAAACTGG

AGAGCACACGTCTGAACTCTTTCCCTACACGACGCTCTTCCGATCTNNNCAACAGATGTTTCCAAACTGG

GCACACGTCTGAACTCTTTCCCTACACGACGCTCTTCCGATCTNNNGTGATAATGTGCCGCGAATTGTA

GCACACGTCTGAACTCTTTCCCTACACGACGCTCTTCCGATCTNNNGTGATAATGTGCCGCGAATTGTA

GAGCACACGTCTGAACTCTTTCCCTACACGACGCTCTTCCGATCTNNNGAGGCTGAGGCAAGAGAATA

GAGCACACGTCTGAACTCTTTCCCTACACGACGCTCTTCCGATCTNNNGAGGCTGAGGCAAGAGAATA

AGCACACGTCTGAACTCTTTCCCTACACGACGCTCTTCCGATCTNNNTTATGAAGCCAGCATCATCCTGA

AGCACACGTCTGAACTCTTTCCCTACACGACGCTCTTCCGATCTNNNTTATGAAGCCAGCATCATCCTGA

GCACACGTCTGAACTCTTTCCCTACACGACGCTCTTCCGATCTNNNAATCTGTAGGACTTGCCAACAGT

GCACACGTCTGAACTCTTTCCCTACACGACGCTCTTCCGATCTNNNAATCTGTAGGACTTGCCAACAGT

AGCACACGTCTGAACTCTTTCCCTACACGACGCTCTTCCGATCTNNNTTTAGTCACAGCCGGTTCCTTC

AGCACACGTCTGAACTCTTTCCCTACACGACGCTCTTCCGATCTNNNTTTAGTCACAGCCGGTTCCTTC

AGCACACGTCTGAACTCTTTCCCTACACGACGCTCTTCCGATCTNNNAGATTGCAGTGAGCTGAGATT

AGCACACGTCTGAACTCTTTCCCTACACGACGCTCTTCCGATCTNNNAGATTGCAGTGAGCTGAGATT

AGAGCACACGTCTGAACTCTTTCCCTACACGACGCTCTTCCGATCTNNNGCAAGGAGGTAAAGAGGGA

AGAGCACACGTCTGAACTCTTTCCCTACACGACGCTCTTCCGATCTNNNGCAAGGAGGTAAAGAGGGA

GCACACGTCTGAACTCTTTCCCTACACGACGCTCTTCCGATCTNNNTCCCTTTCTCCACATCTCTACCACA

GCACACGTCTGAACTCTTTCCCTACACGACGCTCTTCCGATCTNNNTCCCTTTCTCCACATCTCTACCACA

GCACACGTCTGAACTCTTTCCCTACACGACGCTCTTCCGATCTNNNGATGATGCAGGAGGATTGCTTG

GCACACGTCTGAACTCTTTCCCTACACGACGCTCTTCCGATCTNNNGATGATGCAGGAGGATTGCTTG

AGAGCACACGTCTGAACTCTTTCCCTACACGACGCTCTTCCGATCTNNNCCCTTCAGTGGAACAAGCTAT

AGAGCACACGTCTGAACTCTTTCCCTACACGACGCTCTTCCGATCTNNNCCCTTCAGTGGAACAAGCTAT

ACACGTCTGAACTCTTTCCCTACACGACGCTCTTCCGATCTNNNTCCCAGCCTAATGTACCACATTTCAT

ACACGTCTGAACTCTTTCCCTACACGACGCTCTTCCGATCTNNNTCCCAGCCTAATGTACCACATTTCAT

GCACACGTCTGAACTCTTTCCCTACACGACGCTCTTCCGATCTNNNGGAGTTTGCAGTGAACAGACAT

GCACACGTCTGAACTCTTTCCCTACACGACGCTCTTCCGATCTNNNGGAGTTTGCAGTGAACAGACAT

AGAGCACACGTCTGAACTCTTTCCCTACACGACGCTCTTCCGATCTNNNCTAGCAACCTGGGCTGGAGA

AGAGCACACGTCTGAACTCTTTCCCTACACGACGCTCTTCCGATCTNNNCTAGCAACCTGGGCTGGAGA

AGCACACGTCTGAACTCTTTCCCTACACGACGCTCTTCCGATCTNNNGTCAACATGACAAAACCCCATCT

AGCACACGTCTGAACTCTTTCCCTACACGACGCTCTTCCGATCTNNNGTCAACATGACAAAACCCCATCT

AGCACACGTCTGAACTCTTTCCCTACACGACGCTCTTCCGATCTNNNGATCCCCACAGTTCTTCATTGCT

AGCACACGTCTGAACTCTTTCCCTACACGACGCTCTTCCGATCTNNNGATCCCCACAGTTCTTCATTGCT

AGCACACGTCTGAACTCTTTCCCTACACGACGCTCTTCCGATCTNNNATCCCCTTTGCACCTTTACTCAT

AGCACACGTCTGAACTCTTTCCCTACACGACGCTCTTCCGATCTNNNATCCCCTTTGCACCTTTACTCAT

AGCACACGTCTGAACTCTTTCCCTACACGACGCTCTTCCGATCTNNNTGCCCAAGGTCAAGTTTAGATCA

AGCACACGTCTGAACTCTTTCCCTACACGACGCTCTTCCGATCTNNNTGCCCAAGGTCAAGTTTAGATCA
GAGCACACGTCTGAACTCTTTCCCTACACGACGCTCTTCCGATCTNNNCTGTAGTCCCAGCTACTCAGGA
GAGCACACGTCTGAACTCTTTCCCTACACGACGCTCTTCCGATCTNNNCTGTAGTCCCAGCTACTCAGGA
AGCACACGTCTGAACTCTTTCCCTACACGACGCTCTTCCGATCTNNNCAGTCCACATGCTAACGTTGATG
AGCACACGTCTGAACTCTTTCCCTACACGACGCTCTTCCGATCTNNNCAGTCCACATGCTAACGTTGATG
GAGCACACGTCTGAACTCTTTCCCTACACGACGCTCTTCCGATCTNNNCATACTGGAAGGATGTGGGA
GAGCACACGTCTGAACTCTTTCCCTACACGACGCTCTTCCGATCTNNNCATACTGGAAGGATGTGGGA
GAGCACACGTCTGAACTCTTTCCCTACACGACGCTCTTCCGATCTNNNTAAATGGGAGGAAGGAGTGC
GAGCACACGTCTGAACTCTTTCCCTACACGACGCTCTTCCGATCTNNNTAAATGGGAGGAAGGAGTGC
GAGCACACGTCTGAACTCTTTCCCTACACGACGCTCTTCCGATCTNNNCGCCTGTAATCCCAAGCTATTT
GAGCACACGTCTGAACTCTTTCCCTACACGACGCTCTTCCGATCTNNNCGCCTGTAATCCCAAGCTATTT
GAGCACACGTCTGAACTCTTTCCCTACACGACGCTCTTCCGATCTNNNTGGCATGCTCAGAAAGTTTTTG
GAGCACACGTCTGAACTCTTTCCCTACACGACGCTCTTCCGATCTNNNTGGCATGCTCAGAAAGTTTTTG
GAGCACACGTCTGAACTCTTTCCCTACACGACGCTCTTCCGATCTNNNTACACAAGTCTGCTATACGCCA
GAGCACACGTCTGAACTCTTTCCCTACACGACGCTCTTCCGATCTNNNTACACAAGTCTGCTATACGCCA
GAGCACACGTCTGAACTCTTTCCCTACACGACGCTCTTCCGATCTNNNACCATGGAATACTATGCAGCC
GAGCACACGTCTGAACTCTTTCCCTACACGACGCTCTTCCGATCTNNNACCATGGAATACTATGCAGCC
GAGCACACGTCTGAACTCTTTCCCTACACGACGCTCTTCCGATCTNNNGCACTGGAGTCTGAGAAACC
GAGCACACGTCTGAACTCTTTCCCTACACGACGCTCTTCCGATCTNNNGCACTGGAGTCTGAGAAACC
AGCACACGTCTGAACTCTTTCCCTACACGACGCTCTTCCGATCTNNNAGAAATGCAGCTGATTTTTGTG
AGCACACGTCTGAACTCTTTCCCTACACGACGCTCTTCCGATCTNNNAGAAATGCAGCTGATTTTTGTG
GAGCACACGTCTGAACTCTTTCCCTACACGACGCTCTTCCGATCTNNNCAGAAGAATTGCTTGAACCCA
GAGCACACGTCTGAACTCTTTCCCTACACGACGCTCTTCCGATCTNNNCAGAAGAATTGCTTGAACCCA
GCACACGTCTGAACTCTTTCCCTACACGACGCTCTTCCGATCTNNNTGTAGTCCCAGCTACTCAGGAGC
GCACACGTCTGAACTCTTTCCCTACACGACGCTCTTCCGATCTNNNTGTAGTCCCAGCTACTCAGGAGC
AAGAGCACACGTCTGAACTCTTTCCCTACACGACGCTCTTCCGATCTNNNCATTTCAGAGGTTGGCTTTG
AAGAGCACACGTCTGAACTCTTTCCCTACACGACGCTCTTCCGATCTNNNCATTTCAGAGGTTGGCTTTG
GAGCACACGTCTGAACTCTTTCCCTACACGACGCTCTTCCGATCTNNNTGGACTGTTCTACCAGATCCTC
GAGCACACGTCTGAACTCTTTCCCTACACGACGCTCTTCCGATCTNNNTGGACTGTTCTACCAGATCCTC
GAGCACACGTCTGAACTCTTTCCCTACACGACGCTCTTCCGATCTNNNCAGGAAAATGGCATGAACCCA
GAGCACACGTCTGAACTCTTTCCCTACACGACGCTCTTCCGATCTNNNCAGGAAAATGGCATGAACCCA
AGCACACGTCTGAACTCTTTCCCTACACGACGCTCTTCCGATCTNNNCCTGGCATCTTTTTGCTGTATTG
AGCACACGTCTGAACTCTTTCCCTACACGACGCTCTTCCGATCTNNNCCTGGCATCTTTTTGCTGTATTG
AGCACACGTCTGAACTCTTTCCCTACACGACGCTCTTCCGATCTNNNAGTCATATTTCTCTTTCTCTGCCT
AGCACACGTCTGAACTCTTTCCCTACACGACGCTCTTCCGATCTNNNAGTCATATTTCTCTTTCTCTGCCT
GCACACGTCTGAACTCTTTCCCTACACGACGCTCTTCCGATCTNNNCATCCTTGTCTTGTGCTGGTTTTCA
GCACACGTCTGAACTCTTTCCCTACACGACGCTCTTCCGATCTNNNCATCCTTGTCTTGTGCTGGTTTTCA
GAGCACACGTCTGAACTCTTTCCCTACACGACGCTCTTCCGATCTNNNTCACTTGAACCTTGGAGGAGA
GAGCACACGTCTGAACTCTTTCCCTACACGACGCTCTTCCGATCTNNNTCACTTGAACCTTGGAGGAGA
AGCACACGTCTGAACTCTTTCCCTACACGACGCTCTTCCGATCTNNNACACACTTGTAATCCCAGCTACT
AGCACACGTCTGAACTCTTTCCCTACACGACGCTCTTCCGATCTNNNACACACTTGTAATCCCAGCTACT
GAGCACACGTCTGAACTCTTTCCCTACACGACGCTCTTCCGATCTNNNACAAGAACGAAACTCCATCTC
GAGCACACGTCTGAACTCTTTCCCTACACGACGCTCTTCCGATCTNNNACAAGAACGAAACTCCATCTC

GAGCACACGTCTGAACTCTTTCCCTACACGACGCTCTTCCGATCTNNNACAACAAAAGACCCAGTGCAT
GAGCACACGTCTGAACTCTTTCCCTACACGACGCTCTTCCGATCTNNNACAACAAAAGACCCAGTGCAT
AGCACACGTCTGAACTCTTTCCCTACACGACGCTCTTCCGATCTNNNACCTACTCATTTATCACCTTTGCA
AGCACACGTCTGAACTCTTTCCCTACACGACGCTCTTCCGATCTNNNACCTACTCATTTATCACCTTTGCA
GAGCACACGTCTGAACTCTTTCCCTACACGACGCTCTTCCGATCTNNNATTCACAAGGATGGGATGTCA
GAGCACACGTCTGAACTCTTTCCCTACACGACGCTCTTCCGATCTNNNATTCACAAGGATGGGATGTCA
GCACACGTCTGAACTCTTTCCCTACACGACGCTCTTCCGATCTNNNTTCCTCACAGAAAATTCCCTCGTCA
GCACACGTCTGAACTCTTTCCCTACACGACGCTCTTCCGATCTNNNTTCCTCACAGAAAATTCCCTCGTCA
AGAGCACACGTCTGAACTCTTTCCCTACACGACGCTCTTCCGATCTNNNGAGTGGTTTGCTCATTGTCAT
AGAGCACACGTCTGAACTCTTTCCCTACACGACGCTCTTCCGATCTNNNGAGTGGTTTGCTCATTGTCAT
GAGCACACGTCTGAACTCTTTCCCTACACGACGCTCTTCCGATCTNNNTGCCCTATTTCTTACACAGCACT
GAGCACACGTCTGAACTCTTTCCCTACACGACGCTCTTCCGATCTNNNTGCCCTATTTCTTACACAGCACT
AGCACACGTCTGAACTCTTTCCCTACACGACGCTCTTCCGATCTNNNCGTGCAGAATTTCATGTGGACA
AGCACACGTCTGAACTCTTTCCCTACACGACGCTCTTCCGATCTNNNCGTGCAGAATTTCATGTGGACA
GCACACGTCTGAACTCTTTCCCTACACGACGCTCTTCCGATCTNNNATCTGTCCCACCAAACATTAGCTC
GCACACGTCTGAACTCTTTCCCTACACGACGCTCTTCCGATCTNNNATCTGTCCCACCAAACATTAGCTC
AGCACACGTCTGAACTCTTTCCCTACACGACGCTCTTCCGATCTNNNAGGATTTTGCAGAGCATAGGGA
AGCACACGTCTGAACTCTTTCCCTACACGACGCTCTTCCGATCTNNNAGGATTTTGCAGAGCATAGGGA
AGCACACGTCTGAACTCTTTCCCTACACGACGCTCTTCCGATCTNNNAAGGAAGTGTTCAAGGTGGTT
AGCACACGTCTGAACTCTTTCCCTACACGACGCTCTTCCGATCTNNNAAGGAAGTGTTCAAGGTGGTT
AGCACACGTCTGAACTCTTTCCCTACACGACGCTCTTCCGATCTNNNAGTTTCGGTTTTCTGGGATAGG
AGCACACGTCTGAACTCTTTCCCTACACGACGCTCTTCCGATCTNNNAGTTTCGGTTTTCTGGGATAGG
AGCACACGTCTGAACTCTTTCCCTACACGACGCTCTTCCGATCTNNNGGAGACAAGAGTGAGACTCCA
AGCACACGTCTGAACTCTTTCCCTACACGACGCTCTTCCGATCTNNNGGAGACAAGAGTGAGACTCCA
AGAGCACACGTCTGAACTCTTTCCCTACACGACGCTCTTCCGATCTNNNGTTACAGTGAGCTGAGATTG
AGAGCACACGTCTGAACTCTTTCCCTACACGACGCTCTTCCGATCTNNNGTTACAGTGAGCTGAGATTG
AGCACACGTCTGAACTCTTTCCCTACACGACGCTCTTCCGATCTNNNATAACGCCTGAAAGCACGTACTA
AGCACACGTCTGAACTCTTTCCCTACACGACGCTCTTCCGATCTNNNATAACGCCTGAAAGCACGTACTA
AGCACACGTCTGAACTCTTTCCCTACACGACGCTCTTCCGATCTNNNTCTGGGTATGCCTCTGATCATTG
AGCACACGTCTGAACTCTTTCCCTACACGACGCTCTTCCGATCTNNNTCTGGGTATGCCTCTGATCATTG
AGCACACGTCTGAACTCTTTCCCTACACGACGCTCTTCCGATCTNNNCATTTATCTCTACCTGGCCCCTCC
AGCACACGTCTGAACTCTTTCCCTACACGACGCTCTTCCGATCTNNNCATTTATCTCTACCTGGCCCCTCC
AGAGCACACGTCTGAACTCTTTCCCTACACGACGCTCTTCCGATCTNNNCATAAAACTGCCTCCAATGCA
AGAGCACACGTCTGAACTCTTTCCCTACACGACGCTCTTCCGATCTNNNCATAAAACTGCCTCCAATGCA
GAGCACACGTCTGAACTCTTTCCCTACACGACGCTCTTCCGATCTNNNAACATACAGCTGGCTGAGAAG
GAGCACACGTCTGAACTCTTTCCCTACACGACGCTCTTCCGATCTNNNAACATACAGCTGGCTGAGAAG
GAGCACACGTCTGAACTCTTTCCCTACACGACGCTCTTCCGATCTNNNTATGGCAGACTCTGTTCATTG
GAGCACACGTCTGAACTCTTTCCCTACACGACGCTCTTCCGATCTNNNTATGGCAGACTCTGTTCATTG
AGCACACGTCTGAACTCTTTCCCTACACGACGCTCTTCCGATCTNNNAAGTGCTGGGATTATAAGCGTG
AGCACACGTCTGAACTCTTTCCCTACACGACGCTCTTCCGATCTNNNAAGTGCTGGGATTATAAGCGTG
AGCACACGTCTGAACTCTTTCCCTACACGACGCTCTTCCGATCTNNNTTGAACTCTGGACCTCAGGTGAT
AGCACACGTCTGAACTCTTTCCCTACACGACGCTCTTCCGATCTNNNTTGAACTCTGGACCTCAGGTGAT
AGCACACGTCTGAACTCTTTCCCTACACGACGCTCTTCCGATCTNNNTGTTTCCCAAGAACTCACATTGC

GCACACGTCTGAACTCTTTCCCTACACGACGCTCTTCCGATCTNNNTGTTTCCCAAGAACTCACATTGC
AGCACACGTCTGAACTCTTTCCCTACACGACGCTCTTCCGATCTNNNCAGCAGAGAGCCTGAGGAAG
AGCACACGTCTGAACTCTTTCCCTACACGACGCTCTTCCGATCTNNNCAGCAGAGAGCCTGAGGAAG
GAGCACACGTCTGAACTCTTTCCCTACACGACGCTCTTCCGATCTNNNCTTGGGTGACAGAGCAAGA
GAGCACACGTCTGAACTCTTTCCCTACACGACGCTCTTCCGATCTNNNCTTGGGTGACAGAGCAAGA
GCACACGTCTGAACTCTTTCCCTACACGACGCTCTTCCGATCTNNNACCCTGACCTCGAGAATGAATT
GCACACGTCTGAACTCTTTCCCTACACGACGCTCTTCCGATCTNNNACCCTGACCTCGAGAATGAATT
GAGCACACGTCTGAACTCTTTCCCTACACGACGCTCTTCCGATCTNNNATGAACACTGTGAGATAGGT
GAGCACACGTCTGAACTCTTTCCCTACACGACGCTCTTCCGATCTNNNATGAACACTGTGAGATAGGT
GCACACGTCTGAACTCTTTCCCTACACGACGCTCTTCCGATCTNNNGTCACTGGAGAAGCTGAAAAT
GCACACGTCTGAACTCTTTCCCTACACGACGCTCTTCCGATCTNNNGTCACTGGAGAAGCTGAAAAT
GAGCACACGTCTGAACTCTTTCCCTACACGACGCTCTTCCGATCTNNNCTGAGATGAGGCCACTGTAC
GAGCACACGTCTGAACTCTTTCCCTACACGACGCTCTTCCGATCTNNNCTGAGATGAGGCCACTGTAC
GCACACGTCTGAACTCTTTCCCTACACGACGCTCTTCCGATCTNNNTAGTGTCTTTTGATGCACATGG
GCACACGTCTGAACTCTTTCCCTACACGACGCTCTTCCGATCTNNNTAGTGTCTTTTGATGCACATGG
AGCACACGTCTGAACTCTTTCCCTACACGACGCTCTTCCGATCTNNNTTTCATATTCCTGAGGCAGTG
AGCACACGTCTGAACTCTTTCCCTACACGACGCTCTTCCGATCTNNNTTTCATATTCCTGAGGCAGTG
GCACACGTCTGAACTCTTTCCCTACACGACGCTCTTCCGATCTNNNAGGTCACAGCTTCTCATCACAAT
GCACACGTCTGAACTCTTTCCCTACACGACGCTCTTCCGATCTNNNAGGTCACAGCTTCTCATCACAAT
AGCACACGTCTGAACTCTTTCCCTACACGACGCTCTTCCGATCTNNNGCTGTTACAAACTCTGCCTGAA
AGCACACGTCTGAACTCTTTCCCTACACGACGCTCTTCCGATCTNNNGCTGTTACAAACTCTGCCTGAA
AGCACACGTCTGAACTCTTTCCCTACACGACGCTCTTCCGATCTNNNAGACATTGTTCTGTGATCTGC
AGCACACGTCTGAACTCTTTCCCTACACGACGCTCTTCCGATCTNNNAGACATTGTTCTGTGATCTGC
AGCACACGTCTGAACTCTTTCCCTACACGACGCTCTTCCGATCTNNNCAGAGTGAGACCCTGCCTAAA
AGCACACGTCTGAACTCTTTCCCTACACGACGCTCTTCCGATCTNNNCAGAGTGAGACCCTGCCTAAA
GCACACGTCTGAACTCTTTCCCTACACGACGCTCTTCCGATCTNNNTCCCAAGAACCTAGCAAAAGAA
GCACACGTCTGAACTCTTTCCCTACACGACGCTCTTCCGATCTNNNTCCCAAGAACCTAGCAAAAGAA
AGCACACGTCTGAACTCTTTCCCTACACGACGCTCTTCCGATCTNNNCATCCTGGCTAACATGTGAAA
AGCACACGTCTGAACTCTTTCCCTACACGACGCTCTTCCGATCTNNNCATCCTGGCTAACATGTGAAA
GCACACGTCTGAACTCTTTCCCTACACGACGCTCTTCCGATCTNNNCTAGTCCCAAACTCCTGACCTTG
GCACACGTCTGAACTCTTTCCCTACACGACGCTCTTCCGATCTNNNCTAGTCCCAAACTCCTGACCTTG
GAGCACACGTCTGAACTCTTTCCCTACACGACGCTCTTCCGATCTNNNTGAGTGACAGAGTGAGACCC
GAGCACACGTCTGAACTCTTTCCCTACACGACGCTCTTCCGATCTNNNTGAGTGACAGAGTGAGACCC
GCACACGTCTGAACTCTTTCCCTACACGACGCTCTTCCGATCTNNNTGATCAGTCTGGTCTTGAACTCC
GCACACGTCTGAACTCTTTCCCTACACGACGCTCTTCCGATCTNNNTGATCAGTCTGGTCTTGAACTCC
AGCACACGTCTGAACTCTTTCCCTACACGACGCTCTTCCGATCTNNNGATGGTTGACGTCTTCTAAGG
AGCACACGTCTGAACTCTTTCCCTACACGACGCTCTTCCGATCTNNNGATGGTTGACGTCTTCTAAGG
GCACACGTCTGAACTCTTTCCCTACACGACGCTCTTCCGATCTNNNCGCCCTCACTCCTCTGTCCTTCAT
GCACACGTCTGAACTCTTTCCCTACACGACGCTCTTCCGATCTNNNCGCCCTCACTCCTCTGTCCTTCAT
GAGCACACGTCTGAACTCTTTCCCTACACGACGCTCTTCCGATCTNNNGAGAAGAGAGCCCAAGAGA
GAGCACACGTCTGAACTCTTTCCCTACACGACGCTCTTCCGATCTNNNGAGAAGAGAGCCCAAGAGA
GCACACGTCTGAACTCTTTCCCTACACGACGCTCTTCCGATCTNNNGTAGCTCAGTCCTGTAATCCCAG
GCACACGTCTGAACTCTTTCCCTACACGACGCTCTTCCGATCTNNNGTAGCTCAGTCCTGTAATCCCAG

GAGCACACGTCTGAACTCTTTCCCTACACGACGCTCTTCCGATCTNNNTCCATCAGTTGTAGCAAACAT

GAGCACACGTCTGAACTCTTTCCCTACACGACGCTCTTCCGATCTNNNTCCATCAGTTGTAGCAAACAT

GAGCACACGTCTGAACTCTTTCCCTACACGACGCTCTTCCGATCTNNNTCCCAAAGTGCTAGGATTACA

GAGCACACGTCTGAACTCTTTCCCTACACGACGCTCTTCCGATCTNNNTCCCAAAGTGCTAGGATTACA

AGCACACGTCTGAACTCTTTCCCTACACGACGCTCTTCCGATCTNNNTGTCTTCTGTAGGTTGTCTTTCA

AGCACACGTCTGAACTCTTTCCCTACACGACGCTCTTCCGATCTNNNTGTCTTCTGTAGGTTGTCTTTCA

GAGCACACGTCTGAACTCTTTCCCTACACGACGCTCTTCCGATCTNNNTACAAATCACTGTGACTACGG

GAGCACACGTCTGAACTCTTTCCCTACACGACGCTCTTCCGATCTNNNTACAAATCACTGTGACTACGG

AGCACACGTCTGAACTCTTTCCCTACACGACGCTCTTCCGATCTNNNGACCAGCATCATCCTGATACCAA

AGCACACGTCTGAACTCTTTCCCTACACGACGCTCTTCCGATCTNNNGACCAGCATCATCCTGATACCAA

AGCACACGTCTGAACTCTTTCCCTACACGACGCTCTTCCGATCTNNNAACTGACGTTCACACTCTCAGT

AGCACACGTCTGAACTCTTTCCCTACACGACGCTCTTCCGATCTNNNAACTGACGTTCACACTCTCAGT

AGCACACGTCTGAACTCTTTCCCTACACGACGCTCTTCCGATCTNNNTTGTTCAGGCCTTAGCATAGCA

AGCACACGTCTGAACTCTTTCCCTACACGACGCTCTTCCGATCTNNNTTGTTCAGGCCTTAGCATAGCA

CACACGTCTGAACTCTTTCCCTACACGACGCTCTTCCGATCTNNNCTGGCTGAGTTAAGGATCTTCATC

CACACGTCTGAACTCTTTCCCTACACGACGCTCTTCCGATCTNNNCTGGCTGAGTTAAGGATCTTCATC

AGCACACGTCTGAACTCTTTCCCTACACGACGCTCTTCCGATCTNNNAACTTGTGTTTTCCATGCCATCC

AGCACACGTCTGAACTCTTTCCCTACACGACGCTCTTCCGATCTNNNAACTTGTGTTTTCCATGCCATCC

GCACACGTCTGAACTCTTTCCCTACACGACGCTCTTCCGATCTNNNCTCTATTTGGTAGGAGCCAGCAT

GCACACGTCTGAACTCTTTCCCTACACGACGCTCTTCCGATCTNNNCTCTATTTGGTAGGAGCCAGCAT

GAGCACACGTCTGAACTCTTTCCCTACACGACGCTCTTCCGATCTNNNTGAGTCAGGAGAATGGAGTG

GAGCACACGTCTGAACTCTTTCCCTACACGACGCTCTTCCGATCTNNNTGAGTCAGGAGAATGGAGTG

GAGCACACGTCTGAACTCTTTCCCTACACGACGCTCTTCCGATCTNNNGAGGCAGGAAAATCACTTGAA

GAGCACACGTCTGAACTCTTTCCCTACACGACGCTCTTCCGATCTNNNGAGGCAGGAAAATCACTTGAA

GAGCACACGTCTGAACTCTTTCCCTACACGACGCTCTTCCGATCTNNNGGAAGTTGTAGTGAGCCGAG

GAGCACACGTCTGAACTCTTTCCCTACACGACGCTCTTCCGATCTNNNGGAAGTTGTAGTGAGCCGAG

AGCACACGTCTGAACTCTTTCCCTACACGACGCTCTTCCGATCTNNNGAGACTCACTTGAACCTGGGAG

AGCACACGTCTGAACTCTTTCCCTACACGACGCTCTTCCGATCTNNNGAGACTCACTTGAACCTGGGAG

GCACACGTCTGAACTCTTTCCCTACACGACGCTCTTCCGATCTNNNTTGAAAATTCTTAAGCTCATCGCA

GCACACGTCTGAACTCTTTCCCTACACGACGCTCTTCCGATCTNNNTTGAAAATTCTTAAGCTCATCGCA

GAGCACACGTCTGAACTCTTTCCCTACACGACGCTCTTCCGATCTNNNAGACAGGAGAATCACTTGAA

GAGCACACGTCTGAACTCTTTCCCTACACGACGCTCTTCCGATCTNNNAGACAGGAGAATCACTTGAA

AGCACACGTCTGAACTCTTTCCCTACACGACGCTCTTCCGATCTNNNTCCTGTCATCATAATGCTAGCT

AGCACACGTCTGAACTCTTTCCCTACACGACGCTCTTCCGATCTNNNTCCTGTCATCATAATGCTAGCT

AGCACACGTCTGAACTCTTTCCCTACACGACGCTCTTCCGATCTNNNTTGGGGATTGTTAGAAGCCTCT

AGCACACGTCTGAACTCTTTCCCTACACGACGCTCTTCCGATCTNNNTTGGGGATTGTTAGAAGCCTCT

GAGCACACGTCTGAACTCTTTCCCTACACGACGCTCTTCCGATCTNNNGCCTAAAAGGATGGGGAAG

GAGCACACGTCTGAACTCTTTCCCTACACGACGCTCTTCCGATCTNNNGCCTAAAAGGATGGGGAAG

GCACACGTCTGAACTCTTTCCCTACACGACGCTCTTCCGATCTNNNGATCTTTTTGCAACACCACCAGAC

GCACACGTCTGAACTCTTTCCCTACACGACGCTCTTCCGATCTNNNGATCTTTTTGCAACACCACCAGAC

GAGCACACGTCTGAACTCTTTCCCTACACGACGCTCTTCCGATCTNNNGCAGGAGAATTGCTTGAAGA

GAGCACACGTCTGAACTCTTTCCCTACACGACGCTCTTCCGATCTNNNGCAGGAGAATTGCTTGAAGA

AGCACACGTCTGAACTCTTTCCCTACACGACGCTCTTCCGATCTNNNGGTCATGTCACCCTCCTTCTTAG

AGCACACGTCTGAACTCTTTCCCTACACGACGCTCTTCCGATCTNNNGGTCATGTCACCCTCCTTCTTAG
GAGCACACGTCTGAACTCTTTCCCTACACGACGCTCTTCCGATCTNNNCACCTGTAGTCCCAGCTACTTG
GAGCACACGTCTGAACTCTTTCCCTACACGACGCTCTTCCGATCTNNNCACCTGTAGTCCCAGCTACTTG
AGCACACGTCTGAACTCTTTCCCTACACGACGCTCTTCCGATCTNNNATAATCCTAGCACTTTGGGAGG
AGCACACGTCTGAACTCTTTCCCTACACGACGCTCTTCCGATCTNNNATAATCCTAGCACTTTGGGAGG
AGCACACGTCTGAACTCTTTCCCTACACGACGCTCTTCCGATCTNNNGTTGGTATGTCCATTCCCCTTGC
AGCACACGTCTGAACTCTTTCCCTACACGACGCTCTTCCGATCTNNNGTTGGTATGTCCATTCCCCTTGC
AGCACACGTCTGAACTCTTTCCCTACACGACGCTCTTCCGATCTNNNATTTGAAGCTCTGCTGTTTGGTC
AGCACACGTCTGAACTCTTTCCCTACACGACGCTCTTCCGATCTNNNATTTGAAGCTCTGCTGTTTGGTC
GAGCACACGTCTGAACTCTTTCCCTACACGACGCTCTTCCGATCTNNNAGGCTGAGGCAGAATTACTTG
GAGCACACGTCTGAACTCTTTCCCTACACGACGCTCTTCCGATCTNNNAGGCTGAGGCAGAATTACTTG
GAGCACACGTCTGAACTCTTTCCCTACACGACGCTCTTCCGATCTNNNAAGTAACTGGGACCACAGGTA
GAGCACACGTCTGAACTCTTTCCCTACACGACGCTCTTCCGATCTNNNAAGTAACTGGGACCACAGGTA
AGAGCACACGTCTGAACTCTTTCCCTACACGACGCTCTTCCGATCTNNNTCACTTGAACCTGCCTAGATC
AGAGCACACGTCTGAACTCTTTCCCTACACGACGCTCTTCCGATCTNNNTCACTTGAACCTGCCTAGATC
GAGCACACGTCTGAACTCTTTCCCTACACGACGCTCTTCCGATCTNNNCATACACACACACACACACACTC
GAGCACACGTCTGAACTCTTTCCCTACACGACGCTCTTCCGATCTNNNCATACACACACACACACACACTC
AGCACACGTCTGAACTCTTTCCCTACACGACGCTCTTCCGATCTNNNAGCCCAGAAATAGATCCACACAA
AGCACACGTCTGAACTCTTTCCCTACACGACGCTCTTCCGATCTNNNAGCCCAGAAATAGATCCACACAA
AGCACACGTCTGAACTCTTTCCCTACACGACGCTCTTCCGATCTNNNCCGAGTAGCTGGGACTACAGAC
AGCACACGTCTGAACTCTTTCCCTACACGACGCTCTTCCGATCTNNNCCGAGTAGCTGGGACTACAGAC
AGAGCACACGTCTGAACTCTTTCCCTACACGACGCTCTTCCGATCTNNNTGGACTCTGTCAATCAGGACA
AGAGCACACGTCTGAACTCTTTCCCTACACGACGCTCTTCCGATCTNNNTGGACTCTGTCAATCAGGACA
AGCACACGTCTGAACTCTTTCCCTACACGACGCTCTTCCGATCTNNNGTAGCTGGGATTACAGGCATGA
AGCACACGTCTGAACTCTTTCCCTACACGACGCTCTTCCGATCTNNNGTAGCTGGGATTACAGGCATGA
AGCACACGTCTGAACTCTTTCCCTACACGACGCTCTTCCGATCTNNNCCCTGAGACCTAACACACCTAAC
AGCACACGTCTGAACTCTTTCCCTACACGACGCTCTTCCGATCTNNNCCCTGAGACCTAACACACCTAAC
AGCACACGTCTGAACTCTTTCCCTACACGACGCTCTTCCGATCTNNNGGCCAACAGTGAGGAGCTATAA
AGCACACGTCTGAACTCTTTCCCTACACGACGCTCTTCCGATCTNNNGGCCAACAGTGAGGAGCTATAA
AGCACACGTCTGAACTCTTTCCCTACACGACGCTCTTCCGATCTNNNGGAGGTTACAGTGAGCTGAGA
AGCACACGTCTGAACTCTTTCCCTACACGACGCTCTTCCGATCTNNNGGAGGTTACAGTGAGCTGAGA
AGCACACGTCTGAACTCTTTCCCTACACGACGCTCTTCCGATCTNNNACGCCCGGCTAATTTTTCTTTTCC
AGCACACGTCTGAACTCTTTCCCTACACGACGCTCTTCCGATCTNNNACGCCCGGCTAATTTTTCTTTTCC
AGCACACGTCTGAACTCTTTCCCTACACGACGCTCTTCCGATCTNNNCCCATCAGAGATATTGGCCTGTA
AGCACACGTCTGAACTCTTTCCCTACACGACGCTCTTCCGATCTNNNCCCATCAGAGATATTGGCCTGTA
AGCACACGTCTGAACTCTTTCCCTACACGACGCTCTTCCGATCTNNNAAATTTACAGGGCACACTCTTGC
AGCACACGTCTGAACTCTTTCCCTACACGACGCTCTTCCGATCTNNNAAATTTACAGGGCACACTCTTGC
GAGCACACGTCTGAACTCTTTCCCTACACGACGCTCTTCCGATCTNNNGGAGTGAACAGTCTTGTCTGC
GAGCACACGTCTGAACTCTTTCCCTACACGACGCTCTTCCGATCTNNNGGAGTGAACAGTCTTGTCTGC
AGCACACGTCTGAACTCTTTCCCTACACGACGCTCTTCCGATCTNNNATGGACAATTTGGTAAGCACAC
AGCACACGTCTGAACTCTTTCCCTACACGACGCTCTTCCGATCTNNNATGGACAATTTGGTAAGCACAC
AGAGCACACGTCTGAACTCTTTCCCTACACGACGCTCTTCCGATCTNNNTGTAGTCTCAGCTACTCTGGA
AGAGCACACGTCTGAACTCTTTCCCTACACGACGCTCTTCCGATCTNNNTGTAGTCTCAGCTACTCTGGA

GAGCACACGTCTGAACTCTTTCCCTACACGACGCTCTTCCGATCTNNNTTTTTTGACTGTTGTGTAGG
GAGCACACGTCTGAACTCTTTCCCTACACGACGCTCTTCCGATCTNNNTTTTTTGACTGTTGTGTAGG
GCACACGTCTGAACTCTTTCCCTACACGACGCTCTTCCGATCTNNNGGCTTTTGTAAGTAGTGCTGCA
GCACACGTCTGAACTCTTTCCCTACACGACGCTCTTCCGATCTNNNGGCTTTTGTAAGTAGTGCTGCA
AGCACACGTCTGAACTCTTTCCCTACACGACGCTCTTCCGATCTNNNAGGGTCACAGATGGCTCTTTTA
AGCACACGTCTGAACTCTTTCCCTACACGACGCTCTTCCGATCTNNNAGGGTCACAGATGGCTCTTTTA
GAGCACACGTCTGAACTCTTTCCCTACACGACGCTCTTCCGATCTNNNTATCAAGATAGTACAGAGAA
GAGCACACGTCTGAACTCTTTCCCTACACGACGCTCTTCCGATCTNNNTATCAAGATAGTACAGAGAA
AGCACACGTCTGAACTCTTTCCCTACACGACGCTCTTCCGATCTNNNCCAGAGATTAGGATTGCAACTC
AGCACACGTCTGAACTCTTTCCCTACACGACGCTCTTCCGATCTNNNCCAGAGATTAGGATTGCAACTC
AGCACACGTCTGAACTCTTTCCCTACACGACGCTCTTCCGATCTNNNCAACTTTGCTCTTCTCCACGCCA
AGCACACGTCTGAACTCTTTCCCTACACGACGCTCTTCCGATCTNNNCAACTTTGCTCTTCTCCACGCCA
GAGCACACGTCTGAACTCTTTCCCTACACGACGCTCTTCCGATCTNNNGGTTTGATGTGGGTTGGGTT
GAGCACACGTCTGAACTCTTTCCCTACACGACGCTCTTCCGATCTNNNGGTTTGATGTGGGTTGGGTT
AGCACACGTCTGAACTCTTTCCCTACACGACGCTCTTCCGATCTNNNAGGGCAGTTGGGTCATGTTATT
AGCACACGTCTGAACTCTTTCCCTACACGACGCTCTTCCGATCTNNNAGGGCAGTTGGGTCATGTTATT
AGCACACGTCTGAACTCTTTCCCTACACGACGCTCTTCCGATCTNNNACAACACAGAGGATCCCATTTC
AGCACACGTCTGAACTCTTTCCCTACACGACGCTCTTCCGATCTNNNACAACACAGAGGATCCCATTTC
AGCACACGTCTGAACTCTTTCCCTACACGACGCTCTTCCGATCTNNNAGAATTGCTTGAACCTGGGAGT
AGCACACGTCTGAACTCTTTCCCTACACGACGCTCTTCCGATCTNNNAGAATTGCTTGAACCTGGGAGT
AGCACACGTCTGAACTCTTTCCCTACACGACGCTCTTCCGATCTNNNGCTGATCTCTATTCATAGCCTG
AGCACACGTCTGAACTCTTTCCCTACACGACGCTCTTCCGATCTNNNGCTGATCTCTATTCATAGCCTG
GAGCACACGTCTGAACTCTTTCCCTACACGACGCTCTTCCGATCTNNNTGAATTAGCTGTGTGCCACT
GAGCACACGTCTGAACTCTTTCCCTACACGACGCTCTTCCGATCTNNNTGAATTAGCTGTGTGCCACT
AGCACACGTCTGAACTCTTTCCCTACACGACGCTCTTCCGATCTNNNTTTTGAGGTGTAGTACTTCATT
AGCACACGTCTGAACTCTTTCCCTACACGACGCTCTTCCGATCTNNNTTTTGAGGTGTAGTACTTCATT
AGCACACGTCTGAACTCTTTCCCTACACGACGCTCTTCCGATCTNNNAGACTCCGTCTCAAAAAGGGA
AGCACACGTCTGAACTCTTTCCCTACACGACGCTCTTCCGATCTNNNAGACTCCGTCTCAAAAAGGGA
AGCACACGTCTGAACTCTTTCCCTACACGACGCTCTTCCGATCTNNNTGTCGTGACCATTGTGAACAT
AGCACACGTCTGAACTCTTTCCCTACACGACGCTCTTCCGATCTNNNTGTCGTGACCATTGTGAACAT
AGCACACGTCTGAACTCTTTCCCTACACGACGCTCTTCCGATCTNNNTCCTTTGGTTACCCTTCCTTCTG
AGCACACGTCTGAACTCTTTCCCTACACGACGCTCTTCCGATCTNNNTCCTTTGGTTACCCTTCCTTCTG
GAGCACACGTCTGAACTCTTTCCCTACACGACGCTCTTCCGATCTNNNGCAACAGAGTGAGACCTAGG
GAGCACACGTCTGAACTCTTTCCCTACACGACGCTCTTCCGATCTNNNGCAACAGAGTGAGACCTAGG
AGAGCACACGTCTGAACTCTTTCCCTACACGACGCTCTTCCGATCTNNNGGTGACAGAGTGAGACCTT
AGAGCACACGTCTGAACTCTTTCCCTACACGACGCTCTTCCGATCTNNNGGTGACAGAGTGAGACCTT
AGCACACGTCTGAACTCTTTCCCTACACGACGCTCTTCCGATCTNNNGTAATCTGAGGTAAAATGATG
AGCACACGTCTGAACTCTTTCCCTACACGACGCTCTTCCGATCTNNNGTAATCTGAGGTAAAATGATG
AGCACACGTCTGAACTCTTTCCCTACACGACGCTCTTCCGATCTNNNATCTGCTGAGCTAGATCACTTG
AGCACACGTCTGAACTCTTTCCCTACACGACGCTCTTCCGATCTNNNATCTGCTGAGCTAGATCACTTG
GCACACGTCTGAACTCTTTCCCTACACGACGCTCTTCCGATCTNNNTGCCATATTCATGTTCCACCTGTC
GCACACGTCTGAACTCTTTCCCTACACGACGCTCTTCCGATCTNNNTGCCATATTCATGTTCCACCTGTC
GAGCACACGTCTGAACTCTTTCCCTACACGACGCTCTTCCGATCTNNNGTCCCACCAATACCACCAGAT

GAGCACACGTCTGAACTCTTTCCCTACACGACGCTCTTCCGATCTNNNGTCCCCACCAATACCACCAGAT
GAGCACACGTCTGAACTCTTTCCCTACACGACGCTCTTCCGATCTNNNTGTGGCTATTGTAAATGGGAT
GAGCACACGTCTGAACTCTTTCCCTACACGACGCTCTTCCGATCTNNNTGTGGCTATTGTAAATGGGAT
GAGCACACGTCTGAACTCTTTCCCTACACGACGCTCTTCCGATCTNNNTGTCTGTAACATTAACTCGGGC
GAGCACACGTCTGAACTCTTTCCCTACACGACGCTCTTCCGATCTNNNTGTCTGTAACATTAACTCGGGC
GAGCACACGTCTGAACTCTTTCCCTACACGACGCTCTTCCGATCTNNNCAAGACCGGGCCATTGTACTC
GAGCACACGTCTGAACTCTTTCCCTACACGACGCTCTTCCGATCTNNNCAAGACCGGGCCATTGTACTC
AGCACACGTCTGAACTCTTTCCCTACACGACGCTCTTCCGATCTNNNGTAGGAGTAAGGGAAGAGGA
AGCACACGTCTGAACTCTTTCCCTACACGACGCTCTTCCGATCTNNNGTAGGAGTAAGGGAAGAGGA
GAGCACACGTCTGAACTCTTTCCCTACACGACGCTCTTCCGATCTNNNCCTCACTTACCTCAAAGGGTTG
GAGCACACGTCTGAACTCTTTCCCTACACGACGCTCTTCCGATCTNNNCCTCACTTACCTCAAAGGGTTG
AGCACACGTCTGAACTCTTTCCCTACACGACGCTCTTCCGATCTNNNTGGACTAAAATTTCTAACAAAC
AGCACACGTCTGAACTCTTTCCCTACACGACGCTCTTCCGATCTNNNTGGACTAAAATTTCTAACAAAC
AGCACACGTCTGAACTCTTTCCCTACACGACGCTCTTCCGATCTNNNCAGCACTTTAAGTAAAGAATCT
AGCACACGTCTGAACTCTTTCCCTACACGACGCTCTTCCGATCTNNNCAGCACTTTAAGTAAAGAATCT
AGCACACGTCTGAACTCTTTCCCTACACGACGCTCTTCCGATCTNNNGCCAAGGAATGTCGGTTTGTT
AGCACACGTCTGAACTCTTTCCCTACACGACGCTCTTCCGATCTNNNGCCAAGGAATGTCGGTTTGTT
AGCACACGTCTGAACTCTTTCCCTACACGACGCTCTTCCGATCTNNNAAATACAAAAATTAGCCGGGCG
AGCACACGTCTGAACTCTTTCCCTACACGACGCTCTTCCGATCTNNNAAATACAAAAATTAGCCGGGCG
AGCACACGTCTGAACTCTTTCCCTACACGACGCTCTTCCGATCTNNNGATCACGAGGTCAGGAGATTG
AGCACACGTCTGAACTCTTTCCCTACACGACGCTCTTCCGATCTNNNGATCACGAGGTCAGGAGATTG
GAGCACACGTCTGAACTCTTTCCCTACACGACGCTCTTCCGATCTNNNGATGGAGTCTCACTCTGTCACA
GAGCACACGTCTGAACTCTTTCCCTACACGACGCTCTTCCGATCTNNNGATGGAGTCTCACTCTGTCACA
AGAGCACACGTCTGAACTCTTTCCCTACACGACGCTCTTCCGATCTNNNTCTTCTGTTTAAGTCCCAGCCT
AGAGCACACGTCTGAACTCTTTCCCTACACGACGCTCTTCCGATCTNNNTCTTCTGTTTAAGTCCCAGCCT
AGCACACGTCTGAACTCTTTCCCTACACGACGCTCTTCCGATCTNNNGGAAGGAAGGAAGAAAGGAA
AGCACACGTCTGAACTCTTTCCCTACACGACGCTCTTCCGATCTNNNGGAAGGAAGGAAGAAAGGAA
GAGCACACGTCTGAACTCTTTCCCTACACGACGCTCTTCCGATCTNNNTGTAGAAGATTGTTGCCTCTCA
GAGCACACGTCTGAACTCTTTCCCTACACGACGCTCTTCCGATCTNNNTGTAGAAGATTGTTGCCTCTCA
GAGCACACGTCTGAACTCTTTCCCTACACGACGCTCTTCCGATCTNNNCCCAGACAGACTTCAGATTTG
GAGCACACGTCTGAACTCTTTCCCTACACGACGCTCTTCCGATCTNNNCCCAGACAGACTTCAGATTTG
GAGCACACGTCTGAACTCTTTCCCTACACGACGCTCTTCCGATCTNNNGTGGCCTGGGATTTTTTTCTG
GAGCACACGTCTGAACTCTTTCCCTACACGACGCTCTTCCGATCTNNNGTGGCCTGGGATTTTTTTCTG
GCACACGTCTGAACTCTTTCCCTACACGACGCTCTTCCGATCTNNNCTACCCTGTAGCTATGAGCAGTG
GCACACGTCTGAACTCTTTCCCTACACGACGCTCTTCCGATCTNNNCTACCCTGTAGCTATGAGCAGTG
GAGCACACGTCTGAACTCTTTCCCTACACGACGCTCTTCCGATCTNNNCTTGGGTTGAGGGAAGTTCTG
GAGCACACGTCTGAACTCTTTCCCTACACGACGCTCTTCCGATCTNNNCTTGGGTTGAGGGAAGTTCTG
AGAGCACACGTCTGAACTCTTTCCCTACACGACGCTCTTCCGATCTNNNCCAAAGTGCTGGAATTACAG
AGAGCACACGTCTGAACTCTTTCCCTACACGACGCTCTTCCGATCTNNNCCAAAGTGCTGGAATTACAG
GAGCACACGTCTGAACTCTTTCCCTACACGACGCTCTTCCGATCTNNNTCCACTCGTTCCAAAGTGACAT
GAGCACACGTCTGAACTCTTTCCCTACACGACGCTCTTCCGATCTNNNTCCACTCGTTCCAAAGTGACAT
CACACGTCTGAACTCTTTCCCTACACGACGCTCTTCCGATCTNNNCCCACAGAAACTCAGTCCATATCA
CACACGTCTGAACTCTTTCCCTACACGACGCTCTTCCGATCTNNNCCCACAGAAACTCAGTCCATATCA

GAGCACACGTCTGAACTCTTTCCCTACACGACGCTCTTCCGATCTNNNCCGTGGTACCTTCGAGGCTT
GAGCACACGTCTGAACTCTTTCCCTACACGACGCTCTTCCGATCTNNNCCGTGGTACCTTCGAGGCTT
GAGCACACGTCTGAACTCTTTCCCTACACGACGCTCTTCCGATCTNNNCTGTATTCCAGCTACTTGGG
GAGCACACGTCTGAACTCTTTCCCTACACGACGCTCTTCCGATCTNNNCTGTATTCCAGCTACTTGGG
GAGCACACGTCTGAACTCTTTCCCTACACGACGCTCTTCCGATCTNNNCTTTCATGAGGACTGGAGGT
GAGCACACGTCTGAACTCTTTCCCTACACGACGCTCTTCCGATCTNNNCTTTCATGAGGACTGGAGGT
AGCACACGTCTGAACTCTTTCCCTACACGACGCTCTTCCGATCTNNNTCCTCATTTTCCAACTCCTCCAG
AGCACACGTCTGAACTCTTTCCCTACACGACGCTCTTCCGATCTNNNTCCTCATTTTCCAACTCCTCCAG
AGCACACGTCTGAACTCTTTCCCTACACGACGCTCTTCCGATCTNNNGCTGCGGATTGTCATCAAAAA
AGCACACGTCTGAACTCTTTCCCTACACGACGCTCTTCCGATCTNNNGCTGCGGATTGTCATCAAAAA
AGCACACGTCTGAACTCTTTCCCTACACGACGCTCTTCCGATCTNNNGAGAACTGCTTGAACCTGGG
AGCACACGTCTGAACTCTTTCCCTACACGACGCTCTTCCGATCTNNNGAGAACTGCTTGAACCTGGG
GAGCACACGTCTGAACTCTTTCCCTACACGACGCTCTTCCGATCTNNNCACTATCAGGTGCAGAATGT
GAGCACACGTCTGAACTCTTTCCCTACACGACGCTCTTCCGATCTNNNCACTATCAGGTGCAGAATGT
GAGCACACGTCTGAACTCTTTCCCTACACGACGCTCTTCCGATCTNNNGGATTACAGGTATGAGGCAC
GAGCACACGTCTGAACTCTTTCCCTACACGACGCTCTTCCGATCTNNNGGATTACAGGTATGAGGCAC
GAGCACACGTCTGAACTCTTTCCCTACACGACGCTCTTCCGATCTNNNGGACACCCCTAAGCTAAAG
GAGCACACGTCTGAACTCTTTCCCTACACGACGCTCTTCCGATCTNNNGGACACCCCTAAGCTAAAG
AGCACACGTCTGAACTCTTTCCCTACACGACGCTCTTCCGATCTNNNGAGAATCACTCGAACCAGGGA
AGCACACGTCTGAACTCTTTCCCTACACGACGCTCTTCCGATCTNNNGAGAATCACTCGAACCAGGGA
GAGCACACGTCTGAACTCTTTCCCTACACGACGCTCTTCCGATCTNNNACGCCTGTAGTCCCAACTATT
GAGCACACGTCTGAACTCTTTCCCTACACGACGCTCTTCCGATCTNNNACGCCTGTAGTCCCAACTATT
GAGCACACGTCTGAACTCTTTCCCTACACGACGCTCTTCCGATCTNNNGCAAAGACACACATAGGCTC
GAGCACACGTCTGAACTCTTTCCCTACACGACGCTCTTCCGATCTNNNGCAAAGACACACATAGGCTC
AGCACACGTCTGAACTCTTTCCCTACACGACGCTCTTCCGATCTNNNCTTTCTTCCTGTCTGCCTCTGTC
AGCACACGTCTGAACTCTTTCCCTACACGACGCTCTTCCGATCTNNNCTTTCTTCCTGTCTGCCTCTGTC
GCACACGTCTGAACTCTTTCCCTACACGACGCTCTTCCGATCTNNNTGCACCACATCTTCTGTAGAGAC
GCACACGTCTGAACTCTTTCCCTACACGACGCTCTTCCGATCTNNNTGCACCACATCTTCTGTAGAGAC
GCACACGTCTGAACTCTTTCCCTACACGACGCTCTTCCGATCTNNNTCGTTTTTGCCCAATCACCATTC
GCACACGTCTGAACTCTTTCCCTACACGACGCTCTTCCGATCTNNNTCGTTTTTGCCCAATCACCATTC
AGCACACGTCTGAACTCTTTCCCTACACGACGCTCTTCCGATCTNNNCACCGCACAGCCACATATTCTA
AGCACACGTCTGAACTCTTTCCCTACACGACGCTCTTCCGATCTNNNCACCGCACAGCCACATATTCTA
AGCACACGTCTGAACTCTTTCCCTACACGACGCTCTTCCGATCTNNNTTGGTTGTCAAGTTTCATTTGG
AGCACACGTCTGAACTCTTTCCCTACACGACGCTCTTCCGATCTNNNTTGGTTGTCAAGTTTCATTTGG
AGCACACGTCTGAACTCTTTCCCTACACGACGCTCTTCCGATCTNNNGGTACCTCAGCTGGAAATGAA
AGCACACGTCTGAACTCTTTCCCTACACGACGCTCTTCCGATCTNNNGGTACCTCAGCTGGAAATGAA
AGCACACGTCTGAACTCTTTCCCTACACGACGCTCTTCCGATCTNNNCTCCCAAAATACACCTCCCTCA
AGCACACGTCTGAACTCTTTCCCTACACGACGCTCTTCCGATCTNNNCTCCCAAAATACACCTCCCTCA
AGCACACGTCTGAACTCTTTCCCTACACGACGCTCTTCCGATCTNNNTCAGGATTATCCAACTTTACAG
AGCACACGTCTGAACTCTTTCCCTACACGACGCTCTTCCGATCTNNNTCAGGATTATCCAACTTTACAG
GCACACGTCTGAACTCTTTCCCTACACGACGCTCTTCCGATCTNNNGGAAAGGTCTGTTGAGAGGGT
GCACACGTCTGAACTCTTTCCCTACACGACGCTCTTCCGATCTNNNGGAAAGGTCTGTTGAGAGGGT
AGCACACGTCTGAACTCTTTCCCTACACGACGCTCTTCCGATCTNNNGAGCCAGACCCCATCTTTTTTC

AGCACACGTCTGAACTCTTTCCCTACACGACGCTCTTCCGATCTNNNGAGCCAGACCCCATCTTTTTTC
AGCACACGTCTGAACTCTTTCCCTACACGACGCTCTTCCGATCTNNNTTTGGATTATACACACACACAC
AGCACACGTCTGAACTCTTTCCCTACACGACGCTCTTCCGATCTNNNTTTGGATTATACACACACACAC
GAGCACACGTCTGAACTCTTTCCCTACACGACGCTCTTCCGATCTNNNTTCAATCGAATCTCCTGCCTCA
GAGCACACGTCTGAACTCTTTCCCTACACGACGCTCTTCCGATCTNNNTTCAATCGAATCTCCTGCCTCA
AGCACACGTCTGAACTCTTTCCCTACACGACGCTCTTCCGATCTNNNTATGCAGTGTCATGACCATGTC
AGCACACGTCTGAACTCTTTCCCTACACGACGCTCTTCCGATCTNNNTATGCAGTGTCATGACCATGTC
AGCACACGTCTGAACTCTTTCCCTACACGACGCTCTTCCGATCTNNNGAGGTGGAGCTTGCAGTGAG
AGCACACGTCTGAACTCTTTCCCTACACGACGCTCTTCCGATCTNNNGAGGTGGAGCTTGCAGTGAG
AGCACACGTCTGAACTCTTTCCCTACACGACGCTCTTCCGATCTNNNATAGAGATGTGCTCAGGGACT
AGCACACGTCTGAACTCTTTCCCTACACGACGCTCTTCCGATCTNNNATAGAGATGTGCTCAGGGACT
AGCACACGTCTGAACTCTTTCCCTACACGACGCTCTTCCGATCTNNNAATAGGCTAGGCTGCTACCTC
AGCACACGTCTGAACTCTTTCCCTACACGACGCTCTTCCGATCTNNNAATAGGCTAGGCTGCTACCTC
AGCACACGTCTGAACTCTTTCCCTACACGACGCTCTTCCGATCTNNNCACTACCAGCTTCTCTCTTTCCC
AGCACACGTCTGAACTCTTTCCCTACACGACGCTCTTCCGATCTNNNCACTACCAGCTTCTCTCTTTCCC
AGCACACGTCTGAACTCTTTCCCTACACGACGCTCTTCCGATCTNNNTACACATCCTAAATTTGGTAGG
AGCACACGTCTGAACTCTTTCCCTACACGACGCTCTTCCGATCTNNNTACACATCCTAAATTTGGTAGG
AGCACACGTCTGAACTCTTTCCCTACACGACGCTCTTCCGATCTNNNTGATAGGAATGTTTGATAGTG
AGCACACGTCTGAACTCTTTCCCTACACGACGCTCTTCCGATCTNNNTGATAGGAATGTTTGATAGTG
AGCACACGTCTGAACTCTTTCCCTACACGACGCTCTTCCGATCTNNNTGTGCAGAGTGAAAGGTGGT
AGCACACGTCTGAACTCTTTCCCTACACGACGCTCTTCCGATCTNNNTGTGCAGAGTGAAAGGTGGT
AGCACACGTCTGAACTCTTTCCCTACACGACGCTCTTCCGATCTNNNAGTTACAACTGACAGGGAAGG
AGCACACGTCTGAACTCTTTCCCTACACGACGCTCTTCCGATCTNNNAGTTACAACTGACAGGGAAGG
GCACACGTCTGAACTCTTTCCCTACACGACGCTCTTCCGATCTNNNACTGGTATAGTGCTGGTTTGTT
GCACACGTCTGAACTCTTTCCCTACACGACGCTCTTCCGATCTNNNACTGGTATAGTGCTGGTTTGTT
AGCACACGTCTGAACTCTTTCCCTACACGACGCTCTTCCGATCTNNNTGTAGTGGTCAGTCATGGCTT
AGCACACGTCTGAACTCTTTCCCTACACGACGCTCTTCCGATCTNNNTGTAGTGGTCAGTCATGGCTT
GAGCACACGTCTGAACTCTTTCCCTACACGACGCTCTTCCGATCTNNNTGGGCGTTGCTTGTCTCATAA
GAGCACACGTCTGAACTCTTTCCCTACACGACGCTCTTCCGATCTNNNTGGGCGTTGCTTGTCTCATAA
CACACGTCTGAACTCTTTCCCTACACGACGCTCTTCCGATCTNNNTTCCTTCCTTCCTCCATCTCTCTCAT
CACACGTCTGAACTCTTTCCCTACACGACGCTCTTCCGATCTNNNTTCCTTCCTTCCTCCATCTCTCTCAT
GCACACGTCTGAACTCTTTCCCTACACGACGCTCTTCCGATCTNNNCTGCGTTCTTACCAGCATTTTGT
GCACACGTCTGAACTCTTTCCCTACACGACGCTCTTCCGATCTNNNCTGCGTTCTTACCAGCATTTTGT
AGCACACGTCTGAACTCTTTCCCTACACGACGCTCTTCCGATCTNNNGCCAAGATCATGCCATTGTACT
AGCACACGTCTGAACTCTTTCCCTACACGACGCTCTTCCGATCTNNNGCCAAGATCATGCCATTGTACT
GAGCACACGTCTGAACTCTTTCCCTACACGACGCTCTTCCGATCTNNNCTGAGTGTAGAGCTTGGAAA
GAGCACACGTCTGAACTCTTTCCCTACACGACGCTCTTCCGATCTNNNCTGAGTGTAGAGCTTGGAAA
GAGCACACGTCTGAACTCTTTCCCTACACGACGCTCTTCCGATCTNNNATGATAAAAACCAAGATGTT
GAGCACACGTCTGAACTCTTTCCCTACACGACGCTCTTCCGATCTNNNATGATAAAAACCAAGATGTT
GCACACGTCTGAACTCTTTCCCTACACGACGCTCTTCCGATCTNNNGGATTCTTGTTCAAAAGTGGCCA
GCACACGTCTGAACTCTTTCCCTACACGACGCTCTTCCGATCTNNNGGATTCTTGTTCAAAAGTGGCCA
ACACGTCTGAACTCTTTCCCTACACGACGCTCTTCCGATCTNNNAAAAATACAAAAATGAGCCGGCAT
ACACGTCTGAACTCTTTCCCTACACGACGCTCTTCCGATCTNNNAAAAATACAAAAATGAGCCGGCAT

AGCACACGTCTGAACTCTTTCCCTACACGACGCTCTTCCGATCTNNNTCTCTTTCTATGCAACGCTCCTT
AGCACACGTCTGAACTCTTTCCCTACACGACGCTCTTCCGATCTNNNTCTCTTTCTATGCAACGCTCCTT
GAGCACACGTCTGAACTCTTTCCCTACACGACGCTCTTCCGATCTNNNATGTGTGGTTGTAGGAGAT
GAGCACACGTCTGAACTCTTTCCCTACACGACGCTCTTCCGATCTNNNATGTGTGGTTGTAGGAGAT
AGCACACGTCTGAACTCTTTCCCTACACGACGCTCTTCCGATCTNNNACCCTGAAGCCAGTAATCCAA
AGCACACGTCTGAACTCTTTCCCTACACGACGCTCTTCCGATCTNNNACCCTGAAGCCAGTAATCCAA
AGCACACGTCTGAACTCTTTCCCTACACGACGCTCTTCCGATCTNNNAAAGACACCTGTGAATGCTAC
AGCACACGTCTGAACTCTTTCCCTACACGACGCTCTTCCGATCTNNNAAAGACACCTGTGAATGCTAC
AGCACACGTCTGAACTCTTTCCCTACACGACGCTCTTCCGATCTNNNCCGAGGTCTACAAGGCTGGAT
AGCACACGTCTGAACTCTTTCCCTACACGACGCTCTTCCGATCTNNNCCGAGGTCTACAAGGCTGGAT
AGCACACGTCTGAACTCTTTCCCTACACGACGCTCTTCCGATCTNNNGGAGGGACACGGATTTCTTTA
AGCACACGTCTGAACTCTTTCCCTACACGACGCTCTTCCGATCTNNNGGAGGGACACGGATTTCTTTA
AGCACACGTCTGAACTCTTTCCCTACACGACGCTCTTCCGATCTNNNTGTGTCGGTCTAGGAGGTTAT
AGCACACGTCTGAACTCTTTCCCTACACGACGCTCTTCCGATCTNNNTGTGTCGGTCTAGGAGGTTAT
AGCACACGTCTGAACTCTTTCCCTACACGACGCTCTTCCGATCTNNNGGGACATGACAATTGCTTGA
AGCACACGTCTGAACTCTTTCCCTACACGACGCTCTTCCGATCTNNNGGGACATGACAATTGCTTGA
AGCACACGTCTGAACTCTTTCCCTACACGACGCTCTTCCGATCTNNNTGACTCTACCCTATCCATGCTCA
AGCACACGTCTGAACTCTTTCCCTACACGACGCTCTTCCGATCTNNNTGACTCTACCCTATCCATGCTCA
AGCACACGTCTGAACTCTTTCCCTACACGACGCTCTTCCGATCTNNNCAGACCTGCTCTAAAACAAAAG
AGCACACGTCTGAACTCTTTCCCTACACGACGCTCTTCCGATCTNNNCAGACCTGCTCTAAAACAAAAG
GCACACGTCTGAACTCTTTCCCTACACGACGCTCTTCCGATCTNNNACATTCAAACAAGGCAGGTCAA
GCACACGTCTGAACTCTTTCCCTACACGACGCTCTTCCGATCTNNNACATTCAAACAAGGCAGGTCAA
GAGCACACGTCTGAACTCTTTCCCTACACGACGCTCTTCCGATCTNNNGGGCTCGAGATAGAGGTTT
GAGCACACGTCTGAACTCTTTCCCTACACGACGCTCTTCCGATCTNNNGGGCTCGAGATAGAGGTTT
CACACGTCTGAACTCTTTCCCTACACGACGCTCTTCCGATCTNNNTGCTAAAGCGAATCCTGGGCTCA
CACACGTCTGAACTCTTTCCCTACACGACGCTCTTCCGATCTNNNTGCTAAAGCGAATCCTGGGCTCA
AGCACACGTCTGAACTCTTTCCCTACACGACGCTCTTCCGATCTNNNTCGTTCCTTCAGTTCTTCTGGA
AGCACACGTCTGAACTCTTTCCCTACACGACGCTCTTCCGATCTNNNTCGTTCCTTCAGTTCTTCTGGA
AGCACACGTCTGAACTCTTTCCCTACACGACGCTCTTCCGATCTNNNACAACAGACTGCTTGCAATTCT
AGCACACGTCTGAACTCTTTCCCTACACGACGCTCTTCCGATCTNNNACAACAGACTGCTTGCAATTCT
AGCACACGTCTGAACTCTTTCCCTACACGACGCTCTTCCGATCTNNNCACCTGTAATCCCAGCTACTCA
AGCACACGTCTGAACTCTTTCCCTACACGACGCTCTTCCGATCTNNNCACCTGTAATCCCAGCTACTCA
GCACACGTCTGAACTCTTTCCCTACACGACGCTCTTCCGATCTNNNCTTGAACTCCTGACCTCGTGATC
GCACACGTCTGAACTCTTTCCCTACACGACGCTCTTCCGATCTNNNCTTGAACTCCTGACCTCGTGATC
AGCACACGTCTGAACTCTTTCCCTACACGACGCTCTTCCGATCTNNNCTTTGGGAAGGAAGAAGGG
AGCACACGTCTGAACTCTTTCCCTACACGACGCTCTTCCGATCTNNNCTTTGGGAAGGAAGAAGGG
GAGCACACGTCTGAACTCTTTCCCTACACGACGCTCTTCCGATCTNNNCATGGAAAAATTGTGTTCCG
GAGCACACGTCTGAACTCTTTCCCTACACGACGCTCTTCCGATCTNNNCATGGAAAAATTGTGTTCCG
AGCACACGTCTGAACTCTTTCCCTACACGACGCTCTTCCGATCTNNNTCCCATGACTCATGATGGAAT
AGCACACGTCTGAACTCTTTCCCTACACGACGCTCTTCCGATCTNNNTCCCATGACTCATGATGGAAT
AGCACACGTCTGAACTCTTTCCCTACACGACGCTCTTCCGATCTNNNATTCCCCAACTTCACCTTCACAC
AGCACACGTCTGAACTCTTTCCCTACACGACGCTCTTCCGATCTNNNATTCCCCAACTTCACCTTCACAC
GAGCACACGTCTGAACTCTTTCCCTACACGACGCTCTTCCGATCTNNNCATGCTACTACACTCCAGTCT

GAGCACACGTCTGAACTCTTTCCCTACACGACGCTCTTCCGATCTNNNCATGCTACTACACTCCAGTCT

GAGCACACGTCTGAACTCTTTCCCTACACGACGCTCTTCCGATCTNNNGCGACAGAGTGAGACTCCAT

GAGCACACGTCTGAACTCTTTCCCTACACGACGCTCTTCCGATCTNNNGCGACAGAGTGAGACTCCAT

AGCACACGTCTGAACTCTTTCCCTACACGACGCTCTTCCGATCTNNNGGAAAGCCAACTCCTGGATTCA

AGCACACGTCTGAACTCTTTCCCTACACGACGCTCTTCCGATCTNNNGGAAAGCCAACTCCTGGATTCA

GCACACGTCTGAACTCTTTCCCTACACGACGCTCTTCCGATCTNNNTCAATCTCCTGCTTCTCAGCTTCC

GCACACGTCTGAACTCTTTCCCTACACGACGCTCTTCCGATCTNNNTCAATCTCCTGCTTCTCAGCTTCC

AGCACACGTCTGAACTCTTTCCCTACACGACGCTCTTCCGATCTNNNAGGGTGTTTAGTAGCATCTGT

AGCACACGTCTGAACTCTTTCCCTACACGACGCTCTTCCGATCTNNNAGGGTGTTTAGTAGCATCTGT

AGCACACGTCTGAACTCTTTCCCTACACGACGCTCTTCCGATCTNNNTTACATGGTTGGAGCAGGAGA

AGCACACGTCTGAACTCTTTCCCTACACGACGCTCTTCCGATCTNNNTTACATGGTTGGAGCAGGAGA

GCACACGTCTGAACTCTTTCCCTACACGACGCTCTTCCGATCTNNNAGAAAGTGACCAGCCAGTAGAT

GCACACGTCTGAACTCTTTCCCTACACGACGCTCTTCCGATCTNNNAGAAAGTGACCAGCCAGTAGAT

AGCACACGTCTGAACTCTTTCCCTACACGACGCTCTTCCGATCTNNNCTTGGGCTCACTAGGGTTACC

AGCACACGTCTGAACTCTTTCCCTACACGACGCTCTTCCGATCTNNNCTTGGGCTCACTAGGGTTACC

AGCACACGTCTGAACTCTTTCCCTACACGACGCTCTTCCGATCTNNNCTGTAATCTCAGCTACTCGGGG

AGCACACGTCTGAACTCTTTCCCTACACGACGCTCTTCCGATCTNNNCTGTAATCTCAGCTACTCGGGG

GAGCACACGTCTGAACTCTTTCCCTACACGACGCTCTTCCGATCTNNNTGAGTGGATCAGAGAGCTC

GAGCACACGTCTGAACTCTTTCCCTACACGACGCTCTTCCGATCTNNNTGAGTGGATCAGAGAGCTC

GCACACGTCTGAACTCTTTCCCTACACGACGCTCTTCCGATCTNNNCTTAGGTGACACAGTGAGACTC

GCACACGTCTGAACTCTTTCCCTACACGACGCTCTTCCGATCTNNNCTTAGGTGACACAGTGAGACTC

AGCACACGTCTGAACTCTTTCCCTACACGACGCTCTTCCGATCTNNNAAAAAAATTAGCCGGGCGTGAT

AGCACACGTCTGAACTCTTTCCCTACACGACGCTCTTCCGATCTNNNAAAAAAATTAGCCGGGCGTGAT

AGAGCACACGTCTGAACTCTTTCCCTACACGACGCTCTTCCGATCTNNNTCAAATATCACAGCTAGGG

AGAGCACACGTCTGAACTCTTTCCCTACACGACGCTCTTCCGATCTNNNTCAAATATCACAGCTAGGG

AGAGCACACGTCTGAACTCTTTCCCTACACGACGCTCTTCCGATCTNNNCAAGAGAATTGCTTGAACC

AGAGCACACGTCTGAACTCTTTCCCTACACGACGCTCTTCCGATCTNNNCAAGAGAATTGCTTGAACC

GCACACGTCTGAACTCTTTCCCTACACGACGCTCTTCCGATCTNNNGTGAGCCAAGATTGTGCAACTG

GCACACGTCTGAACTCTTTCCCTACACGACGCTCTTCCGATCTNNNGTGAGCCAAGATTGTGCAACTG

GAGCACACGTCTGAACTCTTTCCCTACACGACGCTCTTCCGATCTNNNTGTTGGGGTCACTGATTGTA

GAGCACACGTCTGAACTCTTTCCCTACACGACGCTCTTCCGATCTNNNTGTTGGGGTCACTGATTGTA

AGCACACGTCTGAACTCTTTCCCTACACGACGCTCTTCCGATCTNNNCATGTGTCTTTGAACCCATGCA

AGCACACGTCTGAACTCTTTCCCTACACGACGCTCTTCCGATCTNNNCATGTGTCTTTGAACCCATGCA

AGCACACGTCTGAACTCTTTCCCTACACGACGCTCTTCCGATCTNNNGCAAAAGTACTTGTCTAGCCTC

AGCACACGTCTGAACTCTTTCCCTACACGACGCTCTTCCGATCTNNNGCAAAAGTACTTGTCTAGCCTC

GCACACGTCTGAACTCTTTCCCTACACGACGCTCTTCCGATCTNNNTCGAATTCTACTCAGGGACGGA

GCACACGTCTGAACTCTTTCCCTACACGACGCTCTTCCGATCTNNNTCGAATTCTACTCAGGGACGGA

GCACACGTCTGAACTCTTTCCCTACACGACGCTCTTCCGATCTNNNCACAAGTCCCCTGATACCTGTTT

GCACACGTCTGAACTCTTTCCCTACACGACGCTCTTCCGATCTNNNCACAAGTCCCCTGATACCTGTTT

GAGCACACGTCTGAACTCTTTCCCTACACGACGCTCTTCCGATCTNNNTAATGTGACAGTGGTCTGGG

GAGCACACGTCTGAACTCTTTCCCTACACGACGCTCTTCCGATCTNNNTAATGTGACAGTGGTCTGGG

GCACACGTCTGAACTCTTTCCCTACACGACGCTCTTCCGATCTNNNTCCCATTTAGGAATGACACTGCA

GCACACGTCTGAACTCTTTCCCTACACGACGCTCTTCCGATCTNNNTCCCATTTAGGAATGACACTGCA

GAGCACACGTCTGAACTCTTTCCCTACACGACGCTCTTCCGATCTNNNAAACGGAGCAGTCATTTTGGA

GAGCACACGTCTGAACTCTTTCCCTACACGACGCTCTTCCGATCTNNNAAACGGAGCAGTCATTTTGGA

AGAGCACACGTCTGAACTCTTTCCCTACACGACGCTCTTCCGATCTNNNCTGACATCTGTAATCCCAGCA

AGAGCACACGTCTGAACTCTTTCCCTACACGACGCTCTTCCGATCTNNNCTGACATCTGTAATCCCAGCA

CACACGTCTGAACTCTTTCCCTACACGACGCTCTTCCGATCTNNNCCACTCCCGGCATAATAGTGAAAC

CACACGTCTGAACTCTTTCCCTACACGACGCTCTTCCGATCTNNNCCACTCCCGGCATAATAGTGAAAC

AGCACACGTCTGAACTCTTTCCCTACACGACGCTCTTCCGATCTNNNGGTCTCTATGCACAACAGTGAG

AGCACACGTCTGAACTCTTTCCCTACACGACGCTCTTCCGATCTNNNGGTCTCTATGCACAACAGTGAG

GAGCACACGTCTGAACTCTTTCCCTACACGACGCTCTTCCGATCTNNNCAGCTCCTCTTTGTACATCTGG

GAGCACACGTCTGAACTCTTTCCCTACACGACGCTCTTCCGATCTNNNCAGCTCCTCTTTGTACATCTGG

AGCACACGTCTGAACTCTTTCCCTACACGACGCTCTTCCGATCTNNNCTCCTGAGTAGCTGGGATTACA

AGCACACGTCTGAACTCTTTCCCTACACGACGCTCTTCCGATCTNNNCTCCTGAGTAGCTGGGATTACA

GAGCACACGTCTGAACTCTTTCCCTACACGACGCTCTTCCGATCTNNNGGTAAGGGGGATTTCTGGCTA

GAGCACACGTCTGAACTCTTTCCCTACACGACGCTCTTCCGATCTNNNGGTAAGGGGGATTTCTGGCTA

AGCACACGTCTGAACTCTTTCCCTACACGACGCTCTTCCGATCTNNNAAAGAAAGAGAGAGAGGGGG

AGCACACGTCTGAACTCTTTCCCTACACGACGCTCTTCCGATCTNNNAAAGAAAGAGAGAGAGGGGG

AGCACACGTCTGAACTCTTTCCCTACACGACGCTCTTCCGATCTNNNCAGGTGGCTTTTCAATGATCTG

AGCACACGTCTGAACTCTTTCCCTACACGACGCTCTTCCGATCTNNNCAGGTGGCTTTTCAATGATCTG

GCACACGTCTGAACTCTTTCCCTACACGACGCTCTTCCGATCTNNNACACAAACCAAGCCACCACTATAC

GCACACGTCTGAACTCTTTCCCTACACGACGCTCTTCCGATCTNNNACACAAACCAAGCCACCACTATAC

GAGCACACGTCTGAACTCTTTCCCTACACGACGCTCTTCCGATCTNNNTCACTTGTCATTGTGAGACACC

GAGCACACGTCTGAACTCTTTCCCTACACGACGCTCTTCCGATCTNNNTCACTTGTCATTGTGAGACACC

GCACACGTCTGAACTCTTTCCCTACACGACGCTCTTCCGATCTNNNTCGACCATTCCAGTATCTCCTACT

GCACACGTCTGAACTCTTTCCCTACACGACGCTCTTCCGATCTNNNTCGACCATTCCAGTATCTCCTACT

AGAGCACACGTCTGAACTCTTTCCCTACACGACGCTCTTCCGATCTNNNTCTCCAGGTTACACAGCGTAG

AGAGCACACGTCTGAACTCTTTCCCTACACGACGCTCTTCCGATCTNNNTCTCCAGGTTACACAGCGTAG

AGCACACGTCTGAACTCTTTCCCTACACGACGCTCTTCCGATCTNNNAACAAGGGAGATCATGGTTCCT

AGCACACGTCTGAACTCTTTCCCTACACGACGCTCTTCCGATCTNNNAACAAGGGAGATCATGGTTCCT

GAGCACACGTCTGAACTCTTTCCCTACACGACGCTCTTCCGATCTNNNATCTGCATAGGAAAGTGCTCA

GAGCACACGTCTGAACTCTTTCCCTACACGACGCTCTTCCGATCTNNNATCTGCATAGGAAAGTGCTCA

AGCACACGTCTGAACTCTTTCCCTACACGACGCTCTTCCGATCTNNNGAGACCCATTTTGCTCTCCCTTAC

AGCACACGTCTGAACTCTTTCCCTACACGACGCTCTTCCGATCTNNNGAGACCCATTTTGCTCTCCCTTAC

GCACACGTCTGAACTCTTTCCCTACACGACGCTCTTCCGATCTNNNCTTGCAGTGAGCCAAGATGACCA

GCACACGTCTGAACTCTTTCCCTACACGACGCTCTTCCGATCTNNNCTTGCAGTGAGCCAAGATGACCA

GAGCACACGTCTGAACTCTTTCCCTACACGACGCTCTTCCGATCTNNNTTCTCACTTAGAGCTGTCCTTC

GAGCACACGTCTGAACTCTTTCCCTACACGACGCTCTTCCGATCTNNNTTCTCACTTAGAGCTGTCCTTC

AGCACACGTCTGAACTCTTTCCCTACACGACGCTCTTCCGATCTNNNTGCAACTTTTTCTTCCTTCCTTCC

AGCACACGTCTGAACTCTTTCCCTACACGACGCTCTTCCGATCTNNNTGCAACTTTTTCTTCCTTCCTTCC

AGCACACGTCTGAACTCTTTCCCTACACGACGCTCTTCCGATCTNNNTGCACATGGAAAAGCAAAAGTC

AGCACACGTCTGAACTCTTTCCCTACACGACGCTCTTCCGATCTNNNTGCACATGGAAAAGCAAAAGTC

AGCACACGTCTGAACTCTTTCCCTACACGACGCTCTTCCGATCTNNNAAAAAATTACCCAGGCATGGTG

AGCACACGTCTGAACTCTTTCCCTACACGACGCTCTTCCGATCTNNNAAAAAATTACCCAGGCATGGTG

AGCACACGTCTGAACTCTTTCCCTACACGACGCTCTTCCGATCTNNNATACACCACTTCCACTTTCACGA

AGCACACGTCTGAACTCTTTCCCTACACGACGCTCTTCCGATCTNNNATACACCACTTCCACTTTCACGA
AGCACACGTCTGAACTCTTTCCCTACACGACGCTCTTCCGATCTNNNATTTTGCACTCACTCTGTCTTGC
AGCACACGTCTGAACTCTTTCCCTACACGACGCTCTTCCGATCTNNNATTTTGCACTCACTCTGTCTTGC
AGAGCACACGTCTGAACTCTTTCCCTACACGACGCTCTTCCGATCTNNNTGGAAACAAAACGGACCAA
AGAGCACACGTCTGAACTCTTTCCCTACACGACGCTCTTCCGATCTNNNTGGAAACAAAACGGACCAA
AGCACACGTCTGAACTCTTTCCCTACACGACGCTCTTCCGATCTNNNAGATTGGGGCACTCTACGAAT
AGCACACGTCTGAACTCTTTCCCTACACGACGCTCTTCCGATCTNNNAGATTGGGGCACTCTACGAAT
AGCACACGTCTGAACTCTTTCCCTACACGACGCTCTTCCGATCTNNNCATGATGCTGGCCTCATGAATG
AGCACACGTCTGAACTCTTTCCCTACACGACGCTCTTCCGATCTNNNCATGATGCTGGCCTCATGAATG
AGCACACGTCTGAACTCTTTCCCTACACGACGCTCTTCCGATCTNNNATCGTGTCACCCACAACTTCTCT
AGCACACGTCTGAACTCTTTCCCTACACGACGCTCTTCCGATCTNNNATCGTGTCACCCACAACTTCTCT
GCACACGTCTGAACTCTTTCCCTACACGACGCTCTTCCGATCTNNNAAGATGACGCCTGTGTCCAAAG
GCACACGTCTGAACTCTTTCCCTACACGACGCTCTTCCGATCTNNNAAGATGACGCCTGTGTCCAAAG
GCACACGTCTGAACTCTTTCCCTACACGACGCTCTTCCGATCTNNNCCAAAGAACACAGTAAAGCCCAG
GCACACGTCTGAACTCTTTCCCTACACGACGCTCTTCCGATCTNNNCCAAAGAACACAGTAAAGCCCAG
AAGAGCACACGTCTGAACTCTTTCCCTACACGACGCTCTTCCGATCTNNNTAAAGAGACAGCTGCACT
AAGAGCACACGTCTGAACTCTTTCCCTACACGACGCTCTTCCGATCTNNNTAAAGAGACAGCTGCACT
AGCACACGTCTGAACTCTTTCCCTACACGACGCTCTTCCGATCTNNNTTTTTGAGTCGGTGTCTCACTC
AGCACACGTCTGAACTCTTTCCCTACACGACGCTCTTCCGATCTNNNTTTTTGAGTCGGTGTCTCACTC
GAGCACACGTCTGAACTCTTTCCCTACACGACGCTCTTCCGATCTNNNAGGGTGTTGCTTCCTCTACCAT
GAGCACACGTCTGAACTCTTTCCCTACACGACGCTCTTCCGATCTNNNAGGGTGTTGCTTCCTCTACCAT
AGCACACGTCTGAACTCTTTCCCTACACGACGCTCTTCCGATCTNNNCATAACTGTGTGAGCTGGCTT
AGCACACGTCTGAACTCTTTCCCTACACGACGCTCTTCCGATCTNNNCATAACTGTGTGAGCTGGCTT
AGCACACGTCTGAACTCTTTCCCTACACGACGCTCTTCCGATCTNNNAGCACAAAGGAATTTCTTGGT
AGCACACGTCTGAACTCTTTCCCTACACGACGCTCTTCCGATCTNNNAGCACAAAGGAATTTCTTGGT
GCACACGTCTGAACTCTTTCCCTACACGACGCTCTTCCGATCTNNNTCACAAGAACAGCATGCAGAAA
GCACACGTCTGAACTCTTTCCCTACACGACGCTCTTCCGATCTNNNTCACAAGAACAGCATGCAGAAA
GCACACGTCTGAACTCTTTCCCTACACGACGCTCTTCCGATCTNNNATAGTGAACTCCTGGCCTCTAG
GCACACGTCTGAACTCTTTCCCTACACGACGCTCTTCCGATCTNNNATAGTGAACTCCTGGCCTCTAG
AGCACACGTCTGAACTCTTTCCCTACACGACGCTCTTCCGATCTNNNATTTCACCCATAATCCCAGCAC
AGCACACGTCTGAACTCTTTCCCTACACGACGCTCTTCCGATCTNNNATTTCACCCATAATCCCAGCAC
GAGCACACGTCTGAACTCTTTCCCTACACGACGCTCTTCCGATCTNNNGGACGAGTGGGGTAATAAG
GAGCACACGTCTGAACTCTTTCCCTACACGACGCTCTTCCGATCTNNNGGACGAGTGGGGTAATAAG
AGCACACGTCTGAACTCTTTCCCTACACGACGCTCTTCCGATCTNNNGTCAAGAGGCTGAGGAAGAA
AGCACACGTCTGAACTCTTTCCCTACACGACGCTCTTCCGATCTNNNGTCAAGAGGCTGAGGAAGAA
AGCACACGTCTGAACTCTTTCCCTACACGACGCTCTTCCGATCTNNNAGAGCTAACATTTACTGCATG
AGCACACGTCTGAACTCTTTCCCTACACGACGCTCTTCCGATCTNNNAGAGCTAACATTTACTGCATG
AGCACACGTCTGAACTCTTTCCCTACACGACGCTCTTCCGATCTNNNCACTCAGAATACGGGCACACAT
AGCACACGTCTGAACTCTTTCCCTACACGACGCTCTTCCGATCTNNNCACTCAGAATACGGGCACACAT
AGCACACGTCTGAACTCTTTCCCTACACGACGCTCTTCCGATCTNNNAGAAAAAGGCAGAGGAAAG
AGCACACGTCTGAACTCTTTCCCTACACGACGCTCTTCCGATCTNNNAGAAAAAGGCAGAGGAAAG
AGCACACGTCTGAACTCTTTCCCTACACGACGCTCTTCCGATCTNNNCTCTGAAGAGTGCACATTGTT
AGCACACGTCTGAACTCTTTCCCTACACGACGCTCTTCCGATCTNNNCTCTGAAGAGTGCACATTGTT

AAGAGCACACGTCTGAACTCTTTCCCTACACGACGCTCTTCCGATCTNNNACAAAAATTAGCCAGGTG
AAGAGCACACGTCTGAACTCTTTCCCTACACGACGCTCTTCCGATCTNNNACAAAAATTAGCCAGGTG
GCACACGTCTGAACTCTTTCCCTACACGACGCTCTTCCGATCTNNNAGTGAATGCACCCTTTCTGAGAT
GCACACGTCTGAACTCTTTCCCTACACGACGCTCTTCCGATCTNNNAGTGAATGCACCCTTTCTGAGAT
AGCACACGTCTGAACTCTTTCCCTACACGACGCTCTTCCGATCTNNNGTCCAGCCAGTATTGAGGCTTA
AGCACACGTCTGAACTCTTTCCCTACACGACGCTCTTCCGATCTNNNGTCCAGCCAGTATTGAGGCTTA
AGCACACGTCTGAACTCTTTCCCTACACGACGCTCTTCCGATCTNNNTCCTACCTTTCCCATTTATAAAT
AGCACACGTCTGAACTCTTTCCCTACACGACGCTCTTCCGATCTNNNTCCTACCTTTCCCATTTATAAAT
AGCACACGTCTGAACTCTTTCCCTACACGACGCTCTTCCGATCTNNNTGAGAATCACTTGAACCCAGGA
AGCACACGTCTGAACTCTTTCCCTACACGACGCTCTTCCGATCTNNNTGAGAATCACTTGAACCCAGGA
AGCACACGTCTGAACTCTTTCCCTACACGACGCTCTTCCGATCTNNNCCAAGTAGCTGAGACTACAGG
AGCACACGTCTGAACTCTTTCCCTACACGACGCTCTTCCGATCTNNNCCAAGTAGCTGAGACTACAGG
AGCACACGTCTGAACTCTTTCCCTACACGACGCTCTTCCGATCTNNNTGCATACCTGTAATCCCAGCTA
AGCACACGTCTGAACTCTTTCCCTACACGACGCTCTTCCGATCTNNNTGCATACCTGTAATCCCAGCTA
AGCACACGTCTGAACTCTTTCCCTACACGACGCTCTTCCGATCTNNNCTAGGAGTTTCCTCACAGAGCT
AGCACACGTCTGAACTCTTTCCCTACACGACGCTCTTCCGATCTNNNCTAGGAGTTTCCTCACAGAGCT
GAGCACACGTCTGAACTCTTTCCCTACACGACGCTCTTCCGATCTNNNCAACTCCTGCATTCGACTGTT
GAGCACACGTCTGAACTCTTTCCCTACACGACGCTCTTCCGATCTNNNCAACTCCTGCATTCGACTGTT
GAGCACACGTCTGAACTCTTTCCCTACACGACGCTCTTCCGATCTNNNCTGAGGCAGGAGAATCACTT
GAGCACACGTCTGAACTCTTTCCCTACACGACGCTCTTCCGATCTNNNCTGAGGCAGGAGAATCACTT
AGCACACGTCTGAACTCTTTCCCTACACGACGCTCTTCCGATCTNNNGCCAAACGGGGAAGTTCTATCT
AGCACACGTCTGAACTCTTTCCCTACACGACGCTCTTCCGATCTNNNGCCAAACGGGGAAGTTCTATCT
CACACGTCTGAACTCTTTCCCTACACGACGCTCTTCCGATCTNNNGTGATACTGGAAACACACACAGG
CACACGTCTGAACTCTTTCCCTACACGACGCTCTTCCGATCTNNNGTGATACTGGAAACACACACAGG
GCACACGTCTGAACTCTTTCCCTACACGACGCTCTTCCGATCTNNNCCTCTTACTCATTCATCATCTAAC
GCACACGTCTGAACTCTTTCCCTACACGACGCTCTTCCGATCTNNNCCTCTTACTCATTCATCATCTAAC
GAGCACACGTCTGAACTCTTTCCCTACACGACGCTCTTCCGATCTNNNCTTACACCTGTAATCCCAGCA
GAGCACACGTCTGAACTCTTTCCCTACACGACGCTCTTCCGATCTNNNCTTACACCTGTAATCCCAGCA
AGAGCACACGTCTGAACTCTTTCCCTACACGACGCTCTTCCGATCTNNNCAAAAAAGGAGGCTCAAGC
AGAGCACACGTCTGAACTCTTTCCCTACACGACGCTCTTCCGATCTNNNCAAAAAAGGAGGCTCAAGC
AGCACACGTCTGAACTCTTTCCCTACACGACGCTCTTCCGATCTNNNCGACAAGAGTAAGACCCTGTCT
AGCACACGTCTGAACTCTTTCCCTACACGACGCTCTTCCGATCTNNNCGACAAGAGTAAGACCCTGTCT
GAGCACACGTCTGAACTCTTTCCCTACACGACGCTCTTCCGATCTNNNGTGGTCTGTGTATGCATGCA
GAGCACACGTCTGAACTCTTTCCCTACACGACGCTCTTCCGATCTNNNGTGGTCTGTGTATGCATGCA
AGCACACGTCTGAACTCTTTCCCTACACGACGCTCTTCCGATCTNNNAGCAAAGAGAAAACAAGAGTG
AGCACACGTCTGAACTCTTTCCCTACACGACGCTCTTCCGATCTNNNAGCAAAGAGAAAACAAGAGTG
AGCACACGTCTGAACTCTTTCCCTACACGACGCTCTTCCGATCTNNNTCTCTTCCACCTTGGTCTTCCTA
AGCACACGTCTGAACTCTTTCCCTACACGACGCTCTTCCGATCTNNNTCTCTTCCACCTTGGTCTTCCTA
GCACACGTCTGAACTCTTTCCCTACACGACGCTCTTCCGATCTNNNGATCGTGTCACTATACTCCAGCC
GCACACGTCTGAACTCTTTCCCTACACGACGCTCTTCCGATCTNNNGATCGTGTCACTATACTCCAGCC
AGAGCACACGTCTGAACTCTTTCCCTACACGACGCTCTTCCGATCTNNNAAAGAGAGAGACAGACAG
AGAGCACACGTCTGAACTCTTTCCCTACACGACGCTCTTCCGATCTNNNAAAGAGAGAGACAGACAG
AGCACACGTCTGAACTCTTTCCCTACACGACGCTCTTCCGATCTNNNCATTTAGATCCAGATACCGAAT

AGCACACGTCTGAACTCTTTCCCTACACGACGCTCTTCCGATCTNNNCATTTAGATCCAGATACCGAATA

AGCACACGTCTGAACTCTTTCCCTACACGACGCTCTTCCGATCTNNNCTCTCTCTCTGCTGGTCTTTC

AGCACACGTCTGAACTCTTTCCCTACACGACGCTCTTCCGATCTNNNCTCTCTCTCTGCTGGTCTTTC

GAGCACACGTCTGAACTCTTTCCCTACACGACGCTCTTCCGATCTNNNTCTTTTGTCACTTAGGTTAAA

GAGCACACGTCTGAACTCTTTCCCTACACGACGCTCTTCCGATCTNNNTCTTTTGTCACTTAGGTTAAA

AGCACACGTCTGAACTCTTTCCCTACACGACGCTCTTCCGATCTNNNTGAGGCAGAAGAATTGCTTGA

AGCACACGTCTGAACTCTTTCCCTACACGACGCTCTTCCGATCTNNNTGAGGCAGAAGAATTGCTTGA

GAGCACACGTCTGAACTCTTTCCCTACACGACGCTCTTCCGATCTNNNAATGAGCCGAGATCATGTCA

GAGCACACGTCTGAACTCTTTCCCTACACGACGCTCTTCCGATCTNNNAATGAGCCGAGATCATGTCA

AGCACACGTCTGAACTCTTTCCCTACACGACGCTCTTCCGATCTNNNAGAGTTATCCCTCCTTTTGGCAA

AGCACACGTCTGAACTCTTTCCCTACACGACGCTCTTCCGATCTNNNAGAGTTATCCCTCCTTTTGGCAA

GCACACGTCTGAACTCTTTCCCTACACGACGCTCTTCCGATCTNNNCTGTAGTCCCAGCTACTCATGAG

GCACACGTCTGAACTCTTTCCCTACACGACGCTCTTCCGATCTNNNCTGTAGTCCCAGCTACTCATGAG

GCACACGTCTGAACTCTTTCCCTACACGACGCTCTTCCGATCTNNNGTACCAGTTTTCAAGGGGAATG

GCACACGTCTGAACTCTTTCCCTACACGACGCTCTTCCGATCTNNNGTACCAGTTTTCAAGGGGAATG

AGCACACGTCTGAACTCTTTCCCTACACGACGCTCTTCCGATCTNNNAATGGTCCCTAACACTTCCTCAC

AGCACACGTCTGAACTCTTTCCCTACACGACGCTCTTCCGATCTNNNAATGGTCCCTAACACTTCCTCAC

AGCACACGTCTGAACTCTTTCCCTACACGACGCTCTTCCGATCTNNNTTCTTGCATAACAGCACCTGCA

AGCACACGTCTGAACTCTTTCCCTACACGACGCTCTTCCGATCTNNNTTCTTGCATAACAGCACCTGCA

AGCACACGTCTGAACTCTTTCCCTACACGACGCTCTTCCGATCTNNNATTGGATTGAATAACACTGGC

AGCACACGTCTGAACTCTTTCCCTACACGACGCTCTTCCGATCTNNNATTGGATTGAATAACACTGGC

GCACACGTCTGAACTCTTTCCCTACACGACGCTCTTCCGATCTNNNGCATCTTCTCTCACAGGTCCTAC

GCACACGTCTGAACTCTTTCCCTACACGACGCTCTTCCGATCTNNNGCATCTTCTCTCACAGGTCCTAC

GAGCACACGTCTGAACTCTTTCCCTACACGACGCTCTTCCGATCTNNNTGTCTATGCAGGACTTTGTTC

GAGCACACGTCTGAACTCTTTCCCTACACGACGCTCTTCCGATCTNNNTGTCTATGCAGGACTTTGTTC

GCACACGTCTGAACTCTTTCCCTACACGACGCTCTTCCGATCTNNNTGGGAAGTGAGGAGCGTCTCCAT

GCACACGTCTGAACTCTTTCCCTACACGACGCTCTTCCGATCTNNNTGGGAAGTGAGGAGCGTCTCCAT

GAGCACACGTCTGAACTCTTTCCCTACACGACGCTCTTCCGATCTNNNACTTGGTATACTCACTCCTCCC

GAGCACACGTCTGAACTCTTTCCCTACACGACGCTCTTCCGATCTNNNACTTGGTATACTCACTCCTCCC

AGCACACGTCTGAACTCTTTCCCTACACGACGCTCTTCCGATCTNNNCTCTATTCCTACCACTCATTAAG

AGCACACGTCTGAACTCTTTCCCTACACGACGCTCTTCCGATCTNNNCTCTATTCCTACCACTCATTAAG

GAGCACACGTCTGAACTCTTTCCCTACACGACGCTCTTCCGATCTNNNGAGGTAGGAGAATAGCGTGA

GAGCACACGTCTGAACTCTTTCCCTACACGACGCTCTTCCGATCTNNNGAGGTAGGAGAATAGCGTGA

GAGCACACGTCTGAACTCTTTCCCTACACGACGCTCTTCCGATCTNNNCTGAGGTGGGAGAATCACTT

GAGCACACGTCTGAACTCTTTCCCTACACGACGCTCTTCCGATCTNNNCTGAGGTGGGAGAATCACTT

GCACACGTCTGAACTCTTTCCCTACACGACGCTCTTCCGATCTNNNGTATATGGGGTGGAGTTAGCC

GCACACGTCTGAACTCTTTCCCTACACGACGCTCTTCCGATCTNNNGTATATGGGGTGGAGTTAGCC

GCACACGTCTGAACTCTTTCCCTACACGACGCTCTTCCGATCTNNNATTTAGAAGCCAGTTGAGCCACT

GCACACGTCTGAACTCTTTCCCTACACGACGCTCTTCCGATCTNNNATTTAGAAGCCAGTTGAGCCACT

CACACGTCTGAACTCTTTCCCTACACGACGCTCTTCCGATCTNNNGCTGGAGAATTGCTTGAACCTGC

CACACGTCTGAACTCTTTCCCTACACGACGCTCTTCCGATCTNNNGCTGGAGAATTGCTTGAACCTGC

GAGCACACGTCTGAACTCTTTCCCTACACGACGCTCTTCCGATCTNNNCGCTGTGTTTTGGAAAACTGA

GAGCACACGTCTGAACTCTTTCCCTACACGACGCTCTTCCGATCTNNNCGCTGTGTTTTGGAAAACTGA

GAGCACACGTCTGAACTCTTTCCCTACACGACGCTCTTCCGATCTNNNAAGGAGATGCAAAAACAATGC
GAGCACACGTCTGAACTCTTTCCCTACACGACGCTCTTCCGATCTNNNAAGGAGATGCAAAAACAATGC
AGCACACGTCTGAACTCTTTCCCTACACGACGCTCTTCCGATCTNNNGAGGATCACTTGAACCCAGGA
AGCACACGTCTGAACTCTTTCCCTACACGACGCTCTTCCGATCTNNNGAGGATCACTTGAACCCAGGA
GAGCACACGTCTGAACTCTTTCCCTACACGACGCTCTTCCGATCTNNNATGCCATCTAGGACTTCCATAC
GAGCACACGTCTGAACTCTTTCCCTACACGACGCTCTTCCGATCTNNNATGCCATCTAGGACTTCCATAC
AGCACACGTCTGAACTCTTTCCCTACACGACGCTCTTCCGATCTNNNATATGTGTGTGTGTGTGTGTGT
AGCACACGTCTGAACTCTTTCCCTACACGACGCTCTTCCGATCTNNNATATGTGTGTGTGTGTGTGTGT
GAGCACACGTCTGAACTCTTTCCCTACACGACGCTCTTCCGATCTNNNGACGTTGAGTCTCCAAATTGCC
GAGCACACGTCTGAACTCTTTCCCTACACGACGCTCTTCCGATCTNNNGACGTTGAGTCTCCAAATTGCC
AGAGCACACGTCTGAACTCTTTCCCTACACGACGCTCTTCCGATCTNNNAATACAAAAAATTAGCCGGC
AGAGCACACGTCTGAACTCTTTCCCTACACGACGCTCTTCCGATCTNNNAATACAAAAAATTAGCCGGC
GCACACGTCTGAACTCTTTCCCTACACGACGCTCTTCCGATCTNNNTCTCTCTCTCTCTTTCCCTCCCAT
GCACACGTCTGAACTCTTTCCCTACACGACGCTCTTCCGATCTNNNTCTCTCTCTCTCTTTCCCTCCCAT
GAGCACACGTCTGAACTCTTTCCCTACACGACGCTCTTCCGATCTNNNCCACCACGCCCAACTAATTATTT
GAGCACACGTCTGAACTCTTTCCCTACACGACGCTCTTCCGATCTNNNCCACCACGCCCAACTAATTATTT
AGCACACGTCTGAACTCTTTCCCTACACGACGCTCTTCCGATCTNNNGATCTCGGCTAACTGATACGTC
AGCACACGTCTGAACTCTTTCCCTACACGACGCTCTTCCGATCTNNNGATCTCGGCTAACTGATACGTC
GCACACGTCTGAACTCTTTCCCTACACGACGCTCTTCCGATCTNNNAAAGGCTGTCTTCAAGTGCATTC
GCACACGTCTGAACTCTTTCCCTACACGACGCTCTTCCGATCTNNNAAAGGCTGTCTTCAAGTGCATTC
GAGCACACGTCTGAACTCTTTCCCTACACGACGCTCTTCCGATCTNNNCGACGCCCAGCTAATTTTTGT
GAGCACACGTCTGAACTCTTTCCCTACACGACGCTCTTCCGATCTNNNCGACGCCCAGCTAATTTTTGT
GAGCACACGTCTGAACTCTTTCCCTACACGACGCTCTTCCGATCTNNNAAAAAAAGACCAGTCTGGCCA
GAGCACACGTCTGAACTCTTTCCCTACACGACGCTCTTCCGATCTNNNAAAAAAAGACCAGTCTGGCCA
GCACACGTCTGAACTCTTTCCCTACACGACGCTCTTCCGATCTNNNGAGTGGACTGGTAAAACTTTCAT
GCACACGTCTGAACTCTTTCCCTACACGACGCTCTTCCGATCTNNNGAGTGGACTGGTAAAACTTTCAT
GAGCACACGTCTGAACTCTTTCCCTACACGACGCTCTTCCGATCTNNNAACCTGTAGTTCCAGCTACTTG
GAGCACACGTCTGAACTCTTTCCCTACACGACGCTCTTCCGATCTNNNAACCTGTAGTTCCAGCTACTTG
GAGCACACGTCTGAACTCTTTCCCTACACGACGCTCTTCCGATCTNNNCATGATCGTGCCACTGTACTCC
GAGCACACGTCTGAACTCTTTCCCTACACGACGCTCTTCCGATCTNNNCATGATCGTGCCACTGTACTCC
AGAGCACACGTCTGAACTCTTTCCCTACACGACGCTCTTCCGATCTNNNTAGGGATGCCAAAAGAGCCT
AGAGCACACGTCTGAACTCTTTCCCTACACGACGCTCTTCCGATCTNNNTAGGGATGCCAAAAGAGCCT
GCACACGTCTGAACTCTTTCCCTACACGACGCTCTTCCGATCTNNNAAGAGTACAAGGCAGGTTTAGG
GCACACGTCTGAACTCTTTCCCTACACGACGCTCTTCCGATCTNNNAAGAGTACAAGGCAGGTTTAGG
GCACACGTCTGAACTCTTTCCCTACACGACGCTCTTCCGATCTNNNCGTGTGGTAATCTGAGCTGCATC
GCACACGTCTGAACTCTTTCCCTACACGACGCTCTTCCGATCTNNNCGTGTGGTAATCTGAGCTGCATC
GAGCACACGTCTGAACTCTTTCCCTACACGACGCTCTTCCGATCTNNNCCAGCTATTTGGGAAGATGAG
GAGCACACGTCTGAACTCTTTCCCTACACGACGCTCTTCCGATCTNNNCCAGCTATTTGGGAAGATGAG
AGCACACGTCTGAACTCTTTCCCTACACGACGCTCTTCCGATCTNNNGAAGTTGGAGGAAGCCAGAAA
AGCACACGTCTGAACTCTTTCCCTACACGACGCTCTTCCGATCTNNNGAAGTTGGAGGAAGCCAGAAA
GAAGAGCACACGTCTGAACTCTTTCCCTACACGACGCTCTTCCGATCTNNNCCATTGTGTACCATTGTC
GAAGAGCACACGTCTGAACTCTTTCCCTACACGACGCTCTTCCGATCTNNNCCATTGTGTACCATTGTC
GAGCACACGTCTGAACTCTTTCCCTACACGACGCTCTTCCGATCTNNNCGTTTTGGTGGGAGTTCTCA

GAGCACACGTCTGAACTCTTTCCCTACACGACGCTCTTCCGATCTNNNCGTTTTTGGTGGGAGTTCTCA
AGCACACGTCTGAACTCTTTCCCTACACGACGCTCTTCCGATCTNNNAGTTCATGCATGTAAAGGTGTT
AGCACACGTCTGAACTCTTTCCCTACACGACGCTCTTCCGATCTNNNAGTTCATGCATGTAAAGGTGTT
AGAGCACACGTCTGAACTCTTTCCCTACACGACGCTCTTCCGATCTNNNCAGACATACTTCTCTCCACCCA
AGAGCACACGTCTGAACTCTTTCCCTACACGACGCTCTTCCGATCTNNNCAGACATACTTCTCTCCACCCA
GAGCACACGTCTGAACTCTTTCCCTACACGACGCTCTTCCGATCTNNNCGTGTTAGTCAGGATGGTCTT
GAGCACACGTCTGAACTCTTTCCCTACACGACGCTCTTCCGATCTNNNCGTGTTAGTCAGGATGGTCTT
GAGCACACGTCTGAACTCTTTCCCTACACGACGCTCTTCCGATCTNNNTTCTCTCTCTCTCTCTCTCATG
GAGCACACGTCTGAACTCTTTCCCTACACGACGCTCTTCCGATCTNNNTTCTCTCTCTCTCTCTCTCATG
GCACACGTCTGAACTCTTTCCCTACACGACGCTCTTCCGATCTNNNTACTGACAGACTGGTGCTGGTAA
GCACACGTCTGAACTCTTTCCCTACACGACGCTCTTCCGATCTNNNTACTGACAGACTGGTGCTGGTAA
GAGCACACGTCTGAACTCTTTCCCTACACGACGCTCTTCCGATCTNNNATGTGGAGAACAAGGCAAGA
GAGCACACGTCTGAACTCTTTCCCTACACGACGCTCTTCCGATCTNNNATGTGGAGAACAAGGCAAGA
GAGCACACGTCTGAACTCTTTCCCTACACGACGCTCTTCCGATCTNNNCAGATCTGTGGAAGAAAAAGC
GAGCACACGTCTGAACTCTTTCCCTACACGACGCTCTTCCGATCTNNNCAGATCTGTGGAAGAAAAAGC
GAGCACACGTCTGAACTCTTTCCCTACACGACGCTCTTCCGATCTNNNAAAATCACTTGAGCCTTCTGG
GAGCACACGTCTGAACTCTTTCCCTACACGACGCTCTTCCGATCTNNNAAAATCACTTGAGCCTTCTGG
GAGCACACGTCTGAACTCTTTCCCTACACGACGCTCTTCCGATCTNNNCTGAGGCAGGAGAATCACTTG
GAGCACACGTCTGAACTCTTTCCCTACACGACGCTCTTCCGATCTNNNCTGAGGCAGGAGAATCACTTG
GAGCACACGTCTGAACTCTTTCCCTACACGACGCTCTTCCGATCTNNNTTAGATGGAGTCTCACTGTGT
GAGCACACGTCTGAACTCTTTCCCTACACGACGCTCTTCCGATCTNNNTTAGATGGAGTCTCACTGTGT
AGCACACGTCTGAACTCTTTCCCTACACGACGCTCTTCCGATCTNNNTCTTCGCTAGAGAAAGACAGCA
AGCACACGTCTGAACTCTTTCCCTACACGACGCTCTTCCGATCTNNNTCTTCGCTAGAGAAAGACAGCA
AGCACACGTCTGAACTCTTTCCCTACACGACGCTCTTCCGATCTNNNAGTACCTGCTTTCATTCTAGGG
AGCACACGTCTGAACTCTTTCCCTACACGACGCTCTTCCGATCTNNNAGTACCTGCTTTCATTCTAGGG
AGAGCACACGTCTGAACTCTTTCCCTACACGACGCTCTTCCGATCTNNNGCATGTAGTCCTGGCTATTC
AGAGCACACGTCTGAACTCTTTCCCTACACGACGCTCTTCCGATCTNNNGCATGTAGTCCTGGCTATTC
AGAGCACACGTCTGAACTCTTTCCCTACACGACGCTCTTCCGATCTNNNGAGGATCACTTGAACCTGG
AGAGCACACGTCTGAACTCTTTCCCTACACGACGCTCTTCCGATCTNNNGAGGATCACTTGAACCTGG
GAGCACACGTCTGAACTCTTTCCCTACACGACGCTCTTCCGATCTNNNAGTTAGCCAGGTGTGGTAAT
GAGCACACGTCTGAACTCTTTCCCTACACGACGCTCTTCCGATCTNNNAGTTAGCCAGGTGTGGTAAT
GCACACGTCTGAACTCTTTCCCTACACGACGCTCTTCCGATCTNNNCACCTTCTTCTGTGAAACTGCATCC
GCACACGTCTGAACTCTTTCCCTACACGACGCTCTTCCGATCTNNNCACCTTCTTCTGTGAAACTGCATCC
GAGCACACGTCTGAACTCTTTCCCTACACGACGCTCTTCCGATCTNNNGTCAGGAGTTCTAGACCAAG
GAGCACACGTCTGAACTCTTTCCCTACACGACGCTCTTCCGATCTNNNGTCAGGAGTTCTAGACCAAG
AGCACACGTCTGAACTCTTTCCCTACACGACGCTCTTCCGATCTNNNCACAAGCCCATGCAGATTTCTTC
AGCACACGTCTGAACTCTTTCCCTACACGACGCTCTTCCGATCTNNNCACAAGCCCATGCAGATTTCTTC
AGCACACGTCTGAACTCTTTCCCTACACGACGCTCTTCCGATCTNNNCCTTTACCATCAATCCATCCCCAC
AGCACACGTCTGAACTCTTTCCCTACACGACGCTCTTCCGATCTNNNCCTTTACCATCAATCCATCCCCAC
GCACACGTCTGAACTCTTTCCCTACACGACGCTCTTCCGATCTNNNGAGAGTTGATCTTCTCCCCTCAGC
GCACACGTCTGAACTCTTTCCCTACACGACGCTCTTCCGATCTNNNGAGAGTTGATCTTCTCCCCTCAGC
GAGCACACGTCTGAACTCTTTCCCTACACGACGCTCTTCCGATCTNNNAATCTCTGCCTCCTGGATTCAA
GAGCACACGTCTGAACTCTTTCCCTACACGACGCTCTTCCGATCTNNNAATCTCTGCCTCCTGGATTCAA

GCACACGTCTGAACTCTTTCCCTACACGACGCTCTTCCGATCTNNNCTCAGTATCTCCAGCTTGCCAACC
GCACACGTCTGAACTCTTTCCCTACACGACGCTCTTCCGATCTNNNCTCAGTATCTCCAGCTTGCCAACC
AGAGCACACGTCTGAACTCTTTCCCTACACGACGCTCTTCCGATCTNNNGCTAAGAGAAGAGGGGAC
AGAGCACACGTCTGAACTCTTTCCCTACACGACGCTCTTCCGATCTNNNGCTAAGAGAAGAGGGGAC
GAGCACACGTCTGAACTCTTTCCCTACACGACGCTCTTCCGATCTNNNAGAGACGAGAGAATCACAGA
GAGCACACGTCTGAACTCTTTCCCTACACGACGCTCTTCCGATCTNNNAGAGACGAGAGAATCACAGA
GAGCACACGTCTGAACTCTTTCCCTACACGACGCTCTTCCGATCTNNNCTGCATGCGGCTTTCTTTTTTT
GAGCACACGTCTGAACTCTTTCCCTACACGACGCTCTTCCGATCTNNNCTGCATGCGGCTTTCTTTTTTT
AGAGCACACGTCTGAACTCTTTCCCTACACGACGCTCTTCCGATCTNNNGTTGCAGTGAGATGAGAT
AGAGCACACGTCTGAACTCTTTCCCTACACGACGCTCTTCCGATCTNNNGTTGCAGTGAGATGAGAT
AGCACACGTCTGAACTCTTTCCCTACACGACGCTCTTCCGATCTNNNGGGACTTTCTGTGGACAATCT
AGCACACGTCTGAACTCTTTCCCTACACGACGCTCTTCCGATCTNNNGGGACTTTCTGTGGACAATCT
AGCACACGTCTGAACTCTTTCCCTACACGACGCTCTTCCGATCTNNNCCTGCATTGCTACTCTTTTCTCCC
AGCACACGTCTGAACTCTTTCCCTACACGACGCTCTTCCGATCTNNNCCTGCATTGCTACTCTTTTCTCCC
AGCACACGTCTGAACTCTTTCCCTACACGACGCTCTTCCGATCTNNNTGCTAAATGCCCAAAGTGAAG
AGCACACGTCTGAACTCTTTCCCTACACGACGCTCTTCCGATCTNNNTGCTAAATGCCCAAAGTGAAG
GAGCACACGTCTGAACTCTTTCCCTACACGACGCTCTTCCGATCTNNNCCATTTCCCATGAATGATCAG
GAGCACACGTCTGAACTCTTTCCCTACACGACGCTCTTCCGATCTNNNCCATTTCCCATGAATGATCAG
AGCACACGTCTGAACTCTTTCCCTACACGACGCTCTTCCGATCTNNNGTGGTGTCACCTACTTGGGAGC
AGCACACGTCTGAACTCTTTCCCTACACGACGCTCTTCCGATCTNNNGTGGTGTCACCTACTTGGGAGC
AGCACACGTCTGAACTCTTTCCCTACACGACGCTCTTCCGATCTNNNTGAGAGTCCCTCTTTTCCACTTG
AGCACACGTCTGAACTCTTTCCCTACACGACGCTCTTCCGATCTNNNTGAGAGTCCCTCTTTTCCACTTG
GAGCACACGTCTGAACTCTTTCCCTACACGACGCTCTTCCGATCTNNNCCCTCAATATTGAATGGCCTCC
GAGCACACGTCTGAACTCTTTCCCTACACGACGCTCTTCCGATCTNNNCCCTCAATATTGAATGGCCTCC
AGCACACGTCTGAACTCTTTCCCTACACGACGCTCTTCCGATCTNNNCCTCTTTGTGCCTCTGGTAGAAT
AGCACACGTCTGAACTCTTTCCCTACACGACGCTCTTCCGATCTNNNCCTCTTTGTGCCTCTGGTAGAAT
GCACACGTCTGAACTCTTTCCCTACACGACGCTCTTCCGATCTNNNACCAAATTCTACGAGCAGGACTT
GCACACGTCTGAACTCTTTCCCTACACGACGCTCTTCCGATCTNNNACCAAATTCTACGAGCAGGACTT
GAGCACACGTCTGAACTCTTTCCCTACACGACGCTCTTCCGATCTNNNAAATATGGAACCAGCCCAAAT
GAGCACACGTCTGAACTCTTTCCCTACACGACGCTCTTCCGATCTNNNAAATATGGAACCAGCCCAAAT
AGCACACGTCTGAACTCTTTCCCTACACGACGCTCTTCCGATCTNNNAGGGAAGGCAAATACAGAGTC
AGCACACGTCTGAACTCTTTCCCTACACGACGCTCTTCCGATCTNNNAGGGAAGGCAAATACAGAGTC
GAGCACACGTCTGAACTCTTTCCCTACACGACGCTCTTCCGATCTNNNATTCAAACCAGAGCAGACATC
GAGCACACGTCTGAACTCTTTCCCTACACGACGCTCTTCCGATCTNNNATTCAAACCAGAGCAGACATC
AGAGCACACGTCTGAACTCTTTCCCTACACGACGCTCTTCCGATCTNNNCATGTCTGTAGTCCCAGCTAC
AGAGCACACGTCTGAACTCTTTCCCTACACGACGCTCTTCCGATCTNNNCATGTCTGTAGTCCCAGCTAC
GAGCACACGTCTGAACTCTTTCCCTACACGACGCTCTTCCGATCTNNNAGGAGTAAAGAATTGGGTCA
GAGCACACGTCTGAACTCTTTCCCTACACGACGCTCTTCCGATCTNNNAGGAGTAAAGAATTGGGTCA
AGCACACGTCTGAACTCTTTCCCTACACGACGCTCTTCCGATCTNNNCCAAGATCACACCTCTGCACTCA
AGCACACGTCTGAACTCTTTCCCTACACGACGCTCTTCCGATCTNNNCCAAGATCACACCTCTGCACTCA
GCACACGTCTGAACTCTTTCCCTACACGACGCTCTTCCGATCTNNNTGAGCCTTTGTAAGAGGGCAATT
GCACACGTCTGAACTCTTTCCCTACACGACGCTCTTCCGATCTNNNTGAGCCTTTGTAAGAGGGCAATT
AGAGCACACGTCTGAACTCTTTCCCTACACGACGCTCTTCCGATCTNNNGGGACAAATGAATCCAGATT

AGAGCACACGTCTGAACTCTTTCCCTACACGACGCTCTTCCGATCTNNNGGGACAAATGAATCCAGATT
AGCACACGTCTGAACTCTTTCCCTACACGACGCTCTTCCGATCTNNNCTCCCTAAGCCAGTCAGAGAAAC
AGCACACGTCTGAACTCTTTCCCTACACGACGCTCTTCCGATCTNNNCTCCCTAAGCCAGTCAGAGAAAC
AGAGCACACGTCTGAACTCTTTCCCTACACGACGCTCTTCCGATCTNNNAAAGAAGCGTAATAGCTGG
AGAGCACACGTCTGAACTCTTTCCCTACACGACGCTCTTCCGATCTNNNAAAGAAGCGTAATAGCTGG
GAGCACACGTCTGAACTCTTTCCCTACACGACGCTCTTCCGATCTNNNAGACAGGAGAATTGCTTGAAC
GAGCACACGTCTGAACTCTTTCCCTACACGACGCTCTTCCGATCTNNNAGACAGGAGAATTGCTTGAAC
AGCACACGTCTGAACTCTTTCCCTACACGACGCTCTTCCGATCTNNNCAGCCTGGGCAATAGGAGTAAA
AGCACACGTCTGAACTCTTTCCCTACACGACGCTCTTCCGATCTNNNCAGCCTGGGCAATAGGAGTAAA
AGCACACGTCTGAACTCTTTCCCTACACGACGCTCTTCCGATCTNNNCACCTGTAATCCCAGCTACTCAG
AGCACACGTCTGAACTCTTTCCCTACACGACGCTCTTCCGATCTNNNCACCTGTAATCCCAGCTACTCAG
AGCACACGTCTGAACTCTTTCCCTACACGACGCTCTTCCGATCTNNNAAAACCCTGCTCCATTGTTTGC
AGCACACGTCTGAACTCTTTCCCTACACGACGCTCTTCCGATCTNNNAAAACCCTGCTCCATTGTTTGC
AGCACACGTCTGAACTCTTTCCCTACACGACGCTCTTCCGATCTNNNATCTTTGACCAACATTTCCCCTC
AGCACACGTCTGAACTCTTTCCCTACACGACGCTCTTCCGATCTNNNATCTTTGACCAACATTTCCCCTC
AGCACACGTCTGAACTCTTTCCCTACACGACGCTCTTCCGATCTNNNAAATCCTTTCTCCCCTCATGGTGC
AGCACACGTCTGAACTCTTTCCCTACACGACGCTCTTCCGATCTNNNAAATCCTTTCTCCCCTCATGGTGC
AGCACACGTCTGAACTCTTTCCCTACACGACGCTCTTCCGATCTNNNTCTCTCTCTCTCTCTCTCCCTCA
AGCACACGTCTGAACTCTTTCCCTACACGACGCTCTTCCGATCTNNNTCTCTCTCTCTCTCTCTCCCTCA
AGCACACGTCTGAACTCTTTCCCTACACGACGCTCTTCCGATCTNNNCAAATCTGTGTTTGGTGCATTG
AGCACACGTCTGAACTCTTTCCCTACACGACGCTCTTCCGATCTNNNCAAATCTGTGTTTGGTGCATTG
AAGAGCACACGTCTGAACTCTTTCCCTACACGACGCTCTTCCGATCTNNNTGGACTGAATTACACTACC
AAGAGCACACGTCTGAACTCTTTCCCTACACGACGCTCTTCCGATCTNNNTGGACTGAATTACACTACC
GAGCACACGTCTGAACTCTTTCCCTACACGACGCTCTTCCGATCTNNNTGCTACACTCAAAGTTTTGGCC
GAGCACACGTCTGAACTCTTTCCCTACACGACGCTCTTCCGATCTNNNTGCTACACTCAAAGTTTTGGCC
AGCACACGTCTGAACTCTTTCCCTACACGACGCTCTTCCGATCTNNNCACAGAAGGAACATGTTGACAC
AGCACACGTCTGAACTCTTTCCCTACACGACGCTCTTCCGATCTNNNCACAGAAGGAACATGTTGACAC
AGCACACGTCTGAACTCTTTCCCTACACGACGCTCTTCCGATCTNNNATAATTTACCCCCCAGAAGCTCCC
AGCACACGTCTGAACTCTTTCCCTACACGACGCTCTTCCGATCTNNNATAATTTACCCCCCAGAAGCTCCC
GAGCACACGTCTGAACTCTTTCCCTACACGACGCTCTTCCGATCTNNNGAACCCCACTAGGCTAACAGTA
GAGCACACGTCTGAACTCTTTCCCTACACGACGCTCTTCCGATCTNNNGAACCCCACTAGGCTAACAGTA
AGCACACGTCTGAACTCTTTCCCTACACGACGCTCTTCCGATCTNNNAAAGTGCCCTGAATTCTTTGAGC
AGCACACGTCTGAACTCTTTCCCTACACGACGCTCTTCCGATCTNNNAAAGTGCCCTGAATTCTTTGAGC
AGCACACGTCTGAACTCTTTCCCTACACGACGCTCTTCCGATCTNNNAGACACACATATCATATGCACA
AGCACACGTCTGAACTCTTTCCCTACACGACGCTCTTCCGATCTNNNAGACACACATATCATATGCACA
AGCACACGTCTGAACTCTTTCCCTACACGACGCTCTTCCGATCTNNNTTGAACAGCTTGAGGTACAGGT
AGCACACGTCTGAACTCTTTCCCTACACGACGCTCTTCCGATCTNNNTTGAACAGCTTGAGGTACAGGT
GAGCACACGTCTGAACTCTTTCCCTACACGACGCTCTTCCGATCTNNNAAAAAATTAGCCTGGTGTGGT
GAGCACACGTCTGAACTCTTTCCCTACACGACGCTCTTCCGATCTNNNAAAAAATTAGCCTGGTGTGGT
AGCACACGTCTGAACTCTTTCCCTACACGACGCTCTTCCGATCTNNNGAGGCAAGAGAATCGCTTGAA
AGCACACGTCTGAACTCTTTCCCTACACGACGCTCTTCCGATCTNNNGAGGCAAGAGAATCGCTTGAA
AGCACACGTCTGAACTCTTTCCCTACACGACGCTCTTCCGATCTNNNGAACAACAAGTAGTGCCACAGA
AGCACACGTCTGAACTCTTTCCCTACACGACGCTCTTCCGATCTNNNGAACAACAAGTAGTGCCACAGA

AGCACACGTCTGAACTCTTTCCCTACACGACGCTCTTCCGATCTNNNCATGCAATCTGAGCTCTGAATG
AGCACACGTCTGAACTCTTTCCCTACACGACGCTCTTCCGATCTNNNCATGCAATCTGAGCTCTGAATG
GCACACGTCTGAACTCTTTCCCTACACGACGCTCTTCCGATCTNNNACACCTGTAGTCCCAGCTACTAG
GCACACGTCTGAACTCTTTCCCTACACGACGCTCTTCCGATCTNNNACACCTGTAGTCCCAGCTACTAG
GAGCACACGTCTGAACTCTTTCCCTACACGACGCTCTTCCGATCTNNNAAAGCTGCAGTGAGCCATGA
GAGCACACGTCTGAACTCTTTCCCTACACGACGCTCTTCCGATCTNNNAAAGCTGCAGTGAGCCATGA
AGCACACGTCTGAACTCTTTCCCTACACGACGCTCTTCCGATCTNNNGAGGCAGGAGAATGGTGTGA
AGCACACGTCTGAACTCTTTCCCTACACGACGCTCTTCCGATCTNNNGAGGCAGGAGAATGGTGTGA
GCACACGTCTGAACTCTTTCCCTACACGACGCTCTTCCGATCTNNNTCTCTGCCTGGAATTCTCTTTCC
GCACACGTCTGAACTCTTTCCCTACACGACGCTCTTCCGATCTNNNTCTCTGCCTGGAATTCTCTTTCC
AGCACACGTCTGAACTCTTTCCCTACACGACGCTCTTCCGATCTNNNTACTGCCATGGCTATTAGTGT
AGCACACGTCTGAACTCTTTCCCTACACGACGCTCTTCCGATCTNNNTACTGCCATGGCTATTAGTGT
AGCACACGTCTGAACTCTTTCCCTACACGACGCTCTTCCGATCTNNNAGAATGTCCAAGAAACCCTAC
AGCACACGTCTGAACTCTTTCCCTACACGACGCTCTTCCGATCTNNNAGAATGTCCAAGAAACCCTAC
GAGCACACGTCTGAACTCTTTCCCTACACGACGCTCTTCCGATCTNNNCCCTTCCCGAACAAACTCTAT
GAGCACACGTCTGAACTCTTTCCCTACACGACGCTCTTCCGATCTNNNCCCTTCCCGAACAAACTCTAT
AGCACACGTCTGAACTCTTTCCCTACACGACGCTCTTCCGATCTNNNGCTTGTGTTGGCCCTTTTACTA
AGCACACGTCTGAACTCTTTCCCTACACGACGCTCTTCCGATCTNNNGCTTGTGTTGGCCCTTTTACTA
GAGCACACGTCTGAACTCTTTCCCTACACGACGCTCTTCCGATCTNNNCTGAGGCAGGAGAATCACT
GAGCACACGTCTGAACTCTTTCCCTACACGACGCTCTTCCGATCTNNNCTGAGGCAGGAGAATCACT
AGCACACGTCTGAACTCTTTCCCTACACGACGCTCTTCCGATCTNNNACATGGTGAGAACTCGTTTCT
AGCACACGTCTGAACTCTTTCCCTACACGACGCTCTTCCGATCTNNNACATGGTGAGAACTCGTTTCT
GCACACGTCTGAACTCTTTCCCTACACGACGCTCTTCCGATCTNNNACAAAGTAGCTGAGCCGACTTT
GCACACGTCTGAACTCTTTCCCTACACGACGCTCTTCCGATCTNNNACAAAGTAGCTGAGCCGACTTT
CACGTCTGAACTCTTTCCCTACACGACGCTCTTCCGATCTNNNGAGTACCCCTTCACCTGTTTACCCAT
CACGTCTGAACTCTTTCCCTACACGACGCTCTTCCGATCTNNNGAGTACCCCTTCACCTGTTTACCCAT
AGCACACGTCTGAACTCTTTCCCTACACGACGCTCTTCCGATCTNNNGCTAGAGTGCAGTGATGTGAT
AGCACACGTCTGAACTCTTTCCCTACACGACGCTCTTCCGATCTNNNGCTAGAGTGCAGTGATGTGAT
AGCACACGTCTGAACTCTTTCCCTACACGACGCTCTTCCGATCTNNNCAGCCTAGACTTCTGGACTCAA
AGCACACGTCTGAACTCTTTCCCTACACGACGCTCTTCCGATCTNNNCAGCCTAGACTTCTGGACTCAA
GCACACGTCTGAACTCTTTCCCTACACGACGCTCTTCCGATCTNNNCCTGTAGCATGCCCTCCATATAG
GCACACGTCTGAACTCTTTCCCTACACGACGCTCTTCCGATCTNNNCCTGTAGCATGCCCTCCATATAG
AGCACACGTCTGAACTCTTTCCCTACACGACGCTCTTCCGATCTNNNGTATTCCCAGCTACTTGGTAG
AGCACACGTCTGAACTCTTTCCCTACACGACGCTCTTCCGATCTNNNGTATTCCCAGCTACTTGGTAG
GAGCACACGTCTGAACTCTTTCCCTACACGACGCTCTTCCGATCTNNNTGTGTGGAATCAGAAAGAA
GAGCACACGTCTGAACTCTTTCCCTACACGACGCTCTTCCGATCTNNNTGTGTGGAATCAGAAAGAA
GAGCACACGTCTGAACTCTTTCCCTACACGACGCTCTTCCGATCTNNNGAGATTGAGCCACTGCATTC
GAGCACACGTCTGAACTCTTTCCCTACACGACGCTCTTCCGATCTNNNGAGATTGAGCCACTGCATTC
GCACACGTCTGAACTCTTTCCCTACACGACGCTCTTCCGATCTNNNTTTCGTTAGTCTAGCTAGTGGT
GCACACGTCTGAACTCTTTCCCTACACGACGCTCTTCCGATCTNNNTTTCGTTAGTCTAGCTAGTGGT
GCACACGTCTGAACTCTTTCCCTACACGACGCTCTTCCGATCTNNNGGTCACCATAGGCTACATTTAC
GCACACGTCTGAACTCTTTCCCTACACGACGCTCTTCCGATCTNNNGGTCACCATAGGCTACATTTAC
GCACACGTCTGAACTCTTTCCCTACACGACGCTCTTCCGATCTNNNGAACATGCTCATACACACCCACA

GCACACGTCTGAACTCTTTCCCTACACGACGCTCTTCCGATCTNNNGAACATGCTCATACACACCCACA

AGCACACGTCTGAACTCTTTCCCTACACGACGCTCTTCCGATCTNNNGCAAAGCAAGGTTTGTACTCA

AGCACACGTCTGAACTCTTTCCCTACACGACGCTCTTCCGATCTNNNGCAAAGCAAGGTTTGTACTCA

AGCACACGTCTGAACTCTTTCCCTACACGACGCTCTTCCGATCTNNNCTCAAGGGATGCTAAACAGTC

AGCACACGTCTGAACTCTTTCCCTACACGACGCTCTTCCGATCTNNNCTCAAGGGATGCTAAACAGTC

AGCACACGTCTGAACTCTTTCCCTACACGACGCTCTTCCGATCTNNNAATATTGCGCCACTGCACTCTA

AGCACACGTCTGAACTCTTTCCCTACACGACGCTCTTCCGATCTNNNAATATTGCGCCACTGCACTCTA

GAGCACACGTCTGAACTCTTTCCCTACACGACGCTCTTCCGATCTNNNAGGTTCAGCTGAGACATCAA

GAGCACACGTCTGAACTCTTTCCCTACACGACGCTCTTCCGATCTNNNAGGTTCAGCTGAGACATCAA

AGCACACGTCTGAACTCTTTCCCTACACGACGCTCTTCCGATCTNNNCTCTCTCTCTCACACACACA

AGCACACGTCTGAACTCTTTCCCTACACGACGCTCTTCCGATCTNNNCTCTCTCTCTCACACACACA

GCACACGTCTGAACTCTTTCCCTACACGACGCTCTTCCGATCTNNNATTCTTCACATTCCTCCCTCTGGC

GCACACGTCTGAACTCTTTCCCTACACGACGCTCTTCCGATCTNNNATTCTTCACATTCCTCCCTCTGGC

GCACACGTCTGAACTCTTTCCCTACACGACGCTCTTCCGATCTNNNCACCTGTAATCCCAGCTACTCGC

GCACACGTCTGAACTCTTTCCCTACACGACGCTCTTCCGATCTNNNCACCTGTAATCCCAGCTACTCGC

AGCACACGTCTGAACTCTTTCCCTACACGACGCTCTTCCGATCTNNNTACAAAAAATTAGCCGGGCAA

AGCACACGTCTGAACTCTTTCCCTACACGACGCTCTTCCGATCTNNNTACAAAAAATTAGCCGGGCAA

AGCACACGTCTGAACTCTTTCCCTACACGACGCTCTTCCGATCTNNNAGACTGAGGCACAAGAATCAC

AGCACACGTCTGAACTCTTTCCCTACACGACGCTCTTCCGATCTNNNAGACTGAGGCACAAGAATCAC

GAGCACACGTCTGAACTCTTTCCCTACACGACGCTCTTCCGATCTNNNGAGCTTGCAGTGAGCCAAGA

GAGCACACGTCTGAACTCTTTCCCTACACGACGCTCTTCCGATCTNNNGAGCTTGCAGTGAGCCAAGA

AGCACACGTCTGAACTCTTTCCCTACACGACGCTCTTCCGATCTNNNGACCTGGTTTCTTCAACACCAAG

AGCACACGTCTGAACTCTTTCCCTACACGACGCTCTTCCGATCTNNNGACCTGGTTTCTTCAACACCAAG

GAGCACACGTCTGAACTCTTTCCCTACACGACGCTCTTCCGATCTNNNGAGAATGGCATGAACCCGG

GAGCACACGTCTGAACTCTTTCCCTACACGACGCTCTTCCGATCTNNNGAGAATGGCATGAACCCGG

AGCACACGTCTGAACTCTTTCCCTACACGACGCTCTTCCGATCTNNNCTCTCAGCTACTCGGGACTTT

AGCACACGTCTGAACTCTTTCCCTACACGACGCTCTTCCGATCTNNNCTCTCAGCTACTCGGGACTTT

GAGCACACGTCTGAACTCTTTCCCTACACGACGCTCTTCCGATCTNNNAAACCACATCCACACAGTGAC

GAGCACACGTCTGAACTCTTTCCCTACACGACGCTCTTCCGATCTNNNAAACCACATCCACACAGTGAC

AGCACACGTCTGAACTCTTTCCCTACACGACGCTCTTCCGATCTNNNCATCATGCCAAGGTAAAGAAG

AGCACACGTCTGAACTCTTTCCCTACACGACGCTCTTCCGATCTNNNCATCATGCCAAGGTAAAGAAG

AGCACACGTCTGAACTCTTTCCCTACACGACGCTCTTCCGATCTNNNCTTATAGGGAGGAACTGGCCA

AGCACACGTCTGAACTCTTTCCCTACACGACGCTCTTCCGATCTNNNCTTATAGGGAGGAACTGGCCA

AGAGCACACGTCTGAACTCTTTCCCTACACGACGCTCTTCCGATCTNNNATACTGCCTCTTCTCCATTCT

AGAGCACACGTCTGAACTCTTTCCCTACACGACGCTCTTCCGATCTNNNATACTGCCTCTTCTCCATTCT

AGCACACGTCTGAACTCTTTCCCTACACGACGCTCTTCCGATCTNNNGCTCTTTTTGTCAGACTGTGTC

AGCACACGTCTGAACTCTTTCCCTACACGACGCTCTTCCGATCTNNNGCTCTTTTTGTCAGACTGTGTC

CACACGTCTGAACTCTTTCCCTACACGACGCTCTTCCGATCTNNNCGATGTTGTGCAGGAATTTGGTA

CACACGTCTGAACTCTTTCCCTACACGACGCTCTTCCGATCTNNNCGATGTTGTGCAGGAATTTGGTA

AGCACACGTCTGAACTCTTTCCCTACACGACGCTCTTCCGATCTNNNAGCAGTCAGATATTTGGCCATC

AGCACACGTCTGAACTCTTTCCCTACACGACGCTCTTCCGATCTNNNAGCAGTCAGATATTTGGCCATC

AGCACACGTCTGAACTCTTTCCCTACACGACGCTCTTCCGATCTNNNGCCCAGCCTTTCTTTTTCTTTCTC

GCACACGTCTGAACTCTTTCCCTACACGACGCTCTTCCGATCTNNNGCCCAGCCTTTCTTTTTCTTTCTC

AGCACACGTCTGAACTCTTTCCCTACACGACGCTCTTCCGATCTNNNTTTGAAAGAAATGGTCTCGGCT

AGCACACGTCTGAACTCTTTCCCTACACGACGCTCTTCCGATCTNNNTTTGAAAGAAATGGTCTCGGCT

AGCACACGTCTGAACTCTTTCCCTACACGACGCTCTTCCGATCTNNNTATTCCTAAAAATGGTCCCCCG

AGCACACGTCTGAACTCTTTCCCTACACGACGCTCTTCCGATCTNNNTATTCCTAAAAATGGTCCCCCG

AGCACACGTCTGAACTCTTTCCCTACACGACGCTCTTCCGATCTNNNTCTCAGCTGGATAATGTCCCCA

AGCACACGTCTGAACTCTTTCCCTACACGACGCTCTTCCGATCTNNNTCTCAGCTGGATAATGTCCCCA

AGCACACGTCTGAACTCTTTCCCTACACGACGCTCTTCCGATCTNNNCAGGGCTTCTGTCTTTCTTGAT

AGCACACGTCTGAACTCTTTCCCTACACGACGCTCTTCCGATCTNNNCAGGGCTTCTGTCTTTCTTGAT

AGCACACGTCTGAACTCTTTCCCTACACGACGCTCTTCCGATCTNNNAAAAAAATTAGCTGGGCACGG

AGCACACGTCTGAACTCTTTCCCTACACGACGCTCTTCCGATCTNNNAAAAAAATTAGCTGGGCACGG

AGAGCACACGTCTGAACTCTTTCCCTACACGACGCTCTTCCGATCTNNNGGAGGGACTGAAGATAGC

AGAGCACACGTCTGAACTCTTTCCCTACACGACGCTCTTCCGATCTNNNGGAGGGACTGAAGATAGC

GAGCACACGTCTGAACTCTTTCCCTACACGACGCTCTTCCGATCTNNNTTTCCCATTTGTCCCATGTGTT

GAGCACACGTCTGAACTCTTTCCCTACACGACGCTCTTCCGATCTNNNTTTCCCATTTGTCCCATGTGTT

AGAGCACACGTCTGAACTCTTTCCCTACACGACGCTCTTCCGATCTNNNATCACCACATTCCATTTTCTG

AGAGCACACGTCTGAACTCTTTCCCTACACGACGCTCTTCCGATCTNNNATCACCACATTCCATTTTCTG

GCACACGTCTGAACTCTTTCCCTACACGACGCTCTTCCGATCTNNNCTTTGGCCCTCTGCAAATAAAGC

GCACACGTCTGAACTCTTTCCCTACACGACGCTCTTCCGATCTNNNCTTTGGCCCTCTGCAAATAAAGC

GCACACGTCTGAACTCTTTCCCTACACGACGCTCTTCCGATCTNNNCTGAGGCAGGAGAATCACTTGA

GCACACGTCTGAACTCTTTCCCTACACGACGCTCTTCCGATCTNNNCTGAGGCAGGAGAATCACTTGA

AGCACACGTCTGAACTCTTTCCCTACACGACGCTCTTCCGATCTNNNACCCGAGGAATTCACAGCTAA

AGCACACGTCTGAACTCTTTCCCTACACGACGCTCTTCCGATCTNNNACCCGAGGAATTCACAGCTAA

GCACACGTCTGAACTCTTTCCCTACACGACGCTCTTCCGATCTNNNAGGCTGGTTTCAAGTTTTGGGA

GCACACGTCTGAACTCTTTCCCTACACGACGCTCTTCCGATCTNNNAGGCTGGTTTCAAGTTTTGGGA

GCACACGTCTGAACTCTTTCCCTACACGACGCTCTTCCGATCTNNNGGAAGCAGGAACAAAATGGTG

GCACACGTCTGAACTCTTTCCCTACACGACGCTCTTCCGATCTNNNGGAAGCAGGAACAAAATGGTG

AGCACACGTCTGAACTCTTTCCCTACACGACGCTCTTCCGATCTNNNAGAAATGCAGCTGATTTTTGTC

AGCACACGTCTGAACTCTTTCCCTACACGACGCTCTTCCGATCTNNNAGAAATGCAGCTGATTTTTGTC

GCACACGTCTGAACTCTTTCCCTACACGACGCTCTTCCGATCTNNNGTGGATCATGAGGTCAGGAGA

GCACACGTCTGAACTCTTTCCCTACACGACGCTCTTCCGATCTNNNGTGGATCATGAGGTCAGGAGA

AGCACACGTCTGAACTCTTTCCCTACACGACGCTCTTCCGATCTNNNTGGGTGACAGAGTGAGATTCT

AGCACACGTCTGAACTCTTTCCCTACACGACGCTCTTCCGATCTNNNTGGGTGACAGAGTGAGATTCT

AGCACACGTCTGAACTCTTTCCCTACACGACGCTCTTCCGATCTNNNCTGGGATTACAATTGTGAGCCA

AGCACACGTCTGAACTCTTTCCCTACACGACGCTCTTCCGATCTNNNCTGGGATTACAATTGTGAGCCA

GAGCACACGTCTGAACTCTTTCCCTACACGACGCTCTTCCGATCTNNNATGCCTGTAGTCCTAGCTACT

GAGCACACGTCTGAACTCTTTCCCTACACGACGCTCTTCCGATCTNNNATGCCTGTAGTCCTAGCTACT

AGCACACGTCTGAACTCTTTCCCTACACGACGCTCTTCCGATCTNNNTGGGGACAATAATACACCAAC

AGCACACGTCTGAACTCTTTCCCTACACGACGCTCTTCCGATCTNNNTGGGGACAATAATACACCAAC

AGCACACGTCTGAACTCTTTCCCTACACGACGCTCTTCCGATCTNNNCTTGCAGTGAGCCAAGATCAA

AGCACACGTCTGAACTCTTTCCCTACACGACGCTCTTCCGATCTNNNCTTGCAGTGAGCCAAGATCAA

AGCACACGTCTGAACTCTTTCCCTACACGACGCTCTTCCGATCTNNNTCTAAATGGCCTTCTGACTCCA

AGCACACGTCTGAACTCTTTCCCTACACGACGCTCTTCCGATCTNNNTCTAAATGGCCTTCTGACTCCA

AGCACACGTCTGAACTCTTTCCCTACACGACGCTCTTCCGATCTNNNTGATTGCTTGTCTTACAGAGCC

AGCACACGTCTGAACTCTTTCCCTACACGACGCTCTTCCGATCTNNNTGATTGCTTGTCTTACAGAGCCT
GAGCACACGTCTGAACTCTTTCCCTACACGACGCTCTTCCGATCTNNNCTGAGGCAGGAGAATGACTT
GAGCACACGTCTGAACTCTTTCCCTACACGACGCTCTTCCGATCTNNNCTGAGGCAGGAGAATGACTT
AGCACACGTCTGAACTCTTTCCCTACACGACGCTCTTCCGATCTNNNAAAATGCTGGGATTACACGCAT
AGCACACGTCTGAACTCTTTCCCTACACGACGCTCTTCCGATCTNNNAAAATGCTGGGATTACACGCAT
AGCACACGTCTGAACTCTTTCCCTACACGACGCTCTTCCGATCTNNNCAGGAAAATGGCGTGAACCTGC
AGCACACGTCTGAACTCTTTCCCTACACGACGCTCTTCCGATCTNNNCAGGAAAATGGCGTGAACCTGC
GAGCACACGTCTGAACTCTTTCCCTACACGACGCTCTTCCGATCTNNNAGCTCCAGGAACTCCAGTAAA
GAGCACACGTCTGAACTCTTTCCCTACACGACGCTCTTCCGATCTNNNAGCTCCAGGAACTCCAGTAAA
AGCACACGTCTGAACTCTTTCCCTACACGACGCTCTTCCGATCTNNNATACAAAAATTAGCTGGGCGTG
AGCACACGTCTGAACTCTTTCCCTACACGACGCTCTTCCGATCTNNNATACAAAAATTAGCTGGGCGTG
AGCACACGTCTGAACTCTTTCCCTACACGACGCTCTTCCGATCTNNNAGGTGTCACAGGAGGAGATCTA
AGCACACGTCTGAACTCTTTCCCTACACGACGCTCTTCCGATCTNNNAGGTGTCACAGGAGGAGATCTA
AGCACACGTCTGAACTCTTTCCCTACACGACGCTCTTCCGATCTNNNTGTTGAGGATTTCTGCACCCATA
AGCACACGTCTGAACTCTTTCCCTACACGACGCTCTTCCGATCTNNNTGTTGAGGATTTCTGCACCCATA
GAGCACACGTCTGAACTCTTTCCCTACACGACGCTCTTCCGATCTNNNGCTTTGAATTTTTTCCAAGAC
GAGCACACGTCTGAACTCTTTCCCTACACGACGCTCTTCCGATCTNNNGCTTTGAATTTTTTCCAAGAC
AGCACACGTCTGAACTCTTTCCCTACACGACGCTCTTCCGATCTNNNCAGGAAAATCGCTTGAACCTGG
AGCACACGTCTGAACTCTTTCCCTACACGACGCTCTTCCGATCTNNNCAGGAAAATCGCTTGAACCTGG
AGAGCACACGTCTGAACTCTTTCCCTACACGACGCTCTTCCGATCTNNNTCTGATAGGAACTTGCATT
AGAGCACACGTCTGAACTCTTTCCCTACACGACGCTCTTCCGATCTNNNTCTGATAGGAACTTGCATT
AGCACACGTCTGAACTCTTTCCCTACACGACGCTCTTCCGATCTNNNAAGTAGCTGAGACTACATTGCG
AGCACACGTCTGAACTCTTTCCCTACACGACGCTCTTCCGATCTNNNAAGTAGCTGAGACTACATTGCG
AGCACACGTCTGAACTCTTTCCCTACACGACGCTCTTCCGATCTNNNATGAGCATAGAAGCCCGGATA
AGCACACGTCTGAACTCTTTCCCTACACGACGCTCTTCCGATCTNNNATGAGCATAGAAGCCCGGATA
AGCACACGTCTGAACTCTTTCCCTACACGACGCTCTTCCGATCTNNNGTGTAGTGATGGATGCCTGTA
AGCACACGTCTGAACTCTTTCCCTACACGACGCTCTTCCGATCTNNNGTGTAGTGATGGATGCCTGTA
AGCACACGTCTGAACTCTTTCCCTACACGACGCTCTTCCGATCTNNNCAACATCCATTCTGCAGCCTATG
AGCACACGTCTGAACTCTTTCCCTACACGACGCTCTTCCGATCTNNNCAACATCCATTCTGCAGCCTATG
AGCACACGTCTGAACTCTTTCCCTACACGACGCTCTTCCGATCTNNNAGTGGGGCTATACTTTTTGGT
AGCACACGTCTGAACTCTTTCCCTACACGACGCTCTTCCGATCTNNNAGTGGGGCTATACTTTTTGGT
AGCACACGTCTGAACTCTTTCCCTACACGACGCTCTTCCGATCTNNNACGTTCAACCAGATTTCCAGGT
AGCACACGTCTGAACTCTTTCCCTACACGACGCTCTTCCGATCTNNNACGTTCAACCAGATTTCCAGGT
GCACACGTCTGAACTCTTTCCCTACACGACGCTCTTCCGATCTNNNATATACACAAAAGGAGGGCTGC
GCACACGTCTGAACTCTTTCCCTACACGACGCTCTTCCGATCTNNNATATACACAAAAGGAGGGCTGC
AGCACACGTCTGAACTCTTTCCCTACACGACGCTCTTCCGATCTNNNCAATCTCAGCTCACTGCAATCTC
AGCACACGTCTGAACTCTTTCCCTACACGACGCTCTTCCGATCTNNNCAATCTCAGCTCACTGCAATCTC
GCACACGTCTGAACTCTTTCCCTACACGACGCTCTTCCGATCTNNNGGAACTTACAGTGAGCCGAGAT
GCACACGTCTGAACTCTTTCCCTACACGACGCTCTTCCGATCTNNNGGAACTTACAGTGAGCCGAGAT
GCACACGTCTGAACTCTTTCCCTACACGACGCTCTTCCGATCTNNNCATAGGGCAGTCAGGGTAAACTT
GCACACGTCTGAACTCTTTCCCTACACGACGCTCTTCCGATCTNNNCATAGGGCAGTCAGGGTAAACTT
AGCACACGTCTGAACTCTTTCCCTACACGACGCTCTTCCGATCTNNNGTCTTTGTCCAGGGTCTCATGAT
AGCACACGTCTGAACTCTTTCCCTACACGACGCTCTTCCGATCTNNNGTCTTTGTCCAGGGTCTCATGAT

GCACACGTCTGAACTCTTTCCCTACACGACGCTCTTCCGATCTNNNGTTATGTATCAAGTACCTTTGCCA
GCACACGTCTGAACTCTTTCCCTACACGACGCTCTTCCGATCTNNNGTTATGTATCAAGTACCTTTGCCA
AGCACACGTCTGAACTCTTTCCCTACACGACGCTCTTCCGATCTNNNGATCACGCTACTGCACTCCAACA
AGCACACGTCTGAACTCTTTCCCTACACGACGCTCTTCCGATCTNNNGATCACGCTACTGCACTCCAACA
GAGCACACGTCTGAACTCTTTCCCTACACGACGCTCTTCCGATCTNNNATACAAAAAATGAGCCGGGCA
GAGCACACGTCTGAACTCTTTCCCTACACGACGCTCTTCCGATCTNNNATACAAAAAATGAGCCGGGCA
AGCACACGTCTGAACTCTTTCCCTACACGACGCTCTTCCGATCTNNNTGGTATTTCCTTGTGAACAGTGT
AGCACACGTCTGAACTCTTTCCCTACACGACGCTCTTCCGATCTNNNTGGTATTTCCTTGTGAACAGTGT
GAGCACACGTCTGAACTCTTTCCCTACACGACGCTCTTCCGATCTNNNACACAGTGAAACCCCATCTCTA
GAGCACACGTCTGAACTCTTTCCCTACACGACGCTCTTCCGATCTNNNACACAGTGAAACCCCATCTCTA
GCACACGTCTGAACTCTTTCCCTACACGACGCTCTTCCGATCTNNNCTGTAGTTCCAGCTACTTGGGAG
GCACACGTCTGAACTCTTTCCCTACACGACGCTCTTCCGATCTNNNCTGTAGTTCCAGCTACTTGGGAG
AGCACACGTCTGAACTCTTTCCCTACACGACGCTCTTCCGATCTNNNCCGTCTAAGTATCCTAGGCAAC
AGCACACGTCTGAACTCTTTCCCTACACGACGCTCTTCCGATCTNNNCCGTCTAAGTATCCTAGGCAAC
AGCACACGTCTGAACTCTTTCCCTACACGACGCTCTTCCGATCTNNNTCAAATTCCATGGTCCTCCTCTGC
AGCACACGTCTGAACTCTTTCCCTACACGACGCTCTTCCGATCTNNNTCAAATTCCATGGTCCTCCTCTGC
GAGCACACGTCTGAACTCTTTCCCTACACGACGCTCTTCCGATCTNNNGAGGCTGAGGCAGGAGAATC
GAGCACACGTCTGAACTCTTTCCCTACACGACGCTCTTCCGATCTNNNGAGGCTGAGGCAGGAGAATC
GAGCACACGTCTGAACTCTTTCCCTACACGACGCTCTTCCGATCTNNNGAAGTGAGCTAGCAAACCTG
GAGCACACGTCTGAACTCTTTCCCTACACGACGCTCTTCCGATCTNNNGAAGTGAGCTAGCAAACCTG
AGCACACGTCTGAACTCTTTCCCTACACGACGCTCTTCCGATCTNNNCAACCTAGGTGGTCATGAATGC
AGCACACGTCTGAACTCTTTCCCTACACGACGCTCTTCCGATCTNNNCAACCTAGGTGGTCATGAATGC
AGAGCACACGTCTGAACTCTTTCCCTACACGACGCTCTTCCGATCTNNNGCCAAGATCACATCATTTGCA
AGAGCACACGTCTGAACTCTTTCCCTACACGACGCTCTTCCGATCTNNNGCCAAGATCACATCATTTGCA
AGCACACGTCTGAACTCTTTCCCTACACGACGCTCTTCCGATCTNNNCAAACAAACACACACACACACACC
AGCACACGTCTGAACTCTTTCCCTACACGACGCTCTTCCGATCTNNNCAAACAAACACACACACACACACC
AGCACACGTCTGAACTCTTTCCCTACACGACGCTCTTCCGATCTNNNAGCTTCCTGTGACTTGCAAGAT
AGCACACGTCTGAACTCTTTCCCTACACGACGCTCTTCCGATCTNNNAGCTTCCTGTGACTTGCAAGAT
AGCACACGTCTGAACTCTTTCCCTACACGACGCTCTTCCGATCTNNNCAAGGTGACCTAGAAGCAGTCT
AGCACACGTCTGAACTCTTTCCCTACACGACGCTCTTCCGATCTNNNCAAGGTGACCTAGAAGCAGTCT
AGCACACGTCTGAACTCTTTCCCTACACGACGCTCTTCCGATCTNNNGAGCCGAGATCGCGACATTACAT
AGCACACGTCTGAACTCTTTCCCTACACGACGCTCTTCCGATCTNNNGAGCCGAGATCGCGACATTACAT
GAGCACACGTCTGAACTCTTTCCCTACACGACGCTCTTCCGATCTNNNCCTTGTCAATCACCTCCTGCTT
GAGCACACGTCTGAACTCTTTCCCTACACGACGCTCTTCCGATCTNNNCCTTGTCAATCACCTCCTGCTT
GAGCACACGTCTGAACTCTTTCCCTACACGACGCTCTTCCGATCTNNNGTGTCTCCTGTTACCAACCGAA
GAGCACACGTCTGAACTCTTTCCCTACACGACGCTCTTCCGATCTNNNGTGTCTCCTGTTACCAACCGAA
GAGCACACGTCTGAACTCTTTCCCTACACGACGCTCTTCCGATCTNNNAGTCTCTGCACCTAATACTGCA
GAGCACACGTCTGAACTCTTTCCCTACACGACGCTCTTCCGATCTNNNAGTCTCTGCACCTAATACTGCA
GAGCACACGTCTGAACTCTTTCCCTACACGACGCTCTTCCGATCTNNNTCTCCTTTCTCTCTGAGCAAACA
GAGCACACGTCTGAACTCTTTCCCTACACGACGCTCTTCCGATCTNNNTCTCCTTTCTCTCTGAGCAAACA
GAGCACACGTCTGAACTCTTTCCCTACACGACGCTCTTCCGATCTNNNTGAAGCTCACAAGTCACTTCTC
GAGCACACGTCTGAACTCTTTCCCTACACGACGCTCTTCCGATCTNNNTGAAGCTCACAAGTCACTTCTC
AGCACACGTCTGAACTCTTTCCCTACACGACGCTCTTCCGATCTNNNCCCCTTCTCTCTCTCTCTCTCCA

AGCACACGTCTGAACTCTTTCCCTACACGACGCTCTTCCGATCTNNNCCCCTTCTCTCTCTCTCTCTCTCCA
GAGCACACGTCTGAACTCTTTCCCTACACGACGCTCTTCCGATCTNNNCTCATCCACCAGCTGAACTAGA
GAGCACACGTCTGAACTCTTTCCCTACACGACGCTCTTCCGATCTNNNCTCATCCACCAGCTGAACTAGA
GAGCACACGTCTGAACTCTTTCCCTACACGACGCTCTTCCGATCTNNNACCAGAGAATGTTAACCAGG
GAGCACACGTCTGAACTCTTTCCCTACACGACGCTCTTCCGATCTNNNACCAGAGAATGTTAACCAGG
GAGCACACGTCTGAACTCTTTCCCTACACGACGCTCTTCCGATCTNNNGGCTTCTGCTTCCAAATGTTTC
GAGCACACGTCTGAACTCTTTCCCTACACGACGCTCTTCCGATCTNNNGGCTTCTGCTTCCAAATGTTTC
AGCACACGTCTGAACTCTTTCCCTACACGACGCTCTTCCGATCTNNNGTGTGTGTGTGTGTGATTG
AGCACACGTCTGAACTCTTTCCCTACACGACGCTCTTCCGATCTNNNGTGTGTGTGTGTGTGATTG
AGCACACGTCTGAACTCTTTCCCTACACGACGCTCTTCCGATCTNNNCTGAGACAGGAGAATCGCTTGA
AGCACACGTCTGAACTCTTTCCCTACACGACGCTCTTCCGATCTNNNCTGAGACAGGAGAATCGCTTGA
AGCACACGTCTGAACTCTTTCCCTACACGACGCTCTTCCGATCTNNNCATTGGGAGCTTTGCTATGCTA
AGCACACGTCTGAACTCTTTCCCTACACGACGCTCTTCCGATCTNNNCATTGGGAGCTTTGCTATGCTA
AGCACACGTCTGAACTCTTTCCCTACACGACGCTCTTCCGATCTNNNTCCCCTTGAAAACTAGCACAAG
AGCACACGTCTGAACTCTTTCCCTACACGACGCTCTTCCGATCTNNNTCCCCTTGAAAACTAGCACAAG
AGCACACGTCTGAACTCTTTCCCTACACGACGCTCTTCCGATCTNNNAAGGAAATTGAGGCTCAGAGA
AGCACACGTCTGAACTCTTTCCCTACACGACGCTCTTCCGATCTNNNAAGGAAATTGAGGCTCAGAGA
GCACACGTCTGAACTCTTTCCCTACACGACGCTCTTCCGATCTNNNCTCTACCAGGCAACTTTCCCAAGC
GCACACGTCTGAACTCTTTCCCTACACGACGCTCTTCCGATCTNNNCTCTACCAGGCAACTTTCCCAAGC
GCACACGTCTGAACTCTTTCCCTACACGACGCTCTTCCGATCTNNNCCTCCTCCTAGAAGCCCTCCCATCC
GCACACGTCTGAACTCTTTCCCTACACGACGCTCTTCCGATCTNNNCCTCCTCCTAGAAGCCCTCCCATCC
GCACACGTCTGAACTCTTTCCCTACACGACGCTCTTCCGATCTNNNCCCAGAAACTTAAATATCTGAATC
GCACACGTCTGAACTCTTTCCCTACACGACGCTCTTCCGATCTNNNCCCAGAAACTTAAATATCTGAATC
AGCACACGTCTGAACTCTTTCCCTACACGACGCTCTTCCGATCTNNNAGTAGCTGGGACTATAGGCGTA
AGCACACGTCTGAACTCTTTCCCTACACGACGCTCTTCCGATCTNNNAGTAGCTGGGACTATAGGCGTA
GAGCACACGTCTGAACTCTTTCCCTACACGACGCTCTTCCGATCTNNNGCTGCAGTGAACCATGATCAT
GAGCACACGTCTGAACTCTTTCCCTACACGACGCTCTTCCGATCTNNNGCTGCAGTGAACCATGATCAT
AGCACACGTCTGAACTCTTTCCCTACACGACGCTCTTCCGATCTNNNGCTTCTTTCCTCACTTCCCTACAC
AGCACACGTCTGAACTCTTTCCCTACACGACGCTCTTCCGATCTNNNGCTTCTTTCCTCACTTCCCTACAC
AGCACACGTCTGAACTCTTTCCCTACACGACGCTCTTCCGATCTNNNATATGCCTGTAATCCCAGCTACC
AGCACACGTCTGAACTCTTTCCCTACACGACGCTCTTCCGATCTNNNATATGCCTGTAATCCCAGCTACC
GAGCACACGTCTGAACTCTTTCCCTACACGACGCTCTTCCGATCTNNNATTTCCTGTGGTCCTATGTTC
GAGCACACGTCTGAACTCTTTCCCTACACGACGCTCTTCCGATCTNNNATTTCCTGTGGTCCTATGTTC
GAGCACACGTCTGAACTCTTTCCCTACACGACGCTCTTCCGATCTNNNTCATGTTAGTCCCTTCACACA
GAGCACACGTCTGAACTCTTTCCCTACACGACGCTCTTCCGATCTNNNTCATGTTAGTCCCTTCACACA
AGCACACGTCTGAACTCTTTCCCTACACGACGCTCTTCCGATCTNNNAAGGTGAGAGAGAAGCCAGAT
AGCACACGTCTGAACTCTTTCCCTACACGACGCTCTTCCGATCTNNNAAGGTGAGAGAGAAGCCAGAT
AGCACACGTCTGAACTCTTTCCCTACACGACGCTCTTCCGATCTNNNAGAAAGAAACACCTCGGAGAA
AGCACACGTCTGAACTCTTTCCCTACACGACGCTCTTCCGATCTNNNAGAAAGAAACACCTCGGAGAA
AGCACACGTCTGAACTCTTTCCCTACACGACGCTCTTCCGATCTNNNAAGACCCATCAGTGTGCTGTAT
AGCACACGTCTGAACTCTTTCCCTACACGACGCTCTTCCGATCTNNNAAGACCCATCAGTGTGCTGTAT
AGCACACGTCTGAACTCTTTCCCTACACGACGCTCTTCCGATCTNNNCTCTCTTCCTTCCTTGTCTGCATC
AGCACACGTCTGAACTCTTTCCCTACACGACGCTCTTCCGATCTNNNCTCTCTTCCTTCCTTGTCTGCATC

AGCACACGTCTGAACTCTTTCCCTACACGACGCTCTTCCGATCTNNNGGATCACTTGAGACCAGGAGT
AGCACACGTCTGAACTCTTTCCCTACACGACGCTCTTCCGATCTNNNGGATCACTTGAGACCAGGAGT
GCACACGTCTGAACTCTTTCCCTACACGACGCTCTTCCGATCTNNNACCCCTCTGACCTCTTCTACTACC
GCACACGTCTGAACTCTTTCCCTACACGACGCTCTTCCGATCTNNNACCCCTCTGACCTCTTCTACTACC
GAGCACACGTCTGAACTCTTTCCCTACACGACGCTCTTCCGATCTNNNCAGATCATGAGGTCAGGAGA
GAGCACACGTCTGAACTCTTTCCCTACACGACGCTCTTCCGATCTNNNCAGATCATGAGGTCAGGAGA
AGCACACGTCTGAACTCTTTCCCTACACGACGCTCTTCCGATCTNNNTATAACCCCAGCTACTTGCAAG
AGCACACGTCTGAACTCTTTCCCTACACGACGCTCTTCCGATCTNNNTATAACCCCAGCTACTTGCAAG
GCACACGTCTGAACTCTTTCCCTACACGACGCTCTTCCGATCTNNNGTTTGTATTCCTGCCTCCTCCAGC
GCACACGTCTGAACTCTTTCCCTACACGACGCTCTTCCGATCTNNNGTTTGTATTCCTGCCTCCTCCAGC
GAGCACACGTCTGAACTCTTTCCCTACACGACGCTCTTCCGATCTNNNGAACCAGAACTCCCTCCCTAA
GAGCACACGTCTGAACTCTTTCCCTACACGACGCTCTTCCGATCTNNNGAACCAGAACTCCCTCCCTAA
AGCACACGTCTGAACTCTTTCCCTACACGACGCTCTTCCGATCTNNNGTGACCTCCCTTCTTTCTTTCCT
AGCACACGTCTGAACTCTTTCCCTACACGACGCTCTTCCGATCTNNNGTGACCTCCCTTCTTTCTTTCCT
ACACGTCTGAACTCTTTCCCTACACGACGCTCTTCCGATCTNNNATGTGATTTGTTTGAATTCAGATCA
ACACGTCTGAACTCTTTCCCTACACGACGCTCTTCCGATCTNNNATGTGATTTGTTTGAATTCAGATCA
AGCACACGTCTGAACTCTTTCCCTACACGACGCTCTTCCGATCTNNNCGACCACTTCAAGAGACCTCTA
AGCACACGTCTGAACTCTTTCCCTACACGACGCTCTTCCGATCTNNNCGACCACTTCAAGAGACCTCTA
AGCACACGTCTGAACTCTTTCCCTACACGACGCTCTTCCGATCTNNNGTCATCAACTCATGGCCAATTC
AGCACACGTCTGAACTCTTTCCCTACACGACGCTCTTCCGATCTNNNGTCATCAACTCATGGCCAATTC
AGCACACGTCTGAACTCTTTCCCTACACGACGCTCTTCCGATCTNNNATCATTGGAGATGCTAAGCCA
AGCACACGTCTGAACTCTTTCCCTACACGACGCTCTTCCGATCTNNNATCATTGGAGATGCTAAGCCA
AGCACACGTCTGAACTCTTTCCCTACACGACGCTCTTCCGATCTNNNAAGATCACACCACTGCACTGT
AGCACACGTCTGAACTCTTTCCCTACACGACGCTCTTCCGATCTNNNAAGATCACACCACTGCACTGT
AGCACACGTCTGAACTCTTTCCCTACACGACGCTCTTCCGATCTNNNACAACCTAGAACAGTGCTTAGC
AGCACACGTCTGAACTCTTTCCCTACACGACGCTCTTCCGATCTNNNACAACCTAGAACAGTGCTTAGC
GAGCACACGTCTGAACTCTTTCCCTACACGACGCTCTTCCGATCTNNNATGGAAGGGTGTGTTTGGT
GAGCACACGTCTGAACTCTTTCCCTACACGACGCTCTTCCGATCTNNNATGGAAGGGTGTGTTTGGT
GCACACGTCTGAACTCTTTCCCTACACGACGCTCTTCCGATCTNNNTGAAGGGCATCAATTGTGATTC
GCACACGTCTGAACTCTTTCCCTACACGACGCTCTTCCGATCTNNNTGAAGGGCATCAATTGTGATTC
GCACACGTCTGAACTCTTTCCCTACACGACGCTCTTCCGATCTNNNTCTTTCTTTCCTCCCTTCCTTCTCA
GCACACGTCTGAACTCTTTCCCTACACGACGCTCTTCCGATCTNNNTCTTTCTTTCCTCCCTTCCTTCTCA
GCACACGTCTGAACTCTTTCCCTACACGACGCTCTTCCGATCTNNNGAGGAAGAGAACTGCTTGAAC
GCACACGTCTGAACTCTTTCCCTACACGACGCTCTTCCGATCTNNNGAGGAAGAGAACTGCTTGAAC
AAGAGCACACGTCTGAACTCTTTCCCTACACGACGCTCTTCCGATCTNNNGCAGGAGAATTGCTTGAA
AAGAGCACACGTCTGAACTCTTTCCCTACACGACGCTCTTCCGATCTNNNGCAGGAGAATTGCTTGAA
GAGCACACGTCTGAACTCTTTCCCTACACGACGCTCTTCCGATCTNNNATTGAGTAGGAGTGAGCCA
GAGCACACGTCTGAACTCTTTCCCTACACGACGCTCTTCCGATCTNNNATTGAGTAGGAGTGAGCCA
GCACACGTCTGAACTCTTTCCCTACACGACGCTCTTCCGATCTNNNTGAAGTACAGTGGCACTATCTC
GCACACGTCTGAACTCTTTCCCTACACGACGCTCTTCCGATCTNNNTGAAGTACAGTGGCACTATCTC
GCACACGTCTGAACTCTTTCCCTACACGACGCTCTTCCGATCTNNNGAGAATTGCTTGAACCTGGGAA
GCACACGTCTGAACTCTTTCCCTACACGACGCTCTTCCGATCTNNNGAGAATTGCTTGAACCTGGGAA
CACACGTCTGAACTCTTTCCCTACACGACGCTCTTCCGATCTNNNAACCCTCGAATCTAGCTTACTGTC

GCACACGTCTGAACTCTTTCCCTACACGACGCTCTTCCGATCTNNNAACCCTCGAATCTAGCTTACTGTCA

GAGCACACGTCTGAACTCTTTCCCTACACGACGCTCTTCCGATCTNNNCTACTCTCTGACCCCCAGTGTC

GAGCACACGTCTGAACTCTTTCCCTACACGACGCTCTTCCGATCTNNNCTACTCTCTGACCCCCAGTGTC

AGCACACGTCTGAACTCTTTCCCTACACGACGCTCTTCCGATCTNNNCATGATCACACCACTACACTCCA

AGCACACGTCTGAACTCTTTCCCTACACGACGCTCTTCCGATCTNNNCATGATCACACCACTACACTCCA

GAGCACACGTCTGAACTCTTTCCCTACACGACGCTCTTCCGATCTNNNTAATCCTAGCTACTGGGGAGG

GAGCACACGTCTGAACTCTTTCCCTACACGACGCTCTTCCGATCTNNNTAATCCTAGCTACTGGGGAGG

GAGCACACGTCTGAACTCTTTCCCTACACGACGCTCTTCCGATCTNNNTCAAGAGTCTGAGGGAGGAG

GAGCACACGTCTGAACTCTTTCCCTACACGACGCTCTTCCGATCTNNNTCAAGAGTCTGAGGGAGGAG

GAGCACACGTCTGAACTCTTTCCCTACACGACGCTCTTCCGATCTNNNCTGTAATCCCAGCTACTCAGGA

GAGCACACGTCTGAACTCTTTCCCTACACGACGCTCTTCCGATCTNNNCTGTAATCCCAGCTACTCAGGA

GAGCACACGTCTGAACTCTTTCCCTACACGACGCTCTTCCGATCTNNNCTTGCAGTGAGATGAGATTG

GAGCACACGTCTGAACTCTTTCCCTACACGACGCTCTTCCGATCTNNNCTTGCAGTGAGATGAGATTG

GAGCACACGTCTGAACTCTTTCCCTACACGACGCTCTTCCGATCTNNNGCCAGTCCCACTACTTTACTGA

GAGCACACGTCTGAACTCTTTCCCTACACGACGCTCTTCCGATCTNNNGCCAGTCCCACTACTTTACTGA

AGCACACGTCTGAACTCTTTCCCTACACGACGCTCTTCCGATCTNNNTCAAGCTTTTCTGCCTCATTTGCC

AGCACACGTCTGAACTCTTTCCCTACACGACGCTCTTCCGATCTNNNTCAAGCTTTTCTGCCTCATTTGCC

GAGCACACGTCTGAACTCTTTCCCTACACGACGCTCTTCCGATCTNNNCTGACTGTGCCACTATACTCCA

GAGCACACGTCTGAACTCTTTCCCTACACGACGCTCTTCCGATCTNNNCTGACTGTGCCACTATACTCCA

AGCACACGTCTGAACTCTTTCCCTACACGACGCTCTTCCGATCTNNNCAGACTCTTGAGCCAGGTAAGC

AGCACACGTCTGAACTCTTTCCCTACACGACGCTCTTCCGATCTNNNCAGACTCTTGAGCCAGGTAAGC

GAGCACACGTCTGAACTCTTTCCCTACACGACGCTCTTCCGATCTNNNAACTCCTAGCCTGAAGTGATCT

GAGCACACGTCTGAACTCTTTCCCTACACGACGCTCTTCCGATCTNNNAACTCCTAGCCTGAAGTGATCT

GAGCACACGTCTGAACTCTTTCCCTACACGACGCTCTTCCGATCTNNNGCAGACATCTGTAATCCCAGC

GAGCACACGTCTGAACTCTTTCCCTACACGACGCTCTTCCGATCTNNNGCAGACATCTGTAATCCCAGC

AGCACACGTCTGAACTCTTTCCCTACACGACGCTCTTCCGATCTNNNATATTACATTTCAGAGTTCACCT

AGCACACGTCTGAACTCTTTCCCTACACGACGCTCTTCCGATCTNNNATATTACATTTCAGAGTTCACCT

AGCACACGTCTGAACTCTTTCCCTACACGACGCTCTTCCGATCTNNNCTGTGATCCCAGCTACTCAGGA

AGCACACGTCTGAACTCTTTCCCTACACGACGCTCTTCCGATCTNNNCTGTGATCCCAGCTACTCAGGA

AGCACACGTCTGAACTCTTTCCCTACACGACGCTCTTCCGATCTNNNTGTCACATTTCTCATATTTTCAA

AGCACACGTCTGAACTCTTTCCCTACACGACGCTCTTCCGATCTNNNTGTCACATTTCTCATATTTTCAA

CACACGTCTGAACTCTTTCCCTACACGACGCTCTTCCGATCTNNNCGTCATTGCTTTTGGACTATCATCG

CACACGTCTGAACTCTTTCCCTACACGACGCTCTTCCGATCTNNNCGTCATTGCTTTTGGACTATCATCG

GAGCACACGTCTGAACTCTTTCCCTACACGACGCTCTTCCGATCTNNNACAAACAGCATACTTGAGTTG

GAGCACACGTCTGAACTCTTTCCCTACACGACGCTCTTCCGATCTNNNACAAACAGCATACTTGAGTTG

GAGCACACGTCTGAACTCTTTCCCTACACGACGCTCTTCCGATCTNNNCGGTAAGCTGAGATCATGTCA

GAGCACACGTCTGAACTCTTTCCCTACACGACGCTCTTCCGATCTNNNCGGTAAGCTGAGATCATGTCA

AGAGCACACGTCTGAACTCTTTCCCTACACGACGCTCTTCCGATCTNNNGAGATCACACCATTGCACTTC

AGAGCACACGTCTGAACTCTTTCCCTACACGACGCTCTTCCGATCTNNNGAGATCACACCATTGCACTTC

AGCACACGTCTGAACTCTTTCCCTACACGACGCTCTTCCGATCTNNNATCAGGAGACAGCGTTTTGAGA

AGCACACGTCTGAACTCTTTCCCTACACGACGCTCTTCCGATCTNNNATCAGGAGACAGCGTTTTGAGA

AGCACACGTCTGAACTCTTTCCCTACACGACGCTCTTCCGATCTNNNCCCACAAAGGTAACATCTTGCAA

AGCACACGTCTGAACTCTTTCCCTACACGACGCTCTTCCGATCTNNNCCCACAAAGGTAACATCTTGCAA

AGCACACGTCTGAACTCTTTCCCTACACGACGCTCTTCCGATCTNNNTAGGAGAATCGCTTGAATCCA

AGCACACGTCTGAACTCTTTCCCTACACGACGCTCTTCCGATCTNNNTAGGAGAATCGCTTGAATCCA

ACGTCTGAACTCTTTCCCTACACGACGCTCTTCCGATCTNNNACAGACATTGGCCACCTCTCCATCGGA

ACGTCTGAACTCTTTCCCTACACGACGCTCTTCCGATCTNNNACAGACATTGGCCACCTCTCCATCGGA

AGCACACGTCTGAACTCTTTCCCTACACGACGCTCTTCCGATCTNNNCAGGGCTCTTGTGCACTTAGA

AGCACACGTCTGAACTCTTTCCCTACACGACGCTCTTCCGATCTNNNCAGGGCTCTTGTGCACTTAGA

GAGCACACGTCTGAACTCTTTCCCTACACGACGCTCTTCCGATCTNNNGAGTAGAGTGACTGTGGGA

GAGCACACGTCTGAACTCTTTCCCTACACGACGCTCTTCCGATCTNNNGAGTAGAGTGACTGTGGGA

GCACACGTCTGAACTCTTTCCCTACACGACGCTCTTCCGATCTNNNAAAGGTGTATGTGAAACTTGC

GCACACGTCTGAACTCTTTCCCTACACGACGCTCTTCCGATCTNNNAAAGGTGTATGTGAAACTTGC

GCACACGTCTGAACTCTTTCCCTACACGACGCTCTTCCGATCTNNNAATTGCTAAATAGTGGCATGCC

GCACACGTCTGAACTCTTTCCCTACACGACGCTCTTCCGATCTNNNAATTGCTAAATAGTGGCATGCC

AAGAGCACACGTCTGAACTCTTTCCCTACACGACGCTCTTCCGATCTNNNAACGTCTTCTCTTCCCTCCT

AAGAGCACACGTCTGAACTCTTTCCCTACACGACGCTCTTCCGATCTNNNAACGTCTTCTCTTCCCTCCT

GAGCACACGTCTGAACTCTTTCCCTACACGACGCTCTTCCGATCTNNNTAATTCCAGTACTTTGGGAG

GAGCACACGTCTGAACTCTTTCCCTACACGACGCTCTTCCGATCTNNNTAATTCCAGTACTTTGGGAG

AGCACACGTCTGAACTCTTTCCCTACACGACGCTCTTCCGATCTNNNAGAAAGAGAGAGAGGGAGA

AGCACACGTCTGAACTCTTTCCCTACACGACGCTCTTCCGATCTNNNAGAAAGAGAGAGAGGGAGA

GCACACGTCTGAACTCTTTCCCTACACGACGCTCTTCCGATCTNNNTAAAATACCTGTTTGTGGCCCCA

GCACACGTCTGAACTCTTTCCCTACACGACGCTCTTCCGATCTNNNTAAAATACCTGTTTGTGGCCCCA

GCACACGTCTGAACTCTTTCCCTACACGACGCTCTTCCGATCTNNNTTTCACAGATGCTCTCCTCATCCC

GCACACGTCTGAACTCTTTCCCTACACGACGCTCTTCCGATCTNNNTTTCACAGATGCTCTCCTCATCCC

AGCACACGTCTGAACTCTTTCCCTACACGACGCTCTTCCGATCTNNNCTGGCTGGTATCTGTGATCTC

GAGCACACGTCTGAACTCTTTCCCTACACGACGCTCTTCCGATCTNNNCTGGCTGGTATCTGTGATCTC

AGCACACGTCTGAACTCTTTCCCTACACGACGCTCTTCCGATCTNNNTGTGCCCAGCTTTTTCACTTA

AGCACACGTCTGAACTCTTTCCCTACACGACGCTCTTCCGATCTNNNTGTGCCCAGCTTTTTCACTTA

GCACACGTCTGAACTCTTTCCCTACACGACGCTCTTCCGATCTNNNGTTCCTGTAGTCCCAGCTACTTG

GCACACGTCTGAACTCTTTCCCTACACGACGCTCTTCCGATCTNNNGTTCCTGTAGTCCCAGCTACTTG

AGCACACGTCTGAACTCTTTCCCTACACGACGCTCTTCCGATCTNNNATTGTCTTCTTTTAGGCCCCAG

AGCACACGTCTGAACTCTTTCCCTACACGACGCTCTTCCGATCTNNNATTGTCTTCTTTTAGGCCCCAG

GAGCACACGTCTGAACTCTTTCCCTACACGACGCTCTTCCGATCTNNNGAGAGATCTAGGTGAGGGG

GAGCACACGTCTGAACTCTTTCCCTACACGACGCTCTTCCGATCTNNNGAGAGATCTAGGTGAGGGG

AGCACACGTCTGAACTCTTTCCCTACACGACGCTCTTCCGATCTNNNGCAGGAGAATTGCTTGAACCT

AGCACACGTCTGAACTCTTTCCCTACACGACGCTCTTCCGATCTNNNGCAGGAGAATTGCTTGAACCT

AGCACACGTCTGAACTCTTTCCCTACACGACGCTCTTCCGATCTNNNTCCAATCTGGATGGCTTTGAT

AGCACACGTCTGAACTCTTTCCCTACACGACGCTCTTCCGATCTNNNTCCAATCTGGATGGCTTTGAT

AGAGCACACGTCTGAACTCTTTCCCTACACGACGCTCTTCCGATCTNNNCCGTTTGGGAATTCAAAGA

AGAGCACACGTCTGAACTCTTTCCCTACACGACGCTCTTCCGATCTNNNCCGTTTGGGAATTCAAAGA

AGCACACGTCTGAACTCTTTCCCTACACGACGCTCTTCCGATCTNNNAGAATTGCTTGAATCCTGGAG

AGCACACGTCTGAACTCTTTCCCTACACGACGCTCTTCCGATCTNNNAGAATTGCTTGAATCCTGGAG

GCACACGTCTGAACTCTTTCCCTACACGACGCTCTTCCGATCTNNNCGTGCCCGACTACATTCACATAT

GCACACGTCTGAACTCTTTCCCTACACGACGCTCTTCCGATCTNNNCGTGCCCGACTACATTCACATAT

AGCACACGTCTGAACTCTTTCCCTACACGACGCTCTTCCGATCTNNNAGGCTGAAACAGGAGAATCG

GAGCACACGTCTGAACTCTTTCCCTACACGACGCTCTTCCGATCTNNNAGGCTGAAACAGGAGAATCG
GAGCACACGTCTGAACTCTTTCCCTACACGACGCTCTTCCGATCTNNNAAATTAGTTGGGAATGGTGG
GAGCACACGTCTGAACTCTTTCCCTACACGACGCTCTTCCGATCTNNNAAATTAGTTGGGAATGGTGG
AGCACACGTCTGAACTCTTTCCCTACACGACGCTCTTCCGATCTNNNGAAGGTTGCAATGAGCCAAGA
AGCACACGTCTGAACTCTTTCCCTACACGACGCTCTTCCGATCTNNNGAAGGTTGCAATGAGCCAAGA
GCACACGTCTGAACTCTTTCCCTACACGACGCTCTTCCGATCTNNNGCAAGGGAAAATGGAATCGGTA
GCACACGTCTGAACTCTTTCCCTACACGACGCTCTTCCGATCTNNNGCAAGGGAAAATGGAATCGGTA
AGCACACGTCTGAACTCTTTCCCTACACGACGCTCTTCCGATCTNNNCCACACAAGCTTCTAAATTGGG
AGCACACGTCTGAACTCTTTCCCTACACGACGCTCTTCCGATCTNNNCCACACAAGCTTCTAAATTGGG
AGCACACGTCTGAACTCTTTCCCTACACGACGCTCTTCCGATCTNNNCTGAGGCAGGAGAATCACTTGA
AGCACACGTCTGAACTCTTTCCCTACACGACGCTCTTCCGATCTNNNCTGAGGCAGGAGAATCACTTGA
AGCACACGTCTGAACTCTTTCCCTACACGACGCTCTTCCGATCTNNNAGCAAGCTCCCATCTCTACAAAA
AGCACACGTCTGAACTCTTTCCCTACACGACGCTCTTCCGATCTNNNAGCAAGCTCCCATCTCTACAAAA
AGCACACGTCTGAACTCTTTCCCTACACGACGCTCTTCCGATCTNNNAACATGAGTTTGTCAGTGGAC
AGCACACGTCTGAACTCTTTCCCTACACGACGCTCTTCCGATCTNNNAACATGAGTTTGTCAGTGGAC
GAGCACACGTCTGAACTCTTTCCCTACACGACGCTCTTCCGATCTNNNTCAAGTATGGCCCCATCTGAG
GAGCACACGTCTGAACTCTTTCCCTACACGACGCTCTTCCGATCTNNNTCAAGTATGGCCCCATCTGAG
AGAGCACACGTCTGAACTCTTTCCCTACACGACGCTCTTCCGATCTNNNCTGAGGCAGAGAATTGCTTG
AGAGCACACGTCTGAACTCTTTCCCTACACGACGCTCTTCCGATCTNNNCTGAGGCAGAGAATTGCTTG
GAGCACACGTCTGAACTCTTTCCCTACACGACGCTCTTCCGATCTNNNCTCTGCTATGATCAGGGTATCA
GAGCACACGTCTGAACTCTTTCCCTACACGACGCTCTTCCGATCTNNNCTCTGCTATGATCAGGGTATCA
AGCACACGTCTGAACTCTTTCCCTACACGACGCTCTTCCGATCTNNNGTTCTTGTGGTGCTGGAGAATT
AGCACACGTCTGAACTCTTTCCCTACACGACGCTCTTCCGATCTNNNGTTCTTGTGGTGCTGGAGAATT
GCACACGTCTGAACTCTTTCCCTACACGACGCTCTTCCGATCTNNNGATGGTTTCCAGCTTCATCCATG
GCACACGTCTGAACTCTTTCCCTACACGACGCTCTTCCGATCTNNNGATGGTTTCCAGCTTCATCCATG
AGCACACGTCTGAACTCTTTCCCTACACGACGCTCTTCCGATCTNNNACCTTCCCTCCACTATTGTCCTAC
AGCACACGTCTGAACTCTTTCCCTACACGACGCTCTTCCGATCTNNNACCTTCCCTCCACTATTGTCCTAC
AGCACACGTCTGAACTCTTTCCCTACACGACGCTCTTCCGATCTNNNCAGCAAAACCAGTATCAAGGCT
AGCACACGTCTGAACTCTTTCCCTACACGACGCTCTTCCGATCTNNNCAGCAAAACCAGTATCAAGGCT
AGCACACGTCTGAACTCTTTCCCTACACGACGCTCTTCCGATCTNNNCTTCTTTTCCAAGTCATTCATTAC
AGCACACGTCTGAACTCTTTCCCTACACGACGCTCTTCCGATCTNNNCTTCTTTTCCAAGTCATTCATTAC
GCACACGTCTGAACTCTTTCCCTACACGACGCTCTTCCGATCTNNNTGTAATAATAGTAGAAATAAAGT
GCACACGTCTGAACTCTTTCCCTACACGACGCTCTTCCGATCTNNNTGTAATAATAGTAGAAATAAAGT
AGCACACGTCTGAACTCTTTCCCTACACGACGCTCTTCCGATCTNNNTGTTGGCATACAGTTGTTCATA
AGCACACGTCTGAACTCTTTCCCTACACGACGCTCTTCCGATCTNNNTGTTGGCATACAGTTGTTCATA
GAGCACACGTCTGAACTCTTTCCCTACACGACGCTCTTCCGATCTNNNATTGAAGAGATGGAGCTCAG
GAGCACACGTCTGAACTCTTTCCCTACACGACGCTCTTCCGATCTNNNATTGAAGAGATGGAGCTCAG
AGCACACGTCTGAACTCTTTCCCTACACGACGCTCTTCCGATCTNNNTCAACTGCTCTACCCCACTAAAC
AGCACACGTCTGAACTCTTTCCCTACACGACGCTCTTCCGATCTNNNTCAACTGCTCTACCCCACTAAAC
GAGCACACGTCTGAACTCTTTCCCTACACGACGCTCTTCCGATCTNNNTTGCTATGGATTTGCCAGTCCT
GAGCACACGTCTGAACTCTTTCCCTACACGACGCTCTTCCGATCTNNNTTGCTATGGATTTGCCAGTCCT
AGCACACGTCTGAACTCTTTCCCTACACGACGCTCTTCCGATCTNNNGTCTCTGAGTCCATTCTTGGGT
AGCACACGTCTGAACTCTTTCCCTACACGACGCTCTTCCGATCTNNNGTCTCTGAGTCCATTCTTGGGT

AGCACACGTCTGAACTCTTTCCCTACACGACGCTCTTCCGATCTNNNCAAAAATTAGCCGGGTGAGAT

AGCACACGTCTGAACTCTTTCCCTACACGACGCTCTTCCGATCTNNNCAAAAATTAGCCGGGTGAGAT

AGCACACGTCTGAACTCTTTCCCTACACGACGCTCTTCCGATCTNNNAAAGTGCTGGGATTACAGGCAT

AGCACACGTCTGAACTCTTTCCCTACACGACGCTCTTCCGATCTNNNAAAGTGCTGGGATTACAGGCAT

GCACACGTCTGAACTCTTTCCCTACACGACGCTCTTCCGATCTNNNAAGGACACAGAGCTAAGACATG

GCACACGTCTGAACTCTTTCCCTACACGACGCTCTTCCGATCTNNNAAGGACACAGAGCTAAGACATG

ACACGTCTGAACTCTTTCCCTACACGACGCTCTTCCGATCTNNNTTCTCAATGAGCTGTTGGGTACACA

ACACGTCTGAACTCTTTCCCTACACGACGCTCTTCCGATCTNNNTTCTCAATGAGCTGTTGGGTACACA

GCACACGTCTGAACTCTTTCCCTACACGACGCTCTTCCGATCTNNNTCCCAAAGTGCTGAGAATACAGG

GCACACGTCTGAACTCTTTCCCTACACGACGCTCTTCCGATCTNNNTCCCAAAGTGCTGAGAATACAGG

GAGCACACGTCTGAACTCTTTCCCTACACGACGCTCTTCCGATCTNNNCTGTTTCCTTCCCATTGTGTTG

GAGCACACGTCTGAACTCTTTCCCTACACGACGCTCTTCCGATCTNNNCTGTTTCCTTCCCATTGTGTTG

AGCACACGTCTGAACTCTTTCCCTACACGACGCTCTTCCGATCTNNNGAATGCCGGATGCTTGACTTCT

AGCACACGTCTGAACTCTTTCCCTACACGACGCTCTTCCGATCTNNNGAATGCCGGATGCTTGACTTCT

GAGCACACGTCTGAACTCTTTCCCTACACGACGCTCTTCCGATCTNNNACTCACTCTAAATGCCATGTGT

GAGCACACGTCTGAACTCTTTCCCTACACGACGCTCTTCCGATCTNNNACTCACTCTAAATGCCATGTGT

AGCACACGTCTGAACTCTTTCCCTACACGACGCTCTTCCGATCTNNNTAATTTCCCTATTAGCCAGAAAT

AGCACACGTCTGAACTCTTTCCCTACACGACGCTCTTCCGATCTNNNTAATTTCCCTATTAGCCAGAAAT

AGCACACGTCTGAACTCTTTCCCTACACGACGCTCTTCCGATCTNNNAACACTGGTTACGTGTATCTGG

AGCACACGTCTGAACTCTTTCCCTACACGACGCTCTTCCGATCTNNNAACACTGGTTACGTGTATCTGG

AGCACACGTCTGAACTCTTTCCCTACACGACGCTCTTCCGATCTNNNCTCAAACTCCTGACCTCGTGAT

AGCACACGTCTGAACTCTTTCCCTACACGACGCTCTTCCGATCTNNNCTCAAACTCCTGACCTCGTGAT

GAGCACACGTCTGAACTCTTTCCCTACACGACGCTCTTCCGATCTNNNTCCATTTGTTTGGGAAATCTCT

GAGCACACGTCTGAACTCTTTCCCTACACGACGCTCTTCCGATCTNNNTCCATTTGTTTGGGAAATCTCT

GAGCACACGTCTGAACTCTTTCCCTACACGACGCTCTTCCGATCTNNNCTCTCTCTCTCACACACACAC

GAGCACACGTCTGAACTCTTTCCCTACACGACGCTCTTCCGATCTNNNCTCTCTCTCTCACACACACAC

GAGCACACGTCTGAACTCTTTCCCTACACGACGCTCTTCCGATCTNNNAAGACTGACAGAAGAGCTGT

GAGCACACGTCTGAACTCTTTCCCTACACGACGCTCTTCCGATCTNNNAAGACTGACAGAAGAGCTGT

AGCACACGTCTGAACTCTTTCCCTACACGACGCTCTTCCGATCTNNNACCAAATAACTGTGGGGCTAT

AGCACACGTCTGAACTCTTTCCCTACACGACGCTCTTCCGATCTNNNACCAAATAACTGTGGGGCTAT

AGCACACGTCTGAACTCTTTCCCTACACGACGCTCTTCCGATCTNNNTCAACAACCTGAATTTCCTCCAG

AGCACACGTCTGAACTCTTTCCCTACACGACGCTCTTCCGATCTNNNTCAACAACCTGAATTTCCTCCAG

AGCACACGTCTGAACTCTTTCCCTACACGACGCTCTTCCGATCTNNNTCTGTAATCCCAGCTACTCAGGA

AGCACACGTCTGAACTCTTTCCCTACACGACGCTCTTCCGATCTNNNTCTGTAATCCCAGCTACTCAGGA

GAGCACACGTCTGAACTCTTTCCCTACACGACGCTCTTCCGATCTNNNAAGAGTTTTTTTGGTGGGGG

GAGCACACGTCTGAACTCTTTCCCTACACGACGCTCTTCCGATCTNNNAAGAGTTTTTTTGGTGGGGG

GAGCACACGTCTGAACTCTTTCCCTACACGACGCTCTTCCGATCTNNNGCACATGCATGTTTATAGCAG

GAGCACACGTCTGAACTCTTTCCCTACACGACGCTCTTCCGATCTNNNGCACATGCATGTTTATAGCAG

GAGCACACGTCTGAACTCTTTCCCTACACGACGCTCTTCCGATCTNNNGAGAGAGCAGAGTCGGTAGA

GAGCACACGTCTGAACTCTTTCCCTACACGACGCTCTTCCGATCTNNNGAGAGAGCAGAGTCGGTAGA

GAGCACACGTCTGAACTCTTTCCCTACACGACGCTCTTCCGATCTNNNTGATATGAAAATTGAATAGCC

GAGCACACGTCTGAACTCTTTCCCTACACGACGCTCTTCCGATCTNNNTGATATGAAAATTGAATAGCC

GAGCACACGTCTGAACTCTTTCCCTACACGACGCTCTTCCGATCTNNNCAAGAGAATTGCTTGAACCTG

AGCACACGTCTGAACTCTTTCCCTACACGACGCTCTTCCGATCTNNNCAAGAGAATTGCTTGAACCTG
GAGCACACGTCTGAACTCTTTCCCTACACGACGCTCTTCCGATCTNNNCCACTTGGAGTATTGAGGGA
GAGCACACGTCTGAACTCTTTCCCTACACGACGCTCTTCCGATCTNNNCCACTTGGAGTATTGAGGGA
AGCACACGTCTGAACTCTTTCCCTACACGACGCTCTTCCGATCTNNNCGAATGTAAGAGAACAGGGT
AGCACACGTCTGAACTCTTTCCCTACACGACGCTCTTCCGATCTNNNCGAATGTAAGAGAACAGGGT
AAGAGCACACGTCTGAACTCTTTCCCTACACGACGCTCTTCCGATCTNNNGGGAAAGAGGGAGACAG
AAGAGCACACGTCTGAACTCTTTCCCTACACGACGCTCTTCCGATCTNNNGGGAAAGAGGGAGACAG
AGCACACGTCTGAACTCTTTCCCTACACGACGCTCTTCCGATCTNNNCCACTTCTTGGGTTGAAGCAAT
AGCACACGTCTGAACTCTTTCCCTACACGACGCTCTTCCGATCTNNNCCACTTCTTGGGTTGAAGCAAT
AGCACACGTCTGAACTCTTTCCCTACACGACGCTCTTCCGATCTNNNCTTGAACCTAGGAGGCAGAAG
AGCACACGTCTGAACTCTTTCCCTACACGACGCTCTTCCGATCTNNNCTTGAACCTAGGAGGCAGAAG
GAGCACACGTCTGAACTCTTTCCCTACACGACGCTCTTCCGATCTNNNCAACCCAATGCAAGGAAGCT
GAGCACACGTCTGAACTCTTTCCCTACACGACGCTCTTCCGATCTNNNCAACCCAATGCAAGGAAGCT
GCACACGTCTGAACTCTTTCCCTACACGACGCTCTTCCGATCTNNNTTGAACTCCTAGGCTCCAGTGAT
GCACACGTCTGAACTCTTTCCCTACACGACGCTCTTCCGATCTNNNTTGAACTCCTAGGCTCCAGTGAT
GAGCACACGTCTGAACTCTTTCCCTACACGACGCTCTTCCGATCTNNNAGTGAGCTATGATTGCACCA
GAGCACACGTCTGAACTCTTTCCCTACACGACGCTCTTCCGATCTNNNAGTGAGCTATGATTGCACCA
GAGCACACGTCTGAACTCTTTCCCTACACGACGCTCTTCCGATCTNNNCCCTTTCTTTCTCCCGAACAGA
GAGCACACGTCTGAACTCTTTCCCTACACGACGCTCTTCCGATCTNNNCCCTTTCTTTCTCCCGAACAGA
GAGCACACGTCTGAACTCTTTCCCTACACGACGCTCTTCCGATCTNNNCTGAGTTCAGTGCCTCACAAT
GAGCACACGTCTGAACTCTTTCCCTACACGACGCTCTTCCGATCTNNNCTGAGTTCAGTGCCTCACAAT
GAGCACACGTCTGAACTCTTTCCCTACACGACGCTCTTCCGATCTNNNCTGAGGGAGGAGAATGGTT
GAGCACACGTCTGAACTCTTTCCCTACACGACGCTCTTCCGATCTNNNCTGAGGGAGGAGAATGGTT
GAGCACACGTCTGAACTCTTTCCCTACACGACGCTCTTCCGATCTNNNGAATCTCTTGAAACTGGAAG
GAGCACACGTCTGAACTCTTTCCCTACACGACGCTCTTCCGATCTNNNGAATCTCTTGAAACTGGAAG
GCACACGTCTGAACTCTTTCCCTACACGACGCTCTTCCGATCTNNNGAGATTGCACCACTACACTCCAG
GCACACGTCTGAACTCTTTCCCTACACGACGCTCTTCCGATCTNNNGAGATTGCACCACTACACTCCAG
AGAGCACACGTCTGAACTCTTTCCCTACACGACGCTCTTCCGATCTNNNCATTTGCAGGAATGTGTAG
AGAGCACACGTCTGAACTCTTTCCCTACACGACGCTCTTCCGATCTNNNCATTTGCAGGAATGTGTAG
GAGCACACGTCTGAACTCTTTCCCTACACGACGCTCTTCCGATCTNNNTGGGCAACAGAGCAAGACTC
GAGCACACGTCTGAACTCTTTCCCTACACGACGCTCTTCCGATCTNNNTGGGCAACAGAGCAAGACTC
GAGCACACGTCTGAACTCTTTCCCTACACGACGCTCTTCCGATCTNNNGGTGAGCTGAGATTTATTGT
GAGCACACGTCTGAACTCTTTCCCTACACGACGCTCTTCCGATCTNNNGGTGAGCTGAGATTTATTGT
GAGCACACGTCTGAACTCTTTCCCTACACGACGCTCTTCCGATCTNNNCTGTTTTAATCTGAAAAGAG
GAGCACACGTCTGAACTCTTTCCCTACACGACGCTCTTCCGATCTNNNCTGTTTTAATCTGAAAAGAG
AGAGCACACGTCTGAACTCTTTCCCTACACGACGCTCTTCCGATCTNNNATACGCAATTTGTGTGTGT
AGAGCACACGTCTGAACTCTTTCCCTACACGACGCTCTTCCGATCTNNNATACGCAATTTGTGTGTGT
CACACGTCTGAACTCTTTCCCTACACGACGCTCTTCCGATCTNNNCCTCAGTTATTTGATGGGAGGCTC
CACACGTCTGAACTCTTTCCCTACACGACGCTCTTCCGATCTNNNCCTCAGTTATTTGATGGGAGGCTC
GCACACGTCTGAACTCTTTCCCTACACGACGCTCTTCCGATCTNNNAAAAAGTTGAAACCGTTTTCGT
GCACACGTCTGAACTCTTTCCCTACACGACGCTCTTCCGATCTNNNAAAAAGTTGAAACCGTTTTCGT
GCACACGTCTGAACTCTTTCCCTACACGACGCTCTTCCGATCTNNNTTTCAGGTGACAGAGTGAGACT
GCACACGTCTGAACTCTTTCCCTACACGACGCTCTTCCGATCTNNNTTTCAGGTGACAGAGTGAGACT

AGAGCACACGTCTGAACTCTTTCCCTACACGACGCTCTTCCGATCTNNNGGTGGAGGCTATGAGGGT
AGAGCACACGTCTGAACTCTTTCCCTACACGACGCTCTTCCGATCTNNNGGTGGAGGCTATGAGGGT
CACACGTCTGAACTCTTTCCCTACACGACGCTCTTCCGATCTNNNTCCTCTCTTCTTCCAGAGACTCACA
CACACGTCTGAACTCTTTCCCTACACGACGCTCTTCCGATCTNNNTCCTCTCTTCTTCCAGAGACTCACA
AGCACACGTCTGAACTCTTTCCCTACACGACGCTCTTCCGATCTNNNTTACACACACACACACACACAA
AGCACACGTCTGAACTCTTTCCCTACACGACGCTCTTCCGATCTNNNTTACACACACACACACACACAA
AGCACACGTCTGAACTCTTTCCCTACACGACGCTCTTCCGATCTNNNAAAGAAAGATCAGGGACAGC
AGCACACGTCTGAACTCTTTCCCTACACGACGCTCTTCCGATCTNNNAAAGAAAGATCAGGGACAGC
GCACACGTCTGAACTCTTTCCCTACACGACGCTCTTCCGATCTNNNCAAAGTGCTGGAAAGTACAGG
GCACACGTCTGAACTCTTTCCCTACACGACGCTCTTCCGATCTNNNCAAAGTGCTGGAAAGTACAGG
AAGAGCACACGTCTGAACTCTTTCCCTACACGACGCTCTTCCGATCTNNNATTTCAGCCTCTTGTTCTG
AAGAGCACACGTCTGAACTCTTTCCCTACACGACGCTCTTCCGATCTNNNATTTCAGCCTCTTGTTCTG
GAGCACACGTCTGAACTCTTTCCCTACACGACGCTCTTCCGATCTNNNGGACTGTTAGGAAGGACGT
GAGCACACGTCTGAACTCTTTCCCTACACGACGCTCTTCCGATCTNNNGGACTGTTAGGAAGGACGT
GAGCACACGTCTGAACTCTTTCCCTACACGACGCTCTTCCGATCTNNNGGAGAGTTGCTTTAGTCCCA
GAGCACACGTCTGAACTCTTTCCCTACACGACGCTCTTCCGATCTNNNGGAGAGTTGCTTTAGTCCCA
AGCACACGTCTGAACTCTTTCCCTACACGACGCTCTTCCGATCTNNNCTGGGCAAGAGACTGAGACT
AGCACACGTCTGAACTCTTTCCCTACACGACGCTCTTCCGATCTNNNCTGGGCAAGAGACTGAGACT
AGCACACGTCTGAACTCTTTCCCTACACGACGCTCTTCCGATCTNNNGTGTGAACCAAGGCATACCAA
AGCACACGTCTGAACTCTTTCCCTACACGACGCTCTTCCGATCTNNNGTGTGAACCAAGGCATACCAA
AGCACACGTCTGAACTCTTTCCCTACACGACGCTCTTCCGATCTNNNGTTTTCTGGCATCTTTCTGCCT
AGCACACGTCTGAACTCTTTCCCTACACGACGCTCTTCCGATCTNNNGTTTTCTGGCATCTTTCTGCCT
AGCACACGTCTGAACTCTTTCCCTACACGACGCTCTTCCGATCTNNNCCTAAAACAGTGCATCCCAAGT
AGCACACGTCTGAACTCTTTCCCTACACGACGCTCTTCCGATCTNNNCCTAAAACAGTGCATCCCAAGT
GAGCACACGTCTGAACTCTTTCCCTACACGACGCTCTTCCGATCTNNNGATTCTCCCTCTATCGACACA
GAGCACACGTCTGAACTCTTTCCCTACACGACGCTCTTCCGATCTNNNGATTCTCCCTCTATCGACACA
GAGCACACGTCTGAACTCTTTCCCTACACGACGCTCTTCCGATCTNNNAGACAAGGAAACTAATGATA
GAGCACACGTCTGAACTCTTTCCCTACACGACGCTCTTCCGATCTNNNAGACAAGGAAACTAATGATA
GCACACGTCTGAACTCTTTCCCTACACGACGCTCTTCCGATCTNNNCCAAATTGCTGGGATTACAGG
GCACACGTCTGAACTCTTTCCCTACACGACGCTCTTCCGATCTNNNCCAAATTGCTGGGATTACAGG
GCACACGTCTGAACTCTTTCCCTACACGACGCTCTTCCGATCTNNNCACCCCCAACAAAGATTCCTTCA
GCACACGTCTGAACTCTTTCCCTACACGACGCTCTTCCGATCTNNNCACCCCCAACAAAGATTCCTTCA
AGCACACGTCTGAACTCTTTCCCTACACGACGCTCTTCCGATCTNNNGCGAAAAACTTGCTCCTCTATC
AGCACACGTCTGAACTCTTTCCCTACACGACGCTCTTCCGATCTNNNGCGAAAAACTTGCTCCTCTATC
GCACACGTCTGAACTCTTTCCCTACACGACGCTCTTCCGATCTNNNATAGCAGTAGCAGAAGGACGT
GCACACGTCTGAACTCTTTCCCTACACGACGCTCTTCCGATCTNNNATAGCAGTAGCAGAAGGACGT
GCACACGTCTGAACTCTTTCCCTACACGACGCTCTTCCGATCTNNNGTATCCTTCAGAACCTTTGGCA
GCACACGTCTGAACTCTTTCCCTACACGACGCTCTTCCGATCTNNNGTATCCTTCAGAACCTTTGGCA
GAGCACACGTCTGAACTCTTTCCCTACACGACGCTCTTCCGATCTNNNTAAAAAAACAGGGAGAGG
GAGCACACGTCTGAACTCTTTCCCTACACGACGCTCTTCCGATCTNNNTAAAAAAACAGGGAGAGG
AGCACACGTCTGAACTCTTTCCCTACACGACGCTCTTCCGATCTNNNTACAAGGACAAGAAGCTGAA
AGCACACGTCTGAACTCTTTCCCTACACGACGCTCTTCCGATCTNNNTACAAGGACAAGAAGCTGAA
AGCACACGTCTGAACTCTTTCCCTACACGACGCTCTTCCGATCTNNNAAAACCTGTAAGCCCAGTCTG

AGCACACGTCTGAACTCTTTCCCTACACGACGCTCTTCCGATCTNNNAAAACCTGTAAGCCCAGTCTG
GCACACGTCTGAACTCTTTCCCTACACGACGCTCTTCCGATCTNNNAGTATCCTTCTTTCATAATGAATG
GCACACGTCTGAACTCTTTCCCTACACGACGCTCTTCCGATCTNNNAGTATCCTTCTTTCATAATGAATG
GCACACGTCTGAACTCTTTCCCTACACGACGCTCTTCCGATCTNNNGAGCATTTGTGGGCTATTGTACC
GCACACGTCTGAACTCTTTCCCTACACGACGCTCTTCCGATCTNNNGAGCATTTGTGGGCTATTGTACC
GCACACGTCTGAACTCTTTCCCTACACGACGCTCTTCCGATCTNNNAACTGTACCTTGGCCCCTTTTATC
GCACACGTCTGAACTCTTTCCCTACACGACGCTCTTCCGATCTNNNAACTGTACCTTGGCCCCTTTTATC
AGAGCACACGTCTGAACTCTTTCCCTACACGACGCTCTTCCGATCTNNNGAGGGCACTTCCTTACCTGA
AGAGCACACGTCTGAACTCTTTCCCTACACGACGCTCTTCCGATCTNNNGAGGGCACTTCCTTACCTGA
GCACACGTCTGAACTCTTTCCCTACACGACGCTCTTCCGATCTNNNGCAGTTTCAACTTTCCATTCCCTCA
GCACACGTCTGAACTCTTTCCCTACACGACGCTCTTCCGATCTNNNGCAGTTTCAACTTTCCATTCCCTCA
AGCACACGTCTGAACTCTTTCCCTACACGACGCTCTTCCGATCTNNNGAATCTCTTGAACCCAGGAGG
AGCACACGTCTGAACTCTTTCCCTACACGACGCTCTTCCGATCTNNNGAATCTCTTGAACCCAGGAGG
GAGCACACGTCTGAACTCTTTCCCTACACGACGCTCTTCCGATCTNNNCAGTCACATGGTCTGACCCTA
GAGCACACGTCTGAACTCTTTCCCTACACGACGCTCTTCCGATCTNNNCAGTCACATGGTCTGACCCTA
AGCACACGTCTGAACTCTTTCCCTACACGACGCTCTTCCGATCTNNNCTTCACCATCAAGGATGTCTCCA
AGCACACGTCTGAACTCTTTCCCTACACGACGCTCTTCCGATCTNNNCTTCACCATCAAGGATGTCTCCA
GCACACGTCTGAACTCTTTCCCTACACGACGCTCTTCCGATCTNNNTCTTTTGGCTACCGTCAAACTTG
GCACACGTCTGAACTCTTTCCCTACACGACGCTCTTCCGATCTNNNTCTTTTGGCTACCGTCAAACTTG
GCACACGTCTGAACTCTTTCCCTACACGACGCTCTTCCGATCTNNNGCTCCCAAGTCAAGGATCTGTTT
GCACACGTCTGAACTCTTTCCCTACACGACGCTCTTCCGATCTNNNGCTCCCAAGTCAAGGATCTGTTT
GAGCACACGTCTGAACTCTTTCCCTACACGACGCTCTTCCGATCTNNNTCTGATAGGAACTTGCATTGC
GAGCACACGTCTGAACTCTTTCCCTACACGACGCTCTTCCGATCTNNNTCTGATAGGAACTTGCATTGC
AGCACACGTCTGAACTCTTTCCCTACACGACGCTCTTCCGATCTNNNCTGACTCAAGACTTCTCCAGCT
AGCACACGTCTGAACTCTTTCCCTACACGACGCTCTTCCGATCTNNNCTGACTCAAGACTTCTCCAGCT
GAGCACACGTCTGAACTCTTTCCCTACACGACGCTCTTCCGATCTNNNCCAGGAGTTCAAGGTTACAGT
GAGCACACGTCTGAACTCTTTCCCTACACGACGCTCTTCCGATCTNNNCCAGGAGTTCAAGGTTACAGT
AGCACACGTCTGAACTCTTTCCCTACACGACGCTCTTCCGATCTNNNAAGTGGTCCTCAGCTAAGAAA
AGCACACGTCTGAACTCTTTCCCTACACGACGCTCTTCCGATCTNNNAAGTGGTCCTCAGCTAAGAAA
AGCACACGTCTGAACTCTTTCCCTACACGACGCTCTTCCGATCTNNNCCAGCTAAGAGTGTTTCGGTCA
AGCACACGTCTGAACTCTTTCCCTACACGACGCTCTTCCGATCTNNNCCAGCTAAGAGTGTTTCGGTCA
AGCACACGTCTGAACTCTTTCCCTACACGACGCTCTTCCGATCTNNNGTGATTTCAGCTCACTGCAAACT
AGCACACGTCTGAACTCTTTCCCTACACGACGCTCTTCCGATCTNNNGTGATTTCAGCTCACTGCAAACT
GAGCACACGTCTGAACTCTTTCCCTACACGACGCTCTTCCGATCTNNNGCGACAGAGCAAGATCTTGTC
GAGCACACGTCTGAACTCTTTCCCTACACGACGCTCTTCCGATCTNNNGCGACAGAGCAAGATCTTGTC
GCACACGTCTGAACTCTTTCCCTACACGACGCTCTTCCGATCTNNNCCTTCTCGAGGCCTCATAACTAGC
GCACACGTCTGAACTCTTTCCCTACACGACGCTCTTCCGATCTNNNCCTTCTCGAGGCCTCATAACTAGC
ACACGTCTGAACTCTTTCCCTACACGACGCTCTTCCGATCTNNNTCTGACTGCTCTTTTCACCCATCCATC
ACACGTCTGAACTCTTTCCCTACACGACGCTCTTCCGATCTNNNTCTGACTGCTCTTTTCACCCATCCATC
GCACACGTCTGAACTCTTTCCCTACACGACGCTCTTCCGATCTNNNAGGCACCAAAGACATTATGGGTA
GCACACGTCTGAACTCTTTCCCTACACGACGCTCTTCCGATCTNNNAGGCACCAAAGACATTATGGGTA
AGCACACGTCTGAACTCTTTCCCTACACGACGCTCTTCCGATCTNNNAGGAAAAATACTCAGCCGGTC
AGCACACGTCTGAACTCTTTCCCTACACGACGCTCTTCCGATCTNNNAGGAAAAATACTCAGCCGGTC

AGCACACGTCTGAACTCTTTCCCTACACGACGCTCTTCCGATCTNNNCGTAACAATAATAGTTCCTAAC
AGCACACGTCTGAACTCTTTCCCTACACGACGCTCTTCCGATCTNNNCGTAACAATAATAGTTCCTAAC
GAGCACACGTCTGAACTCTTTCCCTACACGACGCTCTTCCGATCTNNNCCACAAAGCAGAAGCAGGTA
GAGCACACGTCTGAACTCTTTCCCTACACGACGCTCTTCCGATCTNNNCCACAAAGCAGAAGCAGGTA
GCACACGTCTGAACTCTTTCCCTACACGACGCTCTTCCGATCTNNNCCTCTGTGCGTATCTCTATGGAG
GCACACGTCTGAACTCTTTCCCTACACGACGCTCTTCCGATCTNNNCCTCTGTGCGTATCTCTATGGAG
GAGCACACGTCTGAACTCTTTCCCTACACGACGCTCTTCCGATCTNNNGCCCAGCCATCTTTCTTTCTTT
GAGCACACGTCTGAACTCTTTCCCTACACGACGCTCTTCCGATCTNNNGCCCAGCCATCTTTCTTTCTTT
AGCACACGTCTGAACTCTTTCCCTACACGACGCTCTTCCGATCTNNNAATCATGTAGCCCCTCCTGTTTT
AGCACACGTCTGAACTCTTTCCCTACACGACGCTCTTCCGATCTNNNAATCATGTAGCCCCTCCTGTTTT
AGCACACGTCTGAACTCTTTCCCTACACGACGCTCTTCCGATCTNNNTGTTCATCAGACCTTCCCTCCTT
AGCACACGTCTGAACTCTTTCCCTACACGACGCTCTTCCGATCTNNNTGTTCATCAGACCTTCCCTCCTT
GAGCACACGTCTGAACTCTTTCCCTACACGACGCTCTTCCGATCTNNNGGTGGTAGGTCTATAGGCAA
GAGCACACGTCTGAACTCTTTCCCTACACGACGCTCTTCCGATCTNNNGGTGGTAGGTCTATAGGCAA
GAGCACACGTCTGAACTCTTTCCCTACACGACGCTCTTCCGATCTNNNATCCTGCTGCCAAGTACAACT
GAGCACACGTCTGAACTCTTTCCCTACACGACGCTCTTCCGATCTNNNATCCTGCTGCCAAGTACAACT
GAGCACACGTCTGAACTCTTTCCCTACACGACGCTCTTCCGATCTNNNATGTTCACATGTGGCTTAAAC
GAGCACACGTCTGAACTCTTTCCCTACACGACGCTCTTCCGATCTNNNATGTTCACATGTGGCTTAAAC
GCACACGTCTGAACTCTTTCCCTACACGACGCTCTTCCGATCTNNNGTGAGCACTGCAAGATAAAAGC
GCACACGTCTGAACTCTTTCCCTACACGACGCTCTTCCGATCTNNNGTGAGCACTGCAAGATAAAAGC
AGCACACGTCTGAACTCTTTCCCTACACGACGCTCTTCCGATCTNNNGCGCTTCTATAATCCCAGCTACT
AGCACACGTCTGAACTCTTTCCCTACACGACGCTCTTCCGATCTNNNGCGCTTCTATAATCCCAGCTACT
GAGCACACGTCTGAACTCTTTCCCTACACGACGCTCTTCCGATCTNNNCAAGATCATACCACTGCACTC
GAGCACACGTCTGAACTCTTTCCCTACACGACGCTCTTCCGATCTNNNCAAGATCATACCACTGCACTC
AGCACACGTCTGAACTCTTTCCCTACACGACGCTCTTCCGATCTNNNCCCACTGGATCTAGTATGGCA
AGCACACGTCTGAACTCTTTCCCTACACGACGCTCTTCCGATCTNNNCCCACTGGATCTAGTATGGCA
AGCACACGTCTGAACTCTTTCCCTACACGACGCTCTTCCGATCTNNNGCACCACCAAAACTGAGGAAG
AGCACACGTCTGAACTCTTTCCCTACACGACGCTCTTCCGATCTNNNGCACCACCAAAACTGAGGAAG
AGCACACGTCTGAACTCTTTCCCTACACGACGCTCTTCCGATCTNNNTGGGAATTATGACAGGTGCTTG
AGCACACGTCTGAACTCTTTCCCTACACGACGCTCTTCCGATCTNNNTGGGAATTATGACAGGTGCTTG
GCACACGTCTGAACTCTTTCCCTACACGACGCTCTTCCGATCTNNNAGTGCCAGAGTACCACATTCTTT
GCACACGTCTGAACTCTTTCCCTACACGACGCTCTTCCGATCTNNNAGTGCCAGAGTACCACATTCTTT
AGCACACGTCTGAACTCTTTCCCTACACGACGCTCTTCCGATCTNNNACTTTTGGATTTGGAAGCAACC
AGCACACGTCTGAACTCTTTCCCTACACGACGCTCTTCCGATCTNNNACTTTTGGATTTGGAAGCAACC
GAGCACACGTCTGAACTCTTTCCCTACACGACGCTCTTCCGATCTNNNCTCCCAAGCAGCTGAAATTAC
GAGCACACGTCTGAACTCTTTCCCTACACGACGCTCTTCCGATCTNNNCTCCCAAGCAGCTGAAATTAC
GCACACGTCTGAACTCTTTCCCTACACGACGCTCTTCCGATCTNNNTGCTGTCTGTTCTGTGAGTCTA
GCACACGTCTGAACTCTTTCCCTACACGACGCTCTTCCGATCTNNNTGCTGTCTGTTCTGTGAGTCTA
GCACACGTCTGAACTCTTTCCCTACACGACGCTCTTCCGATCTNNNGAACCCAGGAGGCAGAGCTTCA
GCACACGTCTGAACTCTTTCCCTACACGACGCTCTTCCGATCTNNNGAACCCAGGAGGCAGAGCTTCA
AGCACACGTCTGAACTCTTTCCCTACACGACGCTCTTCCGATCTNNNCAGCTGTGTTCTGTTTGGATAG
AGCACACGTCTGAACTCTTTCCCTACACGACGCTCTTCCGATCTNNNCAGCTGTGTTCTGTTTGGATAG
AGCACACGTCTGAACTCTTTCCCTACACGACGCTCTTCCGATCTNNNTCTAATCATGTCACTACGCTCCA

AGCACACGTCTGAACTCTTTCCCTACACGACGCTCTTCCGATCTNNNTCTAATCATGTCACTACGCTCCA
AGCACACGTCTGAACTCTTTCCCTACACGACGCTCTTCCGATCTNNNATGTGTTGGCTCACTCCTGTAA
AGCACACGTCTGAACTCTTTCCCTACACGACGCTCTTCCGATCTNNNATGTGTTGGCTCACTCCTGTAA
GAGCACACGTCTGAACTCTTTCCCTACACGACGCTCTTCCGATCTNNNACAATACTGTATGTCGTCGAG
GAGCACACGTCTGAACTCTTTCCCTACACGACGCTCTTCCGATCTNNNACAATACTGTATGTCGTCGAG
GAGCACACGTCTGAACTCTTTCCCTACACGACGCTCTTCCGATCTNNNCCGTCTTTGTGGCTCTCATCTA
GAGCACACGTCTGAACTCTTTCCCTACACGACGCTCTTCCGATCTNNNCCGTCTTTGTGGCTCTCATCTA
GAGCACACGTCTGAACTCTTTCCCTACACGACGCTCTTCCGATCTNNNCCAGTAGGGAAGAGGGAAGT
GAGCACACGTCTGAACTCTTTCCCTACACGACGCTCTTCCGATCTNNNCCAGTAGGGAAGAGGGAAGT
AGCACACGTCTGAACTCTTTCCCTACACGACGCTCTTCCGATCTNNNTGATTGCAGTGAGTCAAGATCCT
AGCACACGTCTGAACTCTTTCCCTACACGACGCTCTTCCGATCTNNNTGATTGCAGTGAGTCAAGATCCT
AGCACACGTCTGAACTCTTTCCCTACACGACGCTCTTCCGATCTNNNGTCAGAGGAACTGGTACTTCA
AGCACACGTCTGAACTCTTTCCCTACACGACGCTCTTCCGATCTNNNGTCAGAGGAACTGGTACTTCA
AGCACACGTCTGAACTCTTTCCCTACACGACGCTCTTCCGATCTNNNCAATTCTGGAACTTGACCTGCT
AGCACACGTCTGAACTCTTTCCCTACACGACGCTCTTCCGATCTNNNCAATTCTGGAACTTGACCTGCT
AGCACACGTCTGAACTCTTTCCCTACACGACGCTCTTCCGATCTNNNACAGATATTCTCAAAGACAGTG
AGCACACGTCTGAACTCTTTCCCTACACGACGCTCTTCCGATCTNNNACAGATATTCTCAAAGACAGTG
AGCACACGTCTGAACTCTTTCCCTACACGACGCTCTTCCGATCTNNNGCTCTTTGGGGTCTCTCTCTCC
AGCACACGTCTGAACTCTTTCCCTACACGACGCTCTTCCGATCTNNNGCTCTTTGGGGTCTCTCTCTCC
AGCACACGTCTGAACTCTTTCCCTACACGACGCTCTTCCGATCTNNNTGTTGTTGCAAATGACAGATG
AGCACACGTCTGAACTCTTTCCCTACACGACGCTCTTCCGATCTNNNTGTTGTTGCAAATGACAGATG
AGCACACGTCTGAACTCTTTCCCTACACGACGCTCTTCCGATCTNNNGACACTCTGAAGGGTTCATTTG
AGCACACGTCTGAACTCTTTCCCTACACGACGCTCTTCCGATCTNNNGACACTCTGAAGGGTTCATTTG
AGCACACGTCTGAACTCTTTCCCTACACGACGCTCTTCCGATCTNNNGATGGGCACACACAGAGAGATA
AGCACACGTCTGAACTCTTTCCCTACACGACGCTCTTCCGATCTNNNGATGGGCACACACAGAGAGATA
AGCACACGTCTGAACTCTTTCCCTACACGACGCTCTTCCGATCTNNNCTGTAATCCCAGCTACTCAGGAC
AGCACACGTCTGAACTCTTTCCCTACACGACGCTCTTCCGATCTNNNCTGTAATCCCAGCTACTCAGGAC
GAGCACACGTCTGAACTCTTTCCCTACACGACGCTCTTCCGATCTNNNGAGGCAGAAGAATCGCTTGAA
GAGCACACGTCTGAACTCTTTCCCTACACGACGCTCTTCCGATCTNNNGAGGCAGAAGAATCGCTTGAA
GAGCACACGTCTGAACTCTTTCCCTACACGACGCTCTTCCGATCTNNNGGCCCCTTCCTTCCAAGATTTTA
GAGCACACGTCTGAACTCTTTCCCTACACGACGCTCTTCCGATCTNNNGGCCCCTTCCTTCCAAGATTTTA
AGAGCACACGTCTGAACTCTTTCCCTACACGACGCTCTTCCGATCTNNNAAGGTTATGGTGAGCTATG
AGAGCACACGTCTGAACTCTTTCCCTACACGACGCTCTTCCGATCTNNNAAGGTTATGGTGAGCTATG
GAGCACACGTCTGAACTCTTTCCCTACACGACGCTCTTCCGATCTNNNGTTTTGCACTGTGGGAAAAAT
GAGCACACGTCTGAACTCTTTCCCTACACGACGCTCTTCCGATCTNNNGTTTTGCACTGTGGGAAAAAT
GAGCACACGTCTGAACTCTTTCCCTACACGACGCTCTTCCGATCTNNNCTGAAGCAGGAGAATGGTGT
GAGCACACGTCTGAACTCTTTCCCTACACGACGCTCTTCCGATCTNNNCTGAAGCAGGAGAATGGTGT
AGCACACGTCTGAACTCTTTCCCTACACGACGCTCTTCCGATCTNNNAGTGCTGGGATTATAGGCATGA
AGCACACGTCTGAACTCTTTCCCTACACGACGCTCTTCCGATCTNNNAGTGCTGGGATTATAGGCATGA
CACACGTCTGAACTCTTTCCCTACACGACGCTCTTCCGATCTNNNTCAAAGCAGTACTAAAGGGGCAT
CACACGTCTGAACTCTTTCCCTACACGACGCTCTTCCGATCTNNNTCAAAGCAGTACTAAAGGGGCAT
AGAGCACACGTCTGAACTCTTTCCCTACACGACGCTCTTCCGATCTNNNTCACAACTTAAGCTGGTGTCT
AGAGCACACGTCTGAACTCTTTCCCTACACGACGCTCTTCCGATCTNNNTCACAACTTAAGCTGGTGTCT

AGCACACGTCTGAACTCTTTCCCTACACGACGCTCTTCCGATCTNNNCCAGGTTCAAGTGATTCTCTTGC

AGCACACGTCTGAACTCTTTCCCTACACGACGCTCTTCCGATCTNNNCCAGGTTCAAGTGATTCTCTTGC

GAGCACACGTCTGAACTCTTTCCCTACACGACGCTCTTCCGATCTNNNCCCCTGATAGTTACACCTTAGC

GAGCACACGTCTGAACTCTTTCCCTACACGACGCTCTTCCGATCTNNNCCCCTGATAGTTACACCTTAGC

AGAGCACACGTCTGAACTCTTTCCCTACACGACGCTCTTCCGATCTNNNCCACCTCGGGTAGAATTTGAT

AGAGCACACGTCTGAACTCTTTCCCTACACGACGCTCTTCCGATCTNNNCCACCTCGGGTAGAATTTGAT

AGCACACGTCTGAACTCTTTCCCTACACGACGCTCTTCCGATCTNNNTGATCTCTCCATGTTTTCCAGGTC

AGCACACGTCTGAACTCTTTCCCTACACGACGCTCTTCCGATCTNNNTGATCTCTCCATGTTTTCCAGGTC

AGCACACGTCTGAACTCTTTCCCTACACGACGCTCTTCCGATCTNNNTTTGACTTAGGCATGGATTTCCC

AGCACACGTCTGAACTCTTTCCCTACACGACGCTCTTCCGATCTNNNTTTGACTTAGGCATGGATTTCCC

GAGCACACGTCTGAACTCTTTCCCTACACGACGCTCTTCCGATCTNNNCCCAAATGCCCATCAATCAACA

GAGCACACGTCTGAACTCTTTCCCTACACGACGCTCTTCCGATCTNNNCCCAAATGCCCATCAATCAACA

AGCACACGTCTGAACTCTTTCCCTACACGACGCTCTTCCGATCTNNNTGAGAAAACCTATGAATGTAG

AGCACACGTCTGAACTCTTTCCCTACACGACGCTCTTCCGATCTNNNTGAGAAAACCTATGAATGTAG

AGCACACGTCTGAACTCTTTCCCTACACGACGCTCTTCCGATCTNNNCAGGCATTTTTATAGCAGCACG

AGCACACGTCTGAACTCTTTCCCTACACGACGCTCTTCCGATCTNNNCAGGCATTTTTATAGCAGCACG

AGCACACGTCTGAACTCTTTCCCTACACGACGCTCTTCCGATCTNNNAAAATTAGCCAGGCATGATGGT

AGCACACGTCTGAACTCTTTCCCTACACGACGCTCTTCCGATCTNNNAAAATTAGCCAGGCATGATGGT

GCACACGTCTGAACTCTTTCCCTACACGACGCTCTTCCGATCTNNNTACCCACCTCTTTTCCACTTTCCCAT

GCACACGTCTGAACTCTTTCCCTACACGACGCTCTTCCGATCTNNNTACCCACCTCTTTTCCACTTTCCCAT

GAGCACACGTCTGAACTCTTTCCCTACACGACGCTCTTCCGATCTNNNGCAAAGCAACATTGATCCAGC

GAGCACACGTCTGAACTCTTTCCCTACACGACGCTCTTCCGATCTNNNGCAAAGCAACATTGATCCAGC

AGCACACGTCTGAACTCTTTCCCTACACGACGCTCTTCCGATCTNNNGTTTGAGCTGATGGATACCCTG

AGCACACGTCTGAACTCTTTCCCTACACGACGCTCTTCCGATCTNNNGTTTGAGCTGATGGATACCCTG

CACACGTCTGAACTCTTTCCCTACACGACGCTCTTCCGATCTNNNCTCTCCTTCTCCCTTTTTCCTCCCATC

CACACGTCTGAACTCTTTCCCTACACGACGCTCTTCCGATCTNNNCTCTCCTTCTCCCTTTTTCCTCCCATC

AGCACACGTCTGAACTCTTTCCCTACACGACGCTCTTCCGATCTNNNATGGGTACAAAGCATAGTGCTC

AGCACACGTCTGAACTCTTTCCCTACACGACGCTCTTCCGATCTNNNATGGGTACAAAGCATAGTGCTC

GAGCACACGTCTGAACTCTTTCCCTACACGACGCTCTTCCGATCTNNNCAACATGGAGAAACCCAGTCT

GAGCACACGTCTGAACTCTTTCCCTACACGACGCTCTTCCGATCTNNNCAACATGGAGAAACCCAGTCT

AGCACACGTCTGAACTCTTTCCCTACACGACGCTCTTCCGATCTNNNGATCCTGGTGAATTTCTTTGCCA

AGCACACGTCTGAACTCTTTCCCTACACGACGCTCTTCCGATCTNNNGATCCTGGTGAATTTCTTTGCCA

AGCACACGTCTGAACTCTTTCCCTACACGACGCTCTTCCGATCTNNNTGCTGGTCAATTAGAGGTAAGC

AGCACACGTCTGAACTCTTTCCCTACACGACGCTCTTCCGATCTNNNTGCTGGTCAATTAGAGGTAAGC

AGCACACGTCTGAACTCTTTCCCTACACGACGCTCTTCCGATCTNNNCCATACATGTATTGGCCACTTGT

AGCACACGTCTGAACTCTTTCCCTACACGACGCTCTTCCGATCTNNNCCATACATGTATTGGCCACTTGT

AGCACACGTCTGAACTCTTTCCCTACACGACGCTCTTCCGATCTNNNTCGAGTAATGAGTTCAGATGGC

AGCACACGTCTGAACTCTTTCCCTACACGACGCTCTTCCGATCTNNNTCGAGTAATGAGTTCAGATGGC

GAGCACACGTCTGAACTCTTTCCCTACACGACGCTCTTCCGATCTNNNGCATACCTGTAATCCCAGCTAC

GAGCACACGTCTGAACTCTTTCCCTACACGACGCTCTTCCGATCTNNNGCATACCTGTAATCCCAGCTAC

GAGCACACGTCTGAACTCTTTCCCTACACGACGCTCTTCCGATCTNNNCTCACACACACACACATACACA

GAGCACACGTCTGAACTCTTTCCCTACACGACGCTCTTCCGATCTNNNCTCACACACACACACATACACA

GAGCACACGTCTGAACTCTTTCCCTACACGACGCTCTTCCGATCTNNNGCTACTCTGTCTTATCCAGCCTC

AGCACACGTCTGAACTCTTTCCCTACACGACGCTCTTCCGATCTNNNGCTACTCTGTCTTATCCAGCCT
GCACACGTCTGAACTCTTTCCCTACACGACGCTCTTCCGATCTNNNGAGGTTGCAGTGAGCCTAGAT
GCACACGTCTGAACTCTTTCCCTACACGACGCTCTTCCGATCTNNNGAGGTTGCAGTGAGCCTAGAT
GAGCACACGTCTGAACTCTTTCCCTACACGACGCTCTTCCGATCTNNNGCACCTGTAGTCTCAGCTACT
GAGCACACGTCTGAACTCTTTCCCTACACGACGCTCTTCCGATCTNNNGCACCTGTAGTCTCAGCTACT
AGCACACGTCTGAACTCTTTCCCTACACGACGCTCTTCCGATCTNNNGGATCCCTTGAACCTAGGAGT
AGCACACGTCTGAACTCTTTCCCTACACGACGCTCTTCCGATCTNNNGGATCCCTTGAACCTAGGAGT
GCACACGTCTGAACTCTTTCCCTACACGACGCTCTTCCGATCTNNNGTTTGCAGTGAGCTGAGATCAT
GCACACGTCTGAACTCTTTCCCTACACGACGCTCTTCCGATCTNNNGTTTGCAGTGAGCTGAGATCAT
CACACGTCTGAACTCTTTCCCTACACGACGCTCTTCCGATCTNNNCCTGTCTCAAACACACACACACCA
CACACGTCTGAACTCTTTCCCTACACGACGCTCTTCCGATCTNNNCCTGTCTCAAACACACACACACCA
GCACACGTCTGAACTCTTTCCCTACACGACGCTCTTCCGATCTNNNTTTAGCCATGACCACTTATGCCT
GCACACGTCTGAACTCTTTCCCTACACGACGCTCTTCCGATCTNNNTTTAGCCATGACCACTTATGCCT
CACACGTCTGAACTCTTTCCCTACACGACGCTCTTCCGATCTNNNATTCCCATCTCCCTCCCTAGTCCAT
CACACGTCTGAACTCTTTCCCTACACGACGCTCTTCCGATCTNNNATTCCCATCTCCCTCCCTAGTCCAT
CACACGTCTGAACTCTTTCCCTACACGACGCTCTTCCGATCTNNNCATCTTGGATTTCAGTGCCCTTCC
CACACGTCTGAACTCTTTCCCTACACGACGCTCTTCCGATCTNNNCATCTTGGATTTCAGTGCCCTTCC
GAGCACACGTCTGAACTCTTTCCCTACACGACGCTCTTCCGATCTNNNACTTGCCCCCACAATTTTCTTT
GAGCACACGTCTGAACTCTTTCCCTACACGACGCTCTTCCGATCTNNNACTTGCCCCCACAATTTTCTTT
AGCACACGTCTGAACTCTTTCCCTACACGACGCTCTTCCGATCTNNNTTGCAGTGAGCTGAGATCACC
AGCACACGTCTGAACTCTTTCCCTACACGACGCTCTTCCGATCTNNNTTGCAGTGAGCTGAGATCACC
GAGCACACGTCTGAACTCTTTCCCTACACGACGCTCTTCCGATCTNNNTAGGCATGAAGATAGCTGTC
GAGCACACGTCTGAACTCTTTCCCTACACGACGCTCTTCCGATCTNNNTAGGCATGAAGATAGCTGTC
CACACGTCTGAACTCTTTCCCTACACGACGCTCTTCCGATCTNNNCCCGGTCAAGTTTTTTTTCTGGTCA
CACACGTCTGAACTCTTTCCCTACACGACGCTCTTCCGATCTNNNCCCGGTCAAGTTTTTTTTCTGGTCA
AGCACACGTCTGAACTCTTTCCCTACACGACGCTCTTCCGATCTNNNCACAATTCCATAACTCCCCCAC
AGCACACGTCTGAACTCTTTCCCTACACGACGCTCTTCCGATCTNNNCACAATTCCATAACTCCCCCAC
GCACACGTCTGAACTCTTTCCCTACACGACGCTCTTCCGATCTNNNGAACTATCATCCCTGACGCCTAG
GCACACGTCTGAACTCTTTCCCTACACGACGCTCTTCCGATCTNNNGAACTATCATCCCTGACGCCTAG
AGCACACGTCTGAACTCTTTCCCTACACGACGCTCTTCCGATCTNNNTAACCGCTGGACTTTGTTGAA
AGCACACGTCTGAACTCTTTCCCTACACGACGCTCTTCCGATCTNNNTAACCGCTGGACTTTGTTGAA
GCACACGTCTGAACTCTTTCCCTACACGACGCTCTTCCGATCTNNNAGGGAAATGGGGAAAGGGTA
GCACACGTCTGAACTCTTTCCCTACACGACGCTCTTCCGATCTNNNAGGGAAATGGGGAAAGGGTA
AGCACACGTCTGAACTCTTTCCCTACACGACGCTCTTCCGATCTNNNCTGATGCCTGCAAATCCTTGG
AGCACACGTCTGAACTCTTTCCCTACACGACGCTCTTCCGATCTNNNCTGATGCCTGCAAATCCTTGG
GCACACGTCTGAACTCTTTCCCTACACGACGCTCTTCCGATCTNNNCGTCCTGATGATTTTTATTTTTG
GCACACGTCTGAACTCTTTCCCTACACGACGCTCTTCCGATCTNNNCGTCCTGATGATTTTTATTTTTG
GAGCACACGTCTGAACTCTTTCCCTACACGACGCTCTTCCGATCTNNNTTGAGACAGGAGAATGGCA
GAGCACACGTCTGAACTCTTTCCCTACACGACGCTCTTCCGATCTNNNTTGAGACAGGAGAATGGCA
AGCACACGTCTGAACTCTTTCCCTACACGACGCTCTTCCGATCTNNNGAAGGGAGGAAACCACTGTA
AGCACACGTCTGAACTCTTTCCCTACACGACGCTCTTCCGATCTNNNGAAGGGAGGAAACCACTGTA
GCACACGTCTGAACTCTTTCCCTACACGACGCTCTTCCGATCTNNNGCCTCCTCTTCTTCCTCTTCTTCC
GCACACGTCTGAACTCTTTCCCTACACGACGCTCTTCCGATCTNNNGCCTCCTCTTCTTCCTCTTCTTCC

AGCACACGTCTGAACTCTTTCCCTACACGACGCTCTTCCGATCTNNNCTGTGAATCCATCTGGTCCTAG

AGCACACGTCTGAACTCTTTCCCTACACGACGCTCTTCCGATCTNNNCTGTGAATCCATCTGGTCCTAG

GAGCACACGTCTGAACTCTTTCCCTACACGACGCTCTTCCGATCTNNNCCATACTGTGCATCAAATCTG

GAGCACACGTCTGAACTCTTTCCCTACACGACGCTCTTCCGATCTNNNCCATACTGTGCATCAAATCTG

GCACACGTCTGAACTCTTTCCCTACACGACGCTCTTCCGATCTNNNTTTCCTGGGCACATACAACCTC

GCACACGTCTGAACTCTTTCCCTACACGACGCTCTTCCGATCTNNNTTTCCTGGGCACATACAACCTC

AGCACACGTCTGAACTCTTTCCCTACACGACGCTCTTCCGATCTNNNCCACTGATATTTGTAGCTGGG

AGCACACGTCTGAACTCTTTCCCTACACGACGCTCTTCCGATCTNNNCCACTGATATTTGTAGCTGGG

AGCACACGTCTGAACTCTTTCCCTACACGACGCTCTTCCGATCTNNNCTGTCATATCAGCTACTCGGGA

AGCACACGTCTGAACTCTTTCCCTACACGACGCTCTTCCGATCTNNNCTGTCATATCAGCTACTCGGGA

GAGCACACGTCTGAACTCTTTCCCTACACGACGCTCTTCCGATCTNNNGGTGAGGGGGATAGGGAA

GAGCACACGTCTGAACTCTTTCCCTACACGACGCTCTTCCGATCTNNNGGTGAGGGGGATAGGGAA

AGCACACGTCTGAACTCTTTCCCTACACGACGCTCTTCCGATCTNNNCTCTAGCCTGGGCAATGAGAG

AGCACACGTCTGAACTCTTTCCCTACACGACGCTCTTCCGATCTNNNCTCTAGCCTGGGCAATGAGAG

GAGCACACGTCTGAACTCTTTCCCTACACGACGCTCTTCCGATCTNNNTTGGAATGGGAATGCCTACT

GAGCACACGTCTGAACTCTTTCCCTACACGACGCTCTTCCGATCTNNNTTGGAATGGGAATGCCTACT

GAGCACACGTCTGAACTCTTTCCCTACACGACGCTCTTCCGATCTNNNGGTGAAAGAATGAGACCCTG

GAGCACACGTCTGAACTCTTTCCCTACACGACGCTCTTCCGATCTNNNGGTGAAAGAATGAGACCCTG

AGCACACGTCTGAACTCTTTCCCTACACGACGCTCTTCCGATCTNNNCACCTGTAATCCCAGCTACTCAG

AGCACACGTCTGAACTCTTTCCCTACACGACGCTCTTCCGATCTNNNCACCTGTAATCCCAGCTACTCAG

GCACACGTCTGAACTCTTTCCCTACACGACGCTCTTCCGATCTNNNAGGAGGCTAAGGTAGGAGAAT

GCACACGTCTGAACTCTTTCCCTACACGACGCTCTTCCGATCTNNNAGGAGGCTAAGGTAGGAGAAT

AGCACACGTCTGAACTCTTTCCCTACACGACGCTCTTCCGATCTNNNTCCAGGTTACCCAATTTTTTGCC

AGCACACGTCTGAACTCTTTCCCTACACGACGCTCTTCCGATCTNNNTCCAGGTTACCCAATTTTTTGCC

CACGTCTGAACTCTTTCCCTACACGACGCTCTTCCGATCTNNNGATGTTCCTCAGTCCAAAACCATCGG

CACGTCTGAACTCTTTCCCTACACGACGCTCTTCCGATCTNNNGATGTTCCTCAGTCCAAAACCATCGG

GCACACGTCTGAACTCTTTCCCTACACGACGCTCTTCCGATCTNNNCTTTGCCTTTTCTCTGCTCTTGG

GCACACGTCTGAACTCTTTCCCTACACGACGCTCTTCCGATCTNNNCTTTGCCTTTTCTCTGCTCTTGG

GCACACGTCTGAACTCTTTCCCTACACGACGCTCTTCCGATCTNNNTGCTCCCTCAAATCAGTGAAAG

GCACACGTCTGAACTCTTTCCCTACACGACGCTCTTCCGATCTNNNTGCTCCCTCAAATCAGTGAAAG

AGCACACGTCTGAACTCTTTCCCTACACGACGCTCTTCCGATCTNNNTGACAGACTCACACTACCACAT

AGCACACGTCTGAACTCTTTCCCTACACGACGCTCTTCCGATCTNNNTGACAGACTCACACTACCACAT

GAGCACACGTCTGAACTCTTTCCCTACACGACGCTCTTCCGATCTNNNAGCACAAAACAGACTGAGAC

GAGCACACGTCTGAACTCTTTCCCTACACGACGCTCTTCCGATCTNNNAGCACAAAACAGACTGAGAC

AGCACACGTCTGAACTCTTTCCCTACACGACGCTCTTCCGATCTNNNCTGGTGATACCCAGGTGAACA

AGCACACGTCTGAACTCTTTCCCTACACGACGCTCTTCCGATCTNNNCTGGTGATACCCAGGTGAACA

GCACACGTCTGAACTCTTTCCCTACACGACGCTCTTCCGATCTNNNATTATAGACGCCCACCACTACAC

GCACACGTCTGAACTCTTTCCCTACACGACGCTCTTCCGATCTNNNATTATAGACGCCCACCACTACAC

GCACACGTCTGAACTCTTTCCCTACACGACGCTCTTCCGATCTNNNCAGAGCCTGACAAGGATTCAGA

GCACACGTCTGAACTCTTTCCCTACACGACGCTCTTCCGATCTNNNCAGAGCCTGACAAGGATTCAGA

GCACACGTCTGAACTCTTTCCCTACACGACGCTCTTCCGATCTNNNAACACCATCAAGATCTTCCCTCC

GCACACGTCTGAACTCTTTCCCTACACGACGCTCTTCCGATCTNNNAACACCATCAAGATCTTCCCTCC

AGCACACGTCTGAACTCTTTCCCTACACGACGCTCTTCCGATCTNNNATCATAGCTCACTGCAACCTCA

AGCACACGTCTGAACTCTTTCCCTACACGACGCTCTTCCGATCTNNNATCATAGCTCACTGCAACCTCA
AGCACACGTCTGAACTCTTTCCCTACACGACGCTCTTCCGATCTNNNAGTGAAGAGGAATGTGGGCT
AGCACACGTCTGAACTCTTTCCCTACACGACGCTCTTCCGATCTNNNAGTGAAGAGGAATGTGGGCT
GAGCACACGTCTGAACTCTTTCCCTACACGACGCTCTTCCGATCTNNNGGTCATGCACAAATAAATCC
GAGCACACGTCTGAACTCTTTCCCTACACGACGCTCTTCCGATCTNNNGGTCATGCACAAATAAATCC
AGCACACGTCTGAACTCTTTCCCTACACGACGCTCTTCCGATCTNNNCTGTACTTAGTCCTGCGAACAC
AGCACACGTCTGAACTCTTTCCCTACACGACGCTCTTCCGATCTNNNCTGTACTTAGTCCTGCGAACAC
GCACACGTCTGAACTCTTTCCCTACACGACGCTCTTCCGATCTNNNGCCTTTGCTATGACTTTCTGCAT
GCACACGTCTGAACTCTTTCCCTACACGACGCTCTTCCGATCTNNNGCCTTTGCTATGACTTTCTGCAT
AGCACACGTCTGAACTCTTTCCCTACACGACGCTCTTCCGATCTNNNCCTGCTCTTCATCAATCCTCCCT
AGCACACGTCTGAACTCTTTCCCTACACGACGCTCTTCCGATCTNNNCCTGCTCTTCATCAATCCTCCCT
AGCACACGTCTGAACTCTTTCCCTACACGACGCTCTTCCGATCTNNNTATAATTGCACTACAGGCTGGC
AGCACACGTCTGAACTCTTTCCCTACACGACGCTCTTCCGATCTNNNTATAATTGCACTACAGGCTGGC
GAGCACACGTCTGAACTCTTTCCCTACACGACGCTCTTCCGATCTNNNCTCTCCTCCTGTGACCCTTC
GAGCACACGTCTGAACTCTTTCCCTACACGACGCTCTTCCGATCTNNNCTCTCCTCCTGTGACCCTTC
AAGAGCACACGTCTGAACTCTTTCCCTACACGACGCTCTTCCGATCTNNNGAGGACCAACACATTGTC
AAGAGCACACGTCTGAACTCTTTCCCTACACGACGCTCTTCCGATCTNNNGAGGACCAACACATTGTC
AGCACACGTCTGAACTCTTTCCCTACACGACGCTCTTCCGATCTNNNCTGTGCAGCATGATTCCTAAC
AGCACACGTCTGAACTCTTTCCCTACACGACGCTCTTCCGATCTNNNCTGTGCAGCATGATTCCTAAC
AGCACACGTCTGAACTCTTTCCCTACACGACGCTCTTCCGATCTNNNAATATCTGGCTTCGTGTGAGC
AGCACACGTCTGAACTCTTTCCCTACACGACGCTCTTCCGATCTNNNAATATCTGGCTTCGTGTGAGC
CACACGTCTGAACTCTTTCCCTACACGACGCTCTTCCGATCTNNNGCCCTATGGAATTGTAACTTGCCC
CACACGTCTGAACTCTTTCCCTACACGACGCTCTTCCGATCTNNNGCCCTATGGAATTGTAACTTGCCC
GCACACGTCTGAACTCTTTCCCTACACGACGCTCTTCCGATCTNNNCACCTGGAAATTCTCCCTTCAGA
GCACACGTCTGAACTCTTTCCCTACACGACGCTCTTCCGATCTNNNCACCTGGAAATTCTCCCTTCAGA
CACACGTCTGAACTCTTTCCCTACACGACGCTCTTCCGATCTNNNACCTCTCCATCTTTGTTTAGCACTC
CACACGTCTGAACTCTTTCCCTACACGACGCTCTTCCGATCTNNNACCTCTCCATCTTTGTTTAGCACTC
GAGCACACGTCTGAACTCTTTCCCTACACGACGCTCTTCCGATCTNNNAATACAAAAAATTAGCCGGG
GAGCACACGTCTGAACTCTTTCCCTACACGACGCTCTTCCGATCTNNNAATACAAAAAATTAGCCGGG
GAGCACACGTCTGAACTCTTTCCCTACACGACGCTCTTCCGATCTNNNTGAGTCCAGGAATTCAAGAC
GAGCACACGTCTGAACTCTTTCCCTACACGACGCTCTTCCGATCTNNNTGAGTCCAGGAATTCAAGAC
AGCACACGTCTGAACTCTTTCCCTACACGACGCTCTTCCGATCTNNNCAACATGAGACTTGCTGCACAT
AGCACACGTCTGAACTCTTTCCCTACACGACGCTCTTCCGATCTNNNCAACATGAGACTTGCTGCACAT
GCACACGTCTGAACTCTTTCCCTACACGACGCTCTTCCGATCTNNNAACAGGGAGACACATACATCAC
GCACACGTCTGAACTCTTTCCCTACACGACGCTCTTCCGATCTNNNAACAGGGAGACACATACATCAC
AGCACACGTCTGAACTCTTTCCCTACACGACGCTCTTCCGATCTNNNCGGTGATGCCAGTGTAAACTTT
AGCACACGTCTGAACTCTTTCCCTACACGACGCTCTTCCGATCTNNNCGGTGATGCCAGTGTAAACTTT
GCACACGTCTGAACTCTTTCCCTACACGACGCTCTTCCGATCTNNNCTGTGACCTGGCCTCACATGCAT
GCACACGTCTGAACTCTTTCCCTACACGACGCTCTTCCGATCTNNNCTGTGACCTGGCCTCACATGCAT
AGCACACGTCTGAACTCTTTCCCTACACGACGCTCTTCCGATCTNNNACCTTCCCTCCACTATTGTCCTA
AGCACACGTCTGAACTCTTTCCCTACACGACGCTCTTCCGATCTNNNACCTTCCCTCCACTATTGTCCTA
GAGCACACGTCTGAACTCTTTCCCTACACGACGCTCTTCCGATCTNNNGCAGAGTCGGAAAGAATGC
GAGCACACGTCTGAACTCTTTCCCTACACGACGCTCTTCCGATCTNNNGCAGAGTCGGAAAGAATGC

AGCACACGTCTGAACTCTTTCCCTACACGACGCTCTTCCGATCTNNNAGGAAATCAGAGAGAGTGGGA

AGCACACGTCTGAACTCTTTCCCTACACGACGCTCTTCCGATCTNNNAGGAAATCAGAGAGAGTGGGA

AGCACACGTCTGAACTCTTTCCCTACACGACGCTCTTCCGATCTNNNTCTGAGGTGGGGTTTCAGAATT

AGCACACGTCTGAACTCTTTCCCTACACGACGCTCTTCCGATCTNNNTCTGAGGTGGGGTTTCAGAATT

GAGCACACGTCTGAACTCTTTCCCTACACGACGCTCTTCCGATCTNNNTGAATGGATCCCTTCTTTGGA

GAGCACACGTCTGAACTCTTTCCCTACACGACGCTCTTCCGATCTNNNTGAATGGATCCCTTCTTTGGA

GAGCACACGTCTGAACTCTTTCCCTACACGACGCTCTTCCGATCTNNNCCAAGATTGCACCACTACACTT

GAGCACACGTCTGAACTCTTTCCCTACACGACGCTCTTCCGATCTNNNCCAAGATTGCACCACTACACTT

AGCACACGTCTGAACTCTTTCCCTACACGACGCTCTTCCGATCTNNNGTTGCAGTGAGCCGAAATCAT

AGCACACGTCTGAACTCTTTCCCTACACGACGCTCTTCCGATCTNNNGTTGCAGTGAGCCGAAATCAT

AGAGCACACGTCTGAACTCTTTCCCTACACGACGCTCTTCCGATCTNNNGGAAGATTCCGCCTCAAAAA

AGAGCACACGTCTGAACTCTTTCCCTACACGACGCTCTTCCGATCTNNNGGAAGATTCCGCCTCAAAAA

GAGCACACGTCTGAACTCTTTCCCTACACGACGCTCTTCCGATCTNNNTGGGTGACAGTGAGATCCTAT

GAGCACACGTCTGAACTCTTTCCCTACACGACGCTCTTCCGATCTNNNTGGGTGACAGTGAGATCCTAT

GAGCACACGTCTGAACTCTTTCCCTACACGACGCTCTTCCGATCTNNNACAGATTGCTGAGAAGTTGG

GAGCACACGTCTGAACTCTTTCCCTACACGACGCTCTTCCGATCTNNNACAGATTGCTGAGAAGTTGG

AGCACACGTCTGAACTCTTTCCCTACACGACGCTCTTCCGATCTNNNCCTTTGTCTCTCTCTGCATCTGTC

AGCACACGTCTGAACTCTTTCCCTACACGACGCTCTTCCGATCTNNNCCTTTGTCTCTCTCTGCATCTGTC

AGCACACGTCTGAACTCTTTCCCTACACGACGCTCTTCCGATCTNNNTGTCCTTTGCTATGTTGGCAGTA

AGCACACGTCTGAACTCTTTCCCTACACGACGCTCTTCCGATCTNNNTGTCCTTTGCTATGTTGGCAGTA

AGCACACGTCTGAACTCTTTCCCTACACGACGCTCTTCCGATCTNNNACTTGACCTATTGGACTGTTGT

AGCACACGTCTGAACTCTTTCCCTACACGACGCTCTTCCGATCTNNNACTTGACCTATTGGACTGTTGT

GAGCACACGTCTGAACTCTTTCCCTACACGACGCTCTTCCGATCTNNNCACAATTTCGGCTCACTGCAAG

GAGCACACGTCTGAACTCTTTCCCTACACGACGCTCTTCCGATCTNNNCACAATTTCGGCTCACTGCAAG

GAGCACACGTCTGAACTCTTTCCCTACACGACGCTCTTCCGATCTNNNCAATAGCATTTTCATCCCCTGC

GAGCACACGTCTGAACTCTTTCCCTACACGACGCTCTTCCGATCTNNNCAATAGCATTTTCATCCCCTGC

AGCACACGTCTGAACTCTTTCCCTACACGACGCTCTTCCGATCTNNNTAGCTTTCTGTCATATCCTGCCC

AGCACACGTCTGAACTCTTTCCCTACACGACGCTCTTCCGATCTNNNTAGCTTTCTGTCATATCCTGCCC

GAGCACACGTCTGAACTCTTTCCCTACACGACGCTCTTCCGATCTNNNCTCCTGTCTGGGGTATGGAAT

GAGCACACGTCTGAACTCTTTCCCTACACGACGCTCTTCCGATCTNNNCTCCTGTCTGGGGTATGGAAT

AGCACACGTCTGAACTCTTTCCCTACACGACGCTCTTCCGATCTNNNGATGGTCCTAGATGGTCCCATT

AGCACACGTCTGAACTCTTTCCCTACACGACGCTCTTCCGATCTNNNGATGGTCCTAGATGGTCCCATT

GAGCACACGTCTGAACTCTTTCCCTACACGACGCTCTTCCGATCTNNNTAGCACCGCTAGTACTAGTAC

GAGCACACGTCTGAACTCTTTCCCTACACGACGCTCTTCCGATCTNNNTAGCACCGCTAGTACTAGTAC

AGCACACGTCTGAACTCTTTCCCTACACGACGCTCTTCCGATCTNNNCCTGGGCAATTTAGCAAGAAA

AGCACACGTCTGAACTCTTTCCCTACACGACGCTCTTCCGATCTNNNCCTGGGCAATTTAGCAAGAAA

AGCACACGTCTGAACTCTTTCCCTACACGACGCTCTTCCGATCTNNNTTGAGGCAGGAGAATTGCTTGA

AGCACACGTCTGAACTCTTTCCCTACACGACGCTCTTCCGATCTNNNTTGAGGCAGGAGAATTGCTTGA

AGCACACGTCTGAACTCTTTCCCTACACGACGCTCTTCCGATCTNNNTATTCAGAGATTAGTAGGGGC

AGCACACGTCTGAACTCTTTCCCTACACGACGCTCTTCCGATCTNNNTATTCAGAGATTAGTAGGGGC

GAGCACACGTCTGAACTCTTTCCCTACACGACGCTCTTCCGATCTNNNCCTCACATTATATTGATAGAG

GAGCACACGTCTGAACTCTTTCCCTACACGACGCTCTTCCGATCTNNNCCTCACATTATATTGATAGAG

GAGCACACGTCTGAACTCTTTCCCTACACGACGCTCTTCCGATCTNNNATTTACCCTCAGAGCCTGAAG

GAGCACACGTCTGAACTCTTTCCCTACACGACGCTCTTCCGATCTNNNATTTACCCTCAGAGCCTGAAG
AGCACACGTCTGAACTCTTTCCCTACACGACGCTCTTCCGATCTNNNTGCATGGAAGTTCAGGACATAC
AGCACACGTCTGAACTCTTTCCCTACACGACGCTCTTCCGATCTNNNTGCATGGAAGTTCAGGACATAC
GAGCACACGTCTGAACTCTTTCCCTACACGACGCTCTTCCGATCTNNNGTCTCTGTTCAAGTCTCTCTGC
GAGCACACGTCTGAACTCTTTCCCTACACGACGCTCTTCCGATCTNNNGTCTCTGTTCAAGTCTCTCTGC
GAGCACACGTCTGAACTCTTTCCCTACACGACGCTCTTCCGATCTNNNAATACAAGCTACTAACGCCTCC
GAGCACACGTCTGAACTCTTTCCCTACACGACGCTCTTCCGATCTNNNAATACAAGCTACTAACGCCTCC
AGCACACGTCTGAACTCTTTCCCTACACGACGCTCTTCCGATCTNNNAAGAAAAAAGACTGTGCAGGG
AGCACACGTCTGAACTCTTTCCCTACACGACGCTCTTCCGATCTNNNAAGAAAAAAGACTGTGCAGGG
AGCACACGTCTGAACTCTTTCCCTACACGACGCTCTTCCGATCTNNNAAAAAAAAATTGGCCAGGTGTG
AGCACACGTCTGAACTCTTTCCCTACACGACGCTCTTCCGATCTNNNAAAAAAAAATTGGCCAGGTGTG
AGCACACGTCTGAACTCTTTCCCTACACGACGCTCTTCCGATCTNNNAGGTCTGTTGCCTGTTGTGATA
AGCACACGTCTGAACTCTTTCCCTACACGACGCTCTTCCGATCTNNNAGGTCTGTTGCCTGTTGTGATA
AGCACACGTCTGAACTCTTTCCCTACACGACGCTCTTCCGATCTNNNAAGTGTGCATAACCTTAGAGCC
AGCACACGTCTGAACTCTTTCCCTACACGACGCTCTTCCGATCTNNNAAGTGTGCATAACCTTAGAGCC
AGCACACGTCTGAACTCTTTCCCTACACGACGCTCTTCCGATCTNNNGTGCCTCTAATTGCAGCTACTTC
AGCACACGTCTGAACTCTTTCCCTACACGACGCTCTTCCGATCTNNNGTGCCTCTAATTGCAGCTACTTC
GAGCACACGTCTGAACTCTTTCCCTACACGACGCTCTTCCGATCTNNNAAAATGCAGACTATTGGGCCC
GAGCACACGTCTGAACTCTTTCCCTACACGACGCTCTTCCGATCTNNNAAAATGCAGACTATTGGGCCC
AGAGCACACGTCTGAACTCTTTCCCTACACGACGCTCTTCCGATCTNNNAGGAGAATGGCGTGAACCT
AGAGCACACGTCTGAACTCTTTCCCTACACGACGCTCTTCCGATCTNNNAGGAGAATGGCGTGAACCT
AGCACACGTCTGAACTCTTTCCCTACACGACGCTCTTCCGATCTNNNCCATCCAGTTTTCTCCAGTACCAC
AGCACACGTCTGAACTCTTTCCCTACACGACGCTCTTCCGATCTNNNCCATCCAGTTTTCTCCAGTACCAC
GAGCACACGTCTGAACTCTTTCCCTACACGACGCTCTTCCGATCTNNNGTACTAGGGCTTCAAGTCAGC
GAGCACACGTCTGAACTCTTTCCCTACACGACGCTCTTCCGATCTNNNGTACTAGGGCTTCAAGTCAGC
GAGCACACGTCTGAACTCTTTCCCTACACGACGCTCTTCCGATCTNNNTGAGGTCAGGAGTTCGAGAC
GAGCACACGTCTGAACTCTTTCCCTACACGACGCTCTTCCGATCTNNNTGAGGTCAGGAGTTCGAGAC
AGCACACGTCTGAACTCTTTCCCTACACGACGCTCTTCCGATCTNNNAAGGCCCAGCAATTAGGAATA
AGCACACGTCTGAACTCTTTCCCTACACGACGCTCTTCCGATCTNNNAAGGCCCAGCAATTAGGAATA
AGCACACGTCTGAACTCTTTCCCTACACGACGCTCTTCCGATCTNNNATATCCTGACCTCATGATCTGCC
AGCACACGTCTGAACTCTTTCCCTACACGACGCTCTTCCGATCTNNNATATCCTGACCTCATGATCTGCC
GAGCACACGTCTGAACTCTTTCCCTACACGACGCTCTTCCGATCTNNNTACACACATACACCCATACACC
GAGCACACGTCTGAACTCTTTCCCTACACGACGCTCTTCCGATCTNNNTACACACATACACCCATACACC
AGAGCACACGTCTGAACTCTTTCCCTACACGACGCTCTTCCGATCTNNNGGGAGGCTGAGATAGGAG
AGAGCACACGTCTGAACTCTTTCCCTACACGACGCTCTTCCGATCTNNNGGGAGGCTGAGATAGGAG
AAGAGCACACGTCTGAACTCTTTCCCTACACGACGCTCTTCCGATCTNNNCCTCAAAAGGGCAAATCAA
AAGAGCACACGTCTGAACTCTTTCCCTACACGACGCTCTTCCGATCTNNNCCTCAAAAGGGCAAATCAA
AGCACACGTCTGAACTCTTTCCCTACACGACGCTCTTCCGATCTNNNTTTAGGAGATGAGGGGAACAA
AGCACACGTCTGAACTCTTTCCCTACACGACGCTCTTCCGATCTNNNTTTAGGAGATGAGGGGAACAA
GAGCACACGTCTGAACTCTTTCCCTACACGACGCTCTTCCGATCTNNNATACAGGATTGACTAGTGGT
GAGCACACGTCTGAACTCTTTCCCTACACGACGCTCTTCCGATCTNNNATACAGGATTGACTAGTGGT
AGAGCACACGTCTGAACTCTTTCCCTACACGACGCTCTTCCGATCTNNNAGAGAGAGAGAGAGAGAG
AGAGCACACGTCTGAACTCTTTCCCTACACGACGCTCTTCCGATCTNNNAGAGAGAGAGAGAGAGAG

AGCACACGTCTGAACTCTTTCCCTACACGACGCTCTTCCGATCTNNNAGTGAAACACTGGGAGATGT
AGCACACGTCTGAACTCTTTCCCTACACGACGCTCTTCCGATCTNNNAGTGAAACACTGGGAGATGT
AGCACACGTCTGAACTCTTTCCCTACACGACGCTCTTCCGATCTNNNGATTGGGCTGTTGCTGGTATA
AGCACACGTCTGAACTCTTTCCCTACACGACGCTCTTCCGATCTNNNGATTGGGCTGTTGCTGGTATA
CACACGTCTGAACTCTTTCCCTACACGACGCTCTTCCGATCTNNNACCTCAAGTGAGTCTCTTGTAGA
CACACGTCTGAACTCTTTCCCTACACGACGCTCTTCCGATCTNNNACCTCAAGTGAGTCTCTTGTAGA
AGCACACGTCTGAACTCTTTCCCTACACGACGCTCTTCCGATCTNNNCCCCATCAAGATTCAGATCACC
AGCACACGTCTGAACTCTTTCCCTACACGACGCTCTTCCGATCTNNNCCCCATCAAGATTCAGATCACC
GCACACGTCTGAACTCTTTCCCTACACGACGCTCTTCCGATCTNNNATGTCAGAGTCCAGTTGATGCT
GCACACGTCTGAACTCTTTCCCTACACGACGCTCTTCCGATCTNNNATGTCAGAGTCCAGTTGATGCT
AGCACACGTCTGAACTCTTTCCCTACACGACGCTCTTCCGATCTNNNCTTGTTCTGTTGCTGCCTTTCTT
AGCACACGTCTGAACTCTTTCCCTACACGACGCTCTTCCGATCTNNNCTTGTTCTGTTGCTGCCTTTCTT
GCACACGTCTGAACTCTTTCCCTACACGACGCTCTTCCGATCTNNNTGTTGAAGAGCCTTCGTTGAGA
GCACACGTCTGAACTCTTTCCCTACACGACGCTCTTCCGATCTNNNTGTTGAAGAGCCTTCGTTGAGA
AGCACACGTCTGAACTCTTTCCCTACACGACGCTCTTCCGATCTNNNCTCCATCACCTACAGAGTGAC
AGCACACGTCTGAACTCTTTCCCTACACGACGCTCTTCCGATCTNNNCTCCATCACCTACAGAGTGAC
AGCACACGTCTGAACTCTTTCCCTACACGACGCTCTTCCGATCTNNNACATGGCTCACCCCTTTGAATA
AGCACACGTCTGAACTCTTTCCCTACACGACGCTCTTCCGATCTNNNACATGGCTCACCCCTTTGAATA
AGCACACGTCTGAACTCTTTCCCTACACGACGCTCTTCCGATCTNNNGCAAACAGTGATGCAATCCG
AGCACACGTCTGAACTCTTTCCCTACACGACGCTCTTCCGATCTNNNGCAAACAGTGATGCAATCCG
AGCACACGTCTGAACTCTTTCCCTACACGACGCTCTTCCGATCTNNNACACACTTGTAATCCCAGCTAC
AGCACACGTCTGAACTCTTTCCCTACACGACGCTCTTCCGATCTNNNACACACTTGTAATCCCAGCTAC
GAGCACACGTCTGAACTCTTTCCCTACACGACGCTCTTCCGATCTNNNAGGAATAATGGGGAAGAG
GAGCACACGTCTGAACTCTTTCCCTACACGACGCTCTTCCGATCTNNNAGGAATAATGGGGAAGAG
GAGCACACGTCTGAACTCTTTCCCTACACGACGCTCTTCCGATCTNNNACCAGGGAAAGACTCTTGTT
GAGCACACGTCTGAACTCTTTCCCTACACGACGCTCTTCCGATCTNNNACCAGGGAAAGACTCTTGTT
AGAGCACACGTCTGAACTCTTTCCCTACACGACGCTCTTCCGATCTNNNTGCACTCCATCACGACATG
AGAGCACACGTCTGAACTCTTTCCCTACACGACGCTCTTCCGATCTNNNTGCACTCCATCACGACATG
CACACGTCTGAACTCTTTCCCTACACGACGCTCTTCCGATCTNNNTTGGTTTGTAACGAAGTGAACAT
CACACGTCTGAACTCTTTCCCTACACGACGCTCTTCCGATCTNNNTTGGTTTGTAACGAAGTGAACAT
GAGCACACGTCTGAACTCTTTCCCTACACGACGCTCTTCCGATCTNNNAAAAATAGCCGTGTGTGGC
GAGCACACGTCTGAACTCTTTCCCTACACGACGCTCTTCCGATCTNNNAAAAATAGCCGTGTGTGGC
GAGCACACGTCTGAACTCTTTCCCTACACGACGCTCTTCCGATCTNNNAACTGAGATTGTACCACTGC
GAGCACACGTCTGAACTCTTTCCCTACACGACGCTCTTCCGATCTNNNAACTGAGATTGTACCACTGC
GAGCACACGTCTGAACTCTTTCCCTACACGACGCTCTTCCGATCTNNNGCCTGTAATCTCAGCTACTCA
GAGCACACGTCTGAACTCTTTCCCTACACGACGCTCTTCCGATCTNNNGCCTGTAATCTCAGCTACTCA
AGCACACGTCTGAACTCTTTCCCTACACGACGCTCTTCCGATCTNNNCCCTCCCTTTTCTTTGGAAGAG
AGCACACGTCTGAACTCTTTCCCTACACGACGCTCTTCCGATCTNNNCCCTCCCTTTTCTTTGGAAGAG
AGCACACGTCTGAACTCTTTCCCTACACGACGCTCTTCCGATCTNNNCACCTGTAGTCCCAGCTACTTG
AGCACACGTCTGAACTCTTTCCCTACACGACGCTCTTCCGATCTNNNCACCTGTAGTCCCAGCTACTTG
AGCACACGTCTGAACTCTTTCCCTACACGACGCTCTTCCGATCTNNNATGGATGTTGCCACTATCACT
AGCACACGTCTGAACTCTTTCCCTACACGACGCTCTTCCGATCTNNNATGGATGTTGCCACTATCACT
AGCACACGTCTGAACTCTTTCCCTACACGACGCTCTTCCGATCTNNNGGAAGAGGAGCTTGCAGTGA

AGCACACGTCTGAACTCTTTCCCTACACGACGCTCTTCCGATCTNNNGGAAGAGGAGCTTGCAGTGA
AGAGCACACGTCTGAACTCTTTCCCTACACGACGCTCTTCCGATCTNNNCTGGTTGTGGATACTGAG
AGAGCACACGTCTGAACTCTTTCCCTACACGACGCTCTTCCGATCTNNNCTGGTTGTGGATACTGAG
AGCACACGTCTGAACTCTTTCCCTACACGACGCTCTTCCGATCTNNNTTGCCCTAACTAGTCACACAGT
AGCACACGTCTGAACTCTTTCCCTACACGACGCTCTTCCGATCTNNNTTGCCCTAACTAGTCACACAGT
GAGCACACGTCTGAACTCTTTCCCTACACGACGCTCTTCCGATCTNNNCGGAGTAACATGAGGGAGG
GAGCACACGTCTGAACTCTTTCCCTACACGACGCTCTTCCGATCTNNNCGGAGTAACATGAGGGAGG
GAGCACACGTCTGAACTCTTTCCCTACACGACGCTCTTCCGATCTNNNAAATAGATGTGTGAGCTGG
GAGCACACGTCTGAACTCTTTCCCTACACGACGCTCTTCCGATCTNNNAAATAGATGTGTGAGCTGG
AGCACACGTCTGAACTCTTTCCCTACACGACGCTCTTCCGATCTNNNGAGTAAAACGTCAGGAGAGA
AGCACACGTCTGAACTCTTTCCCTACACGACGCTCTTCCGATCTNNNGAGTAAAACGTCAGGAGAGA
GAGCACACGTCTGAACTCTTTCCCTACACGACGCTCTTCCGATCTNNNAATACAAAAAATTAGCCGGG
GAGCACACGTCTGAACTCTTTCCCTACACGACGCTCTTCCGATCTNNNAATACAAAAAATTAGCCGGG
GCACACGTCTGAACTCTTTCCCTACACGACGCTCTTCCGATCTNNNACTCTACAAAGAAATGCATTGTG
GCACACGTCTGAACTCTTTCCCTACACGACGCTCTTCCGATCTNNNACTCTACAAAGAAATGCATTGTG
GAGCACACGTCTGAACTCTTTCCCTACACGACGCTCTTCCGATCTNNNCTTCCCAAGTTCAAGCGATTC
GAGCACACGTCTGAACTCTTTCCCTACACGACGCTCTTCCGATCTNNNCTTCCCAAGTTCAAGCGATTC
GCACACGTCTGAACTCTTTCCCTACACGACGCTCTTCCGATCTNNNGAAGGAGGATCGCTTGAGACTA
GCACACGTCTGAACTCTTTCCCTACACGACGCTCTTCCGATCTNNNGAAGGAGGATCGCTTGAGACTA
AGCACACGTCTGAACTCTTTCCCTACACGACGCTCTTCCGATCTNNNTCTATGAACGCCTAGTTGCTTC
AGCACACGTCTGAACTCTTTCCCTACACGACGCTCTTCCGATCTNNNTCTATGAACGCCTAGTTGCTTC
GCACACGTCTGAACTCTTTCCCTACACGACGCTCTTCCGATCTNNNCTTCTTCTAAGCCCTCCAGACTCC
GCACACGTCTGAACTCTTTCCCTACACGACGCTCTTCCGATCTNNNCTTCTTCTAAGCCCTCCAGACTCC
AGCACACGTCTGAACTCTTTCCCTACACGACGCTCTTCCGATCTNNNGTTCAAGTGATTCTCCTCCCTC
AGCACACGTCTGAACTCTTTCCCTACACGACGCTCTTCCGATCTNNNGTTCAAGTGATTCTCCTCCCTC
AGCACACGTCTGAACTCTTTCCCTACACGACGCTCTTCCGATCTNNNGGCTGATAGAACCAGATCCTG
AGCACACGTCTGAACTCTTTCCCTACACGACGCTCTTCCGATCTNNNGGCTGATAGAACCAGATCCTG
AGCACACGTCTGAACTCTTTCCCTACACGACGCTCTTCCGATCTNNNCTCTAGACTGTAGAATGCCCG
AGCACACGTCTGAACTCTTTCCCTACACGACGCTCTTCCGATCTNNNCTCTAGACTGTAGAATGCCCG
GAGCACACGTCTGAACTCTTTCCCTACACGACGCTCTTCCGATCTNNNTGTAATGTGAGCACAGAGAC
GAGCACACGTCTGAACTCTTTCCCTACACGACGCTCTTCCGATCTNNNTGTAATGTGAGCACAGAGAC
AGCACACGTCTGAACTCTTTCCCTACACGACGCTCTTCCGATCTNNNAGAAGAGGGACCTGTGAAAG
AGCACACGTCTGAACTCTTTCCCTACACGACGCTCTTCCGATCTNNNAGAAGAGGGACCTGTGAAAG
GCACACGTCTGAACTCTTTCCCTACACGACGCTCTTCCGATCTNNNGGACAGCCGAAACCATTCATTAG
GCACACGTCTGAACTCTTTCCCTACACGACGCTCTTCCGATCTNNNGGACAGCCGAAACCATTCATTAG
GCACACGTCTGAACTCTTTCCCTACACGACGCTCTTCCGATCTNNNGCCATTTTCTGCCTCATAGCAT
GCACACGTCTGAACTCTTTCCCTACACGACGCTCTTCCGATCTNNNGCCATTTTCTGCCTCATAGCAT
AGCACACGTCTGAACTCTTTCCCTACACGACGCTCTTCCGATCTNNNGATTTTCTCTCCCCCAAGAAGC
AGCACACGTCTGAACTCTTTCCCTACACGACGCTCTTCCGATCTNNNGATTTTCTCTCCCCCAAGAAGC
AGAGCACACGTCTGAACTCTTTCCCTACACGACGCTCTTCCGATCTNNNCAGCATCTTGGTTCTGCAG
AGAGCACACGTCTGAACTCTTTCCCTACACGACGCTCTTCCGATCTNNNCAGCATCTTGGTTCTGCAG
AGCACACGTCTGAACTCTTTCCCTACACGACGCTCTTCCGATCTNNNGTGTGTGTGTGTGTGTACTT
AGCACACGTCTGAACTCTTTCCCTACACGACGCTCTTCCGATCTNNNGTGTGTGTGTGTGTGTACTT

GAGCACACGTCTGAACTCTTTCCCTACACGACGCTCTTCCGATCTNNNCAAGGCAGACTGTTTGGACAT
GAGCACACGTCTGAACTCTTTCCCTACACGACGCTCTTCCGATCTNNNCAAGGCAGACTGTTTGGACAT
GAGCACACGTCTGAACTCTTTCCCTACACGACGCTCTTCCGATCTNNNTGGGGAAAGTCTCATCTTGCC
GAGCACACGTCTGAACTCTTTCCCTACACGACGCTCTTCCGATCTNNNTGGGGAAAGTCTCATCTTGCC
GAGCACACGTCTGAACTCTTTCCCTACACGACGCTCTTCCGATCTNNNATCCTCAGCAAACTAATGCAG
GAGCACACGTCTGAACTCTTTCCCTACACGACGCTCTTCCGATCTNNNATCCTCAGCAAACTAATGCAG
GAGCACACGTCTGAACTCTTTCCCTACACGACGCTCTTCCGATCTNNNGTAATGAAGTTCTTCTAGTTCA
GAGCACACGTCTGAACTCTTTCCCTACACGACGCTCTTCCGATCTNNNGTAATGAAGTTCTTCTAGTTCA
GAGCACACGTCTGAACTCTTTCCCTACACGACGCTCTTCCGATCTNNNAGGCTTCAGTGAGCTATGATCA
GAGCACACGTCTGAACTCTTTCCCTACACGACGCTCTTCCGATCTNNNAGGCTTCAGTGAGCTATGATCA
GAGCACACGTCTGAACTCTTTCCCTACACGACGCTCTTCCGATCTNNNCCTGGGATGGTCAGCTTATTTG
GAGCACACGTCTGAACTCTTTCCCTACACGACGCTCTTCCGATCTNNNCCTGGGATGGTCAGCTTATTTG
AGCACACGTCTGAACTCTTTCCCTACACGACGCTCTTCCGATCTNNNCTCCGTCTCTAAAAAACAACCCCC
AGCACACGTCTGAACTCTTTCCCTACACGACGCTCTTCCGATCTNNNCTCCGTCTCTAAAAAACAACCCCC
AGCACACGTCTGAACTCTTTCCCTACACGACGCTCTTCCGATCTNNNCAGATACCAGAGGGAAGTGTG
AGCACACGTCTGAACTCTTTCCCTACACGACGCTCTTCCGATCTNNNCAGATACCAGAGGGAAGTGTG
AGCACACGTCTGAACTCTTTCCCTACACGACGCTCTTCCGATCTNNNCTGTCTACAGCCTCCTAAATGCA
AGCACACGTCTGAACTCTTTCCCTACACGACGCTCTTCCGATCTNNNCTGTCTACAGCCTCCTAAATGCA
AGCACACGTCTGAACTCTTTCCCTACACGACGCTCTTCCGATCTNNNCCATACTGTAGCCATTTGAGG
AGCACACGTCTGAACTCTTTCCCTACACGACGCTCTTCCGATCTNNNCCATACTGTAGCCATTTGAGG
GAGCACACGTCTGAACTCTTTCCCTACACGACGCTCTTCCGATCTNNNGGTTCAGGGTGATGTCCAGAG
GAGCACACGTCTGAACTCTTTCCCTACACGACGCTCTTCCGATCTNNNGGTTCAGGGTGATGTCCAGAG
GAGCACACGTCTGAACTCTTTCCCTACACGACGCTCTTCCGATCTNNNTGACATGCACCTATAGTCCCAA
GAGCACACGTCTGAACTCTTTCCCTACACGACGCTCTTCCGATCTNNNTGACATGCACCTATAGTCCCAA
AGCACACGTCTGAACTCTTTCCCTACACGACGCTCTTCCGATCTNNNTAAATGGAGAAATCTTCTTCTG
AGCACACGTCTGAACTCTTTCCCTACACGACGCTCTTCCGATCTNNNTAAATGGAGAAATCTTCTTCTG
AGCACACGTCTGAACTCTTTCCCTACACGACGCTCTTCCGATCTNNNCACCTGTAGTCCCAGCTACTTGC
AGCACACGTCTGAACTCTTTCCCTACACGACGCTCTTCCGATCTNNNCACCTGTAGTCCCAGCTACTTGC
GAGCACACGTCTGAACTCTTTCCCTACACGACGCTCTTCCGATCTNNNGGGTGACAGAGTAAGATTCCC
GAGCACACGTCTGAACTCTTTCCCTACACGACGCTCTTCCGATCTNNNGGGTGACAGAGTAAGATTCCC
AGCACACGTCTGAACTCTTTCCCTACACGACGCTCTTCCGATCTNNNAAAGATGAGAGCAAGAGTCAG
AGCACACGTCTGAACTCTTTCCCTACACGACGCTCTTCCGATCTNNNAAAGATGAGAGCAAGAGTCAG
GAGCACACGTCTGAACTCTTTCCCTACACGACGCTCTTCCGATCTNNNCTGACGACTCTTTTTCAGATCA
GAGCACACGTCTGAACTCTTTCCCTACACGACGCTCTTCCGATCTNNNCTGACGACTCTTTTTCAGATCA
GAGCACACGTCTGAACTCTTTCCCTACACGACGCTCTTCCGATCTNNNAAAAAATTAGCCGGACGTGGT
GAGCACACGTCTGAACTCTTTCCCTACACGACGCTCTTCCGATCTNNNAAAAAATTAGCCGGACGTGGT
GAGCACACGTCTGAACTCTTTCCCTACACGACGCTCTTCCGATCTNNNACCTCTACCTCCTGAGTTCAAC
GAGCACACGTCTGAACTCTTTCCCTACACGACGCTCTTCCGATCTNNNACCTCTACCTCCTGAGTTCAAC
AGCACACGTCTGAACTCTTTCCCTACACGACGCTCTTCCGATCTNNNCCCTACGTTGTAGTCACAAGTGT
AGCACACGTCTGAACTCTTTCCCTACACGACGCTCTTCCGATCTNNNCCCTACGTTGTAGTCACAAGTGT
AGAGCACACGTCTGAACTCTTTCCCTACACGACGCTCTTCCGATCTNNNAACAGCAAAGTCATGGAATC
AGAGCACACGTCTGAACTCTTTCCCTACACGACGCTCTTCCGATCTNNNAACAGCAAAGTCATGGAATC
AGCACACGTCTGAACTCTTTCCCTACACGACGCTCTTCCGATCTNNNAAAATTAGCTCAACTTGGTGGC

AGCACACGTCTGAACTCTTTCCCTACACGACGCTCTTCCGATCTNNNAAAATTAGCTCAACTTGGTGGC
AGCACACGTCTGAACTCTTTCCCTACACGACGCTCTTCCGATCTNNNGCAACAAAATGAGACCCTGTCT
AGCACACGTCTGAACTCTTTCCCTACACGACGCTCTTCCGATCTNNNGCAACAAAATGAGACCCTGTCT
AGCACACGTCTGAACTCTTTCCCTACACGACGCTCTTCCGATCTNNNCATCTTTACATTTTTTTGCAGTTC
AGCACACGTCTGAACTCTTTCCCTACACGACGCTCTTCCGATCTNNNCATCTTTACATTTTTTTGCAGTTC
AGCACACGTCTGAACTCTTTCCCTACACGACGCTCTTCCGATCTNNNGTCTCCTCCTCCAATAAGCTCAGC
AGCACACGTCTGAACTCTTTCCCTACACGACGCTCTTCCGATCTNNNGTCTCCTCCTCCAATAAGCTCAGC
AGCACACGTCTGAACTCTTTCCCTACACGACGCTCTTCCGATCTNNNAAGTGCTGAGATTACAGGCATG
AGCACACGTCTGAACTCTTTCCCTACACGACGCTCTTCCGATCTNNNAAGTGCTGAGATTACAGGCATG
GAGCACACGTCTGAACTCTTTCCCTACACGACGCTCTTCCGATCTNNNTCCAACTCCAGAAGTCTCAACA
GAGCACACGTCTGAACTCTTTCCCTACACGACGCTCTTCCGATCTNNNTCCAACTCCAGAAGTCTCAACA
AGCACACGTCTGAACTCTTTCCCTACACGACGCTCTTCCGATCTNNNACCATGTCCAGCTAAATTTTTGC
AGCACACGTCTGAACTCTTTCCCTACACGACGCTCTTCCGATCTNNNACCATGTCCAGCTAAATTTTTGC
GCACACGTCTGAACTCTTTCCCTACACGACGCTCTTCCGATCTNNNCAGGTTTAGAAGGGGTGTCTGG
GCACACGTCTGAACTCTTTCCCTACACGACGCTCTTCCGATCTNNNCAGGTTTAGAAGGGGTGTCTGG
AGCACACGTCTGAACTCTTTCCCTACACGACGCTCTTCCGATCTNNNGCTGCACATGACAGGATTTCAT
AGCACACGTCTGAACTCTTTCCCTACACGACGCTCTTCCGATCTNNNGCTGCACATGACAGGATTTCAT
GCACACGTCTGAACTCTTTCCCTACACGACGCTCTTCCGATCTNNNTATACCTTGGATGAGCTTCCTGCC
GCACACGTCTGAACTCTTTCCCTACACGACGCTCTTCCGATCTNNNTATACCTTGGATGAGCTTCCTGCC
AGCACACGTCTGAACTCTTTCCCTACACGACGCTCTTCCGATCTNNNGGCATACTTGTTTGTTCCGCAT
AGCACACGTCTGAACTCTTTCCCTACACGACGCTCTTCCGATCTNNNGGCATACTTGTTTGTTCCGCAT
AGAGCACACGTCTGAACTCTTTCCCTACACGACGCTCTTCCGATCTNNNTCGTTCCAGAAATCTCTTGG
AGAGCACACGTCTGAACTCTTTCCCTACACGACGCTCTTCCGATCTNNNTCGTTCCAGAAATCTCTTGG
AGCACACGTCTGAACTCTTTCCCTACACGACGCTCTTCCGATCTNNNCACCATGCACAGCTGATTTTGTA
AGCACACGTCTGAACTCTTTCCCTACACGACGCTCTTCCGATCTNNNCACCATGCACAGCTGATTTTGTA
GAGCACACGTCTGAACTCTTTCCCTACACGACGCTCTTCCGATCTNNNCTGTAATCCCAGCTACTCAGGA
GAGCACACGTCTGAACTCTTTCCCTACACGACGCTCTTCCGATCTNNNCTGTAATCCCAGCTACTCAGGA
GAGCACACGTCTGAACTCTTTCCCTACACGACGCTCTTCCGATCTNNNTGAGATACGCTACTCTCAGGAC
GAGCACACGTCTGAACTCTTTCCCTACACGACGCTCTTCCGATCTNNNTGAGATACGCTACTCTCAGGAC
GAGCACACGTCTGAACTCTTTCCCTACACGACGCTCTTCCGATCTNNNTGTCACCTCGGTGTTATTCTTC
GAGCACACGTCTGAACTCTTTCCCTACACGACGCTCTTCCGATCTNNNTGTCACCTCGGTGTTATTCTTC
AGCACACGTCTGAACTCTTTCCCTACACGACGCTCTTCCGATCTNNNCTTCCATTGACCTCTGTGTCCATC
AGCACACGTCTGAACTCTTTCCCTACACGACGCTCTTCCGATCTNNNCTTCCATTGACCTCTGTGTCCATC
GAGCACACGTCTGAACTCTTTCCCTACACGACGCTCTTCCGATCTNNNCCTTTTCTAAAGGATGAAAAAA
GAGCACACGTCTGAACTCTTTCCCTACACGACGCTCTTCCGATCTNNNCCTTTTCTAAAGGATGAAAAAA
GCACACGTCTGAACTCTTTCCCTACACGACGCTCTTCCGATCTNNNTGTCTTGAGGTGAAAGAGGTTGA
GCACACGTCTGAACTCTTTCCCTACACGACGCTCTTCCGATCTNNNTGTCTTGAGGTGAAAGAGGTTGA
CACACGTCTGAACTCTTTCCCTACACGACGCTCTTCCGATCTNNNCTTTTCAAACATGAGGAGATGCATC
CACACGTCTGAACTCTTTCCCTACACGACGCTCTTCCGATCTNNNCTTTTCAAACATGAGGAGATGCATC
AGCACACGTCTGAACTCTTTCCCTACACGACGCTCTTCCGATCTNNNATCTGGAATGTGTGAGTTTGTG
AGCACACGTCTGAACTCTTTCCCTACACGACGCTCTTCCGATCTNNNATCTGGAATGTGTGAGTTTGTG
AGCACACGTCTGAACTCTTTCCCTACACGACGCTCTTCCGATCTNNNCCCAAATGATGAAGTGATGCAC
AGCACACGTCTGAACTCTTTCCCTACACGACGCTCTTCCGATCTNNNCCCAAATGATGAAGTGATGCAC

AGCACACGTCTGAACTCTTTCCCTACACGACGCTCTTCCGATCTNNNCAGGAAAATCGCTTGAACCCA
AGCACACGTCTGAACTCTTTCCCTACACGACGCTCTTCCGATCTNNNCAGGAAAATCGCTTGAACCCA
AGCACACGTCTGAACTCTTTCCCTACACGACGCTCTTCCGATCTNNNAAGAGTGTGGATCATTGTGGG
AGCACACGTCTGAACTCTTTCCCTACACGACGCTCTTCCGATCTNNNAAGAGTGTGGATCATTGTGGG
AGCACACGTCTGAACTCTTTCCCTACACGACGCTCTTCCGATCTNNNGGGAATGGAGGCTTCATTCTA
AGCACACGTCTGAACTCTTTCCCTACACGACGCTCTTCCGATCTNNNGGGAATGGAGGCTTCATTCTA
GCACACGTCTGAACTCTTTCCCTACACGACGCTCTTCCGATCTNNNGTGTGGTCCCGTGGGTAGAACA
GCACACGTCTGAACTCTTTCCCTACACGACGCTCTTCCGATCTNNNGTGTGGTCCCGTGGGTAGAACA
GAGCACACGTCTGAACTCTTTCCCTACACGACGCTCTTCCGATCTNNNATTTAGGAACCAAGACAGGC
GAGCACACGTCTGAACTCTTTCCCTACACGACGCTCTTCCGATCTNNNATTTAGGAACCAAGACAGGC
GCACACGTCTGAACTCTTTCCCTACACGACGCTCTTCCGATCTNNNCCAAGTTCCTCTTCCTCTTCTCCCA
GCACACGTCTGAACTCTTTCCCTACACGACGCTCTTCCGATCTNNNCCAAGTTCCTCTTCCTCTTCTCCCA
GAGCACACGTCTGAACTCTTTCCCTACACGACGCTCTTCCGATCTNNNCTACAGGATTCTGACCCTCCC
GAGCACACGTCTGAACTCTTTCCCTACACGACGCTCTTCCGATCTNNNCTACAGGATTCTGACCCTCCC
GAAGAGCACACGTCTGAACTCTTTCCCTACACGACGCTCTTCCGATCTNNNCAGACAGTGACCCTTGC
GAAGAGCACACGTCTGAACTCTTTCCCTACACGACGCTCTTCCGATCTNNNCAGACAGTGACCCTTGC
CACACGTCTGAACTCTTTCCCTACACGACGCTCTTCCGATCTNNNCTTGCAGTGAGCCAAGATCATGC
CACACGTCTGAACTCTTTCCCTACACGACGCTCTTCCGATCTNNNCTTGCAGTGAGCCAAGATCATGC
GCACACGTCTGAACTCTTTCCCTACACGACGCTCTTCCGATCTNNNCATTCAAATTTTACAGGATCTGA
GCACACGTCTGAACTCTTTCCCTACACGACGCTCTTCCGATCTNNNCATTCAAATTTTACAGGATCTGA
GAGCACACGTCTGAACTCTTTCCCTACACGACGCTCTTCCGATCTNNNTGATATGACCCTTCGTTTTTTT
GAGCACACGTCTGAACTCTTTCCCTACACGACGCTCTTCCGATCTNNNTGATATGACCCTTCGTTTTTTT
GCACACGTCTGAACTCTTTCCCTACACGACGCTCTTCCGATCTNNNGTAATCCTAGCTACTTGGGAGG
GCACACGTCTGAACTCTTTCCCTACACGACGCTCTTCCGATCTNNNGTAATCCTAGCTACTTGGGAGG
AGCACACGTCTGAACTCTTTCCCTACACGACGCTCTTCCGATCTNNNAGTGTGAATCCTCCAACTGTGT
AGCACACGTCTGAACTCTTTCCCTACACGACGCTCTTCCGATCTNNNAGTGTGAATCCTCCAACTGTGT
AGCACACGTCTGAACTCTTTCCCTACACGACGCTCTTCCGATCTNNNCATGCCATTCTCTGTTCCACAA
AGCACACGTCTGAACTCTTTCCCTACACGACGCTCTTCCGATCTNNNCATGCCATTCTCTGTTCCACAA
AGAGCACACGTCTGAACTCTTTCCCTACACGACGCTCTTCCGATCTNNNCTGACTCCGAAAAACCCATG
AGAGCACACGTCTGAACTCTTTCCCTACACGACGCTCTTCCGATCTNNNCTGACTCCGAAAAACCCATG
AGCACACGTCTGAACTCTTTCCCTACACGACGCTCTTCCGATCTNNNTTGGTCAGGCTAGTCTTGAAC
AGCACACGTCTGAACTCTTTCCCTACACGACGCTCTTCCGATCTNNNTTGGTCAGGCTAGTCTTGAAC
GAGCACACGTCTGAACTCTTTCCCTACACGACGCTCTTCCGATCTNNNTCCTGTTCTTCCTAGAGCTTG
GAGCACACGTCTGAACTCTTTCCCTACACGACGCTCTTCCGATCTNNNTCCTGTTCTTCCTAGAGCTTG
AGCACACGTCTGAACTCTTTCCCTACACGACGCTCTTCCGATCTNNNAATTTCTTTCTTCACAGCGAGC
AGCACACGTCTGAACTCTTTCCCTACACGACGCTCTTCCGATCTNNNAATTTCTTTCTTCACAGCGAGC
AGCACACGTCTGAACTCTTTCCCTACACGACGCTCTTCCGATCTNNNGGGTGTGGAGTATAGCACAGA
AGCACACGTCTGAACTCTTTCCCTACACGACGCTCTTCCGATCTNNNGGGTGTGGAGTATAGCACAGA
GAGCACACGTCTGAACTCTTTCCCTACACGACGCTCTTCCGATCTNNNGTGGAGTTTGCAGTGAGTA
GAGCACACGTCTGAACTCTTTCCCTACACGACGCTCTTCCGATCTNNNGTGGAGTTTGCAGTGAGTA
GAGCACACGTCTGAACTCTTTCCCTACACGACGCTCTTCCGATCTNNNTGCCTGTAATCCCAGCTACTT
GAGCACACGTCTGAACTCTTTCCCTACACGACGCTCTTCCGATCTNNNTGCCTGTAATCCCAGCTACTT
AGCACACGTCTGAACTCTTTCCCTACACGACGCTCTTCCGATCTNNNCTTGCTGGGAGTGGTAAACC

AGCACACGTCTGAACTCTTTCCCTACACGACGCTCTTCCGATCTNNNCTTGCTGGGAGTGGTAAACC
AGCACACGTCTGAACTCTTTCCCTACACGACGCTCTTCCGATCTNNNACAGTCACCACAGAAGAATCA
AGCACACGTCTGAACTCTTTCCCTACACGACGCTCTTCCGATCTNNNACAGTCACCACAGAAGAATCA
AGCACACGTCTGAACTCTTTCCCTACACGACGCTCTTCCGATCTNNNGAGAATGGCATGAACCCAGGA
AGCACACGTCTGAACTCTTTCCCTACACGACGCTCTTCCGATCTNNNGAGAATGGCATGAACCCAGGA
AGCACACGTCTGAACTCTTTCCCTACACGACGCTCTTCCGATCTNNNAAAAATTAGCTTGGAGTGGTG
AGCACACGTCTGAACTCTTTCCCTACACGACGCTCTTCCGATCTNNNAAAAATTAGCTTGGAGTGGTG
AGCACACGTCTGAACTCTTTCCCTACACGACGCTCTTCCGATCTNNNGAGGTTGGCGAAATTTTTCTG
AGCACACGTCTGAACTCTTTCCCTACACGACGCTCTTCCGATCTNNNGAGGTTGGCGAAATTTTTCTG
AGCACACGTCTGAACTCTTTCCCTACACGACGCTCTTCCGATCTNNNCCTGGGAAATAGAACGAGACT
AGCACACGTCTGAACTCTTTCCCTACACGACGCTCTTCCGATCTNNNCCTGGGAAATAGAACGAGACT
GAGCACACGTCTGAACTCTTTCCCTACACGACGCTCTTCCGATCTNNNTATGACAGGGAGCTAACTGA
GAGCACACGTCTGAACTCTTTCCCTACACGACGCTCTTCCGATCTNNNTATGACAGGGAGCTAACTGA
AGCACACGTCTGAACTCTTTCCCTACACGACGCTCTTCCGATCTNNNGGGACCCTGCAAAATATCTAC
AGCACACGTCTGAACTCTTTCCCTACACGACGCTCTTCCGATCTNNNGGGACCCTGCAAAATATCTAC
GAGCACACGTCTGAACTCTTTCCCTACACGACGCTCTTCCGATCTNNNCCTGTAGTCCCAGCTACTTGG
GAGCACACGTCTGAACTCTTTCCCTACACGACGCTCTTCCGATCTNNNCCTGTAGTCCCAGCTACTTGG
GCACACGTCTGAACTCTTTCCCTACACGACGCTCTTCCGATCTNNNAGATTTGTGCATTCATCACCAC
GCACACGTCTGAACTCTTTCCCTACACGACGCTCTTCCGATCTNNNAGATTTGTGCATTCATCACCAC
AGAGCACACGTCTGAACTCTTTCCCTACACGACGCTCTTCCGATCTNNNGTGAAGATTGCAGTGAGC
AGAGCACACGTCTGAACTCTTTCCCTACACGACGCTCTTCCGATCTNNNGTGAAGATTGCAGTGAGC
AGCACACGTCTGAACTCTTTCCCTACACGACGCTCTTCCGATCTNNNATTATTGTTGACCAACGGGGC
AGCACACGTCTGAACTCTTTCCCTACACGACGCTCTTCCGATCTNNNATTATTGTTGACCAACGGGGC
CACACGTCTGAACTCTTTCCCTACACGACGCTCTTCCGATCTNNNTTGTGCTACAAATGGCTCCAAAGC
CACACGTCTGAACTCTTTCCCTACACGACGCTCTTCCGATCTNNNTTGTGCTACAAATGGCTCCAAAGC
GCACACGTCTGAACTCTTTCCCTACACGACGCTCTTCCGATCTNNNCCCAGGAGAAGGTTTTTTGAA
GCACACGTCTGAACTCTTTCCCTACACGACGCTCTTCCGATCTNNNCCCAGGAGAAGGTTTTTTGAA
AGCACACGTCTGAACTCTTTCCCTACACGACGCTCTTCCGATCTNNNGGGAGATTAGAACGTGGACA
AGCACACGTCTGAACTCTTTCCCTACACGACGCTCTTCCGATCTNNNGGGAGATTAGAACGTGGACA
GCACACGTCTGAACTCTTTCCCTACACGACGCTCTTCCGATCTNNNAAGTGCTGAGATTGTAGGCATG
GCACACGTCTGAACTCTTTCCCTACACGACGCTCTTCCGATCTNNNAAGTGCTGAGATTGTAGGCATG
AGCACACGTCTGAACTCTTTCCCTACACGACGCTCTTCCGATCTNNNGATGGTACTACTGCATTCCAG
AGCACACGTCTGAACTCTTTCCCTACACGACGCTCTTCCGATCTNNNGATGGTACTACTGCATTCCAG
GAGCACACGTCTGAACTCTTTCCCTACACGACGCTCTTCCGATCTNNNCCTTCCAGGACTTCCAGAAAT
GAGCACACGTCTGAACTCTTTCCCTACACGACGCTCTTCCGATCTNNNCCTTCCAGGACTTCCAGAAAT
GAGCACACGTCTGAACTCTTTCCCTACACGACGCTCTTCCGATCTNNNGGGGGTGAGGGGATAAAG
GAGCACACGTCTGAACTCTTTCCCTACACGACGCTCTTCCGATCTNNNGGGGGTGAGGGGATAAAG
GCACACGTCTGAACTCTTTCCCTACACGACGCTCTTCCGATCTNNNGGCTCGAATAATTTCTTGCACGA
GCACACGTCTGAACTCTTTCCCTACACGACGCTCTTCCGATCTNNNGGCTCGAATAATTTCTTGCACGA
GCACACGTCTGAACTCTTTCCCTACACGACGCTCTTCCGATCTNNNAATGAAAAAAAAGAGCACCCCC
GCACACGTCTGAACTCTTTCCCTACACGACGCTCTTCCGATCTNNNAATGAAAAAAAAGAGCACCCCC
GCACACGTCTGAACTCTTTCCCTACACGACGCTCTTCCGATCTNNNATAGGAGGGGCTTTTTCACTCA
GCACACGTCTGAACTCTTTCCCTACACGACGCTCTTCCGATCTNNNATAGGAGGGGCTTTTTCACTCA

AGCACACGTCTGAACTCTTTCCCTACACGACGCTCTTCCGATCTNNNCTGTGTTGGCCATCACTTAGTT
AGCACACGTCTGAACTCTTTCCCTACACGACGCTCTTCCGATCTNNNCTGTGTTGGCCATCACTTAGTT
GAGCACACGTCTGAACTCTTTCCCTACACGACGCTCTTCCGATCTNNNTTGAGAGAGAGAGAGAGTGT
GAGCACACGTCTGAACTCTTTCCCTACACGACGCTCTTCCGATCTNNNTTGAGAGAGAGAGAGAGTGT
GAGCACACGTCTGAACTCTTTCCCTACACGACGCTCTTCCGATCTNNNGGCAAGTGGTCCCTTTATGTG
GAGCACACGTCTGAACTCTTTCCCTACACGACGCTCTTCCGATCTNNNGGCAAGTGGTCCCTTTATGTG
GAGCACACGTCTGAACTCTTTCCCTACACGACGCTCTTCCGATCTNNNTCACACCTGTAATCCCAACACT
GAGCACACGTCTGAACTCTTTCCCTACACGACGCTCTTCCGATCTNNNTCACACCTGTAATCCCAACACT
AGAGCACACGTCTGAACTCTTTCCCTACACGACGCTCTTCCGATCTNNNCTGAGGCAGGATAATCGCTT
AGAGCACACGTCTGAACTCTTTCCCTACACGACGCTCTTCCGATCTNNNCTGAGGCAGGATAATCGCTT
GCACACGTCTGAACTCTTTCCCTACACGACGCTCTTCCGATCTNNNGAGCCAGTTGTTCAGTTTCCATG
GCACACGTCTGAACTCTTTCCCTACACGACGCTCTTCCGATCTNNNGAGCCAGTTGTTCAGTTTCCATG
AGCACACGTCTGAACTCTTTCCCTACACGACGCTCTTCCGATCTNNNCTGAGTGACAAAGATTGGCGA
AGCACACGTCTGAACTCTTTCCCTACACGACGCTCTTCCGATCTNNNCTGAGTGACAAAGATTGGCGA
GAGCACACGTCTGAACTCTTTCCCTACACGACGCTCTTCCGATCTNNNTGGTGGGAAGGGAAAGAAA
GAGCACACGTCTGAACTCTTTCCCTACACGACGCTCTTCCGATCTNNNTGGTGGGAAGGGAAAGAAA
GAGCACACGTCTGAACTCTTTCCCTACACGACGCTCTTCCGATCTNNNGGAATCAAGTATGCCCAATGT
GAGCACACGTCTGAACTCTTTCCCTACACGACGCTCTTCCGATCTNNNGGAATCAAGTATGCCCAATGT
AGAGCACACGTCTGAACTCTTTCCCTACACGACGCTCTTCCGATCTNNNAATGAGATACTGTATCCCG
AGAGCACACGTCTGAACTCTTTCCCTACACGACGCTCTTCCGATCTNNNAATGAGATACTGTATCCCG
AGCACACGTCTGAACTCTTTCCCTACACGACGCTCTTCCGATCTNNNCACCTGTAATCCCAGCTACTCA
AGCACACGTCTGAACTCTTTCCCTACACGACGCTCTTCCGATCTNNNCACCTGTAATCCCAGCTACTCA
GCACACGTCTGAACTCTTTCCCTACACGACGCTCTTCCGATCTNNNTTTGGGGTGGGGACACAGTCT
GCACACGTCTGAACTCTTTCCCTACACGACGCTCTTCCGATCTNNNTTTGGGGTGGGGACACAGTCT
GAGCACACGTCTGAACTCTTTCCCTACACGACGCTCTTCCGATCTNNNCCCAGAACAGGCAAAGATAT
GAGCACACGTCTGAACTCTTTCCCTACACGACGCTCTTCCGATCTNNNCCCAGAACAGGCAAAGATAT
GCACACGTCTGAACTCTTTCCCTACACGACGCTCTTCCGATCTNNNCAGTTGTAGAACTCCCTTAGGCA
GCACACGTCTGAACTCTTTCCCTACACGACGCTCTTCCGATCTNNNCAGTTGTAGAACTCCCTTAGGCA
AGCACACGTCTGAACTCTTTCCCTACACGACGCTCTTCCGATCTNNNTTGAGACAGGAGAATCGCTTG
AGCACACGTCTGAACTCTTTCCCTACACGACGCTCTTCCGATCTNNNTTGAGACAGGAGAATCGCTTG
AGCACACGTCTGAACTCTTTCCCTACACGACGCTCTTCCGATCTNNNTGACCCCAGGTTATTTCTGCAT
AGCACACGTCTGAACTCTTTCCCTACACGACGCTCTTCCGATCTNNNTGACCCCAGGTTATTTCTGCAT
GAGCACACGTCTGAACTCTTTCCCTACACGACGCTCTTCCGATCTNNNTACTGTCTCACACTACTGGCTT
GAGCACACGTCTGAACTCTTTCCCTACACGACGCTCTTCCGATCTNNNTACTGTCTCACACTACTGGCTT
AGCACACGTCTGAACTCTTTCCCTACACGACGCTCTTCCGATCTNNNGGGAGGGGTAAGAAAGCAAT
AGCACACGTCTGAACTCTTTCCCTACACGACGCTCTTCCGATCTNNNGGGAGGGGTAAGAAAGCAAT
GCACACGTCTGAACTCTTTCCCTACACGACGCTCTTCCGATCTNNNGGTGACAGAGTGAGATTTGGT
GCACACGTCTGAACTCTTTCCCTACACGACGCTCTTCCGATCTNNNGGTGACAGAGTGAGATTTGGT
AGAGCACACGTCTGAACTCTTTCCCTACACGACGCTCTTCCGATCTNNNGCCTAATGCTCTTCCTTTAT
AGAGCACACGTCTGAACTCTTTCCCTACACGACGCTCTTCCGATCTNNNGCCTAATGCTCTTCCTTTAT
AGCACACGTCTGAACTCTTTCCCTACACGACGCTCTTCCGATCTNNNTGAATCAAAATCGAACCTGGC
AGCACACGTCTGAACTCTTTCCCTACACGACGCTCTTCCGATCTNNNTGAATCAAAATCGAACCTGGC
AGCACACGTCTGAACTCTTTCCCTACACGACGCTCTTCCGATCTNNNAGGCTTTCCGAAGAACTTCAAA

AGCACACGTCTGAACTCTTTCCCTACACGACGCTCTTCCGATCTNNNAGGCTTTCCGAAGAACTTCAAA
AGCACACGTCTGAACTCTTTCCCTACACGACGCTCTTCCGATCTNNNAACACAAAAATTAGCTGGGG
AGCACACGTCTGAACTCTTTCCCTACACGACGCTCTTCCGATCTNNNAACACAAAAATTAGCTGGGG
AGCACACGTCTGAACTCTTTCCCTACACGACGCTCTTCCGATCTNNNAACCAATGGAAAGGACACAA
AGCACACGTCTGAACTCTTTCCCTACACGACGCTCTTCCGATCTNNNAACCAATGGAAAGGACACAA
AGCACACGTCTGAACTCTTTCCCTACACGACGCTCTTCCGATCTNNNAGAGGTTGCAGTATGACTCGA
AGCACACGTCTGAACTCTTTCCCTACACGACGCTCTTCCGATCTNNNAGAGGTTGCAGTATGACTCGA
AGCACACGTCTGAACTCTTTCCCTACACGACGCTCTTCCGATCTNNNAGCAGAGGAAGGAAATGATT
AGCACACGTCTGAACTCTTTCCCTACACGACGCTCTTCCGATCTNNNAGCAGAGGAAGGAAATGATT
AGCACACGTCTGAACTCTTTCCCTACACGACGCTCTTCCGATCTNNNTGTCATGATCTTGGCTCACTGA
AGCACACGTCTGAACTCTTTCCCTACACGACGCTCTTCCGATCTNNNTGTCATGATCTTGGCTCACTGA
AGCACACGTCTGAACTCTTTCCCTACACGACGCTCTTCCGATCTNNNGGATGCTGTCGACAAGTTTCT
AGCACACGTCTGAACTCTTTCCCTACACGACGCTCTTCCGATCTNNNGGATGCTGTCGACAAGTTTCT
AGCACACGTCTGAACTCTTTCCCTACACGACGCTCTTCCGATCTNNNCAGTGGAATGGTGAGCTCTCG
AGCACACGTCTGAACTCTTTCCCTACACGACGCTCTTCCGATCTNNNCAGTGGAATGGTGAGCTCTCG
AGCACACGTCTGAACTCTTTCCCTACACGACGCTCTTCCGATCTNNNGAAGAAGACGAGGGGGAGG
AGCACACGTCTGAACTCTTTCCCTACACGACGCTCTTCCGATCTNNNGAAGAAGACGAGGGGGAGG
AGAGCACACGTCTGAACTCTTTCCCTACACGACGCTCTTCCGATCTNNNGAGAATGGTGTGAACCCAG
AGAGCACACGTCTGAACTCTTTCCCTACACGACGCTCTTCCGATCTNNNGAGAATGGTGTGAACCCAG
AGCACACGTCTGAACTCTTTCCCTACACGACGCTCTTCCGATCTNNNGACTTGTTTCCAAGGCTGACA
AGCACACGTCTGAACTCTTTCCCTACACGACGCTCTTCCGATCTNNNGACTTGTTTCCAAGGCTGACA
AGCACACGTCTGAACTCTTTCCCTACACGACGCTCTTCCGATCTNNNGTAATCCCAGCTACTCAGGCAG
AGCACACGTCTGAACTCTTTCCCTACACGACGCTCTTCCGATCTNNNGTAATCCCAGCTACTCAGGCAG
GCACACGTCTGAACTCTTTCCCTACACGACGCTCTTCCGATCTNNNTCTCTACATTGCTGGCTTTTCCTC
GCACACGTCTGAACTCTTTCCCTACACGACGCTCTTCCGATCTNNNTCTCTACATTGCTGGCTTTTCCTC
AGCACACGTCTGAACTCTTTCCCTACACGACGCTCTTCCGATCTNNNTGTAATCCTAGCTACTCGGGAG
AGCACACGTCTGAACTCTTTCCCTACACGACGCTCTTCCGATCTNNNTGTAATCCTAGCTACTCGGGAG
AGCACACGTCTGAACTCTTTCCCTACACGACGCTCTTCCGATCTNNNAACCTAAGTGCCCATCAACAG
AGCACACGTCTGAACTCTTTCCCTACACGACGCTCTTCCGATCTNNNAACCTAAGTGCCCATCAACAG
AGCACACGTCTGAACTCTTTCCCTACACGACGCTCTTCCGATCTNNNGAGGCAGGAGAATTGCTTGA
AGCACACGTCTGAACTCTTTCCCTACACGACGCTCTTCCGATCTNNNGAGGCAGGAGAATTGCTTGA
GAGCACACGTCTGAACTCTTTCCCTACACGACGCTCTTCCGATCTNNNCTGGACACAGGTAACCAAGA
GAGCACACGTCTGAACTCTTTCCCTACACGACGCTCTTCCGATCTNNNCTGGACACAGGTAACCAAGA
AGCACACGTCTGAACTCTTTCCCTACACGACGCTCTTCCGATCTNNNGCCTTGGGAATGTAGCTTTGTT
AGCACACGTCTGAACTCTTTCCCTACACGACGCTCTTCCGATCTNNNGCCTTGGGAATGTAGCTTTGTT
CACACGTCTGAACTCTTTCCCTACACGACGCTCTTCCGATCTNNNTCTCTTGTTTTCAGCCGTGATC
CACACGTCTGAACTCTTTCCCTACACGACGCTCTTCCGATCTNNNTCTCTTGTTTTCAGCCGTGATC
AGCACACGTCTGAACTCTTTCCCTACACGACGCTCTTCCGATCTNNNTGTCCCTCACAGATGTTTGAGA
AGCACACGTCTGAACTCTTTCCCTACACGACGCTCTTCCGATCTNNNTGTCCCTCACAGATGTTTGAGA
AGCACACGTCTGAACTCTTTCCCTACACGACGCTCTTCCGATCTNNNTAAAAGCAAAAATCCCCAGCC
AGCACACGTCTGAACTCTTTCCCTACACGACGCTCTTCCGATCTNNNTAAAAGCAAAAATCCCCAGCC
CACACGTCTGAACTCTTTCCCTACACGACGCTCTTCCGATCTNNNCTCATTCCTGTAATCCCAGCACTCA
CACACGTCTGAACTCTTTCCCTACACGACGCTCTTCCGATCTNNNCTCATTCCTGTAATCCCAGCACTCA

AGCACACGTCTGAACTCTTTCCCTACACGACGCTCTTCCGATCTNNNCTGAGGCAGGAGAATCACTTG

AGCACACGTCTGAACTCTTTCCCTACACGACGCTCTTCCGATCTNNNCTGAGGCAGGAGAATCACTTG

AGAGCACACGTCTGAACTCTTTCCCTACACGACGCTCTTCCGATCTNNNCACAATGTGCACATGTACCCT

AGAGCACACGTCTGAACTCTTTCCCTACACGACGCTCTTCCGATCTNNNCACAATGTGCACATGTACCCT

GAGCACACGTCTGAACTCTTTCCCTACACGACGCTCTTCCGATCTNNNACCAACCAAGATGACCATCAAC

GAGCACACGTCTGAACTCTTTCCCTACACGACGCTCTTCCGATCTNNNACCAACCAAGATGACCATCAAC

GAGCACACGTCTGAACTCTTTCCCTACACGACGCTCTTCCGATCTNNNCTCACATCCCTAGTCACTCACTG

GAGCACACGTCTGAACTCTTTCCCTACACGACGCTCTTCCGATCTNNNCTCACATCCCTAGTCACTCACTG

AGCACACGTCTGAACTCTTTCCCTACACGACGCTCTTCCGATCTNNNTTTGCAAAAACCTCGCCTCTAAG

AGCACACGTCTGAACTCTTTCCCTACACGACGCTCTTCCGATCTNNNTTTGCAAAAACCTCGCCTCTAAG

AGAGCACACGTCTGAACTCTTTCCCTACACGACGCTCTTCCGATCTNNNGGCACCTCTTCTCAGATTGTC

AGAGCACACGTCTGAACTCTTTCCCTACACGACGCTCTTCCGATCTNNNGGCACCTCTTCTCAGATTGTC

AGAGCACACGTCTGAACTCTTTCCCTACACGACGCTCTTCCGATCTNNNCCTACAGAAGCTACAGAGGA

AGAGCACACGTCTGAACTCTTTCCCTACACGACGCTCTTCCGATCTNNNCCTACAGAAGCTACAGAGGA

GAGCACACGTCTGAACTCTTTCCCTACACGACGCTCTTCCGATCTNNNTTGCTTTCTTTACCTCCTGCTCT

GAGCACACGTCTGAACTCTTTCCCTACACGACGCTCTTCCGATCTNNNTTGCTTTCTTTACCTCCTGCTCT

GAGCACACGTCTGAACTCTTTCCCTACACGACGCTCTTCCGATCTNNNTTAGTCACAAATGCAGCACACA

GAGCACACGTCTGAACTCTTTCCCTACACGACGCTCTTCCGATCTNNNTTAGTCACAAATGCAGCACACA

AGAGCACACGTCTGAACTCTTTCCCTACACGACGCTCTTCCGATCTNNNCACCTGTAGTCCCAGCTACTT

AGAGCACACGTCTGAACTCTTTCCCTACACGACGCTCTTCCGATCTNNNCACCTGTAGTCCCAGCTACTT

AGCACACGTCTGAACTCTTTCCCTACACGACGCTCTTCCGATCTNNNACCAATGAGTTTGCCTTTTCCAG

AGCACACGTCTGAACTCTTTCCCTACACGACGCTCTTCCGATCTNNNACCAATGAGTTTGCCTTTTCCAG

AGCACACGTCTGAACTCTTTCCCTACACGACGCTCTTCCGATCTNNNGGTGCATACCTGTAATCCCATCT

AGCACACGTCTGAACTCTTTCCCTACACGACGCTCTTCCGATCTNNNGGTGCATACCTGTAATCCCATCT

GAGCACACGTCTGAACTCTTTCCCTACACGACGCTCTTCCGATCTNNNCTTCCCAAGTAGCTGGAACTAC

GAGCACACGTCTGAACTCTTTCCCTACACGACGCTCTTCCGATCTNNNCTTCCCAAGTAGCTGGAACTAC

GAGCACACGTCTGAACTCTTTCCCTACACGACGCTCTTCCGATCTNNNAGGATGGAGGGAGAGAAAG

GAGCACACGTCTGAACTCTTTCCCTACACGACGCTCTTCCGATCTNNNAGGATGGAGGGAGAGAAAG

AGAGCACACGTCTGAACTCTTTCCCTACACGACGCTCTTCCGATCTNNNCGGGCTTCTCTTCACTTCTTG

AGAGCACACGTCTGAACTCTTTCCCTACACGACGCTCTTCCGATCTNNNCGGGCTTCTCTTCACTTCTTG

AGAGCACACGTCTGAACTCTTTCCCTACACGACGCTCTTCCGATCTNNNCTGAGAGAGTGGGGAAAAC

AGAGCACACGTCTGAACTCTTTCCCTACACGACGCTCTTCCGATCTNNNCTGAGAGAGTGGGGAAAAC

AGCACACGTCTGAACTCTTTCCCTACACGACGCTCTTCCGATCTNNNCACCAGGAAGAGAAAATACCCC

AGCACACGTCTGAACTCTTTCCCTACACGACGCTCTTCCGATCTNNNCACCAGGAAGAGAAAATACCCC

AGCACACGTCTGAACTCTTTCCCTACACGACGCTCTTCCGATCTNNNCCTCCCAGTCTCATTTTGGAGAA

AGCACACGTCTGAACTCTTTCCCTACACGACGCTCTTCCGATCTNNNCCTCCCAGTCTCATTTTGGAGAA

GAGCACACGTCTGAACTCTTTCCCTACACGACGCTCTTCCGATCTNNNGTGTGCCTATAATCCCAGCTA

GAGCACACGTCTGAACTCTTTCCCTACACGACGCTCTTCCGATCTNNNGTGTGCCTATAATCCCAGCTA

GAGCACACGTCTGAACTCTTTCCCTACACGACGCTCTTCCGATCTNNNCAGAGGTTGCAGTGAACCAAC

GAGCACACGTCTGAACTCTTTCCCTACACGACGCTCTTCCGATCTNNNCAGAGGTTGCAGTGAACCAAC

CACACGTCTGAACTCTTTCCCTACACGACGCTCTTCCGATCTNNNCAAGTTCTACAATTACCACAAGCAT

CACACGTCTGAACTCTTTCCCTACACGACGCTCTTCCGATCTNNNCAAGTTCTACAATTACCACAAGCAT

AGCACACGTCTGAACTCTTTCCCTACACGACGCTCTTCCGATCTNNNCTGAGGCAGGAAGATCACTTGA

AGCACACGTCTGAACTCTTTCCCTACACGACGCTCTTCCGATCTNNNCTGAGGCAGGAAGATCACTTGA
GCACACGTCTGAACTCTTTCCCTACACGACGCTCTTCCGATCTNNNAAGGTGGTCTGAATATGGAACC
GCACACGTCTGAACTCTTTCCCTACACGACGCTCTTCCGATCTNNNAAGGTGGTCTGAATATGGAACC
AGCACACGTCTGAACTCTTTCCCTACACGACGCTCTTCCGATCTNNNCGTTTGGGATCTTAGCACACATC
AGCACACGTCTGAACTCTTTCCCTACACGACGCTCTTCCGATCTNNNCGTTTGGGATCTTAGCACACATC
AGCACACGTCTGAACTCTTTCCCTACACGACGCTCTTCCGATCTNNNAGAACATAAGCTCCATGAGGAC
AGCACACGTCTGAACTCTTTCCCTACACGACGCTCTTCCGATCTNNNAGAACATAAGCTCCATGAGGAC
AAGAGCACACGTCTGAACTCTTTCCCTACACGACGCTCTTCCGATCTNNNGTGGAAGCACAGCCATAT
AAGAGCACACGTCTGAACTCTTTCCCTACACGACGCTCTTCCGATCTNNNGTGGAAGCACAGCCATAT
AGCACACGTCTGAACTCTTTCCCTACACGACGCTCTTCCGATCTNNNCATGATCGTGTCACTGTACTCCA
AGCACACGTCTGAACTCTTTCCCTACACGACGCTCTTCCGATCTNNNCATGATCGTGTCACTGTACTCCA
GCACACGTCTGAACTCTTTCCCTACACGACGCTCTTCCGATCTNNNTGAGCTGAGCTACACAAGTCTTC
GCACACGTCTGAACTCTTTCCCTACACGACGCTCTTCCGATCTNNNTGAGCTGAGCTACACAAGTCTTC
AGCACACGTCTGAACTCTTTCCCTACACGACGCTCTTCCGATCTNNNAAGTGCCTTCCAATTACGAGCCC
AGCACACGTCTGAACTCTTTCCCTACACGACGCTCTTCCGATCTNNNAAGTGCCTTCCAATTACGAGCCC
GAGCACACGTCTGAACTCTTTCCCTACACGACGCTCTTCCGATCTNNNTTCTGTCTCCCTTCCTCTCTTCT
GAGCACACGTCTGAACTCTTTCCCTACACGACGCTCTTCCGATCTNNNTTCTGTCTCCCTTCCTCTCTTCT
AGCACACGTCTGAACTCTTTCCCTACACGACGCTCTTCCGATCTNNNGATCTAGGAGCTGAGTCCTACC
AGCACACGTCTGAACTCTTTCCCTACACGACGCTCTTCCGATCTNNNGATCTAGGAGCTGAGTCCTACC
AGCACACGTCTGAACTCTTTCCCTACACGACGCTCTTCCGATCTNNNATCACTCCAATCTCTGCCTTTGTC
AGCACACGTCTGAACTCTTTCCCTACACGACGCTCTTCCGATCTNNNATCACTCCAATCTCTGCCTTTGTC
CACACGTCTGAACTCTTTCCCTACACGACGCTCTTCCGATCTNNNTCCATTCTGGAGCTTTGTAGCATCC
CACACGTCTGAACTCTTTCCCTACACGACGCTCTTCCGATCTNNNTCCATTCTGGAGCTTTGTAGCATCC
GCACACGTCTGAACTCTTTCCCTACACGACGCTCTTCCGATCTNNNCTGAGTGAGCAGTAACGAAGGT
GCACACGTCTGAACTCTTTCCCTACACGACGCTCTTCCGATCTNNNCTGAGTGAGCAGTAACGAAGGT
AGCACACGTCTGAACTCTTTCCCTACACGACGCTCTTCCGATCTNNNGTAGTGTGATCATGGCTCATTG
AGCACACGTCTGAACTCTTTCCCTACACGACGCTCTTCCGATCTNNNGTAGTGTGATCATGGCTCATTG
AGCACACGTCTGAACTCTTTCCCTACACGACGCTCTTCCGATCTNNNGTGGCCTACTAGGAAGTGTTTG
AGCACACGTCTGAACTCTTTCCCTACACGACGCTCTTCCGATCTNNNGTGGCCTACTAGGAAGTGTTTG
AGAGCACACGTCTGAACTCTTTCCCTACACGACGCTCTTCCGATCTNNNGCACATGCACAGTGGAAAA
AGAGCACACGTCTGAACTCTTTCCCTACACGACGCTCTTCCGATCTNNNGCACATGCACAGTGGAAAA
AGCACACGTCTGAACTCTTTCCCTACACGACGCTCTTCCGATCTNNNAAACAGAGTGAGCCCTTCTTCAA
AGCACACGTCTGAACTCTTTCCCTACACGACGCTCTTCCGATCTNNNAAACAGAGTGAGCCCTTCTTCAA
AGAGCACACGTCTGAACTCTTTCCCTACACGACGCTCTTCCGATCTNNNAAGGAGAAGAGAGAGACC
AGAGCACACGTCTGAACTCTTTCCCTACACGACGCTCTTCCGATCTNNNAAGGAGAAGAGAGAGACC
AGCACACGTCTGAACTCTTTCCCTACACGACGCTCTTCCGATCTNNNCAGACAAAATTTGTTGCTGGTG
AGCACACGTCTGAACTCTTTCCCTACACGACGCTCTTCCGATCTNNNCAGACAAAATTTGTTGCTGGTG
AGCACACGTCTGAACTCTTTCCCTACACGACGCTCTTCCGATCTNNNCTACATGCCCTGTGTGTCTATCCC
AGCACACGTCTGAACTCTTTCCCTACACGACGCTCTTCCGATCTNNNCTACATGCCCTGTGTGTCTATCCC
GAGCACACGTCTGAACTCTTTCCCTACACGACGCTCTTCCGATCTNNNAAGGAGAAGAGGTAGGTCAC
GAGCACACGTCTGAACTCTTTCCCTACACGACGCTCTTCCGATCTNNNAAGGAGAAGAGGTAGGTCAC
GAGCACACGTCTGAACTCTTTCCCTACACGACGCTCTTCCGATCTNNNTGTGTCTGTGATTTAGAGAACT
GAGCACACGTCTGAACTCTTTCCCTACACGACGCTCTTCCGATCTNNNTGTGTCTGTGATTTAGAGAACT

GAGCACACGTCTGAACTCTTTCCCTACACGACGCTCTTCCGATCTNNNGTCTGCTCAGTTCCCGCTAAG
GAGCACACGTCTGAACTCTTTCCCTACACGACGCTCTTCCGATCTNNNGTCTGCTCAGTTCCCGCTAAG
AGCACACGTCTGAACTCTTTCCCTACACGACGCTCTTCCGATCTNNNGCCCTCTTGTGCTGTTGAATAG
AGCACACGTCTGAACTCTTTCCCTACACGACGCTCTTCCGATCTNNNGCCCTCTTGTGCTGTTGAATAG
AGCACACGTCTGAACTCTTTCCCTACACGACGCTCTTCCGATCTNNNGCATCCGGCCTTATTTATGCATT
AGCACACGTCTGAACTCTTTCCCTACACGACGCTCTTCCGATCTNNNGCATCCGGCCTTATTTATGCATT
AGCACACGTCTGAACTCTTTCCCTACACGACGCTCTTCCGATCTNNNGGGACTTTACTTGGTGAGTCT
AGCACACGTCTGAACTCTTTCCCTACACGACGCTCTTCCGATCTNNNGGGACTTTACTTGGTGAGTCT
GCACACGTCTGAACTCTTTCCCTACACGACGCTCTTCCGATCTNNNAGAGTTACTCAACAGGAAACTT
GCACACGTCTGAACTCTTTCCCTACACGACGCTCTTCCGATCTNNNAGAGTTACTCAACAGGAAACTT
AGCACACGTCTGAACTCTTTCCCTACACGACGCTCTTCCGATCTNNNAAGTGGTATGTGACTGCATAG
AGCACACGTCTGAACTCTTTCCCTACACGACGCTCTTCCGATCTNNNAAGTGGTATGTGACTGCATAG
AGCACACGTCTGAACTCTTTCCCTACACGACGCTCTTCCGATCTNNNAAGACCCATCAGTGTGCTGTA
AGCACACGTCTGAACTCTTTCCCTACACGACGCTCTTCCGATCTNNNAAGACCCATCAGTGTGCTGTA
AGCACACGTCTGAACTCTTTCCCTACACGACGCTCTTCCGATCTNNNCAATCTTGGCTCACTGCAATCT
AGCACACGTCTGAACTCTTTCCCTACACGACGCTCTTCCGATCTNNNCAATCTTGGCTCACTGCAATCT
GAGCACACGTCTGAACTCTTTCCCTACACGACGCTCTTCCGATCTNNNATACAGAATCTGGCACTGGGT
GAGCACACGTCTGAACTCTTTCCCTACACGACGCTCTTCCGATCTNNNATACAGAATCTGGCACTGGGT
AGCACACGTCTGAACTCTTTCCCTACACGACGCTCTTCCGATCTNNNGTACAGAAGAAGGAAGCTTGG
AGCACACGTCTGAACTCTTTCCCTACACGACGCTCTTCCGATCTNNNGTACAGAAGAAGGAAGCTTGG
GAGCACACGTCTGAACTCTTTCCCTACACGACGCTCTTCCGATCTNNNGTATCTGTGTGTGTGTGTGTG
GAGCACACGTCTGAACTCTTTCCCTACACGACGCTCTTCCGATCTNNNGTATCTGTGTGTGTGTGTGTG
GAGCACACGTCTGAACTCTTTCCCTACACGACGCTCTTCCGATCTNNNGGAACAGAAGTGAGGGATG
GAGCACACGTCTGAACTCTTTCCCTACACGACGCTCTTCCGATCTNNNGGAACAGAAGTGAGGGATG
AGCACACGTCTGAACTCTTTCCCTACACGACGCTCTTCCGATCTNNNTCAACGGTGGGGAGAATATTA
AGCACACGTCTGAACTCTTTCCCTACACGACGCTCTTCCGATCTNNNTCAACGGTGGGGAGAATATTA
GCACACGTCTGAACTCTTTCCCTACACGACGCTCTTCCGATCTNNNTCAGCTCCAGAATTTCTGCTTGA
GCACACGTCTGAACTCTTTCCCTACACGACGCTCTTCCGATCTNNNTCAGCTCCAGAATTTCTGCTTGA
AGCACACGTCTGAACTCTTTCCCTACACGACGCTCTTCCGATCTNNNCCTCATACTCTGGCCATCAGTTT
AGCACACGTCTGAACTCTTTCCCTACACGACGCTCTTCCGATCTNNNCCTCATACTCTGGCCATCAGTTT
GCACACGTCTGAACTCTTTCCCTACACGACGCTCTTCCGATCTNNNGGAGAAGCAGCACACCTAGTTT
GCACACGTCTGAACTCTTTCCCTACACGACGCTCTTCCGATCTNNNGGAGAAGCAGCACACCTAGTTT
GCACACGTCTGAACTCTTTCCCTACACGACGCTCTTCCGATCTNNNTTCTCCTTCATAGGCTATTTTAGC
GCACACGTCTGAACTCTTTCCCTACACGACGCTCTTCCGATCTNNNTTCTCCTTCATAGGCTATTTTAGC
GAGCACACGTCTGAACTCTTTCCCTACACGACGCTCTTCCGATCTNNNTGAACAGGTAGAGGATGGTG
GAGCACACGTCTGAACTCTTTCCCTACACGACGCTCTTCCGATCTNNNTGAACAGGTAGAGGATGGTG
GAGCACACGTCTGAACTCTTTCCCTACACGACGCTCTTCCGATCTNNNATGAAAAATTAGCCAGGCGT
GAGCACACGTCTGAACTCTTTCCCTACACGACGCTCTTCCGATCTNNNATGAAAAATTAGCCAGGCGT
GAGCACACGTCTGAACTCTTTCCCTACACGACGCTCTTCCGATCTNNNACGAAGATCCAATTCCAAGG
GAGCACACGTCTGAACTCTTTCCCTACACGACGCTCTTCCGATCTNNNACGAAGATCCAATTCCAAGG
AGCACACGTCTGAACTCTTTCCCTACACGACGCTCTTCCGATCTNNNTCTGTGCCTCTTGTTCTCTGATC
AGCACACGTCTGAACTCTTTCCCTACACGACGCTCTTCCGATCTNNNTCTGTGCCTCTTGTTCTCTGATC
AGCACACGTCTGAACTCTTTCCCTACACGACGCTCTTCCGATCTNNNCTAAGATGCAGTCCACAATGC

AGCACACGTCTGAACTCTTTCCCTACACGACGCTCTTCCGATCTNNNCTAAGATGCAGTCCACAATGCA

GAGCACACGTCTGAACTCTTTCCCTACACGACGCTCTTCCGATCTNNNTACATAGCGGTTGTTGATGGC

GAGCACACGTCTGAACTCTTTCCCTACACGACGCTCTTCCGATCTNNNTACATAGCGGTTGTTGATGGC

CACACGTCTGAACTCTTTCCCTACACGACGCTCTTCCGATCTNNNCGACTCTGCTGCTGCTTTTTATCA

CACACGTCTGAACTCTTTCCCTACACGACGCTCTTCCGATCTNNNCGACTCTGCTGCTGCTTTTTATCA

GAGCACACGTCTGAACTCTTTCCCTACACGACGCTCTTCCGATCTNNNACCAATGACGTTCTTCACAGAA

GAGCACACGTCTGAACTCTTTCCCTACACGACGCTCTTCCGATCTNNNACCAATGACGTTCTTCACAGAA

AGCACACGTCTGAACTCTTTCCCTACACGACGCTCTTCCGATCTNNNTACCAGAGCAACACCAATTCAG

AGCACACGTCTGAACTCTTTCCCTACACGACGCTCTTCCGATCTNNNTACCAGAGCAACACCAATTCAG

AGAGCACACGTCTGAACTCTTTCCCTACACGACGCTCTTCCGATCTNNNACTAACCCACTACCTTCCCAG

AGAGCACACGTCTGAACTCTTTCCCTACACGACGCTCTTCCGATCTNNNACTAACCCACTACCTTCCCAG

AGCACACGTCTGAACTCTTTCCCTACACGACGCTCTTCCGATCTNNNGCAGGGGCTTAGTAAAATGAC

AGCACACGTCTGAACTCTTTCCCTACACGACGCTCTTCCGATCTNNNGCAGGGGCTTAGTAAAATGAC

GAGCACACGTCTGAACTCTTTCCCTACACGACGCTCTTCCGATCTNNNTGAGAGACACTTCTGAAGCCA

GAGCACACGTCTGAACTCTTTCCCTACACGACGCTCTTCCGATCTNNNTGAGAGACACTTCTGAAGCCA

AGAGCACACGTCTGAACTCTTTCCCTACACGACGCTCTTCCGATCTNNNAATGAGCTTTGATGCCCTTTC

AGAGCACACGTCTGAACTCTTTCCCTACACGACGCTCTTCCGATCTNNNAATGAGCTTTGATGCCCTTTC

AGCACACGTCTGAACTCTTTCCCTACACGACGCTCTTCCGATCTNNNAGGCTCTGTGATGATTGTGCTTA

AGCACACGTCTGAACTCTTTCCCTACACGACGCTCTTCCGATCTNNNAGGCTCTGTGATGATTGTGCTTA

AGCACACGTCTGAACTCTTTCCCTACACGACGCTCTTCCGATCTNNNTCCCCTATTCCCTTGTGCTTGTAC

AGCACACGTCTGAACTCTTTCCCTACACGACGCTCTTCCGATCTNNNTCCCCTATTCCCTTGTGCTTGTAC

GAGCACACGTCTGAACTCTTTCCCTACACGACGCTCTTCCGATCTNNNGAGAATCACTTGAACCTGGG

GAGCACACGTCTGAACTCTTTCCCTACACGACGCTCTTCCGATCTNNNGAGAATCACTTGAACCTGGG

AAGAGCACACGTCTGAACTCTTTCCCTACACGACGCTCTTCCGATCTNNNAGAATCGCTTGAACCCAGG

AAGAGCACACGTCTGAACTCTTTCCCTACACGACGCTCTTCCGATCTNNNAGAATCGCTTGAACCCAGG

AGCACACGTCTGAACTCTTTCCCTACACGACGCTCTTCCGATCTNNNCTTTTTCTCCCTCTCAGATGCAT

AGCACACGTCTGAACTCTTTCCCTACACGACGCTCTTCCGATCTNNNCTTTTTCTCCCTCTCAGATGCAT

AGAGCACACGTCTGAACTCTTTCCCTACACGACGCTCTTCCGATCTNNNGAGAAGAGATGATTGGGTG

AGAGCACACGTCTGAACTCTTTCCCTACACGACGCTCTTCCGATCTNNNGAGAAGAGATGATTGGGTG

GAGCACACGTCTGAACTCTTTCCCTACACGACGCTCTTCCGATCTNNNCCAAAGTTGATCAGGCCTAGT

GAGCACACGTCTGAACTCTTTCCCTACACGACGCTCTTCCGATCTNNNCCAAAGTTGATCAGGCCTAGT

AGAGCACACGTCTGAACTCTTTCCCTACACGACGCTCTTCCGATCTNNNCTGAGACAGGGGAATTGCTT

AGAGCACACGTCTGAACTCTTTCCCTACACGACGCTCTTCCGATCTNNNCTGAGACAGGGGAATTGCTT

AGCACACGTCTGAACTCTTTCCCTACACGACGCTCTTCCGATCTNNNTGTGTGTGTGTGTGTGAAA

AGCACACGTCTGAACTCTTTCCCTACACGACGCTCTTCCGATCTNNNTGTGTGTGTGTGTGTGAAA

AGCACACGTCTGAACTCTTTCCCTACACGACGCTCTTCCGATCTNNNGCTGATTGTAGTTTTGGGCTGT

AGCACACGTCTGAACTCTTTCCCTACACGACGCTCTTCCGATCTNNNGCTGATTGTAGTTTTGGGCTGT

AGCACACGTCTGAACTCTTTCCCTACACGACGCTCTTCCGATCTNNNATTGCCTGAGCTCAGTAGTTCA

AGCACACGTCTGAACTCTTTCCCTACACGACGCTCTTCCGATCTNNNATTGCCTGAGCTCAGTAGTTCA

GAGCACACGTCTGAACTCTTTCCCTACACGACGCTCTTCCGATCTNNNATGGAGGGCAATGTGCTTTAC

GAGCACACGTCTGAACTCTTTCCCTACACGACGCTCTTCCGATCTNNNATGGAGGGCAATGTGCTTTAC

GAGCACACGTCTGAACTCTTTCCCTACACGACGCTCTTCCGATCTNNNCACACATAGGTTCTTGGCCAAA

GAGCACACGTCTGAACTCTTTCCCTACACGACGCTCTTCCGATCTNNNCACACATAGGTTCTTGGCCAAA

GCACACGTCTGAACTCTTTCCCTACACGACGCTCTTCCGATCTNNNTCTGAGAGATTTTGGTGCACTC

GCACACGTCTGAACTCTTTCCCTACACGACGCTCTTCCGATCTNNNTCTGAGAGATTTTGGTGCACTC

CACACGTCTGAACTCTTTCCCTACACGACGCTCTTCCGATCTNNNAGGAAACACAGGATTCTATGATAT

CACACGTCTGAACTCTTTCCCTACACGACGCTCTTCCGATCTNNNAGGAAACACAGGATTCTATGATAT

GAGCACACGTCTGAACTCTTTCCCTACACGACGCTCTTCCGATCTNNNGGGTGCTGGACATCTCTACT

GAGCACACGTCTGAACTCTTTCCCTACACGACGCTCTTCCGATCTNNNGGGTGCTGGACATCTCTACT

GAGCACACGTCTGAACTCTTTCCCTACACGACGCTCTTCCGATCTNNNTCACCTTTTCTGTCCTTGGTCT

GAGCACACGTCTGAACTCTTTCCCTACACGACGCTCTTCCGATCTNNNTCACCTTTTCTGTCCTTGGTCT

AGCACACGTCTGAACTCTTTCCCTACACGACGCTCTTCCGATCTNNNCCATGTGTTTTGAGTGGCTTCT

AGCACACGTCTGAACTCTTTCCCTACACGACGCTCTTCCGATCTNNNCCATGTGTTTTGAGTGGCTTCT

AGAGCACACGTCTGAACTCTTTCCCTACACGACGCTCTTCCGATCTNNNGATCACAAGGTCAGGAGAT

AGAGCACACGTCTGAACTCTTTCCCTACACGACGCTCTTCCGATCTNNNGATCACAAGGTCAGGAGAT

AGCACACGTCTGAACTCTTTCCCTACACGACGCTCTTCCGATCTNNNGCTTGCAGTGAGCTCAAATTG

AGCACACGTCTGAACTCTTTCCCTACACGACGCTCTTCCGATCTNNNGCTTGCAGTGAGCTCAAATTG

AGCACACGTCTGAACTCTTTCCCTACACGACGCTCTTCCGATCTNNNTGCTTTTACTGAGAAAGAAGC

AGCACACGTCTGAACTCTTTCCCTACACGACGCTCTTCCGATCTNNNTGCTTTTACTGAGAAAGAAGC

AGAGCACACGTCTGAACTCTTTCCCTACACGACGCTCTTCCGATCTNNNGAGAATCAGTTGAACCTGG

AGAGCACACGTCTGAACTCTTTCCCTACACGACGCTCTTCCGATCTNNNGAGAATCAGTTGAACCTGG

AGCACACGTCTGAACTCTTTCCCTACACGACGCTCTTCCGATCTNNNGAGCTTACCTATTATCCTCAGC

AGCACACGTCTGAACTCTTTCCCTACACGACGCTCTTCCGATCTNNNGAGCTTACCTATTATCCTCAGC

GAGCACACGTCTGAACTCTTTCCCTACACGACGCTCTTCCGATCTNNNCCAAATACAGGAAACTCGACT

GAGCACACGTCTGAACTCTTTCCCTACACGACGCTCTTCCGATCTNNNCCAAATACAGGAAACTCGACT

GCACACGTCTGAACTCTTTCCCTACACGACGCTCTTCCGATCTNNNATTTCCCTTCCTTGGAGTTCTGG

GCACACGTCTGAACTCTTTCCCTACACGACGCTCTTCCGATCTNNNATTTCCCTTCCTTGGAGTTCTGG

AGAGCACACGTCTGAACTCTTTCCCTACACGACGCTCTTCCGATCTNNNCACAGCATTCCTCACCAGAA

AGAGCACACGTCTGAACTCTTTCCCTACACGACGCTCTTCCGATCTNNNCACAGCATTCCTCACCAGAA

GAGCACACGTCTGAACTCTTTCCCTACACGACGCTCTTCCGATCTNNNCTGAGGCAGGAGAATCACTT

GAGCACACGTCTGAACTCTTTCCCTACACGACGCTCTTCCGATCTNNNCTGAGGCAGGAGAATCACTT

GCACACGTCTGAACTCTTTCCCTACACGACGCTCTTCCGATCTNNNAGTTAATGGTATGGATGTGCCA

GCACACGTCTGAACTCTTTCCCTACACGACGCTCTTCCGATCTNNNAGTTAATGGTATGGATGTGCCA

AGCACACGTCTGAACTCTTTCCCTACACGACGCTCTTCCGATCTNNNGACTAAGCTCTCTCATGACCCA

AGCACACGTCTGAACTCTTTCCCTACACGACGCTCTTCCGATCTNNNGACTAAGCTCTCTCATGACCCA

GAGCACACGTCTGAACTCTTTCCCTACACGACGCTCTTCCGATCTNNNCTATTGGAGAGGTTGTGGG

GAGCACACGTCTGAACTCTTTCCCTACACGACGCTCTTCCGATCTNNNCTATTGGAGAGGTTGTGGG

GCACACGTCTGAACTCTTTCCCTACACGACGCTCTTCCGATCTNNNGATGCGTTTCTTTGGGACTAGT

GCACACGTCTGAACTCTTTCCCTACACGACGCTCTTCCGATCTNNNGATGCGTTTCTTTGGGACTAGT

GCACACGTCTGAACTCTTTCCCTACACGACGCTCTTCCGATCTNNNAATTCAATCCTGCAGCTTTCCAC

GCACACGTCTGAACTCTTTCCCTACACGACGCTCTTCCGATCTNNNAATTCAATCCTGCAGCTTTCCAC

AGCACACGTCTGAACTCTTTCCCTACACGACGCTCTTCCGATCTNNNTTGCAATTTTCTTCTCAGTGGG

AGCACACGTCTGAACTCTTTCCCTACACGACGCTCTTCCGATCTNNNTTGCAATTTTCTTCTCAGTGGG

GAGCACACGTCTGAACTCTTTCCCTACACGACGCTCTTCCGATCTNNNTGAGATTGTGCCATTGTACTC

GAGCACACGTCTGAACTCTTTCCCTACACGACGCTCTTCCGATCTNNNTGAGATTGTGCCATTGTACTC

AGCACACGTCTGAACTCTTTCCCTACACGACGCTCTTCCGATCTNNNTCCGAGACTATCCTGACCAAC

GAGCACACGTCTGAACTCTTTCCCTACACGACGCTCTTCCGATCTNNNTCCGAGACTATCCTGACCAACC
AGCACACGTCTGAACTCTTTCCCTACACGACGCTCTTCCGATCTNNNGACTGATGCAGGAGAATTGCT
AGCACACGTCTGAACTCTTTCCCTACACGACGCTCTTCCGATCTNNNGACTGATGCAGGAGAATTGCT
AGAGCACACGTCTGAACTCTTTCCCTACACGACGCTCTTCCGATCTNNNGTGTGAGGCATAAATCCTG
AGAGCACACGTCTGAACTCTTTCCCTACACGACGCTCTTCCGATCTNNNGTGTGAGGCATAAATCCTG
AGAGCACACGTCTGAACTCTTTCCCTACACGACGCTCTTCCGATCTNNNCAAGCATCTGATGGCTACAC
AGAGCACACGTCTGAACTCTTTCCCTACACGACGCTCTTCCGATCTNNNCAAGCATCTGATGGCTACAC
AGCACACGTCTGAACTCTTTCCCTACACGACGCTCTTCCGATCTNNNACACAGTGAAACCCTGTCTCTAC
AGCACACGTCTGAACTCTTTCCCTACACGACGCTCTTCCGATCTNNNACACAGTGAAACCCTGTCTCTAC
GAGCACACGTCTGAACTCTTTCCCTACACGACGCTCTTCCGATCTNNNACCCCTCCTCCTTTATTACCCAG
GAGCACACGTCTGAACTCTTTCCCTACACGACGCTCTTCCGATCTNNNACCCCTCCTCCTTTATTACCCAG
AGCACACGTCTGAACTCTTTCCCTACACGACGCTCTTCCGATCTNNNGAGAATGGCATGAACCCAGAA
AGCACACGTCTGAACTCTTTCCCTACACGACGCTCTTCCGATCTNNNGAGAATGGCATGAACCCAGAA
GAGCACACGTCTGAACTCTTTCCCTACACGACGCTCTTCCGATCTNNNACAAAAATTAGCCAGGCGTGA
GAGCACACGTCTGAACTCTTTCCCTACACGACGCTCTTCCGATCTNNNACAAAAATTAGCCAGGCGTGA
AGCACACGTCTGAACTCTTTCCCTACACGACGCTCTTCCGATCTNNNACCCCCACACCCAGATAATTTTTC
AGCACACGTCTGAACTCTTTCCCTACACGACGCTCTTCCGATCTNNNACCCCCACACCCAGATAATTTTTC
AGCACACGTCTGAACTCTTTCCCTACACGACGCTCTTCCGATCTNNNCTAAGGCAGGAGAATCGCTTGA
AGCACACGTCTGAACTCTTTCCCTACACGACGCTCTTCCGATCTNNNCTAAGGCAGGAGAATCGCTTGA
GCACACGTCTGAACTCTTTCCCTACACGACGCTCTTCCGATCTNNNTGCTGATCATCCTGGCTGTATACC
GCACACGTCTGAACTCTTTCCCTACACGACGCTCTTCCGATCTNNNTGCTGATCATCCTGGCTGTATACC
GAGCACACGTCTGAACTCTTTCCCTACACGACGCTCTTCCGATCTNNNCTTTAGTGCCAGCAGACCATT
GAGCACACGTCTGAACTCTTTCCCTACACGACGCTCTTCCGATCTNNNCTTTAGTGCCAGCAGACCATT
AGCACACGTCTGAACTCTTTCCCTACACGACGCTCTTCCGATCTNNNACACCTGTAATCCCAGCTACTTG
AGCACACGTCTGAACTCTTTCCCTACACGACGCTCTTCCGATCTNNNACACCTGTAATCCCAGCTACTTG
GCACACGTCTGAACTCTTTCCCTACACGACGCTCTTCCGATCTNNNTCAGATCACCAAGTATTCTTGTGT
GCACACGTCTGAACTCTTTCCCTACACGACGCTCTTCCGATCTNNNTCAGATCACCAAGTATTCTTGTGT
AGCACACGTCTGAACTCTTTCCCTACACGACGCTCTTCCGATCTNNNCCAGAGAGCAAGGAAGACTGA
AGCACACGTCTGAACTCTTTCCCTACACGACGCTCTTCCGATCTNNNCCAGAGAGCAAGGAAGACTGA
GAGCACACGTCTGAACTCTTTCCCTACACGACGCTCTTCCGATCTNNNGCCAGACCCCACAAATCAAGT
GAGCACACGTCTGAACTCTTTCCCTACACGACGCTCTTCCGATCTNNNGCCAGACCCCACAAATCAAGT
GAGCACACGTCTGAACTCTTTCCCTACACGACGCTCTTCCGATCTNNNAAAACAAAAGTGCATGCCACAC
GAGCACACGTCTGAACTCTTTCCCTACACGACGCTCTTCCGATCTNNNAAAACAAAAGTGCATGCCACAC
AGCACACGTCTGAACTCTTTCCCTACACGACGCTCTTCCGATCTNNNCTTTATCTCCTGGGCTCAAGTGT
AGCACACGTCTGAACTCTTTCCCTACACGACGCTCTTCCGATCTNNNCTTTATCTCCTGGGCTCAAGTGT
AGCACACGTCTGAACTCTTTCCCTACACGACGCTCTTCCGATCTNNNGCCCTGATAAACCCTTCAGACCT
AGCACACGTCTGAACTCTTTCCCTACACGACGCTCTTCCGATCTNNNGCCCTGATAAACCCTTCAGACCT
AGCACACGTCTGAACTCTTTCCCTACACGACGCTCTTCCGATCTNNNACCTCTTCTTCACCAGGCTATTG
AGCACACGTCTGAACTCTTTCCCTACACGACGCTCTTCCGATCTNNNACCTCTTCTTCACCAGGCTATTG
AGCACACGTCTGAACTCTTTCCCTACACGACGCTCTTCCGATCTNNNCATGCCTGTAATCCCAGCTACTC
AGCACACGTCTGAACTCTTTCCCTACACGACGCTCTTCCGATCTNNNCATGCCTGTAATCCCAGCTACTC
GAGCACACGTCTGAACTCTTTCCCTACACGACGCTCTTCCGATCTNNNAAAAAATTAGCCGGGCGTAGT
GAGCACACGTCTGAACTCTTTCCCTACACGACGCTCTTCCGATCTNNNAAAAAATTAGCCGGGCGTAGT

AGCACACGTCTGAACTCTTTCCCTACACGACGCTCTTCCGATCTNNNCAGTAGTGGGACATGGTCAGAA
AGCACACGTCTGAACTCTTTCCCTACACGACGCTCTTCCGATCTNNNCAGTAGTGGGACATGGTCAGAA
AGAGCACACGTCTGAACTCTTTCCCTACACGACGCTCTTCCGATCTNNNGGGCTCTATTCACAAGTGGG
AGAGCACACGTCTGAACTCTTTCCCTACACGACGCTCTTCCGATCTNNNGGGCTCTATTCACAAGTGGG
GAGCACACGTCTGAACTCTTTCCCTACACGACGCTCTTCCGATCTNNNTGTATTCTCCTTACCCCAACCCC
GAGCACACGTCTGAACTCTTTCCCTACACGACGCTCTTCCGATCTNNNTGTATTCTCCTTACCCCAACCCC
AGAGCACACGTCTGAACTCTTTCCCTACACGACGCTCTTCCGATCTNNNCTCTCTCTCTGTGTGTGTGTG
AGAGCACACGTCTGAACTCTTTCCCTACACGACGCTCTTCCGATCTNNNCTCTCTCTCTGTGTGTGTGTG
GAGCACACGTCTGAACTCTTTCCCTACACGACGCTCTTCCGATCTNNNTCCCTTACCTTCTCCCAAGCAAT
GAGCACACGTCTGAACTCTTTCCCTACACGACGCTCTTCCGATCTNNNTCCCTTACCTTCTCCCAAGCAAT
AGAGCACACGTCTGAACTCTTTCCCTACACGACGCTCTTCCGATCTNNNATGGATGGGGATGGGGAG
AGAGCACACGTCTGAACTCTTTCCCTACACGACGCTCTTCCGATCTNNNATGGATGGGGATGGGGAG
AGAGCACACGTCTGAACTCTTTCCCTACACGACGCTCTTCCGATCTNNNGGGCTCTATTCACAAGTGGG
AGAGCACACGTCTGAACTCTTTCCCTACACGACGCTCTTCCGATCTNNNGGGCTCTATTCACAAGTGGG
CACACGTCTGAACTCTTTCCCTACACGACGCTCTTCCGATCTNNNGGGATGGAACTTGACGTGGAACA
CACACGTCTGAACTCTTTCCCTACACGACGCTCTTCCGATCTNNNGGGATGGAACTTGACGTGGAACA
GAGCACACGTCTGAACTCTTTCCCTACACGACGCTCTTCCGATCTNNNAGCAGAACTGAATAGCATGTG
GAGCACACGTCTGAACTCTTTCCCTACACGACGCTCTTCCGATCTNNNAGCAGAACTGAATAGCATGTG
GAGCACACGTCTGAACTCTTTCCCTACACGACGCTCTTCCGATCTNNNCTTCAAAATGCCCTGCTGCTTT
GAGCACACGTCTGAACTCTTTCCCTACACGACGCTCTTCCGATCTNNNCTTCAAAATGCCCTGCTGCTTT
AGCACACGTCTGAACTCTTTCCCTACACGACGCTCTTCCGATCTNNNTAAGAGGTACCTGGTGTCCATG
AGCACACGTCTGAACTCTTTCCCTACACGACGCTCTTCCGATCTNNNTAAGAGGTACCTGGTGTCCATG
AGAGCACACGTCTGAACTCTTTCCCTACACGACGCTCTTCCGATCTNNNCAATGGTACCTCCCGATCTAA
AGAGCACACGTCTGAACTCTTTCCCTACACGACGCTCTTCCGATCTNNNCAATGGTACCTCCCGATCTAA
AGAGCACACGTCTGAACTCTTTCCCTACACGACGCTCTTCCGATCTNNNCCAATATTGACCTCAGCAGAG
AGAGCACACGTCTGAACTCTTTCCCTACACGACGCTCTTCCGATCTNNNCCAATATTGACCTCAGCAGAG
AGCACACGTCTGAACTCTTTCCCTACACGACGCTCTTCCGATCTNNNCTGAGGCAGGAGAATCACTTGA
AGCACACGTCTGAACTCTTTCCCTACACGACGCTCTTCCGATCTNNNCTGAGGCAGGAGAATCACTTGA
GAGCACACGTCTGAACTCTTTCCCTACACGACGCTCTTCCGATCTNNNAAGATTGCACCACTGTACTCC
GAGCACACGTCTGAACTCTTTCCCTACACGACGCTCTTCCGATCTNNNAAGATTGCACCACTGTACTCC
CACGTCTGAACTCTTTCCCTACACGACGCTCTTCCGATCTNNNAGGCTTCATTTCTGTCATTTCATCGGA
CACGTCTGAACTCTTTCCCTACACGACGCTCTTCCGATCTNNNAGGCTTCATTTCTGTCATTTCATCGGA
AGCACACGTCTGAACTCTTTCCCTACACGACGCTCTTCCGATCTNNNAAATCAAGCCAGTGATAGGGTG
AGCACACGTCTGAACTCTTTCCCTACACGACGCTCTTCCGATCTNNNAAATCAAGCCAGTGATAGGGTG
AGCACACGTCTGAACTCTTTCCCTACACGACGCTCTTCCGATCTNNNTCATCAGTGTCAGGAAAGCAAC
AGCACACGTCTGAACTCTTTCCCTACACGACGCTCTTCCGATCTNNNTCATCAGTGTCAGGAAAGCAAC
AGCACACGTCTGAACTCTTTCCCTACACGACGCTCTTCCGATCTNNNATTATACAGACTGCCAGCCTGA
AGCACACGTCTGAACTCTTTCCCTACACGACGCTCTTCCGATCTNNNATTATACAGACTGCCAGCCTGA
AGCACACGTCTGAACTCTTTCCCTACACGACGCTCTTCCGATCTNNNGTAGTGTAGCAGTCAGGGTTC
AGCACACGTCTGAACTCTTTCCCTACACGACGCTCTTCCGATCTNNNGTAGTGTAGCAGTCAGGGTTC
GAGCACACGTCTGAACTCTTTCCCTACACGACGCTCTTCCGATCTNNNCCAAGATTGTACCATTGCACTC
GAGCACACGTCTGAACTCTTTCCCTACACGACGCTCTTCCGATCTNNNCCAAGATTGTACCATTGCACTC
AGCACACGTCTGAACTCTTTCCCTACACGACGCTCTTCCGATCTNNNTGCATCATCTGCATCACACACATG

AGCACACGTCTGAACTCTTTCCCTACACGACGCTCTTCCGATCTNNNTGCATCATCTGCATCACACACAT
GAGCACACGTCTGAACTCTTTCCCTACACGACGCTCTTCCGATCTNNNTTGAAGTAGCCTCTGCAAAAT
GAGCACACGTCTGAACTCTTTCCCTACACGACGCTCTTCCGATCTNNNTTGAAGTAGCCTCTGCAAAAT
GAGCACACGTCTGAACTCTTTCCCTACACGACGCTCTTCCGATCTNNNACCTGTAATCCCAGCTACTTTG
GAGCACACGTCTGAACTCTTTCCCTACACGACGCTCTTCCGATCTNNNACCTGTAATCCCAGCTACTTTG
GAGCACACGTCTGAACTCTTTCCCTACACGACGCTCTTCCGATCTNNNTGAGGTGTGGATTCATCTCAC
GAGCACACGTCTGAACTCTTTCCCTACACGACGCTCTTCCGATCTNNNTGAGGTGTGGATTCATCTCAC
AGCACACGTCTGAACTCTTTCCCTACACGACGCTCTTCCGATCTNNNAGTCCTAGCTAGCAAGAGACT
AGCACACGTCTGAACTCTTTCCCTACACGACGCTCTTCCGATCTNNNAGTCCTAGCTAGCAAGAGACT
AGAGCACACGTCTGAACTCTTTCCCTACACGACGCTCTTCCGATCTNNNACACTCTCTGACTGGCAAGA
AGAGCACACGTCTGAACTCTTTCCCTACACGACGCTCTTCCGATCTNNNACACTCTCTGACTGGCAAGA
AGCACACGTCTGAACTCTTTCCCTACACGACGCTCTTCCGATCTNNNGACCATTGCCATAGACATCGAG
AGCACACGTCTGAACTCTTTCCCTACACGACGCTCTTCCGATCTNNNGACCATTGCCATAGACATCGAG
AGCACACGTCTGAACTCTTTCCCTACACGACGCTCTTCCGATCTNNNTTCGTGCCCTCTTATTGTGGAA
AGCACACGTCTGAACTCTTTCCCTACACGACGCTCTTCCGATCTNNNTTCGTGCCCTCTTATTGTGGAA
GAGCACACGTCTGAACTCTTTCCCTACACGACGCTCTTCCGATCTNNNTCCCTAAATGCAACTAGAAGG
GAGCACACGTCTGAACTCTTTCCCTACACGACGCTCTTCCGATCTNNNTCCCTAAATGCAACTAGAAGG
GAGCACACGTCTGAACTCTTTCCCTACACGACGCTCTTCCGATCTNNNAAAAATTAGCTGGGCATGAT
GAGCACACGTCTGAACTCTTTCCCTACACGACGCTCTTCCGATCTNNNAAAAATTAGCTGGGCATGAT
AGCACACGTCTGAACTCTTTCCCTACACGACGCTCTTCCGATCTNNNCGAATGATGAGAGATTTCCCAC
AGCACACGTCTGAACTCTTTCCCTACACGACGCTCTTCCGATCTNNNCGAATGATGAGAGATTTCCCAC
GAGCACACGTCTGAACTCTTTCCCTACACGACGCTCTTCCGATCTNNNATAGTGTGCTAGTGTGTGTGT
GAGCACACGTCTGAACTCTTTCCCTACACGACGCTCTTCCGATCTNNNATAGTGTGCTAGTGTGTGTGT
GCACACGTCTGAACTCTTTCCCTACACGACGCTCTTCCGATCTNNNTTAATTTGATCACATCTGATTCCC
GCACACGTCTGAACTCTTTCCCTACACGACGCTCTTCCGATCTNNNTTAATTTGATCACATCTGATTCCC
AGCACACGTCTGAACTCTTTCCCTACACGACGCTCTTCCGATCTNNNCATCGGTGTACTGTATCCAGGA
AGCACACGTCTGAACTCTTTCCCTACACGACGCTCTTCCGATCTNNNCATCGGTGTACTGTATCCAGGA
AGCACACGTCTGAACTCTTTCCCTACACGACGCTCTTCCGATCTNNNTCAGTGACACCGTGTGGATAT
AGCACACGTCTGAACTCTTTCCCTACACGACGCTCTTCCGATCTNNNTCAGTGACACCGTGTGGATAT
GAGCACACGTCTGAACTCTTTCCCTACACGACGCTCTTCCGATCTNNNAGGGCTCAGAAGACCTACTC
GAGCACACGTCTGAACTCTTTCCCTACACGACGCTCTTCCGATCTNNNAGGGCTCAGAAGACCTACTC
AGAGCACACGTCTGAACTCTTTCCCTACACGACGCTCTTCCGATCTNNNTGTCTCTCCAAAACCTTAGG
AGAGCACACGTCTGAACTCTTTCCCTACACGACGCTCTTCCGATCTNNNTGTCTCTCCAAAACCTTAGG
GAGCACACGTCTGAACTCTTTCCCTACACGACGCTCTTCCGATCTNNNAGCAGGGAATATGGTATGCA
GAGCACACGTCTGAACTCTTTCCCTACACGACGCTCTTCCGATCTNNNAGCAGGGAATATGGTATGCA
AGCACACGTCTGAACTCTTTCCCTACACGACGCTCTTCCGATCTNNNGGCAGAGCTGTTTACTACTCAC
AGCACACGTCTGAACTCTTTCCCTACACGACGCTCTTCCGATCTNNNGGCAGAGCTGTTTACTACTCAC
GAGCACACGTCTGAACTCTTTCCCTACACGACGCTCTTCCGATCTNNNGTGAAGTCAACCAGCTTGAT
GAGCACACGTCTGAACTCTTTCCCTACACGACGCTCTTCCGATCTNNNGTGAAGTCAACCAGCTTGAT
GAGCACACGTCTGAACTCTTTCCCTACACGACGCTCTTCCGATCTNNNTCAGGTCCTCTTCTAAAGCCA
GAGCACACGTCTGAACTCTTTCCCTACACGACGCTCTTCCGATCTNNNTCAGGTCCTCTTCTAAAGCCA
AGAGCACACGTCTGAACTCTTTCCCTACACGACGCTCTTCCGATCTNNNGAGGCTGAGGCAGGAGAAT
AGAGCACACGTCTGAACTCTTTCCCTACACGACGCTCTTCCGATCTNNNGAGGCTGAGGCAGGAGAAT

CACACGTCTGAACTCTTTCCCTACACGACGCTCTTCCGATCTNNNTTCATAATGGTGCTTTCAAACCAT

CACACGTCTGAACTCTTTCCCTACACGACGCTCTTCCGATCTNNNTTCATAATGGTGCTTTCAAACCAT

AGCACACGTCTGAACTCTTTCCCTACACGACGCTCTTCCGATCTNNNAGAAGGCTGAGACAGGAGAA

AGCACACGTCTGAACTCTTTCCCTACACGACGCTCTTCCGATCTNNNAGAAGGCTGAGACAGGAGAA

GCACACGTCTGAACTCTTTCCCTACACGACGCTCTTCCGATCTNNNTGTGGTACTAGCTACTCAGGAG

GCACACGTCTGAACTCTTTCCCTACACGACGCTCTTCCGATCTNNNTGTGGTACTAGCTACTCAGGAG

AGCACACGTCTGAACTCTTTCCCTACACGACGCTCTTCCGATCTNNNGGTTTAGGGTGGGATCTGAA

AGCACACGTCTGAACTCTTTCCCTACACGACGCTCTTCCGATCTNNNGGTTTAGGGTGGGATCTGAA

AGCACACGTCTGAACTCTTTCCCTACACGACGCTCTTCCGATCTNNNCAGAGTTACAAAGAGGAGCT

AGCACACGTCTGAACTCTTTCCCTACACGACGCTCTTCCGATCTNNNCAGAGTTACAAAGAGGAGCT

GCACACGTCTGAACTCTTTCCCTACACGACGCTCTTCCGATCTNNNCGTGGGATTAGATTGGGAGGT

GCACACGTCTGAACTCTTTCCCTACACGACGCTCTTCCGATCTNNNCGTGGGATTAGATTGGGAGGT

GAGCACACGTCTGAACTCTTTCCCTACACGACGCTCTTCCGATCTNNNTTTTGTAGGTTTGAAGAGG

GAGCACACGTCTGAACTCTTTCCCTACACGACGCTCTTCCGATCTNNNTTTTGTAGGTTTGAAGAGG

AGCACACGTCTGAACTCTTTCCCTACACGACGCTCTTCCGATCTNNNGCAGCCCACAAGAATGAGAG

AGCACACGTCTGAACTCTTTCCCTACACGACGCTCTTCCGATCTNNNGCAGCCCACAAGAATGAGAG

GCACACGTCTGAACTCTTTCCCTACACGACGCTCTTCCGATCTNNNGGAGAATTGCTTGAACTTGGG

GCACACGTCTGAACTCTTTCCCTACACGACGCTCTTCCGATCTNNNGGAGAATTGCTTGAACTTGGG

AGCACACGTCTGAACTCTTTCCCTACACGACGCTCTTCCGATCTNNNTGAGGCAGGAGAATCACTTG

AGCACACGTCTGAACTCTTTCCCTACACGACGCTCTTCCGATCTNNNTGAGGCAGGAGAATCACTTG

AGCACACGTCTGAACTCTTTCCCTACACGACGCTCTTCCGATCTNNNGTGAGCTGAGATTGCACCATT

AGCACACGTCTGAACTCTTTCCCTACACGACGCTCTTCCGATCTNNNGTGAGCTGAGATTGCACCATT

AGCACACGTCTGAACTCTTTCCCTACACGACGCTCTTCCGATCTNNNGGAGGGAATCACCAGTACATG

AGCACACGTCTGAACTCTTTCCCTACACGACGCTCTTCCGATCTNNNGGAGGGAATCACCAGTACATG

AGCACACGTCTGAACTCTTTCCCTACACGACGCTCTTCCGATCTNNNTTTGCCAAGACAGCAACTCTTC

AGCACACGTCTGAACTCTTTCCCTACACGACGCTCTTCCGATCTNNNTTTGCCAAGACAGCAACTCTTC

GAGCACACGTCTGAACTCTTTCCCTACACGACGCTCTTCCGATCTNNNAATAAAGTGTGTGTGTGCA

GAGCACACGTCTGAACTCTTTCCCTACACGACGCTCTTCCGATCTNNNAATAAAGTGTGTGTGTGCA

AGCACACGTCTGAACTCTTTCCCTACACGACGCTCTTCCGATCTNNNTGGGGAAAGTGATCCTGTCAT

AGCACACGTCTGAACTCTTTCCCTACACGACGCTCTTCCGATCTNNNTGGGGAAAGTGATCCTGTCAT

AGCACACGTCTGAACTCTTTCCCTACACGACGCTCTTCCGATCTNNNCAGAGCAAAATGAAAACACGC

AGCACACGTCTGAACTCTTTCCCTACACGACGCTCTTCCGATCTNNNCAGAGCAAAATGAAAACACGC

AGCACACGTCTGAACTCTTTCCCTACACGACGCTCTTCCGATCTNNNTACTATGCTGTCCTCTTCTGCA

AGCACACGTCTGAACTCTTTCCCTACACGACGCTCTTCCGATCTNNNTACTATGCTGTCCTCTTCTGCA

GCACACGTCTGAACTCTTTCCCTACACGACGCTCTTCCGATCTNNNCAGGTACTTGTTAGGGTCTCCT

GCACACGTCTGAACTCTTTCCCTACACGACGCTCTTCCGATCTNNNCAGGTACTTGTTAGGGTCTCCT

GAGCACACGTCTGAACTCTTTCCCTACACGACGCTCTTCCGATCTNNNACGATTTCCTGGATATGAAC

GAGCACACGTCTGAACTCTTTCCCTACACGACGCTCTTCCGATCTNNNACGATTTCCTGGATATGAAC

GAGCACACGTCTGAACTCTTTCCCTACACGACGCTCTTCCGATCTNNNAGATGATCACATTAGACCG

GAGCACACGTCTGAACTCTTTCCCTACACGACGCTCTTCCGATCTNNNAGATGATCACATTAGACCG

AGCACACGTCTGAACTCTTTCCCTACACGACGCTCTTCCGATCTNNNCACACACGCACACACACATATA

AGCACACGTCTGAACTCTTTCCCTACACGACGCTCTTCCGATCTNNNCACACACGCACACACACATATA

AGCACACGTCTGAACTCTTTCCCTACACGACGCTCTTCCGATCTNNNAGTCACATATTCATAGGCCAG

AGCACACGTCTGAACTCTTTCCCTACACGACGCTCTTCCGATCTNNNAGTCACATATTCATAGGCCAGG

AGCACACGTCTGAACTCTTTCCCTACACGACGCTCTTCCGATCTNNNACAGCTTGTATCATTCCTTCTGA

AGCACACGTCTGAACTCTTTCCCTACACGACGCTCTTCCGATCTNNNACAGCTTGTATCATTCCTTCTGA

AGCACACGTCTGAACTCTTTCCCTACACGACGCTCTTCCGATCTNNNGCCTACCCGTTTTCTTATTGAGA

AGCACACGTCTGAACTCTTTCCCTACACGACGCTCTTCCGATCTNNNGCCTACCCGTTTTCTTATTGAGA

GAGCACACGTCTGAACTCTTTCCCTACACGACGCTCTTCCGATCTNNNTATGGTTTCACTCTGTGTCCCT

GAGCACACGTCTGAACTCTTTCCCTACACGACGCTCTTCCGATCTNNNTATGGTTTCACTCTGTGTCCCT

AGCACACGTCTGAACTCTTTCCCTACACGACGCTCTTCCGATCTNNNCTGCTGTCAGCTAAAGATGAGC

AGCACACGTCTGAACTCTTTCCCTACACGACGCTCTTCCGATCTNNNCTGCTGTCAGCTAAAGATGAGC

GAGCACACGTCTGAACTCTTTCCCTACACGACGCTCTTCCGATCTNNNAAATTGTGTCTAGGGCTGAGA

GAGCACACGTCTGAACTCTTTCCCTACACGACGCTCTTCCGATCTNNNAAATTGTGTCTAGGGCTGAGA

AGCACACGTCTGAACTCTTTCCCTACACGACGCTCTTCCGATCTNNNATTCTAGCTACTCAGGAGGCTGA

AGCACACGTCTGAACTCTTTCCCTACACGACGCTCTTCCGATCTNNNATTCTAGCTACTCAGGAGGCTGA

AGAGCACACGTCTGAACTCTTTCCCTACACGACGCTCTTCCGATCTNNNTCCTGACTATAAGCCAGAGAC

AGAGCACACGTCTGAACTCTTTCCCTACACGACGCTCTTCCGATCTNNNTCCTGACTATAAGCCAGAGAC

AGCACACGTCTGAACTCTTTCCCTACACGACGCTCTTCCGATCTNNNGTGCCTGTAATCCCAGCTACTAG

AGCACACGTCTGAACTCTTTCCCTACACGACGCTCTTCCGATCTNNNGTGCCTGTAATCCCAGCTACTAG

AGCACACGTCTGAACTCTTTCCCTACACGACGCTCTTCCGATCTNNNATATTCTTCATCTCAAAAGACAC

AGCACACGTCTGAACTCTTTCCCTACACGACGCTCTTCCGATCTNNNATATTCTTCATCTCAAAAGACAC

AGAGCACACGTCTGAACTCTTTCCCTACACGACGCTCTTCCGATCTNNNTGAGGCAGGAGAATCTCTT

AGAGCACACGTCTGAACTCTTTCCCTACACGACGCTCTTCCGATCTNNNTGAGGCAGGAGAATCTCTT

ACACGTCTGAACTCTTTCCCTACACGACGCTCTTCCGATCTNNNCTGTTCATGTCTTTGCCCACTTTCATC

ACACGTCTGAACTCTTTCCCTACACGACGCTCTTCCGATCTNNNCTGTTCATGTCTTTGCCCACTTTCATC

GAGCACACGTCTGAACTCTTTCCCTACACGACGCTCTTCCGATCTNNNAGAGAAATGACCACTTCTGAC

GAGCACACGTCTGAACTCTTTCCCTACACGACGCTCTTCCGATCTNNNAGAGAAATGACCACTTCTGAC

AGCACACGTCTGAACTCTTTCCCTACACGACGCTCTTCCGATCTNNNAGTGCTGGGATTACAGGTGTAA

AGCACACGTCTGAACTCTTTCCCTACACGACGCTCTTCCGATCTNNNAGTGCTGGGATTACAGGTGTAA

AGAGCACACGTCTGAACTCTTTCCCTACACGACGCTCTTCCGATCTNNNAGCATAAATCAGATGCCACT

AGAGCACACGTCTGAACTCTTTCCCTACACGACGCTCTTCCGATCTNNNAGCATAAATCAGATGCCACT

AGAGCACACGTCTGAACTCTTTCCCTACACGACGCTCTTCCGATCTNNNCACTCAGGGTCTTAGTGAAC

AGAGCACACGTCTGAACTCTTTCCCTACACGACGCTCTTCCGATCTNNNCACTCAGGGTCTTAGTGAAC

GAGCACACGTCTGAACTCTTTCCCTACACGACGCTCTTCCGATCTNNNTATAGTCCCAGCTACTCAGGA

GAGCACACGTCTGAACTCTTTCCCTACACGACGCTCTTCCGATCTNNNTATAGTCCCAGCTACTCAGGA

AGCACACGTCTGAACTCTTTCCCTACACGACGCTCTTCCGATCTNNNATGTATACATGTGCCATGCTGGT

AGCACACGTCTGAACTCTTTCCCTACACGACGCTCTTCCGATCTNNNATGTATACATGTGCCATGCTGGT

GAGCACACGTCTGAACTCTTTCCCTACACGACGCTCTTCCGATCTNNNTGGACACAGTTTTCTCCAACAC

GAGCACACGTCTGAACTCTTTCCCTACACGACGCTCTTCCGATCTNNNTGGACACAGTTTTCTCCAACAC

GAGCACACGTCTGAACTCTTTCCCTACACGACGCTCTTCCGATCTNNNAGGAGAATGGCTTGAACCCG

GAGCACACGTCTGAACTCTTTCCCTACACGACGCTCTTCCGATCTNNNAGGAGAATGGCTTGAACCCG

GAGCACACGTCTGAACTCTTTCCCTACACGACGCTCTTCCGATCTNNNTGAACCAGGAAAGTGGAGCTT

GAGCACACGTCTGAACTCTTTCCCTACACGACGCTCTTCCGATCTNNNTGAACCAGGAAAGTGGAGCTT

GAGCACACGTCTGAACTCTTTCCCTACACGACGCTCTTCCGATCTNNNTGGTATGGGGTAATATGGCT

GAGCACACGTCTGAACTCTTTCCCTACACGACGCTCTTCCGATCTNNNTGGTATGGGGTAATATGGCT

GAGCACACGTCTGAACTCTTTCCCTACACGACGCTCTTCCGATCTNNNCAGTGTCAAAAATGTGGTCAG
GAGCACACGTCTGAACTCTTTCCCTACACGACGCTCTTCCGATCTNNNCAGTGTCAAAAATGTGGTCAG
GAGCACACGTCTGAACTCTTTCCCTACACGACGCTCTTCCGATCTNNNGAAGAATGGTGTGAACCTGGC
GAGCACACGTCTGAACTCTTTCCCTACACGACGCTCTTCCGATCTNNNGAAGAATGGTGTGAACCTGGC
AGAGCACACGTCTGAACTCTTTCCCTACACGACGCTCTTCCGATCTNNNCTCACAATCAACGAGGAAGA
AGAGCACACGTCTGAACTCTTTCCCTACACGACGCTCTTCCGATCTNNNCTCACAATCAACGAGGAAGA
AGCACACGTCTGAACTCTTTCCCTACACGACGCTCTTCCGATCTNNNCACACACACACACACATATGGAC
AGCACACGTCTGAACTCTTTCCCTACACGACGCTCTTCCGATCTNNNCACACACACACACACATATGGAC
GAGCACACGTCTGAACTCTTTCCCTACACGACGCTCTTCCGATCTNNNCCTGCAAAGTTCAGTCTTGACA
GAGCACACGTCTGAACTCTTTCCCTACACGACGCTCTTCCGATCTNNNCCTGCAAAGTTCAGTCTTGACA
AGCACACGTCTGAACTCTTTCCCTACACGACGCTCTTCCGATCTNNNAGTTTGGGTAAGGTATGAGCAC
AGCACACGTCTGAACTCTTTCCCTACACGACGCTCTTCCGATCTNNNAGTTTGGGTAAGGTATGAGCAC
AGCACACGTCTGAACTCTTTCCCTACACGACGCTCTTCCGATCTNNNTCAGATTCCCAAGGCTGAATGAA
AGCACACGTCTGAACTCTTTCCCTACACGACGCTCTTCCGATCTNNNTCAGATTCCCAAGGCTGAATGAA
GAGCACACGTCTGAACTCTTTCCCTACACGACGCTCTTCCGATCTNNNAACCACGGTGCCATTATATGAC
GAGCACACGTCTGAACTCTTTCCCTACACGACGCTCTTCCGATCTNNNAACCACGGTGCCATTATATGAC
GAGCACACGTCTGAACTCTTTCCCTACACGACGCTCTTCCGATCTNNNAAGTTCAACATCAAGATGCCA
GAGCACACGTCTGAACTCTTTCCCTACACGACGCTCTTCCGATCTNNNAAGTTCAACATCAAGATGCCA
GAGCACACGTCTGAACTCTTTCCCTACACGACGCTCTTCCGATCTNNNTCTTGTGAATTGCTTCTAACTG
GAGCACACGTCTGAACTCTTTCCCTACACGACGCTCTTCCGATCTNNNTCTTGTGAATTGCTTCTAACTG
GAGCACACGTCTGAACTCTTTCCCTACACGACGCTCTTCCGATCTNNNTTGTAGTGAGCCTAGATTGTC
GAGCACACGTCTGAACTCTTTCCCTACACGACGCTCTTCCGATCTNNNTTGTAGTGAGCCTAGATTGTC
GAGCACACGTCTGAACTCTTTCCCTACACGACGCTCTTCCGATCTNNNGAGAATCGCTTGAACCCAGG
GAGCACACGTCTGAACTCTTTCCCTACACGACGCTCTTCCGATCTNNNGAGAATCGCTTGAACCCAGG
GAGCACACGTCTGAACTCTTTCCCTACACGACGCTCTTCCGATCTNNNAGAATCACTTCAACCTGGGAG
GAGCACACGTCTGAACTCTTTCCCTACACGACGCTCTTCCGATCTNNNAGAATCACTTCAACCTGGGAG
AGCACACGTCTGAACTCTTTCCCTACACGACGCTCTTCCGATCTNNNAAAATCTCACCACTCCACTCCAAC
AGCACACGTCTGAACTCTTTCCCTACACGACGCTCTTCCGATCTNNNAAAATCTCACCACTCCACTCCAAC
GAGCACACGTCTGAACTCTTTCCCTACACGACGCTCTTCCGATCTNNNAGGGCAGTAGTATACACACAC
GAGCACACGTCTGAACTCTTTCCCTACACGACGCTCTTCCGATCTNNNAGGGCAGTAGTATACACACAC
ACACGTCTGAACTCTTTCCCTACACGACGCTCTTCCGATCTNNNGAACCCCACATTCGTGTCATGTCCAT
ACACGTCTGAACTCTTTCCCTACACGACGCTCTTCCGATCTNNNGAACCCCACATTCGTGTCATGTCCAT
AGCACACGTCTGAACTCTTTCCCTACACGACGCTCTTCCGATCTNNNTACAGCATTGGAGTAGCACTTG
AGCACACGTCTGAACTCTTTCCCTACACGACGCTCTTCCGATCTNNNTACAGCATTGGAGTAGCACTTG
AGCACACGTCTGAACTCTTTCCCTACACGACGCTCTTCCGATCTNNNCATAGATTAGGCATGCCCAAGC
AGCACACGTCTGAACTCTTTCCCTACACGACGCTCTTCCGATCTNNNCATAGATTAGGCATGCCCAAGC
AGCACACGTCTGAACTCTTTCCCTACACGACGCTCTTCCGATCTNNNTGAAATACCACCTAATCTTTCTT
AGCACACGTCTGAACTCTTTCCCTACACGACGCTCTTCCGATCTNNNTGAAATACCACCTAATCTTTCTT
AGCACACGTCTGAACTCTTTCCCTACACGACGCTCTTCCGATCTNNNCTGCTTTTCTGGACCAAACCAATC
AGCACACGTCTGAACTCTTTCCCTACACGACGCTCTTCCGATCTNNNCTGCTTTTCTGGACCAAACCAATC
AGCACACGTCTGAACTCTTTCCCTACACGACGCTCTTCCGATCTNNNAGGACCATATGGATTCACAGCT
AGCACACGTCTGAACTCTTTCCCTACACGACGCTCTTCCGATCTNNNAGGACCATATGGATTCACAGCT
AGAGCACACGTCTGAACTCTTTCCCTACACGACGCTCTTCCGATCTNNNGCAGGAGAATTGCTTGAACT

AGAGCACACGTCTGAACTCTTTCCCTACACGACGCTCTTCCGATCTNNNGCAGGAGAATTGCTTGAAC
AGAGCACACGTCTGAACTCTTTCCCTACACGACGCTCTTCCGATCTNNNTGCTCCCAATACTGGTTCAT
AGAGCACACGTCTGAACTCTTTCCCTACACGACGCTCTTCCGATCTNNNTGCTCCCAATACTGGTTCAT
GAGCACACGTCTGAACTCTTTCCCTACACGACGCTCTTCCGATCTNNNATGATCTCAGGTCACTGCAAG
GAGCACACGTCTGAACTCTTTCCCTACACGACGCTCTTCCGATCTNNNATGATCTCAGGTCACTGCAAG
AGCACACGTCTGAACTCTTTCCCTACACGACGCTCTTCCGATCTNNNGGACACATGAGAGGATGTTCAC
AGCACACGTCTGAACTCTTTCCCTACACGACGCTCTTCCGATCTNNNGGACACATGAGAGGATGTTCAC
AGCACACGTCTGAACTCTTTCCCTACACGACGCTCTTCCGATCTNNNTGATTCCATTACTTTAGCGCTG
AGCACACGTCTGAACTCTTTCCCTACACGACGCTCTTCCGATCTNNNTGATTCCATTACTTTAGCGCTG
AGCACACGTCTGAACTCTTTCCCTACACGACGCTCTTCCGATCTNNNCTTGCAGCTGCCGATATATCACT
AGCACACGTCTGAACTCTTTCCCTACACGACGCTCTTCCGATCTNNNCTTGCAGCTGCCGATATATCACT
CACACGTCTGAACTCTTTCCCTACACGACGCTCTTCCGATCTNNNATAACCTTCCTAGTACCTGAGCATC
CACACGTCTGAACTCTTTCCCTACACGACGCTCTTCCGATCTNNNATAACCTTCCTAGTACCTGAGCATC
GAGCACACGTCTGAACTCTTTCCCTACACGACGCTCTTCCGATCTNNNAACTCCTCAAGAATGGACTCA
GAGCACACGTCTGAACTCTTTCCCTACACGACGCTCTTCCGATCTNNNAACTCCTCAAGAATGGACTCA
AGCACACGTCTGAACTCTTTCCCTACACGACGCTCTTCCGATCTNNNATAGTTTGGCCATTGATGTTGC
AGCACACGTCTGAACTCTTTCCCTACACGACGCTCTTCCGATCTNNNATAGTTTGGCCATTGATGTTGC
AGCACACGTCTGAACTCTTTCCCTACACGACGCTCTTCCGATCTNNNGTAATCTCAGCTACTCAGGAGG
AGCACACGTCTGAACTCTTTCCCTACACGACGCTCTTCCGATCTNNNGTAATCTCAGCTACTCAGGAGG
AGCACACGTCTGAACTCTTTCCCTACACGACGCTCTTCCGATCTNNNCACCATTCCCGGCTAATTTTTGT
AGCACACGTCTGAACTCTTTCCCTACACGACGCTCTTCCGATCTNNNCACCATTCCCGGCTAATTTTTGT
AGCACACGTCTGAACTCTTTCCCTACACGACGCTCTTCCGATCTNNNGCGAAAGGAAGCTTTGCAATTT
AGCACACGTCTGAACTCTTTCCCTACACGACGCTCTTCCGATCTNNNGCGAAAGGAAGCTTTGCAATTT
AGCACACGTCTGAACTCTTTCCCTACACGACGCTCTTCCGATCTNNNTGGTAGTTGGATCTAGATGGC
AGCACACGTCTGAACTCTTTCCCTACACGACGCTCTTCCGATCTNNNTGGTAGTTGGATCTAGATGGC
GAGCACACGTCTGAACTCTTTCCCTACACGACGCTCTTCCGATCTNNNTGGTGGAGTCTTTAGTTTTTT
GAGCACACGTCTGAACTCTTTCCCTACACGACGCTCTTCCGATCTNNNTGGTGGAGTCTTTAGTTTTTT
AGCACACGTCTGAACTCTTTCCCTACACGACGCTCTTCCGATCTNNNAGCCTTGTTCACAGATGAGAAG
AGCACACGTCTGAACTCTTTCCCTACACGACGCTCTTCCGATCTNNNAGCCTTGTTCACAGATGAGAAG
AGCACACGTCTGAACTCTTTCCCTACACGACGCTCTTCCGATCTNNNGCACTTGTAATCCCAGCTACTTG
AGCACACGTCTGAACTCTTTCCCTACACGACGCTCTTCCGATCTNNNGCACTTGTAATCCCAGCTACTTG
AGCACACGTCTGAACTCTTTCCCTACACGACGCTCTTCCGATCTNNNAAATTAAAATCTCTTACCTTCC
AGCACACGTCTGAACTCTTTCCCTACACGACGCTCTTCCGATCTNNNAAATTAAAATCTCTTACCTTCC
AGCACACGTCTGAACTCTTTCCCTACACGACGCTCTTCCGATCTNNNTGTCCTACCCTGTACTGTCACTG
AGCACACGTCTGAACTCTTTCCCTACACGACGCTCTTCCGATCTNNNTGTCCTACCCTGTACTGTCACTG
GCACACGTCTGAACTCTTTCCCTACACGACGCTCTTCCGATCTNNNAAAAAGTGAGGGAAAAGGCTGG
GCACACGTCTGAACTCTTTCCCTACACGACGCTCTTCCGATCTNNNAAAAAGTGAGGGAAAAGGCTGG
GCACACGTCTGAACTCTTTCCCTACACGACGCTCTTCCGATCTNNNGCAGTACCTACCACTATGCCATT
GCACACGTCTGAACTCTTTCCCTACACGACGCTCTTCCGATCTNNNGCAGTACCTACCACTATGCCATT
GCACACGTCTGAACTCTTTCCCTACACGACGCTCTTCCGATCTNNNACATACAAACTCGTGTGCATGTA
GCACACGTCTGAACTCTTTCCCTACACGACGCTCTTCCGATCTNNNACATACAAACTCGTGTGCATGTA
AGCACACGTCTGAACTCTTTCCCTACACGACGCTCTTCCGATCTNNNTAAAGTACTGTCTGCATGGAG
AGCACACGTCTGAACTCTTTCCCTACACGACGCTCTTCCGATCTNNNTAAAGTACTGTCTGCATGGAG

AGCACACGTCTGAACTCTTTCCCTACACGACGCTCTTCCGATCTNNNTTTTTGAGATGGAGTCTTGCC

AGCACACGTCTGAACTCTTTCCCTACACGACGCTCTTCCGATCTNNNTTTTTGAGATGGAGTCTTGCC

AGCACACGTCTGAACTCTTTCCCTACACGACGCTCTTCCGATCTNNNTCCCTCGCCATTGTGAAAAAAAA

AGCACACGTCTGAACTCTTTCCCTACACGACGCTCTTCCGATCTNNNTCCCTCGCCATTGTGAAAAAAAA

AGCACACGTCTGAACTCTTTCCCTACACGACGCTCTTCCGATCTNNNACCACTACGCCAGGCTAATTAT

AGCACACGTCTGAACTCTTTCCCTACACGACGCTCTTCCGATCTNNNACCACTACGCCAGGCTAATTAT

GCACACGTCTGAACTCTTTCCCTACACGACGCTCTTCCGATCTNNNCCATTAGGCTGCACATTTCCAAA

GCACACGTCTGAACTCTTTCCCTACACGACGCTCTTCCGATCTNNNCCATTAGGCTGCACATTTCCAAA

AGCACACGTCTGAACTCTTTCCCTACACGACGCTCTTCCGATCTNNNGAGGTTACAGTGAGCCGAGA

AGCACACGTCTGAACTCTTTCCCTACACGACGCTCTTCCGATCTNNNGAGGTTACAGTGAGCCGAGA

AGCACACGTCTGAACTCTTTCCCTACACGACGCTCTTCCGATCTNNNAACAAAAGTCACTGTTAATTCT

AGCACACGTCTGAACTCTTTCCCTACACGACGCTCTTCCGATCTNNNAACAAAAGTCACTGTTAATTCT

AGCACACGTCTGAACTCTTTCCCTACACGACGCTCTTCCGATCTNNNTTTTTTGTAGACACGGGGTTTC

AGCACACGTCTGAACTCTTTCCCTACACGACGCTCTTCCGATCTNNNTTTTTTGTAGACACGGGGTTTC

GAGCACACGTCTGAACTCTTTCCCTACACGACGCTCTTCCGATCTNNNATTGTGTGATTTCTGCTGCAC

GAGCACACGTCTGAACTCTTTCCCTACACGACGCTCTTCCGATCTNNNATTGTGTGATTTCTGCTGCAC

GCACACGTCTGAACTCTTTCCCTACACGACGCTCTTCCGATCTNNNACTAAGGTTCATCTTGTCCCTGG

GCACACGTCTGAACTCTTTCCCTACACGACGCTCTTCCGATCTNNNACTAAGGTTCATCTTGTCCCTGG

AGCACACGTCTGAACTCTTTCCCTACACGACGCTCTTCCGATCTNNNTGAACACTTGGTGCTTTATTGC

AGCACACGTCTGAACTCTTTCCCTACACGACGCTCTTCCGATCTNNNTGAACACTTGGTGCTTTATTGC

AGCACACGTCTGAACTCTTTCCCTACACGACGCTCTTCCGATCTNNNCCACGCCCAGCTAATTTTTGTA

AGCACACGTCTGAACTCTTTCCCTACACGACGCTCTTCCGATCTNNNCCACGCCCAGCTAATTTTTGTA

AGCACACGTCTGAACTCTTTCCCTACACGACGCTCTTCCGATCTNNNCTCTTCCTTCCTTCCTTCCTTCCC

AGCACACGTCTGAACTCTTTCCCTACACGACGCTCTTCCGATCTNNNCTCTTCCTTCCTTCCTTCCTTCCC

AGAGCACACGTCTGAACTCTTTCCCTACACGACGCTCTTCCGATCTNNNACACCTGTAGTCCCAGCTAC

AGAGCACACGTCTGAACTCTTTCCCTACACGACGCTCTTCCGATCTNNNACACCTGTAGTCCCAGCTAC

GAGCACACGTCTGAACTCTTTCCCTACACGACGCTCTTCCGATCTNNNATGTAAATATGTGTGTGTGT

GAGCACACGTCTGAACTCTTTCCCTACACGACGCTCTTCCGATCTNNNATGTAAATATGTGTGTGTGT

GAGCACACGTCTGAACTCTTTCCCTACACGACGCTCTTCCGATCTNNNGTGAGCGAGCATGAACCTAT

GAGCACACGTCTGAACTCTTTCCCTACACGACGCTCTTCCGATCTNNNGTGAGCGAGCATGAACCTAT

AGCACACGTCTGAACTCTTTCCCTACACGACGCTCTTCCGATCTNNNGAATGCACATACACACACACA

AGCACACGTCTGAACTCTTTCCCTACACGACGCTCTTCCGATCTNNNGAATGCACATACACACACACA

GCACACGTCTGAACTCTTTCCCTACACGACGCTCTTCCGATCTNNNTTTGGCCCAACAGGCTATTTCAT

GCACACGTCTGAACTCTTTCCCTACACGACGCTCTTCCGATCTNNNTTTGGCCCAACAGGCTATTTCAT

AGCACACGTCTGAACTCTTTCCCTACACGACGCTCTTCCGATCTNNNCATCAACACCTTCTGTAGAGCA

AGCACACGTCTGAACTCTTTCCCTACACGACGCTCTTCCGATCTNNNCATCAACACCTTCTGTAGAGCA

GCACACGTCTGAACTCTTTCCCTACACGACGCTCTTCCGATCTNNNTCAGTTCTCATAGTTTTCTGGTG

GCACACGTCTGAACTCTTTCCCTACACGACGCTCTTCCGATCTNNNTCAGTTCTCATAGTTTTCTGGTG

AGCACACGTCTGAACTCTTTCCCTACACGACGCTCTTCCGATCTNNNATATATGGGAGGAAAGGCCAT

AGCACACGTCTGAACTCTTTCCCTACACGACGCTCTTCCGATCTNNNATATATGGGAGGAAAGGCCAT

GCACACGTCTGAACTCTTTCCCTACACGACGCTCTTCCGATCTNNNACGATCTCACCACTGCATTCTAG

GCACACGTCTGAACTCTTTCCCTACACGACGCTCTTCCGATCTNNNACGATCTCACCACTGCATTCTAG

AGCACACGTCTGAACTCTTTCCCTACACGACGCTCTTCCGATCTNNNTAAGGCAGGAGAATCGCTTG

AGCACACGTCTGAACTCTTTCCCTACACGACGCTCTTCCGATCTNNNTAAGGCAGGAGAATCGCTTG
GAGCACACGTCTGAACTCTTTCCCTACACGACGCTCTTCCGATCTNNNGGTTTTGGTTCATGATACGG
GAGCACACGTCTGAACTCTTTCCCTACACGACGCTCTTCCGATCTNNNGGTTTTGGTTCATGATACGG
AGCACACGTCTGAACTCTTTCCCTACACGACGCTCTTCCGATCTNNNAGGGAGCAGTTCCTTCTTTCA
AGCACACGTCTGAACTCTTTCCCTACACGACGCTCTTCCGATCTNNNAGGGAGCAGTTCCTTCTTTCA
AGCACACGTCTGAACTCTTTCCCTACACGACGCTCTTCCGATCTNNNCAATACCCATCTCTATTTAAG
AGCACACGTCTGAACTCTTTCCCTACACGACGCTCTTCCGATCTNNNCAATACCCATCTCTATTTAAG
GCACACGTCTGAACTCTTTCCCTACACGACGCTCTTCCGATCTNNNATAGGAGTGGAGGCTAACTTCC
GCACACGTCTGAACTCTTTCCCTACACGACGCTCTTCCGATCTNNNATAGGAGTGGAGGCTAACTTCC
CACACGTCTGAACTCTTTCCCTACACGACGCTCTTCCGATCTNNNCTGAGGCAGGAGAATCACTTGA
CACACGTCTGAACTCTTTCCCTACACGACGCTCTTCCGATCTNNNCTGAGGCAGGAGAATCACTTGA
GAGCACACGTCTGAACTCTTTCCCTACACGACGCTCTTCCGATCTNNNCAGATCTGACTCTAAAGCCCA
GAGCACACGTCTGAACTCTTTCCCTACACGACGCTCTTCCGATCTNNNCAGATCTGACTCTAAAGCCCA
GCACACGTCTGAACTCTTTCCCTACACGACGCTCTTCCGATCTNNNCTCACCTCAGCATCTCTAGTAGC
GCACACGTCTGAACTCTTTCCCTACACGACGCTCTTCCGATCTNNNCTCACCTCAGCATCTCTAGTAGC
GCACACGTCTGAACTCTTTCCCTACACGACGCTCTTCCGATCTNNNCACAGATGTAGTGGTGAGGGA
GCACACGTCTGAACTCTTTCCCTACACGACGCTCTTCCGATCTNNNCACAGATGTAGTGGTGAGGGA
AGCACACGTCTGAACTCTTTCCCTACACGACGCTCTTCCGATCTNNNGCTAGGAATACAGGCATGTG
AGCACACGTCTGAACTCTTTCCCTACACGACGCTCTTCCGATCTNNNGCTAGGAATACAGGCATGTG
GCACACGTCTGAACTCTTTCCCTACACGACGCTCTTCCGATCTNNNCAGCGATCACAACAGCTTCAGAT
GCACACGTCTGAACTCTTTCCCTACACGACGCTCTTCCGATCTNNNCAGCGATCACAACAGCTTCAGAT
GAGCACACGTCTGAACTCTTTCCCTACACGACGCTCTTCCGATCTNNNGACAGAGCAAGACTCCGTCT
GAGCACACGTCTGAACTCTTTCCCTACACGACGCTCTTCCGATCTNNNGACAGAGCAAGACTCCGTCT
AGCACACGTCTGAACTCTTTCCCTACACGACGCTCTTCCGATCTNNNACCAATTGCTTCATGACATCCTT
AGCACACGTCTGAACTCTTTCCCTACACGACGCTCTTCCGATCTNNNACCAATTGCTTCATGACATCCTT
AGCACACGTCTGAACTCTTTCCCTACACGACGCTCTTCCGATCTNNNAAAAAAATTAGCCAGGCAGTC
AGCACACGTCTGAACTCTTTCCCTACACGACGCTCTTCCGATCTNNNAAAAAAATTAGCCAGGCAGTC
AGCACACGTCTGAACTCTTTCCCTACACGACGCTCTTCCGATCTNNNGAGCAAGTGGGCCAGAGATGC
AGCACACGTCTGAACTCTTTCCCTACACGACGCTCTTCCGATCTNNNGAGCAAGTGGGCCAGAGATGC
AGAGCACACGTCTGAACTCTTTCCCTACACGACGCTCTTCCGATCTNNNGGCACCTATAATCCCAGCTA
AGAGCACACGTCTGAACTCTTTCCCTACACGACGCTCTTCCGATCTNNNGGCACCTATAATCCCAGCTA
GAGCACACGTCTGAACTCTTTCCCTACACGACGCTCTTCCGATCTNNNGATTGCAGTGAACCGAGATT
GAGCACACGTCTGAACTCTTTCCCTACACGACGCTCTTCCGATCTNNNGATTGCAGTGAACCGAGATT
AGCACACGTCTGAACTCTTTCCCTACACGACGCTCTTCCGATCTNNNTGTTTTAGCAATACCACAGCG
AGCACACGTCTGAACTCTTTCCCTACACGACGCTCTTCCGATCTNNNTGTTTTAGCAATACCACAGCG
AGCACACGTCTGAACTCTTTCCCTACACGACGCTCTTCCGATCTNNNCAAGATCACACCACTGTACTC
AGCACACGTCTGAACTCTTTCCCTACACGACGCTCTTCCGATCTNNNCAAGATCACACCACTGTACTC
AGCACACGTCTGAACTCTTTCCCTACACGACGCTCTTCCGATCTNNNGCTTGAAAAATACCTGAGGGC
AGCACACGTCTGAACTCTTTCCCTACACGACGCTCTTCCGATCTNNNGCTTGAAAAATACCTGAGGGC
AGCACACGTCTGAACTCTTTCCCTACACGACGCTCTTCCGATCTNNNACGACTTGTCTTCAACTTTGCT
AGCACACGTCTGAACTCTTTCCCTACACGACGCTCTTCCGATCTNNNACGACTTGTCTTCAACTTTGCT
AGCACACGTCTGAACTCTTTCCCTACACGACGCTCTTCCGATCTNNNTCAGATGAGACTCAGCACATT
AGCACACGTCTGAACTCTTTCCCTACACGACGCTCTTCCGATCTNNNTCAGATGAGACTCAGCACATT

AGCACACGTCTGAACTCTTTCCCTACACGACGCTCTTCCGATCTNNNCCTCAGTTCCTCATTGTCAAGGC
AGCACACGTCTGAACTCTTTCCCTACACGACGCTCTTCCGATCTNNNCCTCAGTTCCTCATTGTCAAGGC
GAGCACACGTCTGAACTCTTTCCCTACACGACGCTCTTCCGATCTNNNTAATTCCAGTACTTTGGGAGG
GAGCACACGTCTGAACTCTTTCCCTACACGACGCTCTTCCGATCTNNNTAATTCCAGTACTTTGGGAGG
GCACACGTCTGAACTCTTTCCCTACACGACGCTCTTCCGATCTNNNCTTCTCCTTGGAAATGGACTGGA
GCACACGTCTGAACTCTTTCCCTACACGACGCTCTTCCGATCTNNNCTTCTCCTTGGAAATGGACTGGA
GAGCACACGTCTGAACTCTTTCCCTACACGACGCTCTTCCGATCTNNNAAGGAACTGGCATTACTGGCA
GAGCACACGTCTGAACTCTTTCCCTACACGACGCTCTTCCGATCTNNNAAGGAACTGGCATTACTGGCA
GAGCACACGTCTGAACTCTTTCCCTACACGACGCTCTTCCGATCTNNNGGGAAAGGAGGTGAAAAAA
GAGCACACGTCTGAACTCTTTCCCTACACGACGCTCTTCCGATCTNNNGGGAAAGGAGGTGAAAAAA
GAGCACACGTCTGAACTCTTTCCCTACACGACGCTCTTCCGATCTNNNTTCCCTTCCCATCTTTTTGGCAG
GAGCACACGTCTGAACTCTTTCCCTACACGACGCTCTTCCGATCTNNNTTCCCTTCCCATCTTTTTGGCAG
AGCACACGTCTGAACTCTTTCCCTACACGACGCTCTTCCGATCTNNNTATTATGACATACTGCTGGGCC
AGCACACGTCTGAACTCTTTCCCTACACGACGCTCTTCCGATCTNNNTATTATGACATACTGCTGGGCC
AGCACACGTCTGAACTCTTTCCCTACACGACGCTCTTCCGATCTNNNGCTGTGTGTAGAGACAGGACTA
AGCACACGTCTGAACTCTTTCCCTACACGACGCTCTTCCGATCTNNNGCTGTGTGTAGAGACAGGACTA
AGCACACGTCTGAACTCTTTCCCTACACGACGCTCTTCCGATCTNNNAGCCTGTAGTTCCAGCTATTCA
AGCACACGTCTGAACTCTTTCCCTACACGACGCTCTTCCGATCTNNNAGCCTGTAGTTCCAGCTATTCA
GAGCACACGTCTGAACTCTTTCCCTACACGACGCTCTTCCGATCTNNNCTGAGGCAGAAGAATTGCTTG
GAGCACACGTCTGAACTCTTTCCCTACACGACGCTCTTCCGATCTNNNCTGAGGCAGAAGAATTGCTTG
GAGCACACGTCTGAACTCTTTCCCTACACGACGCTCTTCCGATCTNNNTCTCAGGGTCTCACTCTGTCAC
GAGCACACGTCTGAACTCTTTCCCTACACGACGCTCTTCCGATCTNNNTCTCAGGGTCTCACTCTGTCAC
GAGCACACGTCTGAACTCTTTCCCTACACGACGCTCTTCCGATCTNNNCTGAGCCAGGAGAATCACTTG
GAGCACACGTCTGAACTCTTTCCCTACACGACGCTCTTCCGATCTNNNCTGAGCCAGGAGAATCACTTG
ACACGTCTGAACTCTTTCCCTACACGACGCTCTTCCGATCTNNNCTATGACAGCGTCACCTGGAGCATCC
ACACGTCTGAACTCTTTCCCTACACGACGCTCTTCCGATCTNNNCTATGACAGCGTCACCTGGAGCATCC
GCACACGTCTGAACTCTTTCCCTACACGACGCTCTTCCGATCTNNNAGCATGTGATGTGTTTCCATTTGT
GCACACGTCTGAACTCTTTCCCTACACGACGCTCTTCCGATCTNNNAGCATGTGATGTGTTTCCATTTGT
GCACACGTCTGAACTCTTTCCCTACACGACGCTCTTCCGATCTNNNATCTTCAGGCCTTTGCTCAACTGC
GCACACGTCTGAACTCTTTCCCTACACGACGCTCTTCCGATCTNNNATCTTCAGGCCTTTGCTCAACTGC
AGCACACGTCTGAACTCTTTCCCTACACGACGCTCTTCCGATCTNNNCCTGCAGTGAGCCGAGATTGCA
AGCACACGTCTGAACTCTTTCCCTACACGACGCTCTTCCGATCTNNNCCTGCAGTGAGCCGAGATTGCA
GAGCACACGTCTGAACTCTTTCCCTACACGACGCTCTTCCGATCTNNNATTGCAGAAAAGGTTGACTGA
GAGCACACGTCTGAACTCTTTCCCTACACGACGCTCTTCCGATCTNNNATTGCAGAAAAGGTTGACTGA
AGCACACGTCTGAACTCTTTCCCTACACGACGCTCTTCCGATCTNNNCCTTTGGTTCCCAGATCCATGTA
AGCACACGTCTGAACTCTTTCCCTACACGACGCTCTTCCGATCTNNNCCTTTGGTTCCCAGATCCATGTA
AGAGCACACGTCTGAACTCTTTCCCTACACGACGCTCTTCCGATCTNNNGAGGCTGAGGCAGGAGAA
AGAGCACACGTCTGAACTCTTTCCCTACACGACGCTCTTCCGATCTNNNGAGGCTGAGGCAGGAGAA
GAGCACACGTCTGAACTCTTTCCCTACACGACGCTCTTCCGATCTNNNCTGTCCAGGTAGGTATGGGA
GAGCACACGTCTGAACTCTTTCCCTACACGACGCTCTTCCGATCTNNNCTGTCCAGGTAGGTATGGGA
AGCACACGTCTGAACTCTTTCCCTACACGACGCTCTTCCGATCTNNNCCCGGCCTTTTTTCTTTCTTTCTC
AGCACACGTCTGAACTCTTTCCCTACACGACGCTCTTCCGATCTNNNCCCGGCCTTTTTTCTTTCTTTCTC
GAGCACACGTCTGAACTCTTTCCCTACACGACGCTCTTCCGATCTNNNGCACTCATAAGTCATCCCCTTC

GAGCACACGTCTGAACTCTTTCCCTACACGACGCTCTTCCGATCTNNNGCACTCATAAGTCATCCCCTTC
AGCACACGTCTGAACTCTTTCCCTACACGACGCTCTTCCGATCTNNNCCATGGTGGAGTTCTAGCTACT
AGCACACGTCTGAACTCTTTCCCTACACGACGCTCTTCCGATCTNNNCCATGGTGGAGTTCTAGCTACT
AGCACACGTCTGAACTCTTTCCCTACACGACGCTCTTCCGATCTNNNTGTACACACGAATGATCCATGG
AGCACACGTCTGAACTCTTTCCCTACACGACGCTCTTCCGATCTNNNTGTACACACGAATGATCCATGG
GAGCACACGTCTGAACTCTTTCCCTACACGACGCTCTTCCGATCTNNNTCCTAAAGTGCTGGGATTACA
GAGCACACGTCTGAACTCTTTCCCTACACGACGCTCTTCCGATCTNNNTCCTAAAGTGCTGGGATTACA
AGCACACGTCTGAACTCTTTCCCTACACGACGCTCTTCCGATCTNNNCTGCAAGAAACAATCACCAGAGT
AGCACACGTCTGAACTCTTTCCCTACACGACGCTCTTCCGATCTNNNCTGCAAGAAACAATCACCAGAGT
AGAGCACACGTCTGAACTCTTTCCCTACACGACGCTCTTCCGATCTNNNACCAAGTGGGAGGACAAAT
AGAGCACACGTCTGAACTCTTTCCCTACACGACGCTCTTCCGATCTNNNACCAAGTGGGAGGACAAAT
AGCACACGTCTGAACTCTTTCCCTACACGACGCTCTTCCGATCTNNNCCTGGGGTTATTGTGAGTTACC
AGCACACGTCTGAACTCTTTCCCTACACGACGCTCTTCCGATCTNNNCCTGGGGTTATTGTGAGTTACC
AGCACACGTCTGAACTCTTTCCCTACACGACGCTCTTCCGATCTNNNTGTATGTTGTCTGGTCAGGAAC
AGCACACGTCTGAACTCTTTCCCTACACGACGCTCTTCCGATCTNNNTGTATGTTGTCTGGTCAGGAAC
GAGCACACGTCTGAACTCTTTCCCTACACGACGCTCTTCCGATCTNNNGCTGGAGAATTGCTTGAACCT
GAGCACACGTCTGAACTCTTTCCCTACACGACGCTCTTCCGATCTNNNGCTGGAGAATTGCTTGAACCT
AGCACACGTCTGAACTCTTTCCCTACACGACGCTCTTCCGATCTNNNGAATCTTGAGACACCTCTCTGG
AGCACACGTCTGAACTCTTTCCCTACACGACGCTCTTCCGATCTNNNGAATCTTGAGACACCTCTCTGG
AGCACACGTCTGAACTCTTTCCCTACACGACGCTCTTCCGATCTNNNTCCTACATGGGAGGGTTTTCTT
AGCACACGTCTGAACTCTTTCCCTACACGACGCTCTTCCGATCTNNNTCCTACATGGGAGGGTTTTCTT
AGCACACGTCTGAACTCTTTCCCTACACGACGCTCTTCCGATCTNNNCTGTAATCCCAGCTACTCAGGAC
AGCACACGTCTGAACTCTTTCCCTACACGACGCTCTTCCGATCTNNNCTGTAATCCCAGCTACTCAGGAC
GAGCACACGTCTGAACTCTTTCCCTACACGACGCTCTTCCGATCTNNNAGTGAACTGTGATTGTGCCAT
GAGCACACGTCTGAACTCTTTCCCTACACGACGCTCTTCCGATCTNNNAGTGAACTGTGATTGTGCCAT
AGCACACGTCTGAACTCTTTCCCTACACGACGCTCTTCCGATCTNNNGCCTCCTAGGTTCAAGTGATTCT
AGCACACGTCTGAACTCTTTCCCTACACGACGCTCTTCCGATCTNNNGCCTCCTAGGTTCAAGTGATTCT
GAGCACACGTCTGAACTCTTTCCCTACACGACGCTCTTCCGATCTNNNTGAGGCAAGAGAATTGCTTGA
GAGCACACGTCTGAACTCTTTCCCTACACGACGCTCTTCCGATCTNNNTGAGGCAAGAGAATTGCTTGA
GAGCACACGTCTGAACTCTTTCCCTACACGACGCTCTTCCGATCTNNNGGAGGTGAAGGTTGCAGTGA
GAGCACACGTCTGAACTCTTTCCCTACACGACGCTCTTCCGATCTNNNGGAGGTGAAGGTTGCAGTGA
GAGCACACGTCTGAACTCTTTCCCTACACGACGCTCTTCCGATCTNNNTGTGGTACTGGCATAACAACA
GAGCACACGTCTGAACTCTTTCCCTACACGACGCTCTTCCGATCTNNNTGTGGTACTGGCATAACAACA
AGCACACGTCTGAACTCTTTCCCTACACGACGCTCTTCCGATCTNNNCACACTCGTGTAATCCCAGCTAC
AGCACACGTCTGAACTCTTTCCCTACACGACGCTCTTCCGATCTNNNCACACTCGTGTAATCCCAGCTAC
AGCACACGTCTGAACTCTTTCCCTACACGACGCTCTTCCGATCTNNNAGCATGCCCACAAGATCTAGAA
AGCACACGTCTGAACTCTTTCCCTACACGACGCTCTTCCGATCTNNNAGCATGCCCACAAGATCTAGAA
GAGCACACGTCTGAACTCTTTCCCTACACGACGCTCTTCCGATCTNNNGAATCATGCCATTACACTCCAG
GAGCACACGTCTGAACTCTTTCCCTACACGACGCTCTTCCGATCTNNNGAATCATGCCATTACACTCCAG
AGCACACGTCTGAACTCTTTCCCTACACGACGCTCTTCCGATCTNNNAGAATTACTTGAACCTGGGAGG
AGCACACGTCTGAACTCTTTCCCTACACGACGCTCTTCCGATCTNNNAGAATTACTTGAACCTGGGAGG
AGCACACGTCTGAACTCTTTCCCTACACGACGCTCTTCCGATCTNNNATACAAAAATTAGCTGGGCGTG
AGCACACGTCTGAACTCTTTCCCTACACGACGCTCTTCCGATCTNNNATACAAAAATTAGCTGGGCGTG

GAGCACACGTCTGAACTCTTTCCCTACACGACGCTCTTCCGATCTNNNTAGTCCCAGCTACTCAGGAGA
GAGCACACGTCTGAACTCTTTCCCTACACGACGCTCTTCCGATCTNNNTAGTCCCAGCTACTCAGGAGA
AGCACACGTCTGAACTCTTTCCCTACACGACGCTCTTCCGATCTNNNTGAGAATCACTTGAACCTGGGA
AGCACACGTCTGAACTCTTTCCCTACACGACGCTCTTCCGATCTNNNTGAGAATCACTTGAACCTGGGA
AGAGCACACGTCTGAACTCTTTCCCTACACGACGCTCTTCCGATCTNNNCTGAGGTGGGAGGATCATT
AGAGCACACGTCTGAACTCTTTCCCTACACGACGCTCTTCCGATCTNNNCTGAGGTGGGAGGATCATT
AGCACACGTCTGAACTCTTTCCCTACACGACGCTCTTCCGATCTNNNCAATGCCCTCTTTGATTCTGCTC
AGCACACGTCTGAACTCTTTCCCTACACGACGCTCTTCCGATCTNNNCAATGCCCTCTTTGATTCTGCTC
GAGCACACGTCTGAACTCTTTCCCTACACGACGCTCTTCCGATCTNNNTCAGTTTTGCCCTAGAGTGAG
GAGCACACGTCTGAACTCTTTCCCTACACGACGCTCTTCCGATCTNNNTCAGTTTTGCCCTAGAGTGAG
AGCACACGTCTGAACTCTTTCCCTACACGACGCTCTTCCGATCTNNNACAGAAATTTTGACTCCTTAGA
AGCACACGTCTGAACTCTTTCCCTACACGACGCTCTTCCGATCTNNNACAGAAATTTTGACTCCTTAGA
GAGCACACGTCTGAACTCTTTCCCTACACGACGCTCTTCCGATCTNNNCGTCTGAGAAGTGAGGAGCC
GAGCACACGTCTGAACTCTTTCCCTACACGACGCTCTTCCGATCTNNNCGTCTGAGAAGTGAGGAGCC
CACACGTCTGAACTCTTTCCCTACACGACGCTCTTCCGATCTNNNGAAAGCTGTTCAGAAACATCCATC
CACACGTCTGAACTCTTTCCCTACACGACGCTCTTCCGATCTNNNGAAAGCTGTTCAGAAACATCCATC
GAGCACACGTCTGAACTCTTTCCCTACACGACGCTCTTCCGATCTNNNAGAGGTTGCAGTGAGTCAAA
GAGCACACGTCTGAACTCTTTCCCTACACGACGCTCTTCCGATCTNNNAGAGGTTGCAGTGAGTCAAA
GAGCACACGTCTGAACTCTTTCCCTACACGACGCTCTTCCGATCTNNNCCTTTGAGCTCAGGAGTTTGA
GAGCACACGTCTGAACTCTTTCCCTACACGACGCTCTTCCGATCTNNNCCTTTGAGCTCAGGAGTTTGA
GAGCACACGTCTGAACTCTTTCCCTACACGACGCTCTTCCGATCTNNNGAGGTTACAGTGAGCCGAGAT
GAGCACACGTCTGAACTCTTTCCCTACACGACGCTCTTCCGATCTNNNGAGGTTACAGTGAGCCGAGAT
AGCACACGTCTGAACTCTTTCCCTACACGACGCTCTTCCGATCTNNNATTACAAACAGGCCAACATGTC
AGCACACGTCTGAACTCTTTCCCTACACGACGCTCTTCCGATCTNNNATTACAAACAGGCCAACATGTC
AGAGCACACGTCTGAACTCTTTCCCTACACGACGCTCTTCCGATCTNNNAATCAGGTTTAGTGGCAAGA
AGAGCACACGTCTGAACTCTTTCCCTACACGACGCTCTTCCGATCTNNNAATCAGGTTTAGTGGCAAGA
CACACGTCTGAACTCTTTCCCTACACGACGCTCTTCCGATCTNNNGCCAAGAGGAAAAGGAGCTAATA
CACACGTCTGAACTCTTTCCCTACACGACGCTCTTCCGATCTNNNGCCAAGAGGAAAAGGAGCTAATA
AGCACACGTCTGAACTCTTTCCCTACACGACGCTCTTCCGATCTNNNTGATGGAGTGAAACCCTTTCTC
AGCACACGTCTGAACTCTTTCCCTACACGACGCTCTTCCGATCTNNNTGATGGAGTGAAACCCTTTCTC
GAGCACACGTCTGAACTCTTTCCCTACACGACGCTCTTCCGATCTNNNCAAATGTGTTTGCGGTTATG
GAGCACACGTCTGAACTCTTTCCCTACACGACGCTCTTCCGATCTNNNCAAATGTGTTTGCGGTTATG
GAGCACACGTCTGAACTCTTTCCCTACACGACGCTCTTCCGATCTNNNGGTTTCTCCTGAGGCCTCTTTA
GAGCACACGTCTGAACTCTTTCCCTACACGACGCTCTTCCGATCTNNNGGTTTCTCCTGAGGCCTCTTTA
AGCACACGTCTGAACTCTTTCCCTACACGACGCTCTTCCGATCTNNNGAGACACTGAGCTAGCTTGAA
AGCACACGTCTGAACTCTTTCCCTACACGACGCTCTTCCGATCTNNNGAGACACTGAGCTAGCTTGAA
AGCACACGTCTGAACTCTTTCCCTACACGACGCTCTTCCGATCTNNNCCACATTGAAGGAGGTGATCCA
AGCACACGTCTGAACTCTTTCCCTACACGACGCTCTTCCGATCTNNNCCACATTGAAGGAGGTGATCCA
GCACACGTCTGAACTCTTTCCCTACACGACGCTCTTCCGATCTNNNCACCCAGCCTCTATCTTTCCTTCCA
GCACACGTCTGAACTCTTTCCCTACACGACGCTCTTCCGATCTNNNCACCCAGCCTCTATCTTTCCTTCCA
GCACACGTCTGAACTCTTTCCCTACACGACGCTCTTCCGATCTNNNGAAGGAGACACCAGTCCAGAAA
GCACACGTCTGAACTCTTTCCCTACACGACGCTCTTCCGATCTNNNGAAGGAGACACCAGTCCAGAAA
AGAGCACACGTCTGAACTCTTTCCCTACACGACGCTCTTCCGATCTNNNGCTTTTGGTCATGTGAAGTT

AGAGCACACGTCTGAACTCTTTCCCTACACGACGCTCTTCCGATCTNNNGCTTTTGGTCATGTGAAGTT
GAGCACACGTCTGAACTCTTTCCCTACACGACGCTCTTCCGATCTNNNTCACATCTGTAATCCCAGCACA
GAGCACACGTCTGAACTCTTTCCCTACACGACGCTCTTCCGATCTNNNTCACATCTGTAATCCCAGCACA
GCACACGTCTGAACTCTTTCCCTACACGACGCTCTTCCGATCTNNNACAAAATGCCTCAGGTCCTTCAG
GCACACGTCTGAACTCTTTCCCTACACGACGCTCTTCCGATCTNNNACAAAATGCCTCAGGTCCTTCAG
GAGCACACGTCTGAACTCTTTCCCTACACGACGCTCTTCCGATCTNNNCTTCCTCCTACAGAATAGGCCA
GAGCACACGTCTGAACTCTTTCCCTACACGACGCTCTTCCGATCTNNNCTTCCTCCTACAGAATAGGCCA
ACACGTCTGAACTCTTTCCCTACACGACGCTCTTCCGATCTNNNGTCTCCTCCCTCCCTTCCTCCATCGGA
ACACGTCTGAACTCTTTCCCTACACGACGCTCTTCCGATCTNNNGTCTCCTCCCTCCCTTCCTCCATCGGA
GAGCACACGTCTGAACTCTTTCCCTACACGACGCTCTTCCGATCTNNNTATCCTTTTTCCTCCCAGCTGA
GAGCACACGTCTGAACTCTTTCCCTACACGACGCTCTTCCGATCTNNNTATCCTTTTTCCTCCCAGCTGA
AGAGCACACGTCTGAACTCTTTCCCTACACGACGCTCTTCCGATCTNNNCAGTTTGTGCTTATGGATGG
AGAGCACACGTCTGAACTCTTTCCCTACACGACGCTCTTCCGATCTNNNCAGTTTGTGCTTATGGATGG
GAGCACACGTCTGAACTCTTTCCCTACACGACGCTCTTCCGATCTNNNGGGGATTCTGTCACAGATG
GAGCACACGTCTGAACTCTTTCCCTACACGACGCTCTTCCGATCTNNNGGGGATTCTGTCACAGATG
AGCACACGTCTGAACTCTTTCCCTACACGACGCTCTTCCGATCTNNNACTGCACTCCAGCCTAAATAACA
AGCACACGTCTGAACTCTTTCCCTACACGACGCTCTTCCGATCTNNNACTGCACTCCAGCCTAAATAACA
GAGCACACGTCTGAACTCTTTCCCTACACGACGCTCTTCCGATCTNNNTGAGGAAAAGGAGGTGCCTA
GAGCACACGTCTGAACTCTTTCCCTACACGACGCTCTTCCGATCTNNNTGAGGAAAAGGAGGTGCCTA
GAGCACACGTCTGAACTCTTTCCCTACACGACGCTCTTCCGATCTNNNACTACAACACTTCCTTCACCGT
GAGCACACGTCTGAACTCTTTCCCTACACGACGCTCTTCCGATCTNNNACTACAACACTTCCTTCACCGT
AGCACACGTCTGAACTCTTTCCCTACACGACGCTCTTCCGATCTNNNAAGTGGGAGTATCATTTGGAC
AGCACACGTCTGAACTCTTTCCCTACACGACGCTCTTCCGATCTNNNAAGTGGGAGTATCATTTGGAC
AGAGCACACGTCTGAACTCTTTCCCTACACGACGCTCTTCCGATCTNNNTCCTCCAGAAGCCATTACTT
AGAGCACACGTCTGAACTCTTTCCCTACACGACGCTCTTCCGATCTNNNTCCTCCAGAAGCCATTACTT
AGAGCACACGTCTGAACTCTTTCCCTACACGACGCTCTTCCGATCTNNNATGGTAGTACACGCCTGTAA
AGAGCACACGTCTGAACTCTTTCCCTACACGACGCTCTTCCGATCTNNNATGGTAGTACACGCCTGTAA
AGCACACGTCTGAACTCTTTCCCTACACGACGCTCTTCCGATCTNNNATTCTGCAGCACTTTTGTCTACT
AGCACACGTCTGAACTCTTTCCCTACACGACGCTCTTCCGATCTNNNATTCTGCAGCACTTTTGTCTACT
GCACACGTCTGAACTCTTTCCCTACACGACGCTCTTCCGATCTNNNAAACTGCTGGGATTACAGGTGTC
GCACACGTCTGAACTCTTTCCCTACACGACGCTCTTCCGATCTNNNAAACTGCTGGGATTACAGGTGTC
GAGCACACGTCTGAACTCTTTCCCTACACGACGCTCTTCCGATCTNNNCAACAAGAGTGAAACACCGTC
GAGCACACGTCTGAACTCTTTCCCTACACGACGCTCTTCCGATCTNNNCAACAAGAGTGAAACACCGTC
AGAGCACACGTCTGAACTCTTTCCCTACACGACGCTCTTCCGATCTNNNCGGAACAGGTACCTTGCTAA
AGAGCACACGTCTGAACTCTTTCCCTACACGACGCTCTTCCGATCTNNNCGGAACAGGTACCTTGCTAA
GAGCACACGTCTGAACTCTTTCCCTACACGACGCTCTTCCGATCTNNNGGGACTTCTGCAAAACCATACC
GAGCACACGTCTGAACTCTTTCCCTACACGACGCTCTTCCGATCTNNNGGGACTTCTGCAAAACCATACC
CACACGTCTGAACTCTTTCCCTACACGACGCTCTTCCGATCTNNNCATTCCTTTGTCAGAGCCACAAGCA
CACACGTCTGAACTCTTTCCCTACACGACGCTCTTCCGATCTNNNCATTCCTTTGTCAGAGCCACAAGCA
GAGCACACGTCTGAACTCTTTCCCTACACGACGCTCTTCCGATCTNNNCCTCTGTCTCTCTCTCCTCTCTC
GAGCACACGTCTGAACTCTTTCCCTACACGACGCTCTTCCGATCTNNNCCTCTGTCTCTCTCTCCTCTCTC
AGCACACGTCTGAACTCTTTCCCTACACGACGCTCTTCCGATCTNNNTGGTGCTGACCTCATGAAGTA
AGCACACGTCTGAACTCTTTCCCTACACGACGCTCTTCCGATCTNNNTGGTGCTGACCTCATGAAGTA

GAGCACACGTCTGAACTCTTTCCCTACACGACGCTCTTCCGATCTNNNAAAAAAACAATGATGGGGTC
GAGCACACGTCTGAACTCTTTCCCTACACGACGCTCTTCCGATCTNNNAAAAAAACAATGATGGGGTC
GCACACGTCTGAACTCTTTCCCTACACGACGCTCTTCCGATCTNNNGTTGCATATTCTTCATTTCATTGC
GCACACGTCTGAACTCTTTCCCTACACGACGCTCTTCCGATCTNNNGTTGCATATTCTTCATTTCATTGC
GAGCACACGTCTGAACTCTTTCCCTACACGACGCTCTTCCGATCTNNNAGCTTAGGATGGTGCAAAGT
GAGCACACGTCTGAACTCTTTCCCTACACGACGCTCTTCCGATCTNNNAGCTTAGGATGGTGCAAAGT
GAGCACACGTCTGAACTCTTTCCCTACACGACGCTCTTCCGATCTNNNTCAATATAGCTGTTTGGGCCAC
GAGCACACGTCTGAACTCTTTCCCTACACGACGCTCTTCCGATCTNNNTCAATATAGCTGTTTGGGCCAC
AGAGCACACGTCTGAACTCTTTCCCTACACGACGCTCTTCCGATCTNNNGGAGGTACAAACAGCAGAG
AGAGCACACGTCTGAACTCTTTCCCTACACGACGCTCTTCCGATCTNNNGGAGGTACAAACAGCAGAG
GAGCACACGTCTGAACTCTTTCCCTACACGACGCTCTTCCGATCTNNNGCCTCTGTGCACCTTTCATTCCA
GAGCACACGTCTGAACTCTTTCCCTACACGACGCTCTTCCGATCTNNNGCCTCTGTGCACCTTTCATTCCA
AGCACACGTCTGAACTCTTTCCCTACACGACGCTCTTCCGATCTNNNCAGAGGAGAGAGGCAGCTTGC
AGCACACGTCTGAACTCTTTCCCTACACGACGCTCTTCCGATCTNNNCAGAGGAGAGAGGCAGCTTGC
AGCACACGTCTGAACTCTTTCCCTACACGACGCTCTTCCGATCTNNNAGTTCACACCTGCAGATTCTAGC
AGCACACGTCTGAACTCTTTCCCTACACGACGCTCTTCCGATCTNNNAGTTCACACCTGCAGATTCTAGC
GCACACGTCTGAACTCTTTCCCTACACGACGCTCTTCCGATCTNNNGAAGAATGCTCAGAGCTTGCAGC
GCACACGTCTGAACTCTTTCCCTACACGACGCTCTTCCGATCTNNNGAAGAATGCTCAGAGCTTGCAGC
AGCACACGTCTGAACTCTTTCCCTACACGACGCTCTTCCGATCTNNNAGTGGCATGATCTCATCTCACTC
AGCACACGTCTGAACTCTTTCCCTACACGACGCTCTTCCGATCTNNNAGTGGCATGATCTCATCTCACTC
GAGCACACGTCTGAACTCTTTCCCTACACGACGCTCTTCCGATCTNNNGGTGGTGCATATCTGTAATCCC
GAGCACACGTCTGAACTCTTTCCCTACACGACGCTCTTCCGATCTNNNGGTGGTGCATATCTGTAATCCC
GAGCACACGTCTGAACTCTTTCCCTACACGACGCTCTTCCGATCTNNNCACCAGTGAATGCTGCAAAAC
GAGCACACGTCTGAACTCTTTCCCTACACGACGCTCTTCCGATCTNNNCACCAGTGAATGCTGCAAAAC
GAGCACACGTCTGAACTCTTTCCCTACACGACGCTCTTCCGATCTNNNCTTGCAGTGAGCTGAGACACG
GAGCACACGTCTGAACTCTTTCCCTACACGACGCTCTTCCGATCTNNNCTTGCAGTGAGCTGAGACACG
GAGCACACGTCTGAACTCTTTCCCTACACGACGCTCTTCCGATCTNNNAAATGTTAATAAGTCCCTTTTG
GAGCACACGTCTGAACTCTTTCCCTACACGACGCTCTTCCGATCTNNNAAATGTTAATAAGTCCCTTTTG
GCACACGTCTGAACTCTTTCCCTACACGACGCTCTTCCGATCTNNNCAAAGACAGAACTTAACCTCTCAT
GCACACGTCTGAACTCTTTCCCTACACGACGCTCTTCCGATCTNNNCAAAGACAGAACTTAACCTCTCAT
GAGCACACGTCTGAACTCTTTCCCTACACGACGCTCTTCCGATCTNNNATTCATGCCCTCCCTGGAATAT
GAGCACACGTCTGAACTCTTTCCCTACACGACGCTCTTCCGATCTNNNATTCATGCCCTCCCTGGAATAT
AGCACACGTCTGAACTCTTTCCCTACACGACGCTCTTCCGATCTNNNTACCGAAGTTGACCACAGGTAA
AGCACACGTCTGAACTCTTTCCCTACACGACGCTCTTCCGATCTNNNTACCGAAGTTGACCACAGGTAA
GAGCACACGTCTGAACTCTTTCCCTACACGACGCTCTTCCGATCTNNNGCGAATGGTGCAACTTTAGAT
GAGCACACGTCTGAACTCTTTCCCTACACGACGCTCTTCCGATCTNNNGCGAATGGTGCAACTTTAGAT
GAGCACACGTCTGAACTCTTTCCCTACACGACGCTCTTCCGATCTNNNTGAGGCAGGAGAATTGCTTG
GAGCACACGTCTGAACTCTTTCCCTACACGACGCTCTTCCGATCTNNNTGAGGCAGGAGAATTGCTTG
GAGCACACGTCTGAACTCTTTCCCTACACGACGCTCTTCCGATCTNNNCTGTAGTTCCAGCTACTTGGGA
GAGCACACGTCTGAACTCTTTCCCTACACGACGCTCTTCCGATCTNNNCTGTAGTTCCAGCTACTTGGGA
GAGCACACGTCTGAACTCTTTCCCTACACGACGCTCTTCCGATCTNNNAGGCTGTAATGAGCTATGATT
GAGCACACGTCTGAACTCTTTCCCTACACGACGCTCTTCCGATCTNNNAGGCTGTAATGAGCTATGATT
GAGCACACGTCTGAACTCTTTCCCTACACGACGCTCTTCCGATCTNNNGTTGTGCTACAGGGGAGCAAA

GAGCACACGTCTGAACTCTTTCCCTACACGACGCTCTTCCGATCTNNNGTTGTGCTACAGGGGAGCAAA
GAGCACACGTCTGAACTCTTTCCCTACACGACGCTCTTCCGATCTNNNTATGAAGGTTAGGCAGTTCCA
GAGCACACGTCTGAACTCTTTCCCTACACGACGCTCTTCCGATCTNNNTATGAAGGTTAGGCAGTTCCA
AGCACACGTCTGAACTCTTTCCCTACACGACGCTCTTCCGATCTNNNGTCTGACTTCATCCAAGGGTTCT
AGCACACGTCTGAACTCTTTCCCTACACGACGCTCTTCCGATCTNNNGTCTGACTTCATCCAAGGGTTCT
AGCACACGTCTGAACTCTTTCCCTACACGACGCTCTTCCGATCTNNNCATCCTAGGCTCAAGTGATCCTC
AGCACACGTCTGAACTCTTTCCCTACACGACGCTCTTCCGATCTNNNCATCCTAGGCTCAAGTGATCCTC
GAGCACACGTCTGAACTCTTTCCCTACACGACGCTCTTCCGATCTNNNACGAGACAAATAGCCACACAC
GAGCACACGTCTGAACTCTTTCCCTACACGACGCTCTTCCGATCTNNNACGAGACAAATAGCCACACAC
GAGCACACGTCTGAACTCTTTCCCTACACGACGCTCTTCCGATCTNNNGTGAGCTGAGATCACACTACT
GAGCACACGTCTGAACTCTTTCCCTACACGACGCTCTTCCGATCTNNNGTGAGCTGAGATCACACTACT
GAGCACACGTCTGAACTCTTTCCCTACACGACGCTCTTCCGATCTNNNCTGAGGCAGGAGAATCACTTG
GAGCACACGTCTGAACTCTTTCCCTACACGACGCTCTTCCGATCTNNNCTGAGGCAGGAGAATCACTTG
GCACACGTCTGAACTCTTTCCCTACACGACGCTCTTCCGATCTNNNGGGTGTAGAATTTTAGGCTGGG
GCACACGTCTGAACTCTTTCCCTACACGACGCTCTTCCGATCTNNNGGGTGTAGAATTTTAGGCTGGG
AGCACACGTCTGAACTCTTTCCCTACACGACGCTCTTCCGATCTNNNCACATACACAGAGAGACACACC
AGCACACGTCTGAACTCTTTCCCTACACGACGCTCTTCCGATCTNNNCACATACACAGAGAGACACACC
GAGCACACGTCTGAACTCTTTCCCTACACGACGCTCTTCCGATCTNNNACACATAGGATTGTTGAGGG
GAGCACACGTCTGAACTCTTTCCCTACACGACGCTCTTCCGATCTNNNACACATAGGATTGTTGAGGG
GAGCACACGTCTGAACTCTTTCCCTACACGACGCTCTTCCGATCTNNNTCTCTCTCTCTCTCTCTCTCTC
GAGCACACGTCTGAACTCTTTCCCTACACGACGCTCTTCCGATCTNNNTCTCTCTCTCTCTCTCTCTCTC
GAGCACACGTCTGAACTCTTTCCCTACACGACGCTCTTCCGATCTNNNACATGGCGAAATCTGTCTCTAC
GAGCACACGTCTGAACTCTTTCCCTACACGACGCTCTTCCGATCTNNNACATGGCGAAATCTGTCTCTAC
AGCACACGTCTGAACTCTTTCCCTACACGACGCTCTTCCGATCTNNNAAGAAACGCATGTCTTCTGTGA
AGCACACGTCTGAACTCTTTCCCTACACGACGCTCTTCCGATCTNNNAAGAAACGCATGTCTTCTGTGA
GAGCACACGTCTGAACTCTTTCCCTACACGACGCTCTTCCGATCTNNNGTAGCTGGAATTACACACATGC
GAGCACACGTCTGAACTCTTTCCCTACACGACGCTCTTCCGATCTNNNGTAGCTGGAATTACACACATGC
GAGCACACGTCTGAACTCTTTCCCTACACGACGCTCTTCCGATCTNNNTGTTTTCAAATTTCCTCTCCGCC
GAGCACACGTCTGAACTCTTTCCCTACACGACGCTCTTCCGATCTNNNTGTTTTCAAATTTCCTCTCCGCC
AGCACACGTCTGAACTCTTTCCCTACACGACGCTCTTCCGATCTNNNGAGAATTGCTTGAATCCGGGAA
AGCACACGTCTGAACTCTTTCCCTACACGACGCTCTTCCGATCTNNNGAGAATTGCTTGAATCCGGGAA
GAGCACACGTCTGAACTCTTTCCCTACACGACGCTCTTCCGATCTNNNATGGTCATGGTGTTCCTATCT
GAGCACACGTCTGAACTCTTTCCCTACACGACGCTCTTCCGATCTNNNATGGTCATGGTGTTCCTATCT
GCACACGTCTGAACTCTTTCCCTACACGACGCTCTTCCGATCTNNNGTCAGTGGTAGTTTGATGGGGA
GCACACGTCTGAACTCTTTCCCTACACGACGCTCTTCCGATCTNNNGTCAGTGGTAGTTTGATGGGGA
AGCACACGTCTGAACTCTTTCCCTACACGACGCTCTTCCGATCTNNNAAGCAACTCTTTACCACCTGGAT
AGCACACGTCTGAACTCTTTCCCTACACGACGCTCTTCCGATCTNNNAAGCAACTCTTTACCACCTGGAT
GAGCACACGTCTGAACTCTTTCCCTACACGACGCTCTTCCGATCTNNNCACCTGTAGTCCCAGCTACTTG
GAGCACACGTCTGAACTCTTTCCCTACACGACGCTCTTCCGATCTNNNCACCTGTAGTCCCAGCTACTTG
GCACACGTCTGAACTCTTTCCCTACACGACGCTCTTCCGATCTNNNCTCCTGAATAGCTGGGACTACAG
GCACACGTCTGAACTCTTTCCCTACACGACGCTCTTCCGATCTNNNCTCCTGAATAGCTGGGACTACAG
GAGCACACGTCTGAACTCTTTCCCTACACGACGCTCTTCCGATCTNNNTTTTTTGCAGGGGGGTTTGAT
GAGCACACGTCTGAACTCTTTCCCTACACGACGCTCTTCCGATCTNNNTTTTTTGCAGGGGGGTTTGAT

AGCACACGTCTGAACTCTTTCCCTACACGACGCTCTTCCGATCTNNNTAAAGGGAGTCATTTGCCAGG
AGCACACGTCTGAACTCTTTCCCTACACGACGCTCTTCCGATCTNNNTAAAGGGAGTCATTTGCCAGG
GCACACGTCTGAACTCTTTCCCTACACGACGCTCTTCCGATCTNNNCAGAACACCTCATCCCCCAAATA
GCACACGTCTGAACTCTTTCCCTACACGACGCTCTTCCGATCTNNNCAGAACACCTCATCCCCCAAATA
AGCACACGTCTGAACTCTTTCCCTACACGACGCTCTTCCGATCTNNNAAGGCTTGTAAGTCATTCACCC
AGCACACGTCTGAACTCTTTCCCTACACGACGCTCTTCCGATCTNNNAAGGCTTGTAAGTCATTCACCC
AGAGCACACGTCTGAACTCTTTCCCTACACGACGCTCTTCCGATCTNNNGAGATTGCGCCATTGCACT
AGAGCACACGTCTGAACTCTTTCCCTACACGACGCTCTTCCGATCTNNNGAGATTGCGCCATTGCACT
GAGCACACGTCTGAACTCTTTCCCTACACGACGCTCTTCCGATCTNNNCTAATATCTGGGGGGGAGA
GAGCACACGTCTGAACTCTTTCCCTACACGACGCTCTTCCGATCTNNNCTAATATCTGGGGGGGAGA
AGCACACGTCTGAACTCTTTCCCTACACGACGCTCTTCCGATCTNNNCAGACTGGAAGACAAGGACT
AGCACACGTCTGAACTCTTTCCCTACACGACGCTCTTCCGATCTNNNCAGACTGGAAGACAAGGACT
AGCACACGTCTGAACTCTTTCCCTACACGACGCTCTTCCGATCTNNNTCTCCCGTAAATTGGTCCGAAT
AGCACACGTCTGAACTCTTTCCCTACACGACGCTCTTCCGATCTNNNTCTCCCGTAAATTGGTCCGAAT
GAGCACACGTCTGAACTCTTTCCCTACACGACGCTCTTCCGATCTNNNAAAAATTAGCCAAGCATCGT
GAGCACACGTCTGAACTCTTTCCCTACACGACGCTCTTCCGATCTNNNAAAAATTAGCCAAGCATCGT
GAGCACACGTCTGAACTCTTTCCCTACACGACGCTCTTCCGATCTNNNCTTGCAGTGAGCCGAGATCG
GAGCACACGTCTGAACTCTTTCCCTACACGACGCTCTTCCGATCTNNNCTTGCAGTGAGCCGAGATCG
CACACGTCTGAACTCTTTCCCTACACGACGCTCTTCCGATCTNNNAGAATCTGACTGCACATCTGGTT
CACACGTCTGAACTCTTTCCCTACACGACGCTCTTCCGATCTNNNAGAATCTGACTGCACATCTGGTT
GCACACGTCTGAACTCTTTCCCTACACGACGCTCTTCCGATCTNNNGAAACTGCCCCCATGATTCAAAT
GCACACGTCTGAACTCTTTCCCTACACGACGCTCTTCCGATCTNNNGAAACTGCCCCCATGATTCAAAT
AGCACACGTCTGAACTCTTTCCCTACACGACGCTCTTCCGATCTNNNGCCTGGCAAAAGAAAAAGTCT
AGCACACGTCTGAACTCTTTCCCTACACGACGCTCTTCCGATCTNNNGCCTGGCAAAAGAAAAAGTCT
GCACACGTCTGAACTCTTTCCCTACACGACGCTCTTCCGATCTNNNTTGAACTTCCAGATGCAAAAGG
GCACACGTCTGAACTCTTTCCCTACACGACGCTCTTCCGATCTNNNTTGAACTTCCAGATGCAAAAGG
GCACACGTCTGAACTCTTTCCCTACACGACGCTCTTCCGATCTNNNGGCTGTAACCAGGACCATCTTT
GCACACGTCTGAACTCTTTCCCTACACGACGCTCTTCCGATCTNNNGGCTGTAACCAGGACCATCTTT
AGCACACGTCTGAACTCTTTCCCTACACGACGCTCTTCCGATCTNNNCTTAGTGATTTTGGGTGCCG
AGCACACGTCTGAACTCTTTCCCTACACGACGCTCTTCCGATCTNNNCTTAGTGATTTTGGGTGCCG
AGCACACGTCTGAACTCTTTCCCTACACGACGCTCTTCCGATCTNNNAAACATGCACACAGAACTTCC
AGCACACGTCTGAACTCTTTCCCTACACGACGCTCTTCCGATCTNNNAAACATGCACACAGAACTTCC
AGCACACGTCTGAACTCTTTCCCTACACGACGCTCTTCCGATCTNNNTCCTTTCCTTTCCTTTCCTTTCT
AGCACACGTCTGAACTCTTTCCCTACACGACGCTCTTCCGATCTNNNTCCTTTCCTTTCCTTTCCTTTCT
GCACACGTCTGAACTCTTTCCCTACACGACGCTCTTCCGATCTNNNACAAAGCAGGACTCTGTCTCAA
GCACACGTCTGAACTCTTTCCCTACACGACGCTCTTCCGATCTNNNACAAAGCAGGACTCTGTCTCAA
GAGCACACGTCTGAACTCTTTCCCTACACGACGCTCTTCCGATCTNNNCTGTAATCCCAGCTACTCAG
GAGCACACGTCTGAACTCTTTCCCTACACGACGCTCTTCCGATCTNNNCTGTAATCCCAGCTACTCAG
AGCACACGTCTGAACTCTTTCCCTACACGACGCTCTTCCGATCTNNNTACTGGGAAGTGAGGAGCCC
AGCACACGTCTGAACTCTTTCCCTACACGACGCTCTTCCGATCTNNNTACTGGGAAGTGAGGAGCCC
AGCACACGTCTGAACTCTTTCCCTACACGACGCTCTTCCGATCTNNNAGAAAAACCCTCATAACCTGC
AGCACACGTCTGAACTCTTTCCCTACACGACGCTCTTCCGATCTNNNAGAAAAACCCTCATAACCTGC
AGCACACGTCTGAACTCTTTCCCTACACGACGCTCTTCCGATCTNNNTCAGGAGAATCACTTGAACCC

AGCACACGTCTGAACTCTTTCCCTACACGACGCTCTTCCGATCTNNNTCAGGAGAATCACTTGAACCC
GAGCACACGTCTGAACTCTTTCCCTACACGACGCTCTTCCGATCTNNNCTCTGTTCTCAGCCACAGG
GAGCACACGTCTGAACTCTTTCCCTACACGACGCTCTTCCGATCTNNNCTCTGTTCTCAGCCACAGG
GAGCACACGTCTGAACTCTTTCCCTACACGACGCTCTTCCGATCTNNNACTGAGTCAATTCAATCAGCA
GAGCACACGTCTGAACTCTTTCCCTACACGACGCTCTTCCGATCTNNNACTGAGTCAATTCAATCAGCA
AGCACACGTCTGAACTCTTTCCCTACACGACGCTCTTCCGATCTNNNTGCTCTGAGGCTGAATTTTCAA
AGCACACGTCTGAACTCTTTCCCTACACGACGCTCTTCCGATCTNNNTGCTCTGAGGCTGAATTTTCAA
AGCACACGTCTGAACTCTTTCCCTACACGACGCTCTTCCGATCTNNNTCTCTGGGTGTAGAGCTTTAGG
AGCACACGTCTGAACTCTTTCCCTACACGACGCTCTTCCGATCTNNNTCTCTGGGTGTAGAGCTTTAGG
AGCACACGTCTGAACTCTTTCCCTACACGACGCTCTTCCGATCTNNNATGCTCAGTCACATGGTTCTCAT
AGCACACGTCTGAACTCTTTCCCTACACGACGCTCTTCCGATCTNNNATGCTCAGTCACATGGTTCTCAT
GAGCACACGTCTGAACTCTTTCCCTACACGACGCTCTTCCGATCTNNNAGTTAGGGTTTTCCAGAGAA
GAGCACACGTCTGAACTCTTTCCCTACACGACGCTCTTCCGATCTNNNAGTTAGGGTTTTCCAGAGAA
AGCACACGTCTGAACTCTTTCCCTACACGACGCTCTTCCGATCTNNNAGGGCATGATCCAGCTTTTCTT
AGCACACGTCTGAACTCTTTCCCTACACGACGCTCTTCCGATCTNNNAGGGCATGATCCAGCTTTTCTT
AGCACACGTCTGAACTCTTTCCCTACACGACGCTCTTCCGATCTNNNTCATCATCTTGATCTTCAAATT
AGCACACGTCTGAACTCTTTCCCTACACGACGCTCTTCCGATCTNNNTCATCATCTTGATCTTCAAATT
GAGCACACGTCTGAACTCTTTCCCTACACGACGCTCTTCCGATCTNNNAGTGAAGTGTGAGTGAGTG
GAGCACACGTCTGAACTCTTTCCCTACACGACGCTCTTCCGATCTNNNAGTGAAGTGTGAGTGAGTG
GAGCACACGTCTGAACTCTTTCCCTACACGACGCTCTTCCGATCTNNNTCACCATGTTGTCTAGACTGG
GAGCACACGTCTGAACTCTTTCCCTACACGACGCTCTTCCGATCTNNNTCACCATGTTGTCTAGACTGG
GAGCACACGTCTGAACTCTTTCCCTACACGACGCTCTTCCGATCTNNNCTGGACAACAAGAGTGAGAC
GAGCACACGTCTGAACTCTTTCCCTACACGACGCTCTTCCGATCTNNNCTGGACAACAAGAGTGAGAC
AGAGCACACGTCTGAACTCTTTCCCTACACGACGCTCTTCCGATCTNNNTATCTGCATCCCCTCTAGTCG
AGAGCACACGTCTGAACTCTTTCCCTACACGACGCTCTTCCGATCTNNNTATCTGCATCCCCTCTAGTCG
GAGCACACGTCTGAACTCTTTCCCTACACGACGCTCTTCCGATCTNNNTAGGAGAATTGCTTGAACCC
GAGCACACGTCTGAACTCTTTCCCTACACGACGCTCTTCCGATCTNNNTAGGAGAATTGCTTGAACCC
GAGCACACGTCTGAACTCTTTCCCTACACGACGCTCTTCCGATCTNNNTTGGCTTCCCTTTTCTCATCTG
GAGCACACGTCTGAACTCTTTCCCTACACGACGCTCTTCCGATCTNNNTTGGCTTCCCTTTTCTCATCTG
AGCACACGTCTGAACTCTTTCCCTACACGACGCTCTTCCGATCTNNNAAATAATCTACCATCCCTCCCGC
AGCACACGTCTGAACTCTTTCCCTACACGACGCTCTTCCGATCTNNNAAATAATCTACCATCCCTCCCGC
GAGCACACGTCTGAACTCTTTCCCTACACGACGCTCTTCCGATCTNNNCTCTCTTGAAGCTGCCCTGAA
GAGCACACGTCTGAACTCTTTCCCTACACGACGCTCTTCCGATCTNNNCTCTCTTGAAGCTGCCCTGAA
GAGCACACGTCTGAACTCTTTCCCTACACGACGCTCTTCCGATCTNNNGAAGAGGAGTAAGTGCCACT
GAGCACACGTCTGAACTCTTTCCCTACACGACGCTCTTCCGATCTNNNGAAGAGGAGTAAGTGCCACT
AGCACACGTCTGAACTCTTTCCCTACACGACGCTCTTCCGATCTNNNTGTTTCTGCTCTCTGGTGAAGA
AGCACACGTCTGAACTCTTTCCCTACACGACGCTCTTCCGATCTNNNTGTTTCTGCTCTCTGGTGAAGA
CACACGTCTGAACTCTTTCCCTACACGACGCTCTTCCGATCTNNNCTCATGCCCTCTCCTCTGCCCATCG
CACACGTCTGAACTCTTTCCCTACACGACGCTCTTCCGATCTNNNCTCATGCCCTCTCCTCTGCCCATCG
GAGCACACGTCTGAACTCTTTCCCTACACGACGCTCTTCCGATCTNNNGAGGTTGCAGTGAGCTGAG
GAGCACACGTCTGAACTCTTTCCCTACACGACGCTCTTCCGATCTNNNGAGGTTGCAGTGAGCTGAG
GAGCACACGTCTGAACTCTTTCCCTACACGACGCTCTTCCGATCTNNNATACAGAGAGACCCTTGTGCA
GAGCACACGTCTGAACTCTTTCCCTACACGACGCTCTTCCGATCTNNNATACAGAGAGACCCTTGTGCA

GAGCACACGTCTGAACTCTTTCCCTACACGACGCTCTTCCGATCTNNNTAGTCTTAGCTACTCAGGAGG

GAGCACACGTCTGAACTCTTTCCCTACACGACGCTCTTCCGATCTNNNTAGTCTTAGCTACTCAGGAGG

AGCACACGTCTGAACTCTTTCCCTACACGACGCTCTTCCGATCTNNNTGTAATCCCAGCTACTCAGGAG

AGCACACGTCTGAACTCTTTCCCTACACGACGCTCTTCCGATCTNNNTGTAATCCCAGCTACTCAGGAG

GAGCACACGTCTGAACTCTTTCCCTACACGACGCTCTTCCGATCTNNNGCCTGTAGTCCCAGCTACTTGG

GAGCACACGTCTGAACTCTTTCCCTACACGACGCTCTTCCGATCTNNNGCCTGTAGTCCCAGCTACTTGG

AGCACACGTCTGAACTCTTTCCCTACACGACGCTCTTCCGATCTNNNTTAGCCTCTGCCCTTCCCTTTAG

AGCACACGTCTGAACTCTTTCCCTACACGACGCTCTTCCGATCTNNNTTAGCCTCTGCCCTTCCCTTTAG

AGAGCACACGTCTGAACTCTTTCCCTACACGACGCTCTTCCGATCTNNNCCCACTAACTCGTCATCTAGCA

AGAGCACACGTCTGAACTCTTTCCCTACACGACGCTCTTCCGATCTNNNCCCACTAACTCGTCATCTAGCA

AGCACACGTCTGAACTCTTTCCCTACACGACGCTCTTCCGATCTNNNTTTATAAAACCGTCAGACCCCATG

AGCACACGTCTGAACTCTTTCCCTACACGACGCTCTTCCGATCTNNNTTTATAAAACCGTCAGACCCCATG

AGCACACGTCTGAACTCTTTCCCTACACGACGCTCTTCCGATCTNNNAAATTCTTCATCATATGTTTGCT

AGCACACGTCTGAACTCTTTCCCTACACGACGCTCTTCCGATCTNNNAAATTCTTCATCATATGTTTGCT

GCACACGTCTGAACTCTTTCCCTACACGACGCTCTTCCGATCTNNNAAAAAAAAGCCACCGCAGTAACA

GCACACGTCTGAACTCTTTCCCTACACGACGCTCTTCCGATCTNNNAAAAAAAAGCCACCGCAGTAACA

ACACGTCTGAACTCTTTCCCTACACGACGCTCTTCCGATCTNNNGAAAAGAACTCAGTACCTGAATCAT

ACACGTCTGAACTCTTTCCCTACACGACGCTCTTCCGATCTNNNGAAAAGAACTCAGTACCTGAATCAT

AGCACACGTCTGAACTCTTTCCCTACACGACGCTCTTCCGATCTNNNGTTGTGGTACTGGAGCAAGAC

AGCACACGTCTGAACTCTTTCCCTACACGACGCTCTTCCGATCTNNNGTTGTGGTACTGGAGCAAGAC

GAGCACACGTCTGAACTCTTTCCCTACACGACGCTCTTCCGATCTNNNTAGGAGAATTGCTTGAACCAG

GAGCACACGTCTGAACTCTTTCCCTACACGACGCTCTTCCGATCTNNNTAGGAGAATTGCTTGAACCAG

GAGCACACGTCTGAACTCTTTCCCTACACGACGCTCTTCCGATCTNNNTATTTAGGGCAAAGGGGCAT

GAGCACACGTCTGAACTCTTTCCCTACACGACGCTCTTCCGATCTNNNTATTTAGGGCAAAGGGGCAT

GAGCACACGTCTGAACTCTTTCCCTACACGACGCTCTTCCGATCTNNNATTTCCACAGACAGAGGATTC

GAGCACACGTCTGAACTCTTTCCCTACACGACGCTCTTCCGATCTNNNATTTCCACAGACAGAGGATTC

GAGCACACGTCTGAACTCTTTCCCTACACGACGCTCTTCCGATCTNNNTGGAGAAGAGGCCAAAAGCA

GAGCACACGTCTGAACTCTTTCCCTACACGACGCTCTTCCGATCTNNNTGGAGAAGAGGCCAAAAGCA

AGCACACGTCTGAACTCTTTCCCTACACGACGCTCTTCCGATCTNNNCCTGTAATCCCAGCTACTCAGG

AGCACACGTCTGAACTCTTTCCCTACACGACGCTCTTCCGATCTNNNCCTGTAATCCCAGCTACTCAGG

GCACACGTCTGAACTCTTTCCCTACACGACGCTCTTCCGATCTNNNGTGCCTATGTCCTCATTCTTTGCCA

GCACACGTCTGAACTCTTTCCCTACACGACGCTCTTCCGATCTNNNGTGCCTATGTCCTCATTCTTTGCCA

AGCACACGTCTGAACTCTTTCCCTACACGACGCTCTTCCGATCTNNNGACTACACACACCACCATATCCA

AGCACACGTCTGAACTCTTTCCCTACACGACGCTCTTCCGATCTNNNGACTACACACACCACCATATCCA

AGAGCACACGTCTGAACTCTTTCCCTACACGACGCTCTTCCGATCTNNNTCATGAGGTCAGGAGATCGA

AGAGCACACGTCTGAACTCTTTCCCTACACGACGCTCTTCCGATCTNNNTCATGAGGTCAGGAGATCGA

GAGCACACGTCTGAACTCTTTCCCTACACGACGCTCTTCCGATCTNNNTATGAGCCCTTGTGAACAATG

GAGCACACGTCTGAACTCTTTCCCTACACGACGCTCTTCCGATCTNNNTATGAGCCCTTGTGAACAATG

AGCACACGTCTGAACTCTTTCCCTACACGACGCTCTTCCGATCTNNNGCAGGTATTGACTGGGTATCAG

AGCACACGTCTGAACTCTTTCCCTACACGACGCTCTTCCGATCTNNNGCAGGTATTGACTGGGTATCAG

GAGCACACGTCTGAACTCTTTCCCTACACGACGCTCTTCCGATCTNNNTGTTTGATCATGGTGTCTGCA

GAGCACACGTCTGAACTCTTTCCCTACACGACGCTCTTCCGATCTNNNTGTTTGATCATGGTGTCTGCA

AGAGCACACGTCTGAACTCTTTCCCTACACGACGCTCTTCCGATCTNNNTGTCTGCTGTGCAGAAGTTTT

GAGCACACGTCTGAACTCTTTCCCTACACGACGCTCTTCCGATCTNNNTGTCTGCTGTGCAGAAGTTT

AGCACACGTCTGAACTCTTTCCCTACACGACGCTCTTCCGATCTNNNTACCATTCTGGGGTCTGAAGG

AGCACACGTCTGAACTCTTTCCCTACACGACGCTCTTCCGATCTNNNTACCATTCTGGGGTCTGAAGG

GAGCACACGTCTGAACTCTTTCCCTACACGACGCTCTTCCGATCTNNNTTTGGATGCTTCCAGGTCAGT

GAGCACACGTCTGAACTCTTTCCCTACACGACGCTCTTCCGATCTNNNTTTGGATGCTTCCAGGTCAGT

GCACACGTCTGAACTCTTTCCCTACACGACGCTCTTCCGATCTNNNCAGAGCTACCCAACCCCTCCCATC

GCACACGTCTGAACTCTTTCCCTACACGACGCTCTTCCGATCTNNNCAGAGCTACCCAACCCCTCCCATC

GAGCACACGTCTGAACTCTTTCCCTACACGACGCTCTTCCGATCTNNNCCCAGATAGCAGGAGTTCAT

GAGCACACGTCTGAACTCTTTCCCTACACGACGCTCTTCCGATCTNNNCCCAGATAGCAGGAGTTCAT

AGCACACGTCTGAACTCTTTCCCTACACGACGCTCTTCCGATCTNNNGGCCTATAGGGTTTCTGCAAAG

AGCACACGTCTGAACTCTTTCCCTACACGACGCTCTTCCGATCTNNNGGCCTATAGGGTTTCTGCAAAG

GCACACGTCTGAACTCTTTCCCTACACGACGCTCTTCCGATCTNNNGCGACCCCCTGCAATAATAGATT

GCACACGTCTGAACTCTTTCCCTACACGACGCTCTTCCGATCTNNNGCGACCCCCTGCAATAATAGATT

GAGCACACGTCTGAACTCTTTCCCTACACGACGCTCTTCCGATCTNNNAAGGGAAGTGTCTTAGTCCC

GAGCACACGTCTGAACTCTTTCCCTACACGACGCTCTTCCGATCTNNNAAGGGAAGTGTCTTAGTCCC

AGCACACGTCTGAACTCTTTCCCTACACGACGCTCTTCCGATCTNNNCAAACTCAATCCTCCCTCTCTCCC

AGCACACGTCTGAACTCTTTCCCTACACGACGCTCTTCCGATCTNNNCAAACTCAATCCTCCCTCTCTCCC

AGCACACGTCTGAACTCTTTCCCTACACGACGCTCTTCCGATCTNNNCAACTGTATCTTTTGCATAAAA

AGCACACGTCTGAACTCTTTCCCTACACGACGCTCTTCCGATCTNNNCAACTGTATCTTTTGCATAAAA

GAGCACACGTCTGAACTCTTTCCCTACACGACGCTCTTCCGATCTNNNAGGCATAGTTGGAAAAGGCT

GAGCACACGTCTGAACTCTTTCCCTACACGACGCTCTTCCGATCTNNNAGGCATAGTTGGAAAAGGCT

AGCACACGTCTGAACTCTTTCCCTACACGACGCTCTTCCGATCTNNNTTGGCCACCAAACTGAATAGTC

AGCACACGTCTGAACTCTTTCCCTACACGACGCTCTTCCGATCTNNNTTGGCCACCAAACTGAATAGTC

GAGCACACGTCTGAACTCTTTCCCTACACGACGCTCTTCCGATCTNNNAGGCCTCTGATTCCTCACTGAT

GAGCACACGTCTGAACTCTTTCCCTACACGACGCTCTTCCGATCTNNNAGGCCTCTGATTCCTCACTGAT

ACGTCTGAACTCTTTCCCTACACGACGCTCTTCCGATCTNNNGGGACACCCAACCCAAATGGCATCGG

ACGTCTGAACTCTTTCCCTACACGACGCTCTTCCGATCTNNNGGGACACCCAACCCAAATGGCATCGG

AGCACACGTCTGAACTCTTTCCCTACACGACGCTCTTCCGATCTNNNGAACTTGCAGTGAGCCGAGAT

AGCACACGTCTGAACTCTTTCCCTACACGACGCTCTTCCGATCTNNNGAACTTGCAGTGAGCCGAGAT

AGCACACGTCTGAACTCTTTCCCTACACGACGCTCTTCCGATCTNNNGTTCCATGTGGCATTCGACCT

AGCACACGTCTGAACTCTTTCCCTACACGACGCTCTTCCGATCTNNNGTTCCATGTGGCATTCGACCT

GCACACGTCTGAACTCTTTCCCTACACGACGCTCTTCCGATCTNNNTAGAAGGGTTGGTGTATGTATG

GCACACGTCTGAACTCTTTCCCTACACGACGCTCTTCCGATCTNNNTAGAAGGGTTGGTGTATGTATG

AGCACACGTCTGAACTCTTTCCCTACACGACGCTCTTCCGATCTNNNACAAAACCAAGTCCAACTAAGC

AGCACACGTCTGAACTCTTTCCCTACACGACGCTCTTCCGATCTNNNACAAAACCAAGTCCAACTAAGC

AGCACACGTCTGAACTCTTTCCCTACACGACGCTCTTCCGATCTNNNACTAGTGGCAATCTGAGAGA

AGCACACGTCTGAACTCTTTCCCTACACGACGCTCTTCCGATCTNNNACTAGTGGCAATCTGAGAGA

GAGCACACGTCTGAACTCTTTCCCTACACGACGCTCTTCCGATCTNNNCTGAGGCAGGAGAATCACTT

GAGCACACGTCTGAACTCTTTCCCTACACGACGCTCTTCCGATCTNNNCTGAGGCAGGAGAATCACTT

GAGCACACGTCTGAACTCTTTCCCTACACGACGCTCTTCCGATCTNNNCAACATAGTGAAACCCCGTCT

GAGCACACGTCTGAACTCTTTCCCTACACGACGCTCTTCCGATCTNNNCAACATAGTGAAACCCCGTCT

GCACACGTCTGAACTCTTTCCCTACACGACGCTCTTCCGATCTNNNGTACTGGGATGACAGGTCTAAG

GCACACGTCTGAACTCTTTCCCTACACGACGCTCTTCCGATCTNNNGTACTGGGATGACAGGTCTAAG

GAGCACACGTCTGAACTCTTTCCCTACACGACGCTCTTCCGATCTNNNCTTGCTGTCGATTGGATTGG
GAGCACACGTCTGAACTCTTTCCCTACACGACGCTCTTCCGATCTNNNCTTGCTGTCGATTGGATTGG
AGCACACGTCTGAACTCTTTCCCTACACGACGCTCTTCCGATCTNNNCTTGCAGTGAGCCAAGATTGC
AGCACACGTCTGAACTCTTTCCCTACACGACGCTCTTCCGATCTNNNCTTGCAGTGAGCCAAGATTGC
CACACGTCTGAACTCTTTCCCTACACGACGCTCTTCCGATCTNNNTCTTTGAAGCCCTTTTCCTTGGC
CACACGTCTGAACTCTTTCCCTACACGACGCTCTTCCGATCTNNNTCTTTGAAGCCCTTTTCCTTGGC
GCACACGTCTGAACTCTTTCCCTACACGACGCTCTTCCGATCTNNNGTATAGAAGGTGTAGACCAGC
GCACACGTCTGAACTCTTTCCCTACACGACGCTCTTCCGATCTNNNGTATAGAAGGTGTAGACCAGC
GCACACGTCTGAACTCTTTCCCTACACGACGCTCTTCCGATCTNNNCATCATCTATCTTCCTGCTGTGG
GCACACGTCTGAACTCTTTCCCTACACGACGCTCTTCCGATCTNNNCATCATCTATCTTCCTGCTGTGG
GAGCACACGTCTGAACTCTTTCCCTACACGACGCTCTTCCGATCTNNNCTGTAATCACAGCTACTGGG
GAGCACACGTCTGAACTCTTTCCCTACACGACGCTCTTCCGATCTNNNCTGTAATCACAGCTACTGGG
GCACACGTCTGAACTCTTTCCCTACACGACGCTCTTCCGATCTNNNCAGATGGTATTTTAAGATTCCTG
GCACACGTCTGAACTCTTTCCCTACACGACGCTCTTCCGATCTNNNCAGATGGTATTTTAAGATTCCTG
GCACACGTCTGAACTCTTTCCCTACACGACGCTCTTCCGATCTNNNACTCCATACACAAGGCCCCTATTC
GCACACGTCTGAACTCTTTCCCTACACGACGCTCTTCCGATCTNNNACTCCATACACAAGGCCCCTATTC
GAGCACACGTCTGAACTCTTTCCCTACACGACGCTCTTCCGATCTNNNTGAGCATGGGATGTTTCCATT
GAGCACACGTCTGAACTCTTTCCCTACACGACGCTCTTCCGATCTNNNTGAGCATGGGATGTTTCCATT
AGCACACGTCTGAACTCTTTCCCTACACGACGCTCTTCCGATCTNNNGCTACCTGGTGGCTTCATCTAT
AGCACACGTCTGAACTCTTTCCCTACACGACGCTCTTCCGATCTNNNGCTACCTGGTGGCTTCATCTAT
AGCACACGTCTGAACTCTTTCCCTACACGACGCTCTTCCGATCTNNNGAGGCTGAGGCAGGAGAATG
AGCACACGTCTGAACTCTTTCCCTACACGACGCTCTTCCGATCTNNNGAGGCTGAGGCAGGAGAATG
AGCACACGTCTGAACTCTTTCCCTACACGACGCTCTTCCGATCTNNNCAGCAGATGTGTCTCATTCAC
AGCACACGTCTGAACTCTTTCCCTACACGACGCTCTTCCGATCTNNNCAGCAGATGTGTCTCATTCAC
GCACACGTCTGAACTCTTTCCCTACACGACGCTCTTCCGATCTNNNGATGCAGGAGAATCGCTTGAAC
GCACACGTCTGAACTCTTTCCCTACACGACGCTCTTCCGATCTNNNGATGCAGGAGAATCGCTTGAAC
AGAGCACACGTCTGAACTCTTTCCCTACACGACGCTCTTCCGATCTNNNTCAGTCATCTGTATAAAATG
AGAGCACACGTCTGAACTCTTTCCCTACACGACGCTCTTCCGATCTNNNTCAGTCATCTGTATAAAATG
GCACACGTCTGAACTCTTTCCCTACACGACGCTCTTCCGATCTNNNATCCTAAAACATGACATGCCCCTC
GCACACGTCTGAACTCTTTCCCTACACGACGCTCTTCCGATCTNNNATCCTAAAACATGACATGCCCCTC
GCACACGTCTGAACTCTTTCCCTACACGACGCTCTTCCGATCTNNNCAGGAGATGGAGATTACGGTG
GCACACGTCTGAACTCTTTCCCTACACGACGCTCTTCCGATCTNNNCAGGAGATGGAGATTACGGTG
AGCACACGTCTGAACTCTTTCCCTACACGACGCTCTTCCGATCTNNNTTCAGCTAGTTCATAGGGGAC
AGCACACGTCTGAACTCTTTCCCTACACGACGCTCTTCCGATCTNNNTTCAGCTAGTTCATAGGGGAC
GAGCACACGTCTGAACTCTTTCCCTACACGACGCTCTTCCGATCTNNNCAGGAGACTAAGGCAGGAG
GAGCACACGTCTGAACTCTTTCCCTACACGACGCTCTTCCGATCTNNNCAGGAGACTAAGGCAGGAG
CACACGTCTGAACTCTTTCCCTACACGACGCTCTTCCGATCTNNNGGGCGATAGAGTAAGACTCCATC
CACACGTCTGAACTCTTTCCCTACACGACGCTCTTCCGATCTNNNGGGCGATAGAGTAAGACTCCATC
AGCACACGTCTGAACTCTTTCCCTACACGACGCTCTTCCGATCTNNNGACTGGTACATAGTAGGTGCTC
AGCACACGTCTGAACTCTTTCCCTACACGACGCTCTTCCGATCTNNNGACTGGTACATAGTAGGTGCTC
GAGCACACGTCTGAACTCTTTCCCTACACGACGCTCTTCCGATCTNNNCCTCTCCATATACTCTGCTCCAT
GAGCACACGTCTGAACTCTTTCCCTACACGACGCTCTTCCGATCTNNNCCTCTCCATATACTCTGCTCCAT
AGCACACGTCTGAACTCTTTCCCTACACGACGCTCTTCCGATCTNNNCCTTTGGTAAGCCTCCTTTGTC

GAGCACACGTCTGAACTCTTTCCCTACACGACGCTCTTCCGATCTNNNCCTTTGGTAAGCCTCCTTTGTC
GAGCACACGTCTGAACTCTTTCCCTACACGACGCTCTTCCGATCTNNNGCCAAAGCTATCCAGAGCAAA
GAGCACACGTCTGAACTCTTTCCCTACACGACGCTCTTCCGATCTNNNGCCAAAGCTATCCAGAGCAAA
GAGCACACGTCTGAACTCTTTCCCTACACGACGCTCTTCCGATCTNNNCAACATGGAGAAACCCTGTCTC
GAGCACACGTCTGAACTCTTTCCCTACACGACGCTCTTCCGATCTNNNCAACATGGAGAAACCCTGTCTC
AGCACACGTCTGAACTCTTTCCCTACACGACGCTCTTCCGATCTNNNGCAAGCATAGCTCTCAAAAGAC
AGCACACGTCTGAACTCTTTCCCTACACGACGCTCTTCCGATCTNNNGCAAGCATAGCTCTCAAAAGAC
AGCACACGTCTGAACTCTTTCCCTACACGACGCTCTTCCGATCTNNNGCACCCGTTAGCTCATCATTAC
AGCACACGTCTGAACTCTTTCCCTACACGACGCTCTTCCGATCTNNNGCACCCGTTAGCTCATCATTAC
AGCACACGTCTGAACTCTTTCCCTACACGACGCTCTTCCGATCTNNNATTGGTTTCAGAACTGCTTGTA
AGCACACGTCTGAACTCTTTCCCTACACGACGCTCTTCCGATCTNNNATTGGTTTCAGAACTGCTTGTA
AGCACACGTCTGAACTCTTTCCCTACACGACGCTCTTCCGATCTNNNGGGTCTCCATCTTGCAGACTATC
AGCACACGTCTGAACTCTTTCCCTACACGACGCTCTTCCGATCTNNNGGGTCTCCATCTTGCAGACTATC
AGCACACGTCTGAACTCTTTCCCTACACGACGCTCTTCCGATCTNNNGTGCCTGTAATCCCAGCTACTCC
AGCACACGTCTGAACTCTTTCCCTACACGACGCTCTTCCGATCTNNNGTGCCTGTAATCCCAGCTACTCC
AGCACACGTCTGAACTCTTTCCCTACACGACGCTCTTCCGATCTNNNCAAGAACAAAACCCCGTCTCAAA
AGCACACGTCTGAACTCTTTCCCTACACGACGCTCTTCCGATCTNNNCAAGAACAAAACCCCGTCTCAAA
AGCACACGTCTGAACTCTTTCCCTACACGACGCTCTTCCGATCTNNNGTACACGGGTCTGGAAAATTTC
AGCACACGTCTGAACTCTTTCCCTACACGACGCTCTTCCGATCTNNNGTACACGGGTCTGGAAAATTTC
GCACACGTCTGAACTCTTTCCCTACACGACGCTCTTCCGATCTNNNGAATCTCCATTGACGTTTGGGTG
GCACACGTCTGAACTCTTTCCCTACACGACGCTCTTCCGATCTNNNGAATCTCCATTGACGTTTGGGTG
GCACACGTCTGAACTCTTTCCCTACACGACGCTCTTCCGATCTNNNAGATGTTGCACCGTATTTCTTTCCA
GCACACGTCTGAACTCTTTCCCTACACGACGCTCTTCCGATCTNNNAGATGTTGCACCGTATTTCTTTCCA
AAGAGCACACGTCTGAACTCTTTCCCTACACGACGCTCTTCCGATCTNNNTTGATCGTGAGTTTGCTCA
AAGAGCACACGTCTGAACTCTTTCCCTACACGACGCTCTTCCGATCTNNNTTGATCGTGAGTTTGCTCA
AGCACACGTCTGAACTCTTTCCCTACACGACGCTCTTCCGATCTNNNATGCAAAGGATGTTCGAAGCAA
AGCACACGTCTGAACTCTTTCCCTACACGACGCTCTTCCGATCTNNNATGCAAAGGATGTTCGAAGCAA
AGCACACGTCTGAACTCTTTCCCTACACGACGCTCTTCCGATCTNNNAGGTAGGAGAATTGCTTGAAC
AGCACACGTCTGAACTCTTTCCCTACACGACGCTCTTCCGATCTNNNAGGTAGGAGAATTGCTTGAAC
GCACACGTCTGAACTCTTTCCCTACACGACGCTCTTCCGATCTNNNAGAATCACAGCCTAATCCATCCAC
GCACACGTCTGAACTCTTTCCCTACACGACGCTCTTCCGATCTNNNAGAATCACAGCCTAATCCATCCAC
AGCACACGTCTGAACTCTTTCCCTACACGACGCTCTTCCGATCTNNNCCTAGTAGGTGTTCAGAAGGTG
AGCACACGTCTGAACTCTTTCCCTACACGACGCTCTTCCGATCTNNNCCTAGTAGGTGTTCAGAAGGTG
GAGCACACGTCTGAACTCTTTCCCTACACGACGCTCTTCCGATCTNNNATGCATTCAATTGCGAGAACA
GAGCACACGTCTGAACTCTTTCCCTACACGACGCTCTTCCGATCTNNNATGCATTCAATTGCGAGAACA
GAGCACACGTCTGAACTCTTTCCCTACACGACGCTCTTCCGATCTNNNCACCTTTGCCTCATTGTGACAA
GAGCACACGTCTGAACTCTTTCCCTACACGACGCTCTTCCGATCTNNNCACCTTTGCCTCATTGTGACAA
GCACACGTCTGAACTCTTTCCCTACACGACGCTCTTCCGATCTNNNGCTGCAGTGAGCTGAGATAGAT
GCACACGTCTGAACTCTTTCCCTACACGACGCTCTTCCGATCTNNNGCTGCAGTGAGCTGAGATAGAT
AGAGCACACGTCTGAACTCTTTCCCTACACGACGCTCTTCCGATCTNNNCACTAGGCAGAGGTTGTAGT
AGAGCACACGTCTGAACTCTTTCCCTACACGACGCTCTTCCGATCTNNNCACTAGGCAGAGGTTGTAGT
GAGCACACGTCTGAACTCTTTCCCTACACGACGCTCTTCCGATCTNNNGCTTGCAGTAAGTGAGATCAT
GAGCACACGTCTGAACTCTTTCCCTACACGACGCTCTTCCGATCTNNNGCTTGCAGTAAGTGAGATCAT

AGAGCACACGTCTGAACTCTTTCCCTACACGACGCTCTTCCGATCTNNNGATTCAAGGGGGCTCTTTCT
AGAGCACACGTCTGAACTCTTTCCCTACACGACGCTCTTCCGATCTNNNGATTCAAGGGGGCTCTTTCT
AGCACACGTCTGAACTCTTTCCCTACACGACGCTCTTCCGATCTNNNTTTCTAGCCTAGCTTGTGACAGC
AGCACACGTCTGAACTCTTTCCCTACACGACGCTCTTCCGATCTNNNTTTCTAGCCTAGCTTGTGACAGC
AGCACACGTCTGAACTCTTTCCCTACACGACGCTCTTCCGATCTNNNCTCAGGAGGATTAGGGAGGAC
AGCACACGTCTGAACTCTTTCCCTACACGACGCTCTTCCGATCTNNNCTCAGGAGGATTAGGGAGGAC
AGCACACGTCTGAACTCTTTCCCTACACGACGCTCTTCCGATCTNNNAAACAGGATGAGGTAAGGCAC
AGCACACGTCTGAACTCTTTCCCTACACGACGCTCTTCCGATCTNNNAAACAGGATGAGGTAAGGCAC
GAGCACACGTCTGAACTCTTTCCCTACACGACGCTCTTCCGATCTNNNCAGTATTCAGTGGAGGTAGC
GAGCACACGTCTGAACTCTTTCCCTACACGACGCTCTTCCGATCTNNNCAGTATTCAGTGGAGGTAGC
GAGCACACGTCTGAACTCTTTCCCTACACGACGCTCTTCCGATCTNNNGGCCCACTGTTCTAAACCTACA
GAGCACACGTCTGAACTCTTTCCCTACACGACGCTCTTCCGATCTNNNGGCCCACTGTTCTAAACCTACA
AGCACACGTCTGAACTCTTTCCCTACACGACGCTCTTCCGATCTNNNTCACTCAGAGAAATACAAAGCT
AGCACACGTCTGAACTCTTTCCCTACACGACGCTCTTCCGATCTNNNTCACTCAGAGAAATACAAAGCT
GCACACGTCTGAACTCTTTCCCTACACGACGCTCTTCCGATCTNNNTTCAAAAATTTACATGGGGCCGG
GCACACGTCTGAACTCTTTCCCTACACGACGCTCTTCCGATCTNNNTTCAAAAATTTACATGGGGCCGG
GAGCACACGTCTGAACTCTTTCCCTACACGACGCTCTTCCGATCTNNNAAGTATGGCTCGTGGTCTACA
GAGCACACGTCTGAACTCTTTCCCTACACGACGCTCTTCCGATCTNNNAAGTATGGCTCGTGGTCTACA
AGCACACGTCTGAACTCTTTCCCTACACGACGCTCTTCCGATCTNNNGCCGGTTAGTCTAAGATCAAGC
AGCACACGTCTGAACTCTTTCCCTACACGACGCTCTTCCGATCTNNNGCCGGTTAGTCTAAGATCAAGC
AGCACACGTCTGAACTCTTTCCCTACACGACGCTCTTCCGATCTNNNAGATTGTGCCACTACACTCTAG
AGCACACGTCTGAACTCTTTCCCTACACGACGCTCTTCCGATCTNNNAGATTGTGCCACTACACTCTAG
AGAGCACACGTCTGAACTCTTTCCCTACACGACGCTCTTCCGATCTNNNGACCTTTCAGAGGTAGGATC
AGAGCACACGTCTGAACTCTTTCCCTACACGACGCTCTTCCGATCTNNNGACCTTTCAGAGGTAGGATC
AGCACACGTCTGAACTCTTTCCCTACACGACGCTCTTCCGATCTNNNTTGGAGGATCACTTGAGTCAAC
AGCACACGTCTGAACTCTTTCCCTACACGACGCTCTTCCGATCTNNNTTGGAGGATCACTTGAGTCAAC
GAGCACACGTCTGAACTCTTTCCCTACACGACGCTCTTCCGATCTNNNAATATGTTTTTTTTCCTTCCCC
GAGCACACGTCTGAACTCTTTCCCTACACGACGCTCTTCCGATCTNNNAATATGTTTTTTTTCCTTCCCC
GAGCACACGTCTGAACTCTTTCCCTACACGACGCTCTTCCGATCTNNNGAGCAAGACCCTGTCTCAAAA
GAGCACACGTCTGAACTCTTTCCCTACACGACGCTCTTCCGATCTNNNGAGCAAGACCCTGTCTCAAAA
CACACGTCTGAACTCTTTCCCTACACGACGCTCTTCCGATCTNNNTGGGGGGTAGGTTTTCATTCAATC
CACACGTCTGAACTCTTTCCCTACACGACGCTCTTCCGATCTNNNTGGGGGGTAGGTTTTCATTCAATC
GCACACGTCTGAACTCTTTCCCTACACGACGCTCTTCCGATCTNNNATGTGCCTGTTTCTACTGGATGT
GCACACGTCTGAACTCTTTCCCTACACGACGCTCTTCCGATCTNNNATGTGCCTGTTTCTACTGGATGT
ACACGTCTGAACTCTTTCCCTACACGACGCTCTTCCGATCTNNNCGACTCCCTCAGCTTGCGAGCATCG
ACACGTCTGAACTCTTTCCCTACACGACGCTCTTCCGATCTNNNCGACTCCCTCAGCTTGCGAGCATCG
AGCACACGTCTGAACTCTTTCCCTACACGACGCTCTTCCGATCTNNNCTGGGACTATAGGCATCTGTCA
AGCACACGTCTGAACTCTTTCCCTACACGACGCTCTTCCGATCTNNNCTGGGACTATAGGCATCTGTCA
GCACACGTCTGAACTCTTTCCCTACACGACGCTCTTCCGATCTNNNCATGCCTGTAGTCCCAGATACTC
GCACACGTCTGAACTCTTTCCCTACACGACGCTCTTCCGATCTNNNCATGCCTGTAGTCCCAGATACTC
GAGCACACGTCTGAACTCTTTCCCTACACGACGCTCTTCCGATCTNNNGTGCCTGTAATCCCAGCTACTT
GAGCACACGTCTGAACTCTTTCCCTACACGACGCTCTTCCGATCTNNNGTGCCTGTAATCCCAGCTACTT
GAGCACACGTCTGAACTCTTTCCCTACACGACGCTCTTCCGATCTNNNCTTGCAGTGAGCCGAGATCAA

AGCACACGTCTGAACTCTTTCCCTACACGACGCTCTTCCGATCTNNNCTTGCAGTGAGCCGAGATCAA
AGCACACGTCTGAACTCTTTCCCTACACGACGCTCTTCCGATCTNNNACCATCACTGAGTGTGGAAAG
AGCACACGTCTGAACTCTTTCCCTACACGACGCTCTTCCGATCTNNNACCATCACTGAGTGTGGAAAG
GAGCACACGTCTGAACTCTTTCCCTACACGACGCTCTTCCGATCTNNNCTTCACTTCTGACTCTTCCCCA
GAGCACACGTCTGAACTCTTTCCCTACACGACGCTCTTCCGATCTNNNCTTCACTTCTGACTCTTCCCCA
AGCACACGTCTGAACTCTTTCCCTACACGACGCTCTTCCGATCTNNNAGAAGGAGAGAGGGAAGAA
AGCACACGTCTGAACTCTTTCCCTACACGACGCTCTTCCGATCTNNNAGAAGGAGAGAGGGAAGAA
GCACACGTCTGAACTCTTTCCCTACACGACGCTCTTCCGATCTNNNGAGATCACACCATTGCACTCCAG
GCACACGTCTGAACTCTTTCCCTACACGACGCTCTTCCGATCTNNNGAGATCACACCATTGCACTCCAG
CACACGTCTGAACTCTTTCCCTACACGACGCTCTTCCGATCTNNNAAACTATACCCGTCACGTTTCCAC
CACACGTCTGAACTCTTTCCCTACACGACGCTCTTCCGATCTNNNAAACTATACCCGTCACGTTTCCAC
AGAGCACACGTCTGAACTCTTTCCCTACACGACGCTCTTCCGATCTNNNCTCACAATGCTTCACATGGG
AGAGCACACGTCTGAACTCTTTCCCTACACGACGCTCTTCCGATCTNNNCTCACAATGCTTCACATGGG
GAGCACACGTCTGAACTCTTTCCCTACACGACGCTCTTCCGATCTNNNCGACAGAGTGAGACTCCGTC
GAGCACACGTCTGAACTCTTTCCCTACACGACGCTCTTCCGATCTNNNCGACAGAGTGAGACTCCGTC
AGCACACGTCTGAACTCTTTCCCTACACGACGCTCTTCCGATCTNNNCTCAAACTCCTGACCTCAAGTG
AGCACACGTCTGAACTCTTTCCCTACACGACGCTCTTCCGATCTNNNCTCAAACTCCTGACCTCAAGTG
AAGAGCACACGTCTGAACTCTTTCCCTACACGACGCTCTTCCGATCTNNNGCTGACAAATCACCATTTT
AAGAGCACACGTCTGAACTCTTTCCCTACACGACGCTCTTCCGATCTNNNGCTGACAAATCACCATTTT
CACACGTCTGAACTCTTTCCCTACACGACGCTCTTCCGATCTNNNTGCACCATTTTCTCTATCAGCATCG
CACACGTCTGAACTCTTTCCCTACACGACGCTCTTCCGATCTNNNTGCACCATTTTCTCTATCAGCATCG
CACACGTCTGAACTCTTTCCCTACACGACGCTCTTCCGATCTNNNTGATGGCCAACTAGTCTAACATGG
CACACGTCTGAACTCTTTCCCTACACGACGCTCTTCCGATCTNNNTGATGGCCAACTAGTCTAACATGG
AGCACACGTCTGAACTCTTTCCCTACACGACGCTCTTCCGATCTNNNGCTGCAAGTCCAAACAGGTAA
AGCACACGTCTGAACTCTTTCCCTACACGACGCTCTTCCGATCTNNNGCTGCAAGTCCAAACAGGTAA
AGCACACGTCTGAACTCTTTCCCTACACGACGCTCTTCCGATCTNNNCCCAGTGAGGAGGAATCTAG
AGCACACGTCTGAACTCTTTCCCTACACGACGCTCTTCCGATCTNNNCCCAGTGAGGAGGAATCTAG
GCACACGTCTGAACTCTTTCCCTACACGACGCTCTTCCGATCTNNNTATAGTCCCAGCTACTCGAGAG
GCACACGTCTGAACTCTTTCCCTACACGACGCTCTTCCGATCTNNNTATAGTCCCAGCTACTCGAGAG
GAGCACACGTCTGAACTCTTTCCCTACACGACGCTCTTCCGATCTNNNGTGGTCCTTTCTACTGCTTGA
GAGCACACGTCTGAACTCTTTCCCTACACGACGCTCTTCCGATCTNNNGTGGTCCTTTCTACTGCTTGA
AGAGCACACGTCTGAACTCTTTCCCTACACGACGCTCTTCCGATCTNNNACAAAAATTATCCAGGCGTG
AGAGCACACGTCTGAACTCTTTCCCTACACGACGCTCTTCCGATCTNNNACAAAAATTATCCAGGCGTG
AGCACACGTCTGAACTCTTTCCCTACACGACGCTCTTCCGATCTNNNCAGGAGAATCACTTGAACCCA
AGCACACGTCTGAACTCTTTCCCTACACGACGCTCTTCCGATCTNNNCAGGAGAATCACTTGAACCCA
GAGCACACGTCTGAACTCTTTCCCTACACGACGCTCTTCCGATCTNNNGGTGGTCGTCTTGTAGTCG
GAGCACACGTCTGAACTCTTTCCCTACACGACGCTCTTCCGATCTNNNGGTGGTCGTCTTGTAGTCG
AGCACACGTCTGAACTCTTTCCCTACACGACGCTCTTCCGATCTNNNGGGAGTGTGAGTTTTACGCTT
AGCACACGTCTGAACTCTTTCCCTACACGACGCTCTTCCGATCTNNNGGGAGTGTGAGTTTTACGCTT
AGCACACGTCTGAACTCTTTCCCTACACGACGCTCTTCCGATCTNNNGAGATGAGGTTTCACCATGTTG
AGCACACGTCTGAACTCTTTCCCTACACGACGCTCTTCCGATCTNNNGAGATGAGGTTTCACCATGTTG
AGCACACGTCTGAACTCTTTCCCTACACGACGCTCTTCCGATCTNNNCGCCTGTTAGTGTCTTTTTCTG
AGCACACGTCTGAACTCTTTCCCTACACGACGCTCTTCCGATCTNNNCGCCTGTTAGTGTCTTTTTCTG

GAGCACACGTCTGAACTCTTTCCCTACACGACGCTCTTCCGATCTNNNCTGTAATTGCAGCTACTGGG
GAGCACACGTCTGAACTCTTTCCCTACACGACGCTCTTCCGATCTNNNCTGTAATTGCAGCTACTGGG
GAGCACACGTCTGAACTCTTTCCCTACACGACGCTCTTCCGATCTNNNAAGAAATGAAGTCCTCAGGG
GAGCACACGTCTGAACTCTTTCCCTACACGACGCTCTTCCGATCTNNNAAGAAATGAAGTCCTCAGGG
GAGCACACGTCTGAACTCTTTCCCTACACGACGCTCTTCCGATCTNNNCTGTAGTCCCAGCTACACAGG
GAGCACACGTCTGAACTCTTTCCCTACACGACGCTCTTCCGATCTNNNCTGTAGTCCCAGCTACACAGG
AGCACACGTCTGAACTCTTTCCCTACACGACGCTCTTCCGATCTNNNAGGGCTCTATCAAACCATGTTG
AGCACACGTCTGAACTCTTTCCCTACACGACGCTCTTCCGATCTNNNAGGGCTCTATCAAACCATGTTG
AGCACACGTCTGAACTCTTTCCCTACACGACGCTCTTCCGATCTNNNTCTCTTGTTTTCAGCCGTGAT
AGCACACGTCTGAACTCTTTCCCTACACGACGCTCTTCCGATCTNNNTCTCTTGTTTTCAGCCGTGAT
AGCACACGTCTGAACTCTTTCCCTACACGACGCTCTTCCGATCTNNNTTGGAGAGTTATGAGCGCCTG
AGCACACGTCTGAACTCTTTCCCTACACGACGCTCTTCCGATCTNNNTTGGAGAGTTATGAGCGCCTG
AGCACACGTCTGAACTCTTTCCCTACACGACGCTCTTCCGATCTNNNGCCCGGCCATGATTATTTCTTT
AGCACACGTCTGAACTCTTTCCCTACACGACGCTCTTCCGATCTNNNGCCCGGCCATGATTATTTCTTT
CACGTCTGAACTCTTTCCCTACACGACGCTCTTCCGATCTNNNAGTGACTCAATGTTCTTGTAGTCCAC
CACGTCTGAACTCTTTCCCTACACGACGCTCTTCCGATCTNNNAGTGACTCAATGTTCTTGTAGTCCAC
GAGCACACGTCTGAACTCTTTCCCTACACGACGCTCTTCCGATCTNNNGAATTGGCTTTTTCCCATGGT
GAGCACACGTCTGAACTCTTTCCCTACACGACGCTCTTCCGATCTNNNGAATTGGCTTTTTCCCATGGT
AGCACACGTCTGAACTCTTTCCCTACACGACGCTCTTCCGATCTNNNGATCAAATTACTGCACTCCAGCC
AGCACACGTCTGAACTCTTTCCCTACACGACGCTCTTCCGATCTNNNGATCAAATTACTGCACTCCAGCC
GAGCACACGTCTGAACTCTTTCCCTACACGACGCTCTTCCGATCTNNNCGTGATCTCAGCTCACTGCAA
GAGCACACGTCTGAACTCTTTCCCTACACGACGCTCTTCCGATCTNNNCGTGATCTCAGCTCACTGCAA
GAGCACACGTCTGAACTCTTTCCCTACACGACGCTCTTCCGATCTNNNAGGATAATGGCCACCAGTTTC
GAGCACACGTCTGAACTCTTTCCCTACACGACGCTCTTCCGATCTNNNAGGATAATGGCCACCAGTTTC
GAGCACACGTCTGAACTCTTTCCCTACACGACGCTCTTCCGATCTNNNGGTACACACCTGTAATCTCAG
GAGCACACGTCTGAACTCTTTCCCTACACGACGCTCTTCCGATCTNNNGGTACACACCTGTAATCTCAG
CACACGTCTGAACTCTTTCCCTACACGACGCTCTTCCGATCTNNNGATCCAGAAGCAGAATAAGAGCAT
CACACGTCTGAACTCTTTCCCTACACGACGCTCTTCCGATCTNNNGATCCAGAAGCAGAATAAGAGCAT
AGCACACGTCTGAACTCTTTCCCTACACGACGCTCTTCCGATCTNNNCTAATGGAGTCGAGGAGCAAC
AGCACACGTCTGAACTCTTTCCCTACACGACGCTCTTCCGATCTNNNCTAATGGAGTCGAGGAGCAAC
GAGCACACGTCTGAACTCTTTCCCTACACGACGCTCTTCCGATCTNNNACCTGTAATCCCATCTACTCG
GAGCACACGTCTGAACTCTTTCCCTACACGACGCTCTTCCGATCTNNNACCTGTAATCCCATCTACTCG
AGCACACGTCTGAACTCTTTCCCTACACGACGCTCTTCCGATCTNNNCTCCCAAGTAGCTGGAGTTACA
AGCACACGTCTGAACTCTTTCCCTACACGACGCTCTTCCGATCTNNNCTCCCAAGTAGCTGGAGTTACA
GAGCACACGTCTGAACTCTTTCCCTACACGACGCTCTTCCGATCTNNNCAACACCAATACACCTCACTCC
GAGCACACGTCTGAACTCTTTCCCTACACGACGCTCTTCCGATCTNNNCAACACCAATACACCTCACTCC
AGCACACGTCTGAACTCTTTCCCTACACGACGCTCTTCCGATCTNNNCCAACTGTTTGCCATGAATTTG
AGCACACGTCTGAACTCTTTCCCTACACGACGCTCTTCCGATCTNNNCCAACTGTTTGCCATGAATTTG
GAGCACACGTCTGAACTCTTTCCCTACACGACGCTCTTCCGATCTNNNGTGTGCCTATAATCCCAGCTA
GAGCACACGTCTGAACTCTTTCCCTACACGACGCTCTTCCGATCTNNNGTGTGCCTATAATCCCAGCTA
GCACACGTCTGAACTCTTTCCCTACACGACGCTCTTCCGATCTNNNTCATAAAGGAATCCATTGGCCAT
GCACACGTCTGAACTCTTTCCCTACACGACGCTCTTCCGATCTNNNTCATAAAGGAATCCATTGGCCAT
GAGCACACGTCTGAACTCTTTCCCTACACGACGCTCTTCCGATCTNNNCAAAGTGCTGGGATTACAGG

GAGCACACGTCTGAACTCTTTCCCTACACGACGCTCTTCCGATCTNNNCAAAGTGCTGGGATTACAGG
GCACACGTCTGAACTCTTTCCCTACACGACGCTCTTCCGATCTNNNTGTGATTCTCTTTGAGCCAAGGA
GCACACGTCTGAACTCTTTCCCTACACGACGCTCTTCCGATCTNNNTGTGATTCTCTTTGAGCCAAGGA
AGCACACGTCTGAACTCTTTCCCTACACGACGCTCTTCCGATCTNNNATGCTGTGGTTTAGGTGATGC
AGCACACGTCTGAACTCTTTCCCTACACGACGCTCTTCCGATCTNNNATGCTGTGGTTTAGGTGATGC
GAGCACACGTCTGAACTCTTTCCCTACACGACGCTCTTCCGATCTNNNTCCCTTGAACCCAGAAGATCA
GAGCACACGTCTGAACTCTTTCCCTACACGACGCTCTTCCGATCTNNNTCCCTTGAACCCAGAAGATCA
GAGCACACGTCTGAACTCTTTCCCTACACGACGCTCTTCCGATCTNNNTGAGTATGAGACCACCTCCTT
GAGCACACGTCTGAACTCTTTCCCTACACGACGCTCTTCCGATCTNNNTGAGTATGAGACCACCTCCTT
GCACACGTCTGAACTCTTTCCCTACACGACGCTCTTCCGATCTNNNCCTTCTGATTTTTGTCCCGCTTC
GCACACGTCTGAACTCTTTCCCTACACGACGCTCTTCCGATCTNNNCCTTCTGATTTTTGTCCCGCTTC
GAGCACACGTCTGAACTCTTTCCCTACACGACGCTCTTCCGATCTNNNGGTCAATGGACTGGAGGTT
GAGCACACGTCTGAACTCTTTCCCTACACGACGCTCTTCCGATCTNNNGGTCAATGGACTGGAGGTT
AGCACACGTCTGAACTCTTTCCCTACACGACGCTCTTCCGATCTNNNTGGGGATGGCATTGAATCTGT
AGCACACGTCTGAACTCTTTCCCTACACGACGCTCTTCCGATCTNNNTGGGGATGGCATTGAATCTGT
GCACACGTCTGAACTCTTTCCCTACACGACGCTCTTCCGATCTNNNAAGTTCCAGGGAAATTGTGAGG
GCACACGTCTGAACTCTTTCCCTACACGACGCTCTTCCGATCTNNNAAGTTCCAGGGAAATTGTGAGG
AGCACACGTCTGAACTCTTTCCCTACACGACGCTCTTCCGATCTNNNTAGGGTCCTGAAGTGAGTGTT
AGCACACGTCTGAACTCTTTCCCTACACGACGCTCTTCCGATCTNNNTAGGGTCCTGAAGTGAGTGTT
AGCACACGTCTGAACTCTTTCCCTACACGACGCTCTTCCGATCTNNNAGTTTGTGTGCAGAAAGATGT
AGCACACGTCTGAACTCTTTCCCTACACGACGCTCTTCCGATCTNNNAGTTTGTGTGCAGAAAGATGT
AGCACACGTCTGAACTCTTTCCCTACACGACGCTCTTCCGATCTNNNGCTAACAGCAGAACTTCCATTG
AGCACACGTCTGAACTCTTTCCCTACACGACGCTCTTCCGATCTNNNGCTAACAGCAGAACTTCCATTG
GAGCACACGTCTGAACTCTTTCCCTACACGACGCTCTTCCGATCTNNNGAGGTTGCAGTGAGCTGAG
GAGCACACGTCTGAACTCTTTCCCTACACGACGCTCTTCCGATCTNNNGAGGTTGCAGTGAGCTGAG
GCACACGTCTGAACTCTTTCCCTACACGACGCTCTTCCGATCTNNNAAGCGCCTGTAGTTTTAGCTACT
GCACACGTCTGAACTCTTTCCCTACACGACGCTCTTCCGATCTNNNAAGCGCCTGTAGTTTTAGCTACT
AGCACACGTCTGAACTCTTTCCCTACACGACGCTCTTCCGATCTNNNATAGGAAGTCCCATGACCAAA
AGCACACGTCTGAACTCTTTCCCTACACGACGCTCTTCCGATCTNNNATAGGAAGTCCCATGACCAAA
AGCACACGTCTGAACTCTTTCCCTACACGACGCTCTTCCGATCTNNNCACCTGTAATCCCAGCATTTTG
AGCACACGTCTGAACTCTTTCCCTACACGACGCTCTTCCGATCTNNNCACCTGTAATCCCAGCATTTTG
AGCACACGTCTGAACTCTTTCCCTACACGACGCTCTTCCGATCTNNNGAGAACCAAGCAGGATAAAAG
AGCACACGTCTGAACTCTTTCCCTACACGACGCTCTTCCGATCTNNNGAGAACCAAGCAGGATAAAAG
AGCACACGTCTGAACTCTTTCCCTACACGACGCTCTTCCGATCTNNNGATTGAGACCATCCTGGCTAA
AGCACACGTCTGAACTCTTTCCCTACACGACGCTCTTCCGATCTNNNGATTGAGACCATCCTGGCTAA
AGCACACGTCTGAACTCTTTCCCTACACGACGCTCTTCCGATCTNNNCACCACACCCAGCTAATTTTTG
AGCACACGTCTGAACTCTTTCCCTACACGACGCTCTTCCGATCTNNNCACCACACCCAGCTAATTTTTG
AGCACACGTCTGAACTCTTTCCCTACACGACGCTCTTCCGATCTNNNGGGACAAGATCGAGACTCCAT
AGCACACGTCTGAACTCTTTCCCTACACGACGCTCTTCCGATCTNNNGGGACAAGATCGAGACTCCAT
GAGCACACGTCTGAACTCTTTCCCTACACGACGCTCTTCCGATCTNNNCTCTGTATGGAATCCACATAC
GAGCACACGTCTGAACTCTTTCCCTACACGACGCTCTTCCGATCTNNNCTCTGTATGGAATCCACATAC
GAGCACACGTCTGAACTCTTTCCCTACACGACGCTCTTCCGATCTNNNGAAAAACAGGGTTGACAAGG
GAGCACACGTCTGAACTCTTTCCCTACACGACGCTCTTCCGATCTNNNGAAAAACAGGGTTGACAAGG

GAGCACACGTCTGAACTCTTTCCCTACACGACGCTCTTCCGATCTNNNAGGGGTGTGGTTCTTTTCTTT
GAGCACACGTCTGAACTCTTTCCCTACACGACGCTCTTCCGATCTNNNAGGGGTGTGGTTCTTTTCTTT
AGCACACGTCTGAACTCTTTCCCTACACGACGCTCTTCCGATCTNNNAGAAGAGAAGCTGAATTGGC
AGCACACGTCTGAACTCTTTCCCTACACGACGCTCTTCCGATCTNNNAGAAGAGAAGCTGAATTGGC
AGCACACGTCTGAACTCTTTCCCTACACGACGCTCTTCCGATCTNNNGTTCTTTGTATTCCCATCCCCTC
AGCACACGTCTGAACTCTTTCCCTACACGACGCTCTTCCGATCTNNNGTTCTTTGTATTCCCATCCCCTC
AGCACACGTCTGAACTCTTTCCCTACACGACGCTCTTCCGATCTNNNGCAACACAACGAGACTCAGTTT
AGCACACGTCTGAACTCTTTCCCTACACGACGCTCTTCCGATCTNNNGCAACACAACGAGACTCAGTTT
AGCACACGTCTGAACTCTTTCCCTACACGACGCTCTTCCGATCTNNNCCTGAGACTGGGTAATTTTGGG
AGCACACGTCTGAACTCTTTCCCTACACGACGCTCTTCCGATCTNNNCCTGAGACTGGGTAATTTTGGG
AGCACACGTCTGAACTCTTTCCCTACACGACGCTCTTCCGATCTNNNTTCCTGAGTACCTGGGACTACA
AGCACACGTCTGAACTCTTTCCCTACACGACGCTCTTCCGATCTNNNTTCCTGAGTACCTGGGACTACA
AGCACACGTCTGAACTCTTTCCCTACACGACGCTCTTCCGATCTNNNAATTGGTGAATTGGAGTCTTCG
AGCACACGTCTGAACTCTTTCCCTACACGACGCTCTTCCGATCTNNNAATTGGTGAATTGGAGTCTTCG
AGCACACGTCTGAACTCTTTCCCTACACGACGCTCTTCCGATCTNNNGAGGTTGCAGTGAGCTGAGA
AGCACACGTCTGAACTCTTTCCCTACACGACGCTCTTCCGATCTNNNGAGGTTGCAGTGAGCTGAGA
GAGCACACGTCTGAACTCTTTCCCTACACGACGCTCTTCCGATCTNNNCCATGAGGGAGAAGAAATCA
GAGCACACGTCTGAACTCTTTCCCTACACGACGCTCTTCCGATCTNNNCCATGAGGGAGAAGAAATCA
AGCACACGTCTGAACTCTTTCCCTACACGACGCTCTTCCGATCTNNNGGACAGACTGACACAGATACA
AGCACACGTCTGAACTCTTTCCCTACACGACGCTCTTCCGATCTNNNGGACAGACTGACACAGATACA
AGCACACGTCTGAACTCTTTCCCTACACGACGCTCTTCCGATCTNNNTATCCTAGCATTTTGGGAGGCA
AGCACACGTCTGAACTCTTTCCCTACACGACGCTCTTCCGATCTNNNTATCCTAGCATTTTGGGAGGCA
AGAGCACACGTCTGAACTCTTTCCCTACACGACGCTCTTCCGATCTNNNGTACAAAAATTAGCCAGTC
AGAGCACACGTCTGAACTCTTTCCCTACACGACGCTCTTCCGATCTNNNGTACAAAAATTAGCCAGTC
AGCACACGTCTGAACTCTTTCCCTACACGACGCTCTTCCGATCTNNNCCACCACACCCTGCTAATTTTTA
AGCACACGTCTGAACTCTTTCCCTACACGACGCTCTTCCGATCTNNNCCACCACACCCTGCTAATTTTTA
AGCACACGTCTGAACTCTTTCCCTACACGACGCTCTTCCGATCTNNNCAACCAGTTCAGCCTGGAGTA
AGCACACGTCTGAACTCTTTCCCTACACGACGCTCTTCCGATCTNNNCAACCAGTTCAGCCTGGAGTA
AGCACACGTCTGAACTCTTTCCCTACACGACGCTCTTCCGATCTNNNCTCCACCGTCTGTGTATCGGTA
AGCACACGTCTGAACTCTTTCCCTACACGACGCTCTTCCGATCTNNNCTCCACCGTCTGTGTATCGGTA
GAGCACACGTCTGAACTCTTTCCCTACACGACGCTCTTCCGATCTNNNCAGAGGTTACAGTGAGCCAA
GAGCACACGTCTGAACTCTTTCCCTACACGACGCTCTTCCGATCTNNNCAGAGGTTACAGTGAGCCAA
AGCACACGTCTGAACTCTTTCCCTACACGACGCTCTTCCGATCTNNNGATTTAGAGACCCATTTGCTGG
AGCACACGTCTGAACTCTTTCCCTACACGACGCTCTTCCGATCTNNNGATTTAGAGACCCATTTGCTGG
AGCACACGTCTGAACTCTTTCCCTACACGACGCTCTTCCGATCTNNNACACCTGTAATCCCAGCTACTT
AGCACACGTCTGAACTCTTTCCCTACACGACGCTCTTCCGATCTNNNACACCTGTAATCCCAGCTACTT
AGCACACGTCTGAACTCTTTCCCTACACGACGCTCTTCCGATCTNNNAAAAAATTAGGCGGGTGTAG
AGCACACGTCTGAACTCTTTCCCTACACGACGCTCTTCCGATCTNNNAAAAAATTAGGCGGGTGTAG
AGCACACGTCTGAACTCTTTCCCTACACGACGCTCTTCCGATCTNNNTGTAATCCCAGCTAGTCAGGA
AGCACACGTCTGAACTCTTTCCCTACACGACGCTCTTCCGATCTNNNTGTAATCCCAGCTAGTCAGGA
AGCACACGTCTGAACTCTTTCCCTACACGACGCTCTTCCGATCTNNNTTCCTTCTTCCTTCCTTCCTTCCCA
AGCACACGTCTGAACTCTTTCCCTACACGACGCTCTTCCGATCTNNNTTCCTTCTTCCTTCCTTCCTTCCCA
AGCACACGTCTGAACTCTTTCCCTACACGACGCTCTTCCGATCTNNNGTAGCGTTTAGAGGAGTCTTC

AGCACACGTCTGAACTCTTTCCCTACACGACGCTCTTCCGATCTNNNGTAGCGTTTAGAGGAGTCTTC
GCACACGTCTGAACTCTTTCCCTACACGACGCTCTTCCGATCTNNNTGACAGGGTTGCAAAGTTGTTT
GCACACGTCTGAACTCTTTCCCTACACGACGCTCTTCCGATCTNNNTGACAGGGTTGCAAAGTTGTTT
AGCACACGTCTGAACTCTTTCCCTACACGACGCTCTTCCGATCTNNNAAGATGTGCACAGGTTATATG
AGCACACGTCTGAACTCTTTCCCTACACGACGCTCTTCCGATCTNNNAAGATGTGCACAGGTTATATG
AGCACACGTCTGAACTCTTTCCCTACACGACGCTCTTCCGATCTNNNAAGGTTGCAAAGGCTGGAATA
AGCACACGTCTGAACTCTTTCCCTACACGACGCTCTTCCGATCTNNNAAGGTTGCAAAGGCTGGAATA
AGAGCACACGTCTGAACTCTTTCCCTACACGACGCTCTTCCGATCTNNNTAGAAAACACCAGGTCTGAG
AGAGCACACGTCTGAACTCTTTCCCTACACGACGCTCTTCCGATCTNNNTAGAAAACACCAGGTCTGAG
GAGCACACGTCTGAACTCTTTCCCTACACGACGCTCTTCCGATCTNNNACGCCTGTAATCCCAGTAATCT
GAGCACACGTCTGAACTCTTTCCCTACACGACGCTCTTCCGATCTNNNACGCCTGTAATCCCAGTAATCT
GCACACGTCTGAACTCTTTCCCTACACGACGCTCTTCCGATCTNNNCATAGGGTTCATCCTCCTGGGAA
GCACACGTCTGAACTCTTTCCCTACACGACGCTCTTCCGATCTNNNCATAGGGTTCATCCTCCTGGGAA
AGAGCACACGTCTGAACTCTTTCCCTACACGACGCTCTTCCGATCTNNNGCAACAAAGGGAGACTCCAT
AGAGCACACGTCTGAACTCTTTCCCTACACGACGCTCTTCCGATCTNNNGCAACAAAGGGAGACTCCAT
GAGCACACGTCTGAACTCTTTCCCTACACGACGCTCTTCCGATCTNNNTCAAACCCATTGCTTCCAATTGT
GAGCACACGTCTGAACTCTTTCCCTACACGACGCTCTTCCGATCTNNNTCAAACCCATTGCTTCCAATTGT
AGCACACGTCTGAACTCTTTCCCTACACGACGCTCTTCCGATCTNNNGAGAATCACTGGAACTCGGGA
AGCACACGTCTGAACTCTTTCCCTACACGACGCTCTTCCGATCTNNNGAGAATCACTGGAACTCGGGA
AGCACACGTCTGAACTCTTTCCCTACACGACGCTCTTCCGATCTNNNCGCCCGGCTTTTCTTTTTTTTTT
AGCACACGTCTGAACTCTTTCCCTACACGACGCTCTTCCGATCTNNNCGCCCGGCTTTTCTTTTTTTTTT
GCACACGTCTGAACTCTTTCCCTACACGACGCTCTTCCGATCTNNNTGGATTCACACTTTTCCCTCTACAC
GCACACGTCTGAACTCTTTCCCTACACGACGCTCTTCCGATCTNNNTGGATTCACACTTTTCCCTCTACAC
AGCACACGTCTGAACTCTTTCCCTACACGACGCTCTTCCGATCTNNNTGTGTGGGTTGGTCTCTTTATC
AGCACACGTCTGAACTCTTTCCCTACACGACGCTCTTCCGATCTNNNTGTGTGGGTTGGTCTCTTTATC
AGAGCACACGTCTGAACTCTTTCCCTACACGACGCTCTTCCGATCTNNNGCATTCCTGTAATCCCAGCTA
AGAGCACACGTCTGAACTCTTTCCCTACACGACGCTCTTCCGATCTNNNGCATTCCTGTAATCCCAGCTA
GCACACGTCTGAACTCTTTCCCTACACGACGCTCTTCCGATCTNNNACGTTTTCAGGGGAAATACTGG
GCACACGTCTGAACTCTTTCCCTACACGACGCTCTTCCGATCTNNNACGTTTTCAGGGGAAATACTGG
GAGCACACGTCTGAACTCTTTCCCTACACGACGCTCTTCCGATCTNNNTTGTCACTGGAATAGATGCA
GAGCACACGTCTGAACTCTTTCCCTACACGACGCTCTTCCGATCTNNNTTGTCACTGGAATAGATGCA
AGCACACGTCTGAACTCTTTCCCTACACGACGCTCTTCCGATCTNNNGATGATGGTAACCAGGTACAT
AGCACACGTCTGAACTCTTTCCCTACACGACGCTCTTCCGATCTNNNGATGATGGTAACCAGGTACAT
GAGCACACGTCTGAACTCTTTCCCTACACGACGCTCTTCCGATCTNNNTGAGTGTGATGTTAGTTTTG
GAGCACACGTCTGAACTCTTTCCCTACACGACGCTCTTCCGATCTNNNTGAGTGTGATGTTAGTTTTG
AGCACACGTCTGAACTCTTTCCCTACACGACGCTCTTCCGATCTNNNGGAGAATCGCTTGAACCTGGA
AGCACACGTCTGAACTCTTTCCCTACACGACGCTCTTCCGATCTNNNGGAGAATCGCTTGAACCTGGA
GAGCACACGTCTGAACTCTTTCCCTACACGACGCTCTTCCGATCTNNNAAGAGCATACCAACCCTACCT
GAGCACACGTCTGAACTCTTTCCCTACACGACGCTCTTCCGATCTNNNAAGAGCATACCAACCCTACCT
AGCACACGTCTGAACTCTTTCCCTACACGACGCTCTTCCGATCTNNNAGAATTGCTTGAATTTGGGAG
AGCACACGTCTGAACTCTTTCCCTACACGACGCTCTTCCGATCTNNNAGAATTGCTTGAATTTGGGAG
GAGCACACGTCTGAACTCTTTCCCTACACGACGCTCTTCCGATCTNNNGTGGTATCGCATTGTGGTTTT
GAGCACACGTCTGAACTCTTTCCCTACACGACGCTCTTCCGATCTNNNGTGGTATCGCATTGTGGTTTT

GCACACGTCTGAACTCTTTCCCTACACGACGCTCTTCCGATCTNNNGAAGCAGAAGAATCACTTGAG

GCACACGTCTGAACTCTTTCCCTACACGACGCTCTTCCGATCTNNNGAAGCAGAAGAATCACTTGAG

AGCACACGTCTGAACTCTTTCCCTACACGACGCTCTTCCGATCTNNNCAACATGGAGAAACCCCGTAT

AGCACACGTCTGAACTCTTTCCCTACACGACGCTCTTCCGATCTNNNCAACATGGAGAAACCCCGTAT

GCACACGTCTGAACTCTTTCCCTACACGACGCTCTTCCGATCTNNNGATGGGGCTCTAGGGACATAG

GCACACGTCTGAACTCTTTCCCTACACGACGCTCTTCCGATCTNNNGATGGGGCTCTAGGGACATAG

CACACGTCTGAACTCTTTCCCTACACGACGCTCTTCCGATCTNNNCCACGAAAACCCTAGTGAATCAC

CACACGTCTGAACTCTTTCCCTACACGACGCTCTTCCGATCTNNNCCACGAAAACCCTAGTGAATCAC

GCACACGTCTGAACTCTTTCCCTACACGACGCTCTTCCGATCTNNNGTGTGACCATCACCTATGTCTG

GCACACGTCTGAACTCTTTCCCTACACGACGCTCTTCCGATCTNNNGTGTGACCATCACCTATGTCTG

AGCACACGTCTGAACTCTTTCCCTACACGACGCTCTTCCGATCTNNNGAGGGGAGACTTCTCAACTCA

AGCACACGTCTGAACTCTTTCCCTACACGACGCTCTTCCGATCTNNNGAGGGGAGACTTCTCAACTCA

GCACACGTCTGAACTCTTTCCCTACACGACGCTCTTCCGATCTNNNAACTCGCTTGATTTTGAATACCA

GCACACGTCTGAACTCTTTCCCTACACGACGCTCTTCCGATCTNNNAACTCGCTTGATTTTGAATACCA

GCACACGTCTGAACTCTTTCCCTACACGACGCTCTTCCGATCTNNNACCAGAGCCAGATGATTTCACTT

GCACACGTCTGAACTCTTTCCCTACACGACGCTCTTCCGATCTNNNACCAGAGCCAGATGATTTCACTT

AGAGCACACGTCTGAACTCTTTCCCTACACGACGCTCTTCCGATCTNNNCTGGGGAGAACTAAGGCA

AGAGCACACGTCTGAACTCTTTCCCTACACGACGCTCTTCCGATCTNNNCTGGGGAGAACTAAGGCA

GCACACGTCTGAACTCTTTCCCTACACGACGCTCTTCCGATCTNNNGCATGCTCACTTGATACAAGGT

GCACACGTCTGAACTCTTTCCCTACACGACGCTCTTCCGATCTNNNGCATGCTCACTTGATACAAGGT

GAGCACACGTCTGAACTCTTTCCCTACACGACGCTCTTCCGATCTNNNTGGGTATCTGGATAGGGTTT

GAGCACACGTCTGAACTCTTTCCCTACACGACGCTCTTCCGATCTNNNTGGGTATCTGGATAGGGTTT

GCACACGTCTGAACTCTTTCCCTACACGACGCTCTTCCGATCTNNNTTTAGTAGAGACGGGGTTTCAC

GCACACGTCTGAACTCTTTCCCTACACGACGCTCTTCCGATCTNNNTTTAGTAGAGACGGGGTTTCAC

GAGCACACGTCTGAACTCTTTCCCTACACGACGCTCTTCCGATCTNNNCCAAACATTCTGTCCTGAAGC

GAGCACACGTCTGAACTCTTTCCCTACACGACGCTCTTCCGATCTNNNCCAAACATTCTGTCCTGAAGC

GCACACGTCTGAACTCTTTCCCTACACGACGCTCTTCCGATCTNNNTCCAAGACAGTAGACTTTGGG

GCACACGTCTGAACTCTTTCCCTACACGACGCTCTTCCGATCTNNNTCCAAGACAGTAGACTTTGGG

CACACGTCTGAACTCTTTCCCTACACGACGCTCTTCCGATCTNNNTCTATCCAGTCTGCCTGTCCTTACA

CACACGTCTGAACTCTTTCCCTACACGACGCTCTTCCGATCTNNNTCTATCCAGTCTGCCTGTCCTTACA

AGCACACGTCTGAACTCTTTCCCTACACGACGCTCTTCCGATCTNNNAGGATTGCTTGAGCCTAGGAA

AGCACACGTCTGAACTCTTTCCCTACACGACGCTCTTCCGATCTNNNAGGATTGCTTGAGCCTAGGAA

GCACACGTCTGAACTCTTTCCCTACACGACGCTCTTCCGATCTNNNTGCTCAGCATTCTCCCATCATAA

GCACACGTCTGAACTCTTTCCCTACACGACGCTCTTCCGATCTNNNTGCTCAGCATTCTCCCATCATAA

GAGCACACGTCTGAACTCTTTCCCTACACGACGCTCTTCCGATCTNNNGTGCTGTGACTCTGTGTTTTC

GAGCACACGTCTGAACTCTTTCCCTACACGACGCTCTTCCGATCTNNNGTGCTGTGACTCTGTGTTTTC

GCACACGTCTGAACTCTTTCCCTACACGACGCTCTTCCGATCTNNNGAGACAAGAGGCGAGGAGAA

GCACACGTCTGAACTCTTTCCCTACACGACGCTCTTCCGATCTNNNGAGACAAGAGGCGAGGAGAA

GAGCACACGTCTGAACTCTTTCCCTACACGACGCTCTTCCGATCTNNNAGGGACAACAAAGAGGAAC

GAGCACACGTCTGAACTCTTTCCCTACACGACGCTCTTCCGATCTNNNAGGGACAACAAAGAGGAAC

AGCACACGTCTGAACTCTTTCCCTACACGACGCTCTTCCGATCTNNNTCCAAAGGTACGTGGCTCATA

AGCACACGTCTGAACTCTTTCCCTACACGACGCTCTTCCGATCTNNNTCCAAAGGTACGTGGCTCATA

GAGCACACGTCTGAACTCTTTCCCTACACGACGCTCTTCCGATCTNNNGTGCCTGTAGTCCCATCTACT

GAGCACACGTCTGAACTCTTTCCCTACACGACGCTCTTCCGATCTNNNGTGCCTGTAGTCCCATCTACTC
GCACACGTCTGAACTCTTTCCCTACACGACGCTCTTCCGATCTNNNCTACCCTGTAGCTATGAGCAGTG
GCACACGTCTGAACTCTTTCCCTACACGACGCTCTTCCGATCTNNNCTACCCTGTAGCTATGAGCAGTG
AAGAGCACACGTCTGAACTCTTTCCCTACACGACGCTCTTCCGATCTNNNTAGCCTGGGTGACAAAGTC
AAGAGCACACGTCTGAACTCTTTCCCTACACGACGCTCTTCCGATCTNNNTAGCCTGGGTGACAAAGTC
GAGCACACGTCTGAACTCTTTCCCTACACGACGCTCTTCCGATCTNNNGAGGCTGAGGCAGGAGAATG
GAGCACACGTCTGAACTCTTTCCCTACACGACGCTCTTCCGATCTNNNGAGGCTGAGGCAGGAGAATG
GCACACGTCTGAACTCTTTCCCTACACGACGCTCTTCCGATCTNNNGGCACAACACTTTCTGCATTGTAC
GCACACGTCTGAACTCTTTCCCTACACGACGCTCTTCCGATCTNNNGGCACAACACTTTCTGCATTGTAC
AAGAGCACACGTCTGAACTCTTTCCCTACACGACGCTCTTCCGATCTNNNTTGAGGTTGCAGTGAGCTA
AAGAGCACACGTCTGAACTCTTTCCCTACACGACGCTCTTCCGATCTNNNTTGAGGTTGCAGTGAGCTA
AGCACACGTCTGAACTCTTTCCCTACACGACGCTCTTCCGATCTNNNTAGTGGCTTAGATAAAAAATCC
AGCACACGTCTGAACTCTTTCCCTACACGACGCTCTTCCGATCTNNNTAGTGGCTTAGATAAAAAATCC
AGAGCACACGTCTGAACTCTTTCCCTACACGACGCTCTTCCGATCTNNNGGGACTATGTCTCTTTCCTCT
AGAGCACACGTCTGAACTCTTTCCCTACACGACGCTCTTCCGATCTNNNGGGACTATGTCTCTTTCCTCT
GAGCACACGTCTGAACTCTTTCCCTACACGACGCTCTTCCGATCTNNNGGAGAATCACTCAAACCTGGC
GAGCACACGTCTGAACTCTTTCCCTACACGACGCTCTTCCGATCTNNNGGAGAATCACTCAAACCTGGC
GCACACGTCTGAACTCTTTCCCTACACGACGCTCTTCCGATCTNNNCTCCCATGGCCTTATTCCTCCATC
GCACACGTCTGAACTCTTTCCCTACACGACGCTCTTCCGATCTNNNCTCCCATGGCCTTATTCCTCCATC
AGAGCACACGTCTGAACTCTTTCCCTACACGACGCTCTTCCGATCTNNNTGTGTCCCTACTTCCATACCTC
AGAGCACACGTCTGAACTCTTTCCCTACACGACGCTCTTCCGATCTNNNTGTGTCCCTACTTCCATACCTC
GAGCACACGTCTGAACTCTTTCCCTACACGACGCTCTTCCGATCTNNNGCATCATGCCAAGTGATAAGA
GAGCACACGTCTGAACTCTTTCCCTACACGACGCTCTTCCGATCTNNNGCATCATGCCAAGTGATAAGA
AGCACACGTCTGAACTCTTTCCCTACACGACGCTCTTCCGATCTNNNTGACACCTGTAGAAGCAGAACT
AGCACACGTCTGAACTCTTTCCCTACACGACGCTCTTCCGATCTNNNTGACACCTGTAGAAGCAGAACT
GCACACGTCTGAACTCTTTCCCTACACGACGCTCTTCCGATCTNNNTTTCAAATAAAACTTGTACTGGGC
GCACACGTCTGAACTCTTTCCCTACACGACGCTCTTCCGATCTNNNTTTCAAATAAAACTTGTACTGGGC
AGAGCACACGTCTGAACTCTTTCCCTACACGACGCTCTTCCGATCTNNNTTTGTGTAGGGGGAAGAA
AGAGCACACGTCTGAACTCTTTCCCTACACGACGCTCTTCCGATCTNNNTTTGTGTAGGGGGAAGAA
CACACGTCTGAACTCTTTCCCTACACGACGCTCTTCCGATCTNNNGAAGATACTTACTGTAGTTCTCACAT
CACACGTCTGAACTCTTTCCCTACACGACGCTCTTCCGATCTNNNGAAGATACTTACTGTAGTTCTCACAT
GAGCACACGTCTGAACTCTTTCCCTACACGACGCTCTTCCGATCTNNNGAATGGGGTTGGGGAGAAT
GAGCACACGTCTGAACTCTTTCCCTACACGACGCTCTTCCGATCTNNNGAATGGGGTTGGGGAGAAT
GAGCACACGTCTGAACTCTTTCCCTACACGACGCTCTTCCGATCTNNNCACACTCATCTCTCCTAGAACCC
GAGCACACGTCTGAACTCTTTCCCTACACGACGCTCTTCCGATCTNNNCACACTCATCTCTCCTAGAACCC
GCACACGTCTGAACTCTTTCCCTACACGACGCTCTTCCGATCTNNNAACTCTTGCCTCCTTTCCTTCAGCA
GCACACGTCTGAACTCTTTCCCTACACGACGCTCTTCCGATCTNNNAACTCTTGCCTCCTTTCCTTCAGCA
GAGCACACGTCTGAACTCTTTCCCTACACGACGCTCTTCCGATCTNNNGTGCCTCTAGTTAGCCATCTTG
GAGCACACGTCTGAACTCTTTCCCTACACGACGCTCTTCCGATCTNNNGTGCCTCTAGTTAGCCATCTTG
GAGCACACGTCTGAACTCTTTCCCTACACGACGCTCTTCCGATCTNNNCTCCTGAGTAGCTGGGATTAC
GAGCACACGTCTGAACTCTTTCCCTACACGACGCTCTTCCGATCTNNNCTCCTGAGTAGCTGGGATTAC
GCACACGTCTGAACTCTTTCCCTACACGACGCTCTTCCGATCTNNNAAAGAAGCTCTTGCCTCACAGATC
GCACACGTCTGAACTCTTTCCCTACACGACGCTCTTCCGATCTNNNAAAGAAGCTCTTGCCTCACAGATC

AGCACACGTCTGAACTCTTTCCCTACACGACGCTCTTCCGATCTNNNGATTGACAGTTCTGTCTCCACCT

AGCACACGTCTGAACTCTTTCCCTACACGACGCTCTTCCGATCTNNNGATTGACAGTTCTGTCTCCACCT

GAGCACACGTCTGAACTCTTTCCCTACACGACGCTCTTCCGATCTNNNCTGTAATCCCAGCTACTCGGAA

GAGCACACGTCTGAACTCTTTCCCTACACGACGCTCTTCCGATCTNNNCTGTAATCCCAGCTACTCGGAA

AGCACACGTCTGAACTCTTTCCCTACACGACGCTCTTCCGATCTNNNGGTGTGAGGGGAAGAAAGAGA

AGCACACGTCTGAACTCTTTCCCTACACGACGCTCTTCCGATCTNNNGGTGTGAGGGGAAGAAAGAGA

GAGCACACGTCTGAACTCTTTCCCTACACGACGCTCTTCCGATCTNNNCTTGAGGTCGGGAGTTCTAG

GAGCACACGTCTGAACTCTTTCCCTACACGACGCTCTTCCGATCTNNNCTTGAGGTCGGGAGTTCTAG

GAAGAGCACACGTCTGAACTCTTTCCCTACACGACGCTCTTCCGATCTNNNACCAAAAGATACAGGGG

GAAGAGCACACGTCTGAACTCTTTCCCTACACGACGCTCTTCCGATCTNNNACCAAAAGATACAGGGG

GAGCACACGTCTGAACTCTTTCCCTACACGACGCTCTTCCGATCTNNNACTGCATTACGTTTCCACCTAC

GAGCACACGTCTGAACTCTTTCCCTACACGACGCTCTTCCGATCTNNNACTGCATTACGTTTCCACCTAC

AGCACACGTCTGAACTCTTTCCCTACACGACGCTCTTCCGATCTNNNACTTATGGGTGAATCATGTTATA

AGCACACGTCTGAACTCTTTCCCTACACGACGCTCTTCCGATCTNNNACTTATGGGTGAATCATGTTATA

GAGCACACGTCTGAACTCTTTCCCTACACGACGCTCTTCCGATCTNNNACAATCATAAGTCACAGGAGG

GAGCACACGTCTGAACTCTTTCCCTACACGACGCTCTTCCGATCTNNNACAATCATAAGTCACAGGAGG

AGCACACGTCTGAACTCTTTCCCTACACGACGCTCTTCCGATCTNNNACCATCTTCAGAAAAATGAGCC

AGCACACGTCTGAACTCTTTCCCTACACGACGCTCTTCCGATCTNNNACCATCTTCAGAAAAATGAGCC

AGCACACGTCTGAACTCTTTCCCTACACGACGCTCTTCCGATCTNNNGCCTGTAGTCCCAGCTACTTGCA

AGCACACGTCTGAACTCTTTCCCTACACGACGCTCTTCCGATCTNNNGCCTGTAGTCCCAGCTACTTGCA

AGCACACGTCTGAACTCTTTCCCTACACGACGCTCTTCCGATCTNNNAGCACAGTACCCCTACAACTTTT

AGCACACGTCTGAACTCTTTCCCTACACGACGCTCTTCCGATCTNNNAGCACAGTACCCCTACAACTTTT

GCACACGTCTGAACTCTTTCCCTACACGACGCTCTTCCGATCTNNNCATCATGCTTCAGTGGTCACATGC

GCACACGTCTGAACTCTTTCCCTACACGACGCTCTTCCGATCTNNNCATCATGCTTCAGTGGTCACATGC

AGCACACGTCTGAACTCTTTCCCTACACGACGCTCTTCCGATCTNNNCTCTCCCTTCCTTTCTTCCTTCCCA

AGCACACGTCTGAACTCTTTCCCTACACGACGCTCTTCCGATCTNNNCTCTCCCTTCCTTTCTTCCTTCCCA

AGCACACGTCTGAACTCTTTCCCTACACGACGCTCTTCCGATCTNNNATTCTCCTGCCTCAGTATCCCAAC

AGCACACGTCTGAACTCTTTCCCTACACGACGCTCTTCCGATCTNNNATTCTCCTGCCTCAGTATCCCAAC

AGCACACGTCTGAACTCTTTCCCTACACGACGCTCTTCCGATCTNNNCTTGAACTCCTGACCTCAAGTGA

AGCACACGTCTGAACTCTTTCCCTACACGACGCTCTTCCGATCTNNNCTTGAACTCCTGACCTCAAGTGA

AGCACACGTCTGAACTCTTTCCCTACACGACGCTCTTCCGATCTNNNGCTCATGTCTGCCTAACTACCTAC

AGCACACGTCTGAACTCTTTCCCTACACGACGCTCTTCCGATCTNNNGCTCATGTCTGCCTAACTACCTAC

GAGCACACGTCTGAACTCTTTCCCTACACGACGCTCTTCCGATCTNNNCACTTGAGGTCAGGAGTTTG

GAGCACACGTCTGAACTCTTTCCCTACACGACGCTCTTCCGATCTNNNCACTTGAGGTCAGGAGTTTG

AGCACACGTCTGAACTCTTTCCCTACACGACGCTCTTCCGATCTNNNCAGAGTCTCTCCTCTCTTCCTCTC

AGCACACGTCTGAACTCTTTCCCTACACGACGCTCTTCCGATCTNNNCAGAGTCTCTCCTCTCTTCCTCTC

GAGCACACGTCTGAACTCTTTCCCTACACGACGCTCTTCCGATCTNNNAGAGTAGCTGGGACTACAGGC

GAGCACACGTCTGAACTCTTTCCCTACACGACGCTCTTCCGATCTNNNAGAGTAGCTGGGACTACAGGC

AGCACACGTCTGAACTCTTTCCCTACACGACGCTCTTCCGATCTNNNCTCCCAGCCTCAAAAGAACAATC

AGCACACGTCTGAACTCTTTCCCTACACGACGCTCTTCCGATCTNNNCTCCCAGCCTCAAAAGAACAATC

GCACACGTCTGAACTCTTTCCCTACACGACGCTCTTCCGATCTNNNTACCAGTCTTGACCCCAAACAATC

GCACACGTCTGAACTCTTTCCCTACACGACGCTCTTCCGATCTNNNTACCAGTCTTGACCCCAAACAATC

GAGCACACGTCTGAACTCTTTCCCTACACGACGCTCTTCCGATCTNNNAAGGCAGTGTGTGATTTAGGC

AGCACACGTCTGAACTCTTTCCCTACACGACGCTCTTCCGATCTNNNAAGGCAGTGTGTGATTTAGG
AGCACACGTCTGAACTCTTTCCCTACACGACGCTCTTCCGATCTNNNTCCCGCAGTCTTATTTATACCCG
AGCACACGTCTGAACTCTTTCCCTACACGACGCTCTTCCGATCTNNNTCCCGCAGTCTTATTTATACCCG
GAGCACACGTCTGAACTCTTTCCCTACACGACGCTCTTCCGATCTNNNGGGACACCAGCAATAGTAAC
GAGCACACGTCTGAACTCTTTCCCTACACGACGCTCTTCCGATCTNNNGGGACACCAGCAATAGTAAC
GAGCACACGTCTGAACTCTTTCCCTACACGACGCTCTTCCGATCTNNNCCAGCAGATTCGGTGATAGA
GAGCACACGTCTGAACTCTTTCCCTACACGACGCTCTTCCGATCTNNNCCAGCAGATTCGGTGATAGA
AGCACACGTCTGAACTCTTTCCCTACACGACGCTCTTCCGATCTNNNTCTCCCTCCATTCCTCTAGTG
AGCACACGTCTGAACTCTTTCCCTACACGACGCTCTTCCGATCTNNNTCTCCCTCCATTCCTCTAGTG
AGCACACGTCTGAACTCTTTCCCTACACGACGCTCTTCCGATCTNNNGTGCCTGTAGTCCCAGCTATTT
AGCACACGTCTGAACTCTTTCCCTACACGACGCTCTTCCGATCTNNNGTGCCTGTAGTCCCAGCTATTT
AGCACACGTCTGAACTCTTTCCCTACACGACGCTCTTCCGATCTNNNCTTGTTTTCTTCCCAGGCATCT
AGCACACGTCTGAACTCTTTCCCTACACGACGCTCTTCCGATCTNNNCTTGTTTTCTTCCCAGGCATCT
GAGCACACGTCTGAACTCTTTCCCTACACGACGCTCTTCCGATCTNNNTATGTGGTCTCAGCTACTCA
GAGCACACGTCTGAACTCTTTCCCTACACGACGCTCTTCCGATCTNNNTATGTGGTCTCAGCTACTCA
CACACGTCTGAACTCTTTCCCTACACGACGCTCTTCCGATCTNNNCTGTGTCCATGTCAGAGCAACAG
CACACGTCTGAACTCTTTCCCTACACGACGCTCTTCCGATCTNNNCTGTGTCCATGTCAGAGCAACAG
AGCACACGTCTGAACTCTTTCCCTACACGACGCTCTTCCGATCTNNNAGCTTTTTGATTTGCTGCTGG
AGCACACGTCTGAACTCTTTCCCTACACGACGCTCTTCCGATCTNNNAGCTTTTTGATTTGCTGCTGG
AGCACACGTCTGAACTCTTTCCCTACACGACGCTCTTCCGATCTNNNGAGCTTGCAATGAGCCAAGAT
AGCACACGTCTGAACTCTTTCCCTACACGACGCTCTTCCGATCTNNNGAGCTTGCAATGAGCCAAGAT
AGCACACGTCTGAACTCTTTCCCTACACGACGCTCTTCCGATCTNNNGACGAAGAATGGTCCCTGTAC
AGCACACGTCTGAACTCTTTCCCTACACGACGCTCTTCCGATCTNNNGACGAAGAATGGTCCCTGTAC
AGCACACGTCTGAACTCTTTCCCTACACGACGCTCTTCCGATCTNNNGTGATCTTGGAGCCCCTAAGA
AGCACACGTCTGAACTCTTTCCCTACACGACGCTCTTCCGATCTNNNGTGATCTTGGAGCCCCTAAGA
AGCACACGTCTGAACTCTTTCCCTACACGACGCTCTTCCGATCTNNNCTCCAGCTGCACAAAAGCAATA
AGCACACGTCTGAACTCTTTCCCTACACGACGCTCTTCCGATCTNNNCTCCAGCTGCACAAAAGCAATA
AGCACACGTCTGAACTCTTTCCCTACACGACGCTCTTCCGATCTNNNATGATGGTGATGATGGTGATG
AGCACACGTCTGAACTCTTTCCCTACACGACGCTCTTCCGATCTNNNATGATGGTGATGATGGTGATG
AGCACACGTCTGAACTCTTTCCCTACACGACGCTCTTCCGATCTNNNAGAGGCCCTCTTACTACAGAG
AGCACACGTCTGAACTCTTTCCCTACACGACGCTCTTCCGATCTNNNAGAGGCCCTCTTACTACAGAG
GAGCACACGTCTGAACTCTTTCCCTACACGACGCTCTTCCGATCTNNNGGCTGAGACAGAAGAATTG
GAGCACACGTCTGAACTCTTTCCCTACACGACGCTCTTCCGATCTNNNGGCTGAGACAGAAGAATTG
AGCACACGTCTGAACTCTTTCCCTACACGACGCTCTTCCGATCTNNNTTGCAAGTAATTACAATGATGA
AGCACACGTCTGAACTCTTTCCCTACACGACGCTCTTCCGATCTNNNTTGCAAGTAATTACAATGATGA
GAGCACACGTCTGAACTCTTTCCCTACACGACGCTCTTCCGATCTNNNTCAATTACCATCCTTCCAGAAC
AGCACACGTCTGAACTCTTTCCCTACACGACGCTCTTCCGATCTNNNTCAATTACCATCCTTCCAGAAC
AGCACACGTCTGAACTCTTTCCCTACACGACGCTCTTCCGATCTNNNCAAGATGGGAGGATCACTAGA
AGCACACGTCTGAACTCTTTCCCTACACGACGCTCTTCCGATCTNNNCAAGATGGGAGGATCACTAGA
AGCACACGTCTGAACTCTTTCCCTACACGACGCTCTTCCGATCTNNNTCCATCCCTTTCCCAAATCTCTG
AGCACACGTCTGAACTCTTTCCCTACACGACGCTCTTCCGATCTNNNTCCATCCCTTTCCCAAATCTCTG
GCACACGTCTGAACTCTTTCCCTACACGACGCTCTTCCGATCTNNNCCATAAGAAGCAAGCACCTAAG
GCACACGTCTGAACTCTTTCCCTACACGACGCTCTTCCGATCTNNNCCATAAGAAGCAAGCACCTAAG

AGCACACGTCTGAACTCTTTCCCTACACGACGCTCTTCCGATCTNNNGCATCCTTGTCTTGTTCCAGTTC
AGCACACGTCTGAACTCTTTCCCTACACGACGCTCTTCCGATCTNNNGCATCCTTGTCTTGTTCCAGTTC
AAGAGCACACGTCTGAACTCTTTCCCTACACGACGCTCTTCCGATCTNNNTGTCTGATGTTGGATGTCC
AAGAGCACACGTCTGAACTCTTTCCCTACACGACGCTCTTCCGATCTNNNTGTCTGATGTTGGATGTCC
AGCACACGTCTGAACTCTTTCCCTACACGACGCTCTTCCGATCTNNNATTACAAACAGGCCAACATGTCC
AGCACACGTCTGAACTCTTTCCCTACACGACGCTCTTCCGATCTNNNATTACAAACAGGCCAACATGTCC
ACACGTCTGAACTCTTTCCCTACACGACGCTCTTCCGATCTNNNGAGATCACCTCGGCCCCTTGCATCGC
ACACGTCTGAACTCTTTCCCTACACGACGCTCTTCCGATCTNNNGAGATCACCTCGGCCCCTTGCATCGC
GCACACGTCTGAACTCTTTCCCTACACGACGCTCTTCCGATCTNNNGGAGACCAGCAGCACAGTATATA
GCACACGTCTGAACTCTTTCCCTACACGACGCTCTTCCGATCTNNNGGAGACCAGCAGCACAGTATATA
GAGCACACGTCTGAACTCTTTCCCTACACGACGCTCTTCCGATCTNNNGTCCAACCTCTCATCACTACCTG
GAGCACACGTCTGAACTCTTTCCCTACACGACGCTCTTCCGATCTNNNGTCCAACCTCTCATCACTACCTG
AGAGCACACGTCTGAACTCTTTCCCTACACGACGCTCTTCCGATCTNNNCTGTAATCCCAGATACTCGGA
AGAGCACACGTCTGAACTCTTTCCCTACACGACGCTCTTCCGATCTNNNCTGTAATCCCAGATACTCGGA
AGCACACGTCTGAACTCTTTCCCTACACGACGCTCTTCCGATCTNNNTAAAGCCCTTTCTTGGCTGCAAT
AGCACACGTCTGAACTCTTTCCCTACACGACGCTCTTCCGATCTNNNTAAAGCCCTTTCTTGGCTGCAAT
AGAGCACACGTCTGAACTCTTTCCCTACACGACGCTCTTCCGATCTNNNGGGGATGGAAGTCATGGTT
AGAGCACACGTCTGAACTCTTTCCCTACACGACGCTCTTCCGATCTNNNGGGGATGGAAGTCATGGTT
AGCACACGTCTGAACTCTTTCCCTACACGACGCTCTTCCGATCTNNNGCCACTGTTTACCAAGACACTGA
AGCACACGTCTGAACTCTTTCCCTACACGACGCTCTTCCGATCTNNNGCCACTGTTTACCAAGACACTGA
AGCACACGTCTGAACTCTTTCCCTACACGACGCTCTTCCGATCTNNNAAACAGGGAGGGGAACATCAT
AGCACACGTCTGAACTCTTTCCCTACACGACGCTCTTCCGATCTNNNAAACAGGGAGGGGAACATCAT
AGAGCACACGTCTGAACTCTTTCCCTACACGACGCTCTTCCGATCTNNNTCACGCCTGTAATCTTAGCAC
AGAGCACACGTCTGAACTCTTTCCCTACACGACGCTCTTCCGATCTNNNTCACGCCTGTAATCTTAGCAC
AGCACACGTCTGAACTCTTTCCCTACACGACGCTCTTCCGATCTNNNTCCAAATTCCACCACAGATTTCAC
AGCACACGTCTGAACTCTTTCCCTACACGACGCTCTTCCGATCTNNNTCCAAATTCCACCACAGATTTCAC
AGCACACGTCTGAACTCTTTCCCTACACGACGCTCTTCCGATCTNNNTCTCGTGACTTTGTCATTGGGA
AGCACACGTCTGAACTCTTTCCCTACACGACGCTCTTCCGATCTNNNTCTCGTGACTTTGTCATTGGGA
AGCACACGTCTGAACTCTTTCCCTACACGACGCTCTTCCGATCTNNNACCCCCTCGACTATAGTTAGCAT
AGCACACGTCTGAACTCTTTCCCTACACGACGCTCTTCCGATCTNNNACCCCCTCGACTATAGTTAGCAT
AGCACACGTCTGAACTCTTTCCCTACACGACGCTCTTCCGATCTNNNGGCCAGCATCATCCTGATACTAA
AGCACACGTCTGAACTCTTTCCCTACACGACGCTCTTCCGATCTNNNGGCCAGCATCATCCTGATACTAA
GAGCACACGTCTGAACTCTTTCCCTACACGACGCTCTTCCGATCTNNNGTGCACATGGCATAAAATCCGT
GAGCACACGTCTGAACTCTTTCCCTACACGACGCTCTTCCGATCTNNNGTGCACATGGCATAAAATCCGT
AGCACACGTCTGAACTCTTTCCCTACACGACGCTCTTCCGATCTNNNTCTCCGACCTTGCATGTTTATCAC
AGCACACGTCTGAACTCTTTCCCTACACGACGCTCTTCCGATCTNNNTCTCCGACCTTGCATGTTTATCAC
AGCACACGTCTGAACTCTTTCCCTACACGACGCTCTTCCGATCTNNNAGTGAAGTCAAAAAGGAGTGA
AGCACACGTCTGAACTCTTTCCCTACACGACGCTCTTCCGATCTNNNAGTGAAGTCAAAAAGGAGTGA
AGCACACGTCTGAACTCTTTCCCTACACGACGCTCTTCCGATCTNNNATACCAAAAACCCAAAGCAAGC
AGCACACGTCTGAACTCTTTCCCTACACGACGCTCTTCCGATCTNNNATACCAAAAACCCAAAGCAAGC
AGCACACGTCTGAACTCTTTCCCTACACGACGCTCTTCCGATCTNNNACATACCTCTGTTCTGGCTCTGT
AGCACACGTCTGAACTCTTTCCCTACACGACGCTCTTCCGATCTNNNACATACCTCTGTTCTGGCTCTGT
AGCACACGTCTGAACTCTTTCCCTACACGACGCTCTTCCGATCTNNNAAGTATAAAAAATCAGCCGGG

AGCACACGTCTGAACTCTTTCCCTACACGACGCTCTTCCGATCTNNNAAGTATAAAAAATCAGCCGGG
AGCACACGTCTGAACTCTTTCCCTACACGACGCTCTTCCGATCTNNNAGGCAGTCTCTAGTTCTCTTCCT
AGCACACGTCTGAACTCTTTCCCTACACGACGCTCTTCCGATCTNNNAGGCAGTCTCTAGTTCTCTTCCT
GCACACGTCTGAACTCTTTCCCTACACGACGCTCTTCCGATCTNNNAAGTTATAAGGAGAGGGGAGG
GCACACGTCTGAACTCTTTCCCTACACGACGCTCTTCCGATCTNNNAAGTTATAAGGAGAGGGGAGG
AGCACACGTCTGAACTCTTTCCCTACACGACGCTCTTCCGATCTNNNGTGGGGGCCAATATTCAACATT
AGCACACGTCTGAACTCTTTCCCTACACGACGCTCTTCCGATCTNNNGTGGGGGCCAATATTCAACATT
AGCACACGTCTGAACTCTTTCCCTACACGACGCTCTTCCGATCTNNNCCACCATAGGAAGGACAAATTG
AGCACACGTCTGAACTCTTTCCCTACACGACGCTCTTCCGATCTNNNCCACCATAGGAAGGACAAATTG
AGCACACGTCTGAACTCTTTCCCTACACGACGCTCTTCCGATCTNNNACATATGTTGGTGGATGCTGTT
AGCACACGTCTGAACTCTTTCCCTACACGACGCTCTTCCGATCTNNNACATATGTTGGTGGATGCTGTT
GAGCACACGTCTGAACTCTTTCCCTACACGACGCTCTTCCGATCTNNNAGAAGTTATTTTGCCTCCCTG
GAGCACACGTCTGAACTCTTTCCCTACACGACGCTCTTCCGATCTNNNAGAAGTTATTTTGCCTCCCTG
GAGCACACGTCTGAACTCTTTCCCTACACGACGCTCTTCCGATCTNNNTGGAGGGGCCTAGAAATATT
GAGCACACGTCTGAACTCTTTCCCTACACGACGCTCTTCCGATCTNNNTGGAGGGGCCTAGAAATATT
GCACACGTCTGAACTCTTTCCCTACACGACGCTCTTCCGATCTNNNACAGGTTGAGCATCCCTTATCTG
GCACACGTCTGAACTCTTTCCCTACACGACGCTCTTCCGATCTNNNACAGGTTGAGCATCCCTTATCTG
AGCACACGTCTGAACTCTTTCCCTACACGACGCTCTTCCGATCTNNNCTTGTACATGAGTTTCATCCCGC
AGCACACGTCTGAACTCTTTCCCTACACGACGCTCTTCCGATCTNNNCTTGTACATGAGTTTCATCCCGC
GCACACGTCTGAACTCTTTCCCTACACGACGCTCTTCCGATCTNNNCCCTGATGGCTAATGATGTTCCTC
GCACACGTCTGAACTCTTTCCCTACACGACGCTCTTCCGATCTNNNCCCTGATGGCTAATGATGTTCCTC
AGCACACGTCTGAACTCTTTCCCTACACGACGCTCTTCCGATCTNNNCAGTGACGGATTAGACGCTAA
AGCACACGTCTGAACTCTTTCCCTACACGACGCTCTTCCGATCTNNNCAGTGACGGATTAGACGCTAA
GAGCACACGTCTGAACTCTTTCCCTACACGACGCTCTTCCGATCTNNNTTGTGTTTTCTCTGTCTCCCACC
GAGCACACGTCTGAACTCTTTCCCTACACGACGCTCTTCCGATCTNNNTTGTGTTTTCTCTGTCTCCCACC
GAGCACACGTCTGAACTCTTTCCCTACACGACGCTCTTCCGATCTNNNTTCAAGCTGAGGCAGGAGAA
GAGCACACGTCTGAACTCTTTCCCTACACGACGCTCTTCCGATCTNNNTTCAAGCTGAGGCAGGAGAA
GCACACGTCTGAACTCTTTCCCTACACGACGCTCTTCCGATCTNNNATAGACTCCAATCGCATTCTTCATC
GCACACGTCTGAACTCTTTCCCTACACGACGCTCTTCCGATCTNNNATAGACTCCAATCGCATTCTTCATC
AGCACACGTCTGAACTCTTTCCCTACACGACGCTCTTCCGATCTNNNCATGGGCAGGTGAAAAGAAGA
AGCACACGTCTGAACTCTTTCCCTACACGACGCTCTTCCGATCTNNNCATGGGCAGGTGAAAAGAAGA
GAGCACACGTCTGAACTCTTTCCCTACACGACGCTCTTCCGATCTNNNTGCTGCGAATGCCATTATTTCT
GAGCACACGTCTGAACTCTTTCCCTACACGACGCTCTTCCGATCTNNNTGCTGCGAATGCCATTATTTCT
AGCACACGTCTGAACTCTTTCCCTACACGACGCTCTTCCGATCTNNNTTTGCAAAGCCAAGAAGTGTG
AGCACACGTCTGAACTCTTTCCCTACACGACGCTCTTCCGATCTNNNTTTGCAAAGCCAAGAAGTGTG
GAGCACACGTCTGAACTCTTTCCCTACACGACGCTCTTCCGATCTNNNCCCCAGAAAAGCACTTGACAA
GAGCACACGTCTGAACTCTTTCCCTACACGACGCTCTTCCGATCTNNNCCCCAGAAAAGCACTTGACAA
GAGCACACGTCTGAACTCTTTCCCTACACGACGCTCTTCCGATCTNNNACTCCTGAAATCTGCTCACCAA
GAGCACACGTCTGAACTCTTTCCCTACACGACGCTCTTCCGATCTNNNACTCCTGAAATCTGCTCACCAA
CACGTCTGAACTCTTTCCCTACACGACGCTCTTCCGATCTNNNTCCTTTGACTAGAGAAAGCTGGCCAT
CACGTCTGAACTCTTTCCCTACACGACGCTCTTCCGATCTNNNTCCTTTGACTAGAGAAAGCTGGCCAT
GCACACGTCTGAACTCTTTCCCTACACGACGCTCTTCCGATCTNNNTTCAAGGTCATTGTTCCAGCAAG
GCACACGTCTGAACTCTTTCCCTACACGACGCTCTTCCGATCTNNNTTCAAGGTCATTGTTCCAGCAAG

AGAGCACACGTCTGAACTCTTTCCCTACACGACGCTCTTCCGATCTNNNTGAGGTAGGAGAATCGCTTC
AGAGCACACGTCTGAACTCTTTCCCTACACGACGCTCTTCCGATCTNNNTGAGGTAGGAGAATCGCTTC
GAGCACACGTCTGAACTCTTTCCCTACACGACGCTCTTCCGATCTNNNTGTACTGCTTACTCTCCAAAGG
GAGCACACGTCTGAACTCTTTCCCTACACGACGCTCTTCCGATCTNNNTGTACTGCTTACTCTCCAAAGG
AGCACACGTCTGAACTCTTTCCCTACACGACGCTCTTCCGATCTNNNCTTCAGCCTGGATGACAAGAGT
AGCACACGTCTGAACTCTTTCCCTACACGACGCTCTTCCGATCTNNNCTTCAGCCTGGATGACAAGAGT
AGCACACGTCTGAACTCTTTCCCTACACGACGCTCTTCCGATCTNNNGCCTGGCCATTATGTAGCTTTT
AGCACACGTCTGAACTCTTTCCCTACACGACGCTCTTCCGATCTNNNGCCTGGCCATTATGTAGCTTTT
AAGAGCACACGTCTGAACTCTTTCCCTACACGACGCTCTTCCGATCTNNNTCGGAAAATATCACCATGC
AAGAGCACACGTCTGAACTCTTTCCCTACACGACGCTCTTCCGATCTNNNTCGGAAAATATCACCATGC
GAGCACACGTCTGAACTCTTTCCCTACACGACGCTCTTCCGATCTNNNGTGCTCTTCTCCTTATAGCCACA
GAGCACACGTCTGAACTCTTTCCCTACACGACGCTCTTCCGATCTNNNGTGCTCTTCTCCTTATAGCCACA
AGAGCACACGTCTGAACTCTTTCCCTACACGACGCTCTTCCGATCTNNNGCCAAGTGTTCGTGAGTTTT
AGAGCACACGTCTGAACTCTTTCCCTACACGACGCTCTTCCGATCTNNNGCCAAGTGTTCGTGAGTTTT
GAGCACACGTCTGAACTCTTTCCCTACACGACGCTCTTCCGATCTNNNGGATGGTCTTGAACTCCTGAC
GAGCACACGTCTGAACTCTTTCCCTACACGACGCTCTTCCGATCTNNNGGATGGTCTTGAACTCCTGAC
AGCACACGTCTGAACTCTTTCCCTACACGACGCTCTTCCGATCTNNNGGGCTACAGTGTAGGACTTTGA
AGCACACGTCTGAACTCTTTCCCTACACGACGCTCTTCCGATCTNNNGGGCTACAGTGTAGGACTTTGA
AGCACACGTCTGAACTCTTTCCCTACACGACGCTCTTCCGATCTNNNCATCCTGGTGGGTACAGAGTTT
AGCACACGTCTGAACTCTTTCCCTACACGACGCTCTTCCGATCTNNNCATCCTGGTGGGTACAGAGTTT
AGAGCACACGTCTGAACTCTTTCCCTACACGACGCTCTTCCGATCTNNNTGGTGGCACAGCTATTCATAA
AGAGCACACGTCTGAACTCTTTCCCTACACGACGCTCTTCCGATCTNNNTGGTGGCACAGCTATTCATAA
GAGCACACGTCTGAACTCTTTCCCTACACGACGCTCTTCCGATCTNNNCATGATCACACCACTGTACTCCA
GAGCACACGTCTGAACTCTTTCCCTACACGACGCTCTTCCGATCTNNNCATGATCACACCACTGTACTCCA
AGCACACGTCTGAACTCTTTCCCTACACGACGCTCTTCCGATCTNNNGGAAGGAGGAGTTTGCAGTG
AGCACACGTCTGAACTCTTTCCCTACACGACGCTCTTCCGATCTNNNGGAAGGAGGAGTTTGCAGTG
AGAGCACACGTCTGAACTCTTTCCCTACACGACGCTCTTCCGATCTNNNCTCTTCCAGAAGCTTCTACCCA
AGAGCACACGTCTGAACTCTTTCCCTACACGACGCTCTTCCGATCTNNNCTCTTCCAGAAGCTTCTACCCA
AGCACACGTCTGAACTCTTTCCCTACACGACGCTCTTCCGATCTNNNCGCACCTGTAATCCTAGCTACTC
AGCACACGTCTGAACTCTTTCCCTACACGACGCTCTTCCGATCTNNNCGCACCTGTAATCCTAGCTACTC
ACACGTCTGAACTCTTTCCCTACACGACGCTCTTCCGATCTNNNCTCCTCACTTCCTAGTAGGGGCCATC
ACACGTCTGAACTCTTTCCCTACACGACGCTCTTCCGATCTNNNCTCCTCACTTCCTAGTAGGGGCCATC
AGAGCACACGTCTGAACTCTTTCCCTACACGACGCTCTTCCGATCTNNNGAGACCTCAGTGATTTGGGC
AGAGCACACGTCTGAACTCTTTCCCTACACGACGCTCTTCCGATCTNNNGAGACCTCAGTGATTTGGGC
GAGCACACGTCTGAACTCTTTCCCTACACGACGCTCTTCCGATCTNNNCAGCCTGGACAACAAGAGAGA
GAGCACACGTCTGAACTCTTTCCCTACACGACGCTCTTCCGATCTNNNCAGCCTGGACAACAAGAGAGA
GAGCACACGTCTGAACTCTTTCCCTACACGACGCTCTTCCGATCTNNNCCAAACACTGTACTGACGTCCA
GAGCACACGTCTGAACTCTTTCCCTACACGACGCTCTTCCGATCTNNNCCAAACACTGTACTGACGTCCA
GCACACGTCTGAACTCTTTCCCTACACGACGCTCTTCCGATCTNNNTGAAGACCCTGTCAGGTATCTCTC
GCACACGTCTGAACTCTTTCCCTACACGACGCTCTTCCGATCTNNNTGAAGACCCTGTCAGGTATCTCTC
GAGCACACGTCTGAACTCTTTCCCTACACGACGCTCTTCCGATCTNNNCACCCACCTTTTGGGTTGTTACT
GAGCACACGTCTGAACTCTTTCCCTACACGACGCTCTTCCGATCTNNNCACCCACCTTTTGGGTTGTTACT
GAGCACACGTCTGAACTCTTTCCCTACACGACGCTCTTCCGATCTNNNGCTTCCCAAGTAGCTAGGACT

GAGCACACGTCTGAACTCTTTCCCTACACGACGCTCTTCCGATCTNNNGCTTCCCAAGTAGCTAGGACT

GAGCACACGTCTGAACTCTTTCCCTACACGACGCTCTTCCGATCTNNNCCAGACTTCTACCTGTGCTCAT

GAGCACACGTCTGAACTCTTTCCCTACACGACGCTCTTCCGATCTNNNCCAGACTTCTACCTGTGCTCAT

AGCACACGTCTGAACTCTTTCCCTACACGACGCTCTTCCGATCTNNNCCCCATGAAGATCCTTAACACA

AGCACACGTCTGAACTCTTTCCCTACACGACGCTCTTCCGATCTNNNCCCCATGAAGATCCTTAACACA

AGAGCACACGTCTGAACTCTTTCCCTACACGACGCTCTTCCGATCTNNNCAACAGCTGGTGTCTTGGTTT

AGAGCACACGTCTGAACTCTTTCCCTACACGACGCTCTTCCGATCTNNNCAACAGCTGGTGTCTTGGTTT

AGCACACGTCTGAACTCTTTCCCTACACGACGCTCTTCCGATCTNNNCGACAGTTCAGCCTTATCCTCTG

AGCACACGTCTGAACTCTTTCCCTACACGACGCTCTTCCGATCTNNNCGACAGTTCAGCCTTATCCTCTG

AGCACACGTCTGAACTCTTTCCCTACACGACGCTCTTCCGATCTNNNCATCTATCACGTCTGGATGCAGA

AGCACACGTCTGAACTCTTTCCCTACACGACGCTCTTCCGATCTNNNCATCTATCACGTCTGGATGCAGA

AGCACACGTCTGAACTCTTTCCCTACACGACGCTCTTCCGATCTNNNATCAGAGACTGCCATCTTGGTAC

AGCACACGTCTGAACTCTTTCCCTACACGACGCTCTTCCGATCTNNNATCAGAGACTGCCATCTTGGTAC

GAGCACACGTCTGAACTCTTTCCCTACACGACGCTCTTCCGATCTNNNTCTCCCTCTACGAACTGCTTTTC

GAGCACACGTCTGAACTCTTTCCCTACACGACGCTCTTCCGATCTNNNTCTCCCTCTACGAACTGCTTTTC

AGCACACGTCTGAACTCTTTCCCTACACGACGCTCTTCCGATCTNNNTCTTTGGGATTTTGAGCCATGTC

AGCACACGTCTGAACTCTTTCCCTACACGACGCTCTTCCGATCTNNNTCTTTGGGATTTTGAGCCATGTC

AGAGCACACGTCTGAACTCTTTCCCTACACGACGCTCTTCCGATCTNNNTGAGGCAGGAGAATCACTTC

AGAGCACACGTCTGAACTCTTTCCCTACACGACGCTCTTCCGATCTNNNTGAGGCAGGAGAATCACTTC

GAGCACACGTCTGAACTCTTTCCCTACACGACGCTCTTCCGATCTNNNCTAACGTGTGTGTTCATGCATC

GAGCACACGTCTGAACTCTTTCCCTACACGACGCTCTTCCGATCTNNNCTAACGTGTGTGTTCATGCATC

AGCACACGTCTGAACTCTTTCCCTACACGACGCTCTTCCGATCTNNNCATTGTGTGTGTGTGTGTGTGT

AGCACACGTCTGAACTCTTTCCCTACACGACGCTCTTCCGATCTNNNCATTGTGTGTGTGTGTGTGTGT

GCACACGTCTGAACTCTTTCCCTACACGACGCTCTTCCGATCTNNNTGTTGAGAAATTGCAGAGTGCTG

GCACACGTCTGAACTCTTTCCCTACACGACGCTCTTCCGATCTNNNTGTTGAGAAATTGCAGAGTGCTG

GAGCACACGTCTGAACTCTTTCCCTACACGACGCTCTTCCGATCTNNNAGACCAAAACTTAGCCCAGCTA

GAGCACACGTCTGAACTCTTTCCCTACACGACGCTCTTCCGATCTNNNAGACCAAAACTTAGCCCAGCTA

GCACACGTCTGAACTCTTTCCCTACACGACGCTCTTCCGATCTNNNTTGATAGTACTCTGTCATTGTCCAT

GCACACGTCTGAACTCTTTCCCTACACGACGCTCTTCCGATCTNNNTTGATAGTACTCTGTCATTGTCCAT

AGAGCACACGTCTGAACTCTTTCCCTACACGACGCTCTTCCGATCTNNNCCAACATGGTGAAACTCCATC

AGAGCACACGTCTGAACTCTTTCCCTACACGACGCTCTTCCGATCTNNNCCAACATGGTGAAACTCCATC

GAGCACACGTCTGAACTCTTTCCCTACACGACGCTCTTCCGATCTNNNCTGAGGCAGGAGAATCACTTC

GAGCACACGTCTGAACTCTTTCCCTACACGACGCTCTTCCGATCTNNNCTGAGGCAGGAGAATCACTTC

GAGCACACGTCTGAACTCTTTCCCTACACGACGCTCTTCCGATCTNNNATTTGGTCAGGAGGGTATGG

GAGCACACGTCTGAACTCTTTCCCTACACGACGCTCTTCCGATCTNNNATTTGGTCAGGAGGGTATGG

AGAGCACACGTCTGAACTCTTTCCCTACACGACGCTCTTCCGATCTNNNCTGAGGCAGGATAACCGTTT

AGAGCACACGTCTGAACTCTTTCCCTACACGACGCTCTTCCGATCTNNNCTGAGGCAGGATAACCGTTT

AGCACACGTCTGAACTCTTTCCCTACACGACGCTCTTCCGATCTNNNGTCAGAAACCGGAAGCCAATTT

AGCACACGTCTGAACTCTTTCCCTACACGACGCTCTTCCGATCTNNNGTCAGAAACCGGAAGCCAATTT

AGAGCACACGTCTGAACTCTTTCCCTACACGACGCTCTTCCGATCTNNNAGCTGGACTTTTAGCCAACT

AGAGCACACGTCTGAACTCTTTCCCTACACGACGCTCTTCCGATCTNNNAGCTGGACTTTTAGCCAACT

AGCACACGTCTGAACTCTTTCCCTACACGACGCTCTTCCGATCTNNNGCAAGTTGCAGTTTACCCATGTA

AGCACACGTCTGAACTCTTTCCCTACACGACGCTCTTCCGATCTNNNGCAAGTTGCAGTTTACCCATGTA

AGCACACGTCTGAACTCTTTCCCTACACGACGCTCTTCCGATCTNNNGAGGTTACAGTGAGCCGAGAT
AGCACACGTCTGAACTCTTTCCCTACACGACGCTCTTCCGATCTNNNGAGGTTACAGTGAGCCGAGAT
GAGCACACGTCTGAACTCTTTCCCTACACGACGCTCTTCCGATCTNNNAGCCAGACATTATTTAGGCGC
GAGCACACGTCTGAACTCTTTCCCTACACGACGCTCTTCCGATCTNNNAGCCAGACATTATTTAGGCGC
AGCACACGTCTGAACTCTTTCCCTACACGACGCTCTTCCGATCTNNNTTTCACTTTGTCCTTGTCTGCCA
AGCACACGTCTGAACTCTTTCCCTACACGACGCTCTTCCGATCTNNNTTTCACTTTGTCCTTGTCTGCCA
AGCACACGTCTGAACTCTTTCCCTACACGACGCTCTTCCGATCTNNNATTGGGGGAGGCTAGGTAGAC
AGCACACGTCTGAACTCTTTCCCTACACGACGCTCTTCCGATCTNNNATTGGGGGAGGCTAGGTAGAC
AGCACACGTCTGAACTCTTTCCCTACACGACGCTCTTCCGATCTNNNACTAGCTGAGACTACAAGTGCA
AGCACACGTCTGAACTCTTTCCCTACACGACGCTCTTCCGATCTNNNACTAGCTGAGACTACAAGTGCA
AGCACACGTCTGAACTCTTTCCCTACACGACGCTCTTCCGATCTNNNTTTCTTCAAGGAGCCCTAGTTCC
AGCACACGTCTGAACTCTTTCCCTACACGACGCTCTTCCGATCTNNNTTTCTTCAAGGAGCCCTAGTTCC
AGCACACGTCTGAACTCTTTCCCTACACGACGCTCTTCCGATCTNNNCTCGAACTTCAGACCTCAAGTGA
AGCACACGTCTGAACTCTTTCCCTACACGACGCTCTTCCGATCTNNNCTCGAACTTCAGACCTCAAGTGA
GCACACGTCTGAACTCTTTCCCTACACGACGCTCTTCCGATCTNNNCGGTACATGGGTAAGAGTGGAT
GCACACGTCTGAACTCTTTCCCTACACGACGCTCTTCCGATCTNNNCGGTACATGGGTAAGAGTGGAT
GAGCACACGTCTGAACTCTTTCCCTACACGACGCTCTTCCGATCTNNNCCATTGATTTCCTGGTGGATTC
GAGCACACGTCTGAACTCTTTCCCTACACGACGCTCTTCCGATCTNNNCCATTGATTTCCTGGTGGATTC
GAGCACACGTCTGAACTCTTTCCCTACACGACGCTCTTCCGATCTNNNGTGTGGACATCAGAGAATCCC
GAGCACACGTCTGAACTCTTTCCCTACACGACGCTCTTCCGATCTNNNGTGTGGACATCAGAGAATCCC
AGCACACGTCTGAACTCTTTCCCTACACGACGCTCTTCCGATCTNNNGATGATCCTAGCACTCCTTCACC
AGCACACGTCTGAACTCTTTCCCTACACGACGCTCTTCCGATCTNNNGATGATCCTAGCACTCCTTCACC
AGAGCACACGTCTGAACTCTTTCCCTACACGACGCTCTTCCGATCTNNNTCTGCCAAGTCAAATTAGGC
AGAGCACACGTCTGAACTCTTTCCCTACACGACGCTCTTCCGATCTNNNTCTGCCAAGTCAAATTAGGC
GAGCACACGTCTGAACTCTTTCCCTACACGACGCTCTTCCGATCTNNNCAGATCACGAGGTCAGGAGAA
GAGCACACGTCTGAACTCTTTCCCTACACGACGCTCTTCCGATCTNNNCAGATCACGAGGTCAGGAGAA
GAGCACACGTCTGAACTCTTTCCCTACACGACGCTCTTCCGATCTNNNTACAAAGTATGAAAGAGGCCC
GAGCACACGTCTGAACTCTTTCCCTACACGACGCTCTTCCGATCTNNNTACAAAGTATGAAAGAGGCCC
AGAGCACACGTCTGAACTCTTTCCCTACACGACGCTCTTCCGATCTNNNGCCATAGAGCGAGACTTCATC
AGAGCACACGTCTGAACTCTTTCCCTACACGACGCTCTTCCGATCTNNNGCCATAGAGCGAGACTTCATC
AGCACACGTCTGAACTCTTTCCCTACACGACGCTCTTCCGATCTNNNCACCATGACCAGCAATCCATTAT
AGCACACGTCTGAACTCTTTCCCTACACGACGCTCTTCCGATCTNNNCACCATGACCAGCAATCCATTAT
GCACACGTCTGAACTCTTTCCCTACACGACGCTCTTCCGATCTNNNGAGAGAGTGAGAGAGAGAGGC
GCACACGTCTGAACTCTTTCCCTACACGACGCTCTTCCGATCTNNNGAGAGAGTGAGAGAGAGAGGC
AGAGCACACGTCTGAACTCTTTCCCTACACGACGCTCTTCCGATCTNNNTGCCACTATCTGATACGGAA
AGAGCACACGTCTGAACTCTTTCCCTACACGACGCTCTTCCGATCTNNNTGCCACTATCTGATACGGAA
GCACACGTCTGAACTCTTTCCCTACACGACGCTCTTCCGATCTNNNGCTTCACAGTAGATAGGTGGCAA
GCACACGTCTGAACTCTTTCCCTACACGACGCTCTTCCGATCTNNNGCTTCACAGTAGATAGGTGGCAA
GAGCACACGTCTGAACTCTTTCCCTACACGACGCTCTTCCGATCTNNNTTAGTAGAGACGGCATTGCAT
GAGCACACGTCTGAACTCTTTCCCTACACGACGCTCTTCCGATCTNNNTTAGTAGAGACGGCATTGCAT
CACACGTCTGAACTCTTTCCCTACACGACGCTCTTCCGATCTNNNTATAGGCAGCCATTCAAAGGTGTC
CACACGTCTGAACTCTTTCCCTACACGACGCTCTTCCGATCTNNNTATAGGCAGCCATTCAAAGGTGTC
GAGCACACGTCTGAACTCTTTCCCTACACGACGCTCTTCCGATCTNNNCCTTGTCTACCTTTCCACTCCTCC

AGCACACGTCTGAACTCTTTCCCTACACGACGCTCTTCCGATCTNNNCCTTGTCTACCTTTCCACTCCTC
GCACACGTCTGAACTCTTTCCCTACACGACGCTCTTCCGATCTNNNATTGGAAGGAGTTGGAAGGAG
GCACACGTCTGAACTCTTTCCCTACACGACGCTCTTCCGATCTNNNATTGGAAGGAGTTGGAAGGAG
GAGCACACGTCTGAACTCTTTCCCTACACGACGCTCTTCCGATCTNNNGGGCCAATTATATCCAGCTC
GAGCACACGTCTGAACTCTTTCCCTACACGACGCTCTTCCGATCTNNNGGGCCAATTATATCCAGCTC
AGCACACGTCTGAACTCTTTCCCTACACGACGCTCTTCCGATCTNNNGCTAGTGGGCTAAGAAAAGA
AGCACACGTCTGAACTCTTTCCCTACACGACGCTCTTCCGATCTNNNGCTAGTGGGCTAAGAAAAGA
GCACACGTCTGAACTCTTTCCCTACACGACGCTCTTCCGATCTNNNCACCACACTGAGTGCCAATATC
GCACACGTCTGAACTCTTTCCCTACACGACGCTCTTCCGATCTNNNCACCACACTGAGTGCCAATATC
AGCACACGTCTGAACTCTTTCCCTACACGACGCTCTTCCGATCTNNNGAACATCAATAACAGAAAGAC
AGCACACGTCTGAACTCTTTCCCTACACGACGCTCTTCCGATCTNNNGAACATCAATAACAGAAAGAC
AGCACACGTCTGAACTCTTTCCCTACACGACGCTCTTCCGATCTNNNGAGGTTGCAATGAGCCAAGA
AGCACACGTCTGAACTCTTTCCCTACACGACGCTCTTCCGATCTNNNGAGGTTGCAATGAGCCAAGA
AGCACACGTCTGAACTCTTTCCCTACACGACGCTCTTCCGATCTNNNCTAGTCGTCCTTCCACACTCTT
AGCACACGTCTGAACTCTTTCCCTACACGACGCTCTTCCGATCTNNNCTAGTCGTCCTTCCACACTCTT
CACACGTCTGAACTCTTTCCCTACACGACGCTCTTCCGATCTNNNCAATGGTGCATTCTTGGAGAAGG
CACACGTCTGAACTCTTTCCCTACACGACGCTCTTCCGATCTNNNCAATGGTGCATTCTTGGAGAAGG
GAGCACACGTCTGAACTCTTTCCCTACACGACGCTCTTCCGATCTNNNTGAGGCACAAGAATCGTTT
GAGCACACGTCTGAACTCTTTCCCTACACGACGCTCTTCCGATCTNNNTGAGGCACAAGAATCGTTT
AGCACACGTCTGAACTCTTTCCCTACACGACGCTCTTCCGATCTNNNTACAAAAAATTAGCCGGGCTT
AGCACACGTCTGAACTCTTTCCCTACACGACGCTCTTCCGATCTNNNTACAAAAAATTAGCCGGGCTT
GCACACGTCTGAACTCTTTCCCTACACGACGCTCTTCCGATCTNNNAATGGAACTGGGGAGAAACACT
GCACACGTCTGAACTCTTTCCCTACACGACGCTCTTCCGATCTNNNAATGGAACTGGGGAGAAACACT
AGCACACGTCTGAACTCTTTCCCTACACGACGCTCTTCCGATCTNNNTGTGGATATTGCTTCACAAGT
AGCACACGTCTGAACTCTTTCCCTACACGACGCTCTTCCGATCTNNNTGTGGATATTGCTTCACAAGT
AGCACACGTCTGAACTCTTTCCCTACACGACGCTCTTCCGATCTNNNAGGTACTTTGATCCCAGCTGA
AGCACACGTCTGAACTCTTTCCCTACACGACGCTCTTCCGATCTNNNAGGTACTTTGATCCCAGCTGA
AGCACACGTCTGAACTCTTTCCCTACACGACGCTCTTCCGATCTNNNATTGTGGTCCCAGCTACTTGG
AGCACACGTCTGAACTCTTTCCCTACACGACGCTCTTCCGATCTNNNATTGTGGTCCCAGCTACTTGG
AGCACACGTCTGAACTCTTTCCCTACACGACGCTCTTCCGATCTNNNCTGAGGCAGGAGAATCACTTG
AGCACACGTCTGAACTCTTTCCCTACACGACGCTCTTCCGATCTNNNCTGAGGCAGGAGAATCACTTG
GAGCACACGTCTGAACTCTTTCCCTACACGACGCTCTTCCGATCTNNNGGAAATCACAAGATTCCTGC
GAGCACACGTCTGAACTCTTTCCCTACACGACGCTCTTCCGATCTNNNGGAAATCACAAGATTCCTGC
GCACACGTCTGAACTCTTTCCCTACACGACGCTCTTCCGATCTNNNTGGCTTTGATGGAACTTTGTTC
GCACACGTCTGAACTCTTTCCCTACACGACGCTCTTCCGATCTNNNTGGCTTTGATGGAACTTTGTTC
GAGCACACGTCTGAACTCTTTCCCTACACGACGCTCTTCCGATCTNNNTCTTAAGCGGCCATGATCCA
GAGCACACGTCTGAACTCTTTCCCTACACGACGCTCTTCCGATCTNNNTCTTAAGCGGCCATGATCCA
GCACACGTCTGAACTCTTTCCCTACACGACGCTCTTCCGATCTNNNGAACTCCTGGCCTCAAGTGATC
GCACACGTCTGAACTCTTTCCCTACACGACGCTCTTCCGATCTNNNGAACTCCTGGCCTCAAGTGATC
AGCACACGTCTGAACTCTTTCCCTACACGACGCTCTTCCGATCTNNNCACCACATCCGGCCAAATATTT
AGCACACGTCTGAACTCTTTCCCTACACGACGCTCTTCCGATCTNNNCACCACATCCGGCCAAATATTT
AGCACACGTCTGAACTCTTTCCCTACACGACGCTCTTCCGATCTNNNTGAGGCAGGAGAATCACTTG
AGCACACGTCTGAACTCTTTCCCTACACGACGCTCTTCCGATCTNNNTGAGGCAGGAGAATCACTTG

GCACACGTCTGAACTCTTTCCCTACACGACGCTCTTCCGATCTNNNTTAGGATGGGAGGATGGTGGA
GCACACGTCTGAACTCTTTCCCTACACGACGCTCTTCCGATCTNNNTTAGGATGGGAGGATGGTGGA
GCACACGTCTGAACTCTTTCCCTACACGACGCTCTTCCGATCTNNNTATAGTCCCAGCTACTCAGGAG
GCACACGTCTGAACTCTTTCCCTACACGACGCTCTTCCGATCTNNNTATAGTCCCAGCTACTCAGGAG
GCACACGTCTGAACTCTTTCCCTACACGACGCTCTTCCGATCTNNNTCAGTCTTGCCAGAAATTTGTCA
GCACACGTCTGAACTCTTTCCCTACACGACGCTCTTCCGATCTNNNTCAGTCTTGCCAGAAATTTGTCA
AGAGCACACGTCTGAACTCTTTCCCTACACGACGCTCTTCCGATCTNNNGATCAGAGTCAGTGGTGT
AGAGCACACGTCTGAACTCTTTCCCTACACGACGCTCTTCCGATCTNNNGATCAGAGTCAGTGGTGT
AGCACACGTCTGAACTCTTTCCCTACACGACGCTCTTCCGATCTNNNCAGTGAGCTGAGATCATACCA
AGCACACGTCTGAACTCTTTCCCTACACGACGCTCTTCCGATCTNNNCAGTGAGCTGAGATCATACCA
GAGCACACGTCTGAACTCTTTCCCTACACGACGCTCTTCCGATCTNNNTGCAGTCCATCCTTTGGTTAT
GAGCACACGTCTGAACTCTTTCCCTACACGACGCTCTTCCGATCTNNNTGCAGTCCATCCTTTGGTTAT
AGCACACGTCTGAACTCTTTCCCTACACGACGCTCTTCCGATCTNNNTCGCAAAAATAAATGCACACCC
AGCACACGTCTGAACTCTTTCCCTACACGACGCTCTTCCGATCTNNNTCGCAAAAATAAATGCACACCC
GAGCACACGTCTGAACTCTTTCCCTACACGACGCTCTTCCGATCTNNNCACTACAACTTGGGTGACAGA
GAGCACACGTCTGAACTCTTTCCCTACACGACGCTCTTCCGATCTNNNCACTACAACTTGGGTGACAGA
GCACACGTCTGAACTCTTTCCCTACACGACGCTCTTCCGATCTNNNATTCTTCTCATCCTCACATGGCAC
GCACACGTCTGAACTCTTTCCCTACACGACGCTCTTCCGATCTNNNATTCTTCTCATCCTCACATGGCAC
AGCACACGTCTGAACTCTTTCCCTACACGACGCTCTTCCGATCTNNNGCAACTGAACACCACATATTCC
AGCACACGTCTGAACTCTTTCCCTACACGACGCTCTTCCGATCTNNNGCAACTGAACACCACATATTCC
GCACACGTCTGAACTCTTTCCCTACACGACGCTCTTCCGATCTNNNGAGGCTGAGGCAGGAGAATGC
GCACACGTCTGAACTCTTTCCCTACACGACGCTCTTCCGATCTNNNGAGGCTGAGGCAGGAGAATGC
GAGCACACGTCTGAACTCTTTCCCTACACGACGCTCTTCCGATCTNNNAAATTTCCCCTTCCTCTCTGTCC
GAGCACACGTCTGAACTCTTTCCCTACACGACGCTCTTCCGATCTNNNAAATTTCCCCTTCCTCTCTGTCC
AGCACACGTCTGAACTCTTTCCCTACACGACGCTCTTCCGATCTNNNACATAACTAACACTCCAGCCTG
AGCACACGTCTGAACTCTTTCCCTACACGACGCTCTTCCGATCTNNNACATAACTAACACTCCAGCCTG
AGCACACGTCTGAACTCTTTCCCTACACGACGCTCTTCCGATCTNNNGCTTTTGATTTCTGTTCGTGTG
AGCACACGTCTGAACTCTTTCCCTACACGACGCTCTTCCGATCTNNNGCTTTTGATTTCTGTTCGTGTG
GAGCACACGTCTGAACTCTTTCCCTACACGACGCTCTTCCGATCTNNNCCCATGAAGACGTCTACAAAT
GAGCACACGTCTGAACTCTTTCCCTACACGACGCTCTTCCGATCTNNNCCCATGAAGACGTCTACAAAT
AGCACACGTCTGAACTCTTTCCCTACACGACGCTCTTCCGATCTNNNTGACTTGTGCTGCAATACACAT
AGCACACGTCTGAACTCTTTCCCTACACGACGCTCTTCCGATCTNNNTGACTTGTGCTGCAATACACAT
GAGCACACGTCTGAACTCTTTCCCTACACGACGCTCTTCCGATCTNNNCAGATCACTTGAGCTCAGGA
GAGCACACGTCTGAACTCTTTCCCTACACGACGCTCTTCCGATCTNNNCAGATCACTTGAGCTCAGGA
AGAGCACACGTCTGAACTCTTTCCCTACACGACGCTCTTCCGATCTNNNGGTACCTCAGATGGAAATG
AGAGCACACGTCTGAACTCTTTCCCTACACGACGCTCTTCCGATCTNNNGGTACCTCAGATGGAAATG
GCACACGTCTGAACTCTTTCCCTACACGACGCTCTTCCGATCTNNNTATGTCCAGTAGTGCTGCTGAA
GCACACGTCTGAACTCTTTCCCTACACGACGCTCTTCCGATCTNNNTATGTCCAGTAGTGCTGCTGAA
GAGCACACGTCTGAACTCTTTCCCTACACGACGCTCTTCCGATCTNNNTGGTCAGCAACTTGAATAGCT
GAGCACACGTCTGAACTCTTTCCCTACACGACGCTCTTCCGATCTNNNTGGTCAGCAACTTGAATAGCT
AGAGCACACGTCTGAACTCTTTCCCTACACGACGCTCTTCCGATCTNNNAGACTCACCTGTCATGTTTC
AGAGCACACGTCTGAACTCTTTCCCTACACGACGCTCTTCCGATCTNNNAGACTCACCTGTCATGTTTC
AGCACACGTCTGAACTCTTTCCCTACACGACGCTCTTCCGATCTNNNAAACAAGCAAAGCACAAGTCC

AGCACACGTCTGAACTCTTTCCCTACACGACGCTCTTCCGATCTNNNAAACAAGCAAAGCACAAGTCC
GAGCACACGTCTGAACTCTTTCCCTACACGACGCTCTTCCGATCTNNNCAAGAAACCTTAGCAACACA
GAGCACACGTCTGAACTCTTTCCCTACACGACGCTCTTCCGATCTNNNCAAGAAACCTTAGCAACACA
AGCACACGTCTGAACTCTTTCCCTACACGACGCTCTTCCGATCTNNNGTTCTGATCTGGTGAAGGAG
AGCACACGTCTGAACTCTTTCCCTACACGACGCTCTTCCGATCTNNNGTTCTGATCTGGTGAAGGAG
AGCACACGTCTGAACTCTTTCCCTACACGACGCTCTTCCGATCTNNNGAGAACAGTATGGGGTTCCT
AGCACACGTCTGAACTCTTTCCCTACACGACGCTCTTCCGATCTNNNGAGAACAGTATGGGGTTCCT
AGCACACGTCTGAACTCTTTCCCTACACGACGCTCTTCCGATCTNNNTACATACTGCACTCCCCTTCAA
AGCACACGTCTGAACTCTTTCCCTACACGACGCTCTTCCGATCTNNNTACATACTGCACTCCCCTTCAA
CACACGTCTGAACTCTTTCCCTACACGACGCTCTTCCGATCTNNNTAGTAGGGACATGTTGGTTTCGC
CACACGTCTGAACTCTTTCCCTACACGACGCTCTTCCGATCTNNNTAGTAGGGACATGTTGGTTTCGC
AGCACACGTCTGAACTCTTTCCCTACACGACGCTCTTCCGATCTNNNGGAGGGAGGTAGGGGGTCA
AGCACACGTCTGAACTCTTTCCCTACACGACGCTCTTCCGATCTNNNGGAGGGAGGTAGGGGGTCA
AGAGCACACGTCTGAACTCTTTCCCTACACGACGCTCTTCCGATCTNNNTCATGGAGGTGAGAAAGA
AGAGCACACGTCTGAACTCTTTCCCTACACGACGCTCTTCCGATCTNNNTCATGGAGGTGAGAAAGA
AGCACACGTCTGAACTCTTTCCCTACACGACGCTCTTCCGATCTNNNTCTGAGTGTGTTTATGTGTGT
AGCACACGTCTGAACTCTTTCCCTACACGACGCTCTTCCGATCTNNNTCTGAGTGTGTTTATGTGTGT
AGCACACGTCTGAACTCTTTCCCTACACGACGCTCTTCCGATCTNNNACGGAAGTATTATATAGCACTT
AGCACACGTCTGAACTCTTTCCCTACACGACGCTCTTCCGATCTNNNACGGAAGTATTATATAGCACTT
GAGCACACGTCTGAACTCTTTCCCTACACGACGCTCTTCCGATCTNNNTTTTAAACTTCTCATATGGT
GAGCACACGTCTGAACTCTTTCCCTACACGACGCTCTTCCGATCTNNNTTTTAAACTTCTCATATGGT
GAGCACACGTCTGAACTCTTTCCCTACACGACGCTCTTCCGATCTNNNAGCAAAGTTAGCCTCACATA
GAGCACACGTCTGAACTCTTTCCCTACACGACGCTCTTCCGATCTNNNAGCAAAGTTAGCCTCACATA
AGCACACGTCTGAACTCTTTCCCTACACGACGCTCTTCCGATCTNNNTCCTTTTGCTGTTTTTGCTGTC
AGCACACGTCTGAACTCTTTCCCTACACGACGCTCTTCCGATCTNNNTCCTTTTGCTGTTTTTGCTGTC
AGCACACGTCTGAACTCTTTCCCTACACGACGCTCTTCCGATCTNNNTGAAATAAAAATCTACAATGG
AGCACACGTCTGAACTCTTTCCCTACACGACGCTCTTCCGATCTNNNTGAAATAAAAATCTACAATGG
AGCACACGTCTGAACTCTTTCCCTACACGACGCTCTTCCGATCTNNNAATGAAGCACTTTAGTTATGG
AGCACACGTCTGAACTCTTTCCCTACACGACGCTCTTCCGATCTNNNAATGAAGCACTTTAGTTATGG
AGCACACGTCTGAACTCTTTCCCTACACGACGCTCTTCCGATCTNNNCTGAACTCACATCCTCCGCAGC
AGCACACGTCTGAACTCTTTCCCTACACGACGCTCTTCCGATCTNNNCTGAACTCACATCCTCCGCAGC
GCACACGTCTGAACTCTTTCCCTACACGACGCTCTTCCGATCTNNNTGCAAAGCTAACATCTTCCAAG
GCACACGTCTGAACTCTTTCCCTACACGACGCTCTTCCGATCTNNNTGCAAAGCTAACATCTTCCAAG
ACACGTCTGAACTCTTTCCCTACACGACGCTCTTCCGATCTNNNACAATGTGTGCTTCAAGTGTCAGCA
ACACGTCTGAACTCTTTCCCTACACGACGCTCTTCCGATCTNNNACAATGTGTGCTTCAAGTGTCAGCA
GAGCACACGTCTGAACTCTTTCCCTACACGACGCTCTTCCGATCTNNNTGAGCTGAGATCACATTACT
GAGCACACGTCTGAACTCTTTCCCTACACGACGCTCTTCCGATCTNNNTGAGCTGAGATCACATTACT
AGCACACGTCTGAACTCTTTCCCTACACGACGCTCTTCCGATCTNNNTTTCTAAACCGGGCTAGAATG
AGCACACGTCTGAACTCTTTCCCTACACGACGCTCTTCCGATCTNNNTTTCTAAACCGGGCTAGAATG
AGCACACGTCTGAACTCTTTCCCTACACGACGCTCTTCCGATCTNNNTGTACCTGTTCACTCACACTTG
AGCACACGTCTGAACTCTTTCCCTACACGACGCTCTTCCGATCTNNNTGTACCTGTTCACTCACACTTG
AGCACACGTCTGAACTCTTTCCCTACACGACGCTCTTCCGATCTNNNGGGGTAAAGCTCCTTGGAAT
AGCACACGTCTGAACTCTTTCCCTACACGACGCTCTTCCGATCTNNNGGGGTAAAGCTCCTTGGAAT

AGCACACGTCTGAACTCTTTCCCTACACGACGCTCTTCCGATCTNNNGCACTCTTGTAATCCCAGCTACT
AGCACACGTCTGAACTCTTTCCCTACACGACGCTCTTCCGATCTNNNGCACTCTTGTAATCCCAGCTACT
AGCACACGTCTGAACTCTTTCCCTACACGACGCTCTTCCGATCTNNNATAAGTATATTAAAGAGACAGC
AGCACACGTCTGAACTCTTTCCCTACACGACGCTCTTCCGATCTNNNATAAGTATATTAAAGAGACAGC
AGCACACGTCTGAACTCTTTCCCTACACGACGCTCTTCCGATCTNNNGGTACATGTGAGTTCCTCCAAG
AGCACACGTCTGAACTCTTTCCCTACACGACGCTCTTCCGATCTNNNGGTACATGTGAGTTCCTCCAAG
GCACACGTCTGAACTCTTTCCCTACACGACGCTCTTCCGATCTNNNCTGTCAGAGAACTTGGTGGCATCA
GCACACGTCTGAACTCTTTCCCTACACGACGCTCTTCCGATCTNNNCTGTCAGAGAACTTGGTGGCATCA
AGCACACGTCTGAACTCTTTCCCTACACGACGCTCTTCCGATCTNNNTAGCATGAAAGCAGCCATAGAC
AGCACACGTCTGAACTCTTTCCCTACACGACGCTCTTCCGATCTNNNTAGCATGAAAGCAGCCATAGAC
GAGCACACGTCTGAACTCTTTCCCTACACGACGCTCTTCCGATCTNNNGCACACTACCACACCCAGATAA
GAGCACACGTCTGAACTCTTTCCCTACACGACGCTCTTCCGATCTNNNGCACACTACCACACCCAGATAA
GCACACGTCTGAACTCTTTCCCTACACGACGCTCTTCCGATCTNNNATCCATCATTCCATTTTTCTTTCCT
GCACACGTCTGAACTCTTTCCCTACACGACGCTCTTCCGATCTNNNATCCATCATTCCATTTTTCTTTCCT
GAGCACACGTCTGAACTCTTTCCCTACACGACGCTCTTCCGATCTNNNTGCCTTGTTCATTCTACTGCTCT
GAGCACACGTCTGAACTCTTTCCCTACACGACGCTCTTCCGATCTNNNTGCCTTGTTCATTCTACTGCTCT
GAGCACACGTCTGAACTCTTTCCCTACACGACGCTCTTCCGATCTNNNTACAAAAGTTAGCCAGGTGTG
GAGCACACGTCTGAACTCTTTCCCTACACGACGCTCTTCCGATCTNNNTACAAAAGTTAGCCAGGTGTG
AGCACACGTCTGAACTCTTTCCCTACACGACGCTCTTCCGATCTNNNTTGCATTTTCCTGCTATTGTGCAC
AGCACACGTCTGAACTCTTTCCCTACACGACGCTCTTCCGATCTNNNTTGCATTTTCCTGCTATTGTGCAC
AGAGCACACGTCTGAACTCTTTCCCTACACGACGCTCTTCCGATCTNNNGCAGTGAGCCAAGATTGCAC
AGAGCACACGTCTGAACTCTTTCCCTACACGACGCTCTTCCGATCTNNNGCAGTGAGCCAAGATTGCAC
GAGCACACGTCTGAACTCTTTCCCTACACGACGCTCTTCCGATCTNNNATCCCTGTAATCCCAGCTACTTG
GAGCACACGTCTGAACTCTTTCCCTACACGACGCTCTTCCGATCTNNNATCCCTGTAATCCCAGCTACTTG
AGCACACGTCTGAACTCTTTCCCTACACGACGCTCTTCCGATCTNNNCTCCACACCGAGTTTTCCCTATTG
AGCACACGTCTGAACTCTTTCCCTACACGACGCTCTTCCGATCTNNNCTCCACACCGAGTTTTCCCTATTG
GCACACGTCTGAACTCTTTCCCTACACGACGCTCTTCCGATCTNNNTGTAATCCCAGCACTTTAGGAGG
GCACACGTCTGAACTCTTTCCCTACACGACGCTCTTCCGATCTNNNTGTAATCCCAGCACTTTAGGAGG
AGCACACGTCTGAACTCTTTCCCTACACGACGCTCTTCCGATCTNNNCCCGGCCCAATATCCTTCTACCAT
AGCACACGTCTGAACTCTTTCCCTACACGACGCTCTTCCGATCTNNNCCCGGCCCAATATCCTTCTACCAT
AGCACACGTCTGAACTCTTTCCCTACACGACGCTCTTCCGATCTNNNACTTCATACCCGGACCACAAATT
AGCACACGTCTGAACTCTTTCCCTACACGACGCTCTTCCGATCTNNNACTTCATACCCGGACCACAAATT
AGCACACGTCTGAACTCTTTCCCTACACGACGCTCTTCCGATCTNNNCTCCTAGCTCTGACTATTCTGCCC
AGCACACGTCTGAACTCTTTCCCTACACGACGCTCTTCCGATCTNNNCTCCTAGCTCTGACTATTCTGCCC
GAGCACACGTCTGAACTCTTTCCCTACACGACGCTCTTCCGATCTNNNAAATACAAAAATTAGCCGGGC
GAGCACACGTCTGAACTCTTTCCCTACACGACGCTCTTCCGATCTNNNAAATACAAAAATTAGCCGGGC
GAGCACACGTCTGAACTCTTTCCCTACACGACGCTCTTCCGATCTNNNTTACCTCTCTCTCAACATTAATA
GAGCACACGTCTGAACTCTTTCCCTACACGACGCTCTTCCGATCTNNNTTACCTCTCTCTCAACATTAATA
AGCACACGTCTGAACTCTTTCCCTACACGACGCTCTTCCGATCTNNNAGATCCTCCACCAGCAAAAAGA
AGCACACGTCTGAACTCTTTCCCTACACGACGCTCTTCCGATCTNNNAGATCCTCCACCAGCAAAAAGA
GAAGAGCACACGTCTGAACTCTTTCCCTACACGACGCTCTTCCGATCTNNNACTAGTTCCATGCATTTT
GAAGAGCACACGTCTGAACTCTTTCCCTACACGACGCTCTTCCGATCTNNNACTAGTTCCATGCATTTT
AGCACACGTCTGAACTCTTTCCCTACACGACGCTCTTCCGATCTNNNTCTGGCATCATCTCCCTTTTCTCC

AGCACACGTCTGAACTCTTTCCCTACACGACGCTCTTCCGATCTNNNTCTGGCATCATCTCCCTTTTCTCC
GAGCACACGTCTGAACTCTTTCCCTACACGACGCTCTTCCGATCTNNNTGATTACTGGCAGGAAATGTG
GAGCACACGTCTGAACTCTTTCCCTACACGACGCTCTTCCGATCTNNNTGATTACTGGCAGGAAATGTG
GCACACGTCTGAACTCTTTCCCTACACGACGCTCTTCCGATCTNNNTCAGTGTAATCCCAGCTACTTGGC
GCACACGTCTGAACTCTTTCCCTACACGACGCTCTTCCGATCTNNNTCAGTGTAATCCCAGCTACTTGGC
AGCACACGTCTGAACTCTTTCCCTACACGACGCTCTTCCGATCTNNNTGTGCAATGATGGTCACTTTTGT
AGCACACGTCTGAACTCTTTCCCTACACGACGCTCTTCCGATCTNNNTGTGCAATGATGGTCACTTTTGT
AGCACACGTCTGAACTCTTTCCCTACACGACGCTCTTCCGATCTNNNCTGATTAGTGGTGGGGCTTTA
AGCACACGTCTGAACTCTTTCCCTACACGACGCTCTTCCGATCTNNNCTGATTAGTGGTGGGGCTTTA
AGCACACGTCTGAACTCTTTCCCTACACGACGCTCTTCCGATCTNNNTCAGACTATTCGTTCAACAGCCA
AGCACACGTCTGAACTCTTTCCCTACACGACGCTCTTCCGATCTNNNTCAGACTATTCGTTCAACAGCCA
AGCACACGTCTGAACTCTTTCCCTACACGACGCTCTTCCGATCTNNNATACTAGCTACTTGGGAGGCTG
AGCACACGTCTGAACTCTTTCCCTACACGACGCTCTTCCGATCTNNNATACTAGCTACTTGGGAGGCTG
AGCACACGTCTGAACTCTTTCCCTACACGACGCTCTTCCGATCTNNNACAGTACTACAGAAGGTCCCCTA
AGCACACGTCTGAACTCTTTCCCTACACGACGCTCTTCCGATCTNNNACAGTACTACAGAAGGTCCCCTA
AGCACACGTCTGAACTCTTTCCCTACACGACGCTCTTCCGATCTNNNACAACACTACCCTCCTCTCTTCCCA
AGCACACGTCTGAACTCTTTCCCTACACGACGCTCTTCCGATCTNNNACAACACTACCCTCCTCTCTTCCCA
AGCACACGTCTGAACTCTTTCCCTACACGACGCTCTTCCGATCTNNNTTTGGGAGGACAGAGACATTCA
AGCACACGTCTGAACTCTTTCCCTACACGACGCTCTTCCGATCTNNNTTTGGGAGGACAGAGACATTCA
AGCACACGTCTGAACTCTTTCCCTACACGACGCTCTTCCGATCTNNNTTCCTGCAGAATAGGATAAGCC
AGCACACGTCTGAACTCTTTCCCTACACGACGCTCTTCCGATCTNNNTTCCTGCAGAATAGGATAAGCC
GCACACGTCTGAACTCTTTCCCTACACGACGCTCTTCCGATCTNNNGACTTTGCAAATGTTGGAGAGCT
GCACACGTCTGAACTCTTTCCCTACACGACGCTCTTCCGATCTNNNGACTTTGCAAATGTTGGAGAGCT
AGCACACGTCTGAACTCTTTCCCTACACGACGCTCTTCCGATCTNNNCCGCGGTGTTGTGTACATCTCCA
AGCACACGTCTGAACTCTTTCCCTACACGACGCTCTTCCGATCTNNNCCGCGGTGTTGTGTACATCTCCA
GAGCACACGTCTGAACTCTTTCCCTACACGACGCTCTTCCGATCTNNNCTTACATTCTCCAAAGTGGCAG
GAGCACACGTCTGAACTCTTTCCCTACACGACGCTCTTCCGATCTNNNCTTACATTCTCCAAAGTGGCAG
GAGCACACGTCTGAACTCTTTCCCTACACGACGCTCTTCCGATCTNNNCAGAACAGCTTGGAATGGCT
GAGCACACGTCTGAACTCTTTCCCTACACGACGCTCTTCCGATCTNNNCAGAACAGCTTGGAATGGCT
AGCACACGTCTGAACTCTTTCCCTACACGACGCTCTTCCGATCTNNNGAAGGTGCCTGTAATCCCAGT
AGCACACGTCTGAACTCTTTCCCTACACGACGCTCTTCCGATCTNNNGAAGGTGCCTGTAATCCCAGT
GAGCACACGTCTGAACTCTTTCCCTACACGACGCTCTTCCGATCTNNNCCCCTCTTAGAAGTTTTTTAAA
GAGCACACGTCTGAACTCTTTCCCTACACGACGCTCTTCCGATCTNNNCCCCTCTTAGAAGTTTTTTAAA
AGCACACGTCTGAACTCTTTCCCTACACGACGCTCTTCCGATCTNNNTGTATGCATGTGACTTGTGTAT
AGCACACGTCTGAACTCTTTCCCTACACGACGCTCTTCCGATCTNNNTGTATGCATGTGACTTGTGTAT
AGCACACGTCTGAACTCTTTCCCTACACGACGCTCTTCCGATCTNNNGTACCAAAAAATCAGCTGGGCA
AGCACACGTCTGAACTCTTTCCCTACACGACGCTCTTCCGATCTNNNGTACCAAAAAATCAGCTGGGCA
AGCACACGTCTGAACTCTTTCCCTACACGACGCTCTTCCGATCTNNNCATTGTGTGTGTCAGTGTGTTCA
AGCACACGTCTGAACTCTTTCCCTACACGACGCTCTTCCGATCTNNNCATTGTGTGTGTCAGTGTGTTCA
AGCACACGTCTGAACTCTTTCCCTACACGACGCTCTTCCGATCTNNNTAAGTAAGGGTAGGAGGCTGC
AGCACACGTCTGAACTCTTTCCCTACACGACGCTCTTCCGATCTNNNTAAGTAAGGGTAGGAGGCTGC
GAGCACACGTCTGAACTCTTTCCCTACACGACGCTCTTCCGATCTNNNGAGAATGGCATGAACCTGGG
GAGCACACGTCTGAACTCTTTCCCTACACGACGCTCTTCCGATCTNNNGAGAATGGCATGAACCTGGG

GCACACGTCTGAACTCTTTCCCTACACGACGCTCTTCCGATCTNNNACCAAAGCAGCAGGAATAGGACA
GCACACGTCTGAACTCTTTCCCTACACGACGCTCTTCCGATCTNNNACCAAAGCAGCAGGAATAGGACA
GAGCACACGTCTGAACTCTTTCCCTACACGACGCTCTTCCGATCTNNNAGGAGGTGGGTGGTAAATA
GAGCACACGTCTGAACTCTTTCCCTACACGACGCTCTTCCGATCTNNNAGGAGGTGGGTGGTAAATA
GCACACGTCTGAACTCTTTCCCTACACGACGCTCTTCCGATCTNNNCTCTCGAGTAGCTGGGACTACAG
GCACACGTCTGAACTCTTTCCCTACACGACGCTCTTCCGATCTNNNCTCTCGAGTAGCTGGGACTACAG
AGCACACGTCTGAACTCTTTCCCTACACGACGCTCTTCCGATCTNNNCCACCCTCAAGTCTCTTCAATCAC
AGCACACGTCTGAACTCTTTCCCTACACGACGCTCTTCCGATCTNNNCCACCCTCAAGTCTCTTCAATCAC
GAGCACACGTCTGAACTCTTTCCCTACACGACGCTCTTCCGATCTNNNGGTATGGTGGCAGGTGTCTA
GAGCACACGTCTGAACTCTTTCCCTACACGACGCTCTTCCGATCTNNNGGTATGGTGGCAGGTGTCTA
AGCACACGTCTGAACTCTTTCCCTACACGACGCTCTTCCGATCTNNNCTGAAGAATTGCTTGAACCTGC
AGCACACGTCTGAACTCTTTCCCTACACGACGCTCTTCCGATCTNNNCTGAAGAATTGCTTGAACCTGC
AGCACACGTCTGAACTCTTTCCCTACACGACGCTCTTCCGATCTNNNTTCAATCTTCCAGGGTGTGTGTC
AGCACACGTCTGAACTCTTTCCCTACACGACGCTCTTCCGATCTNNNTTCAATCTTCCAGGGTGTGTGTC
AGCACACGTCTGAACTCTTTCCCTACACGACGCTCTTCCGATCTNNNATCAAAAGCTGTTGTTCATCCCG
AGCACACGTCTGAACTCTTTCCCTACACGACGCTCTTCCGATCTNNNATCAAAAGCTGTTGTTCATCCCG
AGCACACGTCTGAACTCTTTCCCTACACGACGCTCTTCCGATCTNNNTTCATTTGGTTGCATACACCACC
AGCACACGTCTGAACTCTTTCCCTACACGACGCTCTTCCGATCTNNNTTCATTTGGTTGCATACACCACC
AGCACACGTCTGAACTCTTTCCCTACACGACGCTCTTCCGATCTNNNTTCACACCTTCAGACTCAGACTG
AGCACACGTCTGAACTCTTTCCCTACACGACGCTCTTCCGATCTNNNTTCACACCTTCAGACTCAGACTG
AGCACACGTCTGAACTCTTTCCCTACACGACGCTCTTCCGATCTNNNGGGCTAACCTCTCTTATATAGTG
AGCACACGTCTGAACTCTTTCCCTACACGACGCTCTTCCGATCTNNNGGGCTAACCTCTCTTATATAGTG
AGCACACGTCTGAACTCTTTCCCTACACGACGCTCTTCCGATCTNNNCCAAAGTGCTGGGATTACAGTT
AGCACACGTCTGAACTCTTTCCCTACACGACGCTCTTCCGATCTNNNCCAAAGTGCTGGGATTACAGTT
GAGCACACGTCTGAACTCTTTCCCTACACGACGCTCTTCCGATCTNNNCTGGGATTACAGGAGTGCAT
GAGCACACGTCTGAACTCTTTCCCTACACGACGCTCTTCCGATCTNNNCTGGGATTACAGGAGTGCAT
AGCACACGTCTGAACTCTTTCCCTACACGACGCTCTTCCGATCTNNNCACCTTGCCTGGCCTAGTTATAT
AGCACACGTCTGAACTCTTTCCCTACACGACGCTCTTCCGATCTNNNCACCTTGCCTGGCCTAGTTATAT
AGCACACGTCTGAACTCTTTCCCTACACGACGCTCTTCCGATCTNNNAGGGGAGATTTAGACACACACA
AGCACACGTCTGAACTCTTTCCCTACACGACGCTCTTCCGATCTNNNAGGGGAGATTTAGACACACACA
AGCACACGTCTGAACTCTTTCCCTACACGACGCTCTTCCGATCTNNNATCTGCCAGAAAGGCTTAGAGG
AGCACACGTCTGAACTCTTTCCCTACACGACGCTCTTCCGATCTNNNATCTGCCAGAAAGGCTTAGAGG
AGCACACGTCTGAACTCTTTCCCTACACGACGCTCTTCCGATCTNNNGTGTGTCTGGCCAACATGAATC
AGCACACGTCTGAACTCTTTCCCTACACGACGCTCTTCCGATCTNNNGTGTGTCTGGCCAACATGAATC
AGCACACGTCTGAACTCTTTCCCTACACGACGCTCTTCCGATCTNNNCTTGGTGTCTGAGCTGTACATTG
AGCACACGTCTGAACTCTTTCCCTACACGACGCTCTTCCGATCTNNNCTTGGTGTCTGAGCTGTACATTG
GAGCACACGTCTGAACTCTTTCCCTACACGACGCTCTTCCGATCTNNNCACCTGAGGTCATGAGTTTGA
GAGCACACGTCTGAACTCTTTCCCTACACGACGCTCTTCCGATCTNNNCACCTGAGGTCATGAGTTTGA
GCACACGTCTGAACTCTTTCCCTACACGACGCTCTTCCGATCTNNNAAAGAGAGAGACAGAGAGGGG
GCACACGTCTGAACTCTTTCCCTACACGACGCTCTTCCGATCTNNNAAAGAGAGAGACAGAGAGGGG
GAGCACACGTCTGAACTCTTTCCCTACACGACGCTCTTCCGATCTNNNCAAATGCTAACGAACCACCCCT
GAGCACACGTCTGAACTCTTTCCCTACACGACGCTCTTCCGATCTNNNCAAATGCTAACGAACCACCCCT
GAGCACACGTCTGAACTCTTTCCCTACACGACGCTCTTCCGATCTNNNAGTGACGGTTTGACTTCCTCTT

GAGCACACGTCTGAACTCTTTCCCTACACGACGCTCTTCCGATCTNNNAGTGACGGTTTGACTTCCTCTT
AGCACACGTCTGAACTCTTTCCCTACACGACGCTCTTCCGATCTNNNATTTAGCCAATGAGGAGCCCTA
AGCACACGTCTGAACTCTTTCCCTACACGACGCTCTTCCGATCTNNNATTTAGCCAATGAGGAGCCCTA
GCACACGTCTGAACTCTTTCCCTACACGACGCTCTTCCGATCTNNNACATACAAACTCGTGTGCATGTA
GCACACGTCTGAACTCTTTCCCTACACGACGCTCTTCCGATCTNNNACATACAAACTCGTGTGCATGTA
GCACACGTCTGAACTCTTTCCCTACACGACGCTCTTCCGATCTNNNCACCACACCCAGCTAATTTTTGTC
GCACACGTCTGAACTCTTTCCCTACACGACGCTCTTCCGATCTNNNCACCACACCCAGCTAATTTTTGTC
GAGCACACGTCTGAACTCTTTCCCTACACGACGCTCTTCCGATCTNNNCATGCACCTGTAGTCCCATCTA
GAGCACACGTCTGAACTCTTTCCCTACACGACGCTCTTCCGATCTNNNCATGCACCTGTAGTCCCATCTA
GCACACGTCTGAACTCTTTCCCTACACGACGCTCTTCCGATCTNNNCCACGAAAACCCTAGTGAATCAC
GCACACGTCTGAACTCTTTCCCTACACGACGCTCTTCCGATCTNNNCCACGAAAACCCTAGTGAATCAC
AGCACACGTCTGAACTCTTTCCCTACACGACGCTCTTCCGATCTNNNGCCTAGGCAACAGAGGTAAAC
AGCACACGTCTGAACTCTTTCCCTACACGACGCTCTTCCGATCTNNNGCCTAGGCAACAGAGGTAAAC
AGCACACGTCTGAACTCTTTCCCTACACGACGCTCTTCCGATCTNNNTTGGGAAACTAGGAAGGCATA
AGCACACGTCTGAACTCTTTCCCTACACGACGCTCTTCCGATCTNNNTTGGGAAACTAGGAAGGCATA
AGCACACGTCTGAACTCTTTCCCTACACGACGCTCTTCCGATCTNNNGGAAATAGGGTTTCTTCCAAG
AGCACACGTCTGAACTCTTTCCCTACACGACGCTCTTCCGATCTNNNGGAAATAGGGTTTCTTCCAAG
GAGCACACGTCTGAACTCTTTCCCTACACGACGCTCTTCCGATCTNNNCAGAAGTTTGCTGGAGGAG
GAGCACACGTCTGAACTCTTTCCCTACACGACGCTCTTCCGATCTNNNCAGAAGTTTGCTGGAGGAG
GAGCACACGTCTGAACTCTTTCCCTACACGACGCTCTTCCGATCTNNNAGGGAGGTAGAGGTTGTAG
GAGCACACGTCTGAACTCTTTCCCTACACGACGCTCTTCCGATCTNNNAGGGAGGTAGAGGTTGTAG
GAGCACACGTCTGAACTCTTTCCCTACACGACGCTCTTCCGATCTNNNCTTCTGGTTTCAAGCGATTCTC
GAGCACACGTCTGAACTCTTTCCCTACACGACGCTCTTCCGATCTNNNCTTCTGGTTTCAAGCGATTCTC
GAGCACACGTCTGAACTCTTTCCCTACACGACGCTCTTCCGATCTNNNCTAAAAATTAGCCGGGCATG
GAGCACACGTCTGAACTCTTTCCCTACACGACGCTCTTCCGATCTNNNCTAAAAATTAGCCGGGCATG
AGCACACGTCTGAACTCTTTCCCTACACGACGCTCTTCCGATCTNNNCCAGTTTGGGTAACAGAGTGA
AGCACACGTCTGAACTCTTTCCCTACACGACGCTCTTCCGATCTNNNCCAGTTTGGGTAACAGAGTGA
AGCACACGTCTGAACTCTTTCCCTACACGACGCTCTTCCGATCTNNNTGAGTTCTCATTGGAGGTTCTG
AGCACACGTCTGAACTCTTTCCCTACACGACGCTCTTCCGATCTNNNTGAGTTCTCATTGGAGGTTCTG
GCACACGTCTGAACTCTTTCCCTACACGACGCTCTTCCGATCTNNNGAGGTCAGAAGATCGAGACCAT
GCACACGTCTGAACTCTTTCCCTACACGACGCTCTTCCGATCTNNNGAGGTCAGAAGATCGAGACCAT
GAGCACACGTCTGAACTCTTTCCCTACACGACGCTCTTCCGATCTNNNTCATCTACAGACTCCAACAAG
GAGCACACGTCTGAACTCTTTCCCTACACGACGCTCTTCCGATCTNNNTCATCTACAGACTCCAACAAG
AGCACACGTCTGAACTCTTTCCCTACACGACGCTCTTCCGATCTNNNCCATTGAGGAGAAGGATGGG
AGCACACGTCTGAACTCTTTCCCTACACGACGCTCTTCCGATCTNNNCCATTGAGGAGAAGGATGGG
AGCACACGTCTGAACTCTTTCCCTACACGACGCTCTTCCGATCTNNNCCCATGCCACTCCTATTATCTGT
AGCACACGTCTGAACTCTTTCCCTACACGACGCTCTTCCGATCTNNNCCCATGCCACTCCTATTATCTGT
AGCACACGTCTGAACTCTTTCCCTACACGACGCTCTTCCGATCTNNNTCTCATCCCAGCTACTCAAGGAG
AGCACACGTCTGAACTCTTTCCCTACACGACGCTCTTCCGATCTNNNTCTCATCCCAGCTACTCAAGGAG
GAGCACACGTCTGAACTCTTTCCCTACACGACGCTCTTCCGATCTNNNGTGGCATGGACAAGATAAGG
GAGCACACGTCTGAACTCTTTCCCTACACGACGCTCTTCCGATCTNNNGTGGCATGGACAAGATAAGG
GCACACGTCTGAACTCTTTCCCTACACGACGCTCTTCCGATCTNNNTTCTCTAGCAGGAAGTGGAGAG
GCACACGTCTGAACTCTTTCCCTACACGACGCTCTTCCGATCTNNNTTCTCTAGCAGGAAGTGGAGAG

AGCACACGTCTGAACTCTTTCCCTACACGACGCTCTTCCGATCTNNNTTCTTCAATATTGCCTGGGCTCT
AGCACACGTCTGAACTCTTTCCCTACACGACGCTCTTCCGATCTNNNTTCTTCAATATTGCCTGGGCTCT
AGCACACGTCTGAACTCTTTCCCTACACGACGCTCTTCCGATCTNNNCAAGAGAATTGCTTCAACCTGG
AGCACACGTCTGAACTCTTTCCCTACACGACGCTCTTCCGATCTNNNCAAGAGAATTGCTTCAACCTGG
GCACACGTCTGAACTCTTTCCCTACACGACGCTCTTCCGATCTNNNGAGAAGAGCCAGAGAGAGTATG
GCACACGTCTGAACTCTTTCCCTACACGACGCTCTTCCGATCTNNNGAGAAGAGCCAGAGAGAGTATG
GAGCACACGTCTGAACTCTTTCCCTACACGACGCTCTTCCGATCTNNNTTTGTCACATCCTCAGCAACA
GAGCACACGTCTGAACTCTTTCCCTACACGACGCTCTTCCGATCTNNNTTTGTCACATCCTCAGCAACA
GAGCACACGTCTGAACTCTTTCCCTACACGACGCTCTTCCGATCTNNNCACAGTCCTTCCCATTATTACC
GAGCACACGTCTGAACTCTTTCCCTACACGACGCTCTTCCGATCTNNNCACAGTCCTTCCCATTATTACC
GAGCACACGTCTGAACTCTTTCCCTACACGACGCTCTTCCGATCTNNNGAGCAAGACACCGTCTCAGAA
GAGCACACGTCTGAACTCTTTCCCTACACGACGCTCTTCCGATCTNNNGAGCAAGACACCGTCTCAGAA
AGCACACGTCTGAACTCTTTCCCTACACGACGCTCTTCCGATCTNNNCTGTAGTCTTCAGAACAGCACC
AGCACACGTCTGAACTCTTTCCCTACACGACGCTCTTCCGATCTNNNCTGTAGTCTTCAGAACAGCACC
ACACGTCTGAACTCTTTCCCTACACGACGCTCTTCCGATCTNNNCTCCTTGGCACTCCTTTCCTATGCATC
ACACGTCTGAACTCTTTCCCTACACGACGCTCTTCCGATCTNNNCTCCTTGGCACTCCTTTCCTATGCATC
AGCACACGTCTGAACTCTTTCCCTACACGACGCTCTTCCGATCTNNNGAGAATCGCTTGAACCCAGGA
AGCACACGTCTGAACTCTTTCCCTACACGACGCTCTTCCGATCTNNNGAGAATCGCTTGAACCCAGGA
AGCACACGTCTGAACTCTTTCCCTACACGACGCTCTTCCGATCTNNNCTGTAGTTCCAGCTACTCAGGA
AGCACACGTCTGAACTCTTTCCCTACACGACGCTCTTCCGATCTNNNCTGTAGTTCCAGCTACTCAGGA
AGCACACGTCTGAACTCTTTCCCTACACGACGCTCTTCCGATCTNNNCTGTTACAGGCTCAGACCCCCA
AGCACACGTCTGAACTCTTTCCCTACACGACGCTCTTCCGATCTNNNCTGTTACAGGCTCAGACCCCCA
GAGCACACGTCTGAACTCTTTCCCTACACGACGCTCTTCCGATCTNNNATTAGCCAGGTGTAGAGTCAT
GAGCACACGTCTGAACTCTTTCCCTACACGACGCTCTTCCGATCTNNNATTAGCCAGGTGTAGAGTCAT
AGCACACGTCTGAACTCTTTCCCTACACGACGCTCTTCCGATCTNNNTACAAACATTACCTGGGTGTGC
AGCACACGTCTGAACTCTTTCCCTACACGACGCTCTTCCGATCTNNNTACAAACATTACCTGGGTGTGC
GAGCACACGTCTGAACTCTTTCCCTACACGACGCTCTTCCGATCTNNNTTCTACTTGAGGAGAGGAGA
GAGCACACGTCTGAACTCTTTCCCTACACGACGCTCTTCCGATCTNNNTTCTACTTGAGGAGAGGAGA
AGAGCACACGTCTGAACTCTTTCCCTACACGACGCTCTTCCGATCTNNNCCACACTACAGTCCATTTCCTC
AGAGCACACGTCTGAACTCTTTCCCTACACGACGCTCTTCCGATCTNNNCCACACTACAGTCCATTTCCTC
GAGCACACGTCTGAACTCTTTCCCTACACGACGCTCTTCCGATCTNNNGCTGGCCGCTTTTTTTTTTTTTT
GAGCACACGTCTGAACTCTTTCCCTACACGACGCTCTTCCGATCTNNNGCTGGCCGCTTTTTTTTTTTTTT
GAGCACACGTCTGAACTCTTTCCCTACACGACGCTCTTCCGATCTNNNGCCATGGTGATCTTACTTGAG
GAGCACACGTCTGAACTCTTTCCCTACACGACGCTCTTCCGATCTNNNGCCATGGTGATCTTACTTGAG
GAGCACACGTCTGAACTCTTTCCCTACACGACGCTCTTCCGATCTNNNGCTCCACATTCATGGCAGCTAA
GAGCACACGTCTGAACTCTTTCCCTACACGACGCTCTTCCGATCTNNNGCTCCACATTCATGGCAGCTAA
AGCACACGTCTGAACTCTTTCCCTACACGACGCTCTTCCGATCTNNNGAAGTTGCAGCAAGCTGAGAT
AGCACACGTCTGAACTCTTTCCCTACACGACGCTCTTCCGATCTNNNGAAGTTGCAGCAAGCTGAGAT
AGCACACGTCTGAACTCTTTCCCTACACGACGCTCTTCCGATCTNNNGGGCGACACATACAAAAAGACA
AGCACACGTCTGAACTCTTTCCCTACACGACGCTCTTCCGATCTNNNGGGCGACACATACAAAAAGACA
GCACACGTCTGAACTCTTTCCCTACACGACGCTCTTCCGATCTNNNGGAGTGTGCAGACAGGATTTTTC
GCACACGTCTGAACTCTTTCCCTACACGACGCTCTTCCGATCTNNNGGAGTGTGCAGACAGGATTTTTC
AGAGCACACGTCTGAACTCTTTCCCTACACGACGCTCTTCCGATCTNNNGCAGTGGGAGAAGATGGAA

GAGCACACGTCTGAACTCTTTCCCTACACGACGCTCTTCCGATCTNNNGCAGTGGGAGAAGATGGAA
GCACACGTCTGAACTCTTTCCCTACACGACGCTCTTCCGATCTNNNAGAGTCTCCTGGAAGCTTTCAAG
GCACACGTCTGAACTCTTTCCCTACACGACGCTCTTCCGATCTNNNAGAGTCTCCTGGAAGCTTTCAAG
AGAGCACACGTCTGAACTCTTTCCCTACACGACGCTCTTCCGATCTNNNAGGCTCAGCACAACTTACAC
AGAGCACACGTCTGAACTCTTTCCCTACACGACGCTCTTCCGATCTNNNAGGCTCAGCACAACTTACAC
AGCACACGTCTGAACTCTTTCCCTACACGACGCTCTTCCGATCTNNNCAGGGAAGAGAGAGTGAGAG
AGCACACGTCTGAACTCTTTCCCTACACGACGCTCTTCCGATCTNNNCAGGGAAGAGAGAGTGAGAG
GAGCACACGTCTGAACTCTTTCCCTACACGACGCTCTTCCGATCTNNNCTGTCCTTCAACTATTGGCCAC
GAGCACACGTCTGAACTCTTTCCCTACACGACGCTCTTCCGATCTNNNCTGTCCTTCAACTATTGGCCAC
AGCACACGTCTGAACTCTTTCCCTACACGACGCTCTTCCGATCTNNNAAGCCTATTCCTTCAACCTCTGC
AGCACACGTCTGAACTCTTTCCCTACACGACGCTCTTCCGATCTNNNAAGCCTATTCCTTCAACCTCTGC
GCACACGTCTGAACTCTTTCCCTACACGACGCTCTTCCGATCTNNNTTAGGATGGGAGGATGGTGGA
GCACACGTCTGAACTCTTTCCCTACACGACGCTCTTCCGATCTNNNTTAGGATGGGAGGATGGTGGA
CACACGTCTGAACTCTTTCCCTACACGACGCTCTTCCGATCTNNNTGGAGCTAAGAGTGGCAAGAAT
CACACGTCTGAACTCTTTCCCTACACGACGCTCTTCCGATCTNNNTGGAGCTAAGAGTGGCAAGAAT
GCACACGTCTGAACTCTTTCCCTACACGACGCTCTTCCGATCTNNNTGGTTCTTATGGCAAAAACGTC
GCACACGTCTGAACTCTTTCCCTACACGACGCTCTTCCGATCTNNNTGGTTCTTATGGCAAAAACGTC
AGCACACGTCTGAACTCTTTCCCTACACGACGCTCTTCCGATCTNNNCAATTGAGCCTGTCTGGTTCTC
AGCACACGTCTGAACTCTTTCCCTACACGACGCTCTTCCGATCTNNNCAATTGAGCCTGTCTGGTTCTC
GAGCACACGTCTGAACTCTTTCCCTACACGACGCTCTTCCGATCTNNNCTAGCTCATTGTCCTCCCTGTC
GAGCACACGTCTGAACTCTTTCCCTACACGACGCTCTTCCGATCTNNNCTAGCTCATTGTCCTCCCTGTC
AAGAGCACACGTCTGAACTCTTTCCCTACACGACGCTCTTCCGATCTNNNCTTATTAGGGCTGGCAAA
AAGAGCACACGTCTGAACTCTTTCCCTACACGACGCTCTTCCGATCTNNNCTTATTAGGGCTGGCAAA
GCACACGTCTGAACTCTTTCCCTACACGACGCTCTTCCGATCTNNNACATAGCAGTTCGTGATGTTCCT
GCACACGTCTGAACTCTTTCCCTACACGACGCTCTTCCGATCTNNNACATAGCAGTTCGTGATGTTCCT
GCACACGTCTGAACTCTTTCCCTACACGACGCTCTTCCGATCTNNNTTCTCCTGTGTCAACATCCTGAG
GCACACGTCTGAACTCTTTCCCTACACGACGCTCTTCCGATCTNNNTTCTCCTGTGTCAACATCCTGAG
AGCACACGTCTGAACTCTTTCCCTACACGACGCTCTTCCGATCTNNNCCTCAACCCCTCCTTAGGCTCAT
AGCACACGTCTGAACTCTTTCCCTACACGACGCTCTTCCGATCTNNNCCTCAACCCCTCCTTAGGCTCAT
AGCACACGTCTGAACTCTTTCCCTACACGACGCTCTTCCGATCTNNNCCAGACCCAAACATTTTCCCCTA
AGCACACGTCTGAACTCTTTCCCTACACGACGCTCTTCCGATCTNNNCCAGACCCAAACATTTTCCCCTA
AGCACACGTCTGAACTCTTTCCCTACACGACGCTCTTCCGATCTNNNCACTATGTTGTCCAGGCTGATC
AGCACACGTCTGAACTCTTTCCCTACACGACGCTCTTCCGATCTNNNCACTATGTTGTCCAGGCTGATC
AGCACACGTCTGAACTCTTTCCCTACACGACGCTCTTCCGATCTNNNTAGCTGGGATTATAGGCATGT
AGCACACGTCTGAACTCTTTCCCTACACGACGCTCTTCCGATCTNNNTAGCTGGGATTATAGGCATGT
AGCACACGTCTGAACTCTTTCCCTACACGACGCTCTTCCGATCTNNNCTCCTTCCTTCCTTCCTTCCTTCC
AGCACACGTCTGAACTCTTTCCCTACACGACGCTCTTCCGATCTNNNCTCCTTCCTTCCTTCCTTCCTTCC
AGCACACGTCTGAACTCTTTCCCTACACGACGCTCTTCCGATCTNNNCAAGTAGCTGGGACTATAGGC
AGCACACGTCTGAACTCTTTCCCTACACGACGCTCTTCCGATCTNNNCAAGTAGCTGGGACTATAGGC
CACACGTCTGAACTCTTTCCCTACACGACGCTCTTCCGATCTNNNAAGCCCATAGTACTCCTGCTATGCA
CACACGTCTGAACTCTTTCCCTACACGACGCTCTTCCGATCTNNNAAGCCCATAGTACTCCTGCTATGCA
AGCACACGTCTGAACTCTTTCCCTACACGACGCTCTTCCGATCTNNNCTTTACCTCCCTCCCACTTTACCC
AGCACACGTCTGAACTCTTTCCCTACACGACGCTCTTCCGATCTNNNCTTTACCTCCCTCCCACTTTACCC

AGCACACGTCTGAACTCTTTCCCTACACGACGCTCTTCCGATCTNNNTGGTATTGCCCTATAATAATTC
AGCACACGTCTGAACTCTTTCCCTACACGACGCTCTTCCGATCTNNNTGGTATTGCCCTATAATAATTC
GCACACGTCTGAACTCTTTCCCTACACGACGCTCTTCCGATCTNNNATGCTTGTAATCCCAGCACTTTG
GCACACGTCTGAACTCTTTCCCTACACGACGCTCTTCCGATCTNNNATGCTTGTAATCCCAGCACTTTG
AGAGCACACGTCTGAACTCTTTCCCTACACGACGCTCTTCCGATCTNNNAGACCAGCCCTCTAATGACT
AGAGCACACGTCTGAACTCTTTCCCTACACGACGCTCTTCCGATCTNNNAGACCAGCCCTCTAATGACT
AGCACACGTCTGAACTCTTTCCCTACACGACGCTCTTCCGATCTNNNGACTTTTGAGTGTGGAGCTGA
AGCACACGTCTGAACTCTTTCCCTACACGACGCTCTTCCGATCTNNNGACTTTTGAGTGTGGAGCTGA
GAGCACACGTCTGAACTCTTTCCCTACACGACGCTCTTCCGATCTNNNGAGAATGGCATGAATCGAG
GAGCACACGTCTGAACTCTTTCCCTACACGACGCTCTTCCGATCTNNNGAGAATGGCATGAATCGAG
AGCACACGTCTGAACTCTTTCCCTACACGACGCTCTTCCGATCTNNNAGGCATAAGGGGCATCCATT
AGCACACGTCTGAACTCTTTCCCTACACGACGCTCTTCCGATCTNNNAGGCATAAGGGGCATCCATT
AGCACACGTCTGAACTCTTTCCCTACACGACGCTCTTCCGATCTNNNGGAGTTTGAACTAGCCTGGG
AGCACACGTCTGAACTCTTTCCCTACACGACGCTCTTCCGATCTNNNGGAGTTTGAACTAGCCTGGG
AGAGCACACGTCTGAACTCTTTCCCTACACGACGCTCTTCCGATCTNNNGGTAGTGAGTTGAGATCAT
AGAGCACACGTCTGAACTCTTTCCCTACACGACGCTCTTCCGATCTNNNGGTAGTGAGTTGAGATCAT
CACGTCTGAACTCTTTCCCTACACGACGCTCTTCCGATCTNNNTGGTCTTTACAACCAGTACCAAACAT
CACGTCTGAACTCTTTCCCTACACGACGCTCTTCCGATCTNNNTGGTCTTTACAACCAGTACCAAACAT
AGCACACGTCTGAACTCTTTCCCTACACGACGCTCTTCCGATCTNNNACTCGAAGTGGTTTCTCCCTTT
AGCACACGTCTGAACTCTTTCCCTACACGACGCTCTTCCGATCTNNNACTCGAAGTGGTTTCTCCCTTT
AGCACACGTCTGAACTCTTTCCCTACACGACGCTCTTCCGATCTNNNCTGAGGCAGGAGAATTGCTTG
AGCACACGTCTGAACTCTTTCCCTACACGACGCTCTTCCGATCTNNNCTGAGGCAGGAGAATTGCTTG
AGCACACGTCTGAACTCTTTCCCTACACGACGCTCTTCCGATCTNNNAGAGGTTGCAGTGAGTTGAG
AGCACACGTCTGAACTCTTTCCCTACACGACGCTCTTCCGATCTNNNAGAGGTTGCAGTGAGTTGAG
CACACGTCTGAACTCTTTCCCTACACGACGCTCTTCCGATCTNNNCTCTCCTCTGTCTCTCTCTCTCCCAT
CACACGTCTGAACTCTTTCCCTACACGACGCTCTTCCGATCTNNNCTCTCCTCTGTCTCTCTCTCTCCCAT
AGCACACGTCTGAACTCTTTCCCTACACGACGCTCTTCCGATCTNNNAAGTAGCTGGGATTACAGGCA
AGCACACGTCTGAACTCTTTCCCTACACGACGCTCTTCCGATCTNNNAAGTAGCTGGGATTACAGGCA
CACGTCTGAACTCTTTCCCTACACGACGCTCTTCCGATCTNNNCCATTCCTCTCCTGTGTCCATACCATC
CACGTCTGAACTCTTTCCCTACACGACGCTCTTCCGATCTNNNCCATTCCTCTCCTGTGTCCATACCATC
CACACGTCTGAACTCTTTCCCTACACGACGCTCTTCCGATCTNNNAATTAACTAGTTCCATGCATTTTCA
CACACGTCTGAACTCTTTCCCTACACGACGCTCTTCCGATCTNNNAATTAACTAGTTCCATGCATTTTCA
GCACACGTCTGAACTCTTTCCCTACACGACGCTCTTCCGATCTNNNAACAAATATCTCCACCGTCCGAG
GCACACGTCTGAACTCTTTCCCTACACGACGCTCTTCCGATCTNNNAACAAATATCTCCACCGTCCGAG
AGCACACGTCTGAACTCTTTCCCTACACGACGCTCTTCCGATCTNNNCACTCAGCCTGAACACATCTAC
AGCACACGTCTGAACTCTTTCCCTACACGACGCTCTTCCGATCTNNNCACTCAGCCTGAACACATCTAC
GCACACGTCTGAACTCTTTCCCTACACGACGCTCTTCCGATCTNNNAAGAGCAGAAGAAGCTAGAGG
GCACACGTCTGAACTCTTTCCCTACACGACGCTCTTCCGATCTNNNAAGAGCAGAAGAAGCTAGAGG
GAGCACACGTCTGAACTCTTTCCCTACACGACGCTCTTCCGATCTNNNATCCAGGCTCTTTCCATCTTG
GAGCACACGTCTGAACTCTTTCCCTACACGACGCTCTTCCGATCTNNNATCCAGGCTCTTTCCATCTTG
AGCACACGTCTGAACTCTTTCCCTACACGACGCTCTTCCGATCTNNNCAGAAAGGTCAAATGCTGGA
AGCACACGTCTGAACTCTTTCCCTACACGACGCTCTTCCGATCTNNNCAGAAAGGTCAAATGCTGGA
AGCACACGTCTGAACTCTTTCCCTACACGACGCTCTTCCGATCTNNNAATGCAACACACACTAGCATCA

AGCACACGTCTGAACTCTTTCCCTACACGACGCTCTTCCGATCTNNNAATGCAACACACACTAGCATCA
AGCACACGTCTGAACTCTTTCCCTACACGACGCTCTTCCGATCTNNNTGGGAGGTTGAGGTAGGAGA
AGCACACGTCTGAACTCTTTCCCTACACGACGCTCTTCCGATCTNNNTGGGAGGTTGAGGTAGGAGA
GCACACGTCTGAACTCTTTCCCTACACGACGCTCTTCCGATCTNNNGGCATCCTTGACAAACCAAACTCC
GCACACGTCTGAACTCTTTCCCTACACGACGCTCTTCCGATCTNNNGGCATCCTTGACAAACCAAACTCC
GCACACGTCTGAACTCTTTCCCTACACGACGCTCTTCCGATCTNNNGAGAAGCACCGATCCCAGGGCAT
GCACACGTCTGAACTCTTTCCCTACACGACGCTCTTCCGATCTNNNGAGAAGCACCGATCCCAGGGCAT
AGCACACGTCTGAACTCTTTCCCTACACGACGCTCTTCCGATCTNNNCACATCCTATTGGGCCCTCTTTA
AGCACACGTCTGAACTCTTTCCCTACACGACGCTCTTCCGATCTNNNCACATCCTATTGGGCCCTCTTTA
GAGCACACGTCTGAACTCTTTCCCTACACGACGCTCTTCCGATCTNNNTGACAGAGCAAGAGTCCATTT
GAGCACACGTCTGAACTCTTTCCCTACACGACGCTCTTCCGATCTNNNTGACAGAGCAAGAGTCCATTT
GAGCACACGTCTGAACTCTTTCCCTACACGACGCTCTTCCGATCTNNNACCCTTCTGTCTCTCTCTCTCTC
GAGCACACGTCTGAACTCTTTCCCTACACGACGCTCTTCCGATCTNNNACCCTTCTGTCTCTCTCTCTCTC
AGCACACGTCTGAACTCTTTCCCTACACGACGCTCTTCCGATCTNNNACCACACCCGGCTAATTTTTTTG
AGCACACGTCTGAACTCTTTCCCTACACGACGCTCTTCCGATCTNNNACCACACCCGGCTAATTTTTTTG
AGCACACGTCTGAACTCTTTCCCTACACGACGCTCTTCCGATCTNNNCGGCTTTCAGTGTAGGAGTTAC
AGCACACGTCTGAACTCTTTCCCTACACGACGCTCTTCCGATCTNNNCGGCTTTCAGTGTAGGAGTTAC
GAGCACACGTCTGAACTCTTTCCCTACACGACGCTCTTCCGATCTNNNGCATGAGCCAATTTCTTACCAC
GAGCACACGTCTGAACTCTTTCCCTACACGACGCTCTTCCGATCTNNNGCATGAGCCAATTTCTTACCAC
GAGCACACGTCTGAACTCTTTCCCTACACGACGCTCTTCCGATCTNNNACAATCCGTCTATTGAGGACA
GAGCACACGTCTGAACTCTTTCCCTACACGACGCTCTTCCGATCTNNNACAATCCGTCTATTGAGGACA
AGCACACGTCTGAACTCTTTCCCTACACGACGCTCTTCCGATCTNNNGAAAACCTACCAGGGCAGCTAC
AGCACACGTCTGAACTCTTTCCCTACACGACGCTCTTCCGATCTNNNGAAAACCTACCAGGGCAGCTAC
AGCACACGTCTGAACTCTTTCCCTACACGACGCTCTTCCGATCTNNNCCCAAATGATGAAGTGATGCAG
AGCACACGTCTGAACTCTTTCCCTACACGACGCTCTTCCGATCTNNNCCCAAATGATGAAGTGATGCAG
GAGCACACGTCTGAACTCTTTCCCTACACGACGCTCTTCCGATCTNNNTGGGGAATAGTAGCAGAGGA
GAGCACACGTCTGAACTCTTTCCCTACACGACGCTCTTCCGATCTNNNTGGGGAATAGTAGCAGAGGA
AGCACACGTCTGAACTCTTTCCCTACACGACGCTCTTCCGATCTNNNAGAAGCCAAAGCACACATAGA
AGCACACGTCTGAACTCTTTCCCTACACGACGCTCTTCCGATCTNNNAGAAGCCAAAGCACACATAGA
GAGCACACGTCTGAACTCTTTCCCTACACGACGCTCTTCCGATCTNNNAGTCTCCAATTCCATCCAGGTT
GAGCACACGTCTGAACTCTTTCCCTACACGACGCTCTTCCGATCTNNNAGTCTCCAATTCCATCCAGGTT
GAGCACACGTCTGAACTCTTTCCCTACACGACGCTCTTCCGATCTNNNCCAGTAATGGAGCCTACATACC
GAGCACACGTCTGAACTCTTTCCCTACACGACGCTCTTCCGATCTNNNCCAGTAATGGAGCCTACATACC
GAGCACACGTCTGAACTCTTTCCCTACACGACGCTCTTCCGATCTNNNTGTAAACTCACTCTAGAAAATG
GAGCACACGTCTGAACTCTTTCCCTACACGACGCTCTTCCGATCTNNNTGTAAACTCACTCTAGAAAATG
AGCACACGTCTGAACTCTTTCCCTACACGACGCTCTTCCGATCTNNNAGAAACTCAGCTCCAAAATGGC
AGCACACGTCTGAACTCTTTCCCTACACGACGCTCTTCCGATCTNNNAGAAACTCAGCTCCAAAATGGC
AGAGCACACGTCTGAACTCTTTCCCTACACGACGCTCTTCCGATCTNNNGAGGCAGGAGAATTGTTTGA
AGAGCACACGTCTGAACTCTTTCCCTACACGACGCTCTTCCGATCTNNNGAGGCAGGAGAATTGTTTGA
GAGCACACGTCTGAACTCTTTCCCTACACGACGCTCTTCCGATCTNNNCACCAGGATGTAAGAGGTCC
GAGCACACGTCTGAACTCTTTCCCTACACGACGCTCTTCCGATCTNNNCACCAGGATGTAAGAGGTCC
AGAGCACACGTCTGAACTCTTTCCCTACACGACGCTCTTCCGATCTNNNATTCACTCTGAGACCCTATCC
AGAGCACACGTCTGAACTCTTTCCCTACACGACGCTCTTCCGATCTNNNATTCACTCTGAGACCCTATCC

AGCACACGTCTGAACTCTTTCCCTACACGACGCTCTTCCGATCTNNNTCTGTAATCCCAGCTACTCAGG
AGCACACGTCTGAACTCTTTCCCTACACGACGCTCTTCCGATCTNNNTCTGTAATCCCAGCTACTCAGG
GAGCACACGTCTGAACTCTTTCCCTACACGACGCTCTTCCGATCTNNNCGAAGATTGGCAGCTCAGAT
GAGCACACGTCTGAACTCTTTCCCTACACGACGCTCTTCCGATCTNNNCGAAGATTGGCAGCTCAGAT
ACACGTCTGAACTCTTTCCCTACACGACGCTCTTCCGATCTNNNTTCCCATTTGTTTCATCCGTTCGCAT
ACACGTCTGAACTCTTTCCCTACACGACGCTCTTCCGATCTNNNTTCCCATTTGTTTCATCCGTTCGCAT
GCACACGTCTGAACTCTTTCCCTACACGACGCTCTTCCGATCTNNNCAGGAGAATCGCTTGAACCCAAC
GCACACGTCTGAACTCTTTCCCTACACGACGCTCTTCCGATCTNNNCAGGAGAATCGCTTGAACCCAAC
GAGCACACGTCTGAACTCTTTCCCTACACGACGCTCTTCCGATCTNNNTGGCAGATACTCTTAGGACG
GAGCACACGTCTGAACTCTTTCCCTACACGACGCTCTTCCGATCTNNNTGGCAGATACTCTTAGGACG
AGCACACGTCTGAACTCTTTCCCTACACGACGCTCTTCCGATCTNNNCTGTAATCCCAGCTACTCCAGA
AGCACACGTCTGAACTCTTTCCCTACACGACGCTCTTCCGATCTNNNCTGTAATCCCAGCTACTCCAGA
GCACACGTCTGAACTCTTTCCCTACACGACGCTCTTCCGATCTNNNCCAGGTTCCAAGAGAAACCCTA
GCACACGTCTGAACTCTTTCCCTACACGACGCTCTTCCGATCTNNNCCAGGTTCCAAGAGAAACCCTA
ACGTCTGAACTCTTTCCCTACACGACGCTCTTCCGATCTNNNTCTTTACCCTCTAAGCTGAACATCGGA
ACGTCTGAACTCTTTCCCTACACGACGCTCTTCCGATCTNNNTCTTTACCCTCTAAGCTGAACATCGGA
GCACACGTCTGAACTCTTTCCCTACACGACGCTCTTCCGATCTNNNGACCAAGATGTGGAACAAGTG
GCACACGTCTGAACTCTTTCCCTACACGACGCTCTTCCGATCTNNNGACCAAGATGTGGAACAAGTG
GCACACGTCTGAACTCTTTCCCTACACGACGCTCTTCCGATCTNNNATGAAAGATGTAGAAGCCCAG
GCACACGTCTGAACTCTTTCCCTACACGACGCTCTTCCGATCTNNNATGAAAGATGTAGAAGCCCAG
AGCACACGTCTGAACTCTTTCCCTACACGACGCTCTTCCGATCTNNNCAGAGAAAACAACCGGAGAG
AGCACACGTCTGAACTCTTTCCCTACACGACGCTCTTCCGATCTNNNCAGAGAAAACAACCGGAGAG
AGCACACGTCTGAACTCTTTCCCTACACGACGCTCTTCCGATCTNNNTCCAGATAAACCGAATGCAGT
AGCACACGTCTGAACTCTTTCCCTACACGACGCTCTTCCGATCTNNNTCCAGATAAACCGAATGCAGT
GCACACGTCTGAACTCTTTCCCTACACGACGCTCTTCCGATCTNNNATTTCCTTTCAAAAGAGACAGTC
GCACACGTCTGAACTCTTTCCCTACACGACGCTCTTCCGATCTNNNATTTCCTTTCAAAAGAGACAGTC
AGCACACGTCTGAACTCTTTCCCTACACGACGCTCTTCCGATCTNNNTACTTTCACAACTGTAAGCATA
AGCACACGTCTGAACTCTTTCCCTACACGACGCTCTTCCGATCTNNNTACTTTCACAACTGTAAGCATA
AGCACACGTCTGAACTCTTTCCCTACACGACGCTCTTCCGATCTNNNGTGAATGTGACGGGACAGAC
AGCACACGTCTGAACTCTTTCCCTACACGACGCTCTTCCGATCTNNNGTGAATGTGACGGGACAGAC
GCACACGTCTGAACTCTTTCCCTACACGACGCTCTTCCGATCTNNNTGATTTCTGTGAGAGTTCCTTG
GCACACGTCTGAACTCTTTCCCTACACGACGCTCTTCCGATCTNNNTGATTTCTGTGAGAGTTCCTTG
GAGCACACGTCTGAACTCTTTCCCTACACGACGCTCTTCCGATCTNNNATGCCCCAGAGAGAGGTAGG
GAGCACACGTCTGAACTCTTTCCCTACACGACGCTCTTCCGATCTNNNATGCCCCAGAGAGAGGTAGG
GCACACGTCTGAACTCTTTCCCTACACGACGCTCTTCCGATCTNNNCTCTGCTCAGTTCCTGGACAAAA
GCACACGTCTGAACTCTTTCCCTACACGACGCTCTTCCGATCTNNNCTCTGCTCAGTTCCTGGACAAAA
GAGCACACGTCTGAACTCTTTCCCTACACGACGCTCTTCCGATCTNNNTCCTGGTGGGGTGTTCAATA
GAGCACACGTCTGAACTCTTTCCCTACACGACGCTCTTCCGATCTNNNTCCTGGTGGGGTGTTCAATA
AGCACACGTCTGAACTCTTTCCCTACACGACGCTCTTCCGATCTNNNACCACACAAGACATTTTTCTAC
AGCACACGTCTGAACTCTTTCCCTACACGACGCTCTTCCGATCTNNNACCACACAAGACATTTTTCTAC
AGCACACGTCTGAACTCTTTCCCTACACGACGCTCTTCCGATCTNNNCAGGGTTTCACTGTTTTAGCTG
AGCACACGTCTGAACTCTTTCCCTACACGACGCTCTTCCGATCTNNNCAGGGTTTCACTGTTTTAGCTG
GAGCACACGTCTGAACTCTTTCCCTACACGACGCTCTTCCGATCTNNNTGTGAATATATGCTCAAAGG

GAGCACACGTCTGAACTCTTTCCCTACACGACGCTCTTCCGATCTNNNTGTGAATATATGCTCAAAGG
AGCACACGTCTGAACTCTTTCCCTACACGACGCTCTTCCGATCTNNNATCAGGCATGACATCCTAAGT
AGCACACGTCTGAACTCTTTCCCTACACGACGCTCTTCCGATCTNNNATCAGGCATGACATCCTAAGT
AGCACACGTCTGAACTCTTTCCCTACACGACGCTCTTCCGATCTNNNCTATAGGTTAGGCCCAGGGG
AGCACACGTCTGAACTCTTTCCCTACACGACGCTCTTCCGATCTNNNCTATAGGTTAGGCCCAGGGG
AGCACACGTCTGAACTCTTTCCCTACACGACGCTCTTCCGATCTNNNTATATACACACACACACACACG
AGCACACGTCTGAACTCTTTCCCTACACGACGCTCTTCCGATCTNNNTATATACACACACACACACACG
AGCACACGTCTGAACTCTTTCCCTACACGACGCTCTTCCGATCTNNNCCAACATGGTGAAACTCCATCT
AGCACACGTCTGAACTCTTTCCCTACACGACGCTCTTCCGATCTNNNCCAACATGGTGAAACTCCATCT
AGCACACGTCTGAACTCTTTCCCTACACGACGCTCTTCCGATCTNNNGTTTGATTTGGTGGCTGTTCTG
AGCACACGTCTGAACTCTTTCCCTACACGACGCTCTTCCGATCTNNNGTTTGATTTGGTGGCTGTTCTG
GAGCACACGTCTGAACTCTTTCCCTACACGACGCTCTTCCGATCTNNNGTCTTCCTCCTTCGGAATTCA
GAGCACACGTCTGAACTCTTTCCCTACACGACGCTCTTCCGATCTNNNGTCTTCCTCCTTCGGAATTCA
GAGCACACGTCTGAACTCTTTCCCTACACGACGCTCTTCCGATCTNNNACAGTGAAACCCCGTCTCTAC
GAGCACACGTCTGAACTCTTTCCCTACACGACGCTCTTCCGATCTNNNACAGTGAAACCCCGTCTCTAC
AGCACACGTCTGAACTCTTTCCCTACACGACGCTCTTCCGATCTNNNACCTGATATACCACTGCACAAG
AGCACACGTCTGAACTCTTTCCCTACACGACGCTCTTCCGATCTNNNACCTGATATACCACTGCACAAG
AGCACACGTCTGAACTCTTTCCCTACACGACGCTCTTCCGATCTNNNTTTAGTAGAGACGGGATTTCG
AGCACACGTCTGAACTCTTTCCCTACACGACGCTCTTCCGATCTNNNTTTAGTAGAGACGGGATTTCG
GAGCACACGTCTGAACTCTTTCCCTACACGACGCTCTTCCGATCTNNNTGAAGTGGAGGAGTAGAAAG
GAGCACACGTCTGAACTCTTTCCCTACACGACGCTCTTCCGATCTNNNTGAAGTGGAGGAGTAGAAAG
AGCACACGTCTGAACTCTTTCCCTACACGACGCTCTTCCGATCTNNNGCAGGAGAATTGCTTGAACCTG
AGCACACGTCTGAACTCTTTCCCTACACGACGCTCTTCCGATCTNNNGCAGGAGAATTGCTTGAACCTG
GAGCACACGTCTGAACTCTTTCCCTACACGACGCTCTTCCGATCTNNNTCATGTGTTTCTCTCTGCCTTG
GAGCACACGTCTGAACTCTTTCCCTACACGACGCTCTTCCGATCTNNNTCATGTGTTTCTCTCTGCCTTG
AGCACACGTCTGAACTCTTTCCCTACACGACGCTCTTCCGATCTNNNTGCCCTACTGTCTAGGAGATCA
AGCACACGTCTGAACTCTTTCCCTACACGACGCTCTTCCGATCTNNNTGCCCTACTGTCTAGGAGATCA
GAGCACACGTCTGAACTCTTTCCCTACACGACGCTCTTCCGATCTNNNCCTCCTGAGTAGCTGGGACT
GAGCACACGTCTGAACTCTTTCCCTACACGACGCTCTTCCGATCTNNNCCTCCTGAGTAGCTGGGACT
GAGCACACGTCTGAACTCTTTCCCTACACGACGCTCTTCCGATCTNNNGCGACAAAGCAAGAATCTGT
GAGCACACGTCTGAACTCTTTCCCTACACGACGCTCTTCCGATCTNNNGCGACAAAGCAAGAATCTGT
GAGCACACGTCTGAACTCTTTCCCTACACGACGCTCTTCCGATCTNNNCCTTTTTACACCTCTGCCTTGT
GAGCACACGTCTGAACTCTTTCCCTACACGACGCTCTTCCGATCTNNNCCTTTTTACACCTCTGCCTTGT
AGCACACGTCTGAACTCTTTCCCTACACGACGCTCTTCCGATCTNNNTTCAAGTGATTCTCCTGCCTCAG
AGCACACGTCTGAACTCTTTCCCTACACGACGCTCTTCCGATCTNNNTTCAAGTGATTCTCCTGCCTCAG
GCACACGTCTGAACTCTTTCCCTACACGACGCTCTTCCGATCTNNNGGCAGGAGAATTGCTTGAATCC
GCACACGTCTGAACTCTTTCCCTACACGACGCTCTTCCGATCTNNNGGCAGGAGAATTGCTTGAATCC
GAGCACACGTCTGAACTCTTTCCCTACACGACGCTCTTCCGATCTNNNTGGAAGCCAGGAGAAAGTG
GAGCACACGTCTGAACTCTTTCCCTACACGACGCTCTTCCGATCTNNNTGGAAGCCAGGAGAAAGTG
GAGCACACGTCTGAACTCTTTCCCTACACGACGCTCTTCCGATCTNNNTGAAGACAGGATTGGATTCC
GAGCACACGTCTGAACTCTTTCCCTACACGACGCTCTTCCGATCTNNNTGAAGACAGGATTGGATTCC
AGCACACGTCTGAACTCTTTCCCTACACGACGCTCTTCCGATCTNNNCACCTGTAATCCCAGCTACTTG
AGCACACGTCTGAACTCTTTCCCTACACGACGCTCTTCCGATCTNNNCACCTGTAATCCCAGCTACTTG

AGCACACGTCTGAACTCTTTCCCTACACGACGCTCTTCCGATCTNNNAAATAAGCAATCGGCCAGGT
AGCACACGTCTGAACTCTTTCCCTACACGACGCTCTTCCGATCTNNNAAATAAGCAATCGGCCAGGT
GCACACGTCTGAACTCTTTCCCTACACGACGCTCTTCCGATCTNNNAGGGGGAAATTATTTGTGGTG
GCACACGTCTGAACTCTTTCCCTACACGACGCTCTTCCGATCTNNNAGGGGGAAATTATTTGTGGTG
AGCACACGTCTGAACTCTTTCCCTACACGACGCTCTTCCGATCTNNNTAAGCGCTTGTAATCCCACCTA
AGCACACGTCTGAACTCTTTCCCTACACGACGCTCTTCCGATCTNNNTAAGCGCTTGTAATCCCACCTA
AGCACACGTCTGAACTCTTTCCCTACACGACGCTCTTCCGATCTNNNATGCTCATTGATGTCTGCCAAT
AGCACACGTCTGAACTCTTTCCCTACACGACGCTCTTCCGATCTNNNATGCTCATTGATGTCTGCCAAT
AGCACACGTCTGAACTCTTTCCCTACACGACGCTCTTCCGATCTNNNACTCCTAGTCCCAGCTACTCAG
AGCACACGTCTGAACTCTTTCCCTACACGACGCTCTTCCGATCTNNNACTCCTAGTCCCAGCTACTCAG
GAGCACACGTCTGAACTCTTTCCCTACACGACGCTCTTCCGATCTNNNAGAGAAGGGGTCTGATGAG
GAGCACACGTCTGAACTCTTTCCCTACACGACGCTCTTCCGATCTNNNAGAGAAGGGGTCTGATGAG
GAGCACACGTCTGAACTCTTTCCCTACACGACGCTCTTCCGATCTNNNCGGCATCATTAACTGGGGA
GAGCACACGTCTGAACTCTTTCCCTACACGACGCTCTTCCGATCTNNNCGGCATCATTAACTGGGGA
GCACACGTCTGAACTCTTTCCCTACACGACGCTCTTCCGATCTNNNGGCCACATATTCAGTTTGGTTG
GCACACGTCTGAACTCTTTCCCTACACGACGCTCTTCCGATCTNNNGGCCACATATTCAGTTTGGTTG
AGCACACGTCTGAACTCTTTCCCTACACGACGCTCTTCCGATCTNNNAGGCAGGAGAATCACTTGAA
AGCACACGTCTGAACTCTTTCCCTACACGACGCTCTTCCGATCTNNNAGGCAGGAGAATCACTTGAA
AGCACACGTCTGAACTCTTTCCCTACACGACGCTCTTCCGATCTNNNCATGAGACACAAAGGACTCCC
AGCACACGTCTGAACTCTTTCCCTACACGACGCTCTTCCGATCTNNNCATGAGACACAAAGGACTCCC
AGCACACGTCTGAACTCTTTCCCTACACGACGCTCTTCCGATCTNNNAACTGCAGATCTTGGGACTTCT
AGCACACGTCTGAACTCTTTCCCTACACGACGCTCTTCCGATCTNNNAACTGCAGATCTTGGGACTTCT
CACACGTCTGAACTCTTTCCCTACACGACGCTCTTCCGATCTNNNTTATTCAGTGCTGCAATCCCAACC
CACACGTCTGAACTCTTTCCCTACACGACGCTCTTCCGATCTNNNTTATTCAGTGCTGCAATCCCAACC
AGCACACGTCTGAACTCTTTCCCTACACGACGCTCTTCCGATCTNNNTGCCAGTAATTCCAGCATTTTG
AGCACACGTCTGAACTCTTTCCCTACACGACGCTCTTCCGATCTNNNTGCCAGTAATTCCAGCATTTTG
AGCACACGTCTGAACTCTTTCCCTACACGACGCTCTTCCGATCTNNNGTGTGTGGGTGTGAATGTAT
AGCACACGTCTGAACTCTTTCCCTACACGACGCTCTTCCGATCTNNNGTGTGTGGGTGTGAATGTAT
ACACGTCTGAACTCTTTCCCTACACGACGCTCTTCCGATCTNNNGGAAAAACATTTGCATGGTTATCA
ACACGTCTGAACTCTTTCCCTACACGACGCTCTTCCGATCTNNNGGAAAAACATTTGCATGGTTATCA
GCACACGTCTGAACTCTTTCCCTACACGACGCTCTTCCGATCTNNNAAAAATTAGCCCAGTGTGATGG
GCACACGTCTGAACTCTTTCCCTACACGACGCTCTTCCGATCTNNNAAAAATTAGCCCAGTGTGATGG
CACACGTCTGAACTCTTTCCCTACACGACGCTCTTCCGATCTNNNTTCTGCAGATCACGAAGACACAG
CACACGTCTGAACTCTTTCCCTACACGACGCTCTTCCGATCTNNNTTCTGCAGATCACGAAGACACAG
GCACACGTCTGAACTCTTTCCCTACACGACGCTCTTCCGATCTNNNCTCTGGCTTGAAGTTCTCTCCTT
GCACACGTCTGAACTCTTTCCCTACACGACGCTCTTCCGATCTNNNCTCTGGCTTGAAGTTCTCTCCTT
AGCACACGTCTGAACTCTTTCCCTACACGACGCTCTTCCGATCTNNNATGCCCATGGAAAAGAGAAA
AGCACACGTCTGAACTCTTTCCCTACACGACGCTCTTCCGATCTNNNATGCCCATGGAAAAGAGAAA
AGCACACGTCTGAACTCTTTCCCTACACGACGCTCTTCCGATCTNNNCTGAGGCAGGAGAATCACTT
AGCACACGTCTGAACTCTTTCCCTACACGACGCTCTTCCGATCTNNNCTGAGGCAGGAGAATCACTT
AGCACACGTCTGAACTCTTTCCCTACACGACGCTCTTCCGATCTNNNCTGAGGCAGGAGAATCACTTG
AGCACACGTCTGAACTCTTTCCCTACACGACGCTCTTCCGATCTNNNCTGAGGCAGGAGAATCACTTG
GCACACGTCTGAACTCTTTCCCTACACGACGCTCTTCCGATCTNNNATGGTACACTCAAGGCATTATG

AGCACACGTCTGAACTCTTTCCCTACACGACGCTCTTCCGATCTNNNATGGTACACTCAAGGCATTATG

AGAGCACACGTCTGAACTCTTTCCCTACACGACGCTCTTCCGATCTNNNGGAGAAGACAATGTGTGTG

AGAGCACACGTCTGAACTCTTTCCCTACACGACGCTCTTCCGATCTNNNGGAGAAGACAATGTGTGTG

AGCACACGTCTGAACTCTTTCCCTACACGACGCTCTTCCGATCTNNNAATGTTTGGGCTTCCTCAGAGA

AGCACACGTCTGAACTCTTTCCCTACACGACGCTCTTCCGATCTNNNAATGTTTGGGCTTCCTCAGAGA

GAGCACACGTCTGAACTCTTTCCCTACACGACGCTCTTCCGATCTNNNGCAGAGGTTGTAGTGAGCAA

GAGCACACGTCTGAACTCTTTCCCTACACGACGCTCTTCCGATCTNNNGCAGAGGTTGTAGTGAGCAA

AGCACACGTCTGAACTCTTTCCCTACACGACGCTCTTCCGATCTNNNAACTCATGGCTGTTGGTATAG

AGCACACGTCTGAACTCTTTCCCTACACGACGCTCTTCCGATCTNNNAACTCATGGCTGTTGGTATAG

AGCACACGTCTGAACTCTTTCCCTACACGACGCTCTTCCGATCTNNNCTGTAATTCCAGCTACTCGGGA

AGCACACGTCTGAACTCTTTCCCTACACGACGCTCTTCCGATCTNNNCTGTAATTCCAGCTACTCGGGA

AGCACACGTCTGAACTCTTTCCCTACACGACGCTCTTCCGATCTNNNGCTCTTTGGTTTCAGAAAGTC

AGCACACGTCTGAACTCTTTCCCTACACGACGCTCTTCCGATCTNNNGCTCTTTGGTTTCAGAAAGTC

AGCACACGTCTGAACTCTTTCCCTACACGACGCTCTTCCGATCTNNNGGTGGTTGTGGTGACCTGATA

AGCACACGTCTGAACTCTTTCCCTACACGACGCTCTTCCGATCTNNNGGTGGTTGTGGTGACCTGATA

GAGCACACGTCTGAACTCTTTCCCTACACGACGCTCTTCCGATCTNNNCTAAGGCAGGAGTATCGCTTG

GAGCACACGTCTGAACTCTTTCCCTACACGACGCTCTTCCGATCTNNNCTAAGGCAGGAGTATCGCTTG

GAGCACACGTCTGAACTCTTTCCCTACACGACGCTCTTCCGATCTNNNTGTTTTGGTTTGGTTTTGGGT

GAGCACACGTCTGAACTCTTTCCCTACACGACGCTCTTCCGATCTNNNTGTTTTGGTTTGGTTTTGGGT

GAGCACACGTCTGAACTCTTTCCCTACACGACGCTCTTCCGATCTNNNCTATACTAATGCTCCCTGCCTC

GAGCACACGTCTGAACTCTTTCCCTACACGACGCTCTTCCGATCTNNNCTATACTAATGCTCCCTGCCTC

GCACACGTCTGAACTCTTTCCCTACACGACGCTCTTCCGATCTNNNCTTGAACTCCTGACCTCGTGATC

GCACACGTCTGAACTCTTTCCCTACACGACGCTCTTCCGATCTNNNCTTGAACTCCTGACCTCGTGATC

GAGCACACGTCTGAACTCTTTCCCTACACGACGCTCTTCCGATCTNNNCATTCCCACCAGCAGTGTAAA

GAGCACACGTCTGAACTCTTTCCCTACACGACGCTCTTCCGATCTNNNCATTCCCACCAGCAGTGTAAA

AGCACACGTCTGAACTCTTTCCCTACACGACGCTCTTCCGATCTNNNAGTCCCTGCAGATCTTAGAGA

AGCACACGTCTGAACTCTTTCCCTACACGACGCTCTTCCGATCTNNNAGTCCCTGCAGATCTTAGAGA

GAGCACACGTCTGAACTCTTTCCCTACACGACGCTCTTCCGATCTNNNGCCTTGAGGTTTGCATTCTGG

GAGCACACGTCTGAACTCTTTCCCTACACGACGCTCTTCCGATCTNNNGCCTTGAGGTTTGCATTCTGG

GAGCACACGTCTGAACTCTTTCCCTACACGACGCTCTTCCGATCTNNNGGGGAGAAAGGCAAGTTATT

GAGCACACGTCTGAACTCTTTCCCTACACGACGCTCTTCCGATCTNNNGGGGAGAAAGGCAAGTTATT

AGCACACGTCTGAACTCTTTCCCTACACGACGCTCTTCCGATCTNNNTACAGTAATAGGGATGCGCCA

AGCACACGTCTGAACTCTTTCCCTACACGACGCTCTTCCGATCTNNNTACAGTAATAGGGATGCGCCA

AGCACACGTCTGAACTCTTTCCCTACACGACGCTCTTCCGATCTNNNACCTGTAATCCCAGCTACTTTGG

AGCACACGTCTGAACTCTTTCCCTACACGACGCTCTTCCGATCTNNNACCTGTAATCCCAGCTACTTTGG

AGAGCACACGTCTGAACTCTTTCCCTACACGACGCTCTTCCGATCTNNNGAGGCAGGAGAATTGCTTG

AGAGCACACGTCTGAACTCTTTCCCTACACGACGCTCTTCCGATCTNNNGAGGCAGGAGAATTGCTTG

GAGCACACGTCTGAACTCTTTCCCTACACGACGCTCTTCCGATCTNNNGAGGTTGCAATGAGCCAAGAT

GAGCACACGTCTGAACTCTTTCCCTACACGACGCTCTTCCGATCTNNNGAGGTTGCAATGAGCCAAGAT

AGCACACGTCTGAACTCTTTCCCTACACGACGCTCTTCCGATCTNNNCAAGTCCGAGGCTCTGAGAAG

AGCACACGTCTGAACTCTTTCCCTACACGACGCTCTTCCGATCTNNNCAAGTCCGAGGCTCTGAGAAG

GAGCACACGTCTGAACTCTTTCCCTACACGACGCTCTTCCGATCTNNNGTATTGCCAGCTATTTTGGAG

GAGCACACGTCTGAACTCTTTCCCTACACGACGCTCTTCCGATCTNNNGTATTGCCAGCTATTTTGGAG

AGCACACGTCTGAACTCTTTCCCTACACGACGCTCTTCCGATCTNNNTCAGGTAATGAGAGTCCTCCA

AGCACACGTCTGAACTCTTTCCCTACACGACGCTCTTCCGATCTNNNTCAGGTAATGAGAGTCCTCCA

GCACACGTCTGAACTCTTTCCCTACACGACGCTCTTCCGATCTNNNTTTTCTTGCCTAATTGCCCTGG

GCACACGTCTGAACTCTTTCCCTACACGACGCTCTTCCGATCTNNNTTTTCTTGCCTAATTGCCCTGG

AGCACACGTCTGAACTCTTTCCCTACACGACGCTCTTCCGATCTNNNCAACATGGAGAAACCCTGTCT

AGCACACGTCTGAACTCTTTCCCTACACGACGCTCTTCCGATCTNNNCAACATGGAGAAACCCTGTCT

GAGCACACGTCTGAACTCTTTCCCTACACGACGCTCTTCCGATCTNNNACAAGTTCAGGAAATCGAGA

GAGCACACGTCTGAACTCTTTCCCTACACGACGCTCTTCCGATCTNNNACAAGTTCAGGAAATCGAGA

CACACGTCTGAACTCTTTCCCTACACGACGCTCTTCCGATCTNNNGTCCATGGCCTATTAGGAAGCAT

CACACGTCTGAACTCTTTCCCTACACGACGCTCTTCCGATCTNNNGTCCATGGCCTATTAGGAAGCAT

AGCACACGTCTGAACTCTTTCCCTACACGACGCTCTTCCGATCTNNNTGATATTAATGACCAGTCCAT

AGCACACGTCTGAACTCTTTCCCTACACGACGCTCTTCCGATCTNNNTGATATTAATGACCAGTCCAT

AGCACACGTCTGAACTCTTTCCCTACACGACGCTCTTCCGATCTNNNTAGGCTATTCGATAACCTGCA

AGCACACGTCTGAACTCTTTCCCTACACGACGCTCTTCCGATCTNNNTAGGCTATTCGATAACCTGCA

AGCACACGTCTGAACTCTTTCCCTACACGACGCTCTTCCGATCTNNNGCCAATATGGAGAAACCCTG

AGCACACGTCTGAACTCTTTCCCTACACGACGCTCTTCCGATCTNNNGCCAATATGGAGAAACCCTG

AGCACACGTCTGAACTCTTTCCCTACACGACGCTCTTCCGATCTNNNTCTTTGCCTTTTTGTGCAAGAA

AGCACACGTCTGAACTCTTTCCCTACACGACGCTCTTCCGATCTNNNTCTTTGCCTTTTTGTGCAAGAA

AGCACACGTCTGAACTCTTTCCCTACACGACGCTCTTCCGATCTNNNAACATCAGGAAATGGTCTCTC

AGCACACGTCTGAACTCTTTCCCTACACGACGCTCTTCCGATCTNNNAACATCAGGAAATGGTCTCTC

AGCACACGTCTGAACTCTTTCCCTACACGACGCTCTTCCGATCTNNNAGATAGCTTCTTTCTGTCCAG

AGCACACGTCTGAACTCTTTCCCTACACGACGCTCTTCCGATCTNNNAGATAGCTTCTTTCTGTCCAG

AGCACACGTCTGAACTCTTTCCCTACACGACGCTCTTCCGATCTNNNTCAGCAGTGTTTTTGTAGTTC

AGCACACGTCTGAACTCTTTCCCTACACGACGCTCTTCCGATCTNNNTCAGCAGTGTTTTTGTAGTTC

AGCACACGTCTGAACTCTTTCCCTACACGACGCTCTTCCGATCTNNNGTGCCTGTAATCCCAGCTACT

AGCACACGTCTGAACTCTTTCCCTACACGACGCTCTTCCGATCTNNNGTGCCTGTAATCCCAGCTACT

GCACACGTCTGAACTCTTTCCCTACACGACGCTCTTCCGATCTNNNGGACATGGGGGACTTGAAGAT

GCACACGTCTGAACTCTTTCCCTACACGACGCTCTTCCGATCTNNNGGACATGGGGGACTTGAAGAT

GAGCACACGTCTGAACTCTTTCCCTACACGACGCTCTTCCGATCTNNNATCAGAGTGGATTTGAGCTG

GAGCACACGTCTGAACTCTTTCCCTACACGACGCTCTTCCGATCTNNNATCAGAGTGGATTTGAGCTG

GCACACGTCTGAACTCTTTCCCTACACGACGCTCTTCCGATCTNNNGAGATCATGCCACTGTACCCCCA

GCACACGTCTGAACTCTTTCCCTACACGACGCTCTTCCGATCTNNNGAGATCATGCCACTGTACCCCCA

GCACACGTCTGAACTCTTTCCCTACACGACGCTCTTCCGATCTNNNGTACTTGTCCTCCATGTGTTTG

GCACACGTCTGAACTCTTTCCCTACACGACGCTCTTCCGATCTNNNGTACTTGTCCTCCATGTGTTTG

CACACGTCTGAACTCTTTCCCTACACGACGCTCTTCCGATCTNNNTCTATCCAGTCTGCCTGTCCTTACA

CACACGTCTGAACTCTTTCCCTACACGACGCTCTTCCGATCTNNNTCTATCCAGTCTGCCTGTCCTTACA

AGCACACGTCTGAACTCTTTCCCTACACGACGCTCTTCCGATCTNNNTCTTGCTATGGGACTTGGACTT

AGCACACGTCTGAACTCTTTCCCTACACGACGCTCTTCCGATCTNNNTCTTGCTATGGGACTTGGACTT

GCACACGTCTGAACTCTTTCCCTACACGACGCTCTTCCGATCTNNNCTGTAGTCCCAGCTACTCAGGA

GCACACGTCTGAACTCTTTCCCTACACGACGCTCTTCCGATCTNNNCTGTAGTCCCAGCTACTCAGGA

ACACGTCTGAACTCTTTCCCTACACGACGCTCTTCCGATCTNNNCAAAGGAGAGGAGGAGGAGGAG

ACACGTCTGAACTCTTTCCCTACACGACGCTCTTCCGATCTNNNCAAAGGAGAGGAGGAGGAGGAG

AGCACACGTCTGAACTCTTTCCCTACACGACGCTCTTCCGATCTNNNGGGAGGGAGAGAGAGACTG

AGCACACGTCTGAACTCTTTCCCTACACGACGCTCTTCCGATCTNNNGGGAGGGAGAGAGAGACTGA
AGCACACGTCTGAACTCTTTCCCTACACGACGCTCTTCCGATCTNNNGGTTGCAGTGAGCCATGATTG
AGCACACGTCTGAACTCTTTCCCTACACGACGCTCTTCCGATCTNNNGGTTGCAGTGAGCCATGATTG
GCACACGTCTGAACTCTTTCCCTACACGACGCTCTTCCGATCTNNNGTGCCTCAGATTTTCAATCCCACC
GCACACGTCTGAACTCTTTCCCTACACGACGCTCTTCCGATCTNNNGTGCCTCAGATTTTCAATCCCACC
GAGCACACGTCTGAACTCTTTCCCTACACGACGCTCTTCCGATCTNNNAGAGAGACAGAGAGACAAG
GAGCACACGTCTGAACTCTTTCCCTACACGACGCTCTTCCGATCTNNNAGAGAGACAGAGAGACAAG
AGCACACGTCTGAACTCTTTCCCTACACGACGCTCTTCCGATCTNNNAACTCCTGAAGAGGAGGGGT
AGCACACGTCTGAACTCTTTCCCTACACGACGCTCTTCCGATCTNNNAACTCCTGAAGAGGAGGGGT
AGCACACGTCTGAACTCTTTCCCTACACGACGCTCTTCCGATCTNNNATATTACATTTCAGAGTTCACCT
AGCACACGTCTGAACTCTTTCCCTACACGACGCTCTTCCGATCTNNNATATTACATTTCAGAGTTCACCT
AGCACACGTCTGAACTCTTTCCCTACACGACGCTCTTCCGATCTNNNTCAAGGCTGCAGTGACTTATG
AGCACACGTCTGAACTCTTTCCCTACACGACGCTCTTCCGATCTNNNTCAAGGCTGCAGTGACTTATG
AGCACACGTCTGAACTCTTTCCCTACACGACGCTCTTCCGATCTNNNCATCTGAGGCAGAAACAGAG
AGCACACGTCTGAACTCTTTCCCTACACGACGCTCTTCCGATCTNNNCATCTGAGGCAGAAACAGAG
GAGCACACGTCTGAACTCTTTCCCTACACGACGCTCTTCCGATCTNNNGGTGATTGTATTCTGTGTACT
GAGCACACGTCTGAACTCTTTCCCTACACGACGCTCTTCCGATCTNNNGGTGATTGTATTCTGTGTACT
GAGCACACGTCTGAACTCTTTCCCTACACGACGCTCTTCCGATCTNNNGCAGAATTGTTTCAACCGAG
GAGCACACGTCTGAACTCTTTCCCTACACGACGCTCTTCCGATCTNNNGCAGAATTGTTTCAACCGAG
GAGCACACGTCTGAACTCTTTCCCTACACGACGCTCTTCCGATCTNNNGGTACTGGTACTGGCACAAA
GAGCACACGTCTGAACTCTTTCCCTACACGACGCTCTTCCGATCTNNNGGTACTGGTACTGGCACAAA
ACACGTCTGAACTCTTTCCCTACACGACGCTCTTCCGATCTNNNCTAAATATTGCGGTGATGCTGGGC
ACACGTCTGAACTCTTTCCCTACACGACGCTCTTCCGATCTNNNCTAAATATTGCGGTGATGCTGGGC
AGCACACGTCTGAACTCTTTCCCTACACGACGCTCTTCCGATCTNNNCCTCCAAATGAACCATGGTAGG
AGCACACGTCTGAACTCTTTCCCTACACGACGCTCTTCCGATCTNNNCCTCCAAATGAACCATGGTAGG
GAGCACACGTCTGAACTCTTTCCCTACACGACGCTCTTCCGATCTNNNAGAGAGACAGGCAGAAAAA
GAGCACACGTCTGAACTCTTTCCCTACACGACGCTCTTCCGATCTNNNAGAGAGACAGGCAGAAAAA
GAGCACACGTCTGAACTCTTTCCCTACACGACGCTCTTCCGATCTNNNAAAGGCCCATGGAAACTGAT
GAGCACACGTCTGAACTCTTTCCCTACACGACGCTCTTCCGATCTNNNAAAGGCCCATGGAAACTGAT
AGCACACGTCTGAACTCTTTCCCTACACGACGCTCTTCCGATCTNNNCAACAAGAGTGAAAGTCCGTCT
AGCACACGTCTGAACTCTTTCCCTACACGACGCTCTTCCGATCTNNNCAACAAGAGTGAAAGTCCGTCT
AGCACACGTCTGAACTCTTTCCCTACACGACGCTCTTCCGATCTNNNGTGAGCTATGTTTGTGCCAGT
AGCACACGTCTGAACTCTTTCCCTACACGACGCTCTTCCGATCTNNNGTGAGCTATGTTTGTGCCAGT
GAGCACACGTCTGAACTCTTTCCCTACACGACGCTCTTCCGATCTNNNCAGTGAGCTATGACCATGCT
GAGCACACGTCTGAACTCTTTCCCTACACGACGCTCTTCCGATCTNNNCAGTGAGCTATGACCATGCT
CACACGTCTGAACTCTTTCCCTACACGACGCTCTTCCGATCTNNNACAGGTATAGCAACCAATCAGCA
CACACGTCTGAACTCTTTCCCTACACGACGCTCTTCCGATCTNNNACAGGTATAGCAACCAATCAGCA
AGCACACGTCTGAACTCTTTCCCTACACGACGCTCTTCCGATCTNNNTCCAAGCTGATCTTATGCTGT
AGCACACGTCTGAACTCTTTCCCTACACGACGCTCTTCCGATCTNNNTCCAAGCTGATCTTATGCTGT
AGCACACGTCTGAACTCTTTCCCTACACGACGCTCTTCCGATCTNNNCACCACGCCCAGCTAATTTTTAT
AGCACACGTCTGAACTCTTTCCCTACACGACGCTCTTCCGATCTNNNCACCACGCCCAGCTAATTTTTAT
AGCACACGTCTGAACTCTTTCCCTACACGACGCTCTTCCGATCTNNNCCTAACAGTGAGAAGATTAAG
AGCACACGTCTGAACTCTTTCCCTACACGACGCTCTTCCGATCTNNNCCTAACAGTGAGAAGATTAAG

AGCACACGTCTGAACTCTTTCCCTACACGACGCTCTTCCGATCTNNNTCTCTTCAAGACCCTGCTTTCA
AGCACACGTCTGAACTCTTTCCCTACACGACGCTCTTCCGATCTNNNTCTCTTCAAGACCCTGCTTTCA
CACACGTCTGAACTCTTTCCCTACACGACGCTCTTCCGATCTNNNGTTTCACCATGTTAGCCAGGATG
CACACGTCTGAACTCTTTCCCTACACGACGCTCTTCCGATCTNNNGTTTCACCATGTTAGCCAGGATG
GAGCACACGTCTGAACTCTTTCCCTACACGACGCTCTTCCGATCTNNNACCAAATATCACACCCCTGTT
GAGCACACGTCTGAACTCTTTCCCTACACGACGCTCTTCCGATCTNNNACCAAATATCACACCCCTGTT
AGCACACGTCTGAACTCTTTCCCTACACGACGCTCTTCCGATCTNNNATTGCAACTGGTAATTCAGAG
AGCACACGTCTGAACTCTTTCCCTACACGACGCTCTTCCGATCTNNNATTGCAACTGGTAATTCAGAG
GAGCACACGTCTGAACTCTTTCCCTACACGACGCTCTTCCGATCTNNNGCAAATGTGATGTGGATTTT
GAGCACACGTCTGAACTCTTTCCCTACACGACGCTCTTCCGATCTNNNGCAAATGTGATGTGGATTTT
GCACACGTCTGAACTCTTTCCCTACACGACGCTCTTCCGATCTNNNGCCGCTTCTTCTTCTTCTTCTTCC
GCACACGTCTGAACTCTTTCCCTACACGACGCTCTTCCGATCTNNNGCCGCTTCTTCTTCTTCTTCTTCC
GCACACGTCTGAACTCTTTCCCTACACGACGCTCTTCCGATCTNNNCTGCTGAGAAGCTCAACAACTTC
GCACACGTCTGAACTCTTTCCCTACACGACGCTCTTCCGATCTNNNCTGCTGAGAAGCTCAACAACTTC
AGCACACGTCTGAACTCTTTCCCTACACGACGCTCTTCCGATCTNNNGCAACAGAGTGAGACTCCATC
AGCACACGTCTGAACTCTTTCCCTACACGACGCTCTTCCGATCTNNNGCAACAGAGTGAGACTCCATC
GCACACGTCTGAACTCTTTCCCTACACGACGCTCTTCCGATCTNNNCTCCTTGTCCTCTCCTCTCTGTCC
GCACACGTCTGAACTCTTTCCCTACACGACGCTCTTCCGATCTNNNCTCCTTGTCCTCTCCTCTCTGTCC
GCACACGTCTGAACTCTTTCCCTACACGACGCTCTTCCGATCTNNNGAGGTTGCAGTGAGACAAGATT
GCACACGTCTGAACTCTTTCCCTACACGACGCTCTTCCGATCTNNNGAGGTTGCAGTGAGACAAGATT
AGCACACGTCTGAACTCTTTCCCTACACGACGCTCTTCCGATCTNNNTGCATATCATGCATTTGAAGCC
AGCACACGTCTGAACTCTTTCCCTACACGACGCTCTTCCGATCTNNNTGCATATCATGCATTTGAAGCC
AGCACACGTCTGAACTCTTTCCCTACACGACGCTCTTCCGATCTNNNACTCTGTGCTAGGTTCCGAGT
AGCACACGTCTGAACTCTTTCCCTACACGACGCTCTTCCGATCTNNNACTCTGTGCTAGGTTCCGAGT
GAGCACACGTCTGAACTCTTTCCCTACACGACGCTCTTCCGATCTNNNTGTCTTCTCACTCAGAGGTTG
GAGCACACGTCTGAACTCTTTCCCTACACGACGCTCTTCCGATCTNNNTGTCTTCTCACTCAGAGGTTG
AGCACACGTCTGAACTCTTTCCCTACACGACGCTCTTCCGATCTNNNCTCTGGATGGTCAGATTGTAG
AGCACACGTCTGAACTCTTTCCCTACACGACGCTCTTCCGATCTNNNCTCTGGATGGTCAGATTGTAG
AGCACACGTCTGAACTCTTTCCCTACACGACGCTCTTCCGATCTNNNCTGGTGTCTGGGTGGCTTGTC
AGCACACGTCTGAACTCTTTCCCTACACGACGCTCTTCCGATCTNNNCTGGTGTCTGGGTGGCTTGTC
GCACACGTCTGAACTCTTTCCCTACACGACGCTCTTCCGATCTNNNCCTGGTCCCTGTCATCTCTTTTTC
GCACACGTCTGAACTCTTTCCCTACACGACGCTCTTCCGATCTNNNCCTGGTCCCTGTCATCTCTTTTTC
CACACGTCTGAACTCTTTCCCTACACGACGCTCTTCCGATCTNNNTCAGAACTTACACAGCTCCTCACCA
CACACGTCTGAACTCTTTCCCTACACGACGCTCTTCCGATCTNNNTCAGAACTTACACAGCTCCTCACCA
GCACACGTCTGAACTCTTTCCCTACACGACGCTCTTCCGATCTNNNGGTGATTGGAAGGCCCCTAATA
GCACACGTCTGAACTCTTTCCCTACACGACGCTCTTCCGATCTNNNGGTGATTGGAAGGCCCCTAATA
AGCACACGTCTGAACTCTTTCCCTACACGACGCTCTTCCGATCTNNNAATCCTAATTCCCAGGCCATCT
AGCACACGTCTGAACTCTTTCCCTACACGACGCTCTTCCGATCTNNNAATCCTAATTCCCAGGCCATCT
AGCACACGTCTGAACTCTTTCCCTACACGACGCTCTTCCGATCTNNNATGCCTGTAATACCAGCTACTC
AGCACACGTCTGAACTCTTTCCCTACACGACGCTCTTCCGATCTNNNATGCCTGTAATACCAGCTACTC
AGCACACGTCTGAACTCTTTCCCTACACGACGCTCTTCCGATCTNNNGTTGCAGTGAGCCAAGATCA
AGCACACGTCTGAACTCTTTCCCTACACGACGCTCTTCCGATCTNNNGTTGCAGTGAGCCAAGATCA
GAGCACACGTCTGAACTCTTTCCCTACACGACGCTCTTCCGATCTNNNTGAACTTTTCAGTGTTGCTG

GAGCACACGTCTGAACTCTTTCCCTACACGACGCTCTTCCGATCTNNNTGAACTTTTCAGTGTTGCTGC
GAGCACACGTCTGAACTCTTTCCCTACACGACGCTCTTCCGATCTNNNTTGGTGCTTGGGGAAGATCT
GAGCACACGTCTGAACTCTTTCCCTACACGACGCTCTTCCGATCTNNNTTGGTGCTTGGGGAAGATCT
AGCACACGTCTGAACTCTTTCCCTACACGACGCTCTTCCGATCTNNNGCTGGACATTGAGGAGGCTTTA
AGCACACGTCTGAACTCTTTCCCTACACGACGCTCTTCCGATCTNNNGCTGGACATTGAGGAGGCTTTA
GAGCACACGTCTGAACTCTTTCCCTACACGACGCTCTTCCGATCTNNNCTCATTTCCAAACTGAGCTGCT
GAGCACACGTCTGAACTCTTTCCCTACACGACGCTCTTCCGATCTNNNCTCATTTCCAAACTGAGCTGCT
AGCACACGTCTGAACTCTTTCCCTACACGACGCTCTTCCGATCTNNNGTAATCCCAACTACTCGGCAGG
AGCACACGTCTGAACTCTTTCCCTACACGACGCTCTTCCGATCTNNNGTAATCCCAACTACTCGGCAGG
AGCACACGTCTGAACTCTTTCCCTACACGACGCTCTTCCGATCTNNNGTTGCAGTGAGCCAAGATCATC
AGCACACGTCTGAACTCTTTCCCTACACGACGCTCTTCCGATCTNNNGTTGCAGTGAGCCAAGATCATC
GAGCACACGTCTGAACTCTTTCCCTACACGACGCTCTTCCGATCTNNNAGGGTTGAGAAGAACATTGA
GAGCACACGTCTGAACTCTTTCCCTACACGACGCTCTTCCGATCTNNNAGGGTTGAGAAGAACATTGA
CACGTCTGAACTCTTTCCCTACACGACGCTCTTCCGATCTNNNACTGATTTTGGTACCAAGAGGGACAT
CACGTCTGAACTCTTTCCCTACACGACGCTCTTCCGATCTNNNACTGATTTTGGTACCAAGAGGGACAT
AGAGCACACGTCTGAACTCTTTCCCTACACGACGCTCTTCCGATCTNNNACTCAGGTGCTTTTCGAACTA
AGAGCACACGTCTGAACTCTTTCCCTACACGACGCTCTTCCGATCTNNNACTCAGGTGCTTTTCGAACTA
AGCACACGTCTGAACTCTTTCCCTACACGACGCTCTTCCGATCTNNNGTAATCCCAGCTACTAGTGAGC
AGCACACGTCTGAACTCTTTCCCTACACGACGCTCTTCCGATCTNNNGTAATCCCAGCTACTAGTGAGC
GCACACGTCTGAACTCTTTCCCTACACGACGCTCTTCCGATCTNNNAAGACAAAGAAGAAAAACGGCC
GCACACGTCTGAACTCTTTCCCTACACGACGCTCTTCCGATCTNNNAAGACAAAGAAGAAAAACGGCC
AGCACACGTCTGAACTCTTTCCCTACACGACGCTCTTCCGATCTNNNAAACATTGCCAACTACTCACCCT
AGCACACGTCTGAACTCTTTCCCTACACGACGCTCTTCCGATCTNNNAAACATTGCCAACTACTCACCCT
AGCACACGTCTGAACTCTTTCCCTACACGACGCTCTTCCGATCTNNNCGTAACAATAATAGTTCCTAACA
AGCACACGTCTGAACTCTTTCCCTACACGACGCTCTTCCGATCTNNNCGTAACAATAATAGTTCCTAACA
GCACACGTCTGAACTCTTTCCCTACACGACGCTCTTCCGATCTNNNTCTCAGAATCTGTGACTTGTCTTC
GCACACGTCTGAACTCTTTCCCTACACGACGCTCTTCCGATCTNNNTCTCAGAATCTGTGACTTGTCTTC
AGCACACGTCTGAACTCTTTCCCTACACGACGCTCTTCCGATCTNNNGGGGTACACAGGGTATTTGGT
AGCACACGTCTGAACTCTTTCCCTACACGACGCTCTTCCGATCTNNNGGGGTACACAGGGTATTTGGT
AGCACACGTCTGAACTCTTTCCCTACACGACGCTCTTCCGATCTNNNTTTGTACCCTGCAACTGTCCTTAC
AGCACACGTCTGAACTCTTTCCCTACACGACGCTCTTCCGATCTNNNTTTGTACCCTGCAACTGTCCTTAC
GAGCACACGTCTGAACTCTTTCCCTACACGACGCTCTTCCGATCTNNNGACTGGGCACACACTAGGATC
GAGCACACGTCTGAACTCTTTCCCTACACGACGCTCTTCCGATCTNNNGACTGGGCACACACTAGGATC
GCACACGTCTGAACTCTTTCCCTACACGACGCTCTTCCGATCTNNNGCCTAGGAGCTAGAGGTGAAAT
GCACACGTCTGAACTCTTTCCCTACACGACGCTCTTCCGATCTNNNGCCTAGGAGCTAGAGGTGAAAT
GAGCACACGTCTGAACTCTTTCCCTACACGACGCTCTTCCGATCTNNNCTGAAGTTGCAAAGAGCCATG
GAGCACACGTCTGAACTCTTTCCCTACACGACGCTCTTCCGATCTNNNCTGAAGTTGCAAAGAGCCATG
GAGCACACGTCTGAACTCTTTCCCTACACGACGCTCTTCCGATCTNNNTGATGAAACCCCGTCGTTACT
GAGCACACGTCTGAACTCTTTCCCTACACGACGCTCTTCCGATCTNNNTGATGAAACCCCGTCGTTACT
AGCACACGTCTGAACTCTTTCCCTACACGACGCTCTTCCGATCTNNNGCACAGAACAATGCCCTTCAAT
AGCACACGTCTGAACTCTTTCCCTACACGACGCTCTTCCGATCTNNNGCACAGAACAATGCCCTTCAAT
GCACACGTCTGAACTCTTTCCCTACACGACGCTCTTCCGATCTNNNAGTATGCACACTTTCCTCCACTTC
GCACACGTCTGAACTCTTTCCCTACACGACGCTCTTCCGATCTNNNAGTATGCACACTTTCCTCCACTTC

AGCACACGTCTGAACTCTTTCCCTACACGACGCTCTTCCGATCTNNNGTCCTTCTCACTTGCAGTCACTA
AGCACACGTCTGAACTCTTTCCCTACACGACGCTCTTCCGATCTNNNGTCCTTCTCACTTGCAGTCACTA
AGCACACGTCTGAACTCTTTCCCTACACGACGCTCTTCCGATCTNNNGAGGCTGAGGCAGGAGAATGC
AGCACACGTCTGAACTCTTTCCCTACACGACGCTCTTCCGATCTNNNGAGGCTGAGGCAGGAGAATGC
GAGCACACGTCTGAACTCTTTCCCTACACGACGCTCTTCCGATCTNNNCTGTCTGTGTCTCCTCTTCTGT
GAGCACACGTCTGAACTCTTTCCCTACACGACGCTCTTCCGATCTNNNCTGTCTGTGTCTCCTCTTCTGT
GAGCACACGTCTGAACTCTTTCCCTACACGACGCTCTTCCGATCTNNNTTCCAGGTTTAGTCTTGGAAG
GAGCACACGTCTGAACTCTTTCCCTACACGACGCTCTTCCGATCTNNNTTCCAGGTTTAGTCTTGGAAG
GCACACGTCTGAACTCTTTCCCTACACGACGCTCTTCCGATCTNNNCCTTGGGGAGAAAGCTTCTAGTT
GCACACGTCTGAACTCTTTCCCTACACGACGCTCTTCCGATCTNNNCCTTGGGGAGAAAGCTTCTAGTT
GAGCACACGTCTGAACTCTTTCCCTACACGACGCTCTTCCGATCTNNNCTAAAGTTGAAACAGGCCTTG
GAGCACACGTCTGAACTCTTTCCCTACACGACGCTCTTCCGATCTNNNCTAAAGTTGAAACAGGCCTTG
AGAGCACACGTCTGAACTCTTTCCCTACACGACGCTCTTCCGATCTNNNTCCAGCTAATCCAGAACCAC
AGAGCACACGTCTGAACTCTTTCCCTACACGACGCTCTTCCGATCTNNNTCCAGCTAATCCAGAACCAC
AGCACACGTCTGAACTCTTTCCCTACACGACGCTCTTCCGATCTNNNTCTCTTTACTCACTCTGTCCCCTCA
AGCACACGTCTGAACTCTTTCCCTACACGACGCTCTTCCGATCTNNNTCTCTTTACTCACTCTGTCCCCTCA
GAGCACACGTCTGAACTCTTTCCCTACACGACGCTCTTCCGATCTNNNAGTACACCCAGTTTACCACTTG
GAGCACACGTCTGAACTCTTTCCCTACACGACGCTCTTCCGATCTNNNAGTACACCCAGTTTACCACTTG
GCACACGTCTGAACTCTTTCCCTACACGACGCTCTTCCGATCTNNNGGAAAGTTGGGTTTTGCCAGATT
GCACACGTCTGAACTCTTTCCCTACACGACGCTCTTCCGATCTNNNGGAAAGTTGGGTTTTGCCAGATT
AGCACACGTCTGAACTCTTTCCCTACACGACGCTCTTCCGATCTNNNGATCTCCTGACCTTGTGATCCTC
AGCACACGTCTGAACTCTTTCCCTACACGACGCTCTTCCGATCTNNNGATCTCCTGACCTTGTGATCCTC
AGCACACGTCTGAACTCTTTCCCTACACGACGCTCTTCCGATCTNNNCCAAGTAGGTGGGATTACAGG
AGCACACGTCTGAACTCTTTCCCTACACGACGCTCTTCCGATCTNNNCCAAGTAGGTGGGATTACAGG
AGAGCACACGTCTGAACTCTTTCCCTACACGACGCTCTTCCGATCTNNNTCAGGTGCTAATTGTGCTTG
AGAGCACACGTCTGAACTCTTTCCCTACACGACGCTCTTCCGATCTNNNTCAGGTGCTAATTGTGCTTG
AGAGCACACGTCTGAACTCTTTCCCTACACGACGCTCTTCCGATCTNNNCTGAGGCAGGAGAATGTCTT
AGAGCACACGTCTGAACTCTTTCCCTACACGACGCTCTTCCGATCTNNNCTGAGGCAGGAGAATGTCTT
AGCACACGTCTGAACTCTTTCCCTACACGACGCTCTTCCGATCTNNNGCCTGTAGTCCCAGCTACTTGCA
AGCACACGTCTGAACTCTTTCCCTACACGACGCTCTTCCGATCTNNNGCCTGTAGTCCCAGCTACTTGCA
AGCACACGTCTGAACTCTTTCCCTACACGACGCTCTTCCGATCTNNNGTTGCATCCATGTCTGGTTTTG
AGCACACGTCTGAACTCTTTCCCTACACGACGCTCTTCCGATCTNNNGTTGCATCCATGTCTGGTTTTG
AGCACACGTCTGAACTCTTTCCCTACACGACGCTCTTCCGATCTNNNTTACTTTTGGAATTTTTTGCAAA
AGCACACGTCTGAACTCTTTCCCTACACGACGCTCTTCCGATCTNNNTTACTTTTGGAATTTTTTGCAAA
AGAGCACACGTCTGAACTCTTTCCCTACACGACGCTCTTCCGATCTNNNCGATTCTTCAGGGGGAACAA
AGAGCACACGTCTGAACTCTTTCCCTACACGACGCTCTTCCGATCTNNNCGATTCTTCAGGGGGAACAA
AGCACACGTCTGAACTCTTTCCCTACACGACGCTCTTCCGATCTNNNAGGAAGAAAGGTACAATAGCA
AGCACACGTCTGAACTCTTTCCCTACACGACGCTCTTCCGATCTNNNAGGAAGAAAGGTACAATAGCA
AGCACACGTCTGAACTCTTTCCCTACACGACGCTCTTCCGATCTNNNCAACAGAGTGAGACCCTGTGT
AGCACACGTCTGAACTCTTTCCCTACACGACGCTCTTCCGATCTNNNCAACAGAGTGAGACCCTGTGT
AGCACACGTCTGAACTCTTTCCCTACACGACGCTCTTCCGATCTNNNAGCAAAAGCAGGCATCTTACAT
AGCACACGTCTGAACTCTTTCCCTACACGACGCTCTTCCGATCTNNNAGCAAAAGCAGGCATCTTACAT
GAGCACACGTCTGAACTCTTTCCCTACACGACGCTCTTCCGATCTNNNTGGAGGTCATATGTGAACATG

AGCACACGTCTGAACTCTTTCCCTACACGACGCTCTTCCGATCTNNNTGGAGGTCATATGTGAACATG
GCACACGTCTGAACTCTTTCCCTACACGACGCTCTTCCGATCTNNNGCTTCGGCAGGACAGAAATTA
GCACACGTCTGAACTCTTTCCCTACACGACGCTCTTCCGATCTNNNGCTTCGGCAGGACAGAAATTA
AGCACACGTCTGAACTCTTTCCCTACACGACGCTCTTCCGATCTNNNGGGGTTCCATGAAGAATTCCA
AGCACACGTCTGAACTCTTTCCCTACACGACGCTCTTCCGATCTNNNGGGGTTCCATGAAGAATTCCA
AGCACACGTCTGAACTCTTTCCCTACACGACGCTCTTCCGATCTNNNCTAGGGTTCTCTAGAGGGAC
AGCACACGTCTGAACTCTTTCCCTACACGACGCTCTTCCGATCTNNNCTAGGGTTCTCTAGAGGGAC
AGCACACGTCTGAACTCTTTCCCTACACGACGCTCTTCCGATCTNNNGTAGTTTTGTGTGGGCTGCAT
AGCACACGTCTGAACTCTTTCCCTACACGACGCTCTTCCGATCTNNNGTAGTTTTGTGTGGGCTGCAT
AGCACACGTCTGAACTCTTTCCCTACACGACGCTCTTCCGATCTNNNTCTAGGTCTGATTCTTGGCTTC
AGCACACGTCTGAACTCTTTCCCTACACGACGCTCTTCCGATCTNNNTCTAGGTCTGATTCTTGGCTTC
AGCACACGTCTGAACTCTTTCCCTACACGACGCTCTTCCGATCTNNNCTTCTGGGTAGCTGAGACTAC
AGCACACGTCTGAACTCTTTCCCTACACGACGCTCTTCCGATCTNNNCTTCTGGGTAGCTGAGACTAC
AGCACACGTCTGAACTCTTTCCCTACACGACGCTCTTCCGATCTNNNAAATGCCCACATCGAAAAGCT
AGCACACGTCTGAACTCTTTCCCTACACGACGCTCTTCCGATCTNNNAAATGCCCACATCGAAAAGCT
GAGCACACGTCTGAACTCTTTCCCTACACGACGCTCTTCCGATCTNNNGTACACATTGATTCAGGGGA
GAGCACACGTCTGAACTCTTTCCCTACACGACGCTCTTCCGATCTNNNGTACACATTGATTCAGGGGA
GCACACGTCTGAACTCTTTCCCTACACGACGCTCTTCCGATCTNNNGAGATCACACCATTGCACTCCAG
GCACACGTCTGAACTCTTTCCCTACACGACGCTCTTCCGATCTNNNGAGATCACACCATTGCACTCCAG
AGCACACGTCTGAACTCTTTCCCTACACGACGCTCTTCCGATCTNNNAAAAGCAAACAAAAGGCCCAG
AGCACACGTCTGAACTCTTTCCCTACACGACGCTCTTCCGATCTNNNAAAAGCAAACAAAAGGCCCAG
GCACACGTCTGAACTCTTTCCCTACACGACGCTCTTCCGATCTNNNTTCCTCTCTCGAGCCAGTCTATG
GCACACGTCTGAACTCTTTCCCTACACGACGCTCTTCCGATCTNNNTTCCTCTCTCGAGCCAGTCTATG
GAGCACACGTCTGAACTCTTTCCCTACACGACGCTCTTCCGATCTNNNCCAACAGGCATATGCAAAGA
GAGCACACGTCTGAACTCTTTCCCTACACGACGCTCTTCCGATCTNNNCCAACAGGCATATGCAAAGA
GAGCACACGTCTGAACTCTTTCCCTACACGACGCTCTTCCGATCTNNNTTCCTCATTCCTTATTCCATGC
GAGCACACGTCTGAACTCTTTCCCTACACGACGCTCTTCCGATCTNNNTTCCTCATTCCTTATTCCATGC
AGCACACGTCTGAACTCTTTCCCTACACGACGCTCTTCCGATCTNNNAGAGAACAAGGTCACAGAGCA
AGCACACGTCTGAACTCTTTCCCTACACGACGCTCTTCCGATCTNNNAGAGAACAAGGTCACAGAGCA
GAGCACACGTCTGAACTCTTTCCCTACACGACGCTCTTCCGATCTNNNTCATTCTCCACAACTCTGAAG
GAGCACACGTCTGAACTCTTTCCCTACACGACGCTCTTCCGATCTNNNTCATTCTCCACAACTCTGAAG
GCACACGTCTGAACTCTTTCCCTACACGACGCTCTTCCGATCTNNNGTGGAAGAACCCCCTCATTGTAT
GCACACGTCTGAACTCTTTCCCTACACGACGCTCTTCCGATCTNNNGTGGAAGAACCCCCTCATTGTAT
GAGCACACGTCTGAACTCTTTCCCTACACGACGCTCTTCCGATCTNNNTGAGCTGTCAGGTAAGTGCA
GAGCACACGTCTGAACTCTTTCCCTACACGACGCTCTTCCGATCTNNNTGAGCTGTCAGGTAAGTGCA
AGAGCACACGTCTGAACTCTTTCCCTACACGACGCTCTTCCGATCTNNNGTAATCCCAGCTACTAAGGA
AGAGCACACGTCTGAACTCTTTCCCTACACGACGCTCTTCCGATCTNNNGTAATCCCAGCTACTAAGGA
AGCACACGTCTGAACTCTTTCCCTACACGACGCTCTTCCGATCTNNNAACCACCCCTTAGCACTAATCT
AGCACACGTCTGAACTCTTTCCCTACACGACGCTCTTCCGATCTNNNAACCACCCCTTAGCACTAATCT
GAGCACACGTCTGAACTCTTTCCCTACACGACGCTCTTCCGATCTNNNATCGAGATCAACCTGGACAA
GAGCACACGTCTGAACTCTTTCCCTACACGACGCTCTTCCGATCTNNNATCGAGATCAACCTGGACAA
AGCACACGTCTGAACTCTTTCCCTACACGACGCTCTTCCGATCTNNNGTTCCCGCTGTTCTATTTTCAG
AGCACACGTCTGAACTCTTTCCCTACACGACGCTCTTCCGATCTNNNGTTCCCGCTGTTCTATTTTCAG

GAGCACACGTCTGAACTCTTTCCCTACACGACGCTCTTCCGATCTNNNTACAGTGTGTAGTCTCGTGT
GAGCACACGTCTGAACTCTTTCCCTACACGACGCTCTTCCGATCTNNNTACAGTGTGTAGTCTCGTGT
AGCACACGTCTGAACTCTTTCCCTACACGACGCTCTTCCGATCTNNNTTTCCCAACCCTGGAAAAGAG
AGCACACGTCTGAACTCTTTCCCTACACGACGCTCTTCCGATCTNNNTTTCCCAACCCTGGAAAAGAG
AGCACACGTCTGAACTCTTTCCCTACACGACGCTCTTCCGATCTNNNTCCACATATCTTTCAAGTCTCT
AGCACACGTCTGAACTCTTTCCCTACACGACGCTCTTCCGATCTNNNTCCACATATCTTTCAAGTCTCT
AGCACACGTCTGAACTCTTTCCCTACACGACGCTCTTCCGATCTNNNGAGTGACAGGTGATAGCAAG
AGCACACGTCTGAACTCTTTCCCTACACGACGCTCTTCCGATCTNNNGAGTGACAGGTGATAGCAAG
AGCACACGTCTGAACTCTTTCCCTACACGACGCTCTTCCGATCTNNNCCCAGGAGTTCTAGACCATCCT
AGCACACGTCTGAACTCTTTCCCTACACGACGCTCTTCCGATCTNNNCCCAGGAGTTCTAGACCATCCT
AGCACACGTCTGAACTCTTTCCCTACACGACGCTCTTCCGATCTNNNCCCCTTCACATTGCCTCAACTTA
AGCACACGTCTGAACTCTTTCCCTACACGACGCTCTTCCGATCTNNNCCCCTTCACATTGCCTCAACTTA
AGCACACGTCTGAACTCTTTCCCTACACGACGCTCTTCCGATCTNNNACTTCTCTGTGTGTTTGGGATG
AGCACACGTCTGAACTCTTTCCCTACACGACGCTCTTCCGATCTNNNACTTCTCTGTGTGTTTGGGATG
AGCACACGTCTGAACTCTTTCCCTACACGACGCTCTTCCGATCTNNNGAGGATCTTTCCACGTGGGTA
AGCACACGTCTGAACTCTTTCCCTACACGACGCTCTTCCGATCTNNNGAGGATCTTTCCACGTGGGTA
AGCACACGTCTGAACTCTTTCCCTACACGACGCTCTTCCGATCTNNNTGTGCTTGTAATCCCAGCTACT
AGCACACGTCTGAACTCTTTCCCTACACGACGCTCTTCCGATCTNNNTGTGCTTGTAATCCCAGCTACT
AGCACACGTCTGAACTCTTTCCCTACACGACGCTCTTCCGATCTNNNGCTGGGGTTTAGGTTGTGTTTA
AGCACACGTCTGAACTCTTTCCCTACACGACGCTCTTCCGATCTNNNGCTGGGGTTTAGGTTGTGTTTA
GAGCACACGTCTGAACTCTTTCCCTACACGACGCTCTTCCGATCTNNNACAACTTGGTCCCAAATGGTT
GAGCACACGTCTGAACTCTTTCCCTACACGACGCTCTTCCGATCTNNNACAACTTGGTCCCAAATGGTT
AGCACACGTCTGAACTCTTTCCCTACACGACGCTCTTCCGATCTNNNCACAGATGGTGCTTCCTACAG
AGCACACGTCTGAACTCTTTCCCTACACGACGCTCTTCCGATCTNNNCACAGATGGTGCTTCCTACAG
AGCACACGTCTGAACTCTTTCCCTACACGACGCTCTTCCGATCTNNNCTGTGATCACTCTCAAGACACC
AGCACACGTCTGAACTCTTTCCCTACACGACGCTCTTCCGATCTNNNCTGTGATCACTCTCAAGACACC
AGCACACGTCTGAACTCTTTCCCTACACGACGCTCTTCCGATCTNNNCAAAGAAGCATGTGGAATTCCC
AGCACACGTCTGAACTCTTTCCCTACACGACGCTCTTCCGATCTNNNCAAAGAAGCATGTGGAATTCCC
AGCACACGTCTGAACTCTTTCCCTACACGACGCTCTTCCGATCTNNNGTGGCTGCCTTCAAGAAGATTT
AGCACACGTCTGAACTCTTTCCCTACACGACGCTCTTCCGATCTNNNGTGGCTGCCTTCAAGAAGATTT
GAGCACACGTCTGAACTCTTTCCCTACACGACGCTCTTCCGATCTNNNTAAAGCTAATGCAGACACAC
GAGCACACGTCTGAACTCTTTCCCTACACGACGCTCTTCCGATCTNNNTAAAGCTAATGCAGACACAC
AGCACACGTCTGAACTCTTTCCCTACACGACGCTCTTCCGATCTNNNCCTCCTATCACCTCCTAGCCATAC
AGCACACGTCTGAACTCTTTCCCTACACGACGCTCTTCCGATCTNNNCCTCCTATCACCTCCTAGCCATAC
AGCACACGTCTGAACTCTTTCCCTACACGACGCTCTTCCGATCTNNNTGAGTTAGGGAGGAGTATGTC
AGCACACGTCTGAACTCTTTCCCTACACGACGCTCTTCCGATCTNNNTGAGTTAGGGAGGAGTATGTC
AGCACACGTCTGAACTCTTTCCCTACACGACGCTCTTCCGATCTNNNCACACACACACACACATATGGA
AGCACACGTCTGAACTCTTTCCCTACACGACGCTCTTCCGATCTNNNCACACACACACACACATATGGA
GAGCACACGTCTGAACTCTTTCCCTACACGACGCTCTTCCGATCTNNNTACAAAAATTAGCCGGGCGT
GAGCACACGTCTGAACTCTTTCCCTACACGACGCTCTTCCGATCTNNNTACAAAAATTAGCCGGGCGT
AGCACACGTCTGAACTCTTTCCCTACACGACGCTCTTCCGATCTNNNTTTGTTGAAAATGAAATCTAGT
AGCACACGTCTGAACTCTTTCCCTACACGACGCTCTTCCGATCTNNNTTTGTTGAAAATGAAATCTAGT
AGCACACGTCTGAACTCTTTCCCTACACGACGCTCTTCCGATCTNNNTGCCTCAAGTACAGGGATTAC

AGCACACGTCTGAACTCTTTCCCTACACGACGCTCTTCCGATCTNNNTGCCTCAAGTACAGGGATTACA
GAGCACACGTCTGAACTCTTTCCCTACACGACGCTCTTCCGATCTNNNCTCTAACCACTCTGTCCTTGCTT
GAGCACACGTCTGAACTCTTTCCCTACACGACGCTCTTCCGATCTNNNCTCTAACCACTCTGTCCTTGCTT
AGAGCACACGTCTGAACTCTTTCCCTACACGACGCTCTTCCGATCTNNNGTAGAGGTGGAGATACACC
AGAGCACACGTCTGAACTCTTTCCCTACACGACGCTCTTCCGATCTNNNGTAGAGGTGGAGATACACC
GAGCACACGTCTGAACTCTTTCCCTACACGACGCTCTTCCGATCTNNNAATCCCGGAAGCAAATATGCC
GAGCACACGTCTGAACTCTTTCCCTACACGACGCTCTTCCGATCTNNNAATCCCGGAAGCAAATATGCC
GAGCACACGTCTGAACTCTTTCCCTACACGACGCTCTTCCGATCTNNNTCCAATATGTCAAGTTGCCTTT
GAGCACACGTCTGAACTCTTTCCCTACACGACGCTCTTCCGATCTNNNTCCAATATGTCAAGTTGCCTTT
GAGCACACGTCTGAACTCTTTCCCTACACGACGCTCTTCCGATCTNNNCCCATTTCTTTCACCAACTGAG
GAGCACACGTCTGAACTCTTTCCCTACACGACGCTCTTCCGATCTNNNCCCATTTCTTTCACCAACTGAG
GAGCACACGTCTGAACTCTTTCCCTACACGACGCTCTTCCGATCTNNNAAGACCTGCCACATTAGGGAA
GAGCACACGTCTGAACTCTTTCCCTACACGACGCTCTTCCGATCTNNNAAGACCTGCCACATTAGGGAA
AGCACACGTCTGAACTCTTTCCCTACACGACGCTCTTCCGATCTNNNCTGCTCAGTTACCCAACATGTTT
AGCACACGTCTGAACTCTTTCCCTACACGACGCTCTTCCGATCTNNNCTGCTCAGTTACCCAACATGTTT
AGAGCACACGTCTGAACTCTTTCCCTACACGACGCTCTTCCGATCTNNNCAATCACACTGAAAACCGTC
AGAGCACACGTCTGAACTCTTTCCCTACACGACGCTCTTCCGATCTNNNCAATCACACTGAAAACCGTC
AGCACACGTCTGAACTCTTTCCCTACACGACGCTCTTCCGATCTNNNAGCTCTAAGGCTTCAGGCAATA
AGCACACGTCTGAACTCTTTCCCTACACGACGCTCTTCCGATCTNNNAGCTCTAAGGCTTCAGGCAATA
AGCACACGTCTGAACTCTTTCCCTACACGACGCTCTTCCGATCTNNNTGCCCATTCAGTATAATGTTGG
AGCACACGTCTGAACTCTTTCCCTACACGACGCTCTTCCGATCTNNNTGCCCATTCAGTATAATGTTGG
GCACACGTCTGAACTCTTTCCCTACACGACGCTCTTCCGATCTNNNGATTTACAACCATCCCGGCATTG
GCACACGTCTGAACTCTTTCCCTACACGACGCTCTTCCGATCTNNNGATTTACAACCATCCCGGCATTG
AGCACACGTCTGAACTCTTTCCCTACACGACGCTCTTCCGATCTNNNACCTTTGTGATCTGAACTGGGA
AGCACACGTCTGAACTCTTTCCCTACACGACGCTCTTCCGATCTNNNACCTTTGTGATCTGAACTGGGA
GAGCACACGTCTGAACTCTTTCCCTACACGACGCTCTTCCGATCTNNNGTGGCTCTGTTCCCAAAAAGA
GAGCACACGTCTGAACTCTTTCCCTACACGACGCTCTTCCGATCTNNNGTGGCTCTGTTCCCAAAAAGA
AGCACACGTCTGAACTCTTTCCCTACACGACGCTCTTCCGATCTNNNAGGAGAAAATTTTCAAAAGGCC
AGCACACGTCTGAACTCTTTCCCTACACGACGCTCTTCCGATCTNNNAGGAGAAAATTTTCAAAAGGCC
AGAGCACACGTCTGAACTCTTTCCCTACACGACGCTCTTCCGATCTNNNCAATCTTTCCCCTTTCTCCAGG
AGAGCACACGTCTGAACTCTTTCCCTACACGACGCTCTTCCGATCTNNNCAATCTTTCCCCTTTCTCCAGG
CACACGTCTGAACTCTTTCCCTACACGACGCTCTTCCGATCTNNNCTAGAGCTTTTCCTTTTAAGAGACA
CACACGTCTGAACTCTTTCCCTACACGACGCTCTTCCGATCTNNNCTAGAGCTTTTCCTTTTAAGAGACA
AGCACACGTCTGAACTCTTTCCCTACACGACGCTCTTCCGATCTNNNCGCCTATAATCCCAGCTACTCGC
AGCACACGTCTGAACTCTTTCCCTACACGACGCTCTTCCGATCTNNNCGCCTATAATCCCAGCTACTCGC
GCACACGTCTGAACTCTTTCCCTACACGACGCTCTTCCGATCTNNNGGAACATTCGCCAAAATAGACCAC
GCACACGTCTGAACTCTTTCCCTACACGACGCTCTTCCGATCTNNNGGAACATTCGCCAAAATAGACCAC
AGCACACGTCTGAACTCTTTCCCTACACGACGCTCTTCCGATCTNNNACTTTCCAGACAGGGTTAGGTT
AGCACACGTCTGAACTCTTTCCCTACACGACGCTCTTCCGATCTNNNACTTTCCAGACAGGGTTAGGTT
AGAGCACACGTCTGAACTCTTTCCCTACACGACGCTCTTCCGATCTNNNTTGCGCTTGAGTTTAGTTGC
AGAGCACACGTCTGAACTCTTTCCCTACACGACGCTCTTCCGATCTNNNTTGCGCTTGAGTTTAGTTGC
GAGCACACGTCTGAACTCTTTCCCTACACGACGCTCTTCCGATCTNNNTGTGCAGTGGTGTCTCATTAT
GAGCACACGTCTGAACTCTTTCCCTACACGACGCTCTTCCGATCTNNNTGTGCAGTGGTGTCTCATTAT

GCACACGTCTGAACTCTTTCCCTACACGACGCTCTTCCGATCTNNNTTTCTCACTTGGTGTCTCTGTCC
GCACACGTCTGAACTCTTTCCCTACACGACGCTCTTCCGATCTNNNTTTCTCACTTGGTGTCTCTGTCC
AGCACACGTCTGAACTCTTTCCCTACACGACGCTCTTCCGATCTNNNCATCCTGGCTAACATGGTGAA
AGCACACGTCTGAACTCTTTCCCTACACGACGCTCTTCCGATCTNNNCATCCTGGCTAACATGGTGAA
GAGCACACGTCTGAACTCTTTCCCTACACGACGCTCTTCCGATCTNNNGCCACACCACCTTTCCTTATTT
GAGCACACGTCTGAACTCTTTCCCTACACGACGCTCTTCCGATCTNNNGCCACACCACCTTTCCTTATTT
AGCACACGTCTGAACTCTTTCCCTACACGACGCTCTTCCGATCTNNNAACGATCCTGTAACTCCAAGA
AGCACACGTCTGAACTCTTTCCCTACACGACGCTCTTCCGATCTNNNAACGATCCTGTAACTCCAAGA
AGCACACGTCTGAACTCTTTCCCTACACGACGCTCTTCCGATCTNNNCTGGGTTTCTAGTAAGTAGCC
AGCACACGTCTGAACTCTTTCCCTACACGACGCTCTTCCGATCTNNNCTGGGTTTCTAGTAAGTAGCC
AGCACACGTCTGAACTCTTTCCCTACACGACGCTCTTCCGATCTNNNACATGATGAAACCCCGTCTCTA
AGCACACGTCTGAACTCTTTCCCTACACGACGCTCTTCCGATCTNNNACATGATGAAACCCCGTCTCTA
AGCACACGTCTGAACTCTTTCCCTACACGACGCTCTTCCGATCTNNNATGGAAGAGATAGGAGAGG
AGCACACGTCTGAACTCTTTCCCTACACGACGCTCTTCCGATCTNNNATGGAAGAGATAGGAGAGG
AGCACACGTCTGAACTCTTTCCCTACACGACGCTCTTCCGATCTNNNGTGTCCACGAAGAAACAAATG
AGCACACGTCTGAACTCTTTCCCTACACGACGCTCTTCCGATCTNNNGTGTCCACGAAGAAACAAATG
GCACACGTCTGAACTCTTTCCCTACACGACGCTCTTCCGATCTNNNATTAGGTACTTGGCAAGCACTG
GCACACGTCTGAACTCTTTCCCTACACGACGCTCTTCCGATCTNNNATTAGGTACTTGGCAAGCACTG
GCACACGTCTGAACTCTTTCCCTACACGACGCTCTTCCGATCTNNNTGGGCAACAGAGCAAGACTCCA
GCACACGTCTGAACTCTTTCCCTACACGACGCTCTTCCGATCTNNNTGGGCAACAGAGCAAGACTCCA
GCACACGTCTGAACTCTTTCCCTACACGACGCTCTTCCGATCTNNNTCAAGGATTTCCATAGCACTGC
GCACACGTCTGAACTCTTTCCCTACACGACGCTCTTCCGATCTNNNTCAAGGATTTCCATAGCACTGC
CACACGTCTGAACTCTTTCCCTACACGACGCTCTTCCGATCTNNNTTGGATGGGGATTTGGCAAAAA
CACACGTCTGAACTCTTTCCCTACACGACGCTCTTCCGATCTNNNTTGGATGGGGATTTGGCAAAAA
AGCACACGTCTGAACTCTTTCCCTACACGACGCTCTTCCGATCTNNNATGTGTCCACAATGTCTTGGA
AGCACACGTCTGAACTCTTTCCCTACACGACGCTCTTCCGATCTNNNATGTGTCCACAATGTCTTGGA
GAGCACACGTCTGAACTCTTTCCCTACACGACGCTCTTCCGATCTNNNGAGGCAGGAGAATGGTGT
GAGCACACGTCTGAACTCTTTCCCTACACGACGCTCTTCCGATCTNNNGAGGCAGGAGAATGGTGT
AGCACACGTCTGAACTCTTTCCCTACACGACGCTCTTCCGATCTNNNCCGCGATACCCAAAGTCAATA
AGCACACGTCTGAACTCTTTCCCTACACGACGCTCTTCCGATCTNNNCCGCGATACCCAAAGTCAATA
AGAGCACACGTCTGAACTCTTTCCCTACACGACGCTCTTCCGATCTNNNAAAAATTGGAGCAGCATTG
AGAGCACACGTCTGAACTCTTTCCCTACACGACGCTCTTCCGATCTNNNAAAAATTGGAGCAGCATTG
AGCACACGTCTGAACTCTTTCCCTACACGACGCTCTTCCGATCTNNNTTTAGGAGCTCTGTGTCAGGA
AGCACACGTCTGAACTCTTTCCCTACACGACGCTCTTCCGATCTNNNTTTAGGAGCTCTGTGTCAGGA
GCACACGTCTGAACTCTTTCCCTACACGACGCTCTTCCGATCTNNNGCTTCTGGAGTACCTGGGAATA
GCACACGTCTGAACTCTTTCCCTACACGACGCTCTTCCGATCTNNNGCTTCTGGAGTACCTGGGAATA
AGCACACGTCTGAACTCTTTCCCTACACGACGCTCTTCCGATCTNNNCCCTTCCTCCAAATCTCTGTGA
AGCACACGTCTGAACTCTTTCCCTACACGACGCTCTTCCGATCTNNNCCCTTCCTCCAAATCTCTGTGA
GAGCACACGTCTGAACTCTTTCCCTACACGACGCTCTTCCGATCTNNNTCATGCCTGTAATCTCAGCAC
GAGCACACGTCTGAACTCTTTCCCTACACGACGCTCTTCCGATCTNNNTCATGCCTGTAATCTCAGCAC
GAGCACACGTCTGAACTCTTTCCCTACACGACGCTCTTCCGATCTNNNGGCTGGTCTGAGAAATAAA
GAGCACACGTCTGAACTCTTTCCCTACACGACGCTCTTCCGATCTNNNGGCTGGTCTGAGAAATAAA
AGCACACGTCTGAACTCTTTCCCTACACGACGCTCTTCCGATCTNNNTGCAACTTCTTTTCATTGTGACC

AGCACACGTCTGAACTCTTTCCCTACACGACGCTCTTCCGATCTNNNTGCAACTTCTTTTCATTGTGACC
AGCACACGTCTGAACTCTTTCCCTACACGACGCTCTTCCGATCTNNNGGCCACAAGTCCTTTAGGAG
AGCACACGTCTGAACTCTTTCCCTACACGACGCTCTTCCGATCTNNNGGCCACAAGTCCTTTAGGAG
GCACACGTCTGAACTCTTTCCCTACACGACGCTCTTCCGATCTNNNTCCCTCTGGGACTGACATTATCT
GCACACGTCTGAACTCTTTCCCTACACGACGCTCTTCCGATCTNNNTCCCTCTGGGACTGACATTATCT
GCACACGTCTGAACTCTTTCCCTACACGACGCTCTTCCGATCTNNNGACAGAGCGAGACTCCGTCTCA
GCACACGTCTGAACTCTTTCCCTACACGACGCTCTTCCGATCTNNNGACAGAGCGAGACTCCGTCTCA
AGCACACGTCTGAACTCTTTCCCTACACGACGCTCTTCCGATCTNNNAGTCGGAGGTCTGGTTTTTCT
AGCACACGTCTGAACTCTTTCCCTACACGACGCTCTTCCGATCTNNNAGTCGGAGGTCTGGTTTTTCT
GCACACGTCTGAACTCTTTCCCTACACGACGCTCTTCCGATCTNNNCTTGGCTCATCTTCAAACCGTCT
GCACACGTCTGAACTCTTTCCCTACACGACGCTCTTCCGATCTNNNCTTGGCTCATCTTCAAACCGTCT
AGCACACGTCTGAACTCTTTCCCTACACGACGCTCTTCCGATCTNNNTGAATACCTGGGCTCAAGTGA
AGCACACGTCTGAACTCTTTCCCTACACGACGCTCTTCCGATCTNNNTGAATACCTGGGCTCAAGTGA
GCACACGTCTGAACTCTTTCCCTACACGACGCTCTTCCGATCTNNNGAGGCAGGAGAATGGTGTGAA
GCACACGTCTGAACTCTTTCCCTACACGACGCTCTTCCGATCTNNNGAGGCAGGAGAATGGTGTGAA
AGCACACGTCTGAACTCTTTCCCTACACGACGCTCTTCCGATCTNNNCAATTTTATCTTGAAACCATTCG
AGCACACGTCTGAACTCTTTCCCTACACGACGCTCTTCCGATCTNNNCAATTTTATCTTGAAACCATTCG
GAGCACACGTCTGAACTCTTTCCCTACACGACGCTCTTCCGATCTNNNGTGCCTGTAATCCCAGCTACT
GAGCACACGTCTGAACTCTTTCCCTACACGACGCTCTTCCGATCTNNNGTGCCTGTAATCCCAGCTACT
AGCACACGTCTGAACTCTTTCCCTACACGACGCTCTTCCGATCTNNNTGCACAGTACCCTCTTATAATG
AGCACACGTCTGAACTCTTTCCCTACACGACGCTCTTCCGATCTNNNTGCACAGTACCCTCTTATAATG
AGAGCACACGTCTGAACTCTTTCCCTACACGACGCTCTTCCGATCTNNNTGAAGCAGGGAGGATTGA
AGAGCACACGTCTGAACTCTTTCCCTACACGACGCTCTTCCGATCTNNNTGAAGCAGGGAGGATTGA
AGCACACGTCTGAACTCTTTCCCTACACGACGCTCTTCCGATCTNNNTACAAAAAATTAGCTGGGCG
AGCACACGTCTGAACTCTTTCCCTACACGACGCTCTTCCGATCTNNNTACAAAAAATTAGCTGGGCG
AGCACACGTCTGAACTCTTTCCCTACACGACGCTCTTCCGATCTNNNAAGGTCTCCAAATCGACAGATC
AGCACACGTCTGAACTCTTTCCCTACACGACGCTCTTCCGATCTNNNAAGGTCTCCAAATCGACAGATC
ACACGTCTGAACTCTTTCCCTACACGACGCTCTTCCGATCTNNNTTCTCAATGAGCTGTTGGGCACATC
ACACGTCTGAACTCTTTCCCTACACGACGCTCTTCCGATCTNNNTTCTCAATGAGCTGTTGGGCACATC
AGCACACGTCTGAACTCTTTCCCTACACGACGCTCTTCCGATCTNNNCGGCATTCACATTTGCCATCTT
AGCACACGTCTGAACTCTTTCCCTACACGACGCTCTTCCGATCTNNNCGGCATTCACATTTGCCATCTT
AGCACACGTCTGAACTCTTTCCCTACACGACGCTCTTCCGATCTNNNGATCACAAGGTCAGGAGATCG
AGCACACGTCTGAACTCTTTCCCTACACGACGCTCTTCCGATCTNNNGATCACAAGGTCAGGAGATCG
GCACACGTCTGAACTCTTTCCCTACACGACGCTCTTCCGATCTNNNGCCCAGGGTCAAAATCAAACCA
GCACACGTCTGAACTCTTTCCCTACACGACGCTCTTCCGATCTNNNGCCCAGGGTCAAAATCAAACCA
AGCACACGTCTGAACTCTTTCCCTACACGACGCTCTTCCGATCTNNNTCTTGATACAAGAAAGCTGGC
AGCACACGTCTGAACTCTTTCCCTACACGACGCTCTTCCGATCTNNNTCTTGATACAAGAAAGCTGGC
AGCACACGTCTGAACTCTTTCCCTACACGACGCTCTTCCGATCTNNNCACAAGGTGGTAGTATGGCTT
AGCACACGTCTGAACTCTTTCCCTACACGACGCTCTTCCGATCTNNNCACAAGGTGGTAGTATGGCTT
AGAGCACACGTCTGAACTCTTTCCCTACACGACGCTCTTCCGATCTNNNGCTGAGACAGGAGAATCA
AGAGCACACGTCTGAACTCTTTCCCTACACGACGCTCTTCCGATCTNNNGCTGAGACAGGAGAATCA
GAGCACACGTCTGAACTCTTTCCCTACACGACGCTCTTCCGATCTNNNTGGAATCCTGAACAAGATGC
GAGCACACGTCTGAACTCTTTCCCTACACGACGCTCTTCCGATCTNNNTGGAATCCTGAACAAGATGC

GAGCACACGTCTGAACTCTTTCCCTACACGACGCTCTTCCGATCTNNNGAGAATCAGGCAGGGAGGT
GAGCACACGTCTGAACTCTTTCCCTACACGACGCTCTTCCGATCTNNNGAGAATCAGGCAGGGAGGT
GCACACGTCTGAACTCTTTCCCTACACGACGCTCTTCCGATCTNNNGCCAGGTATGATCTCAGCTACTC
GCACACGTCTGAACTCTTTCCCTACACGACGCTCTTCCGATCTNNNGCCAGGTATGATCTCAGCTACTC
AGCACACGTCTGAACTCTTTCCCTACACGACGCTCTTCCGATCTNNNTGAAATCACTGCCAGGAATCA
AGCACACGTCTGAACTCTTTCCCTACACGACGCTCTTCCGATCTNNNTGAAATCACTGCCAGGAATCA
GAGCACACGTCTGAACTCTTTCCCTACACGACGCTCTTCCGATCTNNNCGGACGATGCTGTTGCTATA
GAGCACACGTCTGAACTCTTTCCCTACACGACGCTCTTCCGATCTNNNCGGACGATGCTGTTGCTATA
GAGCACACGTCTGAACTCTTTCCCTACACGACGCTCTTCCGATCTNNNGGACCCACTGGCTAGAAGAT
GAGCACACGTCTGAACTCTTTCCCTACACGACGCTCTTCCGATCTNNNGGACCCACTGGCTAGAAGAT
AGCACACGTCTGAACTCTTTCCCTACACGACGCTCTTCCGATCTNNNGGCCTCTGGAGTAGCTATTCTT
AGCACACGTCTGAACTCTTTCCCTACACGACGCTCTTCCGATCTNNNGGCCTCTGGAGTAGCTATTCTT
AGCACACGTCTGAACTCTTTCCCTACACGACGCTCTTCCGATCTNNNGTTGGCACATGACTGTAGTCC
AGCACACGTCTGAACTCTTTCCCTACACGACGCTCTTCCGATCTNNNGTTGGCACATGACTGTAGTCC
GAGCACACGTCTGAACTCTTTCCCTACACGACGCTCTTCCGATCTNNNAGGATCCAGTGGCATTTTTTT
GAGCACACGTCTGAACTCTTTCCCTACACGACGCTCTTCCGATCTNNNAGGATCCAGTGGCATTTTTTT
GAGCACACGTCTGAACTCTTTCCCTACACGACGCTCTTCCGATCTNNNGCAGGCTGGAGAGAATGTT
GAGCACACGTCTGAACTCTTTCCCTACACGACGCTCTTCCGATCTNNNGCAGGCTGGAGAGAATGTT
AGCACACGTCTGAACTCTTTCCCTACACGACGCTCTTCCGATCTNNNGATAATGTCAGCATAAGGCCT
AGCACACGTCTGAACTCTTTCCCTACACGACGCTCTTCCGATCTNNNGATAATGTCAGCATAAGGCCT
GAGCACACGTCTGAACTCTTTCCCTACACGACGCTCTTCCGATCTNNNAAAGGTATAGGGATGTGGGT
GAGCACACGTCTGAACTCTTTCCCTACACGACGCTCTTCCGATCTNNNAAAGGTATAGGGATGTGGGT
GCACACGTCTGAACTCTTTCCCTACACGACGCTCTTCCGATCTNNNTGAGGCAGGAGAATCACTTGAA
GCACACGTCTGAACTCTTTCCCTACACGACGCTCTTCCGATCTNNNTGAGGCAGGAGAATCACTTGAA
GAGCACACGTCTGAACTCTTTCCCTACACGACGCTCTTCCGATCTNNNGCTTTGCTTGTCTGGAAAAC
GAGCACACGTCTGAACTCTTTCCCTACACGACGCTCTTCCGATCTNNNGCTTTGCTTGTCTGGAAAAC
AGAGCACACGTCTGAACTCTTTCCCTACACGACGCTCTTCCGATCTNNNAGGAAGAACAACTGGACTT
AGAGCACACGTCTGAACTCTTTCCCTACACGACGCTCTTCCGATCTNNNAGGAAGAACAACTGGACTT
AGCACACGTCTGAACTCTTTCCCTACACGACGCTCTTCCGATCTNNNGGAAAAAGGGTTACAAAAGG
AGCACACGTCTGAACTCTTTCCCTACACGACGCTCTTCCGATCTNNNGGAAAAAGGGTTACAAAAGG
AGCACACGTCTGAACTCTTTCCCTACACGACGCTCTTCCGATCTNNNTGAAAACAGCTGCCTTGAAAC
AGCACACGTCTGAACTCTTTCCCTACACGACGCTCTTCCGATCTNNNTGAAAACAGCTGCCTTGAAAC
ACACGTCTGAACTCTTTCCCTACACGACGCTCTTCCGATCTNNNTCCTTTGACTTAGTCTTGACCCGCAT
ACACGTCTGAACTCTTTCCCTACACGACGCTCTTCCGATCTNNNTCCTTTGACTTAGTCTTGACCCGCAT
GAGCACACGTCTGAACTCTTTCCCTACACGACGCTCTTCCGATCTNNNCTGTAGTCCCAGCTACTCAGG
GAGCACACGTCTGAACTCTTTCCCTACACGACGCTCTTCCGATCTNNNCTGTAGTCCCAGCTACTCAGG
AGCACACGTCTGAACTCTTTCCCTACACGACGCTCTTCCGATCTNNNACACTGAGGACACCTACCGTAG
AGCACACGTCTGAACTCTTTCCCTACACGACGCTCTTCCGATCTNNNACACTGAGGACACCTACCGTAG
GAGCACACGTCTGAACTCTTTCCCTACACGACGCTCTTCCGATCTNNNACACAAAGGGACACCTGTCA
GAGCACACGTCTGAACTCTTTCCCTACACGACGCTCTTCCGATCTNNNACACAAAGGGACACCTGTCA
AGCACACGTCTGAACTCTTTCCCTACACGACGCTCTTCCGATCTNNNTGTAGTTGCAGCTACTTGGGA
AGCACACGTCTGAACTCTTTCCCTACACGACGCTCTTCCGATCTNNNTGTAGTTGCAGCTACTTGGGA
ACACGTCTGAACTCTTTCCCTACACGACGCTCTTCCGATCTNNNGACAAAGCAAAGGTATGAAAGCA

CACACGTCTGAACTCTTTCCCTACACGACGCTCTTCCGATCTNNNGACAAAGCAAAGGTATGAAAGCAT
AGCACACGTCTGAACTCTTTCCCTACACGACGCTCTTCCGATCTNNNGCATCCTCAGCCATGAACATAAA
AGCACACGTCTGAACTCTTTCCCTACACGACGCTCTTCCGATCTNNNGCATCCTCAGCCATGAACATAAA
GAGCACACGTCTGAACTCTTTCCCTACACGACGCTCTTCCGATCTNNNATGAAATGATGCCCCAGATG
GAGCACACGTCTGAACTCTTTCCCTACACGACGCTCTTCCGATCTNNNATGAAATGATGCCCCAGATG
AGCACACGTCTGAACTCTTTCCCTACACGACGCTCTTCCGATCTNNNGCTTGTTTGTTTCTCCCTCTGTT
AGCACACGTCTGAACTCTTTCCCTACACGACGCTCTTCCGATCTNNNGCTTGTTTGTTTCTCCCTCTGTT
AGAGCACACGTCTGAACTCTTTCCCTACACGACGCTCTTCCGATCTNNNCCTAACACTAACTGAATGGG
AGAGCACACGTCTGAACTCTTTCCCTACACGACGCTCTTCCGATCTNNNCCTAACACTAACTGAATGGG
GAGCACACGTCTGAACTCTTTCCCTACACGACGCTCTTCCGATCTNNNCCAAGAGGCCTAGCAAGAGA
GAGCACACGTCTGAACTCTTTCCCTACACGACGCTCTTCCGATCTNNNCCAAGAGGCCTAGCAAGAGA
GAGCACACGTCTGAACTCTTTCCCTACACGACGCTCTTCCGATCTNNNCCCAGATCACACTTCTACACTCC
GAGCACACGTCTGAACTCTTTCCCTACACGACGCTCTTCCGATCTNNNCCCAGATCACACTTCTACACTCC
GAGCACACGTCTGAACTCTTTCCCTACACGACGCTCTTCCGATCTNNNGATGCCTATTTGTTGCTACCAC
GAGCACACGTCTGAACTCTTTCCCTACACGACGCTCTTCCGATCTNNNGATGCCTATTTGTTGCTACCAC
AGCACACGTCTGAACTCTTTCCCTACACGACGCTCTTCCGATCTNNNAGGAGAGAGGAGAAATCTTGC
AGCACACGTCTGAACTCTTTCCCTACACGACGCTCTTCCGATCTNNNAGGAGAGAGGAGAAATCTTGC
AAGAGCACACGTCTGAACTCTTTCCCTACACGACGCTCTTCCGATCTNNNTCTATCTTGAAATGCGTGT
AAGAGCACACGTCTGAACTCTTTCCCTACACGACGCTCTTCCGATCTNNNTCTATCTTGAAATGCGTGT
AGCACACGTCTGAACTCTTTCCCTACACGACGCTCTTCCGATCTNNNATGTTTTTGCATAGGCTTGTCG
AGCACACGTCTGAACTCTTTCCCTACACGACGCTCTTCCGATCTNNNATGTTTTTGCATAGGCTTGTCG
GAGCACACGTCTGAACTCTTTCCCTACACGACGCTCTTCCGATCTNNNGGTGACAAAGGGGAATGGT
GAGCACACGTCTGAACTCTTTCCCTACACGACGCTCTTCCGATCTNNNGGTGACAAAGGGGAATGGT
GAGCACACGTCTGAACTCTTTCCCTACACGACGCTCTTCCGATCTNNNCCAGCCTCCGTCTCAAAAAAA
GAGCACACGTCTGAACTCTTTCCCTACACGACGCTCTTCCGATCTNNNCCAGCCTCCGTCTCAAAAAAA
AGCACACGTCTGAACTCTTTCCCTACACGACGCTCTTCCGATCTNNNTTGTTGTGGTGTGAGAGTAGA
AGCACACGTCTGAACTCTTTCCCTACACGACGCTCTTCCGATCTNNNTTGTTGTGGTGTGAGAGTAGA
GAGCACACGTCTGAACTCTTTCCCTACACGACGCTCTTCCGATCTNNNACATCTTTCAGAACTCAGCTCT
GAGCACACGTCTGAACTCTTTCCCTACACGACGCTCTTCCGATCTNNNACATCTTTCAGAACTCAGCTCT
GAGCACACGTCTGAACTCTTTCCCTACACGACGCTCTTCCGATCTNNNGGCTGAGGTAGGAGAATCAC
GAGCACACGTCTGAACTCTTTCCCTACACGACGCTCTTCCGATCTNNNGGCTGAGGTAGGAGAATCAC
GAGCACACGTCTGAACTCTTTCCCTACACGACGCTCTTCCGATCTNNNAGCAATAGCATTTTCATCCCCT
GAGCACACGTCTGAACTCTTTCCCTACACGACGCTCTTCCGATCTNNNAGCAATAGCATTTTCATCCCCT
GAGCACACGTCTGAACTCTTTCCCTACACGACGCTCTTCCGATCTNNNATGCCTGTAATCCCAGCTACTT
GAGCACACGTCTGAACTCTTTCCCTACACGACGCTCTTCCGATCTNNNATGCCTGTAATCCCAGCTACTT
GAGCACACGTCTGAACTCTTTCCCTACACGACGCTCTTCCGATCTNNNTGTTTCACTGGCAGAAGCAAA
GAGCACACGTCTGAACTCTTTCCCTACACGACGCTCTTCCGATCTNNNTGTTTCACTGGCAGAAGCAAA
AGCACACGTCTGAACTCTTTCCCTACACGACGCTCTTCCGATCTNNNAGTGAGCTGAGATCTCGCTATT
AGCACACGTCTGAACTCTTTCCCTACACGACGCTCTTCCGATCTNNNAGTGAGCTGAGATCTCGCTATT
AGAGCACACGTCTGAACTCTTTCCCTACACGACGCTCTTCCGATCTNNNATGATGAGTATGAGAGCAG
AGAGCACACGTCTGAACTCTTTCCCTACACGACGCTCTTCCGATCTNNNATGATGAGTATGAGAGCAG
GAGCACACGTCTGAACTCTTTCCCTACACGACGCTCTTCCGATCTNNNCAGCACTGTTCACAATAGCCA
GAGCACACGTCTGAACTCTTTCCCTACACGACGCTCTTCCGATCTNNNCAGCACTGTTCACAATAGCCA

AGCACACGTCTGAACTCTTTCCCTACACGACGCTCTTCCGATCTNNNCATTTATGTGAGATGTGGTTTT
AGCACACGTCTGAACTCTTTCCCTACACGACGCTCTTCCGATCTNNNCATTTATGTGAGATGTGGTTTT
GCACACGTCTGAACTCTTTCCCTACACGACGCTCTTCCGATCTNNNGGATGGCATTGAGGGTTTCTTTC
GCACACGTCTGAACTCTTTCCCTACACGACGCTCTTCCGATCTNNNGGATGGCATTGAGGGTTTCTTTC
AGCACACGTCTGAACTCTTTCCCTACACGACGCTCTTCCGATCTNNNCCAAACTCAAGACAATTACTTAT
AGCACACGTCTGAACTCTTTCCCTACACGACGCTCTTCCGATCTNNNCCAAACTCAAGACAATTACTTAT
GAGCACACGTCTGAACTCTTTCCCTACACGACGCTCTTCCGATCTNNNTGTAAAGTTGAAACTACCGCA
GAGCACACGTCTGAACTCTTTCCCTACACGACGCTCTTCCGATCTNNNTGTAAAGTTGAAACTACCGCA
GAGCACACGTCTGAACTCTTTCCCTACACGACGCTCTTCCGATCTNNNGAGAATTGCTTGAACCCAGGA
GAGCACACGTCTGAACTCTTTCCCTACACGACGCTCTTCCGATCTNNNGAGAATTGCTTGAACCCAGGA
AGCACACGTCTGAACTCTTTCCCTACACGACGCTCTTCCGATCTNNNGTTTTGCTGTTTGTTCCTGCCTT
AGCACACGTCTGAACTCTTTCCCTACACGACGCTCTTCCGATCTNNNGTTTTGCTGTTTGTTCCTGCCTT
GAGCACACGTCTGAACTCTTTCCCTACACGACGCTCTTCCGATCTNNNCTCTCAGCTACTCGGGACTTTA
GAGCACACGTCTGAACTCTTTCCCTACACGACGCTCTTCCGATCTNNNCTCTCAGCTACTCGGGACTTTA
AGCACACGTCTGAACTCTTTCCCTACACGACGCTCTTCCGATCTNNNGTGACAGAACCAATCCTTGTGTC
AGCACACGTCTGAACTCTTTCCCTACACGACGCTCTTCCGATCTNNNGTGACAGAACCAATCCTTGTGTC
GAGCACACGTCTGAACTCTTTCCCTACACGACGCTCTTCCGATCTNNNCCAACATCCCCTTATCACACCAC
GAGCACACGTCTGAACTCTTTCCCTACACGACGCTCTTCCGATCTNNNCCAACATCCCCTTATCACACCAC
AGCACACGTCTGAACTCTTTCCCTACACGACGCTCTTCCGATCTNNNTCTTCTCTGGCTGCTTTCAAGAT
AGCACACGTCTGAACTCTTTCCCTACACGACGCTCTTCCGATCTNNNTCTTCTCTGGCTGCTTTCAAGAT
GAGCACACGTCTGAACTCTTTCCCTACACGACGCTCTTCCGATCTNNNGGTGGTGCTTATTTTAAAATT
GAGCACACGTCTGAACTCTTTCCCTACACGACGCTCTTCCGATCTNNNGGTGGTGCTTATTTTAAAATT
GAGCACACGTCTGAACTCTTTCCCTACACGACGCTCTTCCGATCTNNNACTGGAAGAGAAGAAAAGGG
GAGCACACGTCTGAACTCTTTCCCTACACGACGCTCTTCCGATCTNNNACTGGAAGAGAAGAAAAGGG
GAGCACACGTCTGAACTCTTTCCCTACACGACGCTCTTCCGATCTNNNTGCTTGTTTGTTTGTTTGGGA
GAGCACACGTCTGAACTCTTTCCCTACACGACGCTCTTCCGATCTNNNTGCTTGTTTGTTTGTTTGGGA
AGCACACGTCTGAACTCTTTCCCTACACGACGCTCTTCCGATCTNNNTTGCACAATAGGCTGGTTTATG
AGCACACGTCTGAACTCTTTCCCTACACGACGCTCTTCCGATCTNNNTTGCACAATAGGCTGGTTTATG
GCACACGTCTGAACTCTTTCCCTACACGACGCTCTTCCGATCTNNNCAGAAGAACCGCTTTGCTTGAAC
GCACACGTCTGAACTCTTTCCCTACACGACGCTCTTCCGATCTNNNCAGAAGAACCGCTTTGCTTGAAC
GAGCACACGTCTGAACTCTTTCCCTACACGACGCTCTTCCGATCTNNNGACAATACAGCTGGGCAACAT
GAGCACACGTCTGAACTCTTTCCCTACACGACGCTCTTCCGATCTNNNGACAATACAGCTGGGCAACAT
GAGCACACGTCTGAACTCTTTCCCTACACGACGCTCTTCCGATCTNNNCCAGGTTCAAGCAATTCTTCCT
GAGCACACGTCTGAACTCTTTCCCTACACGACGCTCTTCCGATCTNNNCCAGGTTCAAGCAATTCTTCCT
GAGCACACGTCTGAACTCTTTCCCTACACGACGCTCTTCCGATCTNNNACCTTGAGCATTCTGCAACAAC
GAGCACACGTCTGAACTCTTTCCCTACACGACGCTCTTCCGATCTNNNACCTTGAGCATTCTGCAACAAC
AGCACACGTCTGAACTCTTTCCCTACACGACGCTCTTCCGATCTNNNAGATCAGCAAGTGACTGTAGT
AGCACACGTCTGAACTCTTTCCCTACACGACGCTCTTCCGATCTNNNAGATCAGCAAGTGACTGTAGT
ACACGTCTGAACTCTTTCCCTACACGACGCTCTTCCGATCTNNNTAAACTCAGCCTTAGGTCAACCCCAT
ACACGTCTGAACTCTTTCCCTACACGACGCTCTTCCGATCTNNNTAAACTCAGCCTTAGGTCAACCCCAT
GAGCACACGTCTGAACTCTTTCCCTACACGACGCTCTTCCGATCTNNNCAAAAAAAAATTAGCCGGGC
GAGCACACGTCTGAACTCTTTCCCTACACGACGCTCTTCCGATCTNNNCAAAAAAAAATTAGCCGGGC
GAGCACACGTCTGAACTCTTTCCCTACACGACGCTCTTCCGATCTNNNTCACGTTCATCAACCCAATTGT

GAGCACACGTCTGAACTCTTTCCCTACACGACGCTCTTCCGATCTNNNTCACGTTCATCAACCCAATTGT
AAGAGCACACGTCTGAACTCTTTCCCTACACGACGCTCTTCCGATCTNNNGAAAGAGAATTCAGCCAGA
AAGAGCACACGTCTGAACTCTTTCCCTACACGACGCTCTTCCGATCTNNNGAAAGAGAATTCAGCCAGA
AGAGCACACGTCTGAACTCTTTCCCTACACGACGCTCTTCCGATCTNNNGCCAGCAATGGAGATAGAC
AGAGCACACGTCTGAACTCTTTCCCTACACGACGCTCTTCCGATCTNNNGCCAGCAATGGAGATAGAC
GCACACGTCTGAACTCTTTCCCTACACGACGCTCTTCCGATCTNNNCTTCATCTGTATTTATAGCCACTCA
GCACACGTCTGAACTCTTTCCCTACACGACGCTCTTCCGATCTNNNCTTCATCTGTATTTATAGCCACTCA
AGCACACGTCTGAACTCTTTCCCTACACGACGCTCTTCCGATCTNNNCAGTCCCTGGTAGTCAGTTGTAC
AGCACACGTCTGAACTCTTTCCCTACACGACGCTCTTCCGATCTNNNCAGTCCCTGGTAGTCAGTTGTAC
GAGCACACGTCTGAACTCTTTCCCTACACGACGCTCTTCCGATCTNNNGTTTCTTCACAGAGCTACAGGC
GAGCACACGTCTGAACTCTTTCCCTACACGACGCTCTTCCGATCTNNNGTTTCTTCACAGAGCTACAGGC
AGCACACGTCTGAACTCTTTCCCTACACGACGCTCTTCCGATCTNNNGAGGCAGGAGAATGGTGTGAA
AGCACACGTCTGAACTCTTTCCCTACACGACGCTCTTCCGATCTNNNGAGGCAGGAGAATGGTGTGAA
GCACACGTCTGAACTCTTTCCCTACACGACGCTCTTCCGATCTNNNTGCTCTCTTTGTCTTTCTCCCTCCAT
GCACACGTCTGAACTCTTTCCCTACACGACGCTCTTCCGATCTNNNTGCTCTCTTTGTCTTTCTCCCTCCAT
AGCACACGTCTGAACTCTTTCCCTACACGACGCTCTTCCGATCTNNNTTCAAGTGATTCTCCTGCCTCAG
AGCACACGTCTGAACTCTTTCCCTACACGACGCTCTTCCGATCTNNNTTCAAGTGATTCTCCTGCCTCAG
GAGCACACGTCTGAACTCTTTCCCTACACGACGCTCTTCCGATCTNNNTACCACAAAATGACAGCAGCAT
GAGCACACGTCTGAACTCTTTCCCTACACGACGCTCTTCCGATCTNNNTACCACAAAATGACAGCAGCAT
GAGCACACGTCTGAACTCTTTCCCTACACGACGCTCTTCCGATCTNNNTGGGTGACAGAGTGAGAGTC
GAGCACACGTCTGAACTCTTTCCCTACACGACGCTCTTCCGATCTNNNTGGGTGACAGAGTGAGAGTC
AGCACACGTCTGAACTCTTTCCCTACACGACGCTCTTCCGATCTNNNCACTGTGGTGCTCATTTTCAGAC
AGCACACGTCTGAACTCTTTCCCTACACGACGCTCTTCCGATCTNNNCACTGTGGTGCTCATTTTCAGAC
AGCACACGTCTGAACTCTTTCCCTACACGACGCTCTTCCGATCTNNNTCTGTAGCCCATTTTCCTGTGAA
AGCACACGTCTGAACTCTTTCCCTACACGACGCTCTTCCGATCTNNNTCTGTAGCCCATTTTCCTGTGAA
AGCACACGTCTGAACTCTTTCCCTACACGACGCTCTTCCGATCTNNNATGCTGAGTCCAGTTCCCGATAC
AGCACACGTCTGAACTCTTTCCCTACACGACGCTCTTCCGATCTNNNATGCTGAGTCCAGTTCCCGATAC
GAGCACACGTCTGAACTCTTTCCCTACACGACGCTCTTCCGATCTNNNAAGTGATTTGCATGTCATGAT
GAGCACACGTCTGAACTCTTTCCCTACACGACGCTCTTCCGATCTNNNAAGTGATTTGCATGTCATGAT
AAGAGCACACGTCTGAACTCTTTCCCTACACGACGCTCTTCCGATCTNNNTAATCCTAGCTACTGAGGAC
AAGAGCACACGTCTGAACTCTTTCCCTACACGACGCTCTTCCGATCTNNNTAATCCTAGCTACTGAGGAC
AGCACACGTCTGAACTCTTTCCCTACACGACGCTCTTCCGATCTNNNCACACAACTGTACTCTGGTCTGA
AGCACACGTCTGAACTCTTTCCCTACACGACGCTCTTCCGATCTNNNCACACAACTGTACTCTGGTCTGA
GAGCACACGTCTGAACTCTTTCCCTACACGACGCTCTTCCGATCTNNNGCAAATGGCAGGATCTCCTTTT
GAGCACACGTCTGAACTCTTTCCCTACACGACGCTCTTCCGATCTNNNGCAAATGGCAGGATCTCCTTTT
AGCACACGTCTGAACTCTTTCCCTACACGACGCTCTTCCGATCTNNNGTAATCTCAGCTACTCAGGAGG
AGCACACGTCTGAACTCTTTCCCTACACGACGCTCTTCCGATCTNNNGTAATCTCAGCTACTCAGGAGG
AGCACACGTCTGAACTCTTTCCCTACACGACGCTCTTCCGATCTNNNGAGGTTGCAGTGAGCTGAAAT
AGCACACGTCTGAACTCTTTCCCTACACGACGCTCTTCCGATCTNNNGAGGTTGCAGTGAGCTGAAAT
GAGCACACGTCTGAACTCTTTCCCTACACGACGCTCTTCCGATCTNNNCTTGAAAACTTGCAGCCCATGT
GAGCACACGTCTGAACTCTTTCCCTACACGACGCTCTTCCGATCTNNNCTTGAAAACTTGCAGCCCATGT
AGCACACGTCTGAACTCTTTCCCTACACGACGCTCTTCCGATCTNNNCTGTAGTCCCAGCTACTCAGGA
AGCACACGTCTGAACTCTTTCCCTACACGACGCTCTTCCGATCTNNNCTGTAGTCCCAGCTACTCAGGA

GCACACGTCTGAACTCTTTCCCTACACGACGCTCTTCCGATCTNNNTCTTGAGTAAGGCAGTCAAATGC
GCACACGTCTGAACTCTTTCCCTACACGACGCTCTTCCGATCTNNNTCTTGAGTAAGGCAGTCAAATGC
AGCACACGTCTGAACTCTTTCCCTACACGACGCTCTTCCGATCTNNNACTAACATTTCTTACCTGGTTAT
AGCACACGTCTGAACTCTTTCCCTACACGACGCTCTTCCGATCTNNNACTAACATTTCTTACCTGGTTAT
AGCACACGTCTGAACTCTTTCCCTACACGACGCTCTTCCGATCTNNNTCCAAAACCTCATCTCCCTGAAG
AGCACACGTCTGAACTCTTTCCCTACACGACGCTCTTCCGATCTNNNTCCAAAACCTCATCTCCCTGAAG
GAGCACACGTCTGAACTCTTTCCCTACACGACGCTCTTCCGATCTNNNGGGGTCTACTTGAGGGTAAA
GAGCACACGTCTGAACTCTTTCCCTACACGACGCTCTTCCGATCTNNNGGGGTCTACTTGAGGGTAAA
GAGCACACGTCTGAACTCTTTCCCTACACGACGCTCTTCCGATCTNNNCATGCCTGTAATCCCAGCTACT
GAGCACACGTCTGAACTCTTTCCCTACACGACGCTCTTCCGATCTNNNCATGCCTGTAATCCCAGCTACT
GAGCACACGTCTGAACTCTTTCCCTACACGACGCTCTTCCGATCTNNNTGCCAACGTATACCCATTTTGC
GAGCACACGTCTGAACTCTTTCCCTACACGACGCTCTTCCGATCTNNNTGCCAACGTATACCCATTTTGC
AGCACACGTCTGAACTCTTTCCCTACACGACGCTCTTCCGATCTNNNAGCACTACTCAGCAATGAAAGC
AGCACACGTCTGAACTCTTTCCCTACACGACGCTCTTCCGATCTNNNAGCACTACTCAGCAATGAAAGC
GAGCACACGTCTGAACTCTTTCCCTACACGACGCTCTTCCGATCTNNNGTAGAGACAGAGTTTCACCGT
GAGCACACGTCTGAACTCTTTCCCTACACGACGCTCTTCCGATCTNNNGTAGAGACAGAGTTTCACCGT
GAGCACACGTCTGAACTCTTTCCCTACACGACGCTCTTCCGATCTNNNAGGCTCAAACTCTTGACTTCAG
GAGCACACGTCTGAACTCTTTCCCTACACGACGCTCTTCCGATCTNNNAGGCTCAAACTCTTGACTTCAG
GAGCACACGTCTGAACTCTTTCCCTACACGACGCTCTTCCGATCTNNNTGGTTGTATGTCCACAGTCAA
GAGCACACGTCTGAACTCTTTCCCTACACGACGCTCTTCCGATCTNNNTGGTTGTATGTCCACAGTCAA
GCACACGTCTGAACTCTTTCCCTACACGACGCTCTTCCGATCTNNNTGATCTTTACTCTTGGCCACACAC
GCACACGTCTGAACTCTTTCCCTACACGACGCTCTTCCGATCTNNNTGATCTTTACTCTTGGCCACACAC
GAGCACACGTCTGAACTCTTTCCCTACACGACGCTCTTCCGATCTNNNCTACAGCACTGCCATTTCTTCTC
GAGCACACGTCTGAACTCTTTCCCTACACGACGCTCTTCCGATCTNNNCTACAGCACTGCCATTTCTTCTC
AGAGCACACGTCTGAACTCTTTCCCTACACGACGCTCTTCCGATCTNNNAAGATTTAGGTAGCAGACGT
AGAGCACACGTCTGAACTCTTTCCCTACACGACGCTCTTCCGATCTNNNAAGATTTAGGTAGCAGACGT
AGAGCACACGTCTGAACTCTTTCCCTACACGACGCTCTTCCGATCTNNNTGGCAGACTCAACTTCTTAG
AGAGCACACGTCTGAACTCTTTCCCTACACGACGCTCTTCCGATCTNNNTGGCAGACTCAACTTCTTAG
GAGCACACGTCTGAACTCTTTCCCTACACGACGCTCTTCCGATCTNNNAAAAAATTAGCCAGGTGTGG
GAGCACACGTCTGAACTCTTTCCCTACACGACGCTCTTCCGATCTNNNAAAAAATTAGCCAGGTGTGG
GAGCACACGTCTGAACTCTTTCCCTACACGACGCTCTTCCGATCTNNNGGAATAGCACGGAAAGTTGTC
GAGCACACGTCTGAACTCTTTCCCTACACGACGCTCTTCCGATCTNNNGGAATAGCACGGAAAGTTGTC
GAGCACACGTCTGAACTCTTTCCCTACACGACGCTCTTCCGATCTNNNAGCTGCTGTATCTGTTGTAGC
GAGCACACGTCTGAACTCTTTCCCTACACGACGCTCTTCCGATCTNNNAGCTGCTGTATCTGTTGTAGC
AGCACACGTCTGAACTCTTTCCCTACACGACGCTCTTCCGATCTNNNCTTCTCTGGCTACAGCTCGTCCAT
AGCACACGTCTGAACTCTTTCCCTACACGACGCTCTTCCGATCTNNNCTTCTCTGGCTACAGCTCGTCCAT
GCACACGTCTGAACTCTTTCCCTACACGACGCTCTTCCGATCTNNNCAGTGTTGTGCCTCTGAGGATAT
GCACACGTCTGAACTCTTTCCCTACACGACGCTCTTCCGATCTNNNCAGTGTTGTGCCTCTGAGGATAT
GAGCACACGTCTGAACTCTTTCCCTACACGACGCTCTTCCGATCTNNNGAATAGCTTGGATCCGGGAG
GAGCACACGTCTGAACTCTTTCCCTACACGACGCTCTTCCGATCTNNNGAATAGCTTGGATCCGGGAG
AGAGCACACGTCTGAACTCTTTCCCTACACGACGCTCTTCCGATCTNNNCCCAAACTGACCAACGAATCA
AGAGCACACGTCTGAACTCTTTCCCTACACGACGCTCTTCCGATCTNNNCCCAAACTGACCAACGAATCA
GAGCACACGTCTGAACTCTTTCCCTACACGACGCTCTTCCGATCTNNNCATCTCCAGCCTCCTACCCTCCA

GAGCACACGTCTGAACTCTTTCCCTACACGACGCTCTTCCGATCTNNNCATCTCCAGCCTCCTACCCTCCA
AGCACACGTCTGAACTCTTTCCCTACACGACGCTCTTCCGATCTNNNTGGAACTCAGCAAAGAAAGGA
AGCACACGTCTGAACTCTTTCCCTACACGACGCTCTTCCGATCTNNNTGGAACTCAGCAAAGAAAGGA
GAGCACACGTCTGAACTCTTTCCCTACACGACGCTCTTCCGATCTNNNATTATCCATCTGTCCAGCACACA
GAGCACACGTCTGAACTCTTTCCCTACACGACGCTCTTCCGATCTNNNATTATCCATCTGTCCAGCACACA
GAGCACACGTCTGAACTCTTTCCCTACACGACGCTCTTCCGATCTNNNTTACAGGGAATCTTGCATCAC
GAGCACACGTCTGAACTCTTTCCCTACACGACGCTCTTCCGATCTNNNTTACAGGGAATCTTGCATCAC
GCACACGTCTGAACTCTTTCCCTACACGACGCTCTTCCGATCTNNNCTCAAAGTGCTGGGATTACAAGC
GCACACGTCTGAACTCTTTCCCTACACGACGCTCTTCCGATCTNNNCTCAAAGTGCTGGGATTACAAGC
GAGCACACGTCTGAACTCTTTCCCTACACGACGCTCTTCCGATCTNNNATAGCTACCATGCACTACCACA
GAGCACACGTCTGAACTCTTTCCCTACACGACGCTCTTCCGATCTNNNATAGCTACCATGCACTACCACA
GAGCACACGTCTGAACTCTTTCCCTACACGACGCTCTTCCGATCTNNNTCCTGGAAACATACAATCTTCC
GAGCACACGTCTGAACTCTTTCCCTACACGACGCTCTTCCGATCTNNNTCCTGGAAACATACAATCTTCC
AGCACACGTCTGAACTCTTTCCCTACACGACGCTCTTCCGATCTNNNGAGGCAGGAGAATTGCTTGAA
AGCACACGTCTGAACTCTTTCCCTACACGACGCTCTTCCGATCTNNNGAGGCAGGAGAATTGCTTGAA
AGCACACGTCTGAACTCTTTCCCTACACGACGCTCTTCCGATCTNNNCCCAGGTTCAAGCAATTCTCCTA
AGCACACGTCTGAACTCTTTCCCTACACGACGCTCTTCCGATCTNNNCCCAGGTTCAAGCAATTCTCCTA
GAGCACACGTCTGAACTCTTTCCCTACACGACGCTCTTCCGATCTNNNAAAGAAGAGGAAGGGAAGG
GAGCACACGTCTGAACTCTTTCCCTACACGACGCTCTTCCGATCTNNNAAAGAAGAGGAAGGGAAGG
AGCACACGTCTGAACTCTTTCCCTACACGACGCTCTTCCGATCTNNNGATGGTTACACACACACACACAC
AGCACACGTCTGAACTCTTTCCCTACACGACGCTCTTCCGATCTNNNGATGGTTACACACACACACACAC
GCACACGTCTGAACTCTTTCCCTACACGACGCTCTTCCGATCTNNNTGGTAATTCAGATGTGCCCTCAA
GCACACGTCTGAACTCTTTCCCTACACGACGCTCTTCCGATCTNNNTGGTAATTCAGATGTGCCCTCAA
CACACGTCTGAACTCTTTCCCTACACGACGCTCTTCCGATCTNNNCAGTCAACATGTACGTTGCGGCAT
CACACGTCTGAACTCTTTCCCTACACGACGCTCTTCCGATCTNNNCAGTCAACATGTACGTTGCGGCAT
GAGCACACGTCTGAACTCTTTCCCTACACGACGCTCTTCCGATCTNNNAGATGTTGCAATGAGCCAAG
GAGCACACGTCTGAACTCTTTCCCTACACGACGCTCTTCCGATCTNNNAGATGTTGCAATGAGCCAAG
GAGCACACGTCTGAACTCTTTCCCTACACGACGCTCTTCCGATCTNNNCTCATGCCTATGATCGTAGCA
GAGCACACGTCTGAACTCTTTCCCTACACGACGCTCTTCCGATCTNNNCTCATGCCTATGATCGTAGCA
AGCACACGTCTGAACTCTTTCCCTACACGACGCTCTTCCGATCTNNNAAAAAAGGTAGCTGCAATCAG
AGCACACGTCTGAACTCTTTCCCTACACGACGCTCTTCCGATCTNNNAAAAAAGGTAGCTGCAATCAG
AGCACACGTCTGAACTCTTTCCCTACACGACGCTCTTCCGATCTNNNACACTGCCTTTGACATACTGCAT
AGCACACGTCTGAACTCTTTCCCTACACGACGCTCTTCCGATCTNNNACACTGCCTTTGACATACTGCAT
CACACGTCTGAACTCTTTCCCTACACGACGCTCTTCCGATCTNNNTTGCCTGAGAATCCTCCTTTGTTCA
CACACGTCTGAACTCTTTCCCTACACGACGCTCTTCCGATCTNNNTTGCCTGAGAATCCTCCTTTGTTCA
AGCACACGTCTGAACTCTTTCCCTACACGACGCTCTTCCGATCTNNNCTGAGGCAGGAGAATCACTTGA
AGCACACGTCTGAACTCTTTCCCTACACGACGCTCTTCCGATCTNNNCTGAGGCAGGAGAATCACTTGA
GAGCACACGTCTGAACTCTTTCCCTACACGACGCTCTTCCGATCTNNNCTTGCAGTGAGCCGAGATCC
GAGCACACGTCTGAACTCTTTCCCTACACGACGCTCTTCCGATCTNNNCTTGCAGTGAGCCGAGATCC
GCACACGTCTGAACTCTTTCCCTACACGACGCTCTTCCGATCTNNNGTCCTTTCCCATAGCAGCACTCAT
GCACACGTCTGAACTCTTTCCCTACACGACGCTCTTCCGATCTNNNGTCCTTTCCCATAGCAGCACTCAT
GGAAGAGCACACGTCTGAACTCTTTCCCTACACGACGCTCTTCCGATCTNNNGGTGAGCCTGACTGCA
GGAAGAGCACACGTCTGAACTCTTTCCCTACACGACGCTCTTCCGATCTNNNGGTGAGCCTGACTGCA

AGCACACGTCTGAACTCTTTCCCTACACGACGCTCTTCCGATCTNNNGTGGAGCTTATACCTGGATCCT
AGCACACGTCTGAACTCTTTCCCTACACGACGCTCTTCCGATCTNNNGTGGAGCTTATACCTGGATCCT
GAGCACACGTCTGAACTCTTTCCCTACACGACGCTCTTCCGATCTNNNTACATGGGAGTGTTGCATGA
GAGCACACGTCTGAACTCTTTCCCTACACGACGCTCTTCCGATCTNNNTACATGGGAGTGTTGCATGA
GAGCACACGTCTGAACTCTTTCCCTACACGACGCTCTTCCGATCTNNNGAAGAAGAAGGCCAAGAACA
GAGCACACGTCTGAACTCTTTCCCTACACGACGCTCTTCCGATCTNNNGAAGAAGAAGGCCAAGAACA
AGCACACGTCTGAACTCTTTCCCTACACGACGCTCTTCCGATCTNNNCTGAGGCAGGAGAATCACTTGA
AGCACACGTCTGAACTCTTTCCCTACACGACGCTCTTCCGATCTNNNCTGAGGCAGGAGAATCACTTGA
CACACGTCTGAACTCTTTCCCTACACGACGCTCTTCCGATCTNNNCTAGCCTGGACAACACAGTGAGCA
CACACGTCTGAACTCTTTCCCTACACGACGCTCTTCCGATCTNNNCTAGCCTGGACAACACAGTGAGCA
AGCACACGTCTGAACTCTTTCCCTACACGACGCTCTTCCGATCTNNNTATGGGGCACAAACACTACACAT
AGCACACGTCTGAACTCTTTCCCTACACGACGCTCTTCCGATCTNNNTATGGGGCACAAACACTACACAT
ACACGTCTGAACTCTTTCCCTACACGACGCTCTTCCGATCTNNNTCCCTTACAGTCCAAAACACACACATC
ACACGTCTGAACTCTTTCCCTACACGACGCTCTTCCGATCTNNNTCCCTTACAGTCCAAAACACACACATC
GAGCACACGTCTGAACTCTTTCCCTACACGACGCTCTTCCGATCTNNNGGGTTCCGGTCTCTTTATTCTG
GAGCACACGTCTGAACTCTTTCCCTACACGACGCTCTTCCGATCTNNNGGGTTCCGGTCTCTTTATTCTG
AGCACACGTCTGAACTCTTTCCCTACACGACGCTCTTCCGATCTNNNTTACCTGGATTGCAAATGGACCC
AGCACACGTCTGAACTCTTTCCCTACACGACGCTCTTCCGATCTNNNTTACCTGGATTGCAAATGGACCC
ACACGTCTGAACTCTTTCCCTACACGACGCTCTTCCGATCTNNNAGGAAGAATGCGATGTAAAATCATC
ACACGTCTGAACTCTTTCCCTACACGACGCTCTTCCGATCTNNNAGGAAGAATGCGATGTAAAATCATC
AGCACACGTCTGAACTCTTTCCCTACACGACGCTCTTCCGATCTNNNTATTGAACTCCTGGGCTCAAGT
AGCACACGTCTGAACTCTTTCCCTACACGACGCTCTTCCGATCTNNNTATTGAACTCCTGGGCTCAAGT
AGAGCACACGTCTGAACTCTTTCCCTACACGACGCTCTTCCGATCTNNNGAGAGCTGTTGAAAAAGCCT
AGAGCACACGTCTGAACTCTTTCCCTACACGACGCTCTTCCGATCTNNNGAGAGCTGTTGAAAAAGCCT
AGAGCACACGTCTGAACTCTTTCCCTACACGACGCTCTTCCGATCTNNNTTTCACTGCTCAATGCTTCCTC
AGAGCACACGTCTGAACTCTTTCCCTACACGACGCTCTTCCGATCTNNNTTTCACTGCTCAATGCTTCCTC
AGAGCACACGTCTGAACTCTTTCCCTACACGACGCTCTTCCGATCTNNNGAAAATCGCTTGAACTGGGG
AGAGCACACGTCTGAACTCTTTCCCTACACGACGCTCTTCCGATCTNNNGAAAATCGCTTGAACTGGGG
AGCACACGTCTGAACTCTTTCCCTACACGACGCTCTTCCGATCTNNNCCAGTCACGGAATCTTGTTTGTC
AGCACACGTCTGAACTCTTTCCCTACACGACGCTCTTCCGATCTNNNCCAGTCACGGAATCTTGTTTGTC
AGCACACGTCTGAACTCTTTCCCTACACGACGCTCTTCCGATCTNNNGTGGTGGGTGCCTATAATCCTAC
AGCACACGTCTGAACTCTTTCCCTACACGACGCTCTTCCGATCTNNNGTGGTGGGTGCCTATAATCCTAC
AGAGCACACGTCTGAACTCTTTCCCTACACGACGCTCTTCCGATCTNNNTTCCAGGCTTATTAGGCTTCC
AGAGCACACGTCTGAACTCTTTCCCTACACGACGCTCTTCCGATCTNNNTTCCAGGCTTATTAGGCTTCC
GAGCACACGTCTGAACTCTTTCCCTACACGACGCTCTTCCGATCTNNNTTCTGTCTCACTGTCATTCCAGC
GAGCACACGTCTGAACTCTTTCCCTACACGACGCTCTTCCGATCTNNNTTCTGTCTCACTGTCATTCCAGC
GAGCACACGTCTGAACTCTTTCCCTACACGACGCTCTTCCGATCTNNNAAGCCTTTTTATGCATCACCTC
GAGCACACGTCTGAACTCTTTCCCTACACGACGCTCTTCCGATCTNNNAAGCCTTTTTATGCATCACCTC
GAGCACACGTCTGAACTCTTTCCCTACACGACGCTCTTCCGATCTNNNTGGTAAATTTTAGGCCCCACA
GAGCACACGTCTGAACTCTTTCCCTACACGACGCTCTTCCGATCTNNNTGGTAAATTTTAGGCCCCACA
GAGCACACGTCTGAACTCTTTCCCTACACGACGCTCTTCCGATCTNNNGAGCTATAGACTGGAGCTGTT
GAGCACACGTCTGAACTCTTTCCCTACACGACGCTCTTCCGATCTNNNGAGCTATAGACTGGAGCTGTT
GAGCACACGTCTGAACTCTTTCCCTACACGACGCTCTTCCGATCTNNNTTTTCTCCCCTTCTGGAAGACC

GAGCACACGTCTGAACTCTTTCCCTACACGACGCTCTTCCGATCTNNNTTTTCTCCCCTTCTGGAAGACC
AGCACACGTCTGAACTCTTTCCCTACACGACGCTCTTCCGATCTNNNGGACCTTGGTATGGGCAAAAA
AGCACACGTCTGAACTCTTTCCCTACACGACGCTCTTCCGATCTNNNGGACCTTGGTATGGGCAAAAA
GAGCACACGTCTGAACTCTTTCCCTACACGACGCTCTTCCGATCTNNNGCAACGAGCAAAACTCCATCT
GAGCACACGTCTGAACTCTTTCCCTACACGACGCTCTTCCGATCTNNNGCAACGAGCAAAACTCCATCT
GAGCACACGTCTGAACTCTTTCCCTACACGACGCTCTTCCGATCTNNNAGGAAACTCATGCTCAGGTT
GAGCACACGTCTGAACTCTTTCCCTACACGACGCTCTTCCGATCTNNNAGGAAACTCATGCTCAGGTT
AGCACACGTCTGAACTCTTTCCCTACACGACGCTCTTCCGATCTNNNCTCCAACGCAGACTGGTTATTT
AGCACACGTCTGAACTCTTTCCCTACACGACGCTCTTCCGATCTNNNCTCCAACGCAGACTGGTTATTT
CACACGTCTGAACTCTTTCCCTACACGACGCTCTTCCGATCTNNNTCATGAATACTGTGACCTTTCCAT
CACACGTCTGAACTCTTTCCCTACACGACGCTCTTCCGATCTNNNTCATGAATACTGTGACCTTTCCAT
GAGCACACGTCTGAACTCTTTCCCTACACGACGCTCTTCCGATCTNNNCTGAGGCAGGAGAATCACTT
GAGCACACGTCTGAACTCTTTCCCTACACGACGCTCTTCCGATCTNNNCTGAGGCAGGAGAATCACTT
GAGCACACGTCTGAACTCTTTCCCTACACGACGCTCTTCCGATCTNNNCAGTGGAGCTGAATTGCTCAA
GAGCACACGTCTGAACTCTTTCCCTACACGACGCTCTTCCGATCTNNNCAGTGGAGCTGAATTGCTCAA
GCACACGTCTGAACTCTTTCCCTACACGACGCTCTTCCGATCTNNNTTTCTCAGCACCTCTCTCTTTCCCA
GCACACGTCTGAACTCTTTCCCTACACGACGCTCTTCCGATCTNNNTTTCTCAGCACCTCTCTCTTTCCCA
AGCACACGTCTGAACTCTTTCCCTACACGACGCTCTTCCGATCTNNNCTCCTGAGTAGCTGGGACTACA
AGCACACGTCTGAACTCTTTCCCTACACGACGCTCTTCCGATCTNNNCTCCTGAGTAGCTGGGACTACA
AGCACACGTCTGAACTCTTTCCCTACACGACGCTCTTCCGATCTNNNCACACTCTAACATGGGTGACAG
AGCACACGTCTGAACTCTTTCCCTACACGACGCTCTTCCGATCTNNNCACACTCTAACATGGGTGACAG
GAGCACACGTCTGAACTCTTTCCCTACACGACGCTCTTCCGATCTNNNTAAGGCAGGAGAATTGCTTG
GAGCACACGTCTGAACTCTTTCCCTACACGACGCTCTTCCGATCTNNNTAAGGCAGGAGAATTGCTTG
AGCACACGTCTGAACTCTTTCCCTACACGACGCTCTTCCGATCTNNNGTAGGGGAAGATCAGTGTGA
AGCACACGTCTGAACTCTTTCCCTACACGACGCTCTTCCGATCTNNNGTAGGGGAAGATCAGTGTGA
GCACACGTCTGAACTCTTTCCCTACACGACGCTCTTCCGATCTNNNGAGGGTGTATGTGTCCAGGAA
GCACACGTCTGAACTCTTTCCCTACACGACGCTCTTCCGATCTNNNGAGGGTGTATGTGTCCAGGAA
GAGCACACGTCTGAACTCTTTCCCTACACGACGCTCTTCCGATCTNNNATACTGAGAAGGCTGAGGCA
GAGCACACGTCTGAACTCTTTCCCTACACGACGCTCTTCCGATCTNNNATACTGAGAAGGCTGAGGCA
GAGCACACGTCTGAACTCTTTCCCTACACGACGCTCTTCCGATCTNNNATGCTTTGTTTCTTCCTGCAG
GAGCACACGTCTGAACTCTTTCCCTACACGACGCTCTTCCGATCTNNNATGCTTTGTTTCTTCCTGCAG
AGCACACGTCTGAACTCTTTCCCTACACGACGCTCTTCCGATCTNNNAAATGCTTCCATTTGGCAGGAA
AGCACACGTCTGAACTCTTTCCCTACACGACGCTCTTCCGATCTNNNAAATGCTTCCATTTGGCAGGAA
GAGCACACGTCTGAACTCTTTCCCTACACGACGCTCTTCCGATCTNNNCCAGACAAACAAAACCAACTG
GAGCACACGTCTGAACTCTTTCCCTACACGACGCTCTTCCGATCTNNNCCAGACAAACAAAACCAACTG
GCACACGTCTGAACTCTTTCCCTACACGACGCTCTTCCGATCTNNNAAGCATCTTTGATAACCCATTAC
GCACACGTCTGAACTCTTTCCCTACACGACGCTCTTCCGATCTNNNAAGCATCTTTGATAACCCATTAC
AGCACACGTCTGAACTCTTTCCCTACACGACGCTCTTCCGATCTNNNGAAAAGCTGAGGGAAAAGCAG
AGCACACGTCTGAACTCTTTCCCTACACGACGCTCTTCCGATCTNNNGAAAAGCTGAGGGAAAAGCAG
GAGCACACGTCTGAACTCTTTCCCTACACGACGCTCTTCCGATCTNNNTTCTATCCCACACTGCACATTG
GAGCACACGTCTGAACTCTTTCCCTACACGACGCTCTTCCGATCTNNNTTCTATCCCACACTGCACATTG
CACACGTCTGAACTCTTTCCCTACACGACGCTCTTCCGATCTNNNAAAATTATTGAGAACAGAAGCAG
CACACGTCTGAACTCTTTCCCTACACGACGCTCTTCCGATCTNNNAAAATTATTGAGAACAGAAGCAG

GCACACGTCTGAACTCTTTCCCTACACGACGCTCTTCCGATCTNNNTGAAGGGCTCTCAAAATCATCCAC

GCACACGTCTGAACTCTTTCCCTACACGACGCTCTTCCGATCTNNNTGAAGGGCTCTCAAAATCATCCAC

AGCACACGTCTGAACTCTTTCCCTACACGACGCTCTTCCGATCTNNNACGCTAAAGATCCCTACTTAGCA

AGCACACGTCTGAACTCTTTCCCTACACGACGCTCTTCCGATCTNNNACGCTAAAGATCCCTACTTAGCA

GAGCACACGTCTGAACTCTTTCCCTACACGACGCTCTTCCGATCTNNNGTCTCATTCTGTTGTCTGTTGC

GAGCACACGTCTGAACTCTTTCCCTACACGACGCTCTTCCGATCTNNNGTCTCATTCTGTTGTCTGTTGC

GAGCACACGTCTGAACTCTTTCCCTACACGACGCTCTTCCGATCTNNNTCAATTCAGAGAACTGCACGGA

GAGCACACGTCTGAACTCTTTCCCTACACGACGCTCTTCCGATCTNNNTCAATTCAGAGAACTGCACGGA

GAGCACACGTCTGAACTCTTTCCCTACACGACGCTCTTCCGATCTNNNTTAGTCAGGCTGGTCTTGAAC

GAGCACACGTCTGAACTCTTTCCCTACACGACGCTCTTCCGATCTNNNTTAGTCAGGCTGGTCTTGAAC

AGCACACGTCTGAACTCTTTCCCTACACGACGCTCTTCCGATCTNNNGCCATTGCTTGTCTGTAAAGGA

AGCACACGTCTGAACTCTTTCCCTACACGACGCTCTTCCGATCTNNNGCCATTGCTTGTCTGTAAAGGA

AGCACACGTCTGAACTCTTTCCCTACACGACGCTCTTCCGATCTNNNAGCCATGATTACGTCACTGCAT

AGCACACGTCTGAACTCTTTCCCTACACGACGCTCTTCCGATCTNNNAGCCATGATTACGTCACTGCAT

AGCACACGTCTGAACTCTTTCCCTACACGACGCTCTTCCGATCTNNNAGTTGTCTGTGGTATTCACTGC

AGCACACGTCTGAACTCTTTCCCTACACGACGCTCTTCCGATCTNNNAGTTGTCTGTGGTATTCACTGC

GCACACGTCTGAACTCTTTCCCTACACGACGCTCTTCCGATCTNNNTGTAGTGATGTTCCCTTGTTCGTC

GCACACGTCTGAACTCTTTCCCTACACGACGCTCTTCCGATCTNNNTGTAGTGATGTTCCCTTGTTCGTC

AGCACACGTCTGAACTCTTTCCCTACACGACGCTCTTCCGATCTNNNGCCTATAAGGAAGTGTGGACT

AGCACACGTCTGAACTCTTTCCCTACACGACGCTCTTCCGATCTNNNGCCTATAAGGAAGTGTGGACT

AGCACACGTCTGAACTCTTTCCCTACACGACGCTCTTCCGATCTNNNGAGTAACAGGCTTCAGTGGAGA

AGCACACGTCTGAACTCTTTCCCTACACGACGCTCTTCCGATCTNNNGAGTAACAGGCTTCAGTGGAGA

GCACACGTCTGAACTCTTTCCCTACACGACGCTCTTCCGATCTNNNGCCTTTGCACCATGACTCAATAAC

GCACACGTCTGAACTCTTTCCCTACACGACGCTCTTCCGATCTNNNGCCTTTGCACCATGACTCAATAAC

AGCACACGTCTGAACTCTTTCCCTACACGACGCTCTTCCGATCTNNNTCGGACAAAAAATGTGGGGAA

AGCACACGTCTGAACTCTTTCCCTACACGACGCTCTTCCGATCTNNNTCGGACAAAAAATGTGGGGAA

GAGCACACGTCTGAACTCTTTCCCTACACGACGCTCTTCCGATCTNNNGTATTTCTGCAGTCCCAGCTAC

GAGCACACGTCTGAACTCTTTCCCTACACGACGCTCTTCCGATCTNNNGTATTTCTGCAGTCCCAGCTAC

GAGCACACGTCTGAACTCTTTCCCTACACGACGCTCTTCCGATCTNNNCCCCAGAAAGCCTCGTTATATG

GAGCACACGTCTGAACTCTTTCCCTACACGACGCTCTTCCGATCTNNNCCCCAGAAAGCCTCGTTATATG

AGAGCACACGTCTGAACTCTTTCCCTACACGACGCTCTTCCGATCTNNNACTTCAGGATATTTCGCTCTC

AGAGCACACGTCTGAACTCTTTCCCTACACGACGCTCTTCCGATCTNNNACTTCAGGATATTTCGCTCTC

AGCACACGTCTGAACTCTTTCCCTACACGACGCTCTTCCGATCTNNNCACCTGTAGTCCCAGCTACTTGC

AGCACACGTCTGAACTCTTTCCCTACACGACGCTCTTCCGATCTNNNCACCTGTAGTCCCAGCTACTTGC

GAGCACACGTCTGAACTCTTTCCCTACACGACGCTCTTCCGATCTNNNCGATGGCAGGTAAGAGAGAC

GAGCACACGTCTGAACTCTTTCCCTACACGACGCTCTTCCGATCTNNNCGATGGCAGGTAAGAGAGAC

GCACACGTCTGAACTCTTTCCCTACACGACGCTCTTCCGATCTNNNGCAGTGAACCATGATTGCATCAA

GCACACGTCTGAACTCTTTCCCTACACGACGCTCTTCCGATCTNNNGCAGTGAACCATGATTGCATCAA

AGCACACGTCTGAACTCTTTCCCTACACGACGCTCTTCCGATCTNNNGGACCTAAACTCTCAGGCTCAAA

AGCACACGTCTGAACTCTTTCCCTACACGACGCTCTTCCGATCTNNNGGACCTAAACTCTCAGGCTCAAA

GAGCACACGTCTGAACTCTTTCCCTACACGACGCTCTTCCGATCTNNNAGACAATGAATCACCTCCTTGC

GAGCACACGTCTGAACTCTTTCCCTACACGACGCTCTTCCGATCTNNNAGACAATGAATCACCTCCTTGC

AGCACACGTCTGAACTCTTTCCCTACACGACGCTCTTCCGATCTNNNTCTACCCACTCTTTCCTGCTTTCC

AGCACACGTCTGAACTCTTTCCCTACACGACGCTCTTCCGATCTNNNTCTACCCACTCTTTCCTGCTTTCC
GAGCACACGTCTGAACTCTTTCCCTACACGACGCTCTTCCGATCTNNNCCCTGGAGCAAAAGAAATGCA
GAGCACACGTCTGAACTCTTTCCCTACACGACGCTCTTCCGATCTNNNCCCTGGAGCAAAAGAAATGCA
AGCACACGTCTGAACTCTTTCCCTACACGACGCTCTTCCGATCTNNNGAAGACAAACTCTATGACATTTT
AGCACACGTCTGAACTCTTTCCCTACACGACGCTCTTCCGATCTNNNGAAGACAAACTCTATGACATTTT
GAGCACACGTCTGAACTCTTTCCCTACACGACGCTCTTCCGATCTNNNTCTCTTTCCTTCCTTCCTTCCC
GAGCACACGTCTGAACTCTTTCCCTACACGACGCTCTTCCGATCTNNNTCTCTTTCCTTCCTTCCTTCCC
AGAGCACACGTCTGAACTCTTTCCCTACACGACGCTCTTCCGATCTNNNATTACAGTAGTCAGGCATGG
AGAGCACACGTCTGAACTCTTTCCCTACACGACGCTCTTCCGATCTNNNATTACAGTAGTCAGGCATGG
AGCACACGTCTGAACTCTTTCCCTACACGACGCTCTTCCGATCTNNNACGCCCGGCTAATTTTTCTTTTCC
AGCACACGTCTGAACTCTTTCCCTACACGACGCTCTTCCGATCTNNNACGCCCGGCTAATTTTTCTTTTCC
GAGCACACGTCTGAACTCTTTCCCTACACGACGCTCTTCCGATCTNNNATCCACCAACCAAGCCCTATATG
GAGCACACGTCTGAACTCTTTCCCTACACGACGCTCTTCCGATCTNNNATCCACCAACCAAGCCCTATATG
AGCACACGTCTGAACTCTTTCCCTACACGACGCTCTTCCGATCTNNNACCATGTCCAGCTAAATTTTTGC
AGCACACGTCTGAACTCTTTCCCTACACGACGCTCTTCCGATCTNNNACCATGTCCAGCTAAATTTTTGC
AGCACACGTCTGAACTCTTTCCCTACACGACGCTCTTCCGATCTNNNTGAAGCTCACATGGAAGACAG
AGCACACGTCTGAACTCTTTCCCTACACGACGCTCTTCCGATCTNNNTGAAGCTCACATGGAAGACAG
GAGCACACGTCTGAACTCTTTCCCTACACGACGCTCTTCCGATCTNNNCTCTGCTCTTACAGGAGGAACA
GAGCACACGTCTGAACTCTTTCCCTACACGACGCTCTTCCGATCTNNNCTCTGCTCTTACAGGAGGAACA
GAGCACACGTCTGAACTCTTTCCCTACACGACGCTCTTCCGATCTNNNGAGCCTATAATCCCAGCTACTC
GAGCACACGTCTGAACTCTTTCCCTACACGACGCTCTTCCGATCTNNNGAGCCTATAATCCCAGCTACTC
GAGCACACGTCTGAACTCTTTCCCTACACGACGCTCTTCCGATCTNNNCAGGGAGCTAATGTCCAAGGA
GAGCACACGTCTGAACTCTTTCCCTACACGACGCTCTTCCGATCTNNNCAGGGAGCTAATGTCCAAGGA
AGCACACGTCTGAACTCTTTCCCTACACGACGCTCTTCCGATCTNNNCCAGACTACAGTGAACGACTCT
AGCACACGTCTGAACTCTTTCCCTACACGACGCTCTTCCGATCTNNNCCAGACTACAGTGAACGACTCT
AGCACACGTCTGAACTCTTTCCCTACACGACGCTCTTCCGATCTNNNTCTGGTGGTCTAGTGGTTAGG
AGCACACGTCTGAACTCTTTCCCTACACGACGCTCTTCCGATCTNNNTCTGGTGGTCTAGTGGTTAGG
AGCACACGTCTGAACTCTTTCCCTACACGACGCTCTTCCGATCTNNNGGGGTACAAAAGAGCAAAAGC
AGCACACGTCTGAACTCTTTCCCTACACGACGCTCTTCCGATCTNNNGGGGTACAAAAGAGCAAAAGC
GAGCACACGTCTGAACTCTTTCCCTACACGACGCTCTTCCGATCTNNNGGCAAAGGGGTTTAGAGCTTT
GAGCACACGTCTGAACTCTTTCCCTACACGACGCTCTTCCGATCTNNNGGCAAAGGGGTTTAGAGCTTT
GAGCACACGTCTGAACTCTTTCCCTACACGACGCTCTTCCGATCTNNNCCCTACCTCTAAGATGTCTGGC
GAGCACACGTCTGAACTCTTTCCCTACACGACGCTCTTCCGATCTNNNCCCTACCTCTAAGATGTCTGGC
AGCACACGTCTGAACTCTTTCCCTACACGACGCTCTTCCGATCTNNNTCCCGAGTAGCTGGTACTATAGC
AGCACACGTCTGAACTCTTTCCCTACACGACGCTCTTCCGATCTNNNTCCCGAGTAGCTGGTACTATAGC
ACACGTCTGAACTCTTTCCCTACACGACGCTCTTCCGATCTNNNCTAAAAGTTTTGAGCTTCTCAACATG
ACACGTCTGAACTCTTTCCCTACACGACGCTCTTCCGATCTNNNCTAAAAGTTTTGAGCTTCTCAACATG
CACACGTCTGAACTCTTTCCCTACACGACGCTCTTCCGATCTNNNTGTTTACAGGGAATGTAGTTTCCAT
CACACGTCTGAACTCTTTCCCTACACGACGCTCTTCCGATCTNNNTGTTTACAGGGAATGTAGTTTCCAT
AGCACACGTCTGAACTCTTTCCCTACACGACGCTCTTCCGATCTNNNGTTCAAGACCAACATGACCAACA
AGCACACGTCTGAACTCTTTCCCTACACGACGCTCTTCCGATCTNNNGTTCAAGACCAACATGACCAACA
GAGCACACGTCTGAACTCTTTCCCTACACGACGCTCTTCCGATCTNNNCTGAGGCAAGAGAGTTGCTTG
GAGCACACGTCTGAACTCTTTCCCTACACGACGCTCTTCCGATCTNNNCTGAGGCAAGAGAGTTGCTTG

AGCACACGTCTGAACTCTTTCCCTACACGACGCTCTTCCGATCTNNNGCCTTGCGCAAGTACTATGAAA

AGCACACGTCTGAACTCTTTCCCTACACGACGCTCTTCCGATCTNNNGCCTTGCGCAAGTACTATGAAA

CACACGTCTGAACTCTTTCCCTACACGACGCTCTTCCGATCTNNNAAAATTATTGAGAACAGAAGCAG

CACACGTCTGAACTCTTTCCCTACACGACGCTCTTCCGATCTNNNAAAATTATTGAGAACAGAAGCAG

AGCACACGTCTGAACTCTTTCCCTACACGACGCTCTTCCGATCTNNNCAAATGCAGCATGCATGTAGGT

AGCACACGTCTGAACTCTTTCCCTACACGACGCTCTTCCGATCTNNNCAAATGCAGCATGCATGTAGGT

AGCACACGTCTGAACTCTTTCCCTACACGACGCTCTTCCGATCTNNNCTGAGGCAGGAGAATCACTTGA

AGCACACGTCTGAACTCTTTCCCTACACGACGCTCTTCCGATCTNNNCTGAGGCAGGAGAATCACTTGA

GAGCACACGTCTGAACTCTTTCCCTACACGACGCTCTTCCGATCTNNNGAGAATCGCTTGATCCTTGGA

GAGCACACGTCTGAACTCTTTCCCTACACGACGCTCTTCCGATCTNNNGAGAATCGCTTGATCCTTGGA

AGCACACGTCTGAACTCTTTCCCTACACGACGCTCTTCCGATCTNNNTTTTCAGAAAATCACACCAGCCC

AGCACACGTCTGAACTCTTTCCCTACACGACGCTCTTCCGATCTNNNTTTTCAGAAAATCACACCAGCCC

GCACACGTCTGAACTCTTTCCCTACACGACGCTCTTCCGATCTNNNCTGCTCCTGGCCTCCTATTTATTCA

GCACACGTCTGAACTCTTTCCCTACACGACGCTCTTCCGATCTNNNCTGCTCCTGGCCTCCTATTTATTCA

AGAGCACACGTCTGAACTCTTTCCCTACACGACGCTCTTCCGATCTNNNGAAGGGCGAAGTCAGTAGA

AGAGCACACGTCTGAACTCTTTCCCTACACGACGCTCTTCCGATCTNNNGAAGGGCGAAGTCAGTAGA

AGAGCACACGTCTGAACTCTTTCCCTACACGACGCTCTTCCGATCTNNNTGGAAGCCATTAGAGCTGT

AGAGCACACGTCTGAACTCTTTCCCTACACGACGCTCTTCCGATCTNNNTGGAAGCCATTAGAGCTGT

GAGCACACGTCTGAACTCTTTCCCTACACGACGCTCTTCCGATCTNNNTGCTAGAAACAAATCTCCAATG

GAGCACACGTCTGAACTCTTTCCCTACACGACGCTCTTCCGATCTNNNTGCTAGAAACAAATCTCCAATG

GAGCACACGTCTGAACTCTTTCCCTACACGACGCTCTTCCGATCTNNNGGTGACAGAGTGAGACTCCA

GAGCACACGTCTGAACTCTTTCCCTACACGACGCTCTTCCGATCTNNNGGTGACAGAGTGAGACTCCA

GAGCACACGTCTGAACTCTTTCCCTACACGACGCTCTTCCGATCTNNNCCTGATGGGAAGCTATAGAA

GAGCACACGTCTGAACTCTTTCCCTACACGACGCTCTTCCGATCTNNNCCTGATGGGAAGCTATAGAA

CACACGTCTGAACTCTTTCCCTACACGACGCTCTTCCGATCTNNNATCACGAGGTCAGGAGATAGACT

CACACGTCTGAACTCTTTCCCTACACGACGCTCTTCCGATCTNNNATCACGAGGTCAGGAGATAGACT

AGAGCACACGTCTGAACTCTTTCCCTACACGACGCTCTTCCGATCTNNNGACCTTAGAGACATGCTACC

AGAGCACACGTCTGAACTCTTTCCCTACACGACGCTCTTCCGATCTNNNGACCTTAGAGACATGCTACC

AGCACACGTCTGAACTCTTTCCCTACACGACGCTCTTCCGATCTNNNATCGCTGTATTCTTGACTGAGCC

AGCACACGTCTGAACTCTTTCCCTACACGACGCTCTTCCGATCTNNNATCGCTGTATTCTTGACTGAGCC

GAGCACACGTCTGAACTCTTTCCCTACACGACGCTCTTCCGATCTNNNTGAAACAGGGTCTTAGTCTGT

GAGCACACGTCTGAACTCTTTCCCTACACGACGCTCTTCCGATCTNNNTGAAACAGGGTCTTAGTCTGT

AGCACACGTCTGAACTCTTTCCCTACACGACGCTCTTCCGATCTNNNAAGGAAAACTCTGGACAGACA

AGCACACGTCTGAACTCTTTCCCTACACGACGCTCTTCCGATCTNNNAAGGAAAACTCTGGACAGACA

GAGCACACGTCTGAACTCTTTCCCTACACGACGCTCTTCCGATCTNNNCAGATCACTTGAGGTCAGGA

GAGCACACGTCTGAACTCTTTCCCTACACGACGCTCTTCCGATCTNNNCAGATCACTTGAGGTCAGGA

AGAGCACACGTCTGAACTCTTTCCCTACACGACGCTCTTCCGATCTNNNAGCAAAGTTAGCCTCACATA

AGAGCACACGTCTGAACTCTTTCCCTACACGACGCTCTTCCGATCTNNNAGCAAAGTTAGCCTCACATA

AGCACACGTCTGAACTCTTTCCCTACACGACGCTCTTCCGATCTNNNACATGGTGAAATCCCGTCTCTA

AGCACACGTCTGAACTCTTTCCCTACACGACGCTCTTCCGATCTNNNACATGGTGAAATCCCGTCTCTA

AGAGCACACGTCTGAACTCTTTCCCTACACGACGCTCTTCCGATCTNNNCAGCCACACAGTCCAAAACA

AGAGCACACGTCTGAACTCTTTCCCTACACGACGCTCTTCCGATCTNNNCAGCCACACAGTCCAAAACA

GAGCACACGTCTGAACTCTTTCCCTACACGACGCTCTTCCGATCTNNNGTGGAAAGTCAGGAGGAGGT

GAGCACACGTCTGAACTCTTTCCCTACACGACGCTCTTCCGATCTNNNGTGGAAAGTCAGGAGGAGGT
AGCACACGTCTGAACTCTTTCCCTACACGACGCTCTTCCGATCTNNNTGCCTAGGGGAAATTGTAGAA
AGCACACGTCTGAACTCTTTCCCTACACGACGCTCTTCCGATCTNNNTGCCTAGGGGAAATTGTAGAA
GCACACGTCTGAACTCTTTCCCTACACGACGCTCTTCCGATCTNNNGCCTATGGATGTTCAATTGCTCC
GCACACGTCTGAACTCTTTCCCTACACGACGCTCTTCCGATCTNNNGCCTATGGATGTTCAATTGCTCC
AGCACACGTCTGAACTCTTTCCCTACACGACGCTCTTCCGATCTNNNACGGTTGAAGTTACACTGAGC
AGCACACGTCTGAACTCTTTCCCTACACGACGCTCTTCCGATCTNNNACGGTTGAAGTTACACTGAGC
GAGCACACGTCTGAACTCTTTCCCTACACGACGCTCTTCCGATCTNNNAGGAGAATCACTTGAACCCAC
GAGCACACGTCTGAACTCTTTCCCTACACGACGCTCTTCCGATCTNNNAGGAGAATCACTTGAACCCAC
GAGCACACGTCTGAACTCTTTCCCTACACGACGCTCTTCCGATCTNNNCTCCAGTGACCTTTCCTGTGCC
GAGCACACGTCTGAACTCTTTCCCTACACGACGCTCTTCCGATCTNNNCTCCAGTGACCTTTCCTGTGCC
AGCACACGTCTGAACTCTTTCCCTACACGACGCTCTTCCGATCTNNNAGATGTTTATTCGCATCCTTTGT
AGCACACGTCTGAACTCTTTCCCTACACGACGCTCTTCCGATCTNNNAGATGTTTATTCGCATCCTTTGT
AGCACACGTCTGAACTCTTTCCCTACACGACGCTCTTCCGATCTNNNAACAGTCCCCTTCCATATTTCCC
AGCACACGTCTGAACTCTTTCCCTACACGACGCTCTTCCGATCTNNNAACAGTCCCCTTCCATATTTCCC
AGCACACGTCTGAACTCTTTCCCTACACGACGCTCTTCCGATCTNNNGTGAAACAAAAATGATGGCAT
AGCACACGTCTGAACTCTTTCCCTACACGACGCTCTTCCGATCTNNNGTGAAACAAAAATGATGGCAT
GAGCACACGTCTGAACTCTTTCCCTACACGACGCTCTTCCGATCTNNNGAAGCAGGACAATCTCGTGA
GAGCACACGTCTGAACTCTTTCCCTACACGACGCTCTTCCGATCTNNNGAAGCAGGACAATCTCGTGA
AGCACACGTCTGAACTCTTTCCCTACACGACGCTCTTCCGATCTNNNTGAGTGAGACTCTGCCTCAAAA
AGCACACGTCTGAACTCTTTCCCTACACGACGCTCTTCCGATCTNNNTGAGTGAGACTCTGCCTCAAAA
CACACGTCTGAACTCTTTCCCTACACGACGCTCTTCCGATCTNNNAAAGAGGGAGTGACAGTACTGTT
CACACGTCTGAACTCTTTCCCTACACGACGCTCTTCCGATCTNNNAAAGAGGGAGTGACAGTACTGTT
AGAGCACACGTCTGAACTCTTTCCCTACACGACGCTCTTCCGATCTNNNAGACTGCATCTTTCCTCTTGG
AGAGCACACGTCTGAACTCTTTCCCTACACGACGCTCTTCCGATCTNNNAGACTGCATCTTTCCTCTTGG
AGCACACGTCTGAACTCTTTCCCTACACGACGCTCTTCCGATCTNNNCCAGCCATCATCTTGTCCATTTC
AGCACACGTCTGAACTCTTTCCCTACACGACGCTCTTCCGATCTNNNCCAGCCATCATCTTGTCCATTTC
AGCACACGTCTGAACTCTTTCCCTACACGACGCTCTTCCGATCTNNNCAGGAGAATTGCTTGAACCTGG
AGCACACGTCTGAACTCTTTCCCTACACGACGCTCTTCCGATCTNNNCAGGAGAATTGCTTGAACCTGG
AGAGCACACGTCTGAACTCTTTCCCTACACGACGCTCTTCCGATCTNNNAGTTGCCTCTCTGTTGAACTC
AGAGCACACGTCTGAACTCTTTCCCTACACGACGCTCTTCCGATCTNNNAGTTGCCTCTCTGTTGAACTC
GCACACGTCTGAACTCTTTCCCTACACGACGCTCTTCCGATCTNNNCTGGGAAGTGAGGAGCGTCTCCA
GCACACGTCTGAACTCTTTCCCTACACGACGCTCTTCCGATCTNNNCTGGGAAGTGAGGAGCGTCTCCA
AGCACACGTCTGAACTCTTTCCCTACACGACGCTCTTCCGATCTNNNGTAATTCTAGCTACTGGGGAGG
AGCACACGTCTGAACTCTTTCCCTACACGACGCTCTTCCGATCTNNNGTAATTCTAGCTACTGGGGAGG
AGCACACGTCTGAACTCTTTCCCTACACGACGCTCTTCCGATCTNNNTTCAGTGTAGTCCTCAGAAAACA
AGCACACGTCTGAACTCTTTCCCTACACGACGCTCTTCCGATCTNNNTTCAGTGTAGTCCTCAGAAAACA
GAGCACACGTCTGAACTCTTTCCCTACACGACGCTCTTCCGATCTNNNCAGGGACATTGGCCAGAAAA
GAGCACACGTCTGAACTCTTTCCCTACACGACGCTCTTCCGATCTNNNCAGGGACATTGGCCAGAAAA
GCACACGTCTGAACTCTTTCCCTACACGACGCTCTTCCGATCTNNNGTCAAGAAGGTGAGGCTCTCATC
GCACACGTCTGAACTCTTTCCCTACACGACGCTCTTCCGATCTNNNGTCAAGAAGGTGAGGCTCTCATC
AGCACACGTCTGAACTCTTTCCCTACACGACGCTCTTCCGATCTNNNATGGAGAAGTTTCTGAGAAGG
AGCACACGTCTGAACTCTTTCCCTACACGACGCTCTTCCGATCTNNNATGGAGAAGTTTCTGAGAAGG

AGAGCACACGTCTGAACTCTTTCCCTACACGACGCTCTTCCGATCTNNNAGTAGATTGTGAGGGAAG

AGAGCACACGTCTGAACTCTTTCCCTACACGACGCTCTTCCGATCTNNNAGTAGATTGTGAGGGAAG

AGCACACGTCTGAACTCTTTCCCTACACGACGCTCTTCCGATCTNNNAGCCAACAGAACATTGCATTCT

AGCACACGTCTGAACTCTTTCCCTACACGACGCTCTTCCGATCTNNNAGCCAACAGAACATTGCATTCT

GAGCACACGTCTGAACTCTTTCCCTACACGACGCTCTTCCGATCTNNNCTCCAAAACACACACACACACA

GAGCACACGTCTGAACTCTTTCCCTACACGACGCTCTTCCGATCTNNNCTCCAAAACACACACACACACA

AGCACACGTCTGAACTCTTTCCCTACACGACGCTCTTCCGATCTNNNCAAGAGAATTGCTTGAACCTG

AGCACACGTCTGAACTCTTTCCCTACACGACGCTCTTCCGATCTNNNCAAGAGAATTGCTTGAACCTG

AGCACACGTCTGAACTCTTTCCCTACACGACGCTCTTCCGATCTNNNTGTGTCCCAAACTAAATTCCCC

AGCACACGTCTGAACTCTTTCCCTACACGACGCTCTTCCGATCTNNNTGTGTCCCAAACTAAATTCCCC

AAGAGCACACGTCTGAACTCTTTCCCTACACGACGCTCTTCCGATCTNNNGTACCTGATCTGATGACAC

AAGAGCACACGTCTGAACTCTTTCCCTACACGACGCTCTTCCGATCTNNNGTACCTGATCTGATGACAC

AGAGCACACGTCTGAACTCTTTCCCTACACGACGCTCTTCCGATCTNNNATAAAAATGAGCCGGGCAG

AGAGCACACGTCTGAACTCTTTCCCTACACGACGCTCTTCCGATCTNNNATAAAAATGAGCCGGGCAG

AGCACACGTCTGAACTCTTTCCCTACACGACGCTCTTCCGATCTNNNTCAAGAGATTGTCCTGTCTCAG

AGCACACGTCTGAACTCTTTCCCTACACGACGCTCTTCCGATCTNNNTCAAGAGATTGTCCTGTCTCAG

AGCACACGTCTGAACTCTTTCCCTACACGACGCTCTTCCGATCTNNNATGTGGAAATTCAGTTGTGCCA

AGCACACGTCTGAACTCTTTCCCTACACGACGCTCTTCCGATCTNNNATGTGGAAATTCAGTTGTGCCA

AGCACACGTCTGAACTCTTTCCCTACACGACGCTCTTCCGATCTNNNCCGACAGAGTGAGACTCCATCT

AGCACACGTCTGAACTCTTTCCCTACACGACGCTCTTCCGATCTNNNCCGACAGAGTGAGACTCCATCT

AGCACACGTCTGAACTCTTTCCCTACACGACGCTCTTCCGATCTNNNGAGGCTGAGGCAGGAGAATG

AGCACACGTCTGAACTCTTTCCCTACACGACGCTCTTCCGATCTNNNGAGGCTGAGGCAGGAGAATG

AGCACACGTCTGAACTCTTTCCCTACACGACGCTCTTCCGATCTNNNCCACTTCTTGTTTGGTGCTGTAC

AGCACACGTCTGAACTCTTTCCCTACACGACGCTCTTCCGATCTNNNCCACTTCTTGTTTGGTGCTGTAC

AGCACACGTCTGAACTCTTTCCCTACACGACGCTCTTCCGATCTNNNCAACTTTGGGGCATAGTTTGAG

AGCACACGTCTGAACTCTTTCCCTACACGACGCTCTTCCGATCTNNNCAACTTTGGGGCATAGTTTGAG

AGCACACGTCTGAACTCTTTCCCTACACGACGCTCTTCCGATCTNNNTTTATTCTGGGCGTGATTCTGG

AGCACACGTCTGAACTCTTTCCCTACACGACGCTCTTCCGATCTNNNTTTATTCTGGGCGTGATTCTGG

GAGCACACGTCTGAACTCTTTCCCTACACGACGCTCTTCCGATCTNNNGGTCTGGGCAAATGTTTGTT

GAGCACACGTCTGAACTCTTTCCCTACACGACGCTCTTCCGATCTNNNGGTCTGGGCAAATGTTTGTT

AGCACACGTCTGAACTCTTTCCCTACACGACGCTCTTCCGATCTNNNGAGATTGCAGTGAGCCGAGAT

AGCACACGTCTGAACTCTTTCCCTACACGACGCTCTTCCGATCTNNNGAGATTGCAGTGAGCCGAGAT

GAGCACACGTCTGAACTCTTTCCCTACACGACGCTCTTCCGATCTNNNGCAGGAGAATTGCTTGAACCT

GAGCACACGTCTGAACTCTTTCCCTACACGACGCTCTTCCGATCTNNNGCAGGAGAATTGCTTGAACCT

AGCACACGTCTGAACTCTTTCCCTACACGACGCTCTTCCGATCTNNNTTCCATGTGGTTGTAGGACTG

AGCACACGTCTGAACTCTTTCCCTACACGACGCTCTTCCGATCTNNNTTCCATGTGGTTGTAGGACTG

GAGCACACGTCTGAACTCTTTCCCTACACGACGCTCTTCCGATCTNNNATGCATTGCTCAAGTACTCAG

GAGCACACGTCTGAACTCTTTCCCTACACGACGCTCTTCCGATCTNNNATGCATTGCTCAAGTACTCAG

GAGCACACGTCTGAACTCTTTCCCTACACGACGCTCTTCCGATCTNNNTTTGGGCTCTGAATCTACATG

GAGCACACGTCTGAACTCTTTCCCTACACGACGCTCTTCCGATCTNNNTTTGGGCTCTGAATCTACATG

AGCACACGTCTGAACTCTTTCCCTACACGACGCTCTTCCGATCTNNNCCACACCGGGCTAATTTTTGTA

AGCACACGTCTGAACTCTTTCCCTACACGACGCTCTTCCGATCTNNNCCACACCGGGCTAATTTTTGTA

AAGAGCACACGTCTGAACTCTTTCCCTACACGACGCTCTTCCGATCTNNNGACTGTGTTTCTCCCTTCT

AAGAGCACACGTCTGAACTCTTTCCCTACACGACGCTCTTCCGATCTNNNGACTGTGTTTCTCCCTTCTC
GAGCACACGTCTGAACTCTTTCCCTACACGACGCTCTTCCGATCTNNNTAAGGTCAGGAGTTCGAAAC
GAGCACACGTCTGAACTCTTTCCCTACACGACGCTCTTCCGATCTNNNTAAGGTCAGGAGTTCGAAAC
AGCACACGTCTGAACTCTTTCCCTACACGACGCTCTTCCGATCTNNNCTTCTGGGAATATCAGCTGCTG
AGCACACGTCTGAACTCTTTCCCTACACGACGCTCTTCCGATCTNNNCTTCTGGGAATATCAGCTGCTG
AGCACACGTCTGAACTCTTTCCCTACACGACGCTCTTCCGATCTNNNTCATGCCTCCCAAGATTGAATCA
AGCACACGTCTGAACTCTTTCCCTACACGACGCTCTTCCGATCTNNNTCATGCCTCCCAAGATTGAATCA
GAGCACACGTCTGAACTCTTTCCCTACACGACGCTCTTCCGATCTNNNTCAGTACTATGACTCCTACAAC
GAGCACACGTCTGAACTCTTTCCCTACACGACGCTCTTCCGATCTNNNTCAGTACTATGACTCCTACAAC
GAGCACACGTCTGAACTCTTTCCCTACACGACGCTCTTCCGATCTNNNCCGTTTTCAAGAAAATTGCAG
GAGCACACGTCTGAACTCTTTCCCTACACGACGCTCTTCCGATCTNNNCCGTTTTCAAGAAAATTGCAG
GAGCACACGTCTGAACTCTTTCCCTACACGACGCTCTTCCGATCTNNNTACCTGTAGTCCCAGCTACTCA
GAGCACACGTCTGAACTCTTTCCCTACACGACGCTCTTCCGATCTNNNTACCTGTAGTCCCAGCTACTCA
AGCACACGTCTGAACTCTTTCCCTACACGACGCTCTTCCGATCTNNNTCTTTCTCTCTCTCTCTCCCTCCCA
AGCACACGTCTGAACTCTTTCCCTACACGACGCTCTTCCGATCTNNNTCTTTCTCTCTCTCTCTCCCTCCCA
GAGCACACGTCTGAACTCTTTCCCTACACGACGCTCTTCCGATCTNNNGGGTCTCTCTCTCTCTCTCTCTC
GAGCACACGTCTGAACTCTTTCCCTACACGACGCTCTTCCGATCTNNNGGGTCTCTCTCTCTCTCTCTCTC
AAGAGCACACGTCTGAACTCTTTCCCTACACGACGCTCTTCCGATCTNNNTAACCTAGACCAACATGGG
AAGAGCACACGTCTGAACTCTTTCCCTACACGACGCTCTTCCGATCTNNNTAACCTAGACCAACATGGG
AGAGCACACGTCTGAACTCTTTCCCTACACGACGCTCTTCCGATCTNNNATATGTCCTCTGTGATCACGA
AGAGCACACGTCTGAACTCTTTCCCTACACGACGCTCTTCCGATCTNNNATATGTCCTCTGTGATCACGA
GAGCACACGTCTGAACTCTTTCCCTACACGACGCTCTTCCGATCTNNNATTTCTCCCACCTTGCTATAGGA
GAGCACACGTCTGAACTCTTTCCCTACACGACGCTCTTCCGATCTNNNATTTCTCCCACCTTGCTATAGGA
AGCACACGTCTGAACTCTTTCCCTACACGACGCTCTTCCGATCTNNNCATTTCAGCCTGCACAATAGCTA
AGCACACGTCTGAACTCTTTCCCTACACGACGCTCTTCCGATCTNNNCATTTCAGCCTGCACAATAGCTA
AGCACACGTCTGAACTCTTTCCCTACACGACGCTCTTCCGATCTNNNTACCCACAGCTAACAACATACCG
AGCACACGTCTGAACTCTTTCCCTACACGACGCTCTTCCGATCTNNNTACCCACAGCTAACAACATACCG
AGCACACGTCTGAACTCTTTCCCTACACGACGCTCTTCCGATCTNNNGAGGCAGGAGAATTGCTTCAA
AGCACACGTCTGAACTCTTTCCCTACACGACGCTCTTCCGATCTNNNGAGGCAGGAGAATTGCTTCAA
GAGCACACGTCTGAACTCTTTCCCTACACGACGCTCTTCCGATCTNNNGGAGACTAGTAACAACTCTTC
GAGCACACGTCTGAACTCTTTCCCTACACGACGCTCTTCCGATCTNNNGGAGACTAGTAACAACTCTTC
CACGTCTGAACTCTTTCCCTACACGACGCTCTTCCGATCTNNNATTTTGCCATTTCACATCAGCATCGGA
CACGTCTGAACTCTTTCCCTACACGACGCTCTTCCGATCTNNNATTTTGCCATTTCACATCAGCATCGGA
GCACACGTCTGAACTCTTTCCCTACACGACGCTCTTCCGATCTNNNCTTTACATCCACTCTCCCAGCATCAT
GCACACGTCTGAACTCTTTCCCTACACGACGCTCTTCCGATCTNNNCTTTACATCCACTCTCCCAGCATCAT
AGCACACGTCTGAACTCTTTCCCTACACGACGCTCTTCCGATCTNNNCACTCAAGATCCCAAACTCCAGA
AGCACACGTCTGAACTCTTTCCCTACACGACGCTCTTCCGATCTNNNCACTCAAGATCCCAAACTCCAGA
AGAGCACACGTCTGAACTCTTTCCCTACACGACGCTCTTCCGATCTNNNGTTCATGCACTCACTTGACAC
AGAGCACACGTCTGAACTCTTTCCCTACACGACGCTCTTCCGATCTNNNGTTCATGCACTCACTTGACAC
AGCACACGTCTGAACTCTTTCCCTACACGACGCTCTTCCGATCTNNNTAAAGAGTCAAGACCCATCGGT
AGCACACGTCTGAACTCTTTCCCTACACGACGCTCTTCCGATCTNNNTAAAGAGTCAAGACCCATCGGT
AGCACACGTCTGAACTCTTTCCCTACACGACGCTCTTCCGATCTNNNAAGATCACTCCATTGCACTCCAC
AGCACACGTCTGAACTCTTTCCCTACACGACGCTCTTCCGATCTNNNAAGATCACTCCATTGCACTCCAC

AGCACACGTCTGAACTCTTTCCCTACACGACGCTCTTCCGATCTNNNTGGAATCTTCTGTGTTGTCTTT
AGCACACGTCTGAACTCTTTCCCTACACGACGCTCTTCCGATCTNNNTGGAATCTTCTGTGTTGTCTTT
AGCACACGTCTGAACTCTTTCCCTACACGACGCTCTTCCGATCTNNNGGGTTCACTTGGCATGGAATAT
AGCACACGTCTGAACTCTTTCCCTACACGACGCTCTTCCGATCTNNNGGGTTCACTTGGCATGGAATAT
AGAGCACACGTCTGAACTCTTTCCCTACACGACGCTCTTCCGATCTNNNAAGATGGATAGGACTTGAG
AGAGCACACGTCTGAACTCTTTCCCTACACGACGCTCTTCCGATCTNNNAAGATGGATAGGACTTGAG
AGCACACGTCTGAACTCTTTCCCTACACGACGCTCTTCCGATCTNNNACCACGTTCGGCTAATTTTTGT
AGCACACGTCTGAACTCTTTCCCTACACGACGCTCTTCCGATCTNNNACCACGTTCGGCTAATTTTTGT
AGCACACGTCTGAACTCTTTCCCTACACGACGCTCTTCCGATCTNNNCCCTTTTCTTGGCCTGTTATGT
AGCACACGTCTGAACTCTTTCCCTACACGACGCTCTTCCGATCTNNNCCCTTTTCTTGGCCTGTTATGT
AGCACACGTCTGAACTCTTTCCCTACACGACGCTCTTCCGATCTNNNGAGCCAATCATTCTCTTTGCTGG
AGCACACGTCTGAACTCTTTCCCTACACGACGCTCTTCCGATCTNNNGAGCCAATCATTCTCTTTGCTGG
AGCACACGTCTGAACTCTTTCCCTACACGACGCTCTTCCGATCTNNNCTGATCATTTGGCTCTTTTTTCA
AGCACACGTCTGAACTCTTTCCCTACACGACGCTCTTCCGATCTNNNCTGATCATTTGGCTCTTTTTTCA
AGCACACGTCTGAACTCTTTCCCTACACGACGCTCTTCCGATCTNNNAAATCTTCTGTCCCTTGCCATC
AGCACACGTCTGAACTCTTTCCCTACACGACGCTCTTCCGATCTNNNAAATCTTCTGTCCCTTGCCATC
GAGCACACGTCTGAACTCTTTCCCTACACGACGCTCTTCCGATCTNNNACATTTCCTTCTTTTTATGCCT
GAGCACACGTCTGAACTCTTTCCCTACACGACGCTCTTCCGATCTNNNACATTTCCTTCTTTTTATGCCT
AGCACACGTCTGAACTCTTTCCCTACACGACGCTCTTCCGATCTNNNTCTTGGGCCATACGTAACGAT
AGCACACGTCTGAACTCTTTCCCTACACGACGCTCTTCCGATCTNNNTCTTGGGCCATACGTAACGAT
AGCACACGTCTGAACTCTTTCCCTACACGACGCTCTTCCGATCTNNNTTCCTTCCTTCCTTCCTTCCTTCC
AGCACACGTCTGAACTCTTTCCCTACACGACGCTCTTCCGATCTNNNTTCCTTCCTTCCTTCCTTCCTTCC
AGCACACGTCTGAACTCTTTCCCTACACGACGCTCTTCCGATCTNNNCTTGAGCCCAGGAGTTCGAGC
AGCACACGTCTGAACTCTTTCCCTACACGACGCTCTTCCGATCTNNNCTTGAGCCCAGGAGTTCGAGC
CACACGTCTGAACTCTTTCCCTACACGACGCTCTTCCGATCTNNNTTTTATTCCTCCATTTTGCAAACAT
CACACGTCTGAACTCTTTCCCTACACGACGCTCTTCCGATCTNNNTTTTATTCCTCCATTTTGCAAACAT
GCACACGTCTGAACTCTTTCCCTACACGACGCTCTTCCGATCTNNNAACCTTTGTTGAGCATGTACTGC
GCACACGTCTGAACTCTTTCCCTACACGACGCTCTTCCGATCTNNNAACCTTTGTTGAGCATGTACTGC
GCACACGTCTGAACTCTTTCCCTACACGACGCTCTTCCGATCTNNNATCCTGGCTAACAAGGTGAAACT
GCACACGTCTGAACTCTTTCCCTACACGACGCTCTTCCGATCTNNNATCCTGGCTAACAAGGTGAAACT
GAGCACACGTCTGAACTCTTTCCCTACACGACGCTCTTCCGATCTNNNTGGAGAACTAGGGAATGCC
GAGCACACGTCTGAACTCTTTCCCTACACGACGCTCTTCCGATCTNNNTGGAGAACTAGGGAATGCC
GCACACGTCTGAACTCTTTCCCTACACGACGCTCTTCCGATCTNNNACTGATGAATGCTCAGACTGGG
GCACACGTCTGAACTCTTTCCCTACACGACGCTCTTCCGATCTNNNACTGATGAATGCTCAGACTGGG
AGCACACGTCTGAACTCTTTCCCTACACGACGCTCTTCCGATCTNNNGCTGCATAAGTGAAGATTACG
AGCACACGTCTGAACTCTTTCCCTACACGACGCTCTTCCGATCTNNNGCTGCATAAGTGAAGATTACG
AGCACACGTCTGAACTCTTTCCCTACACGACGCTCTTCCGATCTNNNTAGCTTGCATCTTGGATCTTGG
AGCACACGTCTGAACTCTTTCCCTACACGACGCTCTTCCGATCTNNNTAGCTTGCATCTTGGATCTTGG
AGCACACGTCTGAACTCTTTCCCTACACGACGCTCTTCCGATCTNNNACTTGAACCTGGGAGAAAAAG
AGCACACGTCTGAACTCTTTCCCTACACGACGCTCTTCCGATCTNNNACTTGAACCTGGGAGAAAAAG
GAGCACACGTCTGAACTCTTTCCCTACACGACGCTCTTCCGATCTNNNATGCGTGATTTGCTTCATTGT
GAGCACACGTCTGAACTCTTTCCCTACACGACGCTCTTCCGATCTNNNATGCGTGATTTGCTTCATTGT
GAGCACACGTCTGAACTCTTTCCCTACACGACGCTCTTCCGATCTNNNGAGGCTGATACAGGAGAAT

AGAGCACACGTCTGAACTCTTTCCCTACACGACGCTCTTCCGATCTNNNGAGGCTGATACAGGAGAAT

GAGCACACGTCTGAACTCTTTCCCTACACGACGCTCTTCCGATCTNNNGAGATTCTGTCCAGCCTTACG

GAGCACACGTCTGAACTCTTTCCCTACACGACGCTCTTCCGATCTNNNGAGATTCTGTCCAGCCTTACG

GAGCACACGTCTGAACTCTTTCCCTACACGACGCTCTTCCGATCTNNNTCAGAAATTACTGCATGAGCC

GAGCACACGTCTGAACTCTTTCCCTACACGACGCTCTTCCGATCTNNNTCAGAAATTACTGCATGAGCC

GAGCACACGTCTGAACTCTTTCCCTACACGACGCTCTTCCGATCTNNNTCGAGGGTGACTTTCTTCTCT

GAGCACACGTCTGAACTCTTTCCCTACACGACGCTCTTCCGATCTNNNTCGAGGGTGACTTTCTTCTCT

GAGCACACGTCTGAACTCTTTCCCTACACGACGCTCTTCCGATCTNNNGCAGGAGAATTGCTTGAACCT

GAGCACACGTCTGAACTCTTTCCCTACACGACGCTCTTCCGATCTNNNGCAGGAGAATTGCTTGAACCT

AGAGCACACGTCTGAACTCTTTCCCTACACGACGCTCTTCCGATCTNNNACAAAAATTAGCTGGGCATG

AGAGCACACGTCTGAACTCTTTCCCTACACGACGCTCTTCCGATCTNNNACAAAAATTAGCTGGGCATG

AGCACACGTCTGAACTCTTTCCCTACACGACGCTCTTCCGATCTNNNCTAGGCTAAGGGTGTCCTTCACT

AGCACACGTCTGAACTCTTTCCCTACACGACGCTCTTCCGATCTNNNCTAGGCTAAGGGTGTCCTTCACT

AGCACACGTCTGAACTCTTTCCCTACACGACGCTCTTCCGATCTNNNTTCCGAAACATCACTGTTGAAG

AGCACACGTCTGAACTCTTTCCCTACACGACGCTCTTCCGATCTNNNTTCCGAAACATCACTGTTGAAG

AGCACACGTCTGAACTCTTTCCCTACACGACGCTCTTCCGATCTNNNAACATGATACAGTGCTGCAGA

AGCACACGTCTGAACTCTTTCCCTACACGACGCTCTTCCGATCTNNNAACATGATACAGTGCTGCAGA

AGCACACGTCTGAACTCTTTCCCTACACGACGCTCTTCCGATCTNNNCCGAACTACGTCACTCTGTGTA

AGCACACGTCTGAACTCTTTCCCTACACGACGCTCTTCCGATCTNNNCCGAACTACGTCACTCTGTGTA

GAGCACACGTCTGAACTCTTTCCCTACACGACGCTCTTCCGATCTNNNGCAGGAGAATCATTTGAACC

GAGCACACGTCTGAACTCTTTCCCTACACGACGCTCTTCCGATCTNNNGCAGGAGAATCATTTGAACC

AGCACACGTCTGAACTCTTTCCCTACACGACGCTCTTCCGATCTNNNCTCTGGTGATGGATCTGATCA

AGCACACGTCTGAACTCTTTCCCTACACGACGCTCTTCCGATCTNNNCTCTGGTGATGGATCTGATCA

GCACACGTCTGAACTCTTTCCCTACACGACGCTCTTCCGATCTNNNCTGTCTGCAGAGAGAGGCTGCAT

GCACACGTCTGAACTCTTTCCCTACACGACGCTCTTCCGATCTNNNCTGTCTGCAGAGAGAGGCTGCAT

AGCACACGTCTGAACTCTTTCCCTACACGACGCTCTTCCGATCTNNNGCTCATGCCTGTAATCTCAACA

AGCACACGTCTGAACTCTTTCCCTACACGACGCTCTTCCGATCTNNNGCTCATGCCTGTAATCTCAACA

AGCACACGTCTGAACTCTTTCCCTACACGACGCTCTTCCGATCTNNNAATACAAAAAATTAGCCGGGC

AGCACACGTCTGAACTCTTTCCCTACACGACGCTCTTCCGATCTNNNAATACAAAAAATTAGCCGGGC

GAGCACACGTCTGAACTCTTTCCCTACACGACGCTCTTCCGATCTNNNAGATCGAGACCATCATGGCT

GAGCACACGTCTGAACTCTTTCCCTACACGACGCTCTTCCGATCTNNNAGATCGAGACCATCATGGCT

AGCACACGTCTGAACTCTTTCCCTACACGACGCTCTTCCGATCTNNNACCAAAATTTCTGCCATTCCTTT

AGCACACGTCTGAACTCTTTCCCTACACGACGCTCTTCCGATCTNNNACCAAAATTTCTGCCATTCCTTT

GAGCACACGTCTGAACTCTTTCCCTACACGACGCTCTTCCGATCTNNNATATGGCGAAACCCTGTCTCT

GAGCACACGTCTGAACTCTTTCCCTACACGACGCTCTTCCGATCTNNNATATGGCGAAACCCTGTCTCT

AAGAGCACACGTCTGAACTCTTTCCCTACACGACGCTCTTCCGATCTNNNAATTTGCTCAGTTGCAGGG

AAGAGCACACGTCTGAACTCTTTCCCTACACGACGCTCTTCCGATCTNNNAATTTGCTCAGTTGCAGGG

GAGCACACGTCTGAACTCTTTCCCTACACGACGCTCTTCCGATCTNNNCAGTGGATTTGGACTGCAGA

GAGCACACGTCTGAACTCTTTCCCTACACGACGCTCTTCCGATCTNNNCAGTGGATTTGGACTGCAGA

GAGCACACGTCTGAACTCTTTCCCTACACGACGCTCTTCCGATCTNNNCAAGGCAACAGAACAAAACTC

GAGCACACGTCTGAACTCTTTCCCTACACGACGCTCTTCCGATCTNNNCAAGGCAACAGAACAAAACTC

AAGAGCACACGTCTGAACTCTTTCCCTACACGACGCTCTTCCGATCTNNNGAACAATCTGCAGAAGAG

AAGAGCACACGTCTGAACTCTTTCCCTACACGACGCTCTTCCGATCTNNNGAACAATCTGCAGAAGAG

AGCACACGTCTGAACTCTTTCCCTACACGACGCTCTTCCGATCTNNNCGCTATTGGAGGGCTCAGTTA
AGCACACGTCTGAACTCTTTCCCTACACGACGCTCTTCCGATCTNNNCGCTATTGGAGGGCTCAGTTA
GCACACGTCTGAACTCTTTCCCTACACGACGCTCTTCCGATCTNNNGGTTTGTTCCCGAATTGTTTCCT
GCACACGTCTGAACTCTTTCCCTACACGACGCTCTTCCGATCTNNNGGTTTGTTCCCGAATTGTTTCCT
AGCACACGTCTGAACTCTTTCCCTACACGACGCTCTTCCGATCTNNNTGCTGGGTTCAGGCAATTATA
AGCACACGTCTGAACTCTTTCCCTACACGACGCTCTTCCGATCTNNNTGCTGGGTTCAGGCAATTATA
AGCACACGTCTGAACTCTTTCCCTACACGACGCTCTTCCGATCTNNNACATGTACAAGTCTCCCAAC
AGCACACGTCTGAACTCTTTCCCTACACGACGCTCTTCCGATCTNNNACATGTACAAGTCTCCCAAC
CACACGTCTGAACTCTTTCCCTACACGACGCTCTTCCGATCTNNNTCTGGCTGTGTAGAAATGAAAG
CACACGTCTGAACTCTTTCCCTACACGACGCTCTTCCGATCTNNNTCTGGCTGTGTAGAAATGAAAG
AGCACACGTCTGAACTCTTTCCCTACACGACGCTCTTCCGATCTNNNACTCAACCCCTTTCACAAGAGT
AGCACACGTCTGAACTCTTTCCCTACACGACGCTCTTCCGATCTNNNACTCAACCCCTTTCACAAGAGT
AGCACACGTCTGAACTCTTTCCCTACACGACGCTCTTCCGATCTNNNTGCCTGTAATCCCAGATATTCG
AGCACACGTCTGAACTCTTTCCCTACACGACGCTCTTCCGATCTNNNTGCCTGTAATCCCAGATATTCG
GAGCACACGTCTGAACTCTTTCCCTACACGACGCTCTTCCGATCTNNNTAAAGCAGGAGGATCATCTG
GAGCACACGTCTGAACTCTTTCCCTACACGACGCTCTTCCGATCTNNNTAAAGCAGGAGGATCATCTG
GCACACGTCTGAACTCTTTCCCTACACGACGCTCTTCCGATCTNNNCATCCTGGATGACTATGACTCAT
GCACACGTCTGAACTCTTTCCCTACACGACGCTCTTCCGATCTNNNCATCCTGGATGACTATGACTCAT
AGCACACGTCTGAACTCTTTCCCTACACGACGCTCTTCCGATCTNNNGCCAGAGTAAGCCCATTTTT
AGCACACGTCTGAACTCTTTCCCTACACGACGCTCTTCCGATCTNNNGCCAGAGTAAGCCCATTTTT
AGCACACGTCTGAACTCTTTCCCTACACGACGCTCTTCCGATCTNNNAGAGAATCGCTTGAACCTGAG
AGCACACGTCTGAACTCTTTCCCTACACGACGCTCTTCCGATCTNNNAGAGAATCGCTTGAACCTGAG
GCACACGTCTGAACTCTTTCCCTACACGACGCTCTTCCGATCTNNNGCCGGCTTCAAATGTCTTCTTTT
GCACACGTCTGAACTCTTTCCCTACACGACGCTCTTCCGATCTNNNGCCGGCTTCAAATGTCTTCTTTT
CACACGTCTGAACTCTTTCCCTACACGACGCTCTTCCGATCTNNNCCCGGCCTCAATACTGTATTCTTC
CACACGTCTGAACTCTTTCCCTACACGACGCTCTTCCGATCTNNNCCCGGCCTCAATACTGTATTCTTC
AGAGCACACGTCTGAACTCTTTCCCTACACGACGCTCTTCCGATCTNNNCAGAATGATGTCAGGACTT
AGAGCACACGTCTGAACTCTTTCCCTACACGACGCTCTTCCGATCTNNNCAGAATGATGTCAGGACTT
GAGCACACGTCTGAACTCTTTCCCTACACGACGCTCTTCCGATCTNNNCTGTCTTCTACCAGCATCCAT
GAGCACACGTCTGAACTCTTTCCCTACACGACGCTCTTCCGATCTNNNCTGTCTTCTACCAGCATCCAT
GCACACGTCTGAACTCTTTCCCTACACGACGCTCTTCCGATCTNNNGAACTCCTGACCTCATGTGATCC
GCACACGTCTGAACTCTTTCCCTACACGACGCTCTTCCGATCTNNNGAACTCCTGACCTCATGTGATCC
AGCACACGTCTGAACTCTTTCCCTACACGACGCTCTTCCGATCTNNNTACTTACGTCGGCAGACCTTAC
AGCACACGTCTGAACTCTTTCCCTACACGACGCTCTTCCGATCTNNNTACTTACGTCGGCAGACCTTAC
GCACACGTCTGAACTCTTTCCCTACACGACGCTCTTCCGATCTNNNCATATGCACATTATGGCTCCTCG
GCACACGTCTGAACTCTTTCCCTACACGACGCTCTTCCGATCTNNNCATATGCACATTATGGCTCCTCG
AGCACACGTCTGAACTCTTTCCCTACACGACGCTCTTCCGATCTNNNTCAGGATTCTACAGCCTCAGA
AGCACACGTCTGAACTCTTTCCCTACACGACGCTCTTCCGATCTNNNTCAGGATTCTACAGCCTCAGA
AGCACACGTCTGAACTCTTTCCCTACACGACGCTCTTCCGATCTNNNTTGTGGTGACCTGAGATTGTA
AGCACACGTCTGAACTCTTTCCCTACACGACGCTCTTCCGATCTNNNTTGTGGTGACCTGAGATTGTA
AGCACACGTCTGAACTCTTTCCCTACACGACGCTCTTCCGATCTNNNTGGATCCTGAGGTCAAGAGTT
AGCACACGTCTGAACTCTTTCCCTACACGACGCTCTTCCGATCTNNNTGGATCCTGAGGTCAAGAGTT
GCACACGTCTGAACTCTTTCCCTACACGACGCTCTTCCGATCTNNNCTGTGTTCCAGGATGCGTACTA

AGCACACGTCTGAACTCTTTCCCTACACGACGCTCTTCCGATCTNNNCTGTGTTCCAGGATGCGTACTAC

GAGCACACGTCTGAACTCTTTCCCTACACGACGCTCTTCCGATCTNNNATTTGTTAGGGTTCCTCTGGG

GAGCACACGTCTGAACTCTTTCCCTACACGACGCTCTTCCGATCTNNNATTTGTTAGGGTTCCTCTGGG

AGCACACGTCTGAACTCTTTCCCTACACGACGCTCTTCCGATCTNNNTGTGTGTGTGTGTGTATGGGT

AGCACACGTCTGAACTCTTTCCCTACACGACGCTCTTCCGATCTNNNTGTGTGTGTGTGTGTATGGGT

AGCACACGTCTGAACTCTTTCCCTACACGACGCTCTTCCGATCTNNNAGTCTCTGTAGGACAGTATTCC

AGCACACGTCTGAACTCTTTCCCTACACGACGCTCTTCCGATCTNNNAGTCTCTGTAGGACAGTATTCC

GAGCACACGTCTGAACTCTTTCCCTACACGACGCTCTTCCGATCTNNNCCTCACACCTGTTAGGATGAAC

GAGCACACGTCTGAACTCTTTCCCTACACGACGCTCTTCCGATCTNNNCCTCACACCTGTTAGGATGAAC

AGCACACGTCTGAACTCTTTCCCTACACGACGCTCTTCCGATCTNNNAGAAAACGGTAGCATGACAGC

AGCACACGTCTGAACTCTTTCCCTACACGACGCTCTTCCGATCTNNNAGAAAACGGTAGCATGACAGC

GAGCACACGTCTGAACTCTTTCCCTACACGACGCTCTTCCGATCTNNNCTGGGTCTCTTCAACAATGTCC

GAGCACACGTCTGAACTCTTTCCCTACACGACGCTCTTCCGATCTNNNCTGGGTCTCTTCAACAATGTCC

AGCACACGTCTGAACTCTTTCCCTACACGACGCTCTTCCGATCTNNNGGTTGAAGAACTCCCGGGAATA

AGCACACGTCTGAACTCTTTCCCTACACGACGCTCTTCCGATCTNNNGGTTGAAGAACTCCCGGGAATA

GAGCACACGTCTGAACTCTTTCCCTACACGACGCTCTTCCGATCTNNNCTCAAGCCTGTAATCCTAGCAC

GAGCACACGTCTGAACTCTTTCCCTACACGACGCTCTTCCGATCTNNNCTCAAGCCTGTAATCCTAGCAC

AGCACACGTCTGAACTCTTTCCCTACACGACGCTCTTCCGATCTNNNAATGGTACCAGCTCGTCTTTGT

AGCACACGTCTGAACTCTTTCCCTACACGACGCTCTTCCGATCTNNNAATGGTACCAGCTCGTCTTTGT

GAGCACACGTCTGAACTCTTTCCCTACACGACGCTCTTCCGATCTNNNCTGTGATTGCACCACTTTGCTC

GAGCACACGTCTGAACTCTTTCCCTACACGACGCTCTTCCGATCTNNNCTGTGATTGCACCACTTTGCTC

GAGCACACGTCTGAACTCTTTCCCTACACGACGCTCTTCCGATCTNNNTCAGGCCCAAGCTAATGGTTAT

GAGCACACGTCTGAACTCTTTCCCTACACGACGCTCTTCCGATCTNNNTCAGGCCCAAGCTAATGGTTAT

AGCACACGTCTGAACTCTTTCCCTACACGACGCTCTTCCGATCTNNNCTTTGGTCTTTGATGACGGTGA

AGCACACGTCTGAACTCTTTCCCTACACGACGCTCTTCCGATCTNNNCTTTGGTCTTTGATGACGGTGA

AGAGCACACGTCTGAACTCTTTCCCTACACGACGCTCTTCCGATCTNNNCTGTAGTCCCAGCTACTCAGA

AGAGCACACGTCTGAACTCTTTCCCTACACGACGCTCTTCCGATCTNNNCTGTAGTCCCAGCTACTCAGA

AGCACACGTCTGAACTCTTTCCCTACACGACGCTCTTCCGATCTNNNCACTGTCTCCTCTCCATGGTCCC

AGCACACGTCTGAACTCTTTCCCTACACGACGCTCTTCCGATCTNNNCACTGTCTCCTCTCCATGGTCCC

GCACACGTCTGAACTCTTTCCCTACACGACGCTCTTCCGATCTNNNGCAAGATCTGACATCCAGAACCA

GCACACGTCTGAACTCTTTCCCTACACGACGCTCTTCCGATCTNNNGCAAGATCTGACATCCAGAACCA

GAGCACACGTCTGAACTCTTTCCCTACACGACGCTCTTCCGATCTNNNATGGGTAGATCTTGCCATTGT

GAGCACACGTCTGAACTCTTTCCCTACACGACGCTCTTCCGATCTNNNATGGGTAGATCTTGCCATTGT

AGCACACGTCTGAACTCTTTCCCTACACGACGCTCTTCCGATCTNNNAGTGGTGTAATCATGGCTCACT

AGCACACGTCTGAACTCTTTCCCTACACGACGCTCTTCCGATCTNNNAGTGGTGTAATCATGGCTCACT

GAGCACACGTCTGAACTCTTTCCCTACACGACGCTCTTCCGATCTNNNTGAATAGAATGGGAGGCGGA

GAGCACACGTCTGAACTCTTTCCCTACACGACGCTCTTCCGATCTNNNTGAATAGAATGGGAGGCGGA

GAGCACACGTCTGAACTCTTTCCCTACACGACGCTCTTCCGATCTNNNGAAGCAGGAGAATCACTTGA

GAGCACACGTCTGAACTCTTTCCCTACACGACGCTCTTCCGATCTNNNGAAGCAGGAGAATCACTTGA

AGCACACGTCTGAACTCTTTCCCTACACGACGCTCTTCCGATCTNNNTGGCAATGCTGTATTCTTCTCCTC

AGCACACGTCTGAACTCTTTCCCTACACGACGCTCTTCCGATCTNNNTGGCAATGCTGTATTCTTCTCCTC

AGCACACGTCTGAACTCTTTCCCTACACGACGCTCTTCCGATCTNNNTCTGCGAGGGTACTAGTGAGAC

AGCACACGTCTGAACTCTTTCCCTACACGACGCTCTTCCGATCTNNNTCTGCGAGGGTACTAGTGAGAC

AGAGCACACGTCTGAACTCTTTCCCTACACGACGCTCTTCCGATCTNNNTTTGGGAGGCTGAGATAGT

AGAGCACACGTCTGAACTCTTTCCCTACACGACGCTCTTCCGATCTNNNTTTGGGAGGCTGAGATAGT

AGCACACGTCTGAACTCTTTCCCTACACGACGCTCTTCCGATCTNNNTTGTATTCTTAGTAGCGACGGG

AGCACACGTCTGAACTCTTTCCCTACACGACGCTCTTCCGATCTNNNTTGTATTCTTAGTAGCGACGGG

AGCACACGTCTGAACTCTTTCCCTACACGACGCTCTTCCGATCTNNNCCGAGTAGCTGGGACTACAGGC

AGCACACGTCTGAACTCTTTCCCTACACGACGCTCTTCCGATCTNNNCCGAGTAGCTGGGACTACAGGC

GCACACGTCTGAACTCTTTCCCTACACGACGCTCTTCCGATCTNNNTGGAGTCCCAATCACGATTATT

GCACACGTCTGAACTCTTTCCCTACACGACGCTCTTCCGATCTNNNTGGAGTCCCAATCACGATTATT

GAGCACACGTCTGAACTCTTTCCCTACACGACGCTCTTCCGATCTNNNAAAGAATGTTTGTATACAAAC

GAGCACACGTCTGAACTCTTTCCCTACACGACGCTCTTCCGATCTNNNAAAGAATGTTTGTATACAAAC

AGCACACGTCTGAACTCTTTCCCTACACGACGCTCTTCCGATCTNNNCAAAGAGCCAAGCAAGGTAACA

AGCACACGTCTGAACTCTTTCCCTACACGACGCTCTTCCGATCTNNNCAAAGAGCCAAGCAAGGTAACA

CACACGTCTGAACTCTTTCCCTACACGACGCTCTTCCGATCTNNNGTAATCCCAGTTACTCAGGAGGCC

CACACGTCTGAACTCTTTCCCTACACGACGCTCTTCCGATCTNNNGTAATCCCAGTTACTCAGGAGGCC

AGCACACGTCTGAACTCTTTCCCTACACGACGCTCTTCCGATCTNNNAGACAGAAGAATCGCTTGAACC

AGCACACGTCTGAACTCTTTCCCTACACGACGCTCTTCCGATCTNNNAGACAGAAGAATCGCTTGAACC

GCACACGTCTGAACTCTTTCCCTACACGACGCTCTTCCGATCTNNNCCAGGAAGTCCAAAAAGGCATTT

GCACACGTCTGAACTCTTTCCCTACACGACGCTCTTCCGATCTNNNCCAGGAAGTCCAAAAAGGCATTT

AGCACACGTCTGAACTCTTTCCCTACACGACGCTCTTCCGATCTNNNAAGGATGCACAAGTTCCAAGGT

AGCACACGTCTGAACTCTTTCCCTACACGACGCTCTTCCGATCTNNNAAGGATGCACAAGTTCCAAGGT

AGAGCACACGTCTGAACTCTTTCCCTACACGACGCTCTTCCGATCTNNNGCACATGACTGTAACTCCAG

AGAGCACACGTCTGAACTCTTTCCCTACACGACGCTCTTCCGATCTNNNGCACATGACTGTAACTCCAG

GCACACGTCTGAACTCTTTCCCTACACGACGCTCTTCCGATCTNNNCCACCACACCAGGCTAATTTTTCCA

GCACACGTCTGAACTCTTTCCCTACACGACGCTCTTCCGATCTNNNCCACCACACCAGGCTAATTTTTCCA

AGCACACGTCTGAACTCTTTCCCTACACGACGCTCTTCCGATCTNNNCACCTGTAGTCCTGGCTACTTG

AGCACACGTCTGAACTCTTTCCCTACACGACGCTCTTCCGATCTNNNCACCTGTAGTCCTGGCTACTTG

GAGCACACGTCTGAACTCTTTCCCTACACGACGCTCTTCCGATCTNNNATTTGAGCTAAGGAGTACCAC

GAGCACACGTCTGAACTCTTTCCCTACACGACGCTCTTCCGATCTNNNATTTGAGCTAAGGAGTACCAC

AGCACACGTCTGAACTCTTTCCCTACACGACGCTCTTCCGATCTNNNAGCAGTACAAGTGGTTTTTGGT

AGCACACGTCTGAACTCTTTCCCTACACGACGCTCTTCCGATCTNNNAGCAGTACAAGTGGTTTTTGGT

AGCACACGTCTGAACTCTTTCCCTACACGACGCTCTTCCGATCTNNNGTGGCATTATCTGTGCTTACTG

AGCACACGTCTGAACTCTTTCCCTACACGACGCTCTTCCGATCTNNNGTGGCATTATCTGTGCTTACTG

AAGAGCACACGTCTGAACTCTTTCCCTACACGACGCTCTTCCGATCTNNNCCTTCTTATGGAGTGTCTTC

AAGAGCACACGTCTGAACTCTTTCCCTACACGACGCTCTTCCGATCTNNNCCTTCTTATGGAGTGTCTTC

GAGCACACGTCTGAACTCTTTCCCTACACGACGCTCTTCCGATCTNNNTCTGTAATTCGAACTCTGCCAC

GAGCACACGTCTGAACTCTTTCCCTACACGACGCTCTTCCGATCTNNNTCTGTAATTCGAACTCTGCCAC

GAGCACACGTCTGAACTCTTTCCCTACACGACGCTCTTCCGATCTNNNTGCATCCTAAACTGCTGGAGAA

GAGCACACGTCTGAACTCTTTCCCTACACGACGCTCTTCCGATCTNNNTGCATCCTAAACTGCTGGAGAA

GCACACGTCTGAACTCTTTCCCTACACGACGCTCTTCCGATCTNNNCAAGAAGAGCCACCAAGACAATC

GCACACGTCTGAACTCTTTCCCTACACGACGCTCTTCCGATCTNNNCAAGAAGAGCCACCAAGACAATC

AGCACACGTCTGAACTCTTTCCCTACACGACGCTCTTCCGATCTNNNCCTGATTTCAAAGGCTGTGGTT

AGCACACGTCTGAACTCTTTCCCTACACGACGCTCTTCCGATCTNNNCCTGATTTCAAAGGCTGTGGTT

AGCACACGTCTGAACTCTTTCCCTACACGACGCTCTTCCGATCTNNNCAGAACTGTGAGCCAATCAAAC

AGCACACGTCTGAACTCTTTCCCTACACGACGCTCTTCCGATCTNNNCAGAACTGTGAGCCAATCAAA
AGCACACGTCTGAACTCTTTCCCTACACGACGCTCTTCCGATCTNNNCGAGGACCCAAAGAACAACAT
AGCACACGTCTGAACTCTTTCCCTACACGACGCTCTTCCGATCTNNNCGAGGACCCAAAGAACAACAT
GCACACGTCTGAACTCTTTCCCTACACGACGCTCTTCCGATCTNNNCCACGGTGCTCTCATAGATATCA
GCACACGTCTGAACTCTTTCCCTACACGACGCTCTTCCGATCTNNNCCACGGTGCTCTCATAGATATCA
AGCACACGTCTGAACTCTTTCCCTACACGACGCTCTTCCGATCTNNNCCAAAGTGCTGGAATTACAGGT
AGCACACGTCTGAACTCTTTCCCTACACGACGCTCTTCCGATCTNNNCCAAAGTGCTGGAATTACAGGT
AGCACACGTCTGAACTCTTTCCCTACACGACGCTCTTCCGATCTNNNAATGGTAGGGAAGATGGGAG
AGCACACGTCTGAACTCTTTCCCTACACGACGCTCTTCCGATCTNNNAATGGTAGGGAAGATGGGAG
AGCACACGTCTGAACTCTTTCCCTACACGACGCTCTTCCGATCTNNNTCTCACTTCTGCTATGAAACCA
AGCACACGTCTGAACTCTTTCCCTACACGACGCTCTTCCGATCTNNNTCTCACTTCTGCTATGAAACCA
GAGCACACGTCTGAACTCTTTCCCTACACGACGCTCTTCCGATCTNNNAGCCTGAACAACAACAGTGA
GAGCACACGTCTGAACTCTTTCCCTACACGACGCTCTTCCGATCTNNNAGCCTGAACAACAACAGTGA
AGCACACGTCTGAACTCTTTCCCTACACGACGCTCTTCCGATCTNNNCCACCATGCCCAGCTAATTTTTT
AGCACACGTCTGAACTCTTTCCCTACACGACGCTCTTCCGATCTNNNCCACCATGCCCAGCTAATTTTTT
AGAGCACACGTCTGAACTCTTTCCCTACACGACGCTCTTCCGATCTNNNGAGGCATGAGAATCGCTTG
AGAGCACACGTCTGAACTCTTTCCCTACACGACGCTCTTCCGATCTNNNGAGGCATGAGAATCGCTTG
AGCACACGTCTGAACTCTTTCCCTACACGACGCTCTTCCGATCTNNNTTTAATGAAGAAGAGAAGTC
AGCACACGTCTGAACTCTTTCCCTACACGACGCTCTTCCGATCTNNNTTTAATGAAGAAGAGAAGTC
GAGCACACGTCTGAACTCTTTCCCTACACGACGCTCTTCCGATCTNNNGAGCTTGTAGTGAGCCGAG
GAGCACACGTCTGAACTCTTTCCCTACACGACGCTCTTCCGATCTNNNGAGCTTGTAGTGAGCCGAG
AGCACACGTCTGAACTCTTTCCCTACACGACGCTCTTCCGATCTNNNGGTTTTTGGAAGGCTCAGTACA
AGCACACGTCTGAACTCTTTCCCTACACGACGCTCTTCCGATCTNNNGGTTTTTGGAAGGCTCAGTACA
GCACACGTCTGAACTCTTTCCCTACACGACGCTCTTCCGATCTNNNTTGTGCACATGTACCCTAGAACT
GCACACGTCTGAACTCTTTCCCTACACGACGCTCTTCCGATCTNNNTTGTGCACATGTACCCTAGAACT
GAGCACACGTCTGAACTCTTTCCCTACACGACGCTCTTCCGATCTNNNGCAGATGGCTCTCTTTTTCCA
GAGCACACGTCTGAACTCTTTCCCTACACGACGCTCTTCCGATCTNNNGCAGATGGCTCTCTTTTTCCA
AGCACACGTCTGAACTCTTTCCCTACACGACGCTCTTCCGATCTNNNAAGTTCCTTCCTTTCAGCTCTGA
AGCACACGTCTGAACTCTTTCCCTACACGACGCTCTTCCGATCTNNNAAGTTCCTTCCTTTCAGCTCTGA
AGCACACGTCTGAACTCTTTCCCTACACGACGCTCTTCCGATCTNNNGCCCCCTTTGTGTCCTACTTAT
AGCACACGTCTGAACTCTTTCCCTACACGACGCTCTTCCGATCTNNNGCCCCCTTTGTGTCCTACTTAT
GAGCACACGTCTGAACTCTTTCCCTACACGACGCTCTTCCGATCTNNNCGTGAAGTGAAAATTGCATC
GAGCACACGTCTGAACTCTTTCCCTACACGACGCTCTTCCGATCTNNNCGTGAAGTGAAAATTGCATC
AGCACACGTCTGAACTCTTTCCCTACACGACGCTCTTCCGATCTNNNCTAGACCCCCACATTGAGTCAAT
AGCACACGTCTGAACTCTTTCCCTACACGACGCTCTTCCGATCTNNNCTAGACCCCCACATTGAGTCAAT
GAGCACACGTCTGAACTCTTTCCCTACACGACGCTCTTCCGATCTNNNCATGGCGAACACCTAGTTGAA
GAGCACACGTCTGAACTCTTTCCCTACACGACGCTCTTCCGATCTNNNCATGGCGAACACCTAGTTGAA
AGCACACGTCTGAACTCTTTCCCTACACGACGCTCTTCCGATCTNNNCCCACCAACAGTGTAAAAGCAT
AGCACACGTCTGAACTCTTTCCCTACACGACGCTCTTCCGATCTNNNCCCACCAACAGTGTAAAAGCAT
AGCACACGTCTGAACTCTTTCCCTACACGACGCTCTTCCGATCTNNNAAAAAAAGGTACTGGGGAGCT
AGCACACGTCTGAACTCTTTCCCTACACGACGCTCTTCCGATCTNNNAAAAAAAGGTACTGGGGAGCT
AGCACACGTCTGAACTCTTTCCCTACACGACGCTCTTCCGATCTNNNAGCAAATATTGGAAGTGGGG
AGCACACGTCTGAACTCTTTCCCTACACGACGCTCTTCCGATCTNNNAGCAAATATTGGAAGTGGGG

AGCACACGTCTGAACTCTTTCCCTACACGACGCTCTTCCGATCTNNNTGTTTTTCCCAGGAACACTCTCT

AGCACACGTCTGAACTCTTTCCCTACACGACGCTCTTCCGATCTNNNTGTTTTTCCCAGGAACACTCTCT

AGCACACGTCTGAACTCTTTCCCTACACGACGCTCTTCCGATCTNNNGTGTGTACATGTGTGTGTGTGA

AGCACACGTCTGAACTCTTTCCCTACACGACGCTCTTCCGATCTNNNGTGTGTACATGTGTGTGTGTGA

GAGCACACGTCTGAACTCTTTCCCTACACGACGCTCTTCCGATCTNNNCCCAGTGCTAACAGCTATCCTG

GAGCACACGTCTGAACTCTTTCCCTACACGACGCTCTTCCGATCTNNNCCCAGTGCTAACAGCTATCCTG

GAGCACACGTCTGAACTCTTTCCCTACACGACGCTCTTCCGATCTNNNGCCTTCCATGCCAAGAAGATTT

GAGCACACGTCTGAACTCTTTCCCTACACGACGCTCTTCCGATCTNNNGCCTTCCATGCCAAGAAGATTT

GAGCACACGTCTGAACTCTTTCCCTACACGACGCTCTTCCGATCTNNNCCCTGTAAGAAGAAACACCGA

GAGCACACGTCTGAACTCTTTCCCTACACGACGCTCTTCCGATCTNNNCCCTGTAAGAAGAAACACCGA

CACACGTCTGAACTCTTTCCCTACACGACGCTCTTCCGATCTNNNCCCCAATTTCGGCGATTCTGACATC

CACACGTCTGAACTCTTTCCCTACACGACGCTCTTCCGATCTNNNCCCCAATTTCGGCGATTCTGACATC

CACACGTCTGAACTCTTTCCCTACACGACGCTCTTCCGATCTNNNGTTCAAACGACTCTCTGCTTCAGCA

CACACGTCTGAACTCTTTCCCTACACGACGCTCTTCCGATCTNNNGTTCAAACGACTCTCTGCTTCAGCA

GAGCACACGTCTGAACTCTTTCCCTACACGACGCTCTTCCGATCTNNNCTCCCAAAGTGCTGCGATTACA

GAGCACACGTCTGAACTCTTTCCCTACACGACGCTCTTCCGATCTNNNCTCCCAAAGTGCTGCGATTACA

GAGCACACGTCTGAACTCTTTCCCTACACGACGCTCTTCCGATCTNNNATTAGGGACCCACTTACAGAA

GAGCACACGTCTGAACTCTTTCCCTACACGACGCTCTTCCGATCTNNNATTAGGGACCCACTTACAGAA

GAGCACACGTCTGAACTCTTTCCCTACACGACGCTCTTCCGATCTNNNCCAGGGGTCTACATTTTTCCTC

GAGCACACGTCTGAACTCTTTCCCTACACGACGCTCTTCCGATCTNNNCCAGGGGTCTACATTTTTCCTC

GAGCACACGTCTGAACTCTTTCCCTACACGACGCTCTTCCGATCTNNNGTTGCAGTGAGATGAGATTCT

GAGCACACGTCTGAACTCTTTCCCTACACGACGCTCTTCCGATCTNNNGTTGCAGTGAGATGAGATTCT

AGAGCACACGTCTGAACTCTTTCCCTACACGACGCTCTTCCGATCTNNNATACTGATCTCCATCCTCCTG

AGAGCACACGTCTGAACTCTTTCCCTACACGACGCTCTTCCGATCTNNNATACTGATCTCCATCCTCCTG

AGAGCACACGTCTGAACTCTTTCCCTACACGACGCTCTTCCGATCTNNNCAGAGAGAACTGCAGAGAG

AGAGCACACGTCTGAACTCTTTCCCTACACGACGCTCTTCCGATCTNNNCAGAGAGAACTGCAGAGAG

GCACACGTCTGAACTCTTTCCCTACACGACGCTCTTCCGATCTNNNGCTCTCCATTTTCGCTCACACATCA

GCACACGTCTGAACTCTTTCCCTACACGACGCTCTTCCGATCTNNNGCTCTCCATTTTCGCTCACACATCA

AGAGCACACGTCTGAACTCTTTCCCTACACGACGCTCTTCCGATCTNNNGAGGCAGGATAATTGCTTGA

AGAGCACACGTCTGAACTCTTTCCCTACACGACGCTCTTCCGATCTNNNGAGGCAGGATAATTGCTTGA

AGCACACGTCTGAACTCTTTCCCTACACGACGCTCTTCCGATCTNNNACCACCATAATCCCAGCTACTTG

AGCACACGTCTGAACTCTTTCCCTACACGACGCTCTTCCGATCTNNNACCACCATAATCCCAGCTACTTG

AGCACACGTCTGAACTCTTTCCCTACACGACGCTCTTCCGATCTNNNGTGAGCTGTAATTGTGCTACTG

AGCACACGTCTGAACTCTTTCCCTACACGACGCTCTTCCGATCTNNNGTGAGCTGTAATTGTGCTACTG

CACACGTCTGAACTCTTTCCCTACACGACGCTCTTCCGATCTNNNTGTGTCGGAGTTCTAACGATGTTC

CACACGTCTGAACTCTTTCCCTACACGACGCTCTTCCGATCTNNNTGTGTCGGAGTTCTAACGATGTTC

GAGCACACGTCTGAACTCTTTCCCTACACGACGCTCTTCCGATCTNNNAAAACGTTGATGAAGGTCCAG

GAGCACACGTCTGAACTCTTTCCCTACACGACGCTCTTCCGATCTNNNAAAACGTTGATGAAGGTCCAG

GAGCACACGTCTGAACTCTTTCCCTACACGACGCTCTTCCGATCTNNNCTGTAGTCCCAGCTACTTGGA

GAGCACACGTCTGAACTCTTTCCCTACACGACGCTCTTCCGATCTNNNCTGTAGTCCCAGCTACTTGGA

GAGCACACGTCTGAACTCTTTCCCTACACGACGCTCTTCCGATCTNNNCTCGGTGAATCATAAGAGAG

GAGCACACGTCTGAACTCTTTCCCTACACGACGCTCTTCCGATCTNNNCTCGGTGAATCATAAGAGAG

GAGCACACGTCTGAACTCTTTCCCTACACGACGCTCTTCCGATCTNNNATTCAGGGACCTTCTACCTCAG

GAGCACACGTCTGAACTCTTTCCCTACACGACGCTCTTCCGATCTNNNATTCAGGGACCTTCTACCTCAG

GAGCACACGTCTGAACTCTTTCCCTACACGACGCTCTTCCGATCTNNNGGGCTGTAAGGTTTCTAGGCT

GAGCACACGTCTGAACTCTTTCCCTACACGACGCTCTTCCGATCTNNNGGGCTGTAAGGTTTCTAGGCT

AGCACACGTCTGAACTCTTTCCCTACACGACGCTCTTCCGATCTNNNATGGTTTTCTCCTTTTGCTGGG

AGCACACGTCTGAACTCTTTCCCTACACGACGCTCTTCCGATCTNNNATGGTTTTCTCCTTTTGCTGGG

AGCACACGTCTGAACTCTTTCCCTACACGACGCTCTTCCGATCTNNNCTGAGACGTGAGAATCCCTTGA

AGCACACGTCTGAACTCTTTCCCTACACGACGCTCTTCCGATCTNNNCTGAGACGTGAGAATCCCTTGA

GAGCACACGTCTGAACTCTTTCCCTACACGACGCTCTTCCGATCTNNNCTGATTGTCAGCTATGGTCTC

GAGCACACGTCTGAACTCTTTCCCTACACGACGCTCTTCCGATCTNNNCTGATTGTCAGCTATGGTCTC

GAGCACACGTCTGAACTCTTTCCCTACACGACGCTCTTCCGATCTNNNCAAGATTGCTCCATTGCACTTC

GAGCACACGTCTGAACTCTTTCCCTACACGACGCTCTTCCGATCTNNNCAAGATTGCTCCATTGCACTTC

GAGCACACGTCTGAACTCTTTCCCTACACGACGCTCTTCCGATCTNNNTGAAAAGCAGTAGCAGTCAGC

GAGCACACGTCTGAACTCTTTCCCTACACGACGCTCTTCCGATCTNNNTGAAAAGCAGTAGCAGTCAGC

CACACGTCTGAACTCTTTCCCTACACGACGCTCTTCCGATCTNNNTACATCAAGAGGTCTCCTTGCATCG

CACACGTCTGAACTCTTTCCCTACACGACGCTCTTCCGATCTNNNTACATCAAGAGGTCTCCTTGCATCG

AGAGCACACGTCTGAACTCTTTCCCTACACGACGCTCTTCCGATCTNNNTGGTGCTAAGCAGTTGAATT

AGAGCACACGTCTGAACTCTTTCCCTACACGACGCTCTTCCGATCTNNNTGGTGCTAAGCAGTTGAATT

AGCACACGTCTGAACTCTTTCCCTACACGACGCTCTTCCGATCTNNNTCTATGAACTAGTGGGCCTGAG

AGCACACGTCTGAACTCTTTCCCTACACGACGCTCTTCCGATCTNNNTCTATGAACTAGTGGGCCTGAG

GAGCACACGTCTGAACTCTTTCCCTACACGACGCTCTTCCGATCTNNNAATTGGCACCAAGACAGTCAC

GAGCACACGTCTGAACTCTTTCCCTACACGACGCTCTTCCGATCTNNNAATTGGCACCAAGACAGTCAC

GAGCACACGTCTGAACTCTTTCCCTACACGACGCTCTTCCGATCTNNNTCTGTAATCCCAGCTACTCAGG

GAGCACACGTCTGAACTCTTTCCCTACACGACGCTCTTCCGATCTNNNTCTGTAATCCCAGCTACTCAGG

AGAGCACACGTCTGAACTCTTTCCCTACACGACGCTCTTCCGATCTNNNAGCTATTATAGGGGTGTGTC

AGAGCACACGTCTGAACTCTTTCCCTACACGACGCTCTTCCGATCTNNNAGCTATTATAGGGGTGTGTC

GAGCACACGTCTGAACTCTTTCCCTACACGACGCTCTTCCGATCTNNNCCCAAACTCTCAACTTTCCCCTA

GAGCACACGTCTGAACTCTTTCCCTACACGACGCTCTTCCGATCTNNNCCCAAACTCTCAACTTTCCCCTA

AGCACACGTCTGAACTCTTTCCCTACACGACGCTCTTCCGATCTNNNATTTCACACCCACTAGAATGGCT

AGCACACGTCTGAACTCTTTCCCTACACGACGCTCTTCCGATCTNNNATTTCACACCCACTAGAATGGCT

AGCACACGTCTGAACTCTTTCCCTACACGACGCTCTTCCGATCTNNNGCATGGCTTCAGAGCTTGAGTA

AGCACACGTCTGAACTCTTTCCCTACACGACGCTCTTCCGATCTNNNGCATGGCTTCAGAGCTTGAGTA

AGCACACGTCTGAACTCTTTCCCTACACGACGCTCTTCCGATCTNNNGATCAAATTCACAAAGCCTGCCT

AGCACACGTCTGAACTCTTTCCCTACACGACGCTCTTCCGATCTNNNGATCAAATTCACAAAGCCTGCCT

AGAGCACACGTCTGAACTCTTTCCCTACACGACGCTCTTCCGATCTNNNAAAGTGGTTCACCACCTCTCT

AGAGCACACGTCTGAACTCTTTCCCTACACGACGCTCTTCCGATCTNNNAAAGTGGTTCACCACCTCTCT

GAGCACACGTCTGAACTCTTTCCCTACACGACGCTCTTCCGATCTNNNTTCTTCAGGTTCTGCAGCAAGA

GAGCACACGTCTGAACTCTTTCCCTACACGACGCTCTTCCGATCTNNNTTCTTCAGGTTCTGCAGCAAGA

GAGCACACGTCTGAACTCTTTCCCTACACGACGCTCTTCCGATCTNNNAAGAAGTTGAGAGACTGCTT

GAGCACACGTCTGAACTCTTTCCCTACACGACGCTCTTCCGATCTNNNAAGAAGTTGAGAGACTGCTT

GCACACGTCTGAACTCTTTCCCTACACGACGCTCTTCCGATCTNNNGCACATTCAGGAATCCCTAAGCAG

GCACACGTCTGAACTCTTTCCCTACACGACGCTCTTCCGATCTNNNGCACATTCAGGAATCCCTAAGCAG

AGCACACGTCTGAACTCTTTCCCTACACGACGCTCTTCCGATCTNNNTGTAATCCCAGCTACTCAGGAG

AGCACACGTCTGAACTCTTTCCCTACACGACGCTCTTCCGATCTNNNTGTAATCCCAGCTACTCAGGAG

GCACACGTCTGAACTCTTTCCCTACACGACGCTCTTCCGATCTNNNTGAGGCAGGAGAATAGTGTGAA

GCACACGTCTGAACTCTTTCCCTACACGACGCTCTTCCGATCTNNNTGAGGCAGGAGAATAGTGTGAA

AGCACACGTCTGAACTCTTTCCCTACACGACGCTCTTCCGATCTNNNACCTTTGTGAAAAGCTTGGAAC

AGCACACGTCTGAACTCTTTCCCTACACGACGCTCTTCCGATCTNNNACCTTTGTGAAAAGCTTGGAAC

AGCACACGTCTGAACTCTTTCCCTACACGACGCTCTTCCGATCTNNNGAGGTTGCAGTGAGCTGAGATC

AGCACACGTCTGAACTCTTTCCCTACACGACGCTCTTCCGATCTNNNGAGGTTGCAGTGAGCTGAGATC

GAGCACACGTCTGAACTCTTTCCCTACACGACGCTCTTCCGATCTNNNTTGTAGTGAGCTCAGATCGTC

GAGCACACGTCTGAACTCTTTCCCTACACGACGCTCTTCCGATCTNNNTTGTAGTGAGCTCAGATCGTC

GAGCACACGTCTGAACTCTTTCCCTACACGACGCTCTTCCGATCTNNNGTGTGTGTGTGTGTGTGTGTA

GAGCACACGTCTGAACTCTTTCCCTACACGACGCTCTTCCGATCTNNNGTGTGTGTGTGTGTGTGTGTA

GAGCACACGTCTGAACTCTTTCCCTACACGACGCTCTTCCGATCTNNNGACCATGCCTGTTCCCTACATA

GAGCACACGTCTGAACTCTTTCCCTACACGACGCTCTTCCGATCTNNNGACCATGCCTGTTCCCTACATA

GCACACGTCTGAACTCTTTCCCTACACGACGCTCTTCCGATCTNNNTGGGTCTAGGTACTCTCTTCTGGC

GCACACGTCTGAACTCTTTCCCTACACGACGCTCTTCCGATCTNNNTGGGTCTAGGTACTCTCTTCTGGC

GAGCACACGTCTGAACTCTTTCCCTACACGACGCTCTTCCGATCTNNNGGTGGGGAGCAAAATCTTTC

GAGCACACGTCTGAACTCTTTCCCTACACGACGCTCTTCCGATCTNNNGGTGGGGAGCAAAATCTTTC

AGCACACGTCTGAACTCTTTCCCTACACGACGCTCTTCCGATCTNNNGGTGATGGTTGCACAACAATGT

AGCACACGTCTGAACTCTTTCCCTACACGACGCTCTTCCGATCTNNNGGTGATGGTTGCACAACAATGT

GAGCACACGTCTGAACTCTTTCCCTACACGACGCTCTTCCGATCTNNNGAGGTACTGCCTTGGTGATAA

GAGCACACGTCTGAACTCTTTCCCTACACGACGCTCTTCCGATCTNNNGAGGTACTGCCTTGGTGATAA

GAGCACACGTCTGAACTCTTTCCCTACACGACGCTCTTCCGATCTNNNGGGAGAAGTCAGGGAAGCAT

GAGCACACGTCTGAACTCTTTCCCTACACGACGCTCTTCCGATCTNNNGGGAGAAGTCAGGGAAGCAT

GAGCACACGTCTGAACTCTTTCCCTACACGACGCTCTTCCGATCTNNNACCCATAATCCTAGCACTTTGC

GAGCACACGTCTGAACTCTTTCCCTACACGACGCTCTTCCGATCTNNNACCCATAATCCTAGCACTTTGC

AGCACACGTCTGAACTCTTTCCCTACACGACGCTCTTCCGATCTNNNGTGGCTCACTCCTGTAATCCTAC

AGCACACGTCTGAACTCTTTCCCTACACGACGCTCTTCCGATCTNNNGTGGCTCACTCCTGTAATCCTAC

GAGCACACGTCTGAACTCTTTCCCTACACGACGCTCTTCCGATCTNNNAGTATAAGACTGTGTGTATGT

GAGCACACGTCTGAACTCTTTCCCTACACGACGCTCTTCCGATCTNNNAGTATAAGACTGTGTGTATGT

GCACACGTCTGAACTCTTTCCCTACACGACGCTCTTCCGATCTNNNTTTCCAGGGAAGCAGTTTAACATC

GCACACGTCTGAACTCTTTCCCTACACGACGCTCTTCCGATCTNNNTTTCCAGGGAAGCAGTTTAACATC

GAGCACACGTCTGAACTCTTTCCCTACACGACGCTCTTCCGATCTNNNTCGAGACGTTGGACACATTCT

GAGCACACGTCTGAACTCTTTCCCTACACGACGCTCTTCCGATCTNNNTCGAGACGTTGGACACATTCT

GAGCACACGTCTGAACTCTTTCCCTACACGACGCTCTTCCGATCTNNNCTGATGACCCTTGACTGTCTGA

GAGCACACGTCTGAACTCTTTCCCTACACGACGCTCTTCCGATCTNNNCTGATGACCCTTGACTGTCTGA

GAGCACACGTCTGAACTCTTTCCCTACACGACGCTCTTCCGATCTNNNTGCCTGTAATCCCAGCTATTCA

GAGCACACGTCTGAACTCTTTCCCTACACGACGCTCTTCCGATCTNNNTGCCTGTAATCCCAGCTATTCA

AGCACACGTCTGAACTCTTTCCCTACACGACGCTCTTCCGATCTNNNACCCACATTTATAACCCCAGCAT

AGCACACGTCTGAACTCTTTCCCTACACGACGCTCTTCCGATCTNNNACCCACATTTATAACCCCAGCAT

AGAGCACACGTCTGAACTCTTTCCCTACACGACGCTCTTCCGATCTNNNCTACAGTGAGGAAGGCCAGA

AGAGCACACGTCTGAACTCTTTCCCTACACGACGCTCTTCCGATCTNNNCTACAGTGAGGAAGGCCAGA

GCACACGTCTGAACTCTTTCCCTACACGACGCTCTTCCGATCTNNNCTGTAGTACCAGCTACTTGGGAC

GCACACGTCTGAACTCTTTCCCTACACGACGCTCTTCCGATCTNNNCTGTAGTACCAGCTACTTGGGAC

GAGCACACGTCTGAACTCTTTCCCTACACGACGCTCTTCCGATCTNNNAAAATATTAGCGTGACGTGGT

GAGCACACGTCTGAACTCTTTCCCTACACGACGCTCTTCCGATCTNNNAAAATATTAGCGTGACGTGG
AGCACACGTCTGAACTCTTTCCCTACACGACGCTCTTCCGATCTNNNTAGTCACAGCTACTCTAGAGG
AGCACACGTCTGAACTCTTTCCCTACACGACGCTCTTCCGATCTNNNTAGTCACAGCTACTCTAGAGG
AGCACACGTCTGAACTCTTTCCCTACACGACGCTCTTCCGATCTNNNATGTGCTTGTAGTCACAGCTAC
AGCACACGTCTGAACTCTTTCCCTACACGACGCTCTTCCGATCTNNNATGTGCTTGTAGTCACAGCTAC
AGCACACGTCTGAACTCTTTCCCTACACGACGCTCTTCCGATCTNNNTTTGAGAACAGCCTGGGTAACA
AGCACACGTCTGAACTCTTTCCCTACACGACGCTCTTCCGATCTNNNTTTGAGAACAGCCTGGGTAACA
AGCACACGTCTGAACTCTTTCCCTACACGACGCTCTTCCGATCTNNNTGGCATGGCCATTATTTCATCCA
AGCACACGTCTGAACTCTTTCCCTACACGACGCTCTTCCGATCTNNNTGGCATGGCCATTATTTCATCCA
AGCACACGTCTGAACTCTTTCCCTACACGACGCTCTTCCGATCTNNNCCATCTCCCCGTAAGATCTGATG
AGCACACGTCTGAACTCTTTCCCTACACGACGCTCTTCCGATCTNNNCCATCTCCCCGTAAGATCTGATG
AGAGCACACGTCTGAACTCTTTCCCTACACGACGCTCTTCCGATCTNNNATATGTGCCATCAGAACCCTA
AGAGCACACGTCTGAACTCTTTCCCTACACGACGCTCTTCCGATCTNNNATATGTGCCATCAGAACCCTA
AGAGCACACGTCTGAACTCTTTCCCTACACGACGCTCTTCCGATCTNNNGGAGCAGGTAACATTTCTGA
AGAGCACACGTCTGAACTCTTTCCCTACACGACGCTCTTCCGATCTNNNGGAGCAGGTAACATTTCTGA
GAGCACACGTCTGAACTCTTTCCCTACACGACGCTCTTCCGATCTNNNTCACATCCCTCCCTTAGGTAGA
GAGCACACGTCTGAACTCTTTCCCTACACGACGCTCTTCCGATCTNNNTCACATCCCTCCCTTAGGTAGA
AGCACACGTCTGAACTCTTTCCCTACACGACGCTCTTCCGATCTNNNTAGTCACCAGTGGCCAGCTCCA
AGCACACGTCTGAACTCTTTCCCTACACGACGCTCTTCCGATCTNNNTAGTCACCAGTGGCCAGCTCCA
GCACACGTCTGAACTCTTTCCCTACACGACGCTCTTCCGATCTNNNCTCACCCACTGTCCTCTGACTCCAT
GCACACGTCTGAACTCTTTCCCTACACGACGCTCTTCCGATCTNNNCTCACCCACTGTCCTCTGACTCCAT
AGAGCACACGTCTGAACTCTTTCCCTACACGACGCTCTTCCGATCTNNNGGCACCTGTAATCCCAGTTAC
AGAGCACACGTCTGAACTCTTTCCCTACACGACGCTCTTCCGATCTNNNGGCACCTGTAATCCCAGTTAC
GAGCACACGTCTGAACTCTTTCCCTACACGACGCTCTTCCGATCTNNNCAAGAAGGAAAATGTGTGTG
GAGCACACGTCTGAACTCTTTCCCTACACGACGCTCTTCCGATCTNNNCAAGAAGGAAAATGTGTGTG
AGAGCACACGTCTGAACTCTTTCCCTACACGACGCTCTTCCGATCTNNNTTCCAAGTTTCCTCAGATCCC
AGAGCACACGTCTGAACTCTTTCCCTACACGACGCTCTTCCGATCTNNNTTCCAAGTTTCCTCAGATCCC
GAGCACACGTCTGAACTCTTTCCCTACACGACGCTCTTCCGATCTNNNGTCAGGGTTCTCTATGTGTGT
GAGCACACGTCTGAACTCTTTCCCTACACGACGCTCTTCCGATCTNNNGTCAGGGTTCTCTATGTGTGT
GAGCACACGTCTGAACTCTTTCCCTACACGACGCTCTTCCGATCTNNNATATGTGTGTGTGTGTGTGTG
GAGCACACGTCTGAACTCTTTCCCTACACGACGCTCTTCCGATCTNNNATATGTGTGTGTGTGTGTGTG
AGCACACGTCTGAACTCTTTCCCTACACGACGCTCTTCCGATCTNNNGCACTTTTCTAGGTCCTTTTCGT
AGCACACGTCTGAACTCTTTCCCTACACGACGCTCTTCCGATCTNNNGCACTTTTCTAGGTCCTTTTCGT
AGCACACGTCTGAACTCTTTCCCTACACGACGCTCTTCCGATCTNNNCTCAATTGTGGCCAGGATCTGA
AGCACACGTCTGAACTCTTTCCCTACACGACGCTCTTCCGATCTNNNCTCAATTGTGGCCAGGATCTGA
AGCACACGTCTGAACTCTTTCCCTACACGACGCTCTTCCGATCTNNNGAGGATTTCCCCTATGCTGTTCT
AGCACACGTCTGAACTCTTTCCCTACACGACGCTCTTCCGATCTNNNGAGGATTTCCCCTATGCTGTTCT
GAGCACACGTCTGAACTCTTTCCCTACACGACGCTCTTCCGATCTNNNCAGCTACCCTGACCGAAGTGC
GAGCACACGTCTGAACTCTTTCCCTACACGACGCTCTTCCGATCTNNNCAGCTACCCTGACCGAAGTGC
GAGCACACGTCTGAACTCTTTCCCTACACGACGCTCTTCCGATCTNNNCTGTAGTCCCAGCTTCTCAGG
GAGCACACGTCTGAACTCTTTCCCTACACGACGCTCTTCCGATCTNNNCTGTAGTCCCAGCTTCTCAGG
AGCACACGTCTGAACTCTTTCCCTACACGACGCTCTTCCGATCTNNNCGTGGAATCTCGGGGTCTATTA
AGCACACGTCTGAACTCTTTCCCTACACGACGCTCTTCCGATCTNNNCGTGGAATCTCGGGGTCTATTA

GAGCACACGTCTGAACTCTTTCCCTACACGACGCTCTTCCGATCTNNNAGGCAGGAGAATTTCTTGAA
GAGCACACGTCTGAACTCTTTCCCTACACGACGCTCTTCCGATCTNNNAGGCAGGAGAATTTCTTGAA
AGAGCACACGTCTGAACTCTTTCCCTACACGACGCTCTTCCGATCTNNNTTTTGTTGTGGCCATAGTTG
AGAGCACACGTCTGAACTCTTTCCCTACACGACGCTCTTCCGATCTNNNTTTTGTTGTGGCCATAGTTG
CACACGTCTGAACTCTTTCCCTACACGACGCTCTTCCGATCTNNNGAGTACACGTTGAGGGGTTGGCA
CACACGTCTGAACTCTTTCCCTACACGACGCTCTTCCGATCTNNNGAGTACACGTTGAGGGGTTGGCA
GAGCACACGTCTGAACTCTTTCCCTACACGACGCTCTTCCGATCTNNNCAACAAACACTGCCAAAATGCT
GAGCACACGTCTGAACTCTTTCCCTACACGACGCTCTTCCGATCTNNNCAACAAACACTGCCAAAATGCT
AGCACACGTCTGAACTCTTTCCCTACACGACGCTCTTCCGATCTNNNTTTCCTGCTTTGTGTGTGCTTAG
AGCACACGTCTGAACTCTTTCCCTACACGACGCTCTTCCGATCTNNNTTTCCTGCTTTGTGTGTGCTTAG
GAGCACACGTCTGAACTCTTTCCCTACACGACGCTCTTCCGATCTNNNTTTGGAGAGTCTGAACTGAC
GAGCACACGTCTGAACTCTTTCCCTACACGACGCTCTTCCGATCTNNNTTTGGAGAGTCTGAACTGAC
AGCACACGTCTGAACTCTTTCCCTACACGACGCTCTTCCGATCTNNNCAACGGCCTTACATACCAGTGAA
AGCACACGTCTGAACTCTTTCCCTACACGACGCTCTTCCGATCTNNNCAACGGCCTTACATACCAGTGAA
GAGCACACGTCTGAACTCTTTCCCTACACGACGCTCTTCCGATCTNNNATCCAGGAAAGTGTGAAGGA
GAGCACACGTCTGAACTCTTTCCCTACACGACGCTCTTCCGATCTNNNATCCAGGAAAGTGTGAAGGA
AGCACACGTCTGAACTCTTTCCCTACACGACGCTCTTCCGATCTNNNTACGTCGACTTTCCTGTTCTGTT
AGCACACGTCTGAACTCTTTCCCTACACGACGCTCTTCCGATCTNNNTACGTCGACTTTCCTGTTCTGTT
GCACACGTCTGAACTCTTTCCCTACACGACGCTCTTCCGATCTNNNTCCCCAACTCCAATCTCTTTTCCA
GCACACGTCTGAACTCTTTCCCTACACGACGCTCTTCCGATCTNNNTCCCCAACTCCAATCTCTTTTCCA
AGCACACGTCTGAACTCTTTCCCTACACGACGCTCTTCCGATCTNNNAATTTGGGAAATTCACAGCCTG
AGCACACGTCTGAACTCTTTCCCTACACGACGCTCTTCCGATCTNNNAATTTGGGAAATTCACAGCCTG
AGCACACGTCTGAACTCTTTCCCTACACGACGCTCTTCCGATCTNNNGACTTTTTCACAGAAAACAAAAA
AGCACACGTCTGAACTCTTTCCCTACACGACGCTCTTCCGATCTNNNGACTTTTTCACAGAAAACAAAAA
AGAGCACACGTCTGAACTCTTTCCCTACACGACGCTCTTCCGATCTNNNGCCCATTGCTTGAGACATTAG
AGAGCACACGTCTGAACTCTTTCCCTACACGACGCTCTTCCGATCTNNNGCCCATTGCTTGAGACATTAG
GAGCACACGTCTGAACTCTTTCCCTACACGACGCTCTTCCGATCTNNNCAGGTGCCTATAATCCCAGCT
GAGCACACGTCTGAACTCTTTCCCTACACGACGCTCTTCCGATCTNNNCAGGTGCCTATAATCCCAGCT
GAGCACACGTCTGAACTCTTTCCCTACACGACGCTCTTCCGATCTNNNAAAGATCCTCAAGTGGTAAG
GAGCACACGTCTGAACTCTTTCCCTACACGACGCTCTTCCGATCTNNNAAAGATCCTCAAGTGGTAAG
AGCACACGTCTGAACTCTTTCCCTACACGACGCTCTTCCGATCTNNNGAGGCTAAAGCAGGAGAATCA
AGCACACGTCTGAACTCTTTCCCTACACGACGCTCTTCCGATCTNNNGAGGCTAAAGCAGGAGAATCA
AGCACACGTCTGAACTCTTTCCCTACACGACGCTCTTCCGATCTNNNCAATATAGCGCCATTGCACTCAA
AGCACACGTCTGAACTCTTTCCCTACACGACGCTCTTCCGATCTNNNCAATATAGCGCCATTGCACTCAA
GAGCACACGTCTGAACTCTTTCCCTACACGACGCTCTTCCGATCTNNNCCAAAGTCAGCCCAGTGAGA
GAGCACACGTCTGAACTCTTTCCCTACACGACGCTCTTCCGATCTNNNCCAAAGTCAGCCCAGTGAGA
AGAGCACACGTCTGAACTCTTTCCCTACACGACGCTCTTCCGATCTNNNCTGTAATCCCAACTACCCAGG
AGAGCACACGTCTGAACTCTTTCCCTACACGACGCTCTTCCGATCTNNNCTGTAATCCCAACTACCCAGG
GAGCACACGTCTGAACTCTTTCCCTACACGACGCTCTTCCGATCTNNNTTGGTGCTATGGTCTTTCAGA
GAGCACACGTCTGAACTCTTTCCCTACACGACGCTCTTCCGATCTNNNTTGGTGCTATGGTCTTTCAGA
AGCACACGTCTGAACTCTTTCCCTACACGACGCTCTTCCGATCTNNNCCCGGCCAAGACTTTTTCTGCA
AGCACACGTCTGAACTCTTTCCCTACACGACGCTCTTCCGATCTNNNCCCGGCCAAGACTTTTTCTGCA
GAGCACACGTCTGAACTCTTTCCCTACACGACGCTCTTCCGATCTNNNATCAACAGGCTAAGTATGCT

GAGCACACGTCTGAACTCTTTCCCTACACGACGCTCTTCCGATCTNNNATCAACAGGCTAAGTATGCT
AGCACACGTCTGAACTCTTTCCCTACACGACGCTCTTCCGATCTNNNAGTTTTAGAGCTTGTGGGGTC
AGCACACGTCTGAACTCTTTCCCTACACGACGCTCTTCCGATCTNNNAGTTTTAGAGCTTGTGGGGTC
AGCACACGTCTGAACTCTTTCCCTACACGACGCTCTTCCGATCTNNNGATCACGCCATTGCATTCCAG
AGCACACGTCTGAACTCTTTCCCTACACGACGCTCTTCCGATCTNNNGATCACGCCATTGCATTCCAG
AGCACACGTCTGAACTCTTTCCCTACACGACGCTCTTCCGATCTNNNGTGACAGAGTGAGACTCCCT
AGCACACGTCTGAACTCTTTCCCTACACGACGCTCTTCCGATCTNNNGTGACAGAGTGAGACTCCCT
CACGTCTGAACTCTTTCCCTACACGACGCTCTTCCGATCTNNNGAGGAGGGAAGGCAAGAGATGCA
CACGTCTGAACTCTTTCCCTACACGACGCTCTTCCGATCTNNNGAGGAGGGAAGGCAAGAGATGCA
AGCACACGTCTGAACTCTTTCCCTACACGACGCTCTTCCGATCTNNNATATAGCACCTGTGTGTGTGT
AGCACACGTCTGAACTCTTTCCCTACACGACGCTCTTCCGATCTNNNATATAGCACCTGTGTGTGTGT
GAGCACACGTCTGAACTCTTTCCCTACACGACGCTCTTCCGATCTNNNGAGAATTGCTTGAACCCAGG
GAGCACACGTCTGAACTCTTTCCCTACACGACGCTCTTCCGATCTNNNGAGAATTGCTTGAACCCAGG
GCACACGTCTGAACTCTTTCCCTACACGACGCTCTTCCGATCTNNNTAGCCTGGGTGACAAGAGTGA
GCACACGTCTGAACTCTTTCCCTACACGACGCTCTTCCGATCTNNNTAGCCTGGGTGACAAGAGTGA
GCACACGTCTGAACTCTTTCCCTACACGACGCTCTTCCGATCTNNNTAGTGAAATCTCTCAGATTGA
GCACACGTCTGAACTCTTTCCCTACACGACGCTCTTCCGATCTNNNTAGTGAAATCTCTCAGATTGA
GAGCACACGTCTGAACTCTTTCCCTACACGACGCTCTTCCGATCTNNNTGTAATGTTAGCACTTCGGG
GAGCACACGTCTGAACTCTTTCCCTACACGACGCTCTTCCGATCTNNNTGTAATGTTAGCACTTCGGG
AGCACACGTCTGAACTCTTTCCCTACACGACGCTCTTCCGATCTNNNTCTGGGTCTGTTGCATTTCCAT
AGCACACGTCTGAACTCTTTCCCTACACGACGCTCTTCCGATCTNNNTCTGGGTCTGTTGCATTTCCAT
GCACACGTCTGAACTCTTTCCCTACACGACGCTCTTCCGATCTNNNATGCTTCAGAGATGTTCCCTGAG
GCACACGTCTGAACTCTTTCCCTACACGACGCTCTTCCGATCTNNNATGCTTCAGAGATGTTCCCTGAG
GCACACGTCTGAACTCTTTCCCTACACGACGCTCTTCCGATCTNNNAGTGGTTGCTCTAAGGTATCCA
GCACACGTCTGAACTCTTTCCCTACACGACGCTCTTCCGATCTNNNAGTGGTTGCTCTAAGGTATCCA
GCACACGTCTGAACTCTTTCCCTACACGACGCTCTTCCGATCTNNNTATACTCACACACACTTCCAGGCC
GCACACGTCTGAACTCTTTCCCTACACGACGCTCTTCCGATCTNNNTATACTCACACACACTTCCAGGCC
GCACACGTCTGAACTCTTTCCCTACACGACGCTCTTCCGATCTNNNCACCTCCAACATTGTGTGTCAAA
GCACACGTCTGAACTCTTTCCCTACACGACGCTCTTCCGATCTNNNCACCTCCAACATTGTGTGTCAAA
CACGTCTGAACTCTTTCCCTACACGACGCTCTTCCGATCTNNNTTCAAGTGATTCTCCTGCCTCAGCAT
CACGTCTGAACTCTTTCCCTACACGACGCTCTTCCGATCTNNNTTCAAGTGATTCTCCTGCCTCAGCAT
AGAGCACACGTCTGAACTCTTTCCCTACACGACGCTCTTCCGATCTNNNGAGCCCGTATTCTACAAGA
AGAGCACACGTCTGAACTCTTTCCCTACACGACGCTCTTCCGATCTNNNGAGCCCGTATTCTACAAGA
AGCACACGTCTGAACTCTTTCCCTACACGACGCTCTTCCGATCTNNNTCAGGTTTGTGGAAGTGTAA
AGCACACGTCTGAACTCTTTCCCTACACGACGCTCTTCCGATCTNNNTCAGGTTTGTGGAAGTGTAA
AGCACACGTCTGAACTCTTTCCCTACACGACGCTCTTCCGATCTNNNATTCACAAGCCTCATGGTAAC
AGCACACGTCTGAACTCTTTCCCTACACGACGCTCTTCCGATCTNNNATTCACAAGCCTCATGGTAAC
AGCACACGTCTGAACTCTTTCCCTACACGACGCTCTTCCGATCTNNNAGCCAGTGCCATATTGACTTA
AGCACACGTCTGAACTCTTTCCCTACACGACGCTCTTCCGATCTNNNAGCCAGTGCCATATTGACTTA
AGCACACGTCTGAACTCTTTCCCTACACGACGCTCTTCCGATCTNNNCGCTTATAGTCCCAGCTACTCG
AGCACACGTCTGAACTCTTTCCCTACACGACGCTCTTCCGATCTNNNCGCTTATAGTCCCAGCTACTCG
GCACACGTCTGAACTCTTTCCCTACACGACGCTCTTCCGATCTNNNCAGGAAGGAAGGAAGGAAGG
GCACACGTCTGAACTCTTTCCCTACACGACGCTCTTCCGATCTNNNCAGGAAGGAAGGAAGGAAGG

GCACACGTCTGAACTCTTTCCCTACACGACGCTCTTCCGATCTNNNGACCCCAACAATTGTATAAAATA
GCACACGTCTGAACTCTTTCCCTACACGACGCTCTTCCGATCTNNNGACCCCAACAATTGTATAAAATA
GAGCACACGTCTGAACTCTTTCCCTACACGACGCTCTTCCGATCTNNNCGAAGGTTGATCTTGAGAAT
GAGCACACGTCTGAACTCTTTCCCTACACGACGCTCTTCCGATCTNNNCGAAGGTTGATCTTGAGAAT
AGCACACGTCTGAACTCTTTCCCTACACGACGCTCTTCCGATCTNNNGACAGAGCGAGACTGTAGGG
AGCACACGTCTGAACTCTTTCCCTACACGACGCTCTTCCGATCTNNNGACAGAGCGAGACTGTAGGG
GAGCACACGTCTGAACTCTTTCCCTACACGACGCTCTTCCGATCTNNNATGCTCTGGGTTCTGTTTTTT
GAGCACACGTCTGAACTCTTTCCCTACACGACGCTCTTCCGATCTNNNATGCTCTGGGTTCTGTTTTTT
CACACGTCTGAACTCTTTCCCTACACGACGCTCTTCCGATCTNNNAGACATGCAGTTCCAGCTACCCAT
CACACGTCTGAACTCTTTCCCTACACGACGCTCTTCCGATCTNNNAGACATGCAGTTCCAGCTACCCAT
GCACACGTCTGAACTCTTTCCCTACACGACGCTCTTCCGATCTNNNGGATGAATGGAGGACTCAAGAC
GCACACGTCTGAACTCTTTCCCTACACGACGCTCTTCCGATCTNNNGGATGAATGGAGGACTCAAGAC
ACACGTCTGAACTCTTTCCCTACACGACGCTCTTCCGATCTNNNCTCTCTGACGTTTCCCAAGTGAGCA
ACACGTCTGAACTCTTTCCCTACACGACGCTCTTCCGATCTNNNCTCTCTGACGTTTCCCAAGTGAGCA
AGCACACGTCTGAACTCTTTCCCTACACGACGCTCTTCCGATCTNNNCATTGCTGATGATCTTGAGGCT
AGCACACGTCTGAACTCTTTCCCTACACGACGCTCTTCCGATCTNNNCATTGCTGATGATCTTGAGGCT
GCACACGTCTGAACTCTTTCCCTACACGACGCTCTTCCGATCTNNNAGTGAGCTGAGATCATGCCATT
GCACACGTCTGAACTCTTTCCCTACACGACGCTCTTCCGATCTNNNAGTGAGCTGAGATCATGCCATT
GAGCACACGTCTGAACTCTTTCCCTACACGACGCTCTTCCGATCTNNNACATGACATAGCCTGTGTGTA
GAGCACACGTCTGAACTCTTTCCCTACACGACGCTCTTCCGATCTNNNACATGACATAGCCTGTGTGTA
GAGCACACGTCTGAACTCTTTCCCTACACGACGCTCTTCCGATCTNNNCCCAAATAACCGAATGACTCC
GAGCACACGTCTGAACTCTTTCCCTACACGACGCTCTTCCGATCTNNNCCCAAATAACCGAATGACTCC
GAGCACACGTCTGAACTCTTTCCCTACACGACGCTCTTCCGATCTNNNGAGCTGGGGAGATGGATGA
GAGCACACGTCTGAACTCTTTCCCTACACGACGCTCTTCCGATCTNNNGAGCTGGGGAGATGGATGA
GAGCACACGTCTGAACTCTTTCCCTACACGACGCTCTTCCGATCTNNNTGGATGTACTACGGATGTCTC
GAGCACACGTCTGAACTCTTTCCCTACACGACGCTCTTCCGATCTNNNTGGATGTACTACGGATGTCTC
AGCACACGTCTGAACTCTTTCCCTACACGACGCTCTTCCGATCTNNNTCAGCCTCCTGAGTAGTAGCTG
AGCACACGTCTGAACTCTTTCCCTACACGACGCTCTTCCGATCTNNNTCAGCCTCCTGAGTAGTAGCTG
AGCACACGTCTGAACTCTTTCCCTACACGACGCTCTTCCGATCTNNNTGGGCAATAGAACAAGACTCC
AGCACACGTCTGAACTCTTTCCCTACACGACGCTCTTCCGATCTNNNTGGGCAATAGAACAAGACTCC
GAGCACACGTCTGAACTCTTTCCCTACACGACGCTCTTCCGATCTNNNGAAACAGAGTGAGGCTCCGT
GAGCACACGTCTGAACTCTTTCCCTACACGACGCTCTTCCGATCTNNNGAAACAGAGTGAGGCTCCGT
GCACACGTCTGAACTCTTTCCCTACACGACGCTCTTCCGATCTNNNCATCAACAAAAAGGACGTCCAC
GCACACGTCTGAACTCTTTCCCTACACGACGCTCTTCCGATCTNNNCATCAACAAAAAGGACGTCCAC
AGAGCACACGTCTGAACTCTTTCCCTACACGACGCTCTTCCGATCTNNNTGTAGTACCAGCTACTCAG
AGAGCACACGTCTGAACTCTTTCCCTACACGACGCTCTTCCGATCTNNNTGTAGTACCAGCTACTCAG
GCACACGTCTGAACTCTTTCCCTACACGACGCTCTTCCGATCTNNNAGGTAATGCATATTAACAACTT
GCACACGTCTGAACTCTTTCCCTACACGACGCTCTTCCGATCTNNNAGGTAATGCATATTAACAACTT
AGCACACGTCTGAACTCTTTCCCTACACGACGCTCTTCCGATCTNNNAGTGGTCAGAAAGCACAAGTG
AGCACACGTCTGAACTCTTTCCCTACACGACGCTCTTCCGATCTNNNAGTGGTCAGAAAGCACAAGTG
AGCACACGTCTGAACTCTTTCCCTACACGACGCTCTTCCGATCTNNNGGTGCACAGAGCCTAAGTAAC
AGCACACGTCTGAACTCTTTCCCTACACGACGCTCTTCCGATCTNNNGGTGCACAGAGCCTAAGTAAC
GAGCACACGTCTGAACTCTTTCCCTACACGACGCTCTTCCGATCTNNNATAGCTACCGAGAACACACCA

GAGCACACGTCTGAACTCTTTCCCTACACGACGCTCTTCCGATCTNNNATAGCTACCGAGAACACACCA
GCACACGTCTGAACTCTTTCCCTACACGACGCTCTTCCGATCTNNNGGTTAGTTTTCGCAACATACGGT
GCACACGTCTGAACTCTTTCCCTACACGACGCTCTTCCGATCTNNNGGTTAGTTTTCGCAACATACGGT
CACACGTCTGAACTCTTTCCCTACACGACGCTCTTCCGATCTNNNAAAGAGACCCACATGATAGTGTAT
CACACGTCTGAACTCTTTCCCTACACGACGCTCTTCCGATCTNNNAAAGAGACCCACATGATAGTGTAT
GAGCACACGTCTGAACTCTTTCCCTACACGACGCTCTTCCGATCTNNNCGTGGGGATTACAATTCGAG
GAGCACACGTCTGAACTCTTTCCCTACACGACGCTCTTCCGATCTNNNCGTGGGGATTACAATTCGAG
AGCACACGTCTGAACTCTTTCCCTACACGACGCTCTTCCGATCTNNNGAACTTAGCCTTCTGACAATGC
AGCACACGTCTGAACTCTTTCCCTACACGACGCTCTTCCGATCTNNNGAACTTAGCCTTCTGACAATGC
GAGCACACGTCTGAACTCTTTCCCTACACGACGCTCTTCCGATCTNNNTGCCTGTAATTCCAGCTACTC
GAGCACACGTCTGAACTCTTTCCCTACACGACGCTCTTCCGATCTNNNTGCCTGTAATTCCAGCTACTC
GAGCACACGTCTGAACTCTTTCCCTACACGACGCTCTTCCGATCTNNNCACGAGCTTCCCATATGAGAT
GAGCACACGTCTGAACTCTTTCCCTACACGACGCTCTTCCGATCTNNNCACGAGCTTCCCATATGAGAT
AGCACACGTCTGAACTCTTTCCCTACACGACGCTCTTCCGATCTNNNGTTTTCCCCCTACCAGATACACC
AGCACACGTCTGAACTCTTTCCCTACACGACGCTCTTCCGATCTNNNGTTTTCCCCCTACCAGATACACC
GAGCACACGTCTGAACTCTTTCCCTACACGACGCTCTTCCGATCTNNNCTTGAGCAATACAGACAGACC
GAGCACACGTCTGAACTCTTTCCCTACACGACGCTCTTCCGATCTNNNCTTGAGCAATACAGACAGACC
AGCACACGTCTGAACTCTTTCCCTACACGACGCTCTTCCGATCTNNNAAAAAAAAAAATTAGTCGGGCG
AGCACACGTCTGAACTCTTTCCCTACACGACGCTCTTCCGATCTNNNAAAAAAAAAAATTAGTCGGGCG
GCACACGTCTGAACTCTTTCCCTACACGACGCTCTTCCGATCTNNNCCTATCCAGGAGGTGAGGGGC
GCACACGTCTGAACTCTTTCCCTACACGACGCTCTTCCGATCTNNNCCTATCCAGGAGGTGAGGGGC
AGCACACGTCTGAACTCTTTCCCTACACGACGCTCTTCCGATCTNNNAGTGCTGGGATTACAGGTGTA
AGCACACGTCTGAACTCTTTCCCTACACGACGCTCTTCCGATCTNNNAGTGCTGGGATTACAGGTGTA
GAGCACACGTCTGAACTCTTTCCCTACACGACGCTCTTCCGATCTNNNTCAAAGACATGCAAAACTGAG
GAGCACACGTCTGAACTCTTTCCCTACACGACGCTCTTCCGATCTNNNTCAAAGACATGCAAAACTGAG
GAGCACACGTCTGAACTCTTTCCCTACACGACGCTCTTCCGATCTNNNACTTGTTCATCCGTATTGTGG
GAGCACACGTCTGAACTCTTTCCCTACACGACGCTCTTCCGATCTNNNACTTGTTCATCCGTATTGTGG
GAGCACACGTCTGAACTCTTTCCCTACACGACGCTCTTCCGATCTNNNTAATCCCAGCTACTCAGGAGG
GAGCACACGTCTGAACTCTTTCCCTACACGACGCTCTTCCGATCTNNNTAATCCCAGCTACTCAGGAGG
AGCACACGTCTGAACTCTTTCCCTACACGACGCTCTTCCGATCTNNNGAGCCAGTAGGGAACAACTCA
AGCACACGTCTGAACTCTTTCCCTACACGACGCTCTTCCGATCTNNNGAGCCAGTAGGGAACAACTCA
GCACACGTCTGAACTCTTTCCCTACACGACGCTCTTCCGATCTNNNATGTTATGATGCAGCAAGAAGG
GCACACGTCTGAACTCTTTCCCTACACGACGCTCTTCCGATCTNNNATGTTATGATGCAGCAAGAAGG
GAGCACACGTCTGAACTCTTTCCCTACACGACGCTCTTCCGATCTNNNTTTCAGTGAGCCGAGATAGTG
GAGCACACGTCTGAACTCTTTCCCTACACGACGCTCTTCCGATCTNNNTTTCAGTGAGCCGAGATAGTG
GAGCACACGTCTGAACTCTTTCCCTACACGACGCTCTTCCGATCTNNNTCTTGTCTGTCTGTCTGTGTCTC
GAGCACACGTCTGAACTCTTTCCCTACACGACGCTCTTCCGATCTNNNTCTTGTCTGTCTGTCTGTGTCTC
CACGTCTGAACTCTTTCCCTACACGACGCTCTTCCGATCTNNNCTGTTGTTAAACTGTCTTCACCATCGG
CACGTCTGAACTCTTTCCCTACACGACGCTCTTCCGATCTNNNCTGTTGTTAAACTGTCTTCACCATCGG
AGAGCACACGTCTGAACTCTTTCCCTACACGACGCTCTTCCGATCTNNNATGCCTGTAATCCCAGCTACT
AGAGCACACGTCTGAACTCTTTCCCTACACGACGCTCTTCCGATCTNNNATGCCTGTAATCCCAGCTACT
GAGCACACGTCTGAACTCTTTCCCTACACGACGCTCTTCCGATCTNNNACTGTGTCCAAATAAGGCAAA
GAGCACACGTCTGAACTCTTTCCCTACACGACGCTCTTCCGATCTNNNACTGTGTCCAAATAAGGCAAA

GCACACGTCTGAACTCTTTCCCTACACGACGCTCTTCCGATCTNNNCACAACACCAGAGGCTGAAAAA
GCACACGTCTGAACTCTTTCCCTACACGACGCTCTTCCGATCTNNNCACAACACCAGAGGCTGAAAAA
AGCACACGTCTGAACTCTTTCCCTACACGACGCTCTTCCGATCTNNNCTGGCCAGGAGAGAAAAGAA
AGCACACGTCTGAACTCTTTCCCTACACGACGCTCTTCCGATCTNNNCTGGCCAGGAGAGAAAAGAA
AGCACACGTCTGAACTCTTTCCCTACACGACGCTCTTCCGATCTNNNGAGGCTGAGGCAGGAGAATG
AGCACACGTCTGAACTCTTTCCCTACACGACGCTCTTCCGATCTNNNGAGGCTGAGGCAGGAGAATG
GCACACGTCTGAACTCTTTCCCTACACGACGCTCTTCCGATCTNNNACACATCATTTTGTCTCAGCCCTC
GCACACGTCTGAACTCTTTCCCTACACGACGCTCTTCCGATCTNNNACACATCATTTTGTCTCAGCCCTC
AGCACACGTCTGAACTCTTTCCCTACACGACGCTCTTCCGATCTNNNCCCTAGCACAGTGCCTTGTATA
AGCACACGTCTGAACTCTTTCCCTACACGACGCTCTTCCGATCTNNNCCCTAGCACAGTGCCTTGTATA
CACACGTCTGAACTCTTTCCCTACACGACGCTCTTCCGATCTNNNGATTCCCCCGTACAGTTTCAGGCA
CACACGTCTGAACTCTTTCCCTACACGACGCTCTTCCGATCTNNNGATTCCCCCGTACAGTTTCAGGCA
AGCACACGTCTGAACTCTTTCCCTACACGACGCTCTTCCGATCTNNNCCACCTATGAAGTGAGGAGCA
AGCACACGTCTGAACTCTTTCCCTACACGACGCTCTTCCGATCTNNNCCACCTATGAAGTGAGGAGCA
AGCACACGTCTGAACTCTTTCCCTACACGACGCTCTTCCGATCTNNNGTTCTTGTTGGGGAACAATTG
AGCACACGTCTGAACTCTTTCCCTACACGACGCTCTTCCGATCTNNNGTTCTTGTTGGGGAACAATTG
GCACACGTCTGAACTCTTTCCCTACACGACGCTCTTCCGATCTNNNGAGAGAAGCATTGTAGGGGAG
GCACACGTCTGAACTCTTTCCCTACACGACGCTCTTCCGATCTNNNGAGAGAAGCATTGTAGGGGAG
CACACGTCTGAACTCTTTCCCTACACGACGCTCTTCCGATCTNNNATCGTTTTCCGATGGAAGAGGTC
CACACGTCTGAACTCTTTCCCTACACGACGCTCTTCCGATCTNNNATCGTTTTCCGATGGAAGAGGTC
GCACACGTCTGAACTCTTTCCCTACACGACGCTCTTCCGATCTNNNAACTCGCTTGATTTTGAATACCA
GCACACGTCTGAACTCTTTCCCTACACGACGCTCTTCCGATCTNNNAACTCGCTTGATTTTGAATACCA
AGCACACGTCTGAACTCTTTCCCTACACGACGCTCTTCCGATCTNNNAAATACTAGTGAGGGTGGG
AGCACACGTCTGAACTCTTTCCCTACACGACGCTCTTCCGATCTNNNAAATACTAGTGAGGGTGGG
AGAGCACACGTCTGAACTCTTTCCCTACACGACGCTCTTCCGATCTNNNAAAGGCTTTGATAGGTTCT
AGAGCACACGTCTGAACTCTTTCCCTACACGACGCTCTTCCGATCTNNNAAAGGCTTTGATAGGTTCT
AGCACACGTCTGAACTCTTTCCCTACACGACGCTCTTCCGATCTNNNTGTAATCACAGCTACTTGGGA
AGCACACGTCTGAACTCTTTCCCTACACGACGCTCTTCCGATCTNNNTGTAATCACAGCTACTTGGGA
GCACACGTCTGAACTCTTTCCCTACACGACGCTCTTCCGATCTNNNCTTCCTGGGTTCAAGCAATTCT
GCACACGTCTGAACTCTTTCCCTACACGACGCTCTTCCGATCTNNNCTTCCTGGGTTCAAGCAATTCT
AGAGCACACGTCTGAACTCTTTCCCTACACGACGCTCTTCCGATCTNNNGGCAAATAACACATGGTCC
AGAGCACACGTCTGAACTCTTTCCCTACACGACGCTCTTCCGATCTNNNGGCAAATAACACATGGTCC
GCACACGTCTGAACTCTTTCCCTACACGACGCTCTTCCGATCTNNNTCTGTGTGTGTCTGTGTCCTAA
GCACACGTCTGAACTCTTTCCCTACACGACGCTCTTCCGATCTNNNTCTGTGTGTGTCTGTGTCCTAA
AGAGCACACGTCTGAACTCTTTCCCTACACGACGCTCTTCCGATCTNNNCAGACAGGGTGGGTAAAG
AGAGCACACGTCTGAACTCTTTCCCTACACGACGCTCTTCCGATCTNNNCAGACAGGGTGGGTAAAG
GCACACGTCTGAACTCTTTCCCTACACGACGCTCTTCCGATCTNNNTGTTCCTCTCTCTGTCCTCTCATC
GCACACGTCTGAACTCTTTCCCTACACGACGCTCTTCCGATCTNNNTGTTCCTCTCTCTGTCCTCTCATC
GAGCACACGTCTGAACTCTTTCCCTACACGACGCTCTTCCGATCTNNNTAGCTTGAGACCAGACTGG
GAGCACACGTCTGAACTCTTTCCCTACACGACGCTCTTCCGATCTNNNTAGCTTGAGACCAGACTGG
GAGCACACGTCTGAACTCTTTCCCTACACGACGCTCTTCCGATCTNNNTGCTGAAATAAGACAGGTC
GAGCACACGTCTGAACTCTTTCCCTACACGACGCTCTTCCGATCTNNNTGCTGAAATAAGACAGGTC
GAGCACACGTCTGAACTCTTTCCCTACACGACGCTCTTCCGATCTNNNAAAAATAAGCTAGGGGTGG

AGAGCACACGTCTGAACTCTTTCCCTACACGACGCTCTTCCGATCTNNNAAAAATAAGCTAGGGGTGG
GCACACGTCTGAACTCTTTCCCTACACGACGCTCTTCCGATCTNNNTGTTTTGTGGCTGCAATCCTACTC
GCACACGTCTGAACTCTTTCCCTACACGACGCTCTTCCGATCTNNNTGTTTTGTGGCTGCAATCCTACTC
GAGCACACGTCTGAACTCTTTCCCTACACGACGCTCTTCCGATCTNNNACCCGAGGACGAAAACTTGTA
GAGCACACGTCTGAACTCTTTCCCTACACGACGCTCTTCCGATCTNNNACCCGAGGACGAAAACTTGTA
AGCACACGTCTGAACTCTTTCCCTACACGACGCTCTTCCGATCTNNNGGCCCGAGAGCTAAGAATAGTT
AGCACACGTCTGAACTCTTTCCCTACACGACGCTCTTCCGATCTNNNGGCCCGAGAGCTAAGAATAGTT
AGCACACGTCTGAACTCTTTCCCTACACGACGCTCTTCCGATCTNNNGATGTGATTGCTGCTGAAGAT
AGCACACGTCTGAACTCTTTCCCTACACGACGCTCTTCCGATCTNNNGATGTGATTGCTGCTGAAGAT
AGCACACGTCTGAACTCTTTCCCTACACGACGCTCTTCCGATCTNNNGCCTGTGGTGTGTTAGAGAGAA
AGCACACGTCTGAACTCTTTCCCTACACGACGCTCTTCCGATCTNNNGCCTGTGGTGTGTTAGAGAGAA
AGCACACGTCTGAACTCTTTCCCTACACGACGCTCTTCCGATCTNNNCTTCTGGGTTCAGGCGATTCTC
AGCACACGTCTGAACTCTTTCCCTACACGACGCTCTTCCGATCTNNNCTTCTGGGTTCAGGCGATTCTC
AGCACACGTCTGAACTCTTTCCCTACACGACGCTCTTCCGATCTNNNCTCTCTTGGTCCCAGAAATGGA
AGCACACGTCTGAACTCTTTCCCTACACGACGCTCTTCCGATCTNNNCTCTCTTGGTCCCAGAAATGGA
AGCACACGTCTGAACTCTTTCCCTACACGACGCTCTTCCGATCTNNNATATCTGGACAAGGGGAATGC
AGCACACGTCTGAACTCTTTCCCTACACGACGCTCTTCCGATCTNNNATATCTGGACAAGGGGAATGC
AGAGCACACGTCTGAACTCTTTCCCTACACGACGCTCTTCCGATCTNNNAAGAAAGTGGAGTCAGGCTT
AGAGCACACGTCTGAACTCTTTCCCTACACGACGCTCTTCCGATCTNNNAAGAAAGTGGAGTCAGGCTT
GAGCACACGTCTGAACTCTTTCCCTACACGACGCTCTTCCGATCTNNNAGAAATGAAGAGGCAGAGGT
GAGCACACGTCTGAACTCTTTCCCTACACGACGCTCTTCCGATCTNNNAGAAATGAAGAGGCAGAGGT
AGCACACGTCTGAACTCTTTCCCTACACGACGCTCTTCCGATCTNNNTTTGCAAAGGTGGTTTCACTGT
AGCACACGTCTGAACTCTTTCCCTACACGACGCTCTTCCGATCTNNNTTTGCAAAGGTGGTTTCACTGT
GCACACGTCTGAACTCTTTCCCTACACGACGCTCTTCCGATCTNNNCCTCTATTCCTGCTCTCTTCACCCAT
GCACACGTCTGAACTCTTTCCCTACACGACGCTCTTCCGATCTNNNCCTCTATTCCTGCTCTCTTCACCCAT
GAGCACACGTCTGAACTCTTTCCCTACACGACGCTCTTCCGATCTNNNAGTGATTCTTCATCCTCAGCCT
GAGCACACGTCTGAACTCTTTCCCTACACGACGCTCTTCCGATCTNNNAGTGATTCTTCATCCTCAGCCT
GAGCACACGTCTGAACTCTTTCCCTACACGACGCTCTTCCGATCTNNNCATCCCTGTAATCCCAGCTACT
GAGCACACGTCTGAACTCTTTCCCTACACGACGCTCTTCCGATCTNNNCATCCCTGTAATCCCAGCTACT
AGAGCACACGTCTGAACTCTTTCCCTACACGACGCTCTTCCGATCTNNNCTCTGCTTGATGCTGCCTCTAT
AGAGCACACGTCTGAACTCTTTCCCTACACGACGCTCTTCCGATCTNNNCTCTGCTTGATGCTGCCTCTAT
GAGCACACGTCTGAACTCTTTCCCTACACGACGCTCTTCCGATCTNNNTTTCTATCTATGGCTGTGGCT
GAGCACACGTCTGAACTCTTTCCCTACACGACGCTCTTCCGATCTNNNTTTCTATCTATGGCTGTGGCT
AGAGCACACGTCTGAACTCTTTCCCTACACGACGCTCTTCCGATCTNNNCCACGGTTGTCTCAATCTCT
AGAGCACACGTCTGAACTCTTTCCCTACACGACGCTCTTCCGATCTNNNCCACGGTTGTCTCAATCTCT
AGAGCACACGTCTGAACTCTTTCCCTACACGACGCTCTTCCGATCTNNNTTGTTTCCATGGCTTCGTGAT
AGAGCACACGTCTGAACTCTTTCCCTACACGACGCTCTTCCGATCTNNNTTGTTTCCATGGCTTCGTGAT
GAGCACACGTCTGAACTCTTTCCCTACACGACGCTCTTCCGATCTNNNGAGGCTGGAGTGAGCTATGAT
GAGCACACGTCTGAACTCTTTCCCTACACGACGCTCTTCCGATCTNNNGAGGCTGGAGTGAGCTATGAT
GAGCACACGTCTGAACTCTTTCCCTACACGACGCTCTTCCGATCTNNNTTGCTGGAATCTCTCTGCTCTT
GAGCACACGTCTGAACTCTTTCCCTACACGACGCTCTTCCGATCTNNNTTGCTGGAATCTCTCTGCTCTT
AGCACACGTCTGAACTCTTTCCCTACACGACGCTCTTCCGATCTNNNCATGACCTACAGCCTCAGGTTC
AGCACACGTCTGAACTCTTTCCCTACACGACGCTCTTCCGATCTNNNCATGACCTACAGCCTCAGGTTC

GAGCACACGTCTGAACTCTTTCCCTACACGACGCTCTTCCGATCTNNNACTCTAGGAGTAGGAGGTA
GAGCACACGTCTGAACTCTTTCCCTACACGACGCTCTTCCGATCTNNNACTCTAGGAGTAGGAGGTA
GCACACGTCTGAACTCTTTCCCTACACGACGCTCTTCCGATCTNNNGAGAATCTCCGCAAGAAAGGG
GCACACGTCTGAACTCTTTCCCTACACGACGCTCTTCCGATCTNNNGAGAATCTCCGCAAGAAAGGG
AGCACACGTCTGAACTCTTTCCCTACACGACGCTCTTCCGATCTNNNAAGAAAAGCTGTAGGTGTTT
AGCACACGTCTGAACTCTTTCCCTACACGACGCTCTTCCGATCTNNNAAGAAAAGCTGTAGGTGTTT
CACACGTCTGAACTCTTTCCCTACACGACGCTCTTCCGATCTNNNATAGGTTGCAGTGAGCCAAGAT
CACACGTCTGAACTCTTTCCCTACACGACGCTCTTCCGATCTNNNATAGGTTGCAGTGAGCCAAGAT
GCACACGTCTGAACTCTTTCCCTACACGACGCTCTTCCGATCTNNNACTGGTCCAGTTCTCCATTCAAA
GCACACGTCTGAACTCTTTCCCTACACGACGCTCTTCCGATCTNNNACTGGTCCAGTTCTCCATTCAAA
GCACACGTCTGAACTCTTTCCCTACACGACGCTCTTCCGATCTNNNTGAGGGAGGAGAATCACTTGA
GCACACGTCTGAACTCTTTCCCTACACGACGCTCTTCCGATCTNNNTGAGGGAGGAGAATCACTTGA
AGAGCACACGTCTGAACTCTTTCCCTACACGACGCTCTTCCGATCTNNNAGAGATTGCAGTGAGCTG
AGAGCACACGTCTGAACTCTTTCCCTACACGACGCTCTTCCGATCTNNNAGAGATTGCAGTGAGCTG
GCACACGTCTGAACTCTTTCCCTACACGACGCTCTTCCGATCTNNNTGGTAGAAAAGGAAGCAGTCA
GCACACGTCTGAACTCTTTCCCTACACGACGCTCTTCCGATCTNNNTGGTAGAAAAGGAAGCAGTCA
AGCACACGTCTGAACTCTTTCCCTACACGACGCTCTTCCGATCTNNNCTGAGGTGGGAGGATCTCTA
AGCACACGTCTGAACTCTTTCCCTACACGACGCTCTTCCGATCTNNNCTGAGGTGGGAGGATCTCTA
AGCACACGTCTGAACTCTTTCCCTACACGACGCTCTTCCGATCTNNNGGTTACAGTGAGCTGAGATCA
AGCACACGTCTGAACTCTTTCCCTACACGACGCTCTTCCGATCTNNNGGTTACAGTGAGCTGAGATCA
GAGCACACGTCTGAACTCTTTCCCTACACGACGCTCTTCCGATCTNNNTTTTGTGTGTGTGTGTGTGT
GAGCACACGTCTGAACTCTTTCCCTACACGACGCTCTTCCGATCTNNNTTTTGTGTGTGTGTGTGTGT
CACACGTCTGAACTCTTTCCCTACACGACGCTCTTCCGATCTNNNCCTTCTCCAGTATGACCTCATCCCA
CACACGTCTGAACTCTTTCCCTACACGACGCTCTTCCGATCTNNNCCTTCTCCAGTATGACCTCATCCCA
GCACACGTCTGAACTCTTTCCCTACACGACGCTCTTCCGATCTNNNGGACACATAACCAAATTGTGCG
GCACACGTCTGAACTCTTTCCCTACACGACGCTCTTCCGATCTNNNGGACACATAACCAAATTGTGCG
AGCACACGTCTGAACTCTTTCCCTACACGACGCTCTTCCGATCTNNNTCTTTTCCTCTTGAGTAGCCCA
AGCACACGTCTGAACTCTTTCCCTACACGACGCTCTTCCGATCTNNNTCTTTTCCTCTTGAGTAGCCCA
AGCACACGTCTGAACTCTTTCCCTACACGACGCTCTTCCGATCTNNNGAGAATTGCTTGAACTTGGGA
AGCACACGTCTGAACTCTTTCCCTACACGACGCTCTTCCGATCTNNNGAGAATTGCTTGAACTTGGGA
GAGCACACGTCTGAACTCTTTCCCTACACGACGCTCTTCCGATCTNNNAAGCCTGGGTAGGTACACAT
GAGCACACGTCTGAACTCTTTCCCTACACGACGCTCTTCCGATCTNNNAAGCCTGGGTAGGTACACAT
AGCACACGTCTGAACTCTTTCCCTACACGACGCTCTTCCGATCTNNNAAAAAATTAGCCAGGCTTGGT
AGCACACGTCTGAACTCTTTCCCTACACGACGCTCTTCCGATCTNNNAAAAAATTAGCCAGGCTTGGT
GAGCACACGTCTGAACTCTTTCCCTACACGACGCTCTTCCGATCTNNNAATGCTCTCAGGCAAAAAAC
GAGCACACGTCTGAACTCTTTCCCTACACGACGCTCTTCCGATCTNNNAATGCTCTCAGGCAAAAAAC
AGCACACGTCTGAACTCTTTCCCTACACGACGCTCTTCCGATCTNNNAACCAACTCCTCTTCTGCTGGC
AGCACACGTCTGAACTCTTTCCCTACACGACGCTCTTCCGATCTNNNAACCAACTCCTCTTCTGCTGGC
CACACGTCTGAACTCTTTCCCTACACGACGCTCTTCCGATCTNNNCAGGGTTTCACCAAATTAGCCAGC
CACACGTCTGAACTCTTTCCCTACACGACGCTCTTCCGATCTNNNCAGGGTTTCACCAAATTAGCCAGC
AGCACACGTCTGAACTCTTTCCCTACACGACGCTCTTCCGATCTNNNTGCCTGTAGTCCCAGCTATTTG
AGCACACGTCTGAACTCTTTCCCTACACGACGCTCTTCCGATCTNNNTGCCTGTAGTCCCAGCTATTTG
GCACACGTCTGAACTCTTTCCCTACACGACGCTCTTCCGATCTNNNACTGAAACTTTGCTAAACTGTCA

GCACACGTCTGAACTCTTTCCCTACACGACGCTCTTCCGATCTNNNACTGAAACTTTGCTAAACTGTCA
GCACACGTCTGAACTCTTTCCCTACACGACGCTCTTCCGATCTNNNTGTGTACTAAGTGCTGCTCCATC
GCACACGTCTGAACTCTTTCCCTACACGACGCTCTTCCGATCTNNNTGTGTACTAAGTGCTGCTCCATC
AGCACACGTCTGAACTCTTTCCCTACACGACGCTCTTCCGATCTNNNGAAAAGGGAAGAGAAGGCTC
AGCACACGTCTGAACTCTTTCCCTACACGACGCTCTTCCGATCTNNNGAAAAGGGAAGAGAAGGCTC
GCACACGTCTGAACTCTTTCCCTACACGACGCTCTTCCGATCTNNNCTTTCATTGGCAATGTCGGTCTT
GCACACGTCTGAACTCTTTCCCTACACGACGCTCTTCCGATCTNNNCTTTCATTGGCAATGTCGGTCTT
GAGCACACGTCTGAACTCTTTCCCTACACGACGCTCTTCCGATCTNNNAACTCCAACCTGGGCAATAAG
GAGCACACGTCTGAACTCTTTCCCTACACGACGCTCTTCCGATCTNNNAACTCCAACCTGGGCAATAAG
GCACACGTCTGAACTCTTTCCCTACACGACGCTCTTCCGATCTNNNTTGAGCGTCTACTATGTGTCAG
GCACACGTCTGAACTCTTTCCCTACACGACGCTCTTCCGATCTNNNTTGAGCGTCTACTATGTGTCAG
AGCACACGTCTGAACTCTTTCCCTACACGACGCTCTTCCGATCTNNNGTAGTTCCCAGCTACTCAGGAT
AGCACACGTCTGAACTCTTTCCCTACACGACGCTCTTCCGATCTNNNGTAGTTCCCAGCTACTCAGGAT
GAGCACACGTCTGAACTCTTTCCCTACACGACGCTCTTCCGATCTNNNGAAAAAGTGCTGACCCCAGA
GAGCACACGTCTGAACTCTTTCCCTACACGACGCTCTTCCGATCTNNNGAAAAAGTGCTGACCCCAGA
GCACACGTCTGAACTCTTTCCCTACACGACGCTCTTCCGATCTNNNACTCAGACTATGTCAGGGATTG
GCACACGTCTGAACTCTTTCCCTACACGACGCTCTTCCGATCTNNNACTCAGACTATGTCAGGGATTG
AGAGCACACGTCTGAACTCTTTCCCTACACGACGCTCTTCCGATCTNNNGAGACAGGAGAATGGCTAG
AGAGCACACGTCTGAACTCTTTCCCTACACGACGCTCTTCCGATCTNNNGAGACAGGAGAATGGCTAG
CACACGTCTGAACTCTTTCCCTACACGACGCTCTTCCGATCTNNNGTTTGATACAAAATGGGGTGGG
CACACGTCTGAACTCTTTCCCTACACGACGCTCTTCCGATCTNNNGTTTGATACAAAATGGGGTGGG
GCACACGTCTGAACTCTTTCCCTACACGACGCTCTTCCGATCTNNNCCGAGTAGATGGGATTACAGG
GCACACGTCTGAACTCTTTCCCTACACGACGCTCTTCCGATCTNNNCCGAGTAGATGGGATTACAGG
AGCACACGTCTGAACTCTTTCCCTACACGACGCTCTTCCGATCTNNNAGTCTGTAACGTTACTTCTTAA
AGCACACGTCTGAACTCTTTCCCTACACGACGCTCTTCCGATCTNNNAGTCTGTAACGTTACTTCTTAA
AGCACACGTCTGAACTCTTTCCCTACACGACGCTCTTCCGATCTNNNACTGGCCATCACTACTGTACAT
AGCACACGTCTGAACTCTTTCCCTACACGACGCTCTTCCGATCTNNNACTGGCCATCACTACTGTACAT
GCACACGTCTGAACTCTTTCCCTACACGACGCTCTTCCGATCTNNNCAGCAACAAAAACCAGACAAAC
GCACACGTCTGAACTCTTTCCCTACACGACGCTCTTCCGATCTNNNCAGCAACAAAAACCAGACAAAC
AGCACACGTCTGAACTCTTTCCCTACACGACGCTCTTCCGATCTNNNTGTTGTTGTTGACCTTTTGACC
AGCACACGTCTGAACTCTTTCCCTACACGACGCTCTTCCGATCTNNNTGTTGTTGTTGACCTTTTGACC
AGAGCACACGTCTGAACTCTTTCCCTACACGACGCTCTTCCGATCTNNNACGTCACTATTATTCCCCAGT
AGAGCACACGTCTGAACTCTTTCCCTACACGACGCTCTTCCGATCTNNNACGTCACTATTATTCCCCAGT
GCACACGTCTGAACTCTTTCCCTACACGACGCTCTTCCGATCTNNNTTTAGGGTCAATGTTTCCTGGG
GCACACGTCTGAACTCTTTCCCTACACGACGCTCTTCCGATCTNNNTTTAGGGTCAATGTTTCCTGGG
GCACACGTCTGAACTCTTTCCCTACACGACGCTCTTCCGATCTNNNGGTGGGAGACATCCTTTCAGAT
GCACACGTCTGAACTCTTTCCCTACACGACGCTCTTCCGATCTNNNGGTGGGAGACATCCTTTCAGAT
GAGCACACGTCTGAACTCTTTCCCTACACGACGCTCTTCCGATCTNNNCATCCTGGCTAACATGGTGAA
GAGCACACGTCTGAACTCTTTCCCTACACGACGCTCTTCCGATCTNNNCATCCTGGCTAACATGGTGAA
GAGCACACGTCTGAACTCTTTCCCTACACGACGCTCTTCCGATCTNNNCTGTAATCCCAGCTACTCAGG
GAGCACACGTCTGAACTCTTTCCCTACACGACGCTCTTCCGATCTNNNCTGTAATCCCAGCTACTCAGG
GAGCACACGTCTGAACTCTTTCCCTACACGACGCTCTTCCGATCTNNNTTGCAATGAGCTGAGAACAT
GAGCACACGTCTGAACTCTTTCCCTACACGACGCTCTTCCGATCTNNNTTGCAATGAGCTGAGAACAT

GCACACGTCTGAACTCTTTCCCTACACGACGCTCTTCCGATCTNNNCGACATCCGACCTTGTTTCTTTTCA
GCACACGTCTGAACTCTTTCCCTACACGACGCTCTTCCGATCTNNNCGACATCCGACCTTGTTTCTTTTCA
AGCACACGTCTGAACTCTTTCCCTACACGACGCTCTTCCGATCTNNNGATGCAATCATAGCTCACTGCA
AGCACACGTCTGAACTCTTTCCCTACACGACGCTCTTCCGATCTNNNGATGCAATCATAGCTCACTGCA
GAGCACACGTCTGAACTCTTTCCCTACACGACGCTCTTCCGATCTNNNGTGAAGGTACTTTGAGGTGG
GAGCACACGTCTGAACTCTTTCCCTACACGACGCTCTTCCGATCTNNNGTGAAGGTACTTTGAGGTGG
GAGCACACGTCTGAACTCTTTCCCTACACGACGCTCTTCCGATCTNNNTGTGCCCGGCATAATTTTTCTT
GAGCACACGTCTGAACTCTTTCCCTACACGACGCTCTTCCGATCTNNNTGTGCCCGGCATAATTTTTCTT
GAGCACACGTCTGAACTCTTTCCCTACACGACGCTCTTCCGATCTNNNTTTAGTGAAGACGGGGTTTCT
GAGCACACGTCTGAACTCTTTCCCTACACGACGCTCTTCCGATCTNNNTTTAGTGAAGACGGGGTTTCT
AGCACACGTCTGAACTCTTTCCCTACACGACGCTCTTCCGATCTNNNAAAAAATTAGCTGGGCATGGTG
AGCACACGTCTGAACTCTTTCCCTACACGACGCTCTTCCGATCTNNNAAAAAATTAGCTGGGCATGGTG
GCACACGTCTGAACTCTTTCCCTACACGACGCTCTTCCGATCTNNNCCACTCTATAGGTTGCCTGTTCACA
GCACACGTCTGAACTCTTTCCCTACACGACGCTCTTCCGATCTNNNCCACTCTATAGGTTGCCTGTTCACA
GAGCACACGTCTGAACTCTTTCCCTACACGACGCTCTTCCGATCTNNNGGTGATAAAGCGAGACTCAG
GAGCACACGTCTGAACTCTTTCCCTACACGACGCTCTTCCGATCTNNNGGTGATAAAGCGAGACTCAG
GAGCACACGTCTGAACTCTTTCCCTACACGACGCTCTTCCGATCTNNNGTCATCCCTCAAGGTTCTCACT
GAGCACACGTCTGAACTCTTTCCCTACACGACGCTCTTCCGATCTNNNGTCATCCCTCAAGGTTCTCACT
CACACGTCTGAACTCTTTCCCTACACGACGCTCTTCCGATCTNNNTGATAGGAAGCGGTGACAATGAG
CACACGTCTGAACTCTTTCCCTACACGACGCTCTTCCGATCTNNNTGATAGGAAGCGGTGACAATGAG
AGCACACGTCTGAACTCTTTCCCTACACGACGCTCTTCCGATCTNNNGATTGGGGTTGTTTTCAAGGTC
AGCACACGTCTGAACTCTTTCCCTACACGACGCTCTTCCGATCTNNNGATTGGGGTTGTTTTCAAGGTC
GAGCACACGTCTGAACTCTTTCCCTACACGACGCTCTTCCGATCTNNNTGAATGGAGATTACACCACTG
GAGCACACGTCTGAACTCTTTCCCTACACGACGCTCTTCCGATCTNNNTGAATGGAGATTACACCACTG
GAGCACACGTCTGAACTCTTTCCCTACACGACGCTCTTCCGATCTNNNAACCCCTGGTATAGAAAGCAT
GAGCACACGTCTGAACTCTTTCCCTACACGACGCTCTTCCGATCTNNNAACCCCTGGTATAGAAAGCAT
GAGCACACGTCTGAACTCTTTCCCTACACGACGCTCTTCCGATCTNNNGTTGGATCTGGAACCTGGGTA
GAGCACACGTCTGAACTCTTTCCCTACACGACGCTCTTCCGATCTNNNGTTGGATCTGGAACCTGGGTA
GAGCACACGTCTGAACTCTTTCCCTACACGACGCTCTTCCGATCTNNNCAATCTCTGGGTTTTGGGAAC
GAGCACACGTCTGAACTCTTTCCCTACACGACGCTCTTCCGATCTNNNCAATCTCTGGGTTTTGGGAAC
GAGCACACGTCTGAACTCTTTCCCTACACGACGCTCTTCCGATCTNNNCGACAGAGCAAGACATCGTCT
GAGCACACGTCTGAACTCTTTCCCTACACGACGCTCTTCCGATCTNNNCGACAGAGCAAGACATCGTCT
AGCACACGTCTGAACTCTTTCCCTACACGACGCTCTTCCGATCTNNNAGAGACCTTGTCCAATAGCTCT
AGCACACGTCTGAACTCTTTCCCTACACGACGCTCTTCCGATCTNNNAGAGACCTTGTCCAATAGCTCT
GCACACGTCTGAACTCTTTCCCTACACGACGCTCTTCCGATCTNNNTCTCCTCTCTCTCTCCTTCCTTGCAT
GCACACGTCTGAACTCTTTCCCTACACGACGCTCTTCCGATCTNNNTCTCCTCTCTCTCTCCTTCCTTGCAT
GAGCACACGTCTGAACTCTTTCCCTACACGACGCTCTTCCGATCTNNNCTCTCTCCATATCTTGTGCTCC
GAGCACACGTCTGAACTCTTTCCCTACACGACGCTCTTCCGATCTNNNCTCTCTCCATATCTTGTGCTCC
AGCACACGTCTGAACTCTTTCCCTACACGACGCTCTTCCGATCTNNNCCAGCCAGAGGGTATTTTCCAT
AGCACACGTCTGAACTCTTTCCCTACACGACGCTCTTCCGATCTNNNCCAGCCAGAGGGTATTTTCCAT
GCACACGTCTGAACTCTTTCCCTACACGACGCTCTTCCGATCTNNNCAAACATTTGCAGAGGGACCGCA
GCACACGTCTGAACTCTTTCCCTACACGACGCTCTTCCGATCTNNNCAAACATTTGCAGAGGGACCGCA
AGCACACGTCTGAACTCTTTCCCTACACGACGCTCTTCCGATCTNNNACACTCCTCCATGTAACCATGTG

AGCACACGTCTGAACTCTTTCCCTACACGACGCTCTTCCGATCTNNNACACTCCTCCATGTAACCATGTG
AGCACACGTCTGAACTCTTTCCCTACACGACGCTCTTCCGATCTNNNTCCTAACACTATCCCTCCTCCAG
AGCACACGTCTGAACTCTTTCCCTACACGACGCTCTTCCGATCTNNNTCCTAACACTATCCCTCCTCCAG
AAGAGCACACGTCTGAACTCTTTCCCTACACGACGCTCTTCCGATCTNNNAGATTGGAGTAAGCCGAG
AAGAGCACACGTCTGAACTCTTTCCCTACACGACGCTCTTCCGATCTNNNAGATTGGAGTAAGCCGAG
AGCACACGTCTGAACTCTTTCCCTACACGACGCTCTTCCGATCTNNNTTGCTGACCCAGATTCTGTTTT
AGCACACGTCTGAACTCTTTCCCTACACGACGCTCTTCCGATCTNNNTTGCTGACCCAGATTCTGTTTT
AGCACACGTCTGAACTCTTTCCCTACACGACGCTCTTCCGATCTNNNAAACAGACCAATCAGCTCTCTG
AGCACACGTCTGAACTCTTTCCCTACACGACGCTCTTCCGATCTNNNAAACAGACCAATCAGCTCTCTG
AGAGCACACGTCTGAACTCTTTCCCTACACGACGCTCTTCCGATCTNNNACCTACATCCTGTCTCCATCC
AGAGCACACGTCTGAACTCTTTCCCTACACGACGCTCTTCCGATCTNNNACCTACATCCTGTCTCCATCC
GAGCACACGTCTGAACTCTTTCCCTACACGACGCTCTTCCGATCTNNNAACACACACTGCTTCACAAAT
GAGCACACGTCTGAACTCTTTCCCTACACGACGCTCTTCCGATCTNNNAACACACACTGCTTCACAAAT
GAGCACACGTCTGAACTCTTTCCCTACACGACGCTCTTCCGATCTNNNGTTATTCCAGCTACTAGGGAG
GAGCACACGTCTGAACTCTTTCCCTACACGACGCTCTTCCGATCTNNNGTTATTCCAGCTACTAGGGAG
AGAGCACACGTCTGAACTCTTTCCCTACACGACGCTCTTCCGATCTNNNGTTGTCCCATGGCAAACACT
AGAGCACACGTCTGAACTCTTTCCCTACACGACGCTCTTCCGATCTNNNGTTGTCCCATGGCAAACACT
AGCACACGTCTGAACTCTTTCCCTACACGACGCTCTTCCGATCTNNNGAACTGCTCGCGTTGGAAAAT
AGCACACGTCTGAACTCTTTCCCTACACGACGCTCTTCCGATCTNNNGAACTGCTCGCGTTGGAAAAT
GAGCACACGTCTGAACTCTTTCCCTACACGACGCTCTTCCGATCTNNNGAGAATCGTGTGAACCCAGG
GAGCACACGTCTGAACTCTTTCCCTACACGACGCTCTTCCGATCTNNNGAGAATCGTGTGAACCCAGG
AGAGCACACGTCTGAACTCTTTCCCTACACGACGCTCTTCCGATCTNNNGAGAATTGCTTGAACCTGGG
AGAGCACACGTCTGAACTCTTTCCCTACACGACGCTCTTCCGATCTNNNGAGAATTGCTTGAACCTGGG
AGCACACGTCTGAACTCTTTCCCTACACGACGCTCTTCCGATCTNNNTGCCCTTTCCTTCTTCTTCTTCTC
AGCACACGTCTGAACTCTTTCCCTACACGACGCTCTTCCGATCTNNNTGCCCTTTCCTTCTTCTTCTTCTC
GAGCACACGTCTGAACTCTTTCCCTACACGACGCTCTTCCGATCTNNNATTTGCTTAGGTCTAGGAGCA
GAGCACACGTCTGAACTCTTTCCCTACACGACGCTCTTCCGATCTNNNATTTGCTTAGGTCTAGGAGCA
AGCACACGTCTGAACTCTTTCCCTACACGACGCTCTTCCGATCTNNNCCCACCTTAGCTTCCTAAGTAGC
AGCACACGTCTGAACTCTTTCCCTACACGACGCTCTTCCGATCTNNNCCCACCTTAGCTTCCTAAGTAGC
AGCACACGTCTGAACTCTTTCCCTACACGACGCTCTTCCGATCTNNNTGCCCTTCTTTGTCGTTTTTGAT
AGCACACGTCTGAACTCTTTCCCTACACGACGCTCTTCCGATCTNNNTGCCCTTCTTTGTCGTTTTTGAT
AGCACACGTCTGAACTCTTTCCCTACACGACGCTCTTCCGATCTNNNCACTGAAGGAAAAGAGATGCA
AGCACACGTCTGAACTCTTTCCCTACACGACGCTCTTCCGATCTNNNCACTGAAGGAAAAGAGATGCA
GAGCACACGTCTGAACTCTTTCCCTACACGACGCTCTTCCGATCTNNNAAGAACAGCTCCACATCAAAA
GAGCACACGTCTGAACTCTTTCCCTACACGACGCTCTTCCGATCTNNNAAGAACAGCTCCACATCAAAA
GAGCACACGTCTGAACTCTTTCCCTACACGACGCTCTTCCGATCTNNNTGTCTTCCACAAAACTGTTCCCT
GAGCACACGTCTGAACTCTTTCCCTACACGACGCTCTTCCGATCTNNNTGTCTTCCACAAAACTGTTCCCT
AGCACACGTCTGAACTCTTTCCCTACACGACGCTCTTCCGATCTNNNGCTGGCTGCAAAATTTTTCACA
AGCACACGTCTGAACTCTTTCCCTACACGACGCTCTTCCGATCTNNNGCTGGCTGCAAAATTTTTCACA
AGCACACGTCTGAACTCTTTCCCTACACGACGCTCTTCCGATCTNNNGTTGTGGTGACCTGAGATCGTA
AGCACACGTCTGAACTCTTTCCCTACACGACGCTCTTCCGATCTNNNGTTGTGGTGACCTGAGATCGTA
GAGCACACGTCTGAACTCTTTCCCTACACGACGCTCTTCCGATCTNNNGGCCTTCTCCAATAGCTTAGCA
GAGCACACGTCTGAACTCTTTCCCTACACGACGCTCTTCCGATCTNNNGGCCTTCTCCAATAGCTTAGCA

CACACGTCTGAACTCTTTCCCTACACGACGCTCTTCCGATCTNNNTAGTAGAGATGGGGTTTCTCCGTC
CACACGTCTGAACTCTTTCCCTACACGACGCTCTTCCGATCTNNNTAGTAGAGATGGGGTTTCTCCGTC
GAGCACACGTCTGAACTCTTTCCCTACACGACGCTCTTCCGATCTNNNATCTGGGCCTAGAGTTTTCTT
GAGCACACGTCTGAACTCTTTCCCTACACGACGCTCTTCCGATCTNNNATCTGGGCCTAGAGTTTTCTT
GAGCACACGTCTGAACTCTTTCCCTACACGACGCTCTTCCGATCTNNNGAACATATCCTTCTCGCTAGGC
GAGCACACGTCTGAACTCTTTCCCTACACGACGCTCTTCCGATCTNNNGAACATATCCTTCTCGCTAGGC
GAGCACACGTCTGAACTCTTTCCCTACACGACGCTCTTCCGATCTNNNATGCCTGTAATCTCAGCTACTC
GAGCACACGTCTGAACTCTTTCCCTACACGACGCTCTTCCGATCTNNNATGCCTGTAATCTCAGCTACTC
AGCACACGTCTGAACTCTTTCCCTACACGACGCTCTTCCGATCTNNNAGGAATTGAATGGGTAGGGCA
AGCACACGTCTGAACTCTTTCCCTACACGACGCTCTTCCGATCTNNNAGGAATTGAATGGGTAGGGCA
AGCACACGTCTGAACTCTTTCCCTACACGACGCTCTTCCGATCTNNNACAGACTATGAGCGTGAGAATC
AGCACACGTCTGAACTCTTTCCCTACACGACGCTCTTCCGATCTNNNACAGACTATGAGCGTGAGAATC
GAGCACACGTCTGAACTCTTTCCCTACACGACGCTCTTCCGATCTNNNCACCTTTCATTGTGGACCTATC
GAGCACACGTCTGAACTCTTTCCCTACACGACGCTCTTCCGATCTNNNCACCTTTCATTGTGGACCTATC
AGCACACGTCTGAACTCTTTCCCTACACGACGCTCTTCCGATCTNNNTCTCCTAGTCTTCTCTCCCCATTC
AGCACACGTCTGAACTCTTTCCCTACACGACGCTCTTCCGATCTNNNTCTCCTAGTCTTCTCTCCCCATTC
GAGCACACGTCTGAACTCTTTCCCTACACGACGCTCTTCCGATCTNNNACTGGCCTCAAGTTTGTTCTAC
GAGCACACGTCTGAACTCTTTCCCTACACGACGCTCTTCCGATCTNNNACTGGCCTCAAGTTTGTTCTAC
AGCACACGTCTGAACTCTTTCCCTACACGACGCTCTTCCGATCTNNNATATGCTTGAAGGGAAAGGCT
AGCACACGTCTGAACTCTTTCCCTACACGACGCTCTTCCGATCTNNNATATGCTTGAAGGGAAAGGCT
CACACGTCTGAACTCTTTCCCTACACGACGCTCTTCCGATCTNNNCTTCCTCAGCAGGAGATAGACCACA
CACACGTCTGAACTCTTTCCCTACACGACGCTCTTCCGATCTNNNCTTCCTCAGCAGGAGATAGACCACA
AGCACACGTCTGAACTCTTTCCCTACACGACGCTCTTCCGATCTNNNCGCCAATCTACCACTGACCTTAT
AGCACACGTCTGAACTCTTTCCCTACACGACGCTCTTCCGATCTNNNCGCCAATCTACCACTGACCTTAT
AGCACACGTCTGAACTCTTTCCCTACACGACGCTCTTCCGATCTNNNTGTATCTCTGCTCCAGAATGCTC
AGCACACGTCTGAACTCTTTCCCTACACGACGCTCTTCCGATCTNNNTGTATCTCTGCTCCAGAATGCTC
AGCACACGTCTGAACTCTTTCCCTACACGACGCTCTTCCGATCTNNNACCACGCCTGGCTAATTTCTAACA
AGCACACGTCTGAACTCTTTCCCTACACGACGCTCTTCCGATCTNNNACCACGCCTGGCTAATTTCTAACA
GAGCACACGTCTGAACTCTTTCCCTACACGACGCTCTTCCGATCTNNNTGCAGTGAGCTATGATTGTGC
GAGCACACGTCTGAACTCTTTCCCTACACGACGCTCTTCCGATCTNNNTGCAGTGAGCTATGATTGTGC
AGCACACGTCTGAACTCTTTCCCTACACGACGCTCTTCCGATCTNNNTGTAAGTGTGCAGTTCAGTGGT
AGCACACGTCTGAACTCTTTCCCTACACGACGCTCTTCCGATCTNNNTGTAAGTGTGCAGTTCAGTGGT
AGCACACGTCTGAACTCTTTCCCTACACGACGCTCTTCCGATCTNNNATCCCTCTTTCCTTGTCTTTGCTCA
AGCACACGTCTGAACTCTTTCCCTACACGACGCTCTTCCGATCTNNNATCCCTCTTTCCTTGTCTTTGCTCA
AGCACACGTCTGAACTCTTTCCCTACACGACGCTCTTCCGATCTNNNGAGCCAGGAGAATTGTTTGAA
AGCACACGTCTGAACTCTTTCCCTACACGACGCTCTTCCGATCTNNNGAGCCAGGAGAATTGTTTGAA
AGCACACGTCTGAACTCTTTCCCTACACGACGCTCTTCCGATCTNNNACAAACATCCACAGCCTTACCAT
AGCACACGTCTGAACTCTTTCCCTACACGACGCTCTTCCGATCTNNNACAAACATCCACAGCCTTACCAT
AGCACACGTCTGAACTCTTTCCCTACACGACGCTCTTCCGATCTNNNTCCCTTTGCTTATATATCTCTTTA
AGCACACGTCTGAACTCTTTCCCTACACGACGCTCTTCCGATCTNNNTCCCTTTGCTTATATATCTCTTTA
AGCACACGTCTGAACTCTTTCCCTACACGACGCTCTTCCGATCTNNNTGCCTCATTTGACACAACAAAG
AGCACACGTCTGAACTCTTTCCCTACACGACGCTCTTCCGATCTNNNTGCCTCATTTGACACAACAAAG
AGCACACGTCTGAACTCTTTCCCTACACGACGCTCTTCCGATCTNNNACCTTTGAGCTATTGTACAACCT

AGCACACGTCTGAACTCTTTCCCTACACGACGCTCTTCCGATCTNNNACCTTTGAGCTATTGTACAACCT
AGCACACGTCTGAACTCTTTCCCTACACGACGCTCTTCCGATCTNNNCCAAAGTGCTGGAATTACTGGT
AGCACACGTCTGAACTCTTTCCCTACACGACGCTCTTCCGATCTNNNCCAAAGTGCTGGAATTACTGGT
AGCACACGTCTGAACTCTTTCCCTACACGACGCTCTTCCGATCTNNNACAGGAGTTCTACTGGCTTTCT
AGCACACGTCTGAACTCTTTCCCTACACGACGCTCTTCCGATCTNNNACAGGAGTTCTACTGGCTTTCT
GAGCACACGTCTGAACTCTTTCCCTACACGACGCTCTTCCGATCTNNNGCAGAGGTCCTGTAGAGACA
GAGCACACGTCTGAACTCTTTCCCTACACGACGCTCTTCCGATCTNNNGCAGAGGTCCTGTAGAGACA
GAGCACACGTCTGAACTCTTTCCCTACACGACGCTCTTCCGATCTNNNTCCTTTATCCACACTGTAGCCA
GAGCACACGTCTGAACTCTTTCCCTACACGACGCTCTTCCGATCTNNNTCCTTTATCCACACTGTAGCCA
AGCACACGTCTGAACTCTTTCCCTACACGACGCTCTTCCGATCTNNNGGAATCTCTGGTGGAAGCCAC
AGCACACGTCTGAACTCTTTCCCTACACGACGCTCTTCCGATCTNNNGGAATCTCTGGTGGAAGCCAC
AGCACACGTCTGAACTCTTTCCCTACACGACGCTCTTCCGATCTNNNCTGTAGTCCCCGCTACTCAGAA
AGCACACGTCTGAACTCTTTCCCTACACGACGCTCTTCCGATCTNNNCTGTAGTCCCCGCTACTCAGAA
GAGCACACGTCTGAACTCTTTCCCTACACGACGCTCTTCCGATCTNNNCTCTCACACACACATGTTCACA
GAGCACACGTCTGAACTCTTTCCCTACACGACGCTCTTCCGATCTNNNCTCTCACACACACATGTTCACA
AGCACACGTCTGAACTCTTTCCCTACACGACGCTCTTCCGATCTNNNTCCTAAAGTGCTGGGATTACAG
AGCACACGTCTGAACTCTTTCCCTACACGACGCTCTTCCGATCTNNNTCCTAAAGTGCTGGGATTACAG
GCACACGTCTGAACTCTTTCCCTACACGACGCTCTTCCGATCTNNNGCCACCTAGATATTCAATGACCC
GCACACGTCTGAACTCTTTCCCTACACGACGCTCTTCCGATCTNNNGCCACCTAGATATTCAATGACCC
AGAGCACACGTCTGAACTCTTTCCCTACACGACGCTCTTCCGATCTNNNCAAGTGGCCAACGAACATAT
AGAGCACACGTCTGAACTCTTTCCCTACACGACGCTCTTCCGATCTNNNCAAGTGGCCAACGAACATAT
AGCACACGTCTGAACTCTTTCCCTACACGACGCTCTTCCGATCTNNNTGACAACAGATATCTGAAGGCG
AGCACACGTCTGAACTCTTTCCCTACACGACGCTCTTCCGATCTNNNTGACAACAGATATCTGAAGGCG
CACACGTCTGAACTCTTTCCCTACACGACGCTCTTCCGATCTNNNGTCATCTCATGCCCTTTCACTCCCAT
CACACGTCTGAACTCTTTCCCTACACGACGCTCTTCCGATCTNNNGTCATCTCATGCCCTTTCACTCCCAT
GAGCACACGTCTGAACTCTTTCCCTACACGACGCTCTTCCGATCTNNNGCTTCACTACAACCTCTTACCCT
GAGCACACGTCTGAACTCTTTCCCTACACGACGCTCTTCCGATCTNNNGCTTCACTACAACCTCTTACCCT
AGCACACGTCTGAACTCTTTCCCTACACGACGCTCTTCCGATCTNNNTCTTTCCTCCTTTCAACTCCCTCC
AGCACACGTCTGAACTCTTTCCCTACACGACGCTCTTCCGATCTNNNTCTTTCCTCCTTTCAACTCCCTCC
AGCACACGTCTGAACTCTTTCCCTACACGACGCTCTTCCGATCTNNNTGCTAGGATTACAGGCATGAT
AGCACACGTCTGAACTCTTTCCCTACACGACGCTCTTCCGATCTNNNTGCTAGGATTACAGGCATGAT
AGAGCACACGTCTGAACTCTTTCCCTACACGACGCTCTTCCGATCTNNNAATAAGTGTCTCACTGTGCA
AGAGCACACGTCTGAACTCTTTCCCTACACGACGCTCTTCCGATCTNNNAATAAGTGTCTCACTGTGCA
GAGCACACGTCTGAACTCTTTCCCTACACGACGCTCTTCCGATCTNNNTTCATCCGGAATCTATCGGTT
GAGCACACGTCTGAACTCTTTCCCTACACGACGCTCTTCCGATCTNNNTTCATCCGGAATCTATCGGTT
GAGCACACGTCTGAACTCTTTCCCTACACGACGCTCTTCCGATCTNNNCCAACAGTGTATAAGCACTCC
GAGCACACGTCTGAACTCTTTCCCTACACGACGCTCTTCCGATCTNNNCCAACAGTGTATAAGCACTCC
GAGCACACGTCTGAACTCTTTCCCTACACGACGCTCTTCCGATCTNNNACCAAAGCTGTGAAGTTGTCA
GAGCACACGTCTGAACTCTTTCCCTACACGACGCTCTTCCGATCTNNNACCAAAGCTGTGAAGTTGTCA
AGCACACGTCTGAACTCTTTCCCTACACGACGCTCTTCCGATCTNNNTGGTCCAAAGGTTTTGTAAGTG
AGCACACGTCTGAACTCTTTCCCTACACGACGCTCTTCCGATCTNNNTGGTCCAAAGGTTTTGTAAGTG
GAGCACACGTCTGAACTCTTTCCCTACACGACGCTCTTCCGATCTNNNGCAAAGATTCCTGTAGCCGTT
GAGCACACGTCTGAACTCTTTCCCTACACGACGCTCTTCCGATCTNNNGCAAAGATTCCTGTAGCCGTT

AGCACACGTCTGAACTCTTTCCCTACACGACGCTCTTCCGATCTNNNAATTTGTAATGATTTTTCTGTG

AGCACACGTCTGAACTCTTTCCCTACACGACGCTCTTCCGATCTNNNAATTTGTAATGATTTTTCTGTG

AGCACACGTCTGAACTCTTTCCCTACACGACGCTCTTCCGATCTNNNGTAACTGGGACTACAGGCATA

AGCACACGTCTGAACTCTTTCCCTACACGACGCTCTTCCGATCTNNNGTAACTGGGACTACAGGCATA

AGCACACGTCTGAACTCTTTCCCTACACGACGCTCTTCCGATCTNNNAAAGTGGAATCATCACGACTT

AGCACACGTCTGAACTCTTTCCCTACACGACGCTCTTCCGATCTNNNAAAGTGGAATCATCACGACTT

GAGCACACGTCTGAACTCTTTCCCTACACGACGCTCTTCCGATCTNNNAGACATCCATTCCCAACAAA

GAGCACACGTCTGAACTCTTTCCCTACACGACGCTCTTCCGATCTNNNAGACATCCATTCCCAACAAA

AGCACACGTCTGAACTCTTTCCCTACACGACGCTCTTCCGATCTNNNCCAACATGGTGAAAACCTGTC

AGCACACGTCTGAACTCTTTCCCTACACGACGCTCTTCCGATCTNNNCCAACATGGTGAAAACCTGTC

AGCACACGTCTGAACTCTTTCCCTACACGACGCTCTTCCGATCTNNNACCTTGAAAAACCCTCACCTG

AGCACACGTCTGAACTCTTTCCCTACACGACGCTCTTCCGATCTNNNACCTTGAAAAACCCTCACCTG

GCACACGTCTGAACTCTTTCCCTACACGACGCTCTTCCGATCTNNNCTCTTCTCTCCCTTTCTCTGCTGCA

GCACACGTCTGAACTCTTTCCCTACACGACGCTCTTCCGATCTNNNCTCTTCTCTCCCTTTCTCTGCTGCA

AGCACACGTCTGAACTCTTTCCCTACACGACGCTCTTCCGATCTNNNTTCAGATTCTTACTGGCACTG

AGCACACGTCTGAACTCTTTCCCTACACGACGCTCTTCCGATCTNNNTTCAGATTCTTACTGGCACTG

GAGCACACGTCTGAACTCTTTCCCTACACGACGCTCTTCCGATCTNNNCAAAACAGCATGGCACTGGA

GAGCACACGTCTGAACTCTTTCCCTACACGACGCTCTTCCGATCTNNNCAAAACAGCATGGCACTGGA

GAGCACACGTCTGAACTCTTTCCCTACACGACGCTCTTCCGATCTNNNACTCAGGTTGTTGTTGTTGTT

GAGCACACGTCTGAACTCTTTCCCTACACGACGCTCTTCCGATCTNNNACTCAGGTTGTTGTTGTTGTT

AGCACACGTCTGAACTCTTTCCCTACACGACGCTCTTCCGATCTNNNGAGTAGAAACTGCTAAGCCC

AGCACACGTCTGAACTCTTTCCCTACACGACGCTCTTCCGATCTNNNGAGTAGAAACTGCTAAGCCC

AGCACACGTCTGAACTCTTTCCCTACACGACGCTCTTCCGATCTNNNATCTCTTTCTGGCTTTGCAGG

AGCACACGTCTGAACTCTTTCCCTACACGACGCTCTTCCGATCTNNNATCTCTTTCTGGCTTTGCAGG

AGCACACGTCTGAACTCTTTCCCTACACGACGCTCTTCCGATCTNNNAAAGGGATAATGGGGTGTCT

AGCACACGTCTGAACTCTTTCCCTACACGACGCTCTTCCGATCTNNNAAAGGGATAATGGGGTGTCT

GCACACGTCTGAACTCTTTCCCTACACGACGCTCTTCCGATCTNNNAGTGATCTTTCTAGAGCGTGCTC

GCACACGTCTGAACTCTTTCCCTACACGACGCTCTTCCGATCTNNNAGTGATCTTTCTAGAGCGTGCTC

GCACACGTCTGAACTCTTTCCCTACACGACGCTCTTCCGATCTNNNTCAAAATCACTGATTTTCTTCTTC

GCACACGTCTGAACTCTTTCCCTACACGACGCTCTTCCGATCTNNNTCAAAATCACTGATTTTCTTCTTC

GAGCACACGTCTGAACTCTTTCCCTACACGACGCTCTTCCGATCTNNNATTAGCCAGGAGAATCGCTT

GAGCACACGTCTGAACTCTTTCCCTACACGACGCTCTTCCGATCTNNNATTAGCCAGGAGAATCGCTT

AGCACACGTCTGAACTCTTTCCCTACACGACGCTCTTCCGATCTNNNTAGCGAAACCCCATCTCTAGAG

AGCACACGTCTGAACTCTTTCCCTACACGACGCTCTTCCGATCTNNNTAGCGAAACCCCATCTCTAGAG

GAGCACACGTCTGAACTCTTTCCCTACACGACGCTCTTCCGATCTNNNGCCTGTAATCCCAGCTACTTG

GAGCACACGTCTGAACTCTTTCCCTACACGACGCTCTTCCGATCTNNNGCCTGTAATCCCAGCTACTTG

GAGCACACGTCTGAACTCTTTCCCTACACGACGCTCTTCCGATCTNNNCTCACAAGAAGCCTCTCCTG

GAGCACACGTCTGAACTCTTTCCCTACACGACGCTCTTCCGATCTNNNCTCACAAGAAGCCTCTCCTG

AGCACACGTCTGAACTCTTTCCCTACACGACGCTCTTCCGATCTNNNCCAAAATGCTGGGATTACAGG

AGCACACGTCTGAACTCTTTCCCTACACGACGCTCTTCCGATCTNNNCCAAAATGCTGGGATTACAGG

AGCACACGTCTGAACTCTTTCCCTACACGACGCTCTTCCGATCTNNNCTGAGGCAGGAGAATCACTTG

AGCACACGTCTGAACTCTTTCCCTACACGACGCTCTTCCGATCTNNNCTGAGGCAGGAGAATCACTTG

GCACACGTCTGAACTCTTTCCCTACACGACGCTCTTCCGATCTNNNCCTCAGCCTCCTAAAGTTCTGAT

AGCACACGTCTGAACTCTTTCCCTACACGACGCTCTTCCGATCTNNNCCTCAGCCTCCTAAAGTTCTGAT
AGCACACGTCTGAACTCTTTCCCTACACGACGCTCTTCCGATCTNNNAAACGATGTGTTGATGGATGCA
AGCACACGTCTGAACTCTTTCCCTACACGACGCTCTTCCGATCTNNNAAACGATGTGTTGATGGATGCA
AGCACACGTCTGAACTCTTTCCCTACACGACGCTCTTCCGATCTNNNATTCCTCCCTCTCCCTAGTGTTT
AGCACACGTCTGAACTCTTTCCCTACACGACGCTCTTCCGATCTNNNATTCCTCCCTCTCCCTAGTGTTT
GCACACGTCTGAACTCTTTCCCTACACGACGCTCTTCCGATCTNNNAAAAACAGGCATGGAGTTTGCAG
GCACACGTCTGAACTCTTTCCCTACACGACGCTCTTCCGATCTNNNAAAAACAGGCATGGAGTTTGCAG
GAGCACACGTCTGAACTCTTTCCCTACACGACGCTCTTCCGATCTNNNGAGATTGGGTTGTCACTGTGT
GAGCACACGTCTGAACTCTTTCCCTACACGACGCTCTTCCGATCTNNNGAGATTGGGTTGTCACTGTGT
AGCACACGTCTGAACTCTTTCCCTACACGACGCTCTTCCGATCTNNNACATTACTTGGGAGATATGGC
AGCACACGTCTGAACTCTTTCCCTACACGACGCTCTTCCGATCTNNNACATTACTTGGGAGATATGGC
AGCACACGTCTGAACTCTTTCCCTACACGACGCTCTTCCGATCTNNNTCCTCAGGGTCCCCTTTATTCAT
AGCACACGTCTGAACTCTTTCCCTACACGACGCTCTTCCGATCTNNNTCCTCAGGGTCCCCTTTATTCAT
AGCACACGTCTGAACTCTTTCCCTACACGACGCTCTTCCGATCTNNNGCTAAGCTGCAAACTACATAGC
AGCACACGTCTGAACTCTTTCCCTACACGACGCTCTTCCGATCTNNNGCTAAGCTGCAAACTACATAGC
GAGCACACGTCTGAACTCTTTCCCTACACGACGCTCTTCCGATCTNNNTCCTCGTGAACATAGATGCAA
GAGCACACGTCTGAACTCTTTCCCTACACGACGCTCTTCCGATCTNNNTCCTCGTGAACATAGATGCAA
AGAGCACACGTCTGAACTCTTTCCCTACACGACGCTCTTCCGATCTNNNAATGCGACAATAACCAGAA
AGAGCACACGTCTGAACTCTTTCCCTACACGACGCTCTTCCGATCTNNNAATGCGACAATAACCAGAA
GCACACGTCTGAACTCTTTCCCTACACGACGCTCTTCCGATCTNNNTGAAAGGCTTTATCCTGATCAAA
GCACACGTCTGAACTCTTTCCCTACACGACGCTCTTCCGATCTNNNTGAAAGGCTTTATCCTGATCAAA
AGCACACGTCTGAACTCTTTCCCTACACGACGCTCTTCCGATCTNNNAAACAATCCTCTCACATCAGCCT
AGCACACGTCTGAACTCTTTCCCTACACGACGCTCTTCCGATCTNNNAAACAATCCTCTCACATCAGCCT
AGCACACGTCTGAACTCTTTCCCTACACGACGCTCTTCCGATCTNNNAGTCCCCAAAGTTCATTGTGTCT
AGCACACGTCTGAACTCTTTCCCTACACGACGCTCTTCCGATCTNNNAGTCCCCAAAGTTCATTGTGTCT
AGCACACGTCTGAACTCTTTCCCTACACGACGCTCTTCCGATCTNNNTCAAGTTTCAAAAAGCCGCTGA
AGCACACGTCTGAACTCTTTCCCTACACGACGCTCTTCCGATCTNNNTCAAGTTTCAAAAAGCCGCTGA
AGCACACGTCTGAACTCTTTCCCTACACGACGCTCTTCCGATCTNNNGTGATTGTACCACTGCCCTTCA
AGCACACGTCTGAACTCTTTCCCTACACGACGCTCTTCCGATCTNNNGTGATTGTACCACTGCCCTTCA
GAGCACACGTCTGAACTCTTTCCCTACACGACGCTCTTCCGATCTNNNATGCCTGTAATCCCAGCTACTT
GAGCACACGTCTGAACTCTTTCCCTACACGACGCTCTTCCGATCTNNNATGCCTGTAATCCCAGCTACTT
CACACGTCTGAACTCTTTCCCTACACGACGCTCTTCCGATCTNNNTCTCATTCCCTCTCTCCTTCTGTCAT
CACACGTCTGAACTCTTTCCCTACACGACGCTCTTCCGATCTNNNTCTCATTCCCTCTCTCCTTCTGTCAT
AGCACACGTCTGAACTCTTTCCCTACACGACGCTCTTCCGATCTNNNGAGGCTGAGGCAGGAGAATG
AGCACACGTCTGAACTCTTTCCCTACACGACGCTCTTCCGATCTNNNGAGGCTGAGGCAGGAGAATG
AGCACACGTCTGAACTCTTTCCCTACACGACGCTCTTCCGATCTNNNGAGTTTGAGGCTGTAGTGAGA
AGCACACGTCTGAACTCTTTCCCTACACGACGCTCTTCCGATCTNNNGAGTTTGAGGCTGTAGTGAGA
AGCACACGTCTGAACTCTTTCCCTACACGACGCTCTTCCGATCTNNNCCCAGTCTCAGGCAGTTCTTTA
AGCACACGTCTGAACTCTTTCCCTACACGACGCTCTTCCGATCTNNNCCCAGTCTCAGGCAGTTCTTTA
GCACACGTCTGAACTCTTTCCCTACACGACGCTCTTCCGATCTNNNGGCTGAATCCAAAGACAGGTTT
GCACACGTCTGAACTCTTTCCCTACACGACGCTCTTCCGATCTNNNGGCTGAATCCAAAGACAGGTTT
AGCACACGTCTGAACTCTTTCCCTACACGACGCTCTTCCGATCTNNNGAGCTTGCAGTGAGTCGAAAT
AGCACACGTCTGAACTCTTTCCCTACACGACGCTCTTCCGATCTNNNGAGCTTGCAGTGAGTCGAAAT

ACACGTCTGAACTCTTTCCCTACACGACGCTCTTCCGATCTNNNGGGCTTCCCTAAGAGAGCTCTCCAT
ACACGTCTGAACTCTTTCCCTACACGACGCTCTTCCGATCTNNNGGGCTTCCCTAAGAGAGCTCTCCAT
AGCACACGTCTGAACTCTTTCCCTACACGACGCTCTTCCGATCTNNNTGTGTCATTGTTGATTTCCTGG
AGCACACGTCTGAACTCTTTCCCTACACGACGCTCTTCCGATCTNNNTGTGTCATTGTTGATTTCCTGG
GCACACGTCTGAACTCTTTCCCTACACGACGCTCTTCCGATCTNNNCTGAGTAGCTGGGACTACAGGT
GCACACGTCTGAACTCTTTCCCTACACGACGCTCTTCCGATCTNNNCTGAGTAGCTGGGACTACAGGT
GCACACGTCTGAACTCTTTCCCTACACGACGCTCTTCCGATCTNNNCTTTGGACCTCTCCCATGTGTTA
GCACACGTCTGAACTCTTTCCCTACACGACGCTCTTCCGATCTNNNCTTTGGACCTCTCCCATGTGTTA
GCACACGTCTGAACTCTTTCCCTACACGACGCTCTTCCGATCTNNNGAGAGAGGTTGGGGGGTCAG
GCACACGTCTGAACTCTTTCCCTACACGACGCTCTTCCGATCTNNNGAGAGAGGTTGGGGGGTCAG
GCACACGTCTGAACTCTTTCCCTACACGACGCTCTTCCGATCTNNNTCTCTCTGGTTAGCTGTATCCCT
GCACACGTCTGAACTCTTTCCCTACACGACGCTCTTCCGATCTNNNTCTCTCTGGTTAGCTGTATCCCT
AGCACACGTCTGAACTCTTTCCCTACACGACGCTCTTCCGATCTNNNCCTGTCTCAAAAATGCACACAC
AGCACACGTCTGAACTCTTTCCCTACACGACGCTCTTCCGATCTNNNCCTGTCTCAAAAATGCACACAC
CACACGTCTGAACTCTTTCCCTACACGACGCTCTTCCGATCTNNNTGAGTTTCTTGCCAGGCTAGAAT
CACACGTCTGAACTCTTTCCCTACACGACGCTCTTCCGATCTNNNTGAGTTTCTTGCCAGGCTAGAAT
CACACGTCTGAACTCTTTCCCTACACGACGCTCTTCCGATCTNNNCAAGTCTTTCACCCCATTGTCACC
CACACGTCTGAACTCTTTCCCTACACGACGCTCTTCCGATCTNNNCAAGTCTTTCACCCCATTGTCACC
GAGCACACGTCTGAACTCTTTCCCTACACGACGCTCTTCCGATCTNNNTTTGAGAAAGGGTCCCACTTT
GAGCACACGTCTGAACTCTTTCCCTACACGACGCTCTTCCGATCTNNNTTTGAGAAAGGGTCCCACTTT
GAGCACACGTCTGAACTCTTTCCCTACACGACGCTCTTCCGATCTNNNGGAGGCAGAAATTTCAGTGA
GAGCACACGTCTGAACTCTTTCCCTACACGACGCTCTTCCGATCTNNNGGAGGCAGAAATTTCAGTGA
AGAGCACACGTCTGAACTCTTTCCCTACACGACGCTCTTCCGATCTNNNGGCAAGAGAGAGACAGAG
AGAGCACACGTCTGAACTCTTTCCCTACACGACGCTCTTCCGATCTNNNGGCAAGAGAGAGACAGAG
CACGTCTGAACTCTTTCCCTACACGACGCTCTTCCGATCTNNNGCTCCTTTCAAGAATTTTTGCCGCAT
CACGTCTGAACTCTTTCCCTACACGACGCTCTTCCGATCTNNNGCTCCTTTCAAGAATTTTTGCCGCAT
GAGCACACGTCTGAACTCTTTCCCTACACGACGCTCTTCCGATCTNNNTGACAAGAAAAAAATGGGA
GAGCACACGTCTGAACTCTTTCCCTACACGACGCTCTTCCGATCTNNNTGACAAGAAAAAAATGGGA
GAGCACACGTCTGAACTCTTTCCCTACACGACGCTCTTCCGATCTNNNCCCTCTTTAGTCGAATGGTGG
GAGCACACGTCTGAACTCTTTCCCTACACGACGCTCTTCCGATCTNNNCCCTCTTTAGTCGAATGGTGG
GAGCACACGTCTGAACTCTTTCCCTACACGACGCTCTTCCGATCTNNNTTGGGACACAGAAATAGAGC
GAGCACACGTCTGAACTCTTTCCCTACACGACGCTCTTCCGATCTNNNTTGGGACACAGAAATAGAGC
AGCACACGTCTGAACTCTTTCCCTACACGACGCTCTTCCGATCTNNNACCAATGAGGCAGTTTAGACC
AGCACACGTCTGAACTCTTTCCCTACACGACGCTCTTCCGATCTNNNACCAATGAGGCAGTTTAGACC
AGCACACGTCTGAACTCTTTCCCTACACGACGCTCTTCCGATCTNNNAACTAGATCGGAAGCTGAAGT
AGCACACGTCTGAACTCTTTCCCTACACGACGCTCTTCCGATCTNNNAACTAGATCGGAAGCTGAAGT
GAGCACACGTCTGAACTCTTTCCCTACACGACGCTCTTCCGATCTNNNCTCCTGAGTAGCTGGGATTA
GAGCACACGTCTGAACTCTTTCCCTACACGACGCTCTTCCGATCTNNNCTCCTGAGTAGCTGGGATTA
GCACACGTCTGAACTCTTTCCCTACACGACGCTCTTCCGATCTNNNGACATTAAAAGACCTACCAATTG
GCACACGTCTGAACTCTTTCCCTACACGACGCTCTTCCGATCTNNNGACATTAAAAGACCTACCAATTG
GAGCACACGTCTGAACTCTTTCCCTACACGACGCTCTTCCGATCTNNNACCATCAGCAGACATTTCCATC
GAGCACACGTCTGAACTCTTTCCCTACACGACGCTCTTCCGATCTNNNACCATCAGCAGACATTTCCATC
GAGCACACGTCTGAACTCTTTCCCTACACGACGCTCTTCCGATCTNNNTTGAGGCACGAGAATCACTT

AGAGCACACGTCTGAACTCTTTCCCTACACGACGCTCTTCCGATCTNNNTTGAGGCACGAGAATCACTT

GCACACGTCTGAACTCTTTCCCTACACGACGCTCTTCCGATCTNNNGGAATCTGCCCTGTGTCGTCCAT

GCACACGTCTGAACTCTTTCCCTACACGACGCTCTTCCGATCTNNNGGAATCTGCCCTGTGTCGTCCAT

GAGCACACGTCTGAACTCTTTCCCTACACGACGCTCTTCCGATCTNNNGGCCTCCCAAAGTACTGGAAT

GAGCACACGTCTGAACTCTTTCCCTACACGACGCTCTTCCGATCTNNNGGCCTCCCAAAGTACTGGAAT

AGCACACGTCTGAACTCTTTCCCTACACGACGCTCTTCCGATCTNNNATGGGCATTTCCTTTCCCTAACT

AGCACACGTCTGAACTCTTTCCCTACACGACGCTCTTCCGATCTNNNATGGGCATTTCCTTTCCCTAACT

AGCACACGTCTGAACTCTTTCCCTACACGACGCTCTTCCGATCTNNNCTAGAAGAAAACTGAGTGCGT

AGCACACGTCTGAACTCTTTCCCTACACGACGCTCTTCCGATCTNNNCTAGAAGAAAACTGAGTGCGT

AGCACACGTCTGAACTCTTTCCCTACACGACGCTCTTCCGATCTNNNCAGCTGCTAGGGAGATTGAGA

AGCACACGTCTGAACTCTTTCCCTACACGACGCTCTTCCGATCTNNNCAGCTGCTAGGGAGATTGAGA

AGCACACGTCTGAACTCTTTCCCTACACGACGCTCTTCCGATCTNNNGGTTCATTCCTGTAATCCCAGC

AGCACACGTCTGAACTCTTTCCCTACACGACGCTCTTCCGATCTNNNGGTTCATTCCTGTAATCCCAGC

AGCACACGTCTGAACTCTTTCCCTACACGACGCTCTTCCGATCTNNNACATGCTTATAAAAATGTCCAT

AGCACACGTCTGAACTCTTTCCCTACACGACGCTCTTCCGATCTNNNACATGCTTATAAAAATGTCCAT

AGAGCACACGTCTGAACTCTTTCCCTACACGACGCTCTTCCGATCTNNNAAAAAATTAGCCAGGTGTG

AGAGCACACGTCTGAACTCTTTCCCTACACGACGCTCTTCCGATCTNNNAAAAAATTAGCCAGGTGTG

AGCACACGTCTGAACTCTTTCCCTACACGACGCTCTTCCGATCTNNNCATTGTAAACCTTAGGCTGCAG

AGCACACGTCTGAACTCTTTCCCTACACGACGCTCTTCCGATCTNNNCATTGTAAACCTTAGGCTGCAG

GAGCACACGTCTGAACTCTTTCCCTACACGACGCTCTTCCGATCTNNNCCATCTTTCATCCTACTGCCCAT

GAGCACACGTCTGAACTCTTTCCCTACACGACGCTCTTCCGATCTNNNCCATCTTTCATCCTACTGCCCAT

AGCACACGTCTGAACTCTTTCCCTACACGACGCTCTTCCGATCTNNNTACAAAAAAGGCTCCAAGCACT

AGCACACGTCTGAACTCTTTCCCTACACGACGCTCTTCCGATCTNNNTACAAAAAAGGCTCCAAGCACT

AGCACACGTCTGAACTCTTTCCCTACACGACGCTCTTCCGATCTNNNTTGGTTTCTTCTGAGGCCTCTTT

AGCACACGTCTGAACTCTTTCCCTACACGACGCTCTTCCGATCTNNNTTGGTTTCTTCTGAGGCCTCTTT

GCACACGTCTGAACTCTTTCCCTACACGACGCTCTTCCGATCTNNNAAAAACTCACTCCATGCTCACCTC

GCACACGTCTGAACTCTTTCCCTACACGACGCTCTTCCGATCTNNNAAAAACTCACTCCATGCTCACCTC

GAGCACACGTCTGAACTCTTTCCCTACACGACGCTCTTCCGATCTNNNGTGCCTGTAATCCCAGCTACTT

GAGCACACGTCTGAACTCTTTCCCTACACGACGCTCTTCCGATCTNNNGTGCCTGTAATCCCAGCTACTT

GCACACGTCTGAACTCTTTCCCTACACGACGCTCTTCCGATCTNNNTTGAGGAGGGGAGAAGTTACTC

GCACACGTCTGAACTCTTTCCCTACACGACGCTCTTCCGATCTNNNTTGAGGAGGGGAGAAGTTACTC

AGCACACGTCTGAACTCTTTCCCTACACGACGCTCTTCCGATCTNNNCCTTCTAACCTCTTCAGCCCAAT

AGCACACGTCTGAACTCTTTCCCTACACGACGCTCTTCCGATCTNNNCCTTCTAACCTCTTCAGCCCAAT

GAGCACACGTCTGAACTCTTTCCCTACACGACGCTCTTCCGATCTNNNAGCACTTTGTAAGCCTCTACA

GAGCACACGTCTGAACTCTTTCCCTACACGACGCTCTTCCGATCTNNNAGCACTTTGTAAGCCTCTACA

AGAGCACACGTCTGAACTCTTTCCCTACACGACGCTCTTCCGATCTNNNAGGTTTTCCGGCCTTACATTA

AGAGCACACGTCTGAACTCTTTCCCTACACGACGCTCTTCCGATCTNNNAGGTTTTCCGGCCTTACATTA

AGCACACGTCTGAACTCTTTCCCTACACGACGCTCTTCCGATCTNNNTAGCTCAGTTAGGTCCAAGCAA

AGCACACGTCTGAACTCTTTCCCTACACGACGCTCTTCCGATCTNNNTAGCTCAGTTAGGTCCAAGCAA

GCACACGTCTGAACTCTTTCCCTACACGACGCTCTTCCGATCTNNNATATTCCCTTTGTCGTGTCTGGTC

GCACACGTCTGAACTCTTTCCCTACACGACGCTCTTCCGATCTNNNATATTCCCTTTGTCGTGTCTGGTC

GAGCACACGTCTGAACTCTTTCCCTACACGACGCTCTTCCGATCTNNNATAGTCCTGGTGTCATGTTGA

GAGCACACGTCTGAACTCTTTCCCTACACGACGCTCTTCCGATCTNNNATAGTCCTGGTGTCATGTTGA

GAGCACACGTCTGAACTCTTTCCCTACACGACGCTCTTCCGATCTNNNCTTGGTGGTGGCAATGAGAG
GAGCACACGTCTGAACTCTTTCCCTACACGACGCTCTTCCGATCTNNNCTTGGTGGTGGCAATGAGAG
AGCACACGTCTGAACTCTTTCCCTACACGACGCTCTTCCGATCTNNNGAGTCTTGCTCTGTTGCCCAGC
AGCACACGTCTGAACTCTTTCCCTACACGACGCTCTTCCGATCTNNNGAGTCTTGCTCTGTTGCCCAGC
ACACGTCTGAACTCTTTCCCTACACGACGCTCTTCCGATCTNNNGACAGCCGGACCAGGCATACTGCAT
ACACGTCTGAACTCTTTCCCTACACGACGCTCTTCCGATCTNNNGACAGCCGGACCAGGCATACTGCAT
AGCACACGTCTGAACTCTTTCCCTACACGACGCTCTTCCGATCTNNNGACTTGGTGTGTAGGTTCCTTC
AGCACACGTCTGAACTCTTTCCCTACACGACGCTCTTCCGATCTNNNGACTTGGTGTGTAGGTTCCTTC
AGCACACGTCTGAACTCTTTCCCTACACGACGCTCTTCCGATCTNNNCAAGTAGCTGGGACTACAGGC
AGCACACGTCTGAACTCTTTCCCTACACGACGCTCTTCCGATCTNNNCAAGTAGCTGGGACTACAGGC
GAGCACACGTCTGAACTCTTTCCCTACACGACGCTCTTCCGATCTNNNATTGGTGGTGCATGGCTGTAG
GAGCACACGTCTGAACTCTTTCCCTACACGACGCTCTTCCGATCTNNNATTGGTGGTGCATGGCTGTAG
AGAGCACACGTCTGAACTCTTTCCCTACACGACGCTCTTCCGATCTNNNGTAATCCCAGCTACTTAGGA
AGAGCACACGTCTGAACTCTTTCCCTACACGACGCTCTTCCGATCTNNNGTAATCCCAGCTACTTAGGA
GAGCACACGTCTGAACTCTTTCCCTACACGACGCTCTTCCGATCTNNNACAGACTAAGACGAGGGTGT
GAGCACACGTCTGAACTCTTTCCCTACACGACGCTCTTCCGATCTNNNACAGACTAAGACGAGGGTGT
AGCACACGTCTGAACTCTTTCCCTACACGACGCTCTTCCGATCTNNNCAGAAGAATTGCCCAAACCCAG
AGCACACGTCTGAACTCTTTCCCTACACGACGCTCTTCCGATCTNNNCAGAAGAATTGCCCAAACCCAG
GAGCACACGTCTGAACTCTTTCCCTACACGACGCTCTTCCGATCTNNNTACTCAGCCTGTTTTCTAAGC
GAGCACACGTCTGAACTCTTTCCCTACACGACGCTCTTCCGATCTNNNTACTCAGCCTGTTTTCTAAGC
GAGCACACGTCTGAACTCTTTCCCTACACGACGCTCTTCCGATCTNNNGGCAAGACTTTATCATGACAG
GAGCACACGTCTGAACTCTTTCCCTACACGACGCTCTTCCGATCTNNNGGCAAGACTTTATCATGACAG
AGCACACGTCTGAACTCTTTCCCTACACGACGCTCTTCCGATCTNNNTTTAGTAGAGACAGGGTTTCG
AGCACACGTCTGAACTCTTTCCCTACACGACGCTCTTCCGATCTNNNTTTAGTAGAGACAGGGTTTCG
GAGCACACGTCTGAACTCTTTCCCTACACGACGCTCTTCCGATCTNNNCATGCCTGTAATCCCAGCTAC
GAGCACACGTCTGAACTCTTTCCCTACACGACGCTCTTCCGATCTNNNCATGCCTGTAATCCCAGCTAC
AGCACACGTCTGAACTCTTTCCCTACACGACGCTCTTCCGATCTNNNTACAACTGCCCTCTGAGTTCAA
AGCACACGTCTGAACTCTTTCCCTACACGACGCTCTTCCGATCTNNNTACAACTGCCCTCTGAGTTCAA
AGCACACGTCTGAACTCTTTCCCTACACGACGCTCTTCCGATCTNNNCTCCAGTTCTGCCGGCTTCTCAT
AGCACACGTCTGAACTCTTTCCCTACACGACGCTCTTCCGATCTNNNCTCCAGTTCTGCCGGCTTCTCAT
AGAGCACACGTCTGAACTCTTTCCCTACACGACGCTCTTCCGATCTNNNGAGAACAAAAAACCAAACAC
AGAGCACACGTCTGAACTCTTTCCCTACACGACGCTCTTCCGATCTNNNGAGAACAAAAAACCAAACAC
AGAGCACACGTCTGAACTCTTTCCCTACACGACGCTCTTCCGATCTNNNCAGTTGCCATTGAGTAGCAA
AGAGCACACGTCTGAACTCTTTCCCTACACGACGCTCTTCCGATCTNNNCAGTTGCCATTGAGTAGCAA
AGCACACGTCTGAACTCTTTCCCTACACGACGCTCTTCCGATCTNNNAAGAATCTCTTGAACCCAGGAG
AGCACACGTCTGAACTCTTTCCCTACACGACGCTCTTCCGATCTNNNAAGAATCTCTTGAACCCAGGAG
AGCACACGTCTGAACTCTTTCCCTACACGACGCTCTTCCGATCTNNNGTGGGTACTTTAGCAACCTCAG
AGCACACGTCTGAACTCTTTCCCTACACGACGCTCTTCCGATCTNNNGTGGGTACTTTAGCAACCTCAG
GAGCACACGTCTGAACTCTTTCCCTACACGACGCTCTTCCGATCTNNNTCCTTCCTGTGCGGAAATAAG
GAGCACACGTCTGAACTCTTTCCCTACACGACGCTCTTCCGATCTNNNTCCTTCCTGTGCGGAAATAAG
AGAGCACACGTCTGAACTCTTTCCCTACACGACGCTCTTCCGATCTNNNAGAATTGCTTGAATCTGGGA
AGAGCACACGTCTGAACTCTTTCCCTACACGACGCTCTTCCGATCTNNNAGAATTGCTTGAATCTGGGA
AGCACACGTCTGAACTCTTTCCCTACACGACGCTCTTCCGATCTNNNAATGGTAGTTTGTATTTCTGTG

AGCACACGTCTGAACTCTTTCCCTACACGACGCTCTTCCGATCTNNNAATGGTAGTTTGTATTTCTGTG
AGCACACGTCTGAACTCTTTCCCTACACGACGCTCTTCCGATCTNNNCCAAAGTACTGGGATTACAGG
AGCACACGTCTGAACTCTTTCCCTACACGACGCTCTTCCGATCTNNNCCAAAGTACTGGGATTACAGG
GCACACGTCTGAACTCTTTCCCTACACGACGCTCTTCCGATCTNNNCTCCAGAGTCCTCAAGTTCCAGT
GCACACGTCTGAACTCTTTCCCTACACGACGCTCTTCCGATCTNNNCTCCAGAGTCCTCAAGTTCCAGT
AGAGCACACGTCTGAACTCTTTCCCTACACGACGCTCTTCCGATCTNNNGATGAGCATTGCTGGAAGA
AGAGCACACGTCTGAACTCTTTCCCTACACGACGCTCTTCCGATCTNNNGATGAGCATTGCTGGAAGA
CACGTCTGAACTCTTTCCCTACACGACGCTCTTCCGATCTNNNCATCCAATGGAGAGAGGACAACGCA
CACGTCTGAACTCTTTCCCTACACGACGCTCTTCCGATCTNNNCATCCAATGGAGAGAGGACAACGCA
AGCACACGTCTGAACTCTTTCCCTACACGACGCTCTTCCGATCTNNNCCACACTGTCTCTCTCTCTCTC
AGCACACGTCTGAACTCTTTCCCTACACGACGCTCTTCCGATCTNNNCCACACTGTCTCTCTCTCTCTC
AGCACACGTCTGAACTCTTTCCCTACACGACGCTCTTCCGATCTNNNGATTCTAAAAAAGAGGCCGGG
AGCACACGTCTGAACTCTTTCCCTACACGACGCTCTTCCGATCTNNNGATTCTAAAAAAGAGGCCGGG
AGCACACGTCTGAACTCTTTCCCTACACGACGCTCTTCCGATCTNNNCTCCACCTCTCTAAGTACTGGG
AGCACACGTCTGAACTCTTTCCCTACACGACGCTCTTCCGATCTNNNCTCCACCTCTCTAAGTACTGGG
GAGCACACGTCTGAACTCTTTCCCTACACGACGCTCTTCCGATCTNNNCCACACCCAGTAATGTTTGCA
GAGCACACGTCTGAACTCTTTCCCTACACGACGCTCTTCCGATCTNNNCCACACCCAGTAATGTTTGCA
AGAGCACACGTCTGAACTCTTTCCCTACACGACGCTCTTCCGATCTNNNGAGAATCACTTGAACCGGG
AGAGCACACGTCTGAACTCTTTCCCTACACGACGCTCTTCCGATCTNNNGAGAATCACTTGAACCGGG
GAGCACACGTCTGAACTCTTTCCCTACACGACGCTCTTCCGATCTNNNCCACCAGTTCCATTCTTCCCTAT
GAGCACACGTCTGAACTCTTTCCCTACACGACGCTCTTCCGATCTNNNCCACCAGTTCCATTCTTCCCTAT
GAGCACACGTCTGAACTCTTTCCCTACACGACGCTCTTCCGATCTNNNCACCTGTAATCCCAGCTACTC
GAGCACACGTCTGAACTCTTTCCCTACACGACGCTCTTCCGATCTNNNCACCTGTAATCCCAGCTACTC
CACACGTCTGAACTCTTTCCCTACACGACGCTCTTCCGATCTNNNCTAACTATGGACCCGCTGTAGACC
CACACGTCTGAACTCTTTCCCTACACGACGCTCTTCCGATCTNNNCTAACTATGGACCCGCTGTAGACC
AGCACACGTCTGAACTCTTTCCCTACACGACGCTCTTCCGATCTNNNGAGAATTGCTTGAACCCAGGA
AGCACACGTCTGAACTCTTTCCCTACACGACGCTCTTCCGATCTNNNGAGAATTGCTTGAACCCAGGA
GAGCACACGTCTGAACTCTTTCCCTACACGACGCTCTTCCGATCTNNNGAAAACAAAATGCTGAAGAA
GAGCACACGTCTGAACTCTTTCCCTACACGACGCTCTTCCGATCTNNNGAAAACAAAATGCTGAAGAA
AGCACACGTCTGAACTCTTTCCCTACACGACGCTCTTCCGATCTNNNATCTCCTTTGCTTCCCCAAGGC
AGCACACGTCTGAACTCTTTCCCTACACGACGCTCTTCCGATCTNNNATCTCCTTTGCTTCCCCAAGGC
AGCACACGTCTGAACTCTTTCCCTACACGACGCTCTTCCGATCTNNNGAGGCACGAGAATTACTTGAA
AGCACACGTCTGAACTCTTTCCCTACACGACGCTCTTCCGATCTNNNGAGGCACGAGAATTACTTGAA
GCACACGTCTGAACTCTTTCCCTACACGACGCTCTTCCGATCTNNNACCTTCCCTCCACTATTGTCCTAC
GCACACGTCTGAACTCTTTCCCTACACGACGCTCTTCCGATCTNNNACCTTCCCTCCACTATTGTCCTAC
ACACGTCTGAACTCTTTCCCTACACGACGCTCTTCCGATCTNNNTCTTCTATTCCCAGCTTTTTGAGGAC
ACACGTCTGAACTCTTTCCCTACACGACGCTCTTCCGATCTNNNTCTTCTATTCCCAGCTTTTTGAGGAC
GAGCACACGTCTGAACTCTTTCCCTACACGACGCTCTTCCGATCTNNNATCAGAGCCAGTCTCACCTTC
GAGCACACGTCTGAACTCTTTCCCTACACGACGCTCTTCCGATCTNNNATCAGAGCCAGTCTCACCTTC
GAGCACACGTCTGAACTCTTTCCCTACACGACGCTCTTCCGATCTNNNGTGTTAGCCTGAAAGAGCCT
GAGCACACGTCTGAACTCTTTCCCTACACGACGCTCTTCCGATCTNNNGTGTTAGCCTGAAAGAGCCT
GAGCACACGTCTGAACTCTTTCCCTACACGACGCTCTTCCGATCTNNNCTGACATTCCTTTCACCCAGA
GAGCACACGTCTGAACTCTTTCCCTACACGACGCTCTTCCGATCTNNNCTGACATTCCTTTCACCCAGA

GAGCACACGTCTGAACTCTTTCCCTACACGACGCTCTTCCGATCTNNNTCAGAACCTCACACTTACACA
GAGCACACGTCTGAACTCTTTCCCTACACGACGCTCTTCCGATCTNNNTCAGAACCTCACACTTACACA
GAGCACACGTCTGAACTCTTTCCCTACACGACGCTCTTCCGATCTNNNCCATGAGAACAGTATGGGGG
GAGCACACGTCTGAACTCTTTCCCTACACGACGCTCTTCCGATCTNNNCCATGAGAACAGTATGGGGG
AGCACACGTCTGAACTCTTTCCCTACACGACGCTCTTCCGATCTNNNAGGGCCCAGCTAGAAAAACAA
AGCACACGTCTGAACTCTTTCCCTACACGACGCTCTTCCGATCTNNNAGGGCCCAGCTAGAAAAACAA
AGCACACGTCTGAACTCTTTCCCTACACGACGCTCTTCCGATCTNNNGGCAGGAGAATCACTTGAACC
AGCACACGTCTGAACTCTTTCCCTACACGACGCTCTTCCGATCTNNNGGCAGGAGAATCACTTGAACC
AGCACACGTCTGAACTCTTTCCCTACACGACGCTCTTCCGATCTNNNCAGATCACACTACTGCACTCCA
AGCACACGTCTGAACTCTTTCCCTACACGACGCTCTTCCGATCTNNNCAGATCACACTACTGCACTCCA
AGCACACGTCTGAACTCTTTCCCTACACGACGCTCTTCCGATCTNNNGGCAACATAGCAAGACCTCTCT
AGCACACGTCTGAACTCTTTCCCTACACGACGCTCTTCCGATCTNNNGGCAACATAGCAAGACCTCTCT
AGCACACGTCTGAACTCTTTCCCTACACGACGCTCTTCCGATCTNNNATATTGCCCATGCTGGTGTCAA
AGCACACGTCTGAACTCTTTCCCTACACGACGCTCTTCCGATCTNNNATATTGCCCATGCTGGTGTCAA
AGCACACGTCTGAACTCTTTCCCTACACGACGCTCTTCCGATCTNNNACACATGTATACCAGCAAGGG
AGCACACGTCTGAACTCTTTCCCTACACGACGCTCTTCCGATCTNNNACACATGTATACCAGCAAGGG
AGCACACGTCTGAACTCTTTCCCTACACGACGCTCTTCCGATCTNNNATCCACGTGCACATATCTACACA
AGCACACGTCTGAACTCTTTCCCTACACGACGCTCTTCCGATCTNNNATCCACGTGCACATATCTACACA
AGCACACGTCTGAACTCTTTCCCTACACGACGCTCTTCCGATCTNNNGATCTCAAGCAGTCAAGAAAG
AGCACACGTCTGAACTCTTTCCCTACACGACGCTCTTCCGATCTNNNGATCTCAAGCAGTCAAGAAAG
AGCACACGTCTGAACTCTTTCCCTACACGACGCTCTTCCGATCTNNNTGTCCATGAGGTATCCAAACA
AGCACACGTCTGAACTCTTTCCCTACACGACGCTCTTCCGATCTNNNTGTCCATGAGGTATCCAAACA
GCACACGTCTGAACTCTTTCCCTACACGACGCTCTTCCGATCTNNNATGGGACCTGATCTGCATTTCTT
GCACACGTCTGAACTCTTTCCCTACACGACGCTCTTCCGATCTNNNATGGGACCTGATCTGCATTTCTT
GCACACGTCTGAACTCTTTCCCTACACGACGCTCTTCCGATCTNNNTCCTTCAAATGTCCCCATAAGCT
GCACACGTCTGAACTCTTTCCCTACACGACGCTCTTCCGATCTNNNTCCTTCAAATGTCCCCATAAGCT
AAGAGCACACGTCTGAACTCTTTCCCTACACGACGCTCTTCCGATCTNNNGGAAGATTACTTGAGCCT
AAGAGCACACGTCTGAACTCTTTCCCTACACGACGCTCTTCCGATCTNNNGGAAGATTACTTGAGCCT
AGCACACGTCTGAACTCTTTCCCTACACGACGCTCTTCCGATCTNNNGCCAAGATTCCCACTTGCGATA
AGCACACGTCTGAACTCTTTCCCTACACGACGCTCTTCCGATCTNNNGCCAAGATTCCCACTTGCGATA
AGCACACGTCTGAACTCTTTCCCTACACGACGCTCTTCCGATCTNNNCTGTAGTTCCAGTTACTCGGGA
AGCACACGTCTGAACTCTTTCCCTACACGACGCTCTTCCGATCTNNNCTGTAGTTCCAGTTACTCGGGA
GAGCACACGTCTGAACTCTTTCCCTACACGACGCTCTTCCGATCTNNNGTTGTTCCAAAGATCAGTTCC
GAGCACACGTCTGAACTCTTTCCCTACACGACGCTCTTCCGATCTNNNGTTGTTCCAAAGATCAGTTCC
AGCACACGTCTGAACTCTTTCCCTACACGACGCTCTTCCGATCTNNNCTGATTGCAGCTGGAGTAATT
AGCACACGTCTGAACTCTTTCCCTACACGACGCTCTTCCGATCTNNNCTGATTGCAGCTGGAGTAATT
AGCACACGTCTGAACTCTTTCCCTACACGACGCTCTTCCGATCTNNNACACCTGTAATCCCAGCTACAT
AGCACACGTCTGAACTCTTTCCCTACACGACGCTCTTCCGATCTNNNACACCTGTAATCCCAGCTACAT
AGCACACGTCTGAACTCTTTCCCTACACGACGCTCTTCCGATCTNNNGAGGCACGAGAATTACTTGAA
AGCACACGTCTGAACTCTTTCCCTACACGACGCTCTTCCGATCTNNNGAGGCACGAGAATTACTTGAA
AGCACACGTCTGAACTCTTTCCCTACACGACGCTCTTCCGATCTNNNAAAAGGGCACTTACACGATGTT
AGCACACGTCTGAACTCTTTCCCTACACGACGCTCTTCCGATCTNNNAAAAGGGCACTTACACGATGTT
GAGCACACGTCTGAACTCTTTCCCTACACGACGCTCTTCCGATCTNNNCATGAAATTGAATGTGGGGT

GAGCACACGTCTGAACTCTTTCCCTACACGACGCTCTTCCGATCTNNNCATGAAATTGAATGTGGGGT

GAGCACACGTCTGAACTCTTTCCCTACACGACGCTCTTCCGATCTNNNAATTAGCTGAATTTGGTGGCA

GAGCACACGTCTGAACTCTTTCCCTACACGACGCTCTTCCGATCTNNNAATTAGCTGAATTTGGTGGCA

AGCACACGTCTGAACTCTTTCCCTACACGACGCTCTTCCGATCTNNNTCTCAGGAAGGCAAAGTTTC

AGCACACGTCTGAACTCTTTCCCTACACGACGCTCTTCCGATCTNNNTCTCAGGAAGGCAAAGTTTC

GAGCACACGTCTGAACTCTTTCCCTACACGACGCTCTTCCGATCTNNNTTCAATGAGCTGAGATCACAC

GAGCACACGTCTGAACTCTTTCCCTACACGACGCTCTTCCGATCTNNNTTCAATGAGCTGAGATCACAC

AGCACACGTCTGAACTCTTTCCCTACACGACGCTCTTCCGATCTNNNCAATCTTGGCTTACTGCAACCTC

AGCACACGTCTGAACTCTTTCCCTACACGACGCTCTTCCGATCTNNNCAATCTTGGCTTACTGCAACCTC

AGCACACGTCTGAACTCTTTCCCTACACGACGCTCTTCCGATCTNNNGGATCACCTGAGGTCAGAAGT

AGCACACGTCTGAACTCTTTCCCTACACGACGCTCTTCCGATCTNNNGGATCACCTGAGGTCAGAAGT

CACACGTCTGAACTCTTTCCCTACACGACGCTCTTCCGATCTNNNTACTTTTTCTTTGCATTTGTTCCAT

CACACGTCTGAACTCTTTCCCTACACGACGCTCTTCCGATCTNNNTACTTTTTCTTTGCATTTGTTCCAT

AGCACACGTCTGAACTCTTTCCCTACACGACGCTCTTCCGATCTNNNCCTACTCCTCAGCCCTTGGGCAT

AGCACACGTCTGAACTCTTTCCCTACACGACGCTCTTCCGATCTNNNCCTACTCCTCAGCCCTTGGGCAT

GAGCACACGTCTGAACTCTTTCCCTACACGACGCTCTTCCGATCTNNNATGCCTTCTATCACAGTTGTGG

GAGCACACGTCTGAACTCTTTCCCTACACGACGCTCTTCCGATCTNNNATGCCTTCTATCACAGTTGTGG

AGCACACGTCTGAACTCTTTCCCTACACGACGCTCTTCCGATCTNNNGAGCTTACAGTGAGCCGAGAT

AGCACACGTCTGAACTCTTTCCCTACACGACGCTCTTCCGATCTNNNGAGCTTACAGTGAGCCGAGAT

AGCACACGTCTGAACTCTTTCCCTACACGACGCTCTTCCGATCTNNNCACAGAATTAGGTGGGGAAAG

AGCACACGTCTGAACTCTTTCCCTACACGACGCTCTTCCGATCTNNNCACAGAATTAGGTGGGGAAAG

GAGCACACGTCTGAACTCTTTCCCTACACGACGCTCTTCCGATCTNNNTAGAGGTTGCAGTCAGCTGAA

GAGCACACGTCTGAACTCTTTCCCTACACGACGCTCTTCCGATCTNNNTAGAGGTTGCAGTCAGCTGAA

AGAGCACACGTCTGAACTCTTTCCCTACACGACGCTCTTCCGATCTNNNGCCGCACATATTGTATCCTTT

AGAGCACACGTCTGAACTCTTTCCCTACACGACGCTCTTCCGATCTNNNGCCGCACATATTGTATCCTTT

AGCACACGTCTGAACTCTTTCCCTACACGACGCTCTTCCGATCTNNNCGAACTCCTAACCTCAGGTGATC

AGCACACGTCTGAACTCTTTCCCTACACGACGCTCTTCCGATCTNNNCGAACTCCTAACCTCAGGTGATC

GAGCACACGTCTGAACTCTTTCCCTACACGACGCTCTTCCGATCTNNNCACCAGCTCAAAGTATCCCCTA

GAGCACACGTCTGAACTCTTTCCCTACACGACGCTCTTCCGATCTNNNCACCAGCTCAAAGTATCCCCTA

GAGCACACGTCTGAACTCTTTCCCTACACGACGCTCTTCCGATCTNNNGAACCACTGACCTAGACTAAG

GAGCACACGTCTGAACTCTTTCCCTACACGACGCTCTTCCGATCTNNNGAACCACTGACCTAGACTAAG

GAGCACACGTCTGAACTCTTTCCCTACACGACGCTCTTCCGATCTNNNACTCAGCACTTGTAGAACCATG

GAGCACACGTCTGAACTCTTTCCCTACACGACGCTCTTCCGATCTNNNACTCAGCACTTGTAGAACCATG

GAGCACACGTCTGAACTCTTTCCCTACACGACGCTCTTCCGATCTNNNAGATGGGGTTTCTTCATGTTG

GAGCACACGTCTGAACTCTTTCCCTACACGACGCTCTTCCGATCTNNNAGATGGGGTTTCTTCATGTTG

AGAGCACACGTCTGAACTCTTTCCCTACACGACGCTCTTCCGATCTNNNCCCCTAAAATGTTGACAGGG

AGAGCACACGTCTGAACTCTTTCCCTACACGACGCTCTTCCGATCTNNNCCCCTAAAATGTTGACAGGG

AGCACACGTCTGAACTCTTTCCCTACACGACGCTCTTCCGATCTNNNGGGCAAGTTACTTCATTTCTGG

AGCACACGTCTGAACTCTTTCCCTACACGACGCTCTTCCGATCTNNNGGGCAAGTTACTTCATTTCTGG

GAGCACACGTCTGAACTCTTTCCCTACACGACGCTCTTCCGATCTNNNATGTGGTGAGCCTCTCTTTGAT

GAGCACACGTCTGAACTCTTTCCCTACACGACGCTCTTCCGATCTNNNATGTGGTGAGCCTCTCTTTGAT

GAGCACACGTCTGAACTCTTTCCCTACACGACGCTCTTCCGATCTNNNCTCCTACTTGTGTTGCTCTG

GAGCACACGTCTGAACTCTTTCCCTACACGACGCTCTTCCGATCTNNNCTCCTACTTGTGTTGCTCTG

AGCACACGTCTGAACTCTTTCCCTACACGACGCTCTTCCGATCTNNNTGAGTCAAGATCCTTCCATTG
AGCACACGTCTGAACTCTTTCCCTACACGACGCTCTTCCGATCTNNNTGAGTCAAGATCCTTCCATTG
GAGCACACGTCTGAACTCTTTCCCTACACGACGCTCTTCCGATCTNNNGAATGAATCTGAGTTTATCT
GAGCACACGTCTGAACTCTTTCCCTACACGACGCTCTTCCGATCTNNNGAATGAATCTGAGTTTATCT
ACACGTCTGAACTCTTTCCCTACACGACGCTCTTCCGATCTNNNGAAATAGGGGAGGCTTGAGTCTTC
ACACGTCTGAACTCTTTCCCTACACGACGCTCTTCCGATCTNNNGAAATAGGGGAGGCTTGAGTCTTC
CACACGTCTGAACTCTTTCCCTACACGACGCTCTTCCGATCTNNNTTTGACAAAACATCCTGGGCTG
CACACGTCTGAACTCTTTCCCTACACGACGCTCTTCCGATCTNNNTTTGACAAAACATCCTGGGCTG
GAGCACACGTCTGAACTCTTTCCCTACACGACGCTCTTCCGATCTNNNCACCTGTAATCCCTGCTACTC
GAGCACACGTCTGAACTCTTTCCCTACACGACGCTCTTCCGATCTNNNCACCTGTAATCCCTGCTACTC
AGCACACGTCTGAACTCTTTCCCTACACGACGCTCTTCCGATCTNNNGGATTACCAGTATGAGCCACC
AGCACACGTCTGAACTCTTTCCCTACACGACGCTCTTCCGATCTNNNGGATTACCAGTATGAGCCACC
AGCACACGTCTGAACTCTTTCCCTACACGACGCTCTTCCGATCTNNNGTGAGTGCTGAGATCATGCTA
AGCACACGTCTGAACTCTTTCCCTACACGACGCTCTTCCGATCTNNNGTGAGTGCTGAGATCATGCTA
GAGCACACGTCTGAACTCTTTCCCTACACGACGCTCTTCCGATCTNNNGATGAGGTGGTAGCTGCTT
GAGCACACGTCTGAACTCTTTCCCTACACGACGCTCTTCCGATCTNNNGATGAGGTGGTAGCTGCTT
AGCACACGTCTGAACTCTTTCCCTACACGACGCTCTTCCGATCTNNNGATAGTTTGAGCCCAGGAGT
AGCACACGTCTGAACTCTTTCCCTACACGACGCTCTTCCGATCTNNNGATAGTTTGAGCCCAGGAGT
AGCACACGTCTGAACTCTTTCCCTACACGACGCTCTTCCGATCTNNNACCTTCCCTCCACTATTGTCCTA
AGCACACGTCTGAACTCTTTCCCTACACGACGCTCTTCCGATCTNNNACCTTCCCTCCACTATTGTCCTA
AGCACACGTCTGAACTCTTTCCCTACACGACGCTCTTCCGATCTNNNTGTGCTGTAAAGCAAGTTCCA
AGCACACGTCTGAACTCTTTCCCTACACGACGCTCTTCCGATCTNNNTGTGCTGTAAAGCAAGTTCCA
CACACGTCTGAACTCTTTCCCTACACGACGCTCTTCCGATCTNNNTCAAAACACAACTTCTCCTGGAAC
CACACGTCTGAACTCTTTCCCTACACGACGCTCTTCCGATCTNNNTCAAAACACAACTTCTCCTGGAAC
GCACACGTCTGAACTCTTTCCCTACACGACGCTCTTCCGATCTNNNCTACCCAATCCCTCACTGTTCTCC
GCACACGTCTGAACTCTTTCCCTACACGACGCTCTTCCGATCTNNNCTACCCAATCCCTCACTGTTCTCC
GAGCACACGTCTGAACTCTTTCCCTACACGACGCTCTTCCGATCTNNNCTTTCCTGATTGGCTGGCTCA
GAGCACACGTCTGAACTCTTTCCCTACACGACGCTCTTCCGATCTNNNCTTTCCTGATTGGCTGGCTCA
AGCACACGTCTGAACTCTTTCCCTACACGACGCTCTTCCGATCTNNNCTCCCAAGTTCAAGCGATCCTCC
AGCACACGTCTGAACTCTTTCCCTACACGACGCTCTTCCGATCTNNNCTCCCAAGTTCAAGCGATCCTCC
GAGCACACGTCTGAACTCTTTCCCTACACGACGCTCTTCCGATCTNNNTACCACCTGCTGAAGCTCTAT
GAGCACACGTCTGAACTCTTTCCCTACACGACGCTCTTCCGATCTNNNTACCACCTGCTGAAGCTCTAT
GCACACGTCTGAACTCTTTCCCTACACGACGCTCTTCCGATCTNNNTAGTCTTTTATCCCGCCTCCAAC
GCACACGTCTGAACTCTTTCCCTACACGACGCTCTTCCGATCTNNNTAGTCTTTTATCCCGCCTCCAAC
AGCACACGTCTGAACTCTTTCCCTACACGACGCTCTTCCGATCTNNNCCTAGTCTGTTGTCTGGCTGA
AGCACACGTCTGAACTCTTTCCCTACACGACGCTCTTCCGATCTNNNCCTAGTCTGTTGTCTGGCTGA
GAGCACACGTCTGAACTCTTTCCCTACACGACGCTCTTCCGATCTNNNAGCATTGGAGAGTTGCACTT
GAGCACACGTCTGAACTCTTTCCCTACACGACGCTCTTCCGATCTNNNAGCATTGGAGAGTTGCACTT
GAGCACACGTCTGAACTCTTTCCCTACACGACGCTCTTCCGATCTNNNTACAAGCGTCAGGATTCTGG
GAGCACACGTCTGAACTCTTTCCCTACACGACGCTCTTCCGATCTNNNTACAAGCGTCAGGATTCTGG
GCACACGTCTGAACTCTTTCCCTACACGACGCTCTTCCGATCTNNNGTTCATGCCATTCTCCTGCTTCCA
GCACACGTCTGAACTCTTTCCCTACACGACGCTCTTCCGATCTNNNGTTCATGCCATTCTCCTGCTTCCA
GCACACGTCTGAACTCTTTCCCTACACGACGCTCTTCCGATCTNNNTTGAGACAGGAGAATCGCTTGA

GCACACGTCTGAACTCTTTCCCTACACGACGCTCTTCCGATCTNNNTTGAGACAGGAGAATCGCTTGA

AGCACACGTCTGAACTCTTTCCCTACACGACGCTCTTCCGATCTNNNGCCTGTAATTGTAGTGCTTTG

AGCACACGTCTGAACTCTTTCCCTACACGACGCTCTTCCGATCTNNNGCCTGTAATTGTAGTGCTTTG

CACACGTCTGAACTCTTTCCCTACACGACGCTCTTCCGATCTNNNGTCCTGTGTCTCTTCCTTCAGAGC

CACACGTCTGAACTCTTTCCCTACACGACGCTCTTCCGATCTNNNGTCCTGTGTCTCTTCCTTCAGAGC

GAGCACACGTCTGAACTCTTTCCCTACACGACGCTCTTCCGATCTNNNAATGGTGCAATATTGGCTCA

GAGCACACGTCTGAACTCTTTCCCTACACGACGCTCTTCCGATCTNNNAATGGTGCAATATTGGCTCA

GCACACGTCTGAACTCTTTCCCTACACGACGCTCTTCCGATCTNNNGACACCCCTACTCTTAGAATGG

GCACACGTCTGAACTCTTTCCCTACACGACGCTCTTCCGATCTNNNGACACCCCTACTCTTAGAATGG

GCACACGTCTGAACTCTTTCCCTACACGACGCTCTTCCGATCTNNNTTGCCTATGCTTGCTGTTCTTAC

GCACACGTCTGAACTCTTTCCCTACACGACGCTCTTCCGATCTNNNTTGCCTATGCTTGCTGTTCTTAC

GCACACGTCTGAACTCTTTCCCTACACGACGCTCTTCCGATCTNNNCAGGCAGATTACTTGAGGTCAG

GCACACGTCTGAACTCTTTCCCTACACGACGCTCTTCCGATCTNNNCAGGCAGATTACTTGAGGTCAG

AGCACACGTCTGAACTCTTTCCCTACACGACGCTCTTCCGATCTNNNAATCCTAGCTATTCAGGAGCC

AGCACACGTCTGAACTCTTTCCCTACACGACGCTCTTCCGATCTNNNAATCCTAGCTATTCAGGAGCC

AGCACACGTCTGAACTCTTTCCCTACACGACGCTCTTCCGATCTNNNTTAATTTTGGTTCCACAGAAT

AGCACACGTCTGAACTCTTTCCCTACACGACGCTCTTCCGATCTNNNTTAATTTTGGTTCCACAGAAT

AAGAGCACACGTCTGAACTCTTTCCCTACACGACGCTCTTCCGATCTNNNTGTAACAAGACTGATGG

AAGAGCACACGTCTGAACTCTTTCCCTACACGACGCTCTTCCGATCTNNNTGTAACAAGACTGATGG

GCACACGTCTGAACTCTTTCCCTACACGACGCTCTTCCGATCTNNNGGACGTCAGTCAGAAAGGATG

GCACACGTCTGAACTCTTTCCCTACACGACGCTCTTCCGATCTNNNGGACGTCAGTCAGAAAGGATG

GCACACGTCTGAACTCTTTCCCTACACGACGCTCTTCCGATCTNNNCAGAAGAGTCATGTTCCACTCCC

GCACACGTCTGAACTCTTTCCCTACACGACGCTCTTCCGATCTNNNCAGAAGAGTCATGTTCCACTCCC

AGCACACGTCTGAACTCTTTCCCTACACGACGCTCTTCCGATCTNNNCAACAACAGAGTGAGACCCTGT

AGCACACGTCTGAACTCTTTCCCTACACGACGCTCTTCCGATCTNNNCAACAACAGAGTGAGACCCTGT

GCACACGTCTGAACTCTTTCCCTACACGACGCTCTTCCGATCTNNNACCAATTTTGAGTTATGTGTGC

GCACACGTCTGAACTCTTTCCCTACACGACGCTCTTCCGATCTNNNACCAATTTTGAGTTATGTGTGC

AGCACACGTCTGAACTCTTTCCCTACACGACGCTCTTCCGATCTNNNGCTTGGCAGTGAACTGAATTCT

AGCACACGTCTGAACTCTTTCCCTACACGACGCTCTTCCGATCTNNNGCTTGGCAGTGAACTGAATTCT

AGCACACGTCTGAACTCTTTCCCTACACGACGCTCTTCCGATCTNNNTGTTGCATGCCTGTTCTTTTTC

AGCACACGTCTGAACTCTTTCCCTACACGACGCTCTTCCGATCTNNNTGTTGCATGCCTGTTCTTTTTC

AGCACACGTCTGAACTCTTTCCCTACACGACGCTCTTCCGATCTNNNGTGAACTCCTGGTCTCAAGTG

AGCACACGTCTGAACTCTTTCCCTACACGACGCTCTTCCGATCTNNNGTGAACTCCTGGTCTCAAGTG

AGCACACGTCTGAACTCTTTCCCTACACGACGCTCTTCCGATCTNNNAAGGTTCCTGACAACTCATTCC

AGCACACGTCTGAACTCTTTCCCTACACGACGCTCTTCCGATCTNNNAAGGTTCCTGACAACTCATTCC

AGCACACGTCTGAACTCTTTCCCTACACGACGCTCTTCCGATCTNNNGAGGCTCAGCTTCATCATCAG

AGCACACGTCTGAACTCTTTCCCTACACGACGCTCTTCCGATCTNNNGAGGCTCAGCTTCATCATCAG

AGCACACGTCTGAACTCTTTCCCTACACGACGCTCTTCCGATCTNNNATTTATCCCTGGTGGCTCCAAA

AGCACACGTCTGAACTCTTTCCCTACACGACGCTCTTCCGATCTNNNATTTATCCCTGGTGGCTCCAAA

GAGCACACGTCTGAACTCTTTCCCTACACGACGCTCTTCCGATCTNNNTGGGCGACAGAGCTAAACTC

GAGCACACGTCTGAACTCTTTCCCTACACGACGCTCTTCCGATCTNNNTGGGCGACAGAGCTAAACTC

GAGCACACGTCTGAACTCTTTCCCTACACGACGCTCTTCCGATCTNNNGAGGTTGCAGTGAGCCAAG

GAGCACACGTCTGAACTCTTTCCCTACACGACGCTCTTCCGATCTNNNGAGGTTGCAGTGAGCCAAG

GCACACGTCTGAACTCTTTCCCTACACGACGCTCTTCCGATCTNNNGAATGTTGCAGTGAGCTGAGAT
GCACACGTCTGAACTCTTTCCCTACACGACGCTCTTCCGATCTNNNGAATGTTGCAGTGAGCTGAGAT
AGCACACGTCTGAACTCTTTCCCTACACGACGCTCTTCCGATCTNNNTTGAAATGCCAGCCATGAAAGT
AGCACACGTCTGAACTCTTTCCCTACACGACGCTCTTCCGATCTNNNTTGAAATGCCAGCCATGAAAGT
AGAGCACACGTCTGAACTCTTTCCCTACACGACGCTCTTCCGATCTNNNAGGTGACTTGTTGCCTGAA
AGAGCACACGTCTGAACTCTTTCCCTACACGACGCTCTTCCGATCTNNNAGGTGACTTGTTGCCTGAA
AGCACACGTCTGAACTCTTTCCCTACACGACGCTCTTCCGATCTNNNCTCACTCTCACTCTCTCCTCTCCC
AGCACACGTCTGAACTCTTTCCCTACACGACGCTCTTCCGATCTNNNCTCACTCTCACTCTCTCCTCTCCC
AGAGCACACGTCTGAACTCTTTCCCTACACGACGCTCTTCCGATCTNNNTTGAGGAATGCAGCAGTCTA
AGAGCACACGTCTGAACTCTTTCCCTACACGACGCTCTTCCGATCTNNNTTGAGGAATGCAGCAGTCTA
AGCACACGTCTGAACTCTTTCCCTACACGACGCTCTTCCGATCTNNNACACAGCAGTTTTTCCACCAAAT
AGCACACGTCTGAACTCTTTCCCTACACGACGCTCTTCCGATCTNNNACACAGCAGTTTTTCCACCAAAT
AGCACACGTCTGAACTCTTTCCCTACACGACGCTCTTCCGATCTNNNACCTCAAATCAAAAACACACAAC
AGCACACGTCTGAACTCTTTCCCTACACGACGCTCTTCCGATCTNNNACCTCAAATCAAAAACACACAAC
AGCACACGTCTGAACTCTTTCCCTACACGACGCTCTTCCGATCTNNNAAAGGGGGAAATAGAGAGCCA
AGCACACGTCTGAACTCTTTCCCTACACGACGCTCTTCCGATCTNNNAAAGGGGGAAATAGAGAGCCA
AGCACACGTCTGAACTCTTTCCCTACACGACGCTCTTCCGATCTNNNGAGGCTGAGGCAGGAGAATGC
AGCACACGTCTGAACTCTTTCCCTACACGACGCTCTTCCGATCTNNNGAGGCTGAGGCAGGAGAATGC
GAGCACACGTCTGAACTCTTTCCCTACACGACGCTCTTCCGATCTNNNATAGGCTTTTCAGCTTGCTTTG
GAGCACACGTCTGAACTCTTTCCCTACACGACGCTCTTCCGATCTNNNATAGGCTTTTCAGCTTGCTTTG
AGCACACGTCTGAACTCTTTCCCTACACGACGCTCTTCCGATCTNNNAGCCTTCACACCACATCCAGCAT
AGCACACGTCTGAACTCTTTCCCTACACGACGCTCTTCCGATCTNNNAGCCTTCACACCACATCCAGCAT
AGCACACGTCTGAACTCTTTCCCTACACGACGCTCTTCCGATCTNNNGGGCCTACAAAACAACTATTCCC
AGCACACGTCTGAACTCTTTCCCTACACGACGCTCTTCCGATCTNNNGGGCCTACAAAACAACTATTCCC
GAGCACACGTCTGAACTCTTTCCCTACACGACGCTCTTCCGATCTNNNTGGAACAGGTGGAAAGATG
GAGCACACGTCTGAACTCTTTCCCTACACGACGCTCTTCCGATCTNNNTGGAACAGGTGGAAAGATG
AGCACACGTCTGAACTCTTTCCCTACACGACGCTCTTCCGATCTNNNAGCAGAGCCTGATGATGTAAA
AGCACACGTCTGAACTCTTTCCCTACACGACGCTCTTCCGATCTNNNAGCAGAGCCTGATGATGTAAA
GAGCACACGTCTGAACTCTTTCCCTACACGACGCTCTTCCGATCTNNNCCTTCTAGGAGGTGCTTCTCAT
GAGCACACGTCTGAACTCTTTCCCTACACGACGCTCTTCCGATCTNNNCCTTCTAGGAGGTGCTTCTCAT
GAGCACACGTCTGAACTCTTTCCCTACACGACGCTCTTCCGATCTNNNCTTGCAGAGAGCCGAGATCG
GAGCACACGTCTGAACTCTTTCCCTACACGACGCTCTTCCGATCTNNNCTTGCAGAGAGCCGAGATCG
AGCACACGTCTGAACTCTTTCCCTACACGACGCTCTTCCGATCTNNNCACAAGAAAGATGTGCCCTCAT
AGCACACGTCTGAACTCTTTCCCTACACGACGCTCTTCCGATCTNNNCACAAGAAAGATGTGCCCTCAT
AGAGCACACGTCTGAACTCTTTCCCTACACGACGCTCTTCCGATCTNNNCATGTTGGATCACGCCTGTA
AGAGCACACGTCTGAACTCTTTCCCTACACGACGCTCTTCCGATCTNNNCATGTTGGATCACGCCTGTA
GAGCACACGTCTGAACTCTTTCCCTACACGACGCTCTTCCGATCTNNNTGTGAGATGTTTGCAGTTTGT
GAGCACACGTCTGAACTCTTTCCCTACACGACGCTCTTCCGATCTNNNTGTGAGATGTTTGCAGTTTGT
AGCACACGTCTGAACTCTTTCCCTACACGACGCTCTTCCGATCTNNNATGAAATGCCCCAAGGTATCTG
AGCACACGTCTGAACTCTTTCCCTACACGACGCTCTTCCGATCTNNNATGAAATGCCCCAAGGTATCTG
CACACGTCTGAACTCTTTCCCTACACGACGCTCTTCCGATCTNNNATTTAGCCACTCTAGGAAATGTTT
CACACGTCTGAACTCTTTCCCTACACGACGCTCTTCCGATCTNNNATTTAGCCACTCTAGGAAATGTTT
AGAGCACACGTCTGAACTCTTTCCCTACACGACGCTCTTCCGATCTNNNCAAAAATTAGCAGGGTGTG

GAGCACACGTCTGAACTCTTTCCCTACACGACGCTCTTCCGATCTNNNCAAAAATTAGCAGGGTGTG
GCACACGTCTGAACTCTTTCCCTACACGACGCTCTTCCGATCTNNNCTGAGGCAGGAGAATCACTTGA
GCACACGTCTGAACTCTTTCCCTACACGACGCTCTTCCGATCTNNNCTGAGGCAGGAGAATCACTTGA
GAGCACACGTCTGAACTCTTTCCCTACACGACGCTCTTCCGATCTNNNAGAAAGCCTCTGTGACATTG
GAGCACACGTCTGAACTCTTTCCCTACACGACGCTCTTCCGATCTNNNAGAAAGCCTCTGTGACATTG
GAGCACACGTCTGAACTCTTTCCCTACACGACGCTCTTCCGATCTNNNGAGGTTGCAGTGAGCCAAG
GAGCACACGTCTGAACTCTTTCCCTACACGACGCTCTTCCGATCTNNNGAGGTTGCAGTGAGCCAAG
GAGCACACGTCTGAACTCTTTCCCTACACGACGCTCTTCCGATCTNNNGGGAGGTTGTAGGATATCA
GAGCACACGTCTGAACTCTTTCCCTACACGACGCTCTTCCGATCTNNNGGGAGGTTGTAGGATATCA
GCACACGTCTGAACTCTTTCCCTACACGACGCTCTTCCGATCTNNNATTTTCTGGGTGAGAGACGAA
GCACACGTCTGAACTCTTTCCCTACACGACGCTCTTCCGATCTNNNATTTTCTGGGTGAGAGACGAA
GAAGAGCACACGTCTGAACTCTTTCCCTACACGACGCTCTTCCGATCTNNNACAATAAGGATGACTG
GAAGAGCACACGTCTGAACTCTTTCCCTACACGACGCTCTTCCGATCTNNNACAATAAGGATGACTG
AGCACACGTCTGAACTCTTTCCCTACACGACGCTCTTCCGATCTNNNCTCCTGAGTAGCTGGGATTAC
AGCACACGTCTGAACTCTTTCCCTACACGACGCTCTTCCGATCTNNNCTCCTGAGTAGCTGGGATTAC
GCACACGTCTGAACTCTTTCCCTACACGACGCTCTTCCGATCTNNNGAGAATACCTTGAACCTGGGAG
GCACACGTCTGAACTCTTTCCCTACACGACGCTCTTCCGATCTNNNGAGAATACCTTGAACCTGGGAG
AGCACACGTCTGAACTCTTTCCCTACACGACGCTCTTCCGATCTNNNAGTTCTTCCAAAGAGAGTGCT
AGCACACGTCTGAACTCTTTCCCTACACGACGCTCTTCCGATCTNNNAGTTCTTCCAAAGAGAGTGCT
GCACACGTCTGAACTCTTTCCCTACACGACGCTCTTCCGATCTNNNTTCCTCAAGAAGGCTTGATTTG
GCACACGTCTGAACTCTTTCCCTACACGACGCTCTTCCGATCTNNNTTCCTCAAGAAGGCTTGATTTG
GCACACGTCTGAACTCTTTCCCTACACGACGCTCTTCCGATCTNNNACCTGTTGTAGAGATGGGGTT
GCACACGTCTGAACTCTTTCCCTACACGACGCTCTTCCGATCTNNNACCTGTTGTAGAGATGGGGTT
AGCACACGTCTGAACTCTTTCCCTACACGACGCTCTTCCGATCTNNNAACCCATATTTCTCTCTGCCTGG
AGCACACGTCTGAACTCTTTCCCTACACGACGCTCTTCCGATCTNNNAACCCATATTTCTCTCTGCCTGG
AGCACACGTCTGAACTCTTTCCCTACACGACGCTCTTCCGATCTNNNCTGTCTCCTTTGTCTTGGTAGC
AGCACACGTCTGAACTCTTTCCCTACACGACGCTCTTCCGATCTNNNCTGTCTCCTTTGTCTTGGTAGC
CACACGTCTGAACTCTTTCCCTACACGACGCTCTTCCGATCTNNNCCACGAAAACCCTAGTGAATCAC
CACACGTCTGAACTCTTTCCCTACACGACGCTCTTCCGATCTNNNCCACGAAAACCCTAGTGAATCAC
GCACACGTCTGAACTCTTTCCCTACACGACGCTCTTCCGATCTNNNTGCAATAAACATACGCATGCAG
GCACACGTCTGAACTCTTTCCCTACACGACGCTCTTCCGATCTNNNTGCAATAAACATACGCATGCAG
GCACACGTCTGAACTCTTTCCCTACACGACGCTCTTCCGATCTNNNCTCCTCCTTCCTTTCTCTCTC
GCACACGTCTGAACTCTTTCCCTACACGACGCTCTTCCGATCTNNNCTCCTCCTTCCTTTCTCTCTC
GAGCACACGTCTGAACTCTTTCCCTACACGACGCTCTTCCGATCTNNNCACTCACCTTTACTTTGCTCTG
GAGCACACGTCTGAACTCTTTCCCTACACGACGCTCTTCCGATCTNNNCACTCACCTTTACTTTGCTCTG
GAGCACACGTCTGAACTCTTTCCCTACACGACGCTCTTCCGATCTNNNGCAAGAGGTAGCACAAAACA
GAGCACACGTCTGAACTCTTTCCCTACACGACGCTCTTCCGATCTNNNGCAAGAGGTAGCACAAAACA
CACGTCTGAACTCTTTCCCTACACGACGCTCTTCCGATCTNNNCTTCTCTCTCCATGATTACTGCATCG
CACGTCTGAACTCTTTCCCTACACGACGCTCTTCCGATCTNNNCTTCTCTCTCCATGATTACTGCATCG
AGCACACGTCTGAACTCTTTCCCTACACGACGCTCTTCCGATCTNNNGCCTGTAGTCCCAGCTACTTG
AGCACACGTCTGAACTCTTTCCCTACACGACGCTCTTCCGATCTNNNGCCTGTAGTCCCAGCTACTTG
GAGCACACGTCTGAACTCTTTCCCTACACGACGCTCTTCCGATCTNNNCTCAATCTGTCTCTGCTTCTG
GAGCACACGTCTGAACTCTTTCCCTACACGACGCTCTTCCGATCTNNNCTCAATCTGTCTCTGCTTCTG

GAGCACACGTCTGAACTCTTTCCCTACACGACGCTCTTCCGATCTNNNTAAGTGGAACGTTAGCCTTG

GAGCACACGTCTGAACTCTTTCCCTACACGACGCTCTTCCGATCTNNNTAAGTGGAACGTTAGCCTTG

ACACGTCTGAACTCTTTCCCTACACGACGCTCTTCCGATCTNNNCGATCTCCTGACCTCATGATCTGCAT

ACACGTCTGAACTCTTTCCCTACACGACGCTCTTCCGATCTNNNCGATCTCCTGACCTCATGATCTGCAT

AGCACACGTCTGAACTCTTTCCCTACACGACGCTCTTCCGATCTNNNGCCTCATGTCAGGACTTTGAA

AGCACACGTCTGAACTCTTTCCCTACACGACGCTCTTCCGATCTNNNGCCTCATGTCAGGACTTTGAA

GCACACGTCTGAACTCTTTCCCTACACGACGCTCTTCCGATCTNNNGACCTCCAAACAAAAAGGAAGG

GCACACGTCTGAACTCTTTCCCTACACGACGCTCTTCCGATCTNNNGACCTCCAAACAAAAAGGAAGG

AGAGCACACGTCTGAACTCTTTCCCTACACGACGCTCTTCCGATCTNNNCTTTTCTTTCAGGCTTCCTAT

AGAGCACACGTCTGAACTCTTTCCCTACACGACGCTCTTCCGATCTNNNCTTTTCTTTCAGGCTTCCTAT

GAGCACACGTCTGAACTCTTTCCCTACACGACGCTCTTCCGATCTNNNTAGGGATCTTAGCTCACTGCA

GAGCACACGTCTGAACTCTTTCCCTACACGACGCTCTTCCGATCTNNNTAGGGATCTTAGCTCACTGCA

AGCACACGTCTGAACTCTTTCCCTACACGACGCTCTTCCGATCTNNNTCCCTTTCCATTTCCTTTTCCCTC

AGCACACGTCTGAACTCTTTCCCTACACGACGCTCTTCCGATCTNNNTCCCTTTCCATTTCCTTTTCCCTC

GCACACGTCTGAACTCTTTCCCTACACGACGCTCTTCCGATCTNNNGTTGCAGTGAGCCAAGATCATG

GCACACGTCTGAACTCTTTCCCTACACGACGCTCTTCCGATCTNNNGTTGCAGTGAGCCAAGATCATG

AGCACACGTCTGAACTCTTTCCCTACACGACGCTCTTCCGATCTNNNGACTAGAATTCACCAGGCTGG

AGCACACGTCTGAACTCTTTCCCTACACGACGCTCTTCCGATCTNNNGACTAGAATTCACCAGGCTGG

AGCACACGTCTGAACTCTTTCCCTACACGACGCTCTTCCGATCTNNNGAGAATCGTTTGAACCCAAGA

AGCACACGTCTGAACTCTTTCCCTACACGACGCTCTTCCGATCTNNNGAGAATCGTTTGAACCCAAGA

AGCACACGTCTGAACTCTTTCCCTACACGACGCTCTTCCGATCTNNNCTGTAATCCCAGCTACTCAGGA

AGCACACGTCTGAACTCTTTCCCTACACGACGCTCTTCCGATCTNNNCTGTAATCCCAGCTACTCAGGA

AGCACACGTCTGAACTCTTTCCCTACACGACGCTCTTCCGATCTNNNGAGCCACAAAATTCAGCTCTA

AGCACACGTCTGAACTCTTTCCCTACACGACGCTCTTCCGATCTNNNGAGCCACAAAATTCAGCTCTA

GAGCACACGTCTGAACTCTTTCCCTACACGACGCTCTTCCGATCTNNNGAGGCTGAGGCAGGAGAAT

GAGCACACGTCTGAACTCTTTCCCTACACGACGCTCTTCCGATCTNNNGAGGCTGAGGCAGGAGAAT

ACACGTCTGAACTCTTTCCCTACACGACGCTCTTCCGATCTNNNTGTTTACTGTAAGACAAAAAACACA

ACACGTCTGAACTCTTTCCCTACACGACGCTCTTCCGATCTNNNTGTTTACTGTAAGACAAAAAACACA

AGCACACGTCTGAACTCTTTCCCTACACGACGCTCTTCCGATCTNNNAATTACCGTGAGAGTGAGGG

AGCACACGTCTGAACTCTTTCCCTACACGACGCTCTTCCGATCTNNNAATTACCGTGAGAGTGAGGG

GAGCACACGTCTGAACTCTTTCCCTACACGACGCTCTTCCGATCTNNNGTTGGGACCAGTGACTTCTA

GAGCACACGTCTGAACTCTTTCCCTACACGACGCTCTTCCGATCTNNNGTTGGGACCAGTGACTTCTA

ACACGTCTGAACTCTTTCCCTACACGACGCTCTTCCGATCTNNNCCGCCAATATTTAGGCTTTGCAGCA

ACACGTCTGAACTCTTTCCCTACACGACGCTCTTCCGATCTNNNCCGCCAATATTTAGGCTTTGCAGCA

AGCACACGTCTGAACTCTTTCCCTACACGACGCTCTTCCGATCTNNNGTTGGTGCTAACATCCCTGTTT

AGCACACGTCTGAACTCTTTCCCTACACGACGCTCTTCCGATCTNNNGTTGGTGCTAACATCCCTGTTT

AGCACACGTCTGAACTCTTTCCCTACACGACGCTCTTCCGATCTNNNTCTACCAATGTACCTCCAGGAA

AGCACACGTCTGAACTCTTTCCCTACACGACGCTCTTCCGATCTNNNTCTACCAATGTACCTCCAGGAA

AGCACACGTCTGAACTCTTTCCCTACACGACGCTCTTCCGATCTNNNAGGTTGCAGTGAGCTGAGAT

AGCACACGTCTGAACTCTTTCCCTACACGACGCTCTTCCGATCTNNNAGGTTGCAGTGAGCTGAGAT

AGCACACGTCTGAACTCTTTCCCTACACGACGCTCTTCCGATCTNNNCAGATCTGGGAGCTCCTGTCCA

AGCACACGTCTGAACTCTTTCCCTACACGACGCTCTTCCGATCTNNNCAGATCTGGGAGCTCCTGTCCA

AGCACACGTCTGAACTCTTTCCCTACACGACGCTCTTCCGATCTNNNTAATGCTTTCACATTTCCAGCG

AGCACACGTCTGAACTCTTTCCCTACACGACGCTCTTCCGATCTNNNTAATGCTTTCACATTTCCAGCGG

GCACACGTCTGAACTCTTTCCCTACACGACGCTCTTCCGATCTNNNTTACTAAGCTCTTTCCCTGTGCCC

GCACACGTCTGAACTCTTTCCCTACACGACGCTCTTCCGATCTNNNTTACTAAGCTCTTTCCCTGTGCCC

AGCACACGTCTGAACTCTTTCCCTACACGACGCTCTTCCGATCTNNNGATCTCCTGACCTTGTGATCCTC

AGCACACGTCTGAACTCTTTCCCTACACGACGCTCTTCCGATCTNNNGATCTCCTGACCTTGTGATCCTC

AGCACACGTCTGAACTCTTTCCCTACACGACGCTCTTCCGATCTNNNTCTCTTCGTTTGGGAGTAATGC

AGCACACGTCTGAACTCTTTCCCTACACGACGCTCTTCCGATCTNNNTCTCTTCGTTTGGGAGTAATGC

GAGCACACGTCTGAACTCTTTCCCTACACGACGCTCTTCCGATCTNNNTAAATACCCTTCTCGGTGTGAC

GAGCACACGTCTGAACTCTTTCCCTACACGACGCTCTTCCGATCTNNNTAAATACCCTTCTCGGTGTGAC

AGAGCACACGTCTGAACTCTTTCCCTACACGACGCTCTTCCGATCTNNNGACACTTATTTTTTCTTTGTA

AGAGCACACGTCTGAACTCTTTCCCTACACGACGCTCTTCCGATCTNNNGACACTTATTTTTTCTTTGTA

AGCACACGTCTGAACTCTTTCCCTACACGACGCTCTTCCGATCTNNNGTAAATAGTCTCAGCTCCTCGG

AGCACACGTCTGAACTCTTTCCCTACACGACGCTCTTCCGATCTNNNGTAAATAGTCTCAGCTCCTCGG

AGCACACGTCTGAACTCTTTCCCTACACGACGCTCTTCCGATCTNNNGCATTCTATTCCTGTTCCATGGC

AGCACACGTCTGAACTCTTTCCCTACACGACGCTCTTCCGATCTNNNGCATTCTATTCCTGTTCCATGGC

AGCACACGTCTGAACTCTTTCCCTACACGACGCTCTTCCGATCTNNNGAAGGTAGGTGGAAGAGAGT

AGCACACGTCTGAACTCTTTCCCTACACGACGCTCTTCCGATCTNNNGAAGGTAGGTGGAAGAGAGT

CACACGTCTGAACTCTTTCCCTACACGACGCTCTTCCGATCTNNNACTGAAATGCCCCTAATGGATCAC

CACACGTCTGAACTCTTTCCCTACACGACGCTCTTCCGATCTNNNACTGAAATGCCCCTAATGGATCAC

GCACACGTCTGAACTCTTTCCCTACACGACGCTCTTCCGATCTNNNCTGTGCCATCTGATGTCCATTTCC

GCACACGTCTGAACTCTTTCCCTACACGACGCTCTTCCGATCTNNNCTGTGCCATCTGATGTCCATTTCC

GAGCACACGTCTGAACTCTTTCCCTACACGACGCTCTTCCGATCTNNNGGCAAAAGTGATGGGATGAC

GAGCACACGTCTGAACTCTTTCCCTACACGACGCTCTTCCGATCTNNNGGCAAAAGTGATGGGATGAC

GCACACGTCTGAACTCTTTCCCTACACGACGCTCTTCCGATCTNNNTCCATCTCACAACGGTATGAGTT

GCACACGTCTGAACTCTTTCCCTACACGACGCTCTTCCGATCTNNNTCCATCTCACAACGGTATGAGTT

GAGCACACGTCTGAACTCTTTCCCTACACGACGCTCTTCCGATCTNNNCCCCAATATTCATTGAGCAAC

GAGCACACGTCTGAACTCTTTCCCTACACGACGCTCTTCCGATCTNNNCCCCAATATTCATTGAGCAAC

GAGCACACGTCTGAACTCTTTCCCTACACGACGCTCTTCCGATCTNNNCACGCCCGGCTAATTATTGAA

GAGCACACGTCTGAACTCTTTCCCTACACGACGCTCTTCCGATCTNNNCACGCCCGGCTAATTATTGAA

GAGCACACGTCTGAACTCTTTCCCTACACGACGCTCTTCCGATCTNNNAATAGTTTGCCTGTAGGTTGC

GAGCACACGTCTGAACTCTTTCCCTACACGACGCTCTTCCGATCTNNNAATAGTTTGCCTGTAGGTTGC

GCACACGTCTGAACTCTTTCCCTACACGACGCTCTTCCGATCTNNNAGTGCTACAATAAACATATGCAT

GCACACGTCTGAACTCTTTCCCTACACGACGCTCTTCCGATCTNNNAGTGCTACAATAAACATATGCAT

AGAGCACACGTCTGAACTCTTTCCCTACACGACGCTCTTCCGATCTNNNCAGTTAGGGTGAAGGAGAC

AGAGCACACGTCTGAACTCTTTCCCTACACGACGCTCTTCCGATCTNNNCAGTTAGGGTGAAGGAGAC

AGAGCACACGTCTGAACTCTTTCCCTACACGACGCTCTTCCGATCTNNNGCTACCTCCTATTCTGCACTT

AGAGCACACGTCTGAACTCTTTCCCTACACGACGCTCTTCCGATCTNNNGCTACCTCCTATTCTGCACTT

AGAGCACACGTCTGAACTCTTTCCCTACACGACGCTCTTCCGATCTNNNGATGTGTAAAGAATATACA

AGAGCACACGTCTGAACTCTTTCCCTACACGACGCTCTTCCGATCTNNNGATGTGTAAAGAATATACA

GAGCACACGTCTGAACTCTTTCCCTACACGACGCTCTTCCGATCTNNNTGTTTTTGGCAAAAGCTCTTC

GAGCACACGTCTGAACTCTTTCCCTACACGACGCTCTTCCGATCTNNNTGTTTTTGGCAAAAGCTCTTC

AGCACACGTCTGAACTCTTTCCCTACACGACGCTCTTCCGATCTNNNCTGTTTTGTGCTGCTGTAACAG

AGCACACGTCTGAACTCTTTCCCTACACGACGCTCTTCCGATCTNNNCTGTTTTGTGCTGCTGTAACAG

GCACACGTCTGAACTCTTTCCCTACACGACGCTCTTCCGATCTNNNCTAGCTCTCATTGCTGTCTTTGCC
GCACACGTCTGAACTCTTTCCCTACACGACGCTCTTCCGATCTNNNCTAGCTCTCATTGCTGTCTTTGCC
GAGCACACGTCTGAACTCTTTCCCTACACGACGCTCTTCCGATCTNNNTGTGTGTCTGTAATCCCAGCT
GAGCACACGTCTGAACTCTTTCCCTACACGACGCTCTTCCGATCTNNNTGTGTGTCTGTAATCCCAGCT
AGCACACGTCTGAACTCTTTCCCTACACGACGCTCTTCCGATCTNNNCTTTGAAACTGTGTGCCATTTG
AGCACACGTCTGAACTCTTTCCCTACACGACGCTCTTCCGATCTNNNCTTTGAAACTGTGTGCCATTTG
AGCACACGTCTGAACTCTTTCCCTACACGACGCTCTTCCGATCTNNNGATTTCTGCTTCTCTGGTGCTTC
AGCACACGTCTGAACTCTTTCCCTACACGACGCTCTTCCGATCTNNNGATTTCTGCTTCTCTGGTGCTTC
AGCACACGTCTGAACTCTTTCCCTACACGACGCTCTTCCGATCTNNNGCCCACTTTCAGCACTTCTATTC
AGCACACGTCTGAACTCTTTCCCTACACGACGCTCTTCCGATCTNNNGCCCACTTTCAGCACTTCTATTC
AGCACACGTCTGAACTCTTTCCCTACACGACGCTCTTCCGATCTNNNTCTTCCAATCTGTGAACCCACTT
AGCACACGTCTGAACTCTTTCCCTACACGACGCTCTTCCGATCTNNNTCTTCCAATCTGTGAACCCACTT
GAGCACACGTCTGAACTCTTTCCCTACACGACGCTCTTCCGATCTNNNACCTGTAGTCCCAGCTACTCG
GAGCACACGTCTGAACTCTTTCCCTACACGACGCTCTTCCGATCTNNNACCTGTAGTCCCAGCTACTCG
AGCACACGTCTGAACTCTTTCCCTACACGACGCTCTTCCGATCTNNNTCCTTCTTTCCTATCTTCCTGTCT
AGCACACGTCTGAACTCTTTCCCTACACGACGCTCTTCCGATCTNNNTCCTTCTTTCCTATCTTCCTGTCT
GAGCACACGTCTGAACTCTTTCCCTACACGACGCTCTTCCGATCTNNNTGCAATGACCCATGATCACACT
GAGCACACGTCTGAACTCTTTCCCTACACGACGCTCTTCCGATCTNNNTGCAATGACCCATGATCACACT
AGAGCACACGTCTGAACTCTTTCCCTACACGACGCTCTTCCGATCTNNNGGAGCACAGAAAGACAGAA
AGAGCACACGTCTGAACTCTTTCCCTACACGACGCTCTTCCGATCTNNNGGAGCACAGAAAGACAGAA
GCACACGTCTGAACTCTTTCCCTACACGACGCTCTTCCGATCTNNNATTGACCCTAGGAAATGAGGTCC
GCACACGTCTGAACTCTTTCCCTACACGACGCTCTTCCGATCTNNNATTGACCCTAGGAAATGAGGTCC
AGCACACGTCTGAACTCTTTCCCTACACGACGCTCTTCCGATCTNNNCTGACATATCATTGTTACCAAAG
AGCACACGTCTGAACTCTTTCCCTACACGACGCTCTTCCGATCTNNNCTGACATATCATTGTTACCAAAG
AAGAGCACACGTCTGAACTCTTTCCCTACACGACGCTCTTCCGATCTNNNGGTAAAAAAATCAGGTTGG
AAGAGCACACGTCTGAACTCTTTCCCTACACGACGCTCTTCCGATCTNNNGGTAAAAAAATCAGGTTGG
GAGCACACGTCTGAACTCTTTCCCTACACGACGCTCTTCCGATCTNNNACTGAGTGTAAAGGTGTCCTT
GAGCACACGTCTGAACTCTTTCCCTACACGACGCTCTTCCGATCTNNNACTGAGTGTAAAGGTGTCCTT
AGCACACGTCTGAACTCTTTCCCTACACGACGCTCTTCCGATCTNNNATACTCGCAAGTGACAGAACTG
AGCACACGTCTGAACTCTTTCCCTACACGACGCTCTTCCGATCTNNNATACTCGCAAGTGACAGAACTG
AGCACACGTCTGAACTCTTTCCCTACACGACGCTCTTCCGATCTNNNCCAGCAACAAAAGATGGGATTC
AGCACACGTCTGAACTCTTTCCCTACACGACGCTCTTCCGATCTNNNCCAGCAACAAAAGATGGGATTC
AGAGCACACGTCTGAACTCTTTCCCTACACGACGCTCTTCCGATCTNNNTGTAGTCCTAGCTACTCAGGA
AGAGCACACGTCTGAACTCTTTCCCTACACGACGCTCTTCCGATCTNNNTGTAGTCCTAGCTACTCAGGA
GAGCACACGTCTGAACTCTTTCCCTACACGACGCTCTTCCGATCTNNNTGAAACCCTGTCTCTACTGCAA
GAGCACACGTCTGAACTCTTTCCCTACACGACGCTCTTCCGATCTNNNTGAAACCCTGTCTCTACTGCAA
GCACACGTCTGAACTCTTTCCCTACACGACGCTCTTCCGATCTNNNAGCACCTTCATGATGGAATTCCT
GCACACGTCTGAACTCTTTCCCTACACGACGCTCTTCCGATCTNNNAGCACCTTCATGATGGAATTCCT
AGCACACGTCTGAACTCTTTCCCTACACGACGCTCTTCCGATCTNNNCAATCTCACAGAGCTATGCTTG
AGCACACGTCTGAACTCTTTCCCTACACGACGCTCTTCCGATCTNNNCAATCTCACAGAGCTATGCTTG
AGAGCACACGTCTGAACTCTTTCCCTACACGACGCTCTTCCGATCTNNNTACTCTCATTGTAGGTCCAAA
AGAGCACACGTCTGAACTCTTTCCCTACACGACGCTCTTCCGATCTNNNTACTCTCATTGTAGGTCCAAA
GAGCACACGTCTGAACTCTTTCCCTACACGACGCTCTTCCGATCTNNNTTGCTCTGATTTTACTGACTGG

GAGCACACGTCTGAACTCTTTCCCTACACGACGCTCTTCCGATCTNNNTTGCTCTGATTTTACTGACTGG
AGCACACGTCTGAACTCTTTCCCTACACGACGCTCTTCCGATCTNNNTCTCCCTCTCTCTCTCTCTCA
AGCACACGTCTGAACTCTTTCCCTACACGACGCTCTTCCGATCTNNNTCTCCCTCTCTCTCTCTCTCA
GAGCACACGTCTGAACTCTTTCCCTACACGACGCTCTTCCGATCTNNNGCTCCTAACAAGAAAAGATGG
GAGCACACGTCTGAACTCTTTCCCTACACGACGCTCTTCCGATCTNNNGCTCCTAACAAGAAAAGATGG
AGCACACGTCTGAACTCTTTCCCTACACGACGCTCTTCCGATCTNNNATACTGCCATGCCCAGCTAATTT
AGCACACGTCTGAACTCTTTCCCTACACGACGCTCTTCCGATCTNNNATACTGCCATGCCCAGCTAATTT
AGCACACGTCTGAACTCTTTCCCTACACGACGCTCTTCCGATCTNNNCTGAGGCAGGAGAATCGTTTG
AGCACACGTCTGAACTCTTTCCCTACACGACGCTCTTCCGATCTNNNCTGAGGCAGGAGAATCGTTTG
GAGCACACGTCTGAACTCTTTCCCTACACGACGCTCTTCCGATCTNNNGGAGAGGCAAGAACTCCTGA
GAGCACACGTCTGAACTCTTTCCCTACACGACGCTCTTCCGATCTNNNGGAGAGGCAAGAACTCCTGA
AGCACACGTCTGAACTCTTTCCCTACACGACGCTCTTCCGATCTNNNGAGATCACGCCACTGCACTCCAT
AGCACACGTCTGAACTCTTTCCCTACACGACGCTCTTCCGATCTNNNGAGATCACGCCACTGCACTCCAT
GAGCACACGTCTGAACTCTTTCCCTACACGACGCTCTTCCGATCTNNNGGAGTTTTACTTTGTTGCCCA
GAGCACACGTCTGAACTCTTTCCCTACACGACGCTCTTCCGATCTNNNGGAGTTTTACTTTGTTGCCCA
AGCACACGTCTGAACTCTTTCCCTACACGACGCTCTTCCGATCTNNNAGCTGCAATGTAAACAGAAAA
AGCACACGTCTGAACTCTTTCCCTACACGACGCTCTTCCGATCTNNNAGCTGCAATGTAAACAGAAAA
GAGCACACGTCTGAACTCTTTCCCTACACGACGCTCTTCCGATCTNNNATGTGGGAATTCTTTGTTGC
GAGCACACGTCTGAACTCTTTCCCTACACGACGCTCTTCCGATCTNNNATGTGGGAATTCTTTGTTGC
GAGCACACGTCTGAACTCTTTCCCTACACGACGCTCTTCCGATCTNNNTCTTTCTGGACAGACTGGGA
GAGCACACGTCTGAACTCTTTCCCTACACGACGCTCTTCCGATCTNNNTCTTTCTGGACAGACTGGGA
GAGCACACGTCTGAACTCTTTCCCTACACGACGCTCTTCCGATCTNNNGAATCGCTTGAACCAGGGAGA
GAGCACACGTCTGAACTCTTTCCCTACACGACGCTCTTCCGATCTNNNGAATCGCTTGAACCAGGGAGA
AGCACACGTCTGAACTCTTTCCCTACACGACGCTCTTCCGATCTNNNAGACAGGAGAATTGCTTGAAC
AGCACACGTCTGAACTCTTTCCCTACACGACGCTCTTCCGATCTNNNAGACAGGAGAATTGCTTGAAC
GAGCACACGTCTGAACTCTTTCCCTACACGACGCTCTTCCGATCTNNNGACTCTTTCATTTGGCCAAACG
GAGCACACGTCTGAACTCTTTCCCTACACGACGCTCTTCCGATCTNNNGACTCTTTCATTTGGCCAAACG
GAGCACACGTCTGAACTCTTTCCCTACACGACGCTCTTCCGATCTNNNATTCCATCCAGGTTGCTGAGA
GAGCACACGTCTGAACTCTTTCCCTACACGACGCTCTTCCGATCTNNNATTCCATCCAGGTTGCTGAGA
AGCACACGTCTGAACTCTTTCCCTACACGACGCTCTTCCGATCTNNNCTCAAGTGATTCTCCCTCCTCAG
AGCACACGTCTGAACTCTTTCCCTACACGACGCTCTTCCGATCTNNNCTCAAGTGATTCTCCCTCCTCAG
GCACACGTCTGAACTCTTTCCCTACACGACGCTCTTCCGATCTNNNTGGTTCTCTTCAGCTCCAACATTCA
GCACACGTCTGAACTCTTTCCCTACACGACGCTCTTCCGATCTNNNTGGTTCTCTTCAGCTCCAACATTCA
AGCACACGTCTGAACTCTTTCCCTACACGACGCTCTTCCGATCTNNNCTCCACCTCCCAGATTCAAATGA
AGCACACGTCTGAACTCTTTCCCTACACGACGCTCTTCCGATCTNNNCTCCACCTCCCAGATTCAAATGA
GAGCACACGTCTGAACTCTTTCCCTACACGACGCTCTTCCGATCTNNNAAGCTTCTCTATGCCCTTGTT
GAGCACACGTCTGAACTCTTTCCCTACACGACGCTCTTCCGATCTNNNAAGCTTCTCTATGCCCTTGTT
GAGCACACGTCTGAACTCTTTCCCTACACGACGCTCTTCCGATCTNNNAGTTTCCATGTGACAATTGTG
GAGCACACGTCTGAACTCTTTCCCTACACGACGCTCTTCCGATCTNNNAGTTTCCATGTGACAATTGTG
AGCACACGTCTGAACTCTTTCCCTACACGACGCTCTTCCGATCTNNNGACAGGGAGAGAGAATGCAA
AGCACACGTCTGAACTCTTTCCCTACACGACGCTCTTCCGATCTNNNGACAGGGAGAGAGAATGCAA
GAGCACACGTCTGAACTCTTTCCCTACACGACGCTCTTCCGATCTNNNTTGGAGTATACCCAGGCAACA
GAGCACACGTCTGAACTCTTTCCCTACACGACGCTCTTCCGATCTNNNTTGGAGTATACCCAGGCAACA

AGCACACGTCTGAACTCTTTCCCTACACGACGCTCTTCCGATCTNNNGGAGATTCACCTGCAGGAATA
AGCACACGTCTGAACTCTTTCCCTACACGACGCTCTTCCGATCTNNNGGAGATTCACCTGCAGGAATA
GAGCACACGTCTGAACTCTTTCCCTACACGACGCTCTTCCGATCTNNNCTGAAAATCGATTTCCAGGCT
GAGCACACGTCTGAACTCTTTCCCTACACGACGCTCTTCCGATCTNNNCTGAAAATCGATTTCCAGGCT
GAGCACACGTCTGAACTCTTTCCCTACACGACGCTCTTCCGATCTNNNTGTCTCTGTCTCTGTCTGTCTC
GAGCACACGTCTGAACTCTTTCCCTACACGACGCTCTTCCGATCTNNNTGTCTCTGTCTCTGTCTGTCTC
GCACACGTCTGAACTCTTTCCCTACACGACGCTCTTCCGATCTNNNTGTGTGTGAAAGGGAAAACGA
GCACACGTCTGAACTCTTTCCCTACACGACGCTCTTCCGATCTNNNTGTGTGTGAAAGGGAAAACGA
AGCACACGTCTGAACTCTTTCCCTACACGACGCTCTTCCGATCTNNNTATCATCTTGGAGGGGTGTC
AGCACACGTCTGAACTCTTTCCCTACACGACGCTCTTCCGATCTNNNTATCATCTTGGAGGGGTGTC
AAGAGCACACGTCTGAACTCTTTCCCTACACGACGCTCTTCCGATCTNNNTGAGTGATAAGCCAGACT
AAGAGCACACGTCTGAACTCTTTCCCTACACGACGCTCTTCCGATCTNNNTGAGTGATAAGCCAGACT
AGCACACGTCTGAACTCTTTCCCTACACGACGCTCTTCCGATCTNNNTCATTCATTCATTCCCCCCACA
AGCACACGTCTGAACTCTTTCCCTACACGACGCTCTTCCGATCTNNNTCATTCATTCATTCCCCCCACA
AGCACACGTCTGAACTCTTTCCCTACACGACGCTCTTCCGATCTNNNGGCATCTGTAGTCCCAACTAG
AGCACACGTCTGAACTCTTTCCCTACACGACGCTCTTCCGATCTNNNGGCATCTGTAGTCCCAACTAG
AGCACACGTCTGAACTCTTTCCCTACACGACGCTCTTCCGATCTNNNGACCTCTAAGCACAGATGAGT
AGCACACGTCTGAACTCTTTCCCTACACGACGCTCTTCCGATCTNNNGACCTCTAAGCACAGATGAGT
AGCACACGTCTGAACTCTTTCCCTACACGACGCTCTTCCGATCTNNNAATATGCAGCAACTGAGTTGA
AGCACACGTCTGAACTCTTTCCCTACACGACGCTCTTCCGATCTNNNAATATGCAGCAACTGAGTTGA
AGCACACGTCTGAACTCTTTCCCTACACGACGCTCTTCCGATCTNNNGAGAATCTCTTGAACCCAGGA
AGCACACGTCTGAACTCTTTCCCTACACGACGCTCTTCCGATCTNNNGAGAATCTCTTGAACCCAGGA
AGCACACGTCTGAACTCTTTCCCTACACGACGCTCTTCCGATCTNNNAAACTTAGACTGCAGTGTTCC
AGCACACGTCTGAACTCTTTCCCTACACGACGCTCTTCCGATCTNNNAAACTTAGACTGCAGTGTTCC
AGCACACGTCTGAACTCTTTCCCTACACGACGCTCTTCCGATCTNNNAATGCCATACACCACTTTTGTC
AGCACACGTCTGAACTCTTTCCCTACACGACGCTCTTCCGATCTNNNAATGCCATACACCACTTTTGTC
CACGTCTGAACTCTTTCCCTACACGACGCTCTTCCGATCTNNNTCCCAAAGTGCTAGGATTACAGGCA
CACGTCTGAACTCTTTCCCTACACGACGCTCTTCCGATCTNNNTCCCAAAGTGCTAGGATTACAGGCA
AGCACACGTCTGAACTCTTTCCCTACACGACGCTCTTCCGATCTNNNGAAGAATGGTGTGAACCCAG
AGCACACGTCTGAACTCTTTCCCTACACGACGCTCTTCCGATCTNNNGAAGAATGGTGTGAACCCAG
AGCACACGTCTGAACTCTTTCCCTACACGACGCTCTTCCGATCTNNNCTCTTGTGTGTGTTGCTGTGT
AGCACACGTCTGAACTCTTTCCCTACACGACGCTCTTCCGATCTNNNCTCTTGTGTGTGTTGCTGTGT
AGCACACGTCTGAACTCTTTCCCTACACGACGCTCTTCCGATCTNNNAGTGAGCACCAATTGTTTGTC
AGCACACGTCTGAACTCTTTCCCTACACGACGCTCTTCCGATCTNNNAGTGAGCACCAATTGTTTGTC
GCACACGTCTGAACTCTTTCCCTACACGACGCTCTTCCGATCTNNNGATCACAAGGTCAGGAGATCGA
GCACACGTCTGAACTCTTTCCCTACACGACGCTCTTCCGATCTNNNGATCACAAGGTCAGGAGATCGA
AGCACACGTCTGAACTCTTTCCCTACACGACGCTCTTCCGATCTNNNGTTTTCACAAGAACTTCCTGGG
AGCACACGTCTGAACTCTTTCCCTACACGACGCTCTTCCGATCTNNNGTTTTCACAAGAACTTCCTGGG
GAGCACACGTCTGAACTCTTTCCCTACACGACGCTCTTCCGATCTNNNTGCCTGTGCTTTTGTTATCCA
GAGCACACGTCTGAACTCTTTCCCTACACGACGCTCTTCCGATCTNNNTGCCTGTGCTTTTGTTATCCA
AGCACACGTCTGAACTCTTTCCCTACACGACGCTCTTCCGATCTNNNAGTGAGCAGAAATAGTGCCAC
AGCACACGTCTGAACTCTTTCCCTACACGACGCTCTTCCGATCTNNNAGTGAGCAGAAATAGTGCCAC
GAGCACACGTCTGAACTCTTTCCCTACACGACGCTCTTCCGATCTNNNAGGAAGGTGAAGTGAACAA

GAGCACACGTCTGAACTCTTTCCCTACACGACGCTCTTCCGATCTNNNAGGAAGGTGAAGTGAACAA
GAGCACACGTCTGAACTCTTTCCCTACACGACGCTCTTCCGATCTNNNTCTTTTTTCTTTTAGGTTCTG
GAGCACACGTCTGAACTCTTTCCCTACACGACGCTCTTCCGATCTNNNTCTTTTTTCTTTTAGGTTCTG
AGCACACGTCTGAACTCTTTCCCTACACGACGCTCTTCCGATCTNNNGTTCAAGCGATTCTCCTGCATC
AGCACACGTCTGAACTCTTTCCCTACACGACGCTCTTCCGATCTNNNGTTCAAGCGATTCTCCTGCATC
GAGCACACGTCTGAACTCTTTCCCTACACGACGCTCTTCCGATCTNNNCCACGCCCGGATAATTTTTG
GAGCACACGTCTGAACTCTTTCCCTACACGACGCTCTTCCGATCTNNNCCACGCCCGGATAATTTTTG
GAGCACACGTCTGAACTCTTTCCCTACACGACGCTCTTCCGATCTNNNGTACATTTTTGTGCCCGTGTG
GAGCACACGTCTGAACTCTTTCCCTACACGACGCTCTTCCGATCTNNNGTACATTTTTGTGCCCGTGTG
GAGCACACGTCTGAACTCTTTCCCTACACGACGCTCTTCCGATCTNNNTCTTTGCCAGCCAGAGTATCA
GAGCACACGTCTGAACTCTTTCCCTACACGACGCTCTTCCGATCTNNNTCTTTGCCAGCCAGAGTATCA
GAGCACACGTCTGAACTCTTTCCCTACACGACGCTCTTCCGATCTNNNCTTAGGTCACGAACCCTACAG
GAGCACACGTCTGAACTCTTTCCCTACACGACGCTCTTCCGATCTNNNCTTAGGTCACGAACCCTACAG
AGCACACGTCTGAACTCTTTCCCTACACGACGCTCTTCCGATCTNNNTCACTTTATCACATGCACATGG
AGCACACGTCTGAACTCTTTCCCTACACGACGCTCTTCCGATCTNNNTCACTTTATCACATGCACATGG
GAGCACACGTCTGAACTCTTTCCCTACACGACGCTCTTCCGATCTNNNATGCCTGTAATCCCAGCTACT
GAGCACACGTCTGAACTCTTTCCCTACACGACGCTCTTCCGATCTNNNATGCCTGTAATCCCAGCTACT
AGCACACGTCTGAACTCTTTCCCTACACGACGCTCTTCCGATCTNNNCTGCTCTAAAGGACCCTACTGA
AGCACACGTCTGAACTCTTTCCCTACACGACGCTCTTCCGATCTNNNCTGCTCTAAAGGACCCTACTGA
GCACACGTCTGAACTCTTTCCCTACACGACGCTCTTCCGATCTNNNCTGTCAGCATAACCTGAGGAACT
GCACACGTCTGAACTCTTTCCCTACACGACGCTCTTCCGATCTNNNCTGTCAGCATAACCTGAGGAACT
GCACACGTCTGAACTCTTTCCCTACACGACGCTCTTCCGATCTNNNCTGTAATCCCAGCTACTCAGGAG
GCACACGTCTGAACTCTTTCCCTACACGACGCTCTTCCGATCTNNNCTGTAATCCCAGCTACTCAGGAG
GAGCACACGTCTGAACTCTTTCCCTACACGACGCTCTTCCGATCTNNNTGCTCTACTTTTTTGCTCCAG
GAGCACACGTCTGAACTCTTTCCCTACACGACGCTCTTCCGATCTNNNTGCTCTACTTTTTTGCTCCAG
GAGCACACGTCTGAACTCTTTCCCTACACGACGCTCTTCCGATCTNNNCAGAGGTTGTAGTGAGCCAA
GAGCACACGTCTGAACTCTTTCCCTACACGACGCTCTTCCGATCTNNNCAGAGGTTGTAGTGAGCCAA
GCACACGTCTGAACTCTTTCCCTACACGACGCTCTTCCGATCTNNNTCAAAAGCTCTGCAGGTGATTTG
GCACACGTCTGAACTCTTTCCCTACACGACGCTCTTCCGATCTNNNTCAAAAGCTCTGCAGGTGATTTG
GCACACGTCTGAACTCTTTCCCTACACGACGCTCTTCCGATCTNNNCTGAGGCAGGAGAATCACTTGA
GCACACGTCTGAACTCTTTCCCTACACGACGCTCTTCCGATCTNNNCTGAGGCAGGAGAATCACTTGA
AGCACACGTCTGAACTCTTTCCCTACACGACGCTCTTCCGATCTNNNGTACCATCCCAGGCGGTCTTCA
AGCACACGTCTGAACTCTTTCCCTACACGACGCTCTTCCGATCTNNNGTACCATCCCAGGCGGTCTTCA
AGCACACGTCTGAACTCTTTCCCTACACGACGCTCTTCCGATCTNNNTTTAGTAGAGACGGGGTTTCA
AGCACACGTCTGAACTCTTTCCCTACACGACGCTCTTCCGATCTNNNTTTAGTAGAGACGGGGTTTCA
AGCACACGTCTGAACTCTTTCCCTACACGACGCTCTTCCGATCTNNNAAGGAGGTGACAAGTGCTATG
AGCACACGTCTGAACTCTTTCCCTACACGACGCTCTTCCGATCTNNNAAGGAGGTGACAAGTGCTATG
GAGCACACGTCTGAACTCTTTCCCTACACGACGCTCTTCCGATCTNNNTCCCTCTGTACATGTCTATCC
GAGCACACGTCTGAACTCTTTCCCTACACGACGCTCTTCCGATCTNNNTCCCTCTGTACATGTCTATCC
GAGCACACGTCTGAACTCTTTCCCTACACGACGCTCTTCCGATCTNNNTCATTCCATGCATCAGACTGT
GAGCACACGTCTGAACTCTTTCCCTACACGACGCTCTTCCGATCTNNNTCATTCCATGCATCAGACTGT
GAGCACACGTCTGAACTCTTTCCCTACACGACGCTCTTCCGATCTNNNCTGAGACAGGAGAATCGCTT
GAGCACACGTCTGAACTCTTTCCCTACACGACGCTCTTCCGATCTNNNCTGAGACAGGAGAATCGCTT

AGCACACGTCTGAACTCTTTCCCTACACGACGCTCTTCCGATCTNNNGTTACATTGCTGCACACATCTG
AGCACACGTCTGAACTCTTTCCCTACACGACGCTCTTCCGATCTNNNGTTACATTGCTGCACACATCTG
GAGCACACGTCTGAACTCTTTCCCTACACGACGCTCTTCCGATCTNNNGAATTGCTTGAACCCGTGAG
GAGCACACGTCTGAACTCTTTCCCTACACGACGCTCTTCCGATCTNNNGAATTGCTTGAACCCGTGAG
GAGCACACGTCTGAACTCTTTCCCTACACGACGCTCTTCCGATCTNNNTGGAACATTGCATGGTTTAT
GAGCACACGTCTGAACTCTTTCCCTACACGACGCTCTTCCGATCTNNNTGGAACATTGCATGGTTTAT
GAGCACACGTCTGAACTCTTTCCCTACACGACGCTCTTCCGATCTNNNTGGGCTGTACTTTCACAACTT
GAGCACACGTCTGAACTCTTTCCCTACACGACGCTCTTCCGATCTNNNTGGGCTGTACTTTCACAACTT
AGCACACGTCTGAACTCTTTCCCTACACGACGCTCTTCCGATCTNNNCTAAAACACTCAGGTCTCGGG
AGCACACGTCTGAACTCTTTCCCTACACGACGCTCTTCCGATCTNNNCTAAAACACTCAGGTCTCGGG
GAGCACACGTCTGAACTCTTTCCCTACACGACGCTCTTCCGATCTNNNCATGGTGAAACTCTGCCTCTA
GAGCACACGTCTGAACTCTTTCCCTACACGACGCTCTTCCGATCTNNNCATGGTGAAACTCTGCCTCTA
GCACACGTCTGAACTCTTTCCCTACACGACGCTCTTCCGATCTNNNTGGTTGATGGTGTTGCTCAATT
GCACACGTCTGAACTCTTTCCCTACACGACGCTCTTCCGATCTNNNTGGTTGATGGTGTTGCTCAATT
AGAGCACACGTCTGAACTCTTTCCCTACACGACGCTCTTCCGATCTNNNACCTTGTCAAAAAAAAAAA
AGAGCACACGTCTGAACTCTTTCCCTACACGACGCTCTTCCGATCTNNNACCTTGTCAAAAAAAAAAA
GAGCACACGTCTGAACTCTTTCCCTACACGACGCTCTTCCGATCTNNNCAAGCCTTTGTAATGGTGAG
GAGCACACGTCTGAACTCTTTCCCTACACGACGCTCTTCCGATCTNNNCAAGCCTTTGTAATGGTGAG
AGCACACGTCTGAACTCTTTCCCTACACGACGCTCTTCCGATCTNNNGTCCATACTTACCTAATTTTGA
AGCACACGTCTGAACTCTTTCCCTACACGACGCTCTTCCGATCTNNNGTCCATACTTACCTAATTTTGA
AGCACACGTCTGAACTCTTTCCCTACACGACGCTCTTCCGATCTNNNGTTGTGTGACTTGTCTGTGTG
AGCACACGTCTGAACTCTTTCCCTACACGACGCTCTTCCGATCTNNNGTTGTGTGACTTGTCTGTGTG
AGCACACGTCTGAACTCTTTCCCTACACGACGCTCTTCCGATCTNNNAGAATTGCTTGAACCAGGGAG
AGCACACGTCTGAACTCTTTCCCTACACGACGCTCTTCCGATCTNNNAGAATTGCTTGAACCAGGGAG
CACACGTCTGAACTCTTTCCCTACACGACGCTCTTCCGATCTNNNGCAAATGCTCCTCATTTCCTCCTCA
CACACGTCTGAACTCTTTCCCTACACGACGCTCTTCCGATCTNNNGCAAATGCTCCTCATTTCCTCCTCA
GAGCACACGTCTGAACTCTTTCCCTACACGACGCTCTTCCGATCTNNNTGCCATTACTAAGCTGTTTCG
GAGCACACGTCTGAACTCTTTCCCTACACGACGCTCTTCCGATCTNNNTGCCATTACTAAGCTGTTTCG
AGCACACGTCTGAACTCTTTCCCTACACGACGCTCTTCCGATCTNNNCTGAGGCAGGAGAATTGCTT
AGCACACGTCTGAACTCTTTCCCTACACGACGCTCTTCCGATCTNNNCTGAGGCAGGAGAATTGCTT
AGCACACGTCTGAACTCTTTCCCTACACGACGCTCTTCCGATCTNNNCAAATTTGCACTGCCATCAGAG
AGCACACGTCTGAACTCTTTCCCTACACGACGCTCTTCCGATCTNNNCAAATTTGCACTGCCATCAGAG
GAGCACACGTCTGAACTCTTTCCCTACACGACGCTCTTCCGATCTNNNTAATGCAGCATCCTGTTTGGT
GAGCACACGTCTGAACTCTTTCCCTACACGACGCTCTTCCGATCTNNNTAATGCAGCATCCTGTTTGGT
AGCACACGTCTGAACTCTTTCCCTACACGACGCTCTTCCGATCTNNNGAGACTCGACAGCACAGAGA
AGCACACGTCTGAACTCTTTCCCTACACGACGCTCTTCCGATCTNNNGAGACTCGACAGCACAGAGA
GCACACGTCTGAACTCTTTCCCTACACGACGCTCTTCCGATCTNNNATCCCTGCTTTCTTCTACTCTGC
GCACACGTCTGAACTCTTTCCCTACACGACGCTCTTCCGATCTNNNATCCCTGCTTTCTTCTACTCTGC
AGCACACGTCTGAACTCTTTCCCTACACGACGCTCTTCCGATCTNNNACAAATTTCACAGAGCCTGGTT
AGCACACGTCTGAACTCTTTCCCTACACGACGCTCTTCCGATCTNNNACAAATTTCACAGAGCCTGGTT
GCACACGTCTGAACTCTTTCCCTACACGACGCTCTTCCGATCTNNNGTTAGAGGTGTGTCTCCTGGAA
GCACACGTCTGAACTCTTTCCCTACACGACGCTCTTCCGATCTNNNGTTAGAGGTGTGTCTCCTGGAA
AGCACACGTCTGAACTCTTTCCCTACACGACGCTCTTCCGATCTNNNAAAATAAATGTCGGTGGACCA

AGCACACGTCTGAACTCTTTCCCTACACGACGCTCTTCCGATCTNNNAAAATAAATGTCGGTGGACCA
GCACACGTCTGAACTCTTTCCCTACACGACGCTCTTCCGATCTNNNCTGTAATCCCAGCTACTCCAGAG
GCACACGTCTGAACTCTTTCCCTACACGACGCTCTTCCGATCTNNNCTGTAATCCCAGCTACTCCAGAG
GAGCACACGTCTGAACTCTTTCCCTACACGACGCTCTTCCGATCTNNNGTATGTGAGGTCGCAATTGT
GAGCACACGTCTGAACTCTTTCCCTACACGACGCTCTTCCGATCTNNNGTATGTGAGGTCGCAATTGT
GCACACGTCTGAACTCTTTCCCTACACGACGCTCTTCCGATCTNNNCAGCCTATGAATTCCCTGAGCAG
GCACACGTCTGAACTCTTTCCCTACACGACGCTCTTCCGATCTNNNCAGCCTATGAATTCCCTGAGCAG
GAGCACACGTCTGAACTCTTTCCCTACACGACGCTCTTCCGATCTNNNAGACAAACCATATCCCTGCCA
GAGCACACGTCTGAACTCTTTCCCTACACGACGCTCTTCCGATCTNNNAGACAAACCATATCCCTGCCA
GCACACGTCTGAACTCTTTCCCTACACGACGCTCTTCCGATCTNNNAGGTCTTGAGTCTTTGGCTTCA
GCACACGTCTGAACTCTTTCCCTACACGACGCTCTTCCGATCTNNNAGGTCTTGAGTCTTTGGCTTCA
GAGCACACGTCTGAACTCTTTCCCTACACGACGCTCTTCCGATCTNNNTTCAAGACCAGCCTGAGCAA
GAGCACACGTCTGAACTCTTTCCCTACACGACGCTCTTCCGATCTNNNTTCAAGACCAGCCTGAGCAA
CACACGTCTGAACTCTTTCCCTACACGACGCTCTTCCGATCTNNNTGCTAAGCATGGGGAGAGAATTT
CACACGTCTGAACTCTTTCCCTACACGACGCTCTTCCGATCTNNNTGCTAAGCATGGGGAGAGAATTT
GCACACGTCTGAACTCTTTCCCTACACGACGCTCTTCCGATCTNNNGATCAGAGCCTAAGAGACCTGT
GCACACGTCTGAACTCTTTCCCTACACGACGCTCTTCCGATCTNNNGATCAGAGCCTAAGAGACCTGT
CACACGTCTGAACTCTTTCCCTACACGACGCTCTTCCGATCTNNNATCACTTTCCCACTCACCATCTCCA
CACACGTCTGAACTCTTTCCCTACACGACGCTCTTCCGATCTNNNATCACTTTCCCACTCACCATCTCCA
AGCACACGTCTGAACTCTTTCCCTACACGACGCTCTTCCGATCTNNNAGAATCATTTGAGCCCAGGAG
AGCACACGTCTGAACTCTTTCCCTACACGACGCTCTTCCGATCTNNNAGAATCATTTGAGCCCAGGAG
GAGCACACGTCTGAACTCTTTCCCTACACGACGCTCTTCCGATCTNNNTGCCCGGCCCATATTATGTA
GAGCACACGTCTGAACTCTTTCCCTACACGACGCTCTTCCGATCTNNNTGCCCGGCCCATATTATGTA
GCACACGTCTGAACTCTTTCCCTACACGACGCTCTTCCGATCTNNNCCACAGGTGGCAAGAACTCTTA
GCACACGTCTGAACTCTTTCCCTACACGACGCTCTTCCGATCTNNNCCACAGGTGGCAAGAACTCTTA
AGCACACGTCTGAACTCTTTCCCTACACGACGCTCTTCCGATCTNNNATTTCTGAGTCTGCAGTGGGA
AGCACACGTCTGAACTCTTTCCCTACACGACGCTCTTCCGATCTNNNATTTCTGAGTCTGCAGTGGGA
GAGCACACGTCTGAACTCTTTCCCTACACGACGCTCTTCCGATCTNNNCAGGAGAATTGCTTGAAACC
GAGCACACGTCTGAACTCTTTCCCTACACGACGCTCTTCCGATCTNNNCAGGAGAATTGCTTGAAACC
AGCACACGTCTGAACTCTTTCCCTACACGACGCTCTTCCGATCTNNNTCTGGAGAGGTTGTCCTTTTCT
AGCACACGTCTGAACTCTTTCCCTACACGACGCTCTTCCGATCTNNNTCTGGAGAGGTTGTCCTTTTCT
AGCACACGTCTGAACTCTTTCCCTACACGACGCTCTTCCGATCTNNNTTTGCCCCTTTCCATGGTAAAAG
AGCACACGTCTGAACTCTTTCCCTACACGACGCTCTTCCGATCTNNNTTTGCCCCTTTCCATGGTAAAAG
AGCACACGTCTGAACTCTTTCCCTACACGACGCTCTTCCGATCTNNNGCAAACCCATAACAAATGTTGC
AGCACACGTCTGAACTCTTTCCCTACACGACGCTCTTCCGATCTNNNGCAAACCCATAACAAATGTTGC
GAGCACACGTCTGAACTCTTTCCCTACACGACGCTCTTCCGATCTNNNTGTTGTTCTACCGTCAAATCG
GAGCACACGTCTGAACTCTTTCCCTACACGACGCTCTTCCGATCTNNNTGTTGTTCTACCGTCAAATCG
AGCACACGTCTGAACTCTTTCCCTACACGACGCTCTTCCGATCTNNNCCTGCTTGACATGTGTGACCAT
AGCACACGTCTGAACTCTTTCCCTACACGACGCTCTTCCGATCTNNNCCTGCTTGACATGTGTGACCAT
GCACACGTCTGAACTCTTTCCCTACACGACGCTCTTCCGATCTNNNGGCCTCTATTCCTACTGTCATGG
GCACACGTCTGAACTCTTTCCCTACACGACGCTCTTCCGATCTNNNGGCCTCTATTCCTACTGTCATGG
AGCACACGTCTGAACTCTTTCCCTACACGACGCTCTTCCGATCTNNNTGGCAGCTGCAATAGACAAAA
AGCACACGTCTGAACTCTTTCCCTACACGACGCTCTTCCGATCTNNNTGGCAGCTGCAATAGACAAAA

AGCACACGTCTGAACTCTTTCCCTACACGACGCTCTTCCGATCTNNNAGATCGCACCTTACCTCAACAAT
AGCACACGTCTGAACTCTTTCCCTACACGACGCTCTTCCGATCTNNNAGATCGCACCTTACCTCAACAAT
AGCACACGTCTGAACTCTTTCCCTACACGACGCTCTTCCGATCTNNNGTAGAGTCAAGGTTTCACCGTC
AGCACACGTCTGAACTCTTTCCCTACACGACGCTCTTCCGATCTNNNGTAGAGTCAAGGTTTCACCGTC
AGCACACGTCTGAACTCTTTCCCTACACGACGCTCTTCCGATCTNNNGCTAGGAAACAAGTGCCTCAAT
AGCACACGTCTGAACTCTTTCCCTACACGACGCTCTTCCGATCTNNNGCTAGGAAACAAGTGCCTCAAT
GAGCACACGTCTGAACTCTTTCCCTACACGACGCTCTTCCGATCTNNNCTGTAGTCCCAGCTACTCAGG
GAGCACACGTCTGAACTCTTTCCCTACACGACGCTCTTCCGATCTNNNCTGTAGTCCCAGCTACTCAGG
AGCACACGTCTGAACTCTTTCCCTACACGACGCTCTTCCGATCTNNNAGAATTGCTTGAACCTGGGAG
AGCACACGTCTGAACTCTTTCCCTACACGACGCTCTTCCGATCTNNNAGAATTGCTTGAACCTGGGAG
AGCACACGTCTGAACTCTTTCCCTACACGACGCTCTTCCGATCTNNNTTGTGAATAGTGCTGCAATGGA
AGCACACGTCTGAACTCTTTCCCTACACGACGCTCTTCCGATCTNNNTTGTGAATAGTGCTGCAATGGA
AGCACACGTCTGAACTCTTTCCCTACACGACGCTCTTCCGATCTNNNATACAAATTCTTTATTTCCTCCCC
AGCACACGTCTGAACTCTTTCCCTACACGACGCTCTTCCGATCTNNNATACAAATTCTTTATTTCCTCCCC
AGCACACGTCTGAACTCTTTCCCTACACGACGCTCTTCCGATCTNNNGTCATCCTAGCACTTTGGGAGG
AGCACACGTCTGAACTCTTTCCCTACACGACGCTCTTCCGATCTNNNGTCATCCTAGCACTTTGGGAGG
GAGCACACGTCTGAACTCTTTCCCTACACGACGCTCTTCCGATCTNNNGGTTGCTGGGGATATTGAGA
GAGCACACGTCTGAACTCTTTCCCTACACGACGCTCTTCCGATCTNNNGGTTGCTGGGGATATTGAGA
AGCACACGTCTGAACTCTTTCCCTACACGACGCTCTTCCGATCTNNNAGATCATCATAAAGGTCTTCGTT
AGCACACGTCTGAACTCTTTCCCTACACGACGCTCTTCCGATCTNNNAGATCATCATAAAGGTCTTCGTT
GAGCACACGTCTGAACTCTTTCCCTACACGACGCTCTTCCGATCTNNNTTGTCTGGATTTGAGCCATAC
GAGCACACGTCTGAACTCTTTCCCTACACGACGCTCTTCCGATCTNNNTTGTCTGGATTTGAGCCATAC
GAGCACACGTCTGAACTCTTTCCCTACACGACGCTCTTCCGATCTNNNGGAGATCAAGGCTGCAGTGA
GAGCACACGTCTGAACTCTTTCCCTACACGACGCTCTTCCGATCTNNNGGAGATCAAGGCTGCAGTGA
AGAGCACACGTCTGAACTCTTTCCCTACACGACGCTCTTCCGATCTNNNATACCAGCTGAAGCAGATAA
AGAGCACACGTCTGAACTCTTTCCCTACACGACGCTCTTCCGATCTNNNATACCAGCTGAAGCAGATAA
AGCACACGTCTGAACTCTTTCCCTACACGACGCTCTTCCGATCTNNNGAGAATCACTTGAACCAGGAGG
AGCACACGTCTGAACTCTTTCCCTACACGACGCTCTTCCGATCTNNNGAGAATCACTTGAACCAGGAGG
GCACACGTCTGAACTCTTTCCCTACACGACGCTCTTCCGATCTNNNTGTGCTATGGATTGGTGGAAAGA
GCACACGTCTGAACTCTTTCCCTACACGACGCTCTTCCGATCTNNNTGTGCTATGGATTGGTGGAAAGA
AGCACACGTCTGAACTCTTTCCCTACACGACGCTCTTCCGATCTNNNTTCAAAAGTAGGACACACCACC
AGCACACGTCTGAACTCTTTCCCTACACGACGCTCTTCCGATCTNNNTTCAAAAGTAGGACACACCACC
GAGCACACGTCTGAACTCTTTCCCTACACGACGCTCTTCCGATCTNNNGCCATCCCTGTGTGTGAGTTC
GAGCACACGTCTGAACTCTTTCCCTACACGACGCTCTTCCGATCTNNNGCCATCCCTGTGTGTGAGTTC
AGCACACGTCTGAACTCTTTCCCTACACGACGCTCTTCCGATCTNNNAAACTGGCTGAATGAGACAGAG
AGCACACGTCTGAACTCTTTCCCTACACGACGCTCTTCCGATCTNNNAAACTGGCTGAATGAGACAGAG
AGCACACGTCTGAACTCTTTCCCTACACGACGCTCTTCCGATCTNNNCTTGCAGTGAGCCAAGATTGCC
AGCACACGTCTGAACTCTTTCCCTACACGACGCTCTTCCGATCTNNNCTTGCAGTGAGCCAAGATTGCC
AGCACACGTCTGAACTCTTTCCCTACACGACGCTCTTCCGATCTNNNTCGCTGTAGATCTGACGTACACT
AGCACACGTCTGAACTCTTTCCCTACACGACGCTCTTCCGATCTNNNTCGCTGTAGATCTGACGTACACT
AGCACACGTCTGAACTCTTTCCCTACACGACGCTCTTCCGATCTNNNGGTGCCTGTAATCCCAACTACTC
AGCACACGTCTGAACTCTTTCCCTACACGACGCTCTTCCGATCTNNNGGTGCCTGTAATCCCAACTACTC
AGAGCACACGTCTGAACTCTTTCCCTACACGACGCTCTTCCGATCTNNNGCGACAAAGTGAGACTCCAT

AGAGCACACGTCTGAACTCTTTCCCTACACGACGCTCTTCCGATCTNNNGCGACAAAGTGAGACTCCA
GAGCACACGTCTGAACTCTTTCCCTACACGACGCTCTTCCGATCTNNNCACTATTCACTTGGCCCACTCT
GAGCACACGTCTGAACTCTTTCCCTACACGACGCTCTTCCGATCTNNNCACTATTCACTTGGCCCACTCT
AGCACACGTCTGAACTCTTTCCCTACACGACGCTCTTCCGATCTNNNCTCTAGGGATTTCTTCTCCTGG
AGCACACGTCTGAACTCTTTCCCTACACGACGCTCTTCCGATCTNNNCTCTAGGGATTTCTTCTCCTGG
AGCACACGTCTGAACTCTTTCCCTACACGACGCTCTTCCGATCTNNNTGTTGCACCCTATCAACTCATCA
AGCACACGTCTGAACTCTTTCCCTACACGACGCTCTTCCGATCTNNNTGTTGCACCCTATCAACTCATCA
AGCACACGTCTGAACTCTTTCCCTACACGACGCTCTTCCGATCTNNNACGGTTTCAAACAATGTGCTG
AGCACACGTCTGAACTCTTTCCCTACACGACGCTCTTCCGATCTNNNACGGTTTCAAACAATGTGCTG
GAGCACACGTCTGAACTCTTTCCCTACACGACGCTCTTCCGATCTNNNCCATCTGCAAAGACGGACAA
GAGCACACGTCTGAACTCTTTCCCTACACGACGCTCTTCCGATCTNNNCCATCTGCAAAGACGGACAA
GAGCACACGTCTGAACTCTTTCCCTACACGACGCTCTTCCGATCTNNNCAGGAGAATGGTGTGAACGT
GAGCACACGTCTGAACTCTTTCCCTACACGACGCTCTTCCGATCTNNNCAGGAGAATGGTGTGAACGT
GAGCACACGTCTGAACTCTTTCCCTACACGACGCTCTTCCGATCTNNNACAAAACAAAAAACCCACCA
GAGCACACGTCTGAACTCTTTCCCTACACGACGCTCTTCCGATCTNNNACAAAACAAAAAACCCACCA
GCACACGTCTGAACTCTTTCCCTACACGACGCTCTTCCGATCTNNNCCAGATCAAAGACAACATCGGT
GCACACGTCTGAACTCTTTCCCTACACGACGCTCTTCCGATCTNNNCCAGATCAAAGACAACATCGGT
GAGCACACGTCTGAACTCTTTCCCTACACGACGCTCTTCCGATCTNNNGTAGAAAGGAGAGGTTTGG
GAGCACACGTCTGAACTCTTTCCCTACACGACGCTCTTCCGATCTNNNGTAGAAAGGAGAGGTTTGG
GCACACGTCTGAACTCTTTCCCTACACGACGCTCTTCCGATCTNNNTAAAAGGCGTGAAATAGTAACG
GCACACGTCTGAACTCTTTCCCTACACGACGCTCTTCCGATCTNNNTAAAAGGCGTGAAATAGTAACG
AGCACACGTCTGAACTCTTTCCCTACACGACGCTCTTCCGATCTNNNGGATCACTTGAGGTCAGGAGT
AGCACACGTCTGAACTCTTTCCCTACACGACGCTCTTCCGATCTNNNGGATCACTTGAGGTCAGGAGT
GCACACGTCTGAACTCTTTCCCTACACGACGCTCTTCCGATCTNNNCTCCAGAGTCCTCAAGTTCCAGT
GCACACGTCTGAACTCTTTCCCTACACGACGCTCTTCCGATCTNNNCTCCAGAGTCCTCAAGTTCCAGT
GCACACGTCTGAACTCTTTCCCTACACGACGCTCTTCCGATCTNNNTTCCACTACTTTGTCTCTGACCCC
GCACACGTCTGAACTCTTTCCCTACACGACGCTCTTCCGATCTNNNTTCCACTACTTTGTCTCTGACCCC
AGCACACGTCTGAACTCTTTCCCTACACGACGCTCTTCCGATCTNNNGGAGCAGGAGAGTTTCAGCTT
AGCACACGTCTGAACTCTTTCCCTACACGACGCTCTTCCGATCTNNNGGAGCAGGAGAGTTTCAGCTT
GAGCACACGTCTGAACTCTTTCCCTACACGACGCTCTTCCGATCTNNNGAGGCTGAGGCAGGAGAAT
GAGCACACGTCTGAACTCTTTCCCTACACGACGCTCTTCCGATCTNNNGAGGCTGAGGCAGGAGAAT
GAGCACACGTCTGAACTCTTTCCCTACACGACGCTCTTCCGATCTNNNAAAACTGAAGCATCAAAGCCA
GAGCACACGTCTGAACTCTTTCCCTACACGACGCTCTTCCGATCTNNNAAAACTGAAGCATCAAAGCCA
AGCACACGTCTGAACTCTTTCCCTACACGACGCTCTTCCGATCTNNNACTGCTCCATCATCTTGGGATT
AGCACACGTCTGAACTCTTTCCCTACACGACGCTCTTCCGATCTNNNACTGCTCCATCATCTTGGGATT
GAGCACACGTCTGAACTCTTTCCCTACACGACGCTCTTCCGATCTNNNCAAGGAACAGTTGGAGACC
GAGCACACGTCTGAACTCTTTCCCTACACGACGCTCTTCCGATCTNNNCAAGGAACAGTTGGAGACC
GAGCACACGTCTGAACTCTTTCCCTACACGACGCTCTTCCGATCTNNNCTCCTTGCCTTCTGCCATAATT
GAGCACACGTCTGAACTCTTTCCCTACACGACGCTCTTCCGATCTNNNCTCCTTGCCTTCTGCCATAATT
GAGCACACGTCTGAACTCTTTCCCTACACGACGCTCTTCCGATCTNNNCCACTCTCGAAAATAGTTTGG
GAGCACACGTCTGAACTCTTTCCCTACACGACGCTCTTCCGATCTNNNCCACTCTCGAAAATAGTTTGG
GAGCACACGTCTGAACTCTTTCCCTACACGACGCTCTTCCGATCTNNNCACTCAAAAGGTATCCGTGG
GAGCACACGTCTGAACTCTTTCCCTACACGACGCTCTTCCGATCTNNNCACTCAAAAGGTATCCGTGG

AGCACACGTCTGAACTCTTTCCCTACACGACGCTCTTCCGATCTNNNCGGCAAAAAGTTTCCTTCTAGC
AGCACACGTCTGAACTCTTTCCCTACACGACGCTCTTCCGATCTNNNCGGCAAAAAGTTTCCTTCTAGC
GCACACGTCTGAACTCTTTCCCTACACGACGCTCTTCCGATCTNNNAACCACTAGTCTGCTCTCTTTGGC
GCACACGTCTGAACTCTTTCCCTACACGACGCTCTTCCGATCTNNNAACCACTAGTCTGCTCTCTTTGGC
GAGCACACGTCTGAACTCTTTCCCTACACGACGCTCTTCCGATCTNNNCTGAGGCAGGAGAATCACTT
GAGCACACGTCTGAACTCTTTCCCTACACGACGCTCTTCCGATCTNNNCTGAGGCAGGAGAATCACTT
ACACGTCTGAACTCTTTCCCTACACGACGCTCTTCCGATCTNNNTGGGATTGTGTGGAATCTCTAGACA
ACACGTCTGAACTCTTTCCCTACACGACGCTCTTCCGATCTNNNTGGGATTGTGTGGAATCTCTAGACA
GAGCACACGTCTGAACTCTTTCCCTACACGACGCTCTTCCGATCTNNNTGCACTGGTTTTTGATCTTGC
GAGCACACGTCTGAACTCTTTCCCTACACGACGCTCTTCCGATCTNNNTGCACTGGTTTTTGATCTTGC
GAGCACACGTCTGAACTCTTTCCCTACACGACGCTCTTCCGATCTNNNCTCTAACCCATGAAGACACAG
GAGCACACGTCTGAACTCTTTCCCTACACGACGCTCTTCCGATCTNNNCTCTAACCCATGAAGACACAG
GCACACGTCTGAACTCTTTCCCTACACGACGCTCTTCCGATCTNNNGCACTGTGCATACATAAGCCAAA
GCACACGTCTGAACTCTTTCCCTACACGACGCTCTTCCGATCTNNNGCACTGTGCATACATAAGCCAAA
AGCACACGTCTGAACTCTTTCCCTACACGACGCTCTTCCGATCTNNNGCCTAGTGAAGCATGTGTAGG
AGCACACGTCTGAACTCTTTCCCTACACGACGCTCTTCCGATCTNNNGCCTAGTGAAGCATGTGTAGG
AGAGCACACGTCTGAACTCTTTCCCTACACGACGCTCTTCCGATCTNNNGATTATCGATTGAGCCCAG
AGAGCACACGTCTGAACTCTTTCCCTACACGACGCTCTTCCGATCTNNNGATTATCGATTGAGCCCAG
GAGCACACGTCTGAACTCTTTCCCTACACGACGCTCTTCCGATCTNNNAATGTTGTCCTGCAGGGTCA
GAGCACACGTCTGAACTCTTTCCCTACACGACGCTCTTCCGATCTNNNAATGTTGTCCTGCAGGGTCA
GAGCACACGTCTGAACTCTTTCCCTACACGACGCTCTTCCGATCTNNNCCTCCTTAGGTATATTCGGG
GAGCACACGTCTGAACTCTTTCCCTACACGACGCTCTTCCGATCTNNNCCTCCTTAGGTATATTCGGG
AGCACACGTCTGAACTCTTTCCCTACACGACGCTCTTCCGATCTNNNGCAGTAGAAACGGGATCTGA
AGCACACGTCTGAACTCTTTCCCTACACGACGCTCTTCCGATCTNNNGCAGTAGAAACGGGATCTGA
GAGCACACGTCTGAACTCTTTCCCTACACGACGCTCTTCCGATCTNNNCACAAACCACTCCCAGAGAAC
GAGCACACGTCTGAACTCTTTCCCTACACGACGCTCTTCCGATCTNNNCACAAACCACTCCCAGAGAAC
GCACACGTCTGAACTCTTTCCCTACACGACGCTCTTCCGATCTNNNGTCTCTTGAATCCTCATCAGCAT
GCACACGTCTGAACTCTTTCCCTACACGACGCTCTTCCGATCTNNNGTCTCTTGAATCCTCATCAGCAT
AGAGCACACGTCTGAACTCTTTCCCTACACGACGCTCTTCCGATCTNNNCTGTAATCACAGCTACTCGG
AGAGCACACGTCTGAACTCTTTCCCTACACGACGCTCTTCCGATCTNNNCTGTAATCACAGCTACTCGG
GAGCACACGTCTGAACTCTTTCCCTACACGACGCTCTTCCGATCTNNNGAGAATCGTTTGAACCCAGG
GAGCACACGTCTGAACTCTTTCCCTACACGACGCTCTTCCGATCTNNNGAGAATCGTTTGAACCCAGG
GAGCACACGTCTGAACTCTTTCCCTACACGACGCTCTTCCGATCTNNNTTTATTCTGGGGGAGGTAAT
GAGCACACGTCTGAACTCTTTCCCTACACGACGCTCTTCCGATCTNNNTTTATTCTGGGGGAGGTAAT
CACGTCTGAACTCTTTCCCTACACGACGCTCTTCCGATCTNNNTGGCTGCCACTTGTATGTTTTTTCAT
CACGTCTGAACTCTTTCCCTACACGACGCTCTTCCGATCTNNNTGGCTGCCACTTGTATGTTTTTTCAT
GCACACGTCTGAACTCTTTCCCTACACGACGCTCTTCCGATCTNNNTGTAGAAGTACTGTTATGGCTG
GCACACGTCTGAACTCTTTCCCTACACGACGCTCTTCCGATCTNNNTGTAGAAGTACTGTTATGGCTG
AGCACACGTCTGAACTCTTTCCCTACACGACGCTCTTCCGATCTNNNAAAAATGCAATGCCTGGATCC
AGCACACGTCTGAACTCTTTCCCTACACGACGCTCTTCCGATCTNNNAAAAATGCAATGCCTGGATCC
GAGCACACGTCTGAACTCTTTCCCTACACGACGCTCTTCCGATCTNNNAGAAGTGTGCAGACAGCTCT
GAGCACACGTCTGAACTCTTTCCCTACACGACGCTCTTCCGATCTNNNAGAAGTGTGCAGACAGCTCT
GCACACGTCTGAACTCTTTCCCTACACGACGCTCTTCCGATCTNNNCCACCACCAAACAGTGACCATAA

AGCACACGTCTGAACTCTTTCCCTACACGACGCTCTTCCGATCTNNNCCACCACCAAACAGTGACCATAA
AGCACACGTCTGAACTCTTTCCCTACACGACGCTCTTCCGATCTNNNCATCTGTAAATCCAGCTGCTTGG
AGCACACGTCTGAACTCTTTCCCTACACGACGCTCTTCCGATCTNNNCATCTGTAAATCCAGCTGCTTGG
AGCACACGTCTGAACTCTTTCCCTACACGACGCTCTTCCGATCTNNNCAGTTTTGAGCTCCACTGCATTC
AGCACACGTCTGAACTCTTTCCCTACACGACGCTCTTCCGATCTNNNCAGTTTTGAGCTCCACTGCATTC
AGCACACGTCTGAACTCTTTCCCTACACGACGCTCTTCCGATCTNNNATTCATAGCCACTCTGATTCCCC
AGCACACGTCTGAACTCTTTCCCTACACGACGCTCTTCCGATCTNNNATTCATAGCCACTCTGATTCCCC
AGAGCACACGTCTGAACTCTTTCCCTACACGACGCTCTTCCGATCTNNNGCGTATTAAGGTTTGAGAAT
AGAGCACACGTCTGAACTCTTTCCCTACACGACGCTCTTCCGATCTNNNGCGTATTAAGGTTTGAGAAT
AGCACACGTCTGAACTCTTTCCCTACACGACGCTCTTCCGATCTNNNCACCTGTAATCCCAGCTACTTGG
AGCACACGTCTGAACTCTTTCCCTACACGACGCTCTTCCGATCTNNNCACCTGTAATCCCAGCTACTTGG
AGCACACGTCTGAACTCTTTCCCTACACGACGCTCTTCCGATCTNNNCTGTAGTCCCAGCTACTCAAGA
AGCACACGTCTGAACTCTTTCCCTACACGACGCTCTTCCGATCTNNNCTGTAGTCCCAGCTACTCAAGA
AGCACACGTCTGAACTCTTTCCCTACACGACGCTCTTCCGATCTNNNTACACTGCATAGCCTTTCCAACT
AGCACACGTCTGAACTCTTTCCCTACACGACGCTCTTCCGATCTNNNTACACTGCATAGCCTTTCCAACT
GCACACGTCTGAACTCTTTCCCTACACGACGCTCTTCCGATCTNNNCGGCCTTTGTTCAGTCTACATTG
GCACACGTCTGAACTCTTTCCCTACACGACGCTCTTCCGATCTNNNCGGCCTTTGTTCAGTCTACATTG
AGCACACGTCTGAACTCTTTCCCTACACGACGCTCTTCCGATCTNNNTCCTAAAGTGCTGGGATTACAG
AGCACACGTCTGAACTCTTTCCCTACACGACGCTCTTCCGATCTNNNTCCTAAAGTGCTGGGATTACAG
AGCACACGTCTGAACTCTTTCCCTACACGACGCTCTTCCGATCTNNNAGGCCCATATATTCAGGCTTGA
AGCACACGTCTGAACTCTTTCCCTACACGACGCTCTTCCGATCTNNNAGGCCCATATATTCAGGCTTGA
AGCACACGTCTGAACTCTTTCCCTACACGACGCTCTTCCGATCTNNNACCTTGGAGTCTGAAAGCCTTA
AGCACACGTCTGAACTCTTTCCCTACACGACGCTCTTCCGATCTNNNACCTTGGAGTCTGAAAGCCTTA
AGCACACGTCTGAACTCTTTCCCTACACGACGCTCTTCCGATCTNNNTCAAAGATATGTGTGTCCGTGT
AGCACACGTCTGAACTCTTTCCCTACACGACGCTCTTCCGATCTNNNTCAAAGATATGTGTGTCCGTGT
GAGCACACGTCTGAACTCTTTCCCTACACGACGCTCTTCCGATCTNNNCTTAGGTGGCTGAGGCAGAAT
GAGCACACGTCTGAACTCTTTCCCTACACGACGCTCTTCCGATCTNNNCTTAGGTGGCTGAGGCAGAAT
AGCACACGTCTGAACTCTTTCCCTACACGACGCTCTTCCGATCTNNNCTTTCACGGGGAAGTTGAGGA
AGCACACGTCTGAACTCTTTCCCTACACGACGCTCTTCCGATCTNNNCTTTCACGGGGAAGTTGAGGA
GAGCACACGTCTGAACTCTTTCCCTACACGACGCTCTTCCGATCTNNNGCCATGAATCAAGGAATGTAG
GAGCACACGTCTGAACTCTTTCCCTACACGACGCTCTTCCGATCTNNNGCCATGAATCAAGGAATGTAG
GAGCACACGTCTGAACTCTTTCCCTACACGACGCTCTTCCGATCTNNNTGAGTGAGAAATACTTCCTG
GAGCACACGTCTGAACTCTTTCCCTACACGACGCTCTTCCGATCTNNNTGAGTGAGAAATACTTCCTG
AGCACACGTCTGAACTCTTTCCCTACACGACGCTCTTCCGATCTNNNCCTCAAACAATCCTCCTGTCTCA
AGCACACGTCTGAACTCTTTCCCTACACGACGCTCTTCCGATCTNNNCCTCAAACAATCCTCCTGTCTCA
AGCACACGTCTGAACTCTTTCCCTACACGACGCTCTTCCGATCTNNNAGCCATTCCACCAGCTTACATAT
AGCACACGTCTGAACTCTTTCCCTACACGACGCTCTTCCGATCTNNNAGCCATTCCACCAGCTTACATAT
AGAGCACACGTCTGAACTCTTTCCCTACACGACGCTCTTCCGATCTNNNGGGGGTGTTGGTGGTATTA
AGAGCACACGTCTGAACTCTTTCCCTACACGACGCTCTTCCGATCTNNNGGGGGTGTTGGTGGTATTA
GAGCACACGTCTGAACTCTTTCCCTACACGACGCTCTTCCGATCTNNNTAATTTCTCTCCTGTCTTGCCC
GAGCACACGTCTGAACTCTTTCCCTACACGACGCTCTTCCGATCTNNNTAATTTCTCTCCTGTCTTGCCC
GAGCACACGTCTGAACTCTTTCCCTACACGACGCTCTTCCGATCTNNNGCACACATTTCTAAACCCGTCA
GAGCACACGTCTGAACTCTTTCCCTACACGACGCTCTTCCGATCTNNNGCACACATTTCTAAACCCGTCA

AGCACACGTCTGAACTCTTTCCCTACACGACGCTCTTCCGATCTNNNTGAGGCCCAAGATTCATTTTC
AGCACACGTCTGAACTCTTTCCCTACACGACGCTCTTCCGATCTNNNTGAGGCCCAAGATTCATTTTC
AGCACACGTCTGAACTCTTTCCCTACACGACGCTCTTCCGATCTNNNTTAGCTGCTGGATCTCACTTG
AGCACACGTCTGAACTCTTTCCCTACACGACGCTCTTCCGATCTNNNTTAGCTGCTGGATCTCACTTG
AGCACACGTCTGAACTCTTTCCCTACACGACGCTCTTCCGATCTNNNCCTCATTTCTGCAGTTGGTCA
AGCACACGTCTGAACTCTTTCCCTACACGACGCTCTTCCGATCTNNNCCTCATTTCTGCAGTTGGTCA
AGCACACGTCTGAACTCTTTCCCTACACGACGCTCTTCCGATCTNNNTTTCCCCCTATTTTCTCCAGTG
AGCACACGTCTGAACTCTTTCCCTACACGACGCTCTTCCGATCTNNNTTTCCCCCTATTTTCTCCAGTG
AGCACACGTCTGAACTCTTTCCCTACACGACGCTCTTCCGATCTNNNAAAGCCAAAATAAGCACAAG
AGCACACGTCTGAACTCTTTCCCTACACGACGCTCTTCCGATCTNNNAAAGCCAAAATAAGCACAAG
AGCACACGTCTGAACTCTTTCCCTACACGACGCTCTTCCGATCTNNNGGCAACAAGAGGAAAACTGT
AGCACACGTCTGAACTCTTTCCCTACACGACGCTCTTCCGATCTNNNGGCAACAAGAGGAAAACTGT
AGCACACGTCTGAACTCTTTCCCTACACGACGCTCTTCCGATCTNNNAAGAAAAAAGTGCCACAGGG
AGCACACGTCTGAACTCTTTCCCTACACGACGCTCTTCCGATCTNNNAAGAAAAAAGTGCCACAGGG
GAGCACACGTCTGAACTCTTTCCCTACACGACGCTCTTCCGATCTNNNCTCACCTCTAATCCCAGCA
GAGCACACGTCTGAACTCTTTCCCTACACGACGCTCTTCCGATCTNNNCTCACCTCTAATCCCAGCA
AGCACACGTCTGAACTCTTTCCCTACACGACGCTCTTCCGATCTNNNGGGACTCCTCACAGTTTAGTT
AGCACACGTCTGAACTCTTTCCCTACACGACGCTCTTCCGATCTNNNGGGACTCCTCACAGTTTAGTT
AGCACACGTCTGAACTCTTTCCCTACACGACGCTCTTCCGATCTNNNAAAAAGACACCTCCTGGCATAG
AGCACACGTCTGAACTCTTTCCCTACACGACGCTCTTCCGATCTNNNAAAAAGACACCTCCTGGCATAG
AGCACACGTCTGAACTCTTTCCCTACACGACGCTCTTCCGATCTNNNACAACAGGGCATCCAGAAAT
AGCACACGTCTGAACTCTTTCCCTACACGACGCTCTTCCGATCTNNNACAACAGGGCATCCAGAAAT
AGCACACGTCTGAACTCTTTCCCTACACGACGCTCTTCCGATCTNNNGACAGAGCGAGACTCCGTCT
AGCACACGTCTGAACTCTTTCCCTACACGACGCTCTTCCGATCTNNNGACAGAGCGAGACTCCGTCT
AGCACACGTCTGAACTCTTTCCCTACACGACGCTCTTCCGATCTNNNTCCCGAGAGTAGGTGGAATT
AGCACACGTCTGAACTCTTTCCCTACACGACGCTCTTCCGATCTNNNTCCCGAGAGTAGGTGGAATT
AGCACACGTCTGAACTCTTTCCCTACACGACGCTCTTCCGATCTNNNGTTGCAGTGAGCTGAGATCTT
AGCACACGTCTGAACTCTTTCCCTACACGACGCTCTTCCGATCTNNNGTTGCAGTGAGCTGAGATCTT
AGCACACGTCTGAACTCTTTCCCTACACGACGCTCTTCCGATCTNNNACGCCACTTTACTTAGGACACA
AGCACACGTCTGAACTCTTTCCCTACACGACGCTCTTCCGATCTNNNACGCCACTTTACTTAGGACACA
GCACACGTCTGAACTCTTTCCCTACACGACGCTCTTCCGATCTNNNGTTGACTGCTGCTCTGTAGAGA
GCACACGTCTGAACTCTTTCCCTACACGACGCTCTTCCGATCTNNNGTTGACTGCTGCTCTGTAGAGA
GAGCACACGTCTGAACTCTTTCCCTACACGACGCTCTTCCGATCTNNNTCTGCTTAGTTTCCTCCAACCT
GAGCACACGTCTGAACTCTTTCCCTACACGACGCTCTTCCGATCTNNNTCTGCTTAGTTTCCTCCAACCT
CACACGTCTGAACTCTTTCCCTACACGACGCTCTTCCGATCTNNNGATTCCTCTAACCTGCCATCCATC
CACACGTCTGAACTCTTTCCCTACACGACGCTCTTCCGATCTNNNGATTCCTCTAACCTGCCATCCATC
GCACACGTCTGAACTCTTTCCCTACACGACGCTCTTCCGATCTNNNCTGTAATCCCAGCTACTCAGGAG
GCACACGTCTGAACTCTTTCCCTACACGACGCTCTTCCGATCTNNNCTGTAATCCCAGCTACTCAGGAG
GAGCACACGTCTGAACTCTTTCCCTACACGACGCTCTTCCGATCTNNNAGTTGAAGGGATTCACTCTT
GAGCACACGTCTGAACTCTTTCCCTACACGACGCTCTTCCGATCTNNNAGTTGAAGGGATTCACTCTT
AGCACACGTCTGAACTCTTTCCCTACACGACGCTCTTCCGATCTNNNGGCTGAAAACTAGCCTTGGA
AGCACACGTCTGAACTCTTTCCCTACACGACGCTCTTCCGATCTNNNGGCTGAAAACTAGCCTTGGA
GAGCACACGTCTGAACTCTTTCCCTACACGACGCTCTTCCGATCTNNNGAATCGCTTGAACCCAGGA

GAGCACACGTCTGAACTCTTTCCCTACACGACGCTCTTCCGATCTNNNGAATCGCTTGAACCCAGGA
AGAGCACACGTCTGAACTCTTTCCCTACACGACGCTCTTCCGATCTNNNCACCACTGAGCTCCAGCCT
AGAGCACACGTCTGAACTCTTTCCCTACACGACGCTCTTCCGATCTNNNCACCACTGAGCTCCAGCCT
AGCACACGTCTGAACTCTTTCCCTACACGACGCTCTTCCGATCTNNNCTAGGTGTGTTCCTAGGGGA
AGCACACGTCTGAACTCTTTCCCTACACGACGCTCTTCCGATCTNNNCTAGGTGTGTTCCTAGGGGA
GAGCACACGTCTGAACTCTTTCCCTACACGACGCTCTTCCGATCTNNNTGACAAAGACATACCCGAGA
GAGCACACGTCTGAACTCTTTCCCTACACGACGCTCTTCCGATCTNNNTGACAAAGACATACCCGAGA
GCACACGTCTGAACTCTTTCCCTACACGACGCTCTTCCGATCTNNNTGTTTGTACCAGCTATCTCGAG
GCACACGTCTGAACTCTTTCCCTACACGACGCTCTTCCGATCTNNNTGTTTGTACCAGCTATCTCGAG
AGCACACGTCTGAACTCTTTCCCTACACGACGCTCTTCCGATCTNNNGTGCCTATAACCCCAGCTACTTG
AGCACACGTCTGAACTCTTTCCCTACACGACGCTCTTCCGATCTNNNGTGCCTATAACCCCAGCTACTTG
AGCACACGTCTGAACTCTTTCCCTACACGACGCTCTTCCGATCTNNNACCTGACATGTGGAAAGAGTC
AGCACACGTCTGAACTCTTTCCCTACACGACGCTCTTCCGATCTNNNACCTGACATGTGGAAAGAGTC
AGCACACGTCTGAACTCTTTCCCTACACGACGCTCTTCCGATCTNNNGCTCCCACAATGCCTGAAATAA
AGCACACGTCTGAACTCTTTCCCTACACGACGCTCTTCCGATCTNNNGCTCCCACAATGCCTGAAATAA
AGCACACGTCTGAACTCTTTCCCTACACGACGCTCTTCCGATCTNNNCAGGAGAATCACTTGAACCCA
AGCACACGTCTGAACTCTTTCCCTACACGACGCTCTTCCGATCTNNNCAGGAGAATCACTTGAACCCA
GAGCACACGTCTGAACTCTTTCCCTACACGACGCTCTTCCGATCTNNNGGGAGCTTTCTGTCCTGTATC
GAGCACACGTCTGAACTCTTTCCCTACACGACGCTCTTCCGATCTNNNGGGAGCTTTCTGTCCTGTATC
CACACGTCTGAACTCTTTCCCTACACGACGCTCTTCCGATCTNNNTCTCAGCTTCCTTGTGCTTAGAGC
CACACGTCTGAACTCTTTCCCTACACGACGCTCTTCCGATCTNNNTCTCAGCTTCCTTGTGCTTAGAGC
AGCACACGTCTGAACTCTTTCCCTACACGACGCTCTTCCGATCTNNNTGGACTGTCCAAATAAACGTAT
AGCACACGTCTGAACTCTTTCCCTACACGACGCTCTTCCGATCTNNNTGGACTGTCCAAATAAACGTAT
AGCACACGTCTGAACTCTTTCCCTACACGACGCTCTTCCGATCTNNNAGGAGGCTGAGACAGGAGAA
AGCACACGTCTGAACTCTTTCCCTACACGACGCTCTTCCGATCTNNNAGGAGGCTGAGACAGGAGAA
GAGCACACGTCTGAACTCTTTCCCTACACGACGCTCTTCCGATCTNNNGAGATAGCACCACTACACTCC
GAGCACACGTCTGAACTCTTTCCCTACACGACGCTCTTCCGATCTNNNGAGATAGCACCACTACACTCC
AGCACACGTCTGAACTCTTTCCCTACACGACGCTCTTCCGATCTNNNCCTTGTATCACATCAGGGGCAT
AGCACACGTCTGAACTCTTTCCCTACACGACGCTCTTCCGATCTNNNCCTTGTATCACATCAGGGGCAT
GAGCACACGTCTGAACTCTTTCCCTACACGACGCTCTTCCGATCTNNNGTTCATCCATGTTGTTGCAAA
GAGCACACGTCTGAACTCTTTCCCTACACGACGCTCTTCCGATCTNNNGTTCATCCATGTTGTTGCAAA
GAGCACACGTCTGAACTCTTTCCCTACACGACGCTCTTCCGATCTNNNTAGGGTCATCAAAAGGGTTC
GAGCACACGTCTGAACTCTTTCCCTACACGACGCTCTTCCGATCTNNNTAGGGTCATCAAAAGGGTTC
GCACACGTCTGAACTCTTTCCCTACACGACGCTCTTCCGATCTNNNGCACATCCAGCTGTTAGTCTTG
GCACACGTCTGAACTCTTTCCCTACACGACGCTCTTCCGATCTNNNGCACATCCAGCTGTTAGTCTTG
GCACACGTCTGAACTCTTTCCCTACACGACGCTCTTCCGATCTNNNCAGAGCTTGCAGTGAGTCGAG
GCACACGTCTGAACTCTTTCCCTACACGACGCTCTTCCGATCTNNNCAGAGCTTGCAGTGAGTCGAG
GCACACGTCTGAACTCTTTCCCTACACGACGCTCTTCCGATCTNNNACACAGCAAATATGGTCACAAG
GCACACGTCTGAACTCTTTCCCTACACGACGCTCTTCCGATCTNNNACACAGCAAATATGGTCACAAG
GCACACGTCTGAACTCTTTCCCTACACGACGCTCTTCCGATCTNNNCTCCTACAGAGCAGTTCACTGG
GCACACGTCTGAACTCTTTCCCTACACGACGCTCTTCCGATCTNNNCTCCTACAGAGCAGTTCACTGG
GCACACGTCTGAACTCTTTCCCTACACGACGCTCTTCCGATCTNNNTCTTGGATAAGCCTGATGTAAT
GCACACGTCTGAACTCTTTCCCTACACGACGCTCTTCCGATCTNNNTCTTGGATAAGCCTGATGTAAT

GAGCACACGTCTGAACTCTTTCCCTACACGACGCTCTTCCGATCTNNNGTGCTTCTGGAAACCTAAGGG
GAGCACACGTCTGAACTCTTTCCCTACACGACGCTCTTCCGATCTNNNGTGCTTCTGGAAACCTAAGGG
GAGCACACGTCTGAACTCTTTCCCTACACGACGCTCTTCCGATCTNNNTGCCCAGCTAGTGTGTATGTG
GAGCACACGTCTGAACTCTTTCCCTACACGACGCTCTTCCGATCTNNNTGCCCAGCTAGTGTGTATGTG
CACACGTCTGAACTCTTTCCCTACACGACGCTCTTCCGATCTNNNCTGATTTCTGATCCAGCACCATCG
CACACGTCTGAACTCTTTCCCTACACGACGCTCTTCCGATCTNNNCTGATTTCTGATCCAGCACCATCG
AGCACACGTCTGAACTCTTTCCCTACACGACGCTCTTCCGATCTNNNAGTAAGCCAAGATTGTGCCATT
AGCACACGTCTGAACTCTTTCCCTACACGACGCTCTTCCGATCTNNNAGTAAGCCAAGATTGTGCCATT
GAGCACACGTCTGAACTCTTTCCCTACACGACGCTCTTCCGATCTNNNGTTGCAGTGAGCCGAGATCA
GAGCACACGTCTGAACTCTTTCCCTACACGACGCTCTTCCGATCTNNNGTTGCAGTGAGCCGAGATCA
AGAGCACACGTCTGAACTCTTTCCCTACACGACGCTCTTCCGATCTNNNCAGATCTCTACTGCCACCAAA
AGAGCACACGTCTGAACTCTTTCCCTACACGACGCTCTTCCGATCTNNNCAGATCTCTACTGCCACCAAA
AGCACACGTCTGAACTCTTTCCCTACACGACGCTCTTCCGATCTNNNAAAATAACACTGAGCCATGCAG
AGCACACGTCTGAACTCTTTCCCTACACGACGCTCTTCCGATCTNNNAAAATAACACTGAGCCATGCAG
GAGCACACGTCTGAACTCTTTCCCTACACGACGCTCTTCCGATCTNNNAATACAAAAAATTAGCCGGGC
GAGCACACGTCTGAACTCTTTCCCTACACGACGCTCTTCCGATCTNNNAATACAAAAAATTAGCCGGGC
AGCACACGTCTGAACTCTTTCCCTACACGACGCTCTTCCGATCTNNNTTTCCTTCTGTTATCCCTCCCTCC
AGCACACGTCTGAACTCTTTCCCTACACGACGCTCTTCCGATCTNNNTTTCCTTCTGTTATCCCTCCCTCC
GAGCACACGTCTGAACTCTTTCCCTACACGACGCTCTTCCGATCTNNNCACATCATTGCTGGAGCCTTA
GAGCACACGTCTGAACTCTTTCCCTACACGACGCTCTTCCGATCTNNNCACATCATTGCTGGAGCCTTA
AGCACACGTCTGAACTCTTTCCCTACACGACGCTCTTCCGATCTNNNTGCCTACTAACTTGGACCCATT
AGCACACGTCTGAACTCTTTCCCTACACGACGCTCTTCCGATCTNNNTGCCTACTAACTTGGACCCATT
AGAGCACACGTCTGAACTCTTTCCCTACACGACGCTCTTCCGATCTNNNAATGATAGACCCCTGGAGC
AGAGCACACGTCTGAACTCTTTCCCTACACGACGCTCTTCCGATCTNNNAATGATAGACCCCTGGAGC
AGCACACGTCTGAACTCTTTCCCTACACGACGCTCTTCCGATCTNNNCACCCCCAGGATTCTTACAGAA
AGCACACGTCTGAACTCTTTCCCTACACGACGCTCTTCCGATCTNNNCACCCCCAGGATTCTTACAGAA
AGCACACGTCTGAACTCTTTCCCTACACGACGCTCTTCCGATCTNNNGAGTTTGAGGCTGTAGTGAGC
AGCACACGTCTGAACTCTTTCCCTACACGACGCTCTTCCGATCTNNNGAGTTTGAGGCTGTAGTGAGC
GAGCACACGTCTGAACTCTTTCCCTACACGACGCTCTTCCGATCTNNNCCCTAAACCTTGATAGGCACAG
GAGCACACGTCTGAACTCTTTCCCTACACGACGCTCTTCCGATCTNNNCCCTAAACCTTGATAGGCACAG
AGAGCACACGTCTGAACTCTTTCCCTACACGACGCTCTTCCGATCTNNNAATTACTGCAGCAACACATCT
AGAGCACACGTCTGAACTCTTTCCCTACACGACGCTCTTCCGATCTNNNAATTACTGCAGCAACACATCT
AGCACACGTCTGAACTCTTTCCCTACACGACGCTCTTCCGATCTNNNGGACATAAGGAGGGACAAAGA
AGCACACGTCTGAACTCTTTCCCTACACGACGCTCTTCCGATCTNNNGGACATAAGGAGGGACAAAGA
GAGCACACGTCTGAACTCTTTCCCTACACGACGCTCTTCCGATCTNNNTCTACTGGACTGATGGGAAA
GAGCACACGTCTGAACTCTTTCCCTACACGACGCTCTTCCGATCTNNNTCTACTGGACTGATGGGAAA
AGCACACGTCTGAACTCTTTCCCTACACGACGCTCTTCCGATCTNNNCTTTGTGGATGTTGCAGGGGA
AGCACACGTCTGAACTCTTTCCCTACACGACGCTCTTCCGATCTNNNCTTTGTGGATGTTGCAGGGGA
GAGCACACGTCTGAACTCTTTCCCTACACGACGCTCTTCCGATCTNNNTTTTCTAGGGAACTGCTGTCA
GAGCACACGTCTGAACTCTTTCCCTACACGACGCTCTTCCGATCTNNNTTTTCTAGGGAACTGCTGTCA
AGAGCACACGTCTGAACTCTTTCCCTACACGACGCTCTTCCGATCTNNNCTGGCAGAACAAAAAGAGA
AGAGCACACGTCTGAACTCTTTCCCTACACGACGCTCTTCCGATCTNNNCTGGCAGAACAAAAAGAGA
GAGCACACGTCTGAACTCTTTCCCTACACGACGCTCTTCCGATCTNNNGACACAACAGAATCGCTTGA

GAGCACACGTCTGAACTCTTTCCCTACACGACGCTCTTCCGATCTNNNGACACAACAGAATCGCTTGA
CACGTCTGAACTCTTTCCCTACACGACGCTCTTCCGATCTNNNTCATGGGATACATCATGACATTTTGA
CACGTCTGAACTCTTTCCCTACACGACGCTCTTCCGATCTNNNTCATGGGATACATCATGACATTTTGA
CACACGTCTGAACTCTTTCCCTACACGACGCTCTTCCGATCTNNNCCTACAGGTGTTAGCCGGAACCAT
CACACGTCTGAACTCTTTCCCTACACGACGCTCTTCCGATCTNNNCCTACAGGTGTTAGCCGGAACCAT
GAGCACACGTCTGAACTCTTTCCCTACACGACGCTCTTCCGATCTNNNTGTGTAAAGGAAATGGTGT
GAGCACACGTCTGAACTCTTTCCCTACACGACGCTCTTCCGATCTNNNTGTGTAAAGGAAATGGTGT
AGCACACGTCTGAACTCTTTCCCTACACGACGCTCTTCCGATCTNNNGGCCTCTGGAGTAGCTATTCTT
AGCACACGTCTGAACTCTTTCCCTACACGACGCTCTTCCGATCTNNNGGCCTCTGGAGTAGCTATTCTT
GAGCACACGTCTGAACTCTTTCCCTACACGACGCTCTTCCGATCTNNNCCTCAGCAAAGTCTCAGGAT
GAGCACACGTCTGAACTCTTTCCCTACACGACGCTCTTCCGATCTNNNCCTCAGCAAAGTCTCAGGAT
AGCACACGTCTGAACTCTTTCCCTACACGACGCTCTTCCGATCTNNNTTTCTCCATGACACTTCCCACA
AGCACACGTCTGAACTCTTTCCCTACACGACGCTCTTCCGATCTNNNTTTCTCCATGACACTTCCCACA
AGCACACGTCTGAACTCTTTCCCTACACGACGCTCTTCCGATCTNNNCCACGGTAGAGTTGCAGATAC
AGCACACGTCTGAACTCTTTCCCTACACGACGCTCTTCCGATCTNNNCCACGGTAGAGTTGCAGATAC
GCACACGTCTGAACTCTTTCCCTACACGACGCTCTTCCGATCTNNNCGCCTCTATTGGTTTTGTGCTTA
GCACACGTCTGAACTCTTTCCCTACACGACGCTCTTCCGATCTNNNCGCCTCTATTGGTTTTGTGCTTA
AGCACACGTCTGAACTCTTTCCCTACACGACGCTCTTCCGATCTNNNCGGCAAAAAGTTTCCTTCTAGC
AGCACACGTCTGAACTCTTTCCCTACACGACGCTCTTCCGATCTNNNCGGCAAAAAGTTTCCTTCTAGC
GCACACGTCTGAACTCTTTCCCTACACGACGCTCTTCCGATCTNNNTCACCCAGCCCCTTATGATTTTC
GCACACGTCTGAACTCTTTCCCTACACGACGCTCTTCCGATCTNNNTCACCCAGCCCCTTATGATTTTC
AGCACACGTCTGAACTCTTTCCCTACACGACGCTCTTCCGATCTNNNAAGGAAAACTCTGGACAGACA
AGCACACGTCTGAACTCTTTCCCTACACGACGCTCTTCCGATCTNNNAAGGAAAACTCTGGACAGACA
AGCACACGTCTGAACTCTTTCCCTACACGACGCTCTTCCGATCTNNNCGCTGGTTGTTGGCAGAATAT
AGCACACGTCTGAACTCTTTCCCTACACGACGCTCTTCCGATCTNNNCGCTGGTTGTTGGCAGAATAT
GAGCACACGTCTGAACTCTTTCCCTACACGACGCTCTTCCGATCTNNNGCAGGAGAATCGTTTGAACT
GAGCACACGTCTGAACTCTTTCCCTACACGACGCTCTTCCGATCTNNNGCAGGAGAATCGTTTGAACT
GCACACGTCTGAACTCTTTCCCTACACGACGCTCTTCCGATCTNNNTAAGTTCACAGTCATAGTGGGC
GCACACGTCTGAACTCTTTCCCTACACGACGCTCTTCCGATCTNNNTAAGTTCACAGTCATAGTGGGC
GAGCACACGTCTGAACTCTTTCCCTACACGACGCTCTTCCGATCTNNNGTCAGTCCTTAGATTCAGAG
GAGCACACGTCTGAACTCTTTCCCTACACGACGCTCTTCCGATCTNNNGTCAGTCCTTAGATTCAGAG
AGCACACGTCTGAACTCTTTCCCTACACGACGCTCTTCCGATCTNNNGGGACTCATCGTCGATGTTCTT
AGCACACGTCTGAACTCTTTCCCTACACGACGCTCTTCCGATCTNNNGGGACTCATCGTCGATGTTCTT
AGCACACGTCTGAACTCTTTCCCTACACGACGCTCTTCCGATCTNNNGACAGGTGAATACAGACAGC
AGCACACGTCTGAACTCTTTCCCTACACGACGCTCTTCCGATCTNNNGACAGGTGAATACAGACAGC
GAGCACACGTCTGAACTCTTTCCCTACACGACGCTCTTCCGATCTNNNCACTGGATTGCAAGTAGGCT
GAGCACACGTCTGAACTCTTTCCCTACACGACGCTCTTCCGATCTNNNCACTGGATTGCAAGTAGGCT
GAGCACACGTCTGAACTCTTTCCCTACACGACGCTCTTCCGATCTNNNATGTACCACCATCCCCAGAAA
GAGCACACGTCTGAACTCTTTCCCTACACGACGCTCTTCCGATCTNNNATGTACCACCATCCCCAGAAA
AGCACACGTCTGAACTCTTTCCCTACACGACGCTCTTCCGATCTNNNCAAGATGGTGCTACTGTACTC
AGCACACGTCTGAACTCTTTCCCTACACGACGCTCTTCCGATCTNNNCAAGATGGTGCTACTGTACTC
AGCACACGTCTGAACTCTTTCCCTACACGACGCTCTTCCGATCTNNNCTCAAGCCCTCCACTAATGAGA
AGCACACGTCTGAACTCTTTCCCTACACGACGCTCTTCCGATCTNNNCTCAAGCCCTCCACTAATGAGA

AGCACACGTCTGAACTCTTTCCCTACACGACGCTCTTCCGATCTNNNGGATTTTGGGCTGCTCTTTTCA
AGCACACGTCTGAACTCTTTCCCTACACGACGCTCTTCCGATCTNNNGGATTTTGGGCTGCTCTTTTCA
AGCACACGTCTGAACTCTTTCCCTACACGACGCTCTTCCGATCTNNNAGGGACCCCAATTATTGAAGG
AGCACACGTCTGAACTCTTTCCCTACACGACGCTCTTCCGATCTNNNAGGGACCCCAATTATTGAAGG
AGCACACGTCTGAACTCTTTCCCTACACGACGCTCTTCCGATCTNNNCATCTTGCTTGTTTCTTCCAGCA
AGCACACGTCTGAACTCTTTCCCTACACGACGCTCTTCCGATCTNNNCATCTTGCTTGTTTCTTCCAGCA
GAGCACACGTCTGAACTCTTTCCCTACACGACGCTCTTCCGATCTNNNGGATTTTCAGAACCAACTGGC
GAGCACACGTCTGAACTCTTTCCCTACACGACGCTCTTCCGATCTNNNGGATTTTCAGAACCAACTGGC
AGCACACGTCTGAACTCTTTCCCTACACGACGCTCTTCCGATCTNNNGAATCACTTGAAGAACCTCGGC
AGCACACGTCTGAACTCTTTCCCTACACGACGCTCTTCCGATCTNNNGAATCACTTGAAGAACCTCGGC
AGCACACGTCTGAACTCTTTCCCTACACGACGCTCTTCCGATCTNNNCACAATAGCTCACGCCTGTAATC
AGCACACGTCTGAACTCTTTCCCTACACGACGCTCTTCCGATCTNNNCACAATAGCTCACGCCTGTAATC
AGCACACGTCTGAACTCTTTCCCTACACGACGCTCTTCCGATCTNNNCAAATGGTGGTAAGGCTGGGA
AGCACACGTCTGAACTCTTTCCCTACACGACGCTCTTCCGATCTNNNCAAATGGTGGTAAGGCTGGGA
GAGCACACGTCTGAACTCTTTCCCTACACGACGCTCTTCCGATCTNNNTTTGCCTTGTGAGGTTTCTGT
GAGCACACGTCTGAACTCTTTCCCTACACGACGCTCTTCCGATCTNNNTTTGCCTTGTGAGGTTTCTGT
AAGAGCACACGTCTGAACTCTTTCCCTACACGACGCTCTTCCGATCTNNNACATACGGCTATACGACAA
AAGAGCACACGTCTGAACTCTTTCCCTACACGACGCTCTTCCGATCTNNNACATACGGCTATACGACAA
AGCACACGTCTGAACTCTTTCCCTACACGACGCTCTTCCGATCTNNNATGTGCAGGTTGGTTACATAGC
AGCACACGTCTGAACTCTTTCCCTACACGACGCTCTTCCGATCTNNNATGTGCAGGTTGGTTACATAGC
AGCACACGTCTGAACTCTTTCCCTACACGACGCTCTTCCGATCTNNNTATAAGGCATGTTCCTCCGTCTC
AGCACACGTCTGAACTCTTTCCCTACACGACGCTCTTCCGATCTNNNTATAAGGCATGTTCCTCCGTCTC
GCACACGTCTGAACTCTTTCCCTACACGACGCTCTTCCGATCTNNNCAGCCATGTTGAAATTCCATCCTC
GCACACGTCTGAACTCTTTCCCTACACGACGCTCTTCCGATCTNNNCAGCCATGTTGAAATTCCATCCTC
AGCACACGTCTGAACTCTTTCCCTACACGACGCTCTTCCGATCTNNNGTTTGCAGACAGCCACTTCTCTA
AGCACACGTCTGAACTCTTTCCCTACACGACGCTCTTCCGATCTNNNGTTTGCAGACAGCCACTTCTCTA
GAGCACACGTCTGAACTCTTTCCCTACACGACGCTCTTCCGATCTNNNACACACACACACACACACAAAT
GAGCACACGTCTGAACTCTTTCCCTACACGACGCTCTTCCGATCTNNNACACACACACACACACACAAAT
GCACACGTCTGAACTCTTTCCCTACACGACGCTCTTCCGATCTNNNACCTCTCTGTGGGAAGAATAACTC
GCACACGTCTGAACTCTTTCCCTACACGACGCTCTTCCGATCTNNNACCTCTCTGTGGGAAGAATAACTC
GCACACGTCTGAACTCTTTCCCTACACGACGCTCTTCCGATCTNNNCTTCTGGATTCCACTCCTTCTCCCA
GCACACGTCTGAACTCTTTCCCTACACGACGCTCTTCCGATCTNNNCTTCTGGATTCCACTCCTTCTCCCA
AGCACACGTCTGAACTCTTTCCCTACACGACGCTCTTCCGATCTNNNCTGTTCAGGGTTTATAGGAGGC
AGCACACGTCTGAACTCTTTCCCTACACGACGCTCTTCCGATCTNNNCTGTTCAGGGTTTATAGGAGGC
GCACACGTCTGAACTCTTTCCCTACACGACGCTCTTCCGATCTNNNTACTCAGGAGACTAAGGGGAGA
GCACACGTCTGAACTCTTTCCCTACACGACGCTCTTCCGATCTNNNTACTCAGGAGACTAAGGGGAGA
GAGCACACGTCTGAACTCTTTCCCTACACGACGCTCTTCCGATCTNNNAAGTGCATCTGTGGAGTGAT
GAGCACACGTCTGAACTCTTTCCCTACACGACGCTCTTCCGATCTNNNAAGTGCATCTGTGGAGTGAT
AGCACACGTCTGAACTCTTTCCCTACACGACGCTCTTCCGATCTNNNGGATCACGAGGTCAGAAGATC
AGCACACGTCTGAACTCTTTCCCTACACGACGCTCTTCCGATCTNNNGGATCACGAGGTCAGAAGATC
GCACACGTCTGAACTCTTTCCCTACACGACGCTCTTCCGATCTNNNCAGTCCAAATCCAACTGAGGAGA
GCACACGTCTGAACTCTTTCCCTACACGACGCTCTTCCGATCTNNNCAGTCCAAATCCAACTGAGGAGA
GAGCACACGTCTGAACTCTTTCCCTACACGACGCTCTTCCGATCTNNNCACCTGTAATCCCAGCTACTCA

GAGCACACGTCTGAACTCTTTCCCTACACGACGCTCTTCCGATCTNNNCACCTGTAATCCCAGCTACTCA
AGCACACGTCTGAACTCTTTCCCTACACGACGCTCTTCCGATCTNNNATTCCACAGGTCCCTGTTCTCTT
AGCACACGTCTGAACTCTTTCCCTACACGACGCTCTTCCGATCTNNNATTCCACAGGTCCCTGTTCTCTT
GAGCACACGTCTGAACTCTTTCCCTACACGACGCTCTTCCGATCTNNNAGCTGAAGCCATTATGTTACC
GAGCACACGTCTGAACTCTTTCCCTACACGACGCTCTTCCGATCTNNNAGCTGAAGCCATTATGTTACC
AGCACACGTCTGAACTCTTTCCCTACACGACGCTCTTCCGATCTNNNGTCTTTTCCAGCTTCTAGAGGC
AGCACACGTCTGAACTCTTTCCCTACACGACGCTCTTCCGATCTNNNGTCTTTTCCAGCTTCTAGAGGC
AGAGCACACGTCTGAACTCTTTCCCTACACGACGCTCTTCCGATCTNNNTGTGGATCTTGTTGAGTGTG
AGAGCACACGTCTGAACTCTTTCCCTACACGACGCTCTTCCGATCTNNNTGTGGATCTTGTTGAGTGTG
AGCACACGTCTGAACTCTTTCCCTACACGACGCTCTTCCGATCTNNNCTCATCTGCTCTGCTCAGTAGA
AGCACACGTCTGAACTCTTTCCCTACACGACGCTCTTCCGATCTNNNCTCATCTGCTCTGCTCAGTAGA
GAGCACACGTCTGAACTCTTTCCCTACACGACGCTCTTCCGATCTNNNGGCTCACATCTGTAATCCTAG
GAGCACACGTCTGAACTCTTTCCCTACACGACGCTCTTCCGATCTNNNGGCTCACATCTGTAATCCTAG
AGCACACGTCTGAACTCTTTCCCTACACGACGCTCTTCCGATCTNNNAGGTTCAAGCCATTCTCATGTC
AGCACACGTCTGAACTCTTTCCCTACACGACGCTCTTCCGATCTNNNAGGTTCAAGCCATTCTCATGTC
GAGCACACGTCTGAACTCTTTCCCTACACGACGCTCTTCCGATCTNNNCTTGAACTCCTGACCTCATGTG
GAGCACACGTCTGAACTCTTTCCCTACACGACGCTCTTCCGATCTNNNCTTGAACTCCTGACCTCATGTG
AGAGCACACGTCTGAACTCTTTCCCTACACGACGCTCTTCCGATCTNNNTCCATAAAGCCCTCTTGGTCT
AGAGCACACGTCTGAACTCTTTCCCTACACGACGCTCTTCCGATCTNNNTCCATAAAGCCCTCTTGGTCT
AGCACACGTCTGAACTCTTTCCCTACACGACGCTCTTCCGATCTNNNAGAGCTGGTGATGTGTGATGT
AGCACACGTCTGAACTCTTTCCCTACACGACGCTCTTCCGATCTNNNAGAGCTGGTGATGTGTGATGT
GAGCACACGTCTGAACTCTTTCCCTACACGACGCTCTTCCGATCTNNNATTCAGAGGCACAATAGGAG
GAGCACACGTCTGAACTCTTTCCCTACACGACGCTCTTCCGATCTNNNATTCAGAGGCACAATAGGAG
AGCACACGTCTGAACTCTTTCCCTACACGACGCTCTTCCGATCTNNNTGAGGCAGAAGAATCATCTGA
AGCACACGTCTGAACTCTTTCCCTACACGACGCTCTTCCGATCTNNNTGAGGCAGAAGAATCATCTGA
GAGCACACGTCTGAACTCTTTCCCTACACGACGCTCTTCCGATCTNNNGGTCGCACTTTCCAATTCTCTC
GAGCACACGTCTGAACTCTTTCCCTACACGACGCTCTTCCGATCTNNNGGTCGCACTTTCCAATTCTCTC
AGCACACGTCTGAACTCTTTCCCTACACGACGCTCTTCCGATCTNNNAGCCAGTTATCCCACCATCATTT
AGCACACGTCTGAACTCTTTCCCTACACGACGCTCTTCCGATCTNNNAGCCAGTTATCCCACCATCATTT
AGAGCACACGTCTGAACTCTTTCCCTACACGACGCTCTTCCGATCTNNNGGAGCCCTTCAATTCTATCA
AGAGCACACGTCTGAACTCTTTCCCTACACGACGCTCTTCCGATCTNNNGGAGCCCTTCAATTCTATCA
GAGCACACGTCTGAACTCTTTCCCTACACGACGCTCTTCCGATCTNNNTAATATGAGGCACAGCGTGAT
GAGCACACGTCTGAACTCTTTCCCTACACGACGCTCTTCCGATCTNNNTAATATGAGGCACAGCGTGAT
GAGCACACGTCTGAACTCTTTCCCTACACGACGCTCTTCCGATCTNNNAAAGGCAAACCCAACAGGAAA
GAGCACACGTCTGAACTCTTTCCCTACACGACGCTCTTCCGATCTNNNAAAGGCAAACCCAACAGGAAA
AGCACACGTCTGAACTCTTTCCCTACACGACGCTCTTCCGATCTNNNGGCAAGGATCTAGAAGCAAGA
AGCACACGTCTGAACTCTTTCCCTACACGACGCTCTTCCGATCTNNNGGCAAGGATCTAGAAGCAAGA
AGCACACGTCTGAACTCTTTCCCTACACGACGCTCTTCCGATCTNNNCCTCCTGAGGTACAACTTCCATC
AGCACACGTCTGAACTCTTTCCCTACACGACGCTCTTCCGATCTNNNCCTCCTGAGGTACAACTTCCATC
AGCACACGTCTGAACTCTTTCCCTACACGACGCTCTTCCGATCTNNNCACTTGAGCTCAGGAGTTTGAC
AGCACACGTCTGAACTCTTTCCCTACACGACGCTCTTCCGATCTNNNCACTTGAGCTCAGGAGTTTGAC
CACACGTCTGAACTCTTTCCCTACACGACGCTCTTCCGATCTNNNACACCACCAGCTTGTCATATCTTCA
CACACGTCTGAACTCTTTCCCTACACGACGCTCTTCCGATCTNNNACACCACCAGCTTGTCATATCTTCA

AGAGCACACGTCTGAACTCTTTCCCTACACGACGCTCTTCCGATCTNNNCATTACTAGGGGAGGATTGT
AGAGCACACGTCTGAACTCTTTCCCTACACGACGCTCTTCCGATCTNNNCATTACTAGGGGAGGATTGT
AGCACACGTCTGAACTCTTTCCCTACACGACGCTCTTCCGATCTNNNGCTAAGCTGCAAACTACATAGCC
AGCACACGTCTGAACTCTTTCCCTACACGACGCTCTTCCGATCTNNNGCTAAGCTGCAAACTACATAGCC
AGCACACGTCTGAACTCTTTCCCTACACGACGCTCTTCCGATCTNNNGGGTGAACCTGATAACCAATCT
AGCACACGTCTGAACTCTTTCCCTACACGACGCTCTTCCGATCTNNNGGGTGAACCTGATAACCAATCT
AGCACACGTCTGAACTCTTTCCCTACACGACGCTCTTCCGATCTNNNTCCTCCTCCTCCTCTTCTTCTTCCA
AGCACACGTCTGAACTCTTTCCCTACACGACGCTCTTCCGATCTNNNTCCTCCTCCTCCTCTTCTTCTTCCA
AGCACACGTCTGAACTCTTTCCCTACACGACGCTCTTCCGATCTNNNACCAAGGGCTAGGAATCTTCTTC
AGCACACGTCTGAACTCTTTCCCTACACGACGCTCTTCCGATCTNNNACCAAGGGCTAGGAATCTTCTTC
AGCACACGTCTGAACTCTTTCCCTACACGACGCTCTTCCGATCTNNNGGAGACAAGAATGAAACGCCA
AGCACACGTCTGAACTCTTTCCCTACACGACGCTCTTCCGATCTNNNGGAGACAAGAATGAAACGCCA
AGCACACGTCTGAACTCTTTCCCTACACGACGCTCTTCCGATCTNNNCCATCACAGCTTCTGTTTCCAGA
AGCACACGTCTGAACTCTTTCCCTACACGACGCTCTTCCGATCTNNNCCATCACAGCTTCTGTTTCCAGA
GAGCACACGTCTGAACTCTTTCCCTACACGACGCTCTTCCGATCTNNNGGGCGACAATCATCAGGTTA
GAGCACACGTCTGAACTCTTTCCCTACACGACGCTCTTCCGATCTNNNGGGCGACAATCATCAGGTTA
GAGCACACGTCTGAACTCTTTCCCTACACGACGCTCTTCCGATCTNNNATCCAAAAATTAGCTGGGCAT
GAGCACACGTCTGAACTCTTTCCCTACACGACGCTCTTCCGATCTNNNATCCAAAAATTAGCTGGGCAT
AGCACACGTCTGAACTCTTTCCCTACACGACGCTCTTCCGATCTNNNGACTCAGTTCCCACATCCTAAGC
AGCACACGTCTGAACTCTTTCCCTACACGACGCTCTTCCGATCTNNNGACTCAGTTCCCACATCCTAAGC
GAGCACACGTCTGAACTCTTTCCCTACACGACGCTCTTCCGATCTNNNGCCAGCATCACCAGCAAAAGA
GAGCACACGTCTGAACTCTTTCCCTACACGACGCTCTTCCGATCTNNNGCCAGCATCACCAGCAAAAGA
AGCACACGTCTGAACTCTTTCCCTACACGACGCTCTTCCGATCTNNNTGCACTGGGTTCTTCAACATTTT
AGCACACGTCTGAACTCTTTCCCTACACGACGCTCTTCCGATCTNNNTGCACTGGGTTCTTCAACATTTT
GAGCACACGTCTGAACTCTTTCCCTACACGACGCTCTTCCGATCTNNNGCTTTCTGGCCAACATGAATCT
GAGCACACGTCTGAACTCTTTCCCTACACGACGCTCTTCCGATCTNNNGCTTTCTGGCCAACATGAATCT
AGCACACGTCTGAACTCTTTCCCTACACGACGCTCTTCCGATCTNNNCTTTAGGGAGGCTTTTAAAAAC
AGCACACGTCTGAACTCTTTCCCTACACGACGCTCTTCCGATCTNNNCTTTAGGGAGGCTTTTAAAAAC
GAGCACACGTCTGAACTCTTTCCCTACACGACGCTCTTCCGATCTNNNCTGTGTCAGCAGAGTGACTGC
GAGCACACGTCTGAACTCTTTCCCTACACGACGCTCTTCCGATCTNNNCTGTGTCAGCAGAGTGACTGC
CACACGTCTGAACTCTTTCCCTACACGACGCTCTTCCGATCTNNNCCACCATCATCATAATCACCCATCGG
CACACGTCTGAACTCTTTCCCTACACGACGCTCTTCCGATCTNNNCCACCATCATCATAATCACCCATCGG
GAGCACACGTCTGAACTCTTTCCCTACACGACGCTCTTCCGATCTNNNTCTAGGAACCTCAAGGCAAAC
GAGCACACGTCTGAACTCTTTCCCTACACGACGCTCTTCCGATCTNNNTCTAGGAACCTCAAGGCAAAC
GAGCACACGTCTGAACTCTTTCCCTACACGACGCTCTTCCGATCTNNNATCAAGCAATGCACAGAGACA
GAGCACACGTCTGAACTCTTTCCCTACACGACGCTCTTCCGATCTNNNATCAAGCAATGCACAGAGACA
GAGCACACGTCTGAACTCTTTCCCTACACGACGCTCTTCCGATCTNNNCCATGTGGAGATGATGCATAT
GAGCACACGTCTGAACTCTTTCCCTACACGACGCTCTTCCGATCTNNNCCATGTGGAGATGATGCATAT
AGCACACGTCTGAACTCTTTCCCTACACGACGCTCTTCCGATCTNNNGGTCCACCTCATTCTCTACAGTGC
AGCACACGTCTGAACTCTTTCCCTACACGACGCTCTTCCGATCTNNNGGTCCACCTCATTCTCTACAGTGC
AGAGCACACGTCTGAACTCTTTCCCTACACGACGCTCTTCCGATCTNNNTTTGCCAAACAGAGCTCTGAA
AGAGCACACGTCTGAACTCTTTCCCTACACGACGCTCTTCCGATCTNNNTTTGCCAAACAGAGCTCTGAA
AGCACACGTCTGAACTCTTTCCCTACACGACGCTCTTCCGATCTNNNATTTGCAAGGGACAAACAACCA

AGCACACGTCTGAACTCTTTCCCTACACGACGCTCTTCCGATCTNNNATTTGCAAGGGACAAACAACCA
AGCACACGTCTGAACTCTTTCCCTACACGACGCTCTTCCGATCTNNNCCAGCAGTGTAGAAGTGTTCTC
AGCACACGTCTGAACTCTTTCCCTACACGACGCTCTTCCGATCTNNNCCAGCAGTGTAGAAGTGTTCTC
GCACACGTCTGAACTCTTTCCCTACACGACGCTCTTCCGATCTNNNCCATTAGCTCTGCTCTTCCCTAGCA
GCACACGTCTGAACTCTTTCCCTACACGACGCTCTTCCGATCTNNNCCATTAGCTCTGCTCTTCCCTAGCA
GAGCACACGTCTGAACTCTTTCCCTACACGACGCTCTTCCGATCTNNNAGGAAGGAGAAGGACCTAT
GAGCACACGTCTGAACTCTTTCCCTACACGACGCTCTTCCGATCTNNNAGGAAGGAGAAGGACCTAT
GAGCACACGTCTGAACTCTTTCCCTACACGACGCTCTTCCGATCTNNNGTGGAGAGGCAGGAAAATA
GAGCACACGTCTGAACTCTTTCCCTACACGACGCTCTTCCGATCTNNNGTGGAGAGGCAGGAAAATA
GAGCACACGTCTGAACTCTTTCCCTACACGACGCTCTTCCGATCTNNNGCTTGTTGGGCAGGTATACT
GAGCACACGTCTGAACTCTTTCCCTACACGACGCTCTTCCGATCTNNNGCTTGTTGGGCAGGTATACT
GAGCACACGTCTGAACTCTTTCCCTACACGACGCTCTTCCGATCTNNNGAGGTTGCAGTGAGCCATAAT
GAGCACACGTCTGAACTCTTTCCCTACACGACGCTCTTCCGATCTNNNGAGGTTGCAGTGAGCCATAAT
GAGCACACGTCTGAACTCTTTCCCTACACGACGCTCTTCCGATCTNNNCCTTAGCACCAGCCAGAAAAT
GAGCACACGTCTGAACTCTTTCCCTACACGACGCTCTTCCGATCTNNNCCTTAGCACCAGCCAGAAAAT
AGCACACGTCTGAACTCTTTCCCTACACGACGCTCTTCCGATCTNNNTTCTCCAAGAGAGGGTCCCTAA
AGCACACGTCTGAACTCTTTCCCTACACGACGCTCTTCCGATCTNNNTTCTCCAAGAGAGGGTCCCTAA
AGCACACGTCTGAACTCTTTCCCTACACGACGCTCTTCCGATCTNNNTATTTTGGAAAAACAAACGGAT
AGCACACGTCTGAACTCTTTCCCTACACGACGCTCTTCCGATCTNNNTATTTTGGAAAAACAAACGGAT
GAGCACACGTCTGAACTCTTTCCCTACACGACGCTCTTCCGATCTNNNAGTGATAAAAATCAGGCCCA
GAGCACACGTCTGAACTCTTTCCCTACACGACGCTCTTCCGATCTNNNAGTGATAAAAATCAGGCCCA
ACACGTCTGAACTCTTTCCCTACACGACGCTCTTCCGATCTNNNCATGTAATTGTCAGCACGGGATCCAT
ACACGTCTGAACTCTTTCCCTACACGACGCTCTTCCGATCTNNNCATGTAATTGTCAGCACGGGATCCAT
GAGCACACGTCTGAACTCTTTCCCTACACGACGCTCTTCCGATCTNNNACTCAGAATTGCTTTGGCTGT
GAGCACACGTCTGAACTCTTTCCCTACACGACGCTCTTCCGATCTNNNACTCAGAATTGCTTTGGCTGT
GAGCACACGTCTGAACTCTTTCCCTACACGACGCTCTTCCGATCTNNNTGCAGAGGGCTGAAAGAAAA
GAGCACACGTCTGAACTCTTTCCCTACACGACGCTCTTCCGATCTNNNTGCAGAGGGCTGAAAGAAAA
AGCACACGTCTGAACTCTTTCCCTACACGACGCTCTTCCGATCTNNNAATTCGTGGTATGCCCGTTTTTT
AGCACACGTCTGAACTCTTTCCCTACACGACGCTCTTCCGATCTNNNAATTCGTGGTATGCCCGTTTTTT
AGCACACGTCTGAACTCTTTCCCTACACGACGCTCTTCCGATCTNNNGTCTTCAATCCCCAATCCAGTCT
AGCACACGTCTGAACTCTTTCCCTACACGACGCTCTTCCGATCTNNNGTCTTCAATCCCCAATCCAGTCT
GAGCACACGTCTGAACTCTTTCCCTACACGACGCTCTTCCGATCTNNNATGGGTTGTCTCTAACAGCACT
GAGCACACGTCTGAACTCTTTCCCTACACGACGCTCTTCCGATCTNNNATGGGTTGTCTCTAACAGCACT
CACACGTCTGAACTCTTTCCCTACACGACGCTCTTCCGATCTNNNGTTGATCTCCTGACCTCGTGATCCA
CACACGTCTGAACTCTTTCCCTACACGACGCTCTTCCGATCTNNNGTTGATCTCCTGACCTCGTGATCCA
AGCACACGTCTGAACTCTTTCCCTACACGACGCTCTTCCGATCTNNNGAGGCTGAGGCAGGAGAATG
AGCACACGTCTGAACTCTTTCCCTACACGACGCTCTTCCGATCTNNNGAGGCTGAGGCAGGAGAATG
ACACGTCTGAACTCTTTCCCTACACGACGCTCTTCCGATCTNNNCAAAACCCTTTCCTTCTTCCCAGCATC
ACACGTCTGAACTCTTTCCCTACACGACGCTCTTCCGATCTNNNCAAAACCCTTTCCTTCTTCCCAGCATC
AGAGCACACGTCTGAACTCTTTCCCTACACGACGCTCTTCCGATCTNNNCTACATCAGTATCATCACTTG
AGAGCACACGTCTGAACTCTTTCCCTACACGACGCTCTTCCGATCTNNNCTACATCAGTATCATCACTTG
AGCACACGTCTGAACTCTTTCCCTACACGACGCTCTTCCGATCTNNNGTGTATATGCATGTGTGTGTG
AGCACACGTCTGAACTCTTTCCCTACACGACGCTCTTCCGATCTNNNGTGTATATGCATGTGTGTGTG

AGCACACGTCTGAACTCTTTCCCTACACGACGCTCTTCCGATCTNNNGACAATTTCACTTTCCCAAGGG

AGCACACGTCTGAACTCTTTCCCTACACGACGCTCTTCCGATCTNNNGACAATTTCACTTTCCCAAGGG

AGCACACGTCTGAACTCTTTCCCTACACGACGCTCTTCCGATCTNNNTGTTGTTGCAAATGACAGATG

AGCACACGTCTGAACTCTTTCCCTACACGACGCTCTTCCGATCTNNNTGTTGTTGCAAATGACAGATG

GCACACGTCTGAACTCTTTCCCTACACGACGCTCTTCCGATCTNNNCCAAAAGCATCTGATCAGTGTGG

GCACACGTCTGAACTCTTTCCCTACACGACGCTCTTCCGATCTNNNCCAAAAGCATCTGATCAGTGTGG

AGCACACGTCTGAACTCTTTCCCTACACGACGCTCTTCCGATCTNNNTGTTAACACAAGATTTTTGTAT

AGCACACGTCTGAACTCTTTCCCTACACGACGCTCTTCCGATCTNNNTGTTAACACAAGATTTTTGTAT

AGCACACGTCTGAACTCTTTCCCTACACGACGCTCTTCCGATCTNNNGCCCTACCTATGGTGTTAGGAT

AGCACACGTCTGAACTCTTTCCCTACACGACGCTCTTCCGATCTNNNGCCCTACCTATGGTGTTAGGAT

GAGCACACGTCTGAACTCTTTCCCTACACGACGCTCTTCCGATCTNNNTTCCTCAAGGTGTGCAAAGTT

GAGCACACGTCTGAACTCTTTCCCTACACGACGCTCTTCCGATCTNNNTTCCTCAAGGTGTGCAAAGTT

CACACGTCTGAACTCTTTCCCTACACGACGCTCTTCCGATCTNNNAAATTAGCCAAGCATGATGGTGG

CACACGTCTGAACTCTTTCCCTACACGACGCTCTTCCGATCTNNNAAATTAGCCAAGCATGATGGTGG

CACACGTCTGAACTCTTTCCCTACACGACGCTCTTCCGATCTNNNCTGTGATTCCAGCTACTCAGGAGC

CACACGTCTGAACTCTTTCCCTACACGACGCTCTTCCGATCTNNNCTGTGATTCCAGCTACTCAGGAGC

GAGCACACGTCTGAACTCTTTCCCTACACGACGCTCTTCCGATCTNNNCCTCTGAATTGGGAGCACAGT

GAGCACACGTCTGAACTCTTTCCCTACACGACGCTCTTCCGATCTNNNCCTCTGAATTGGGAGCACAGT

CACACGTCTGAACTCTTTCCCTACACGACGCTCTTCCGATCTNNNTCCTTCTTTCCCTCTCTCCTTCCCATC

CACACGTCTGAACTCTTTCCCTACACGACGCTCTTCCGATCTNNNTCCTTCTTTCCCTCTCTCCTTCCCATC

AGCACACGTCTGAACTCTTTCCCTACACGACGCTCTTCCGATCTNNNTTACACACACACACACACACAC

AGCACACGTCTGAACTCTTTCCCTACACGACGCTCTTCCGATCTNNNTTACACACACACACACACACAC

GAGCACACGTCTGAACTCTTTCCCTACACGACGCTCTTCCGATCTNNNTATTCCCTCTCTCCCTCCTCATC

GAGCACACGTCTGAACTCTTTCCCTACACGACGCTCTTCCGATCTNNNTATTCCCTCTCTCCCTCCTCATC

GAGCACACGTCTGAACTCTTTCCCTACACGACGCTCTTCCGATCTNNNGTAGAGGCGTGAGTTCAAAGA

GAGCACACGTCTGAACTCTTTCCCTACACGACGCTCTTCCGATCTNNNGTAGAGGCGTGAGTTCAAAGA

GAGCACACGTCTGAACTCTTTCCCTACACGACGCTCTTCCGATCTNNNAAGCAAAGGTTGAACTGGCT

GAGCACACGTCTGAACTCTTTCCCTACACGACGCTCTTCCGATCTNNNAAGCAAAGGTTGAACTGGCT

GAGCACACGTCTGAACTCTTTCCCTACACGACGCTCTTCCGATCTNNNTGTAATGCATCTGTGTTCCAG

GAGCACACGTCTGAACTCTTTCCCTACACGACGCTCTTCCGATCTNNNTGTAATGCATCTGTGTTCCAG

GCACACGTCTGAACTCTTTCCCTACACGACGCTCTTCCGATCTNNNGAGCCTACCTTTTCTTCTGAGGG

GCACACGTCTGAACTCTTTCCCTACACGACGCTCTTCCGATCTNNNGAGCCTACCTTTTCTTCTGAGGG

GAGCACACGTCTGAACTCTTTCCCTACACGACGCTCTTCCGATCTNNNCTGTAATCCCAGCTACTCAGG

GAGCACACGTCTGAACTCTTTCCCTACACGACGCTCTTCCGATCTNNNCTGTAATCCCAGCTACTCAGG

AGAGCACACGTCTGAACTCTTTCCCTACACGACGCTCTTCCGATCTNNNGACAAAGAGTCAAGCCTTGA

AGAGCACACGTCTGAACTCTTTCCCTACACGACGCTCTTCCGATCTNNNGACAAAGAGTCAAGCCTTGA

CACACGTCTGAACTCTTTCCCTACACGACGCTCTTCCGATCTNNNAGCTCAACTCTGGTGTGAATGATC

CACACGTCTGAACTCTTTCCCTACACGACGCTCTTCCGATCTNNNAGCTCAACTCTGGTGTGAATGATC

AGCACACGTCTGAACTCTTTCCCTACACGACGCTCTTCCGATCTNNNAAAGTGCTGGGATTAGAGGAC

AGCACACGTCTGAACTCTTTCCCTACACGACGCTCTTCCGATCTNNNAAAGTGCTGGGATTAGAGGAC

AGCACACGTCTGAACTCTTTCCCTACACGACGCTCTTCCGATCTNNNCCTGAGCCTTGTAAGTTGAGACA

AGCACACGTCTGAACTCTTTCCCTACACGACGCTCTTCCGATCTNNNCCTGAGCCTTGTAAGTTGAGACA

GAGCACACGTCTGAACTCTTTCCCTACACGACGCTCTTCCGATCTNNNATGAGCCCTTAGAGAACCTTT

GAGCACACGTCTGAACTCTTTCCCTACACGACGCTCTTCCGATCTNNNATGAGCCCTTAGAGAACCTTT

AGCACACGTCTGAACTCTTTCCCTACACGACGCTCTTCCGATCTNNNGCCAAAAATACATGTGAGGTG

AGCACACGTCTGAACTCTTTCCCTACACGACGCTCTTCCGATCTNNNGCCAAAAATACATGTGAGGTG

AGCACACGTCTGAACTCTTTCCCTACACGACGCTCTTCCGATCTNNNGTAACAAAACACCACCTGTTCC

AGCACACGTCTGAACTCTTTCCCTACACGACGCTCTTCCGATCTNNNGTAACAAAACACCACCTGTTCC

AGAGCACACGTCTGAACTCTTTCCCTACACGACGCTCTTCCGATCTNNNGAGAAACTGACAGAGAGG

AGAGCACACGTCTGAACTCTTTCCCTACACGACGCTCTTCCGATCTNNNGAGAAACTGACAGAGAGG

AGCACACGTCTGAACTCTTTCCCTACACGACGCTCTTCCGATCTNNNGAAAAAGCACAGGGTGACATC

AGCACACGTCTGAACTCTTTCCCTACACGACGCTCTTCCGATCTNNNGAAAAAGCACAGGGTGACATC

AGCACACGTCTGAACTCTTTCCCTACACGACGCTCTTCCGATCTNNNACCCTTTGGAACACTCTTCTCTG

AGCACACGTCTGAACTCTTTCCCTACACGACGCTCTTCCGATCTNNNACCCTTTGGAACACTCTTCTCTG

GCACACGTCTGAACTCTTTCCCTACACGACGCTCTTCCGATCTNNNCACTCAAGCTGGACCATCAAATCC

GCACACGTCTGAACTCTTTCCCTACACGACGCTCTTCCGATCTNNNCACTCAAGCTGGACCATCAAATCC

AGCACACGTCTGAACTCTTTCCCTACACGACGCTCTTCCGATCTNNNATAGAACTGCACAAAACAGCTT

AGCACACGTCTGAACTCTTTCCCTACACGACGCTCTTCCGATCTNNNATAGAACTGCACAAAACAGCTT

GAGCACACGTCTGAACTCTTTCCCTACACGACGCTCTTCCGATCTNNNTCATGCCTGTAATCTCAGCAC

GAGCACACGTCTGAACTCTTTCCCTACACGACGCTCTTCCGATCTNNNTCATGCCTGTAATCTCAGCAC

GAGCACACGTCTGAACTCTTTCCCTACACGACGCTCTTCCGATCTNNNACAATTCTACTGACTCAGCCT

GAGCACACGTCTGAACTCTTTCCCTACACGACGCTCTTCCGATCTNNNACAATTCTACTGACTCAGCCT

GCACACGTCTGAACTCTTTCCCTACACGACGCTCTTCCGATCTNNNTGTAGTCCTAGCTACTCAGGAG

GCACACGTCTGAACTCTTTCCCTACACGACGCTCTTCCGATCTNNNTGTAGTCCTAGCTACTCAGGAG

GAGCACACGTCTGAACTCTTTCCCTACACGACGCTCTTCCGATCTNNNAGAATCACTTGAATCCAGGA

GAGCACACGTCTGAACTCTTTCCCTACACGACGCTCTTCCGATCTNNNAGAATCACTTGAATCCAGGA

GCACACGTCTGAACTCTTTCCCTACACGACGCTCTTCCGATCTNNNTTTCAGGCTGCAAATTTTCCCAA

GCACACGTCTGAACTCTTTCCCTACACGACGCTCTTCCGATCTNNNTTTCAGGCTGCAAATTTTCCCAA

GAGCACACGTCTGAACTCTTTCCCTACACGACGCTCTTCCGATCTNNNCTCATCTTTGTGGGCTTCTACC

GAGCACACGTCTGAACTCTTTCCCTACACGACGCTCTTCCGATCTNNNCTCATCTTTGTGGGCTTCTACC

GAGCACACGTCTGAACTCTTTCCCTACACGACGCTCTTCCGATCTNNNTTTGTGAGCACCAGAATCTGT

GAGCACACGTCTGAACTCTTTCCCTACACGACGCTCTTCCGATCTNNNTTTGTGAGCACCAGAATCTGT

GCACACGTCTGAACTCTTTCCCTACACGACGCTCTTCCGATCTNNNAGCTGAGAACATAGGCATGAACT

GCACACGTCTGAACTCTTTCCCTACACGACGCTCTTCCGATCTNNNAGCTGAGAACATAGGCATGAACT

GAGCACACGTCTGAACTCTTTCCCTACACGACGCTCTTCCGATCTNNNTGGGGTTGTTTGTGAAACTT

GAGCACACGTCTGAACTCTTTCCCTACACGACGCTCTTCCGATCTNNNTGGGGTTGTTTGTGAAACTT

GCACACGTCTGAACTCTTTCCCTACACGACGCTCTTCCGATCTNNNTCTTCTCCTTCTCCTTCTCCTTCTC

GCACACGTCTGAACTCTTTCCCTACACGACGCTCTTCCGATCTNNNTCTTCTCCTTCTCCTTCTCCTTCTC

GAGCACACGTCTGAACTCTTTCCCTACACGACGCTCTTCCGATCTNNNCGTAATCCCAGCTACTCAGGA

GAGCACACGTCTGAACTCTTTCCCTACACGACGCTCTTCCGATCTNNNCGTAATCCCAGCTACTCAGGA

GAGCACACGTCTGAACTCTTTCCCTACACGACGCTCTTCCGATCTNNNCTCTTTTGTCTCTCCCTCTCTGC

GAGCACACGTCTGAACTCTTTCCCTACACGACGCTCTTCCGATCTNNNCTCTTTTGTCTCTCCCTCTCTGC

GCACACGTCTGAACTCTTTCCCTACACGACGCTCTTCCGATCTNNNCCGCCATTTTCTTTTCCTCATCACA

GCACACGTCTGAACTCTTTCCCTACACGACGCTCTTCCGATCTNNNCCGCCATTTTCTTTTCCTCATCACA

AGCACACGTCTGAACTCTTTCCCTACACGACGCTCTTCCGATCTNNNTGCAAAGAGTATAGGGAAGT

AGCACACGTCTGAACTCTTTCCCTACACGACGCTCTTCCGATCTNNNTGCAAAGAGTATAGGGAAGT

AGCACACGTCTGAACTCTTTCCCTACACGACGCTCTTCCGATCTNNNTAACCTAGACCAACATGGGCTG
AGCACACGTCTGAACTCTTTCCCTACACGACGCTCTTCCGATCTNNNTAACCTAGACCAACATGGGCTG
CACACGTCTGAACTCTTTCCCTACACGACGCTCTTCCGATCTNNNGGCCCTTCCTTTAGTTCAAGTCCCA
CACACGTCTGAACTCTTTCCCTACACGACGCTCTTCCGATCTNNNGGCCCTTCCTTTAGTTCAAGTCCCA
AGCACACGTCTGAACTCTTTCCCTACACGACGCTCTTCCGATCTNNNGGCAGCAAGAATGACTTTGAA
AGCACACGTCTGAACTCTTTCCCTACACGACGCTCTTCCGATCTNNNGGCAGCAAGAATGACTTTGAA
GCACACGTCTGAACTCTTTCCCTACACGACGCTCTTCCGATCTNNNTTTAGCAAGGTCTCTGTGCAGC
GCACACGTCTGAACTCTTTCCCTACACGACGCTCTTCCGATCTNNNTTTAGCAAGGTCTCTGTGCAGC
CACACGTCTGAACTCTTTCCCTACACGACGCTCTTCCGATCTNNNATTTCTCCCTAAGGCACCCAGATCAT
CACACGTCTGAACTCTTTCCCTACACGACGCTCTTCCGATCTNNNATTTCTCCCTAAGGCACCCAGATCAT
AGCACACGTCTGAACTCTTTCCCTACACGACGCTCTTCCGATCTNNNGAGGGAGGAGAACCACTTGAA
AGCACACGTCTGAACTCTTTCCCTACACGACGCTCTTCCGATCTNNNGAGGGAGGAGAACCACTTGAA
GAGCACACGTCTGAACTCTTTCCCTACACGACGCTCTTCCGATCTNNNGAGGGCTCCTTGTTTTTCTTCA
GAGCACACGTCTGAACTCTTTCCCTACACGACGCTCTTCCGATCTNNNGAGGGCTCCTTGTTTTTCTTCA
AGCACACGTCTGAACTCTTTCCCTACACGACGCTCTTCCGATCTNNNGCTTGCAGACTGTAACCCTTTTT
AGCACACGTCTGAACTCTTTCCCTACACGACGCTCTTCCGATCTNNNGCTTGCAGACTGTAACCCTTTTT
AGCACACGTCTGAACTCTTTCCCTACACGACGCTCTTCCGATCTNNNAAAATTAGCTGGGTTTGGTGG
AGCACACGTCTGAACTCTTTCCCTACACGACGCTCTTCCGATCTNNNAAAATTAGCTGGGTTTGGTGG
AGCACACGTCTGAACTCTTTCCCTACACGACGCTCTTCCGATCTNNNATAGGAGGGCAGGGGTGAAAA
AGCACACGTCTGAACTCTTTCCCTACACGACGCTCTTCCGATCTNNNATAGGAGGGCAGGGGTGAAAA
GCACACGTCTGAACTCTTTCCCTACACGACGCTCTTCCGATCTNNNCTATGTTTTGTGGAAAGAGAAG
GCACACGTCTGAACTCTTTCCCTACACGACGCTCTTCCGATCTNNNCTATGTTTTGTGGAAAGAGAAG
GAGCACACGTCTGAACTCTTTCCCTACACGACGCTCTTCCGATCTNNNTTGCCTTGTATACGTGTGTGT
GAGCACACGTCTGAACTCTTTCCCTACACGACGCTCTTCCGATCTNNNTTGCCTTGTATACGTGTGTGT
GAAGAGCACACGTCTGAACTCTTTCCCTACACGACGCTCTTCCGATCTNNNGCAAACTGAAGCAAATAT
GAAGAGCACACGTCTGAACTCTTTCCCTACACGACGCTCTTCCGATCTNNNGCAAACTGAAGCAAATAT
GCACACGTCTGAACTCTTTCCCTACACGACGCTCTTCCGATCTNNNGGCATCCCTGAACAAAATTGTGA
GCACACGTCTGAACTCTTTCCCTACACGACGCTCTTCCGATCTNNNGGCATCCCTGAACAAAATTGTGA
GAGCACACGTCTGAACTCTTTCCCTACACGACGCTCTTCCGATCTNNNAGTGCTGGGATTATAGGCATG
GAGCACACGTCTGAACTCTTTCCCTACACGACGCTCTTCCGATCTNNNAGTGCTGGGATTATAGGCATG
CACGTCTGAACTCTTTCCCTACACGACGCTCTTCCGATCTNNNTGACCTAGCAATCCCACTACAAGCATG
CACGTCTGAACTCTTTCCCTACACGACGCTCTTCCGATCTNNNTGACCTAGCAATCCCACTACAAGCATG
AGCACACGTCTGAACTCTTTCCCTACACGACGCTCTTCCGATCTNNNAAAAAATTAGCTGGGTGTGGTG
AGCACACGTCTGAACTCTTTCCCTACACGACGCTCTTCCGATCTNNNAAAAAATTAGCTGGGTGTGGTG
GCACACGTCTGAACTCTTTCCCTACACGACGCTCTTCCGATCTNNNGTCTTCTCCCTACAGGACCTTGCAT
GCACACGTCTGAACTCTTTCCCTACACGACGCTCTTCCGATCTNNNGTCTTCTCCCTACAGGACCTTGCAT
AGCACACGTCTGAACTCTTTCCCTACACGACGCTCTTCCGATCTNNNCTTGCCATTCACAACAGTTGCA
AGCACACGTCTGAACTCTTTCCCTACACGACGCTCTTCCGATCTNNNCTTGCCATTCACAACAGTTGCA
AGCACACGTCTGAACTCTTTCCCTACACGACGCTCTTCCGATCTNNNAGTTTATTGTGAAAACATGATG
AGCACACGTCTGAACTCTTTCCCTACACGACGCTCTTCCGATCTNNNAGTTTATTGTGAAAACATGATG
AGCACACGTCTGAACTCTTTCCCTACACGACGCTCTTCCGATCTNNNTAAACGGGCAGAATACATGAG
AGCACACGTCTGAACTCTTTCCCTACACGACGCTCTTCCGATCTNNNTAAACGGGCAGAATACATGAG
GCACACGTCTGAACTCTTTCCCTACACGACGCTCTTCCGATCTNNNCTGTAATCCCAGCTACTCGGAAG

GCACACGTCTGAACTCTTTCCCTACACGACGCTCTTCCGATCTNNNCTGTAATCCCAGCTACTCGGAAG
AGCACACGTCTGAACTCTTTCCCTACACGACGCTCTTCCGATCTNNNTCCAAAAATTCTAACGTGAGG
AGCACACGTCTGAACTCTTTCCCTACACGACGCTCTTCCGATCTNNNTCCAAAAATTCTAACGTGAGG
AGCACACGTCTGAACTCTTTCCCTACACGACGCTCTTCCGATCTNNNCAAATGCTCACAGGTGACAG
AGCACACGTCTGAACTCTTTCCCTACACGACGCTCTTCCGATCTNNNCAAATGCTCACAGGTGACAG
AGCACACGTCTGAACTCTTTCCCTACACGACGCTCTTCCGATCTNNNTTCAGACAGCCACAAATGTCT
AGCACACGTCTGAACTCTTTCCCTACACGACGCTCTTCCGATCTNNNTTCAGACAGCCACAAATGTCT
AGCACACGTCTGAACTCTTTCCCTACACGACGCTCTTCCGATCTNNNCCACATAGGAGTAGCCTGACT
AGCACACGTCTGAACTCTTTCCCTACACGACGCTCTTCCGATCTNNNCCACATAGGAGTAGCCTGACT
AGCACACGTCTGAACTCTTTCCCTACACGACGCTCTTCCGATCTNNNAGAAACTGGCAAAGGTGTTT
AGCACACGTCTGAACTCTTTCCCTACACGACGCTCTTCCGATCTNNNAGAAACTGGCAAAGGTGTTT
AGCACACGTCTGAACTCTTTCCCTACACGACGCTCTTCCGATCTNNNCTCTTCTCAAGTACACCCACC
AGCACACGTCTGAACTCTTTCCCTACACGACGCTCTTCCGATCTNNNCTCTTCTCAAGTACACCCACC
AGCACACGTCTGAACTCTTTCCCTACACGACGCTCTTCCGATCTNNNAGGTTACAGTGAGCTGAGATC
AGCACACGTCTGAACTCTTTCCCTACACGACGCTCTTCCGATCTNNNAGGTTACAGTGAGCTGAGATC
GCACACGTCTGAACTCTTTCCCTACACGACGCTCTTCCGATCTNNNGAATCTAGGAGATGGCAGAGG
GCACACGTCTGAACTCTTTCCCTACACGACGCTCTTCCGATCTNNNGAATCTAGGAGATGGCAGAGG
GAGCACACGTCTGAACTCTTTCCCTACACGACGCTCTTCCGATCTNNNTTTGCGTGTAACCTATGCACA
GAGCACACGTCTGAACTCTTTCCCTACACGACGCTCTTCCGATCTNNNTTTGCGTGTAACCTATGCACA
AGCACACGTCTGAACTCTTTCCCTACACGACGCTCTTCCGATCTNNNCAGAGGCTGCTGTCATGTCTG
AGCACACGTCTGAACTCTTTCCCTACACGACGCTCTTCCGATCTNNNCAGAGGCTGCTGTCATGTCTG
AGCACACGTCTGAACTCTTTCCCTACACGACGCTCTTCCGATCTNNNATGAGGGGAGGCTGTTGTTAT
AGCACACGTCTGAACTCTTTCCCTACACGACGCTCTTCCGATCTNNNATGAGGGGAGGCTGTTGTTAT
AGCACACGTCTGAACTCTTTCCCTACACGACGCTCTTCCGATCTNNNGGAGTACACATACATAGCCACC
AGCACACGTCTGAACTCTTTCCCTACACGACGCTCTTCCGATCTNNNGGAGTACACATACATAGCCACC
GAGCACACGTCTGAACTCTTTCCCTACACGACGCTCTTCCGATCTNNNGATCTTTGACAAAGGGGCA
GAGCACACGTCTGAACTCTTTCCCTACACGACGCTCTTCCGATCTNNNGATCTTTGACAAAGGGGCA
GCACACGTCTGAACTCTTTCCCTACACGACGCTCTTCCGATCTNNNGGAACTTGAGTGGATGTCTTCCT
GCACACGTCTGAACTCTTTCCCTACACGACGCTCTTCCGATCTNNNGGAACTTGAGTGGATGTCTTCCT
AGCACACGTCTGAACTCTTTCCCTACACGACGCTCTTCCGATCTNNNAGCCTTCCAAGAAGCTGAAA
AGCACACGTCTGAACTCTTTCCCTACACGACGCTCTTCCGATCTNNNAGCCTTCCAAGAAGCTGAAA
GCACACGTCTGAACTCTTTCCCTACACGACGCTCTTCCGATCTNNNGCCTGTAGTCCCAGCTACTTGCA
GCACACGTCTGAACTCTTTCCCTACACGACGCTCTTCCGATCTNNNGCCTGTAGTCCCAGCTACTTGCA
AGCACACGTCTGAACTCTTTCCCTACACGACGCTCTTCCGATCTNNNATTGGGGTATTGCTATTCCCAG
AGCACACGTCTGAACTCTTTCCCTACACGACGCTCTTCCGATCTNNNATTGGGGTATTGCTATTCCCAG
AGCACACGTCTGAACTCTTTCCCTACACGACGCTCTTCCGATCTNNNCACTTGAATCAGGAGATGGAG
AGCACACGTCTGAACTCTTTCCCTACACGACGCTCTTCCGATCTNNNCACTTGAATCAGGAGATGGAG
GCACACGTCTGAACTCTTTCCCTACACGACGCTCTTCCGATCTNNNTCCAGTAAATCTCCCACCCAAAG
GCACACGTCTGAACTCTTTCCCTACACGACGCTCTTCCGATCTNNNTCCAGTAAATCTCCCACCCAAAG
GAGCACACGTCTGAACTCTTTCCCTACACGACGCTCTTCCGATCTNNNTTGCACCTGTAATCCCAGCTA
GAGCACACGTCTGAACTCTTTCCCTACACGACGCTCTTCCGATCTNNNTTGCACCTGTAATCCCAGCTA
AGCACACGTCTGAACTCTTTCCCTACACGACGCTCTTCCGATCTNNNCCGCCCTGAAAAATCTGTTTCT
AGCACACGTCTGAACTCTTTCCCTACACGACGCTCTTCCGATCTNNNCCGCCCTGAAAAATCTGTTTCT

AGCACACGTCTGAACTCTTTCCCTACACGACGCTCTTCCGATCTNNNAGGTGGATTCACAGCTGAATT

AGCACACGTCTGAACTCTTTCCCTACACGACGCTCTTCCGATCTNNNAGGTGGATTCACAGCTGAATT

GAGCACACGTCTGAACTCTTTCCCTACACGACGCTCTTCCGATCTNNNACACTACCAAGATTCCAGAG

GAGCACACGTCTGAACTCTTTCCCTACACGACGCTCTTCCGATCTNNNACACTACCAAGATTCCAGAG

AGCACACGTCTGAACTCTTTCCCTACACGACGCTCTTCCGATCTNNNCTGGGAAGTCAAGGCTTTAGT

AGCACACGTCTGAACTCTTTCCCTACACGACGCTCTTCCGATCTNNNCTGGGAAGTCAAGGCTTTAGT

GCACACGTCTGAACTCTTTCCCTACACGACGCTCTTCCGATCTNNNGGAATCTGCCCTGTGTCGTCCAT

GCACACGTCTGAACTCTTTCCCTACACGACGCTCTTCCGATCTNNNGGAATCTGCCCTGTGTCGTCCAT

GAGCACACGTCTGAACTCTTTCCCTACACGACGCTCTTCCGATCTNNNCCAGCCCAATCCTTCCTTTCTA

GAGCACACGTCTGAACTCTTTCCCTACACGACGCTCTTCCGATCTNNNCCAGCCCAATCCTTCCTTTCTA

AGCACACGTCTGAACTCTTTCCCTACACGACGCTCTTCCGATCTNNNGAGAATCGCTTGAACCCTGGA

AGCACACGTCTGAACTCTTTCCCTACACGACGCTCTTCCGATCTNNNGAGAATCGCTTGAACCCTGGA

AGCACACGTCTGAACTCTTTCCCTACACGACGCTCTTCCGATCTNNNCTGAGGCAGGAGAATCACTTG

AGCACACGTCTGAACTCTTTCCCTACACGACGCTCTTCCGATCTNNNCTGAGGCAGGAGAATCACTTG

AGAGCACACGTCTGAACTCTTTCCCTACACGACGCTCTTCCGATCTNNNTCATGAGAGGGTGAGGTA

AGAGCACACGTCTGAACTCTTTCCCTACACGACGCTCTTCCGATCTNNNTCATGAGAGGGTGAGGTA

AGCACACGTCTGAACTCTTTCCCTACACGACGCTCTTCCGATCTNNNGGATCACGAGGTCAGAAGATC

AGCACACGTCTGAACTCTTTCCCTACACGACGCTCTTCCGATCTNNNGGATCACGAGGTCAGAAGATC

GAGCACACGTCTGAACTCTTTCCCTACACGACGCTCTTCCGATCTNNNGAAGGCAGAGGTTACAGTG

GAGCACACGTCTGAACTCTTTCCCTACACGACGCTCTTCCGATCTNNNGAAGGCAGAGGTTACAGTG

AGCACACGTCTGAACTCTTTCCCTACACGACGCTCTTCCGATCTNNNAATCCCAGCTACTTGGAAGTCT

AGCACACGTCTGAACTCTTTCCCTACACGACGCTCTTCCGATCTNNNAATCCCAGCTACTTGGAAGTCT

AGCACACGTCTGAACTCTTTCCCTACACGACGCTCTTCCGATCTNNNCTCCTTTTATCGTCTCCCTACGG

AGCACACGTCTGAACTCTTTCCCTACACGACGCTCTTCCGATCTNNNCTCCTTTTATCGTCTCCCTACGG

GAGCACACGTCTGAACTCTTTCCCTACACGACGCTCTTCCGATCTNNNCACCAGACAGGCCCATTTTTC

GAGCACACGTCTGAACTCTTTCCCTACACGACGCTCTTCCGATCTNNNCACCAGACAGGCCCATTTTTC

GAGCACACGTCTGAACTCTTTCCCTACACGACGCTCTTCCGATCTNNNCTCTGAACAGACCAATAGCG

GAGCACACGTCTGAACTCTTTCCCTACACGACGCTCTTCCGATCTNNNCTCTGAACAGACCAATAGCG

AGCACACGTCTGAACTCTTTCCCTACACGACGCTCTTCCGATCTNNNACACATGCAATTGACCATCACA

AGCACACGTCTGAACTCTTTCCCTACACGACGCTCTTCCGATCTNNNACACATGCAATTGACCATCACA

AGCACACGTCTGAACTCTTTCCCTACACGACGCTCTTCCGATCTNNNGAATGCTAGAGATCCTTGTCCC

AGCACACGTCTGAACTCTTTCCCTACACGACGCTCTTCCGATCTNNNGAATGCTAGAGATCCTTGTCCC

GCACACGTCTGAACTCTTTCCCTACACGACGCTCTTCCGATCTNNNGGATGGTCTCAATTCCCCGATA

GCACACGTCTGAACTCTTTCCCTACACGACGCTCTTCCGATCTNNNGGATGGTCTCAATTCCCCGATA

AGCACACGTCTGAACTCTTTCCCTACACGACGCTCTTCCGATCTNNNAACTGAGATTGTACTCCAGCCT

AGCACACGTCTGAACTCTTTCCCTACACGACGCTCTTCCGATCTNNNAACTGAGATTGTACTCCAGCCT

GAGCACACGTCTGAACTCTTTCCCTACACGACGCTCTTCCGATCTNNNGCGACAGCAGAGAGAGACTC

GAGCACACGTCTGAACTCTTTCCCTACACGACGCTCTTCCGATCTNNNGCGACAGCAGAGAGAGACTC

AGAGCACACGTCTGAACTCTTTCCCTACACGACGCTCTTCCGATCTNNNGTACCAGCTGTAACATCGA

AGAGCACACGTCTGAACTCTTTCCCTACACGACGCTCTTCCGATCTNNNGTACCAGCTGTAACATCGA

CACACGTCTGAACTCTTTCCCTACACGACGCTCTTCCGATCTNNNACTCTTTCTAGACAGGCGGAATGC

CACACGTCTGAACTCTTTCCCTACACGACGCTCTTCCGATCTNNNACTCTTTCTAGACAGGCGGAATGC

GCACACGTCTGAACTCTTTCCCTACACGACGCTCTTCCGATCTNNNTTTGCAGAAGTGAATCTGGGGT

GCACACGTCTGAACTCTTTCCCTACACGACGCTCTTCCGATCTNNNTTTGCAGAAGTGAATCTGGGGTA
AGCACACGTCTGAACTCTTTCCCTACACGACGCTCTTCCGATCTNNNCACCTTAGCCTCCTGAGTAACTG
AGCACACGTCTGAACTCTTTCCCTACACGACGCTCTTCCGATCTNNNCACCTTAGCCTCCTGAGTAACTG
AGCACACGTCTGAACTCTTTCCCTACACGACGCTCTTCCGATCTNNNCAGCAGTGGTGTTCTCTAGACT
AGCACACGTCTGAACTCTTTCCCTACACGACGCTCTTCCGATCTNNNCAGCAGTGGTGTTCTCTAGACT
GCACACGTCTGAACTCTTTCCCTACACGACGCTCTTCCGATCTNNNTCCTCTTGGTGATGGGGTTGTAT
GCACACGTCTGAACTCTTTCCCTACACGACGCTCTTCCGATCTNNNTCCTCTTGGTGATGGGGTTGTAT
GAGCACACGTCTGAACTCTTTCCCTACACGACGCTCTTCCGATCTNNNTGGGAGAGCCAGAGGAAATA
GAGCACACGTCTGAACTCTTTCCCTACACGACGCTCTTCCGATCTNNNTGGGAGAGCCAGAGGAAATA
AGAGCACACGTCTGAACTCTTTCCCTACACGACGCTCTTCCGATCTNNNCTGAGGCAGAGAATTGCTTG
AGAGCACACGTCTGAACTCTTTCCCTACACGACGCTCTTCCGATCTNNNCTGAGGCAGAGAATTGCTTG
GAGCACACGTCTGAACTCTTTCCCTACACGACGCTCTTCCGATCTNNNGACACCGTTGTTAGGTGGCTA
GAGCACACGTCTGAACTCTTTCCCTACACGACGCTCTTCCGATCTNNNGACACCGTTGTTAGGTGGCTA
CACACGTCTGAACTCTTTCCCTACACGACGCTCTTCCGATCTNNNCTCATTACGTTTTAGATGGGATTCA
CACACGTCTGAACTCTTTCCCTACACGACGCTCTTCCGATCTNNNCTCATTACGTTTTAGATGGGATTCA
GAGCACACGTCTGAACTCTTTCCCTACACGACGCTCTTCCGATCTNNNTGCTTCTTTGTGGCTGAGATTT
GAGCACACGTCTGAACTCTTTCCCTACACGACGCTCTTCCGATCTNNNTGCTTCTTTGTGGCTGAGATTT
GAGCACACGTCTGAACTCTTTCCCTACACGACGCTCTTCCGATCTNNNTCTGTGTCCGGTTTGTATGACA
GAGCACACGTCTGAACTCTTTCCCTACACGACGCTCTTCCGATCTNNNTCTGTGTCCGGTTTGTATGACA
AGAGCACACGTCTGAACTCTTTCCCTACACGACGCTCTTCCGATCTNNNTGCAGATCACAGGGGATATG
AGAGCACACGTCTGAACTCTTTCCCTACACGACGCTCTTCCGATCTNNNTGCAGATCACAGGGGATATG
AGCACACGTCTGAACTCTTTCCCTACACGACGCTCTTCCGATCTNNNAGGCAAGAGAATTGCTTGAACA
AGCACACGTCTGAACTCTTTCCCTACACGACGCTCTTCCGATCTNNNAGGCAAGAGAATTGCTTGAACA
GAGCACACGTCTGAACTCTTTCCCTACACGACGCTCTTCCGATCTNNNGAAGGGCCCAGGAGAGTAAC
GAGCACACGTCTGAACTCTTTCCCTACACGACGCTCTTCCGATCTNNNGAAGGGCCCAGGAGAGTAAC
AGAGCACACGTCTGAACTCTTTCCCTACACGACGCTCTTCCGATCTNNNCAGATATGGCATTCAAGGAA
AGAGCACACGTCTGAACTCTTTCCCTACACGACGCTCTTCCGATCTNNNCAGATATGGCATTCAAGGAA
AGCACACGTCTGAACTCTTTCCCTACACGACGCTCTTCCGATCTNNNGAAGCAGGAGAATCGCTTGAA
AGCACACGTCTGAACTCTTTCCCTACACGACGCTCTTCCGATCTNNNGAAGCAGGAGAATCGCTTGAA
GAGCACACGTCTGAACTCTTTCCCTACACGACGCTCTTCCGATCTNNNTTGATAGGACTGTGAGTGTGC
GAGCACACGTCTGAACTCTTTCCCTACACGACGCTCTTCCGATCTNNNTTGATAGGACTGTGAGTGTGC
GAGCACACGTCTGAACTCTTTCCCTACACGACGCTCTTCCGATCTNNNGTATATGGCTCCACACAAATG
GAGCACACGTCTGAACTCTTTCCCTACACGACGCTCTTCCGATCTNNNGTATATGGCTCCACACAAATG
GAGCACACGTCTGAACTCTTTCCCTACACGACGCTCTTCCGATCTNNNTTTCACCTCCACTTCCTGAGTA
GAGCACACGTCTGAACTCTTTCCCTACACGACGCTCTTCCGATCTNNNTTTCACCTCCACTTCCTGAGTA
AGCACACGTCTGAACTCTTTCCCTACACGACGCTCTTCCGATCTNNNCTAGAGCAACACTTGTCAAAGG
AGCACACGTCTGAACTCTTTCCCTACACGACGCTCTTCCGATCTNNNCTAGAGCAACACTTGTCAAAGG
GAGCACACGTCTGAACTCTTTCCCTACACGACGCTCTTCCGATCTNNNTATGGTTTTTGCAGTCCACAGA
GAGCACACGTCTGAACTCTTTCCCTACACGACGCTCTTCCGATCTNNNTATGGTTTTTGCAGTCCACAGA
AGCACACGTCTGAACTCTTTCCCTACACGACGCTCTTCCGATCTNNNAATGGTGTTTGGCTTTCTTTGGC
AGCACACGTCTGAACTCTTTCCCTACACGACGCTCTTCCGATCTNNNAATGGTGTTTGGCTTTCTTTGGC
AGCACACGTCTGAACTCTTTCCCTACACGACGCTCTTCCGATCTNNNCAAGCAAAAGTTGGGGTGTAG
AGCACACGTCTGAACTCTTTCCCTACACGACGCTCTTCCGATCTNNNCAAGCAAAAGTTGGGGTGTAG

AGCACACGTCTGAACTCTTTCCCTACACGACGCTCTTCCGATCTNNNAGTGCGTTATGAACCCAGCTAT
AGCACACGTCTGAACTCTTTCCCTACACGACGCTCTTCCGATCTNNNAGTGCGTTATGAACCCAGCTAT
GAGCACACGTCTGAACTCTTTCCCTACACGACGCTCTTCCGATCTNNNTCCAGTTATATGCCCCAATGC
GAGCACACGTCTGAACTCTTTCCCTACACGACGCTCTTCCGATCTNNNTCCAGTTATATGCCCCAATGC
AGCACACGTCTGAACTCTTTCCCTACACGACGCTCTTCCGATCTNNNTATAAGTAAAAGTGTGGGCAT
AGCACACGTCTGAACTCTTTCCCTACACGACGCTCTTCCGATCTNNNTATAAGTAAAAGTGTGGGCAT
AGCACACGTCTGAACTCTTTCCCTACACGACGCTCTTCCGATCTNNNAATGGGCAGACTGGAGTAAA
AGCACACGTCTGAACTCTTTCCCTACACGACGCTCTTCCGATCTNNNAATGGGCAGACTGGAGTAAA
GAGCACACGTCTGAACTCTTTCCCTACACGACGCTCTTCCGATCTNNNCAGGCAGTTTAAATCTCGTT
GAGCACACGTCTGAACTCTTTCCCTACACGACGCTCTTCCGATCTNNNCAGGCAGTTTAAATCTCGTT
AGCACACGTCTGAACTCTTTCCCTACACGACGCTCTTCCGATCTNNNCGACAAGAGCAAAAACTACGT
AGCACACGTCTGAACTCTTTCCCTACACGACGCTCTTCCGATCTNNNCGACAAGAGCAAAAACTACGT
AGCACACGTCTGAACTCTTTCCCTACACGACGCTCTTCCGATCTNNNGGTTGCCATGCAGATGTCTAA
AGCACACGTCTGAACTCTTTCCCTACACGACGCTCTTCCGATCTNNNGGTTGCCATGCAGATGTCTAA
AAGAGCACACGTCTGAACTCTTTCCCTACACGACGCTCTTCCGATCTNNNGAGGATGAGGCAGGAGA
AAGAGCACACGTCTGAACTCTTTCCCTACACGACGCTCTTCCGATCTNNNGAGGATGAGGCAGGAGA
AGCACACGTCTGAACTCTTTCCCTACACGACGCTCTTCCGATCTNNNTCAACCTCTTAGGCTCAAGCAAT
AGCACACGTCTGAACTCTTTCCCTACACGACGCTCTTCCGATCTNNNTCAACCTCTTAGGCTCAAGCAAT
AGCACACGTCTGAACTCTTTCCCTACACGACGCTCTTCCGATCTNNNCTCCTGAGTAGCTGGGACTACA
AGCACACGTCTGAACTCTTTCCCTACACGACGCTCTTCCGATCTNNNCTCCTGAGTAGCTGGGACTACA
AAGAGCACACGTCTGAACTCTTTCCCTACACGACGCTCTTCCGATCTNNNCCCGAGAGTTTTATGTTGC
AAGAGCACACGTCTGAACTCTTTCCCTACACGACGCTCTTCCGATCTNNNCCCGAGAGTTTTATGTTGC
AGCACACGTCTGAACTCTTTCCCTACACGACGCTCTTCCGATCTNNNCAACCCCTTGATCCCCTTGAATA
AGCACACGTCTGAACTCTTTCCCTACACGACGCTCTTCCGATCTNNNCAACCCCTTGATCCCCTTGAATA
GAGCACACGTCTGAACTCTTTCCCTACACGACGCTCTTCCGATCTNNNCCATTCTCCTGCCTTAGTCTCT
GAGCACACGTCTGAACTCTTTCCCTACACGACGCTCTTCCGATCTNNNCCATTCTCCTGCCTTAGTCTCT
AGCACACGTCTGAACTCTTTCCCTACACGACGCTCTTCCGATCTNNNACGAAAAAGATACTTGCACACG
AGCACACGTCTGAACTCTTTCCCTACACGACGCTCTTCCGATCTNNNACGAAAAAGATACTTGCACACG
GAGCACACGTCTGAACTCTTTCCCTACACGACGCTCTTCCGATCTNNNTGGTCAGGATGTCAAGACTC
GAGCACACGTCTGAACTCTTTCCCTACACGACGCTCTTCCGATCTNNNTGGTCAGGATGTCAAGACTC
GCACACGTCTGAACTCTTTCCCTACACGACGCTCTTCCGATCTNNNGGAATCTGCCCTGTGTCGTCCAT
GCACACGTCTGAACTCTTTCCCTACACGACGCTCTTCCGATCTNNNGGAATCTGCCCTGTGTCGTCCAT
AGCACACGTCTGAACTCTTTCCCTACACGACGCTCTTCCGATCTNNNATCATGCCATTGTACTCCATCCA
AGCACACGTCTGAACTCTTTCCCTACACGACGCTCTTCCGATCTNNNATCATGCCATTGTACTCCATCCA
AGCACACGTCTGAACTCTTTCCCTACACGACGCTCTTCCGATCTNNNCACCTCTAACTCGGGTCTGTCCA
AGCACACGTCTGAACTCTTTCCCTACACGACGCTCTTCCGATCTNNNCACCTCTAACTCGGGTCTGTCCA
GAGCACACGTCTGAACTCTTTCCCTACACGACGCTCTTCCGATCTNNNGATGCATGAGGTCACAACAGA
GAGCACACGTCTGAACTCTTTCCCTACACGACGCTCTTCCGATCTNNNGATGCATGAGGTCACAACAGA
AGCACACGTCTGAACTCTTTCCCTACACGACGCTCTTCCGATCTNNNGGTGAAAGAGTGAGACCCTGT
AGCACACGTCTGAACTCTTTCCCTACACGACGCTCTTCCGATCTNNNGGTGAAAGAGTGAGACCCTGT
AGCACACGTCTGAACTCTTTCCCTACACGACGCTCTTCCGATCTNNNAGTTTGAGAATCTCCAGCACAGT
AGCACACGTCTGAACTCTTTCCCTACACGACGCTCTTCCGATCTNNNAGTTTGAGAATCTCCAGCACAGT
AGAGCACACGTCTGAACTCTTTCCCTACACGACGCTCTTCCGATCTNNNACGCTTTCATTACTGTCTGCCT

GAGCACACGTCTGAACTCTTTCCCTACACGACGCTCTTCCGATCTNNNACGCTTTCATTACTGTCTGCC
AGCACACGTCTGAACTCTTTCCCTACACGACGCTCTTCCGATCTNNNAGGAGAATGGCATGAACCCTG
AGCACACGTCTGAACTCTTTCCCTACACGACGCTCTTCCGATCTNNNAGGAGAATGGCATGAACCCTG
GAGCACACGTCTGAACTCTTTCCCTACACGACGCTCTTCCGATCTNNNGGCAGGAGAATCACTTGAA
GAGCACACGTCTGAACTCTTTCCCTACACGACGCTCTTCCGATCTNNNGGCAGGAGAATCACTTGAA
GCACACGTCTGAACTCTTTCCCTACACGACGCTCTTCCGATCTNNNGAATCATCCAGCCTCAAATGTCG
GCACACGTCTGAACTCTTTCCCTACACGACGCTCTTCCGATCTNNNGAATCATCCAGCCTCAAATGTCG
AGCACACGTCTGAACTCTTTCCCTACACGACGCTCTTCCGATCTNNNGTGGTAGGCTAGAGTTCTGCT
AGCACACGTCTGAACTCTTTCCCTACACGACGCTCTTCCGATCTNNNGTGGTAGGCTAGAGTTCTGCT
GCACACGTCTGAACTCTTTCCCTACACGACGCTCTTCCGATCTNNNGGCTGGGATTGATTAAGGGTG
GCACACGTCTGAACTCTTTCCCTACACGACGCTCTTCCGATCTNNNGGCTGGGATTGATTAAGGGTG
GCACACGTCTGAACTCTTTCCCTACACGACGCTCTTCCGATCTNNNTTATTTAGCACAGTCCTCCACCCC
GCACACGTCTGAACTCTTTCCCTACACGACGCTCTTCCGATCTNNNTTATTTAGCACAGTCCTCCACCCC
GAGCACACGTCTGAACTCTTTCCCTACACGACGCTCTTCCGATCTNNNAACTCCTGACCTCAAGTGATC
GAGCACACGTCTGAACTCTTTCCCTACACGACGCTCTTCCGATCTNNNAACTCCTGACCTCAAGTGATC
ACACGTCTGAACTCTTTCCCTACACGACGCTCTTCCGATCTNNNCCATATGCCATTCCAGTGATGGACA
ACACGTCTGAACTCTTTCCCTACACGACGCTCTTCCGATCTNNNCCATATGCCATTCCAGTGATGGACA
AGAGCACACGTCTGAACTCTTTCCCTACACGACGCTCTTCCGATCTNNNGAGGCTGAGGCAGGAGA
AGAGCACACGTCTGAACTCTTTCCCTACACGACGCTCTTCCGATCTNNNGAGGCTGAGGCAGGAGA
AGCACACGTCTGAACTCTTTCCCTACACGACGCTCTTCCGATCTNNNGTGGTGTGATCTTGAATCACT
AGCACACGTCTGAACTCTTTCCCTACACGACGCTCTTCCGATCTNNNGTGGTGTGATCTTGAATCACT
GAGCACACGTCTGAACTCTTTCCCTACACGACGCTCTTCCGATCTNNNTACCCATGACCTCTTGTTCTG
GAGCACACGTCTGAACTCTTTCCCTACACGACGCTCTTCCGATCTNNNTACCCATGACCTCTTGTTCTG
GCACACGTCTGAACTCTTTCCCTACACGACGCTCTTCCGATCTNNNATAGAAATGCCCCAATTCCTGGT
GCACACGTCTGAACTCTTTCCCTACACGACGCTCTTCCGATCTNNNATAGAAATGCCCCAATTCCTGGT
GCACACGTCTGAACTCTTTCCCTACACGACGCTCTTCCGATCTNNNAGAACTTTGCTGCTGTTTTGGT
GCACACGTCTGAACTCTTTCCCTACACGACGCTCTTCCGATCTNNNAGAACTTTGCTGCTGTTTTGGT
AGCACACGTCTGAACTCTTTCCCTACACGACGCTCTTCCGATCTNNNTTGACTGTGAAGTCCTCAGGT
AGCACACGTCTGAACTCTTTCCCTACACGACGCTCTTCCGATCTNNNTTGACTGTGAAGTCCTCAGGT
AGCACACGTCTGAACTCTTTCCCTACACGACGCTCTTCCGATCTNNNAAGCTACTTAGGTGGGTGAA
AGCACACGTCTGAACTCTTTCCCTACACGACGCTCTTCCGATCTNNNAAGCTACTTAGGTGGGTGAA
GCACACGTCTGAACTCTTTCCCTACACGACGCTCTTCCGATCTNNNAGAATGTGCTGATGCCTGTAG
GCACACGTCTGAACTCTTTCCCTACACGACGCTCTTCCGATCTNNNAGAATGTGCTGATGCCTGTAG
GCACACGTCTGAACTCTTTCCCTACACGACGCTCTTCCGATCTNNNTAAAGAGGGTTGGAGAATTGG
GCACACGTCTGAACTCTTTCCCTACACGACGCTCTTCCGATCTNNNTAAAGAGGGTTGGAGAATTGG
AGCACACGTCTGAACTCTTTCCCTACACGACGCTCTTCCGATCTNNNCTCATGGTGGTTTTGACTTGCA
AGCACACGTCTGAACTCTTTCCCTACACGACGCTCTTCCGATCTNNNCTCATGGTGGTTTTGACTTGCA
AGCACACGTCTGAACTCTTTCCCTACACGACGCTCTTCCGATCTNNNGCTGTGGAGAAAAAGGAATG
AGCACACGTCTGAACTCTTTCCCTACACGACGCTCTTCCGATCTNNNGCTGTGGAGAAAAAGGAATG
GAGCACACGTCTGAACTCTTTCCCTACACGACGCTCTTCCGATCTNNNGAGAAGAATGATCAAGGTG
GAGCACACGTCTGAACTCTTTCCCTACACGACGCTCTTCCGATCTNNNGAGAAGAATGATCAAGGTG
GCACACGTCTGAACTCTTTCCCTACACGACGCTCTTCCGATCTNNNGCCCATCTAGATTTGGAAGGG
GCACACGTCTGAACTCTTTCCCTACACGACGCTCTTCCGATCTNNNGCCCATCTAGATTTGGAAGGG

AGAGCACACGTCTGAACTCTTTCCCTACACGACGCTCTTCCGATCTNNNCCAGGCTAGGGTTTTTCTCAT
AGAGCACACGTCTGAACTCTTTCCCTACACGACGCTCTTCCGATCTNNNCCAGGCTAGGGTTTTTCTCAT
AGCACACGTCTGAACTCTTTCCCTACACGACGCTCTTCCGATCTNNNGAGGTTGCAGTGAGCTGAGAT
AGCACACGTCTGAACTCTTTCCCTACACGACGCTCTTCCGATCTNNNGAGGTTGCAGTGAGCTGAGAT
AGCACACGTCTGAACTCTTTCCCTACACGACGCTCTTCCGATCTNNNGGCAGGGTCAATATTCTGTACC
AGCACACGTCTGAACTCTTTCCCTACACGACGCTCTTCCGATCTNNNGGCAGGGTCAATATTCTGTACC
GAGCACACGTCTGAACTCTTTCCCTACACGACGCTCTTCCGATCTNNNTCAAAACCACAATGCGATACCA
GAGCACACGTCTGAACTCTTTCCCTACACGACGCTCTTCCGATCTNNNTCAAAACCACAATGCGATACCA
GAGCACACGTCTGAACTCTTTCCCTACACGACGCTCTTCCGATCTNNNGCTACATGTACGTGTTGAAAG
GAGCACACGTCTGAACTCTTTCCCTACACGACGCTCTTCCGATCTNNNGCTACATGTACGTGTTGAAAG
AGCACACGTCTGAACTCTTTCCCTACACGACGCTCTTCCGATCTNNNGAGTTCTGTTGCAGTAATGCCAC
AGCACACGTCTGAACTCTTTCCCTACACGACGCTCTTCCGATCTNNNGAGTTCTGTTGCAGTAATGCCAC
AGCACACGTCTGAACTCTTTCCCTACACGACGCTCTTCCGATCTNNNCCCAAATGATGAAGTGATGCAG
AGCACACGTCTGAACTCTTTCCCTACACGACGCTCTTCCGATCTNNNCCCAAATGATGAAGTGATGCAG
GAGCACACGTCTGAACTCTTTCCCTACACGACGCTCTTCCGATCTNNNTTTGAGATGTGGTTTCGCTCTT
GAGCACACGTCTGAACTCTTTCCCTACACGACGCTCTTCCGATCTNNNTTTGAGATGTGGTTTCGCTCTT
AGCACACGTCTGAACTCTTTCCCTACACGACGCTCTTCCGATCTNNNTAATCAAAGTCAGCAATTTATG
AGCACACGTCTGAACTCTTTCCCTACACGACGCTCTTCCGATCTNNNTAATCAAAGTCAGCAATTTATG
AGAGCACACGTCTGAACTCTTTCCCTACACGACGCTCTTCCGATCTNNNGGAGAATCACTTGAACCCAG
AGAGCACACGTCTGAACTCTTTCCCTACACGACGCTCTTCCGATCTNNNGGAGAATCACTTGAACCCAG
AGAGCACACGTCTGAACTCTTTCCCTACACGACGCTCTTCCGATCTNNNGACTGTAATCCCAGCTACTCA
AGAGCACACGTCTGAACTCTTTCCCTACACGACGCTCTTCCGATCTNNNGACTGTAATCCCAGCTACTCA
AGAGCACACGTCTGAACTCTTTCCCTACACGACGCTCTTCCGATCTNNNAAGTCACCCAAACAGCAAAG
AGAGCACACGTCTGAACTCTTTCCCTACACGACGCTCTTCCGATCTNNNAAGTCACCCAAACAGCAAAG
AGAGCACACGTCTGAACTCTTTCCCTACACGACGCTCTTCCGATCTNNNCATGTCATAGGCTGCTGTTTT
AGAGCACACGTCTGAACTCTTTCCCTACACGACGCTCTTCCGATCTNNNCATGTCATAGGCTGCTGTTTT
GAGCACACGTCTGAACTCTTTCCCTACACGACGCTCTTCCGATCTNNNCACCTCTCCAGATCAAGCTCTA
GAGCACACGTCTGAACTCTTTCCCTACACGACGCTCTTCCGATCTNNNCACCTCTCCAGATCAAGCTCTA
GAGCACACGTCTGAACTCTTTCCCTACACGACGCTCTTCCGATCTNNNATAGCAAACACATTGGCACAG
GAGCACACGTCTGAACTCTTTCCCTACACGACGCTCTTCCGATCTNNNATAGCAAACACATTGGCACAG
GAGCACACGTCTGAACTCTTTCCCTACACGACGCTCTTCCGATCTNNNTTAGAAAAGGCCTAACCACTCC
GAGCACACGTCTGAACTCTTTCCCTACACGACGCTCTTCCGATCTNNNTTAGAAAAGGCCTAACCACTCC
AGCACACGTCTGAACTCTTTCCCTACACGACGCTCTTCCGATCTNNNGTACTTTCTCTCTCTCCCCCTCTC
AGCACACGTCTGAACTCTTTCCCTACACGACGCTCTTCCGATCTNNNGTACTTTCTCTCTCTCCCCCTCTC
GAGCACACGTCTGAACTCTTTCCCTACACGACGCTCTTCCGATCTNNNGAGTCGGGAATGTTCCTCACT
GAGCACACGTCTGAACTCTTTCCCTACACGACGCTCTTCCGATCTNNNGAGTCGGGAATGTTCCTCACT
AGAGCACACGTCTGAACTCTTTCCCTACACGACGCTCTTCCGATCTNNNAGACAGAGAGATGGAGGG
AGAGCACACGTCTGAACTCTTTCCCTACACGACGCTCTTCCGATCTNNNAGACAGAGAGATGGAGGG
AGAGCACACGTCTGAACTCTTTCCCTACACGACGCTCTTCCGATCTNNNGGAATGCGAGAAAACAGGC
AGAGCACACGTCTGAACTCTTTCCCTACACGACGCTCTTCCGATCTNNNGGAATGCGAGAAAACAGGC
GAGCACACGTCTGAACTCTTTCCCTACACGACGCTCTTCCGATCTNNNTGAGAGTATTTACCTTTTGATG
GAGCACACGTCTGAACTCTTTCCCTACACGACGCTCTTCCGATCTNNNTGAGAGTATTTACCTTTTGATG
AGAGCACACGTCTGAACTCTTTCCCTACACGACGCTCTTCCGATCTNNNCAGCTGGACATTGAGGAGG

AGAGCACACGTCTGAACTCTTTCCCTACACGACGCTCTTCCGATCTNNNCAGCTGGACATTGAGGAGG
GAGCACACGTCTGAACTCTTTCCCTACACGACGCTCTTCCGATCTNNNACGTTACACAGAAACCTAGACC
GAGCACACGTCTGAACTCTTTCCCTACACGACGCTCTTCCGATCTNNNACGTTACACAGAAACCTAGACC
AGCACACGTCTGAACTCTTTCCCTACACGACGCTCTTCCGATCTNNNCAGGTTGCTGTGAATGCCATTA
AGCACACGTCTGAACTCTTTCCCTACACGACGCTCTTCCGATCTNNNCAGGTTGCTGTGAATGCCATTA
GCACACGTCTGAACTCTTTCCCTACACGACGCTCTTCCGATCTNNNTAAACTTCCTTCCCCTGCTCCTCCA
GCACACGTCTGAACTCTTTCCCTACACGACGCTCTTCCGATCTNNNTAAACTTCCTTCCCCTGCTCCTCCA
AGCACACGTCTGAACTCTTTCCCTACACGACGCTCTTCCGATCTNNNTTCCACTTGGTCTTCAGAGTCAG
AGCACACGTCTGAACTCTTTCCCTACACGACGCTCTTCCGATCTNNNTTCCACTTGGTCTTCAGAGTCAG
AGCACACGTCTGAACTCTTTCCCTACACGACGCTCTTCCGATCTNNNGCTGAAGCATAAGAATCGCTTG
AGCACACGTCTGAACTCTTTCCCTACACGACGCTCTTCCGATCTNNNGCTGAAGCATAAGAATCGCTTG
AGCACACGTCTGAACTCTTTCCCTACACGACGCTCTTCCGATCTNNNTGAAGTCTAAAAGCCATTCTGCT
AGCACACGTCTGAACTCTTTCCCTACACGACGCTCTTCCGATCTNNNTGAAGTCTAAAAGCCATTCTGCT
AGCACACGTCTGAACTCTTTCCCTACACGACGCTCTTCCGATCTNNNCTGAGCCAAGGGAAAATCCTAG
AGCACACGTCTGAACTCTTTCCCTACACGACGCTCTTCCGATCTNNNCTGAGCCAAGGGAAAATCCTAG
GAGCACACGTCTGAACTCTTTCCCTACACGACGCTCTTCCGATCTNNNTGACATTCCCAACTAGGATCGG
GAGCACACGTCTGAACTCTTTCCCTACACGACGCTCTTCCGATCTNNNTGACATTCCCAACTAGGATCGG
GAGCACACGTCTGAACTCTTTCCCTACACGACGCTCTTCCGATCTNNNGTGTCTAATCGAAGTGTCTGCC
GAGCACACGTCTGAACTCTTTCCCTACACGACGCTCTTCCGATCTNNNGTGTCTAATCGAAGTGTCTGCC
AGAGCACACGTCTGAACTCTTTCCCTACACGACGCTCTTCCGATCTNNNCGCGAGACGGAGGTTGCAAT
AGAGCACACGTCTGAACTCTTTCCCTACACGACGCTCTTCCGATCTNNNCGCGAGACGGAGGTTGCAAT
GAGCACACGTCTGAACTCTTTCCCTACACGACGCTCTTCCGATCTNNNGGAGAAGCTTCCTTCCTGGTT
GAGCACACGTCTGAACTCTTTCCCTACACGACGCTCTTCCGATCTNNNGGAGAAGCTTCCTTCCTGGTT
AGCACACGTCTGAACTCTTTCCCTACACGACGCTCTTCCGATCTNNNCTGTGGTTTTTGCAGACCCATA
AGCACACGTCTGAACTCTTTCCCTACACGACGCTCTTCCGATCTNNNCTGTGGTTTTTGCAGACCCATA
GAGCACACGTCTGAACTCTTTCCCTACACGACGCTCTTCCGATCTNNNGAAAACCCTGTAAAACCACCCA
GAGCACACGTCTGAACTCTTTCCCTACACGACGCTCTTCCGATCTNNNGAAAACCCTGTAAAACCACCCA
GCACACGTCTGAACTCTTTCCCTACACGACGCTCTTCCGATCTNNNCTTTTCTGAAGCCTGAAAAAACAT
GCACACGTCTGAACTCTTTCCCTACACGACGCTCTTCCGATCTNNNCTTTTCTGAAGCCTGAAAAAACAT
AGAGCACACGTCTGAACTCTTTCCCTACACGACGCTCTTCCGATCTNNNGAGTCTCACTCTGTCACCCAG
AGAGCACACGTCTGAACTCTTTCCCTACACGACGCTCTTCCGATCTNNNGAGTCTCACTCTGTCACCCAG
AGCACACGTCTGAACTCTTTCCCTACACGACGCTCTTCCGATCTNNNATGCCTGTAATCCCAGCTACTTC
AGCACACGTCTGAACTCTTTCCCTACACGACGCTCTTCCGATCTNNNATGCCTGTAATCCCAGCTACTTC
AGCACACGTCTGAACTCTTTCCCTACACGACGCTCTTCCGATCTNNNTGCATACTACTACTGCCTGGAAC
AGCACACGTCTGAACTCTTTCCCTACACGACGCTCTTCCGATCTNNNTGCATACTACTACTGCCTGGAAC
GCACACGTCTGAACTCTTTCCCTACACGACGCTCTTCCGATCTNNNGGCTCGGTCCTCTCAAGTGGCATC
GCACACGTCTGAACTCTTTCCCTACACGACGCTCTTCCGATCTNNNGGCTCGGTCCTCTCAAGTGGCATC
AGCACACGTCTGAACTCTTTCCCTACACGACGCTCTTCCGATCTNNNTACATCAAGTTCTTGGAGGGAC
AGCACACGTCTGAACTCTTTCCCTACACGACGCTCTTCCGATCTNNNTACATCAAGTTCTTGGAGGGAC
GAGCACACGTCTGAACTCTTTCCCTACACGACGCTCTTCCGATCTNNNTATTTCTTTCAGCTTGGGCCTCT
GAGCACACGTCTGAACTCTTTCCCTACACGACGCTCTTCCGATCTNNNTATTTCTTTCAGCTTGGGCCTCT
GAGCACACGTCTGAACTCTTTCCCTACACGACGCTCTTCCGATCTNNNAAGTCTTTATAGTCAGCCCCAG
GAGCACACGTCTGAACTCTTTCCCTACACGACGCTCTTCCGATCTNNNAAGTCTTTATAGTCAGCCCCAG

AGCACACGTCTGAACTCTTTCCCTACACGACGCTCTTCCGATCTNNNGATCAAGACCATCCTGGCTAAC

AGCACACGTCTGAACTCTTTCCCTACACGACGCTCTTCCGATCTNNNGATCAAGACCATCCTGGCTAAC

AGCACACGTCTGAACTCTTTCCCTACACGACGCTCTTCCGATCTNNNAAGAAAAAGTGGATGGAGGT

AGCACACGTCTGAACTCTTTCCCTACACGACGCTCTTCCGATCTNNNAAGAAAAAGTGGATGGAGGT

GCACACGTCTGAACTCTTTCCCTACACGACGCTCTTCCGATCTNNNGCCACCACTCAGCTAATTTTT

GCACACGTCTGAACTCTTTCCCTACACGACGCTCTTCCGATCTNNNGCCACCACTCAGCTAATTTTT

AGCACACGTCTGAACTCTTTCCCTACACGACGCTCTTCCGATCTNNNATGCCTGTAATCCCAGCTACTT

AGCACACGTCTGAACTCTTTCCCTACACGACGCTCTTCCGATCTNNNATGCCTGTAATCCCAGCTACTT

AGCACACGTCTGAACTCTTTCCCTACACGACGCTCTTCCGATCTNNNGTTGCATGCCACTAAGTTCGTT

AGCACACGTCTGAACTCTTTCCCTACACGACGCTCTTCCGATCTNNNGTTGCATGCCACTAAGTTCGTT

AGCACACGTCTGAACTCTTTCCCTACACGACGCTCTTCCGATCTNNNGTAATCCTAGCTACTTGGGAG

AGCACACGTCTGAACTCTTTCCCTACACGACGCTCTTCCGATCTNNNGTAATCCTAGCTACTTGGGAG

GCACACGTCTGAACTCTTTCCCTACACGACGCTCTTCCGATCTNNNTCCACATTTTGCCATTGTCAGACC

GCACACGTCTGAACTCTTTCCCTACACGACGCTCTTCCGATCTNNNTCCACATTTTGCCATTGTCAGACC

AGCACACGTCTGAACTCTTTCCCTACACGACGCTCTTCCGATCTNNNGCATGAGAATTGCTTGAACCT

AGCACACGTCTGAACTCTTTCCCTACACGACGCTCTTCCGATCTNNNGCATGAGAATTGCTTGAACCT

AGAGCACACGTCTGAACTCTTTCCCTACACGACGCTCTTCCGATCTNNNAGAATGGCATGAACCTGG

AGAGCACACGTCTGAACTCTTTCCCTACACGACGCTCTTCCGATCTNNNAGAATGGCATGAACCTGG

GCACACGTCTGAACTCTTTCCCTACACGACGCTCTTCCGATCTNNNTTACACTGCATTTACTCCACCCAC

GCACACGTCTGAACTCTTTCCCTACACGACGCTCTTCCGATCTNNNTTACACTGCATTTACTCCACCCAC

GCACACGTCTGAACTCTTTCCCTACACGACGCTCTTCCGATCTNNNGGAGACAAAAAAATGGTGGGA

GCACACGTCTGAACTCTTTCCCTACACGACGCTCTTCCGATCTNNNGGAGACAAAAAAATGGTGGGA

AGCACACGTCTGAACTCTTTCCCTACACGACGCTCTTCCGATCTNNNGAGACAGGAGAATTGCTTGA

AGCACACGTCTGAACTCTTTCCCTACACGACGCTCTTCCGATCTNNNGAGACAGGAGAATTGCTTGA

AGCACACGTCTGAACTCTTTCCCTACACGACGCTCTTCCGATCTNNNATTTGTTACTCTGTGTGTGTG

AGCACACGTCTGAACTCTTTCCCTACACGACGCTCTTCCGATCTNNNATTTGTTACTCTGTGTGTGTG

ACACGTCTGAACTCTTTCCCTACACGACGCTCTTCCGATCTNNNCACTGTTTTCGGACCTTCCATGCAT

ACACGTCTGAACTCTTTCCCTACACGACGCTCTTCCGATCTNNNCACTGTTTTCGGACCTTCCATGCAT

GCACACGTCTGAACTCTTTCCCTACACGACGCTCTTCCGATCTNNNTCCAGAGTAGCTAGGACTACAG

GCACACGTCTGAACTCTTTCCCTACACGACGCTCTTCCGATCTNNNTCCAGAGTAGCTAGGACTACAG

GCACACGTCTGAACTCTTTCCCTACACGACGCTCTTCCGATCTNNNCATTCAAGGAGCATGCAAAGTG

GCACACGTCTGAACTCTTTCCCTACACGACGCTCTTCCGATCTNNNCATTCAAGGAGCATGCAAAGTG

GAGCACACGTCTGAACTCTTTCCCTACACGACGCTCTTCCGATCTNNNACCAAATATCACACCCCTGTT

GAGCACACGTCTGAACTCTTTCCCTACACGACGCTCTTCCGATCTNNNACCAAATATCACACCCCTGTT

AAGAGCACACGTCTGAACTCTTTCCCTACACGACGCTCTTCCGATCTNNNGTCTTTCTCAGACCCCCTCA

AAGAGCACACGTCTGAACTCTTTCCCTACACGACGCTCTTCCGATCTNNNGTCTTTCTCAGACCCCCTCA

AGCACACGTCTGAACTCTTTCCCTACACGACGCTCTTCCGATCTNNNCAGGATTTCTTTGTTTCTGTCC

AGCACACGTCTGAACTCTTTCCCTACACGACGCTCTTCCGATCTNNNCAGGATTTCTTTGTTTCTGTCC

AGCACACGTCTGAACTCTTTCCCTACACGACGCTCTTCCGATCTNNNGAACAAAGGATCCCAGTGAAA

AGCACACGTCTGAACTCTTTCCCTACACGACGCTCTTCCGATCTNNNGAACAAAGGATCCCAGTGAAA

AGCACACGTCTGAACTCTTTCCCTACACGACGCTCTTCCGATCTNNNCTTCATGTCTGACCTAGCACAG

AGCACACGTCTGAACTCTTTCCCTACACGACGCTCTTCCGATCTNNNCTTCATGTCTGACCTAGCACAG

AGCACACGTCTGAACTCTTTCCCTACACGACGCTCTTCCGATCTNNNGAGAGGAGCTAGAGCCAAAG

AGCACACGTCTGAACTCTTTCCCTACACGACGCTCTTCCGATCTNNNGAGAGGAGCTAGAGCCAAAG
AGCACACGTCTGAACTCTTTCCCTACACGACGCTCTTCCGATCTNNNGCTGAAATCTTGGTGGATTGT
AGCACACGTCTGAACTCTTTCCCTACACGACGCTCTTCCGATCTNNNGCTGAAATCTTGGTGGATTGT
AGCACACGTCTGAACTCTTTCCCTACACGACGCTCTTCCGATCTNNNGGGTTCAAGTGATTTTTGTG
AGCACACGTCTGAACTCTTTCCCTACACGACGCTCTTCCGATCTNNNGGGTTCAAGTGATTTTTGTG
AGCACACGTCTGAACTCTTTCCCTACACGACGCTCTTCCGATCTNNNCTTAGTGGTAGGGGAGGCAA
AGCACACGTCTGAACTCTTTCCCTACACGACGCTCTTCCGATCTNNNCTTAGTGGTAGGGGAGGCAA
AGCACACGTCTGAACTCTTTCCCTACACGACGCTCTTCCGATCTNNNATCCATTCAAAAACAGCTCAGG
AGCACACGTCTGAACTCTTTCCCTACACGACGCTCTTCCGATCTNNNATCCATTCAAAAACAGCTCAGG
GAGCACACGTCTGAACTCTTTCCCTACACGACGCTCTTCCGATCTNNNCAGTCACTCTGGCTATGTGA
GAGCACACGTCTGAACTCTTTCCCTACACGACGCTCTTCCGATCTNNNCAGTCACTCTGGCTATGTGA
GAGCACACGTCTGAACTCTTTCCCTACACGACGCTCTTCCGATCTNNNTGAGGAGGGTCTCAGAAGG
GAGCACACGTCTGAACTCTTTCCCTACACGACGCTCTTCCGATCTNNNTGAGGAGGGTCTCAGAAGG
GAGCACACGTCTGAACTCTTTCCCTACACGACGCTCTTCCGATCTNNNTTGAGAAAAGACTTTCCCAG
GAGCACACGTCTGAACTCTTTCCCTACACGACGCTCTTCCGATCTNNNTTGAGAAAAGACTTTCCCAG
GAGCACACGTCTGAACTCTTTCCCTACACGACGCTCTTCCGATCTNNNACTGGCAAGGAACTATATTG
GAGCACACGTCTGAACTCTTTCCCTACACGACGCTCTTCCGATCTNNNACTGGCAAGGAACTATATTG
CACACGTCTGAACTCTTTCCCTACACGACGCTCTTCCGATCTNNNGAAGCAGGAGAATCACTTGAACC
CACACGTCTGAACTCTTTCCCTACACGACGCTCTTCCGATCTNNNGAAGCAGGAGAATCACTTGAACC
AGCACACGTCTGAACTCTTTCCCTACACGACGCTCTTCCGATCTNNNAAAATTATCTGTGTGTGGCGG
AGCACACGTCTGAACTCTTTCCCTACACGACGCTCTTCCGATCTNNNAAAATTATCTGTGTGTGGCGG
GAGCACACGTCTGAACTCTTTCCCTACACGACGCTCTTCCGATCTNNNGAGAATTGCTTGAACCCAGG
GAGCACACGTCTGAACTCTTTCCCTACACGACGCTCTTCCGATCTNNNGAGAATTGCTTGAACCCAGG
GAGCACACGTCTGAACTCTTTCCCTACACGACGCTCTTCCGATCTNNNCTGTGTTTGAGAAAGGCTAG
GAGCACACGTCTGAACTCTTTCCCTACACGACGCTCTTCCGATCTNNNCTGTGTTTGAGAAAGGCTAG
AGCACACGTCTGAACTCTTTCCCTACACGACGCTCTTCCGATCTNNNGACTCACAGGTCAAAGCTAAA
AGCACACGTCTGAACTCTTTCCCTACACGACGCTCTTCCGATCTNNNGACTCACAGGTCAAAGCTAAA
AGCACACGTCTGAACTCTTTCCCTACACGACGCTCTTCCGATCTNNNGAATATCCTGTCCCCAACAACC
AGCACACGTCTGAACTCTTTCCCTACACGACGCTCTTCCGATCTNNNGAATATCCTGTCCCCAACAACC
GCACACGTCTGAACTCTTTCCCTACACGACGCTCTTCCGATCTNNNGGATCACTTGAGGTCAGGAGTT
GCACACGTCTGAACTCTTTCCCTACACGACGCTCTTCCGATCTNNNGGATCACTTGAGGTCAGGAGTT
GCACACGTCTGAACTCTTTCCCTACACGACGCTCTTCCGATCTNNNCTGAGGCAGGAGAATCACTTGA
GCACACGTCTGAACTCTTTCCCTACACGACGCTCTTCCGATCTNNNCTGAGGCAGGAGAATCACTTGA
AGCACACGTCTGAACTCTTTCCCTACACGACGCTCTTCCGATCTNNNGCCTCCTCCTACACAACATTGAT
AGCACACGTCTGAACTCTTTCCCTACACGACGCTCTTCCGATCTNNNGCCTCCTCCTACACAACATTGAT
AGCACACGTCTGAACTCTTTCCCTACACGACGCTCTTCCGATCTNNNAATTGAACCTTTAAGTATGAG
AGCACACGTCTGAACTCTTTCCCTACACGACGCTCTTCCGATCTNNNAATTGAACCTTTAAGTATGAG
AGCACACGTCTGAACTCTTTCCCTACACGACGCTCTTCCGATCTNNNATTGCTTGAACATAGGAGGC
AGCACACGTCTGAACTCTTTCCCTACACGACGCTCTTCCGATCTNNNATTGCTTGAACATAGGAGGC
GAGCACACGTCTGAACTCTTTCCCTACACGACGCTCTTCCGATCTNNNCCAACATGGTGAAATCCCATC
GAGCACACGTCTGAACTCTTTCCCTACACGACGCTCTTCCGATCTNNNCCAACATGGTGAAATCCCATC
AGCACACGTCTGAACTCTTTCCCTACACGACGCTCTTCCGATCTNNNAAAGTGTGTGTGTGTGTGTG
AGCACACGTCTGAACTCTTTCCCTACACGACGCTCTTCCGATCTNNNAAAGTGTGTGTGTGTGTGTG

GAGCACACGTCTGAACTCTTTCCCTACACGACGCTCTTCCGATCTNNNCCACTTGGGTTAGAGTGAACC
GAGCACACGTCTGAACTCTTTCCCTACACGACGCTCTTCCGATCTNNNCCACTTGGGTTAGAGTGAACC
GAGCACACGTCTGAACTCTTTCCCTACACGACGCTCTTCCGATCTNNNGGTGAAAACTCATTTGTGGTC
GAGCACACGTCTGAACTCTTTCCCTACACGACGCTCTTCCGATCTNNNGGTGAAAACTCATTTGTGGTC
AGCACACGTCTGAACTCTTTCCCTACACGACGCTCTTCCGATCTNNNATATTTCCCCGCATCTTTCCTGAC
AGCACACGTCTGAACTCTTTCCCTACACGACGCTCTTCCGATCTNNNATATTTCCCCGCATCTTTCCTGAC
GCACACGTCTGAACTCTTTCCCTACACGACGCTCTTCCGATCTNNNAGGAATGGCCATACTTCCTCACTC
GCACACGTCTGAACTCTTTCCCTACACGACGCTCTTCCGATCTNNNAGGAATGGCCATACTTCCTCACTC
GAGCACACGTCTGAACTCTTTCCCTACACGACGCTCTTCCGATCTNNNTGGTTCAAAAGCGATTCTTGCA
GAGCACACGTCTGAACTCTTTCCCTACACGACGCTCTTCCGATCTNNNTGGTTCAAAAGCGATTCTTGCA
GAGCACACGTCTGAACTCTTTCCCTACACGACGCTCTTCCGATCTNNNTTTGTGCCTTCATCAAGCAAG
GAGCACACGTCTGAACTCTTTCCCTACACGACGCTCTTCCGATCTNNNTTTGTGCCTTCATCAAGCAAG
AGCACACGTCTGAACTCTTTCCCTACACGACGCTCTTCCGATCTNNNACTCAGGGCTTCTAACAACTTCC
AGCACACGTCTGAACTCTTTCCCTACACGACGCTCTTCCGATCTNNNACTCAGGGCTTCTAACAACTTCC
GAGCACACGTCTGAACTCTTTCCCTACACGACGCTCTTCCGATCTNNNCAACAAGAGTGAGACTCCACCT
GAGCACACGTCTGAACTCTTTCCCTACACGACGCTCTTCCGATCTNNNCAACAAGAGTGAGACTCCACCT
AGCACACGTCTGAACTCTTTCCCTACACGACGCTCTTCCGATCTNNNCCAAAATCCAAAGTGCTGGGAA
AGCACACGTCTGAACTCTTTCCCTACACGACGCTCTTCCGATCTNNNCCAAAATCCAAAGTGCTGGGAA
GAGCACACGTCTGAACTCTTTCCCTACACGACGCTCTTCCGATCTNNNAGAGACACAAAAACCGTTCAAA
GAGCACACGTCTGAACTCTTTCCCTACACGACGCTCTTCCGATCTNNNAGAGACACAAAAACCGTTCAAA
GAGCACACGTCTGAACTCTTTCCCTACACGACGCTCTTCCGATCTNNNTGCATGACTTGTTTTATTGCAC
GAGCACACGTCTGAACTCTTTCCCTACACGACGCTCTTCCGATCTNNNTGCATGACTTGTTTTATTGCAC
AGCACACGTCTGAACTCTTTCCCTACACGACGCTCTTCCGATCTNNNCCATTTGTTTGTGCTACCCACAG
AGCACACGTCTGAACTCTTTCCCTACACGACGCTCTTCCGATCTNNNCCATTTGTTTGTGCTACCCACAG
GAGCACACGTCTGAACTCTTTCCCTACACGACGCTCTTCCGATCTNNNCCATTTCGTGGCCTCTTCCATTA
GAGCACACGTCTGAACTCTTTCCCTACACGACGCTCTTCCGATCTNNNCCATTTCGTGGCCTCTTCCATTA
AGCACACGTCTGAACTCTTTCCCTACACGACGCTCTTCCGATCTNNNGGGTGACAGAGTAAGACTCCA
AGCACACGTCTGAACTCTTTCCCTACACGACGCTCTTCCGATCTNNNGGGTGACAGAGTAAGACTCCA
GAGCACACGTCTGAACTCTTTCCCTACACGACGCTCTTCCGATCTNNNCTAAAGTCACAACTCCTCTGCC
GAGCACACGTCTGAACTCTTTCCCTACACGACGCTCTTCCGATCTNNNCTAAAGTCACAACTCCTCTGCC
GAGCACACGTCTGAACTCTTTCCCTACACGACGCTCTTCCGATCTNNNTTGCAGACTCATAGAGGTACTC
GAGCACACGTCTGAACTCTTTCCCTACACGACGCTCTTCCGATCTNNNTTGCAGACTCATAGAGGTACTC
AGCACACGTCTGAACTCTTTCCCTACACGACGCTCTTCCGATCTNNNGAACATTCCATGTGCACTTGTGA
AGCACACGTCTGAACTCTTTCCCTACACGACGCTCTTCCGATCTNNNGAACATTCCATGTGCACTTGTGA
GAGCACACGTCTGAACTCTTTCCCTACACGACGCTCTTCCGATCTNNNGACCCATGATGGAACCCATGA
GAGCACACGTCTGAACTCTTTCCCTACACGACGCTCTTCCGATCTNNNGACCCATGATGGAACCCATGA
GAGCACACGTCTGAACTCTTTCCCTACACGACGCTCTTCCGATCTNNNCTCATGCCTGTAATCCTAGCAG
GAGCACACGTCTGAACTCTTTCCCTACACGACGCTCTTCCGATCTNNNCTCATGCCTGTAATCCTAGCAG
AGCACACGTCTGAACTCTTTCCCTACACGACGCTCTTCCGATCTNNNTAGGGCCAATCATTTTGCTTTG
AGCACACGTCTGAACTCTTTCCCTACACGACGCTCTTCCGATCTNNNTAGGGCCAATCATTTTGCTTTG
AGCACACGTCTGAACTCTTTCCCTACACGACGCTCTTCCGATCTNNNCTTAAAATCTCCAAATCAGAGAG
AGCACACGTCTGAACTCTTTCCCTACACGACGCTCTTCCGATCTNNNCTTAAAATCTCCAAATCAGAGAG
AGCACACGTCTGAACTCTTTCCCTACACGACGCTCTTCCGATCTNNNATCTTCAGGTCATTTGTCTGGGG

AGCACACGTCTGAACTCTTTCCCTACACGACGCTCTTCCGATCTNNNATCTTCAGGTCATTTGTCTGGG
GAGCACACGTCTGAACTCTTTCCCTACACGACGCTCTTCCGATCTNNNAACATTCGTGAACCATTGGCA
GAGCACACGTCTGAACTCTTTCCCTACACGACGCTCTTCCGATCTNNNAACATTCGTGAACCATTGGCA
AGCACACGTCTGAACTCTTTCCCTACACGACGCTCTTCCGATCTNNNTAGTTCACTGAAGCCTTGACCT
AGCACACGTCTGAACTCTTTCCCTACACGACGCTCTTCCGATCTNNNTAGTTCACTGAAGCCTTGACCT
AGCACACGTCTGAACTCTTTCCCTACACGACGCTCTTCCGATCTNNNTCCTAGAGATTGGAAAACGGCT
AGCACACGTCTGAACTCTTTCCCTACACGACGCTCTTCCGATCTNNNTCCTAGAGATTGGAAAACGGCT
AGCACACGTCTGAACTCTTTCCCTACACGACGCTCTTCCGATCTNNNGGATTCCCCAGTAATCAAACTG
AGCACACGTCTGAACTCTTTCCCTACACGACGCTCTTCCGATCTNNNGGATTCCCCAGTAATCAAACTG
AGCACACGTCTGAACTCTTTCCCTACACGACGCTCTTCCGATCTNNNCCCCCCAAGAAAAGATGAGAGA
AGCACACGTCTGAACTCTTTCCCTACACGACGCTCTTCCGATCTNNNCCCCCCAAGAAAAGATGAGAGA
AGCACACGTCTGAACTCTTTCCCTACACGACGCTCTTCCGATCTNNNCATGATCTCGGCTCAATTCAACC
AGCACACGTCTGAACTCTTTCCCTACACGACGCTCTTCCGATCTNNNCATGATCTCGGCTCAATTCAACC
AGCACACGTCTGAACTCTTTCCCTACACGACGCTCTTCCGATCTNNNCTACTCTACAGCAGTGGGGTAA
AGCACACGTCTGAACTCTTTCCCTACACGACGCTCTTCCGATCTNNNCTACTCTACAGCAGTGGGGTAA
GAGCACACGTCTGAACTCTTTCCCTACACGACGCTCTTCCGATCTNNNATGCCTGTAATCCCAGCTACTT
GAGCACACGTCTGAACTCTTTCCCTACACGACGCTCTTCCGATCTNNNATGCCTGTAATCCCAGCTACTT
GCACACGTCTGAACTCTTTCCCTACACGACGCTCTTCCGATCTNNNGCAAGACTTGAAGAACACACACA
GCACACGTCTGAACTCTTTCCCTACACGACGCTCTTCCGATCTNNNGCAAGACTTGAAGAACACACACA
GAGCACACGTCTGAACTCTTTCCCTACACGACGCTCTTCCGATCTNNNAGAATCACTTGAGCCTGGGA
GAGCACACGTCTGAACTCTTTCCCTACACGACGCTCTTCCGATCTNNNAGAATCACTTGAGCCTGGGA
GAGCACACGTCTGAACTCTTTCCCTACACGACGCTCTTCCGATCTNNNACGGAGATGCCATCAATCAAG
GAGCACACGTCTGAACTCTTTCCCTACACGACGCTCTTCCGATCTNNNACGGAGATGCCATCAATCAAG
AGCACACGTCTGAACTCTTTCCCTACACGACGCTCTTCCGATCTNNNTGGCAAGAATACCACAGAAGTA
AGCACACGTCTGAACTCTTTCCCTACACGACGCTCTTCCGATCTNNNTGGCAAGAATACCACAGAAGTA
AGCACACGTCTGAACTCTTTCCCTACACGACGCTCTTCCGATCTNNNCCGTCTAAGTATCCTAGGCAACC
AGCACACGTCTGAACTCTTTCCCTACACGACGCTCTTCCGATCTNNNCCGTCTAAGTATCCTAGGCAACC
AGCACACGTCTGAACTCTTTCCCTACACGACGCTCTTCCGATCTNNNGAGGGAAGGCAGGATGAGAA
AGCACACGTCTGAACTCTTTCCCTACACGACGCTCTTCCGATCTNNNGAGGGAAGGCAGGATGAGAA
AGCACACGTCTGAACTCTTTCCCTACACGACGCTCTTCCGATCTNNNTTTGTCTTGGTTTGAAGATGGC
AGCACACGTCTGAACTCTTTCCCTACACGACGCTCTTCCGATCTNNNTTTGTCTTGGTTTGAAGATGGC
GAGCACACGTCTGAACTCTTTCCCTACACGACGCTCTTCCGATCTNNNCTCGGCCTCCCAAAACTTTTTG
GAGCACACGTCTGAACTCTTTCCCTACACGACGCTCTTCCGATCTNNNCTCGGCCTCCCAAAACTTTTTG
AGCACACGTCTGAACTCTTTCCCTACACGACGCTCTTCCGATCTNNNAATCCTTCCACCAAAACAGGCTA
AGCACACGTCTGAACTCTTTCCCTACACGACGCTCTTCCGATCTNNNAATCCTTCCACCAAAACAGGCTA
AGAGCACACGTCTGAACTCTTTCCCTACACGACGCTCTTCCGATCTNNNCCAAACTGTGCCCTAACCTTT
AGAGCACACGTCTGAACTCTTTCCCTACACGACGCTCTTCCGATCTNNNCCAAACTGTGCCCTAACCTTT
AGCACACGTCTGAACTCTTTCCCTACACGACGCTCTTCCGATCTNNNTGGTTTCGATCTCTTGACCTTGT
AGCACACGTCTGAACTCTTTCCCTACACGACGCTCTTCCGATCTNNNTGGTTTCGATCTCTTGACCTTGT
AGCACACGTCTGAACTCTTTCCCTACACGACGCTCTTCCGATCTNNNTGCTCGTAACATTCTCTCAGTTCA
AGCACACGTCTGAACTCTTTCCCTACACGACGCTCTTCCGATCTNNNTGCTCGTAACATTCTCTCAGTTCA
GAGCACACGTCTGAACTCTTTCCCTACACGACGCTCTTCCGATCTNNNATAGCCCATAGGGTTGTGTGA
GAGCACACGTCTGAACTCTTTCCCTACACGACGCTCTTCCGATCTNNNATAGCCCATAGGGTTGTGTGA

GAGCACACGTCTGAACTCTTTCCCTACACGACGCTCTTCCGATCTNNNCCAGACTGGGCAACATAGTTA
GAGCACACGTCTGAACTCTTTCCCTACACGACGCTCTTCCGATCTNNNCCAGACTGGGCAACATAGTTA
GAGCACACGTCTGAACTCTTTCCCTACACGACGCTCTTCCGATCTNNNGGTTTGTGTCTTGGCTTCTACA
GAGCACACGTCTGAACTCTTTCCCTACACGACGCTCTTCCGATCTNNNGGTTTGTGTCTTGGCTTCTACA
AGCACACGTCTGAACTCTTTCCCTACACGACGCTCTTCCGATCTNNNTGTTGAAGCCACATGTCCAAAAT
AGCACACGTCTGAACTCTTTCCCTACACGACGCTCTTCCGATCTNNNTGTTGAAGCCACATGTCCAAAAT
AGCACACGTCTGAACTCTTTCCCTACACGACGCTCTTCCGATCTNNNGCCTGTAGTCCCAGCTACTTGCA
AGCACACGTCTGAACTCTTTCCCTACACGACGCTCTTCCGATCTNNNGCCTGTAGTCCCAGCTACTTGCA
GCACACGTCTGAACTCTTTCCCTACACGACGCTCTTCCGATCTNNNAGCACCGTTTCTTCTTTCTCTGTCA
GCACACGTCTGAACTCTTTCCCTACACGACGCTCTTCCGATCTNNNAGCACCGTTTCTTCTTTCTCTGTCA
AGCACACGTCTGAACTCTTTCCCTACACGACGCTCTTCCGATCTNNNCAGGTCCCATATCCTGCATCTTC
AGCACACGTCTGAACTCTTTCCCTACACGACGCTCTTCCGATCTNNNCAGGTCCCATATCCTGCATCTTC
AGCACACGTCTGAACTCTTTCCCTACACGACGCTCTTCCGATCTNNNACCATGAACCTGACTTTCCTTGA
AGCACACGTCTGAACTCTTTCCCTACACGACGCTCTTCCGATCTNNNACCATGAACCTGACTTTCCTTGA
GAGCACACGTCTGAACTCTTTCCCTACACGACGCTCTTCCGATCTNNNGTTAGCTGTTCAGTGTGCCTT
GAGCACACGTCTGAACTCTTTCCCTACACGACGCTCTTCCGATCTNNNGTTAGCTGTTCAGTGTGCCTT
AGCACACGTCTGAACTCTTTCCCTACACGACGCTCTTCCGATCTNNNTGTGCCTTGCAGAGAATACTTCT
AGCACACGTCTGAACTCTTTCCCTACACGACGCTCTTCCGATCTNNNTGTGCCTTGCAGAGAATACTTCT
AGCACACGTCTGAACTCTTTCCCTACACGACGCTCTTCCGATCTNNNCTTCCACAACCCTACTCAACACA
AGCACACGTCTGAACTCTTTCCCTACACGACGCTCTTCCGATCTNNNCTTCCACAACCCTACTCAACACA
GAGCACACGTCTGAACTCTTTCCCTACACGACGCTCTTCCGATCTNNNCTTCCAAAGTGCTGGGATAACA
GAGCACACGTCTGAACTCTTTCCCTACACGACGCTCTTCCGATCTNNNCTTCCAAAGTGCTGGGATAACA
AGCACACGTCTGAACTCTTTCCCTACACGACGCTCTTCCGATCTNNNGATCCACTTGCCTCACTGTTAGA
AGCACACGTCTGAACTCTTTCCCTACACGACGCTCTTCCGATCTNNNGATCCACTTGCCTCACTGTTAGA
GAGCACACGTCTGAACTCTTTCCCTACACGACGCTCTTCCGATCTNNNCCCATCATGGAGAGATGATC
GAGCACACGTCTGAACTCTTTCCCTACACGACGCTCTTCCGATCTNNNCCCATCATGGAGAGATGATC
AGCACACGTCTGAACTCTTTCCCTACACGACGCTCTTCCGATCTNNNCCATCTCACACCAATCAGAATGC
AGCACACGTCTGAACTCTTTCCCTACACGACGCTCTTCCGATCTNNNCCATCTCACACCAATCAGAATGC
ACACGTCTGAACTCTTTCCCTACACGACGCTCTTCCGATCTNNNCTCCCAACAGGACCCTGACTCATCGG
ACACGTCTGAACTCTTTCCCTACACGACGCTCTTCCGATCTNNNCTCCCAACAGGACCCTGACTCATCGG
AGCACACGTCTGAACTCTTTCCCTACACGACGCTCTTCCGATCTNNNGCTTAACGGAACTTATAAAACA
AGCACACGTCTGAACTCTTTCCCTACACGACGCTCTTCCGATCTNNNGCTTAACGGAACTTATAAAACA
GAGCACACGTCTGAACTCTTTCCCTACACGACGCTCTTCCGATCTNNNTCTGTGTGTGTGAGTGCATAT
GAGCACACGTCTGAACTCTTTCCCTACACGACGCTCTTCCGATCTNNNTCTGTGTGTGTGAGTGCATAT
AGCACACGTCTGAACTCTTTCCCTACACGACGCTCTTCCGATCTNNNACATAGTTTTGAGCCCCACAGAT
AGCACACGTCTGAACTCTTTCCCTACACGACGCTCTTCCGATCTNNNACATAGTTTTGAGCCCCACAGAT
GCACACGTCTGAACTCTTTCCCTACACGACGCTCTTCCGATCTNNNCCGCTCTCATCACTTTCCAACTTCA
GCACACGTCTGAACTCTTTCCCTACACGACGCTCTTCCGATCTNNNCCGCTCTCATCACTTTCCAACTTCA
GCACACGTCTGAACTCTTTCCCTACACGACGCTCTTCCGATCTNNNAATTCTTGGTCTTCTGTATCTTTCA
GCACACGTCTGAACTCTTTCCCTACACGACGCTCTTCCGATCTNNNAATTCTTGGTCTTCTGTATCTTTCA
AGCACACGTCTGAACTCTTTCCCTACACGACGCTCTTCCGATCTNNNCAACCGGCCCAAAATGTACTCCA
AGCACACGTCTGAACTCTTTCCCTACACGACGCTCTTCCGATCTNNNCAACCGGCCCAAAATGTACTCCA
GAGCACACGTCTGAACTCTTTCCCTACACGACGCTCTTCCGATCTNNNATACAACAACAACAGGCAAG

GAGCACACGTCTGAACTCTTTCCCTACACGACGCTCTTCCGATCTNNNATACAACAACAACAGGCAAG

AGCACACGTCTGAACTCTTTCCCTACACGACGCTCTTCCGATCTNNNAGGAGAATGGCGTGAAACCAG

AGCACACGTCTGAACTCTTTCCCTACACGACGCTCTTCCGATCTNNNAGGAGAATGGCGTGAAACCAG

AGCACACGTCTGAACTCTTTCCCTACACGACGCTCTTCCGATCTNNNGAAGAGGACAGATAGCCTACA

AGCACACGTCTGAACTCTTTCCCTACACGACGCTCTTCCGATCTNNNGAAGAGGACAGATAGCCTACA

GAGCACACGTCTGAACTCTTTCCCTACACGACGCTCTTCCGATCTNNNGGCCATCCTATCTCCACAAGA

GAGCACACGTCTGAACTCTTTCCCTACACGACGCTCTTCCGATCTNNNGGCCATCCTATCTCCACAAGA

CACACGTCTGAACTCTTTCCCTACACGACGCTCTTCCGATCTNNNCTGAATAGCGGATAGTTGTCCATC

CACACGTCTGAACTCTTTCCCTACACGACGCTCTTCCGATCTNNNCTGAATAGCGGATAGTTGTCCATC

GCACACGTCTGAACTCTTTCCCTACACGACGCTCTTCCGATCTNNNCCTTTATTCCCTTTCGTCCTCCTCA

GCACACGTCTGAACTCTTTCCCTACACGACGCTCTTCCGATCTNNNCCTTTATTCCCTTTCGTCCTCCTCA

CACACGTCTGAACTCTTTCCCTACACGACGCTCTTCCGATCTNNNGATGAAGACATCCCAGGAGCTAG

CACACGTCTGAACTCTTTCCCTACACGACGCTCTTCCGATCTNNNGATGAAGACATCCCAGGAGCTAG

AGCACACGTCTGAACTCTTTCCCTACACGACGCTCTTCCGATCTNNNTCACAGATATCATCCCCCACTCC

AGCACACGTCTGAACTCTTTCCCTACACGACGCTCTTCCGATCTNNNTCACAGATATCATCCCCCACTCC

AGCACACGTCTGAACTCTTTCCCTACACGACGCTCTTCCGATCTNNNCAGGACAATGGCATGAACTCA

AGCACACGTCTGAACTCTTTCCCTACACGACGCTCTTCCGATCTNNNCAGGACAATGGCATGAACTCA

GAGCACACGTCTGAACTCTTTCCCTACACGACGCTCTTCCGATCTNNNGCAGGAGAATTGCTTGATCC

GAGCACACGTCTGAACTCTTTCCCTACACGACGCTCTTCCGATCTNNNGCAGGAGAATTGCTTGATCC

GAGCACACGTCTGAACTCTTTCCCTACACGACGCTCTTCCGATCTNNNCCTCACACCAGGTCCTTTATTC

GAGCACACGTCTGAACTCTTTCCCTACACGACGCTCTTCCGATCTNNNCCTCACACCAGGTCCTTTATTC

GCACACGTCTGAACTCTTTCCCTACACGACGCTCTTCCGATCTNNNTGGGTGACAGAGTGAGATTTTT

GCACACGTCTGAACTCTTTCCCTACACGACGCTCTTCCGATCTNNNTGGGTGACAGAGTGAGATTTTT

AGCACACGTCTGAACTCTTTCCCTACACGACGCTCTTCCGATCTNNNAATGTAGCAGACCTCCAGAACA

AGCACACGTCTGAACTCTTTCCCTACACGACGCTCTTCCGATCTNNNAATGTAGCAGACCTCCAGAACA

AGCACACGTCTGAACTCTTTCCCTACACGACGCTCTTCCGATCTNNNCTGCTTCCTTTCACTTTCAGGTG

AGCACACGTCTGAACTCTTTCCCTACACGACGCTCTTCCGATCTNNNCTGCTTCCTTTCACTTTCAGGTG

AGAGCACACGTCTGAACTCTTTCCCTACACGACGCTCTTCCGATCTNNNCAGAGGTAGCAGTGAGTGA

AGAGCACACGTCTGAACTCTTTCCCTACACGACGCTCTTCCGATCTNNNCAGAGGTAGCAGTGAGTGA

GAGCACACGTCTGAACTCTTTCCCTACACGACGCTCTTCCGATCTNNNGCGTTATGCACAAAATTTCCC

GAGCACACGTCTGAACTCTTTCCCTACACGACGCTCTTCCGATCTNNNGCGTTATGCACAAAATTTCCC

AGCACACGTCTGAACTCTTTCCCTACACGACGCTCTTCCGATCTNNNAGGTTGAGCATGAGATTTTCTT

AGCACACGTCTGAACTCTTTCCCTACACGACGCTCTTCCGATCTNNNAGGTTGAGCATGAGATTTTCTT

GAGCACACGTCTGAACTCTTTCCCTACACGACGCTCTTCCGATCTNNNTGCAATCTCAGCTACTTCAGAG

GAGCACACGTCTGAACTCTTTCCCTACACGACGCTCTTCCGATCTNNNTGCAATCTCAGCTACTTCAGAG

AGCACACGTCTGAACTCTTTCCCTACACGACGCTCTTCCGATCTNNNCAGTCTCCTGAGTAGCTAGGAC

AGCACACGTCTGAACTCTTTCCCTACACGACGCTCTTCCGATCTNNNCAGTCTCCTGAGTAGCTAGGAC

GAGCACACGTCTGAACTCTTTCCCTACACGACGCTCTTCCGATCTNNNAACTTGTAGGAAGGAAGCTG

GAGCACACGTCTGAACTCTTTCCCTACACGACGCTCTTCCGATCTNNNAACTTGTAGGAAGGAAGCTG

AGCACACGTCTGAACTCTTTCCCTACACGACGCTCTTCCGATCTNNNCTCTGAGGAGCGAGGGACTGC

AGCACACGTCTGAACTCTTTCCCTACACGACGCTCTTCCGATCTNNNCTCTGAGGAGCGAGGGACTGC

AGCACACGTCTGAACTCTTTCCCTACACGACGCTCTTCCGATCTNNNCAGCTGTCTGAAACACACTCAA

AGCACACGTCTGAACTCTTTCCCTACACGACGCTCTTCCGATCTNNNCAGCTGTCTGAAACACACTCAA

GAGCACACGTCTGAACTCTTTCCCTACACGACGCTCTTCCGATCTNNNAGCACCATATTTTCCTCAGGG

GAGCACACGTCTGAACTCTTTCCCTACACGACGCTCTTCCGATCTNNNAGCACCATATTTTCCTCAGGG

AGCACACGTCTGAACTCTTTCCCTACACGACGCTCTTCCGATCTNNNTTCATGGTAAGTGCCCTCCTCTA

AGCACACGTCTGAACTCTTTCCCTACACGACGCTCTTCCGATCTNNNTTCATGGTAAGTGCCCTCCTCTA

AGCACACGTCTGAACTCTTTCCCTACACGACGCTCTTCCGATCTNNNAGACAGGAGAATCACTTGAACC

AGCACACGTCTGAACTCTTTCCCTACACGACGCTCTTCCGATCTNNNAGACAGGAGAATCACTTGAACC

GAGCACACGTCTGAACTCTTTCCCTACACGACGCTCTTCCGATCTNNNGCCATTTGTGGAGCCATATA

GAGCACACGTCTGAACTCTTTCCCTACACGACGCTCTTCCGATCTNNNGCCATTTGTGGAGCCATATA

AGCACACGTCTGAACTCTTTCCCTACACGACGCTCTTCCGATCTNNNGGCTCAGGTCATTTCAGTAGGA

AGCACACGTCTGAACTCTTTCCCTACACGACGCTCTTCCGATCTNNNGGCTCAGGTCATTTCAGTAGGA

AGCACACGTCTGAACTCTTTCCCTACACGACGCTCTTCCGATCTNNNATCTACACACTGTCTCCAGCTTG

AGCACACGTCTGAACTCTTTCCCTACACGACGCTCTTCCGATCTNNNATCTACACACTGTCTCCAGCTTG

AGCACACGTCTGAACTCTTTCCCTACACGACGCTCTTCCGATCTNNNGAACTGTGATTGTGCCACTACA

AGCACACGTCTGAACTCTTTCCCTACACGACGCTCTTCCGATCTNNNGAACTGTGATTGTGCCACTACA

ACACGTCTGAACTCTTTCCCTACACGACGCTCTTCCGATCTNNNCCAGTTTTCTAGACACCTGTGGACAT

ACACGTCTGAACTCTTTCCCTACACGACGCTCTTCCGATCTNNNCCAGTTTTCTAGACACCTGTGGACAT

AGCACACGTCTGAACTCTTTCCCTACACGACGCTCTTCCGATCTNNNCCAAGTTGGTCACACAATAGCCT

AGCACACGTCTGAACTCTTTCCCTACACGACGCTCTTCCGATCTNNNCCAAGTTGGTCACACAATAGCCT

AGCACACGTCTGAACTCTTTCCCTACACGACGCTCTTCCGATCTNNNTTCTTCCTTTCCTTTCCTCTCCCCA

AGCACACGTCTGAACTCTTTCCCTACACGACGCTCTTCCGATCTNNNTTCTTCCTTTCCTTTCCTCTCCCCA

AGCACACGTCTGAACTCTTTCCCTACACGACGCTCTTCCGATCTNNNGGCTTTTTGCACTAACAGCAACT

AGCACACGTCTGAACTCTTTCCCTACACGACGCTCTTCCGATCTNNNGGCTTTTTGCACTAACAGCAACT

AGCACACGTCTGAACTCTTTCCCTACACGACGCTCTTCCGATCTNNNGGATTCTCTAAAGGCTCAGTGC

AGCACACGTCTGAACTCTTTCCCTACACGACGCTCTTCCGATCTNNNGGATTCTCTAAAGGCTCAGTGC

GAGCACACGTCTGAACTCTTTCCCTACACGACGCTCTTCCGATCTNNNCATTCATCCTACTCCTGAGCCTC

GAGCACACGTCTGAACTCTTTCCCTACACGACGCTCTTCCGATCTNNNCATTCATCCTACTCCTGAGCCTC

AGCACACGTCTGAACTCTTTCCCTACACGACGCTCTTCCGATCTNNNTGTAGTCTGTAGGTTGCAGATG

AGCACACGTCTGAACTCTTTCCCTACACGACGCTCTTCCGATCTNNNTGTAGTCTGTAGGTTGCAGATG

GAGCACACGTCTGAACTCTTTCCCTACACGACGCTCTTCCGATCTNNNGACTGGAGGAGCAATGGGAA

GAGCACACGTCTGAACTCTTTCCCTACACGACGCTCTTCCGATCTNNNGACTGGAGGAGCAATGGGAA

AGCACACGTCTGAACTCTTTCCCTACACGACGCTCTTCCGATCTNNNAGTGTCTATACTGTGGCTGGTTC

AGCACACGTCTGAACTCTTTCCCTACACGACGCTCTTCCGATCTNNNAGTGTCTATACTGTGGCTGGTTC

AGCACACGTCTGAACTCTTTCCCTACACGACGCTCTTCCGATCTNNNACAGAAACAGTGATAAAACAAA

AGCACACGTCTGAACTCTTTCCCTACACGACGCTCTTCCGATCTNNNACAGAAACAGTGATAAAACAAA

AGCACACGTCTGAACTCTTTCCCTACACGACGCTCTTCCGATCTNNNCAAACAGCTGGAAGACGTACAT

AGCACACGTCTGAACTCTTTCCCTACACGACGCTCTTCCGATCTNNNCAAACAGCTGGAAGACGTACAT

AGCACACGTCTGAACTCTTTCCCTACACGACGCTCTTCCGATCTNNNGAGCTTGCAGTGAGCCAAGATC

AGCACACGTCTGAACTCTTTCCCTACACGACGCTCTTCCGATCTNNNGAGCTTGCAGTGAGCCAAGATC

AGCACACGTCTGAACTCTTTCCCTACACGACGCTCTTCCGATCTNNNAATTGCCCATCAACCAATGAGT

AGCACACGTCTGAACTCTTTCCCTACACGACGCTCTTCCGATCTNNNAATTGCCCATCAACCAATGAGT

GCACACGTCTGAACTCTTTCCCTACACGACGCTCTTCCGATCTNNNGAAAACAAAACATGCTCGAAGGC

GCACACGTCTGAACTCTTTCCCTACACGACGCTCTTCCGATCTNNNGAAAACAAAACATGCTCGAAGGC

GAGCACACGTCTGAACTCTTTCCCTACACGACGCTCTTCCGATCTNNNACGCCTGGCTAACATTTGTATT

GAGCACACGTCTGAACTCTTTCCCTACACGACGCTCTTCCGATCTNNNACGCCTGGCTAACATTTGTAT
GCACACGTCTGAACTCTTTCCCTACACGACGCTCTTCCGATCTNNNTGCGTATCTGTAATCCCAGCTAC
GCACACGTCTGAACTCTTTCCCTACACGACGCTCTTCCGATCTNNNTGCGTATCTGTAATCCCAGCTAC
AGCACACGTCTGAACTCTTTCCCTACACGACGCTCTTCCGATCTNNNCAGCTGCTCAGAATCTGTAGA
AGCACACGTCTGAACTCTTTCCCTACACGACGCTCTTCCGATCTNNNCAGCTGCTCAGAATCTGTAGA
GCACACGTCTGAACTCTTTCCCTACACGACGCTCTTCCGATCTNNNGACGAAGCCCCACATTTATTTG
GCACACGTCTGAACTCTTTCCCTACACGACGCTCTTCCGATCTNNNGACGAAGCCCCACATTTATTTG
AGCACACGTCTGAACTCTTTCCCTACACGACGCTCTTCCGATCTNNNACCTGAGCGATCTGTAACATCT
AGCACACGTCTGAACTCTTTCCCTACACGACGCTCTTCCGATCTNNNACCTGAGCGATCTGTAACATCT
AGCACACGTCTGAACTCTTTCCCTACACGACGCTCTTCCGATCTNNNCAAGATCGCCCTACTGCACTCC
AGCACACGTCTGAACTCTTTCCCTACACGACGCTCTTCCGATCTNNNCAAGATCGCCCTACTGCACTCC
GAGCACACGTCTGAACTCTTTCCCTACACGACGCTCTTCCGATCTNNNAACATAGGAGATGGAGGTT
GAGCACACGTCTGAACTCTTTCCCTACACGACGCTCTTCCGATCTNNNAACATAGGAGATGGAGGTT
AGCACACGTCTGAACTCTTTCCCTACACGACGCTCTTCCGATCTNNNAGTAGCTGGGACTATAGGTG
AGCACACGTCTGAACTCTTTCCCTACACGACGCTCTTCCGATCTNNNAGTAGCTGGGACTATAGGTG
AGCACACGTCTGAACTCTTTCCCTACACGACGCTCTTCCGATCTNNNGCAGATGGGAAAAGGTTTCA
AGCACACGTCTGAACTCTTTCCCTACACGACGCTCTTCCGATCTNNNGCAGATGGGAAAAGGTTTCA
GCACACGTCTGAACTCTTTCCCTACACGACGCTCTTCCGATCTNNNAGAAAAGGAGAGTCGGCTATG
GCACACGTCTGAACTCTTTCCCTACACGACGCTCTTCCGATCTNNNAGAAAAGGAGAGTCGGCTATG
AGCACACGTCTGAACTCTTTCCCTACACGACGCTCTTCCGATCTNNNCGTAATCCCAGCTACTCAGGA
AGCACACGTCTGAACTCTTTCCCTACACGACGCTCTTCCGATCTNNNCGTAATCCCAGCTACTCAGGA
GCACACGTCTGAACTCTTTCCCTACACGACGCTCTTCCGATCTNNNAATGATGCTCCCAGGTACACTTT
GCACACGTCTGAACTCTTTCCCTACACGACGCTCTTCCGATCTNNNAATGATGCTCCCAGGTACACTTT
AGCACACGTCTGAACTCTTTCCCTACACGACGCTCTTCCGATCTNNNAGTAGCACTGCTATACCAA
AGCACACGTCTGAACTCTTTCCCTACACGACGCTCTTCCGATCTNNNAGTAGCACTGCTATACCAA
AGCACACGTCTGAACTCTTTCCCTACACGACGCTCTTCCGATCTNNNGAGATCACACCATTGCATTCCA
AGCACACGTCTGAACTCTTTCCCTACACGACGCTCTTCCGATCTNNNGAGATCACACCATTGCATTCCA
AGCACACGTCTGAACTCTTTCCCTACACGACGCTCTTCCGATCTNNNGTAAATAGTCTCAGCTCCTCG
AGCACACGTCTGAACTCTTTCCCTACACGACGCTCTTCCGATCTNNNGTAAATAGTCTCAGCTCCTCG
GAGCACACGTCTGAACTCTTTCCCTACACGACGCTCTTCCGATCTNNNGATCGTGCCACTACACTCCAG
GAGCACACGTCTGAACTCTTTCCCTACACGACGCTCTTCCGATCTNNNGATCGTGCCACTACACTCCAG
GCACACGTCTGAACTCTTTCCCTACACGACGCTCTTCCGATCTNNNGATAATGGCGTGATCCTGGGA
GCACACGTCTGAACTCTTTCCCTACACGACGCTCTTCCGATCTNNNGATAATGGCGTGATCCTGGGA
GAGCACACGTCTGAACTCTTTCCCTACACGACGCTCTTCCGATCTNNNTCCTCTGCTTTGGTTACGTGA
GAGCACACGTCTGAACTCTTTCCCTACACGACGCTCTTCCGATCTNNNTCCTCTGCTTTGGTTACGTGA
AGCACACGTCTGAACTCTTTCCCTACACGACGCTCTTCCGATCTNNNACTGTTTCTAGCCTTCAGGTGA
AGCACACGTCTGAACTCTTTCCCTACACGACGCTCTTCCGATCTNNNACTGTTTCTAGCCTTCAGGTGA
AGCACACGTCTGAACTCTTTCCCTACACGACGCTCTTCCGATCTNNNAATAAATTGCAAGAAAGGCC
AGCACACGTCTGAACTCTTTCCCTACACGACGCTCTTCCGATCTNNNAATAAATTGCAAGAAAGGCC
AGAGCACACGTCTGAACTCTTTCCCTACACGACGCTCTTCCGATCTNNNAGATTGTTCTGAAGAAGCC
AGAGCACACGTCTGAACTCTTTCCCTACACGACGCTCTTCCGATCTNNNAGATTGTTCTGAAGAAGCC
GCACACGTCTGAACTCTTTCCCTACACGACGCTCTTCCGATCTNNNCCCATACGTTTTGGTATGCTGT
GCACACGTCTGAACTCTTTCCCTACACGACGCTCTTCCGATCTNNNCCCATACGTTTTGGTATGCTGT

CACGTCTGAACTCTTTCCCTACACGACGCTCTTCCGATCTNNNACGTTGTGCATATGTACCCTAGACAT
CACGTCTGAACTCTTTCCCTACACGACGCTCTTCCGATCTNNNACGTTGTGCATATGTACCCTAGACAT
AGCACACGTCTGAACTCTTTCCCTACACGACGCTCTTCCGATCTNNNCTGAGGGAGGATAACTGCTT
AGCACACGTCTGAACTCTTTCCCTACACGACGCTCTTCCGATCTNNNCTGAGGGAGGATAACTGCTT
AGAGCACACGTCTGAACTCTTTCCCTACACGACGCTCTTCCGATCTNNNGAGGAGAAGAGAGGAGA
AGAGCACACGTCTGAACTCTTTCCCTACACGACGCTCTTCCGATCTNNNGAGGAGAAGAGAGGAGA
GAGCACACGTCTGAACTCTTTCCCTACACGACGCTCTTCCGATCTNNNCCTGTTATTCCCTGCCCATTCT
GAGCACACGTCTGAACTCTTTCCCTACACGACGCTCTTCCGATCTNNNCCTGTTATTCCCTGCCCATTCT
AGCACACGTCTGAACTCTTTCCCTACACGACGCTCTTCCGATCTNNNTTCAAGTGATAGAAATGGCC
AGCACACGTCTGAACTCTTTCCCTACACGACGCTCTTCCGATCTNNNTTCAAGTGATAGAAATGGCC
GAGCACACGTCTGAACTCTTTCCCTACACGACGCTCTTCCGATCTNNNGTCAGGAAACTACTGGGACT
GAGCACACGTCTGAACTCTTTCCCTACACGACGCTCTTCCGATCTNNNGTCAGGAAACTACTGGGACT
GAAGAGCACACGTCTGAACTCTTTCCCTACACGACGCTCTTCCGATCTNNNGTTGCAGTGAACCGAG
GAAGAGCACACGTCTGAACTCTTTCCCTACACGACGCTCTTCCGATCTNNNGTTGCAGTGAACCGAG
GAGCACACGTCTGAACTCTTTCCCTACACGACGCTCTTCCGATCTNNNGCCCAAGAACTGTGCTAAGA
GAGCACACGTCTGAACTCTTTCCCTACACGACGCTCTTCCGATCTNNNGCCCAAGAACTGTGCTAAGA
AGCACACGTCTGAACTCTTTCCCTACACGACGCTCTTCCGATCTNNNTCAACCACTTTCCGTTTTCCTGT
AGCACACGTCTGAACTCTTTCCCTACACGACGCTCTTCCGATCTNNNTCAACCACTTTCCGTTTTCCTGT
GCACACGTCTGAACTCTTTCCCTACACGACGCTCTTCCGATCTNNNCCACCACACCAGGCTAATTTTTCC
GCACACGTCTGAACTCTTTCCCTACACGACGCTCTTCCGATCTNNNCCACCACACCAGGCTAATTTTTCC
AGAGCACACGTCTGAACTCTTTCCCTACACGACGCTCTTCCGATCTNNNGCTACAGTCCTCCTATCAGA
AGAGCACACGTCTGAACTCTTTCCCTACACGACGCTCTTCCGATCTNNNGCTACAGTCCTCCTATCAGA
GAGCACACGTCTGAACTCTTTCCCTACACGACGCTCTTCCGATCTNNNTTGCAAAGATATGGAGCCAA
GAGCACACGTCTGAACTCTTTCCCTACACGACGCTCTTCCGATCTNNNTTGCAAAGATATGGAGCCAA
AGCACACGTCTGAACTCTTTCCCTACACGACGCTCTTCCGATCTNNNCTAGCACCTTTTCCCACAGTTT
AGCACACGTCTGAACTCTTTCCCTACACGACGCTCTTCCGATCTNNNCTAGCACCTTTTCCCACAGTTT
AGCACACGTCTGAACTCTTTCCCTACACGACGCTCTTCCGATCTNNNACAAGAGAGTAGATACGTCA
AGCACACGTCTGAACTCTTTCCCTACACGACGCTCTTCCGATCTNNNACAAGAGAGTAGATACGTCA
AGCACACGTCTGAACTCTTTCCCTACACGACGCTCTTCCGATCTNNNAATAAACTGCAGGGAGGACA
AGCACACGTCTGAACTCTTTCCCTACACGACGCTCTTCCGATCTNNNAATAAACTGCAGGGAGGACA
AGCACACGTCTGAACTCTTTCCCTACACGACGCTCTTCCGATCTNNNGTAATCCCAGCTACTTAGGAG
AGCACACGTCTGAACTCTTTCCCTACACGACGCTCTTCCGATCTNNNGTAATCCCAGCTACTTAGGAG
GCACACGTCTGAACTCTTTCCCTACACGACGCTCTTCCGATCTNNNCTTGTTAGCTCGCTGGCTCTTTA
GCACACGTCTGAACTCTTTCCCTACACGACGCTCTTCCGATCTNNNCTTGTTAGCTCGCTGGCTCTTTA
GAGCACACGTCTGAACTCTTTCCCTACACGACGCTCTTCCGATCTNNNTGTGCATGTGTGTATCATTTT
GAGCACACGTCTGAACTCTTTCCCTACACGACGCTCTTCCGATCTNNNTGTGCATGTGTGTATCATTTT
AGCACACGTCTGAACTCTTTCCCTACACGACGCTCTTCCGATCTNNNTGCTGTTTTGGTTACTGTAGCT
AGCACACGTCTGAACTCTTTCCCTACACGACGCTCTTCCGATCTNNNTGCTGTTTTGGTTACTGTAGCT
AGCACACGTCTGAACTCTTTCCCTACACGACGCTCTTCCGATCTNNNAGATCACACCACTGAACTCTA
AGCACACGTCTGAACTCTTTCCCTACACGACGCTCTTCCGATCTNNNAGATCACACCACTGAACTCTA
GAGCACACGTCTGAACTCTTTCCCTACACGACGCTCTTCCGATCTNNNGTTGCAGTGAGCCAAGATTG
GAGCACACGTCTGAACTCTTTCCCTACACGACGCTCTTCCGATCTNNNGTTGCAGTGAGCCAAGATTG
ACACGTCTGAACTCTTTCCCTACACGACGCTCTTCCGATCTNNNCCAGCCATTTGGGGAAAGATTACC

CACACGTCTGAACTCTTTCCCTACACGACGCTCTTCCGATCTNNNCCAGCCATTTGGGGAAAGATTACC
AGCACACGTCTGAACTCTTTCCCTACACGACGCTCTTCCGATCTNNNTCCTGCCTTCCTTCCTTTCTTAGC
AGCACACGTCTGAACTCTTTCCCTACACGACGCTCTTCCGATCTNNNTCCTGCCTTCCTTCCTTTCTTAGC
GAGCACACGTCTGAACTCTTTCCCTACACGACGCTCTTCCGATCTNNNCAGCAGGCTGTGTAGACTTC
GAGCACACGTCTGAACTCTTTCCCTACACGACGCTCTTCCGATCTNNNCAGCAGGCTGTGTAGACTTC
GCACACGTCTGAACTCTTTCCCTACACGACGCTCTTCCGATCTNNNAGTATCTGGGACTACAGGTTCCC
GCACACGTCTGAACTCTTTCCCTACACGACGCTCTTCCGATCTNNNAGTATCTGGGACTACAGGTTCCC
GCACACGTCTGAACTCTTTCCCTACACGACGCTCTTCCGATCTNNNCTAAGGATAATGGCCTCCACTCCC
GCACACGTCTGAACTCTTTCCCTACACGACGCTCTTCCGATCTNNNCTAAGGATAATGGCCTCCACTCCC
GAGCACACGTCTGAACTCTTTCCCTACACGACGCTCTTCCGATCTNNNACTCTTATCACCCAGGCTAGA
GAGCACACGTCTGAACTCTTTCCCTACACGACGCTCTTCCGATCTNNNACTCTTATCACCCAGGCTAGA
AGCACACGTCTGAACTCTTTCCCTACACGACGCTCTTCCGATCTNNNGCCTTCTGAGTAGCTGGGATTA
AGCACACGTCTGAACTCTTTCCCTACACGACGCTCTTCCGATCTNNNGCCTTCTGAGTAGCTGGGATTA
GAGCACACGTCTGAACTCTTTCCCTACACGACGCTCTTCCGATCTNNNGCTATTCTTATGGGCTGTGCC
GAGCACACGTCTGAACTCTTTCCCTACACGACGCTCTTCCGATCTNNNGCTATTCTTATGGGCTGTGCC
GCACACGTCTGAACTCTTTCCCTACACGACGCTCTTCCGATCTNNNAAGGAGGTACCACTAGTCAACA
GCACACGTCTGAACTCTTTCCCTACACGACGCTCTTCCGATCTNNNAAGGAGGTACCACTAGTCAACA
AGAGCACACGTCTGAACTCTTTCCCTACACGACGCTCTTCCGATCTNNNAGCTACTCAGGAAAAAGGG
AGAGCACACGTCTGAACTCTTTCCCTACACGACGCTCTTCCGATCTNNNAGCTACTCAGGAAAAAGGG
GAGCACACGTCTGAACTCTTTCCCTACACGACGCTCTTCCGATCTNNNTGCAGTAAGATATGATCACGC
GAGCACACGTCTGAACTCTTTCCCTACACGACGCTCTTCCGATCTNNNTGCAGTAAGATATGATCACGC
GAGCACACGTCTGAACTCTTTCCCTACACGACGCTCTTCCGATCTNNNAGTTGCAGTAAGCTGAGATC
GAGCACACGTCTGAACTCTTTCCCTACACGACGCTCTTCCGATCTNNNAGTTGCAGTAAGCTGAGATC
AGCACACGTCTGAACTCTTTCCCTACACGACGCTCTTCCGATCTNNNAGGGGCTTCTTCTTTTTTCTTCC
AGCACACGTCTGAACTCTTTCCCTACACGACGCTCTTCCGATCTNNNAGGGGCTTCTTCTTTTTTCTTCC
AGCACACGTCTGAACTCTTTCCCTACACGACGCTCTTCCGATCTNNNTTTGTTTCTCTTTGGTGGAGGG
AGCACACGTCTGAACTCTTTCCCTACACGACGCTCTTCCGATCTNNNTTTGTTTCTCTTTGGTGGAGGG
AGCACACGTCTGAACTCTTTCCCTACACGACGCTCTTCCGATCTNNNGGACCAAACTTTGAGGTAGGT
AGCACACGTCTGAACTCTTTCCCTACACGACGCTCTTCCGATCTNNNGGACCAAACTTTGAGGTAGGT
GCACACGTCTGAACTCTTTCCCTACACGACGCTCTTCCGATCTNNNCACATCTGACCCAAAATGCCATTC
GCACACGTCTGAACTCTTTCCCTACACGACGCTCTTCCGATCTNNNCACATCTGACCCAAAATGCCATTC
GCACACGTCTGAACTCTTTCCCTACACGACGCTCTTCCGATCTNNNGCTCTGTTTTCTCTCGCAGAATC
GCACACGTCTGAACTCTTTCCCTACACGACGCTCTTCCGATCTNNNGCTCTGTTTTCTCTCGCAGAATC
GAGCACACGTCTGAACTCTTTCCCTACACGACGCTCTTCCGATCTNNNAGTTCATGCTGCAGTAGGAA
GAGCACACGTCTGAACTCTTTCCCTACACGACGCTCTTCCGATCTNNNAGTTCATGCTGCAGTAGGAA
GCACACGTCTGAACTCTTTCCCTACACGACGCTCTTCCGATCTNNNTGTTTTCTCCTACTATGGCCTGTC
GCACACGTCTGAACTCTTTCCCTACACGACGCTCTTCCGATCTNNNTGTTTTCTCCTACTATGGCCTGTC
GAGCACACGTCTGAACTCTTTCCCTACACGACGCTCTTCCGATCTNNNGTGTGGTGGTATTTCTGAGG
GAGCACACGTCTGAACTCTTTCCCTACACGACGCTCTTCCGATCTNNNGTGTGGTGGTATTTCTGAGG
GAGCACACGTCTGAACTCTTTCCCTACACGACGCTCTTCCGATCTNNNTTTCAGGAGAGGGGAGGAA
GAGCACACGTCTGAACTCTTTCCCTACACGACGCTCTTCCGATCTNNNTTTCAGGAGAGGGGAGGAA
GCACACGTCTGAACTCTTTCCCTACACGACGCTCTTCCGATCTNNNGTGAGTTGATGGGTACAGGTCT
GCACACGTCTGAACTCTTTCCCTACACGACGCTCTTCCGATCTNNNGTGAGTTGATGGGTACAGGTCT

GAGCACACGTCTGAACTCTTTCCCTACACGACGCTCTTCCGATCTNNNACGTAAAATGCAGATTGCCAA

GAGCACACGTCTGAACTCTTTCCCTACACGACGCTCTTCCGATCTNNNACGTAAAATGCAGATTGCCAA

AGCACACGTCTGAACTCTTTCCCTACACGACGCTCTTCCGATCTNNNCCCTCTCAAAGTGCTGGGATTAT

AGCACACGTCTGAACTCTTTCCCTACACGACGCTCTTCCGATCTNNNCCCTCTCAAAGTGCTGGGATTAT

GAGCACACGTCTGAACTCTTTCCCTACACGACGCTCTTCCGATCTNNNATACAAAAATTAGCTGGGCGT

GAGCACACGTCTGAACTCTTTCCCTACACGACGCTCTTCCGATCTNNNATACAAAAATTAGCTGGGCGT

GAGCACACGTCTGAACTCTTTCCCTACACGACGCTCTTCCGATCTNNNCCTCGTCCTCCTCCATCCTAGCA

GAGCACACGTCTGAACTCTTTCCCTACACGACGCTCTTCCGATCTNNNCCTCGTCCTCCTCCATCCTAGCA

GCACACGTCTGAACTCTTTCCCTACACGACGCTCTTCCGATCTNNNTTCCAAACAGGGACACATAACCT

GCACACGTCTGAACTCTTTCCCTACACGACGCTCTTCCGATCTNNNTTCCAAACAGGGACACATAACCT

AGCACACGTCTGAACTCTTTCCCTACACGACGCTCTTCCGATCTNNNTGGATCACTTGAACCTATGAGG

AGCACACGTCTGAACTCTTTCCCTACACGACGCTCTTCCGATCTNNNTGGATCACTTGAACCTATGAGG

AGCACACGTCTGAACTCTTTCCCTACACGACGCTCTTCCGATCTNNNTCCAGAGGTGGCTAGAGTTGA

AGCACACGTCTGAACTCTTTCCCTACACGACGCTCTTCCGATCTNNNTCCAGAGGTGGCTAGAGTTGA

CACACGTCTGAACTCTTTCCCTACACGACGCTCTTCCGATCTNNNGATACCAATGCTAGCCTGAAGGAC

CACACGTCTGAACTCTTTCCCTACACGACGCTCTTCCGATCTNNNGATACCAATGCTAGCCTGAAGGAC

GAGCACACGTCTGAACTCTTTCCCTACACGACGCTCTTCCGATCTNNNCTTCCCTCTCCCCCTTGATGTAG

GAGCACACGTCTGAACTCTTTCCCTACACGACGCTCTTCCGATCTNNNCTTCCCTCTCCCCCTTGATGTAG

AGCACACGTCTGAACTCTTTCCCTACACGACGCTCTTCCGATCTNNNAAAATAGCCCAAACATGTGGA

AGCACACGTCTGAACTCTTTCCCTACACGACGCTCTTCCGATCTNNNAAAATAGCCCAAACATGTGGA

AGCACACGTCTGAACTCTTTCCCTACACGACGCTCTTCCGATCTNNNGAGCTGAGATCATGTCATTGCA

AGCACACGTCTGAACTCTTTCCCTACACGACGCTCTTCCGATCTNNNGAGCTGAGATCATGTCATTGCA

CACACGTCTGAACTCTTTCCCTACACGACGCTCTTCCGATCTNNNCCAACTCTGTCTGGCTTCCTAGCAT

CACACGTCTGAACTCTTTCCCTACACGACGCTCTTCCGATCTNNNCCAACTCTGTCTGGCTTCCTAGCAT

AGCACACGTCTGAACTCTTTCCCTACACGACGCTCTTCCGATCTNNNTCCTTGTCCCTTCTTCTCCCTTTCA

AGCACACGTCTGAACTCTTTCCCTACACGACGCTCTTCCGATCTNNNTCCTTGTCCCTTCTTCTCCCTTTCA

AAGAGCACACGTCTGAACTCTTTCCCTACACGACGCTCTTCCGATCTNNNGAGGCTGAGGCAGGAGA

AAGAGCACACGTCTGAACTCTTTCCCTACACGACGCTCTTCCGATCTNNNGAGGCTGAGGCAGGAGA

GAAGAGCACACGTCTGAACTCTTTCCCTACACGACGCTCTTCCGATCTNNNGCCCTGTCTGGAAGGAAT

GAAGAGCACACGTCTGAACTCTTTCCCTACACGACGCTCTTCCGATCTNNNGCCCTGTCTGGAAGGAAT

AGAGCACACGTCTGAACTCTTTCCCTACACGACGCTCTTCCGATCTNNNAAGGAGTGGTGTGCTTGGT

AGAGCACACGTCTGAACTCTTTCCCTACACGACGCTCTTCCGATCTNNNAAGGAGTGGTGTGCTTGGT

GAGCACACGTCTGAACTCTTTCCCTACACGACGCTCTTCCGATCTNNNTCCCGAAATTGAACCAGGAAG

GAGCACACGTCTGAACTCTTTCCCTACACGACGCTCTTCCGATCTNNNTCCCGAAATTGAACCAGGAAG

AGCACACGTCTGAACTCTTTCCCTACACGACGCTCTTCCGATCTNNNCCCTGGTTTGTCTTTGGTTGAT

AGCACACGTCTGAACTCTTTCCCTACACGACGCTCTTCCGATCTNNNCCCTGGTTTGTCTTTGGTTGAT

GCACACGTCTGAACTCTTTCCCTACACGACGCTCTTCCGATCTNNNAGAGATTCACAGACTGGAACTG

GCACACGTCTGAACTCTTTCCCTACACGACGCTCTTCCGATCTNNNAGAGATTCACAGACTGGAACTG

GAGCACACGTCTGAACTCTTTCCCTACACGACGCTCTTCCGATCTNNNATCCCATCCAGCTGATACTAG

GAGCACACGTCTGAACTCTTTCCCTACACGACGCTCTTCCGATCTNNNATCCCATCCAGCTGATACTAG

GAGCACACGTCTGAACTCTTTCCCTACACGACGCTCTTCCGATCTNNNAAAAAGAAATTTTGCGGGAG

GAGCACACGTCTGAACTCTTTCCCTACACGACGCTCTTCCGATCTNNNAAAAAGAAATTTTGCGGGAG

ACACGTCTGAACTCTTTCCCTACACGACGCTCTTCCGATCTNNNTCCCTTTACCTCTCTTACGTTGCCAT

ACACGTCTGAACTCTTTCCCTACACGACGCTCTTCCGATCTNNNTCCCTTTACCTCTCTTACGTTGCCATC
GAGCACACGTCTGAACTCTTTCCCTACACGACGCTCTTCCGATCTNNNCACCTGTAGTCCCAGCTATTCA
GAGCACACGTCTGAACTCTTTCCCTACACGACGCTCTTCCGATCTNNNCACCTGTAGTCCCAGCTATTCA
GAGCACACGTCTGAACTCTTTCCCTACACGACGCTCTTCCGATCTNNNTTACCTGGATCCACTGAAAGC
GAGCACACGTCTGAACTCTTTCCCTACACGACGCTCTTCCGATCTNNNTTACCTGGATCCACTGAAAGC
GAGCACACGTCTGAACTCTTTCCCTACACGACGCTCTTCCGATCTNNNGTGGACTAAGGCAAAAGGT
GAGCACACGTCTGAACTCTTTCCCTACACGACGCTCTTCCGATCTNNNGTGGACTAAGGCAAAAGGT
GAGCACACGTCTGAACTCTTTCCCTACACGACGCTCTTCCGATCTNNNCAGAGGTTGTAGTGAGCTGT
GAGCACACGTCTGAACTCTTTCCCTACACGACGCTCTTCCGATCTNNNCAGAGGTTGTAGTGAGCTGT
GAGCACACGTCTGAACTCTTTCCCTACACGACGCTCTTCCGATCTNNNCAGGATATTACTCTGTCACCCC
GAGCACACGTCTGAACTCTTTCCCTACACGACGCTCTTCCGATCTNNNCAGGATATTACTCTGTCACCCC
AGCACACGTCTGAACTCTTTCCCTACACGACGCTCTTCCGATCTNNNTGGAAAACACAGGTAGTTTGA
AGCACACGTCTGAACTCTTTCCCTACACGACGCTCTTCCGATCTNNNTGGAAAACACAGGTAGTTTGA
GAGCACACGTCTGAACTCTTTCCCTACACGACGCTCTTCCGATCTNNNATATGTGTGTGTGTGTGTGTC
GAGCACACGTCTGAACTCTTTCCCTACACGACGCTCTTCCGATCTNNNATATGTGTGTGTGTGTGTGTC
GAGCACACGTCTGAACTCTTTCCCTACACGACGCTCTTCCGATCTNNNCCATTAGCTCTGGGGACACTTA
GAGCACACGTCTGAACTCTTTCCCTACACGACGCTCTTCCGATCTNNNCCATTAGCTCTGGGGACACTTA
GCACACGTCTGAACTCTTTCCCTACACGACGCTCTTCCGATCTNNNGAGAGCACATCCCAGGGTATCCA
GCACACGTCTGAACTCTTTCCCTACACGACGCTCTTCCGATCTNNNGAGAGCACATCCCAGGGTATCCA
AGCACACGTCTGAACTCTTTCCCTACACGACGCTCTTCCGATCTNNNCTCCCTTCAGTGTCTGCTAGAA
AGCACACGTCTGAACTCTTTCCCTACACGACGCTCTTCCGATCTNNNCTCCCTTCAGTGTCTGCTAGAA
AGCACACGTCTGAACTCTTTCCCTACACGACGCTCTTCCGATCTNNNCTACTGACTGGGTAGACGCTAT
AGCACACGTCTGAACTCTTTCCCTACACGACGCTCTTCCGATCTNNNCTACTGACTGGGTAGACGCTAT
AGCACACGTCTGAACTCTTTCCCTACACGACGCTCTTCCGATCTNNNCAGGGGATATGAGTCTAGCAC
AGCACACGTCTGAACTCTTTCCCTACACGACGCTCTTCCGATCTNNNCAGGGGATATGAGTCTAGCAC
AGCACACGTCTGAACTCTTTCCCTACACGACGCTCTTCCGATCTNNNGTGCATGTCCACTCTACAAATG
AGCACACGTCTGAACTCTTTCCCTACACGACGCTCTTCCGATCTNNNGTGCATGTCCACTCTACAAATG
AGCACACGTCTGAACTCTTTCCCTACACGACGCTCTTCCGATCTNNNCTCCTGAGTAGCTGGGATTACA
AGCACACGTCTGAACTCTTTCCCTACACGACGCTCTTCCGATCTNNNCTCCTGAGTAGCTGGGATTACA
AAGAGCACACGTCTGAACTCTTTCCCTACACGACGCTCTTCCGATCTNNNCAAGATTGTGTCACGGCAC
AAGAGCACACGTCTGAACTCTTTCCCTACACGACGCTCTTCCGATCTNNNCAAGATTGTGTCACGGCAC
GCACACGTCTGAACTCTTTCCCTACACGACGCTCTTCCGATCTNNNTAGAATCCGTGAATCCCTGAAGC
GCACACGTCTGAACTCTTTCCCTACACGACGCTCTTCCGATCTNNNTAGAATCCGTGAATCCCTGAAGC
AGCACACGTCTGAACTCTTTCCCTACACGACGCTCTTCCGATCTNNNCTGAGGCAGGAAAATTGCTTG
AGCACACGTCTGAACTCTTTCCCTACACGACGCTCTTCCGATCTNNNCTGAGGCAGGAAAATTGCTTG
GAGCACACGTCTGAACTCTTTCCCTACACGACGCTCTTCCGATCTNNNTTGCAAGATCCAGGCAAAAGT
GAGCACACGTCTGAACTCTTTCCCTACACGACGCTCTTCCGATCTNNNTTGCAAGATCCAGGCAAAAGT
AGCACACGTCTGAACTCTTTCCCTACACGACGCTCTTCCGATCTNNNAGGTTGAGCATGAGATTTTCTT
AGCACACGTCTGAACTCTTTCCCTACACGACGCTCTTCCGATCTNNNAGGTTGAGCATGAGATTTTCTT
AGCACACGTCTGAACTCTTTCCCTACACGACGCTCTTCCGATCTNNNACTCTCAAAACGTGTCCAGAGAA
AGCACACGTCTGAACTCTTTCCCTACACGACGCTCTTCCGATCTNNNACTCTCAAAACGTGTCCAGAGAA
GAGCACACGTCTGAACTCTTTCCCTACACGACGCTCTTCCGATCTNNNAAACTCCTAGGCTCAAGTGAT
GAGCACACGTCTGAACTCTTTCCCTACACGACGCTCTTCCGATCTNNNAAACTCCTAGGCTCAAGTGAT

GCACACGTCTGAACTCTTTCCCTACACGACGCTCTTCCGATCTNNNCAAGTGTCAAGGTTTCCCACATC
GCACACGTCTGAACTCTTTCCCTACACGACGCTCTTCCGATCTNNNCAAGTGTCAAGGTTTCCCACATC
AGCACACGTCTGAACTCTTTCCCTACACGACGCTCTTCCGATCTNNNGGAGAATTGCTTGAACTTGGT
AGCACACGTCTGAACTCTTTCCCTACACGACGCTCTTCCGATCTNNNGGAGAATTGCTTGAACTTGGT
GCACACGTCTGAACTCTTTCCCTACACGACGCTCTTCCGATCTNNNAGAATTAGGAAATCGGTGGTA
GCACACGTCTGAACTCTTTCCCTACACGACGCTCTTCCGATCTNNNAGAATTAGGAAATCGGTGGTA
AGCACACGTCTGAACTCTTTCCCTACACGACGCTCTTCCGATCTNNNACAGCAAGAAGATATCCAAGG
AGCACACGTCTGAACTCTTTCCCTACACGACGCTCTTCCGATCTNNNACAGCAAGAAGATATCCAAGG
GAGCACACGTCTGAACTCTTTCCCTACACGACGCTCTTCCGATCTNNNGAAGCACTGCTCTAGACAGTT
GAGCACACGTCTGAACTCTTTCCCTACACGACGCTCTTCCGATCTNNNGAAGCACTGCTCTAGACAGTT
AGCACACGTCTGAACTCTTTCCCTACACGACGCTCTTCCGATCTNNNATACAGGTGCATCCTACCACAT
AGCACACGTCTGAACTCTTTCCCTACACGACGCTCTTCCGATCTNNNATACAGGTGCATCCTACCACAT
GCACACGTCTGAACTCTTTCCCTACACGACGCTCTTCCGATCTNNNTGCCAGTGATACTCTAGGTGAG
GCACACGTCTGAACTCTTTCCCTACACGACGCTCTTCCGATCTNNNTGCCAGTGATACTCTAGGTGAG
GCACACGTCTGAACTCTTTCCCTACACGACGCTCTTCCGATCTNNNTCCTTGGACCTCTCATCACCTATC
GCACACGTCTGAACTCTTTCCCTACACGACGCTCTTCCGATCTNNNTCCTTGGACCTCTCATCACCTATC
GCACACGTCTGAACTCTTTCCCTACACGACGCTCTTCCGATCTNNNTGTCACTGGATAATACCTGCAA
GCACACGTCTGAACTCTTTCCCTACACGACGCTCTTCCGATCTNNNTGTCACTGGATAATACCTGCAA
AGCACACGTCTGAACTCTTTCCCTACACGACGCTCTTCCGATCTNNNGACAAGGGGAATGCTATCACT
AGCACACGTCTGAACTCTTTCCCTACACGACGCTCTTCCGATCTNNNGACAAGGGGAATGCTATCACT
GCACACGTCTGAACTCTTTCCCTACACGACGCTCTTCCGATCTNNNATCTGGAGGCAGGTATAAAAC
GCACACGTCTGAACTCTTTCCCTACACGACGCTCTTCCGATCTNNNATCTGGAGGCAGGTATAAAAC
GAGCACACGTCTGAACTCTTTCCCTACACGACGCTCTTCCGATCTNNNGGACACCACTCTTGTCTCAGA
GAGCACACGTCTGAACTCTTTCCCTACACGACGCTCTTCCGATCTNNNGGACACCACTCTTGTCTCAGA
AGCACACGTCTGAACTCTTTCCCTACACGACGCTCTTCCGATCTNNNGCTTTATGCTTTCAGCCAGAGA
AGCACACGTCTGAACTCTTTCCCTACACGACGCTCTTCCGATCTNNNGCTTTATGCTTTCAGCCAGAGA
GCACACGTCTGAACTCTTTCCCTACACGACGCTCTTCCGATCTNNNCTGAGACAGGAGAGTCGTTTGA
GCACACGTCTGAACTCTTTCCCTACACGACGCTCTTCCGATCTNNNCTGAGACAGGAGAGTCGTTTGA
CACACGTCTGAACTCTTTCCCTACACGACGCTCTTCCGATCTNNNCACCTCGTCTTCTTGAATCCAGAC
CACACGTCTGAACTCTTTCCCTACACGACGCTCTTCCGATCTNNNCACCTCGTCTTCTTGAATCCAGAC
GAGCACACGTCTGAACTCTTTCCCTACACGACGCTCTTCCGATCTNNNCCCTACCTCTGTTGCAGTGTA
GAGCACACGTCTGAACTCTTTCCCTACACGACGCTCTTCCGATCTNNNCCCTACCTCTGTTGCAGTGTA
AAGAGCACACGTCTGAACTCTTTCCCTACACGACGCTCTTCCGATCTNNNGAGGCTAAGGTGGAAGA
AAGAGCACACGTCTGAACTCTTTCCCTACACGACGCTCTTCCGATCTNNNGAGGCTAAGGTGGAAGA
GAGCACACGTCTGAACTCTTTCCCTACACGACGCTCTTCCGATCTNNNTGGGAGACTGAGACACGAAA
GAGCACACGTCTGAACTCTTTCCCTACACGACGCTCTTCCGATCTNNNTGGGAGACTGAGACACGAAA
AGCACACGTCTGAACTCTTTCCCTACACGACGCTCTTCCGATCTNNNAACACTTCCTTTGTTTCTGGCA
AGCACACGTCTGAACTCTTTCCCTACACGACGCTCTTCCGATCTNNNAACACTTCCTTTGTTTCTGGCA
AGCACACGTCTGAACTCTTTCCCTACACGACGCTCTTCCGATCTNNNAAATAGAGTGAGCCATGATTG
AGCACACGTCTGAACTCTTTCCCTACACGACGCTCTTCCGATCTNNNAAATAGAGTGAGCCATGATTG
CACACGTCTGAACTCTTTCCCTACACGACGCTCTTCCGATCTNNNTAAATACAATGGGGAGAGGGGG
CACACGTCTGAACTCTTTCCCTACACGACGCTCTTCCGATCTNNNTAAATACAATGGGGAGAGGGGG
AGCACACGTCTGAACTCTTTCCCTACACGACGCTCTTCCGATCTNNNAAGACTTACTTGTGGGGTTTG

AGCACACGTCTGAACTCTTTCCCTACACGACGCTCTTCCGATCTNNNAAGACTTACTTGTGGGGTTTG

AGCACACGTCTGAACTCTTTCCCTACACGACGCTCTTCCGATCTNNNGGAAGCTGATGTGGGAGGAT

AGCACACGTCTGAACTCTTTCCCTACACGACGCTCTTCCGATCTNNNGGAAGCTGATGTGGGAGGAT

GAGCACACGTCTGAACTCTTTCCCTACACGACGCTCTTCCGATCTNNNTTTTCTACCATGGCACCAGT

GAGCACACGTCTGAACTCTTTCCCTACACGACGCTCTTCCGATCTNNNTTTTCTACCATGGCACCAGT

AGCACACGTCTGAACTCTTTCCCTACACGACGCTCTTCCGATCTNNNTGGGAACAGTAGTTTCAAGTG

AGCACACGTCTGAACTCTTTCCCTACACGACGCTCTTCCGATCTNNNTGGGAACAGTAGTTTCAAGTG

AGCACACGTCTGAACTCTTTCCCTACACGACGCTCTTCCGATCTNNNACAGGAATCCATGACTGTGAC

AGCACACGTCTGAACTCTTTCCCTACACGACGCTCTTCCGATCTNNNACAGGAATCCATGACTGTGAC

GAGCACACGTCTGAACTCTTTCCCTACACGACGCTCTTCCGATCTNNNGCACACCAGTACAGAGGAAC

GAGCACACGTCTGAACTCTTTCCCTACACGACGCTCTTCCGATCTNNNGCACACCAGTACAGAGGAAC

AGCACACGTCTGAACTCTTTCCCTACACGACGCTCTTCCGATCTNNNCTGAGACAGGAGAATCGCTTG

AGCACACGTCTGAACTCTTTCCCTACACGACGCTCTTCCGATCTNNNCTGAGACAGGAGAATCGCTTG

GCACACGTCTGAACTCTTTCCCTACACGACGCTCTTCCGATCTNNNAGAAGATCTTGTAGCCAAAGCC

GCACACGTCTGAACTCTTTCCCTACACGACGCTCTTCCGATCTNNNAGAAGATCTTGTAGCCAAAGCC

CACGTCTGAACTCTTTCCCTACACGACGCTCTTCCGATCTNNNCTCAGACAGCTGGTTCTGCGCATCG

CACGTCTGAACTCTTTCCCTACACGACGCTCTTCCGATCTNNNCTCAGACAGCTGGTTCTGCGCATCG

AGAGCACACGTCTGAACTCTTTCCCTACACGACGCTCTTCCGATCTNNNTACGTGAGACTTGGAATTG

AGAGCACACGTCTGAACTCTTTCCCTACACGACGCTCTTCCGATCTNNNTACGTGAGACTTGGAATTG

AGCACACGTCTGAACTCTTTCCCTACACGACGCTCTTCCGATCTNNNTTTCAGGTACACCAATCAGAC

AGCACACGTCTGAACTCTTTCCCTACACGACGCTCTTCCGATCTNNNTTTCAGGTACACCAATCAGAC

AGCACACGTCTGAACTCTTTCCCTACACGACGCTCTTCCGATCTNNNTTAGGACACATTTTACCAGGC

AGCACACGTCTGAACTCTTTCCCTACACGACGCTCTTCCGATCTNNNTTAGGACACATTTTACCAGGC

AGCACACGTCTGAACTCTTTCCCTACACGACGCTCTTCCGATCTNNNTAAGCAGGTTAGGGTAGCCAG

AGCACACGTCTGAACTCTTTCCCTACACGACGCTCTTCCGATCTNNNTAAGCAGGTTAGGGTAGCCAG

AGCACACGTCTGAACTCTTTCCCTACACGACGCTCTTCCGATCTNNNTGAGTGCATTGATATCAAGCCC

AGCACACGTCTGAACTCTTTCCCTACACGACGCTCTTCCGATCTNNNTGAGTGCATTGATATCAAGCCC

AGCACACGTCTGAACTCTTTCCCTACACGACGCTCTTCCGATCTNNNCTGTGCTTGACACACATGACGT

AGCACACGTCTGAACTCTTTCCCTACACGACGCTCTTCCGATCTNNNCTGTGCTTGACACACATGACGT

GCACACGTCTGAACTCTTTCCCTACACGACGCTCTTCCGATCTNNNGGATGTCTGGACGCTATAGTTG

GCACACGTCTGAACTCTTTCCCTACACGACGCTCTTCCGATCTNNNGGATGTCTGGACGCTATAGTTG

AGCACACGTCTGAACTCTTTCCCTACACGACGCTCTTCCGATCTNNNCATGATCAATTCCAGCTTCCTG

AGCACACGTCTGAACTCTTTCCCTACACGACGCTCTTCCGATCTNNNCATGATCAATTCCAGCTTCCTG

GAGCACACGTCTGAACTCTTTCCCTACACGACGCTCTTCCGATCTNNNCATAGGTTGCAGTGAGTGGA

GAGCACACGTCTGAACTCTTTCCCTACACGACGCTCTTCCGATCTNNNCATAGGTTGCAGTGAGTGGA

CACACGTCTGAACTCTTTCCCTACACGACGCTCTTCCGATCTNNNCTGAACAAGGAGTCCGGAATACA

CACACGTCTGAACTCTTTCCCTACACGACGCTCTTCCGATCTNNNCTGAACAAGGAGTCCGGAATACA

GAGCACACGTCTGAACTCTTTCCCTACACGACGCTCTTCCGATCTNNNAGGTATGAAAGTGGAAGCC

GAGCACACGTCTGAACTCTTTCCCTACACGACGCTCTTCCGATCTNNNAGGTATGAAAGTGGAAGCC

CACACGTCTGAACTCTTTCCCTACACGACGCTCTTCCGATCTNNNTCCCTCTTTGTCTCTCTGTCTCTCA

CACACGTCTGAACTCTTTCCCTACACGACGCTCTTCCGATCTNNNTCCCTCTTTGTCTCTCTGTCTCTCA

GAGCACACGTCTGAACTCTTTCCCTACACGACGCTCTTCCGATCTNNNATCAGAATTAAGACTAGAGA

GAGCACACGTCTGAACTCTTTCCCTACACGACGCTCTTCCGATCTNNNATCAGAATTAAGACTAGAGA

GAGCACACGTCTGAACTCTTTCCCTACACGACGCTCTTCCGATCTNNNTCAATCACGTACTGTCTGCA
GAGCACACGTCTGAACTCTTTCCCTACACGACGCTCTTCCGATCTNNNTCAATCACGTACTGTCTGCA
GCACACGTCTGAACTCTTTCCCTACACGACGCTCTTCCGATCTNNNTCAACATCCTGAGCTCAAGTGA
GCACACGTCTGAACTCTTTCCCTACACGACGCTCTTCCGATCTNNNTCAACATCCTGAGCTCAAGTGA
AGCACACGTCTGAACTCTTTCCCTACACGACGCTCTTCCGATCTNNNTGTGCTTGTAATCCCAGCTACT
AGCACACGTCTGAACTCTTTCCCTACACGACGCTCTTCCGATCTNNNTGTGCTTGTAATCCCAGCTACT
AGCACACGTCTGAACTCTTTCCCTACACGACGCTCTTCCGATCTNNNATGATGTAGCTGTCTTCTGCA
AGCACACGTCTGAACTCTTTCCCTACACGACGCTCTTCCGATCTNNNATGATGTAGCTGTCTTCTGCA
GAGCACACGTCTGAACTCTTTCCCTACACGACGCTCTTCCGATCTNNNCCAGGAGATAAGCCCAAAGT
GAGCACACGTCTGAACTCTTTCCCTACACGACGCTCTTCCGATCTNNNCCAGGAGATAAGCCCAAAGT
AAGAGCACACGTCTGAACTCTTTCCCTACACGACGCTCTTCCGATCTNNNCTTGAATCCAGGAGGCAG
AAGAGCACACGTCTGAACTCTTTCCCTACACGACGCTCTTCCGATCTNNNCTTGAATCCAGGAGGCAG
AGCACACGTCTGAACTCTTTCCCTACACGACGCTCTTCCGATCTNNNCTGTAGTCCCAGCTACTCAGG
AGCACACGTCTGAACTCTTTCCCTACACGACGCTCTTCCGATCTNNNCTGTAGTCCCAGCTACTCAGG
GAGCACACGTCTGAACTCTTTCCCTACACGACGCTCTTCCGATCTNNNAGTCCAGGTTGAATTGGCTT
GAGCACACGTCTGAACTCTTTCCCTACACGACGCTCTTCCGATCTNNNAGTCCAGGTTGAATTGGCTT
AGCACACGTCTGAACTCTTTCCCTACACGACGCTCTTCCGATCTNNNGATTGCAGTAAGCCAAGATCA
AGCACACGTCTGAACTCTTTCCCTACACGACGCTCTTCCGATCTNNNGATTGCAGTAAGCCAAGATCA
AGCACACGTCTGAACTCTTTCCCTACACGACGCTCTTCCGATCTNNNGTGCATGCACATTTCTCTGAG
AGCACACGTCTGAACTCTTTCCCTACACGACGCTCTTCCGATCTNNNGTGCATGCACATTTCTCTGAG
AGCACACGTCTGAACTCTTTCCCTACACGACGCTCTTCCGATCTNNNCACACTCTACTGTACTGACGCC
AGCACACGTCTGAACTCTTTCCCTACACGACGCTCTTCCGATCTNNNCACACTCTACTGTACTGACGCC
AGCACACGTCTGAACTCTTTCCCTACACGACGCTCTTCCGATCTNNNTTGTTCTTCAGGCAGAGTTCTG
AGCACACGTCTGAACTCTTTCCCTACACGACGCTCTTCCGATCTNNNTTGTTCTTCAGGCAGAGTTCTG
GAGCACACGTCTGAACTCTTTCCCTACACGACGCTCTTCCGATCTNNNCAACATCGGACTTGGCAATG
GAGCACACGTCTGAACTCTTTCCCTACACGACGCTCTTCCGATCTNNNCAACATCGGACTTGGCAATG
GCACACGTCTGAACTCTTTCCCTACACGACGCTCTTCCGATCTNNNCAGGGTGTTCACCTTGCTTTCCA
GCACACGTCTGAACTCTTTCCCTACACGACGCTCTTCCGATCTNNNCAGGGTGTTCACCTTGCTTTCCA
GCACACGTCTGAACTCTTTCCCTACACGACGCTCTTCCGATCTNNNCGAGAGGGAGGAGGGAAGTA
GCACACGTCTGAACTCTTTCCCTACACGACGCTCTTCCGATCTNNNCGAGAGGGAGGAGGGAAGTA
AGCACACGTCTGAACTCTTTCCCTACACGACGCTCTTCCGATCTNNNGTAATTTTCGGGTTTGGTGTG
AGCACACGTCTGAACTCTTTCCCTACACGACGCTCTTCCGATCTNNNGTAATTTTCGGGTTTGGTGTG
GAGCACACGTCTGAACTCTTTCCCTACACGACGCTCTTCCGATCTNNNGAGGTCCCAGCTACTCAGGA
GAGCACACGTCTGAACTCTTTCCCTACACGACGCTCTTCCGATCTNNNGAGGTCCCAGCTACTCAGGA
AGCACACGTCTGAACTCTTTCCCTACACGACGCTCTTCCGATCTNNNGGGTACATGAACCTACACGTG
AGCACACGTCTGAACTCTTTCCCTACACGACGCTCTTCCGATCTNNNGGGTACATGAACCTACACGTG
GCACACGTCTGAACTCTTTCCCTACACGACGCTCTTCCGATCTNNNCCCAAGAGTCAGGAGATTATGG
GCACACGTCTGAACTCTTTCCCTACACGACGCTCTTCCGATCTNNNCCCAAGAGTCAGGAGATTATGG
AGCACACGTCTGAACTCTTTCCCTACACGACGCTCTTCCGATCTNNNACTGTGTCTAGTTAGTGCATG
AGCACACGTCTGAACTCTTTCCCTACACGACGCTCTTCCGATCTNNNACTGTGTCTAGTTAGTGCATG
AGCACACGTCTGAACTCTTTCCCTACACGACGCTCTTCCGATCTNNNCTTCCCGGGTTCAAGCAATTAT
AGCACACGTCTGAACTCTTTCCCTACACGACGCTCTTCCGATCTNNNCTTCCCGGGTTCAAGCAATTAT
GAGCACACGTCTGAACTCTTTCCCTACACGACGCTCTTCCGATCTNNNTGTCCAACAGGTAATGGCAT

GAGCACACGTCTGAACTCTTTCCCTACACGACGCTCTTCCGATCTNNNTGTCCAACAGGTAATGGCAT
AGCACACGTCTGAACTCTTTCCCTACACGACGCTCTTCCGATCTNNNAGCCACCATGGTAGGAAATCTT
AGCACACGTCTGAACTCTTTCCCTACACGACGCTCTTCCGATCTNNNAGCCACCATGGTAGGAAATCTT
GCACACGTCTGAACTCTTTCCCTACACGACGCTCTTCCGATCTNNNGTTTGACTTCGGAGCTGAGAAA
GCACACGTCTGAACTCTTTCCCTACACGACGCTCTTCCGATCTNNNGTTTGACTTCGGAGCTGAGAAA
GCACACGTCTGAACTCTTTCCCTACACGACGCTCTTCCGATCTNNNCTGAAATCCCAGCTACTCAGGAG
GCACACGTCTGAACTCTTTCCCTACACGACGCTCTTCCGATCTNNNCTGAAATCCCAGCTACTCAGGAG
GAGCACACGTCTGAACTCTTTCCCTACACGACGCTCTTCCGATCTNNNGTGGCTGAGAAACCTCAAAAG
GAGCACACGTCTGAACTCTTTCCCTACACGACGCTCTTCCGATCTNNNGTGGCTGAGAAACCTCAAAAG
AGCACACGTCTGAACTCTTTCCCTACACGACGCTCTTCCGATCTNNNTGGACCAATCAGCACTCTGTAA
AGCACACGTCTGAACTCTTTCCCTACACGACGCTCTTCCGATCTNNNTGGACCAATCAGCACTCTGTAA
AAGAGCACACGTCTGAACTCTTTCCCTACACGACGCTCTTCCGATCTNNNACAATGGACCACTATTCAG
AAGAGCACACGTCTGAACTCTTTCCCTACACGACGCTCTTCCGATCTNNNACAATGGACCACTATTCAG
AAGAGCACACGTCTGAACTCTTTCCCTACACGACGCTCTTCCGATCTNNNTGCACAGACTCAAGAAACA
AAGAGCACACGTCTGAACTCTTTCCCTACACGACGCTCTTCCGATCTNNNTGCACAGACTCAAGAAACA
GCACACGTCTGAACTCTTTCCCTACACGACGCTCTTCCGATCTNNNAGGGTGCCTCTTTTTCCATCTAG
GCACACGTCTGAACTCTTTCCCTACACGACGCTCTTCCGATCTNNNAGGGTGCCTCTTTTTCCATCTAG
AGAGCACACGTCTGAACTCTTTCCCTACACGACGCTCTTCCGATCTNNNCTGAGGCAGGAGAATCACT
AGAGCACACGTCTGAACTCTTTCCCTACACGACGCTCTTCCGATCTNNNCTGAGGCAGGAGAATCACT
AGAGCACACGTCTGAACTCTTTCCCTACACGACGCTCTTCCGATCTNNNAATACAAAAAATTAGCCGG
AGAGCACACGTCTGAACTCTTTCCCTACACGACGCTCTTCCGATCTNNNAATACAAAAAATTAGCCGG
GCACACGTCTGAACTCTTTCCCTACACGACGCTCTTCCGATCTNNNATAATGAGGTTTTTAACTTGTGC
GCACACGTCTGAACTCTTTCCCTACACGACGCTCTTCCGATCTNNNATAATGAGGTTTTTAACTTGTGC
AGCACACGTCTGAACTCTTTCCCTACACGACGCTCTTCCGATCTNNNAAGAATAGCTTGAACCCAGGA
AGCACACGTCTGAACTCTTTCCCTACACGACGCTCTTCCGATCTNNNAAGAATAGCTTGAACCCAGGA
AGCACACGTCTGAACTCTTTCCCTACACGACGCTCTTCCGATCTNNNTCACGCCTGTAATCTTAGCACT
AGCACACGTCTGAACTCTTTCCCTACACGACGCTCTTCCGATCTNNNTCACGCCTGTAATCTTAGCACT
GAGCACACGTCTGAACTCTTTCCCTACACGACGCTCTTCCGATCTNNNGGATCACTTGAGGTCAGGAG
GAGCACACGTCTGAACTCTTTCCCTACACGACGCTCTTCCGATCTNNNGGATCACTTGAGGTCAGGAG
AGCACACGTCTGAACTCTTTCCCTACACGACGCTCTTCCGATCTNNNGGTGACAGAGAGAGACACCAT
AGCACACGTCTGAACTCTTTCCCTACACGACGCTCTTCCGATCTNNNGGTGACAGAGAGAGACACCAT
CACACGTCTGAACTCTTTCCCTACACGACGCTCTTCCGATCTNNNGTGGTAGAGCATATACGCCACTCC
CACACGTCTGAACTCTTTCCCTACACGACGCTCTTCCGATCTNNNGTGGTAGAGCATATACGCCACTCC
GAGCACACGTCTGAACTCTTTCCCTACACGACGCTCTTCCGATCTNNNGAGGCAGGAAAATCACTTGA
GAGCACACGTCTGAACTCTTTCCCTACACGACGCTCTTCCGATCTNNNGAGGCAGGAAAATCACTTGA
GAGCACACGTCTGAACTCTTTCCCTACACGACGCTCTTCCGATCTNNNCTGTAATCACAGCTACTCGGG
GAGCACACGTCTGAACTCTTTCCCTACACGACGCTCTTCCGATCTNNNCTGTAATCACAGCTACTCGGG
AGCACACGTCTGAACTCTTTCCCTACACGACGCTCTTCCGATCTNNNGAATGTGGAAGAGACTGTGAC
AGCACACGTCTGAACTCTTTCCCTACACGACGCTCTTCCGATCTNNNGAATGTGGAAGAGACTGTGAC
AGCACACGTCTGAACTCTTTCCCTACACGACGCTCTTCCGATCTNNNTCCAAGTGCAAGAAACAAAGCA
AGCACACGTCTGAACTCTTTCCCTACACGACGCTCTTCCGATCTNNNTCCAAGTGCAAGAAACAAAGCA
GCACACGTCTGAACTCTTTCCCTACACGACGCTCTTCCGATCTNNNCCATGATTCTAGCCCTCAGAGTC
GCACACGTCTGAACTCTTTCCCTACACGACGCTCTTCCGATCTNNNCCATGATTCTAGCCCTCAGAGTC

AGCACACGTCTGAACTCTTTCCCTACACGACGCTCTTCCGATCTNNNACCCCTTGAACCTAAAAGTTG
AGCACACGTCTGAACTCTTTCCCTACACGACGCTCTTCCGATCTNNNACCCCTTGAACCTAAAAGTTG
GCACACGTCTGAACTCTTTCCCTACACGACGCTCTTCCGATCTNNNTGTTCAACTTGTCCTTCTTTCACT
GCACACGTCTGAACTCTTTCCCTACACGACGCTCTTCCGATCTNNNTGTTCAACTTGTCCTTCTTTCACT
AGCACACGTCTGAACTCTTTCCCTACACGACGCTCTTCCGATCTNNNGAGATGAAGGAACCTCCACTCT
AGCACACGTCTGAACTCTTTCCCTACACGACGCTCTTCCGATCTNNNGAGATGAAGGAACCTCCACTCT
GAGCACACGTCTGAACTCTTTCCCTACACGACGCTCTTCCGATCTNNNGTAAATAGTCTCAGCTCCTCG
GAGCACACGTCTGAACTCTTTCCCTACACGACGCTCTTCCGATCTNNNGTAAATAGTCTCAGCTCCTCG
GAGCACACGTCTGAACTCTTTCCCTACACGACGCTCTTCCGATCTNNNGATCTGTCCTGTCTGCGGATT
GAGCACACGTCTGAACTCTTTCCCTACACGACGCTCTTCCGATCTNNNGATCTGTCCTGTCTGCGGATT
GAGCACACGTCTGAACTCTTTCCCTACACGACGCTCTTCCGATCTNNNCGCCTGTAATCCCAGCTACTA
GAGCACACGTCTGAACTCTTTCCCTACACGACGCTCTTCCGATCTNNNCGCCTGTAATCCCAGCTACTA
AGAGCACACGTCTGAACTCTTTCCCTACACGACGCTCTTCCGATCTNNNAAGAGTCAACCTCAAGCTA
AGAGCACACGTCTGAACTCTTTCCCTACACGACGCTCTTCCGATCTNNNAAGAGTCAACCTCAAGCTA
GCACACGTCTGAACTCTTTCCCTACACGACGCTCTTCCGATCTNNNGCAAGTGACTGAGATTGTCCCT
GCACACGTCTGAACTCTTTCCCTACACGACGCTCTTCCGATCTNNNGCAAGTGACTGAGATTGTCCCT
AGAGCACACGTCTGAACTCTTTCCCTACACGACGCTCTTCCGATCTNNNCTGTAGTCTCAGCTACTCGG
AGAGCACACGTCTGAACTCTTTCCCTACACGACGCTCTTCCGATCTNNNCTGTAGTCTCAGCTACTCGG
GAGCACACGTCTGAACTCTTTCCCTACACGACGCTCTTCCGATCTNNNGGGAGTGAGCTTTCTCTTGT
GAGCACACGTCTGAACTCTTTCCCTACACGACGCTCTTCCGATCTNNNGGGAGTGAGCTTTCTCTTGT
AGCACACGTCTGAACTCTTTCCCTACACGACGCTCTTCCGATCTNNNGTTGCAGTGAGCTGAGATTGC
AGCACACGTCTGAACTCTTTCCCTACACGACGCTCTTCCGATCTNNNGTTGCAGTGAGCTGAGATTGC
AGCACACGTCTGAACTCTTTCCCTACACGACGCTCTTCCGATCTNNNCTGTGATCCCAGCATTTTGGAA
AGCACACGTCTGAACTCTTTCCCTACACGACGCTCTTCCGATCTNNNCTGTGATCCCAGCATTTTGGAA
AGCACACGTCTGAACTCTTTCCCTACACGACGCTCTTCCGATCTNNNCCTGTCTCTGGTGCTTTTCTAG
AGCACACGTCTGAACTCTTTCCCTACACGACGCTCTTCCGATCTNNNCCTGTCTCTGGTGCTTTTCTAG
AGAGCACACGTCTGAACTCTTTCCCTACACGACGCTCTTCCGATCTNNNCACAATGTGCACATGTACCC
AGAGCACACGTCTGAACTCTTTCCCTACACGACGCTCTTCCGATCTNNNCACAATGTGCACATGTACCC
AGCACACGTCTGAACTCTTTCCCTACACGACGCTCTTCCGATCTNNNAGGCAAAAAATCAGCAAGCAT
AGCACACGTCTGAACTCTTTCCCTACACGACGCTCTTCCGATCTNNNAGGCAAAAAATCAGCAAGCAT
AGCACACGTCTGAACTCTTTCCCTACACGACGCTCTTCCGATCTNNNGGAGACAGGAGAATCGCTTGA
AGCACACGTCTGAACTCTTTCCCTACACGACGCTCTTCCGATCTNNNGGAGACAGGAGAATCGCTTGA
AAGAGCACACGTCTGAACTCTTTCCCTACACGACGCTCTTCCGATCTNNNAAGAAACTGAGAATTCTG
AAGAGCACACGTCTGAACTCTTTCCCTACACGACGCTCTTCCGATCTNNNAAGAAACTGAGAATTCTG
GAGCACACGTCTGAACTCTTTCCCTACACGACGCTCTTCCGATCTNNNAGAAAATCCTCCTTCAGCTTTC
GAGCACACGTCTGAACTCTTTCCCTACACGACGCTCTTCCGATCTNNNAGAAAATCCTCCTTCAGCTTTC
GCACACGTCTGAACTCTTTCCCTACACGACGCTCTTCCGATCTNNNACCACGCCTGGCTAATTTTTGTA
GCACACGTCTGAACTCTTTCCCTACACGACGCTCTTCCGATCTNNNACCACGCCTGGCTAATTTTTGTA
AGCACACGTCTGAACTCTTTCCCTACACGACGCTCTTCCGATCTNNNGAACCTGCATTGTGACTGCTG
AGCACACGTCTGAACTCTTTCCCTACACGACGCTCTTCCGATCTNNNGAACCTGCATTGTGACTGCTG
AGCACACGTCTGAACTCTTTCCCTACACGACGCTCTTCCGATCTNNNGAGGTTGCAGTGAGCCAAAAT
AGCACACGTCTGAACTCTTTCCCTACACGACGCTCTTCCGATCTNNNGAGGTTGCAGTGAGCCAAAAT
AGAGCACACGTCTGAACTCTTTCCCTACACGACGCTCTTCCGATCTNNNTTGTTTCTACAGCTTGTGAT

AGAGCACACGTCTGAACTCTTTCCCTACACGACGCTCTTCCGATCTNNNTTGTTTCTACAGCTTGTGAT
AGCACACGTCTGAACTCTTTCCCTACACGACGCTCTTCCGATCTNNNAAAATTATTCTGGCGTGGTGGT
AGCACACGTCTGAACTCTTTCCCTACACGACGCTCTTCCGATCTNNNAAAATTATTCTGGCGTGGTGGT
AGCACACGTCTGAACTCTTTCCCTACACGACGCTCTTCCGATCTNNNACTGGAGAAAAACTAAGTCCTT
AGCACACGTCTGAACTCTTTCCCTACACGACGCTCTTCCGATCTNNNACTGGAGAAAAACTAAGTCCTT
AGCACACGTCTGAACTCTTTCCCTACACGACGCTCTTCCGATCTNNNATGCTCTCCCAAGCTTATCTTCC
AGCACACGTCTGAACTCTTTCCCTACACGACGCTCTTCCGATCTNNNATGCTCTCCCAAGCTTATCTTCC
GAGCACACGTCTGAACTCTTTCCCTACACGACGCTCTTCCGATCTNNNGATTACAGTGCCTGGCCATAG
GAGCACACGTCTGAACTCTTTCCCTACACGACGCTCTTCCGATCTNNNGATTACAGTGCCTGGCCATAG
GAGCACACGTCTGAACTCTTTCCCTACACGACGCTCTTCCGATCTNNNTGCACCCATCAACTTGTCATCT
GAGCACACGTCTGAACTCTTTCCCTACACGACGCTCTTCCGATCTNNNTGCACCCATCAACTTGTCATCT
GAGCACACGTCTGAACTCTTTCCCTACACGACGCTCTTCCGATCTNNNTACAGCAAGTGTGTGTTTGTG
GAGCACACGTCTGAACTCTTTCCCTACACGACGCTCTTCCGATCTNNNTACAGCAAGTGTGTGTTTGTG
GAGCACACGTCTGAACTCTTTCCCTACACGACGCTCTTCCGATCTNNNGACAGTTCTCTCTTCAGACAG
GAGCACACGTCTGAACTCTTTCCCTACACGACGCTCTTCCGATCTNNNGACAGTTCTCTCTTCAGACAG
AGCACACGTCTGAACTCTTTCCCTACACGACGCTCTTCCGATCTNNNCTGTTGAACAACCATGTCTGGAC
AGCACACGTCTGAACTCTTTCCCTACACGACGCTCTTCCGATCTNNNCTGTTGAACAACCATGTCTGGAC
AGCACACGTCTGAACTCTTTCCCTACACGACGCTCTTCCGATCTNNNCACTTAGTTTTACGACCTTGGG
AGCACACGTCTGAACTCTTTCCCTACACGACGCTCTTCCGATCTNNNCACTTAGTTTTACGACCTTGGG
GAGCACACGTCTGAACTCTTTCCCTACACGACGCTCTTCCGATCTNNNTGATACAAGGGTGTACAACTG
GAGCACACGTCTGAACTCTTTCCCTACACGACGCTCTTCCGATCTNNNTGATACAAGGGTGTACAACTG
AGCACACGTCTGAACTCTTTCCCTACACGACGCTCTTCCGATCTNNNCTGAGGCAGGAGAATCACTTCA
AGCACACGTCTGAACTCTTTCCCTACACGACGCTCTTCCGATCTNNNCTGAGGCAGGAGAATCACTTCA
GCACACGTCTGAACTCTTTCCCTACACGACGCTCTTCCGATCTNNNTCCTTCCTTCCTTCCTTCCTTTCCAT
GCACACGTCTGAACTCTTTCCCTACACGACGCTCTTCCGATCTNNNTCCTTCCTTCCTTCCTTCCTTTCCAT
AGCACACGTCTGAACTCTTTCCCTACACGACGCTCTTCCGATCTNNNTAGTTCTCTCATGAAAGGCCCAC
AGCACACGTCTGAACTCTTTCCCTACACGACGCTCTTCCGATCTNNNTAGTTCTCTCATGAAAGGCCCAC
GAGCACACGTCTGAACTCTTTCCCTACACGACGCTCTTCCGATCTNNNAATCCACAGACCAAGAGATTCC
GAGCACACGTCTGAACTCTTTCCCTACACGACGCTCTTCCGATCTNNNAATCCACAGACCAAGAGATTCC
GAGCACACGTCTGAACTCTTTCCCTACACGACGCTCTTCCGATCTNNNAACCAACAACAGGTTCATCTTC
GAGCACACGTCTGAACTCTTTCCCTACACGACGCTCTTCCGATCTNNNAACCAACAACAGGTTCATCTTC
GAGCACACGTCTGAACTCTTTCCCTACACGACGCTCTTCCGATCTNNNCAACAAACACTGCCAAAATGCT
GAGCACACGTCTGAACTCTTTCCCTACACGACGCTCTTCCGATCTNNNCAACAAACACTGCCAAAATGCT
CACGTCTGAACTCTTTCCCTACACGACGCTCTTCCGATCTNNNGGGAGAGAGACCCAGGAAAATGCAT
CACGTCTGAACTCTTTCCCTACACGACGCTCTTCCGATCTNNNGGGAGAGAGACCCAGGAAAATGCAT
AGCACACGTCTGAACTCTTTCCCTACACGACGCTCTTCCGATCTNNNCCTCCCAAAGTGCTGTGATTAC
AGCACACGTCTGAACTCTTTCCCTACACGACGCTCTTCCGATCTNNNCCTCCCAAAGTGCTGTGATTAC
AGAGCACACGTCTGAACTCTTTCCCTACACGACGCTCTTCCGATCTNNNCCTAAAGACAGCAAGAGAGC
AGAGCACACGTCTGAACTCTTTCCCTACACGACGCTCTTCCGATCTNNNCCTAAAGACAGCAAGAGAGC
AGCACACGTCTGAACTCTTTCCCTACACGACGCTCTTCCGATCTNNNGTCTTCTCAAATACCAAGCTGCC
AGCACACGTCTGAACTCTTTCCCTACACGACGCTCTTCCGATCTNNNGTCTTCTCAAATACCAAGCTGCC
AGAGCACACGTCTGAACTCTTTCCCTACACGACGCTCTTCCGATCTNNNTCTGAGGAGGTGTTGGAGTA
AGAGCACACGTCTGAACTCTTTCCCTACACGACGCTCTTCCGATCTNNNTCTGAGGAGGTGTTGGAGTA

AGCACACGTCTGAACTCTTTCCCTACACGACGCTCTTCCGATCTNNNTGCTTATCAGCCATCTTTCCACT

AGCACACGTCTGAACTCTTTCCCTACACGACGCTCTTCCGATCTNNNTGCTTATCAGCCATCTTTCCACT

GAGCACACGTCTGAACTCTTTCCCTACACGACGCTCTTCCGATCTNNNTAGGTTGCATGAGCTGAGAT

GAGCACACGTCTGAACTCTTTCCCTACACGACGCTCTTCCGATCTNNNTAGGTTGCATGAGCTGAGAT

AGCACACGTCTGAACTCTTTCCCTACACGACGCTCTTCCGATCTNNNGATTGAGACCATCCTGGCTAA

AGCACACGTCTGAACTCTTTCCCTACACGACGCTCTTCCGATCTNNNGATTGAGACCATCCTGGCTAA

GAGCACACGTCTGAACTCTTTCCCTACACGACGCTCTTCCGATCTNNNTGAGATGGAACCTCACTCTGT

GAGCACACGTCTGAACTCTTTCCCTACACGACGCTCTTCCGATCTNNNTGAGATGGAACCTCACTCTGT

AGCACACGTCTGAACTCTTTCCCTACACGACGCTCTTCCGATCTNNNCAGAGCAACTTTCATTCCTGCAA

AGCACACGTCTGAACTCTTTCCCTACACGACGCTCTTCCGATCTNNNCAGAGCAACTTTCATTCCTGCAA

GAGCACACGTCTGAACTCTTTCCCTACACGACGCTCTTCCGATCTNNNTGTCTTGAACCCAGGACAGTT

GAGCACACGTCTGAACTCTTTCCCTACACGACGCTCTTCCGATCTNNNTGTCTTGAACCCAGGACAGTT

GCACACGTCTGAACTCTTTCCCTACACGACGCTCTTCCGATCTNNNTTGACTGGAGATGGAGGTCAAA

GCACACGTCTGAACTCTTTCCCTACACGACGCTCTTCCGATCTNNNTTGACTGGAGATGGAGGTCAAA

GCACACGTCTGAACTCTTTCCCTACACGACGCTCTTCCGATCTNNNCCCGTCATTATGATGTTAGCTGG

GCACACGTCTGAACTCTTTCCCTACACGACGCTCTTCCGATCTNNNCCCGTCATTATGATGTTAGCTGG

GCACACGTCTGAACTCTTTCCCTACACGACGCTCTTCCGATCTNNNATCAAAGTTCTGAGACTTAATCTC

GCACACGTCTGAACTCTTTCCCTACACGACGCTCTTCCGATCTNNNATCAAAGTTCTGAGACTTAATCTC

GAGCACACGTCTGAACTCTTTCCCTACACGACGCTCTTCCGATCTNNNTCCTCCTTTCCCCTTACTCTCA

GAGCACACGTCTGAACTCTTTCCCTACACGACGCTCTTCCGATCTNNNTCCTCCTTTCCCCTTACTCTCA

AGAGCACACGTCTGAACTCTTTCCCTACACGACGCTCTTCCGATCTNNNTTTGCCTAGTATTGCTTTGG

AGAGCACACGTCTGAACTCTTTCCCTACACGACGCTCTTCCGATCTNNNTTTGCCTAGTATTGCTTTGG

AGCACACGTCTGAACTCTTTCCCTACACGACGCTCTTCCGATCTNNNGGTTACTCCATAGGCAGAGCA

AGCACACGTCTGAACTCTTTCCCTACACGACGCTCTTCCGATCTNNNGGTTACTCCATAGGCAGAGCA

AGCACACGTCTGAACTCTTTCCCTACACGACGCTCTTCCGATCTNNNTCCCTTTTTCTTTCTCTCTCTCTC

AGCACACGTCTGAACTCTTTCCCTACACGACGCTCTTCCGATCTNNNTCCCTTTTTCTTTCTCTCTCTCTC

GAGCACACGTCTGAACTCTTTCCCTACACGACGCTCTTCCGATCTNNNTTGGGCCACAGTGATAAAG

GAGCACACGTCTGAACTCTTTCCCTACACGACGCTCTTCCGATCTNNNTTGGGCCACAGTGATAAAG

AGCACACGTCTGAACTCTTTCCCTACACGACGCTCTTCCGATCTNNNTGTGCATACTGTGATCTCACCT

AGCACACGTCTGAACTCTTTCCCTACACGACGCTCTTCCGATCTNNNTGTGCATACTGTGATCTCACCT

GCACACGTCTGAACTCTTTCCCTACACGACGCTCTTCCGATCTNNNAACTCTTGACCTCAGGTGATCTG

GCACACGTCTGAACTCTTTCCCTACACGACGCTCTTCCGATCTNNNAACTCTTGACCTCAGGTGATCTG

AGCACACGTCTGAACTCTTTCCCTACACGACGCTCTTCCGATCTNNNCTCATGCCTATAATCCCAGCAC

AGCACACGTCTGAACTCTTTCCCTACACGACGCTCTTCCGATCTNNNCTCATGCCTATAATCCCAGCAC

AGCACACGTCTGAACTCTTTCCCTACACGACGCTCTTCCGATCTNNNTCAGACCCACCATGTCATTCTTT

AGCACACGTCTGAACTCTTTCCCTACACGACGCTCTTCCGATCTNNNTCAGACCCACCATGTCATTCTTT

AGCACACGTCTGAACTCTTTCCCTACACGACGCTCTTCCGATCTNNNGAGACCAGCCACTCCCAAATGC

AGCACACGTCTGAACTCTTTCCCTACACGACGCTCTTCCGATCTNNNGAGACCAGCCACTCCCAAATGC

AGCACACGTCTGAACTCTTTCCCTACACGACGCTCTTCCGATCTNNNAAAGTGCTGGAATTACATGCG

AGCACACGTCTGAACTCTTTCCCTACACGACGCTCTTCCGATCTNNNAAAGTGCTGGAATTACATGCG

GAGCACACGTCTGAACTCTTTCCCTACACGACGCTCTTCCGATCTNNNTTAATTGACACTTGATGGCCC

GAGCACACGTCTGAACTCTTTCCCTACACGACGCTCTTCCGATCTNNNTTAATTGACACTTGATGGCCC

AGCACACGTCTGAACTCTTTCCCTACACGACGCTCTTCCGATCTNNNTGGCTGACGTGATATTCAAGCT

AGCACACGTCTGAACTCTTTCCCTACACGACGCTCTTCCGATCTNNNTGGCTGACGTGATATTCAAGCT
AGCACACGTCTGAACTCTTTCCCTACACGACGCTCTTCCGATCTNNNCTTAGCCAGGGCAATCAGAAA
AGCACACGTCTGAACTCTTTCCCTACACGACGCTCTTCCGATCTNNNCTTAGCCAGGGCAATCAGAAA
GAGCACACGTCTGAACTCTTTCCCTACACGACGCTCTTCCGATCTNNNCATATGGTTATCTTGGAATAT
GAGCACACGTCTGAACTCTTTCCCTACACGACGCTCTTCCGATCTNNNCATATGGTTATCTTGGAATAT
AGCACACGTCTGAACTCTTTCCCTACACGACGCTCTTCCGATCTNNNGAAGGTTACAGTGAGCCGAG
AGCACACGTCTGAACTCTTTCCCTACACGACGCTCTTCCGATCTNNNGAAGGTTACAGTGAGCCGAG
AGCACACGTCTGAACTCTTTCCCTACACGACGCTCTTCCGATCTNNNAGTGAAATTCCGCCTCAAAAAC
AGCACACGTCTGAACTCTTTCCCTACACGACGCTCTTCCGATCTNNNAGTGAAATTCCGCCTCAAAAAC
GAGCACACGTCTGAACTCTTTCCCTACACGACGCTCTTCCGATCTNNNGATGTGACAGTGAATGCAAA
GAGCACACGTCTGAACTCTTTCCCTACACGACGCTCTTCCGATCTNNNGATGTGACAGTGAATGCAAA
AGCACACGTCTGAACTCTTTCCCTACACGACGCTCTTCCGATCTNNNGCGAAAGTCATGGGTTTGATC
AGCACACGTCTGAACTCTTTCCCTACACGACGCTCTTCCGATCTNNNGCGAAAGTCATGGGTTTGATC
AGCACACGTCTGAACTCTTTCCCTACACGACGCTCTTCCGATCTNNNGTTTTTCAGGTAAATGGTCTTA
AGCACACGTCTGAACTCTTTCCCTACACGACGCTCTTCCGATCTNNNGTTTTTCAGGTAAATGGTCTTA
GAGCACACGTCTGAACTCTTTCCCTACACGACGCTCTTCCGATCTNNNGGAAGACAGCATCCTTTCACC
GAGCACACGTCTGAACTCTTTCCCTACACGACGCTCTTCCGATCTNNNGGAAGACAGCATCCTTTCACC
AGCACACGTCTGAACTCTTTCCCTACACGACGCTCTTCCGATCTNNNGAGGTTGCAGTGAGCTGAGAT
AGCACACGTCTGAACTCTTTCCCTACACGACGCTCTTCCGATCTNNNGAGGTTGCAGTGAGCTGAGAT
AGCACACGTCTGAACTCTTTCCCTACACGACGCTCTTCCGATCTNNNTGTCCCTGATCTAGATGAGTGG
AGCACACGTCTGAACTCTTTCCCTACACGACGCTCTTCCGATCTNNNTGTCCCTGATCTAGATGAGTGG
AGCACACGTCTGAACTCTTTCCCTACACGACGCTCTTCCGATCTNNNACCTATTTGTAGCTGTCACAGG
AGCACACGTCTGAACTCTTTCCCTACACGACGCTCTTCCGATCTNNNACCTATTTGTAGCTGTCACAGG
AGAGCACACGTCTGAACTCTTTCCCTACACGACGCTCTTCCGATCTNNNGCCTCAGGTGGTGAGATTA
AGAGCACACGTCTGAACTCTTTCCCTACACGACGCTCTTCCGATCTNNNGCCTCAGGTGGTGAGATTA
GAGCACACGTCTGAACTCTTTCCCTACACGACGCTCTTCCGATCTNNNATCCAATCCCGGATCCTACAGT
GAGCACACGTCTGAACTCTTTCCCTACACGACGCTCTTCCGATCTNNNATCCAATCCCGGATCCTACAGT
AGCACACGTCTGAACTCTTTCCCTACACGACGCTCTTCCGATCTNNNGTGAGCCAAGATCATGCCATTG
AGCACACGTCTGAACTCTTTCCCTACACGACGCTCTTCCGATCTNNNGTGAGCCAAGATCATGCCATTG
GCACACGTCTGAACTCTTTCCCTACACGACGCTCTTCCGATCTNNNGCAGATTGGGAACAGAGGAGT
GCACACGTCTGAACTCTTTCCCTACACGACGCTCTTCCGATCTNNNGCAGATTGGGAACAGAGGAGT
AGAGCACACGTCTGAACTCTTTCCCTACACGACGCTCTTCCGATCTNNNACAGGAGCCACACTATCTTC
AGAGCACACGTCTGAACTCTTTCCCTACACGACGCTCTTCCGATCTNNNACAGGAGCCACACTATCTTC
AGCACACGTCTGAACTCTTTCCCTACACGACGCTCTTCCGATCTNNNGCTTGTGCTTGACTGATTCTGA
AGCACACGTCTGAACTCTTTCCCTACACGACGCTCTTCCGATCTNNNGCTTGTGCTTGACTGATTCTGA
GCACACGTCTGAACTCTTTCCCTACACGACGCTCTTCCGATCTNNNAGTAAAAGTCCCAGAGGAATGC
GCACACGTCTGAACTCTTTCCCTACACGACGCTCTTCCGATCTNNNAGTAAAAGTCCCAGAGGAATGC
GAGCACACGTCTGAACTCTTTCCCTACACGACGCTCTTCCGATCTNNNAGACTGAGGCAGGAGAATTC
GAGCACACGTCTGAACTCTTTCCCTACACGACGCTCTTCCGATCTNNNAGACTGAGGCAGGAGAATTC
GAGCACACGTCTGAACTCTTTCCCTACACGACGCTCTTCCGATCTNNNCAAAATGAGGGAGGCAATA
GAGCACACGTCTGAACTCTTTCCCTACACGACGCTCTTCCGATCTNNNCAAAATGAGGGAGGCAATA
AGAGCACACGTCTGAACTCTTTCCCTACACGACGCTCTTCCGATCTNNNCAGATAGAGACAGAGGAAG
AGAGCACACGTCTGAACTCTTTCCCTACACGACGCTCTTCCGATCTNNNCAGATAGAGACAGAGGAAG

AGCACACGTCTGAACTCTTTCCCTACACGACGCTCTTCCGATCTNNNATTCTGCCTTTCTGATTCACCC
AGCACACGTCTGAACTCTTTCCCTACACGACGCTCTTCCGATCTNNNATTCTGCCTTTCTGATTCACCC
AGCACACGTCTGAACTCTTTCCCTACACGACGCTCTTCCGATCTNNNGCAAGAGAATCACTTGAATCCG
AGCACACGTCTGAACTCTTTCCCTACACGACGCTCTTCCGATCTNNNGCAAGAGAATCACTTGAATCCG
GCACACGTCTGAACTCTTTCCCTACACGACGCTCTTCCGATCTNNNTGGGAATCTCCATTATGCAATG
GCACACGTCTGAACTCTTTCCCTACACGACGCTCTTCCGATCTNNNTGGGAATCTCCATTATGCAATG
GCACACGTCTGAACTCTTTCCCTACACGACGCTCTTCCGATCTNNNCCAGCTCAGTGGTTAGATTGAG
GCACACGTCTGAACTCTTTCCCTACACGACGCTCTTCCGATCTNNNCCAGCTCAGTGGTTAGATTGAG
GCACACGTCTGAACTCTTTCCCTACACGACGCTCTTCCGATCTNNNAATCCATCCTGGTTTCAGATACAT
GCACACGTCTGAACTCTTTCCCTACACGACGCTCTTCCGATCTNNNAATCCATCCTGGTTTCAGATACAT
AGCACACGTCTGAACTCTTTCCCTACACGACGCTCTTCCGATCTNNNAGGAAACCTGAATCTCAGTTT
AGCACACGTCTGAACTCTTTCCCTACACGACGCTCTTCCGATCTNNNAGGAAACCTGAATCTCAGTTT
ACACGTCTGAACTCTTTCCCTACACGACGCTCTTCCGATCTNNNTTTATGCGAGCCATTGATGACTGCA
ACACGTCTGAACTCTTTCCCTACACGACGCTCTTCCGATCTNNNTTTATGCGAGCCATTGATGACTGCA
AGCACACGTCTGAACTCTTTCCCTACACGACGCTCTTCCGATCTNNNTTACTATGACTTGCTTCAGGGG
AGCACACGTCTGAACTCTTTCCCTACACGACGCTCTTCCGATCTNNNTTACTATGACTTGCTTCAGGGG
GAGCACACGTCTGAACTCTTTCCCTACACGACGCTCTTCCGATCTNNNCAATGGGTGAAGGCTTTGAA
GAGCACACGTCTGAACTCTTTCCCTACACGACGCTCTTCCGATCTNNNCAATGGGTGAAGGCTTTGAA
GAGCACACGTCTGAACTCTTTCCCTACACGACGCTCTTCCGATCTNNNTGAAGTCAGGAGTTTGAGAC
GAGCACACGTCTGAACTCTTTCCCTACACGACGCTCTTCCGATCTNNNTGAAGTCAGGAGTTTGAGAC
GAGCACACGTCTGAACTCTTTCCCTACACGACGCTCTTCCGATCTNNNAGGAGAACTGCTTGAACCCA
GAGCACACGTCTGAACTCTTTCCCTACACGACGCTCTTCCGATCTNNNAGGAGAACTGCTTGAACCCA
AGCACACGTCTGAACTCTTTCCCTACACGACGCTCTTCCGATCTNNNGGTTGCAGTAAGCCAAGATCA
AGCACACGTCTGAACTCTTTCCCTACACGACGCTCTTCCGATCTNNNGGTTGCAGTAAGCCAAGATCA
AGCACACGTCTGAACTCTTTCCCTACACGACGCTCTTCCGATCTNNNACTAAAGCATGGAGAAGTCCC
AGCACACGTCTGAACTCTTTCCCTACACGACGCTCTTCCGATCTNNNACTAAAGCATGGAGAAGTCCC
AGCACACGTCTGAACTCTTTCCCTACACGACGCTCTTCCGATCTNNNGCAACATGGTGAGACTCCATCT
AGCACACGTCTGAACTCTTTCCCTACACGACGCTCTTCCGATCTNNNGCAACATGGTGAGACTCCATCT
AGCACACGTCTGAACTCTTTCCCTACACGACGCTCTTCCGATCTNNNCACAATCTTGGCTCACTGCAACC
AGCACACGTCTGAACTCTTTCCCTACACGACGCTCTTCCGATCTNNNCACAATCTTGGCTCACTGCAACC
AGCACACGTCTGAACTCTTTCCCTACACGACGCTCTTCCGATCTNNNAGCCAGTTATTTCAGCACCTTT
AGCACACGTCTGAACTCTTTCCCTACACGACGCTCTTCCGATCTNNNAGCCAGTTATTTCAGCACCTTT
AGCACACGTCTGAACTCTTTCCCTACACGACGCTCTTCCGATCTNNNCAGGGTCTGTATTGGTGACTG
AGCACACGTCTGAACTCTTTCCCTACACGACGCTCTTCCGATCTNNNCAGGGTCTGTATTGGTGACTG
GAGCACACGTCTGAACTCTTTCCCTACACGACGCTCTTCCGATCTNNNAAAAGGCAGAGGCATTGGA
GAGCACACGTCTGAACTCTTTCCCTACACGACGCTCTTCCGATCTNNNAAAAGGCAGAGGCATTGGA
AGCACACGTCTGAACTCTTTCCCTACACGACGCTCTTCCGATCTNNNGCAGGAGAATTGCTTGAACCTG
AGCACACGTCTGAACTCTTTCCCTACACGACGCTCTTCCGATCTNNNGCAGGAGAATTGCTTGAACCTG
AGCACACGTCTGAACTCTTTCCCTACACGACGCTCTTCCGATCTNNNGGCTTTGACTTGAAGCTTGGA
AGCACACGTCTGAACTCTTTCCCTACACGACGCTCTTCCGATCTNNNGGCTTTGACTTGAAGCTTGGA
GAGCACACGTCTGAACTCTTTCCCTACACGACGCTCTTCCGATCTNNNACTCATATATGTCTGGCAGCA
GAGCACACGTCTGAACTCTTTCCCTACACGACGCTCTTCCGATCTNNNACTCATATATGTCTGGCAGCA
AGCACACGTCTGAACTCTTTCCCTACACGACGCTCTTCCGATCTNNNGTGTGTGTGTGTGTGTGT

AGCACACGTCTGAACTCTTTCCCTACACGACGCTCTTCCGATCTNNNGTGTGTGTGTGTGTGTGTGT
ACACGTCTGAACTCTTTCCCTACACGACGCTCTTCCGATCTNNNTGCATATCATTTATATGGTATATCA
ACACGTCTGAACTCTTTCCCTACACGACGCTCTTCCGATCTNNNTGCATATCATTTATATGGTATATCA
GCACACGTCTGAACTCTTTCCCTACACGACGCTCTTCCGATCTNNNAGTCTTGCTCTGTTGCCCAGCAT
GCACACGTCTGAACTCTTTCCCTACACGACGCTCTTCCGATCTNNNAGTCTTGCTCTGTTGCCCAGCAT
AGCACACGTCTGAACTCTTTCCCTACACGACGCTCTTCCGATCTNNNCAACTCCTGACCTCGTGATCTA
AGCACACGTCTGAACTCTTTCCCTACACGACGCTCTTCCGATCTNNNCAACTCCTGACCTCGTGATCTA
GCACACGTCTGAACTCTTTCCCTACACGACGCTCTTCCGATCTNNNTTAGAGGTATCACAACAGAGGC
GCACACGTCTGAACTCTTTCCCTACACGACGCTCTTCCGATCTNNNTTAGAGGTATCACAACAGAGGC
AGCACACGTCTGAACTCTTTCCCTACACGACGCTCTTCCGATCTNNNAGAAGATCGTTCAAAGCTAGG
AGCACACGTCTGAACTCTTTCCCTACACGACGCTCTTCCGATCTNNNAGAAGATCGTTCAAAGCTAGG
GAGCACACGTCTGAACTCTTTCCCTACACGACGCTCTTCCGATCTNNNAAAACAGGGACTACTAGCAC
GAGCACACGTCTGAACTCTTTCCCTACACGACGCTCTTCCGATCTNNNAAAACAGGGACTACTAGCAC
CACACGTCTGAACTCTTTCCCTACACGACGCTCTTCCGATCTNNNAAATACCATAGTCTGGGGGTGG
CACACGTCTGAACTCTTTCCCTACACGACGCTCTTCCGATCTNNNAAATACCATAGTCTGGGGGTGG
AGCACACGTCTGAACTCTTTCCCTACACGACGCTCTTCCGATCTNNNTTCCTTTGCTTCCTCCTCCTTTTC
AGCACACGTCTGAACTCTTTCCCTACACGACGCTCTTCCGATCTNNNTTCCTTTGCTTCCTCCTCCTTTTC
AGAGCACACGTCTGAACTCTTTCCCTACACGACGCTCTTCCGATCTNNNCTGTGACACAGTAGACTAG
AGAGCACACGTCTGAACTCTTTCCCTACACGACGCTCTTCCGATCTNNNCTGTGACACAGTAGACTAG
GCACACGTCTGAACTCTTTCCCTACACGACGCTCTTCCGATCTNNNACTTGTTGGCCATTTGTATGTCT
GCACACGTCTGAACTCTTTCCCTACACGACGCTCTTCCGATCTNNNACTTGTTGGCCATTTGTATGTCT
AGCACACGTCTGAACTCTTTCCCTACACGACGCTCTTCCGATCTNNNGTCAAAACCCTCCTGTCCATTTG
AGCACACGTCTGAACTCTTTCCCTACACGACGCTCTTCCGATCTNNNGTCAAAACCCTCCTGTCCATTTG
GCACACGTCTGAACTCTTTCCCTACACGACGCTCTTCCGATCTNNNCTCCACTGTGCTGCGAAAGTCAT
GCACACGTCTGAACTCTTTCCCTACACGACGCTCTTCCGATCTNNNCTCCACTGTGCTGCGAAAGTCAT
GAGCACACGTCTGAACTCTTTCCCTACACGACGCTCTTCCGATCTNNNCTCATGTCTGTAATCCCAGCA
GAGCACACGTCTGAACTCTTTCCCTACACGACGCTCTTCCGATCTNNNCTCATGTCTGTAATCCCAGCA
GAAGAGCACACGTCTGAACTCTTTCCCTACACGACGCTCTTCCGATCTNNNCTGATCCAGATCGGGA
GAAGAGCACACGTCTGAACTCTTTCCCTACACGACGCTCTTCCGATCTNNNCTGATCCAGATCGGGA
GAGCACACGTCTGAACTCTTTCCCTACACGACGCTCTTCCGATCTNNNTTGAACCTGGGAGATGGAG
GAGCACACGTCTGAACTCTTTCCCTACACGACGCTCTTCCGATCTNNNTTGAACCTGGGAGATGGAG
AGCACACGTCTGAACTCTTTCCCTACACGACGCTCTTCCGATCTNNNTGTGTGTGTGTGTGTATA
AGCACACGTCTGAACTCTTTCCCTACACGACGCTCTTCCGATCTNNNTGTGTGTGTGTGTGTATA
GCACACGTCTGAACTCTTTCCCTACACGACGCTCTTCCGATCTNNNAGTCTTTTGGTGAACCTGTGTC
GCACACGTCTGAACTCTTTCCCTACACGACGCTCTTCCGATCTNNNAGTCTTTTGGTGAACCTGTGTC
AGCACACGTCTGAACTCTTTCCCTACACGACGCTCTTCCGATCTNNNTTGATTTAGCGCCAATACTGCA
AGCACACGTCTGAACTCTTTCCCTACACGACGCTCTTCCGATCTNNNTTGATTTAGCGCCAATACTGCA
AGCACACGTCTGAACTCTTTCCCTACACGACGCTCTTCCGATCTNNNCTGAGGTGGGAGGATCATTTG
AGCACACGTCTGAACTCTTTCCCTACACGACGCTCTTCCGATCTNNNCTGAGGTGGGAGGATCATTTG
GCACACGTCTGAACTCTTTCCCTACACGACGCTCTTCCGATCTNNNTTCTGCTTTAGGAGAAACCCCAA
GCACACGTCTGAACTCTTTCCCTACACGACGCTCTTCCGATCTNNNTTCTGCTTTAGGAGAAACCCCAA
AGCACACGTCTGAACTCTTTCCCTACACGACGCTCTTCCGATCTNNNACCACACTGTTTTCATTACTGT
AGCACACGTCTGAACTCTTTCCCTACACGACGCTCTTCCGATCTNNNACCACACTGTTTTCATTACTGT

CACGTCTGAACTCTTTCCCTACACGACGCTCTTCCGATCTNNNCTTGATGGGGATGGCATTGAATCCA
CACGTCTGAACTCTTTCCCTACACGACGCTCTTCCGATCTNNNCTTGATGGGGATGGCATTGAATCCA
GAGCACACGTCTGAACTCTTTCCCTACACGACGCTCTTCCGATCTNNNGCCCTTGTGTTCTACTCACTG
GAGCACACGTCTGAACTCTTTCCCTACACGACGCTCTTCCGATCTNNNGCCCTTGTGTTCTACTCACTG
AAGAGCACACGTCTGAACTCTTTCCCTACACGACGCTCTTCCGATCTNNNGAGAATCGCTTGAATCTG
AAGAGCACACGTCTGAACTCTTTCCCTACACGACGCTCTTCCGATCTNNNGAGAATCGCTTGAATCTG
AGCACACGTCTGAACTCTTTCCCTACACGACGCTCTTCCGATCTNNNAAAGCTACCACCATAAAGCTCG
AGCACACGTCTGAACTCTTTCCCTACACGACGCTCTTCCGATCTNNNAAAGCTACCACCATAAAGCTCG
AGCACACGTCTGAACTCTTTCCCTACACGACGCTCTTCCGATCTNNNTATAATCCCAGCTACTAGGGGG
AGCACACGTCTGAACTCTTTCCCTACACGACGCTCTTCCGATCTNNNTATAATCCCAGCTACTAGGGGG
AGCACACGTCTGAACTCTTTCCCTACACGACGCTCTTCCGATCTNNNATCTCTGCAGTGTGGATTGAA
AGCACACGTCTGAACTCTTTCCCTACACGACGCTCTTCCGATCTNNNATCTCTGCAGTGTGGATTGAA
AGCACACGTCTGAACTCTTTCCCTACACGACGCTCTTCCGATCTNNNGCAACAGAGCAAGACTTCATCT
AGCACACGTCTGAACTCTTTCCCTACACGACGCTCTTCCGATCTNNNGCAACAGAGCAAGACTTCATCT
GCACACGTCTGAACTCTTTCCCTACACGACGCTCTTCCGATCTNNNGTTGAGGTTTGAGGAAGGAGA
GCACACGTCTGAACTCTTTCCCTACACGACGCTCTTCCGATCTNNNGTTGAGGTTTGAGGAAGGAGA
CACACGTCTGAACTCTTTCCCTACACGACGCTCTTCCGATCTNNNGTTGTTTCAGGTGGGAGGGTAAA
CACACGTCTGAACTCTTTCCCTACACGACGCTCTTCCGATCTNNNGTTGTTTCAGGTGGGAGGGTAAA
AGCACACGTCTGAACTCTTTCCCTACACGACGCTCTTCCGATCTNNNGTGTTTGAGGCAAGTCAGTAA
AGCACACGTCTGAACTCTTTCCCTACACGACGCTCTTCCGATCTNNNGTGTTTGAGGCAAGTCAGTAA
AGCACACGTCTGAACTCTTTCCCTACACGACGCTCTTCCGATCTNNNTACCATCTCAACAAGTCAATTC
AGCACACGTCTGAACTCTTTCCCTACACGACGCTCTTCCGATCTNNNTACCATCTCAACAAGTCAATTC
AGCACACGTCTGAACTCTTTCCCTACACGACGCTCTTCCGATCTNNNACATTTCTGCCCTAGGTCACTTT
AGCACACGTCTGAACTCTTTCCCTACACGACGCTCTTCCGATCTNNNACATTTCTGCCCTAGGTCACTTT
AGCACACGTCTGAACTCTTTCCCTACACGACGCTCTTCCGATCTNNNCATGTGTTTTCTTGGCTGCCTA
AGCACACGTCTGAACTCTTTCCCTACACGACGCTCTTCCGATCTNNNCATGTGTTTTCTTGGCTGCCTA
CACGTCTGAACTCTTTCCCTACACGACGCTCTTCCGATCTNNNGTCTCCTCCCTTCTCACCCCCATCGGAC
CACGTCTGAACTCTTTCCCTACACGACGCTCTTCCGATCTNNNGTCTCCTCCCTTCTCACCCCCATCGGAC
AGCACACGTCTGAACTCTTTCCCTACACGACGCTCTTCCGATCTNNNGCTTGCAATGAGCCGAGATTG
AGCACACGTCTGAACTCTTTCCCTACACGACGCTCTTCCGATCTNNNGCTTGCAATGAGCCGAGATTG
GAGCACACGTCTGAACTCTTTCCCTACACGACGCTCTTCCGATCTNNNAGTCCCAGAAGAATTGAGCA
GAGCACACGTCTGAACTCTTTCCCTACACGACGCTCTTCCGATCTNNNAGTCCCAGAAGAATTGAGCA
AGCACACGTCTGAACTCTTTCCCTACACGACGCTCTTCCGATCTNNNTTCTCCCAGGGGAACATACTGT
AGCACACGTCTGAACTCTTTCCCTACACGACGCTCTTCCGATCTNNNTTCTCCCAGGGGAACATACTGT
GAGCACACGTCTGAACTCTTTCCCTACACGACGCTCTTCCGATCTNNNGGCTGAGGTAGGAAAATTG
GAGCACACGTCTGAACTCTTTCCCTACACGACGCTCTTCCGATCTNNNGGCTGAGGTAGGAAAATTG
GAGCACACGTCTGAACTCTTTCCCTACACGACGCTCTTCCGATCTNNNAAAAAAAAAAATTAGCCGGGC
GAGCACACGTCTGAACTCTTTCCCTACACGACGCTCTTCCGATCTNNNAAAAAAAAAAATTAGCCGGGC
CACACGTCTGAACTCTTTCCCTACACGACGCTCTTCCGATCTNNNGGATGTCGTAACAACAGCTGTTTC
CACACGTCTGAACTCTTTCCCTACACGACGCTCTTCCGATCTNNNGGATGTCGTAACAACAGCTGTTTC
AGCACACGTCTGAACTCTTTCCCTACACGACGCTCTTCCGATCTNNNTGGCAAGACATAAACTTGTAG
AGCACACGTCTGAACTCTTTCCCTACACGACGCTCTTCCGATCTNNNTGGCAAGACATAAACTTGTAG
AGCACACGTCTGAACTCTTTCCCTACACGACGCTCTTCCGATCTNNNAGAGGGTTTCAGTCAACTTCCT

AGCACACGTCTGAACTCTTTCCCTACACGACGCTCTTCCGATCTNNNAGAGGGTTTCAGTCAACTTCCT
AGCACACGTCTGAACTCTTTCCCTACACGACGCTCTTCCGATCTNNNTGAAGGTTGCAGTGAGCTAAG
AGCACACGTCTGAACTCTTTCCCTACACGACGCTCTTCCGATCTNNNTGAAGGTTGCAGTGAGCTAAG
AGCACACGTCTGAACTCTTTCCCTACACGACGCTCTTCCGATCTNNNCGTTCATGGGTTAGAAGATAG
AGCACACGTCTGAACTCTTTCCCTACACGACGCTCTTCCGATCTNNNCGTTCATGGGTTAGAAGATAG
AGCACACGTCTGAACTCTTTCCCTACACGACGCTCTTCCGATCTNNNGACAACAGAGTGAGACTCCGT
AGCACACGTCTGAACTCTTTCCCTACACGACGCTCTTCCGATCTNNNGACAACAGAGTGAGACTCCGT
AGCACACGTCTGAACTCTTTCCCTACACGACGCTCTTCCGATCTNNNCAAAGAGGAGGTTCAATGTCT
AGCACACGTCTGAACTCTTTCCCTACACGACGCTCTTCCGATCTNNNCAAAGAGGAGGTTCAATGTCT
AGCACACGTCTGAACTCTTTCCCTACACGACGCTCTTCCGATCTNNNTCCTCTCCTCTCTCTCTCTCTCTCA
AGCACACGTCTGAACTCTTTCCCTACACGACGCTCTTCCGATCTNNNTCCTCTCCTCTCTCTCTCTCTCTCA
AGCACACGTCTGAACTCTTTCCCTACACGACGCTCTTCCGATCTNNNGAGGCTGAGGCAGGAGAAT
AGCACACGTCTGAACTCTTTCCCTACACGACGCTCTTCCGATCTNNNGAGGCTGAGGCAGGAGAAT
AGCACACGTCTGAACTCTTTCCCTACACGACGCTCTTCCGATCTNNNAGGAGAATCGCTTTGGAAACT
AGCACACGTCTGAACTCTTTCCCTACACGACGCTCTTCCGATCTNNNAGGAGAATCGCTTTGGAAACT
AGCACACGTCTGAACTCTTTCCCTACACGACGCTCTTCCGATCTNNNTGAGAATCACTTGAACCCAGG
AGCACACGTCTGAACTCTTTCCCTACACGACGCTCTTCCGATCTNNNTGAGAATCACTTGAACCCAGG
AGCACACGTCTGAACTCTTTCCCTACACGACGCTCTTCCGATCTNNNACATGGTGAAATCCCGTCTCTA
AGCACACGTCTGAACTCTTTCCCTACACGACGCTCTTCCGATCTNNNACATGGTGAAATCCCGTCTCTA
AGAGCACACGTCTGAACTCTTTCCCTACACGACGCTCTTCCGATCTNNNTGGCACTACTCAAAAAGCCA
AGAGCACACGTCTGAACTCTTTCCCTACACGACGCTCTTCCGATCTNNNTGGCACTACTCAAAAAGCCA
GAGCACACGTCTGAACTCTTTCCCTACACGACGCTCTTCCGATCTNNNACCATGAAGTGCATTCCTGT
GAGCACACGTCTGAACTCTTTCCCTACACGACGCTCTTCCGATCTNNNACCATGAAGTGCATTCCTGT
AGCACACGTCTGAACTCTTTCCCTACACGACGCTCTTCCGATCTNNNTCTCTCTCTCTCTCTTTCTCTC
AGCACACGTCTGAACTCTTTCCCTACACGACGCTCTTCCGATCTNNNTCTCTCTCTCTCTCTTTCTCTC
AGCACACGTCTGAACTCTTTCCCTACACGACGCTCTTCCGATCTNNNAAAAAATTAGCCGGGTGTGGT
AGCACACGTCTGAACTCTTTCCCTACACGACGCTCTTCCGATCTNNNAAAAAATTAGCCGGGTGTGGT
GCACACGTCTGAACTCTTTCCCTACACGACGCTCTTCCGATCTNNNACCTTCTCTGCAGGAATGTCAAT
GCACACGTCTGAACTCTTTCCCTACACGACGCTCTTCCGATCTNNNACCTTCTCTGCAGGAATGTCAAT
AGCACACGTCTGAACTCTTTCCCTACACGACGCTCTTCCGATCTNNNATGTGTAAGACAGCAGGATCCA
AGCACACGTCTGAACTCTTTCCCTACACGACGCTCTTCCGATCTNNNATGTGTAAGACAGCAGGATCCA
AGCACACGTCTGAACTCTTTCCCTACACGACGCTCTTCCGATCTNNNTAAAGGGAGTCATTTGCCAGG
AGCACACGTCTGAACTCTTTCCCTACACGACGCTCTTCCGATCTNNNTAAAGGGAGTCATTTGCCAGG
GAGCACACGTCTGAACTCTTTCCCTACACGACGCTCTTCCGATCTNNNACTTGAGAGATGAGATGTAA
GAGCACACGTCTGAACTCTTTCCCTACACGACGCTCTTCCGATCTNNNACTTGAGAGATGAGATGTAA
CACACGTCTGAACTCTTTCCCTACACGACGCTCTTCCGATCTNNNGAAATTAATTAGAAGAGTTGCCAC
CACACGTCTGAACTCTTTCCCTACACGACGCTCTTCCGATCTNNNGAAATTAATTAGAAGAGTTGCCAC
AGCACACGTCTGAACTCTTTCCCTACACGACGCTCTTCCGATCTNNNTCCTGACCTCAGATGAGATGAG
AGCACACGTCTGAACTCTTTCCCTACACGACGCTCTTCCGATCTNNNTCCTGACCTCAGATGAGATGAG
GAGCACACGTCTGAACTCTTTCCCTACACGACGCTCTTCCGATCTNNNACGGTTTTCAGCTACATTCCAC
GAGCACACGTCTGAACTCTTTCCCTACACGACGCTCTTCCGATCTNNNACGGTTTTCAGCTACATTCCAC
GCACACGTCTGAACTCTTTCCCTACACGACGCTCTTCCGATCTNNNTCCTTCTCTTGCTCCAAAAAGGT
GCACACGTCTGAACTCTTTCCCTACACGACGCTCTTCCGATCTNNNTCCTTCTCTTGCTCCAAAAAGGT

AGCACACGTCTGAACTCTTTCCCTACACGACGCTCTTCCGATCTNNNGACTTCTCTGATGGTGAGGA
AGCACACGTCTGAACTCTTTCCCTACACGACGCTCTTCCGATCTNNNGACTTCTCTGATGGTGAGGA
AGCACACGTCTGAACTCTTTCCCTACACGACGCTCTTCCGATCTNNNGCCCAGGAGCCCTAAATAAAT
AGCACACGTCTGAACTCTTTCCCTACACGACGCTCTTCCGATCTNNNGCCCAGGAGCCCTAAATAAAT
GCACACGTCTGAACTCTTTCCCTACACGACGCTCTTCCGATCTNNNGCTTGCAATGAGCCGAGATGC
GCACACGTCTGAACTCTTTCCCTACACGACGCTCTTCCGATCTNNNGCTTGCAATGAGCCGAGATGC
GAGCACACGTCTGAACTCTTTCCCTACACGACGCTCTTCCGATCTNNNATGAAGAAATGGGGCCACTT
GAGCACACGTCTGAACTCTTTCCCTACACGACGCTCTTCCGATCTNNNATGAAGAAATGGGGCCACTT
GCACACGTCTGAACTCTTTCCCTACACGACGCTCTTCCGATCTNNNCACGCTCTCTTTCTAGTTCCCTT
GCACACGTCTGAACTCTTTCCCTACACGACGCTCTTCCGATCTNNNCACGCTCTCTTTCTAGTTCCCTT
GAGCACACGTCTGAACTCTTTCCCTACACGACGCTCTTCCGATCTNNNTCCACTCGTTCCAAAGTGACA
GAGCACACGTCTGAACTCTTTCCCTACACGACGCTCTTCCGATCTNNNTCCACTCGTTCCAAAGTGACA
GAGCACACGTCTGAACTCTTTCCCTACACGACGCTCTTCCGATCTNNNGCCCTAGCTACTTTGAAGTCT
GAGCACACGTCTGAACTCTTTCCCTACACGACGCTCTTCCGATCTNNNGCCCTAGCTACTTTGAAGTCT
GCACACGTCTGAACTCTTTCCCTACACGACGCTCTTCCGATCTNNNTTAGTGGGATCCCTGTGTTGAA
GCACACGTCTGAACTCTTTCCCTACACGACGCTCTTCCGATCTNNNTTAGTGGGATCCCTGTGTTGAA
GCACACGTCTGAACTCTTTCCCTACACGACGCTCTTCCGATCTNNNCTGGGCAACAAGAGCAAAACTC
GCACACGTCTGAACTCTTTCCCTACACGACGCTCTTCCGATCTNNNCTGGGCAACAAGAGCAAAACTC
GCACACGTCTGAACTCTTTCCCTACACGACGCTCTTCCGATCTNNNAAGCCTGCAAAATGATGAGTG
GCACACGTCTGAACTCTTTCCCTACACGACGCTCTTCCGATCTNNNAAGCCTGCAAAATGATGAGTG
GCACACGTCTGAACTCTTTCCCTACACGACGCTCTTCCGATCTNNNCCCCATCAAGTGAACTCTGGAA
GCACACGTCTGAACTCTTTCCCTACACGACGCTCTTCCGATCTNNNCCCCATCAAGTGAACTCTGGAA
GCACACGTCTGAACTCTTTCCCTACACGACGCTCTTCCGATCTNNNACATTGCCTACTTCTTCTGCGAC
GCACACGTCTGAACTCTTTCCCTACACGACGCTCTTCCGATCTNNNACATTGCCTACTTCTTCTGCGAC
GCACACGTCTGAACTCTTTCCCTACACGACGCTCTTCCGATCTNNNCACACCTGGCCTCTTTTTTTCTTC
GCACACGTCTGAACTCTTTCCCTACACGACGCTCTTCCGATCTNNNCACACCTGGCCTCTTTTTTTCTTC
AGCACACGTCTGAACTCTTTCCCTACACGACGCTCTTCCGATCTNNNGGGAGGAAAACCAGGGAAT
AGCACACGTCTGAACTCTTTCCCTACACGACGCTCTTCCGATCTNNNGGGAGGAAAACCAGGGAAT
AGCACACGTCTGAACTCTTTCCCTACACGACGCTCTTCCGATCTNNNCTCCTGGGTTCAAGCGATTCT
AGCACACGTCTGAACTCTTTCCCTACACGACGCTCTTCCGATCTNNNCTCCTGGGTTCAAGCGATTCT
AGCACACGTCTGAACTCTTTCCCTACACGACGCTCTTCCGATCTNNNGGTGTTTAGGGGTGCTTGTTG
AGCACACGTCTGAACTCTTTCCCTACACGACGCTCTTCCGATCTNNNGGTGTTTAGGGGTGCTTGTTG
AAGAGCACACGTCTGAACTCTTTCCCTACACGACGCTCTTCCGATCTNNNCTTGTAGTGAGCTGAGAT
AAGAGCACACGTCTGAACTCTTTCCCTACACGACGCTCTTCCGATCTNNNCTTGTAGTGAGCTGAGAT
GCACACGTCTGAACTCTTTCCCTACACGACGCTCTTCCGATCTNNNAGGTTCAAGTGATTCTTCTGCCT
GCACACGTCTGAACTCTTTCCCTACACGACGCTCTTCCGATCTNNNAGGTTCAAGTGATTCTTCTGCCT
GAGCACACGTCTGAACTCTTTCCCTACACGACGCTCTTCCGATCTNNNTATACAAACAAAGCTTCAAC
GAGCACACGTCTGAACTCTTTCCCTACACGACGCTCTTCCGATCTNNNTATACAAACAAAGCTTCAAC
GCACACGTCTGAACTCTTTCCCTACACGACGCTCTTCCGATCTNNNCTGAGATGGGAGGATCATTGA
GCACACGTCTGAACTCTTTCCCTACACGACGCTCTTCCGATCTNNNCTGAGATGGGAGGATCATTGA
AGCACACGTCTGAACTCTTTCCCTACACGACGCTCTTCCGATCTNNNTGGTATCAGGGTTATACTGG
AGCACACGTCTGAACTCTTTCCCTACACGACGCTCTTCCGATCTNNNTGGTATCAGGGTTATACTGG
GCACACGTCTGAACTCTTTCCCTACACGACGCTCTTCCGATCTNNNCTGGTTCCATTCGCAGAGGTAT

AGCACACGTCTGAACTCTTTCCCTACACGACGCTCTTCCGATCTNNNCTGGTTCCATTCGCAGAGGTATA
AGCACACGTCTGAACTCTTTCCCTACACGACGCTCTTCCGATCTNNNAAAGTCAGTGCTGGTGAAGTG
AGCACACGTCTGAACTCTTTCCCTACACGACGCTCTTCCGATCTNNNAAAGTCAGTGCTGGTGAAGTG
GAGCACACGTCTGAACTCTTTCCCTACACGACGCTCTTCCGATCTNNNCGCATCGCATGTTAGGTTTATC
GAGCACACGTCTGAACTCTTTCCCTACACGACGCTCTTCCGATCTNNNCGCATCGCATGTTAGGTTTATC
GAGCACACGTCTGAACTCTTTCCCTACACGACGCTCTTCCGATCTNNNACACATAGCTCAAACATCCTGT
GAGCACACGTCTGAACTCTTTCCCTACACGACGCTCTTCCGATCTNNNACACATAGCTCAAACATCCTGT
AGCACACGTCTGAACTCTTTCCCTACACGACGCTCTTCCGATCTNNNAAGTAGCTGGGATTACAGGCAC
AGCACACGTCTGAACTCTTTCCCTACACGACGCTCTTCCGATCTNNNAAGTAGCTGGGATTACAGGCAC
AGCACACGTCTGAACTCTTTCCCTACACGACGCTCTTCCGATCTNNNAAGAGAGGGAAGGGTAGTTG
AGCACACGTCTGAACTCTTTCCCTACACGACGCTCTTCCGATCTNNNAAGAGAGGGAAGGGTAGTTG
GAGCACACGTCTGAACTCTTTCCCTACACGACGCTCTTCCGATCTNNNAGCTTGTCTGTGAGGAATTGT
GAGCACACGTCTGAACTCTTTCCCTACACGACGCTCTTCCGATCTNNNAGCTTGTCTGTGAGGAATTGT
AGCACACGTCTGAACTCTTTCCCTACACGACGCTCTTCCGATCTNNNGAGAATGGAGTGAACCTGAGA
AGCACACGTCTGAACTCTTTCCCTACACGACGCTCTTCCGATCTNNNGAGAATGGAGTGAACCTGAGA
GAGCACACGTCTGAACTCTTTCCCTACACGACGCTCTTCCGATCTNNNAAGTTGGTTGTGACTGGAGA
GAGCACACGTCTGAACTCTTTCCCTACACGACGCTCTTCCGATCTNNNAAGTTGGTTGTGACTGGAGA
CACACGTCTGAACTCTTTCCCTACACGACGCTCTTCCGATCTNNNCTGGGGTCTGTGGTCCTGAAGCAT
CACACGTCTGAACTCTTTCCCTACACGACGCTCTTCCGATCTNNNCTGGGGTCTGTGGTCCTGAAGCAT
AGCACACGTCTGAACTCTTTCCCTACACGACGCTCTTCCGATCTNNNGCACTCCAACAAAGAGAACAAC
AGCACACGTCTGAACTCTTTCCCTACACGACGCTCTTCCGATCTNNNGCACTCCAACAAAGAGAACAAC
AGCACACGTCTGAACTCTTTCCCTACACGACGCTCTTCCGATCTNNNCAATATCCTTCAAAAACTTCAGAC
AGCACACGTCTGAACTCTTTCCCTACACGACGCTCTTCCGATCTNNNCAATATCCTTCAAAAACTTCAGAC
AGAGCACACGTCTGAACTCTTTCCCTACACGACGCTCTTCCGATCTNNNAGTCTCAGTGGACTCATTTGA
AGAGCACACGTCTGAACTCTTTCCCTACACGACGCTCTTCCGATCTNNNAGTCTCAGTGGACTCATTTGA
AGCACACGTCTGAACTCTTTCCCTACACGACGCTCTTCCGATCTNNNACTCTTGATGCCCAAAACAACA
AGCACACGTCTGAACTCTTTCCCTACACGACGCTCTTCCGATCTNNNACTCTTGATGCCCAAAACAACA
AGCACACGTCTGAACTCTTTCCCTACACGACGCTCTTCCGATCTNNNCTTCACCTTCCATCTAGTCTTCCA
AGCACACGTCTGAACTCTTTCCCTACACGACGCTCTTCCGATCTNNNCTTCACCTTCCATCTAGTCTTCCA
AGCACACGTCTGAACTCTTTCCCTACACGACGCTCTTCCGATCTNNNGGCATATGGGGGCATAAAAAGT
AGCACACGTCTGAACTCTTTCCCTACACGACGCTCTTCCGATCTNNNGGCATATGGGGGCATAAAAAGT
AGCACACGTCTGAACTCTTTCCCTACACGACGCTCTTCCGATCTNNNTACTCTGTGTACACGCCATTTGAC
AGCACACGTCTGAACTCTTTCCCTACACGACGCTCTTCCGATCTNNNTACTCTGTGTACACGCCATTTGAC
AGCACACGTCTGAACTCTTTCCCTACACGACGCTCTTCCGATCTNNNGAACTCTTGACCTCAGGTGATC
AGCACACGTCTGAACTCTTTCCCTACACGACGCTCTTCCGATCTNNNGAACTCTTGACCTCAGGTGATC
AGCACACGTCTGAACTCTTTCCCTACACGACGCTCTTCCGATCTNNNTCTCCCGTGATTTCAAACTTTGC
AGCACACGTCTGAACTCTTTCCCTACACGACGCTCTTCCGATCTNNNTCTCCCGTGATTTCAAACTTTGC
AGCACACGTCTGAACTCTTTCCCTACACGACGCTCTTCCGATCTNNNTGCCTACAGATGAAGAATTGTC
AGCACACGTCTGAACTCTTTCCCTACACGACGCTCTTCCGATCTNNNTGCCTACAGATGAAGAATTGTC
GAGCACACGTCTGAACTCTTTCCCTACACGACGCTCTTCCGATCTNNNGTAGGTGACATCCCATACCCAT
GAGCACACGTCTGAACTCTTTCCCTACACGACGCTCTTCCGATCTNNNGTAGGTGACATCCCATACCCAT
GAGCACACGTCTGAACTCTTTCCCTACACGACGCTCTTCCGATCTNNNTTAGCTCACTTAATGACGTATC
GAGCACACGTCTGAACTCTTTCCCTACACGACGCTCTTCCGATCTNNNTTAGCTCACTTAATGACGTATC

GCACACGTCTGAACTCTTTCCCTACACGACGCTCTTCCGATCTNNNCCACATGCTCTCTTGAACCTACT
GCACACGTCTGAACTCTTTCCCTACACGACGCTCTTCCGATCTNNNCCACATGCTCTCTTGAACCTACT
AGCACACGTCTGAACTCTTTCCCTACACGACGCTCTTCCGATCTNNNCCTTTGGTCATAACATTTTCATT
AGCACACGTCTGAACTCTTTCCCTACACGACGCTCTTCCGATCTNNNCCTTTGGTCATAACATTTTCATT
AGAGCACACGTCTGAACTCTTTCCCTACACGACGCTCTTCCGATCTNNNGTGTGCAATTGGGTATGTT
AGAGCACACGTCTGAACTCTTTCCCTACACGACGCTCTTCCGATCTNNNGTGTGCAATTGGGTATGTT
AGCACACGTCTGAACTCTTTCCCTACACGACGCTCTTCCGATCTNNNTCGCTGTGATAATGAGGTGTC
AGCACACGTCTGAACTCTTTCCCTACACGACGCTCTTCCGATCTNNNTCGCTGTGATAATGAGGTGTC
GCACACGTCTGAACTCTTTCCCTACACGACGCTCTTCCGATCTNNNCTATGGGAAGGAAACCTCGAG
GCACACGTCTGAACTCTTTCCCTACACGACGCTCTTCCGATCTNNNCTATGGGAAGGAAACCTCGAG
AGCACACGTCTGAACTCTTTCCCTACACGACGCTCTTCCGATCTNNNACTGCCAAGAATAAAAGGAGC
AGCACACGTCTGAACTCTTTCCCTACACGACGCTCTTCCGATCTNNNACTGCCAAGAATAAAAGGAGC
AGCACACGTCTGAACTCTTTCCCTACACGACGCTCTTCCGATCTNNNCAGGAAAACTGCTTGAACCCA
AGCACACGTCTGAACTCTTTCCCTACACGACGCTCTTCCGATCTNNNCAGGAAAACTGCTTGAACCCA
AGCACACGTCTGAACTCTTTCCCTACACGACGCTCTTCCGATCTNNNCTGCAACCATCTCCTATCCTACC
AGCACACGTCTGAACTCTTTCCCTACACGACGCTCTTCCGATCTNNNCTGCAACCATCTCCTATCCTACC
GAGCACACGTCTGAACTCTTTCCCTACACGACGCTCTTCCGATCTNNNTGAGGCAGGAGAATCTCTTG
GAGCACACGTCTGAACTCTTTCCCTACACGACGCTCTTCCGATCTNNNTGAGGCAGGAGAATCTCTTG
AGCACACGTCTGAACTCTTTCCCTACACGACGCTCTTCCGATCTNNNACCCTTGGTAGGCCTTTGTATT
AGCACACGTCTGAACTCTTTCCCTACACGACGCTCTTCCGATCTNNNACCCTTGGTAGGCCTTTGTATT
AGAGCACACGTCTGAACTCTTTCCCTACACGACGCTCTTCCGATCTNNNCAGGACCCATCAGTGTGTTG
AGAGCACACGTCTGAACTCTTTCCCTACACGACGCTCTTCCGATCTNNNCAGGACCCATCAGTGTGTTG
AGCACACGTCTGAACTCTTTCCCTACACGACGCTCTTCCGATCTNNNCTTGAGTCACCCCTCTTCCAAC
AGCACACGTCTGAACTCTTTCCCTACACGACGCTCTTCCGATCTNNNCTTGAGTCACCCCTCTTCCAAC
GCACACGTCTGAACTCTTTCCCTACACGACGCTCTTCCGATCTNNNTTTTGAGTTTGTCTGTTCCTGGA
GCACACGTCTGAACTCTTTCCCTACACGACGCTCTTCCGATCTNNNTTTTGAGTTTGTCTGTTCCTGGA
ACACGTCTGAACTCTTTCCCTACACGACGCTCTTCCGATCTNNNGCATGTTTACTGAAAGACATGGGTC
ACACGTCTGAACTCTTTCCCTACACGACGCTCTTCCGATCTNNNGCATGTTTACTGAAAGACATGGGTC
AGCACACGTCTGAACTCTTTCCCTACACGACGCTCTTCCGATCTNNNTCATTCAATTGTGTCTTTCCAGG
AGCACACGTCTGAACTCTTTCCCTACACGACGCTCTTCCGATCTNNNTCATTCAATTGTGTCTTTCCAGG
AGCACACGTCTGAACTCTTTCCCTACACGACGCTCTTCCGATCTNNNGCAACAAAAGCGAGATTCTGT
AGCACACGTCTGAACTCTTTCCCTACACGACGCTCTTCCGATCTNNNGCAACAAAAGCGAGATTCTGT
CACACGTCTGAACTCTTTCCCTACACGACGCTCTTCCGATCTNNNCCATGTTTACATTCCCTTTATTTCA
CACACGTCTGAACTCTTTCCCTACACGACGCTCTTCCGATCTNNNCCATGTTTACATTCCCTTTATTTCA
CACACGTCTGAACTCTTTCCCTACACGACGCTCTTCCGATCTNNNGCTCCTGGCCAGTTTTCACTATAC
CACACGTCTGAACTCTTTCCCTACACGACGCTCTTCCGATCTNNNGCTCCTGGCCAGTTTTCACTATAC
AGCACACGTCTGAACTCTTTCCCTACACGACGCTCTTCCGATCTNNNTGTCAGTGTACCTCAGTGGAA
AGCACACGTCTGAACTCTTTCCCTACACGACGCTCTTCCGATCTNNNTGTCAGTGTACCTCAGTGGAA
GAGCACACGTCTGAACTCTTTCCCTACACGACGCTCTTCCGATCTNNNGAATCTGATATGACCCTTCG
GAGCACACGTCTGAACTCTTTCCCTACACGACGCTCTTCCGATCTNNNGAATCTGATATGACCCTTCG
AGCACACGTCTGAACTCTTTCCCTACACGACGCTCTTCCGATCTNNNTCCACAACTTGCGTGCAATTATT
AGCACACGTCTGAACTCTTTCCCTACACGACGCTCTTCCGATCTNNNTCCACAACTTGCGTGCAATTATT
AGCACACGTCTGAACTCTTTCCCTACACGACGCTCTTCCGATCTNNNGGGCCATTTTCATCAGAAGAC

AGCACACGTCTGAACTCTTTCCCTACACGACGCTCTTCCGATCTNNNGGGCCATTTTCATCAGAAGAC
CACACGTCTGAACTCTTTCCCTACACGACGCTCTTCCGATCTNNNTCATCTACAAGGTTGAGGTTGCT
CACACGTCTGAACTCTTTCCCTACACGACGCTCTTCCGATCTNNNTCATCTACAAGGTTGAGGTTGCT
AGCACACGTCTGAACTCTTTCCCTACACGACGCTCTTCCGATCTNNNATTGCAGTGAGCTATGATTGT
AGCACACGTCTGAACTCTTTCCCTACACGACGCTCTTCCGATCTNNNATTGCAGTGAGCTATGATTGT
AGCACACGTCTGAACTCTTTCCCTACACGACGCTCTTCCGATCTNNNAGGCTGGAGGAACATCATGA
AGCACACGTCTGAACTCTTTCCCTACACGACGCTCTTCCGATCTNNNAGGCTGGAGGAACATCATGA
GCACACGTCTGAACTCTTTCCCTACACGACGCTCTTCCGATCTNNNTGTCCGGAAAGGGTGATTCTTT
GCACACGTCTGAACTCTTTCCCTACACGACGCTCTTCCGATCTNNNTGTCCGGAAAGGGTGATTCTTT
CACACGTCTGAACTCTTTCCCTACACGACGCTCTTCCGATCTNNNTACTGGGAAGTGAGGAGCCCCAT
CACACGTCTGAACTCTTTCCCTACACGACGCTCTTCCGATCTNNNTACTGGGAAGTGAGGAGCCCCAT
GAGCACACGTCTGAACTCTTTCCCTACACGACGCTCTTCCGATCTNNNCTAGTGCTGCAATTCCCTGTT
GAGCACACGTCTGAACTCTTTCCCTACACGACGCTCTTCCGATCTNNNCTAGTGCTGCAATTCCCTGTT
GCACACGTCTGAACTCTTTCCCTACACGACGCTCTTCCGATCTNNNCTGAGAGAAACTGCTGGGGAG
GCACACGTCTGAACTCTTTCCCTACACGACGCTCTTCCGATCTNNNCTGAGAGAAACTGCTGGGGAG
GCACACGTCTGAACTCTTTCCCTACACGACGCTCTTCCGATCTNNNTGCTGGGATTACAGGCATGAG
GCACACGTCTGAACTCTTTCCCTACACGACGCTCTTCCGATCTNNNTGCTGGGATTACAGGCATGAG
AGCACACGTCTGAACTCTTTCCCTACACGACGCTCTTCCGATCTNNNTTCTTGTTGCCGAGATTGGAG
AGCACACGTCTGAACTCTTTCCCTACACGACGCTCTTCCGATCTNNNTTCTTGTTGCCGAGATTGGAG
AGAGCACACGTCTGAACTCTTTCCCTACACGACGCTCTTCCGATCTNNNCTGTAGTCCCAGCTACTCA
AGAGCACACGTCTGAACTCTTTCCCTACACGACGCTCTTCCGATCTNNNCTGTAGTCCCAGCTACTCA
AGCACACGTCTGAACTCTTTCCCTACACGACGCTCTTCCGATCTNNNGCTGCTATTTGAGAAACTGTG
AGCACACGTCTGAACTCTTTCCCTACACGACGCTCTTCCGATCTNNNGCTGCTATTTGAGAAACTGTG
CACGTCTGAACTCTTTCCCTACACGACGCTCTTCCGATCTNNNCCATGGTGTATTTGTGCCACATTCAT
CACGTCTGAACTCTTTCCCTACACGACGCTCTTCCGATCTNNNCCATGGTGTATTTGTGCCACATTCAT
GAGCACACGTCTGAACTCTTTCCCTACACGACGCTCTTCCGATCTNNNTTGGCCATGTTTTCTTCAAGT
GAGCACACGTCTGAACTCTTTCCCTACACGACGCTCTTCCGATCTNNNTTGGCCATGTTTTCTTCAAGT
AGCACACGTCTGAACTCTTTCCCTACACGACGCTCTTCCGATCTNNNTGAGGTACTTGGCTTTGAAGG
AGCACACGTCTGAACTCTTTCCCTACACGACGCTCTTCCGATCTNNNTGAGGTACTTGGCTTTGAAGG
AGCACACGTCTGAACTCTTTCCCTACACGACGCTCTTCCGATCTNNNCCCAAACACTGGGAGACAGAT
AGCACACGTCTGAACTCTTTCCCTACACGACGCTCTTCCGATCTNNNCCCAAACACTGGGAGACAGAT
GCACACGTCTGAACTCTTTCCCTACACGACGCTCTTCCGATCTNNNCCAAAGTGCTGGGATTACATGC
GCACACGTCTGAACTCTTTCCCTACACGACGCTCTTCCGATCTNNNCCAAAGTGCTGGGATTACATGC
AGCACACGTCTGAACTCTTTCCCTACACGACGCTCTTCCGATCTNNNTTCGCTGTCATATTCCTGGCTA
AGCACACGTCTGAACTCTTTCCCTACACGACGCTCTTCCGATCTNNNTTCGCTGTCATATTCCTGGCTA
AGCACACGTCTGAACTCTTTCCCTACACGACGCTCTTCCGATCTNNNAGATTTATCTTGGCCTTGCTG
AGCACACGTCTGAACTCTTTCCCTACACGACGCTCTTCCGATCTNNNAGATTTATCTTGGCCTTGCTG
AGCACACGTCTGAACTCTTTCCCTACACGACGCTCTTCCGATCTNNNACCATCACCTACAAGGAAACCT
AGCACACGTCTGAACTCTTTCCCTACACGACGCTCTTCCGATCTNNNACCATCACCTACAAGGAAACCT
AGCACACGTCTGAACTCTTTCCCTACACGACGCTCTTCCGATCTNNNAAACAAGCACTGGCCTTGAG
AGCACACGTCTGAACTCTTTCCCTACACGACGCTCTTCCGATCTNNNAAACAAGCACTGGCCTTGAG
CACACGTCTGAACTCTTTCCCTACACGACGCTCTTCCGATCTNNNAAGGGAGTGCTTAGATCATGGT
CACACGTCTGAACTCTTTCCCTACACGACGCTCTTCCGATCTNNNAAGGGAGTGCTTAGATCATGGT

GCACACGTCTGAACTCTTTCCCTACACGACGCTCTTCCGATCTNNNAGGATCATTCAGGCATTTGCTCA

GCACACGTCTGAACTCTTTCCCTACACGACGCTCTTCCGATCTNNNAGGATCATTCAGGCATTTGCTCA

AGCACACGTCTGAACTCTTTCCCTACACGACGCTCTTCCGATCTNNNGAGGCAGGAGAATTGTTTGAA

AGCACACGTCTGAACTCTTTCCCTACACGACGCTCTTCCGATCTNNNGAGGCAGGAGAATTGTTTGAA

GAGCACACGTCTGAACTCTTTCCCTACACGACGCTCTTCCGATCTNNNCCCACAACAGAGTCATAGGTT

GAGCACACGTCTGAACTCTTTCCCTACACGACGCTCTTCCGATCTNNNCCCACAACAGAGTCATAGGTT

AGCACACGTCTGAACTCTTTCCCTACACGACGCTCTTCCGATCTNNNAGAATTGCTTGAATTTGGGAG

AGCACACGTCTGAACTCTTTCCCTACACGACGCTCTTCCGATCTNNNAGAATTGCTTGAATTTGGGAG

AGCACACGTCTGAACTCTTTCCCTACACGACGCTCTTCCGATCTNNNAGAGCCTTTGTCACATTCCTGAT

AGCACACGTCTGAACTCTTTCCCTACACGACGCTCTTCCGATCTNNNAGAGCCTTTGTCACATTCCTGAT

GAGCACACGTCTGAACTCTTTCCCTACACGACGCTCTTCCGATCTNNNCCTGATTGTTGGTGGGATCAA

GAGCACACGTCTGAACTCTTTCCCTACACGACGCTCTTCCGATCTNNNCCTGATTGTTGGTGGGATCAA

AGCACACGTCTGAACTCTTTCCCTACACGACGCTCTTCCGATCTNNNCTGTAATCCCAGCTACTCGGAA

AGCACACGTCTGAACTCTTTCCCTACACGACGCTCTTCCGATCTNNNCTGTAATCCCAGCTACTCGGAA

AGCACACGTCTGAACTCTTTCCCTACACGACGCTCTTCCGATCTNNNCACAGACAGAGTCACATAAGCCT

AGCACACGTCTGAACTCTTTCCCTACACGACGCTCTTCCGATCTNNNCACAGACAGAGTCACATAAGCCT

GAGCACACGTCTGAACTCTTTCCCTACACGACGCTCTTCCGATCTNNNGAGGCAGGAGAATCGCTTGA

GAGCACACGTCTGAACTCTTTCCCTACACGACGCTCTTCCGATCTNNNGAGGCAGGAGAATCGCTTGA

GAGCACACGTCTGAACTCTTTCCCTACACGACGCTCTTCCGATCTNNNGAACTACAGCATAAGGAACGC

GAGCACACGTCTGAACTCTTTCCCTACACGACGCTCTTCCGATCTNNNGAACTACAGCATAAGGAACGC

GAGCACACGTCTGAACTCTTTCCCTACACGACGCTCTTCCGATCTNNNGACAGGTCTTTTTCTGCCACT

GAGCACACGTCTGAACTCTTTCCCTACACGACGCTCTTCCGATCTNNNGACAGGTCTTTTTCTGCCACT

AGCACACGTCTGAACTCTTTCCCTACACGACGCTCTTCCGATCTNNNCCACGCCTGGCTAATTTTTGTATC

AGCACACGTCTGAACTCTTTCCCTACACGACGCTCTTCCGATCTNNNCCACGCCTGGCTAATTTTTGTATC

AGCACACGTCTGAACTCTTTCCCTACACGACGCTCTTCCGATCTNNNCCTCTTTCTTCTCCCTCTCCTTCCA

AGCACACGTCTGAACTCTTTCCCTACACGACGCTCTTCCGATCTNNNCCTCTTTCTTCTCCCTCTCCTTCCA

AGCACACGTCTGAACTCTTTCCCTACACGACGCTCTTCCGATCTNNNAGGAGCTTGTAGTGAGCCTAGA

AGCACACGTCTGAACTCTTTCCCTACACGACGCTCTTCCGATCTNNNAGGAGCTTGTAGTGAGCCTAGA

AGAGCACACGTCTGAACTCTTTCCCTACACGACGCTCTTCCGATCTNNNGACAGAGCAAGACTCCGTCTC

AGAGCACACGTCTGAACTCTTTCCCTACACGACGCTCTTCCGATCTNNNGACAGAGCAAGACTCCGTCTC

GAGCACACGTCTGAACTCTTTCCCTACACGACGCTCTTCCGATCTNNNTGGTGAACAGTGACAGTTTGA

GAGCACACGTCTGAACTCTTTCCCTACACGACGCTCTTCCGATCTNNNTGGTGAACAGTGACAGTTTGA

GAGCACACGTCTGAACTCTTTCCCTACACGACGCTCTTCCGATCTNNNGCAACATAGTGAGACCCTGTC

GAGCACACGTCTGAACTCTTTCCCTACACGACGCTCTTCCGATCTNNNGCAACATAGTGAGACCCTGTC

AGCACACGTCTGAACTCTTTCCCTACACGACGCTCTTCCGATCTNNNACGGAAGTCTCGCTCTGTTGCAT

AGCACACGTCTGAACTCTTTCCCTACACGACGCTCTTCCGATCTNNNACGGAAGTCTCGCTCTGTTGCAT

AGCACACGTCTGAACTCTTTCCCTACACGACGCTCTTCCGATCTNNNGCACACCCACCAAGTTCTCTAAT

AGCACACGTCTGAACTCTTTCCCTACACGACGCTCTTCCGATCTNNNGCACACCCACCAAGTTCTCTAAT

AGCACACGTCTGAACTCTTTCCCTACACGACGCTCTTCCGATCTNNNGTAATCCCCAGGTTATCTCAGGC

AGCACACGTCTGAACTCTTTCCCTACACGACGCTCTTCCGATCTNNNGTAATCCCCAGGTTATCTCAGGC

GCACACGTCTGAACTCTTTCCCTACACGACGCTCTTCCGATCTNNNTGTATCCCATCCCAACCCAATACC

GCACACGTCTGAACTCTTTCCCTACACGACGCTCTTCCGATCTNNNTGTATCCCATCCCAACCCAATACC

AGCACACGTCTGAACTCTTTCCCTACACGACGCTCTTCCGATCTNNNAGAATCACTTGAATCCAGGAG

AGCACACGTCTGAACTCTTTCCCTACACGACGCTCTTCCGATCTNNNAGAATCACTTGAATCCAGGAG
AGCACACGTCTGAACTCTTTCCCTACACGACGCTCTTCCGATCTNNNAGGGGTGCACCAAAATTTTCAA
AGCACACGTCTGAACTCTTTCCCTACACGACGCTCTTCCGATCTNNNAGGGGTGCACCAAAATTTTCAA
AGCACACGTCTGAACTCTTTCCCTACACGACGCTCTTCCGATCTNNNTGTCACCACTCTAAACATAGGG
AGCACACGTCTGAACTCTTTCCCTACACGACGCTCTTCCGATCTNNNTGTCACCACTCTAAACATAGGG
GAGCACACGTCTGAACTCTTTCCCTACACGACGCTCTTCCGATCTNNNGTGGTTGCTTTCTCATGATGCT
GAGCACACGTCTGAACTCTTTCCCTACACGACGCTCTTCCGATCTNNNGTGGTTGCTTTCTCATGATGCT
AGAGCACACGTCTGAACTCTTTCCCTACACGACGCTCTTCCGATCTNNNGTCAAAATCAATTGCTGAAG
AGAGCACACGTCTGAACTCTTTCCCTACACGACGCTCTTCCGATCTNNNGTCAAAATCAATTGCTGAAG
AGCACACGTCTGAACTCTTTCCCTACACGACGCTCTTCCGATCTNNNTGCCACTTGGTGTCTCCATTTA
AGCACACGTCTGAACTCTTTCCCTACACGACGCTCTTCCGATCTNNNTGCCACTTGGTGTCTCCATTTA
GAGCACACGTCTGAACTCTTTCCCTACACGACGCTCTTCCGATCTNNNCTTGATGCTCACCAGTACTCCA
GAGCACACGTCTGAACTCTTTCCCTACACGACGCTCTTCCGATCTNNNCTTGATGCTCACCAGTACTCCA
GAGCACACGTCTGAACTCTTTCCCTACACGACGCTCTTCCGATCTNNNTTCTTTTTTTTAGTCGCCCCC
GAGCACACGTCTGAACTCTTTCCCTACACGACGCTCTTCCGATCTNNNTTCTTTTTTTTAGTCGCCCCC
GAGCACACGTCTGAACTCTTTCCCTACACGACGCTCTTCCGATCTNNNACCACTTGTGAGCACTTTTCTT
GAGCACACGTCTGAACTCTTTCCCTACACGACGCTCTTCCGATCTNNNACCACTTGTGAGCACTTTTCTT
AGCACACGTCTGAACTCTTTCCCTACACGACGCTCTTCCGATCTNNNTACATGCCTGTAATCCCAGCTA
AGCACACGTCTGAACTCTTTCCCTACACGACGCTCTTCCGATCTNNNTACATGCCTGTAATCCCAGCTA
GAGCACACGTCTGAACTCTTTCCCTACACGACGCTCTTCCGATCTNNNCACACACTCTAGACATTCACCC
GAGCACACGTCTGAACTCTTTCCCTACACGACGCTCTTCCGATCTNNNCACACACTCTAGACATTCACCC
AGCACACGTCTGAACTCTTTCCCTACACGACGCTCTTCCGATCTNNNTTCTTTCCTCTGTGAGCAAAAGT
AGCACACGTCTGAACTCTTTCCCTACACGACGCTCTTCCGATCTNNNTTCTTTCCTCTGTGAGCAAAAGT
GAGCACACGTCTGAACTCTTTCCCTACACGACGCTCTTCCGATCTNNNCCTGTAATCTCACCTACTGGG
GAGCACACGTCTGAACTCTTTCCCTACACGACGCTCTTCCGATCTNNNCCTGTAATCTCACCTACTGGG
AAGAGCACACGTCTGAACTCTTTCCCTACACGACGCTCTTCCGATCTNNNATGAACTTGTCTTCCCGTCG
AAGAGCACACGTCTGAACTCTTTCCCTACACGACGCTCTTCCGATCTNNNATGAACTTGTCTTCCCGTCG
AGCACACGTCTGAACTCTTTCCCTACACGACGCTCTTCCGATCTNNNTGAGAATCATTTTGACAACCAC
AGCACACGTCTGAACTCTTTCCCTACACGACGCTCTTCCGATCTNNNTGAGAATCATTTTGACAACCAC
GAGCACACGTCTGAACTCTTTCCCTACACGACGCTCTTCCGATCTNNNCAAATCCTGGACACTGTGGTA
GAGCACACGTCTGAACTCTTTCCCTACACGACGCTCTTCCGATCTNNNCAAATCCTGGACACTGTGGTA
GAGCACACGTCTGAACTCTTTCCCTACACGACGCTCTTCCGATCTNNNCCACTAACAGCCACTGTATCCT
GAGCACACGTCTGAACTCTTTCCCTACACGACGCTCTTCCGATCTNNNCCACTAACAGCCACTGTATCCT
AGCACACGTCTGAACTCTTTCCCTACACGACGCTCTTCCGATCTNNNGTATTTTTCGTATTTCTTGCAAA
AGCACACGTCTGAACTCTTTCCCTACACGACGCTCTTCCGATCTNNNGTATTTTTCGTATTTCTTGCAAA
GAGCACACGTCTGAACTCTTTCCCTACACGACGCTCTTCCGATCTNNNCTCTTCTTACAACTGCCTGCCT
GAGCACACGTCTGAACTCTTTCCCTACACGACGCTCTTCCGATCTNNNCTCTTCTTACAACTGCCTGCCT
GAGCACACGTCTGAACTCTTTCCCTACACGACGCTCTTCCGATCTNNNACCTGAACTTGAGGAAAACTG
GAGCACACGTCTGAACTCTTTCCCTACACGACGCTCTTCCGATCTNNNACCTGAACTTGAGGAAAACTG
AGCACACGTCTGAACTCTTTCCCTACACGACGCTCTTCCGATCTNNNCATGGCGTAGGTAGGGACTAT
AGCACACGTCTGAACTCTTTCCCTACACGACGCTCTTCCGATCTNNNCATGGCGTAGGTAGGGACTAT
GAGCACACGTCTGAACTCTTTCCCTACACGACGCTCTTCCGATCTNNNGGGTGGAAAATAAGGTCAGG
GAGCACACGTCTGAACTCTTTCCCTACACGACGCTCTTCCGATCTNNNGGGTGGAAAATAAGGTCAGG

GAGCACACGTCTGAACTCTTTCCCTACACGACGCTCTTCCGATCTNNNAATTCTCACTCCCATCCCATGT

GAGCACACGTCTGAACTCTTTCCCTACACGACGCTCTTCCGATCTNNNAATTCTCACTCCCATCCCATGT

AGCACACGTCTGAACTCTTTCCCTACACGACGCTCTTCCGATCTNNNACCATCAAGTAGTTGTGTGAC

AGCACACGTCTGAACTCTTTCCCTACACGACGCTCTTCCGATCTNNNACCATCAAGTAGTTGTGTGAC

AGAGCACACGTCTGAACTCTTTCCCTACACGACGCTCTTCCGATCTNNNGAGGAAAACTGCTTTTGTC

AGAGCACACGTCTGAACTCTTTCCCTACACGACGCTCTTCCGATCTNNNGAGGAAAACTGCTTTTGTC

AGCACACGTCTGAACTCTTTCCCTACACGACGCTCTTCCGATCTNNNTTTAGGAGCACCATTCATCAGC

AGCACACGTCTGAACTCTTTCCCTACACGACGCTCTTCCGATCTNNNTTTAGGAGCACCATTCATCAGC

GAGCACACGTCTGAACTCTTTCCCTACACGACGCTCTTCCGATCTNNNATGTCAGAAGGAGTAGGAG

GAGCACACGTCTGAACTCTTTCCCTACACGACGCTCTTCCGATCTNNNATGTCAGAAGGAGTAGGAG

GCACACGTCTGAACTCTTTCCCTACACGACGCTCTTCCGATCTNNNGTTTAAATTCTCCACAGATTGTAC

GCACACGTCTGAACTCTTTCCCTACACGACGCTCTTCCGATCTNNNGTTTAAATTCTCCACAGATTGTAC

AAGAGCACACGTCTGAACTCTTTCCCTACACGACGCTCTTCCGATCTNNNTCAGTGTTTTGAAGAACAT

AAGAGCACACGTCTGAACTCTTTCCCTACACGACGCTCTTCCGATCTNNNTCAGTGTTTTGAAGAACAT

GCACACGTCTGAACTCTTTCCCTACACGACGCTCTTCCGATCTNNNTTTCTTTGAGACAGGGTCTTGCT

GCACACGTCTGAACTCTTTCCCTACACGACGCTCTTCCGATCTNNNTTTCTTTGAGACAGGGTCTTGCT

AGCACACGTCTGAACTCTTTCCCTACACGACGCTCTTCCGATCTNNNGTGGTAAGTCATGTATGTTGC

AGCACACGTCTGAACTCTTTCCCTACACGACGCTCTTCCGATCTNNNGTGGTAAGTCATGTATGTTGC

AGCACACGTCTGAACTCTTTCCCTACACGACGCTCTTCCGATCTNNNAGACTATTCCCAAGCACGTCTT

AGCACACGTCTGAACTCTTTCCCTACACGACGCTCTTCCGATCTNNNAGACTATTCCCAAGCACGTCTT

GAGCACACGTCTGAACTCTTTCCCTACACGACGCTCTTCCGATCTNNNGTTGGGTGTGTACATGTGTG

GAGCACACGTCTGAACTCTTTCCCTACACGACGCTCTTCCGATCTNNNGTTGGGTGTGTACATGTGTG

AGCACACGTCTGAACTCTTTCCCTACACGACGCTCTTCCGATCTNNNAACTGTTCTGTCTCATGACTGC

AGCACACGTCTGAACTCTTTCCCTACACGACGCTCTTCCGATCTNNNAACTGTTCTGTCTCATGACTGC

CACACGTCTGAACTCTTTCCCTACACGACGCTCTTCCGATCTNNNGGATTGAGGTGAATCAGTCCAG

CACACGTCTGAACTCTTTCCCTACACGACGCTCTTCCGATCTNNNGGATTGAGGTGAATCAGTCCAG

GCACACGTCTGAACTCTTTCCCTACACGACGCTCTTCCGATCTNNNCCCAGAAATAGACCCACAGGAA

GCACACGTCTGAACTCTTTCCCTACACGACGCTCTTCCGATCTNNNCCCAGAAATAGACCCACAGGAA

AGCACACGTCTGAACTCTTTCCCTACACGACGCTCTTCCGATCTNNNCATGCCTGTAATCCCAGCTACT

AGCACACGTCTGAACTCTTTCCCTACACGACGCTCTTCCGATCTNNNCATGCCTGTAATCCCAGCTACT

GCACACGTCTGAACTCTTTCCCTACACGACGCTCTTCCGATCTNNNACTCCTTTCTCCATAATGTGCTTC

GCACACGTCTGAACTCTTTCCCTACACGACGCTCTTCCGATCTNNNACTCCTTTCTCCATAATGTGCTTC

GAGCACACGTCTGAACTCTTTCCCTACACGACGCTCTTCCGATCTNNNTTTGTATTGGATTGACTGCTG

GAGCACACGTCTGAACTCTTTCCCTACACGACGCTCTTCCGATCTNNNTTTGTATTGGATTGACTGCTG

AGCACACGTCTGAACTCTTTCCCTACACGACGCTCTTCCGATCTNNNAGGCAGGAGAATCATTTGAAC

AGCACACGTCTGAACTCTTTCCCTACACGACGCTCTTCCGATCTNNNAGGCAGGAGAATCATTTGAAC

GAGCACACGTCTGAACTCTTTCCCTACACGACGCTCTTCCGATCTNNNGTTTTGTTTCAAGATTCCCCC

GAGCACACGTCTGAACTCTTTCCCTACACGACGCTCTTCCGATCTNNNGTTTTGTTTCAAGATTCCCCC

GAGCACACGTCTGAACTCTTTCCCTACACGACGCTCTTCCGATCTNNNAAGGTCACTGAAGCTTCCCT

GAGCACACGTCTGAACTCTTTCCCTACACGACGCTCTTCCGATCTNNNAAGGTCACTGAAGCTTCCCT

AGCACACGTCTGAACTCTTTCCCTACACGACGCTCTTCCGATCTNNNTTGAAGAGGATCCCACCTTCCT

AGCACACGTCTGAACTCTTTCCCTACACGACGCTCTTCCGATCTNNNTTGAAGAGGATCCCACCTTCCT

GAGCACACGTCTGAACTCTTTCCCTACACGACGCTCTTCCGATCTNNNCCAAAGTGGGAGAATTGCTT

GAGCACACGTCTGAACTCTTTCCCTACACGACGCTCTTCCGATCTNNNCCAAAGTGGGAGAATTGCTT

GAAGAGCACACGTCTGAACTCTTTCCCTACACGACGCTCTTCCGATCTNNNCAATGCTCTGTGTGGCA

GAAGAGCACACGTCTGAACTCTTTCCCTACACGACGCTCTTCCGATCTNNNCAATGCTCTGTGTGGCA

AGCACACGTCTGAACTCTTTCCCTACACGACGCTCTTCCGATCTNNNATCACTTGAACTTGTGAGGCA

AGCACACGTCTGAACTCTTTCCCTACACGACGCTCTTCCGATCTNNNATCACTTGAACTTGTGAGGCA

GCACACGTCTGAACTCTTTCCCTACACGACGCTCTTCCGATCTNNNTCTCTGTCTTTCCTCCTCTGTCAT

GCACACGTCTGAACTCTTTCCCTACACGACGCTCTTCCGATCTNNNTCTCTGTCTTTCCTCCTCTGTCAT

AGCACACGTCTGAACTCTTTCCCTACACGACGCTCTTCCGATCTNNNGCAACAGAGCGAGATTCCATCT

AGCACACGTCTGAACTCTTTCCCTACACGACGCTCTTCCGATCTNNNGCAACAGAGCGAGATTCCATCT

AGCACACGTCTGAACTCTTTCCCTACACGACGCTCTTCCGATCTNNNTGACATTGAGCCTTCTTTCTCTC

AGCACACGTCTGAACTCTTTCCCTACACGACGCTCTTCCGATCTNNNTGACATTGAGCCTTCTTTCTCTC

AGCACACGTCTGAACTCTTTCCCTACACGACGCTCTTCCGATCTNNNACACAAAAATTAGCTGGGGTG

AGCACACGTCTGAACTCTTTCCCTACACGACGCTCTTCCGATCTNNNACACAAAAATTAGCTGGGGTG

GCACACGTCTGAACTCTTTCCCTACACGACGCTCTTCCGATCTNNNGTCTACCATTTTGATAGCTTAGC

GCACACGTCTGAACTCTTTCCCTACACGACGCTCTTCCGATCTNNNGTCTACCATTTTGATAGCTTAGC

GCACACGTCTGAACTCTTTCCCTACACGACGCTCTTCCGATCTNNNTTTCAGCTCCTATATCATTTTTGCA

GCACACGTCTGAACTCTTTCCCTACACGACGCTCTTCCGATCTNNNTTTCAGCTCCTATATCATTTTTGCA

GCACACGTCTGAACTCTTTCCCTACACGACGCTCTTCCGATCTNNNGCAGGAGAATTTCTTGAACCCA

GCACACGTCTGAACTCTTTCCCTACACGACGCTCTTCCGATCTNNNGCAGGAGAATTTCTTGAACCCA

AGCACACGTCTGAACTCTTTCCCTACACGACGCTCTTCCGATCTNNNGTCTCAAGCTCCTGACCAAGTA

AGCACACGTCTGAACTCTTTCCCTACACGACGCTCTTCCGATCTNNNGTCTCAAGCTCCTGACCAAGTA

AGCACACGTCTGAACTCTTTCCCTACACGACGCTCTTCCGATCTNNNGCAGAGACACAAATCCAAACCA

AGCACACGTCTGAACTCTTTCCCTACACGACGCTCTTCCGATCTNNNGCAGAGACACAAATCCAAACCA

AGCACACGTCTGAACTCTTTCCCTACACGACGCTCTTCCGATCTNNNATATCCAATTTCATATACTGGCT

AGCACACGTCTGAACTCTTTCCCTACACGACGCTCTTCCGATCTNNNATATCCAATTTCATATACTGGCT

GCACACGTCTGAACTCTTTCCCTACACGACGCTCTTCCGATCTNNNGAGTAGTTGGCCTTCCCTTTGTC

GCACACGTCTGAACTCTTTCCCTACACGACGCTCTTCCGATCTNNNGAGTAGTTGGCCTTCCCTTTGTC

GAGCACACGTCTGAACTCTTTCCCTACACGACGCTCTTCCGATCTNNNAACAAAAAGAAGCCAGAGG

GAGCACACGTCTGAACTCTTTCCCTACACGACGCTCTTCCGATCTNNNAACAAAAAGAAGCCAGAGG

AGCACACGTCTGAACTCTTTCCCTACACGACGCTCTTCCGATCTNNNAAAAATTAGCTGTGCGTGGTG

AGCACACGTCTGAACTCTTTCCCTACACGACGCTCTTCCGATCTNNNAAAAATTAGCTGTGCGTGGTG

AGAGCACACGTCTGAACTCTTTCCCTACACGACGCTCTTCCGATCTNNNGCACACATCTCTAATCCCAG

AGAGCACACGTCTGAACTCTTTCCCTACACGACGCTCTTCCGATCTNNNGCACACATCTCTAATCCCAG

GAGCACACGTCTGAACTCTTTCCCTACACGACGCTCTTCCGATCTNNNTGTGATCCTAGCACTTTGGGA

GAGCACACGTCTGAACTCTTTCCCTACACGACGCTCTTCCGATCTNNNTGTGATCCTAGCACTTTGGGA

AGCACACGTCTGAACTCTTTCCCTACACGACGCTCTTCCGATCTNNNGTGACAGAAAAAGACTTCGCC

AGCACACGTCTGAACTCTTTCCCTACACGACGCTCTTCCGATCTNNNGTGACAGAAAAAGACTTCGCC

AGCACACGTCTGAACTCTTTCCCTACACGACGCTCTTCCGATCTNNNAAAAAAATTAGCTGGGCGTGG

AGCACACGTCTGAACTCTTTCCCTACACGACGCTCTTCCGATCTNNNAAAAAAATTAGCTGGGCGTGG

AGCACACGTCTGAACTCTTTCCCTACACGACGCTCTTCCGATCTNNNTCCTATTTTCATTAGGGCAGTC

AGCACACGTCTGAACTCTTTCCCTACACGACGCTCTTCCGATCTNNNTCCTATTTTCATTAGGGCAGTC

AGCACACGTCTGAACTCTTTCCCTACACGACGCTCTTCCGATCTNNNCTCATGCCTGTAATCCCAGCTA

AGCACACGTCTGAACTCTTTCCCTACACGACGCTCTTCCGATCTNNNCTCATGCCTGTAATCCCAGCTA

GCACACGTCTGAACTCTTTCCCTACACGACGCTCTTCCGATCTNNNCTCTTGAACTCCTGACCTTGTGA
GCACACGTCTGAACTCTTTCCCTACACGACGCTCTTCCGATCTNNNCTCTTGAACTCCTGACCTTGTGA
AGCACACGTCTGAACTCTTTCCCTACACGACGCTCTTCCGATCTNNNCTGTGAATCTATCTGGTCCTG
AGCACACGTCTGAACTCTTTCCCTACACGACGCTCTTCCGATCTNNNCTGTGAATCTATCTGGTCCTG
GCACACGTCTGAACTCTTTCCCTACACGACGCTCTTCCGATCTNNNTTTGTTGGATTTTTCACAAAA
GCACACGTCTGAACTCTTTCCCTACACGACGCTCTTCCGATCTNNNTTTGTTGGATTTTTCACAAAA
AGCACACGTCTGAACTCTTTCCCTACACGACGCTCTTCCGATCTNNNTGTCTCAAGTTTGGTGATTTC
AGCACACGTCTGAACTCTTTCCCTACACGACGCTCTTCCGATCTNNNTGTCTCAAGTTTGGTGATTTC
GCACACGTCTGAACTCTTTCCCTACACGACGCTCTTCCGATCTNNNTTTGTGCCACAAACATTCCACTTC
GCACACGTCTGAACTCTTTCCCTACACGACGCTCTTCCGATCTNNNTTTGTGCCACAAACATTCCACTTC
GCACACGTCTGAACTCTTTCCCTACACGACGCTCTTCCGATCTNNNGAGTTTGGGCAGAAGGTTGTT
GCACACGTCTGAACTCTTTCCCTACACGACGCTCTTCCGATCTNNNGAGTTTGGGCAGAAGGTTGTT
AGCACACGTCTGAACTCTTTCCCTACACGACGCTCTTCCGATCTNNNGTTGCACTTTCCTTCCTTGTTG
AGCACACGTCTGAACTCTTTCCCTACACGACGCTCTTCCGATCTNNNGTTGCACTTTCCTTCCTTGTTG
AGCACACGTCTGAACTCTTTCCCTACACGACGCTCTTCCGATCTNNNGGGAGAGGCAAAAGGTGAG
AGCACACGTCTGAACTCTTTCCCTACACGACGCTCTTCCGATCTNNNGGGAGAGGCAAAAGGTGAG
GCACACGTCTGAACTCTTTCCCTACACGACGCTCTTCCGATCTNNNCTGTAATCCCAGCTACTCAGGAG
GCACACGTCTGAACTCTTTCCCTACACGACGCTCTTCCGATCTNNNCTGTAATCCCAGCTACTCAGGAG
GCACACGTCTGAACTCTTTCCCTACACGACGCTCTTCCGATCTNNNGACCCTCCTGGAAACTCACCTCAT
GCACACGTCTGAACTCTTTCCCTACACGACGCTCTTCCGATCTNNNGACCCTCCTGGAAACTCACCTCAT
AGCACACGTCTGAACTCTTTCCCTACACGACGCTCTTCCGATCTNNNGGAATATTTCTGTGCTTGCAG
AGCACACGTCTGAACTCTTTCCCTACACGACGCTCTTCCGATCTNNNGGAATATTTCTGTGCTTGCAG
GCACACGTCTGAACTCTTTCCCTACACGACGCTCTTCCGATCTNNNCTCATCTTCTAGCTCCTTCTTCAT
GCACACGTCTGAACTCTTTCCCTACACGACGCTCTTCCGATCTNNNCTCATCTTCTAGCTCCTTCTTCAT
AGCACACGTCTGAACTCTTTCCCTACACGACGCTCTTCCGATCTNNNGTTTCCTGAGTAGCTGGGACT
AGCACACGTCTGAACTCTTTCCCTACACGACGCTCTTCCGATCTNNNGTTTCCTGAGTAGCTGGGACT
AGCACACGTCTGAACTCTTTCCCTACACGACGCTCTTCCGATCTNNNCCCTCTACATTGATGCCAAAA
AGCACACGTCTGAACTCTTTCCCTACACGACGCTCTTCCGATCTNNNCCCTCTACATTGATGCCAAAA
GAGCACACGTCTGAACTCTTTCCCTACACGACGCTCTTCCGATCTNNNGAGGCTGCAGTGAGCTATG
GAGCACACGTCTGAACTCTTTCCCTACACGACGCTCTTCCGATCTNNNGAGGCTGCAGTGAGCTATG
ACACGTCTGAACTCTTTCCCTACACGACGCTCTTCCGATCTNNNTCCACACCCCCAGGATTCTTCATCG
ACACGTCTGAACTCTTTCCCTACACGACGCTCTTCCGATCTNNNTCCACACCCCCAGGATTCTTCATCG
GCACACGTCTGAACTCTTTCCCTACACGACGCTCTTCCGATCTNNNACATGGTGAAACTCCGTCTCTA
GCACACGTCTGAACTCTTTCCCTACACGACGCTCTTCCGATCTNNNACATGGTGAAACTCCGTCTCTA
GCACACGTCTGAACTCTTTCCCTACACGACGCTCTTCCGATCTNNNTGGTTAGTGCCTTGTTTTGGTT
GCACACGTCTGAACTCTTTCCCTACACGACGCTCTTCCGATCTNNNTGGTTAGTGCCTTGTTTTGGTT
AGCACACGTCTGAACTCTTTCCCTACACGACGCTCTTCCGATCTNNNCTCTATTGCCAAGCGTTCTGAA
AGCACACGTCTGAACTCTTTCCCTACACGACGCTCTTCCGATCTNNNCTCTATTGCCAAGCGTTCTGAA
GCACACGTCTGAACTCTTTCCCTACACGACGCTCTTCCGATCTNNNATGTGTGGGTCTTCTACGTCAA
GCACACGTCTGAACTCTTTCCCTACACGACGCTCTTCCGATCTNNNATGTGTGGGTCTTCTACGTCAA
GAGCACACGTCTGAACTCTTTCCCTACACGACGCTCTTCCGATCTNNNACACACCTTCTATGTACCCACC
GAGCACACGTCTGAACTCTTTCCCTACACGACGCTCTTCCGATCTNNNACACACCTTCTATGTACCCACC
AAGAGCACACGTCTGAACTCTTTCCCTACACGACGCTCTTCCGATCTNNNACATAATGTAGACCGCAA

AAGAGCACACGTCTGAACTCTTTCCCTACACGACGCTCTTCCGATCTNNNACATAATGTAGACCGCAA
AGCACACGTCTGAACTCTTTCCCTACACGACGCTCTTCCGATCTNNNTGCTAAACAGAAAAGAGAAAG
AGCACACGTCTGAACTCTTTCCCTACACGACGCTCTTCCGATCTNNNTGCTAAACAGAAAAGAGAAAG
GCACACGTCTGAACTCTTTCCCTACACGACGCTCTTCCGATCTNNNTTGGAGGTCAGGTTAGTCAAC
GCACACGTCTGAACTCTTTCCCTACACGACGCTCTTCCGATCTNNNTTGGAGGTCAGGTTAGTCAAC
CACACGTCTGAACTCTTTCCCTACACGACGCTCTTCCGATCTNNNCAGGGTTTGGGAACAGAAGACT
CACACGTCTGAACTCTTTCCCTACACGACGCTCTTCCGATCTNNNCAGGGTTTGGGAACAGAAGACT
AGCACACGTCTGAACTCTTTCCCTACACGACGCTCTTCCGATCTNNNTTGAGATGGAGTCTTGCTCTG
AGCACACGTCTGAACTCTTTCCCTACACGACGCTCTTCCGATCTNNNTTGAGATGGAGTCTTGCTCTG
AGCACACGTCTGAACTCTTTCCCTACACGACGCTCTTCCGATCTNNNGTCTCTCCAGCCAGAAAGAGT
AGCACACGTCTGAACTCTTTCCCTACACGACGCTCTTCCGATCTNNNGTCTCTCCAGCCAGAAAGAGT
AGCACACGTCTGAACTCTTTCCCTACACGACGCTCTTCCGATCTNNNTTTAATGAAGAAGAGAAGTC
AGCACACGTCTGAACTCTTTCCCTACACGACGCTCTTCCGATCTNNNTTTAATGAAGAAGAGAAGTC
GAGCACACGTCTGAACTCTTTCCCTACACGACGCTCTTCCGATCTNNNTCAGAGATACTCCATGGGTG
GAGCACACGTCTGAACTCTTTCCCTACACGACGCTCTTCCGATCTNNNTCAGAGATACTCCATGGGTG
GAGCACACGTCTGAACTCTTTCCCTACACGACGCTCTTCCGATCTNNNAAACAATTCTCCTGTCTCAGC
GAGCACACGTCTGAACTCTTTCCCTACACGACGCTCTTCCGATCTNNNAAACAATTCTCCTGTCTCAGC
AGCACACGTCTGAACTCTTTCCCTACACGACGCTCTTCCGATCTNNNACATTGAGAAAAGGAGGAA
AGCACACGTCTGAACTCTTTCCCTACACGACGCTCTTCCGATCTNNNACATTGAGAAAAGGAGGAA
AGCACACGTCTGAACTCTTTCCCTACACGACGCTCTTCCGATCTNNNCTGAAGCCTTCTGTTCACATGA
AGCACACGTCTGAACTCTTTCCCTACACGACGCTCTTCCGATCTNNNCTGAAGCCTTCTGTTCACATGA
GAGCACACGTCTGAACTCTTTCCCTACACGACGCTCTTCCGATCTNNNGTATCCCCAGCTACTAAAGA
GAGCACACGTCTGAACTCTTTCCCTACACGACGCTCTTCCGATCTNNNGTATCCCCAGCTACTAAAGA
GCACACGTCTGAACTCTTTCCCTACACGACGCTCTTCCGATCTNNNTGGGTGACAGAGTGAGACTCCC
GCACACGTCTGAACTCTTTCCCTACACGACGCTCTTCCGATCTNNNTGGGTGACAGAGTGAGACTCCC
GCACACGTCTGAACTCTTTCCCTACACGACGCTCTTCCGATCTNNNCAAGACTCAGCAGCAAATCTCTC
GCACACGTCTGAACTCTTTCCCTACACGACGCTCTTCCGATCTNNNCAAGACTCAGCAGCAAATCTCTC
AGCACACGTCTGAACTCTTTCCCTACACGACGCTCTTCCGATCTNNNTTCTCCCCTGTTCCTCTAAGGAT
AGCACACGTCTGAACTCTTTCCCTACACGACGCTCTTCCGATCTNNNTTCTCCCCTGTTCCTCTAAGGAT
GCACACGTCTGAACTCTTTCCCTACACGACGCTCTTCCGATCTNNNAATCATGTGGTGATCTCTTGCAC
GCACACGTCTGAACTCTTTCCCTACACGACGCTCTTCCGATCTNNNAATCATGTGGTGATCTCTTGCAC
GCACACGTCTGAACTCTTTCCCTACACGACGCTCTTCCGATCTNNNGGAGAATCACTTGAACCTGGG
GCACACGTCTGAACTCTTTCCCTACACGACGCTCTTCCGATCTNNNGGAGAATCACTTGAACCTGGG
AGCACACGTCTGAACTCTTTCCCTACACGACGCTCTTCCGATCTNNNTTTGTTGCTCTATCTTGCAGC
AGCACACGTCTGAACTCTTTCCCTACACGACGCTCTTCCGATCTNNNTTTGTTGCTCTATCTTGCAGC
AGAGCACACGTCTGAACTCTTTCCCTACACGACGCTCTTCCGATCTNNNGAGAGAGAGAGAGAGGG
AGAGCACACGTCTGAACTCTTTCCCTACACGACGCTCTTCCGATCTNNNGAGAGAGAGAGAGAGGG
AGCACACGTCTGAACTCTTTCCCTACACGACGCTCTTCCGATCTNNNATCAAAATCCTACCTTTACTCTT
AGCACACGTCTGAACTCTTTCCCTACACGACGCTCTTCCGATCTNNNATCAAAATCCTACCTTTACTCTT
GCACACGTCTGAACTCTTTCCCTACACGACGCTCTTCCGATCTNNNTGTTGATATGGGTGCCTTACAG
GCACACGTCTGAACTCTTTCCCTACACGACGCTCTTCCGATCTNNNTGTTGATATGGGTGCCTTACAG
AGCACACGTCTGAACTCTTTCCCTACACGACGCTCTTCCGATCTNNNAGGCTGAGGCAGAATTACTT
AGCACACGTCTGAACTCTTTCCCTACACGACGCTCTTCCGATCTNNNAGGCTGAGGCAGAATTACTT

AGCACACGTCTGAACTCTTTCCCTACACGACGCTCTTCCGATCTNNNCCGGCTGATTTCTGGGTTGGCA

AGCACACGTCTGAACTCTTTCCCTACACGACGCTCTTCCGATCTNNNCCGGCTGATTTCTGGGTTGGCA

AGAGCACACGTCTGAACTCTTTCCCTACACGACGCTCTTCCGATCTNNNTGAGGCAGGAGAATACTGT

AGAGCACACGTCTGAACTCTTTCCCTACACGACGCTCTTCCGATCTNNNTGAGGCAGGAGAATACTGT

AGAGCACACGTCTGAACTCTTTCCCTACACGACGCTCTTCCGATCTNNNTTATTCCCACTTGCCAACTCT

AGAGCACACGTCTGAACTCTTTCCCTACACGACGCTCTTCCGATCTNNNTTATTCCCACTTGCCAACTCT

GAGCACACGTCTGAACTCTTTCCCTACACGACGCTCTTCCGATCTNNNGAAACATGGTGAGACCCCTGT

GAGCACACGTCTGAACTCTTTCCCTACACGACGCTCTTCCGATCTNNNGAAACATGGTGAGACCCCTGT

GCACACGTCTGAACTCTTTCCCTACACGACGCTCTTCCGATCTNNNTAATACCCACTTCCTCAACAGCCC

GCACACGTCTGAACTCTTTCCCTACACGACGCTCTTCCGATCTNNNTAATACCCACTTCCTCAACAGCCC

GAGCACACGTCTGAACTCTTTCCCTACACGACGCTCTTCCGATCTNNNACTTAGCTTTTGAGGCTCATCC

GAGCACACGTCTGAACTCTTTCCCTACACGACGCTCTTCCGATCTNNNACTTAGCTTTTGAGGCTCATCC

GCACACGTCTGAACTCTTTCCCTACACGACGCTCTTCCGATCTNNNACACTGAGATGCTTCTTGTGCTA

GCACACGTCTGAACTCTTTCCCTACACGACGCTCTTCCGATCTNNNACACTGAGATGCTTCTTGTGCTA

AGAGCACACGTCTGAACTCTTTCCCTACACGACGCTCTTCCGATCTNNNGGACAGAAAACGAAACACC

AGAGCACACGTCTGAACTCTTTCCCTACACGACGCTCTTCCGATCTNNNGGACAGAAAACGAAACACC

GAGCACACGTCTGAACTCTTTCCCTACACGACGCTCTTCCGATCTNNNGGTGGATCACTAGGTCAGG

GAGCACACGTCTGAACTCTTTCCCTACACGACGCTCTTCCGATCTNNNGGTGGATCACTAGGTCAGG

GCACACGTCTGAACTCTTTCCCTACACGACGCTCTTCCGATCTNNNGGGCAACAAGAGCAAAACTCAA

GCACACGTCTGAACTCTTTCCCTACACGACGCTCTTCCGATCTNNNGGGCAACAAGAGCAAAACTCAA

AGCACACGTCTGAACTCTTTCCCTACACGACGCTCTTCCGATCTNNNTGTGCATCCACATACATAGGGA

AGCACACGTCTGAACTCTTTCCCTACACGACGCTCTTCCGATCTNNNTGTGCATCCACATACATAGGGA

GAGCACACGTCTGAACTCTTTCCCTACACGACGCTCTTCCGATCTNNNAGGAGAATGGCATGAACCCG

GAGCACACGTCTGAACTCTTTCCCTACACGACGCTCTTCCGATCTNNNAGGAGAATGGCATGAACCCG

GAGCACACGTCTGAACTCTTTCCCTACACGACGCTCTTCCGATCTNNNCTGGAGTGTCAGCCATTTTTT

GAGCACACGTCTGAACTCTTTCCCTACACGACGCTCTTCCGATCTNNNCTGGAGTGTCAGCCATTTTTT

GAGCACACGTCTGAACTCTTTCCCTACACGACGCTCTTCCGATCTNNNGAACTTTGCACCAGCTTCTACA

GAGCACACGTCTGAACTCTTTCCCTACACGACGCTCTTCCGATCTNNNGAACTTTGCACCAGCTTCTACA

AGCACACGTCTGAACTCTTTCCCTACACGACGCTCTTCCGATCTNNNACAAAAATTAGCCAGGTGTGG

AGCACACGTCTGAACTCTTTCCCTACACGACGCTCTTCCGATCTNNNACAAAAATTAGCCAGGTGTGG

GAGCACACGTCTGAACTCTTTCCCTACACGACGCTCTTCCGATCTNNNTATCCCTGGGAGAAAGCCAG

GAGCACACGTCTGAACTCTTTCCCTACACGACGCTCTTCCGATCTNNNTATCCCTGGGAGAAAGCCAG

GAGCACACGTCTGAACTCTTTCCCTACACGACGCTCTTCCGATCTNNNTGAGGGGCATGTATGAGTGG

GAGCACACGTCTGAACTCTTTCCCTACACGACGCTCTTCCGATCTNNNTGAGGGGCATGTATGAGTGG

GAGCACACGTCTGAACTCTTTCCCTACACGACGCTCTTCCGATCTNNNGAGGCAGGAGAATAGCGTG

GAGCACACGTCTGAACTCTTTCCCTACACGACGCTCTTCCGATCTNNNGAGGCAGGAGAATAGCGTG

GCACACGTCTGAACTCTTTCCCTACACGACGCTCTTCCGATCTNNNCCTAGTCGTCCTCTCCTCGAGCA

GCACACGTCTGAACTCTTTCCCTACACGACGCTCTTCCGATCTNNNCCTAGTCGTCCTCTCCTCGAGCA

AGCACACGTCTGAACTCTTTCCCTACACGACGCTCTTCCGATCTNNNGTGAAGCTTTCCCAGGTGTTTT

AGCACACGTCTGAACTCTTTCCCTACACGACGCTCTTCCGATCTNNNGTGAAGCTTTCCCAGGTGTTTT

GCACACGTCTGAACTCTTTCCCTACACGACGCTCTTCCGATCTNNNTATTTTCTCCCAGTCTGCAGCTT

GCACACGTCTGAACTCTTTCCCTACACGACGCTCTTCCGATCTNNNTATTTTCTCCCAGTCTGCAGCTT

GAGCACACGTCTGAACTCTTTCCCTACACGACGCTCTTCCGATCTNNNAGCAATTCTTCTACCTCAGCCT

GAGCACACGTCTGAACTCTTTCCCTACACGACGCTCTTCCGATCTNNNAGCAATTCTTCTACCTCAGCCT

AGCACACGTCTGAACTCTTTCCCTACACGACGCTCTTCCGATCTNNNCGATGCCCGAAACAGGTCAGCA

AGCACACGTCTGAACTCTTTCCCTACACGACGCTCTTCCGATCTNNNCGATGCCCGAAACAGGTCAGCA

AGCACACGTCTGAACTCTTTCCCTACACGACGCTCTTCCGATCTNNNGGTGACAGAGTGAGACTCCTTC

AGCACACGTCTGAACTCTTTCCCTACACGACGCTCTTCCGATCTNNNGGTGACAGAGTGAGACTCCTTC

GAGCACACGTCTGAACTCTTTCCCTACACGACGCTCTTCCGATCTNNNATCATGTTCTGGTCCTTCCTTG

GAGCACACGTCTGAACTCTTTCCCTACACGACGCTCTTCCGATCTNNNATCATGTTCTGGTCCTTCCTTG

ACACGTCTGAACTCTTTCCCTACACGACGCTCTTCCGATCTNNNGCGTGGAACAGTTATTACAGCTGCA

ACACGTCTGAACTCTTTCCCTACACGACGCTCTTCCGATCTNNNGCGTGGAACAGTTATTACAGCTGCA

GCACACGTCTGAACTCTTTCCCTACACGACGCTCTTCCGATCTNNNTGCTTCTAATTCACCTTGTGCATC

GCACACGTCTGAACTCTTTCCCTACACGACGCTCTTCCGATCTNNNTGCTTCTAATTCACCTTGTGCATC

GAGCACACGTCTGAACTCTTTCCCTACACGACGCTCTTCCGATCTNNNTGGTCCTGGAATTTTTCACTGC

GAGCACACGTCTGAACTCTTTCCCTACACGACGCTCTTCCGATCTNNNTGGTCCTGGAATTTTTCACTGC

GAGCACACGTCTGAACTCTTTCCCTACACGACGCTCTTCCGATCTNNNAGGATTGCAAACCCTAGTCTCT

GAGCACACGTCTGAACTCTTTCCCTACACGACGCTCTTCCGATCTNNNAGGATTGCAAACCCTAGTCTCT

AGCACACGTCTGAACTCTTTCCCTACACGACGCTCTTCCGATCTNNNCTTGCAATACTTGTGGTGACTG

AGCACACGTCTGAACTCTTTCCCTACACGACGCTCTTCCGATCTNNNCTTGCAATACTTGTGGTGACTG

GAGCACACGTCTGAACTCTTTCCCTACACGACGCTCTTCCGATCTNNNCCCAACCAGCTTGTAGCCTAAT

GAGCACACGTCTGAACTCTTTCCCTACACGACGCTCTTCCGATCTNNNCCCAACCAGCTTGTAGCCTAAT

GCACACGTCTGAACTCTTTCCCTACACGACGCTCTTCCGATCTNNNTTTGTACTGCTGAATGTGAAGGC

GCACACGTCTGAACTCTTTCCCTACACGACGCTCTTCCGATCTNNNTTTGTACTGCTGAATGTGAAGGC

GAGCACACGTCTGAACTCTTTCCCTACACGACGCTCTTCCGATCTNNNTGTGCTTTGGCATTTATACAGC

GAGCACACGTCTGAACTCTTTCCCTACACGACGCTCTTCCGATCTNNNTGTGCTTTGGCATTTATACAGC

AGCACACGTCTGAACTCTTTCCCTACACGACGCTCTTCCGATCTNNNTTTCCAACTTGGTTGCATTCTCCC

AGCACACGTCTGAACTCTTTCCCTACACGACGCTCTTCCGATCTNNNTTTCCAACTTGGTTGCATTCTCCC

AGAGCACACGTCTGAACTCTTTCCCTACACGACGCTCTTCCGATCTNNNGCAGGAAAACGGTGTGAACC

AGAGCACACGTCTGAACTCTTTCCCTACACGACGCTCTTCCGATCTNNNGCAGGAAAACGGTGTGAACC

GAGCACACGTCTGAACTCTTTCCCTACACGACGCTCTTCCGATCTNNNAGAAATGGGAGGATCACTTG

GAGCACACGTCTGAACTCTTTCCCTACACGACGCTCTTCCGATCTNNNAGAAATGGGAGGATCACTTG

ACACGTCTGAACTCTTTCCCTACACGACGCTCTTCCGATCTNNNTCTAATCCCACCACACAGAGGTACAT

ACACGTCTGAACTCTTTCCCTACACGACGCTCTTCCGATCTNNNTCTAATCCCACCACACAGAGGTACAT

AGCACACGTCTGAACTCTTTCCCTACACGACGCTCTTCCGATCTNNNCTCCTTTCCCAACACAGAGGGCA

AGCACACGTCTGAACTCTTTCCCTACACGACGCTCTTCCGATCTNNNCTCCTTTCCCAACACAGAGGGCA

GAGCACACGTCTGAACTCTTTCCCTACACGACGCTCTTCCGATCTNNNGAGGACATTGACTGGGGAGT

GAGCACACGTCTGAACTCTTTCCCTACACGACGCTCTTCCGATCTNNNGAGGACATTGACTGGGGAGT

GAGCACACGTCTGAACTCTTTCCCTACACGACGCTCTTCCGATCTNNNGCTGAGACAGGAGAATCACTT

GAGCACACGTCTGAACTCTTTCCCTACACGACGCTCTTCCGATCTNNNGCTGAGACAGGAGAATCACTT

AGCACACGTCTGAACTCTTTCCCTACACGACGCTCTTCCGATCTNNNCTGTCGTCTAGCAGAGAACCTTT

AGCACACGTCTGAACTCTTTCCCTACACGACGCTCTTCCGATCTNNNCTGTCGTCTAGCAGAGAACCTTT

AGCACACGTCTGAACTCTTTCCCTACACGACGCTCTTCCGATCTNNNTGTAGGACACCGTCATGGTACT

AGCACACGTCTGAACTCTTTCCCTACACGACGCTCTTCCGATCTNNNTGTAGGACACCGTCATGGTACT

GAGCACACGTCTGAACTCTTTCCCTACACGACGCTCTTCCGATCTNNNTTTTACGGATGAGGAAAGGG

GAGCACACGTCTGAACTCTTTCCCTACACGACGCTCTTCCGATCTNNNTTTTACGGATGAGGAAAGGG

GAGCACACGTCTGAACTCTTTCCCTACACGACGCTCTTCCGATCTNNNCTAGAGTGCAGTGGTACGATC

GAGCACACGTCTGAACTCTTTCCCTACACGACGCTCTTCCGATCTNNNCTAGAGTGCAGTGGTACGATC

GAGCACACGTCTGAACTCTTTCCCTACACGACGCTCTTCCGATCTNNNTTGCTGTGACTTCTGTTGGCT

GAGCACACGTCTGAACTCTTTCCCTACACGACGCTCTTCCGATCTNNNTTGCTGTGACTTCTGTTGGCT

GAGCACACGTCTGAACTCTTTCCCTACACGACGCTCTTCCGATCTNNNGTGAGGTGGTGATGGTTTTG

GAGCACACGTCTGAACTCTTTCCCTACACGACGCTCTTCCGATCTNNNGTGAGGTGGTGATGGTTTTG

GAGCACACGTCTGAACTCTTTCCCTACACGACGCTCTTCCGATCTNNNAAAAACATGGTGGCTTCTGTC

GAGCACACGTCTGAACTCTTTCCCTACACGACGCTCTTCCGATCTNNNAAAAACATGGTGGCTTCTGTC

GAGCACACGTCTGAACTCTTTCCCTACACGACGCTCTTCCGATCTNNNTCTGTGCCTCCATCAAAATACC

GAGCACACGTCTGAACTCTTTCCCTACACGACGCTCTTCCGATCTNNNTCTGTGCCTCCATCAAAATACC

AGCACACGTCTGAACTCTTTCCCTACACGACGCTCTTCCGATCTNNNATCCATTGCTGGTGAACTAGTGT

AGCACACGTCTGAACTCTTTCCCTACACGACGCTCTTCCGATCTNNNATCCATTGCTGGTGAACTAGTGT

AGAGCACACGTCTGAACTCTTTCCCTACACGACGCTCTTCCGATCTNNNGCCCCTTTGAACTCACCTCTTT

AGAGCACACGTCTGAACTCTTTCCCTACACGACGCTCTTCCGATCTNNNGCCCCTTTGAACTCACCTCTTT

GAGCACACGTCTGAACTCTTTCCCTACACGACGCTCTTCCGATCTNNNTCAGTTTCTACATCAGGACAG

GAGCACACGTCTGAACTCTTTCCCTACACGACGCTCTTCCGATCTNNNTCAGTTTCTACATCAGGACAG

CACACGTCTGAACTCTTTCCCTACACGACGCTCTTCCGATCTNNNCTCCCCTCATCCCACTAAGACATCG

CACACGTCTGAACTCTTTCCCTACACGACGCTCTTCCGATCTNNNCTCCCCTCATCCCACTAAGACATCG

GAGCACACGTCTGAACTCTTTCCCTACACGACGCTCTTCCGATCTNNNACGTGTAGTTCTGGTTACTTGC

GAGCACACGTCTGAACTCTTTCCCTACACGACGCTCTTCCGATCTNNNACGTGTAGTTCTGGTTACTTGC

GCACACGTCTGAACTCTTTCCCTACACGACGCTCTTCCGATCTNNNAGCCATATTTTTGGGAGGCAGAC

GCACACGTCTGAACTCTTTCCCTACACGACGCTCTTCCGATCTNNNAGCCATATTTTTGGGAGGCAGAC

GAGCACACGTCTGAACTCTTTCCCTACACGACGCTCTTCCGATCTNNNTGCTTTATCCTTAGGTTCCTCT

GAGCACACGTCTGAACTCTTTCCCTACACGACGCTCTTCCGATCTNNNTGCTTTATCCTTAGGTTCCTCT

AGCACACGTCTGAACTCTTTCCCTACACGACGCTCTTCCGATCTNNNAATGAGTAGGTCTTTCTCTGGC

AGCACACGTCTGAACTCTTTCCCTACACGACGCTCTTCCGATCTNNNAATGAGTAGGTCTTTCTCTGGC

GCACACGTCTGAACTCTTTCCCTACACGACGCTCTTCCGATCTNNNAATGCAGAAGGGGTACTCTCAAC

GCACACGTCTGAACTCTTTCCCTACACGACGCTCTTCCGATCTNNNAATGCAGAAGGGGTACTCTCAAC

GCACACGTCTGAACTCTTTCCCTACACGACGCTCTTCCGATCTNNNAAAAACCAAGCTGTGAGTGGATC

GCACACGTCTGAACTCTTTCCCTACACGACGCTCTTCCGATCTNNNAAAAACCAAGCTGTGAGTGGATC

AGCACACGTCTGAACTCTTTCCCTACACGACGCTCTTCCGATCTNNNGGTTTCTGTGAGCCAAGATTGT

AGCACACGTCTGAACTCTTTCCCTACACGACGCTCTTCCGATCTNNNGGTTTCTGTGAGCCAAGATTGT

GAGCACACGTCTGAACTCTTTCCCTACACGACGCTCTTCCGATCTNNNGAAAACAGAGGTTGGGTAGT

GAGCACACGTCTGAACTCTTTCCCTACACGACGCTCTTCCGATCTNNNGAAAACAGAGGTTGGGTAGT

AGAGCACACGTCTGAACTCTTTCCCTACACGACGCTCTTCCGATCTNNNTTGAGGCAGGAGAATTGCTT

AGAGCACACGTCTGAACTCTTTCCCTACACGACGCTCTTCCGATCTNNNTTGAGGCAGGAGAATTGCTT

GAGCACACGTCTGAACTCTTTCCCTACACGACGCTCTTCCGATCTNNNCTGAGGCATGAGAATTGCTTG

GAGCACACGTCTGAACTCTTTCCCTACACGACGCTCTTCCGATCTNNNCTGAGGCATGAGAATTGCTTG

GAGCACACGTCTGAACTCTTTCCCTACACGACGCTCTTCCGATCTNNNGGAAGGGGAGAGAGAGAGA

GAGCACACGTCTGAACTCTTTCCCTACACGACGCTCTTCCGATCTNNNGGAAGGGGAGAGAGAGAGA

GCACACGTCTGAACTCTTTCCCTACACGACGCTCTTCCGATCTNNNTGGGTAAGAATTCCACACATCCAC

GCACACGTCTGAACTCTTTCCCTACACGACGCTCTTCCGATCTNNNTGGGTAAGAATTCCACACATCCAC

GAGCACACGTCTGAACTCTTTCCCTACACGACGCTCTTCCGATCTNNNCGTTTATTTGGACCTCACGGAA

AGCACACGTCTGAACTCTTTCCCTACACGACGCTCTTCCGATCTNNNCGTTTATTGGACCTCACGGA

AGCACACGTCTGAACTCTTTCCCTACACGACGCTCTTCCGATCTNNNGGGTCTTCCTTAAAAATTGGG

AGCACACGTCTGAACTCTTTCCCTACACGACGCTCTTCCGATCTNNNGGGTCTTCCTTAAAAATTGGG

AGCACACGTCTGAACTCTTTCCCTACACGACGCTCTTCCGATCTNNNTGAGCAGACATCTCCCTTTACA

AGCACACGTCTGAACTCTTTCCCTACACGACGCTCTTCCGATCTNNNTGAGCAGACATCTCCCTTTACA

AGCACACGTCTGAACTCTTTCCCTACACGACGCTCTTCCGATCTNNNCTGGGGTCACATATAGCTCAGT

AGCACACGTCTGAACTCTTTCCCTACACGACGCTCTTCCGATCTNNNCTGGGGTCACATATAGCTCAGT

AGCACACGTCTGAACTCTTTCCCTACACGACGCTCTTCCGATCTNNNCAAAATCCCTGCTACATTTGCC

AGCACACGTCTGAACTCTTTCCCTACACGACGCTCTTCCGATCTNNNCAAAATCCCTGCTACATTTGCC

GCACACGTCTGAACTCTTTCCCTACACGACGCTCTTCCGATCTNNNCATCCCTTCGGCTCTGGCTCCATC

GCACACGTCTGAACTCTTTCCCTACACGACGCTCTTCCGATCTNNNCATCCCTTCGGCTCTGGCTCCATC

AGCACACGTCTGAACTCTTTCCCTACACGACGCTCTTCCGATCTNNNTTGTGGCTCAAACATTTGGCTT

AGCACACGTCTGAACTCTTTCCCTACACGACGCTCTTCCGATCTNNNTTGTGGCTCAAACATTTGGCTT

GAGCACACGTCTGAACTCTTTCCCTACACGACGCTCTTCCGATCTNNNGAGAATCGTTTGAACCCAGG

GAGCACACGTCTGAACTCTTTCCCTACACGACGCTCTTCCGATCTNNNGAGAATCGTTTGAACCCAGG

AGCACACGTCTGAACTCTTTCCCTACACGACGCTCTTCCGATCTNNNTTGCCATTTGTCTGGTGATTCA

AGCACACGTCTGAACTCTTTCCCTACACGACGCTCTTCCGATCTNNNTTGCCATTTGTCTGGTGATTCA

GAGCACACGTCTGAACTCTTTCCCTACACGACGCTCTTCCGATCTNNNTTTTCTCAAAGCTCCACTAGG

GAGCACACGTCTGAACTCTTTCCCTACACGACGCTCTTCCGATCTNNNTTTTCTCAAAGCTCCACTAGG

ACACGTCTGAACTCTTTCCCTACACGACGCTCTTCCGATCTNNNTATCAGGGAGACTTTGAAGGCACC

ACACGTCTGAACTCTTTCCCTACACGACGCTCTTCCGATCTNNNTATCAGGGAGACTTTGAAGGCACC

GCACACGTCTGAACTCTTTCCCTACACGACGCTCTTCCGATCTNNNAAAAATTAGCCAGACGTGGTAG

GCACACGTCTGAACTCTTTCCCTACACGACGCTCTTCCGATCTNNNAAAAATTAGCCAGACGTGGTAG

GCACACGTCTGAACTCTTTCCCTACACGACGCTCTTCCGATCTNNNCCAGTGCACGTGACCTTATTTTC

GCACACGTCTGAACTCTTTCCCTACACGACGCTCTTCCGATCTNNNCCAGTGCACGTGACCTTATTTTC

GAGCACACGTCTGAACTCTTTCCCTACACGACGCTCTTCCGATCTNNNCTAGGCGTGCGCTAGAGGAA

GAGCACACGTCTGAACTCTTTCCCTACACGACGCTCTTCCGATCTNNNCTAGGCGTGCGCTAGAGGAA

AGCACACGTCTGAACTCTTTCCCTACACGACGCTCTTCCGATCTNNNCTGCTGAACATGGATTGCTGA

AGCACACGTCTGAACTCTTTCCCTACACGACGCTCTTCCGATCTNNNCTGCTGAACATGGATTGCTGA

AGAGCACACGTCTGAACTCTTTCCCTACACGACGCTCTTCCGATCTNNNTTTGGTGGACAGTTGGTGT

AGAGCACACGTCTGAACTCTTTCCCTACACGACGCTCTTCCGATCTNNNTTTGGTGGACAGTTGGTGT

AGCACACGTCTGAACTCTTTCCCTACACGACGCTCTTCCGATCTNNNGGATAATTCACCAGAGAGCTG

AGCACACGTCTGAACTCTTTCCCTACACGACGCTCTTCCGATCTNNNGGATAATTCACCAGAGAGCTG

GAGCACACGTCTGAACTCTTTCCCTACACGACGCTCTTCCGATCTNNNCAAGAGAATTGCTTGAACCCA

GAGCACACGTCTGAACTCTTTCCCTACACGACGCTCTTCCGATCTNNNCAAGAGAATTGCTTGAACCCA

GAGCACACGTCTGAACTCTTTCCCTACACGACGCTCTTCCGATCTNNNTGTGAGCCTCTGTGAAAATCT

GAGCACACGTCTGAACTCTTTCCCTACACGACGCTCTTCCGATCTNNNTGTGAGCCTCTGTGAAAATCT

AGCACACGTCTGAACTCTTTCCCTACACGACGCTCTTCCGATCTNNNGCAGAGGTTATAGTGAGCCA

AGCACACGTCTGAACTCTTTCCCTACACGACGCTCTTCCGATCTNNNGCAGAGGTTATAGTGAGCCA

GAGCACACGTCTGAACTCTTTCCCTACACGACGCTCTTCCGATCTNNNCTTGCAATGAGCTGAGATCA

GAGCACACGTCTGAACTCTTTCCCTACACGACGCTCTTCCGATCTNNNCTTGCAATGAGCTGAGATCA

GCACACGTCTGAACTCTTTCCCTACACGACGCTCTTCCGATCTNNNATAGCAAGAACAGAGAGTGAC

GCACACGTCTGAACTCTTTCCCTACACGACGCTCTTCCGATCTNNNATAGCAAGAACAGAGAGTGAC

GAGCACACGTCTGAACTCTTTCCCTACACGACGCTCTTCCGATCTNNNGAGAGACTGAGGCAAGAGA

GAGCACACGTCTGAACTCTTTCCCTACACGACGCTCTTCCGATCTNNNGAGAGACTGAGGCAAGAGA

GAGCACACGTCTGAACTCTTTCCCTACACGACGCTCTTCCGATCTNNNTGATATTCAAGGTGGTGGGA

GAGCACACGTCTGAACTCTTTCCCTACACGACGCTCTTCCGATCTNNNTGATATTCAAGGTGGTGGGA

AGCACACGTCTGAACTCTTTCCCTACACGACGCTCTTCCGATCTNNNTGCACATTTGTATCTCCTGGGT

AGCACACGTCTGAACTCTTTCCCTACACGACGCTCTTCCGATCTNNNTGCACATTTGTATCTCCTGGGT

GAGCACACGTCTGAACTCTTTCCCTACACGACGCTCTTCCGATCTNNNCTTTGATCCACAAATCGGCCT

GAGCACACGTCTGAACTCTTTCCCTACACGACGCTCTTCCGATCTNNNCTTTGATCCACAAATCGGCCT

CACACGTCTGAACTCTTTCCCTACACGACGCTCTTCCGATCTNNNTCCAACCAGATAGGCCCACTATAC

CACACGTCTGAACTCTTTCCCTACACGACGCTCTTCCGATCTNNNTCCAACCAGATAGGCCCACTATAC

ACACGTCTGAACTCTTTCCCTACACGACGCTCTTCCGATCTNNNATATATGTGCCATCTTTCTACATCAT

ACACGTCTGAACTCTTTCCCTACACGACGCTCTTCCGATCTNNNATATATGTGCCATCTTTCTACATCAT

AGCACACGTCTGAACTCTTTCCCTACACGACGCTCTTCCGATCTNNNCCACGCCCAGCTAATTTTTGTA

AGCACACGTCTGAACTCTTTCCCTACACGACGCTCTTCCGATCTNNNCCACGCCCAGCTAATTTTTGTA

GCACACGTCTGAACTCTTTCCCTACACGACGCTCTTCCGATCTNNNTCCCAGAAAGAAGTCTGTTTCTC

GCACACGTCTGAACTCTTTCCCTACACGACGCTCTTCCGATCTNNNTCCCAGAAAGAAGTCTGTTTCTC

GAGCACACGTCTGAACTCTTTCCCTACACGACGCTCTTCCGATCTNNNTCTACACACATACACACACAC

GAGCACACGTCTGAACTCTTTCCCTACACGACGCTCTTCCGATCTNNNTCTACACACATACACACACAC

GAGCACACGTCTGAACTCTTTCCCTACACGACGCTCTTCCGATCTNNNGCCTGTAGTCCCAGCTATTTG

GAGCACACGTCTGAACTCTTTCCCTACACGACGCTCTTCCGATCTNNNGCCTGTAGTCCCAGCTATTTG

AGCACACGTCTGAACTCTTTCCCTACACGACGCTCTTCCGATCTNNNGAAGGTTGCAATGAGCCAAGA

AGCACACGTCTGAACTCTTTCCCTACACGACGCTCTTCCGATCTNNNGAAGGTTGCAATGAGCCAAGA

GAGCACACGTCTGAACTCTTTCCCTACACGACGCTCTTCCGATCTNNNATGACCTTCCTTGTCTCTCTCT

GAGCACACGTCTGAACTCTTTCCCTACACGACGCTCTTCCGATCTNNNATGACCTTCCTTGTCTCTCTCT

GCACACGTCTGAACTCTTTCCCTACACGACGCTCTTCCGATCTNNNCTCCCAGGTTCAAGTGATTCTCA

GCACACGTCTGAACTCTTTCCCTACACGACGCTCTTCCGATCTNNNCTCCCAGGTTCAAGTGATTCTCA

CACACGTCTGAACTCTTTCCCTACACGACGCTCTTCCGATCTNNNAAGATAAAAGGAAACCTGACTGC

CACACGTCTGAACTCTTTCCCTACACGACGCTCTTCCGATCTNNNAAGATAAAAGGAAACCTGACTGC

AGCACACGTCTGAACTCTTTCCCTACACGACGCTCTTCCGATCTNNNACCTTCCCTCCACTATTGTCCTAC

AGCACACGTCTGAACTCTTTCCCTACACGACGCTCTTCCGATCTNNNACCTTCCCTCCACTATTGTCCTAC

GCACACGTCTGAACTCTTTCCCTACACGACGCTCTTCCGATCTNNNAAAGTGGAACAGGTATGCCGT

GCACACGTCTGAACTCTTTCCCTACACGACGCTCTTCCGATCTNNNAAAGTGGAACAGGTATGCCGT

CACGTCTGAACTCTTTCCCTACACGACGCTCTTCCGATCTNNNTGCCAAAGAAACCAAGCTCTTTTCAT

CACGTCTGAACTCTTTCCCTACACGACGCTCTTCCGATCTNNNTGCCAAAGAAACCAAGCTCTTTTCAT

AGAGCACACGTCTGAACTCTTTCCCTACACGACGCTCTTCCGATCTNNNGTTGATAGTCCTGGGAGCC

AGAGCACACGTCTGAACTCTTTCCCTACACGACGCTCTTCCGATCTNNNGTTGATAGTCCTGGGAGCC

AGCACACGTCTGAACTCTTTCCCTACACGACGCTCTTCCGATCTNNNACCTTTGATGGAAATTTGTGT

AGCACACGTCTGAACTCTTTCCCTACACGACGCTCTTCCGATCTNNNACCTTTGATGGAAATTTGTGT

GAGCACACGTCTGAACTCTTTCCCTACACGACGCTCTTCCGATCTNNNAGTCTCAGTGGACTCATTTGA

GAGCACACGTCTGAACTCTTTCCCTACACGACGCTCTTCCGATCTNNNAGTCTCAGTGGACTCATTTGA

CACACGTCTGAACTCTTTCCCTACACGACGCTCTTCCGATCTNNNGCACATGCCTATAATCCCAGCTAC

CACACGTCTGAACTCTTTCCCTACACGACGCTCTTCCGATCTNNNGCACATGCCTATAATCCCAGCTAC

GAGCACACGTCTGAACTCTTTCCCTACACGACGCTCTTCCGATCTNNNCACAACTGTAGTCCCAGCTAC

GAGCACACGTCTGAACTCTTTCCCTACACGACGCTCTTCCGATCTNNNCACAACTGTAGTCCCAGCTAC
AGCACACGTCTGAACTCTTTCCCTACACGACGCTCTTCCGATCTNNNGGGACTTTACTTGGTGAGTCT
AGCACACGTCTGAACTCTTTCCCTACACGACGCTCTTCCGATCTNNNGGGACTTTACTTGGTGAGTCT
GCACACGTCTGAACTCTTTCCCTACACGACGCTCTTCCGATCTNNNATCAAGATAACTAGCCCCAAGG
GCACACGTCTGAACTCTTTCCCTACACGACGCTCTTCCGATCTNNNATCAAGATAACTAGCCCCAAGG
AGCACACGTCTGAACTCTTTCCCTACACGACGCTCTTCCGATCTNNNTGCTTTGTATTTGGCCAAGAC
AGCACACGTCTGAACTCTTTCCCTACACGACGCTCTTCCGATCTNNNTGCTTTGTATTTGGCCAAGAC
ACACGTCTGAACTCTTTCCCTACACGACGCTCTTCCGATCTNNNTTGAAAAAAGGCCCTTCCTCATCCA
ACACGTCTGAACTCTTTCCCTACACGACGCTCTTCCGATCTNNNTTGAAAAAAGGCCCTTCCTCATCCA
AGCACACGTCTGAACTCTTTCCCTACACGACGCTCTTCCGATCTNNNAATCCCCTACTTCTCACTTTTG
AGCACACGTCTGAACTCTTTCCCTACACGACGCTCTTCCGATCTNNNAATCCCCTACTTCTCACTTTTG
AGCACACGTCTGAACTCTTTCCCTACACGACGCTCTTCCGATCTNNNATCATAACCCAGGCTTTCAACC
AGCACACGTCTGAACTCTTTCCCTACACGACGCTCTTCCGATCTNNNATCATAACCCAGGCTTTCAACC
AGCACACGTCTGAACTCTTTCCCTACACGACGCTCTTCCGATCTNNNCTTGGTAGCACTCACCTGTAGT
AGCACACGTCTGAACTCTTTCCCTACACGACGCTCTTCCGATCTNNNCTTGGTAGCACTCACCTGTAGT
GCACACGTCTGAACTCTTTCCCTACACGACGCTCTTCCGATCTNNNTGCAGCTTTCTTGGCTTCTAGTT
GCACACGTCTGAACTCTTTCCCTACACGACGCTCTTCCGATCTNNNTGCAGCTTTCTTGGCTTCTAGTT
AGCACACGTCTGAACTCTTTCCCTACACGACGCTCTTCCGATCTNNNTTGAGAGAGAAGAGGCAGGT
AGCACACGTCTGAACTCTTTCCCTACACGACGCTCTTCCGATCTNNNTTGAGAGAGAAGAGGCAGGT
AGCACACGTCTGAACTCTTTCCCTACACGACGCTCTTCCGATCTNNNGCAGGAGAATCACTTGAACCT
AGCACACGTCTGAACTCTTTCCCTACACGACGCTCTTCCGATCTNNNGCAGGAGAATCACTTGAACCT
GCACACGTCTGAACTCTTTCCCTACACGACGCTCTTCCGATCTNNNAGCTACTCACACATTAGAAGCT
GCACACGTCTGAACTCTTTCCCTACACGACGCTCTTCCGATCTNNNAGCTACTCACACATTAGAAGCT
AGCACACGTCTGAACTCTTTCCCTACACGACGCTCTTCCGATCTNNNGAACTTGCAGTGAGCCAAGA
AGCACACGTCTGAACTCTTTCCCTACACGACGCTCTTCCGATCTNNNGAACTTGCAGTGAGCCAAGA
AGCACACGTCTGAACTCTTTCCCTACACGACGCTCTTCCGATCTNNNTTCAAGAAATAAACCTTCGGCC
AGCACACGTCTGAACTCTTTCCCTACACGACGCTCTTCCGATCTNNNTTCAAGAAATAAACCTTCGGCC
GAGCACACGTCTGAACTCTTTCCCTACACGACGCTCTTCCGATCTNNNGCACACTTACTTGGCACTTCA
GAGCACACGTCTGAACTCTTTCCCTACACGACGCTCTTCCGATCTNNNGCACACTTACTTGGCACTTCA
AGCACACGTCTGAACTCTTTCCCTACACGACGCTCTTCCGATCTNNNGAGTGGTTCCCGGAAACTAAG
AGCACACGTCTGAACTCTTTCCCTACACGACGCTCTTCCGATCTNNNGAGTGGTTCCCGGAAACTAAG
AGCACACGTCTGAACTCTTTCCCTACACGACGCTCTTCCGATCTNNNCATCTTGGACCATTGGACTCTC
AGCACACGTCTGAACTCTTTCCCTACACGACGCTCTTCCGATCTNNNCATCTTGGACCATTGGACTCTC
AGCACACGTCTGAACTCTTTCCCTACACGACGCTCTTCCGATCTNNNCTTCTAGCCCCTTTTACACCCTT
AGCACACGTCTGAACTCTTTCCCTACACGACGCTCTTCCGATCTNNNCTTCTAGCCCCTTTTACACCCTT
GAGCACACGTCTGAACTCTTTCCCTACACGACGCTCTTCCGATCTNNNCCTAAAGACAGCAAGAGAG
GAGCACACGTCTGAACTCTTTCCCTACACGACGCTCTTCCGATCTNNNCCTAAAGACAGCAAGAGAG
GAGCACACGTCTGAACTCTTTCCCTACACGACGCTCTTCCGATCTNNNTTGGGCTACTGATTTGGGGA
GAGCACACGTCTGAACTCTTTCCCTACACGACGCTCTTCCGATCTNNNTTGGGCTACTGATTTGGGGA
GCACACGTCTGAACTCTTTCCCTACACGACGCTCTTCCGATCTNNNCCTCCCTTTAGCCACAGTATCAA
GCACACGTCTGAACTCTTTCCCTACACGACGCTCTTCCGATCTNNNCCTCCCTTTAGCCACAGTATCAA
GCACACGTCTGAACTCTTTCCCTACACGACGCTCTTCCGATCTNNNAACTCTTGACCTCAGGTGATCTG
GCACACGTCTGAACTCTTTCCCTACACGACGCTCTTCCGATCTNNNAACTCTTGACCTCAGGTGATCTG

AGCACACGTCTGAACTCTTTCCCTACACGACGCTCTTCCGATCTNNNAGGAGAATGGCATGAACCCG
AGCACACGTCTGAACTCTTTCCCTACACGACGCTCTTCCGATCTNNNAGGAGAATGGCATGAACCCG
AGCACACGTCTGAACTCTTTCCCTACACGACGCTCTTCCGATCTNNNTCAGTTACTGTGTGTGGCCTA
AGCACACGTCTGAACTCTTTCCCTACACGACGCTCTTCCGATCTNNNTCAGTTACTGTGTGTGGCCTA
AGCACACGTCTGAACTCTTTCCCTACACGACGCTCTTCCGATCTNNNATTCTTGATCTTCCAGACCTGC
AGCACACGTCTGAACTCTTTCCCTACACGACGCTCTTCCGATCTNNNATTCTTGATCTTCCAGACCTGC
AGCACACGTCTGAACTCTTTCCCTACACGACGCTCTTCCGATCTNNNTGTGGATGTCCTTTTTCACTCC
AGCACACGTCTGAACTCTTTCCCTACACGACGCTCTTCCGATCTNNNTGTGGATGTCCTTTTTCACTCC
AGCACACGTCTGAACTCTTTCCCTACACGACGCTCTTCCGATCTNNNCGGCTCTCTCCTCGTCTTTTAAA
AGCACACGTCTGAACTCTTTCCCTACACGACGCTCTTCCGATCTNNNCGGCTCTCTCCTCGTCTTTTAAA
AGCACACGTCTGAACTCTTTCCCTACACGACGCTCTTCCGATCTNNNATCCTCTACAGACAAGCAAATG
AGCACACGTCTGAACTCTTTCCCTACACGACGCTCTTCCGATCTNNNATCCTCTACAGACAAGCAAATG
GCACACGTCTGAACTCTTTCCCTACACGACGCTCTTCCGATCTNNNCAAGAACAAGCAAACAAGCCAA
GCACACGTCTGAACTCTTTCCCTACACGACGCTCTTCCGATCTNNNCAAGAACAAGCAAACAAGCCAA
AGCACACGTCTGAACTCTTTCCCTACACGACGCTCTTCCGATCTNNNATTCAGACATTGCTTTGTGGT
AGCACACGTCTGAACTCTTTCCCTACACGACGCTCTTCCGATCTNNNATTCAGACATTGCTTTGTGGT
AGCACACGTCTGAACTCTTTCCCTACACGACGCTCTTCCGATCTNNNAGAATTGCTTGAACCCAGGAG
AGCACACGTCTGAACTCTTTCCCTACACGACGCTCTTCCGATCTNNNAGAATTGCTTGAACCCAGGAG
GAGCACACGTCTGAACTCTTTCCCTACACGACGCTCTTCCGATCTNNNCCACTCTCTAACACTTCTGTG
GAGCACACGTCTGAACTCTTTCCCTACACGACGCTCTTCCGATCTNNNCCACTCTCTAACACTTCTGTG
AGCACACGTCTGAACTCTTTCCCTACACGACGCTCTTCCGATCTNNNTGTAATCCCAGCTTCTTGTGAG
AGCACACGTCTGAACTCTTTCCCTACACGACGCTCTTCCGATCTNNNTGTAATCCCAGCTTCTTGTGAG
AGCACACGTCTGAACTCTTTCCCTACACGACGCTCTTCCGATCTNNNGAGATTGCAGTGAGCCGAGAT
AGCACACGTCTGAACTCTTTCCCTACACGACGCTCTTCCGATCTNNNGAGATTGCAGTGAGCCGAGAT
GCACACGTCTGAACTCTTTCCCTACACGACGCTCTTCCGATCTNNNTTTAGAGGGAAAACGGTGATG
GCACACGTCTGAACTCTTTCCCTACACGACGCTCTTCCGATCTNNNTTTAGAGGGAAAACGGTGATG
AGCACACGTCTGAACTCTTTCCCTACACGACGCTCTTCCGATCTNNNAAAGAATTTTGCCAGCCCAAA
AGCACACGTCTGAACTCTTTCCCTACACGACGCTCTTCCGATCTNNNAAAGAATTTTGCCAGCCCAAA
GAGCACACGTCTGAACTCTTTCCCTACACGACGCTCTTCCGATCTNNNAAAGTGCTGGGATTACGGAC
GAGCACACGTCTGAACTCTTTCCCTACACGACGCTCTTCCGATCTNNNAAAGTGCTGGGATTACGGAC
CACACGTCTGAACTCTTTCCCTACACGACGCTCTTCCGATCTNNNGCAAACAATTCCAACAGGGGTAT
CACACGTCTGAACTCTTTCCCTACACGACGCTCTTCCGATCTNNNGCAAACAATTCCAACAGGGGTAT
AGCACACGTCTGAACTCTTTCCCTACACGACGCTCTTCCGATCTNNNTCAAGAGGTCTCCTTGTCCTCA
AGCACACGTCTGAACTCTTTCCCTACACGACGCTCTTCCGATCTNNNTCAAGAGGTCTCCTTGTCCTCA
GAGCACACGTCTGAACTCTTTCCCTACACGACGCTCTTCCGATCTNNNACAAAAGGGCTTATTGGAA
GAGCACACGTCTGAACTCTTTCCCTACACGACGCTCTTCCGATCTNNNACAAAAGGGCTTATTGGAA
CACACGTCTGAACTCTTTCCCTACACGACGCTCTTCCGATCTNNNATCCCTGACCTTTGCTACCATTTCA
CACACGTCTGAACTCTTTCCCTACACGACGCTCTTCCGATCTNNNATCCCTGACCTTTGCTACCATTTCA
GAGCACACGTCTGAACTCTTTCCCTACACGACGCTCTTCCGATCTNNNGGGAGGGATAGCATTGGG
GAGCACACGTCTGAACTCTTTCCCTACACGACGCTCTTCCGATCTNNNGGGAGGGATAGCATTGGG
AGCACACGTCTGAACTCTTTCCCTACACGACGCTCTTCCGATCTNNNCCTTGTCATTTTGGGGTCATTC
AGCACACGTCTGAACTCTTTCCCTACACGACGCTCTTCCGATCTNNNCCTTGTCATTTTGGGGTCATTC
AGCACACGTCTGAACTCTTTCCCTACACGACGCTCTTCCGATCTNNNTTGACTGAACCCACATTGTACC

AGCACACGTCTGAACTCTTTCCCTACACGACGCTCTTCCGATCTNNNTTGACTGAACCCACATTGTACC
AGCACACGTCTGAACTCTTTCCCTACACGACGCTCTTCCGATCTNNNTACCAGTGATCAAATTCAGCGG
AGCACACGTCTGAACTCTTTCCCTACACGACGCTCTTCCGATCTNNNTACCAGTGATCAAATTCAGCGG
AGCACACGTCTGAACTCTTTCCCTACACGACGCTCTTCCGATCTNNNTGCTGGGATTACAGGATTACA
AGCACACGTCTGAACTCTTTCCCTACACGACGCTCTTCCGATCTNNNTGCTGGGATTACAGGATTACA
GCACACGTCTGAACTCTTTCCCTACACGACGCTCTTCCGATCTNNNGAGGCTGAGGCAGGAGAATGC
GCACACGTCTGAACTCTTTCCCTACACGACGCTCTTCCGATCTNNNGAGGCTGAGGCAGGAGAATGC
CACGTCTGAACTCTTTCCCTACACGACGCTCTTCCGATCTNNNTTCTATTTCTGTGGGGTCAGTGGCAT
CACGTCTGAACTCTTTCCCTACACGACGCTCTTCCGATCTNNNTTCTATTTCTGTGGGGTCAGTGGCAT
AGCACACGTCTGAACTCTTTCCCTACACGACGCTCTTCCGATCTNNNCCAACATGGTGAAACTGCATC
AGCACACGTCTGAACTCTTTCCCTACACGACGCTCTTCCGATCTNNNCCAACATGGTGAAACTGCATC
AGCACACGTCTGAACTCTTTCCCTACACGACGCTCTTCCGATCTNNNTGGGCATCTTTAGGTATTTCGC
AGCACACGTCTGAACTCTTTCCCTACACGACGCTCTTCCGATCTNNNTGGGCATCTTTAGGTATTTCGC
AGCACACGTCTGAACTCTTTCCCTACACGACGCTCTTCCGATCTNNNTCCCTCATGTTCTGTCAGTTAG
AGCACACGTCTGAACTCTTTCCCTACACGACGCTCTTCCGATCTNNNTCCCTCATGTTCTGTCAGTTAG
AGCACACGTCTGAACTCTTTCCCTACACGACGCTCTTCCGATCTNNNGCAGGAGAATCATTTGAACCC
AGCACACGTCTGAACTCTTTCCCTACACGACGCTCTTCCGATCTNNNGCAGGAGAATCATTTGAACCC
AGCACACGTCTGAACTCTTTCCCTACACGACGCTCTTCCGATCTNNNCATCTGAGAAGTGAGGAGCCC
AGCACACGTCTGAACTCTTTCCCTACACGACGCTCTTCCGATCTNNNCATCTGAGAAGTGAGGAGCCC
AGCACACGTCTGAACTCTTTCCCTACACGACGCTCTTCCGATCTNNNGGGGGAAGCAGATTTATCAAT
AGCACACGTCTGAACTCTTTCCCTACACGACGCTCTTCCGATCTNNNGGGGGAAGCAGATTTATCAAT
GCACACGTCTGAACTCTTTCCCTACACGACGCTCTTCCGATCTNNNCCGGTTAGCTCAGTTGGTAAGA
GCACACGTCTGAACTCTTTCCCTACACGACGCTCTTCCGATCTNNNCCGGTTAGCTCAGTTGGTAAGA
AGCACACGTCTGAACTCTTTCCCTACACGACGCTCTTCCGATCTNNNTTTGGGTTGGTTCCAAGACTTT
AGCACACGTCTGAACTCTTTCCCTACACGACGCTCTTCCGATCTNNNTTTGGGTTGGTTCCAAGACTTT
AGCACACGTCTGAACTCTTTCCCTACACGACGCTCTTCCGATCTNNNCTTATGAACTCTCCAGATTGGC
AGCACACGTCTGAACTCTTTCCCTACACGACGCTCTTCCGATCTNNNCTTATGAACTCTCCAGATTGGC
AGAGCACACGTCTGAACTCTTTCCCTACACGACGCTCTTCCGATCTNNNGCCAGTTGAGGAAGTAGAA
AGAGCACACGTCTGAACTCTTTCCCTACACGACGCTCTTCCGATCTNNNGCCAGTTGAGGAAGTAGAA
GCACACGTCTGAACTCTTTCCCTACACGACGCTCTTCCGATCTNNNTCTGATTTACATAGTTTTAAGGC
GCACACGTCTGAACTCTTTCCCTACACGACGCTCTTCCGATCTNNNTCTGATTTACATAGTTTTAAGGC
AGCACACGTCTGAACTCTTTCCCTACACGACGCTCTTCCGATCTNNNCGCATTTTTCAGAAGCACTTCCT
AGCACACGTCTGAACTCTTTCCCTACACGACGCTCTTCCGATCTNNNCGCATTTTTCAGAAGCACTTCCT
AGCACACGTCTGAACTCTTTCCCTACACGACGCTCTTCCGATCTNNNATGGTTTCTGTGGAGTCTAGC
AGCACACGTCTGAACTCTTTCCCTACACGACGCTCTTCCGATCTNNNATGGTTTCTGTGGAGTCTAGC
GAGCACACGTCTGAACTCTTTCCCTACACGACGCTCTTCCGATCTNNNTTTGGTTACTACAGCTTTGCAT
GAGCACACGTCTGAACTCTTTCCCTACACGACGCTCTTCCGATCTNNNTTTGGTTACTACAGCTTTGCAT
GCACACGTCTGAACTCTTTCCCTACACGACGCTCTTCCGATCTNNNTTTTTCCTTCCCTTCCCTTGTCTCA
GCACACGTCTGAACTCTTTCCCTACACGACGCTCTTCCGATCTNNNTTTTTCCTTCCCTTCCCTTGTCTCA
AGCACACGTCTGAACTCTTTCCCTACACGACGCTCTTCCGATCTNNNCCAGGACCTAGTGGAATCATG
AGCACACGTCTGAACTCTTTCCCTACACGACGCTCTTCCGATCTNNNCCAGGACCTAGTGGAATCATG
AGCACACGTCTGAACTCTTTCCCTACACGACGCTCTTCCGATCTNNNTGAGGCAGGAAAGTCACTTG
AGCACACGTCTGAACTCTTTCCCTACACGACGCTCTTCCGATCTNNNTGAGGCAGGAAAGTCACTTG

GAGCACACGTCTGAACTCTTTCCCTACACGACGCTCTTCCGATCTNNNTAAGCCCTCCAAACTGTTCAACT
GAGCACACGTCTGAACTCTTTCCCTACACGACGCTCTTCCGATCTNNNTAAGCCCTCCAAACTGTTCAACT
AGCACACGTCTGAACTCTTTCCCTACACGACGCTCTTCCGATCTNNNAGTAGAAACGGGATTTTGCCAT
AGCACACGTCTGAACTCTTTCCCTACACGACGCTCTTCCGATCTNNNAGTAGAAACGGGATTTTGCCAT
AGCACACGTCTGAACTCTTTCCCTACACGACGCTCTTCCGATCTNNNCTTTGCTTCCTTCCTTGCATGTC
AGCACACGTCTGAACTCTTTCCCTACACGACGCTCTTCCGATCTNNNCTTTGCTTCCTTCCTTGCATGTC
AGCACACGTCTGAACTCTTTCCCTACACGACGCTCTTCCGATCTNNNTGAGCTTAGGAGTTCAAGACCA
AGCACACGTCTGAACTCTTTCCCTACACGACGCTCTTCCGATCTNNNTGAGCTTAGGAGTTCAAGACCA
AGCACACGTCTGAACTCTTTCCCTACACGACGCTCTTCCGATCTNNNTCCATCCATATCTGAGTGACTCT
AGCACACGTCTGAACTCTTTCCCTACACGACGCTCTTCCGATCTNNNTCCATCCATATCTGAGTGACTCT
AGCACACGTCTGAACTCTTTCCCTACACGACGCTCTTCCGATCTNNNCAAAGATTAGTTGGGTGTGGTC
AGCACACGTCTGAACTCTTTCCCTACACGACGCTCTTCCGATCTNNNCAAAGATTAGTTGGGTGTGGTC
GAGCACACGTCTGAACTCTTTCCCTACACGACGCTCTTCCGATCTNNNGGAATCACTTGAACCTGGGAG
GAGCACACGTCTGAACTCTTTCCCTACACGACGCTCTTCCGATCTNNNGGAATCACTTGAACCTGGGAG
AGCACACGTCTGAACTCTTTCCCTACACGACGCTCTTCCGATCTNNNTTCTAAAGCTTCAGCCTCCACTTC
AGCACACGTCTGAACTCTTTCCCTACACGACGCTCTTCCGATCTNNNTTCTAAAGCTTCAGCCTCCACTTC
CACACGTCTGAACTCTTTCCCTACACGACGCTCTTCCGATCTNNNAATGGTTGTCCTGTTTTCGTTTGCA
CACACGTCTGAACTCTTTCCCTACACGACGCTCTTCCGATCTNNNAATGGTTGTCCTGTTTTCGTTTGCA
AGCACACGTCTGAACTCTTTCCCTACACGACGCTCTTCCGATCTNNNATTTGAACAGGCTCTGAGGTTC
AGCACACGTCTGAACTCTTTCCCTACACGACGCTCTTCCGATCTNNNATTTGAACAGGCTCTGAGGTTC
AGCACACGTCTGAACTCTTTCCCTACACGACGCTCTTCCGATCTNNNGTGTGCACATTTAGTTCAGCTGA
AGCACACGTCTGAACTCTTTCCCTACACGACGCTCTTCCGATCTNNNGTGTGCACATTTAGTTCAGCTGA
GCACACGTCTGAACTCTTTCCCTACACGACGCTCTTCCGATCTNNNCTGAGGCAGGAGAATCACTTGAA
GCACACGTCTGAACTCTTTCCCTACACGACGCTCTTCCGATCTNNNCTGAGGCAGGAGAATCACTTGAA
GAGCACACGTCTGAACTCTTTCCCTACACGACGCTCTTCCGATCTNNNGTCATACTACTAACGGGGCAC
GAGCACACGTCTGAACTCTTTCCCTACACGACGCTCTTCCGATCTNNNGTCATACTACTAACGGGGCAC
AGAGCACACGTCTGAACTCTTTCCCTACACGACGCTCTTCCGATCTNNNGCAGTAACGAGGATGAGGA
AGAGCACACGTCTGAACTCTTTCCCTACACGACGCTCTTCCGATCTNNNGCAGTAACGAGGATGAGGA
GAGCACACGTCTGAACTCTTTCCCTACACGACGCTCTTCCGATCTNNNTCTGATCACTCTCACAGGCTAG
GAGCACACGTCTGAACTCTTTCCCTACACGACGCTCTTCCGATCTNNNTCTGATCACTCTCACAGGCTAG
AGCACACGTCTGAACTCTTTCCCTACACGACGCTCTTCCGATCTNNNTTGAAGTGATTCTCCTGCCTCAC
AGCACACGTCTGAACTCTTTCCCTACACGACGCTCTTCCGATCTNNNTTGAAGTGATTCTCCTGCCTCAC
GAGCACACGTCTGAACTCTTTCCCTACACGACGCTCTTCCGATCTNNNTCATTCATTCATTCCCCCACAT
GAGCACACGTCTGAACTCTTTCCCTACACGACGCTCTTCCGATCTNNNTCATTCATTCATTCCCCCACAT
GAGCACACGTCTGAACTCTTTCCCTACACGACGCTCTTCCGATCTNNNGAGAATCACTTGAACCCGAGA
GAGCACACGTCTGAACTCTTTCCCTACACGACGCTCTTCCGATCTNNNGAGAATCACTTGAACCCGAGA
CACACGTCTGAACTCTTTCCCTACACGACGCTCTTCCGATCTNNNTATACGTTTATACCTTCCGCGCTCAT
CACACGTCTGAACTCTTTCCCTACACGACGCTCTTCCGATCTNNNTATACGTTTATACCTTCCGCGCTCAT
AGCACACGTCTGAACTCTTTCCCTACACGACGCTCTTCCGATCTNNNAATTACCACTTCCCTTCCAGACCC
AGCACACGTCTGAACTCTTTCCCTACACGACGCTCTTCCGATCTNNNAATTACCACTTCCCTTCCAGACCC
GAGCACACGTCTGAACTCTTTCCCTACACGACGCTCTTCCGATCTNNNGCAATCGAATCTACCCCACATT
GAGCACACGTCTGAACTCTTTCCCTACACGACGCTCTTCCGATCTNNNGCAATCGAATCTACCCCACATT
GAGCACACGTCTGAACTCTTTCCCTACACGACGCTCTTCCGATCTNNNCAAGATCATGCCACTGTACTC

GAGCACACGTCTGAACTCTTTCCCTACACGACGCTCTTCCGATCTNNNCAAGATCATGCCACTGTACTC
GAGCACACGTCTGAACTCTTTCCCTACACGACGCTCTTCCGATCTNNNGGGCGACAGAGTGAGATTCT
GAGCACACGTCTGAACTCTTTCCCTACACGACGCTCTTCCGATCTNNNGGGCGACAGAGTGAGATTCT
AGCACACGTCTGAACTCTTTCCCTACACGACGCTCTTCCGATCTNNNTGCCCTAAAGTCTCAGTTCACT
AGCACACGTCTGAACTCTTTCCCTACACGACGCTCTTCCGATCTNNNTGCCCTAAAGTCTCAGTTCACT
GAGCACACGTCTGAACTCTTTCCCTACACGACGCTCTTCCGATCTNNNCACGAGAATTGCTTGAACTG
GAGCACACGTCTGAACTCTTTCCCTACACGACGCTCTTCCGATCTNNNCACGAGAATTGCTTGAACTG
GAGCACACGTCTGAACTCTTTCCCTACACGACGCTCTTCCGATCTNNNGAGATTACGCTACTGCACTCC
GAGCACACGTCTGAACTCTTTCCCTACACGACGCTCTTCCGATCTNNNGAGATTACGCTACTGCACTCC
ACACGTCTGAACTCTTTCCCTACACGACGCTCTTCCGATCTNNNATCAACTCTTCTCTGGTTCTCGGCA
ACACGTCTGAACTCTTTCCCTACACGACGCTCTTCCGATCTNNNATCAACTCTTCTCTGGTTCTCGGCA
AGCACACGTCTGAACTCTTTCCCTACACGACGCTCTTCCGATCTNNNTTCAAGCAATTCTCCTGCCTCA
AGCACACGTCTGAACTCTTTCCCTACACGACGCTCTTCCGATCTNNNTTCAAGCAATTCTCCTGCCTCA
GCACACGTCTGAACTCTTTCCCTACACGACGCTCTTCCGATCTNNNAAAATTAGCCAGGTGTTGTGG
GCACACGTCTGAACTCTTTCCCTACACGACGCTCTTCCGATCTNNNAAAATTAGCCAGGTGTTGTGG
GAGCACACGTCTGAACTCTTTCCCTACACGACGCTCTTCCGATCTNNNCCAGACAGAAGCTGAGGAA
GAGCACACGTCTGAACTCTTTCCCTACACGACGCTCTTCCGATCTNNNCCAGACAGAAGCTGAGGAA
GAGCACACGTCTGAACTCTTTCCCTACACGACGCTCTTCCGATCTNNNAATTCCTAGGATGGGTACAG
GAGCACACGTCTGAACTCTTTCCCTACACGACGCTCTTCCGATCTNNNAATTCCTAGGATGGGTACAG
GCACACGTCTGAACTCTTTCCCTACACGACGCTCTTCCGATCTNNNGCTCCCTTTGCAGTGGTAAATTT
GCACACGTCTGAACTCTTTCCCTACACGACGCTCTTCCGATCTNNNGCTCCCTTTGCAGTGGTAAATTT
AGCACACGTCTGAACTCTTTCCCTACACGACGCTCTTCCGATCTNNNTCTTGTACATGGGAAGAGAG
AGCACACGTCTGAACTCTTTCCCTACACGACGCTCTTCCGATCTNNNTCTTGTACATGGGAAGAGAG
AGCACACGTCTGAACTCTTTCCCTACACGACGCTCTTCCGATCTNNNTTTAGTTGTCATGTCCCCTTGG
AGCACACGTCTGAACTCTTTCCCTACACGACGCTCTTCCGATCTNNNTTTAGTTGTCATGTCCCCTTGG
AGCACACGTCTGAACTCTTTCCCTACACGACGCTCTTCCGATCTNNNGAACAATAATGGAAATGTTA
AGCACACGTCTGAACTCTTTCCCTACACGACGCTCTTCCGATCTNNNGAACAATAATGGAAATGTTA
AGCACACGTCTGAACTCTTTCCCTACACGACGCTCTTCCGATCTNNNCTCAAGCGATCCTCTTTCTCAC
AGCACACGTCTGAACTCTTTCCCTACACGACGCTCTTCCGATCTNNNCTCAAGCGATCCTCTTTCTCAC
AGCACACGTCTGAACTCTTTCCCTACACGACGCTCTTCCGATCTNNNTAAATCTAGATATCAGCTGAT
AGCACACGTCTGAACTCTTTCCCTACACGACGCTCTTCCGATCTNNNTAAATCTAGATATCAGCTGAT
GCACACGTCTGAACTCTTTCCCTACACGACGCTCTTCCGATCTNNNTGCTTTCTCTTTTGTTTGTGTCAC
GCACACGTCTGAACTCTTTCCCTACACGACGCTCTTCCGATCTNNNTGCTTTCTCTTTTGTTTGTGTCAC
GCACACGTCTGAACTCTTTCCCTACACGACGCTCTTCCGATCTNNNAGAAAATCCAGTTTCTAGGGGC
GCACACGTCTGAACTCTTTCCCTACACGACGCTCTTCCGATCTNNNAGAAAATCCAGTTTCTAGGGGC
AGCACACGTCTGAACTCTTTCCCTACACGACGCTCTTCCGATCTNNNCTATGTAGCAGGCACAGGAGA
AGCACACGTCTGAACTCTTTCCCTACACGACGCTCTTCCGATCTNNNCTATGTAGCAGGCACAGGAGA
GCACACGTCTGAACTCTTTCCCTACACGACGCTCTTCCGATCTNNNTCTCTACATTGCTGGCTTTTCCT
GCACACGTCTGAACTCTTTCCCTACACGACGCTCTTCCGATCTNNNTCTCTACATTGCTGGCTTTTCCT
AAGAGCACACGTCTGAACTCTTTCCCTACACGACGCTCTTCCGATCTNNNTTTGGCATTGGTCTGACT
AAGAGCACACGTCTGAACTCTTTCCCTACACGACGCTCTTCCGATCTNNNTTTGGCATTGGTCTGACT
AGCACACGTCTGAACTCTTTCCCTACACGACGCTCTTCCGATCTNNNGAGAATCACTTGAACCCAGG
AGCACACGTCTGAACTCTTTCCCTACACGACGCTCTTCCGATCTNNNGAGAATCACTTGAACCCAGG

GCACACGTCTGAACTCTTTCCCTACACGACGCTCTTCCGATCTNNNATGATGATGGGCTCCTGTAATC
GCACACGTCTGAACTCTTTCCCTACACGACGCTCTTCCGATCTNNNATGATGATGGGCTCCTGTAATC
AGCACACGTCTGAACTCTTTCCCTACACGACGCTCTTCCGATCTNNNTAGAAAGGCCTTAGGACAAA
AGCACACGTCTGAACTCTTTCCCTACACGACGCTCTTCCGATCTNNNTAGAAAGGCCTTAGGACAAA
GCACACGTCTGAACTCTTTCCCTACACGACGCTCTTCCGATCTNNNTAACAAGCCGCTCTCCCTAATTT
GCACACGTCTGAACTCTTTCCCTACACGACGCTCTTCCGATCTNNNTAACAAGCCGCTCTCCCTAATTT
GCACACGTCTGAACTCTTTCCCTACACGACGCTCTTCCGATCTNNNTCAAACTTGATCCTACAGCCAG
GCACACGTCTGAACTCTTTCCCTACACGACGCTCTTCCGATCTNNNTCAAACTTGATCCTACAGCCAG
AGCACACGTCTGAACTCTTTCCCTACACGACGCTCTTCCGATCTNNNTACAGCCTGGCTAGTTGAAAG
AGCACACGTCTGAACTCTTTCCCTACACGACGCTCTTCCGATCTNNNTACAGCCTGGCTAGTTGAAAG
AAGAGCACACGTCTGAACTCTTTCCCTACACGACGCTCTTCCGATCTNNNGAGATCACACCACTGCAT
AAGAGCACACGTCTGAACTCTTTCCCTACACGACGCTCTTCCGATCTNNNGAGATCACACCACTGCAT
AGCACACGTCTGAACTCTTTCCCTACACGACGCTCTTCCGATCTNNNAGCCTGGCTAATATGGTGAAA
AGCACACGTCTGAACTCTTTCCCTACACGACGCTCTTCCGATCTNNNAGCCTGGCTAATATGGTGAAA
GCACACGTCTGAACTCTTTCCCTACACGACGCTCTTCCGATCTNNNCTGGGTTCAAGTGATTCTTCTG
GCACACGTCTGAACTCTTTCCCTACACGACGCTCTTCCGATCTNNNCTGGGTTCAAGTGATTCTTCTG
GCACACGTCTGAACTCTTTCCCTACACGACGCTCTTCCGATCTNNNCCACCACGTCTGGCTATTTTTTTC
GCACACGTCTGAACTCTTTCCCTACACGACGCTCTTCCGATCTNNNCCACCACGTCTGGCTATTTTTTTC
GAGCACACGTCTGAACTCTTTCCCTACACGACGCTCTTCCGATCTNNNCAGGAAAATGGCGTGAACCT
GAGCACACGTCTGAACTCTTTCCCTACACGACGCTCTTCCGATCTNNNCAGGAAAATGGCGTGAACCT
GAGCACACGTCTGAACTCTTTCCCTACACGACGCTCTTCCGATCTNNNTGCTCATGCTTTTGGTGTCAT
GAGCACACGTCTGAACTCTTTCCCTACACGACGCTCTTCCGATCTNNNTGCTCATGCTTTTGGTGTCAT
GCACACGTCTGAACTCTTTCCCTACACGACGCTCTTCCGATCTNNNGCCTGTAGTCCCAGCTACTTGCA
GCACACGTCTGAACTCTTTCCCTACACGACGCTCTTCCGATCTNNNGCCTGTAGTCCCAGCTACTTGCA
AGCACACGTCTGAACTCTTTCCCTACACGACGCTCTTCCGATCTNNNGTGAGCTATGTTTGTGCCAGT
AGCACACGTCTGAACTCTTTCCCTACACGACGCTCTTCCGATCTNNNGTGAGCTATGTTTGTGCCAGT
GCACACGTCTGAACTCTTTCCCTACACGACGCTCTTCCGATCTNNNACACCATGGTATACTGCAGCTA
GCACACGTCTGAACTCTTTCCCTACACGACGCTCTTCCGATCTNNNACACCATGGTATACTGCAGCTA
GAGCACACGTCTGAACTCTTTCCCTACACGACGCTCTTCCGATCTNNNACAGGAGTACTTTTGACGCTC
GAGCACACGTCTGAACTCTTTCCCTACACGACGCTCTTCCGATCTNNNACAGGAGTACTTTTGACGCTC
GCACACGTCTGAACTCTTTCCCTACACGACGCTCTTCCGATCTNNNATCATGAGGTCAGGAGATCGAG
GCACACGTCTGAACTCTTTCCCTACACGACGCTCTTCCGATCTNNNATCATGAGGTCAGGAGATCGAG
AGCACACGTCTGAACTCTTTCCCTACACGACGCTCTTCCGATCTNNNGAATTTTGAAGTGGTGGCAGG
AGCACACGTCTGAACTCTTTCCCTACACGACGCTCTTCCGATCTNNNGAATTTTGAAGTGGTGGCAGG
GAGCACACGTCTGAACTCTTTCCCTACACGACGCTCTTCCGATCTNNNCTGTAGTTTCAGCTACTCGGG
GAGCACACGTCTGAACTCTTTCCCTACACGACGCTCTTCCGATCTNNNCTGTAGTTTCAGCTACTCGGG
GCACACGTCTGAACTCTTTCCCTACACGACGCTCTTCCGATCTNNNCTTTGCCTACCAGAAACGTAGTG
GCACACGTCTGAACTCTTTCCCTACACGACGCTCTTCCGATCTNNNCTTTGCCTACCAGAAACGTAGTG
GAGCACACGTCTGAACTCTTTCCCTACACGACGCTCTTCCGATCTNNNAGCCATCACATGATAAACTCC
GAGCACACGTCTGAACTCTTTCCCTACACGACGCTCTTCCGATCTNNNAGCCATCACATGATAAACTCC
AGCACACGTCTGAACTCTTTCCCTACACGACGCTCTTCCGATCTNNNGAGGGGCCAGAAGGAAAATG
AGCACACGTCTGAACTCTTTCCCTACACGACGCTCTTCCGATCTNNNGAGGGGCCAGAAGGAAAATG
GCACACGTCTGAACTCTTTCCCTACACGACGCTCTTCCGATCTNNNAAGCACCATTCATTTCATTTTCAT

GCACACGTCTGAACTCTTTCCCTACACGACGCTCTTCCGATCTNNNAAGCACCATTCATTTCATTTTCAT

GCACACGTCTGAACTCTTTCCCTACACGACGCTCTTCCGATCTNNNGACCACTAAGAGAAGGAAAGC

GCACACGTCTGAACTCTTTCCCTACACGACGCTCTTCCGATCTNNNGACCACTAAGAGAAGGAAAGC

AGAGCACACGTCTGAACTCTTTCCCTACACGACGCTCTTCCGATCTNNNACTAGAACCTGGGCGAGA

AGAGCACACGTCTGAACTCTTTCCCTACACGACGCTCTTCCGATCTNNNACTAGAACCTGGGCGAGA

GAGCACACGTCTGAACTCTTTCCCTACACGACGCTCTTCCGATCTNNNTCTGTAATCCCAGCTACTCAG

GAGCACACGTCTGAACTCTTTCCCTACACGACGCTCTTCCGATCTNNNTCTGTAATCCCAGCTACTCAG

AGCACACGTCTGAACTCTTTCCCTACACGACGCTCTTCCGATCTNNNAACTCCTGACCTCAAGTGATCT

AGCACACGTCTGAACTCTTTCCCTACACGACGCTCTTCCGATCTNNNAACTCCTGACCTCAAGTGATCT

AGCACACGTCTGAACTCTTTCCCTACACGACGCTCTTCCGATCTNNNAGCACTTTGAGAAATTTGAGG

AGCACACGTCTGAACTCTTTCCCTACACGACGCTCTTCCGATCTNNNAGCACTTTGAGAAATTTGAGG

GCACACGTCTGAACTCTTTCCCTACACGACGCTCTTCCGATCTNNNGGCATCACCAGGAGGATATGA

GCACACGTCTGAACTCTTTCCCTACACGACGCTCTTCCGATCTNNNGGCATCACCAGGAGGATATGA

CACACGTCTGAACTCTTTCCCTACACGACGCTCTTCCGATCTNNNTTTTTTTCTTGTAGGCTGGGCACC

CACACGTCTGAACTCTTTCCCTACACGACGCTCTTCCGATCTNNNTTTTTTTCTTGTAGGCTGGGCACC

AGCACACGTCTGAACTCTTTCCCTACACGACGCTCTTCCGATCTNNNGAGGCAGGAGAATCGCTTGA

AGCACACGTCTGAACTCTTTCCCTACACGACGCTCTTCCGATCTNNNGAGGCAGGAGAATCGCTTGA

AGCACACGTCTGAACTCTTTCCCTACACGACGCTCTTCCGATCTNNNTGTTTGAGCCCAAAAGTTTGA

AGCACACGTCTGAACTCTTTCCCTACACGACGCTCTTCCGATCTNNNTGTTTGAGCCCAAAAGTTTGA

GAGCACACGTCTGAACTCTTTCCCTACACGACGCTCTTCCGATCTNNNTACCGTTCCGAGGTAACCTAA

GAGCACACGTCTGAACTCTTTCCCTACACGACGCTCTTCCGATCTNNNTACCGTTCCGAGGTAACCTAA

AGCACACGTCTGAACTCTTTCCCTACACGACGCTCTTCCGATCTNNNTGAAGTGGGAGTACTGTTTGA

AGCACACGTCTGAACTCTTTCCCTACACGACGCTCTTCCGATCTNNNTGAAGTGGGAGTACTGTTTGA

AGCACACGTCTGAACTCTTTCCCTACACGACGCTCTTCCGATCTNNNTGAGGTGTGAGTGATACTTTTG

AGCACACGTCTGAACTCTTTCCCTACACGACGCTCTTCCGATCTNNNTGAGGTGTGAGTGATACTTTTG

GAGCACACGTCTGAACTCTTTCCCTACACGACGCTCTTCCGATCTNNNACACCTTTCATGATGAAAACA

GAGCACACGTCTGAACTCTTTCCCTACACGACGCTCTTCCGATCTNNNACACCTTTCATGATGAAAACA

GCACACGTCTGAACTCTTTCCCTACACGACGCTCTTCCGATCTNNNTCTCTGGAGTGCAGACTTCAAA

GCACACGTCTGAACTCTTTCCCTACACGACGCTCTTCCGATCTNNNTCTCTGGAGTGCAGACTTCAAA

AGCACACGTCTGAACTCTTTCCCTACACGACGCTCTTCCGATCTNNNTGAGGCAGGAGAATCTCTTGA

AGCACACGTCTGAACTCTTTCCCTACACGACGCTCTTCCGATCTNNNTGAGGCAGGAGAATCTCTTGA

AGCACACGTCTGAACTCTTTCCCTACACGACGCTCTTCCGATCTNNNCACTCCATGTTCAGTGCTCTTTG

AGCACACGTCTGAACTCTTTCCCTACACGACGCTCTTCCGATCTNNNCACTCCATGTTCAGTGCTCTTTG

GAGCACACGTCTGAACTCTTTCCCTACACGACGCTCTTCCGATCTNNNCCATAATCCCAGCTACTCGGG

GAGCACACGTCTGAACTCTTTCCCTACACGACGCTCTTCCGATCTNNNCCATAATCCCAGCTACTCGGG

AGCACACGTCTGAACTCTTTCCCTACACGACGCTCTTCCGATCTNNNCCCACCCATCCATCATTCATCTA

AGCACACGTCTGAACTCTTTCCCTACACGACGCTCTTCCGATCTNNNCCCACCCATCCATCATTCATCTA

AGCACACGTCTGAACTCTTTCCCTACACGACGCTCTTCCGATCTNNNCATCCTGGCTAACATGGTGAAA

AGCACACGTCTGAACTCTTTCCCTACACGACGCTCTTCCGATCTNNNCATCCTGGCTAACATGGTGAAA

AGCACACGTCTGAACTCTTTCCCTACACGACGCTCTTCCGATCTNNNAGCTGAGATACTGGCAATTGG

AGCACACGTCTGAACTCTTTCCCTACACGACGCTCTTCCGATCTNNNAGCTGAGATACTGGCAATTGG

AGCACACGTCTGAACTCTTTCCCTACACGACGCTCTTCCGATCTNNNACAAAAATTAGTCGGGCATGG

AGCACACGTCTGAACTCTTTCCCTACACGACGCTCTTCCGATCTNNNACAAAAATTAGTCGGGCATGG

AGCACACGTCTGAACTCTTTCCCTACACGACGCTCTTCCGATCTNNNAATCACTTGAACTTGGGAGGA

AGCACACGTCTGAACTCTTTCCCTACACGACGCTCTTCCGATCTNNNAATCACTTGAACTTGGGAGGA

GAGCACACGTCTGAACTCTTTCCCTACACGACGCTCTTCCGATCTNNNTCCACATTCAATTCATGACGGT

GAGCACACGTCTGAACTCTTTCCCTACACGACGCTCTTCCGATCTNNNTCCACATTCAATTCATGACGGT

AGCACACGTCTGAACTCTTTCCCTACACGACGCTCTTCCGATCTNNNGGAACCAGTGTACCTCTCAACA

AGCACACGTCTGAACTCTTTCCCTACACGACGCTCTTCCGATCTNNNGGAACCAGTGTACCTCTCAACA

AGCACACGTCTGAACTCTTTCCCTACACGACGCTCTTCCGATCTNNNGAGCTTGTAGTGAGCCGAGAT

AGCACACGTCTGAACTCTTTCCCTACACGACGCTCTTCCGATCTNNNGAGCTTGTAGTGAGCCGAGAT

GAGCACACGTCTGAACTCTTTCCCTACACGACGCTCTTCCGATCTNNNCCCAAAATTTCACTGACACAC

GAGCACACGTCTGAACTCTTTCCCTACACGACGCTCTTCCGATCTNNNCCCAAAATTTCACTGACACAC

AGCACACGTCTGAACTCTTTCCCTACACGACGCTCTTCCGATCTNNNATACTCATCTACAGGAGGGGA

AGCACACGTCTGAACTCTTTCCCTACACGACGCTCTTCCGATCTNNNATACTCATCTACAGGAGGGGA

AGCACACGTCTGAACTCTTTCCCTACACGACGCTCTTCCGATCTNNNATAGGTTGCTGTGAGCCATGAT

AGCACACGTCTGAACTCTTTCCCTACACGACGCTCTTCCGATCTNNNATAGGTTGCTGTGAGCCATGAT

CACACGTCTGAACTCTTTCCCTACACGACGCTCTTCCGATCTNNNCAGCTCTGGTGTTGTCGTGGCATC

CACACGTCTGAACTCTTTCCCTACACGACGCTCTTCCGATCTNNNCAGCTCTGGTGTTGTCGTGGCATC

GAGCACACGTCTGAACTCTTTCCCTACACGACGCTCTTCCGATCTNNNATTCATGGGGTGTTCAGCCTA

GAGCACACGTCTGAACTCTTTCCCTACACGACGCTCTTCCGATCTNNNATTCATGGGGTGTTCAGCCTA

GAGCACACGTCTGAACTCTTTCCCTACACGACGCTCTTCCGATCTNNNCATATTGAGCCCCTAGTCTGT

GAGCACACGTCTGAACTCTTTCCCTACACGACGCTCTTCCGATCTNNNCATATTGAGCCCCTAGTCTGT

GAGCACACGTCTGAACTCTTTCCCTACACGACGCTCTTCCGATCTNNNCTGAGGCAGGAGAATTGCTA

GAGCACACGTCTGAACTCTTTCCCTACACGACGCTCTTCCGATCTNNNCTGAGGCAGGAGAATTGCTA

GAGCACACGTCTGAACTCTTTCCCTACACGACGCTCTTCCGATCTNNNCTCCTATTTCCCTAGGGCCAAA

GAGCACACGTCTGAACTCTTTCCCTACACGACGCTCTTCCGATCTNNNCTCCTATTTCCCTAGGGCCAAA

AGCACACGTCTGAACTCTTTCCCTACACGACGCTCTTCCGATCTNNNGCCTCTGTAATAGCTGGGACTA

AGCACACGTCTGAACTCTTTCCCTACACGACGCTCTTCCGATCTNNNGCCTCTGTAATAGCTGGGACTA

AGCACACGTCTGAACTCTTTCCCTACACGACGCTCTTCCGATCTNNNGCAGTTCTTCCACTTTAGCTTCC

AGCACACGTCTGAACTCTTTCCCTACACGACGCTCTTCCGATCTNNNGCAGTTCTTCCACTTTAGCTTCC

GAGCACACGTCTGAACTCTTTCCCTACACGACGCTCTTCCGATCTNNNACACATGGGAATGGGTAACTT

GAGCACACGTCTGAACTCTTTCCCTACACGACGCTCTTCCGATCTNNNACACATGGGAATGGGTAACTT

GAGCACACGTCTGAACTCTTTCCCTACACGACGCTCTTCCGATCTNNNTGGACTAGGTTGTGAGTTCCA

GAGCACACGTCTGAACTCTTTCCCTACACGACGCTCTTCCGATCTNNNTGGACTAGGTTGTGAGTTCCA

GAGCACACGTCTGAACTCTTTCCCTACACGACGCTCTTCCGATCTNNNTTGACTTTCCCACAGATCTTGG

GAGCACACGTCTGAACTCTTTCCCTACACGACGCTCTTCCGATCTNNNTTGACTTTCCCACAGATCTTGG

GAGCACACGTCTGAACTCTTTCCCTACACGACGCTCTTCCGATCTNNNTGGCTGGATATGAAATTCTG

GAGCACACGTCTGAACTCTTTCCCTACACGACGCTCTTCCGATCTNNNTGGCTGGATATGAAATTCTG

AGCACACGTCTGAACTCTTTCCCTACACGACGCTCTTCCGATCTNNNACAGTGGCTGAGTTGAACAAA

AGCACACGTCTGAACTCTTTCCCTACACGACGCTCTTCCGATCTNNNACAGTGGCTGAGTTGAACAAA

GAGCACACGTCTGAACTCTTTCCCTACACGACGCTCTTCCGATCTNNNGGCTCACTCCTGTAATTCTAG

GAGCACACGTCTGAACTCTTTCCCTACACGACGCTCTTCCGATCTNNNGGCTCACTCCTGTAATTCTAG

AGCACACGTCTGAACTCTTTCCCTACACGACGCTCTTCCGATCTNNNCCTTCTGTTAGAAGATAAATTG

AGCACACGTCTGAACTCTTTCCCTACACGACGCTCTTCCGATCTNNNCCTTCTGTTAGAAGATAAATTG

GAGCACACGTCTGAACTCTTTCCCTACACGACGCTCTTCCGATCTNNNGTAATCCCAGCTACTCAAGAG

GAGCACACGTCTGAACTCTTTCCCTACACGACGCTCTTCCGATCTNNNGTAATCCCAGCTACTCAAGAG
AGCACACGTCTGAACTCTTTCCCTACACGACGCTCTTCCGATCTNNNAGTACCCTCTCATACTGTCCTGC
AGCACACGTCTGAACTCTTTCCCTACACGACGCTCTTCCGATCTNNNAGTACCCTCTCATACTGTCCTGC
GAGCACACGTCTGAACTCTTTCCCTACACGACGCTCTTCCGATCTNNNCTGAGCTGTGATCACACCACT
GAGCACACGTCTGAACTCTTTCCCTACACGACGCTCTTCCGATCTNNNCTGAGCTGTGATCACACCACT
GCACACGTCTGAACTCTTTCCCTACACGACGCTCTTCCGATCTNNNCTCTTGTGGAGAAGGTTGGCTA
GCACACGTCTGAACTCTTTCCCTACACGACGCTCTTCCGATCTNNNCTCTTGTGGAGAAGGTTGGCTA
GAGCACACGTCTGAACTCTTTCCCTACACGACGCTCTTCCGATCTNNNCAAGGAGGATGGCAGATTTT
GAGCACACGTCTGAACTCTTTCCCTACACGACGCTCTTCCGATCTNNNCAAGGAGGATGGCAGATTTT
GAGCACACGTCTGAACTCTTTCCCTACACGACGCTCTTCCGATCTNNNCTGCTGGTTGCTTGAGAAGA
GAGCACACGTCTGAACTCTTTCCCTACACGACGCTCTTCCGATCTNNNCTGCTGGTTGCTTGAGAAGA
GCACACGTCTGAACTCTTTCCCTACACGACGCTCTTCCGATCTNNNCTTGAAAGATTCAGGCTGATGG
GCACACGTCTGAACTCTTTCCCTACACGACGCTCTTCCGATCTNNNCTTGAAAGATTCAGGCTGATGG
AGCACACGTCTGAACTCTTTCCCTACACGACGCTCTTCCGATCTNNNGCAATTTACGTAGATGATTTTC
AGCACACGTCTGAACTCTTTCCCTACACGACGCTCTTCCGATCTNNNGCAATTTACGTAGATGATTTTC
GAGCACACGTCTGAACTCTTTCCCTACACGACGCTCTTCCGATCTNNNAACATGGCGAAATCCCATCTC
GAGCACACGTCTGAACTCTTTCCCTACACGACGCTCTTCCGATCTNNNAACATGGCGAAATCCCATCTC
GAGCACACGTCTGAACTCTTTCCCTACACGACGCTCTTCCGATCTNNNCATCTTTAGCATCCATTCCCTT
GAGCACACGTCTGAACTCTTTCCCTACACGACGCTCTTCCGATCTNNNCATCTTTAGCATCCATTCCCTT
AGCACACGTCTGAACTCTTTCCCTACACGACGCTCTTCCGATCTNNNGAGGCTGAGGCAGGAGAATG
AGCACACGTCTGAACTCTTTCCCTACACGACGCTCTTCCGATCTNNNGAGGCTGAGGCAGGAGAATG
GCACACGTCTGAACTCTTTCCCTACACGACGCTCTTCCGATCTNNNCCACTGCAACACACTGCTATAAG
GCACACGTCTGAACTCTTTCCCTACACGACGCTCTTCCGATCTNNNCCACTGCAACACACTGCTATAAG
AGCACACGTCTGAACTCTTTCCCTACACGACGCTCTTCCGATCTNNNCTGAAAGTGCAGAGGAAAGCT
AGCACACGTCTGAACTCTTTCCCTACACGACGCTCTTCCGATCTNNNCTGAAAGTGCAGAGGAAAGCT
AGCACACGTCTGAACTCTTTCCCTACACGACGCTCTTCCGATCTNNNCCAAACACCGCATATTCTCACT
AGCACACGTCTGAACTCTTTCCCTACACGACGCTCTTCCGATCTNNNCCAAACACCGCATATTCTCACT
AGCACACGTCTGAACTCTTTCCCTACACGACGCTCTTCCGATCTNNNGTATTGGCTAGGAATGGCTA
AGCACACGTCTGAACTCTTTCCCTACACGACGCTCTTCCGATCTNNNGTATTGGCTAGGAATGGCTA
GAGCACACGTCTGAACTCTTTCCCTACACGACGCTCTTCCGATCTNNNCTGAGGCAGGATAATCGCTT
GAGCACACGTCTGAACTCTTTCCCTACACGACGCTCTTCCGATCTNNNCTGAGGCAGGATAATCGCTT
AGCACACGTCTGAACTCTTTCCCTACACGACGCTCTTCCGATCTNNNCTCCAGACGTTTTAGGAATTCG
AGCACACGTCTGAACTCTTTCCCTACACGACGCTCTTCCGATCTNNNCTCCAGACGTTTTAGGAATTCG
AGCACACGTCTGAACTCTTTCCCTACACGACGCTCTTCCGATCTNNNATTGATGGAAGGGGCAAGAT
AGCACACGTCTGAACTCTTTCCCTACACGACGCTCTTCCGATCTNNNATTGATGGAAGGGGCAAGAT
AGCACACGTCTGAACTCTTTCCCTACACGACGCTCTTCCGATCTNNNAAACACTATTCAGCACAACCAG
AGCACACGTCTGAACTCTTTCCCTACACGACGCTCTTCCGATCTNNNAAACACTATTCAGCACAACCAG
AGAGCACACGTCTGAACTCTTTCCCTACACGACGCTCTTCCGATCTNNNATTCACTCAGAAAAACGCTG
AGAGCACACGTCTGAACTCTTTCCCTACACGACGCTCTTCCGATCTNNNATTCACTCAGAAAAACGCTG
AGCACACGTCTGAACTCTTTCCCTACACGACGCTCTTCCGATCTNNNTTTCTCAGCCCCACATGTATTCA
AGCACACGTCTGAACTCTTTCCCTACACGACGCTCTTCCGATCTNNNTTTCTCAGCCCCACATGTATTCA
GAGCACACGTCTGAACTCTTTCCCTACACGACGCTCTTCCGATCTNNNCTGTAATCCCAGCTACTCGGA
GAGCACACGTCTGAACTCTTTCCCTACACGACGCTCTTCCGATCTNNNCTGTAATCCCAGCTACTCGGA

AGCACACGTCTGAACTCTTTCCCTACACGACGCTCTTCCGATCTNNNGCCTGTAATCCCAACACTTTGA
AGCACACGTCTGAACTCTTTCCCTACACGACGCTCTTCCGATCTNNNGCCTGTAATCCCAACACTTTGA
AGCACACGTCTGAACTCTTTCCCTACACGACGCTCTTCCGATCTNNNTTGTAGAAAGGGTTGGCAGA
AGCACACGTCTGAACTCTTTCCCTACACGACGCTCTTCCGATCTNNNTTGTAGAAAGGGTTGGCAGA
GAGCACACGTCTGAACTCTTTCCCTACACGACGCTCTTCCGATCTNNNAAGTAGCTGGGACTACAAGT
GAGCACACGTCTGAACTCTTTCCCTACACGACGCTCTTCCGATCTNNNAAGTAGCTGGGACTACAAGT
CACACGTCTGAACTCTTTCCCTACACGACGCTCTTCCGATCTNNNCTTCCTTCTCCCAGCACCTCCATCGG
CACACGTCTGAACTCTTTCCCTACACGACGCTCTTCCGATCTNNNCTTCCTTCTCCCAGCACCTCCATCGG
AGCACACGTCTGAACTCTTTCCCTACACGACGCTCTTCCGATCTNNNTACTACAAACTCACAGCAGCCA
AGCACACGTCTGAACTCTTTCCCTACACGACGCTCTTCCGATCTNNNTACTACAAACTCACAGCAGCCA
ACACGTCTGAACTCTTTCCCTACACGACGCTCTTCCGATCTNNNCATGGTTTGGCTCTCTGTTTGTCCA
ACACGTCTGAACTCTTTCCCTACACGACGCTCTTCCGATCTNNNCATGGTTTGGCTCTCTGTTTGTCCA
GAGCACACGTCTGAACTCTTTCCCTACACGACGCTCTTCCGATCTNNNATCTCTTGAGCCTAGGAAGT
GAGCACACGTCTGAACTCTTTCCCTACACGACGCTCTTCCGATCTNNNATCTCTTGAGCCTAGGAAGT
AGCACACGTCTGAACTCTTTCCCTACACGACGCTCTTCCGATCTNNNTGAGGCAGGAGAATCACTTGA
AGCACACGTCTGAACTCTTTCCCTACACGACGCTCTTCCGATCTNNNTGAGGCAGGAGAATCACTTGA
GCACACGTCTGAACTCTTTCCCTACACGACGCTCTTCCGATCTNNNGGCTCCAGGTCAGAGTCTAAAT
GCACACGTCTGAACTCTTTCCCTACACGACGCTCTTCCGATCTNNNGGCTCCAGGTCAGAGTCTAAAT
ACACGTCTGAACTCTTTCCCTACACGACGCTCTTCCGATCTNNNGATGGGGTTGTCATGCTCGATCAT
ACACGTCTGAACTCTTTCCCTACACGACGCTCTTCCGATCTNNNGATGGGGTTGTCATGCTCGATCAT
AGCACACGTCTGAACTCTTTCCCTACACGACGCTCTTCCGATCTNNNCACATGCCTATAGTCCCAGCTA
AGCACACGTCTGAACTCTTTCCCTACACGACGCTCTTCCGATCTNNNCACATGCCTATAGTCCCAGCTA
GCACACGTCTGAACTCTTTCCCTACACGACGCTCTTCCGATCTNNNAGCCCAGTTCCATATTGAGGTT
GCACACGTCTGAACTCTTTCCCTACACGACGCTCTTCCGATCTNNNAGCCCAGTTCCATATTGAGGTT
CACACGTCTGAACTCTTTCCCTACACGACGCTCTTCCGATCTNNNCTATGGCACTCATTCTGTCCAGTC
CACACGTCTGAACTCTTTCCCTACACGACGCTCTTCCGATCTNNNCTATGGCACTCATTCTGTCCAGTC
GAGCACACGTCTGAACTCTTTCCCTACACGACGCTCTTCCGATCTNNNTACATAGGCAGCTTTGGATG
GAGCACACGTCTGAACTCTTTCCCTACACGACGCTCTTCCGATCTNNNTACATAGGCAGCTTTGGATG
GAGCACACGTCTGAACTCTTTCCCTACACGACGCTCTTCCGATCTNNNCGAGTTGAAAAAGGGAAAG
GAGCACACGTCTGAACTCTTTCCCTACACGACGCTCTTCCGATCTNNNCGAGTTGAAAAAGGGAAAG
GCACACGTCTGAACTCTTTCCCTACACGACGCTCTTCCGATCTNNNCGAGTACAAGTTCTTCTGGACG
GCACACGTCTGAACTCTTTCCCTACACGACGCTCTTCCGATCTNNNCGAGTACAAGTTCTTCTGGACG
AGCACACGTCTGAACTCTTTCCCTACACGACGCTCTTCCGATCTNNNGGAGAATGACAGAGATATCA
AGCACACGTCTGAACTCTTTCCCTACACGACGCTCTTCCGATCTNNNGGAGAATGACAGAGATATCA
GCACACGTCTGAACTCTTTCCCTACACGACGCTCTTCCGATCTNNNCTCGAACTTCTGACTTCAGGTG
GCACACGTCTGAACTCTTTCCCTACACGACGCTCTTCCGATCTNNNCTCGAACTTCTGACTTCAGGTG
AGCACACGTCTGAACTCTTTCCCTACACGACGCTCTTCCGATCTNNNGCATTCCTTGTCATGTGCTTTCT
AGCACACGTCTGAACTCTTTCCCTACACGACGCTCTTCCGATCTNNNGCATTCCTTGTCATGTGCTTTCT
AGCACACGTCTGAACTCTTTCCCTACACGACGCTCTTCCGATCTNNNTGGTATTCTGTTGTACTGTGG
AGCACACGTCTGAACTCTTTCCCTACACGACGCTCTTCCGATCTNNNTGGTATTCTGTTGTACTGTGG
GAGCACACGTCTGAACTCTTTCCCTACACGACGCTCTTCCGATCTNNNGAGGGGTATCATGTTTGGTT
GAGCACACGTCTGAACTCTTTCCCTACACGACGCTCTTCCGATCTNNNGAGGGGTATCATGTTTGGTT
AGCACACGTCTGAACTCTTTCCCTACACGACGCTCTTCCGATCTNNNCTCCTGAGGGGGAATTTGGTT

GAGCACACGTCTGAACTCTTTCCCTACACGACGCTCTTCCGATCTNNNCTCCTGAGGGGGAATTTGGTT

GCACACGTCTGAACTCTTTCCCTACACGACGCTCTTCCGATCTNNNGTTTGTCAAGAAATGGTTGCAC

GCACACGTCTGAACTCTTTCCCTACACGACGCTCTTCCGATCTNNNGTTTGTCAAGAAATGGTTGCAC

AGCACACGTCTGAACTCTTTCCCTACACGACGCTCTTCCGATCTNNNTCGATCTTCGAGCCAAAGATG

AGCACACGTCTGAACTCTTTCCCTACACGACGCTCTTCCGATCTNNNTCGATCTTCGAGCCAAAGATG

AGCACACGTCTGAACTCTTTCCCTACACGACGCTCTTCCGATCTNNNTATCAGAGAAGGGTTTCAGAA

AGCACACGTCTGAACTCTTTCCCTACACGACGCTCTTCCGATCTNNNTATCAGAGAAGGGTTTCAGAA

GCACACGTCTGAACTCTTTCCCTACACGACGCTCTTCCGATCTNNNAGCTAGCCAAAATTGCACCATTG

GCACACGTCTGAACTCTTTCCCTACACGACGCTCTTCCGATCTNNNAGCTAGCCAAAATTGCACCATTG

AGAGCACACGTCTGAACTCTTTCCCTACACGACGCTCTTCCGATCTNNNCTCCCTCTTTCTCTCACCT

AGAGCACACGTCTGAACTCTTTCCCTACACGACGCTCTTCCGATCTNNNCTCCCTCTTTCTCTCACCT

AGCACACGTCTGAACTCTTTCCCTACACGACGCTCTTCCGATCTNNNATTTGTTCCTTCTTGCCATGCTCC

AGCACACGTCTGAACTCTTTCCCTACACGACGCTCTTCCGATCTNNNATTTGTTCCTTCTTGCCATGCTCC

AGCACACGTCTGAACTCTTTCCCTACACGACGCTCTTCCGATCTNNNAACCAGTACATGAGGCAATCCA

AGCACACGTCTGAACTCTTTCCCTACACGACGCTCTTCCGATCTNNNAACCAGTACATGAGGCAATCCA

GAGCACACGTCTGAACTCTTTCCCTACACGACGCTCTTCCGATCTNNNAGTATGCTGAGACTGTCACTT

GAGCACACGTCTGAACTCTTTCCCTACACGACGCTCTTCCGATCTNNNAGTATGCTGAGACTGTCACTT

GAGCACACGTCTGAACTCTTTCCCTACACGACGCTCTTCCGATCTNNNATACCTAGTTCTGCTTTCCCTG

GAGCACACGTCTGAACTCTTTCCCTACACGACGCTCTTCCGATCTNNNATACCTAGTTCTGCTTTCCCTG

AGCACACGTCTGAACTCTTTCCCTACACGACGCTCTTCCGATCTNNNCAGGAGTTCAAGACCAGCCTAT

AGCACACGTCTGAACTCTTTCCCTACACGACGCTCTTCCGATCTNNNCAGGAGTTCAAGACCAGCCTAT

GAGCACACGTCTGAACTCTTTCCCTACACGACGCTCTTCCGATCTNNNCTAAGGCAGGAGGATCACTTC

GAGCACACGTCTGAACTCTTTCCCTACACGACGCTCTTCCGATCTNNNCTAAGGCAGGAGGATCACTTC

GCACACGTCTGAACTCTTTCCCTACACGACGCTCTTCCGATCTNNNCTCCCACACCCCAAAAGACCATCC

GCACACGTCTGAACTCTTTCCCTACACGACGCTCTTCCGATCTNNNCTCCCACACCCCAAAAGACCATCC

GAGCACACGTCTGAACTCTTTCCCTACACGACGCTCTTCCGATCTNNNAGCTCCCAGTGATATCAACACA

GAGCACACGTCTGAACTCTTTCCCTACACGACGCTCTTCCGATCTNNNAGCTCCCAGTGATATCAACACA

AGCACACGTCTGAACTCTTTCCCTACACGACGCTCTTCCGATCTNNNCTCCAGGCACTCGGTTCTTTCCAT

AGCACACGTCTGAACTCTTTCCCTACACGACGCTCTTCCGATCTNNNCTCCAGGCACTCGGTTCTTTCCAT

AGCACACGTCTGAACTCTTTCCCTACACGACGCTCTTCCGATCTNNNCTGCCTCAGCCTCCCAAATAGCA

AGCACACGTCTGAACTCTTTCCCTACACGACGCTCTTCCGATCTNNNCTGCCTCAGCCTCCCAAATAGCA

GCACACGTCTGAACTCTTTCCCTACACGACGCTCTTCCGATCTNNNGGCAGAGTGTAAATCCCAAATC

GCACACGTCTGAACTCTTTCCCTACACGACGCTCTTCCGATCTNNNGGCAGAGTGTAAATCCCAAATC

GAGCACACGTCTGAACTCTTTCCCTACACGACGCTCTTCCGATCTNNNACAATCTTCACAATCCAGCCTCT

GAGCACACGTCTGAACTCTTTCCCTACACGACGCTCTTCCGATCTNNNACAATCTTCACAATCCAGCCTCT

AGCACACGTCTGAACTCTTTCCCTACACGACGCTCTTCCGATCTNNNCAAGAGGTTGAAGCAGCAGTTA

AGCACACGTCTGAACTCTTTCCCTACACGACGCTCTTCCGATCTNNNCAAGAGGTTGAAGCAGCAGTTA

GAGCACACGTCTGAACTCTTTCCCTACACGACGCTCTTCCGATCTNNNGCTAACTTTGTGTGTGTGTGTG

GAGCACACGTCTGAACTCTTTCCCTACACGACGCTCTTCCGATCTNNNGCTAACTTTGTGTGTGTGTGTG

GAGCACACGTCTGAACTCTTTCCCTACACGACGCTCTTCCGATCTNNNAGGCAGAAGAATCACTTGAAC

GAGCACACGTCTGAACTCTTTCCCTACACGACGCTCTTCCGATCTNNNAGGCAGAAGAATCACTTGAAC

AGCACACGTCTGAACTCTTTCCCTACACGACGCTCTTCCGATCTNNNAGTCAAGCTAGACTGCTGCTAT

AGCACACGTCTGAACTCTTTCCCTACACGACGCTCTTCCGATCTNNNAGTCAAGCTAGACTGCTGCTAT

CACACGTCTGAACTCTTTCCCTACACGACGCTCTTCCGATCTNNNCACTGCTACCTGGTGGCTGGCATC
CACACGTCTGAACTCTTTCCCTACACGACGCTCTTCCGATCTNNNCACTGCTACCTGGTGGCTGGCATC
GAGCACACGTCTGAACTCTTTCCCTACACGACGCTCTTCCGATCTNNNGCCACATGCTGCTGAAAATG
GAGCACACGTCTGAACTCTTTCCCTACACGACGCTCTTCCGATCTNNNGCCACATGCTGCTGAAAATG
GCACACGTCTGAACTCTTTCCCTACACGACGCTCTTCCGATCTNNNGTGTTGGGAGGGATAACAGAC
GCACACGTCTGAACTCTTTCCCTACACGACGCTCTTCCGATCTNNNGTGTTGGGAGGGATAACAGAC
GCACACGTCTGAACTCTTTCCCTACACGACGCTCTTCCGATCTNNNCTGACTTGAGTGCAGTGATCTCT
GCACACGTCTGAACTCTTTCCCTACACGACGCTCTTCCGATCTNNNCTGACTTGAGTGCAGTGATCTCT
AAGAGCACACGTCTGAACTCTTTCCCTACACGACGCTCTTCCGATCTNNNAGACAAGGGAATCACTTG
AAGAGCACACGTCTGAACTCTTTCCCTACACGACGCTCTTCCGATCTNNNAGACAAGGGAATCACTTG
AGCACACGTCTGAACTCTTTCCCTACACGACGCTCTTCCGATCTNNNTTAGTAGAGATGGGGTTTCAC
AGCACACGTCTGAACTCTTTCCCTACACGACGCTCTTCCGATCTNNNTTAGTAGAGATGGGGTTTCAC
CACACGTCTGAACTCTTTCCCTACACGACGCTCTTCCGATCTNNNACTTACAAGGGATGTGAAGGACC
CACACGTCTGAACTCTTTCCCTACACGACGCTCTTCCGATCTNNNACTTACAAGGGATGTGAAGGACC
AGCACACGTCTGAACTCTTTCCCTACACGACGCTCTTCCGATCTNNNCTAGGCGCCAGTTCTCTTGAG
AGCACACGTCTGAACTCTTTCCCTACACGACGCTCTTCCGATCTNNNCTAGGCGCCAGTTCTCTTGAG
GAGCACACGTCTGAACTCTTTCCCTACACGACGCTCTTCCGATCTNNNTGATAAAGGGGATATCACCA
GAGCACACGTCTGAACTCTTTCCCTACACGACGCTCTTCCGATCTNNNTGATAAAGGGGATATCACCA
AGCACACGTCTGAACTCTTTCCCTACACGACGCTCTTCCGATCTNNNCAACAACGCAGAGCCTCAAATA
AGCACACGTCTGAACTCTTTCCCTACACGACGCTCTTCCGATCTNNNCAACAACGCAGAGCCTCAAATA
GAGCACACGTCTGAACTCTTTCCCTACACGACGCTCTTCCGATCTNNNGAAGTTGCAGTGAGCTGAG
GAGCACACGTCTGAACTCTTTCCCTACACGACGCTCTTCCGATCTNNNGAAGTTGCAGTGAGCTGAG
AGCACACGTCTGAACTCTTTCCCTACACGACGCTCTTCCGATCTNNNGTTCCTGTAGTCCCAGCTACTTG
AGCACACGTCTGAACTCTTTCCCTACACGACGCTCTTCCGATCTNNNGTTCCTGTAGTCCCAGCTACTTG
AGCACACGTCTGAACTCTTTCCCTACACGACGCTCTTCCGATCTNNNCTAACCTCAAGTGATCCACCCTC
AGCACACGTCTGAACTCTTTCCCTACACGACGCTCTTCCGATCTNNNCTAACCTCAAGTGATCCACCCTC
GCACACGTCTGAACTCTTTCCCTACACGACGCTCTTCCGATCTNNNTCATGCATGTAATCCCCACACTT
GCACACGTCTGAACTCTTTCCCTACACGACGCTCTTCCGATCTNNNTCATGCATGTAATCCCCACACTT
GAGCACACGTCTGAACTCTTTCCCTACACGACGCTCTTCCGATCTNNNGTTACTTTGGGCAAGACATT
GAGCACACGTCTGAACTCTTTCCCTACACGACGCTCTTCCGATCTNNNGTTACTTTGGGCAAGACATT
AGCACACGTCTGAACTCTTTCCCTACACGACGCTCTTCCGATCTNNNCTCCCGAGTAGCTGGAATTACA
AGCACACGTCTGAACTCTTTCCCTACACGACGCTCTTCCGATCTNNNCTCCCGAGTAGCTGGAATTACA
GAGCACACGTCTGAACTCTTTCCCTACACGACGCTCTTCCGATCTNNNCAGATCACGAGGTCAGGAGA
GAGCACACGTCTGAACTCTTTCCCTACACGACGCTCTTCCGATCTNNNCAGATCACGAGGTCAGGAGA
GCACACGTCTGAACTCTTTCCCTACACGACGCTCTTCCGATCTNNNATCTTTTTGCTGATGGAGGGTC
GCACACGTCTGAACTCTTTCCCTACACGACGCTCTTCCGATCTNNNATCTTTTTGCTGATGGAGGGTC
AGAGCACACGTCTGAACTCTTTCCCTACACGACGCTCTTCCGATCTNNNGATGTAGGACTCGTGGTTT
AGAGCACACGTCTGAACTCTTTCCCTACACGACGCTCTTCCGATCTNNNGATGTAGGACTCGTGGTTT
GAGCACACGTCTGAACTCTTTCCCTACACGACGCTCTTCCGATCTNNNTAATCCTGGAGTAGAGTGAG
GAGCACACGTCTGAACTCTTTCCCTACACGACGCTCTTCCGATCTNNNTAATCCTGGAGTAGAGTGAG
GAGCACACGTCTGAACTCTTTCCCTACACGACGCTCTTCCGATCTNNNTGGGTGACAGAGCAAGACTC
GAGCACACGTCTGAACTCTTTCCCTACACGACGCTCTTCCGATCTNNNTGGGTGACAGAGCAAGACTC
AGCACACGTCTGAACTCTTTCCCTACACGACGCTCTTCCGATCTNNNTCAACTTATCTTTCAACACCCAG

AGCACACGTCTGAACTCTTTCCCTACACGACGCTCTTCCGATCTNNNTCAACTTATCTTTCAACACCCAG
AGCACACGTCTGAACTCTTTCCCTACACGACGCTCTTCCGATCTNNNGGCTCATTATCCCGGGATAAC
AGCACACGTCTGAACTCTTTCCCTACACGACGCTCTTCCGATCTNNNGGCTCATTATCCCGGGATAAC
AGCACACGTCTGAACTCTTTCCCTACACGACGCTCTTCCGATCTNNNGAAAGGCCTCTGGTCAAGAC
AGCACACGTCTGAACTCTTTCCCTACACGACGCTCTTCCGATCTNNNGAAAGGCCTCTGGTCAAGAC
AGCACACGTCTGAACTCTTTCCCTACACGACGCTCTTCCGATCTNNNTTCCGATGTGAAGGCTTATTT
AGCACACGTCTGAACTCTTTCCCTACACGACGCTCTTCCGATCTNNNTTCCGATGTGAAGGCTTATTT
AGCACACGTCTGAACTCTTTCCCTACACGACGCTCTTCCGATCTNNNATGTGACTCACTCTTGACTTCC
AGCACACGTCTGAACTCTTTCCCTACACGACGCTCTTCCGATCTNNNATGTGACTCACTCTTGACTTCC
AGCACACGTCTGAACTCTTTCCCTACACGACGCTCTTCCGATCTNNNATTGGTATCAGGCTCCAAGAA
AGCACACGTCTGAACTCTTTCCCTACACGACGCTCTTCCGATCTNNNATTGGTATCAGGCTCCAAGAA
GCACACGTCTGAACTCTTTCCCTACACGACGCTCTTCCGATCTNNNTTTGTGTGTCAGCATAATAATT
GCACACGTCTGAACTCTTTCCCTACACGACGCTCTTCCGATCTNNNTTTGTGTGTCAGCATAATAATT
AGCACACGTCTGAACTCTTTCCCTACACGACGCTCTTCCGATCTNNNTGCAGCTGTGAAAAGAAATG
AGCACACGTCTGAACTCTTTCCCTACACGACGCTCTTCCGATCTNNNTGCAGCTGTGAAAAGAAATG
CACACGTCTGAACTCTTTCCCTACACGACGCTCTTCCGATCTNNNACACGCTTGTAATCGTAGCTACT
CACACGTCTGAACTCTTTCCCTACACGACGCTCTTCCGATCTNNNACACGCTTGTAATCGTAGCTACT
AGCACACGTCTGAACTCTTTCCCTACACGACGCTCTTCCGATCTNNNAGTACGTGTCATACCGTCTTG
AGCACACGTCTGAACTCTTTCCCTACACGACGCTCTTCCGATCTNNNAGTACGTGTCATACCGTCTTG
AGCACACGTCTGAACTCTTTCCCTACACGACGCTCTTCCGATCTNNNATTTTTGGAGCATGTGTGTG
AGCACACGTCTGAACTCTTTCCCTACACGACGCTCTTCCGATCTNNNATTTTTGGAGCATGTGTGTG
GAGCACACGTCTGAACTCTTTCCCTACACGACGCTCTTCCGATCTNNNGAGAGTGCAGTCATCTCTCC
GAGCACACGTCTGAACTCTTTCCCTACACGACGCTCTTCCGATCTNNNGAGAGTGCAGTCATCTCTCC
GCACACGTCTGAACTCTTTCCCTACACGACGCTCTTCCGATCTNNNAGCTAGATACAGAGTGCCCATT
GCACACGTCTGAACTCTTTCCCTACACGACGCTCTTCCGATCTNNNAGCTAGATACAGAGTGCCCATT
GCACACGTCTGAACTCTTTCCCTACACGACGCTCTTCCGATCTNNNTACAGTTCATCTTGTGGAGGGG
GCACACGTCTGAACTCTTTCCCTACACGACGCTCTTCCGATCTNNNTACAGTTCATCTTGTGGAGGGG
GCACACGTCTGAACTCTTTCCCTACACGACGCTCTTCCGATCTNNNGTGGTGCTCATTCCCTGATATCT
GCACACGTCTGAACTCTTTCCCTACACGACGCTCTTCCGATCTNNNGTGGTGCTCATTCCCTGATATCT
AGCACACGTCTGAACTCTTTCCCTACACGACGCTCTTCCGATCTNNNACCCCCTGGTTTCTTGTCATTT
AGCACACGTCTGAACTCTTTCCCTACACGACGCTCTTCCGATCTNNNACCCCCTGGTTTCTTGTCATTT
GAGCACACGTCTGAACTCTTTCCCTACACGACGCTCTTCCGATCTNNNCCCTCCTTTCCTAGCTGTCTC
GAGCACACGTCTGAACTCTTTCCCTACACGACGCTCTTCCGATCTNNNCCCTCCTTTCCTAGCTGTCTC
AGCACACGTCTGAACTCTTTCCCTACACGACGCTCTTCCGATCTNNNTTCAGTGAGTTGAGATTGCA
AGCACACGTCTGAACTCTTTCCCTACACGACGCTCTTCCGATCTNNNTTCAGTGAGTTGAGATTGCA
AGCACACGTCTGAACTCTTTCCCTACACGACGCTCTTCCGATCTNNNTGTCCCAAATGGTCTGATTTTG
AGCACACGTCTGAACTCTTTCCCTACACGACGCTCTTCCGATCTNNNTGTCCCAAATGGTCTGATTTTG
GCACACGTCTGAACTCTTTCCCTACACGACGCTCTTCCGATCTNNNCTAAGGGCAGGAGAATCACTTG
GCACACGTCTGAACTCTTTCCCTACACGACGCTCTTCCGATCTNNNCTAAGGGCAGGAGAATCACTTG
AGCACACGTCTGAACTCTTTCCCTACACGACGCTCTTCCGATCTNNNAGTCAATTGTGTGTGTGTGTG
AGCACACGTCTGAACTCTTTCCCTACACGACGCTCTTCCGATCTNNNAGTCAATTGTGTGTGTGTGTG
AGCACACGTCTGAACTCTTTCCCTACACGACGCTCTTCCGATCTNNNCATCTGAATGGACTTCCTCCT
AGCACACGTCTGAACTCTTTCCCTACACGACGCTCTTCCGATCTNNNCATCTGAATGGACTTCCTCCT

GAGCACACGTCTGAACTCTTTCCCTACACGACGCTCTTCCGATCTNNNCTGTCTTAGTAATGGGGTGGA
GAGCACACGTCTGAACTCTTTCCCTACACGACGCTCTTCCGATCTNNNCTGTCTTAGTAATGGGGTGGA
AGCACACGTCTGAACTCTTTCCCTACACGACGCTCTTCCGATCTNNNACCACGCCTGGCTAATTTTTTTG
AGCACACGTCTGAACTCTTTCCCTACACGACGCTCTTCCGATCTNNNACCACGCCTGGCTAATTTTTTTG
AGCACACGTCTGAACTCTTTCCCTACACGACGCTCTTCCGATCTNNNTTCACAGAGCATTGGAGACTGA
AGCACACGTCTGAACTCTTTCCCTACACGACGCTCTTCCGATCTNNNTTCACAGAGCATTGGAGACTGA
GAGCACACGTCTGAACTCTTTCCCTACACGACGCTCTTCCGATCTNNNGCCCAGCCTCCATTCTTTTTAT
GAGCACACGTCTGAACTCTTTCCCTACACGACGCTCTTCCGATCTNNNGCCCAGCCTCCATTCTTTTTAT
GAGCACACGTCTGAACTCTTTCCCTACACGACGCTCTTCCGATCTNNNCCTTTCTTCCAATACTATTGAG
GAGCACACGTCTGAACTCTTTCCCTACACGACGCTCTTCCGATCTNNNCCTTTCTTCCAATACTATTGAG
AGCACACGTCTGAACTCTTTCCCTACACGACGCTCTTCCGATCTNNNAATTCTTTGCATGGGCTGTTGG
AGCACACGTCTGAACTCTTTCCCTACACGACGCTCTTCCGATCTNNNAATTCTTTGCATGGGCTGTTGG
GAGCACACGTCTGAACTCTTTCCCTACACGACGCTCTTCCGATCTNNNTTCTCACCTAGATCCACCAAAG
GAGCACACGTCTGAACTCTTTCCCTACACGACGCTCTTCCGATCTNNNTTCTCACCTAGATCCACCAAAG
AGCACACGTCTGAACTCTTTCCCTACACGACGCTCTTCCGATCTNNNTTTGTCTTGCACCTTAGGTACCA
AGCACACGTCTGAACTCTTTCCCTACACGACGCTCTTCCGATCTNNNTTTGTCTTGCACCTTAGGTACCA
AGCACACGTCTGAACTCTTTCCCTACACGACGCTCTTCCGATCTNNNGTAACTCCCTATAGGCACAAGC
AGCACACGTCTGAACTCTTTCCCTACACGACGCTCTTCCGATCTNNNGTAACTCCCTATAGGCACAAGC
AGCACACGTCTGAACTCTTTCCCTACACGACGCTCTTCCGATCTNNNCACATACACACATGTCCACACAC
AGCACACGTCTGAACTCTTTCCCTACACGACGCTCTTCCGATCTNNNCACATACACACATGTCCACACAC
GAGCACACGTCTGAACTCTTTCCCTACACGACGCTCTTCCGATCTNNNACTTCGCACACACACACACATA
GAGCACACGTCTGAACTCTTTCCCTACACGACGCTCTTCCGATCTNNNACTTCGCACACACACACACATA
AGAGCACACGTCTGAACTCTTTCCCTACACGACGCTCTTCCGATCTNNNACTGTGGACATCTTGCTGTAG
AGAGCACACGTCTGAACTCTTTCCCTACACGACGCTCTTCCGATCTNNNACTGTGGACATCTTGCTGTAG
AGCACACGTCTGAACTCTTTCCCTACACGACGCTCTTCCGATCTNNNGTTTGCAGTGAGCCAAGATCAT
AGCACACGTCTGAACTCTTTCCCTACACGACGCTCTTCCGATCTNNNGTTTGCAGTGAGCCAAGATCAT
AGCACACGTCTGAACTCTTTCCCTACACGACGCTCTTCCGATCTNNNATCTAGTTCCCGTTAGCTCAACG
AGCACACGTCTGAACTCTTTCCCTACACGACGCTCTTCCGATCTNNNATCTAGTTCCCGTTAGCTCAACG
GAGCACACGTCTGAACTCTTTCCCTACACGACGCTCTTCCGATCTNNNTCATCTGGAATCTGTCGGTTCG
GAGCACACGTCTGAACTCTTTCCCTACACGACGCTCTTCCGATCTNNNTCATCTGGAATCTGTCGGTTCG
GAAGAGCACACGTCTGAACTCTTTCCCTACACGACGCTCTTCCGATCTNNNGAGGCTAGACCTTCTCTG
GAAGAGCACACGTCTGAACTCTTTCCCTACACGACGCTCTTCCGATCTNNNGAGGCTAGACCTTCTCTG
GAGCACACGTCTGAACTCTTTCCCTACACGACGCTCTTCCGATCTNNNTCAAGATGAATGTGCTGTTTA
GAGCACACGTCTGAACTCTTTCCCTACACGACGCTCTTCCGATCTNNNTCAAGATGAATGTGCTGTTTA
GAGCACACGTCTGAACTCTTTCCCTACACGACGCTCTTCCGATCTNNNGGACACTGAAAGGGACTGAAG
GAGCACACGTCTGAACTCTTTCCCTACACGACGCTCTTCCGATCTNNNGGACACTGAAAGGGACTGAAG
AGCACACGTCTGAACTCTTTCCCTACACGACGCTCTTCCGATCTNNNTTACAAGGTGCGTATTCTCTGCT
AGCACACGTCTGAACTCTTTCCCTACACGACGCTCTTCCGATCTNNNTTACAAGGTGCGTATTCTCTGCT
CACACGTCTGAACTCTTTCCCTACACGACGCTCTTCCGATCTNNNTCCTTTCCTCCCTCCTCCTATTTCATG
CACACGTCTGAACTCTTTCCCTACACGACGCTCTTCCGATCTNNNTCCTTTCCTCCCTCCTCCTATTTCATG
GAGCACACGTCTGAACTCTTTCCCTACACGACGCTCTTCCGATCTNNNAAGGAGTTCTGGTGTCACTGA
GAGCACACGTCTGAACTCTTTCCCTACACGACGCTCTTCCGATCTNNNAAGGAGTTCTGGTGTCACTGA
AGCACACGTCTGAACTCTTTCCCTACACGACGCTCTTCCGATCTNNNGGAAGTCCCCTGAGATGTGAA

AGCACACGTCTGAACTCTTTCCCTACACGACGCTCTTCCGATCTNNNGGAAGTCCCCTGAGATGTGAA
AGCACACGTCTGAACTCTTTCCCTACACGACGCTCTTCCGATCTNNNAGGAGGATGAAGCAGGAGA
AGCACACGTCTGAACTCTTTCCCTACACGACGCTCTTCCGATCTNNNAGGAGGATGAAGCAGGAGA
AGCACACGTCTGAACTCTTTCCCTACACGACGCTCTTCCGATCTNNNTACAAACTACTCAGGAGGCTC
AGCACACGTCTGAACTCTTTCCCTACACGACGCTCTTCCGATCTNNNTACAAACTACTCAGGAGGCTC
AGCACACGTCTGAACTCTTTCCCTACACGACGCTCTTCCGATCTNNNGACAATTTCACTTTCCCAAGGG
AGCACACGTCTGAACTCTTTCCCTACACGACGCTCTTCCGATCTNNNGACAATTTCACTTTCCCAAGGG
AGCACACGTCTGAACTCTTTCCCTACACGACGCTCTTCCGATCTNNNTCATAGGTGAAAGGGTCTGAA
AGCACACGTCTGAACTCTTTCCCTACACGACGCTCTTCCGATCTNNNTCATAGGTGAAAGGGTCTGAA
GAGCACACGTCTGAACTCTTTCCCTACACGACGCTCTTCCGATCTNNNCACAGGCATTTCATAGGGTT
GAGCACACGTCTGAACTCTTTCCCTACACGACGCTCTTCCGATCTNNNCACAGGCATTTCATAGGGTT
GAGCACACGTCTGAACTCTTTCCCTACACGACGCTCTTCCGATCTNNNTTCTGTGATTCACATCTGAGG
GAGCACACGTCTGAACTCTTTCCCTACACGACGCTCTTCCGATCTNNNTTCTGTGATTCACATCTGAGG
AGCACACGTCTGAACTCTTTCCCTACACGACGCTCTTCCGATCTNNNTTACCTCATTTCCCCCACCCTTT
AGCACACGTCTGAACTCTTTCCCTACACGACGCTCTTCCGATCTNNNTTACCTCATTTCCCCCACCCTTT
GAGCACACGTCTGAACTCTTTCCCTACACGACGCTCTTCCGATCTNNNCTGAAGCAGGAGAATGGCA
GAGCACACGTCTGAACTCTTTCCCTACACGACGCTCTTCCGATCTNNNCTGAAGCAGGAGAATGGCA
GAGCACACGTCTGAACTCTTTCCCTACACGACGCTCTTCCGATCTNNNCTGGAGGCTGAAACAGGAG
GAGCACACGTCTGAACTCTTTCCCTACACGACGCTCTTCCGATCTNNNCTGGAGGCTGAAACAGGAG
AGCACACGTCTGAACTCTTTCCCTACACGACGCTCTTCCGATCTNNNTAATCAACATACGCATGGGAC
AGCACACGTCTGAACTCTTTCCCTACACGACGCTCTTCCGATCTNNNTAATCAACATACGCATGGGAC
AGCACACGTCTGAACTCTTTCCCTACACGACGCTCTTCCGATCTNNNACTCCAGGCTCTGTCTCAAAAA
AGCACACGTCTGAACTCTTTCCCTACACGACGCTCTTCCGATCTNNNACTCCAGGCTCTGTCTCAAAAA
AGCACACGTCTGAACTCTTTCCCTACACGACGCTCTTCCGATCTNNNTACAAAAAATTAGCCGGGCAT
AGCACACGTCTGAACTCTTTCCCTACACGACGCTCTTCCGATCTNNNTACAAAAAATTAGCCGGGCAT
CACACGTCTGAACTCTTTCCCTACACGACGCTCTTCCGATCTNNNCTGGAATTTCTGACTTGGTGTGG
CACACGTCTGAACTCTTTCCCTACACGACGCTCTTCCGATCTNNNCTGGAATTTCTGACTTGGTGTGG
AGCACACGTCTGAACTCTTTCCCTACACGACGCTCTTCCGATCTNNNGGCCGTCTTCTCCTTATCTTACA
AGCACACGTCTGAACTCTTTCCCTACACGACGCTCTTCCGATCTNNNGGCCGTCTTCTCCTTATCTTACA
GCACACGTCTGAACTCTTTCCCTACACGACGCTCTTCCGATCTNNNGAGAAACCCGGAGTCGTGAAAC
GCACACGTCTGAACTCTTTCCCTACACGACGCTCTTCCGATCTNNNGAGAAACCCGGAGTCGTGAAAC
GAGCACACGTCTGAACTCTTTCCCTACACGACGCTCTTCCGATCTNNNGTGCCTGAGCTGAAAGATGA
GAGCACACGTCTGAACTCTTTCCCTACACGACGCTCTTCCGATCTNNNGTGCCTGAGCTGAAAGATGA
GAGCACACGTCTGAACTCTTTCCCTACACGACGCTCTTCCGATCTNNNTGAGTCCGAAAAGAGAGTCA
GAGCACACGTCTGAACTCTTTCCCTACACGACGCTCTTCCGATCTNNNTGAGTCCGAAAAGAGAGTCA
AGCACACGTCTGAACTCTTTCCCTACACGACGCTCTTCCGATCTNNNTCACTTTATCACCCAGGATAGC
AGCACACGTCTGAACTCTTTCCCTACACGACGCTCTTCCGATCTNNNTCACTTTATCACCCAGGATAGC
GAGCACACGTCTGAACTCTTTCCCTACACGACGCTCTTCCGATCTNNNATGTGGAACACAGAAGTTTC
GAGCACACGTCTGAACTCTTTCCCTACACGACGCTCTTCCGATCTNNNATGTGGAACACAGAAGTTTC
GCACACGTCTGAACTCTTTCCCTACACGACGCTCTTCCGATCTNNNCCGGCATTACAGGCATGAGCCAT
GCACACGTCTGAACTCTTTCCCTACACGACGCTCTTCCGATCTNNNCCGGCATTACAGGCATGAGCCAT
AGCACACGTCTGAACTCTTTCCCTACACGACGCTCTTCCGATCTNNNAAAAACGAAATGAAGCTCCCA
AGCACACGTCTGAACTCTTTCCCTACACGACGCTCTTCCGATCTNNNAAAAACGAAATGAAGCTCCCA

CACACGTCTGAACTCTTTCCCTACACGACGCTCTTCCGATCTNNNGAGTCCAGAGATCGAGGCCGCATC

CACACGTCTGAACTCTTTCCCTACACGACGCTCTTCCGATCTNNNGAGTCCAGAGATCGAGGCCGCATC

AGCACACGTCTGAACTCTTTCCCTACACGACGCTCTTCCGATCTNNNCATGTGGCTTTAGCACCAATACC

AGCACACGTCTGAACTCTTTCCCTACACGACGCTCTTCCGATCTNNNCATGTGGCTTTAGCACCAATACC

AGAGCACACGTCTGAACTCTTTCCCTACACGACGCTCTTCCGATCTNNNCCTATGTTCGCCATCTAGGA

AGAGCACACGTCTGAACTCTTTCCCTACACGACGCTCTTCCGATCTNNNCCTATGTTCGCCATCTAGGA

GAGCACACGTCTGAACTCTTTCCCTACACGACGCTCTTCCGATCTNNNGCAATTATTCACAACTGGGGC

GAGCACACGTCTGAACTCTTTCCCTACACGACGCTCTTCCGATCTNNNGCAATTATTCACAACTGGGGC

GAGCACACGTCTGAACTCTTTCCCTACACGACGCTCTTCCGATCTNNNAAGAAATGGGTCCTCTTGAGC

GAGCACACGTCTGAACTCTTTCCCTACACGACGCTCTTCCGATCTNNNAAGAAATGGGTCCTCTTGAGC

GAGCACACGTCTGAACTCTTTCCCTACACGACGCTCTTCCGATCTNNNAAGGCTGCAGTGAGTTGATT

GAGCACACGTCTGAACTCTTTCCCTACACGACGCTCTTCCGATCTNNNAAGGCTGCAGTGAGTTGATT

AGCACACGTCTGAACTCTTTCCCTACACGACGCTCTTCCGATCTNNNCACAATCATAGCTCACTGCAACC

AGCACACGTCTGAACTCTTTCCCTACACGACGCTCTTCCGATCTNNNCACAATCATAGCTCACTGCAACC

GCACACGTCTGAACTCTTTCCCTACACGACGCTCTTCCGATCTNNNGGAAGCTGAGGTAGGAGAATCA

GCACACGTCTGAACTCTTTCCCTACACGACGCTCTTCCGATCTNNNGGAAGCTGAGGTAGGAGAATCA

GAGCACACGTCTGAACTCTTTCCCTACACGACGCTCTTCCGATCTNNNTTCAGGCATTTCCCCTCATGAC

GAGCACACGTCTGAACTCTTTCCCTACACGACGCTCTTCCGATCTNNNTTCAGGCATTTCCCCTCATGAC

AGAGCACACGTCTGAACTCTTTCCCTACACGACGCTCTTCCGATCTNNNGACTCCATCTCTCCCTCTCTCT

AGAGCACACGTCTGAACTCTTTCCCTACACGACGCTCTTCCGATCTNNNGACTCCATCTCTCCCTCTCTCT

CACACGTCTGAACTCTTTCCCTACACGACGCTCTTCCGATCTNNNCTCCGTCTCAAAAAAAACAAAACAT

CACACGTCTGAACTCTTTCCCTACACGACGCTCTTCCGATCTNNNCTCCGTCTCAAAAAAAACAAAACAT

GCACACGTCTGAACTCTTTCCCTACACGACGCTCTTCCGATCTNNNAAAAACCCTAGGCCACACTGATTC

GCACACGTCTGAACTCTTTCCCTACACGACGCTCTTCCGATCTNNNAAAAACCCTAGGCCACACTGATTC

GAGCACACGTCTGAACTCTTTCCCTACACGACGCTCTTCCGATCTNNNACACATGAGGGCTAAAGATG

GAGCACACGTCTGAACTCTTTCCCTACACGACGCTCTTCCGATCTNNNACACATGAGGGCTAAAGATG

AGCACACGTCTGAACTCTTTCCCTACACGACGCTCTTCCGATCTNNNACCCTCACAAGATTGGTCTTTCA

AGCACACGTCTGAACTCTTTCCCTACACGACGCTCTTCCGATCTNNNACCCTCACAAGATTGGTCTTTCA

AGCACACGTCTGAACTCTTTCCCTACACGACGCTCTTCCGATCTNNNTTGGGTAGACATCAGGTTGTAG

AGCACACGTCTGAACTCTTTCCCTACACGACGCTCTTCCGATCTNNNTTGGGTAGACATCAGGTTGTAG

GAGCACACGTCTGAACTCTTTCCCTACACGACGCTCTTCCGATCTNNNGGCAAGAGAGTGAGACTCCAT

GAGCACACGTCTGAACTCTTTCCCTACACGACGCTCTTCCGATCTNNNGGCAAGAGAGTGAGACTCCAT

GAGCACACGTCTGAACTCTTTCCCTACACGACGCTCTTCCGATCTNNNCTTCCCAAACTGATTAGTGCTG

GAGCACACGTCTGAACTCTTTCCCTACACGACGCTCTTCCGATCTNNNCTTCCCAAACTGATTAGTGCTG

GAGCACACGTCTGAACTCTTTCCCTACACGACGCTCTTCCGATCTNNNCGTTACTAACTAGAGGGGAGC

GAGCACACGTCTGAACTCTTTCCCTACACGACGCTCTTCCGATCTNNNCGTTACTAACTAGAGGGGAGC

GAGCACACGTCTGAACTCTTTCCCTACACGACGCTCTTCCGATCTNNNATGTCTGTAATCCCAGCTACCTC

GAGCACACGTCTGAACTCTTTCCCTACACGACGCTCTTCCGATCTNNNATGTCTGTAATCCCAGCTACCTC

AGAGCACACGTCTGAACTCTTTCCCTACACGACGCTCTTCCGATCTNNNCTGAGGCAGGAGAATCACT

AGAGCACACGTCTGAACTCTTTCCCTACACGACGCTCTTCCGATCTNNNCTGAGGCAGGAGAATCACT

AGCACACGTCTGAACTCTTTCCCTACACGACGCTCTTCCGATCTNNNGCTTTCCTTCACCCAAGACATTCC

AGCACACGTCTGAACTCTTTCCCTACACGACGCTCTTCCGATCTNNNGCTTTCCTTCACCCAAGACATTCC

GAGCACACGTCTGAACTCTTTCCCTACACGACGCTCTTCCGATCTNNNGGCAGAGCTTATGAAAAGGCA

AGCACACGTCTGAACTCTTTCCCTACACGACGCTCTTCCGATCTNNNGGCAGAGCTTATGAAAAGGC
AAGAGCACACGTCTGAACTCTTTCCCTACACGACGCTCTTCCGATCTNNNAAAAATTAGCTGAGTGT
AAGAGCACACGTCTGAACTCTTTCCCTACACGACGCTCTTCCGATCTNNNAAAAATTAGCTGAGTGT
GCACACGTCTGAACTCTTTCCCTACACGACGCTCTTCCGATCTNNNCTCAGTCAGCCAGTACCACGCAT
GCACACGTCTGAACTCTTTCCCTACACGACGCTCTTCCGATCTNNNCTCAGTCAGCCAGTACCACGCAT
AGCACACGTCTGAACTCTTTCCCTACACGACGCTCTTCCGATCTNNNCATGGTGGTAGGTGCCTGTAA
AGCACACGTCTGAACTCTTTCCCTACACGACGCTCTTCCGATCTNNNCATGGTGGTAGGTGCCTGTAA
AGCACACGTCTGAACTCTTTCCCTACACGACGCTCTTCCGATCTNNNGAATTTGGCTGTGAATCCCTC
AGCACACGTCTGAACTCTTTCCCTACACGACGCTCTTCCGATCTNNNGAATTTGGCTGTGAATCCCTC
AGCACACGTCTGAACTCTTTCCCTACACGACGCTCTTCCGATCTNNNGCAACAGAGTGAGAGTCCATC
AGCACACGTCTGAACTCTTTCCCTACACGACGCTCTTCCGATCTNNNGCAACAGAGTGAGAGTCCATC
AGAGCACACGTCTGAACTCTTTCCCTACACGACGCTCTTCCGATCTNNNGTTAGGACGATGGAACCAA
AGAGCACACGTCTGAACTCTTTCCCTACACGACGCTCTTCCGATCTNNNGTTAGGACGATGGAACCAA
GCACACGTCTGAACTCTTTCCCTACACGACGCTCTTCCGATCTNNNGAGGCAAGAGAATGGGTGTG
GCACACGTCTGAACTCTTTCCCTACACGACGCTCTTCCGATCTNNNGAGGCAAGAGAATGGGTGTG
AGCACACGTCTGAACTCTTTCCCTACACGACGCTCTTCCGATCTNNNCCCATCTATCCAGAAGCTGAT
AGCACACGTCTGAACTCTTTCCCTACACGACGCTCTTCCGATCTNNNCCCATCTATCCAGAAGCTGAT
GAGCACACGTCTGAACTCTTTCCCTACACGACGCTCTTCCGATCTNNNGCTGATGATGAGTCTCCTCCT
GAGCACACGTCTGAACTCTTTCCCTACACGACGCTCTTCCGATCTNNNGCTGATGATGAGTCTCCTCCT
GCACACGTCTGAACTCTTTCCCTACACGACGCTCTTCCGATCTNNNTACAAAAAATTAGCCGGGCGTG
GCACACGTCTGAACTCTTTCCCTACACGACGCTCTTCCGATCTNNNTACAAAAAATTAGCCGGGCGTG
CACACGTCTGAACTCTTTCCCTACACGACGCTCTTCCGATCTNNNGTTTGACAATAATCTCACAAAACC
CACACGTCTGAACTCTTTCCCTACACGACGCTCTTCCGATCTNNNGTTTGACAATAATCTCACAAAACC
CACACGTCTGAACTCTTTCCCTACACGACGCTCTTCCGATCTNNNCCTCCTAAGGGATTGATTGGCATC
CACACGTCTGAACTCTTTCCCTACACGACGCTCTTCCGATCTNNNCCTCCTAAGGGATTGATTGGCATC
GCACACGTCTGAACTCTTTCCCTACACGACGCTCTTCCGATCTNNNGTTCTGCCTTTAGCTGGTGACAA
GCACACGTCTGAACTCTTTCCCTACACGACGCTCTTCCGATCTNNNGTTCTGCCTTTAGCTGGTGACAA
GCACACGTCTGAACTCTTTCCCTACACGACGCTCTTCCGATCTNNNTGCACAGACCTATGAAGACTGA
GCACACGTCTGAACTCTTTCCCTACACGACGCTCTTCCGATCTNNNTGCACAGACCTATGAAGACTGA
GCACACGTCTGAACTCTTTCCCTACACGACGCTCTTCCGATCTNNNATCCTTCTCAGGTTGAATTGGCT
GCACACGTCTGAACTCTTTCCCTACACGACGCTCTTCCGATCTNNNATCCTTCTCAGGTTGAATTGGCT
GAGCACACGTCTGAACTCTTTCCCTACACGACGCTCTTCCGATCTNNNCTGTGAAGGTGACTGTAGC
GAGCACACGTCTGAACTCTTTCCCTACACGACGCTCTTCCGATCTNNNCTGTGAAGGTGACTGTAGC
CACACGTCTGAACTCTTTCCCTACACGACGCTCTTCCGATCTNNNCATGGGATGTGCTTCCATTTGTT
CACACGTCTGAACTCTTTCCCTACACGACGCTCTTCCGATCTNNNCATGGGATGTGCTTCCATTTGTT
AGCACACGTCTGAACTCTTTCCCTACACGACGCTCTTCCGATCTNNNTAACACAGTGAAACCCTGTCT
AGCACACGTCTGAACTCTTTCCCTACACGACGCTCTTCCGATCTNNNTAACACAGTGAAACCCTGTCT
AGCACACGTCTGAACTCTTTCCCTACACGACGCTCTTCCGATCTNNNTGCCTTATTTGCAATGTTTGG
AGCACACGTCTGAACTCTTTCCCTACACGACGCTCTTCCGATCTNNNTGCCTTATTTGCAATGTTTGG
GCACACGTCTGAACTCTTTCCCTACACGACGCTCTTCCGATCTNNNCTAAGGCAGGAGAATCGCTTGA
GCACACGTCTGAACTCTTTCCCTACACGACGCTCTTCCGATCTNNNCTAAGGCAGGAGAATCGCTTGA
GAGCACACGTCTGAACTCTTTCCCTACACGACGCTCTTCCGATCTNNNTATATGCTGTTGCCATGGG
GAGCACACGTCTGAACTCTTTCCCTACACGACGCTCTTCCGATCTNNNTATATGCTGTTGCCATGGG

GAGCACACGTCTGAACTCTTTCCCTACACGACGCTCTTCCGATCTNNNAGATTCAGGACGCATTCTACC
GAGCACACGTCTGAACTCTTTCCCTACACGACGCTCTTCCGATCTNNNAGATTCAGGACGCATTCTACC
GAGCACACGTCTGAACTCTTTCCCTACACGACGCTCTTCCGATCTNNNGAGGCTGAGGCAGGAGAAT
GAGCACACGTCTGAACTCTTTCCCTACACGACGCTCTTCCGATCTNNNGAGGCTGAGGCAGGAGAAT
AGCACACGTCTGAACTCTTTCCCTACACGACGCTCTTCCGATCTNNNTGTCCACGGACCAACATAGAG
AGCACACGTCTGAACTCTTTCCCTACACGACGCTCTTCCGATCTNNNTGTCCACGGACCAACATAGAG
GCACACGTCTGAACTCTTTCCCTACACGACGCTCTTCCGATCTNNNCCAAATGCTGTCGTTTTTGCCTC
GCACACGTCTGAACTCTTTCCCTACACGACGCTCTTCCGATCTNNNCCAAATGCTGTCGTTTTTGCCTC
GAGCACACGTCTGAACTCTTTCCCTACACGACGCTCTTCCGATCTNNNAAAAAAAAAAGTGTGGGGA
GAGCACACGTCTGAACTCTTTCCCTACACGACGCTCTTCCGATCTNNNAAAAAAAAAAGTGTGGGGA
CACACGTCTGAACTCTTTCCCTACACGACGCTCTTCCGATCTNNNTGAAAATTAAGTGCTATGGTAAG
CACACGTCTGAACTCTTTCCCTACACGACGCTCTTCCGATCTNNNTGAAAATTAAGTGCTATGGTAAG
AGCACACGTCTGAACTCTTTCCCTACACGACGCTCTTCCGATCTNNNAATCAGAGTTCAGTCCGCAGT
AGCACACGTCTGAACTCTTTCCCTACACGACGCTCTTCCGATCTNNNAATCAGAGTTCAGTCCGCAGT
GCACACGTCTGAACTCTTTCCCTACACGACGCTCTTCCGATCTNNNTAGAAAGGGTGGATTTGGTGG
GCACACGTCTGAACTCTTTCCCTACACGACGCTCTTCCGATCTNNNTAGAAAGGGTGGATTTGGTGG
AGCACACGTCTGAACTCTTTCCCTACACGACGCTCTTCCGATCTNNNACAAAAATAAGCCAGGTGTGG
AGCACACGTCTGAACTCTTTCCCTACACGACGCTCTTCCGATCTNNNACAAAAATAAGCCAGGTGTGG
GCACACGTCTGAACTCTTTCCCTACACGACGCTCTTCCGATCTNNNTCAAAGAATGTCTCCTCCCTGTT
GCACACGTCTGAACTCTTTCCCTACACGACGCTCTTCCGATCTNNNTCAAAGAATGTCTCCTCCCTGTT
AGCACACGTCTGAACTCTTTCCCTACACGACGCTCTTCCGATCTNNNGATCAAGACCATCCTGGCTAAC
AGCACACGTCTGAACTCTTTCCCTACACGACGCTCTTCCGATCTNNNGATCAAGACCATCCTGGCTAAC
GAGCACACGTCTGAACTCTTTCCCTACACGACGCTCTTCCGATCTNNNACATGTATTCACACTTCACGT
GAGCACACGTCTGAACTCTTTCCCTACACGACGCTCTTCCGATCTNNNACATGTATTCACACTTCACGT
GCACACGTCTGAACTCTTTCCCTACACGACGCTCTTCCGATCTNNNAAACTCCTGACCTCAAGTGATCC
GCACACGTCTGAACTCTTTCCCTACACGACGCTCTTCCGATCTNNNAAACTCCTGACCTCAAGTGATCC
AGCACACGTCTGAACTCTTTCCCTACACGACGCTCTTCCGATCTNNNTGAGGCAGAAGAATTGCTCAA
AGCACACGTCTGAACTCTTTCCCTACACGACGCTCTTCCGATCTNNNTGAGGCAGAAGAATTGCTCAA
AGCACACGTCTGAACTCTTTCCCTACACGACGCTCTTCCGATCTNNNGCTGAGGCAGGAGAAAAGTA
AGCACACGTCTGAACTCTTTCCCTACACGACGCTCTTCCGATCTNNNGCTGAGGCAGGAGAAAAGTA
GCACACGTCTGAACTCTTTCCCTACACGACGCTCTTCCGATCTNNNATACAAAAATTAGCCTGGCGTGG
GCACACGTCTGAACTCTTTCCCTACACGACGCTCTTCCGATCTNNNATACAAAAATTAGCCTGGCGTGG
AGCACACGTCTGAACTCTTTCCCTACACGACGCTCTTCCGATCTNNNATGCTATCCATCCGAGTCATCA
AGCACACGTCTGAACTCTTTCCCTACACGACGCTCTTCCGATCTNNNATGCTATCCATCCGAGTCATCA
GAGCACACGTCTGAACTCTTTCCCTACACGACGCTCTTCCGATCTNNNCATCCACTCATTGATTGATGG
GAGCACACGTCTGAACTCTTTCCCTACACGACGCTCTTCCGATCTNNNCATCCACTCATTGATTGATGG
AGCACACGTCTGAACTCTTTCCCTACACGACGCTCTTCCGATCTNNNGCGGGTAGATCACAAGTTCAG
AGCACACGTCTGAACTCTTTCCCTACACGACGCTCTTCCGATCTNNNGCGGGTAGATCACAAGTTCAG
AGCACACGTCTGAACTCTTTCCCTACACGACGCTCTTCCGATCTNNNCTGAGTAACTGGGATTACAGG
AGCACACGTCTGAACTCTTTCCCTACACGACGCTCTTCCGATCTNNNCTGAGTAACTGGGATTACAGG
GCACACGTCTGAACTCTTTCCCTACACGACGCTCTTCCGATCTNNNATATTTGTTTGACAAGGGACCG
GCACACGTCTGAACTCTTTCCCTACACGACGCTCTTCCGATCTNNNATATTTGTTTGACAAGGGACCG
GAGCACACGTCTGAACTCTTTCCCTACACGACGCTCTTCCGATCTNNNTTGGAGTATGAGGCAGCAG

GAGCACACGTCTGAACTCTTTCCCTACACGACGCTCTTCCGATCTNNNTTGGAGTATGAGGCAGCAG
GAGCACACGTCTGAACTCTTTCCCTACACGACGCTCTTCCGATCTNNNCCTTGCCAGCTAAGCTTTCTTT
GAGCACACGTCTGAACTCTTTCCCTACACGACGCTCTTCCGATCTNNNCCTTGCCAGCTAAGCTTTCTTT
GAGCACACGTCTGAACTCTTTCCCTACACGACGCTCTTCCGATCTNNNTCTGCCAGAGAGAGATTGTG
GAGCACACGTCTGAACTCTTTCCCTACACGACGCTCTTCCGATCTNNNTCTGCCAGAGAGAGATTGTG
GCACACGTCTGAACTCTTTCCCTACACGACGCTCTTCCGATCTNNNGGAATCTGCCCTGTGTCGTCCAT
GCACACGTCTGAACTCTTTCCCTACACGACGCTCTTCCGATCTNNNGGAATCTGCCCTGTGTCGTCCAT
GAGCACACGTCTGAACTCTTTCCCTACACGACGCTCTTCCGATCTNNNTGTAGTCCCAGCTACTCAGGA
GAGCACACGTCTGAACTCTTTCCCTACACGACGCTCTTCCGATCTNNNTGTAGTCCCAGCTACTCAGGA
GCACACGTCTGAACTCTTTCCCTACACGACGCTCTTCCGATCTNNNAGGAAAGCTGAGAAGCATGAGT
GCACACGTCTGAACTCTTTCCCTACACGACGCTCTTCCGATCTNNNAGGAAAGCTGAGAAGCATGAGT
GCACACGTCTGAACTCTTTCCCTACACGACGCTCTTCCGATCTNNNTTGTCTGTCAATTTGTAGTGCCA
GCACACGTCTGAACTCTTTCCCTACACGACGCTCTTCCGATCTNNNTTGTCTGTCAATTTGTAGTGCCA
GAGCACACGTCTGAACTCTTTCCCTACACGACGCTCTTCCGATCTNNNAATGGCTAAGATGACACAAG
GAGCACACGTCTGAACTCTTTCCCTACACGACGCTCTTCCGATCTNNNAATGGCTAAGATGACACAAG
GAGCACACGTCTGAACTCTTTCCCTACACGACGCTCTTCCGATCTNNNTGATGCCAGAAGCACAACTT
GAGCACACGTCTGAACTCTTTCCCTACACGACGCTCTTCCGATCTNNNTGATGCCAGAAGCACAACTT
AGCACACGTCTGAACTCTTTCCCTACACGACGCTCTTCCGATCTNNNTGTAAGGAACTAGCTGCATGG
AGCACACGTCTGAACTCTTTCCCTACACGACGCTCTTCCGATCTNNNTGTAAGGAACTAGCTGCATGG
AGCACACGTCTGAACTCTTTCCCTACACGACGCTCTTCCGATCTNNNGTAATCTCAGCTACTCAGGAG
AGCACACGTCTGAACTCTTTCCCTACACGACGCTCTTCCGATCTNNNGTAATCTCAGCTACTCAGGAG
GAGCACACGTCTGAACTCTTTCCCTACACGACGCTCTTCCGATCTNNNACGCCTGTAATCCCAAAACTTT
GAGCACACGTCTGAACTCTTTCCCTACACGACGCTCTTCCGATCTNNNACGCCTGTAATCCCAAAACTTT
AGCACACGTCTGAACTCTTTCCCTACACGACGCTCTTCCGATCTNNNAAGGCTGTGTGACTTTCCAGTA
AGCACACGTCTGAACTCTTTCCCTACACGACGCTCTTCCGATCTNNNAAGGCTGTGTGACTTTCCAGTA
AAGAGCACACGTCTGAACTCTTTCCCTACACGACGCTCTTCCGATCTNNNTCACTACTTGGGAGTCTG
AAGAGCACACGTCTGAACTCTTTCCCTACACGACGCTCTTCCGATCTNNNTCACTACTTGGGAGTCTG
AGCACACGTCTGAACTCTTTCCCTACACGACGCTCTTCCGATCTNNNCTGAGACAGGAGAATGGCTTG
AGCACACGTCTGAACTCTTTCCCTACACGACGCTCTTCCGATCTNNNCTGAGACAGGAGAATGGCTTG
GCACACGTCTGAACTCTTTCCCTACACGACGCTCTTCCGATCTNNNCTGAGGCAGGAGAATCACTTGA
GCACACGTCTGAACTCTTTCCCTACACGACGCTCTTCCGATCTNNNCTGAGGCAGGAGAATCACTTGA
AGCACACGTCTGAACTCTTTCCCTACACGACGCTCTTCCGATCTNNNTGAACTGGACCAATACTCTCTG
AGCACACGTCTGAACTCTTTCCCTACACGACGCTCTTCCGATCTNNNTGAACTGGACCAATACTCTCTG
GCACACGTCTGAACTCTTTCCCTACACGACGCTCTTCCGATCTNNNGCCTTCCTCACTTAGCTTAGCTTC
GCACACGTCTGAACTCTTTCCCTACACGACGCTCTTCCGATCTNNNGCCTTCCTCACTTAGCTTAGCTTC
GCACACGTCTGAACTCTTTCCCTACACGACGCTCTTCCGATCTNNNCCCGGCCTTTTTCCTTTTTTCTTC
GCACACGTCTGAACTCTTTCCCTACACGACGCTCTTCCGATCTNNNCCCGGCCTTTTTCCTTTTTTCTTC
AGCACACGTCTGAACTCTTTCCCTACACGACGCTCTTCCGATCTNNNCTCCACGCATTCCTTACCTATGA
AGCACACGTCTGAACTCTTTCCCTACACGACGCTCTTCCGATCTNNNCTCCACGCATTCCTTACCTATGA
AGCACACGTCTGAACTCTTTCCCTACACGACGCTCTTCCGATCTNNNGCACCGAACCTACAGTCACTATT
AGCACACGTCTGAACTCTTTCCCTACACGACGCTCTTCCGATCTNNNGCACCGAACCTACAGTCACTATT
GCACACGTCTGAACTCTTTCCCTACACGACGCTCTTCCGATCTNNNACCACACCCAGCTAATTTTGGTA
GCACACGTCTGAACTCTTTCCCTACACGACGCTCTTCCGATCTNNNACCACACCCAGCTAATTTTGGTA

AGCACACGTCTGAACTCTTTCCCTACACGACGCTCTTCCGATCTNNNGACTGGGAGGCCTTTGATACT
AGCACACGTCTGAACTCTTTCCCTACACGACGCTCTTCCGATCTNNNGACTGGGAGGCCTTTGATACT
GAGCACACGTCTGAACTCTTTCCCTACACGACGCTCTTCCGATCTNNNAAAAACCCACTGCAAACACT
GAGCACACGTCTGAACTCTTTCCCTACACGACGCTCTTCCGATCTNNNAAAAACCCACTGCAAACACT
AGCACACGTCTGAACTCTTTCCCTACACGACGCTCTTCCGATCTNNNGATTACAGACGTGTACCACAC
AGCACACGTCTGAACTCTTTCCCTACACGACGCTCTTCCGATCTNNNGATTACAGACGTGTACCACAC
CACACGTCTGAACTCTTTCCCTACACGACGCTCTTCCGATCTNNNCTGTTTGAAAGCTTCCAGAAGGG
CACACGTCTGAACTCTTTCCCTACACGACGCTCTTCCGATCTNNNCTGTTTGAAAGCTTCCAGAAGGG
AGCACACGTCTGAACTCTTTCCCTACACGACGCTCTTCCGATCTNNNCTGAGAACAAGGAACCACATT
AGCACACGTCTGAACTCTTTCCCTACACGACGCTCTTCCGATCTNNNCTGAGAACAAGGAACCACATT
AGCACACGTCTGAACTCTTTCCCTACACGACGCTCTTCCGATCTNNNACCCATGATGGAACCAATGAT
AGCACACGTCTGAACTCTTTCCCTACACGACGCTCTTCCGATCTNNNACCCATGATGGAACCAATGAT
AGCACACGTCTGAACTCTTTCCCTACACGACGCTCTTCCGATCTNNNTTTTCTTCCTCCTCTCTGCTGAC
AGCACACGTCTGAACTCTTTCCCTACACGACGCTCTTCCGATCTNNNTTTTCTTCCTCCTCTCTGCTGAC
CACACGTCTGAACTCTTTCCCTACACGACGCTCTTCCGATCTNNNCTTTTGGGGTTTCTTGGACATCCC
CACACGTCTGAACTCTTTCCCTACACGACGCTCTTCCGATCTNNNCTTTTGGGGTTTCTTGGACATCCC
GCACACGTCTGAACTCTTTCCCTACACGACGCTCTTCCGATCTNNNTCAAGGCCCCGTCTCTACAAAAG
GCACACGTCTGAACTCTTTCCCTACACGACGCTCTTCCGATCTNNNTCAAGGCCCCGTCTCTACAAAAG
GAGCACACGTCTGAACTCTTTCCCTACACGACGCTCTTCCGATCTNNNCTTCCTGTTCCTCTGCCTTTTT
GAGCACACGTCTGAACTCTTTCCCTACACGACGCTCTTCCGATCTNNNCTTCCTGTTCCTCTGCCTTTTT
GCACACGTCTGAACTCTTTCCCTACACGACGCTCTTCCGATCTNNNGACTAGTGATAGGGGTGTGTG
GCACACGTCTGAACTCTTTCCCTACACGACGCTCTTCCGATCTNNNGACTAGTGATAGGGGTGTGTG
AGCACACGTCTGAACTCTTTCCCTACACGACGCTCTTCCGATCTNNNCGCTTCCCTGTTGTATCCTGTCA
AGCACACGTCTGAACTCTTTCCCTACACGACGCTCTTCCGATCTNNNCGCTTCCCTGTTGTATCCTGTCA
ACACGTCTGAACTCTTTCCCTACACGACGCTCTTCCGATCTNNNTGAAGAAGATGTGGATAACTACCA
ACACGTCTGAACTCTTTCCCTACACGACGCTCTTCCGATCTNNNTGAAGAAGATGTGGATAACTACCA
AGCACACGTCTGAACTCTTTCCCTACACGACGCTCTTCCGATCTNNNGTTCAAAACCAACCTGGGCAA
AGCACACGTCTGAACTCTTTCCCTACACGACGCTCTTCCGATCTNNNGTTCAAAACCAACCTGGGCAA
AGCACACGTCTGAACTCTTTCCCTACACGACGCTCTTCCGATCTNNNGGTGGATAGTGTGCAATAGG
AGCACACGTCTGAACTCTTTCCCTACACGACGCTCTTCCGATCTNNNGGTGGATAGTGTGCAATAGG
AGCACACGTCTGAACTCTTTCCCTACACGACGCTCTTCCGATCTNNNAGAACTCTCCAAACTGCTTTCC
AGCACACGTCTGAACTCTTTCCCTACACGACGCTCTTCCGATCTNNNAGAACTCTCCAAACTGCTTTCC
GAGCACACGTCTGAACTCTTTCCCTACACGACGCTCTTCCGATCTNNNTGATTATTTGAAAAGCTTCGC
GAGCACACGTCTGAACTCTTTCCCTACACGACGCTCTTCCGATCTNNNTGATTATTTGAAAAGCTTCGC
AGCACACGTCTGAACTCTTTCCCTACACGACGCTCTTCCGATCTNNNGGAGGCTGAGACAGGAGAAT
AGCACACGTCTGAACTCTTTCCCTACACGACGCTCTTCCGATCTNNNGGAGGCTGAGACAGGAGAAT
GAGCACACGTCTGAACTCTTTCCCTACACGACGCTCTTCCGATCTNNNTTTGTGGTGTGTATGTGTGT
GAGCACACGTCTGAACTCTTTCCCTACACGACGCTCTTCCGATCTNNNTTTGTGGTGTGTATGTGTGT
CACACGTCTGAACTCTTTCCCTACACGACGCTCTTCCGATCTNNNTACTGTACAAATGCATAACCTTTC
CACACGTCTGAACTCTTTCCCTACACGACGCTCTTCCGATCTNNNTACTGTACAAATGCATAACCTTTC
AGCACACGTCTGAACTCTTTCCCTACACGACGCTCTTCCGATCTNNNTACAGTCCCACCAAAAGTGTAG
AGCACACGTCTGAACTCTTTCCCTACACGACGCTCTTCCGATCTNNNTACAGTCCCACCAAAAGTGTAG
GCACACGTCTGAACTCTTTCCCTACACGACGCTCTTCCGATCTNNNGTTTCTCACTTTAGCCAATTCCAT

GCACACGTCTGAACTCTTTCCCTACACGACGCTCTTCCGATCTNNNGTTTCTCACTTTAGCCAATTCCATC
GCACACGTCTGAACTCTTTCCCTACACGACGCTCTTCCGATCTNNNCTCCAGAGACAGGATCAGCTTGCA
GCACACGTCTGAACTCTTTCCCTACACGACGCTCTTCCGATCTNNNCTCCAGAGACAGGATCAGCTTGCA
GAGCACACGTCTGAACTCTTTCCCTACACGACGCTCTTCCGATCTNNNCCTTTATTCCTTCTCTTCTTTTCC
GAGCACACGTCTGAACTCTTTCCCTACACGACGCTCTTCCGATCTNNNCCTTTATTCCTTCTCTTCTTTTCC
GCACACGTCTGAACTCTTTCCCTACACGACGCTCTTCCGATCTNNNCTGAGACAGGAGAATCGCTTGAA
GCACACGTCTGAACTCTTTCCCTACACGACGCTCTTCCGATCTNNNCTGAGACAGGAGAATCGCTTGAA
AGCACACGTCTGAACTCTTTCCCTACACGACGCTCTTCCGATCTNNNTCTTTATGTGCCCATGAGACCA
AGCACACGTCTGAACTCTTTCCCTACACGACGCTCTTCCGATCTNNNTCTTTATGTGCCCATGAGACCA
AGCACACGTCTGAACTCTTTCCCTACACGACGCTCTTCCGATCTNNNAAGCATGAAGAGTCCCAGAACT
AGCACACGTCTGAACTCTTTCCCTACACGACGCTCTTCCGATCTNNNAAGCATGAAGAGTCCCAGAACT
GAGCACACGTCTGAACTCTTTCCCTACACGACGCTCTTCCGATCTNNNAGGTTCAAGTGATTCTTCTGC
GAGCACACGTCTGAACTCTTTCCCTACACGACGCTCTTCCGATCTNNNAGGTTCAAGTGATTCTTCTGC
AGCACACGTCTGAACTCTTTCCCTACACGACGCTCTTCCGATCTNNNAGCCTTTGTTTATCTGGGAAGG
AGCACACGTCTGAACTCTTTCCCTACACGACGCTCTTCCGATCTNNNAGCCTTTGTTTATCTGGGAAGG
AGCACACGTCTGAACTCTTTCCCTACACGACGCTCTTCCGATCTNNNATGCCTGGTACCAATATGTCTG
AGCACACGTCTGAACTCTTTCCCTACACGACGCTCTTCCGATCTNNNATGCCTGGTACCAATATGTCTG
CACACGTCTGAACTCTTTCCCTACACGACGCTCTTCCGATCTNNNGTTAATGAATCCGGTAGAAGATCA
CACACGTCTGAACTCTTTCCCTACACGACGCTCTTCCGATCTNNNGTTAATGAATCCGGTAGAAGATCA
GAGCACACGTCTGAACTCTTTCCCTACACGACGCTCTTCCGATCTNNNTGAGGCTAGCATGATTCTGAT
GAGCACACGTCTGAACTCTTTCCCTACACGACGCTCTTCCGATCTNNNTGAGGCTAGCATGATTCTGAT
GAGCACACGTCTGAACTCTTTCCCTACACGACGCTCTTCCGATCTNNNAAACGATCCTTCTTCTTCAGCCT
GAGCACACGTCTGAACTCTTTCCCTACACGACGCTCTTCCGATCTNNNAAACGATCCTTCTTCTTCAGCCT
GAGCACACGTCTGAACTCTTTCCCTACACGACGCTCTTCCGATCTNNNGAGAACTCTGATGTTTGTAAA
GAGCACACGTCTGAACTCTTTCCCTACACGACGCTCTTCCGATCTNNNGAGAACTCTGATGTTTGTAAA
AGCACACGTCTGAACTCTTTCCCTACACGACGCTCTTCCGATCTNNNATATGTGTGTGTGTGTGTGTGT
AGCACACGTCTGAACTCTTTCCCTACACGACGCTCTTCCGATCTNNNATATGTGTGTGTGTGTGTGTGT
GAGCACACGTCTGAACTCTTTCCCTACACGACGCTCTTCCGATCTNNNGATGCTGTCCCATCAGACGTA
GAGCACACGTCTGAACTCTTTCCCTACACGACGCTCTTCCGATCTNNNGATGCTGTCCCATCAGACGTA
AGCACACGTCTGAACTCTTTCCCTACACGACGCTCTTCCGATCTNNNGTGAAACCCCGTCTCTACTAAAA
AGCACACGTCTGAACTCTTTCCCTACACGACGCTCTTCCGATCTNNNGTGAAACCCCGTCTCTACTAAAA
AGCACACGTCTGAACTCTTTCCCTACACGACGCTCTTCCGATCTNNNGGATCAATCAGAAGGCAATGG
AGCACACGTCTGAACTCTTTCCCTACACGACGCTCTTCCGATCTNNNGGATCAATCAGAAGGCAATGG
AGCACACGTCTGAACTCTTTCCCTACACGACGCTCTTCCGATCTNNNTGTGCTTTGTGACATGATGGTG
AGCACACGTCTGAACTCTTTCCCTACACGACGCTCTTCCGATCTNNNTGTGCTTTGTGACATGATGGTG
GAGCACACGTCTGAACTCTTTCCCTACACGACGCTCTTCCGATCTNNNGAGGTTACAGTGAGCCGAGAT
GAGCACACGTCTGAACTCTTTCCCTACACGACGCTCTTCCGATCTNNNGAGGTTACAGTGAGCCGAGAT
AGCACACGTCTGAACTCTTTCCCTACACGACGCTCTTCCGATCTNNNACGGAGCTAAAACATCCATGTC
AGCACACGTCTGAACTCTTTCCCTACACGACGCTCTTCCGATCTNNNACGGAGCTAAAACATCCATGTC
GAGCACACGTCTGAACTCTTTCCCTACACGACGCTCTTCCGATCTNNNGCGTCTCTAGGATATCGACAG
GAGCACACGTCTGAACTCTTTCCCTACACGACGCTCTTCCGATCTNNNGCGTCTCTAGGATATCGACAG
GAGCACACGTCTGAACTCTTTCCCTACACGACGCTCTTCCGATCTNNNTATCATGCGTATTGGCCTGTAC
GAGCACACGTCTGAACTCTTTCCCTACACGACGCTCTTCCGATCTNNNTATCATGCGTATTGGCCTGTAC

GAGCACACGTCTGAACTCTTTCCCTACACGACGCTCTTCCGATCTNNNGGTGAGATGAGTGAAACTCT
GAGCACACGTCTGAACTCTTTCCCTACACGACGCTCTTCCGATCTNNNGGTGAGATGAGTGAAACTCT
GAGCACACGTCTGAACTCTTTCCCTACACGACGCTCTTCCGATCTNNNCATGGTGGCACGCTCTTATA
GAGCACACGTCTGAACTCTTTCCCTACACGACGCTCTTCCGATCTNNNCATGGTGGCACGCTCTTATA
GAGCACACGTCTGAACTCTTTCCCTACACGACGCTCTTCCGATCTNNNGGACTTGGACTTTCAGAACTG
GAGCACACGTCTGAACTCTTTCCCTACACGACGCTCTTCCGATCTNNNGGACTTGGACTTTCAGAACTG
GAGCACACGTCTGAACTCTTTCCCTACACGACGCTCTTCCGATCTNNNAGGGAAGACAATCCTTCTGT
GAGCACACGTCTGAACTCTTTCCCTACACGACGCTCTTCCGATCTNNNAGGGAAGACAATCCTTCTGT
AGAGCACACGTCTGAACTCTTTCCCTACACGACGCTCTTCCGATCTNNNCTGTAGTCCCAGCTACTTGT
AGAGCACACGTCTGAACTCTTTCCCTACACGACGCTCTTCCGATCTNNNCTGTAGTCCCAGCTACTTGT
AGCACACGTCTGAACTCTTTCCCTACACGACGCTCTTCCGATCTNNNATTACACTCAGATGCTTCCCTCC
AGCACACGTCTGAACTCTTTCCCTACACGACGCTCTTCCGATCTNNNATTACACTCAGATGCTTCCCTCC
AGCACACGTCTGAACTCTTTCCCTACACGACGCTCTTCCGATCTNNNACTAGAACAGTAAGAAGCAGC
AGCACACGTCTGAACTCTTTCCCTACACGACGCTCTTCCGATCTNNNACTAGAACAGTAAGAAGCAGC
AGCACACGTCTGAACTCTTTCCCTACACGACGCTCTTCCGATCTNNNACCATGCTCTCAGACTTCCTAAC
AGCACACGTCTGAACTCTTTCCCTACACGACGCTCTTCCGATCTNNNACCATGCTCTCAGACTTCCTAAC
AGCACACGTCTGAACTCTTTCCCTACACGACGCTCTTCCGATCTNNNCACACATATTGTCATGAGGCCA
AGCACACGTCTGAACTCTTTCCCTACACGACGCTCTTCCGATCTNNNCACACATATTGTCATGAGGCCA
GAGCACACGTCTGAACTCTTTCCCTACACGACGCTCTTCCGATCTNNNCTCAAACGATCCTCTTGCCTC
GAGCACACGTCTGAACTCTTTCCCTACACGACGCTCTTCCGATCTNNNCTCAAACGATCCTCTTGCCTC
AGCACACGTCTGAACTCTTTCCCTACACGACGCTCTTCCGATCTNNNCAAGGCCAAGTAGATACAGTC
AGCACACGTCTGAACTCTTTCCCTACACGACGCTCTTCCGATCTNNNCAAGGCCAAGTAGATACAGTC
GAGCACACGTCTGAACTCTTTCCCTACACGACGCTCTTCCGATCTNNNCTGTTCTGTACCTTGACTGTG
GAGCACACGTCTGAACTCTTTCCCTACACGACGCTCTTCCGATCTNNNCTGTTCTGTACCTTGACTGTG
GAGCACACGTCTGAACTCTTTCCCTACACGACGCTCTTCCGATCTNNNAGCCAAGCAGGAAACAAAGA
GAGCACACGTCTGAACTCTTTCCCTACACGACGCTCTTCCGATCTNNNAGCCAAGCAGGAAACAAAGA
GAGCACACGTCTGAACTCTTTCCCTACACGACGCTCTTCCGATCTNNNGAGATTGCAGTGAGCCAAGA
GAGCACACGTCTGAACTCTTTCCCTACACGACGCTCTTCCGATCTNNNGAGATTGCAGTGAGCCAAGA
GCACACGTCTGAACTCTTTCCCTACACGACGCTCTTCCGATCTNNNTGATGTAGAAAGTTAAAGTGTT
GCACACGTCTGAACTCTTTCCCTACACGACGCTCTTCCGATCTNNNTGATGTAGAAAGTTAAAGTGTT
GCACACGTCTGAACTCTTTCCCTACACGACGCTCTTCCGATCTNNNCCGAGTAGCTGAGATTACAGGT
GCACACGTCTGAACTCTTTCCCTACACGACGCTCTTCCGATCTNNNCCGAGTAGCTGAGATTACAGGT
GAGCACACGTCTGAACTCTTTCCCTACACGACGCTCTTCCGATCTNNNCCTAGCTTCAAGTGACTCAGG
GAGCACACGTCTGAACTCTTTCCCTACACGACGCTCTTCCGATCTNNNCCTAGCTTCAAGTGACTCAGG
AGCACACGTCTGAACTCTTTCCCTACACGACGCTCTTCCGATCTNNNAACAGAATAGATGCTTGCTTG
AGCACACGTCTGAACTCTTTCCCTACACGACGCTCTTCCGATCTNNNAACAGAATAGATGCTTGCTTG
GAGCACACGTCTGAACTCTTTCCCTACACGACGCTCTTCCGATCTNNNATTGAGTTGCTTGCCGCTATT
GAGCACACGTCTGAACTCTTTCCCTACACGACGCTCTTCCGATCTNNNATTGAGTTGCTTGCCGCTATT
AGAGCACACGTCTGAACTCTTTCCCTACACGACGCTCTTCCGATCTNNNGTGTGTGTATTTGCCTTGCA
AGAGCACACGTCTGAACTCTTTCCCTACACGACGCTCTTCCGATCTNNNGTGTGTGTATTTGCCTTGCA
AGCACACGTCTGAACTCTTTCCCTACACGACGCTCTTCCGATCTNNNCTAATCCTTCTGATTTGCGCACC
AGCACACGTCTGAACTCTTTCCCTACACGACGCTCTTCCGATCTNNNCTAATCCTTCTGATTTGCGCACC
GAGCACACGTCTGAACTCTTTCCCTACACGACGCTCTTCCGATCTNNNGCCTCTGTGCACCTTTCATTCC

AGCACACGTCTGAACTCTTTCCCTACACGACGCTCTTCCGATCTNNNGCCTCTGTGCACCTTTCATTCC
GCACACGTCTGAACTCTTTCCCTACACGACGCTCTTCCGATCTNNNCTCATGCCTATAATCCAAGCCCT
GCACACGTCTGAACTCTTTCCCTACACGACGCTCTTCCGATCTNNNCTCATGCCTATAATCCAAGCCCT
AAGAGCACACGTCTGAACTCTTTCCCTACACGACGCTCTTCCGATCTNNNACTTCTCAACTTCCAGACT
AAGAGCACACGTCTGAACTCTTTCCCTACACGACGCTCTTCCGATCTNNNACTTCTCAACTTCCAGACT
GAGCACACGTCTGAACTCTTTCCCTACACGACGCTCTTCCGATCTNNNCCAGCCTGGGCAATATGATA
GAGCACACGTCTGAACTCTTTCCCTACACGACGCTCTTCCGATCTNNNCCAGCCTGGGCAATATGATA
AGCACACGTCTGAACTCTTTCCCTACACGACGCTCTTCCGATCTNNNGCTGAGATGGTACCACTACA
AGCACACGTCTGAACTCTTTCCCTACACGACGCTCTTCCGATCTNNNGCTGAGATGGTACCACTACA
GCACACGTCTGAACTCTTTCCCTACACGACGCTCTTCCGATCTNNNTGCCATTTTCTTAGACCCAAAGC
GCACACGTCTGAACTCTTTCCCTACACGACGCTCTTCCGATCTNNNTGCCATTTTCTTAGACCCAAAGC
GCACACGTCTGAACTCTTTCCCTACACGACGCTCTTCCGATCTNNNCAGTAAATTTCCATCTGCTGCCC
GCACACGTCTGAACTCTTTCCCTACACGACGCTCTTCCGATCTNNNCAGTAAATTTCCATCTGCTGCCC
AGCACACGTCTGAACTCTTTCCCTACACGACGCTCTTCCGATCTNNNGAGAATGGTGTGAACCTGGG
AGCACACGTCTGAACTCTTTCCCTACACGACGCTCTTCCGATCTNNNGAGAATGGTGTGAACCTGGG
AGAGCACACGTCTGAACTCTTTCCCTACACGACGCTCTTCCGATCTNNNGATGCCTGTAATCCCAGCTA
AGAGCACACGTCTGAACTCTTTCCCTACACGACGCTCTTCCGATCTNNNGATGCCTGTAATCCCAGCTA
GAGCACACGTCTGAACTCTTTCCCTACACGACGCTCTTCCGATCTNNNCCTACCCTCTTCCTGTCTCCTA
GAGCACACGTCTGAACTCTTTCCCTACACGACGCTCTTCCGATCTNNNCCTACCCTCTTCCTGTCTCCTA
GCACACGTCTGAACTCTTTCCCTACACGACGCTCTTCCGATCTNNNACAAGGGATAGCATTGTGGTG
GCACACGTCTGAACTCTTTCCCTACACGACGCTCTTCCGATCTNNNACAAGGGATAGCATTGTGGTG
AGCACACGTCTGAACTCTTTCCCTACACGACGCTCTTCCGATCTNNNTGGAGACAGGAGAATTGCTTG
AGCACACGTCTGAACTCTTTCCCTACACGACGCTCTTCCGATCTNNNTGGAGACAGGAGAATTGCTTG
AGCACACGTCTGAACTCTTTCCCTACACGACGCTCTTCCGATCTNNNAGCAGAGGGAGAGAGATAG
AGCACACGTCTGAACTCTTTCCCTACACGACGCTCTTCCGATCTNNNAGCAGAGGGAGAGAGATAG
GCACACGTCTGAACTCTTTCCCTACACGACGCTCTTCCGATCTNNNGGCTGTAGGTCTCTGTGTTCTT
GCACACGTCTGAACTCTTTCCCTACACGACGCTCTTCCGATCTNNNGGCTGTAGGTCTCTGTGTTCTT
GCACACGTCTGAACTCTTTCCCTACACGACGCTCTTCCGATCTNNNTACGACAGAGTGAGACTCCATC
GCACACGTCTGAACTCTTTCCCTACACGACGCTCTTCCGATCTNNNTACGACAGAGTGAGACTCCATC
GAGCACACGTCTGAACTCTTTCCCTACACGACGCTCTTCCGATCTNNNTCTTACAAAGTACAGTGGG
GAGCACACGTCTGAACTCTTTCCCTACACGACGCTCTTCCGATCTNNNTCTTACAAAGTACAGTGGG
AGCACACGTCTGAACTCTTTCCCTACACGACGCTCTTCCGATCTNNNTTCTGTAGAAGCAGCATCTCAC
AGCACACGTCTGAACTCTTTCCCTACACGACGCTCTTCCGATCTNNNTTCTGTAGAAGCAGCATCTCAC
AGCACACGTCTGAACTCTTTCCCTACACGACGCTCTTCCGATCTNNNCAGAGGTTACAGTGAGCTGA
AGCACACGTCTGAACTCTTTCCCTACACGACGCTCTTCCGATCTNNNCAGAGGTTACAGTGAGCTGA
GCACACGTCTGAACTCTTTCCCTACACGACGCTCTTCCGATCTNNNCCTGAGAGGAATTAGTCGTCCA
GCACACGTCTGAACTCTTTCCCTACACGACGCTCTTCCGATCTNNNCCTGAGAGGAATTAGTCGTCCA
GCACACGTCTGAACTCTTTCCCTACACGACGCTCTTCCGATCTNNNTCCGTAGTGTTAGGGTTGCTTT
GCACACGTCTGAACTCTTTCCCTACACGACGCTCTTCCGATCTNNNTCCGTAGTGTTAGGGTTGCTTT
GCACACGTCTGAACTCTTTCCCTACACGACGCTCTTCCGATCTNNNTTTAGAACCAAGGACACTCCCA
GCACACGTCTGAACTCTTTCCCTACACGACGCTCTTCCGATCTNNNTTTAGAACCAAGGACACTCCCA
GAGCACACGTCTGAACTCTTTCCCTACACGACGCTCTTCCGATCTNNNCCAAACACCGCATATTCTCAC
GAGCACACGTCTGAACTCTTTCCCTACACGACGCTCTTCCGATCTNNNCCAAACACCGCATATTCTCAC

AGCACACGTCTGAACTCTTTCCCTACACGACGCTCTTCCGATCTNNNCACATGCACGATGCTCTTATCTG

AGCACACGTCTGAACTCTTTCCCTACACGACGCTCTTCCGATCTNNNCACATGCACGATGCTCTTATCTG

GAGCACACGTCTGAACTCTTTCCCTACACGACGCTCTTCCGATCTNNNAAGGCCAATCTGATACAGAC

GAGCACACGTCTGAACTCTTTCCCTACACGACGCTCTTCCGATCTNNNAAGGCCAATCTGATACAGAC

AGCACACGTCTGAACTCTTTCCCTACACGACGCTCTTCCGATCTNNNAAAGGAAGCTAGGGTCTTTGC

AGCACACGTCTGAACTCTTTCCCTACACGACGCTCTTCCGATCTNNNAAAGGAAGCTAGGGTCTTTGC

AGCACACGTCTGAACTCTTTCCCTACACGACGCTCTTCCGATCTNNNTCAATGGAACACCCTCCTCCAA

AGCACACGTCTGAACTCTTTCCCTACACGACGCTCTTCCGATCTNNNTCAATGGAACACCCTCCTCCAA

CACACGTCTGAACTCTTTCCCTACACGACGCTCTTCCGATCTNNNTTAAGCCAGTGTTAGGGCATGTG

CACACGTCTGAACTCTTTCCCTACACGACGCTCTTCCGATCTNNNTTAAGCCAGTGTTAGGGCATGTG

AGCACACGTCTGAACTCTTTCCCTACACGACGCTCTTCCGATCTNNNTCTGTTTTGTGAGCAGTTCTCC

AGCACACGTCTGAACTCTTTCCCTACACGACGCTCTTCCGATCTNNNTCTGTTTTGTGAGCAGTTCTCC

GAGCACACGTCTGAACTCTTTCCCTACACGACGCTCTTCCGATCTNNNCCAATCACCACCCAATAAAGCT

GAGCACACGTCTGAACTCTTTCCCTACACGACGCTCTTCCGATCTNNNCCAATCACCACCCAATAAAGCT

GAGCACACGTCTGAACTCTTTCCCTACACGACGCTCTTCCGATCTNNNGTGTGTTTTTTTTTTTGGGGG

GAGCACACGTCTGAACTCTTTCCCTACACGACGCTCTTCCGATCTNNNGTGTGTTTTTTTTTTTGGGGG

AGCACACGTCTGAACTCTTTCCCTACACGACGCTCTTCCGATCTNNNCTTCAGGATAAGCCCACTCTCT

AGCACACGTCTGAACTCTTTCCCTACACGACGCTCTTCCGATCTNNNCTTCAGGATAAGCCCACTCTCT

AGCACACGTCTGAACTCTTTCCCTACACGACGCTCTTCCGATCTNNNTGTTCAGGCAGACACTAAGCTA

AGCACACGTCTGAACTCTTTCCCTACACGACGCTCTTCCGATCTNNNTGTTCAGGCAGACACTAAGCTA

GAGCACACGTCTGAACTCTTTCCCTACACGACGCTCTTCCGATCTNNNTTGGTAGGTTGTACATGTCCA

GAGCACACGTCTGAACTCTTTCCCTACACGACGCTCTTCCGATCTNNNTTGGTAGGTTGTACATGTCCA

AGCACACGTCTGAACTCTTTCCCTACACGACGCTCTTCCGATCTNNNATTGTGCGATGGAGTCTCACT

AGCACACGTCTGAACTCTTTCCCTACACGACGCTCTTCCGATCTNNNATTGTGCGATGGAGTCTCACT

GAGCACACGTCTGAACTCTTTCCCTACACGACGCTCTTCCGATCTNNNTTGTGTGTGTTTGTGGTGAG

GAGCACACGTCTGAACTCTTTCCCTACACGACGCTCTTCCGATCTNNNTTGTGTGTGTTTGTGGTGAG

AGCACACGTCTGAACTCTTTCCCTACACGACGCTCTTCCGATCTNNNAGTGAACCATGATAGTGCCAC

AGCACACGTCTGAACTCTTTCCCTACACGACGCTCTTCCGATCTNNNAGTGAACCATGATAGTGCCAC

AGCACACGTCTGAACTCTTTCCCTACACGACGCTCTTCCGATCTNNNCAAGTGATTCTCTTGCTTCAGC

AGCACACGTCTGAACTCTTTCCCTACACGACGCTCTTCCGATCTNNNCAAGTGATTCTCTTGCTTCAGC

GAGCACACGTCTGAACTCTTTCCCTACACGACGCTCTTCCGATCTNNNCGGTGAAAGTCCGTCTCTACT

GAGCACACGTCTGAACTCTTTCCCTACACGACGCTCTTCCGATCTNNNCGGTGAAAGTCCGTCTCTACT

GCACACGTCTGAACTCTTTCCCTACACGACGCTCTTCCGATCTNNNCCGTCTGAGAAGTGAGGAGCC

GCACACGTCTGAACTCTTTCCCTACACGACGCTCTTCCGATCTNNNCCGTCTGAGAAGTGAGGAGCC

AGCACACGTCTGAACTCTTTCCCTACACGACGCTCTTCCGATCTNNNCAATAGAGCTTTCTGGCAACTG

AGCACACGTCTGAACTCTTTCCCTACACGACGCTCTTCCGATCTNNNCAATAGAGCTTTCTGGCAACTG

AGCACACGTCTGAACTCTTTCCCTACACGACGCTCTTCCGATCTNNNGCTTCTCTTTTTCCCCAGGAGTA

AGCACACGTCTGAACTCTTTCCCTACACGACGCTCTTCCGATCTNNNGCTTCTCTTTTTCCCCAGGAGTA

GAGCACACGTCTGAACTCTTTCCCTACACGACGCTCTTCCGATCTNNNCTGAGCGGGCAAAAAATGAG

GAGCACACGTCTGAACTCTTTCCCTACACGACGCTCTTCCGATCTNNNCTGAGCGGGCAAAAAATGAG

GAGCACACGTCTGAACTCTTTCCCTACACGACGCTCTTCCGATCTNNNCACCGCACAGCCACATATTCT

GAGCACACGTCTGAACTCTTTCCCTACACGACGCTCTTCCGATCTNNNCACCGCACAGCCACATATTCT

AGCACACGTCTGAACTCTTTCCCTACACGACGCTCTTCCGATCTNNNGAGATCGTACCACTGCACTCCC

AGCACACGTCTGAACTCTTTCCCTACACGACGCTCTTCCGATCTNNNGAGATCGTACCACTGCACTCCC

AGCACACGTCTGAACTCTTTCCCTACACGACGCTCTTCCGATCTNNNCAGGTTCAAGAGATCCTCCCAC

AGCACACGTCTGAACTCTTTCCCTACACGACGCTCTTCCGATCTNNNCAGGTTCAAGAGATCCTCCCAC

GCACACGTCTGAACTCTTTCCCTACACGACGCTCTTCCGATCTNNNAAATGGACTGCCCTTTACCTAGC

GCACACGTCTGAACTCTTTCCCTACACGACGCTCTTCCGATCTNNNAAATGGACTGCCCTTTACCTAGC

AGAGCACACGTCTGAACTCTTTCCCTACACGACGCTCTTCCGATCTNNNGTGTTGTGCAGATTGAATG

AGAGCACACGTCTGAACTCTTTCCCTACACGACGCTCTTCCGATCTNNNGTGTTGTGCAGATTGAATG

AGCACACGTCTGAACTCTTTCCCTACACGACGCTCTTCCGATCTNNNATCCTTAGGCCTCTGGAGACAT

AGCACACGTCTGAACTCTTTCCCTACACGACGCTCTTCCGATCTNNNATCCTTAGGCCTCTGGAGACAT

AGCACACGTCTGAACTCTTTCCCTACACGACGCTCTTCCGATCTNNNTTCCTTCCTTCCTTCCTTCCTTCC

AGCACACGTCTGAACTCTTTCCCTACACGACGCTCTTCCGATCTNNNTTCCTTCCTTCCTTCCTTCCTTCC

AGAGCACACGTCTGAACTCTTTCCCTACACGACGCTCTTCCGATCTNNNAGAACATGCCTGGTGCATA

AGAGCACACGTCTGAACTCTTTCCCTACACGACGCTCTTCCGATCTNNNAGAACATGCCTGGTGCATA

AGCACACGTCTGAACTCTTTCCCTACACGACGCTCTTCCGATCTNNNCCACCACACCCAGCTAATTTTTC

AGCACACGTCTGAACTCTTTCCCTACACGACGCTCTTCCGATCTNNNCCACCACACCCAGCTAATTTTTC

AGCACACGTCTGAACTCTTTCCCTACACGACGCTCTTCCGATCTNNNGAGATGATAGCCAGGTGTGG

AGCACACGTCTGAACTCTTTCCCTACACGACGCTCTTCCGATCTNNNGAGATGATAGCCAGGTGTGG

GCACACGTCTGAACTCTTTCCCTACACGACGCTCTTCCGATCTNNNTGGTCTTTACACTGCTTTTGTGA

GCACACGTCTGAACTCTTTCCCTACACGACGCTCTTCCGATCTNNNTGGTCTTTACACTGCTTTTGTGA

AGCACACGTCTGAACTCTTTCCCTACACGACGCTCTTCCGATCTNNNTGCATGGGTATGGCTTTTATGT

AGCACACGTCTGAACTCTTTCCCTACACGACGCTCTTCCGATCTNNNTGCATGGGTATGGCTTTTATGT

GAGCACACGTCTGAACTCTTTCCCTACACGACGCTCTTCCGATCTNNNAAATGGGGAAGCCTAGGAA

GAGCACACGTCTGAACTCTTTCCCTACACGACGCTCTTCCGATCTNNNAAATGGGGAAGCCTAGGAA

CACACGTCTGAACTCTTTCCCTACACGACGCTCTTCCGATCTNNNAGGTGGTTCCTGAACAACAATTTC

CACACGTCTGAACTCTTTCCCTACACGACGCTCTTCCGATCTNNNAGGTGGTTCCTGAACAACAATTTC

AGCACACGTCTGAACTCTTTCCCTACACGACGCTCTTCCGATCTNNNAAATTAGCTAGGCATCCTGGTG

AGCACACGTCTGAACTCTTTCCCTACACGACGCTCTTCCGATCTNNNAAATTAGCTAGGCATCCTGGTG

AGCACACGTCTGAACTCTTTCCCTACACGACGCTCTTCCGATCTNNNATCTCTGCTCACTGTAAGCTCT

AGCACACGTCTGAACTCTTTCCCTACACGACGCTCTTCCGATCTNNNATCTCTGCTCACTGTAAGCTCT

GAGCACACGTCTGAACTCTTTCCCTACACGACGCTCTTCCGATCTNNNCTGAGATCGTGCCACTGTACT

GAGCACACGTCTGAACTCTTTCCCTACACGACGCTCTTCCGATCTNNNCTGAGATCGTGCCACTGTACT

GAGCACACGTCTGAACTCTTTCCCTACACGACGCTCTTCCGATCTNNNTATCCTACTGAGGCTTTCTGG

GAGCACACGTCTGAACTCTTTCCCTACACGACGCTCTTCCGATCTNNNTATCCTACTGAGGCTTTCTGG

AGCACACGTCTGAACTCTTTCCCTACACGACGCTCTTCCGATCTNNNTTTTCTGTGAATCATGAGCCTG

AGCACACGTCTGAACTCTTTCCCTACACGACGCTCTTCCGATCTNNNTTTTCTGTGAATCATGAGCCTG

GCACACGTCTGAACTCTTTCCCTACACGACGCTCTTCCGATCTNNNACACCTGACAATGTCCAGTTCAA

GCACACGTCTGAACTCTTTCCCTACACGACGCTCTTCCGATCTNNNACACCTGACAATGTCCAGTTCAA

AGCACACGTCTGAACTCTTTCCCTACACGACGCTCTTCCGATCTNNNCTGTACCTGAAGCAGACAGAG

AGCACACGTCTGAACTCTTTCCCTACACGACGCTCTTCCGATCTNNNCTGTACCTGAAGCAGACAGAG

AGCACACGTCTGAACTCTTTCCCTACACGACGCTCTTCCGATCTNNNGGTTTCTGTCTTTACCAAGGTG

AGCACACGTCTGAACTCTTTCCCTACACGACGCTCTTCCGATCTNNNGGTTTCTGTCTTTACCAAGGTG

GAGCACACGTCTGAACTCTTTCCCTACACGACGCTCTTCCGATCTNNNCACTTTTCGAGGAGTCGTCTC

GAGCACACGTCTGAACTCTTTCCCTACACGACGCTCTTCCGATCTNNNCACTTTTCGAGGAGTCGTCTC

AGCACACGTCTGAACTCTTTCCCTACACGACGCTCTTCCGATCTNNNAGAATTGCTTGAACCCAGGAG
AGCACACGTCTGAACTCTTTCCCTACACGACGCTCTTCCGATCTNNNAGAATTGCTTGAACCCAGGAG
AGCACACGTCTGAACTCTTTCCCTACACGACGCTCTTCCGATCTNNNAGTGTAAGAGTGTGTGTGTG
AGCACACGTCTGAACTCTTTCCCTACACGACGCTCTTCCGATCTNNNAGTGTAAGAGTGTGTGTGTG
AGCACACGTCTGAACTCTTTCCCTACACGACGCTCTTCCGATCTNNNTGAAAGTGGCTTTATGCATGTC
AGCACACGTCTGAACTCTTTCCCTACACGACGCTCTTCCGATCTNNNTGAAAGTGGCTTTATGCATGTC
AGCACACGTCTGAACTCTTTCCCTACACGACGCTCTTCCGATCTNNNTGTGGTGTTTCTAGTGCTCATC
AGCACACGTCTGAACTCTTTCCCTACACGACGCTCTTCCGATCTNNNTGTGGTGTTTCTAGTGCTCATC
AGCACACGTCTGAACTCTTTCCCTACACGACGCTCTTCCGATCTNNNTCAAACCAGGGAGTATTTTGC
AGCACACGTCTGAACTCTTTCCCTACACGACGCTCTTCCGATCTNNNTCAAACCAGGGAGTATTTTGC
AGCACACGTCTGAACTCTTTCCCTACACGACGCTCTTCCGATCTNNNTTACCATGTTGGCCAGGATAG
AGCACACGTCTGAACTCTTTCCCTACACGACGCTCTTCCGATCTNNNTTACCATGTTGGCCAGGATAG
GAGCACACGTCTGAACTCTTTCCCTACACGACGCTCTTCCGATCTNNNTATTAGGGGGTGGGGACTT
GAGCACACGTCTGAACTCTTTCCCTACACGACGCTCTTCCGATCTNNNTATTAGGGGGTGGGGACTT
GAGCACACGTCTGAACTCTTTCCCTACACGACGCTCTTCCGATCTNNNATGTCGTTTTGATGGCAAGA
GAGCACACGTCTGAACTCTTTCCCTACACGACGCTCTTCCGATCTNNNATGTCGTTTTGATGGCAAGA
AGCACACGTCTGAACTCTTTCCCTACACGACGCTCTTCCGATCTNNNCCTGGGTGACACAATGAGACT
AGCACACGTCTGAACTCTTTCCCTACACGACGCTCTTCCGATCTNNNCCTGGGTGACACAATGAGACT
GAGCACACGTCTGAACTCTTTCCCTACACGACGCTCTTCCGATCTNNNACATTGCATGTTCTCACTTTG
GAGCACACGTCTGAACTCTTTCCCTACACGACGCTCTTCCGATCTNNNACATTGCATGTTCTCACTTTG
AGCACACGTCTGAACTCTTTCCCTACACGACGCTCTTCCGATCTNNNTCTTAGAAAGGCTTTCCTAATG
AGCACACGTCTGAACTCTTTCCCTACACGACGCTCTTCCGATCTNNNTCTTAGAAAGGCTTTCCTAATG
GCACACGTCTGAACTCTTTCCCTACACGACGCTCTTCCGATCTNNNCCAACCCAGCATCTTTCTAGTCC
GCACACGTCTGAACTCTTTCCCTACACGACGCTCTTCCGATCTNNNCCAACCCAGCATCTTTCTAGTCC
GAGCACACGTCTGAACTCTTTCCCTACACGACGCTCTTCCGATCTNNNTTGAGGTCAGCAGTTCAAGA
GAGCACACGTCTGAACTCTTTCCCTACACGACGCTCTTCCGATCTNNNTTGAGGTCAGCAGTTCAAGA
CACGTCTGAACTCTTTCCCTACACGACGCTCTTCCGATCTNNNTAAGGCTGAATTTCCCAGGAAGGCA
CACGTCTGAACTCTTTCCCTACACGACGCTCTTCCGATCTNNNTAAGGCTGAATTTCCCAGGAAGGCA
AGCACACGTCTGAACTCTTTCCCTACACGACGCTCTTCCGATCTNNNCCCAACAAAACAAGCACATCAA
AGCACACGTCTGAACTCTTTCCCTACACGACGCTCTTCCGATCTNNNCCCAACAAAACAAGCACATCAA
AGCACACGTCTGAACTCTTTCCCTACACGACGCTCTTCCGATCTNNNAGAAACCACAGAGATATCAGC
AGCACACGTCTGAACTCTTTCCCTACACGACGCTCTTCCGATCTNNNAGAAACCACAGAGATATCAGC
GAGCACACGTCTGAACTCTTTCCCTACACGACGCTCTTCCGATCTNNNATACAAAAAGTTAGCCAGGC
GAGCACACGTCTGAACTCTTTCCCTACACGACGCTCTTCCGATCTNNNATACAAAAAGTTAGCCAGGC
GAGCACACGTCTGAACTCTTTCCCTACACGACGCTCTTCCGATCTNNNTGTGTGTGTGTCTGTGCTTGT
GAGCACACGTCTGAACTCTTTCCCTACACGACGCTCTTCCGATCTNNNTGTGTGTGTGTCTGTGCTTGT
GAGCACACGTCTGAACTCTTTCCCTACACGACGCTCTTCCGATCTNNNTAAATGCCAACCAAAGCCAC
GAGCACACGTCTGAACTCTTTCCCTACACGACGCTCTTCCGATCTNNNTAAATGCCAACCAAAGCCAC
AGCACACGTCTGAACTCTTTCCCTACACGACGCTCTTCCGATCTNNNCTGATACTGGGCCTCATGCAAT
AGCACACGTCTGAACTCTTTCCCTACACGACGCTCTTCCGATCTNNNCTGATACTGGGCCTCATGCAAT
AGCACACGTCTGAACTCTTTCCCTACACGACGCTCTTCCGATCTNNNTAGCAAAGTAGGCAGCTCTTG
AGCACACGTCTGAACTCTTTCCCTACACGACGCTCTTCCGATCTNNNTAGCAAAGTAGGCAGCTCTTG
AGCACACGTCTGAACTCTTTCCCTACACGACGCTCTTCCGATCTNNNTCTCCAGACTGCTTCTAGGAT

GAGCACACGTCTGAACTCTTTCCCTACACGACGCTCTTCCGATCTNNNTCTCCAGACTGCTTCTAGGATG
GCACACGTCTGAACTCTTTCCCTACACGACGCTCTTCCGATCTNNNTTGACTTCCCGGCCATTTAGTAACA
GCACACGTCTGAACTCTTTCCCTACACGACGCTCTTCCGATCTNNNTTGACTTCCCGGCCATTTAGTAACA
GAGCACACGTCTGAACTCTTTCCCTACACGACGCTCTTCCGATCTNNNTTGCATTACGTCACGCTACCCT
GAGCACACGTCTGAACTCTTTCCCTACACGACGCTCTTCCGATCTNNNTTGCATTACGTCACGCTACCCT
GAGCACACGTCTGAACTCTTTCCCTACACGACGCTCTTCCGATCTNNNACAGTGGTGCTTGTGTTACCT
GAGCACACGTCTGAACTCTTTCCCTACACGACGCTCTTCCGATCTNNNACAGTGGTGCTTGTGTTACCT
AGCACACGTCTGAACTCTTTCCCTACACGACGCTCTTCCGATCTNNNGGCAACAGAAGAAGACTCCATG
AGCACACGTCTGAACTCTTTCCCTACACGACGCTCTTCCGATCTNNNGGCAACAGAAGAAGACTCCATG
AGCACACGTCTGAACTCTTTCCCTACACGACGCTCTTCCGATCTNNNCAGGGGAAAGAGATGTTTGGC
AGCACACGTCTGAACTCTTTCCCTACACGACGCTCTTCCGATCTNNNCAGGGGAAAGAGATGTTTGGC
GAGCACACGTCTGAACTCTTTCCCTACACGACGCTCTTCCGATCTNNNTCTTCCACTCTGGTTACTCAAG
GAGCACACGTCTGAACTCTTTCCCTACACGACGCTCTTCCGATCTNNNTCTTCCACTCTGGTTACTCAAG
AGCACACGTCTGAACTCTTTCCCTACACGACGCTCTTCCGATCTNNNAGGTTGCAGTGAACCAAGATA
AGCACACGTCTGAACTCTTTCCCTACACGACGCTCTTCCGATCTNNNAGGTTGCAGTGAACCAAGATA
GAGCACACGTCTGAACTCTTTCCCTACACGACGCTCTTCCGATCTNNNGACTGTACCCTTGACCTTCTGA
GAGCACACGTCTGAACTCTTTCCCTACACGACGCTCTTCCGATCTNNNGACTGTACCCTTGACCTTCTGA
AGCACACGTCTGAACTCTTTCCCTACACGACGCTCTTCCGATCTNNNGAGAGGTAGAGCTTGCAGTGA
AGCACACGTCTGAACTCTTTCCCTACACGACGCTCTTCCGATCTNNNGAGAGGTAGAGCTTGCAGTGA
GAGCACACGTCTGAACTCTTTCCCTACACGACGCTCTTCCGATCTNNNTTTCAAATGGGAAACACTCAGC
GAGCACACGTCTGAACTCTTTCCCTACACGACGCTCTTCCGATCTNNNTTTCAAATGGGAAACACTCAGC
AGCACACGTCTGAACTCTTTCCCTACACGACGCTCTTCCGATCTNNNAGAGAAACAGAACCAACAGGAC
AGCACACGTCTGAACTCTTTCCCTACACGACGCTCTTCCGATCTNNNAGAGAAACAGAACCAACAGGAC
GAGCACACGTCTGAACTCTTTCCCTACACGACGCTCTTCCGATCTNNNTCACCCTCTCACTTCAAAAAGCT
GAGCACACGTCTGAACTCTTTCCCTACACGACGCTCTTCCGATCTNNNTCACCCTCTCACTTCAAAAAGCT
GAGCACACGTCTGAACTCTTTCCCTACACGACGCTCTTCCGATCTNNNGAGACTTGTTGCCATGGAGAC
GAGCACACGTCTGAACTCTTTCCCTACACGACGCTCTTCCGATCTNNNGAGACTTGTTGCCATGGAGAC
AGCACACGTCTGAACTCTTTCCCTACACGACGCTCTTCCGATCTNNNGAAGCCCTAGGAGATGAATCCA
AGCACACGTCTGAACTCTTTCCCTACACGACGCTCTTCCGATCTNNNGAAGCCCTAGGAGATGAATCCA
GAGCACACGTCTGAACTCTTTCCCTACACGACGCTCTTCCGATCTNNNTTACTTAGGAGGCTGAGGTG
GAGCACACGTCTGAACTCTTTCCCTACACGACGCTCTTCCGATCTNNNTTACTTAGGAGGCTGAGGTG
AGCACACGTCTGAACTCTTTCCCTACACGACGCTCTTCCGATCTNNNGAACTCCCGACTTCAGTTGATCT
AGCACACGTCTGAACTCTTTCCCTACACGACGCTCTTCCGATCTNNNGAACTCCCGACTTCAGTTGATCT
AGCACACGTCTGAACTCTTTCCCTACACGACGCTCTTCCGATCTNNNCTCATTCTCTTGCTCCTCGACTTC
AGCACACGTCTGAACTCTTTCCCTACACGACGCTCTTCCGATCTNNNCTCATTCTCTTGCTCCTCGACTTC
GAGCACACGTCTGAACTCTTTCCCTACACGACGCTCTTCCGATCTNNNTATGAAGAGCCTGGGATGAG
GAGCACACGTCTGAACTCTTTCCCTACACGACGCTCTTCCGATCTNNNTATGAAGAGCCTGGGATGAG
GAGCACACGTCTGAACTCTTTCCCTACACGACGCTCTTCCGATCTNNNCCAATTGCAAACAGATCAGGT
GAGCACACGTCTGAACTCTTTCCCTACACGACGCTCTTCCGATCTNNNCCAATTGCAAACAGATCAGGT
AGAGCACACGTCTGAACTCTTTCCCTACACGACGCTCTTCCGATCTNNNGAGGCAAGAGAATCGCTTGA
AGAGCACACGTCTGAACTCTTTCCCTACACGACGCTCTTCCGATCTNNNGAGGCAAGAGAATCGCTTGA
AGCACACGTCTGAACTCTTTCCCTACACGACGCTCTTCCGATCTNNNAGCAGTGTATAAGTGGTGCCTT
AGCACACGTCTGAACTCTTTCCCTACACGACGCTCTTCCGATCTNNNAGCAGTGTATAAGTGGTGCCTT

CACACGTCTGAACTCTTTCCCTACACGACGCTCTTCCGATCTNNNGGTGTTTGAAGAGTTTTGTCCATCC
CACACGTCTGAACTCTTTCCCTACACGACGCTCTTCCGATCTNNNGGTGTTTGAAGAGTTTTGTCCATCC
AGCACACGTCTGAACTCTTTCCCTACACGACGCTCTTCCGATCTNNNCAGGGAAATCAGCTGAGACTCA
AGCACACGTCTGAACTCTTTCCCTACACGACGCTCTTCCGATCTNNNCAGGGAAATCAGCTGAGACTCA
AGCACACGTCTGAACTCTTTCCCTACACGACGCTCTTCCGATCTNNNGTGTGTTTGTGTGTGTGTGT
AGCACACGTCTGAACTCTTTCCCTACACGACGCTCTTCCGATCTNNNGTGTGTTTGTGTGTGTGTGT
AGCACACGTCTGAACTCTTTCCCTACACGACGCTCTTCCGATCTNNNCGTCTCTGCACTCCAGACTGCAT
AGCACACGTCTGAACTCTTTCCCTACACGACGCTCTTCCGATCTNNNCGTCTCTGCACTCCAGACTGCAT
GAGCACACGTCTGAACTCTTTCCCTACACGACGCTCTTCCGATCTNNNACTCCTCAGTACAACGTTGAG
GAGCACACGTCTGAACTCTTTCCCTACACGACGCTCTTCCGATCTNNNACTCCTCAGTACAACGTTGAG
AGAGCACACGTCTGAACTCTTTCCCTACACGACGCTCTTCCGATCTNNNCATGTTGGCCTAAGGACTATC
AGAGCACACGTCTGAACTCTTTCCCTACACGACGCTCTTCCGATCTNNNCATGTTGGCCTAAGGACTATC
GAGCACACGTCTGAACTCTTTCCCTACACGACGCTCTTCCGATCTNNNAGGACTTCAACTCAACAAAGA
GAGCACACGTCTGAACTCTTTCCCTACACGACGCTCTTCCGATCTNNNAGGACTTCAACTCAACAAAGA
ACACGTCTGAACTCTTTCCCTACACGACGCTCTTCCGATCTNNNGCGTACTTTGTCTTCTTGTAACATCG
ACACGTCTGAACTCTTTCCCTACACGACGCTCTTCCGATCTNNNGCGTACTTTGTCTTCTTGTAACATCG
AGCACACGTCTGAACTCTTTCCCTACACGACGCTCTTCCGATCTNNNGCTCCAATCTCCCTCTCTCTCTCC
AGCACACGTCTGAACTCTTTCCCTACACGACGCTCTTCCGATCTNNNGCTCCAATCTCCCTCTCTCTCTCC
AGCACACGTCTGAACTCTTTCCCTACACGACGCTCTTCCGATCTNNNGCCTATAATCCCAGCTACTCAGG
AGCACACGTCTGAACTCTTTCCCTACACGACGCTCTTCCGATCTNNNGCCTATAATCCCAGCTACTCAGG
AGCACACGTCTGAACTCTTTCCCTACACGACGCTCTTCCGATCTNNNCTGAGGCAGGAGAATCACTTGA
AGCACACGTCTGAACTCTTTCCCTACACGACGCTCTTCCGATCTNNNCTGAGGCAGGAGAATCACTTGA
GAGCACACGTCTGAACTCTTTCCCTACACGACGCTCTTCCGATCTNNNTACACGTGTGGCGGTACTTAC
GAGCACACGTCTGAACTCTTTCCCTACACGACGCTCTTCCGATCTNNNTACACGTGTGGCGGTACTTAC
GAGCACACGTCTGAACTCTTTCCCTACACGACGCTCTTCCGATCTNNNTGGTGGCATATAGTAAGCAG
GAGCACACGTCTGAACTCTTTCCCTACACGACGCTCTTCCGATCTNNNTGGTGGCATATAGTAAGCAG
AGCACACGTCTGAACTCTTTCCCTACACGACGCTCTTCCGATCTNNNTATCCACCAGTGCCCAATGTTTA
AGCACACGTCTGAACTCTTTCCCTACACGACGCTCTTCCGATCTNNNTATCCACCAGTGCCCAATGTTTA
AGCACACGTCTGAACTCTTTCCCTACACGACGCTCTTCCGATCTNNNCTGATGTCAGGTCCTCCACAGCA
AGCACACGTCTGAACTCTTTCCCTACACGACGCTCTTCCGATCTNNNCTGATGTCAGGTCCTCCACAGCA
CACACGTCTGAACTCTTTCCCTACACGACGCTCTTCCGATCTNNNATGAGACATGTCAGCAACTCTGTCA
CACACGTCTGAACTCTTTCCCTACACGACGCTCTTCCGATCTNNNATGAGACATGTCAGCAACTCTGTCA
GAGCACACGTCTGAACTCTTTCCCTACACGACGCTCTTCCGATCTNNNACAGGACCACAGGCTTCTCTAC
GAGCACACGTCTGAACTCTTTCCCTACACGACGCTCTTCCGATCTNNNACAGGACCACAGGCTTCTCTAC
AGCACACGTCTGAACTCTTTCCCTACACGACGCTCTTCCGATCTNNNAGACGAGACGTACCATTTTTTG
AGCACACGTCTGAACTCTTTCCCTACACGACGCTCTTCCGATCTNNNAGACGAGACGTACCATTTTTTG
AGCACACGTCTGAACTCTTTCCCTACACGACGCTCTTCCGATCTNNNCCAGCTGTTCTTTGTACATCTGT
AGCACACGTCTGAACTCTTTCCCTACACGACGCTCTTCCGATCTNNNCCAGCTGTTCTTTGTACATCTGT
GCACACGTCTGAACTCTTTCCCTACACGACGCTCTTCCGATCTNNNTGAGCTGAGCTACACAAGTCTTC
GCACACGTCTGAACTCTTTCCCTACACGACGCTCTTCCGATCTNNNTGAGCTGAGCTACACAAGTCTTC
AGAGCACACGTCTGAACTCTTTCCCTACACGACGCTCTTCCGATCTNNNCTAAGGCAGGAGGATTGCTT
AGAGCACACGTCTGAACTCTTTCCCTACACGACGCTCTTCCGATCTNNNCTAAGGCAGGAGGATTGCTT
AGCACACGTCTGAACTCTTTCCCTACACGACGCTCTTCCGATCTNNNCTCTTGTCCCCCTTTCCTGTACCA

AGCACACGTCTGAACTCTTTCCCTACACGACGCTCTTCCGATCTNNNCTCTTGTCCCCCTTTCCTGTACCA
AGCACACGTCTGAACTCTTTCCCTACACGACGCTCTTCCGATCTNNNGTCTGCACACATAATTCACGCT
AGCACACGTCTGAACTCTTTCCCTACACGACGCTCTTCCGATCTNNNGTCTGCACACATAATTCACGCT
AGCACACGTCTGAACTCTTTCCCTACACGACGCTCTTCCGATCTNNNTGGGACTAGAGGTACACACAC
AGCACACGTCTGAACTCTTTCCCTACACGACGCTCTTCCGATCTNNNTGGGACTAGAGGTACACACAC
ACACGTCTGAACTCTTTCCCTACACGACGCTCTTCCGATCTNNNATTGTGTGTCTCATGATGTTCATCG
ACACGTCTGAACTCTTTCCCTACACGACGCTCTTCCGATCTNNNATTGTGTGTCTCATGATGTTCATCG
AGCACACGTCTGAACTCTTTCCCTACACGACGCTCTTCCGATCTNNNAGGGCAGACTCCATCTCAAAAA
AGCACACGTCTGAACTCTTTCCCTACACGACGCTCTTCCGATCTNNNAGGGCAGACTCCATCTCAAAAA
GAGCACACGTCTGAACTCTTTCCCTACACGACGCTCTTCCGATCTNNNTAGAGTACTTAGGGGGAAG
GAGCACACGTCTGAACTCTTTCCCTACACGACGCTCTTCCGATCTNNNTAGAGTACTTAGGGGGAAG
CACACGTCTGAACTCTTTCCCTACACGACGCTCTTCCGATCTNNNCCATCTGGTTCCTTCTCATCTGGCA
CACACGTCTGAACTCTTTCCCTACACGACGCTCTTCCGATCTNNNCCATCTGGTTCCTTCTCATCTGGCA
GCACACGTCTGAACTCTTTCCCTACACGACGCTCTTCCGATCTNNNAATACAAAAAATTAGCCGGGCG
GCACACGTCTGAACTCTTTCCCTACACGACGCTCTTCCGATCTNNNAATACAAAAAATTAGCCGGGCG
GAGCACACGTCTGAACTCTTTCCCTACACGACGCTCTTCCGATCTNNNCCTGGACAATATGGAGGAAC
GAGCACACGTCTGAACTCTTTCCCTACACGACGCTCTTCCGATCTNNNCCTGGACAATATGGAGGAAC
AGCACACGTCTGAACTCTTTCCCTACACGACGCTCTTCCGATCTNNNGCTGAAAATAGTCCGATGAGT
AGCACACGTCTGAACTCTTTCCCTACACGACGCTCTTCCGATCTNNNGCTGAAAATAGTCCGATGAGT
CACACGTCTGAACTCTTTCCCTACACGACGCTCTTCCGATCTNNNCACAACGACCACATCCTCATAGACA
CACACGTCTGAACTCTTTCCCTACACGACGCTCTTCCGATCTNNNCACAACGACCACATCCTCATAGACA
GAGCACACGTCTGAACTCTTTCCCTACACGACGCTCTTCCGATCTNNNTAAATCTTGTGCCTTGGGGAT
GAGCACACGTCTGAACTCTTTCCCTACACGACGCTCTTCCGATCTNNNTAAATCTTGTGCCTTGGGGAT
AGCACACGTCTGAACTCTTTCCCTACACGACGCTCTTCCGATCTNNNCCTCCTTCTAGCCCTTGTTTTCT
AGCACACGTCTGAACTCTTTCCCTACACGACGCTCTTCCGATCTNNNCCTCCTTCTAGCCCTTGTTTTCT
GCACACGTCTGAACTCTTTCCCTACACGACGCTCTTCCGATCTNNNCTGAAATTGGTCACTGGGTTGT
GCACACGTCTGAACTCTTTCCCTACACGACGCTCTTCCGATCTNNNCTGAAATTGGTCACTGGGTTGT
AGCACACGTCTGAACTCTTTCCCTACACGACGCTCTTCCGATCTNNNCATTCCCAGGGAGCAGAAAAT
AGCACACGTCTGAACTCTTTCCCTACACGACGCTCTTCCGATCTNNNCATTCCCAGGGAGCAGAAAAT
AGCACACGTCTGAACTCTTTCCCTACACGACGCTCTTCCGATCTNNNTCCATTCACTAGCTTCCCCTCAA
AGCACACGTCTGAACTCTTTCCCTACACGACGCTCTTCCGATCTNNNTCCATTCACTAGCTTCCCCTCAA
AGCACACGTCTGAACTCTTTCCCTACACGACGCTCTTCCGATCTNNNTGACCATCTACTATGTGTCAGT
AGCACACGTCTGAACTCTTTCCCTACACGACGCTCTTCCGATCTNNNTGACCATCTACTATGTGTCAGT
GAGCACACGTCTGAACTCTTTCCCTACACGACGCTCTTCCGATCTNNNCTCCCAGTGGTAGCCTAGTCT
GAGCACACGTCTGAACTCTTTCCCTACACGACGCTCTTCCGATCTNNNCTCCCAGTGGTAGCCTAGTCT
CACACGTCTGAACTCTTTCCCTACACGACGCTCTTCCGATCTNNNCGGCGATGCTGGAATTCTTGCAT
CACACGTCTGAACTCTTTCCCTACACGACGCTCTTCCGATCTNNNCGGCGATGCTGGAATTCTTGCAT
GCACACGTCTGAACTCTTTCCCTACACGACGCTCTTCCGATCTNNNCTGAGGCAGGAGAATCACTTGA
GCACACGTCTGAACTCTTTCCCTACACGACGCTCTTCCGATCTNNNCTGAGGCAGGAGAATCACTTGA
GCACACGTCTGAACTCTTTCCCTACACGACGCTCTTCCGATCTNNNATCTAGTCTTCTCACCTGTCCCAC
GCACACGTCTGAACTCTTTCCCTACACGACGCTCTTCCGATCTNNNATCTAGTCTTCTCACCTGTCCCAC
AGCACACGTCTGAACTCTTTCCCTACACGACGCTCTTCCGATCTNNNACGTTTATTGTAGGTTAGGCG
AGCACACGTCTGAACTCTTTCCCTACACGACGCTCTTCCGATCTNNNACGTTTATTGTAGGTTAGGCG

AGCACACGTCTGAACTCTTTCCCTACACGACGCTCTTCCGATCTNNNTGGTCATAAGGGTTAGCTTTG
AGCACACGTCTGAACTCTTTCCCTACACGACGCTCTTCCGATCTNNNTGGTCATAAGGGTTAGCTTTG
AGCACACGTCTGAACTCTTTCCCTACACGACGCTCTTCCGATCTNNNGATCACTTAGGGTTAGGAGCT
AGCACACGTCTGAACTCTTTCCCTACACGACGCTCTTCCGATCTNNNGATCACTTAGGGTTAGGAGCT
AGCACACGTCTGAACTCTTTCCCTACACGACGCTCTTCCGATCTNNNATCCAGGTAGCAGTGAAAGTCT
AGCACACGTCTGAACTCTTTCCCTACACGACGCTCTTCCGATCTNNNATCCAGGTAGCAGTGAAAGTCT
AGAGCACACGTCTGAACTCTTTCCCTACACGACGCTCTTCCGATCTNNNGGCTTGACCTCTCTGAAGA
AGAGCACACGTCTGAACTCTTTCCCTACACGACGCTCTTCCGATCTNNNGGCTTGACCTCTCTGAAGA
GCACACGTCTGAACTCTTTCCCTACACGACGCTCTTCCGATCTNNNTGCAGAATTCCAGAACTTCCTGA
GCACACGTCTGAACTCTTTCCCTACACGACGCTCTTCCGATCTNNNTGCAGAATTCCAGAACTTCCTGA
GCACACGTCTGAACTCTTTCCCTACACGACGCTCTTCCGATCTNNNAGGAAATGATGTTCAGGCCTCT
GCACACGTCTGAACTCTTTCCCTACACGACGCTCTTCCGATCTNNNAGGAAATGATGTTCAGGCCTCT
AGCACACGTCTGAACTCTTTCCCTACACGACGCTCTTCCGATCTNNNCAAACACAGCTCACTAAAGCCT
AGCACACGTCTGAACTCTTTCCCTACACGACGCTCTTCCGATCTNNNCAAACACAGCTCACTAAAGCCT
GCACACGTCTGAACTCTTTCCCTACACGACGCTCTTCCGATCTNNNGACCATGACTGTGTGAGGAAGA
GCACACGTCTGAACTCTTTCCCTACACGACGCTCTTCCGATCTNNNGACCATGACTGTGTGAGGAAGA
AGCACACGTCTGAACTCTTTCCCTACACGACGCTCTTCCGATCTNNNCAGCTACTTAGGAGGTGAAG
AGCACACGTCTGAACTCTTTCCCTACACGACGCTCTTCCGATCTNNNCAGCTACTTAGGAGGTGAAG
GAGCACACGTCTGAACTCTTTCCCTACACGACGCTCTTCCGATCTNNNACCTGTAATCCTAGCACTTTG
GAGCACACGTCTGAACTCTTTCCCTACACGACGCTCTTCCGATCTNNNACCTGTAATCCTAGCACTTTG
AGCACACGTCTGAACTCTTTCCCTACACGACGCTCTTCCGATCTNNNCTGAGGCAGGAGAATCACTTG
AGCACACGTCTGAACTCTTTCCCTACACGACGCTCTTCCGATCTNNNCTGAGGCAGGAGAATCACTTG
AGCACACGTCTGAACTCTTTCCCTACACGACGCTCTTCCGATCTNNNATTTTCTTGTAGACGGCTGGA
AGCACACGTCTGAACTCTTTCCCTACACGACGCTCTTCCGATCTNNNATTTTCTTGTAGACGGCTGGA
AGCACACGTCTGAACTCTTTCCCTACACGACGCTCTTCCGATCTNNNCTCTGAACTTTTTCCTTGCTGGG
AGCACACGTCTGAACTCTTTCCCTACACGACGCTCTTCCGATCTNNNCTCTGAACTTTTTCCTTGCTGGG
AGCACACGTCTGAACTCTTTCCCTACACGACGCTCTTCCGATCTNNNAAGGTGTGTGCCTTTTACTGTG
AGCACACGTCTGAACTCTTTCCCTACACGACGCTCTTCCGATCTNNNAAGGTGTGTGCCTTTTACTGTG
GAGCACACGTCTGAACTCTTTCCCTACACGACGCTCTTCCGATCTNNNGTCTTGTGCTGGTCCATTTCT
GAGCACACGTCTGAACTCTTTCCCTACACGACGCTCTTCCGATCTNNNGTCTTGTGCTGGTCCATTTCT
AGCACACGTCTGAACTCTTTCCCTACACGACGCTCTTCCGATCTNNNTCTCTGCGATATCAGGAGACTC
AGCACACGTCTGAACTCTTTCCCTACACGACGCTCTTCCGATCTNNNTCTCTGCGATATCAGGAGACTC
GCACACGTCTGAACTCTTTCCCTACACGACGCTCTTCCGATCTNNNGTCAGTGATCTCTGAGGCTTCTT
GCACACGTCTGAACTCTTTCCCTACACGACGCTCTTCCGATCTNNNGTCAGTGATCTCTGAGGCTTCTT
AGCACACGTCTGAACTCTTTCCCTACACGACGCTCTTCCGATCTNNNAACCAGAAGTATCGCTTGAAC
AGCACACGTCTGAACTCTTTCCCTACACGACGCTCTTCCGATCTNNNAACCAGAAGTATCGCTTGAAC
GAGCACACGTCTGAACTCTTTCCCTACACGACGCTCTTCCGATCTNNNAGCTTTGGCATTGAGAAATG
GAGCACACGTCTGAACTCTTTCCCTACACGACGCTCTTCCGATCTNNNAGCTTTGGCATTGAGAAATG
AGCACACGTCTGAACTCTTTCCCTACACGACGCTCTTCCGATCTNNNACAATGGCTCCTCTTATGGCTTC
AGCACACGTCTGAACTCTTTCCCTACACGACGCTCTTCCGATCTNNNACAATGGCTCCTCTTATGGCTTC
AGCACACGTCTGAACTCTTTCCCTACACGACGCTCTTCCGATCTNNNTTGACCTCCTGAGTAGCTAGG
AGCACACGTCTGAACTCTTTCCCTACACGACGCTCTTCCGATCTNNNTTGACCTCCTGAGTAGCTAGG
AGCACACGTCTGAACTCTTTCCCTACACGACGCTCTTCCGATCTNNNAGGCAGTTTCACAGATAGCTT

AGCACACGTCTGAACTCTTTCCCTACACGACGCTCTTCCGATCTNNNAGGCAGTTTCACAGATAGCTT

GCACACGTCTGAACTCTTTCCCTACACGACGCTCTTCCGATCTNNNCCACAAAAGCCAATTCTCTCCTG

GCACACGTCTGAACTCTTTCCCTACACGACGCTCTTCCGATCTNNNCCACAAAAGCCAATTCTCTCCTG

AGCACACGTCTGAACTCTTTCCCTACACGACGCTCTTCCGATCTNNNTACAAAGAACCCCTGCAAGATG

AGCACACGTCTGAACTCTTTCCCTACACGACGCTCTTCCGATCTNNNTACAAAGAACCCCTGCAAGATG

GAGCACACGTCTGAACTCTTTCCCTACACGACGCTCTTCCGATCTNNNTGAGAGACCTTGACCCTGAA

GAGCACACGTCTGAACTCTTTCCCTACACGACGCTCTTCCGATCTNNNTGAGAGACCTTGACCCTGAA

GAGCACACGTCTGAACTCTTTCCCTACACGACGCTCTTCCGATCTNNNAACCTGTAAGTCTCTGCTTG

GAGCACACGTCTGAACTCTTTCCCTACACGACGCTCTTCCGATCTNNNAACCTGTAAGTCTCTGCTTG

AGCACACGTCTGAACTCTTTCCCTACACGACGCTCTTCCGATCTNNNTGAGCCAAGATTGTGCCACTC

AGCACACGTCTGAACTCTTTCCCTACACGACGCTCTTCCGATCTNNNTGAGCCAAGATTGTGCCACTC

AGAGCACACGTCTGAACTCTTTCCCTACACGACGCTCTTCCGATCTNNNCACTCAGGAGTTGAATAG

AGAGCACACGTCTGAACTCTTTCCCTACACGACGCTCTTCCGATCTNNNCACTCAGGAGTTGAATAG

AGCACACGTCTGAACTCTTTCCCTACACGACGCTCTTCCGATCTNNNCCCCGACGACGTGGACTACC

AGCACACGTCTGAACTCTTTCCCTACACGACGCTCTTCCGATCTNNNCCCCGACGACGTGGACTACC

AGCACACGTCTGAACTCTTTCCCTACACGACGCTCTTCCGATCTNNNCTAAGGCAGGGGAATCACTTG

AGCACACGTCTGAACTCTTTCCCTACACGACGCTCTTCCGATCTNNNCTAAGGCAGGGGAATCACTTG

AGCACACGTCTGAACTCTTTCCCTACACGACGCTCTTCCGATCTNNNCCTGGAGTTTCCCCAATGTTTT

AGCACACGTCTGAACTCTTTCCCTACACGACGCTCTTCCGATCTNNNCCTGGAGTTTCCCCAATGTTTT

AGCACACGTCTGAACTCTTTCCCTACACGACGCTCTTCCGATCTNNNCAATTTGTCAAGGGGTGAGTG

AGCACACGTCTGAACTCTTTCCCTACACGACGCTCTTCCGATCTNNNCAATTTGTCAAGGGGTGAGTG

AGCACACGTCTGAACTCTTTCCCTACACGACGCTCTTCCGATCTNNNTAGCCTAGACTCCTTCTCTCCAC

AGCACACGTCTGAACTCTTTCCCTACACGACGCTCTTCCGATCTNNNTAGCCTAGACTCCTTCTCTCCAC

GAGCACACGTCTGAACTCTTTCCCTACACGACGCTCTTCCGATCTNNNGATTGCTTGAGTCCAGGATT

GAGCACACGTCTGAACTCTTTCCCTACACGACGCTCTTCCGATCTNNNGATTGCTTGAGTCCAGGATT

GCACACGTCTGAACTCTTTCCCTACACGACGCTCTTCCGATCTNNNAGGGAGGAAGGACTGCATTTG

GCACACGTCTGAACTCTTTCCCTACACGACGCTCTTCCGATCTNNNAGGGAGGAAGGACTGCATTTG

GAGCACACGTCTGAACTCTTTCCCTACACGACGCTCTTCCGATCTNNNGACCTTTGATCCAGGCTCCTT

GAGCACACGTCTGAACTCTTTCCCTACACGACGCTCTTCCGATCTNNNGACCTTTGATCCAGGCTCCTT

AGCACACGTCTGAACTCTTTCCCTACACGACGCTCTTCCGATCTNNNCCACCACACCCTGCTAATTTTTC

AGCACACGTCTGAACTCTTTCCCTACACGACGCTCTTCCGATCTNNNCCACCACACCCTGCTAATTTTTC

AGCACACGTCTGAACTCTTTCCCTACACGACGCTCTTCCGATCTNNNTCAGGCTACAGGTAGGGAAA

AGCACACGTCTGAACTCTTTCCCTACACGACGCTCTTCCGATCTNNNTCAGGCTACAGGTAGGGAAA

GAGCACACGTCTGAACTCTTTCCCTACACGACGCTCTTCCGATCTNNNAGTAGACAGAGATTGCAGTG

GAGCACACGTCTGAACTCTTTCCCTACACGACGCTCTTCCGATCTNNNAGTAGACAGAGATTGCAGTG

GCACACGTCTGAACTCTTTCCCTACACGACGCTCTTCCGATCTNNNAGGAGAATTGTTTGAATCCGGG

GCACACGTCTGAACTCTTTCCCTACACGACGCTCTTCCGATCTNNNAGGAGAATTGTTTGAATCCGGG

AGCACACGTCTGAACTCTTTCCCTACACGACGCTCTTCCGATCTNNNCTCATCCCTGAGTGAGGTTTA

AGCACACGTCTGAACTCTTTCCCTACACGACGCTCTTCCGATCTNNNCTCATCCCTGAGTGAGGTTTA

AGCACACGTCTGAACTCTTTCCCTACACGACGCTCTTCCGATCTNNNAGCCCAATCACAGAAACCAGAA

AGCACACGTCTGAACTCTTTCCCTACACGACGCTCTTCCGATCTNNNAGCCCAATCACAGAAACCAGAA

AGCACACGTCTGAACTCTTTCCCTACACGACGCTCTTCCGATCTNNNCTCTGCCCCCAATTTAGAGTGA

AGCACACGTCTGAACTCTTTCCCTACACGACGCTCTTCCGATCTNNNCTCTGCCCCCAATTTAGAGTGA

GCACACGTCTGAACTCTTTCCCTACACGACGCTCTTCCGATCTNNNATCCTGGAAACCCTGGTTCTTGCA

GCACACGTCTGAACTCTTTCCCTACACGACGCTCTTCCGATCTNNNATCCTGGAAACCCTGGTTCTTGCA

AGCACACGTCTGAACTCTTTCCCTACACGACGCTCTTCCGATCTNNNCCCGTCCTCCAAAACTTTTTTC

AGCACACGTCTGAACTCTTTCCCTACACGACGCTCTTCCGATCTNNNCCCGTCCTCCAAAACTTTTTTC

CACACGTCTGAACTCTTTCCCTACACGACGCTCTTCCGATCTNNNGGGATGACTTCTTAGACCCCTTGCA

CACACGTCTGAACTCTTTCCCTACACGACGCTCTTCCGATCTNNNGGGATGACTTCTTAGACCCCTTGCA

AGCACACGTCTGAACTCTTTCCCTACACGACGCTCTTCCGATCTNNNTGAGTCAGGAAGAAGTACCTCC

AGCACACGTCTGAACTCTTTCCCTACACGACGCTCTTCCGATCTNNNTGAGTCAGGAAGAAGTACCTCC

AGCACACGTCTGAACTCTTTCCCTACACGACGCTCTTCCGATCTNNNTTTGTTCCTTTTCAGTGCTGAGT

AGCACACGTCTGAACTCTTTCCCTACACGACGCTCTTCCGATCTNNNTTTGTTCCTTTTCAGTGCTGAGT

AGCACACGTCTGAACTCTTTCCCTACACGACGCTCTTCCGATCTNNNGTTGAGGCAGAAGGATTGCTT

AGCACACGTCTGAACTCTTTCCCTACACGACGCTCTTCCGATCTNNNGTTGAGGCAGAAGGATTGCTT

AGCACACGTCTGAACTCTTTCCCTACACGACGCTCTTCCGATCTNNNCTACCATAGTCCTCAAGGTGCTC

AGCACACGTCTGAACTCTTTCCCTACACGACGCTCTTCCGATCTNNNCTACCATAGTCCTCAAGGTGCTC

AGAGCACACGTCTGAACTCTTTCCCTACACGACGCTCTTCCGATCTNNNTATAGTCCCAGCTACTCAGGA

AGAGCACACGTCTGAACTCTTTCCCTACACGACGCTCTTCCGATCTNNNTATAGTCCCAGCTACTCAGGA

GAGCACACGTCTGAACTCTTTCCCTACACGACGCTCTTCCGATCTNNNGTCCTGTTGAGAGAAATCCCA

GAGCACACGTCTGAACTCTTTCCCTACACGACGCTCTTCCGATCTNNNGTCCTGTTGAGAGAAATCCCA

GAGCACACGTCTGAACTCTTTCCCTACACGACGCTCTTCCGATCTNNNACAGCTCCTGGATTTCTTGATC

GAGCACACGTCTGAACTCTTTCCCTACACGACGCTCTTCCGATCTNNNACAGCTCCTGGATTTCTTGATC

GAGCACACGTCTGAACTCTTTCCCTACACGACGCTCTTCCGATCTNNNCAGGAAAATCGCTTGAACCTC

GAGCACACGTCTGAACTCTTTCCCTACACGACGCTCTTCCGATCTNNNCAGGAAAATCGCTTGAACCTC

GAGCACACGTCTGAACTCTTTCCCTACACGACGCTCTTCCGATCTNNNCGATGCTCTGTGGATTAGGG

GAGCACACGTCTGAACTCTTTCCCTACACGACGCTCTTCCGATCTNNNCGATGCTCTGTGGATTAGGG

CACACGTCTGAACTCTTTCCCTACACGACGCTCTTCCGATCTNNNACTGTTGGTGTTGTCTATGAGCAC

CACACGTCTGAACTCTTTCCCTACACGACGCTCTTCCGATCTNNNACTGTTGGTGTTGTCTATGAGCAC

GCACACGTCTGAACTCTTTCCCTACACGACGCTCTTCCGATCTNNNTCCCAAAGTGCTGAGATTACAGG

GCACACGTCTGAACTCTTTCCCTACACGACGCTCTTCCGATCTNNNTCCCAAAGTGCTGAGATTACAGG

AGCACACGTCTGAACTCTTTCCCTACACGACGCTCTTCCGATCTNNNACAGAAGAAATAACTGGCTGGC

AGCACACGTCTGAACTCTTTCCCTACACGACGCTCTTCCGATCTNNNACAGAAGAAATAACTGGCTGGC

AGCACACGTCTGAACTCTTTCCCTACACGACGCTCTTCCGATCTNNNCAACATTGGTGGACAGTTACAC

AGCACACGTCTGAACTCTTTCCCTACACGACGCTCTTCCGATCTNNNCAACATTGGTGGACAGTTACAC

GAGCACACGTCTGAACTCTTTCCCTACACGACGCTCTTCCGATCTNNNATTGCTTGAGCCTAGGAGTTCA

GAGCACACGTCTGAACTCTTTCCCTACACGACGCTCTTCCGATCTNNNATTGCTTGAGCCTAGGAGTTCA

AGCACACGTCTGAACTCTTTCCCTACACGACGCTCTTCCGATCTNNNATTCTCCTGCCTCAGTATCCCAAC

AGCACACGTCTGAACTCTTTCCCTACACGACGCTCTTCCGATCTNNNATTCTCCTGCCTCAGTATCCCAAC

AGCACACGTCTGAACTCTTTCCCTACACGACGCTCTTCCGATCTNNNCAATCCTAGACATTTCCCCCCAA

AGCACACGTCTGAACTCTTTCCCTACACGACGCTCTTCCGATCTNNNCAATCCTAGACATTTCCCCCCAA

GAGCACACGTCTGAACTCTTTCCCTACACGACGCTCTTCCGATCTNNNTCTGTAACCCTCTAACCATTGCC

GAGCACACGTCTGAACTCTTTCCCTACACGACGCTCTTCCGATCTNNNTCTGTAACCCTCTAACCATTGCC

AGCACACGTCTGAACTCTTTCCCTACACGACGCTCTTCCGATCTNNNAGTGAGCAAAGATCATGCCATT

AGCACACGTCTGAACTCTTTCCCTACACGACGCTCTTCCGATCTNNNAGTGAGCAAAGATCATGCCATT

CACACGTCTGAACTCTTTCCCTACACGACGCTCTTCCGATCTNNNAGCGGAATTTTCTCTCCAGACACCA

CACACGTCTGAACTCTTTCCCTACACGACGCTCTTCCGATCTNNNAGCGGAATTTTCTCTCCAGACACCA
AGCACACGTCTGAACTCTTTCCCTACACGACGCTCTTCCGATCTNNNTGTGAGGTAGGTAGAGTTTGG
AGCACACGTCTGAACTCTTTCCCTACACGACGCTCTTCCGATCTNNNTGTGAGGTAGGTAGAGTTTGG
GAGCACACGTCTGAACTCTTTCCCTACACGACGCTCTTCCGATCTNNNTTGGAAAAGCACCTTCTTTCAC
GAGCACACGTCTGAACTCTTTCCCTACACGACGCTCTTCCGATCTNNNTTGGAAAAGCACCTTCTTTCAC
AGCACACGTCTGAACTCTTTCCCTACACGACGCTCTTCCGATCTNNNAGTGGTGGCAATTACAGGACTA
AGCACACGTCTGAACTCTTTCCCTACACGACGCTCTTCCGATCTNNNAGTGGTGGCAATTACAGGACTA
AGCACACGTCTGAACTCTTTCCCTACACGACGCTCTTCCGATCTNNNCTGGCTTACACTTTGCAGTCTTC
AGCACACGTCTGAACTCTTTCCCTACACGACGCTCTTCCGATCTNNNCTGGCTTACACTTTGCAGTCTTC
GAGCACACGTCTGAACTCTTTCCCTACACGACGCTCTTCCGATCTNNNTACCCAGGAAGTAAGAGGAG
GAGCACACGTCTGAACTCTTTCCCTACACGACGCTCTTCCGATCTNNNTACCCAGGAAGTAAGAGGAG
AGAGCACACGTCTGAACTCTTTCCCTACACGACGCTCTTCCGATCTNNNCGTTCTCCATCAAGGATATAC
AGAGCACACGTCTGAACTCTTTCCCTACACGACGCTCTTCCGATCTNNNCGTTCTCCATCAAGGATATAC
GAGCACACGTCTGAACTCTTTCCCTACACGACGCTCTTCCGATCTNNNGAGGTTGCAGTGAGCTGAGAT
GAGCACACGTCTGAACTCTTTCCCTACACGACGCTCTTCCGATCTNNNGAGGTTGCAGTGAGCTGAGAT
AGAGCACACGTCTGAACTCTTTCCCTACACGACGCTCTTCCGATCTNNNCCTCTCCTATTTCCGATGGTT
AGAGCACACGTCTGAACTCTTTCCCTACACGACGCTCTTCCGATCTNNNCCTCTCCTATTTCCGATGGTT
AGCACACGTCTGAACTCTTTCCCTACACGACGCTCTTCCGATCTNNNATACTCATCACAAGCCTTGGAG
AGCACACGTCTGAACTCTTTCCCTACACGACGCTCTTCCGATCTNNNATACTCATCACAAGCCTTGGAG
AGCACACGTCTGAACTCTTTCCCTACACGACGCTCTTCCGATCTNNNTTTCCTCATTTTTGCCCAACTGCC
AGCACACGTCTGAACTCTTTCCCTACACGACGCTCTTCCGATCTNNNTTTCCTCATTTTTGCCCAACTGCC
GAGCACACGTCTGAACTCTTTCCCTACACGACGCTCTTCCGATCTNNNTAGTCCCAGCTACTCAGGAGT
GAGCACACGTCTGAACTCTTTCCCTACACGACGCTCTTCCGATCTNNNTAGTCCCAGCTACTCAGGAGT
AGCACACGTCTGAACTCTTTCCCTACACGACGCTCTTCCGATCTNNNATTGCAGTGAGTGAGCTGAGA
AGCACACGTCTGAACTCTTTCCCTACACGACGCTCTTCCGATCTNNNATTGCAGTGAGTGAGCTGAGA
AGCACACGTCTGAACTCTTTCCCTACACGACGCTCTTCCGATCTNNNTGGCTTCTGTCACTTAGCATCATC
AGCACACGTCTGAACTCTTTCCCTACACGACGCTCTTCCGATCTNNNTGGCTTCTGTCACTTAGCATCATC
GCACACGTCTGAACTCTTTCCCTACACGACGCTCTTCCGATCTNNNTCCTGAGTAGCTGGGACTATAGC
GCACACGTCTGAACTCTTTCCCTACACGACGCTCTTCCGATCTNNNTCCTGAGTAGCTGGGACTATAGC
AGCACACGTCTGAACTCTTTCCCTACACGACGCTCTTCCGATCTNNNCTGTCATGCTCCAGTCACAGTTT
AGCACACGTCTGAACTCTTTCCCTACACGACGCTCTTCCGATCTNNNCTGTCATGCTCCAGTCACAGTTT
GAGCACACGTCTGAACTCTTTCCCTACACGACGCTCTTCCGATCTNNNGGAGTTTGCAATGAGCCAAG
GAGCACACGTCTGAACTCTTTCCCTACACGACGCTCTTCCGATCTNNNGGAGTTTGCAATGAGCCAAG
GAGCACACGTCTGAACTCTTTCCCTACACGACGCTCTTCCGATCTNNNCCTGTAGCAAGGCAAATTTAG
GAGCACACGTCTGAACTCTTTCCCTACACGACGCTCTTCCGATCTNNNCCTGTAGCAAGGCAAATTTAG
GAGCACACGTCTGAACTCTTTCCCTACACGACGCTCTTCCGATCTNNNGCCTGTAGTCCCAGCTACTTGC
GAGCACACGTCTGAACTCTTTCCCTACACGACGCTCTTCCGATCTNNNGCCTGTAGTCCCAGCTACTTGC
AGCACACGTCTGAACTCTTTCCCTACACGACGCTCTTCCGATCTNNNGAGGCAGGAGAATTGCTTGAA
AGCACACGTCTGAACTCTTTCCCTACACGACGCTCTTCCGATCTNNNGAGGCAGGAGAATTGCTTGAA
AGCACACGTCTGAACTCTTTCCCTACACGACGCTCTTCCGATCTNNNTGACTTTCTCCTCACCTCCATCAC
AGCACACGTCTGAACTCTTTCCCTACACGACGCTCTTCCGATCTNNNTGACTTTCTCCTCACCTCCATCAC
AGCACACGTCTGAACTCTTTCCCTACACGACGCTCTTCCGATCTNNNACCTATAGGATCCTCTCCCTCCTC
AGCACACGTCTGAACTCTTTCCCTACACGACGCTCTTCCGATCTNNNACCTATAGGATCCTCTCCCTCCTC

AGCACACGTCTGAACTCTTTCCCTACACGACGCTCTTCCGATCTNNNCTGAGGCAGGAGAATCTCTTG
AGCACACGTCTGAACTCTTTCCCTACACGACGCTCTTCCGATCTNNNCTGAGGCAGGAGAATCTCTTG
GAGCACACGTCTGAACTCTTTCCCTACACGACGCTCTTCCGATCTNNNACATATATGGGTATGGGAGT
GAGCACACGTCTGAACTCTTTCCCTACACGACGCTCTTCCGATCTNNNACATATATGGGTATGGGAGT
AGAGCACACGTCTGAACTCTTTCCCTACACGACGCTCTTCCGATCTNNNTGACGTCCAAATTAGTGCCT
AGAGCACACGTCTGAACTCTTTCCCTACACGACGCTCTTCCGATCTNNNTGACGTCCAAATTAGTGCCT
AGCACACGTCTGAACTCTTTCCCTACACGACGCTCTTCCGATCTNNNTTTCCTCTTCAAAGCATCAGGG
AGCACACGTCTGAACTCTTTCCCTACACGACGCTCTTCCGATCTNNNTTTCCTCTTCAAAGCATCAGGG
GAGCACACGTCTGAACTCTTTCCCTACACGACGCTCTTCCGATCTNNNAGAGGATTGTCCCATGATAAA
GAGCACACGTCTGAACTCTTTCCCTACACGACGCTCTTCCGATCTNNNAGAGGATTGTCCCATGATAAA
GAGCACACGTCTGAACTCTTTCCCTACACGACGCTCTTCCGATCTNNNGAAATAGGTCTCGCTGAAGG
GAGCACACGTCTGAACTCTTTCCCTACACGACGCTCTTCCGATCTNNNGAAATAGGTCTCGCTGAAGG
AGCACACGTCTGAACTCTTTCCCTACACGACGCTCTTCCGATCTNNNACGTACTTATCAGCTCACGTTC
AGCACACGTCTGAACTCTTTCCCTACACGACGCTCTTCCGATCTNNNACGTACTTATCAGCTCACGTTC
AGCACACGTCTGAACTCTTTCCCTACACGACGCTCTTCCGATCTNNNCAAAAATTAGCTGGGTGTGGT
AGCACACGTCTGAACTCTTTCCCTACACGACGCTCTTCCGATCTNNNCAAAAATTAGCTGGGTGTGGT
GAGCACACGTCTGAACTCTTTCCCTACACGACGCTCTTCCGATCTNNNTTTGTATTTCTCCCACCCAACC
GAGCACACGTCTGAACTCTTTCCCTACACGACGCTCTTCCGATCTNNNTTTGTATTTCTCCCACCCAACC
GAGCACACGTCTGAACTCTTTCCCTACACGACGCTCTTCCGATCTNNNGAAATGGTAGCCAATTCTGG
GAGCACACGTCTGAACTCTTTCCCTACACGACGCTCTTCCGATCTNNNGAAATGGTAGCCAATTCTGG
AGCACACGTCTGAACTCTTTCCCTACACGACGCTCTTCCGATCTNNNCATGCACCCTTTAGTCCCAGATA
AGCACACGTCTGAACTCTTTCCCTACACGACGCTCTTCCGATCTNNNCATGCACCCTTTAGTCCCAGATA
AGCACACGTCTGAACTCTTTCCCTACACGACGCTCTTCCGATCTNNNGCCACCACAGTCAGCTAATTTTT
AGCACACGTCTGAACTCTTTCCCTACACGACGCTCTTCCGATCTNNNGCCACCACAGTCAGCTAATTTTT
GAGCACACGTCTGAACTCTTTCCCTACACGACGCTCTTCCGATCTNNNGTAGGAGACTGAGTCTGGAG
GAGCACACGTCTGAACTCTTTCCCTACACGACGCTCTTCCGATCTNNNGTAGGAGACTGAGTCTGGAG
GAGCACACGTCTGAACTCTTTCCCTACACGACGCTCTTCCGATCTNNNAAGCAATGTACTCGAAGGAG
GAGCACACGTCTGAACTCTTTCCCTACACGACGCTCTTCCGATCTNNNAAGCAATGTACTCGAAGGAG
AGCACACGTCTGAACTCTTTCCCTACACGACGCTCTTCCGATCTNNNCTGGGTGACAAGAGCAAAACT
AGCACACGTCTGAACTCTTTCCCTACACGACGCTCTTCCGATCTNNNCTGGGTGACAAGAGCAAAACT
GCACACGTCTGAACTCTTTCCCTACACGACGCTCTTCCGATCTNNNGTGAAATTAACTGTTAGGAGAA
GCACACGTCTGAACTCTTTCCCTACACGACGCTCTTCCGATCTNNNGTGAAATTAACTGTTAGGAGAA
GAGCACACGTCTGAACTCTTTCCCTACACGACGCTCTTCCGATCTNNNCTGCAGTGAGCCAAAACCAT
GAGCACACGTCTGAACTCTTTCCCTACACGACGCTCTTCCGATCTNNNCTGCAGTGAGCCAAAACCAT
GAGCACACGTCTGAACTCTTTCCCTACACGACGCTCTTCCGATCTNNNGGTCAAGGAAGTGGTATGG
GAGCACACGTCTGAACTCTTTCCCTACACGACGCTCTTCCGATCTNNNGGTCAAGGAAGTGGTATGG
GAGCACACGTCTGAACTCTTTCCCTACACGACGCTCTTCCGATCTNNNCATCCATGATACGTGTATGTG
GAGCACACGTCTGAACTCTTTCCCTACACGACGCTCTTCCGATCTNNNCATCCATGATACGTGTATGTG
AGCACACGTCTGAACTCTTTCCCTACACGACGCTCTTCCGATCTNNNGGGTGGACACAATCTTCTCGAG
AGCACACGTCTGAACTCTTTCCCTACACGACGCTCTTCCGATCTNNNGGGTGGACACAATCTTCTCGAG
AGCACACGTCTGAACTCTTTCCCTACACGACGCTCTTCCGATCTNNNCTTGTTGCTGCATCCTCTGGAG
AGCACACGTCTGAACTCTTTCCCTACACGACGCTCTTCCGATCTNNNCTTGTTGCTGCATCCTCTGGAG
GAGCACACGTCTGAACTCTTTCCCTACACGACGCTCTTCCGATCTNNNGCATGGACCTTATTGGCTCTT

GAGCACACGTCTGAACTCTTTCCCTACACGACGCTCTTCCGATCTNNNGCATGGACCTTATTGGCTCTT
AGCACACGTCTGAACTCTTTCCCTACACGACGCTCTTCCGATCTNNNCGGCCCCCAAAGATTCTTATTTC
AGCACACGTCTGAACTCTTTCCCTACACGACGCTCTTCCGATCTNNNCGGCCCCCAAAGATTCTTATTTC
GAGCACACGTCTGAACTCTTTCCCTACACGACGCTCTTCCGATCTNNNCTTTCGATGCAAAAACGAAGCA
GAGCACACGTCTGAACTCTTTCCCTACACGACGCTCTTCCGATCTNNNCTTTCGATGCAAAAACGAAGCA
GAGCACACGTCTGAACTCTTTCCCTACACGACGCTCTTCCGATCTNNNAAATGAGATGAAACCGTGGA
GAGCACACGTCTGAACTCTTTCCCTACACGACGCTCTTCCGATCTNNNAAATGAGATGAAACCGTGGA
AGAGCACACGTCTGAACTCTTTCCCTACACGACGCTCTTCCGATCTNNNAGAATCACTTGAACCAAGGA
AGAGCACACGTCTGAACTCTTTCCCTACACGACGCTCTTCCGATCTNNNAGAATCACTTGAACCAAGGA
AGCACACGTCTGAACTCTTTCCCTACACGACGCTCTTCCGATCTNNNAATTAGCCAGAGATGATGGTGC
AGCACACGTCTGAACTCTTTCCCTACACGACGCTCTTCCGATCTNNNAATTAGCCAGAGATGATGGTGC
AGCACACGTCTGAACTCTTTCCCTACACGACGCTCTTCCGATCTNNNTTCCCCTCTGAATGATCTCTCTC
AGCACACGTCTGAACTCTTTCCCTACACGACGCTCTTCCGATCTNNNTTCCCCTCTGAATGATCTCTCTC
AGCACACGTCTGAACTCTTTCCCTACACGACGCTCTTCCGATCTNNNATCAGTTGAAAGACCCTTGGGT
AGCACACGTCTGAACTCTTTCCCTACACGACGCTCTTCCGATCTNNNATCAGTTGAAAGACCCTTGGGT
ACACGTCTGAACTCTTTCCCTACACGACGCTCTTCCGATCTNNNTCCCAGCCTAATGTACCACATTTCATC
ACACGTCTGAACTCTTTCCCTACACGACGCTCTTCCGATCTNNNTCCCAGCCTAATGTACCACATTTCATC
AGCACACGTCTGAACTCTTTCCCTACACGACGCTCTTCCGATCTNNNGGCTTGCTTCAGTTCCTTCAAAT
AGCACACGTCTGAACTCTTTCCCTACACGACGCTCTTCCGATCTNNNGGCTTGCTTCAGTTCCTTCAAAT
GAGCACACGTCTGAACTCTTTCCCTACACGACGCTCTTCCGATCTNNNTGCTTGGTACCAGCTTGAACTA
GAGCACACGTCTGAACTCTTTCCCTACACGACGCTCTTCCGATCTNNNTGCTTGGTACCAGCTTGAACTA
AGCACACGTCTGAACTCTTTCCCTACACGACGCTCTTCCGATCTNNNAAATTTTAGCCTGCTGGAGAGC
AGCACACGTCTGAACTCTTTCCCTACACGACGCTCTTCCGATCTNNNAAATTTTAGCCTGCTGGAGAGC
GAGCACACGTCTGAACTCTTTCCCTACACGACGCTCTTCCGATCTNNNACACATGTAATCCCAGCTACTC
GAGCACACGTCTGAACTCTTTCCCTACACGACGCTCTTCCGATCTNNNACACATGTAATCCCAGCTACTC
AGCACACGTCTGAACTCTTTCCCTACACGACGCTCTTCCGATCTNNNTCTCCACACTTCAACAATTGTGC
AGCACACGTCTGAACTCTTTCCCTACACGACGCTCTTCCGATCTNNNTCTCCACACTTCAACAATTGTGC
AGCACACGTCTGAACTCTTTCCCTACACGACGCTCTTCCGATCTNNNCACCAGGTTGGAGATCCTCTTCC
AGCACACGTCTGAACTCTTTCCCTACACGACGCTCTTCCGATCTNNNCACCAGGTTGGAGATCCTCTTCC
GAGCACACGTCTGAACTCTTTCCCTACACGACGCTCTTCCGATCTNNNGGTATTTGTCCTAGTGCTCTCC
GAGCACACGTCTGAACTCTTTCCCTACACGACGCTCTTCCGATCTNNNGGTATTTGTCCTAGTGCTCTCC
AGCACACGTCTGAACTCTTTCCCTACACGACGCTCTTCCGATCTNNNAAAGAAAACAAGGACATTTGC
AGCACACGTCTGAACTCTTTCCCTACACGACGCTCTTCCGATCTNNNAAAGAAAACAAGGACATTTGC
GAGCACACGTCTGAACTCTTTCCCTACACGACGCTCTTCCGATCTNNNAGTAGAATGATGGTTGCCAG
GAGCACACGTCTGAACTCTTTCCCTACACGACGCTCTTCCGATCTNNNAGTAGAATGATGGTTGCCAG
AGCACACGTCTGAACTCTTTCCCTACACGACGCTCTTCCGATCTNNNTTTGAAGGCTTTGTGTTGACCT
AGCACACGTCTGAACTCTTTCCCTACACGACGCTCTTCCGATCTNNNTTTGAAGGCTTTGTGTTGACCT
GCACACGTCTGAACTCTTTCCCTACACGACGCTCTTCCGATCTNNNCCACGAAAACCCTAGTGAATCACC
GCACACGTCTGAACTCTTTCCCTACACGACGCTCTTCCGATCTNNNCCACGAAAACCCTAGTGAATCACC
AAGAGCACACGTCTGAACTCTTTCCCTACACGACGCTCTTCCGATCTNNNTGCATTTTGTAAGCACCTTC
AAGAGCACACGTCTGAACTCTTTCCCTACACGACGCTCTTCCGATCTNNNTGCATTTTGTAAGCACCTTC
AGCACACGTCTGAACTCTTTCCCTACACGACGCTCTTCCGATCTNNNGCTTCATGCAACTGCGTAAAAT
AGCACACGTCTGAACTCTTTCCCTACACGACGCTCTTCCGATCTNNNGCTTCATGCAACTGCGTAAAAT

AGAGCACACGTCTGAACTCTTTCCCTACACGACGCTCTTCCGATCTNNNCAAAAAATTAGCCAGGTTT
AGAGCACACGTCTGAACTCTTTCCCTACACGACGCTCTTCCGATCTNNNCAAAAAATTAGCCAGGTTT
GCACACGTCTGAACTCTTTCCCTACACGACGCTCTTCCGATCTNNNAGGATTTAGACGGGGATGAAG
GCACACGTCTGAACTCTTTCCCTACACGACGCTCTTCCGATCTNNNAGGATTTAGACGGGGATGAAG
AGCACACGTCTGAACTCTTTCCCTACACGACGCTCTTCCGATCTNNNCTCCAACGCAGACTGGTTATTT
AGCACACGTCTGAACTCTTTCCCTACACGACGCTCTTCCGATCTNNNCTCCAACGCAGACTGGTTATTT
AGAGCACACGTCTGAACTCTTTCCCTACACGACGCTCTTCCGATCTNNNTGATGTACTGGACTGTTGT
AGAGCACACGTCTGAACTCTTTCCCTACACGACGCTCTTCCGATCTNNNTGATGTACTGGACTGTTGT
GAGCACACGTCTGAACTCTTTCCCTACACGACGCTCTTCCGATCTNNNGAAAAGAGGTCCTAACTGGG
GAGCACACGTCTGAACTCTTTCCCTACACGACGCTCTTCCGATCTNNNGAAAAGAGGTCCTAACTGGG
GAGCACACGTCTGAACTCTTTCCCTACACGACGCTCTTCCGATCTNNNGTAATCCCTGCTACTCAAGA
GAGCACACGTCTGAACTCTTTCCCTACACGACGCTCTTCCGATCTNNNGTAATCCCTGCTACTCAAGA
GCACACGTCTGAACTCTTTCCCTACACGACGCTCTTCCGATCTNNNCTCCTGAGTAGCTGGAACTACA
GCACACGTCTGAACTCTTTCCCTACACGACGCTCTTCCGATCTNNNCTCCTGAGTAGCTGGAACTACA
GAGCACACGTCTGAACTCTTTCCCTACACGACGCTCTTCCGATCTNNNCTGAGGCAGGAGAATTGCTT
GAGCACACGTCTGAACTCTTTCCCTACACGACGCTCTTCCGATCTNNNCTGAGGCAGGAGAATTGCTT
GCACACGTCTGAACTCTTTCCCTACACGACGCTCTTCCGATCTNNNTTTCAAGACATTTGTGTAGTTGC
GCACACGTCTGAACTCTTTCCCTACACGACGCTCTTCCGATCTNNNTTTCAAGACATTTGTGTAGTTGC
AGCACACGTCTGAACTCTTTCCCTACACGACGCTCTTCCGATCTNNNCTCATCTCTTCTCTTTTCCTCCAT
AGCACACGTCTGAACTCTTTCCCTACACGACGCTCTTCCGATCTNNNCTCATCTCTTCTCTTTTCCTCCAT
AGCACACGTCTGAACTCTTTCCCTACACGACGCTCTTCCGATCTNNNATCCTGGAGATGTCTTTTGCC
AGCACACGTCTGAACTCTTTCCCTACACGACGCTCTTCCGATCTNNNATCCTGGAGATGTCTTTTGCC
CACACGTCTGAACTCTTTCCCTACACGACGCTCTTCCGATCTNNNTATGCACCTCCTGTAATCTGAGCC
CACACGTCTGAACTCTTTCCCTACACGACGCTCTTCCGATCTNNNTATGCACCTCCTGTAATCTGAGCC
CACACGTCTGAACTCTTTCCCTACACGACGCTCTTCCGATCTNNNATCACTCTGTTCTGTGCCTGATTC
CACACGTCTGAACTCTTTCCCTACACGACGCTCTTCCGATCTNNNATCACTCTGTTCTGTGCCTGATTC
AGCACACGTCTGAACTCTTTCCCTACACGACGCTCTTCCGATCTNNNTACATGGCTTGGTAGGGATG
AGCACACGTCTGAACTCTTTCCCTACACGACGCTCTTCCGATCTNNNTACATGGCTTGGTAGGGATG
GAGCACACGTCTGAACTCTTTCCCTACACGACGCTCTTCCGATCTNNNCTGTAATCCCAGCAACTTGG
GAGCACACGTCTGAACTCTTTCCCTACACGACGCTCTTCCGATCTNNNCTGTAATCCCAGCAACTTGG
AGCACACGTCTGAACTCTTTCCCTACACGACGCTCTTCCGATCTNNNGCAATGACAGCAAAACTACGT
AGCACACGTCTGAACTCTTTCCCTACACGACGCTCTTCCGATCTNNNGCAATGACAGCAAAACTACGT
GCACACGTCTGAACTCTTTCCCTACACGACGCTCTTCCGATCTNNNTCTGTGTGCCTCTGCCTCTCCAT
GCACACGTCTGAACTCTTTCCCTACACGACGCTCTTCCGATCTNNNTCTGTGTGCCTCTGCCTCTCCAT
GAGCACACGTCTGAACTCTTTCCCTACACGACGCTCTTCCGATCTNNNGACTGGTAAAAGGCTTTTCC
GAGCACACGTCTGAACTCTTTCCCTACACGACGCTCTTCCGATCTNNNGACTGGTAAAAGGCTTTTCC
GCACACGTCTGAACTCTTTCCCTACACGACGCTCTTCCGATCTNNNTTCCAGCTATTTGGGAGACTGA
GCACACGTCTGAACTCTTTCCCTACACGACGCTCTTCCGATCTNNNTTCCAGCTATTTGGGAGACTGA
GCACACGTCTGAACTCTTTCCCTACACGACGCTCTTCCGATCTNNNGCTTTCACTATTGCCTCCTCTCT
GCACACGTCTGAACTCTTTCCCTACACGACGCTCTTCCGATCTNNNGCTTTCACTATTGCCTCCTCTCT
AGCACACGTCTGAACTCTTTCCCTACACGACGCTCTTCCGATCTNNNGTAGTGAGTTGTGAGTCCTCC
AGCACACGTCTGAACTCTTTCCCTACACGACGCTCTTCCGATCTNNNGTAGTGAGTTGTGAGTCCTCC
AGCACACGTCTGAACTCTTTCCCTACACGACGCTCTTCCGATCTNNNAGGCATTGAGGAAACAGATG

AGCACACGTCTGAACTCTTTCCCTACACGACGCTCTTCCGATCTNNNAGGCATTGAGGAAACAGATGT

AGCACACGTCTGAACTCTTTCCCTACACGACGCTCTTCCGATCTNNNACGTACCAGTAACCATACCCAAC

AGCACACGTCTGAACTCTTTCCCTACACGACGCTCTTCCGATCTNNNACGTACCAGTAACCATACCCAAC

AGCACACGTCTGAACTCTTTCCCTACACGACGCTCTTCCGATCTNNNCACCACCATACTCCATAGGACAC

AGCACACGTCTGAACTCTTTCCCTACACGACGCTCTTCCGATCTNNNCACCACCATACTCCATAGGACAC

AGCACACGTCTGAACTCTTTCCCTACACGACGCTCTTCCGATCTNNNAGGCAAGTCAGTCACTTCAGTT

AGCACACGTCTGAACTCTTTCCCTACACGACGCTCTTCCGATCTNNNAGGCAAGTCAGTCACTTCAGTT

AGAGCACACGTCTGAACTCTTTCCCTACACGACGCTCTTCCGATCTNNNTGTGACCTTCTGTTGTGAAC

AGAGCACACGTCTGAACTCTTTCCCTACACGACGCTCTTCCGATCTNNNTGTGACCTTCTGTTGTGAAC

GAGCACACGTCTGAACTCTTTCCCTACACGACGCTCTTCCGATCTNNNGAAAGAGAAAGAGGTTCCCC

GAGCACACGTCTGAACTCTTTCCCTACACGACGCTCTTCCGATCTNNNGAAAGAGAAAGAGGTTCCCC

CACACGTCTGAACTCTTTCCCTACACGACGCTCTTCCGATCTNNNCCAGTATGACAAGGTATGATAAGC

CACACGTCTGAACTCTTTCCCTACACGACGCTCTTCCGATCTNNNCCAGTATGACAAGGTATGATAAGC

GAGCACACGTCTGAACTCTTTCCCTACACGACGCTCTTCCGATCTNNNAAGCACACATAAGGAGGATG

GAGCACACGTCTGAACTCTTTCCCTACACGACGCTCTTCCGATCTNNNAAGCACACATAAGGAGGATG

GCACACGTCTGAACTCTTTCCCTACACGACGCTCTTCCGATCTNNNACTGTTTTTGATTGTGCAGCTGA

GCACACGTCTGAACTCTTTCCCTACACGACGCTCTTCCGATCTNNNACTGTTTTTGATTGTGCAGCTGA

AGAGCACACGTCTGAACTCTTTCCCTACACGACGCTCTTCCGATCTNNNTGAGGCAGGAGTATCACTTC

AGAGCACACGTCTGAACTCTTTCCCTACACGACGCTCTTCCGATCTNNNTGAGGCAGGAGTATCACTTC

GCACACGTCTGAACTCTTTCCCTACACGACGCTCTTCCGATCTNNNGTCCAGATGAAGCTCCCAGAATG

GCACACGTCTGAACTCTTTCCCTACACGACGCTCTTCCGATCTNNNGTCCAGATGAAGCTCCCAGAATG

AGCACACGTCTGAACTCTTTCCCTACACGACGCTCTTCCGATCTNNNAGAGCAGTGTGTCCTCATCTTTT

AGCACACGTCTGAACTCTTTCCCTACACGACGCTCTTCCGATCTNNNAGAGCAGTGTGTCCTCATCTTTT

AGAGCACACGTCTGAACTCTTTCCCTACACGACGCTCTTCCGATCTNNNCTAAGGCACGAGAATTGCTT

AGAGCACACGTCTGAACTCTTTCCCTACACGACGCTCTTCCGATCTNNNCTAAGGCACGAGAATTGCTT

AGCACACGTCTGAACTCTTTCCCTACACGACGCTCTTCCGATCTNNNAAGAATCCTATGGAGAAGGCT

AGCACACGTCTGAACTCTTTCCCTACACGACGCTCTTCCGATCTNNNAAGAATCCTATGGAGAAGGCT

AGAGCACACGTCTGAACTCTTTCCCTACACGACGCTCTTCCGATCTNNNGTCATCAAAGCCGAATCCGA

AGAGCACACGTCTGAACTCTTTCCCTACACGACGCTCTTCCGATCTNNNGTCATCAAAGCCGAATCCGA

GAGCACACGTCTGAACTCTTTCCCTACACGACGCTCTTCCGATCTNNNACATGCATACACTTCTGTCTTCA

GAGCACACGTCTGAACTCTTTCCCTACACGACGCTCTTCCGATCTNNNACATGCATACACTTCTGTCTTCA

AGAGCACACGTCTGAACTCTTTCCCTACACGACGCTCTTCCGATCTNNNATAACATTAAGCCAATTTTCT

AGAGCACACGTCTGAACTCTTTCCCTACACGACGCTCTTCCGATCTNNNATAACATTAAGCCAATTTTCT

GAGCACACGTCTGAACTCTTTCCCTACACGACGCTCTTCCGATCTNNNGCCTATAATCCCAGCTACTCAG

GAGCACACGTCTGAACTCTTTCCCTACACGACGCTCTTCCGATCTNNNGCCTATAATCCCAGCTACTCAG

GAGCACACGTCTGAACTCTTTCCCTACACGACGCTCTTCCGATCTNNNTGAGGTTGCAGTGAACTATGG

GAGCACACGTCTGAACTCTTTCCCTACACGACGCTCTTCCGATCTNNNTGAGGTTGCAGTGAACTATGG

AGAGCACACGTCTGAACTCTTTCCCTACACGACGCTCTTCCGATCTNNNAGCACCATTGAGAGATCTGG

AGAGCACACGTCTGAACTCTTTCCCTACACGACGCTCTTCCGATCTNNNAGCACCATTGAGAGATCTGG

AGCACACGTCTGAACTCTTTCCCTACACGACGCTCTTCCGATCTNNNCTCATCTCCCCTGCTCTCTTTTTC

AGCACACGTCTGAACTCTTTCCCTACACGACGCTCTTCCGATCTNNNCTCATCTCCCCTGCTCTCTTTTTC

GAGCACACGTCTGAACTCTTTCCCTACACGACGCTCTTCCGATCTNNNTATTGTTCTGTGGGACGAGGG

GAGCACACGTCTGAACTCTTTCCCTACACGACGCTCTTCCGATCTNNNTATTGTTCTGTGGGACGAGGG

ACACGTCTGAACTCTTTCCCTACACGACGCTCTTCCGATCTNNNCAGGGTTAGGGTTGGAAGACCCAT
ACACGTCTGAACTCTTTCCCTACACGACGCTCTTCCGATCTNNNCAGGGTTAGGGTTGGAAGACCCAT
AGCACACGTCTGAACTCTTTCCCTACACGACGCTCTTCCGATCTNNNGGCAACAGAGTGAGAATCCA
AGCACACGTCTGAACTCTTTCCCTACACGACGCTCTTCCGATCTNNNGGCAACAGAGTGAGAATCCA
AGCACACGTCTGAACTCTTTCCCTACACGACGCTCTTCCGATCTNNNAGCCAGCATCATTCTGATACCA
AGCACACGTCTGAACTCTTTCCCTACACGACGCTCTTCCGATCTNNNAGCCAGCATCATTCTGATACCA
GCACACGTCTGAACTCTTTCCCTACACGACGCTCTTCCGATCTNNNCTGAGGCAGGAGAATAGCTTGA
GCACACGTCTGAACTCTTTCCCTACACGACGCTCTTCCGATCTNNNCTGAGGCAGGAGAATAGCTTGA
AGCACACGTCTGAACTCTTTCCCTACACGACGCTCTTCCGATCTNNNCCTCTCATCCAGGCTGAACTTT
AGCACACGTCTGAACTCTTTCCCTACACGACGCTCTTCCGATCTNNNCCTCTCATCCAGGCTGAACTTT
AGCACACGTCTGAACTCTTTCCCTACACGACGCTCTTCCGATCTNNNGAGCTTGCAGTGAGAGGAGA
AGCACACGTCTGAACTCTTTCCCTACACGACGCTCTTCCGATCTNNNGAGCTTGCAGTGAGAGGAGA
AGCACACGTCTGAACTCTTTCCCTACACGACGCTCTTCCGATCTNNNACAGGAGAATAGCTTGAACCA
AGCACACGTCTGAACTCTTTCCCTACACGACGCTCTTCCGATCTNNNACAGGAGAATAGCTTGAACCA
AGCACACGTCTGAACTCTTTCCCTACACGACGCTCTTCCGATCTNNNGGCAACAGAGTAAGACACTG
AGCACACGTCTGAACTCTTTCCCTACACGACGCTCTTCCGATCTNNNGGCAACAGAGTAAGACACTG
AGCACACGTCTGAACTCTTTCCCTACACGACGCTCTTCCGATCTNNNGGCTGAGGTAGGAGGATTTCT
AGCACACGTCTGAACTCTTTCCCTACACGACGCTCTTCCGATCTNNNGGCTGAGGTAGGAGGATTTCT
AGCACACGTCTGAACTCTTTCCCTACACGACGCTCTTCCGATCTNNNACTATAAGCTGTGGCATCTGG
AGCACACGTCTGAACTCTTTCCCTACACGACGCTCTTCCGATCTNNNACTATAAGCTGTGGCATCTGG
AGAGCACACGTCTGAACTCTTTCCCTACACGACGCTCTTCCGATCTNNNGTGAGGAGAGTTGTGGGG
AGAGCACACGTCTGAACTCTTTCCCTACACGACGCTCTTCCGATCTNNNGTGAGGAGAGTTGTGGGG
GCACACGTCTGAACTCTTTCCCTACACGACGCTCTTCCGATCTNNNCACATTGGTTGCTGTAGTCCAAG
GCACACGTCTGAACTCTTTCCCTACACGACGCTCTTCCGATCTNNNCACATTGGTTGCTGTAGTCCAAG
GCACACGTCTGAACTCTTTCCCTACACGACGCTCTTCCGATCTNNNTATGAACAATAACGTGAGATCC
GCACACGTCTGAACTCTTTCCCTACACGACGCTCTTCCGATCTNNNTATGAACAATAACGTGAGATCC
AGCACACGTCTGAACTCTTTCCCTACACGACGCTCTTCCGATCTNNNCTCACAGTCTCTGCTTCAGTCAT
AGCACACGTCTGAACTCTTTCCCTACACGACGCTCTTCCGATCTNNNCTCACAGTCTCTGCTTCAGTCAT
AGCACACGTCTGAACTCTTTCCCTACACGACGCTCTTCCGATCTNNNTGGCAGCTTCCTATTGCAATTT
AGCACACGTCTGAACTCTTTCCCTACACGACGCTCTTCCGATCTNNNTGGCAGCTTCCTATTGCAATTT
CACACGTCTGAACTCTTTCCCTACACGACGCTCTTCCGATCTNNNGAAAGCAGCCAATCAACAGCTCAT
CACACGTCTGAACTCTTTCCCTACACGACGCTCTTCCGATCTNNNGAAAGCAGCCAATCAACAGCTCAT
AGCACACGTCTGAACTCTTTCCCTACACGACGCTCTTCCGATCTNNNACACACATACACATTTTTTTCTG
AGCACACGTCTGAACTCTTTCCCTACACGACGCTCTTCCGATCTNNNACACACATACACATTTTTTTCTG
AGCACACGTCTGAACTCTTTCCCTACACGACGCTCTTCCGATCTNNNGAACAGAATTACAGGAAGTTT
AGCACACGTCTGAACTCTTTCCCTACACGACGCTCTTCCGATCTNNNGAACAGAATTACAGGAAGTTT
AGCACACGTCTGAACTCTTTCCCTACACGACGCTCTTCCGATCTNNNCTGGGCAACAAAGATTCCATC
AGCACACGTCTGAACTCTTTCCCTACACGACGCTCTTCCGATCTNNNCTGGGCAACAAAGATTCCATC
AGCACACGTCTGAACTCTTTCCCTACACGACGCTCTTCCGATCTNNNCCAAGAAAGGACTCACATCCA
AGCACACGTCTGAACTCTTTCCCTACACGACGCTCTTCCGATCTNNNCCAAGAAAGGACTCACATCCA
AGCACACGTCTGAACTCTTTCCCTACACGACGCTCTTCCGATCTNNNTCTTGTAGTCCCAGCTACTCTG
AGCACACGTCTGAACTCTTTCCCTACACGACGCTCTTCCGATCTNNNTCTTGTAGTCCCAGCTACTCTG
AGCACACGTCTGAACTCTTTCCCTACACGACGCTCTTCCGATCTNNNGAGGATTGCTTGAACCCAGG

AGCACACGTCTGAACTCTTTCCCTACACGACGCTCTTCCGATCTNNNGAGGATTGCTTGAACCCAGG

AGCACACGTCTGAACTCTTTCCCTACACGACGCTCTTCCGATCTNNNCAAAAAATTAGCCAGGCGTGG

AGCACACGTCTGAACTCTTTCCCTACACGACGCTCTTCCGATCTNNNCAAAAAATTAGCCAGGCGTGG

AAGAGCACACGTCTGAACTCTTTCCCTACACGACGCTCTTCCGATCTNNNATCCTTGTGCAGAAATGG

AAGAGCACACGTCTGAACTCTTTCCCTACACGACGCTCTTCCGATCTNNNATCCTTGTGCAGAAATGG

GAGCACACGTCTGAACTCTTTCCCTACACGACGCTCTTCCGATCTNNNGGCTGGTCTGAGAAATAAA

GAGCACACGTCTGAACTCTTTCCCTACACGACGCTCTTCCGATCTNNNGGCTGGTCTGAGAAATAAA

CACACGTCTGAACTCTTTCCCTACACGACGCTCTTCCGATCTNNNCCAGCCTCCAGAATTGTGAAAAA

CACACGTCTGAACTCTTTCCCTACACGACGCTCTTCCGATCTNNNCCAGCCTCCAGAATTGTGAAAAA

GCACACGTCTGAACTCTTTCCCTACACGACGCTCTTCCGATCTNNNAATTCTAGACATTGGTCTCGCC

GCACACGTCTGAACTCTTTCCCTACACGACGCTCTTCCGATCTNNNAATTCTAGACATTGGTCTCGCC

AGCACACGTCTGAACTCTTTCCCTACACGACGCTCTTCCGATCTNNNGCAGGGAGCCAATCAACATAT

AGCACACGTCTGAACTCTTTCCCTACACGACGCTCTTCCGATCTNNNGCAGGGAGCCAATCAACATAT

AGCACACGTCTGAACTCTTTCCCTACACGACGCTCTTCCGATCTNNNAATCACTTGTACCTGGGAAGT

AGCACACGTCTGAACTCTTTCCCTACACGACGCTCTTCCGATCTNNNAATCACTTGTACCTGGGAAGT

AGCACACGTCTGAACTCTTTCCCTACACGACGCTCTTCCGATCTNNNGTGAATCACTTGAGGTCAGGA

AGCACACGTCTGAACTCTTTCCCTACACGACGCTCTTCCGATCTNNNGTGAATCACTTGAGGTCAGGA

GAGCACACGTCTGAACTCTTTCCCTACACGACGCTCTTCCGATCTNNNGAGGCTGAGGCAGGAGAA

GAGCACACGTCTGAACTCTTTCCCTACACGACGCTCTTCCGATCTNNNGAGGCTGAGGCAGGAGAA

AGCACACGTCTGAACTCTTTCCCTACACGACGCTCTTCCGATCTNNNTTTTGTGTGTGTGTGTGTGTG

AGCACACGTCTGAACTCTTTCCCTACACGACGCTCTTCCGATCTNNNTTTTGTGTGTGTGTGTGTGTG

AGCACACGTCTGAACTCTTTCCCTACACGACGCTCTTCCGATCTNNNGTAAGCCGAGATCATGACATT

AGCACACGTCTGAACTCTTTCCCTACACGACGCTCTTCCGATCTNNNGTAAGCCGAGATCATGACATT

AGCACACGTCTGAACTCTTTCCCTACACGACGCTCTTCCGATCTNNNGACCAAATGAGATGCAAAGCC

AGCACACGTCTGAACTCTTTCCCTACACGACGCTCTTCCGATCTNNNGACCAAATGAGATGCAAAGCC

GCACACGTCTGAACTCTTTCCCTACACGACGCTCTTCCGATCTNNNCCTCCATCAGGGTTCTTTGTTTC

GCACACGTCTGAACTCTTTCCCTACACGACGCTCTTCCGATCTNNNCCTCCATCAGGGTTCTTTGTTTC

AGCACACGTCTGAACTCTTTCCCTACACGACGCTCTTCCGATCTNNNCCTCCTTCTTGCTTCTTTTCAGCC

AGCACACGTCTGAACTCTTTCCCTACACGACGCTCTTCCGATCTNNNCCTCCTTCTTGCTTCTTTTCAGCC

AGCACACGTCTGAACTCTTTCCCTACACGACGCTCTTCCGATCTNNNTACAGATGTAAGCCACGAGCCT

AGCACACGTCTGAACTCTTTCCCTACACGACGCTCTTCCGATCTNNNTACAGATGTAAGCCACGAGCCT

GAGCACACGTCTGAACTCTTTCCCTACACGACGCTCTTCCGATCTNNNACTCTTCTGCTACACCCTCCTA

GAGCACACGTCTGAACTCTTTCCCTACACGACGCTCTTCCGATCTNNNACTCTTCTGCTACACCCTCCTA

GAGCACACGTCTGAACTCTTTCCCTACACGACGCTCTTCCGATCTNNNTTTTAGGGTAAAAGTGGCC

GAGCACACGTCTGAACTCTTTCCCTACACGACGCTCTTCCGATCTNNNTTTTAGGGTAAAAGTGGCC

AGCACACGTCTGAACTCTTTCCCTACACGACGCTCTTCCGATCTNNNAAAAAATTAGACGGGTGTGGT

AGCACACGTCTGAACTCTTTCCCTACACGACGCTCTTCCGATCTNNNAAAAAATTAGACGGGTGTGGT

AGCACACGTCTGAACTCTTTCCCTACACGACGCTCTTCCGATCTNNNCCATCACGCCGAGCTAATTTTTT

AGCACACGTCTGAACTCTTTCCCTACACGACGCTCTTCCGATCTNNNCCATCACGCCGAGCTAATTTTTT

AGCACACGTCTGAACTCTTTCCCTACACGACGCTCTTCCGATCTNNNGGCTGTCTCATGGAGAAGAG

AGCACACGTCTGAACTCTTTCCCTACACGACGCTCTTCCGATCTNNNGGCTGTCTCATGGAGAAGAG

AGCACACGTCTGAACTCTTTCCCTACACGACGCTCTTCCGATCTNNNTTCATCATAGGCTTCAAAGGG

AGCACACGTCTGAACTCTTTCCCTACACGACGCTCTTCCGATCTNNNTTCATCATAGGCTTCAAAGGG

AGCACACGTCTGAACTCTTTCCCTACACGACGCTCTTCCGATCTNNNTCCAAGTCAAATGCTGCAATTC
AGCACACGTCTGAACTCTTTCCCTACACGACGCTCTTCCGATCTNNNTCCAAGTCAAATGCTGCAATTC
GAGCACACGTCTGAACTCTTTCCCTACACGACGCTCTTCCGATCTNNNAGGACTTTGACTTGCATCTTG
GAGCACACGTCTGAACTCTTTCCCTACACGACGCTCTTCCGATCTNNNAGGACTTTGACTTGCATCTTG
GAGCACACGTCTGAACTCTTTCCCTACACGACGCTCTTCCGATCTNNNGCTTGTGTTGCTTTTTGGACA
GAGCACACGTCTGAACTCTTTCCCTACACGACGCTCTTCCGATCTNNNGCTTGTGTTGCTTTTTGGACA
AGCACACGTCTGAACTCTTTCCCTACACGACGCTCTTCCGATCTNNNCAGGAAGGGAAAACCAATGG
AGCACACGTCTGAACTCTTTCCCTACACGACGCTCTTCCGATCTNNNCAGGAAGGGAAAACCAATGG
AGCACACGTCTGAACTCTTTCCCTACACGACGCTCTTCCGATCTNNNGAGACAGGATTTGAGAGCAGA
AGCACACGTCTGAACTCTTTCCCTACACGACGCTCTTCCGATCTNNNGAGACAGGATTTGAGAGCAGA
CACACGTCTGAACTCTTTCCCTACACGACGCTCTTCCGATCTNNNGATGTTAACACAAATGAAGTCCC
CACACGTCTGAACTCTTTCCCTACACGACGCTCTTCCGATCTNNNGATGTTAACACAAATGAAGTCCC
GAGCACACGTCTGAACTCTTTCCCTACACGACGCTCTTCCGATCTNNNAGACCAAGTCTAGGCTTGATT
GAGCACACGTCTGAACTCTTTCCCTACACGACGCTCTTCCGATCTNNNAGACCAAGTCTAGGCTTGATT
GAGCACACGTCTGAACTCTTTCCCTACACGACGCTCTTCCGATCTNNNAACAGGATTTCAAAGGCAGA
GAGCACACGTCTGAACTCTTTCCCTACACGACGCTCTTCCGATCTNNNAACAGGATTTCAAAGGCAGA
AGCACACGTCTGAACTCTTTCCCTACACGACGCTCTTCCGATCTNNNGAGGTTACAGCGAGCTATGAT
AGCACACGTCTGAACTCTTTCCCTACACGACGCTCTTCCGATCTNNNGAGGTTACAGCGAGCTATGAT
GAGCACACGTCTGAACTCTTTCCCTACACGACGCTCTTCCGATCTNNNCTCAGGTTTCACGCATGTGTG
GAGCACACGTCTGAACTCTTTCCCTACACGACGCTCTTCCGATCTNNNCTCAGGTTTCACGCATGTGTG
AGCACACGTCTGAACTCTTTCCCTACACGACGCTCTTCCGATCTNNNCTGACTCTGTCTAAGGTCCAAC
AGCACACGTCTGAACTCTTTCCCTACACGACGCTCTTCCGATCTNNNCTGACTCTGTCTAAGGTCCAAC
GAGCACACGTCTGAACTCTTTCCCTACACGACGCTCTTCCGATCTNNNCACCTATAATCCCAGCTACTCG
GAGCACACGTCTGAACTCTTTCCCTACACGACGCTCTTCCGATCTNNNCACCTATAATCCCAGCTACTCG
AGAGCACACGTCTGAACTCTTTCCCTACACGACGCTCTTCCGATCTNNNGACAGAGCCAGACTCCGTC
AGAGCACACGTCTGAACTCTTTCCCTACACGACGCTCTTCCGATCTNNNGACAGAGCCAGACTCCGTC
AGCACACGTCTGAACTCTTTCCCTACACGACGCTCTTCCGATCTNNNCTGTAATCCCAGCACTTTGGAA
AGCACACGTCTGAACTCTTTCCCTACACGACGCTCTTCCGATCTNNNCTGTAATCCCAGCACTTTGGAA
AGCACACGTCTGAACTCTTTCCCTACACGACGCTCTTCCGATCTNNNAGAAAGGTTTGCTTCTGATGCT
AGCACACGTCTGAACTCTTTCCCTACACGACGCTCTTCCGATCTNNNAGAAAGGTTTGCTTCTGATGCT
GAGCACACGTCTGAACTCTTTCCCTACACGACGCTCTTCCGATCTNNNCCAAAGTGACCACATAGATGG
GAGCACACGTCTGAACTCTTTCCCTACACGACGCTCTTCCGATCTNNNCCAAAGTGACCACATAGATGG
AGAGCACACGTCTGAACTCTTTCCCTACACGACGCTCTTCCGATCTNNNCTGTAGTCCCAGCTACTTGG
AGAGCACACGTCTGAACTCTTTCCCTACACGACGCTCTTCCGATCTNNNCTGTAGTCCCAGCTACTTGG
AGCACACGTCTGAACTCTTTCCCTACACGACGCTCTTCCGATCTNNNTTTTTCTTTCTCAGCCTTGCCAGC
AGCACACGTCTGAACTCTTTCCCTACACGACGCTCTTCCGATCTNNNTTTTTCTTTCTCAGCCTTGCCAGC
GAGCACACGTCTGAACTCTTTCCCTACACGACGCTCTTCCGATCTNNNGCACACAGAAAGCACTCAGT
GAGCACACGTCTGAACTCTTTCCCTACACGACGCTCTTCCGATCTNNNGCACACAGAAAGCACTCAGT
AGCACACGTCTGAACTCTTTCCCTACACGACGCTCTTCCGATCTNNNCGGACATTTGGAATCCCAGCTA
AGCACACGTCTGAACTCTTTCCCTACACGACGCTCTTCCGATCTNNNCGGACATTTGGAATCCCAGCTA
GCACACGTCTGAACTCTTTCCCTACACGACGCTCTTCCGATCTNNNAGGAAAGCAAGAGAAAAGTGT
GCACACGTCTGAACTCTTTCCCTACACGACGCTCTTCCGATCTNNNAGGAAAGCAAGAGAAAAGTGT
GAGCACACGTCTGAACTCTTTCCCTACACGACGCTCTTCCGATCTNNNGGGAGGCTGAGGTACAAGA

AGCACACGTCTGAACTCTTTCCCTACACGACGCTCTTCCGATCTNNNGGGAGGCTGAGGTACAAGA

GCACACGTCTGAACTCTTTCCCTACACGACGCTCTTCCGATCTNNNAGAAATGTCAGGCTCAAACCAG

GCACACGTCTGAACTCTTTCCCTACACGACGCTCTTCCGATCTNNNAGAAATGTCAGGCTCAAACCAG

GCACACGTCTGAACTCTTTCCCTACACGACGCTCTTCCGATCTNNNAGTCAGGAGAATCACTTGAACC

GCACACGTCTGAACTCTTTCCCTACACGACGCTCTTCCGATCTNNNAGTCAGGAGAATCACTTGAACC

GAGCACACGTCTGAACTCTTTCCCTACACGACGCTCTTCCGATCTNNNTCAGGAGGCTGAGGTAGGA

GAGCACACGTCTGAACTCTTTCCCTACACGACGCTCTTCCGATCTNNNTCAGGAGGCTGAGGTAGGA

GCACACGTCTGAACTCTTTCCCTACACGACGCTCTTCCGATCTNNNCTGGACAGCACTCCTTTATGTCT

GCACACGTCTGAACTCTTTCCCTACACGACGCTCTTCCGATCTNNNCTGGACAGCACTCCTTTATGTCT

GAGCACACGTCTGAACTCTTTCCCTACACGACGCTCTTCCGATCTNNNGAGCTTGCAGTGAGCCAAG

GAGCACACGTCTGAACTCTTTCCCTACACGACGCTCTTCCGATCTNNNGAGCTTGCAGTGAGCCAAG

AGCACACGTCTGAACTCTTTCCCTACACGACGCTCTTCCGATCTNNNAGATTGTGGTGAGCTGAGAT

AGCACACGTCTGAACTCTTTCCCTACACGACGCTCTTCCGATCTNNNAGATTGTGGTGAGCTGAGAT

ACACGTCTGAACTCTTTCCCTACACGACGCTCTTCCGATCTNNNGTGTTCCTAGCAAAACTGTGCATC

ACACGTCTGAACTCTTTCCCTACACGACGCTCTTCCGATCTNNNGTGTTCCTAGCAAAACTGTGCATC

GCACACGTCTGAACTCTTTCCCTACACGACGCTCTTCCGATCTNNNATGACTTAGCCAGGTGTTGAA

GCACACGTCTGAACTCTTTCCCTACACGACGCTCTTCCGATCTNNNATGACTTAGCCAGGTGTTGAA

AGCACACGTCTGAACTCTTTCCCTACACGACGCTCTTCCGATCTNNNAAGAAAGAGCATGTGCAAAG

AGCACACGTCTGAACTCTTTCCCTACACGACGCTCTTCCGATCTNNNAAGAAAGAGCATGTGCAAAG

AGCACACGTCTGAACTCTTTCCCTACACGACGCTCTTCCGATCTNNNAATTTGCAGTAAGCCAAGTTCC

AGCACACGTCTGAACTCTTTCCCTACACGACGCTCTTCCGATCTNNNAATTTGCAGTAAGCCAAGTTCC

GCACACGTCTGAACTCTTTCCCTACACGACGCTCTTCCGATCTNNNGAGGCAGGAGAATTGCTTGAA

GCACACGTCTGAACTCTTTCCCTACACGACGCTCTTCCGATCTNNNGAGGCAGGAGAATTGCTTGAA

GCACACGTCTGAACTCTTTCCCTACACGACGCTCTTCCGATCTNNNGTGACAATTGGCGAGAGAGAG

GCACACGTCTGAACTCTTTCCCTACACGACGCTCTTCCGATCTNNNGTGACAATTGGCGAGAGAGAG

AGCACACGTCTGAACTCTTTCCCTACACGACGCTCTTCCGATCTNNNACAGAAGGTCTCCCTGAAAGT

AGCACACGTCTGAACTCTTTCCCTACACGACGCTCTTCCGATCTNNNACAGAAGGTCTCCCTGAAAGT

AGCACACGTCTGAACTCTTTCCCTACACGACGCTCTTCCGATCTNNNTATGAGTACGTGTGTGTGTGT

AGCACACGTCTGAACTCTTTCCCTACACGACGCTCTTCCGATCTNNNTATGAGTACGTGTGTGTGTGT

AGCACACGTCTGAACTCTTTCCCTACACGACGCTCTTCCGATCTNNNAGAATGAGGGATGAGGGAT

AGCACACGTCTGAACTCTTTCCCTACACGACGCTCTTCCGATCTNNNAGAATGAGGGATGAGGGAT

AGCACACGTCTGAACTCTTTCCCTACACGACGCTCTTCCGATCTNNNGCCTGAGCAATGCAACAGATTT

AGCACACGTCTGAACTCTTTCCCTACACGACGCTCTTCCGATCTNNNGCCTGAGCAATGCAACAGATTT

GCACACGTCTGAACTCTTTCCCTACACGACGCTCTTCCGATCTNNNCCAGAGACAATCACCACACTAC

GCACACGTCTGAACTCTTTCCCTACACGACGCTCTTCCGATCTNNNCCAGAGACAATCACCACACTAC

GAGCACACGTCTGAACTCTTTCCCTACACGACGCTCTTCCGATCTNNNGTGAAACCCTGTCTCCACTAA

GAGCACACGTCTGAACTCTTTCCCTACACGACGCTCTTCCGATCTNNNGTGAAACCCTGTCTCCACTAA

AGCACACGTCTGAACTCTTTCCCTACACGACGCTCTTCCGATCTNNNAACATCCTGAGTAGCTGGGAC

AGCACACGTCTGAACTCTTTCCCTACACGACGCTCTTCCGATCTNNNAACATCCTGAGTAGCTGGGAC

AGCACACGTCTGAACTCTTTCCCTACACGACGCTCTTCCGATCTNNNACTGGTTGAAAGAGTTCACCG

AGCACACGTCTGAACTCTTTCCCTACACGACGCTCTTCCGATCTNNNACTGGTTGAAAGAGTTCACCG

GAGCACACGTCTGAACTCTTTCCCTACACGACGCTCTTCCGATCTNNNCTGGGTGTCATCCAGGTATT

GAGCACACGTCTGAACTCTTTCCCTACACGACGCTCTTCCGATCTNNNCTGGGTGTCATCCAGGTATT

GAGCACACGTCTGAACTCTTTCCCTACACGACGCTCTTCCGATCTNNNCAGAATCCAGAGCATAGCACT
GAGCACACGTCTGAACTCTTTCCCTACACGACGCTCTTCCGATCTNNNCAGAATCCAGAGCATAGCACT
GAGCACACGTCTGAACTCTTTCCCTACACGACGCTCTTCCGATCTNNNGGGCAACAAGAGTGAAATT
GAGCACACGTCTGAACTCTTTCCCTACACGACGCTCTTCCGATCTNNNGGGCAACAAGAGTGAAATT
GAGCACACGTCTGAACTCTTTCCCTACACGACGCTCTTCCGATCTNNNGAATCTCTTGAAACTGGAAG
GAGCACACGTCTGAACTCTTTCCCTACACGACGCTCTTCCGATCTNNNGAATCTCTTGAAACTGGAAG
GCACACGTCTGAACTCTTTCCCTACACGACGCTCTTCCGATCTNNNGAGAATCCCTTGAACCCAGGAG
GCACACGTCTGAACTCTTTCCCTACACGACGCTCTTCCGATCTNNNGAGAATCCCTTGAACCCAGGAG
AGCACACGTCTGAACTCTTTCCCTACACGACGCTCTTCCGATCTNNNGATCTACCCCATTGGCTCTTTG
AGCACACGTCTGAACTCTTTCCCTACACGACGCTCTTCCGATCTNNNGATCTACCCCATTGGCTCTTTG
AGCACACGTCTGAACTCTTTCCCTACACGACGCTCTTCCGATCTNNNCATTACAAGTCTTGCTGGTGT
AGCACACGTCTGAACTCTTTCCCTACACGACGCTCTTCCGATCTNNNCATTACAAGTCTTGCTGGTGT
AGCACACGTCTGAACTCTTTCCCTACACGACGCTCTTCCGATCTNNNTATCCTATCCCTTCCCCCAACTTC
AGCACACGTCTGAACTCTTTCCCTACACGACGCTCTTCCGATCTNNNTATCCTATCCCTTCCCCCAACTTC
GAGCACACGTCTGAACTCTTTCCCTACACGACGCTCTTCCGATCTNNNCATGAGTGTCAACCCCAGAAT
GAGCACACGTCTGAACTCTTTCCCTACACGACGCTCTTCCGATCTNNNCATGAGTGTCAACCCCAGAAT
AGCACACGTCTGAACTCTTTCCCTACACGACGCTCTTCCGATCTNNNCTGAGGCAGGAGAATTGGTTG
AGCACACGTCTGAACTCTTTCCCTACACGACGCTCTTCCGATCTNNNCTGAGGCAGGAGAATTGGTTG
AGCACACGTCTGAACTCTTTCCCTACACGACGCTCTTCCGATCTNNNGCATTGGCATCTTTTTGTTCGT
AGCACACGTCTGAACTCTTTCCCTACACGACGCTCTTCCGATCTNNNGCATTGGCATCTTTTTGTTCGT
AGCACACGTCTGAACTCTTTCCCTACACGACGCTCTTCCGATCTNNNGAATCCAGGCATTCAGGACCA
AGCACACGTCTGAACTCTTTCCCTACACGACGCTCTTCCGATCTNNNGAATCCAGGCATTCAGGACCA
AGCACACGTCTGAACTCTTTCCCTACACGACGCTCTTCCGATCTNNNCATGCCTCCTCTTCTTCCTTGAA
AGCACACGTCTGAACTCTTTCCCTACACGACGCTCTTCCGATCTNNNCATGCCTCCTCTTCTTCCTTGAA
GAGCACACGTCTGAACTCTTTCCCTACACGACGCTCTTCCGATCTNNNACATTTGTGTGTGTGTGTGT
GAGCACACGTCTGAACTCTTTCCCTACACGACGCTCTTCCGATCTNNNACATTTGTGTGTGTGTGTGT
AGCACACGTCTGAACTCTTTCCCTACACGACGCTCTTCCGATCTNNNATATCAGTGTTCTCCCTTGCAG
AGCACACGTCTGAACTCTTTCCCTACACGACGCTCTTCCGATCTNNNATATCAGTGTTCTCCCTTGCAG
AGCACACGTCTGAACTCTTTCCCTACACGACGCTCTTCCGATCTNNNGATAAAAGTTGCAAGGCTTCC
AGCACACGTCTGAACTCTTTCCCTACACGACGCTCTTCCGATCTNNNGATAAAAGTTGCAAGGCTTCC
GCACACGTCTGAACTCTTTCCCTACACGACGCTCTTCCGATCTNNNTGTTGTCAACAAAACATTCCTCAC
GCACACGTCTGAACTCTTTCCCTACACGACGCTCTTCCGATCTNNNTGTTGTCAACAAAACATTCCTCAC
GCACACGTCTGAACTCTTTCCCTACACGACGCTCTTCCGATCTNNNTCAACAATTGTCTGTTCATGGCC
GCACACGTCTGAACTCTTTCCCTACACGACGCTCTTCCGATCTNNNTCAACAATTGTCTGTTCATGGCC
AGCACACGTCTGAACTCTTTCCCTACACGACGCTCTTCCGATCTNNNATTCCCCAACTTCACCTTCACAC
AGCACACGTCTGAACTCTTTCCCTACACGACGCTCTTCCGATCTNNNATTCCCCAACTTCACCTTCACAC
AGCACACGTCTGAACTCTTTCCCTACACGACGCTCTTCCGATCTNNNTATACCAGTTCAGGAGGACAG
AGCACACGTCTGAACTCTTTCCCTACACGACGCTCTTCCGATCTNNNTATACCAGTTCAGGAGGACAG
AGCACACGTCTGAACTCTTTCCCTACACGACGCTCTTCCGATCTNNNCACCCAGAAGATGAGACTATG
AGCACACGTCTGAACTCTTTCCCTACACGACGCTCTTCCGATCTNNNCACCCAGAAGATGAGACTATG
AGCACACGTCTGAACTCTTTCCCTACACGACGCTCTTCCGATCTNNNGGCAACAACTCTCTCCTCCAAT
AGCACACGTCTGAACTCTTTCCCTACACGACGCTCTTCCGATCTNNNGGCAACAACTCTCTCCTCCAAT
GAGCACACGTCTGAACTCTTTCCCTACACGACGCTCTTCCGATCTNNNGGCTCTATTCTTTCTCACAGG

GAGCACACGTCTGAACTCTTTCCCTACACGACGCTCTTCCGATCTNNNGGCTCTATTCTTTCTCACAGGC

AGCACACGTCTGAACTCTTTCCCTACACGACGCTCTTCCGATCTNNNCGGTCTCACCTTGGCTAAGTAT

AGCACACGTCTGAACTCTTTCCCTACACGACGCTCTTCCGATCTNNNCGGTCTCACCTTGGCTAAGTAT

AGAGCACACGTCTGAACTCTTTCCCTACACGACGCTCTTCCGATCTNNNCTTCATCTAGTGCAGGGTCCT

AGAGCACACGTCTGAACTCTTTCCCTACACGACGCTCTTCCGATCTNNNCTTCATCTAGTGCAGGGTCCT

AGCACACGTCTGAACTCTTTCCCTACACGACGCTCTTCCGATCTNNNGCCGTCAAGAGAAAGGATTGT

AGCACACGTCTGAACTCTTTCCCTACACGACGCTCTTCCGATCTNNNGCCGTCAAGAGAAAGGATTGT

AGCACACGTCTGAACTCTTTCCCTACACGACGCTCTTCCGATCTNNNTTTTGTGCAAGCAACATCTCCC

AGCACACGTCTGAACTCTTTCCCTACACGACGCTCTTCCGATCTNNNTTTTGTGCAAGCAACATCTCCC

AGCACACGTCTGAACTCTTTCCCTACACGACGCTCTTCCGATCTNNNTGTTTGATCTTCTCTGTGGGCT

AGCACACGTCTGAACTCTTTCCCTACACGACGCTCTTCCGATCTNNNTGTTTGATCTTCTCTGTGGGCT

GAGCACACGTCTGAACTCTTTCCCTACACGACGCTCTTCCGATCTNNNCTTCTAGCTGAAACCATCACCC

GAGCACACGTCTGAACTCTTTCCCTACACGACGCTCTTCCGATCTNNNCTTCTAGCTGAAACCATCACCC

AGCACACGTCTGAACTCTTTCCCTACACGACGCTCTTCCGATCTNNNAAAAAATTAGCCAGGCTTGGTC

AGCACACGTCTGAACTCTTTCCCTACACGACGCTCTTCCGATCTNNNAAAAAATTAGCCAGGCTTGGTC

GCACACGTCTGAACTCTTTCCCTACACGACGCTCTTCCGATCTNNNGGAGTTACAAACCAGTGCTCCAT

GCACACGTCTGAACTCTTTCCCTACACGACGCTCTTCCGATCTNNNGGAGTTACAAACCAGTGCTCCAT

GCACACGTCTGAACTCTTTCCCTACACGACGCTCTTCCGATCTNNNCTGTTTTCAGGTTCATTCTGCAT

GCACACGTCTGAACTCTTTCCCTACACGACGCTCTTCCGATCTNNNCTGTTTTCAGGTTCATTCTGCAT

GAGCACACGTCTGAACTCTTTCCCTACACGACGCTCTTCCGATCTNNNAATTCAGGGAGCACAAATCAA

GAGCACACGTCTGAACTCTTTCCCTACACGACGCTCTTCCGATCTNNNAATTCAGGGAGCACAAATCAA

GAGCACACGTCTGAACTCTTTCCCTACACGACGCTCTTCCGATCTNNNTCATACCCTCCGCCAGTATAGA

GAGCACACGTCTGAACTCTTTCCCTACACGACGCTCTTCCGATCTNNNTCATACCCTCCGCCAGTATAGA

GAGCACACGTCTGAACTCTTTCCCTACACGACGCTCTTCCGATCTNNNCATGGTGGTGTGCATCTGTAA

GAGCACACGTCTGAACTCTTTCCCTACACGACGCTCTTCCGATCTNNNCATGGTGGTGTGCATCTGTAA

GAGCACACGTCTGAACTCTTTCCCTACACGACGCTCTTCCGATCTNNNATCCATGTTGCTGCAAATGAC

GAGCACACGTCTGAACTCTTTCCCTACACGACGCTCTTCCGATCTNNNATCCATGTTGCTGCAAATGAC

GAGCACACGTCTGAACTCTTTCCCTACACGACGCTCTTCCGATCTNNNAGATCAAGACCATCCTGACCA

GAGCACACGTCTGAACTCTTTCCCTACACGACGCTCTTCCGATCTNNNAGATCAAGACCATCCTGACCA

GAGCACACGTCTGAACTCTTTCCCTACACGACGCTCTTCCGATCTNNNACTTAGAGATTCTAGCCAGAG

GAGCACACGTCTGAACTCTTTCCCTACACGACGCTCTTCCGATCTNNNACTTAGAGATTCTAGCCAGAG

AGCACACGTCTGAACTCTTTCCCTACACGACGCTCTTCCGATCTNNNCATCCTCCCTTGGAAGTGTCTAG

AGCACACGTCTGAACTCTTTCCCTACACGACGCTCTTCCGATCTNNNCATCCTCCCTTGGAAGTGTCTAG

AGCACACGTCTGAACTCTTTCCCTACACGACGCTCTTCCGATCTNNNCTTCAATGCCACCTCGTTCCTCAT

AGCACACGTCTGAACTCTTTCCCTACACGACGCTCTTCCGATCTNNNCTTCAATGCCACCTCGTTCCTCAT

AAGAGCACACGTCTGAACTCTTTCCCTACACGACGCTCTTCCGATCTNNNGTATTCTTTGTTCCATTGCC

AAGAGCACACGTCTGAACTCTTTCCCTACACGACGCTCTTCCGATCTNNNGTATTCTTTGTTCCATTGCC

AGCACACGTCTGAACTCTTTCCCTACACGACGCTCTTCCGATCTNNNCTTGCAGGGATTTCATATTAAAA

AGCACACGTCTGAACTCTTTCCCTACACGACGCTCTTCCGATCTNNNCTTGCAGGGATTTCATATTAAAA

GAGCACACGTCTGAACTCTTTCCCTACACGACGCTCTTCCGATCTNNNATTTAGGAATGAGGAAGCTGC

GAGCACACGTCTGAACTCTTTCCCTACACGACGCTCTTCCGATCTNNNATTTAGGAATGAGGAAGCTGC

GAGCACACGTCTGAACTCTTTCCCTACACGACGCTCTTCCGATCTNNNTTACACAGGCAGGGAAATCTC

GAGCACACGTCTGAACTCTTTCCCTACACGACGCTCTTCCGATCTNNNTTACACAGGCAGGGAAATCTC

GAGCACACGTCTGAACTCTTTCCCTACACGACGCTCTTCCGATCTNNNTCCAGGTTTAGGAGGGGTCTC
GAGCACACGTCTGAACTCTTTCCCTACACGACGCTCTTCCGATCTNNNTCCAGGTTTAGGAGGGGTCTC
AGCACACGTCTGAACTCTTTCCCTACACGACGCTCTTCCGATCTNNNGAAGGTTGCAGTGAGCTGAA
AGCACACGTCTGAACTCTTTCCCTACACGACGCTCTTCCGATCTNNNGAAGGTTGCAGTGAGCTGAA
AGCACACGTCTGAACTCTTTCCCTACACGACGCTCTTCCGATCTNNNCTTGGAGAATGAATTGCTGTG
AGCACACGTCTGAACTCTTTCCCTACACGACGCTCTTCCGATCTNNNCTTGGAGAATGAATTGCTGTG
GAGCACACGTCTGAACTCTTTCCCTACACGACGCTCTTCCGATCTNNNAAATGTTCTTACCACCCCTCTG
GAGCACACGTCTGAACTCTTTCCCTACACGACGCTCTTCCGATCTNNNAAATGTTCTTACCACCCCTCTG
GAGCACACGTCTGAACTCTTTCCCTACACGACGCTCTTCCGATCTNNNTGCCTGGTTTGTCCACTTCTA
GAGCACACGTCTGAACTCTTTCCCTACACGACGCTCTTCCGATCTNNNTGCCTGGTTTGTCCACTTCTA
CACACGTCTGAACTCTTTCCCTACACGACGCTCTTCCGATCTNNNTGTGGATGAAGGTAAGCACGAAC
CACACGTCTGAACTCTTTCCCTACACGACGCTCTTCCGATCTNNNTGTGGATGAAGGTAAGCACGAAC
AGCACACGTCTGAACTCTTTCCCTACACGACGCTCTTCCGATCTNNNAAAGCAACTTGTGTTTCCTTTTG
AGCACACGTCTGAACTCTTTCCCTACACGACGCTCTTCCGATCTNNNAAAGCAACTTGTGTTTCCTTTTG
GAGCACACGTCTGAACTCTTTCCCTACACGACGCTCTTCCGATCTNNNATAGTGTATTGTATGCTGAAG
GAGCACACGTCTGAACTCTTTCCCTACACGACGCTCTTCCGATCTNNNATAGTGTATTGTATGCTGAAG
GCACACGTCTGAACTCTTTCCCTACACGACGCTCTTCCGATCTNNNTCTCTTCTTTCCCTGACAAGTGG
GCACACGTCTGAACTCTTTCCCTACACGACGCTCTTCCGATCTNNNTCTCTTCTTTCCCTGACAAGTGG
GAGCACACGTCTGAACTCTTTCCCTACACGACGCTCTTCCGATCTNNNCAGTAGTTCCAGCTGCTCATG
GAGCACACGTCTGAACTCTTTCCCTACACGACGCTCTTCCGATCTNNNCAGTAGTTCCAGCTGCTCATG
GAGCACACGTCTGAACTCTTTCCCTACACGACGCTCTTCCGATCTNNNCCACGCCCAGCTAATTTTTGT
GAGCACACGTCTGAACTCTTTCCCTACACGACGCTCTTCCGATCTNNNCCACGCCCAGCTAATTTTTGT
AGAGCACACGTCTGAACTCTTTCCCTACACGACGCTCTTCCGATCTNNNGAGAATAGCTTGAACCCAG
AGAGCACACGTCTGAACTCTTTCCCTACACGACGCTCTTCCGATCTNNNGAGAATAGCTTGAACCCAG
GAGCACACGTCTGAACTCTTTCCCTACACGACGCTCTTCCGATCTNNNCTTCTCCACAAGCCAGTATTCC
GAGCACACGTCTGAACTCTTTCCCTACACGACGCTCTTCCGATCTNNNCTTCTCCACAAGCCAGTATTCC
AAGAGCACACGTCTGAACTCTTTCCCTACACGACGCTCTTCCGATCTNNNCGTCTCAACAGAAGAAAG
AAGAGCACACGTCTGAACTCTTTCCCTACACGACGCTCTTCCGATCTNNNCGTCTCAACAGAAGAAAG
AGCACACGTCTGAACTCTTTCCCTACACGACGCTCTTCCGATCTNNNCTGTAGTCCCAGCTACTCAGGA
AGCACACGTCTGAACTCTTTCCCTACACGACGCTCTTCCGATCTNNNCTGTAGTCCCAGCTACTCAGGA
GAGCACACGTCTGAACTCTTTCCCTACACGACGCTCTTCCGATCTNNNCCGAGTTTGGTGATTGCGAAT
GAGCACACGTCTGAACTCTTTCCCTACACGACGCTCTTCCGATCTNNNCCGAGTTTGGTGATTGCGAAT
AAGAGCACACGTCTGAACTCTTTCCCTACACGACGCTCTTCCGATCTNNNTCTGCCATCACCTCAGCTTT
AAGAGCACACGTCTGAACTCTTTCCCTACACGACGCTCTTCCGATCTNNNTCTGCCATCACCTCAGCTTT
GAGCACACGTCTGAACTCTTTCCCTACACGACGCTCTTCCGATCTNNNGCCTTCTACATGCCTTGTGGTT
GAGCACACGTCTGAACTCTTTCCCTACACGACGCTCTTCCGATCTNNNGCCTTCTACATGCCTTGTGGTT
GAGCACACGTCTGAACTCTTTCCCTACACGACGCTCTTCCGATCTNNNGAGGCAGGAGAATCGCTTG
GAGCACACGTCTGAACTCTTTCCCTACACGACGCTCTTCCGATCTNNNGAGGCAGGAGAATCGCTTG
GAGCACACGTCTGAACTCTTTCCCTACACGACGCTCTTCCGATCTNNNGCAGATTAGAGAGCAACGTC
GAGCACACGTCTGAACTCTTTCCCTACACGACGCTCTTCCGATCTNNNGCAGATTAGAGAGCAACGTC
AGCACACGTCTGAACTCTTTCCCTACACGACGCTCTTCCGATCTNNNCACCCAAAATAGTGATGAGGA
AGCACACGTCTGAACTCTTTCCCTACACGACGCTCTTCCGATCTNNNCACCCAAAATAGTGATGAGGA
GAGCACACGTCTGAACTCTTTCCCTACACGACGCTCTTCCGATCTNNNGGGAAACTACAACAGCACAA

GAGCACACGTCTGAACTCTTTCCCTACACGACGCTCTTCCGATCTNNNGGGAAACTACAACAGCACAAT
AGAGCACACGTCTGAACTCTTTCCCTACACGACGCTCTTCCGATCTNNNATAAGCTCTGTAAGGCCTGAA
AGAGCACACGTCTGAACTCTTTCCCTACACGACGCTCTTCCGATCTNNNATAAGCTCTGTAAGGCCTGAA
GAGCACACGTCTGAACTCTTTCCCTACACGACGCTCTTCCGATCTNNNCTGATCCCCTTCAAGAACATGC
GAGCACACGTCTGAACTCTTTCCCTACACGACGCTCTTCCGATCTNNNCTGATCCCCTTCAAGAACATGC
AGCACACGTCTGAACTCTTTCCCTACACGACGCTCTTCCGATCTNNNCGCTTCTCTCTCTCTCTCTCTCA
AGCACACGTCTGAACTCTTTCCCTACACGACGCTCTTCCGATCTNNNCGCTTCTCTCTCTCTCTCTCTCA
GAGCACACGTCTGAACTCTTTCCCTACACGACGCTCTTCCGATCTNNNCACTTACACATACCTTTGGGGC
GAGCACACGTCTGAACTCTTTCCCTACACGACGCTCTTCCGATCTNNNCACTTACACATACCTTTGGGGC
AGCACACGTCTGAACTCTTTCCCTACACGACGCTCTTCCGATCTNNNTGAAACCAAATTATGCGACCAG
AGCACACGTCTGAACTCTTTCCCTACACGACGCTCTTCCGATCTNNNTGAAACCAAATTATGCGACCAG
AGCACACGTCTGAACTCTTTCCCTACACGACGCTCTTCCGATCTNNNATGTGTTGGCTCACTCCTGTAA
AGCACACGTCTGAACTCTTTCCCTACACGACGCTCTTCCGATCTNNNATGTGTTGGCTCACTCCTGTAA
GCACACGTCTGAACTCTTTCCCTACACGACGCTCTTCCGATCTNNNTTCTGAAACTGAAGAGTCAAACAT
GCACACGTCTGAACTCTTTCCCTACACGACGCTCTTCCGATCTNNNTTCTGAAACTGAAGAGTCAAACAT
GAGCACACGTCTGAACTCTTTCCCTACACGACGCTCTTCCGATCTNNNAGGTAGCAACAGGAGAGAGC
GAGCACACGTCTGAACTCTTTCCCTACACGACGCTCTTCCGATCTNNNAGGTAGCAACAGGAGAGAGC
GAGCACACGTCTGAACTCTTTCCCTACACGACGCTCTTCCGATCTNNNAAGGACAAACCAAACTGAAAC
GAGCACACGTCTGAACTCTTTCCCTACACGACGCTCTTCCGATCTNNNAAGGACAAACCAAACTGAAAC
AGCACACGTCTGAACTCTTTCCCTACACGACGCTCTTCCGATCTNNNGAGGCAGGAGAATTGCTTGAAT
AGCACACGTCTGAACTCTTTCCCTACACGACGCTCTTCCGATCTNNNGAGGCAGGAGAATTGCTTGAAT
GAGCACACGTCTGAACTCTTTCCCTACACGACGCTCTTCCGATCTNNNTCTCTATCCAGAATCCTCTCCCC
GAGCACACGTCTGAACTCTTTCCCTACACGACGCTCTTCCGATCTNNNTCTCTATCCAGAATCCTCTCCCC
GCACACGTCTGAACTCTTTCCCTACACGACGCTCTTCCGATCTNNNTCCAACATCTTGTTGCCATTGGAC
GCACACGTCTGAACTCTTTCCCTACACGACGCTCTTCCGATCTNNNTCCAACATCTTGTTGCCATTGGAC
GAGCACACGTCTGAACTCTTTCCCTACACGACGCTCTTCCGATCTNNNGCCTGTAGTCCCAGCTACTTGG
GAGCACACGTCTGAACTCTTTCCCTACACGACGCTCTTCCGATCTNNNGCCTGTAGTCCCAGCTACTTGG
GAGCACACGTCTGAACTCTTTCCCTACACGACGCTCTTCCGATCTNNNTGGGAGTGATGCAACATTAGG
GAGCACACGTCTGAACTCTTTCCCTACACGACGCTCTTCCGATCTNNNTGGGAGTGATGCAACATTAGG
AGCACACGTCTGAACTCTTTCCCTACACGACGCTCTTCCGATCTNNNAAAGGGAGAGAGAGAAGGGA
AGCACACGTCTGAACTCTTTCCCTACACGACGCTCTTCCGATCTNNNAAAGGGAGAGAGAGAAGGGA
AAGAGCACACGTCTGAACTCTTTCCCTACACGACGCTCTTCCGATCTNNNCTGTCGATCCAATCACCTCT
AAGAGCACACGTCTGAACTCTTTCCCTACACGACGCTCTTCCGATCTNNNCTGTCGATCCAATCACCTCT
AGCACACGTCTGAACTCTTTCCCTACACGACGCTCTTCCGATCTNNNGAGGTTACAGTGAGCCAAGAT
AGCACACGTCTGAACTCTTTCCCTACACGACGCTCTTCCGATCTNNNGAGGTTACAGTGAGCCAAGAT
GAGCACACGTCTGAACTCTTTCCCTACACGACGCTCTTCCGATCTNNNGGCCACATATTCAGTTTGGTTC
GAGCACACGTCTGAACTCTTTCCCTACACGACGCTCTTCCGATCTNNNGGCCACATATTCAGTTTGGTTC
AGCACACGTCTGAACTCTTTCCCTACACGACGCTCTTCCGATCTNNNGGACTTTCCTTTGTAGTGTTGCC
AGCACACGTCTGAACTCTTTCCCTACACGACGCTCTTCCGATCTNNNGGACTTTCCTTTGTAGTGTTGCC
GAGCACACGTCTGAACTCTTTCCCTACACGACGCTCTTCCGATCTNNNGGCACCTGTAATCCTAGCTACT
GAGCACACGTCTGAACTCTTTCCCTACACGACGCTCTTCCGATCTNNNGGCACCTGTAATCCTAGCTACT
GCACACGTCTGAACTCTTTCCCTACACGACGCTCTTCCGATCTNNNACAAGAAACCAAGAAGCTTGCTC
GCACACGTCTGAACTCTTTCCCTACACGACGCTCTTCCGATCTNNNACAAGAAACCAAGAAGCTTGCTC

AGCACACGTCTGAACTCTTTCCCTACACGACGCTCTTCCGATCTNNNCTTCCACTGCAAGGAGAGTTA

AGCACACGTCTGAACTCTTTCCCTACACGACGCTCTTCCGATCTNNNCTTCCACTGCAAGGAGAGTTA

AGCACACGTCTGAACTCTTTCCCTACACGACGCTCTTCCGATCTNNNAAGCTTGTGGCACAGAGTAA

AGCACACGTCTGAACTCTTTCCCTACACGACGCTCTTCCGATCTNNNAAGCTTGTGGCACAGAGTAA

AGCACACGTCTGAACTCTTTCCCTACACGACGCTCTTCCGATCTNNNTCCCCCAGAACCCTTCTATACT

AGCACACGTCTGAACTCTTTCCCTACACGACGCTCTTCCGATCTNNNTCCCCCAGAACCCTTCTATACT

AGCACACGTCTGAACTCTTTCCCTACACGACGCTCTTCCGATCTNNNGTTTTCTGCTGAAAGGTCCAC

AGCACACGTCTGAACTCTTTCCCTACACGACGCTCTTCCGATCTNNNGTTTTCTGCTGAAAGGTCCAC

GCACACGTCTGAACTCTTTCCCTACACGACGCTCTTCCGATCTNNNGAATGACTGAAGTATCAACATT

GCACACGTCTGAACTCTTTCCCTACACGACGCTCTTCCGATCTNNNGAATGACTGAAGTATCAACATT

AGCACACGTCTGAACTCTTTCCCTACACGACGCTCTTCCGATCTNNNCTGAACTAGCTGCAAGACGTT

AGCACACGTCTGAACTCTTTCCCTACACGACGCTCTTCCGATCTNNNCTGAACTAGCTGCAAGACGTT

GCACACGTCTGAACTCTTTCCCTACACGACGCTCTTCCGATCTNNNAACTAGTCTCTGTGCCAGAAAG

GCACACGTCTGAACTCTTTCCCTACACGACGCTCTTCCGATCTNNNAACTAGTCTCTGTGCCAGAAAG

AGCACACGTCTGAACTCTTTCCCTACACGACGCTCTTCCGATCTNNNACATCAACAAACATAGGAGTG

AGCACACGTCTGAACTCTTTCCCTACACGACGCTCTTCCGATCTNNNACATCAACAAACATAGGAGTG

AGCACACGTCTGAACTCTTTCCCTACACGACGCTCTTCCGATCTNNNAGCAGCAAATTCCATCAGCATT

AGCACACGTCTGAACTCTTTCCCTACACGACGCTCTTCCGATCTNNNAGCAGCAAATTCCATCAGCATT

AGAGCACACGTCTGAACTCTTTCCCTACACGACGCTCTTCCGATCTNNNGAGAAGGAAGGAAGGAA

AGAGCACACGTCTGAACTCTTTCCCTACACGACGCTCTTCCGATCTNNNGAGAAGGAAGGAAGGAA

GCACACGTCTGAACTCTTTCCCTACACGACGCTCTTCCGATCTNNNCTCTCTCTCCTGTCTCTCTGTCA

GCACACGTCTGAACTCTTTCCCTACACGACGCTCTTCCGATCTNNNCTCTCTCTCCTGTCTCTCTGTCA

GAGCACACGTCTGAACTCTTTCCCTACACGACGCTCTTCCGATCTNNNCTGTAATCCCAGCTACTCGGT

GAGCACACGTCTGAACTCTTTCCCTACACGACGCTCTTCCGATCTNNNCTGTAATCCCAGCTACTCGGT

GCACACGTCTGAACTCTTTCCCTACACGACGCTCTTCCGATCTNNNCATCTGTGGTCCCAGCTACTTAG

GCACACGTCTGAACTCTTTCCCTACACGACGCTCTTCCGATCTNNNCATCTGTGGTCCCAGCTACTTAG

AGCACACGTCTGAACTCTTTCCCTACACGACGCTCTTCCGATCTNNNCTGGAAACTATCAAACACAAC

AGCACACGTCTGAACTCTTTCCCTACACGACGCTCTTCCGATCTNNNCTGGAAACTATCAAACACAAC

AGCACACGTCTGAACTCTTTCCCTACACGACGCTCTTCCGATCTNNNCAACCCCAAAGCCTGAGATAG

AGCACACGTCTGAACTCTTTCCCTACACGACGCTCTTCCGATCTNNNCAACCCCAAAGCCTGAGATAG

AGCACACGTCTGAACTCTTTCCCTACACGACGCTCTTCCGATCTNNNGGTCTCTTACAGCATCAGCATT

AGCACACGTCTGAACTCTTTCCCTACACGACGCTCTTCCGATCTNNNGGTCTCTTACAGCATCAGCATT

GCACACGTCTGAACTCTTTCCCTACACGACGCTCTTCCGATCTNNNAATTCCAGCTCCTGCACATCTAT

GCACACGTCTGAACTCTTTCCCTACACGACGCTCTTCCGATCTNNNAATTCCAGCTCCTGCACATCTAT

GAGCACACGTCTGAACTCTTTCCCTACACGACGCTCTTCCGATCTNNNTTTAGCTGAAATTTGCCACAG

GAGCACACGTCTGAACTCTTTCCCTACACGACGCTCTTCCGATCTNNNTTTAGCTGAAATTTGCCACAG

GCACACGTCTGAACTCTTTCCCTACACGACGCTCTTCCGATCTNNNGCTCCCCAGTTATTCCTATGGTC

GCACACGTCTGAACTCTTTCCCTACACGACGCTCTTCCGATCTNNNGCTCCCCAGTTATTCCTATGGTC

GCACACGTCTGAACTCTTTCCCTACACGACGCTCTTCCGATCTNNNGACTTTCCTTTTCTGCCTGGAAC

GCACACGTCTGAACTCTTTCCCTACACGACGCTCTTCCGATCTNNNGACTTTCCTTTTCTGCCTGGAAC

AGAGCACACGTCTGAACTCTTTCCCTACACGACGCTCTTCCGATCTNNNAGGCTGAGTCACGAGAATC

AGAGCACACGTCTGAACTCTTTCCCTACACGACGCTCTTCCGATCTNNNAGGCTGAGTCACGAGAATC

CACACGTCTGAACTCTTTCCCTACACGACGCTCTTCCGATCTNNNCATCAACTCCACTGACCTGGTAAC

CACACGTCTGAACTCTTTCCCTACACGACGCTCTTCCGATCTNNNCATCAACTCCACTGACCTGGTAAC

GAGCACACGTCTGAACTCTTTCCCTACACGACGCTCTTCCGATCTNNNTTCTGTTTTGCAAAGTCTGTG

GAGCACACGTCTGAACTCTTTCCCTACACGACGCTCTTCCGATCTNNNTTCTGTTTTGCAAAGTCTGTG

GAGCACACGTCTGAACTCTTTCCCTACACGACGCTCTTCCGATCTNNNGAGGCTGAGGCAGGAGAAT

GAGCACACGTCTGAACTCTTTCCCTACACGACGCTCTTCCGATCTNNNGAGGCTGAGGCAGGAGAAT

GCACACGTCTGAACTCTTTCCCTACACGACGCTCTTCCGATCTNNNTGATGATCTGTTCTGCCTCCTTC

GCACACGTCTGAACTCTTTCCCTACACGACGCTCTTCCGATCTNNNTGATGATCTGTTCTGCCTCCTTC

GAGCACACGTCTGAACTCTTTCCCTACACGACGCTCTTCCGATCTNNNGATTCAAGTGATTCCCCTGCC

GAGCACACGTCTGAACTCTTTCCCTACACGACGCTCTTCCGATCTNNNGATTCAAGTGATTCCCCTGCC

AGCACACGTCTGAACTCTTTCCCTACACGACGCTCTTCCGATCTNNNAATGCGGAAAGGTCCTCAAGA

AGCACACGTCTGAACTCTTTCCCTACACGACGCTCTTCCGATCTNNNAATGCGGAAAGGTCCTCAAGA

CACACGTCTGAACTCTTTCCCTACACGACGCTCTTCCGATCTNNNTATAACACATCTACTGAAAAAGCC

CACACGTCTGAACTCTTTCCCTACACGACGCTCTTCCGATCTNNNTATAACACATCTACTGAAAAAGCC

AGCACACGTCTGAACTCTTTCCCTACACGACGCTCTTCCGATCTNNNACAGAGTAAGACCTCGTCTCA

AGCACACGTCTGAACTCTTTCCCTACACGACGCTCTTCCGATCTNNNACAGAGTAAGACCTCGTCTCA

GCACACGTCTGAACTCTTTCCCTACACGACGCTCTTCCGATCTNNNCAACAGGGTATAAGCATTCCCC

GCACACGTCTGAACTCTTTCCCTACACGACGCTCTTCCGATCTNNNCAACAGGGTATAAGCATTCCCC

GCACACGTCTGAACTCTTTCCCTACACGACGCTCTTCCGATCTNNNCTCCTCGGCTCAAGGAAAGTAC

GCACACGTCTGAACTCTTTCCCTACACGACGCTCTTCCGATCTNNNCTCCTCGGCTCAAGGAAAGTAC

AGCACACGTCTGAACTCTTTCCCTACACGACGCTCTTCCGATCTNNNCCATCCCTCAGTACACTCAGAA

AGCACACGTCTGAACTCTTTCCCTACACGACGCTCTTCCGATCTNNNCCATCCCTCAGTACACTCAGAA

GAGCACACGTCTGAACTCTTTCCCTACACGACGCTCTTCCGATCTNNNAGTTTCTCCTCATTTTTGCCAG

GAGCACACGTCTGAACTCTTTCCCTACACGACGCTCTTCCGATCTNNNAGTTTCTCCTCATTTTTGCCAG

AGCACACGTCTGAACTCTTTCCCTACACGACGCTCTTCCGATCTNNNAACACCTCAAGGAAAGGTCAA

AGCACACGTCTGAACTCTTTCCCTACACGACGCTCTTCCGATCTNNNAACACCTCAAGGAAAGGTCAA

GAGCACACGTCTGAACTCTTTCCCTACACGACGCTCTTCCGATCTNNNGAGAATCCCTTGAACCCAGG

GAGCACACGTCTGAACTCTTTCCCTACACGACGCTCTTCCGATCTNNNGAGAATCCCTTGAACCCAGG

AGCACACGTCTGAACTCTTTCCCTACACGACGCTCTTCCGATCTNNNCCAGGGCTAGAACTTCCATCT

AGCACACGTCTGAACTCTTTCCCTACACGACGCTCTTCCGATCTNNNCCAGGGCTAGAACTTCCATCT

GAAGAGCACACGTCTGAACTCTTTCCCTACACGACGCTCTTCCGATCTNNNCAGCACTGAGCCGAGAT

GAAGAGCACACGTCTGAACTCTTTCCCTACACGACGCTCTTCCGATCTNNNCAGCACTGAGCCGAGAT

ACACGTCTGAACTCTTTCCCTACACGACGCTCTTCCGATCTNNNGAGAAACACGTCCTCAAGCCCCATC

ACACGTCTGAACTCTTTCCCTACACGACGCTCTTCCGATCTNNNGAGAAACACGTCCTCAAGCCCCATC

GAGCACACGTCTGAACTCTTTCCCTACACGACGCTCTTCCGATCTNNNGTCTCACTCTGTCACCCATACT

GAGCACACGTCTGAACTCTTTCCCTACACGACGCTCTTCCGATCTNNNGTCTCACTCTGTCACCCATACT

GCACACGTCTGAACTCTTTCCCTACACGACGCTCTTCCGATCTNNNCTCCCAGGTTCAAGCGATTCTCC

GCACACGTCTGAACTCTTTCCCTACACGACGCTCTTCCGATCTNNNCTCCCAGGTTCAAGCGATTCTCC

GCACACGTCTGAACTCTTTCCCTACACGACGCTCTTCCGATCTNNNTTTGTCTTCCAGACTCTTCCCTGC

GCACACGTCTGAACTCTTTCCCTACACGACGCTCTTCCGATCTNNNTTTGTCTTCCAGACTCTTCCCTGC

GAGCACACGTCTGAACTCTTTCCCTACACGACGCTCTTCCGATCTNNNGTTTTCCCCAGGTGGTTCTTT

GAGCACACGTCTGAACTCTTTCCCTACACGACGCTCTTCCGATCTNNNGTTTTCCCCAGGTGGTTCTTT

CACACGTCTGAACTCTTTCCCTACACGACGCTCTTCCGATCTNNNACATAATTGGGTCTTTGTCATATT

CACACGTCTGAACTCTTTCCCTACACGACGCTCTTCCGATCTNNNACATAATTGGGTCTTTGTCATATT

AGCACACGTCTGAACTCTTTCCCTACACGACGCTCTTCCGATCTNNNTAATCACTTGAACCCAGGAAGC
AGCACACGTCTGAACTCTTTCCCTACACGACGCTCTTCCGATCTNNNTAATCACTTGAACCCAGGAAGC
AGAGCACACGTCTGAACTCTTTCCCTACACGACGCTCTTCCGATCTNNNTGCATGTTCATAGCAGCACA
AGAGCACACGTCTGAACTCTTTCCCTACACGACGCTCTTCCGATCTNNNTGCATGTTCATAGCAGCACA
AGCACACGTCTGAACTCTTTCCCTACACGACGCTCTTCCGATCTNNNTTCCAAAGTGCTGGGATTACA
AGCACACGTCTGAACTCTTTCCCTACACGACGCTCTTCCGATCTNNNTTCCAAAGTGCTGGGATTACA
AGCACACGTCTGAACTCTTTCCCTACACGACGCTCTTCCGATCTNNNAACTGCTATCTACTGCTTAACCA
AGCACACGTCTGAACTCTTTCCCTACACGACGCTCTTCCGATCTNNNAACTGCTATCTACTGCTTAACCA
AGCACACGTCTGAACTCTTTCCCTACACGACGCTCTTCCGATCTNNNGAACTTAGAGGGAGAGGAGA
AGCACACGTCTGAACTCTTTCCCTACACGACGCTCTTCCGATCTNNNGAACTTAGAGGGAGAGGAGA
AGCACACGTCTGAACTCTTTCCCTACACGACGCTCTTCCGATCTNNNCTAAGAGTTCCCAGTGGATTG
AGCACACGTCTGAACTCTTTCCCTACACGACGCTCTTCCGATCTNNNCTAAGAGTTCCCAGTGGATTG
ACACGTCTGAACTCTTTCCCTACACGACGCTCTTCCGATCTNNNTGGAAGAAACTTCAGGCAAACAAC
ACACGTCTGAACTCTTTCCCTACACGACGCTCTTCCGATCTNNNTGGAAGAAACTTCAGGCAAACAAC
AGCACACGTCTGAACTCTTTCCCTACACGACGCTCTTCCGATCTNNNTCAGAACCTACTGCAACCTCTT
AGCACACGTCTGAACTCTTTCCCTACACGACGCTCTTCCGATCTNNNTCAGAACCTACTGCAACCTCTT
GAGCACACGTCTGAACTCTTTCCCTACACGACGCTCTTCCGATCTNNNGGGTGACAGTGTGAGACTCT
GAGCACACGTCTGAACTCTTTCCCTACACGACGCTCTTCCGATCTNNNGGGTGACAGTGTGAGACTCT
GCACACGTCTGAACTCTTTCCCTACACGACGCTCTTCCGATCTNNNCCCAAATAGTTTTTGGCAAAGGC
GCACACGTCTGAACTCTTTCCCTACACGACGCTCTTCCGATCTNNNCCCAAATAGTTTTTGGCAAAGGC
AGCACACGTCTGAACTCTTTCCCTACACGACGCTCTTCCGATCTNNNGTGACCTCGATGCTAAATAGC
AGCACACGTCTGAACTCTTTCCCTACACGACGCTCTTCCGATCTNNNGTGACCTCGATGCTAAATAGC
GAGCACACGTCTGAACTCTTTCCCTACACGACGCTCTTCCGATCTNNNCTGTAGTCCCAGCTACTTTGG
GAGCACACGTCTGAACTCTTTCCCTACACGACGCTCTTCCGATCTNNNCTGTAGTCCCAGCTACTTTGG
AGCACACGTCTGAACTCTTTCCCTACACGACGCTCTTCCGATCTNNNCTCAATTGTTTTGAGCAGCAGG
AGCACACGTCTGAACTCTTTCCCTACACGACGCTCTTCCGATCTNNNCTCAATTGTTTTGAGCAGCAGG
GAGCACACGTCTGAACTCTTTCCCTACACGACGCTCTTCCGATCTNNNCAAACACTTCACACACACACA
GAGCACACGTCTGAACTCTTTCCCTACACGACGCTCTTCCGATCTNNNCAAACACTTCACACACACACA
GAGCACACGTCTGAACTCTTTCCCTACACGACGCTCTTCCGATCTNNNGCATAGAGCTCGAGGTTGAT
GAGCACACGTCTGAACTCTTTCCCTACACGACGCTCTTCCGATCTNNNGCATAGAGCTCGAGGTTGAT
GCACACGTCTGAACTCTTTCCCTACACGACGCTCTTCCGATCTNNNTTATATGGTGGTGAAGTCTGGG
GCACACGTCTGAACTCTTTCCCTACACGACGCTCTTCCGATCTNNNTTATATGGTGGTGAAGTCTGGG
AGCACACGTCTGAACTCTTTCCCTACACGACGCTCTTCCGATCTNNNTCTGATGATGGGCTCTGTATT
AGCACACGTCTGAACTCTTTCCCTACACGACGCTCTTCCGATCTNNNTCTGATGATGGGCTCTGTATT
AGCACACGTCTGAACTCTTTCCCTACACGACGCTCTTCCGATCTNNNAAGAGACACCAACCACAAGATG
AGCACACGTCTGAACTCTTTCCCTACACGACGCTCTTCCGATCTNNNAAGAGACACCAACCACAAGATG
AGCACACGTCTGAACTCTTTCCCTACACGACGCTCTTCCGATCTNNNTTTGATTCTCACTCTGTTGCCCT
AGCACACGTCTGAACTCTTTCCCTACACGACGCTCTTCCGATCTNNNTTTGATTCTCACTCTGTTGCCCT
AGCACACGTCTGAACTCTTTCCCTACACGACGCTCTTCCGATCTNNNGTACTTCTGTAGTGTCCAGCCA
AGCACACGTCTGAACTCTTTCCCTACACGACGCTCTTCCGATCTNNNGTACTTCTGTAGTGTCCAGCCA
AGCACACGTCTGAACTCTTTCCCTACACGACGCTCTTCCGATCTNNNGGAAACCACCACATCTACCTGA
AGCACACGTCTGAACTCTTTCCCTACACGACGCTCTTCCGATCTNNNGGAAACCACCACATCTACCTGA
AGCACACGTCTGAACTCTTTCCCTACACGACGCTCTTCCGATCTNNNAACTGTCTTTTCACGTCCATTGC

AGCACACGTCTGAACTCTTTCCCTACACGACGCTCTTCCGATCTNNNAACTGTCTTTTCACGTCCATTGC
GCACACGTCTGAACTCTTTCCCTACACGACGCTCTTCCGATCTNNNGGGTGATCTGTGTGTCCTTACAT
GCACACGTCTGAACTCTTTCCCTACACGACGCTCTTCCGATCTNNNGGGTGATCTGTGTGTCCTTACAT
GAGCACACGTCTGAACTCTTTCCCTACACGACGCTCTTCCGATCTNNNCAGGAGAATTGCTTGAACCAA
GAGCACACGTCTGAACTCTTTCCCTACACGACGCTCTTCCGATCTNNNCAGGAGAATTGCTTGAACCAA
GAGCACACGTCTGAACTCTTTCCCTACACGACGCTCTTCCGATCTNNNGATATGTGGATGATGATTTTC
GAGCACACGTCTGAACTCTTTCCCTACACGACGCTCTTCCGATCTNNNGATATGTGGATGATGATTTTC
GAGCACACGTCTGAACTCTTTCCCTACACGACGCTCTTCCGATCTNNNTGTGTGTGTGCCTATGTGTGT
GAGCACACGTCTGAACTCTTTCCCTACACGACGCTCTTCCGATCTNNNTGTGTGTGTGCCTATGTGTGT
AGCACACGTCTGAACTCTTTCCCTACACGACGCTCTTCCGATCTNNNGAAATCCTTGGGAGGTGGACA
AGCACACGTCTGAACTCTTTCCCTACACGACGCTCTTCCGATCTNNNGAAATCCTTGGGAGGTGGACA
AGCACACGTCTGAACTCTTTCCCTACACGACGCTCTTCCGATCTNNNTTTTAGCTCCCTCTTATGTGGG
AGCACACGTCTGAACTCTTTCCCTACACGACGCTCTTCCGATCTNNNTTTTAGCTCCCTCTTATGTGGG
GAGCACACGTCTGAACTCTTTCCCTACACGACGCTCTTCCGATCTNNNCACACCAGTTTTCTTCCTGCTC
GAGCACACGTCTGAACTCTTTCCCTACACGACGCTCTTCCGATCTNNNCACACCAGTTTTCTTCCTGCTC
AGCACACGTCTGAACTCTTTCCCTACACGACGCTCTTCCGATCTNNNATATTACATTTCAGAGTTCACCT
AGCACACGTCTGAACTCTTTCCCTACACGACGCTCTTCCGATCTNNNATATTACATTTCAGAGTTCACCT
GAGCACACGTCTGAACTCTTTCCCTACACGACGCTCTTCCGATCTNNNAGAATTGCTTGAATCTGGGA
GAGCACACGTCTGAACTCTTTCCCTACACGACGCTCTTCCGATCTNNNAGAATTGCTTGAATCTGGGA
GAGCACACGTCTGAACTCTTTCCCTACACGACGCTCTTCCGATCTNNNCTGGATGAGGTTGGAGTCCAT
GAGCACACGTCTGAACTCTTTCCCTACACGACGCTCTTCCGATCTNNNCTGGATGAGGTTGGAGTCCAT
AGCACACGTCTGAACTCTTTCCCTACACGACGCTCTTCCGATCTNNNCCTTTTTTTTTTTTTGCGGGGG
AGCACACGTCTGAACTCTTTCCCTACACGACGCTCTTCCGATCTNNNCCTTTTTTTTTTTTTGCGGGGG
AGAGCACACGTCTGAACTCTTTCCCTACACGACGCTCTTCCGATCTNNNCTAGGGATTTCAATGCTGCA
AGAGCACACGTCTGAACTCTTTCCCTACACGACGCTCTTCCGATCTNNNCTAGGGATTTCAATGCTGCA
GAGCACACGTCTGAACTCTTTCCCTACACGACGCTCTTCCGATCTNNNAATTCCATCCAAGTTGCTGCAA
GAGCACACGTCTGAACTCTTTCCCTACACGACGCTCTTCCGATCTNNNAATTCCATCCAAGTTGCTGCAA
GAGCACACGTCTGAACTCTTTCCCTACACGACGCTCTTCCGATCTNNNTCCATCCCATCAGGACAGACTT
GAGCACACGTCTGAACTCTTTCCCTACACGACGCTCTTCCGATCTNNNTCCATCCCATCAGGACAGACTT
GAGCACACGTCTGAACTCTTTCCCTACACGACGCTCTTCCGATCTNNNCATGCCTGTAATCCCAGCTAC
GAGCACACGTCTGAACTCTTTCCCTACACGACGCTCTTCCGATCTNNNCATGCCTGTAATCCCAGCTAC
GAGCACACGTCTGAACTCTTTCCCTACACGACGCTCTTCCGATCTNNNGGCAAAATGAGGAGTGGCAT
GAGCACACGTCTGAACTCTTTCCCTACACGACGCTCTTCCGATCTNNNGGCAAAATGAGGAGTGGCAT
AGCACACGTCTGAACTCTTTCCCTACACGACGCTCTTCCGATCTNNNATATAGGGAGGTGAAGGGGA
AGCACACGTCTGAACTCTTTCCCTACACGACGCTCTTCCGATCTNNNATATAGGGAGGTGAAGGGGA
AGCACACGTCTGAACTCTTTCCCTACACGACGCTCTTCCGATCTNNNATGATGGTGATGATGGTGATG
AGCACACGTCTGAACTCTTTCCCTACACGACGCTCTTCCGATCTNNNATGATGGTGATGATGGTGATG
GAGCACACGTCTGAACTCTTTCCCTACACGACGCTCTTCCGATCTNNNAATGTGCTGGGATTATAGGG
GAGCACACGTCTGAACTCTTTCCCTACACGACGCTCTTCCGATCTNNNAATGTGCTGGGATTATAGGG
AGCACACGTCTGAACTCTTTCCCTACACGACGCTCTTCCGATCTNNNGACCTTTCCCCCTCTATGACTTCC
AGCACACGTCTGAACTCTTTCCCTACACGACGCTCTTCCGATCTNNNGACCTTTCCCCCTCTATGACTTCC
GAGCACACGTCTGAACTCTTTCCCTACACGACGCTCTTCCGATCTNNNTAAAAATTATGTTAGACCTGG
GAGCACACGTCTGAACTCTTTCCCTACACGACGCTCTTCCGATCTNNNTAAAAATTATGTTAGACCTGG

GCACACGTCTGAACTCTTTCCCTACACGACGCTCTTCCGATCTNNNCACAGCACAGAATCCTACCTCAT

GCACACGTCTGAACTCTTTCCCTACACGACGCTCTTCCGATCTNNNCACAGCACAGAATCCTACCTCAT

GCACACGTCTGAACTCTTTCCCTACACGACGCTCTTCCGATCTNNNGGACAGCTCACCTGATTGTTCTA

GCACACGTCTGAACTCTTTCCCTACACGACGCTCTTCCGATCTNNNGGACAGCTCACCTGATTGTTCTA

AGCACACGTCTGAACTCTTTCCCTACACGACGCTCTTCCGATCTNNNCCAGAAGGGAGTGACATGAC

AGCACACGTCTGAACTCTTTCCCTACACGACGCTCTTCCGATCTNNNCCAGAAGGGAGTGACATGAC

AGCACACGTCTGAACTCTTTCCCTACACGACGCTCTTCCGATCTNNNTCATTGTCTTGATCTGTCTTG

AGCACACGTCTGAACTCTTTCCCTACACGACGCTCTTCCGATCTNNNTCATTGTCTTGATCTGTCTTG

GAGCACACGTCTGAACTCTTTCCCTACACGACGCTCTTCCGATCTNNNGAGCCCAGGACTTTTAGGTT

GAGCACACGTCTGAACTCTTTCCCTACACGACGCTCTTCCGATCTNNNGAGCCCAGGACTTTTAGGTT

AGCACACGTCTGAACTCTTTCCCTACACGACGCTCTTCCGATCTNNNATGGTGTCAGTGCAAAGTCTA

AGCACACGTCTGAACTCTTTCCCTACACGACGCTCTTCCGATCTNNNATGGTGTCAGTGCAAAGTCTA

ACACGTCTGAACTCTTTCCCTACACGACGCTCTTCCGATCTNNNCTCCTGAAATATCAACCCATGCATCG

ACACGTCTGAACTCTTTCCCTACACGACGCTCTTCCGATCTNNNCTCCTGAAATATCAACCCATGCATCG

AGCACACGTCTGAACTCTTTCCCTACACGACGCTCTTCCGATCTNNNCGATTTGAGTCCTGGAAGCTG

AGCACACGTCTGAACTCTTTCCCTACACGACGCTCTTCCGATCTNNNCGATTTGAGTCCTGGAAGCTG

AGCACACGTCTGAACTCTTTCCCTACACGACGCTCTTCCGATCTNNNAAAAAATTAGCTGGCCATGGT

AGCACACGTCTGAACTCTTTCCCTACACGACGCTCTTCCGATCTNNNAAAAAATTAGCTGGCCATGGT

AGCACACGTCTGAACTCTTTCCCTACACGACGCTCTTCCGATCTNNNACGAGAATCACTTGAACATGG

AGCACACGTCTGAACTCTTTCCCTACACGACGCTCTTCCGATCTNNNACGAGAATCACTTGAACATGG

GAGCACACGTCTGAACTCTTTCCCTACACGACGCTCTTCCGATCTNNNGGGGACAAAGAAAGACTCC

GAGCACACGTCTGAACTCTTTCCCTACACGACGCTCTTCCGATCTNNNGGGGACAAAGAAAGACTCC

GAGCACACGTCTGAACTCTTTCCCTACACGACGCTCTTCCGATCTNNNCATATTGGCTAGGCTGGTCT

GAGCACACGTCTGAACTCTTTCCCTACACGACGCTCTTCCGATCTNNNCATATTGGCTAGGCTGGTCT

AGCACACGTCTGAACTCTTTCCCTACACGACGCTCTTCCGATCTNNNGGACGTGTAGGGCCAATATA

AGCACACGTCTGAACTCTTTCCCTACACGACGCTCTTCCGATCTNNNGGACGTGTAGGGCCAATATA

AGCACACGTCTGAACTCTTTCCCTACACGACGCTCTTCCGATCTNNNTAGTGTCTACTGCTTGTTGGG

AGCACACGTCTGAACTCTTTCCCTACACGACGCTCTTCCGATCTNNNTAGTGTCTACTGCTTGTTGGG

GCACACGTCTGAACTCTTTCCCTACACGACGCTCTTCCGATCTNNNCACGTCCGTGAGATGTTACCTTT

GCACACGTCTGAACTCTTTCCCTACACGACGCTCTTCCGATCTNNNCACGTCCGTGAGATGTTACCTTT

AGCACACGTCTGAACTCTTTCCCTACACGACGCTCTTCCGATCTNNNTTGACTCTAGGTAAGCCTCTT

AGCACACGTCTGAACTCTTTCCCTACACGACGCTCTTCCGATCTNNNTTGACTCTAGGTAAGCCTCTT

GCACACGTCTGAACTCTTTCCCTACACGACGCTCTTCCGATCTNNNCACTCGTGACTAAACAGGCACA

GCACACGTCTGAACTCTTTCCCTACACGACGCTCTTCCGATCTNNNCACTCGTGACTAAACAGGCACA

CACACGTCTGAACTCTTTCCCTACACGACGCTCTTCCGATCTNNNCCCGTTACCCCCATTTTTCTTTTCA

CACACGTCTGAACTCTTTCCCTACACGACGCTCTTCCGATCTNNNCCCGTTACCCCCATTTTTCTTTTCA

GAGCACACGTCTGAACTCTTTCCCTACACGACGCTCTTCCGATCTNNNTTTAAATGCTCCCGGACCATC

GAGCACACGTCTGAACTCTTTCCCTACACGACGCTCTTCCGATCTNNNTTTAAATGCTCCCGGACCATC

AGCACACGTCTGAACTCTTTCCCTACACGACGCTCTTCCGATCTNNNCAGTGAGCAGAGGTCATGCC

AGCACACGTCTGAACTCTTTCCCTACACGACGCTCTTCCGATCTNNNCAGTGAGCAGAGGTCATGCC

AGCACACGTCTGAACTCTTTCCCTACACGACGCTCTTCCGATCTNNNGATCTCCATCCCCACTAACACTG

AGCACACGTCTGAACTCTTTCCCTACACGACGCTCTTCCGATCTNNNGATCTCCATCCCCACTAACACTG

CACACGTCTGAACTCTTTCCCTACACGACGCTCTTCCGATCTNNNTACAAAAAATTAGCCGCGTGTGG

CACACGTCTGAACTCTTTCCCTACACGACGCTCTTCCGATCTNNNTACAAAAAATTAGCCGCGTGTGG
GAGCACACGTCTGAACTCTTTCCCTACACGACGCTCTTCCGATCTNNNTACTGGCCAACAGAGTTTGG
GAGCACACGTCTGAACTCTTTCCCTACACGACGCTCTTCCGATCTNNNTACTGGCCAACAGAGTTTGG
AGCACACGTCTGAACTCTTTCCCTACACGACGCTCTTCCGATCTNNNCCAGCTGTCACCTCCTTTTTTTT
AGCACACGTCTGAACTCTTTCCCTACACGACGCTCTTCCGATCTNNNCCAGCTGTCACCTCCTTTTTTTT
AGCACACGTCTGAACTCTTTCCCTACACGACGCTCTTCCGATCTNNNCTTCAGCCTGGATGACAAGAGT
AGCACACGTCTGAACTCTTTCCCTACACGACGCTCTTCCGATCTNNNCTTCAGCCTGGATGACAAGAGT
GAGCACACGTCTGAACTCTTTCCCTACACGACGCTCTTCCGATCTNNNCAGACAATGGCCATGACCAAA
GAGCACACGTCTGAACTCTTTCCCTACACGACGCTCTTCCGATCTNNNCAGACAATGGCCATGACCAAA
GAGCACACGTCTGAACTCTTTCCCTACACGACGCTCTTCCGATCTNNNCTCTTGACCACTCGACTTCCTAC
GAGCACACGTCTGAACTCTTTCCCTACACGACGCTCTTCCGATCTNNNCTCTTGACCACTCGACTTCCTAC
AGCACACGTCTGAACTCTTTCCCTACACGACGCTCTTCCGATCTNNNCTTGCAGTGAGCCAAGATTGT
AGCACACGTCTGAACTCTTTCCCTACACGACGCTCTTCCGATCTNNNCTTGCAGTGAGCCAAGATTGT
AGCACACGTCTGAACTCTTTCCCTACACGACGCTCTTCCGATCTNNNTCCTGTTTTGTTGTTGTTGTTTC
AGCACACGTCTGAACTCTTTCCCTACACGACGCTCTTCCGATCTNNNTCCTGTTTTGTTGTTGTTGTTTC
GAGCACACGTCTGAACTCTTTCCCTACACGACGCTCTTCCGATCTNNNGTACTCCTGAGCGCTCATATTG
GAGCACACGTCTGAACTCTTTCCCTACACGACGCTCTTCCGATCTNNNGTACTCCTGAGCGCTCATATTG
AGAGCACACGTCTGAACTCTTTCCCTACACGACGCTCTTCCGATCTNNNGCAGGAGAATTTCTTGAATC
AGAGCACACGTCTGAACTCTTTCCCTACACGACGCTCTTCCGATCTNNNGCAGGAGAATTTCTTGAATC
GAGCACACGTCTGAACTCTTTCCCTACACGACGCTCTTCCGATCTNNNCTGTGCTCTAAGTCTGGTGTA
GAGCACACGTCTGAACTCTTTCCCTACACGACGCTCTTCCGATCTNNNCTGTGCTCTAAGTCTGGTGTA
GAGCACACGTCTGAACTCTTTCCCTACACGACGCTCTTCCGATCTNNNTGGAAAGAACCACAATGTTCA
GAGCACACGTCTGAACTCTTTCCCTACACGACGCTCTTCCGATCTNNNTGGAAAGAACCACAATGTTCA
GAGCACACGTCTGAACTCTTTCCCTACACGACGCTCTTCCGATCTNNNAGGGAGTAGAGAACAAAGG
GAGCACACGTCTGAACTCTTTCCCTACACGACGCTCTTCCGATCTNNNAGGGAGTAGAGAACAAAGG
GAGCACACGTCTGAACTCTTTCCCTACACGACGCTCTTCCGATCTNNNGAGATGGCATCACTGTACTCC
GAGCACACGTCTGAACTCTTTCCCTACACGACGCTCTTCCGATCTNNNGAGATGGCATCACTGTACTCC
GCACACGTCTGAACTCTTTCCCTACACGACGCTCTTCCGATCTNNNCTACTATGTGCTCCTATGCACCC
GCACACGTCTGAACTCTTTCCCTACACGACGCTCTTCCGATCTNNNCTACTATGTGCTCCTATGCACCC
AGCACACGTCTGAACTCTTTCCCTACACGACGCTCTTCCGATCTNNNTGTTTCTAGGACCCATGTTGAG
AGCACACGTCTGAACTCTTTCCCTACACGACGCTCTTCCGATCTNNNTGTTTCTAGGACCCATGTTGAG
GAGCACACGTCTGAACTCTTTCCCTACACGACGCTCTTCCGATCTNNNGCATCCAACAAGGTAAAATCC
GAGCACACGTCTGAACTCTTTCCCTACACGACGCTCTTCCGATCTNNNGCATCCAACAAGGTAAAATCC
GAGCACACGTCTGAACTCTTTCCCTACACGACGCTCTTCCGATCTNNNTGTCCTCATAGTACAAGTACAG
GAGCACACGTCTGAACTCTTTCCCTACACGACGCTCTTCCGATCTNNNTGTCCTCATAGTACAAGTACAG
AGCACACGTCTGAACTCTTTCCCTACACGACGCTCTTCCGATCTNNNCTGTAGTCCCAGCTACACAGGA
AGCACACGTCTGAACTCTTTCCCTACACGACGCTCTTCCGATCTNNNCTGTAGTCCCAGCTACACAGGA
AGCACACGTCTGAACTCTTTCCCTACACGACGCTCTTCCGATCTNNNAGTTTTCAACCTATTGCTAACTG
AGCACACGTCTGAACTCTTTCCCTACACGACGCTCTTCCGATCTNNNAGTTTTCAACCTATTGCTAACTG
AGCACACGTCTGAACTCTTTCCCTACACGACGCTCTTCCGATCTNNNTGAGAACTCCACCCCTATCACTC
AGCACACGTCTGAACTCTTTCCCTACACGACGCTCTTCCGATCTNNNTGAGAACTCCACCCCTATCACTC
AGAGCACACGTCTGAACTCTTTCCCTACACGACGCTCTTCCGATCTNNNCTGGACAACATGGAGAAAC
AGAGCACACGTCTGAACTCTTTCCCTACACGACGCTCTTCCGATCTNNNCTGGACAACATGGAGAAAC

GCACACGTCTGAACTCTTTCCCTACACGACGCTCTTCCGATCTNNNAGATTTCTGTTTGGGCTTGTGT
GCACACGTCTGAACTCTTTCCCTACACGACGCTCTTCCGATCTNNNAGATTTCTGTTTGGGCTTGTGT
GCACACGTCTGAACTCTTTCCCTACACGACGCTCTTCCGATCTNNNTACTGCAATGAAAAATATTCCT
GCACACGTCTGAACTCTTTCCCTACACGACGCTCTTCCGATCTNNNTACTGCAATGAAAAATATTCCT
ACACGTCTGAACTCTTTCCCTACACGACGCTCTTCCGATCTNNNTTAGTCAGGCTGGTCTTGAACTCC
ACACGTCTGAACTCTTTCCCTACACGACGCTCTTCCGATCTNNNTTAGTCAGGCTGGTCTTGAACTCC
GCACACGTCTGAACTCTTTCCCTACACGACGCTCTTCCGATCTNNNCACACCTGTAGTCTCAGCTACTC
GCACACGTCTGAACTCTTTCCCTACACGACGCTCTTCCGATCTNNNCACACCTGTAGTCTCAGCTACTC
GCACACGTCTGAACTCTTTCCCTACACGACGCTCTTCCGATCTNNNCAAAGGCCAGAGTTTCAGGAAA
GCACACGTCTGAACTCTTTCCCTACACGACGCTCTTCCGATCTNNNCAAAGGCCAGAGTTTCAGGAAA
GCACACGTCTGAACTCTTTCCCTACACGACGCTCTTCCGATCTNNNATTCCAAGCAAAGGAGGAGACA
GCACACGTCTGAACTCTTTCCCTACACGACGCTCTTCCGATCTNNNATTCCAAGCAAAGGAGGAGACA
AGCACACGTCTGAACTCTTTCCCTACACGACGCTCTTCCGATCTNNNCTCGCCTATGGTAGCTTCTCAA
AGCACACGTCTGAACTCTTTCCCTACACGACGCTCTTCCGATCTNNNCTCGCCTATGGTAGCTTCTCAA
GCACACGTCTGAACTCTTTCCCTACACGACGCTCTTCCGATCTNNNAAGACAAAGAAGAAAAACGGC
GCACACGTCTGAACTCTTTCCCTACACGACGCTCTTCCGATCTNNNAAGACAAAGAAGAAAAACGGC
AGCACACGTCTGAACTCTTTCCCTACACGACGCTCTTCCGATCTNNNATACAAAATTACCCAGGCGTG
AGCACACGTCTGAACTCTTTCCCTACACGACGCTCTTCCGATCTNNNATACAAAATTACCCAGGCGTG
GAGCACACGTCTGAACTCTTTCCCTACACGACGCTCTTCCGATCTNNNCTGTAGTCCCAGCTACTCAG
GAGCACACGTCTGAACTCTTTCCCTACACGACGCTCTTCCGATCTNNNCTGTAGTCCCAGCTACTCAG
GCACACGTCTGAACTCTTTCCCTACACGACGCTCTTCCGATCTNNNTACACGTTGAATTGAGGGGGA
GCACACGTCTGAACTCTTTCCCTACACGACGCTCTTCCGATCTNNNTACACGTTGAATTGAGGGGGA
GCACACGTCTGAACTCTTTCCCTACACGACGCTCTTCCGATCTNNNATGGGTTTGGGTTTGATTTGC
GCACACGTCTGAACTCTTTCCCTACACGACGCTCTTCCGATCTNNNATGGGTTTGGGTTTGATTTGC
GAGCACACGTCTGAACTCTTTCCCTACACGACGCTCTTCCGATCTNNNCTGAGGCAGGAGAATCACTT
GAGCACACGTCTGAACTCTTTCCCTACACGACGCTCTTCCGATCTNNNCTGAGGCAGGAGAATCACTT
GAGCACACGTCTGAACTCTTTCCCTACACGACGCTCTTCCGATCTNNNTTGCAATGGTTAAGGTGAA
GAGCACACGTCTGAACTCTTTCCCTACACGACGCTCTTCCGATCTNNNTTGCAATGGTTAAGGTGAA
AGCACACGTCTGAACTCTTTCCCTACACGACGCTCTTCCGATCTNNNCTTCAAGTTTTCCATGATCTTT
AGCACACGTCTGAACTCTTTCCCTACACGACGCTCTTCCGATCTNNNCTTCAAGTTTTCCATGATCTTT
GCACACGTCTGAACTCTTTCCCTACACGACGCTCTTCCGATCTNNNTCACCGGCTGTATTTTCACAAAC
GCACACGTCTGAACTCTTTCCCTACACGACGCTCTTCCGATCTNNNTCACCGGCTGTATTTTCACAAAC
AGAGCACACGTCTGAACTCTTTCCCTACACGACGCTCTTCCGATCTNNNAGGATAAAAGCGAGTGCC
AGAGCACACGTCTGAACTCTTTCCCTACACGACGCTCTTCCGATCTNNNAGGATAAAAGCGAGTGCC
GAGCACACGTCTGAACTCTTTCCCTACACGACGCTCTTCCGATCTNNNTTTACATATCATCTCGTCCAAT
GAGCACACGTCTGAACTCTTTCCCTACACGACGCTCTTCCGATCTNNNTTTACATATCATCTCGTCCAAT
GAGCACACGTCTGAACTCTTTCCCTACACGACGCTCTTCCGATCTNNNTGAAATTATCCCCAAGCCCAC
GAGCACACGTCTGAACTCTTTCCCTACACGACGCTCTTCCGATCTNNNTGAAATTATCCCCAAGCCCAC
CACACGTCTGAACTCTTTCCCTACACGACGCTCTTCCGATCTNNNGAGGCTGAAGCAAGAGAATCTC
CACACGTCTGAACTCTTTCCCTACACGACGCTCTTCCGATCTNNNGAGGCTGAAGCAAGAGAATCTC
AGCACACGTCTGAACTCTTTCCCTACACGACGCTCTTCCGATCTNNNGATCTGCATCCTGGAAACCCTG
AGCACACGTCTGAACTCTTTCCCTACACGACGCTCTTCCGATCTNNNGATCTGCATCCTGGAAACCCTG
GCACACGTCTGAACTCTTTCCCTACACGACGCTCTTCCGATCTNNNCAGAACAGGATTTACCAGTCCC

GCACACGTCTGAACTCTTTCCCTACACGACGCTCTTCCGATCTNNNCAGAACAGGATTTACCAGTCCC
AGCACACGTCTGAACTCTTTCCCTACACGACGCTCTTCCGATCTNNNCACACCTGTAGTCCCACTTACT
AGCACACGTCTGAACTCTTTCCCTACACGACGCTCTTCCGATCTNNNCACACCTGTAGTCCCACTTACT
GAGCACACGTCTGAACTCTTTCCCTACACGACGCTCTTCCGATCTNNNTGCACTTGTAATCTCGGCTAC
GAGCACACGTCTGAACTCTTTCCCTACACGACGCTCTTCCGATCTNNNTGCACTTGTAATCTCGGCTAC
AGCACACGTCTGAACTCTTTCCCTACACGACGCTCTTCCGATCTNNNCAGTGCCTTAGTTGAGACTCA
AGCACACGTCTGAACTCTTTCCCTACACGACGCTCTTCCGATCTNNNCAGTGCCTTAGTTGAGACTCA
GCACACGTCTGAACTCTTTCCCTACACGACGCTCTTCCGATCTNNNCAGAACAGAACCAATAGGGGG
GCACACGTCTGAACTCTTTCCCTACACGACGCTCTTCCGATCTNNNCAGAACAGAACCAATAGGGGG
GCACACGTCTGAACTCTTTCCCTACACGACGCTCTTCCGATCTNNNTAAAAAGGGCCTTGGGATTGT
GCACACGTCTGAACTCTTTCCCTACACGACGCTCTTCCGATCTNNNTAAAAAGGGCCTTGGGATTGT
AGCACACGTCTGAACTCTTTCCCTACACGACGCTCTTCCGATCTNNNGCCTCAGTTTGTCCATCTGTAG
AGCACACGTCTGAACTCTTTCCCTACACGACGCTCTTCCGATCTNNNGCCTCAGTTTGTCCATCTGTAG
GCACACGTCTGAACTCTTTCCCTACACGACGCTCTTCCGATCTNNNTCCCATTATAAGCAACACCATGA
GCACACGTCTGAACTCTTTCCCTACACGACGCTCTTCCGATCTNNNTCCCATTATAAGCAACACCATGA
GCACACGTCTGAACTCTTTCCCTACACGACGCTCTTCCGATCTNNNAAAGCAGCTAGTACTTCTCACC
GCACACGTCTGAACTCTTTCCCTACACGACGCTCTTCCGATCTNNNAAAGCAGCTAGTACTTCTCACC
AGCACACGTCTGAACTCTTTCCCTACACGACGCTCTTCCGATCTNNNGGAGCAGAGAAGTTACAGTG
AGCACACGTCTGAACTCTTTCCCTACACGACGCTCTTCCGATCTNNNGGAGCAGAGAAGTTACAGTG
AGCACACGTCTGAACTCTTTCCCTACACGACGCTCTTCCGATCTNNNTACGTATATTCCCACCACGCATT
AGCACACGTCTGAACTCTTTCCCTACACGACGCTCTTCCGATCTNNNTACGTATATTCCCACCACGCATT
AGCACACGTCTGAACTCTTTCCCTACACGACGCTCTTCCGATCTNNNTACCTGGCACACAGTGTTGATT
AGCACACGTCTGAACTCTTTCCCTACACGACGCTCTTCCGATCTNNNTACCTGGCACACAGTGTTGATT
GCACACGTCTGAACTCTTTCCCTACACGACGCTCTTCCGATCTNNNGGGTAACAGAGCGAGACTTCGC
GCACACGTCTGAACTCTTTCCCTACACGACGCTCTTCCGATCTNNNGGGTAACAGAGCGAGACTTCGC
GAGCACACGTCTGAACTCTTTCCCTACACGACGCTCTTCCGATCTNNNTTTGACCACGACATTTGATTG
GAGCACACGTCTGAACTCTTTCCCTACACGACGCTCTTCCGATCTNNNTTTGACCACGACATTTGATTG
GAGCACACGTCTGAACTCTTTCCCTACACGACGCTCTTCCGATCTNNNTGTGGAATCGTGTTGGAGAA
GAGCACACGTCTGAACTCTTTCCCTACACGACGCTCTTCCGATCTNNNTGTGGAATCGTGTTGGAGAA
GCACACGTCTGAACTCTTTCCCTACACGACGCTCTTCCGATCTNNNAGACAGGAGAATCGCTTGAAC
GCACACGTCTGAACTCTTTCCCTACACGACGCTCTTCCGATCTNNNAGACAGGAGAATCGCTTGAAC
GCACACGTCTGAACTCTTTCCCTACACGACGCTCTTCCGATCTNNNTTCCAGGAACGTAACATTCATCT
GCACACGTCTGAACTCTTTCCCTACACGACGCTCTTCCGATCTNNNTTCCAGGAACGTAACATTCATCT
AGCACACGTCTGAACTCTTTCCCTACACGACGCTCTTCCGATCTNNNGCACACACACACTTTCTAATCTC
AGCACACGTCTGAACTCTTTCCCTACACGACGCTCTTCCGATCTNNNGCACACACACACTTTCTAATCTC
AGCACACGTCTGAACTCTTTCCCTACACGACGCTCTTCCGATCTNNNCCAACATGGTGAAACTGCATC
AGCACACGTCTGAACTCTTTCCCTACACGACGCTCTTCCGATCTNNNCCAACATGGTGAAACTGCATC
GCACACGTCTGAACTCTTTCCCTACACGACGCTCTTCCGATCTNNNAAGTATGTGTGGGGAGACAG
GCACACGTCTGAACTCTTTCCCTACACGACGCTCTTCCGATCTNNNAAGTATGTGTGGGGAGACAG
AGCACACGTCTGAACTCTTTCCCTACACGACGCTCTTCCGATCTNNNACTCACCTCTAAGAGTGGG
AGCACACGTCTGAACTCTTTCCCTACACGACGCTCTTCCGATCTNNNACTCACCTCTAAGAGTGGG
CACACGTCTGAACTCTTTCCCTACACGACGCTCTTCCGATCTNNNCTACTGCCTTTTTTTTTTTTTCTCAT
CACACGTCTGAACTCTTTCCCTACACGACGCTCTTCCGATCTNNNCTACTGCCTTTTTTTTTTTTTCTCAT

AGCACACGTCTGAACTCTTTCCCTACACGACGCTCTTCCGATCTNNNCAACGAATCCAGGAGTTTGTTC
AGCACACGTCTGAACTCTTTCCCTACACGACGCTCTTCCGATCTNNNCAACGAATCCAGGAGTTTGTTC
GCACACGTCTGAACTCTTTCCCTACACGACGCTCTTCCGATCTNNNGGCTAAGTCCTGTCTCTGTAGTG
GCACACGTCTGAACTCTTTCCCTACACGACGCTCTTCCGATCTNNNGGCTAAGTCCTGTCTCTGTAGTG
GCACACGTCTGAACTCTTTCCCTACACGACGCTCTTCCGATCTNNNCAGGGAGTCTGGAAGGAGGTA
GCACACGTCTGAACTCTTTCCCTACACGACGCTCTTCCGATCTNNNCAGGGAGTCTGGAAGGAGGTA
AGCACACGTCTGAACTCTTTCCCTACACGACGCTCTTCCGATCTNNNTGAGGGAGGAGAATCACTTGA
AGCACACGTCTGAACTCTTTCCCTACACGACGCTCTTCCGATCTNNNTGAGGGAGGAGAATCACTTGA
GAGCACACGTCTGAACTCTTTCCCTACACGACGCTCTTCCGATCTNNNTACCCAATGACCTTTGGCAAG
GAGCACACGTCTGAACTCTTTCCCTACACGACGCTCTTCCGATCTNNNTACCCAATGACCTTTGGCAAG
AGCACACGTCTGAACTCTTTCCCTACACGACGCTCTTCCGATCTNNNTCCCTGTCTACTGTTGTGATTG
AGCACACGTCTGAACTCTTTCCCTACACGACGCTCTTCCGATCTNNNTCCCTGTCTACTGTTGTGATTG
GCACACGTCTGAACTCTTTCCCTACACGACGCTCTTCCGATCTNNNCTCGGAGGGTCTCAGTTTCAAA
GCACACGTCTGAACTCTTTCCCTACACGACGCTCTTCCGATCTNNNCTCGGAGGGTCTCAGTTTCAAA
GAGCACACGTCTGAACTCTTTCCCTACACGACGCTCTTCCGATCTNNNATTTGACTCACGTTCCCCACTA
GAGCACACGTCTGAACTCTTTCCCTACACGACGCTCTTCCGATCTNNNATTTGACTCACGTTCCCCACTA
AGCACACGTCTGAACTCTTTCCCTACACGACGCTCTTCCGATCTNNNCCCAGTAGCTACTACAGGCATG
AGCACACGTCTGAACTCTTTCCCTACACGACGCTCTTCCGATCTNNNCCCAGTAGCTACTACAGGCATG
GAGCACACGTCTGAACTCTTTCCCTACACGACGCTCTTCCGATCTNNNTTGGAGATTTGGTTGAGAGA
GAGCACACGTCTGAACTCTTTCCCTACACGACGCTCTTCCGATCTNNNTTGGAGATTTGGTTGAGAGA
GCACACGTCTGAACTCTTTCCCTACACGACGCTCTTCCGATCTNNNATTTACACACACACGCACACAAA
GCACACGTCTGAACTCTTTCCCTACACGACGCTCTTCCGATCTNNNATTTACACACACACGCACACAAA
AGAGCACACGTCTGAACTCTTTCCCTACACGACGCTCTTCCGATCTNNNACATGGCAATATCATGCTTTG
AGAGCACACGTCTGAACTCTTTCCCTACACGACGCTCTTCCGATCTNNNACATGGCAATATCATGCTTTG
GAGCACACGTCTGAACTCTTTCCCTACACGACGCTCTTCCGATCTNNNCCATCAGCTTTGTGCTAGGTG
GAGCACACGTCTGAACTCTTTCCCTACACGACGCTCTTCCGATCTNNNCCATCAGCTTTGTGCTAGGTG
AGCACACGTCTGAACTCTTTCCCTACACGACGCTCTTCCGATCTNNNTCTTTCTGCCACACCTTACTCAG
AGCACACGTCTGAACTCTTTCCCTACACGACGCTCTTCCGATCTNNNTCTTTCTGCCACACCTTACTCAG
GCACACGTCTGAACTCTTTCCCTACACGACGCTCTTCCGATCTNNNCTGAAGCAAAGGCTGGAGCAAT
GCACACGTCTGAACTCTTTCCCTACACGACGCTCTTCCGATCTNNNCTGAAGCAAAGGCTGGAGCAAT
GAGCACACGTCTGAACTCTTTCCCTACACGACGCTCTTCCGATCTNNNTGACACAGAGACATGAAGTA
GAGCACACGTCTGAACTCTTTCCCTACACGACGCTCTTCCGATCTNNNTGACACAGAGACATGAAGTA
GAGCACACGTCTGAACTCTTTCCCTACACGACGCTCTTCCGATCTNNNCAAGAGCATGCTGACAAGTTC
GAGCACACGTCTGAACTCTTTCCCTACACGACGCTCTTCCGATCTNNNCAAGAGCATGCTGACAAGTTC
GCACACGTCTGAACTCTTTCCCTACACGACGCTCTTCCGATCTNNNGCCATTTGCCTACATTTTAGCAG
GCACACGTCTGAACTCTTTCCCTACACGACGCTCTTCCGATCTNNNGCCATTTGCCTACATTTTAGCAG
AGCACACGTCTGAACTCTTTCCCTACACGACGCTCTTCCGATCTNNNTCCTTTTGGTACCTTCCTCAGTG
AGCACACGTCTGAACTCTTTCCCTACACGACGCTCTTCCGATCTNNNTCCTTTTGGTACCTTCCTCAGTG
GAGCACACGTCTGAACTCTTTCCCTACACGACGCTCTTCCGATCTNNNAAAATGTGTTGCAATGGGGG
GAGCACACGTCTGAACTCTTTCCCTACACGACGCTCTTCCGATCTNNNAAAATGTGTTGCAATGGGGG
AGCACACGTCTGAACTCTTTCCCTACACGACGCTCTTCCGATCTNNNGGCTCGAAACTCCATCTCAAAA
AGCACACGTCTGAACTCTTTCCCTACACGACGCTCTTCCGATCTNNNGGCTCGAAACTCCATCTCAAAA
AGCACACGTCTGAACTCTTTCCCTACACGACGCTCTTCCGATCTNNNTTTGGTTTCAGACCACCACTAG

AGCACACGTCTGAACTCTTTCCCTACACGACGCTCTTCCGATCTNNNTTTGGTTTCAGACCACCACTAG
GAGCACACGTCTGAACTCTTTCCCTACACGACGCTCTTCCGATCTNNNGCTGTAGTAAGCCATGTTTG
GAGCACACGTCTGAACTCTTTCCCTACACGACGCTCTTCCGATCTNNNGCTGTAGTAAGCCATGTTTG
AGCACACGTCTGAACTCTTTCCCTACACGACGCTCTTCCGATCTNNNGGAACATCTCTCTCTCCAGTCA
AGCACACGTCTGAACTCTTTCCCTACACGACGCTCTTCCGATCTNNNGGAACATCTCTCTCTCCAGTCA
GAGCACACGTCTGAACTCTTTCCCTACACGACGCTCTTCCGATCTNNNGATCACAAGGTCAGGAGATC
GAGCACACGTCTGAACTCTTTCCCTACACGACGCTCTTCCGATCTNNNGATCACAAGGTCAGGAGATC
GAGCACACGTCTGAACTCTTTCCCTACACGACGCTCTTCCGATCTNNNGCCTGTAATCCCAACACTTTGA
GAGCACACGTCTGAACTCTTTCCCTACACGACGCTCTTCCGATCTNNNGCCTGTAATCCCAACACTTTGA
AGCACACGTCTGAACTCTTTCCCTACACGACGCTCTTCCGATCTNNNGGATTACAGGTATGAGCCACC
AGCACACGTCTGAACTCTTTCCCTACACGACGCTCTTCCGATCTNNNGGATTACAGGTATGAGCCACC
GAGCACACGTCTGAACTCTTTCCCTACACGACGCTCTTCCGATCTNNNGGCCTACCAAAGTTCTGGGAT
GAGCACACGTCTGAACTCTTTCCCTACACGACGCTCTTCCGATCTNNNGGCCTACCAAAGTTCTGGGAT
AGCACACGTCTGAACTCTTTCCCTACACGACGCTCTTCCGATCTNNNACACCTCAGATATATTTCTTCAT
AGCACACGTCTGAACTCTTTCCCTACACGACGCTCTTCCGATCTNNNACACCTCAGATATATTTCTTCAT
GAGCACACGTCTGAACTCTTTCCCTACACGACGCTCTTCCGATCTNNNAAAGACGTTTTATGAGGCTG
GAGCACACGTCTGAACTCTTTCCCTACACGACGCTCTTCCGATCTNNNAAAGACGTTTTATGAGGCTG
GAGCACACGTCTGAACTCTTTCCCTACACGACGCTCTTCCGATCTNNNCTTTGGCAAAATAGAAGGAG
GAGCACACGTCTGAACTCTTTCCCTACACGACGCTCTTCCGATCTNNNCTTTGGCAAAATAGAAGGAG
GAGCACACGTCTGAACTCTTTCCCTACACGACGCTCTTCCGATCTNNNAAGCAGGGGTCAGATCATAC
GAGCACACGTCTGAACTCTTTCCCTACACGACGCTCTTCCGATCTNNNAAGCAGGGGTCAGATCATAC
AGCACACGTCTGAACTCTTTCCCTACACGACGCTCTTCCGATCTNNNTAGAAAACAGAAGAATGGCGG
AGCACACGTCTGAACTCTTTCCCTACACGACGCTCTTCCGATCTNNNTAGAAAACAGAAGAATGGCGG
AGCACACGTCTGAACTCTTTCCCTACACGACGCTCTTCCGATCTNNNACAATTCTGTGATCTCCAGACA
AGCACACGTCTGAACTCTTTCCCTACACGACGCTCTTCCGATCTNNNACAATTCTGTGATCTCCAGACA
GCACACGTCTGAACTCTTTCCCTACACGACGCTCTTCCGATCTNNNGCATAAACTGAAAGGCATGGAC
GCACACGTCTGAACTCTTTCCCTACACGACGCTCTTCCGATCTNNNGCATAAACTGAAAGGCATGGAC
GCACACGTCTGAACTCTTTCCCTACACGACGCTCTTCCGATCTNNNCTGAGGCAGGAGGATTTCTTGAA
GCACACGTCTGAACTCTTTCCCTACACGACGCTCTTCCGATCTNNNCTGAGGCAGGAGGATTTCTTGAA
GAGCACACGTCTGAACTCTTTCCCTACACGACGCTCTTCCGATCTNNNCTCATGCCTATAATCCCAGCAC
GAGCACACGTCTGAACTCTTTCCCTACACGACGCTCTTCCGATCTNNNCTCATGCCTATAATCCCAGCAC
GAGCACACGTCTGAACTCTTTCCCTACACGACGCTCTTCCGATCTNNNCACCTGGCCCCTGCAATATATA
GAGCACACGTCTGAACTCTTTCCCTACACGACGCTCTTCCGATCTNNNCACCTGGCCCCTGCAATATATA
GAGCACACGTCTGAACTCTTTCCCTACACGACGCTCTTCCGATCTNNNAATACAAAAAATTAGCCGGG
GAGCACACGTCTGAACTCTTTCCCTACACGACGCTCTTCCGATCTNNNAATACAAAAAATTAGCCGGG
ACACGTCTGAACTCTTTCCCTACACGACGCTCTTCCGATCTNNNCAACTGTCTTCAAACGGCTTAGGCAT
ACACGTCTGAACTCTTTCCCTACACGACGCTCTTCCGATCTNNNCAACTGTCTTCAAACGGCTTAGGCAT
GAGCACACGTCTGAACTCTTTCCCTACACGACGCTCTTCCGATCTNNNCCTGGGCAACAAGAGTGAAA
GAGCACACGTCTGAACTCTTTCCCTACACGACGCTCTTCCGATCTNNNCCTGGGCAACAAGAGTGAAA
GAGCACACGTCTGAACTCTTTCCCTACACGACGCTCTTCCGATCTNNNGAGTCTGGTGAGTTGGGAAA
GAGCACACGTCTGAACTCTTTCCCTACACGACGCTCTTCCGATCTNNNGAGTCTGGTGAGTTGGGAAA
GAGCACACGTCTGAACTCTTTCCCTACACGACGCTCTTCCGATCTNNNGCAGGAGAATTGCTTGAACCT
GAGCACACGTCTGAACTCTTTCCCTACACGACGCTCTTCCGATCTNNNGCAGGAGAATTGCTTGAACCT

GAGCACACGTCTGAACTCTTTCCCTACACGACGCTCTTCCGATCTNNNGGAGGACCAACTTTCACATG

GAGCACACGTCTGAACTCTTTCCCTACACGACGCTCTTCCGATCTNNNGGAGGACCAACTTTCACATG

GAGCACACGTCTGAACTCTTTCCCTACACGACGCTCTTCCGATCTNNNAGCAAAGAAGGCCAAGAGA

GAGCACACGTCTGAACTCTTTCCCTACACGACGCTCTTCCGATCTNNNAGCAAAGAAGGCCAAGAGA

AGCACACGTCTGAACTCTTTCCCTACACGACGCTCTTCCGATCTNNNGCCAATTTGCAGGAAATATAG

AGCACACGTCTGAACTCTTTCCCTACACGACGCTCTTCCGATCTNNNGCCAATTTGCAGGAAATATAG

GAGCACACGTCTGAACTCTTTCCCTACACGACGCTCTTCCGATCTNNNGAGAGTCCCTGACACATGTC

GAGCACACGTCTGAACTCTTTCCCTACACGACGCTCTTCCGATCTNNNGAGAGTCCCTGACACATGTC

AGCACACGTCTGAACTCTTTCCCTACACGACGCTCTTCCGATCTNNNAGGGTGTCTTCAATTTTTTGG

AGCACACGTCTGAACTCTTTCCCTACACGACGCTCTTCCGATCTNNNAGGGTGTCTTCAATTTTTTGG

AGCACACGTCTGAACTCTTTCCCTACACGACGCTCTTCCGATCTNNNAGAGATTGCAGTGAGCTGAGA

AGCACACGTCTGAACTCTTTCCCTACACGACGCTCTTCCGATCTNNNAGAGATTGCAGTGAGCTGAGA

AGCACACGTCTGAACTCTTTCCCTACACGACGCTCTTCCGATCTNNNTGAGACCAGGAATTCAAGAC

AGCACACGTCTGAACTCTTTCCCTACACGACGCTCTTCCGATCTNNNTGAGACCAGGAATTCAAGAC

GCACACGTCTGAACTCTTTCCCTACACGACGCTCTTCCGATCTNNNAAGTCTGTCCCACATCCTCTCTCC

GCACACGTCTGAACTCTTTCCCTACACGACGCTCTTCCGATCTNNNAAGTCTGTCCCACATCCTCTCTCC

CACACGTCTGAACTCTTTCCCTACACGACGCTCTTCCGATCTNNNCCCCCTAGGAAGTGGATTTGGCAT

CACACGTCTGAACTCTTTCCCTACACGACGCTCTTCCGATCTNNNCCCCCTAGGAAGTGGATTTGGCAT

AGCACACGTCTGAACTCTTTCCCTACACGACGCTCTTCCGATCTNNNGAGTACAGGAATGGAAGGAG

AGCACACGTCTGAACTCTTTCCCTACACGACGCTCTTCCGATCTNNNGAGTACAGGAATGGAAGGAG

GCACACGTCTGAACTCTTTCCCTACACGACGCTCTTCCGATCTNNNTTTCTCAGCACCTCTCTCTTTCCCA

GCACACGTCTGAACTCTTTCCCTACACGACGCTCTTCCGATCTNNNTTTCTCAGCACCTCTCTCTTTCCCA

GCACACGTCTGAACTCTTTCCCTACACGACGCTCTTCCGATCTNNNGGGAATCTGCCCTGTGTCATCCA

GCACACGTCTGAACTCTTTCCCTACACGACGCTCTTCCGATCTNNNGGGAATCTGCCCTGTGTCATCCA

GAGCACACGTCTGAACTCTTTCCCTACACGACGCTCTTCCGATCTNNNCTGAGGCAGGAGAATTGCTT

GAGCACACGTCTGAACTCTTTCCCTACACGACGCTCTTCCGATCTNNNCTGAGGCAGGAGAATTGCTT

AGCACACGTCTGAACTCTTTCCCTACACGACGCTCTTCCGATCTNNNCCCTGCCCTCTTACATTTTCTGA

AGCACACGTCTGAACTCTTTCCCTACACGACGCTCTTCCGATCTNNNCCCTGCCCTCTTACATTTTCTGA

AGAGCACACGTCTGAACTCTTTCCCTACACGACGCTCTTCCGATCTNNNTCAACTCTGACTAGCCACAC

AGAGCACACGTCTGAACTCTTTCCCTACACGACGCTCTTCCGATCTNNNTCAACTCTGACTAGCCACAC

AGCACACGTCTGAACTCTTTCCCTACACGACGCTCTTCCGATCTNNNTTTCTCCAGAAAGGCAGCTCTT

AGCACACGTCTGAACTCTTTCCCTACACGACGCTCTTCCGATCTNNNTTTCTCCAGAAAGGCAGCTCTT

AGCACACGTCTGAACTCTTTCCCTACACGACGCTCTTCCGATCTNNNGACAACAAGGGTGTTACACAT

AGCACACGTCTGAACTCTTTCCCTACACGACGCTCTTCCGATCTNNNGACAACAAGGGTGTTACACAT

AAGAGCACACGTCTGAACTCTTTCCCTACACGACGCTCTTCCGATCTNNNTGAAATACCTAGTGGGGA

AAGAGCACACGTCTGAACTCTTTCCCTACACGACGCTCTTCCGATCTNNNTGAAATACCTAGTGGGGA

GAGCACACGTCTGAACTCTTTCCCTACACGACGCTCTTCCGATCTNNNTCCCTACATGCATCAGTACAT

GAGCACACGTCTGAACTCTTTCCCTACACGACGCTCTTCCGATCTNNNTCCCTACATGCATCAGTACAT

AGCACACGTCTGAACTCTTTCCCTACACGACGCTCTTCCGATCTNNNTCACTATTCCTGGCTTGAGTTCC

AGCACACGTCTGAACTCTTTCCCTACACGACGCTCTTCCGATCTNNNTCACTATTCCTGGCTTGAGTTCC

AGCACACGTCTGAACTCTTTCCCTACACGACGCTCTTCCGATCTNNNATGCCTGTAATCCCAGTTACTC

AGCACACGTCTGAACTCTTTCCCTACACGACGCTCTTCCGATCTNNNATGCCTGTAATCCCAGTTACTC

AGCACACGTCTGAACTCTTTCCCTACACGACGCTCTTCCGATCTNNNAATACAAAATTAGCTGGGCGT

AGCACACGTCTGAACTCTTTCCCTACACGACGCTCTTCCGATCTNNNAATACAAAATTAGCTGGGCGT
AGCACACGTCTGAACTCTTTCCCTACACGACGCTCTTCCGATCTNNNCCCTATCTACCTGGCCATTCAT
AGCACACGTCTGAACTCTTTCCCTACACGACGCTCTTCCGATCTNNNCCCTATCTACCTGGCCATTCAT
GCACACGTCTGAACTCTTTCCCTACACGACGCTCTTCCGATCTNNNGTTGAGCTGAGATCCTGCCATTA
GCACACGTCTGAACTCTTTCCCTACACGACGCTCTTCCGATCTNNNGTTGAGCTGAGATCCTGCCATTA
AGCACACGTCTGAACTCTTTCCCTACACGACGCTCTTCCGATCTNNNCAACATGATGAAACCCCCTCTCT
AGCACACGTCTGAACTCTTTCCCTACACGACGCTCTTCCGATCTNNNCAACATGATGAAACCCCCTCTCT
GAGCACACGTCTGAACTCTTTCCCTACACGACGCTCTTCCGATCTNNNACAAAAATTACCCAGTGTGG
GAGCACACGTCTGAACTCTTTCCCTACACGACGCTCTTCCGATCTNNNACAAAAATTACCCAGTGTGG
ACACGTCTGAACTCTTTCCCTACACGACGCTCTTCCGATCTNNNCAATCCATGTTCATAAAGAGGAACA
ACACGTCTGAACTCTTTCCCTACACGACGCTCTTCCGATCTNNNCAATCCATGTTCATAAAGAGGAACA
CACACGTCTGAACTCTTTCCCTACACGACGCTCTTCCGATCTNNNTGTAATCCCAGCTACTCAGGAGA
CACACGTCTGAACTCTTTCCCTACACGACGCTCTTCCGATCTNNNTGTAATCCCAGCTACTCAGGAGA
AGCACACGTCTGAACTCTTTCCCTACACGACGCTCTTCCGATCTNNNACCTGTAGTCCCAGCTACTCGC
AGCACACGTCTGAACTCTTTCCCTACACGACGCTCTTCCGATCTNNNACCTGTAGTCCCAGCTACTCGC
AGAGCACACGTCTGAACTCTTTCCCTACACGACGCTCTTCCGATCTNNNCCTGGGCAAAAAGAGCAAA
AGAGCACACGTCTGAACTCTTTCCCTACACGACGCTCTTCCGATCTNNNCCTGGGCAAAAAGAGCAAA
AGCACACGTCTGAACTCTTTCCCTACACGACGCTCTTCCGATCTNNNACACACACATACATGCATTTCC
AGCACACGTCTGAACTCTTTCCCTACACGACGCTCTTCCGATCTNNNACACACACATACATGCATTTCC
AGCACACGTCTGAACTCTTTCCCTACACGACGCTCTTCCGATCTNNNGTAATCCCAGCTACTTAGGAG
AGCACACGTCTGAACTCTTTCCCTACACGACGCTCTTCCGATCTNNNGTAATCCCAGCTACTTAGGAG
AGCACACGTCTGAACTCTTTCCCTACACGACGCTCTTCCGATCTNNNCCCAAATTTGAGGACTGTGAC
AGCACACGTCTGAACTCTTTCCCTACACGACGCTCTTCCGATCTNNNCCCAAATTTGAGGACTGTGAC
GCACACGTCTGAACTCTTTCCCTACACGACGCTCTTCCGATCTNNNTTCTGGCTGTTCCTATGGCTAAA
GCACACGTCTGAACTCTTTCCCTACACGACGCTCTTCCGATCTNNNTTCTGGCTGTTCCTATGGCTAAA
AGCACACGTCTGAACTCTTTCCCTACACGACGCTCTTCCGATCTNNNATGGACTGTACCCATTTGCATC
AGCACACGTCTGAACTCTTTCCCTACACGACGCTCTTCCGATCTNNNATGGACTGTACCCATTTGCATC
GAGCACACGTCTGAACTCTTTCCCTACACGACGCTCTTCCGATCTNNNGAATCATCACTTGAACCGGG
GAGCACACGTCTGAACTCTTTCCCTACACGACGCTCTTCCGATCTNNNGAATCATCACTTGAACCGGG
GCACACGTCTGAACTCTTTCCCTACACGACGCTCTTCCGATCTNNNTAGTACTCAAGCAGCCCTTTCAG
GCACACGTCTGAACTCTTTCCCTACACGACGCTCTTCCGATCTNNNTAGTACTCAAGCAGCCCTTTCAG
GAGCACACGTCTGAACTCTTTCCCTACACGACGCTCTTCCGATCTNNNTCATGCCAGTAATCTCAGCAC
GAGCACACGTCTGAACTCTTTCCCTACACGACGCTCTTCCGATCTNNNTCATGCCAGTAATCTCAGCAC
AGCACACGTCTGAACTCTTTCCCTACACGACGCTCTTCCGATCTNNNTCTACAGCAGTGCTCACTTTTC
AGCACACGTCTGAACTCTTTCCCTACACGACGCTCTTCCGATCTNNNTCTACAGCAGTGCTCACTTTTC
GAGCACACGTCTGAACTCTTTCCCTACACGACGCTCTTCCGATCTNNNTAGTTTCGGAATGTGTGTGT
GAGCACACGTCTGAACTCTTTCCCTACACGACGCTCTTCCGATCTNNNTAGTTTCGGAATGTGTGTGT
AGCACACGTCTGAACTCTTTCCCTACACGACGCTCTTCCGATCTNNNATAGACACAATTCTATCCTAAG
AGCACACGTCTGAACTCTTTCCCTACACGACGCTCTTCCGATCTNNNATAGACACAATTCTATCCTAAG
AGCACACGTCTGAACTCTTTCCCTACACGACGCTCTTCCGATCTNNNTCCTCCTGCACATAAAAACATCT
AGCACACGTCTGAACTCTTTCCCTACACGACGCTCTTCCGATCTNNNTCCTCCTGCACATAAAAACATCT
AGCACACGTCTGAACTCTTTCCCTACACGACGCTCTTCCGATCTNNNCAGATGTCCTCTGCTTTTCATGT
AGCACACGTCTGAACTCTTTCCCTACACGACGCTCTTCCGATCTNNNCAGATGTCCTCTGCTTTTCATGT

AGCACACGTCTGAACTCTTTCCCTACACGACGCTCTTCCGATCTNNNAGGATTCAAGTACAGCCTCCTTG
AGCACACGTCTGAACTCTTTCCCTACACGACGCTCTTCCGATCTNNNAGGATTCAAGTACAGCCTCCTTG
GCACACGTCTGAACTCTTTCCCTACACGACGCTCTTCCGATCTNNNCCTCTATTCCTGCTCTCTTCACCCAT
GCACACGTCTGAACTCTTTCCCTACACGACGCTCTTCCGATCTNNNCCTCTATTCCTGCTCTCTTCACCCAT
AGAGCACACGTCTGAACTCTTTCCCTACACGACGCTCTTCCGATCTNNNAGCAGGATGAGAAGTGAAC
AGAGCACACGTCTGAACTCTTTCCCTACACGACGCTCTTCCGATCTNNNAGCAGGATGAGAAGTGAAC
AGCACACGTCTGAACTCTTTCCCTACACGACGCTCTTCCGATCTNNNCAGAGCAGCTGGAACTACAGG
AGCACACGTCTGAACTCTTTCCCTACACGACGCTCTTCCGATCTNNNCAGAGCAGCTGGAACTACAGG
GAGCACACGTCTGAACTCTTTCCCTACACGACGCTCTTCCGATCTNNNGACTTCCTCCTTAGTCAGAGCT
GAGCACACGTCTGAACTCTTTCCCTACACGACGCTCTTCCGATCTNNNGACTTCCTCCTTAGTCAGAGCT
GAGCACACGTCTGAACTCTTTCCCTACACGACGCTCTTCCGATCTNNNAGACAAGGAAACTAATGATA
GAGCACACGTCTGAACTCTTTCCCTACACGACGCTCTTCCGATCTNNNAGACAAGGAAACTAATGATA
AGCACACGTCTGAACTCTTTCCCTACACGACGCTCTTCCGATCTNNNATGGAAGGTAGTCCCATAGGTC
AGCACACGTCTGAACTCTTTCCCTACACGACGCTCTTCCGATCTNNNATGGAAGGTAGTCCCATAGGTC
AGCACACGTCTGAACTCTTTCCCTACACGACGCTCTTCCGATCTNNNTGGCATAGCTGTGATTATGTGAT
AGCACACGTCTGAACTCTTTCCCTACACGACGCTCTTCCGATCTNNNTGGCATAGCTGTGATTATGTGAT
AGCACACGTCTGAACTCTTTCCCTACACGACGCTCTTCCGATCTNNNGAGAGGAATGTAAGGCTTGCCA
AGCACACGTCTGAACTCTTTCCCTACACGACGCTCTTCCGATCTNNNGAGAGGAATGTAAGGCTTGCCA
AGCACACGTCTGAACTCTTTCCCTACACGACGCTCTTCCGATCTNNNCTGAGTAGGTGGGATTACAGGT
AGCACACGTCTGAACTCTTTCCCTACACGACGCTCTTCCGATCTNNNCTGAGTAGGTGGGATTACAGGT
GAGCACACGTCTGAACTCTTTCCCTACACGACGCTCTTCCGATCTNNNCACCTGTAGTCCCAGCTACTTG
GAGCACACGTCTGAACTCTTTCCCTACACGACGCTCTTCCGATCTNNNCACCTGTAGTCCCAGCTACTTG
GAGCACACGTCTGAACTCTTTCCCTACACGACGCTCTTCCGATCTNNNAGCTGTTCTTTGGAGGTTTGA
GAGCACACGTCTGAACTCTTTCCCTACACGACGCTCTTCCGATCTNNNAGCTGTTCTTTGGAGGTTTGA
AGCACACGTCTGAACTCTTTCCCTACACGACGCTCTTCCGATCTNNNACCCTTTAGCAAACAGGTCAGAA
AGCACACGTCTGAACTCTTTCCCTACACGACGCTCTTCCGATCTNNNACCCTTTAGCAAACAGGTCAGAA
AGCACACGTCTGAACTCTTTCCCTACACGACGCTCTTCCGATCTNNNGGACCAGGTGGAGGTAATTGA
AGCACACGTCTGAACTCTTTCCCTACACGACGCTCTTCCGATCTNNNGGACCAGGTGGAGGTAATTGA
AGAGCACACGTCTGAACTCTTTCCCTACACGACGCTCTTCCGATCTNNNTATTCATTCTGGGCTTCCCTT
AGAGCACACGTCTGAACTCTTTCCCTACACGACGCTCTTCCGATCTNNNTATTCATTCTGGGCTTCCCTT
GAGCACACGTCTGAACTCTTTCCCTACACGACGCTCTTCCGATCTNNNAAAAAAAAAATAGCCGCGTGT
GAGCACACGTCTGAACTCTTTCCCTACACGACGCTCTTCCGATCTNNNAAAAAAAAAATAGCCGCGTGT
AGAGCACACGTCTGAACTCTTTCCCTACACGACGCTCTTCCGATCTNNNTGGACTTGGCAGAAGAATC
AGAGCACACGTCTGAACTCTTTCCCTACACGACGCTCTTCCGATCTNNNTGGACTTGGCAGAAGAATC
GAGCACACGTCTGAACTCTTTCCCTACACGACGCTCTTCCGATCTNNNATCTATCTACGTGTGCACCGTA
GAGCACACGTCTGAACTCTTTCCCTACACGACGCTCTTCCGATCTNNNATCTATCTACGTGTGCACCGTA
AGCACACGTCTGAACTCTTTCCCTACACGACGCTCTTCCGATCTNNNCTGGGTTCAGGTGATTTTTCCA
AGCACACGTCTGAACTCTTTCCCTACACGACGCTCTTCCGATCTNNNCTGGGTTCAGGTGATTTTTCCA
AGCACACGTCTGAACTCTTTCCCTACACGACGCTCTTCCGATCTNNNCTGTAATCCCAGCTACTCAGGAG
AGCACACGTCTGAACTCTTTCCCTACACGACGCTCTTCCGATCTNNNCTGTAATCCCAGCTACTCAGGAG
GCACACGTCTGAACTCTTTCCCTACACGACGCTCTTCCGATCTNNNCAATCTCAGTTCACTGCAAGCTCCA
GCACACGTCTGAACTCTTTCCCTACACGACGCTCTTCCGATCTNNNCAATCTCAGTTCACTGCAAGCTCCA
GAGCACACGTCTGAACTCTTTCCCTACACGACGCTCTTCCGATCTNNNAGGGTGATCATAGCTTACTGC

GAGCACACGTCTGAACTCTTTCCCTACACGACGCTCTTCCGATCTNNNAGGGTGATCATAGCTTACTGC
AGCACACGTCTGAACTCTTTCCCTACACGACGCTCTTCCGATCTNNNGGTTGCAGTGAGCTGTGATTG
AGCACACGTCTGAACTCTTTCCCTACACGACGCTCTTCCGATCTNNNGGTTGCAGTGAGCTGTGATTG
AGCACACGTCTGAACTCTTTCCCTACACGACGCTCTTCCGATCTNNNCAGAGAATCGGATTCCCCAGG
AGCACACGTCTGAACTCTTTCCCTACACGACGCTCTTCCGATCTNNNCAGAGAATCGGATTCCCCAGG
AGCACACGTCTGAACTCTTTCCCTACACGACGCTCTTCCGATCTNNNCAGAAGTTGCAGTGAGCTGAAA
AGCACACGTCTGAACTCTTTCCCTACACGACGCTCTTCCGATCTNNNCAGAAGTTGCAGTGAGCTGAAA
GAGCACACGTCTGAACTCTTTCCCTACACGACGCTCTTCCGATCTNNNTTTCTCCCCTGAAAGATGGA
GAGCACACGTCTGAACTCTTTCCCTACACGACGCTCTTCCGATCTNNNTTTCTCCCCTGAAAGATGGA
AGAGCACACGTCTGAACTCTTTCCCTACACGACGCTCTTCCGATCTNNNGAGTTTGAGACCAGCCTAGA
AGAGCACACGTCTGAACTCTTTCCCTACACGACGCTCTTCCGATCTNNNGAGTTTGAGACCAGCCTAGA
AGCACACGTCTGAACTCTTTCCCTACACGACGCTCTTCCGATCTNNNGAGAATCGCTTGATCCTGGGA
AGCACACGTCTGAACTCTTTCCCTACACGACGCTCTTCCGATCTNNNGAGAATCGCTTGATCCTGGGA
GAGCACACGTCTGAACTCTTTCCCTACACGACGCTCTTCCGATCTNNNCACTTCTCCCCAACTTTCTCTGT
GAGCACACGTCTGAACTCTTTCCCTACACGACGCTCTTCCGATCTNNNCACTTCTCCCCAACTTTCTCTGT
AGAGCACACGTCTGAACTCTTTCCCTACACGACGCTCTTCCGATCTNNNAGTGCTAGGATTACAGGCAT
AGAGCACACGTCTGAACTCTTTCCCTACACGACGCTCTTCCGATCTNNNAGTGCTAGGATTACAGGCAT
AGAGCACACGTCTGAACTCTTTCCCTACACGACGCTCTTCCGATCTNNNGAGACCCGTCAGAGCTCTCA
AGAGCACACGTCTGAACTCTTTCCCTACACGACGCTCTTCCGATCTNNNGAGACCCGTCAGAGCTCTCA
GAGCACACGTCTGAACTCTTTCCCTACACGACGCTCTTCCGATCTNNNAATTTCTTCCACCAGGCAACCT
GAGCACACGTCTGAACTCTTTCCCTACACGACGCTCTTCCGATCTNNNAATTTCTTCCACCAGGCAACCT
GAGCACACGTCTGAACTCTTTCCCTACACGACGCTCTTCCGATCTNNNTGAGGCAGGAGAATCTCTTGA
GAGCACACGTCTGAACTCTTTCCCTACACGACGCTCTTCCGATCTNNNTGAGGCAGGAGAATCTCTTGA
AGCACACGTCTGAACTCTTTCCCTACACGACGCTCTTCCGATCTNNNCCAATCAAAAGTCAACATGGCGA
AGCACACGTCTGAACTCTTTCCCTACACGACGCTCTTCCGATCTNNNCCAATCAAAAGTCAACATGGCGA
GAGCACACGTCTGAACTCTTTCCCTACACGACGCTCTTCCGATCTNNNTACAGATGTGCACCACTATGCT
GAGCACACGTCTGAACTCTTTCCCTACACGACGCTCTTCCGATCTNNNTACAGATGTGCACCACTATGCT
AGAGCACACGTCTGAACTCTTTCCCTACACGACGCTCTTCCGATCTNNNAGGTAGGCATTGGTATGTG
AGAGCACACGTCTGAACTCTTTCCCTACACGACGCTCTTCCGATCTNNNAGGTAGGCATTGGTATGTG
AGCACACGTCTGAACTCTTTCCCTACACGACGCTCTTCCGATCTNNNGGGTGGAGCTAAAGACAGATG
AGCACACGTCTGAACTCTTTCCCTACACGACGCTCTTCCGATCTNNNGGGTGGAGCTAAAGACAGATG
GAGCACACGTCTGAACTCTTTCCCTACACGACGCTCTTCCGATCTNNNCTGAGGCAGGAGAATCACTTG
GAGCACACGTCTGAACTCTTTCCCTACACGACGCTCTTCCGATCTNNNCTGAGGCAGGAGAATCACTTG
GAGCACACGTCTGAACTCTTTCCCTACACGACGCTCTTCCGATCTNNNTGAGGTAGGAGAATCGCTTGA
GAGCACACGTCTGAACTCTTTCCCTACACGACGCTCTTCCGATCTNNNTGAGGTAGGAGAATCGCTTGA
AGCACACGTCTGAACTCTTTCCCTACACGACGCTCTTCCGATCTNNNAAAACCTAAAACCGTGCAATGCT
AGCACACGTCTGAACTCTTTCCCTACACGACGCTCTTCCGATCTNNNAAAACCTAAAACCGTGCAATGCT
AGCACACGTCTGAACTCTTTCCCTACACGACGCTCTTCCGATCTNNNTATGATGTCTGTGTGGAAAGC
AGCACACGTCTGAACTCTTTCCCTACACGACGCTCTTCCGATCTNNNTATGATGTCTGTGTGGAAAGC
GAGCACACGTCTGAACTCTTTCCCTACACGACGCTCTTCCGATCTNNNGTAAGTCCGCAGTCAACATCA
GAGCACACGTCTGAACTCTTTCCCTACACGACGCTCTTCCGATCTNNNGTAAGTCCGCAGTCAACATCA
AGCACACGTCTGAACTCTTTCCCTACACGACGCTCTTCCGATCTNNNTAAAACCCTCAGACTCCACATCC
AGCACACGTCTGAACTCTTTCCCTACACGACGCTCTTCCGATCTNNNTAAAACCCTCAGACTCCACATCC

AGCACACGTCTGAACTCTTTCCCTACACGACGCTCTTCCGATCTNNNAATAAATGTGATTTGCCTTCTAG
AGCACACGTCTGAACTCTTTCCCTACACGACGCTCTTCCGATCTNNNAATAAATGTGATTTGCCTTCTAG
AGCACACGTCTGAACTCTTTCCCTACACGACGCTCTTCCGATCTNNNGTTGGTCCTTGAAATAAAGGCC
AGCACACGTCTGAACTCTTTCCCTACACGACGCTCTTCCGATCTNNNGTTGGTCCTTGAAATAAAGGCC
AGCACACGTCTGAACTCTTTCCCTACACGACGCTCTTCCGATCTNNNCTGGGTGCAAACAGTTGAGAA
AGCACACGTCTGAACTCTTTCCCTACACGACGCTCTTCCGATCTNNNCTGGGTGCAAACAGTTGAGAA
GAGCACACGTCTGAACTCTTTCCCTACACGACGCTCTTCCGATCTNNNTGGGTGTGTGTGTGTATATGT
GAGCACACGTCTGAACTCTTTCCCTACACGACGCTCTTCCGATCTNNNTGGGTGTGTGTGTGTATATGT
CACACGTCTGAACTCTTTCCCTACACGACGCTCTTCCGATCTNNNGCCGTTAGGAGTATTTGAATCATC
CACACGTCTGAACTCTTTCCCTACACGACGCTCTTCCGATCTNNNGCCGTTAGGAGTATTTGAATCATC
AGCACACGTCTGAACTCTTTCCCTACACGACGCTCTTCCGATCTNNNTGCCAAGAAATAAGGTCCTCTC
AGCACACGTCTGAACTCTTTCCCTACACGACGCTCTTCCGATCTNNNTGCCAAGAAATAAGGTCCTCTC
AGCACACGTCTGAACTCTTTCCCTACACGACGCTCTTCCGATCTNNNTCTTGAAATCCTGAGCTCGTGAT
AGCACACGTCTGAACTCTTTCCCTACACGACGCTCTTCCGATCTNNNTCTTGAAATCCTGAGCTCGTGAT
AGAGCACACGTCTGAACTCTTTCCCTACACGACGCTCTTCCGATCTNNNATGGCTAATTCTAGAACTGG
AGAGCACACGTCTGAACTCTTTCCCTACACGACGCTCTTCCGATCTNNNATGGCTAATTCTAGAACTGG
AGCACACGTCTGAACTCTTTCCCTACACGACGCTCTTCCGATCTNNNCTGGTGACGATGTGAAGAAAA
AGCACACGTCTGAACTCTTTCCCTACACGACGCTCTTCCGATCTNNNCTGGTGACGATGTGAAGAAAA
AGCACACGTCTGAACTCTTTCCCTACACGACGCTCTTCCGATCTNNNACTGGAATATCTGCATGCAGGA
AGCACACGTCTGAACTCTTTCCCTACACGACGCTCTTCCGATCTNNNACTGGAATATCTGCATGCAGGA
AGAGCACACGTCTGAACTCTTTCCCTACACGACGCTCTTCCGATCTNNNGAGGGATTTAGAAGCTCCAG
AGAGCACACGTCTGAACTCTTTCCCTACACGACGCTCTTCCGATCTNNNGAGGGATTTAGAAGCTCCAG
GAGCACACGTCTGAACTCTTTCCCTACACGACGCTCTTCCGATCTNNNAGAATTGCTTGAACCTGGGAG
GAGCACACGTCTGAACTCTTTCCCTACACGACGCTCTTCCGATCTNNNAGAATTGCTTGAACCTGGGAG
GAGCACACGTCTGAACTCTTTCCCTACACGACGCTCTTCCGATCTNNNCCTTTTCATCATGCCCCATGTAC
GAGCACACGTCTGAACTCTTTCCCTACACGACGCTCTTCCGATCTNNNCCTTTTCATCATGCCCCATGTAC
AGAGCACACGTCTGAACTCTTTCCCTACACGACGCTCTTCCGATCTNNNGAGGTTGCAGTGAGCTGAGA
AGAGCACACGTCTGAACTCTTTCCCTACACGACGCTCTTCCGATCTNNNGAGGTTGCAGTGAGCTGAGA
ACACGTCTGAACTCTTTCCCTACACGACGCTCTTCCGATCTNNNCTTTTCTCTGTGCACTCGCTCCATCGC
ACACGTCTGAACTCTTTCCCTACACGACGCTCTTCCGATCTNNNCTTTTCTCTGTGCACTCGCTCCATCGC
GAGCACACGTCTGAACTCTTTCCCTACACGACGCTCTTCCGATCTNNNGTAGACTCACACAAAGCAACA
GAGCACACGTCTGAACTCTTTCCCTACACGACGCTCTTCCGATCTNNNGTAGACTCACACAAAGCAACA
GAGCACACGTCTGAACTCTTTCCCTACACGACGCTCTTCCGATCTNNNACCCTGTACCCCTAAATATTTCA
GAGCACACGTCTGAACTCTTTCCCTACACGACGCTCTTCCGATCTNNNACCCTGTACCCCTAAATATTTCA
AGCACACGTCTGAACTCTTTCCCTACACGACGCTCTTCCGATCTNNNCCACTGCACCTGAGAAAGATAC
AGCACACGTCTGAACTCTTTCCCTACACGACGCTCTTCCGATCTNNNCCACTGCACCTGAGAAAGATAC
AGCACACGTCTGAACTCTTTCCCTACACGACGCTCTTCCGATCTNNNCTAGATAACTTGGCATCCTGGCT
AGCACACGTCTGAACTCTTTCCCTACACGACGCTCTTCCGATCTNNNCTAGATAACTTGGCATCCTGGCT
GAGCACACGTCTGAACTCTTTCCCTACACGACGCTCTTCCGATCTNNNCCCATAGGATTTCTTGTTGCTC
GAGCACACGTCTGAACTCTTTCCCTACACGACGCTCTTCCGATCTNNNCCCATAGGATTTCTTGTTGCTC
GAGCACACGTCTGAACTCTTTCCCTACACGACGCTCTTCCGATCTNNNTTCCGTAGCAATTGCACCATTT
GAGCACACGTCTGAACTCTTTCCCTACACGACGCTCTTCCGATCTNNNTTCCGTAGCAATTGCACCATTT
GAGCACACGTCTGAACTCTTTCCCTACACGACGCTCTTCCGATCTNNNATGGAAAGACGGGTTTCATG

GAGCACACGTCTGAACTCTTTCCCTACACGACGCTCTTCCGATCTNNNATGGAAAGACGGGTTTCATG
GAGCACACGTCTGAACTCTTTCCCTACACGACGCTCTTCCGATCTNNNTTGCACAGCTGGAGCAAGAT
GAGCACACGTCTGAACTCTTTCCCTACACGACGCTCTTCCGATCTNNNTTGCACAGCTGGAGCAAGAT
GAGCACACGTCTGAACTCTTTCCCTACACGACGCTCTTCCGATCTNNNTGGTTTTCCATTCCTGAGTTA
GAGCACACGTCTGAACTCTTTCCCTACACGACGCTCTTCCGATCTNNNTGGTTTTCCATTCCTGAGTTA
GCACACGTCTGAACTCTTTCCCTACACGACGCTCTTCCGATCTNNNCAGTGGCCACTTCAGGAAAAAT
GCACACGTCTGAACTCTTTCCCTACACGACGCTCTTCCGATCTNNNCAGTGGCCACTTCAGGAAAAAT
AGCACACGTCTGAACTCTTTCCCTACACGACGCTCTTCCGATCTNNNCTGCCAAGTTGACTAAAAGGGT
AGCACACGTCTGAACTCTTTCCCTACACGACGCTCTTCCGATCTNNNCTGCCAAGTTGACTAAAAGGGT
AGAGCACACGTCTGAACTCTTTCCCTACACGACGCTCTTCCGATCTNNNCAAGGAGGCATACATCTTGC
AGAGCACACGTCTGAACTCTTTCCCTACACGACGCTCTTCCGATCTNNNCAAGGAGGCATACATCTTGC
AGCACACGTCTGAACTCTTTCCCTACACGACGCTCTTCCGATCTNNNAATCAGAGCCACTGGATTGGA
AGCACACGTCTGAACTCTTTCCCTACACGACGCTCTTCCGATCTNNNAATCAGAGCCACTGGATTGGA
AGCACACGTCTGAACTCTTTCCCTACACGACGCTCTTCCGATCTNNNTTATACACAAAGCAGCCCCTAC
AGCACACGTCTGAACTCTTTCCCTACACGACGCTCTTCCGATCTNNNTTATACACAAAGCAGCCCCTAC
AGCACACGTCTGAACTCTTTCCCTACACGACGCTCTTCCGATCTNNNTCTTGCAGAGATCTTTCACCTC
AGCACACGTCTGAACTCTTTCCCTACACGACGCTCTTCCGATCTNNNTCTTGCAGAGATCTTTCACCTC
GAGCACACGTCTGAACTCTTTCCCTACACGACGCTCTTCCGATCTNNNAACAGAGCTCGCTTAGATGG
GAGCACACGTCTGAACTCTTTCCCTACACGACGCTCTTCCGATCTNNNAACAGAGCTCGCTTAGATGG
GAGCACACGTCTGAACTCTTTCCCTACACGACGCTCTTCCGATCTNNNCTGAGTAGCTGGAACCATAGG
GAGCACACGTCTGAACTCTTTCCCTACACGACGCTCTTCCGATCTNNNCTGAGTAGCTGGAACCATAGG
GAGCACACGTCTGAACTCTTTCCCTACACGACGCTCTTCCGATCTNNNCAATCTGGAGAAGAATTTTGA
GAGCACACGTCTGAACTCTTTCCCTACACGACGCTCTTCCGATCTNNNCAATCTGGAGAAGAATTTTGA
AGAGCACACGTCTGAACTCTTTCCCTACACGACGCTCTTCCGATCTNNNGAGAATCACTTGAACCCAGG
AGAGCACACGTCTGAACTCTTTCCCTACACGACGCTCTTCCGATCTNNNGAGAATCACTTGAACCCAGG
AGCACACGTCTGAACTCTTTCCCTACACGACGCTCTTCCGATCTNNNATCACTTGAAGCCAGGAGTTTG
AGCACACGTCTGAACTCTTTCCCTACACGACGCTCTTCCGATCTNNNATCACTTGAAGCCAGGAGTTTG
GAGCACACGTCTGAACTCTTTCCCTACACGACGCTCTTCCGATCTNNNACTGCTTACTGTTCATGTAGC
GAGCACACGTCTGAACTCTTTCCCTACACGACGCTCTTCCGATCTNNNACTGCTTACTGTTCATGTAGC
GAGCACACGTCTGAACTCTTTCCCTACACGACGCTCTTCCGATCTNNNTCCTTCCTGAGTAGCTGGAAC
GAGCACACGTCTGAACTCTTTCCCTACACGACGCTCTTCCGATCTNNNTCCTTCCTGAGTAGCTGGAAC
AGCACACGTCTGAACTCTTTCCCTACACGACGCTCTTCCGATCTNNNTCATTGCCTCAGAGGAACTTTG
AGCACACGTCTGAACTCTTTCCCTACACGACGCTCTTCCGATCTNNNTCATTGCCTCAGAGGAACTTTG
CACACGTCTGAACTCTTTCCCTACACGACGCTCTTCCGATCTNNNGAGGGTGGGCTTTCAGAACTAGC
CACACGTCTGAACTCTTTCCCTACACGACGCTCTTCCGATCTNNNGAGGGTGGGCTTTCAGAACTAGC
GAGCACACGTCTGAACTCTTTCCCTACACGACGCTCTTCCGATCTNNNGGGCCTAACTCCCATTTTCTCT
GAGCACACGTCTGAACTCTTTCCCTACACGACGCTCTTCCGATCTNNNGGGCCTAACTCCCATTTTCTCT
GAGCACACGTCTGAACTCTTTCCCTACACGACGCTCTTCCGATCTNNNTAGTAGCCCAGTCTGGTCTTT
GAGCACACGTCTGAACTCTTTCCCTACACGACGCTCTTCCGATCTNNNTAGTAGCCCAGTCTGGTCTTT
GAGCACACGTCTGAACTCTTTCCCTACACGACGCTCTTCCGATCTNNNTCACTACTTGGGAGTCTGCTG
GAGCACACGTCTGAACTCTTTCCCTACACGACGCTCTTCCGATCTNNNTCACTACTTGGGAGTCTGCTG
AGAGCACACGTCTGAACTCTTTCCCTACACGACGCTCTTCCGATCTNNNGCCTACCTACCTCCTCAGGAT
AGAGCACACGTCTGAACTCTTTCCCTACACGACGCTCTTCCGATCTNNNGCCTACCTACCTCCTCAGGAT

AGCACACGTCTGAACTCTTTCCCTACACGACGCTCTTCCGATCTNNNCCAATTCATACTGGGGACTCCA
AGCACACGTCTGAACTCTTTCCCTACACGACGCTCTTCCGATCTNNNCCAATTCATACTGGGGACTCCA
AGCACACGTCTGAACTCTTTCCCTACACGACGCTCTTCCGATCTNNNTTCAAGCGATTCTCCTGCCTTAG
AGCACACGTCTGAACTCTTTCCCTACACGACGCTCTTCCGATCTNNNTTCAAGCGATTCTCCTGCCTTAG
AGAGCACACGTCTGAACTCTTTCCCTACACGACGCTCTTCCGATCTNNNAAAAATTAGCCTGGCATGGT
AGAGCACACGTCTGAACTCTTTCCCTACACGACGCTCTTCCGATCTNNNAAAAATTAGCCTGGCATGGT
GAGCACACGTCTGAACTCTTTCCCTACACGACGCTCTTCCGATCTNNNGAATGAGCTTGGAGTTAGTC
GAGCACACGTCTGAACTCTTTCCCTACACGACGCTCTTCCGATCTNNNGAATGAGCTTGGAGTTAGTC
AGCACACGTCTGAACTCTTTCCCTACACGACGCTCTTCCGATCTNNNGTCACCTGTAATCCCAGCTACT
AGCACACGTCTGAACTCTTTCCCTACACGACGCTCTTCCGATCTNNNGTCACCTGTAATCCCAGCTACT
GCACACGTCTGAACTCTTTCCCTACACGACGCTCTTCCGATCTNNNTTTGTCCTCAGAAGACACGGTC
GCACACGTCTGAACTCTTTCCCTACACGACGCTCTTCCGATCTNNNTTTGTCCTCAGAAGACACGGTC
AGCACACGTCTGAACTCTTTCCCTACACGACGCTCTTCCGATCTNNNTCTTCATCGCTACTGTGAAAGCT
AGCACACGTCTGAACTCTTTCCCTACACGACGCTCTTCCGATCTNNNTCTTCATCGCTACTGTGAAAGCT
AGCACACGTCTGAACTCTTTCCCTACACGACGCTCTTCCGATCTNNNCCAGCATTCAGTCAGAAACACC
AGCACACGTCTGAACTCTTTCCCTACACGACGCTCTTCCGATCTNNNCCAGCATTCAGTCAGAAACACC
AGCACACGTCTGAACTCTTTCCCTACACGACGCTCTTCCGATCTNNNTTGAATTCCTGGGCTCATGTGA
AGCACACGTCTGAACTCTTTCCCTACACGACGCTCTTCCGATCTNNNTTGAATTCCTGGGCTCATGTGA
ACACGTCTGAACTCTTTCCCTACACGACGCTCTTCCGATCTNNNAAATCCCTGTTGCTGTGTGTTCTCAT
ACACGTCTGAACTCTTTCCCTACACGACGCTCTTCCGATCTNNNAAATCCCTGTTGCTGTGTGTTCTCAT
GAGCACACGTCTGAACTCTTTCCCTACACGACGCTCTTCCGATCTNNNTTCGAGACCAACCTGACCAAC
GAGCACACGTCTGAACTCTTTCCCTACACGACGCTCTTCCGATCTNNNTTCGAGACCAACCTGACCAAC
GAGCACACGTCTGAACTCTTTCCCTACACGACGCTCTTCCGATCTNNNGAGGTTGCAGTGAGCCAAGA
GAGCACACGTCTGAACTCTTTCCCTACACGACGCTCTTCCGATCTNNNGAGGTTGCAGTGAGCCAAGA
AGCACACGTCTGAACTCTTTCCCTACACGACGCTCTTCCGATCTNNNATGTCATTTAGCGTCTTGGAGG
AGCACACGTCTGAACTCTTTCCCTACACGACGCTCTTCCGATCTNNNATGTCATTTAGCGTCTTGGAGG
GAGCACACGTCTGAACTCTTTCCCTACACGACGCTCTTCCGATCTNNNCATGGGATGTGTTTGGTTGGT
GAGCACACGTCTGAACTCTTTCCCTACACGACGCTCTTCCGATCTNNNCATGGGATGTGTTTGGTTGGT
GAGCACACGTCTGAACTCTTTCCCTACACGACGCTCTTCCGATCTNNNAAGTTGACGCTTGTTTGCTTAC
GAGCACACGTCTGAACTCTTTCCCTACACGACGCTCTTCCGATCTNNNAAGTTGACGCTTGTTTGCTTAC
GCACACGTCTGAACTCTTTCCCTACACGACGCTCTTCCGATCTNNNGCTGTGATCCAGTAGAAGGTAC
GCACACGTCTGAACTCTTTCCCTACACGACGCTCTTCCGATCTNNNGCTGTGATCCAGTAGAAGGTAC
CACACGTCTGAACTCTTTCCCTACACGACGCTCTTCCGATCTNNNCTGCAGATACCGGATGGAGGCAT
CACACGTCTGAACTCTTTCCCTACACGACGCTCTTCCGATCTNNNCTGCAGATACCGGATGGAGGCAT
AGCACACGTCTGAACTCTTTCCCTACACGACGCTCTTCCGATCTNNNCTGTAGTCCCAGCTACTCAGGA
AGCACACGTCTGAACTCTTTCCCTACACGACGCTCTTCCGATCTNNNCTGTAGTCCCAGCTACTCAGGA
GAGCACACGTCTGAACTCTTTCCCTACACGACGCTCTTCCGATCTNNNTACGTGAACAAAGTCAAGGCA
GAGCACACGTCTGAACTCTTTCCCTACACGACGCTCTTCCGATCTNNNTACGTGAACAAAGTCAAGGCA
GAGCACACGTCTGAACTCTTTCCCTACACGACGCTCTTCCGATCTNNNTGAGAGAGGAGAATGGCATC
GAGCACACGTCTGAACTCTTTCCCTACACGACGCTCTTCCGATCTNNNTGAGAGAGGAGAATGGCATC
GAGCACACGTCTGAACTCTTTCCCTACACGACGCTCTTCCGATCTNNNGGCAAGAGAGTGAGACTCCAT
GAGCACACGTCTGAACTCTTTCCCTACACGACGCTCTTCCGATCTNNNGGCAAGAGAGTGAGACTCCAT
GAGCACACGTCTGAACTCTTTCCCTACACGACGCTCTTCCGATCTNNNTAGTATTAGACACTGATGGAT

GAGCACACGTCTGAACTCTTTCCCTACACGACGCTCTTCCGATCTNNNTAGTATTAGACACTGATGGAT
GAGCACACGTCTGAACTCTTTCCCTACACGACGCTCTTCCGATCTNNNTCACAAGGTCAAGAGATGGA
GAGCACACGTCTGAACTCTTTCCCTACACGACGCTCTTCCGATCTNNNTCACAAGGTCAAGAGATGGA
AGAGCACACGTCTGAACTCTTTCCCTACACGACGCTCTTCCGATCTNNNGAGAATCTCTTGAACCTGGC
AGAGCACACGTCTGAACTCTTTCCCTACACGACGCTCTTCCGATCTNNNGAGAATCTCTTGAACCTGGC
GAGCACACGTCTGAACTCTTTCCCTACACGACGCTCTTCCGATCTNNNTTGTATCCCCATTGCTGGTGTT
GAGCACACGTCTGAACTCTTTCCCTACACGACGCTCTTCCGATCTNNNTTGTATCCCCATTGCTGGTGTT
AGCACACGTCTGAACTCTTTCCCTACACGACGCTCTTCCGATCTNNNGTTTTCACTTAGGGCCAAGGTT
AGCACACGTCTGAACTCTTTCCCTACACGACGCTCTTCCGATCTNNNGTTTTCACTTAGGGCCAAGGTT
AGCACACGTCTGAACTCTTTCCCTACACGACGCTCTTCCGATCTNNNGAGCAAGACTCCGTCTCAAAAA
AGCACACGTCTGAACTCTTTCCCTACACGACGCTCTTCCGATCTNNNGAGCAAGACTCCGTCTCAAAAA
AGAGCACACGTCTGAACTCTTTCCCTACACGACGCTCTTCCGATCTNNNTCTTGGTTTTGATGGCTGTT
AGAGCACACGTCTGAACTCTTTCCCTACACGACGCTCTTCCGATCTNNNTCTTGGTTTTGATGGCTGTT
AGCACACGTCTGAACTCTTTCCCTACACGACGCTCTTCCGATCTNNNTCTACATCCCCTCAAACACTTGGC
AGCACACGTCTGAACTCTTTCCCTACACGACGCTCTTCCGATCTNNNTCTACATCCCCTCAAACACTTGGC
GCACACGTCTGAACTCTTTCCCTACACGACGCTCTTCCGATCTNNNCTTCTGAAAGCTTTGCCTTCACAC
GCACACGTCTGAACTCTTTCCCTACACGACGCTCTTCCGATCTNNNCTTCTGAAAGCTTTGCCTTCACAC
AGCACACGTCTGAACTCTTTCCCTACACGACGCTCTTCCGATCTNNNCAAAAGTGAACAGAGATGCAG
AGCACACGTCTGAACTCTTTCCCTACACGACGCTCTTCCGATCTNNNCAAAAGTGAACAGAGATGCAG
AGCACACGTCTGAACTCTTTCCCTACACGACGCTCTTCCGATCTNNNTCCTTCTTCTCCTTCTCCTTCCTCA
AGCACACGTCTGAACTCTTTCCCTACACGACGCTCTTCCGATCTNNNTCCTTCTTCTCCTTCTCCTTCCTCA
AGCACACGTCTGAACTCTTTCCCTACACGACGCTCTTCCGATCTNNNTCATCCTGTTCCTCTTTGCTGAA
AGCACACGTCTGAACTCTTTCCCTACACGACGCTCTTCCGATCTNNNTCATCCTGTTCCTCTTTGCTGAA
GAGCACACGTCTGAACTCTTTCCCTACACGACGCTCTTCCGATCTNNNACCCCTGGCTTGAAAATTTGG
GAGCACACGTCTGAACTCTTTCCCTACACGACGCTCTTCCGATCTNNNACCCCTGGCTTGAAAATTTGG
GAGCACACGTCTGAACTCTTTCCCTACACGACGCTCTTCCGATCTNNNTTTTTCAAGAACCGATGCAG
GAGCACACGTCTGAACTCTTTCCCTACACGACGCTCTTCCGATCTNNNTTTTTCAAGAACCGATGCAG
AGCACACGTCTGAACTCTTTCCCTACACGACGCTCTTCCGATCTNNNGTGGGTCCCTTTCTAAAATGTG
AGCACACGTCTGAACTCTTTCCCTACACGACGCTCTTCCGATCTNNNGTGGGTCCCTTTCTAAAATGTG
AGCACACGTCTGAACTCTTTCCCTACACGACGCTCTTCCGATCTNNNCACTGGGGTGGGAAACAATACT
AGCACACGTCTGAACTCTTTCCCTACACGACGCTCTTCCGATCTNNNCACTGGGGTGGGAAACAATACT
AGCACACGTCTGAACTCTTTCCCTACACGACGCTCTTCCGATCTNNNATGCCTGCAATCCTAGCTACTCC
AGCACACGTCTGAACTCTTTCCCTACACGACGCTCTTCCGATCTNNNATGCCTGCAATCCTAGCTACTCC
AGCACACGTCTGAACTCTTTCCCTACACGACGCTCTTCCGATCTNNNCTGAGACACATCCTCCACTAAGG
AGCACACGTCTGAACTCTTTCCCTACACGACGCTCTTCCGATCTNNNCTGAGACACATCCTCCACTAAGG
GAGCACACGTCTGAACTCTTTCCCTACACGACGCTCTTCCGATCTNNNGCAGGAGAATTGCTTGAACTC
GAGCACACGTCTGAACTCTTTCCCTACACGACGCTCTTCCGATCTNNNGCAGGAGAATTGCTTGAACTC
GAGCACACGTCTGAACTCTTTCCCTACACGACGCTCTTCCGATCTNNNTTGTAGGTCCCGTTCATGAAA
GAGCACACGTCTGAACTCTTTCCCTACACGACGCTCTTCCGATCTNNNTTGTAGGTCCCGTTCATGAAA
AGCACACGTCTGAACTCTTTCCCTACACGACGCTCTTCCGATCTNNNGAGTGAGCTCATGGCAACTCTA
AGCACACGTCTGAACTCTTTCCCTACACGACGCTCTTCCGATCTNNNGAGTGAGCTCATGGCAACTCTA
AGCACACGTCTGAACTCTTTCCCTACACGACGCTCTTCCGATCTNNNGAAGATCATGAGGTCAGGGGT
AGCACACGTCTGAACTCTTTCCCTACACGACGCTCTTCCGATCTNNNGAAGATCATGAGGTCAGGGGT

AGCACACGTCTGAACTCTTTCCCTACACGACGCTCTTCCGATCTNNNAAGAACACCGTTACCCAATGA
AGCACACGTCTGAACTCTTTCCCTACACGACGCTCTTCCGATCTNNNAAGAACACCGTTACCCAATGA
GCACACGTCTGAACTCTTTCCCTACACGACGCTCTTCCGATCTNNNGTCTCTCTCTCCCCTCTTTCTCA
GCACACGTCTGAACTCTTTCCCTACACGACGCTCTTCCGATCTNNNGTCTCTCTCTCCCCTCTTTCTCA
GAGCACACGTCTGAACTCTTTCCCTACACGACGCTCTTCCGATCTNNNTTCCGGGCTTAGAAAGTTAG
GAGCACACGTCTGAACTCTTTCCCTACACGACGCTCTTCCGATCTNNNTTCCGGGCTTAGAAAGTTAG
AGCACACGTCTGAACTCTTTCCCTACACGACGCTCTTCCGATCTNNNAACGTAAACTGGGTCTTATG
AGCACACGTCTGAACTCTTTCCCTACACGACGCTCTTCCGATCTNNNAACGTAAACTGGGTCTTATG
AGCACACGTCTGAACTCTTTCCCTACACGACGCTCTTCCGATCTNNNAAAAAATTAGCCGGGCGTAGT
AGCACACGTCTGAACTCTTTCCCTACACGACGCTCTTCCGATCTNNNAAAAAATTAGCCGGGCGTAGT
AGCACACGTCTGAACTCTTTCCCTACACGACGCTCTTCCGATCTNNNGAGACAGGAGGATGGCTTG
AGCACACGTCTGAACTCTTTCCCTACACGACGCTCTTCCGATCTNNNGAGACAGGAGGATGGCTTG
GAGCACACGTCTGAACTCTTTCCCTACACGACGCTCTTCCGATCTNNNCGACAAGAACAAAACTCCGT
GAGCACACGTCTGAACTCTTTCCCTACACGACGCTCTTCCGATCTNNNCGACAAGAACAAAACTCCGT
AGCACACGTCTGAACTCTTTCCCTACACGACGCTCTTCCGATCTNNNTCAGGAGGTTGAGGTAGGAG
AGCACACGTCTGAACTCTTTCCCTACACGACGCTCTTCCGATCTNNNTCAGGAGGTTGAGGTAGGAG
AGCACACGTCTGAACTCTTTCCCTACACGACGCTCTTCCGATCTNNNTGTTTCTAGGACCCATGTTGAG
AGCACACGTCTGAACTCTTTCCCTACACGACGCTCTTCCGATCTNNNTGTTTCTAGGACCCATGTTGAG
AGCACACGTCTGAACTCTTTCCCTACACGACGCTCTTCCGATCTNNNAGCAACAGAACAAGACTCCAT
AGCACACGTCTGAACTCTTTCCCTACACGACGCTCTTCCGATCTNNNAGCAACAGAACAAGACTCCAT
ACACGTCTGAACTCTTTCCCTACACGACGCTCTTCCGATCTNNNGGAAAAACCAGAGACCACTCACTC
ACACGTCTGAACTCTTTCCCTACACGACGCTCTTCCGATCTNNNGGAAAAACCAGAGACCACTCACTC
AGCACACGTCTGAACTCTTTCCCTACACGACGCTCTTCCGATCTNNNTTCCTATCAGGCTAGCATTCTG
AGCACACGTCTGAACTCTTTCCCTACACGACGCTCTTCCGATCTNNNTTCCTATCAGGCTAGCATTCTG
AGCACACGTCTGAACTCTTTCCCTACACGACGCTCTTCCGATCTNNNCATTGAGTGGGGATTAGCACT
AGCACACGTCTGAACTCTTTCCCTACACGACGCTCTTCCGATCTNNNCATTGAGTGGGGATTAGCACT
GAGCACACGTCTGAACTCTTTCCCTACACGACGCTCTTCCGATCTNNNACCAGCATCTAGTGGGTAGT
GAGCACACGTCTGAACTCTTTCCCTACACGACGCTCTTCCGATCTNNNACCAGCATCTAGTGGGTAGT
AGCACACGTCTGAACTCTTTCCCTACACGACGCTCTTCCGATCTNNNTATGTTGAATTCATTTTCAGGT
AGCACACGTCTGAACTCTTTCCCTACACGACGCTCTTCCGATCTNNNTATGTTGAATTCATTTTCAGGT
GCACACGTCTGAACTCTTTCCCTACACGACGCTCTTCCGATCTNNNTGTAGAAGCAGAAGTTGTGGA
GCACACGTCTGAACTCTTTCCCTACACGACGCTCTTCCGATCTNNNTGTAGAAGCAGAAGTTGTGGA
AGCACACGTCTGAACTCTTTCCCTACACGACGCTCTTCCGATCTNNNGTGCACCTGTAATCTCAGCTAC
AGCACACGTCTGAACTCTTTCCCTACACGACGCTCTTCCGATCTNNNGTGCACCTGTAATCTCAGCTAC
GAGCACACGTCTGAACTCTTTCCCTACACGACGCTCTTCCGATCTNNNGACATGAAAGCTGAGCCAGA
GAGCACACGTCTGAACTCTTTCCCTACACGACGCTCTTCCGATCTNNNGACATGAAAGCTGAGCCAGA
AGCACACGTCTGAACTCTTTCCCTACACGACGCTCTTCCGATCTNNNAAGTATACTCAGTCAAGCAGC
AGCACACGTCTGAACTCTTTCCCTACACGACGCTCTTCCGATCTNNNAAGTATACTCAGTCAAGCAGC
AGCACACGTCTGAACTCTTTCCCTACACGACGCTCTTCCGATCTNNNGGATGACGGACCATTCTATCC
AGCACACGTCTGAACTCTTTCCCTACACGACGCTCTTCCGATCTNNNGGATGACGGACCATTCTATCC
AGCACACGTCTGAACTCTTTCCCTACACGACGCTCTTCCGATCTNNNGTGACTCCTCCCTCACTCATTTT
AGCACACGTCTGAACTCTTTCCCTACACGACGCTCTTCCGATCTNNNGTGACTCCTCCCTCACTCATTTT
GAGCACACGTCTGAACTCTTTCCCTACACGACGCTCTTCCGATCTNNNTACACAGCTCTTTGATGAGG

GAGCACACGTCTGAACTCTTTCCCTACACGACGCTCTTCCGATCTNNNTACACAGCTCTTTGATGAGGA
AGCACACGTCTGAACTCTTTCCCTACACGACGCTCTTCCGATCTNNNGCACCCCAGTTTCCAAAATTTGT
AGCACACGTCTGAACTCTTTCCCTACACGACGCTCTTCCGATCTNNNGCACCCCAGTTTCCAAAATTTGT
AGCACACGTCTGAACTCTTTCCCTACACGACGCTCTTCCGATCTNNNTCCTCAATTTCTGCCTCCATCTCC
AGCACACGTCTGAACTCTTTCCCTACACGACGCTCTTCCGATCTNNNTCCTCAATTTCTGCCTCCATCTCC
AGCACACGTCTGAACTCTTTCCCTACACGACGCTCTTCCGATCTNNNGCCTGTAATCTCAGCACTTTGTC
AGCACACGTCTGAACTCTTTCCCTACACGACGCTCTTCCGATCTNNNGCCTGTAATCTCAGCACTTTGTC
AGAGCACACGTCTGAACTCTTTCCCTACACGACGCTCTTCCGATCTNNNGAGGCAGGATAATTGCTTG
AGAGCACACGTCTGAACTCTTTCCCTACACGACGCTCTTCCGATCTNNNGAGGCAGGATAATTGCTTG
AGCACACGTCTGAACTCTTTCCCTACACGACGCTCTTCCGATCTNNNAGCATATCTGCGACTACCTAAG
AGCACACGTCTGAACTCTTTCCCTACACGACGCTCTTCCGATCTNNNAGCATATCTGCGACTACCTAAG
AGAGCACACGTCTGAACTCTTTCCCTACACGACGCTCTTCCGATCTNNNTGTCTTTCTCTCTTCCCCTTTC
AGAGCACACGTCTGAACTCTTTCCCTACACGACGCTCTTCCGATCTNNNTGTCTTTCTCTCTTCCCCTTTC
GCACACGTCTGAACTCTTTCCCTACACGACGCTCTTCCGATCTNNNGAGTAAGTCTATCATCAATTTGG
GCACACGTCTGAACTCTTTCCCTACACGACGCTCTTCCGATCTNNNGAGTAAGTCTATCATCAATTTGG
GAGCACACGTCTGAACTCTTTCCCTACACGACGCTCTTCCGATCTNNNAAGCACCTGTAATCCCAGCTAC
GAGCACACGTCTGAACTCTTTCCCTACACGACGCTCTTCCGATCTNNNAAGCACCTGTAATCCCAGCTAC
AGCACACGTCTGAACTCTTTCCCTACACGACGCTCTTCCGATCTNNNGCTTCTAAAACACCATTTAAAAT
AGCACACGTCTGAACTCTTTCCCTACACGACGCTCTTCCGATCTNNNGCTTCTAAAACACCATTTAAAAT
AGCACACGTCTGAACTCTTTCCCTACACGACGCTCTTCCGATCTNNNCTGCCTCTCAGTTCAAGCAATTC
AGCACACGTCTGAACTCTTTCCCTACACGACGCTCTTCCGATCTNNNCTGCCTCTCAGTTCAAGCAATTC
AGCACACGTCTGAACTCTTTCCCTACACGACGCTCTTCCGATCTNNNCCACGCCCGACTAATTTTTGTAT
AGCACACGTCTGAACTCTTTCCCTACACGACGCTCTTCCGATCTNNNCCACGCCCGACTAATTTTTGTAT
GAGCACACGTCTGAACTCTTTCCCTACACGACGCTCTTCCGATCTNNNAGTCTCAAGTGTCACCAGATT
GAGCACACGTCTGAACTCTTTCCCTACACGACGCTCTTCCGATCTNNNAGTCTCAAGTGTCACCAGATT
GAGCACACGTCTGAACTCTTTCCCTACACGACGCTCTTCCGATCTNNNAAAATGCAGACAGACACACAC
GAGCACACGTCTGAACTCTTTCCCTACACGACGCTCTTCCGATCTNNNAAAATGCAGACAGACACACAC
AGAGCACACGTCTGAACTCTTTCCCTACACGACGCTCTTCCGATCTNNNCCCACTAACTCGTCATCTAGCA
AGAGCACACGTCTGAACTCTTTCCCTACACGACGCTCTTCCGATCTNNNCCCACTAACTCGTCATCTAGCA
AGCACACGTCTGAACTCTTTCCCTACACGACGCTCTTCCGATCTNNNTGGGACATCTAGGTACCTCAACT
AGCACACGTCTGAACTCTTTCCCTACACGACGCTCTTCCGATCTNNNTGGGACATCTAGGTACCTCAACT
AGCACACGTCTGAACTCTTTCCCTACACGACGCTCTTCCGATCTNNNTGTAATCCCAGCTACTCAGGAG
AGCACACGTCTGAACTCTTTCCCTACACGACGCTCTTCCGATCTNNNTGTAATCCCAGCTACTCAGGAG
GAGCACACGTCTGAACTCTTTCCCTACACGACGCTCTTCCGATCTNNNACGAAAATCAGGCAAGGACAA
GAGCACACGTCTGAACTCTTTCCCTACACGACGCTCTTCCGATCTNNNACGAAAATCAGGCAAGGACAA
AGCACACGTCTGAACTCTTTCCCTACACGACGCTCTTCCGATCTNNNCTTGCATCCAGGTGGTCTATCAA
AGCACACGTCTGAACTCTTTCCCTACACGACGCTCTTCCGATCTNNNCTTGCATCCAGGTGGTCTATCAA
GAGCACACGTCTGAACTCTTTCCCTACACGACGCTCTTCCGATCTNNNCTTGTCACATCTTCTCTGGCTCT
GAGCACACGTCTGAACTCTTTCCCTACACGACGCTCTTCCGATCTNNNCTTGTCACATCTTCTCTGGCTCT
AGAGCACACGTCTGAACTCTTTCCCTACACGACGCTCTTCCGATCTNNNTCATTCTTAGGGGATCCATGC
AGAGCACACGTCTGAACTCTTTCCCTACACGACGCTCTTCCGATCTNNNTCATTCTTAGGGGATCCATGC
GAGCACACGTCTGAACTCTTTCCCTACACGACGCTCTTCCGATCTNNNAAATGCCCATCAATCAACACGT
GAGCACACGTCTGAACTCTTTCCCTACACGACGCTCTTCCGATCTNNNAAATGCCCATCAATCAACACGT

GAGCACACGTCTGAACTCTTTCCCTACACGACGCTCTTCCGATCTNNNACTGACACACACATACACACAC
GAGCACACGTCTGAACTCTTTCCCTACACGACGCTCTTCCGATCTNNNACTGACACACACATACACACAC
GAGCACACGTCTGAACTCTTTCCCTACACGACGCTCTTCCGATCTNNNAGGTATGTGGCCTCATTTCTCA
GAGCACACGTCTGAACTCTTTCCCTACACGACGCTCTTCCGATCTNNNAGGTATGTGGCCTCATTTCTCA
AGCACACGTCTGAACTCTTTCCCTACACGACGCTCTTCCGATCTNNNAAAAATTATTGGGTGTGGGG
AGCACACGTCTGAACTCTTTCCCTACACGACGCTCTTCCGATCTNNNAAAAATTATTGGGTGTGGGG
AGCACACGTCTGAACTCTTTCCCTACACGACGCTCTTCCGATCTNNNCCTTTCTCCTTTCCCTTTTCCCTCA
AGCACACGTCTGAACTCTTTCCCTACACGACGCTCTTCCGATCTNNNCCTTTCTCCTTTCCCTTTTCCCTCA
AGCACACGTCTGAACTCTTTCCCTACACGACGCTCTTCCGATCTNNNTCTCTCCAACTCCACTAGTAGCAC
AGCACACGTCTGAACTCTTTCCCTACACGACGCTCTTCCGATCTNNNTCTCTCCAACTCCACTAGTAGCAC
GAGCACACGTCTGAACTCTTTCCCTACACGACGCTCTTCCGATCTNNNCCAGGCTGCAGATATTTTCCTT
GAGCACACGTCTGAACTCTTTCCCTACACGACGCTCTTCCGATCTNNNCCAGGCTGCAGATATTTTCCTT
AGCACACGTCTGAACTCTTTCCCTACACGACGCTCTTCCGATCTNNNTTTTTAGAATATGTTATGCTGG
AGCACACGTCTGAACTCTTTCCCTACACGACGCTCTTCCGATCTNNNTTTTTAGAATATGTTATGCTGG
AGCACACGTCTGAACTCTTTCCCTACACGACGCTCTTCCGATCTNNNACCAACAGTAACCAAGCTGAGA
AGCACACGTCTGAACTCTTTCCCTACACGACGCTCTTCCGATCTNNNACCAACAGTAACCAAGCTGAGA
AGCACACGTCTGAACTCTTTCCCTACACGACGCTCTTCCGATCTNNNCACACACACACTCATTTTCCCTCCA
AGCACACGTCTGAACTCTTTCCCTACACGACGCTCTTCCGATCTNNNCACACACACACTCATTTTCCCTCCA
AGCACACGTCTGAACTCTTTCCCTACACGACGCTCTTCCGATCTNNNTGTATGCCTGTAATCCCAGCTAC
AGCACACGTCTGAACTCTTTCCCTACACGACGCTCTTCCGATCTNNNTGTATGCCTGTAATCCCAGCTAC
GAGCACACGTCTGAACTCTTTCCCTACACGACGCTCTTCCGATCTNNNGATCACACAGATCCAGCCATT
GAGCACACGTCTGAACTCTTTCCCTACACGACGCTCTTCCGATCTNNNGATCACACAGATCCAGCCATT
GAGCACACGTCTGAACTCTTTCCCTACACGACGCTCTTCCGATCTNNNGGACTTTGACACCTAAGGACCA
GAGCACACGTCTGAACTCTTTCCCTACACGACGCTCTTCCGATCTNNNGGACTTTGACACCTAAGGACCA
AGCACACGTCTGAACTCTTTCCCTACACGACGCTCTTCCGATCTNNNTTGTCCACATGTTGTCTGTAGGT
AGCACACGTCTGAACTCTTTCCCTACACGACGCTCTTCCGATCTNNNTTGTCCACATGTTGTCTGTAGGT
AGCACACGTCTGAACTCTTTCCCTACACGACGCTCTTCCGATCTNNNGCACTCAGTAATCGTTTGGGAA
AGCACACGTCTGAACTCTTTCCCTACACGACGCTCTTCCGATCTNNNGCACTCAGTAATCGTTTGGGAA
GAGCACACGTCTGAACTCTTTCCCTACACGACGCTCTTCCGATCTNNNAGGAGTTCCCAGAGAAGAAAA
GAGCACACGTCTGAACTCTTTCCCTACACGACGCTCTTCCGATCTNNNAGGAGTTCCCAGAGAAGAAAA
GCACACGTCTGAACTCTTTCCCTACACGACGCTCTTCCGATCTNNNCTTATTGAAGCTCTGTCCTCCATC
GCACACGTCTGAACTCTTTCCCTACACGACGCTCTTCCGATCTNNNCTTATTGAAGCTCTGTCCTCCATC
GAGCACACGTCTGAACTCTTTCCCTACACGACGCTCTTCCGATCTNNNACCAGGAAAGTGGAAAATGG
GAGCACACGTCTGAACTCTTTCCCTACACGACGCTCTTCCGATCTNNNACCAGGAAAGTGGAAAATGG
GAGCACACGTCTGAACTCTTTCCCTACACGACGCTCTTCCGATCTNNNCACAGGACAGAGGGAATCAAT
GAGCACACGTCTGAACTCTTTCCCTACACGACGCTCTTCCGATCTNNNCACAGGACAGAGGGAATCAAT
AGCACACGTCTGAACTCTTTCCCTACACGACGCTCTTCCGATCTNNNAGTAATACAATGTTCTTGTCATG
AGCACACGTCTGAACTCTTTCCCTACACGACGCTCTTCCGATCTNNNAGTAATACAATGTTCTTGTCATG
AGCACACGTCTGAACTCTTTCCCTACACGACGCTCTTCCGATCTNNNAGATTGTCCATTTAGTCGGGGT
AGCACACGTCTGAACTCTTTCCCTACACGACGCTCTTCCGATCTNNNAGATTGTCCATTTAGTCGGGGT
GAGCACACGTCTGAACTCTTTCCCTACACGACGCTCTTCCGATCTNNNCTGTCATCCCAGCTATTCAGG
GAGCACACGTCTGAACTCTTTCCCTACACGACGCTCTTCCGATCTNNNCTGTCATCCCAGCTATTCAGG
AGCACACGTCTGAACTCTTTCCCTACACGACGCTCTTCCGATCTNNNCTCCAAAGTAGCTGGGACTACAC

GCACACGTCTGAACTCTTTCCCTACACGACGCTCTTCCGATCTNNNCTCCAAAGTAGCTGGGACTACA
CACACGTCTGAACTCTTTCCCTACACGACGCTCTTCCGATCTNNNTAAGTTGATGGAGGAAGTAAAT
CACACGTCTGAACTCTTTCCCTACACGACGCTCTTCCGATCTNNNTAAGTTGATGGAGGAAGTAAAT
GCACACGTCTGAACTCTTTCCCTACACGACGCTCTTCCGATCTNNNGTAGGGCATATTTACATCTTTG
GCACACGTCTGAACTCTTTCCCTACACGACGCTCTTCCGATCTNNNGTAGGGCATATTTACATCTTTG
AGCACACGTCTGAACTCTTTCCCTACACGACGCTCTTCCGATCTNNNTATTCTAGAGGCCTGATGTCCC
AGCACACGTCTGAACTCTTTCCCTACACGACGCTCTTCCGATCTNNNTATTCTAGAGGCCTGATGTCCC
GAGCACACGTCTGAACTCTTTCCCTACACGACGCTCTTCCGATCTNNNTTCTGACCTCAAGTAATCCAC
GAGCACACGTCTGAACTCTTTCCCTACACGACGCTCTTCCGATCTNNNTTCTGACCTCAAGTAATCCAC
GCACACGTCTGAACTCTTTCCCTACACGACGCTCTTCCGATCTNNNTGATGAGTGTTGAAGGGCTAG
GCACACGTCTGAACTCTTTCCCTACACGACGCTCTTCCGATCTNNNTGATGAGTGTTGAAGGGCTAG
AGAGCACACGTCTGAACTCTTTCCCTACACGACGCTCTTCCGATCTNNNGAGAGGGGAATGAGAATG
AGAGCACACGTCTGAACTCTTTCCCTACACGACGCTCTTCCGATCTNNNGAGAGGGGAATGAGAATG
CACACGTCTGAACTCTTTCCCTACACGACGCTCTTCCGATCTNNNAGTAATCACTCAGGAGCAGGTTG
CACACGTCTGAACTCTTTCCCTACACGACGCTCTTCCGATCTNNNAGTAATCACTCAGGAGCAGGTTG
GAGCACACGTCTGAACTCTTTCCCTACACGACGCTCTTCCGATCTNNNGCAATCCCCAAAGGTACAGG
GAGCACACGTCTGAACTCTTTCCCTACACGACGCTCTTCCGATCTNNNGCAATCCCCAAAGGTACAGG
GAGCACACGTCTGAACTCTTTCCCTACACGACGCTCTTCCGATCTNNNGATGAGGTAAGGAAGGGT
GAGCACACGTCTGAACTCTTTCCCTACACGACGCTCTTCCGATCTNNNGATGAGGTAAGGAAGGGT
GAGCACACGTCTGAACTCTTTCCCTACACGACGCTCTTCCGATCTNNNTAGCATTACAGGTTCACTCCA
GAGCACACGTCTGAACTCTTTCCCTACACGACGCTCTTCCGATCTNNNTAGCATTACAGGTTCACTCCA
GCACACGTCTGAACTCTTTCCCTACACGACGCTCTTCCGATCTNNNTGCCCGTCTAAGAGGCTATTTTC
GCACACGTCTGAACTCTTTCCCTACACGACGCTCTTCCGATCTNNNTGCCCGTCTAAGAGGCTATTTTC
AGCACACGTCTGAACTCTTTCCCTACACGACGCTCTTCCGATCTNNNTATAGTCCCAGCTACTCAGGA
AGCACACGTCTGAACTCTTTCCCTACACGACGCTCTTCCGATCTNNNTATAGTCCCAGCTACTCAGGA
AGCACACGTCTGAACTCTTTCCCTACACGACGCTCTTCCGATCTNNNCTGTAATCCCAGCTACTCAGG
AGCACACGTCTGAACTCTTTCCCTACACGACGCTCTTCCGATCTNNNCTGTAATCCCAGCTACTCAGG
GAGCACACGTCTGAACTCTTTCCCTACACGACGCTCTTCCGATCTNNNCAACACTGTGTCTTCCACCAA
GAGCACACGTCTGAACTCTTTCCCTACACGACGCTCTTCCGATCTNNNCAACACTGTGTCTTCCACCAA
GCACACGTCTGAACTCTTTCCCTACACGACGCTCTTCCGATCTNNNTGTTGTTGTTGTTGTTGTTGTTT
GCACACGTCTGAACTCTTTCCCTACACGACGCTCTTCCGATCTNNNTGTTGTTGTTGTTGTTGTTGTTT
AGCACACGTCTGAACTCTTTCCCTACACGACGCTCTTCCGATCTNNNACACACACACATATACACACAC
AGCACACGTCTGAACTCTTTCCCTACACGACGCTCTTCCGATCTNNNACACACACACATATACACACAC
AGCACACGTCTGAACTCTTTCCCTACACGACGCTCTTCCGATCTNNNTGAGGCAGAAGAATTGCTTGA
AGCACACGTCTGAACTCTTTCCCTACACGACGCTCTTCCGATCTNNNTGAGGCAGAAGAATTGCTTGA
AGCACACGTCTGAACTCTTTCCCTACACGACGCTCTTCCGATCTNNNGGTGGGGCTCTAAACTTATGTC
AGCACACGTCTGAACTCTTTCCCTACACGACGCTCTTCCGATCTNNNGGTGGGGCTCTAAACTTATGTC
AGCACACGTCTGAACTCTTTCCCTACACGACGCTCTTCCGATCTNNNCAGATGACTTTATCTAACACCTG
AGCACACGTCTGAACTCTTTCCCTACACGACGCTCTTCCGATCTNNNCAGATGACTTTATCTAACACCTG
GCACACGTCTGAACTCTTTCCCTACACGACGCTCTTCCGATCTNNNCTTGAATGAGTTTACACGGCTG
GCACACGTCTGAACTCTTTCCCTACACGACGCTCTTCCGATCTNNNCTTGAATGAGTTTACACGGCTG
AGCACACGTCTGAACTCTTTCCCTACACGACGCTCTTCCGATCTNNNCTGTAGCTGGTATTACAGGCAC
AGCACACGTCTGAACTCTTTCCCTACACGACGCTCTTCCGATCTNNNCTGTAGCTGGTATTACAGGCAC

AGCACACGTCTGAACTCTTTCCCTACACGACGCTCTTCCGATCTNNNTTCAAGCGATTCTCCTGTCTCAG

AGCACACGTCTGAACTCTTTCCCTACACGACGCTCTTCCGATCTNNNTTCAAGCGATTCTCCTGTCTCAG

GAGCACACGTCTGAACTCTTTCCCTACACGACGCTCTTCCGATCTNNNTTTGTTGCTCAAAGCCAGGTA

GAGCACACGTCTGAACTCTTTCCCTACACGACGCTCTTCCGATCTNNNTTTGTTGCTCAAAGCCAGGTA

GCACACGTCTGAACTCTTTCCCTACACGACGCTCTTCCGATCTNNNTCATCCCTCCTCCTACCCATAAACAT

GCACACGTCTGAACTCTTTCCCTACACGACGCTCTTCCGATCTNNNTCATCCCTCCTCCTACCCATAAACAT

AGCACACGTCTGAACTCTTTCCCTACACGACGCTCTTCCGATCTNNNATACAAAAAATTAGCCGGGCGT

AGCACACGTCTGAACTCTTTCCCTACACGACGCTCTTCCGATCTNNNATACAAAAAATTAGCCGGGCGT

GAGCACACGTCTGAACTCTTTCCCTACACGACGCTCTTCCGATCTNNNTGTACTCTAACAACCTGTAGG

GAGCACACGTCTGAACTCTTTCCCTACACGACGCTCTTCCGATCTNNNTGTACTCTAACAACCTGTAGG

GAGCACACGTCTGAACTCTTTCCCTACACGACGCTCTTCCGATCTNNNAACAATCTCTGTGTCGTCTGTC

GAGCACACGTCTGAACTCTTTCCCTACACGACGCTCTTCCGATCTNNNAACAATCTCTGTGTCGTCTGTC

GAGCACACGTCTGAACTCTTTCCCTACACGACGCTCTTCCGATCTNNNGAGAATCACTTGAGCCCAGGA

GAGCACACGTCTGAACTCTTTCCCTACACGACGCTCTTCCGATCTNNNGAGAATCACTTGAGCCCAGGA

AGCACACGTCTGAACTCTTTCCCTACACGACGCTCTTCCGATCTNNNCTTGTGCAACTAAGGGAAAGTG

AGCACACGTCTGAACTCTTTCCCTACACGACGCTCTTCCGATCTNNNCTTGTGCAACTAAGGGAAAGTG

AGCACACGTCTGAACTCTTTCCCTACACGACGCTCTTCCGATCTNNNGAGAGAGCGAGAGAGAGAGA

AGCACACGTCTGAACTCTTTCCCTACACGACGCTCTTCCGATCTNNNGAGAGAGCGAGAGAGAGAGA

AGCACACGTCTGAACTCTTTCCCTACACGACGCTCTTCCGATCTNNNTGAAGGGATGGGGATGGTAG

AGCACACGTCTGAACTCTTTCCCTACACGACGCTCTTCCGATCTNNNTGAAGGGATGGGGATGGTAG

GAGCACACGTCTGAACTCTTTCCCTACACGACGCTCTTCCGATCTNNNAAAAATTAGCCAGGTGTGGTG

GAGCACACGTCTGAACTCTTTCCCTACACGACGCTCTTCCGATCTNNNAAAAATTAGCCAGGTGTGGTG

GAGCACACGTCTGAACTCTTTCCCTACACGACGCTCTTCCGATCTNNNGGTGCCGACAAGTCCTCTAAT

GAGCACACGTCTGAACTCTTTCCCTACACGACGCTCTTCCGATCTNNNGGTGCCGACAAGTCCTCTAAT

GCACACGTCTGAACTCTTTCCCTACACGACGCTCTTCCGATCTNNNACATTGCTCAGGCTAGTCTTGAA

GCACACGTCTGAACTCTTTCCCTACACGACGCTCTTCCGATCTNNNACATTGCTCAGGCTAGTCTTGAA

GAGCACACGTCTGAACTCTTTCCCTACACGACGCTCTTCCGATCTNNNTACTTGAGCCTAGGAGGTAGA

GAGCACACGTCTGAACTCTTTCCCTACACGACGCTCTTCCGATCTNNNTACTTGAGCCTAGGAGGTAGA

GAGCACACGTCTGAACTCTTTCCCTACACGACGCTCTTCCGATCTNNNGAGAATGGCATGAACCTGGG

GAGCACACGTCTGAACTCTTTCCCTACACGACGCTCTTCCGATCTNNNGAGAATGGCATGAACCTGGG

GCACACGTCTGAACTCTTTCCCTACACGACGCTCTTCCGATCTNNNACTGTACCACCATCCACTACAACCA

GCACACGTCTGAACTCTTTCCCTACACGACGCTCTTCCGATCTNNNACTGTACCACCATCCACTACAACCA

AGCACACGTCTGAACTCTTTCCCTACACGACGCTCTTCCGATCTNNNCTGAGGCAGAGAATTGCTTGA

AGCACACGTCTGAACTCTTTCCCTACACGACGCTCTTCCGATCTNNNCTGAGGCAGAGAATTGCTTGA

AGCACACGTCTGAACTCTTTCCCTACACGACGCTCTTCCGATCTNNNCCAATTCTATACAGGTTGCTGC

AGCACACGTCTGAACTCTTTCCCTACACGACGCTCTTCCGATCTNNNCCAATTCTATACAGGTTGCTGC

AGCACACGTCTGAACTCTTTCCCTACACGACGCTCTTCCGATCTNNNTATCACACAGTCTTTGGGTCCAG

AGCACACGTCTGAACTCTTTCCCTACACGACGCTCTTCCGATCTNNNTATCACACAGTCTTTGGGTCCAG

GAGCACACGTCTGAACTCTTTCCCTACACGACGCTCTTCCGATCTNNNAGTCAGGAGAATTGCTTGAA

GAGCACACGTCTGAACTCTTTCCCTACACGACGCTCTTCCGATCTNNNAGTCAGGAGAATTGCTTGAA

GCACACGTCTGAACTCTTTCCCTACACGACGCTCTTCCGATCTNNNGGAAGTGACCCTTTCCATAAGGA

GCACACGTCTGAACTCTTTCCCTACACGACGCTCTTCCGATCTNNNGGAAGTGACCCTTTCCATAAGGA

AGCACACGTCTGAACTCTTTCCCTACACGACGCTCTTCCGATCTNNNCCTCTGTGTGAAATTTTGGGGT

AGCACACGTCTGAACTCTTTCCCTACACGACGCTCTTCCGATCTNNNCCTCTGTGTGAAATTTTGGGGT
AGCACACGTCTGAACTCTTTCCCTACACGACGCTCTTCCGATCTNNNCACCTGTAATCCCAGCTACTCAC
AGCACACGTCTGAACTCTTTCCCTACACGACGCTCTTCCGATCTNNNCACCTGTAATCCCAGCTACTCAC
AGCACACGTCTGAACTCTTTCCCTACACGACGCTCTTCCGATCTNNNCAGGAGACAGGGTTTTGAGATC
AGCACACGTCTGAACTCTTTCCCTACACGACGCTCTTCCGATCTNNNCAGGAGACAGGGTTTTGAGATC
AGCACACGTCTGAACTCTTTCCCTACACGACGCTCTTCCGATCTNNNTCACTGTGGGAAAGGAAGAAA
AGCACACGTCTGAACTCTTTCCCTACACGACGCTCTTCCGATCTNNNTCACTGTGGGAAAGGAAGAAA
AGCACACGTCTGAACTCTTTCCCTACACGACGCTCTTCCGATCTNNNTAAGGCAGCAGAATCACTTGAA
AGCACACGTCTGAACTCTTTCCCTACACGACGCTCTTCCGATCTNNNTAAGGCAGCAGAATCACTTGAA
AGCACACGTCTGAACTCTTTCCCTACACGACGCTCTTCCGATCTNNNACACTTGTAATCCCAGCTACTCG
AGCACACGTCTGAACTCTTTCCCTACACGACGCTCTTCCGATCTNNNACACTTGTAATCCCAGCTACTCG
GCACACGTCTGAACTCTTTCCCTACACGACGCTCTTCCGATCTNNNGATTAAAGATGGAGGCTCTGCCC
GCACACGTCTGAACTCTTTCCCTACACGACGCTCTTCCGATCTNNNGATTAAAGATGGAGGCTCTGCCC
GCACACGTCTGAACTCTTTCCCTACACGACGCTCTTCCGATCTNNNTCAACTCATTCTCGGTAAAGGTCA
GCACACGTCTGAACTCTTTCCCTACACGACGCTCTTCCGATCTNNNTCAACTCATTCTCGGTAAAGGTCA
AGAGCACACGTCTGAACTCTTTCCCTACACGACGCTCTTCCGATCTNNNTCAAGGCTGCAGTGAGTTAT
AGAGCACACGTCTGAACTCTTTCCCTACACGACGCTCTTCCGATCTNNNTCAAGGCTGCAGTGAGTTAT
AGCACACGTCTGAACTCTTTCCCTACACGACGCTCTTCCGATCTNNNGTAGATCAGTGCTGTTCGGAAC
AGCACACGTCTGAACTCTTTCCCTACACGACGCTCTTCCGATCTNNNGTAGATCAGTGCTGTTCGGAAC
GAGCACACGTCTGAACTCTTTCCCTACACGACGCTCTTCCGATCTNNNCTGGAACTCCTGACCTCATGA
GAGCACACGTCTGAACTCTTTCCCTACACGACGCTCTTCCGATCTNNNCTGGAACTCCTGACCTCATGA
GAGCACACGTCTGAACTCTTTCCCTACACGACGCTCTTCCGATCTNNNAGGAGTTCAAGACCAGCCTAG
GAGCACACGTCTGAACTCTTTCCCTACACGACGCTCTTCCGATCTNNNAGGAGTTCAAGACCAGCCTAG
CACACGTCTGAACTCTTTCCCTACACGACGCTCTTCCGATCTNNNGACAGAAGGAGAAGATGGAGCAT
CACACGTCTGAACTCTTTCCCTACACGACGCTCTTCCGATCTNNNGACAGAAGGAGAAGATGGAGCAT
GAGCACACGTCTGAACTCTTTCCCTACACGACGCTCTTCCGATCTNNNACTGAACTCAACTCCTCAGCAA
GAGCACACGTCTGAACTCTTTCCCTACACGACGCTCTTCCGATCTNNNACTGAACTCAACTCCTCAGCAA
AGAGCACACGTCTGAACTCTTTCCCTACACGACGCTCTTCCGATCTNNNCTTGTCCCAAGTCTCCTGAGA
AGAGCACACGTCTGAACTCTTTCCCTACACGACGCTCTTCCGATCTNNNCTTGTCCCAAGTCTCCTGAGA
GAGCACACGTCTGAACTCTTTCCCTACACGACGCTCTTCCGATCTNNNTACAAAAATTAGCGGGGTGTC
GAGCACACGTCTGAACTCTTTCCCTACACGACGCTCTTCCGATCTNNNTACAAAAATTAGCGGGGTGTC
GAGCACACGTCTGAACTCTTTCCCTACACGACGCTCTTCCGATCTNNNGTTTCCTTTTGACAGACTGTTC
GAGCACACGTCTGAACTCTTTCCCTACACGACGCTCTTCCGATCTNNNGTTTCCTTTTGACAGACTGTTC
AGCACACGTCTGAACTCTTTCCCTACACGACGCTCTTCCGATCTNNNTCATCCCTTCACTCACAAATGCTC
AGCACACGTCTGAACTCTTTCCCTACACGACGCTCTTCCGATCTNNNTCATCCCTTCACTCACAAATGCTC
AGCACACGTCTGAACTCTTTCCCTACACGACGCTCTTCCGATCTNNNGGTGCCTATAATCCCAGCTACTC
AGCACACGTCTGAACTCTTTCCCTACACGACGCTCTTCCGATCTNNNGGTGCCTATAATCCCAGCTACTC
GAGCACACGTCTGAACTCTTTCCCTACACGACGCTCTTCCGATCTNNNTTCTCTGCTGAATTGTTCTTGC
GAGCACACGTCTGAACTCTTTCCCTACACGACGCTCTTCCGATCTNNNTTCTCTGCTGAATTGTTCTTGC
AGCACACGTCTGAACTCTTTCCCTACACGACGCTCTTCCGATCTNNNTGTTTTGTATAGTGGCAGCAGC
AGCACACGTCTGAACTCTTTCCCTACACGACGCTCTTCCGATCTNNNTGTTTTGTATAGTGGCAGCAGC
AGCACACGTCTGAACTCTTTCCCTACACGACGCTCTTCCGATCTNNNTTAGTCTTTGGCCTGGTTTACTC
AGCACACGTCTGAACTCTTTCCCTACACGACGCTCTTCCGATCTNNNTTAGTCTTTGGCCTGGTTTACTC

GAGCACACGTCTGAACTCTTTCCCTACACGACGCTCTTCCGATCTNNNCAGGAGTTCGAGACTATACT
GAGCACACGTCTGAACTCTTTCCCTACACGACGCTCTTCCGATCTNNNCAGGAGTTCGAGACTATACT
AGCACACGTCTGAACTCTTTCCCTACACGACGCTCTTCCGATCTNNNCCTGTTCCCTCAATAACCCTCTTC
AGCACACGTCTGAACTCTTTCCCTACACGACGCTCTTCCGATCTNNNCCTGTTCCCTCAATAACCCTCTTC
GCACACGTCTGAACTCTTTCCCTACACGACGCTCTTCCGATCTNNNCTATTGATTGGACAGCTTGAATC
GCACACGTCTGAACTCTTTCCCTACACGACGCTCTTCCGATCTNNNCTATTGATTGGACAGCTTGAATC
AGCACACGTCTGAACTCTTTCCCTACACGACGCTCTTCCGATCTNNNTTTCCGTGAAACCTAGTGCTGA
AGCACACGTCTGAACTCTTTCCCTACACGACGCTCTTCCGATCTNNNTTTCCGTGAAACCTAGTGCTGA
GAGCACACGTCTGAACTCTTTCCCTACACGACGCTCTTCCGATCTNNNGAAGGATACAGTGGGGGAT
GAGCACACGTCTGAACTCTTTCCCTACACGACGCTCTTCCGATCTNNNGAAGGATACAGTGGGGGAT
GAGCACACGTCTGAACTCTTTCCCTACACGACGCTCTTCCGATCTNNNTGTTTTTATCCGGAAACTGGT
GAGCACACGTCTGAACTCTTTCCCTACACGACGCTCTTCCGATCTNNNTGTTTTTATCCGGAAACTGGT
AGCACACGTCTGAACTCTTTCCCTACACGACGCTCTTCCGATCTNNNGTTTGCTGCACCCATAAACTCAT
AGCACACGTCTGAACTCTTTCCCTACACGACGCTCTTCCGATCTNNNGTTTGCTGCACCCATAAACTCAT
AAGAGCACACGTCTGAACTCTTTCCCTACACGACGCTCTTCCGATCTNNNGATTCACAAACATCCCCCTT
AAGAGCACACGTCTGAACTCTTTCCCTACACGACGCTCTTCCGATCTNNNGATTCACAAACATCCCCCTT
GCACACGTCTGAACTCTTTCCCTACACGACGCTCTTCCGATCTNNNAAGTCATGCAGAGTTCCACCATT
GCACACGTCTGAACTCTTTCCCTACACGACGCTCTTCCGATCTNNNAAGTCATGCAGAGTTCCACCATT
CACACGTCTGAACTCTTTCCCTACACGACGCTCTTCCGATCTNNNAGCCTCATAAAACAAGAGCTCCAC
CACACGTCTGAACTCTTTCCCTACACGACGCTCTTCCGATCTNNNAGCCTCATAAAACAAGAGCTCCAC
AGAGCACACGTCTGAACTCTTTCCCTACACGACGCTCTTCCGATCTNNNTCCTATGGGATACAGCAAA
AGAGCACACGTCTGAACTCTTTCCCTACACGACGCTCTTCCGATCTNNNTCCTATGGGATACAGCAAA
CACACGTCTGAACTCTTTCCCTACACGACGCTCTTCCGATCTNNNCAATCGCCAGTCAGACACTCACAT
CACACGTCTGAACTCTTTCCCTACACGACGCTCTTCCGATCTNNNCAATCGCCAGTCAGACACTCACAT
AGCACACGTCTGAACTCTTTCCCTACACGACGCTCTTCCGATCTNNNGTGATTCTCTTGAGTAGCTGG
AGCACACGTCTGAACTCTTTCCCTACACGACGCTCTTCCGATCTNNNGTGATTCTCTTGAGTAGCTGG
GAGCACACGTCTGAACTCTTTCCCTACACGACGCTCTTCCGATCTNNNCAGGAGTTTGAGACCAGCCT
GAGCACACGTCTGAACTCTTTCCCTACACGACGCTCTTCCGATCTNNNCAGGAGTTTGAGACCAGCCT
AGCACACGTCTGAACTCTTTCCCTACACGACGCTCTTCCGATCTNNNGTAAATAGTCTCAGCTCCTCGG
AGCACACGTCTGAACTCTTTCCCTACACGACGCTCTTCCGATCTNNNGTAAATAGTCTCAGCTCCTCGG
GAGCACACGTCTGAACTCTTTCCCTACACGACGCTCTTCCGATCTNNNACACAGTCATCTGCTATTAGT
GAGCACACGTCTGAACTCTTTCCCTACACGACGCTCTTCCGATCTNNNACACAGTCATCTGCTATTAGT
AGCACACGTCTGAACTCTTTCCCTACACGACGCTCTTCCGATCTNNNCTTCCTTCCTTCCTTCCTCCCTCC
AGCACACGTCTGAACTCTTTCCCTACACGACGCTCTTCCGATCTNNNCTTCCTTCCTTCCTTCCTCCCTCC
AGCACACGTCTGAACTCTTTCCCTACACGACGCTCTTCCGATCTNNNCCACACCAAATGCTTCTTCAGAA
AGCACACGTCTGAACTCTTTCCCTACACGACGCTCTTCCGATCTNNNCCACACCAAATGCTTCTTCAGAA
AGCACACGTCTGAACTCTTTCCCTACACGACGCTCTTCCGATCTNNNTGACTTGTTCATGGTGGAGAA
AGCACACGTCTGAACTCTTTCCCTACACGACGCTCTTCCGATCTNNNTGACTTGTTCATGGTGGAGAA
GCACACGTCTGAACTCTTTCCCTACACGACGCTCTTCCGATCTNNNAGTAAGCAGAGATCACACCACTG
GCACACGTCTGAACTCTTTCCCTACACGACGCTCTTCCGATCTNNNAGTAAGCAGAGATCACACCACTG
GAGCACACGTCTGAACTCTTTCCCTACACGACGCTCTTCCGATCTNNNGAGTTTGCAGTGAGCCGAG
GAGCACACGTCTGAACTCTTTCCCTACACGACGCTCTTCCGATCTNNNGAGTTTGCAGTGAGCCGAG
GCACACGTCTGAACTCTTTCCCTACACGACGCTCTTCCGATCTNNNTTTCATTGTCTGTGTGCATCACC

GCACACGTCTGAACTCTTTCCCTACACGACGCTCTTCCGATCTNNNTTTCATTGTCTGTGTGCATCACC
AGCACACGTCTGAACTCTTTCCCTACACGACGCTCTTCCGATCTNNNACTGGTGGTATTTGGATTGCT
AGCACACGTCTGAACTCTTTCCCTACACGACGCTCTTCCGATCTNNNACTGGTGGTATTTGGATTGCT
GAGCACACGTCTGAACTCTTTCCCTACACGACGCTCTTCCGATCTNNNGGGGTTTGGCAGAGATGTG
GAGCACACGTCTGAACTCTTTCCCTACACGACGCTCTTCCGATCTNNNGGGGTTTGGCAGAGATGTG
AGCACACGTCTGAACTCTTTCCCTACACGACGCTCTTCCGATCTNNNACTACAGACATGCATCACCAAG
AGCACACGTCTGAACTCTTTCCCTACACGACGCTCTTCCGATCTNNNACTACAGACATGCATCACCAAG
AGCACACGTCTGAACTCTTTCCCTACACGACGCTCTTCCGATCTNNNTCCTAAGTAGCTGAGACCACAG
AGCACACGTCTGAACTCTTTCCCTACACGACGCTCTTCCGATCTNNNTCCTAAGTAGCTGAGACCACAG
AGCACACGTCTGAACTCTTTCCCTACACGACGCTCTTCCGATCTNNNCAGCCAAGGTTGTGATGTTAC
AGCACACGTCTGAACTCTTTCCCTACACGACGCTCTTCCGATCTNNNCAGCCAAGGTTGTGATGTTAC
AGCACACGTCTGAACTCTTTCCCTACACGACGCTCTTCCGATCTNNNAGAATCACTTCAATCCAGGAG
AGCACACGTCTGAACTCTTTCCCTACACGACGCTCTTCCGATCTNNNAGAATCACTTCAATCCAGGAG
AGCACACGTCTGAACTCTTTCCCTACACGACGCTCTTCCGATCTNNNAGGTTGAGGCAGGAGAATCA
AGCACACGTCTGAACTCTTTCCCTACACGACGCTCTTCCGATCTNNNAGGTTGAGGCAGGAGAATCA
AGCACACGTCTGAACTCTTTCCCTACACGACGCTCTTCCGATCTNNNGGCTAACACGGTAAACCTAGT
AGCACACGTCTGAACTCTTTCCCTACACGACGCTCTTCCGATCTNNNGGCTAACACGGTAAACCTAGT
GAGCACACGTCTGAACTCTTTCCCTACACGACGCTCTTCCGATCTNNNTTTGTGCCCTGCTAAAAATCT
GAGCACACGTCTGAACTCTTTCCCTACACGACGCTCTTCCGATCTNNNTTTGTGCCCTGCTAAAAATCT
AGCACACGTCTGAACTCTTTCCCTACACGACGCTCTTCCGATCTNNNACTTTCTGTTTGCCCAATTGGAT
AGCACACGTCTGAACTCTTTCCCTACACGACGCTCTTCCGATCTNNNACTTTCTGTTTGCCCAATTGGAT
GCACACGTCTGAACTCTTTCCCTACACGACGCTCTTCCGATCTNNNTGGTTTCAAACTCCTGACCTCAG
GCACACGTCTGAACTCTTTCCCTACACGACGCTCTTCCGATCTNNNTGGTTTCAAACTCCTGACCTCAG
AGCACACGTCTGAACTCTTTCCCTACACGACGCTCTTCCGATCTNNNAGAATTGCTTCAACCTGGGAG
AGCACACGTCTGAACTCTTTCCCTACACGACGCTCTTCCGATCTNNNAGAATTGCTTCAACCTGGGAG
AGCACACGTCTGAACTCTTTCCCTACACGACGCTCTTCCGATCTNNNAGAGAACCACTTGAACCCAGA
AGCACACGTCTGAACTCTTTCCCTACACGACGCTCTTCCGATCTNNNAGAGAACCACTTGAACCCAGA
GAGCACACGTCTGAACTCTTTCCCTACACGACGCTCTTCCGATCTNNNCCCACCACCTTCATTTCTAGCT
GAGCACACGTCTGAACTCTTTCCCTACACGACGCTCTTCCGATCTNNNCCCACCACCTTCATTTCTAGCT
AGCACACGTCTGAACTCTTTCCCTACACGACGCTCTTCCGATCTNNNATGAGATCTCAGTTCACTGCAC
AGCACACGTCTGAACTCTTTCCCTACACGACGCTCTTCCGATCTNNNATGAGATCTCAGTTCACTGCAC
GCACACGTCTGAACTCTTTCCCTACACGACGCTCTTCCGATCTNNNTCCTTCCTTCCTTCCTTCCTTTCCA
GCACACGTCTGAACTCTTTCCCTACACGACGCTCTTCCGATCTNNNTCCTTCCTTCCTTCCTTCCTTTCCA
AGCACACGTCTGAACTCTTTCCCTACACGACGCTCTTCCGATCTNNNCAGGGAGTTTCATTTTTATCTC
AGCACACGTCTGAACTCTTTCCCTACACGACGCTCTTCCGATCTNNNCAGGGAGTTTCATTTTTATCTC
GCACACGTCTGAACTCTTTCCCTACACGACGCTCTTCCGATCTNNNCTCTTGAACTCCTGACCTTGTGA
GCACACGTCTGAACTCTTTCCCTACACGACGCTCTTCCGATCTNNNCTCTTGAACTCCTGACCTTGTGA
GCACACGTCTGAACTCTTTCCCTACACGACGCTCTTCCGATCTNNNCCAAGAAGATCAGAGTTCTGCC
GCACACGTCTGAACTCTTTCCCTACACGACGCTCTTCCGATCTNNNCCAAGAAGATCAGAGTTCTGCC
AGCACACGTCTGAACTCTTTCCCTACACGACGCTCTTCCGATCTNNNACCACACCTGGCTAAGTTTTG
AGCACACGTCTGAACTCTTTCCCTACACGACGCTCTTCCGATCTNNNACCACACCTGGCTAAGTTTTG
AGCACACGTCTGAACTCTTTCCCTACACGACGCTCTTCCGATCTNNNCACAATGCGATACCATCTCACA
AGCACACGTCTGAACTCTTTCCCTACACGACGCTCTTCCGATCTNNNCACAATGCGATACCATCTCACA

AGCACACGTCTGAACTCTTTCCCTACACGACGCTCTTCCGATCTNNNGAGGTCATGGTAGTGTGAGG
AGCACACGTCTGAACTCTTTCCCTACACGACGCTCTTCCGATCTNNNGAGGTCATGGTAGTGTGAGG
AGCACACGTCTGAACTCTTTCCCTACACGACGCTCTTCCGATCTNNNAGGCTGAAGCAAGAGAACGG
AGCACACGTCTGAACTCTTTCCCTACACGACGCTCTTCCGATCTNNNAGGCTGAAGCAAGAGAACGG
AGCACACGTCTGAACTCTTTCCCTACACGACGCTCTTCCGATCTNNNCAAACCAAGGTAAGTGAAGGT
AGCACACGTCTGAACTCTTTCCCTACACGACGCTCTTCCGATCTNNNCAAACCAAGGTAAGTGAAGGT
AGCACACGTCTGAACTCTTTCCCTACACGACGCTCTTCCGATCTNNNATTTCAGGTAGAGGAAGCCAC
AGCACACGTCTGAACTCTTTCCCTACACGACGCTCTTCCGATCTNNNATTTCAGGTAGAGGAAGCCAC
GAGCACACGTCTGAACTCTTTCCCTACACGACGCTCTTCCGATCTNNNCTGCCCCGCCATTTTTTTTTTT
GAGCACACGTCTGAACTCTTTCCCTACACGACGCTCTTCCGATCTNNNCTGCCCCGCCATTTTTTTTTTT
GCACACGTCTGAACTCTTTCCCTACACGACGCTCTTCCGATCTNNNATCCTCGAGCATCCTTTCTTCAGC
GCACACGTCTGAACTCTTTCCCTACACGACGCTCTTCCGATCTNNNATCCTCGAGCATCCTTTCTTCAGC
AGCACACGTCTGAACTCTTTCCCTACACGACGCTCTTCCGATCTNNNTGGCTGTGTAGTATTCCATGG
AGCACACGTCTGAACTCTTTCCCTACACGACGCTCTTCCGATCTNNNTGGCTGTGTAGTATTCCATGG
AGCACACGTCTGAACTCTTTCCCTACACGACGCTCTTCCGATCTNNNTTTCACAGCCTTAGCCATTGTTG
AGCACACGTCTGAACTCTTTCCCTACACGACGCTCTTCCGATCTNNNTTTCACAGCCTTAGCCATTGTTG
AGCACACGTCTGAACTCTTTCCCTACACGACGCTCTTCCGATCTNNNCTAGGCGACAGAGCAAGACT
AGCACACGTCTGAACTCTTTCCCTACACGACGCTCTTCCGATCTNNNCTAGGCGACAGAGCAAGACT
AGCACACGTCTGAACTCTTTCCCTACACGACGCTCTTCCGATCTNNNCCCTGGTTTGCAATTCATCTAC
AGCACACGTCTGAACTCTTTCCCTACACGACGCTCTTCCGATCTNNNCCCTGGTTTGCAATTCATCTAC
AGCACACGTCTGAACTCTTTCCCTACACGACGCTCTTCCGATCTNNNAGACCCTGTCTCAGAAAAACAC
AGCACACGTCTGAACTCTTTCCCTACACGACGCTCTTCCGATCTNNNAGACCCTGTCTCAGAAAAACAC
AGAGCACACGTCTGAACTCTTTCCCTACACGACGCTCTTCCGATCTNNNCTTGGGAGGCTAGGTTGAA
AGAGCACACGTCTGAACTCTTTCCCTACACGACGCTCTTCCGATCTNNNCTTGGGAGGCTAGGTTGAA
GAGCACACGTCTGAACTCTTTCCCTACACGACGCTCTTCCGATCTNNNGAGACAGGAGAATGGCGTG
GAGCACACGTCTGAACTCTTTCCCTACACGACGCTCTTCCGATCTNNNGAGACAGGAGAATGGCGTG
GAGCACACGTCTGAACTCTTTCCCTACACGACGCTCTTCCGATCTNNNGTATGTGTACAGGCATGTGT
GAGCACACGTCTGAACTCTTTCCCTACACGACGCTCTTCCGATCTNNNGTATGTGTACAGGCATGTGT
AGAGCACACGTCTGAACTCTTTCCCTACACGACGCTCTTCCGATCTNNNACGTCCTAGGAATGGTGAT
AGAGCACACGTCTGAACTCTTTCCCTACACGACGCTCTTCCGATCTNNNACGTCCTAGGAATGGTGAT
GAGCACACGTCTGAACTCTTTCCCTACACGACGCTCTTCCGATCTNNNAGTGTGGTAACCTATTGCTGC
GAGCACACGTCTGAACTCTTTCCCTACACGACGCTCTTCCGATCTNNNAGTGTGGTAACCTATTGCTGC
AGCACACGTCTGAACTCTTTCCCTACACGACGCTCTTCCGATCTNNNAATTTCACTCCCTAGTTCAGGC
AGCACACGTCTGAACTCTTTCCCTACACGACGCTCTTCCGATCTNNNAATTTCACTCCCTAGTTCAGGC
AGAGCACACGTCTGAACTCTTTCCCTACACGACGCTCTTCCGATCTNNNTCAGATAGGGGGAACACAG
AGAGCACACGTCTGAACTCTTTCCCTACACGACGCTCTTCCGATCTNNNTCAGATAGGGGGAACACAG
AGCACACGTCTGAACTCTTTCCCTACACGACGCTCTTCCGATCTNNNGCCTTTGCTTTCTTTGCTTTCA
AGCACACGTCTGAACTCTTTCCCTACACGACGCTCTTCCGATCTNNNGCCTTTGCTTTCTTTGCTTTCA
GAGCACACGTCTGAACTCTTTCCCTACACGACGCTCTTCCGATCTNNNGCATATGAACACACACCCATC
GAGCACACGTCTGAACTCTTTCCCTACACGACGCTCTTCCGATCTNNNGCATATGAACACACACCCATC
GAGCACACGTCTGAACTCTTTCCCTACACGACGCTCTTCCGATCTNNNGCAACTTAGCAAGACCTTGTC
GAGCACACGTCTGAACTCTTTCCCTACACGACGCTCTTCCGATCTNNNGCAACTTAGCAAGACCTTGTC
AGCACACGTCTGAACTCTTTCCCTACACGACGCTCTTCCGATCTNNNCGTGGATGACAGATCTGACCT

AGCACACGTCTGAACTCTTTCCCTACACGACGCTCTTCCGATCTNNNCGTGGATGACAGATCTGACCT

GCACACGTCTGAACTCTTTCCCTACACGACGCTCTTCCGATCTNNNCTGAGGCAGGAGAATTGCTTG

GCACACGTCTGAACTCTTTCCCTACACGACGCTCTTCCGATCTNNNCTGAGGCAGGAGAATTGCTTG

AGCACACGTCTGAACTCTTTCCCTACACGACGCTCTTCCGATCTNNNGAAAGATCATGGCTCTGGGA

AGCACACGTCTGAACTCTTTCCCTACACGACGCTCTTCCGATCTNNNGAAAGATCATGGCTCTGGGA

AGCACACGTCTGAACTCTTTCCCTACACGACGCTCTTCCGATCTNNNTTGTGGAACAGGGAGCTGAG

AGCACACGTCTGAACTCTTTCCCTACACGACGCTCTTCCGATCTNNNTTGTGGAACAGGGAGCTGAG

GCACACGTCTGAACTCTTTCCCTACACGACGCTCTTCCGATCTNNNTCCTCCTCCTTCTTCTCCTTCTTCA

GCACACGTCTGAACTCTTTCCCTACACGACGCTCTTCCGATCTNNNTCCTCCTCCTTCTTCTCCTTCTTCA

AGAGCACACGTCTGAACTCTTTCCCTACACGACGCTCTTCCGATCTNNNCTGCCTGTTTTCTGAATCAC

AGAGCACACGTCTGAACTCTTTCCCTACACGACGCTCTTCCGATCTNNNCTGCCTGTTTTCTGAATCAC

AGCACACGTCTGAACTCTTTCCCTACACGACGCTCTTCCGATCTNNNGAGTGCAATGGTATGATCTTG

AGCACACGTCTGAACTCTTTCCCTACACGACGCTCTTCCGATCTNNNGAGTGCAATGGTATGATCTTG

GCACACGTCTGAACTCTTTCCCTACACGACGCTCTTCCGATCTNNNCTGAGAGTACACCACTGAACTC

GCACACGTCTGAACTCTTTCCCTACACGACGCTCTTCCGATCTNNNCTGAGAGTACACCACTGAACTC

AGAGCACACGTCTGAACTCTTTCCCTACACGACGCTCTTCCGATCTNNNCTACTAGGCCCTACACACAC

AGAGCACACGTCTGAACTCTTTCCCTACACGACGCTCTTCCGATCTNNNCTACTAGGCCCTACACACAC

AGCACACGTCTGAACTCTTTCCCTACACGACGCTCTTCCGATCTNNNTGGGGCTTAGATATCTACCCA

GAGCACACGTCTGAACTCTTTCCCTACACGACGCTCTTCCGATCTNNNTGGGGCTTAGATATCTACCCA

AGCACACGTCTGAACTCTTTCCCTACACGACGCTCTTCCGATCTNNNGTCTTTAGGAGACACACACA

GAGCACACGTCTGAACTCTTTCCCTACACGACGCTCTTCCGATCTNNNGTCTTTAGGAGACACACACA

GCACACGTCTGAACTCTTTCCCTACACGACGCTCTTCCGATCTNNNCTGAGGCAGGAGAATCACTTGA

GCACACGTCTGAACTCTTTCCCTACACGACGCTCTTCCGATCTNNNCTGAGGCAGGAGAATCACTTGA

AGCACACGTCTGAACTCTTTCCCTACACGACGCTCTTCCGATCTNNNGATCACCTGAGGTCAGCAGTT

AGCACACGTCTGAACTCTTTCCCTACACGACGCTCTTCCGATCTNNNGATCACCTGAGGTCAGCAGTT

GCACACGTCTGAACTCTTTCCCTACACGACGCTCTTCCGATCTNNNGATGAACCCGATTCGTCTGTTG

GCACACGTCTGAACTCTTTCCCTACACGACGCTCTTCCGATCTNNNGATGAACCCGATTCGTCTGTTG

CACACGTCTGAACTCTTTCCCTACACGACGCTCTTCCGATCTNNNAGAAAGACTTGTGCCTTGGAAGA

CACACGTCTGAACTCTTTCCCTACACGACGCTCTTCCGATCTNNNAGAAAGACTTGTGCCTTGGAAGA

GCACACGTCTGAACTCTTTCCCTACACGACGCTCTTCCGATCTNNNGCATGACAGTAGAAGAGCTGC

GCACACGTCTGAACTCTTTCCCTACACGACGCTCTTCCGATCTNNNGCATGACAGTAGAAGAGCTGC

AGCACACGTCTGAACTCTTTCCCTACACGACGCTCTTCCGATCTNNNTGAGGCAGGAGAATGGTATG

AGCACACGTCTGAACTCTTTCCCTACACGACGCTCTTCCGATCTNNNTGAGGCAGGAGAATGGTATG

AGCACACGTCTGAACTCTTTCCCTACACGACGCTCTTCCGATCTNNNATTGCTTGAGCCTAGGAGTTT

AGCACACGTCTGAACTCTTTCCCTACACGACGCTCTTCCGATCTNNNATTGCTTGAGCCTAGGAGTTT

GAGCACACGTCTGAACTCTTTCCCTACACGACGCTCTTCCGATCTNNNCTGTAGTCCCAGCTACTCGAG

GAGCACACGTCTGAACTCTTTCCCTACACGACGCTCTTCCGATCTNNNCTGTAGTCCCAGCTACTCGAG

AGCACACGTCTGAACTCTTTCCCTACACGACGCTCTTCCGATCTNNNGTAATAGTCTTCCAGGGCAGA

AGCACACGTCTGAACTCTTTCCCTACACGACGCTCTTCCGATCTNNNGTAATAGTCTTCCAGGGCAGA

AAGAGCACACGTCTGAACTCTTTCCCTACACGACGCTCTTCCGATCTNNNGCGACACCCTGTCCCTCTA

AAGAGCACACGTCTGAACTCTTTCCCTACACGACGCTCTTCCGATCTNNNGCGACACCCTGTCCCTCTA

GCACACGTCTGAACTCTTTCCCTACACGACGCTCTTCCGATCTNNNAAAAATGAGCCTTTCTCCTCCCC

GCACACGTCTGAACTCTTTCCCTACACGACGCTCTTCCGATCTNNNAAAAATGAGCCTTTCTCCTCCCC

CACACGTCTGAACTCTTTCCCTACACGACGCTCTTCCGATCTNNNCTCAAATTCTCTTGCTTTTCTATCA

CACACGTCTGAACTCTTTCCCTACACGACGCTCTTCCGATCTNNNCTCAAATTCTCTTGCTTTTCTATCA

AGCACACGTCTGAACTCTTTCCCTACACGACGCTCTTCCGATCTNNNATTACAAGCATGAGCCATGTG

AGCACACGTCTGAACTCTTTCCCTACACGACGCTCTTCCGATCTNNNATTACAAGCATGAGCCATGTG

AGCACACGTCTGAACTCTTTCCCTACACGACGCTCTTCCGATCTNNNCTGAGGCAGGAGAATCACTT

AGCACACGTCTGAACTCTTTCCCTACACGACGCTCTTCCGATCTNNNCTGAGGCAGGAGAATCACTT

ACACGTCTGAACTCTTTCCCTACACGACGCTCTTCCGATCTNNNAGAATTTTAGGTTGGTGGGCCTTC

ACACGTCTGAACTCTTTCCCTACACGACGCTCTTCCGATCTNNNAGAATTTTAGGTTGGTGGGCCTTC

GAGCACACGTCTGAACTCTTTCCCTACACGACGCTCTTCCGATCTNNNTATGACAGGGAGCTAACTGA

GAGCACACGTCTGAACTCTTTCCCTACACGACGCTCTTCCGATCTNNNTATGACAGGGAGCTAACTGA

GCACACGTCTGAACTCTTTCCCTACACGACGCTCTTCCGATCTNNNGATGAGTAAGGTCGGTTGTGT

GCACACGTCTGAACTCTTTCCCTACACGACGCTCTTCCGATCTNNNGATGAGTAAGGTCGGTTGTGT

GAGCACACGTCTGAACTCTTTCCCTACACGACGCTCTTCCGATCTNNNTCAACCTCGCCAAAAATCAAA

GAGCACACGTCTGAACTCTTTCCCTACACGACGCTCTTCCGATCTNNNTCAACCTCGCCAAAAATCAAA

GAGCACACGTCTGAACTCTTTCCCTACACGACGCTCTTCCGATCTNNNGTGTGTGAAGGGATCAGG

GAGCACACGTCTGAACTCTTTCCCTACACGACGCTCTTCCGATCTNNNGTGTGTGAAGGGATCAGG

AGCACACGTCTGAACTCTTTCCCTACACGACGCTCTTCCGATCTNNNTGGAAGATGAGAAAACACTT

AGCACACGTCTGAACTCTTTCCCTACACGACGCTCTTCCGATCTNNNTGGAAGATGAGAAAACACTT

AGCACACGTCTGAACTCTTTCCCTACACGACGCTCTTCCGATCTNNNTGCTTGTAATCCCAGCTACTC

AGCACACGTCTGAACTCTTTCCCTACACGACGCTCTTCCGATCTNNNTGCTTGTAATCCCAGCTACTC

AGCACACGTCTGAACTCTTTCCCTACACGACGCTCTTCCGATCTNNNGAGTTTGAGCAACATAGCGAG

AGCACACGTCTGAACTCTTTCCCTACACGACGCTCTTCCGATCTNNNGAGTTTGAGCAACATAGCGAG

GCACACGTCTGAACTCTTTCCCTACACGACGCTCTTCCGATCTNNNCCTCTTTTCTTCCTAGCCTGGTTC

GCACACGTCTGAACTCTTTCCCTACACGACGCTCTTCCGATCTNNNCCTCTTTTCTTCCTAGCCTGGTTC

AGCACACGTCTGAACTCTTTCCCTACACGACGCTCTTCCGATCTNNNAAAAAATTAGCCGGACATGG

AGCACACGTCTGAACTCTTTCCCTACACGACGCTCTTCCGATCTNNNAAAAAATTAGCCGGACATGG

GCACACGTCTGAACTCTTTCCCTACACGACGCTCTTCCGATCTNNNAGTGACAGATTCATTCGCACTCC

GCACACGTCTGAACTCTTTCCCTACACGACGCTCTTCCGATCTNNNAGTGACAGATTCATTCGCACTCC

GCACACGTCTGAACTCTTTCCCTACACGACGCTCTTCCGATCTNNNCCCGCAGTTTTTTGTACAGACAT

GCACACGTCTGAACTCTTTCCCTACACGACGCTCTTCCGATCTNNNCCCGCAGTTTTTTGTACAGACAT

AGCACACGTCTGAACTCTTTCCCTACACGACGCTCTTCCGATCTNNNTCATGCTACCTTGCTGAACTCAT

AGCACACGTCTGAACTCTTTCCCTACACGACGCTCTTCCGATCTNNNTCATGCTACCTTGCTGAACTCAT

AGCACACGTCTGAACTCTTTCCCTACACGACGCTCTTCCGATCTNNNTTTCAGCCTGAAGAACTCCCTT

AGCACACGTCTGAACTCTTTCCCTACACGACGCTCTTCCGATCTNNNTTTCAGCCTGAAGAACTCCCTT

CACACGTCTGAACTCTTTCCCTACACGACGCTCTTCCGATCTNNNGGACTGAACCCGAGGATTCACCA

CACACGTCTGAACTCTTTCCCTACACGACGCTCTTCCGATCTNNNGGACTGAACCCGAGGATTCACCA

AGAGCACACGTCTGAACTCTTTCCCTACACGACGCTCTTCCGATCTNNNGGAATGGGTTTGAGGTGG

AGAGCACACGTCTGAACTCTTTCCCTACACGACGCTCTTCCGATCTNNNGGAATGGGTTTGAGGTGG

AGCACACGTCTGAACTCTTTCCCTACACGACGCTCTTCCGATCTNNNGGGGATGAGAGAGAGAGAA

AGCACACGTCTGAACTCTTTCCCTACACGACGCTCTTCCGATCTNNNGGGGATGAGAGAGAGAGAA

GCACACGTCTGAACTCTTTCCCTACACGACGCTCTTCCGATCTNNNTGAGGTAGGAGAATGGCATGA

GCACACGTCTGAACTCTTTCCCTACACGACGCTCTTCCGATCTNNNTGAGGTAGGAGAATGGCATGA

GCACACGTCTGAACTCTTTCCCTACACGACGCTCTTCCGATCTNNNCTCCTCCACAAACAAAGCAAAC

AGCACACGTCTGAACTCTTTCCCTACACGACGCTCTTCCGATCTNNNCTCCTCCACAAACAAAGCAAACA
GAGCACACGTCTGAACTCTTTCCCTACACGACGCTCTTCCGATCTNNNAATTGTCCCATAAGCATCTCCC
GAGCACACGTCTGAACTCTTTCCCTACACGACGCTCTTCCGATCTNNNAATTGTCCCATAAGCATCTCCC
GAGCACACGTCTGAACTCTTTCCCTACACGACGCTCTTCCGATCTNNNTACAAAAAATTAGCCAGGCGA
GAGCACACGTCTGAACTCTTTCCCTACACGACGCTCTTCCGATCTNNNTACAAAAAATTAGCCAGGCGA
GAGCACACGTCTGAACTCTTTCCCTACACGACGCTCTTCCGATCTNNNCACAGAAAATGCTTCTCTCTGC
GAGCACACGTCTGAACTCTTTCCCTACACGACGCTCTTCCGATCTNNNCACAGAAAATGCTTCTCTCTGC
AGCACACGTCTGAACTCTTTCCCTACACGACGCTCTTCCGATCTNNNGTCTCCCTGTCTCTCTCTTCTCTCA
AGCACACGTCTGAACTCTTTCCCTACACGACGCTCTTCCGATCTNNNGTCTCCCTGTCTCTCTCTTCTCTCA
AGAGCACACGTCTGAACTCTTTCCCTACACGACGCTCTTCCGATCTNNNGTGAAAACAGGAGGTTGGA
AGAGCACACGTCTGAACTCTTTCCCTACACGACGCTCTTCCGATCTNNNGTGAAAACAGGAGGTTGGA
GAGCACACGTCTGAACTCTTTCCCTACACGACGCTCTTCCGATCTNNNCACTAGCAATTTTCATCCAGCG
GAGCACACGTCTGAACTCTTTCCCTACACGACGCTCTTCCGATCTNNNCACTAGCAATTTTCATCCAGCG
AGCACACGTCTGAACTCTTTCCCTACACGACGCTCTTCCGATCTNNNCTGGGCAACAAAAGAGAAACTC
AGCACACGTCTGAACTCTTTCCCTACACGACGCTCTTCCGATCTNNNCTGGGCAACAAAAGAGAAACTC
AGCACACGTCTGAACTCTTTCCCTACACGACGCTCTTCCGATCTNNNAGAGCTTTCAGTGAGCCAAGAT
AGCACACGTCTGAACTCTTTCCCTACACGACGCTCTTCCGATCTNNNAGAGCTTTCAGTGAGCCAAGAT
AAGAGCACACGTCTGAACTCTTTCCCTACACGACGCTCTTCCGATCTNNNCAATCTCGGCTCACTACAAC
AAGAGCACACGTCTGAACTCTTTCCCTACACGACGCTCTTCCGATCTNNNCAATCTCGGCTCACTACAAC
AAGAGCACACGTCTGAACTCTTTCCCTACACGACGCTCTTCCGATCTNNNCGCACCTATAGTCCTTGCTA
AAGAGCACACGTCTGAACTCTTTCCCTACACGACGCTCTTCCGATCTNNNCGCACCTATAGTCCTTGCTA
AGCACACGTCTGAACTCTTTCCCTACACGACGCTCTTCCGATCTNNNCCCTCCTCCTATACCTCTCAACACA
AGCACACGTCTGAACTCTTTCCCTACACGACGCTCTTCCGATCTNNNCCCTCCTCCTATACCTCTCAACACA
GAGCACACGTCTGAACTCTTTCCCTACACGACGCTCTTCCGATCTNNNGCCTTTGCATGTGTCTGTGTT
GAGCACACGTCTGAACTCTTTCCCTACACGACGCTCTTCCGATCTNNNGCCTTTGCATGTGTCTGTGTT
GCACACGTCTGAACTCTTTCCCTACACGACGCTCTTCCGATCTNNNCCAGTCTGTTTTTCAGCAAAGGTC
GCACACGTCTGAACTCTTTCCCTACACGACGCTCTTCCGATCTNNNCCAGTCTGTTTTTCAGCAAAGGTC
AGCACACGTCTGAACTCTTTCCCTACACGACGCTCTTCCGATCTNNNAAAAGAGGGACTGACAGGGTT
AGCACACGTCTGAACTCTTTCCCTACACGACGCTCTTCCGATCTNNNAAAAGAGGGACTGACAGGGTT
AGCACACGTCTGAACTCTTTCCCTACACGACGCTCTTCCGATCTNNNAAGGAGAATCACTTGAACCTGG
AGCACACGTCTGAACTCTTTCCCTACACGACGCTCTTCCGATCTNNNAAGGAGAATCACTTGAACCTGG
AGAGCACACGTCTGAACTCTTTCCCTACACGACGCTCTTCCGATCTNNNAGCACATCATGGGACTTCTC
AGAGCACACGTCTGAACTCTTTCCCTACACGACGCTCTTCCGATCTNNNAGCACATCATGGGACTTCTC
GAGCACACGTCTGAACTCTTTCCCTACACGACGCTCTTCCGATCTNNNTCACCTGGGTATAGCCTCAAGT
GAGCACACGTCTGAACTCTTTCCCTACACGACGCTCTTCCGATCTNNNTCACCTGGGTATAGCCTCAAGT
CACACGTCTGAACTCTTTCCCTACACGACGCTCTTCCGATCTNNNTGATTGGTTATTCAACAGATATGCA
CACACGTCTGAACTCTTTCCCTACACGACGCTCTTCCGATCTNNNTGATTGGTTATTCAACAGATATGCA
AGCACACGTCTGAACTCTTTCCCTACACGACGCTCTTCCGATCTNNNTAATCTCAGCACTTTGGGAAGC
AGCACACGTCTGAACTCTTTCCCTACACGACGCTCTTCCGATCTNNNTAATCTCAGCACTTTGGGAAGC
AGCACACGTCTGAACTCTTTCCCTACACGACGCTCTTCCGATCTNNNTATGTTCCCCTTTCTCTGCAACAC
AGCACACGTCTGAACTCTTTCCCTACACGACGCTCTTCCGATCTNNNTATGTTCCCCTTTCTCTGCAACAC
GAGCACACGTCTGAACTCTTTCCCTACACGACGCTCTTCCGATCTNNNGCAGGATAATCGTTTGAACCC
GAGCACACGTCTGAACTCTTTCCCTACACGACGCTCTTCCGATCTNNNGCAGGATAATCGTTTGAACCC

GAGCACACGTCTGAACTCTTTCCCTACACGACGCTCTTCCGATCTNNNCTGAGGTGGGAGGATTACT
GAGCACACGTCTGAACTCTTTCCCTACACGACGCTCTTCCGATCTNNNCTGAGGTGGGAGGATTACT
AGCACACGTCTGAACTCTTTCCCTACACGACGCTCTTCCGATCTNNNTTGGTGTACAGTTCTGGTTGT
AGCACACGTCTGAACTCTTTCCCTACACGACGCTCTTCCGATCTNNNTTGGTGTACAGTTCTGGTTGT
AGCACACGTCTGAACTCTTTCCCTACACGACGCTCTTCCGATCTNNNAGTGGCATGATCACAGTTCAC
AGCACACGTCTGAACTCTTTCCCTACACGACGCTCTTCCGATCTNNNAGTGGCATGATCACAGTTCAC
AGCACACGTCTGAACTCTTTCCCTACACGACGCTCTTCCGATCTNNNGCAGTTCTATCATGGTTAAGA
AGCACACGTCTGAACTCTTTCCCTACACGACGCTCTTCCGATCTNNNGCAGTTCTATCATGGTTAAGA
GAGCACACGTCTGAACTCTTTCCCTACACGACGCTCTTCCGATCTNNNTATTTCCCAACGCTGTTCTAG
GAGCACACGTCTGAACTCTTTCCCTACACGACGCTCTTCCGATCTNNNTATTTCCCAACGCTGTTCTAG
GAGCACACGTCTGAACTCTTTCCCTACACGACGCTCTTCCGATCTNNNGGTGACAGAGAGAGATTCC
GAGCACACGTCTGAACTCTTTCCCTACACGACGCTCTTCCGATCTNNNGGTGACAGAGAGAGATTCC
GCACACGTCTGAACTCTTTCCCTACACGACGCTCTTCCGATCTNNNCTCTAACCTCACGTTTCTCCACAC
GCACACGTCTGAACTCTTTCCCTACACGACGCTCTTCCGATCTNNNCTCTAACCTCACGTTTCTCCACAC
AGCACACGTCTGAACTCTTTCCCTACACGACGCTCTTCCGATCTNNNGTAACATGTGAGCTGGAGAG
AGCACACGTCTGAACTCTTTCCCTACACGACGCTCTTCCGATCTNNNGTAACATGTGAGCTGGAGAG
GCACACGTCTGAACTCTTTCCCTACACGACGCTCTTCCGATCTNNNTGCTCTTCCACCTTCTCTTTACCC
GCACACGTCTGAACTCTTTCCCTACACGACGCTCTTCCGATCTNNNTGCTCTTCCACCTTCTCTTTACCC
GAGCACACGTCTGAACTCTTTCCCTACACGACGCTCTTCCGATCTNNNCCTGCTTTCTCAGGTTTGTTTT
GAGCACACGTCTGAACTCTTTCCCTACACGACGCTCTTCCGATCTNNNCCTGCTTTCTCAGGTTTGTTTT
AGCACACGTCTGAACTCTTTCCCTACACGACGCTCTTCCGATCTNNNCCTGACCTCAAGTGATTTGCTT
AGCACACGTCTGAACTCTTTCCCTACACGACGCTCTTCCGATCTNNNCCTGACCTCAAGTGATTTGCTT
AGCACACGTCTGAACTCTTTCCCTACACGACGCTCTTCCGATCTNNNCGAGAATTGTTCCATGATAAAT
AGCACACGTCTGAACTCTTTCCCTACACGACGCTCTTCCGATCTNNNCGAGAATTGTTCCATGATAAAT
AGCACACGTCTGAACTCTTTCCCTACACGACGCTCTTCCGATCTNNNCTGTCACATTTCATAGAAACTG
AGCACACGTCTGAACTCTTTCCCTACACGACGCTCTTCCGATCTNNNCTGTCACATTTCATAGAAACTG
GAGCACACGTCTGAACTCTTTCCCTACACGACGCTCTTCCGATCTNNNCTGAGGCAGGAGAATCACTT
GAGCACACGTCTGAACTCTTTCCCTACACGACGCTCTTCCGATCTNNNCTGAGGCAGGAGAATCACTT
AGCACACGTCTGAACTCTTTCCCTACACGACGCTCTTCCGATCTNNNGTTTACCCTGGTAATGATGCT
AGCACACGTCTGAACTCTTTCCCTACACGACGCTCTTCCGATCTNNNGTTTACCCTGGTAATGATGCT
AGCACACGTCTGAACTCTTTCCCTACACGACGCTCTTCCGATCTNNNGTTCTTCAGATCAAGGTCCCCA
AGCACACGTCTGAACTCTTTCCCTACACGACGCTCTTCCGATCTNNNGTTCTTCAGATCAAGGTCCCCA
AGCACACGTCTGAACTCTTTCCCTACACGACGCTCTTCCGATCTNNNACAAACGTGCAGGTGTCTTTAT
AGCACACGTCTGAACTCTTTCCCTACACGACGCTCTTCCGATCTNNNACAAACGTGCAGGTGTCTTTAT
AGCACACGTCTGAACTCTTTCCCTACACGACGCTCTTCCGATCTNNNGATTCCCAGTGCTAGAGCTGA
AGCACACGTCTGAACTCTTTCCCTACACGACGCTCTTCCGATCTNNNGATTCCCAGTGCTAGAGCTGA
AGCACACGTCTGAACTCTTTCCCTACACGACGCTCTTCCGATCTNNNAGGCTCTTGAAGATGGTGTTC
AGCACACGTCTGAACTCTTTCCCTACACGACGCTCTTCCGATCTNNNAGGCTCTTGAAGATGGTGTTC
CACACGTCTGAACTCTTTCCCTACACGACGCTCTTCCGATCTNNNTGGGTGACAGAGCGAGACTCCAT
CACACGTCTGAACTCTTTCCCTACACGACGCTCTTCCGATCTNNNTGGGTGACAGAGCGAGACTCCAT
GAGCACACGTCTGAACTCTTTCCCTACACGACGCTCTTCCGATCTNNNAATAGGTTACAGCGAGCCAA
GAGCACACGTCTGAACTCTTTCCCTACACGACGCTCTTCCGATCTNNNAATAGGTTACAGCGAGCCAA
AGCACACGTCTGAACTCTTTCCCTACACGACGCTCTTCCGATCTNNNTGCATCTCTGTTTGTCAGTCAC

AGCACACGTCTGAACTCTTTCCCTACACGACGCTCTTCCGATCTNNNTGCATCTCTGTTTGTCAGTCAC
AGCACACGTCTGAACTCTTTCCCTACACGACGCTCTTCCGATCTNNNTCATTTGCTTCAGAGGATGGCT
AGCACACGTCTGAACTCTTTCCCTACACGACGCTCTTCCGATCTNNNTCATTTGCTTCAGAGGATGGCT
GCACACGTCTGAACTCTTTCCCTACACGACGCTCTTCCGATCTNNNGACAGACCATTGAGACAGAGG
GCACACGTCTGAACTCTTTCCCTACACGACGCTCTTCCGATCTNNNGACAGACCATTGAGACAGAGG
GAGCACACGTCTGAACTCTTTCCCTACACGACGCTCTTCCGATCTNNNCTCTCAGTGTCTTGATCAGGC
GAGCACACGTCTGAACTCTTTCCCTACACGACGCTCTTCCGATCTNNNCTCTCAGTGTCTTGATCAGGC
GAGCACACGTCTGAACTCTTTCCCTACACGACGCTCTTCCGATCTNNNTGCTGTTCTCCTAACCAATGT
GAGCACACGTCTGAACTCTTTCCCTACACGACGCTCTTCCGATCTNNNTGCTGTTCTCCTAACCAATGT
AGCACACGTCTGAACTCTTTCCCTACACGACGCTCTTCCGATCTNNNGAACTAGCTCCTGCATAGCACT
AGCACACGTCTGAACTCTTTCCCTACACGACGCTCTTCCGATCTNNNGAACTAGCTCCTGCATAGCACT
AGCACACGTCTGAACTCTTTCCCTACACGACGCTCTTCCGATCTNNNCGACAGTGAGATCCTGTGTCA
AGCACACGTCTGAACTCTTTCCCTACACGACGCTCTTCCGATCTNNNCGACAGTGAGATCCTGTGTCA
AGCACACGTCTGAACTCTTTCCCTACACGACGCTCTTCCGATCTNNNGGTGCCTATAATCCCAGCTACT
AGCACACGTCTGAACTCTTTCCCTACACGACGCTCTTCCGATCTNNNGGTGCCTATAATCCCAGCTACT
AGCACACGTCTGAACTCTTTCCCTACACGACGCTCTTCCGATCTNNNGGGAGGGATAGCATTGGGA
AGCACACGTCTGAACTCTTTCCCTACACGACGCTCTTCCGATCTNNNGGGAGGGATAGCATTGGGA
GAGCACACGTCTGAACTCTTTCCCTACACGACGCTCTTCCGATCTNNNTACTGTTCACACAGCATCTT
GAGCACACGTCTGAACTCTTTCCCTACACGACGCTCTTCCGATCTNNNTACTGTTCACACAGCATCTT
GAGCACACGTCTGAACTCTTTCCCTACACGACGCTCTTCCGATCTNNNCACCATCCATCTTCATTCCAGG
GAGCACACGTCTGAACTCTTTCCCTACACGACGCTCTTCCGATCTNNNCACCATCCATCTTCATTCCAGG
AGCACACGTCTGAACTCTTTCCCTACACGACGCTCTTCCGATCTNNNAGTGAGCCGAGATCTCATCAT
AGCACACGTCTGAACTCTTTCCCTACACGACGCTCTTCCGATCTNNNAGTGAGCCGAGATCTCATCAT
AGCACACGTCTGAACTCTTTCCCTACACGACGCTCTTCCGATCTNNNGTCTGTTCATGTACTTTGCCCA
AGCACACGTCTGAACTCTTTCCCTACACGACGCTCTTCCGATCTNNNGTCTGTTCATGTACTTTGCCCA
CACGTCTGAACTCTTTCCCTACACGACGCTCTTCCGATCTNNNCCCGTCTACGAGAAGTGCATGCATC
CACGTCTGAACTCTTTCCCTACACGACGCTCTTCCGATCTNNNCCCGTCTACGAGAAGTGCATGCATC
GCACACGTCTGAACTCTTTCCCTACACGACGCTCTTCCGATCTNNNTTTATAGCTGAGTAGTGCTCCG
GCACACGTCTGAACTCTTTCCCTACACGACGCTCTTCCGATCTNNNTTTATAGCTGAGTAGTGCTCCG
AGCACACGTCTGAACTCTTTCCCTACACGACGCTCTTCCGATCTNNNTAAAGACCAAAGGCACTCTTC
AGCACACGTCTGAACTCTTTCCCTACACGACGCTCTTCCGATCTNNNTAAAGACCAAAGGCACTCTTC
AGCACACGTCTGAACTCTTTCCCTACACGACGCTCTTCCGATCTNNNCGAGGAGTGGGGTAATTCCT
AGCACACGTCTGAACTCTTTCCCTACACGACGCTCTTCCGATCTNNNCGAGGAGTGGGGTAATTCCT
AGCACACGTCTGAACTCTTTCCCTACACGACGCTCTTCCGATCTNNNGATACAGATGCTACTTGACTTA
AGCACACGTCTGAACTCTTTCCCTACACGACGCTCTTCCGATCTNNNGATACAGATGCTACTTGACTTA
AGCACACGTCTGAACTCTTTCCCTACACGACGCTCTTCCGATCTNNNTGATGAAATTCTGGTGCACAC
AGCACACGTCTGAACTCTTTCCCTACACGACGCTCTTCCGATCTNNNTGATGAAATTCTGGTGCACAC
GAGCACACGTCTGAACTCTTTCCCTACACGACGCTCTTCCGATCTNNNATCCTTTTCTACCTCTGCTCC
GAGCACACGTCTGAACTCTTTCCCTACACGACGCTCTTCCGATCTNNNATCCTTTTCTACCTCTGCTCC
GCACACGTCTGAACTCTTTCCCTACACGACGCTCTTCCGATCTNNNACACTCGCACTCACATAAATCCT
GCACACGTCTGAACTCTTTCCCTACACGACGCTCTTCCGATCTNNNACACTCGCACTCACATAAATCCT
GCACACGTCTGAACTCTTTCCCTACACGACGCTCTTCCGATCTNNNTGCTTTGCTTTTACTTGGTGTCT
GCACACGTCTGAACTCTTTCCCTACACGACGCTCTTCCGATCTNNNTGCTTTGCTTTTACTTGGTGTCT

GAGCACACGTCTGAACTCTTTCCCTACACGACGCTCTTCCGATCTNNNAGGCTGTGTTAGATCAGGTTG
GAGCACACGTCTGAACTCTTTCCCTACACGACGCTCTTCCGATCTNNNAGGCTGTGTTAGATCAGGTTG
AGCACACGTCTGAACTCTTTCCCTACACGACGCTCTTCCGATCTNNNCTGAGGCAGGAGAATTGCTTGA
AGCACACGTCTGAACTCTTTCCCTACACGACGCTCTTCCGATCTNNNCTGAGGCAGGAGAATTGCTTGA
GAGCACACGTCTGAACTCTTTCCCTACACGACGCTCTTCCGATCTNNNATCGTTTTCTGTTACAGGAGGC
GAGCACACGTCTGAACTCTTTCCCTACACGACGCTCTTCCGATCTNNNATCGTTTTCTGTTACAGGAGGC
AGCACACGTCTGAACTCTTTCCCTACACGACGCTCTTCCGATCTNNNATGCCTGTAATCCCAGCTACTTG
AGCACACGTCTGAACTCTTTCCCTACACGACGCTCTTCCGATCTNNNATGCCTGTAATCCCAGCTACTTG
GAGCACACGTCTGAACTCTTTCCCTACACGACGCTCTTCCGATCTNNNTAAGAAATTGGTGGCTAGAAT
GAGCACACGTCTGAACTCTTTCCCTACACGACGCTCTTCCGATCTNNNTAAGAAATTGGTGGCTAGAAT
GCACACGTCTGAACTCTTTCCCTACACGACGCTCTTCCGATCTNNNGAGGAGGGCACTAAGGACTAGC
GCACACGTCTGAACTCTTTCCCTACACGACGCTCTTCCGATCTNNNGAGGAGGGCACTAAGGACTAGC
AGCACACGTCTGAACTCTTTCCCTACACGACGCTCTTCCGATCTNNNGGTGAAAAAAGGAAAGGCAGC
AGCACACGTCTGAACTCTTTCCCTACACGACGCTCTTCCGATCTNNNGGTGAAAAAAGGAAAGGCAGC
GAGCACACGTCTGAACTCTTTCCCTACACGACGCTCTTCCGATCTNNNCCTGGCAATCACAGTGTGAAA
GAGCACACGTCTGAACTCTTTCCCTACACGACGCTCTTCCGATCTNNNCCTGGCAATCACAGTGTGAAA
AGAGCACACGTCTGAACTCTTTCCCTACACGACGCTCTTCCGATCTNNNGCCTGTAATCCTAGCTACTCA
AGAGCACACGTCTGAACTCTTTCCCTACACGACGCTCTTCCGATCTNNNGCCTGTAATCCTAGCTACTCA
GCACACGTCTGAACTCTTTCCCTACACGACGCTCTTCCGATCTNNNTGTCTTAGAACTTCCTCGACACCC
GCACACGTCTGAACTCTTTCCCTACACGACGCTCTTCCGATCTNNNTGTCTTAGAACTTCCTCGACACCC
AGAGCACACGTCTGAACTCTTTCCCTACACGACGCTCTTCCGATCTNNNTCTACACACATACACACACAC
AGAGCACACGTCTGAACTCTTTCCCTACACGACGCTCTTCCGATCTNNNTCTACACACATACACACACAC
AGAGCACACGTCTGAACTCTTTCCCTACACGACGCTCTTCCGATCTNNNCCAGGTATTCAGGAGGCATA
AGAGCACACGTCTGAACTCTTTCCCTACACGACGCTCTTCCGATCTNNNCCAGGTATTCAGGAGGCATA
AGCACACGTCTGAACTCTTTCCCTACACGACGCTCTTCCGATCTNNNGTAGGTCGGGAACTGCTAAAGT
AGCACACGTCTGAACTCTTTCCCTACACGACGCTCTTCCGATCTNNNGTAGGTCGGGAACTGCTAAAGT
GAGCACACGTCTGAACTCTTTCCCTACACGACGCTCTTCCGATCTNNNCTGGTCTGTTACAGTCGGACA
GAGCACACGTCTGAACTCTTTCCCTACACGACGCTCTTCCGATCTNNNCTGGTCTGTTACAGTCGGACA
GAGCACACGTCTGAACTCTTTCCCTACACGACGCTCTTCCGATCTNNNAAACTCCTGACCTCAAGTGAT
GAGCACACGTCTGAACTCTTTCCCTACACGACGCTCTTCCGATCTNNNAAACTCCTGACCTCAAGTGAT
GCACACGTCTGAACTCTTTCCCTACACGACGCTCTTCCGATCTNNNGGTTTGAAGAGGTGTCTTGCTGA
GCACACGTCTGAACTCTTTCCCTACACGACGCTCTTCCGATCTNNNGGTTTGAAGAGGTGTCTTGCTGA
AGAGCACACGTCTGAACTCTTTCCCTACACGACGCTCTTCCGATCTNNNTGTTCTTTTGCAGTCAGTATC
AGAGCACACGTCTGAACTCTTTCCCTACACGACGCTCTTCCGATCTNNNTGTTCTTTTGCAGTCAGTATC
GAGCACACGTCTGAACTCTTTCCCTACACGACGCTCTTCCGATCTNNNAGTTGCCTCTAGATGCTCTTTC
GAGCACACGTCTGAACTCTTTCCCTACACGACGCTCTTCCGATCTNNNAGTTGCCTCTAGATGCTCTTTC
AGAGCACACGTCTGAACTCTTTCCCTACACGACGCTCTTCCGATCTNNNCTCTCCAGGGGTTCGCTCTTA
AGAGCACACGTCTGAACTCTTTCCCTACACGACGCTCTTCCGATCTNNNCTCTCCAGGGGTTCGCTCTTA
GAGCACACGTCTGAACTCTTTCCCTACACGACGCTCTTCCGATCTNNNGGGTGGGACTTTTCCAGAAAA
GAGCACACGTCTGAACTCTTTCCCTACACGACGCTCTTCCGATCTNNNGGGTGGGACTTTTCCAGAAAA
GAGCACACGTCTGAACTCTTTCCCTACACGACGCTCTTCCGATCTNNNAGACTGTAGTCCCAGCTACTCA
GAGCACACGTCTGAACTCTTTCCCTACACGACGCTCTTCCGATCTNNNAGACTGTAGTCCCAGCTACTCA
AGCACACGTCTGAACTCTTTCCCTACACGACGCTCTTCCGATCTNNNCACCCGGCACTATAAACGTTTAT

AGCACACGTCTGAACTCTTTCCCTACACGACGCTCTTCCGATCTNNNCACCCGGCACTATAAACGTTTAT

AGCACACGTCTGAACTCTTTCCCTACACGACGCTCTTCCGATCTNNNACAGTCATTCTACTCATCACTTG

AGCACACGTCTGAACTCTTTCCCTACACGACGCTCTTCCGATCTNNNACAGTCATTCTACTCATCACTTG

AGCACACGTCTGAACTCTTTCCCTACACGACGCTCTTCCGATCTNNNGAGTGTTCCACCTTTTGGCTTTT

AGCACACGTCTGAACTCTTTCCCTACACGACGCTCTTCCGATCTNNNGAGTGTTCCACCTTTTGGCTTTT

GCACACGTCTGAACTCTTTCCCTACACGACGCTCTTCCGATCTNNNTAACCTAGACCAACATGGGCTGA

GCACACGTCTGAACTCTTTCCCTACACGACGCTCTTCCGATCTNNNTAACCTAGACCAACATGGGCTGA

AGCACACGTCTGAACTCTTTCCCTACACGACGCTCTTCCGATCTNNNCAGCTTCAGCAAGAAACAGACA

AGCACACGTCTGAACTCTTTCCCTACACGACGCTCTTCCGATCTNNNCAGCTTCAGCAAGAAACAGACA

GAGCACACGTCTGAACTCTTTCCCTACACGACGCTCTTCCGATCTNNNGTTGCAGTGAGCTGAGATCAT

GAGCACACGTCTGAACTCTTTCCCTACACGACGCTCTTCCGATCTNNNGTTGCAGTGAGCTGAGATCAT

GAGCACACGTCTGAACTCTTTCCCTACACGACGCTCTTCCGATCTNNNGAATGCCAAAGACTCACTTTC

GAGCACACGTCTGAACTCTTTCCCTACACGACGCTCTTCCGATCTNNNGAATGCCAAAGACTCACTTTC

AGCACACGTCTGAACTCTTTCCCTACACGACGCTCTTCCGATCTNNNGACCTCATTTTGCCTGAATCACC

AGCACACGTCTGAACTCTTTCCCTACACGACGCTCTTCCGATCTNNNGACCTCATTTTGCCTGAATCACC

AGCACACGTCTGAACTCTTTCCCTACACGACGCTCTTCCGATCTNNNGTGTCATTTTTCATTCCCTTAACA

AGCACACGTCTGAACTCTTTCCCTACACGACGCTCTTCCGATCTNNNGTGTCATTTTTCATTCCCTTAACA

GAGCACACGTCTGAACTCTTTCCCTACACGACGCTCTTCCGATCTNNNTTTCATTGTCTGTGTGCATCAC

GAGCACACGTCTGAACTCTTTCCCTACACGACGCTCTTCCGATCTNNNTTTCATTGTCTGTGTGCATCAC

AAGAGCACACGTCTGAACTCTTTCCCTACACGACGCTCTTCCGATCTNNNACCCAAGTCTCCTAATTGTC

AAGAGCACACGTCTGAACTCTTTCCCTACACGACGCTCTTCCGATCTNNNACCCAAGTCTCCTAATTGTC

GAGCACACGTCTGAACTCTTTCCCTACACGACGCTCTTCCGATCTNNNGGCATCTGTTCTTGTTTGCACA

GAGCACACGTCTGAACTCTTTCCCTACACGACGCTCTTCCGATCTNNNGGCATCTGTTCTTGTTTGCACA

CACACGTCTGAACTCTTTCCCTACACGACGCTCTTCCGATCTNNNGATCTTCCCCCTGCAGGTTCCATCG

CACACGTCTGAACTCTTTCCCTACACGACGCTCTTCCGATCTNNNGATCTTCCCCCTGCAGGTTCCATCG

GCACACGTCTGAACTCTTTCCCTACACGACGCTCTTCCGATCTNNNGAGGCTGAGGCAGGAGAATGCA

GCACACGTCTGAACTCTTTCCCTACACGACGCTCTTCCGATCTNNNGAGGCTGAGGCAGGAGAATGCA

AGCACACGTCTGAACTCTTTCCCTACACGACGCTCTTCCGATCTNNNATTGCATCAGACTTGAGACACC

AGCACACGTCTGAACTCTTTCCCTACACGACGCTCTTCCGATCTNNNATTGCATCAGACTTGAGACACC

AGCACACGTCTGAACTCTTTCCCTACACGACGCTCTTCCGATCTNNNCAGAGGAGTCGTCTATGCCAAT

AGCACACGTCTGAACTCTTTCCCTACACGACGCTCTTCCGATCTNNNCAGAGGAGTCGTCTATGCCAAT

GAGCACACGTCTGAACTCTTTCCCTACACGACGCTCTTCCGATCTNNNATACCTGCAGGACGATCTTCTT

GAGCACACGTCTGAACTCTTTCCCTACACGACGCTCTTCCGATCTNNNATACCTGCAGGACGATCTTCTT

GCACACGTCTGAACTCTTTCCCTACACGACGCTCTTCCGATCTNNNGTAGTCTCAAGCAGATGTTAATGC

GCACACGTCTGAACTCTTTCCCTACACGACGCTCTTCCGATCTNNNGTAGTCTCAAGCAGATGTTAATGC

AGAGCACACGTCTGAACTCTTTCCCTACACGACGCTCTTCCGATCTNNNTCTGAACATCTATCGAGTGTG

AGAGCACACGTCTGAACTCTTTCCCTACACGACGCTCTTCCGATCTNNNTCTGAACATCTATCGAGTGTG

GAGCACACGTCTGAACTCTTTCCCTACACGACGCTCTTCCGATCTNNNGCCCTCAACATGGTTGGTTCTA

GAGCACACGTCTGAACTCTTTCCCTACACGACGCTCTTCCGATCTNNNGCCCTCAACATGGTTGGTTCTA

AGCACACGTCTGAACTCTTTCCCTACACGACGCTCTTCCGATCTNNNAGTCACAGTTCTCTAGCACTTGC

AGCACACGTCTGAACTCTTTCCCTACACGACGCTCTTCCGATCTNNNAGTCACAGTTCTCTAGCACTTGC

GAGCACACGTCTGAACTCTTTCCCTACACGACGCTCTTCCGATCTNNNTTCCACATTTGCATGTGCTCTC

GAGCACACGTCTGAACTCTTTCCCTACACGACGCTCTTCCGATCTNNNTTCCACATTTGCATGTGCTCTC

GAGCACACGTCTGAACTCTTTCCCTACACGACGCTCTTCCGATCTNNNCAAGGATGTTCCCAGTCCAC
GAGCACACGTCTGAACTCTTTCCCTACACGACGCTCTTCCGATCTNNNCAAGGATGTTCCCAGTCCAC
GCACACGTCTGAACTCTTTCCCTACACGACGCTCTTCCGATCTNNNGCAGAAACTTTACAAGCCAGGA
GCACACGTCTGAACTCTTTCCCTACACGACGCTCTTCCGATCTNNNGCAGAAACTTTACAAGCCAGGA
AGCACACGTCTGAACTCTTTCCCTACACGACGCTCTTCCGATCTNNNTGCAACAGGAATGAATTTGGC
AGCACACGTCTGAACTCTTTCCCTACACGACGCTCTTCCGATCTNNNTGCAACAGGAATGAATTTGGC
GAGCACACGTCTGAACTCTTTCCCTACACGACGCTCTTCCGATCTNNNCACACCTAAGACTCTGCTACC
GAGCACACGTCTGAACTCTTTCCCTACACGACGCTCTTCCGATCTNNNCACACCTAAGACTCTGCTACC
GAGCACACGTCTGAACTCTTTCCCTACACGACGCTCTTCCGATCTNNNGGAAGATAGCTTGAGTCCAG
GAGCACACGTCTGAACTCTTTCCCTACACGACGCTCTTCCGATCTNNNGGAAGATAGCTTGAGTCCAG
GAGCACACGTCTGAACTCTTTCCCTACACGACGCTCTTCCGATCTNNNGTAATCTCAGCTACTCAGGAG
GAGCACACGTCTGAACTCTTTCCCTACACGACGCTCTTCCGATCTNNNGTAATCTCAGCTACTCAGGAG
GCACACGTCTGAACTCTTTCCCTACACGACGCTCTTCCGATCTNNNCATTTCAACAACCACAGCAGACT
GCACACGTCTGAACTCTTTCCCTACACGACGCTCTTCCGATCTNNNCATTTCAACAACCACAGCAGACT
AGCACACGTCTGAACTCTTTCCCTACACGACGCTCTTCCGATCTNNNGTGTCACTCTTGTCCCATAGAG
AGCACACGTCTGAACTCTTTCCCTACACGACGCTCTTCCGATCTNNNGTGTCACTCTTGTCCCATAGAG
GAGCACACGTCTGAACTCTTTCCCTACACGACGCTCTTCCGATCTNNNGACCTAGACAGGACTGTGGA
GAGCACACGTCTGAACTCTTTCCCTACACGACGCTCTTCCGATCTNNNGACCTAGACAGGACTGTGGA
GAGCACACGTCTGAACTCTTTCCCTACACGACGCTCTTCCGATCTNNNAAGTTGGAGTGCAATCCTCTT
GAGCACACGTCTGAACTCTTTCCCTACACGACGCTCTTCCGATCTNNNAAGTTGGAGTGCAATCCTCTT
GAGCACACGTCTGAACTCTTTCCCTACACGACGCTCTTCCGATCTNNNAGACACACACTGTCTAATAGG
GAGCACACGTCTGAACTCTTTCCCTACACGACGCTCTTCCGATCTNNNAGACACACACTGTCTAATAGG
GCACACGTCTGAACTCTTTCCCTACACGACGCTCTTCCGATCTNNNGAAAGAAAGAAAGAAAGGGG
GCACACGTCTGAACTCTTTCCCTACACGACGCTCTTCCGATCTNNNGAAAGAAAGAAAGAAAGGGG
AGCACACGTCTGAACTCTTTCCCTACACGACGCTCTTCCGATCTNNNACTGTTGACTCTGCTTCTCATGT
AGCACACGTCTGAACTCTTTCCCTACACGACGCTCTTCCGATCTNNNACTGTTGACTCTGCTTCTCATGT
AGCACACGTCTGAACTCTTTCCCTACACGACGCTCTTCCGATCTNNNGCAACAAGAGTGAAAATCTGC
AGCACACGTCTGAACTCTTTCCCTACACGACGCTCTTCCGATCTNNNGCAACAAGAGTGAAAATCTGC
GCACACGTCTGAACTCTTTCCCTACACGACGCTCTTCCGATCTNNNCTCCACCCAAAAGGAGCAAAAT
GCACACGTCTGAACTCTTTCCCTACACGACGCTCTTCCGATCTNNNCTCCACCCAAAAGGAGCAAAAT
GAGCACACGTCTGAACTCTTTCCCTACACGACGCTCTTCCGATCTNNNGAAGGAGGATGGACATCAC
GAGCACACGTCTGAACTCTTTCCCTACACGACGCTCTTCCGATCTNNNGAAGGAGGATGGACATCAC
GAGCACACGTCTGAACTCTTTCCCTACACGACGCTCTTCCGATCTNNNTGTAGTGTTTCAATATAATG
GAGCACACGTCTGAACTCTTTCCCTACACGACGCTCTTCCGATCTNNNTGTAGTGTTTCAATATAATG
GAGCACACGTCTGAACTCTTTCCCTACACGACGCTCTTCCGATCTNNNGTGCCTGTGCAAAAATGTTC
GAGCACACGTCTGAACTCTTTCCCTACACGACGCTCTTCCGATCTNNNGTGCCTGTGCAAAAATGTTC
GAGCACACGTCTGAACTCTTTCCCTACACGACGCTCTTCCGATCTNNNGAAATATTAAGAAGTCTCCCT
GAGCACACGTCTGAACTCTTTCCCTACACGACGCTCTTCCGATCTNNNGAAATATTAAGAAGTCTCCCT
GAGCACACGTCTGAACTCTTTCCCTACACGACGCTCTTCCGATCTNNNAAAGGGGGAAACGTGTACAT
GAGCACACGTCTGAACTCTTTCCCTACACGACGCTCTTCCGATCTNNNAAAGGGGGAAACGTGTACAT
AGCACACGTCTGAACTCTTTCCCTACACGACGCTCTTCCGATCTNNNGATAATCGGATCATGGAGGCA
AGCACACGTCTGAACTCTTTCCCTACACGACGCTCTTCCGATCTNNNGATAATCGGATCATGGAGGCA
AGCACACGTCTGAACTCTTTCCCTACACGACGCTCTTCCGATCTNNNTAATCACTTGAATCCAGGAGG

AGCACACGTCTGAACTCTTTCCCTACACGACGCTCTTCCGATCTNNNTAATCACTTGAATCCAGGAGGC
AGAGCACACGTCTGAACTCTTTCCCTACACGACGCTCTTCCGATCTNNNGTGGCAGGTACCTGTAATC
AGAGCACACGTCTGAACTCTTTCCCTACACGACGCTCTTCCGATCTNNNGTGGCAGGTACCTGTAATC
GCACACGTCTGAACTCTTTCCCTACACGACGCTCTTCCGATCTNNNCACCCATAATCCCACCTACTCGCAT
GCACACGTCTGAACTCTTTCCCTACACGACGCTCTTCCGATCTNNNCACCCATAATCCCACCTACTCGCAT
GAGCACACGTCTGAACTCTTTCCCTACACGACGCTCTTCCGATCTNNNTTGAGGAATGCAGCAGTCTA
GAGCACACGTCTGAACTCTTTCCCTACACGACGCTCTTCCGATCTNNNTTGAGGAATGCAGCAGTCTA
AGCACACGTCTGAACTCTTTCCCTACACGACGCTCTTCCGATCTNNNTGGAGACCCACCTTAAAGTTTG
AGCACACGTCTGAACTCTTTCCCTACACGACGCTCTTCCGATCTNNNTGGAGACCCACCTTAAAGTTTG
GAGCACACGTCTGAACTCTTTCCCTACACGACGCTCTTCCGATCTNNNTGTCATCCCAGTTACACAGAAC
GAGCACACGTCTGAACTCTTTCCCTACACGACGCTCTTCCGATCTNNNTGTCATCCCAGTTACACAGAAC
GCACACGTCTGAACTCTTTCCCTACACGACGCTCTTCCGATCTNNNGTGGTGGGAACTCATAATGACC
GCACACGTCTGAACTCTTTCCCTACACGACGCTCTTCCGATCTNNNGTGGTGGGAACTCATAATGACC
GAGCACACGTCTGAACTCTTTCCCTACACGACGCTCTTCCGATCTNNNTTAGGAAATGTGCCCCAAATT
GAGCACACGTCTGAACTCTTTCCCTACACGACGCTCTTCCGATCTNNNTTAGGAAATGTGCCCCAAATT
GAGCACACGTCTGAACTCTTTCCCTACACGACGCTCTTCCGATCTNNNAGTGAAGTGTGAGTGAGTGT
GAGCACACGTCTGAACTCTTTCCCTACACGACGCTCTTCCGATCTNNNAGTGAAGTGTGAGTGAGTGT
GCACACGTCTGAACTCTTTCCCTACACGACGCTCTTCCGATCTNNNCAGGGAAATGACACTGGGAGCC
GCACACGTCTGAACTCTTTCCCTACACGACGCTCTTCCGATCTNNNCAGGGAAATGACACTGGGAGCC
GAGCACACGTCTGAACTCTTTCCCTACACGACGCTCTTCCGATCTNNNATCTCCATCCAACTGAGTCATG
GAGCACACGTCTGAACTCTTTCCCTACACGACGCTCTTCCGATCTNNNATCTCCATCCAACTGAGTCATG
AGCACACGTCTGAACTCTTTCCCTACACGACGCTCTTCCGATCTNNNAAACGCCTTTATTTGCATGACCA
AGCACACGTCTGAACTCTTTCCCTACACGACGCTCTTCCGATCTNNNAAACGCCTTTATTTGCATGACCA
GAGCACACGTCTGAACTCTTTCCCTACACGACGCTCTTCCGATCTNNNGAGAACTACTTGAACCTGGG
GAGCACACGTCTGAACTCTTTCCCTACACGACGCTCTTCCGATCTNNNGAGAACTACTTGAACCTGGG
GAGCACACGTCTGAACTCTTTCCCTACACGACGCTCTTCCGATCTNNNGCTGTAAAATAGCAGGCAGT
GAGCACACGTCTGAACTCTTTCCCTACACGACGCTCTTCCGATCTNNNGCTGTAAAATAGCAGGCAGT
GCACACGTCTGAACTCTTTCCCTACACGACGCTCTTCCGATCTNNNGCATTCCTTGGAAATCCGCTTTAC
GCACACGTCTGAACTCTTTCCCTACACGACGCTCTTCCGATCTNNNGCATTCCTTGGAAATCCGCTTTAC
GAGCACACGTCTGAACTCTTTCCCTACACGACGCTCTTCCGATCTNNNTAGCCCTGAGTCTCAGTGAAA
GAGCACACGTCTGAACTCTTTCCCTACACGACGCTCTTCCGATCTNNNTAGCCCTGAGTCTCAGTGAAA
AGCACACGTCTGAACTCTTTCCCTACACGACGCTCTTCCGATCTNNNAAGGGGAAGAAAGGCATGTC
AGCACACGTCTGAACTCTTTCCCTACACGACGCTCTTCCGATCTNNNAAGGGGAAGAAAGGCATGTC
AGCACACGTCTGAACTCTTTCCCTACACGACGCTCTTCCGATCTNNNTGACTTGGAGGCACTGTTTTTC
AGCACACGTCTGAACTCTTTCCCTACACGACGCTCTTCCGATCTNNNTGACTTGGAGGCACTGTTTTTC
AGAGCACACGTCTGAACTCTTTCCCTACACGACGCTCTTCCGATCTNNNGAGAATCGCTTCAACCCAGG
AGAGCACACGTCTGAACTCTTTCCCTACACGACGCTCTTCCGATCTNNNGAGAATCGCTTCAACCCAGG
AGAGCACACGTCTGAACTCTTTCCCTACACGACGCTCTTCCGATCTNNNTGCACATGTATCCCAGAACG
AGAGCACACGTCTGAACTCTTTCCCTACACGACGCTCTTCCGATCTNNNTGCACATGTATCCCAGAACG
GCACACGTCTGAACTCTTTCCCTACACGACGCTCTTCCGATCTNNNGGGGCTACAATCCAAGATGAGAT
GCACACGTCTGAACTCTTTCCCTACACGACGCTCTTCCGATCTNNNGGGGCTACAATCCAAGATGAGAT
AGCACACGTCTGAACTCTTTCCCTACACGACGCTCTTCCGATCTNNNCCTTGAAACTTAGCACAAAGAT
AGCACACGTCTGAACTCTTTCCCTACACGACGCTCTTCCGATCTNNNCCTTGAAACTTAGCACAAAGAT

AGCACACGTCTGAACTCTTTCCCTACACGACGCTCTTCCGATCTNNNAATTCCTCAGCACACTCCAATCT

AGCACACGTCTGAACTCTTTCCCTACACGACGCTCTTCCGATCTNNNAATTCCTCAGCACACTCCAATCT

GCACACGTCTGAACTCTTTCCCTACACGACGCTCTTCCGATCTNNNGAGTCTGAGGCAGGAGAATGG

GCACACGTCTGAACTCTTTCCCTACACGACGCTCTTCCGATCTNNNGAGTCTGAGGCAGGAGAATGG

AGAGCACACGTCTGAACTCTTTCCCTACACGACGCTCTTCCGATCTNNNCCTAGGAGACAGAGCAAGA

AGAGCACACGTCTGAACTCTTTCCCTACACGACGCTCTTCCGATCTNNNCCTAGGAGACAGAGCAAGA

GCACACGTCTGAACTCTTTCCCTACACGACGCTCTTCCGATCTNNNTGCCTCTGAGCTTAGACTTTGTT

GCACACGTCTGAACTCTTTCCCTACACGACGCTCTTCCGATCTNNNTGCCTCTGAGCTTAGACTTTGTT

GAGCACACGTCTGAACTCTTTCCCTACACGACGCTCTTCCGATCTNNNCTGGACAACATAGCAAAACCC

GAGCACACGTCTGAACTCTTTCCCTACACGACGCTCTTCCGATCTNNNCTGGACAACATAGCAAAACCC

AGCACACGTCTGAACTCTTTCCCTACACGACGCTCTTCCGATCTNNNGAGCTTGCAGTGAGTCGAGAT

AGCACACGTCTGAACTCTTTCCCTACACGACGCTCTTCCGATCTNNNGAGCTTGCAGTGAGTCGAGAT

AGAGCACACGTCTGAACTCTTTCCCTACACGACGCTCTTCCGATCTNNNTTACCTAAGCGAGTGGAGTT

AGAGCACACGTCTGAACTCTTTCCCTACACGACGCTCTTCCGATCTNNNTTACCTAAGCGAGTGGAGTT

GAGCACACGTCTGAACTCTTTCCCTACACGACGCTCTTCCGATCTNNNGAGTTTGCAGTGAGCCGAGAT

GAGCACACGTCTGAACTCTTTCCCTACACGACGCTCTTCCGATCTNNNGAGTTTGCAGTGAGCCGAGAT

GAGCACACGTCTGAACTCTTTCCCTACACGACGCTCTTCCGATCTNNNTCCAACCCAGCATCTTTCTAGT

GAGCACACGTCTGAACTCTTTCCCTACACGACGCTCTTCCGATCTNNNTCCAACCCAGCATCTTTCTAGT

GAGCACACGTCTGAACTCTTTCCCTACACGACGCTCTTCCGATCTNNNATGGCTGCATAGTATTCCATG

GAGCACACGTCTGAACTCTTTCCCTACACGACGCTCTTCCGATCTNNNATGGCTGCATAGTATTCCATG

AGAGCACACGTCTGAACTCTTTCCCTACACGACGCTCTTCCGATCTNNNCTTGCAGTGAGCCAAGATTA

AGAGCACACGTCTGAACTCTTTCCCTACACGACGCTCTTCCGATCTNNNCTTGCAGTGAGCCAAGATTA

AGCACACGTCTGAACTCTTTCCCTACACGACGCTCTTCCGATCTNNNCATCCACACATTGTATTTCTTAC

AGCACACGTCTGAACTCTTTCCCTACACGACGCTCTTCCGATCTNNNCATCCACACATTGTATTTCTTAC

GAGCACACGTCTGAACTCTTTCCCTACACGACGCTCTTCCGATCTNNNGAGAATGGCATGAACCCCAGA

GAGCACACGTCTGAACTCTTTCCCTACACGACGCTCTTCCGATCTNNNGAGAATGGCATGAACCCCAGA

GAGCACACGTCTGAACTCTTTCCCTACACGACGCTCTTCCGATCTNNNCCTGATCCTCTTTCTTCTCCCAC

GAGCACACGTCTGAACTCTTTCCCTACACGACGCTCTTCCGATCTNNNCCTGATCCTCTTTCTTCTCCCAC

AGCACACGTCTGAACTCTTTCCCTACACGACGCTCTTCCGATCTNNNTGGGAGATAGAGTGAGACGT

AGCACACGTCTGAACTCTTTCCCTACACGACGCTCTTCCGATCTNNNTGGGAGATAGAGTGAGACGT

AGCACACGTCTGAACTCTTTCCCTACACGACGCTCTTCCGATCTNNNGATTGCTTGAACATGGGAGGAT

AGCACACGTCTGAACTCTTTCCCTACACGACGCTCTTCCGATCTNNNGATTGCTTGAACATGGGAGGAT

GAGCACACGTCTGAACTCTTTCCCTACACGACGCTCTTCCGATCTNNNAGATGCTAACTTTGAGCCCTA

GAGCACACGTCTGAACTCTTTCCCTACACGACGCTCTTCCGATCTNNNAGATGCTAACTTTGAGCCCTA

GCACACGTCTGAACTCTTTCCCTACACGACGCTCTTCCGATCTNNNTCCTCTGTCTCTCTCTCCTTCTGCAT

GCACACGTCTGAACTCTTTCCCTACACGACGCTCTTCCGATCTNNNTCCTCTGTCTCTCTCTCCTTCTGCAT

AGCACACGTCTGAACTCTTTCCCTACACGACGCTCTTCCGATCTNNNGTGAGCCAAGATCATGCCATTG

AGCACACGTCTGAACTCTTTCCCTACACGACGCTCTTCCGATCTNNNGTGAGCCAAGATCATGCCATTG

AGCACACGTCTGAACTCTTTCCCTACACGACGCTCTTCCGATCTNNNTTGAGTGGTAGGATTCAAGGG

AGCACACGTCTGAACTCTTTCCCTACACGACGCTCTTCCGATCTNNNTTGAGTGGTAGGATTCAAGGG

AGCACACGTCTGAACTCTTTCCCTACACGACGCTCTTCCGATCTNNNCACCTGTAGTCCCAGCTACTTGC

AGCACACGTCTGAACTCTTTCCCTACACGACGCTCTTCCGATCTNNNCACCTGTAGTCCCAGCTACTTGC

AGAGCACACGTCTGAACTCTTTCCCTACACGACGCTCTTCCGATCTNNNAAGGTTGCAGTGAGCTAAGA

GAGCACACGTCTGAACTCTTTCCCTACACGACGCTCTTCCGATCTNNNAAGGTTGCAGTGAGCTAAG
AGCACACGTCTGAACTCTTTCCCTACACGACGCTCTTCCGATCTNNNATGTGTGTCCTAGAGATCCTGG
AGCACACGTCTGAACTCTTTCCCTACACGACGCTCTTCCGATCTNNNATGTGTGTCCTAGAGATCCTGG
GCACACGTCTGAACTCTTTCCCTACACGACGCTCTTCCGATCTNNNCATGGCGAGACCCTATGTTTAC
GCACACGTCTGAACTCTTTCCCTACACGACGCTCTTCCGATCTNNNCATGGCGAGACCCTATGTTTAC
AGCACACGTCTGAACTCTTTCCCTACACGACGCTCTTCCGATCTNNNAGATTGTGCTGCTGTAAACAT
AGCACACGTCTGAACTCTTTCCCTACACGACGCTCTTCCGATCTNNNAGATTGTGCTGCTGTAAACAT
GCACACGTCTGAACTCTTTCCCTACACGACGCTCTTCCGATCTNNNTGGTTCTATCACAACTCACTGCA
GCACACGTCTGAACTCTTTCCCTACACGACGCTCTTCCGATCTNNNTGGTTCTATCACAACTCACTGCA
GAGCACACGTCTGAACTCTTTCCCTACACGACGCTCTTCCGATCTNNNAGAGGTTGCAATGAGCTGA
GAGCACACGTCTGAACTCTTTCCCTACACGACGCTCTTCCGATCTNNNAGAGGTTGCAATGAGCTGA
AGCACACGTCTGAACTCTTTCCCTACACGACGCTCTTCCGATCTNNNCGACATCAGGAAAAGCACTTCT
AGCACACGTCTGAACTCTTTCCCTACACGACGCTCTTCCGATCTNNNCGACATCAGGAAAAGCACTTCT
AGCACACGTCTGAACTCTTTCCCTACACGACGCTCTTCCGATCTNNNTCCTGTCTTCTATTCCCTTCCCT
AGCACACGTCTGAACTCTTTCCCTACACGACGCTCTTCCGATCTNNNTCCTGTCTTCTATTCCCTTCCCT
CACACGTCTGAACTCTTTCCCTACACGACGCTCTTCCGATCTNNNCCTGGCCTATGTACCCATTTCTTC
CACACGTCTGAACTCTTTCCCTACACGACGCTCTTCCGATCTNNNCCTGGCCTATGTACCCATTTCTTC
AGCACACGTCTGAACTCTTTCCCTACACGACGCTCTTCCGATCTNNNGATCACGAGGTCAGGAGTTCA
AGCACACGTCTGAACTCTTTCCCTACACGACGCTCTTCCGATCTNNNGATCACGAGGTCAGGAGTTCA
GCACACGTCTGAACTCTTTCCCTACACGACGCTCTTCCGATCTNNNGACTGTAGGCACATGCTACCAT
GCACACGTCTGAACTCTTTCCCTACACGACGCTCTTCCGATCTNNNGACTGTAGGCACATGCTACCAT
AGCACACGTCTGAACTCTTTCCCTACACGACGCTCTTCCGATCTNNNTATGATGAGCATAGTGAGCAC
AGCACACGTCTGAACTCTTTCCCTACACGACGCTCTTCCGATCTNNNTATGATGAGCATAGTGAGCAC
GCACACGTCTGAACTCTTTCCCTACACGACGCTCTTCCGATCTNNNGCTAATCACCAACCATGCCACTA
GCACACGTCTGAACTCTTTCCCTACACGACGCTCTTCCGATCTNNNGCTAATCACCAACCATGCCACTA
AGCACACGTCTGAACTCTTTCCCTACACGACGCTCTTCCGATCTNNNGCTGAGGTAGGATAACTGCTT
AGCACACGTCTGAACTCTTTCCCTACACGACGCTCTTCCGATCTNNNGCTGAGGTAGGATAACTGCTT
GAGCACACGTCTGAACTCTTTCCCTACACGACGCTCTTCCGATCTNNNAGCTTGTACGAAGAACTGAG
GAGCACACGTCTGAACTCTTTCCCTACACGACGCTCTTCCGATCTNNNAGCTTGTACGAAGAACTGAG
CACGTCTGAACTCTTTCCCTACACGACGCTCTTCCGATCTNNNTTGTACATGTGAAGTGCTTTGCACA
CACGTCTGAACTCTTTCCCTACACGACGCTCTTCCGATCTNNNTTGTACATGTGAAGTGCTTTGCACA
AGCACACGTCTGAACTCTTTCCCTACACGACGCTCTTCCGATCTNNNACCCTCATCCATTTTACAGGCA
AGCACACGTCTGAACTCTTTCCCTACACGACGCTCTTCCGATCTNNNACCCTCATCCATTTTACAGGCA
AGCACACGTCTGAACTCTTTCCCTACACGACGCTCTTCCGATCTNNNTGCCCATGAACTTCATAGACT
AGCACACGTCTGAACTCTTTCCCTACACGACGCTCTTCCGATCTNNNTGCCCATGAACTTCATAGACT
CACACGTCTGAACTCTTTCCCTACACGACGCTCTTCCGATCTNNNGAATTTCAGGTATTAAAGGAAC
CACACGTCTGAACTCTTTCCCTACACGACGCTCTTCCGATCTNNNGAATTTCAGGTATTAAAGGAAC
GCACACGTCTGAACTCTTTCCCTACACGACGCTCTTCCGATCTNNNTTTTGTTCCAGTTCTCAGGGGA
GCACACGTCTGAACTCTTTCCCTACACGACGCTCTTCCGATCTNNNTTTTGTTCCAGTTCTCAGGGGA
AGAGCACACGTCTGAACTCTTTCCCTACACGACGCTCTTCCGATCTNNNGAGAATCACTTGAACCTGG
AGAGCACACGTCTGAACTCTTTCCCTACACGACGCTCTTCCGATCTNNNGAGAATCACTTGAACCTGG
GCACACGTCTGAACTCTTTCCCTACACGACGCTCTTCCGATCTNNNAAGTAGCTGGGATTACAGGCA
GCACACGTCTGAACTCTTTCCCTACACGACGCTCTTCCGATCTNNNAAGTAGCTGGGATTACAGGCA

GCACACGTCTGAACTCTTTCCCTACACGACGCTCTTCCGATCTNNNGCTCCTCTTCTTCTTCCCCATAGCA

GCACACGTCTGAACTCTTTCCCTACACGACGCTCTTCCGATCTNNNGCTCCTCTTCTTCTTCCCCATAGCA

AGAGCACACGTCTGAACTCTTTCCCTACACGACGCTCTTCCGATCTNNNGGATTGCTTAGAGCTCAGGA

AGAGCACACGTCTGAACTCTTTCCCTACACGACGCTCTTCCGATCTNNNGGATTGCTTAGAGCTCAGGA

GAGCACACGTCTGAACTCTTTCCCTACACGACGCTCTTCCGATCTNNNAAAAAAAAAATTAGCCGGGC

GAGCACACGTCTGAACTCTTTCCCTACACGACGCTCTTCCGATCTNNNAAAAAAAAAATTAGCCGGGC

GAAGAGCACACGTCTGAACTCTTTCCCTACACGACGCTCTTCCGATCTNNNGAGCTTGCAGTGAGCCTA

GAAGAGCACACGTCTGAACTCTTTCCCTACACGACGCTCTTCCGATCTNNNGAGCTTGCAGTGAGCCTA

AGCACACGTCTGAACTCTTTCCCTACACGACGCTCTTCCGATCTNNNATCACACCTCACTGATGAAACCA

AGCACACGTCTGAACTCTTTCCCTACACGACGCTCTTCCGATCTNNNATCACACCTCACTGATGAAACCA

AGAGCACACGTCTGAACTCTTTCCCTACACGACGCTCTTCCGATCTNNNAACCTTGGAACTGACACCGA

AGAGCACACGTCTGAACTCTTTCCCTACACGACGCTCTTCCGATCTNNNAACCTTGGAACTGACACCGA

GAGCACACGTCTGAACTCTTTCCCTACACGACGCTCTTCCGATCTNNNCACCCCTTCCAAGATTGACAAA

GAGCACACGTCTGAACTCTTTCCCTACACGACGCTCTTCCGATCTNNNCACCCCTTCCAAGATTGACAAA

AGCACACGTCTGAACTCTTTCCCTACACGACGCTCTTCCGATCTNNNCTGAGGCAGGAGAATCACTTG

AGCACACGTCTGAACTCTTTCCCTACACGACGCTCTTCCGATCTNNNCTGAGGCAGGAGAATCACTTG

AGAGCACACGTCTGAACTCTTTCCCTACACGACGCTCTTCCGATCTNNNTGTTGACTCTTCTACTCTCTCG

AGAGCACACGTCTGAACTCTTTCCCTACACGACGCTCTTCCGATCTNNNTGTTGACTCTTCTACTCTCTCG

AGCACACGTCTGAACTCTTTCCCTACACGACGCTCTTCCGATCTNNNCCCTCAGTCTTGTTCCCACATTA

AGCACACGTCTGAACTCTTTCCCTACACGACGCTCTTCCGATCTNNNCCCTCAGTCTTGTTCCCACATTA

AGCACACGTCTGAACTCTTTCCCTACACGACGCTCTTCCGATCTNNNGAACTCCTGAACTCAAGGGATC

AGCACACGTCTGAACTCTTTCCCTACACGACGCTCTTCCGATCTNNNGAACTCCTGAACTCAAGGGATC

AGCACACGTCTGAACTCTTTCCCTACACGACGCTCTTCCGATCTNNNCTTTTCCTAGGCTCCAGGTGTT

AGCACACGTCTGAACTCTTTCCCTACACGACGCTCTTCCGATCTNNNCTTTTCCTAGGCTCCAGGTGTT

AGCACACGTCTGAACTCTTTCCCTACACGACGCTCTTCCGATCTNNNCAGAGTTTTTGCGCTGGTTTTT

AGCACACGTCTGAACTCTTTCCCTACACGACGCTCTTCCGATCTNNNCAGAGTTTTTGCGCTGGTTTTT

AAGAGCACACGTCTGAACTCTTTCCCTACACGACGCTCTTCCGATCTNNNGTGAGCCAAGATTGTGCCA

AAGAGCACACGTCTGAACTCTTTCCCTACACGACGCTCTTCCGATCTNNNGTGAGCCAAGATTGTGCCA

AGAGCACACGTCTGAACTCTTTCCCTACACGACGCTCTTCCGATCTNNNCCCTGAAGTTTGGGTGACAG

AGAGCACACGTCTGAACTCTTTCCCTACACGACGCTCTTCCGATCTNNNCCCTGAAGTTTGGGTGACAG

GAGCACACGTCTGAACTCTTTCCCTACACGACGCTCTTCCGATCTNNNAGTGGTCAGATGTCAGCAGAA

GAGCACACGTCTGAACTCTTTCCCTACACGACGCTCTTCCGATCTNNNAGTGGTCAGATGTCAGCAGAA

AAGAGCACACGTCTGAACTCTTTCCCTACACGACGCTCTTCCGATCTNNNCGTCTCAACAGAAGAAAGG

AAGAGCACACGTCTGAACTCTTTCCCTACACGACGCTCTTCCGATCTNNNCGTCTCAACAGAAGAAAGG

AGAGCACACGTCTGAACTCTTTCCCTACACGACGCTCTTCCGATCTNNNTGACCCAGGATTTCTACTCAG

AGAGCACACGTCTGAACTCTTTCCCTACACGACGCTCTTCCGATCTNNNTGACCCAGGATTTCTACTCAG

AGAGCACACGTCTGAACTCTTTCCCTACACGACGCTCTTCCGATCTNNNGGGTAGCAAAGAATTATGC

AGAGCACACGTCTGAACTCTTTCCCTACACGACGCTCTTCCGATCTNNNGGGTAGCAAAGAATTATGC

AGCACACGTCTGAACTCTTTCCCTACACGACGCTCTTCCGATCTNNNTTGTGACAGAACTCACCCACTCC

AGCACACGTCTGAACTCTTTCCCTACACGACGCTCTTCCGATCTNNNTTGTGACAGAACTCACCCACTCC

AGCACACGTCTGAACTCTTTCCCTACACGACGCTCTTCCGATCTNNNGCAGTATTTGAAACACATGGC

AGCACACGTCTGAACTCTTTCCCTACACGACGCTCTTCCGATCTNNNGCAGTATTTGAAACACATGGC

AGAGCACACGTCTGAACTCTTTCCCTACACGACGCTCTTCCGATCTNNNTTTCCAATGATAGTAGGAAA

GAGCACACGTCTGAACTCTTTCCCTACACGACGCTCTTCCGATCTNNNTTTCCAATGATAGTAGGAAA
AGCACACGTCTGAACTCTTTCCCTACACGACGCTCTTCCGATCTNNNGGTTCAAGCAATTCTTCTGCCT
AGCACACGTCTGAACTCTTTCCCTACACGACGCTCTTCCGATCTNNNGGTTCAAGCAATTCTTCTGCCT
CACACGTCTGAACTCTTTCCCTACACGACGCTCTTCCGATCTNNNGAGTATCTAGTAATTTGGAAGGT
CACACGTCTGAACTCTTTCCCTACACGACGCTCTTCCGATCTNNNGAGTATCTAGTAATTTGGAAGGT
GCACACGTCTGAACTCTTTCCCTACACGACGCTCTTCCGATCTNNNAGCTAGAGGCACTAGGTTCTG
GCACACGTCTGAACTCTTTCCCTACACGACGCTCTTCCGATCTNNNAGCTAGAGGCACTAGGTTCTG
GCACACGTCTGAACTCTTTCCCTACACGACGCTCTTCCGATCTNNNCTAGAAACACACAATGGCCCCTC
GCACACGTCTGAACTCTTTCCCTACACGACGCTCTTCCGATCTNNNCTAGAAACACACAATGGCCCCTC
AGCACACGTCTGAACTCTTTCCCTACACGACGCTCTTCCGATCTNNNTGAGTGACAAGGGCGAAACTT
AGCACACGTCTGAACTCTTTCCCTACACGACGCTCTTCCGATCTNNNTGAGTGACAAGGGCGAAACTT
AGCACACGTCTGAACTCTTTCCCTACACGACGCTCTTCCGATCTNNNGAGGTCTCACTTCCTAGCTTG
AGCACACGTCTGAACTCTTTCCCTACACGACGCTCTTCCGATCTNNNGAGGTCTCACTTCCTAGCTTG
AGAGCACACGTCTGAACTCTTTCCCTACACGACGCTCTTCCGATCTNNNACTATCCTTCCTGCCCTAGA
AGAGCACACGTCTGAACTCTTTCCCTACACGACGCTCTTCCGATCTNNNACTATCCTTCCTGCCCTAGA
AGCACACGTCTGAACTCTTTCCCTACACGACGCTCTTCCGATCTNNNGCGCCTATAATCCCAGTTACTC
AGCACACGTCTGAACTCTTTCCCTACACGACGCTCTTCCGATCTNNNGCGCCTATAATCCCAGTTACTC
GAGCACACGTCTGAACTCTTTCCCTACACGACGCTCTTCCGATCTNNNATTTTCACATCTGCCAACCAA
GAGCACACGTCTGAACTCTTTCCCTACACGACGCTCTTCCGATCTNNNATTTTCACATCTGCCAACCAA
GAGCACACGTCTGAACTCTTTCCCTACACGACGCTCTTCCGATCTNNNGTAATTTCAGGGCAAGGCA
GAGCACACGTCTGAACTCTTTCCCTACACGACGCTCTTCCGATCTNNNGTAATTTCAGGGCAAGGCA
AGCACACGTCTGAACTCTTTCCCTACACGACGCTCTTCCGATCTNNNTTGACATACCTCAGCACACAC
AGCACACGTCTGAACTCTTTCCCTACACGACGCTCTTCCGATCTNNNTTGACATACCTCAGCACACAC
AGCACACGTCTGAACTCTTTCCCTACACGACGCTCTTCCGATCTNNNAAATAAGTTAGTTGGGGGTG
AGCACACGTCTGAACTCTTTCCCTACACGACGCTCTTCCGATCTNNNAAATAAGTTAGTTGGGGGTG
ACACGTCTGAACTCTTTCCCTACACGACGCTCTTCCGATCTNNNCTGAAGAAATAAAGTTCTTTTTTTT
ACACGTCTGAACTCTTTCCCTACACGACGCTCTTCCGATCTNNNCTGAAGAAATAAAGTTCTTTTTTTT
GCACACGTCTGAACTCTTTCCCTACACGACGCTCTTCCGATCTNNNCTTGCGTTTCATGTAGTCCATCC
GCACACGTCTGAACTCTTTCCCTACACGACGCTCTTCCGATCTNNNCTTGCGTTTCATGTAGTCCATCC
AGCACACGTCTGAACTCTTTCCCTACACGACGCTCTTCCGATCTNNNTGCCAAAATATCCTAGGTGCCT
AGCACACGTCTGAACTCTTTCCCTACACGACGCTCTTCCGATCTNNNTGCCAAAATATCCTAGGTGCCT
CACACGTCTGAACTCTTTCCCTACACGACGCTCTTCCGATCTNNNTCCCTCTCCATCTCTCTTTGTCTCAT
CACACGTCTGAACTCTTTCCCTACACGACGCTCTTCCGATCTNNNTCCCTCTCCATCTCTCTTTGTCTCAT
GAGCACACGTCTGAACTCTTTCCCTACACGACGCTCTTCCGATCTNNNGGGGACTAGTGGATCATCG
GAGCACACGTCTGAACTCTTTCCCTACACGACGCTCTTCCGATCTNNNGGGGACTAGTGGATCATCG
AGCACACGTCTGAACTCTTTCCCTACACGACGCTCTTCCGATCTNNNTTGTCTAGCTAACTGCAACAGC
AGCACACGTCTGAACTCTTTCCCTACACGACGCTCTTCCGATCTNNNTTGTCTAGCTAACTGCAACAGC
AGCACACGTCTGAACTCTTTCCCTACACGACGCTCTTCCGATCTNNNTGATGTTTGCCTTGGTTCTTTG
AGCACACGTCTGAACTCTTTCCCTACACGACGCTCTTCCGATCTNNNTGATGTTTGCCTTGGTTCTTTG
AGCACACGTCTGAACTCTTTCCCTACACGACGCTCTTCCGATCTNNNTGAGTCACAGAGAGAGACTG
AGCACACGTCTGAACTCTTTCCCTACACGACGCTCTTCCGATCTNNNTGAGTCACAGAGAGAGACTG
AGCACACGTCTGAACTCTTTCCCTACACGACGCTCTTCCGATCTNNNTTGAGGATTTTCGCAGCAATG
AGCACACGTCTGAACTCTTTCCCTACACGACGCTCTTCCGATCTNNNTTGAGGATTTTCGCAGCAATG

GAGCACACGTCTGAACTCTTTCCCTACACGACGCTCTTCCGATCTNNNGAGGCAGGAAAATCACTTG

GAGCACACGTCTGAACTCTTTCCCTACACGACGCTCTTCCGATCTNNNGAGGCAGGAAAATCACTTG

GCACACGTCTGAACTCTTTCCCTACACGACGCTCTTCCGATCTNNNCTTGCAGTGAGCTGAAATCATG

GCACACGTCTGAACTCTTTCCCTACACGACGCTCTTCCGATCTNNNCTTGCAGTGAGCTGAAATCATG

AGCACACGTCTGAACTCTTTCCCTACACGACGCTCTTCCGATCTNNNGGTCCATAAGCCCCAAGTAAA

AGCACACGTCTGAACTCTTTCCCTACACGACGCTCTTCCGATCTNNNGGTCCATAAGCCCCAAGTAAA

AGCACACGTCTGAACTCTTTCCCTACACGACGCTCTTCCGATCTNNNATTGCCCTTCTGTTTACTGCA

AGCACACGTCTGAACTCTTTCCCTACACGACGCTCTTCCGATCTNNNATTGCCCTTCTGTTTACTGCA

GAGCACACGTCTGAACTCTTTCCCTACACGACGCTCTTCCGATCTNNNCATGTTTATCTGGTGCAGGT

GAGCACACGTCTGAACTCTTTCCCTACACGACGCTCTTCCGATCTNNNCATGTTTATCTGGTGCAGGT

AGCACACGTCTGAACTCTTTCCCTACACGACGCTCTTCCGATCTNNNAGGGAGAGAGAGAGAGAG

AGCACACGTCTGAACTCTTTCCCTACACGACGCTCTTCCGATCTNNNAGGGAGAGAGAGAGAGAG

GCACACGTCTGAACTCTTTCCCTACACGACGCTCTTCCGATCTNNNGAGAAGGGTGAATGACAGAG

GCACACGTCTGAACTCTTTCCCTACACGACGCTCTTCCGATCTNNNGAGAAGGGTGAATGACAGAG

AGCACACGTCTGAACTCTTTCCCTACACGACGCTCTTCCGATCTNNNAGAAAACTGCACTTCAAGCTTG

AGCACACGTCTGAACTCTTTCCCTACACGACGCTCTTCCGATCTNNNAGAAAACTGCACTTCAAGCTTG

CACACGTCTGAACTCTTTCCCTACACGACGCTCTTCCGATCTNNNCCACCACACCAGGCTAATTTTTCC

CACACGTCTGAACTCTTTCCCTACACGACGCTCTTCCGATCTNNNCCACCACACCAGGCTAATTTTTCC

AGCACACGTCTGAACTCTTTCCCTACACGACGCTCTTCCGATCTNNNGGGACAGAGTGAGACTCCATC

AGCACACGTCTGAACTCTTTCCCTACACGACGCTCTTCCGATCTNNNGGGACAGAGTGAGACTCCATC

AGCACACGTCTGAACTCTTTCCCTACACGACGCTCTTCCGATCTNNNGAGAACACATAGACTCAGGGT

AGCACACGTCTGAACTCTTTCCCTACACGACGCTCTTCCGATCTNNNGAGAACACATAGACTCAGGGT

AGCACACGTCTGAACTCTTTCCCTACACGACGCTCTTCCGATCTNNNGCTTCTGCTCACTCAGACCTAA

AGCACACGTCTGAACTCTTTCCCTACACGACGCTCTTCCGATCTNNNGCTTCTGCTCACTCAGACCTAA

CACACGTCTGAACTCTTTCCCTACACGACGCTCTTCCGATCTNNNGGTTGCAGTGAGCTGATATTGT

CACACGTCTGAACTCTTTCCCTACACGACGCTCTTCCGATCTNNNGGTTGCAGTGAGCTGATATTGT

AGCACACGTCTGAACTCTTTCCCTACACGACGCTCTTCCGATCTNNNTGGATCACAAGGTCAGGAAAT

AGCACACGTCTGAACTCTTTCCCTACACGACGCTCTTCCGATCTNNNTGGATCACAAGGTCAGGAAAT

GCACACGTCTGAACTCTTTCCCTACACGACGCTCTTCCGATCTNNNCCTCATCCCCTGAGTATGTGGA

GCACACGTCTGAACTCTTTCCCTACACGACGCTCTTCCGATCTNNNCCTCATCCCCTGAGTATGTGGA

CACACGTCTGAACTCTTTCCCTACACGACGCTCTTCCGATCTNNNTGAACAGTGGATGTCGCTTTTTCC

CACACGTCTGAACTCTTTCCCTACACGACGCTCTTCCGATCTNNNTGAACAGTGGATGTCGCTTTTTCC

GCACACGTCTGAACTCTTTCCCTACACGACGCTCTTCCGATCTNNNGAGGCAGGAGAATGGTGTGAA

GCACACGTCTGAACTCTTTCCCTACACGACGCTCTTCCGATCTNNNGAGGCAGGAGAATGGTGTGAA

GCACACGTCTGAACTCTTTCCCTACACGACGCTCTTCCGATCTNNNTTTATATCCCTGTAGAACTGAAG

GCACACGTCTGAACTCTTTCCCTACACGACGCTCTTCCGATCTNNNTTTATATCCCTGTAGAACTGAAG

GCACACGTCTGAACTCTTTCCCTACACGACGCTCTTCCGATCTNNNGTGACAGAGTGAGATTCCATC

GCACACGTCTGAACTCTTTCCCTACACGACGCTCTTCCGATCTNNNGTGACAGAGTGAGATTCCATC

AGCACACGTCTGAACTCTTTCCCTACACGACGCTCTTCCGATCTNNNGACCCGCATTGATAATCAGAG

AGCACACGTCTGAACTCTTTCCCTACACGACGCTCTTCCGATCTNNNGACCCGCATTGATAATCAGAG

GCACACGTCTGAACTCTTTCCCTACACGACGCTCTTCCGATCTNNNGCTCTTGCCCTCTTCTCTGGTTA

GCACACGTCTGAACTCTTTCCCTACACGACGCTCTTCCGATCTNNNGCTCTTGCCCTCTTCTCTGGTTA

AGCACACGTCTGAACTCTTTCCCTACACGACGCTCTTCCGATCTNNNTCAAGAGTCTGAGGGAGGAG

AGCACACGTCTGAACTCTTTCCCTACACGACGCTCTTCCGATCTNNNTCAAGAGTCTGAGGGAGGAG
AGCACACGTCTGAACTCTTTCCCTACACGACGCTCTTCCGATCTNNNGTGGTGCAATCCTAGCTCAGTA
AGCACACGTCTGAACTCTTTCCCTACACGACGCTCTTCCGATCTNNNGTGGTGCAATCCTAGCTCAGTA
GCACACGTCTGAACTCTTTCCCTACACGACGCTCTTCCGATCTNNNGCCTCCCAAAGTGCTGAGATTAT
GCACACGTCTGAACTCTTTCCCTACACGACGCTCTTCCGATCTNNNGCCTCCCAAAGTGCTGAGATTAT
GAGCACACGTCTGAACTCTTTCCCTACACGACGCTCTTCCGATCTNNNACATTTGTCTGTGTGGGGAG
GAGCACACGTCTGAACTCTTTCCCTACACGACGCTCTTCCGATCTNNNACATTTGTCTGTGTGGGGAG
GCACACGTCTGAACTCTTTCCCTACACGACGCTCTTCCGATCTNNNATTGGTGGTGGGGAGTAGAAT
GCACACGTCTGAACTCTTTCCCTACACGACGCTCTTCCGATCTNNNATTGGTGGTGGGGAGTAGAAT
AGCACACGTCTGAACTCTTTCCCTACACGACGCTCTTCCGATCTNNNCATGCAGATACACATGTACACA
AGCACACGTCTGAACTCTTTCCCTACACGACGCTCTTCCGATCTNNNCATGCAGATACACATGTACACA
ACACGTCTGAACTCTTTCCCTACACGACGCTCTTCCGATCTNNNAAACATTTTCAGACCTATGGAAACA
ACACGTCTGAACTCTTTCCCTACACGACGCTCTTCCGATCTNNNAAACATTTTCAGACCTATGGAAACA
GCACACGTCTGAACTCTTTCCCTACACGACGCTCTTCCGATCTNNNCACCTGTAGTCCCAGCTACTTGC
GCACACGTCTGAACTCTTTCCCTACACGACGCTCTTCCGATCTNNNCACCTGTAGTCCCAGCTACTTGC
GAGCACACGTCTGAACTCTTTCCCTACACGACGCTCTTCCGATCTNNNACCAAAGACCGCATATTCTCA
GAGCACACGTCTGAACTCTTTCCCTACACGACGCTCTTCCGATCTNNNACCAAAGACCGCATATTCTCA
GCACACGTCTGAACTCTTTCCCTACACGACGCTCTTCCGATCTNNNCTCTGGCTACACATCTCCTGTCCA
GCACACGTCTGAACTCTTTCCCTACACGACGCTCTTCCGATCTNNNCTCTGGCTACACATCTCCTGTCCA
GCACACGTCTGAACTCTTTCCCTACACGACGCTCTTCCGATCTNNNCTGTTGTGGGAGGATTGCTTGA
GCACACGTCTGAACTCTTTCCCTACACGACGCTCTTCCGATCTNNNCTGTTGTGGGAGGATTGCTTGA
GCACACGTCTGAACTCTTTCCCTACACGACGCTCTTCCGATCTNNNCAAACACAGTGAAACCCCATCT
GCACACGTCTGAACTCTTTCCCTACACGACGCTCTTCCGATCTNNNCAAACACAGTGAAACCCCATCT
GAGCACACGTCTGAACTCTTTCCCTACACGACGCTCTTCCGATCTNNNAGAGATTGCAGTGAGCTGA
GAGCACACGTCTGAACTCTTTCCCTACACGACGCTCTTCCGATCTNNNAGAGATTGCAGTGAGCTGA
GAGCACACGTCTGAACTCTTTCCCTACACGACGCTCTTCCGATCTNNNAGTAGCTGCCCTTAGAAAGT
GAGCACACGTCTGAACTCTTTCCCTACACGACGCTCTTCCGATCTNNNAGTAGCTGCCCTTAGAAAGT
AGCACACGTCTGAACTCTTTCCCTACACGACGCTCTTCCGATCTNNNTGAGTGACAAAGTGAGACACT
AGCACACGTCTGAACTCTTTCCCTACACGACGCTCTTCCGATCTNNNTGAGTGACAAAGTGAGACACT
GCACACGTCTGAACTCTTTCCCTACACGACGCTCTTCCGATCTNNNGAGGTGAGCGTCTATCCCTGGC
GCACACGTCTGAACTCTTTCCCTACACGACGCTCTTCCGATCTNNNGAGGTGAGCGTCTATCCCTGGC
GAGCACACGTCTGAACTCTTTCCCTACACGACGCTCTTCCGATCTNNNGCTATGAAGAGGGCTCAGG
GAGCACACGTCTGAACTCTTTCCCTACACGACGCTCTTCCGATCTNNNGCTATGAAGAGGGCTCAGG
GCACACGTCTGAACTCTTTCCCTACACGACGCTCTTCCGATCTNNNGAGAATCTCTTGAACTGGGGA
GCACACGTCTGAACTCTTTCCCTACACGACGCTCTTCCGATCTNNNGAGAATCTCTTGAACTGGGGA
GAGCACACGTCTGAACTCTTTCCCTACACGACGCTCTTCCGATCTNNNCTGTAATCCCAGCTACTCAGG
GAGCACACGTCTGAACTCTTTCCCTACACGACGCTCTTCCGATCTNNNCTGTAATCCCAGCTACTCAGG
AGCACACGTCTGAACTCTTTCCCTACACGACGCTCTTCCGATCTNNNTCACATTGACATTCCGTTTGAC
AGCACACGTCTGAACTCTTTCCCTACACGACGCTCTTCCGATCTNNNTCACATTGACATTCCGTTTGAC
AGCACACGTCTGAACTCTTTCCCTACACGACGCTCTTCCGATCTNNNGAAAGTAGGTAGGGTGGAAG
AGCACACGTCTGAACTCTTTCCCTACACGACGCTCTTCCGATCTNNNGAAAGTAGGTAGGGTGGAAG
GAGCACACGTCTGAACTCTTTCCCTACACGACGCTCTTCCGATCTNNNGTCACGTTTGTGTGTGTGTG
GAGCACACGTCTGAACTCTTTCCCTACACGACGCTCTTCCGATCTNNNGTCACGTTTGTGTGTGTGTG

AGCACACGTCTGAACTCTTTCCCTACACGACGCTCTTCCGATCTNNNGACCTTTGATCCAGGCTCCTT
AGCACACGTCTGAACTCTTTCCCTACACGACGCTCTTCCGATCTNNNGACCTTTGATCCAGGCTCCTT
AGCACACGTCTGAACTCTTTCCCTACACGACGCTCTTCCGATCTNNNACCAAAACTGGCAAATGGAG
AGCACACGTCTGAACTCTTTCCCTACACGACGCTCTTCCGATCTNNNACCAAAACTGGCAAATGGAG
GCACACGTCTGAACTCTTTCCCTACACGACGCTCTTCCGATCTNNNTTAGTCCCTGCCCAATCTCATTC
GCACACGTCTGAACTCTTTCCCTACACGACGCTCTTCCGATCTNNNTTAGTCCCTGCCCAATCTCATTC
AGCACACGTCTGAACTCTTTCCCTACACGACGCTCTTCCGATCTNNNTGGTACATACACTCACACCAAC
AGCACACGTCTGAACTCTTTCCCTACACGACGCTCTTCCGATCTNNNTGGTACATACACTCACACCAAC
GCACACGTCTGAACTCTTTCCCTACACGACGCTCTTCCGATCTNNNATTTTTAGAAAGCTTCCGGCCAG
GCACACGTCTGAACTCTTTCCCTACACGACGCTCTTCCGATCTNNNATTTTTAGAAAGCTTCCGGCCAG
AGCACACGTCTGAACTCTTTCCCTACACGACGCTCTTCCGATCTNNNCCCAAATGATGAAGTGATGCA
AGCACACGTCTGAACTCTTTCCCTACACGACGCTCTTCCGATCTNNNCCCAAATGATGAAGTGATGCA
CACACGTCTGAACTCTTTCCCTACACGACGCTCTTCCGATCTNNNCATGCAGTTGATACACGAGCATT
CACACGTCTGAACTCTTTCCCTACACGACGCTCTTCCGATCTNNNCATGCAGTTGATACACGAGCATT
GCACACGTCTGAACTCTTTCCCTACACGACGCTCTTCCGATCTNNNAATACAATAAAAATTAACACCTG
GCACACGTCTGAACTCTTTCCCTACACGACGCTCTTCCGATCTNNNAATACAATAAAAATTAACACCTG
AGCACACGTCTGAACTCTTTCCCTACACGACGCTCTTCCGATCTNNNGTTTCAATTGTGTGTGTGTGCA
AGCACACGTCTGAACTCTTTCCCTACACGACGCTCTTCCGATCTNNNGTTTCAATTGTGTGTGTGTGCA
GAGCACACGTCTGAACTCTTTCCCTACACGACGCTCTTCCGATCTNNNCACCTGTAGTCCCAGCTACTT
GAGCACACGTCTGAACTCTTTCCCTACACGACGCTCTTCCGATCTNNNCACCTGTAGTCCCAGCTACTT
AGCACACGTCTGAACTCTTTCCCTACACGACGCTCTTCCGATCTNNNCACGATGATCCCCTTTGCTTTTG
AGCACACGTCTGAACTCTTTCCCTACACGACGCTCTTCCGATCTNNNCACGATGATCCCCTTTGCTTTTG
GCACACGTCTGAACTCTTTCCCTACACGACGCTCTTCCGATCTNNNATAAATTCTGTGACACTGGGGC
GCACACGTCTGAACTCTTTCCCTACACGACGCTCTTCCGATCTNNNATAAATTCTGTGACACTGGGGC
AGCACACGTCTGAACTCTTTCCCTACACGACGCTCTTCCGATCTNNNGAATAACTGGGACTACAGGC
AGCACACGTCTGAACTCTTTCCCTACACGACGCTCTTCCGATCTNNNGAATAACTGGGACTACAGGC
AGCACACGTCTGAACTCTTTCCCTACACGACGCTCTTCCGATCTNNNAGCCAGTTGACACAAAAGTCT
AGCACACGTCTGAACTCTTTCCCTACACGACGCTCTTCCGATCTNNNAGCCAGTTGACACAAAAGTCT
AGCACACGTCTGAACTCTTTCCCTACACGACGCTCTTCCGATCTNNNTTCGAAGTCGGGATCTAGGTC
AGCACACGTCTGAACTCTTTCCCTACACGACGCTCTTCCGATCTNNNTTCGAAGTCGGGATCTAGGTC
AGCACACGTCTGAACTCTTTCCCTACACGACGCTCTTCCGATCTNNNTTGAAGGGGGTTAGAAGAGA
AGCACACGTCTGAACTCTTTCCCTACACGACGCTCTTCCGATCTNNNTTGAAGGGGGTTAGAAGAGA
AGCACACGTCTGAACTCTTTCCCTACACGACGCTCTTCCGATCTNNNCACCAAGAAACAGAACCAGCA
AGCACACGTCTGAACTCTTTCCCTACACGACGCTCTTCCGATCTNNNCACCAAGAAACAGAACCAGCA
AGCACACGTCTGAACTCTTTCCCTACACGACGCTCTTCCGATCTNNNCTGAGGCAGGAGAATCGTTTG
AGCACACGTCTGAACTCTTTCCCTACACGACGCTCTTCCGATCTNNNCTGAGGCAGGAGAATCGTTTG
AGCACACGTCTGAACTCTTTCCCTACACGACGCTCTTCCGATCTNNNTCAGGTACTCCCGGGCCTAGCA
AGCACACGTCTGAACTCTTTCCCTACACGACGCTCTTCCGATCTNNNTCAGGTACTCCCGGGCCTAGCA
AGCACACGTCTGAACTCTTTCCCTACACGACGCTCTTCCGATCTNNNACTGTTTATGAAGATGAAACTT
AGCACACGTCTGAACTCTTTCCCTACACGACGCTCTTCCGATCTNNNACTGTTTATGAAGATGAAACTT
AGCACACGTCTGAACTCTTTCCCTACACGACGCTCTTCCGATCTNNNCTTGCAGTGAGCCAAGATTGC
AGCACACGTCTGAACTCTTTCCCTACACGACGCTCTTCCGATCTNNNCTTGCAGTGAGCCAAGATTGC
AGCACACGTCTGAACTCTTTCCCTACACGACGCTCTTCCGATCTNNNTGCTGAGTCTCCTCTGGAATC

GAGCACACGTCTGAACTCTTTCCCTACACGACGCTCTTCCGATCTNNNTGCTGAGTCTCCTCTGGAATCT
AGCACACGTCTGAACTCTTTCCCTACACGACGCTCTTCCGATCTNNNTCCTTGTTCTACTCAAGTGCACA
AGCACACGTCTGAACTCTTTCCCTACACGACGCTCTTCCGATCTNNNTCCTTGTTCTACTCAAGTGCACA
AGCACACGTCTGAACTCTTTCCCTACACGACGCTCTTCCGATCTNNNAAAATGTGTGTGTGTGTGTGT
AGCACACGTCTGAACTCTTTCCCTACACGACGCTCTTCCGATCTNNNAAAATGTGTGTGTGTGTGTGT
GAGCACACGTCTGAACTCTTTCCCTACACGACGCTCTTCCGATCTNNNAAACATTCTCCAGGGTACAGC
GAGCACACGTCTGAACTCTTTCCCTACACGACGCTCTTCCGATCTNNNAAACATTCTCCAGGGTACAGC
GAGCACACGTCTGAACTCTTTCCCTACACGACGCTCTTCCGATCTNNNGCTACTCTATTCCCAGCTACTC
GAGCACACGTCTGAACTCTTTCCCTACACGACGCTCTTCCGATCTNNNGCTACTCTATTCCCAGCTACTC
GAGCACACGTCTGAACTCTTTCCCTACACGACGCTCTTCCGATCTNNNCTTGAGCCCAGAGATTTGAGA
GAGCACACGTCTGAACTCTTTCCCTACACGACGCTCTTCCGATCTNNNCTTGAGCCCAGAGATTTGAGA
AGCACACGTCTGAACTCTTTCCCTACACGACGCTCTTCCGATCTNNNTTAGAAAACAAAAGCCAATGTA
AGCACACGTCTGAACTCTTTCCCTACACGACGCTCTTCCGATCTNNNTTAGAAAACAAAAGCCAATGTA
GAGCACACGTCTGAACTCTTTCCCTACACGACGCTCTTCCGATCTNNNCTCAATCTGTCTCTGCTTCTGC
GAGCACACGTCTGAACTCTTTCCCTACACGACGCTCTTCCGATCTNNNCTCAATCTGTCTCTGCTTCTGC
GCACACGTCTGAACTCTTTCCCTACACGACGCTCTTCCGATCTNNNAGAAGGTTCAATGACCTCCCAGA
GCACACGTCTGAACTCTTTCCCTACACGACGCTCTTCCGATCTNNNAGAAGGTTCAATGACCTCCCAGA
ACACGTCTGAACTCTTTCCCTACACGACGCTCTTCCGATCTNNNCACATGTGTATGTCCTGTCTCATCGG
ACACGTCTGAACTCTTTCCCTACACGACGCTCTTCCGATCTNNNCACATGTGTATGTCCTGTCTCATCGG
GAGCACACGTCTGAACTCTTTCCCTACACGACGCTCTTCCGATCTNNNGGTGTCTTGGACCAGGATGAT
GAGCACACGTCTGAACTCTTTCCCTACACGACGCTCTTCCGATCTNNNGGTGTCTTGGACCAGGATGAT
AGCACACGTCTGAACTCTTTCCCTACACGACGCTCTTCCGATCTNNNTCTTCCTCCTCCTCCTCTTCTTCCA
AGCACACGTCTGAACTCTTTCCCTACACGACGCTCTTCCGATCTNNNTCTTCCTCCTCCTCCTCTTCTTCCA
GAGCACACGTCTGAACTCTTTCCCTACACGACGCTCTTCCGATCTNNNAGTGCCTGCATTCCCTTAGGAC
GAGCACACGTCTGAACTCTTTCCCTACACGACGCTCTTCCGATCTNNNAGTGCCTGCATTCCCTTAGGAC
GAGCACACGTCTGAACTCTTTCCCTACACGACGCTCTTCCGATCTNNNTTACCACCCAGGATGATTGTGA
GAGCACACGTCTGAACTCTTTCCCTACACGACGCTCTTCCGATCTNNNTTACCACCCAGGATGATTGTGA
AGCACACGTCTGAACTCTTTCCCTACACGACGCTCTTCCGATCTNNNTGTCTATTCTCCACAGGCTTTGC
AGCACACGTCTGAACTCTTTCCCTACACGACGCTCTTCCGATCTNNNTGTCTATTCTCCACAGGCTTTGC
CACACGTCTGAACTCTTTCCCTACACGACGCTCTTCCGATCTNNNCTAGAGCTTTTCCTTTTAAGAGACA
CACACGTCTGAACTCTTTCCCTACACGACGCTCTTCCGATCTNNNCTAGAGCTTTTCCTTTTAAGAGACA
GCACACGTCTGAACTCTTTCCCTACACGACGCTCTTCCGATCTNNNACCCAAGAAATGCAAAATCTGAG
GCACACGTCTGAACTCTTTCCCTACACGACGCTCTTCCGATCTNNNACCCAAGAAATGCAAAATCTGAG
GAGCACACGTCTGAACTCTTTCCCTACACGACGCTCTTCCGATCTNNNACTGGCCTTTGAAATATCTGG
GAGCACACGTCTGAACTCTTTCCCTACACGACGCTCTTCCGATCTNNNACTGGCCTTTGAAATATCTGG
AGCACACGTCTGAACTCTTTCCCTACACGACGCTCTTCCGATCTNNNATTTCCTTGCCAACTAGTAAAAA
AGCACACGTCTGAACTCTTTCCCTACACGACGCTCTTCCGATCTNNNATTTCCTTGCCAACTAGTAAAAA
AGCACACGTCTGAACTCTTTCCCTACACGACGCTCTTCCGATCTNNNTATCTTTGTCAGCCACGTGTACA
AGCACACGTCTGAACTCTTTCCCTACACGACGCTCTTCCGATCTNNNTATCTTTGTCAGCCACGTGTACA
AGCACACGTCTGAACTCTTTCCCTACACGACGCTCTTCCGATCTNNNTGTGGGGCACTGAGTATACAA
AGCACACGTCTGAACTCTTTCCCTACACGACGCTCTTCCGATCTNNNTGTGGGGCACTGAGTATACAA
GCACACGTCTGAACTCTTTCCCTACACGACGCTCTTCCGATCTNNNGCCATGGTGATCTTACTTGAGGG
GCACACGTCTGAACTCTTTCCCTACACGACGCTCTTCCGATCTNNNGCCATGGTGATCTTACTTGAGGG

AGCACACGTCTGAACTCTTTCCCTACACGACGCTCTTCCGATCTNNNTGTTCCTATTACCACCAAGCCAA
AGCACACGTCTGAACTCTTTCCCTACACGACGCTCTTCCGATCTNNNTGTTCCTATTACCACCAAGCCAA
AGAGCACACGTCTGAACTCTTTCCCTACACGACGCTCTTCCGATCTNNNCACTGTTGTAAGCCCTCTCA
AGAGCACACGTCTGAACTCTTTCCCTACACGACGCTCTTCCGATCTNNNCACTGTTGTAAGCCCTCTCA
GAGCACACGTCTGAACTCTTTCCCTACACGACGCTCTTCCGATCTNNNGAGGTGGGAGAATTGCTTGA
GAGCACACGTCTGAACTCTTTCCCTACACGACGCTCTTCCGATCTNNNGAGGTGGGAGAATTGCTTGA
GAGCACACGTCTGAACTCTTTCCCTACACGACGCTCTTCCGATCTNNNGGGCGAAAGAGTGAGATTTT
GAGCACACGTCTGAACTCTTTCCCTACACGACGCTCTTCCGATCTNNNGGGCGAAAGAGTGAGATTTT
GAGCACACGTCTGAACTCTTTCCCTACACGACGCTCTTCCGATCTNNNAAAAAATTAGCCAGGCGTGGT
GAGCACACGTCTGAACTCTTTCCCTACACGACGCTCTTCCGATCTNNNAAAAAATTAGCCAGGCGTGGT
GAGCACACGTCTGAACTCTTTCCCTACACGACGCTCTTCCGATCTNNNTTCAGTCTACTTTGGGTCTGTC
GAGCACACGTCTGAACTCTTTCCCTACACGACGCTCTTCCGATCTNNNTTCAGTCTACTTTGGGTCTGTC
AGCACACGTCTGAACTCTTTCCCTACACGACGCTCTTCCGATCTNNNTCTTTACTTTCTTTTTAACAGGCC
AGCACACGTCTGAACTCTTTCCCTACACGACGCTCTTCCGATCTNNNTCTTTACTTTCTTTTTAACAGGCC
GAGCACACGTCTGAACTCTTTCCCTACACGACGCTCTTCCGATCTNNNGCCTCTGAAAACAACAGAACA
GAGCACACGTCTGAACTCTTTCCCTACACGACGCTCTTCCGATCTNNNGCCTCTGAAAACAACAGAACA
AGCACACGTCTGAACTCTTTCCCTACACGACGCTCTTCCGATCTNNNGGTTTCTGACCCTAAAGTGCCTT
AGCACACGTCTGAACTCTTTCCCTACACGACGCTCTTCCGATCTNNNGGTTTCTGACCCTAAAGTGCCTT
AGCACACGTCTGAACTCTTTCCCTACACGACGCTCTTCCGATCTNNNGTATGCCATCATGCCCAGCTAAC
AGCACACGTCTGAACTCTTTCCCTACACGACGCTCTTCCGATCTNNNGTATGCCATCATGCCCAGCTAAC
AGCACACGTCTGAACTCTTTCCCTACACGACGCTCTTCCGATCTNNNCACATACACCTTCCCAAGACTGA
AGCACACGTCTGAACTCTTTCCCTACACGACGCTCTTCCGATCTNNNCACATACACCTTCCCAAGACTGA
GAGCACACGTCTGAACTCTTTCCCTACACGACGCTCTTCCGATCTNNNGTATATGGCTCCACACAAATG
GAGCACACGTCTGAACTCTTTCCCTACACGACGCTCTTCCGATCTNNNGTATATGGCTCCACACAAATG
GAGCACACGTCTGAACTCTTTCCCTACACGACGCTCTTCCGATCTNNNCACCTACTTCCACTTTCCCTTCA
GAGCACACGTCTGAACTCTTTCCCTACACGACGCTCTTCCGATCTNNNCACCTACTTCCACTTTCCCTTCA
AGAGCACACGTCTGAACTCTTTCCCTACACGACGCTCTTCCGATCTNNNCTATCTCAGCTCACTGCAACC
AGAGCACACGTCTGAACTCTTTCCCTACACGACGCTCTTCCGATCTNNNCTATCTCAGCTCACTGCAACC
AGCACACGTCTGAACTCTTTCCCTACACGACGCTCTTCCGATCTNNNAATTTACACGCCATCATTACCGTC
AGCACACGTCTGAACTCTTTCCCTACACGACGCTCTTCCGATCTNNNAATTTACACGCCATCATTACCGTC
AGCACACGTCTGAACTCTTTCCCTACACGACGCTCTTCCGATCTNNNACAAAGCTGGGCTATAAAAACA
AGCACACGTCTGAACTCTTTCCCTACACGACGCTCTTCCGATCTNNNACAAAGCTGGGCTATAAAAACA
AGAGCACACGTCTGAACTCTTTCCCTACACGACGCTCTTCCGATCTNNNAAAATTATTTGGGCATGGTG
AGAGCACACGTCTGAACTCTTTCCCTACACGACGCTCTTCCGATCTNNNAAAATTATTTGGGCATGGTG
AGCACACGTCTGAACTCTTTCCCTACACGACGCTCTTCCGATCTNNNGTGGAGATCCTGCCAATTCACT
AGCACACGTCTGAACTCTTTCCCTACACGACGCTCTTCCGATCTNNNGTGGAGATCCTGCCAATTCACT
AGCACACGTCTGAACTCTTTCCCTACACGACGCTCTTCCGATCTNNNATCTCCCTTTTGTTTTACAAGTCA
AGCACACGTCTGAACTCTTTCCCTACACGACGCTCTTCCGATCTNNNATCTCCCTTTTGTTTTACAAGTCA
AGCACACGTCTGAACTCTTTCCCTACACGACGCTCTTCCGATCTNNNATTAACAATCGGGGAAAAATAA
AGCACACGTCTGAACTCTTTCCCTACACGACGCTCTTCCGATCTNNNATTAACAATCGGGGAAAAATAA
AGCACACGTCTGAACTCTTTCCCTACACGACGCTCTTCCGATCTNNNAAAAGTGGGGAGAGTAGCAGA
AGCACACGTCTGAACTCTTTCCCTACACGACGCTCTTCCGATCTNNNAAAAGTGGGGAGAGTAGCAGA
AGAGCACACGTCTGAACTCTTTCCCTACACGACGCTCTTCCGATCTNNNATCAGAGAGTGGAGAAAG

AGAGCACACGTCTGAACTCTTTCCCTACACGACGCTCTTCCGATCTNNNATCAGAGAGTGGAGAAAG
GCACACGTCTGAACTCTTTCCCTACACGACGCTCTTCCGATCTNNNCCCTTTTATGACCCCAGATGGA
GCACACGTCTGAACTCTTTCCCTACACGACGCTCTTCCGATCTNNNCCCTTTTATGACCCCAGATGGA
AGAGCACACGTCTGAACTCTTTCCCTACACGACGCTCTTCCGATCTNNNAAAATTTGTACCAGGAATG
AGAGCACACGTCTGAACTCTTTCCCTACACGACGCTCTTCCGATCTNNNAAAATTTGTACCAGGAATG
GCACACGTCTGAACTCTTTCCCTACACGACGCTCTTCCGATCTNNNTCCTTGCCTACATCCTATCTCCTC
GCACACGTCTGAACTCTTTCCCTACACGACGCTCTTCCGATCTNNNTCCTTGCCTACATCCTATCTCCTC
AGCACACGTCTGAACTCTTTCCCTACACGACGCTCTTCCGATCTNNNATGAGCTAGATAGTCAACCCC
AGCACACGTCTGAACTCTTTCCCTACACGACGCTCTTCCGATCTNNNATGAGCTAGATAGTCAACCCC
GAGCACACGTCTGAACTCTTTCCCTACACGACGCTCTTCCGATCTNNNCTGGCTTGAATGATGCATCT
GAGCACACGTCTGAACTCTTTCCCTACACGACGCTCTTCCGATCTNNNCTGGCTTGAATGATGCATCT
GAGCACACGTCTGAACTCTTTCCCTACACGACGCTCTTCCGATCTNNNAGAAGCAGCAGGATAGTGA
GAGCACACGTCTGAACTCTTTCCCTACACGACGCTCTTCCGATCTNNNAGAAGCAGCAGGATAGTGA
GAGCACACGTCTGAACTCTTTCCCTACACGACGCTCTTCCGATCTNNNAGGAGTGAATGGTTCTGTC
GAGCACACGTCTGAACTCTTTCCCTACACGACGCTCTTCCGATCTNNNAGGAGTGAATGGTTCTGTC
AGCACACGTCTGAACTCTTTCCCTACACGACGCTCTTCCGATCTNNNGATCTCAGCCTCATTCCTCCT
AGCACACGTCTGAACTCTTTCCCTACACGACGCTCTTCCGATCTNNNGATCTCAGCCTCATTCCTCCT
AGCACACGTCTGAACTCTTTCCCTACACGACGCTCTTCCGATCTNNNGGAAGTCTAAGATCATGGTG
AGCACACGTCTGAACTCTTTCCCTACACGACGCTCTTCCGATCTNNNGGAAGTCTAAGATCATGGTG
GCACACGTCTGAACTCTTTCCCTACACGACGCTCTTCCGATCTNNNTATGCTTAGGTCTCTTGGTGAG
GCACACGTCTGAACTCTTTCCCTACACGACGCTCTTCCGATCTNNNTATGCTTAGGTCTCTTGGTGAG
GCACACGTCTGAACTCTTTCCCTACACGACGCTCTTCCGATCTNNNCAATTGCACTTTTCAGCTCCAGA
GCACACGTCTGAACTCTTTCCCTACACGACGCTCTTCCGATCTNNNCAATTGCACTTTTCAGCTCCAGA
GAGCACACGTCTGAACTCTTTCCCTACACGACGCTCTTCCGATCTNNNAGCTGACAGAAACAAAATG
GAGCACACGTCTGAACTCTTTCCCTACACGACGCTCTTCCGATCTNNNAGCTGACAGAAACAAAATG
GCACACGTCTGAACTCTTTCCCTACACGACGCTCTTCCGATCTNNNTCAGTGAGCACTCATTCCCTAC
GCACACGTCTGAACTCTTTCCCTACACGACGCTCTTCCGATCTNNNTCAGTGAGCACTCATTCCCTAC
AGCACACGTCTGAACTCTTTCCCTACACGACGCTCTTCCGATCTNNNCTGCATTCCCATATTGCTCTGT
AGCACACGTCTGAACTCTTTCCCTACACGACGCTCTTCCGATCTNNNCTGCATTCCCATATTGCTCTGT
AGCACACGTCTGAACTCTTTCCCTACACGACGCTCTTCCGATCTNNNGACAGAGCAAGCTCCTGTTTC
AGCACACGTCTGAACTCTTTCCCTACACGACGCTCTTCCGATCTNNNGACAGAGCAAGCTCCTGTTTC
AGCACACGTCTGAACTCTTTCCCTACACGACGCTCTTCCGATCTNNNTTTTGAGGGAGATGTGAACTC
AGCACACGTCTGAACTCTTTCCCTACACGACGCTCTTCCGATCTNNNTTTTGAGGGAGATGTGAACTC
GAGCACACGTCTGAACTCTTTCCCTACACGACGCTCTTCCGATCTNNNATGGGATTGTGGGGATATG
GAGCACACGTCTGAACTCTTTCCCTACACGACGCTCTTCCGATCTNNNATGGGATTGTGGGGATATG
GCACACGTCTGAACTCTTTCCCTACACGACGCTCTTCCGATCTNNNTAGTCAGGAGGCTGAAGTTGA
GCACACGTCTGAACTCTTTCCCTACACGACGCTCTTCCGATCTNNNTAGTCAGGAGGCTGAAGTTGA
AGCACACGTCTGAACTCTTTCCCTACACGACGCTCTTCCGATCTNNNCTTGAGCCCAGGAAGTTGAGG
AGCACACGTCTGAACTCTTTCCCTACACGACGCTCTTCCGATCTNNNCTTGAGCCCAGGAAGTTGAGG
GCACACGTCTGAACTCTTTCCCTACACGACGCTCTTCCGATCTNNNCTGTAGTCCCAGCTATTCAGGA
GCACACGTCTGAACTCTTTCCCTACACGACGCTCTTCCGATCTNNNCTGTAGTCCCAGCTATTCAGGA
AGCACACGTCTGAACTCTTTCCCTACACGACGCTCTTCCGATCTNNNCTTGAACCCAGGAGACAGAGT
AGCACACGTCTGAACTCTTTCCCTACACGACGCTCTTCCGATCTNNNCTTGAACCCAGGAGACAGAGT

AGCACACGTCTGAACTCTTTCCCTACACGACGCTCTTCCGATCTNNNTGGTGTGAGAGTGTCATGAAG

AGCACACGTCTGAACTCTTTCCCTACACGACGCTCTTCCGATCTNNNTGGTGTGAGAGTGTCATGAAG

GCACACGTCTGAACTCTTTCCCTACACGACGCTCTTCCGATCTNNNATCCTCACCATCTGGCTCTTCAAC

GCACACGTCTGAACTCTTTCCCTACACGACGCTCTTCCGATCTNNNATCCTCACCATCTGGCTCTTCAAC

GAGCACACGTCTGAACTCTTTCCCTACACGACGCTCTTCCGATCTNNNTCAAGTAATTCTCCAGCCTCA

GAGCACACGTCTGAACTCTTTCCCTACACGACGCTCTTCCGATCTNNNTCAAGTAATTCTCCAGCCTCA

AGCACACGTCTGAACTCTTTCCCTACACGACGCTCTTCCGATCTNNNGATTTCAGACTGACAGGGACA

AGCACACGTCTGAACTCTTTCCCTACACGACGCTCTTCCGATCTNNNGATTTCAGACTGACAGGGACA

GAGCACACGTCTGAACTCTTTCCCTACACGACGCTCTTCCGATCTNNNCTGAGGTGGCAGGATTACCT

GAGCACACGTCTGAACTCTTTCCCTACACGACGCTCTTCCGATCTNNNCTGAGGTGGCAGGATTACCT

AGCACACGTCTGAACTCTTTCCCTACACGACGCTCTTCCGATCTNNNTTTCCAGGCTTAGACACTACCC

AGCACACGTCTGAACTCTTTCCCTACACGACGCTCTTCCGATCTNNNTTTCCAGGCTTAGACACTACCC

GCACACGTCTGAACTCTTTCCCTACACGACGCTCTTCCGATCTNNNGACTTCTGAGAGGCCAATGTCA

GCACACGTCTGAACTCTTTCCCTACACGACGCTCTTCCGATCTNNNGACTTCTGAGAGGCCAATGTCA

AGCACACGTCTGAACTCTTTCCCTACACGACGCTCTTCCGATCTNNNAGTTTTGTTACAAACACTGCC

AGCACACGTCTGAACTCTTTCCCTACACGACGCTCTTCCGATCTNNNAGTTTTGTTACAAACACTGCC

GAGCACACGTCTGAACTCTTTCCCTACACGACGCTCTTCCGATCTNNNTCCTTATCATGCCTAGCACTGA

GAGCACACGTCTGAACTCTTTCCCTACACGACGCTCTTCCGATCTNNNTCCTTATCATGCCTAGCACTGA

AGCACACGTCTGAACTCTTTCCCTACACGACGCTCTTCCGATCTNNNTTGCAAACCTAATAGGCTTCCCT

AGCACACGTCTGAACTCTTTCCCTACACGACGCTCTTCCGATCTNNNTTGCAAACCTAATAGGCTTCCCT

AGCACACGTCTGAACTCTTTCCCTACACGACGCTCTTCCGATCTNNNTCTGAATGGGAAGTGGTATAG

AGCACACGTCTGAACTCTTTCCCTACACGACGCTCTTCCGATCTNNNTCTGAATGGGAAGTGGTATAG

AGCACACGTCTGAACTCTTTCCCTACACGACGCTCTTCCGATCTNNNATCTCAGCACTTTGGAAGGACA

AGCACACGTCTGAACTCTTTCCCTACACGACGCTCTTCCGATCTNNNATCTCAGCACTTTGGAAGGACA

GAGCACACGTCTGAACTCTTTCCCTACACGACGCTCTTCCGATCTNNNTGCTACATAGTGAAACCTTGC

GAGCACACGTCTGAACTCTTTCCCTACACGACGCTCTTCCGATCTNNNTGCTACATAGTGAAACCTTGC

AGAGCACACGTCTGAACTCTTTCCCTACACGACGCTCTTCCGATCTNNNTTCTGGTGTCTAGACTCAGC

AGAGCACACGTCTGAACTCTTTCCCTACACGACGCTCTTCCGATCTNNNTTCTGGTGTCTAGACTCAGC

GAGCACACGTCTGAACTCTTTCCCTACACGACGCTCTTCCGATCTNNNGTAGAAAGGAGAGGTTTGGG

GAGCACACGTCTGAACTCTTTCCCTACACGACGCTCTTCCGATCTNNNGTAGAAAGGAGAGGTTTGGG

GCACACGTCTGAACTCTTTCCCTACACGACGCTCTTCCGATCTNNNTGATGTTGAGCACCTCTTCATGT

GCACACGTCTGAACTCTTTCCCTACACGACGCTCTTCCGATCTNNNTGATGTTGAGCACCTCTTCATGT

GAGCACACGTCTGAACTCTTTCCCTACACGACGCTCTTCCGATCTNNNGTAGAGTTTTGCCCTTTGTGG

GAGCACACGTCTGAACTCTTTCCCTACACGACGCTCTTCCGATCTNNNGTAGAGTTTTGCCCTTTGTGG

AGCACACGTCTGAACTCTTTCCCTACACGACGCTCTTCCGATCTNNNGCTGTTTTAACTTTATTGTGAT

AGCACACGTCTGAACTCTTTCCCTACACGACGCTCTTCCGATCTNNNGCTGTTTTAACTTTATTGTGAT

GCACACGTCTGAACTCTTTCCCTACACGACGCTCTTCCGATCTNNNCCTGTAGTCCCAGCTACTAGGGC

GCACACGTCTGAACTCTTTCCCTACACGACGCTCTTCCGATCTNNNCCTGTAGTCCCAGCTACTAGGGC

GAGCACACGTCTGAACTCTTTCCCTACACGACGCTCTTCCGATCTNNNTGAGGTAGGAGAATCGCTTGA

GAGCACACGTCTGAACTCTTTCCCTACACGACGCTCTTCCGATCTNNNTGAGGTAGGAGAATCGCTTGA

AGCACACGTCTGAACTCTTTCCCTACACGACGCTCTTCCGATCTNNNAAGCAAAGAGTCGATCATGAG

AGCACACGTCTGAACTCTTTCCCTACACGACGCTCTTCCGATCTNNNAAGCAAAGAGTCGATCATGAG

AGCACACGTCTGAACTCTTTCCCTACACGACGCTCTTCCGATCTNNNAAGCATTGGGATTACAGGTGT

AGCACACGTCTGAACTCTTTCCCTACACGACGCTCTTCCGATCTNNNAAGCATTGGGATTACAGGTGT
GCACACGTCTGAACTCTTTCCCTACACGACGCTCTTCCGATCTNNNGCCTGTAGTCCCAGCTACTTGC
GCACACGTCTGAACTCTTTCCCTACACGACGCTCTTCCGATCTNNNGCCTGTAGTCCCAGCTACTTGC
GAGCACACGTCTGAACTCTTTCCCTACACGACGCTCTTCCGATCTNNNCCATGTAGACACTCTTCAGGG
GAGCACACGTCTGAACTCTTTCCCTACACGACGCTCTTCCGATCTNNNCCATGTAGACACTCTTCAGGG
CACACGTCTGAACTCTTTCCCTACACGACGCTCTTCCGATCTNNNTCCCTCTCTCTCTCTCTCTCTCAT
CACACGTCTGAACTCTTTCCCTACACGACGCTCTTCCGATCTNNNTCCCTCTCTCTCTCTCTCTCTCAT
CACACGTCTGAACTCTTTCCCTACACGACGCTCTTCCGATCTNNNCTGACACATACAATCAGTTTTTAC
CACACGTCTGAACTCTTTCCCTACACGACGCTCTTCCGATCTNNNCTGACACATACAATCAGTTTTTAC
AGCACACGTCTGAACTCTTTCCCTACACGACGCTCTTCCGATCTNNNGATCACAAGGTCAGGAGATC
AGCACACGTCTGAACTCTTTCCCTACACGACGCTCTTCCGATCTNNNGATCACAAGGTCAGGAGATC
AGCACACGTCTGAACTCTTTCCCTACACGACGCTCTTCCGATCTNNNTAACAATGGTGGAAAGGGAG
AGCACACGTCTGAACTCTTTCCCTACACGACGCTCTTCCGATCTNNNTAACAATGGTGGAAAGGGAG
AGCACACGTCTGAACTCTTTCCCTACACGACGCTCTTCCGATCTNNNATGTGGCAATATCATTGAAG
AGCACACGTCTGAACTCTTTCCCTACACGACGCTCTTCCGATCTNNNATGTGGCAATATCATTGAAG
AGCACACGTCTGAACTCTTTCCCTACACGACGCTCTTCCGATCTNNNTTATAGAGGAACGTAATAGTC
AGCACACGTCTGAACTCTTTCCCTACACGACGCTCTTCCGATCTNNNTTATAGAGGAACGTAATAGTC
AGCACACGTCTGAACTCTTTCCCTACACGACGCTCTTCCGATCTNNNGGCCCCTCTTGTGTGAAATAA
AGCACACGTCTGAACTCTTTCCCTACACGACGCTCTTCCGATCTNNNGGCCCCTCTTGTGTGAAATAA
AAGAGCACACGTCTGAACTCTTTCCCTACACGACGCTCTTCCGATCTNNNTGTAATTCCAGCTACTTGG
AAGAGCACACGTCTGAACTCTTTCCCTACACGACGCTCTTCCGATCTNNNTGTAATTCCAGCTACTTGG
AGCACACGTCTGAACTCTTTCCCTACACGACGCTCTTCCGATCTNNNTATTTTCCAGACCCTGCACTTGA
GCACACGTCTGAACTCTTTCCCTACACGACGCTCTTCCGATCTNNNTATTTTCCAGACCCTGCACTTGA
GCACACGTCTGAACTCTTTCCCTACACGACGCTCTTCCGATCTNNNCTTCTGGGTTCAGGCGATTCTC
GCACACGTCTGAACTCTTTCCCTACACGACGCTCTTCCGATCTNNNCTTCTGGGTTCAGGCGATTCTC
GCACACGTCTGAACTCTTTCCCTACACGACGCTCTTCCGATCTNNNGTGCCTGTAATCCCAGCTACTCC
GCACACGTCTGAACTCTTTCCCTACACGACGCTCTTCCGATCTNNNGTGCCTGTAATCCCAGCTACTCC
AGCACACGTCTGAACTCTTTCCCTACACGACGCTCTTCCGATCTNNNGGTGAACAAGACCCATAATCC
AGCACACGTCTGAACTCTTTCCCTACACGACGCTCTTCCGATCTNNNGGTGAACAAGACCCATAATCC
GCACACGTCTGAACTCTTTCCCTACACGACGCTCTTCCGATCTNNNCAAAGGTCTCGAAGAGGATTG
GCACACGTCTGAACTCTTTCCCTACACGACGCTCTTCCGATCTNNNCAAAGGTCTCGAAGAGGATTG
AGCACACGTCTGAACTCTTTCCCTACACGACGCTCTTCCGATCTNNNACAGAAAGAACATGCTGAATT
AGCACACGTCTGAACTCTTTCCCTACACGACGCTCTTCCGATCTNNNACAGAAAGAACATGCTGAATT
GCACACGTCTGAACTCTTTCCCTACACGACGCTCTTCCGATCTNNNTTGAGGTAGGGGAATTGCTTG
GCACACGTCTGAACTCTTTCCCTACACGACGCTCTTCCGATCTNNNTTGAGGTAGGGGAATTGCTTG
AGCACACGTCTGAACTCTTTCCCTACACGACGCTCTTCCGATCTNNNATGGAGAAACCCCGTCTCTACT
AGCACACGTCTGAACTCTTTCCCTACACGACGCTCTTCCGATCTNNNATGGAGAAACCCCGTCTCTACT
AGCACACGTCTGAACTCTTTCCCTACACGACGCTCTTCCGATCTNNNGCATCTGTAATCCCAGCTACT
AGCACACGTCTGAACTCTTTCCCTACACGACGCTCTTCCGATCTNNNGCATCTGTAATCCCAGCTACT
GAGCACACGTCTGAACTCTTTCCCTACACGACGCTCTTCCGATCTNNNCCTCTAACTCTGTCCAAGTTG
GAGCACACGTCTGAACTCTTTCCCTACACGACGCTCTTCCGATCTNNNCCTCTAACTCTGTCCAAGTTG
GAGCACACGTCTGAACTCTTTCCCTACACGACGCTCTTCCGATCTNNNCTGAGGCAGGAGAATCACTT
GAGCACACGTCTGAACTCTTTCCCTACACGACGCTCTTCCGATCTNNNCTGAGGCAGGAGAATCACTT

GAGCACACGTCTGAACTCTTTCCCTACACGACGCTCTTCCGATCTNNNGGTACTAAGACTAATGCAGCT
GAGCACACGTCTGAACTCTTTCCCTACACGACGCTCTTCCGATCTNNNGGTACTAAGACTAATGCAGCT
AGCACACGTCTGAACTCTTTCCCTACACGACGCTCTTCCGATCTNNNTCTGGTCCTCATCTTCATTCTGC
AGCACACGTCTGAACTCTTTCCCTACACGACGCTCTTCCGATCTNNNTCTGGTCCTCATCTTCATTCTGC
GAGCACACGTCTGAACTCTTTCCCTACACGACGCTCTTCCGATCTNNNCGATTCTTCAGGGGGAACAA
GAGCACACGTCTGAACTCTTTCCCTACACGACGCTCTTCCGATCTNNNCGATTCTTCAGGGGGAACAA
AGCACACGTCTGAACTCTTTCCCTACACGACGCTCTTCCGATCTNNNCGCTTGTTTTTGTCAGCTTTGTC
AGCACACGTCTGAACTCTTTCCCTACACGACGCTCTTCCGATCTNNNCGCTTGTTTTTGTCAGCTTTGTC
GAGCACACGTCTGAACTCTTTCCCTACACGACGCTCTTCCGATCTNNNGAGGCTGAGGCAGGAGAATG
GAGCACACGTCTGAACTCTTTCCCTACACGACGCTCTTCCGATCTNNNGAGGCTGAGGCAGGAGAATG
GCACACGTCTGAACTCTTTCCCTACACGACGCTCTTCCGATCTNNNATCATACTGCTAAGAATTCAAAC
GCACACGTCTGAACTCTTTCCCTACACGACGCTCTTCCGATCTNNNATCATACTGCTAAGAATTCAAAC
AGCACACGTCTGAACTCTTTCCCTACACGACGCTCTTCCGATCTNNNGTTTCACCATGTTAGCCAGGAT
AGCACACGTCTGAACTCTTTCCCTACACGACGCTCTTCCGATCTNNNGTTTCACCATGTTAGCCAGGAT
GAGCACACGTCTGAACTCTTTCCCTACACGACGCTCTTCCGATCTNNNAGGAGAATTGCTTGAACCTGG
GAGCACACGTCTGAACTCTTTCCCTACACGACGCTCTTCCGATCTNNNAGGAGAATTGCTTGAACCTGG
AGCACACGTCTGAACTCTTTCCCTACACGACGCTCTTCCGATCTNNNCTGGGATCCAATACCTTACGCTG
AGCACACGTCTGAACTCTTTCCCTACACGACGCTCTTCCGATCTNNNCTGGGATCCAATACCTTACGCTG
AGCACACGTCTGAACTCTTTCCCTACACGACGCTCTTCCGATCTNNNAGGCCAGCATAATTTGATACCC
AGCACACGTCTGAACTCTTTCCCTACACGACGCTCTTCCGATCTNNNAGGCCAGCATAATTTGATACCC
AGCACACGTCTGAACTCTTTCCCTACACGACGCTCTTCCGATCTNNNCCTGTATCATAGTGCCTGGAAC
AGCACACGTCTGAACTCTTTCCCTACACGACGCTCTTCCGATCTNNNCCTGTATCATAGTGCCTGGAAC
AGCACACGTCTGAACTCTTTCCCTACACGACGCTCTTCCGATCTNNNTGCACACTGTGGATTCGTAAGA
AGCACACGTCTGAACTCTTTCCCTACACGACGCTCTTCCGATCTNNNTGCACACTGTGGATTCGTAAGA
AGCACACGTCTGAACTCTTTCCCTACACGACGCTCTTCCGATCTNNNGGGAGAATCATTTGAACCCAGC
AGCACACGTCTGAACTCTTTCCCTACACGACGCTCTTCCGATCTNNNGGGAGAATCATTTGAACCCAGC
AGCACACGTCTGAACTCTTTCCCTACACGACGCTCTTCCGATCTNNNAGGCCTTCTTTGTTATCAGCTGA
AGCACACGTCTGAACTCTTTCCCTACACGACGCTCTTCCGATCTNNNAGGCCTTCTTTGTTATCAGCTGA
GAGCACACGTCTGAACTCTTTCCCTACACGACGCTCTTCCGATCTNNNTGTGGGCTTTCTTTTCTCTGTC
GAGCACACGTCTGAACTCTTTCCCTACACGACGCTCTTCCGATCTNNNTGTGGGCTTTCTTTTCTCTGTC
GAGCACACGTCTGAACTCTTTCCCTACACGACGCTCTTCCGATCTNNNGCTGCACAGTATTCCATGGAG
GAGCACACGTCTGAACTCTTTCCCTACACGACGCTCTTCCGATCTNNNGCTGCACAGTATTCCATGGAG
AGCACACGTCTGAACTCTTTCCCTACACGACGCTCTTCCGATCTNNNAGTCACTTCTGCAGGATTTAGCA
AGCACACGTCTGAACTCTTTCCCTACACGACGCTCTTCCGATCTNNNAGTCACTTCTGCAGGATTTAGCA
GAGCACACGTCTGAACTCTTTCCCTACACGACGCTCTTCCGATCTNNNGAACGGAGGCCCAGAGAGAT
GAGCACACGTCTGAACTCTTTCCCTACACGACGCTCTTCCGATCTNNNGAACGGAGGCCCAGAGAGAT
AGCACACGTCTGAACTCTTTCCCTACACGACGCTCTTCCGATCTNNNTACTGATGGTGGGGCGTATATT
AGCACACGTCTGAACTCTTTCCCTACACGACGCTCTTCCGATCTNNNTACTGATGGTGGGGCGTATATT
GAGCACACGTCTGAACTCTTTCCCTACACGACGCTCTTCCGATCTNNNTGAGATGCTGAGATTGTGTGT
GAGCACACGTCTGAACTCTTTCCCTACACGACGCTCTTCCGATCTNNNTGAGATGCTGAGATTGTGTGT
AGCACACGTCTGAACTCTTTCCCTACACGACGCTCTTCCGATCTNNNGGAGGTTGTAGTGAGCTGAGA
AGCACACGTCTGAACTCTTTCCCTACACGACGCTCTTCCGATCTNNNGGAGGTTGTAGTGAGCTGAGA
CACACGTCTGAACTCTTTCCCTACACGACGCTCTTCCGATCTNNNCACAACGACCACATCCTCATAGACA

CACACGTCTGAACTCTTTCCCTACACGACGCTCTTCCGATCTNNNCACAACGACCACATCCTCATAGACA
AGCACACGTCTGAACTCTTTCCCTACACGACGCTCTTCCGATCTNNNGTTGATTTATACCCTGATAGAGT
AGCACACGTCTGAACTCTTTCCCTACACGACGCTCTTCCGATCTNNNGTTGATTTATACCCTGATAGAGT
AGAGCACACGTCTGAACTCTTTCCCTACACGACGCTCTTCCGATCTNNNTACAAATGAATCTGCTCTCCC
AGAGCACACGTCTGAACTCTTTCCCTACACGACGCTCTTCCGATCTNNNTACAAATGAATCTGCTCTCCC
GAGCACACGTCTGAACTCTTTCCCTACACGACGCTCTTCCGATCTNNNTCCCTAACTGTGGATCCTACTG
GAGCACACGTCTGAACTCTTTCCCTACACGACGCTCTTCCGATCTNNNTCCCTAACTGTGGATCCTACTG
AGCACACGTCTGAACTCTTTCCCTACACGACGCTCTTCCGATCTNNNCTCAGATTGGGTGGTAATCTAC
AGCACACGTCTGAACTCTTTCCCTACACGACGCTCTTCCGATCTNNNCTCAGATTGGGTGGTAATCTAC
GAGCACACGTCTGAACTCTTTCCCTACACGACGCTCTTCCGATCTNNNAACTACTCGGGAAGTTGAGGT
GAGCACACGTCTGAACTCTTTCCCTACACGACGCTCTTCCGATCTNNNAACTACTCGGGAAGTTGAGGT
GCACACGTCTGAACTCTTTCCCTACACGACGCTCTTCCGATCTNNNTTGCATTTCTCTAACGATTTGCACA
GCACACGTCTGAACTCTTTCCCTACACGACGCTCTTCCGATCTNNNTTGCATTTCTCTAACGATTTGCACA
GAGCACACGTCTGAACTCTTTCCCTACACGACGCTCTTCCGATCTNNNGGGTCCATTCAGATTCAGACA
GAGCACACGTCTGAACTCTTTCCCTACACGACGCTCTTCCGATCTNNNGGGTCCATTCAGATTCAGACA
GAGCACACGTCTGAACTCTTTCCCTACACGACGCTCTTCCGATCTNNNGGTGGCAGATACTCTGTTTCTC
GAGCACACGTCTGAACTCTTTCCCTACACGACGCTCTTCCGATCTNNNGGTGGCAGATACTCTGTTTCTC
GAGCACACGTCTGAACTCTTTCCCTACACGACGCTCTTCCGATCTNNNAGACACACAGACATACACACAC
GAGCACACGTCTGAACTCTTTCCCTACACGACGCTCTTCCGATCTNNNAGACACACAGACATACACACAC
AGCACACGTCTGAACTCTTTCCCTACACGACGCTCTTCCGATCTNNNCTACAACTGTTACCAAGCAGCTC
AGCACACGTCTGAACTCTTTCCCTACACGACGCTCTTCCGATCTNNNCTACAACTGTTACCAAGCAGCTC
GAGCACACGTCTGAACTCTTTCCCTACACGACGCTCTTCCGATCTNNNGAGGCTGAAATGAGAGGATCA
GAGCACACGTCTGAACTCTTTCCCTACACGACGCTCTTCCGATCTNNNGAGGCTGAAATGAGAGGATCA
AGCACACGTCTGAACTCTTTCCCTACACGACGCTCTTCCGATCTNNNTACAGTGAGCCGAGATCATGCT
AGCACACGTCTGAACTCTTTCCCTACACGACGCTCTTCCGATCTNNNTACAGTGAGCCGAGATCATGCT
GAGCACACGTCTGAACTCTTTCCCTACACGACGCTCTTCCGATCTNNNATGTGATTGTGTCATCCCTACC
GAGCACACGTCTGAACTCTTTCCCTACACGACGCTCTTCCGATCTNNNATGTGATTGTGTCATCCCTACC
GAGCACACGTCTGAACTCTTTCCCTACACGACGCTCTTCCGATCTNNNGCAGTAATGCAGATCAATGAC
GAGCACACGTCTGAACTCTTTCCCTACACGACGCTCTTCCGATCTNNNGCAGTAATGCAGATCAATGAC
AGCACACGTCTGAACTCTTTCCCTACACGACGCTCTTCCGATCTNNNGGGACAAAAAGGGTGTGATCT
AGCACACGTCTGAACTCTTTCCCTACACGACGCTCTTCCGATCTNNNGGGACAAAAAGGGTGTGATCT
AGAGCACACGTCTGAACTCTTTCCCTACACGACGCTCTTCCGATCTNNNTGTTACTCCAAGTTCTCCACT
AGAGCACACGTCTGAACTCTTTCCCTACACGACGCTCTTCCGATCTNNNTGTTACTCCAAGTTCTCCACT
AGCACACGTCTGAACTCTTTCCCTACACGACGCTCTTCCGATCTNNNTTTTTGGAACTGGAGATGTTCAT
AGCACACGTCTGAACTCTTTCCCTACACGACGCTCTTCCGATCTNNNTTTTTGGAACTGGAGATGTTCAT
GAGCACACGTCTGAACTCTTTCCCTACACGACGCTCTTCCGATCTNNNAATATTGGCTCTCTACCAGGTC
GAGCACACGTCTGAACTCTTTCCCTACACGACGCTCTTCCGATCTNNNAATATTGGCTCTCTACCAGGTC
GAGCACACGTCTGAACTCTTTCCCTACACGACGCTCTTCCGATCTNNNTTTGTTTGGTTGGTTGGTTGG
GAGCACACGTCTGAACTCTTTCCCTACACGACGCTCTTCCGATCTNNNTTTGTTTGGTTGGTTGGTTGG
CACACGTCTGAACTCTTTCCCTACACGACGCTCTTCCGATCTNNNAGGCTGTCTCTTCTCTTCTCTCTCAT
CACACGTCTGAACTCTTTCCCTACACGACGCTCTTCCGATCTNNNAGGCTGTCTCTTCTCTTCTCTCTCAT
GAGCACACGTCTGAACTCTTTCCCTACACGACGCTCTTCCGATCTNNNGCTCATTATGCTAGGAGTCTC
GAGCACACGTCTGAACTCTTTCCCTACACGACGCTCTTCCGATCTNNNGCTCATTATGCTAGGAGTCTC

GAGCACACGTCTGAACTCTTTCCCTACACGACGCTCTTCCGATCTNNNTGCAGTGAGAGTTGGAAGATT

GAGCACACGTCTGAACTCTTTCCCTACACGACGCTCTTCCGATCTNNNTGCAGTGAGAGTTGGAAGATT

GAGCACACGTCTGAACTCTTTCCCTACACGACGCTCTTCCGATCTNNNAGAATCTCTTGAATCAGGGAG

GAGCACACGTCTGAACTCTTTCCCTACACGACGCTCTTCCGATCTNNNAGAATCTCTTGAATCAGGGAG

AGCACACGTCTGAACTCTTTCCCTACACGACGCTCTTCCGATCTNNNCTAGAGTACAGTGTGCGATCTC

AGCACACGTCTGAACTCTTTCCCTACACGACGCTCTTCCGATCTNNNCTAGAGTACAGTGTGCGATCTC

GAGCACACGTCTGAACTCTTTCCCTACACGACGCTCTTCCGATCTNNNTGTAGTCTCAGCTACTCGAGA

GAGCACACGTCTGAACTCTTTCCCTACACGACGCTCTTCCGATCTNNNTGTAGTCTCAGCTACTCGAGA

AGCACACGTCTGAACTCTTTCCCTACACGACGCTCTTCCGATCTNNNCCAATCTCAAAGGTTACAACGTC

AGCACACGTCTGAACTCTTTCCCTACACGACGCTCTTCCGATCTNNNCCAATCTCAAAGGTTACAACGTC

GCACACGTCTGAACTCTTTCCCTACACGACGCTCTTCCGATCTNNNGAGATGGCAGAAACGGAGAGAA

GCACACGTCTGAACTCTTTCCCTACACGACGCTCTTCCGATCTNNNGAGATGGCAGAAACGGAGAGAA

GAGCACACGTCTGAACTCTTTCCCTACACGACGCTCTTCCGATCTNNNAAAAATTAGCCGGTGTGATGG

GAGCACACGTCTGAACTCTTTCCCTACACGACGCTCTTCCGATCTNNNAAAAATTAGCCGGTGTGATGG

AGCACACGTCTGAACTCTTTCCCTACACGACGCTCTTCCGATCTNNNGCCTGTTTTCTAAAGAGCTTCC

AGCACACGTCTGAACTCTTTCCCTACACGACGCTCTTCCGATCTNNNGCCTGTTTTCTAAAGAGCTTCC

GAGCACACGTCTGAACTCTTTCCCTACACGACGCTCTTCCGATCTNNNAAGAGAGTCTTCCTTTGCGAC

GAGCACACGTCTGAACTCTTTCCCTACACGACGCTCTTCCGATCTNNNAAGAGAGTCTTCCTTTGCGAC

AGCACACGTCTGAACTCTTTCCCTACACGACGCTCTTCCGATCTNNNGGCATTAGCTAGAACTGCAATG

AGCACACGTCTGAACTCTTTCCCTACACGACGCTCTTCCGATCTNNNGGCATTAGCTAGAACTGCAATG

AAGAGCACACGTCTGAACTCTTTCCCTACACGACGCTCTTCCGATCTNNNCACTCCAGGCTCGGTGACAC

AAGAGCACACGTCTGAACTCTTTCCCTACACGACGCTCTTCCGATCTNNNCACTCCAGGCTCGGTGACAC

AGCACACGTCTGAACTCTTTCCCTACACGACGCTCTTCCGATCTNNNGTACACCAACATGCCTGGCTAAC

AGCACACGTCTGAACTCTTTCCCTACACGACGCTCTTCCGATCTNNNGTACACCAACATGCCTGGCTAAC

AGCACACGTCTGAACTCTTTCCCTACACGACGCTCTTCCGATCTNNNGAAATTGATTGAACCTGGGAGC

AGCACACGTCTGAACTCTTTCCCTACACGACGCTCTTCCGATCTNNNGAAATTGATTGAACCTGGGAGC

AGCACACGTCTGAACTCTTTCCCTACACGACGCTCTTCCGATCTNNNCTGCCTTTTTGCACTTGTTGTTGC

AGCACACGTCTGAACTCTTTCCCTACACGACGCTCTTCCGATCTNNNCTGCCTTTTTGCACTTGTTGTTGC

AGCACACGTCTGAACTCTTTCCCTACACGACGCTCTTCCGATCTNNNACCACGTCCAGCCAAAAAAAAAC

AGCACACGTCTGAACTCTTTCCCTACACGACGCTCTTCCGATCTNNNACCACGTCCAGCCAAAAAAAAAC

AGCACACGTCTGAACTCTTTCCCTACACGACGCTCTTCCGATCTNNNCCTCCTCAAGAAGCATCCCTTGCA

AGCACACGTCTGAACTCTTTCCCTACACGACGCTCTTCCGATCTNNNCCTCCTCAAGAAGCATCCCTTGCA

AGCACACGTCTGAACTCTTTCCCTACACGACGCTCTTCCGATCTNNNTCTGCCTACACCATCCAATCTTC

AGCACACGTCTGAACTCTTTCCCTACACGACGCTCTTCCGATCTNNNTCTGCCTACACCATCCAATCTTC

GCACACGTCTGAACTCTTTCCCTACACGACGCTCTTCCGATCTNNNGCCCTTGTTCAGCAATTACCTTTCA

GCACACGTCTGAACTCTTTCCCTACACGACGCTCTTCCGATCTNNNGCCCTTGTTCAGCAATTACCTTTCA

GAGCACACGTCTGAACTCTTTCCCTACACGACGCTCTTCCGATCTNNNTGTCAGAGAACATCTCACTGT

GAGCACACGTCTGAACTCTTTCCCTACACGACGCTCTTCCGATCTNNNTGTCAGAGAACATCTCACTGT

AGCACACGTCTGAACTCTTTCCCTACACGACGCTCTTCCGATCTNNNGACAAATATGCCATGGTGAAG

AGCACACGTCTGAACTCTTTCCCTACACGACGCTCTTCCGATCTNNNGACAAATATGCCATGGTGAAG

AGCACACGTCTGAACTCTTTCCCTACACGACGCTCTTCCGATCTNNNTTCCATGGTGTATATGTGCCAC

AGCACACGTCTGAACTCTTTCCCTACACGACGCTCTTCCGATCTNNNTTCCATGGTGTATATGTGCCAC

GCACACGTCTGAACTCTTTCCCTACACGACGCTCTTCCGATCTNNNATCAAAGTTCTGAGACTTAATCTC

GCACACGTCTGAACTCTTTCCCTACACGACGCTCTTCCGATCTNNNATCAAAGTTCTGAGACTTAATCTC
GCACACGTCTGAACTCTTTCCCTACACGACGCTCTTCCGATCTNNNGAATCTGCCATTGGTGCATACAG
GCACACGTCTGAACTCTTTCCCTACACGACGCTCTTCCGATCTNNNGAATCTGCCATTGGTGCATACAG
AGCACACGTCTGAACTCTTTCCCTACACGACGCTCTTCCGATCTNNNGCCTCCCAAAGTGCTTGAATTAC
AGCACACGTCTGAACTCTTTCCCTACACGACGCTCTTCCGATCTNNNGCCTCCCAAAGTGCTTGAATTAC
AAGAGCACACGTCTGAACTCTTTCCCTACACGACGCTCTTCCGATCTNNNCTGTAGTCCCAGCTACTCAG
AAGAGCACACGTCTGAACTCTTTCCCTACACGACGCTCTTCCGATCTNNNCTGTAGTCCCAGCTACTCAG
AGCACACGTCTGAACTCTTTCCCTACACGACGCTCTTCCGATCTNNNTATAGATGTGTGCCACCATGCCC
AGCACACGTCTGAACTCTTTCCCTACACGACGCTCTTCCGATCTNNNTATAGATGTGTGCCACCATGCCC
AGCACACGTCTGAACTCTTTCCCTACACGACGCTCTTCCGATCTNNNCTGGTCTCCAACTCTTGACCTCC
AGCACACGTCTGAACTCTTTCCCTACACGACGCTCTTCCGATCTNNNCTGGTCTCCAACTCTTGACCTCC
AGCACACGTCTGAACTCTTTCCCTACACGACGCTCTTCCGATCTNNNGTCCAAAACTGCTGCCTCTAAT
AGCACACGTCTGAACTCTTTCCCTACACGACGCTCTTCCGATCTNNNGTCCAAAACTGCTGCCTCTAAT
ACACGTCTGAACTCTTTCCCTACACGACGCTCTTCCGATCTNNNGGCTCCTCACTTCCCAGTAGGCATCG
ACACGTCTGAACTCTTTCCCTACACGACGCTCTTCCGATCTNNNGGCTCCTCACTTCCCAGTAGGCATCG
GAGCACACGTCTGAACTCTTTCCCTACACGACGCTCTTCCGATCTNNNCTTTCTGTTCTTCGTTCATTCC
GAGCACACGTCTGAACTCTTTCCCTACACGACGCTCTTCCGATCTNNNCTTTCTGTTCTTCGTTCATTCC
GAGCACACGTCTGAACTCTTTCCCTACACGACGCTCTTCCGATCTNNNAGTTTCACTATGTTGGCCAGG
GAGCACACGTCTGAACTCTTTCCCTACACGACGCTCTTCCGATCTNNNAGTTTCACTATGTTGGCCAGG
AGAGCACACGTCTGAACTCTTTCCCTACACGACGCTCTTCCGATCTNNNCTCATACTCCCTCCCTCCCTTC
AGAGCACACGTCTGAACTCTTTCCCTACACGACGCTCTTCCGATCTNNNCTCATACTCCCTCCCTCCCTTC
AGAGCACACGTCTGAACTCTTTCCCTACACGACGCTCTTCCGATCTNNNCACAAAGATGTCCATGTAGC
AGAGCACACGTCTGAACTCTTTCCCTACACGACGCTCTTCCGATCTNNNCACAAAGATGTCCATGTAGC
GAGCACACGTCTGAACTCTTTCCCTACACGACGCTCTTCCGATCTNNNTCTCTTGAACCCTGGAGTTTGA
GAGCACACGTCTGAACTCTTTCCCTACACGACGCTCTTCCGATCTNNNTCTCTTGAACCCTGGAGTTTGA
CACACGTCTGAACTCTTTCCCTACACGACGCTCTTCCGATCTNNNAGTACCTCTCTCTTTCCCCTCTCCAT
CACACGTCTGAACTCTTTCCCTACACGACGCTCTTCCGATCTNNNAGTACCTCTCTCTTTCCCCTCTCCAT
AGCACACGTCTGAACTCTTTCCCTACACGACGCTCTTCCGATCTNNNAATCATAGCTCCCTGTAGCCTT
AGCACACGTCTGAACTCTTTCCCTACACGACGCTCTTCCGATCTNNNAATCATAGCTCCCTGTAGCCTT
GAGCACACGTCTGAACTCTTTCCCTACACGACGCTCTTCCGATCTNNNAAGTACAAAAAAATTAGCCGG
GAGCACACGTCTGAACTCTTTCCCTACACGACGCTCTTCCGATCTNNNAAGTACAAAAAAATTAGCCGG
GCACACGTCTGAACTCTTTCCCTACACGACGCTCTTCCGATCTNNNTCGTCATCTTCACAGTTGGTCTTC
GCACACGTCTGAACTCTTTCCCTACACGACGCTCTTCCGATCTNNNTCGTCATCTTCACAGTTGGTCTTC
GCACACGTCTGAACTCTTTCCCTACACGACGCTCTTCCGATCTNNNCTTACCCACACCCCATCCAATTACA
GCACACGTCTGAACTCTTTCCCTACACGACGCTCTTCCGATCTNNNCTTACCCACACCCCATCCAATTACA
GAGCACACGTCTGAACTCTTTCCCTACACGACGCTCTTCCGATCTNNNGGCAACAGAGTGAGACTCCA
GAGCACACGTCTGAACTCTTTCCCTACACGACGCTCTTCCGATCTNNNGGCAACAGAGTGAGACTCCA
GAGCACACGTCTGAACTCTTTCCCTACACGACGCTCTTCCGATCTNNNCGTGGAGATAAACTGGGGAT
GAGCACACGTCTGAACTCTTTCCCTACACGACGCTCTTCCGATCTNNNCGTGGAGATAAACTGGGGAT
AGCACACGTCTGAACTCTTTCCCTACACGACGCTCTTCCGATCTNNNCCCTTTCATCAACCTGGGATCA
AGCACACGTCTGAACTCTTTCCCTACACGACGCTCTTCCGATCTNNNCCCTTTCATCAACCTGGGATCA
GAGCACACGTCTGAACTCTTTCCCTACACGACGCTCTTCCGATCTNNNCTGTAATCCTGGCTACTTGGG
GAGCACACGTCTGAACTCTTTCCCTACACGACGCTCTTCCGATCTNNNCTGTAATCCTGGCTACTTGGG

AGCACACGTCTGAACTCTTTCCCTACACGACGCTCTTCCGATCTNNNCACCTCCACACCTGTATAAATGC
AGCACACGTCTGAACTCTTTCCCTACACGACGCTCTTCCGATCTNNNCACCTCCACACCTGTATAAATGC
AGCACACGTCTGAACTCTTTCCCTACACGACGCTCTTCCGATCTNNNCAGAAGAATTGCTTGAACCTGC
AGCACACGTCTGAACTCTTTCCCTACACGACGCTCTTCCGATCTNNNCAGAAGAATTGCTTGAACCTGC
AGCACACGTCTGAACTCTTTCCCTACACGACGCTCTTCCGATCTNNNTTGTTGTGCAGAAGCTTTTAGC
AGCACACGTCTGAACTCTTTCCCTACACGACGCTCTTCCGATCTNNNTTGTTGTGCAGAAGCTTTTAGC
GCACACGTCTGAACTCTTTCCCTACACGACGCTCTTCCGATCTNNNTGTTGAAGAAATTCTTGAGAACC
GCACACGTCTGAACTCTTTCCCTACACGACGCTCTTCCGATCTNNNTGTTGAAGAAATTCTTGAGAACC
AGCACACGTCTGAACTCTTTCCCTACACGACGCTCTTCCGATCTNNNAGTTTGTGAGGGGTGGTGATT
AGCACACGTCTGAACTCTTTCCCTACACGACGCTCTTCCGATCTNNNAGTTTGTGAGGGGTGGTGATT
AGCACACGTCTGAACTCTTTCCCTACACGACGCTCTTCCGATCTNNNTGCTGGATCAAATGGTGGTTCTA
AGCACACGTCTGAACTCTTTCCCTACACGACGCTCTTCCGATCTNNNTGCTGGATCAAATGGTGGTTCTA
GAGCACACGTCTGAACTCTTTCCCTACACGACGCTCTTCCGATCTNNNACTCTCATGAATGGAATCGGT
GAGCACACGTCTGAACTCTTTCCCTACACGACGCTCTTCCGATCTNNNACTCTCATGAATGGAATCGGT
GAGCACACGTCTGAACTCTTTCCCTACACGACGCTCTTCCGATCTNNNCAGAATTGCTTGAGCCCAGAT
GAGCACACGTCTGAACTCTTTCCCTACACGACGCTCTTCCGATCTNNNCAGAATTGCTTGAGCCCAGAT
GAGCACACGTCTGAACTCTTTCCCTACACGACGCTCTTCCGATCTNNNGAGGCAGGAGAATCGCTCAAA
GAGCACACGTCTGAACTCTTTCCCTACACGACGCTCTTCCGATCTNNNGAGGCAGGAGAATCGCTCAAA
GAGCACACGTCTGAACTCTTTCCCTACACGACGCTCTTCCGATCTNNNTGCCTGTAATACCAGCTACTTC
GAGCACACGTCTGAACTCTTTCCCTACACGACGCTCTTCCGATCTNNNTGCCTGTAATACCAGCTACTTC
AGCACACGTCTGAACTCTTTCCCTACACGACGCTCTTCCGATCTNNNAATACACCTTATCACACAATGAC
AGCACACGTCTGAACTCTTTCCCTACACGACGCTCTTCCGATCTNNNAATACACCTTATCACACAATGAC
AGCACACGTCTGAACTCTTTCCCTACACGACGCTCTTCCGATCTNNNAGGCAAGTTCCTTTCAGCTATG
AGCACACGTCTGAACTCTTTCCCTACACGACGCTCTTCCGATCTNNNAGGCAAGTTCCTTTCAGCTATG
AGCACACGTCTGAACTCTTTCCCTACACGACGCTCTTCCGATCTNNNTCCCATTGGTAGGACCTAGAAG
AGCACACGTCTGAACTCTTTCCCTACACGACGCTCTTCCGATCTNNNTCCCATTGGTAGGACCTAGAAG
GAGCACACGTCTGAACTCTTTCCCTACACGACGCTCTTCCGATCTNNNCCTGGCTCTACTCCCGATTTCT
GAGCACACGTCTGAACTCTTTCCCTACACGACGCTCTTCCGATCTNNNCCTGGCTCTACTCCCGATTTCT
GAGCACACGTCTGAACTCTTTCCCTACACGACGCTCTTCCGATCTNNNATGGTGCCTTCATGATGGGAT
GAGCACACGTCTGAACTCTTTCCCTACACGACGCTCTTCCGATCTNNNATGGTGCCTTCATGATGGGAT
AGCACACGTCTGAACTCTTTCCCTACACGACGCTCTTCCGATCTNNNCTGAGGCAGAAGAATGGTGTC
AGCACACGTCTGAACTCTTTCCCTACACGACGCTCTTCCGATCTNNNCTGAGGCAGAAGAATGGTGTC
AGCACACGTCTGAACTCTTTCCCTACACGACGCTCTTCCGATCTNNNCACCAACCTTCATTTTCTGGTCC
AGCACACGTCTGAACTCTTTCCCTACACGACGCTCTTCCGATCTNNNCACCAACCTTCATTTTCTGGTCC
GAGCACACGTCTGAACTCTTTCCCTACACGACGCTCTTCCGATCTNNNAAAAAAATTAGCCGGGCGTA
GAGCACACGTCTGAACTCTTTCCCTACACGACGCTCTTCCGATCTNNNAAAAAAATTAGCCGGGCGTA
AGCACACGTCTGAACTCTTTCCCTACACGACGCTCTTCCGATCTNNNTGTGAGCCCAAAATGGAGAAA
AGCACACGTCTGAACTCTTTCCCTACACGACGCTCTTCCGATCTNNNTGTGAGCCCAAAATGGAGAAA
GCACACGTCTGAACTCTTTCCCTACACGACGCTCTTCCGATCTNNNCACAGAGAGGTTACATTGGTGG
GCACACGTCTGAACTCTTTCCCTACACGACGCTCTTCCGATCTNNNCACAGAGAGGTTACATTGGTGG
AGCACACGTCTGAACTCTTTCCCTACACGACGCTCTTCCGATCTNNNTGGTGAAGGTAGATCCAAACGT
AGCACACGTCTGAACTCTTTCCCTACACGACGCTCTTCCGATCTNNNTGGTGAAGGTAGATCCAAACGT
GCACACGTCTGAACTCTTTCCCTACACGACGCTCTTCCGATCTNNNTCCTGTTGAAGACTTGGTGGAAA

GCACACGTCTGAACTCTTTCCCTACACGACGCTCTTCCGATCTNNNTCCTGTTGAAGACTTGGTGGAAA
GAGCACACGTCTGAACTCTTTCCCTACACGACGCTCTTCCGATCTNNNATTGGTGCCCCCTTTTTTATTG
GAGCACACGTCTGAACTCTTTCCCTACACGACGCTCTTCCGATCTNNNATTGGTGCCCCCTTTTTTATTG
GCACACGTCTGAACTCTTTCCCTACACGACGCTCTTCCGATCTNNNACTTGAGGATTGCAAACTGGTTG
GCACACGTCTGAACTCTTTCCCTACACGACGCTCTTCCGATCTNNNACTTGAGGATTGCAAACTGGTTG
AGAGCACACGTCTGAACTCTTTCCCTACACGACGCTCTTCCGATCTNNNGATTCAGGGCAATGGATGT
AGAGCACACGTCTGAACTCTTTCCCTACACGACGCTCTTCCGATCTNNNGATTCAGGGCAATGGATGT
GAGCACACGTCTGAACTCTTTCCCTACACGACGCTCTTCCGATCTNNNGCCAACTGTACCTTGAAGTCA
GAGCACACGTCTGAACTCTTTCCCTACACGACGCTCTTCCGATCTNNNGCCAACTGTACCTTGAAGTCA
GAGCACACGTCTGAACTCTTTCCCTACACGACGCTCTTCCGATCTNNNCATCCCACGTGACTATCCCTA
GAGCACACGTCTGAACTCTTTCCCTACACGACGCTCTTCCGATCTNNNCATCCCACGTGACTATCCCTA
GCACACGTCTGAACTCTTTCCCTACACGACGCTCTTCCGATCTNNNAGCATTCACACAGAAAACAGAGC
GCACACGTCTGAACTCTTTCCCTACACGACGCTCTTCCGATCTNNNAGCATTCACACAGAAAACAGAGC
GAGCACACGTCTGAACTCTTTCCCTACACGACGCTCTTCCGATCTNNNTGTTCCAGCTAAAGATGCTTC
GAGCACACGTCTGAACTCTTTCCCTACACGACGCTCTTCCGATCTNNNTGTTCCAGCTAAAGATGCTTC
AGCACACGTCTGAACTCTTTCCCTACACGACGCTCTTCCGATCTNNNAGTCCTAGCTACTCTGGAGACT
AGCACACGTCTGAACTCTTTCCCTACACGACGCTCTTCCGATCTNNNAGTCCTAGCTACTCTGGAGACT
GAGCACACGTCTGAACTCTTTCCCTACACGACGCTCTTCCGATCTNNNCTCATGCCTATAATCCCTGCAC
GAGCACACGTCTGAACTCTTTCCCTACACGACGCTCTTCCGATCTNNNCTCATGCCTATAATCCCTGCAC
GAGCACACGTCTGAACTCTTTCCCTACACGACGCTCTTCCGATCTNNNCAAGATCATACCTCTGCACTC
GAGCACACGTCTGAACTCTTTCCCTACACGACGCTCTTCCGATCTNNNCAAGATCATACCTCTGCACTC
AGCACACGTCTGAACTCTTTCCCTACACGACGCTCTTCCGATCTNNNTTCAATCCTTGCTGCTGTGATTG
AGCACACGTCTGAACTCTTTCCCTACACGACGCTCTTCCGATCTNNNTTCAATCCTTGCTGCTGTGATTG
AGAGCACACGTCTGAACTCTTTCCCTACACGACGCTCTTCCGATCTNNNTTACTTGGTGCAAAAGTCCA
AGAGCACACGTCTGAACTCTTTCCCTACACGACGCTCTTCCGATCTNNNTTACTTGGTGCAAAAGTCCA
GCACACGTCTGAACTCTTTCCCTACACGACGCTCTTCCGATCTNNNGGAAGTGACCCTTTCCATAAGGA
GCACACGTCTGAACTCTTTCCCTACACGACGCTCTTCCGATCTNNNGGAAGTGACCCTTTCCATAAGGA
AGCACACGTCTGAACTCTTTCCCTACACGACGCTCTTCCGATCTNNNACACACATGCACATACAAGCAA
AGCACACGTCTGAACTCTTTCCCTACACGACGCTCTTCCGATCTNNNACACACATGCACATACAAGCAA
AGCACACGTCTGAACTCTTTCCCTACACGACGCTCTTCCGATCTNNNCAACTGGAAACATGTGATCTG
AGCACACGTCTGAACTCTTTCCCTACACGACGCTCTTCCGATCTNNNCAACTGGAAACATGTGATCTG
ACACGTCTGAACTCTTTCCCTACACGACGCTCTTCCGATCTNNNGAAGATACTTACTGTAGTTCTCACA
ACACGTCTGAACTCTTTCCCTACACGACGCTCTTCCGATCTNNNGAAGATACTTACTGTAGTTCTCACA
AGAGCACACGTCTGAACTCTTTCCCTACACGACGCTCTTCCGATCTNNNAGGAAAGGCAGAATATAGG
AGAGCACACGTCTGAACTCTTTCCCTACACGACGCTCTTCCGATCTNNNAGGAAAGGCAGAATATAGG
AGCACACGTCTGAACTCTTTCCCTACACGACGCTCTTCCGATCTNNNTCCATTCACTAGCTTCCCCTCAA
AGCACACGTCTGAACTCTTTCCCTACACGACGCTCTTCCGATCTNNNTCCATTCACTAGCTTCCCCTCAA
GAGCACACGTCTGAACTCTTTCCCTACACGACGCTCTTCCGATCTNNNCCAACATGGTGAAACTCTGTCA
GAGCACACGTCTGAACTCTTTCCCTACACGACGCTCTTCCGATCTNNNCCAACATGGTGAAACTCTGTCA
AGCACACGTCTGAACTCTTTCCCTACACGACGCTCTTCCGATCTNNNGTAGTCCCAGCTACTCGGAGA
AGCACACGTCTGAACTCTTTCCCTACACGACGCTCTTCCGATCTNNNGTAGTCCCAGCTACTCGGAGA
GAGCACACGTCTGAACTCTTTCCCTACACGACGCTCTTCCGATCTNNNACAGTGAAACCCGGTCTCTAC
GAGCACACGTCTGAACTCTTTCCCTACACGACGCTCTTCCGATCTNNNACAGTGAAACCCGGTCTCTAC

AGCACACGTCTGAACTCTTTCCCTACACGACGCTCTTCCGATCTNNNGCATTCATCATATTTTCTATTT
AGCACACGTCTGAACTCTTTCCCTACACGACGCTCTTCCGATCTNNNGCATTCATCATATTTTCTATTT
GAGCACACGTCTGAACTCTTTCCCTACACGACGCTCTTCCGATCTNNNCCCAGCCAATCCTTTCTTCTCT
GAGCACACGTCTGAACTCTTTCCCTACACGACGCTCTTCCGATCTNNNCCCAGCCAATCCTTTCTTCTCT
GCACACGTCTGAACTCTTTCCCTACACGACGCTCTTCCGATCTNNNGGTGGTTCATGCCTATAATCCC
GCACACGTCTGAACTCTTTCCCTACACGACGCTCTTCCGATCTNNNGGTGGTTCATGCCTATAATCCC
AGCACACGTCTGAACTCTTTCCCTACACGACGCTCTTCCGATCTNNNTACTTAGGAGGCTGAGGTAG
AGCACACGTCTGAACTCTTTCCCTACACGACGCTCTTCCGATCTNNNTACTTAGGAGGCTGAGGTAG
GAGCACACGTCTGAACTCTTTCCCTACACGACGCTCTTCCGATCTNNNTTGCCTTTCCTGTCTTCATCTC
GAGCACACGTCTGAACTCTTTCCCTACACGACGCTCTTCCGATCTNNNTTGCCTTTCCTGTCTTCATCTC
GAGCACACGTCTGAACTCTTTCCCTACACGACGCTCTTCCGATCTNNNTCATCTAGGAGTGGGGAGA
GAGCACACGTCTGAACTCTTTCCCTACACGACGCTCTTCCGATCTNNNTCATCTAGGAGTGGGGAGA
AGCACACGTCTGAACTCTTTCCCTACACGACGCTCTTCCGATCTNNNCTGGTAAAGTGTATGGATGG
AGCACACGTCTGAACTCTTTCCCTACACGACGCTCTTCCGATCTNNNCTGGTAAAGTGTATGGATGG
GCACACGTCTGAACTCTTTCCCTACACGACGCTCTTCCGATCTNNNTATCCTGTGCATGGCTCAGTCA
GCACACGTCTGAACTCTTTCCCTACACGACGCTCTTCCGATCTNNNTATCCTGTGCATGGCTCAGTCA
AGCACACGTCTGAACTCTTTCCCTACACGACGCTCTTCCGATCTNNNAAAGTACTGGGATTACAGGC
AGCACACGTCTGAACTCTTTCCCTACACGACGCTCTTCCGATCTNNNAAAGTACTGGGATTACAGGC
AGCACACGTCTGAACTCTTTCCCTACACGACGCTCTTCCGATCTNNNTTCAGATTTACTTCCGACAGG
AGCACACGTCTGAACTCTTTCCCTACACGACGCTCTTCCGATCTNNNTTCAGATTTACTTCCGACAGG
AGCACACGTCTGAACTCTTTCCCTACACGACGCTCTTCCGATCTNNNCCATATGATCCAGCAATCCCA
AGCACACGTCTGAACTCTTTCCCTACACGACGCTCTTCCGATCTNNNCCATATGATCCAGCAATCCCA
AGCACACGTCTGAACTCTTTCCCTACACGACGCTCTTCCGATCTNNNCTCCCTGTGTGAAGCATCAAA
AGCACACGTCTGAACTCTTTCCCTACACGACGCTCTTCCGATCTNNNCTCCCTGTGTGAAGCATCAAA
GCACACGTCTGAACTCTTTCCCTACACGACGCTCTTCCGATCTNNNTTCTACTCCTGAGAGAGGGTG
GCACACGTCTGAACTCTTTCCCTACACGACGCTCTTCCGATCTNNNTTCTACTCCTGAGAGAGGGTG
GCACACGTCTGAACTCTTTCCCTACACGACGCTCTTCCGATCTNNNTCCATCTCAGTCTCTTCCTTCCTC
GCACACGTCTGAACTCTTTCCCTACACGACGCTCTTCCGATCTNNNTCCATCTCAGTCTCTTCCTTCCTC
AGCACACGTCTGAACTCTTTCCCTACACGACGCTCTTCCGATCTNNNCCCAGATTCCCTTCTACAGTTC
AGCACACGTCTGAACTCTTTCCCTACACGACGCTCTTCCGATCTNNNCCCAGATTCCCTTCTACAGTTC
AGCACACGTCTGAACTCTTTCCCTACACGACGCTCTTCCGATCTNNNCACAGAAAGCTCTCCAGGTCT
AGCACACGTCTGAACTCTTTCCCTACACGACGCTCTTCCGATCTNNNCACAGAAAGCTCTCCAGGTCT
GCACACGTCTGAACTCTTTCCCTACACGACGCTCTTCCGATCTNNNTCTTCTTTCAGCTGACTCTGTCC
GCACACGTCTGAACTCTTTCCCTACACGACGCTCTTCCGATCTNNNTCTTCTTTCAGCTGACTCTGTCC
GCACACGTCTGAACTCTTTCCCTACACGACGCTCTTCCGATCTNNNCACAGTGTATTTGCATGCAGGA
GCACACGTCTGAACTCTTTCCCTACACGACGCTCTTCCGATCTNNNCACAGTGTATTTGCATGCAGGA
AGAGCACACGTCTGAACTCTTTCCCTACACGACGCTCTTCCGATCTNNNCTGAGGCAGGAAAATTGCT
AGAGCACACGTCTGAACTCTTTCCCTACACGACGCTCTTCCGATCTNNNCTGAGGCAGGAAAATTGCT
GCACACGTCTGAACTCTTTCCCTACACGACGCTCTTCCGATCTNNNGTTCCTCCAACTGTGTCCCATTAC
GCACACGTCTGAACTCTTTCCCTACACGACGCTCTTCCGATCTNNNGTTCCTCCAACTGTGTCCCATTAC
AGCACACGTCTGAACTCTTTCCCTACACGACGCTCTTCCGATCTNNNTGCTGGTAAGGATGTGGAGA
AGCACACGTCTGAACTCTTTCCCTACACGACGCTCTTCCGATCTNNNTGCTGGTAAGGATGTGGAGA
CACACGTCTGAACTCTTTCCCTACACGACGCTCTTCCGATCTNNNTGCATTCTTATACATAGCTAATAT

CACACGTCTGAACTCTTTCCCTACACGACGCTCTTCCGATCTNNNTGCATTCTTATACATAGCTAATATG
CACACGTCTGAACTCTTTCCCTACACGACGCTCTTCCGATCTNNNGAACTCCCAACCTCAGGTGATCCA
CACACGTCTGAACTCTTTCCCTACACGACGCTCTTCCGATCTNNNGAACTCCCAACCTCAGGTGATCCA
AGCACACGTCTGAACTCTTTCCCTACACGACGCTCTTCCGATCTNNNCTGAGGCAGGAGAATAGCTTG
AGCACACGTCTGAACTCTTTCCCTACACGACGCTCTTCCGATCTNNNCTGAGGCAGGAGAATAGCTTG
AGCACACGTCTGAACTCTTTCCCTACACGACGCTCTTCCGATCTNNNACAGAGAAACAGAACCAATG
AGCACACGTCTGAACTCTTTCCCTACACGACGCTCTTCCGATCTNNNACAGAGAAACAGAACCAATG
GAGCACACGTCTGAACTCTTTCCCTACACGACGCTCTTCCGATCTNNNTGTGTAGTGGGACCAATGAG
GAGCACACGTCTGAACTCTTTCCCTACACGACGCTCTTCCGATCTNNNTGTGTAGTGGGACCAATGAG
GCACACGTCTGAACTCTTTCCCTACACGACGCTCTTCCGATCTNNNCTGTAGTCCTAGCTACTCGGGA
GCACACGTCTGAACTCTTTCCCTACACGACGCTCTTCCGATCTNNNCTGTAGTCCTAGCTACTCGGGA
GCACACGTCTGAACTCTTTCCCTACACGACGCTCTTCCGATCTNNNTGGATTCAGCTAGCTCAACTTGT
GCACACGTCTGAACTCTTTCCCTACACGACGCTCTTCCGATCTNNNTGGATTCAGCTAGCTCAACTTGT
AGCACACGTCTGAACTCTTTCCCTACACGACGCTCTTCCGATCTNNNCCAAATGTTTTTATGATCTCTG
AGCACACGTCTGAACTCTTTCCCTACACGACGCTCTTCCGATCTNNNCCAAATGTTTTTATGATCTCTG
GAGCACACGTCTGAACTCTTTCCCTACACGACGCTCTTCCGATCTNNNATTTTGGGCAGCTTTCAGTG
GAGCACACGTCTGAACTCTTTCCCTACACGACGCTCTTCCGATCTNNNATTTTGGGCAGCTTTCAGTG
GCACACGTCTGAACTCTTTCCCTACACGACGCTCTTCCGATCTNNNCGGGGGTCTTATGAACTACAGT
GCACACGTCTGAACTCTTTCCCTACACGACGCTCTTCCGATCTNNNCGGGGGTCTTATGAACTACAGT
GCACACGTCTGAACTCTTTCCCTACACGACGCTCTTCCGATCTNNNGCACAATGTCCAATGTCAACTG
GCACACGTCTGAACTCTTTCCCTACACGACGCTCTTCCGATCTNNNGCACAATGTCCAATGTCAACTG
AGCACACGTCTGAACTCTTTCCCTACACGACGCTCTTCCGATCTNNNGAACAAGTAACAGCTGGTCAC
AGCACACGTCTGAACTCTTTCCCTACACGACGCTCTTCCGATCTNNNGAACAAGTAACAGCTGGTCAC
GAGCACACGTCTGAACTCTTTCCCTACACGACGCTCTTCCGATCTNNNGTAGTCTCAGCTACTCAGGA
GAGCACACGTCTGAACTCTTTCCCTACACGACGCTCTTCCGATCTNNNGTAGTCTCAGCTACTCAGGA
GAGCACACGTCTGAACTCTTTCCCTACACGACGCTCTTCCGATCTNNNACACACTGGTAGTCTCTGTA
GAGCACACGTCTGAACTCTTTCCCTACACGACGCTCTTCCGATCTNNNACACACTGGTAGTCTCTGTA
AGAGCACACGTCTGAACTCTTTCCCTACACGACGCTCTTCCGATCTNNNAGAAACACACGGAGGACAT
AGAGCACACGTCTGAACTCTTTCCCTACACGACGCTCTTCCGATCTNNNAGAAACACACGGAGGACAT
AAGAGCACACGTCTGAACTCTTTCCCTACACGACGCTCTTCCGATCTNNNGCCTAGGGGAAGAGAGT
AAGAGCACACGTCTGAACTCTTTCCCTACACGACGCTCTTCCGATCTNNNGCCTAGGGGAAGAGAGT
AGCACACGTCTGAACTCTTTCCCTACACGACGCTCTTCCGATCTNNNGGAGACAGAGTGAGACTCCAT
AGCACACGTCTGAACTCTTTCCCTACACGACGCTCTTCCGATCTNNNGGAGACAGAGTGAGACTCCAT
AGCACACGTCTGAACTCTTTCCCTACACGACGCTCTTCCGATCTNNNACCAGCCTAAGCAACATGAGA
AGCACACGTCTGAACTCTTTCCCTACACGACGCTCTTCCGATCTNNNACCAGCCTAAGCAACATGAGA
GCACACGTCTGAACTCTTTCCCTACACGACGCTCTTCCGATCTNNNGTTATGTGGCCATCTGTTATCAT
GCACACGTCTGAACTCTTTCCCTACACGACGCTCTTCCGATCTNNNGTTATGTGGCCATCTGTTATCAT
GAGCACACGTCTGAACTCTTTCCCTACACGACGCTCTTCCGATCTNNNTGTTAGTGATTATTCAGCAA
GAGCACACGTCTGAACTCTTTCCCTACACGACGCTCTTCCGATCTNNNTGTTAGTGATTATTCAGCAA
GAGCACACGTCTGAACTCTTTCCCTACACGACGCTCTTCCGATCTNNNGGATTTCCCGTCCTTCCGAAT
GAGCACACGTCTGAACTCTTTCCCTACACGACGCTCTTCCGATCTNNNGGATTTCCCGTCCTTCCGAAT
GCACACGTCTGAACTCTTTCCCTACACGACGCTCTTCCGATCTNNNAGGCTCATGAGATCTGATGGAT
GCACACGTCTGAACTCTTTCCCTACACGACGCTCTTCCGATCTNNNAGGCTCATGAGATCTGATGGAT

AGCACACGTCTGAACTCTTTCCCTACACGACGCTCTTCCGATCTNNNGAGGCTGAGGCAGGAGAATG

AGCACACGTCTGAACTCTTTCCCTACACGACGCTCTTCCGATCTNNNGAGGCTGAGGCAGGAGAATG

ACACGTCTGAACTCTTTCCCTACACGACGCTCTTCCGATCTNNNGTGGCTTTCACTGTAAATTTGCATC

ACACGTCTGAACTCTTTCCCTACACGACGCTCTTCCGATCTNNNGTGGCTTTCACTGTAAATTTGCATC

AGCACACGTCTGAACTCTTTCCCTACACGACGCTCTTCCGATCTNNNCCCAGGTATTGAGTCCAGAAT

AGCACACGTCTGAACTCTTTCCCTACACGACGCTCTTCCGATCTNNNCCCAGGTATTGAGTCCAGAAT

AGCACACGTCTGAACTCTTTCCCTACACGACGCTCTTCCGATCTNNNGGAGGTTGCAGTGAGCTTAG

AGCACACGTCTGAACTCTTTCCCTACACGACGCTCTTCCGATCTNNNGGAGGTTGCAGTGAGCTTAG

AGCACACGTCTGAACTCTTTCCCTACACGACGCTCTTCCGATCTNNNAGGAGAATGGCTTGGCTTGAA

AGCACACGTCTGAACTCTTTCCCTACACGACGCTCTTCCGATCTNNNAGGAGAATGGCTTGGCTTGAA

AGCACACGTCTGAACTCTTTCCCTACACGACGCTCTTCCGATCTNNNCTGAGGCATGAGAATCCTTCG

AGCACACGTCTGAACTCTTTCCCTACACGACGCTCTTCCGATCTNNNCTGAGGCATGAGAATCCTTCG

GCACACGTCTGAACTCTTTCCCTACACGACGCTCTTCCGATCTNNNGAATTTCTGAATGATGAAAACA

GCACACGTCTGAACTCTTTCCCTACACGACGCTCTTCCGATCTNNNGAATTTCTGAATGATGAAAACA

GAGCACACGTCTGAACTCTTTCCCTACACGACGCTCTTCCGATCTNNNGAGAACAGCGGATGGATAC

GAGCACACGTCTGAACTCTTTCCCTACACGACGCTCTTCCGATCTNNNGAGAACAGCGGATGGATAC

AGCACACGTCTGAACTCTTTCCCTACACGACGCTCTTCCGATCTNNNGAGGCTGAGGCAGGAGAATG

AGCACACGTCTGAACTCTTTCCCTACACGACGCTCTTCCGATCTNNNGAGGCTGAGGCAGGAGAATG

AGCACACGTCTGAACTCTTTCCCTACACGACGCTCTTCCGATCTNNNCAAAAAATTAGCCGGGCGTAG

AGCACACGTCTGAACTCTTTCCCTACACGACGCTCTTCCGATCTNNNCAAAAAATTAGCCGGGCGTAG

AGCACACGTCTGAACTCTTTCCCTACACGACGCTCTTCCGATCTNNNTGTTTGTCTCCTCTTCATTGTCG

AGCACACGTCTGAACTCTTTCCCTACACGACGCTCTTCCGATCTNNNTGTTTGTCTCCTCTTCATTGTCG

AGCACACGTCTGAACTCTTTCCCTACACGACGCTCTTCCGATCTNNNCCACCACACCTGGCTAATTTTT

AGCACACGTCTGAACTCTTTCCCTACACGACGCTCTTCCGATCTNNNCCACCACACCTGGCTAATTTTT

GAGCACACGTCTGAACTCTTTCCCTACACGACGCTCTTCCGATCTNNNTCTCTTGTGACTCAAAACACT

GAGCACACGTCTGAACTCTTTCCCTACACGACGCTCTTCCGATCTNNNTCTCTTGTGACTCAAAACACT

GAGCACACGTCTGAACTCTTTCCCTACACGACGCTCTTCCGATCTNNNTGCTTTCAATAACGTGGGCTT

GAGCACACGTCTGAACTCTTTCCCTACACGACGCTCTTCCGATCTNNNTGCTTTCAATAACGTGGGCTT

GAGCACACGTCTGAACTCTTTCCCTACACGACGCTCTTCCGATCTNNNAAAAAGCTGGGATTGTGTGT

GAGCACACGTCTGAACTCTTTCCCTACACGACGCTCTTCCGATCTNNNAAAAAGCTGGGATTGTGTGT

AGCACACGTCTGAACTCTTTCCCTACACGACGCTCTTCCGATCTNNNCTTCTACGACTGTGTCATAGCA

AGCACACGTCTGAACTCTTTCCCTACACGACGCTCTTCCGATCTNNNCTTCTACGACTGTGTCATAGCA

GCACACGTCTGAACTCTTTCCCTACACGACGCTCTTCCGATCTNNNAAACAGGCCAATTTTTTGTGGA

GCACACGTCTGAACTCTTTCCCTACACGACGCTCTTCCGATCTNNNAAACAGGCCAATTTTTTGTGGA

AGCACACGTCTGAACTCTTTCCCTACACGACGCTCTTCCGATCTNNNACAGTGTATATGTGTGTGTGT

AGCACACGTCTGAACTCTTTCCCTACACGACGCTCTTCCGATCTNNNACAGTGTATATGTGTGTGTGT

GCACACGTCTGAACTCTTTCCCTACACGACGCTCTTCCGATCTNNNTTTGCTCCAAGCTATTAAAATAG

GCACACGTCTGAACTCTTTCCCTACACGACGCTCTTCCGATCTNNNTTTGCTCCAAGCTATTAAAATAG

GAGCACACGTCTGAACTCTTTCCCTACACGACGCTCTTCCGATCTNNNGAGCCACATACGTAGTCTGGA

GAGCACACGTCTGAACTCTTTCCCTACACGACGCTCTTCCGATCTNNNGAGCCACATACGTAGTCTGGA

AGCACACGTCTGAACTCTTTCCCTACACGACGCTCTTCCGATCTNNNAGAATCACTTGAACCCTGGAG

AGCACACGTCTGAACTCTTTCCCTACACGACGCTCTTCCGATCTNNNAGAATCACTTGAACCCTGGAG

AGCACACGTCTGAACTCTTTCCCTACACGACGCTCTTCCGATCTNNNTCCTGAACCCTAGTGATTCTCC

AGCACACGTCTGAACTCTTTCCCTACACGACGCTCTTCCGATCTNNNTCCTGAACCCTAGTGATTCTCCT
AGCACACGTCTGAACTCTTTCCCTACACGACGCTCTTCCGATCTNNNGTGGACATTTGGGAGGGTATCT
AGCACACGTCTGAACTCTTTCCCTACACGACGCTCTTCCGATCTNNNGTGGACATTTGGGAGGGTATCT
GAGCACACGTCTGAACTCTTTCCCTACACGACGCTCTTCCGATCTNNNTATTTGAGAGGCTGAGGTGC
GAGCACACGTCTGAACTCTTTCCCTACACGACGCTCTTCCGATCTNNNTATTTGAGAGGCTGAGGTGC
AGCACACGTCTGAACTCTTTCCCTACACGACGCTCTTCCGATCTNNNAACAAGGCAAGGATGTTTGTCT
AGCACACGTCTGAACTCTTTCCCTACACGACGCTCTTCCGATCTNNNAACAAGGCAAGGATGTTTGTCT
AGAGCACACGTCTGAACTCTTTCCCTACACGACGCTCTTCCGATCTNNNGGAGAATCACGTGAACCTGC
AGAGCACACGTCTGAACTCTTTCCCTACACGACGCTCTTCCGATCTNNNGGAGAATCACGTGAACCTGC
AGAGCACACGTCTGAACTCTTTCCCTACACGACGCTCTTCCGATCTNNNCCATGGCCAAGACCTCAGTG
AGAGCACACGTCTGAACTCTTTCCCTACACGACGCTCTTCCGATCTNNNCCATGGCCAAGACCTCAGTG
GCACACGTCTGAACTCTTTCCCTACACGACGCTCTTCCGATCTNNNAAGATTGCTCATGGCCAGTGTTGC
GCACACGTCTGAACTCTTTCCCTACACGACGCTCTTCCGATCTNNNAAGATTGCTCATGGCCAGTGTTGC
GCACACGTCTGAACTCTTTCCCTACACGACGCTCTTCCGATCTNNNGACAGAAGGGAGCAAGTGTTTT
GCACACGTCTGAACTCTTTCCCTACACGACGCTCTTCCGATCTNNNGACAGAAGGGAGCAAGTGTTTT
GAGCACACGTCTGAACTCTTTCCCTACACGACGCTCTTCCGATCTNNNGAAACCCTACAAGCCAGAAGA
GAGCACACGTCTGAACTCTTTCCCTACACGACGCTCTTCCGATCTNNNGAAACCCTACAAGCCAGAAGA
AGCACACGTCTGAACTCTTTCCCTACACGACGCTCTTCCGATCTNNNACTCCAACCTAAGTGACAGAGT
AGCACACGTCTGAACTCTTTCCCTACACGACGCTCTTCCGATCTNNNACTCCAACCTAAGTGACAGAGT
GAGCACACGTCTGAACTCTTTCCCTACACGACGCTCTTCCGATCTNNNCGGGGGAGAGGAGAATAAGA
GAGCACACGTCTGAACTCTTTCCCTACACGACGCTCTTCCGATCTNNNCGGGGGAGAGGAGAATAAGA
AGCACACGTCTGAACTCTTTCCCTACACGACGCTCTTCCGATCTNNNCCTCCCTCCTTCTCTCTTTTCTTCA
AGCACACGTCTGAACTCTTTCCCTACACGACGCTCTTCCGATCTNNNCCTCCCTCCTTCTCTCTTTTCTTCA
GCACACGTCTGAACTCTTTCCCTACACGACGCTCTTCCGATCTNNNTATTTTATCTTTAGGGCTTAGGTC
GCACACGTCTGAACTCTTTCCCTACACGACGCTCTTCCGATCTNNNTATTTTATCTTTAGGGCTTAGGTC
GAGCACACGTCTGAACTCTTTCCCTACACGACGCTCTTCCGATCTNNNGGAAGGCCCTGTCTCAAAAAA
GAGCACACGTCTGAACTCTTTCCCTACACGACGCTCTTCCGATCTNNNGGAAGGCCCTGTCTCAAAAAA
AGCACACGTCTGAACTCTTTCCCTACACGACGCTCTTCCGATCTNNNACTCATGTCTCTGTCATTGTCCC
AGCACACGTCTGAACTCTTTCCCTACACGACGCTCTTCCGATCTNNNACTCATGTCTCTGTCATTGTCCC
AGCACACGTCTGAACTCTTTCCCTACACGACGCTCTTCCGATCTNNNGTGACAATTAGCCAAAGGTACCC
AGCACACGTCTGAACTCTTTCCCTACACGACGCTCTTCCGATCTNNNGTGACAATTAGCCAAAGGTACCC
AGCACACGTCTGAACTCTTTCCCTACACGACGCTCTTCCGATCTNNNCACCTGTAATCCTAGCACTCTGG
AGCACACGTCTGAACTCTTTCCCTACACGACGCTCTTCCGATCTNNNCACCTGTAATCCTAGCACTCTGG
AGCACACGTCTGAACTCTTTCCCTACACGACGCTCTTCCGATCTNNNAGACACAGGAAGCAAACGATC
AGCACACGTCTGAACTCTTTCCCTACACGACGCTCTTCCGATCTNNNAGACACAGGAAGCAAACGATC
GAGCACACGTCTGAACTCTTTCCCTACACGACGCTCTTCCGATCTNNNTGTTGATCCTAATCCCAAACTC
GAGCACACGTCTGAACTCTTTCCCTACACGACGCTCTTCCGATCTNNNTGTTGATCCTAATCCCAAACTC
AAGAGCACACGTCTGAACTCTTTCCCTACACGACGCTCTTCCGATCTNNNTAGCAAGGTTCAGAAGCAT
AAGAGCACACGTCTGAACTCTTTCCCTACACGACGCTCTTCCGATCTNNNTAGCAAGGTTCAGAAGCAT
GAGCACACGTCTGAACTCTTTCCCTACACGACGCTCTTCCGATCTNNNAAAAAATTAGCTGGGTGTGG
GAGCACACGTCTGAACTCTTTCCCTACACGACGCTCTTCCGATCTNNNAAAAAATTAGCTGGGTGTGG
GAGCACACGTCTGAACTCTTTCCCTACACGACGCTCTTCCGATCTNNNACATCGTCCTCAACATTGACCT
GAGCACACGTCTGAACTCTTTCCCTACACGACGCTCTTCCGATCTNNNACATCGTCCTCAACATTGACCT

AGAGCACACGTCTGAACTCTTTCCCTACACGACGCTCTTCCGATCTNNNCCCATGTCCAAGGATAAAAG
AGAGCACACGTCTGAACTCTTTCCCTACACGACGCTCTTCCGATCTNNNCCCATGTCCAAGGATAAAAG
AGCACACGTCTGAACTCTTTCCCTACACGACGCTCTTCCGATCTNNNGTTCTCATCTCGTCCTCTCCATCC
AGCACACGTCTGAACTCTTTCCCTACACGACGCTCTTCCGATCTNNNGTTCTCATCTCGTCCTCTCCATCC
GCACACGTCTGAACTCTTTCCCTACACGACGCTCTTCCGATCTNNNGAGGCTGAGGCAGGAGAATGCA
GCACACGTCTGAACTCTTTCCCTACACGACGCTCTTCCGATCTNNNGAGGCTGAGGCAGGAGAATGCA
AGCACACGTCTGAACTCTTTCCCTACACGACGCTCTTCCGATCTNNNTACAAAAAATTAGCCGGGCGTA
AGCACACGTCTGAACTCTTTCCCTACACGACGCTCTTCCGATCTNNNTACAAAAAATTAGCCGGGCGTA
AGAGCACACGTCTGAACTCTTTCCCTACACGACGCTCTTCCGATCTNNNATTGAGCCGAGATCAACCCAT
AGAGCACACGTCTGAACTCTTTCCCTACACGACGCTCTTCCGATCTNNNATTGAGCCGAGATCAACCCAT
AGAGCACACGTCTGAACTCTTTCCCTACACGACGCTCTTCCGATCTNNNCAGCAACTTGGATGAAGCTA
AGAGCACACGTCTGAACTCTTTCCCTACACGACGCTCTTCCGATCTNNNCAGCAACTTGGATGAAGCTA
AGCACACGTCTGAACTCTTTCCCTACACGACGCTCTTCCGATCTNNNTCTGGAATGATACACCACTCCAA
AGCACACGTCTGAACTCTTTCCCTACACGACGCTCTTCCGATCTNNNTCTGGAATGATACACCACTCCAA
GAGCACACGTCTGAACTCTTTCCCTACACGACGCTCTTCCGATCTNNNCAGTTATCTCAGAGCCCTGTCT
GAGCACACGTCTGAACTCTTTCCCTACACGACGCTCTTCCGATCTNNNCAGTTATCTCAGAGCCCTGTCT
AGCACACGTCTGAACTCTTTCCCTACACGACGCTCTTCCGATCTNNNCTGAGATGGGAGGATCACTTGA
AGCACACGTCTGAACTCTTTCCCTACACGACGCTCTTCCGATCTNNNCTGAGATGGGAGGATCACTTGA
GAGCACACGTCTGAACTCTTTCCCTACACGACGCTCTTCCGATCTNNNCTATACAGGCATGGTTGCCTC
GAGCACACGTCTGAACTCTTTCCCTACACGACGCTCTTCCGATCTNNNCTATACAGGCATGGTTGCCTC
AGCACACGTCTGAACTCTTTCCCTACACGACGCTCTTCCGATCTNNNGACTTTTCATACCCTGCTCTCCAC
AGCACACGTCTGAACTCTTTCCCTACACGACGCTCTTCCGATCTNNNGACTTTTCATACCCTGCTCTCCAC
AGAGCACACGTCTGAACTCTTTCCCTACACGACGCTCTTCCGATCTNNNTCGTCCTTACTAACTTCATTGC
AGAGCACACGTCTGAACTCTTTCCCTACACGACGCTCTTCCGATCTNNNTCGTCCTTACTAACTTCATTGC
GAGCACACGTCTGAACTCTTTCCCTACACGACGCTCTTCCGATCTNNNATTGGTTAGGGTTCCTCATTG
GAGCACACGTCTGAACTCTTTCCCTACACGACGCTCTTCCGATCTNNNATTGGTTAGGGTTCCTCATTG
AGCACACGTCTGAACTCTTTCCCTACACGACGCTCTTCCGATCTNNNAACAATTCGACATCCCCCACTAA
AGCACACGTCTGAACTCTTTCCCTACACGACGCTCTTCCGATCTNNNAACAATTCGACATCCCCCACTAA
GAGCACACGTCTGAACTCTTTCCCTACACGACGCTCTTCCGATCTNNNATCTCTTGCACTTTCAGCAGCT
GAGCACACGTCTGAACTCTTTCCCTACACGACGCTCTTCCGATCTNNNATCTCTTGCACTTTCAGCAGCT
AGCACACGTCTGAACTCTTTCCCTACACGACGCTCTTCCGATCTNNNACAGCAAGTATTTTAAGTTTTGT
AGCACACGTCTGAACTCTTTCCCTACACGACGCTCTTCCGATCTNNNACAGCAAGTATTTTAAGTTTTGT
AGAGCACACGTCTGAACTCTTTCCCTACACGACGCTCTTCCGATCTNNNTACACGAGAGAGAGAGAGGA
AGAGCACACGTCTGAACTCTTTCCCTACACGACGCTCTTCCGATCTNNNTACACGAGAGAGAGAGAGGA
CACACGTCTGAACTCTTTCCCTACACGACGCTCTTCCGATCTNNNTTTCTTCGCCCTCTTAGAACACACA
CACACGTCTGAACTCTTTCCCTACACGACGCTCTTCCGATCTNNNTTTCTTCGCCCTCTTAGAACACACA
GAGCACACGTCTGAACTCTTTCCCTACACGACGCTCTTCCGATCTNNNGTCTCTAGGGCTTGACAATGT
GAGCACACGTCTGAACTCTTTCCCTACACGACGCTCTTCCGATCTNNNGTCTCTAGGGCTTGACAATGT
AGCACACGTCTGAACTCTTTCCCTACACGACGCTCTTCCGATCTNNNCGTGTTGCTTTGTGTGTTCCATA
AGCACACGTCTGAACTCTTTCCCTACACGACGCTCTTCCGATCTNNNCGTGTTGCTTTGTGTGTTCCATA
GAGCACACGTCTGAACTCTTTCCCTACACGACGCTCTTCCGATCTNNNAAAATGTGTGTGTGTGTGTG
GAGCACACGTCTGAACTCTTTCCCTACACGACGCTCTTCCGATCTNNNAAAATGTGTGTGTGTGTGTG
GAGCACACGTCTGAACTCTTTCCCTACACGACGCTCTTCCGATCTNNNAGAAGTACAGTGGCATGATC

GAGCACACGTCTGAACTCTTTCCCTACACGACGCTCTTCCGATCTNNNAGAAGTACAGTGGCATGATCT

GAGCACACGTCTGAACTCTTTCCCTACACGACGCTCTTCCGATCTNNNGGGTCACAAACACCAACTTTGA

GAGCACACGTCTGAACTCTTTCCCTACACGACGCTCTTCCGATCTNNNGGGTCACAAACACCAACTTTGA

GAGCACACGTCTGAACTCTTTCCCTACACGACGCTCTTCCGATCTNNNAAATGGCCACAGCAACTACAG

GAGCACACGTCTGAACTCTTTCCCTACACGACGCTCTTCCGATCTNNNAAATGGCCACAGCAACTACAG

GAGCACACGTCTGAACTCTTTCCCTACACGACGCTCTTCCGATCTNNNGTAATTCTTTTTGTGGCCGGG

GAGCACACGTCTGAACTCTTTCCCTACACGACGCTCTTCCGATCTNNNGTAATTCTTTTTGTGGCCGGG

ACACGTCTGAACTCTTTCCCTACACGACGCTCTTCCGATCTNNNGGGATCAACCAGCACGTTTTTATCAT

ACACGTCTGAACTCTTTCCCTACACGACGCTCTTCCGATCTNNNGGGATCAACCAGCACGTTTTTATCAT

AGCACACGTCTGAACTCTTTCCCTACACGACGCTCTTCCGATCTNNNGAGGTTGCAGTGAACCAAGAT

AGCACACGTCTGAACTCTTTCCCTACACGACGCTCTTCCGATCTNNNGAGGTTGCAGTGAACCAAGAT

AGCACACGTCTGAACTCTTTCCCTACACGACGCTCTTCCGATCTNNNCTTACACTTACAGTCCCAGCACT

AGCACACGTCTGAACTCTTTCCCTACACGACGCTCTTCCGATCTNNNCTTACACTTACAGTCCCAGCACT

GAGCACACGTCTGAACTCTTTCCCTACACGACGCTCTTCCGATCTNNNAAAAAATTAGCCGGATGTGGT

GAGCACACGTCTGAACTCTTTCCCTACACGACGCTCTTCCGATCTNNNAAAAAATTAGCCGGATGTGGT

GCACACGTCTGAACTCTTTCCCTACACGACGCTCTTCCGATCTNNNCAAGACCCATACACGATTCAGCAC

GCACACGTCTGAACTCTTTCCCTACACGACGCTCTTCCGATCTNNNCAAGACCCATACACGATTCAGCAC

AGCACACGTCTGAACTCTTTCCCTACACGACGCTCTTCCGATCTNNNGGCAATTAGTTGAAAGAGTTAT

AGCACACGTCTGAACTCTTTCCCTACACGACGCTCTTCCGATCTNNNGGCAATTAGTTGAAAGAGTTAT

AGCACACGTCTGAACTCTTTCCCTACACGACGCTCTTCCGATCTNNNCAAGCAGTGACAAAAGAACCAC

AGCACACGTCTGAACTCTTTCCCTACACGACGCTCTTCCGATCTNNNCAAGCAGTGACAAAAGAACCAC

AGAGCACACGTCTGAACTCTTTCCCTACACGACGCTCTTCCGATCTNNNGATACCAACAGACCTGGGAG

AGAGCACACGTCTGAACTCTTTCCCTACACGACGCTCTTCCGATCTNNNGATACCAACAGACCTGGGAG

AGCACACGTCTGAACTCTTTCCCTACACGACGCTCTTCCGATCTNNNGAGCCTGCAGAGTGTGTAGAG

AGCACACGTCTGAACTCTTTCCCTACACGACGCTCTTCCGATCTNNNGAGCCTGCAGAGTGTGTAGAG

GAGCACACGTCTGAACTCTTTCCCTACACGACGCTCTTCCGATCTNNNAAAAAGCACTCTCCCAAACAT

GAGCACACGTCTGAACTCTTTCCCTACACGACGCTCTTCCGATCTNNNAAAAAGCACTCTCCCAAACAT

AGAGCACACGTCTGAACTCTTTCCCTACACGACGCTCTTCCGATCTNNNGTCTACCCTGCTGGATGAAC

AGAGCACACGTCTGAACTCTTTCCCTACACGACGCTCTTCCGATCTNNNGTCTACCCTGCTGGATGAAC

GAGCACACGTCTGAACTCTTTCCCTACACGACGCTCTTCCGATCTNNNGCAGAGCTTATAATGAGCGG

GAGCACACGTCTGAACTCTTTCCCTACACGACGCTCTTCCGATCTNNNGCAGAGCTTATAATGAGCGG

GAGCACACGTCTGAACTCTTTCCCTACACGACGCTCTTCCGATCTNNNGCACTGTGATGTATTATTTTCA

GAGCACACGTCTGAACTCTTTCCCTACACGACGCTCTTCCGATCTNNNGCACTGTGATGTATTATTTTCA

AGCACACGTCTGAACTCTTTCCCTACACGACGCTCTTCCGATCTNNNAAGATTCAAACTCTGAGGTTGG

AGCACACGTCTGAACTCTTTCCCTACACGACGCTCTTCCGATCTNNNAAGATTCAAACTCTGAGGTTGG

CACACGTCTGAACTCTTTCCCTACACGACGCTCTTCCGATCTNNNCTAGAATAACCCCAAGGCTCCTCCAT

CACACGTCTGAACTCTTTCCCTACACGACGCTCTTCCGATCTNNNCTAGAATAACCCCAAGGCTCCTCCAT

AGCACACGTCTGAACTCTTTCCCTACACGACGCTCTTCCGATCTNNNTTCAAGCAATTCTCCCACCTCAGC

AGCACACGTCTGAACTCTTTCCCTACACGACGCTCTTCCGATCTNNNTTCAAGCAATTCTCCCACCTCAGC

AGAGCACACGTCTGAACTCTTTCCCTACACGACGCTCTTCCGATCTNNNTTTAGCATGCATGGATTTTGC

AGAGCACACGTCTGAACTCTTTCCCTACACGACGCTCTTCCGATCTNNNTTTAGCATGCATGGATTTTGC

GAGCACACGTCTGAACTCTTTCCCTACACGACGCTCTTCCGATCTNNNCCGATCTCTAGAGTGTTGGTTC

GAGCACACGTCTGAACTCTTTCCCTACACGACGCTCTTCCGATCTNNNCCGATCTCTAGAGTGTTGGTTC

GAGCACACGTCTGAACTCTTTCCCTACACGACGCTCTTCCGATCTNNNCAGAGAAAGAAAGGGAGAG
GAGCACACGTCTGAACTCTTTCCCTACACGACGCTCTTCCGATCTNNNCAGAGAAAGAAAGGGAGAG
AGAGCACACGTCTGAACTCTTTCCCTACACGACGCTCTTCCGATCTNNNGAGAATGGCGTGAACTCAG
AGAGCACACGTCTGAACTCTTTCCCTACACGACGCTCTTCCGATCTNNNGAGAATGGCGTGAACTCAG
GCACACGTCTGAACTCTTTCCCTACACGACGCTCTTCCGATCTNNNATGCAGCCAGTTAGTGATAGAG
GCACACGTCTGAACTCTTTCCCTACACGACGCTCTTCCGATCTNNNATGCAGCCAGTTAGTGATAGAG
GCACACGTCTGAACTCTTTCCCTACACGACGCTCTTCCGATCTNNNCAACCCTTTGCATCTCTCTCTCTC
GCACACGTCTGAACTCTTTCCCTACACGACGCTCTTCCGATCTNNNCAACCCTTTGCATCTCTCTCTCTC
GCACACGTCTGAACTCTTTCCCTACACGACGCTCTTCCGATCTNNNACTACTTACTTCTGCATCAGACAA
GCACACGTCTGAACTCTTTCCCTACACGACGCTCTTCCGATCTNNNACTACTTACTTCTGCATCAGACAA
AGCACACGTCTGAACTCTTTCCCTACACGACGCTCTTCCGATCTNNNACTTGGTGAAACTCCGTCTCTA
AGCACACGTCTGAACTCTTTCCCTACACGACGCTCTTCCGATCTNNNACTTGGTGAAACTCCGTCTCTA
AGCACACGTCTGAACTCTTTCCCTACACGACGCTCTTCCGATCTNNNTTTTGAGAGAGTTTCACGCCA
AGCACACGTCTGAACTCTTTCCCTACACGACGCTCTTCCGATCTNNNTTTTGAGAGAGTTTCACGCCA
AGCACACGTCTGAACTCTTTCCCTACACGACGCTCTTCCGATCTNNNTAGTTTCAGAATTGAATTGCAC
AGCACACGTCTGAACTCTTTCCCTACACGACGCTCTTCCGATCTNNNTAGTTTCAGAATTGAATTGCAC
GCACACGTCTGAACTCTTTCCCTACACGACGCTCTTCCGATCTNNNTCAAGGCAGGAGGATTGTTTG
GCACACGTCTGAACTCTTTCCCTACACGACGCTCTTCCGATCTNNNTCAAGGCAGGAGGATTGTTTG
GAGCACACGTCTGAACTCTTTCCCTACACGACGCTCTTCCGATCTNNNATCCACCAACCAAGCCCTATA
GAGCACACGTCTGAACTCTTTCCCTACACGACGCTCTTCCGATCTNNNATCCACCAACCAAGCCCTATA
GAGCACACGTCTGAACTCTTTCCCTACACGACGCTCTTCCGATCTNNNAAACACTGAGGAAGATTGA
GAGCACACGTCTGAACTCTTTCCCTACACGACGCTCTTCCGATCTNNNAAACACTGAGGAAGATTGA
AGCACACGTCTGAACTCTTTCCCTACACGACGCTCTTCCGATCTNNNGAATTGCTTGAACCCAGTAGT
AGCACACGTCTGAACTCTTTCCCTACACGACGCTCTTCCGATCTNNNGAATTGCTTGAACCCAGTAGT
GCACACGTCTGAACTCTTTCCCTACACGACGCTCTTCCGATCTNNNCCACCATGCCCTGCTAATTTTTC
GCACACGTCTGAACTCTTTCCCTACACGACGCTCTTCCGATCTNNNCCACCATGCCCTGCTAATTTTTC
GCACACGTCTGAACTCTTTCCCTACACGACGCTCTTCCGATCTNNNAAGATCAAGGTGCCAGCATCTA
GCACACGTCTGAACTCTTTCCCTACACGACGCTCTTCCGATCTNNNAAGATCAAGGTGCCAGCATCTA
GCACACGTCTGAACTCTTTCCCTACACGACGCTCTTCCGATCTNNNGCTGCAAAGGACATGATCTTGT
GCACACGTCTGAACTCTTTCCCTACACGACGCTCTTCCGATCTNNNGCTGCAAAGGACATGATCTTGT
CACACGTCTGAACTCTTTCCCTACACGACGCTCTTCCGATCTNNNAGGGAGGCACTATCACAAAATGA
CACACGTCTGAACTCTTTCCCTACACGACGCTCTTCCGATCTNNNAGGGAGGCACTATCACAAAATGA
AGCACACGTCTGAACTCTTTCCCTACACGACGCTCTTCCGATCTNNNCCCAGCTGGTGGAAAATATGT
AGCACACGTCTGAACTCTTTCCCTACACGACGCTCTTCCGATCTNNNCCCAGCTGGTGGAAAATATGT
AGCACACGTCTGAACTCTTTCCCTACACGACGCTCTTCCGATCTNNNAAGGTTGCAAAGGCTGGAATA
AGCACACGTCTGAACTCTTTCCCTACACGACGCTCTTCCGATCTNNNAAGGTTGCAAAGGCTGGAATA
AGCACACGTCTGAACTCTTTCCCTACACGACGCTCTTCCGATCTNNNTACCTTAGTCCCAGCTACTCCT
AGCACACGTCTGAACTCTTTCCCTACACGACGCTCTTCCGATCTNNNTACCTTAGTCCCAGCTACTCCT
GCACACGTCTGAACTCTTTCCCTACACGACGCTCTTCCGATCTNNNAAAAAGGAGAGGGGACACTTT
GCACACGTCTGAACTCTTTCCCTACACGACGCTCTTCCGATCTNNNAAAAAGGAGAGGGGACACTTT
GCACACGTCTGAACTCTTTCCCTACACGACGCTCTTCCGATCTNNNATGTTCCTTTAGCACCTGTCCAAC
GCACACGTCTGAACTCTTTCCCTACACGACGCTCTTCCGATCTNNNATGTTCCTTTAGCACCTGTCCAAC
AGCACACGTCTGAACTCTTTCCCTACACGACGCTCTTCCGATCTNNNTGTGTGTGTGTGTGTGTGTAT

GAGCACACGTCTGAACTCTTTCCCTACACGACGCTCTTCCGATCTNNNTGTGTGTGTGTGTGTGTGTAT
AGCACACGTCTGAACTCTTTCCCTACACGACGCTCTTCCGATCTNNNGTGAAGGCCTGAACTTTTGTCTT
AGCACACGTCTGAACTCTTTCCCTACACGACGCTCTTCCGATCTNNNGTGAAGGCCTGAACTTTTGTCTT
AGCACACGTCTGAACTCTTTCCCTACACGACGCTCTTCCGATCTNNNGCAACAGCTACATGGCACAAAA
AGCACACGTCTGAACTCTTTCCCTACACGACGCTCTTCCGATCTNNNGCAACAGCTACATGGCACAAAA
GAGCACACGTCTGAACTCTTTCCCTACACGACGCTCTTCCGATCTNNNCTAGGTGGGGGTATGAATTGG
GAGCACACGTCTGAACTCTTTCCCTACACGACGCTCTTCCGATCTNNNCTAGGTGGGGGTATGAATTGG
GAGCACACGTCTGAACTCTTTCCCTACACGACGCTCTTCCGATCTNNNCTAGGAAACACACACACACACA
GAGCACACGTCTGAACTCTTTCCCTACACGACGCTCTTCCGATCTNNNCTAGGAAACACACACACACACA
AGCACACGTCTGAACTCTTTCCCTACACGACGCTCTTCCGATCTNNNTCCAGGACACTAAGCAAAACAT
AGCACACGTCTGAACTCTTTCCCTACACGACGCTCTTCCGATCTNNNTCCAGGACACTAAGCAAAACAT
AGCACACGTCTGAACTCTTTCCCTACACGACGCTCTTCCGATCTNNNGATCAGTGCATTCCCACTTTAGC
AGCACACGTCTGAACTCTTTCCCTACACGACGCTCTTCCGATCTNNNGATCAGTGCATTCCCACTTTAGC
GCACACGTCTGAACTCTTTCCCTACACGACGCTCTTCCGATCTNNNGCAACTTCAGGGAAGCATTTTCT
GCACACGTCTGAACTCTTTCCCTACACGACGCTCTTCCGATCTNNNGCAACTTCAGGGAAGCATTTTCT
GAGCACACGTCTGAACTCTTTCCCTACACGACGCTCTTCCGATCTNNNCCTGTCTCTGGTGCTTTTCTAGT
GAGCACACGTCTGAACTCTTTCCCTACACGACGCTCTTCCGATCTNNNCCTGTCTCTGGTGCTTTTCTAGT
GCACACGTCTGAACTCTTTCCCTACACGACGCTCTTCCGATCTNNNTGTGCTTTTGGTGTCAAACTGAA
GCACACGTCTGAACTCTTTCCCTACACGACGCTCTTCCGATCTNNNTGTGCTTTTGGTGTCAAACTGAA
AGCACACGTCTGAACTCTTTCCCTACACGACGCTCTTCCGATCTNNNGCAGGCAAAGAGTAAGCATTCA
AGCACACGTCTGAACTCTTTCCCTACACGACGCTCTTCCGATCTNNNGCAGGCAAAGAGTAAGCATTCA
AGCACACGTCTGAACTCTTTCCCTACACGACGCTCTTCCGATCTNNNTGTACCTGCAATGAGGTTTTG
AGCACACGTCTGAACTCTTTCCCTACACGACGCTCTTCCGATCTNNNTGTACCTGCAATGAGGTTTTG
GAGCACACGTCTGAACTCTTTCCCTACACGACGCTCTTCCGATCTNNNGAAGCTGTAGTGAGCAGAGAT
GAGCACACGTCTGAACTCTTTCCCTACACGACGCTCTTCCGATCTNNNGAAGCTGTAGTGAGCAGAGAT
AGCACACGTCTGAACTCTTTCCCTACACGACGCTCTTCCGATCTNNNAAGCAGAGGCAGAAGAATAGC
AGCACACGTCTGAACTCTTTCCCTACACGACGCTCTTCCGATCTNNNAAGCAGAGGCAGAAGAATAGC
AGAGCACACGTCTGAACTCTTTCCCTACACGACGCTCTTCCGATCTNNNGAGAATCACTTGAACCCAGG
AGAGCACACGTCTGAACTCTTTCCCTACACGACGCTCTTCCGATCTNNNGAGAATCACTTGAACCCAGG
AGCACACGTCTGAACTCTTTCCCTACACGACGCTCTTCCGATCTNNNTGAAGTGGGTTTCTCCATTGCTA
AGCACACGTCTGAACTCTTTCCCTACACGACGCTCTTCCGATCTNNNTGAAGTGGGTTTCTCCATTGCTA
AGAGCACACGTCTGAACTCTTTCCCTACACGACGCTCTTCCGATCTNNNTCGTGCTGCGTGTATTATTT
AGAGCACACGTCTGAACTCTTTCCCTACACGACGCTCTTCCGATCTNNNTCGTGCTGCGTGTATTATTT
AGCACACGTCTGAACTCTTTCCCTACACGACGCTCTTCCGATCTNNNCAAAAATTTGCTGCACATGGTG
AGCACACGTCTGAACTCTTTCCCTACACGACGCTCTTCCGATCTNNNCAAAAATTTGCTGCACATGGTG
GAGCACACGTCTGAACTCTTTCCCTACACGACGCTCTTCCGATCTNNNTCTGACACATATTTCCGATGCC
GAGCACACGTCTGAACTCTTTCCCTACACGACGCTCTTCCGATCTNNNTCTGACACATATTTCCGATGCC
GAGCACACGTCTGAACTCTTTCCCTACACGACGCTCTTCCGATCTNNNGGCTCTAGAGGTGGGTGAAT
GAGCACACGTCTGAACTCTTTCCCTACACGACGCTCTTCCGATCTNNNGGCTCTAGAGGTGGGTGAAT
AGCACACGTCTGAACTCTTTCCCTACACGACGCTCTTCCGATCTNNNACCAAATGACAACTAAAATCCGC
AGCACACGTCTGAACTCTTTCCCTACACGACGCTCTTCCGATCTNNNACCAAATGACAACTAAAATCCGC
AGCACACGTCTGAACTCTTTCCCTACACGACGCTCTTCCGATCTNNNAGTGAACAGGCAACCTACAGAA
AGCACACGTCTGAACTCTTTCCCTACACGACGCTCTTCCGATCTNNNAGTGAACAGGCAACCTACAGAA

AGCACACGTCTGAACTCTTTCCCTACACGACGCTCTTCCGATCTNNNGGTTGAAAAGCACTGTGGTACA
AGCACACGTCTGAACTCTTTCCCTACACGACGCTCTTCCGATCTNNNGGTTGAAAAGCACTGTGGTACA
AGAGCACACGTCTGAACTCTTTCCCTACACGACGCTCTTCCGATCTNNNAAGGAAGGAACGAAGGAAT
AGAGCACACGTCTGAACTCTTTCCCTACACGACGCTCTTCCGATCTNNNAAGGAAGGAACGAAGGAAT
AGCACACGTCTGAACTCTTTCCCTACACGACGCTCTTCCGATCTNNNGTATCAACGTGGTCATCTGGGT
AGCACACGTCTGAACTCTTTCCCTACACGACGCTCTTCCGATCTNNNGTATCAACGTGGTCATCTGGGT
AGCACACGTCTGAACTCTTTCCCTACACGACGCTCTTCCGATCTNNNTGGTGGTAACTTGATGTATAAA
AGCACACGTCTGAACTCTTTCCCTACACGACGCTCTTCCGATCTNNNTGGTGGTAACTTGATGTATAAA
GAGCACACGTCTGAACTCTTTCCCTACACGACGCTCTTCCGATCTNNNTGGGTTGAGATGATGGGAAA
GAGCACACGTCTGAACTCTTTCCCTACACGACGCTCTTCCGATCTNNNTGGGTTGAGATGATGGGAAA
AGCACACGTCTGAACTCTTTCCCTACACGACGCTCTTCCGATCTNNNCCTGAGGGTAAGGGTCAAGAT
AGCACACGTCTGAACTCTTTCCCTACACGACGCTCTTCCGATCTNNNCCTGAGGGTAAGGGTCAAGAT
GAGCACACGTCTGAACTCTTTCCCTACACGACGCTCTTCCGATCTNNNACAAGAATCTTGAACCTGGGA
GAGCACACGTCTGAACTCTTTCCCTACACGACGCTCTTCCGATCTNNNACAAGAATCTTGAACCTGGGA
CACACGTCTGAACTCTTTCCCTACACGACGCTCTTCCGATCTNNNCTTCCTGCTTTAGAATCCCTCATCGG
CACACGTCTGAACTCTTTCCCTACACGACGCTCTTCCGATCTNNNCTTCCTGCTTTAGAATCCCTCATCGG
AGCACACGTCTGAACTCTTTCCCTACACGACGCTCTTCCGATCTNNNTCACCAGCAAATCTGATCCTTCA
AGCACACGTCTGAACTCTTTCCCTACACGACGCTCTTCCGATCTNNNTCACCAGCAAATCTGATCCTTCA
GAGCACACGTCTGAACTCTTTCCCTACACGACGCTCTTCCGATCTNNNATTTTTCCCACTCAACTGCTG
GAGCACACGTCTGAACTCTTTCCCTACACGACGCTCTTCCGATCTNNNATTTTTCCCACTCAACTGCTG
AGAGCACACGTCTGAACTCTTTCCCTACACGACGCTCTTCCGATCTNNNCTGTAATCTCAGCTACTCGG
AGAGCACACGTCTGAACTCTTTCCCTACACGACGCTCTTCCGATCTNNNCTGTAATCTCAGCTACTCGG
CACACGTCTGAACTCTTTCCCTACACGACGCTCTTCCGATCTNNNAGTGTAGATAGATGAAACCTTTAC
CACACGTCTGAACTCTTTCCCTACACGACGCTCTTCCGATCTNNNAGTGTAGATAGATGAAACCTTTAC
GAGCACACGTCTGAACTCTTTCCCTACACGACGCTCTTCCGATCTNNNTACACTCCTGGGATAAATGTG
GAGCACACGTCTGAACTCTTTCCCTACACGACGCTCTTCCGATCTNNNTACACTCCTGGGATAAATGTG
GAGCACACGTCTGAACTCTTTCCCTACACGACGCTCTTCCGATCTNNNGGCGACAGTATGAGGATCCAT
GAGCACACGTCTGAACTCTTTCCCTACACGACGCTCTTCCGATCTNNNGGCGACAGTATGAGGATCCAT
AGCACACGTCTGAACTCTTTCCCTACACGACGCTCTTCCGATCTNNNCACTTGAGGTCAGGAGTTCAA
AGCACACGTCTGAACTCTTTCCCTACACGACGCTCTTCCGATCTNNNCACTTGAGGTCAGGAGTTCAA
GAGCACACGTCTGAACTCTTTCCCTACACGACGCTCTTCCGATCTNNNAGTTTGAGACTGGGCAACAT
GAGCACACGTCTGAACTCTTTCCCTACACGACGCTCTTCCGATCTNNNAGTTTGAGACTGGGCAACAT
GAGCACACGTCTGAACTCTTTCCCTACACGACGCTCTTCCGATCTNNNAAATCCACAGACCTTTCCTCCT
GAGCACACGTCTGAACTCTTTCCCTACACGACGCTCTTCCGATCTNNNAAATCCACAGACCTTTCCTCCT
AGCACACGTCTGAACTCTTTCCCTACACGACGCTCTTCCGATCTNNNTCCACTGTCTGTTCAAATGTCCA
AGCACACGTCTGAACTCTTTCCCTACACGACGCTCTTCCGATCTNNNTCCACTGTCTGTTCAAATGTCCA
AGCACACGTCTGAACTCTTTCCCTACACGACGCTCTTCCGATCTNNNCTGTAAAAATGACCTCCCTTCCAT
AGCACACGTCTGAACTCTTTCCCTACACGACGCTCTTCCGATCTNNNCTGTAAAAATGACCTCCCTTCCAT
GCACACGTCTGAACTCTTTCCCTACACGACGCTCTTCCGATCTNNNTGAGCCTTTGTAAGAGGGCAATT
GCACACGTCTGAACTCTTTCCCTACACGACGCTCTTCCGATCTNNNTGAGCCTTTGTAAGAGGGCAATT
GAGCACACGTCTGAACTCTTTCCCTACACGACGCTCTTCCGATCTNNNTCATTCATTCATTCCCCCCACAT
GAGCACACGTCTGAACTCTTTCCCTACACGACGCTCTTCCGATCTNNNTCATTCATTCATTCCCCCCACAT
AGCACACGTCTGAACTCTTTCCCTACACGACGCTCTTCCGATCTNNNTAGTCCCAGCTACTCAGAAGACT

GCACACGTCTGAACTCTTTCCCTACACGACGCTCTTCCGATCTNNNTAGTCCCAGCTACTCAGAAGACT

GAGCACACGTCTGAACTCTTTCCCTACACGACGCTCTTCCGATCTNNNCATGTGAAGGCTTTGGAGGA

GAGCACACGTCTGAACTCTTTCCCTACACGACGCTCTTCCGATCTNNNCATGTGAAGGCTTTGGAGGA

GCACACGTCTGAACTCTTTCCCTACACGACGCTCTTCCGATCTNNNCGGTGAAAGCTCGTCTCTACTAA

GCACACGTCTGAACTCTTTCCCTACACGACGCTCTTCCGATCTNNNCGGTGAAAGCTCGTCTCTACTAA

GCACACGTCTGAACTCTTTCCCTACACGACGCTCTTCCGATCTNNNCACCACCACCAAGTGTTCCTACCA

GCACACGTCTGAACTCTTTCCCTACACGACGCTCTTCCGATCTNNNCACCACCACCAAGTGTTCCTACCA

AGCACACGTCTGAACTCTTTCCCTACACGACGCTCTTCCGATCTNNNGCAAATTGCTGTTCATTGCCTC

AGCACACGTCTGAACTCTTTCCCTACACGACGCTCTTCCGATCTNNNGCAAATTGCTGTTCATTGCCTC

AGCACACGTCTGAACTCTTTCCCTACACGACGCTCTTCCGATCTNNNTACTGGGAAGTGAGGAGCCC

AGCACACGTCTGAACTCTTTCCCTACACGACGCTCTTCCGATCTNNNTACTGGGAAGTGAGGAGCCC

GAGCACACGTCTGAACTCTTTCCCTACACGACGCTCTTCCGATCTNNNGAGGAAAGAAGGAAGGAAG

GAGCACACGTCTGAACTCTTTCCCTACACGACGCTCTTCCGATCTNNNGAGGAAAGAAGGAAGGAAG

AGCACACGTCTGAACTCTTTCCCTACACGACGCTCTTCCGATCTNNNGACAGAGTAACACTCCGTCTCG

AGCACACGTCTGAACTCTTTCCCTACACGACGCTCTTCCGATCTNNNGACAGAGTAACACTCCGTCTCG

GAGCACACGTCTGAACTCTTTCCCTACACGACGCTCTTCCGATCTNNNAAAGTTGCAGTGAGCTGAGA

GAGCACACGTCTGAACTCTTTCCCTACACGACGCTCTTCCGATCTNNNAAAGTTGCAGTGAGCTGAGA

AGCACACGTCTGAACTCTTTCCCTACACGACGCTCTTCCGATCTNNNAGTTGTGCCAGGTGAAATTCAA

AGCACACGTCTGAACTCTTTCCCTACACGACGCTCTTCCGATCTNNNAGTTGTGCCAGGTGAAATTCAA

AGCACACGTCTGAACTCTTTCCCTACACGACGCTCTTCCGATCTNNNTGTTTTAGAAGCTGCTCCCTTTG

AGCACACGTCTGAACTCTTTCCCTACACGACGCTCTTCCGATCTNNNTGTTTTAGAAGCTGCTCCCTTTG

GAGCACACGTCTGAACTCTTTCCCTACACGACGCTCTTCCGATCTNNNATAGCAGCTCTCCTTCCAGTTT

GAGCACACGTCTGAACTCTTTCCCTACACGACGCTCTTCCGATCTNNNATAGCAGCTCTCCTTCCAGTTT

AGAGCACACGTCTGAACTCTTTCCCTACACGACGCTCTTCCGATCTNNNGAAGCGAGAAAGGTCTGA

AGAGCACACGTCTGAACTCTTTCCCTACACGACGCTCTTCCGATCTNNNGAAGCGAGAAAGGTCTGA

AGAGCACACGTCTGAACTCTTTCCCTACACGACGCTCTTCCGATCTNNNGCTACTTCTTTGATTTCTGGA

AGAGCACACGTCTGAACTCTTTCCCTACACGACGCTCTTCCGATCTNNNGCTACTTCTTTGATTTCTGGA

GAGCACACGTCTGAACTCTTTCCCTACACGACGCTCTTCCGATCTNNNAGGAACCTTGGAGATAGTTG

GAGCACACGTCTGAACTCTTTCCCTACACGACGCTCTTCCGATCTNNNAGGAACCTTGGAGATAGTTG

GCACACGTCTGAACTCTTTCCCTACACGACGCTCTTCCGATCTNNNTCGTACAGAGGCATCTTACTGTG

GCACACGTCTGAACTCTTTCCCTACACGACGCTCTTCCGATCTNNNTCGTACAGAGGCATCTTACTGTG

AGCACACGTCTGAACTCTTTCCCTACACGACGCTCTTCCGATCTNNNCAGCTATTCTGGAGGCTAAGGT

AGCACACGTCTGAACTCTTTCCCTACACGACGCTCTTCCGATCTNNNCAGCTATTCTGGAGGCTAAGGT

GAGCACACGTCTGAACTCTTTCCCTACACGACGCTCTTCCGATCTNNNCCTTCAGGAAGACTCACTTGCT

GAGCACACGTCTGAACTCTTTCCCTACACGACGCTCTTCCGATCTNNNCCTTCAGGAAGACTCACTTGCT

GCACACGTCTGAACTCTTTCCCTACACGACGCTCTTCCGATCTNNNAGCTGAGGTGATACACTCTCTCT

GCACACGTCTGAACTCTTTCCCTACACGACGCTCTTCCGATCTNNNAGCTGAGGTGATACACTCTCTCT

AGCACACGTCTGAACTCTTTCCCTACACGACGCTCTTCCGATCTNNNGTGAACGCCAAGAATGAGTTCT

AGCACACGTCTGAACTCTTTCCCTACACGACGCTCTTCCGATCTNNNGTGAACGCCAAGAATGAGTTCT

GAGCACACGTCTGAACTCTTTCCCTACACGACGCTCTTCCGATCTNNNAGGCAAGAGCAAACAAATTCG

GAGCACACGTCTGAACTCTTTCCCTACACGACGCTCTTCCGATCTNNNAGGCAAGAGCAAACAAATTCG

GAGCACACGTCTGAACTCTTTCCCTACACGACGCTCTTCCGATCTNNNCTCAGCCCTCACCCTGGTTTCC

GAGCACACGTCTGAACTCTTTCCCTACACGACGCTCTTCCGATCTNNNCTCAGCCCTCACCCTGGTTTCC

AGCACACGTCTGAACTCTTTCCCTACACGACGCTCTTCCGATCTNNNTGAGGTAGGAGAATTGCCTG
AGCACACGTCTGAACTCTTTCCCTACACGACGCTCTTCCGATCTNNNTGAGGTAGGAGAATTGCCTG
AGCACACGTCTGAACTCTTTCCCTACACGACGCTCTTCCGATCTNNNCAGAGACATTGACACGTAAG
AGCACACGTCTGAACTCTTTCCCTACACGACGCTCTTCCGATCTNNNCAGAGACATTGACACGTAAG
GAGCACACGTCTGAACTCTTTCCCTACACGACGCTCTTCCGATCTNNNGATGCAAGAGGATTGCTTGA
GAGCACACGTCTGAACTCTTTCCCTACACGACGCTCTTCCGATCTNNNGATGCAAGAGGATTGCTTGA
GCACACGTCTGAACTCTTTCCCTACACGACGCTCTTCCGATCTNNNTTCAGCTAAAGACTCTGTGACC
GCACACGTCTGAACTCTTTCCCTACACGACGCTCTTCCGATCTNNNTTCAGCTAAAGACTCTGTGACC
GAGCACACGTCTGAACTCTTTCCCTACACGACGCTCTTCCGATCTNNNATCTTCTCCAAGATTTTCGGC
GAGCACACGTCTGAACTCTTTCCCTACACGACGCTCTTCCGATCTNNNATCTTCTCCAAGATTTTCGGC
GCACACGTCTGAACTCTTTCCCTACACGACGCTCTTCCGATCTNNNGGCAGGAGAATCACTTGAACCT
GCACACGTCTGAACTCTTTCCCTACACGACGCTCTTCCGATCTNNNGGCAGGAGAATCACTTGAACCT
AGCACACGTCTGAACTCTTTCCCTACACGACGCTCTTCCGATCTNNNGAGGCAGAAGAATTGCTTGT
AGCACACGTCTGAACTCTTTCCCTACACGACGCTCTTCCGATCTNNNGAGGCAGAAGAATTGCTTGT
GAGCACACGTCTGAACTCTTTCCCTACACGACGCTCTTCCGATCTNNNATCTTTCCCAGATTGGCTTGG
GAGCACACGTCTGAACTCTTTCCCTACACGACGCTCTTCCGATCTNNNATCTTTCCCAGATTGGCTTGG
GAGCACACGTCTGAACTCTTTCCCTACACGACGCTCTTCCGATCTNNNGGCTAAGGCAGGAAAATCTC
GAGCACACGTCTGAACTCTTTCCCTACACGACGCTCTTCCGATCTNNNGGCTAAGGCAGGAAAATCTC
AGCACACGTCTGAACTCTTTCCCTACACGACGCTCTTCCGATCTNNNTGAGGCAGAAGAATTGCTTG
AGCACACGTCTGAACTCTTTCCCTACACGACGCTCTTCCGATCTNNNTGAGGCAGAAGAATTGCTTG
AGCACACGTCTGAACTCTTTCCCTACACGACGCTCTTCCGATCTNNNAGTCAACAAATTGGTTTACTG
AGCACACGTCTGAACTCTTTCCCTACACGACGCTCTTCCGATCTNNNAGTCAACAAATTGGTTTACTG
AGCACACGTCTGAACTCTTTCCCTACACGACGCTCTTCCGATCTNNNCGTTATCCTGGAGGCATAGCAT
AGCACACGTCTGAACTCTTTCCCTACACGACGCTCTTCCGATCTNNNCGTTATCCTGGAGGCATAGCAT
GCACACGTCTGAACTCTTTCCCTACACGACGCTCTTCCGATCTNNNGAAGCAGAAGAATCGCTGGAA
GCACACGTCTGAACTCTTTCCCTACACGACGCTCTTCCGATCTNNNGAAGCAGAAGAATCGCTGGAA
GCACACGTCTGAACTCTTTCCCTACACGACGCTCTTCCGATCTNNNTCATACCTAAAAAACCCAGGCC
GCACACGTCTGAACTCTTTCCCTACACGACGCTCTTCCGATCTNNNTCATACCTAAAAAACCCAGGCC
AGCACACGTCTGAACTCTTTCCCTACACGACGCTCTTCCGATCTNNNTGTTCCATTGATCTATGTGTGT
AGCACACGTCTGAACTCTTTCCCTACACGACGCTCTTCCGATCTNNNTGTTCCATTGATCTATGTGTGT
GCACACGTCTGAACTCTTTCCCTACACGACGCTCTTCCGATCTNNNTGCCAGACTAGCGATATATGAG
GCACACGTCTGAACTCTTTCCCTACACGACGCTCTTCCGATCTNNNTGCCAGACTAGCGATATATGAG
GCACACGTCTGAACTCTTTCCCTACACGACGCTCTTCCGATCTNNNCTGTAGTCTCAGCTACTTGGGA
GCACACGTCTGAACTCTTTCCCTACACGACGCTCTTCCGATCTNNNCTGTAGTCTCAGCTACTTGGGA
GAGCACACGTCTGAACTCTTTCCCTACACGACGCTCTTCCGATCTNNNTTGATCCCAATGTCTTCATCA
GAGCACACGTCTGAACTCTTTCCCTACACGACGCTCTTCCGATCTNNNTTGATCCCAATGTCTTCATCA
GCACACGTCTGAACTCTTTCCCTACACGACGCTCTTCCGATCTNNNCATCTCCATAGTAGCCCCCTTTCC
GCACACGTCTGAACTCTTTCCCTACACGACGCTCTTCCGATCTNNNCATCTCCATAGTAGCCCCCTTTCC
AGCACACGTCTGAACTCTTTCCCTACACGACGCTCTTCCGATCTNNNTGATGCAGGAGAATTGCTTGA
AGCACACGTCTGAACTCTTTCCCTACACGACGCTCTTCCGATCTNNNTGATGCAGGAGAATTGCTTGA
GCACACGTCTGAACTCTTTCCCTACACGACGCTCTTCCGATCTNNNTCATTGTAGCTATAGACTATTGG
GCACACGTCTGAACTCTTTCCCTACACGACGCTCTTCCGATCTNNNTCATTGTAGCTATAGACTATTGG
GAGCACACGTCTGAACTCTTTCCCTACACGACGCTCTTCCGATCTNNNTCATTCATTCATTCCCCCACAT

AGCACACGTCTGAACTCTTTCCCTACACGACGCTCTTCCGATCTNNNTCATTCATTCATTCCCCCACAT
GAGCACACGTCTGAACTCTTTCCCTACACGACGCTCTTCCGATCTNNNAGACAAAACTTGTGCCTGAA
AGCACACGTCTGAACTCTTTCCCTACACGACGCTCTTCCGATCTNNNAGACAAAACTTGTGCCTGAA
AGCACACGTCTGAACTCTTTCCCTACACGACGCTCTTCCGATCTNNNTCTGGGTATAAGTCCTCTGTTG
AGCACACGTCTGAACTCTTTCCCTACACGACGCTCTTCCGATCTNNNTCTGGGTATAAGTCCTCTGTTG
GCACACGTCTGAACTCTTTCCCTACACGACGCTCTTCCGATCTNNNACATGGCATCTCATTGTGGTTTG
GCACACGTCTGAACTCTTTCCCTACACGACGCTCTTCCGATCTNNNACATGGCATCTCATTGTGGTTTG
GAGCACACGTCTGAACTCTTTCCCTACACGACGCTCTTCCGATCTNNNGAGAATTACTTGAGCCCAG
GAGCACACGTCTGAACTCTTTCCCTACACGACGCTCTTCCGATCTNNNGAGAATTACTTGAGCCCAG
AGCACACGTCTGAACTCTTTCCCTACACGACGCTCTTCCGATCTNNNACACACATACACACACATACAC
AGCACACGTCTGAACTCTTTCCCTACACGACGCTCTTCCGATCTNNNACACACATACACACACATACAC
AGCACACGTCTGAACTCTTTCCCTACACGACGCTCTTCCGATCTNNNTAGGTTGCATGAGCTGAGAT
AGCACACGTCTGAACTCTTTCCCTACACGACGCTCTTCCGATCTNNNTAGGTTGCATGAGCTGAGAT
GCACACGTCTGAACTCTTTCCCTACACGACGCTCTTCCGATCTNNNCTAGTGCTGGGGAACTTTGAGA
GCACACGTCTGAACTCTTTCCCTACACGACGCTCTTCCGATCTNNNCTAGTGCTGGGGAACTTTGAGA
GAGCACACGTCTGAACTCTTTCCCTACACGACGCTCTTCCGATCTNNNAAATGAGTCCAGTTCCTCCA
GAGCACACGTCTGAACTCTTTCCCTACACGACGCTCTTCCGATCTNNNAAATGAGTCCAGTTCCTCCA
GAGCACACGTCTGAACTCTTTCCCTACACGACGCTCTTCCGATCTNNNGGAGGTAGAGGCCACATGA
GAGCACACGTCTGAACTCTTTCCCTACACGACGCTCTTCCGATCTNNNGGAGGTAGAGGCCACATGA
GCACACGTCTGAACTCTTTCCCTACACGACGCTCTTCCGATCTNNNCTTCCAATAGAGGCTTTGGCTT
GCACACGTCTGAACTCTTTCCCTACACGACGCTCTTCCGATCTNNNCTTCCAATAGAGGCTTTGGCTT
GAGCACACGTCTGAACTCTTTCCCTACACGACGCTCTTCCGATCTNNNGCGAAACCCGGTTTCTACTA
GAGCACACGTCTGAACTCTTTCCCTACACGACGCTCTTCCGATCTNNNGCGAAACCCGGTTTCTACTA
AGCACACGTCTGAACTCTTTCCCTACACGACGCTCTTCCGATCTNNNTAGGAAGCTAAATTCAGCCC
AGCACACGTCTGAACTCTTTCCCTACACGACGCTCTTCCGATCTNNNTAGGAAGCTAAATTCAGCCC
AGAGCACACGTCTGAACTCTTTCCCTACACGACGCTCTTCCGATCTNNNAAAAAAACATCTGCCCACAC
AGAGCACACGTCTGAACTCTTTCCCTACACGACGCTCTTCCGATCTNNNAAAAAAACATCTGCCCACAC
GCACACGTCTGAACTCTTTCCCTACACGACGCTCTTCCGATCTNNNCATGTGTCTGTAGTCCCAGCTA
GCACACGTCTGAACTCTTTCCCTACACGACGCTCTTCCGATCTNNNCATGTGTCTGTAGTCCCAGCTA
GCACACGTCTGAACTCTTTCCCTACACGACGCTCTTCCGATCTNNNGCAGAGTTTACAGTGAGCTGA
GCACACGTCTGAACTCTTTCCCTACACGACGCTCTTCCGATCTNNNGCAGAGTTTACAGTGAGCTGA
GAGCACACGTCTGAACTCTTTCCCTACACGACGCTCTTCCGATCTNNNAGGAATGGTCTCGGAAAAA
GAGCACACGTCTGAACTCTTTCCCTACACGACGCTCTTCCGATCTNNNAGGAATGGTCTCGGAAAAA
AGCACACGTCTGAACTCTTTCCCTACACGACGCTCTTCCGATCTNNNCACAGACATCCCTGAGCTGAGC
AGCACACGTCTGAACTCTTTCCCTACACGACGCTCTTCCGATCTNNNCACAGACATCCCTGAGCTGAGC
GAGCACACGTCTGAACTCTTTCCCTACACGACGCTCTTCCGATCTNNNATCTCCAGGTAGAGAGTGT
GAGCACACGTCTGAACTCTTTCCCTACACGACGCTCTTCCGATCTNNNATCTCCAGGTAGAGAGTGT
GAGCACACGTCTGAACTCTTTCCCTACACGACGCTCTTCCGATCTNNNGAACTTGAGCAGGAAGTCG
GAGCACACGTCTGAACTCTTTCCCTACACGACGCTCTTCCGATCTNNNGAACTTGAGCAGGAAGTCG
AGCACACGTCTGAACTCTTTCCCTACACGACGCTCTTCCGATCTNNNCTGGGCAACATAGTAAGACCC
AGCACACGTCTGAACTCTTTCCCTACACGACGCTCTTCCGATCTNNNCTGGGCAACATAGTAAGACCC
GAGCACACGTCTGAACTCTTTCCCTACACGACGCTCTTCCGATCTNNNGAGGCAGGAGAATCGCTTG
GAGCACACGTCTGAACTCTTTCCCTACACGACGCTCTTCCGATCTNNNGAGGCAGGAGAATCGCTTG

AGCACACGTCTGAACTCTTTCCCTACACGACGCTCTTCCGATCTNNNGCACTGGGTTCAGAATTTCCTCT
AGCACACGTCTGAACTCTTTCCCTACACGACGCTCTTCCGATCTNNNGCACTGGGTTCAGAATTTCCTCT
AGCACACGTCTGAACTCTTTCCCTACACGACGCTCTTCCGATCTNNNTTTGGCCTTTCTCACCAGAGTA
AGCACACGTCTGAACTCTTTCCCTACACGACGCTCTTCCGATCTNNNTTTGGCCTTTCTCACCAGAGTA
AGCACACGTCTGAACTCTTTCCCTACACGACGCTCTTCCGATCTNNNTTCTTGCCTTCTCCTTTATCCCCA
AGCACACGTCTGAACTCTTTCCCTACACGACGCTCTTCCGATCTNNNTTCTTGCCTTCTCCTTTATCCCCA
GAGCACACGTCTGAACTCTTTCCCTACACGACGCTCTTCCGATCTNNNTCTACATGCTATCAAGGGCCA
GAGCACACGTCTGAACTCTTTCCCTACACGACGCTCTTCCGATCTNNNTCTACATGCTATCAAGGGCCA
GAGCACACGTCTGAACTCTTTCCCTACACGACGCTCTTCCGATCTNNNGGGATGAGAGTTCCTAAGCT
GAGCACACGTCTGAACTCTTTCCCTACACGACGCTCTTCCGATCTNNNGGGATGAGAGTTCCTAAGCT
AGCACACGTCTGAACTCTTTCCCTACACGACGCTCTTCCGATCTNNNCCTGTTTTGCTCACCCTCTTTTCC
AGCACACGTCTGAACTCTTTCCCTACACGACGCTCTTCCGATCTNNNCCTGTTTTGCTCACCCTCTTTTCC
AGAGCACACGTCTGAACTCTTTCCCTACACGACGCTCTTCCGATCTNNNGTGGCAGGAAGTGGAGAA
AGAGCACACGTCTGAACTCTTTCCCTACACGACGCTCTTCCGATCTNNNGTGGCAGGAAGTGGAGAA
GAGCACACGTCTGAACTCTTTCCCTACACGACGCTCTTCCGATCTNNNTCCGAATATGCTTGGCCTTGA
GAGCACACGTCTGAACTCTTTCCCTACACGACGCTCTTCCGATCTNNNTCCGAATATGCTTGGCCTTGA
AGCACACGTCTGAACTCTTTCCCTACACGACGCTCTTCCGATCTNNNCAGATTCCCAAGCAAGGGAGAA
AGCACACGTCTGAACTCTTTCCCTACACGACGCTCTTCCGATCTNNNCAGATTCCCAAGCAAGGGAGAA
AGCACACGTCTGAACTCTTTCCCTACACGACGCTCTTCCGATCTNNNCTTGGCTTCTAGTTTCCTTTCATC
AGCACACGTCTGAACTCTTTCCCTACACGACGCTCTTCCGATCTNNNCTTGGCTTCTAGTTTCCTTTCATC
GAGCACACGTCTGAACTCTTTCCCTACACGACGCTCTTCCGATCTNNNTGCGCCTGGCTAATTTTTGTAA
GAGCACACGTCTGAACTCTTTCCCTACACGACGCTCTTCCGATCTNNNTGCGCCTGGCTAATTTTTGTAA
GAGCACACGTCTGAACTCTTTCCCTACACGACGCTCTTCCGATCTNNNTTTTTTTCCTAGACGTGTGCCA
GAGCACACGTCTGAACTCTTTCCCTACACGACGCTCTTCCGATCTNNNTTTTTTTCCTAGACGTGTGCCA
AGCACACGTCTGAACTCTTTCCCTACACGACGCTCTTCCGATCTNNNGAAGTACAGGTGTGAACCACTC
AGCACACGTCTGAACTCTTTCCCTACACGACGCTCTTCCGATCTNNNGAAGTACAGGTGTGAACCACTC
GAGCACACGTCTGAACTCTTTCCCTACACGACGCTCTTCCGATCTNNNCCCAGAACTTTGAGAGGTTGA
GAGCACACGTCTGAACTCTTTCCCTACACGACGCTCTTCCGATCTNNNCCCAGAACTTTGAGAGGTTGA
GAGCACACGTCTGAACTCTTTCCCTACACGACGCTCTTCCGATCTNNNTGACCATACTGTCAAAGGCAA
GAGCACACGTCTGAACTCTTTCCCTACACGACGCTCTTCCGATCTNNNTGACCATACTGTCAAAGGCAA
GAGCACACGTCTGAACTCTTTCCCTACACGACGCTCTTCCGATCTNNNAGAGGTTGCAGTGAGTCAAG
GAGCACACGTCTGAACTCTTTCCCTACACGACGCTCTTCCGATCTNNNAGAGGTTGCAGTGAGTCAAG
GAGCACACGTCTGAACTCTTTCCCTACACGACGCTCTTCCGATCTNNNCACGATGATGTCATAAACAGC
GAGCACACGTCTGAACTCTTTCCCTACACGACGCTCTTCCGATCTNNNCACGATGATGTCATAAACAGC
AGAGCACACGTCTGAACTCTTTCCCTACACGACGCTCTTCCGATCTNNNTATCAGAGAAGCCCGATTCT
AGAGCACACGTCTGAACTCTTTCCCTACACGACGCTCTTCCGATCTNNNTATCAGAGAAGCCCGATTCT
GCACACGTCTGAACTCTTTCCCTACACGACGCTCTTCCGATCTNNNCCAGAACCTCTGGACTCTACACA
GCACACGTCTGAACTCTTTCCCTACACGACGCTCTTCCGATCTNNNCCAGAACCTCTGGACTCTACACA
AAGAGCACACGTCTGAACTCTTTCCCTACACGACGCTCTTCCGATCTNNNCCCGAGAGTTTTATGTTGC
AAGAGCACACGTCTGAACTCTTTCCCTACACGACGCTCTTCCGATCTNNNCCCGAGAGTTTTATGTTGC
GCACACGTCTGAACTCTTTCCCTACACGACGCTCTTCCGATCTNNNAAGTTCCTTTGTGCTTCCTGTGTC
GCACACGTCTGAACTCTTTCCCTACACGACGCTCTTCCGATCTNNNAAGTTCCTTTGTGCTTCCTGTGTC
AGCACACGTCTGAACTCTTTCCCTACACGACGCTCTTCCGATCTNNNATTTTCATGCTGCTCTTTTGGCA

GCACACGTCTGAACTCTTTCCCTACACGACGCTCTTCCGATCTNNNATTTCATGCTGCTCTTTTGGCA
GCACACGTCTGAACTCTTTCCCTACACGACGCTCTTCCGATCTNNNTCCTCAGTGACGTTCATCTTGTT
GCACACGTCTGAACTCTTTCCCTACACGACGCTCTTCCGATCTNNNTCCTCAGTGACGTTCATCTTGTT
AGCACACGTCTGAACTCTTTCCCTACACGACGCTCTTCCGATCTNNNGTCCTACTCATTGCTGGTGGA
AGCACACGTCTGAACTCTTTCCCTACACGACGCTCTTCCGATCTNNNGTCCTACTCATTGCTGGTGGA
CACACGTCTGAACTCTTTCCCTACACGACGCTCTTCCGATCTNNNTGGGTGACAGAGCGAGACTCCAT
CACACGTCTGAACTCTTTCCCTACACGACGCTCTTCCGATCTNNNTGGGTGACAGAGCGAGACTCCAT
GCACACGTCTGAACTCTTTCCCTACACGACGCTCTTCCGATCTNNNTCCATGTCTCTCTCTCTCTGTGTC
GCACACGTCTGAACTCTTTCCCTACACGACGCTCTTCCGATCTNNNTCCATGTCTCTCTCTCTCTGTGTC
AGAGCACACGTCTGAACTCTTTCCCTACACGACGCTCTTCCGATCTNNNTGGAGCAAGATGGCAGATA
AGAGCACACGTCTGAACTCTTTCCCTACACGACGCTCTTCCGATCTNNNTGGAGCAAGATGGCAGATA
GAGCACACGTCTGAACTCTTTCCCTACACGACGCTCTTCCGATCTNNNAAATTCCCACAGAAGGGAAT
GAGCACACGTCTGAACTCTTTCCCTACACGACGCTCTTCCGATCTNNNAAATTCCCACAGAAGGGAAT
GCACACGTCTGAACTCTTTCCCTACACGACGCTCTTCCGATCTNNNGCTACGTGCTCTCCTACCACCAT
GCACACGTCTGAACTCTTTCCCTACACGACGCTCTTCCGATCTNNNGCTACGTGCTCTCCTACCACCAT
AGCACACGTCTGAACTCTTTCCCTACACGACGCTCTTCCGATCTNNNCACATCAGGTGCGGTGAAGTC
AGCACACGTCTGAACTCTTTCCCTACACGACGCTCTTCCGATCTNNNCACATCAGGTGCGGTGAAGTC
GCACACGTCTGAACTCTTTCCCTACACGACGCTCTTCCGATCTNNNGGCTACACAAACTGCTATATGG
GCACACGTCTGAACTCTTTCCCTACACGACGCTCTTCCGATCTNNNGGCTACACAAACTGCTATATGG
GAGCACACGTCTGAACTCTTTCCCTACACGACGCTCTTCCGATCTNNNTGCAAGCCTAGTTAGGTGTTG
GAGCACACGTCTGAACTCTTTCCCTACACGACGCTCTTCCGATCTNNNTGCAAGCCTAGTTAGGTGTTG
AGCACACGTCTGAACTCTTTCCCTACACGACGCTCTTCCGATCTNNNCTGCCATGGTTGGTAACCCTAT
AGCACACGTCTGAACTCTTTCCCTACACGACGCTCTTCCGATCTNNNCTGCCATGGTTGGTAACCCTAT
AGCACACGTCTGAACTCTTTCCCTACACGACGCTCTTCCGATCTNNNTCTGTAATCCCAGCTACTCAGG
AGCACACGTCTGAACTCTTTCCCTACACGACGCTCTTCCGATCTNNNTCTGTAATCCCAGCTACTCAGG
AGCACACGTCTGAACTCTTTCCCTACACGACGCTCTTCCGATCTNNNACAAAGCAAAAACCCCATCTCTG
AGCACACGTCTGAACTCTTTCCCTACACGACGCTCTTCCGATCTNNNACAAAGCAAAAACCCCATCTCTG
AGCACACGTCTGAACTCTTTCCCTACACGACGCTCTTCCGATCTNNNGCCTTCTGACAACACTAATGCT
AGCACACGTCTGAACTCTTTCCCTACACGACGCTCTTCCGATCTNNNGCCTTCTGACAACACTAATGCT
GAGCACACGTCTGAACTCTTTCCCTACACGACGCTCTTCCGATCTNNNGGTTTGTATCCATTGCTGGT
GAGCACACGTCTGAACTCTTTCCCTACACGACGCTCTTCCGATCTNNNGGTTTGTATCCATTGCTGGT
AGCACACGTCTGAACTCTTTCCCTACACGACGCTCTTCCGATCTNNNTAGACGGGAGGTTTTATGATG
AGCACACGTCTGAACTCTTTCCCTACACGACGCTCTTCCGATCTNNNTAGACGGGAGGTTTTATGATG
GCACACGTCTGAACTCTTTCCCTACACGACGCTCTTCCGATCTNNNCGAAGGATTTGAAGCACCTTACC
GCACACGTCTGAACTCTTTCCCTACACGACGCTCTTCCGATCTNNNCGAAGGATTTGAAGCACCTTACC
AGCACACGTCTGAACTCTTTCCCTACACGACGCTCTTCCGATCTNNNCATACACACACACACACACACAA
AGCACACGTCTGAACTCTTTCCCTACACGACGCTCTTCCGATCTNNNCATACACACACACACACACACAA
GAGCACACGTCTGAACTCTTTCCCTACACGACGCTCTTCCGATCTNNNTAGCCCCTTTACCTTGATGCA
GAGCACACGTCTGAACTCTTTCCCTACACGACGCTCTTCCGATCTNNNTAGCCCCTTTACCTTGATGCA
GCACACGTCTGAACTCTTTCCCTACACGACGCTCTTCCGATCTNNNCAGTTACTAGGAAGGCTGAGGT
GCACACGTCTGAACTCTTTCCCTACACGACGCTCTTCCGATCTNNNCAGTTACTAGGAAGGCTGAGGT
AGCACACGTCTGAACTCTTTCCCTACACGACGCTCTTCCGATCTNNNGAGGCAGGAGAATTGTTTGA
AGCACACGTCTGAACTCTTTCCCTACACGACGCTCTTCCGATCTNNNGAGGCAGGAGAATTGTTTGA

GAGCACACGTCTGAACTCTTTCCCTACACGACGCTCTTCCGATCTNNNCTGTAATGCTAGCTACTCGGG
GAGCACACGTCTGAACTCTTTCCCTACACGACGCTCTTCCGATCTNNNCTGTAATGCTAGCTACTCGGG
AGAGCACACGTCTGAACTCTTTCCCTACACGACGCTCTTCCGATCTNNNAGCACTAGGTGTTACATTCT
AGAGCACACGTCTGAACTCTTTCCCTACACGACGCTCTTCCGATCTNNNAGCACTAGGTGTTACATTCT
AGCACACGTCTGAACTCTTTCCCTACACGACGCTCTTCCGATCTNNNGTCTGTGAAATCATACAGTATTT
AGCACACGTCTGAACTCTTTCCCTACACGACGCTCTTCCGATCTNNNGTCTGTGAAATCATACAGTATTT
GAGCACACGTCTGAACTCTTTCCCTACACGACGCTCTTCCGATCTNNNAAGGCCTTTAGGTGCTTACAT
GAGCACACGTCTGAACTCTTTCCCTACACGACGCTCTTCCGATCTNNNAAGGCCTTTAGGTGCTTACAT
GAGCACACGTCTGAACTCTTTCCCTACACGACGCTCTTCCGATCTNNNATGTGAAATATGGGAGGCTG
GAGCACACGTCTGAACTCTTTCCCTACACGACGCTCTTCCGATCTNNNATGTGAAATATGGGAGGCTG
GAGCACACGTCTGAACTCTTTCCCTACACGACGCTCTTCCGATCTNNNGCTGAGGCAGGATAATCTCT
GAGCACACGTCTGAACTCTTTCCCTACACGACGCTCTTCCGATCTNNNGCTGAGGCAGGATAATCTCT
GAGCACACGTCTGAACTCTTTCCCTACACGACGCTCTTCCGATCTNNNTTACCTCATTCTCCCACTTCAGC
GAGCACACGTCTGAACTCTTTCCCTACACGACGCTCTTCCGATCTNNNTTACCTCATTCTCCCACTTCAGC
GCACACGTCTGAACTCTTTCCCTACACGACGCTCTTCCGATCTNNNACTTTTCTTTCTCTCTTTCTTTCCAT
GCACACGTCTGAACTCTTTCCCTACACGACGCTCTTCCGATCTNNNACTTTTCTTTCTCTCTTTCTTTCCAT
CACACGTCTGAACTCTTTCCCTACACGACGCTCTTCCGATCTNNNCTATCTCTCAACATGTGCTGCATCG
CACACGTCTGAACTCTTTCCCTACACGACGCTCTTCCGATCTNNNCTATCTCTCAACATGTGCTGCATCG
GAGCACACGTCTGAACTCTTTCCCTACACGACGCTCTTCCGATCTNNNGCAGAAAGAATTAACAGACAT
GAGCACACGTCTGAACTCTTTCCCTACACGACGCTCTTCCGATCTNNNGCAGAAAGAATTAACAGACAT
AGCACACGTCTGAACTCTTTCCCTACACGACGCTCTTCCGATCTNNNGTCACCTGTATGGATACCAGCA
AGCACACGTCTGAACTCTTTCCCTACACGACGCTCTTCCGATCTNNNGTCACCTGTATGGATACCAGCA
AGAGCACACGTCTGAACTCTTTCCCTACACGACGCTCTTCCGATCTNNNCACCTGTAATCCCAGCTACTT
AGAGCACACGTCTGAACTCTTTCCCTACACGACGCTCTTCCGATCTNNNCACCTGTAATCCCAGCTACTT
CACGTCTGAACTCTTTCCCTACACGACGCTCTTCCGATCTNNNCCGTCTCCTCCCTTCTCACCATCGGAC
CACGTCTGAACTCTTTCCCTACACGACGCTCTTCCGATCTNNNCCGTCTCCTCCCTTCTCACCATCGGAC
AGCACACGTCTGAACTCTTTCCCTACACGACGCTCTTCCGATCTNNNCCATTGTACTCTAGCCTGGACA
AGCACACGTCTGAACTCTTTCCCTACACGACGCTCTTCCGATCTNNNCCATTGTACTCTAGCCTGGACA
AGCACACGTCTGAACTCTTTCCCTACACGACGCTCTTCCGATCTNNNACCAAGAAGGGTGCTGAGTTT
AGCACACGTCTGAACTCTTTCCCTACACGACGCTCTTCCGATCTNNNACCAAGAAGGGTGCTGAGTTT
CACACGTCTGAACTCTTTCCCTACACGACGCTCTTCCGATCTNNNACGCCCGGCTAATTTTTGTATTTCA
CACACGTCTGAACTCTTTCCCTACACGACGCTCTTCCGATCTNNNACGCCCGGCTAATTTTTGTATTTCA
GAGCACACGTCTGAACTCTTTCCCTACACGACGCTCTTCCGATCTNNNTCAAAGCCTTCGGTTTCAAAA
GAGCACACGTCTGAACTCTTTCCCTACACGACGCTCTTCCGATCTNNNTCAAAGCCTTCGGTTTCAAAA
AGCACACGTCTGAACTCTTTCCCTACACGACGCTCTTCCGATCTNNNCAGTGACCTGAGATCCTACCAA
AGCACACGTCTGAACTCTTTCCCTACACGACGCTCTTCCGATCTNNNCAGTGACCTGAGATCCTACCAA
GAGCACACGTCTGAACTCTTTCCCTACACGACGCTCTTCCGATCTNNNGGAGAAGGGAAATCAGCCAA
GAGCACACGTCTGAACTCTTTCCCTACACGACGCTCTTCCGATCTNNNGGAGAAGGGAAATCAGCCAA
GAGCACACGTCTGAACTCTTTCCCTACACGACGCTCTTCCGATCTNNNGGAGGTTGTAGTGAGCCAAG
GAGCACACGTCTGAACTCTTTCCCTACACGACGCTCTTCCGATCTNNNGGAGGTTGTAGTGAGCCAAG
AGAGCACACGTCTGAACTCTTTCCCTACACGACGCTCTTCCGATCTNNNTCCAGGCCCTTTTGTCTGTCC
AGAGCACACGTCTGAACTCTTTCCCTACACGACGCTCTTCCGATCTNNNTCCAGGCCCTTTTGTCTGTCC
GCACACGTCTGAACTCTTTCCCTACACGACGCTCTTCCGATCTNNNCTCTCCAGGTGGCCACAGTAGCA

GCACACGTCTGAACTCTTTCCCTACACGACGCTCTTCCGATCTNNNCTCTCCAGGTGGCCACAGTAGCAT
GCACACGTCTGAACTCTTTCCCTACACGACGCTCTTCCGATCTNNNAGTCAAATGGAGGCATCTAGGTG
GCACACGTCTGAACTCTTTCCCTACACGACGCTCTTCCGATCTNNNAGTCAAATGGAGGCATCTAGGTG
AGCACACGTCTGAACTCTTTCCCTACACGACGCTCTTCCGATCTNNNTTCTCTGAAGGTTGGACTGTGA
AGCACACGTCTGAACTCTTTCCCTACACGACGCTCTTCCGATCTNNNTTCTCTGAAGGTTGGACTGTGA
GAGCACACGTCTGAACTCTTTCCCTACACGACGCTCTTCCGATCTNNNCCAATGACTACAACACCTCTCC
GAGCACACGTCTGAACTCTTTCCCTACACGACGCTCTTCCGATCTNNNCCAATGACTACAACACCTCTCC
GAGCACACGTCTGAACTCTTTCCCTACACGACGCTCTTCCGATCTNNNAGTTCTGTGCTTGCAAAAGTG
GAGCACACGTCTGAACTCTTTCCCTACACGACGCTCTTCCGATCTNNNAGTTCTGTGCTTGCAAAAGTG
GCACACGTCTGAACTCTTTCCCTACACGACGCTCTTCCGATCTNNNATTTCATCCTAGGGGTGCAGCAT
GCACACGTCTGAACTCTTTCCCTACACGACGCTCTTCCGATCTNNNATTTCATCCTAGGGGTGCAGCAT
GAGCACACGTCTGAACTCTTTCCCTACACGACGCTCTTCCGATCTNNNTACAAAATTAGCTGGGCATGG
GAGCACACGTCTGAACTCTTTCCCTACACGACGCTCTTCCGATCTNNNTACAAAATTAGCTGGGCATGG
GCACACGTCTGAACTCTTTCCCTACACGACGCTCTTCCGATCTNNNATTATGCATGAGCTTCCAACCAGC
GCACACGTCTGAACTCTTTCCCTACACGACGCTCTTCCGATCTNNNATTATGCATGAGCTTCCAACCAGC
GCACACGTCTGAACTCTTTCCCTACACGACGCTCTTCCGATCTNNNCCTCTCTTTCTCTCCATTGCCTTCAT
GCACACGTCTGAACTCTTTCCCTACACGACGCTCTTCCGATCTNNNCCTCTCTTTCTCTCCATTGCCTTCAT
AGCACACGTCTGAACTCTTTCCCTACACGACGCTCTTCCGATCTNNNCCTGGCAAATTGGGATTGAAA
AGCACACGTCTGAACTCTTTCCCTACACGACGCTCTTCCGATCTNNNCCTGGCAAATTGGGATTGAAA
AGAGCACACGTCTGAACTCTTTCCCTACACGACGCTCTTCCGATCTNNNCACGTGTAGTCCCAGCTACTC
AGAGCACACGTCTGAACTCTTTCCCTACACGACGCTCTTCCGATCTNNNCACGTGTAGTCCCAGCTACTC
GCACACGTCTGAACTCTTTCCCTACACGACGCTCTTCCGATCTNNNATTCTGTACCCCCACAACTTACCCA
GCACACGTCTGAACTCTTTCCCTACACGACGCTCTTCCGATCTNNNATTCTGTACCCCCACAACTTACCCA
AGCACACGTCTGAACTCTTTCCCTACACGACGCTCTTCCGATCTNNNTCAGACATCAATGCAAAGGGAG
AGCACACGTCTGAACTCTTTCCCTACACGACGCTCTTCCGATCTNNNTCAGACATCAATGCAAAGGGAG
GAGCACACGTCTGAACTCTTTCCCTACACGACGCTCTTCCGATCTNNNGGAGTGAATGCCTTATAAGGC
GAGCACACGTCTGAACTCTTTCCCTACACGACGCTCTTCCGATCTNNNGGAGTGAATGCCTTATAAGGC
GAGCACACGTCTGAACTCTTTCCCTACACGACGCTCTTCCGATCTNNNGCAGGAGAATTGCTTGAACCT
GAGCACACGTCTGAACTCTTTCCCTACACGACGCTCTTCCGATCTNNNGCAGGAGAATTGCTTGAACCT
AGCACACGTCTGAACTCTTTCCCTACACGACGCTCTTCCGATCTNNNTTGGAACTTCTTGAAGCCTCCTT
AGCACACGTCTGAACTCTTTCCCTACACGACGCTCTTCCGATCTNNNTTGGAACTTCTTGAAGCCTCCTT
AGAGCACACGTCTGAACTCTTTCCCTACACGACGCTCTTCCGATCTNNNTGGCATTTGGAATTGCACTTC
AGAGCACACGTCTGAACTCTTTCCCTACACGACGCTCTTCCGATCTNNNTGGCATTTGGAATTGCACTTC
AGAGCACACGTCTGAACTCTTTCCCTACACGACGCTCTTCCGATCTNNNCTCAACGAAAGCCTCAAAAA
AGAGCACACGTCTGAACTCTTTCCCTACACGACGCTCTTCCGATCTNNNCTCAACGAAAGCCTCAAAAA
AGCACACGTCTGAACTCTTTCCCTACACGACGCTCTTCCGATCTNNNTGTGTGTGTGTGTGGAGAT
AGCACACGTCTGAACTCTTTCCCTACACGACGCTCTTCCGATCTNNNTGTGTGTGTGTGTGGAGAT
AGCACACGTCTGAACTCTTTCCCTACACGACGCTCTTCCGATCTNNNTCACTTGAGTCCAGGAGTTCAA
AGCACACGTCTGAACTCTTTCCCTACACGACGCTCTTCCGATCTNNNTCACTTGAGTCCAGGAGTTCAA
AGAGCACACGTCTGAACTCTTTCCCTACACGACGCTCTTCCGATCTNNNATGGTCTTGATCTCTTGACCT
AGAGCACACGTCTGAACTCTTTCCCTACACGACGCTCTTCCGATCTNNNATGGTCTTGATCTCTTGACCT
GAGCACACGTCTGAACTCTTTCCCTACACGACGCTCTTCCGATCTNNNTTCATCAAACAGCCAGGAGTG
GAGCACACGTCTGAACTCTTTCCCTACACGACGCTCTTCCGATCTNNNTTCATCAAACAGCCAGGAGTG

CACACGTCTGAACTCTTTCCCTACACGACGCTCTTCCGATCTNNNATGGCCCGTCCTGAATAACATAAC
CACACGTCTGAACTCTTTCCCTACACGACGCTCTTCCGATCTNNNATGGCCCGTCCTGAATAACATAAC
GAGCACACGTCTGAACTCTTTCCCTACACGACGCTCTTCCGATCTNNNTGATACCCATTGAAGGATGCA
GAGCACACGTCTGAACTCTTTCCCTACACGACGCTCTTCCGATCTNNNTGATACCCATTGAAGGATGCA
AGCACACGTCTGAACTCTTTCCCTACACGACGCTCTTCCGATCTNNNCCCATAGAACATGGAATTGGG
AGCACACGTCTGAACTCTTTCCCTACACGACGCTCTTCCGATCTNNNCCCATAGAACATGGAATTGGG
AGCACACGTCTGAACTCTTTCCCTACACGACGCTCTTCCGATCTNNNCACGCCCAGCTAACTTTTTGTAT
AGCACACGTCTGAACTCTTTCCCTACACGACGCTCTTCCGATCTNNNCACGCCCAGCTAACTTTTTGTAT
CACACGTCTGAACTCTTTCCCTACACGACGCTCTTCCGATCTNNNTTTTTTGTTTTAAGATAGCTTGCCA
CACACGTCTGAACTCTTTCCCTACACGACGCTCTTCCGATCTNNNTTTTTTGTTTTAAGATAGCTTGCCA
GCACACGTCTGAACTCTTTCCCTACACGACGCTCTTCCGATCTNNNTCCTGTTTTGTCTTACGTCTTGAC
GCACACGTCTGAACTCTTTCCCTACACGACGCTCTTCCGATCTNNNTCCTGTTTTGTCTTACGTCTTGAC
GCACACGTCTGAACTCTTTCCCTACACGACGCTCTTCCGATCTNNNGAGAGAGAGAGACAGGAGAGA
GCACACGTCTGAACTCTTTCCCTACACGACGCTCTTCCGATCTNNNGAGAGAGAGAGACAGGAGAGA
AGCACACGTCTGAACTCTTTCCCTACACGACGCTCTTCCGATCTNNNGAGGCATGAGAATCGCTTGAA
AGCACACGTCTGAACTCTTTCCCTACACGACGCTCTTCCGATCTNNNGAGGCATGAGAATCGCTTGAA
GAGCACACGTCTGAACTCTTTCCCTACACGACGCTCTTCCGATCTNNNAGCCTGGGAGACATAGTTAGA
GAGCACACGTCTGAACTCTTTCCCTACACGACGCTCTTCCGATCTNNNAGCCTGGGAGACATAGTTAGA
AGCACACGTCTGAACTCTTTCCCTACACGACGCTCTTCCGATCTNNNATTGGCAACAAACCATATGTGC
AGCACACGTCTGAACTCTTTCCCTACACGACGCTCTTCCGATCTNNNATTGGCAACAAACCATATGTGC
GAGCACACGTCTGAACTCTTTCCCTACACGACGCTCTTCCGATCTNNNATTTGCAACCCAGTAAAAGCAC
GAGCACACGTCTGAACTCTTTCCCTACACGACGCTCTTCCGATCTNNNATTTGCAACCCAGTAAAAGCAC
AGCACACGTCTGAACTCTTTCCCTACACGACGCTCTTCCGATCTNNNTACTTCTGGGAGAAGCGCTTGA
AGCACACGTCTGAACTCTTTCCCTACACGACGCTCTTCCGATCTNNNTACTTCTGGGAGAAGCGCTTGA
AGCACACGTCTGAACTCTTTCCCTACACGACGCTCTTCCGATCTNNNGGAGACTGAGATGGGAGAATC
AGCACACGTCTGAACTCTTTCCCTACACGACGCTCTTCCGATCTNNNGGAGACTGAGATGGGAGAATC
AGAGCACACGTCTGAACTCTTTCCCTACACGACGCTCTTCCGATCTNNNGGAGAATCACTTGAACCTGC
AGAGCACACGTCTGAACTCTTTCCCTACACGACGCTCTTCCGATCTNNNGGAGAATCACTTGAACCTGC
GAGCACACGTCTGAACTCTTTCCCTACACGACGCTCTTCCGATCTNNNCAGTTTCCATGTAGACCCCTTG
GAGCACACGTCTGAACTCTTTCCCTACACGACGCTCTTCCGATCTNNNCAGTTTCCATGTAGACCCCTTG
GAGCACACGTCTGAACTCTTTCCCTACACGACGCTCTTCCGATCTNNNATCCTGTTCTCTCTCAGCCAAAC
GAGCACACGTCTGAACTCTTTCCCTACACGACGCTCTTCCGATCTNNNATCCTGTTCTCTCTCAGCCAAAC
GAGCACACGTCTGAACTCTTTCCCTACACGACGCTCTTCCGATCTNNNTAAGGTGGAAGGATTGCTTGA
GAGCACACGTCTGAACTCTTTCCCTACACGACGCTCTTCCGATCTNNNTAAGGTGGAAGGATTGCTTGA
AGAGCACACGTCTGAACTCTTTCCCTACACGACGCTCTTCCGATCTNNNTGGAGGATGCAATGAGCTAT
AGAGCACACGTCTGAACTCTTTCCCTACACGACGCTCTTCCGATCTNNNTGGAGGATGCAATGAGCTAT
GAGCACACGTCTGAACTCTTTCCCTACACGACGCTCTTCCGATCTNNNTTCAGGAATGTTGTCCATCCCA
GAGCACACGTCTGAACTCTTTCCCTACACGACGCTCTTCCGATCTNNNTTCAGGAATGTTGTCCATCCCA
AGCACACGTCTGAACTCTTTCCCTACACGACGCTCTTCCGATCTNNNCTTTCATGGGCTGGTGTTTAGTC
AGCACACGTCTGAACTCTTTCCCTACACGACGCTCTTCCGATCTNNNCTTTCATGGGCTGGTGTTTAGTC
ACACGTCTGAACTCTTTCCCTACACGACGCTCTTCCGATCTNNNCACATCTGACCCAAATGCCATTTCAT
ACACGTCTGAACTCTTTCCCTACACGACGCTCTTCCGATCTNNNCACATCTGACCCAAATGCCATTTCAT
GCACACGTCTGAACTCTTTCCCTACACGACGCTCTTCCGATCTNNNGTTATGTATCAAGTACCTTTGCCA

GCACACGTCTGAACTCTTTCCCTACACGACGCTCTTCCGATCTNNNGTTATGTATCAAGTACCTTTGCCA

AGAGCACACGTCTGAACTCTTTCCCTACACGACGCTCTTCCGATCTNNNAAGATGAGAGAGAGGCAGG

AGAGCACACGTCTGAACTCTTTCCCTACACGACGCTCTTCCGATCTNNNAAGATGAGAGAGAGGCAGG

GAGCACACGTCTGAACTCTTTCCCTACACGACGCTCTTCCGATCTNNNTGCTCAGATCATTGGAACACTC

GAGCACACGTCTGAACTCTTTCCCTACACGACGCTCTTCCGATCTNNNTGCTCAGATCATTGGAACACTC

GAGCACACGTCTGAACTCTTTCCCTACACGACGCTCTTCCGATCTNNNATCTATCCATCTGACAAAGGG

GAGCACACGTCTGAACTCTTTCCCTACACGACGCTCTTCCGATCTNNNATCTATCCATCTGACAAAGGG

GAGCACACGTCTGAACTCTTTCCCTACACGACGCTCTTCCGATCTNNNACTCCCACCTTACCCCTTCTAGC

GAGCACACGTCTGAACTCTTTCCCTACACGACGCTCTTCCGATCTNNNACTCCCACCTTACCCCTTCTAGC

AGAGCACACGTCTGAACTCTTTCCCTACACGACGCTCTTCCGATCTNNNAGAGCTGTGGTTCTGTCTAA

AGAGCACACGTCTGAACTCTTTCCCTACACGACGCTCTTCCGATCTNNNAGAGCTGTGGTTCTGTCTAA

AGCACACGTCTGAACTCTTTCCCTACACGACGCTCTTCCGATCTNNNCAGGCTGGTATGCTCAGTGATA

AGCACACGTCTGAACTCTTTCCCTACACGACGCTCTTCCGATCTNNNCAGGCTGGTATGCTCAGTGATA

AGCACACGTCTGAACTCTTTCCCTACACGACGCTCTTCCGATCTNNNTGAGGCAGGAGAATCACTTGAA

AGCACACGTCTGAACTCTTTCCCTACACGACGCTCTTCCGATCTNNNTGAGGCAGGAGAATCACTTGAA

AGCACACGTCTGAACTCTTTCCCTACACGACGCTCTTCCGATCTNNNTGTCAAGTCTTCTCTTGCCATGA

AGCACACGTCTGAACTCTTTCCCTACACGACGCTCTTCCGATCTNNNTGTCAAGTCTTCTCTTGCCATGA

GAGCACACGTCTGAACTCTTTCCCTACACGACGCTCTTCCGATCTNNNTTCACAACCAATGGTGAATGA

GAGCACACGTCTGAACTCTTTCCCTACACGACGCTCTTCCGATCTNNNTTCACAACCAATGGTGAATGA

GCACACGTCTGAACTCTTTCCCTACACGACGCTCTTCCGATCTNNNAGCTTACCTGTCCTCTCATTTGGCA

GCACACGTCTGAACTCTTTCCCTACACGACGCTCTTCCGATCTNNNAGCTTACCTGTCCTCTCATTTGGCA

GAGCACACGTCTGAACTCTTTCCCTACACGACGCTCTTCCGATCTNNNAGTGTAAAATCCTACATCCTTG

GAGCACACGTCTGAACTCTTTCCCTACACGACGCTCTTCCGATCTNNNAGTGTAAAATCCTACATCCTTG

GAGCACACGTCTGAACTCTTTCCCTACACGACGCTCTTCCGATCTNNNAGGGAATCTGTCTCTCACTCTC

GAGCACACGTCTGAACTCTTTCCCTACACGACGCTCTTCCGATCTNNNAGGGAATCTGTCTCTCACTCTC

GAGCACACGTCTGAACTCTTTCCCTACACGACGCTCTTCCGATCTNNNCACTGCTCTCGTCTTCAACAAA

GAGCACACGTCTGAACTCTTTCCCTACACGACGCTCTTCCGATCTNNNCACTGCTCTCGTCTTCAACAAA

GAGCACACGTCTGAACTCTTTCCCTACACGACGCTCTTCCGATCTNNNAATGAGCAGAGATCCAGCCA

GAGCACACGTCTGAACTCTTTCCCTACACGACGCTCTTCCGATCTNNNAATGAGCAGAGATCCAGCCA

AGAGCACACGTCTGAACTCTTTCCCTACACGACGCTCTTCCGATCTNNNACAGAATGAGGCCTTGTCTG

AGAGCACACGTCTGAACTCTTTCCCTACACGACGCTCTTCCGATCTNNNACAGAATGAGGCCTTGTCTG

GAGCACACGTCTGAACTCTTTCCCTACACGACGCTCTTCCGATCTNNNGACTACAGACACACAACACACA

GAGCACACGTCTGAACTCTTTCCCTACACGACGCTCTTCCGATCTNNNGACTACAGACACACAACACACA

GCACACGTCTGAACTCTTTCCCTACACGACGCTCTTCCGATCTNNNTAGTAGAGATGGGGTTTGCCAT

GCACACGTCTGAACTCTTTCCCTACACGACGCTCTTCCGATCTNNNTAGTAGAGATGGGGTTTGCCAT

AGAGCACACGTCTGAACTCTTTCCCTACACGACGCTCTTCCGATCTNNNTGGCTCTCCCAGAGTCTACAT

AGAGCACACGTCTGAACTCTTTCCCTACACGACGCTCTTCCGATCTNNNTGGCTCTCCCAGAGTCTACAT

AGCACACGTCTGAACTCTTTCCCTACACGACGCTCTTCCGATCTNNNCAAGAACAAAACCCCGTCTCAAA

AGCACACGTCTGAACTCTTTCCCTACACGACGCTCTTCCGATCTNNNCAAGAACAAAACCCCGTCTCAAA

ACACGTCTGAACTCTTTCCCTACACGACGCTCTTCCGATCTNNNTAGATGCCGGAATTTCCCTTGAACAT

ACACGTCTGAACTCTTTCCCTACACGACGCTCTTCCGATCTNNNTAGATGCCGGAATTTCCCTTGAACAT

GAGCACACGTCTGAACTCTTTCCCTACACGACGCTCTTCCGATCTNNNGAGGCTGAGGCAGGAGAATG

GAGCACACGTCTGAACTCTTTCCCTACACGACGCTCTTCCGATCTNNNGAGGCTGAGGCAGGAGAATG

AGCACACGTCTGAACTCTTTCCCTACACGACGCTCTTCCGATCTNNNGATTCACGCCATTCTTCTGCCC
AGCACACGTCTGAACTCTTTCCCTACACGACGCTCTTCCGATCTNNNGATTCACGCCATTCTTCTGCCC
AGCACACGTCTGAACTCTTTCCCTACACGACGCTCTTCCGATCTNNNTGGTACATGCCTATACTTCCAG
AGCACACGTCTGAACTCTTTCCCTACACGACGCTCTTCCGATCTNNNTGGTACATGCCTATACTTCCAG
AGCACACGTCTGAACTCTTTCCCTACACGACGCTCTTCCGATCTNNNCTTGTCAATACCGCAAACTCCA
AGCACACGTCTGAACTCTTTCCCTACACGACGCTCTTCCGATCTNNNCTTGTCAATACCGCAAACTCCA
AGCACACGTCTGAACTCTTTCCCTACACGACGCTCTTCCGATCTNNNCCTCCTCTTTTCTGGGAATGTGT
AGCACACGTCTGAACTCTTTCCCTACACGACGCTCTTCCGATCTNNNCCTCCTCTTTTCTGGGAATGTGT
GAGCACACGTCTGAACTCTTTCCCTACACGACGCTCTTCCGATCTNNNGGACTCAAAAGCATGCAAAC
GAGCACACGTCTGAACTCTTTCCCTACACGACGCTCTTCCGATCTNNNGGACTCAAAAGCATGCAAAC
AGCACACGTCTGAACTCTTTCCCTACACGACGCTCTTCCGATCTNNNAAGCAGCATCTCTTATTGCAGT
AGCACACGTCTGAACTCTTTCCCTACACGACGCTCTTCCGATCTNNNAAGCAGCATCTCTTATTGCAGT
AGCACACGTCTGAACTCTTTCCCTACACGACGCTCTTCCGATCTNNNTGGGAGAATCACTTGAACTCA
AGCACACGTCTGAACTCTTTCCCTACACGACGCTCTTCCGATCTNNNTGGGAGAATCACTTGAACTCA
GCACACGTCTGAACTCTTTCCCTACACGACGCTCTTCCGATCTNNNTCACTGTGGATGGAGAATGATG
GCACACGTCTGAACTCTTTCCCTACACGACGCTCTTCCGATCTNNNTCACTGTGGATGGAGAATGATG
AGCACACGTCTGAACTCTTTCCCTACACGACGCTCTTCCGATCTNNNTTTATACAGGGCTGAGGCTTTG
AGCACACGTCTGAACTCTTTCCCTACACGACGCTCTTCCGATCTNNNTTTATACAGGGCTGAGGCTTTG
AGCACACGTCTGAACTCTTTCCCTACACGACGCTCTTCCGATCTNNNCCCAAATGATGAAGTGATGCA
AGCACACGTCTGAACTCTTTCCCTACACGACGCTCTTCCGATCTNNNCCCAAATGATGAAGTGATGCA
CACGTCTGAACTCTTTCCCTACACGACGCTCTTCCGATCTNNNTGCAAGTTCACCAGCTTATGTTGCAT
CACGTCTGAACTCTTTCCCTACACGACGCTCTTCCGATCTNNNTGCAAGTTCACCAGCTTATGTTGCAT
AGCACACGTCTGAACTCTTTCCCTACACGACGCTCTTCCGATCTNNNATGAGGTCGATAAAAACCCCT
AGCACACGTCTGAACTCTTTCCCTACACGACGCTCTTCCGATCTNNNATGAGGTCGATAAAAACCCCT
ACACGTCTGAACTCTTTCCCTACACGACGCTCTTCCGATCTNNNCATAAAGGGTGTGATCTAGAGCAT
ACACGTCTGAACTCTTTCCCTACACGACGCTCTTCCGATCTNNNCATAAAGGGTGTGATCTAGAGCAT
AGCACACGTCTGAACTCTTTCCCTACACGACGCTCTTCCGATCTNNNGAGAGGCTGAGGCACAAGAA
AGCACACGTCTGAACTCTTTCCCTACACGACGCTCTTCCGATCTNNNGAGAGGCTGAGGCACAAGAA
GAGCACACGTCTGAACTCTTTCCCTACACGACGCTCTTCCGATCTNNNTTGGGAGTGCCATTAGTTCA
GAGCACACGTCTGAACTCTTTCCCTACACGACGCTCTTCCGATCTNNNTTGGGAGTGCCATTAGTTCA
GAGCACACGTCTGAACTCTTTCCCTACACGACGCTCTTCCGATCTNNNTCCCATTCTTCAAAGCTGACT
GAGCACACGTCTGAACTCTTTCCCTACACGACGCTCTTCCGATCTNNNTCCCATTCTTCAAAGCTGACT
GCACACGTCTGAACTCTTTCCCTACACGACGCTCTTCCGATCTNNNCTGTAAAAGGGCCTATGGGAG
GCACACGTCTGAACTCTTTCCCTACACGACGCTCTTCCGATCTNNNCTGTAAAAGGGCCTATGGGAG
GCACACGTCTGAACTCTTTCCCTACACGACGCTCTTCCGATCTNNNTTGTCTTTGTGGGTTGCCTTTTT
GCACACGTCTGAACTCTTTCCCTACACGACGCTCTTCCGATCTNNNTTGTCTTTGTGGGTTGCCTTTTT
GCACACGTCTGAACTCTTTCCCTACACGACGCTCTTCCGATCTNNNCCATCTAGCCTAGCCCCCATTTAC
GCACACGTCTGAACTCTTTCCCTACACGACGCTCTTCCGATCTNNNCCATCTAGCCTAGCCCCCATTTAC
GCACACGTCTGAACTCTTTCCCTACACGACGCTCTTCCGATCTNNNAGTAGCTAGATGTATGAGACCA
GCACACGTCTGAACTCTTTCCCTACACGACGCTCTTCCGATCTNNNAGTAGCTAGATGTATGAGACCA
GAGCACACGTCTGAACTCTTTCCCTACACGACGCTCTTCCGATCTNNNTGCTTGAATCCATGACATCCT
GAGCACACGTCTGAACTCTTTCCCTACACGACGCTCTTCCGATCTNNNTGCTTGAATCCATGACATCCT
GCACACGTCTGAACTCTTTCCCTACACGACGCTCTTCCGATCTNNNCTCTCGCTCCTCCTCCTCAGCATCG

GCACACGTCTGAACTCTTTCCCTACACGACGCTCTTCCGATCTNNNCTCTCGCTCCTCCTCCTCAGCATCG
GAGCACACGTCTGAACTCTTTCCCTACACGACGCTCTTCCGATCTNNNATGCCTGTAATCCCAGCATTTT
GAGCACACGTCTGAACTCTTTCCCTACACGACGCTCTTCCGATCTNNNATGCCTGTAATCCCAGCATTTT
GGAAGAGCACACGTCTGAACTCTTTCCCTACACGACGCTCTTCCGATCTNNNAAGAGAATTGCTTGAA
GGAAGAGCACACGTCTGAACTCTTTCCCTACACGACGCTCTTCCGATCTNNNAAGAGAATTGCTTGAA
AGCACACGTCTGAACTCTTTCCCTACACGACGCTCTTCCGATCTNNNCACTGCTTTCACTAACTGTTGG
AGCACACGTCTGAACTCTTTCCCTACACGACGCTCTTCCGATCTNNNCACTGCTTTCACTAACTGTTGG
GAGCACACGTCTGAACTCTTTCCCTACACGACGCTCTTCCGATCTNNNAAAAAAATTAGCCAGGCGTGG
GAGCACACGTCTGAACTCTTTCCCTACACGACGCTCTTCCGATCTNNNAAAAAAATTAGCCAGGCGTGG
AGCACACGTCTGAACTCTTTCCCTACACGACGCTCTTCCGATCTNNNGCTACCACAATTCAACACAGCA
AGCACACGTCTGAACTCTTTCCCTACACGACGCTCTTCCGATCTNNNGCTACCACAATTCAACACAGCA
GAGCACACGTCTGAACTCTTTCCCTACACGACGCTCTTCCGATCTNNNAAGGAAAAGCCAGTACAGCAT
GAGCACACGTCTGAACTCTTTCCCTACACGACGCTCTTCCGATCTNNNAAGGAAAAGCCAGTACAGCAT
AGCACACGTCTGAACTCTTTCCCTACACGACGCTCTTCCGATCTNNNTTCCAGAAACTATTATCAGCAT
AGCACACGTCTGAACTCTTTCCCTACACGACGCTCTTCCGATCTNNNTTCCAGAAACTATTATCAGCAT
GAGCACACGTCTGAACTCTTTCCCTACACGACGCTCTTCCGATCTNNNTGTAATCCCAGCTACTTGGAA
GAGCACACGTCTGAACTCTTTCCCTACACGACGCTCTTCCGATCTNNNTGTAATCCCAGCTACTTGGAA
AGCACACGTCTGAACTCTTTCCCTACACGACGCTCTTCCGATCTNNNGGGAGTTGAACAATGAGAACA
AGCACACGTCTGAACTCTTTCCCTACACGACGCTCTTCCGATCTNNNGGGAGTTGAACAATGAGAACA
GAGCACACGTCTGAACTCTTTCCCTACACGACGCTCTTCCGATCTNNNGCCTGGGTGATGAAACCATC
GAGCACACGTCTGAACTCTTTCCCTACACGACGCTCTTCCGATCTNNNGCCTGGGTGATGAAACCATC
GAGCACACGTCTGAACTCTTTCCCTACACGACGCTCTTCCGATCTNNNTACAAAACTTAGCTGGGCTTG
GAGCACACGTCTGAACTCTTTCCCTACACGACGCTCTTCCGATCTNNNTACAAAACTTAGCTGGGCTTG
AGCACACGTCTGAACTCTTTCCCTACACGACGCTCTTCCGATCTNNNGCAAGGGATCCAATGATGTGA
AGCACACGTCTGAACTCTTTCCCTACACGACGCTCTTCCGATCTNNNGCAAGGGATCCAATGATGTGA
GAGCACACGTCTGAACTCTTTCCCTACACGACGCTCTTCCGATCTNNNTTATTTGCAGAACCATTCAGCG
GAGCACACGTCTGAACTCTTTCCCTACACGACGCTCTTCCGATCTNNNTTATTTGCAGAACCATTCAGCG
AGCACACGTCTGAACTCTTTCCCTACACGACGCTCTTCCGATCTNNNTAGAAAGGAGACAGCCGAATT
AGCACACGTCTGAACTCTTTCCCTACACGACGCTCTTCCGATCTNNNTAGAAAGGAGACAGCCGAATT
GCACACGTCTGAACTCTTTCCCTACACGACGCTCTTCCGATCTNNNTATCCGTGTAGACTTGGGAGGAT
GCACACGTCTGAACTCTTTCCCTACACGACGCTCTTCCGATCTNNNTATCCGTGTAGACTTGGGAGGAT
ACACGTCTGAACTCTTTCCCTACACGACGCTCTTCCGATCTNNNTCTCAATGAGCTGTTGGGTACACCA
ACACGTCTGAACTCTTTCCCTACACGACGCTCTTCCGATCTNNNTCTCAATGAGCTGTTGGGTACACCA
AGCACACGTCTGAACTCTTTCCCTACACGACGCTCTTCCGATCTNNNAAAATTAGCTGGGTGTAGTGG
AGCACACGTCTGAACTCTTTCCCTACACGACGCTCTTCCGATCTNNNAAAATTAGCTGGGTGTAGTGG
AGAGCACACGTCTGAACTCTTTCCCTACACGACGCTCTTCCGATCTNNNAGTGGGGGAAGGAGATGTT
AGAGCACACGTCTGAACTCTTTCCCTACACGACGCTCTTCCGATCTNNNAGTGGGGGAAGGAGATGTT
AGCACACGTCTGAACTCTTTCCCTACACGACGCTCTTCCGATCTNNNTAGTGATTCAGATAGTAGTTGT
AGCACACGTCTGAACTCTTTCCCTACACGACGCTCTTCCGATCTNNNTAGTGATTCAGATAGTAGTTGT
GAGCACACGTCTGAACTCTTTCCCTACACGACGCTCTTCCGATCTNNNCGTTTCCTTTAACCACATAATT
GAGCACACGTCTGAACTCTTTCCCTACACGACGCTCTTCCGATCTNNNCGTTTCCTTTAACCACATAATT
GAGCACACGTCTGAACTCTTTCCCTACACGACGCTCTTCCGATCTNNNGAGCCAGTCTCTCTTTTCTCCT
GAGCACACGTCTGAACTCTTTCCCTACACGACGCTCTTCCGATCTNNNGAGCCAGTCTCTCTTTTCTCCT

GCACACGTCTGAACTCTTTCCCTACACGACGCTCTTCCGATCTNNNGGAATCTGCCCTGTGTCGTCCAT

GCACACGTCTGAACTCTTTCCCTACACGACGCTCTTCCGATCTNNNGGAATCTGCCCTGTGTCGTCCAT

GAGCACACGTCTGAACTCTTTCCCTACACGACGCTCTTCCGATCTNNNCTTCAGTTCCTAAGGTGCCTC

GAGCACACGTCTGAACTCTTTCCCTACACGACGCTCTTCCGATCTNNNCTTCAGTTCCTAAGGTGCCTC

GCACACGTCTGAACTCTTTCCCTACACGACGCTCTTCCGATCTNNNTGGGACTGACTTGTGGTAATCT

GCACACGTCTGAACTCTTTCCCTACACGACGCTCTTCCGATCTNNNTGGGACTGACTTGTGGTAATCT

AGCACACGTCTGAACTCTTTCCCTACACGACGCTCTTCCGATCTNNNGATCTGAGACAGGAGAATCGC

AGCACACGTCTGAACTCTTTCCCTACACGACGCTCTTCCGATCTNNNGATCTGAGACAGGAGAATCGC

AGCACACGTCTGAACTCTTTCCCTACACGACGCTCTTCCGATCTNNNCTCACCTTGAAACCTGGCAGA

AGCACACGTCTGAACTCTTTCCCTACACGACGCTCTTCCGATCTNNNCTCACCTTGAAACCTGGCAGA

AGCACACGTCTGAACTCTTTCCCTACACGACGCTCTTCCGATCTNNNCTCCTGAGTAGCTGGGACTACA

AGCACACGTCTGAACTCTTTCCCTACACGACGCTCTTCCGATCTNNNCTCCTGAGTAGCTGGGACTACA

GAGCACACGTCTGAACTCTTTCCCTACACGACGCTCTTCCGATCTNNNATGAGCCTTAGGACAATGCT

GAGCACACGTCTGAACTCTTTCCCTACACGACGCTCTTCCGATCTNNNATGAGCCTTAGGACAATGCT

AGCACACGTCTGAACTCTTTCCCTACACGACGCTCTTCCGATCTNNNTCATAAAGTTGTGATCTAAATG

AGCACACGTCTGAACTCTTTCCCTACACGACGCTCTTCCGATCTNNNTCATAAAGTTGTGATCTAAATG

AGCACACGTCTGAACTCTTTCCCTACACGACGCTCTTCCGATCTNNNCATACACCCATGTGCTCTCTCC

AGCACACGTCTGAACTCTTTCCCTACACGACGCTCTTCCGATCTNNNCATACACCCATGTGCTCTCTCC

ACACGTCTGAACTCTTTCCCTACACGACGCTCTTCCGATCTNNNTTCCTCTGGAGCCCACTCTCCATCGG

ACACGTCTGAACTCTTTCCCTACACGACGCTCTTCCGATCTNNNTTCCTCTGGAGCCCACTCTCCATCGG

ACACGTCTGAACTCTTTCCCTACACGACGCTCTTCCGATCTNNNAGTGAGATCAAGAGGAAGACCATC

ACACGTCTGAACTCTTTCCCTACACGACGCTCTTCCGATCTNNNAGTGAGATCAAGAGGAAGACCATC

AGCACACGTCTGAACTCTTTCCCTACACGACGCTCTTCCGATCTNNNGTTCAAGCGATTCTCTTGCCTCC

AGCACACGTCTGAACTCTTTCCCTACACGACGCTCTTCCGATCTNNNGTTCAAGCGATTCTCTTGCCTCC

GAGCACACGTCTGAACTCTTTCCCTACACGACGCTCTTCCGATCTNNNGCCTGTAGTCCCAGCTACTCT

GAGCACACGTCTGAACTCTTTCCCTACACGACGCTCTTCCGATCTNNNGCCTGTAGTCCCAGCTACTCT

AGCACACGTCTGAACTCTTTCCCTACACGACGCTCTTCCGATCTNNNTTTTCTCCTCCTTCCAGTTACCG

AGCACACGTCTGAACTCTTTCCCTACACGACGCTCTTCCGATCTNNNTTTTCTCCTCCTTCCAGTTACCG

GCACACGTCTGAACTCTTTCCCTACACGACGCTCTTCCGATCTNNNGCCTGTAGTCCCAGCTACTTGCA

GCACACGTCTGAACTCTTTCCCTACACGACGCTCTTCCGATCTNNNGCCTGTAGTCCCAGCTACTTGCA

AGCACACGTCTGAACTCTTTCCCTACACGACGCTCTTCCGATCTNNNTTTTCTCCTCGACTCCCTCTTG

AGCACACGTCTGAACTCTTTCCCTACACGACGCTCTTCCGATCTNNNTTTTCTCCTCGACTCCCTCTTG

GAGCACACGTCTGAACTCTTTCCCTACACGACGCTCTTCCGATCTNNNCACACACACACACACACACAT

GAGCACACGTCTGAACTCTTTCCCTACACGACGCTCTTCCGATCTNNNCACACACACACACACACACAT

GCACACGTCTGAACTCTTTCCCTACACGACGCTCTTCCGATCTNNNAACTTGGAATCCAAAGTGCTGTG

GCACACGTCTGAACTCTTTCCCTACACGACGCTCTTCCGATCTNNNAACTTGGAATCCAAAGTGCTGTG

AGCACACGTCTGAACTCTTTCCCTACACGACGCTCTTCCGATCTNNNCTTTAGGTGAGAAGGGTTTCGG

AGCACACGTCTGAACTCTTTCCCTACACGACGCTCTTCCGATCTNNNCTTTAGGTGAGAAGGGTTTCGG

AGAGCACACGTCTGAACTCTTTCCCTACACGACGCTCTTCCGATCTNNNTCTGTAATCCCAGCTACTCAG

AGAGCACACGTCTGAACTCTTTCCCTACACGACGCTCTTCCGATCTNNNTCTGTAATCCCAGCTACTCAG

GCACACGTCTGAACTCTTTCCCTACACGACGCTCTTCCGATCTNNNTCTGGATCATCTCCAGCTCCATT

GCACACGTCTGAACTCTTTCCCTACACGACGCTCTTCCGATCTNNNTCTGGATCATCTCCAGCTCCATT

AGAGCACACGTCTGAACTCTTTCCCTACACGACGCTCTTCCGATCTNNNCACCATGTTTGTGAAGTAGG

GAGCACACGTCTGAACTCTTTCCCTACACGACGCTCTTCCGATCTNNNCACCATGTTTGTGAAGTAGG
GCACACGTCTGAACTCTTTCCCTACACGACGCTCTTCCGATCTNNNCAAGTCCCTTCCCCTTTCCATAAC
GCACACGTCTGAACTCTTTCCCTACACGACGCTCTTCCGATCTNNNCAAGTCCCTTCCCCTTTCCATAAC
GCACACGTCTGAACTCTTTCCCTACACGACGCTCTTCCGATCTNNNATGGATTGGGAAGAACTCTGCT
GCACACGTCTGAACTCTTTCCCTACACGACGCTCTTCCGATCTNNNATGGATTGGGAAGAACTCTGCT
AGCACACGTCTGAACTCTTTCCCTACACGACGCTCTTCCGATCTNNNTACAGATGAAAAGACTGAGGC
AGCACACGTCTGAACTCTTTCCCTACACGACGCTCTTCCGATCTNNNTACAGATGAAAAGACTGAGGC
AGCACACGTCTGAACTCTTTCCCTACACGACGCTCTTCCGATCTNNNGGGCAATATGGCAAGATCAC
AGCACACGTCTGAACTCTTTCCCTACACGACGCTCTTCCGATCTNNNGGGCAATATGGCAAGATCAC
AGCACACGTCTGAACTCTTTCCCTACACGACGCTCTTCCGATCTNNNGCACATGCCTATAATCCCAGCT
AGCACACGTCTGAACTCTTTCCCTACACGACGCTCTTCCGATCTNNNGCACATGCCTATAATCCCAGCT
AGCACACGTCTGAACTCTTTCCCTACACGACGCTCTTCCGATCTNNNAAGGTCCCAAGTTGGATGATCT
AGCACACGTCTGAACTCTTTCCCTACACGACGCTCTTCCGATCTNNNAAGGTCCCAAGTTGGATGATCT
GCACACGTCTGAACTCTTTCCCTACACGACGCTCTTCCGATCTNNNGTTCAGTTTTGTCCATCAAGGCA
GCACACGTCTGAACTCTTTCCCTACACGACGCTCTTCCGATCTNNNGTTCAGTTTTGTCCATCAAGGCA
GAGCACACGTCTGAACTCTTTCCCTACACGACGCTCTTCCGATCTNNNTCAAGCAATTCTCCTGCCTTA
GAGCACACGTCTGAACTCTTTCCCTACACGACGCTCTTCCGATCTNNNTCAAGCAATTCTCCTGCCTTA
GCACACGTCTGAACTCTTTCCCTACACGACGCTCTTCCGATCTNNNTGAGAGCAGAGACATCTTTCCT
GCACACGTCTGAACTCTTTCCCTACACGACGCTCTTCCGATCTNNNTGAGAGCAGAGACATCTTTCCT
GCACACGTCTGAACTCTTTCCCTACACGACGCTCTTCCGATCTNNNATGTCAGGAGGCTTTCACAGTAC
GCACACGTCTGAACTCTTTCCCTACACGACGCTCTTCCGATCTNNNATGTCAGGAGGCTTTCACAGTAC
AGCACACGTCTGAACTCTTTCCCTACACGACGCTCTTCCGATCTNNNTAGCTCAGTACACATGCACACA
AGCACACGTCTGAACTCTTTCCCTACACGACGCTCTTCCGATCTNNNTAGCTCAGTACACATGCACACA
AGCACACGTCTGAACTCTTTCCCTACACGACGCTCTTCCGATCTNNNCTGAGACAGGAGAATGGCAT
AGCACACGTCTGAACTCTTTCCCTACACGACGCTCTTCCGATCTNNNCTGAGACAGGAGAATGGCAT
GCACACGTCTGAACTCTTTCCCTACACGACGCTCTTCCGATCTNNNCCATCTCCACATACCTCTTCCTGC
GCACACGTCTGAACTCTTTCCCTACACGACGCTCTTCCGATCTNNNCCATCTCCACATACCTCTTCCTGC
AGCACACGTCTGAACTCTTTCCCTACACGACGCTCTTCCGATCTNNNATGAGATGATTTGCCACTATG
AGCACACGTCTGAACTCTTTCCCTACACGACGCTCTTCCGATCTNNNATGAGATGATTTGCCACTATG
GCACACGTCTGAACTCTTTCCCTACACGACGCTCTTCCGATCTNNNGATTTTGGCTCACTGCAACCTCC
GCACACGTCTGAACTCTTTCCCTACACGACGCTCTTCCGATCTNNNGATTTTGGCTCACTGCAACCTCC
GCACACGTCTGAACTCTTTCCCTACACGACGCTCTTCCGATCTNNNGTTCTGTACATCTTGACAAGTGC
GCACACGTCTGAACTCTTTCCCTACACGACGCTCTTCCGATCTNNNGTTCTGTACATCTTGACAAGTGC
AGCACACGTCTGAACTCTTTCCCTACACGACGCTCTTCCGATCTNNNCATGCCTGTAATCCCAGCTACTC
AGCACACGTCTGAACTCTTTCCCTACACGACGCTCTTCCGATCTNNNCATGCCTGTAATCCCAGCTACTC
AGCACACGTCTGAACTCTTTCCCTACACGACGCTCTTCCGATCTNNNGGATCAGTGTCTTGACTTTTA
AGCACACGTCTGAACTCTTTCCCTACACGACGCTCTTCCGATCTNNNGGATCAGTGTCTTGACTTTTA
GAGCACACGTCTGAACTCTTTCCCTACACGACGCTCTTCCGATCTNNNAAGCTAAGGCACAAGAATCG
GAGCACACGTCTGAACTCTTTCCCTACACGACGCTCTTCCGATCTNNNAAGCTAAGGCACAAGAATCG
AGCACACGTCTGAACTCTTTCCCTACACGACGCTCTTCCGATCTNNNCACATCTTCGCCAGCATTTGTAA
AGCACACGTCTGAACTCTTTCCCTACACGACGCTCTTCCGATCTNNNCACATCTTCGCCAGCATTTGTAA
AGCACACGTCTGAACTCTTTCCCTACACGACGCTCTTCCGATCTNNNTGCTATTGTTCTTTTTCCTGTGT
AGCACACGTCTGAACTCTTTCCCTACACGACGCTCTTCCGATCTNNNTGCTATTGTTCTTTTTCCTGTGT

AGCACACGTCTGAACTCTTTCCCTACACGACGCTCTTCCGATCTNNNATGGTTGCTCTTCCTTTTACCCT
AGCACACGTCTGAACTCTTTCCCTACACGACGCTCTTCCGATCTNNNATGGTTGCTCTTCCTTTTACCCT
GCACACGTCTGAACTCTTTCCCTACACGACGCTCTTCCGATCTNNNAGTCCAGTTATTTGTTCGGGACA
GCACACGTCTGAACTCTTTCCCTACACGACGCTCTTCCGATCTNNNAGTCCAGTTATTTGTTCGGGACA
GAGCACACGTCTGAACTCTTTCCCTACACGACGCTCTTCCGATCTNNNCTTGAACTCCTGAACTTGCGA
GAGCACACGTCTGAACTCTTTCCCTACACGACGCTCTTCCGATCTNNNCTTGAACTCCTGAACTTGCGA
GAGCACACGTCTGAACTCTTTCCCTACACGACGCTCTTCCGATCTNNNTGAATCCTCCAGCCTTGTTCTT
GAGCACACGTCTGAACTCTTTCCCTACACGACGCTCTTCCGATCTNNNTGAATCCTCCAGCCTTGTTCTT
GCACACGTCTGAACTCTTTCCCTACACGACGCTCTTCCGATCTNNNACTTGTGAAGTAGACTCTGCTG
GCACACGTCTGAACTCTTTCCCTACACGACGCTCTTCCGATCTNNNACTTGTGAAGTAGACTCTGCTG
AGCACACGTCTGAACTCTTTCCCTACACGACGCTCTTCCGATCTNNNTCTCCCAGCTTTATATGCCCATC
AGCACACGTCTGAACTCTTTCCCTACACGACGCTCTTCCGATCTNNNTCTCCCAGCTTTATATGCCCATC
AGCACACGTCTGAACTCTTTCCCTACACGACGCTCTTCCGATCTNNNTTCAATGTGTGTGTCTATGCA
AGCACACGTCTGAACTCTTTCCCTACACGACGCTCTTCCGATCTNNNTTCAATGTGTGTGTCTATGCA
AGCACACGTCTGAACTCTTTCCCTACACGACGCTCTTCCGATCTNNNTCTAAAGACCACCTAGTCCCAG
AGCACACGTCTGAACTCTTTCCCTACACGACGCTCTTCCGATCTNNNTCTAAAGACCACCTAGTCCCAG
GCACACGTCTGAACTCTTTCCCTACACGACGCTCTTCCGATCTNNNGAGAAAGGACTAAACCCTCAGC
GCACACGTCTGAACTCTTTCCCTACACGACGCTCTTCCGATCTNNNGAGAAAGGACTAAACCCTCAGC
AGCACACGTCTGAACTCTTTCCCTACACGACGCTCTTCCGATCTNNNACCATTTGAATCACCAACACC
AGCACACGTCTGAACTCTTTCCCTACACGACGCTCTTCCGATCTNNNACCATTTGAATCACCAACACC
GAGCACACGTCTGAACTCTTTCCCTACACGACGCTCTTCCGATCTNNNGCAGTGGAAGGGTGTTTTTT
GAGCACACGTCTGAACTCTTTCCCTACACGACGCTCTTCCGATCTNNNGCAGTGGAAGGGTGTTTTTT
AGCACACGTCTGAACTCTTTCCCTACACGACGCTCTTCCGATCTNNNGCTGAGGGAAGAAGAAGAG
AGCACACGTCTGAACTCTTTCCCTACACGACGCTCTTCCGATCTNNNGCTGAGGGAAGAAGAAGAG
AGCACACGTCTGAACTCTTTCCCTACACGACGCTCTTCCGATCTNNNGCCCATTCCATTTGGCCTAATTA
AGCACACGTCTGAACTCTTTCCCTACACGACGCTCTTCCGATCTNNNGCCCATTCCATTTGGCCTAATTA
AGCACACGTCTGAACTCTTTCCCTACACGACGCTCTTCCGATCTNNNGGGACAGAGTGAGACTCCATC
AGCACACGTCTGAACTCTTTCCCTACACGACGCTCTTCCGATCTNNNGGGACAGAGTGAGACTCCATC
AGCACACGTCTGAACTCTTTCCCTACACGACGCTCTTCCGATCTNNNTCCCAAACTTCTCAGAAGTTTTC
AGCACACGTCTGAACTCTTTCCCTACACGACGCTCTTCCGATCTNNNTCCCAAACTTCTCAGAAGTTTTC
AGCACACGTCTGAACTCTTTCCCTACACGACGCTCTTCCGATCTNNNTAAGAGAGTCAGGTGCTCTTC
AGCACACGTCTGAACTCTTTCCCTACACGACGCTCTTCCGATCTNNNTAAGAGAGTCAGGTGCTCTTC
GAGCACACGTCTGAACTCTTTCCCTACACGACGCTCTTCCGATCTNNNAGGAGGCTGAGACAGAAGA
GAGCACACGTCTGAACTCTTTCCCTACACGACGCTCTTCCGATCTNNNAGGAGGCTGAGACAGAAGA
AGCACACGTCTGAACTCTTTCCCTACACGACGCTCTTCCGATCTNNNAGGCAGGAGAATCATTTGAAC
AGCACACGTCTGAACTCTTTCCCTACACGACGCTCTTCCGATCTNNNAGGCAGGAGAATCATTTGAAC
GAGCACACGTCTGAACTCTTTCCCTACACGACGCTCTTCCGATCTNNNAGGGCAAAACCTTAGCTACAG
GAGCACACGTCTGAACTCTTTCCCTACACGACGCTCTTCCGATCTNNNAGGGCAAAACCTTAGCTACAG
AGCACACGTCTGAACTCTTTCCCTACACGACGCTCTTCCGATCTNNNATGCAGGAGAAATTGCTTGAA
AGCACACGTCTGAACTCTTTCCCTACACGACGCTCTTCCGATCTNNNATGCAGGAGAAATTGCTTGAA
AGAGCACACGTCTGAACTCTTTCCCTACACGACGCTCTTCCGATCTNNNGCAGATCACCTAAGGTCAG
AGAGCACACGTCTGAACTCTTTCCCTACACGACGCTCTTCCGATCTNNNGCAGATCACCTAAGGTCAG
GCACACGTCTGAACTCTTTCCCTACACGACGCTCTTCCGATCTNNNCACTGTCTGCTGTGAAGGTCTA

AGCACACGTCTGAACTCTTTCCCTACACGACGCTCTTCCGATCTNNNCACTGTCTGCTGTGAAGGTCTA
GAGCACACGTCTGAACTCTTTCCCTACACGACGCTCTTCCGATCTNNNGCGTGTGTGTGTGTATGTTT
GAGCACACGTCTGAACTCTTTCCCTACACGACGCTCTTCCGATCTNNNGCGTGTGTGTGTGTATGTTT
GAGCACACGTCTGAACTCTTTCCCTACACGACGCTCTTCCGATCTNNNCCCCTAAAAATGTGTCCGTTCT
GAGCACACGTCTGAACTCTTTCCCTACACGACGCTCTTCCGATCTNNNCCCCTAAAAATGTGTCCGTTCT
AGCACACGTCTGAACTCTTTCCCTACACGACGCTCTTCCGATCTNNNCCTAGGAGGTAGAGTTTGCAG
AGCACACGTCTGAACTCTTTCCCTACACGACGCTCTTCCGATCTNNNCCTAGGAGGTAGAGTTTGCAG
AGCACACGTCTGAACTCTTTCCCTACACGACGCTCTTCCGATCTNNNTCTCTCCCTCTCTCTCTCTC
AGCACACGTCTGAACTCTTTCCCTACACGACGCTCTTCCGATCTNNNTCTCTCCCTCTCTCTCTCTC
GAGCACACGTCTGAACTCTTTCCCTACACGACGCTCTTCCGATCTNNNGGAGCCTGTGGAGATGCTATA
GAGCACACGTCTGAACTCTTTCCCTACACGACGCTCTTCCGATCTNNNGGAGCCTGTGGAGATGCTATA
AGAGCACACGTCTGAACTCTTTCCCTACACGACGCTCTTCCGATCTNNNGTATAGAAGCGCGAAGATG
AGAGCACACGTCTGAACTCTTTCCCTACACGACGCTCTTCCGATCTNNNGTATAGAAGCGCGAAGATG
AGAGCACACGTCTGAACTCTTTCCCTACACGACGCTCTTCCGATCTNNNGGAACTAATCAGCAGGATG
AGAGCACACGTCTGAACTCTTTCCCTACACGACGCTCTTCCGATCTNNNGGAACTAATCAGCAGGATG
CACACGTCTGAACTCTTTCCCTACACGACGCTCTTCCGATCTNNNACAGGGGAAACTACCAAACACTTC
CACACGTCTGAACTCTTTCCCTACACGACGCTCTTCCGATCTNNNACAGGGGAAACTACCAAACACTTC
AGAGCACACGTCTGAACTCTTTCCCTACACGACGCTCTTCCGATCTNNNGTCAGCGCATTCTTACCCTCT
AGAGCACACGTCTGAACTCTTTCCCTACACGACGCTCTTCCGATCTNNNGTCAGCGCATTCTTACCCTCT
GAGCACACGTCTGAACTCTTTCCCTACACGACGCTCTTCCGATCTNNNAATACAAGAGAACGAGAGGT
GAGCACACGTCTGAACTCTTTCCCTACACGACGCTCTTCCGATCTNNNAATACAAGAGAACGAGAGGT
GAGCACACGTCTGAACTCTTTCCCTACACGACGCTCTTCCGATCTNNNCACGTTCTCCTTGGAAGAGAG
GAGCACACGTCTGAACTCTTTCCCTACACGACGCTCTTCCGATCTNNNCACGTTCTCCTTGGAAGAGAG
AGCACACGTCTGAACTCTTTCCCTACACGACGCTCTTCCGATCTNNNAAAAAATTAGCCAGGTGAGGT
AGCACACGTCTGAACTCTTTCCCTACACGACGCTCTTCCGATCTNNNAAAAAATTAGCCAGGTGAGGT
AGCACACGTCTGAACTCTTTCCCTACACGACGCTCTTCCGATCTNNNTACACATAAGCACACCAGGCTAA
AGCACACGTCTGAACTCTTTCCCTACACGACGCTCTTCCGATCTNNNTACACATAAGCACACCAGGCTAA
GCACACGTCTGAACTCTTTCCCTACACGACGCTCTTCCGATCTNNNAGGCCTCAGAGACGTAAACAAA
GCACACGTCTGAACTCTTTCCCTACACGACGCTCTTCCGATCTNNNAGGCCTCAGAGACGTAAACAAA
GCACACGTCTGAACTCTTTCCCTACACGACGCTCTTCCGATCTNNNTTGGGTCCCTATATCTGTGAGGA
GCACACGTCTGAACTCTTTCCCTACACGACGCTCTTCCGATCTNNNTTGGGTCCCTATATCTGTGAGGA
AGCACACGTCTGAACTCTTTCCCTACACGACGCTCTTCCGATCTNNNATGATCTCGGCTCACTGCAAGC
AGCACACGTCTGAACTCTTTCCCTACACGACGCTCTTCCGATCTNNNATGATCTCGGCTCACTGCAAGC
GAGCACACGTCTGAACTCTTTCCCTACACGACGCTCTTCCGATCTNNNGCCATCTAATTAGCACCCCCAT
GAGCACACGTCTGAACTCTTTCCCTACACGACGCTCTTCCGATCTNNNGCCATCTAATTAGCACCCCCAT
AGCACACGTCTGAACTCTTTCCCTACACGACGCTCTTCCGATCTNNNTTTGGAAGCTGGTATCAAGGG
AGCACACGTCTGAACTCTTTCCCTACACGACGCTCTTCCGATCTNNNTTTGGAAGCTGGTATCAAGGG
AGCACACGTCTGAACTCTTTCCCTACACGACGCTCTTCCGATCTNNNAAAGTGCTGGGATTACAGGCA
AGCACACGTCTGAACTCTTTCCCTACACGACGCTCTTCCGATCTNNNAAAGTGCTGGGATTACAGGCA
AGCACACGTCTGAACTCTTTCCCTACACGACGCTCTTCCGATCTNNNGAGGCTGAGGCAGGAGAATG
AGCACACGTCTGAACTCTTTCCCTACACGACGCTCTTCCGATCTNNNGAGGCTGAGGCAGGAGAATG
CACACGTCTGAACTCTTTCCCTACACGACGCTCTTCCGATCTNNNTTTAGTAGAGACGGGGTTTCACCC
CACACGTCTGAACTCTTTCCCTACACGACGCTCTTCCGATCTNNNTTTAGTAGAGACGGGGTTTCACCC

GAGCACACGTCTGAACTCTTTCCCTACACGACGCTCTTCCGATCTNNNGATGCCTGTAATCCCAGCTACT
GAGCACACGTCTGAACTCTTTCCCTACACGACGCTCTTCCGATCTNNNGATGCCTGTAATCCCAGCTACT
AGCACACGTCTGAACTCTTTCCCTACACGACGCTCTTCCGATCTNNNCGTGGCCAACAAGCATATTGTA
AGCACACGTCTGAACTCTTTCCCTACACGACGCTCTTCCGATCTNNNCGTGGCCAACAAGCATATTGTA
AGCACACGTCTGAACTCTTTCCCTACACGACGCTCTTCCGATCTNNNTTTCATGGGCTTTGTCTTTGGTG
AGCACACGTCTGAACTCTTTCCCTACACGACGCTCTTCCGATCTNNNTTTCATGGGCTTTGTCTTTGGTG
AGCACACGTCTGAACTCTTTCCCTACACGACGCTCTTCCGATCTNNNCTAGAGTGGCCAAGTAACATG
AGCACACGTCTGAACTCTTTCCCTACACGACGCTCTTCCGATCTNNNCTAGAGTGGCCAAGTAACATG
AGCACACGTCTGAACTCTTTCCCTACACGACGCTCTTCCGATCTNNNCCCTTTTCCCTTACACCAGATCAC
AGCACACGTCTGAACTCTTTCCCTACACGACGCTCTTCCGATCTNNNCCCTTTTCCCTTACACCAGATCAC
AGAGCACACGTCTGAACTCTTTCCCTACACGACGCTCTTCCGATCTNNNTCAAGACCATCCTGGCTAACA
AGAGCACACGTCTGAACTCTTTCCCTACACGACGCTCTTCCGATCTNNNTCAAGACCATCCTGGCTAACA
GAGCACACGTCTGAACTCTTTCCCTACACGACGCTCTTCCGATCTNNNCTGAGGCAGGAAAATTGCTTG
GAGCACACGTCTGAACTCTTTCCCTACACGACGCTCTTCCGATCTNNNCTGAGGCAGGAAAATTGCTTG
AGCACACGTCTGAACTCTTTCCCTACACGACGCTCTTCCGATCTNNNGGAGCTATGGCACAGATACTGC
AGCACACGTCTGAACTCTTTCCCTACACGACGCTCTTCCGATCTNNNGGAGCTATGGCACAGATACTGC
GAGCACACGTCTGAACTCTTTCCCTACACGACGCTCTTCCGATCTNNNGAGGCTGAGGCAAGAGAATT
GAGCACACGTCTGAACTCTTTCCCTACACGACGCTCTTCCGATCTNNNGAGGCTGAGGCAAGAGAATT
AGAGCACACGTCTGAACTCTTTCCCTACACGACGCTCTTCCGATCTNNNACAAAAATTAGCTGGGTGTC
AGAGCACACGTCTGAACTCTTTCCCTACACGACGCTCTTCCGATCTNNNACAAAAATTAGCTGGGTGTC
CACACGTCTGAACTCTTTCCCTACACGACGCTCTTCCGATCTNNNTTTATTTGTAATTTTTGCTGCTGCAT
CACACGTCTGAACTCTTTCCCTACACGACGCTCTTCCGATCTNNNTTTATTTGTAATTTTTGCTGCTGCAT
AGCACACGTCTGAACTCTTTCCCTACACGACGCTCTTCCGATCTNNNAAACCTCCACATCCTCTCCTTTCC
AGCACACGTCTGAACTCTTTCCCTACACGACGCTCTTCCGATCTNNNAAACCTCCACATCCTCTCCTTTCC
GAGCACACGTCTGAACTCTTTCCCTACACGACGCTCTTCCGATCTNNNCAAGCCTCCAGATCTAGATAG
GAGCACACGTCTGAACTCTTTCCCTACACGACGCTCTTCCGATCTNNNCAAGCCTCCAGATCTAGATAG
AGCACACGTCTGAACTCTTTCCCTACACGACGCTCTTCCGATCTNNNGCAGGAGAATTGCTTGAACCTC
AGCACACGTCTGAACTCTTTCCCTACACGACGCTCTTCCGATCTNNNGCAGGAGAATTGCTTGAACCTC
GAGCACACGTCTGAACTCTTTCCCTACACGACGCTCTTCCGATCTNNNTGTCATTACACTGCCATACGTC
GAGCACACGTCTGAACTCTTTCCCTACACGACGCTCTTCCGATCTNNNTGTCATTACACTGCCATACGTC
AGCACACGTCTGAACTCTTTCCCTACACGACGCTCTTCCGATCTNNNGGTATAGCCTGTGGGAAGGAA
AGCACACGTCTGAACTCTTTCCCTACACGACGCTCTTCCGATCTNNNGGTATAGCCTGTGGGAAGGAA
AGCACACGTCTGAACTCTTTCCCTACACGACGCTCTTCCGATCTNNNCAAAGTGTGGAAGCACTTAGGC
AGCACACGTCTGAACTCTTTCCCTACACGACGCTCTTCCGATCTNNNCAAAGTGTGGAAGCACTTAGGC
GAGCACACGTCTGAACTCTTTCCCTACACGACGCTCTTCCGATCTNNNCTCTTCTTTCGCCTCTCAGAACT
GAGCACACGTCTGAACTCTTTCCCTACACGACGCTCTTCCGATCTNNNCTCTTCTTTCGCCTCTCAGAACT
AGCACACGTCTGAACTCTTTCCCTACACGACGCTCTTCCGATCTNNNCTCTAGTCCCAGCTACTCAGGA
AGCACACGTCTGAACTCTTTCCCTACACGACGCTCTTCCGATCTNNNCTCTAGTCCCAGCTACTCAGGA
AGCACACGTCTGAACTCTTTCCCTACACGACGCTCTTCCGATCTNNNCACCAGTCCATCCTCTGTATCAC
AGCACACGTCTGAACTCTTTCCCTACACGACGCTCTTCCGATCTNNNCACCAGTCCATCCTCTGTATCAC
GAGCACACGTCTGAACTCTTTCCCTACACGACGCTCTTCCGATCTNNNCTGGCTTCAGACCTCACTCAAT
GAGCACACGTCTGAACTCTTTCCCTACACGACGCTCTTCCGATCTNNNCTGGCTTCAGACCTCACTCAAT
AGCACACGTCTGAACTCTTTCCCTACACGACGCTCTTCCGATCTNNNGGAATGTGGTTTTACTGCTGGT

AGCACACGTCTGAACTCTTTCCCTACACGACGCTCTTCCGATCTNNNGGAATGTGGTTTTACTGCTGG
AGCACACGTCTGAACTCTTTCCCTACACGACGCTCTTCCGATCTNNNGATCCGGTGCTGTAGATGATG
AGCACACGTCTGAACTCTTTCCCTACACGACGCTCTTCCGATCTNNNGATCCGGTGCTGTAGATGATG
GCACACGTCTGAACTCTTTCCCTACACGACGCTCTTCCGATCTNNNGAGAGGGAGAGGGAGAGCTC
GCACACGTCTGAACTCTTTCCCTACACGACGCTCTTCCGATCTNNNGAGAGGGAGAGGGAGAGCTC
AGCACACGTCTGAACTCTTTCCCTACACGACGCTCTTCCGATCTNNNGAGGTTGCAATGAGCCAAGA
AGCACACGTCTGAACTCTTTCCCTACACGACGCTCTTCCGATCTNNNGAGGTTGCAATGAGCCAAGA
GAGCACACGTCTGAACTCTTTCCCTACACGACGCTCTTCCGATCTNNNTTCTCCAGAGAAACACACACA
GAGCACACGTCTGAACTCTTTCCCTACACGACGCTCTTCCGATCTNNNTTCTCCAGAGAAACACACACA
GCACACGTCTGAACTCTTTCCCTACACGACGCTCTTCCGATCTNNNCCCATCCAGTAGTTCTCTTTCC
GCACACGTCTGAACTCTTTCCCTACACGACGCTCTTCCGATCTNNNCCCATCCAGTAGTTCTCTTTCC
AGCACACGTCTGAACTCTTTCCCTACACGACGCTCTTCCGATCTNNNCTGCAGATCTTGGGACTACTCA
AGCACACGTCTGAACTCTTTCCCTACACGACGCTCTTCCGATCTNNNCTGCAGATCTTGGGACTACTCA
AGAGCACACGTCTGAACTCTTTCCCTACACGACGCTCTTCCGATCTNNNAGGAGTTTGTTCAGAGCAC
AGAGCACACGTCTGAACTCTTTCCCTACACGACGCTCTTCCGATCTNNNAGGAGTTTGTTCAGAGCAC
AGCACACGTCTGAACTCTTTCCCTACACGACGCTCTTCCGATCTNNNCCCTCAGCAAATGTTTGAGATC
AGCACACGTCTGAACTCTTTCCCTACACGACGCTCTTCCGATCTNNNCCCTCAGCAAATGTTTGAGATC
GAGCACACGTCTGAACTCTTTCCCTACACGACGCTCTTCCGATCTNNNTGAGGCAGAAGAATTGCTTG
GAGCACACGTCTGAACTCTTTCCCTACACGACGCTCTTCCGATCTNNNTGAGGCAGAAGAATTGCTTG
AGAGCACACGTCTGAACTCTTTCCCTACACGACGCTCTTCCGATCTNNNCATGCTGGCCTTTACTGAAT
AGAGCACACGTCTGAACTCTTTCCCTACACGACGCTCTTCCGATCTNNNCATGCTGGCCTTTACTGAAT
AGCACACGTCTGAACTCTTTCCCTACACGACGCTCTTCCGATCTNNNCATGTGATACAGGACCAGGGT
AGCACACGTCTGAACTCTTTCCCTACACGACGCTCTTCCGATCTNNNCATGTGATACAGGACCAGGGT
GAGCACACGTCTGAACTCTTTCCCTACACGACGCTCTTCCGATCTNNNAGGCAGGAGAATCATTTGAA
GAGCACACGTCTGAACTCTTTCCCTACACGACGCTCTTCCGATCTNNNAGGCAGGAGAATCATTTGAA
AGCACACGTCTGAACTCTTTCCCTACACGACGCTCTTCCGATCTNNNTGAGTGTCATACATTGGTCCCC
AGCACACGTCTGAACTCTTTCCCTACACGACGCTCTTCCGATCTNNNTGAGTGTCATACATTGGTCCCC
GAGCACACGTCTGAACTCTTTCCCTACACGACGCTCTTCCGATCTNNNGTGTGCAACCTGATATTCTT
GAGCACACGTCTGAACTCTTTCCCTACACGACGCTCTTCCGATCTNNNGTGTGCAACCTGATATTCTT
CACGTCTGAACTCTTTCCCTACACGACGCTCTTCCGATCTNNNGTCTCCTCCCTTCTCACCCCCATCGGAC
CACGTCTGAACTCTTTCCCTACACGACGCTCTTCCGATCTNNNGTCTCCTCCCTTCTCACCCCCATCGGAC
AGAGCACACGTCTGAACTCTTTCCCTACACGACGCTCTTCCGATCTNNNGAGAAGGCCCTGCTTTCAG
AGAGCACACGTCTGAACTCTTTCCCTACACGACGCTCTTCCGATCTNNNGAGAAGGCCCTGCTTTCAG
GCACACGTCTGAACTCTTTCCCTACACGACGCTCTTCCGATCTNNNTTTCCTTAATATTGGCATGAATT
GCACACGTCTGAACTCTTTCCCTACACGACGCTCTTCCGATCTNNNTTTCCTTAATATTGGCATGAATT
AGAGCACACGTCTGAACTCTTTCCCTACACGACGCTCTTCCGATCTNNNGTAGAAAAAGCTTGGATGG
AGAGCACACGTCTGAACTCTTTCCCTACACGACGCTCTTCCGATCTNNNGTAGAAAAAGCTTGGATGG
CACACGTCTGAACTCTTTCCCTACACGACGCTCTTCCGATCTNNNGGAGAGAGAAATGCAAATCGC
CACACGTCTGAACTCTTTCCCTACACGACGCTCTTCCGATCTNNNGGAGAGAGAAATGCAAATCGC
GAGCACACGTCTGAACTCTTTCCCTACACGACGCTCTTCCGATCTNNNAGGCAAGAGTATCAGTTGAG
GAGCACACGTCTGAACTCTTTCCCTACACGACGCTCTTCCGATCTNNNAGGCAAGAGTATCAGTTGAG
GAGCACACGTCTGAACTCTTTCCCTACACGACGCTCTTCCGATCTNNNACCCCAGAAAGGTAACACATT
GAGCACACGTCTGAACTCTTTCCCTACACGACGCTCTTCCGATCTNNNACCCCAGAAAGGTAACACATT

AGCACACGTCTGAACTCTTTCCCTACACGACGCTCTTCCGATCTNNNACAGATACTCACTTTCTCCACCA
AGCACACGTCTGAACTCTTTCCCTACACGACGCTCTTCCGATCTNNNACAGATACTCACTTTCTCCACCA
GCACACGTCTGAACTCTTTCCCTACACGACGCTCTTCCGATCTNNNCTTCTTTGAGTTCTACCGCCTCCC
GCACACGTCTGAACTCTTTCCCTACACGACGCTCTTCCGATCTNNNCTTCTTTGAGTTCTACCGCCTCCC
GAGCACACGTCTGAACTCTTTCCCTACACGACGCTCTTCCGATCTNNNGAGAATGGCATGAACCCAGG
GAGCACACGTCTGAACTCTTTCCCTACACGACGCTCTTCCGATCTNNNGAGAATGGCATGAACCCAGG
AGCACACGTCTGAACTCTTTCCCTACACGACGCTCTTCCGATCTNNNCAGTGAGCTGAGATCATACCAC
AGCACACGTCTGAACTCTTTCCCTACACGACGCTCTTCCGATCTNNNCAGTGAGCTGAGATCATACCAC
AGCACACGTCTGAACTCTTTCCCTACACGACGCTCTTCCGATCTNNNGTCAAGAGATCGAGACCATCCT
AGCACACGTCTGAACTCTTTCCCTACACGACGCTCTTCCGATCTNNNGTCAAGAGATCGAGACCATCCT
AGAGCACACGTCTGAACTCTTTCCCTACACGACGCTCTTCCGATCTNNNGCTACTCTGGAGGCTGAAG
AGAGCACACGTCTGAACTCTTTCCCTACACGACGCTCTTCCGATCTNNNGCTACTCTGGAGGCTGAAG
AGAGCACACGTCTGAACTCTTTCCCTACACGACGCTCTTCCGATCTNNNGAGACCTGCTTCGTCAGTG
AGAGCACACGTCTGAACTCTTTCCCTACACGACGCTCTTCCGATCTNNNGAGACCTGCTTCGTCAGTG
GCACACGTCTGAACTCTTTCCCTACACGACGCTCTTCCGATCTNNNTCTGAACTTTTCCACTTTAGAGCA
GCACACGTCTGAACTCTTTCCCTACACGACGCTCTTCCGATCTNNNTCTGAACTTTTCCACTTTAGAGCA
GAGCACACGTCTGAACTCTTTCCCTACACGACGCTCTTCCGATCTNNNATCCCTAAGACAGAGGCAAAG
GAGCACACGTCTGAACTCTTTCCCTACACGACGCTCTTCCGATCTNNNATCCCTAAGACAGAGGCAAAG
AGCACACGTCTGAACTCTTTCCCTACACGACGCTCTTCCGATCTNNNGTTGCAGTGAGCTGAGATTGAG
AGCACACGTCTGAACTCTTTCCCTACACGACGCTCTTCCGATCTNNNGTTGCAGTGAGCTGAGATTGAG
AGCACACGTCTGAACTCTTTCCCTACACGACGCTCTTCCGATCTNNNTGCACCATAGTAGTTTTGGGGA
AGCACACGTCTGAACTCTTTCCCTACACGACGCTCTTCCGATCTNNNTGCACCATAGTAGTTTTGGGGA
AGCACACGTCTGAACTCTTTCCCTACACGACGCTCTTCCGATCTNNNGCCTGGACAACAAGAGTGAAA
AGCACACGTCTGAACTCTTTCCCTACACGACGCTCTTCCGATCTNNNGCCTGGACAACAAGAGTGAAA
CACACGTCTGAACTCTTTCCCTACACGACGCTCTTCCGATCTNNNCTCATAGCACTCAGGGCACTTACCA
CACACGTCTGAACTCTTTCCCTACACGACGCTCTTCCGATCTNNNCTCATAGCACTCAGGGCACTTACCA
GAGCACACGTCTGAACTCTTTCCCTACACGACGCTCTTCCGATCTNNNGAGAGAGCAAGGTGGGAACA
GAGCACACGTCTGAACTCTTTCCCTACACGACGCTCTTCCGATCTNNNGAGAGAGCAAGGTGGGAACA
AGCACACGTCTGAACTCTTTCCCTACACGACGCTCTTCCGATCTNNNTTTTTTTTTTTTTTAACAGGCAG
AGCACACGTCTGAACTCTTTCCCTACACGACGCTCTTCCGATCTNNNTTTTTTTTTTTTTTAACAGGCAG
GAGCACACGTCTGAACTCTTTCCCTACACGACGCTCTTCCGATCTNNNGAAGTACCAGTAGGAACACAG
GAGCACACGTCTGAACTCTTTCCCTACACGACGCTCTTCCGATCTNNNGAAGTACCAGTAGGAACACAG
AGCACACGTCTGAACTCTTTCCCTACACGACGCTCTTCCGATCTNNNGCCGAGAGTTGGACTATTGATG
AGCACACGTCTGAACTCTTTCCCTACACGACGCTCTTCCGATCTNNNGCCGAGAGTTGGACTATTGATG
GAGCACACGTCTGAACTCTTTCCCTACACGACGCTCTTCCGATCTNNNCTGGGTCAGGAAGCTCTTTAT
GAGCACACGTCTGAACTCTTTCCCTACACGACGCTCTTCCGATCTNNNCTGGGTCAGGAAGCTCTTTAT
AGCACACGTCTGAACTCTTTCCCTACACGACGCTCTTCCGATCTNNNGAACAAAGCCAGAGTCATCACA
AGCACACGTCTGAACTCTTTCCCTACACGACGCTCTTCCGATCTNNNGAACAAAGCCAGAGTCATCACA
GAGCACACGTCTGAACTCTTTCCCTACACGACGCTCTTCCGATCTNNNGCAAAAGAGGAAGGAACAGA
GAGCACACGTCTGAACTCTTTCCCTACACGACGCTCTTCCGATCTNNNGCAAAAGAGGAAGGAACAGA
GAGCACACGTCTGAACTCTTTCCCTACACGACGCTCTTCCGATCTNNNCTCCTGAGTAGCTGGACGACA
GAGCACACGTCTGAACTCTTTCCCTACACGACGCTCTTCCGATCTNNNCTCCTGAGTAGCTGGACGACA
GAGCACACGTCTGAACTCTTTCCCTACACGACGCTCTTCCGATCTNNNATCTGACTTCCCACTGAGAGA

GAGCACACGTCTGAACTCTTTCCCTACACGACGCTCTTCCGATCTNNNATCTGACTTCCCACTGAGAG
AGCACACGTCTGAACTCTTTCCCTACACGACGCTCTTCCGATCTNNNGATGTACCTATGGTGCTAGTG
AGCACACGTCTGAACTCTTTCCCTACACGACGCTCTTCCGATCTNNNGATGTACCTATGGTGCTAGTG
AGCACACGTCTGAACTCTTTCCCTACACGACGCTCTTCCGATCTNNNTTACCCAAAAGGAATGCATGCA
AGCACACGTCTGAACTCTTTCCCTACACGACGCTCTTCCGATCTNNNTTACCCAAAAGGAATGCATGCA
AGCACACGTCTGAACTCTTTCCCTACACGACGCTCTTCCGATCTNNNTTTCTTTGTCCTCCTCTCCCACC
AGCACACGTCTGAACTCTTTCCCTACACGACGCTCTTCCGATCTNNNTTTCTTTGTCCTCCTCTCCCACC
GCACACGTCTGAACTCTTTCCCTACACGACGCTCTTCCGATCTNNNGACAGAGCAAGACTACGTCTCA
GCACACGTCTGAACTCTTTCCCTACACGACGCTCTTCCGATCTNNNGACAGAGCAAGACTACGTCTCA
AGCACACGTCTGAACTCTTTCCCTACACGACGCTCTTCCGATCTNNNATATTACATTTCAGAGTTCACCT
AGCACACGTCTGAACTCTTTCCCTACACGACGCTCTTCCGATCTNNNATATTACATTTCAGAGTTCACCT
AGCACACGTCTGAACTCTTTCCCTACACGACGCTCTTCCGATCTNNNGAAAGACAGTTGACGTGTCA
AGCACACGTCTGAACTCTTTCCCTACACGACGCTCTTCCGATCTNNNGAAAGACAGTTGACGTGTCA
AGCACACGTCTGAACTCTTTCCCTACACGACGCTCTTCCGATCTNNNTATAGGTGTGAGCCATATTGC
AGCACACGTCTGAACTCTTTCCCTACACGACGCTCTTCCGATCTNNNTATAGGTGTGAGCCATATTGC
GCACACGTCTGAACTCTTTCCCTACACGACGCTCTTCCGATCTNNNAGACCAGTTTTCTTGCGGAAAT
GCACACGTCTGAACTCTTTCCCTACACGACGCTCTTCCGATCTNNNAGACCAGTTTTCTTGCGGAAAT
ACACGTCTGAACTCTTTCCCTACACGACGCTCTTCCGATCTNNNTGATCAGATAGGAGCACAAGCAT
ACACGTCTGAACTCTTTCCCTACACGACGCTCTTCCGATCTNNNTGATCAGATAGGAGCACAAGCAT
CACACGTCTGAACTCTTTCCCTACACGACGCTCTTCCGATCTNNNCCTCAGCTACTCCTCAGGCACATCG
CACACGTCTGAACTCTTTCCCTACACGACGCTCTTCCGATCTNNNCCTCAGCTACTCCTCAGGCACATCG
GAGCACACGTCTGAACTCTTTCCCTACACGACGCTCTTCCGATCTNNNTGACAGGAAAGTTCGATTTC
GAGCACACGTCTGAACTCTTTCCCTACACGACGCTCTTCCGATCTNNNTGACAGGAAAGTTCGATTTC
AGCACACGTCTGAACTCTTTCCCTACACGACGCTCTTCCGATCTNNNACAGCTTACCTTCTACATGTCT
AGCACACGTCTGAACTCTTTCCCTACACGACGCTCTTCCGATCTNNNACAGCTTACCTTCTACATGTCT
GCACACGTCTGAACTCTTTCCCTACACGACGCTCTTCCGATCTNNNCATCTTGCCCAACCCCATTATTCC
GCACACGTCTGAACTCTTTCCCTACACGACGCTCTTCCGATCTNNNCATCTTGCCCAACCCCATTATTCC
AGCACACGTCTGAACTCTTTCCCTACACGACGCTCTTCCGATCTNNNCTGAGGCAGAAGAATCACTTG
AGCACACGTCTGAACTCTTTCCCTACACGACGCTCTTCCGATCTNNNCTGAGGCAGAAGAATCACTTG
AGCACACGTCTGAACTCTTTCCCTACACGACGCTCTTCCGATCTNNNCCCAGGTTCAAGCAATTCTTCT
AGCACACGTCTGAACTCTTTCCCTACACGACGCTCTTCCGATCTNNNCCCAGGTTCAAGCAATTCTTCT
AGCACACGTCTGAACTCTTTCCCTACACGACGCTCTTCCGATCTNNNGTGGCTATCTTGTTTGCCCTTT
AGCACACGTCTGAACTCTTTCCCTACACGACGCTCTTCCGATCTNNNGTGGCTATCTTGTTTGCCCTTT
AGCACACGTCTGAACTCTTTCCCTACACGACGCTCTTCCGATCTNNNATCATCTTAGGCAGAAGAACG
AGCACACGTCTGAACTCTTTCCCTACACGACGCTCTTCCGATCTNNNATCATCTTAGGCAGAAGAACG
GCACACGTCTGAACTCTTTCCCTACACGACGCTCTTCCGATCTNNNATATGGGGTCTCACTCTGCTGTC
GCACACGTCTGAACTCTTTCCCTACACGACGCTCTTCCGATCTNNNATATGGGGTCTCACTCTGCTGTC
GAGCACACGTCTGAACTCTTTCCCTACACGACGCTCTTCCGATCTNNNTTTGTTATTGCCCAGACTTGT
GAGCACACGTCTGAACTCTTTCCCTACACGACGCTCTTCCGATCTNNNTTTGTTATTGCCCAGACTTGT
AGCACACGTCTGAACTCTTTCCCTACACGACGCTCTTCCGATCTNNNAGGTGATGTTGTAAGACAGG
AGCACACGTCTGAACTCTTTCCCTACACGACGCTCTTCCGATCTNNNAGGTGATGTTGTAAGACAGG
GCACACGTCTGAACTCTTTCCCTACACGACGCTCTTCCGATCTNNNGGTCTCCATGTTCAGTGGATTC
GCACACGTCTGAACTCTTTCCCTACACGACGCTCTTCCGATCTNNNGGTCTCCATGTTCAGTGGATTC

AGCACACGTCTGAACTCTTTCCCTACACGACGCTCTTCCGATCTNNNATGTGCTTCCCCAAAATAGTGA
AGCACACGTCTGAACTCTTTCCCTACACGACGCTCTTCCGATCTNNNATGTGCTTCCCCAAAATAGTGA
GAGCACACGTCTGAACTCTTTCCCTACACGACGCTCTTCCGATCTNNNTGTAATCCCAGCTACTCAGGA
GAGCACACGTCTGAACTCTTTCCCTACACGACGCTCTTCCGATCTNNNTGTAATCCCAGCTACTCAGGA
GCACACGTCTGAACTCTTTCCCTACACGACGCTCTTCCGATCTNNNCTCAGTTTGGTTGAAAATCCCGG
GCACACGTCTGAACTCTTTCCCTACACGACGCTCTTCCGATCTNNNCTCAGTTTGGTTGAAAATCCCGG
GAGCACACGTCTGAACTCTTTCCCTACACGACGCTCTTCCGATCTNNNAGATTGCTCTTTGCCAATTTAC
GAGCACACGTCTGAACTCTTTCCCTACACGACGCTCTTCCGATCTNNNAGATTGCTCTTTGCCAATTTAC
AGCACACGTCTGAACTCTTTCCCTACACGACGCTCTTCCGATCTNNNAGTGAGCCAAGATCATACCACT
AGCACACGTCTGAACTCTTTCCCTACACGACGCTCTTCCGATCTNNNAGTGAGCCAAGATCATACCACT
GCACACGTCTGAACTCTTTCCCTACACGACGCTCTTCCGATCTNNNGATGTGAAGGGATCTGCAGAGT
GCACACGTCTGAACTCTTTCCCTACACGACGCTCTTCCGATCTNNNGATGTGAAGGGATCTGCAGAGT
GCACACGTCTGAACTCTTTCCCTACACGACGCTCTTCCGATCTNNNGGCTAGTCTTGAACTGCTGATCT
GCACACGTCTGAACTCTTTCCCTACACGACGCTCTTCCGATCTNNNGGCTAGTCTTGAACTGCTGATCT
CACACGTCTGAACTCTTTCCCTACACGACGCTCTTCCGATCTNNNCAGGCTGGTTTTGAATTCCTGATC
CACACGTCTGAACTCTTTCCCTACACGACGCTCTTCCGATCTNNNCAGGCTGGTTTTGAATTCCTGATC
AGAGCACACGTCTGAACTCTTTCCCTACACGACGCTCTTCCGATCTNNNGGGAGAGGAAAAAAGAG
AGAGCACACGTCTGAACTCTTTCCCTACACGACGCTCTTCCGATCTNNNGGGAGAGGAAAAAAGAG
AGCACACGTCTGAACTCTTTCCCTACACGACGCTCTTCCGATCTNNNTTCTCCTTACCCTGCTCTGTTTC
AGCACACGTCTGAACTCTTTCCCTACACGACGCTCTTCCGATCTNNNTTCTCCTTACCCTGCTCTGTTTC
GAGCACACGTCTGAACTCTTTCCCTACACGACGCTCTTCCGATCTNNNCACCTGTAGTCCCAGCTATTCA
GAGCACACGTCTGAACTCTTTCCCTACACGACGCTCTTCCGATCTNNNCACCTGTAGTCCCAGCTATTCA
AGCACACGTCTGAACTCTTTCCCTACACGACGCTCTTCCGATCTNNNATTCGCTAGGAGCTGTTTGAA
AGCACACGTCTGAACTCTTTCCCTACACGACGCTCTTCCGATCTNNNATTCGCTAGGAGCTGTTTGAA
AGCACACGTCTGAACTCTTTCCCTACACGACGCTCTTCCGATCTNNNTGAACCCAGGAGGCAAAGTTTC
AGCACACGTCTGAACTCTTTCCCTACACGACGCTCTTCCGATCTNNNTGAACCCAGGAGGCAAAGTTTC
GAGCACACGTCTGAACTCTTTCCCTACACGACGCTCTTCCGATCTNNNTATCTCTCACTCTTCTGGGGGT
GAGCACACGTCTGAACTCTTTCCCTACACGACGCTCTTCCGATCTNNNTATCTCTCACTCTTCTGGGGGT
AGCACACGTCTGAACTCTTTCCCTACACGACGCTCTTCCGATCTNNNGATTTAGCTCACTTCTGCTGCA
AGCACACGTCTGAACTCTTTCCCTACACGACGCTCTTCCGATCTNNNGATTTAGCTCACTTCTGCTGCA
AGCACACGTCTGAACTCTTTCCCTACACGACGCTCTTCCGATCTNNNAACCTAAGTGCCCATCAACAGA
AGCACACGTCTGAACTCTTTCCCTACACGACGCTCTTCCGATCTNNNAACCTAAGTGCCCATCAACAGA
AGCACACGTCTGAACTCTTTCCCTACACGACGCTCTTCCGATCTNNNTGATAGCAGAGAGTGTGTGT
AGCACACGTCTGAACTCTTTCCCTACACGACGCTCTTCCGATCTNNNTGATAGCAGAGAGTGTGTGT
GAGCACACGTCTGAACTCTTTCCCTACACGACGCTCTTCCGATCTNNNGCTTTCCTCACTATTCCAGTT
GAGCACACGTCTGAACTCTTTCCCTACACGACGCTCTTCCGATCTNNNGCTTTCCTCACTATTCCAGTT
GAGCACACGTCTGAACTCTTTCCCTACACGACGCTCTTCCGATCTNNNCTGGCCAGAGAAGTTTTTGAC
GAGCACACGTCTGAACTCTTTCCCTACACGACGCTCTTCCGATCTNNNCTGGCCAGAGAAGTTTTTGAC
AGCACACGTCTGAACTCTTTCCCTACACGACGCTCTTCCGATCTNNNTATGATTTTTGCCCTGTCCGAGA
AGCACACGTCTGAACTCTTTCCCTACACGACGCTCTTCCGATCTNNNTATGATTTTTGCCCTGTCCGAGA
GAGCACACGTCTGAACTCTTTCCCTACACGACGCTCTTCCGATCTNNNCAAGGGTAAGTCTCACAATC
GAGCACACGTCTGAACTCTTTCCCTACACGACGCTCTTCCGATCTNNNCAAGGGTAAGTCTCACAATC
AGCACACGTCTGAACTCTTTCCCTACACGACGCTCTTCCGATCTNNNTTTACGCCATTATCCTGCCTCAC

AGCACACGTCTGAACTCTTTCCCTACACGACGCTCTTCCGATCTNNNTTTACGCCATTATCCTGCCTCA
AGCACACGTCTGAACTCTTTCCCTACACGACGCTCTTCCGATCTNNNAGAGTCCCATCTCAGTTACCTG
AGCACACGTCTGAACTCTTTCCCTACACGACGCTCTTCCGATCTNNNAGAGTCCCATCTCAGTTACCTG
GAGCACACGTCTGAACTCTTTCCCTACACGACGCTCTTCCGATCTNNNCACAGAGCAAGTTCCAGAAT
GAGCACACGTCTGAACTCTTTCCCTACACGACGCTCTTCCGATCTNNNCACAGAGCAAGTTCCAGAAT
CACACGTCTGAACTCTTTCCCTACACGACGCTCTTCCGATCTNNNGGGGAGAAACTCAGTGTCTTCTC
CACACGTCTGAACTCTTTCCCTACACGACGCTCTTCCGATCTNNNGGGGAGAAACTCAGTGTCTTCTC
AGCACACGTCTGAACTCTTTCCCTACACGACGCTCTTCCGATCTNNNCCCAGGACTGTGAGCTAGAAC
AGCACACGTCTGAACTCTTTCCCTACACGACGCTCTTCCGATCTNNNCCCAGGACTGTGAGCTAGAAC
AGCACACGTCTGAACTCTTTCCCTACACGACGCTCTTCCGATCTNNNGATTACAGGCATGTGTCACCAT
AGCACACGTCTGAACTCTTTCCCTACACGACGCTCTTCCGATCTNNNGATTACAGGCATGTGTCACCAT
AGCACACGTCTGAACTCTTTCCCTACACGACGCTCTTCCGATCTNNNAAGTTGGTCTTGCTCCCTGAG
AGCACACGTCTGAACTCTTTCCCTACACGACGCTCTTCCGATCTNNNAAGTTGGTCTTGCTCCCTGAG
GAGCACACGTCTGAACTCTTTCCCTACACGACGCTCTTCCGATCTNNNGAGACGGATTGGAGGTTGC
GAGCACACGTCTGAACTCTTTCCCTACACGACGCTCTTCCGATCTNNNGAGACGGATTGGAGGTTGC
AGCACACGTCTGAACTCTTTCCCTACACGACGCTCTTCCGATCTNNNCACAGTTCCTGCTTCAATGGA
AGCACACGTCTGAACTCTTTCCCTACACGACGCTCTTCCGATCTNNNCACAGTTCCTGCTTCAATGGA
CACACGTCTGAACTCTTTCCCTACACGACGCTCTTCCGATCTNNNCTTCACATAAAGGAACATGGTTC
CACACGTCTGAACTCTTTCCCTACACGACGCTCTTCCGATCTNNNCTTCACATAAAGGAACATGGTTC
GCACACGTCTGAACTCTTTCCCTACACGACGCTCTTCCGATCTNNNGGAGGTTGAGGTAGGAGAATC
GCACACGTCTGAACTCTTTCCCTACACGACGCTCTTCCGATCTNNNGGAGGTTGAGGTAGGAGAATC
GAGCACACGTCTGAACTCTTTCCCTACACGACGCTCTTCCGATCTNNNTCTCCATCCAAGAAGTCACC
GAGCACACGTCTGAACTCTTTCCCTACACGACGCTCTTCCGATCTNNNTCTCCATCCAAGAAGTCACC
GAGCACACGTCTGAACTCTTTCCCTACACGACGCTCTTCCGATCTNNNTGAGCTGAGATTGTTCCACT
GAGCACACGTCTGAACTCTTTCCCTACACGACGCTCTTCCGATCTNNNTGAGCTGAGATTGTTCCACT
GCACACGTCTGAACTCTTTCCCTACACGACGCTCTTCCGATCTNNNGATGTTTCCTCTCCTCGTGGTAC
GCACACGTCTGAACTCTTTCCCTACACGACGCTCTTCCGATCTNNNGATGTTTCCTCTCCTCGTGGTAC
GCACACGTCTGAACTCTTTCCCTACACGACGCTCTTCCGATCTNNNAGCCTGGCTAATTTCAAAAACG
GCACACGTCTGAACTCTTTCCCTACACGACGCTCTTCCGATCTNNNAGCCTGGCTAATTTCAAAAACG
AGCACACGTCTGAACTCTTTCCCTACACGACGCTCTTCCGATCTNNNGCACTCAATTTCCACCAGAATC
AGCACACGTCTGAACTCTTTCCCTACACGACGCTCTTCCGATCTNNNGCACTCAATTTCCACCAGAATC
AGCACACGTCTGAACTCTTTCCCTACACGACGCTCTTCCGATCTNNNATTACCCAGGTGAAGGAATTG
AGCACACGTCTGAACTCTTTCCCTACACGACGCTCTTCCGATCTNNNATTACCCAGGTGAAGGAATTG
AGCACACGTCTGAACTCTTTCCCTACACGACGCTCTTCCGATCTNNNAGAATTGCTTGAATCCAGGAG
AGCACACGTCTGAACTCTTTCCCTACACGACGCTCTTCCGATCTNNNAGAATTGCTTGAATCCAGGAG
AGCACACGTCTGAACTCTTTCCCTACACGACGCTCTTCCGATCTNNNTACTGGGAAGTGAGGAGCCC
AGCACACGTCTGAACTCTTTCCCTACACGACGCTCTTCCGATCTNNNTACTGGGAAGTGAGGAGCCC
AGCACACGTCTGAACTCTTTCCCTACACGACGCTCTTCCGATCTNNNCAAACAATCATGTAATAGACA
AGCACACGTCTGAACTCTTTCCCTACACGACGCTCTTCCGATCTNNNCAAACAATCATGTAATAGACA
AGCACACGTCTGAACTCTTTCCCTACACGACGCTCTTCCGATCTNNNGTTCCTGGGAAGGTATAGCA
AGCACACGTCTGAACTCTTTCCCTACACGACGCTCTTCCGATCTNNNGTTCCTGGGAAGGTATAGCA
AGCACACGTCTGAACTCTTTCCCTACACGACGCTCTTCCGATCTNNNTGAGTGCTAACTGTCAAGTGT
AGCACACGTCTGAACTCTTTCCCTACACGACGCTCTTCCGATCTNNNTGAGTGCTAACTGTCAAGTGT

AGCACACGTCTGAACTCTTTCCCTACACGACGCTCTTCCGATCTNNNTGGGAAGAAGTGGGAGGAA
AGCACACGTCTGAACTCTTTCCCTACACGACGCTCTTCCGATCTNNNTGGGAAGAAGTGGGAGGAA
AGCACACGTCTGAACTCTTTCCCTACACGACGCTCTTCCGATCTNNNTGAGGTTGAAAAACCCTGCTC
AGCACACGTCTGAACTCTTTCCCTACACGACGCTCTTCCGATCTNNNTGAGGTTGAAAAACCCTGCTC
AGCACACGTCTGAACTCTTTCCCTACACGACGCTCTTCCGATCTNNNGAGGGGAAAGCCTCCGTTTTA
AGCACACGTCTGAACTCTTTCCCTACACGACGCTCTTCCGATCTNNNGAGGGGAAAGCCTCCGTTTTA
AGAGCACACGTCTGAACTCTTTCCCTACACGACGCTCTTCCGATCTNNNATGCTCTGCTCAATAACAGG
AGAGCACACGTCTGAACTCTTTCCCTACACGACGCTCTTCCGATCTNNNATGCTCTGCTCAATAACAGG
GAGCACACGTCTGAACTCTTTCCCTACACGACGCTCTTCCGATCTNNNCATGTACAGGGCCACTTCTAG
GAGCACACGTCTGAACTCTTTCCCTACACGACGCTCTTCCGATCTNNNCATGTACAGGGCCACTTCTAG
AGCACACGTCTGAACTCTTTCCCTACACGACGCTCTTCCGATCTNNNGTGGATCACGAGGTTAGGAG
AGCACACGTCTGAACTCTTTCCCTACACGACGCTCTTCCGATCTNNNGTGGATCACGAGGTTAGGAG
GCACACGTCTGAACTCTTTCCCTACACGACGCTCTTCCGATCTNNNCAGAGTTTACCTCCTAGGGAAG
GCACACGTCTGAACTCTTTCCCTACACGACGCTCTTCCGATCTNNNCAGAGTTTACCTCCTAGGGAAG
AGCACACGTCTGAACTCTTTCCCTACACGACGCTCTTCCGATCTNNNGGCACCTGTAATACCAGCTAC
AGCACACGTCTGAACTCTTTCCCTACACGACGCTCTTCCGATCTNNNGGCACCTGTAATACCAGCTAC
AGCACACGTCTGAACTCTTTCCCTACACGACGCTCTTCCGATCTNNNGAGGCAGTAGAATTGCTTGAA
AGCACACGTCTGAACTCTTTCCCTACACGACGCTCTTCCGATCTNNNGAGGCAGTAGAATTGCTTGAA
GCACACGTCTGAACTCTTTCCCTACACGACGCTCTTCCGATCTNNNCCACCATGCCCAGCTAATATTTG
GCACACGTCTGAACTCTTTCCCTACACGACGCTCTTCCGATCTNNNCCACCATGCCCAGCTAATATTTG
GAGCACACGTCTGAACTCTTTCCCTACACGACGCTCTTCCGATCTNNNCTGAGGTTGTAGTGAGCCAA
GAGCACACGTCTGAACTCTTTCCCTACACGACGCTCTTCCGATCTNNNCTGAGGTTGTAGTGAGCCAA
AGCACACGTCTGAACTCTTTCCCTACACGACGCTCTTCCGATCTNNNGGACACTTCTCTTACCTGCCAA
AGCACACGTCTGAACTCTTTCCCTACACGACGCTCTTCCGATCTNNNGGACACTTCTCTTACCTGCCAA
AGCACACGTCTGAACTCTTTCCCTACACGACGCTCTTCCGATCTNNNCAATGCATGAGTGTGCTGTGT
AGCACACGTCTGAACTCTTTCCCTACACGACGCTCTTCCGATCTNNNCAATGCATGAGTGTGCTGTGT
GCACACGTCTGAACTCTTTCCCTACACGACGCTCTTCCGATCTNNNGGCCTCCTAAATTGCTGGGATTA
GCACACGTCTGAACTCTTTCCCTACACGACGCTCTTCCGATCTNNNGGCCTCCTAAATTGCTGGGATTA
AGCACACGTCTGAACTCTTTCCCTACACGACGCTCTTCCGATCTNNNATGCCTGTAATCCCAGCTACTT
AGCACACGTCTGAACTCTTTCCCTACACGACGCTCTTCCGATCTNNNATGCCTGTAATCCCAGCTACTT
AGCACACGTCTGAACTCTTTCCCTACACGACGCTCTTCCGATCTNNNGCTGGGACTACATATGAACACC
AGCACACGTCTGAACTCTTTCCCTACACGACGCTCTTCCGATCTNNNGCTGGGACTACATATGAACACC
AGCACACGTCTGAACTCTTTCCCTACACGACGCTCTTCCGATCTNNNAAGCAGAAGCATCAGAGTCA
AGCACACGTCTGAACTCTTTCCCTACACGACGCTCTTCCGATCTNNNAAGCAGAAGCATCAGAGTCA
AGCACACGTCTGAACTCTTTCCCTACACGACGCTCTTCCGATCTNNNGAACTGAGATCCCACCACTGT
AGCACACGTCTGAACTCTTTCCCTACACGACGCTCTTCCGATCTNNNGAACTGAGATCCCACCACTGT
GAGCACACGTCTGAACTCTTTCCCTACACGACGCTCTTCCGATCTNNNTGCTTTCCTATCACTTCTGCC
GAGCACACGTCTGAACTCTTTCCCTACACGACGCTCTTCCGATCTNNNTGCTTTCCTATCACTTCTGCC
AGCACACGTCTGAACTCTTTCCCTACACGACGCTCTTCCGATCTNNNAGTTACCTGCCTTCCATTGTCAT
AGCACACGTCTGAACTCTTTCCCTACACGACGCTCTTCCGATCTNNNAGTTACCTGCCTTCCATTGTCAT
AGCACACGTCTGAACTCTTTCCCTACACGACGCTCTTCCGATCTNNNTCATGGTGGCTTCATGAGATCT
AGCACACGTCTGAACTCTTTCCCTACACGACGCTCTTCCGATCTNNNTCATGGTGGCTTCATGAGATCT
AGCACACGTCTGAACTCTTTCCCTACACGACGCTCTTCCGATCTNNNGAGATTGTGCCACTGAACTCC

AGCACACGTCTGAACTCTTTCCCTACACGACGCTCTTCCGATCTNNNGAGATTGTGCCACTGAACTCC
AGCACACGTCTGAACTCTTTCCCTACACGACGCTCTTCCGATCTNNNGGGCTTGCTTTTGAGATTTCTG
AGCACACGTCTGAACTCTTTCCCTACACGACGCTCTTCCGATCTNNNGGGCTTGCTTTTGAGATTTCTG
AGAGCACACGTCTGAACTCTTTCCCTACACGACGCTCTTCCGATCTNNNAGGGAATTACAGCTGCCGA
AGAGCACACGTCTGAACTCTTTCCCTACACGACGCTCTTCCGATCTNNNAGGGAATTACAGCTGCCGA
AGCACACGTCTGAACTCTTTCCCTACACGACGCTCTTCCGATCTNNNTGCATGAGCTAGGTCTTCAAA
AGCACACGTCTGAACTCTTTCCCTACACGACGCTCTTCCGATCTNNNTGCATGAGCTAGGTCTTCAAA
GCACACGTCTGAACTCTTTCCCTACACGACGCTCTTCCGATCTNNNCTTTTCCCCTTCCTTCCTTCCTTCA
GCACACGTCTGAACTCTTTCCCTACACGACGCTCTTCCGATCTNNNCTTTTCCCCTTCCTTCCTTCCTTCA
AGCACACGTCTGAACTCTTTCCCTACACGACGCTCTTCCGATCTNNNCACATGTAGTCCCAGCTACTCA
AGCACACGTCTGAACTCTTTCCCTACACGACGCTCTTCCGATCTNNNCACATGTAGTCCCAGCTACTCA
AGCACACGTCTGAACTCTTTCCCTACACGACGCTCTTCCGATCTNNNGTCCTCTTGCCCCCATCTTAGC
AGCACACGTCTGAACTCTTTCCCTACACGACGCTCTTCCGATCTNNNGTCCTCTTGCCCCCATCTTAGC
AGCACACGTCTGAACTCTTTCCCTACACGACGCTCTTCCGATCTNNNGTGGATCACTTGAGGTCAGGA
AGCACACGTCTGAACTCTTTCCCTACACGACGCTCTTCCGATCTNNNGTGGATCACTTGAGGTCAGGA
AGCACACGTCTGAACTCTTTCCCTACACGACGCTCTTCCGATCTNNNAAATACAAAACGTTAGCCGGGT
AGCACACGTCTGAACTCTTTCCCTACACGACGCTCTTCCGATCTNNNAAATACAAAACGTTAGCCGGGT
AGCACACGTCTGAACTCTTTCCCTACACGACGCTCTTCCGATCTNNNAGTAGCTAGGATTACAAGGCA
AGCACACGTCTGAACTCTTTCCCTACACGACGCTCTTCCGATCTNNNAGTAGCTAGGATTACAAGGCA
AGCACACGTCTGAACTCTTTCCCTACACGACGCTCTTCCGATCTNNNTGTCCAGCTTTCTCTTTCTCTCCC
AGCACACGTCTGAACTCTTTCCCTACACGACGCTCTTCCGATCTNNNTGTCCAGCTTTCTCTTTCTCTCCC
CACACGTCTGAACTCTTTCCCTACACGACGCTCTTCCGATCTNNNAACGAGGACATTTGCATCCTTTGC
CACACGTCTGAACTCTTTCCCTACACGACGCTCTTCCGATCTNNNAACGAGGACATTTGCATCCTTTGC
GAGCACACGTCTGAACTCTTTCCCTACACGACGCTCTTCCGATCTNNNCACATTTAATCCATAACTCCT
GAGCACACGTCTGAACTCTTTCCCTACACGACGCTCTTCCGATCTNNNCACATTTAATCCATAACTCCT
AGCACACGTCTGAACTCTTTCCCTACACGACGCTCTTCCGATCTNNNTAACCTCCTTTGTCCTGAATCC
AGCACACGTCTGAACTCTTTCCCTACACGACGCTCTTCCGATCTNNNTAACCTCCTTTGTCCTGAATCC
AGCACACGTCTGAACTCTTTCCCTACACGACGCTCTTCCGATCTNNNGCTTTCTTGACTCAGGTGTGAT
AGCACACGTCTGAACTCTTTCCCTACACGACGCTCTTCCGATCTNNNGCTTTCTTGACTCAGGTGTGAT
AGCACACGTCTGAACTCTTTCCCTACACGACGCTCTTCCGATCTNNNAAAAGAATTAGCCAGACGTGG
AGCACACGTCTGAACTCTTTCCCTACACGACGCTCTTCCGATCTNNNAAAAGAATTAGCCAGACGTGG
GAGCACACGTCTGAACTCTTTCCCTACACGACGCTCTTCCGATCTNNNCAAGAGAAAAGCGAACTCCC
GAGCACACGTCTGAACTCTTTCCCTACACGACGCTCTTCCGATCTNNNCAAGAGAAAAGCGAACTCCC
CACACGTCTGAACTCTTTCCCTACACGACGCTCTTCCGATCTNNNCTTAGTTACCTTTTTCCATGTTTCA
CACACGTCTGAACTCTTTCCCTACACGACGCTCTTCCGATCTNNNCTTAGTTACCTTTTTCCATGTTTCA
GAGCACACGTCTGAACTCTTTCCCTACACGACGCTCTTCCGATCTNNNGTGATGGCACGCACTTATAAT
GAGCACACGTCTGAACTCTTTCCCTACACGACGCTCTTCCGATCTNNNGTGATGGCACGCACTTATAAT
GAGCACACGTCTGAACTCTTTCCCTACACGACGCTCTTCCGATCTNNNTGGACTATTCAGTGGATATC
GAGCACACGTCTGAACTCTTTCCCTACACGACGCTCTTCCGATCTNNNTGGACTATTCAGTGGATATC
AGCACACGTCTGAACTCTTTCCCTACACGACGCTCTTCCGATCTNNNACCACGCCCAGCTATTTTTTTT
AGCACACGTCTGAACTCTTTCCCTACACGACGCTCTTCCGATCTNNNACCACGCCCAGCTATTTTTTTT
GCACACGTCTGAACTCTTTCCCTACACGACGCTCTTCCGATCTNNNCAACAGAGCACATTTTAGGGGG
GCACACGTCTGAACTCTTTCCCTACACGACGCTCTTCCGATCTNNNCAACAGAGCACATTTTAGGGGG

AGCACACGTCTGAACTCTTTCCCTACACGACGCTCTTCCGATCTNNNAAAAAATTAGCCAGGCGTGGT
AGCACACGTCTGAACTCTTTCCCTACACGACGCTCTTCCGATCTNNNAAAAAATTAGCCAGGCGTGGT
AGCACACGTCTGAACTCTTTCCCTACACGACGCTCTTCCGATCTNNNAGAGGGGAAGCAATTCACTGA
AGCACACGTCTGAACTCTTTCCCTACACGACGCTCTTCCGATCTNNNAGAGGGGAAGCAATTCACTGA
GAGCACACGTCTGAACTCTTTCCCTACACGACGCTCTTCCGATCTNNNCCTCCTCTTGGGAACTGTAAC
GAGCACACGTCTGAACTCTTTCCCTACACGACGCTCTTCCGATCTNNNCCTCCTCTTGGGAACTGTAAC
AGCACACGTCTGAACTCTTTCCCTACACGACGCTCTTCCGATCTNNNGTTCCTCTTTTGGTGTGCATTCT
AGCACACGTCTGAACTCTTTCCCTACACGACGCTCTTCCGATCTNNNGTTCCTCTTTTGGTGTGCATTCT
AGCACACGTCTGAACTCTTTCCCTACACGACGCTCTTCCGATCTNNNCAAGTTGTACCCCAAACACCTT
AGCACACGTCTGAACTCTTTCCCTACACGACGCTCTTCCGATCTNNNCAAGTTGTACCCCAAACACCTT
GCACACGTCTGAACTCTTTCCCTACACGACGCTCTTCCGATCTNNNTCCATCTGTCTCTCTCTTCCACTCA
GCACACGTCTGAACTCTTTCCCTACACGACGCTCTTCCGATCTNNNTCCATCTGTCTCTCTCTTCCACTCA
AGCACACGTCTGAACTCTTTCCCTACACGACGCTCTTCCGATCTNNNAGACCTCATGGATGACCTCTA
AGCACACGTCTGAACTCTTTCCCTACACGACGCTCTTCCGATCTNNNAGACCTCATGGATGACCTCTA
GCACACGTCTGAACTCTTTCCCTACACGACGCTCTTCCGATCTNNNAAGCCAATATCTCACCAAGGTGT
GCACACGTCTGAACTCTTTCCCTACACGACGCTCTTCCGATCTNNNAAGCCAATATCTCACCAAGGTGT
AGCACACGTCTGAACTCTTTCCCTACACGACGCTCTTCCGATCTNNNCATCCTCCTCAATACCGACATGT
AGCACACGTCTGAACTCTTTCCCTACACGACGCTCTTCCGATCTNNNCATCCTCCTCAATACCGACATGT
GAGCACACGTCTGAACTCTTTCCCTACACGACGCTCTTCCGATCTNNNTCAGAGGCTAATCAGTCCTGT
GAGCACACGTCTGAACTCTTTCCCTACACGACGCTCTTCCGATCTNNNTCAGAGGCTAATCAGTCCTGT
GCACACGTCTGAACTCTTTCCCTACACGACGCTCTTCCGATCTNNNTCCTCTGCAGTCCAATACTCCTAC
GCACACGTCTGAACTCTTTCCCTACACGACGCTCTTCCGATCTNNNTCCTCTGCAGTCCAATACTCCTAC
AGAGCACACGTCTGAACTCTTTCCCTACACGACGCTCTTCCGATCTNNNTGAGAAAGAGAGAGCGAG
AGAGCACACGTCTGAACTCTTTCCCTACACGACGCTCTTCCGATCTNNNTGAGAAAGAGAGAGCGAG
GAGCACACGTCTGAACTCTTTCCCTACACGACGCTCTTCCGATCTNNNGGGTCTCTGTTAGATAGGC
GAGCACACGTCTGAACTCTTTCCCTACACGACGCTCTTCCGATCTNNNGGGTCTCTGTTAGATAGGC
GAGCACACGTCTGAACTCTTTCCCTACACGACGCTCTTCCGATCTNNNTTTGCGGAGATCAAGTGTT
GAGCACACGTCTGAACTCTTTCCCTACACGACGCTCTTCCGATCTNNNTTTGCGGAGATCAAGTGTT
AGCACACGTCTGAACTCTTTCCCTACACGACGCTCTTCCGATCTNNNACATAGCAAGACCTCATCTCTG
AGCACACGTCTGAACTCTTTCCCTACACGACGCTCTTCCGATCTNNNACATAGCAAGACCTCATCTCTG
GAGCACACGTCTGAACTCTTTCCCTACACGACGCTCTTCCGATCTNNNGTGTTTCGCTCTCATGCTGAA
GAGCACACGTCTGAACTCTTTCCCTACACGACGCTCTTCCGATCTNNNGTGTTTCGCTCTCATGCTGAA
AGAGCACACGTCTGAACTCTTTCCCTACACGACGCTCTTCCGATCTNNNAGAGCAGACTCCACCTCAAA
AGAGCACACGTCTGAACTCTTTCCCTACACGACGCTCTTCCGATCTNNNAGAGCAGACTCCACCTCAAA
GAGCACACGTCTGAACTCTTTCCCTACACGACGCTCTTCCGATCTNNNTTTCTCTCTCACACACACACAC
GAGCACACGTCTGAACTCTTTCCCTACACGACGCTCTTCCGATCTNNNTTTCTCTCTCACACACACACAC
AGCACACGTCTGAACTCTTTCCCTACACGACGCTCTTCCGATCTNNNCTGTAGTCCCAGCTACTCAGG
AGCACACGTCTGAACTCTTTCCCTACACGACGCTCTTCCGATCTNNNCTGTAGTCCCAGCTACTCAGG
GAGCACACGTCTGAACTCTTTCCCTACACGACGCTCTTCCGATCTNNNGAGATCGAGTCACTGCACTC
GAGCACACGTCTGAACTCTTTCCCTACACGACGCTCTTCCGATCTNNNGAGATCGAGTCACTGCACTC
AGCACACGTCTGAACTCTTTCCCTACACGACGCTCTTCCGATCTNNNAAAGTCTCACTCTGTCATCCAG
AGCACACGTCTGAACTCTTTCCCTACACGACGCTCTTCCGATCTNNNAAAGTCTCACTCTGTCATCCAG
AGCACACGTCTGAACTCTTTCCCTACACGACGCTCTTCCGATCTNNNCAGCAAAATGCCTTCAGACTTG

AGCACACGTCTGAACTCTTTCCCTACACGACGCTCTTCCGATCTNNNCAGCAAAATGCCTTCAGACTTG
AGCACACGTCTGAACTCTTTCCCTACACGACGCTCTTCCGATCTNNNTCAGCCTTCTGAGTAGTTGGA
AGCACACGTCTGAACTCTTTCCCTACACGACGCTCTTCCGATCTNNNTCAGCCTTCTGAGTAGTTGGA
GCACACGTCTGAACTCTTTCCCTACACGACGCTCTTCCGATCTNNNGGAGAAGTGTGTCCTATCTTCG
GCACACGTCTGAACTCTTTCCCTACACGACGCTCTTCCGATCTNNNGGAGAAGTGTGTCCTATCTTCG
GAGCACACGTCTGAACTCTTTCCCTACACGACGCTCTTCCGATCTNNNGGATGAGGAGGGAGAGAG
GAGCACACGTCTGAACTCTTTCCCTACACGACGCTCTTCCGATCTNNNGGATGAGGAGGGAGAGAG
GAGCACACGTCTGAACTCTTTCCCTACACGACGCTCTTCCGATCTNNNGTGCAGACTACACAGGTAG
GAGCACACGTCTGAACTCTTTCCCTACACGACGCTCTTCCGATCTNNNGTGCAGACTACACAGGTAG
GAGCACACGTCTGAACTCTTTCCCTACACGACGCTCTTCCGATCTNNNGGCTGAGACAGGAGAATCT
GAGCACACGTCTGAACTCTTTCCCTACACGACGCTCTTCCGATCTNNNGGCTGAGACAGGAGAATCT
AGCACACGTCTGAACTCTTTCCCTACACGACGCTCTTCCGATCTNNNTTACCATATTGCCCATGCTGGT
AGCACACGTCTGAACTCTTTCCCTACACGACGCTCTTCCGATCTNNNTTACCATATTGCCCATGCTGGT
AGCACACGTCTGAACTCTTTCCCTACACGACGCTCTTCCGATCTNNNCACCCACTGCTCTAGATTTGG
AGCACACGTCTGAACTCTTTCCCTACACGACGCTCTTCCGATCTNNNCACCCACTGCTCTAGATTTGG
AGCACACGTCTGAACTCTTTCCCTACACGACGCTCTTCCGATCTNNNGGTGTAATAGGAGAACCAG
AGCACACGTCTGAACTCTTTCCCTACACGACGCTCTTCCGATCTNNNGGTGTAATAGGAGAACCAG
CACACGTCTGAACTCTTTCCCTACACGACGCTCTTCCGATCTNNNTAGTCAGCTCATTGATGTCCGTCC
CACACGTCTGAACTCTTTCCCTACACGACGCTCTTCCGATCTNNNTAGTCAGCTCATTGATGTCCGTCC
AGCACACGTCTGAACTCTTTCCCTACACGACGCTCTTCCGATCTNNNCTGAGGCAGGAGAATTGCTT
AGCACACGTCTGAACTCTTTCCCTACACGACGCTCTTCCGATCTNNNCTGAGGCAGGAGAATTGCTT
GAGCACACGTCTGAACTCTTTCCCTACACGACGCTCTTCCGATCTNNNTCCATCAGGTTTCTTGTGTTT
GAGCACACGTCTGAACTCTTTCCCTACACGACGCTCTTCCGATCTNNNTCCATCAGGTTTCTTGTGTTT
AGCACACGTCTGAACTCTTTCCCTACACGACGCTCTTCCGATCTNNNGAGGCTGAGGCAGGAGAATG
AGCACACGTCTGAACTCTTTCCCTACACGACGCTCTTCCGATCTNNNGAGGCTGAGGCAGGAGAATG
AGCACACGTCTGAACTCTTTCCCTACACGACGCTCTTCCGATCTNNNGAGTAGAAAGGAGAGGTTTG
AGCACACGTCTGAACTCTTTCCCTACACGACGCTCTTCCGATCTNNNGAGTAGAAAGGAGAGGTTTG
AGCACACGTCTGAACTCTTTCCCTACACGACGCTCTTCCGATCTNNNGACAGAGCAAGACTCCGTCTC
AGCACACGTCTGAACTCTTTCCCTACACGACGCTCTTCCGATCTNNNGACAGAGCAAGACTCCGTCTC
GCACACGTCTGAACTCTTTCCCTACACGACGCTCTTCCGATCTNNNCAGGAGATTGAGACCACAGTG
GCACACGTCTGAACTCTTTCCCTACACGACGCTCTTCCGATCTNNNCAGGAGATTGAGACCACAGTG
AGCACACGTCTGAACTCTTTCCCTACACGACGCTCTTCCGATCTNNNGTTTGTGGTATTCTCTAATGGC
AGCACACGTCTGAACTCTTTCCCTACACGACGCTCTTCCGATCTNNNGTTTGTGGTATTCTCTAATGGC
GCACACGTCTGAACTCTTTCCCTACACGACGCTCTTCCGATCTNNNCAACCAGTTTGTACCCAGGGATA
GCACACGTCTGAACTCTTTCCCTACACGACGCTCTTCCGATCTNNNCAACCAGTTTGTACCCAGGGATA
GCACACGTCTGAACTCTTTCCCTACACGACGCTCTTCCGATCTNNNGAGATCAAATCATGCCTCAGTG
GCACACGTCTGAACTCTTTCCCTACACGACGCTCTTCCGATCTNNNGAGATCAAATCATGCCTCAGTG
GAGCACACGTCTGAACTCTTTCCCTACACGACGCTCTTCCGATCTNNNCCTGTCACCTCCCAGTATGGA
GAGCACACGTCTGAACTCTTTCCCTACACGACGCTCTTCCGATCTNNNCCTGTCACCTCCCAGTATGGA
GCACACGTCTGAACTCTTTCCCTACACGACGCTCTTCCGATCTNNNCCTCCTTCTTGCTTCTTTTCAGCC
GCACACGTCTGAACTCTTTCCCTACACGACGCTCTTCCGATCTNNNCCTCCTTCTTGCTTCTTTTCAGCC
AGCACACGTCTGAACTCTTTCCCTACACGACGCTCTTCCGATCTNNNCAGTTTGGTAAGTTTCTCAAT
AGCACACGTCTGAACTCTTTCCCTACACGACGCTCTTCCGATCTNNNCAGTTTGGTAAGTTTCTCAAT

AGCACACGTCTGAACTCTTTCCCTACACGACGCTCTTCCGATCTNNNTACCACAAAATGACAGCAGCA
AGCACACGTCTGAACTCTTTCCCTACACGACGCTCTTCCGATCTNNNTACCACAAAATGACAGCAGCA
GAGCACACGTCTGAACTCTTTCCCTACACGACGCTCTTCCGATCTNNNCTGTAGTTCCAGCTACTTGG
GAGCACACGTCTGAACTCTTTCCCTACACGACGCTCTTCCGATCTNNNCTGTAGTTCCAGCTACTTGG
GAGCACACGTCTGAACTCTTTCCCTACACGACGCTCTTCCGATCTNNNCTGAGGCAGGAGAATTGCTT
GAGCACACGTCTGAACTCTTTCCCTACACGACGCTCTTCCGATCTNNNCTGAGGCAGGAGAATTGCTT
AGAGCACACGTCTGAACTCTTTCCCTACACGACGCTCTTCCGATCTNNNAGATGGAGGATGGAAAGA
AGAGCACACGTCTGAACTCTTTCCCTACACGACGCTCTTCCGATCTNNNAGATGGAGGATGGAAAGA
AAGAGCACACGTCTGAACTCTTTCCCTACACGACGCTCTTCCGATCTNNNATGAGAATTGCTTGAGCCT
AAGAGCACACGTCTGAACTCTTTCCCTACACGACGCTCTTCCGATCTNNNATGAGAATTGCTTGAGCCT
AGCACACGTCTGAACTCTTTCCCTACACGACGCTCTTCCGATCTNNNGATTGAGACCATCCTGGCTAA
AGCACACGTCTGAACTCTTTCCCTACACGACGCTCTTCCGATCTNNNGATTGAGACCATCCTGGCTAA
AGAGCACACGTCTGAACTCTTTCCCTACACGACGCTCTTCCGATCTNNNAGAAATGGGCATGTAAGGT
AGAGCACACGTCTGAACTCTTTCCCTACACGACGCTCTTCCGATCTNNNAGAAATGGGCATGTAAGGT
GCACACGTCTGAACTCTTTCCCTACACGACGCTCTTCCGATCTNNNAGCCAGACCCTGTCTCTAAAAAG
GCACACGTCTGAACTCTTTCCCTACACGACGCTCTTCCGATCTNNNAGCCAGACCCTGTCTCTAAAAAG
AGAGCACACGTCTGAACTCTTTCCCTACACGACGCTCTTCCGATCTNNNCTTCTTGCCTTCCTTCCTTGTT
AGAGCACACGTCTGAACTCTTTCCCTACACGACGCTCTTCCGATCTNNNCTTCTTGCCTTCCTTCCTTGTT
AGCACACGTCTGAACTCTTTCCCTACACGACGCTCTTCCGATCTNNNTGGGGGTCTGTTTACATGGAT
AGCACACGTCTGAACTCTTTCCCTACACGACGCTCTTCCGATCTNNNTGGGGGTCTGTTTACATGGAT
GAGCACACGTCTGAACTCTTTCCCTACACGACGCTCTTCCGATCTNNNTTCAATCCATTTGCGTCTTTGG
GAGCACACGTCTGAACTCTTTCCCTACACGACGCTCTTCCGATCTNNNTTCAATCCATTTGCGTCTTTGG
AGCACACGTCTGAACTCTTTCCCTACACGACGCTCTTCCGATCTNNNATAATGCCAGGTGGTGATCAG
AGCACACGTCTGAACTCTTTCCCTACACGACGCTCTTCCGATCTNNNATAATGCCAGGTGGTGATCAG
GCACACGTCTGAACTCTTTCCCTACACGACGCTCTTCCGATCTNNNTCCCCTCTCCTCTCTTTCTTTCTCAT
GCACACGTCTGAACTCTTTCCCTACACGACGCTCTTCCGATCTNNNTCCCCTCTCCTCTCTTTCTTTCTCAT
GCACACGTCTGAACTCTTTCCCTACACGACGCTCTTCCGATCTNNNCCTGACTACGGCAGCAACTTATA
GCACACGTCTGAACTCTTTCCCTACACGACGCTCTTCCGATCTNNNCCTGACTACGGCAGCAACTTATA
AGCACACGTCTGAACTCTTTCCCTACACGACGCTCTTCCGATCTNNNCTCTATTATCCCTGAGGCAACC
AGCACACGTCTGAACTCTTTCCCTACACGACGCTCTTCCGATCTNNNCTCTATTATCCCTGAGGCAACC
AGCACACGTCTGAACTCTTTCCCTACACGACGCTCTTCCGATCTNNNGCCTGTAGTCCCAGCTACTTGC
AGCACACGTCTGAACTCTTTCCCTACACGACGCTCTTCCGATCTNNNGCCTGTAGTCCCAGCTACTTGC
AGCACACGTCTGAACTCTTTCCCTACACGACGCTCTTCCGATCTNNNGAGTTCGAGACCAGCTTGAGTA
AGCACACGTCTGAACTCTTTCCCTACACGACGCTCTTCCGATCTNNNGAGTTCGAGACCAGCTTGAGTA
AGCACACGTCTGAACTCTTTCCCTACACGACGCTCTTCCGATCTNNNGGATTGAGGTGAATCAGTCCA
AGCACACGTCTGAACTCTTTCCCTACACGACGCTCTTCCGATCTNNNGGATTGAGGTGAATCAGTCCA
CACACGTCTGAACTCTTTCCCTACACGACGCTCTTCCGATCTNNNGAGAGAGGGTCACAGAGGACAG
CACACGTCTGAACTCTTTCCCTACACGACGCTCTTCCGATCTNNNGAGAGAGGGTCACAGAGGACAG
AGCACACGTCTGAACTCTTTCCCTACACGACGCTCTTCCGATCTNNNTAACAACACACACACACACACA
AGCACACGTCTGAACTCTTTCCCTACACGACGCTCTTCCGATCTNNNTAACAACACACACACACACACA
AGCACACGTCTGAACTCTTTCCCTACACGACGCTCTTCCGATCTNNNCACACACACATACACACACACA
AGCACACGTCTGAACTCTTTCCCTACACGACGCTCTTCCGATCTNNNCACACACACATACACACACACA
GAGCACACGTCTGAACTCTTTCCCTACACGACGCTCTTCCGATCTNNNGAACCAATCCAAATGCCCATC

GAGCACACGTCTGAACTCTTTCCCTACACGACGCTCTTCCGATCTNNNGAACCAATCCAAATGCCCATC
GAGCACACGTCTGAACTCTTTCCCTACACGACGCTCTTCCGATCTNNNGAGGCTGAGGCAGGAGAAT
GAGCACACGTCTGAACTCTTTCCCTACACGACGCTCTTCCGATCTNNNGAGGCTGAGGCAGGAGAAT
GAGCACACGTCTGAACTCTTTCCCTACACGACGCTCTTCCGATCTNNNTGGATTGCCTAGGTATACAC
GAGCACACGTCTGAACTCTTTCCCTACACGACGCTCTTCCGATCTNNNTGGATTGCCTAGGTATACAC
GAGCACACGTCTGAACTCTTTCCCTACACGACGCTCTTCCGATCTNNNTGTCAACAAAGCACAGTTTG
GAGCACACGTCTGAACTCTTTCCCTACACGACGCTCTTCCGATCTNNNTGTCAACAAAGCACAGTTTG
AGCACACGTCTGAACTCTTTCCCTACACGACGCTCTTCCGATCTNNNTGCCACCTTTCACCTTATACTGG
AGCACACGTCTGAACTCTTTCCCTACACGACGCTCTTCCGATCTNNNTGCCACCTTTCACCTTATACTGG
AGCACACGTCTGAACTCTTTCCCTACACGACGCTCTTCCGATCTNNNTCTGAAAAATAGCAGGCCAGA
AGCACACGTCTGAACTCTTTCCCTACACGACGCTCTTCCGATCTNNNTCTGAAAAATAGCAGGCCAGA
AGCACACGTCTGAACTCTTTCCCTACACGACGCTCTTCCGATCTNNNAAGAATAGCTTGAACCCAGGAG
AGCACACGTCTGAACTCTTTCCCTACACGACGCTCTTCCGATCTNNNAAGAATAGCTTGAACCCAGGAG
GCACACGTCTGAACTCTTTCCCTACACGACGCTCTTCCGATCTNNNCTGTAATCCCAGCTACTCCAGAG
GCACACGTCTGAACTCTTTCCCTACACGACGCTCTTCCGATCTNNNCTGTAATCCCAGCTACTCCAGAG
GAGCACACGTCTGAACTCTTTCCCTACACGACGCTCTTCCGATCTNNNATATACAGGCAGGACACACA
GAGCACACGTCTGAACTCTTTCCCTACACGACGCTCTTCCGATCTNNNATATACAGGCAGGACACACA
AGCACACGTCTGAACTCTTTCCCTACACGACGCTCTTCCGATCTNNNTATCCTGTTGATTTGGGGTGG
AGCACACGTCTGAACTCTTTCCCTACACGACGCTCTTCCGATCTNNNTATCCTGTTGATTTGGGGTGG
AGCACACGTCTGAACTCTTTCCCTACACGACGCTCTTCCGATCTNNNCTTGACAACAGACAAAGCAAG
AGCACACGTCTGAACTCTTTCCCTACACGACGCTCTTCCGATCTNNNCTTGACAACAGACAAAGCAAG
AGCACACGTCTGAACTCTTTCCCTACACGACGCTCTTCCGATCTNNNGAGGTTACAGTGAGCTGAGAT
AGCACACGTCTGAACTCTTTCCCTACACGACGCTCTTCCGATCTNNNGAGGTTACAGTGAGCTGAGAT
GAGCACACGTCTGAACTCTTTCCCTACACGACGCTCTTCCGATCTNNNCCTGACTTCATTTTAGGCACTG
GAGCACACGTCTGAACTCTTTCCCTACACGACGCTCTTCCGATCTNNNCCTGACTTCATTTTAGGCACTG
AGCACACGTCTGAACTCTTTCCCTACACGACGCTCTTCCGATCTNNNTGCATTTGTTTGCTCTTGCTTCT
AGCACACGTCTGAACTCTTTCCCTACACGACGCTCTTCCGATCTNNNTGCATTTGTTTGCTCTTGCTTCT
GCACACGTCTGAACTCTTTCCCTACACGACGCTCTTCCGATCTNNNCAGGATGGCATTTCTTAGGGAC
GCACACGTCTGAACTCTTTCCCTACACGACGCTCTTCCGATCTNNNCAGGATGGCATTTCTTAGGGAC
GAGCACACGTCTGAACTCTTTCCCTACACGACGCTCTTCCGATCTNNNAGTACACTCCTGCCCAAATAC
GAGCACACGTCTGAACTCTTTCCCTACACGACGCTCTTCCGATCTNNNAGTACACTCCTGCCCAAATAC
GCACACGTCTGAACTCTTTCCCTACACGACGCTCTTCCGATCTNNNCTGGGTGTTCTGCTCTTCTGTAT
GCACACGTCTGAACTCTTTCCCTACACGACGCTCTTCCGATCTNNNCTGGGTGTTCTGCTCTTCTGTAT
AGCACACGTCTGAACTCTTTCCCTACACGACGCTCTTCCGATCTNNNGGGAGAATCATTTGAACCTGG
AGCACACGTCTGAACTCTTTCCCTACACGACGCTCTTCCGATCTNNNGGGAGAATCATTTGAACCTGG
CACACGTCTGAACTCTTTCCCTACACGACGCTCTTCCGATCTNNNTAGTCATAGTTACTCGGGAGGCT
CACACGTCTGAACTCTTTCCCTACACGACGCTCTTCCGATCTNNNTAGTCATAGTTACTCGGGAGGCT
AGCACACGTCTGAACTCTTTCCCTACACGACGCTCTTCCGATCTNNNAATGAATGTGGGGCTGAAAAG
AGCACACGTCTGAACTCTTTCCCTACACGACGCTCTTCCGATCTNNNAATGAATGTGGGGCTGAAAAG
GCACACGTCTGAACTCTTTCCCTACACGACGCTCTTCCGATCTNNNTTGTCATATTTGCGTACCCACCC
GCACACGTCTGAACTCTTTCCCTACACGACGCTCTTCCGATCTNNNTTGTCATATTTGCGTACCCACCC
GAGCACACGTCTGAACTCTTTCCCTACACGACGCTCTTCCGATCTNNNGGATGCTCTACTCTAATTGG
GAGCACACGTCTGAACTCTTTCCCTACACGACGCTCTTCCGATCTNNNGGATGCTCTACTCTAATTGG

GAGCACACGTCTGAACTCTTTCCCTACACGACGCTCTTCCGATCTNNNGGCCAGGAGAGACTTTATT
GAGCACACGTCTGAACTCTTTCCCTACACGACGCTCTTCCGATCTNNNGGCCAGGAGAGACTTTATT
GCACACGTCTGAACTCTTTCCCTACACGACGCTCTTCCGATCTNNNCACCAGTTTCTCTGCTTTCCACC
GCACACGTCTGAACTCTTTCCCTACACGACGCTCTTCCGATCTNNNCACCAGTTTCTCTGCTTTCCACC
GCACACGTCTGAACTCTTTCCCTACACGACGCTCTTCCGATCTNNNGGGCGACAGAGAAAAACTTCA
GCACACGTCTGAACTCTTTCCCTACACGACGCTCTTCCGATCTNNNGGGCGACAGAGAAAAACTTCA
GCACACGTCTGAACTCTTTCCCTACACGACGCTCTTCCGATCTNNNCTCATGTCCAAGGGCAAAAGAA
GCACACGTCTGAACTCTTTCCCTACACGACGCTCTTCCGATCTNNNCTCATGTCCAAGGGCAAAAGAA
AGCACACGTCTGAACTCTTTCCCTACACGACGCTCTTCCGATCTNNNGGAGGTTACAGTGAGCTGAG
AGCACACGTCTGAACTCTTTCCCTACACGACGCTCTTCCGATCTNNNGGAGGTTACAGTGAGCTGAG
GAGCACACGTCTGAACTCTTTCCCTACACGACGCTCTTCCGATCTNNNATCTGTAGTCCCAGCTACTC
GAGCACACGTCTGAACTCTTTCCCTACACGACGCTCTTCCGATCTNNNATCTGTAGTCCCAGCTACTC
GAGCACACGTCTGAACTCTTTCCCTACACGACGCTCTTCCGATCTNNNTCTTGAGTCCAGGAGTTTGA
GAGCACACGTCTGAACTCTTTCCCTACACGACGCTCTTCCGATCTNNNTCTTGAGTCCAGGAGTTTGA
GAGCACACGTCTGAACTCTTTCCCTACACGACGCTCTTCCGATCTNNNCCAGCAACTAAGTGACTTGC
GAGCACACGTCTGAACTCTTTCCCTACACGACGCTCTTCCGATCTNNNCCAGCAACTAAGTGACTTGC
AGCACACGTCTGAACTCTTTCCCTACACGACGCTCTTCCGATCTNNNGTGAGGCTGGAGAATTCCTTG
AGCACACGTCTGAACTCTTTCCCTACACGACGCTCTTCCGATCTNNNGTGAGGCTGGAGAATTCCTTG
AGCACACGTCTGAACTCTTTCCCTACACGACGCTCTTCCGATCTNNNAGGTCTGAGAGAAGGAGAAG
AGCACACGTCTGAACTCTTTCCCTACACGACGCTCTTCCGATCTNNNAGGTCTGAGAGAAGGAGAAG
GAGCACACGTCTGAACTCTTTCCCTACACGACGCTCTTCCGATCTNNNACATTTTGGCATGATTTTGCA
GAGCACACGTCTGAACTCTTTCCCTACACGACGCTCTTCCGATCTNNNACATTTTGGCATGATTTTGCA
AGCACACGTCTGAACTCTTTCCCTACACGACGCTCTTCCGATCTNNNTGACTGTAATCCCAGCTACTT
AGCACACGTCTGAACTCTTTCCCTACACGACGCTCTTCCGATCTNNNTGACTGTAATCCCAGCTACTT
GCACACGTCTGAACTCTTTCCCTACACGACGCTCTTCCGATCTNNNCTGGGCAAGAAAGTGAGACAC
GCACACGTCTGAACTCTTTCCCTACACGACGCTCTTCCGATCTNNNCTGGGCAAGAAAGTGAGACAC
AGCACACGTCTGAACTCTTTCCCTACACGACGCTCTTCCGATCTNNNTTCCTGAAATGTGACACATGC
AGCACACGTCTGAACTCTTTCCCTACACGACGCTCTTCCGATCTNNNTTCCTGAAATGTGACACATGC
AGCACACGTCTGAACTCTTTCCCTACACGACGCTCTTCCGATCTNNNCATCTGCTGTCTTTTCTTTGCCA
AGCACACGTCTGAACTCTTTCCCTACACGACGCTCTTCCGATCTNNNCATCTGCTGTCTTTTCTTTGCCA
AGCACACGTCTGAACTCTTTCCCTACACGACGCTCTTCCGATCTNNNAGGAGAGTAAAAAGGAGACA
AGCACACGTCTGAACTCTTTCCCTACACGACGCTCTTCCGATCTNNNAGGAGAGTAAAAAGGAGACA
AGCACACGTCTGAACTCTTTCCCTACACGACGCTCTTCCGATCTNNNCTCAGTCTCCCAAGTAGCTGAA
AGCACACGTCTGAACTCTTTCCCTACACGACGCTCTTCCGATCTNNNCTCAGTCTCCCAAGTAGCTGAA
GAGCACACGTCTGAACTCTTTCCCTACACGACGCTCTTCCGATCTNNNTGAGTTGAGATCTCACCAAT
GAGCACACGTCTGAACTCTTTCCCTACACGACGCTCTTCCGATCTNNNTGAGTTGAGATCTCACCAAT
CACACGTCTGAACTCTTTCCCTACACGACGCTCTTCCGATCTNNNCTGACCTCGGATGGTGAGCTCAT
CACACGTCTGAACTCTTTCCCTACACGACGCTCTTCCGATCTNNNCTGACCTCGGATGGTGAGCTCAT
GAGCACACGTCTGAACTCTTTCCCTACACGACGCTCTTCCGATCTNNNATCAGGAAAAGTGTCCTCTA
GAGCACACGTCTGAACTCTTTCCCTACACGACGCTCTTCCGATCTNNNATCAGGAAAAGTGTCCTCTA
GCACACGTCTGAACTCTTTCCCTACACGACGCTCTTCCGATCTNNNGCTGCCTATCCTAGAGCTTTTCT
GCACACGTCTGAACTCTTTCCCTACACGACGCTCTTCCGATCTNNNGCTGCCTATCCTAGAGCTTTTCT
GCACACGTCTGAACTCTTTCCCTACACGACGCTCTTCCGATCTNNNCCTATAGTCTCAGCTGCTTGGA

AGCACACGTCTGAACTCTTTCCCTACACGACGCTCTTCCGATCTNNNCCTATAGTCTCAGCTGCTTGGAC
AGCACACGTCTGAACTCTTTCCCTACACGACGCTCTTCCGATCTNNNCGACGAAGAAGAGGAGGATGA
AGCACACGTCTGAACTCTTTCCCTACACGACGCTCTTCCGATCTNNNCGACGAAGAAGAGGAGGATGA
GAGCACACGTCTGAACTCTTTCCCTACACGACGCTCTTCCGATCTNNNACTGGTTTCTGTGCATTTGAAG
GAGCACACGTCTGAACTCTTTCCCTACACGACGCTCTTCCGATCTNNNACTGGTTTCTGTGCATTTGAAG
AGCACACGTCTGAACTCTTTCCCTACACGACGCTCTTCCGATCTNNNACATACTGAGGATATGTGGCAG
AGCACACGTCTGAACTCTTTCCCTACACGACGCTCTTCCGATCTNNNACATACTGAGGATATGTGGCAG
AGCACACGTCTGAACTCTTTCCCTACACGACGCTCTTCCGATCTNNNCCTTGTGATCTGGTCTCCATCTT
AGCACACGTCTGAACTCTTTCCCTACACGACGCTCTTCCGATCTNNNCCTTGTGATCTGGTCTCCATCTT
GCACACGTCTGAACTCTTTCCCTACACGACGCTCTTCCGATCTNNNATCAAAGTTCTGAGACTTAATCTC
GCACACGTCTGAACTCTTTCCCTACACGACGCTCTTCCGATCTNNNATCAAAGTTCTGAGACTTAATCTC
AGCACACGTCTGAACTCTTTCCCTACACGACGCTCTTCCGATCTNNNCGGTTTGTGGACAACATGTACA
AGCACACGTCTGAACTCTTTCCCTACACGACGCTCTTCCGATCTNNNCGGTTTGTGGACAACATGTACA
AGCACACGTCTGAACTCTTTCCCTACACGACGCTCTTCCGATCTNNNCAGACACCTCTCTAGACCTTGTC
AGCACACGTCTGAACTCTTTCCCTACACGACGCTCTTCCGATCTNNNCAGACACCTCTCTAGACCTTGTC
GAGCACACGTCTGAACTCTTTCCCTACACGACGCTCTTCCGATCTNNNGAATCACTTGAACCTGGGAGT
GAGCACACGTCTGAACTCTTTCCCTACACGACGCTCTTCCGATCTNNNGAATCACTTGAACCTGGGAGT
GAGCACACGTCTGAACTCTTTCCCTACACGACGCTCTTCCGATCTNNNCCTCCTTCAACATGTTCCCATTT
GAGCACACGTCTGAACTCTTTCCCTACACGACGCTCTTCCGATCTNNNCCTCCTTCAACATGTTCCCATTT
GAGCACACGTCTGAACTCTTTCCCTACACGACGCTCTTCCGATCTNNNCGTTGTTCAGTGGGGGACTTT
GAGCACACGTCTGAACTCTTTCCCTACACGACGCTCTTCCGATCTNNNCGTTGTTCAGTGGGGGACTTT
AGCACACGTCTGAACTCTTTCCCTACACGACGCTCTTCCGATCTNNNTCTGTTCCTGAAATGAACGACCA
AGCACACGTCTGAACTCTTTCCCTACACGACGCTCTTCCGATCTNNNTCTGTTCCTGAAATGAACGACCA
GCACACGTCTGAACTCTTTCCCTACACGACGCTCTTCCGATCTNNNAAAAACTCACTCCATGCTCACCTCA
GCACACGTCTGAACTCTTTCCCTACACGACGCTCTTCCGATCTNNNAAAAACTCACTCCATGCTCACCTCA
GAGCACACGTCTGAACTCTTTCCCTACACGACGCTCTTCCGATCTNNNTTGTCTATGTCCTAAGTGCTGC
GAGCACACGTCTGAACTCTTTCCCTACACGACGCTCTTCCGATCTNNNTTGTCTATGTCCTAAGTGCTGC
GAGCACACGTCTGAACTCTTTCCCTACACGACGCTCTTCCGATCTNNNGGGGAGGTGTGTAGCTATTTT
GAGCACACGTCTGAACTCTTTCCCTACACGACGCTCTTCCGATCTNNNGGGGAGGTGTGTAGCTATTTT
AGCACACGTCTGAACTCTTTCCCTACACGACGCTCTTCCGATCTNNNTGCTATTGTGAATTGTGCTGCA
AGCACACGTCTGAACTCTTTCCCTACACGACGCTCTTCCGATCTNNNTGCTATTGTGAATTGTGCTGCA
AGCACACGTCTGAACTCTTTCCCTACACGACGCTCTTCCGATCTNNNTGGGTGACACATCAAGTCTCAAT
AGCACACGTCTGAACTCTTTCCCTACACGACGCTCTTCCGATCTNNNTGGGTGACACATCAAGTCTCAAT
AGCACACGTCTGAACTCTTTCCCTACACGACGCTCTTCCGATCTNNNGCAGATTTTAGCATGGAACCTG
AGCACACGTCTGAACTCTTTCCCTACACGACGCTCTTCCGATCTNNNGCAGATTTTAGCATGGAACCTG
AGCACACGTCTGAACTCTTTCCCTACACGACGCTCTTCCGATCTNNNGACTCCTCCAAAAAGCTCCTACA
AGCACACGTCTGAACTCTTTCCCTACACGACGCTCTTCCGATCTNNNGACTCCTCCAAAAAGCTCCTACA
GAGCACACGTCTGAACTCTTTCCCTACACGACGCTCTTCCGATCTNNNCCATCACCTTCAAAAGTTTCCT
GAGCACACGTCTGAACTCTTTCCCTACACGACGCTCTTCCGATCTNNNCCATCACCTTCAAAAGTTTCCT
GAGCACACGTCTGAACTCTTTCCCTACACGACGCTCTTCCGATCTNNNCTGTAATCCCAGCTTCTCAGGA
GAGCACACGTCTGAACTCTTTCCCTACACGACGCTCTTCCGATCTNNNCTGTAATCCCAGCTTCTCAGGA
GAGCACACGTCTGAACTCTTTCCCTACACGACGCTCTTCCGATCTNNNACAGGGAGATAGGACTTGAA
GAGCACACGTCTGAACTCTTTCCCTACACGACGCTCTTCCGATCTNNNACAGGGAGATAGGACTTGAA

AGCACACGTCTGAACTCTTTCCCTACACGACGCTCTTCCGATCTNNNTGAGGTGAATTTCCACCCTTA
AGCACACGTCTGAACTCTTTCCCTACACGACGCTCTTCCGATCTNNNTGAGGTGAATTTCCACCCTTA
AGCACACGTCTGAACTCTTTCCCTACACGACGCTCTTCCGATCTNNNCAAGGGTCATTTATGGTTGCA
AGCACACGTCTGAACTCTTTCCCTACACGACGCTCTTCCGATCTNNNCAAGGGTCATTTATGGTTGCA
AGCACACGTCTGAACTCTTTCCCTACACGACGCTCTTCCGATCTNNNCCTCATCCCCTGAGTATGTGG
AGCACACGTCTGAACTCTTTCCCTACACGACGCTCTTCCGATCTNNNCCTCATCCCCTGAGTATGTGG
AGCACACGTCTGAACTCTTTCCCTACACGACGCTCTTCCGATCTNNNGCCTAGGCAAAAGAACAAGAC
AGCACACGTCTGAACTCTTTCCCTACACGACGCTCTTCCGATCTNNNGCCTAGGCAAAAGAACAAGAC
AGCACACGTCTGAACTCTTTCCCTACACGACGCTCTTCCGATCTNNNTCTACGAGGAAACGAAGGTGA
AGCACACGTCTGAACTCTTTCCCTACACGACGCTCTTCCGATCTNNNTCTACGAGGAAACGAAGGTGA
AGCACACGTCTGAACTCTTTCCCTACACGACGCTCTTCCGATCTNNNTTCACACCATTCTCCTGCCTCCA
AGCACACGTCTGAACTCTTTCCCTACACGACGCTCTTCCGATCTNNNTTCACACCATTCTCCTGCCTCCA
GAGCACACGTCTGAACTCTTTCCCTACACGACGCTCTTCCGATCTNNNAGGACACCTGTTTTTCCTCAT
GAGCACACGTCTGAACTCTTTCCCTACACGACGCTCTTCCGATCTNNNAGGACACCTGTTTTTCCTCAT
AGCACACGTCTGAACTCTTTCCCTACACGACGCTCTTCCGATCTNNNTGAGCTGAGACCACACCATTG
AGCACACGTCTGAACTCTTTCCCTACACGACGCTCTTCCGATCTNNNTGAGCTGAGACCACACCATTG
ACACGTCTGAACTCTTTCCCTACACGACGCTCTTCCGATCTNNNCCTTCAGGCATTCTTACAGGGTGCA
ACACGTCTGAACTCTTTCCCTACACGACGCTCTTCCGATCTNNNCCTTCAGGCATTCTTACAGGGTGCA
CACACGTCTGAACTCTTTCCCTACACGACGCTCTTCCGATCTNNNTCCACCCCTTGTTTGGCATATAAC
CACACGTCTGAACTCTTTCCCTACACGACGCTCTTCCGATCTNNNTCCACCCCTTGTTTGGCATATAAC
GCACACGTCTGAACTCTTTCCCTACACGACGCTCTTCCGATCTNNNGGTCTCAAACTCCTGACCTCAGC
GCACACGTCTGAACTCTTTCCCTACACGACGCTCTTCCGATCTNNNGGTCTCAAACTCCTGACCTCAGC
GAGCACACGTCTGAACTCTTTCCCTACACGACGCTCTTCCGATCTNNNTTGCCACAACAGCAGTATAG
GAGCACACGTCTGAACTCTTTCCCTACACGACGCTCTTCCGATCTNNNTTGCCACAACAGCAGTATAG
GCACACGTCTGAACTCTTTCCCTACACGACGCTCTTCCGATCTNNNGCCCAAGCTACAGAAGCATTTTT
GCACACGTCTGAACTCTTTCCCTACACGACGCTCTTCCGATCTNNNGCCCAAGCTACAGAAGCATTTTT
AGCACACGTCTGAACTCTTTCCCTACACGACGCTCTTCCGATCTNNNCACTGTGAGATGGTAGCTGCTT
AGCACACGTCTGAACTCTTTCCCTACACGACGCTCTTCCGATCTNNNCACTGTGAGATGGTAGCTGCTT
AGCACACGTCTGAACTCTTTCCCTACACGACGCTCTTCCGATCTNNNTCATTCACCACCGTTCTACTAGC
AGCACACGTCTGAACTCTTTCCCTACACGACGCTCTTCCGATCTNNNTCATTCACCACCGTTCTACTAGC
AGCACACGTCTGAACTCTTTCCCTACACGACGCTCTTCCGATCTNNNAACAAAACTCTGGTTCTCTCTG
AGCACACGTCTGAACTCTTTCCCTACACGACGCTCTTCCGATCTNNNAACAAAACTCTGGTTCTCTCTG
GAGCACACGTCTGAACTCTTTCCCTACACGACGCTCTTCCGATCTNNNGATCACAAGGTCAGGAGATC
GAGCACACGTCTGAACTCTTTCCCTACACGACGCTCTTCCGATCTNNNGATCACAAGGTCAGGAGATC
AGCACACGTCTGAACTCTTTCCCTACACGACGCTCTTCCGATCTNNNGAGTCAGTGAGTGGTGAGTG
AGCACACGTCTGAACTCTTTCCCTACACGACGCTCTTCCGATCTNNNGAGTCAGTGAGTGGTGAGTG
AGCACACGTCTGAACTCTTTCCCTACACGACGCTCTTCCGATCTNNNGGGGAGCCCTATCTGTTTCTT
AGCACACGTCTGAACTCTTTCCCTACACGACGCTCTTCCGATCTNNNGGGGAGCCCTATCTGTTTCTT
GCACACGTCTGAACTCTTTCCCTACACGACGCTCTTCCGATCTNNNTTCTTAGGTCCAACAGGAAACCC
GCACACGTCTGAACTCTTTCCCTACACGACGCTCTTCCGATCTNNNTTCTTAGGTCCAACAGGAAACCC
AGCACACGTCTGAACTCTTTCCCTACACGACGCTCTTCCGATCTNNNTTCTTCTTCTTCCTCCTCCTCCTC
AGCACACGTCTGAACTCTTTCCCTACACGACGCTCTTCCGATCTNNNTTCTTCTTCTTCCTCCTCCTCCTC
GAGCACACGTCTGAACTCTTTCCCTACACGACGCTCTTCCGATCTNNNGGTGCTACTGTTTATGGCAA

GAGCACACGTCTGAACTCTTTCCCTACACGACGCTCTTCCGATCTNNNGGTGCTACTGTTTATGGCAA

AGCACACGTCTGAACTCTTTCCCTACACGACGCTCTTCCGATCTNNNGCACACACATGTTTATAGCAG

AGCACACGTCTGAACTCTTTCCCTACACGACGCTCTTCCGATCTNNNGCACACACATGTTTATAGCAG

GCACACGTCTGAACTCTTTCCCTACACGACGCTCTTCCGATCTNNNTTAGGATGGGAGGATGGTGGA

GCACACGTCTGAACTCTTTCCCTACACGACGCTCTTCCGATCTNNNTTAGGATGGGAGGATGGTGGA

AGCACACGTCTGAACTCTTTCCCTACACGACGCTCTTCCGATCTNNNTGAGAAAAGCGTGGTTTTTCA

AGCACACGTCTGAACTCTTTCCCTACACGACGCTCTTCCGATCTNNNTGAGAAAAGCGTGGTTTTTCA

AAGAGCACACGTCTGAACTCTTTCCCTACACGACGCTCTTCCGATCTNNNGAGACAGGAGAATGGCG

AAGAGCACACGTCTGAACTCTTTCCCTACACGACGCTCTTCCGATCTNNNGAGACAGGAGAATGGCG

GAGCACACGTCTGAACTCTTTCCCTACACGACGCTCTTCCGATCTNNNAAAATGGTGTCTCCTGCCTG

GAGCACACGTCTGAACTCTTTCCCTACACGACGCTCTTCCGATCTNNNAAAATGGTGTCTCCTGCCTG

AGCACACGTCTGAACTCTTTCCCTACACGACGCTCTTCCGATCTNNNCTCACTTGGATGGTCCTAAAG

AGCACACGTCTGAACTCTTTCCCTACACGACGCTCTTCCGATCTNNNCTCACTTGGATGGTCCTAAAG

GAGCACACGTCTGAACTCTTTCCCTACACGACGCTCTTCCGATCTNNNTTCCTGAAGTATGGTACCTT

GAGCACACGTCTGAACTCTTTCCCTACACGACGCTCTTCCGATCTNNNTTCCTGAAGTATGGTACCTT

GCACACGTCTGAACTCTTTCCCTACACGACGCTCTTCCGATCTNNNAGATGGTCAGATTCCCCCTTTTC

GCACACGTCTGAACTCTTTCCCTACACGACGCTCTTCCGATCTNNNAGATGGTCAGATTCCCCCTTTTC

AGCACACGTCTGAACTCTTTCCCTACACGACGCTCTTCCGATCTNNNTTGGCTGTATGTGAGATAGCT

AGCACACGTCTGAACTCTTTCCCTACACGACGCTCTTCCGATCTNNNTTGGCTGTATGTGAGATAGCT

GAGCACACGTCTGAACTCTTTCCCTACACGACGCTCTTCCGATCTNNNTGAGTTTCTAAGCTACTGGCA

GAGCACACGTCTGAACTCTTTCCCTACACGACGCTCTTCCGATCTNNNTGAGTTTCTAAGCTACTGGCA

AGCACACGTCTGAACTCTTTCCCTACACGACGCTCTTCCGATCTNNNGATCGAGACCATCCTGGCTAA

AGCACACGTCTGAACTCTTTCCCTACACGACGCTCTTCCGATCTNNNGATCGAGACCATCCTGGCTAA

GCACACGTCTGAACTCTTTCCCTACACGACGCTCTTCCGATCTNNNCATGCCTGTAATCCCAGCTACTCC

GCACACGTCTGAACTCTTTCCCTACACGACGCTCTTCCGATCTNNNCATGCCTGTAATCCCAGCTACTCC

AGCACACGTCTGAACTCTTTCCCTACACGACGCTCTTCCGATCTNNNCTTTGCTCCTCTGCTCTGAGAT

AGCACACGTCTGAACTCTTTCCCTACACGACGCTCTTCCGATCTNNNCTTTGCTCCTCTGCTCTGAGAT

AGCACACGTCTGAACTCTTTCCCTACACGACGCTCTTCCGATCTNNNAGCCCCCACTATAACTTGTGTT

AGCACACGTCTGAACTCTTTCCCTACACGACGCTCTTCCGATCTNNNAGCCCCCACTATAACTTGTGTT

GCACACGTCTGAACTCTTTCCCTACACGACGCTCTTCCGATCTNNNTCCAAGGATATCAAAGGAGGA

GCACACGTCTGAACTCTTTCCCTACACGACGCTCTTCCGATCTNNNTCCAAGGATATCAAAGGAGGA

AGCACACGTCTGAACTCTTTCCCTACACGACGCTCTTCCGATCTNNNTGACAGAAAGGACAAACTCCC

AGCACACGTCTGAACTCTTTCCCTACACGACGCTCTTCCGATCTNNNTGACAGAAAGGACAAACTCCC

AGCACACGTCTGAACTCTTTCCCTACACGACGCTCTTCCGATCTNNNGGTGTCCATTTCTCCAAACACA

AGCACACGTCTGAACTCTTTCCCTACACGACGCTCTTCCGATCTNNNGGTGTCCATTTCTCCAAACACA

GAGCACACGTCTGAACTCTTTCCCTACACGACGCTCTTCCGATCTNNNCTGAGGCAGGAGAATCACTT

GAGCACACGTCTGAACTCTTTCCCTACACGACGCTCTTCCGATCTNNNCTGAGGCAGGAGAATCACTT

AGCACACGTCTGAACTCTTTCCCTACACGACGCTCTTCCGATCTNNNGTGAGAAGCAAACTGTTCTCT

AGCACACGTCTGAACTCTTTCCCTACACGACGCTCTTCCGATCTNNNGTGAGAAGCAAACTGTTCTCT

GAGCACACGTCTGAACTCTTTCCCTACACGACGCTCTTCCGATCTNNNCTTCCAAGTAGCTGGGACTA

GAGCACACGTCTGAACTCTTTCCCTACACGACGCTCTTCCGATCTNNNCTTCCAAGTAGCTGGGACTA

AGCACACGTCTGAACTCTTTCCCTACACGACGCTCTTCCGATCTNNNCGAATGTTAGAATGGGGAAA

AGCACACGTCTGAACTCTTTCCCTACACGACGCTCTTCCGATCTNNNCGAATGTTAGAATGGGGAAA

GAGCACACGTCTGAACTCTTTCCCTACACGACGCTCTTCCGATCTNNNGGTTCAAGTGATTTTCCTGCC

GAGCACACGTCTGAACTCTTTCCCTACACGACGCTCTTCCGATCTNNNGGTTCAAGTGATTTTCCTGCC

CACACGTCTGAACTCTTTCCCTACACGACGCTCTTCCGATCTNNNAAACAAGTGACTTCTGCGAATGG

CACACGTCTGAACTCTTTCCCTACACGACGCTCTTCCGATCTNNNAAACAAGTGACTTCTGCGAATGG

AGCACACGTCTGAACTCTTTCCCTACACGACGCTCTTCCGATCTNNNACAGCACTCCTTCTGGATAAAC

AGCACACGTCTGAACTCTTTCCCTACACGACGCTCTTCCGATCTNNNACAGCACTCCTTCTGGATAAAC

AGCACACGTCTGAACTCTTTCCCTACACGACGCTCTTCCGATCTNNNGGAAGGATCCATGTCGGTAA

AGCACACGTCTGAACTCTTTCCCTACACGACGCTCTTCCGATCTNNNGGAAGGATCCATGTCGGTAA

ACACGTCTGAACTCTTTCCCTACACGACGCTCTTCCGATCTNNNAGAGGAGACAGAGAGAGATCAGG

ACACGTCTGAACTCTTTCCCTACACGACGCTCTTCCGATCTNNNAGAGGAGACAGAGAGAGATCAGG

GCACACGTCTGAACTCTTTCCCTACACGACGCTCTTCCGATCTNNNGTGATCCCAGACACTCAGAGAA

GCACACGTCTGAACTCTTTCCCTACACGACGCTCTTCCGATCTNNNGTGATCCCAGACACTCAGAGAA

GCACACGTCTGAACTCTTTCCCTACACGACGCTCTTCCGATCTNNNCCAATGTCAGCCTTCTTCAGCGC

GCACACGTCTGAACTCTTTCCCTACACGACGCTCTTCCGATCTNNNCCAATGTCAGCCTTCTTCAGCGC

AGCACACGTCTGAACTCTTTCCCTACACGACGCTCTTCCGATCTNNNCCCGGCAGCTGTTTCTACTATA

AGCACACGTCTGAACTCTTTCCCTACACGACGCTCTTCCGATCTNNNCCCGGCAGCTGTTTCTACTATA

AGCACACGTCTGAACTCTTTCCCTACACGACGCTCTTCCGATCTNNNCTGGCTTCAAGCAATTCTCCTGC

AGCACACGTCTGAACTCTTTCCCTACACGACGCTCTTCCGATCTNNNCTGGCTTCAAGCAATTCTCCTGC

AGCACACGTCTGAACTCTTTCCCTACACGACGCTCTTCCGATCTNNNGATACAAGTGGAGGAGTAGC

AGCACACGTCTGAACTCTTTCCCTACACGACGCTCTTCCGATCTNNNGATACAAGTGGAGGAGTAGC

GCACACGTCTGAACTCTTTCCCTACACGACGCTCTTCCGATCTNNNACATCTTACAAGTACAGCTGCCA

GCACACGTCTGAACTCTTTCCCTACACGACGCTCTTCCGATCTNNNACATCTTACAAGTACAGCTGCCA

GAGCACACGTCTGAACTCTTTCCCTACACGACGCTCTTCCGATCTNNNGCCAATATGGAGAAACCCTG

GAGCACACGTCTGAACTCTTTCCCTACACGACGCTCTTCCGATCTNNNGCCAATATGGAGAAACCCTG

GAGCACACGTCTGAACTCTTTCCCTACACGACGCTCTTCCGATCTNNNGTGTAGTTCCAGCTACTTGG

GAGCACACGTCTGAACTCTTTCCCTACACGACGCTCTTCCGATCTNNNGTGTAGTTCCAGCTACTTGG

GAGCACACGTCTGAACTCTTTCCCTACACGACGCTCTTCCGATCTNNNTCTCTGGAGATCCATACCACC

GAGCACACGTCTGAACTCTTTCCCTACACGACGCTCTTCCGATCTNNNTCTCTGGAGATCCATACCACC

GAGCACACGTCTGAACTCTTTCCCTACACGACGCTCTTCCGATCTNNNTCAAACCGGAGCATCATTTCA

GAGCACACGTCTGAACTCTTTCCCTACACGACGCTCTTCCGATCTNNNTCAAACCGGAGCATCATTTCA

AGCACACGTCTGAACTCTTTCCCTACACGACGCTCTTCCGATCTNNNCAATGGAGGGGAACTGAAGAC

AGCACACGTCTGAACTCTTTCCCTACACGACGCTCTTCCGATCTNNNCAATGGAGGGGAACTGAAGAC

GCACACGTCTGAACTCTTTCCCTACACGACGCTCTTCCGATCTNNNGTCCCTGGAAGATGCTGTTTTC

GCACACGTCTGAACTCTTTCCCTACACGACGCTCTTCCGATCTNNNGTCCCTGGAAGATGCTGTTTTC

AGCACACGTCTGAACTCTTTCCCTACACGACGCTCTTCCGATCTNNNTTTACACTCCCAAACTCCAGTG

AGCACACGTCTGAACTCTTTCCCTACACGACGCTCTTCCGATCTNNNTTTACACTCCCAAACTCCAGTG

GAGCACACGTCTGAACTCTTTCCCTACACGACGCTCTTCCGATCTNNNAAGCATTGGGATTACAGGT

GAGCACACGTCTGAACTCTTTCCCTACACGACGCTCTTCCGATCTNNNAAGCATTGGGATTACAGGT

AGCACACGTCTGAACTCTTTCCCTACACGACGCTCTTCCGATCTNNNTGTGAACTCTGTTACGCATCAG

AGCACACGTCTGAACTCTTTCCCTACACGACGCTCTTCCGATCTNNNTGTGAACTCTGTTACGCATCAG

GAGCACACGTCTGAACTCTTTCCCTACACGACGCTCTTCCGATCTNNNAAAGGTAGAGCCATACCTCC

GAGCACACGTCTGAACTCTTTCCCTACACGACGCTCTTCCGATCTNNNAAAGGTAGAGCCATACCTCC

AGCACACGTCTGAACTCTTTCCCTACACGACGCTCTTCCGATCTNNNGGCAGATCATTTGAGGTCAGG

AGCACACGTCTGAACTCTTTCCCTACACGACGCTCTTCCGATCTNNNGGCAGATCATTTGAGGTCAGG
GAGCACACGTCTGAACTCTTTCCCTACACGACGCTCTTCCGATCTNNNTTGGGAACTATCTTTAGGGG
GAGCACACGTCTGAACTCTTTCCCTACACGACGCTCTTCCGATCTNNNTTGGGAACTATCTTTAGGGG
AGCACACGTCTGAACTCTTTCCCTACACGACGCTCTTCCGATCTNNNGAGGATTCATCTCACTCCTCCA
AGCACACGTCTGAACTCTTTCCCTACACGACGCTCTTCCGATCTNNNGAGGATTCATCTCACTCCTCCA
CACACGTCTGAACTCTTTCCCTACACGACGCTCTTCCGATCTNNNAGGCAATACAATTCCTTTAATTTCA
CACACGTCTGAACTCTTTCCCTACACGACGCTCTTCCGATCTNNNAGGCAATACAATTCCTTTAATTTCA
GCACACGTCTGAACTCTTTCCCTACACGACGCTCTTCCGATCTNNNGCAGAAGGCTTGTTTTATTGCA
GCACACGTCTGAACTCTTTCCCTACACGACGCTCTTCCGATCTNNNGCAGAAGGCTTGTTTTATTGCA
AGCACACGTCTGAACTCTTTCCCTACACGACGCTCTTCCGATCTNNNCTAACCCTGTCCTGGAAAAAG
AGCACACGTCTGAACTCTTTCCCTACACGACGCTCTTCCGATCTNNNCTAACCCTGTCCTGGAAAAAG
GCACACGTCTGAACTCTTTCCCTACACGACGCTCTTCCGATCTNNNTAGAGTGTAGATGCATCAAAATC
GCACACGTCTGAACTCTTTCCCTACACGACGCTCTTCCGATCTNNNTAGAGTGTAGATGCATCAAAATC
GAGCACACGTCTGAACTCTTTCCCTACACGACGCTCTTCCGATCTNNNGATGGACAGGAGATGTGGAT
GAGCACACGTCTGAACTCTTTCCCTACACGACGCTCTTCCGATCTNNNGATGGACAGGAGATGTGGAT
AGCACACGTCTGAACTCTTTCCCTACACGACGCTCTTCCGATCTNNNAGCCCTTTCAGTGGTTCTTGAT
AGCACACGTCTGAACTCTTTCCCTACACGACGCTCTTCCGATCTNNNAGCCCTTTCAGTGGTTCTTGAT
AGAGCACACGTCTGAACTCTTTCCCTACACGACGCTCTTCCGATCTNNNATCCCCCTATTGCCTTTTCTC
AGAGCACACGTCTGAACTCTTTCCCTACACGACGCTCTTCCGATCTNNNATCCCCCTATTGCCTTTTCTC
CACACGTCTGAACTCTTTCCCTACACGACGCTCTTCCGATCTNNNCTCTTTGCTGCACATTATCCATCGG
CACACGTCTGAACTCTTTCCCTACACGACGCTCTTCCGATCTNNNCTCTTTGCTGCACATTATCCATCGG
AGCACACGTCTGAACTCTTTCCCTACACGACGCTCTTCCGATCTNNNTTCCATGTGAACTTGACTTTTTC
AGCACACGTCTGAACTCTTTCCCTACACGACGCTCTTCCGATCTNNNTTCCATGTGAACTTGACTTTTTC
GAGCACACGTCTGAACTCTTTCCCTACACGACGCTCTTCCGATCTNNNTACAAACACACACACACACACA
GAGCACACGTCTGAACTCTTTCCCTACACGACGCTCTTCCGATCTNNNTACAAACACACACACACACACA
AGCACACGTCTGAACTCTTTCCCTACACGACGCTCTTCCGATCTNNNATATCTGCCCAGTGCCATCTTCT
AGCACACGTCTGAACTCTTTCCCTACACGACGCTCTTCCGATCTNNNATATCTGCCCAGTGCCATCTTCT
AGCACACGTCTGAACTCTTTCCCTACACGACGCTCTTCCGATCTNNNGGTACTTGACCTCAATCCACCA
AGCACACGTCTGAACTCTTTCCCTACACGACGCTCTTCCGATCTNNNGGTACTTGACCTCAATCCACCA
AAGAGCACACGTCTGAACTCTTTCCCTACACGACGCTCTTCCGATCTNNNGGACAAGTCCCAACTCTATC
AAGAGCACACGTCTGAACTCTTTCCCTACACGACGCTCTTCCGATCTNNNGGACAAGTCCCAACTCTATC
AGCACACGTCTGAACTCTTTCCCTACACGACGCTCTTCCGATCTNNNATGGCCAAATGCATGACAGAA
AGCACACGTCTGAACTCTTTCCCTACACGACGCTCTTCCGATCTNNNATGGCCAAATGCATGACAGAA
AGCACACGTCTGAACTCTTTCCCTACACGACGCTCTTCCGATCTNNNCTCCTAGGTTCAAGCGATTCTCC
AGCACACGTCTGAACTCTTTCCCTACACGACGCTCTTCCGATCTNNNCTCCTAGGTTCAAGCGATTCTCC
AGAGCACACGTCTGAACTCTTTCCCTACACGACGCTCTTCCGATCTNNNGGAGTTTGCAGTGAGCTGA
AGAGCACACGTCTGAACTCTTTCCCTACACGACGCTCTTCCGATCTNNNGGAGTTTGCAGTGAGCTGA
GAGCACACGTCTGAACTCTTTCCCTACACGACGCTCTTCCGATCTNNNGTCCTTACAAGACGGGGAGT
GAGCACACGTCTGAACTCTTTCCCTACACGACGCTCTTCCGATCTNNNGTCCTTACAAGACGGGGAGT
GAGCACACGTCTGAACTCTTTCCCTACACGACGCTCTTCCGATCTNNNATCCTGGGCTTTTTCCTCTTCA
GAGCACACGTCTGAACTCTTTCCCTACACGACGCTCTTCCGATCTNNNATCCTGGGCTTTTTCCTCTTCA
GCACACGTCTGAACTCTTTCCCTACACGACGCTCTTCCGATCTNNNCTCTTCCTACCTCTCTGACCACA
GCACACGTCTGAACTCTTTCCCTACACGACGCTCTTCCGATCTNNNCTCTTCCTACCTCTCTGACCACA

GAGCACACGTCTGAACTCTTTCCCTACACGACGCTCTTCCGATCTNNNCCAGGAAGAAACTGAATCCC
GAGCACACGTCTGAACTCTTTCCCTACACGACGCTCTTCCGATCTNNNCCAGGAAGAAACTGAATCCC
AGCACACGTCTGAACTCTTTCCCTACACGACGCTCTTCCGATCTNNNTAGAGGATAATTGCTTGAAC
AGCACACGTCTGAACTCTTTCCCTACACGACGCTCTTCCGATCTNNNTAGAGGATAATTGCTTGAAC
CACACGTCTGAACTCTTTCCCTACACGACGCTCTTCCGATCTNNNAACTCTAACTCCCCATGGAGTCC
CACACGTCTGAACTCTTTCCCTACACGACGCTCTTCCGATCTNNNAACTCTAACTCCCCATGGAGTCC
AGCACACGTCTGAACTCTTTCCCTACACGACGCTCTTCCGATCTNNNCTGAGGCAGGAGAATCACTT
AGCACACGTCTGAACTCTTTCCCTACACGACGCTCTTCCGATCTNNNCTGAGGCAGGAGAATCACTT
AGCACACGTCTGAACTCTTTCCCTACACGACGCTCTTCCGATCTNNNATACAAAAAAATTAGCCGGGC
AGCACACGTCTGAACTCTTTCCCTACACGACGCTCTTCCGATCTNNNATACAAAAAAATTAGCCGGGC
GCACACGTCTGAACTCTTTCCCTACACGACGCTCTTCCGATCTNNNTAGCTATGTGACCTTGAGCAGA
GCACACGTCTGAACTCTTTCCCTACACGACGCTCTTCCGATCTNNNTAGCTATGTGACCTTGAGCAGA
GCACACGTCTGAACTCTTTCCCTACACGACGCTCTTCCGATCTNNNCTTTGGGAAGGAGGAGACAG
GCACACGTCTGAACTCTTTCCCTACACGACGCTCTTCCGATCTNNNCTTTGGGAAGGAGGAGACAG
GCACACGTCTGAACTCTTTCCCTACACGACGCTCTTCCGATCTNNNACTCCTGCAATTCTGACTCTGAG
GCACACGTCTGAACTCTTTCCCTACACGACGCTCTTCCGATCTNNNACTCCTGCAATTCTGACTCTGAG
AGCACACGTCTGAACTCTTTCCCTACACGACGCTCTTCCGATCTNNNCTGGTGTTTGTTCATCCTGTC
AGCACACGTCTGAACTCTTTCCCTACACGACGCTCTTCCGATCTNNNCTGGTGTTTGTTCATCCTGTC
AGCACACGTCTGAACTCTTTCCCTACACGACGCTCTTCCGATCTNNNAACATAGAGTGATCCTGTGTG
AGCACACGTCTGAACTCTTTCCCTACACGACGCTCTTCCGATCTNNNAACATAGAGTGATCCTGTGTG
GCACACGTCTGAACTCTTTCCCTACACGACGCTCTTCCGATCTNNNTCTGACTTGCAGCACTATTGAG
GCACACGTCTGAACTCTTTCCCTACACGACGCTCTTCCGATCTNNNTCTGACTTGCAGCACTATTGAG
GAGCACACGTCTGAACTCTTTCCCTACACGACGCTCTTCCGATCTNNNGTGTGATACTGACCAGCAAA
GAGCACACGTCTGAACTCTTTCCCTACACGACGCTCTTCCGATCTNNNGTGTGATACTGACCAGCAAA
GCACACGTCTGAACTCTTTCCCTACACGACGCTCTTCCGATCTNNNAGAGGTGAAGATGGGGGTAA
GCACACGTCTGAACTCTTTCCCTACACGACGCTCTTCCGATCTNNNAGAGGTGAAGATGGGGGTAA
CACACGTCTGAACTCTTTCCCTACACGACGCTCTTCCGATCTNNNACTGGCTTCGTTCACTTAGCATAC
CACACGTCTGAACTCTTTCCCTACACGACGCTCTTCCGATCTNNNACTGGCTTCGTTCACTTAGCATAC
AGCACACGTCTGAACTCTTTCCCTACACGACGCTCTTCCGATCTNNNATTGAGGCCATCCTGAATAGC
AGCACACGTCTGAACTCTTTCCCTACACGACGCTCTTCCGATCTNNNATTGAGGCCATCCTGAATAGC
AGCACACGTCTGAACTCTTTCCCTACACGACGCTCTTCCGATCTNNNTACAAAAATGAGCTGGGAGT
AGCACACGTCTGAACTCTTTCCCTACACGACGCTCTTCCGATCTNNNTACAAAAATGAGCTGGGAGT
CACACGTCTGAACTCTTTCCCTACACGACGCTCTTCCGATCTNNNTTGTGGGAAGGAAACCAATGAGA
CACACGTCTGAACTCTTTCCCTACACGACGCTCTTCCGATCTNNNTTGTGGGAAGGAAACCAATGAGA
GAGCACACGTCTGAACTCTTTCCCTACACGACGCTCTTCCGATCTNNNCTGGAGAATCACTTGAACCT
GAGCACACGTCTGAACTCTTTCCCTACACGACGCTCTTCCGATCTNNNCTGGAGAATCACTTGAACCT
GCACACGTCTGAACTCTTTCCCTACACGACGCTCTTCCGATCTNNNAAAAGAGGTGAGGCCCATGAG
GCACACGTCTGAACTCTTTCCCTACACGACGCTCTTCCGATCTNNNAAAAGAGGTGAGGCCCATGAG
AGCACACGTCTGAACTCTTTCCCTACACGACGCTCTTCCGATCTNNNAACTTCAGAGAATTAGAGGCC
AGCACACGTCTGAACTCTTTCCCTACACGACGCTCTTCCGATCTNNNAACTTCAGAGAATTAGAGGCC
GCACACGTCTGAACTCTTTCCCTACACGACGCTCTTCCGATCTNNNAGGCAGGAGAATTACTTGAAC
GCACACGTCTGAACTCTTTCCCTACACGACGCTCTTCCGATCTNNNAGGCAGGAGAATTACTTGAAC
AGAGCACACGTCTGAACTCTTTCCCTACACGACGCTCTTCCGATCTNNNCACCCACATTCAAACCATAG

AGAGCACACGTCTGAACTCTTTCCCTACACGACGCTCTTCCGATCTNNNCACCCACATTCAAACCATAG
GAGCACACGTCTGAACTCTTTCCCTACACGACGCTCTTCCGATCTNNNCCGATCTCTAGAGTGTTGGTT
GAGCACACGTCTGAACTCTTTCCCTACACGACGCTCTTCCGATCTNNNCCGATCTCTAGAGTGTTGGTT
AGCACACGTCTGAACTCTTTCCCTACACGACGCTCTTCCGATCTNNNAGGCAAGGGTATCAAAGAGTC
AGCACACGTCTGAACTCTTTCCCTACACGACGCTCTTCCGATCTNNNAGGCAAGGGTATCAAAGAGTC
AGAGCACACGTCTGAACTCTTTCCCTACACGACGCTCTTCCGATCTNNNGGAGGTTGTGATGAGCTGA
AGAGCACACGTCTGAACTCTTTCCCTACACGACGCTCTTCCGATCTNNNGGAGGTTGTGATGAGCTGA
AGCACACGTCTGAACTCTTTCCCTACACGACGCTCTTCCGATCTNNNAGCACTCCAAATATCCTCTTGCA
AGCACACGTCTGAACTCTTTCCCTACACGACGCTCTTCCGATCTNNNAGCACTCCAAATATCCTCTTGCA
AGCACACGTCTGAACTCTTTCCCTACACGACGCTCTTCCGATCTNNNGGAAATGGGGTCGACTTTTGA
AGCACACGTCTGAACTCTTTCCCTACACGACGCTCTTCCGATCTNNNGGAAATGGGGTCGACTTTTGA
AGCACACGTCTGAACTCTTTCCCTACACGACGCTCTTCCGATCTNNNCTCTTCTGTGATCTGGTCCCTTT
AGCACACGTCTGAACTCTTTCCCTACACGACGCTCTTCCGATCTNNNCTCTTCTGTGATCTGGTCCCTTT
GAGCACACGTCTGAACTCTTTCCCTACACGACGCTCTTCCGATCTNNNATTACCTCCCACTGGGTTCTTT
GAGCACACGTCTGAACTCTTTCCCTACACGACGCTCTTCCGATCTNNNATTACCTCCCACTGGGTTCTTT
GCACACGTCTGAACTCTTTCCCTACACGACGCTCTTCCGATCTNNNATTGATTTCACTTATTACCTTGG
GCACACGTCTGAACTCTTTCCCTACACGACGCTCTTCCGATCTNNNATTGATTTCACTTATTACCTTGG
AGAGCACACGTCTGAACTCTTTCCCTACACGACGCTCTTCCGATCTNNNAGAGAAAGAAGGTACAGTG
AGAGCACACGTCTGAACTCTTTCCCTACACGACGCTCTTCCGATCTNNNAGAGAAAGAAGGTACAGTG
GCACACGTCTGAACTCTTTCCCTACACGACGCTCTTCCGATCTNNNGAAGTTGGAGAACTCATTGGGC
GCACACGTCTGAACTCTTTCCCTACACGACGCTCTTCCGATCTNNNGAAGTTGGAGAACTCATTGGGC
AGAGCACACGTCTGAACTCTTTCCCTACACGACGCTCTTCCGATCTNNNCTCACGCCTGTAATCCCAATA
AGAGCACACGTCTGAACTCTTTCCCTACACGACGCTCTTCCGATCTNNNCTCACGCCTGTAATCCCAATA
GAGCACACGTCTGAACTCTTTCCCTACACGACGCTCTTCCGATCTNNNCATTGCATTACAGCCTGGTTGA
GAGCACACGTCTGAACTCTTTCCCTACACGACGCTCTTCCGATCTNNNCATTGCATTACAGCCTGGTTGA
AGCACACGTCTGAACTCTTTCCCTACACGACGCTCTTCCGATCTNNNTGTTTGTGTGTGTGAGACAGAG
AGCACACGTCTGAACTCTTTCCCTACACGACGCTCTTCCGATCTNNNTGTTTGTGTGTGTGAGACAGAG
AGCACACGTCTGAACTCTTTCCCTACACGACGCTCTTCCGATCTNNNCACAGCACTATTCACAACAGCA
AGCACACGTCTGAACTCTTTCCCTACACGACGCTCTTCCGATCTNNNCACAGCACTATTCACAACAGCA
GAGCACACGTCTGAACTCTTTCCCTACACGACGCTCTTCCGATCTNNNCCAAACTCTTCTCAATCCCTGGA
GAGCACACGTCTGAACTCTTTCCCTACACGACGCTCTTCCGATCTNNNCCAAACTCTTCTCAATCCCTGGA
GAGCACACGTCTGAACTCTTTCCCTACACGACGCTCTTCCGATCTNNNGTTGGGGGTATTGAGAGAGT
GAGCACACGTCTGAACTCTTTCCCTACACGACGCTCTTCCGATCTNNNGTTGGGGGTATTGAGAGAGT
AGCACACGTCTGAACTCTTTCCCTACACGACGCTCTTCCGATCTNNNGACCCCAATTCTCTCTCCTCAAG
AGCACACGTCTGAACTCTTTCCCTACACGACGCTCTTCCGATCTNNNGACCCCAATTCTCTCTCCTCAAG
AGAGCACACGTCTGAACTCTTTCCCTACACGACGCTCTTCCGATCTNNNCTGTGTTCAGGAACCTGGT
AGAGCACACGTCTGAACTCTTTCCCTACACGACGCTCTTCCGATCTNNNCTGTGTTCAGGAACCTGGT
AGCACACGTCTGAACTCTTTCCCTACACGACGCTCTTCCGATCTNNNTATGCAATAACTCTGGAAGCAG
AGCACACGTCTGAACTCTTTCCCTACACGACGCTCTTCCGATCTNNNTATGCAATAACTCTGGAAGCAG
AGCACACGTCTGAACTCTTTCCCTACACGACGCTCTTCCGATCTNNNCAGGAGAATTGCTTGAACACAG
AGCACACGTCTGAACTCTTTCCCTACACGACGCTCTTCCGATCTNNNCAGGAGAATTGCTTGAACACAG
AGCACACGTCTGAACTCTTTCCCTACACGACGCTCTTCCGATCTNNNCATGGGCGGAAGCTTTTTGAAA
AGCACACGTCTGAACTCTTTCCCTACACGACGCTCTTCCGATCTNNNCATGGGCGGAAGCTTTTTGAAA

GCACACGTCTGAACTCTTTCCCTACACGACGCTCTTCCGATCTNNNAAACTCCTGACCTCAAGTGATCC
GCACACGTCTGAACTCTTTCCCTACACGACGCTCTTCCGATCTNNNAAACTCCTGACCTCAAGTGATCC
AGCACACGTCTGAACTCTTTCCCTACACGACGCTCTTCCGATCTNNNGGACATGTTTAGGGAGGTGTA
AGCACACGTCTGAACTCTTTCCCTACACGACGCTCTTCCGATCTNNNGGACATGTTTAGGGAGGTGTA
AGCACACGTCTGAACTCTTTCCCTACACGACGCTCTTCCGATCTNNNTGGACAGGATAATTGCAAGGT
AGCACACGTCTGAACTCTTTCCCTACACGACGCTCTTCCGATCTNNNTGGACAGGATAATTGCAAGGT
GCACACGTCTGAACTCTTTCCCTACACGACGCTCTTCCGATCTNNNCTTGCAGTGAGCCGAGATTGCA
GCACACGTCTGAACTCTTTCCCTACACGACGCTCTTCCGATCTNNNCTTGCAGTGAGCCGAGATTGCA
GAGCACACGTCTGAACTCTTTCCCTACACGACGCTCTTCCGATCTNNNCATGGTGGCTCATGCTTGTA
GAGCACACGTCTGAACTCTTTCCCTACACGACGCTCTTCCGATCTNNNCATGGTGGCTCATGCTTGTA
ACACGTCTGAACTCTTTCCCTACACGACGCTCTTCCGATCTNNNGGCTCCTCACTTCCCAGTAGGCATC
ACACGTCTGAACTCTTTCCCTACACGACGCTCTTCCGATCTNNNGGCTCCTCACTTCCCAGTAGGCATC
AGCACACGTCTGAACTCTTTCCCTACACGACGCTCTTCCGATCTNNNCTGAGGCAGGAGAATCACTTG
AGCACACGTCTGAACTCTTTCCCTACACGACGCTCTTCCGATCTNNNCTGAGGCAGGAGAATCACTTG
AGCACACGTCTGAACTCTTTCCCTACACGACGCTCTTCCGATCTNNNCACAGACATAAACCTCCTCCTG
AGCACACGTCTGAACTCTTTCCCTACACGACGCTCTTCCGATCTNNNCACAGACATAAACCTCCTCCTG
AGCACACGTCTGAACTCTTTCCCTACACGACGCTCTTCCGATCTNNNACCTGAGGTCAAGAGTTCAAG
AGCACACGTCTGAACTCTTTCCCTACACGACGCTCTTCCGATCTNNNACCTGAGGTCAAGAGTTCAAG
AGCACACGTCTGAACTCTTTCCCTACACGACGCTCTTCCGATCTNNNACAGCATTCTAGACAGTTCCTCT
AGCACACGTCTGAACTCTTTCCCTACACGACGCTCTTCCGATCTNNNACAGCATTCTAGACAGTTCCTCT
GCACACGTCTGAACTCTTTCCCTACACGACGCTCTTCCGATCTNNNTCTGTGTGTATTTCGCTCTATGC
GCACACGTCTGAACTCTTTCCCTACACGACGCTCTTCCGATCTNNNTCTGTGTGTATTTCGCTCTATGC
AGCACACGTCTGAACTCTTTCCCTACACGACGCTCTTCCGATCTNNNTATGGGAAAGCACAACACATC
AGCACACGTCTGAACTCTTTCCCTACACGACGCTCTTCCGATCTNNNTATGGGAAAGCACAACACATC
AGCACACGTCTGAACTCTTTCCCTACACGACGCTCTTCCGATCTNNNGAGATCATGCCACTACACTCCA
AGCACACGTCTGAACTCTTTCCCTACACGACGCTCTTCCGATCTNNNGAGATCATGCCACTACACTCCA
AGCACACGTCTGAACTCTTTCCCTACACGACGCTCTTCCGATCTNNNTTCCATTCATGTGTGTGTGTGT
AGCACACGTCTGAACTCTTTCCCTACACGACGCTCTTCCGATCTNNNTTCCATTCATGTGTGTGTGTGT
AGAGCACACGTCTGAACTCTTTCCCTACACGACGCTCTTCCGATCTNNNGCAACATGGATGAATCACG
AGAGCACACGTCTGAACTCTTTCCCTACACGACGCTCTTCCGATCTNNNGCAACATGGATGAATCACG
AGCACACGTCTGAACTCTTTCCCTACACGACGCTCTTCCGATCTNNNCTTGCAGTGAGCCGAGATCAC
AGCACACGTCTGAACTCTTTCCCTACACGACGCTCTTCCGATCTNNNCTTGCAGTGAGCCGAGATCAC
GAGCACACGTCTGAACTCTTTCCCTACACGACGCTCTTCCGATCTNNNTTCGTCTGGAGGGAAAGAG
GAGCACACGTCTGAACTCTTTCCCTACACGACGCTCTTCCGATCTNNNTTCGTCTGGAGGGAAAGAG
GAGCACACGTCTGAACTCTTTCCCTACACGACGCTCTTCCGATCTNNNGCCTCAGTGAGCCATTATTTT
GAGCACACGTCTGAACTCTTTCCCTACACGACGCTCTTCCGATCTNNNGCCTCAGTGAGCCATTATTTT
AGCACACGTCTGAACTCTTTCCCTACACGACGCTCTTCCGATCTNNNACTGCTCAAAAGGAACTCAGT
AGCACACGTCTGAACTCTTTCCCTACACGACGCTCTTCCGATCTNNNACTGCTCAAAAGGAACTCAGT
GAGCACACGTCTGAACTCTTTCCCTACACGACGCTCTTCCGATCTNNNTGCTGGAGGCAAATCACATA
GAGCACACGTCTGAACTCTTTCCCTACACGACGCTCTTCCGATCTNNNTGCTGGAGGCAAATCACATA
AGCACACGTCTGAACTCTTTCCCTACACGACGCTCTTCCGATCTNNNCCCATGAACAACTTCCATGAAC
AGCACACGTCTGAACTCTTTCCCTACACGACGCTCTTCCGATCTNNNCCCATGAACAACTTCCATGAAC
AGCACACGTCTGAACTCTTTCCCTACACGACGCTCTTCCGATCTNNNGCTTTCTCTTGTGTACCAAACC

AGCACACGTCTGAACTCTTTCCCTACACGACGCTCTTCCGATCTNNNGCTTTCTCTTGTGTACCAAACC
GCACACGTCTGAACTCTTTCCCTACACGACGCTCTTCCGATCTNNNCTTGAGCTAAGATGCACTGAAG
GCACACGTCTGAACTCTTTCCCTACACGACGCTCTTCCGATCTNNNCTTGAGCTAAGATGCACTGAAG
AGCACACGTCTGAACTCTTTCCCTACACGACGCTCTTCCGATCTNNNGGGTGATGCAAGAGAATCAA
AGCACACGTCTGAACTCTTTCCCTACACGACGCTCTTCCGATCTNNNGGGTGATGCAAGAGAATCAA
CACACGTCTGAACTCTTTCCCTACACGACGCTCTTCCGATCTNNNTGTCCATTCTCTTGGGAGCACTAC
CACACGTCTGAACTCTTTCCCTACACGACGCTCTTCCGATCTNNNTGTCCATTCTCTTGGGAGCACTAC
GAGCACACGTCTGAACTCTTTCCCTACACGACGCTCTTCCGATCTNNNTGGATCAATAGTGTGTGCCT
GAGCACACGTCTGAACTCTTTCCCTACACGACGCTCTTCCGATCTNNNTGGATCAATAGTGTGTGCCT
GAGCACACGTCTGAACTCTTTCCCTACACGACGCTCTTCCGATCTNNNTAATCTCCCTTCACTGCAAGC
GAGCACACGTCTGAACTCTTTCCCTACACGACGCTCTTCCGATCTNNNTAATCTCCCTTCACTGCAAGC
GAGCACACGTCTGAACTCTTTCCCTACACGACGCTCTTCCGATCTNNNACATCTTACTTTGCCTCTTGG
GAGCACACGTCTGAACTCTTTCCCTACACGACGCTCTTCCGATCTNNNACATCTTACTTTGCCTCTTGG
GAGCACACGTCTGAACTCTTTCCCTACACGACGCTCTTCCGATCTNNNCCCTAGGACTCCTCAGAAGA
GAGCACACGTCTGAACTCTTTCCCTACACGACGCTCTTCCGATCTNNNCCCTAGGACTCCTCAGAAGA
GAGCACACGTCTGAACTCTTTCCCTACACGACGCTCTTCCGATCTNNNAAGCCCCCTAAAGATGGAGA
GAGCACACGTCTGAACTCTTTCCCTACACGACGCTCTTCCGATCTNNNAAGCCCCCTAAAGATGGAGA
GCACACGTCTGAACTCTTTCCCTACACGACGCTCTTCCGATCTNNNATCTTCATACTCTGCAGGCAACC
GCACACGTCTGAACTCTTTCCCTACACGACGCTCTTCCGATCTNNNATCTTCATACTCTGCAGGCAACC
AGCACACGTCTGAACTCTTTCCCTACACGACGCTCTTCCGATCTNNNAGTTTGAGGCTGCAGTAAGCT
AGCACACGTCTGAACTCTTTCCCTACACGACGCTCTTCCGATCTNNNAGTTTGAGGCTGCAGTAAGCT
GCACACGTCTGAACTCTTTCCCTACACGACGCTCTTCCGATCTNNNTTATTTGTGTTTGCTTCCCCCCC
GCACACGTCTGAACTCTTTCCCTACACGACGCTCTTCCGATCTNNNTTATTTGTGTTTGCTTCCCCCCC
AGCACACGTCTGAACTCTTTCCCTACACGACGCTCTTCCGATCTNNNGTGACAGAACAAGACTCCGTCT
AGCACACGTCTGAACTCTTTCCCTACACGACGCTCTTCCGATCTNNNGTGACAGAACAAGACTCCGTCT
GAGCACACGTCTGAACTCTTTCCCTACACGACGCTCTTCCGATCTNNNGTTGCAGTGAGCTGAGATC
GAGCACACGTCTGAACTCTTTCCCTACACGACGCTCTTCCGATCTNNNGTTGCAGTGAGCTGAGATC
GCACACGTCTGAACTCTTTCCCTACACGACGCTCTTCCGATCTNNNACTTCAGTCTCCCAAAGTAGCTG
GCACACGTCTGAACTCTTTCCCTACACGACGCTCTTCCGATCTNNNACTTCAGTCTCCCAAAGTAGCTG
AGCACACGTCTGAACTCTTTCCCTACACGACGCTCTTCCGATCTNNNCAGATGGTTGCTGTTTCACTC
AGCACACGTCTGAACTCTTTCCCTACACGACGCTCTTCCGATCTNNNCAGATGGTTGCTGTTTCACTC
AGCACACGTCTGAACTCTTTCCCTACACGACGCTCTTCCGATCTNNNCACACACACACACATACACA
AGCACACGTCTGAACTCTTTCCCTACACGACGCTCTTCCGATCTNNNCACACACACACACATACACA
GAGCACACGTCTGAACTCTTTCCCTACACGACGCTCTTCCGATCTNNNTCCACTGCTCTGGCTTAGTGA
GAGCACACGTCTGAACTCTTTCCCTACACGACGCTCTTCCGATCTNNNTCCACTGCTCTGGCTTAGTGA
AGCACACGTCTGAACTCTTTCCCTACACGACGCTCTTCCGATCTNNNTTGTAGTGAGCCAAAATCATG
AGCACACGTCTGAACTCTTTCCCTACACGACGCTCTTCCGATCTNNNTTGTAGTGAGCCAAAATCATG
GAGCACACGTCTGAACTCTTTCCCTACACGACGCTCTTCCGATCTNNNAGACCCAGTCTCCCAAAAGA
GAGCACACGTCTGAACTCTTTCCCTACACGACGCTCTTCCGATCTNNNAGACCCAGTCTCCCAAAAGA
AGAGCACACGTCTGAACTCTTTCCCTACACGACGCTCTTCCGATCTNNNAGCAACACTAGACATGGTCT
AGAGCACACGTCTGAACTCTTTCCCTACACGACGCTCTTCCGATCTNNNAGCAACACTAGACATGGTCT
AGCACACGTCTGAACTCTTTCCCTACACGACGCTCTTCCGATCTNNNTGTCCTGGGCCACACATAAAA
AGCACACGTCTGAACTCTTTCCCTACACGACGCTCTTCCGATCTNNNTGTCCTGGGCCACACATAAAA

AGCACACGTCTGAACTCTTTCCCTACACGACGCTCTTCCGATCTNNNTCCTCAAGGAACATATGGCCAA
AGCACACGTCTGAACTCTTTCCCTACACGACGCTCTTCCGATCTNNNTCCTCAAGGAACATATGGCCAA
AGCACACGTCTGAACTCTTTCCCTACACGACGCTCTTCCGATCTNNNGTAATCACAGCTACTCGGGAG
AGCACACGTCTGAACTCTTTCCCTACACGACGCTCTTCCGATCTNNNGTAATCACAGCTACTCGGGAG
GCACACGTCTGAACTCTTTCCCTACACGACGCTCTTCCGATCTNNNATCACCACACCCAGCTAATTTG
GCACACGTCTGAACTCTTTCCCTACACGACGCTCTTCCGATCTNNNATCACCACACCCAGCTAATTTG
AGCACACGTCTGAACTCTTTCCCTACACGACGCTCTTCCGATCTNNNCCAGAAGCTAAGGCAGGAGAA
AGCACACGTCTGAACTCTTTCCCTACACGACGCTCTTCCGATCTNNNCCAGAAGCTAAGGCAGGAGAA
GAGCACACGTCTGAACTCTTTCCCTACACGACGCTCTTCCGATCTNNNCTTGGGTGACAGAGTAAGAC
GAGCACACGTCTGAACTCTTTCCCTACACGACGCTCTTCCGATCTNNNCTTGGGTGACAGAGTAAGAC
AGCACACGTCTGAACTCTTTCCCTACACGACGCTCTTCCGATCTNNNGTGATTATACCAACGTAGGGC
AGCACACGTCTGAACTCTTTCCCTACACGACGCTCTTCCGATCTNNNGTGATTATACCAACGTAGGGC
AGCACACGTCTGAACTCTTTCCCTACACGACGCTCTTCCGATCTNNNTGTTTTGTGAAACACTGGCTTC
AGCACACGTCTGAACTCTTTCCCTACACGACGCTCTTCCGATCTNNNTGTTTTGTGAAACACTGGCTTC
AGCACACGTCTGAACTCTTTCCCTACACGACGCTCTTCCGATCTNNNAGTCTGTCTCTCACTGCCATTCT
AGCACACGTCTGAACTCTTTCCCTACACGACGCTCTTCCGATCTNNNAGTCTGTCTCTCACTGCCATTCT
AGCACACGTCTGAACTCTTTCCCTACACGACGCTCTTCCGATCTNNNGAGCTTACAGTGAGCCGAGAT
AGCACACGTCTGAACTCTTTCCCTACACGACGCTCTTCCGATCTNNNGAGCTTACAGTGAGCCGAGAT
AGCACACGTCTGAACTCTTTCCCTACACGACGCTCTTCCGATCTNNNAGAATCGCTTGAACCTGGGAG
AGCACACGTCTGAACTCTTTCCCTACACGACGCTCTTCCGATCTNNNAGAATCGCTTGAACCTGGGAG
GAGCACACGTCTGAACTCTTTCCCTACACGACGCTCTTCCGATCTNNNGAGGCAGGAGAATCGCTTGA
GAGCACACGTCTGAACTCTTTCCCTACACGACGCTCTTCCGATCTNNNGAGGCAGGAGAATCGCTTGA
AGCACACGTCTGAACTCTTTCCCTACACGACGCTCTTCCGATCTNNNGCTCCAAGAAGAACTCAGTCTG
AGCACACGTCTGAACTCTTTCCCTACACGACGCTCTTCCGATCTNNNGCTCCAAGAAGAACTCAGTCTG
AGCACACGTCTGAACTCTTTCCCTACACGACGCTCTTCCGATCTNNNATGTTGCTTTTCTGATTATTTCT
AGCACACGTCTGAACTCTTTCCCTACACGACGCTCTTCCGATCTNNNATGTTGCTTTTCTGATTATTTCT
CACACGTCTGAACTCTTTCCCTACACGACGCTCTTCCGATCTNNNAGAATGAGCTGTCACCATAGCATC
CACACGTCTGAACTCTTTCCCTACACGACGCTCTTCCGATCTNNNAGAATGAGCTGTCACCATAGCATC
GCACACGTCTGAACTCTTTCCCTACACGACGCTCTTCCGATCTNNNAAGAATCACTTGAACCTGGGAG
GCACACGTCTGAACTCTTTCCCTACACGACGCTCTTCCGATCTNNNAAGAATCACTTGAACCTGGGAG
GAGCACACGTCTGAACTCTTTCCCTACACGACGCTCTTCCGATCTNNNGAGGGACAGAGACAGAAAG
GAGCACACGTCTGAACTCTTTCCCTACACGACGCTCTTCCGATCTNNNGAGGGACAGAGACAGAAAG
AGCACACGTCTGAACTCTTTCCCTACACGACGCTCTTCCGATCTNNNGTCATCCTAGCTACTCAGAAGG
AGCACACGTCTGAACTCTTTCCCTACACGACGCTCTTCCGATCTNNNGTCATCCTAGCTACTCAGAAGG
GAGCACACGTCTGAACTCTTTCCCTACACGACGCTCTTCCGATCTNNNGTCTTTGGCATTCACACTGGA
GAGCACACGTCTGAACTCTTTCCCTACACGACGCTCTTCCGATCTNNNGTCTTTGGCATTCACACTGGA
AGCACACGTCTGAACTCTTTCCCTACACGACGCTCTTCCGATCTNNNTATCCCAGGTTCAAGCAGTTCT
AGCACACGTCTGAACTCTTTCCCTACACGACGCTCTTCCGATCTNNNTATCCCAGGTTCAAGCAGTTCT
GAGCACACGTCTGAACTCTTTCCCTACACGACGCTCTTCCGATCTNNNACAGTACAGGAAGCAAGAGT
GAGCACACGTCTGAACTCTTTCCCTACACGACGCTCTTCCGATCTNNNACAGTACAGGAAGCAAGAGT
AGCACACGTCTGAACTCTTTCCCTACACGACGCTCTTCCGATCTNNNCACCATGCCTGCTGATTTTTGT
AGCACACGTCTGAACTCTTTCCCTACACGACGCTCTTCCGATCTNNNCACCATGCCTGCTGATTTTTGT
GAGCACACGTCTGAACTCTTTCCCTACACGACGCTCTTCCGATCTNNNTTTTTACGGCAGCCTGTTTAG

GAGCACACGTCTGAACTCTTTCCCTACACGACGCTCTTCCGATCTNNNTTTTTACGGCAGCCTGTTTAG
GAGCACACGTCTGAACTCTTTCCCTACACGACGCTCTTCCGATCTNNNGTGATCAAACCACTGCACTCT
GAGCACACGTCTGAACTCTTTCCCTACACGACGCTCTTCCGATCTNNNGTGATCAAACCACTGCACTCT
AGCACACGTCTGAACTCTTTCCCTACACGACGCTCTTCCGATCTNNNATTGAGAAGGCTGCAGTTAG
AGCACACGTCTGAACTCTTTCCCTACACGACGCTCTTCCGATCTNNNATTGAGAAGGCTGCAGTTAG
AGCACACGTCTGAACTCTTTCCCTACACGACGCTCTTCCGATCTNNNTAGTGGCTGCATAGTCTCCTTT
AGCACACGTCTGAACTCTTTCCCTACACGACGCTCTTCCGATCTNNNTAGTGGCTGCATAGTCTCCTTT
AGCACACGTCTGAACTCTTTCCCTACACGACGCTCTTCCGATCTNNNATTTGCTCCTACCCTCACTTGTT
AGCACACGTCTGAACTCTTTCCCTACACGACGCTCTTCCGATCTNNNATTTGCTCCTACCCTCACTTGTT
AGCACACGTCTGAACTCTTTCCCTACACGACGCTCTTCCGATCTNNNCTTACCCTTGGCACTACTCACTT
AGCACACGTCTGAACTCTTTCCCTACACGACGCTCTTCCGATCTNNNCTTACCCTTGGCACTACTCACTT
AGCACACGTCTGAACTCTTTCCCTACACGACGCTCTTCCGATCTNNNTTTCCAGTTTGCACAGGAAGA
AGCACACGTCTGAACTCTTTCCCTACACGACGCTCTTCCGATCTNNNTTTCCAGTTTGCACAGGAAGA
AGCACACGTCTGAACTCTTTCCCTACACGACGCTCTTCCGATCTNNNTAATGAGACATGATCGTGCCA
AGCACACGTCTGAACTCTTTCCCTACACGACGCTCTTCCGATCTNNNTAATGAGACATGATCGTGCCA
GAGCACACGTCTGAACTCTTTCCCTACACGACGCTCTTCCGATCTNNNGGCAACAGCATCTGGACTAT
GAGCACACGTCTGAACTCTTTCCCTACACGACGCTCTTCCGATCTNNNGGCAACAGCATCTGGACTAT
GAGCACACGTCTGAACTCTTTCCCTACACGACGCTCTTCCGATCTNNNAATGCTGTTCTCATGATCGCA
GAGCACACGTCTGAACTCTTTCCCTACACGACGCTCTTCCGATCTNNNAATGCTGTTCTCATGATCGCA
CACACGTCTGAACTCTTTCCCTACACGACGCTCTTCCGATCTNNNCATTTTTATTGTTTCCGGAATGC
CACACGTCTGAACTCTTTCCCTACACGACGCTCTTCCGATCTNNNCATTTTTATTGTTTCCGGAATGC
GAGCACACGTCTGAACTCTTTCCCTACACGACGCTCTTCCGATCTNNNATACACCTGCTGTCCTTCACTT
GAGCACACGTCTGAACTCTTTCCCTACACGACGCTCTTCCGATCTNNNATACACCTGCTGTCCTTCACTT
GAGCACACGTCTGAACTCTTTCCCTACACGACGCTCTTCCGATCTNNNATTGGGTTGCTTGCCTTTTTC
GAGCACACGTCTGAACTCTTTCCCTACACGACGCTCTTCCGATCTNNNATTGGGTTGCTTGCCTTTTTC
AGCACACGTCTGAACTCTTTCCCTACACGACGCTCTTCCGATCTNNNTCCTAACACTTTGGGAGGTCAA
AGCACACGTCTGAACTCTTTCCCTACACGACGCTCTTCCGATCTNNNTCCTAACACTTTGGGAGGTCAA
AGCACACGTCTGAACTCTTTCCCTACACGACGCTCTTCCGATCTNNNTCAACCCCTCTTAGAGCAGTTT
AGCACACGTCTGAACTCTTTCCCTACACGACGCTCTTCCGATCTNNNTCAACCCCTCTTAGAGCAGTTT
AGCACACGTCTGAACTCTTTCCCTACACGACGCTCTTCCGATCTNNNTGATGGCTTTGTTTGGTGGATT
AGCACACGTCTGAACTCTTTCCCTACACGACGCTCTTCCGATCTNNNTGATGGCTTTGTTTGGTGGATT
GAGCACACGTCTGAACTCTTTCCCTACACGACGCTCTTCCGATCTNNNAGGAAGGCAAGTTTTTCTGC
GAGCACACGTCTGAACTCTTTCCCTACACGACGCTCTTCCGATCTNNNAGGAAGGCAAGTTTTTCTGC
AGAGCACACGTCTGAACTCTTTCCCTACACGACGCTCTTCCGATCTNNNGAGAATGGCATGAACCTGG
AGAGCACACGTCTGAACTCTTTCCCTACACGACGCTCTTCCGATCTNNNGAGAATGGCATGAACCTGG
GAGCACACGTCTGAACTCTTTCCCTACACGACGCTCTTCCGATCTNNNGACAAAATGAGACTCCACCTC
GAGCACACGTCTGAACTCTTTCCCTACACGACGCTCTTCCGATCTNNNGACAAAATGAGACTCCACCTC
AGCACACGTCTGAACTCTTTCCCTACACGACGCTCTTCCGATCTNNNAGTGATTCTTCTGTCTCAGCCTC
AGCACACGTCTGAACTCTTTCCCTACACGACGCTCTTCCGATCTNNNAGTGATTCTTCTGTCTCAGCCTC
GAGCACACGTCTGAACTCTTTCCCTACACGACGCTCTTCCGATCTNNNCTTTGTTGCCTCTCATGTCCAA
GAGCACACGTCTGAACTCTTTCCCTACACGACGCTCTTCCGATCTNNNCTTTGTTGCCTCTCATGTCCAA
GCACACGTCTGAACTCTTTCCCTACACGACGCTCTTCCGATCTNNNAGGCCCACATCATTCATTTCCAT
GCACACGTCTGAACTCTTTCCCTACACGACGCTCTTCCGATCTNNNAGGCCCACATCATTCATTTCCAT

AGCACACGTCTGAACTCTTTCCCTACACGACGCTCTTCCGATCTNNNTGTGTGCATATCTCTGTGTCCA

AGCACACGTCTGAACTCTTTCCCTACACGACGCTCTTCCGATCTNNNTGTGTGCATATCTCTGTGTCCA

GCACACGTCTGAACTCTTTCCCTACACGACGCTCTTCCGATCTNNNACTTAGGATTGGAGATAGCGC

GCACACGTCTGAACTCTTTCCCTACACGACGCTCTTCCGATCTNNNACTTAGGATTGGAGATAGCGC

AGCACACGTCTGAACTCTTTCCCTACACGACGCTCTTCCGATCTNNNGATCTCAGCTCACTGCAAACTC

AGCACACGTCTGAACTCTTTCCCTACACGACGCTCTTCCGATCTNNNGATCTCAGCTCACTGCAAACTC

GAGCACACGTCTGAACTCTTTCCCTACACGACGCTCTTCCGATCTNNNACGGCTACTTTTCCACAAAA

GAGCACACGTCTGAACTCTTTCCCTACACGACGCTCTTCCGATCTNNNACGGCTACTTTTCCACAAAA

GAGCACACGTCTGAACTCTTTCCCTACACGACGCTCTTCCGATCTNNNCAACACTGCACTTCATTCAGA

GAGCACACGTCTGAACTCTTTCCCTACACGACGCTCTTCCGATCTNNNCAACACTGCACTTCATTCAGA

AGCACACGTCTGAACTCTTTCCCTACACGACGCTCTTCCGATCTNNNCAGTTCCTTCTGGAGGCTCTA

AGCACACGTCTGAACTCTTTCCCTACACGACGCTCTTCCGATCTNNNCAGTTCCTTCTGGAGGCTCTA

GAGCACACGTCTGAACTCTTTCCCTACACGACGCTCTTCCGATCTNNNGGCTTTGTATTTTGCCCACTT

GAGCACACGTCTGAACTCTTTCCCTACACGACGCTCTTCCGATCTNNNGGCTTTGTATTTTGCCCACTT

AGCACACGTCTGAACTCTTTCCCTACACGACGCTCTTCCGATCTNNNAAACAATTATGACACACCCTGC

AGCACACGTCTGAACTCTTTCCCTACACGACGCTCTTCCGATCTNNNAAACAATTATGACACACCCTGC

GCACACGTCTGAACTCTTTCCCTACACGACGCTCTTCCGATCTNNNGCTCGGAGAAGTAGAGCGCACA

GCACACGTCTGAACTCTTTCCCTACACGACGCTCTTCCGATCTNNNGCTCGGAGAAGTAGAGCGCACA

ACACGTCTGAACTCTTTCCCTACACGACGCTCTTCCGATCTNNNGTAGGCAGCTTTCCTCCTTTCTCCAT

ACACGTCTGAACTCTTTCCCTACACGACGCTCTTCCGATCTNNNGTAGGCAGCTTTCCTCCTTTCTCCAT

GAGCACACGTCTGAACTCTTTCCCTACACGACGCTCTTCCGATCTNNNCCCTTTTTGCTTCTCAGAGA

GAGCACACGTCTGAACTCTTTCCCTACACGACGCTCTTCCGATCTNNNCCCTTTTTGCTTCTCAGAGA

GCACACGTCTGAACTCTTTCCCTACACGACGCTCTTCCGATCTNNNAGAAGTACGAAACAGCAAACA

GCACACGTCTGAACTCTTTCCCTACACGACGCTCTTCCGATCTNNNAGAAGTACGAAACAGCAAACA

AGAGCACACGTCTGAACTCTTTCCCTACACGACGCTCTTCCGATCTNNNTCCCTTCTCTTTGAACATGA

AGAGCACACGTCTGAACTCTTTCCCTACACGACGCTCTTCCGATCTNNNTCCCTTCTCTTTGAACATGA

GAGCACACGTCTGAACTCTTTCCCTACACGACGCTCTTCCGATCTNNNAGTGAACTGAAATCGTGCCA

GAGCACACGTCTGAACTCTTTCCCTACACGACGCTCTTCCGATCTNNNAGTGAACTGAAATCGTGCCA

GCACACGTCTGAACTCTTTCCCTACACGACGCTCTTCCGATCTNNNGTTGCTCCAATACAGCCTCATTC

GCACACGTCTGAACTCTTTCCCTACACGACGCTCTTCCGATCTNNNGTTGCTCCAATACAGCCTCATTC

AGCACACGTCTGAACTCTTTCCCTACACGACGCTCTTCCGATCTNNNCTTGTCTGGTAAGCTGGAGGA

AGCACACGTCTGAACTCTTTCCCTACACGACGCTCTTCCGATCTNNNCTTGTCTGGTAAGCTGGAGGA

AGCACACGTCTGAACTCTTTCCCTACACGACGCTCTTCCGATCTNNNAGAGGTTGCAGTGAGTTGAG

AGCACACGTCTGAACTCTTTCCCTACACGACGCTCTTCCGATCTNNNAGAGGTTGCAGTGAGTTGAG

GAGCACACGTCTGAACTCTTTCCCTACACGACGCTCTTCCGATCTNNNAATAAATTCAGCCAGGCACA

GAGCACACGTCTGAACTCTTTCCCTACACGACGCTCTTCCGATCTNNNAATAAATTCAGCCAGGCACA

GCACACGTCTGAACTCTTTCCCTACACGACGCTCTTCCGATCTNNNAGCCATCTTCAACAATGAAAGC

GCACACGTCTGAACTCTTTCCCTACACGACGCTCTTCCGATCTNNNAGCCATCTTCAACAATGAAAGC

AGCACACGTCTGAACTCTTTCCCTACACGACGCTCTTCCGATCTNNNATAACACCTACATCTGCCACCT

AGCACACGTCTGAACTCTTTCCCTACACGACGCTCTTCCGATCTNNNATAACACCTACATCTGCCACCT

AGCACACGTCTGAACTCTTTCCCTACACGACGCTCTTCCGATCTNNNGGTGGGAATGTACAATGGCA

AGCACACGTCTGAACTCTTTCCCTACACGACGCTCTTCCGATCTNNNGGTGGGAATGTACAATGGCA

GCACACGTCTGAACTCTTTCCCTACACGACGCTCTTCCGATCTNNNCTTGGAGGCCACTATAATCTCCC

GCACACGTCTGAACTCTTTCCCTACACGACGCTCTTCCGATCTNNNCTTGGAGGCCACTATAATCTCCC
AGCACACGTCTGAACTCTTTCCCTACACGACGCTCTTCCGATCTNNNGAGATCAAGACCATCCTGGCA
AGCACACGTCTGAACTCTTTCCCTACACGACGCTCTTCCGATCTNNNGAGATCAAGACCATCCTGGCA
GCACACGTCTGAACTCTTTCCCTACACGACGCTCTTCCGATCTNNNAATGGGTCATCATAAAGCCAGG
GCACACGTCTGAACTCTTTCCCTACACGACGCTCTTCCGATCTNNNAATGGGTCATCATAAAGCCAGG
GAGCACACGTCTGAACTCTTTCCCTACACGACGCTCTTCCGATCTNNNTGATGACGCAATCTTAGCTCA
GAGCACACGTCTGAACTCTTTCCCTACACGACGCTCTTCCGATCTNNNTGATGACGCAATCTTAGCTCA
AGCACACGTCTGAACTCTTTCCCTACACGACGCTCTTCCGATCTNNNGGAACCACAAAAGACCCAGA
AGCACACGTCTGAACTCTTTCCCTACACGACGCTCTTCCGATCTNNNGGAACCACAAAAGACCCAGA
GCACACGTCTGAACTCTTTCCCTACACGACGCTCTTCCGATCTNNNAGCTGGGATTACAGGTGCATG
GCACACGTCTGAACTCTTTCCCTACACGACGCTCTTCCGATCTNNNAGCTGGGATTACAGGTGCATG
GAGCACACGTCTGAACTCTTTCCCTACACGACGCTCTTCCGATCTNNNGGAAGATCAGTTTCGGTGCA
GAGCACACGTCTGAACTCTTTCCCTACACGACGCTCTTCCGATCTNNNGGAAGATCAGTTTCGGTGCA
GCACACGTCTGAACTCTTTCCCTACACGACGCTCTTCCGATCTNNNTACAAAAATTAGCCAGGCGTGG
GCACACGTCTGAACTCTTTCCCTACACGACGCTCTTCCGATCTNNNTACAAAAATTAGCCAGGCGTGG
AGCACACGTCTGAACTCTTTCCCTACACGACGCTCTTCCGATCTNNNTTCAACAAAGGGATTCACTCTC
AGCACACGTCTGAACTCTTTCCCTACACGACGCTCTTCCGATCTNNNTTCAACAAAGGGATTCACTCTC
AGCACACGTCTGAACTCTTTCCCTACACGACGCTCTTCCGATCTNNNCTCTGTCTCTCTCTCTCCAC
AGCACACGTCTGAACTCTTTCCCTACACGACGCTCTTCCGATCTNNNCTCTGTCTCTCTCTCTCCAC
AGCACACGTCTGAACTCTTTCCCTACACGACGCTCTTCCGATCTNNNGATCAGAGTGAACAGGCAACC
AGCACACGTCTGAACTCTTTCCCTACACGACGCTCTTCCGATCTNNNGATCAGAGTGAACAGGCAACC
GCACACGTCTGAACTCTTTCCCTACACGACGCTCTTCCGATCTNNNGCCAGCTGGATTCTTTGTTTTCT
GCACACGTCTGAACTCTTTCCCTACACGACGCTCTTCCGATCTNNNGCCAGCTGGATTCTTTGTTTTCT
GAGCACACGTCTGAACTCTTTCCCTACACGACGCTCTTCCGATCTNNNCGATGCTCTGTGGATTAGGG
GAGCACACGTCTGAACTCTTTCCCTACACGACGCTCTTCCGATCTNNNCGATGCTCTGTGGATTAGGG
AGCACACGTCTGAACTCTTTCCCTACACGACGCTCTTCCGATCTNNNGGAGGCTGTTGGCTTCTGTAT
AGCACACGTCTGAACTCTTTCCCTACACGACGCTCTTCCGATCTNNNGGAGGCTGTTGGCTTCTGTAT
AGCACACGTCTGAACTCTTTCCCTACACGACGCTCTTCCGATCTNNNTTCGAGGCCAGCTGAGAGAG
AGCACACGTCTGAACTCTTTCCCTACACGACGCTCTTCCGATCTNNNTTCGAGGCCAGCTGAGAGAG
GCACACGTCTGAACTCTTTCCCTACACGACGCTCTTCCGATCTNNNGCTTCTCAGCCACAGTCAGTCAT
GCACACGTCTGAACTCTTTCCCTACACGACGCTCTTCCGATCTNNNGCTTCTCAGCCACAGTCAGTCAT
CACACGTCTGAACTCTTTCCCTACACGACGCTCTTCCGATCTNNNGGAGCTTTTCTACCCAGAACTCAC
CACACGTCTGAACTCTTTCCCTACACGACGCTCTTCCGATCTNNNGGAGCTTTTCTACCCAGAACTCAC
GCACACGTCTGAACTCTTTCCCTACACGACGCTCTTCCGATCTNNNTGTTTGTGTGTGTGAGACAGAG
GCACACGTCTGAACTCTTTCCCTACACGACGCTCTTCCGATCTNNNTGTTTGTGTGTGTGAGACAGAG
CACACGTCTGAACTCTTTCCCTACACGACGCTCTTCCGATCTNNNGCTGATATCTGCATTTGGCCATA
CACACGTCTGAACTCTTTCCCTACACGACGCTCTTCCGATCTNNNGCTGATATCTGCATTTGGCCATA
GAGCACACGTCTGAACTCTTTCCCTACACGACGCTCTTCCGATCTNNNGAAGCGAGAGGAATGTGAC
GAGCACACGTCTGAACTCTTTCCCTACACGACGCTCTTCCGATCTNNNGAAGCGAGAGGAATGTGAC
AGCACACGTCTGAACTCTTTCCCTACACGACGCTCTTCCGATCTNNNACCACGCCCACCTAATTTTTTT
AGCACACGTCTGAACTCTTTCCCTACACGACGCTCTTCCGATCTNNNACCACGCCCACCTAATTTTTTT
GCACACGTCTGAACTCTTTCCCTACACGACGCTCTTCCGATCTNNNTCCCATTTAGGAATGACACTGCA
GCACACGTCTGAACTCTTTCCCTACACGACGCTCTTCCGATCTNNNTCCCATTTAGGAATGACACTGCA

AGCACACGTCTGAACTCTTTCCCTACACGACGCTCTTCCGATCTNNNGCTCAGGTCATCTTCCTCTTTC
AGCACACGTCTGAACTCTTTCCCTACACGACGCTCTTCCGATCTNNNGCTCAGGTCATCTTCCTCTTTC
GAGCACACGTCTGAACTCTTTCCCTACACGACGCTCTTCCGATCTNNNGAGGCAGGAGACTCTCTTGA
GAGCACACGTCTGAACTCTTTCCCTACACGACGCTCTTCCGATCTNNNGAGGCAGGAGACTCTCTTGA
GAGCACACGTCTGAACTCTTTCCCTACACGACGCTCTTCCGATCTNNNGACATTCTGATGGAGGAGG
GAGCACACGTCTGAACTCTTTCCCTACACGACGCTCTTCCGATCTNNNGACATTCTGATGGAGGAGG
GCACACGTCTGAACTCTTTCCCTACACGACGCTCTTCCGATCTNNNGAGGTGGAATAATTGCTTGAG
GCACACGTCTGAACTCTTTCCCTACACGACGCTCTTCCGATCTNNNGAGGTGGAATAATTGCTTGAG
GAGCACACGTCTGAACTCTTTCCCTACACGACGCTCTTCCGATCTNNNCTCAGCCTCCCAAGTAGTGTG
GAGCACACGTCTGAACTCTTTCCCTACACGACGCTCTTCCGATCTNNNCTCAGCCTCCCAAGTAGTGTG
AGCACACGTCTGAACTCTTTCCCTACACGACGCTCTTCCGATCTNNNCGGAAGGACTGGACAGACA
AGCACACGTCTGAACTCTTTCCCTACACGACGCTCTTCCGATCTNNNCGGAAGGACTGGACAGACA
AGCACACGTCTGAACTCTTTCCCTACACGACGCTCTTCCGATCTNNNGGCCTGAGTCTTTCATCAAGAG
AGCACACGTCTGAACTCTTTCCCTACACGACGCTCTTCCGATCTNNNGGCCTGAGTCTTTCATCAAGAG
AGCACACGTCTGAACTCTTTCCCTACACGACGCTCTTCCGATCTNNNCTGAGGCAGGAGAATCACTTG
AGCACACGTCTGAACTCTTTCCCTACACGACGCTCTTCCGATCTNNNCTGAGGCAGGAGAATCACTTG
AGCACACGTCTGAACTCTTTCCCTACACGACGCTCTTCCGATCTNNNCCAGAGTCATCCAAATTGTCAC
AGCACACGTCTGAACTCTTTCCCTACACGACGCTCTTCCGATCTNNNCCAGAGTCATCCAAATTGTCAC
AGCACACGTCTGAACTCTTTCCCTACACGACGCTCTTCCGATCTNNNAAGGTCAGCAGGGGAGAAA
AGCACACGTCTGAACTCTTTCCCTACACGACGCTCTTCCGATCTNNNAAGGTCAGCAGGGGAGAAA
AGCACACGTCTGAACTCTTTCCCTACACGACGCTCTTCCGATCTNNNAGCAGATAAAAGCAAGCTGG
AGCACACGTCTGAACTCTTTCCCTACACGACGCTCTTCCGATCTNNNAGCAGATAAAAGCAAGCTGG
GAGCACACGTCTGAACTCTTTCCCTACACGACGCTCTTCCGATCTNNNTGAGCTGAGATAGGGTATT
GAGCACACGTCTGAACTCTTTCCCTACACGACGCTCTTCCGATCTNNNTGAGCTGAGATAGGGTATT
AGCACACGTCTGAACTCTTTCCCTACACGACGCTCTTCCGATCTNNNTTCAGCTACATCATCTTAGGCC
AGCACACGTCTGAACTCTTTCCCTACACGACGCTCTTCCGATCTNNNTTCAGCTACATCATCTTAGGCC
GAGCACACGTCTGAACTCTTTCCCTACACGACGCTCTTCCGATCTNNNAAAAATTAGCTGGGCGTAA
GAGCACACGTCTGAACTCTTTCCCTACACGACGCTCTTCCGATCTNNNAAAAATTAGCTGGGCGTAA
AGCACACGTCTGAACTCTTTCCCTACACGACGCTCTTCCGATCTNNNGAGGTGCAGAGGCTTCTCAAG
AGCACACGTCTGAACTCTTTCCCTACACGACGCTCTTCCGATCTNNNGAGGTGCAGAGGCTTCTCAAG
AGCACACGTCTGAACTCTTTCCCTACACGACGCTCTTCCGATCTNNNTAAGGTAAAAAGCAAACGGCC
AGCACACGTCTGAACTCTTTCCCTACACGACGCTCTTCCGATCTNNNTAAGGTAAAAAGCAAACGGCC
GAGCACACGTCTGAACTCTTTCCCTACACGACGCTCTTCCGATCTNNNGGCACCTATAATCCCAGCTAC
GAGCACACGTCTGAACTCTTTCCCTACACGACGCTCTTCCGATCTNNNGGCACCTATAATCCCAGCTAC
GAGCACACGTCTGAACTCTTTCCCTACACGACGCTCTTCCGATCTNNNCATCCTCTTTCCGGATCCCAA
GAGCACACGTCTGAACTCTTTCCCTACACGACGCTCTTCCGATCTNNNCATCCTCTTTCCGGATCCCAA
AGCACACGTCTGAACTCTTTCCCTACACGACGCTCTTCCGATCTNNNTTAACTCAGAGAAGTTTCCATT
AGCACACGTCTGAACTCTTTCCCTACACGACGCTCTTCCGATCTNNNTTAACTCAGAGAAGTTTCCATT
AGCACACGTCTGAACTCTTTCCCTACACGACGCTCTTCCGATCTNNNGACACAATCTGCCAAAACTCAC
AGCACACGTCTGAACTCTTTCCCTACACGACGCTCTTCCGATCTNNNGACACAATCTGCCAAAACTCAC
GCACACGTCTGAACTCTTTCCCTACACGACGCTCTTCCGATCTNNNGGAATCCATGTGTCTTGCTCCTT
GCACACGTCTGAACTCTTTCCCTACACGACGCTCTTCCGATCTNNNGGAATCCATGTGTCTTGCTCCTT
AGCACACGTCTGAACTCTTTCCCTACACGACGCTCTTCCGATCTNNNCATCACGAGGTCAGGAGATCG

AGCACACGTCTGAACTCTTTCCCTACACGACGCTCTTCCGATCTNNNCATCACGAGGTCAGGAGATCG
AGAGCACACGTCTGAACTCTTTCCCTACACGACGCTCTTCCGATCTNNNCCTCCAGTTTTCCATGTACCT
AGAGCACACGTCTGAACTCTTTCCCTACACGACGCTCTTCCGATCTNNNCCTCCAGTTTTCCATGTACCT
AGCACACGTCTGAACTCTTTCCCTACACGACGCTCTTCCGATCTNNNTTTGTAGCGAGTTGAGATTGT
AGCACACGTCTGAACTCTTTCCCTACACGACGCTCTTCCGATCTNNNTTTGTAGCGAGTTGAGATTGT
AGCACACGTCTGAACTCTTTCCCTACACGACGCTCTTCCGATCTNNNACAATGAGAGTCTGTGGCATT
AGCACACGTCTGAACTCTTTCCCTACACGACGCTCTTCCGATCTNNNACAATGAGAGTCTGTGGCATT
GAGCACACGTCTGAACTCTTTCCCTACACGACGCTCTTCCGATCTNNNGGAATCACTTGAACCTGGGA
GAGCACACGTCTGAACTCTTTCCCTACACGACGCTCTTCCGATCTNNNGGAATCACTTGAACCTGGGA
GCACACGTCTGAACTCTTTCCCTACACGACGCTCTTCCGATCTNNNTCTCAAATTGCAAAGGAGGCAT
GCACACGTCTGAACTCTTTCCCTACACGACGCTCTTCCGATCTNNNTCTCAAATTGCAAAGGAGGCAT
AGCACACGTCTGAACTCTTTCCCTACACGACGCTCTTCCGATCTNNNCTGAGGCAGGAGAATCACTT
AGCACACGTCTGAACTCTTTCCCTACACGACGCTCTTCCGATCTNNNCTGAGGCAGGAGAATCACTT
AGCACACGTCTGAACTCTTTCCCTACACGACGCTCTTCCGATCTNNNATTTGAAAGGCACAGCATTC
AGCACACGTCTGAACTCTTTCCCTACACGACGCTCTTCCGATCTNNNATTTGAAAGGCACAGCATTC
AGCACACGTCTGAACTCTTTCCCTACACGACGCTCTTCCGATCTNNNTAGCAGATAGCACTGCAGTTT
AGCACACGTCTGAACTCTTTCCCTACACGACGCTCTTCCGATCTNNNTAGCAGATAGCACTGCAGTTT
AGCACACGTCTGAACTCTTTCCCTACACGACGCTCTTCCGATCTNNNGTTACAGCAGCCTGAATGAA
AGCACACGTCTGAACTCTTTCCCTACACGACGCTCTTCCGATCTNNNGTTACAGCAGCCTGAATGAA
AGCACACGTCTGAACTCTTTCCCTACACGACGCTCTTCCGATCTNNNATTTAGTTCCATGTGGCAAG
AGCACACGTCTGAACTCTTTCCCTACACGACGCTCTTCCGATCTNNNATTTAGTTCCATGTGGCAAG
GCACACGTCTGAACTCTTTCCCTACACGACGCTCTTCCGATCTNNNAAGAACAGACCATGAGCAGAGG
GCACACGTCTGAACTCTTTCCCTACACGACGCTCTTCCGATCTNNNAAGAACAGACCATGAGCAGAGG
GCACACGTCTGAACTCTTTCCCTACACGACGCTCTTCCGATCTNNNCAGGTTCAAGCAATTCTCCTGC
GCACACGTCTGAACTCTTTCCCTACACGACGCTCTTCCGATCTNNNCAGGTTCAAGCAATTCTCCTGC
GCACACGTCTGAACTCTTTCCCTACACGACGCTCTTCCGATCTNNNACTAGAACCCAGGCTTTTTGAC
GCACACGTCTGAACTCTTTCCCTACACGACGCTCTTCCGATCTNNNACTAGAACCCAGGCTTTTTGAC
AGCACACGTCTGAACTCTTTCCCTACACGACGCTCTTCCGATCTNNNCAGTAAGTTCAGGCAACAGCT
AGCACACGTCTGAACTCTTTCCCTACACGACGCTCTTCCGATCTNNNCAGTAAGTTCAGGCAACAGCT
AGCACACGTCTGAACTCTTTCCCTACACGACGCTCTTCCGATCTNNNTACAACTGTGGGGGTAAAAC
AGCACACGTCTGAACTCTTTCCCTACACGACGCTCTTCCGATCTNNNTACAACTGTGGGGGTAAAAC
GAGCACACGTCTGAACTCTTTCCCTACACGACGCTCTTCCGATCTNNNCCCTTTGCAACACCATACAAT
GAGCACACGTCTGAACTCTTTCCCTACACGACGCTCTTCCGATCTNNNCCCTTTGCAACACCATACAAT
GAGCACACGTCTGAACTCTTTCCCTACACGACGCTCTTCCGATCTNNNAAAAAAAAAAAGTTAGCCGG
GAGCACACGTCTGAACTCTTTCCCTACACGACGCTCTTCCGATCTNNNAAAAAAAAAAAGTTAGCCGG
AGCACACGTCTGAACTCTTTCCCTACACGACGCTCTTCCGATCTNNNGCTTGGCCAAATTAGTCCCATT
AGCACACGTCTGAACTCTTTCCCTACACGACGCTCTTCCGATCTNNNGCTTGGCCAAATTAGTCCCATT
AGCACACGTCTGAACTCTTTCCCTACACGACGCTCTTCCGATCTNNNGGTGATGGGTAAATGACTTT
AGCACACGTCTGAACTCTTTCCCTACACGACGCTCTTCCGATCTNNNGGTGATGGGTAAATGACTTT
AGCACACGTCTGAACTCTTTCCCTACACGACGCTCTTCCGATCTNNNGGAGACCCTGTCACCAAAAAAA
AGCACACGTCTGAACTCTTTCCCTACACGACGCTCTTCCGATCTNNNGGAGACCCTGTCACCAAAAAAA
AGCACACGTCTGAACTCTTTCCCTACACGACGCTCTTCCGATCTNNNAAGTTTTGGTCACACTCTCACC
AGCACACGTCTGAACTCTTTCCCTACACGACGCTCTTCCGATCTNNNAAGTTTTGGTCACACTCTCACC

AGCACACGTCTGAACTCTTTCCCTACACGACGCTCTTCCGATCTNNNGGCTGTATCCAATAGCATTTGC
AGCACACGTCTGAACTCTTTCCCTACACGACGCTCTTCCGATCTNNNGGCTGTATCCAATAGCATTTGC
GAGCACACGTCTGAACTCTTTCCCTACACGACGCTCTTCCGATCTNNNGAGAATGGCATGAACCTGGG
GAGCACACGTCTGAACTCTTTCCCTACACGACGCTCTTCCGATCTNNNGAGAATGGCATGAACCTGGG
CACACGTCTGAACTCTTTCCCTACACGACGCTCTTCCGATCTNNNTCTCTCTGTCCTTCCTTCCTCTTCAT
CACACGTCTGAACTCTTTCCCTACACGACGCTCTTCCGATCTNNNTCTCTCTGTCCTTCCTTCCTCTTCAT
GAGCACACGTCTGAACTCTTTCCCTACACGACGCTCTTCCGATCTNNNAGCAAGATGTGTGAATGGG
GAGCACACGTCTGAACTCTTTCCCTACACGACGCTCTTCCGATCTNNNAGCAAGATGTGTGAATGGG
GAGCACACGTCTGAACTCTTTCCCTACACGACGCTCTTCCGATCTNNNCCTGGCTCTACTCCCGATTTCT
GAGCACACGTCTGAACTCTTTCCCTACACGACGCTCTTCCGATCTNNNCCTGGCTCTACTCCCGATTTCT
AGAGCACACGTCTGAACTCTTTCCCTACACGACGCTCTTCCGATCTNNNGATTGTGTGAGTGTGTGAG
AGAGCACACGTCTGAACTCTTTCCCTACACGACGCTCTTCCGATCTNNNGATTGTGTGAGTGTGTGAG
AGCACACGTCTGAACTCTTTCCCTACACGACGCTCTTCCGATCTNNNTGTCTTCCTTCCTCCAACTCTTGC
AGCACACGTCTGAACTCTTTCCCTACACGACGCTCTTCCGATCTNNNTGTCTTCCTTCCTCCAACTCTTGC
GAGCACACGTCTGAACTCTTTCCCTACACGACGCTCTTCCGATCTNNNGTCCAGAACCCAGTTACTGTG
GAGCACACGTCTGAACTCTTTCCCTACACGACGCTCTTCCGATCTNNNGTCCAGAACCCAGTTACTGTG
GCACACGTCTGAACTCTTTCCCTACACGACGCTCTTCCGATCTNNNATCTATGTGGAAGTGCAGCAAG
GCACACGTCTGAACTCTTTCCCTACACGACGCTCTTCCGATCTNNNATCTATGTGGAAGTGCAGCAAG
AGCACACGTCTGAACTCTTTCCCTACACGACGCTCTTCCGATCTNNNTGTCTGCAGTTCTAGGCTTGTA
AGCACACGTCTGAACTCTTTCCCTACACGACGCTCTTCCGATCTNNNTGTCTGCAGTTCTAGGCTTGTA
AGCACACGTCTGAACTCTTTCCCTACACGACGCTCTTCCGATCTNNNGCAGTGGTGTAATCACAGACC
AGCACACGTCTGAACTCTTTCCCTACACGACGCTCTTCCGATCTNNNGCAGTGGTGTAATCACAGACC
AGCACACGTCTGAACTCTTTCCCTACACGACGCTCTTCCGATCTNNNGCTGGAAAGATACTGCCCAATT
AGCACACGTCTGAACTCTTTCCCTACACGACGCTCTTCCGATCTNNNGCTGGAAAGATACTGCCCAATT
GAAGAGCACACGTCTGAACTCTTTCCCTACACGACGCTCTTCCGATCTNNNACTGCACTTGCACTTCAT
GAAGAGCACACGTCTGAACTCTTTCCCTACACGACGCTCTTCCGATCTNNNACTGCACTTGCACTTCAT
GAGCACACGTCTGAACTCTTTCCCTACACGACGCTCTTCCGATCTNNNATGTGCATTGCCTCATGGAA
GAGCACACGTCTGAACTCTTTCCCTACACGACGCTCTTCCGATCTNNNATGTGCATTGCCTCATGGAA
AGCACACGTCTGAACTCTTTCCCTACACGACGCTCTTCCGATCTNNNACCAGAATCACTTGAACCCACA
AGCACACGTCTGAACTCTTTCCCTACACGACGCTCTTCCGATCTNNNACCAGAATCACTTGAACCCACA
GAGCACACGTCTGAACTCTTTCCCTACACGACGCTCTTCCGATCTNNNAGACAAGATCTGGCTCTGTCA
GAGCACACGTCTGAACTCTTTCCCTACACGACGCTCTTCCGATCTNNNAGACAAGATCTGGCTCTGTCA
AGCACACGTCTGAACTCTTTCCCTACACGACGCTCTTCCGATCTNNNGATTTTAAGTGTCACAGGTAAA
AGCACACGTCTGAACTCTTTCCCTACACGACGCTCTTCCGATCTNNNGATTTTAAGTGTCACAGGTAAA
AGCACACGTCTGAACTCTTTCCCTACACGACGCTCTTCCGATCTNNNCTTCTCAGCCTCCATAAGTGCAT
AGCACACGTCTGAACTCTTTCCCTACACGACGCTCTTCCGATCTNNNCTTCTCAGCCTCCATAAGTGCAT
GAGCACACGTCTGAACTCTTTCCCTACACGACGCTCTTCCGATCTNNNAGAAGAGCAGGCAGTAAACC
GAGCACACGTCTGAACTCTTTCCCTACACGACGCTCTTCCGATCTNNNAGAAGAGCAGGCAGTAAACC
AGCACACGTCTGAACTCTTTCCCTACACGACGCTCTTCCGATCTNNNGATCTCAGCTCACTGTAACCTC
AGCACACGTCTGAACTCTTTCCCTACACGACGCTCTTCCGATCTNNNGATCTCAGCTCACTGTAACCTC
GAGCACACGTCTGAACTCTTTCCCTACACGACGCTCTTCCGATCTNNNGAGAATTGCTTGAACCCAGG
GAGCACACGTCTGAACTCTTTCCCTACACGACGCTCTTCCGATCTNNNGAGAATTGCTTGAACCCAGG
AGCACACGTCTGAACTCTTTCCCTACACGACGCTCTTCCGATCTNNNAAGAATTTCCAGAGAGTTGTAT

AGCACACGTCTGAACTCTTTCCCTACACGACGCTCTTCCGATCTNNNAAGAATTTCCAGAGAGTTGTAT
GAGCACACGTCTGAACTCTTTCCCTACACGACGCTCTTCCGATCTNNNGGTGCTGACATTGCTAGTATT
GAGCACACGTCTGAACTCTTTCCCTACACGACGCTCTTCCGATCTNNNGGTGCTGACATTGCTAGTATT
ACACGTCTGAACTCTTTCCCTACACGACGCTCTTCCGATCTNNNATGTCCAATAAACACGGTGCATCCAT
ACACGTCTGAACTCTTTCCCTACACGACGCTCTTCCGATCTNNNATGTCCAATAAACACGGTGCATCCAT
GCACACGTCTGAACTCTTTCCCTACACGACGCTCTTCCGATCTNNNTAGGAGAAAACTTGAGCCCAGGA
GCACACGTCTGAACTCTTTCCCTACACGACGCTCTTCCGATCTNNNTAGGAGAAAACTTGAGCCCAGGA
AGCACACGTCTGAACTCTTTCCCTACACGACGCTCTTCCGATCTNNNGCTCACAGCTATAATCCCAGCAT
AGCACACGTCTGAACTCTTTCCCTACACGACGCTCTTCCGATCTNNNGCTCACAGCTATAATCCCAGCAT
GAGCACACGTCTGAACTCTTTCCCTACACGACGCTCTTCCGATCTNNNGCCTACACAGTAAGAGCAAGA
GAGCACACGTCTGAACTCTTTCCCTACACGACGCTCTTCCGATCTNNNGCCTACACAGTAAGAGCAAGA
AGCACACGTCTGAACTCTTTCCCTACACGACGCTCTTCCGATCTNNNCCAAGTAGGTGGGATTACAGG
AGCACACGTCTGAACTCTTTCCCTACACGACGCTCTTCCGATCTNNNCCAAGTAGGTGGGATTACAGG
AGCACACGTCTGAACTCTTTCCCTACACGACGCTCTTCCGATCTNNNTCTGCAGTAACTCTTTTCTGTTT
AGCACACGTCTGAACTCTTTCCCTACACGACGCTCTTCCGATCTNNNTCTGCAGTAACTCTTTTCTGTTT
AGCACACGTCTGAACTCTTTCCCTACACGACGCTCTTCCGATCTNNNTCTAAGAAGTCCGGGATTTGAC
AGCACACGTCTGAACTCTTTCCCTACACGACGCTCTTCCGATCTNNNTCTAAGAAGTCCGGGATTTGAC
AGCACACGTCTGAACTCTTTCCCTACACGACGCTCTTCCGATCTNNNAAAGTGAAAAAGCCCCACAACA
AGCACACGTCTGAACTCTTTCCCTACACGACGCTCTTCCGATCTNNNAAAGTGAAAAAGCCCCACAACA
GAGCACACGTCTGAACTCTTTCCCTACACGACGCTCTTCCGATCTNNNACCCAGTCAGATAGGATGGCT
GAGCACACGTCTGAACTCTTTCCCTACACGACGCTCTTCCGATCTNNNACCCAGTCAGATAGGATGGCT
GAGCACACGTCTGAACTCTTTCCCTACACGACGCTCTTCCGATCTNNNCTGTAATCCCAGCTACTCAGG
GAGCACACGTCTGAACTCTTTCCCTACACGACGCTCTTCCGATCTNNNCTGTAATCCCAGCTACTCAGG
AGCACACGTCTGAACTCTTTCCCTACACGACGCTCTTCCGATCTNNNCAGGAGAATTGCTTGAACCAGG
AGCACACGTCTGAACTCTTTCCCTACACGACGCTCTTCCGATCTNNNCAGGAGAATTGCTTGAACCAGG
GAGCACACGTCTGAACTCTTTCCCTACACGACGCTCTTCCGATCTNNNCAAAAAAATTAGCCGTGCGTG
GAGCACACGTCTGAACTCTTTCCCTACACGACGCTCTTCCGATCTNNNCAAAAAAATTAGCCGTGCGTG
GAGCACACGTCTGAACTCTTTCCCTACACGACGCTCTTCCGATCTNNNTGAAAGAGCAGGCATCCCTAA
GAGCACACGTCTGAACTCTTTCCCTACACGACGCTCTTCCGATCTNNNTGAAAGAGCAGGCATCCCTAA
AGCACACGTCTGAACTCTTTCCCTACACGACGCTCTTCCGATCTNNNCCATTCCTCCCTCCACTCAGGCAT
AGCACACGTCTGAACTCTTTCCCTACACGACGCTCTTCCGATCTNNNCCATTCCTCCCTCCACTCAGGCAT
GAGCACACGTCTGAACTCTTTCCCTACACGACGCTCTTCCGATCTNNNGGGAATTTAGGAGCCCCATAC
GAGCACACGTCTGAACTCTTTCCCTACACGACGCTCTTCCGATCTNNNGGGAATTTAGGAGCCCCATAC
GAGCACACGTCTGAACTCTTTCCCTACACGACGCTCTTCCGATCTNNNCGTTCCTGTCACTTCCATAGGA
GAGCACACGTCTGAACTCTTTCCCTACACGACGCTCTTCCGATCTNNNCGTTCCTGTCACTTCCATAGGA
GAGCACACGTCTGAACTCTTTCCCTACACGACGCTCTTCCGATCTNNNAACACTAGCAGGAAAGCAAAA
GAGCACACGTCTGAACTCTTTCCCTACACGACGCTCTTCCGATCTNNNAACACTAGCAGGAAAGCAAAA
GCACACGTCTGAACTCTTTCCCTACACGACGCTCTTCCGATCTNNNTACATGCCTGAGTTTGTGTGTCTC
GCACACGTCTGAACTCTTTCCCTACACGACGCTCTTCCGATCTNNNTACATGCCTGAGTTTGTGTGTCTC
AGAGCACACGTCTGAACTCTTTCCCTACACGACGCTCTTCCGATCTNNNTGTGTGTGTGTGTGTGTAT
AGAGCACACGTCTGAACTCTTTCCCTACACGACGCTCTTCCGATCTNNNTGTGTGTGTGTGTGTGTAT
AGCACACGTCTGAACTCTTTCCCTACACGACGCTCTTCCGATCTNNNTACCCTATTCAACGAGATGAGC
AGCACACGTCTGAACTCTTTCCCTACACGACGCTCTTCCGATCTNNNTACCCTATTCAACGAGATGAGC

AGCACACGTCTGAACTCTTTCCCTACACGACGCTCTTCCGATCTNNNAAAAAGAGAAACCAAGGGGG
AGCACACGTCTGAACTCTTTCCCTACACGACGCTCTTCCGATCTNNNAAAAAGAGAAACCAAGGGGG
GAGCACACGTCTGAACTCTTTCCCTACACGACGCTCTTCCGATCTNNNCTGAGGCAGGAGAATCACTTC
GAGCACACGTCTGAACTCTTTCCCTACACGACGCTCTTCCGATCTNNNCTGAGGCAGGAGAATCACTTC
AGCACACGTCTGAACTCTTTCCCTACACGACGCTCTTCCGATCTNNNAAACCTTTCCTGGGGAATCACCC
AGCACACGTCTGAACTCTTTCCCTACACGACGCTCTTCCGATCTNNNAAACCTTTCCTGGGGAATCACCC
GAGCACACGTCTGAACTCTTTCCCTACACGACGCTCTTCCGATCTNNNGCAACAACAGCAAAACTCTGAG
GAGCACACGTCTGAACTCTTTCCCTACACGACGCTCTTCCGATCTNNNGCAACAACAGCAAAACTCTGAG
GAGCACACGTCTGAACTCTTTCCCTACACGACGCTCTTCCGATCTNNNACTGTGCCATCTCATTGTCTCT
GAGCACACGTCTGAACTCTTTCCCTACACGACGCTCTTCCGATCTNNNACTGTGCCATCTCATTGTCTCT
AGCACACGTCTGAACTCTTTCCCTACACGACGCTCTTCCGATCTNNNTCGTTGATTAGAGAGCCTGAAG
AGCACACGTCTGAACTCTTTCCCTACACGACGCTCTTCCGATCTNNNTCGTTGATTAGAGAGCCTGAAG
GAGCACACGTCTGAACTCTTTCCCTACACGACGCTCTTCCGATCTNNNTGTGATGCAGGTTCCAAATTG
GAGCACACGTCTGAACTCTTTCCCTACACGACGCTCTTCCGATCTNNNTGTGATGCAGGTTCCAAATTG
GAGCACACGTCTGAACTCTTTCCCTACACGACGCTCTTCCGATCTNNNTGTAATTCCAGCTACTTGAGG
GAGCACACGTCTGAACTCTTTCCCTACACGACGCTCTTCCGATCTNNNTGTAATTCCAGCTACTTGAGG
AGCACACGTCTGAACTCTTTCCCTACACGACGCTCTTCCGATCTNNNGACCGAGAATATGAGCTCTAGC
AGCACACGTCTGAACTCTTTCCCTACACGACGCTCTTCCGATCTNNNGACCGAGAATATGAGCTCTAGC
AGCACACGTCTGAACTCTTTCCCTACACGACGCTCTTCCGATCTNNNCTCTAACTCACGACCACCATCTCC
AGCACACGTCTGAACTCTTTCCCTACACGACGCTCTTCCGATCTNNNCTCTAACTCACGACCACCATCTCC
AGAGCACACGTCTGAACTCTTTCCCTACACGACGCTCTTCCGATCTNNNTGAGCATCGTTAAGGCCCAC
AGAGCACACGTCTGAACTCTTTCCCTACACGACGCTCTTCCGATCTNNNTGAGCATCGTTAAGGCCCAC
AGCACACGTCTGAACTCTTTCCCTACACGACGCTCTTCCGATCTNNNCATAGCAAGACCCCATCTCTACC
AGCACACGTCTGAACTCTTTCCCTACACGACGCTCTTCCGATCTNNNCATAGCAAGACCCCATCTCTACC
GAGCACACGTCTGAACTCTTTCCCTACACGACGCTCTTCCGATCTNNNGTCTAGGGGACAGAGAGAGA
GAGCACACGTCTGAACTCTTTCCCTACACGACGCTCTTCCGATCTNNNGTCTAGGGGACAGAGAGAGA
ACACGTCTGAACTCTTTCCCTACACGACGCTCTTCCGATCTNNNATCCAAGAATATGCCCTTTCCATCGG
ACACGTCTGAACTCTTTCCCTACACGACGCTCTTCCGATCTNNNATCCAAGAATATGCCCTTTCCATCGG
AGCACACGTCTGAACTCTTTCCCTACACGACGCTCTTCCGATCTNNNTAAACTGTGTCTTTTGTAGTTTC
AGCACACGTCTGAACTCTTTCCCTACACGACGCTCTTCCGATCTNNNTAAACTGTGTCTTTTGTAGTTTC
AGAGCACACGTCTGAACTCTTTCCCTACACGACGCTCTTCCGATCTNNNCGTGCACATGTACTCCTTGAA
AGAGCACACGTCTGAACTCTTTCCCTACACGACGCTCTTCCGATCTNNNCGTGCACATGTACTCCTTGAA
GAGCACACGTCTGAACTCTTTCCCTACACGACGCTCTTCCGATCTNNNTTAGCAAAGGACTTGGCACAG
GAGCACACGTCTGAACTCTTTCCCTACACGACGCTCTTCCGATCTNNNTTAGCAAAGGACTTGGCACAG
GAGCACACGTCTGAACTCTTTCCCTACACGACGCTCTTCCGATCTNNNGTGATGCCTTCAGCTTTGTTCT
GAGCACACGTCTGAACTCTTTCCCTACACGACGCTCTTCCGATCTNNNGTGATGCCTTCAGCTTTGTTCT
GAGCACACGTCTGAACTCTTTCCCTACACGACGCTCTTCCGATCTNNNCGTCCTACTTGGCAGGAGATTT
GAGCACACGTCTGAACTCTTTCCCTACACGACGCTCTTCCGATCTNNNCGTCCTACTTGGCAGGAGATTT
AGCACACGTCTGAACTCTTTCCCTACACGACGCTCTTCCGATCTNNNCTCTAAATTATCCAGACTGAAG
AGCACACGTCTGAACTCTTTCCCTACACGACGCTCTTCCGATCTNNNCTCTAAATTATCCAGACTGAAG
AGCACACGTCTGAACTCTTTCCCTACACGACGCTCTTCCGATCTNNNCTCCAACGCAGACTGGTTATTT
AGCACACGTCTGAACTCTTTCCCTACACGACGCTCTTCCGATCTNNNCTCCAACGCAGACTGGTTATTT
AGCACACGTCTGAACTCTTTCCCTACACGACGCTCTTCCGATCTNNNGAGGCTGCCATGAAGATCTCTA

AGCACACGTCTGAACTCTTTCCCTACACGACGCTCTTCCGATCTNNNGAGGCTGCCATGAAGATCTCTA
AGAGCACACGTCTGAACTCTTTCCCTACACGACGCTCTTCCGATCTNNNCCACAAAATGAGAAAGGCCT
AGAGCACACGTCTGAACTCTTTCCCTACACGACGCTCTTCCGATCTNNNCCACAAAATGAGAAAGGCCT
GAGCACACGTCTGAACTCTTTCCCTACACGACGCTCTTCCGATCTNNNGGATAAAAGGGCACTAGGTC
GAGCACACGTCTGAACTCTTTCCCTACACGACGCTCTTCCGATCTNNNGGATAAAAGGGCACTAGGTC
GAGCACACGTCTGAACTCTTTCCCTACACGACGCTCTTCCGATCTNNNCCCTCATGTTCTTAGTGGGTAC
GAGCACACGTCTGAACTCTTTCCCTACACGACGCTCTTCCGATCTNNNCCCTCATGTTCTTAGTGGGTAC
CACACGTCTGAACTCTTTCCCTACACGACGCTCTTCCGATCTNNNTGTTTTGTTGTTGTTGTTGTTGTCA
CACACGTCTGAACTCTTTCCCTACACGACGCTCTTCCGATCTNNNTGTTTTGTTGTTGTTGTTGTTGTCA
GAGCACACGTCTGAACTCTTTCCCTACACGACGCTCTTCCGATCTNNNTTGTGACTAGCTGTGTGTGTA
GAGCACACGTCTGAACTCTTTCCCTACACGACGCTCTTCCGATCTNNNTTGTGACTAGCTGTGTGTGTA
GAGCACACGTCTGAACTCTTTCCCTACACGACGCTCTTCCGATCTNNNAGACTAGGCTTATCAGCTGCT
GAGCACACGTCTGAACTCTTTCCCTACACGACGCTCTTCCGATCTNNNAGACTAGGCTTATCAGCTGCT
CACACGTCTGAACTCTTTCCCTACACGACGCTCTTCCGATCTNNNCCAGAGGTGGGACTACAATCTTGC
CACACGTCTGAACTCTTTCCCTACACGACGCTCTTCCGATCTNNNCCAGAGGTGGGACTACAATCTTGC
AGAGCACACGTCTGAACTCTTTCCCTACACGACGCTCTTCCGATCTNNNCATCTGTGTGGAGGTTACTG
AGAGCACACGTCTGAACTCTTTCCCTACACGACGCTCTTCCGATCTNNNCATCTGTGTGGAGGTTACTG
AGAGCACACGTCTGAACTCTTTCCCTACACGACGCTCTTCCGATCTNNNCTTGCAGTGAGCCGAGATCG
AGAGCACACGTCTGAACTCTTTCCCTACACGACGCTCTTCCGATCTNNNCTTGCAGTGAGCCGAGATCG
GAGCACACGTCTGAACTCTTTCCCTACACGACGCTCTTCCGATCTNNNCAAGCAGCTGGGATTACAAG
GAGCACACGTCTGAACTCTTTCCCTACACGACGCTCTTCCGATCTNNNCAAGCAGCTGGGATTACAAG
GAGCACACGTCTGAACTCTTTCCCTACACGACGCTCTTCCGATCTNNNCTGGAAGACCCTGGCAATGTTA
GAGCACACGTCTGAACTCTTTCCCTACACGACGCTCTTCCGATCTNNNCTGGAAGACCCTGGCAATGTTA
AGCACACGTCTGAACTCTTTCCCTACACGACGCTCTTCCGATCTNNNTTGCGTCTGAATCCTGTAACACA
AGCACACGTCTGAACTCTTTCCCTACACGACGCTCTTCCGATCTNNNTTGCGTCTGAATCCTGTAACACA
AGCACACGTCTGAACTCTTTCCCTACACGACGCTCTTCCGATCTNNNGATATGTTTCATTTGGGGCCAC
AGCACACGTCTGAACTCTTTCCCTACACGACGCTCTTCCGATCTNNNGATATGTTTCATTTGGGGCCAC
GAGCACACGTCTGAACTCTTTCCCTACACGACGCTCTTCCGATCTNNNCTGTAATCCCAGCTACTCAGGA
GAGCACACGTCTGAACTCTTTCCCTACACGACGCTCTTCCGATCTNNNCTGTAATCCCAGCTACTCAGGA
AGCACACGTCTGAACTCTTTCCCTACACGACGCTCTTCCGATCTNNNTAATGCAGTGGCTTGATAATGG
AGCACACGTCTGAACTCTTTCCCTACACGACGCTCTTCCGATCTNNNTAATGCAGTGGCTTGATAATGG
AGCACACGTCTGAACTCTTTCCCTACACGACGCTCTTCCGATCTNNNTGGTGCCACCTATCACATGAAT
AGCACACGTCTGAACTCTTTCCCTACACGACGCTCTTCCGATCTNNNTGGTGCCACCTATCACATGAAT
GCACACGTCTGAACTCTTTCCCTACACGACGCTCTTCCGATCTNNNAGAGGACTCTCTCTCCTTGATGAC
GCACACGTCTGAACTCTTTCCCTACACGACGCTCTTCCGATCTNNNAGAGGACTCTCTCTCCTTGATGAC
GAGCACACGTCTGAACTCTTTCCCTACACGACGCTCTTCCGATCTNNNCTGAGGCAGGAGAATCACTTG
GAGCACACGTCTGAACTCTTTCCCTACACGACGCTCTTCCGATCTNNNCTGAGGCAGGAGAATCACTTG
AGCACACGTCTGAACTCTTTCCCTACACGACGCTCTTCCGATCTNNNCAGTCTGGGCAACATAGTGAGA
AGCACACGTCTGAACTCTTTCCCTACACGACGCTCTTCCGATCTNNNCAGTCTGGGCAACATAGTGAGA
GAGCACACGTCTGAACTCTTTCCCTACACGACGCTCTTCCGATCTNNNGATCTGGAACACGACAAAGA
GAGCACACGTCTGAACTCTTTCCCTACACGACGCTCTTCCGATCTNNNGATCTGGAACACGACAAAGA
AGCACACGTCTGAACTCTTTCCCTACACGACGCTCTTCCGATCTNNNGCCGAGATCAAGCCACTATACT
AGCACACGTCTGAACTCTTTCCCTACACGACGCTCTTCCGATCTNNNGCCGAGATCAAGCCACTATACT

GAGCACACGTCTGAACTCTTTCCCTACACGACGCTCTTCCGATCTNNNGGGCGACAGAGTGAGATTCT
GAGCACACGTCTGAACTCTTTCCCTACACGACGCTCTTCCGATCTNNNGGGCGACAGAGTGAGATTCT
AGCACACGTCTGAACTCTTTCCCTACACGACGCTCTTCCGATCTNNNACAAAGTGGATGTAAGGCAAT
AGCACACGTCTGAACTCTTTCCCTACACGACGCTCTTCCGATCTNNNACAAAGTGGATGTAAGGCAAT
CACGTCTGAACTCTTTCCCTACACGACGCTCTTCCGATCTNNNGGAAGCAAACAAACGAGACAGAACAT
CACGTCTGAACTCTTTCCCTACACGACGCTCTTCCGATCTNNNGGAAGCAAACAAACGAGACAGAACAT
GAGCACACGTCTGAACTCTTTCCCTACACGACGCTCTTCCGATCTNNNCAGCTGCTCAGAATCTGTAGAC
GAGCACACGTCTGAACTCTTTCCCTACACGACGCTCTTCCGATCTNNNCAGCTGCTCAGAATCTGTAGAC
AGCACACGTCTGAACTCTTTCCCTACACGACGCTCTTCCGATCTNNNCAACCCTCCCGACTTCTTCTAAC
AGCACACGTCTGAACTCTTTCCCTACACGACGCTCTTCCGATCTNNNCAACCCTCCCGACTTCTTCTAAC
AGCACACGTCTGAACTCTTTCCCTACACGACGCTCTTCCGATCTNNNCCTGAGCGAAAGAGTGAGACT
AGCACACGTCTGAACTCTTTCCCTACACGACGCTCTTCCGATCTNNNCCTGAGCGAAAGAGTGAGACT
GAGCACACGTCTGAACTCTTTCCCTACACGACGCTCTTCCGATCTNNNCTGCTCAGCTTCTCTCTCTCATT
GAGCACACGTCTGAACTCTTTCCCTACACGACGCTCTTCCGATCTNNNCTGCTCAGCTTCTCTCTCTCATT
GCACACGTCTGAACTCTTTCCCTACACGACGCTCTTCCGATCTNNNCATCCATCTCCTTTGCTTTGCATC
GCACACGTCTGAACTCTTTCCCTACACGACGCTCTTCCGATCTNNNCATCCATCTCCTTTGCTTTGCATC
GAGCACACGTCTGAACTCTTTCCCTACACGACGCTCTTCCGATCTNNNTTGCTGCAAATGCCATTACTTC
GAGCACACGTCTGAACTCTTTCCCTACACGACGCTCTTCCGATCTNNNTTGCTGCAAATGCCATTACTTC
AGAGCACACGTCTGAACTCTTTCCCTACACGACGCTCTTCCGATCTNNNGGATGCCATGTCTGGAAATA
AGAGCACACGTCTGAACTCTTTCCCTACACGACGCTCTTCCGATCTNNNGGATGCCATGTCTGGAAATA
AGAGCACACGTCTGAACTCTTTCCCTACACGACGCTCTTCCGATCTNNNAGATGTCACCAGTCATCCGA
AGAGCACACGTCTGAACTCTTTCCCTACACGACGCTCTTCCGATCTNNNAGATGTCACCAGTCATCCGA
AGAGCACACGTCTGAACTCTTTCCCTACACGACGCTCTTCCGATCTNNNAGGCAAGAGAATCACTTGAA
AGAGCACACGTCTGAACTCTTTCCCTACACGACGCTCTTCCGATCTNNNAGGCAAGAGAATCACTTGAA
GAGCACACGTCTGAACTCTTTCCCTACACGACGCTCTTCCGATCTNNNGAGAAATAAGGAAGCAAGCA
GAGCACACGTCTGAACTCTTTCCCTACACGACGCTCTTCCGATCTNNNGAGAAATAAGGAAGCAAGCA
AGCACACGTCTGAACTCTTTCCCTACACGACGCTCTTCCGATCTNNNTCCCCATGATCTCTCTCTCTCTCC
AGCACACGTCTGAACTCTTTCCCTACACGACGCTCTTCCGATCTNNNTCCCCATGATCTCTCTCTCTCTCC
GCACACGTCTGAACTCTTTCCCTACACGACGCTCTTCCGATCTNNNGTCGTAGGACAAAGGGCCCCCATG
GCACACGTCTGAACTCTTTCCCTACACGACGCTCTTCCGATCTNNNGTCGTAGGACAAAGGGCCCCCATG
GAGCACACGTCTGAACTCTTTCCCTACACGACGCTCTTCCGATCTNNNGTAGCAGAACAAGACCCTGTC
GAGCACACGTCTGAACTCTTTCCCTACACGACGCTCTTCCGATCTNNNGTAGCAGAACAAGACCCTGTC
GAGCACACGTCTGAACTCTTTCCCTACACGACGCTCTTCCGATCTNNNGGTCAGGAGATCGAGACTATC
GAGCACACGTCTGAACTCTTTCCCTACACGACGCTCTTCCGATCTNNNGGTCAGGAGATCGAGACTATC
GCACACGTCTGAACTCTTTCCCTACACGACGCTCTTCCGATCTNNNTTCAAGCTATTCTCCTGCCTCAGC
GCACACGTCTGAACTCTTTCCCTACACGACGCTCTTCCGATCTNNNTTCAAGCTATTCTCCTGCCTCAGC
GAGCACACGTCTGAACTCTTTCCCTACACGACGCTCTTCCGATCTNNNAGCATTGGTACCAGCTCTTCTT
GAGCACACGTCTGAACTCTTTCCCTACACGACGCTCTTCCGATCTNNNAGCATTGGTACCAGCTCTTCTT
AGAGCACACGTCTGAACTCTTTCCCTACACGACGCTCTTCCGATCTNNNCCTAGGTGGAGTGGCAAAGT
AGAGCACACGTCTGAACTCTTTCCCTACACGACGCTCTTCCGATCTNNNCCTAGGTGGAGTGGCAAAGT
GAGCACACGTCTGAACTCTTTCCCTACACGACGCTCTTCCGATCTNNNAAAAAATTAGCAGGGCATGG
GAGCACACGTCTGAACTCTTTCCCTACACGACGCTCTTCCGATCTNNNAAAAAATTAGCAGGGCATGG
GAGCACACGTCTGAACTCTTTCCCTACACGACGCTCTTCCGATCTNNNCGGATCATTTGAGGTCAGGA

GAGCACACGTCTGAACTCTTTCCCTACACGACGCTCTTCCGATCTNNNCGGATCATTTGAGGTCAGGA
GAGCACACGTCTGAACTCTTTCCCTACACGACGCTCTTCCGATCTNNNCTCAGGGCAGCTCACTCTCTA
GAGCACACGTCTGAACTCTTTCCCTACACGACGCTCTTCCGATCTNNNCTCAGGGCAGCTCACTCTCTA
GCACACGTCTGAACTCTTTCCCTACACGACGCTCTTCCGATCTNNNTGAGTCACAGAGAGTTTCAGTG
GCACACGTCTGAACTCTTTCCCTACACGACGCTCTTCCGATCTNNNTGAGTCACAGAGAGTTTCAGTG
AGCACACGTCTGAACTCTTTCCCTACACGACGCTCTTCCGATCTNNNCTGTTACACACTGACAGACCAC
AGCACACGTCTGAACTCTTTCCCTACACGACGCTCTTCCGATCTNNNCTGTTACACACTGACAGACCAC
AGCACACGTCTGAACTCTTTCCCTACACGACGCTCTTCCGATCTNNNTTTTCTGTCTCTCTGTCTCCCCC
AGCACACGTCTGAACTCTTTCCCTACACGACGCTCTTCCGATCTNNNTTTTCTGTCTCTCTGTCTCCCCC
GCACACGTCTGAACTCTTTCCCTACACGACGCTCTTCCGATCTNNNGCTGTTCAGTTGCTCTTGCTTTC
GCACACGTCTGAACTCTTTCCCTACACGACGCTCTTCCGATCTNNNGCTGTTCAGTTGCTCTTGCTTTC
GAGCACACGTCTGAACTCTTTCCCTACACGACGCTCTTCCGATCTNNNTGACTGAGACAGGGAATGA
GAGCACACGTCTGAACTCTTTCCCTACACGACGCTCTTCCGATCTNNNTGACTGAGACAGGGAATGA
GAGCACACGTCTGAACTCTTTCCCTACACGACGCTCTTCCGATCTNNNGTATCAGTGGACCTAGCATG
GAGCACACGTCTGAACTCTTTCCCTACACGACGCTCTTCCGATCTNNNGTATCAGTGGACCTAGCATG
AGCACACGTCTGAACTCTTTCCCTACACGACGCTCTTCCGATCTNNNATCCCTTGACCTATCAATGCTG
AGCACACGTCTGAACTCTTTCCCTACACGACGCTCTTCCGATCTNNNATCCCTTGACCTATCAATGCTG
AGCACACGTCTGAACTCTTTCCCTACACGACGCTCTTCCGATCTNNNAGCCAACAGTGCCATATAGTG
AGCACACGTCTGAACTCTTTCCCTACACGACGCTCTTCCGATCTNNNAGCCAACAGTGCCATATAGTG
CACACGTCTGAACTCTTTCCCTACACGACGCTCTTCCGATCTNNNTCTAACTACTCCCCACCTCTCTCCA
CACACGTCTGAACTCTTTCCCTACACGACGCTCTTCCGATCTNNNTCTAACTACTCCCCACCTCTCTCCA
AGCACACGTCTGAACTCTTTCCCTACACGACGCTCTTCCGATCTNNNCATAAACAGAGGTCTCAGGAG
AGCACACGTCTGAACTCTTTCCCTACACGACGCTCTTCCGATCTNNNCATAAACAGAGGTCTCAGGAG
AGCACACGTCTGAACTCTTTCCCTACACGACGCTCTTCCGATCTNNNACATGGGTGACAGAATGAGA
AGCACACGTCTGAACTCTTTCCCTACACGACGCTCTTCCGATCTNNNACATGGGTGACAGAATGAGA
AGCACACGTCTGAACTCTTTCCCTACACGACGCTCTTCCGATCTNNNCTGAGGCAGGAGAATCACTTG
AGCACACGTCTGAACTCTTTCCCTACACGACGCTCTTCCGATCTNNNCTGAGGCAGGAGAATCACTTG
GCACACGTCTGAACTCTTTCCCTACACGACGCTCTTCCGATCTNNNAGAGATTGCAGTGAGACGAGA
GCACACGTCTGAACTCTTTCCCTACACGACGCTCTTCCGATCTNNNAGAGATTGCAGTGAGACGAGA
AGCACACGTCTGAACTCTTTCCCTACACGACGCTCTTCCGATCTNNNACCTAAGCTTGATGATGGCAA
AGCACACGTCTGAACTCTTTCCCTACACGACGCTCTTCCGATCTNNNACCTAAGCTTGATGATGGCAA
GAGCACACGTCTGAACTCTTTCCCTACACGACGCTCTTCCGATCTNNNAATCAGGAAGAATCACCTCA
GAGCACACGTCTGAACTCTTTCCCTACACGACGCTCTTCCGATCTNNNAATCAGGAAGAATCACCTCA
AGCACACGTCTGAACTCTTTCCCTACACGACGCTCTTCCGATCTNNNTATTTGGAAAATATTGCATGT
AGCACACGTCTGAACTCTTTCCCTACACGACGCTCTTCCGATCTNNNTATTTGGAAAATATTGCATGT
AGCACACGTCTGAACTCTTTCCCTACACGACGCTCTTCCGATCTNNNTGCTTCAGTTTTTCCCCCAAAAG
AGCACACGTCTGAACTCTTTCCCTACACGACGCTCTTCCGATCTNNNTGCTTCAGTTTTTCCCCCAAAAG
GCACACGTCTGAACTCTTTCCCTACACGACGCTCTTCCGATCTNNNTTAGGATGGGAGGATGGTGGA
GCACACGTCTGAACTCTTTCCCTACACGACGCTCTTCCGATCTNNNTTAGGATGGGAGGATGGTGGA
AGAGCACACGTCTGAACTCTTTCCCTACACGACGCTCTTCCGATCTNNNCCTACTAAATCCTGCCCATC
AGAGCACACGTCTGAACTCTTTCCCTACACGACGCTCTTCCGATCTNNNCCTACTAAATCCTGCCCATC
GCACACGTCTGAACTCTTTCCCTACACGACGCTCTTCCGATCTNNNGGTGTGAGCTCTGAATCCAGTT
GCACACGTCTGAACTCTTTCCCTACACGACGCTCTTCCGATCTNNNGGTGTGAGCTCTGAATCCAGTT

GAGCACACGTCTGAACTCTTTCCCTACACGACGCTCTTCCGATCTNNNACCACACCTTCTTGATTACTG

GAGCACACGTCTGAACTCTTTCCCTACACGACGCTCTTCCGATCTNNNACCACACCTTCTTGATTACTG

GAGCACACGTCTGAACTCTTTCCCTACACGACGCTCTTCCGATCTNNNTCTTGCCTGATTGCTCTAGCT

GAGCACACGTCTGAACTCTTTCCCTACACGACGCTCTTCCGATCTNNNTCTTGCCTGATTGCTCTAGCT

GCACACGTCTGAACTCTTTCCCTACACGACGCTCTTCCGATCTNNNGAGACAGGAGAATTGCTTGAA

GCACACGTCTGAACTCTTTCCCTACACGACGCTCTTCCGATCTNNNGAGACAGGAGAATTGCTTGAA

GCACACGTCTGAACTCTTTCCCTACACGACGCTCTTCCGATCTNNNCAGACTTTCCCTACAGCTCTTGT

GCACACGTCTGAACTCTTTCCCTACACGACGCTCTTCCGATCTNNNCAGACTTTCCCTACAGCTCTTGT

AGCACACGTCTGAACTCTTTCCCTACACGACGCTCTTCCGATCTNNNTGTGCTTTGCTTCTATTGGCTT

AGCACACGTCTGAACTCTTTCCCTACACGACGCTCTTCCGATCTNNNTGTGCTTTGCTTCTATTGGCTT

GCACACGTCTGAACTCTTTCCCTACACGACGCTCTTCCGATCTNNNTTCTGTATCCAGTGTAACAAAAA

GCACACGTCTGAACTCTTTCCCTACACGACGCTCTTCCGATCTNNNTTCTGTATCCAGTGTAACAAAAA

AGCACACGTCTGAACTCTTTCCCTACACGACGCTCTTCCGATCTNNNCTGTAGTCCCAGCTACTTTGG

AGCACACGTCTGAACTCTTTCCCTACACGACGCTCTTCCGATCTNNNCTGTAGTCCCAGCTACTTTGG

AGCACACGTCTGAACTCTTTCCCTACACGACGCTCTTCCGATCTNNNGCCTGTAGTCCCAGCTACTTGC

AGCACACGTCTGAACTCTTTCCCTACACGACGCTCTTCCGATCTNNNGCCTGTAGTCCCAGCTACTTGC

AGCACACGTCTGAACTCTTTCCCTACACGACGCTCTTCCGATCTNNNTTATTGTTTTCATGCTGGGTGC

AGCACACGTCTGAACTCTTTCCCTACACGACGCTCTTCCGATCTNNNTTATTGTTTTCATGCTGGGTGC

GAGCACACGTCTGAACTCTTTCCCTACACGACGCTCTTCCGATCTNNNTTGATTCTACCCATCCATGAG

GAGCACACGTCTGAACTCTTTCCCTACACGACGCTCTTCCGATCTNNNTTGATTCTACCCATCCATGAG

CACGTCTGAACTCTTTCCCTACACGACGCTCTTCCGATCTNNNTCCGATTTTATTCATTTGAGCCTTCTC

CACGTCTGAACTCTTTCCCTACACGACGCTCTTCCGATCTNNNTCCGATTTTATTCATTTGAGCCTTCTC

GAGCACACGTCTGAACTCTTTCCCTACACGACGCTCTTCCGATCTNNNCACCAGTGACACTTTCTATTT

GAGCACACGTCTGAACTCTTTCCCTACACGACGCTCTTCCGATCTNNNCACCAGTGACACTTTCTATTT

GAGCACACGTCTGAACTCTTTCCCTACACGACGCTCTTCCGATCTNNNACAAAAAACAACACGACTGT

GAGCACACGTCTGAACTCTTTCCCTACACGACGCTCTTCCGATCTNNNACAAAAAACAACACGACTGT

AGCACACGTCTGAACTCTTTCCCTACACGACGCTCTTCCGATCTNNNATACAGAGGTCTTGGTTCCAGT

AGCACACGTCTGAACTCTTTCCCTACACGACGCTCTTCCGATCTNNNATACAGAGGTCTTGGTTCCAGT

AGCACACGTCTGAACTCTTTCCCTACACGACGCTCTTCCGATCTNNNAGTACGAGGTGCTTAGGAAG

AGCACACGTCTGAACTCTTTCCCTACACGACGCTCTTCCGATCTNNNAGTACGAGGTGCTTAGGAAG

GAGCACACGTCTGAACTCTTTCCCTACACGACGCTCTTCCGATCTNNNCCTCTGTGAGGGGGTTATTT

GAGCACACGTCTGAACTCTTTCCCTACACGACGCTCTTCCGATCTNNNCCTCTGTGAGGGGGTTATTT

GCACACGTCTGAACTCTTTCCCTACACGACGCTCTTCCGATCTNNNCTCTCCTTGTCTTCTTTCCTGGTC

GCACACGTCTGAACTCTTTCCCTACACGACGCTCTTCCGATCTNNNCTCTCCTTGTCTTCTTTCCTGGTC

GCACACGTCTGAACTCTTTCCCTACACGACGCTCTTCCGATCTNNNGTACACCAACATGCCTGGCTAA

GCACACGTCTGAACTCTTTCCCTACACGACGCTCTTCCGATCTNNNGTACACCAACATGCCTGGCTAA

AGCACACGTCTGAACTCTTTCCCTACACGACGCTCTTCCGATCTNNNTCACCCTACCCATAACTTCATG

AGCACACGTCTGAACTCTTTCCCTACACGACGCTCTTCCGATCTNNNTCACCCTACCCATAACTTCATG

AGCACACGTCTGAACTCTTTCCCTACACGACGCTCTTCCGATCTNNNTGTTGGTCAGACTAGTCTCGA

AGCACACGTCTGAACTCTTTCCCTACACGACGCTCTTCCGATCTNNNTGTTGGTCAGACTAGTCTCGA

AGCACACGTCTGAACTCTTTCCCTACACGACGCTCTTCCGATCTNNNTCAAACTCCTGATCTCAAGCGA

AGCACACGTCTGAACTCTTTCCCTACACGACGCTCTTCCGATCTNNNTCAAACTCCTGATCTCAAGCGA

CACACGTCTGAACTCTTTCCCTACACGACGCTCTTCCGATCTNNNTTACCCTTACTCCAACACTGCAACA

CACACGTCTGAACTCTTTCCCTACACGACGCTCTTCCGATCTNNNTTACCCTTACTCCAACACTGCAACA
GAGCACACGTCTGAACTCTTTCCCTACACGACGCTCTTCCGATCTNNNGAGCCCATGCTCTAAGTAGTC
GAGCACACGTCTGAACTCTTTCCCTACACGACGCTCTTCCGATCTNNNGAGCCCATGCTCTAAGTAGTC
GAGCACACGTCTGAACTCTTTCCCTACACGACGCTCTTCCGATCTNNNGCCTAGGTCGAGTAGGAATG
GAGCACACGTCTGAACTCTTTCCCTACACGACGCTCTTCCGATCTNNNGCCTAGGTCGAGTAGGAATG
AGCACACGTCTGAACTCTTTCCCTACACGACGCTCTTCCGATCTNNNATGGCTGCATAGTATTCCATGG
AGCACACGTCTGAACTCTTTCCCTACACGACGCTCTTCCGATCTNNNATGGCTGCATAGTATTCCATGG
AGCACACGTCTGAACTCTTTCCCTACACGACGCTCTTCCGATCTNNNCTGTAATCCCAGCTACTCAGGA
AGCACACGTCTGAACTCTTTCCCTACACGACGCTCTTCCGATCTNNNCTGTAATCCCAGCTACTCAGGA
AGCACACGTCTGAACTCTTTCCCTACACGACGCTCTTCCGATCTNNNCAACGGCTGGGAGGTCTAACA
AGCACACGTCTGAACTCTTTCCCTACACGACGCTCTTCCGATCTNNNCAACGGCTGGGAGGTCTAACA
GAGCACACGTCTGAACTCTTTCCCTACACGACGCTCTTCCGATCTNNNATCTCCCAGGTTCAAGCAATTC
GAGCACACGTCTGAACTCTTTCCCTACACGACGCTCTTCCGATCTNNNATCTCCCAGGTTCAAGCAATTC
GAGCACACGTCTGAACTCTTTCCCTACACGACGCTCTTCCGATCTNNNTGAAATCCAAAACACTTCTGGC
GAGCACACGTCTGAACTCTTTCCCTACACGACGCTCTTCCGATCTNNNTGAAATCCAAAACACTTCTGGC
GAGCACACGTCTGAACTCTTTCCCTACACGACGCTCTTCCGATCTNNNTTCTGTTGTGGCACCTTCCTAAT
GAGCACACGTCTGAACTCTTTCCCTACACGACGCTCTTCCGATCTNNNTTCTGTTGTGGCACCTTCCTAAT
GAGCACACGTCTGAACTCTTTCCCTACACGACGCTCTTCCGATCTNNNTCACACCTGTAATCCCAACACT
GAGCACACGTCTGAACTCTTTCCCTACACGACGCTCTTCCGATCTNNNTCACACCTGTAATCCCAACACT
GAGCACACGTCTGAACTCTTTCCCTACACGACGCTCTTCCGATCTNNNGCATCTGTAGTCCCAGCTACTC
GAGCACACGTCTGAACTCTTTCCCTACACGACGCTCTTCCGATCTNNNGCATCTGTAGTCCCAGCTACTC
GCACACGTCTGAACTCTTTCCCTACACGACGCTCTTCCGATCTNNNCATTGCCCCTGGTATTCCCACCATC
GCACACGTCTGAACTCTTTCCCTACACGACGCTCTTCCGATCTNNNCATTGCCCCTGGTATTCCCACCATC
GAGCACACGTCTGAACTCTTTCCCTACACGACGCTCTTCCGATCTNNNGTGTTGCGAACAGACAAAAAC
GAGCACACGTCTGAACTCTTTCCCTACACGACGCTCTTCCGATCTNNNGTGTTGCGAACAGACAAAAAC
GAGCACACGTCTGAACTCTTTCCCTACACGACGCTCTTCCGATCTNNNAGCTTGCAGTGAGTCGAGATC
GAGCACACGTCTGAACTCTTTCCCTACACGACGCTCTTCCGATCTNNNAGCTTGCAGTGAGTCGAGATC
AGCACACGTCTGAACTCTTTCCCTACACGACGCTCTTCCGATCTNNNACTCCTACTTTTTCCAGCACGAT
AGCACACGTCTGAACTCTTTCCCTACACGACGCTCTTCCGATCTNNNACTCCTACTTTTTCCAGCACGAT
GAGCACACGTCTGAACTCTTTCCCTACACGACGCTCTTCCGATCTNNNCCATACTTGGCAGTTTTCTTTT
GAGCACACGTCTGAACTCTTTCCCTACACGACGCTCTTCCGATCTNNNCCATACTTGGCAGTTTTCTTTT
ACACGTCTGAACTCTTTCCCTACACGACGCTCTTCCGATCTNNNGGACCCTGAACTCGGAGCAGCATCC
ACACGTCTGAACTCTTTCCCTACACGACGCTCTTCCGATCTNNNGGACCCTGAACTCGGAGCAGCATCC
GCACACGTCTGAACTCTTTCCCTACACGACGCTCTTCCGATCTNNNCTGAGGCAGGAGAATTGCTTGA
GCACACGTCTGAACTCTTTCCCTACACGACGCTCTTCCGATCTNNNCTGAGGCAGGAGAATTGCTTGA
AGCACACGTCTGAACTCTTTCCCTACACGACGCTCTTCCGATCTNNNAATCATCTGGTCCAGCTTCACTT
AGCACACGTCTGAACTCTTTCCCTACACGACGCTCTTCCGATCTNNNAATCATCTGGTCCAGCTTCACTT
GAGCACACGTCTGAACTCTTTCCCTACACGACGCTCTTCCGATCTNNNGGTAGATCCACCTTTCCAACCT
GAGCACACGTCTGAACTCTTTCCCTACACGACGCTCTTCCGATCTNNNGGTAGATCCACCTTTCCAACCT
GCACACGTCTGAACTCTTTCCCTACACGACGCTCTTCCGATCTNNNAGCCATAGTTGAAGGTTTAGGT
GCACACGTCTGAACTCTTTCCCTACACGACGCTCTTCCGATCTNNNAGCCATAGTTGAAGGTTTAGGT
AGAGCACACGTCTGAACTCTTTCCCTACACGACGCTCTTCCGATCTNNNTTCAAGAGCAGAAAACAAAC
AGAGCACACGTCTGAACTCTTTCCCTACACGACGCTCTTCCGATCTNNNTTCAAGAGCAGAAAACAAAC

AGCACACGTCTGAACTCTTTCCCTACACGACGCTCTTCCGATCTNNNCCTGTCCCATTGCTTTTTCTCTGC
AGCACACGTCTGAACTCTTTCCCTACACGACGCTCTTCCGATCTNNNCCTGTCCCATTGCTTTTTCTCTGC
AGCACACGTCTGAACTCTTTCCCTACACGACGCTCTTCCGATCTNNNACGTTTGTTGGCCATTTGTGTA
AGCACACGTCTGAACTCTTTCCCTACACGACGCTCTTCCGATCTNNNACGTTTGTTGGCCATTTGTGTA
GAGCACACGTCTGAACTCTTTCCCTACACGACGCTCTTCCGATCTNNNAGATACTTAGGGCTGTTGTGT
GAGCACACGTCTGAACTCTTTCCCTACACGACGCTCTTCCGATCTNNNAGATACTTAGGGCTGTTGTGT
GAGCACACGTCTGAACTCTTTCCCTACACGACGCTCTTCCGATCTNNNAGAGAATCACTTGAACCCAGC
GAGCACACGTCTGAACTCTTTCCCTACACGACGCTCTTCCGATCTNNNAGAGAATCACTTGAACCCAGC
GAGCACACGTCTGAACTCTTTCCCTACACGACGCTCTTCCGATCTNNNACAACAGGAATGACAGCTAAC
GAGCACACGTCTGAACTCTTTCCCTACACGACGCTCTTCCGATCTNNNACAACAGGAATGACAGCTAAC
AGAGCACACGTCTGAACTCTTTCCCTACACGACGCTCTTCCGATCTNNNGAATTGCTTGAACCTGGGAA
AGAGCACACGTCTGAACTCTTTCCCTACACGACGCTCTTCCGATCTNNNGAATTGCTTGAACCTGGGAA
GAGCACACGTCTGAACTCTTTCCCTACACGACGCTCTTCCGATCTNNNGTGTGGGGGAGTGTATTAGC
GAGCACACGTCTGAACTCTTTCCCTACACGACGCTCTTCCGATCTNNNGTGTGGGGGAGTGTATTAGC
AGCACACGTCTGAACTCTTTCCCTACACGACGCTCTTCCGATCTNNNGTAATCCCAGCTACTTAGGAGC
AGCACACGTCTGAACTCTTTCCCTACACGACGCTCTTCCGATCTNNNGTAATCCCAGCTACTTAGGAGC
GAGCACACGTCTGAACTCTTTCCCTACACGACGCTCTTCCGATCTNNNAGAAAGGTTAAGGTTGGCAGT
GAGCACACGTCTGAACTCTTTCCCTACACGACGCTCTTCCGATCTNNNAGAAAGGTTAAGGTTGGCAGT
AGCACACGTCTGAACTCTTTCCCTACACGACGCTCTTCCGATCTNNNCTGAGGCAGAGAATTGCTTGAA
AGCACACGTCTGAACTCTTTCCCTACACGACGCTCTTCCGATCTNNNCTGAGGCAGAGAATTGCTTGAA
AGAGCACACGTCTGAACTCTTTCCCTACACGACGCTCTTCCGATCTNNNACCTTTTCATCAACAAGCAGT
AGAGCACACGTCTGAACTCTTTCCCTACACGACGCTCTTCCGATCTNNNACCTTTTCATCAACAAGCAGT
AGAGCACACGTCTGAACTCTTTCCCTACACGACGCTCTTCCGATCTNNNACATCATCCAACTAGCAGCA
AGAGCACACGTCTGAACTCTTTCCCTACACGACGCTCTTCCGATCTNNNACATCATCCAACTAGCAGCA
GAGCACACGTCTGAACTCTTTCCCTACACGACGCTCTTCCGATCTNNNGCTCACACCTGTAACCCTAGTA
GAGCACACGTCTGAACTCTTTCCCTACACGACGCTCTTCCGATCTNNNGCTCACACCTGTAACCCTAGTA
GAGCACACGTCTGAACTCTTTCCCTACACGACGCTCTTCCGATCTNNNAGCAACTGGAGATCTGAGAAT
GAGCACACGTCTGAACTCTTTCCCTACACGACGCTCTTCCGATCTNNNAGCAACTGGAGATCTGAGAAT
AGCACACGTCTGAACTCTTTCCCTACACGACGCTCTTCCGATCTNNNGCCACTTCCTCTTCCTCATGTAAC
AGCACACGTCTGAACTCTTTCCCTACACGACGCTCTTCCGATCTNNNGCCACTTCCTCTTCCTCATGTAAC
AGCACACGTCTGAACTCTTTCCCTACACGACGCTCTTCCGATCTNNNTAGGGAGATGGGTTTGAGGGT
AGCACACGTCTGAACTCTTTCCCTACACGACGCTCTTCCGATCTNNNTAGGGAGATGGGTTTGAGGGT
AGCACACGTCTGAACTCTTTCCCTACACGACGCTCTTCCGATCTNNNAAAAGCTGACAGTGGCCTTTAT
AGCACACGTCTGAACTCTTTCCCTACACGACGCTCTTCCGATCTNNNAAAAGCTGACAGTGGCCTTTAT
AGCACACGTCTGAACTCTTTCCCTACACGACGCTCTTCCGATCTNNNCTCTTGCCCTGGCTTCTAAGAGCA
AGCACACGTCTGAACTCTTTCCCTACACGACGCTCTTCCGATCTNNNCTCTTGCCCTGGCTTCTAAGAGCA
AGCACACGTCTGAACTCTTTCCCTACACGACGCTCTTCCGATCTNNNAAACACTGCAACTCCTAGGTGAC
AGCACACGTCTGAACTCTTTCCCTACACGACGCTCTTCCGATCTNNNAAACACTGCAACTCCTAGGTGAC
GAGCACACGTCTGAACTCTTTCCCTACACGACGCTCTTCCGATCTNNNATTGCCAGACTGAGTGTAGAA
GAGCACACGTCTGAACTCTTTCCCTACACGACGCTCTTCCGATCTNNNATTGCCAGACTGAGTGTAGAA
GAGCACACGTCTGAACTCTTTCCCTACACGACGCTCTTCCGATCTNNNAGACCATCATCAGCACGAACTA
GAGCACACGTCTGAACTCTTTCCCTACACGACGCTCTTCCGATCTNNNAGACCATCATCAGCACGAACTA
AGCACACGTCTGAACTCTTTCCCTACACGACGCTCTTCCGATCTNNNCTTCCTTCCTTCCTTCCTTCCTCCA

AGCACACGTCTGAACTCTTTCCCTACACGACGCTCTTCCGATCTNNNCTTCCTTCCTTCCTTCCTTCCTCC
AGCACACGTCTGAACTCTTTCCCTACACGACGCTCTTCCGATCTNNNGTGCACATGTACCCTAGAACTG
AGCACACGTCTGAACTCTTTCCCTACACGACGCTCTTCCGATCTNNNGTGCACATGTACCCTAGAACTG
AGCACACGTCTGAACTCTTTCCCTACACGACGCTCTTCCGATCTNNNAGCTGTTGTACTGTATTGGTTT
AGCACACGTCTGAACTCTTTCCCTACACGACGCTCTTCCGATCTNNNAGCTGTTGTACTGTATTGGTTT
AGCACACGTCTGAACTCTTTCCCTACACGACGCTCTTCCGATCTNNNTGCAACCATATATGCTGGACTG
AGCACACGTCTGAACTCTTTCCCTACACGACGCTCTTCCGATCTNNNTGCAACCATATATGCTGGACTG
GAGCACACGTCTGAACTCTTTCCCTACACGACGCTCTTCCGATCTNNNATTCAGTTTCTCTCGGTGTTCC
GAGCACACGTCTGAACTCTTTCCCTACACGACGCTCTTCCGATCTNNNATTCAGTTTCTCTCGGTGTTCC
GAGCACACGTCTGAACTCTTTCCCTACACGACGCTCTTCCGATCTNNNAGGAAGCCTGGAAATGAAGA
GAGCACACGTCTGAACTCTTTCCCTACACGACGCTCTTCCGATCTNNNAGGAAGCCTGGAAATGAAGA
GAGCACACGTCTGAACTCTTTCCCTACACGACGCTCTTCCGATCTNNNCAGTGTAGTGTGCTCCTCTTC
GAGCACACGTCTGAACTCTTTCCCTACACGACGCTCTTCCGATCTNNNCAGTGTAGTGTGCTCCTCTTC
GAGCACACGTCTGAACTCTTTCCCTACACGACGCTCTTCCGATCTNNNTGCAGCTCTTAACTGCTGGAA
GAGCACACGTCTGAACTCTTTCCCTACACGACGCTCTTCCGATCTNNNTGCAGCTCTTAACTGCTGGAA
AGCACACGTCTGAACTCTTTCCCTACACGACGCTCTTCCGATCTNNNTTCATGCCATTCTCCTGCCTTAG
AGCACACGTCTGAACTCTTTCCCTACACGACGCTCTTCCGATCTNNNTTCATGCCATTCTCCTGCCTTAG
AGCACACGTCTGAACTCTTTCCCTACACGACGCTCTTCCGATCTNNNCCCAAGGTCTGTTGCTGAAACT
AGCACACGTCTGAACTCTTTCCCTACACGACGCTCTTCCGATCTNNNCCCAAGGTCTGTTGCTGAAACT
AGAGCACACGTCTGAACTCTTTCCCTACACGACGCTCTTCCGATCTNNNGAGAATCGCTTGAAACCAG
AGAGCACACGTCTGAACTCTTTCCCTACACGACGCTCTTCCGATCTNNNGAGAATCGCTTGAAACCAG
GAGCACACGTCTGAACTCTTTCCCTACACGACGCTCTTCCGATCTNNNCGCCTATAATCCCAGCTAGTC
GAGCACACGTCTGAACTCTTTCCCTACACGACGCTCTTCCGATCTNNNCGCCTATAATCCCAGCTAGTC
AGCACACGTCTGAACTCTTTCCCTACACGACGCTCTTCCGATCTNNNCAACTCTCCCATGGAAACCCAT
AGCACACGTCTGAACTCTTTCCCTACACGACGCTCTTCCGATCTNNNCAACTCTCCCATGGAAACCCAT
GAGCACACGTCTGAACTCTTTCCCTACACGACGCTCTTCCGATCTNNNCTGTTGTTGCAGTGTCCAGTG
GAGCACACGTCTGAACTCTTTCCCTACACGACGCTCTTCCGATCTNNNCTGTTGTTGCAGTGTCCAGTG
AGCACACGTCTGAACTCTTTCCCTACACGACGCTCTTCCGATCTNNNTATAATGTGGGGTTGCACTGT
AGCACACGTCTGAACTCTTTCCCTACACGACGCTCTTCCGATCTNNNTATAATGTGGGGTTGCACTGT
GCACACGTCTGAACTCTTTCCCTACACGACGCTCTTCCGATCTNNNGGACGGTTTCCCAAGGAATGTA
GCACACGTCTGAACTCTTTCCCTACACGACGCTCTTCCGATCTNNNGGACGGTTTCCCAAGGAATGTA
CACACGTCTGAACTCTTTCCCTACACGACGCTCTTCCGATCTNNNTGCCCATGACCCTCTTCTAGATACA
CACACGTCTGAACTCTTTCCCTACACGACGCTCTTCCGATCTNNNTGCCCATGACCCTCTTCTAGATACA
AGCACACGTCTGAACTCTTTCCCTACACGACGCTCTTCCGATCTNNNCCACATCCTTGATTGTTTGCCTT
AGCACACGTCTGAACTCTTTCCCTACACGACGCTCTTCCGATCTNNNCCACATCCTTGATTGTTTGCCTT
GCACACGTCTGAACTCTTTCCCTACACGACGCTCTTCCGATCTNNNAGAAATCCATACCCAGACACGTC
GCACACGTCTGAACTCTTTCCCTACACGACGCTCTTCCGATCTNNNAGAAATCCATACCCAGACACGTC
GAGCACACGTCTGAACTCTTTCCCTACACGACGCTCTTCCGATCTNNNGTCGACTGTTGGAGGAAGAT
GAGCACACGTCTGAACTCTTTCCCTACACGACGCTCTTCCGATCTNNNGTCGACTGTTGGAGGAAGAT
GAGCACACGTCTGAACTCTTTCCCTACACGACGCTCTTCCGATCTNNNGAAGTCTGACTGTCAGTGGA
GAGCACACGTCTGAACTCTTTCCCTACACGACGCTCTTCCGATCTNNNGAAGTCTGACTGTCAGTGGA
GAGCACACGTCTGAACTCTTTCCCTACACGACGCTCTTCCGATCTNNNCTTCTGAGGCCTAAAGGAACT
GAGCACACGTCTGAACTCTTTCCCTACACGACGCTCTTCCGATCTNNNCTTCTGAGGCCTAAAGGAACT

CACACGTCTGAACTCTTTCCCTACACGACGCTCTTCCGATCTNNNTGATCTCAGCTTACTGCAACCTCC
CACACGTCTGAACTCTTTCCCTACACGACGCTCTTCCGATCTNNNTGATCTCAGCTTACTGCAACCTCC
GCACACGTCTGAACTCTTTCCCTACACGACGCTCTTCCGATCTNNNGTTCAAGAGAGTCTCCTGCCTCC
GCACACGTCTGAACTCTTTCCCTACACGACGCTCTTCCGATCTNNNGTTCAAGAGAGTCTCCTGCCTCC
AGCACACGTCTGAACTCTTTCCCTACACGACGCTCTTCCGATCTNNNGCTCCAAACACAGACACATAC
AGCACACGTCTGAACTCTTTCCCTACACGACGCTCTTCCGATCTNNNGCTCCAAACACAGACACATAC
GCACACGTCTGAACTCTTTCCCTACACGACGCTCTTCCGATCTNNNCCTTATCTCCTAGGCATTGACCCC
GCACACGTCTGAACTCTTTCCCTACACGACGCTCTTCCGATCTNNNCCTTATCTCCTAGGCATTGACCCC
AGCACACGTCTGAACTCTTTCCCTACACGACGCTCTTCCGATCTNNNGGCCCCATTACCCCATTATTTCT
AGCACACGTCTGAACTCTTTCCCTACACGACGCTCTTCCGATCTNNNGGCCCCATTACCCCATTATTTCT
GAGCACACGTCTGAACTCTTTCCCTACACGACGCTCTTCCGATCTNNNAAAGATACCCATTCACCCTCT
GAGCACACGTCTGAACTCTTTCCCTACACGACGCTCTTCCGATCTNNNAAAGATACCCATTCACCCTCT
AGCACACGTCTGAACTCTTTCCCTACACGACGCTCTTCCGATCTNNNTTAGGGCTGGGTTGGAGATTA
AGCACACGTCTGAACTCTTTCCCTACACGACGCTCTTCCGATCTNNNTTAGGGCTGGGTTGGAGATTA
AGCACACGTCTGAACTCTTTCCCTACACGACGCTCTTCCGATCTNNNACCAGACTGGGCAACATAGTA
AGCACACGTCTGAACTCTTTCCCTACACGACGCTCTTCCGATCTNNNACCAGACTGGGCAACATAGTA
AGCACACGTCTGAACTCTTTCCCTACACGACGCTCTTCCGATCTNNNCAAAAAATTAGCTGGGCATGT
AGCACACGTCTGAACTCTTTCCCTACACGACGCTCTTCCGATCTNNNCAAAAAATTAGCTGGGCATGT
GCACACGTCTGAACTCTTTCCCTACACGACGCTCTTCCGATCTNNNGAGCAAAACCCTGTCAGAAAG
GCACACGTCTGAACTCTTTCCCTACACGACGCTCTTCCGATCTNNNGAGCAAAACCCTGTCAGAAAG
AGCACACGTCTGAACTCTTTCCCTACACGACGCTCTTCCGATCTNNNCACTACACACTGATACACACC
AGCACACGTCTGAACTCTTTCCCTACACGACGCTCTTCCGATCTNNNCACTACACACTGATACACACC
AGCACACGTCTGAACTCTTTCCCTACACGACGCTCTTCCGATCTNNNTGTTTCCCTGCAAGACGAATTT
AGCACACGTCTGAACTCTTTCCCTACACGACGCTCTTCCGATCTNNNTGTTTCCCTGCAAGACGAATTT
AGCACACGTCTGAACTCTTTCCCTACACGACGCTCTTCCGATCTNNNAAAAAGAAGTGACAGCTTGGT
AGCACACGTCTGAACTCTTTCCCTACACGACGCTCTTCCGATCTNNNAAAAAGAAGTGACAGCTTGGT
GCACACGTCTGAACTCTTTCCCTACACGACGCTCTTCCGATCTNNNCAGTCCTCCTAACCACTGTGAAG
GCACACGTCTGAACTCTTTCCCTACACGACGCTCTTCCGATCTNNNCAGTCCTCCTAACCACTGTGAAG
AGCACACGTCTGAACTCTTTCCCTACACGACGCTCTTCCGATCTNNNTGTTGCAATGTATTCTGATATA
AGCACACGTCTGAACTCTTTCCCTACACGACGCTCTTCCGATCTNNNTGTTGCAATGTATTCTGATATA
GAGCACACGTCTGAACTCTTTCCCTACACGACGCTCTTCCGATCTNNNAAAAAATTAGATGGGCGTG
GAGCACACGTCTGAACTCTTTCCCTACACGACGCTCTTCCGATCTNNNAAAAAATTAGATGGGCGTG
GCACACGTCTGAACTCTTTCCCTACACGACGCTCTTCCGATCTNNNACTAAAACTCAGCGATACTGCC
GCACACGTCTGAACTCTTTCCCTACACGACGCTCTTCCGATCTNNNACTAAAACTCAGCGATACTGCC
GAGCACACGTCTGAACTCTTTCCCTACACGACGCTCTTCCGATCTNNNGTCCCAGCTGAGGTAAGAG
GAGCACACGTCTGAACTCTTTCCCTACACGACGCTCTTCCGATCTNNNGTCCCAGCTGAGGTAAGAG
GCACACGTCTGAACTCTTTCCCTACACGACGCTCTTCCGATCTNNNGCTCCTGGACTGTCTAAAGGCT
GCACACGTCTGAACTCTTTCCCTACACGACGCTCTTCCGATCTNNNGCTCCTGGACTGTCTAAAGGCT
AGCACACGTCTGAACTCTTTCCCTACACGACGCTCTTCCGATCTNNNTCCATTACTTCTCCCTGGAACT
AGCACACGTCTGAACTCTTTCCCTACACGACGCTCTTCCGATCTNNNTCCATTACTTCTCCCTGGAACT
AGCACACGTCTGAACTCTTTCCCTACACGACGCTCTTCCGATCTNNNTGAAGATTGTGGTGAGCTGA
AGCACACGTCTGAACTCTTTCCCTACACGACGCTCTTCCGATCTNNNTGAAGATTGTGGTGAGCTGA
AGCACACGTCTGAACTCTTTCCCTACACGACGCTCTTCCGATCTNNNACTGAGTCAATTCAATCAGCA

AGCACACGTCTGAACTCTTTCCCTACACGACGCTCTTCCGATCTNNNACTGAGTCAATTCAATCAGCA
GAGCACACGTCTGAACTCTTTCCCTACACGACGCTCTTCCGATCTNNNGGCCACAAAGAAAGAGCAC
GAGCACACGTCTGAACTCTTTCCCTACACGACGCTCTTCCGATCTNNNGGCCACAAAGAAAGAGCAC
AGCACACGTCTGAACTCTTTCCCTACACGACGCTCTTCCGATCTNNNTCCTGAGTAGCTGGGATTACA
AGCACACGTCTGAACTCTTTCCCTACACGACGCTCTTCCGATCTNNNTCCTGAGTAGCTGGGATTACA
AGCACACGTCTGAACTCTTTCCCTACACGACGCTCTTCCGATCTNNNCAATACATGGCATGGCATAGGT
AGCACACGTCTGAACTCTTTCCCTACACGACGCTCTTCCGATCTNNNCAATACATGGCATGGCATAGGT
AGCACACGTCTGAACTCTTTCCCTACACGACGCTCTTCCGATCTNNNCCCATTACTGACAGTCAAGGA
AGCACACGTCTGAACTCTTTCCCTACACGACGCTCTTCCGATCTNNNCCCATTACTGACAGTCAAGGA
GCACACGTCTGAACTCTTTCCCTACACGACGCTCTTCCGATCTNNNCCCTCAGCCAAAGATGGAAAAG
GCACACGTCTGAACTCTTTCCCTACACGACGCTCTTCCGATCTNNNCCCTCAGCCAAAGATGGAAAAG
AGCACACGTCTGAACTCTTTCCCTACACGACGCTCTTCCGATCTNNNAGGAGTGAAGTGGCATAATGT
AGCACACGTCTGAACTCTTTCCCTACACGACGCTCTTCCGATCTNNNAGGAGTGAAGTGGCATAATGT
AGCACACGTCTGAACTCTTTCCCTACACGACGCTCTTCCGATCTNNNGAGGCTGAGGCAGGAGAATG
AGCACACGTCTGAACTCTTTCCCTACACGACGCTCTTCCGATCTNNNGAGGCTGAGGCAGGAGAATG
GCACACGTCTGAACTCTTTCCCTACACGACGCTCTTCCGATCTNNNCCCAGTGTATATTCTTGGCACT
GCACACGTCTGAACTCTTTCCCTACACGACGCTCTTCCGATCTNNNCCCAGTGTATATTCTTGGCACT
GCACACGTCTGAACTCTTTCCCTACACGACGCTCTTCCGATCTNNNGGACTCTGTCAATCAGGCAATC
GCACACGTCTGAACTCTTTCCCTACACGACGCTCTTCCGATCTNNNGGACTCTGTCAATCAGGCAATC
AGCACACGTCTGAACTCTTTCCCTACACGACGCTCTTCCGATCTNNNAGTGGTCTGCAACTAAACCTG
AGCACACGTCTGAACTCTTTCCCTACACGACGCTCTTCCGATCTNNNAGTGGTCTGCAACTAAACCTG
AGCACACGTCTGAACTCTTTCCCTACACGACGCTCTTCCGATCTNNNTTACCCCACTGATCACTCCTAGT
AGCACACGTCTGAACTCTTTCCCTACACGACGCTCTTCCGATCTNNNTTACCCCACTGATCACTCCTAGT
GCACACGTCTGAACTCTTTCCCTACACGACGCTCTTCCGATCTNNNTATACTCTCAGCCTTCCCTTCCAC
GCACACGTCTGAACTCTTTCCCTACACGACGCTCTTCCGATCTNNNTATACTCTCAGCCTTCCCTTCCAC
ACACGTCTGAACTCTTTCCCTACACGACGCTCTTCCGATCTNNNTCACAAGCCATAAAAGCTGAAGCCA
ACACGTCTGAACTCTTTCCCTACACGACGCTCTTCCGATCTNNNTCACAAGCCATAAAAGCTGAAGCCA
AGCACACGTCTGAACTCTTTCCCTACACGACGCTCTTCCGATCTNNNTCAAAATGCTGGAATTACAGG
AGCACACGTCTGAACTCTTTCCCTACACGACGCTCTTCCGATCTNNNTCAAAATGCTGGAATTACAGG
GCACACGTCTGAACTCTTTCCCTACACGACGCTCTTCCGATCTNNNTGCCTTTAGGGTCTTCTCTTCTG
GCACACGTCTGAACTCTTTCCCTACACGACGCTCTTCCGATCTNNNTGCCTTTAGGGTCTTCTCTTCTG
GAGCACACGTCTGAACTCTTTCCCTACACGACGCTCTTCCGATCTNNNCCCACTAACTCGTCATCTAGC
GAGCACACGTCTGAACTCTTTCCCTACACGACGCTCTTCCGATCTNNNCCCACTAACTCGTCATCTAGC
GAGCACACGTCTGAACTCTTTCCCTACACGACGCTCTTCCGATCTNNNGAAGTCAGGAGTTCGAGAC
GAGCACACGTCTGAACTCTTTCCCTACACGACGCTCTTCCGATCTNNNGAAGTCAGGAGTTCGAGAC
GCACACGTCTGAACTCTTTCCCTACACGACGCTCTTCCGATCTNNNACGCCCGGTTAGTTTTTTTTGTT
GCACACGTCTGAACTCTTTCCCTACACGACGCTCTTCCGATCTNNNACGCCCGGTTAGTTTTTTTTGTT
AAGAGCACACGTCTGAACTCTTTCCCTACACGACGCTCTTCCGATCTNNNCCACTCTTTCTTTCTCCAG
AAGAGCACACGTCTGAACTCTTTCCCTACACGACGCTCTTCCGATCTNNNCCACTCTTTCTTTCTCCAG
GCACACGTCTGAACTCTTTCCCTACACGACGCTCTTCCGATCTNNNCATTTGCCTTCCACATTTGGGTC
GCACACGTCTGAACTCTTTCCCTACACGACGCTCTTCCGATCTNNNCATTTGCCTTCCACATTTGGGTC
GCACACGTCTGAACTCTTTCCCTACACGACGCTCTTCCGATCTNNNTTATTTGTGTTTGCTTCCCCCCC
GCACACGTCTGAACTCTTTCCCTACACGACGCTCTTCCGATCTNNNTTATTTGTGTTTGCTTCCCCCCC

GAGCACACGTCTGAACTCTTTCCCTACACGACGCTCTTCCGATCTNNNTGTTCTTGAGCAAATCCAAAC
GAGCACACGTCTGAACTCTTTCCCTACACGACGCTCTTCCGATCTNNNTGTTCTTGAGCAAATCCAAAC
AGCACACGTCTGAACTCTTTCCCTACACGACGCTCTTCCGATCTNNNACATCCATTGACATGAGTTGGT
AGCACACGTCTGAACTCTTTCCCTACACGACGCTCTTCCGATCTNNNACATCCATTGACATGAGTTGGT
AGCACACGTCTGAACTCTTTCCCTACACGACGCTCTTCCGATCTNNNCTTCTAGGAGATGGTGGTCCCT
AGCACACGTCTGAACTCTTTCCCTACACGACGCTCTTCCGATCTNNNCTTCTAGGAGATGGTGGTCCCT
GAGCACACGTCTGAACTCTTTCCCTACACGACGCTCTTCCGATCTNNNTGGGAACTTACAACAGGGACA
GAGCACACGTCTGAACTCTTTCCCTACACGACGCTCTTCCGATCTNNNTGGGAACTTACAACAGGGACA
GAGCACACGTCTGAACTCTTTCCCTACACGACGCTCTTCCGATCTNNNTCAAATTGGGCGACAGAGTAA
GAGCACACGTCTGAACTCTTTCCCTACACGACGCTCTTCCGATCTNNNTCAAATTGGGCGACAGAGTAA
AGCACACGTCTGAACTCTTTCCCTACACGACGCTCTTCCGATCTNNNCTGTAATCCCAGCTACTCAGGA
AGCACACGTCTGAACTCTTTCCCTACACGACGCTCTTCCGATCTNNNCTGTAATCCCAGCTACTCAGGA
GAGCACACGTCTGAACTCTTTCCCTACACGACGCTCTTCCGATCTNNNCTGTAGTCCCAGCTACTCAGGA
GAGCACACGTCTGAACTCTTTCCCTACACGACGCTCTTCCGATCTNNNCTGTAGTCCCAGCTACTCAGGA
AGCACACGTCTGAACTCTTTCCCTACACGACGCTCTTCCGATCTNNNTGCAGACATTGGAAGGTCATCA
AGCACACGTCTGAACTCTTTCCCTACACGACGCTCTTCCGATCTNNNTGCAGACATTGGAAGGTCATCA
AGCACACGTCTGAACTCTTTCCCTACACGACGCTCTTCCGATCTNNNGGCATGTCTTCCTCAGATGGTTA
AGCACACGTCTGAACTCTTTCCCTACACGACGCTCTTCCGATCTNNNGGCATGTCTTCCTCAGATGGTTA
GCACACGTCTGAACTCTTTCCCTACACGACGCTCTTCCGATCTNNNAGAAACCCAAAATCTGTTTTCCAC
GCACACGTCTGAACTCTTTCCCTACACGACGCTCTTCCGATCTNNNAGAAACCCAAAATCTGTTTTCCAC
AGCACACGTCTGAACTCTTTCCCTACACGACGCTCTTCCGATCTNNNGCAGCAGCGTAACTTTTCTTTTTC
AGCACACGTCTGAACTCTTTCCCTACACGACGCTCTTCCGATCTNNNGCAGCAGCGTAACTTTTCTTTTTC
GAGCACACGTCTGAACTCTTTCCCTACACGACGCTCTTCCGATCTNNNGACTGCAGTGATGTTTCTGTG
GAGCACACGTCTGAACTCTTTCCCTACACGACGCTCTTCCGATCTNNNGACTGCAGTGATGTTTCTGTG
AGCACACGTCTGAACTCTTTCCCTACACGACGCTCTTCCGATCTNNNCACATCTGTAGTCCCAGCTACTC
AGCACACGTCTGAACTCTTTCCCTACACGACGCTCTTCCGATCTNNNCACATCTGTAGTCCCAGCTACTC
GAGCACACGTCTGAACTCTTTCCCTACACGACGCTCTTCCGATCTNNNGTGGTACTTGTGGGGATTCTC
GAGCACACGTCTGAACTCTTTCCCTACACGACGCTCTTCCGATCTNNNGTGGTACTTGTGGGGATTCTC
GAGCACACGTCTGAACTCTTTCCCTACACGACGCTCTTCCGATCTNNNCAAAAATTAGCTGGGGATGGT
GAGCACACGTCTGAACTCTTTCCCTACACGACGCTCTTCCGATCTNNNCAAAAATTAGCTGGGGATGGT
GAGCACACGTCTGAACTCTTTCCCTACACGACGCTCTTCCGATCTNNNCCAATGTGGATGGATTGCTTG
GAGCACACGTCTGAACTCTTTCCCTACACGACGCTCTTCCGATCTNNNCCAATGTGGATGGATTGCTTG
AGCACACGTCTGAACTCTTTCCCTACACGACGCTCTTCCGATCTNNNGGAAGGGAATGGGGAGTAGA
AGCACACGTCTGAACTCTTTCCCTACACGACGCTCTTCCGATCTNNNGGAAGGGAATGGGGAGTAGA
GAGCACACGTCTGAACTCTTTCCCTACACGACGCTCTTCCGATCTNNNTGCCACCAGTCTCTCTTTTTTTC
GAGCACACGTCTGAACTCTTTCCCTACACGACGCTCTTCCGATCTNNNTGCCACCAGTCTCTCTTTTTTTC
ACACGTCTGAACTCTTTCCCTACACGACGCTCTTCCGATCTNNNATCAATGTACAAAAGCCAGCAGCCAT
ACACGTCTGAACTCTTTCCCTACACGACGCTCTTCCGATCTNNNATCAATGTACAAAAGCCAGCAGCCAT
AGCACACGTCTGAACTCTTTCCCTACACGACGCTCTTCCGATCTNNNTTTCCCATTCCTGTTTTGCCTTGC
AGCACACGTCTGAACTCTTTCCCTACACGACGCTCTTCCGATCTNNNTTTCCCATTCCTGTTTTGCCTTGC
AGCACACGTCTGAACTCTTTCCCTACACGACGCTCTTCCGATCTNNNGCAACAAAGCGAGACTCTGTCT
AGCACACGTCTGAACTCTTTCCCTACACGACGCTCTTCCGATCTNNNGCAACAAAGCGAGACTCTGTCT
AGCACACGTCTGAACTCTTTCCCTACACGACGCTCTTCCGATCTNNNGAAACATCACACATCTAAGATT

AGCACACGTCTGAACTCTTTCCCTACACGACGCTCTTCCGATCTNNNGAAACATCACACATCTAAGATT
AGCACACGTCTGAACTCTTTCCCTACACGACGCTCTTCCGATCTNNNTTTGGTGTATGTGTGTGTGTGT
AGCACACGTCTGAACTCTTTCCCTACACGACGCTCTTCCGATCTNNNTTTGGTGTATGTGTGTGTGTGT
AGCACACGTCTGAACTCTTTCCCTACACGACGCTCTTCCGATCTNNNCCCCTTCCTAACAGCATGAGATT
AGCACACGTCTGAACTCTTTCCCTACACGACGCTCTTCCGATCTNNNCCCCTTCCTAACAGCATGAGATT
AGCACACGTCTGAACTCTTTCCCTACACGACGCTCTTCCGATCTNNNAGACTGGACTGGAAACTGAAG
AGCACACGTCTGAACTCTTTCCCTACACGACGCTCTTCCGATCTNNNAGACTGGACTGGAAACTGAAG
GCACACGTCTGAACTCTTTCCCTACACGACGCTCTTCCGATCTNNNTATACACACACACACACACACAC
GCACACGTCTGAACTCTTTCCCTACACGACGCTCTTCCGATCTNNNTATACACACACACACACACACAC
AGCACACGTCTGAACTCTTTCCCTACACGACGCTCTTCCGATCTNNNATCTTGGCTCACTACAACCTCTG
AGCACACGTCTGAACTCTTTCCCTACACGACGCTCTTCCGATCTNNNATCTTGGCTCACTACAACCTCTG
GAGCACACGTCTGAACTCTTTCCCTACACGACGCTCTTCCGATCTNNNGTGGACTCAGGCAGATAGAC
GAGCACACGTCTGAACTCTTTCCCTACACGACGCTCTTCCGATCTNNNGTGGACTCAGGCAGATAGAC
GAGCACACGTCTGAACTCTTTCCCTACACGACGCTCTTCCGATCTNNNATTGCTTGAGCCCAGGAATT
GAGCACACGTCTGAACTCTTTCCCTACACGACGCTCTTCCGATCTNNNATTGCTTGAGCCCAGGAATT
GAGCACACGTCTGAACTCTTTCCCTACACGACGCTCTTCCGATCTNNNGAGATGTCAGGAGACTTTTA
GAGCACACGTCTGAACTCTTTCCCTACACGACGCTCTTCCGATCTNNNGAGATGTCAGGAGACTTTTA
GCACACGTCTGAACTCTTTCCCTACACGACGCTCTTCCGATCTNNNGACCAATTATGCCAGCTGTGTA
GCACACGTCTGAACTCTTTCCCTACACGACGCTCTTCCGATCTNNNGACCAATTATGCCAGCTGTGTA
GAGCACACGTCTGAACTCTTTCCCTACACGACGCTCTTCCGATCTNNNGATCTACCAAGCAAATGGACA
GAGCACACGTCTGAACTCTTTCCCTACACGACGCTCTTCCGATCTNNNGATCTACCAAGCAAATGGACA
AGCACACGTCTGAACTCTTTCCCTACACGACGCTCTTCCGATCTNNNAGCATGCTAAGCTGTTACAGAA
AGCACACGTCTGAACTCTTTCCCTACACGACGCTCTTCCGATCTNNNAGCATGCTAAGCTGTTACAGAA
GAGCACACGTCTGAACTCTTTCCCTACACGACGCTCTTCCGATCTNNNCAAGATCGTGTCATTGCACTC
GAGCACACGTCTGAACTCTTTCCCTACACGACGCTCTTCCGATCTNNNCAAGATCGTGTCATTGCACTC
AGCACACGTCTGAACTCTTTCCCTACACGACGCTCTTCCGATCTNNNGTGCCTGTAATCCCAGCTACTT
AGCACACGTCTGAACTCTTTCCCTACACGACGCTCTTCCGATCTNNNGTGCCTGTAATCCCAGCTACTT
AGCACACGTCTGAACTCTTTCCCTACACGACGCTCTTCCGATCTNNNTGTCGATTTTCTGACCTGGTGA
AGCACACGTCTGAACTCTTTCCCTACACGACGCTCTTCCGATCTNNNTGTCGATTTTCTGACCTGGTGA
GAGCACACGTCTGAACTCTTTCCCTACACGACGCTCTTCCGATCTNNNGTATGACTTTTCCCCACTTCCCT
GAGCACACGTCTGAACTCTTTCCCTACACGACGCTCTTCCGATCTNNNGTATGACTTTTCCCCACTTCCCT
AGCACACGTCTGAACTCTTTCCCTACACGACGCTCTTCCGATCTNNNGTTGTAGTGAGCCAAGATTGA
AGCACACGTCTGAACTCTTTCCCTACACGACGCTCTTCCGATCTNNNGTTGTAGTGAGCCAAGATTGA
AGCACACGTCTGAACTCTTTCCCTACACGACGCTCTTCCGATCTNNNTAACTAGGACCCTGAGAGGTGT
AGCACACGTCTGAACTCTTTCCCTACACGACGCTCTTCCGATCTNNNTAACTAGGACCCTGAGAGGTGT
AGCACACGTCTGAACTCTTTCCCTACACGACGCTCTTCCGATCTNNNTTCCACACTCCTCAGATTGTTCCC
AGCACACGTCTGAACTCTTTCCCTACACGACGCTCTTCCGATCTNNNTTCCACACTCCTCAGATTGTTCCC
GAGCACACGTCTGAACTCTTTCCCTACACGACGCTCTTCCGATCTNNNGCCTGGACGACATAGTGAAAC
GAGCACACGTCTGAACTCTTTCCCTACACGACGCTCTTCCGATCTNNNGCCTGGACGACATAGTGAAAC
GAGCACACGTCTGAACTCTTTCCCTACACGACGCTCTTCCGATCTNNNTTCAAGTACCGCCTTCTCTACAC
GAGCACACGTCTGAACTCTTTCCCTACACGACGCTCTTCCGATCTNNNTTCAAGTACCGCCTTCTCTACAC
AGCACACGTCTGAACTCTTTCCCTACACGACGCTCTTCCGATCTNNNTTGACAGTTTGGGCTTCAACGT
AGCACACGTCTGAACTCTTTCCCTACACGACGCTCTTCCGATCTNNNTTGACAGTTTGGGCTTCAACGT

GAGCACACGTCTGAACTCTTTCCCTACACGACGCTCTTCCGATCTNNNAAGCAGAGATTGAGGACACT
GAGCACACGTCTGAACTCTTTCCCTACACGACGCTCTTCCGATCTNNNAAGCAGAGATTGAGGACACT
AGCACACGTCTGAACTCTTTCCCTACACGACGCTCTTCCGATCTNNNCAACCTTAGGCAGTAAAAGGCA
AGCACACGTCTGAACTCTTTCCCTACACGACGCTCTTCCGATCTNNNCAACCTTAGGCAGTAAAAGGCA
CACACGTCTGAACTCTTTCCCTACACGACGCTCTTCCGATCTNNNCACAACGACCACATCCTCATAGACA
CACACGTCTGAACTCTTTCCCTACACGACGCTCTTCCGATCTNNNCACAACGACCACATCCTCATAGACA
AGCACACGTCTGAACTCTTTCCCTACACGACGCTCTTCCGATCTNNNAGAGGGAACAGTAATTGAGG
AGCACACGTCTGAACTCTTTCCCTACACGACGCTCTTCCGATCTNNNAGAGGGAACAGTAATTGAGG
GCACACGTCTGAACTCTTTCCCTACACGACGCTCTTCCGATCTNNNGCACATGCTGTTGGAAAAATGA
GCACACGTCTGAACTCTTTCCCTACACGACGCTCTTCCGATCTNNNGCACATGCTGTTGGAAAAATGA
AGCACACGTCTGAACTCTTTCCCTACACGACGCTCTTCCGATCTNNNAATGGAAAACCATTGCACGTG
AGCACACGTCTGAACTCTTTCCCTACACGACGCTCTTCCGATCTNNNAATGGAAAACCATTGCACGTG
AAGAGCACACGTCTGAACTCTTTCCCTACACGACGCTCTTCCGATCTNNNACAGTAACCTTTTTGCTGT
AAGAGCACACGTCTGAACTCTTTCCCTACACGACGCTCTTCCGATCTNNNACAGTAACCTTTTTGCTGT
AGCACACGTCTGAACTCTTTCCCTACACGACGCTCTTCCGATCTNNNCTCATATTCCCTAGGAACCACCC
AGCACACGTCTGAACTCTTTCCCTACACGACGCTCTTCCGATCTNNNCTCATATTCCCTAGGAACCACCC
GCACACGTCTGAACTCTTTCCCTACACGACGCTCTTCCGATCTNNNTGGATGTACTACGGATGTCTCCC
GCACACGTCTGAACTCTTTCCCTACACGACGCTCTTCCGATCTNNNTGGATGTACTACGGATGTCTCCC
GAGCACACGTCTGAACTCTTTCCCTACACGACGCTCTTCCGATCTNNNCACCTCTTCAATTTTACTTTTT
GAGCACACGTCTGAACTCTTTCCCTACACGACGCTCTTCCGATCTNNNCACCTCTTCAATTTTACTTTTT
GAGCACACGTCTGAACTCTTTCCCTACACGACGCTCTTCCGATCTNNNCTGTTTTTGGAGTCTTTTCTTA
GAGCACACGTCTGAACTCTTTCCCTACACGACGCTCTTCCGATCTNNNCTGTTTTTGGAGTCTTTTCTTA
AGCACACGTCTGAACTCTTTCCCTACACGACGCTCTTCCGATCTNNNGTCACATTCTGAGGTTCCTGGA
AGCACACGTCTGAACTCTTTCCCTACACGACGCTCTTCCGATCTNNNGTCACATTCTGAGGTTCCTGGA
GAGCACACGTCTGAACTCTTTCCCTACACGACGCTCTTCCGATCTNNNCCAAAGTGCTGGGATTACAG
GAGCACACGTCTGAACTCTTTCCCTACACGACGCTCTTCCGATCTNNNCCAAAGTGCTGGGATTACAG
AGCACACGTCTGAACTCTTTCCCTACACGACGCTCTTCCGATCTNNNACAGATTTGGAGGATGGGAAT
AGCACACGTCTGAACTCTTTCCCTACACGACGCTCTTCCGATCTNNNACAGATTTGGAGGATGGGAAT
CACACGTCTGAACTCTTTCCCTACACGACGCTCTTCCGATCTNNNTCTGCGTGCCAAAATAGTCTTTGC
CACACGTCTGAACTCTTTCCCTACACGACGCTCTTCCGATCTNNNTCTGCGTGCCAAAATAGTCTTTGC
AGCACACGTCTGAACTCTTTCCCTACACGACGCTCTTCCGATCTNNNTTGCTGCAAATTCTAGAAGCCT
AGCACACGTCTGAACTCTTTCCCTACACGACGCTCTTCCGATCTNNNTTGCTGCAAATTCTAGAAGCCT
GCACACGTCTGAACTCTTTCCCTACACGACGCTCTTCCGATCTNNNAGAACTAATTGGGAGAGAGCA
GCACACGTCTGAACTCTTTCCCTACACGACGCTCTTCCGATCTNNNAGAACTAATTGGGAGAGAGCA
GAGCACACGTCTGAACTCTTTCCCTACACGACGCTCTTCCGATCTNNNTTAGTCTTTAGGCAGATGGA
GAGCACACGTCTGAACTCTTTCCCTACACGACGCTCTTCCGATCTNNNTTAGTCTTTAGGCAGATGGA
GAGCACACGTCTGAACTCTTTCCCTACACGACGCTCTTCCGATCTNNNCCCCACCCTCCACTTTAACATT
GAGCACACGTCTGAACTCTTTCCCTACACGACGCTCTTCCGATCTNNNCCCCACCCTCCACTTTAACATT
GCACACGTCTGAACTCTTTCCCTACACGACGCTCTTCCGATCTNNNAGGTTGAACACAGTGTTTTTCTC
GCACACGTCTGAACTCTTTCCCTACACGACGCTCTTCCGATCTNNNAGGTTGAACACAGTGTTTTTCTC
GAGCACACGTCTGAACTCTTTCCCTACACGACGCTCTTCCGATCTNNNCTGTCACTTCAGTGGACAGAT
GAGCACACGTCTGAACTCTTTCCCTACACGACGCTCTTCCGATCTNNNCTGTCACTTCAGTGGACAGAT
GAGCACACGTCTGAACTCTTTCCCTACACGACGCTCTTCCGATCTNNNCTCTGTCTCCCGAATTCAAGT

GAGCACACGTCTGAACTCTTTCCCTACACGACGCTCTTCCGATCTNNNCTCTGTCTCCCGAATTCAAGT
AGCACACGTCTGAACTCTTTCCCTACACGACGCTCTTCCGATCTNNNGCTCACAGCTTTTGACACAGTC
AGCACACGTCTGAACTCTTTCCCTACACGACGCTCTTCCGATCTNNNGCTCACAGCTTTTGACACAGTC
AGCACACGTCTGAACTCTTTCCCTACACGACGCTCTTCCGATCTNNNTCTGATGTTCTTAGGATTCCTT
AGCACACGTCTGAACTCTTTCCCTACACGACGCTCTTCCGATCTNNNTCTGATGTTCTTAGGATTCCTT
AGCACACGTCTGAACTCTTTCCCTACACGACGCTCTTCCGATCTNNNAGGTTCAAGCCATTCTCATGTC
AGCACACGTCTGAACTCTTTCCCTACACGACGCTCTTCCGATCTNNNAGGTTCAAGCCATTCTCATGTC
AGCACACGTCTGAACTCTTTCCCTACACGACGCTCTTCCGATCTNNNTCCAGAAGAGGCAAATTCATG
AGCACACGTCTGAACTCTTTCCCTACACGACGCTCTTCCGATCTNNNTCCAGAAGAGGCAAATTCATG
GAGCACACGTCTGAACTCTTTCCCTACACGACGCTCTTCCGATCTNNNCTGAGGCAGGAGAATCACTT
GAGCACACGTCTGAACTCTTTCCCTACACGACGCTCTTCCGATCTNNNCTGAGGCAGGAGAATCACTT
GAGCACACGTCTGAACTCTTTCCCTACACGACGCTCTTCCGATCTNNNTGTGCTATGACCTTCTTTACCT
GAGCACACGTCTGAACTCTTTCCCTACACGACGCTCTTCCGATCTNNNTGTGCTATGACCTTCTTTACCT
GAGCACACGTCTGAACTCTTTCCCTACACGACGCTCTTCCGATCTNNNTTAGTCTCTGTGATGCCCCTTT
GAGCACACGTCTGAACTCTTTCCCTACACGACGCTCTTCCGATCTNNNTTAGTCTCTGTGATGCCCCTTT
AGCACACGTCTGAACTCTTTCCCTACACGACGCTCTTCCGATCTNNNCCCTTTGGGCTCAGAAAAAAC
AGCACACGTCTGAACTCTTTCCCTACACGACGCTCTTCCGATCTNNNCCCTTTGGGCTCAGAAAAAAC
AGAGCACACGTCTGAACTCTTTCCCTACACGACGCTCTTCCGATCTNNNGCATGCACCTGTAGTCTCAA
AGAGCACACGTCTGAACTCTTTCCCTACACGACGCTCTTCCGATCTNNNGCATGCACCTGTAGTCTCAA
GAGCACACGTCTGAACTCTTTCCCTACACGACGCTCTTCCGATCTNNNCGTGCCCAGCCTCATAGATATT
GAGCACACGTCTGAACTCTTTCCCTACACGACGCTCTTCCGATCTNNNCGTGCCCAGCCTCATAGATATT
AGCACACGTCTGAACTCTTTCCCTACACGACGCTCTTCCGATCTNNNCAGAGCCTGCTACTTCTACATGT
AGCACACGTCTGAACTCTTTCCCTACACGACGCTCTTCCGATCTNNNCAGAGCCTGCTACTTCTACATGT
AGCACACGTCTGAACTCTTTCCCTACACGACGCTCTTCCGATCTNNNCCAAGCAATCCTCCTACCTCAGC
AGCACACGTCTGAACTCTTTCCCTACACGACGCTCTTCCGATCTNNNCCAAGCAATCCTCCTACCTCAGC
GAGCACACGTCTGAACTCTTTCCCTACACGACGCTCTTCCGATCTNNNGACAGGTTAGAATTCAGACCC
GAGCACACGTCTGAACTCTTTCCCTACACGACGCTCTTCCGATCTNNNGACAGGTTAGAATTCAGACCC
GAGCACACGTCTGAACTCTTTCCCTACACGACGCTCTTCCGATCTNNNCATACAAACCCAAAACCTCCAG
GAGCACACGTCTGAACTCTTTCCCTACACGACGCTCTTCCGATCTNNNCATACAAACCCAAAACCTCCAG
GAGCACACGTCTGAACTCTTTCCCTACACGACGCTCTTCCGATCTNNNTGCCACTACCCTTCTAGTTCCT
GAGCACACGTCTGAACTCTTTCCCTACACGACGCTCTTCCGATCTNNNTGCCACTACCCTTCTAGTTCCT
AGAGCACACGTCTGAACTCTTTCCCTACACGACGCTCTTCCGATCTNNNGGCAACAAGAGCGAAAATCC
AGAGCACACGTCTGAACTCTTTCCCTACACGACGCTCTTCCGATCTNNNGGCAACAAGAGCGAAAATCC
GAGCACACGTCTGAACTCTTTCCCTACACGACGCTCTTCCGATCTNNNCTGGTAGCTGTGCCCAATAAA
GAGCACACGTCTGAACTCTTTCCCTACACGACGCTCTTCCGATCTNNNCTGGTAGCTGTGCCCAATAAA
AGCACACGTCTGAACTCTTTCCCTACACGACGCTCTTCCGATCTNNNCAGTGAGCTATGATCATGCCA
AGCACACGTCTGAACTCTTTCCCTACACGACGCTCTTCCGATCTNNNCAGTGAGCTATGATCATGCCA
GAGCACACGTCTGAACTCTTTCCCTACACGACGCTCTTCCGATCTNNNAGAGTTTGCAGTGAGCCAAG
GAGCACACGTCTGAACTCTTTCCCTACACGACGCTCTTCCGATCTNNNAGAGTTTGCAGTGAGCCAAG
GAGCACACGTCTGAACTCTTTCCCTACACGACGCTCTTCCGATCTNNNCGGTTCTTCCTGATGCAGACT
GAGCACACGTCTGAACTCTTTCCCTACACGACGCTCTTCCGATCTNNNCGGTTCTTCCTGATGCAGACT
AGAGCACACGTCTGAACTCTTTCCCTACACGACGCTCTTCCGATCTNNNAGCATGATTACTTCTATGGCT
AGAGCACACGTCTGAACTCTTTCCCTACACGACGCTCTTCCGATCTNNNAGCATGATTACTTCTATGGCT

AGCACACGTCTGAACTCTTTCCCTACACGACGCTCTTCCGATCTNNNCACCAGGAAGAGAAAATACCC
AGCACACGTCTGAACTCTTTCCCTACACGACGCTCTTCCGATCTNNNCACCAGGAAGAGAAAATACCC
AGCACACGTCTGAACTCTTTCCCTACACGACGCTCTTCCGATCTNNNAGACATACTTAGCAGCTGATG
AGCACACGTCTGAACTCTTTCCCTACACGACGCTCTTCCGATCTNNNAGACATACTTAGCAGCTGATG
AGCACACGTCTGAACTCTTTCCCTACACGACGCTCTTCCGATCTNNNAAGAATCACTTGAACCTGGGA
AGCACACGTCTGAACTCTTTCCCTACACGACGCTCTTCCGATCTNNNAAGAATCACTTGAACCTGGGA
GCACACGTCTGAACTCTTTCCCTACACGACGCTCTTCCGATCTNNNGTCACCTGTAATCCCAGCTACTC
GCACACGTCTGAACTCTTTCCCTACACGACGCTCTTCCGATCTNNNGTCACCTGTAATCCCAGCTACTC
GCACACGTCTGAACTCTTTCCCTACACGACGCTCTTCCGATCTNNNAGACAGGAGAATTGCTTGAACC
GCACACGTCTGAACTCTTTCCCTACACGACGCTCTTCCGATCTNNNAGACAGGAGAATTGCTTGAACC
GAGCACACGTCTGAACTCTTTCCCTACACGACGCTCTTCCGATCTNNNGGTAGATGACATCGGTGGA
GAGCACACGTCTGAACTCTTTCCCTACACGACGCTCTTCCGATCTNNNGGTAGATGACATCGGTGGA
AGCACACGTCTGAACTCTTTCCCTACACGACGCTCTTCCGATCTNNNGAAGGGCAGTACCTACAGAC
AGCACACGTCTGAACTCTTTCCCTACACGACGCTCTTCCGATCTNNNGAAGGGCAGTACCTACAGAC
GAGCACACGTCTGAACTCTTTCCCTACACGACGCTCTTCCGATCTNNNGCACAAATGAAACATCTCTCC
GAGCACACGTCTGAACTCTTTCCCTACACGACGCTCTTCCGATCTNNNGCACAAATGAAACATCTCTCC
GCACACGTCTGAACTCTTTCCCTACACGACGCTCTTCCGATCTNNNTTGTGCACCTTTCATTCTTGGTC
GCACACGTCTGAACTCTTTCCCTACACGACGCTCTTCCGATCTNNNTTGTGCACCTTTCATTCTTGGTC
AGCACACGTCTGAACTCTTTCCCTACACGACGCTCTTCCGATCTNNNAGAATCACTTGAACCTAGGAG
AGCACACGTCTGAACTCTTTCCCTACACGACGCTCTTCCGATCTNNNAGAATCACTTGAACCTAGGAG
GAGCACACGTCTGAACTCTTTCCCTACACGACGCTCTTCCGATCTNNNAGGAGAATCGCTTGAACCCG
GAGCACACGTCTGAACTCTTTCCCTACACGACGCTCTTCCGATCTNNNAGGAGAATCGCTTGAACCCG
AGCACACGTCTGAACTCTTTCCCTACACGACGCTCTTCCGATCTNNNTGTGCAGACCAAGGAATTGA
AGCACACGTCTGAACTCTTTCCCTACACGACGCTCTTCCGATCTNNNTGTGCAGACCAAGGAATTGA
AGCACACGTCTGAACTCTTTCCCTACACGACGCTCTTCCGATCTNNNTGTAGTCCCAGCTACTCTAGA
AGCACACGTCTGAACTCTTTCCCTACACGACGCTCTTCCGATCTNNNTGTAGTCCCAGCTACTCTAGA
GAGCACACGTCTGAACTCTTTCCCTACACGACGCTCTTCCGATCTNNNAGACAGGAGAATGGCGTGA
GAGCACACGTCTGAACTCTTTCCCTACACGACGCTCTTCCGATCTNNNAGACAGGAGAATGGCGTGA
AGAGCACACGTCTGAACTCTTTCCCTACACGACGCTCTTCCGATCTNNNCAGCTAATGACCACAACCAG
AGAGCACACGTCTGAACTCTTTCCCTACACGACGCTCTTCCGATCTNNNCAGCTAATGACCACAACCAG
AGCACACGTCTGAACTCTTTCCCTACACGACGCTCTTCCGATCTNNNGCTGGAATTACAGGCATAAGC
AGCACACGTCTGAACTCTTTCCCTACACGACGCTCTTCCGATCTNNNGCTGGAATTACAGGCATAAGC
GCACACGTCTGAACTCTTTCCCTACACGACGCTCTTCCGATCTNNNGACCTCTCAACATGTGCCATTTC
GCACACGTCTGAACTCTTTCCCTACACGACGCTCTTCCGATCTNNNGACCTCTCAACATGTGCCATTTC
AGCACACGTCTGAACTCTTTCCCTACACGACGCTCTTCCGATCTNNNTGCATACCTGTAATCCCAGCTA
AGCACACGTCTGAACTCTTTCCCTACACGACGCTCTTCCGATCTNNNTGCATACCTGTAATCCCAGCTA
GCACACGTCTGAACTCTTTCCCTACACGACGCTCTTCCGATCTNNNCCTAGGACATGCCTTCCAAAGA
GCACACGTCTGAACTCTTTCCCTACACGACGCTCTTCCGATCTNNNCCTAGGACATGCCTTCCAAAGA
GCACACGTCTGAACTCTTTCCCTACACGACGCTCTTCCGATCTNNNATGATCTTGCCCCACCCTTATTT
GCACACGTCTGAACTCTTTCCCTACACGACGCTCTTCCGATCTNNNATGATCTTGCCCCACCCTTATTT
AGCACACGTCTGAACTCTTTCCCTACACGACGCTCTTCCGATCTNNNATCCAAATACAAGGAGGTGA
AGCACACGTCTGAACTCTTTCCCTACACGACGCTCTTCCGATCTNNNATCCAAATACAAGGAGGTGA
AGCACACGTCTGAACTCTTTCCCTACACGACGCTCTTCCGATCTNNNATACACACACGTATAATACACA

AGCACACGTCTGAACTCTTTCCCTACACGACGCTCTTCCGATCTNNNATACACACACGTATAATACACA
GAGCACACGTCTGAACTCTTTCCCTACACGACGCTCTTCCGATCTNNNCTGGAGGAAACCTGCAAAA
GAGCACACGTCTGAACTCTTTCCCTACACGACGCTCTTCCGATCTNNNCTGGAGGAAACCTGCAAAA
AGCACACGTCTGAACTCTTTCCCTACACGACGCTCTTCCGATCTNNNGCAGGTTTGGGAAAGAGGAG
AGCACACGTCTGAACTCTTTCCCTACACGACGCTCTTCCGATCTNNNGCAGGTTTGGGAAAGAGGAG
AGCACACGTCTGAACTCTTTCCCTACACGACGCTCTTCCGATCTNNNGCACCCCTATCCCAGTAATAGT
AGCACACGTCTGAACTCTTTCCCTACACGACGCTCTTCCGATCTNNNGCACCCCTATCCCAGTAATAGT
AGCACACGTCTGAACTCTTTCCCTACACGACGCTCTTCCGATCTNNNACCTAAAGTGCCCCAACATTCT
AGCACACGTCTGAACTCTTTCCCTACACGACGCTCTTCCGATCTNNNACCTAAAGTGCCCCAACATTCT
AGCACACGTCTGAACTCTTTCCCTACACGACGCTCTTCCGATCTNNNACTATGGGTGGATAATGAAG
AGCACACGTCTGAACTCTTTCCCTACACGACGCTCTTCCGATCTNNNACTATGGGTGGATAATGAAG
GCACACGTCTGAACTCTTTCCCTACACGACGCTCTTCCGATCTNNNCAAGTCCTAGCCCAATGTTTGTG
GCACACGTCTGAACTCTTTCCCTACACGACGCTCTTCCGATCTNNNCAAGTCCTAGCCCAATGTTTGTG
AGCACACGTCTGAACTCTTTCCCTACACGACGCTCTTCCGATCTNNNGCAAAGAAGCAGTCGTAGATC
AGCACACGTCTGAACTCTTTCCCTACACGACGCTCTTCCGATCTNNNGCAAAGAAGCAGTCGTAGATC
GCACACGTCTGAACTCTTTCCCTACACGACGCTCTTCCGATCTNNNAGTTGGGACCAGGGATATACC
GCACACGTCTGAACTCTTTCCCTACACGACGCTCTTCCGATCTNNNAGTTGGGACCAGGGATATACC
GCACACGTCTGAACTCTTTCCCTACACGACGCTCTTCCGATCTNNNATTCTCTAAAATAACCAAAGCAG
GCACACGTCTGAACTCTTTCCCTACACGACGCTCTTCCGATCTNNNATTCTCTAAAATAACCAAAGCAG
GCACACGTCTGAACTCTTTCCCTACACGACGCTCTTCCGATCTNNNATTTGATCTTGCCGTTTTCCGTTC
GCACACGTCTGAACTCTTTCCCTACACGACGCTCTTCCGATCTNNNATTTGATCTTGCCGTTTTCCGTTC
AGCACACGTCTGAACTCTTTCCCTACACGACGCTCTTCCGATCTNNNCGACAGAGTGTGATCATGTCA
AGCACACGTCTGAACTCTTTCCCTACACGACGCTCTTCCGATCTNNNCGACAGAGTGTGATCATGTCA
AGCACACGTCTGAACTCTTTCCCTACACGACGCTCTTCCGATCTNNNTCATGTCCTGACTTAGCATGC
AGCACACGTCTGAACTCTTTCCCTACACGACGCTCTTCCGATCTNNNTCATGTCCTGACTTAGCATGC
CACACGTCTGAACTCTTTCCCTACACGACGCTCTTCCGATCTNNNACTGGTTGGCTAATACAGGAAAG
CACACGTCTGAACTCTTTCCCTACACGACGCTCTTCCGATCTNNNACTGGTTGGCTAATACAGGAAAG
GCACACGTCTGAACTCTTTCCCTACACGACGCTCTTCCGATCTNNNAAAGCTGCTTCCACATTTTCAGG
GCACACGTCTGAACTCTTTCCCTACACGACGCTCTTCCGATCTNNNAAAGCTGCTTCCACATTTTCAGG
CACACGTCTGAACTCTTTCCCTACACGACGCTCTTCCGATCTNNNGTCGTAGGACAAAGGGCCCCCAT
CACACGTCTGAACTCTTTCCCTACACGACGCTCTTCCGATCTNNNGTCGTAGGACAAAGGGCCCCCAT
AGCACACGTCTGAACTCTTTCCCTACACGACGCTCTTCCGATCTNNNTTCACCCAGTTTTCCCCAATAGT
AGCACACGTCTGAACTCTTTCCCTACACGACGCTCTTCCGATCTNNNTTCACCCAGTTTTCCCCAATAGT
GCACACGTCTGAACTCTTTCCCTACACGACGCTCTTCCGATCTNNNGAGGGGAAAAGATTGCTTGAA
GCACACGTCTGAACTCTTTCCCTACACGACGCTCTTCCGATCTNNNGAGGGGAAAAGATTGCTTGAA
AGCACACGTCTGAACTCTTTCCCTACACGACGCTCTTCCGATCTNNNTTATTCCATGCTTTGGCCATTG
AGCACACGTCTGAACTCTTTCCCTACACGACGCTCTTCCGATCTNNNTTATTCCATGCTTTGGCCATTG
AGCACACGTCTGAACTCTTTCCCTACACGACGCTCTTCCGATCTNNNCTGGGAAACTAAGTCACGTGA
AGCACACGTCTGAACTCTTTCCCTACACGACGCTCTTCCGATCTNNNCTGGGAAACTAAGTCACGTGA
AGCACACGTCTGAACTCTTTCCCTACACGACGCTCTTCCGATCTNNNACAACACAGAGGATCCCATTTC
AGCACACGTCTGAACTCTTTCCCTACACGACGCTCTTCCGATCTNNNACAACACAGAGGATCCCATTTC
AGCACACGTCTGAACTCTTTCCCTACACGACGCTCTTCCGATCTNNNGAGGCAGAGGTAGGAAAATC
AGCACACGTCTGAACTCTTTCCCTACACGACGCTCTTCCGATCTNNNGAGGCAGAGGTAGGAAAATC

GAGCACACGTCTGAACTCTTTCCCTACACGACGCTCTTCCGATCTNNNGCTCTGTATCAAACAAAGGCA
GAGCACACGTCTGAACTCTTTCCCTACACGACGCTCTTCCGATCTNNNGCTCTGTATCAAACAAAGGCA
AGCACACGTCTGAACTCTTTCCCTACACGACGCTCTTCCGATCTNNNAAGAGTCAAGACCCATCAGTGT
AGCACACGTCTGAACTCTTTCCCTACACGACGCTCTTCCGATCTNNNAAGAGTCAAGACCCATCAGTGT
GCACACGTCTGAACTCTTTCCCTACACGACGCTCTTCCGATCTNNNCCAAGAAAGAAAACCTGCAGGA
GCACACGTCTGAACTCTTTCCCTACACGACGCTCTTCCGATCTNNNCCAAGAAAGAAAACCTGCAGGA
AGCACACGTCTGAACTCTTTCCCTACACGACGCTCTTCCGATCTNNNGGTCTGGATATCTTGGGGTTT
AGCACACGTCTGAACTCTTTCCCTACACGACGCTCTTCCGATCTNNNGGTCTGGATATCTTGGGGTTT
GAGCACACGTCTGAACTCTTTCCCTACACGACGCTCTTCCGATCTNNNTTCTGGCAATCTCTGTCTCTGT
GAGCACACGTCTGAACTCTTTCCCTACACGACGCTCTTCCGATCTNNNTTCTGGCAATCTCTGTCTCTGT
AGCACACGTCTGAACTCTTTCCCTACACGACGCTCTTCCGATCTNNNAGCAAAAGCAGTACTAAGTGG
AGCACACGTCTGAACTCTTTCCCTACACGACGCTCTTCCGATCTNNNAGCAAAAGCAGTACTAAGTGG
AGCACACGTCTGAACTCTTTCCCTACACGACGCTCTTCCGATCTNNNTTTGGAGACTGAGGTAGGAG
AGCACACGTCTGAACTCTTTCCCTACACGACGCTCTTCCGATCTNNNTTTGGAGACTGAGGTAGGAG
AGCACACGTCTGAACTCTTTCCCTACACGACGCTCTTCCGATCTNNNTGGGCAACAAAACAAAACTC
AGCACACGTCTGAACTCTTTCCCTACACGACGCTCTTCCGATCTNNNTGGGCAACAAAACAAAACTC
GCACACGTCTGAACTCTTTCCCTACACGACGCTCTTCCGATCTNNNACTGTATCATAGGCCTCCTCCTA
GCACACGTCTGAACTCTTTCCCTACACGACGCTCTTCCGATCTNNNACTGTATCATAGGCCTCCTCCTA
AGCACACGTCTGAACTCTTTCCCTACACGACGCTCTTCCGATCTNNNTTGAAAATATAGTCCCCTGCCC
AGCACACGTCTGAACTCTTTCCCTACACGACGCTCTTCCGATCTNNNTTGAAAATATAGTCCCCTGCCC
GAGCACACGTCTGAACTCTTTCCCTACACGACGCTCTTCCGATCTNNNTGAAATCATTGAACATGGTA
GAGCACACGTCTGAACTCTTTCCCTACACGACGCTCTTCCGATCTNNNTGAAATCATTGAACATGGTA
AGCACACGTCTGAACTCTTTCCCTACACGACGCTCTTCCGATCTNNNCTCCCCAGTAGCTAGGACTACA
AGCACACGTCTGAACTCTTTCCCTACACGACGCTCTTCCGATCTNNNCTCCCCAGTAGCTAGGACTACA
GAGCACACGTCTGAACTCTTTCCCTACACGACGCTCTTCCGATCTNNNTATGGCTCATTGTAGGCTCAC
GAGCACACGTCTGAACTCTTTCCCTACACGACGCTCTTCCGATCTNNNTATGGCTCATTGTAGGCTCAC
GAGCACACGTCTGAACTCTTTCCCTACACGACGCTCTTCCGATCTNNNACCATCATGCCTGGCAATTTTT
GAGCACACGTCTGAACTCTTTCCCTACACGACGCTCTTCCGATCTNNNACCATCATGCCTGGCAATTTTT
GCACACGTCTGAACTCTTTCCCTACACGACGCTCTTCCGATCTNNNGCTGATGGAGATCAATAGAGCC
GCACACGTCTGAACTCTTTCCCTACACGACGCTCTTCCGATCTNNNGCTGATGGAGATCAATAGAGCC
AGCACACGTCTGAACTCTTTCCCTACACGACGCTCTTCCGATCTNNNAGTGAGCCGAGATTGTGTCAT
AGCACACGTCTGAACTCTTTCCCTACACGACGCTCTTCCGATCTNNNAGTGAGCCGAGATTGTGTCAT
AGCACACGTCTGAACTCTTTCCCTACACGACGCTCTTCCGATCTNNNTGTTGTTGTCAAATGACTGCAG
AGCACACGTCTGAACTCTTTCCCTACACGACGCTCTTCCGATCTNNNTGTTGTTGTCAAATGACTGCAG
GAGCACACGTCTGAACTCTTTCCCTACACGACGCTCTTCCGATCTNNNACGAAACTACTTAGGAAAGCC
GAGCACACGTCTGAACTCTTTCCCTACACGACGCTCTTCCGATCTNNNACGAAACTACTTAGGAAAGCC
ACACGTCTGAACTCTTTCCCTACACGACGCTCTTCCGATCTNNNGAAGATACTTACTGTAGTTCTCACA
ACACGTCTGAACTCTTTCCCTACACGACGCTCTTCCGATCTNNNGAAGATACTTACTGTAGTTCTCACA
GAGCACACGTCTGAACTCTTTCCCTACACGACGCTCTTCCGATCTNNNCCTCTGGGGCTCAAATGTTTC
GAGCACACGTCTGAACTCTTTCCCTACACGACGCTCTTCCGATCTNNNCCTCTGGGGCTCAAATGTTTC
GAGCACACGTCTGAACTCTTTCCCTACACGACGCTCTTCCGATCTNNNCCCATGTCCATTTCCTGATCTC
GAGCACACGTCTGAACTCTTTCCCTACACGACGCTCTTCCGATCTNNNCCCATGTCCATTTCCTGATCTC
AGCACACGTCTGAACTCTTTCCCTACACGACGCTCTTCCGATCTNNNGCGGGGTCAGATAATCAGGTA

AGCACACGTCTGAACTCTTTCCCTACACGACGCTCTTCCGATCTNNNGCGGGGTCAGATAATCAGGTA
GAGCACACGTCTGAACTCTTTCCCTACACGACGCTCTTCCGATCTNNNCATTGTGGACCAGGTACTCCTA
GAGCACACGTCTGAACTCTTTCCCTACACGACGCTCTTCCGATCTNNNCATTGTGGACCAGGTACTCCTA
AGCACACGTCTGAACTCTTTCCCTACACGACGCTCTTCCGATCTNNNTCCACAATTGCATGTCAGTTCCT
AGCACACGTCTGAACTCTTTCCCTACACGACGCTCTTCCGATCTNNNTCCACAATTGCATGTCAGTTCCT
GAGCACACGTCTGAACTCTTTCCCTACACGACGCTCTTCCGATCTNNNACTCTGCTTCACTCACCATTCAA
GAGCACACGTCTGAACTCTTTCCCTACACGACGCTCTTCCGATCTNNNACTCTGCTTCACTCACCATTCAA
AGCACACGTCTGAACTCTTTCCCTACACGACGCTCTTCCGATCTNNNGAGAATTCAGATCTCCCTGGGT
AGCACACGTCTGAACTCTTTCCCTACACGACGCTCTTCCGATCTNNNGAGAATTCAGATCTCCCTGGGT
GAGCACACGTCTGAACTCTTTCCCTACACGACGCTCTTCCGATCTNNNAAAAAAAATTAGCTGGGCGTG
GAGCACACGTCTGAACTCTTTCCCTACACGACGCTCTTCCGATCTNNNAAAAAAAATTAGCTGGGCGTG
AGCACACGTCTGAACTCTTTCCCTACACGACGCTCTTCCGATCTNNNGCAGAACCCACATCCTTGTTTACG
AGCACACGTCTGAACTCTTTCCCTACACGACGCTCTTCCGATCTNNNGCAGAACCCACATCCTTGTTTACG
GAGCACACGTCTGAACTCTTTCCCTACACGACGCTCTTCCGATCTNNNGGAAATAAAACAACAGTCGCG
GAGCACACGTCTGAACTCTTTCCCTACACGACGCTCTTCCGATCTNNNGGAAATAAAACAACAGTCGCG
AGAGCACACGTCTGAACTCTTTCCCTACACGACGCTCTTCCGATCTNNNACAAAAATTAGCTGGGTGTG
AGAGCACACGTCTGAACTCTTTCCCTACACGACGCTCTTCCGATCTNNNACAAAAATTAGCTGGGTGTG
GAAGAGCACACGTCTGAACTCTTTCCCTACACGACGCTCTTCCGATCTNNNGTAGTCCCAGCTACTCCG
GAAGAGCACACGTCTGAACTCTTTCCCTACACGACGCTCTTCCGATCTNNNGTAGTCCCAGCTACTCCG
GAGCACACGTCTGAACTCTTTCCCTACACGACGCTCTTCCGATCTNNNTGTTCAAGGCTCTGCACAATTT
GAGCACACGTCTGAACTCTTTCCCTACACGACGCTCTTCCGATCTNNNTGTTCAAGGCTCTGCACAATTT
AGAGCACACGTCTGAACTCTTTCCCTACACGACGCTCTTCCGATCTNNNTATGCTATCGACCGAACATTG
AGAGCACACGTCTGAACTCTTTCCCTACACGACGCTCTTCCGATCTNNNTATGCTATCGACCGAACATTG
GAGCACACGTCTGAACTCTTTCCCTACACGACGCTCTTCCGATCTNNNTTGCTGACCAGAAGGAGTAG
GAGCACACGTCTGAACTCTTTCCCTACACGACGCTCTTCCGATCTNNNTTGCTGACCAGAAGGAGTAG
GAGCACACGTCTGAACTCTTTCCCTACACGACGCTCTTCCGATCTNNNTTCAGCCTGAGCATCTTCTGA
GAGCACACGTCTGAACTCTTTCCCTACACGACGCTCTTCCGATCTNNNTTCAGCCTGAGCATCTTCTGA
AGCACACGTCTGAACTCTTTCCCTACACGACGCTCTTCCGATCTNNNACTTTCAAAATGTTGCCTTGGG
AGCACACGTCTGAACTCTTTCCCTACACGACGCTCTTCCGATCTNNNACTTTCAAAATGTTGCCTTGGG
CACACGTCTGAACTCTTTCCCTACACGACGCTCTTCCGATCTNNNGATTCCACTGACTCAGAAAGCATC
CACACGTCTGAACTCTTTCCCTACACGACGCTCTTCCGATCTNNNGATTCCACTGACTCAGAAAGCATC
GAGCACACGTCTGAACTCTTTCCCTACACGACGCTCTTCCGATCTNNNAATGTAGAAGAAACGCTGTCC
GAGCACACGTCTGAACTCTTTCCCTACACGACGCTCTTCCGATCTNNNAATGTAGAAGAAACGCTGTCC
AGCACACGTCTGAACTCTTTCCCTACACGACGCTCTTCCGATCTNNNAAATTCCCTCAGCTGTAGGAGT
AGCACACGTCTGAACTCTTTCCCTACACGACGCTCTTCCGATCTNNNAAATTCCCTCAGCTGTAGGAGT
GAGCACACGTCTGAACTCTTTCCCTACACGACGCTCTTCCGATCTNNNTATCTGATTGGGTGTTGTCAG
GAGCACACGTCTGAACTCTTTCCCTACACGACGCTCTTCCGATCTNNNTATCTGATTGGGTGTTGTCAG
AGCACACGTCTGAACTCTTTCCCTACACGACGCTCTTCCGATCTNNNTTCTCCTAATGCTATGCCTTCCCG
AGCACACGTCTGAACTCTTTCCCTACACGACGCTCTTCCGATCTNNNTTCTCCTAATGCTATGCCTTCCCG
GAGCACACGTCTGAACTCTTTCCCTACACGACGCTCTTCCGATCTNNNAGAATTACTTGAACCCAGGAG
GAGCACACGTCTGAACTCTTTCCCTACACGACGCTCTTCCGATCTNNNAGAATTACTTGAACCCAGGAG
AGCACACGTCTGAACTCTTTCCCTACACGACGCTCTTCCGATCTNNNACAATTCTGGGAGAGCATCAAG
AGCACACGTCTGAACTCTTTCCCTACACGACGCTCTTCCGATCTNNNACAATTCTGGGAGAGCATCAAG

AGCACACGTCTGAACTCTTTCCCTACACGACGCTCTTCCGATCTNNNGGGAGAAGGGAGAAATAGA
AGCACACGTCTGAACTCTTTCCCTACACGACGCTCTTCCGATCTNNNGGGAGAAGGGAGAAATAGA
AGCACACGTCTGAACTCTTTCCCTACACGACGCTCTTCCGATCTNNNCAGCCCTGACTCTGTTCCATAT
AGCACACGTCTGAACTCTTTCCCTACACGACGCTCTTCCGATCTNNNCAGCCCTGACTCTGTTCCATAT
AGCACACGTCTGAACTCTTTCCCTACACGACGCTCTTCCGATCTNNNTCCTACAGCGGTTTGTAATGGA
AGCACACGTCTGAACTCTTTCCCTACACGACGCTCTTCCGATCTNNNTCCTACAGCGGTTTGTAATGGA
AGCACACGTCTGAACTCTTTCCCTACACGACGCTCTTCCGATCTNNNGCAATAGCATGATCTCGGTTCA
AGCACACGTCTGAACTCTTTCCCTACACGACGCTCTTCCGATCTNNNGCAATAGCATGATCTCGGTTCA
AGCACACGTCTGAACTCTTTCCCTACACGACGCTCTTCCGATCTNNNGCCTCAAGCTGAGTTCTCTTAC
AGCACACGTCTGAACTCTTTCCCTACACGACGCTCTTCCGATCTNNNGCCTCAAGCTGAGTTCTCTTAC
AGCACACGTCTGAACTCTTTCCCTACACGACGCTCTTCCGATCTNNNAAGAAAGAGCATGTGCAAAG
AGCACACGTCTGAACTCTTTCCCTACACGACGCTCTTCCGATCTNNNAAGAAAGAGCATGTGCAAAG
AGCACACGTCTGAACTCTTTCCCTACACGACGCTCTTCCGATCTNNNAGAGCAGAAGACTGACTGTCA
AGCACACGTCTGAACTCTTTCCCTACACGACGCTCTTCCGATCTNNNAGAGCAGAAGACTGACTGTCA
AGCACACGTCTGAACTCTTTCCCTACACGACGCTCTTCCGATCTNNNCCCAAATGATGAAGTGATGCA
AGCACACGTCTGAACTCTTTCCCTACACGACGCTCTTCCGATCTNNNCCCAAATGATGAAGTGATGCA
AGCACACGTCTGAACTCTTTCCCTACACGACGCTCTTCCGATCTNNNTGAGGAAGGAGAATGGTGTG
AGCACACGTCTGAACTCTTTCCCTACACGACGCTCTTCCGATCTNNNTGAGGAAGGAGAATGGTGTG
AGCACACGTCTGAACTCTTTCCCTACACGACGCTCTTCCGATCTNNNCTTGCCCAGCTCATCCTATATCA
AGCACACGTCTGAACTCTTTCCCTACACGACGCTCTTCCGATCTNNNCTTGCCCAGCTCATCCTATATCA
AGCACACGTCTGAACTCTTTCCCTACACGACGCTCTTCCGATCTNNNCTGAACCAGGAGAGTCACTTC
AGCACACGTCTGAACTCTTTCCCTACACGACGCTCTTCCGATCTNNNCTGAACCAGGAGAGTCACTTC
AGCACACGTCTGAACTCTTTCCCTACACGACGCTCTTCCGATCTNNNACTAGAGTAATGGGGCAGAG
AGCACACGTCTGAACTCTTTCCCTACACGACGCTCTTCCGATCTNNNACTAGAGTAATGGGGCAGAG
AGCACACGTCTGAACTCTTTCCCTACACGACGCTCTTCCGATCTNNNGAATGGTGTGAACCTGGGAA
AGCACACGTCTGAACTCTTTCCCTACACGACGCTCTTCCGATCTNNNGAATGGTGTGAACCTGGGAA
AGCACACGTCTGAACTCTTTCCCTACACGACGCTCTTCCGATCTNNNACATACTCACACAAGCTCAGGA
AGCACACGTCTGAACTCTTTCCCTACACGACGCTCTTCCGATCTNNNACATACTCACACAAGCTCAGGA
GCACACGTCTGAACTCTTTCCCTACACGACGCTCTTCCGATCTNNNGGATCAGTGCTGGATGTTTGTT
GCACACGTCTGAACTCTTTCCCTACACGACGCTCTTCCGATCTNNNGGATCAGTGCTGGATGTTTGTT
AGAGCACACGTCTGAACTCTTTCCCTACACGACGCTCTTCCGATCTNNNACATAGGAGAGAGACCCCT
AGAGCACACGTCTGAACTCTTTCCCTACACGACGCTCTTCCGATCTNNNACATAGGAGAGAGACCCCT
AGCACACGTCTGAACTCTTTCCCTACACGACGCTCTTCCGATCTNNNATCCTCCCACACCCTGAGTAAT
AGCACACGTCTGAACTCTTTCCCTACACGACGCTCTTCCGATCTNNNATCCTCCCACACCCTGAGTAAT
AGAGCACACGTCTGAACTCTTTCCCTACACGACGCTCTTCCGATCTNNNACACCTGTAATCCCAGCACT
AGAGCACACGTCTGAACTCTTTCCCTACACGACGCTCTTCCGATCTNNNACACCTGTAATCCCAGCACT
GAGCACACGTCTGAACTCTTTCCCTACACGACGCTCTTCCGATCTNNNGAGGCAGGAAAATCACTTGA
GAGCACACGTCTGAACTCTTTCCCTACACGACGCTCTTCCGATCTNNNGAGGCAGGAAAATCACTTGA
GCACACGTCTGAACTCTTTCCCTACACGACGCTCTTCCGATCTNNNTCCATGCCACTTTCTTTTCTGGT
GCACACGTCTGAACTCTTTCCCTACACGACGCTCTTCCGATCTNNNTCCATGCCACTTTCTTTTCTGGT
GCACACGTCTGAACTCTTTCCCTACACGACGCTCTTCCGATCTNNNGTTCAAAGTCCTTCCCTTTCCAA
GCACACGTCTGAACTCTTTCCCTACACGACGCTCTTCCGATCTNNNGTTCAAAGTCCTTCCCTTTCCAA
AGCACACGTCTGAACTCTTTCCCTACACGACGCTCTTCCGATCTNNNACATCTTTATGGTATAACTGTG

AGCACACGTCTGAACTCTTTCCCTACACGACGCTCTTCCGATCTNNNACATCTTTATGGTATAACTGTG
AGCACACGTCTGAACTCTTTCCCTACACGACGCTCTTCCGATCTNNNGACTCAGATACCTCCAACAGCA
AGCACACGTCTGAACTCTTTCCCTACACGACGCTCTTCCGATCTNNNGACTCAGATACCTCCAACAGCA
AGCACACGTCTGAACTCTTTCCCTACACGACGCTCTTCCGATCTNNNAGGGAGAGAGAGATTGAAGA
AGCACACGTCTGAACTCTTTCCCTACACGACGCTCTTCCGATCTNNNAGGGAGAGAGAGATTGAAGA
AGCACACGTCTGAACTCTTTCCCTACACGACGCTCTTCCGATCTNNNTCTTCCTTTTTCCATTGTGTGCA
AGCACACGTCTGAACTCTTTCCCTACACGACGCTCTTCCGATCTNNNTCTTCCTTTTTCCATTGTGTGCA
GCACACGTCTGAACTCTTTCCCTACACGACGCTCTTCCGATCTNNNACTCTACACACACATACACACACA
GCACACGTCTGAACTCTTTCCCTACACGACGCTCTTCCGATCTNNNACTCTACACACACATACACACACA
GAGCACACGTCTGAACTCTTTCCCTACACGACGCTCTTCCGATCTNNNCCCAGCCTCCTTCATTTTCTGT
GAGCACACGTCTGAACTCTTTCCCTACACGACGCTCTTCCGATCTNNNCCCAGCCTCCTTCATTTTCTGT
AGCACACGTCTGAACTCTTTCCCTACACGACGCTCTTCCGATCTNNNAGAAGGGATCTCGCTATGTTTC
AGCACACGTCTGAACTCTTTCCCTACACGACGCTCTTCCGATCTNNNAGAAGGGATCTCGCTATGTTTC
GAGCACACGTCTGAACTCTTTCCCTACACGACGCTCTTCCGATCTNNNTGTTCACAAGTGACATGGCA
GAGCACACGTCTGAACTCTTTCCCTACACGACGCTCTTCCGATCTNNNTGTTCACAAGTGACATGGCA
AGCACACGTCTGAACTCTTTCCCTACACGACGCTCTTCCGATCTNNNTTGAGGCAGGAGAATAGCTTG
AGCACACGTCTGAACTCTTTCCCTACACGACGCTCTTCCGATCTNNNTTGAGGCAGGAGAATAGCTTG
AGCACACGTCTGAACTCTTTCCCTACACGACGCTCTTCCGATCTNNNTTGGATACCTATACAGATCCAT
AGCACACGTCTGAACTCTTTCCCTACACGACGCTCTTCCGATCTNNNTTGGATACCTATACAGATCCAT
AGCACACGTCTGAACTCTTTCCCTACACGACGCTCTTCCGATCTNNNTATTCTCCAAACAACACTCCCCA
AGCACACGTCTGAACTCTTTCCCTACACGACGCTCTTCCGATCTNNNTATTCTCCAAACAACACTCCCCA
CACACGTCTGAACTCTTTCCCTACACGACGCTCTTCCGATCTNNNGGACTTTGCCTACTCCTCACCATCG
CACACGTCTGAACTCTTTCCCTACACGACGCTCTTCCGATCTNNNGGACTTTGCCTACTCCTCACCATCG
AGCACACGTCTGAACTCTTTCCCTACACGACGCTCTTCCGATCTNNNGATAATCCAAAGGCTTCCCCAG
AGCACACGTCTGAACTCTTTCCCTACACGACGCTCTTCCGATCTNNNGATAATCCAAAGGCTTCCCCAG
AGCACACGTCTGAACTCTTTCCCTACACGACGCTCTTCCGATCTNNNGTCCTTGTGGAGATCTTTCACCA
AGCACACGTCTGAACTCTTTCCCTACACGACGCTCTTCCGATCTNNNGTCCTTGTGGAGATCTTTCACCA
AGCACACGTCTGAACTCTTTCCCTACACGACGCTCTTCCGATCTNNNCCAGGGTCAGCTACATAGTTT
AGCACACGTCTGAACTCTTTCCCTACACGACGCTCTTCCGATCTNNNCCAGGGTCAGCTACATAGTTT
GCACACGTCTGAACTCTTTCCCTACACGACGCTCTTCCGATCTNNNGTGAAGAGACTGAGAGGTCAA
GCACACGTCTGAACTCTTTCCCTACACGACGCTCTTCCGATCTNNNGTGAAGAGACTGAGAGGTCAA
AGAGCACACGTCTGAACTCTTTCCCTACACGACGCTCTTCCGATCTNNNGTACACTGACTGGACCACTA
AGAGCACACGTCTGAACTCTTTCCCTACACGACGCTCTTCCGATCTNNNGTACACTGACTGGACCACTA
AGCACACGTCTGAACTCTTTCCCTACACGACGCTCTTCCGATCTNNNGTTTTCAGGGTCACTAGTGAG
AGCACACGTCTGAACTCTTTCCCTACACGACGCTCTTCCGATCTNNNGTTTTCAGGGTCACTAGTGAG
GCACACGTCTGAACTCTTTCCCTACACGACGCTCTTCCGATCTNNNAGCATCCTTTGGCAATGTAATGG
GCACACGTCTGAACTCTTTCCCTACACGACGCTCTTCCGATCTNNNAGCATCCTTTGGCAATGTAATGG
AGCACACGTCTGAACTCTTTCCCTACACGACGCTCTTCCGATCTNNNTATAATCCTGGCACTTTGGGA
AGCACACGTCTGAACTCTTTCCCTACACGACGCTCTTCCGATCTNNNTATAATCCTGGCACTTTGGGA
AGCACACGTCTGAACTCTTTCCCTACACGACGCTCTTCCGATCTNNNCCCAGGATTGTAATGCCTCCTA
AGCACACGTCTGAACTCTTTCCCTACACGACGCTCTTCCGATCTNNNCCCAGGATTGTAATGCCTCCTA
AAGAGCACACGTCTGAACTCTTTCCCTACACGACGCTCTTCCGATCTNNNCACCTGTAATCCCAGCTACT
AAGAGCACACGTCTGAACTCTTTCCCTACACGACGCTCTTCCGATCTNNNCACCTGTAATCCCAGCTACT

AGCACACGTCTGAACTCTTTCCCTACACGACGCTCTTCCGATCTNNNATAAGCATTCCCTTTCCTCTGCA
AGCACACGTCTGAACTCTTTCCCTACACGACGCTCTTCCGATCTNNNATAAGCATTCCCTTTCCTCTGCA
GCACACGTCTGAACTCTTTCCCTACACGACGCTCTTCCGATCTNNNAGTTCTGGGATTACAGACGTGAG
GCACACGTCTGAACTCTTTCCCTACACGACGCTCTTCCGATCTNNNAGTTCTGGGATTACAGACGTGAG
AGAGCACACGTCTGAACTCTTTCCCTACACGACGCTCTTCCGATCTNNNTGCTAATGACACTGATATCCT
AGAGCACACGTCTGAACTCTTTCCCTACACGACGCTCTTCCGATCTNNNTGCTAATGACACTGATATCCT
GCACACGTCTGAACTCTTTCCCTACACGACGCTCTTCCGATCTNNNTGCCCTCTAGTTACATTGGCTTTCA
GCACACGTCTGAACTCTTTCCCTACACGACGCTCTTCCGATCTNNNTGCCCTCTAGTTACATTGGCTTTCA
AGCACACGTCTGAACTCTTTCCCTACACGACGCTCTTCCGATCTNNNAACATGTAGCTCATTTTCTAGGT
AGCACACGTCTGAACTCTTTCCCTACACGACGCTCTTCCGATCTNNNAACATGTAGCTCATTTTCTAGGT
GAGCACACGTCTGAACTCTTTCCCTACACGACGCTCTTCCGATCTNNNAGTGCTGGGATTATAGGCATG
GAGCACACGTCTGAACTCTTTCCCTACACGACGCTCTTCCGATCTNNNAGTGCTGGGATTATAGGCATG
GAGCACACGTCTGAACTCTTTCCCTACACGACGCTCTTCCGATCTNNNTGAACTCTTGACTTGCAGTTTT
GAGCACACGTCTGAACTCTTTCCCTACACGACGCTCTTCCGATCTNNNTGAACTCTTGACTTGCAGTTTT
AGAGCACACGTCTGAACTCTTTCCCTACACGACGCTCTTCCGATCTNNNAGTTGCTAGAATGTGCTTAG
AGAGCACACGTCTGAACTCTTTCCCTACACGACGCTCTTCCGATCTNNNAGTTGCTAGAATGTGCTTAG
GAGCACACGTCTGAACTCTTTCCCTACACGACGCTCTTCCGATCTNNNTCAGAGCTTCTAGATGCAACTG
GAGCACACGTCTGAACTCTTTCCCTACACGACGCTCTTCCGATCTNNNTCAGAGCTTCTAGATGCAACTG
AGCACACGTCTGAACTCTTTCCCTACACGACGCTCTTCCGATCTNNNGAGGCAGGAGAATTGCTTGAAT
AGCACACGTCTGAACTCTTTCCCTACACGACGCTCTTCCGATCTNNNGAGGCAGGAGAATTGCTTGAAT
AGCACACGTCTGAACTCTTTCCCTACACGACGCTCTTCCGATCTNNNAGCACTTCAAACAATTTGCGTTT
AGCACACGTCTGAACTCTTTCCCTACACGACGCTCTTCCGATCTNNNAGCACTTCAAACAATTTGCGTTT
GAGCACACGTCTGAACTCTTTCCCTACACGACGCTCTTCCGATCTNNNGAATGTGTGTGTGTGGAATG
GAGCACACGTCTGAACTCTTTCCCTACACGACGCTCTTCCGATCTNNNGAATGTGTGTGTGTGGAATG
AGAGCACACGTCTGAACTCTTTCCCTACACGACGCTCTTCCGATCTNNNTGAGGCAGGAGAATTGCTTG
AGAGCACACGTCTGAACTCTTTCCCTACACGACGCTCTTCCGATCTNNNTGAGGCAGGAGAATTGCTTG
AGAGCACACGTCTGAACTCTTTCCCTACACGACGCTCTTCCGATCTNNNAAACATGCTTGTAGTCCCAGG
AGAGCACACGTCTGAACTCTTTCCCTACACGACGCTCTTCCGATCTNNNAAACATGCTTGTAGTCCCAGG
AGCACACGTCTGAACTCTTTCCCTACACGACGCTCTTCCGATCTNNNCTTTGTTGCTGCATCTTCTGGAG
AGCACACGTCTGAACTCTTTCCCTACACGACGCTCTTCCGATCTNNNCTTTGTTGCTGCATCTTCTGGAG
GAGCACACGTCTGAACTCTTTCCCTACACGACGCTCTTCCGATCTNNNGTAAATAGTCTCAGCTCCTCGG
GAGCACACGTCTGAACTCTTTCCCTACACGACGCTCTTCCGATCTNNNGTAAATAGTCTCAGCTCCTCGG
AGAGCACACGTCTGAACTCTTTCCCTACACGACGCTCTTCCGATCTNNNCTTGGGCAACAGAGAAAGA
AGAGCACACGTCTGAACTCTTTCCCTACACGACGCTCTTCCGATCTNNNCTTGGGCAACAGAGAAAGA
GAGCACACGTCTGAACTCTTTCCCTACACGACGCTCTTCCGATCTNNNCTTGCAGTGAGCCGAGATCGC
GAGCACACGTCTGAACTCTTTCCCTACACGACGCTCTTCCGATCTNNNCTTGCAGTGAGCCGAGATCGC
AGCACACGTCTGAACTCTTTCCCTACACGACGCTCTTCCGATCTNNNGGCAAACGCATCAATTTTTCAG
AGCACACGTCTGAACTCTTTCCCTACACGACGCTCTTCCGATCTNNNGGCAAACGCATCAATTTTTCAG
GAGCACACGTCTGAACTCTTTCCCTACACGACGCTCTTCCGATCTNNNGGAACCAACCCAACTGTCTATG
GAGCACACGTCTGAACTCTTTCCCTACACGACGCTCTTCCGATCTNNNGGAACCAACCCAACTGTCTATG
AGCACACGTCTGAACTCTTTCCCTACACGACGCTCTTCCGATCTNNNGCCCCTGGTGTAGAGGAAAAT
AGCACACGTCTGAACTCTTTCCCTACACGACGCTCTTCCGATCTNNNGCCCCTGGTGTAGAGGAAAAT
GAGCACACGTCTGAACTCTTTCCCTACACGACGCTCTTCCGATCTNNNATCCACCAACCAAGCCCTATATG

AGCACACGTCTGAACTCTTTCCCTACACGACGCTCTTCCGATCTNNNATCCACCAACCAAGCCCTATAT
AGCACACGTCTGAACTCTTTCCCTACACGACGCTCTTCCGATCTNNNCTAGGATTTTGCCTCAGTGGT
AGCACACGTCTGAACTCTTTCCCTACACGACGCTCTTCCGATCTNNNCTAGGATTTTGCCTCAGTGGT
GAGCACACGTCTGAACTCTTTCCCTACACGACGCTCTTCCGATCTNNNAGGGAATAAGGGAGAAGGA
GAGCACACGTCTGAACTCTTTCCCTACACGACGCTCTTCCGATCTNNNAGGGAATAAGGGAGAAGGA
AGCACACGTCTGAACTCTTTCCCTACACGACGCTCTTCCGATCTNNNATACCACAGCTTGCTTTCAAA
AGCACACGTCTGAACTCTTTCCCTACACGACGCTCTTCCGATCTNNNATACCACAGCTTGCTTTCAAA
AGCACACGTCTGAACTCTTTCCCTACACGACGCTCTTCCGATCTNNNTGCCATTGCACAAAAGTATTG
AGCACACGTCTGAACTCTTTCCCTACACGACGCTCTTCCGATCTNNNTGCCATTGCACAAAAGTATTG
AGCACACGTCTGAACTCTTTCCCTACACGACGCTCTTCCGATCTNNNAGTGAGCCAAGATTCCATCACT
AGCACACGTCTGAACTCTTTCCCTACACGACGCTCTTCCGATCTNNNAGTGAGCCAAGATTCCATCACT
AAGAGCACACGTCTGAACTCTTTCCCTACACGACGCTCTTCCGATCTNNNGAAAGAGAATTCAGCCAG
AAGAGCACACGTCTGAACTCTTTCCCTACACGACGCTCTTCCGATCTNNNGAAAGAGAATTCAGCCAG
AGCACACGTCTGAACTCTTTCCCTACACGACGCTCTTCCGATCTNNNCAGGTCATGGTTAGGTAAGAG
AGCACACGTCTGAACTCTTTCCCTACACGACGCTCTTCCGATCTNNNCAGGTCATGGTTAGGTAAGAG
AGCACACGTCTGAACTCTTTCCCTACACGACGCTCTTCCGATCTNNNGCTAATATGGAAGCCCTGCA
AGCACACGTCTGAACTCTTTCCCTACACGACGCTCTTCCGATCTNNNGCTAATATGGAAGCCCTGCA
CACACGTCTGAACTCTTTCCCTACACGACGCTCTTCCGATCTNNNCCCAACACACACACACCTATGAGC
CACACGTCTGAACTCTTTCCCTACACGACGCTCTTCCGATCTNNNCCCAACACACACACACCTATGAGC
GCACACGTCTGAACTCTTTCCCTACACGACGCTCTTCCGATCTNNNTTTGACTATTAGCCGCCAAGACT
GCACACGTCTGAACTCTTTCCCTACACGACGCTCTTCCGATCTNNNTTTGACTATTAGCCGCCAAGACT
AGCACACGTCTGAACTCTTTCCCTACACGACGCTCTTCCGATCTNNNCCTGGTTGCATTCATGGAGAT
AGCACACGTCTGAACTCTTTCCCTACACGACGCTCTTCCGATCTNNNCCTGGTTGCATTCATGGAGAT
GCACACGTCTGAACTCTTTCCCTACACGACGCTCTTCCGATCTNNNGTACCTGTAATCCCAGCTACTCG
GCACACGTCTGAACTCTTTCCCTACACGACGCTCTTCCGATCTNNNGTACCTGTAATCCCAGCTACTCG
GAGCACACGTCTGAACTCTTTCCCTACACGACGCTCTTCCGATCTNNNAGCTTTCTCTTGCTGGTCTAC
GAGCACACGTCTGAACTCTTTCCCTACACGACGCTCTTCCGATCTNNNAGCTTTCTCTTGCTGGTCTAC
AGCACACGTCTGAACTCTTTCCCTACACGACGCTCTTCCGATCTNNNATCCCCTGCAGAACCACAAATA
AGCACACGTCTGAACTCTTTCCCTACACGACGCTCTTCCGATCTNNNATCCCCTGCAGAACCACAAATA
AGAGCACACGTCTGAACTCTTTCCCTACACGACGCTCTTCCGATCTNNNAAATTACCTGTGTTCTGGA
AGAGCACACGTCTGAACTCTTTCCCTACACGACGCTCTTCCGATCTNNNAAATTACCTGTGTTCTGGA
GCACACGTCTGAACTCTTTCCCTACACGACGCTCTTCCGATCTNNNTCTTCAATATGGCGCATTGAACC
GCACACGTCTGAACTCTTTCCCTACACGACGCTCTTCCGATCTNNNTCTTCAATATGGCGCATTGAACC
GCACACGTCTGAACTCTTTCCCTACACGACGCTCTTCCGATCTNNNATTAATCCTTCACAAAGTCTTTGC
GCACACGTCTGAACTCTTTCCCTACACGACGCTCTTCCGATCTNNNATTAATCCTTCACAAAGTCTTTGC
AGCACACGTCTGAACTCTTTCCCTACACGACGCTCTTCCGATCTNNNCAGAAATTTTTCATCCCTGGGC
AGCACACGTCTGAACTCTTTCCCTACACGACGCTCTTCCGATCTNNNCAGAAATTTTTCATCCCTGGGC
AGCACACGTCTGAACTCTTTCCCTACACGACGCTCTTCCGATCTNNNCCATGTTGCAAGTGATTCTCC
AGCACACGTCTGAACTCTTTCCCTACACGACGCTCTTCCGATCTNNNCCATGTTGCAAGTGATTCTCC
AGCACACGTCTGAACTCTTTCCCTACACGACGCTCTTCCGATCTNNNAAGAGCCAAATCAGGAACAA
AGCACACGTCTGAACTCTTTCCCTACACGACGCTCTTCCGATCTNNNAAGAGCCAAATCAGGAACAA
CACACGTCTGAACTCTTTCCCTACACGACGCTCTTCCGATCTNNNTGTGAGTGGCTCTCTACTTGTGA
CACACGTCTGAACTCTTTCCCTACACGACGCTCTTCCGATCTNNNTGTGAGTGGCTCTCTACTTGTGA

GCACACGTCTGAACTCTTTCCCTACACGACGCTCTTCCGATCTNNNAAACAGGCTGTAACCTGGTCTT

GCACACGTCTGAACTCTTTCCCTACACGACGCTCTTCCGATCTNNNAAACAGGCTGTAACCTGGTCTT

GAGCACACGTCTGAACTCTTTCCCTACACGACGCTCTTCCGATCTNNNGGTGCCTATAGTTCCAGCTAC

GAGCACACGTCTGAACTCTTTCCCTACACGACGCTCTTCCGATCTNNNGGTGCCTATAGTTCCAGCTAC

AGCACACGTCTGAACTCTTTCCCTACACGACGCTCTTCCGATCTNNNTCACAAGGTCAGGAGTTCAA

AGCACACGTCTGAACTCTTTCCCTACACGACGCTCTTCCGATCTNNNTCACAAGGTCAGGAGTTCAA

CACACGTCTGAACTCTTTCCCTACACGACGCTCTTCCGATCTNNNGTTTGGTTTCTGTTTGTCGGACA

CACACGTCTGAACTCTTTCCCTACACGACGCTCTTCCGATCTNNNGTTTGGTTTCTGTTTGTCGGACA

GCACACGTCTGAACTCTTTCCCTACACGACGCTCTTCCGATCTNNNTGCACTCCAGCCTCAAAAAAAA

GCACACGTCTGAACTCTTTCCCTACACGACGCTCTTCCGATCTNNNTGCACTCCAGCCTCAAAAAAAA

GCACACGTCTGAACTCTTTCCCTACACGACGCTCTTCCGATCTNNNTTCAGTGAGCCATGATTGTACC

GCACACGTCTGAACTCTTTCCCTACACGACGCTCTTCCGATCTNNNTTCAGTGAGCCATGATTGTACC

AGCACACGTCTGAACTCTTTCCCTACACGACGCTCTTCCGATCTNNNTCACTTGAGCTCAGGAGTTTG

AGCACACGTCTGAACTCTTTCCCTACACGACGCTCTTCCGATCTNNNTCACTTGAGCTCAGGAGTTTG

AGCACACGTCTGAACTCTTTCCCTACACGACGCTCTTCCGATCTNNNCAGGCACACACACAGGTATCTA

AGCACACGTCTGAACTCTTTCCCTACACGACGCTCTTCCGATCTNNNCAGGCACACACACAGGTATCTA

GCACACGTCTGAACTCTTTCCCTACACGACGCTCTTCCGATCTNNNGAGCCAAGATCGCACCGTTGCA

GCACACGTCTGAACTCTTTCCCTACACGACGCTCTTCCGATCTNNNGAGCCAAGATCGCACCGTTGCA

AGCACACGTCTGAACTCTTTCCCTACACGACGCTCTTCCGATCTNNNTAGAAGTTGGAAGCAGGAGT

AGCACACGTCTGAACTCTTTCCCTACACGACGCTCTTCCGATCTNNNTAGAAGTTGGAAGCAGGAGT

ACACGTCTGAACTCTTTCCCTACACGACGCTCTTCCGATCTNNNCACTGTTTTCGGACCTTCCATGCAT

ACACGTCTGAACTCTTTCCCTACACGACGCTCTTCCGATCTNNNCACTGTTTTCGGACCTTCCATGCAT

GAGCACACGTCTGAACTCTTTCCCTACACGACGCTCTTCCGATCTNNNGCGATTGGTACAGCCTACTTT

GAGCACACGTCTGAACTCTTTCCCTACACGACGCTCTTCCGATCTNNNGCGATTGGTACAGCCTACTTT

GCACACGTCTGAACTCTTTCCCTACACGACGCTCTTCCGATCTNNNTGTGCTGCTGGATTTTGTTTGAT

GCACACGTCTGAACTCTTTCCCTACACGACGCTCTTCCGATCTNNNTGTGCTGCTGGATTTTGTTTGAT

GCACACGTCTGAACTCTTTCCCTACACGACGCTCTTCCGATCTNNNTTGTATCTCCAACCTAGCTCCCC

GCACACGTCTGAACTCTTTCCCTACACGACGCTCTTCCGATCTNNNTTGTATCTCCAACCTAGCTCCCC

GCACACGTCTGAACTCTTTCCCTACACGACGCTCTTCCGATCTNNNATACATTTGTCTTAGGTTTGGA

GCACACGTCTGAACTCTTTCCCTACACGACGCTCTTCCGATCTNNNATACATTTGTCTTAGGTTTGGA

GCACACGTCTGAACTCTTTCCCTACACGACGCTCTTCCGATCTNNNCCATCATAGTTCCATGGAGGGT

GCACACGTCTGAACTCTTTCCCTACACGACGCTCTTCCGATCTNNNCCATCATAGTTCCATGGAGGGT

GAGCACACGTCTGAACTCTTTCCCTACACGACGCTCTTCCGATCTNNNGTGGCTCTTACTTGAGAATG

GAGCACACGTCTGAACTCTTTCCCTACACGACGCTCTTCCGATCTNNNGTGGCTCTTACTTGAGAATG

AGCACACGTCTGAACTCTTTCCCTACACGACGCTCTTCCGATCTNNNGTGAGTTTCACTTGCTCTGCA

AGCACACGTCTGAACTCTTTCCCTACACGACGCTCTTCCGATCTNNNGTGAGTTTCACTTGCTCTGCA

GCACACGTCTGAACTCTTTCCCTACACGACGCTCTTCCGATCTNNNCAGTTACATGGAAAAACGGCCA

GCACACGTCTGAACTCTTTCCCTACACGACGCTCTTCCGATCTNNNCAGTTACATGGAAAAACGGCCA

ACACGTCTGAACTCTTTCCCTACACGACGCTCTTCCGATCTNNNCTCATTACGTTTTAGATGGGATTCA

ACACGTCTGAACTCTTTCCCTACACGACGCTCTTCCGATCTNNNCTCATTACGTTTTAGATGGGATTCA

AGCACACGTCTGAACTCTTTCCCTACACGACGCTCTTCCGATCTNNNAAGGAATTGTGTGTGTGTGT

AGCACACGTCTGAACTCTTTCCCTACACGACGCTCTTCCGATCTNNNAAGGAATTGTGTGTGTGTGT

AGCACACGTCTGAACTCTTTCCCTACACGACGCTCTTCCGATCTNNNCTGTAATCCCAGCTACTCAGG

GAGCACACGTCTGAACTCTTTCCCTACACGACGCTCTTCCGATCTNNNCTGTAATCCCAGCTACTCAGGA
GAGCACACGTCTGAACTCTTTCCCTACACGACGCTCTTCCGATCTNNNGAGGAAAGAGAAACCCACCCC
GAGCACACGTCTGAACTCTTTCCCTACACGACGCTCTTCCGATCTNNNGAGGAAAGAGAAACCCACCCC
AGCACACGTCTGAACTCTTTCCCTACACGACGCTCTTCCGATCTNNNGTGCTGTATTCAGGAGACCCAT
AGCACACGTCTGAACTCTTTCCCTACACGACGCTCTTCCGATCTNNNGTGCTGTATTCAGGAGACCCAT
AGCACACGTCTGAACTCTTTCCCTACACGACGCTCTTCCGATCTNNNCCCATTTCATGTGCACAAAGCTT
AGCACACGTCTGAACTCTTTCCCTACACGACGCTCTTCCGATCTNNNCCCATTTCATGTGCACAAAGCTT
AGCACACGTCTGAACTCTTTCCCTACACGACGCTCTTCCGATCTNNNCTGAGGCAGGAGAATCACTTGA
AGCACACGTCTGAACTCTTTCCCTACACGACGCTCTTCCGATCTNNNCTGAGGCAGGAGAATCACTTGA
GAGCACACGTCTGAACTCTTTCCCTACACGACGCTCTTCCGATCTNNNTAATCCCAGATACTCAGGAGG
GAGCACACGTCTGAACTCTTTCCCTACACGACGCTCTTCCGATCTNNNTAATCCCAGATACTCAGGAGG
GAGCACACGTCTGAACTCTTTCCCTACACGACGCTCTTCCGATCTNNNCAAGATGCCAAGTTTGTCTCTG
GAGCACACGTCTGAACTCTTTCCCTACACGACGCTCTTCCGATCTNNNCAAGATGCCAAGTTTGTCTCTG
GCACACGTCTGAACTCTTTCCCTACACGACGCTCTTCCGATCTNNNGTTGTAGATTTTGGTCTTGGCCCC
GCACACGTCTGAACTCTTTCCCTACACGACGCTCTTCCGATCTNNNGTTGTAGATTTTGGTCTTGGCCCC
AGCACACGTCTGAACTCTTTCCCTACACGACGCTCTTCCGATCTNNNCCAGGAGTTTGAGGTTACAGT
AGCACACGTCTGAACTCTTTCCCTACACGACGCTCTTCCGATCTNNNCCAGGAGTTTGAGGTTACAGT
AGAGCACACGTCTGAACTCTTTCCCTACACGACGCTCTTCCGATCTNNNGAGGCTGAGGCAGGAGAAT
AGAGCACACGTCTGAACTCTTTCCCTACACGACGCTCTTCCGATCTNNNGAGGCTGAGGCAGGAGAAT
GAGCACACGTCTGAACTCTTTCCCTACACGACGCTCTTCCGATCTNNNTGTTTTCTGATCGGGGAAGTC
GAGCACACGTCTGAACTCTTTCCCTACACGACGCTCTTCCGATCTNNNTGTTTTCTGATCGGGGAAGTC
GCACACGTCTGAACTCTTTCCCTACACGACGCTCTTCCGATCTNNNCCCCTCAAACCATGCAATTTGTCCA
GCACACGTCTGAACTCTTTCCCTACACGACGCTCTTCCGATCTNNNCCCCTCAAACCATGCAATTTGTCCA
CACACGTCTGAACTCTTTCCCTACACGACGCTCTTCCGATCTNNNCACAACGACCACATCCTCATAGACA
CACACGTCTGAACTCTTTCCCTACACGACGCTCTTCCGATCTNNNCACAACGACCACATCCTCATAGACA
GAGCACACGTCTGAACTCTTTCCCTACACGACGCTCTTCCGATCTNNNCAGTGTGAAGAGGTCATGGA
GAGCACACGTCTGAACTCTTTCCCTACACGACGCTCTTCCGATCTNNNCAGTGTGAAGAGGTCATGGA
AGCACACGTCTGAACTCTTTCCCTACACGACGCTCTTCCGATCTNNNAAAGAGCAGAAGGTGACCTTA
AGCACACGTCTGAACTCTTTCCCTACACGACGCTCTTCCGATCTNNNAAAGAGCAGAAGGTGACCTTA
GAGCACACGTCTGAACTCTTTCCCTACACGACGCTCTTCCGATCTNNNTCCCCCTGTAAAGTCACTAGTG
GAGCACACGTCTGAACTCTTTCCCTACACGACGCTCTTCCGATCTNNNTCCCCCTGTAAAGTCACTAGTG
AGCACACGTCTGAACTCTTTCCCTACACGACGCTCTTCCGATCTNNNTGAGGGATACACTGCAGATCA
AGCACACGTCTGAACTCTTTCCCTACACGACGCTCTTCCGATCTNNNTGAGGGATACACTGCAGATCA
GAGCACACGTCTGAACTCTTTCCCTACACGACGCTCTTCCGATCTNNNAAAAAATTAGCTGGGCATGGT
GAGCACACGTCTGAACTCTTTCCCTACACGACGCTCTTCCGATCTNNNAAAAAATTAGCTGGGCATGGT
GAGCACACGTCTGAACTCTTTCCCTACACGACGCTCTTCCGATCTNNNAACTCCTGCTTCAACAAGCCTT
GAGCACACGTCTGAACTCTTTCCCTACACGACGCTCTTCCGATCTNNNAACTCCTGCTTCAACAAGCCTT
AGCACACGTCTGAACTCTTTCCCTACACGACGCTCTTCCGATCTNNNGCCTGTAGTCCCAGCTACTTGCA
AGCACACGTCTGAACTCTTTCCCTACACGACGCTCTTCCGATCTNNNGCCTGTAGTCCCAGCTACTTGCA
AGAGCACACGTCTGAACTCTTTCCCTACACGACGCTCTTCCGATCTNNNAGAAGTTCCAGTGAGCCAAG
AGAGCACACGTCTGAACTCTTTCCCTACACGACGCTCTTCCGATCTNNNAGAAGTTCCAGTGAGCCAAG
GAGCACACGTCTGAACTCTTTCCCTACACGACGCTCTTCCGATCTNNNGACCCTAGTTCAAGGACCAGA
GAGCACACGTCTGAACTCTTTCCCTACACGACGCTCTTCCGATCTNNNGACCCTAGTTCAAGGACCAGA

GAGCACACGTCTGAACTCTTTCCCTACACGACGCTCTTCCGATCTNNNGCCATAGAGCTTGGATCCCTA
GAGCACACGTCTGAACTCTTTCCCTACACGACGCTCTTCCGATCTNNNGCCATAGAGCTTGGATCCCTA
AGCACACGTCTGAACTCTTTCCCTACACGACGCTCTTCCGATCTNNNTTTGACCCATGATGAGACGGAAT
AGCACACGTCTGAACTCTTTCCCTACACGACGCTCTTCCGATCTNNNTTTGACCCATGATGAGACGGAAT
AGCACACGTCTGAACTCTTTCCCTACACGACGCTCTTCCGATCTNNNCTTCAGCTTCTTTCATTGACCCGC
AGCACACGTCTGAACTCTTTCCCTACACGACGCTCTTCCGATCTNNNCTTCAGCTTCTTTCATTGACCCGC
AGCACACGTCTGAACTCTTTCCCTACACGACGCTCTTCCGATCTNNNCTGGGCATGGTCTTCAAAATCA
AGCACACGTCTGAACTCTTTCCCTACACGACGCTCTTCCGATCTNNNCTGGGCATGGTCTTCAAAATCA
AGCACACGTCTGAACTCTTTCCCTACACGACGCTCTTCCGATCTNNNGGATCACTTGAGCCCAGGAATT
AGCACACGTCTGAACTCTTTCCCTACACGACGCTCTTCCGATCTNNNGGATCACTTGAGCCCAGGAATT
AGCACACGTCTGAACTCTTTCCCTACACGACGCTCTTCCGATCTNNNCCAGGTTCATGCCATTCTCCTCA
AGCACACGTCTGAACTCTTTCCCTACACGACGCTCTTCCGATCTNNNCCAGGTTCATGCCATTCTCCTCA
AGCACACGTCTGAACTCTTTCCCTACACGACGCTCTTCCGATCTNNNTCCTCTTACCAGATCTCTCGTGA
AGCACACGTCTGAACTCTTTCCCTACACGACGCTCTTCCGATCTNNNTCCTCTTACCAGATCTCTCGTGA
AGCACACGTCTGAACTCTTTCCCTACACGACGCTCTTCCGATCTNNNTGACCAACATAGTGAAACCCCA
AGCACACGTCTGAACTCTTTCCCTACACGACGCTCTTCCGATCTNNNTGACCAACATAGTGAAACCCCA
AGCACACGTCTGAACTCTTTCCCTACACGACGCTCTTCCGATCTNNNCCACTGCTCTCTACCATTCTACC
AGCACACGTCTGAACTCTTTCCCTACACGACGCTCTTCCGATCTNNNCCACTGCTCTCTACCATTCTACC
AGCACACGTCTGAACTCTTTCCCTACACGACGCTCTTCCGATCTNNNTTCCATGCTTTGGTTCTTCTCCTC
AGCACACGTCTGAACTCTTTCCCTACACGACGCTCTTCCGATCTNNNTTCCATGCTTTGGTTCTTCTCCTC
GCACACGTCTGAACTCTTTCCCTACACGACGCTCTTCCGATCTNNNGGTCCATTCTGCAATCTGTGATG
GCACACGTCTGAACTCTTTCCCTACACGACGCTCTTCCGATCTNNNGGTCCATTCTGCAATCTGTGATG
GAGCACACGTCTGAACTCTTTCCCTACACGACGCTCTTCCGATCTNNNTCCTAAGCCCAGGAATTTGAG
GAGCACACGTCTGAACTCTTTCCCTACACGACGCTCTTCCGATCTNNNTCCTAAGCCCAGGAATTTGAG
GAGCACACGTCTGAACTCTTTCCCTACACGACGCTCTTCCGATCTNNNAATTTATCATTTTATCCTGCCA
GAGCACACGTCTGAACTCTTTCCCTACACGACGCTCTTCCGATCTNNNAATTTATCATTTTATCCTGCCA
GAGCACACGTCTGAACTCTTTCCCTACACGACGCTCTTCCGATCTNNNAAGGCAGATAGTGTTGAGTC
GAGCACACGTCTGAACTCTTTCCCTACACGACGCTCTTCCGATCTNNNAAGGCAGATAGTGTTGAGTC
AGCACACGTCTGAACTCTTTCCCTACACGACGCTCTTCCGATCTNNNGTAATCCCAGCTACTAAGGAGG
AGCACACGTCTGAACTCTTTCCCTACACGACGCTCTTCCGATCTNNNGTAATCCCAGCTACTAAGGAGG
GAGCACACGTCTGAACTCTTTCCCTACACGACGCTCTTCCGATCTNNNGGATCACTTGAGTCTGGGAA
GAGCACACGTCTGAACTCTTTCCCTACACGACGCTCTTCCGATCTNNNGGATCACTTGAGTCTGGGAA
GAGCACACGTCTGAACTCTTTCCCTACACGACGCTCTTCCGATCTNNNTTCGTATGGGCAGACAAGGAA
GAGCACACGTCTGAACTCTTTCCCTACACGACGCTCTTCCGATCTNNNTTCGTATGGGCAGACAAGGAA
GAGCACACGTCTGAACTCTTTCCCTACACGACGCTCTTCCGATCTNNNGGAAACACACAACTTGCCAAGA
GAGCACACGTCTGAACTCTTTCCCTACACGACGCTCTTCCGATCTNNNGGAAACACACAACTTGCCAAGA
AGCACACGTCTGAACTCTTTCCCTACACGACGCTCTTCCGATCTNNNATGCCCTATTTGCTCAAGTCTGA
AGCACACGTCTGAACTCTTTCCCTACACGACGCTCTTCCGATCTNNNATGCCCTATTTGCTCAAGTCTGA
GAGCACACGTCTGAACTCTTTCCCTACACGACGCTCTTCCGATCTNNNCAAAGTGCTGGGATTACAGG
GAGCACACGTCTGAACTCTTTCCCTACACGACGCTCTTCCGATCTNNNCAAAGTGCTGGGATTACAGG
GAGCACACGTCTGAACTCTTTCCCTACACGACGCTCTTCCGATCTNNNACAAAGTACTGACATGTGGCT
GAGCACACGTCTGAACTCTTTCCCTACACGACGCTCTTCCGATCTNNNACAAAGTACTGACATGTGGCT
GCACACGTCTGAACTCTTTCCCTACACGACGCTCTTCCGATCTNNNACAGGCTCACACTACTACTCTCTCA

GCACACGTCTGAACTCTTTCCCTACACGACGCTCTTCCGATCTNNNACAGGCTCACACTACTACTCTCA
GAGCACACGTCTGAACTCTTTCCCTACACGACGCTCTTCCGATCTNNNGCTTCTCCGTGTGATATCAGGA
GAGCACACGTCTGAACTCTTTCCCTACACGACGCTCTTCCGATCTNNNGCTTCTCCGTGTGATATCAGGA
GAGCACACGTCTGAACTCTTTCCCTACACGACGCTCTTCCGATCTNNNGTTGCAGTGACCCAAGATCAT
GAGCACACGTCTGAACTCTTTCCCTACACGACGCTCTTCCGATCTNNNGTTGCAGTGACCCAAGATCAT
GAGCACACGTCTGAACTCTTTCCCTACACGACGCTCTTCCGATCTNNNGATCACACTATTGCACTTGAG
GAGCACACGTCTGAACTCTTTCCCTACACGACGCTCTTCCGATCTNNNGATCACACTATTGCACTTGAG
GCACACGTCTGAACTCTTTCCCTACACGACGCTCTTCCGATCTNNNGGATTACAGGTGTGAGCTACCAT
GCACACGTCTGAACTCTTTCCCTACACGACGCTCTTCCGATCTNNNGGATTACAGGTGTGAGCTACCAT
AGCACACGTCTGAACTCTTTCCCTACACGACGCTCTTCCGATCTNNNCATAAGTGCAAGTGTGTACCT
AGCACACGTCTGAACTCTTTCCCTACACGACGCTCTTCCGATCTNNNCATAAGTGCAAGTGTGTACCT
AGCACACGTCTGAACTCTTTCCCTACACGACGCTCTTCCGATCTNNNTCACGATGACAGTGAAGACCA
AGCACACGTCTGAACTCTTTCCCTACACGACGCTCTTCCGATCTNNNTCACGATGACAGTGAAGACCA
GAGCACACGTCTGAACTCTTTCCCTACACGACGCTCTTCCGATCTNNNTACAAAAAATTAGCCGGGCAT
GAGCACACGTCTGAACTCTTTCCCTACACGACGCTCTTCCGATCTNNNTACAAAAAATTAGCCGGGCAT
CACACGTCTGAACTCTTTCCCTACACGACGCTCTTCCGATCTNNNCCACGTTTAGAAGCGAGGGAGCAT
CACACGTCTGAACTCTTTCCCTACACGACGCTCTTCCGATCTNNNCCACGTTTAGAAGCGAGGGAGCAT
GAGCACACGTCTGAACTCTTTCCCTACACGACGCTCTTCCGATCTNNNTGTCTAGTTCCAGACCCAGAGA
GAGCACACGTCTGAACTCTTTCCCTACACGACGCTCTTCCGATCTNNNTGTCTAGTTCCAGACCCAGAGA
GAGCACACGTCTGAACTCTTTCCCTACACGACGCTCTTCCGATCTNNNGATGTGGCACAGTTCATTCA
GAGCACACGTCTGAACTCTTTCCCTACACGACGCTCTTCCGATCTNNNGATGTGGCACAGTTCATTCA
AGCACACGTCTGAACTCTTTCCCTACACGACGCTCTTCCGATCTNNNATCCCACCAGCAATGTATGTAG
AGCACACGTCTGAACTCTTTCCCTACACGACGCTCTTCCGATCTNNNATCCCACCAGCAATGTATGTAG
GAGCACACGTCTGAACTCTTTCCCTACACGACGCTCTTCCGATCTNNNGGGTAGGAAAAGGGTAGGA
GAGCACACGTCTGAACTCTTTCCCTACACGACGCTCTTCCGATCTNNNGGGTAGGAAAAGGGTAGGA
AGAGCACACGTCTGAACTCTTTCCCTACACGACGCTCTTCCGATCTNNNCTCAGGAGGCTGTAGCAGA
AGAGCACACGTCTGAACTCTTTCCCTACACGACGCTCTTCCGATCTNNNCTCAGGAGGCTGTAGCAGA
AGCACACGTCTGAACTCTTTCCCTACACGACGCTCTTCCGATCTNNNGGACAAGAACTTGGGGAGACA
AGCACACGTCTGAACTCTTTCCCTACACGACGCTCTTCCGATCTNNNGGACAAGAACTTGGGGAGACA
GCACACGTCTGAACTCTTTCCCTACACGACGCTCTTCCGATCTNNNCTGAGGAGGAGGGGAGAGCGCA
GCACACGTCTGAACTCTTTCCCTACACGACGCTCTTCCGATCTNNNCTGAGGAGGAGGGGAGAGCGCA
AGCACACGTCTGAACTCTTTCCCTACACGACGCTCTTCCGATCTNNNAAGTTTACACCATTGCACTCCAT
AGCACACGTCTGAACTCTTTCCCTACACGACGCTCTTCCGATCTNNNAAGTTTACACCATTGCACTCCAT
AGCACACGTCTGAACTCTTTCCCTACACGACGCTCTTCCGATCTNNNATGCCAGGAGAAAGTGTAAGT
AGCACACGTCTGAACTCTTTCCCTACACGACGCTCTTCCGATCTNNNATGCCAGGAGAAAGTGTAAGT
GAGCACACGTCTGAACTCTTTCCCTACACGACGCTCTTCCGATCTNNNAGATGGAGTTTCGCTATTGTT
GAGCACACGTCTGAACTCTTTCCCTACACGACGCTCTTCCGATCTNNNAGATGGAGTTTCGCTATTGTT
AGCACACGTCTGAACTCTTTCCCTACACGACGCTCTTCCGATCTNNNGCTGTTTTTCCTTCTCCTTTCCCA
AGCACACGTCTGAACTCTTTCCCTACACGACGCTCTTCCGATCTNNNGCTGTTTTTCCTTCTCCTTTCCCA
AGCACACGTCTGAACTCTTTCCCTACACGACGCTCTTCCGATCTNNNGACGGTGTCTCTTCAGCTTTCTA
AGCACACGTCTGAACTCTTTCCCTACACGACGCTCTTCCGATCTNNNGACGGTGTCTCTTCAGCTTTCTA
GAGCACACGTCTGAACTCTTTCCCTACACGACGCTCTTCCGATCTNNNTGAGGCAGGAGAATCACTTGT
GAGCACACGTCTGAACTCTTTCCCTACACGACGCTCTTCCGATCTNNNTGAGGCAGGAGAATCACTTGT

GAGCACACGTCTGAACTCTTTCCCTACACGACGCTCTTCCGATCTNNNGGAAGTATTAGGCTGCCTCT
GAGCACACGTCTGAACTCTTTCCCTACACGACGCTCTTCCGATCTNNNGGAAGTATTAGGCTGCCTCT
AGCACACGTCTGAACTCTTTCCCTACACGACGCTCTTCCGATCTNNNCACGTTGTGCACATGTAACCT
AGCACACGTCTGAACTCTTTCCCTACACGACGCTCTTCCGATCTNNNCACGTTGTGCACATGTAACCT
CACACGTCTGAACTCTTTCCCTACACGACGCTCTTCCGATCTNNNGCAGAAGCTAAAACTCGGTGTAC
CACACGTCTGAACTCTTTCCCTACACGACGCTCTTCCGATCTNNNGCAGAAGCTAAAACTCGGTGTAC
ACACGTCTGAACTCTTTCCCTACACGACGCTCTTCCGATCTNNNCCCTTCCCTACAGCTCCTCTCATCGG
ACACGTCTGAACTCTTTCCCTACACGACGCTCTTCCGATCTNNNCCCTTCCCTACAGCTCCTCTCATCGG
GAGCACACGTCTGAACTCTTTCCCTACACGACGCTCTTCCGATCTNNNTGTAAATGACGTGTTGGTTCT
GAGCACACGTCTGAACTCTTTCCCTACACGACGCTCTTCCGATCTNNNTGTAAATGACGTGTTGGTTCT
GCACACGTCTGAACTCTTTCCCTACACGACGCTCTTCCGATCTNNNCAAGAGGAAGTCTCCACACACA
GCACACGTCTGAACTCTTTCCCTACACGACGCTCTTCCGATCTNNNCAAGAGGAAGTCTCCACACACA
AGCACACGTCTGAACTCTTTCCCTACACGACGCTCTTCCGATCTNNNAATCCATCTCACTGGGGGAAA
AGCACACGTCTGAACTCTTTCCCTACACGACGCTCTTCCGATCTNNNAATCCATCTCACTGGGGGAAA
AGCACACGTCTGAACTCTTTCCCTACACGACGCTCTTCCGATCTNNNATCACAGGAAGTCACCCTTGA
AGCACACGTCTGAACTCTTTCCCTACACGACGCTCTTCCGATCTNNNATCACAGGAAGTCACCCTTGA
GCACACGTCTGAACTCTTTCCCTACACGACGCTCTTCCGATCTNNNGGCCTTCAGGAACAGTTTCTTTT
GCACACGTCTGAACTCTTTCCCTACACGACGCTCTTCCGATCTNNNGGCCTTCAGGAACAGTTTCTTTT
AGCACACGTCTGAACTCTTTCCCTACACGACGCTCTTCCGATCTNNNCTCCTGCTGTTCCTGGTCTGCAT
AGCACACGTCTGAACTCTTTCCCTACACGACGCTCTTCCGATCTNNNCTCCTGCTGTTCCTGGTCTGCAT
AGCACACGTCTGAACTCTTTCCCTACACGACGCTCTTCCGATCTNNNTCTCCTCTCTCTCCTCTCTCT
AGCACACGTCTGAACTCTTTCCCTACACGACGCTCTTCCGATCTNNNTCTCCTCTCTCTCCTCTCTCT
GCACACGTCTGAACTCTTTCCCTACACGACGCTCTTCCGATCTNNNCTCAGGCAGGAGAATCACTTG
GCACACGTCTGAACTCTTTCCCTACACGACGCTCTTCCGATCTNNNCTCAGGCAGGAGAATCACTTG
GAGCACACGTCTGAACTCTTTCCCTACACGACGCTCTTCCGATCTNNNTAAGGACAGTCTCAATGGT
GAGCACACGTCTGAACTCTTTCCCTACACGACGCTCTTCCGATCTNNNTAAGGACAGTCTCAATGGT
AGCACACGTCTGAACTCTTTCCCTACACGACGCTCTTCCGATCTNNNCCCTAAGCCCTTCTTCAAGACA
AGCACACGTCTGAACTCTTTCCCTACACGACGCTCTTCCGATCTNNNCCCTAAGCCCTTCTTCAAGACA
AGCACACGTCTGAACTCTTTCCCTACACGACGCTCTTCCGATCTNNNTCATTCAGGAGCCATGTTCAGA
AGCACACGTCTGAACTCTTTCCCTACACGACGCTCTTCCGATCTNNNTCATTCAGGAGCCATGTTCAGA
GAGCACACGTCTGAACTCTTTCCCTACACGACGCTCTTCCGATCTNNNAAGAATTGCTTGAACCTGGG
GAGCACACGTCTGAACTCTTTCCCTACACGACGCTCTTCCGATCTNNNAAGAATTGCTTGAACCTGGG
AGCACACGTCTGAACTCTTTCCCTACACGACGCTCTTCCGATCTNNNAGAAACAAACCAACACCTATG
AGCACACGTCTGAACTCTTTCCCTACACGACGCTCTTCCGATCTNNNAGAAACAAACCAACACCTATG
GCACACGTCTGAACTCTTTCCCTACACGACGCTCTTCCGATCTNNNCTAAGAGGAGGAGCAGGGGG
GCACACGTCTGAACTCTTTCCCTACACGACGCTCTTCCGATCTNNNCTAAGAGGAGGAGCAGGGGG
GAGCACACGTCTGAACTCTTTCCCTACACGACGCTCTTCCGATCTNNNGCTCACAATTGCTAAGTGAGG
GAGCACACGTCTGAACTCTTTCCCTACACGACGCTCTTCCGATCTNNNGCTCACAATTGCTAAGTGAGG
GCACACGTCTGAACTCTTTCCCTACACGACGCTCTTCCGATCTNNNAATTGGAAATTCAGAAGTAGAA
GCACACGTCTGAACTCTTTCCCTACACGACGCTCTTCCGATCTNNNAATTGGAAATTCAGAAGTAGAA
GAGCACACGTCTGAACTCTTTCCCTACACGACGCTCTTCCGATCTNNNGTTGCAGTGAGCTGTGATCAT
GAGCACACGTCTGAACTCTTTCCCTACACGACGCTCTTCCGATCTNNNGTTGCAGTGAGCTGTGATCAT
AGCACACGTCTGAACTCTTTCCCTACACGACGCTCTTCCGATCTNNNGGCGATTTTATTCTTCAGCTTG

AGCACACGTCTGAACTCTTTCCCTACACGACGCTCTTCCGATCTNNNGGCGATTTTATTCTTCAGCTTG
AGCACACGTCTGAACTCTTTCCCTACACGACGCTCTTCCGATCTNNNCTGTGCTGCTTCTTTGAAGTTG
AGCACACGTCTGAACTCTTTCCCTACACGACGCTCTTCCGATCTNNNCTGTGCTGCTTCTTTGAAGTTG
AGCACACGTCTGAACTCTTTCCCTACACGACGCTCTTCCGATCTNNNCAAGAGCAAAACACCGTCTGA
AGCACACGTCTGAACTCTTTCCCTACACGACGCTCTTCCGATCTNNNCAAGAGCAAAACACCGTCTGA
AGAGCACACGTCTGAACTCTTTCCCTACACGACGCTCTTCCGATCTNNNTGAGGCAGGAGAATCACTT
AGAGCACACGTCTGAACTCTTTCCCTACACGACGCTCTTCCGATCTNNNTGAGGCAGGAGAATCACTT
GCACACGTCTGAACTCTTTCCCTACACGACGCTCTTCCGATCTNNNAGGATGATGCTGGCCTCATAAA
GCACACGTCTGAACTCTTTCCCTACACGACGCTCTTCCGATCTNNNAGGATGATGCTGGCCTCATAAA
GCACACGTCTGAACTCTTTCCCTACACGACGCTCTTCCGATCTNNNTTCAGCAGAGAAGTTCTACCAG
GCACACGTCTGAACTCTTTCCCTACACGACGCTCTTCCGATCTNNNTTCAGCAGAGAAGTTCTACCAG
AGCACACGTCTGAACTCTTTCCCTACACGACGCTCTTCCGATCTNNNAAAGTGCTGGAATTACATGCG
AGCACACGTCTGAACTCTTTCCCTACACGACGCTCTTCCGATCTNNNAAAGTGCTGGAATTACATGCG
GCACACGTCTGAACTCTTTCCCTACACGACGCTCTTCCGATCTNNNGTTCTGTGTGTCTGTATGTGTG
GCACACGTCTGAACTCTTTCCCTACACGACGCTCTTCCGATCTNNNGTTCTGTGTGTCTGTATGTGTG
GAGCACACGTCTGAACTCTTTCCCTACACGACGCTCTTCCGATCTNNNTCCTAACAGAACATCAAGAGG
GAGCACACGTCTGAACTCTTTCCCTACACGACGCTCTTCCGATCTNNNTCCTAACAGAACATCAAGAGG
ACACGTCTGAACTCTTTCCCTACACGACGCTCTTCCGATCTNNNTGACTTTGTGGTTACTGATGTCAC
ACACGTCTGAACTCTTTCCCTACACGACGCTCTTCCGATCTNNNTGACTTTGTGGTTACTGATGTCAC
AGCACACGTCTGAACTCTTTCCCTACACGACGCTCTTCCGATCTNNNAGCTTCTGGGTACACATGAGT
AGCACACGTCTGAACTCTTTCCCTACACGACGCTCTTCCGATCTNNNAGCTTCTGGGTACACATGAGT
GCACACGTCTGAACTCTTTCCCTACACGACGCTCTTCCGATCTNNNCAGTGTGTCTTGGCAGTGTTTA
GCACACGTCTGAACTCTTTCCCTACACGACGCTCTTCCGATCTNNNCAGTGTGTCTTGGCAGTGTTTA
AGAGCACACGTCTGAACTCTTTCCCTACACGACGCTCTTCCGATCTNNNACGTGTTTATCTGCTGACCT
AGAGCACACGTCTGAACTCTTTCCCTACACGACGCTCTTCCGATCTNNNACGTGTTTATCTGCTGACCT
GCACACGTCTGAACTCTTTCCCTACACGACGCTCTTCCGATCTNNNCTGTAGTCCCAGCTACTCAGGAG
GCACACGTCTGAACTCTTTCCCTACACGACGCTCTTCCGATCTNNNCTGTAGTCCCAGCTACTCAGGAG
AGCACACGTCTGAACTCTTTCCCTACACGACGCTCTTCCGATCTNNNGGGTCTGGATTGAGACTCCTT
AGCACACGTCTGAACTCTTTCCCTACACGACGCTCTTCCGATCTNNNGGGTCTGGATTGAGACTCCTT
GCACACGTCTGAACTCTTTCCCTACACGACGCTCTTCCGATCTNNNTCACTCGAATCTACACGGGATG
GCACACGTCTGAACTCTTTCCCTACACGACGCTCTTCCGATCTNNNTCACTCGAATCTACACGGGATG
AGCACACGTCTGAACTCTTTCCCTACACGACGCTCTTCCGATCTNNNTCCTTCCTTCCTTCCTCTCTTTCC
AGCACACGTCTGAACTCTTTCCCTACACGACGCTCTTCCGATCTNNNTCCTTCCTTCCTTCCTCTCTTTCC
AGCACACGTCTGAACTCTTTCCCTACACGACGCTCTTCCGATCTNNNAAAGCCAACAAGCAACATGTC
AGCACACGTCTGAACTCTTTCCCTACACGACGCTCTTCCGATCTNNNAAAGCCAACAAGCAACATGTC
GCACACGTCTGAACTCTTTCCCTACACGACGCTCTTCCGATCTNNNTACCTTTGCCCATAAGCATCTGT
GCACACGTCTGAACTCTTTCCCTACACGACGCTCTTCCGATCTNNNTACCTTTGCCCATAAGCATCTGT
GAGCACACGTCTGAACTCTTTCCCTACACGACGCTCTTCCGATCTNNNGGGTCTGTAATCCCACCACTTT
GAGCACACGTCTGAACTCTTTCCCTACACGACGCTCTTCCGATCTNNNGGGTCTGTAATCCCACCACTTT
GAGCACACGTCTGAACTCTTTCCCTACACGACGCTCTTCCGATCTNNNGCAGACTTGCCACAAACTTTC
GAGCACACGTCTGAACTCTTTCCCTACACGACGCTCTTCCGATCTNNNGCAGACTTGCCACAAACTTTC
AGCACACGTCTGAACTCTTTCCCTACACGACGCTCTTCCGATCTNNNAGCATGAGATTCGTAGTGTTCC
AGCACACGTCTGAACTCTTTCCCTACACGACGCTCTTCCGATCTNNNAGCATGAGATTCGTAGTGTTCC

AGCACACGTCTGAACTCTTTCCCTACACGACGCTCTTCCGATCTNNNTTTGTTCGGCATCATCTTGCTT

AGCACACGTCTGAACTCTTTCCCTACACGACGCTCTTCCGATCTNNNTTTGTTCGGCATCATCTTGCTT

AGCACACGTCTGAACTCTTTCCCTACACGACGCTCTTCCGATCTNNNCACAGGCAGATGCATCTTTTCTT

AGCACACGTCTGAACTCTTTCCCTACACGACGCTCTTCCGATCTNNNCACAGGCAGATGCATCTTTTCTT

AGAGCACACGTCTGAACTCTTTCCCTACACGACGCTCTTCCGATCTNNNGGTACCATTTTCTTCAAATCT

AGAGCACACGTCTGAACTCTTTCCCTACACGACGCTCTTCCGATCTNNNGGTACCATTTTCTTCAAATCT

GAGCACACGTCTGAACTCTTTCCCTACACGACGCTCTTCCGATCTNNNGTGCCTGTAATCCCAGCTACTT

GAGCACACGTCTGAACTCTTTCCCTACACGACGCTCTTCCGATCTNNNGTGCCTGTAATCCCAGCTACTT

GCACACGTCTGAACTCTTTCCCTACACGACGCTCTTCCGATCTNNNGAGAATGGTGTGAACCCAGGAG

GCACACGTCTGAACTCTTTCCCTACACGACGCTCTTCCGATCTNNNGAGAATGGTGTGAACCCAGGAG

GAGCACACGTCTGAACTCTTTCCCTACACGACGCTCTTCCGATCTNNNACCAGCGATGTTCTTCACAAAA

GAGCACACGTCTGAACTCTTTCCCTACACGACGCTCTTCCGATCTNNNACCAGCGATGTTCTTCACAAAA

AGCACACGTCTGAACTCTTTCCCTACACGACGCTCTTCCGATCTNNNGGAGAGTCAATTTGTATGCAGC

AGCACACGTCTGAACTCTTTCCCTACACGACGCTCTTCCGATCTNNNGGAGAGTCAATTTGTATGCAGC

CACACGTCTGAACTCTTTCCCTACACGACGCTCTTCCGATCTNNNGTCTTGTGGTGGTGGGGATCCATC

CACACGTCTGAACTCTTTCCCTACACGACGCTCTTCCGATCTNNNGTCTTGTGGTGGTGGGGATCCATC

AGAGCACACGTCTGAACTCTTTCCCTACACGACGCTCTTCCGATCTNNNGATAGAGTTCCTGGCTAGGC

AGAGCACACGTCTGAACTCTTTCCCTACACGACGCTCTTCCGATCTNNNGATAGAGTTCCTGGCTAGGC

AGCACACGTCTGAACTCTTTCCCTACACGACGCTCTTCCGATCTNNNCTATTGCACACTTAGTATGTTAC

AGCACACGTCTGAACTCTTTCCCTACACGACGCTCTTCCGATCTNNNCTATTGCACACTTAGTATGTTAC

GAGCACACGTCTGAACTCTTTCCCTACACGACGCTCTTCCGATCTNNNCTTCTAGCAACAGCAACAGTAC

GAGCACACGTCTGAACTCTTTCCCTACACGACGCTCTTCCGATCTNNNCTTCTAGCAACAGCAACAGTAC

GAGCACACGTCTGAACTCTTTCCCTACACGACGCTCTTCCGATCTNNNCTCTTGTGTGTGTTGCTGTGT

GAGCACACGTCTGAACTCTTTCCCTACACGACGCTCTTCCGATCTNNNCTCTTGTGTGTGTTGCTGTGT

GAGCACACGTCTGAACTCTTTCCCTACACGACGCTCTTCCGATCTNNNGAGCAAGACTCCGTCTCAAGA

GAGCACACGTCTGAACTCTTTCCCTACACGACGCTCTTCCGATCTNNNGAGCAAGACTCCGTCTCAAGA

GAGCACACGTCTGAACTCTTTCCCTACACGACGCTCTTCCGATCTNNNGGCTGTCCATTGTCTTCAGAG

GAGCACACGTCTGAACTCTTTCCCTACACGACGCTCTTCCGATCTNNNGGCTGTCCATTGTCTTCAGAG

GAGCACACGTCTGAACTCTTTCCCTACACGACGCTCTTCCGATCTNNNAAAAAATTAGCTGGGTGTGG

GAGCACACGTCTGAACTCTTTCCCTACACGACGCTCTTCCGATCTNNNAAAAAATTAGCTGGGTGTGG

AGCACACGTCTGAACTCTTTCCCTACACGACGCTCTTCCGATCTNNNCCAACATGCCTGGCTAATTTTTT

AGCACACGTCTGAACTCTTTCCCTACACGACGCTCTTCCGATCTNNNCCAACATGCCTGGCTAATTTTTT

CACACGTCTGAACTCTTTCCCTACACGACGCTCTTCCGATCTNNNCATCTGAGAAGTGAGGAGCCCCAT

CACACGTCTGAACTCTTTCCCTACACGACGCTCTTCCGATCTNNNCATCTGAGAAGTGAGGAGCCCCAT

GAGCACACGTCTGAACTCTTTCCCTACACGACGCTCTTCCGATCTNNNTATTTGACCTGTCCATCTCTGC

GAGCACACGTCTGAACTCTTTCCCTACACGACGCTCTTCCGATCTNNNTATTTGACCTGTCCATCTCTGC

GAGCACACGTCTGAACTCTTTCCCTACACGACGCTCTTCCGATCTNNNGTAATCCCAGCTACTTAGGAG

GAGCACACGTCTGAACTCTTTCCCTACACGACGCTCTTCCGATCTNNNGTAATCCCAGCTACTTAGGAG

GAGCACACGTCTGAACTCTTTCCCTACACGACGCTCTTCCGATCTNNNGGATTGACTGAGCTGATCTGC

GAGCACACGTCTGAACTCTTTCCCTACACGACGCTCTTCCGATCTNNNGGATTGACTGAGCTGATCTGC

GAGCACACGTCTGAACTCTTTCCCTACACGACGCTCTTCCGATCTNNNTTAGTCAGGGTTCTCTAGACG

GAGCACACGTCTGAACTCTTTCCCTACACGACGCTCTTCCGATCTNNNTTAGTCAGGGTTCTCTAGACG

GAGCACACGTCTGAACTCTTTCCCTACACGACGCTCTTCCGATCTNNNATCTCTCCTACATTGGCAAGCA

AGCACACGTCTGAACTCTTTCCCTACACGACGCTCTTCCGATCTNNNATCTCTCCTACATTGGCAAGCA
AGCACACGTCTGAACTCTTTCCCTACACGACGCTCTTCCGATCTNNNCACTTGAGGTCAGGAGTTCA
AGCACACGTCTGAACTCTTTCCCTACACGACGCTCTTCCGATCTNNNCACTTGAGGTCAGGAGTTCA
GCACACGTCTGAACTCTTTCCCTACACGACGCTCTTCCGATCTNNNTCTAACTCTGCCTCCATCCTTTGC
GCACACGTCTGAACTCTTTCCCTACACGACGCTCTTCCGATCTNNNTCTAACTCTGCCTCCATCCTTTGC
GCACACGTCTGAACTCTTTCCCTACACGACGCTCTTCCGATCTNNNCTAAGAGGAGGAGCAGGGGG
GCACACGTCTGAACTCTTTCCCTACACGACGCTCTTCCGATCTNNNCTAAGAGGAGGAGCAGGGGG
GCACACGTCTGAACTCTTTCCCTACACGACGCTCTTCCGATCTNNNATCCCAACTATTTCAGAGGCTGG
GCACACGTCTGAACTCTTTCCCTACACGACGCTCTTCCGATCTNNNATCCCAACTATTTCAGAGGCTGG
GCACACGTCTGAACTCTTTCCCTACACGACGCTCTTCCGATCTNNNCCTCATAAAGAATGCCAGCCTGT
GCACACGTCTGAACTCTTTCCCTACACGACGCTCTTCCGATCTNNNCCTCATAAAGAATGCCAGCCTGT
GCACACGTCTGAACTCTTTCCCTACACGACGCTCTTCCGATCTNNNCTTTGCTTTGCTTTGCTTTGCTT
GCACACGTCTGAACTCTTTCCCTACACGACGCTCTTCCGATCTNNNCTTTGCTTTGCTTTGCTTTGCTT
AGCACACGTCTGAACTCTTTCCCTACACGACGCTCTTCCGATCTNNNCTTTCCGGTTCACACCATTCTCC
AGCACACGTCTGAACTCTTTCCCTACACGACGCTCTTCCGATCTNNNCTTTCCGGTTCACACCATTCTCC
GCACACGTCTGAACTCTTTCCCTACACGACGCTCTTCCGATCTNNNGGACGTTTCAGTGGAATTTTGG
GCACACGTCTGAACTCTTTCCCTACACGACGCTCTTCCGATCTNNNGGACGTTTCAGTGGAATTTTGG
CACACGTCTGAACTCTTTCCCTACACGACGCTCTTCCGATCTNNNGTCAGGTTGCTGTATCTGGGGCA
CACACGTCTGAACTCTTTCCCTACACGACGCTCTTCCGATCTNNNGTCAGGTTGCTGTATCTGGGGCA
AGCACACGTCTGAACTCTTTCCCTACACGACGCTCTTCCGATCTNNNTGAGCTGTTTCCAAGTTTTGT
AGCACACGTCTGAACTCTTTCCCTACACGACGCTCTTCCGATCTNNNTGAGCTGTTTCCAAGTTTTGT
GAGCACACGTCTGAACTCTTTCCCTACACGACGCTCTTCCGATCTNNNAGGAGAAAGTGGAGATGG
GAGCACACGTCTGAACTCTTTCCCTACACGACGCTCTTCCGATCTNNNAGGAGAAAGTGGAGATGG
GAGCACACGTCTGAACTCTTTCCCTACACGACGCTCTTCCGATCTNNNGATGAGGCAGGAGAATCAC
GAGCACACGTCTGAACTCTTTCCCTACACGACGCTCTTCCGATCTNNNGATGAGGCAGGAGAATCAC
GCACACGTCTGAACTCTTTCCCTACACGACGCTCTTCCGATCTNNNTATGAGAGTTCAGGTTGCTCCA
GCACACGTCTGAACTCTTTCCCTACACGACGCTCTTCCGATCTNNNTATGAGAGTTCAGGTTGCTCCA
AGCACACGTCTGAACTCTTTCCCTACACGACGCTCTTCCGATCTNNNAAGTGCAAGAAACACCACATT
AGCACACGTCTGAACTCTTTCCCTACACGACGCTCTTCCGATCTNNNAAGTGCAAGAAACACCACATT
AGCACACGTCTGAACTCTTTCCCTACACGACGCTCTTCCGATCTNNNAAGATTGTGCCACTTCACTCC
AGCACACGTCTGAACTCTTTCCCTACACGACGCTCTTCCGATCTNNNAAGATTGTGCCACTTCACTCC
AGCACACGTCTGAACTCTTTCCCTACACGACGCTCTTCCGATCTNNNGCTTCCTGGTCATTCTCTCTCTT
AGCACACGTCTGAACTCTTTCCCTACACGACGCTCTTCCGATCTNNNGCTTCCTGGTCATTCTCTCTCTT
AGCACACGTCTGAACTCTTTCCCTACACGACGCTCTTCCGATCTNNNCTGGAGGTCCTGCAGGTAATA
AGCACACGTCTGAACTCTTTCCCTACACGACGCTCTTCCGATCTNNNCTGGAGGTCCTGCAGGTAATA
AGCACACGTCTGAACTCTTTCCCTACACGACGCTCTTCCGATCTNNNTAGGAACTTGTCTTTCGGGGA
AGCACACGTCTGAACTCTTTCCCTACACGACGCTCTTCCGATCTNNNTAGGAACTTGTCTTTCGGGGA
AGCACACGTCTGAACTCTTTCCCTACACGACGCTCTTCCGATCTNNNAAAAAACTCTGGCACCATCACT
AGCACACGTCTGAACTCTTTCCCTACACGACGCTCTTCCGATCTNNNAAAAAACTCTGGCACCATCACT
AGCACACGTCTGAACTCTTTCCCTACACGACGCTCTTCCGATCTNNNCTCTGCTTCCCATCTCATCCTAC
AGCACACGTCTGAACTCTTTCCCTACACGACGCTCTTCCGATCTNNNCTCTGCTTCCCATCTCATCCTAC
GCACACGTCTGAACTCTTTCCCTACACGACGCTCTTCCGATCTNNNTGGTGGATTTGAATTGTGTGTG
GCACACGTCTGAACTCTTTCCCTACACGACGCTCTTCCGATCTNNNTGGTGGATTTGAATTGTGTGTG

AGCACACGTCTGAACTCTTTCCCTACACGACGCTCTTCCGATCTNNNGAGATTGCAGTGAGCCAAGAT

AGCACACGTCTGAACTCTTTCCCTACACGACGCTCTTCCGATCTNNNGAGATTGCAGTGAGCCAAGAT

AGCACACGTCTGAACTCTTTCCCTACACGACGCTCTTCCGATCTNNNGGGTTCCTCAGAGAAGGAGAA

AGCACACGTCTGAACTCTTTCCCTACACGACGCTCTTCCGATCTNNNGGGTTCCTCAGAGAAGGAGAA

GCACACGTCTGAACTCTTTCCCTACACGACGCTCTTCCGATCTNNNGTACTTGCTTTTGTGGTTGCAT

GCACACGTCTGAACTCTTTCCCTACACGACGCTCTTCCGATCTNNNGTACTTGCTTTTGTGGTTGCAT

AGCACACGTCTGAACTCTTTCCCTACACGACGCTCTTCCGATCTNNNTGGTCTCAAACACCTAACCTCA

AGCACACGTCTGAACTCTTTCCCTACACGACGCTCTTCCGATCTNNNTGGTCTCAAACACCTAACCTCA

AGCACACGTCTGAACTCTTTCCCTACACGACGCTCTTCCGATCTNNNAGTGCCAGTACATAAACTTCA

AGCACACGTCTGAACTCTTTCCCTACACGACGCTCTTCCGATCTNNNAGTGCCAGTACATAAACTTCA

AGAGCACACGTCTGAACTCTTTCCCTACACGACGCTCTTCCGATCTNNNACCTGGCTAATGAGGTTCA

AGAGCACACGTCTGAACTCTTTCCCTACACGACGCTCTTCCGATCTNNNACCTGGCTAATGAGGTTCA

AGCACACGTCTGAACTCTTTCCCTACACGACGCTCTTCCGATCTNNNGATCACGAGTTCAGGAGATCG

AGCACACGTCTGAACTCTTTCCCTACACGACGCTCTTCCGATCTNNNGATCACGAGTTCAGGAGATCG

GAGCACACGTCTGAACTCTTTCCCTACACGACGCTCTTCCGATCTNNNGGCCATAATCTGCCTTTGTCT

GAGCACACGTCTGAACTCTTTCCCTACACGACGCTCTTCCGATCTNNNGGCCATAATCTGCCTTTGTCT

AGCACACGTCTGAACTCTTTCCCTACACGACGCTCTTCCGATCTNNNGTGTTCTGGGAAAAGCAGTTG

AGCACACGTCTGAACTCTTTCCCTACACGACGCTCTTCCGATCTNNNGTGTTCTGGGAAAAGCAGTTG

GAGCACACGTCTGAACTCTTTCCCTACACGACGCTCTTCCGATCTNNNTGTGGAAACAAAACCACCCTT

GAGCACACGTCTGAACTCTTTCCCTACACGACGCTCTTCCGATCTNNNTGTGGAAACAAAACCACCCTT

GAGCACACGTCTGAACTCTTTCCCTACACGACGCTCTTCCGATCTNNNCACTTGAGACCAGGAGTTTG

GAGCACACGTCTGAACTCTTTCCCTACACGACGCTCTTCCGATCTNNNCACTTGAGACCAGGAGTTTG

AGCACACGTCTGAACTCTTTCCCTACACGACGCTCTTCCGATCTNNNAGCTAGCCCTGTTGACACAAAT

AGCACACGTCTGAACTCTTTCCCTACACGACGCTCTTCCGATCTNNNAGCTAGCCCTGTTGACACAAAT

GCACACGTCTGAACTCTTTCCCTACACGACGCTCTTCCGATCTNNNGCTGAGGCAAGAAGAATCACTT

GCACACGTCTGAACTCTTTCCCTACACGACGCTCTTCCGATCTNNNGCTGAGGCAAGAAGAATCACTT

GAGCACACGTCTGAACTCTTTCCCTACACGACGCTCTTCCGATCTNNNAAAATGTGTTTGGGGGCTTA

GAGCACACGTCTGAACTCTTTCCCTACACGACGCTCTTCCGATCTNNNAAAATGTGTTTGGGGGCTTA

AGCACACGTCTGAACTCTTTCCCTACACGACGCTCTTCCGATCTNNNATCTGGACTTCACTTCTGTGCTT

AGCACACGTCTGAACTCTTTCCCTACACGACGCTCTTCCGATCTNNNATCTGGACTTCACTTCTGTGCTT

AGCACACGTCTGAACTCTTTCCCTACACGACGCTCTTCCGATCTNNNGCCCTGGAGTCTTTGTTCTTCTA

AGCACACGTCTGAACTCTTTCCCTACACGACGCTCTTCCGATCTNNNGCCCTGGAGTCTTTGTTCTTCTA

AGCACACGTCTGAACTCTTTCCCTACACGACGCTCTTCCGATCTNNNAGGCTCAATTGTTTTGAGCAG

AGCACACGTCTGAACTCTTTCCCTACACGACGCTCTTCCGATCTNNNAGGCTCAATTGTTTTGAGCAG

GAGCACACGTCTGAACTCTTTCCCTACACGACGCTCTTCCGATCTNNNACAAGGCAGGAGAATAGCTT

GAGCACACGTCTGAACTCTTTCCCTACACGACGCTCTTCCGATCTNNNACAAGGCAGGAGAATAGCTT

AGAGCACACGTCTGAACTCTTTCCCTACACGACGCTCTTCCGATCTNNNTAGATAAACGCCAGATGCCT

AGAGCACACGTCTGAACTCTTTCCCTACACGACGCTCTTCCGATCTNNNTAGATAAACGCCAGATGCCT

GAGCACACGTCTGAACTCTTTCCCTACACGACGCTCTTCCGATCTNNNTATGATCATGCCTGTGACTAG

GAGCACACGTCTGAACTCTTTCCCTACACGACGCTCTTCCGATCTNNNTATGATCATGCCTGTGACTAG

GAGCACACGTCTGAACTCTTTCCCTACACGACGCTCTTCCGATCTNNNCCAAGGACAGTTTTAAGATT

GAGCACACGTCTGAACTCTTTCCCTACACGACGCTCTTCCGATCTNNNCCAAGGACAGTTTTAAGATT

GAGCACACGTCTGAACTCTTTCCCTACACGACGCTCTTCCGATCTNNNTCTTACAGTCCTGAGGTTGAT

GAGCACACGTCTGAACTCTTTCCCTACACGACGCTCTTCCGATCTNNNTCTTACAGTCCTGAGGTTGAT
AGCACACGTCTGAACTCTTTCCCTACACGACGCTCTTCCGATCTNNNGTTGTAGTATCCTTTCCCCGTGT
AGCACACGTCTGAACTCTTTCCCTACACGACGCTCTTCCGATCTNNNGTTGTAGTATCCTTTCCCCGTGT
GCACACGTCTGAACTCTTTCCCTACACGACGCTCTTCCGATCTNNNGACCAGAGAGGGCTTGGAAAGC
GCACACGTCTGAACTCTTTCCCTACACGACGCTCTTCCGATCTNNNGACCAGAGAGGGCTTGGAAAGC
AGCACACGTCTGAACTCTTTCCCTACACGACGCTCTTCCGATCTNNNTGGTTGAGACTTTCGGGAATAC
AGCACACGTCTGAACTCTTTCCCTACACGACGCTCTTCCGATCTNNNTGGTTGAGACTTTCGGGAATAC
AGCACACGTCTGAACTCTTTCCCTACACGACGCTCTTCCGATCTNNNGCTGAAAAGTGGAAAGAGTGC
AGCACACGTCTGAACTCTTTCCCTACACGACGCTCTTCCGATCTNNNGCTGAAAAGTGGAAAGAGTGC
GAGCACACGTCTGAACTCTTTCCCTACACGACGCTCTTCCGATCTNNNCACCTGAGGAGATACACACA
GAGCACACGTCTGAACTCTTTCCCTACACGACGCTCTTCCGATCTNNNCACCTGAGGAGATACACACA
AGCACACGTCTGAACTCTTTCCCTACACGACGCTCTTCCGATCTNNNTTACAGATCATCAGTATTTGGA
AGCACACGTCTGAACTCTTTCCCTACACGACGCTCTTCCGATCTNNNTTACAGATCATCAGTATTTGGA
GCACACGTCTGAACTCTTTCCCTACACGACGCTCTTCCGATCTNNNCCAGGAGTAAGAAAAATTGGGC
GCACACGTCTGAACTCTTTCCCTACACGACGCTCTTCCGATCTNNNCCAGGAGTAAGAAAAATTGGGC
GAGCACACGTCTGAACTCTTTCCCTACACGACGCTCTTCCGATCTNNNCATGGAGAAATTGTGGGGCT
GAGCACACGTCTGAACTCTTTCCCTACACGACGCTCTTCCGATCTNNNCATGGAGAAATTGTGGGGCT
GAGCACACGTCTGAACTCTTTCCCTACACGACGCTCTTCCGATCTNNNCCCAGCACTTACACATTCTCCTT
GAGCACACGTCTGAACTCTTTCCCTACACGACGCTCTTCCGATCTNNNCCCAGCACTTACACATTCTCCTT
GAGCACACGTCTGAACTCTTTCCCTACACGACGCTCTTCCGATCTNNNGTGACATACCCTTCTCATCACG
GAGCACACGTCTGAACTCTTTCCCTACACGACGCTCTTCCGATCTNNNGTGACATACCCTTCTCATCACG
AGCACACGTCTGAACTCTTTCCCTACACGACGCTCTTCCGATCTNNNGAGAATCACTTGAACCTGGGA
AGCACACGTCTGAACTCTTTCCCTACACGACGCTCTTCCGATCTNNNGAGAATCACTTGAACCTGGGA
AGCACACGTCTGAACTCTTTCCCTACACGACGCTCTTCCGATCTNNNCTGTAGTTCCAGCTACTTGGGA
AGCACACGTCTGAACTCTTTCCCTACACGACGCTCTTCCGATCTNNNCTGTAGTTCCAGCTACTTGGGA
GAGCACACGTCTGAACTCTTTCCCTACACGACGCTCTTCCGATCTNNNCAGCACACACACACACAAAAAA
GAGCACACGTCTGAACTCTTTCCCTACACGACGCTCTTCCGATCTNNNCAGCACACACACACACAAAAAA
AGAGCACACGTCTGAACTCTTTCCCTACACGACGCTCTTCCGATCTNNNGAATCACTTGAACCCAGGAG
AGAGCACACGTCTGAACTCTTTCCCTACACGACGCTCTTCCGATCTNNNGAATCACTTGAACCCAGGAG
GAGCACACGTCTGAACTCTTTCCCTACACGACGCTCTTCCGATCTNNNGTGATCCTTAGTGGGGATTG
GAGCACACGTCTGAACTCTTTCCCTACACGACGCTCTTCCGATCTNNNGTGATCCTTAGTGGGGATTG
GCACACGTCTGAACTCTTTCCCTACACGACGCTCTTCCGATCTNNNACACATGACTGTTCTGAGATGTC
GCACACGTCTGAACTCTTTCCCTACACGACGCTCTTCCGATCTNNNACACATGACTGTTCTGAGATGTC
AGCACACGTCTGAACTCTTTCCCTACACGACGCTCTTCCGATCTNNNAATTCCTGGACACATACACCCTC
AGCACACGTCTGAACTCTTTCCCTACACGACGCTCTTCCGATCTNNNAATTCCTGGACACATACACCCTC
CACACGTCTGAACTCTTTCCCTACACGACGCTCTTCCGATCTNNNCCTCCAGCTCCACCATGACGCATCGC
CACACGTCTGAACTCTTTCCCTACACGACGCTCTTCCGATCTNNNCCTCCAGCTCCACCATGACGCATCGC
GAGCACACGTCTGAACTCTTTCCCTACACGACGCTCTTCCGATCTNNNCACACCCTACTACACCACTTACC
GAGCACACGTCTGAACTCTTTCCCTACACGACGCTCTTCCGATCTNNNCACACCCTACTACACCACTTACC
GAGCACACGTCTGAACTCTTTCCCTACACGACGCTCTTCCGATCTNNNCTCAAGTAGTGCTCCCCTCAAA
GAGCACACGTCTGAACTCTTTCCCTACACGACGCTCTTCCGATCTNNNCTCAAGTAGTGCTCCCCTCAAA
AGCACACGTCTGAACTCTTTCCCTACACGACGCTCTTCCGATCTNNNCAGCCACGAAGTCCTTCTTTTTT
AGCACACGTCTGAACTCTTTCCCTACACGACGCTCTTCCGATCTNNNCAGCCACGAAGTCCTTCTTTTTT

GAGCACACGTCTGAACTCTTTCCCTACACGACGCTCTTCCGATCTNNNCTAATCAGGAAATTTGTGGCC

GAGCACACGTCTGAACTCTTTCCCTACACGACGCTCTTCCGATCTNNNCTAATCAGGAAATTTGTGGCC

GAGCACACGTCTGAACTCTTTCCCTACACGACGCTCTTCCGATCTNNNTCTAGCCCCTCCATGATTTATGT

GAGCACACGTCTGAACTCTTTCCCTACACGACGCTCTTCCGATCTNNNTCTAGCCCCTCCATGATTTATGT

GAGCACACGTCTGAACTCTTTCCCTACACGACGCTCTTCCGATCTNNNCTGTAGTCCCAGCTACTCAGG

GAGCACACGTCTGAACTCTTTCCCTACACGACGCTCTTCCGATCTNNNCTGTAGTCCCAGCTACTCAGG

GCACACGTCTGAACTCTTTCCCTACACGACGCTCTTCCGATCTNNNCTGAGGCAGGAGAATCACTTGA

GCACACGTCTGAACTCTTTCCCTACACGACGCTCTTCCGATCTNNNCTGAGGCAGGAGAATCACTTGA

AGCACACGTCTGAACTCTTTCCCTACACGACGCTCTTCCGATCTNNNGAGTAGCTGAGACTACAGGTG

AGCACACGTCTGAACTCTTTCCCTACACGACGCTCTTCCGATCTNNNGAGTAGCTGAGACTACAGGTG

GAGCACACGTCTGAACTCTTTCCCTACACGACGCTCTTCCGATCTNNNCTGTAGTCCCAGCTACTCAGGA

GAGCACACGTCTGAACTCTTTCCCTACACGACGCTCTTCCGATCTNNNCTGTAGTCCCAGCTACTCAGGA

GAGCACACGTCTGAACTCTTTCCCTACACGACGCTCTTCCGATCTNNNCTGTAGTCCCAGCTACTCAGGA

GAGCACACGTCTGAACTCTTTCCCTACACGACGCTCTTCCGATCTNNNCTGTAGTCCCAGCTACTCAGGA

GAGCACACGTCTGAACTCTTTCCCTACACGACGCTCTTCCGATCTNNNAGATAGAACTCTGGGACTGA

GAGCACACGTCTGAACTCTTTCCCTACACGACGCTCTTCCGATCTNNNAGATAGAACTCTGGGACTGA

AGCACACGTCTGAACTCTTTCCCTACACGACGCTCTTCCGATCTNNNTTGCAGTGTCATTAGGAAGGC

AGCACACGTCTGAACTCTTTCCCTACACGACGCTCTTCCGATCTNNNTTGCAGTGTCATTAGGAAGGC

AGCACACGTCTGAACTCTTTCCCTACACGACGCTCTTCCGATCTNNNGGTGCTTTGATGAAACTTGCTC

AGCACACGTCTGAACTCTTTCCCTACACGACGCTCTTCCGATCTNNNGGTGCTTTGATGAAACTTGCTC

GAGCACACGTCTGAACTCTTTCCCTACACGACGCTCTTCCGATCTNNNTCCTTGGTCTCCCTACAGGAAA

GAGCACACGTCTGAACTCTTTCCCTACACGACGCTCTTCCGATCTNNNTCCTTGGTCTCCCTACAGGAAA

AGCACACGTCTGAACTCTTTCCCTACACGACGCTCTTCCGATCTNNNGCTGGATTTTATCTCCTCCCACA

AGCACACGTCTGAACTCTTTCCCTACACGACGCTCTTCCGATCTNNNGCTGGATTTTATCTCCTCCCACA

GCACACGTCTGAACTCTTTCCCTACACGACGCTCTTCCGATCTNNNAGAAAGTAGAATGGTGGTTGCCA

GCACACGTCTGAACTCTTTCCCTACACGACGCTCTTCCGATCTNNNAGAAAGTAGAATGGTGGTTGCCA

AGCACACGTCTGAACTCTTTCCCTACACGACGCTCTTCCGATCTNNNTGAGCAACAAAGTGAGACTCCA

AGCACACGTCTGAACTCTTTCCCTACACGACGCTCTTCCGATCTNNNTGAGCAACAAAGTGAGACTCCA

GAGCACACGTCTGAACTCTTTCCCTACACGACGCTCTTCCGATCTNNNTATTTTGCAGCTCATAGACAAT

GAGCACACGTCTGAACTCTTTCCCTACACGACGCTCTTCCGATCTNNNTATTTTGCAGCTCATAGACAAT

AGCACACGTCTGAACTCTTTCCCTACACGACGCTCTTCCGATCTNNNGACAGAGCAAGACTCCGTCTCCA

AGCACACGTCTGAACTCTTTCCCTACACGACGCTCTTCCGATCTNNNGACAGAGCAAGACTCCGTCTCCA

AGCACACGTCTGAACTCTTTCCCTACACGACGCTCTTCCGATCTNNNGGCATTTACAGATCATCCAGCC

AGCACACGTCTGAACTCTTTCCCTACACGACGCTCTTCCGATCTNNNGGCATTTACAGATCATCCAGCC

GAGCACACGTCTGAACTCTTTCCCTACACGACGCTCTTCCGATCTNNNCGAGTTCGTCCACTGTTACAGT

GAGCACACGTCTGAACTCTTTCCCTACACGACGCTCTTCCGATCTNNNCGAGTTCGTCCACTGTTACAGT

AGCACACGTCTGAACTCTTTCCCTACACGACGCTCTTCCGATCTNNNGGCTAAATGTGGGTGTGGAAA

AGCACACGTCTGAACTCTTTCCCTACACGACGCTCTTCCGATCTNNNGGCTAAATGTGGGTGTGGAAA

AGCACACGTCTGAACTCTTTCCCTACACGACGCTCTTCCGATCTNNNACCTCTGTACCAATTAGCTGTCC

AGCACACGTCTGAACTCTTTCCCTACACGACGCTCTTCCGATCTNNNACCTCTGTACCAATTAGCTGTCC

GAGCACACGTCTGAACTCTTTCCCTACACGACGCTCTTCCGATCTNNNGTGGGAGAATCTCTTGAACCCA

GAGCACACGTCTGAACTCTTTCCCTACACGACGCTCTTCCGATCTNNNGTGGGAGAATCTCTTGAACCCA

AGCACACGTCTGAACTCTTTCCCTACACGACGCTCTTCCGATCTNNNGGAGGCTGAAAGTAGAACTGA

AGCACACGTCTGAACTCTTTCCCTACACGACGCTCTTCCGATCTNNNGGAGGCTGAAAGTAGAACTGA

GAGCACACGTCTGAACTCTTTCCCTACACGACGCTCTTCCGATCTNNNGAGGATTGCTTGAGCCTGAG

GAGCACACGTCTGAACTCTTTCCCTACACGACGCTCTTCCGATCTNNNGAGGATTGCTTGAGCCTGAG

AGCACACGTCTGAACTCTTTCCCTACACGACGCTCTTCCGATCTNNNCTGAGGCAGGAGAATCACTTG

AGCACACGTCTGAACTCTTTCCCTACACGACGCTCTTCCGATCTNNNCTGAGGCAGGAGAATCACTTG

AGCACACGTCTGAACTCTTTCCCTACACGACGCTCTTCCGATCTNNNACGGCCGTGTAGATGATGAAG

AGCACACGTCTGAACTCTTTCCCTACACGACGCTCTTCCGATCTNNNACGGCCGTGTAGATGATGAAG

ACACGTCTGAACTCTTTCCCTACACGACGCTCTTCCGATCTNNNATGCATCTTTCCCCCTTCTCTTTCATC

ACACGTCTGAACTCTTTCCCTACACGACGCTCTTCCGATCTNNNATGCATCTTTCCCCCTTCTCTTTCATC

GAGCACACGTCTGAACTCTTTCCCTACACGACGCTCTTCCGATCTNNNTGCAAGGGGTAGACTGTGA

GAGCACACGTCTGAACTCTTTCCCTACACGACGCTCTTCCGATCTNNNTGCAAGGGGTAGACTGTGA

GAGCACACGTCTGAACTCTTTCCCTACACGACGCTCTTCCGATCTNNNGATGGGTTATTTGCTTGGTT

GAGCACACGTCTGAACTCTTTCCCTACACGACGCTCTTCCGATCTNNNGATGGGTTATTTGCTTGGTT

AGAGCACACGTCTGAACTCTTTCCCTACACGACGCTCTTCCGATCTNNNATGGTTTCATGATGCTCCAG

AGAGCACACGTCTGAACTCTTTCCCTACACGACGCTCTTCCGATCTNNNATGGTTTCATGATGCTCCAG

GAGCACACGTCTGAACTCTTTCCCTACACGACGCTCTTCCGATCTNNNGGACTTTCTGACCTGAGTTTC

GAGCACACGTCTGAACTCTTTCCCTACACGACGCTCTTCCGATCTNNNGGACTTTCTGACCTGAGTTTC

GAGCACACGTCTGAACTCTTTCCCTACACGACGCTCTTCCGATCTNNNCTTGCTGATGTTCTGTCTGGT

GAGCACACGTCTGAACTCTTTCCCTACACGACGCTCTTCCGATCTNNNCTTGCTGATGTTCTGTCTGGT

GCACACGTCTGAACTCTTTCCCTACACGACGCTCTTCCGATCTNNNTTTTTCCATGTTGTTCAGGCTGG

GCACACGTCTGAACTCTTTCCCTACACGACGCTCTTCCGATCTNNNTTTTTCCATGTTGTTCAGGCTGG

GCACACGTCTGAACTCTTTCCCTACACGACGCTCTTCCGATCTNNNGAGGCTGAGGCAGGAGAATGC

GCACACGTCTGAACTCTTTCCCTACACGACGCTCTTCCGATCTNNNGAGGCTGAGGCAGGAGAATGC

AGCACACGTCTGAACTCTTTCCCTACACGACGCTCTTCCGATCTNNNCCAGGCAAGGAAGAAGAAAA

AGCACACGTCTGAACTCTTTCCCTACACGACGCTCTTCCGATCTNNNCCAGGCAAGGAAGAAGAAAA

AGCACACGTCTGAACTCTTTCCCTACACGACGCTCTTCCGATCTNNNCCAATCTCAGTAAGTTCAGGG

AGCACACGTCTGAACTCTTTCCCTACACGACGCTCTTCCGATCTNNNCCAATCTCAGTAAGTTCAGGG

GAGCACACGTCTGAACTCTTTCCCTACACGACGCTCTTCCGATCTNNNCTGCTTGGCATACCTTTCCTTC

GAGCACACGTCTGAACTCTTTCCCTACACGACGCTCTTCCGATCTNNNCTGCTTGGCATACCTTTCCTTC

CACACGTCTGAACTCTTTCCCTACACGACGCTCTTCCGATCTNNNAGATGATGATGCAGAATCCACCAC

CACACGTCTGAACTCTTTCCCTACACGACGCTCTTCCGATCTNNNAGATGATGATGCAGAATCCACCAC

GAGCACACGTCTGAACTCTTTCCCTACACGACGCTCTTCCGATCTNNNGTCAGGATTGTTGTGGTTTC

GAGCACACGTCTGAACTCTTTCCCTACACGACGCTCTTCCGATCTNNNGTCAGGATTGTTGTGGTTTC

AGCACACGTCTGAACTCTTTCCCTACACGACGCTCTTCCGATCTNNNTTACCAAGTTAGTCAGACCTGG

AGCACACGTCTGAACTCTTTCCCTACACGACGCTCTTCCGATCTNNNTTACCAAGTTAGTCAGACCTGG

AGAGCACACGTCTGAACTCTTTCCCTACACGACGCTCTTCCGATCTNNNATAGACAAGAGGCACACAG

AGAGCACACGTCTGAACTCTTTCCCTACACGACGCTCTTCCGATCTNNNATAGACAAGAGGCACACAG

AGCACACGTCTGAACTCTTTCCCTACACGACGCTCTTCCGATCTNNNTTCCTCTTCTTCCTCTTCCTCCTC

AGCACACGTCTGAACTCTTTCCCTACACGACGCTCTTCCGATCTNNNTTCCTCTTCTTCCTCTTCCTCCTC

AGCACACGTCTGAACTCTTTCCCTACACGACGCTCTTCCGATCTNNNGAGAACTGGGTCTTGAAGCCT

AGCACACGTCTGAACTCTTTCCCTACACGACGCTCTTCCGATCTNNNGAGAACTGGGTCTTGAAGCCT

AGCACACGTCTGAACTCTTTCCCTACACGACGCTCTTCCGATCTNNNTGGGGAAAAATAAGTCCTACA

AGCACACGTCTGAACTCTTTCCCTACACGACGCTCTTCCGATCTNNNTGGGGAAAAATAAGTCCTACA

AGCACACGTCTGAACTCTTTCCCTACACGACGCTCTTCCGATCTNNNTAAGCCTTTTTGATGTGCTGCT
AGCACACGTCTGAACTCTTTCCCTACACGACGCTCTTCCGATCTNNNTAAGCCTTTTTGATGTGCTGCT
GAGCACACGTCTGAACTCTTTCCCTACACGACGCTCTTCCGATCTNNNAGATTGCACCACTACACTCTA
GAGCACACGTCTGAACTCTTTCCCTACACGACGCTCTTCCGATCTNNNAGATTGCACCACTACACTCTA
AGCACACGTCTGAACTCTTTCCCTACACGACGCTCTTCCGATCTNNNTCAACCTGTCTAAACCAGCATCA
AGCACACGTCTGAACTCTTTCCCTACACGACGCTCTTCCGATCTNNNTCAACCTGTCTAAACCAGCATCA
AGCACACGTCTGAACTCTTTCCCTACACGACGCTCTTCCGATCTNNNCTAGACAACACAGCAAAACCCA
AGCACACGTCTGAACTCTTTCCCTACACGACGCTCTTCCGATCTNNNCTAGACAACACAGCAAAACCCA
CACACGTCTGAACTCTTTCCCTACACGACGCTCTTCCGATCTNNNTCTTCCTGTCTCTCTTTCCCTCCCAT
CACACGTCTGAACTCTTTCCCTACACGACGCTCTTCCGATCTNNNTCTTCCTGTCTCTCTTTCCCTCCCAT
AGCACACGTCTGAACTCTTTCCCTACACGACGCTCTTCCGATCTNNNCTGGGAGTTGTAAAAATCTTTT
AGCACACGTCTGAACTCTTTCCCTACACGACGCTCTTCCGATCTNNNCTGGGAGTTGTAAAAATCTTTT
AGAGCACACGTCTGAACTCTTTCCCTACACGACGCTCTTCCGATCTNNNTCCTGAGTTGTTCCACTGTCT
AGAGCACACGTCTGAACTCTTTCCCTACACGACGCTCTTCCGATCTNNNTCCTGAGTTGTTCCACTGTCT
AGAGCACACGTCTGAACTCTTTCCCTACACGACGCTCTTCCGATCTNNNTGTCTCTGGAAGCCTTTACT
AGAGCACACGTCTGAACTCTTTCCCTACACGACGCTCTTCCGATCTNNNTGTCTCTGGAAGCCTTTACT
GCACACGTCTGAACTCTTTCCCTACACGACGCTCTTCCGATCTNNNTTTGCAGGGTCCTAACTCAATCGC
GCACACGTCTGAACTCTTTCCCTACACGACGCTCTTCCGATCTNNNTTTGCAGGGTCCTAACTCAATCGC
GAGCACACGTCTGAACTCTTTCCCTACACGACGCTCTTCCGATCTNNNCACTCCAACCTCTGCTTGTTTTT
GAGCACACGTCTGAACTCTTTCCCTACACGACGCTCTTCCGATCTNNNCACTCCAACCTCTGCTTGTTTTT
GAGCACACGTCTGAACTCTTTCCCTACACGACGCTCTTCCGATCTNNNACTAGCATTCTCCACAGAAGC
GAGCACACGTCTGAACTCTTTCCCTACACGACGCTCTTCCGATCTNNNACTAGCATTCTCCACAGAAGC
GCACACGTCTGAACTCTTTCCCTACACGACGCTCTTCCGATCTNNNGATTTACTGAATACATCCATGAC
GCACACGTCTGAACTCTTTCCCTACACGACGCTCTTCCGATCTNNNGATTTACTGAATACATCCATGAC
GCACACGTCTGAACTCTTTCCCTACACGACGCTCTTCCGATCTNNNAATACCTGGGATCCATTGCCTCT
GCACACGTCTGAACTCTTTCCCTACACGACGCTCTTCCGATCTNNNAATACCTGGGATCCATTGCCTCT
GCACACGTCTGAACTCTTTCCCTACACGACGCTCTTCCGATCTNNNGACAACACTACAGCAAGGACCTA
GCACACGTCTGAACTCTTTCCCTACACGACGCTCTTCCGATCTNNNGACAACACTACAGCAAGGACCTA
AGCACACGTCTGAACTCTTTCCCTACACGACGCTCTTCCGATCTNNNAGAGATGGTTGACCTTTGTCT
AGCACACGTCTGAACTCTTTCCCTACACGACGCTCTTCCGATCTNNNAGAGATGGTTGACCTTTGTCT
AGAGCACACGTCTGAACTCTTTCCCTACACGACGCTCTTCCGATCTNNNGTGACAGAGTGAGACTCC
AGAGCACACGTCTGAACTCTTTCCCTACACGACGCTCTTCCGATCTNNNGTGACAGAGTGAGACTCC
GAGCACACGTCTGAACTCTTTCCCTACACGACGCTCTTCCGATCTNNNACCTGTAATCCTAGCACTTTGG
GAGCACACGTCTGAACTCTTTCCCTACACGACGCTCTTCCGATCTNNNACCTGTAATCCTAGCACTTTGG
GAGCACACGTCTGAACTCTTTCCCTACACGACGCTCTTCCGATCTNNNTTGGTGAGGTTGACGTACTTA
GAGCACACGTCTGAACTCTTTCCCTACACGACGCTCTTCCGATCTNNNTTGGTGAGGTTGACGTACTTA
GAGCACACGTCTGAACTCTTTCCCTACACGACGCTCTTCCGATCTNNNATGTCTGTAATCCCAGCTACT
GAGCACACGTCTGAACTCTTTCCCTACACGACGCTCTTCCGATCTNNNATGTCTGTAATCCCAGCTACT
AGCACACGTCTGAACTCTTTCCCTACACGACGCTCTTCCGATCTNNNCAGACCTTGCCCGTGATGAGCA
AGCACACGTCTGAACTCTTTCCCTACACGACGCTCTTCCGATCTNNNCAGACCTTGCCCGTGATGAGCA
GAGCACACGTCTGAACTCTTTCCCTACACGACGCTCTTCCGATCTNNNTAATCCCAGCTACTCAGGAGG
GAGCACACGTCTGAACTCTTTCCCTACACGACGCTCTTCCGATCTNNNTAATCCCAGCTACTCAGGAGG
AGCACACGTCTGAACTCTTTCCCTACACGACGCTCTTCCGATCTNNNGTAATCAGGGAAAATGCTCTG

AGCACACGTCTGAACTCTTTCCCTACACGACGCTCTTCCGATCTNNNGTAATCAGGGAAAATGCTCTG
AGCACACGTCTGAACTCTTTCCCTACACGACGCTCTTCCGATCTNNNCCTCACCCTTCCAAAGTGAACT
AGCACACGTCTGAACTCTTTCCCTACACGACGCTCTTCCGATCTNNNCCTCACCCTTCCAAAGTGAACT
AGCACACGTCTGAACTCTTTCCCTACACGACGCTCTTCCGATCTNNNCGTAATCCCAGCTACTCAGGAG
AGCACACGTCTGAACTCTTTCCCTACACGACGCTCTTCCGATCTNNNCGTAATCCCAGCTACTCAGGAG
GCACACGTCTGAACTCTTTCCCTACACGACGCTCTTCCGATCTNNNGAACCACGATGTCCTCTCTTGAG
GCACACGTCTGAACTCTTTCCCTACACGACGCTCTTCCGATCTNNNGAACCACGATGTCCTCTCTTGAG
GAGCACACGTCTGAACTCTTTCCCTACACGACGCTCTTCCGATCTNNNAGCCAGCTAGTCATTGATGTT
GAGCACACGTCTGAACTCTTTCCCTACACGACGCTCTTCCGATCTNNNAGCCAGCTAGTCATTGATGTT
GCACACGTCTGAACTCTTTCCCTACACGACGCTCTTCCGATCTNNNGCTAGCACTTGTTCTTTTCCAGG
GCACACGTCTGAACTCTTTCCCTACACGACGCTCTTCCGATCTNNNGCTAGCACTTGTTCTTTTCCAGG
CACACGTCTGAACTCTTTCCCTACACGACGCTCTTCCGATCTNNNTTTCGGTTCCATCTGTGCCTTAGCA
CACACGTCTGAACTCTTTCCCTACACGACGCTCTTCCGATCTNNNTTTCGGTTCCATCTGTGCCTTAGCA
GCACACGTCTGAACTCTTTCCCTACACGACGCTCTTCCGATCTNNNACACCTGTAATCCCAGCTACTTG
GCACACGTCTGAACTCTTTCCCTACACGACGCTCTTCCGATCTNNNACACCTGTAATCCCAGCTACTTG
GAGCACACGTCTGAACTCTTTCCCTACACGACGCTCTTCCGATCTNNNTCAGGTAGAGAACTGCTTGA
GAGCACACGTCTGAACTCTTTCCCTACACGACGCTCTTCCGATCTNNNTCAGGTAGAGAACTGCTTGA
AGCACACGTCTGAACTCTTTCCCTACACGACGCTCTTCCGATCTNNNTTTGGTTACTACAGCTTTGCAT
AGCACACGTCTGAACTCTTTCCCTACACGACGCTCTTCCGATCTNNNTTTGGTTACTACAGCTTTGCAT
AGCACACGTCTGAACTCTTTCCCTACACGACGCTCTTCCGATCTNNNATATGTTCTAGGGTTGGCTGC
AGCACACGTCTGAACTCTTTCCCTACACGACGCTCTTCCGATCTNNNATATGTTCTAGGGTTGGCTGC
GAGCACACGTCTGAACTCTTTCCCTACACGACGCTCTTCCGATCTNNNGAAAAACCCTGCTAGGTGGG
GAGCACACGTCTGAACTCTTTCCCTACACGACGCTCTTCCGATCTNNNGAAAAACCCTGCTAGGTGGG
GCACACGTCTGAACTCTTTCCCTACACGACGCTCTTCCGATCTNNNGAGGTGGGAGGATCTTTTGAG
GCACACGTCTGAACTCTTTCCCTACACGACGCTCTTCCGATCTNNNGAGGTGGGAGGATCTTTTGAG
AGCACACGTCTGAACTCTTTCCCTACACGACGCTCTTCCGATCTNNNCCCACTCCAGACCAATACAATC
AGCACACGTCTGAACTCTTTCCCTACACGACGCTCTTCCGATCTNNNCCCACTCCAGACCAATACAATC
AGCACACGTCTGAACTCTTTCCCTACACGACGCTCTTCCGATCTNNNGAGGCAGGAAAATGGTGTGA
AGCACACGTCTGAACTCTTTCCCTACACGACGCTCTTCCGATCTNNNGAGGCAGGAAAATGGTGTGA
GAGCACACGTCTGAACTCTTTCCCTACACGACGCTCTTCCGATCTNNNCGAGGAGAAAGATCAGATC
GAGCACACGTCTGAACTCTTTCCCTACACGACGCTCTTCCGATCTNNNCGAGGAGAAAGATCAGATC
AGAGCACACGTCTGAACTCTTTCCCTACACGACGCTCTTCCGATCTNNNATGTGTACCTGCCCTAAACA
AGAGCACACGTCTGAACTCTTTCCCTACACGACGCTCTTCCGATCTNNNATGTGTACCTGCCCTAAACA
GCACACGTCTGAACTCTTTCCCTACACGACGCTCTTCCGATCTNNNTCTGCATGTCCAAAGTCTGCTAT
GCACACGTCTGAACTCTTTCCCTACACGACGCTCTTCCGATCTNNNTCTGCATGTCCAAAGTCTGCTAT
GCACACGTCTGAACTCTTTCCCTACACGACGCTCTTCCGATCTNNNGCTGAGGTAGGAGAATTGCTT
GCACACGTCTGAACTCTTTCCCTACACGACGCTCTTCCGATCTNNNGCTGAGGTAGGAGAATTGCTT
GAGCACACGTCTGAACTCTTTCCCTACACGACGCTCTTCCGATCTNNNTATGGAGAAGCACACATGAA
GAGCACACGTCTGAACTCTTTCCCTACACGACGCTCTTCCGATCTNNNTATGGAGAAGCACACATGAA
AGCACACGTCTGAACTCTTTCCCTACACGACGCTCTTCCGATCTNNNAACCCACAGCCAATCTTGTCTAT
AGCACACGTCTGAACTCTTTCCCTACACGACGCTCTTCCGATCTNNNAACCCACAGCCAATCTTGTCTAT
AGCACACGTCTGAACTCTTTCCCTACACGACGCTCTTCCGATCTNNNCTCAGCAGACTTGTCTCCTCTAC
AGCACACGTCTGAACTCTTTCCCTACACGACGCTCTTCCGATCTNNNCTCAGCAGACTTGTCTCCTCTAC

AGCACACGTCTGAACTCTTTCCCTACACGACGCTCTTCCGATCTNNNGAGCAAATGGTGGAGAGGTAG
AGCACACGTCTGAACTCTTTCCCTACACGACGCTCTTCCGATCTNNNGAGCAAATGGTGGAGAGGTAG
AGCACACGTCTGAACTCTTTCCCTACACGACGCTCTTCCGATCTNNNAAAAAGAGTAGGTAGCGGGTT
AGCACACGTCTGAACTCTTTCCCTACACGACGCTCTTCCGATCTNNNAAAAAGAGTAGGTAGCGGGTT
AGCACACGTCTGAACTCTTTCCCTACACGACGCTCTTCCGATCTNNNCCTTCGAACATCGGACTCCATAT
AGCACACGTCTGAACTCTTTCCCTACACGACGCTCTTCCGATCTNNNCCTTCGAACATCGGACTCCATAT
GAGCACACGTCTGAACTCTTTCCCTACACGACGCTCTTCCGATCTNNNGGTTGCAGTGAGCCAAGATA
GAGCACACGTCTGAACTCTTTCCCTACACGACGCTCTTCCGATCTNNNGGTTGCAGTGAGCCAAGATA
AGCACACGTCTGAACTCTTTCCCTACACGACGCTCTTCCGATCTNNNAAAACAACACTGCCCACCATAA
AGCACACGTCTGAACTCTTTCCCTACACGACGCTCTTCCGATCTNNNAAAACAACACTGCCCACCATAA
GAGCACACGTCTGAACTCTTTCCCTACACGACGCTCTTCCGATCTNNNTCCTCTGGAAGTCAGCTAACAA
GAGCACACGTCTGAACTCTTTCCCTACACGACGCTCTTCCGATCTNNNTCCTCTGGAAGTCAGCTAACAA
GAGCACACGTCTGAACTCTTTCCCTACACGACGCTCTTCCGATCTNNNTACCTTTAGTTGGCTCAGGGT
GAGCACACGTCTGAACTCTTTCCCTACACGACGCTCTTCCGATCTNNNTACCTTTAGTTGGCTCAGGGT
GAGCACACGTCTGAACTCTTTCCCTACACGACGCTCTTCCGATCTNNNACATGGTGAAATCCCGTCTCT
GAGCACACGTCTGAACTCTTTCCCTACACGACGCTCTTCCGATCTNNNACATGGTGAAATCCCGTCTCT
GAGCACACGTCTGAACTCTTTCCCTACACGACGCTCTTCCGATCTNNNAAAAATTAGTTGGGTGTGGTG
GAGCACACGTCTGAACTCTTTCCCTACACGACGCTCTTCCGATCTNNNAAAAATTAGTTGGGTGTGGTG
AGCACACGTCTGAACTCTTTCCCTACACGACGCTCTTCCGATCTNNNCAGGAAGCAGTAGAAGTCCAG
AGCACACGTCTGAACTCTTTCCCTACACGACGCTCTTCCGATCTNNNCAGGAAGCAGTAGAAGTCCAG
AGCACACGTCTGAACTCTTTCCCTACACGACGCTCTTCCGATCTNNNGTAGTACATGCCTGTAGTCCCA
AGCACACGTCTGAACTCTTTCCCTACACGACGCTCTTCCGATCTNNNGTAGTACATGCCTGTAGTCCCA
AGCACACGTCTGAACTCTTTCCCTACACGACGCTCTTCCGATCTNNNCTGTAGTGTCCGCTACTCAGGA
AGCACACGTCTGAACTCTTTCCCTACACGACGCTCTTCCGATCTNNNCTGTAGTGTCCGCTACTCAGGA
GAGCACACGTCTGAACTCTTTCCCTACACGACGCTCTTCCGATCTNNNTCATCTATTGCCTGAGAGCACA
GAGCACACGTCTGAACTCTTTCCCTACACGACGCTCTTCCGATCTNNNTCATCTATTGCCTGAGAGCACA
GAGCACACGTCTGAACTCTTTCCCTACACGACGCTCTTCCGATCTNNNGAGGCTGAGGCAGGAGAATG
GAGCACACGTCTGAACTCTTTCCCTACACGACGCTCTTCCGATCTNNNGAGGCTGAGGCAGGAGAATG
GAGCACACGTCTGAACTCTTTCCCTACACGACGCTCTTCCGATCTNNNCCTGTAGTCCCAGCTACTTGGG
GAGCACACGTCTGAACTCTTTCCCTACACGACGCTCTTCCGATCTNNNCCTGTAGTCCCAGCTACTTGGG
GAGCACACGTCTGAACTCTTTCCCTACACGACGCTCTTCCGATCTNNNCTTGAACTCCTGACCTCAAGTG
GAGCACACGTCTGAACTCTTTCCCTACACGACGCTCTTCCGATCTNNNCTTGAACTCCTGACCTCAAGTG
AGCACACGTCTGAACTCTTTCCCTACACGACGCTCTTCCGATCTNNNGCCCGGCCTAATTTTTTTTTGTG
AGCACACGTCTGAACTCTTTCCCTACACGACGCTCTTCCGATCTNNNGCCCGGCCTAATTTTTTTTTGTG
AGCACACGTCTGAACTCTTTCCCTACACGACGCTCTTCCGATCTNNNATGCTCCTATGTGTCTGATCTGG
AGCACACGTCTGAACTCTTTCCCTACACGACGCTCTTCCGATCTNNNATGCTCCTATGTGTCTGATCTGG
AGCACACGTCTGAACTCTTTCCCTACACGACGCTCTTCCGATCTNNNAGGCAACAAATACCATCAGTTGT
AGCACACGTCTGAACTCTTTCCCTACACGACGCTCTTCCGATCTNNNAGGCAACAAATACCATCAGTTGT
AGCACACGTCTGAACTCTTTCCCTACACGACGCTCTTCCGATCTNNNTTGCGTCTGAATCCTGTAACACA
AGCACACGTCTGAACTCTTTCCCTACACGACGCTCTTCCGATCTNNNTTGCGTCTGAATCCTGTAACACA
AGCACACGTCTGAACTCTTTCCCTACACGACGCTCTTCCGATCTNNNTGAACCGCCATTGTATGTTATTT
AGCACACGTCTGAACTCTTTCCCTACACGACGCTCTTCCGATCTNNNTGAACCGCCATTGTATGTTATTT
GCACACGTCTGAACTCTTTCCCTACACGACGCTCTTCCGATCTNNNCTTTTGCATGTTTTGTAGAGATCA

CACACGTCTGAACTCTTTCCCTACACGACGCTCTTCCGATCTNNNCTTTGCATGTTTTGTAGAGATCA
AGCACACGTCTGAACTCTTTCCCTACACGACGCTCTTCCGATCTNNNGAGAATTGCTTGAACCTGGG
AGCACACGTCTGAACTCTTTCCCTACACGACGCTCTTCCGATCTNNNGAGAATTGCTTGAACCTGGG
AGCACACGTCTGAACTCTTTCCCTACACGACGCTCTTCCGATCTNNNAGGAGAATGGCATGAACCCGC
AGCACACGTCTGAACTCTTTCCCTACACGACGCTCTTCCGATCTNNNAGGAGAATGGCATGAACCCGC
GCACACGTCTGAACTCTTTCCCTACACGACGCTCTTCCGATCTNNNAAGCTATGTGCAGATTGAATGC
GCACACGTCTGAACTCTTTCCCTACACGACGCTCTTCCGATCTNNNAAGCTATGTGCAGATTGAATGC
GAGCACACGTCTGAACTCTTTCCCTACACGACGCTCTTCCGATCTNNNCTGGCCCAGAATTATTGCTCA
GAGCACACGTCTGAACTCTTTCCCTACACGACGCTCTTCCGATCTNNNCTGGCCCAGAATTATTGCTCA
GCACACGTCTGAACTCTTTCCCTACACGACGCTCTTCCGATCTNNNAACAGGTGGGGATTACAATTC
GCACACGTCTGAACTCTTTCCCTACACGACGCTCTTCCGATCTNNNAACAGGTGGGGATTACAATTC
GCACACGTCTGAACTCTTTCCCTACACGACGCTCTTCCGATCTNNNATTTAACAACAAGCATGAAAAA
GCACACGTCTGAACTCTTTCCCTACACGACGCTCTTCCGATCTNNNATTTAACAACAAGCATGAAAAA
GCACACGTCTGAACTCTTTCCCTACACGACGCTCTTCCGATCTNNNCACCATGGCTTTCAGAATCAGA
GCACACGTCTGAACTCTTTCCCTACACGACGCTCTTCCGATCTNNNCACCATGGCTTTCAGAATCAGA
CACACGTCTGAACTCTTTCCCTACACGACGCTCTTCCGATCTNNNCATCCTAGCCAACATGGTGAAAC
CACACGTCTGAACTCTTTCCCTACACGACGCTCTTCCGATCTNNNCATCCTAGCCAACATGGTGAAAC
GCACACGTCTGAACTCTTTCCCTACACGACGCTCTTCCGATCTNNNACATGCATGGACCTCCAGAGTT
GCACACGTCTGAACTCTTTCCCTACACGACGCTCTTCCGATCTNNNACATGCATGGACCTCCAGAGTT
AGCACACGTCTGAACTCTTTCCCTACACGACGCTCTTCCGATCTNNNTACAGACAACGGACGGTTTAG
AGCACACGTCTGAACTCTTTCCCTACACGACGCTCTTCCGATCTNNNTACAGACAACGGACGGTTTAG
AGCACACGTCTGAACTCTTTCCCTACACGACGCTCTTCCGATCTNNNAATGCAGGAGAATTGCTTGAA
AGCACACGTCTGAACTCTTTCCCTACACGACGCTCTTCCGATCTNNNAATGCAGGAGAATTGCTTGAA
GCACACGTCTGAACTCTTTCCCTACACGACGCTCTTCCGATCTNNNGGGAAATTCTAGCCCAGTGATC
GCACACGTCTGAACTCTTTCCCTACACGACGCTCTTCCGATCTNNNGGGAAATTCTAGCCCAGTGATC
CACACGTCTGAACTCTTTCCCTACACGACGCTCTTCCGATCTNNNAATGAAAAGGTACCCCTATTATTC
CACACGTCTGAACTCTTTCCCTACACGACGCTCTTCCGATCTNNNAATGAAAAGGTACCCCTATTATTC
GCACACGTCTGAACTCTTTCCCTACACGACGCTCTTCCGATCTNNNAGACAGAGTCTCACTGTTTCAC
GCACACGTCTGAACTCTTTCCCTACACGACGCTCTTCCGATCTNNNAGACAGAGTCTCACTGTTTCAC
GAGCACACGTCTGAACTCTTTCCCTACACGACGCTCTTCCGATCTNNNTGAGTGACAGAGCAAGACT
GAGCACACGTCTGAACTCTTTCCCTACACGACGCTCTTCCGATCTNNNTGAGTGACAGAGCAAGACT
GAGCACACGTCTGAACTCTTTCCCTACACGACGCTCTTCCGATCTNNNATATCCCCAAAGAACCACAG
GAGCACACGTCTGAACTCTTTCCCTACACGACGCTCTTCCGATCTNNNATATCCCCAAAGAACCACAG
AGCACACGTCTGAACTCTTTCCCTACACGACGCTCTTCCGATCTNNNGCAAATGACTCTTTTACAGGG
AGCACACGTCTGAACTCTTTCCCTACACGACGCTCTTCCGATCTNNNGCAAATGACTCTTTTACAGGG
GAGCACACGTCTGAACTCTTTCCCTACACGACGCTCTTCCGATCTNNNTCGGAATCAGTTGCAGTAAC
GAGCACACGTCTGAACTCTTTCCCTACACGACGCTCTTCCGATCTNNNTCGGAATCAGTTGCAGTAAC
GCACACGTCTGAACTCTTTCCCTACACGACGCTCTTCCGATCTNNNGCAGAGATGGAAAGGGAACTC
GCACACGTCTGAACTCTTTCCCTACACGACGCTCTTCCGATCTNNNGCAGAGATGGAAAGGGAACTC
AGCACACGTCTGAACTCTTTCCCTACACGACGCTCTTCCGATCTNNNTCCTGGGCTTAGGTGTTATGAA
AGCACACGTCTGAACTCTTTCCCTACACGACGCTCTTCCGATCTNNNTCCTGGGCTTAGGTGTTATGAA
AGCACACGTCTGAACTCTTTCCCTACACGACGCTCTTCCGATCTNNNATTTTTTCTACTCTTCACTTGT
AGCACACGTCTGAACTCTTTCCCTACACGACGCTCTTCCGATCTNNNATTTTTTCTACTCTTCACTTGT

AGCACACGTCTGAACTCTTTCCCTACACGACGCTCTTCCGATCTNNNTCATGCTCTGATTAGATAATCT

AGCACACGTCTGAACTCTTTCCCTACACGACGCTCTTCCGATCTNNNTCATGCTCTGATTAGATAATCT

AGAGCACACGTCTGAACTCTTTCCCTACACGACGCTCTTCCGATCTNNNACACCTGTAATCCCAGCTAC

AGAGCACACGTCTGAACTCTTTCCCTACACGACGCTCTTCCGATCTNNNACACCTGTAATCCCAGCTAC

AGCACACGTCTGAACTCTTTCCCTACACGACGCTCTTCCGATCTNNNGGGCTTCCTTAGAATACTGCAC

AGCACACGTCTGAACTCTTTCCCTACACGACGCTCTTCCGATCTNNNGGGCTTCCTTAGAATACTGCAC

GAGCACACGTCTGAACTCTTTCCCTACACGACGCTCTTCCGATCTNNNGTAGTCCAGCTACTAGGGAGG

GAGCACACGTCTGAACTCTTTCCCTACACGACGCTCTTCCGATCTNNNGTAGTCCAGCTACTAGGGAGG

GAGCACACGTCTGAACTCTTTCCCTACACGACGCTCTTCCGATCTNNNGATCACGAGGTCAGGAGATC

GAGCACACGTCTGAACTCTTTCCCTACACGACGCTCTTCCGATCTNNNGATCACGAGGTCAGGAGATC

GAGCACACGTCTGAACTCTTTCCCTACACGACGCTCTTCCGATCTNNNCGACTTGCCTAAGACCTGAGTT

GAGCACACGTCTGAACTCTTTCCCTACACGACGCTCTTCCGATCTNNNCGACTTGCCTAAGACCTGAGTT

AGAGCACACGTCTGAACTCTTTCCCTACACGACGCTCTTCCGATCTNNNACAGGAGAATCGTTTGTAC

AGAGCACACGTCTGAACTCTTTCCCTACACGACGCTCTTCCGATCTNNNACAGGAGAATCGTTTGTAC

GAGCACACGTCTGAACTCTTTCCCTACACGACGCTCTTCCGATCTNNNGAGATCCTTGAGTGTGGGTA

GAGCACACGTCTGAACTCTTTCCCTACACGACGCTCTTCCGATCTNNNGAGATCCTTGAGTGTGGGTA

GAGCACACGTCTGAACTCTTTCCCTACACGACGCTCTTCCGATCTNNNGAGAATCGCTTGAACTCAGGA

GAGCACACGTCTGAACTCTTTCCCTACACGACGCTCTTCCGATCTNNNGAGAATCGCTTGAACTCAGGA

AGCACACGTCTGAACTCTTTCCCTACACGACGCTCTTCCGATCTNNNACCTTCCCTCCACTATTGTCCTA

AGCACACGTCTGAACTCTTTCCCTACACGACGCTCTTCCGATCTNNNACCTTCCCTCCACTATTGTCCTA

AGCACACGTCTGAACTCTTTCCCTACACGACGCTCTTCCGATCTNNNTACCAGTGAATTTTTGCAATAAT

AGCACACGTCTGAACTCTTTCCCTACACGACGCTCTTCCGATCTNNNTACCAGTGAATTTTTGCAATAAT

GAGCACACGTCTGAACTCTTTCCCTACACGACGCTCTTCCGATCTNNNTACTGTAATAAGCAGCTGCATC

GAGCACACGTCTGAACTCTTTCCCTACACGACGCTCTTCCGATCTNNNTACTGTAATAAGCAGCTGCATC

GAGCACACGTCTGAACTCTTTCCCTACACGACGCTCTTCCGATCTNNNCTTGAGACCAGACTGACCAAC

GAGCACACGTCTGAACTCTTTCCCTACACGACGCTCTTCCGATCTNNNCTTGAGACCAGACTGACCAAC

GAGCACACGTCTGAACTCTTTCCCTACACGACGCTCTTCCGATCTNNNCGTGACTTCCATAACACATCAG

GAGCACACGTCTGAACTCTTTCCCTACACGACGCTCTTCCGATCTNNNCGTGACTTCCATAACACATCAG

AGCACACGTCTGAACTCTTTCCCTACACGACGCTCTTCCGATCTNNNAGAGCAAGACCCCATCTCAAAA

AGCACACGTCTGAACTCTTTCCCTACACGACGCTCTTCCGATCTNNNAGAGCAAGACCCCATCTCAAAA

CACACGTCTGAACTCTTTCCCTACACGACGCTCTTCCGATCTNNNATACCCCGACTCGATGGTAGAATCA

CACACGTCTGAACTCTTTCCCTACACGACGCTCTTCCGATCTNNNATACCCCGACTCGATGGTAGAATCA

AGCACACGTCTGAACTCTTTCCCTACACGACGCTCTTCCGATCTNNNGAGTTACCACACTGGAGAGAT

AGCACACGTCTGAACTCTTTCCCTACACGACGCTCTTCCGATCTNNNGAGTTACCACACTGGAGAGAT

AGCACACGTCTGAACTCTTTCCCTACACGACGCTCTTCCGATCTNNNTCCAGGCTTTACTCTCTCTCTCTC

AGCACACGTCTGAACTCTTTCCCTACACGACGCTCTTCCGATCTNNNTCCAGGCTTTACTCTCTCTCTCTC

GAGCACACGTCTGAACTCTTTCCCTACACGACGCTCTTCCGATCTNNNATAGCTTATTTCCCCCTCTGCA

GAGCACACGTCTGAACTCTTTCCCTACACGACGCTCTTCCGATCTNNNATAGCTTATTTCCCCCTCTGCA

GAGCACACGTCTGAACTCTTTCCCTACACGACGCTCTTCCGATCTNNNGCAGGAGAATTGCTTGAACCT

GAGCACACGTCTGAACTCTTTCCCTACACGACGCTCTTCCGATCTNNNGCAGGAGAATTGCTTGAACCT

GAGCACACGTCTGAACTCTTTCCCTACACGACGCTCTTCCGATCTNNNGCTCAGGAAATCTACCCACTTC

GAGCACACGTCTGAACTCTTTCCCTACACGACGCTCTTCCGATCTNNNGCTCAGGAAATCTACCCACTTC

AGCACACGTCTGAACTCTTTCCCTACACGACGCTCTTCCGATCTNNNCTCCTGGCTTTGTCCTACCTTTCC

AGCACACGTCTGAACTCTTTCCCTACACGACGCTCTTCCGATCTNNNCTCCTGGCTTTGTCCTACCTTTCC
GAGCACACGTCTGAACTCTTTCCCTACACGACGCTCTTCCGATCTNNNATGAACTCTGGCAAGTAGAG
GAGCACACGTCTGAACTCTTTCCCTACACGACGCTCTTCCGATCTNNNATGAACTCTGGCAAGTAGAG
AGCACACGTCTGAACTCTTTCCCTACACGACGCTCTTCCGATCTNNNAGAATCACTTGAATGCAGGAG
AGCACACGTCTGAACTCTTTCCCTACACGACGCTCTTCCGATCTNNNAGAATCACTTGAATGCAGGAG
AAGAGCACACGTCTGAACTCTTTCCCTACACGACGCTCTTCCGATCTNNNCCCGAGAGTTTTATGTTGC
AAGAGCACACGTCTGAACTCTTTCCCTACACGACGCTCTTCCGATCTNNNCCCGAGAGTTTTATGTTGC
GAGCACACGTCTGAACTCTTTCCCTACACGACGCTCTTCCGATCTNNNCCTTTGCTGCTTTTCTCCATGA
GAGCACACGTCTGAACTCTTTCCCTACACGACGCTCTTCCGATCTNNNCCTTTGCTGCTTTTCTCCATGA
AGCACACGTCTGAACTCTTTCCCTACACGACGCTCTTCCGATCTNNNTTTCTCACATGCCTCTTCCTACCC
AGCACACGTCTGAACTCTTTCCCTACACGACGCTCTTCCGATCTNNNTTTCTCACATGCCTCTTCCTACCC
GCACACGTCTGAACTCTTTCCCTACACGACGCTCTTCCGATCTNNNACACACCTACCACCTTTCGAAGCA
GCACACGTCTGAACTCTTTCCCTACACGACGCTCTTCCGATCTNNNACACACCTACCACCTTTCGAAGCA
GCACACGTCTGAACTCTTTCCCTACACGACGCTCTTCCGATCTNNNGCTGTACAATTGTTGGCAAGGA
GCACACGTCTGAACTCTTTCCCTACACGACGCTCTTCCGATCTNNNGCTGTACAATTGTTGGCAAGGA
AGCACACGTCTGAACTCTTTCCCTACACGACGCTCTTCCGATCTNNNGCCTCTGGGGACTTTGACTTT
AGCACACGTCTGAACTCTTTCCCTACACGACGCTCTTCCGATCTNNNGCCTCTGGGGACTTTGACTTT
AGCACACGTCTGAACTCTTTCCCTACACGACGCTCTTCCGATCTNNNGTAAAGCACTTTTCTGCCTCAGG
AGCACACGTCTGAACTCTTTCCCTACACGACGCTCTTCCGATCTNNNGTAAAGCACTTTTCTGCCTCAGG
AGCACACGTCTGAACTCTTTCCCTACACGACGCTCTTCCGATCTNNNCTCTGGGAATGAAACCCTGTC
AGCACACGTCTGAACTCTTTCCCTACACGACGCTCTTCCGATCTNNNCTCTGGGAATGAAACCCTGTC
AGCACACGTCTGAACTCTTTCCCTACACGACGCTCTTCCGATCTNNNAAAGTGCTGGGATTACAGGT
AGCACACGTCTGAACTCTTTCCCTACACGACGCTCTTCCGATCTNNNAAAGTGCTGGGATTACAGGT
AGCACACGTCTGAACTCTTTCCCTACACGACGCTCTTCCGATCTNNNGCTTCTCAAAGTGCTGGGATTA
AGCACACGTCTGAACTCTTTCCCTACACGACGCTCTTCCGATCTNNNGCTTCTCAAAGTGCTGGGATTA
AGCACACGTCTGAACTCTTTCCCTACACGACGCTCTTCCGATCTNNNTATGTCCTGCTTGGGAAAGTTG
AGCACACGTCTGAACTCTTTCCCTACACGACGCTCTTCCGATCTNNNTATGTCCTGCTTGGGAAAGTTG
AGCACACGTCTGAACTCTTTCCCTACACGACGCTCTTCCGATCTNNNTTTGTTCATGTCCAGGCTTCAT
AGCACACGTCTGAACTCTTTCCCTACACGACGCTCTTCCGATCTNNNTTTGTTCATGTCCAGGCTTCAT
GAAGAGCACACGTCTGAACTCTTTCCCTACACGACGCTCTTCCGATCTNNNGAGGCTGAGGCAGGAG
GAAGAGCACACGTCTGAACTCTTTCCCTACACGACGCTCTTCCGATCTNNNGAGGCTGAGGCAGGAG
AGAGCACACGTCTGAACTCTTTCCCTACACGACGCTCTTCCGATCTNNNAGTAAGCCAATTAGAACATG
AGAGCACACGTCTGAACTCTTTCCCTACACGACGCTCTTCCGATCTNNNAGTAAGCCAATTAGAACATG
AGAGCACACGTCTGAACTCTTTCCCTACACGACGCTCTTCCGATCTNNNGCTTTGGTCTCACAATGTA
AGAGCACACGTCTGAACTCTTTCCCTACACGACGCTCTTCCGATCTNNNGCTTTGGTCTCACAATGTA
AGCACACGTCTGAACTCTTTCCCTACACGACGCTCTTCCGATCTNNNAGCCCCAGTAGAAATGTTCTCT
AGCACACGTCTGAACTCTTTCCCTACACGACGCTCTTCCGATCTNNNAGCCCCAGTAGAAATGTTCTCT
GAGCACACGTCTGAACTCTTTCCCTACACGACGCTCTTCCGATCTNNNGAATCTACCTGAAAGCCCTTT
GAGCACACGTCTGAACTCTTTCCCTACACGACGCTCTTCCGATCTNNNGAATCTACCTGAAAGCCCTTT
GAGCACACGTCTGAACTCTTTCCCTACACGACGCTCTTCCGATCTNNNACCTAAGCCTTGGAGAAGTG
GAGCACACGTCTGAACTCTTTCCCTACACGACGCTCTTCCGATCTNNNACCTAAGCCTTGGAGAAGTG
GCACACGTCTGAACTCTTTCCCTACACGACGCTCTTCCGATCTNNNGACGGGGAAATACTGGGTAGA
GCACACGTCTGAACTCTTTCCCTACACGACGCTCTTCCGATCTNNNGACGGGGAAATACTGGGTAGA

AGCACACGTCTGAACTCTTTCCCTACACGACGCTCTTCCGATCTNNNCAAGAGAATTGCTTGAACCTG

AGCACACGTCTGAACTCTTTCCCTACACGACGCTCTTCCGATCTNNNCAAGAGAATTGCTTGAACCTG

AGCACACGTCTGAACTCTTTCCCTACACGACGCTCTTCCGATCTNNNATCTGCCTCCCCTTGATCTTTTT

AGCACACGTCTGAACTCTTTCCCTACACGACGCTCTTCCGATCTNNNATCTGCCTCCCCTTGATCTTTTT

GAGCACACGTCTGAACTCTTTCCCTACACGACGCTCTTCCGATCTNNNATCCTGTCATTATGATGCTGG

GAGCACACGTCTGAACTCTTTCCCTACACGACGCTCTTCCGATCTNNNATCCTGTCATTATGATGCTGG

GCACACGTCTGAACTCTTTCCCTACACGACGCTCTTCCGATCTNNNTCATAGCTCACTGTAGCCTTGAC

GCACACGTCTGAACTCTTTCCCTACACGACGCTCTTCCGATCTNNNTCATAGCTCACTGTAGCCTTGAC

AGCACACGTCTGAACTCTTTCCCTACACGACGCTCTTCCGATCTNNNCAAGAATTGCTTCAATCCGGG

AGCACACGTCTGAACTCTTTCCCTACACGACGCTCTTCCGATCTNNNCAAGAATTGCTTCAATCCGGG

AGCACACGTCTGAACTCTTTCCCTACACGACGCTCTTCCGATCTNNNAAGGCATATGCATTAGAGAG

AGCACACGTCTGAACTCTTTCCCTACACGACGCTCTTCCGATCTNNNAAGGCATATGCATTAGAGAG

GAGCACACGTCTGAACTCTTTCCCTACACGACGCTCTTCCGATCTNNNACAGAAAGCATATTGGTGGT

GAGCACACGTCTGAACTCTTTCCCTACACGACGCTCTTCCGATCTNNNACAGAAAGCATATTGGTGGT

CACACGTCTGAACTCTTTCCCTACACGACGCTCTTCCGATCTNNNCTCTGGAGTGAGAGCTTGGTGCA

CACACGTCTGAACTCTTTCCCTACACGACGCTCTTCCGATCTNNNCTCTGGAGTGAGAGCTTGGTGCA

AGCACACGTCTGAACTCTTTCCCTACACGACGCTCTTCCGATCTNNNAGAATTACTTGAACTCGGGAG

AGCACACGTCTGAACTCTTTCCCTACACGACGCTCTTCCGATCTNNNAGAATTACTTGAACTCGGGAG

AGCACACGTCTGAACTCTTTCCCTACACGACGCTCTTCCGATCTNNNTCAGAGACTAGTATTGCAACA

AGCACACGTCTGAACTCTTTCCCTACACGACGCTCTTCCGATCTNNNTCAGAGACTAGTATTGCAACA

GAGCACACGTCTGAACTCTTTCCCTACACGACGCTCTTCCGATCTNNNGAAGTCTTACAGGCCAGGAG

GAGCACACGTCTGAACTCTTTCCCTACACGACGCTCTTCCGATCTNNNGAAGTCTTACAGGCCAGGAG

GCACACGTCTGAACTCTTTCCCTACACGACGCTCTTCCGATCTNNNCTTCAGGATCTTGAGGCACTGA

GCACACGTCTGAACTCTTTCCCTACACGACGCTCTTCCGATCTNNNCTTCAGGATCTTGAGGCACTGA

CACGTCTGAACTCTTTCCCTACACGACGCTCTTCCGATCTNNNCCTTACCAAAAAGCAGCCAGACCATC

CACGTCTGAACTCTTTCCCTACACGACGCTCTTCCGATCTNNNCCTTACCAAAAAGCAGCCAGACCATC

GAGCACACGTCTGAACTCTTTCCCTACACGACGCTCTTCCGATCTNNNAATTCCTGGACACATACACCC

GAGCACACGTCTGAACTCTTTCCCTACACGACGCTCTTCCGATCTNNNAATTCCTGGACACATACACCC

GCACACGTCTGAACTCTTTCCCTACACGACGCTCTTCCGATCTNNNCACATTCTGTGCCTAAAGTGCTG

GCACACGTCTGAACTCTTTCCCTACACGACGCTCTTCCGATCTNNNCACATTCTGTGCCTAAAGTGCTG

GAGCACACGTCTGAACTCTTTCCCTACACGACGCTCTTCCGATCTNNNCGCCTATAGTCCCAGCTACTTG

GAGCACACGTCTGAACTCTTTCCCTACACGACGCTCTTCCGATCTNNNCGCCTATAGTCCCAGCTACTTG

GAGCACACGTCTGAACTCTTTCCCTACACGACGCTCTTCCGATCTNNNGAGGCTGAGGCAGGAGAAT

GAGCACACGTCTGAACTCTTTCCCTACACGACGCTCTTCCGATCTNNNGAGGCTGAGGCAGGAGAAT

AGCACACGTCTGAACTCTTTCCCTACACGACGCTCTTCCGATCTNNNAGAATTACTTGAACCTGGGAG

AGCACACGTCTGAACTCTTTCCCTACACGACGCTCTTCCGATCTNNNAGAATTACTTGAACCTGGGAG

GAGCACACGTCTGAACTCTTTCCCTACACGACGCTCTTCCGATCTNNNCTAGATGAGCAGCTCTGTTA

GAGCACACGTCTGAACTCTTTCCCTACACGACGCTCTTCCGATCTNNNCTAGATGAGCAGCTCTGTTA

AGCACACGTCTGAACTCTTTCCCTACACGACGCTCTTCCGATCTNNNAGAGTTGTACAGTCATCACTAC

AGCACACGTCTGAACTCTTTCCCTACACGACGCTCTTCCGATCTNNNAGAGTTGTACAGTCATCACTAC

GAGCACACGTCTGAACTCTTTCCCTACACGACGCTCTTCCGATCTNNNTGAGACCAGCCTGATGAATAT

GAGCACACGTCTGAACTCTTTCCCTACACGACGCTCTTCCGATCTNNNTGAGACCAGCCTGATGAATAT

AGCACACGTCTGAACTCTTTCCCTACACGACGCTCTTCCGATCTNNNCTGTAATCCCAGCTACTCAGGA

AGCACACGTCTGAACTCTTTCCCTACACGACGCTCTTCCGATCTNNNCTGTAATCCCAGCTACTCAGGA
AGCACACGTCTGAACTCTTTCCCTACACGACGCTCTTCCGATCTNNNCATGAGAACTGAAACTGCCCAA
AGCACACGTCTGAACTCTTTCCCTACACGACGCTCTTCCGATCTNNNCATGAGAACTGAAACTGCCCAA
GAGCACACGTCTGAACTCTTTCCCTACACGACGCTCTTCCGATCTNNNAGGATGGGGAGTAAGAAAA
GAGCACACGTCTGAACTCTTTCCCTACACGACGCTCTTCCGATCTNNNAGGATGGGGAGTAAGAAAA
AGCACACGTCTGAACTCTTTCCCTACACGACGCTCTTCCGATCTNNNGGACAGACCCATAGATCTGGA
AGCACACGTCTGAACTCTTTCCCTACACGACGCTCTTCCGATCTNNNGGACAGACCCATAGATCTGGA
GAGCACACGTCTGAACTCTTTCCCTACACGACGCTCTTCCGATCTNNNTCCTTCCTTCCTTCCTTCCTTC
GAGCACACGTCTGAACTCTTTCCCTACACGACGCTCTTCCGATCTNNNTCCTTCCTTCCTTCCTTCCTTC
CACACGTCTGAACTCTTTCCCTACACGACGCTCTTCCGATCTNNNCCTAACCCAGAACACCAGGAGCAT
CACACGTCTGAACTCTTTCCCTACACGACGCTCTTCCGATCTNNNCCTAACCCAGAACACCAGGAGCAT
GAGCACACGTCTGAACTCTTTCCCTACACGACGCTCTTCCGATCTNNNTCACCACCATCCAATTTTCGTT
GAGCACACGTCTGAACTCTTTCCCTACACGACGCTCTTCCGATCTNNNTCACCACCATCCAATTTTCGTT
GAGCACACGTCTGAACTCTTTCCCTACACGACGCTCTTCCGATCTNNNCAACTACTTGGGAGGCTGAGG
GAGCACACGTCTGAACTCTTTCCCTACACGACGCTCTTCCGATCTNNNCAACTACTTGGGAGGCTGAGG
AGCACACGTCTGAACTCTTTCCCTACACGACGCTCTTCCGATCTNNNTCATGCTCTGATTAGATAATCT
AGCACACGTCTGAACTCTTTCCCTACACGACGCTCTTCCGATCTNNNTCATGCTCTGATTAGATAATCT
GAGCACACGTCTGAACTCTTTCCCTACACGACGCTCTTCCGATCTNNNTGGGATGTCACTCTTGTTATG
GAGCACACGTCTGAACTCTTTCCCTACACGACGCTCTTCCGATCTNNNTGGGATGTCACTCTTGTTATG
AGAGCACACGTCTGAACTCTTTCCCTACACGACGCTCTTCCGATCTNNNACATGCTAAGCTTTGTGATCA
AGAGCACACGTCTGAACTCTTTCCCTACACGACGCTCTTCCGATCTNNNACATGCTAAGCTTTGTGATCA
GAGCACACGTCTGAACTCTTTCCCTACACGACGCTCTTCCGATCTNNNCCAAGAGGTAGAAACTGCAG
GAGCACACGTCTGAACTCTTTCCCTACACGACGCTCTTCCGATCTNNNCCAAGAGGTAGAAACTGCAG
GAGCACACGTCTGAACTCTTTCCCTACACGACGCTCTTCCGATCTNNNAGAAATGCCCGATTTACCAAGA
GAGCACACGTCTGAACTCTTTCCCTACACGACGCTCTTCCGATCTNNNAGAAATGCCCGATTTACCAAGA
GAGCACACGTCTGAACTCTTTCCCTACACGACGCTCTTCCGATCTNNNAAGTTAGGGCCGAACTTGAT
GAGCACACGTCTGAACTCTTTCCCTACACGACGCTCTTCCGATCTNNNAAGTTAGGGCCGAACTTGAT
GCACACGTCTGAACTCTTTCCCTACACGACGCTCTTCCGATCTNNNTGTTCTTGCTCTGACTCTCTCTCCA
GCACACGTCTGAACTCTTTCCCTACACGACGCTCTTCCGATCTNNNTGTTCTTGCTCTGACTCTCTCTCCA
ACACGTCTGAACTCTTTCCCTACACGACGCTCTTCCGATCTNNNCTCAATGAGAATTGACCCATCATCGG
ACACGTCTGAACTCTTTCCCTACACGACGCTCTTCCGATCTNNNCTCAATGAGAATTGACCCATCATCGG
AGCACACGTCTGAACTCTTTCCCTACACGACGCTCTTCCGATCTNNNGAGAATGGCGTGAATGTGGA
AGCACACGTCTGAACTCTTTCCCTACACGACGCTCTTCCGATCTNNNGAGAATGGCGTGAATGTGGA
GAGCACACGTCTGAACTCTTTCCCTACACGACGCTCTTCCGATCTNNNACTTTTATTTTTGTCTAGGATG
GAGCACACGTCTGAACTCTTTCCCTACACGACGCTCTTCCGATCTNNNACTTTTATTTTTGTCTAGGATG
GAGCACACGTCTGAACTCTTTCCCTACACGACGCTCTTCCGATCTNNNCATCCTCGCTAACATGGTGAA
GAGCACACGTCTGAACTCTTTCCCTACACGACGCTCTTCCGATCTNNNCATCCTCGCTAACATGGTGAA
GAGCACACGTCTGAACTCTTTCCCTACACGACGCTCTTCCGATCTNNNGAGAATGGTGTGAACCCAGG
GAGCACACGTCTGAACTCTTTCCCTACACGACGCTCTTCCGATCTNNNGAGAATGGTGTGAACCCAGG
AAGAGCACACGTCTGAACTCTTTCCCTACACGACGCTCTTCCGATCTNNNAACTACTCGGGAAGTTGA
AAGAGCACACGTCTGAACTCTTTCCCTACACGACGCTCTTCCGATCTNNNAACTACTCGGGAAGTTGA
GAGCACACGTCTGAACTCTTTCCCTACACGACGCTCTTCCGATCTNNNTCAGGTGGTTTCCATTTGGTA
GAGCACACGTCTGAACTCTTTCCCTACACGACGCTCTTCCGATCTNNNTCAGGTGGTTTCCATTTGGTA

CACGTCTGAACTCTTTCCCTACACGACGCTCTTCCGATCTNNNGGACACAGCCAAACCATATCAAACAT

CACGTCTGAACTCTTTCCCTACACGACGCTCTTCCGATCTNNNGGACACAGCCAAACCATATCAAACAT

AGCACACGTCTGAACTCTTTCCCTACACGACGCTCTTCCGATCTNNNACGTGGTGAAATTAAACAAA

AGCACACGTCTGAACTCTTTCCCTACACGACGCTCTTCCGATCTNNNACGTGGTGAAATTAAACAAA

AGCACACGTCTGAACTCTTTCCCTACACGACGCTCTTCCGATCTNNNCACCTGTAATCCCAGCTACTCA

AGCACACGTCTGAACTCTTTCCCTACACGACGCTCTTCCGATCTNNNCACCTGTAATCCCAGCTACTCA

AGCACACGTCTGAACTCTTTCCCTACACGACGCTCTTCCGATCTNNNCTTCAGTGTGCTCTGATGAACA

AGCACACGTCTGAACTCTTTCCCTACACGACGCTCTTCCGATCTNNNCTTCAGTGTGCTCTGATGAACA

GAGCACACGTCTGAACTCTTTCCCTACACGACGCTCTTCCGATCTNNNAAAAAATTAGCCGGGCGTA

GAGCACACGTCTGAACTCTTTCCCTACACGACGCTCTTCCGATCTNNNAAAAAATTAGCCGGGCGTA

AGCACACGTCTGAACTCTTTCCCTACACGACGCTCTTCCGATCTNNNCCTCCTATGATGGTGGCATCTT

AGCACACGTCTGAACTCTTTCCCTACACGACGCTCTTCCGATCTNNNCCTCCTATGATGGTGGCATCTT

AGCACACGTCTGAACTCTTTCCCTACACGACGCTCTTCCGATCTNNNCCATCATTTTTTCCACTGCCTGT

AGCACACGTCTGAACTCTTTCCCTACACGACGCTCTTCCGATCTNNNCCATCATTTTTTCCACTGCCTGT

GCACACGTCTGAACTCTTTCCCTACACGACGCTCTTCCGATCTNNNGAATGCAGGCATGATAGACCCT

GCACACGTCTGAACTCTTTCCCTACACGACGCTCTTCCGATCTNNNGAATGCAGGCATGATAGACCCT

GAGCACACGTCTGAACTCTTTCCCTACACGACGCTCTTCCGATCTNNNCCTCAGGAAACCCCTTTAGAT

GAGCACACGTCTGAACTCTTTCCCTACACGACGCTCTTCCGATCTNNNCCTCAGGAAACCCCTTTAGAT

GAGCACACGTCTGAACTCTTTCCCTACACGACGCTCTTCCGATCTNNNCTTGCAGTGAGCCGAGATCC

GAGCACACGTCTGAACTCTTTCCCTACACGACGCTCTTCCGATCTNNNCTTGCAGTGAGCCGAGATCC

GCACACGTCTGAACTCTTTCCCTACACGACGCTCTTCCGATCTNNNTGAGTCACAGTTCTCAAGGGAT

GCACACGTCTGAACTCTTTCCCTACACGACGCTCTTCCGATCTNNNTGAGTCACAGTTCTCAAGGGAT

AGCACACGTCTGAACTCTTTCCCTACACGACGCTCTTCCGATCTNNNTAGATCATGCCTGTAATCCCAG

AGCACACGTCTGAACTCTTTCCCTACACGACGCTCTTCCGATCTNNNTAGATCATGCCTGTAATCCCAG

AGCACACGTCTGAACTCTTTCCCTACACGACGCTCTTCCGATCTNNNGTATTTCTGCAGTCCCAGCTAC

AGCACACGTCTGAACTCTTTCCCTACACGACGCTCTTCCGATCTNNNGTATTTCTGCAGTCCCAGCTAC

AGCACACGTCTGAACTCTTTCCCTACACGACGCTCTTCCGATCTNNNCCTCCTCCTCCTTCTTCTTCCTCA

AGCACACGTCTGAACTCTTTCCCTACACGACGCTCTTCCGATCTNNNCCTCCTCCTCCTTCTTCTTCCTCA

GAGCACACGTCTGAACTCTTTCCCTACACGACGCTCTTCCGATCTNNNAAACCTACCCTGAAAGAACA

GAGCACACGTCTGAACTCTTTCCCTACACGACGCTCTTCCGATCTNNNAAACCTACCCTGAAAGAACA

GCACACGTCTGAACTCTTTCCCTACACGACGCTCTTCCGATCTNNNTGGTGTGTGTATGTGTGTGTG

GCACACGTCTGAACTCTTTCCCTACACGACGCTCTTCCGATCTNNNTGGTGTGTGTATGTGTGTGTG

AGCACACGTCTGAACTCTTTCCCTACACGACGCTCTTCCGATCTNNNGATGGTCTTGATCTCCTGACC

AGCACACGTCTGAACTCTTTCCCTACACGACGCTCTTCCGATCTNNNGATGGTCTTGATCTCCTGACC

AGCACACGTCTGAACTCTTTCCCTACACGACGCTCTTCCGATCTNNNTTCCAAAAAACTGAGGAGGA

AGCACACGTCTGAACTCTTTCCCTACACGACGCTCTTCCGATCTNNNTTCCAAAAAACTGAGGAGGA

AGCACACGTCTGAACTCTTTCCCTACACGACGCTCTTCCGATCTNNNACGTTGGCATTGAGTAAGTTT

AGCACACGTCTGAACTCTTTCCCTACACGACGCTCTTCCGATCTNNNACGTTGGCATTGAGTAAGTTT

GAGCACACGTCTGAACTCTTTCCCTACACGACGCTCTTCCGATCTNNNGGTGCCGACAAGTCCTCTAAT

GAGCACACGTCTGAACTCTTTCCCTACACGACGCTCTTCCGATCTNNNGGTGCCGACAAGTCCTCTAAT

AGCACACGTCTGAACTCTTTCCCTACACGACGCTCTTCCGATCTNNNTTGAAGAACGGCTGCCTTTATT

AGCACACGTCTGAACTCTTTCCCTACACGACGCTCTTCCGATCTNNNTTGAAGAACGGCTGCCTTTATT

ACACGTCTGAACTCTTTCCCTACACGACGCTCTTCCGATCTNNNTTTTCTGTCGCTTTAATCTCATTCAT

ACACGTCTGAACTCTTTCCCTACACGACGCTCTTCCGATCTNNNTTTTCTGTCGCTTTAATCTCATTCAT
GCACACGTCTGAACTCTTTCCCTACACGACGCTCTTCCGATCTNNNTCTTTGGGATGAGTGCTCAACT
GCACACGTCTGAACTCTTTCCCTACACGACGCTCTTCCGATCTNNNTCTTTGGGATGAGTGCTCAACT
AGAGCACACGTCTGAACTCTTTCCCTACACGACGCTCTTCCGATCTNNNGTGCATGTATGTGTGTGTA
AGAGCACACGTCTGAACTCTTTCCCTACACGACGCTCTTCCGATCTNNNGTGCATGTATGTGTGTGTA
AGCACACGTCTGAACTCTTTCCCTACACGACGCTCTTCCGATCTNNNTAGATAGTTCCCTTTGCACACC
AGCACACGTCTGAACTCTTTCCCTACACGACGCTCTTCCGATCTNNNTAGATAGTTCCCTTTGCACACC
AGCACACGTCTGAACTCTTTCCCTACACGACGCTCTTCCGATCTNNNTCGTATGAGAACATGAGTGG
AGCACACGTCTGAACTCTTTCCCTACACGACGCTCTTCCGATCTNNNTCGTATGAGAACATGAGTGG
AGCACACGTCTGAACTCTTTCCCTACACGACGCTCTTCCGATCTNNNATACAAAAATTAGCCGGGCAT
AGCACACGTCTGAACTCTTTCCCTACACGACGCTCTTCCGATCTNNNATACAAAAATTAGCCGGGCAT
AGCACACGTCTGAACTCTTTCCCTACACGACGCTCTTCCGATCTNNNAAGCAACCAAGAAAAATCTGG
AGCACACGTCTGAACTCTTTCCCTACACGACGCTCTTCCGATCTNNNAAGCAACCAAGAAAAATCTGG
GCACACGTCTGAACTCTTTCCCTACACGACGCTCTTCCGATCTNNNATATTCATTGCACCAAAATGTAT
GCACACGTCTGAACTCTTTCCCTACACGACGCTCTTCCGATCTNNNATATTCATTGCACCAAAATGTAT
GAGCACACGTCTGAACTCTTTCCCTACACGACGCTCTTCCGATCTNNNCAAAAGAGCCAGACCCAGTC
GAGCACACGTCTGAACTCTTTCCCTACACGACGCTCTTCCGATCTNNNCAAAAGAGCCAGACCCAGTC
AGCACACGTCTGAACTCTTTCCCTACACGACGCTCTTCCGATCTNNNCGTGGACCTGTAATCCTAGCTA
AGCACACGTCTGAACTCTTTCCCTACACGACGCTCTTCCGATCTNNNCGTGGACCTGTAATCCTAGCTA
GCACACGTCTGAACTCTTTCCCTACACGACGCTCTTCCGATCTNNNCCATCTGGTGAAACTGGTTTGT
GCACACGTCTGAACTCTTTCCCTACACGACGCTCTTCCGATCTNNNCCATCTGGTGAAACTGGTTTGT
GCACACGTCTGAACTCTTTCCCTACACGACGCTCTTCCGATCTNNNTTTTGAGATGGAGTCCTGCTCT
GCACACGTCTGAACTCTTTCCCTACACGACGCTCTTCCGATCTNNNTTTTGAGATGGAGTCCTGCTCT
AGCACACGTCTGAACTCTTTCCCTACACGACGCTCTTCCGATCTNNNAAACTGTGTGTGTTTGTGTGT
AGCACACGTCTGAACTCTTTCCCTACACGACGCTCTTCCGATCTNNNAAACTGTGTGTGTTTGTGTGT
AGCACACGTCTGAACTCTTTCCCTACACGACGCTCTTCCGATCTNNNGAGGCAGGAAAATCACTTGA
AGCACACGTCTGAACTCTTTCCCTACACGACGCTCTTCCGATCTNNNGAGGCAGGAAAATCACTTGA
GAGCACACGTCTGAACTCTTTCCCTACACGACGCTCTTCCGATCTNNNTCACTTGAACTCGAAGGACA
GAGCACACGTCTGAACTCTTTCCCTACACGACGCTCTTCCGATCTNNNTCACTTGAACTCGAAGGACA
GCACACGTCTGAACTCTTTCCCTACACGACGCTCTTCCGATCTNNNTCCTCATTCTTTCTTCTGCTGCTC
GCACACGTCTGAACTCTTTCCCTACACGACGCTCTTCCGATCTNNNTCCTCATTCTTTCTTCTGCTGCTC
GAGCACACGTCTGAACTCTTTCCCTACACGACGCTCTTCCGATCTNNNGCTGTCTGCATCTTGGTAAA
GAGCACACGTCTGAACTCTTTCCCTACACGACGCTCTTCCGATCTNNNGCTGTCTGCATCTTGGTAAA
GCACACGTCTGAACTCTTTCCCTACACGACGCTCTTCCGATCTNNNAGGTTGGGTCTGAGCACAAGC
GCACACGTCTGAACTCTTTCCCTACACGACGCTCTTCCGATCTNNNAGGTTGGGTCTGAGCACAAGC
GCACACGTCTGAACTCTTTCCCTACACGACGCTCTTCCGATCTNNNACTGTAACCTCGAACTCCTAGG
GCACACGTCTGAACTCTTTCCCTACACGACGCTCTTCCGATCTNNNACTGTAACCTCGAACTCCTAGG
GCACACGTCTGAACTCTTTCCCTACACGACGCTCTTCCGATCTNNNTTGAGAGAGAGAAAGAGCACG
GCACACGTCTGAACTCTTTCCCTACACGACGCTCTTCCGATCTNNNTTGAGAGAGAGAAAGAGCACG
GCACACGTCTGAACTCTTTCCCTACACGACGCTCTTCCGATCTNNNAAAAAATTAGCCAGGTGTGGT
GCACACGTCTGAACTCTTTCCCTACACGACGCTCTTCCGATCTNNNAAAAAATTAGCCAGGTGTGGT
CACACGTCTGAACTCTTTCCCTACACGACGCTCTTCCGATCTNNNCAGCCTGGGTGATAGAGTAAGA
CACACGTCTGAACTCTTTCCCTACACGACGCTCTTCCGATCTNNNCAGCCTGGGTGATAGAGTAAGA

GAGCACACGTCTGAACTCTTTCCCTACACGACGCTCTTCCGATCTNNNAAGCATGAAGGAGATTTCCA
GAGCACACGTCTGAACTCTTTCCCTACACGACGCTCTTCCGATCTNNNAAGCATGAAGGAGATTTCCA
GCACACGTCTGAACTCTTTCCCTACACGACGCTCTTCCGATCTNNNTGAGTGTAGACTGTTGGAGAAG
GCACACGTCTGAACTCTTTCCCTACACGACGCTCTTCCGATCTNNNTGAGTGTAGACTGTTGGAGAAG
GAGCACACGTCTGAACTCTTTCCCTACACGACGCTCTTCCGATCTNNNGTAATCCCAGCTATACAGGA
GAGCACACGTCTGAACTCTTTCCCTACACGACGCTCTTCCGATCTNNNGTAATCCCAGCTATACAGGA
AGCACACGTCTGAACTCTTTCCCTACACGACGCTCTTCCGATCTNNNAGCAAAGATAGAGGGGTGTC
AGCACACGTCTGAACTCTTTCCCTACACGACGCTCTTCCGATCTNNNAGCAAAGATAGAGGGGTGTC
GCACACGTCTGAACTCTTTCCCTACACGACGCTCTTCCGATCTNNNTGTTCCAACATTTGAAATGCTTC
GCACACGTCTGAACTCTTTCCCTACACGACGCTCTTCCGATCTNNNTGTTCCAACATTTGAAATGCTTC
AGAGCACACGTCTGAACTCTTTCCCTACACGACGCTCTTCCGATCTNNNCCAGGGTGAACATCCTCTGA
AGAGCACACGTCTGAACTCTTTCCCTACACGACGCTCTTCCGATCTNNNCCAGGGTGAACATCCTCTGA
GCACACGTCTGAACTCTTTCCCTACACGACGCTCTTCCGATCTNNNTTGCTGCATCAGTATCCATCAGA
GCACACGTCTGAACTCTTTCCCTACACGACGCTCTTCCGATCTNNNTTGCTGCATCAGTATCCATCAGA
AGCACACGTCTGAACTCTTTCCCTACACGACGCTCTTCCGATCTNNNCACACAGCTGAACAAATGTCC
AGCACACGTCTGAACTCTTTCCCTACACGACGCTCTTCCGATCTNNNCACACAGCTGAACAAATGTCC
AGCACACGTCTGAACTCTTTCCCTACACGACGCTCTTCCGATCTNNNCTCTGCTGTTTTTCTTGTCCGA
AGCACACGTCTGAACTCTTTCCCTACACGACGCTCTTCCGATCTNNNCTCTGCTGTTTTTCTTGTCCGA
AGCACACGTCTGAACTCTTTCCCTACACGACGCTCTTCCGATCTNNNCAGTGAGACCCCTACAACTTCTT
AGCACACGTCTGAACTCTTTCCCTACACGACGCTCTTCCGATCTNNNCAGTGAGACCCCTACAACTTCTT
AGCACACGTCTGAACTCTTTCCCTACACGACGCTCTTCCGATCTNNNATCATACCACTACACTCCCTCCT
AGCACACGTCTGAACTCTTTCCCTACACGACGCTCTTCCGATCTNNNATCATACCACTACACTCCCTCCT
AGCACACGTCTGAACTCTTTCCCTACACGACGCTCTTCCGATCTNNNCTGTAATCCAAGCTACTCGGG
AGCACACGTCTGAACTCTTTCCCTACACGACGCTCTTCCGATCTNNNCTGTAATCCAAGCTACTCGGG
GAGCACACGTCTGAACTCTTTCCCTACACGACGCTCTTCCGATCTNNNGCAGGAGAATTGCTTGAACC
GAGCACACGTCTGAACTCTTTCCCTACACGACGCTCTTCCGATCTNNNGCAGGAGAATTGCTTGAACC
AGCACACGTCTGAACTCTTTCCCTACACGACGCTCTTCCGATCTNNNCTGCTGTGTGAAAATGCTGTT
AGCACACGTCTGAACTCTTTCCCTACACGACGCTCTTCCGATCTNNNCTGCTGTGTGAAAATGCTGTT
AGCACACGTCTGAACTCTTTCCCTACACGACGCTCTTCCGATCTNNNTGGTTTTTTTGTTGGTGGTG
AGCACACGTCTGAACTCTTTCCCTACACGACGCTCTTCCGATCTNNNTGGTTTTTTTGTTGGTGGTG
AGCACACGTCTGAACTCTTTCCCTACACGACGCTCTTCCGATCTNNNCACCTGTAGTGCCAGCTACTC
AGCACACGTCTGAACTCTTTCCCTACACGACGCTCTTCCGATCTNNNCACCTGTAGTGCCAGCTACTC
AGCACACGTCTGAACTCTTTCCCTACACGACGCTCTTCCGATCTNNNAATGGCACCATCACAGTTCACT
AGCACACGTCTGAACTCTTTCCCTACACGACGCTCTTCCGATCTNNNAATGGCACCATCACAGTTCACT
AGCACACGTCTGAACTCTTTCCCTACACGACGCTCTTCCGATCTNNNGATCTTGGCTCACTACAAGCT
AGCACACGTCTGAACTCTTTCCCTACACGACGCTCTTCCGATCTNNNGATCTTGGCTCACTACAAGCT
AGCACACGTCTGAACTCTTTCCCTACACGACGCTCTTCCGATCTNNNGAGAGCTCACCATTACACTCCA
AGCACACGTCTGAACTCTTTCCCTACACGACGCTCTTCCGATCTNNNGAGAGCTCACCATTACACTCCA
GCACACGTCTGAACTCTTTCCCTACACGACGCTCTTCCGATCTNNNTAGTACTAAGTTGGAGGCAGG
GCACACGTCTGAACTCTTTCCCTACACGACGCTCTTCCGATCTNNNTAGTACTAAGTTGGAGGCAGG
AGCACACGTCTGAACTCTTTCCCTACACGACGCTCTTCCGATCTNNNTCACTTGAGGTCAGAAGTTCG
AGCACACGTCTGAACTCTTTCCCTACACGACGCTCTTCCGATCTNNNTCACTTGAGGTCAGAAGTTCG
AGAGCACACGTCTGAACTCTTTCCCTACACGACGCTCTTCCGATCTNNNTAGGGAAAACACCACAAAA

AGAGCACACGTCTGAACTCTTTCCCTACACGACGCTCTTCCGATCTNNNTAGGGAAAACACCACAAAA
AGCACACGTCTGAACTCTTTCCCTACACGACGCTCTTCCGATCTNNNATACATCTCTCCATGGGATGTG
AGCACACGTCTGAACTCTTTCCCTACACGACGCTCTTCCGATCTNNNATACATCTCTCCATGGGATGTG
AGCACACGTCTGAACTCTTTCCCTACACGACGCTCTTCCGATCTNNNCTTTATGGTGGGGAGGAAAGG
AGCACACGTCTGAACTCTTTCCCTACACGACGCTCTTCCGATCTNNNCTTTATGGTGGGGAGGAAAGG
AGCACACGTCTGAACTCTTTCCCTACACGACGCTCTTCCGATCTNNNATTTCGAGAAAGAGCCCTGAG
AGCACACGTCTGAACTCTTTCCCTACACGACGCTCTTCCGATCTNNNATTTCGAGAAAGAGCCCTGAG
GAGCACACGTCTGAACTCTTTCCCTACACGACGCTCTTCCGATCTNNNCCCACCTCATCTTCTCAAGTAG
GAGCACACGTCTGAACTCTTTCCCTACACGACGCTCTTCCGATCTNNNCCCACCTCATCTTCTCAAGTAG
GAGCACACGTCTGAACTCTTTCCCTACACGACGCTCTTCCGATCTNNNGCACCAGGCATTATCAAAGTG
GAGCACACGTCTGAACTCTTTCCCTACACGACGCTCTTCCGATCTNNNGCACCAGGCATTATCAAAGTG
GAGCACACGTCTGAACTCTTTCCCTACACGACGCTCTTCCGATCTNNNGAATCGCTTGAACCTGGGAAA
GAGCACACGTCTGAACTCTTTCCCTACACGACGCTCTTCCGATCTNNNGAATCGCTTGAACCTGGGAAA
GAGCACACGTCTGAACTCTTTCCCTACACGACGCTCTTCCGATCTNNNATTTTCTCTCCCACACACTCCAG
GAGCACACGTCTGAACTCTTTCCCTACACGACGCTCTTCCGATCTNNNATTTTCTCTCCCACACACTCCAG
GAGCACACGTCTGAACTCTTTCCCTACACGACGCTCTTCCGATCTNNNCATTGCAGTGAGCTCAGATCA
GAGCACACGTCTGAACTCTTTCCCTACACGACGCTCTTCCGATCTNNNCATTGCAGTGAGCTCAGATCA
AGCACACGTCTGAACTCTTTCCCTACACGACGCTCTTCCGATCTNNNAGTCACTTCTGCAGGATTTAGCA
AGCACACGTCTGAACTCTTTCCCTACACGACGCTCTTCCGATCTNNNAGTCACTTCTGCAGGATTTAGCA
AGAGCACACGTCTGAACTCTTTCCCTACACGACGCTCTTCCGATCTNNNTACAAAAAATTAGCTGGGCG
AGAGCACACGTCTGAACTCTTTCCCTACACGACGCTCTTCCGATCTNNNTACAAAAAATTAGCTGGGCG
AGCACACGTCTGAACTCTTTCCCTACACGACGCTCTTCCGATCTNNNTCTGGAGTAGACATCCTCCTCTG
AGCACACGTCTGAACTCTTTCCCTACACGACGCTCTTCCGATCTNNNTCTGGAGTAGACATCCTCCTCTG
AGCACACGTCTGAACTCTTTCCCTACACGACGCTCTTCCGATCTNNNTTCTGTCCCTTCCAAAAGCACAA
AGCACACGTCTGAACTCTTTCCCTACACGACGCTCTTCCGATCTNNNTTCTGTCCCTTCCAAAAGCACAA
AGCACACGTCTGAACTCTTTCCCTACACGACGCTCTTCCGATCTNNNATGAACCCCTAAAGCTGACAGA
AGCACACGTCTGAACTCTTTCCCTACACGACGCTCTTCCGATCTNNNATGAACCCCTAAAGCTGACAGA
AGCACACGTCTGAACTCTTTCCCTACACGACGCTCTTCCGATCTNNNAGAAAGGTTTGCTTCTGATGCT
AGCACACGTCTGAACTCTTTCCCTACACGACGCTCTTCCGATCTNNNAGAAAGGTTTGCTTCTGATGCT
AGCACACGTCTGAACTCTTTCCCTACACGACGCTCTTCCGATCTNNNACCTCCCAATTCACTGAGGATTC
AGCACACGTCTGAACTCTTTCCCTACACGACGCTCTTCCGATCTNNNACCTCCCAATTCACTGAGGATTC
GAGCACACGTCTGAACTCTTTCCCTACACGACGCTCTTCCGATCTNNNTTTCACACCATTCTCCTGCCTCC
GAGCACACGTCTGAACTCTTTCCCTACACGACGCTCTTCCGATCTNNNTTTCACACCATTCTCCTGCCTCC
AGCACACGTCTGAACTCTTTCCCTACACGACGCTCTTCCGATCTNNNTTCTGGTAAGAGTCATGTGAGC
AGCACACGTCTGAACTCTTTCCCTACACGACGCTCTTCCGATCTNNNTTCTGGTAAGAGTCATGTGAGC
GAGCACACGTCTGAACTCTTTCCCTACACGACGCTCTTCCGATCTNNNGTTTGAAACTGAGAGGTCAAG
GAGCACACGTCTGAACTCTTTCCCTACACGACGCTCTTCCGATCTNNNGTTTGAAACTGAGAGGTCAAG
GAGCACACGTCTGAACTCTTTCCCTACACGACGCTCTTCCGATCTNNNTTACCCACTGTGGATTTCACCA
GAGCACACGTCTGAACTCTTTCCCTACACGACGCTCTTCCGATCTNNNTTACCCACTGTGGATTTCACCA
CACACGTCTGAACTCTTTCCCTACACGACGCTCTTCCGATCTNNNTGTGACTCTCATCCCCGTTTTAGCAT
CACACGTCTGAACTCTTTCCCTACACGACGCTCTTCCGATCTNNNTGTGACTCTCATCCCCGTTTTAGCAT
AGCACACGTCTGAACTCTTTCCCTACACGACGCTCTTCCGATCTNNNGAGAATCGCTTGAACCTGGGAC
AGCACACGTCTGAACTCTTTCCCTACACGACGCTCTTCCGATCTNNNGAGAATCGCTTGAACCTGGGAC

CACACGTCTGAACTCTTTCCCTACACGACGCTCTTCCGATCTNNNTAGTATCAGAAAAGCTTAAGGCC
CACACGTCTGAACTCTTTCCCTACACGACGCTCTTCCGATCTNNNTAGTATCAGAAAAGCTTAAGGCC
AGCACACGTCTGAACTCTTTCCCTACACGACGCTCTTCCGATCTNNNATTCTGTCTGGACTCTACTTTCA
AGCACACGTCTGAACTCTTTCCCTACACGACGCTCTTCCGATCTNNNATTCTGTCTGGACTCTACTTTCA
AGCACACGTCTGAACTCTTTCCCTACACGACGCTCTTCCGATCTNNNCTGGTCTTGAACTTCTGGACTC
AGCACACGTCTGAACTCTTTCCCTACACGACGCTCTTCCGATCTNNNCTGGTCTTGAACTTCTGGACTC
AGCACACGTCTGAACTCTTTCCCTACACGACGCTCTTCCGATCTNNNCAGCCTGTGAGGTTTAGCTCA
AGCACACGTCTGAACTCTTTCCCTACACGACGCTCTTCCGATCTNNNCAGCCTGTGAGGTTTAGCTCA
AGCACACGTCTGAACTCTTTCCCTACACGACGCTCTTCCGATCTNNNGTCGGGACACAAATCAAAACC
AGCACACGTCTGAACTCTTTCCCTACACGACGCTCTTCCGATCTNNNGTCGGGACACAAATCAAAACC
GAGCACACGTCTGAACTCTTTCCCTACACGACGCTCTTCCGATCTNNNGCTGAGATTACAGGAGCACC
GAGCACACGTCTGAACTCTTTCCCTACACGACGCTCTTCCGATCTNNNGCTGAGATTACAGGAGCACC
GAGCACACGTCTGAACTCTTTCCCTACACGACGCTCTTCCGATCTNNNGAGAATGGCATGAACCTGG
GAGCACACGTCTGAACTCTTTCCCTACACGACGCTCTTCCGATCTNNNGAGAATGGCATGAACCTGG
AGCACACGTCTGAACTCTTTCCCTACACGACGCTCTTCCGATCTNNNGGAGGTTGAAGTGAGCTGA
AGCACACGTCTGAACTCTTTCCCTACACGACGCTCTTCCGATCTNNNGGAGGTTGAAGTGAGCTGA
AGCACACGTCTGAACTCTTTCCCTACACGACGCTCTTCCGATCTNNNGAGGTTGCAGTGAGCTGAG
AGCACACGTCTGAACTCTTTCCCTACACGACGCTCTTCCGATCTNNNGAGGTTGCAGTGAGCTGAG
AGCACACGTCTGAACTCTTTCCCTACACGACGCTCTTCCGATCTNNNTCTGGTTTTCTGAACTATCGGG
AGCACACGTCTGAACTCTTTCCCTACACGACGCTCTTCCGATCTNNNTCTGGTTTTCTGAACTATCGGG
AGCACACGTCTGAACTCTTTCCCTACACGACGCTCTTCCGATCTNNNTCGATTTGAGCTCAGAAGTTC
AGCACACGTCTGAACTCTTTCCCTACACGACGCTCTTCCGATCTNNNTCGATTTGAGCTCAGAAGTTC
GCACACGTCTGAACTCTTTCCCTACACGACGCTCTTCCGATCTNNNTTCTGTTGTGCCTAGTACACTGT
GCACACGTCTGAACTCTTTCCCTACACGACGCTCTTCCGATCTNNNTTCTGTTGTGCCTAGTACACTGT
AGCACACGTCTGAACTCTTTCCCTACACGACGCTCTTCCGATCTNNNGAATCTGTGAGCTCAGAGGGA
AGCACACGTCTGAACTCTTTCCCTACACGACGCTCTTCCGATCTNNNGAATCTGTGAGCTCAGAGGGA
AGCACACGTCTGAACTCTTTCCCTACACGACGCTCTTCCGATCTNNNTGTGTTTGTGTACATGCAAGC
AGCACACGTCTGAACTCTTTCCCTACACGACGCTCTTCCGATCTNNNTGTGTTTGTGTACATGCAAGC
AGCACACGTCTGAACTCTTTCCCTACACGACGCTCTTCCGATCTNNNTCAAAATGAGGGTGTCTGCTG
AGCACACGTCTGAACTCTTTCCCTACACGACGCTCTTCCGATCTNNNTCAAAATGAGGGTGTCTGCTG
CACACGTCTGAACTCTTTCCCTACACGACGCTCTTCCGATCTNNNCCTCCTCTCCTCCTCTCAGGCATCGG
CACACGTCTGAACTCTTTCCCTACACGACGCTCTTCCGATCTNNNCCTCCTCTCCTCCTCTCAGGCATCGG
AGCACACGTCTGAACTCTTTCCCTACACGACGCTCTTCCGATCTNNNGAGAGACCCTCAGATGCACAC
AGCACACGTCTGAACTCTTTCCCTACACGACGCTCTTCCGATCTNNNGAGAGACCCTCAGATGCACAC
AGCACACGTCTGAACTCTTTCCCTACACGACGCTCTTCCGATCTNNNCTCAATCCCTTTCGCAATAGCT
AGCACACGTCTGAACTCTTTCCCTACACGACGCTCTTCCGATCTNNNCTCAATCCCTTTCGCAATAGCT
GAGCACACGTCTGAACTCTTTCCCTACACGACGCTCTTCCGATCTNNNTACACTCTCACACTGTCCTGG
GAGCACACGTCTGAACTCTTTCCCTACACGACGCTCTTCCGATCTNNNTACACTCTCACACTGTCCTGG
AGCACACGTCTGAACTCTTTCCCTACACGACGCTCTTCCGATCTNNNTTGCAGACTTGTCTATCCCACT
AGCACACGTCTGAACTCTTTCCCTACACGACGCTCTTCCGATCTNNNTTGCAGACTTGTCTATCCCACT
CACACGTCTGAACTCTTTCCCTACACGACGCTCTTCCGATCTNNNGTGATACTGGAAACACACACAGG
CACACGTCTGAACTCTTTCCCTACACGACGCTCTTCCGATCTNNNGTGATACTGGAAACACACACAGG
GAGCACACGTCTGAACTCTTTCCCTACACGACGCTCTTCCGATCTNNNAGAGGTAGGGGAACAACA

GAGCACACGTCTGAACTCTTTCCCTACACGACGCTCTTCCGATCTNNNAGAGGTAGGGGAACAACA
AGAGCACACGTCTGAACTCTTTCCCTACACGACGCTCTTCCGATCTNNNGGCTGAGCTTCTCAGTGCAT
AGAGCACACGTCTGAACTCTTTCCCTACACGACGCTCTTCCGATCTNNNGGCTGAGCTTCTCAGTGCAT
GCACACGTCTGAACTCTTTCCCTACACGACGCTCTTCCGATCTNNNGATGGAGGTGGAGGTCCTAGA
GCACACGTCTGAACTCTTTCCCTACACGACGCTCTTCCGATCTNNNGATGGAGGTGGAGGTCCTAGA
GCACACGTCTGAACTCTTTCCCTACACGACGCTCTTCCGATCTNNNCTAAGGATAATGGCCTCCACCT
GCACACGTCTGAACTCTTTCCCTACACGACGCTCTTCCGATCTNNNCTAAGGATAATGGCCTCCACCT
ACACGTCTGAACTCTTTCCCTACACGACGCTCTTCCGATCTNNNCTGAGGGGGTCGTCAGTACCCATC
ACACGTCTGAACTCTTTCCCTACACGACGCTCTTCCGATCTNNNCTGAGGGGGTCGTCAGTACCCATC
GAGCACACGTCTGAACTCTTTCCCTACACGACGCTCTTCCGATCTNNNATCAACCTAAGTGCCCATCAA
GAGCACACGTCTGAACTCTTTCCCTACACGACGCTCTTCCGATCTNNNATCAACCTAAGTGCCCATCAA
CACACGTCTGAACTCTTTCCCTACACGACGCTCTTCCGATCTNNNTATCCGTGTAGACTTGGGAGGA
CACACGTCTGAACTCTTTCCCTACACGACGCTCTTCCGATCTNNNTATCCGTGTAGACTTGGGAGGA
AGCACACGTCTGAACTCTTTCCCTACACGACGCTCTTCCGATCTNNNAGTGTTGTGAGTGTTGGATG
AGCACACGTCTGAACTCTTTCCCTACACGACGCTCTTCCGATCTNNNAGTGTTGTGAGTGTTGGATG
GCACACGTCTGAACTCTTTCCCTACACGACGCTCTTCCGATCTNNNTGAAACCCTTGTCTATTGCTGGT
GCACACGTCTGAACTCTTTCCCTACACGACGCTCTTCCGATCTNNNTGAAACCCTTGTCTATTGCTGGT
GCACACGTCTGAACTCTTTCCCTACACGACGCTCTTCCGATCTNNNTCAGGACAATCCCAAGGTGAAA
GCACACGTCTGAACTCTTTCCCTACACGACGCTCTTCCGATCTNNNTCAGGACAATCCCAAGGTGAAA
GCACACGTCTGAACTCTTTCCCTACACGACGCTCTTCCGATCTNNNTAGGTATACACATGCCATGGTG
GCACACGTCTGAACTCTTTCCCTACACGACGCTCTTCCGATCTNNNTAGGTATACACATGCCATGGTG
GAGCACACGTCTGAACTCTTTCCCTACACGACGCTCTTCCGATCTNNNCTCTATTGTGAAGGAGAGGT
GAGCACACGTCTGAACTCTTTCCCTACACGACGCTCTTCCGATCTNNNCTCTATTGTGAAGGAGAGGT
GCACACGTCTGAACTCTTTCCCTACACGACGCTCTTCCGATCTNNNTGGAATTCACCAGCAATTCACAT
GCACACGTCTGAACTCTTTCCCTACACGACGCTCTTCCGATCTNNNTGGAATTCACCAGCAATTCACAT
GCACACGTCTGAACTCTTTCCCTACACGACGCTCTTCCGATCTNNNGGTCTTCCCTATGGAGTGGATA
GCACACGTCTGAACTCTTTCCCTACACGACGCTCTTCCGATCTNNNGGTCTTCCCTATGGAGTGGATA
AGCACACGTCTGAACTCTTTCCCTACACGACGCTCTTCCGATCTNNNCATCCTGGCTAACATGGTGAA
AGCACACGTCTGAACTCTTTCCCTACACGACGCTCTTCCGATCTNNNCATCCTGGCTAACATGGTGAA
GAGCACACGTCTGAACTCTTTCCCTACACGACGCTCTTCCGATCTNNNGGGTCATGGGAAAAGAATA
GAGCACACGTCTGAACTCTTTCCCTACACGACGCTCTTCCGATCTNNNGGGTCATGGGAAAAGAATA
GAGCACACGTCTGAACTCTTTCCCTACACGACGCTCTTCCGATCTNNNCTCATCTCATTCCTCCCTCAGT
GAGCACACGTCTGAACTCTTTCCCTACACGACGCTCTTCCGATCTNNNCTCATCTCATTCCTCCCTCAGT
AGCACACGTCTGAACTCTTTCCCTACACGACGCTCTTCCGATCTNNNGCAGGAGAATCACTTGAACCT
AGCACACGTCTGAACTCTTTCCCTACACGACGCTCTTCCGATCTNNNGCAGGAGAATCACTTGAACCT
AGCACACGTCTGAACTCTTTCCCTACACGACGCTCTTCCGATCTNNNAGAGAGGAAATAGCTGGGTG
AGCACACGTCTGAACTCTTTCCCTACACGACGCTCTTCCGATCTNNNAGAGAGGAAATAGCTGGGTG
GCACACGTCTGAACTCTTTCCCTACACGACGCTCTTCCGATCTNNNAGGAAACTCGTGACCCTAATTC
GCACACGTCTGAACTCTTTCCCTACACGACGCTCTTCCGATCTNNNAGGAAACTCGTGACCCTAATTC
AGCACACGTCTGAACTCTTTCCCTACACGACGCTCTTCCGATCTNNNACAGTGAAACCCCGTCTCTACT
AGCACACGTCTGAACTCTTTCCCTACACGACGCTCTTCCGATCTNNNACAGTGAAACCCCGTCTCTACT
GCACACGTCTGAACTCTTTCCCTACACGACGCTCTTCCGATCTNNNTGTGTACTTCTATCGGTGAGCAC
GCACACGTCTGAACTCTTTCCCTACACGACGCTCTTCCGATCTNNNTGTGTACTTCTATCGGTGAGCAC

GAGCACACGTCTGAACTCTTTCCCTACACGACGCTCTTCCGATCTNNNCTGTAATTCCAGCTACTCGGG

GAGCACACGTCTGAACTCTTTCCCTACACGACGCTCTTCCGATCTNNNCTGTAATTCCAGCTACTCGGG

AGCACACGTCTGAACTCTTTCCCTACACGACGCTCTTCCGATCTNNNATGACCTGGCTGTTCTTAGTGA

AGCACACGTCTGAACTCTTTCCCTACACGACGCTCTTCCGATCTNNNATGACCTGGCTGTTCTTAGTGA

AGCACACGTCTGAACTCTTTCCCTACACGACGCTCTTCCGATCTNNNTCCTGATCACATCAACAGGGAT

AGCACACGTCTGAACTCTTTCCCTACACGACGCTCTTCCGATCTNNNTCCTGATCACATCAACAGGGAT

AGCACACGTCTGAACTCTTTCCCTACACGACGCTCTTCCGATCTNNNCTCACACTTGGGAAACTTGGGA

AGCACACGTCTGAACTCTTTCCCTACACGACGCTCTTCCGATCTNNNCTCACACTTGGGAAACTTGGGA

ACACGTCTGAACTCTTTCCCTACACGACGCTCTTCCGATCTNNNGGCTCTGGTAACACCACTTTATCAT

ACACGTCTGAACTCTTTCCCTACACGACGCTCTTCCGATCTNNNGGCTCTGGTAACACCACTTTATCAT

GAGCACACGTCTGAACTCTTTCCCTACACGACGCTCTTCCGATCTNNNCTGAGGCAGGAGAATCACTT

GAGCACACGTCTGAACTCTTTCCCTACACGACGCTCTTCCGATCTNNNCTGAGGCAGGAGAATCACTT

CACACGTCTGAACTCTTTCCCTACACGACGCTCTTCCGATCTNNNAGCCTGGTTTTTAAATGGGCATC

CACACGTCTGAACTCTTTCCCTACACGACGCTCTTCCGATCTNNNAGCCTGGTTTTTAAATGGGCATC

AGCACACGTCTGAACTCTTTCCCTACACGACGCTCTTCCGATCTNNNCTATCTTGGCTCACTGCAAGCTC

AGCACACGTCTGAACTCTTTCCCTACACGACGCTCTTCCGATCTNNNCTATCTTGGCTCACTGCAAGCTC

AGCACACGTCTGAACTCTTTCCCTACACGACGCTCTTCCGATCTNNNTCAGGAGATTTGCTTGAACCCT

AGCACACGTCTGAACTCTTTCCCTACACGACGCTCTTCCGATCTNNNTCAGGAGATTTGCTTGAACCCT

GAGCACACGTCTGAACTCTTTCCCTACACGACGCTCTTCCGATCTNNNTCGACCTGCATGCTATAGAGT

GAGCACACGTCTGAACTCTTTCCCTACACGACGCTCTTCCGATCTNNNTCGACCTGCATGCTATAGAGT

AGCACACGTCTGAACTCTTTCCCTACACGACGCTCTTCCGATCTNNNACATCCCTTTGACACTGGTTCTT

AGCACACGTCTGAACTCTTTCCCTACACGACGCTCTTCCGATCTNNNACATCCCTTTGACACTGGTTCTT

GAGCACACGTCTGAACTCTTTCCCTACACGACGCTCTTCCGATCTNNNAACCTGTGCTCACAATCACTAC

GAGCACACGTCTGAACTCTTTCCCTACACGACGCTCTTCCGATCTNNNAACCTGTGCTCACAATCACTAC

GCACACGTCTGAACTCTTTCCCTACACGACGCTCTTCCGATCTNNNTGAAACAACTGCTTCTGGATCCT

GCACACGTCTGAACTCTTTCCCTACACGACGCTCTTCCGATCTNNNTGAAACAACTGCTTCTGGATCCT

AGAGCACACGTCTGAACTCTTTCCCTACACGACGCTCTTCCGATCTNNNGTTGTAGTGAGCTGAGATT

AGAGCACACGTCTGAACTCTTTCCCTACACGACGCTCTTCCGATCTNNNGTTGTAGTGAGCTGAGATT

GAGCACACGTCTGAACTCTTTCCCTACACGACGCTCTTCCGATCTNNNGATTCCTGGCTGGGGTCTGTC

GAGCACACGTCTGAACTCTTTCCCTACACGACGCTCTTCCGATCTNNNGATTCCTGGCTGGGGTCTGTC

AGCACACGTCTGAACTCTTTCCCTACACGACGCTCTTCCGATCTNNNACTGCAAGATGAAGATGAGCT

AGCACACGTCTGAACTCTTTCCCTACACGACGCTCTTCCGATCTNNNACTGCAAGATGAAGATGAGCT

AGCACACGTCTGAACTCTTTCCCTACACGACGCTCTTCCGATCTNNNCCAGATATCAGTGGGTCTCAA

AGCACACGTCTGAACTCTTTCCCTACACGACGCTCTTCCGATCTNNNCCAGATATCAGTGGGTCTCAA

GCACACGTCTGAACTCTTTCCCTACACGACGCTCTTCCGATCTNNNTGAATCCTTGTCTTCTACCCCTGC

GCACACGTCTGAACTCTTTCCCTACACGACGCTCTTCCGATCTNNNTGAATCCTTGTCTTCTACCCCTGC

AGCACACGTCTGAACTCTTTCCCTACACGACGCTCTTCCGATCTNNNTGCCCTTCTTTGTCGTTTTTGAT

AGCACACGTCTGAACTCTTTCCCTACACGACGCTCTTCCGATCTNNNTGCCCTTCTTTGTCGTTTTTGAT

GAGCACACGTCTGAACTCTTTCCCTACACGACGCTCTTCCGATCTNNNAAAGCCACATACCTACAGCCT

GAGCACACGTCTGAACTCTTTCCCTACACGACGCTCTTCCGATCTNNNAAAGCCACATACCTACAGCCT

GAGCACACGTCTGAACTCTTTCCCTACACGACGCTCTTCCGATCTNNNTACAAAAAATTAGCCGGGTG

GAGCACACGTCTGAACTCTTTCCCTACACGACGCTCTTCCGATCTNNNTACAAAAAATTAGCCGGGTG

GAGCACACGTCTGAACTCTTTCCCTACACGACGCTCTTCCGATCTNNNGAGAAAAAGGGGAAGAAA

GAGCACACGTCTGAACTCTTTCCCTACACGACGCTCTTCCGATCTNNNGAGAAAAAGGGGAAGAAAA
GAGCACACGTCTGAACTCTTTCCCTACACGACGCTCTTCCGATCTNNNTCATTTGAGCCAATTCTCCCTG
GAGCACACGTCTGAACTCTTTCCCTACACGACGCTCTTCCGATCTNNNTCATTTGAGCCAATTCTCCCTG
GAGCACACGTCTGAACTCTTTCCCTACACGACGCTCTTCCGATCTNNNCACGGGATCAAACACATCAAA
GAGCACACGTCTGAACTCTTTCCCTACACGACGCTCTTCCGATCTNNNCACGGGATCAAACACATCAAA
GAGCACACGTCTGAACTCTTTCCCTACACGACGCTCTTCCGATCTNNNCCGGTTTTTTGTGCCCTATGTA
GAGCACACGTCTGAACTCTTTCCCTACACGACGCTCTTCCGATCTNNNCCGGTTTTTTGTGCCCTATGTA
AGCACACGTCTGAACTCTTTCCCTACACGACGCTCTTCCGATCTNNNGTGTGTCTGTGTGTCTCAAATG
AGCACACGTCTGAACTCTTTCCCTACACGACGCTCTTCCGATCTNNNGTGTGTCTGTGTGTCTCAAATG
GAGCACACGTCTGAACTCTTTCCCTACACGACGCTCTTCCGATCTNNNACGCCTATAGTCCCAGCTATTT
GAGCACACGTCTGAACTCTTTCCCTACACGACGCTCTTCCGATCTNNNACGCCTATAGTCCCAGCTATTT
GAGCACACGTCTGAACTCTTTCCCTACACGACGCTCTTCCGATCTNNNTAAGGAAGGGTGAGGGGTAT
GAGCACACGTCTGAACTCTTTCCCTACACGACGCTCTTCCGATCTNNNTAAGGAAGGGTGAGGGGTAT
GAGCACACGTCTGAACTCTTTCCCTACACGACGCTCTTCCGATCTNNNCCAGCTAATTTGTGTGTGTGT
GAGCACACGTCTGAACTCTTTCCCTACACGACGCTCTTCCGATCTNNNCCAGCTAATTTGTGTGTGTGT
GAGCACACGTCTGAACTCTTTCCCTACACGACGCTCTTCCGATCTNNNCTGGTTGTGCCACTCAAGAAAT
GAGCACACGTCTGAACTCTTTCCCTACACGACGCTCTTCCGATCTNNNCTGGTTGTGCCACTCAAGAAAT
GAGCACACGTCTGAACTCTTTCCCTACACGACGCTCTTCCGATCTNNNGGAGGAAGGGATGGAAAAA
GAGCACACGTCTGAACTCTTTCCCTACACGACGCTCTTCCGATCTNNNGGAGGAAGGGATGGAAAAA
AGCACACGTCTGAACTCTTTCCCTACACGACGCTCTTCCGATCTNNNCCCCAAAAGAGGACAGATTGTA
AGCACACGTCTGAACTCTTTCCCTACACGACGCTCTTCCGATCTNNNCCCCAAAAGAGGACAGATTGTA
AGAGCACACGTCTGAACTCTTTCCCTACACGACGCTCTTCCGATCTNNNGAGGTTGCAGTGAGCTAAG
AGAGCACACGTCTGAACTCTTTCCCTACACGACGCTCTTCCGATCTNNNGAGGTTGCAGTGAGCTAAG
AGCACACGTCTGAACTCTTTCCCTACACGACGCTCTTCCGATCTNNNACAATCCCACCATTCAGGCATAA
AGCACACGTCTGAACTCTTTCCCTACACGACGCTCTTCCGATCTNNNACAATCCCACCATTCAGGCATAA
GCACACGTCTGAACTCTTTCCCTACACGACGCTCTTCCGATCTNNNGCACAAATGGTCTGTGATTCCAA
GCACACGTCTGAACTCTTTCCCTACACGACGCTCTTCCGATCTNNNGCACAAATGGTCTGTGATTCCAA
GAGCACACGTCTGAACTCTTTCCCTACACGACGCTCTTCCGATCTNNNCACCTGTAATCCCAGCTACTCT
GAGCACACGTCTGAACTCTTTCCCTACACGACGCTCTTCCGATCTNNNCACCTGTAATCCCAGCTACTCT
AGAGCACACGTCTGAACTCTTTCCCTACACGACGCTCTTCCGATCTNNNTTGGGCTCCTTGAAGTTACCT
AGAGCACACGTCTGAACTCTTTCCCTACACGACGCTCTTCCGATCTNNNTTGGGCTCCTTGAAGTTACCT
GAGCACACGTCTGAACTCTTTCCCTACACGACGCTCTTCCGATCTNNNTTGGACAGTACTGCTTTGGTTT
GAGCACACGTCTGAACTCTTTCCCTACACGACGCTCTTCCGATCTNNNTTGGACAGTACTGCTTTGGTTT
CACACGTCTGAACTCTTTCCCTACACGACGCTCTTCCGATCTNNNATTGAAATGATGGTGCTCTGTGCC
CACACGTCTGAACTCTTTCCCTACACGACGCTCTTCCGATCTNNNATTGAAATGATGGTGCTCTGTGCC
AAGAGCACACGTCTGAACTCTTTCCCTACACGACGCTCTTCCGATCTNNNCAAGATCACACCACTACACT
AAGAGCACACGTCTGAACTCTTTCCCTACACGACGCTCTTCCGATCTNNNCAAGATCACACCACTACACT
GAGCACACGTCTGAACTCTTTCCCTACACGACGCTCTTCCGATCTNNNCACTTTGGGAGGTTCTCTAAGC
GAGCACACGTCTGAACTCTTTCCCTACACGACGCTCTTCCGATCTNNNCACTTTGGGAGGTTCTCTAAGC
AGCACACGTCTGAACTCTTTCCCTACACGACGCTCTTCCGATCTNNNGTTGCAGTGAGCCAAGGTTGCA
AGCACACGTCTGAACTCTTTCCCTACACGACGCTCTTCCGATCTNNNGTTGCAGTGAGCCAAGGTTGCA
ACACGTCTGAACTCTTTCCCTACACGACGCTCTTCCGATCTNNNCTCTCAATGAGCTGCTGGGTACCATG
ACACGTCTGAACTCTTTCCCTACACGACGCTCTTCCGATCTNNNCTCTCAATGAGCTGCTGGGTACCATG

GCACACGTCTGAACTCTTTCCCTACACGACGCTCTTCCGATCTNNNGTACTCCCAGCTGTTTGCAAGGC
GCACACGTCTGAACTCTTTCCCTACACGACGCTCTTCCGATCTNNNGTACTCCCAGCTGTTTGCAAGGC
AAGAGCACACGTCTGAACTCTTTCCCTACACGACGCTCTTCCGATCTNNNAATAGCTGTGTGTGGTG
AAGAGCACACGTCTGAACTCTTTCCCTACACGACGCTCTTCCGATCTNNNAATAGCTGTGTGTGGTG
GCACACGTCTGAACTCTTTCCCTACACGACGCTCTTCCGATCTNNNCAGCCTTTTCCTTACCATCCCTAC
GCACACGTCTGAACTCTTTCCCTACACGACGCTCTTCCGATCTNNNCAGCCTTTTCCTTACCATCCCTAC
AGCACACGTCTGAACTCTTTCCCTACACGACGCTCTTCCGATCTNNNCAAGGCCCTGTGAAGAAAAG
AGCACACGTCTGAACTCTTTCCCTACACGACGCTCTTCCGATCTNNNCAAGGCCCTGTGAAGAAAAG
GAGCACACGTCTGAACTCTTTCCCTACACGACGCTCTTCCGATCTNNNACCTTCTCAACATACCTCTGC
GAGCACACGTCTGAACTCTTTCCCTACACGACGCTCTTCCGATCTNNNACCTTCTCAACATACCTCTGC
AGCACACGTCTGAACTCTTTCCCTACACGACGCTCTTCCGATCTNNNCAGTTTTCTGCCTCTTTGGGAA
AGCACACGTCTGAACTCTTTCCCTACACGACGCTCTTCCGATCTNNNCAGTTTTCTGCCTCTTTGGGAA
AGCACACGTCTGAACTCTTTCCCTACACGACGCTCTTCCGATCTNNNTTACTTGGTGCCATCGAAGTCT
AGCACACGTCTGAACTCTTTCCCTACACGACGCTCTTCCGATCTNNNTTACTTGGTGCCATCGAAGTCT
CACACGTCTGAACTCTTTCCCTACACGACGCTCTTCCGATCTNNNGTGTAGAGTGGAGCTGAGGATG
CACACGTCTGAACTCTTTCCCTACACGACGCTCTTCCGATCTNNNGTGTAGAGTGGAGCTGAGGATG
GAGCACACGTCTGAACTCTTTCCCTACACGACGCTCTTCCGATCTNNNGGCCCAACTCTGTCAAAAGA
GAGCACACGTCTGAACTCTTTCCCTACACGACGCTCTTCCGATCTNNNGGCCCAACTCTGTCAAAAGA
CACACGTCTGAACTCTTTCCCTACACGACGCTCTTCCGATCTNNNACACACCACAATTACAGTGTTGTC
CACACGTCTGAACTCTTTCCCTACACGACGCTCTTCCGATCTNNNACACACCACAATTACAGTGTTGTC
AGAGCACACGTCTGAACTCTTTCCCTACACGACGCTCTTCCGATCTNNNAAAGAGTGCAGGGAGGAG
AGAGCACACGTCTGAACTCTTTCCCTACACGACGCTCTTCCGATCTNNNAAAGAGTGCAGGGAGGAG
AGCACACGTCTGAACTCTTTCCCTACACGACGCTCTTCCGATCTNNNGGACGAATGAATGGCAGAGA
AGCACACGTCTGAACTCTTTCCCTACACGACGCTCTTCCGATCTNNNGGACGAATGAATGGCAGAGA
AGCACACGTCTGAACTCTTTCCCTACACGACGCTCTTCCGATCTNNNGCTGTTTACCCTGGTCTCTTTTG
AGCACACGTCTGAACTCTTTCCCTACACGACGCTCTTCCGATCTNNNGCTGTTTACCCTGGTCTCTTTTG
AGCACACGTCTGAACTCTTTCCCTACACGACGCTCTTCCGATCTNNNTTGCACGTAGTTGCAGAATTCA
AGCACACGTCTGAACTCTTTCCCTACACGACGCTCTTCCGATCTNNNTTGCACGTAGTTGCAGAATTCA
AGCACACGTCTGAACTCTTTCCCTACACGACGCTCTTCCGATCTNNNAATCACTTGAACTTGGAGGGC
AGCACACGTCTGAACTCTTTCCCTACACGACGCTCTTCCGATCTNNNAATCACTTGAACTTGGAGGGC
AGAGCACACGTCTGAACTCTTTCCCTACACGACGCTCTTCCGATCTNNNCTCACCTGTAATCCCAGCACT
AGAGCACACGTCTGAACTCTTTCCCTACACGACGCTCTTCCGATCTNNNCTCACCTGTAATCCCAGCACT
AGCACACGTCTGAACTCTTTCCCTACACGACGCTCTTCCGATCTNNNGAGGAGGAGGGAGGGAGGA
AGCACACGTCTGAACTCTTTCCCTACACGACGCTCTTCCGATCTNNNGAGGAGGAGGGAGGGAGGA
AGCACACGTCTGAACTCTTTCCCTACACGACGCTCTTCCGATCTNNNACAGTGGAGCAAACAACCAAA
AGCACACGTCTGAACTCTTTCCCTACACGACGCTCTTCCGATCTNNNACAGTGGAGCAAACAACCAAA
GAGCACACGTCTGAACTCTTTCCCTACACGACGCTCTTCCGATCTNNNCAGGAATGGAAAACCAAACA
GAGCACACGTCTGAACTCTTTCCCTACACGACGCTCTTCCGATCTNNNCAGGAATGGAAAACCAAACA
GAGCACACGTCTGAACTCTTTCCCTACACGACGCTCTTCCGATCTNNNCAGTGTGTGCCATTCTGTAAT
GAGCACACGTCTGAACTCTTTCCCTACACGACGCTCTTCCGATCTNNNCAGTGTGTGCCATTCTGTAAT
AGCACACGTCTGAACTCTTTCCCTACACGACGCTCTTCCGATCTNNNAATGTGTTTTAGGAGCCTCTG
AGCACACGTCTGAACTCTTTCCCTACACGACGCTCTTCCGATCTNNNAATGTGTTTTAGGAGCCTCTG
GAGCACACGTCTGAACTCTTTCCCTACACGACGCTCTTCCGATCTNNNCAAACAAACAGTCCTTCCCTG

GAGCACACGTCTGAACTCTTTCCCTACACGACGCTCTTCCGATCTNNNCAAACAAACAGTCCTTCCCTG
GAGCACACGTCTGAACTCTTTCCCTACACGACGCTCTTCCGATCTNNNATTGATCGATGGCCACTTAG
GAGCACACGTCTGAACTCTTTCCCTACACGACGCTCTTCCGATCTNNNATTGATCGATGGCCACTTAG
GCACACGTCTGAACTCTTTCCCTACACGACGCTCTTCCGATCTNNNGCTTTCCCTCTAGCATAGCATCTC
GCACACGTCTGAACTCTTTCCCTACACGACGCTCTTCCGATCTNNNGCTTTCCCTCTAGCATAGCATCTC
GAGCACACGTCTGAACTCTTTCCCTACACGACGCTCTTCCGATCTNNNTTTGGGGATGCTGCCAGAA
GAGCACACGTCTGAACTCTTTCCCTACACGACGCTCTTCCGATCTNNNTTTGGGGATGCTGCCAGAA
GCACACGTCTGAACTCTTTCCCTACACGACGCTCTTCCGATCTNNNTGTCATTTTTGGACTTTAGAGGA
GCACACGTCTGAACTCTTTCCCTACACGACGCTCTTCCGATCTNNNTGTCATTTTTGGACTTTAGAGGA
GCACACGTCTGAACTCTTTCCCTACACGACGCTCTTCCGATCTNNNGCATTGCTGGATCAAATGCTAGT
GCACACGTCTGAACTCTTTCCCTACACGACGCTCTTCCGATCTNNNGCATTGCTGGATCAAATGCTAGT
GCACACGTCTGAACTCTTTCCCTACACGACGCTCTTCCGATCTNNNCTGAGCTAACACATCATTGACAT
GCACACGTCTGAACTCTTTCCCTACACGACGCTCTTCCGATCTNNNCTGAGCTAACACATCATTGACAT
AGCACACGTCTGAACTCTTTCCCTACACGACGCTCTTCCGATCTNNNTTTGTGCATTGAGGTAGGCTG
AGCACACGTCTGAACTCTTTCCCTACACGACGCTCTTCCGATCTNNNTTTGTGCATTGAGGTAGGCTG
GAGCACACGTCTGAACTCTTTCCCTACACGACGCTCTTCCGATCTNNNTGAGTCACAGAGAGAGACTG
GAGCACACGTCTGAACTCTTTCCCTACACGACGCTCTTCCGATCTNNNTGAGTCACAGAGAGAGACTG
GCACACGTCTGAACTCTTTCCCTACACGACGCTCTTCCGATCTNNNTACCCTTCCTTCAGCAAATCTCCC
GCACACGTCTGAACTCTTTCCCTACACGACGCTCTTCCGATCTNNNTACCCTTCCTTCAGCAAATCTCCC
GCACACGTCTGAACTCTTTCCCTACACGACGCTCTTCCGATCTNNNATGATGATCTCTCTCTCCCTCCCC
GCACACGTCTGAACTCTTTCCCTACACGACGCTCTTCCGATCTNNNATGATGATCTCTCTCTCCCTCCCC
AGCACACGTCTGAACTCTTTCCCTACACGACGCTCTTCCGATCTNNNCTGGGCAACAAGAGCAAAACTC
AGCACACGTCTGAACTCTTTCCCTACACGACGCTCTTCCGATCTNNNCTGGGCAACAAGAGCAAAACTC
AGAGCACACGTCTGAACTCTTTCCCTACACGACGCTCTTCCGATCTNNNATGCCTGGTTCATGCTTTTT
AGAGCACACGTCTGAACTCTTTCCCTACACGACGCTCTTCCGATCTNNNATGCCTGGTTCATGCTTTTT
GAGCACACGTCTGAACTCTTTCCCTACACGACGCTCTTCCGATCTNNNGGGGAAGGTTGTGGAAGTCT
GAGCACACGTCTGAACTCTTTCCCTACACGACGCTCTTCCGATCTNNNGGGGAAGGTTGTGGAAGTCT
GAGCACACGTCTGAACTCTTTCCCTACACGACGCTCTTCCGATCTNNNAAACAGAACCCCAGGGACCT
GAGCACACGTCTGAACTCTTTCCCTACACGACGCTCTTCCGATCTNNNAAACAGAACCCCAGGGACCT
CACACGTCTGAACTCTTTCCCTACACGACGCTCTTCCGATCTNNNATTAGGAAGGCCCCATTTCTCTCCA
CACACGTCTGAACTCTTTCCCTACACGACGCTCTTCCGATCTNNNATTAGGAAGGCCCCATTTCTCTCCA
AGCACACGTCTGAACTCTTTCCCTACACGACGCTCTTCCGATCTNNNTCCAGGCAAGACCATGATAACA
AGCACACGTCTGAACTCTTTCCCTACACGACGCTCTTCCGATCTNNNTCCAGGCAAGACCATGATAACA
AGCACACGTCTGAACTCTTTCCCTACACGACGCTCTTCCGATCTNNNCCTATGACCCATGATGAAACCC
AGCACACGTCTGAACTCTTTCCCTACACGACGCTCTTCCGATCTNNNCCTATGACCCATGATGAAACCC
GCACACGTCTGAACTCTTTCCCTACACGACGCTCTTCCGATCTNNNTGCAAGCCTTATGGTTACATCAA
GCACACGTCTGAACTCTTTCCCTACACGACGCTCTTCCGATCTNNNTGCAAGCCTTATGGTTACATCAA
AGCACACGTCTGAACTCTTTCCCTACACGACGCTCTTCCGATCTNNNGGCAAGTGACAGTGAGGAAGA
AGCACACGTCTGAACTCTTTCCCTACACGACGCTCTTCCGATCTNNNGGCAAGTGACAGTGAGGAAGA
AGCACACGTCTGAACTCTTTCCCTACACGACGCTCTTCCGATCTNNNGCATGAAGACAAGCAGTAACTC
AGCACACGTCTGAACTCTTTCCCTACACGACGCTCTTCCGATCTNNNGCATGAAGACAAGCAGTAACTC
AGAGCACACGTCTGAACTCTTTCCCTACACGACGCTCTTCCGATCTNNNCACTATCTCCTCGTACTTTGG
AGAGCACACGTCTGAACTCTTTCCCTACACGACGCTCTTCCGATCTNNNCACTATCTCCTCGTACTTTGG

GAGCACACGTCTGAACTCTTTCCCTACACGACGCTCTTCCGATCTNNNCGGGATGAACCATGACCTCAG

GAGCACACGTCTGAACTCTTTCCCTACACGACGCTCTTCCGATCTNNNCGGGATGAACCATGACCTCAG

AGAGCACACGTCTGAACTCTTTCCCTACACGACGCTCTTCCGATCTNNNCATCCTGGCTAACACAGTGA

AGAGCACACGTCTGAACTCTTTCCCTACACGACGCTCTTCCGATCTNNNCATCCTGGCTAACACAGTGA

AGCACACGTCTGAACTCTTTCCCTACACGACGCTCTTCCGATCTNNNTTCTCTGGGTCAATCAGGTGATC

AGCACACGTCTGAACTCTTTCCCTACACGACGCTCTTCCGATCTNNNTTCTCTGGGTCAATCAGGTGATC

GAGCACACGTCTGAACTCTTTCCCTACACGACGCTCTTCCGATCTNNNAATCTCAACTCTTTGGGAGGA

GAGCACACGTCTGAACTCTTTCCCTACACGACGCTCTTCCGATCTNNNAATCTCAACTCTTTGGGAGGA

GAGCACACGTCTGAACTCTTTCCCTACACGACGCTCTTCCGATCTNNNGAGACAAGGTTTCACGATGTT

GAGCACACGTCTGAACTCTTTCCCTACACGACGCTCTTCCGATCTNNNGAGACAAGGTTTCACGATGTT

AGCACACGTCTGAACTCTTTCCCTACACGACGCTCTTCCGATCTNNNGCAGTGGATCAATCACAGTTCA

AGCACACGTCTGAACTCTTTCCCTACACGACGCTCTTCCGATCTNNNGCAGTGGATCAATCACAGTTCA

CACACGTCTGAACTCTTTCCCTACACGACGCTCTTCCGATCTNNNCTCCTCTCCTCAGCTCTCCTCCATCG

CACACGTCTGAACTCTTTCCCTACACGACGCTCTTCCGATCTNNNCTCCTCTCCTCAGCTCTCCTCCATCG

GCACACGTCTGAACTCTTTCCCTACACGACGCTCTTCCGATCTNNNCTGGAGTGTCCACTACTCTATGC

GCACACGTCTGAACTCTTTCCCTACACGACGCTCTTCCGATCTNNNCTGGAGTGTCCACTACTCTATGC

AGCACACGTCTGAACTCTTTCCCTACACGACGCTCTTCCGATCTNNNTGCACATTGTTCCACCTATACGA

AGCACACGTCTGAACTCTTTCCCTACACGACGCTCTTCCGATCTNNNTGCACATTGTTCCACCTATACGA

AGCACACGTCTGAACTCTTTCCCTACACGACGCTCTTCCGATCTNNNCTTGCAGTGAGCTGAGATTGTC

AGCACACGTCTGAACTCTTTCCCTACACGACGCTCTTCCGATCTNNNCTTGCAGTGAGCTGAGATTGTC

AGCACACGTCTGAACTCTTTCCCTACACGACGCTCTTCCGATCTNNNGAGACAGGAGAATTGCTTGAA

AGCACACGTCTGAACTCTTTCCCTACACGACGCTCTTCCGATCTNNNGAGACAGGAGAATTGCTTGAA

GAGCACACGTCTGAACTCTTTCCCTACACGACGCTCTTCCGATCTNNNCCCCAGAATGTTTACTGAGAC

GAGCACACGTCTGAACTCTTTCCCTACACGACGCTCTTCCGATCTNNNCCCCAGAATGTTTACTGAGAC

GAGCACACGTCTGAACTCTTTCCCTACACGACGCTCTTCCGATCTNNNAGCCGATCCTTCTTTTGATACCC

GAGCACACGTCTGAACTCTTTCCCTACACGACGCTCTTCCGATCTNNNAGCCGATCCTTCTTTTGATACCC

AGCACACGTCTGAACTCTTTCCCTACACGACGCTCTTCCGATCTNNNTTGACCTCCCAGACATTATCGAG

AGCACACGTCTGAACTCTTTCCCTACACGACGCTCTTCCGATCTNNNTTGACCTCCCAGACATTATCGAG

GAGCACACGTCTGAACTCTTTCCCTACACGACGCTCTTCCGATCTNNNAGGACCTACAACACCAACATCT

GAGCACACGTCTGAACTCTTTCCCTACACGACGCTCTTCCGATCTNNNAGGACCTACAACACCAACATCT

AGCACACGTCTGAACTCTTTCCCTACACGACGCTCTTCCGATCTNNNAGTGTGAGAACAGACTCAACAC

AGCACACGTCTGAACTCTTTCCCTACACGACGCTCTTCCGATCTNNNAGTGTGAGAACAGACTCAACAC

AGCACACGTCTGAACTCTTTCCCTACACGACGCTCTTCCGATCTNNNAGGCTCCATACCTAGAACACAG

AGCACACGTCTGAACTCTTTCCCTACACGACGCTCTTCCGATCTNNNAGGCTCCATACCTAGAACACAG

GAGCACACGTCTGAACTCTTTCCCTACACGACGCTCTTCCGATCTNNNCAAGAAAGGGCAAATGCTTCA

GAGCACACGTCTGAACTCTTTCCCTACACGACGCTCTTCCGATCTNNNCAAGAAAGGGCAAATGCTTCA

GAGCACACGTCTGAACTCTTTCCCTACACGACGCTCTTCCGATCTNNNAGAAAGGTAAAGAGGAGCAG

GAGCACACGTCTGAACTCTTTCCCTACACGACGCTCTTCCGATCTNNNAGAAAGGTAAAGAGGAGCAG

AGAGCACACGTCTGAACTCTTTCCCTACACGACGCTCTTCCGATCTNNNCTGTAGTCCCAGCTACTCGGC

AGAGCACACGTCTGAACTCTTTCCCTACACGACGCTCTTCCGATCTNNNCTGTAGTCCCAGCTACTCGGC

AGCACACGTCTGAACTCTTTCCCTACACGACGCTCTTCCGATCTNNNTTCCTCCCTCTTATCCACATCCTC

AGCACACGTCTGAACTCTTTCCCTACACGACGCTCTTCCGATCTNNNTTCCTCCCTCTTATCCACATCCTC

AGCACACGTCTGAACTCTTTCCCTACACGACGCTCTTCCGATCTNNNCATGTAATGTGGGACTTCTGCAC

GCACACGTCTGAACTCTTTCCCTACACGACGCTCTTCCGATCTNNNCATGTAATGTGGGACTTCTGCA
AGCACACGTCTGAACTCTTTCCCTACACGACGCTCTTCCGATCTNNNAGTAAGCCATGATCACACCACT
AGCACACGTCTGAACTCTTTCCCTACACGACGCTCTTCCGATCTNNNAGTAAGCCATGATCACACCACT
AGCACACGTCTGAACTCTTTCCCTACACGACGCTCTTCCGATCTNNNTGCACAACTGTAGTCCCAGATA
AGCACACGTCTGAACTCTTTCCCTACACGACGCTCTTCCGATCTNNNTGCACAACTGTAGTCCCAGATA
GAGCACACGTCTGAACTCTTTCCCTACACGACGCTCTTCCGATCTNNNGAAGTCCAGGATGCAGAGC
GAGCACACGTCTGAACTCTTTCCCTACACGACGCTCTTCCGATCTNNNGAAGTCCAGGATGCAGAGC
AGCACACGTCTGAACTCTTTCCCTACACGACGCTCTTCCGATCTNNNTTTGTGGCATTGTAACAACTCC
AGCACACGTCTGAACTCTTTCCCTACACGACGCTCTTCCGATCTNNNTTTGTGGCATTGTAACAACTCC
AGCACACGTCTGAACTCTTTCCCTACACGACGCTCTTCCGATCTNNNTGAATGCTTGCTAGAGAGGT
AGCACACGTCTGAACTCTTTCCCTACACGACGCTCTTCCGATCTNNNTGAATGCTTGCTAGAGAGGT
GAGCACACGTCTGAACTCTTTCCCTACACGACGCTCTTCCGATCTNNNGAGGCTGAGGCAGGAGAAT
GAGCACACGTCTGAACTCTTTCCCTACACGACGCTCTTCCGATCTNNNGAGGCTGAGGCAGGAGAAT
AGCACACGTCTGAACTCTTTCCCTACACGACGCTCTTCCGATCTNNNATTTCAGCCTCCCAAGTATCTG
AGCACACGTCTGAACTCTTTCCCTACACGACGCTCTTCCGATCTNNNATTTCAGCCTCCCAAGTATCTG
AGCACACGTCTGAACTCTTTCCCTACACGACGCTCTTCCGATCTNNNTCTCAGAGCTTGAAGACCAGT
AGCACACGTCTGAACTCTTTCCCTACACGACGCTCTTCCGATCTNNNTCTCAGAGCTTGAAGACCAGT
AGCACACGTCTGAACTCTTTCCCTACACGACGCTCTTCCGATCTNNNGAGAATCGCTTGAACCTGAG
AGCACACGTCTGAACTCTTTCCCTACACGACGCTCTTCCGATCTNNNGAGAATCGCTTGAACCTGAG
AGCACACGTCTGAACTCTTTCCCTACACGACGCTCTTCCGATCTNNNTGCTGCAAAATATTGACTGTG
AGCACACGTCTGAACTCTTTCCCTACACGACGCTCTTCCGATCTNNNTGCTGCAAAATATTGACTGTG
GCACACGTCTGAACTCTTTCCCTACACGACGCTCTTCCGATCTNNNGGTTGTCAGTTGATGTCTCCAGA
GCACACGTCTGAACTCTTTCCCTACACGACGCTCTTCCGATCTNNNGGTTGTCAGTTGATGTCTCCAGA
AGCACACGTCTGAACTCTTTCCCTACACGACGCTCTTCCGATCTNNNCTTGCTTTCTCTCTCTTGTGCA
AGCACACGTCTGAACTCTTTCCCTACACGACGCTCTTCCGATCTNNNCTTGCTTTCTCTCTCTTGTGCA
AGCACACGTCTGAACTCTTTCCCTACACGACGCTCTTCCGATCTNNNCTTGAACTCCTGGACTCAAGTG
AGCACACGTCTGAACTCTTTCCCTACACGACGCTCTTCCGATCTNNNCTTGAACTCCTGGACTCAAGTG
AGCACACGTCTGAACTCTTTCCCTACACGACGCTCTTCCGATCTNNNATATGTGGCTTTCAGGTCTGG
AGCACACGTCTGAACTCTTTCCCTACACGACGCTCTTCCGATCTNNNATATGTGGCTTTCAGGTCTGG
GAGCACACGTCTGAACTCTTTCCCTACACGACGCTCTTCCGATCTNNNGCAGAAAAGGTAGGTTCCA
GAGCACACGTCTGAACTCTTTCCCTACACGACGCTCTTCCGATCTNNNGCAGAAAAGGTAGGTTCCA
CACGTCTGAACTCTTTCCCTACACGACGCTCTTCCGATCTNNNATGTCTCACGGTTTGCAACATTTCAT
CACGTCTGAACTCTTTCCCTACACGACGCTCTTCCGATCTNNNATGTCTCACGGTTTGCAACATTTCAT
AGCACACGTCTGAACTCTTTCCCTACACGACGCTCTTCCGATCTNNNCATGCCTGTAATCCCAGCTACT
AGCACACGTCTGAACTCTTTCCCTACACGACGCTCTTCCGATCTNNNCATGCCTGTAATCCCAGCTACT
AGCACACGTCTGAACTCTTTCCCTACACGACGCTCTTCCGATCTNNNCTCAAAAAAAAAAGGTGGGCA
AGCACACGTCTGAACTCTTTCCCTACACGACGCTCTTCCGATCTNNNCTCAAAAAAAAAAGGTGGGCA
GCACACGTCTGAACTCTTTCCCTACACGACGCTCTTCCGATCTNNNAAAGGTCACTTTGGGGCAAAAT
GCACACGTCTGAACTCTTTCCCTACACGACGCTCTTCCGATCTNNNAAAGGTCACTTTGGGGCAAAAT
AGCACACGTCTGAACTCTTTCCCTACACGACGCTCTTCCGATCTNNNGCATGCTATCAGAGATCCCCA
AGCACACGTCTGAACTCTTTCCCTACACGACGCTCTTCCGATCTNNNGCATGCTATCAGAGATCCCCA
GCACACGTCTGAACTCTTTCCCTACACGACGCTCTTCCGATCTNNNATTTCCTCCAGAAGGCCAGAATT
GCACACGTCTGAACTCTTTCCCTACACGACGCTCTTCCGATCTNNNATTTCCTCCAGAAGGCCAGAATT

AGCACACGTCTGAACTCTTTCCCTACACGACGCTCTTCCGATCTNNNGAGGGTGAGGCAGGAGAAT

AGCACACGTCTGAACTCTTTCCCTACACGACGCTCTTCCGATCTNNNGAGGGTGAGGCAGGAGAAT

AGCACACGTCTGAACTCTTTCCCTACACGACGCTCTTCCGATCTNNNCCAGGGCAGGTAAGTCAAACA

AGCACACGTCTGAACTCTTTCCCTACACGACGCTCTTCCGATCTNNNCCAGGGCAGGTAAGTCAAACA

GCACACGTCTGAACTCTTTCCCTACACGACGCTCTTCCGATCTNNNCCTTCCCCTTCTCTCCTTACTACA

GCACACGTCTGAACTCTTTCCCTACACGACGCTCTTCCGATCTNNNCCTTCCCCTTCTCTCCTTACTACA

GAGCACACGTCTGAACTCTTTCCCTACACGACGCTCTTCCGATCTNNNGGGCAGCAACTATATCCACAT

GAGCACACGTCTGAACTCTTTCCCTACACGACGCTCTTCCGATCTNNNGGGCAGCAACTATATCCACAT

GCACACGTCTGAACTCTTTCCCTACACGACGCTCTTCCGATCTNNNATGCACGGCTTTTCTTTGTAGAG

GCACACGTCTGAACTCTTTCCCTACACGACGCTCTTCCGATCTNNNATGCACGGCTTTTCTTTGTAGAG

AGCACACGTCTGAACTCTTTCCCTACACGACGCTCTTCCGATCTNNNTAGGTGGTAACCAGGAAGTG

AGCACACGTCTGAACTCTTTCCCTACACGACGCTCTTCCGATCTNNNTAGGTGGTAACCAGGAAGTG

GAGCACACGTCTGAACTCTTTCCCTACACGACGCTCTTCCGATCTNNNAAGCAAGACCCAAATACATG

GAGCACACGTCTGAACTCTTTCCCTACACGACGCTCTTCCGATCTNNNAAGCAAGACCCAAATACATG

AGCACACGTCTGAACTCTTTCCCTACACGACGCTCTTCCGATCTNNNATGTACCACATGGACCATGAA

AGCACACGTCTGAACTCTTTCCCTACACGACGCTCTTCCGATCTNNNATGTACCACATGGACCATGAA

GAGCACACGTCTGAACTCTTTCCCTACACGACGCTCTTCCGATCTNNNCCTGGACACTGGCAATATGG

GAGCACACGTCTGAACTCTTTCCCTACACGACGCTCTTCCGATCTNNNCCTGGACACTGGCAATATGG

AGCACACGTCTGAACTCTTTCCCTACACGACGCTCTTCCGATCTNNNTAATTGTCCCATCTCTCCACTGC

AGCACACGTCTGAACTCTTTCCCTACACGACGCTCTTCCGATCTNNNTAATTGTCCCATCTCTCCACTGC

AGCACACGTCTGAACTCTTTCCCTACACGACGCTCTTCCGATCTNNNGGTCTCAAACTCCTGACCTAAG

AGCACACGTCTGAACTCTTTCCCTACACGACGCTCTTCCGATCTNNNGGTCTCAAACTCCTGACCTAAG

AGCACACGTCTGAACTCTTTCCCTACACGACGCTCTTCCGATCTNNNTCTCAACCAGGAGAATTGTAG

AGCACACGTCTGAACTCTTTCCCTACACGACGCTCTTCCGATCTNNNTCTCAACCAGGAGAATTGTAG

GCACACGTCTGAACTCTTTCCCTACACGACGCTCTTCCGATCTNNNAACGAATACTGCTCATGGTTGC

GCACACGTCTGAACTCTTTCCCTACACGACGCTCTTCCGATCTNNNAACGAATACTGCTCATGGTTGC

AGCACACGTCTGAACTCTTTCCCTACACGACGCTCTTCCGATCTNNNCAGGCAATCTTGTCCAGCAATA

AGCACACGTCTGAACTCTTTCCCTACACGACGCTCTTCCGATCTNNNCAGGCAATCTTGTCCAGCAATA

AGCACACGTCTGAACTCTTTCCCTACACGACGCTCTTCCGATCTNNNTCCCTCTAAAGCTTAGATGGTA

AGCACACGTCTGAACTCTTTCCCTACACGACGCTCTTCCGATCTNNNTCCCTCTAAAGCTTAGATGGTA

AGCACACGTCTGAACTCTTTCCCTACACGACGCTCTTCCGATCTNNNTGAGGGAGAAGCCAAAAGAT

AGCACACGTCTGAACTCTTTCCCTACACGACGCTCTTCCGATCTNNNTGAGGGAGAAGCCAAAAGAT

CACACGTCTGAACTCTTTCCCTACACGACGCTCTTCCGATCTNNNACCTATTCCACTGGACTGATCCTCA

CACACGTCTGAACTCTTTCCCTACACGACGCTCTTCCGATCTNNNACCTATTCCACTGGACTGATCCTCA

AGCACACGTCTGAACTCTTTCCCTACACGACGCTCTTCCGATCTNNNCTTGTTGAGAGTGGCTGTGAT

AGCACACGTCTGAACTCTTTCCCTACACGACGCTCTTCCGATCTNNNCTTGTTGAGAGTGGCTGTGAT

GAGCACACGTCTGAACTCTTTCCCTACACGACGCTCTTCCGATCTNNNAAACACAACAAACTCAAGCC

GAGCACACGTCTGAACTCTTTCCCTACACGACGCTCTTCCGATCTNNNAAACACAACAAACTCAAGCC

AGAGCACACGTCTGAACTCTTTCCCTACACGACGCTCTTCCGATCTNNNGCAACAGAGCAAGACTTCA

AGAGCACACGTCTGAACTCTTTCCCTACACGACGCTCTTCCGATCTNNNGCAACAGAGCAAGACTTCA

GCACACGTCTGAACTCTTTCCCTACACGACGCTCTTCCGATCTNNNGAGCAACACTCATTCTCCCATGA

GCACACGTCTGAACTCTTTCCCTACACGACGCTCTTCCGATCTNNNGAGCAACACTCATTCTCCCATGA

AGCACACGTCTGAACTCTTTCCCTACACGACGCTCTTCCGATCTNNNGGTATTTGTCCTAGTGCTCTC

GAGCACACGTCTGAACTCTTTCCCTACACGACGCTCTTCCGATCTNNNGGTATTTGTCCTAGTGCTCTC
GAGCACACGTCTGAACTCTTTCCCTACACGACGCTCTTCCGATCTNNNCTCCTGTTTCCCTTCCTTTCCC
GAGCACACGTCTGAACTCTTTCCCTACACGACGCTCTTCCGATCTNNNCTCCTGTTTCCCTTCCTTTCCC
GAGCACACGTCTGAACTCTTTCCCTACACGACGCTCTTCCGATCTNNNGTTGTCCGATTCAATGCAAAT
GAGCACACGTCTGAACTCTTTCCCTACACGACGCTCTTCCGATCTNNNGTTGTCCGATTCAATGCAAAT
AGCACACGTCTGAACTCTTTCCCTACACGACGCTCTTCCGATCTNNNGATATCTTCCCAGCAAAATAATC
AGCACACGTCTGAACTCTTTCCCTACACGACGCTCTTCCGATCTNNNGATATCTTCCCAGCAAAATAATC
GAGCACACGTCTGAACTCTTTCCCTACACGACGCTCTTCCGATCTNNNTGTCAGGGACAAAATTGGACT
GAGCACACGTCTGAACTCTTTCCCTACACGACGCTCTTCCGATCTNNNTGTCAGGGACAAAATTGGACT
GAGCACACGTCTGAACTCTTTCCCTACACGACGCTCTTCCGATCTNNNATTGCTTGAGCCTAGGAGTTC
GAGCACACGTCTGAACTCTTTCCCTACACGACGCTCTTCCGATCTNNNATTGCTTGAGCCTAGGAGTTC
GAGCACACGTCTGAACTCTTTCCCTACACGACGCTCTTCCGATCTNNNAGTATGGAGTAGGTGCTTCTC
GAGCACACGTCTGAACTCTTTCCCTACACGACGCTCTTCCGATCTNNNAGTATGGAGTAGGTGCTTCTC
AGCACACGTCTGAACTCTTTCCCTACACGACGCTCTTCCGATCTNNNAGACAGGAGAATCACTTGAACC
AGCACACGTCTGAACTCTTTCCCTACACGACGCTCTTCCGATCTNNNAGACAGGAGAATCACTTGAACC
AGCACACGTCTGAACTCTTTCCCTACACGACGCTCTTCCGATCTNNNCCTGCCATCATGTTCAGCTCATAC
AGCACACGTCTGAACTCTTTCCCTACACGACGCTCTTCCGATCTNNNCCTGCCATCATGTTCAGCTCATAC
GAGCACACGTCTGAACTCTTTCCCTACACGACGCTCTTCCGATCTNNNTTGTTTTCCACAGTGACTGTCCT
GAGCACACGTCTGAACTCTTTCCCTACACGACGCTCTTCCGATCTNNNTTGTTTTCCACAGTGACTGTCCT
AGCACACGTCTGAACTCTTTCCCTACACGACGCTCTTCCGATCTNNNGAGGTTGTAGTGAGCCGAGAT
AGCACACGTCTGAACTCTTTCCCTACACGACGCTCTTCCGATCTNNNGAGGTTGTAGTGAGCCGAGAT
GAGCACACGTCTGAACTCTTTCCCTACACGACGCTCTTCCGATCTNNNAAATACAAAAATTAGCCGGGC
GAGCACACGTCTGAACTCTTTCCCTACACGACGCTCTTCCGATCTNNNAAATACAAAAATTAGCCGGGC
GAGCACACGTCTGAACTCTTTCCCTACACGACGCTCTTCCGATCTNNNCTGTAGTCCCAGCTACCCTACTC
GAGCACACGTCTGAACTCTTTCCCTACACGACGCTCTTCCGATCTNNNCTGTAGTCCCAGCTACCCTACTC
CACACGTCTGAACTCTTTCCCTACACGACGCTCTTCCGATCTNNNCCAGAATTCAGTGAAGGCCAATGC
CACACGTCTGAACTCTTTCCCTACACGACGCTCTTCCGATCTNNNCCAGAATTCAGTGAAGGCCAATGC
GAGCACACGTCTGAACTCTTTCCCTACACGACGCTCTTCCGATCTNNNAGTTCCTTTGGCTTAGGGATGT
GAGCACACGTCTGAACTCTTTCCCTACACGACGCTCTTCCGATCTNNNAGTTCCTTTGGCTTAGGGATGT
AGCACACGTCTGAACTCTTTCCCTACACGACGCTCTTCCGATCTNNNACATACGTATACATGTGCCATGC
AGCACACGTCTGAACTCTTTCCCTACACGACGCTCTTCCGATCTNNNACATACGTATACATGTGCCATGC
CACACGTCTGAACTCTTTCCCTACACGACGCTCTTCCGATCTNNNTGAGGCAGGAGAAGTACTTGAAC
CACACGTCTGAACTCTTTCCCTACACGACGCTCTTCCGATCTNNNTGAGGCAGGAGAAGTACTTGAAC
AGCACACGTCTGAACTCTTTCCCTACACGACGCTCTTCCGATCTNNNAGAATCACTTGAACTCAGGAGC
AGCACACGTCTGAACTCTTTCCCTACACGACGCTCTTCCGATCTNNNAGAATCACTTGAACTCAGGAGC
GAGCACACGTCTGAACTCTTTCCCTACACGACGCTCTTCCGATCTNNNGTTCTAACCACTGCCCTATCCG
GAGCACACGTCTGAACTCTTTCCCTACACGACGCTCTTCCGATCTNNNGTTCTAACCACTGCCCTATCCG
GCACACGTCTGAACTCTTTCCCTACACGACGCTCTTCCGATCTNNNGTTTGACAATAATCTCACAAAACCA
GCACACGTCTGAACTCTTTCCCTACACGACGCTCTTCCGATCTNNNGTTTGACAATAATCTCACAAAACCA
AGCACACGTCTGAACTCTTTCCCTACACGACGCTCTTCCGATCTNNNCTCTTCCCTGACCTTGGCTTCCA
AGCACACGTCTGAACTCTTTCCCTACACGACGCTCTTCCGATCTNNNCTCTTCCCTGACCTTGGCTTCCA
AGCACACGTCTGAACTCTTTCCCTACACGACGCTCTTCCGATCTNNNTCCAAAGAGTGTCAGTCTTGAG
AGCACACGTCTGAACTCTTTCCCTACACGACGCTCTTCCGATCTNNNTCCAAAGAGTGTCAGTCTTGAG

GCACACGTCTGAACTCTTTCCCTACACGACGCTCTTCCGATCTNNNGAGACAGGCTCTCATTCTGATGCC

GCACACGTCTGAACTCTTTCCCTACACGACGCTCTTCCGATCTNNNGAGACAGGCTCTCATTCTGATGCC

GAGCACACGTCTGAACTCTTTCCCTACACGACGCTCTTCCGATCTNNNGAACTCACTGAGCCAGAACAC

GAGCACACGTCTGAACTCTTTCCCTACACGACGCTCTTCCGATCTNNNGAACTCACTGAGCCAGAACAC

GAGCACACGTCTGAACTCTTTCCCTACACGACGCTCTTCCGATCTNNNCCAAAATGCTGGGATTACAGC

GAGCACACGTCTGAACTCTTTCCCTACACGACGCTCTTCCGATCTNNNCCAAAATGCTGGGATTACAGC

AGCACACGTCTGAACTCTTTCCCTACACGACGCTCTTCCGATCTNNNTACACACATACGTACAAGGCAC

AGCACACGTCTGAACTCTTTCCCTACACGACGCTCTTCCGATCTNNNTACACACATACGTACAAGGCAC

GAGCACACGTCTGAACTCTTTCCCTACACGACGCTCTTCCGATCTNNNATGCCTGTAGTCCCAGCTACT

GAGCACACGTCTGAACTCTTTCCCTACACGACGCTCTTCCGATCTNNNATGCCTGTAGTCCCAGCTACT

AGAGCACACGTCTGAACTCTTTCCCTACACGACGCTCTTCCGATCTNNNCAAGCTCTTGCTAGTGAAGG

AGAGCACACGTCTGAACTCTTTCCCTACACGACGCTCTTCCGATCTNNNCAAGCTCTTGCTAGTGAAGG

AGAGCACACGTCTGAACTCTTTCCCTACACGACGCTCTTCCGATCTNNNATGCCTGTAATCCCAGCTAC

AGAGCACACGTCTGAACTCTTTCCCTACACGACGCTCTTCCGATCTNNNATGCCTGTAATCCCAGCTAC

GAGCACACGTCTGAACTCTTTCCCTACACGACGCTCTTCCGATCTNNNTGTCTCTGTAGCCTCTTGTTCA

GAGCACACGTCTGAACTCTTTCCCTACACGACGCTCTTCCGATCTNNNTGTCTCTGTAGCCTCTTGTTCA

AGCACACGTCTGAACTCTTTCCCTACACGACGCTCTTCCGATCTNNNGAAGGCCTTATACGTCACAAAG

AGCACACGTCTGAACTCTTTCCCTACACGACGCTCTTCCGATCTNNNGAAGGCCTTATACGTCACAAAG

AGCACACGTCTGAACTCTTTCCCTACACGACGCTCTTCCGATCTNNNGGAGACTTGGAGTGTGTCTGA

AGCACACGTCTGAACTCTTTCCCTACACGACGCTCTTCCGATCTNNNGGAGACTTGGAGTGTGTCTGA

GAGCACACGTCTGAACTCTTTCCCTACACGACGCTCTTCCGATCTNNNGCATATGCATGTTTATAGCAG

GAGCACACGTCTGAACTCTTTCCCTACACGACGCTCTTCCGATCTNNNGCATATGCATGTTTATAGCAG

GCACACGTCTGAACTCTTTCCCTACACGACGCTCTTCCGATCTNNNTGTTCTTTGCTTCTATCCCACCCCA

GCACACGTCTGAACTCTTTCCCTACACGACGCTCTTCCGATCTNNNTGTTCTTTGCTTCTATCCCACCCCA

GAGCACACGTCTGAACTCTTTCCCTACACGACGCTCTTCCGATCTNNNTCATTCATTCATTCCCCCCACAT

GAGCACACGTCTGAACTCTTTCCCTACACGACGCTCTTCCGATCTNNNTCATTCATTCATTCCCCCCACAT

GAGCACACGTCTGAACTCTTTCCCTACACGACGCTCTTCCGATCTNNNTCTTGGTGCTGTAGTAACTGC

GAGCACACGTCTGAACTCTTTCCCTACACGACGCTCTTCCGATCTNNNTCTTGGTGCTGTAGTAACTGC

GCACACGTCTGAACTCTTTCCCTACACGACGCTCTTCCGATCTNNNGGTGGGGAGAGAGAAAAAGAG

GCACACGTCTGAACTCTTTCCCTACACGACGCTCTTCCGATCTNNNGGTGGGGAGAGAGAAAAAGAG

GAGCACACGTCTGAACTCTTTCCCTACACGACGCTCTTCCGATCTNNNCCGAACATTCCTTTTCTGGCAT

GAGCACACGTCTGAACTCTTTCCCTACACGACGCTCTTCCGATCTNNNCCGAACATTCCTTTTCTGGCAT

AGCACACGTCTGAACTCTTTCCCTACACGACGCTCTTCCGATCTNNNAACTTTCTGCCCAGTTTTCCTGT

AGCACACGTCTGAACTCTTTCCCTACACGACGCTCTTCCGATCTNNNAACTTTCTGCCCAGTTTTCCTGT

AGCACACGTCTGAACTCTTTCCCTACACGACGCTCTTCCGATCTNNNTCCATCTACACCTTTTTGGAAGTC

AGCACACGTCTGAACTCTTTCCCTACACGACGCTCTTCCGATCTNNNTCCATCTACACCTTTTTGGAAGTC

AGCACACGTCTGAACTCTTTCCCTACACGACGCTCTTCCGATCTNNNTGGGATAGTCAAGATTCTGGC

AGCACACGTCTGAACTCTTTCCCTACACGACGCTCTTCCGATCTNNNTGGGATAGTCAAGATTCTGGC

GCACACGTCTGAACTCTTTCCCTACACGACGCTCTTCCGATCTNNNCCATCATTTGGCAGCACTGACCA

GCACACGTCTGAACTCTTTCCCTACACGACGCTCTTCCGATCTNNNCCATCATTTGGCAGCACTGACCA

GAGCACACGTCTGAACTCTTTCCCTACACGACGCTCTTCCGATCTNNNTGCAGTGAGCTATGATCATGC

GAGCACACGTCTGAACTCTTTCCCTACACGACGCTCTTCCGATCTNNNTGCAGTGAGCTATGATCATGC

GAGCACACGTCTGAACTCTTTCCCTACACGACGCTCTTCCGATCTNNNCTGTAGTCCCAGCTACTCAGGA

GAGCACACGTCTGAACTCTTTCCCTACACGACGCTCTTCCGATCTNNNCTGTAGTCCCAGCTACTCAGGA
AGCACACGTCTGAACTCTTTCCCTACACGACGCTCTTCCGATCTNNNAACAGTCATTTTAGCCAGTGATT
AGCACACGTCTGAACTCTTTCCCTACACGACGCTCTTCCGATCTNNNAACAGTCATTTTAGCCAGTGATT
AGAGCACACGTCTGAACTCTTTCCCTACACGACGCTCTTCCGATCTNNNAGTGAGCCTAGATAATGCCA
AGAGCACACGTCTGAACTCTTTCCCTACACGACGCTCTTCCGATCTNNNAGTGAGCCTAGATAATGCCA
AGCACACGTCTGAACTCTTTCCCTACACGACGCTCTTCCGATCTNNNATGTTATTGTCACAATAAGCAT
AGCACACGTCTGAACTCTTTCCCTACACGACGCTCTTCCGATCTNNNATGTTATTGTCACAATAAGCAT
AGCACACGTCTGAACTCTTTCCCTACACGACGCTCTTCCGATCTNNNACATATTGCTCATGGGAGAAGC
AGCACACGTCTGAACTCTTTCCCTACACGACGCTCTTCCGATCTNNNACATATTGCTCATGGGAGAAGC
AGCACACGTCTGAACTCTTTCCCTACACGACGCTCTTCCGATCTNNNTATGACAAAAAGAAGGGTGTT
AGCACACGTCTGAACTCTTTCCCTACACGACGCTCTTCCGATCTNNNTATGACAAAAAGAAGGGTGTT
GAGCACACGTCTGAACTCTTTCCCTACACGACGCTCTTCCGATCTNNNTGAAACCATTCCAGTGTAAAA
GAGCACACGTCTGAACTCTTTCCCTACACGACGCTCTTCCGATCTNNNTGAAACCATTCCAGTGTAAAA
GCACACGTCTGAACTCTTTCCCTACACGACGCTCTTCCGATCTNNNTCACTCTCCCTCTCTTTCTCTGTCA
GCACACGTCTGAACTCTTTCCCTACACGACGCTCTTCCGATCTNNNTCACTCTCCCTCTCTTTCTCTGTCA
GAGCACACGTCTGAACTCTTTCCCTACACGACGCTCTTCCGATCTNNNAGCAGAACTTTGGGGCCTAA
GAGCACACGTCTGAACTCTTTCCCTACACGACGCTCTTCCGATCTNNNAGCAGAACTTTGGGGCCTAA
AGAGCACACGTCTGAACTCTTTCCCTACACGACGCTCTTCCGATCTNNNTTGTCCTTTTTCTGGCCGTTT
AGAGCACACGTCTGAACTCTTTCCCTACACGACGCTCTTCCGATCTNNNTTGTCCTTTTTCTGGCCGTTT
GAGCACACGTCTGAACTCTTTCCCTACACGACGCTCTTCCGATCTNNNTGAAGTCAGGAGTTTGAGACC
GAGCACACGTCTGAACTCTTTCCCTACACGACGCTCTTCCGATCTNNNTGAAGTCAGGAGTTTGAGACC
GAGCACACGTCTGAACTCTTTCCCTACACGACGCTCTTCCGATCTNNNTAGCTAGACACACACCACAAA
GAGCACACGTCTGAACTCTTTCCCTACACGACGCTCTTCCGATCTNNNTAGCTAGACACACACCACAAA
GAGCACACGTCTGAACTCTTTCCCTACACGACGCTCTTCCGATCTNNNCTGTAATCCCAGCTACTCAGGA
GAGCACACGTCTGAACTCTTTCCCTACACGACGCTCTTCCGATCTNNNCTGTAATCCCAGCTACTCAGGA
AGAGCACACGTCTGAACTCTTTCCCTACACGACGCTCTTCCGATCTNNNCTTTGGCTTTTCCTGGCTTCT
AGAGCACACGTCTGAACTCTTTCCCTACACGACGCTCTTCCGATCTNNNCTTTGGCTTTTCCTGGCTTCT
AGCACACGTCTGAACTCTTTCCCTACACGACGCTCTTCCGATCTNNNATAGGTTCTTGCTGTCACTCAG
AGCACACGTCTGAACTCTTTCCCTACACGACGCTCTTCCGATCTNNNATAGGTTCTTGCTGTCACTCAG
AGCACACGTCTGAACTCTTTCCCTACACGACGCTCTTCCGATCTNNNTGTTAACAACATGAACTATCTTC
AGCACACGTCTGAACTCTTTCCCTACACGACGCTCTTCCGATCTNNNTGTTAACAACATGAACTATCTTC
GAGCACACGTCTGAACTCTTTCCCTACACGACGCTCTTCCGATCTNNNGAGGCAGGAGAATGGTGTGA
GAGCACACGTCTGAACTCTTTCCCTACACGACGCTCTTCCGATCTNNNGAGGCAGGAGAATGGTGTGA
GAGCACACGTCTGAACTCTTTCCCTACACGACGCTCTTCCGATCTNNNCCAAATGCCCATTGATCAGTC
GAGCACACGTCTGAACTCTTTCCCTACACGACGCTCTTCCGATCTNNNCCAAATGCCCATTGATCAGTC
GCACACGTCTGAACTCTTTCCCTACACGACGCTCTTCCGATCTNNNTGCTTTGCACACATTATCTCCCACA
GCACACGTCTGAACTCTTTCCCTACACGACGCTCTTCCGATCTNNNTGCTTTGCACACATTATCTCCCACA
AGCACACGTCTGAACTCTTTCCCTACACGACGCTCTTCCGATCTNNNACGTGGCTGAAGATCTTGATCT
AGCACACGTCTGAACTCTTTCCCTACACGACGCTCTTCCGATCTNNNACGTGGCTGAAGATCTTGATCT
GAGCACACGTCTGAACTCTTTCCCTACACGACGCTCTTCCGATCTNNNACTAATAGAAATAGTCACTTG
GAGCACACGTCTGAACTCTTTCCCTACACGACGCTCTTCCGATCTNNNACTAATAGAAATAGTCACTTG
GCACACGTCTGAACTCTTTCCCTACACGACGCTCTTCCGATCTNNNGAGGATGACTTGAGCCTTGGAG
GCACACGTCTGAACTCTTTCCCTACACGACGCTCTTCCGATCTNNNGAGGATGACTTGAGCCTTGGAG

AGAGCACACGTCTGAACTCTTTCCCTACACGACGCTCTTCCGATCTNNNTAATGGTTACAGTGGGTCTT
AGAGCACACGTCTGAACTCTTTCCCTACACGACGCTCTTCCGATCTNNNTAATGGTTACAGTGGGTCTT
AGCACACGTCTGAACTCTTTCCCTACACGACGCTCTTCCGATCTNNNCAGACACACACTCACCGAAGAG
AGCACACGTCTGAACTCTTTCCCTACACGACGCTCTTCCGATCTNNNCAGACACACACTCACCGAAGAG
AGCACACGTCTGAACTCTTTCCCTACACGACGCTCTTCCGATCTNNNAGTGATCTGCTGAGGATGTATG
AGCACACGTCTGAACTCTTTCCCTACACGACGCTCTTCCGATCTNNNAGTGATCTGCTGAGGATGTATG
GAGCACACGTCTGAACTCTTTCCCTACACGACGCTCTTCCGATCTNNNGTGCCTGTAATCCCAGCTACTT
GAGCACACGTCTGAACTCTTTCCCTACACGACGCTCTTCCGATCTNNNGTGCCTGTAATCCCAGCTACTT
AGCACACGTCTGAACTCTTTCCCTACACGACGCTCTTCCGATCTNNNGGGAACTGTTCTTGAACCTACC
AGCACACGTCTGAACTCTTTCCCTACACGACGCTCTTCCGATCTNNNGGGAACTGTTCTTGAACCTACC
GAGCACACGTCTGAACTCTTTCCCTACACGACGCTCTTCCGATCTNNNAGAGGTGATACTCTCCACAG
GAGCACACGTCTGAACTCTTTCCCTACACGACGCTCTTCCGATCTNNNAGAGGTGATACTCTCCACAG
GAGCACACGTCTGAACTCTTTCCCTACACGACGCTCTTCCGATCTNNNGAGATGGGGCTTTTGACTTT
GAGCACACGTCTGAACTCTTTCCCTACACGACGCTCTTCCGATCTNNNGAGATGGGGCTTTTGACTTT
AGCACACGTCTGAACTCTTTCCCTACACGACGCTCTTCCGATCTNNNCTTCCTTCCTTCCTTCCTTCCTCCA
AGCACACGTCTGAACTCTTTCCCTACACGACGCTCTTCCGATCTNNNCTTCCTTCCTTCCTTCCTTCCTCCA
GAGCACACGTCTGAACTCTTTCCCTACACGACGCTCTTCCGATCTNNNAAGCTTTTCCATAGTCGACCAC
GAGCACACGTCTGAACTCTTTCCCTACACGACGCTCTTCCGATCTNNNAAGCTTTTCCATAGTCGACCAC
AGCACACGTCTGAACTCTTTCCCTACACGACGCTCTTCCGATCTNNNGTAATCCCAGCTACTTAGGAG
AGCACACGTCTGAACTCTTTCCCTACACGACGCTCTTCCGATCTNNNGTAATCCCAGCTACTTAGGAG
AGAGCACACGTCTGAACTCTTTCCCTACACGACGCTCTTCCGATCTNNNCTGTAATCCCAGCTACTCAG
AGAGCACACGTCTGAACTCTTTCCCTACACGACGCTCTTCCGATCTNNNCTGTAATCCCAGCTACTCAG
CACACGTCTGAACTCTTTCCCTACACGACGCTCTTCCGATCTNNNATACTTTTCACACTCTTCCTCCATCG
CACACGTCTGAACTCTTTCCCTACACGACGCTCTTCCGATCTNNNATACTTTTCACACTCTTCCTCCATCG
AGCACACGTCTGAACTCTTTCCCTACACGACGCTCTTCCGATCTNNNTGCTGCCCAAGTTCAAGTTATT
AGCACACGTCTGAACTCTTTCCCTACACGACGCTCTTCCGATCTNNNTGCTGCCCAAGTTCAAGTTATT
GAGCACACGTCTGAACTCTTTCCCTACACGACGCTCTTCCGATCTNNNAAGTATATCTGTTTTTGAAGCA
GAGCACACGTCTGAACTCTTTCCCTACACGACGCTCTTCCGATCTNNNAAGTATATCTGTTTTTGAAGCA
AGCACACGTCTGAACTCTTTCCCTACACGACGCTCTTCCGATCTNNNTTGACTCTGAATTCTTCCCCTGG
AGCACACGTCTGAACTCTTTCCCTACACGACGCTCTTCCGATCTNNNTTGACTCTGAATTCTTCCCCTGG
GAGCACACGTCTGAACTCTTTCCCTACACGACGCTCTTCCGATCTNNNGTGGATCACCTGAGTTCAGAA
GAGCACACGTCTGAACTCTTTCCCTACACGACGCTCTTCCGATCTNNNGTGGATCACCTGAGTTCAGAA
AGAGCACACGTCTGAACTCTTTCCCTACACGACGCTCTTCCGATCTNNNCACTTTCCAGGTTCCAAGCAA
AGAGCACACGTCTGAACTCTTTCCCTACACGACGCTCTTCCGATCTNNNCACTTTCCAGGTTCCAAGCAA
AGCACACGTCTGAACTCTTTCCCTACACGACGCTCTTCCGATCTNNNACAATATCCTTCCTCGACAACGT
AGCACACGTCTGAACTCTTTCCCTACACGACGCTCTTCCGATCTNNNACAATATCCTTCCTCGACAACGT
GAGCACACGTCTGAACTCTTTCCCTACACGACGCTCTTCCGATCTNNNCTGAGGTGGGAAGATCGATT
GAGCACACGTCTGAACTCTTTCCCTACACGACGCTCTTCCGATCTNNNCTGAGGTGGGAAGATCGATT
AGCACACGTCTGAACTCTTTCCCTACACGACGCTCTTCCGATCTNNNGGATGTCAGCAAAATAAGCAT
AGCACACGTCTGAACTCTTTCCCTACACGACGCTCTTCCGATCTNNNGGATGTCAGCAAAATAAGCAT
GCACACGTCTGAACTCTTTCCCTACACGACGCTCTTCCGATCTNNNATAGAGACAGGCACCCAAACTTG
GCACACGTCTGAACTCTTTCCCTACACGACGCTCTTCCGATCTNNNATAGAGACAGGCACCCAAACTTG
AGCACACGTCTGAACTCTTTCCCTACACGACGCTCTTCCGATCTNNNTCTGGCCAGTGCATCTACATTA

AGCACACGTCTGAACTCTTTCCCTACACGACGCTCTTCCGATCTNNNTCTGGCCAGTGCATCTACATTT
GCACACGTCTGAACTCTTTCCCTACACGACGCTCTTCCGATCTNNNAGGACAATGAACTCTTTAGAAT
GCACACGTCTGAACTCTTTCCCTACACGACGCTCTTCCGATCTNNNAGGACAATGAACTCTTTAGAAT
AGCACACGTCTGAACTCTTTCCCTACACGACGCTCTTCCGATCTNNNTCCCAAGCAAAAAACACAGTTG
AGCACACGTCTGAACTCTTTCCCTACACGACGCTCTTCCGATCTNNNTCCCAAGCAAAAAACACAGTTG
CACACGTCTGAACTCTTTCCCTACACGACGCTCTTCCGATCTNNNAAGAGAACTGGAGTGGCTTTGAT
CACACGTCTGAACTCTTTCCCTACACGACGCTCTTCCGATCTNNNAAGAGAACTGGAGTGGCTTTGAT
AGCACACGTCTGAACTCTTTCCCTACACGACGCTCTTCCGATCTNNNCCCAACCATAAGTAGACCAAGC
AGCACACGTCTGAACTCTTTCCCTACACGACGCTCTTCCGATCTNNNCCCAACCATAAGTAGACCAAGC
GCACACGTCTGAACTCTTTCCCTACACGACGCTCTTCCGATCTNNNGTTTGCAGTGAGCTGAGATCAT
GCACACGTCTGAACTCTTTCCCTACACGACGCTCTTCCGATCTNNNGTTTGCAGTGAGCTGAGATCAT
AGCACACGTCTGAACTCTTTCCCTACACGACGCTCTTCCGATCTNNNATCTTGGCTCACTGCAAGCTCC
AGCACACGTCTGAACTCTTTCCCTACACGACGCTCTTCCGATCTNNNATCTTGGCTCACTGCAAGCTCC
AGCACACGTCTGAACTCTTTCCCTACACGACGCTCTTCCGATCTNNNGGGCAACAGAGTAAAACCTTG
AGCACACGTCTGAACTCTTTCCCTACACGACGCTCTTCCGATCTNNNGGGCAACAGAGTAAAACCTTG
AGCACACGTCTGAACTCTTTCCCTACACGACGCTCTTCCGATCTNNNATTCACCTAAGGCATTCCACCA
AGCACACGTCTGAACTCTTTCCCTACACGACGCTCTTCCGATCTNNNATTCACCTAAGGCATTCCACCA
CACACGTCTGAACTCTTTCCCTACACGACGCTCTTCCGATCTNNNCTTCCCCTATCCCCCAACTTCTCAT
CACACGTCTGAACTCTTTCCCTACACGACGCTCTTCCGATCTNNNCTTCCCCTATCCCCCAACTTCTCAT
GAGCACACGTCTGAACTCTTTCCCTACACGACGCTCTTCCGATCTNNNCTTAGTGCTGTTGTTCCTTTG
GAGCACACGTCTGAACTCTTTCCCTACACGACGCTCTTCCGATCTNNNCTTAGTGCTGTTGTTCCTTTG
AGCACACGTCTGAACTCTTTCCCTACACGACGCTCTTCCGATCTNNNAAGAAATCTCAGAGACTCAGC
AGCACACGTCTGAACTCTTTCCCTACACGACGCTCTTCCGATCTNNNAAGAAATCTCAGAGACTCAGC
GAGCACACGTCTGAACTCTTTCCCTACACGACGCTCTTCCGATCTNNNCTACTGTTATGGTACTGTCCC
GAGCACACGTCTGAACTCTTTCCCTACACGACGCTCTTCCGATCTNNNCTACTGTTATGGTACTGTCCC
GCACACGTCTGAACTCTTTCCCTACACGACGCTCTTCCGATCTNNNCACTAACCAAGGTTATCTGCACA
GCACACGTCTGAACTCTTTCCCTACACGACGCTCTTCCGATCTNNNCACTAACCAAGGTTATCTGCACA
GAGCACACGTCTGAACTCTTTCCCTACACGACGCTCTTCCGATCTNNNTCTATCACCTAAAAGGGAAG
GAGCACACGTCTGAACTCTTTCCCTACACGACGCTCTTCCGATCTNNNTCTATCACCTAAAAGGGAAG
GCACACGTCTGAACTCTTTCCCTACACGACGCTCTTCCGATCTNNNATTAAGCATTAACTCATCAACAA
GCACACGTCTGAACTCTTTCCCTACACGACGCTCTTCCGATCTNNNATTAAGCATTAACTCATCAACAA
GCACACGTCTGAACTCTTTCCCTACACGACGCTCTTCCGATCTNNNAACTTTGCTCTCTAGGCTCATCC
GCACACGTCTGAACTCTTTCCCTACACGACGCTCTTCCGATCTNNNAACTTTGCTCTCTAGGCTCATCC
GAGCACACGTCTGAACTCTTTCCCTACACGACGCTCTTCCGATCTNNNCCAGCCTAGGCAATATAGCA
GAGCACACGTCTGAACTCTTTCCCTACACGACGCTCTTCCGATCTNNNCCAGCCTAGGCAATATAGCA
AGCACACGTCTGAACTCTTTCCCTACACGACGCTCTTCCGATCTNNNGAAATCCTCCAGATGCAGCGA
AGCACACGTCTGAACTCTTTCCCTACACGACGCTCTTCCGATCTNNNGAAATCCTCCAGATGCAGCGA
AGAGCACACGTCTGAACTCTTTCCCTACACGACGCTCTTCCGATCTNNNGGATGGATGGATGAATGG
AGAGCACACGTCTGAACTCTTTCCCTACACGACGCTCTTCCGATCTNNNGGATGGATGGATGAATGG
AGCACACGTCTGAACTCTTTCCCTACACGACGCTCTTCCGATCTNNNTTCCCCATAAAGGTGCTAGAAG
AGCACACGTCTGAACTCTTTCCCTACACGACGCTCTTCCGATCTNNNTTCCCCATAAAGGTGCTAGAAG
GAGCACACGTCTGAACTCTTTCCCTACACGACGCTCTTCCGATCTNNNACTAATGGCAACCAAAACAG
GAGCACACGTCTGAACTCTTTCCCTACACGACGCTCTTCCGATCTNNNACTAATGGCAACCAAAACAG

AGCACACGTCTGAACTCTTTCCCTACACGACGCTCTTCCGATCTNNNCTATGGAATCTTCGGGATCCAG
AGCACACGTCTGAACTCTTTCCCTACACGACGCTCTTCCGATCTNNNCTATGGAATCTTCGGGATCCAG
AGCACACGTCTGAACTCTTTCCCTACACGACGCTCTTCCGATCTNNNGAGAACCCAGACAGAAAGCTA
AGCACACGTCTGAACTCTTTCCCTACACGACGCTCTTCCGATCTNNNGAGAACCCAGACAGAAAGCTA
GAGCACACGTCTGAACTCTTTCCCTACACGACGCTCTTCCGATCTNNNATCCTGGAGATGTCTTTGCC
GAGCACACGTCTGAACTCTTTCCCTACACGACGCTCTTCCGATCTNNNATCCTGGAGATGTCTTTGCC
AGCACACGTCTGAACTCTTTCCCTACACGACGCTCTTCCGATCTNNNGGAGGCCAGCATTCATACATT
AGCACACGTCTGAACTCTTTCCCTACACGACGCTCTTCCGATCTNNNGGAGGCCAGCATTCATACATT
AGCACACGTCTGAACTCTTTCCCTACACGACGCTCTTCCGATCTNNNTCCCTGCAACTTTGACAGTAGTT
AGCACACGTCTGAACTCTTTCCCTACACGACGCTCTTCCGATCTNNNTCCCTGCAACTTTGACAGTAGTT
AGCACACGTCTGAACTCTTTCCCTACACGACGCTCTTCCGATCTNNNTGGCACATTTCCATCTGTTTTTC
AGCACACGTCTGAACTCTTTCCCTACACGACGCTCTTCCGATCTNNNTGGCACATTTCCATCTGTTTTTC
AGCACACGTCTGAACTCTTTCCCTACACGACGCTCTTCCGATCTNNNGTCTACATACCTGGCTAGCTTC
AGCACACGTCTGAACTCTTTCCCTACACGACGCTCTTCCGATCTNNNGTCTACATACCTGGCTAGCTTC
AGCACACGTCTGAACTCTTTCCCTACACGACGCTCTTCCGATCTNNNGAGTTTTCCTCCAGCAAAGCCA
AGCACACGTCTGAACTCTTTCCCTACACGACGCTCTTCCGATCTNNNGAGTTTTCCTCCAGCAAAGCCA
ACACGTCTGAACTCTTTCCCTACACGACGCTCTTCCGATCTNNNGAGAAAGAGGGGGTATCCTTGGA
ACACGTCTGAACTCTTTCCCTACACGACGCTCTTCCGATCTNNNGAGAAAGAGGGGGTATCCTTGGA
AGAGCACACGTCTGAACTCTTTCCCTACACGACGCTCTTCCGATCTNNNCAGATCATGAGGTCAGGAG
AGAGCACACGTCTGAACTCTTTCCCTACACGACGCTCTTCCGATCTNNNCAGATCATGAGGTCAGGAG
AGCACACGTCTGAACTCTTTCCCTACACGACGCTCTTCCGATCTNNNGGTGACTTGTCCATGTTCAACA
AGCACACGTCTGAACTCTTTCCCTACACGACGCTCTTCCGATCTNNNGGTGACTTGTCCATGTTCAACA
GCACACGTCTGAACTCTTTCCCTACACGACGCTCTTCCGATCTNNNTGGATGCCTGTAATCCCAACTAC
GCACACGTCTGAACTCTTTCCCTACACGACGCTCTTCCGATCTNNNTGGATGCCTGTAATCCCAACTAC
GAGCACACGTCTGAACTCTTTCCCTACACGACGCTCTTCCGATCTNNNACTGTGTCACTGTCACTACAC
GAGCACACGTCTGAACTCTTTCCCTACACGACGCTCTTCCGATCTNNNACTGTGTCACTGTCACTACAC
GAGCACACGTCTGAACTCTTTCCCTACACGACGCTCTTCCGATCTNNNTTTTGCCCTCCAGGAGACATT
GAGCACACGTCTGAACTCTTTCCCTACACGACGCTCTTCCGATCTNNNTTTTGCCCTCCAGGAGACATT
AGCACACGTCTGAACTCTTTCCCTACACGACGCTCTTCCGATCTNNNTCAAGGTCTCCTGCTAGGTTTT
AGCACACGTCTGAACTCTTTCCCTACACGACGCTCTTCCGATCTNNNTCAAGGTCTCCTGCTAGGTTTT
GAGCACACGTCTGAACTCTTTCCCTACACGACGCTCTTCCGATCTNNNGAAGCTGTTTGGTTGCTCCTA
GAGCACACGTCTGAACTCTTTCCCTACACGACGCTCTTCCGATCTNNNGAAGCTGTTTGGTTGCTCCTA
GAGCACACGTCTGAACTCTTTCCCTACACGACGCTCTTCCGATCTNNNTTCCAAACCGTTCTGTGTTGT
GAGCACACGTCTGAACTCTTTCCCTACACGACGCTCTTCCGATCTNNNTTCCAAACCGTTCTGTGTTGT
GAGCACACGTCTGAACTCTTTCCCTACACGACGCTCTTCCGATCTNNNCAGAAGAATGGCGTGAACCT
GAGCACACGTCTGAACTCTTTCCCTACACGACGCTCTTCCGATCTNNNCAGAAGAATGGCGTGAACCT
AGCACACGTCTGAACTCTTTCCCTACACGACGCTCTTCCGATCTNNNATGCCTGTAATCCCAGCTACTT
AGCACACGTCTGAACTCTTTCCCTACACGACGCTCTTCCGATCTNNNATGCCTGTAATCCCAGCTACTT
GAGCACACGTCTGAACTCTTTCCCTACACGACGCTCTTCCGATCTNNNATCCACCAACCAAGCCCTATA
GAGCACACGTCTGAACTCTTTCCCTACACGACGCTCTTCCGATCTNNNATCCACCAACCAAGCCCTATA
AGCACACGTCTGAACTCTTTCCCTACACGACGCTCTTCCGATCTNNNTCACAACACTGAACTGGGAAG
AGCACACGTCTGAACTCTTTCCCTACACGACGCTCTTCCGATCTNNNTCACAACACTGAACTGGGAAG
AGCACACGTCTGAACTCTTTCCCTACACGACGCTCTTCCGATCTNNNGAGCCTGACAAGTCTTATGGG

AGCACACGTCTGAACTCTTTCCCTACACGACGCTCTTCCGATCTNNNGAGCCTGACAAGTCTTATGGG
AGAGCACACGTCTGAACTCTTTCCCTACACGACGCTCTTCCGATCTNNNACAGGTGCTCAACAAGTTT
AGAGCACACGTCTGAACTCTTTCCCTACACGACGCTCTTCCGATCTNNNACAGGTGCTCAACAAGTTT
AGCACACGTCTGAACTCTTTCCCTACACGACGCTCTTCCGATCTNNNCACCTTAGCCTCCTAAGTAGCT
AGCACACGTCTGAACTCTTTCCCTACACGACGCTCTTCCGATCTNNNCACCTTAGCCTCCTAAGTAGCT
AGCACACGTCTGAACTCTTTCCCTACACGACGCTCTTCCGATCTNNNCTGTAATCCCAGCTACTCAGGA
AGCACACGTCTGAACTCTTTCCCTACACGACGCTCTTCCGATCTNNNCTGTAATCCCAGCTACTCAGGA
AGCACACGTCTGAACTCTTTCCCTACACGACGCTCTTCCGATCTNNNAGGGATAATTGCCTGAACCTGA
AGCACACGTCTGAACTCTTTCCCTACACGACGCTCTTCCGATCTNNNAGGGATAATTGCCTGAACCTGA
ACACGTCTGAACTCTTTCCCTACACGACGCTCTTCCGATCTNNNGTAAAGAAACACACGGTCTGCATC
ACACGTCTGAACTCTTTCCCTACACGACGCTCTTCCGATCTNNNGTAAAGAAACACACGGTCTGCATC
AGAGCACACGTCTGAACTCTTTCCCTACACGACGCTCTTCCGATCTNNNTGTTGCCTTCTCCTTCTAAGT
AGAGCACACGTCTGAACTCTTTCCCTACACGACGCTCTTCCGATCTNNNTGTTGCCTTCTCCTTCTAAGT
AGCACACGTCTGAACTCTTTCCCTACACGACGCTCTTCCGATCTNNNTACTTGTCTCTGACCAGACCTG
AGCACACGTCTGAACTCTTTCCCTACACGACGCTCTTCCGATCTNNNTACTTGTCTCTGACCAGACCTG
CACACGTCTGAACTCTTTCCCTACACGACGCTCTTCCGATCTNNNCTGTGGATTACATGGACCTGCCTT
CACACGTCTGAACTCTTTCCCTACACGACGCTCTTCCGATCTNNNCTGTGGATTACATGGACCTGCCTT
AGCACACGTCTGAACTCTTTCCCTACACGACGCTCTTCCGATCTNNNAACCCCAAGACCATCTACACAA
AGCACACGTCTGAACTCTTTCCCTACACGACGCTCTTCCGATCTNNNAACCCCAAGACCATCTACACAA
AGCACACGTCTGAACTCTTTCCCTACACGACGCTCTTCCGATCTNNNCGTTACGAAATATATCCTGTTT
AGCACACGTCTGAACTCTTTCCCTACACGACGCTCTTCCGATCTNNNCGTTACGAAATATATCCTGTTT
GCACACGTCTGAACTCTTTCCCTACACGACGCTCTTCCGATCTNNNTCATAAAGGAATCCATTGGCCAT
GCACACGTCTGAACTCTTTCCCTACACGACGCTCTTCCGATCTNNNTCATAAAGGAATCCATTGGCCAT
GAGCACACGTCTGAACTCTTTCCCTACACGACGCTCTTCCGATCTNNNTCACAGATTCCATTAACTAGC
GAGCACACGTCTGAACTCTTTCCCTACACGACGCTCTTCCGATCTNNNTCACAGATTCCATTAACTAGC
AGCACACGTCTGAACTCTTTCCCTACACGACGCTCTTCCGATCTNNNAGCCTGCACAGTTATTAGGTT
AGCACACGTCTGAACTCTTTCCCTACACGACGCTCTTCCGATCTNNNAGCCTGCACAGTTATTAGGTT
GCACACGTCTGAACTCTTTCCCTACACGACGCTCTTCCGATCTNNNGAAGTGAGGAGCCACTCTGCCA
GCACACGTCTGAACTCTTTCCCTACACGACGCTCTTCCGATCTNNNGAAGTGAGGAGCCACTCTGCCA
AGAGCACACGTCTGAACTCTTTCCCTACACGACGCTCTTCCGATCTNNNACCTTGTCAAAAAAAAAAA
AGAGCACACGTCTGAACTCTTTCCCTACACGACGCTCTTCCGATCTNNNACCTTGTCAAAAAAAAAAA
GCACACGTCTGAACTCTTTCCCTACACGACGCTCTTCCGATCTNNNCTTGTATTTTTCAGGCTATTTTCC
GCACACGTCTGAACTCTTTCCCTACACGACGCTCTTCCGATCTNNNCTTGTATTTTTCAGGCTATTTTCC
GAGCACACGTCTGAACTCTTTCCCTACACGACGCTCTTCCGATCTNNNCATGCCTGTAATCCCAGCTAC
GAGCACACGTCTGAACTCTTTCCCTACACGACGCTCTTCCGATCTNNNCATGCCTGTAATCCCAGCTAC
GCACACGTCTGAACTCTTTCCCTACACGACGCTCTTCCGATCTNNNCGCCTCACATTCTTTTGTCCTTTC
GCACACGTCTGAACTCTTTCCCTACACGACGCTCTTCCGATCTNNNCGCCTCACATTCTTTTGTCCTTTC
GCACACGTCTGAACTCTTTCCCTACACGACGCTCTTCCGATCTNNNGTTGCTTGCTGGAAAAGACATG
GCACACGTCTGAACTCTTTCCCTACACGACGCTCTTCCGATCTNNNGTTGCTTGCTGGAAAAGACATG
GCACACGTCTGAACTCTTTCCCTACACGACGCTCTTCCGATCTNNNACCCTGTAACTGTGATAACTGTG
GCACACGTCTGAACTCTTTCCCTACACGACGCTCTTCCGATCTNNNACCCTGTAACTGTGATAACTGTG
AGCACACGTCTGAACTCTTTCCCTACACGACGCTCTTCCGATCTNNNAGGAATGTCACTTAGCTGCTTC
AGCACACGTCTGAACTCTTTCCCTACACGACGCTCTTCCGATCTNNNAGGAATGTCACTTAGCTGCTTC

CACGTCTGAACTCTTTCCCTACACGACGCTCTTCCGATCTNNNTGTTTACCTGCCACCATTAGTGTCAT
CACGTCTGAACTCTTTCCCTACACGACGCTCTTCCGATCTNNNTGTTTACCTGCCACCATTAGTGTCAT
GCACACGTCTGAACTCTTTCCCTACACGACGCTCTTCCGATCTNNNGGTTACAGTGACCAAAAACCCA
GCACACGTCTGAACTCTTTCCCTACACGACGCTCTTCCGATCTNNNGGTTACAGTGACCAAAAACCCA
GAGCACACGTCTGAACTCTTTCCCTACACGACGCTCTTCCGATCTNNNCCTTATTGTCAAGGCCTCTCA
GAGCACACGTCTGAACTCTTTCCCTACACGACGCTCTTCCGATCTNNNCCTTATTGTCAAGGCCTCTCA
AGCACACGTCTGAACTCTTTCCCTACACGACGCTCTTCCGATCTNNNATAAGATTCCATCTCACCTGGC
AGCACACGTCTGAACTCTTTCCCTACACGACGCTCTTCCGATCTNNNATAAGATTCCATCTCACCTGGC
CACACGTCTGAACTCTTTCCCTACACGACGCTCTTCCGATCTNNNTTGTTTGCCATTCTCTGTAGGGAC
CACACGTCTGAACTCTTTCCCTACACGACGCTCTTCCGATCTNNNTTGTTTGCCATTCTCTGTAGGGAC
AGCACACGTCTGAACTCTTTCCCTACACGACGCTCTTCCGATCTNNNGATTGAGACCATCCTGGCTAA
AGCACACGTCTGAACTCTTTCCCTACACGACGCTCTTCCGATCTNNNGATTGAGACCATCCTGGCTAA
GAGCACACGTCTGAACTCTTTCCCTACACGACGCTCTTCCGATCTNNNGTGATCGATGACTCCCAGTT
GAGCACACGTCTGAACTCTTTCCCTACACGACGCTCTTCCGATCTNNNGTGATCGATGACTCCCAGTT
GCACACGTCTGAACTCTTTCCCTACACGACGCTCTTCCGATCTNNNTGTAATCTCAGCACTTTGGGAGG
GCACACGTCTGAACTCTTTCCCTACACGACGCTCTTCCGATCTNNNTGTAATCTCAGCACTTTGGGAGG
AGCACACGTCTGAACTCTTTCCCTACACGACGCTCTTCCGATCTNNNTTGCACCAAGCGAATAGAAGT
AGCACACGTCTGAACTCTTTCCCTACACGACGCTCTTCCGATCTNNNTTGCACCAAGCGAATAGAAGT
GAGCACACGTCTGAACTCTTTCCCTACACGACGCTCTTCCGATCTNNNCTATGTGGGTAAAGGCACCA
GAGCACACGTCTGAACTCTTTCCCTACACGACGCTCTTCCGATCTNNNCTATGTGGGTAAAGGCACCA
AGCACACGTCTGAACTCTTTCCCTACACGACGCTCTTCCGATCTNNNCATTTTCCCTCCAGGTGATAGC
AGCACACGTCTGAACTCTTTCCCTACACGACGCTCTTCCGATCTNNNCATTTTCCCTCCAGGTGATAGC
AGCACACGTCTGAACTCTTTCCCTACACGACGCTCTTCCGATCTNNNTTGGGATTACAGGTATGAGCC
AGCACACGTCTGAACTCTTTCCCTACACGACGCTCTTCCGATCTNNNTTGGGATTACAGGTATGAGCC
AGCACACGTCTGAACTCTTTCCCTACACGACGCTCTTCCGATCTNNNTACAAAATTAGGCCAGGTGTG
AGCACACGTCTGAACTCTTTCCCTACACGACGCTCTTCCGATCTNNNTACAAAATTAGGCCAGGTGTG
CACACGTCTGAACTCTTTCCCTACACGACGCTCTTCCGATCTNNNATCCCCTAGGCTTTGAGTTGATCC
CACACGTCTGAACTCTTTCCCTACACGACGCTCTTCCGATCTNNNATCCCCTAGGCTTTGAGTTGATCC
GCACACGTCTGAACTCTTTCCCTACACGACGCTCTTCCGATCTNNNGAGCCATGTTTCCCTAAAAGCA
GCACACGTCTGAACTCTTTCCCTACACGACGCTCTTCCGATCTNNNGAGCCATGTTTCCCTAAAAGCA
AGCACACGTCTGAACTCTTTCCCTACACGACGCTCTTCCGATCTNNNTAGAGTCCTAAATCTCGAACA
AGCACACGTCTGAACTCTTTCCCTACACGACGCTCTTCCGATCTNNNTAGAGTCCTAAATCTCGAACA
GAGCACACGTCTGAACTCTTTCCCTACACGACGCTCTTCCGATCTNNNCCCAGCCACTAAAAGTATCCT
GAGCACACGTCTGAACTCTTTCCCTACACGACGCTCTTCCGATCTNNNCCCAGCCACTAAAAGTATCCT
GAGCACACGTCTGAACTCTTTCCCTACACGACGCTCTTCCGATCTNNNCAAAGTGAAGACCCTGAGTG
GAGCACACGTCTGAACTCTTTCCCTACACGACGCTCTTCCGATCTNNNCAAAGTGAAGACCCTGAGTG
GCACACGTCTGAACTCTTTCCCTACACGACGCTCTTCCGATCTNNNTGACATTTCAGTCAATGGCAGA
GCACACGTCTGAACTCTTTCCCTACACGACGCTCTTCCGATCTNNNTGACATTTCAGTCAATGGCAGA
GCACACGTCTGAACTCTTTCCCTACACGACGCTCTTCCGATCTNNNTGCTCACCTTGACTTTCCTCTTTC
GCACACGTCTGAACTCTTTCCCTACACGACGCTCTTCCGATCTNNNTGCTCACCTTGACTTTCCTCTTTC
AGCACACGTCTGAACTCTTTCCCTACACGACGCTCTTCCGATCTNNNCGACAAAGCGAGACTCTGTCTC
AGCACACGTCTGAACTCTTTCCCTACACGACGCTCTTCCGATCTNNNCGACAAAGCGAGACTCTGTCTC
AGCACACGTCTGAACTCTTTCCCTACACGACGCTCTTCCGATCTNNNGACAAAGTCTCTCTCTGTCACC

AGCACACGTCTGAACTCTTTCCCTACACGACGCTCTTCCGATCTNNNGACAAAGTCTCTCTCTGTCACC
AGCACACGTCTGAACTCTTTCCCTACACGACGCTCTTCCGATCTNNNGGGTCTCAAAAGGGAGATTAC
AGCACACGTCTGAACTCTTTCCCTACACGACGCTCTTCCGATCTNNNGGGTCTCAAAAGGGAGATTAC
GAGCACACGTCTGAACTCTTTCCCTACACGACGCTCTTCCGATCTNNNACAAAAATTAGCTGGGCATGG
GAGCACACGTCTGAACTCTTTCCCTACACGACGCTCTTCCGATCTNNNACAAAAATTAGCTGGGCATGG
GAGCACACGTCTGAACTCTTTCCCTACACGACGCTCTTCCGATCTNNNATACGACCAATCCTCCTGCAA
GAGCACACGTCTGAACTCTTTCCCTACACGACGCTCTTCCGATCTNNNATACGACCAATCCTCCTGCAA
AGCACACGTCTGAACTCTTTCCCTACACGACGCTCTTCCGATCTNNNTGAGGTAGCTGTGGCAATTTC
AGCACACGTCTGAACTCTTTCCCTACACGACGCTCTTCCGATCTNNNTGAGGTAGCTGTGGCAATTTC
GAGCACACGTCTGAACTCTTTCCCTACACGACGCTCTTCCGATCTNNNAACATCATACACCACACCAGA
GAGCACACGTCTGAACTCTTTCCCTACACGACGCTCTTCCGATCTNNNAACATCATACACCACACCAGA
GCACACGTCTGAACTCTTTCCCTACACGACGCTCTTCCGATCTNNNTATGTGGTGTGAATGTGGGGTA
GCACACGTCTGAACTCTTTCCCTACACGACGCTCTTCCGATCTNNNTATGTGGTGTGAATGTGGGGTA
GCACACGTCTGAACTCTTTCCCTACACGACGCTCTTCCGATCTNNNTTGTGGTCTCAGCTATTTGGGAG
GCACACGTCTGAACTCTTTCCCTACACGACGCTCTTCCGATCTNNNTTGTGGTCTCAGCTATTTGGGAG
AGCACACGTCTGAACTCTTTCCCTACACGACGCTCTTCCGATCTNNNCAGAATCCTGACAGGCTCTGAG
AGCACACGTCTGAACTCTTTCCCTACACGACGCTCTTCCGATCTNNNCAGAATCCTGACAGGCTCTGAG
AGCACACGTCTGAACTCTTTCCCTACACGACGCTCTTCCGATCTNNNTGTGTCTGATGGGCAATCTTT
AGCACACGTCTGAACTCTTTCCCTACACGACGCTCTTCCGATCTNNNTGTGTCTGATGGGCAATCTTT
AGAGCACACGTCTGAACTCTTTCCCTACACGACGCTCTTCCGATCTNNNCCATCTGTTGTAGCAAACAT
AGAGCACACGTCTGAACTCTTTCCCTACACGACGCTCTTCCGATCTNNNCCATCTGTTGTAGCAAACAT
AGCACACGTCTGAACTCTTTCCCTACACGACGCTCTTCCGATCTNNNCTCGTGACATAGGGAGACTGT
AGCACACGTCTGAACTCTTTCCCTACACGACGCTCTTCCGATCTNNNCTCGTGACATAGGGAGACTGT
AGCACACGTCTGAACTCTTTCCCTACACGACGCTCTTCCGATCTNNNAGACTGTTGTTAGTACCCAACC
AGCACACGTCTGAACTCTTTCCCTACACGACGCTCTTCCGATCTNNNAGACTGTTGTTAGTACCCAACC
GAGCACACGTCTGAACTCTTTCCCTACACGACGCTCTTCCGATCTNNNCTTGCAGTGAGCTGAGATCA
GAGCACACGTCTGAACTCTTTCCCTACACGACGCTCTTCCGATCTNNNCTTGCAGTGAGCTGAGATCA
AGCACACGTCTGAACTCTTTCCCTACACGACGCTCTTCCGATCTNNNAAAAATTAGCCGGGTGTAGT
AGCACACGTCTGAACTCTTTCCCTACACGACGCTCTTCCGATCTNNNAAAAATTAGCCGGGTGTAGT
GAGCACACGTCTGAACTCTTTCCCTACACGACGCTCTTCCGATCTNNNAAATACGAAATTAGCCAGG
GAGCACACGTCTGAACTCTTTCCCTACACGACGCTCTTCCGATCTNNNAAATACGAAATTAGCCAGG
AGCACACGTCTGAACTCTTTCCCTACACGACGCTCTTCCGATCTNNNACTTAAAGTGGAAATTACAAA
AGCACACGTCTGAACTCTTTCCCTACACGACGCTCTTCCGATCTNNNACTTAAAGTGGAAATTACAAA
GCACACGTCTGAACTCTTTCCCTACACGACGCTCTTCCGATCTNNNTTTTGGAATACGCCAGGAACAT
GCACACGTCTGAACTCTTTCCCTACACGACGCTCTTCCGATCTNNNTTTTGGAATACGCCAGGAACAT
AGCACACGTCTGAACTCTTTCCCTACACGACGCTCTTCCGATCTNNNCTCTACTCCCCATCACTCCCAGC
AGCACACGTCTGAACTCTTTCCCTACACGACGCTCTTCCGATCTNNNCTCTACTCCCCATCACTCCCAGC
AAGAGCACACGTCTGAACTCTTTCCCTACACGACGCTCTTCCGATCTNNNAGACGAGTGACTTCCCTG
AAGAGCACACGTCTGAACTCTTTCCCTACACGACGCTCTTCCGATCTNNNAGACGAGTGACTTCCCTG
GCACACGTCTGAACTCTTTCCCTACACGACGCTCTTCCGATCTNNNTTTCAGGTCTATACAGCACACCT
GCACACGTCTGAACTCTTTCCCTACACGACGCTCTTCCGATCTNNNTTTCAGGTCTATACAGCACACCT
AAGAGCACACGTCTGAACTCTTTCCCTACACGACGCTCTTCCGATCTNNNGAGGCTGCAGTGAGCTAA
AAGAGCACACGTCTGAACTCTTTCCCTACACGACGCTCTTCCGATCTNNNGAGGCTGCAGTGAGCTAA

AGCACACGTCTGAACTCTTTCCCTACACGACGCTCTTCCGATCTNNNTGAGGAGAGAAATGTTTGGG
AGCACACGTCTGAACTCTTTCCCTACACGACGCTCTTCCGATCTNNNTGAGGAGAGAAATGTTTGGG
GAGCACACGTCTGAACTCTTTCCCTACACGACGCTCTTCCGATCTNNNACTTGAGGTCAGGAGTTTGA
GAGCACACGTCTGAACTCTTTCCCTACACGACGCTCTTCCGATCTNNNACTTGAGGTCAGGAGTTTGA
GAGCACACGTCTGAACTCTTTCCCTACACGACGCTCTTCCGATCTNNNTGAGCATCTACTATGTGTCAG
GAGCACACGTCTGAACTCTTTCCCTACACGACGCTCTTCCGATCTNNNTGAGCATCTACTATGTGTCAG
AGCACACGTCTGAACTCTTTCCCTACACGACGCTCTTCCGATCTNNNGGTTGCAGTGAGCCAAGATT
AGCACACGTCTGAACTCTTTCCCTACACGACGCTCTTCCGATCTNNNGGTTGCAGTGAGCCAAGATT
AGCACACGTCTGAACTCTTTCCCTACACGACGCTCTTCCGATCTNNNTCAGACTTGTATACTGGTTGC
AGCACACGTCTGAACTCTTTCCCTACACGACGCTCTTCCGATCTNNNTCAGACTTGTATACTGGTTGC
AGCACACGTCTGAACTCTTTCCCTACACGACGCTCTTCCGATCTNNNGTGGTAGGCACCTATAATCCC
AGCACACGTCTGAACTCTTTCCCTACACGACGCTCTTCCGATCTNNNGTGGTAGGCACCTATAATCCC
AGCACACGTCTGAACTCTTTCCCTACACGACGCTCTTCCGATCTNNNTGGATCCTCTCCTTATTCCTTAG
AGCACACGTCTGAACTCTTTCCCTACACGACGCTCTTCCGATCTNNNTGGATCCTCTCCTTATTCCTTAG
AGCACACGTCTGAACTCTTTCCCTACACGACGCTCTTCCGATCTNNNCTGTGGTTGCACTTGATGAAC
AGCACACGTCTGAACTCTTTCCCTACACGACGCTCTTCCGATCTNNNCTGTGGTTGCACTTGATGAAC
AGCACACGTCTGAACTCTTTCCCTACACGACGCTCTTCCGATCTNNNCTGTCACCTCTCAATTTGAGCCT
AGCACACGTCTGAACTCTTTCCCTACACGACGCTCTTCCGATCTNNNCTGTCACCTCTCAATTTGAGCCT
AGCACACGTCTGAACTCTTTCCCTACACGACGCTCTTCCGATCTNNNGATGGTCTCGATCTCTTGACCT
AGCACACGTCTGAACTCTTTCCCTACACGACGCTCTTCCGATCTNNNGATGGTCTCGATCTCTTGACCT
AGCACACGTCTGAACTCTTTCCCTACACGACGCTCTTCCGATCTNNNCTCACGCCTATAATCCCATCACT
AGCACACGTCTGAACTCTTTCCCTACACGACGCTCTTCCGATCTNNNCTCACGCCTATAATCCCATCACT
AGCACACGTCTGAACTCTTTCCCTACACGACGCTCTTCCGATCTNNNCTGAGGTGGGAGAATCACTTG
AGCACACGTCTGAACTCTTTCCCTACACGACGCTCTTCCGATCTNNNCTGAGGTGGGAGAATCACTTG
GAGCACACGTCTGAACTCTTTCCCTACACGACGCTCTTCCGATCTNNNCTAGCCAGACAGAGAGAAA
GAGCACACGTCTGAACTCTTTCCCTACACGACGCTCTTCCGATCTNNNCTAGCCAGACAGAGAGAAA
GCACACGTCTGAACTCTTTCCCTACACGACGCTCTTCCGATCTNNNAGTAGTGCATTGTACCAAGCAT
GCACACGTCTGAACTCTTTCCCTACACGACGCTCTTCCGATCTNNNAGTAGTGCATTGTACCAAGCAT
AGCACACGTCTGAACTCTTTCCCTACACGACGCTCTTCCGATCTNNNGGAAGAGAAACTGGCCTGAG
AGCACACGTCTGAACTCTTTCCCTACACGACGCTCTTCCGATCTNNNGGAAGAGAAACTGGCCTGAG
AGCACACGTCTGAACTCTTTCCCTACACGACGCTCTTCCGATCTNNNATATTACATTTCAGAGTTCACC
AGCACACGTCTGAACTCTTTCCCTACACGACGCTCTTCCGATCTNNNATATTACATTTCAGAGTTCACC
AGCACACGTCTGAACTCTTTCCCTACACGACGCTCTTCCGATCTNNNCAGAGTTTGCAATGAGCCAAG
AGCACACGTCTGAACTCTTTCCCTACACGACGCTCTTCCGATCTNNNCAGAGTTTGCAATGAGCCAAG
GCACACGTCTGAACTCTTTCCCTACACGACGCTCTTCCGATCTNNNATGCCTGTAATCCCAGCTACTTG
GCACACGTCTGAACTCTTTCCCTACACGACGCTCTTCCGATCTNNNATGCCTGTAATCCCAGCTACTTG
AGCACACGTCTGAACTCTTTCCCTACACGACGCTCTTCCGATCTNNNATTCACAGAACCTTCCAGCAT
AGCACACGTCTGAACTCTTTCCCTACACGACGCTCTTCCGATCTNNNATTCACAGAACCTTCCAGCAT
GAGCACACGTCTGAACTCTTTCCCTACACGACGCTCTTCCGATCTNNNTTATTTCTCCTCCCTCCCCATC
GAGCACACGTCTGAACTCTTTCCCTACACGACGCTCTTCCGATCTNNNTTATTTCTCCTCCCTCCCCATC
GAGCACACGTCTGAACTCTTTCCCTACACGACGCTCTTCCGATCTNNNACATTGGGGCAGAGAATGT
GAGCACACGTCTGAACTCTTTCCCTACACGACGCTCTTCCGATCTNNNACATTGGGGCAGAGAATGT
GAGCACACGTCTGAACTCTTTCCCTACACGACGCTCTTCCGATCTNNNACTCCAAGTTTTCTCCGAAGA

GAGCACACGTCTGAACTCTTTCCCTACACGACGCTCTTCCGATCTNNNACTCCAAGTTTTCTCCGAAGA
GCACACGTCTGAACTCTTTCCCTACACGACGCTCTTCCGATCTNNNGAAAAGAGGCAGGGCTTTGTG
GCACACGTCTGAACTCTTTCCCTACACGACGCTCTTCCGATCTNNNGAAAAGAGGCAGGGCTTTGTG
GAGCACACGTCTGAACTCTTTCCCTACACGACGCTCTTCCGATCTNNNAAATCTTCTCTTCCCCACTG
GAGCACACGTCTGAACTCTTTCCCTACACGACGCTCTTCCGATCTNNNAAATCTTCTCTTCCCCACTG
GCACACGTCTGAACTCTTTCCCTACACGACGCTCTTCCGATCTNNNCCAACAAGTATTTGCTGAGGT
GCACACGTCTGAACTCTTTCCCTACACGACGCTCTTCCGATCTNNNCCAACAAGTATTTGCTGAGGT
AGCACACGTCTGAACTCTTTCCCTACACGACGCTCTTCCGATCTNNNGCCTTACAGATACACAGTTTC
AGCACACGTCTGAACTCTTTCCCTACACGACGCTCTTCCGATCTNNNGCCTTACAGATACACAGTTTC
AGCACACGTCTGAACTCTTTCCCTACACGACGCTCTTCCGATCTNNNAAACAGGAGTGACTTATGCAG
AGCACACGTCTGAACTCTTTCCCTACACGACGCTCTTCCGATCTNNNAAACAGGAGTGACTTATGCAG
GCACACGTCTGAACTCTTTCCCTACACGACGCTCTTCCGATCTNNNTCCAAAGTCCATGTTCTTTCCCTC
GCACACGTCTGAACTCTTTCCCTACACGACGCTCTTCCGATCTNNNTCCAAAGTCCATGTTCTTTCCCTC
GCACACGTCTGAACTCTTTCCCTACACGACGCTCTTCCGATCTNNNGCATGTTTGCCTTACTCTCTCCT
GCACACGTCTGAACTCTTTCCCTACACGACGCTCTTCCGATCTNNNGCATGTTTGCCTTACTCTCTCCT
GAGCACACGTCTGAACTCTTTCCCTACACGACGCTCTTCCGATCTNNNAATACAAAAAATTAGCCGG
GAGCACACGTCTGAACTCTTTCCCTACACGACGCTCTTCCGATCTNNNAATACAAAAAATTAGCCGG
AGCACACGTCTGAACTCTTTCCCTACACGACGCTCTTCCGATCTNNNGGGTATTATAGCTTTGCTGGA
AGCACACGTCTGAACTCTTTCCCTACACGACGCTCTTCCGATCTNNNGGGTATTATAGCTTTGCTGGA
AGCACACGTCTGAACTCTTTCCCTACACGACGCTCTTCCGATCTNNNATTTCTGAGGGCTCTGTTCTGT
AGCACACGTCTGAACTCTTTCCCTACACGACGCTCTTCCGATCTNNNATTTCTGAGGGCTCTGTTCTGT
AGCACACGTCTGAACTCTTTCCCTACACGACGCTCTTCCGATCTNNNGTAAGCCGAGATTGTACCATT
AGCACACGTCTGAACTCTTTCCCTACACGACGCTCTTCCGATCTNNNGTAAGCCGAGATTGTACCATT
AGCACACGTCTGAACTCTTTCCCTACACGACGCTCTTCCGATCTNNNAGCACAATTCACAACTGCAAA
AGCACACGTCTGAACTCTTTCCCTACACGACGCTCTTCCGATCTNNNAGCACAATTCACAACTGCAAA
AGCACACGTCTGAACTCTTTCCCTACACGACGCTCTTCCGATCTNNNCTGAGGCAGGAGAATCACTTG
AGCACACGTCTGAACTCTTTCCCTACACGACGCTCTTCCGATCTNNNCTGAGGCAGGAGAATCACTTG
AGCACACGTCTGAACTCTTTCCCTACACGACGCTCTTCCGATCTNNNACTACATAGTCAGGCAAGGTC
AGCACACGTCTGAACTCTTTCCCTACACGACGCTCTTCCGATCTNNNACTACATAGTCAGGCAAGGTC
AGCACACGTCTGAACTCTTTCCCTACACGACGCTCTTCCGATCTNNNGTGCCTGTAGTCCCAGTTACTC
AGCACACGTCTGAACTCTTTCCCTACACGACGCTCTTCCGATCTNNNGTGCCTGTAGTCCCAGTTACTC
AGCACACGTCTGAACTCTTTCCCTACACGACGCTCTTCCGATCTNNNGAGCTCCAAAAGGAAGACAGG
AGCACACGTCTGAACTCTTTCCCTACACGACGCTCTTCCGATCTNNNGAGCTCCAAAAGGAAGACAGG
AGCACACGTCTGAACTCTTTCCCTACACGACGCTCTTCCGATCTNNNGCAGGCATCTCCTTTTTTTGA
AGCACACGTCTGAACTCTTTCCCTACACGACGCTCTTCCGATCTNNNGCAGGCATCTCCTTTTTTTGA
AGCACACGTCTGAACTCTTTCCCTACACGACGCTCTTCCGATCTNNNTTCCCATCTCAGCCTCCTTAGTA
AGCACACGTCTGAACTCTTTCCCTACACGACGCTCTTCCGATCTNNNTTCCCATCTCAGCCTCCTTAGTA
ACACGTCTGAACTCTTTCCCTACACGACGCTCTTCCGATCTNNNCATCGAGGTTCACAGTTATGCATC
ACACGTCTGAACTCTTTCCCTACACGACGCTCTTCCGATCTNNNCATCGAGGTTCACAGTTATGCATC
AGCACACGTCTGAACTCTTTCCCTACACGACGCTCTTCCGATCTNNNATGAGACAGGAGAATCGCTTG
AGCACACGTCTGAACTCTTTCCCTACACGACGCTCTTCCGATCTNNNATGAGACAGGAGAATCGCTTG
AGCACACGTCTGAACTCTTTCCCTACACGACGCTCTTCCGATCTNNNTCTTTTGCATTTGCTGAGGACT
AGCACACGTCTGAACTCTTTCCCTACACGACGCTCTTCCGATCTNNNTCTTTTGCATTTGCTGAGGACT

AGCACACGTCTGAACTCTTTCCCTACACGACGCTCTTCCGATCTNNNTGGGGAGGGAGTATATGGGA
AGCACACGTCTGAACTCTTTCCCTACACGACGCTCTTCCGATCTNNNTGGGGAGGGAGTATATGGGA
AGAGCACACGTCTGAACTCTTTCCCTACACGACGCTCTTCCGATCTNNNCTGCTGTCCATCAGTGTCCA
AGAGCACACGTCTGAACTCTTTCCCTACACGACGCTCTTCCGATCTNNNCTGCTGTCCATCAGTGTCCA
AGCACACGTCTGAACTCTTTCCCTACACGACGCTCTTCCGATCTNNNCCCAATCCTCAATGTCACACAA
AGCACACGTCTGAACTCTTTCCCTACACGACGCTCTTCCGATCTNNNCCCAATCCTCAATGTCACACAA
GAGCACACGTCTGAACTCTTTCCCTACACGACGCTCTTCCGATCTNNNCAGAGGGAGAGAGAAGGAT
GAGCACACGTCTGAACTCTTTCCCTACACGACGCTCTTCCGATCTNNNCAGAGGGAGAGAGAAGGAT
AGCACACGTCTGAACTCTTTCCCTACACGACGCTCTTCCGATCTNNNTTGAGCTCAGGAATTTGAGAC
AGCACACGTCTGAACTCTTTCCCTACACGACGCTCTTCCGATCTNNNTTGAGCTCAGGAATTTGAGAC
AGCACACGTCTGAACTCTTTCCCTACACGACGCTCTTCCGATCTNNNGGACTCTGAGGTTGGCTAAGA
AGCACACGTCTGAACTCTTTCCCTACACGACGCTCTTCCGATCTNNNGGACTCTGAGGTTGGCTAAGA
GCACACGTCTGAACTCTTTCCCTACACGACGCTCTTCCGATCTNNNTGAAAAAGCAAGGCTAGGGGA
GCACACGTCTGAACTCTTTCCCTACACGACGCTCTTCCGATCTNNNTGAAAAAGCAAGGCTAGGGGA
GAGCACACGTCTGAACTCTTTCCCTACACGACGCTCTTCCGATCTNNNTGGTATGTGACTGTAATCCCA
GAGCACACGTCTGAACTCTTTCCCTACACGACGCTCTTCCGATCTNNNTGGTATGTGACTGTAATCCCA
CACACGTCTGAACTCTTTCCCTACACGACGCTCTTCCGATCTNNNTTCTGGAAAAGCTGGAAAAGCAT
CACACGTCTGAACTCTTTCCCTACACGACGCTCTTCCGATCTNNNTTCTGGAAAAGCTGGAAAAGCAT
GCACACGTCTGAACTCTTTCCCTACACGACGCTCTTCCGATCTNNNTGTTTTACGGTTACAAGGGCACA
GCACACGTCTGAACTCTTTCCCTACACGACGCTCTTCCGATCTNNNTGTTTTACGGTTACAAGGGCACA
AGCACACGTCTGAACTCTTTCCCTACACGACGCTCTTCCGATCTNNNTGAGGTAGGAGAATCGCTTG
AGCACACGTCTGAACTCTTTCCCTACACGACGCTCTTCCGATCTNNNTGAGGTAGGAGAATCGCTTG
GAGCACACGTCTGAACTCTTTCCCTACACGACGCTCTTCCGATCTNNNTGTGCTGTGTTTTGTTTTGGA
GAGCACACGTCTGAACTCTTTCCCTACACGACGCTCTTCCGATCTNNNTGTGCTGTGTTTTGTTTTGGA
AGCACACGTCTGAACTCTTTCCCTACACGACGCTCTTCCGATCTNNNATGCCTGTAATCCCAGCATTTT
AGCACACGTCTGAACTCTTTCCCTACACGACGCTCTTCCGATCTNNNATGCCTGTAATCCCAGCATTTT
AGCACACGTCTGAACTCTTTCCCTACACGACGCTCTTCCGATCTNNNGAGTTGCATGTTTCAGTGAGCT
AGCACACGTCTGAACTCTTTCCCTACACGACGCTCTTCCGATCTNNNGAGTTGCATGTTTCAGTGAGCT
GAGCACACGTCTGAACTCTTTCCCTACACGACGCTCTTCCGATCTNNNGAACATACAGAGAAGCAAC
GAGCACACGTCTGAACTCTTTCCCTACACGACGCTCTTCCGATCTNNNGAACATACAGAGAAGCAAC
GAGCACACGTCTGAACTCTTTCCCTACACGACGCTCTTCCGATCTNNNGGCCTCGAGGTCTTCATTCTTT
GAGCACACGTCTGAACTCTTTCCCTACACGACGCTCTTCCGATCTNNNGGCCTCGAGGTCTTCATTCTTT
GAGCACACGTCTGAACTCTTTCCCTACACGACGCTCTTCCGATCTNNNCCTGGACAACAAGAGCAAAA
GAGCACACGTCTGAACTCTTTCCCTACACGACGCTCTTCCGATCTNNNCCTGGACAACAAGAGCAAAA
GAGCACACGTCTGAACTCTTTCCCTACACGACGCTCTTCCGATCTNNNAGTAGCTGAGAATACAGGCA
GAGCACACGTCTGAACTCTTTCCCTACACGACGCTCTTCCGATCTNNNAGTAGCTGAGAATACAGGCA
GAGCACACGTCTGAACTCTTTCCCTACACGACGCTCTTCCGATCTNNNCAACATGGCAAAACCCAATCT
GAGCACACGTCTGAACTCTTTCCCTACACGACGCTCTTCCGATCTNNNCAACATGGCAAAACCCAATCT
GAGCACACGTCTGAACTCTTTCCCTACACGACGCTCTTCCGATCTNNNGCCTGTAGTCCCAGCTACTTG
GAGCACACGTCTGAACTCTTTCCCTACACGACGCTCTTCCGATCTNNNGCCTGTAGTCCCAGCTACTTG
GCACACGTCTGAACTCTTTCCCTACACGACGCTCTTCCGATCTNNNGGCCTCCTAAATTGCTGGGATTA
GCACACGTCTGAACTCTTTCCCTACACGACGCTCTTCCGATCTNNNGGCCTCCTAAATTGCTGGGATTA
AGCACACGTCTGAACTCTTTCCCTACACGACGCTCTTCCGATCTNNNAATCACTATGAAGTACTAGGT

AGCACACGTCTGAACTCTTTCCCTACACGACGCTCTTCCGATCTNNNAATCACTATGAAGTACTAGGT
GAGCACACGTCTGAACTCTTTCCCTACACGACGCTCTTCCGATCTNNNCCCTGCCTTCAACATTTCACAT
GAGCACACGTCTGAACTCTTTCCCTACACGACGCTCTTCCGATCTNNNCCCTGCCTTCAACATTTCACAT
AGAGCACACGTCTGAACTCTTTCCCTACACGACGCTCTTCCGATCTNNNCCAAAGAATTCTGAACCAAC
AGAGCACACGTCTGAACTCTTTCCCTACACGACGCTCTTCCGATCTNNNCCAAAGAATTCTGAACCAAC
GCACACGTCTGAACTCTTTCCCTACACGACGCTCTTCCGATCTNNNGGTCCCTCTAAGACTTGCAGAA
GCACACGTCTGAACTCTTTCCCTACACGACGCTCTTCCGATCTNNNGGTCCCTCTAAGACTTGCAGAA
GAGCACACGTCTGAACTCTTTCCCTACACGACGCTCTTCCGATCTNNNTAAGTGATCATATTTACCACC
GAGCACACGTCTGAACTCTTTCCCTACACGACGCTCTTCCGATCTNNNTAAGTGATCATATTTACCACC
GAGCACACGTCTGAACTCTTTCCCTACACGACGCTCTTCCGATCTNNNAGGCAGTGACTAAATAAGGA
GAGCACACGTCTGAACTCTTTCCCTACACGACGCTCTTCCGATCTNNNAGGCAGTGACTAAATAAGGA
GAGCACACGTCTGAACTCTTTCCCTACACGACGCTCTTCCGATCTNNNTTGAGTCTATACTACGTGCC
GAGCACACGTCTGAACTCTTTCCCTACACGACGCTCTTCCGATCTNNNTTGAGTCTATACTACGTGCC
AGCACACGTCTGAACTCTTTCCCTACACGACGCTCTTCCGATCTNNNGTCTCTGTCACGTGAATCCCTA
AGCACACGTCTGAACTCTTTCCCTACACGACGCTCTTCCGATCTNNNGTCTCTGTCACGTGAATCCCTA
GCACACGTCTGAACTCTTTCCCTACACGACGCTCTTCCGATCTNNNAATCCCACATACTCGCTACTTGG
GCACACGTCTGAACTCTTTCCCTACACGACGCTCTTCCGATCTNNNAATCCCACATACTCGCTACTTGG
AGCACACGTCTGAACTCTTTCCCTACACGACGCTCTTCCGATCTNNNGGCTAAAACAGTGAAACCTCG
AGCACACGTCTGAACTCTTTCCCTACACGACGCTCTTCCGATCTNNNGGCTAAAACAGTGAAACCTCG
AGCACACGTCTGAACTCTTTCCCTACACGACGCTCTTCCGATCTNNNGGCTGAAATGGGAGGCTCTA
AGCACACGTCTGAACTCTTTCCCTACACGACGCTCTTCCGATCTNNNGGCTGAAATGGGAGGCTCTA
AGCACACGTCTGAACTCTTTCCCTACACGACGCTCTTCCGATCTNNNTTTCCTTCTGTTATCCCTCCCTCC
AGCACACGTCTGAACTCTTTCCCTACACGACGCTCTTCCGATCTNNNTTTCCTTCTGTTATCCCTCCCTCC
GAGCACACGTCTGAACTCTTTCCCTACACGACGCTCTTCCGATCTNNNAAAAAAATTAGCCGGGTGT
GAGCACACGTCTGAACTCTTTCCCTACACGACGCTCTTCCGATCTNNNAAAAAAATTAGCCGGGTGT
AGCACACGTCTGAACTCTTTCCCTACACGACGCTCTTCCGATCTNNNTCTGCTCTTGATGGACTTTTGG
AGCACACGTCTGAACTCTTTCCCTACACGACGCTCTTCCGATCTNNNTCTGCTCTTGATGGACTTTTGG
CACACGTCTGAACTCTTTCCCTACACGACGCTCTTCCGATCTNNNGCTGAGGACATCTCACTTTCCTTCA
CACACGTCTGAACTCTTTCCCTACACGACGCTCTTCCGATCTNNNGCTGAGGACATCTCACTTTCCTTCA
GCACACGTCTGAACTCTTTCCCTACACGACGCTCTTCCGATCTNNNGCCAGAGAATCGTTTGAACTCT
GCACACGTCTGAACTCTTTCCCTACACGACGCTCTTCCGATCTNNNGCCAGAGAATCGTTTGAACTCT
GCACACGTCTGAACTCTTTCCCTACACGACGCTCTTCCGATCTNNNTTTCCCTGTTTCTGACCACAACCC
GCACACGTCTGAACTCTTTCCCTACACGACGCTCTTCCGATCTNNNTTTCCCTGTTTCTGACCACAACCC
GAGCACACGTCTGAACTCTTTCCCTACACGACGCTCTTCCGATCTNNNAAATCTCTTCCAGTGAGTCAC
GAGCACACGTCTGAACTCTTTCCCTACACGACGCTCTTCCGATCTNNNAAATCTCTTCCAGTGAGTCAC
AGCACACGTCTGAACTCTTTCCCTACACGACGCTCTTCCGATCTNNNGAGACAGGAGAATGGCATGG
AGCACACGTCTGAACTCTTTCCCTACACGACGCTCTTCCGATCTNNNGAGACAGGAGAATGGCATGG
AGAGCACACGTCTGAACTCTTTCCCTACACGACGCTCTTCCGATCTNNNGGGTGACAGAATGAGACTC
AGAGCACACGTCTGAACTCTTTCCCTACACGACGCTCTTCCGATCTNNNGGGTGACAGAATGAGACTC
CACACGTCTGAACTCTTTCCCTACACGACGCTCTTCCGATCTNNNTCTGAATCCCTCTTTGTGTGTCCCA
CACACGTCTGAACTCTTTCCCTACACGACGCTCTTCCGATCTNNNTCTGAATCCCTCTTTGTGTGTCCCA
AGCACACGTCTGAACTCTTTCCCTACACGACGCTCTTCCGATCTNNNGAGGCTGAGGCAGGAGAATG
AGCACACGTCTGAACTCTTTCCCTACACGACGCTCTTCCGATCTNNNGAGGCTGAGGCAGGAGAATG

AGCACACGTCTGAACTCTTTCCCTACACGACGCTCTTCCGATCTNNNGGTGAAACTCCGTCTCTACTG
AGCACACGTCTGAACTCTTTCCCTACACGACGCTCTTCCGATCTNNNGGTGAAACTCCGTCTCTACTG
AGCACACGTCTGAACTCTTTCCCTACACGACGCTCTTCCGATCTNNNCATGGGCCACTCAGTTTCAAT
AGCACACGTCTGAACTCTTTCCCTACACGACGCTCTTCCGATCTNNNCATGGGCCACTCAGTTTCAAT
GCACACGTCTGAACTCTTTCCCTACACGACGCTCTTCCGATCTNNNCCCATCTCTCAGCTGGAATATC
GCACACGTCTGAACTCTTTCCCTACACGACGCTCTTCCGATCTNNNCCCATCTCTCAGCTGGAATATC
AGCACACGTCTGAACTCTTTCCCTACACGACGCTCTTCCGATCTNNNTGGGTGACAGAGCAAGACTCC
AGCACACGTCTGAACTCTTTCCCTACACGACGCTCTTCCGATCTNNNTGGGTGACAGAGCAAGACTCC
GCACACGTCTGAACTCTTTCCCTACACGACGCTCTTCCGATCTNNNCAGTTGGTCAGGTTGCTATTGT
GCACACGTCTGAACTCTTTCCCTACACGACGCTCTTCCGATCTNNNCAGTTGGTCAGGTTGCTATTGT
AGCACACGTCTGAACTCTTTCCCTACACGACGCTCTTCCGATCTNNNTGAAACCCCGTCTGTACTGAA
AGCACACGTCTGAACTCTTTCCCTACACGACGCTCTTCCGATCTNNNTGAAACCCCGTCTGTACTGAA
GCACACGTCTGAACTCTTTCCCTACACGACGCTCTTCCGATCTNNNTTCATGTTTTCAGTGTTTGGGCA
GCACACGTCTGAACTCTTTCCCTACACGACGCTCTTCCGATCTNNNTTCATGTTTTCAGTGTTTGGGCA
AGCACACGTCTGAACTCTTTCCCTACACGACGCTCTTCCGATCTNNNGAAGCAGGAGAATCGCTTGAA
AGCACACGTCTGAACTCTTTCCCTACACGACGCTCTTCCGATCTNNNGAAGCAGGAGAATCGCTTGAA
AGCACACGTCTGAACTCTTTCCCTACACGACGCTCTTCCGATCTNNNTTGACTTCATTTCCCATCCCTG
AGCACACGTCTGAACTCTTTCCCTACACGACGCTCTTCCGATCTNNNTTGACTTCATTTCCCATCCCTG
AGAGCACACGTCTGAACTCTTTCCCTACACGACGCTCTTCCGATCTNNNATCATCCCCAGTTCCCAGATC
AGAGCACACGTCTGAACTCTTTCCCTACACGACGCTCTTCCGATCTNNNATCATCCCCAGTTCCCAGATC
AGCACACGTCTGAACTCTTTCCCTACACGACGCTCTTCCGATCTNNNAGGAGTTTGAAACTATCCTGG
AGCACACGTCTGAACTCTTTCCCTACACGACGCTCTTCCGATCTNNNAGGAGTTTGAAACTATCCTGG
AGCACACGTCTGAACTCTTTCCCTACACGACGCTCTTCCGATCTNNNCATTGCCTGTGTTCTCCAGATTC
AGCACACGTCTGAACTCTTTCCCTACACGACGCTCTTCCGATCTNNNCATTGCCTGTGTTCTCCAGATTC
AGCACACGTCTGAACTCTTTCCCTACACGACGCTCTTCCGATCTNNNGAGGCTGAGGCAGGAGAATG
AGCACACGTCTGAACTCTTTCCCTACACGACGCTCTTCCGATCTNNNGAGGCTGAGGCAGGAGAATG
GAGCACACGTCTGAACTCTTTCCCTACACGACGCTCTTCCGATCTNNNGAGTTGCTGGGACTACAGG
GAGCACACGTCTGAACTCTTTCCCTACACGACGCTCTTCCGATCTNNNGAGTTGCTGGGACTACAGG
CACACGTCTGAACTCTTTCCCTACACGACGCTCTTCCGATCTNNNCTTTTTGCCATGGGATTTCCATC
CACACGTCTGAACTCTTTCCCTACACGACGCTCTTCCGATCTNNNCTTTTTGCCATGGGATTTCCATC
GCACACGTCTGAACTCTTTCCCTACACGACGCTCTTCCGATCTNNNATGGTTGCTGGGTCATATGGTA
GCACACGTCTGAACTCTTTCCCTACACGACGCTCTTCCGATCTNNNATGGTTGCTGGGTCATATGGTA
GAGCACACGTCTGAACTCTTTCCCTACACGACGCTCTTCCGATCTNNNAAAGAGTCATTGAGAACAGC
GAGCACACGTCTGAACTCTTTCCCTACACGACGCTCTTCCGATCTNNNAAAGAGTCATTGAGAACAGC
CACACGTCTGAACTCTTTCCCTACACGACGCTCTTCCGATCTNNNAATTCTTTTGGTTTTTGTCTAGGCA
CACACGTCTGAACTCTTTCCCTACACGACGCTCTTCCGATCTNNNAATTCTTTTGGTTTTTGTCTAGGCA
AGCACACGTCTGAACTCTTTCCCTACACGACGCTCTTCCGATCTNNNGAAGGGACAGTTGCCAGAAAA
AGCACACGTCTGAACTCTTTCCCTACACGACGCTCTTCCGATCTNNNGAAGGGACAGTTGCCAGAAAA
GCACACGTCTGAACTCTTTCCCTACACGACGCTCTTCCGATCTNNNTGCCTGTAATCTCAGCTACTTGG
GCACACGTCTGAACTCTTTCCCTACACGACGCTCTTCCGATCTNNNTGCCTGTAATCTCAGCTACTTGG
AGCACACGTCTGAACTCTTTCCCTACACGACGCTCTTCCGATCTNNNACCCTGTAAAGCCATATTGCTCT
AGCACACGTCTGAACTCTTTCCCTACACGACGCTCTTCCGATCTNNNACCCTGTAAAGCCATATTGCTCT
AGAGCACACGTCTGAACTCTTTCCCTACACGACGCTCTTCCGATCTNNNGTTGCAGTGAGCTGAGATC

AGAGCACACGTCTGAACTCTTTCCCTACACGACGCTCTTCCGATCTNNNGTTGCAGTGAGCTGAGATCA
GAGCACACGTCTGAACTCTTTCCCTACACGACGCTCTTCCGATCTNNNCTTTGCCCAGGCTGTAATGTA
GAGCACACGTCTGAACTCTTTCCCTACACGACGCTCTTCCGATCTNNNCTTTGCCCAGGCTGTAATGTA
AGCACACGTCTGAACTCTTTCCCTACACGACGCTCTTCCGATCTNNNACCTGTATTAAGAATCTGTATTC
AGCACACGTCTGAACTCTTTCCCTACACGACGCTCTTCCGATCTNNNACCTGTATTAAGAATCTGTATTC
AGCACACGTCTGAACTCTTTCCCTACACGACGCTCTTCCGATCTNNNGCTTCCTGGATAGCTGGGATTA
AGCACACGTCTGAACTCTTTCCCTACACGACGCTCTTCCGATCTNNNGCTTCCTGGATAGCTGGGATTA
AGCACACGTCTGAACTCTTTCCCTACACGACGCTCTTCCGATCTNNNAAACATGGTGGGCACTTCAGAA
AGCACACGTCTGAACTCTTTCCCTACACGACGCTCTTCCGATCTNNNAAACATGGTGGGCACTTCAGAA
GCACACGTCTGAACTCTTTCCCTACACGACGCTCTTCCGATCTNNNTACTCACCCCTACTCTCCTATGGCA
GCACACGTCTGAACTCTTTCCCTACACGACGCTCTTCCGATCTNNNTACTCACCCCTACTCTCCTATGGCA
AGAGCACACGTCTGAACTCTTTCCCTACACGACGCTCTTCCGATCTNNNCTGAGATTGCACCACTGTACT
AGAGCACACGTCTGAACTCTTTCCCTACACGACGCTCTTCCGATCTNNNCTGAGATTGCACCACTGTACT
AGCACACGTCTGAACTCTTTCCCTACACGACGCTCTTCCGATCTNNNAGAGATTGGGAAGGCAGAATC
AGCACACGTCTGAACTCTTTCCCTACACGACGCTCTTCCGATCTNNNAGAGATTGGGAAGGCAGAATC
AGCACACGTCTGAACTCTTTCCCTACACGACGCTCTTCCGATCTNNNAGAACCCTTACGTTCACTGACT
AGCACACGTCTGAACTCTTTCCCTACACGACGCTCTTCCGATCTNNNAGAACCCTTACGTTCACTGACT
GAGCACACGTCTGAACTCTTTCCCTACACGACGCTCTTCCGATCTNNNCGTCAGGTACATAAGAAGCAC
GAGCACACGTCTGAACTCTTTCCCTACACGACGCTCTTCCGATCTNNNCGTCAGGTACATAAGAAGCAC
GAGCACACGTCTGAACTCTTTCCCTACACGACGCTCTTCCGATCTNNNGAGGTTTCTCTCTGTGTCTCTC
GAGCACACGTCTGAACTCTTTCCCTACACGACGCTCTTCCGATCTNNNGAGGTTTCTCTCTGTGTCTCTC
GCACACGTCTGAACTCTTTCCCTACACGACGCTCTTCCGATCTNNNTTTAGCAGCAGCAACAGAAGAA
GCACACGTCTGAACTCTTTCCCTACACGACGCTCTTCCGATCTNNNTTTAGCAGCAGCAACAGAAGAA
GAGCACACGTCTGAACTCTTTCCCTACACGACGCTCTTCCGATCTNNNGAAACTCTGGCTCTGGAATAC
GAGCACACGTCTGAACTCTTTCCCTACACGACGCTCTTCCGATCTNNNGAAACTCTGGCTCTGGAATAC
AGCACACGTCTGAACTCTTTCCCTACACGACGCTCTTCCGATCTNNNCACGTTGTGCACATGTACCATA
AGCACACGTCTGAACTCTTTCCCTACACGACGCTCTTCCGATCTNNNCACGTTGTGCACATGTACCATA
GAGCACACGTCTGAACTCTTTCCCTACACGACGCTCTTCCGATCTNNNTTGGAAAGTAACAACCCCAGA
GAGCACACGTCTGAACTCTTTCCCTACACGACGCTCTTCCGATCTNNNTTGGAAAGTAACAACCCCAGA
AGCACACGTCTGAACTCTTTCCCTACACGACGCTCTTCCGATCTNNNGTGCTTCAGTTTCATCTCACAAC
AGCACACGTCTGAACTCTTTCCCTACACGACGCTCTTCCGATCTNNNGTGCTTCAGTTTCATCTCACAAC
GAGCACACGTCTGAACTCTTTCCCTACACGACGCTCTTCCGATCTNNNTCATGCTGTCAAACTCTGGTGA
GAGCACACGTCTGAACTCTTTCCCTACACGACGCTCTTCCGATCTNNNTCATGCTGTCAAACTCTGGTGA
GAGCACACGTCTGAACTCTTTCCCTACACGACGCTCTTCCGATCTNNNATCTTTCTCAGAAGCGAGCTC
GAGCACACGTCTGAACTCTTTCCCTACACGACGCTCTTCCGATCTNNNATCTTTCTCAGAAGCGAGCTC
AGCACACGTCTGAACTCTTTCCCTACACGACGCTCTTCCGATCTNNNCTGTGTTCATGCTACTGGACTC
AGCACACGTCTGAACTCTTTCCCTACACGACGCTCTTCCGATCTNNNCTGTGTTCATGCTACTGGACTC
AGCACACGTCTGAACTCTTTCCCTACACGACGCTCTTCCGATCTNNNTTTTTTTGCTTTTGCCTGCCAAA
AGCACACGTCTGAACTCTTTCCCTACACGACGCTCTTCCGATCTNNNTTTTTTTGCTTTTGCCTGCCAAA
AGCACACGTCTGAACTCTTTCCCTACACGACGCTCTTCCGATCTNNNTGCAAAAGAACCTCAATCAAAC
AGCACACGTCTGAACTCTTTCCCTACACGACGCTCTTCCGATCTNNNTGCAAAAGAACCTCAATCAAAC
AGCACACGTCTGAACTCTTTCCCTACACGACGCTCTTCCGATCTNNNGATAGTGAATCAGGCAGGGTGA
AGCACACGTCTGAACTCTTTCCCTACACGACGCTCTTCCGATCTNNNGATAGTGAATCAGGCAGGGTGA

AGCACACGTCTGAACTCTTTCCCTACACGACGCTCTTCCGATCTNNNGACTTGGCTGAGGACTCAAGT
AGCACACGTCTGAACTCTTTCCCTACACGACGCTCTTCCGATCTNNNGACTTGGCTGAGGACTCAAGT
AGCACACGTCTGAACTCTTTCCCTACACGACGCTCTTCCGATCTNNNTGATCCCCACTTTACATTGCCAT
AGCACACGTCTGAACTCTTTCCCTACACGACGCTCTTCCGATCTNNNTGATCCCCACTTTACATTGCCAT
AGCACACGTCTGAACTCTTTCCCTACACGACGCTCTTCCGATCTNNNGTTCACCATCGATGCAAAGGA
AGCACACGTCTGAACTCTTTCCCTACACGACGCTCTTCCGATCTNNNGTTCACCATCGATGCAAAGGA
AGCACACGTCTGAACTCTTTCCCTACACGACGCTCTTCCGATCTNNNAAGGGTGTTAGTGTCCATTCA
AGCACACGTCTGAACTCTTTCCCTACACGACGCTCTTCCGATCTNNNAAGGGTGTTAGTGTCCATTCA
GCACACGTCTGAACTCTTTCCCTACACGACGCTCTTCCGATCTNNNTCTCTGAAGAGGCTCCAGATAG
GCACACGTCTGAACTCTTTCCCTACACGACGCTCTTCCGATCTNNNTCTCTGAAGAGGCTCCAGATAG
AGCACACGTCTGAACTCTTTCCCTACACGACGCTCTTCCGATCTNNNATCAAAGCTCACTGTAGCCTCA
AGCACACGTCTGAACTCTTTCCCTACACGACGCTCTTCCGATCTNNNATCAAAGCTCACTGTAGCCTCA
AGCACACGTCTGAACTCTTTCCCTACACGACGCTCTTCCGATCTNNNCTATACACTTTCCTGCGTGTG
AGCACACGTCTGAACTCTTTCCCTACACGACGCTCTTCCGATCTNNNCTATACACTTTCCTGCGTGTG
GAGCACACGTCTGAACTCTTTCCCTACACGACGCTCTTCCGATCTNNNGGAGCAAAAGAAATGCAACT
GAGCACACGTCTGAACTCTTTCCCTACACGACGCTCTTCCGATCTNNNGGAGCAAAAGAAATGCAACT
GAGCACACGTCTGAACTCTTTCCCTACACGACGCTCTTCCGATCTNNNATGTTCTCACTCTCTTCTACG
GAGCACACGTCTGAACTCTTTCCCTACACGACGCTCTTCCGATCTNNNATGTTCTCACTCTCTTCTACG
AGAGCACACGTCTGAACTCTTTCCCTACACGACGCTCTTCCGATCTNNNAGTCAGAATGCAAGAGAA
AGAGCACACGTCTGAACTCTTTCCCTACACGACGCTCTTCCGATCTNNNAGTCAGAATGCAAGAGAA
AGCACACGTCTGAACTCTTTCCCTACACGACGCTCTTCCGATCTNNNATCTCTTTAACCCTGCCCCAGA
AGCACACGTCTGAACTCTTTCCCTACACGACGCTCTTCCGATCTNNNATCTCTTTAACCCTGCCCCAGA
GCACACGTCTGAACTCTTTCCCTACACGACGCTCTTCCGATCTNNNGAGACTGAGGCAGGAGAATGG
GCACACGTCTGAACTCTTTCCCTACACGACGCTCTTCCGATCTNNNGAGACTGAGGCAGGAGAATGG
AGCACACGTCTGAACTCTTTCCCTACACGACGCTCTTCCGATCTNNNTGTTTTTCAAGGAATTGGCCTG
AGCACACGTCTGAACTCTTTCCCTACACGACGCTCTTCCGATCTNNNTGTTTTTCAAGGAATTGGCCTG
GCACACGTCTGAACTCTTTCCCTACACGACGCTCTTCCGATCTNNNATATCTTGAGCTCAGGAGTTCA
GCACACGTCTGAACTCTTTCCCTACACGACGCTCTTCCGATCTNNNATATCTTGAGCTCAGGAGTTCA
AGCACACGTCTGAACTCTTTCCCTACACGACGCTCTTCCGATCTNNNCTCCTTCTCCAAGCCGAGTCCA
AGCACACGTCTGAACTCTTTCCCTACACGACGCTCTTCCGATCTNNNCTCCTTCTCCAAGCCGAGTCCA
AGCACACGTCTGAACTCTTTCCCTACACGACGCTCTTCCGATCTNNNGCAAAACCTGGTCTATGCCAA
AGCACACGTCTGAACTCTTTCCCTACACGACGCTCTTCCGATCTNNNGCAAAACCTGGTCTATGCCAA
AGCACACGTCTGAACTCTTTCCCTACACGACGCTCTTCCGATCTNNNATACCAAAAATTAGCCGAGC
AGCACACGTCTGAACTCTTTCCCTACACGACGCTCTTCCGATCTNNNATACCAAAAATTAGCCGAGC
AGCACACGTCTGAACTCTTTCCCTACACGACGCTCTTCCGATCTNNNACCTGTAAATTGGGATTGCTCC
AGCACACGTCTGAACTCTTTCCCTACACGACGCTCTTCCGATCTNNNACCTGTAAATTGGGATTGCTCC
AGCACACGTCTGAACTCTTTCCCTACACGACGCTCTTCCGATCTNNNTCAAGTGGAAGGGGCATACT
AGCACACGTCTGAACTCTTTCCCTACACGACGCTCTTCCGATCTNNNTCAAGTGGAAGGGGCATACT
AGCACACGTCTGAACTCTTTCCCTACACGACGCTCTTCCGATCTNNNGGGGAGTCCTAAGACAGACA
AGCACACGTCTGAACTCTTTCCCTACACGACGCTCTTCCGATCTNNNGGGGAGTCCTAAGACAGACA
GAGCACACGTCTGAACTCTTTCCCTACACGACGCTCTTCCGATCTNNNCTGGGATTACAAGGCGTGA
GAGCACACGTCTGAACTCTTTCCCTACACGACGCTCTTCCGATCTNNNCTGGGATTACAAGGCGTGA
AGCACACGTCTGAACTCTTTCCCTACACGACGCTCTTCCGATCTNNNGTCTGCGAAGGGACCGGAAG

AGCACACGTCTGAACTCTTTCCCTACACGACGCTCTTCCGATCTNNNGTCTGCGAAGGGACCGGAAG
AGCACACGTCTGAACTCTTTCCCTACACGACGCTCTTCCGATCTNNNAATGGAGTCTCTCTCTCTCTGT
AGCACACGTCTGAACTCTTTCCCTACACGACGCTCTTCCGATCTNNNAATGGAGTCTCTCTCTCTCTGT
AGCACACGTCTGAACTCTTTCCCTACACGACGCTCTTCCGATCTNNNGCACATGCCTGTAATCCTAGCT
AGCACACGTCTGAACTCTTTCCCTACACGACGCTCTTCCGATCTNNNGCACATGCCTGTAATCCTAGCT
GAGCACACGTCTGAACTCTTTCCCTACACGACGCTCTTCCGATCTNNNGAGACTGAGGAAGAAGAAT
GAGCACACGTCTGAACTCTTTCCCTACACGACGCTCTTCCGATCTNNNGAGACTGAGGAAGAAGAAT
GAGCACACGTCTGAACTCTTTCCCTACACGACGCTCTTCCGATCTNNNTCTACAACAGCCACCTTCCTA
GAGCACACGTCTGAACTCTTTCCCTACACGACGCTCTTCCGATCTNNNTCTACAACAGCCACCTTCCTA
GAGCACACGTCTGAACTCTTTCCCTACACGACGCTCTTCCGATCTNNNGGGACACAGACAGGAATGGA
GAGCACACGTCTGAACTCTTTCCCTACACGACGCTCTTCCGATCTNNNGGGACACAGACAGGAATGGA
AGCACACGTCTGAACTCTTTCCCTACACGACGCTCTTCCGATCTNNNGGAATCCGTTAGCAGAGCCTGC
AGCACACGTCTGAACTCTTTCCCTACACGACGCTCTTCCGATCTNNNGGAATCCGTTAGCAGAGCCTGC
GAGCACACGTCTGAACTCTTTCCCTACACGACGCTCTTCCGATCTNNNTATAATGGCTCCTGCTGCTCTT
GAGCACACGTCTGAACTCTTTCCCTACACGACGCTCTTCCGATCTNNNTATAATGGCTCCTGCTGCTCTT
AGCACACGTCTGAACTCTTTCCCTACACGACGCTCTTCCGATCTNNNGAGCACTGGAGAAGATGAGA
AGCACACGTCTGAACTCTTTCCCTACACGACGCTCTTCCGATCTNNNGAGCACTGGAGAAGATGAGA
AGCACACGTCTGAACTCTTTCCCTACACGACGCTCTTCCGATCTNNNGAGGCTGAAGCAGGAGAAAG
AGCACACGTCTGAACTCTTTCCCTACACGACGCTCTTCCGATCTNNNGAGGCTGAAGCAGGAGAAAG
GAGCACACGTCTGAACTCTTTCCCTACACGACGCTCTTCCGATCTNNNTGCCTCATTTTAACCCCTTTTCC
GAGCACACGTCTGAACTCTTTCCCTACACGACGCTCTTCCGATCTNNNTGCCTCATTTTAACCCCTTTTCC
AGAGCACACGTCTGAACTCTTTCCCTACACGACGCTCTTCCGATCTNNNGGTCCATTGTATGAGGGTG
AGAGCACACGTCTGAACTCTTTCCCTACACGACGCTCTTCCGATCTNNNGGTCCATTGTATGAGGGTG
AGCACACGTCTGAACTCTTTCCCTACACGACGCTCTTCCGATCTNNNACTTCGGTGTGTCCAGAATAGT
AGCACACGTCTGAACTCTTTCCCTACACGACGCTCTTCCGATCTNNNACTTCGGTGTGTCCAGAATAGT
GAGCACACGTCTGAACTCTTTCCCTACACGACGCTCTTCCGATCTNNNCGAGATCACACCATTGTACTC
GAGCACACGTCTGAACTCTTTCCCTACACGACGCTCTTCCGATCTNNNCGAGATCACACCATTGTACTC
AGCACACGTCTGAACTCTTTCCCTACACGACGCTCTTCCGATCTNNNTGCCAAATAATGTGGTTGATGG
AGCACACGTCTGAACTCTTTCCCTACACGACGCTCTTCCGATCTNNNTGCCAAATAATGTGGTTGATGG
AGAGCACACGTCTGAACTCTTTCCCTACACGACGCTCTTCCGATCTNNNAGAAAGTAAGTTGCTTTGCA
AGAGCACACGTCTGAACTCTTTCCCTACACGACGCTCTTCCGATCTNNNAGAAAGTAAGTTGCTTTGCA
GAGCACACGTCTGAACTCTTTCCCTACACGACGCTCTTCCGATCTNNNCAGAATCTCACTCTGTCACCCA
GAGCACACGTCTGAACTCTTTCCCTACACGACGCTCTTCCGATCTNNNCAGAATCTCACTCTGTCACCCA
GAGCACACGTCTGAACTCTTTCCCTACACGACGCTCTTCCGATCTNNNCTTTGTGAAAGGAAGAAGGC
GAGCACACGTCTGAACTCTTTCCCTACACGACGCTCTTCCGATCTNNNCTTTGTGAAAGGAAGAAGGC
AGAGCACACGTCTGAACTCTTTCCCTACACGACGCTCTTCCGATCTNNNGGCCTTGTTTCTCTCCCAGAT
AGAGCACACGTCTGAACTCTTTCCCTACACGACGCTCTTCCGATCTNNNGGCCTTGTTTCTCTCCCAGAT
GCACACGTCTGAACTCTTTCCCTACACGACGCTCTTCCGATCTNNNTTCACGCCACTATACTCCAATGTC
GCACACGTCTGAACTCTTTCCCTACACGACGCTCTTCCGATCTNNNTTCACGCCACTATACTCCAATGTC
GAGCACACGTCTGAACTCTTTCCCTACACGACGCTCTTCCGATCTNNNTTGTGTTCCAGAGTTCAAGAC
GAGCACACGTCTGAACTCTTTCCCTACACGACGCTCTTCCGATCTNNNTTGTGTTCCAGAGTTCAAGAC
GAGCACACGTCTGAACTCTTTCCCTACACGACGCTCTTCCGATCTNNNGAGACTGGGAAGAAAGAGA
GAGCACACGTCTGAACTCTTTCCCTACACGACGCTCTTCCGATCTNNNGAGACTGGGAAGAAAGAGA

AGCACACGTCTGAACTCTTTCCCTACACGACGCTCTTCCGATCTNNNGAGATGGTAGTGAGCCAAGA
AGCACACGTCTGAACTCTTTCCCTACACGACGCTCTTCCGATCTNNNGAGATGGTAGTGAGCCAAGA
AGCACACGTCTGAACTCTTTCCCTACACGACGCTCTTCCGATCTNNNTATAGAAAGCGGAGACACCC
AGCACACGTCTGAACTCTTTCCCTACACGACGCTCTTCCGATCTNNNTATAGAAAGCGGAGACACCC
GCACACGTCTGAACTCTTTCCCTACACGACGCTCTTCCGATCTNNNTCTTTCTGCTGACTTTGGGCTTA
GCACACGTCTGAACTCTTTCCCTACACGACGCTCTTCCGATCTNNNTCTTTCTGCTGACTTTGGGCTTA
GAGCACACGTCTGAACTCTTTCCCTACACGACGCTCTTCCGATCTNNNCTGCTCAGAGTACTGTCACAG
GAGCACACGTCTGAACTCTTTCCCTACACGACGCTCTTCCGATCTNNNCTGCTCAGAGTACTGTCACAG
AGCACACGTCTGAACTCTTTCCCTACACGACGCTCTTCCGATCTNNNTAAGGCAGACTTTCAGAGACC
AGCACACGTCTGAACTCTTTCCCTACACGACGCTCTTCCGATCTNNNTAAGGCAGACTTTCAGAGACC
AGCACACGTCTGAACTCTTTCCCTACACGACGCTCTTCCGATCTNNNTGAGGTAGGAGAATGGCTTG
AGCACACGTCTGAACTCTTTCCCTACACGACGCTCTTCCGATCTNNNTGAGGTAGGAGAATGGCTTG
GAGCACACGTCTGAACTCTTTCCCTACACGACGCTCTTCCGATCTNNNCTCTTTACCCAATGCCCTCAGA
GAGCACACGTCTGAACTCTTTCCCTACACGACGCTCTTCCGATCTNNNCTCTTTACCCAATGCCCTCAGA
AGCACACGTCTGAACTCTTTCCCTACACGACGCTCTTCCGATCTNNNAAAAATCCTGCTTTTTCCTCTGC
AGCACACGTCTGAACTCTTTCCCTACACGACGCTCTTCCGATCTNNNAAAAATCCTGCTTTTTCCTCTGC
AAGAGCACACGTCTGAACTCTTTCCCTACACGACGCTCTTCCGATCTNNNTCCCTCAGGTAGCATCCTA
AAGAGCACACGTCTGAACTCTTTCCCTACACGACGCTCTTCCGATCTNNNTCCCTCAGGTAGCATCCTA
GAGCACACGTCTGAACTCTTTCCCTACACGACGCTCTTCCGATCTNNNCTGAGACAGGAGAATGGCAT
GAGCACACGTCTGAACTCTTTCCCTACACGACGCTCTTCCGATCTNNNCTGAGACAGGAGAATGGCAT
GCACACGTCTGAACTCTTTCCCTACACGACGCTCTTCCGATCTNNNCGCTACTGGGAAGTGAGGAGC
GCACACGTCTGAACTCTTTCCCTACACGACGCTCTTCCGATCTNNNCGCTACTGGGAAGTGAGGAGC
AGCACACGTCTGAACTCTTTCCCTACACGACGCTCTTCCGATCTNNNTTATTTGATCATTTGTTTTTGGC
AGCACACGTCTGAACTCTTTCCCTACACGACGCTCTTCCGATCTNNNTTATTTGATCATTTGTTTTTGGC
GAGCACACGTCTGAACTCTTTCCCTACACGACGCTCTTCCGATCTNNNACCTACTTACCACTTCTGCATC
GAGCACACGTCTGAACTCTTTCCCTACACGACGCTCTTCCGATCTNNNACCTACTTACCACTTCTGCATC
AGCACACGTCTGAACTCTTTCCCTACACGACGCTCTTCCGATCTNNNGAAGCAGGAGAATCACTTGA
AGCACACGTCTGAACTCTTTCCCTACACGACGCTCTTCCGATCTNNNGAAGCAGGAGAATCACTTGA
GCACACGTCTGAACTCTTTCCCTACACGACGCTCTTCCGATCTNNNGTAAGCTTTGCAAACCATCCAGT
GCACACGTCTGAACTCTTTCCCTACACGACGCTCTTCCGATCTNNNGTAAGCTTTGCAAACCATCCAGT
GAGCACACGTCTGAACTCTTTCCCTACACGACGCTCTTCCGATCTNNNTCCTTTCTTCCCCTGTAGCAAG
GAGCACACGTCTGAACTCTTTCCCTACACGACGCTCTTCCGATCTNNNTCCTTTCTTCCCCTGTAGCAAG
GAGCACACGTCTGAACTCTTTCCCTACACGACGCTCTTCCGATCTNNNATTGATATCTCTCTCACACGC
GAGCACACGTCTGAACTCTTTCCCTACACGACGCTCTTCCGATCTNNNATTGATATCTCTCTCACACGC
GAGCACACGTCTGAACTCTTTCCCTACACGACGCTCTTCCGATCTNNNCGCACATACGTGTCAGAATTT
GAGCACACGTCTGAACTCTTTCCCTACACGACGCTCTTCCGATCTNNNCGCACATACGTGTCAGAATTT
AGCACACGTCTGAACTCTTTCCCTACACGACGCTCTTCCGATCTNNNGTCCCCTGTGAGATGAGCTATT
AGCACACGTCTGAACTCTTTCCCTACACGACGCTCTTCCGATCTNNNGTCCCCTGTGAGATGAGCTATT
GAGCACACGTCTGAACTCTTTCCCTACACGACGCTCTTCCGATCTNNNTGAAAGTAAGAGTGAGAGG
GAGCACACGTCTGAACTCTTTCCCTACACGACGCTCTTCCGATCTNNNTGAAAGTAAGAGTGAGAGG
GAGCACACGTCTGAACTCTTTCCCTACACGACGCTCTTCCGATCTNNNTCTGCTTTTCCACAGTGTCAGT
GAGCACACGTCTGAACTCTTTCCCTACACGACGCTCTTCCGATCTNNNTCTGCTTTTCCACAGTGTCAGT
GCACACGTCTGAACTCTTTCCCTACACGACGCTCTTCCGATCTNNNCTGAGAGCTGCTCTAAAGTGAC

GCACACGTCTGAACTCTTTCCCTACACGACGCTCTTCCGATCTNNNCTGAGAGCTGCTCTAAAGTGAC
AGCACACGTCTGAACTCTTTCCCTACACGACGCTCTTCCGATCTNNNGAGAATTGCTTGAACCCTGGA
AGCACACGTCTGAACTCTTTCCCTACACGACGCTCTTCCGATCTNNNGAGAATTGCTTGAACCCTGGA
AGCACACGTCTGAACTCTTTCCCTACACGACGCTCTTCCGATCTNNNCCCCTCTCATGCATATCCAGAA
AGCACACGTCTGAACTCTTTCCCTACACGACGCTCTTCCGATCTNNNCCCCTCTCATGCATATCCAGAA
AGCACACGTCTGAACTCTTTCCCTACACGACGCTCTTCCGATCTNNNATAGTGAAAAAGTGGCCAG
AGCACACGTCTGAACTCTTTCCCTACACGACGCTCTTCCGATCTNNNATAGTGAAAAAGTGGCCAG
GAGCACACGTCTGAACTCTTTCCCTACACGACGCTCTTCCGATCTNNNGCACTAGAACAGCTTGAACT
GAGCACACGTCTGAACTCTTTCCCTACACGACGCTCTTCCGATCTNNNGCACTAGAACAGCTTGAACT
GAGCACACGTCTGAACTCTTTCCCTACACGACGCTCTTCCGATCTNNNTAGGTAGTAGGGAGAGAG
GAGCACACGTCTGAACTCTTTCCCTACACGACGCTCTTCCGATCTNNNTAGGTAGTAGGGAGAGAG
AGCACACGTCTGAACTCTTTCCCTACACGACGCTCTTCCGATCTNNNTAGTTCATTCTCCTGTTGGCAC
AGCACACGTCTGAACTCTTTCCCTACACGACGCTCTTCCGATCTNNNTAGTTCATTCTCCTGTTGGCAC
GCACACGTCTGAACTCTTTCCCTACACGACGCTCTTCCGATCTNNNAAACATCCATCTGCTTTTTGGG
GCACACGTCTGAACTCTTTCCCTACACGACGCTCTTCCGATCTNNNAAACATCCATCTGCTTTTTGGG
AGCACACGTCTGAACTCTTTCCCTACACGACGCTCTTCCGATCTNNNAATACCAAAACCAGGAAGGG
AGCACACGTCTGAACTCTTTCCCTACACGACGCTCTTCCGATCTNNNAATACCAAAACCAGGAAGGG
AAGAGCACACGTCTGAACTCTTTCCCTACACGACGCTCTTCCGATCTNNNCCAAAGGAGACTCAGGG
AAGAGCACACGTCTGAACTCTTTCCCTACACGACGCTCTTCCGATCTNNNCCAAAGGAGACTCAGGG
AGCACACGTCTGAACTCTTTCCCTACACGACGCTCTTCCGATCTNNNAGTGGAGCTTGAATACAAACC
AGCACACGTCTGAACTCTTTCCCTACACGACGCTCTTCCGATCTNNNAGTGGAGCTTGAATACAAACC
GAGCACACGTCTGAACTCTTTCCCTACACGACGCTCTTCCGATCTNNNGACAGCGATAAAGACAAGA
GAGCACACGTCTGAACTCTTTCCCTACACGACGCTCTTCCGATCTNNNGACAGCGATAAAGACAAGA
GAGCACACGTCTGAACTCTTTCCCTACACGACGCTCTTCCGATCTNNNGGAGGCAAGGTCCAACATAA
GAGCACACGTCTGAACTCTTTCCCTACACGACGCTCTTCCGATCTNNNGGAGGCAAGGTCCAACATAA
GAGCACACGTCTGAACTCTTTCCCTACACGACGCTCTTCCGATCTNNNAGGACATGCCCTATAGGTTA
GAGCACACGTCTGAACTCTTTCCCTACACGACGCTCTTCCGATCTNNNAGGACATGCCCTATAGGTTA
GAGCACACGTCTGAACTCTTTCCCTACACGACGCTCTTCCGATCTNNNTGAGGTAGGAGAATCGCTTG
GAGCACACGTCTGAACTCTTTCCCTACACGACGCTCTTCCGATCTNNNTGAGGTAGGAGAATCGCTTG
GCACACGTCTGAACTCTTTCCCTACACGACGCTCTTCCGATCTNNNTCAGTTATACCCTGCCCTTCAACC
GCACACGTCTGAACTCTTTCCCTACACGACGCTCTTCCGATCTNNNTCAGTTATACCCTGCCCTTCAACC
GCACACGTCTGAACTCTTTCCCTACACGACGCTCTTCCGATCTNNNTAATTCACCCCTTCCCAACATGG
GCACACGTCTGAACTCTTTCCCTACACGACGCTCTTCCGATCTNNNTAATTCACCCCTTCCCAACATGG
AGCACACGTCTGAACTCTTTCCCTACACGACGCTCTTCCGATCTNNNACCATGAGCAAGAATCAGTGT
AGCACACGTCTGAACTCTTTCCCTACACGACGCTCTTCCGATCTNNNACCATGAGCAAGAATCAGTGT
AGAGCACACGTCTGAACTCTTTCCCTACACGACGCTCTTCCGATCTNNNGAACTCCAGTGTAGTGCTC
AGAGCACACGTCTGAACTCTTTCCCTACACGACGCTCTTCCGATCTNNNGAACTCCAGTGTAGTGCTC
GCACACGTCTGAACTCTTTCCCTACACGACGCTCTTCCGATCTNNNGGATAACAGGTGTGAACCACT
GCACACGTCTGAACTCTTTCCCTACACGACGCTCTTCCGATCTNNNGGATAACAGGTGTGAACCACT
GCACACGTCTGAACTCTTTCCCTACACGACGCTCTTCCGATCTNNNGTATGCAGTACTGGCCTGAGAA
GCACACGTCTGAACTCTTTCCCTACACGACGCTCTTCCGATCTNNNGTATGCAGTACTGGCCTGAGAA
GCACACGTCTGAACTCTTTCCCTACACGACGCTCTTCCGATCTNNNTTGCCCATGGAGAGGTATTTCT
GCACACGTCTGAACTCTTTCCCTACACGACGCTCTTCCGATCTNNNTTGCCCATGGAGAGGTATTTCT

GAGCACACGTCTGAACTCTTTCCCTACACGACGCTCTTCCGATCTNNNTTCAGAAGGAGGCCTAATTTG
GAGCACACGTCTGAACTCTTTCCCTACACGACGCTCTTCCGATCTNNNTTCAGAAGGAGGCCTAATTTG
AGCACACGTCTGAACTCTTTCCCTACACGACGCTCTTCCGATCTNNNCACACAGACAATGCTCACAGTTT
AGCACACGTCTGAACTCTTTCCCTACACGACGCTCTTCCGATCTNNNCACACAGACAATGCTCACAGTTT
GAGCACACGTCTGAACTCTTTCCCTACACGACGCTCTTCCGATCTNNNCTTGACGTAGGAAACCCAGGT
GAGCACACGTCTGAACTCTTTCCCTACACGACGCTCTTCCGATCTNNNCTTGACGTAGGAAACCCAGGT
GAGCACACGTCTGAACTCTTTCCCTACACGACGCTCTTCCGATCTNNNTTTCTATCTATGGCTGTGGCTG
GAGCACACGTCTGAACTCTTTCCCTACACGACGCTCTTCCGATCTNNNTTTCTATCTATGGCTGTGGCTG
AGAGCACACGTCTGAACTCTTTCCCTACACGACGCTCTTCCGATCTNNNGGAAACATCAAGCTGATCTCT
AGAGCACACGTCTGAACTCTTTCCCTACACGACGCTCTTCCGATCTNNNGGAAACATCAAGCTGATCTCT
AGAGCACACGTCTGAACTCTTTCCCTACACGACGCTCTTCCGATCTNNNTGCCTCATCATTTACCTCAGCA
AGAGCACACGTCTGAACTCTTTCCCTACACGACGCTCTTCCGATCTNNNTGCCTCATCATTTACCTCAGCA
GAGCACACGTCTGAACTCTTTCCCTACACGACGCTCTTCCGATCTNNNACTCTTTGGTTGCTGAACTTTG
GAGCACACGTCTGAACTCTTTCCCTACACGACGCTCTTCCGATCTNNNACTCTTTGGTTGCTGAACTTTG
GAGCACACGTCTGAACTCTTTCCCTACACGACGCTCTTCCGATCTNNNGAATGTCCATTTTTCTCAGCCC
GAGCACACGTCTGAACTCTTTCCCTACACGACGCTCTTCCGATCTNNNGAATGTCCATTTTTCTCAGCCC
AGCACACGTCTGAACTCTTTCCCTACACGACGCTCTTCCGATCTNNNAAGCCTGTAGTCCTAGCTACTCA
AGCACACGTCTGAACTCTTTCCCTACACGACGCTCTTCCGATCTNNNAAGCCTGTAGTCCTAGCTACTCA
GAGCACACGTCTGAACTCTTTCCCTACACGACGCTCTTCCGATCTNNNTCACCTGCATCCAATGAGTTAC
GAGCACACGTCTGAACTCTTTCCCTACACGACGCTCTTCCGATCTNNNTCACCTGCATCCAATGAGTTAC
AAGAGCACACGTCTGAACTCTTTCCCTACACGACGCTCTTCCGATCTNNNACAAAAATTAGCCAGGCTT
AAGAGCACACGTCTGAACTCTTTCCCTACACGACGCTCTTCCGATCTNNNACAAAAATTAGCCAGGCTT
GAGCACACGTCTGAACTCTTTCCCTACACGACGCTCTTCCGATCTNNNTGTAATCCCAGCTACTTGGAAG
GAGCACACGTCTGAACTCTTTCCCTACACGACGCTCTTCCGATCTNNNTGTAATCCCAGCTACTTGGAAG
GAGCACACGTCTGAACTCTTTCCCTACACGACGCTCTTCCGATCTNNNCCCTTGTTGCAGATGAAGTAT
GAGCACACGTCTGAACTCTTTCCCTACACGACGCTCTTCCGATCTNNNCCCTTGTTGCAGATGAAGTAT
AGCACACGTCTGAACTCTTTCCCTACACGACGCTCTTCCGATCTNNNCCAGTCTGGGTTTTAGGAGCTT
AGCACACGTCTGAACTCTTTCCCTACACGACGCTCTTCCGATCTNNNCCAGTCTGGGTTTTAGGAGCTT
AGAGCACACGTCTGAACTCTTTCCCTACACGACGCTCTTCCGATCTNNNAGTGAAACCCTGAAAAGAAA
AGAGCACACGTCTGAACTCTTTCCCTACACGACGCTCTTCCGATCTNNNAGTGAAACCCTGAAAAGAAA
GAGCACACGTCTGAACTCTTTCCCTACACGACGCTCTTCCGATCTNNNTGGGTTCCTAGGAAACTCACA
GAGCACACGTCTGAACTCTTTCCCTACACGACGCTCTTCCGATCTNNNTGGGTTCCTAGGAAACTCACA
AGCACACGTCTGAACTCTTTCCCTACACGACGCTCTTCCGATCTNNNTCAATGCCTCTATTCCAGCCTTTG
AGCACACGTCTGAACTCTTTCCCTACACGACGCTCTTCCGATCTNNNTCAATGCCTCTATTCCAGCCTTTG
GAGCACACGTCTGAACTCTTTCCCTACACGACGCTCTTCCGATCTNNNGGATCAGTTGTGAGATCAAGG
GAGCACACGTCTGAACTCTTTCCCTACACGACGCTCTTCCGATCTNNNGGATCAGTTGTGAGATCAAGG
AGCACACGTCTGAACTCTTTCCCTACACGACGCTCTTCCGATCTNNNCAAAAAAAATTAGCCGGGCATG
AGCACACGTCTGAACTCTTTCCCTACACGACGCTCTTCCGATCTNNNCAAAAAAAATTAGCCGGGCATG
GAGCACACGTCTGAACTCTTTCCCTACACGACGCTCTTCCGATCTNNNGCAAGCAATGAAAACGGCAAA
GAGCACACGTCTGAACTCTTTCCCTACACGACGCTCTTCCGATCTNNNGCAAGCAATGAAAACGGCAAA
GAGCACACGTCTGAACTCTTTCCCTACACGACGCTCTTCCGATCTNNNTGCAGGGAGTCATATTTGGTT
GAGCACACGTCTGAACTCTTTCCCTACACGACGCTCTTCCGATCTNNNTGCAGGGAGTCATATTTGGTT
GAGCACACGTCTGAACTCTTTCCCTACACGACGCTCTTCCGATCTNNNCTATGATTCAAGGGGAGCTCT

GAGCACACGTCTGAACTCTTTCCCTACACGACGCTCTTCCGATCTNNNCTATGATTCAAGGGGAGCTCT
GAGCACACGTCTGAACTCTTTCCCTACACGACGCTCTTCCGATCTNNNACAATCATAAGCCACTGCACTT
GAGCACACGTCTGAACTCTTTCCCTACACGACGCTCTTCCGATCTNNNACAATCATAAGCCACTGCACTT
AAGAGCACACGTCTGAACTCTTTCCCTACACGACGCTCTTCCGATCTNNNAGAGATTGCAGTGAGCCA
AAGAGCACACGTCTGAACTCTTTCCCTACACGACGCTCTTCCGATCTNNNAGAGATTGCAGTGAGCCA
CACACGTCTGAACTCTTTCCCTACACGACGCTCTTCCGATCTNNNAGCAAGAGATTCATTACAAAAGAG
CACACGTCTGAACTCTTTCCCTACACGACGCTCTTCCGATCTNNNAGCAAGAGATTCATTACAAAAGAG
AGAGCACACGTCTGAACTCTTTCCCTACACGACGCTCTTCCGATCTNNNAACAATTTCCCTAACCACCAC
AGAGCACACGTCTGAACTCTTTCCCTACACGACGCTCTTCCGATCTNNNAACAATTTCCCTAACCACCAC
AGCACACGTCTGAACTCTTTCCCTACACGACGCTCTTCCGATCTNNNGGCAATAAGAGTGAAGTCCGT
AGCACACGTCTGAACTCTTTCCCTACACGACGCTCTTCCGATCTNNNGGCAATAAGAGTGAAGTCCGT
AGCACACGTCTGAACTCTTTCCCTACACGACGCTCTTCCGATCTNNNTCCTTCTTTCCTTCCTTCCTTCCCA
AGCACACGTCTGAACTCTTTCCCTACACGACGCTCTTCCGATCTNNNTCCTTCTTTCCTTCCTTCCTTCCCA
AGCACACGTCTGAACTCTTTCCCTACACGACGCTCTTCCGATCTNNNTTTGATTGCTCTGGCTAGGACTT
AGCACACGTCTGAACTCTTTCCCTACACGACGCTCTTCCGATCTNNNTTTGATTGCTCTGGCTAGGACTT
GAGCACACGTCTGAACTCTTTCCCTACACGACGCTCTTCCGATCTNNNTTGTAATCTCAGCTACGTGGG
GAGCACACGTCTGAACTCTTTCCCTACACGACGCTCTTCCGATCTNNNTTGTAATCTCAGCTACGTGGG
GAGCACACGTCTGAACTCTTTCCCTACACGACGCTCTTCCGATCTNNNTCAGACAGACGGTGCCTACTA
GAGCACACGTCTGAACTCTTTCCCTACACGACGCTCTTCCGATCTNNNTCAGACAGACGGTGCCTACTA
GAGCACACGTCTGAACTCTTTCCCTACACGACGCTCTTCCGATCTNNNCTGGAGCAAGATTATCTAATA
GAGCACACGTCTGAACTCTTTCCCTACACGACGCTCTTCCGATCTNNNCTGGAGCAAGATTATCTAATA
AGCACACGTCTGAACTCTTTCCCTACACGACGCTCTTCCGATCTNNNACTGTGAGATGTGTGTGCATA
AGCACACGTCTGAACTCTTTCCCTACACGACGCTCTTCCGATCTNNNACTGTGAGATGTGTGTGCATA
GCACACGTCTGAACTCTTTCCCTACACGACGCTCTTCCGATCTNNNTCTAGGACCTGTTGCTTCTAGGA
GCACACGTCTGAACTCTTTCCCTACACGACGCTCTTCCGATCTNNNTCTAGGACCTGTTGCTTCTAGGA
GCACACGTCTGAACTCTTTCCCTACACGACGCTCTTCCGATCTNNNTCCTACCCCAACAAAACAGAACA
GCACACGTCTGAACTCTTTCCCTACACGACGCTCTTCCGATCTNNNTCCTACCCCAACAAAACAGAACA
GAGCACACGTCTGAACTCTTTCCCTACACGACGCTCTTCCGATCTNNNCCTCCCAGGTTCAAGTAGTTC
GAGCACACGTCTGAACTCTTTCCCTACACGACGCTCTTCCGATCTNNNCCTCCCAGGTTCAAGTAGTTC
GAGCACACGTCTGAACTCTTTCCCTACACGACGCTCTTCCGATCTNNNTTAGCATCTCCCCTCTCTGATC
GAGCACACGTCTGAACTCTTTCCCTACACGACGCTCTTCCGATCTNNNTTAGCATCTCCCCTCTCTGATC
AGAGCACACGTCTGAACTCTTTCCCTACACGACGCTCTTCCGATCTNNNTGCCTTCTCTCTGTCTCTGTCT
AGAGCACACGTCTGAACTCTTTCCCTACACGACGCTCTTCCGATCTNNNTGCCTTCTCTCTGTCTCTGTCT
AGCACACGTCTGAACTCTTTCCCTACACGACGCTCTTCCGATCTNNNGATGGCAAGGTCATTTTCCAACA
AGCACACGTCTGAACTCTTTCCCTACACGACGCTCTTCCGATCTNNNGATGGCAAGGTCATTTTCCAACA
AGCACACGTCTGAACTCTTTCCCTACACGACGCTCTTCCGATCTNNNTTTAGTACAGACGGGGTTTCAC
AGCACACGTCTGAACTCTTTCCCTACACGACGCTCTTCCGATCTNNNTTTAGTACAGACGGGGTTTCAC
AGCACACGTCTGAACTCTTTCCCTACACGACGCTCTTCCGATCTNNNAACAAAATTAGCTGGGTGTGG
AGCACACGTCTGAACTCTTTCCCTACACGACGCTCTTCCGATCTNNNAACAAAATTAGCTGGGTGTGG
GAGCACACGTCTGAACTCTTTCCCTACACGACGCTCTTCCGATCTNNNAATAAAGGTGTTGGCTGCTTC
GAGCACACGTCTGAACTCTTTCCCTACACGACGCTCTTCCGATCTNNNAATAAAGGTGTTGGCTGCTTC
AGAGCACACGTCTGAACTCTTTCCCTACACGACGCTCTTCCGATCTNNNAACTATTTGGGAGGTTGAG
AGAGCACACGTCTGAACTCTTTCCCTACACGACGCTCTTCCGATCTNNNAACTATTTGGGAGGTTGAG

AGCACACGTCTGAACTCTTTCCCTACACGACGCTCTTCCGATCTNNNAAACACAAAAATTAGCTGGGC
AGCACACGTCTGAACTCTTTCCCTACACGACGCTCTTCCGATCTNNNAAACACAAAAATTAGCTGGGC
AGCACACGTCTGAACTCTTTCCCTACACGACGCTCTTCCGATCTNNNCTGCAGTGGATATTTCACTCCC
AGCACACGTCTGAACTCTTTCCCTACACGACGCTCTTCCGATCTNNNCTGCAGTGGATATTTCACTCCC
AGCACACGTCTGAACTCTTTCCCTACACGACGCTCTTCCGATCTNNNGAGTATGGCCTGAACCCAGAA
AGCACACGTCTGAACTCTTTCCCTACACGACGCTCTTCCGATCTNNNGAGTATGGCCTGAACCCAGAA
GAGCACACGTCTGAACTCTTTCCCTACACGACGCTCTTCCGATCTNNNGGAGACAGAGTGAGACTCCA
GAGCACACGTCTGAACTCTTTCCCTACACGACGCTCTTCCGATCTNNNGGAGACAGAGTGAGACTCCA
AGCACACGTCTGAACTCTTTCCCTACACGACGCTCTTCCGATCTNNNGAGGCTGAGGCAGGAGAATG
AGCACACGTCTGAACTCTTTCCCTACACGACGCTCTTCCGATCTNNNGAGGCTGAGGCAGGAGAATG
AGCACACGTCTGAACTCTTTCCCTACACGACGCTCTTCCGATCTNNNTGAGTCCAGAAAGGTATGCCA
AGCACACGTCTGAACTCTTTCCCTACACGACGCTCTTCCGATCTNNNTGAGTCCAGAAAGGTATGCCA
AGCACACGTCTGAACTCTTTCCCTACACGACGCTCTTCCGATCTNNNGTTCAACCATTGTGGAAGTCA
AGCACACGTCTGAACTCTTTCCCTACACGACGCTCTTCCGATCTNNNGTTCAACCATTGTGGAAGTCA
GAGCACACGTCTGAACTCTTTCCCTACACGACGCTCTTCCGATCTNNNTCCCTTGTTTCTGCCTCTTTCTT
GAGCACACGTCTGAACTCTTTCCCTACACGACGCTCTTCCGATCTNNNTCCCTTGTTTCTGCCTCTTTCTT
GAGCACACGTCTGAACTCTTTCCCTACACGACGCTCTTCCGATCTNNNCTGAGTCAGGAGAATCGCTT
GAGCACACGTCTGAACTCTTTCCCTACACGACGCTCTTCCGATCTNNNCTGAGTCAGGAGAATCGCTT
GAGCACACGTCTGAACTCTTTCCCTACACGACGCTCTTCCGATCTNNNGGTCTCAAAAAATCCTCCCTC
GAGCACACGTCTGAACTCTTTCCCTACACGACGCTCTTCCGATCTNNNGGTCTCAAAAAATCCTCCCTC
AGCACACGTCTGAACTCTTTCCCTACACGACGCTCTTCCGATCTNNNTGGGATTTGAGGATTTCCAGA
AGCACACGTCTGAACTCTTTCCCTACACGACGCTCTTCCGATCTNNNTGGGATTTGAGGATTTCCAGA
AGCACACGTCTGAACTCTTTCCCTACACGACGCTCTTCCGATCTNNNCTGTAGTCCCAGCTTCTCAGG
AGCACACGTCTGAACTCTTTCCCTACACGACGCTCTTCCGATCTNNNCTGTAGTCCCAGCTTCTCAGG
GAGCACACGTCTGAACTCTTTCCCTACACGACGCTCTTCCGATCTNNNGGCTCCCTGAATTCAGAGAT
GAGCACACGTCTGAACTCTTTCCCTACACGACGCTCTTCCGATCTNNNGGCTCCCTGAATTCAGAGAT
AGCACACGTCTGAACTCTTTCCCTACACGACGCTCTTCCGATCTNNNCTATCATAGCTCACTGCAGGCT
AGCACACGTCTGAACTCTTTCCCTACACGACGCTCTTCCGATCTNNNCTATCATAGCTCACTGCAGGCT
AGCACACGTCTGAACTCTTTCCCTACACGACGCTCTTCCGATCTNNNGAAAAGAGGGGAGCGGAGTA
AGCACACGTCTGAACTCTTTCCCTACACGACGCTCTTCCGATCTNNNGAAAAGAGGGGAGCGGAGTA
AGCACACGTCTGAACTCTTTCCCTACACGACGCTCTTCCGATCTNNNAGAGTCTCAAATGCCAGGATG
AGCACACGTCTGAACTCTTTCCCTACACGACGCTCTTCCGATCTNNNAGAGTCTCAAATGCCAGGATG
AGCACACGTCTGAACTCTTTCCCTACACGACGCTCTTCCGATCTNNNGAGTCTCGCTCTGTCACTCAG
AGCACACGTCTGAACTCTTTCCCTACACGACGCTCTTCCGATCTNNNGAGTCTCGCTCTGTCACTCAG
GCACACGTCTGAACTCTTTCCCTACACGACGCTCTTCCGATCTNNNAAGCATCTTTGATAACCCATTAC
GCACACGTCTGAACTCTTTCCCTACACGACGCTCTTCCGATCTNNNAAGCATCTTTGATAACCCATTAC
AGCACACGTCTGAACTCTTTCCCTACACGACGCTCTTCCGATCTNNNGCCTGTAATCCCATCTACTCTGG
AGCACACGTCTGAACTCTTTCCCTACACGACGCTCTTCCGATCTNNNGCCTGTAATCCCATCTACTCTGG
AGCACACGTCTGAACTCTTTCCCTACACGACGCTCTTCCGATCTNNNAAGTGTGTGTGTGTTTGTGTC
AGCACACGTCTGAACTCTTTCCCTACACGACGCTCTTCCGATCTNNNAAGTGTGTGTGTGTTTGTGTC
AGCACACGTCTGAACTCTTTCCCTACACGACGCTCTTCCGATCTNNNAGTGTGGTAGGATGCTGTGAT
AGCACACGTCTGAACTCTTTCCCTACACGACGCTCTTCCGATCTNNNAGTGTGGTAGGATGCTGTGAT
GAGCACACGTCTGAACTCTTTCCCTACACGACGCTCTTCCGATCTNNNTTGGCAGTACTCACTGTAGG

GAGCACACGTCTGAACTCTTTCCCTACACGACGCTCTTCCGATCTNNNTTGGCAGTACTCACTGTAGG
GCACACGTCTGAACTCTTTCCCTACACGACGCTCTTCCGATCTNNNAGGAGGACAAATAAGAAGTG
GCACACGTCTGAACTCTTTCCCTACACGACGCTCTTCCGATCTNNNAGGAGGACAAATAAGAAGTG
CACACGTCTGAACTCTTTCCCTACACGACGCTCTTCCGATCTNNNCGCCTTCAAGGATAAGTACAAG
CACACGTCTGAACTCTTTCCCTACACGACGCTCTTCCGATCTNNNCGCCTTCAAGGATAAGTACAAG
AGAGCACACGTCTGAACTCTTTCCCTACACGACGCTCTTCCGATCTNNNTGAGGCAGGAGAATCTCTT
AGAGCACACGTCTGAACTCTTTCCCTACACGACGCTCTTCCGATCTNNNTGAGGCAGGAGAATCTCTT
AGCACACGTCTGAACTCTTTCCCTACACGACGCTCTTCCGATCTNNNTTCAAATTAGAGGGGGTTGAG
AGCACACGTCTGAACTCTTTCCCTACACGACGCTCTTCCGATCTNNNTTCAAATTAGAGGGGGTTGAG
AGCACACGTCTGAACTCTTTCCCTACACGACGCTCTTCCGATCTNNNAGTTTCTCTCCCTGTATTGGCT
AGCACACGTCTGAACTCTTTCCCTACACGACGCTCTTCCGATCTNNNAGTTTCTCTCCCTGTATTGGCT
AGCACACGTCTGAACTCTTTCCCTACACGACGCTCTTCCGATCTNNNTATGACACTTTGCTTGGCCTAC
AGCACACGTCTGAACTCTTTCCCTACACGACGCTCTTCCGATCTNNNTATGACACTTTGCTTGGCCTAC
AGCACACGTCTGAACTCTTTCCCTACACGACGCTCTTCCGATCTNNNGCTGACACCGGTAAGATATCC
AGCACACGTCTGAACTCTTTCCCTACACGACGCTCTTCCGATCTNNNGCTGACACCGGTAAGATATCC
AGCACACGTCTGAACTCTTTCCCTACACGACGCTCTTCCGATCTNNNGAAGCCTGCAAAAAAAAAGG
AGCACACGTCTGAACTCTTTCCCTACACGACGCTCTTCCGATCTNNNGAAGCCTGCAAAAAAAAAGG
CACACGTCTGAACTCTTTCCCTACACGACGCTCTTCCGATCTNNNAACTCACTCCAAAGACCAGGTGC
CACACGTCTGAACTCTTTCCCTACACGACGCTCTTCCGATCTNNNAACTCACTCCAAAGACCAGGTGC
AGCACACGTCTGAACTCTTTCCCTACACGACGCTCTTCCGATCTNNNTCATACACCTTGCCTCTGTTGA
AGCACACGTCTGAACTCTTTCCCTACACGACGCTCTTCCGATCTNNNTCATACACCTTGCCTCTGTTGA
GCACACGTCTGAACTCTTTCCCTACACGACGCTCTTCCGATCTNNNGATCACGCCATTGCACTCCAGC
GCACACGTCTGAACTCTTTCCCTACACGACGCTCTTCCGATCTNNNGATCACGCCATTGCACTCCAGC
GCACACGTCTGAACTCTTTCCCTACACGACGCTCTTCCGATCTNNNTTTTTTCCTGAAGCTGGGTCTCA
GCACACGTCTGAACTCTTTCCCTACACGACGCTCTTCCGATCTNNNTTTTTTCCTGAAGCTGGGTCTCA
AGCACACGTCTGAACTCTTTCCCTACACGACGCTCTTCCGATCTNNNGAAGTTTGCAGTGAGCCGAAA
AGCACACGTCTGAACTCTTTCCCTACACGACGCTCTTCCGATCTNNNGAAGTTTGCAGTGAGCCGAAA
AGCACACGTCTGAACTCTTTCCCTACACGACGCTCTTCCGATCTNNNGACGAACAAGGATGGGAGG
AGCACACGTCTGAACTCTTTCCCTACACGACGCTCTTCCGATCTNNNGACGAACAAGGATGGGAGG
AGCACACGTCTGAACTCTTTCCCTACACGACGCTCTTCCGATCTNNNCCTATTACCCATGATGGAACCC
AGCACACGTCTGAACTCTTTCCCTACACGACGCTCTTCCGATCTNNNCCTATTACCCATGATGGAACCC
GAGCACACGTCTGAACTCTTTCCCTACACGACGCTCTTCCGATCTNNNACCCATCAACCCATCATCTACA
GAGCACACGTCTGAACTCTTTCCCTACACGACGCTCTTCCGATCTNNNACCCATCAACCCATCATCTACA
AGCACACGTCTGAACTCTTTCCCTACACGACGCTCTTCCGATCTNNNGAATGCTCTGTGTAAGCCAA
AGCACACGTCTGAACTCTTTCCCTACACGACGCTCTTCCGATCTNNNGAATGCTCTGTGTAAGCCAA
GAGCACACGTCTGAACTCTTTCCCTACACGACGCTCTTCCGATCTNNNATACAAAAATTAGCAGGGCG
GAGCACACGTCTGAACTCTTTCCCTACACGACGCTCTTCCGATCTNNNATACAAAAATTAGCAGGGCG
AGCACACGTCTGAACTCTTTCCCTACACGACGCTCTTCCGATCTNNNCCCAATCCCTACTTCTGTTTCCT
AGCACACGTCTGAACTCTTTCCCTACACGACGCTCTTCCGATCTNNNCCCAATCCCTACTTCTGTTTCCT
AGCACACGTCTGAACTCTTTCCCTACACGACGCTCTTCCGATCTNNNTCTACCTGGAATTCAAAAGCCT
AGCACACGTCTGAACTCTTTCCCTACACGACGCTCTTCCGATCTNNNTCTACCTGGAATTCAAAAGCCT
AGCACACGTCTGAACTCTTTCCCTACACGACGCTCTTCCGATCTNNNCCGAGGAGGTGGATGTTATA
AGCACACGTCTGAACTCTTTCCCTACACGACGCTCTTCCGATCTNNNCCGAGGAGGTGGATGTTATA

AGCACACGTCTGAACTCTTTCCCTACACGACGCTCTTCCGATCTNNNGAATGCCTGTAATCCCAGCTAC
AGCACACGTCTGAACTCTTTCCCTACACGACGCTCTTCCGATCTNNNGAATGCCTGTAATCCCAGCTAC
GAGCACACGTCTGAACTCTTTCCCTACACGACGCTCTTCCGATCTNNNATATTTGTGTATCTCCAATAG
GAGCACACGTCTGAACTCTTTCCCTACACGACGCTCTTCCGATCTNNNATATTTGTGTATCTCCAATAG
GAGCACACGTCTGAACTCTTTCCCTACACGACGCTCTTCCGATCTNNNACTGTGATGCCCCATCCATAT
GAGCACACGTCTGAACTCTTTCCCTACACGACGCTCTTCCGATCTNNNACTGTGATGCCCCATCCATAT
GAGCACACGTCTGAACTCTTTCCCTACACGACGCTCTTCCGATCTNNNCCAAATGCCATCAATCAACAG
GAGCACACGTCTGAACTCTTTCCCTACACGACGCTCTTCCGATCTNNNCCAAATGCCATCAATCAACAG
AGCACACGTCTGAACTCTTTCCCTACACGACGCTCTTCCGATCTNNNGTGTCATTTGTTCTCCTAGGGG
AGCACACGTCTGAACTCTTTCCCTACACGACGCTCTTCCGATCTNNNGTGTCATTTGTTCTCCTAGGGG
GAGCACACGTCTGAACTCTTTCCCTACACGACGCTCTTCCGATCTNNNGCAGTGACCTATGATTGCATC
GAGCACACGTCTGAACTCTTTCCCTACACGACGCTCTTCCGATCTNNNGCAGTGACCTATGATTGCATC
GCACACGTCTGAACTCTTTCCCTACACGACGCTCTTCCGATCTNNNGAATGCAGAGTTTGCTTGACAG
GCACACGTCTGAACTCTTTCCCTACACGACGCTCTTCCGATCTNNNGAATGCAGAGTTTGCTTGACAG
AGCACACGTCTGAACTCTTTCCCTACACGACGCTCTTCCGATCTNNNGTTCACACCATTGTCTTGCCTCC
AGCACACGTCTGAACTCTTTCCCTACACGACGCTCTTCCGATCTNNNGTTCACACCATTGTCTTGCCTCC
GAGCACACGTCTGAACTCTTTCCCTACACGACGCTCTTCCGATCTNNNTGGCCATAGAAACAGTAATGG
GAGCACACGTCTGAACTCTTTCCCTACACGACGCTCTTCCGATCTNNNTGGCCATAGAAACAGTAATGG
GAGCACACGTCTGAACTCTTTCCCTACACGACGCTCTTCCGATCTNNNTTTGTCAAAACGCGGTAAAT
GAGCACACGTCTGAACTCTTTCCCTACACGACGCTCTTCCGATCTNNNTTTGTCAAAACGCGGTAAAT
AGCACACGTCTGAACTCTTTCCCTACACGACGCTCTTCCGATCTNNNCTGTAATTCCAGCTACTCGGGA
AGCACACGTCTGAACTCTTTCCCTACACGACGCTCTTCCGATCTNNNCTGTAATTCCAGCTACTCGGGA
GAGCACACGTCTGAACTCTTTCCCTACACGACGCTCTTCCGATCTNNNAAGAATGAGATCAGGGGGA
GAGCACACGTCTGAACTCTTTCCCTACACGACGCTCTTCCGATCTNNNAAGAATGAGATCAGGGGGA
ACACGTCTGAACTCTTTCCCTACACGACGCTCTTCCGATCTNNNCTGTTCCTCGCTCTTCTCCATCCATCG
ACACGTCTGAACTCTTTCCCTACACGACGCTCTTCCGATCTNNNCTGTTCCTCGCTCTTCTCCATCCATCG
GAGCACACGTCTGAACTCTTTCCCTACACGACGCTCTTCCGATCTNNNATGCTTTTTAACGTTTTCTTTTT
GAGCACACGTCTGAACTCTTTCCCTACACGACGCTCTTCCGATCTNNNATGCTTTTTAACGTTTTCTTTTT
AGCACACGTCTGAACTCTTTCCCTACACGACGCTCTTCCGATCTNNNGAGGTTGCAGTGAGTCGAGAT
AGCACACGTCTGAACTCTTTCCCTACACGACGCTCTTCCGATCTNNNGAGGTTGCAGTGAGTCGAGAT
AGCACACGTCTGAACTCTTTCCCTACACGACGCTCTTCCGATCTNNNGAGTCTCCTGGATGCCTATTCT
AGCACACGTCTGAACTCTTTCCCTACACGACGCTCTTCCGATCTNNNGAGTCTCCTGGATGCCTATTCT
GAGCACACGTCTGAACTCTTTCCCTACACGACGCTCTTCCGATCTNNNCTGAGGCAGGAGAATCACTT
GAGCACACGTCTGAACTCTTTCCCTACACGACGCTCTTCCGATCTNNNCTGAGGCAGGAGAATCACTT
GAGCACACGTCTGAACTCTTTCCCTACACGACGCTCTTCCGATCTNNNAAAAAATTAGCCAGGCACGG
GAGCACACGTCTGAACTCTTTCCCTACACGACGCTCTTCCGATCTNNNAAAAAATTAGCCAGGCACGG
GAGCACACGTCTGAACTCTTTCCCTACACGACGCTCTTCCGATCTNNNTGGAGTACAGTGGTGTGATCT
GAGCACACGTCTGAACTCTTTCCCTACACGACGCTCTTCCGATCTNNNTGGAGTACAGTGGTGTGATCT
AGCACACGTCTGAACTCTTTCCCTACACGACGCTCTTCCGATCTNNNCACTGGCCTTCTGTTTCTCTCTC
AGCACACGTCTGAACTCTTTCCCTACACGACGCTCTTCCGATCTNNNCACTGGCCTTCTGTTTCTCTCTC
AGCACACGTCTGAACTCTTTCCCTACACGACGCTCTTCCGATCTNNNGCCACCACACCCAGCTATATATA
AGCACACGTCTGAACTCTTTCCCTACACGACGCTCTTCCGATCTNNNGCCACCACACCCAGCTATATATA
GAGCACACGTCTGAACTCTTTCCCTACACGACGCTCTTCCGATCTNNNAATCACTTGAACCTGAGAGGC

AGCACACGTCTGAACTCTTTCCCTACACGACGCTCTTCCGATCTNNNAATCACTTGAACCTGAGAGGC
GCACACGTCTGAACTCTTTCCCTACACGACGCTCTTCCGATCTNNNATGTATGTGTGTGTGTGTGTGT
GCACACGTCTGAACTCTTTCCCTACACGACGCTCTTCCGATCTNNNATGTATGTGTGTGTGTGTGTGT
AGCACACGTCTGAACTCTTTCCCTACACGACGCTCTTCCGATCTNNNAAGAGCCAACAGAATTTGCTG
AGCACACGTCTGAACTCTTTCCCTACACGACGCTCTTCCGATCTNNNAAGAGCCAACAGAATTTGCTG
AGCACACGTCTGAACTCTTTCCCTACACGACGCTCTTCCGATCTNNNCATGCCTGTAATCCCAGCTACTA
AGCACACGTCTGAACTCTTTCCCTACACGACGCTCTTCCGATCTNNNCATGCCTGTAATCCCAGCTACTA
GAGCACACGTCTGAACTCTTTCCCTACACGACGCTCTTCCGATCTNNNCTGTAGTCACAGCTACTCAG
GAGCACACGTCTGAACTCTTTCCCTACACGACGCTCTTCCGATCTNNNCTGTAGTCACAGCTACTCAG
AGCACACGTCTGAACTCTTTCCCTACACGACGCTCTTCCGATCTNNNACGTGGTAAAACCCTGTCTCT
AGCACACGTCTGAACTCTTTCCCTACACGACGCTCTTCCGATCTNNNACGTGGTAAAACCCTGTCTCT
CACACGTCTGAACTCTTTCCCTACACGACGCTCTTCCGATCTNNNAGACATTGCTTTGCATCACTAGA
CACACGTCTGAACTCTTTCCCTACACGACGCTCTTCCGATCTNNNAGACATTGCTTTGCATCACTAGA
AGCACACGTCTGAACTCTTTCCCTACACGACGCTCTTCCGATCTNNNTAGTTTACAGTCTGTTCCGGG
AGCACACGTCTGAACTCTTTCCCTACACGACGCTCTTCCGATCTNNNTAGTTTACAGTCTGTTCCGGG
AGCACACGTCTGAACTCTTTCCCTACACGACGCTCTTCCGATCTNNNCTGCCTGGAAGCTTCTTCTCTAA
AGCACACGTCTGAACTCTTTCCCTACACGACGCTCTTCCGATCTNNNCTGCCTGGAAGCTTCTTCTCTAA
GCACACGTCTGAACTCTTTCCCTACACGACGCTCTTCCGATCTNNNCCACACCTCTCGCATCCTAATATC
GCACACGTCTGAACTCTTTCCCTACACGACGCTCTTCCGATCTNNNCCACACCTCTCGCATCCTAATATC
AGCACACGTCTGAACTCTTTCCCTACACGACGCTCTTCCGATCTNNNGAGGAAAGAAGGGAAGGGA
AGCACACGTCTGAACTCTTTCCCTACACGACGCTCTTCCGATCTNNNGAGGAAAGAAGGGAAGGGA
AGCACACGTCTGAACTCTTTCCCTACACGACGCTCTTCCGATCTNNNATGTTTCTGTTCCTAGGTGAA
AGCACACGTCTGAACTCTTTCCCTACACGACGCTCTTCCGATCTNNNATGTTTCTGTTCCTAGGTGAA
AGCACACGTCTGAACTCTTTCCCTACACGACGCTCTTCCGATCTNNNCCCAGAGCACAGTCAATTTTTG
AGCACACGTCTGAACTCTTTCCCTACACGACGCTCTTCCGATCTNNNCCCAGAGCACAGTCAATTTTTG
CACACGTCTGAACTCTTTCCCTACACGACGCTCTTCCGATCTNNNAGAGATGGAGTTCACAGACACT
CACACGTCTGAACTCTTTCCCTACACGACGCTCTTCCGATCTNNNAGAGATGGAGTTCACAGACACT
GCACACGTCTGAACTCTTTCCCTACACGACGCTCTTCCGATCTNNNTGCCTCTAGGTTTTCCAGTTTGT
GCACACGTCTGAACTCTTTCCCTACACGACGCTCTTCCGATCTNNNTGCCTCTAGGTTTTCCAGTTTGT
AGCACACGTCTGAACTCTTTCCCTACACGACGCTCTTCCGATCTNNNCAAAATCGCTTGAACTGAGG
AGCACACGTCTGAACTCTTTCCCTACACGACGCTCTTCCGATCTNNNCAAAATCGCTTGAACTGAGG
GCACACGTCTGAACTCTTTCCCTACACGACGCTCTTCCGATCTNNNGGGGACAAGAGTGAGACTTCT
GCACACGTCTGAACTCTTTCCCTACACGACGCTCTTCCGATCTNNNGGGGACAAGAGTGAGACTTCT
GCACACGTCTGAACTCTTTCCCTACACGACGCTCTTCCGATCTNNNGCAGTAAATTGCTAAAGGAGTT
GCACACGTCTGAACTCTTTCCCTACACGACGCTCTTCCGATCTNNNGCAGTAAATTGCTAAAGGAGTT
GAGCACACGTCTGAACTCTTTCCCTACACGACGCTCTTCCGATCTNNNCCAGTGAGAATTTCCTTTGA
GAGCACACGTCTGAACTCTTTCCCTACACGACGCTCTTCCGATCTNNNCCAGTGAGAATTTCCTTTGA
GCACACGTCTGAACTCTTTCCCTACACGACGCTCTTCCGATCTNNNATGATCCAATCATCTCCCACCAG
GCACACGTCTGAACTCTTTCCCTACACGACGCTCTTCCGATCTNNNATGATCCAATCATCTCCCACCAG
GCACACGTCTGAACTCTTTCCCTACACGACGCTCTTCCGATCTNNNATGGCAGAGAGGAGAAAAAG
GCACACGTCTGAACTCTTTCCCTACACGACGCTCTTCCGATCTNNNATGGCAGAGAGGAGAAAAAG
AGCACACGTCTGAACTCTTTCCCTACACGACGCTCTTCCGATCTNNNCTCAGTTACCTTTCACTCCCTC
AGCACACGTCTGAACTCTTTCCCTACACGACGCTCTTCCGATCTNNNCTCAGTTACCTTTCACTCCCTC

AGCACACGTCTGAACTCTTTCCCTACACGACGCTCTTCCGATCTNNNGAAACAGATGTTGCAGTGAGC
AGCACACGTCTGAACTCTTTCCCTACACGACGCTCTTCCGATCTNNNGAAACAGATGTTGCAGTGAGC
GAGCACACGTCTGAACTCTTTCCCTACACGACGCTCTTCCGATCTNNNATAGCTACATATGGCCATTGA
GAGCACACGTCTGAACTCTTTCCCTACACGACGCTCTTCCGATCTNNNATAGCTACATATGGCCATTGA
GCACACGTCTGAACTCTTTCCCTACACGACGCTCTTCCGATCTNNNCAGGAGGTGGAAGTTGGAGTTA
GCACACGTCTGAACTCTTTCCCTACACGACGCTCTTCCGATCTNNNCAGGAGGTGGAAGTTGGAGTTA
AGCACACGTCTGAACTCTTTCCCTACACGACGCTCTTCCGATCTNNNGTTCAAGCGATTCTCATGCCTCC
AGCACACGTCTGAACTCTTTCCCTACACGACGCTCTTCCGATCTNNNGTTCAAGCGATTCTCATGCCTCC
AGCACACGTCTGAACTCTTTCCCTACACGACGCTCTTCCGATCTNNNTCTCCAGCAGAATCTTACTGTGT
AGCACACGTCTGAACTCTTTCCCTACACGACGCTCTTCCGATCTNNNTCTCCAGCAGAATCTTACTGTGT
AGCACACGTCTGAACTCTTTCCCTACACGACGCTCTTCCGATCTNNNCTAGTCCCTCCTCCCTATGAGTC
AGCACACGTCTGAACTCTTTCCCTACACGACGCTCTTCCGATCTNNNCTAGTCCCTCCTCCCTATGAGTC
AGCACACGTCTGAACTCTTTCCCTACACGACGCTCTTCCGATCTNNNCAATAGGAAGTGAGGGGTGA
AGCACACGTCTGAACTCTTTCCCTACACGACGCTCTTCCGATCTNNNCAATAGGAAGTGAGGGGTGA
GAGCACACGTCTGAACTCTTTCCCTACACGACGCTCTTCCGATCTNNNGTGTGCCTGTAATCTCAGCTAC
GAGCACACGTCTGAACTCTTTCCCTACACGACGCTCTTCCGATCTNNNGTGTGCCTGTAATCTCAGCTAC
AGCACACGTCTGAACTCTTTCCCTACACGACGCTCTTCCGATCTNNNGACCATGATGCTCGACTAGGAT
AGCACACGTCTGAACTCTTTCCCTACACGACGCTCTTCCGATCTNNNGACCATGATGCTCGACTAGGAT
GCACACGTCTGAACTCTTTCCCTACACGACGCTCTTCCGATCTNNNATTACCTTTCTGCTCAGGAGCAT
GCACACGTCTGAACTCTTTCCCTACACGACGCTCTTCCGATCTNNNATTACCTTTCTGCTCAGGAGCAT
AGCACACGTCTGAACTCTTTCCCTACACGACGCTCTTCCGATCTNNNACTCTTTACCCGGATTCACCAAT
AGCACACGTCTGAACTCTTTCCCTACACGACGCTCTTCCGATCTNNNACTCTTTACCCGGATTCACCAAT
AGCACACGTCTGAACTCTTTCCCTACACGACGCTCTTCCGATCTNNNCGCCCGGCTAATTCTTGTATTTT
AGCACACGTCTGAACTCTTTCCCTACACGACGCTCTTCCGATCTNNNCGCCCGGCTAATTCTTGTATTTT
AGAGCACACGTCTGAACTCTTTCCCTACACGACGCTCTTCCGATCTNNNACTGATCCTCATGCACTCTGT
AGAGCACACGTCTGAACTCTTTCCCTACACGACGCTCTTCCGATCTNNNACTGATCCTCATGCACTCTGT
AGCACACGTCTGAACTCTTTCCCTACACGACGCTCTTCCGATCTNNNCACCTCTCTTCCCACTTGTTCAG
AGCACACGTCTGAACTCTTTCCCTACACGACGCTCTTCCGATCTNNNCACCTCTCTTCCCACTTGTTCAG
AGCACACGTCTGAACTCTTTCCCTACACGACGCTCTTCCGATCTNNNCTCTGGTCCCAAGCATTTTGTTC
AGCACACGTCTGAACTCTTTCCCTACACGACGCTCTTCCGATCTNNNCTCTGGTCCCAAGCATTTTGTTC
AGCACACGTCTGAACTCTTTCCCTACACGACGCTCTTCCGATCTNNNACGTGTGAATTGGGGTAGGAA
AGCACACGTCTGAACTCTTTCCCTACACGACGCTCTTCCGATCTNNNACGTGTGAATTGGGGTAGGAA
GAGCACACGTCTGAACTCTTTCCCTACACGACGCTCTTCCGATCTNNNCTGAGGCAGGAGAATCACTT
GAGCACACGTCTGAACTCTTTCCCTACACGACGCTCTTCCGATCTNNNCTGAGGCAGGAGAATCACTT
AGAGCACACGTCTGAACTCTTTCCCTACACGACGCTCTTCCGATCTNNNTGAAATATGACCGTGACCGA
AGAGCACACGTCTGAACTCTTTCCCTACACGACGCTCTTCCGATCTNNNTGAAATATGACCGTGACCGA
AGCACACGTCTGAACTCTTTCCCTACACGACGCTCTTCCGATCTNNNAATCCCAGCACTTACAGAGGTTG
AGCACACGTCTGAACTCTTTCCCTACACGACGCTCTTCCGATCTNNNAATCCCAGCACTTACAGAGGTTG
GCACACGTCTGAACTCTTTCCCTACACGACGCTCTTCCGATCTNNNTTGTGGTGATGATTGCACAAGT
GCACACGTCTGAACTCTTTCCCTACACGACGCTCTTCCGATCTNNNTTGTGGTGATGATTGCACAAGT
AGCACACGTCTGAACTCTTTCCCTACACGACGCTCTTCCGATCTNNNCAAGAATTTGAGGCTGCAGTGA
AGCACACGTCTGAACTCTTTCCCTACACGACGCTCTTCCGATCTNNNCAAGAATTTGAGGCTGCAGTGA
GAGCACACGTCTGAACTCTTTCCCTACACGACGCTCTTCCGATCTNNNTCCCAGGTTTGAGTGGTATTC

AGCACACGTCTGAACTCTTTCCCTACACGACGCTCTTCCGATCTNNNTCCCAGGTTTGAGTGGTATTC
CACACGTCTGAACTCTTTCCCTACACGACGCTCTTCCGATCTNNNCAGGTTCTCCCTAGCCTACCTCCAT
CACACGTCTGAACTCTTTCCCTACACGACGCTCTTCCGATCTNNNCAGGTTCTCCCTAGCCTACCTCCAT
GAGCACACGTCTGAACTCTTTCCCTACACGACGCTCTTCCGATCTNNNCTGTCTGTCTGTCTGTCTGTC
GAGCACACGTCTGAACTCTTTCCCTACACGACGCTCTTCCGATCTNNNCTGTCTGTCTGTCTGTCTGTC
CACACGTCTGAACTCTTTCCCTACACGACGCTCTTCCGATCTNNNGGCGGCATCTCAGGTTAGTCCAT
CACACGTCTGAACTCTTTCCCTACACGACGCTCTTCCGATCTNNNGGCGGCATCTCAGGTTAGTCCAT
AGCACACGTCTGAACTCTTTCCCTACACGACGCTCTTCCGATCTNNNGAGAATGGCTTGAACCCAGA
AGCACACGTCTGAACTCTTTCCCTACACGACGCTCTTCCGATCTNNNGAGAATGGCTTGAACCCAGA
GAGCACACGTCTGAACTCTTTCCCTACACGACGCTCTTCCGATCTNNNCTCAAATGGAGCCTGAAAAC
GAGCACACGTCTGAACTCTTTCCCTACACGACGCTCTTCCGATCTNNNCTCAAATGGAGCCTGAAAAC
AGCACACGTCTGAACTCTTTCCCTACACGACGCTCTTCCGATCTNNNACACACAAACTTACATGGACA
AGCACACGTCTGAACTCTTTCCCTACACGACGCTCTTCCGATCTNNNACACACAAACTTACATGGACA
GAGCACACGTCTGAACTCTTTCCCTACACGACGCTCTTCCGATCTNNNCATGCCTTATGGCGTTCAGA
GAGCACACGTCTGAACTCTTTCCCTACACGACGCTCTTCCGATCTNNNCATGCCTTATGGCGTTCAGA
GCACACGTCTGAACTCTTTCCCTACACGACGCTCTTCCGATCTNNNACCACCAGTTCAGTACCATCTGT
GCACACGTCTGAACTCTTTCCCTACACGACGCTCTTCCGATCTNNNACCACCAGTTCAGTACCATCTGT
AGCACACGTCTGAACTCTTTCCCTACACGACGCTCTTCCGATCTNNNCGCATACCTGTAATCCCAGCTA
AGCACACGTCTGAACTCTTTCCCTACACGACGCTCTTCCGATCTNNNCGCATACCTGTAATCCCAGCTA
AGCACACGTCTGAACTCTTTCCCTACACGACGCTCTTCCGATCTNNNGCAATGCCTCCTCCTCATGTTA
AGCACACGTCTGAACTCTTTCCCTACACGACGCTCTTCCGATCTNNNGCAATGCCTCCTCCTCATGTTA
AGCACACGTCTGAACTCTTTCCCTACACGACGCTCTTCCGATCTNNNGAGAATGGCATGAACCCAGG
AGCACACGTCTGAACTCTTTCCCTACACGACGCTCTTCCGATCTNNNGAGAATGGCATGAACCCAGG
AGCACACGTCTGAACTCTTTCCCTACACGACGCTCTTCCGATCTNNNGGTGCCTGTAATCCCAACTACT
AGCACACGTCTGAACTCTTTCCCTACACGACGCTCTTCCGATCTNNNGGTGCCTGTAATCCCAACTACT
CACGTCTGAACTCTTTCCCTACACGACGCTCTTCCGATCTNNNCTCATAGTCCCAAATCCAGCTGTCAT
CACGTCTGAACTCTTTCCCTACACGACGCTCTTCCGATCTNNNCTCATAGTCCCAAATCCAGCTGTCAT
AGCACACGTCTGAACTCTTTCCCTACACGACGCTCTTCCGATCTNNNGTAAATAGTCTCAGCTCCTCG
AGCACACGTCTGAACTCTTTCCCTACACGACGCTCTTCCGATCTNNNGTAAATAGTCTCAGCTCCTCG
AGCACACGTCTGAACTCTTTCCCTACACGACGCTCTTCCGATCTNNNGATAATCCCTTGAACCCAGGA
AGCACACGTCTGAACTCTTTCCCTACACGACGCTCTTCCGATCTNNNGATAATCCCTTGAACCCAGGA
GAGCACACGTCTGAACTCTTTCCCTACACGACGCTCTTCCGATCTNNNAAATAAATAATCCTGGGCTG
GAGCACACGTCTGAACTCTTTCCCTACACGACGCTCTTCCGATCTNNNAAATAAATAATCCTGGGCTG
CACACGTCTGAACTCTTTCCCTACACGACGCTCTTCCGATCTNNNCAAGTCAGAAAGCAGATTGGTG
CACACGTCTGAACTCTTTCCCTACACGACGCTCTTCCGATCTNNNCAAGTCAGAAAGCAGATTGGTG
AGCACACGTCTGAACTCTTTCCCTACACGACGCTCTTCCGATCTNNNGATCATGGCCCTAATCCCAACC
AGCACACGTCTGAACTCTTTCCCTACACGACGCTCTTCCGATCTNNNGATCATGGCCCTAATCCCAACC
GCACACGTCTGAACTCTTTCCCTACACGACGCTCTTCCGATCTNNNAACATGGTGAAAACCCGTCTCT
GCACACGTCTGAACTCTTTCCCTACACGACGCTCTTCCGATCTNNNAACATGGTGAAAACCCGTCTCT
AGCACACGTCTGAACTCTTTCCCTACACGACGCTCTTCCGATCTNNNCATGCCTGTAATCCCAGCTACT
AGCACACGTCTGAACTCTTTCCCTACACGACGCTCTTCCGATCTNNNCATGCCTGTAATCCCAGCTACT
AGCACACGTCTGAACTCTTTCCCTACACGACGCTCTTCCGATCTNNNGAGGCTGAGGCAGGAGAATG
AGCACACGTCTGAACTCTTTCCCTACACGACGCTCTTCCGATCTNNNGAGGCTGAGGCAGGAGAATG

GAGCACACGTCTGAACTCTTTCCCTACACGACGCTCTTCCGATCTNNNCCCAAATGCTCAGCAATCAATG
GAGCACACGTCTGAACTCTTTCCCTACACGACGCTCTTCCGATCTNNNCCCAAATGCTCAGCAATCAATG
GAGCACACGTCTGAACTCTTTCCCTACACGACGCTCTTCCGATCTNNNCCTCAGGAAACCCCTTTAGAT
GAGCACACGTCTGAACTCTTTCCCTACACGACGCTCTTCCGATCTNNNCCTCAGGAAACCCCTTTAGAT
AGCACACGTCTGAACTCTTTCCCTACACGACGCTCTTCCGATCTNNNCTGAGATCCCACCACTGTACTTC
AGCACACGTCTGAACTCTTTCCCTACACGACGCTCTTCCGATCTNNNCTGAGATCCCACCACTGTACTTC
GAGCACACGTCTGAACTCTTTCCCTACACGACGCTCTTCCGATCTNNNTGGGGTGGAGAGAGTAGTAT
GAGCACACGTCTGAACTCTTTCCCTACACGACGCTCTTCCGATCTNNNTGGGGTGGAGAGAGTAGTAT
GAGCACACGTCTGAACTCTTTCCCTACACGACGCTCTTCCGATCTNNNTGCAAATTTTCTGCCTGATCCT
GAGCACACGTCTGAACTCTTTCCCTACACGACGCTCTTCCGATCTNNNTGCAAATTTTCTGCCTGATCCT
AGCACACGTCTGAACTCTTTCCCTACACGACGCTCTTCCGATCTNNNTCAAGCAATCCTCCTACCTCAAC
AGCACACGTCTGAACTCTTTCCCTACACGACGCTCTTCCGATCTNNNTCAAGCAATCCTCCTACCTCAAC
AGCACACGTCTGAACTCTTTCCCTACACGACGCTCTTCCGATCTNNNGAGGAAATGGGCTGAGAGAGA
AGCACACGTCTGAACTCTTTCCCTACACGACGCTCTTCCGATCTNNNGAGGAAATGGGCTGAGAGAGA
ACACGTCTGAACTCTTTCCCTACACGACGCTCTTCCGATCTNNNCTGTGACCACGGCATCTGAGCATCG
ACACGTCTGAACTCTTTCCCTACACGACGCTCTTCCGATCTNNNCTGTGACCACGGCATCTGAGCATCG
GAGCACACGTCTGAACTCTTTCCCTACACGACGCTCTTCCGATCTNNNTCCTGGCTTGTAATGGTGGA
GAGCACACGTCTGAACTCTTTCCCTACACGACGCTCTTCCGATCTNNNTCCTGGCTTGTAATGGTGGA
GAGCACACGTCTGAACTCTTTCCCTACACGACGCTCTTCCGATCTNNNCAACATCTTGGGACCCTATCTC
GAGCACACGTCTGAACTCTTTCCCTACACGACGCTCTTCCGATCTNNNCAACATCTTGGGACCCTATCTC
AGAGCACACGTCTGAACTCTTTCCCTACACGACGCTCTTCCGATCTNNNGGAAGAATCACTTGAACCTC
AGAGCACACGTCTGAACTCTTTCCCTACACGACGCTCTTCCGATCTNNNGGAAGAATCACTTGAACCTC
GAGCACACGTCTGAACTCTTTCCCTACACGACGCTCTTCCGATCTNNNCACACTATGTAGCTCATCCTGC
GAGCACACGTCTGAACTCTTTCCCTACACGACGCTCTTCCGATCTNNNCACACTATGTAGCTCATCCTGC
AGAGCACACGTCTGAACTCTTTCCCTACACGACGCTCTTCCGATCTNNNCAGGAGTTTGAGATCACAG
AGAGCACACGTCTGAACTCTTTCCCTACACGACGCTCTTCCGATCTNNNCAGGAGTTTGAGATCACAG
GAGCACACGTCTGAACTCTTTCCCTACACGACGCTCTTCCGATCTNNNGGAGGTTGTAGTGAGCTGA
GAGCACACGTCTGAACTCTTTCCCTACACGACGCTCTTCCGATCTNNNGGAGGTTGTAGTGAGCTGA
GAGCACACGTCTGAACTCTTTCCCTACACGACGCTCTTCCGATCTNNNTTCTCTTTGGCCTTCATGTAGC
GAGCACACGTCTGAACTCTTTCCCTACACGACGCTCTTCCGATCTNNNTTCTCTTTGGCCTTCATGTAGC
AGAGCACACGTCTGAACTCTTTCCCTACACGACGCTCTTCCGATCTNNNGGTAAATAGGTGGGACAA
AGAGCACACGTCTGAACTCTTTCCCTACACGACGCTCTTCCGATCTNNNGGTAAATAGGTGGGACAA
GCACACGTCTGAACTCTTTCCCTACACGACGCTCTTCCGATCTNNNAGACAAGCTGACTCTGCCTACTT
GCACACGTCTGAACTCTTTCCCTACACGACGCTCTTCCGATCTNNNAGACAAGCTGACTCTGCCTACTT
AGCACACGTCTGAACTCTTTCCCTACACGACGCTCTTCCGATCTNNNCTGGAAGGTTGTTATCCTCCAG
AGCACACGTCTGAACTCTTTCCCTACACGACGCTCTTCCGATCTNNNCTGGAAGGTTGTTATCCTCCAG
AGCACACGTCTGAACTCTTTCCCTACACGACGCTCTTCCGATCTNNNGGCAGAGACACAGCAAAAGA
AGCACACGTCTGAACTCTTTCCCTACACGACGCTCTTCCGATCTNNNGGCAGAGACACAGCAAAAGA
AGCACACGTCTGAACTCTTTCCCTACACGACGCTCTTCCGATCTNNNGGCTGCATAGTTTTCCATGGTT
AGCACACGTCTGAACTCTTTCCCTACACGACGCTCTTCCGATCTNNNGGCTGCATAGTTTTCCATGGTT
GAGCACACGTCTGAACTCTTTCCCTACACGACGCTCTTCCGATCTNNNCACCACACCAGGCTAATTTTTG
GAGCACACGTCTGAACTCTTTCCCTACACGACGCTCTTCCGATCTNNNCACCACACCAGGCTAATTTTTG
AGCACACGTCTGAACTCTTTCCCTACACGACGCTCTTCCGATCTNNNCAACAGGCAAAGACAACAGAG

AGCACACGTCTGAACTCTTTCCCTACACGACGCTCTTCCGATCTNNNCAACAGGCAAAGACAACAGAG
CACACGTCTGAACTCTTTCCCTACACGACGCTCTTCCGATCTNNNTTCTCAGTTGTCAGGAAACTCCGC
CACACGTCTGAACTCTTTCCCTACACGACGCTCTTCCGATCTNNNTTCTCAGTTGTCAGGAAACTCCGC
GCACACGTCTGAACTCTTTCCCTACACGACGCTCTTCCGATCTNNNGTGGCTACACACTTCTCATCTCA
GCACACGTCTGAACTCTTTCCCTACACGACGCTCTTCCGATCTNNNGTGGCTACACACTTCTCATCTCA
AGCACACGTCTGAACTCTTTCCCTACACGACGCTCTTCCGATCTNNNCAGGAGAATCACTTGAACCTG
AGCACACGTCTGAACTCTTTCCCTACACGACGCTCTTCCGATCTNNNCAGGAGAATCACTTGAACCTG
GAGCACACGTCTGAACTCTTTCCCTACACGACGCTCTTCCGATCTNNNAAAGGTAAACGTTGCTTGCT
GAGCACACGTCTGAACTCTTTCCCTACACGACGCTCTTCCGATCTNNNAAAGGTAAACGTTGCTTGCT
GCACACGTCTGAACTCTTTCCCTACACGACGCTCTTCCGATCTNNNTCCTCACCTTTCTCTCCCCTATTCA
GCACACGTCTGAACTCTTTCCCTACACGACGCTCTTCCGATCTNNNTCCTCACCTTTCTCTCCCCTATTCA
AGAGCACACGTCTGAACTCTTTCCCTACACGACGCTCTTCCGATCTNNNTAGGTTGAGTTGATTGAGC
AGAGCACACGTCTGAACTCTTTCCCTACACGACGCTCTTCCGATCTNNNTAGGTTGAGTTGATTGAGC
AGCACACGTCTGAACTCTTTCCCTACACGACGCTCTTCCGATCTNNNGAGAATGGCATGAACCTGGG
AGCACACGTCTGAACTCTTTCCCTACACGACGCTCTTCCGATCTNNNGAGAATGGCATGAACCTGGG
GAGCACACGTCTGAACTCTTTCCCTACACGACGCTCTTCCGATCTNNNCTGAAATAACGCCACTGCACT
GAGCACACGTCTGAACTCTTTCCCTACACGACGCTCTTCCGATCTNNNCTGAAATAACGCCACTGCACT
GAGCACACGTCTGAACTCTTTCCCTACACGACGCTCTTCCGATCTNNNTCCTTTGAGCCATAAGAAACA
GAGCACACGTCTGAACTCTTTCCCTACACGACGCTCTTCCGATCTNNNTCCTTTGAGCCATAAGAAACA
GCACACGTCTGAACTCTTTCCCTACACGACGCTCTTCCGATCTNNNCCTGATGCCCTATTGCAGATCTT
GCACACGTCTGAACTCTTTCCCTACACGACGCTCTTCCGATCTNNNCCTGATGCCCTATTGCAGATCTT
AGCACACGTCTGAACTCTTTCCCTACACGACGCTCTTCCGATCTNNNAACAAGGGGGTCTCTTTCTGAT
AGCACACGTCTGAACTCTTTCCCTACACGACGCTCTTCCGATCTNNNAACAAGGGGGTCTCTTTCTGAT
GAGCACACGTCTGAACTCTTTCCCTACACGACGCTCTTCCGATCTNNNCTCCTGAGTAGCTGGGACTA
GAGCACACGTCTGAACTCTTTCCCTACACGACGCTCTTCCGATCTNNNCTCCTGAGTAGCTGGGACTA
GCACACGTCTGAACTCTTTCCCTACACGACGCTCTTCCGATCTNNNCTTCACTCTTATCTGTTACGTAAC
GCACACGTCTGAACTCTTTCCCTACACGACGCTCTTCCGATCTNNNCTTCACTCTTATCTGTTACGTAAC
AGCACACGTCTGAACTCTTTCCCTACACGACGCTCTTCCGATCTNNNGCAACAGGAATGAAACTTCGT
AGCACACGTCTGAACTCTTTCCCTACACGACGCTCTTCCGATCTNNNGCAACAGGAATGAAACTTCGT
GCACACGTCTGAACTCTTTCCCTACACGACGCTCTTCCGATCTNNNCACATACGCTCAGTTCAGCTGC
GCACACGTCTGAACTCTTTCCCTACACGACGCTCTTCCGATCTNNNCACATACGCTCAGTTCAGCTGC
ACACGTCTGAACTCTTTCCCTACACGACGCTCTTCCGATCTNNNTTCTGTGAGAGCTTTTATGACACTC
ACACGTCTGAACTCTTTCCCTACACGACGCTCTTCCGATCTNNNTTCTGTGAGAGCTTTTATGACACTC
GCACACGTCTGAACTCTTTCCCTACACGACGCTCTTCCGATCTNNNCTTGAACTCCTGACCTCAAGTG
GCACACGTCTGAACTCTTTCCCTACACGACGCTCTTCCGATCTNNNCTTGAACTCCTGACCTCAAGTG
GAGCACACGTCTGAACTCTTTCCCTACACGACGCTCTTCCGATCTNNNGTGTGTGTGTGTGTGTGTG
GAGCACACGTCTGAACTCTTTCCCTACACGACGCTCTTCCGATCTNNNGTGTGTGTGTGTGTGTGTG
GCACACGTCTGAACTCTTTCCCTACACGACGCTCTTCCGATCTNNNTCCCTCTCCCCCTACTTTCTTTCCA
GCACACGTCTGAACTCTTTCCCTACACGACGCTCTTCCGATCTNNNTCCCTCTCCCCCTACTTTCTTTCCA
GCACACGTCTGAACTCTTTCCCTACACGACGCTCTTCCGATCTNNNAGTTTTCTTCCCAGGAAACCCAG
GCACACGTCTGAACTCTTTCCCTACACGACGCTCTTCCGATCTNNNAGTTTTCTTCCCAGGAAACCCAG
GCACACGTCTGAACTCTTTCCCTACACGACGCTCTTCCGATCTNNNGGTGACATAAGAGAGACCCTG
GCACACGTCTGAACTCTTTCCCTACACGACGCTCTTCCGATCTNNNGGTGACATAAGAGAGACCCTG

AGCACACGTCTGAACTCTTTCCCTACACGACGCTCTTCCGATCTNNNTTTAGTAGAGACGGGGTTTCA
AGCACACGTCTGAACTCTTTCCCTACACGACGCTCTTCCGATCTNNNTTTAGTAGAGACGGGGTTTCA
GAGCACACGTCTGAACTCTTTCCCTACACGACGCTCTTCCGATCTNNNGAGCTTGCAGTGAGTCGAGA
GAGCACACGTCTGAACTCTTTCCCTACACGACGCTCTTCCGATCTNNNGAGCTTGCAGTGAGTCGAGA
AGCACACGTCTGAACTCTTTCCCTACACGACGCTCTTCCGATCTNNNTTCACACCATTCTCCTGCCTCCAT
AGCACACGTCTGAACTCTTTCCCTACACGACGCTCTTCCGATCTNNNTTCACACCATTCTCCTGCCTCCAT
GCACACGTCTGAACTCTTTCCCTACACGACGCTCTTCCGATCTNNNTTCCAAAGTGCTGGGATTACAGA
GCACACGTCTGAACTCTTTCCCTACACGACGCTCTTCCGATCTNNNTTCCAAAGTGCTGGGATTACAGA
AGCACACGTCTGAACTCTTTCCCTACACGACGCTCTTCCGATCTNNNATGGTCTCAATCTCCTGACTTCG
AGCACACGTCTGAACTCTTTCCCTACACGACGCTCTTCCGATCTNNNATGGTCTCAATCTCCTGACTTCG
GAGCACACGTCTGAACTCTTTCCCTACACGACGCTCTTCCGATCTNNNGAGGCTGAGGCAGGAGAAT
GAGCACACGTCTGAACTCTTTCCCTACACGACGCTCTTCCGATCTNNNGAGGCTGAGGCAGGAGAAT
GAGCACACGTCTGAACTCTTTCCCTACACGACGCTCTTCCGATCTNNNCCATGTCCTTACCAATCGACCA
GAGCACACGTCTGAACTCTTTCCCTACACGACGCTCTTCCGATCTNNNCCATGTCCTTACCAATCGACCA
AGAGCACACGTCTGAACTCTTTCCCTACACGACGCTCTTCCGATCTNNNCTGCTTCATACGGTGAGACA
AGAGCACACGTCTGAACTCTTTCCCTACACGACGCTCTTCCGATCTNNNCTGCTTCATACGGTGAGACA
GAGCACACGTCTGAACTCTTTCCCTACACGACGCTCTTCCGATCTNNNGAGAAGCAGTTACTCTAGAG
GAGCACACGTCTGAACTCTTTCCCTACACGACGCTCTTCCGATCTNNNGAGAAGCAGTTACTCTAGAG
AGCACACGTCTGAACTCTTTCCCTACACGACGCTCTTCCGATCTNNNATTTGGTACTATGCCAAGGAAG
AGCACACGTCTGAACTCTTTCCCTACACGACGCTCTTCCGATCTNNNATTTGGTACTATGCCAAGGAAG
AGCACACGTCTGAACTCTTTCCCTACACGACGCTCTTCCGATCTNNNTCTTAGCTTAGGTCCAGGAAG
AGCACACGTCTGAACTCTTTCCCTACACGACGCTCTTCCGATCTNNNTCTTAGCTTAGGTCCAGGAAG
GAGCACACGTCTGAACTCTTTCCCTACACGACGCTCTTCCGATCTNNNAGTAGCTGAGATTACAAGTG
GAGCACACGTCTGAACTCTTTCCCTACACGACGCTCTTCCGATCTNNNAGTAGCTGAGATTACAAGTG
AGCACACGTCTGAACTCTTTCCCTACACGACGCTCTTCCGATCTNNNGCCTGAATTTGGGAAGAGAAA
AGCACACGTCTGAACTCTTTCCCTACACGACGCTCTTCCGATCTNNNGCCTGAATTTGGGAAGAGAAA
GAGCACACGTCTGAACTCTTTCCCTACACGACGCTCTTCCGATCTNNNGATAGCTTGAGCCTAGGAGTT
GAGCACACGTCTGAACTCTTTCCCTACACGACGCTCTTCCGATCTNNNGATAGCTTGAGCCTAGGAGTT
GAGCACACGTCTGAACTCTTTCCCTACACGACGCTCTTCCGATCTNNNATACGCTTCTGATATGGCAGG
GAGCACACGTCTGAACTCTTTCCCTACACGACGCTCTTCCGATCTNNNATACGCTTCTGATATGGCAGG
AGCACACGTCTGAACTCTTTCCCTACACGACGCTCTTCCGATCTNNNCCACCAATTCCGGACACAATAT
AGCACACGTCTGAACTCTTTCCCTACACGACGCTCTTCCGATCTNNNCCACCAATTCCGGACACAATAT
GAGCACACGTCTGAACTCTTTCCCTACACGACGCTCTTCCGATCTNNNTCATGCCTGTAATCCCAACACT
GAGCACACGTCTGAACTCTTTCCCTACACGACGCTCTTCCGATCTNNNTCATGCCTGTAATCCCAACACT
GAGCACACGTCTGAACTCTTTCCCTACACGACGCTCTTCCGATCTNNNAAAAAAATGTGTCCAGAGCCA
GAGCACACGTCTGAACTCTTTCCCTACACGACGCTCTTCCGATCTNNNAAAAAAATGTGTCCAGAGCCA
AGAGCACACGTCTGAACTCTTTCCCTACACGACGCTCTTCCGATCTNNNGATTGAAAGAAGCCAGGCC
AGAGCACACGTCTGAACTCTTTCCCTACACGACGCTCTTCCGATCTNNNGATTGAAAGAAGCCAGGCC
AGCACACGTCTGAACTCTTTCCCTACACGACGCTCTTCCGATCTNNNCTCAAGTGATCCTCCCATCTCAG
AGCACACGTCTGAACTCTTTCCCTACACGACGCTCTTCCGATCTNNNCTCAAGTGATCCTCCCATCTCAG
AGAGCACACGTCTGAACTCTTTCCCTACACGACGCTCTTCCGATCTNNNCAACAAAGCGAAACTCTGTCT
AGAGCACACGTCTGAACTCTTTCCCTACACGACGCTCTTCCGATCTNNNCAACAAAGCGAAACTCTGTCT
AGCACACGTCTGAACTCTTTCCCTACACGACGCTCTTCCGATCTNNNTGACTTGCCACAGGTCATAATG

AGCACACGTCTGAACTCTTTCCCTACACGACGCTCTTCCGATCTNNNTGACTTGCCACAGGTCATAATG
AGAGCACACGTCTGAACTCTTTCCCTACACGACGCTCTTCCGATCTNNNCTACTCATGAGGCAGGAGAA
AGAGCACACGTCTGAACTCTTTCCCTACACGACGCTCTTCCGATCTNNNCTACTCATGAGGCAGGAGAA
GAGCACACGTCTGAACTCTTTCCCTACACGACGCTCTTCCGATCTNNNGCCCGAGATTTGCATTTCTAAC
GAGCACACGTCTGAACTCTTTCCCTACACGACGCTCTTCCGATCTNNNGCCCGAGATTTGCATTTCTAAC
AGCACACGTCTGAACTCTTTCCCTACACGACGCTCTTCCGATCTNNNGCTGTACCAAGTGATAGGGTG
AGCACACGTCTGAACTCTTTCCCTACACGACGCTCTTCCGATCTNNNGCTGTACCAAGTGATAGGGTG
AGCACACGTCTGAACTCTTTCCCTACACGACGCTCTTCCGATCTNNNGAATCACCAGAGTCCTACAGCA
AGCACACGTCTGAACTCTTTCCCTACACGACGCTCTTCCGATCTNNNGAATCACCAGAGTCCTACAGCA
AGCACACGTCTGAACTCTTTCCCTACACGACGCTCTTCCGATCTNNNCCTGGGAAATAGAACGAGACT
AGCACACGTCTGAACTCTTTCCCTACACGACGCTCTTCCGATCTNNNCCTGGGAAATAGAACGAGACT
AGAGCACACGTCTGAACTCTTTCCCTACACGACGCTCTTCCGATCTNNNTTAGACACAAAAGGGCTGA
AGAGCACACGTCTGAACTCTTTCCCTACACGACGCTCTTCCGATCTNNNTTAGACACAAAAGGGCTGA
AGCACACGTCTGAACTCTTTCCCTACACGACGCTCTTCCGATCTNNNCTCCGAGAAGAACCAGATGCTA
AGCACACGTCTGAACTCTTTCCCTACACGACGCTCTTCCGATCTNNNCTCCGAGAAGAACCAGATGCTA
GAGCACACGTCTGAACTCTTTCCCTACACGACGCTCTTCCGATCTNNNACACAATAACACAGCACTCAG
GAGCACACGTCTGAACTCTTTCCCTACACGACGCTCTTCCGATCTNNNACACAATAACACAGCACTCAG
GAGCACACGTCTGAACTCTTTCCCTACACGACGCTCTTCCGATCTNNNGAGCTTGCAGTGAGCCAAATA
GAGCACACGTCTGAACTCTTTCCCTACACGACGCTCTTCCGATCTNNNGAGCTTGCAGTGAGCCAAATA
AGCACACGTCTGAACTCTTTCCCTACACGACGCTCTTCCGATCTNNNAGGGTCTTACCATATTGCCCATG
AGCACACGTCTGAACTCTTTCCCTACACGACGCTCTTCCGATCTNNNAGGGTCTTACCATATTGCCCATG
AGCACACGTCTGAACTCTTTCCCTACACGACGCTCTTCCGATCTNNNGAGCAAGACCCCATCTCAAAAA
AGCACACGTCTGAACTCTTTCCCTACACGACGCTCTTCCGATCTNNNGAGCAAGACCCCATCTCAAAAA
GAGCACACGTCTGAACTCTTTCCCTACACGACGCTCTTCCGATCTNNNATGACTCTCAAAGCTGACTTCC
GAGCACACGTCTGAACTCTTTCCCTACACGACGCTCTTCCGATCTNNNATGACTCTCAAAGCTGACTTCC
AGCACACGTCTGAACTCTTTCCCTACACGACGCTCTTCCGATCTNNNACGCCACCATACCTGTCTAATTTC
AGCACACGTCTGAACTCTTTCCCTACACGACGCTCTTCCGATCTNNNACGCCACCATACCTGTCTAATTTC
GCACACGTCTGAACTCTTTCCCTACACGACGCTCTTCCGATCTNNNCTGAGATCACACCACTGCACTCCA
GCACACGTCTGAACTCTTTCCCTACACGACGCTCTTCCGATCTNNNCTGAGATCACACCACTGCACTCCA
GAGCACACGTCTGAACTCTTTCCCTACACGACGCTCTTCCGATCTNNNACAGTTTGGGTGTCCTAATCCA
GAGCACACGTCTGAACTCTTTCCCTACACGACGCTCTTCCGATCTNNNACAGTTTGGGTGTCCTAATCCA
AGCACACGTCTGAACTCTTTCCCTACACGACGCTCTTCCGATCTNNNTCTGTGACCTGTAACTTTGCTGA
AGCACACGTCTGAACTCTTTCCCTACACGACGCTCTTCCGATCTNNNTCTGTGACCTGTAACTTTGCTGA
GAGCACACGTCTGAACTCTTTCCCTACACGACGCTCTTCCGATCTNNNGTAGAGACAGAGTTTCACCGT
GAGCACACGTCTGAACTCTTTCCCTACACGACGCTCTTCCGATCTNNNGTAGAGACAGAGTTTCACCGT
AGCACACGTCTGAACTCTTTCCCTACACGACGCTCTTCCGATCTNNNGCCCCACCCAAGATAATGTTTTT
AGCACACGTCTGAACTCTTTCCCTACACGACGCTCTTCCGATCTNNNGCCCCACCCAAGATAATGTTTTT
GCACACGTCTGAACTCTTTCCCTACACGACGCTCTTCCGATCTNNNACATCAAACACCTACTTCTGGCTCA
GCACACGTCTGAACTCTTTCCCTACACGACGCTCTTCCGATCTNNNACATCAAACACCTACTTCTGGCTCA
AGCACACGTCTGAACTCTTTCCCTACACGACGCTCTTCCGATCTNNNAGGAGAATGGCGTGAACCTGCA
AGCACACGTCTGAACTCTTTCCCTACACGACGCTCTTCCGATCTNNNAGGAGAATGGCGTGAACCTGCA
GAGCACACGTCTGAACTCTTTCCCTACACGACGCTCTTCCGATCTNNNCAGGAGAATGGCATGGACCTC
GAGCACACGTCTGAACTCTTTCCCTACACGACGCTCTTCCGATCTNNNCAGGAGAATGGCATGGACCTC

GAGCACACGTCTGAACTCTTTCCCTACACGACGCTCTTCCGATCTNNNCCAGCCTGGCAGAAGAGAAAT
GAGCACACGTCTGAACTCTTTCCCTACACGACGCTCTTCCGATCTNNNCCAGCCTGGCAGAAGAGAAAT
AGAGCACACGTCTGAACTCTTTCCCTACACGACGCTCTTCCGATCTNNNAGGACAGTGACTGCCATTTG
AGAGCACACGTCTGAACTCTTTCCCTACACGACGCTCTTCCGATCTNNNAGGACAGTGACTGCCATTTG
GAGCACACGTCTGAACTCTTTCCCTACACGACGCTCTTCCGATCTNNNCTCTGAACCCCAAGAGAAGAG
GAGCACACGTCTGAACTCTTTCCCTACACGACGCTCTTCCGATCTNNNCTCTGAACCCCAAGAGAAGAG
ACACGTCTGAACTCTTTCCCTACACGACGCTCTTCCGATCTNNNCACCAGCATGTGACTGTAAAGTGCAT
ACACGTCTGAACTCTTTCCCTACACGACGCTCTTCCGATCTNNNCACCAGCATGTGACTGTAAAGTGCAT
AGAGCACACGTCTGAACTCTTTCCCTACACGACGCTCTTCCGATCTNNNCCAACAGGATATGGTTGCGT
AGAGCACACGTCTGAACTCTTTCCCTACACGACGCTCTTCCGATCTNNNCCAACAGGATATGGTTGCGT
GCACACGTCTGAACTCTTTCCCTACACGACGCTCTTCCGATCTNNNGCACTACCACTCCTGGCTAATTTC
GCACACGTCTGAACTCTTTCCCTACACGACGCTCTTCCGATCTNNNGCACTACCACTCCTGGCTAATTTC
GAGCACACGTCTGAACTCTTTCCCTACACGACGCTCTTCCGATCTNNNCAGGAGAATTGCTTGAACTTG
GAGCACACGTCTGAACTCTTTCCCTACACGACGCTCTTCCGATCTNNNCAGGAGAATTGCTTGAACTTG
GAGCACACGTCTGAACTCTTTCCCTACACGACGCTCTTCCGATCTNNNCTAAACAGCCCAGCTATGTCTC
GAGCACACGTCTGAACTCTTTCCCTACACGACGCTCTTCCGATCTNNNCTAAACAGCCCAGCTATGTCTC
CACACGTCTGAACTCTTTCCCTACACGACGCTCTTCCGATCTNNNATCAATTCGCTCTCAGACTCGAACAT
CACACGTCTGAACTCTTTCCCTACACGACGCTCTTCCGATCTNNNATCAATTCGCTCTCAGACTCGAACAT
GAGCACACGTCTGAACTCTTTCCCTACACGACGCTCTTCCGATCTNNNCACCTGTAATCCCAGCTACTCA
GAGCACACGTCTGAACTCTTTCCCTACACGACGCTCTTCCGATCTNNNCACCTGTAATCCCAGCTACTCA
AGCACACGTCTGAACTCTTTCCCTACACGACGCTCTTCCGATCTNNNTTTCTACTTCTCGTCCTCCTGCCA
AGCACACGTCTGAACTCTTTCCCTACACGACGCTCTTCCGATCTNNNTTTCTACTTCTCGTCCTCCTGCCA
GAGCACACGTCTGAACTCTTTCCCTACACGACGCTCTTCCGATCTNNNTATTTCCCTTTGTTCAGCTGTG
GAGCACACGTCTGAACTCTTTCCCTACACGACGCTCTTCCGATCTNNNTATTTCCCTTTGTTCAGCTGTG
AGCACACGTCTGAACTCTTTCCCTACACGACGCTCTTCCGATCTNNNGAGGTTGCAGTGAGCTGAGAT
AGCACACGTCTGAACTCTTTCCCTACACGACGCTCTTCCGATCTNNNGAGGTTGCAGTGAGCTGAGAT
GAGCACACGTCTGAACTCTTTCCCTACACGACGCTCTTCCGATCTNNNTACCAGTTTTCCCGTTTCCTGAA
GAGCACACGTCTGAACTCTTTCCCTACACGACGCTCTTCCGATCTNNNTACCAGTTTTCCCGTTTCCTGAA
AGCACACGTCTGAACTCTTTCCCTACACGACGCTCTTCCGATCTNNNATACAAAAATTAGCTGGGCGTG
AGCACACGTCTGAACTCTTTCCCTACACGACGCTCTTCCGATCTNNNATACAAAAATTAGCTGGGCGTG
AGCACACGTCTGAACTCTTTCCCTACACGACGCTCTTCCGATCTNNNCTAGCTTTGGGGGTTGGTTTTT
AGCACACGTCTGAACTCTTTCCCTACACGACGCTCTTCCGATCTNNNCTAGCTTTGGGGGTTGGTTTTT
GAGCACACGTCTGAACTCTTTCCCTACACGACGCTCTTCCGATCTNNNTAAAATGAGTTGGGGAGGAG
GAGCACACGTCTGAACTCTTTCCCTACACGACGCTCTTCCGATCTNNNTAAAATGAGTTGGGGAGGAG
GAGCACACGTCTGAACTCTTTCCCTACACGACGCTCTTCCGATCTNNNGGACAACAGAGTGAGACTCCA
GAGCACACGTCTGAACTCTTTCCCTACACGACGCTCTTCCGATCTNNNGGACAACAGAGTGAGACTCCA
AGCACACGTCTGAACTCTTTCCCTACACGACGCTCTTCCGATCTNNNCATGTCTGCATTTTTGGTCAGGT
AGCACACGTCTGAACTCTTTCCCTACACGACGCTCTTCCGATCTNNNCATGTCTGCATTTTTGGTCAGGT
GAGCACACGTCTGAACTCTTTCCCTACACGACGCTCTTCCGATCTNNNGCATGCTTAGAGTGCTTTGGA
GAGCACACGTCTGAACTCTTTCCCTACACGACGCTCTTCCGATCTNNNGCATGCTTAGAGTGCTTTGGA
AGCACACGTCTGAACTCTTTCCCTACACGACGCTCTTCCGATCTNNNAAGACATGCCGGAGTGTTGTA
AGCACACGTCTGAACTCTTTCCCTACACGACGCTCTTCCGATCTNNNAAGACATGCCGGAGTGTTGTA
GAGCACACGTCTGAACTCTTTCCCTACACGACGCTCTTCCGATCTNNNGCCTGTAGTCCCAGCTACTTGC

AGCACACGTCTGAACTCTTTCCCTACACGACGCTCTTCCGATCTNNNGCCTGTAGTCCCAGCTACTTG
AGCACACGTCTGAACTCTTTCCCTACACGACGCTCTTCCGATCTNNNTTAGGCTCTTTCTCCATCACAC
AGCACACGTCTGAACTCTTTCCCTACACGACGCTCTTCCGATCTNNNTTAGGCTCTTTCTCCATCACAC
AGCACACGTCTGAACTCTTTCCCTACACGACGCTCTTCCGATCTNNNGCAGGAGAATTGCTTGAACC
AGCACACGTCTGAACTCTTTCCCTACACGACGCTCTTCCGATCTNNNGCAGGAGAATTGCTTGAACC
GCACACGTCTGAACTCTTTCCCTACACGACGCTCTTCCGATCTNNNGAGGCCAAGGGAACATGTGAT
GCACACGTCTGAACTCTTTCCCTACACGACGCTCTTCCGATCTNNNGAGGCCAAGGGAACATGTGAT
AGCACACGTCTGAACTCTTTCCCTACACGACGCTCTTCCGATCTNNNCTTGTGTGTGAGTGTGTGTG
AGCACACGTCTGAACTCTTTCCCTACACGACGCTCTTCCGATCTNNNCTTGTGTGTGAGTGTGTGTG
AGCACACGTCTGAACTCTTTCCCTACACGACGCTCTTCCGATCTNNNGGCACTAAGCTGTTGCCATTT
AGCACACGTCTGAACTCTTTCCCTACACGACGCTCTTCCGATCTNNNGGCACTAAGCTGTTGCCATTT
GAGCACACGTCTGAACTCTTTCCCTACACGACGCTCTTCCGATCTNNNACCACAGGAAAAACGGAGTA
GAGCACACGTCTGAACTCTTTCCCTACACGACGCTCTTCCGATCTNNNACCACAGGAAAAACGGAGTA
GCACACGTCTGAACTCTTTCCCTACACGACGCTCTTCCGATCTNNNATCCCATTTTTTTGTGGTAATA
GCACACGTCTGAACTCTTTCCCTACACGACGCTCTTCCGATCTNNNATCCCATTTTTTTGTGGTAATA
GAGCACACGTCTGAACTCTTTCCCTACACGACGCTCTTCCGATCTNNNACATCAAGATAGCCACAGG
GAGCACACGTCTGAACTCTTTCCCTACACGACGCTCTTCCGATCTNNNACATCAAGATAGCCACAGG
AGCACACGTCTGAACTCTTTCCCTACACGACGCTCTTCCGATCTNNNCACCTGTAATCCCAGCTACTCA
AGCACACGTCTGAACTCTTTCCCTACACGACGCTCTTCCGATCTNNNCACCTGTAATCCCAGCTACTCA
GCACACGTCTGAACTCTTTCCCTACACGACGCTCTTCCGATCTNNNTGTAATCCTAGCACTTTGGGAG
GCACACGTCTGAACTCTTTCCCTACACGACGCTCTTCCGATCTNNNTGTAATCCTAGCACTTTGGGAG
GAGCACACGTCTGAACTCTTTCCCTACACGACGCTCTTCCGATCTNNNTATTCATCTGCTCTCCCTTTT
GAGCACACGTCTGAACTCTTTCCCTACACGACGCTCTTCCGATCTNNNTATTCATCTGCTCTCCCTTTT
AGCACACGTCTGAACTCTTTCCCTACACGACGCTCTTCCGATCTNNNATCTTAGCTGGGGTGATGTTG
AGCACACGTCTGAACTCTTTCCCTACACGACGCTCTTCCGATCTNNNATCTTAGCTGGGGTGATGTTG
AGCACACGTCTGAACTCTTTCCCTACACGACGCTCTTCCGATCTNNNACCTCCACCTTCCTGAACACAT
AGCACACGTCTGAACTCTTTCCCTACACGACGCTCTTCCGATCTNNNACCTCCACCTTCCTGAACACAT
AGCACACGTCTGAACTCTTTCCCTACACGACGCTCTTCCGATCTNNNTTTATGACAGAACCAATTTCA
AGCACACGTCTGAACTCTTTCCCTACACGACGCTCTTCCGATCTNNNTTTATGACAGAACCAATTTCA
GCACACGTCTGAACTCTTTCCCTACACGACGCTCTTCCGATCTNNNCACTGAATGTCTGCTGCCTATTG
GCACACGTCTGAACTCTTTCCCTACACGACGCTCTTCCGATCTNNNCACTGAATGTCTGCTGCCTATTG
AGCACACGTCTGAACTCTTTCCCTACACGACGCTCTTCCGATCTNNNTATAGTCCCAGCTACTCAGGA
AGCACACGTCTGAACTCTTTCCCTACACGACGCTCTTCCGATCTNNNTATAGTCCCAGCTACTCAGGA
GAGCACACGTCTGAACTCTTTCCCTACACGACGCTCTTCCGATCTNNNTCTTCTCTCCTAGGACCTTGTC
GAGCACACGTCTGAACTCTTTCCCTACACGACGCTCTTCCGATCTNNNTCTTCTCTCCTAGGACCTTGTC
AGCACACGTCTGAACTCTTTCCCTACACGACGCTCTTCCGATCTNNNAATTTCTGACCCCACACCTTCTT
AGCACACGTCTGAACTCTTTCCCTACACGACGCTCTTCCGATCTNNNAATTTCTGACCCCACACCTTCTT
AGCACACGTCTGAACTCTTTCCCTACACGACGCTCTTCCGATCTNNNCTGTAGTTCCAGCTATTCGGG
AGCACACGTCTGAACTCTTTCCCTACACGACGCTCTTCCGATCTNNNCTGTAGTTCCAGCTATTCGGG
GAGCACACGTCTGAACTCTTTCCCTACACGACGCTCTTCCGATCTNNNCATGAGGTCGAGGCTGTAG
GAGCACACGTCTGAACTCTTTCCCTACACGACGCTCTTCCGATCTNNNCATGAGGTCGAGGCTGTAG
GCACACGTCTGAACTCTTTCCCTACACGACGCTCTTCCGATCTNNNCTCCTGGCCTTGACCTTCAGCATC
GCACACGTCTGAACTCTTTCCCTACACGACGCTCTTCCGATCTNNNCTCCTGGCCTTGACCTTCAGCATC

AGCACACGTCTGAACTCTTTCCCTACACGACGCTCTTCCGATCTNNNTAGCCAAGGTCTAACCCCAAAT
AGCACACGTCTGAACTCTTTCCCTACACGACGCTCTTCCGATCTNNNTAGCCAAGGTCTAACCCCAAAT
AGCACACGTCTGAACTCTTTCCCTACACGACGCTCTTCCGATCTNNNTGCCACTGATAGGAACAATGTG
AGCACACGTCTGAACTCTTTCCCTACACGACGCTCTTCCGATCTNNNTGCCACTGATAGGAACAATGTG
AGCACACGTCTGAACTCTTTCCCTACACGACGCTCTTCCGATCTNNNTCAATTCGTGGACCAGATGATC
AGCACACGTCTGAACTCTTTCCCTACACGACGCTCTTCCGATCTNNNTCAATTCGTGGACCAGATGATC
AGCACACGTCTGAACTCTTTCCCTACACGACGCTCTTCCGATCTNNNCTGAGGCCAGAGAATTGCTTGA
AGCACACGTCTGAACTCTTTCCCTACACGACGCTCTTCCGATCTNNNCTGAGGCCAGAGAATTGCTTGA
AGCACACGTCTGAACTCTTTCCCTACACGACGCTCTTCCGATCTNNNGAACCTCAAAGGCCCTTATCCTC
AGCACACGTCTGAACTCTTTCCCTACACGACGCTCTTCCGATCTNNNGAACCTCAAAGGCCCTTATCCTC
GAGCACACGTCTGAACTCTTTCCCTACACGACGCTCTTCCGATCTNNNCAGAAAGGTGGGAAGACTTG
GAGCACACGTCTGAACTCTTTCCCTACACGACGCTCTTCCGATCTNNNCAGAAAGGTGGGAAGACTTG
AGCACACGTCTGAACTCTTTCCCTACACGACGCTCTTCCGATCTNNNGAACTCTAACCTGGGCAAGAA
AGCACACGTCTGAACTCTTTCCCTACACGACGCTCTTCCGATCTNNNGAACTCTAACCTGGGCAAGAA
AGAGCACACGTCTGAACTCTTTCCCTACACGACGCTCTTCCGATCTNNNAGAATCAGTAGCCCAAGAGC
AGAGCACACGTCTGAACTCTTTCCCTACACGACGCTCTTCCGATCTNNNAGAATCAGTAGCCCAAGAGC
AGCACACGTCTGAACTCTTTCCCTACACGACGCTCTTCCGATCTNNNTCTGTCCCATCTTTGCATTTGTT
AGCACACGTCTGAACTCTTTCCCTACACGACGCTCTTCCGATCTNNNTCTGTCCCATCTTTGCATTTGTT
AGCACACGTCTGAACTCTTTCCCTACACGACGCTCTTCCGATCTNNNTTTACTGTCACACATGGGCACAA
AGCACACGTCTGAACTCTTTCCCTACACGACGCTCTTCCGATCTNNNTTTACTGTCACACATGGGCACAA
AGCACACGTCTGAACTCTTTCCCTACACGACGCTCTTCCGATCTNNNGGCTTCAGTTTGTGTTCGTAGT
AGCACACGTCTGAACTCTTTCCCTACACGACGCTCTTCCGATCTNNNGGCTTCAGTTTGTGTTCGTAGT
AGCACACGTCTGAACTCTTTCCCTACACGACGCTCTTCCGATCTNNNCAAGCTACATCTCCTAGCCTAG
AGCACACGTCTGAACTCTTTCCCTACACGACGCTCTTCCGATCTNNNCAAGCTACATCTCCTAGCCTAG
AGCACACGTCTGAACTCTTTCCCTACACGACGCTCTTCCGATCTNNNAGGTTGCAGTGAGCTAAAATC
AGCACACGTCTGAACTCTTTCCCTACACGACGCTCTTCCGATCTNNNAGGTTGCAGTGAGCTAAAATC
GCACACGTCTGAACTCTTTCCCTACACGACGCTCTTCCGATCTNNNATAAATCATCTAAGCCATCCTCCA
GCACACGTCTGAACTCTTTCCCTACACGACGCTCTTCCGATCTNNNATAAATCATCTAAGCCATCCTCCA
AGCACACGTCTGAACTCTTTCCCTACACGACGCTCTTCCGATCTNNNTCCTCTTCCTAATAAGCCTGGGA
AGCACACGTCTGAACTCTTTCCCTACACGACGCTCTTCCGATCTNNNTCCTCTTCCTAATAAGCCTGGGA
GCACACGTCTGAACTCTTTCCCTACACGACGCTCTTCCGATCTNNNATCAAAAAGCCCAGGCAAAGATG
GCACACGTCTGAACTCTTTCCCTACACGACGCTCTTCCGATCTNNNATCAAAAAGCCCAGGCAAAGATG
AGCACACGTCTGAACTCTTTCCCTACACGACGCTCTTCCGATCTNNNTTCCCTGAAGCGAGTACACATTG
AGCACACGTCTGAACTCTTTCCCTACACGACGCTCTTCCGATCTNNNTTCCCTGAAGCGAGTACACATTG
GAGCACACGTCTGAACTCTTTCCCTACACGACGCTCTTCCGATCTNNNCTTTCGAGGAGGAAGATGGC
GAGCACACGTCTGAACTCTTTCCCTACACGACGCTCTTCCGATCTNNNCTTTCGAGGAGGAAGATGGC
GAGCACACGTCTGAACTCTTTCCCTACACGACGCTCTTCCGATCTNNNTTGACAGTGTGTTCACAGAGA
GAGCACACGTCTGAACTCTTTCCCTACACGACGCTCTTCCGATCTNNNTTGACAGTGTGTTCACAGAGA
AGCACACGTCTGAACTCTTTCCCTACACGACGCTCTTCCGATCTNNNCTGTAATCCCAGCTACTCAGGAC
AGCACACGTCTGAACTCTTTCCCTACACGACGCTCTTCCGATCTNNNCTGTAATCCCAGCTACTCAGGAC
GAGCACACGTCTGAACTCTTTCCCTACACGACGCTCTTCCGATCTNNNATAGTGGACTGGCTTATGCTC
GAGCACACGTCTGAACTCTTTCCCTACACGACGCTCTTCCGATCTNNNATAGTGGACTGGCTTATGCTC
AGCACACGTCTGAACTCTTTCCCTACACGACGCTCTTCCGATCTNNNTAGTTGTGATGCTATGGTGCTC

AGCACACGTCTGAACTCTTTCCCTACACGACGCTCTTCCGATCTNNNTAGTTGTGATGCTATGGTGCT
GAGCACACGTCTGAACTCTTTCCCTACACGACGCTCTTCCGATCTNNNAGAGAGATGCAGTGATTTG
GAGCACACGTCTGAACTCTTTCCCTACACGACGCTCTTCCGATCTNNNAGAGAGATGCAGTGATTTG
GAGCACACGTCTGAACTCTTTCCCTACACGACGCTCTTCCGATCTNNNAGGGGATAGATGAGATGAC
GAGCACACGTCTGAACTCTTTCCCTACACGACGCTCTTCCGATCTNNNAGGGGATAGATGAGATGAC
AGCACACGTCTGAACTCTTTCCCTACACGACGCTCTTCCGATCTNNNGGCTCATGATATGGCTACAGCT
AGCACACGTCTGAACTCTTTCCCTACACGACGCTCTTCCGATCTNNNGGCTCATGATATGGCTACAGCT
GCACACGTCTGAACTCTTTCCCTACACGACGCTCTTCCGATCTNNNAGGTTATTGACAGGCAATGGAC
GCACACGTCTGAACTCTTTCCCTACACGACGCTCTTCCGATCTNNNAGGTTATTGACAGGCAATGGAC
AGCACACGTCTGAACTCTTTCCCTACACGACGCTCTTCCGATCTNNNATGTCACAAAAAAAAGGCCTTG
AGCACACGTCTGAACTCTTTCCCTACACGACGCTCTTCCGATCTNNNATGTCACAAAAAAAAGGCCTTG
GAGCACACGTCTGAACTCTTTCCCTACACGACGCTCTTCCGATCTNNNACTCTAGCCTGGGTGATAAGA
GAGCACACGTCTGAACTCTTTCCCTACACGACGCTCTTCCGATCTNNNACTCTAGCCTGGGTGATAAGA
GCACACGTCTGAACTCTTTCCCTACACGACGCTCTTCCGATCTNNNTTGTTGTTGTTGTTGAGACGGA
GCACACGTCTGAACTCTTTCCCTACACGACGCTCTTCCGATCTNNNTTGTTGTTGTTGTTGAGACGGA
AGCACACGTCTGAACTCTTTCCCTACACGACGCTCTTCCGATCTNNNAGTAAGCTGAGATCACCAC
AGCACACGTCTGAACTCTTTCCCTACACGACGCTCTTCCGATCTNNNAGTAAGCTGAGATCACCAC
AGCACACGTCTGAACTCTTTCCCTACACGACGCTCTTCCGATCTNNNAGACAGGAGAATCACTTGAAC
AGCACACGTCTGAACTCTTTCCCTACACGACGCTCTTCCGATCTNNNAGACAGGAGAATCACTTGAAC
AGCACACGTCTGAACTCTTTCCCTACACGACGCTCTTCCGATCTNNNACCTACCACCTAGATTTCACGA
AGCACACGTCTGAACTCTTTCCCTACACGACGCTCTTCCGATCTNNNACCTACCACCTAGATTTCACGA
AGCACACGTCTGAACTCTTTCCCTACACGACGCTCTTCCGATCTNNNATGGTCCCCATAGCACATATTC
AGCACACGTCTGAACTCTTTCCCTACACGACGCTCTTCCGATCTNNNATGGTCCCCATAGCACATATTC
GCACACGTCTGAACTCTTTCCCTACACGACGCTCTTCCGATCTNNNTCCACACCTTCTCCAAAATCTGTC
GCACACGTCTGAACTCTTTCCCTACACGACGCTCTTCCGATCTNNNTCCACACCTTCTCCAAAATCTGTC
AGCACACGTCTGAACTCTTTCCCTACACGACGCTCTTCCGATCTNNNTTCAGCAGTTTTCACCACGAAT
AGCACACGTCTGAACTCTTTCCCTACACGACGCTCTTCCGATCTNNNTTCAGCAGTTTTCACCACGAAT
AGCACACGTCTGAACTCTTTCCCTACACGACGCTCTTCCGATCTNNNAGAATTTCCTTTCAGGCAGGCT
AGCACACGTCTGAACTCTTTCCCTACACGACGCTCTTCCGATCTNNNAGAATTTCCTTTCAGGCAGGCT
GAGCACACGTCTGAACTCTTTCCCTACACGACGCTCTTCCGATCTNNNGAGATCATGCCACTACACTCC
GAGCACACGTCTGAACTCTTTCCCTACACGACGCTCTTCCGATCTNNNGAGATCATGCCACTACACTCC
AGCACACGTCTGAACTCTTTCCCTACACGACGCTCTTCCGATCTNNNTGCAATGTGGAGTAAGCCTAA
AGCACACGTCTGAACTCTTTCCCTACACGACGCTCTTCCGATCTNNNTGCAATGTGGAGTAAGCCTAA
GCACACGTCTGAACTCTTTCCCTACACGACGCTCTTCCGATCTNNNTACCAGCGATTTCTCTCACCATCC
GCACACGTCTGAACTCTTTCCCTACACGACGCTCTTCCGATCTNNNTACCAGCGATTTCTCTCACCATCC
AGCACACGTCTGAACTCTTTCCCTACACGACGCTCTTCCGATCTNNNAAAGCTATTTTGTGATTGTTTT
AGCACACGTCTGAACTCTTTCCCTACACGACGCTCTTCCGATCTNNNAAAGCTATTTTGTGATTGTTTT
AGAGCACACGTCTGAACTCTTTCCCTACACGACGCTCTTCCGATCTNNNTAGTCCTAGCTACTTGGAA
AGAGCACACGTCTGAACTCTTTCCCTACACGACGCTCTTCCGATCTNNNTAGTCCTAGCTACTTGGAA
AGCACACGTCTGAACTCTTTCCCTACACGACGCTCTTCCGATCTNNNATTCCATCCCCTCTGTCACTCCC
AGCACACGTCTGAACTCTTTCCCTACACGACGCTCTTCCGATCTNNNATTCCATCCCCTCTGTCACTCCC
AGAGCACACGTCTGAACTCTTTCCCTACACGACGCTCTTCCGATCTNNNTCAGTTCTCCTGTTTCCTTG
AGAGCACACGTCTGAACTCTTTCCCTACACGACGCTCTTCCGATCTNNNTCAGTTCTCCTGTTTCCTTG

AGCACACGTCTGAACTCTTTCCCTACACGACGCTCTTCCGATCTNNNCCCATTTACCAGGAGCTTCTCA
AGCACACGTCTGAACTCTTTCCCTACACGACGCTCTTCCGATCTNNNCCCATTTACCAGGAGCTTCTCA
AGCACACGTCTGAACTCTTTCCCTACACGACGCTCTTCCGATCTNNNGCCAGGTGTAGAAGGCTTCAT
AGCACACGTCTGAACTCTTTCCCTACACGACGCTCTTCCGATCTNNNGCCAGGTGTAGAAGGCTTCAT
AGCACACGTCTGAACTCTTTCCCTACACGACGCTCTTCCGATCTNNNGGAGATTGAGACAACCTGGCT
AGCACACGTCTGAACTCTTTCCCTACACGACGCTCTTCCGATCTNNNGGAGATTGAGACAACCTGGCT
GCACACGTCTGAACTCTTTCCCTACACGACGCTCTTCCGATCTNNNTATTAGCATCTTGGTGGGAGGG
GCACACGTCTGAACTCTTTCCCTACACGACGCTCTTCCGATCTNNNTATTAGCATCTTGGTGGGAGGG
GCACACGTCTGAACTCTTTCCCTACACGACGCTCTTCCGATCTNNNGATTGGCCAACTGAAGAACTGT
GCACACGTCTGAACTCTTTCCCTACACGACGCTCTTCCGATCTNNNGATTGGCCAACTGAAGAACTGT
ACACGTCTGAACTCTTTCCCTACACGACGCTCTTCCGATCTNNNCTCATTACGTTTAGATGGGATTCA
ACACGTCTGAACTCTTTCCCTACACGACGCTCTTCCGATCTNNNCTCATTACGTTTAGATGGGATTCA
AGCACACGTCTGAACTCTTTCCCTACACGACGCTCTTCCGATCTNNNTGTAGTCCTAGTTACTCGGGA
AGCACACGTCTGAACTCTTTCCCTACACGACGCTCTTCCGATCTNNNTGTAGTCCTAGTTACTCGGGA
AGCACACGTCTGAACTCTTTCCCTACACGACGCTCTTCCGATCTNNNTGTAGTCCTAGTTACTCGGGA
AGCACACGTCTGAACTCTTTCCCTACACGACGCTCTTCCGATCTNNNTGTAGTCCTAGTTACTCGGGA
AGCACACGTCTGAACTCTTTCCCTACACGACGCTCTTCCGATCTNNNTGTAGTCCTAGTTACTCGGGA
GCACACGTCTGAACTCTTTCCCTACACGACGCTCTTCCGATCTNNNGTCTCAAACTCCTGACCTCATGA
GCACACGTCTGAACTCTTTCCCTACACGACGCTCTTCCGATCTNNNGTCTCAAACTCCTGACCTCATGA
GCACACGTCTGAACTCTTTCCCTACACGACGCTCTTCCGATCTNNNGTCTCAAACTCCTGACCTCATGA
GCACACGTCTGAACTCTTTCCCTACACGACGCTCTTCCGATCTNNNGTCTCAAACTCCTGACCTCATGA
GCACACGTCTGAACTCTTTCCCTACACGACGCTCTTCCGATCTNNNGTCTCAAACTCCTGACCTCATGA
AGCACACGTCTGAACTCTTTCCCTACACGACGCTCTTCCGATCTNNNTCTTGACTCACTGCAACCTCTA
AGCACACGTCTGAACTCTTTCCCTACACGACGCTCTTCCGATCTNNNTCTTGACTCACTGCAACCTCTA
AGCACACGTCTGAACTCTTTCCCTACACGACGCTCTTCCGATCTNNNTCTTGACTCACTGCAACCTCTA
AGCACACGTCTGAACTCTTTCCCTACACGACGCTCTTCCGATCTNNNTCTTGACTCACTGCAACCTCTA
AGCACACGTCTGAACTCTTTCCCTACACGACGCTCTTCCGATCTNNNTCTTGACTCACTGCAACCTCTA
AGCACACGTCTGAACTCTTTCCCTACACGACGCTCTTCCGATCTNNNATTTCTGCAGCCAGCTTGAAT
AGCACACGTCTGAACTCTTTCCCTACACGACGCTCTTCCGATCTNNNATTTCTGCAGCCAGCTTGAAT
AGCACACGTCTGAACTCTTTCCCTACACGACGCTCTTCCGATCTNNNATTTCTGCAGCCAGCTTGAAT
AGCACACGTCTGAACTCTTTCCCTACACGACGCTCTTCCGATCTNNNATTTCTGCAGCCAGCTTGAAT
AGCACACGTCTGAACTCTTTCCCTACACGACGCTCTTCCGATCTNNNATTTCTGCAGCCAGCTTGAAT
AGCACACGTCTGAACTCTTTCCCTACACGACGCTCTTCCGATCTNNNCCTATCCTCAGTTTTCCTGCCTT
AGCACACGTCTGAACTCTTTCCCTACACGACGCTCTTCCGATCTNNNCCTATCCTCAGTTTTCCTGCCTT
AGCACACGTCTGAACTCTTTCCCTACACGACGCTCTTCCGATCTNNNCCTATCCTCAGTTTTCCTGCCTT
AGCACACGTCTGAACTCTTTCCCTACACGACGCTCTTCCGATCTNNNCCTATCCTCAGTTTTCCTGCCTT
AGCACACGTCTGAACTCTTTCCCTACACGACGCTCTTCCGATCTNNNCCTATCCTCAGTTTTCCTGCCTT
AGCACACGTCTGAACTCTTTCCCTACACGACGCTCTTCCGATCTNNNTGAGGCAGGAGAATCACTTGT
AGCACACGTCTGAACTCTTTCCCTACACGACGCTCTTCCGATCTNNNTGAGGCAGGAGAATCACTTGT
AGCACACGTCTGAACTCTTTCCCTACACGACGCTCTTCCGATCTNNNTGAGGCAGGAGAATCACTTGT
AGCACACGTCTGAACTCTTTCCCTACACGACGCTCTTCCGATCTNNNTGAGGCAGGAGAATCACTTGT
AGCACACGTCTGAACTCTTTCCCTACACGACGCTCTTCCGATCTNNNTGAGGCAGGAGAATCACTTGT
GAGCACACGTCTGAACTCTTTCCCTACACGACGCTCTTCCGATCTNNNGGTGCCTGTAATCCCAGCTA

GAGCACACGTCTGAACTCTTTCCCTACACGACGCTCTTCCGATCTNNNGGTGCCTGTAATCCCAGCTA
GAGCACACGTCTGAACTCTTTCCCTACACGACGCTCTTCCGATCTNNNGGTGCCTGTAATCCCAGCTA
GAGCACACGTCTGAACTCTTTCCCTACACGACGCTCTTCCGATCTNNNGGTGCCTGTAATCCCAGCTA
GAGCACACGTCTGAACTCTTTCCCTACACGACGCTCTTCCGATCTNNNGGTGCCTGTAATCCCAGCTA
GCACACGTCTGAACTCTTTCCCTACACGACGCTCTTCCGATCTNNNAGAAACTCCTACAATACAGGGC
GCACACGTCTGAACTCTTTCCCTACACGACGCTCTTCCGATCTNNNAGAAACTCCTACAATACAGGGC
GCACACGTCTGAACTCTTTCCCTACACGACGCTCTTCCGATCTNNNAGAAACTCCTACAATACAGGGC
GCACACGTCTGAACTCTTTCCCTACACGACGCTCTTCCGATCTNNNAGAAACTCCTACAATACAGGGC
GCACACGTCTGAACTCTTTCCCTACACGACGCTCTTCCGATCTNNNAGAAACTCCTACAATACAGGGC
AGCACACGTCTGAACTCTTTCCCTACACGACGCTCTTCCGATCTNNNTGTAGTCCCAGCTATTCAGGA
AGCACACGTCTGAACTCTTTCCCTACACGACGCTCTTCCGATCTNNNTGTAGTCCCAGCTATTCAGGA
AGCACACGTCTGAACTCTTTCCCTACACGACGCTCTTCCGATCTNNNTGTAGTCCCAGCTATTCAGGA
AGCACACGTCTGAACTCTTTCCCTACACGACGCTCTTCCGATCTNNNTGTAGTCCCAGCTATTCAGGA
AGCACACGTCTGAACTCTTTCCCTACACGACGCTCTTCCGATCTNNNTGTAGTCCCAGCTATTCAGGA
GAGCACACGTCTGAACTCTTTCCCTACACGACGCTCTTCCGATCTNNNTACTAGGGTGACTGAGGTG
GAGCACACGTCTGAACTCTTTCCCTACACGACGCTCTTCCGATCTNNNTACTAGGGTGACTGAGGTG
GAGCACACGTCTGAACTCTTTCCCTACACGACGCTCTTCCGATCTNNNTACTAGGGTGACTGAGGTG
GAGCACACGTCTGAACTCTTTCCCTACACGACGCTCTTCCGATCTNNNTACTAGGGTGACTGAGGTG
GAGCACACGTCTGAACTCTTTCCCTACACGACGCTCTTCCGATCTNNNTACTAGGGTGACTGAGGTG
AGCACACGTCTGAACTCTTTCCCTACACGACGCTCTTCCGATCTNNNCGCAACAGAAGTCTAAATCCTG
AGCACACGTCTGAACTCTTTCCCTACACGACGCTCTTCCGATCTNNNCGCAACAGAAGTCTAAATCCTG
AGCACACGTCTGAACTCTTTCCCTACACGACGCTCTTCCGATCTNNNCGCAACAGAAGTCTAAATCCTG
AGCACACGTCTGAACTCTTTCCCTACACGACGCTCTTCCGATCTNNNCGCAACAGAAGTCTAAATCCTG
AGCACACGTCTGAACTCTTTCCCTACACGACGCTCTTCCGATCTNNNCGCAACAGAAGTCTAAATCCTG
AGCACACGTCTGAACTCTTTCCCTACACGACGCTCTTCCGATCTNNNGGTCATTGCACTCCAAACAGA
AGCACACGTCTGAACTCTTTCCCTACACGACGCTCTTCCGATCTNNNGGTCATTGCACTCCAAACAGA
AGCACACGTCTGAACTCTTTCCCTACACGACGCTCTTCCGATCTNNNGGTCATTGCACTCCAAACAGA
AGCACACGTCTGAACTCTTTCCCTACACGACGCTCTTCCGATCTNNNGGTCATTGCACTCCAAACAGA
AGCACACGTCTGAACTCTTTCCCTACACGACGCTCTTCCGATCTNNNGGTCATTGCACTCCAAACAGA
AGCACACGTCTGAACTCTTTCCCTACACGACGCTCTTCCGATCTNNNATGCAACATCCAAAGCCACTTTT
AGCACACGTCTGAACTCTTTCCCTACACGACGCTCTTCCGATCTNNNATGCAACATCCAAAGCCACTTTT
AGCACACGTCTGAACTCTTTCCCTACACGACGCTCTTCCGATCTNNNATGCAACATCCAAAGCCACTTTT
AGCACACGTCTGAACTCTTTCCCTACACGACGCTCTTCCGATCTNNNATGCAACATCCAAAGCCACTTTT
AGCACACGTCTGAACTCTTTCCCTACACGACGCTCTTCCGATCTNNNATGCAACATCCAAAGCCACTTTT
GCACACGTCTGAACTCTTTCCCTACACGACGCTCTTCCGATCTNNNATATCCCCAAGAACTTGCACAGT
GCACACGTCTGAACTCTTTCCCTACACGACGCTCTTCCGATCTNNNATATCCCCAAGAACTTGCACAGT
GCACACGTCTGAACTCTTTCCCTACACGACGCTCTTCCGATCTNNNATATCCCCAAGAACTTGCACAGT
GCACACGTCTGAACTCTTTCCCTACACGACGCTCTTCCGATCTNNNATATCCCCAAGAACTTGCACAGT
GCACACGTCTGAACTCTTTCCCTACACGACGCTCTTCCGATCTNNNATATCCCCAAGAACTTGCACAGT
AGCACACGTCTGAACTCTTTCCCTACACGACGCTCTTCCGATCTNNNCTGGCTTACACTTGGAGATAG
AGCACACGTCTGAACTCTTTCCCTACACGACGCTCTTCCGATCTNNNCTGGCTTACACTTGGAGATAG
AGCACACGTCTGAACTCTTTCCCTACACGACGCTCTTCCGATCTNNNCTGGCTTACACTTGGAGATAG
AGCACACGTCTGAACTCTTTCCCTACACGACGCTCTTCCGATCTNNNCTGGCTTACACTTGGAGATAG

AGCACACGTCTGAACTCTTTCCCTACACGACGCTCTTCCGATCTNNNCTGGCTTACACTTGGAGATAG
AGCACACGTCTGAACTCTTTCCCTACACGACGCTCTTCCGATCTNNNTGAGAAGGAGGAAAGGGAA
AGCACACGTCTGAACTCTTTCCCTACACGACGCTCTTCCGATCTNNNTGAGAAGGAGGAAAGGGAA
AGCACACGTCTGAACTCTTTCCCTACACGACGCTCTTCCGATCTNNNTGAGAAGGAGGAAAGGGAA
AGCACACGTCTGAACTCTTTCCCTACACGACGCTCTTCCGATCTNNNTGAGAAGGAGGAAAGGGAA
AGCACACGTCTGAACTCTTTCCCTACACGACGCTCTTCCGATCTNNNTGAGAAGGAGGAAAGGGAA
AGCACACGTCTGAACTCTTTCCCTACACGACGCTCTTCCGATCTNNNTTCAAGTGATTCTCCTGCCTCAG
AGCACACGTCTGAACTCTTTCCCTACACGACGCTCTTCCGATCTNNNTTCAAGTGATTCTCCTGCCTCAG
AGCACACGTCTGAACTCTTTCCCTACACGACGCTCTTCCGATCTNNNTTCAAGTGATTCTCCTGCCTCAG
AGCACACGTCTGAACTCTTTCCCTACACGACGCTCTTCCGATCTNNNTTCAAGTGATTCTCCTGCCTCAG
AGCACACGTCTGAACTCTTTCCCTACACGACGCTCTTCCGATCTNNNTTCAAGTGATTCTCCTGCCTCAG
AGCACACGTCTGAACTCTTTCCCTACACGACGCTCTTCCGATCTNNNCACACACACAAAGACACTCAGA
AGCACACGTCTGAACTCTTTCCCTACACGACGCTCTTCCGATCTNNNCACACACACAAAGACACTCAGA
AGCACACGTCTGAACTCTTTCCCTACACGACGCTCTTCCGATCTNNNCACACACACAAAGACACTCAGA
AGCACACGTCTGAACTCTTTCCCTACACGACGCTCTTCCGATCTNNNCACACACACAAAGACACTCAGA
AGCACACGTCTGAACTCTTTCCCTACACGACGCTCTTCCGATCTNNNCACACACACAAAGACACTCAGA
GAGCACACGTCTGAACTCTTTCCCTACACGACGCTCTTCCGATCTNNNCATGACCACTTTCCACTGTTT
GAGCACACGTCTGAACTCTTTCCCTACACGACGCTCTTCCGATCTNNNCATGACCACTTTCCACTGTTT
GAGCACACGTCTGAACTCTTTCCCTACACGACGCTCTTCCGATCTNNNCATGACCACTTTCCACTGTTT
GAGCACACGTCTGAACTCTTTCCCTACACGACGCTCTTCCGATCTNNNCATGACCACTTTCCACTGTTT
GAGCACACGTCTGAACTCTTTCCCTACACGACGCTCTTCCGATCTNNNCATGACCACTTTCCACTGTTT
AGAGCACACGTCTGAACTCTTTCCCTACACGACGCTCTTCCGATCTNNNACAAAATTAGTTGGGCATG
AGAGCACACGTCTGAACTCTTTCCCTACACGACGCTCTTCCGATCTNNNACAAAATTAGTTGGGCATG
AGAGCACACGTCTGAACTCTTTCCCTACACGACGCTCTTCCGATCTNNNACAAAATTAGTTGGGCATG
AGAGCACACGTCTGAACTCTTTCCCTACACGACGCTCTTCCGATCTNNNACAAAATTAGTTGGGCATG
AGAGCACACGTCTGAACTCTTTCCCTACACGACGCTCTTCCGATCTNNNACAAAATTAGTTGGGCATG
AGCACACGTCTGAACTCTTTCCCTACACGACGCTCTTCCGATCTNNNCACTTGAGGTCAGGAGTTCAA
AGCACACGTCTGAACTCTTTCCCTACACGACGCTCTTCCGATCTNNNCACTTGAGGTCAGGAGTTCAA
AGCACACGTCTGAACTCTTTCCCTACACGACGCTCTTCCGATCTNNNCACTTGAGGTCAGGAGTTCAA
AGCACACGTCTGAACTCTTTCCCTACACGACGCTCTTCCGATCTNNNCACTTGAGGTCAGGAGTTCAA
AGCACACGTCTGAACTCTTTCCCTACACGACGCTCTTCCGATCTNNNCACTTGAGGTCAGGAGTTCAA
GAGCACACGTCTGAACTCTTTCCCTACACGACGCTCTTCCGATCTNNNACTGCATGTCAGAATTCATCCC
GAGCACACGTCTGAACTCTTTCCCTACACGACGCTCTTCCGATCTNNNACTGCATGTCAGAATTCATCCC
GAGCACACGTCTGAACTCTTTCCCTACACGACGCTCTTCCGATCTNNNACTGCATGTCAGAATTCATCCC
GAGCACACGTCTGAACTCTTTCCCTACACGACGCTCTTCCGATCTNNNACTGCATGTCAGAATTCATCCC
GAGCACACGTCTGAACTCTTTCCCTACACGACGCTCTTCCGATCTNNNACTGCATGTCAGAATTCATCCC
AGCACACGTCTGAACTCTTTCCCTACACGACGCTCTTCCGATCTNNNAAATAACTTCTCAGGCTGGGTG
AGCACACGTCTGAACTCTTTCCCTACACGACGCTCTTCCGATCTNNNAAATAACTTCTCAGGCTGGGTG
AGCACACGTCTGAACTCTTTCCCTACACGACGCTCTTCCGATCTNNNAAATAACTTCTCAGGCTGGGTG
AGCACACGTCTGAACTCTTTCCCTACACGACGCTCTTCCGATCTNNNAAATAACTTCTCAGGCTGGGTG
AGCACACGTCTGAACTCTTTCCCTACACGACGCTCTTCCGATCTNNNAAATAACTTCTCAGGCTGGGTG
GCACACGTCTGAACTCTTTCCCTACACGACGCTCTTCCGATCTNNNTGTCTATCATCTATCCCTGTCAAC
GCACACGTCTGAACTCTTTCCCTACACGACGCTCTTCCGATCTNNNTGTCTATCATCTATCCCTGTCAAC

GCACACGTCTGAACTCTTTCCCTACACGACGCTCTTCCGATCTNNNTGTCTATCATCTATCCCTGTCAAC
GCACACGTCTGAACTCTTTCCCTACACGACGCTCTTCCGATCTNNNTGTCTATCATCTATCCCTGTCAAC
GCACACGTCTGAACTCTTTCCCTACACGACGCTCTTCCGATCTNNNTGTCTATCATCTATCCCTGTCAAC
GAGCACACGTCTGAACTCTTTCCCTACACGACGCTCTTCCGATCTNNNGCTGCTCTCAAACTCCTGAGTC
GAGCACACGTCTGAACTCTTTCCCTACACGACGCTCTTCCGATCTNNNGCTGCTCTCAAACTCCTGAGTC
GAGCACACGTCTGAACTCTTTCCCTACACGACGCTCTTCCGATCTNNNGCTGCTCTCAAACTCCTGAGTC
GAGCACACGTCTGAACTCTTTCCCTACACGACGCTCTTCCGATCTNNNGCTGCTCTCAAACTCCTGAGTC
GAGCACACGTCTGAACTCTTTCCCTACACGACGCTCTTCCGATCTNNNGCTGCTCTCAAACTCCTGAGTC
GAGCACACGTCTGAACTCTTTCCCTACACGACGCTCTTCCGATCTNNNGAAATAAAAGGTCAGGGCTCA
GAGCACACGTCTGAACTCTTTCCCTACACGACGCTCTTCCGATCTNNNGAAATAAAAGGTCAGGGCTCA
GAGCACACGTCTGAACTCTTTCCCTACACGACGCTCTTCCGATCTNNNGAAATAAAAGGTCAGGGCTCA
GAGCACACGTCTGAACTCTTTCCCTACACGACGCTCTTCCGATCTNNNGAAATAAAAGGTCAGGGCTCA
GAGCACACGTCTGAACTCTTTCCCTACACGACGCTCTTCCGATCTNNNGAAATAAAAGGTCAGGGCTCA
GAGCACACGTCTGAACTCTTTCCCTACACGACGCTCTTCCGATCTNNNCAATCTCAGCTCACTGCAAGCC
GAGCACACGTCTGAACTCTTTCCCTACACGACGCTCTTCCGATCTNNNCAATCTCAGCTCACTGCAAGCC
GAGCACACGTCTGAACTCTTTCCCTACACGACGCTCTTCCGATCTNNNCAATCTCAGCTCACTGCAAGCC
GAGCACACGTCTGAACTCTTTCCCTACACGACGCTCTTCCGATCTNNNCAATCTCAGCTCACTGCAAGCC
GAGCACACGTCTGAACTCTTTCCCTACACGACGCTCTTCCGATCTNNNCAATCTCAGCTCACTGCAAGCC
GAGCACACGTCTGAACTCTTTCCCTACACGACGCTCTTCCGATCTNNNGAGGCAGGAAAATCACTTGA
GAGCACACGTCTGAACTCTTTCCCTACACGACGCTCTTCCGATCTNNNGAGGCAGGAAAATCACTTGA
GAGCACACGTCTGAACTCTTTCCCTACACGACGCTCTTCCGATCTNNNGAGGCAGGAAAATCACTTGA
GAGCACACGTCTGAACTCTTTCCCTACACGACGCTCTTCCGATCTNNNGAGGCAGGAAAATCACTTGA
GAGCACACGTCTGAACTCTTTCCCTACACGACGCTCTTCCGATCTNNNGAGGCAGGAAAATCACTTGA
AGCACACGTCTGAACTCTTTCCCTACACGACGCTCTTCCGATCTNNNGCTTCCATGTGTGCCAAGTAAA
AGCACACGTCTGAACTCTTTCCCTACACGACGCTCTTCCGATCTNNNGCTTCCATGTGTGCCAAGTAAA
AGCACACGTCTGAACTCTTTCCCTACACGACGCTCTTCCGATCTNNNGCTTCCATGTGTGCCAAGTAAA
AGCACACGTCTGAACTCTTTCCCTACACGACGCTCTTCCGATCTNNNGCTTCCATGTGTGCCAAGTAAA
AGCACACGTCTGAACTCTTTCCCTACACGACGCTCTTCCGATCTNNNGCTTCCATGTGTGCCAAGTAAA
AGCACACGTCTGAACTCTTTCCCTACACGACGCTCTTCCGATCTNNNTTTTGTTTTAAATACCGTTCAGA
AGCACACGTCTGAACTCTTTCCCTACACGACGCTCTTCCGATCTNNNTTTTGTTTTAAATACCGTTCAGA
AGCACACGTCTGAACTCTTTCCCTACACGACGCTCTTCCGATCTNNNTTTTGTTTTAAATACCGTTCAGA
AGCACACGTCTGAACTCTTTCCCTACACGACGCTCTTCCGATCTNNNTTTTGTTTTAAATACCGTTCAGA
AGCACACGTCTGAACTCTTTCCCTACACGACGCTCTTCCGATCTNNNTTTTGTTTTAAATACCGTTCAGA
AGCACACGTCTGAACTCTTTCCCTACACGACGCTCTTCCGATCTNNNCAGGGAGATGGGAGGTTTTTT
AGCACACGTCTGAACTCTTTCCCTACACGACGCTCTTCCGATCTNNNCAGGGAGATGGGAGGTTTTTT
AGCACACGTCTGAACTCTTTCCCTACACGACGCTCTTCCGATCTNNNCAGGGAGATGGGAGGTTTTTT
AGCACACGTCTGAACTCTTTCCCTACACGACGCTCTTCCGATCTNNNCAGGGAGATGGGAGGTTTTTT
AGCACACGTCTGAACTCTTTCCCTACACGACGCTCTTCCGATCTNNNCAGGGAGATGGGAGGTTTTTT
AGCACACGTCTGAACTCTTTCCCTACACGACGCTCTTCCGATCTNNNAATTCTTTCAATTTTTCCAGAGA
AGCACACGTCTGAACTCTTTCCCTACACGACGCTCTTCCGATCTNNNAATTCTTTCAATTTTTCCAGAGA
AGCACACGTCTGAACTCTTTCCCTACACGACGCTCTTCCGATCTNNNAATTCTTTCAATTTTTCCAGAGA
AGCACACGTCTGAACTCTTTCCCTACACGACGCTCTTCCGATCTNNNAATTCTTTCAATTTTTCCAGAGA
AGCACACGTCTGAACTCTTTCCCTACACGACGCTCTTCCGATCTNNNAATTCTTTCAATTTTTCCAGAGA

GCACACGTCTGAACTCTTTCCCTACACGACGCTCTTCCGATCTNNNTGTGTGTGTATGTGTGTGTA

GCACACGTCTGAACTCTTTCCCTACACGACGCTCTTCCGATCTNNNTGTGTGTGTATGTGTGTGTA

GCACACGTCTGAACTCTTTCCCTACACGACGCTCTTCCGATCTNNNTGTGTGTGTATGTGTGTGTA

GCACACGTCTGAACTCTTTCCCTACACGACGCTCTTCCGATCTNNNTGTGTGTGTATGTGTGTGTA

GCACACGTCTGAACTCTTTCCCTACACGACGCTCTTCCGATCTNNNTGTGTGTGTATGTGTGTGTA

GAGCACACGTCTGAACTCTTTCCCTACACGACGCTCTTCCGATCTNNNAGGAGAATCTTTTGAACCTG

GAGCACACGTCTGAACTCTTTCCCTACACGACGCTCTTCCGATCTNNNAGGAGAATCTTTTGAACCTG

GAGCACACGTCTGAACTCTTTCCCTACACGACGCTCTTCCGATCTNNNAGGAGAATCTTTTGAACCTG

GAGCACACGTCTGAACTCTTTCCCTACACGACGCTCTTCCGATCTNNNAGGAGAATCTTTTGAACCTG

GAGCACACGTCTGAACTCTTTCCCTACACGACGCTCTTCCGATCTNNNAGGAGAATCTTTTGAACCTG

GAGCACACGTCTGAACTCTTTCCCTACACGACGCTCTTCCGATCTNNNTACACACCTGTAGTCCCAGCT

GAGCACACGTCTGAACTCTTTCCCTACACGACGCTCTTCCGATCTNNNTACACACCTGTAGTCCCAGCT

GAGCACACGTCTGAACTCTTTCCCTACACGACGCTCTTCCGATCTNNNTACACACCTGTAGTCCCAGCT

GAGCACACGTCTGAACTCTTTCCCTACACGACGCTCTTCCGATCTNNNTACACACCTGTAGTCCCAGCT

GAGCACACGTCTGAACTCTTTCCCTACACGACGCTCTTCCGATCTNNNTACACACCTGTAGTCCCAGCT

GCACACGTCTGAACTCTTTCCCTACACGACGCTCTTCCGATCTNNNCTACAGCAACCAAAACAGCATGA

GCACACGTCTGAACTCTTTCCCTACACGACGCTCTTCCGATCTNNNCTACAGCAACCAAAACAGCATGA

GCACACGTCTGAACTCTTTCCCTACACGACGCTCTTCCGATCTNNNCTACAGCAACCAAAACAGCATGA

GCACACGTCTGAACTCTTTCCCTACACGACGCTCTTCCGATCTNNNCTACAGCAACCAAAACAGCATGA

GCACACGTCTGAACTCTTTCCCTACACGACGCTCTTCCGATCTNNNCTACAGCAACCAAAACAGCATGA

AGCACACGTCTGAACTCTTTCCCTACACGACGCTCTTCCGATCTNNNTGGACAGACATGGGCTTCAAT

AGCACACGTCTGAACTCTTTCCCTACACGACGCTCTTCCGATCTNNNTGGACAGACATGGGCTTCAAT

AGCACACGTCTGAACTCTTTCCCTACACGACGCTCTTCCGATCTNNNTGGACAGACATGGGCTTCAAT

AGCACACGTCTGAACTCTTTCCCTACACGACGCTCTTCCGATCTNNNTGGACAGACATGGGCTTCAAT

AGCACACGTCTGAACTCTTTCCCTACACGACGCTCTTCCGATCTNNNTGGACAGACATGGGCTTCAAT

AGCACACGTCTGAACTCTTTCCCTACACGACGCTCTTCCGATCTNNNGTAACACAGCAAAACCCAGAC

AGCACACGTCTGAACTCTTTCCCTACACGACGCTCTTCCGATCTNNNGTAACACAGCAAAACCCAGAC

AGCACACGTCTGAACTCTTTCCCTACACGACGCTCTTCCGATCTNNNGTAACACAGCAAAACCCAGAC

AGCACACGTCTGAACTCTTTCCCTACACGACGCTCTTCCGATCTNNNGTAACACAGCAAAACCCAGAC

AGCACACGTCTGAACTCTTTCCCTACACGACGCTCTTCCGATCTNNNGTAACACAGCAAAACCCAGAC

AGCACACGTCTGAACTCTTTCCCTACACGACGCTCTTCCGATCTNNNCTGAGGCAGGAGAATCACTT

AGCACACGTCTGAACTCTTTCCCTACACGACGCTCTTCCGATCTNNNCTGAGGCAGGAGAATCACTT

AGCACACGTCTGAACTCTTTCCCTACACGACGCTCTTCCGATCTNNNCTGAGGCAGGAGAATCACTT

AGCACACGTCTGAACTCTTTCCCTACACGACGCTCTTCCGATCTNNNCTGAGGCAGGAGAATCACTT

AGCACACGTCTGAACTCTTTCCCTACACGACGCTCTTCCGATCTNNNCTGAGGCAGGAGAATCACTT

AGCACACGTCTGAACTCTTTCCCTACACGACGCTCTTCCGATCTNNNTTGTAGTGAGCTGAGATCGTG

AGCACACGTCTGAACTCTTTCCCTACACGACGCTCTTCCGATCTNNNTTGTAGTGAGCTGAGATCGTG

AGCACACGTCTGAACTCTTTCCCTACACGACGCTCTTCCGATCTNNNTTGTAGTGAGCTGAGATCGTG

AGCACACGTCTGAACTCTTTCCCTACACGACGCTCTTCCGATCTNNNTTGTAGTGAGCTGAGATCGTG

AGCACACGTCTGAACTCTTTCCCTACACGACGCTCTTCCGATCTNNNTTGTAGTGAGCTGAGATCGTG

AGCACACGTCTGAACTCTTTCCCTACACGACGCTCTTCCGATCTNNNGACAATCTCCGGAATCTCTGA

AGCACACGTCTGAACTCTTTCCCTACACGACGCTCTTCCGATCTNNNGACAATCTCCGGAATCTCTGA

AGCACACGTCTGAACTCTTTCCCTACACGACGCTCTTCCGATCTNNNGACAATCTCCGGAATCTCTGA

AGCACACGTCTGAACTCTTTCCCTACACGACGCTCTTCCGATCTNNNGACAATCTCCGGAATCTCTGA
AGCACACGTCTGAACTCTTTCCCTACACGACGCTCTTCCGATCTNNNGACAATCTCCGGAATCTCTGA
GAGCACACGTCTGAACTCTTTCCCTACACGACGCTCTTCCGATCTNNNAGATGATGTAGGAGTGTGT
GAGCACACGTCTGAACTCTTTCCCTACACGACGCTCTTCCGATCTNNNAGATGATGTAGGAGTGTGT
GAGCACACGTCTGAACTCTTTCCCTACACGACGCTCTTCCGATCTNNNAGATGATGTAGGAGTGTGT
GAGCACACGTCTGAACTCTTTCCCTACACGACGCTCTTCCGATCTNNNAGATGATGTAGGAGTGTGT
GAGCACACGTCTGAACTCTTTCCCTACACGACGCTCTTCCGATCTNNNAGATGATGTAGGAGTGTGT
AGAGCACACGTCTGAACTCTTTCCCTACACGACGCTCTTCCGATCTNNNCTCCTTGTTGCCTAGTCTTG
AGAGCACACGTCTGAACTCTTTCCCTACACGACGCTCTTCCGATCTNNNCTCCTTGTTGCCTAGTCTTG
AGAGCACACGTCTGAACTCTTTCCCTACACGACGCTCTTCCGATCTNNNCTCCTTGTTGCCTAGTCTTG
AGAGCACACGTCTGAACTCTTTCCCTACACGACGCTCTTCCGATCTNNNCTCCTTGTTGCCTAGTCTTG
AGAGCACACGTCTGAACTCTTTCCCTACACGACGCTCTTCCGATCTNNNCTCCTTGTTGCCTAGTCTTG
AGCACACGTCTGAACTCTTTCCCTACACGACGCTCTTCCGATCTNNNAACTAGAATGTGATGTGGGT
AGCACACGTCTGAACTCTTTCCCTACACGACGCTCTTCCGATCTNNNAACTAGAATGTGATGTGGGT
AGCACACGTCTGAACTCTTTCCCTACACGACGCTCTTCCGATCTNNNAACTAGAATGTGATGTGGGT
AGCACACGTCTGAACTCTTTCCCTACACGACGCTCTTCCGATCTNNNAACTAGAATGTGATGTGGGT
AGCACACGTCTGAACTCTTTCCCTACACGACGCTCTTCCGATCTNNNAACTAGAATGTGATGTGGGT
GCACACGTCTGAACTCTTTCCCTACACGACGCTCTTCCGATCTNNNCTGGTGTGAGTAATTTTTTTCTA
GCACACGTCTGAACTCTTTCCCTACACGACGCTCTTCCGATCTNNNCTGGTGTGAGTAATTTTTTTCTA
GCACACGTCTGAACTCTTTCCCTACACGACGCTCTTCCGATCTNNNCTGGTGTGAGTAATTTTTTTCTA
GCACACGTCTGAACTCTTTCCCTACACGACGCTCTTCCGATCTNNNCTGGTGTGAGTAATTTTTTTCTA
GCACACGTCTGAACTCTTTCCCTACACGACGCTCTTCCGATCTNNNCTGGTGTGAGTAATTTTTTTCTA
GAGCACACGTCTGAACTCTTTCCCTACACGACGCTCTTCCGATCTNNNATCCAGGTTGCTGTGAATGC
GAGCACACGTCTGAACTCTTTCCCTACACGACGCTCTTCCGATCTNNNATCCAGGTTGCTGTGAATGC
GAGCACACGTCTGAACTCTTTCCCTACACGACGCTCTTCCGATCTNNNATCCAGGTTGCTGTGAATGC
GAGCACACGTCTGAACTCTTTCCCTACACGACGCTCTTCCGATCTNNNATCCAGGTTGCTGTGAATGC
GAGCACACGTCTGAACTCTTTCCCTACACGACGCTCTTCCGATCTNNNATCCAGGTTGCTGTGAATGC
AGCACACGTCTGAACTCTTTCCCTACACGACGCTCTTCCGATCTNNNACCACACCTAAAAGAAACACC
AGCACACGTCTGAACTCTTTCCCTACACGACGCTCTTCCGATCTNNNACCACACCTAAAAGAAACACC
AGCACACGTCTGAACTCTTTCCCTACACGACGCTCTTCCGATCTNNNACCACACCTAAAAGAAACACC
AGCACACGTCTGAACTCTTTCCCTACACGACGCTCTTCCGATCTNNNACCACACCTAAAAGAAACACC
AGCACACGTCTGAACTCTTTCCCTACACGACGCTCTTCCGATCTNNNACCACACCTAAAAGAAACACC
AGCACACGTCTGAACTCTTTCCCTACACGACGCTCTTCCGATCTNNNCAAAAAAAGGTCCCACATAGC
AGCACACGTCTGAACTCTTTCCCTACACGACGCTCTTCCGATCTNNNCAAAAAAAGGTCCCACATAGC
AGCACACGTCTGAACTCTTTCCCTACACGACGCTCTTCCGATCTNNNCAAAAAAAGGTCCCACATAGC
AGCACACGTCTGAACTCTTTCCCTACACGACGCTCTTCCGATCTNNNCAAAAAAAGGTCCCACATAGC
AGCACACGTCTGAACTCTTTCCCTACACGACGCTCTTCCGATCTNNNCAAAAAAAGGTCCCACATAGC
GAGCACACGTCTGAACTCTTTCCCTACACGACGCTCTTCCGATCTNNNAGAAGTGTGTGTGTGTGTG
GAGCACACGTCTGAACTCTTTCCCTACACGACGCTCTTCCGATCTNNNAGAAGTGTGTGTGTGTGTG
GAGCACACGTCTGAACTCTTTCCCTACACGACGCTCTTCCGATCTNNNAGAAGTGTGTGTGTGTGTG
GAGCACACGTCTGAACTCTTTCCCTACACGACGCTCTTCCGATCTNNNAGAAGTGTGTGTGTGTGTG
GAGCACACGTCTGAACTCTTTCCCTACACGACGCTCTTCCGATCTNNNAGAAGTGTGTGTGTGTGTG
GAGCACACGTCTGAACTCTTTCCCTACACGACGCTCTTCCGATCTNNNAGTGAAACCCCGTCTCTACT

GAGCACACGTCTGAACTCTTTCCCTACACGACGCTCTTCCGATCTNNNAGTGAAACCCCGTCTCTACT
GAGCACACGTCTGAACTCTTTCCCTACACGACGCTCTTCCGATCTNNNAGTGAAACCCCGTCTCTACT
GAGCACACGTCTGAACTCTTTCCCTACACGACGCTCTTCCGATCTNNNAGTGAAACCCCGTCTCTACT
GAGCACACGTCTGAACTCTTTCCCTACACGACGCTCTTCCGATCTNNNAGTGAAACCCCGTCTCTACT
AGCACACGTCTGAACTCTTTCCCTACACGACGCTCTTCCGATCTNNNCCTTTTCTTCTGTCTGTTGAGC
AGCACACGTCTGAACTCTTTCCCTACACGACGCTCTTCCGATCTNNNCCTTTTCTTCTGTCTGTTGAGC
AGCACACGTCTGAACTCTTTCCCTACACGACGCTCTTCCGATCTNNNCCTTTTCTTCTGTCTGTTGAGC
AGCACACGTCTGAACTCTTTCCCTACACGACGCTCTTCCGATCTNNNCCTTTTCTTCTGTCTGTTGAGC
AGCACACGTCTGAACTCTTTCCCTACACGACGCTCTTCCGATCTNNNCCTTTTCTTCTGTCTGTTGAGC
GCACACGTCTGAACTCTTTCCCTACACGACGCTCTTCCGATCTNNNAGTCCCTCACTACATTAGAGTCT
GCACACGTCTGAACTCTTTCCCTACACGACGCTCTTCCGATCTNNNAGTCCCTCACTACATTAGAGTCT
GCACACGTCTGAACTCTTTCCCTACACGACGCTCTTCCGATCTNNNAGTCCCTCACTACATTAGAGTCT
GCACACGTCTGAACTCTTTCCCTACACGACGCTCTTCCGATCTNNNAGTCCCTCACTACATTAGAGTCT
GCACACGTCTGAACTCTTTCCCTACACGACGCTCTTCCGATCTNNNAGTCCCTCACTACATTAGAGTCT
GAGCACACGTCTGAACTCTTTCCCTACACGACGCTCTTCCGATCTNNNGTATATACCCACAGGCACCTC
GAGCACACGTCTGAACTCTTTCCCTACACGACGCTCTTCCGATCTNNNGTATATACCCACAGGCACCTC
GAGCACACGTCTGAACTCTTTCCCTACACGACGCTCTTCCGATCTNNNGTATATACCCACAGGCACCTC
GAGCACACGTCTGAACTCTTTCCCTACACGACGCTCTTCCGATCTNNNGTATATACCCACAGGCACCTC
GAGCACACGTCTGAACTCTTTCCCTACACGACGCTCTTCCGATCTNNNGTATATACCCACAGGCACCTC
GAGCACACGTCTGAACTCTTTCCCTACACGACGCTCTTCCGATCTNNNTGATATGACCCTTCGTTTTTTT
GAGCACACGTCTGAACTCTTTCCCTACACGACGCTCTTCCGATCTNNNTGATATGACCCTTCGTTTTTTT
GAGCACACGTCTGAACTCTTTCCCTACACGACGCTCTTCCGATCTNNNTGATATGACCCTTCGTTTTTTT
GAGCACACGTCTGAACTCTTTCCCTACACGACGCTCTTCCGATCTNNNTGATATGACCCTTCGTTTTTTT
GAGCACACGTCTGAACTCTTTCCCTACACGACGCTCTTCCGATCTNNNTGATATGACCCTTCGTTTTTTT
AGCACACGTCTGAACTCTTTCCCTACACGACGCTCTTCCGATCTNNNTCTTCTTGTCTTTGCAGTGTGT
AGCACACGTCTGAACTCTTTCCCTACACGACGCTCTTCCGATCTNNNTCTTCTTGTCTTTGCAGTGTGT
AGCACACGTCTGAACTCTTTCCCTACACGACGCTCTTCCGATCTNNNTCTTCTTGTCTTTGCAGTGTGT
AGCACACGTCTGAACTCTTTCCCTACACGACGCTCTTCCGATCTNNNTCTTCTTGTCTTTGCAGTGTGT
AGCACACGTCTGAACTCTTTCCCTACACGACGCTCTTCCGATCTNNNTCTTCTTGTCTTTGCAGTGTGT
GCACACGTCTGAACTCTTTCCCTACACGACGCTCTTCCGATCTNNNTAGGCAGATTCTCGAAACAGAG
GCACACGTCTGAACTCTTTCCCTACACGACGCTCTTCCGATCTNNNTAGGCAGATTCTCGAAACAGAG
GCACACGTCTGAACTCTTTCCCTACACGACGCTCTTCCGATCTNNNTAGGCAGATTCTCGAAACAGAG
GCACACGTCTGAACTCTTTCCCTACACGACGCTCTTCCGATCTNNNTAGGCAGATTCTCGAAACAGAG
GCACACGTCTGAACTCTTTCCCTACACGACGCTCTTCCGATCTNNNTAGGCAGATTCTCGAAACAGAG
AGCACACGTCTGAACTCTTTCCCTACACGACGCTCTTCCGATCTNNNCATCCAAACACCATCAGGCTAA
AGCACACGTCTGAACTCTTTCCCTACACGACGCTCTTCCGATCTNNNCATCCAAACACCATCAGGCTAA
AGCACACGTCTGAACTCTTTCCCTACACGACGCTCTTCCGATCTNNNCATCCAAACACCATCAGGCTAA
AGCACACGTCTGAACTCTTTCCCTACACGACGCTCTTCCGATCTNNNCATCCAAACACCATCAGGCTAA
AGCACACGTCTGAACTCTTTCCCTACACGACGCTCTTCCGATCTNNNCATCCAAACACCATCAGGCTAA
GCACACGTCTGAACTCTTTCCCTACACGACGCTCTTCCGATCTNNNCTTCCCATAACCTCTGCCTTTCTC
GCACACGTCTGAACTCTTTCCCTACACGACGCTCTTCCGATCTNNNCTTCCCATAACCTCTGCCTTTCTC
GCACACGTCTGAACTCTTTCCCTACACGACGCTCTTCCGATCTNNNCTTCCCATAACCTCTGCCTTTCTC
GCACACGTCTGAACTCTTTCCCTACACGACGCTCTTCCGATCTNNNCTTCCCATAACCTCTGCCTTTCTC

AGCACACGTCTGAACTCTTTCCCTACACGACGCTCTTCCGATCTNNNCTTCCCATAACCTCTGCCTTTCTCA
GAGCACACGTCTGAACTCTTTCCCTACACGACGCTCTTCCGATCTNNNTGAAATCCAGTCCAGTCTGGG
GAGCACACGTCTGAACTCTTTCCCTACACGACGCTCTTCCGATCTNNNTGAAATCCAGTCCAGTCTGGG
GAGCACACGTCTGAACTCTTTCCCTACACGACGCTCTTCCGATCTNNNTGAAATCCAGTCCAGTCTGGG
GAGCACACGTCTGAACTCTTTCCCTACACGACGCTCTTCCGATCTNNNTGAAATCCAGTCCAGTCTGGG
GAGCACACGTCTGAACTCTTTCCCTACACGACGCTCTTCCGATCTNNNTGAAATCCAGTCCAGTCTGGG
GAGCACACGTCTGAACTCTTTCCCTACACGACGCTCTTCCGATCTNNNAATCCCAGCTACTTGAGAGGT
GAGCACACGTCTGAACTCTTTCCCTACACGACGCTCTTCCGATCTNNNAATCCCAGCTACTTGAGAGGT
GAGCACACGTCTGAACTCTTTCCCTACACGACGCTCTTCCGATCTNNNAATCCCAGCTACTTGAGAGGT
GAGCACACGTCTGAACTCTTTCCCTACACGACGCTCTTCCGATCTNNNAATCCCAGCTACTTGAGAGGT
GAGCACACGTCTGAACTCTTTCCCTACACGACGCTCTTCCGATCTNNNAATCCCAGCTACTTGAGAGGT
AGCACACGTCTGAACTCTTTCCCTACACGACGCTCTTCCGATCTNNNGGCAAGACCCTGTCTCTACAAA
AGCACACGTCTGAACTCTTTCCCTACACGACGCTCTTCCGATCTNNNGGCAAGACCCTGTCTCTACAAA
AGCACACGTCTGAACTCTTTCCCTACACGACGCTCTTCCGATCTNNNGGCAAGACCCTGTCTCTACAAA
AGCACACGTCTGAACTCTTTCCCTACACGACGCTCTTCCGATCTNNNGGCAAGACCCTGTCTCTACAAA
AGCACACGTCTGAACTCTTTCCCTACACGACGCTCTTCCGATCTNNNGGCAAGACCCTGTCTCTACAAA
AGCACACGTCTGAACTCTTTCCCTACACGACGCTCTTCCGATCTNNNCTCAAGTGATCCTCCTACCTCAG
AGCACACGTCTGAACTCTTTCCCTACACGACGCTCTTCCGATCTNNNCTCAAGTGATCCTCCTACCTCAG
AGCACACGTCTGAACTCTTTCCCTACACGACGCTCTTCCGATCTNNNCTCAAGTGATCCTCCTACCTCAG
AGCACACGTCTGAACTCTTTCCCTACACGACGCTCTTCCGATCTNNNCTCAAGTGATCCTCCTACCTCAG
AGCACACGTCTGAACTCTTTCCCTACACGACGCTCTTCCGATCTNNNCTCAAGTGATCCTCCTACCTCAG
ACACGTCTGAACTCTTTCCCTACACGACGCTCTTCCGATCTNNNGGATGGTCTTCTTCTTCTGACCTCATG
ACACGTCTGAACTCTTTCCCTACACGACGCTCTTCCGATCTNNNGGATGGTCTTCTTCTTCTGACCTCATG
ACACGTCTGAACTCTTTCCCTACACGACGCTCTTCCGATCTNNNGGATGGTCTTCTTCTTCTGACCTCATG
ACACGTCTGAACTCTTTCCCTACACGACGCTCTTCCGATCTNNNGGATGGTCTTCTTCTTCTGACCTCATG
ACACGTCTGAACTCTTTCCCTACACGACGCTCTTCCGATCTNNNGGATGGTCTTCTTCTTCTGACCTCATG
AGAGCACACGTCTGAACTCTTTCCCTACACGACGCTCTTCCGATCTNNNATCACTTGGGTATGCACTGA
AGAGCACACGTCTGAACTCTTTCCCTACACGACGCTCTTCCGATCTNNNATCACTTGGGTATGCACTGA
AGAGCACACGTCTGAACTCTTTCCCTACACGACGCTCTTCCGATCTNNNATCACTTGGGTATGCACTGA
AGAGCACACGTCTGAACTCTTTCCCTACACGACGCTCTTCCGATCTNNNATCACTTGGGTATGCACTGA
AGAGCACACGTCTGAACTCTTTCCCTACACGACGCTCTTCCGATCTNNNATCACTTGGGTATGCACTGA
AAGAGCACACGTCTGAACTCTTTCCCTACACGACGCTCTTCCGATCTNNNCCAGTTTGATCTGGAAGGC
AAGAGCACACGTCTGAACTCTTTCCCTACACGACGCTCTTCCGATCTNNNCCAGTTTGATCTGGAAGGC
AAGAGCACACGTCTGAACTCTTTCCCTACACGACGCTCTTCCGATCTNNNCCAGTTTGATCTGGAAGGC
AAGAGCACACGTCTGAACTCTTTCCCTACACGACGCTCTTCCGATCTNNNCCAGTTTGATCTGGAAGGC
AAGAGCACACGTCTGAACTCTTTCCCTACACGACGCTCTTCCGATCTNNNCCAGTTTGATCTGGAAGGC
AGCACACGTCTGAACTCTTTCCCTACACGACGCTCTTCCGATCTNNNTCTGTATCTTTGGTCACTCCCTTC
AGCACACGTCTGAACTCTTTCCCTACACGACGCTCTTCCGATCTNNNTCTGTATCTTTGGTCACTCCCTTC
AGCACACGTCTGAACTCTTTCCCTACACGACGCTCTTCCGATCTNNNTCTGTATCTTTGGTCACTCCCTTC
AGCACACGTCTGAACTCTTTCCCTACACGACGCTCTTCCGATCTNNNTCTGTATCTTTGGTCACTCCCTTC
AGCACACGTCTGAACTCTTTCCCTACACGACGCTCTTCCGATCTNNNTCTGTATCTTTGGTCACTCCCTTC
AAGAGCACACGTCTGAACTCTTTCCCTACACGACGCTCTTCCGATCTNNNAGTAGTCTCAGCTACTCGG
AAGAGCACACGTCTGAACTCTTTCCCTACACGACGCTCTTCCGATCTNNNAGTAGTCTCAGCTACTCGG

AAGAGCACACGTCTGAACTCTTTCCCTACACGACGCTCTTCCGATCTNNNAGTAGTCTCAGCTACTCGG
AAGAGCACACGTCTGAACTCTTTCCCTACACGACGCTCTTCCGATCTNNNAGTAGTCTCAGCTACTCGG
AAGAGCACACGTCTGAACTCTTTCCCTACACGACGCTCTTCCGATCTNNNAGTAGTCTCAGCTACTCGG
GAGCACACGTCTGAACTCTTTCCCTACACGACGCTCTTCCGATCTNNNCCCGCCTTTATGGACCAAACTA
GAGCACACGTCTGAACTCTTTCCCTACACGACGCTCTTCCGATCTNNNCCCGCCTTTATGGACCAAACTA
GAGCACACGTCTGAACTCTTTCCCTACACGACGCTCTTCCGATCTNNNCCCGCCTTTATGGACCAAACTA
GAGCACACGTCTGAACTCTTTCCCTACACGACGCTCTTCCGATCTNNNCCCGCCTTTATGGACCAAACTA
GAGCACACGTCTGAACTCTTTCCCTACACGACGCTCTTCCGATCTNNNCCCGCCTTTATGGACCAAACTA
AGCACACGTCTGAACTCTTTCCCTACACGACGCTCTTCCGATCTNNNTGTATCGCTGATAAGGAAGAG
AGCACACGTCTGAACTCTTTCCCTACACGACGCTCTTCCGATCTNNNTGTATCGCTGATAAGGAAGAG
AGCACACGTCTGAACTCTTTCCCTACACGACGCTCTTCCGATCTNNNTGTATCGCTGATAAGGAAGAG
AGCACACGTCTGAACTCTTTCCCTACACGACGCTCTTCCGATCTNNNTGTATCGCTGATAAGGAAGAG
AGCACACGTCTGAACTCTTTCCCTACACGACGCTCTTCCGATCTNNNTGTATCGCTGATAAGGAAGAG
AGCACACGTCTGAACTCTTTCCCTACACGACGCTCTTCCGATCTNNNGCCCTCAACAAGATGTTTTGCCC
AGCACACGTCTGAACTCTTTCCCTACACGACGCTCTTCCGATCTNNNGCCCTCAACAAGATGTTTTGCCC
AGCACACGTCTGAACTCTTTCCCTACACGACGCTCTTCCGATCTNNNGCCCTCAACAAGATGTTTTGCCC
AGCACACGTCTGAACTCTTTCCCTACACGACGCTCTTCCGATCTNNNGCCCTCAACAAGATGTTTTGCCC
AGCACACGTCTGAACTCTTTCCCTACACGACGCTCTTCCGATCTNNNGCCCTCAACAAGATGTTTTGCCC
GAGCACACGTCTGAACTCTTTCCCTACACGACGCTCTTCCGATCTNNNGCCTTATTTAAGCTAAATATGC
GAGCACACGTCTGAACTCTTTCCCTACACGACGCTCTTCCGATCTNNNGCCTTATTTAAGCTAAATATGC
GAGCACACGTCTGAACTCTTTCCCTACACGACGCTCTTCCGATCTNNNGCCTTATTTAAGCTAAATATGC
GAGCACACGTCTGAACTCTTTCCCTACACGACGCTCTTCCGATCTNNNGCCTTATTTAAGCTAAATATGC
GAGCACACGTCTGAACTCTTTCCCTACACGACGCTCTTCCGATCTNNNGCCTTATTTAAGCTAAATATGC
AGCACACGTCTGAACTCTTTCCCTACACGACGCTCTTCCGATCTNNNCAATGGATCTCAGGCTCCACTG
AGCACACGTCTGAACTCTTTCCCTACACGACGCTCTTCCGATCTNNNCAATGGATCTCAGGCTCCACTG
AGCACACGTCTGAACTCTTTCCCTACACGACGCTCTTCCGATCTNNNCAATGGATCTCAGGCTCCACTG
AGCACACGTCTGAACTCTTTCCCTACACGACGCTCTTCCGATCTNNNCAATGGATCTCAGGCTCCACTG
AGCACACGTCTGAACTCTTTCCCTACACGACGCTCTTCCGATCTNNNCAATGGATCTCAGGCTCCACTG
GAGCACACGTCTGAACTCTTTCCCTACACGACGCTCTTCCGATCTNNNATTTAGGAACCAAGACAGGCT
GAGCACACGTCTGAACTCTTTCCCTACACGACGCTCTTCCGATCTNNNATTTAGGAACCAAGACAGGCT
GAGCACACGTCTGAACTCTTTCCCTACACGACGCTCTTCCGATCTNNNATTTAGGAACCAAGACAGGCT
GAGCACACGTCTGAACTCTTTCCCTACACGACGCTCTTCCGATCTNNNATTTAGGAACCAAGACAGGCT
GAGCACACGTCTGAACTCTTTCCCTACACGACGCTCTTCCGATCTNNNATTTAGGAACCAAGACAGGCT
GAGCACACGTCTGAACTCTTTCCCTACACGACGCTCTTCCGATCTNNNCTGTAGTCCCAGCTACTCAGG
GAGCACACGTCTGAACTCTTTCCCTACACGACGCTCTTCCGATCTNNNCTGTAGTCCCAGCTACTCAGG
GAGCACACGTCTGAACTCTTTCCCTACACGACGCTCTTCCGATCTNNNCTGTAGTCCCAGCTACTCAGG
GAGCACACGTCTGAACTCTTTCCCTACACGACGCTCTTCCGATCTNNNCTGTAGTCCCAGCTACTCAGG
GAGCACACGTCTGAACTCTTTCCCTACACGACGCTCTTCCGATCTNNNCTGTAGTCCCAGCTACTCAGG
AGAGCACACGTCTGAACTCTTTCCCTACACGACGCTCTTCCGATCTNNNTGCAATGCCATTCACTTTTTCC
AGAGCACACGTCTGAACTCTTTCCCTACACGACGCTCTTCCGATCTNNNTGCAATGCCATTCACTTTTTCC
AGAGCACACGTCTGAACTCTTTCCCTACACGACGCTCTTCCGATCTNNNTGCAATGCCATTCACTTTTTCC
AGAGCACACGTCTGAACTCTTTCCCTACACGACGCTCTTCCGATCTNNNTGCAATGCCATTCACTTTTTCC
AGAGCACACGTCTGAACTCTTTCCCTACACGACGCTCTTCCGATCTNNNTGCAATGCCATTCACTTTTTCC

AGCACACGTCTGAACTCTTTCCCTACACGACGCTCTTCCGATCTNNNAAAAAATTAGCCAGGTGTGGT
AGCACACGTCTGAACTCTTTCCCTACACGACGCTCTTCCGATCTNNNAAAAAATTAGCCAGGTGTGGT
AGCACACGTCTGAACTCTTTCCCTACACGACGCTCTTCCGATCTNNNAAAAAATTAGCCAGGTGTGGT
AGCACACGTCTGAACTCTTTCCCTACACGACGCTCTTCCGATCTNNNAAAAAATTAGCCAGGTGTGGT
AGCACACGTCTGAACTCTTTCCCTACACGACGCTCTTCCGATCTNNNAAAAAATTAGCCAGGTGTGGT
AGCACACGTCTGAACTCTTTCCCTACACGACGCTCTTCCGATCTNNNGTACAGTTGTCCCTTGGTATCC
AGCACACGTCTGAACTCTTTCCCTACACGACGCTCTTCCGATCTNNNGTACAGTTGTCCCTTGGTATCC
AGCACACGTCTGAACTCTTTCCCTACACGACGCTCTTCCGATCTNNNGTACAGTTGTCCCTTGGTATCC
AGCACACGTCTGAACTCTTTCCCTACACGACGCTCTTCCGATCTNNNGTACAGTTGTCCCTTGGTATCC
AGCACACGTCTGAACTCTTTCCCTACACGACGCTCTTCCGATCTNNNGTACAGTTGTCCCTTGGTATCC
AGCACACGTCTGAACTCTTTCCCTACACGACGCTCTTCCGATCTNNNGGACAGACTATCTAGGCTTGT
AGCACACGTCTGAACTCTTTCCCTACACGACGCTCTTCCGATCTNNNGGACAGACTATCTAGGCTTGT
AGCACACGTCTGAACTCTTTCCCTACACGACGCTCTTCCGATCTNNNGGACAGACTATCTAGGCTTGT
AGCACACGTCTGAACTCTTTCCCTACACGACGCTCTTCCGATCTNNNGGACAGACTATCTAGGCTTGT
AGCACACGTCTGAACTCTTTCCCTACACGACGCTCTTCCGATCTNNNGGACAGACTATCTAGGCTTGT
AGCACACGTCTGAACTCTTTCCCTACACGACGCTCTTCCGATCTNNNGGCAACAGAGTGAGATTCCAT
AGCACACGTCTGAACTCTTTCCCTACACGACGCTCTTCCGATCTNNNGGCAACAGAGTGAGATTCCAT
AGCACACGTCTGAACTCTTTCCCTACACGACGCTCTTCCGATCTNNNGGCAACAGAGTGAGATTCCAT
AGCACACGTCTGAACTCTTTCCCTACACGACGCTCTTCCGATCTNNNGGCAACAGAGTGAGATTCCAT
AGCACACGTCTGAACTCTTTCCCTACACGACGCTCTTCCGATCTNNNGGCAACAGAGTGAGATTCCAT
AGAGCACACGTCTGAACTCTTTCCCTACACGACGCTCTTCCGATCTNNNACTCAAGGCGTTATCATTTC
AGAGCACACGTCTGAACTCTTTCCCTACACGACGCTCTTCCGATCTNNNACTCAAGGCGTTATCATTTC
AGAGCACACGTCTGAACTCTTTCCCTACACGACGCTCTTCCGATCTNNNACTCAAGGCGTTATCATTTC
AGAGCACACGTCTGAACTCTTTCCCTACACGACGCTCTTCCGATCTNNNACTCAAGGCGTTATCATTTC
AGAGCACACGTCTGAACTCTTTCCCTACACGACGCTCTTCCGATCTNNNACTCAAGGCGTTATCATTTC
GAGCACACGTCTGAACTCTTTCCCTACACGACGCTCTTCCGATCTNNNTGTAGTCCCAGCTACTCAGGA
GAGCACACGTCTGAACTCTTTCCCTACACGACGCTCTTCCGATCTNNNTGTAGTCCCAGCTACTCAGGA
GAGCACACGTCTGAACTCTTTCCCTACACGACGCTCTTCCGATCTNNNTGTAGTCCCAGCTACTCAGGA
GAGCACACGTCTGAACTCTTTCCCTACACGACGCTCTTCCGATCTNNNTGTAGTCCCAGCTACTCAGGA
GAGCACACGTCTGAACTCTTTCCCTACACGACGCTCTTCCGATCTNNNTGTAGTCCCAGCTACTCAGGA
AGCACACGTCTGAACTCTTTCCCTACACGACGCTCTTCCGATCTNNNTTTTGGAGAGTCTCTTCCTTGG
AGCACACGTCTGAACTCTTTCCCTACACGACGCTCTTCCGATCTNNNTTTTGGAGAGTCTCTTCCTTGG
AGCACACGTCTGAACTCTTTCCCTACACGACGCTCTTCCGATCTNNNTTTTGGAGAGTCTCTTCCTTGG
AGCACACGTCTGAACTCTTTCCCTACACGACGCTCTTCCGATCTNNNTTTTGGAGAGTCTCTTCCTTGG
AGCACACGTCTGAACTCTTTCCCTACACGACGCTCTTCCGATCTNNNTTTTGGAGAGTCTCTTCCTTGG
GAGCACACGTCTGAACTCTTTCCCTACACGACGCTCTTCCGATCTNNNAGATGAACATTCCCATTCCTGG
GAGCACACGTCTGAACTCTTTCCCTACACGACGCTCTTCCGATCTNNNAGATGAACATTCCCATTCCTGG
GAGCACACGTCTGAACTCTTTCCCTACACGACGCTCTTCCGATCTNNNAGATGAACATTCCCATTCCTGG
GAGCACACGTCTGAACTCTTTCCCTACACGACGCTCTTCCGATCTNNNAGATGAACATTCCCATTCCTGG
GAGCACACGTCTGAACTCTTTCCCTACACGACGCTCTTCCGATCTNNNAGATGAACATTCCCATTCCTGG
AGCACACGTCTGAACTCTTTCCCTACACGACGCTCTTCCGATCTNNNCTTCTTTGACATGTGGTCCTGC
AGCACACGTCTGAACTCTTTCCCTACACGACGCTCTTCCGATCTNNNCTTCTTTGACATGTGGTCCTGC
AGCACACGTCTGAACTCTTTCCCTACACGACGCTCTTCCGATCTNNNCTTCTTTGACATGTGGTCCTGC

AGCACACGTCTGAACTCTTTCCCTACACGACGCTCTTCCGATCTNNNCTTCTTTGACATGTGGTCCTGC
AGCACACGTCTGAACTCTTTCCCTACACGACGCTCTTCCGATCTNNNCTTCTTTGACATGTGGTCCTGC
AGCACACGTCTGAACTCTTTCCCTACACGACGCTCTTCCGATCTNNNTGCTGTCCCTCCATGACTAAAA
AGCACACGTCTGAACTCTTTCCCTACACGACGCTCTTCCGATCTNNNTGCTGTCCCTCCATGACTAAAA
AGCACACGTCTGAACTCTTTCCCTACACGACGCTCTTCCGATCTNNNTGCTGTCCCTCCATGACTAAAA
AGCACACGTCTGAACTCTTTCCCTACACGACGCTCTTCCGATCTNNNTGCTGTCCCTCCATGACTAAAA
AGCACACGTCTGAACTCTTTCCCTACACGACGCTCTTCCGATCTNNNTGCTGTCCCTCCATGACTAAAA
AGCACACGTCTGAACTCTTTCCCTACACGACGCTCTTCCGATCTNNNCTTGAGTGGGTGAGAGAAAGA
AGCACACGTCTGAACTCTTTCCCTACACGACGCTCTTCCGATCTNNNCTTGAGTGGGTGAGAGAAAGA
AGCACACGTCTGAACTCTTTCCCTACACGACGCTCTTCCGATCTNNNCTTGAGTGGGTGAGAGAAAGA
AGCACACGTCTGAACTCTTTCCCTACACGACGCTCTTCCGATCTNNNCTTGAGTGGGTGAGAGAAAGA
AGCACACGTCTGAACTCTTTCCCTACACGACGCTCTTCCGATCTNNNCTTGAGTGGGTGAGAGAAAGA
GAGCACACGTCTGAACTCTTTCCCTACACGACGCTCTTCCGATCTNNNCATATGGGACTATGTGGAGG
GAGCACACGTCTGAACTCTTTCCCTACACGACGCTCTTCCGATCTNNNCATATGGGACTATGTGGAGG
GAGCACACGTCTGAACTCTTTCCCTACACGACGCTCTTCCGATCTNNNCATATGGGACTATGTGGAGG
GAGCACACGTCTGAACTCTTTCCCTACACGACGCTCTTCCGATCTNNNCATATGGGACTATGTGGAGG
GAGCACACGTCTGAACTCTTTCCCTACACGACGCTCTTCCGATCTNNNCATATGGGACTATGTGGAGG
AGCACACGTCTGAACTCTTTCCCTACACGACGCTCTTCCGATCTNNNCTGCCGACATAGTCAGAGTTAC
AGCACACGTCTGAACTCTTTCCCTACACGACGCTCTTCCGATCTNNNCTGCCGACATAGTCAGAGTTAC
AGCACACGTCTGAACTCTTTCCCTACACGACGCTCTTCCGATCTNNNCTGCCGACATAGTCAGAGTTAC
AGCACACGTCTGAACTCTTTCCCTACACGACGCTCTTCCGATCTNNNCTGCCGACATAGTCAGAGTTAC
AGCACACGTCTGAACTCTTTCCCTACACGACGCTCTTCCGATCTNNNCTGCCGACATAGTCAGAGTTAC
AGCACACGTCTGAACTCTTTCCCTACACGACGCTCTTCCGATCTNNNAACTCCTGACCTCAAGTGATCTG
AGCACACGTCTGAACTCTTTCCCTACACGACGCTCTTCCGATCTNNNAACTCCTGACCTCAAGTGATCTG
AGCACACGTCTGAACTCTTTCCCTACACGACGCTCTTCCGATCTNNNAACTCCTGACCTCAAGTGATCTG
AGCACACGTCTGAACTCTTTCCCTACACGACGCTCTTCCGATCTNNNAACTCCTGACCTCAAGTGATCTG
AGCACACGTCTGAACTCTTTCCCTACACGACGCTCTTCCGATCTNNNAACTCCTGACCTCAAGTGATCTG
AGCACACGTCTGAACTCTTTCCCTACACGACGCTCTTCCGATCTNNNTTTGTTGAAAATGAAATCTAGT
AGCACACGTCTGAACTCTTTCCCTACACGACGCTCTTCCGATCTNNNTTTGTTGAAAATGAAATCTAGT
AGCACACGTCTGAACTCTTTCCCTACACGACGCTCTTCCGATCTNNNTTTGTTGAAAATGAAATCTAGT
AGCACACGTCTGAACTCTTTCCCTACACGACGCTCTTCCGATCTNNNTTTGTTGAAAATGAAATCTAGT
AGCACACGTCTGAACTCTTTCCCTACACGACGCTCTTCCGATCTNNNTTTGTTGAAAATGAAATCTAGT
GAGCACACGTCTGAACTCTTTCCCTACACGACGCTCTTCCGATCTNNNGGTAGTGACACACTCCCTTGA
GAGCACACGTCTGAACTCTTTCCCTACACGACGCTCTTCCGATCTNNNGGTAGTGACACACTCCCTTGA
GAGCACACGTCTGAACTCTTTCCCTACACGACGCTCTTCCGATCTNNNGGTAGTGACACACTCCCTTGA
GAGCACACGTCTGAACTCTTTCCCTACACGACGCTCTTCCGATCTNNNGGTAGTGACACACTCCCTTGA
GAGCACACGTCTGAACTCTTTCCCTACACGACGCTCTTCCGATCTNNNGGTAGTGACACACTCCCTTGA
AGCACACGTCTGAACTCTTTCCCTACACGACGCTCTTCCGATCTNNNGACATGGCAAAACCCCATCTCT
AGCACACGTCTGAACTCTTTCCCTACACGACGCTCTTCCGATCTNNNGACATGGCAAAACCCCATCTCT
AGCACACGTCTGAACTCTTTCCCTACACGACGCTCTTCCGATCTNNNGACATGGCAAAACCCCATCTCT
AGCACACGTCTGAACTCTTTCCCTACACGACGCTCTTCCGATCTNNNGACATGGCAAAACCCCATCTCT
AGCACACGTCTGAACTCTTTCCCTACACGACGCTCTTCCGATCTNNNGACATGGCAAAACCCCATCTCT
GAGCACACGTCTGAACTCTTTCCCTACACGACGCTCTTCCGATCTNNNCAACCTCAGAGCTTCCCTAATA

GAGCACACGTCTGAACTCTTTCCCTACACGACGCTCTTCCGATCTNNNCAACCTCAGAGCTTCCCTAATA
GAGCACACGTCTGAACTCTTTCCCTACACGACGCTCTTCCGATCTNNNCAACCTCAGAGCTTCCCTAATA
GAGCACACGTCTGAACTCTTTCCCTACACGACGCTCTTCCGATCTNNNCAACCTCAGAGCTTCCCTAATA
GAGCACACGTCTGAACTCTTTCCCTACACGACGCTCTTCCGATCTNNNCAACCTCAGAGCTTCCCTAATA
AGCACACGTCTGAACTCTTTCCCTACACGACGCTCTTCCGATCTNNNAAGCACATCAGACTAAAACACG
AGCACACGTCTGAACTCTTTCCCTACACGACGCTCTTCCGATCTNNNAAGCACATCAGACTAAAACACG
AGCACACGTCTGAACTCTTTCCCTACACGACGCTCTTCCGATCTNNNAAGCACATCAGACTAAAACACG
AGCACACGTCTGAACTCTTTCCCTACACGACGCTCTTCCGATCTNNNAAGCACATCAGACTAAAACACG
AGCACACGTCTGAACTCTTTCCCTACACGACGCTCTTCCGATCTNNNAAGCACATCAGACTAAAACACG
AGCACACGTCTGAACTCTTTCCCTACACGACGCTCTTCCGATCTNNNATCCCCACGAAACATATTCCAT
AGCACACGTCTGAACTCTTTCCCTACACGACGCTCTTCCGATCTNNNATCCCCACGAAACATATTCCAT
AGCACACGTCTGAACTCTTTCCCTACACGACGCTCTTCCGATCTNNNATCCCCACGAAACATATTCCAT
AGCACACGTCTGAACTCTTTCCCTACACGACGCTCTTCCGATCTNNNATCCCCACGAAACATATTCCAT
AGCACACGTCTGAACTCTTTCCCTACACGACGCTCTTCCGATCTNNNATCCCCACGAAACATATTCCAT
GAGCACACGTCTGAACTCTTTCCCTACACGACGCTCTTCCGATCTNNNAAAAGAAATAAGACACCTCCC
GAGCACACGTCTGAACTCTTTCCCTACACGACGCTCTTCCGATCTNNNAAAAGAAATAAGACACCTCCC
GAGCACACGTCTGAACTCTTTCCCTACACGACGCTCTTCCGATCTNNNAAAAGAAATAAGACACCTCCC
GAGCACACGTCTGAACTCTTTCCCTACACGACGCTCTTCCGATCTNNNAAAAGAAATAAGACACCTCCC
GAGCACACGTCTGAACTCTTTCCCTACACGACGCTCTTCCGATCTNNNAAAAGAAATAAGACACCTCCC
GAGCACACGTCTGAACTCTTTCCCTACACGACGCTCTTCCGATCTNNNCTGAGGCAGGAGAATTGCTT
GAGCACACGTCTGAACTCTTTCCCTACACGACGCTCTTCCGATCTNNNCTGAGGCAGGAGAATTGCTT
GAGCACACGTCTGAACTCTTTCCCTACACGACGCTCTTCCGATCTNNNCTGAGGCAGGAGAATTGCTT
GAGCACACGTCTGAACTCTTTCCCTACACGACGCTCTTCCGATCTNNNCTGAGGCAGGAGAATTGCTT
GAGCACACGTCTGAACTCTTTCCCTACACGACGCTCTTCCGATCTNNNCTGAGGCAGGAGAATTGCTT
GCACACGTCTGAACTCTTTCCCTACACGACGCTCTTCCGATCTNNNCTAAAAGCCTAGACGTCACCACT
GCACACGTCTGAACTCTTTCCCTACACGACGCTCTTCCGATCTNNNCTAAAAGCCTAGACGTCACCACT
GCACACGTCTGAACTCTTTCCCTACACGACGCTCTTCCGATCTNNNCTAAAAGCCTAGACGTCACCACT
GCACACGTCTGAACTCTTTCCCTACACGACGCTCTTCCGATCTNNNCTAAAAGCCTAGACGTCACCACT
GCACACGTCTGAACTCTTTCCCTACACGACGCTCTTCCGATCTNNNCTAAAAGCCTAGACGTCACCACT
AGCACACGTCTGAACTCTTTCCCTACACGACGCTCTTCCGATCTNNNATGGTGGCTTGCAGGTATATCA
AGCACACGTCTGAACTCTTTCCCTACACGACGCTCTTCCGATCTNNNATGGTGGCTTGCAGGTATATCA
AGCACACGTCTGAACTCTTTCCCTACACGACGCTCTTCCGATCTNNNATGGTGGCTTGCAGGTATATCA
AGCACACGTCTGAACTCTTTCCCTACACGACGCTCTTCCGATCTNNNATGGTGGCTTGCAGGTATATCA
AGCACACGTCTGAACTCTTTCCCTACACGACGCTCTTCCGATCTNNNATGGTGGCTTGCAGGTATATCA
AGCACACGTCTGAACTCTTTCCCTACACGACGCTCTTCCGATCTNNNAGTATACATGATGGAAAAGAG
AGCACACGTCTGAACTCTTTCCCTACACGACGCTCTTCCGATCTNNNAGTATACATGATGGAAAAGAG
AGCACACGTCTGAACTCTTTCCCTACACGACGCTCTTCCGATCTNNNAGTATACATGATGGAAAAGAG
AGCACACGTCTGAACTCTTTCCCTACACGACGCTCTTCCGATCTNNNAGTATACATGATGGAAAAGAG
AGCACACGTCTGAACTCTTTCCCTACACGACGCTCTTCCGATCTNNNAGTATACATGATGGAAAAGAG
GCACACGTCTGAACTCTTTCCCTACACGACGCTCTTCCGATCTNNNGCTGAGATCATGCCATTGAACT
GCACACGTCTGAACTCTTTCCCTACACGACGCTCTTCCGATCTNNNGCTGAGATCATGCCATTGAACT
GCACACGTCTGAACTCTTTCCCTACACGACGCTCTTCCGATCTNNNGCTGAGATCATGCCATTGAACT
GCACACGTCTGAACTCTTTCCCTACACGACGCTCTTCCGATCTNNNGCTGAGATCATGCCATTGAACT

GCACACGTCTGAACTCTTTCCCTACACGACGCTCTTCCGATCTNNNGCTGAGATCATGCCATTGAACT
GCACACGTCTGAACTCTTTCCCTACACGACGCTCTTCCGATCTNNNGAGAAGATGGCTGGAGGACGC
GCACACGTCTGAACTCTTTCCCTACACGACGCTCTTCCGATCTNNNGAGAAGATGGCTGGAGGACGC
GCACACGTCTGAACTCTTTCCCTACACGACGCTCTTCCGATCTNNNGAGAAGATGGCTGGAGGACGC
GCACACGTCTGAACTCTTTCCCTACACGACGCTCTTCCGATCTNNNGAGAAGATGGCTGGAGGACGC
GCACACGTCTGAACTCTTTCCCTACACGACGCTCTTCCGATCTNNNGAGAAGATGGCTGGAGGACGC
GCACACGTCTGAACTCTTTCCCTACACGACGCTCTTCCGATCTNNNAATCACTTGAACCTAGGAGGCA
GCACACGTCTGAACTCTTTCCCTACACGACGCTCTTCCGATCTNNNAATCACTTGAACCTAGGAGGCA
GCACACGTCTGAACTCTTTCCCTACACGACGCTCTTCCGATCTNNNAATCACTTGAACCTAGGAGGCA
GCACACGTCTGAACTCTTTCCCTACACGACGCTCTTCCGATCTNNNAATCACTTGAACCTAGGAGGCA
GCACACGTCTGAACTCTTTCCCTACACGACGCTCTTCCGATCTNNNAATCACTTGAACCTAGGAGGCA
GCACACGTCTGAACTCTTTCCCTACACGACGCTCTTCCGATCTNNNGGTGCATACTGGTAATTCCAAG
GCACACGTCTGAACTCTTTCCCTACACGACGCTCTTCCGATCTNNNGGTGCATACTGGTAATTCCAAG
GCACACGTCTGAACTCTTTCCCTACACGACGCTCTTCCGATCTNNNGGTGCATACTGGTAATTCCAAG
GCACACGTCTGAACTCTTTCCCTACACGACGCTCTTCCGATCTNNNGGTGCATACTGGTAATTCCAAG
GCACACGTCTGAACTCTTTCCCTACACGACGCTCTTCCGATCTNNNGGTGCATACTGGTAATTCCAAG
AGCACACGTCTGAACTCTTTCCCTACACGACGCTCTTCCGATCTNNNCTCAGAGTTAGAGGTCTGTGC
AGCACACGTCTGAACTCTTTCCCTACACGACGCTCTTCCGATCTNNNCTCAGAGTTAGAGGTCTGTGC
AGCACACGTCTGAACTCTTTCCCTACACGACGCTCTTCCGATCTNNNCTCAGAGTTAGAGGTCTGTGC
AGCACACGTCTGAACTCTTTCCCTACACGACGCTCTTCCGATCTNNNCTCAGAGTTAGAGGTCTGTGC
AGCACACGTCTGAACTCTTTCCCTACACGACGCTCTTCCGATCTNNNCTCAGAGTTAGAGGTCTGTGC
GAGCACACGTCTGAACTCTTTCCCTACACGACGCTCTTCCGATCTNNNTGTAATCTCAGCACTTTGGG
GAGCACACGTCTGAACTCTTTCCCTACACGACGCTCTTCCGATCTNNNTGTAATCTCAGCACTTTGGG
GAGCACACGTCTGAACTCTTTCCCTACACGACGCTCTTCCGATCTNNNTGTAATCTCAGCACTTTGGG
GAGCACACGTCTGAACTCTTTCCCTACACGACGCTCTTCCGATCTNNNTGTAATCTCAGCACTTTGGG
GAGCACACGTCTGAACTCTTTCCCTACACGACGCTCTTCCGATCTNNNTGTAATCTCAGCACTTTGGG
AGCACACGTCTGAACTCTTTCCCTACACGACGCTCTTCCGATCTNNNCAGGAATGGTACCAGCTCTTCT
AGCACACGTCTGAACTCTTTCCCTACACGACGCTCTTCCGATCTNNNCAGGAATGGTACCAGCTCTTCT
AGCACACGTCTGAACTCTTTCCCTACACGACGCTCTTCCGATCTNNNCAGGAATGGTACCAGCTCTTCT
AGCACACGTCTGAACTCTTTCCCTACACGACGCTCTTCCGATCTNNNCAGGAATGGTACCAGCTCTTCT
AGCACACGTCTGAACTCTTTCCCTACACGACGCTCTTCCGATCTNNNCAGGAATGGTACCAGCTCTTCT
AGCACACGTCTGAACTCTTTCCCTACACGACGCTCTTCCGATCTNNNATCTGAACTGCAGTCCTATAG
AGCACACGTCTGAACTCTTTCCCTACACGACGCTCTTCCGATCTNNNATCTGAACTGCAGTCCTATAG
AGCACACGTCTGAACTCTTTCCCTACACGACGCTCTTCCGATCTNNNATCTGAACTGCAGTCCTATAG
AGCACACGTCTGAACTCTTTCCCTACACGACGCTCTTCCGATCTNNNATCTGAACTGCAGTCCTATAG
AGCACACGTCTGAACTCTTTCCCTACACGACGCTCTTCCGATCTNNNATCTGAACTGCAGTCCTATAG
ACACGTCTGAACTCTTTCCCTACACGACGCTCTTCCGATCTNNNTCCACTTAAAACCAACACCCAGGCA
ACACGTCTGAACTCTTTCCCTACACGACGCTCTTCCGATCTNNNTCCACTTAAAACCAACACCCAGGCA
ACACGTCTGAACTCTTTCCCTACACGACGCTCTTCCGATCTNNNTCCACTTAAAACCAACACCCAGGCA
ACACGTCTGAACTCTTTCCCTACACGACGCTCTTCCGATCTNNNTCCACTTAAAACCAACACCCAGGCA
ACACGTCTGAACTCTTTCCCTACACGACGCTCTTCCGATCTNNNTCCACTTAAAACCAACACCCAGGCA
ACACGTCTGAACTCTTTCCCTACACGACGCTCTTCCGATCTNNNAACTTCACCACCATCGACGTGCATCG
ACACGTCTGAACTCTTTCCCTACACGACGCTCTTCCGATCTNNNAACTTCACCACCATCGACGTGCATCG

ACACGTCTGAACTCTTTCCCTACACGACGCTCTTCCGATCTNNNAACTTCACCACCATCGACGTGCATCG

ACACGTCTGAACTCTTTCCCTACACGACGCTCTTCCGATCTNNNAACTTCACCACCATCGACGTGCATCG

ACACGTCTGAACTCTTTCCCTACACGACGCTCTTCCGATCTNNNAACTTCACCACCATCGACGTGCATCG

AGCACACGTCTGAACTCTTTCCCTACACGACGCTCTTCCGATCTNNNTATGTTTGTTTATGTGCTGTTTA

AGCACACGTCTGAACTCTTTCCCTACACGACGCTCTTCCGATCTNNNTATGTTTGTTTATGTGCTGTTTA

AGCACACGTCTGAACTCTTTCCCTACACGACGCTCTTCCGATCTNNNTATGTTTGTTTATGTGCTGTTTA

AGCACACGTCTGAACTCTTTCCCTACACGACGCTCTTCCGATCTNNNTATGTTTGTTTATGTGCTGTTTA

AGCACACGTCTGAACTCTTTCCCTACACGACGCTCTTCCGATCTNNNTATGTTTGTTTATGTGCTGTTTA

AGCACACGTCTGAACTCTTTCCCTACACGACGCTCTTCCGATCTNNNACTAAGGTTCATCTTGTCCCTGG

AGCACACGTCTGAACTCTTTCCCTACACGACGCTCTTCCGATCTNNNACTAAGGTTCATCTTGTCCCTGG

AGCACACGTCTGAACTCTTTCCCTACACGACGCTCTTCCGATCTNNNACTAAGGTTCATCTTGTCCCTGG

AGCACACGTCTGAACTCTTTCCCTACACGACGCTCTTCCGATCTNNNACTAAGGTTCATCTTGTCCCTGG

AGCACACGTCTGAACTCTTTCCCTACACGACGCTCTTCCGATCTNNNACTAAGGTTCATCTTGTCCCTGG

AGAGCACACGTCTGAACTCTTTCCCTACACGACGCTCTTCCGATCTNNNCCAGCTACTTTGGAGGATGA

AGAGCACACGTCTGAACTCTTTCCCTACACGACGCTCTTCCGATCTNNNCCAGCTACTTTGGAGGATGA

AGAGCACACGTCTGAACTCTTTCCCTACACGACGCTCTTCCGATCTNNNCCAGCTACTTTGGAGGATGA

AGAGCACACGTCTGAACTCTTTCCCTACACGACGCTCTTCCGATCTNNNCCAGCTACTTTGGAGGATGA

AGAGCACACGTCTGAACTCTTTCCCTACACGACGCTCTTCCGATCTNNNCCAGCTACTTTGGAGGATGA

AGCACACGTCTGAACTCTTTCCCTACACGACGCTCTTCCGATCTNNNTCAAGTGATTCTCGAGCCTCATC

AGCACACGTCTGAACTCTTTCCCTACACGACGCTCTTCCGATCTNNNTCAAGTGATTCTCGAGCCTCATC

AGCACACGTCTGAACTCTTTCCCTACACGACGCTCTTCCGATCTNNNTCAAGTGATTCTCGAGCCTCATC

AGCACACGTCTGAACTCTTTCCCTACACGACGCTCTTCCGATCTNNNTCAAGTGATTCTCGAGCCTCATC

AGCACACGTCTGAACTCTTTCCCTACACGACGCTCTTCCGATCTNNNTCAAGTGATTCTCGAGCCTCATC

GCACACGTCTGAACTCTTTCCCTACACGACGCTCTTCCGATCTNNNTGTGTTCATGCCTTGATACCCTGCA

GCACACGTCTGAACTCTTTCCCTACACGACGCTCTTCCGATCTNNNTGTGTTCATGCCTTGATACCCTGCA

GCACACGTCTGAACTCTTTCCCTACACGACGCTCTTCCGATCTNNNTGTGTTCATGCCTTGATACCCTGCA

GCACACGTCTGAACTCTTTCCCTACACGACGCTCTTCCGATCTNNNTGTGTTCATGCCTTGATACCCTGCA

GCACACGTCTGAACTCTTTCCCTACACGACGCTCTTCCGATCTNNNTGTGTTCATGCCTTGATACCCTGCA

AGCACACGTCTGAACTCTTTCCCTACACGACGCTCTTCCGATCTNNNTTGTTTTCAAGCACGAAACTGGC

AGCACACGTCTGAACTCTTTCCCTACACGACGCTCTTCCGATCTNNNTTGTTTTCAAGCACGAAACTGGC

AGCACACGTCTGAACTCTTTCCCTACACGACGCTCTTCCGATCTNNNTTGTTTTCAAGCACGAAACTGGC

AGCACACGTCTGAACTCTTTCCCTACACGACGCTCTTCCGATCTNNNTTGTTTTCAAGCACGAAACTGGC

AGCACACGTCTGAACTCTTTCCCTACACGACGCTCTTCCGATCTNNNTTGTTTTCAAGCACGAAACTGGC

AGCACACGTCTGAACTCTTTCCCTACACGACGCTCTTCCGATCTNNNCCATTAGGGGCAGAGCAGAAT

AGCACACGTCTGAACTCTTTCCCTACACGACGCTCTTCCGATCTNNNCCATTAGGGGCAGAGCAGAAT

AGCACACGTCTGAACTCTTTCCCTACACGACGCTCTTCCGATCTNNNCCATTAGGGGCAGAGCAGAAT

AGCACACGTCTGAACTCTTTCCCTACACGACGCTCTTCCGATCTNNNCCATTAGGGGCAGAGCAGAAT

AGCACACGTCTGAACTCTTTCCCTACACGACGCTCTTCCGATCTNNNCCATTAGGGGCAGAGCAGAAT

GAGCACACGTCTGAACTCTTTCCCTACACGACGCTCTTCCGATCTNNNCCTCTAGAAGAAACACCAAGC

GAGCACACGTCTGAACTCTTTCCCTACACGACGCTCTTCCGATCTNNNCCTCTAGAAGAAACACCAAGC

GAGCACACGTCTGAACTCTTTCCCTACACGACGCTCTTCCGATCTNNNCCTCTAGAAGAAACACCAAGC

GAGCACACGTCTGAACTCTTTCCCTACACGACGCTCTTCCGATCTNNNCCTCTAGAAGAAACACCAAGC

GAGCACACGTCTGAACTCTTTCCCTACACGACGCTCTTCCGATCTNNNCCTCTAGAAGAAACACCAAGC

GCACACGTCTGAACTCTTTCCCTACACGACGCTCTTCCGATCTNNNTCTGTAGAGGAGAGGAGAAGCA

GCACACGTCTGAACTCTTTCCCTACACGACGCTCTTCCGATCTNNNTCTGTAGAGGAGAGGAGAAGCA

GCACACGTCTGAACTCTTTCCCTACACGACGCTCTTCCGATCTNNNTCTGTAGAGGAGAGGAGAAGCA

GCACACGTCTGAACTCTTTCCCTACACGACGCTCTTCCGATCTNNNTCTGTAGAGGAGAGGAGAAGCA

GCACACGTCTGAACTCTTTCCCTACACGACGCTCTTCCGATCTNNNTCTGTAGAGGAGAGGAGAAGCA

GAGCACACGTCTGAACTCTTTCCCTACACGACGCTCTTCCGATCTNNNTGGTACCAGCATCTGGATCAT

GAGCACACGTCTGAACTCTTTCCCTACACGACGCTCTTCCGATCTNNNTGGTACCAGCATCTGGATCAT

GAGCACACGTCTGAACTCTTTCCCTACACGACGCTCTTCCGATCTNNNTGGTACCAGCATCTGGATCAT

GAGCACACGTCTGAACTCTTTCCCTACACGACGCTCTTCCGATCTNNNTGGTACCAGCATCTGGATCAT

GAGCACACGTCTGAACTCTTTCCCTACACGACGCTCTTCCGATCTNNNTGGTACCAGCATCTGGATCAT

GAGCACACGTCTGAACTCTTTCCCTACACGACGCTCTTCCGATCTNNNCCAAATCAGGCTATTTTTTGGC

GAGCACACGTCTGAACTCTTTCCCTACACGACGCTCTTCCGATCTNNNCCAAATCAGGCTATTTTTTGGC

GAGCACACGTCTGAACTCTTTCCCTACACGACGCTCTTCCGATCTNNNCCAAATCAGGCTATTTTTTGGC

GAGCACACGTCTGAACTCTTTCCCTACACGACGCTCTTCCGATCTNNNCCAAATCAGGCTATTTTTTGGC

GAGCACACGTCTGAACTCTTTCCCTACACGACGCTCTTCCGATCTNNNCCAAATCAGGCTATTTTTTGGC

GAGCACACGTCTGAACTCTTTCCCTACACGACGCTCTTCCGATCTNNNATTCAAACATGGGTGACTGCA

GAGCACACGTCTGAACTCTTTCCCTACACGACGCTCTTCCGATCTNNNATTCAAACATGGGTGACTGCA

GAGCACACGTCTGAACTCTTTCCCTACACGACGCTCTTCCGATCTNNNATTCAAACATGGGTGACTGCA

GAGCACACGTCTGAACTCTTTCCCTACACGACGCTCTTCCGATCTNNNATTCAAACATGGGTGACTGCA

GAGCACACGTCTGAACTCTTTCCCTACACGACGCTCTTCCGATCTNNNATTCAAACATGGGTGACTGCA

AGCACACGTCTGAACTCTTTCCCTACACGACGCTCTTCCGATCTNNNAGATTCTGCTTGGTTGTAGCTGA

AGCACACGTCTGAACTCTTTCCCTACACGACGCTCTTCCGATCTNNNAGATTCTGCTTGGTTGTAGCTGA

AGCACACGTCTGAACTCTTTCCCTACACGACGCTCTTCCGATCTNNNAGATTCTGCTTGGTTGTAGCTGA

AGCACACGTCTGAACTCTTTCCCTACACGACGCTCTTCCGATCTNNNAGATTCTGCTTGGTTGTAGCTGA

AGCACACGTCTGAACTCTTTCCCTACACGACGCTCTTCCGATCTNNNAGATTCTGCTTGGTTGTAGCTGA

AGCACACGTCTGAACTCTTTCCCTACACGACGCTCTTCCGATCTNNNCCATGAGGACAGTAGCAATCCC

AGCACACGTCTGAACTCTTTCCCTACACGACGCTCTTCCGATCTNNNCCATGAGGACAGTAGCAATCCC

AGCACACGTCTGAACTCTTTCCCTACACGACGCTCTTCCGATCTNNNCCATGAGGACAGTAGCAATCCC

AGCACACGTCTGAACTCTTTCCCTACACGACGCTCTTCCGATCTNNNCCATGAGGACAGTAGCAATCCC

AGCACACGTCTGAACTCTTTCCCTACACGACGCTCTTCCGATCTNNNCCATGAGGACAGTAGCAATCCC

AGCACACGTCTGAACTCTTTCCCTACACGACGCTCTTCCGATCTNNNCTCCAACGCAGACTGGTTATTT

AGCACACGTCTGAACTCTTTCCCTACACGACGCTCTTCCGATCTNNNCTCCAACGCAGACTGGTTATTT

AGCACACGTCTGAACTCTTTCCCTACACGACGCTCTTCCGATCTNNNCTCCAACGCAGACTGGTTATTT

AGCACACGTCTGAACTCTTTCCCTACACGACGCTCTTCCGATCTNNNCTCCAACGCAGACTGGTTATTT

AGCACACGTCTGAACTCTTTCCCTACACGACGCTCTTCCGATCTNNNCTCCAACGCAGACTGGTTATTT

AGCACACGTCTGAACTCTTTCCCTACACGACGCTCTTCCGATCTNNNGGGAACATAGGCCCTAATACTC

AGCACACGTCTGAACTCTTTCCCTACACGACGCTCTTCCGATCTNNNGGGAACATAGGCCCTAATACTC

AGCACACGTCTGAACTCTTTCCCTACACGACGCTCTTCCGATCTNNNGGGAACATAGGCCCTAATACTC

AGCACACGTCTGAACTCTTTCCCTACACGACGCTCTTCCGATCTNNNGGGAACATAGGCCCTAATACTC

AGCACACGTCTGAACTCTTTCCCTACACGACGCTCTTCCGATCTNNNGGGAACATAGGCCCTAATACTC

GCACACGTCTGAACTCTTTCCCTACACGACGCTCTTCCGATCTNNNAGTTACTTCGTTTCCAGTTGGGAC

GCACACGTCTGAACTCTTTCCCTACACGACGCTCTTCCGATCTNNNAGTTACTTCGTTTCCAGTTGGGAC

GCACACGTCTGAACTCTTTCCCTACACGACGCTCTTCCGATCTNNNAGTTACTTCGTTTCCAGTTGGGAC

CACACGTCTGAACTCTTTCCCTACACGACGCTCTTCCGATCTNNNAGTTACTTCGTTTCCAGTTGGGA
CACACGTCTGAACTCTTTCCCTACACGACGCTCTTCCGATCTNNNAGTTACTTCGTTTCCAGTTGGGA
AGCACACGTCTGAACTCTTTCCCTACACGACGCTCTTCCGATCTNNNCATGGGGAGACATCATCTGTA
AGCACACGTCTGAACTCTTTCCCTACACGACGCTCTTCCGATCTNNNCATGGGGAGACATCATCTGTA
AGCACACGTCTGAACTCTTTCCCTACACGACGCTCTTCCGATCTNNNCATGGGGAGACATCATCTGTA
AGCACACGTCTGAACTCTTTCCCTACACGACGCTCTTCCGATCTNNNCATGGGGAGACATCATCTGTA
AGCACACGTCTGAACTCTTTCCCTACACGACGCTCTTCCGATCTNNNCATGGGGAGACATCATCTGTA
ACACGTCTGAACTCTTTCCCTACACGACGCTCTTCCGATCTNNNTGACACCTTCTTTCTTCCTAGGCCAT
ACACGTCTGAACTCTTTCCCTACACGACGCTCTTCCGATCTNNNTGACACCTTCTTTCTTCCTAGGCCAT
ACACGTCTGAACTCTTTCCCTACACGACGCTCTTCCGATCTNNNTGACACCTTCTTTCTTCCTAGGCCAT
ACACGTCTGAACTCTTTCCCTACACGACGCTCTTCCGATCTNNNTGACACCTTCTTTCTTCCTAGGCCAT
ACACGTCTGAACTCTTTCCCTACACGACGCTCTTCCGATCTNNNTGACACCTTCTTTCTTCCTAGGCCAT
GCACACGTCTGAACTCTTTCCCTACACGACGCTCTTCCGATCTNNNCTGGCTTGTCCTACAGATCAACT
GCACACGTCTGAACTCTTTCCCTACACGACGCTCTTCCGATCTNNNCTGGCTTGTCCTACAGATCAACT
GCACACGTCTGAACTCTTTCCCTACACGACGCTCTTCCGATCTNNNCTGGCTTGTCCTACAGATCAACT
GCACACGTCTGAACTCTTTCCCTACACGACGCTCTTCCGATCTNNNCTGGCTTGTCCTACAGATCAACT
GCACACGTCTGAACTCTTTCCCTACACGACGCTCTTCCGATCTNNNCTGGCTTGTCCTACAGATCAACT
AGCACACGTCTGAACTCTTTCCCTACACGACGCTCTTCCGATCTNNNCCACACTGCTTTTCACAATGGT
AGCACACGTCTGAACTCTTTCCCTACACGACGCTCTTCCGATCTNNNCCACACTGCTTTTCACAATGGT
AGCACACGTCTGAACTCTTTCCCTACACGACGCTCTTCCGATCTNNNCCACACTGCTTTTCACAATGGT
AGCACACGTCTGAACTCTTTCCCTACACGACGCTCTTCCGATCTNNNCCACACTGCTTTTCACAATGGT
AGCACACGTCTGAACTCTTTCCCTACACGACGCTCTTCCGATCTNNNCCACACTGCTTTTCACAATGGT
AGCACACGTCTGAACTCTTTCCCTACACGACGCTCTTCCGATCTNNNACTTGGATGTCTGTTTTCTCC
AGCACACGTCTGAACTCTTTCCCTACACGACGCTCTTCCGATCTNNNACTTGGATGTCTGTTTTCTCC
AGCACACGTCTGAACTCTTTCCCTACACGACGCTCTTCCGATCTNNNACTTGGATGTCTGTTTTCTCC
AGCACACGTCTGAACTCTTTCCCTACACGACGCTCTTCCGATCTNNNACTTGGATGTCTGTTTTCTCC
AGCACACGTCTGAACTCTTTCCCTACACGACGCTCTTCCGATCTNNNACTTGGATGTCTGTTTTCTCC
GAGCACACGTCTGAACTCTTTCCCTACACGACGCTCTTCCGATCTNNNCAAACAGCAAGAGGGAAAT
GAGCACACGTCTGAACTCTTTCCCTACACGACGCTCTTCCGATCTNNNCAAACAGCAAGAGGGAAAT
GAGCACACGTCTGAACTCTTTCCCTACACGACGCTCTTCCGATCTNNNCAAACAGCAAGAGGGAAAT
GAGCACACGTCTGAACTCTTTCCCTACACGACGCTCTTCCGATCTNNNCAAACAGCAAGAGGGAAAT
GAGCACACGTCTGAACTCTTTCCCTACACGACGCTCTTCCGATCTNNNCAAACAGCAAGAGGGAAAT
GCACACGTCTGAACTCTTTCCCTACACGACGCTCTTCCGATCTNNNTCGAATCTCTCATTCCTCAAGCA
GCACACGTCTGAACTCTTTCCCTACACGACGCTCTTCCGATCTNNNTCGAATCTCTCATTCCTCAAGCA
GCACACGTCTGAACTCTTTCCCTACACGACGCTCTTCCGATCTNNNTCGAATCTCTCATTCCTCAAGCA
GCACACGTCTGAACTCTTTCCCTACACGACGCTCTTCCGATCTNNNTCGAATCTCTCATTCCTCAAGCA
GCACACGTCTGAACTCTTTCCCTACACGACGCTCTTCCGATCTNNNTCGAATCTCTCATTCCTCAAGCA
AGCACACGTCTGAACTCTTTCCCTACACGACGCTCTTCCGATCTNNNCTGGAGTGTCTCTTACCACAA
AGCACACGTCTGAACTCTTTCCCTACACGACGCTCTTCCGATCTNNNCTGGAGTGTCTCTTACCACAA
AGCACACGTCTGAACTCTTTCCCTACACGACGCTCTTCCGATCTNNNCTGGAGTGTCTCTTACCACAA
AGCACACGTCTGAACTCTTTCCCTACACGACGCTCTTCCGATCTNNNCTGGAGTGTCTCTTACCACAA
AGCACACGTCTGAACTCTTTCCCTACACGACGCTCTTCCGATCTNNNCTGGAGTGTCTCTTACCACAA
AGCACACGTCTGAACTCTTTCCCTACACGACGCTCTTCCGATCTNNNACAGCTGTGTCTTTGTGTTTG

GAGCACACGTCTGAACTCTTTCCCTACACGACGCTCTTCCGATCTNNNACAGCTGTGTCTTTGTGTTTGT
GAGCACACGTCTGAACTCTTTCCCTACACGACGCTCTTCCGATCTNNNACAGCTGTGTCTTTGTGTTTGT
GAGCACACGTCTGAACTCTTTCCCTACACGACGCTCTTCCGATCTNNNACAGCTGTGTCTTTGTGTTTGT
GAGCACACGTCTGAACTCTTTCCCTACACGACGCTCTTCCGATCTNNNACAGCTGTGTCTTTGTGTTTGT
AGCACACGTCTGAACTCTTTCCCTACACGACGCTCTTCCGATCTNNNTAGACTCCGCATTGAAAAGACC
AGCACACGTCTGAACTCTTTCCCTACACGACGCTCTTCCGATCTNNNTAGACTCCGCATTGAAAAGACC
AGCACACGTCTGAACTCTTTCCCTACACGACGCTCTTCCGATCTNNNTAGACTCCGCATTGAAAAGACC
AGCACACGTCTGAACTCTTTCCCTACACGACGCTCTTCCGATCTNNNTAGACTCCGCATTGAAAAGACC
AGCACACGTCTGAACTCTTTCCCTACACGACGCTCTTCCGATCTNNNTAGACTCCGCATTGAAAAGACC
GAGCACACGTCTGAACTCTTTCCCTACACGACGCTCTTCCGATCTNNNAAGTCTCTCTCTCTCTCTCCC
GAGCACACGTCTGAACTCTTTCCCTACACGACGCTCTTCCGATCTNNNAAGTCTCTCTCTCTCTCTCCC
GAGCACACGTCTGAACTCTTTCCCTACACGACGCTCTTCCGATCTNNNAAGTCTCTCTCTCTCTCTCCC
GAGCACACGTCTGAACTCTTTCCCTACACGACGCTCTTCCGATCTNNNAAGTCTCTCTCTCTCTCTCCC
GAGCACACGTCTGAACTCTTTCCCTACACGACGCTCTTCCGATCTNNNAAGTCTCTCTCTCTCTCTCCC
GGAAGAGCACACGTCTGAACTCTTTCCCTACACGACGCTCTTCCGATCTNNNTCTGACAGAAGCGATGC
GGAAGAGCACACGTCTGAACTCTTTCCCTACACGACGCTCTTCCGATCTNNNTCTGACAGAAGCGATGC
GGAAGAGCACACGTCTGAACTCTTTCCCTACACGACGCTCTTCCGATCTNNNTCTGACAGAAGCGATGC
GGAAGAGCACACGTCTGAACTCTTTCCCTACACGACGCTCTTCCGATCTNNNTCTGACAGAAGCGATGC
GGAAGAGCACACGTCTGAACTCTTTCCCTACACGACGCTCTTCCGATCTNNNTCTGACAGAAGCGATGC
AGCACACGTCTGAACTCTTTCCCTACACGACGCTCTTCCGATCTNNNTTAGCCTTGTGAGTAGTTGGGA
AGCACACGTCTGAACTCTTTCCCTACACGACGCTCTTCCGATCTNNNTTAGCCTTGTGAGTAGTTGGGA
AGCACACGTCTGAACTCTTTCCCTACACGACGCTCTTCCGATCTNNNTTAGCCTTGTGAGTAGTTGGGA
AGCACACGTCTGAACTCTTTCCCTACACGACGCTCTTCCGATCTNNNTTAGCCTTGTGAGTAGTTGGGA
AGCACACGTCTGAACTCTTTCCCTACACGACGCTCTTCCGATCTNNNTTAGCCTTGTGAGTAGTTGGGA
GAGCACACGTCTGAACTCTTTCCCTACACGACGCTCTTCCGATCTNNNGCCAATGGCGTAACTATTCTG
GAGCACACGTCTGAACTCTTTCCCTACACGACGCTCTTCCGATCTNNNGCCAATGGCGTAACTATTCTG
GAGCACACGTCTGAACTCTTTCCCTACACGACGCTCTTCCGATCTNNNGCCAATGGCGTAACTATTCTG
GAGCACACGTCTGAACTCTTTCCCTACACGACGCTCTTCCGATCTNNNGCCAATGGCGTAACTATTCTG
GAGCACACGTCTGAACTCTTTCCCTACACGACGCTCTTCCGATCTNNNGCCAATGGCGTAACTATTCTG
AGAGCACACGTCTGAACTCTTTCCCTACACGACGCTCTTCCGATCTNNNCTAACTACCCAGGAGGCTGA
AGAGCACACGTCTGAACTCTTTCCCTACACGACGCTCTTCCGATCTNNNCTAACTACCCAGGAGGCTGA
AGAGCACACGTCTGAACTCTTTCCCTACACGACGCTCTTCCGATCTNNNCTAACTACCCAGGAGGCTGA
AGAGCACACGTCTGAACTCTTTCCCTACACGACGCTCTTCCGATCTNNNCTAACTACCCAGGAGGCTGA
AGAGCACACGTCTGAACTCTTTCCCTACACGACGCTCTTCCGATCTNNNCTAACTACCCAGGAGGCTGA
AGCACACGTCTGAACTCTTTCCCTACACGACGCTCTTCCGATCTNNNATTATTTGCTTCATTTCTCTGTTC
AGCACACGTCTGAACTCTTTCCCTACACGACGCTCTTCCGATCTNNNATTATTTGCTTCATTTCTCTGTTC
AGCACACGTCTGAACTCTTTCCCTACACGACGCTCTTCCGATCTNNNATTATTTGCTTCATTTCTCTGTTC
AGCACACGTCTGAACTCTTTCCCTACACGACGCTCTTCCGATCTNNNATTATTTGCTTCATTTCTCTGTTC
AGCACACGTCTGAACTCTTTCCCTACACGACGCTCTTCCGATCTNNNATTATTTGCTTCATTTCTCTGTTC
GAGCACACGTCTGAACTCTTTCCCTACACGACGCTCTTCCGATCTNNNCTTCTTTTCTCCTAGGTGGCTC
GAGCACACGTCTGAACTCTTTCCCTACACGACGCTCTTCCGATCTNNNCTTCTTTTCTCCTAGGTGGCTC
GAGCACACGTCTGAACTCTTTCCCTACACGACGCTCTTCCGATCTNNNCTTCTTTTCTCCTAGGTGGCTC
GAGCACACGTCTGAACTCTTTCCCTACACGACGCTCTTCCGATCTNNNCTTCTTTTCTCCTAGGTGGCTC

GAGCACACGTCTGAACTCTTTCCCTACACGACGCTCTTCCGATCTNNNCTTCTTTTCTCCTAGGTGGCT
GCACACGTCTGAACTCTTTCCCTACACGACGCTCTTCCGATCTNNNGAGGCAGATGGGGGTGTCAGC
GCACACGTCTGAACTCTTTCCCTACACGACGCTCTTCCGATCTNNNGAGGCAGATGGGGGTGTCAGC
GCACACGTCTGAACTCTTTCCCTACACGACGCTCTTCCGATCTNNNGAGGCAGATGGGGGTGTCAGC
GCACACGTCTGAACTCTTTCCCTACACGACGCTCTTCCGATCTNNNGAGGCAGATGGGGGTGTCAGC
GCACACGTCTGAACTCTTTCCCTACACGACGCTCTTCCGATCTNNNGAGGCAGATGGGGGTGTCAGC
GCACACGTCTGAACTCTTTCCCTACACGACGCTCTTCCGATCTNNNGATGGGCTGGGGTTTCTCTCCA
GCACACGTCTGAACTCTTTCCCTACACGACGCTCTTCCGATCTNNNGATGGGCTGGGGTTTCTCTCCA
GCACACGTCTGAACTCTTTCCCTACACGACGCTCTTCCGATCTNNNGATGGGCTGGGGTTTCTCTCCA
GCACACGTCTGAACTCTTTCCCTACACGACGCTCTTCCGATCTNNNGATGGGCTGGGGTTTCTCTCCA
GCACACGTCTGAACTCTTTCCCTACACGACGCTCTTCCGATCTNNNGATGGGCTGGGGTTTCTCTCCA
AGCACACGTCTGAACTCTTTCCCTACACGACGCTCTTCCGATCTNNNCGCAGTCACTAACTTACCCTCAT
AGCACACGTCTGAACTCTTTCCCTACACGACGCTCTTCCGATCTNNNCGCAGTCACTAACTTACCCTCAT
AGCACACGTCTGAACTCTTTCCCTACACGACGCTCTTCCGATCTNNNCGCAGTCACTAACTTACCCTCAT
AGCACACGTCTGAACTCTTTCCCTACACGACGCTCTTCCGATCTNNNCGCAGTCACTAACTTACCCTCAT
AGCACACGTCTGAACTCTTTCCCTACACGACGCTCTTCCGATCTNNNCGCAGTCACTAACTTACCCTCAT
AGCACACGTCTGAACTCTTTCCCTACACGACGCTCTTCCGATCTNNNCAATGCCCTCTTTGATTCTGCTC
AGCACACGTCTGAACTCTTTCCCTACACGACGCTCTTCCGATCTNNNCAATGCCCTCTTTGATTCTGCTC
AGCACACGTCTGAACTCTTTCCCTACACGACGCTCTTCCGATCTNNNCAATGCCCTCTTTGATTCTGCTC
AGCACACGTCTGAACTCTTTCCCTACACGACGCTCTTCCGATCTNNNCAATGCCCTCTTTGATTCTGCTC
AGCACACGTCTGAACTCTTTCCCTACACGACGCTCTTCCGATCTNNNCAATGCCCTCTTTGATTCTGCTC
AGCACACGTCTGAACTCTTTCCCTACACGACGCTCTTCCGATCTNNNGGTGTGTAAGCAGCCTAGGTT
AGCACACGTCTGAACTCTTTCCCTACACGACGCTCTTCCGATCTNNNGGTGTGTAAGCAGCCTAGGTT
AGCACACGTCTGAACTCTTTCCCTACACGACGCTCTTCCGATCTNNNGGTGTGTAAGCAGCCTAGGTT
AGCACACGTCTGAACTCTTTCCCTACACGACGCTCTTCCGATCTNNNGGTGTGTAAGCAGCCTAGGTT
AGCACACGTCTGAACTCTTTCCCTACACGACGCTCTTCCGATCTNNNGGTGTGTAAGCAGCCTAGGTT
AGCACACGTCTGAACTCTTTCCCTACACGACGCTCTTCCGATCTNNNCTCAAGCAGCCCTCTATACTAG
AGCACACGTCTGAACTCTTTCCCTACACGACGCTCTTCCGATCTNNNCTCAAGCAGCCCTCTATACTAG
AGCACACGTCTGAACTCTTTCCCTACACGACGCTCTTCCGATCTNNNCTCAAGCAGCCCTCTATACTAG
AGCACACGTCTGAACTCTTTCCCTACACGACGCTCTTCCGATCTNNNCTCAAGCAGCCCTCTATACTAG
AGCACACGTCTGAACTCTTTCCCTACACGACGCTCTTCCGATCTNNNCTCAAGCAGCCCTCTATACTAG
GAGCACACGTCTGAACTCTTTCCCTACACGACGCTCTTCCGATCTNNNGGAAGGGCAGGCCTATTCTA
GAGCACACGTCTGAACTCTTTCCCTACACGACGCTCTTCCGATCTNNNGGAAGGGCAGGCCTATTCTA
GAGCACACGTCTGAACTCTTTCCCTACACGACGCTCTTCCGATCTNNNGGAAGGGCAGGCCTATTCTA
GAGCACACGTCTGAACTCTTTCCCTACACGACGCTCTTCCGATCTNNNGGAAGGGCAGGCCTATTCTA
GAGCACACGTCTGAACTCTTTCCCTACACGACGCTCTTCCGATCTNNNGGAAGGGCAGGCCTATTCTA
AGCACACGTCTGAACTCTTTCCCTACACGACGCTCTTCCGATCTNNNCTCTGGGACGCTAGGATTTCT
AGCACACGTCTGAACTCTTTCCCTACACGACGCTCTTCCGATCTNNNCTCTGGGACGCTAGGATTTCT
AGCACACGTCTGAACTCTTTCCCTACACGACGCTCTTCCGATCTNNNCTCTGGGACGCTAGGATTTCT
AGCACACGTCTGAACTCTTTCCCTACACGACGCTCTTCCGATCTNNNCTCTGGGACGCTAGGATTTCT
AGCACACGTCTGAACTCTTTCCCTACACGACGCTCTTCCGATCTNNNCTCTGGGACGCTAGGATTTCT
AGCACACGTCTGAACTCTTTCCCTACACGACGCTCTTCCGATCTNNNAGTCCGTCTCATGGAATTGAA
AGCACACGTCTGAACTCTTTCCCTACACGACGCTCTTCCGATCTNNNAGTCCGTCTCATGGAATTGAA

GAGCACACGTCTGAACTCTTTCCCTACACGACGCTCTTCCGATCTNNNAGTCCGTCTCATGGAATTGAA
GAGCACACGTCTGAACTCTTTCCCTACACGACGCTCTTCCGATCTNNNAGTCCGTCTCATGGAATTGAA
GAGCACACGTCTGAACTCTTTCCCTACACGACGCTCTTCCGATCTNNNAGTCCGTCTCATGGAATTGAA
GAGCACACGTCTGAACTCTTTCCCTACACGACGCTCTTCCGATCTNNNATTCAAAAAATTAGCCGGGCG
GAGCACACGTCTGAACTCTTTCCCTACACGACGCTCTTCCGATCTNNNATTCAAAAAATTAGCCGGGCG
GAGCACACGTCTGAACTCTTTCCCTACACGACGCTCTTCCGATCTNNNATTCAAAAAATTAGCCGGGCG
GAGCACACGTCTGAACTCTTTCCCTACACGACGCTCTTCCGATCTNNNATTCAAAAAATTAGCCGGGCG
GAGCACACGTCTGAACTCTTTCCCTACACGACGCTCTTCCGATCTNNNATTCAAAAAATTAGCCGGGCG
GAGCACACGTCTGAACTCTTTCCCTACACGACGCTCTTCCGATCTNNNTTACTATTGTTCGCAGGAGTC
GAGCACACGTCTGAACTCTTTCCCTACACGACGCTCTTCCGATCTNNNTTACTATTGTTCGCAGGAGTC
GAGCACACGTCTGAACTCTTTCCCTACACGACGCTCTTCCGATCTNNNTTACTATTGTTCGCAGGAGTC
GAGCACACGTCTGAACTCTTTCCCTACACGACGCTCTTCCGATCTNNNTTACTATTGTTCGCAGGAGTC
GAGCACACGTCTGAACTCTTTCCCTACACGACGCTCTTCCGATCTNNNTTACTATTGTTCGCAGGAGTC
AGAGCACACGTCTGAACTCTTTCCCTACACGACGCTCTTCCGATCTNNNTGGTAGTTTGCTGTGGCTCT
AGAGCACACGTCTGAACTCTTTCCCTACACGACGCTCTTCCGATCTNNNTGGTAGTTTGCTGTGGCTCT
AGAGCACACGTCTGAACTCTTTCCCTACACGACGCTCTTCCGATCTNNNTGGTAGTTTGCTGTGGCTCT
AGAGCACACGTCTGAACTCTTTCCCTACACGACGCTCTTCCGATCTNNNTGGTAGTTTGCTGTGGCTCT
AGAGCACACGTCTGAACTCTTTCCCTACACGACGCTCTTCCGATCTNNNTGGTAGTTTGCTGTGGCTCT
AGCACACGTCTGAACTCTTTCCCTACACGACGCTCTTCCGATCTNNNCTGGTGGAGAAGGACACAACTA
AGCACACGTCTGAACTCTTTCCCTACACGACGCTCTTCCGATCTNNNCTGGTGGAGAAGGACACAACTA
AGCACACGTCTGAACTCTTTCCCTACACGACGCTCTTCCGATCTNNNCTGGTGGAGAAGGACACAACTA
AGCACACGTCTGAACTCTTTCCCTACACGACGCTCTTCCGATCTNNNCTGGTGGAGAAGGACACAACTA
AGCACACGTCTGAACTCTTTCCCTACACGACGCTCTTCCGATCTNNNCTGGTGGAGAAGGACACAACTA
AGCACACGTCTGAACTCTTTCCCTACACGACGCTCTTCCGATCTNNNCATCTTGCAAAAGCTGCAGTAC
AGCACACGTCTGAACTCTTTCCCTACACGACGCTCTTCCGATCTNNNCATCTTGCAAAAGCTGCAGTAC
AGCACACGTCTGAACTCTTTCCCTACACGACGCTCTTCCGATCTNNNCATCTTGCAAAAGCTGCAGTAC
AGCACACGTCTGAACTCTTTCCCTACACGACGCTCTTCCGATCTNNNCATCTTGCAAAAGCTGCAGTAC
AGCACACGTCTGAACTCTTTCCCTACACGACGCTCTTCCGATCTNNNCATCTTGCAAAAGCTGCAGTAC
GAGCACACGTCTGAACTCTTTCCCTACACGACGCTCTTCCGATCTNNNACTTATGGTTTTGAGCAGAGG
GAGCACACGTCTGAACTCTTTCCCTACACGACGCTCTTCCGATCTNNNACTTATGGTTTTGAGCAGAGG
GAGCACACGTCTGAACTCTTTCCCTACACGACGCTCTTCCGATCTNNNACTTATGGTTTTGAGCAGAGG
GAGCACACGTCTGAACTCTTTCCCTACACGACGCTCTTCCGATCTNNNACTTATGGTTTTGAGCAGAGG
GAGCACACGTCTGAACTCTTTCCCTACACGACGCTCTTCCGATCTNNNACTTATGGTTTTGAGCAGAGG
AGCACACGTCTGAACTCTTTCCCTACACGACGCTCTTCCGATCTNNNTCCTTTCTCTATCTCTCTTTCGCC
AGCACACGTCTGAACTCTTTCCCTACACGACGCTCTTCCGATCTNNNTCCTTTCTCTATCTCTCTTTCGCC
AGCACACGTCTGAACTCTTTCCCTACACGACGCTCTTCCGATCTNNNTCCTTTCTCTATCTCTCTTTCGCC
AGCACACGTCTGAACTCTTTCCCTACACGACGCTCTTCCGATCTNNNTCCTTTCTCTATCTCTCTTTCGCC
AGCACACGTCTGAACTCTTTCCCTACACGACGCTCTTCCGATCTNNNTCCTTTCTCTATCTCTCTTTCGCC
AGCACACGTCTGAACTCTTTCCCTACACGACGCTCTTCCGATCTNNNGTCCCTGTGATGTCTGAGAACTT
AGCACACGTCTGAACTCTTTCCCTACACGACGCTCTTCCGATCTNNNGTCCCTGTGATGTCTGAGAACTT
AGCACACGTCTGAACTCTTTCCCTACACGACGCTCTTCCGATCTNNNGTCCCTGTGATGTCTGAGAACTT
AGCACACGTCTGAACTCTTTCCCTACACGACGCTCTTCCGATCTNNNGTCCCTGTGATGTCTGAGAACTT
AGCACACGTCTGAACTCTTTCCCTACACGACGCTCTTCCGATCTNNNGTCCCTGTGATGTCTGAGAACTT

GAGCACACGTCTGAACTCTTTCCCTACACGACGCTCTTCCGATCTNNNAACTATCTGAGGTGGTATGC
GAGCACACGTCTGAACTCTTTCCCTACACGACGCTCTTCCGATCTNNNAACTATCTGAGGTGGTATGC
GAGCACACGTCTGAACTCTTTCCCTACACGACGCTCTTCCGATCTNNNAACTATCTGAGGTGGTATGC
GAGCACACGTCTGAACTCTTTCCCTACACGACGCTCTTCCGATCTNNNAACTATCTGAGGTGGTATGC
GAGCACACGTCTGAACTCTTTCCCTACACGACGCTCTTCCGATCTNNNAACTATCTGAGGTGGTATGC
AGCACACGTCTGAACTCTTTCCCTACACGACGCTCTTCCGATCTNNNGATCTCAGCTCACTGCAACCTC
AGCACACGTCTGAACTCTTTCCCTACACGACGCTCTTCCGATCTNNNGATCTCAGCTCACTGCAACCTC
AGCACACGTCTGAACTCTTTCCCTACACGACGCTCTTCCGATCTNNNGATCTCAGCTCACTGCAACCTC
AGCACACGTCTGAACTCTTTCCCTACACGACGCTCTTCCGATCTNNNGATCTCAGCTCACTGCAACCTC
AGCACACGTCTGAACTCTTTCCCTACACGACGCTCTTCCGATCTNNNGATCTCAGCTCACTGCAACCTC
GAGCACACGTCTGAACTCTTTCCCTACACGACGCTCTTCCGATCTNNNACCCCGTCTCTACTAAGAAGA
GAGCACACGTCTGAACTCTTTCCCTACACGACGCTCTTCCGATCTNNNACCCCGTCTCTACTAAGAAGA
GAGCACACGTCTGAACTCTTTCCCTACACGACGCTCTTCCGATCTNNNACCCCGTCTCTACTAAGAAGA
GAGCACACGTCTGAACTCTTTCCCTACACGACGCTCTTCCGATCTNNNACCCCGTCTCTACTAAGAAGA
GAGCACACGTCTGAACTCTTTCCCTACACGACGCTCTTCCGATCTNNNACCCCGTCTCTACTAAGAAGA
AGCACACGTCTGAACTCTTTCCCTACACGACGCTCTTCCGATCTNNNTGTTATTCCAGCTACTTGGGA
AGCACACGTCTGAACTCTTTCCCTACACGACGCTCTTCCGATCTNNNTGTTATTCCAGCTACTTGGGA
AGCACACGTCTGAACTCTTTCCCTACACGACGCTCTTCCGATCTNNNTGTTATTCCAGCTACTTGGGA
AGCACACGTCTGAACTCTTTCCCTACACGACGCTCTTCCGATCTNNNTGTTATTCCAGCTACTTGGGA
AGCACACGTCTGAACTCTTTCCCTACACGACGCTCTTCCGATCTNNNTGTTATTCCAGCTACTTGGGA
GCACACGTCTGAACTCTTTCCCTACACGACGCTCTTCCGATCTNNNTTCCAAATAGCTGGGACTACAG
GCACACGTCTGAACTCTTTCCCTACACGACGCTCTTCCGATCTNNNTTCCAAATAGCTGGGACTACAG
GCACACGTCTGAACTCTTTCCCTACACGACGCTCTTCCGATCTNNNTTCCAAATAGCTGGGACTACAG
GCACACGTCTGAACTCTTTCCCTACACGACGCTCTTCCGATCTNNNTTCCAAATAGCTGGGACTACAG
GCACACGTCTGAACTCTTTCCCTACACGACGCTCTTCCGATCTNNNTTCCAAATAGCTGGGACTACAG
AGCACACGTCTGAACTCTTTCCCTACACGACGCTCTTCCGATCTNNNGTGTGTTCTTTTTTTCTTTTCCC
AGCACACGTCTGAACTCTTTCCCTACACGACGCTCTTCCGATCTNNNGTGTGTTCTTTTTTTCTTTTCCC
AGCACACGTCTGAACTCTTTCCCTACACGACGCTCTTCCGATCTNNNGTGTGTTCTTTTTTTCTTTTCCC
AGCACACGTCTGAACTCTTTCCCTACACGACGCTCTTCCGATCTNNNGTGTGTTCTTTTTTTCTTTTCCC
AGCACACGTCTGAACTCTTTCCCTACACGACGCTCTTCCGATCTNNNGTGTGTTCTTTTTTTCTTTTCCC
GAGCACACGTCTGAACTCTTTCCCTACACGACGCTCTTCCGATCTNNNTCTTGACCTGTGAAAGCAGT
GAGCACACGTCTGAACTCTTTCCCTACACGACGCTCTTCCGATCTNNNTCTTGACCTGTGAAAGCAGT
GAGCACACGTCTGAACTCTTTCCCTACACGACGCTCTTCCGATCTNNNTCTTGACCTGTGAAAGCAGT
GAGCACACGTCTGAACTCTTTCCCTACACGACGCTCTTCCGATCTNNNTCTTGACCTGTGAAAGCAGT
GAGCACACGTCTGAACTCTTTCCCTACACGACGCTCTTCCGATCTNNNTCTTGACCTGTGAAAGCAGT
AGCACACGTCTGAACTCTTTCCCTACACGACGCTCTTCCGATCTNNNCCTGTCTTCCTCCTCCAACTTTC
AGCACACGTCTGAACTCTTTCCCTACACGACGCTCTTCCGATCTNNNCCTGTCTTCCTCCTCCAACTTTC
AGCACACGTCTGAACTCTTTCCCTACACGACGCTCTTCCGATCTNNNCCTGTCTTCCTCCTCCAACTTTC
AGCACACGTCTGAACTCTTTCCCTACACGACGCTCTTCCGATCTNNNCCTGTCTTCCTCCTCCAACTTTC
AGCACACGTCTGAACTCTTTCCCTACACGACGCTCTTCCGATCTNNNCCTGTCTTCCTCCTCCAACTTTC
GCACACGTCTGAACTCTTTCCCTACACGACGCTCTTCCGATCTNNNCTTCCTGTTCATCTCCTGCTACAC
GCACACGTCTGAACTCTTTCCCTACACGACGCTCTTCCGATCTNNNCTTCCTGTTCATCTCCTGCTACAC
GCACACGTCTGAACTCTTTCCCTACACGACGCTCTTCCGATCTNNNCTTCCTGTTCATCTCCTGCTACAC

AGCACACGTCTGAACTCTTTCCCTACACGACGCTCTTCCGATCTNNNCTTCCTGTTCATCTCCTGCTACAC
AGCACACGTCTGAACTCTTTCCCTACACGACGCTCTTCCGATCTNNNCTTCCTGTTCATCTCCTGCTACAC
AGCACACGTCTGAACTCTTTCCCTACACGACGCTCTTCCGATCTNNNTAGACACTTCACAAAAAGTAGA
AGCACACGTCTGAACTCTTTCCCTACACGACGCTCTTCCGATCTNNNTAGACACTTCACAAAAAGTAGA
AGCACACGTCTGAACTCTTTCCCTACACGACGCTCTTCCGATCTNNNTAGACACTTCACAAAAAGTAGA
AGCACACGTCTGAACTCTTTCCCTACACGACGCTCTTCCGATCTNNNTAGACACTTCACAAAAAGTAGA
AGCACACGTCTGAACTCTTTCCCTACACGACGCTCTTCCGATCTNNNTAGACACTTCACAAAAAGTAGA
AGCACACGTCTGAACTCTTTCCCTACACGACGCTCTTCCGATCTNNNTCAGCATTTAGTCAGTTGCATG
AGCACACGTCTGAACTCTTTCCCTACACGACGCTCTTCCGATCTNNNTCAGCATTTAGTCAGTTGCATG
AGCACACGTCTGAACTCTTTCCCTACACGACGCTCTTCCGATCTNNNTCAGCATTTAGTCAGTTGCATG
AGCACACGTCTGAACTCTTTCCCTACACGACGCTCTTCCGATCTNNNTCAGCATTTAGTCAGTTGCATG
AGCACACGTCTGAACTCTTTCCCTACACGACGCTCTTCCGATCTNNNTCAGCATTTAGTCAGTTGCATG
AGCACACGTCTGAACTCTTTCCCTACACGACGCTCTTCCGATCTNNNGCAGAGACACAAATCCAAACCA
AGCACACGTCTGAACTCTTTCCCTACACGACGCTCTTCCGATCTNNNGCAGAGACACAAATCCAAACCA
AGCACACGTCTGAACTCTTTCCCTACACGACGCTCTTCCGATCTNNNGCAGAGACACAAATCCAAACCA
AGCACACGTCTGAACTCTTTCCCTACACGACGCTCTTCCGATCTNNNGCAGAGACACAAATCCAAACCA
AGCACACGTCTGAACTCTTTCCCTACACGACGCTCTTCCGATCTNNNGCAGAGACACAAATCCAAACCA
AGCACACGTCTGAACTCTTTCCCTACACGACGCTCTTCCGATCTNNNGAGGCTGAGGCAGGAGAATG
AGCACACGTCTGAACTCTTTCCCTACACGACGCTCTTCCGATCTNNNGAGGCTGAGGCAGGAGAATG
AGCACACGTCTGAACTCTTTCCCTACACGACGCTCTTCCGATCTNNNGAGGCTGAGGCAGGAGAATG
AGCACACGTCTGAACTCTTTCCCTACACGACGCTCTTCCGATCTNNNGAGGCTGAGGCAGGAGAATG
AGCACACGTCTGAACTCTTTCCCTACACGACGCTCTTCCGATCTNNNGAGGCTGAGGCAGGAGAATG
GCACACGTCTGAACTCTTTCCCTACACGACGCTCTTCCGATCTNNNCTGAGGTGGGAGAATCACTTGA
GCACACGTCTGAACTCTTTCCCTACACGACGCTCTTCCGATCTNNNCTGAGGTGGGAGAATCACTTGA
GCACACGTCTGAACTCTTTCCCTACACGACGCTCTTCCGATCTNNNCTGAGGTGGGAGAATCACTTGA
GCACACGTCTGAACTCTTTCCCTACACGACGCTCTTCCGATCTNNNCTGAGGTGGGAGAATCACTTGA
GCACACGTCTGAACTCTTTCCCTACACGACGCTCTTCCGATCTNNNCTGAGGTGGGAGAATCACTTGA
ACACGTCTGAACTCTTTCCCTACACGACGCTCTTCCGATCTNNNGTACCTAAATCTGGATCACTCCATCG
ACACGTCTGAACTCTTTCCCTACACGACGCTCTTCCGATCTNNNGTACCTAAATCTGGATCACTCCATCG
ACACGTCTGAACTCTTTCCCTACACGACGCTCTTCCGATCTNNNGTACCTAAATCTGGATCACTCCATCG
ACACGTCTGAACTCTTTCCCTACACGACGCTCTTCCGATCTNNNGTACCTAAATCTGGATCACTCCATCG
ACACGTCTGAACTCTTTCCCTACACGACGCTCTTCCGATCTNNNGTACCTAAATCTGGATCACTCCATCG
AGCACACGTCTGAACTCTTTCCCTACACGACGCTCTTCCGATCTNNNAAGGAATTGTGTGTGTGTGTG
AGCACACGTCTGAACTCTTTCCCTACACGACGCTCTTCCGATCTNNNAAGGAATTGTGTGTGTGTGTG
AGCACACGTCTGAACTCTTTCCCTACACGACGCTCTTCCGATCTNNNAAGGAATTGTGTGTGTGTGTG
AGCACACGTCTGAACTCTTTCCCTACACGACGCTCTTCCGATCTNNNAAGGAATTGTGTGTGTGTGTG
AGCACACGTCTGAACTCTTTCCCTACACGACGCTCTTCCGATCTNNNAAGGAATTGTGTGTGTGTGTG
AGCACACGTCTGAACTCTTTCCCTACACGACGCTCTTCCGATCTNNNGCTTTTGCTCTGCTATCTCCAG
AGCACACGTCTGAACTCTTTCCCTACACGACGCTCTTCCGATCTNNNGCTTTTGCTCTGCTATCTCCAG
AGCACACGTCTGAACTCTTTCCCTACACGACGCTCTTCCGATCTNNNGCTTTTGCTCTGCTATCTCCAG
AGCACACGTCTGAACTCTTTCCCTACACGACGCTCTTCCGATCTNNNGCTTTTGCTCTGCTATCTCCAG
AGCACACGTCTGAACTCTTTCCCTACACGACGCTCTTCCGATCTNNNGCTTTTGCTCTGCTATCTCCAG
AGCACACGTCTGAACTCTTTCCCTACACGACGCTCTTCCGATCTNNNTGAGAAAAACAATTCCTCAGCCA

GCACACGTCTGAACTCTTTCCCTACACGACGCTCTTCCGATCTNNNTGAGAAAAACAATTCCTCAGCC
GCACACGTCTGAACTCTTTCCCTACACGACGCTCTTCCGATCTNNNTGAGAAAAACAATTCCTCAGCC
GCACACGTCTGAACTCTTTCCCTACACGACGCTCTTCCGATCTNNNTGAGAAAAACAATTCCTCAGCC
GCACACGTCTGAACTCTTTCCCTACACGACGCTCTTCCGATCTNNNTGAGAAAAACAATTCCTCAGCC
AGCACACGTCTGAACTCTTTCCCTACACGACGCTCTTCCGATCTNNNGGTAAGAAAGGGGGAAAGA
AGCACACGTCTGAACTCTTTCCCTACACGACGCTCTTCCGATCTNNNGGTAAGAAAGGGGGAAAGA
AGCACACGTCTGAACTCTTTCCCTACACGACGCTCTTCCGATCTNNNGGTAAGAAAGGGGGAAAGA
AGCACACGTCTGAACTCTTTCCCTACACGACGCTCTTCCGATCTNNNGGTAAGAAAGGGGGAAAGA
AGCACACGTCTGAACTCTTTCCCTACACGACGCTCTTCCGATCTNNNGGTAAGAAAGGGGGAAAGA
AGCACACGTCTGAACTCTTTCCCTACACGACGCTCTTCCGATCTNNNCTAAATCCCAACAATATCTCCTT
AGCACACGTCTGAACTCTTTCCCTACACGACGCTCTTCCGATCTNNNCTAAATCCCAACAATATCTCCTT
AGCACACGTCTGAACTCTTTCCCTACACGACGCTCTTCCGATCTNNNCTAAATCCCAACAATATCTCCTT
AGCACACGTCTGAACTCTTTCCCTACACGACGCTCTTCCGATCTNNNCTAAATCCCAACAATATCTCCTT
AGCACACGTCTGAACTCTTTCCCTACACGACGCTCTTCCGATCTNNNCTAAATCCCAACAATATCTCCTT
GCACACGTCTGAACTCTTTCCCTACACGACGCTCTTCCGATCTNNNCCTACTGTGGCCAATCTTCATCT
GCACACGTCTGAACTCTTTCCCTACACGACGCTCTTCCGATCTNNNCCTACTGTGGCCAATCTTCATCT
GCACACGTCTGAACTCTTTCCCTACACGACGCTCTTCCGATCTNNNCCTACTGTGGCCAATCTTCATCT
GCACACGTCTGAACTCTTTCCCTACACGACGCTCTTCCGATCTNNNCCTACTGTGGCCAATCTTCATCT
GCACACGTCTGAACTCTTTCCCTACACGACGCTCTTCCGATCTNNNCCTACTGTGGCCAATCTTCATCT
AAGAGCACACGTCTGAACTCTTTCCCTACACGACGCTCTTCCGATCTNNNTATTCATCCCTTGGTGGA
AAGAGCACACGTCTGAACTCTTTCCCTACACGACGCTCTTCCGATCTNNNTATTCATCCCTTGGTGGA
AAGAGCACACGTCTGAACTCTTTCCCTACACGACGCTCTTCCGATCTNNNTATTCATCCCTTGGTGGA
AAGAGCACACGTCTGAACTCTTTCCCTACACGACGCTCTTCCGATCTNNNTATTCATCCCTTGGTGGA
AAGAGCACACGTCTGAACTCTTTCCCTACACGACGCTCTTCCGATCTNNNTATTCATCCCTTGGTGGA
GCACACGTCTGAACTCTTTCCCTACACGACGCTCTTCCGATCTNNNTAGCTGGGATTACAGGCACTAT
GCACACGTCTGAACTCTTTCCCTACACGACGCTCTTCCGATCTNNNTAGCTGGGATTACAGGCACTAT
GCACACGTCTGAACTCTTTCCCTACACGACGCTCTTCCGATCTNNNTAGCTGGGATTACAGGCACTAT
GCACACGTCTGAACTCTTTCCCTACACGACGCTCTTCCGATCTNNNTAGCTGGGATTACAGGCACTAT
GCACACGTCTGAACTCTTTCCCTACACGACGCTCTTCCGATCTNNNTAGCTGGGATTACAGGCACTAT
AGAGCACACGTCTGAACTCTTTCCCTACACGACGCTCTTCCGATCTNNNGTGAGTCGAGATCATGCC
AGAGCACACGTCTGAACTCTTTCCCTACACGACGCTCTTCCGATCTNNNGTGAGTCGAGATCATGCC
AGAGCACACGTCTGAACTCTTTCCCTACACGACGCTCTTCCGATCTNNNGTGAGTCGAGATCATGCC
AGAGCACACGTCTGAACTCTTTCCCTACACGACGCTCTTCCGATCTNNNGTGAGTCGAGATCATGCC
AGAGCACACGTCTGAACTCTTTCCCTACACGACGCTCTTCCGATCTNNNGTGAGTCGAGATCATGCC
AGCACACGTCTGAACTCTTTCCCTACACGACGCTCTTCCGATCTNNNCTCTAGGGATTTCTTCTCCTGG
AGCACACGTCTGAACTCTTTCCCTACACGACGCTCTTCCGATCTNNNCTCTAGGGATTTCTTCTCCTGG
AGCACACGTCTGAACTCTTTCCCTACACGACGCTCTTCCGATCTNNNCTCTAGGGATTTCTTCTCCTGG
AGCACACGTCTGAACTCTTTCCCTACACGACGCTCTTCCGATCTNNNCTCTAGGGATTTCTTCTCCTGG
AGCACACGTCTGAACTCTTTCCCTACACGACGCTCTTCCGATCTNNNCTCTAGGGATTTCTTCTCCTGG
GCACACGTCTGAACTCTTTCCCTACACGACGCTCTTCCGATCTNNNTGCCTGTAGTCCCAGCTACTCCA
GCACACGTCTGAACTCTTTCCCTACACGACGCTCTTCCGATCTNNNTGCCTGTAGTCCCAGCTACTCCA
GCACACGTCTGAACTCTTTCCCTACACGACGCTCTTCCGATCTNNNTGCCTGTAGTCCCAGCTACTCCA
GCACACGTCTGAACTCTTTCCCTACACGACGCTCTTCCGATCTNNNTGCCTGTAGTCCCAGCTACTCCA

AGCACACGTCTGAACTCTTTCCCTACACGACGCTCTTCCGATCTNNNTGCCTGTAGTCCCAGCTACTCCA
AGCACACGTCTGAACTCTTTCCCTACACGACGCTCTTCCGATCTNNNCAGGAGAATCGCTTGAACCAAG
AGCACACGTCTGAACTCTTTCCCTACACGACGCTCTTCCGATCTNNNCAGGAGAATCGCTTGAACCAAG
AGCACACGTCTGAACTCTTTCCCTACACGACGCTCTTCCGATCTNNNCAGGAGAATCGCTTGAACCAAG
AGCACACGTCTGAACTCTTTCCCTACACGACGCTCTTCCGATCTNNNCAGGAGAATCGCTTGAACCAAG
AGCACACGTCTGAACTCTTTCCCTACACGACGCTCTTCCGATCTNNNCAGGAGAATCGCTTGAACCAAG
AGCACACGTCTGAACTCTTTCCCTACACGACGCTCTTCCGATCTNNNCTTATTTAAGAAATTGCCACAC
AGCACACGTCTGAACTCTTTCCCTACACGACGCTCTTCCGATCTNNNCTTATTTAAGAAATTGCCACAC
AGCACACGTCTGAACTCTTTCCCTACACGACGCTCTTCCGATCTNNNCTTATTTAAGAAATTGCCACAC
AGCACACGTCTGAACTCTTTCCCTACACGACGCTCTTCCGATCTNNNCTTATTTAAGAAATTGCCACAC
AGCACACGTCTGAACTCTTTCCCTACACGACGCTCTTCCGATCTNNNCTTATTTAAGAAATTGCCACAC
AGCACACGTCTGAACTCTTTCCCTACACGACGCTCTTCCGATCTNNNGTTTGGCAAGCTTTTGTTTCTGG
AGCACACGTCTGAACTCTTTCCCTACACGACGCTCTTCCGATCTNNNGTTTGGCAAGCTTTTGTTTCTGG
AGCACACGTCTGAACTCTTTCCCTACACGACGCTCTTCCGATCTNNNGTTTGGCAAGCTTTTGTTTCTGG
AGCACACGTCTGAACTCTTTCCCTACACGACGCTCTTCCGATCTNNNGTTTGGCAAGCTTTTGTTTCTGG
AGCACACGTCTGAACTCTTTCCCTACACGACGCTCTTCCGATCTNNNGTTTGGCAAGCTTTTGTTTCTGG
GCACACGTCTGAACTCTTTCCCTACACGACGCTCTTCCGATCTNNNAGATATAGTTCAGGCAAACTAGC
GCACACGTCTGAACTCTTTCCCTACACGACGCTCTTCCGATCTNNNAGATATAGTTCAGGCAAACTAGC
GCACACGTCTGAACTCTTTCCCTACACGACGCTCTTCCGATCTNNNAGATATAGTTCAGGCAAACTAGC
GCACACGTCTGAACTCTTTCCCTACACGACGCTCTTCCGATCTNNNAGATATAGTTCAGGCAAACTAGC
GCACACGTCTGAACTCTTTCCCTACACGACGCTCTTCCGATCTNNNAGATATAGTTCAGGCAAACTAGC
GAGCACACGTCTGAACTCTTTCCCTACACGACGCTCTTCCGATCTNNNTGGAGATTTCAGTGAGCCAAG
GAGCACACGTCTGAACTCTTTCCCTACACGACGCTCTTCCGATCTNNNTGGAGATTTCAGTGAGCCAAG
GAGCACACGTCTGAACTCTTTCCCTACACGACGCTCTTCCGATCTNNNTGGAGATTTCAGTGAGCCAAG
GAGCACACGTCTGAACTCTTTCCCTACACGACGCTCTTCCGATCTNNNTGGAGATTTCAGTGAGCCAAG
GAGCACACGTCTGAACTCTTTCCCTACACGACGCTCTTCCGATCTNNNTGGAGATTTCAGTGAGCCAAG
AGCACACGTCTGAACTCTTTCCCTACACGACGCTCTTCCGATCTNNNGAAAGTGCAGGCTATATAACTT
AGCACACGTCTGAACTCTTTCCCTACACGACGCTCTTCCGATCTNNNGAAAGTGCAGGCTATATAACTT
AGCACACGTCTGAACTCTTTCCCTACACGACGCTCTTCCGATCTNNNGAAAGTGCAGGCTATATAACTT
AGCACACGTCTGAACTCTTTCCCTACACGACGCTCTTCCGATCTNNNGAAAGTGCAGGCTATATAACTT
AGCACACGTCTGAACTCTTTCCCTACACGACGCTCTTCCGATCTNNNGAAAGTGCAGGCTATATAACTT
AGAGCACACGTCTGAACTCTTTCCCTACACGACGCTCTTCCGATCTNNNTTCAGGATCACACAGCAAGT
AGAGCACACGTCTGAACTCTTTCCCTACACGACGCTCTTCCGATCTNNNTTCAGGATCACACAGCAAGT
AGAGCACACGTCTGAACTCTTTCCCTACACGACGCTCTTCCGATCTNNNTTCAGGATCACACAGCAAGT
AGAGCACACGTCTGAACTCTTTCCCTACACGACGCTCTTCCGATCTNNNTTCAGGATCACACAGCAAGT
AGAGCACACGTCTGAACTCTTTCCCTACACGACGCTCTTCCGATCTNNNTTCAGGATCACACAGCAAGT
GAGCACACGTCTGAACTCTTTCCCTACACGACGCTCTTCCGATCTNNNAGGGAGGAATGTTGAACCAA
GAGCACACGTCTGAACTCTTTCCCTACACGACGCTCTTCCGATCTNNNAGGGAGGAATGTTGAACCAA
GAGCACACGTCTGAACTCTTTCCCTACACGACGCTCTTCCGATCTNNNAGGGAGGAATGTTGAACCAA
GAGCACACGTCTGAACTCTTTCCCTACACGACGCTCTTCCGATCTNNNAGGGAGGAATGTTGAACCAA
GAGCACACGTCTGAACTCTTTCCCTACACGACGCTCTTCCGATCTNNNAGGGAGGAATGTTGAACCAA
CACACGTCTGAACTCTTTCCCTACACGACGCTCTTCCGATCTNNNAAGGAAGTCTTGAATTTGGCCATC
CACACGTCTGAACTCTTTCCCTACACGACGCTCTTCCGATCTNNNAAGGAAGTCTTGAATTTGGCCATC

CACACGTCTGAACTCTTTCCCTACACGACGCTCTTCCGATCTNNNAAGGAAGTCTTGAATTTGGCCATC
CACACGTCTGAACTCTTTCCCTACACGACGCTCTTCCGATCTNNNAAGGAAGTCTTGAATTTGGCCATC
CACACGTCTGAACTCTTTCCCTACACGACGCTCTTCCGATCTNNNAAGGAAGTCTTGAATTTGGCCATC
GCACACGTCTGAACTCTTTCCCTACACGACGCTCTTCCGATCTNNNATCCAAAGTCTTCCTGATGGCTTC
GCACACGTCTGAACTCTTTCCCTACACGACGCTCTTCCGATCTNNNATCCAAAGTCTTCCTGATGGCTTC
GCACACGTCTGAACTCTTTCCCTACACGACGCTCTTCCGATCTNNNATCCAAAGTCTTCCTGATGGCTTC
GCACACGTCTGAACTCTTTCCCTACACGACGCTCTTCCGATCTNNNATCCAAAGTCTTCCTGATGGCTTC
GCACACGTCTGAACTCTTTCCCTACACGACGCTCTTCCGATCTNNNATCCAAAGTCTTCCTGATGGCTTC
CACACGTCTGAACTCTTTCCCTACACGACGCTCTTCCGATCTNNNAACTGTTGGTTATTTGTGTTATTCA
CACACGTCTGAACTCTTTCCCTACACGACGCTCTTCCGATCTNNNAACTGTTGGTTATTTGTGTTATTCA
CACACGTCTGAACTCTTTCCCTACACGACGCTCTTCCGATCTNNNAACTGTTGGTTATTTGTGTTATTCA
CACACGTCTGAACTCTTTCCCTACACGACGCTCTTCCGATCTNNNAACTGTTGGTTATTTGTGTTATTCA
CACACGTCTGAACTCTTTCCCTACACGACGCTCTTCCGATCTNNNAACTGTTGGTTATTTGTGTTATTCA
AGCACACGTCTGAACTCTTTCCCTACACGACGCTCTTCCGATCTNNNCTTGCAGTGAGCCGAGATCTCC
AGCACACGTCTGAACTCTTTCCCTACACGACGCTCTTCCGATCTNNNCTTGCAGTGAGCCGAGATCTCC
AGCACACGTCTGAACTCTTTCCCTACACGACGCTCTTCCGATCTNNNCTTGCAGTGAGCCGAGATCTCC
AGCACACGTCTGAACTCTTTCCCTACACGACGCTCTTCCGATCTNNNCTTGCAGTGAGCCGAGATCTCC
AGCACACGTCTGAACTCTTTCCCTACACGACGCTCTTCCGATCTNNNCTTGCAGTGAGCCGAGATCTCC
AGCACACGTCTGAACTCTTTCCCTACACGACGCTCTTCCGATCTNNNGCCCGGCCTAATATTCTTTTCTTC
AGCACACGTCTGAACTCTTTCCCTACACGACGCTCTTCCGATCTNNNGCCCGGCCTAATATTCTTTTCTTC
AGCACACGTCTGAACTCTTTCCCTACACGACGCTCTTCCGATCTNNNGCCCGGCCTAATATTCTTTTCTTC
AGCACACGTCTGAACTCTTTCCCTACACGACGCTCTTCCGATCTNNNGCCCGGCCTAATATTCTTTTCTTC
AGCACACGTCTGAACTCTTTCCCTACACGACGCTCTTCCGATCTNNNGCCCGGCCTAATATTCTTTTCTTC
AGCACACGTCTGAACTCTTTCCCTACACGACGCTCTTCCGATCTNNNACTACTTCTACTACTACTACACAC
AGCACACGTCTGAACTCTTTCCCTACACGACGCTCTTCCGATCTNNNACTACTTCTACTACTACTACACAC
AGCACACGTCTGAACTCTTTCCCTACACGACGCTCTTCCGATCTNNNACTACTTCTACTACTACTACACAC
AGCACACGTCTGAACTCTTTCCCTACACGACGCTCTTCCGATCTNNNACTACTTCTACTACTACTACACAC
AGCACACGTCTGAACTCTTTCCCTACACGACGCTCTTCCGATCTNNNACTACTTCTACTACTACTACACAC
GAGCACACGTCTGAACTCTTTCCCTACACGACGCTCTTCCGATCTNNNTGGGACCATTTTTGGGAGTCT
GAGCACACGTCTGAACTCTTTCCCTACACGACGCTCTTCCGATCTNNNTGGGACCATTTTTGGGAGTCT
GAGCACACGTCTGAACTCTTTCCCTACACGACGCTCTTCCGATCTNNNTGGGACCATTTTTGGGAGTCT
GAGCACACGTCTGAACTCTTTCCCTACACGACGCTCTTCCGATCTNNNTGGGACCATTTTTGGGAGTCT
GAGCACACGTCTGAACTCTTTCCCTACACGACGCTCTTCCGATCTNNNTGGGACCATTTTTGGGAGTCT
GAGCACACGTCTGAACTCTTTCCCTACACGACGCTCTTCCGATCTNNNCCACAGCTTCTCTGATGGTAA
GAGCACACGTCTGAACTCTTTCCCTACACGACGCTCTTCCGATCTNNNCCACAGCTTCTCTGATGGTAA
GAGCACACGTCTGAACTCTTTCCCTACACGACGCTCTTCCGATCTNNNCCACAGCTTCTCTGATGGTAA
GAGCACACGTCTGAACTCTTTCCCTACACGACGCTCTTCCGATCTNNNCCACAGCTTCTCTGATGGTAA
GAGCACACGTCTGAACTCTTTCCCTACACGACGCTCTTCCGATCTNNNCCACAGCTTCTCTGATGGTAA
GAGCACACGTCTGAACTCTTTCCCTACACGACGCTCTTCCGATCTNNNTGGGGAGGAAAACAAAAGCA
GAGCACACGTCTGAACTCTTTCCCTACACGACGCTCTTCCGATCTNNNTGGGGAGGAAAACAAAAGCA
GAGCACACGTCTGAACTCTTTCCCTACACGACGCTCTTCCGATCTNNNTGGGGAGGAAAACAAAAGCA
GAGCACACGTCTGAACTCTTTCCCTACACGACGCTCTTCCGATCTNNNTGGGGAGGAAAACAAAAGCA
GAGCACACGTCTGAACTCTTTCCCTACACGACGCTCTTCCGATCTNNNTGGGGAGGAAAACAAAAGCA

AGCACACGTCTGAACTCTTTCCCTACACGACGCTCTTCCGATCTNNNCCTTTACCCGCCTCAGAAGTAAT

AGCACACGTCTGAACTCTTTCCCTACACGACGCTCTTCCGATCTNNNCCTTTACCCGCCTCAGAAGTAAT

AGCACACGTCTGAACTCTTTCCCTACACGACGCTCTTCCGATCTNNNCCTTTACCCGCCTCAGAAGTAAT

AGCACACGTCTGAACTCTTTCCCTACACGACGCTCTTCCGATCTNNNCCTTTACCCGCCTCAGAAGTAAT

AGCACACGTCTGAACTCTTTCCCTACACGACGCTCTTCCGATCTNNNCCTTTACCCGCCTCAGAAGTAAT

AGAGCACACGTCTGAACTCTTTCCCTACACGACGCTCTTCCGATCTNNNGGGTAACAAGAGCGAAACT

AGAGCACACGTCTGAACTCTTTCCCTACACGACGCTCTTCCGATCTNNNGGGTAACAAGAGCGAAACT

AGAGCACACGTCTGAACTCTTTCCCTACACGACGCTCTTCCGATCTNNNGGGTAACAAGAGCGAAACT

AGAGCACACGTCTGAACTCTTTCCCTACACGACGCTCTTCCGATCTNNNGGGTAACAAGAGCGAAACT

AGAGCACACGTCTGAACTCTTTCCCTACACGACGCTCTTCCGATCTNNNGGGTAACAAGAGCGAAACT

ACACGTCTGAACTCTTTCCCTACACGACGCTCTTCCGATCTNNNCACCTGCTCCTCTCCCCTAGCATCGGA

ACACGTCTGAACTCTTTCCCTACACGACGCTCTTCCGATCTNNNCACCTGCTCCTCTCCCCTAGCATCGGA

ACACGTCTGAACTCTTTCCCTACACGACGCTCTTCCGATCTNNNCACCTGCTCCTCTCCCCTAGCATCGGA

ACACGTCTGAACTCTTTCCCTACACGACGCTCTTCCGATCTNNNCACCTGCTCCTCTCCCCTAGCATCGGA

ACACGTCTGAACTCTTTCCCTACACGACGCTCTTCCGATCTNNNCACCTGCTCCTCTCCCCTAGCATCGGA

GAGCACACGTCTGAACTCTTTCCCTACACGACGCTCTTCCGATCTNNNGGCTAAGGCACAAGAATCAC

GAGCACACGTCTGAACTCTTTCCCTACACGACGCTCTTCCGATCTNNNGGCTAAGGCACAAGAATCAC

GAGCACACGTCTGAACTCTTTCCCTACACGACGCTCTTCCGATCTNNNGGCTAAGGCACAAGAATCAC

GAGCACACGTCTGAACTCTTTCCCTACACGACGCTCTTCCGATCTNNNGGCTAAGGCACAAGAATCAC

GAGCACACGTCTGAACTCTTTCCCTACACGACGCTCTTCCGATCTNNNGGCTAAGGCACAAGAATCAC

GAGCACACGTCTGAACTCTTTCCCTACACGACGCTCTTCCGATCTNNNGTCCAGCTACTTGAAAGTCT

GAGCACACGTCTGAACTCTTTCCCTACACGACGCTCTTCCGATCTNNNGTCCAGCTACTTGAAAGTCT

GAGCACACGTCTGAACTCTTTCCCTACACGACGCTCTTCCGATCTNNNGTCCAGCTACTTGAAAGTCT

GAGCACACGTCTGAACTCTTTCCCTACACGACGCTCTTCCGATCTNNNGTCCAGCTACTTGAAAGTCT

GAGCACACGTCTGAACTCTTTCCCTACACGACGCTCTTCCGATCTNNNGTCCAGCTACTTGAAAGTCT

AGCACACGTCTGAACTCTTTCCCTACACGACGCTCTTCCGATCTNNNCATTCTGATACTAGTCCTATTCC

AGCACACGTCTGAACTCTTTCCCTACACGACGCTCTTCCGATCTNNNCATTCTGATACTAGTCCTATTCC

AGCACACGTCTGAACTCTTTCCCTACACGACGCTCTTCCGATCTNNNCATTCTGATACTAGTCCTATTCC

AGCACACGTCTGAACTCTTTCCCTACACGACGCTCTTCCGATCTNNNCATTCTGATACTAGTCCTATTCC

AGCACACGTCTGAACTCTTTCCCTACACGACGCTCTTCCGATCTNNNCATTCTGATACTAGTCCTATTCC

AGCACACGTCTGAACTCTTTCCCTACACGACGCTCTTCCGATCTNNNTTCAACTTCCTAGACAAAGACTG

AGCACACGTCTGAACTCTTTCCCTACACGACGCTCTTCCGATCTNNNTTCAACTTCCTAGACAAAGACTG

AGCACACGTCTGAACTCTTTCCCTACACGACGCTCTTCCGATCTNNNTTCAACTTCCTAGACAAAGACTG

AGCACACGTCTGAACTCTTTCCCTACACGACGCTCTTCCGATCTNNNTTCAACTTCCTAGACAAAGACTG

AGCACACGTCTGAACTCTTTCCCTACACGACGCTCTTCCGATCTNNNTTCAACTTCCTAGACAAAGACTG

AGCACACGTCTGAACTCTTTCCCTACACGACGCTCTTCCGATCTNNNGATATCCTCTATTTTGCTGGGG

AGCACACGTCTGAACTCTTTCCCTACACGACGCTCTTCCGATCTNNNGATATCCTCTATTTTGCTGGGG

AGCACACGTCTGAACTCTTTCCCTACACGACGCTCTTCCGATCTNNNGATATCCTCTATTTTGCTGGGG

AGCACACGTCTGAACTCTTTCCCTACACGACGCTCTTCCGATCTNNNGATATCCTCTATTTTGCTGGGG

AGCACACGTCTGAACTCTTTCCCTACACGACGCTCTTCCGATCTNNNGATATCCTCTATTTTGCTGGGG

GAGCACACGTCTGAACTCTTTCCCTACACGACGCTCTTCCGATCTNNNATGCCTGTAATCCCAGCTCTTA

GAGCACACGTCTGAACTCTTTCCCTACACGACGCTCTTCCGATCTNNNATGCCTGTAATCCCAGCTCTTA

GAGCACACGTCTGAACTCTTTCCCTACACGACGCTCTTCCGATCTNNNATGCCTGTAATCCCAGCTCTTA

GAGCACACGTCTGAACTCTTTCCCTACACGACGCTCTTCCGATCTNNNATGCCTGTAATCCCAGCTCTTA
GAGCACACGTCTGAACTCTTTCCCTACACGACGCTCTTCCGATCTNNNATGCCTGTAATCCCAGCTCTTA
AGCACACGTCTGAACTCTTTCCCTACACGACGCTCTTCCGATCTNNNACCAGAATTCACATGGTGGAA
AGCACACGTCTGAACTCTTTCCCTACACGACGCTCTTCCGATCTNNNACCAGAATTCACATGGTGGAA
AGCACACGTCTGAACTCTTTCCCTACACGACGCTCTTCCGATCTNNNACCAGAATTCACATGGTGGAA
AGCACACGTCTGAACTCTTTCCCTACACGACGCTCTTCCGATCTNNNACCAGAATTCACATGGTGGAA
AGCACACGTCTGAACTCTTTCCCTACACGACGCTCTTCCGATCTNNNACCAGAATTCACATGGTGGAA
AGCACACGTCTGAACTCTTTCCCTACACGACGCTCTTCCGATCTNNNGGGGTTGGAAGCATCTTCTGTA
AGCACACGTCTGAACTCTTTCCCTACACGACGCTCTTCCGATCTNNNGGGGTTGGAAGCATCTTCTGTA
AGCACACGTCTGAACTCTTTCCCTACACGACGCTCTTCCGATCTNNNGGGGTTGGAAGCATCTTCTGTA
AGCACACGTCTGAACTCTTTCCCTACACGACGCTCTTCCGATCTNNNGGGGTTGGAAGCATCTTCTGTA
AGCACACGTCTGAACTCTTTCCCTACACGACGCTCTTCCGATCTNNNGGGGTTGGAAGCATCTTCTGTA
AGCACACGTCTGAACTCTTTCCCTACACGACGCTCTTCCGATCTNNNAAGTAGCTGGGATTACAGGCA
AGCACACGTCTGAACTCTTTCCCTACACGACGCTCTTCCGATCTNNNAAGTAGCTGGGATTACAGGCA
AGCACACGTCTGAACTCTTTCCCTACACGACGCTCTTCCGATCTNNNAAGTAGCTGGGATTACAGGCA
AGCACACGTCTGAACTCTTTCCCTACACGACGCTCTTCCGATCTNNNAAGTAGCTGGGATTACAGGCA
AGCACACGTCTGAACTCTTTCCCTACACGACGCTCTTCCGATCTNNNAAGTAGCTGGGATTACAGGCA
AGAGCACACGTCTGAACTCTTTCCCTACACGACGCTCTTCCGATCTNNNGTAAAGGAGTTAGTAGAAA
AGAGCACACGTCTGAACTCTTTCCCTACACGACGCTCTTCCGATCTNNNGTAAAGGAGTTAGTAGAAA
AGAGCACACGTCTGAACTCTTTCCCTACACGACGCTCTTCCGATCTNNNGTAAAGGAGTTAGTAGAAA
AGAGCACACGTCTGAACTCTTTCCCTACACGACGCTCTTCCGATCTNNNGTAAAGGAGTTAGTAGAAA
AGAGCACACGTCTGAACTCTTTCCCTACACGACGCTCTTCCGATCTNNNGTAAAGGAGTTAGTAGAAA
GAGCACACGTCTGAACTCTTTCCCTACACGACGCTCTTCCGATCTNNNCAGAGAAGTTAGGCTTGAAG
GAGCACACGTCTGAACTCTTTCCCTACACGACGCTCTTCCGATCTNNNCAGAGAAGTTAGGCTTGAAG
GAGCACACGTCTGAACTCTTTCCCTACACGACGCTCTTCCGATCTNNNCAGAGAAGTTAGGCTTGAAG
GAGCACACGTCTGAACTCTTTCCCTACACGACGCTCTTCCGATCTNNNCAGAGAAGTTAGGCTTGAAG
GAGCACACGTCTGAACTCTTTCCCTACACGACGCTCTTCCGATCTNNNCAGAGAAGTTAGGCTTGAAG
AGAGCACACGTCTGAACTCTTTCCCTACACGACGCTCTTCCGATCTNNNGGACTTCTGGCTGCAGGTGT
AGAGCACACGTCTGAACTCTTTCCCTACACGACGCTCTTCCGATCTNNNGGACTTCTGGCTGCAGGTGT
AGAGCACACGTCTGAACTCTTTCCCTACACGACGCTCTTCCGATCTNNNGGACTTCTGGCTGCAGGTGT
AGAGCACACGTCTGAACTCTTTCCCTACACGACGCTCTTCCGATCTNNNGGACTTCTGGCTGCAGGTGT
AGAGCACACGTCTGAACTCTTTCCCTACACGACGCTCTTCCGATCTNNNGGACTTCTGGCTGCAGGTGT
GAGCACACGTCTGAACTCTTTCCCTACACGACGCTCTTCCGATCTNNNCTAAGGCAGGAGAATCAGCTT
GAGCACACGTCTGAACTCTTTCCCTACACGACGCTCTTCCGATCTNNNCTAAGGCAGGAGAATCAGCTT
GAGCACACGTCTGAACTCTTTCCCTACACGACGCTCTTCCGATCTNNNCTAAGGCAGGAGAATCAGCTT
GAGCACACGTCTGAACTCTTTCCCTACACGACGCTCTTCCGATCTNNNCTAAGGCAGGAGAATCAGCTT
GAGCACACGTCTGAACTCTTTCCCTACACGACGCTCTTCCGATCTNNNCTAAGGCAGGAGAATCAGCTT
AGAGCACACGTCTGAACTCTTTCCCTACACGACGCTCTTCCGATCTNNNGAAGAATTGCTTGAATCCGG
AGAGCACACGTCTGAACTCTTTCCCTACACGACGCTCTTCCGATCTNNNGAAGAATTGCTTGAATCCGG
AGAGCACACGTCTGAACTCTTTCCCTACACGACGCTCTTCCGATCTNNNGAAGAATTGCTTGAATCCGG
AGAGCACACGTCTGAACTCTTTCCCTACACGACGCTCTTCCGATCTNNNGAAGAATTGCTTGAATCCGG
AGAGCACACGTCTGAACTCTTTCCCTACACGACGCTCTTCCGATCTNNNGAAGAATTGCTTGAATCCGG
GAGCACACGTCTGAACTCTTTCCCTACACGACGCTCTTCCGATCTNNNTTGAGGCAGGAGTATCACTT

GAGCACACGTCTGAACTCTTTCCCTACACGACGCTCTTCCGATCTNNNTTGAGGCAGGAGTATCACTTC
GAGCACACGTCTGAACTCTTTCCCTACACGACGCTCTTCCGATCTNNNTTGAGGCAGGAGTATCACTTC
GAGCACACGTCTGAACTCTTTCCCTACACGACGCTCTTCCGATCTNNNTTGAGGCAGGAGTATCACTTC
GAGCACACGTCTGAACTCTTTCCCTACACGACGCTCTTCCGATCTNNNTTGAGGCAGGAGTATCACTTC
GCACACGTCTGAACTCTTTCCCTACACGACGCTCTTCCGATCTNNNTGGGGAAGTTCTCCTGGATGATA
GCACACGTCTGAACTCTTTCCCTACACGACGCTCTTCCGATCTNNNTGGGGAAGTTCTCCTGGATGATA
GCACACGTCTGAACTCTTTCCCTACACGACGCTCTTCCGATCTNNNTGGGGAAGTTCTCCTGGATGATA
GCACACGTCTGAACTCTTTCCCTACACGACGCTCTTCCGATCTNNNTGGGGAAGTTCTCCTGGATGATA
GCACACGTCTGAACTCTTTCCCTACACGACGCTCTTCCGATCTNNNTGGGGAAGTTCTCCTGGATGATA
AGCACACGTCTGAACTCTTTCCCTACACGACGCTCTTCCGATCTNNNCTGCTATCCACGCTCTGAAAGAA
AGCACACGTCTGAACTCTTTCCCTACACGACGCTCTTCCGATCTNNNCTGCTATCCACGCTCTGAAAGAA
AGCACACGTCTGAACTCTTTCCCTACACGACGCTCTTCCGATCTNNNCTGCTATCCACGCTCTGAAAGAA
AGCACACGTCTGAACTCTTTCCCTACACGACGCTCTTCCGATCTNNNCTGCTATCCACGCTCTGAAAGAA
AGCACACGTCTGAACTCTTTCCCTACACGACGCTCTTCCGATCTNNNCTGCTATCCACGCTCTGAAAGAA
AGCACACGTCTGAACTCTTTCCCTACACGACGCTCTTCCGATCTNNNCGCTGTCCTTTGATAACTCCCTCC
AGCACACGTCTGAACTCTTTCCCTACACGACGCTCTTCCGATCTNNNCGCTGTCCTTTGATAACTCCCTCC
AGCACACGTCTGAACTCTTTCCCTACACGACGCTCTTCCGATCTNNNCGCTGTCCTTTGATAACTCCCTCC
AGCACACGTCTGAACTCTTTCCCTACACGACGCTCTTCCGATCTNNNCGCTGTCCTTTGATAACTCCCTCC
AGCACACGTCTGAACTCTTTCCCTACACGACGCTCTTCCGATCTNNNCGCTGTCCTTTGATAACTCCCTCC
AGCACACGTCTGAACTCTTTCCCTACACGACGCTCTTCCGATCTNNNCTGTAATCCCAGCACTTTGGAA
AGCACACGTCTGAACTCTTTCCCTACACGACGCTCTTCCGATCTNNNCTGTAATCCCAGCACTTTGGAA
AGCACACGTCTGAACTCTTTCCCTACACGACGCTCTTCCGATCTNNNCTGTAATCCCAGCACTTTGGAA
AGCACACGTCTGAACTCTTTCCCTACACGACGCTCTTCCGATCTNNNCTGTAATCCCAGCACTTTGGAA
AGCACACGTCTGAACTCTTTCCCTACACGACGCTCTTCCGATCTNNNCTGTAATCCCAGCACTTTGGAA
AGCACACGTCTGAACTCTTTCCCTACACGACGCTCTTCCGATCTNNNATGGATCAGTGAAGACCAGAAC
AGCACACGTCTGAACTCTTTCCCTACACGACGCTCTTCCGATCTNNNATGGATCAGTGAAGACCAGAAC
AGCACACGTCTGAACTCTTTCCCTACACGACGCTCTTCCGATCTNNNATGGATCAGTGAAGACCAGAAC
AGCACACGTCTGAACTCTTTCCCTACACGACGCTCTTCCGATCTNNNATGGATCAGTGAAGACCAGAAC
AGCACACGTCTGAACTCTTTCCCTACACGACGCTCTTCCGATCTNNNATGGATCAGTGAAGACCAGAAC
GAGCACACGTCTGAACTCTTTCCCTACACGACGCTCTTCCGATCTNNNCACATCCAGTCTCTCTCTGTTG
GAGCACACGTCTGAACTCTTTCCCTACACGACGCTCTTCCGATCTNNNCACATCCAGTCTCTCTCTGTTG
GAGCACACGTCTGAACTCTTTCCCTACACGACGCTCTTCCGATCTNNNCACATCCAGTCTCTCTCTGTTG
GAGCACACGTCTGAACTCTTTCCCTACACGACGCTCTTCCGATCTNNNCACATCCAGTCTCTCTCTGTTG
GAGCACACGTCTGAACTCTTTCCCTACACGACGCTCTTCCGATCTNNNCACATCCAGTCTCTCTCTGTTG
AGCACACGTCTGAACTCTTTCCCTACACGACGCTCTTCCGATCTNNNCTCTTTCTGGTTCTCACTGGCTC
AGCACACGTCTGAACTCTTTCCCTACACGACGCTCTTCCGATCTNNNCTCTTTCTGGTTCTCACTGGCTC
AGCACACGTCTGAACTCTTTCCCTACACGACGCTCTTCCGATCTNNNCTCTTTCTGGTTCTCACTGGCTC
AGCACACGTCTGAACTCTTTCCCTACACGACGCTCTTCCGATCTNNNCTCTTTCTGGTTCTCACTGGCTC
AGCACACGTCTGAACTCTTTCCCTACACGACGCTCTTCCGATCTNNNCTCTTTCTGGTTCTCACTGGCTC
AGCACACGTCTGAACTCTTTCCCTACACGACGCTCTTCCGATCTNNNGAGAATCGCTTGAACCCAGGAG
AGCACACGTCTGAACTCTTTCCCTACACGACGCTCTTCCGATCTNNNGAGAATCGCTTGAACCCAGGAG
AGCACACGTCTGAACTCTTTCCCTACACGACGCTCTTCCGATCTNNNGAGAATCGCTTGAACCCAGGAG
AGCACACGTCTGAACTCTTTCCCTACACGACGCTCTTCCGATCTNNNGAGAATCGCTTGAACCCAGGAG

GCACACGTCTGAACTCTTTCCCTACACGACGCTCTTCCGATCTNNNGAGAATCGCTTGAACCCAGGA
AGCACACGTCTGAACTCTTTCCCTACACGACGCTCTTCCGATCTNNNAAAGTCATTCCATATCCACATG
AGCACACGTCTGAACTCTTTCCCTACACGACGCTCTTCCGATCTNNNAAAGTCATTCCATATCCACATG
AGCACACGTCTGAACTCTTTCCCTACACGACGCTCTTCCGATCTNNNAAAGTCATTCCATATCCACATG
AGCACACGTCTGAACTCTTTCCCTACACGACGCTCTTCCGATCTNNNAAAGTCATTCCATATCCACATG
AGCACACGTCTGAACTCTTTCCCTACACGACGCTCTTCCGATCTNNNAAAGTCATTCCATATCCACATG
GCACACGTCTGAACTCTTTCCCTACACGACGCTCTTCCGATCTNNNATCATACTGCTAAGAATTCAAAC
GCACACGTCTGAACTCTTTCCCTACACGACGCTCTTCCGATCTNNNATCATACTGCTAAGAATTCAAAC
GCACACGTCTGAACTCTTTCCCTACACGACGCTCTTCCGATCTNNNATCATACTGCTAAGAATTCAAAC
GCACACGTCTGAACTCTTTCCCTACACGACGCTCTTCCGATCTNNNATCATACTGCTAAGAATTCAAAC
GCACACGTCTGAACTCTTTCCCTACACGACGCTCTTCCGATCTNNNATCATACTGCTAAGAATTCAAAC
AGCACACGTCTGAACTCTTTCCCTACACGACGCTCTTCCGATCTNNNATATCTGTTGATGTCCACCAG
AGCACACGTCTGAACTCTTTCCCTACACGACGCTCTTCCGATCTNNNATATCTGTTGATGTCCACCAG
AGCACACGTCTGAACTCTTTCCCTACACGACGCTCTTCCGATCTNNNATATCTGTTGATGTCCACCAG
AGCACACGTCTGAACTCTTTCCCTACACGACGCTCTTCCGATCTNNNATATCTGTTGATGTCCACCAG
AGCACACGTCTGAACTCTTTCCCTACACGACGCTCTTCCGATCTNNNATATCTGTTGATGTCCACCAG
CACACGTCTGAACTCTTTCCCTACACGACGCTCTTCCGATCTNNNATCTCTCTGAACTGTCACAACCCC
CACACGTCTGAACTCTTTCCCTACACGACGCTCTTCCGATCTNNNATCTCTCTGAACTGTCACAACCCC
CACACGTCTGAACTCTTTCCCTACACGACGCTCTTCCGATCTNNNATCTCTCTGAACTGTCACAACCCC
CACACGTCTGAACTCTTTCCCTACACGACGCTCTTCCGATCTNNNATCTCTCTGAACTGTCACAACCCC
CACACGTCTGAACTCTTTCCCTACACGACGCTCTTCCGATCTNNNATCTCTCTGAACTGTCACAACCCC
AGCACACGTCTGAACTCTTTCCCTACACGACGCTCTTCCGATCTNNNGAGACTGAGGCAGGGGAATT
AGCACACGTCTGAACTCTTTCCCTACACGACGCTCTTCCGATCTNNNGAGACTGAGGCAGGGGAATT
AGCACACGTCTGAACTCTTTCCCTACACGACGCTCTTCCGATCTNNNGAGACTGAGGCAGGGGAATT
AGCACACGTCTGAACTCTTTCCCTACACGACGCTCTTCCGATCTNNNGAGACTGAGGCAGGGGAATT
AGCACACGTCTGAACTCTTTCCCTACACGACGCTCTTCCGATCTNNNGAGACTGAGGCAGGGGAATT
GCACACGTCTGAACTCTTTCCCTACACGACGCTCTTCCGATCTNNNAGTGCTGGGATTACAGGTATG
GCACACGTCTGAACTCTTTCCCTACACGACGCTCTTCCGATCTNNNAGTGCTGGGATTACAGGTATG
GCACACGTCTGAACTCTTTCCCTACACGACGCTCTTCCGATCTNNNAGTGCTGGGATTACAGGTATG
GCACACGTCTGAACTCTTTCCCTACACGACGCTCTTCCGATCTNNNAGTGCTGGGATTACAGGTATG
GCACACGTCTGAACTCTTTCCCTACACGACGCTCTTCCGATCTNNNAGTGCTGGGATTACAGGTATG
GCACACGTCTGAACTCTTTCCCTACACGACGCTCTTCCGATCTNNNCTCTCTCTCTGTGTGTGTGTG
GCACACGTCTGAACTCTTTCCCTACACGACGCTCTTCCGATCTNNNCTCTCTCTCTGTGTGTGTGTG
GCACACGTCTGAACTCTTTCCCTACACGACGCTCTTCCGATCTNNNCTCTCTCTCTGTGTGTGTGTG
GCACACGTCTGAACTCTTTCCCTACACGACGCTCTTCCGATCTNNNCTCTCTCTCTGTGTGTGTGTG
GCACACGTCTGAACTCTTTCCCTACACGACGCTCTTCCGATCTNNNCTCTCTCTCTGTGTGTGTGTG
GAGCACACGTCTGAACTCTTTCCCTACACGACGCTCTTCCGATCTNNNATCCATGAGGGGCATTTCTT
GAGCACACGTCTGAACTCTTTCCCTACACGACGCTCTTCCGATCTNNNATCCATGAGGGGCATTTCTT
GAGCACACGTCTGAACTCTTTCCCTACACGACGCTCTTCCGATCTNNNATCCATGAGGGGCATTTCTT
GAGCACACGTCTGAACTCTTTCCCTACACGACGCTCTTCCGATCTNNNATCCATGAGGGGCATTTCTT
GAGCACACGTCTGAACTCTTTCCCTACACGACGCTCTTCCGATCTNNNATCCATGAGGGGCATTTCTT
AGCACACGTCTGAACTCTTTCCCTACACGACGCTCTTCCGATCTNNNGTAATCCCAGCTACTGTAGAG
AGCACACGTCTGAACTCTTTCCCTACACGACGCTCTTCCGATCTNNNGTAATCCCAGCTACTGTAGAG

GAGCACACGTCTGAACTCTTTCCCTACACGACGCTCTTCCGATCTNNNGTAATCCCAGCTACTGTAGAG
GAGCACACGTCTGAACTCTTTCCCTACACGACGCTCTTCCGATCTNNNGTAATCCCAGCTACTGTAGAG
GAGCACACGTCTGAACTCTTTCCCTACACGACGCTCTTCCGATCTNNNGTAATCCCAGCTACTGTAGAG
AGCACACGTCTGAACTCTTTCCCTACACGACGCTCTTCCGATCTNNNGCATGAGCTATACTTTGTGCCA
AGCACACGTCTGAACTCTTTCCCTACACGACGCTCTTCCGATCTNNNGCATGAGCTATACTTTGTGCCA
AGCACACGTCTGAACTCTTTCCCTACACGACGCTCTTCCGATCTNNNGCATGAGCTATACTTTGTGCCA
AGCACACGTCTGAACTCTTTCCCTACACGACGCTCTTCCGATCTNNNGCATGAGCTATACTTTGTGCCA
AGCACACGTCTGAACTCTTTCCCTACACGACGCTCTTCCGATCTNNNGCATGAGCTATACTTTGTGCCA
GAGCACACGTCTGAACTCTTTCCCTACACGACGCTCTTCCGATCTNNNTGGTACTGCAGAACTTCTACA
GAGCACACGTCTGAACTCTTTCCCTACACGACGCTCTTCCGATCTNNNTGGTACTGCAGAACTTCTACA
GAGCACACGTCTGAACTCTTTCCCTACACGACGCTCTTCCGATCTNNNTGGTACTGCAGAACTTCTACA
GAGCACACGTCTGAACTCTTTCCCTACACGACGCTCTTCCGATCTNNNTGGTACTGCAGAACTTCTACA
GAGCACACGTCTGAACTCTTTCCCTACACGACGCTCTTCCGATCTNNNTGGTACTGCAGAACTTCTACA
GCACACGTCTGAACTCTTTCCCTACACGACGCTCTTCCGATCTNNNTGTATTAGCAGGGTGAAGGAAT
GCACACGTCTGAACTCTTTCCCTACACGACGCTCTTCCGATCTNNNTGTATTAGCAGGGTGAAGGAAT
GCACACGTCTGAACTCTTTCCCTACACGACGCTCTTCCGATCTNNNTGTATTAGCAGGGTGAAGGAAT
GCACACGTCTGAACTCTTTCCCTACACGACGCTCTTCCGATCTNNNTGTATTAGCAGGGTGAAGGAAT
GCACACGTCTGAACTCTTTCCCTACACGACGCTCTTCCGATCTNNNTGTATTAGCAGGGTGAAGGAAT
AGCACACGTCTGAACTCTTTCCCTACACGACGCTCTTCCGATCTNNNCGAAGTGCTGACAATGATGTGA
AGCACACGTCTGAACTCTTTCCCTACACGACGCTCTTCCGATCTNNNCGAAGTGCTGACAATGATGTGA
AGCACACGTCTGAACTCTTTCCCTACACGACGCTCTTCCGATCTNNNCGAAGTGCTGACAATGATGTGA
AGCACACGTCTGAACTCTTTCCCTACACGACGCTCTTCCGATCTNNNCGAAGTGCTGACAATGATGTGA
AGCACACGTCTGAACTCTTTCCCTACACGACGCTCTTCCGATCTNNNCGAAGTGCTGACAATGATGTGA
AGCACACGTCTGAACTCTTTCCCTACACGACGCTCTTCCGATCTNNNACACCCAGCTAATTTTTAAAAA
AGCACACGTCTGAACTCTTTCCCTACACGACGCTCTTCCGATCTNNNACACCCAGCTAATTTTTAAAAA
AGCACACGTCTGAACTCTTTCCCTACACGACGCTCTTCCGATCTNNNACACCCAGCTAATTTTTAAAAA
AGCACACGTCTGAACTCTTTCCCTACACGACGCTCTTCCGATCTNNNACACCCAGCTAATTTTTAAAAA
AGCACACGTCTGAACTCTTTCCCTACACGACGCTCTTCCGATCTNNNACACCCAGCTAATTTTTAAAAA
GAGCACACGTCTGAACTCTTTCCCTACACGACGCTCTTCCGATCTNNNAGGTCATGTGTGAATGCCACT
GAGCACACGTCTGAACTCTTTCCCTACACGACGCTCTTCCGATCTNNNAGGTCATGTGTGAATGCCACT
GAGCACACGTCTGAACTCTTTCCCTACACGACGCTCTTCCGATCTNNNAGGTCATGTGTGAATGCCACT
GAGCACACGTCTGAACTCTTTCCCTACACGACGCTCTTCCGATCTNNNAGGTCATGTGTGAATGCCACT
GAGCACACGTCTGAACTCTTTCCCTACACGACGCTCTTCCGATCTNNNAGGTCATGTGTGAATGCCACT
AGCACACGTCTGAACTCTTTCCCTACACGACGCTCTTCCGATCTNNNGAGGCAGGAGAATGGTGTGAA
AGCACACGTCTGAACTCTTTCCCTACACGACGCTCTTCCGATCTNNNGAGGCAGGAGAATGGTGTGAA
AGCACACGTCTGAACTCTTTCCCTACACGACGCTCTTCCGATCTNNNGAGGCAGGAGAATGGTGTGAA
AGCACACGTCTGAACTCTTTCCCTACACGACGCTCTTCCGATCTNNNGAGGCAGGAGAATGGTGTGAA
AGCACACGTCTGAACTCTTTCCCTACACGACGCTCTTCCGATCTNNNGAGGCAGGAGAATGGTGTGAA
GAGCACACGTCTGAACTCTTTCCCTACACGACGCTCTTCCGATCTNNNTCCGTCTCAACAACAACAACAA
GAGCACACGTCTGAACTCTTTCCCTACACGACGCTCTTCCGATCTNNNTCCGTCTCAACAACAACAACAA
GAGCACACGTCTGAACTCTTTCCCTACACGACGCTCTTCCGATCTNNNTCCGTCTCAACAACAACAACAA
GAGCACACGTCTGAACTCTTTCCCTACACGACGCTCTTCCGATCTNNNTCCGTCTCAACAACAACAACAA
GAGCACACGTCTGAACTCTTTCCCTACACGACGCTCTTCCGATCTNNNTCCGTCTCAACAACAACAACAA

GCACACGTCTGAACTCTTTCCCTACACGACGCTCTTCCGATCTNNNGGCCCAAAGTGGTTTTATGACC
GCACACGTCTGAACTCTTTCCCTACACGACGCTCTTCCGATCTNNNGGCCCAAAGTGGTTTTATGACC
GCACACGTCTGAACTCTTTCCCTACACGACGCTCTTCCGATCTNNNGGCCCAAAGTGGTTTTATGACC
GCACACGTCTGAACTCTTTCCCTACACGACGCTCTTCCGATCTNNNGGCCCAAAGTGGTTTTATGACC
GCACACGTCTGAACTCTTTCCCTACACGACGCTCTTCCGATCTNNNGGCCCAAAGTGGTTTTATGACC
AGCACACGTCTGAACTCTTTCCCTACACGACGCTCTTCCGATCTNNNCACTGAGACAGGTGCATCATC
AGCACACGTCTGAACTCTTTCCCTACACGACGCTCTTCCGATCTNNNCACTGAGACAGGTGCATCATC
AGCACACGTCTGAACTCTTTCCCTACACGACGCTCTTCCGATCTNNNCACTGAGACAGGTGCATCATC
AGCACACGTCTGAACTCTTTCCCTACACGACGCTCTTCCGATCTNNNCACTGAGACAGGTGCATCATC
AGCACACGTCTGAACTCTTTCCCTACACGACGCTCTTCCGATCTNNNCACTGAGACAGGTGCATCATC
AGCACACGTCTGAACTCTTTCCCTACACGACGCTCTTCCGATCTNNNTCCTGTAAAGAGTAGGGGAT
AGCACACGTCTGAACTCTTTCCCTACACGACGCTCTTCCGATCTNNNTCCTGTAAAGAGTAGGGGAT
AGCACACGTCTGAACTCTTTCCCTACACGACGCTCTTCCGATCTNNNTCCTGTAAAGAGTAGGGGAT
AGCACACGTCTGAACTCTTTCCCTACACGACGCTCTTCCGATCTNNNTCCTGTAAAGAGTAGGGGAT
AGCACACGTCTGAACTCTTTCCCTACACGACGCTCTTCCGATCTNNNTCCTGTAAAGAGTAGGGGAT
GCACACGTCTGAACTCTTTCCCTACACGACGCTCTTCCGATCTNNNTCATGGTTGAATCATTGTGGTT
GCACACGTCTGAACTCTTTCCCTACACGACGCTCTTCCGATCTNNNTCATGGTTGAATCATTGTGGTT
GCACACGTCTGAACTCTTTCCCTACACGACGCTCTTCCGATCTNNNTCATGGTTGAATCATTGTGGTT
GCACACGTCTGAACTCTTTCCCTACACGACGCTCTTCCGATCTNNNTCATGGTTGAATCATTGTGGTT
GCACACGTCTGAACTCTTTCCCTACACGACGCTCTTCCGATCTNNNTCATGGTTGAATCATTGTGGTT
GCACACGTCTGAACTCTTTCCCTACACGACGCTCTTCCGATCTNNNCAAAAAGGCTCATGGAGTGCT
GCACACGTCTGAACTCTTTCCCTACACGACGCTCTTCCGATCTNNNCAAAAAGGCTCATGGAGTGCT
GCACACGTCTGAACTCTTTCCCTACACGACGCTCTTCCGATCTNNNCAAAAAGGCTCATGGAGTGCT
GCACACGTCTGAACTCTTTCCCTACACGACGCTCTTCCGATCTNNNCAAAAAGGCTCATGGAGTGCT
GCACACGTCTGAACTCTTTCCCTACACGACGCTCTTCCGATCTNNNCAAAAAGGCTCATGGAGTGCT
GAGCACACGTCTGAACTCTTTCCCTACACGACGCTCTTCCGATCTNNNACCACACTAAGAAAGAAAGG
GAGCACACGTCTGAACTCTTTCCCTACACGACGCTCTTCCGATCTNNNACCACACTAAGAAAGAAAGG
GAGCACACGTCTGAACTCTTTCCCTACACGACGCTCTTCCGATCTNNNACCACACTAAGAAAGAAAGG
GAGCACACGTCTGAACTCTTTCCCTACACGACGCTCTTCCGATCTNNNACCACACTAAGAAAGAAAGG
GAGCACACGTCTGAACTCTTTCCCTACACGACGCTCTTCCGATCTNNNACCACACTAAGAAAGAAAGG
AGCACACGTCTGAACTCTTTCCCTACACGACGCTCTTCCGATCTNNNCTCAGCAACAGTTGTGACAGT
AGCACACGTCTGAACTCTTTCCCTACACGACGCTCTTCCGATCTNNNCTCAGCAACAGTTGTGACAGT
AGCACACGTCTGAACTCTTTCCCTACACGACGCTCTTCCGATCTNNNCTCAGCAACAGTTGTGACAGT
AGCACACGTCTGAACTCTTTCCCTACACGACGCTCTTCCGATCTNNNCTCAGCAACAGTTGTGACAGT
AGCACACGTCTGAACTCTTTCCCTACACGACGCTCTTCCGATCTNNNCTCAGCAACAGTTGTGACAGT
AGCACACGTCTGAACTCTTTCCCTACACGACGCTCTTCCGATCTNNNATGAAGTTTTAGTGTGGTCTA
AGCACACGTCTGAACTCTTTCCCTACACGACGCTCTTCCGATCTNNNATGAAGTTTTAGTGTGGTCTA
AGCACACGTCTGAACTCTTTCCCTACACGACGCTCTTCCGATCTNNNATGAAGTTTTAGTGTGGTCTA
AGCACACGTCTGAACTCTTTCCCTACACGACGCTCTTCCGATCTNNNATGAAGTTTTAGTGTGGTCTA
AGCACACGTCTGAACTCTTTCCCTACACGACGCTCTTCCGATCTNNNATGAAGTTTTAGTGTGGTCTA
AGCACACGTCTGAACTCTTTCCCTACACGACGCTCTTCCGATCTNNNCTGTAATCCCAGCTACTCAGGA
AGCACACGTCTGAACTCTTTCCCTACACGACGCTCTTCCGATCTNNNCTGTAATCCCAGCTACTCAGGA
AGCACACGTCTGAACTCTTTCCCTACACGACGCTCTTCCGATCTNNNCTGTAATCCCAGCTACTCAGGA

AGCACACGTCTGAACTCTTTCCCTACACGACGCTCTTCCGATCTNNNCTGTAATCCCAGCTACTCAGGA
AGCACACGTCTGAACTCTTTCCCTACACGACGCTCTTCCGATCTNNNCTGTAATCCCAGCTACTCAGGA
AGAGCACACGTCTGAACTCTTTCCCTACACGACGCTCTTCCGATCTNNNCCTGTTGCATGGTTTTGTTT
AGAGCACACGTCTGAACTCTTTCCCTACACGACGCTCTTCCGATCTNNNCCTGTTGCATGGTTTTGTTT
AGAGCACACGTCTGAACTCTTTCCCTACACGACGCTCTTCCGATCTNNNCCTGTTGCATGGTTTTGTTT
AGAGCACACGTCTGAACTCTTTCCCTACACGACGCTCTTCCGATCTNNNCCTGTTGCATGGTTTTGTTT
AGAGCACACGTCTGAACTCTTTCCCTACACGACGCTCTTCCGATCTNNNCCTGTTGCATGGTTTTGTTT
AGCACACGTCTGAACTCTTTCCCTACACGACGCTCTTCCGATCTNNNCAGCAGTGTGAGGACTGACTA
AGCACACGTCTGAACTCTTTCCCTACACGACGCTCTTCCGATCTNNNCAGCAGTGTGAGGACTGACTA
AGCACACGTCTGAACTCTTTCCCTACACGACGCTCTTCCGATCTNNNCAGCAGTGTGAGGACTGACTA
AGCACACGTCTGAACTCTTTCCCTACACGACGCTCTTCCGATCTNNNCAGCAGTGTGAGGACTGACTA
AGCACACGTCTGAACTCTTTCCCTACACGACGCTCTTCCGATCTNNNCAGCAGTGTGAGGACTGACTA
AGCACACGTCTGAACTCTTTCCCTACACGACGCTCTTCCGATCTNNNTAAGATGGCTTAGCTGTGAGT
AGCACACGTCTGAACTCTTTCCCTACACGACGCTCTTCCGATCTNNNTAAGATGGCTTAGCTGTGAGT
AGCACACGTCTGAACTCTTTCCCTACACGACGCTCTTCCGATCTNNNTAAGATGGCTTAGCTGTGAGT
AGCACACGTCTGAACTCTTTCCCTACACGACGCTCTTCCGATCTNNNTAAGATGGCTTAGCTGTGAGT
AGCACACGTCTGAACTCTTTCCCTACACGACGCTCTTCCGATCTNNNTAAGATGGCTTAGCTGTGAGT
GAGCACACGTCTGAACTCTTTCCCTACACGACGCTCTTCCGATCTNNNCAGGAGAATTGCTTGAACCTT
GAGCACACGTCTGAACTCTTTCCCTACACGACGCTCTTCCGATCTNNNCAGGAGAATTGCTTGAACCTT
GAGCACACGTCTGAACTCTTTCCCTACACGACGCTCTTCCGATCTNNNCAGGAGAATTGCTTGAACCTT
GAGCACACGTCTGAACTCTTTCCCTACACGACGCTCTTCCGATCTNNNCAGGAGAATTGCTTGAACCTT
GAGCACACGTCTGAACTCTTTCCCTACACGACGCTCTTCCGATCTNNNCAGGAGAATTGCTTGAACCTT
AGAGCACACGTCTGAACTCTTTCCCTACACGACGCTCTTCCGATCTNNNGAGGTTGCAGTGAGCCAAG
AGAGCACACGTCTGAACTCTTTCCCTACACGACGCTCTTCCGATCTNNNGAGGTTGCAGTGAGCCAAG
AGAGCACACGTCTGAACTCTTTCCCTACACGACGCTCTTCCGATCTNNNGAGGTTGCAGTGAGCCAAG
AGAGCACACGTCTGAACTCTTTCCCTACACGACGCTCTTCCGATCTNNNGAGGTTGCAGTGAGCCAAG
AGAGCACACGTCTGAACTCTTTCCCTACACGACGCTCTTCCGATCTNNNGAGGTTGCAGTGAGCCAAG
AGAGCACACGTCTGAACTCTTTCCCTACACGACGCTCTTCCGATCTNNNTTTAGTAGAGACGGGGTTT
AGAGCACACGTCTGAACTCTTTCCCTACACGACGCTCTTCCGATCTNNNTTTAGTAGAGACGGGGTTT
AGAGCACACGTCTGAACTCTTTCCCTACACGACGCTCTTCCGATCTNNNTTTAGTAGAGACGGGGTTT
AGAGCACACGTCTGAACTCTTTCCCTACACGACGCTCTTCCGATCTNNNTTTAGTAGAGACGGGGTTT
AGAGCACACGTCTGAACTCTTTCCCTACACGACGCTCTTCCGATCTNNNTTTAGTAGAGACGGGGTTT
AGCACACGTCTGAACTCTTTCCCTACACGACGCTCTTCCGATCTNNNAAAATTAGCCAGGCATGTTGG
AGCACACGTCTGAACTCTTTCCCTACACGACGCTCTTCCGATCTNNNAAAATTAGCCAGGCATGTTGG
AGCACACGTCTGAACTCTTTCCCTACACGACGCTCTTCCGATCTNNNAAAATTAGCCAGGCATGTTGG
AGCACACGTCTGAACTCTTTCCCTACACGACGCTCTTCCGATCTNNNAAAATTAGCCAGGCATGTTGG
AGCACACGTCTGAACTCTTTCCCTACACGACGCTCTTCCGATCTNNNAAAATTAGCCAGGCATGTTGG
AGCACACGTCTGAACTCTTTCCCTACACGACGCTCTTCCGATCTNNNTGTAATCCCAGCTACTCAGGAG
AGCACACGTCTGAACTCTTTCCCTACACGACGCTCTTCCGATCTNNNTGTAATCCCAGCTACTCAGGAG
AGCACACGTCTGAACTCTTTCCCTACACGACGCTCTTCCGATCTNNNTGTAATCCCAGCTACTCAGGAG
AGCACACGTCTGAACTCTTTCCCTACACGACGCTCTTCCGATCTNNNTGTAATCCCAGCTACTCAGGAG
AGCACACGTCTGAACTCTTTCCCTACACGACGCTCTTCCGATCTNNNTGTAATCCCAGCTACTCAGGAG
AGCACACGTCTGAACTCTTTCCCTACACGACGCTCTTCCGATCTNNNTGCTCATCAAGTGTCTGGAGAA

GCACACGTCTGAACTCTTTCCCTACACGACGCTCTTCCGATCTNNNTGCTCATCAAGTGTCTGGAGAA
GCACACGTCTGAACTCTTTCCCTACACGACGCTCTTCCGATCTNNNTGCTCATCAAGTGTCTGGAGAA
GCACACGTCTGAACTCTTTCCCTACACGACGCTCTTCCGATCTNNNTGCTCATCAAGTGTCTGGAGAA
GCACACGTCTGAACTCTTTCCCTACACGACGCTCTTCCGATCTNNNTGCTCATCAAGTGTCTGGAGAA
AGCACACGTCTGAACTCTTTCCCTACACGACGCTCTTCCGATCTNNNTGCTCAACATGACCAATCATCA
AGCACACGTCTGAACTCTTTCCCTACACGACGCTCTTCCGATCTNNNTGCTCAACATGACCAATCATCA
AGCACACGTCTGAACTCTTTCCCTACACGACGCTCTTCCGATCTNNNTGCTCAACATGACCAATCATCA
AGCACACGTCTGAACTCTTTCCCTACACGACGCTCTTCCGATCTNNNTGCTCAACATGACCAATCATCA
AGCACACGTCTGAACTCTTTCCCTACACGACGCTCTTCCGATCTNNNTGCTCAACATGACCAATCATCA
AGCACACGTCTGAACTCTTTCCCTACACGACGCTCTTCCGATCTNNNTGTCACCATTGCTGCCAATTAT
AGCACACGTCTGAACTCTTTCCCTACACGACGCTCTTCCGATCTNNNTGTCACCATTGCTGCCAATTAT
AGCACACGTCTGAACTCTTTCCCTACACGACGCTCTTCCGATCTNNNTGTCACCATTGCTGCCAATTAT
AGCACACGTCTGAACTCTTTCCCTACACGACGCTCTTCCGATCTNNNTGTCACCATTGCTGCCAATTAT
AGCACACGTCTGAACTCTTTCCCTACACGACGCTCTTCCGATCTNNNTGTCACCATTGCTGCCAATTAT
GAGCACACGTCTGAACTCTTTCCCTACACGACGCTCTTCCGATCTNNNTACAAAAATTAGCCAGGCGT
GAGCACACGTCTGAACTCTTTCCCTACACGACGCTCTTCCGATCTNNNTACAAAAATTAGCCAGGCGT
GAGCACACGTCTGAACTCTTTCCCTACACGACGCTCTTCCGATCTNNNTACAAAAATTAGCCAGGCGT
GAGCACACGTCTGAACTCTTTCCCTACACGACGCTCTTCCGATCTNNNTACAAAAATTAGCCAGGCGT
GAGCACACGTCTGAACTCTTTCCCTACACGACGCTCTTCCGATCTNNNTACAAAAATTAGCCAGGCGT
GCACACGTCTGAACTCTTTCCCTACACGACGCTCTTCCGATCTNNNATAGAGAACATTTCCAGAATAC
GCACACGTCTGAACTCTTTCCCTACACGACGCTCTTCCGATCTNNNATAGAGAACATTTCCAGAATAC
GCACACGTCTGAACTCTTTCCCTACACGACGCTCTTCCGATCTNNNATAGAGAACATTTCCAGAATAC
GCACACGTCTGAACTCTTTCCCTACACGACGCTCTTCCGATCTNNNATAGAGAACATTTCCAGAATAC
GCACACGTCTGAACTCTTTCCCTACACGACGCTCTTCCGATCTNNNATAGAGAACATTTCCAGAATAC
GCACACGTCTGAACTCTTTCCCTACACGACGCTCTTCCGATCTNNNGGTACCCCTCTCCAATTATTTTG
GCACACGTCTGAACTCTTTCCCTACACGACGCTCTTCCGATCTNNNGGTACCCCTCTCCAATTATTTTG
GCACACGTCTGAACTCTTTCCCTACACGACGCTCTTCCGATCTNNNGGTACCCCTCTCCAATTATTTTG
GCACACGTCTGAACTCTTTCCCTACACGACGCTCTTCCGATCTNNNGGTACCCCTCTCCAATTATTTTG
GCACACGTCTGAACTCTTTCCCTACACGACGCTCTTCCGATCTNNNGGTACCCCTCTCCAATTATTTTG
AGCACACGTCTGAACTCTTTCCCTACACGACGCTCTTCCGATCTNNNTCTTGTTGCAAATGACTGGATC
AGCACACGTCTGAACTCTTTCCCTACACGACGCTCTTCCGATCTNNNTCTTGTTGCAAATGACTGGATC
AGCACACGTCTGAACTCTTTCCCTACACGACGCTCTTCCGATCTNNNTCTTGTTGCAAATGACTGGATC
AGCACACGTCTGAACTCTTTCCCTACACGACGCTCTTCCGATCTNNNTCTTGTTGCAAATGACTGGATC
AGCACACGTCTGAACTCTTTCCCTACACGACGCTCTTCCGATCTNNNTCTTGTTGCAAATGACTGGATC
GCACACGTCTGAACTCTTTCCCTACACGACGCTCTTCCGATCTNNNCTGCTCCATTTTCTACCCAGAC
GCACACGTCTGAACTCTTTCCCTACACGACGCTCTTCCGATCTNNNCTGCTCCATTTTCTACCCAGAC
GCACACGTCTGAACTCTTTCCCTACACGACGCTCTTCCGATCTNNNCTGCTCCATTTTCTACCCAGAC
GCACACGTCTGAACTCTTTCCCTACACGACGCTCTTCCGATCTNNNCTGCTCCATTTTCTACCCAGAC
GCACACGTCTGAACTCTTTCCCTACACGACGCTCTTCCGATCTNNNCTGCTCCATTTTCTACCCAGAC
GAGCACACGTCTGAACTCTTTCCCTACACGACGCTCTTCCGATCTNNNAACTTCAGGTTCCCAAAAGT
GAGCACACGTCTGAACTCTTTCCCTACACGACGCTCTTCCGATCTNNNAACTTCAGGTTCCCAAAAGT
GAGCACACGTCTGAACTCTTTCCCTACACGACGCTCTTCCGATCTNNNAACTTCAGGTTCCCAAAAGT
GAGCACACGTCTGAACTCTTTCCCTACACGACGCTCTTCCGATCTNNNAACTTCAGGTTCCCAAAAGT

GAGCACACGTCTGAACTCTTTCCCTACACGACGCTCTTCCGATCTNNNAACTTCAGGTTCCCAAAAGT
GCACACGTCTGAACTCTTTCCCTACACGACGCTCTTCCGATCTNNNATACCTATATTGTGCTTTAAGGT
GCACACGTCTGAACTCTTTCCCTACACGACGCTCTTCCGATCTNNNATACCTATATTGTGCTTTAAGGT
GCACACGTCTGAACTCTTTCCCTACACGACGCTCTTCCGATCTNNNATACCTATATTGTGCTTTAAGGT
GCACACGTCTGAACTCTTTCCCTACACGACGCTCTTCCGATCTNNNATACCTATATTGTGCTTTAAGGT
GCACACGTCTGAACTCTTTCCCTACACGACGCTCTTCCGATCTNNNATACCTATATTGTGCTTTAAGGT
AGCACACGTCTGAACTCTTTCCCTACACGACGCTCTTCCGATCTNNNGTAATCTCAGCTACTCAGGAG
AGCACACGTCTGAACTCTTTCCCTACACGACGCTCTTCCGATCTNNNGTAATCTCAGCTACTCAGGAG
AGCACACGTCTGAACTCTTTCCCTACACGACGCTCTTCCGATCTNNNGTAATCTCAGCTACTCAGGAG
AGCACACGTCTGAACTCTTTCCCTACACGACGCTCTTCCGATCTNNNGTAATCTCAGCTACTCAGGAG
AGCACACGTCTGAACTCTTTCCCTACACGACGCTCTTCCGATCTNNNGTAATCTCAGCTACTCAGGAG
GAGCACACGTCTGAACTCTTTCCCTACACGACGCTCTTCCGATCTNNNAGGGGGAACGAAATGAAG
GAGCACACGTCTGAACTCTTTCCCTACACGACGCTCTTCCGATCTNNNAGGGGGAACGAAATGAAG
GAGCACACGTCTGAACTCTTTCCCTACACGACGCTCTTCCGATCTNNNAGGGGGAACGAAATGAAG
GAGCACACGTCTGAACTCTTTCCCTACACGACGCTCTTCCGATCTNNNAGGGGGAACGAAATGAAG
GAGCACACGTCTGAACTCTTTCCCTACACGACGCTCTTCCGATCTNNNAGGGGGAACGAAATGAAG
GCACACGTCTGAACTCTTTCCCTACACGACGCTCTTCCGATCTNNNCAGCATGACACTCAAAAGCTGA
GCACACGTCTGAACTCTTTCCCTACACGACGCTCTTCCGATCTNNNCAGCATGACACTCAAAAGCTGA
GCACACGTCTGAACTCTTTCCCTACACGACGCTCTTCCGATCTNNNCAGCATGACACTCAAAAGCTGA
GCACACGTCTGAACTCTTTCCCTACACGACGCTCTTCCGATCTNNNCAGCATGACACTCAAAAGCTGA
GCACACGTCTGAACTCTTTCCCTACACGACGCTCTTCCGATCTNNNCAGCATGACACTCAAAAGCTGA
GCACACGTCTGAACTCTTTCCCTACACGACGCTCTTCCGATCTNNNCATTTCAACAACCACAGCAGACT
GCACACGTCTGAACTCTTTCCCTACACGACGCTCTTCCGATCTNNNCATTTCAACAACCACAGCAGACT
GCACACGTCTGAACTCTTTCCCTACACGACGCTCTTCCGATCTNNNCATTTCAACAACCACAGCAGACT
GCACACGTCTGAACTCTTTCCCTACACGACGCTCTTCCGATCTNNNCATTTCAACAACCACAGCAGACT
GCACACGTCTGAACTCTTTCCCTACACGACGCTCTTCCGATCTNNNCATTTCAACAACCACAGCAGACT
AGCACACGTCTGAACTCTTTCCCTACACGACGCTCTTCCGATCTNNNCTACAGTGAGCCATGGTTGCA
AGCACACGTCTGAACTCTTTCCCTACACGACGCTCTTCCGATCTNNNCTACAGTGAGCCATGGTTGCA
AGCACACGTCTGAACTCTTTCCCTACACGACGCTCTTCCGATCTNNNCTACAGTGAGCCATGGTTGCA
AGCACACGTCTGAACTCTTTCCCTACACGACGCTCTTCCGATCTNNNCTACAGTGAGCCATGGTTGCA
AGCACACGTCTGAACTCTTTCCCTACACGACGCTCTTCCGATCTNNNCTACAGTGAGCCATGGTTGCA
AGCACACGTCTGAACTCTTTCCCTACACGACGCTCTTCCGATCTNNNTGAGGCACAAGAACCATTTGA
AGCACACGTCTGAACTCTTTCCCTACACGACGCTCTTCCGATCTNNNTGAGGCACAAGAACCATTTGA
AGCACACGTCTGAACTCTTTCCCTACACGACGCTCTTCCGATCTNNNTGAGGCACAAGAACCATTTGA
AGCACACGTCTGAACTCTTTCCCTACACGACGCTCTTCCGATCTNNNTGAGGCACAAGAACCATTTGA
AGCACACGTCTGAACTCTTTCCCTACACGACGCTCTTCCGATCTNNNTGAGGCACAAGAACCATTTGA
GCACACGTCTGAACTCTTTCCCTACACGACGCTCTTCCGATCTNNNCGATGCACTGATTTCTGTGTGAA
GCACACGTCTGAACTCTTTCCCTACACGACGCTCTTCCGATCTNNNCGATGCACTGATTTCTGTGTGAA
GCACACGTCTGAACTCTTTCCCTACACGACGCTCTTCCGATCTNNNCGATGCACTGATTTCTGTGTGAA
GCACACGTCTGAACTCTTTCCCTACACGACGCTCTTCCGATCTNNNCGATGCACTGATTTCTGTGTGAA
GCACACGTCTGAACTCTTTCCCTACACGACGCTCTTCCGATCTNNNCGATGCACTGATTTCTGTGTGAA
GAGCACACGTCTGAACTCTTTCCCTACACGACGCTCTTCCGATCTNNNAAAGATCACATGTTCCAGAG
GAGCACACGTCTGAACTCTTTCCCTACACGACGCTCTTCCGATCTNNNAAAGATCACATGTTCCAGAG

GAGCACACGTCTGAACTCTTTCCCTACACGACGCTCTTCCGATCTNNNAAAGATCACATGTTCCAGAG
GAGCACACGTCTGAACTCTTTCCCTACACGACGCTCTTCCGATCTNNNAAAGATCACATGTTCCAGAG
GAGCACACGTCTGAACTCTTTCCCTACACGACGCTCTTCCGATCTNNNAAAGATCACATGTTCCAGAG
GCACACGTCTGAACTCTTTCCCTACACGACGCTCTTCCGATCTNNNCAAGTGACTGTAGTCCCAGCTAC
GCACACGTCTGAACTCTTTCCCTACACGACGCTCTTCCGATCTNNNCAAGTGACTGTAGTCCCAGCTAC
GCACACGTCTGAACTCTTTCCCTACACGACGCTCTTCCGATCTNNNCAAGTGACTGTAGTCCCAGCTAC
GCACACGTCTGAACTCTTTCCCTACACGACGCTCTTCCGATCTNNNCAAGTGACTGTAGTCCCAGCTAC
GCACACGTCTGAACTCTTTCCCTACACGACGCTCTTCCGATCTNNNCAAGTGACTGTAGTCCCAGCTAC
ACACGTCTGAACTCTTTCCCTACACGACGCTCTTCCGATCTNNNCTGTTCCAGCCAAAGACTTCATCCA
ACACGTCTGAACTCTTTCCCTACACGACGCTCTTCCGATCTNNNCTGTTCCAGCCAAAGACTTCATCCA
ACACGTCTGAACTCTTTCCCTACACGACGCTCTTCCGATCTNNNCTGTTCCAGCCAAAGACTTCATCCA
ACACGTCTGAACTCTTTCCCTACACGACGCTCTTCCGATCTNNNCTGTTCCAGCCAAAGACTTCATCCA
ACACGTCTGAACTCTTTCCCTACACGACGCTCTTCCGATCTNNNCTGTTCCAGCCAAAGACTTCATCCA
AGCACACGTCTGAACTCTTTCCCTACACGACGCTCTTCCGATCTNNNTAGGAAAATCACTGGAACCCT
AGCACACGTCTGAACTCTTTCCCTACACGACGCTCTTCCGATCTNNNTAGGAAAATCACTGGAACCCT
AGCACACGTCTGAACTCTTTCCCTACACGACGCTCTTCCGATCTNNNTAGGAAAATCACTGGAACCCT
AGCACACGTCTGAACTCTTTCCCTACACGACGCTCTTCCGATCTNNNTAGGAAAATCACTGGAACCCT
AGCACACGTCTGAACTCTTTCCCTACACGACGCTCTTCCGATCTNNNTAGGAAAATCACTGGAACCCT
GCACACGTCTGAACTCTTTCCCTACACGACGCTCTTCCGATCTNNNTGTTATTTATTTTACTTGCATGTG
GCACACGTCTGAACTCTTTCCCTACACGACGCTCTTCCGATCTNNNTGTTATTTATTTTACTTGCATGTG
GCACACGTCTGAACTCTTTCCCTACACGACGCTCTTCCGATCTNNNTGTTATTTATTTTACTTGCATGTG
GCACACGTCTGAACTCTTTCCCTACACGACGCTCTTCCGATCTNNNTGTTATTTATTTTACTTGCATGTG
GCACACGTCTGAACTCTTTCCCTACACGACGCTCTTCCGATCTNNNTGTTATTTATTTTACTTGCATGTG
GCACACGTCTGAACTCTTTCCCTACACGACGCTCTTCCGATCTNNNAGTGCTTTTTCCTTCTGAAGCATC
GCACACGTCTGAACTCTTTCCCTACACGACGCTCTTCCGATCTNNNAGTGCTTTTTCCTTCTGAAGCATC
GCACACGTCTGAACTCTTTCCCTACACGACGCTCTTCCGATCTNNNAGTGCTTTTTCCTTCTGAAGCATC
GCACACGTCTGAACTCTTTCCCTACACGACGCTCTTCCGATCTNNNAGTGCTTTTTCCTTCTGAAGCATC
GCACACGTCTGAACTCTTTCCCTACACGACGCTCTTCCGATCTNNNAGTGCTTTTTCCTTCTGAAGCATC
GAGCACACGTCTGAACTCTTTCCCTACACGACGCTCTTCCGATCTNNNAAAAAACTTAGCCAGGTGTG
GAGCACACGTCTGAACTCTTTCCCTACACGACGCTCTTCCGATCTNNNAAAAAACTTAGCCAGGTGTG
GAGCACACGTCTGAACTCTTTCCCTACACGACGCTCTTCCGATCTNNNAAAAAACTTAGCCAGGTGTG
GAGCACACGTCTGAACTCTTTCCCTACACGACGCTCTTCCGATCTNNNAAAAAACTTAGCCAGGTGTG
GAGCACACGTCTGAACTCTTTCCCTACACGACGCTCTTCCGATCTNNNAAAAAACTTAGCCAGGTGTG
GCACACGTCTGAACTCTTTCCCTACACGACGCTCTTCCGATCTNNNCATAATCAGTCCCAGCAGTACC
GCACACGTCTGAACTCTTTCCCTACACGACGCTCTTCCGATCTNNNCATAATCAGTCCCAGCAGTACC
GCACACGTCTGAACTCTTTCCCTACACGACGCTCTTCCGATCTNNNCATAATCAGTCCCAGCAGTACC
GCACACGTCTGAACTCTTTCCCTACACGACGCTCTTCCGATCTNNNCATAATCAGTCCCAGCAGTACC
GCACACGTCTGAACTCTTTCCCTACACGACGCTCTTCCGATCTNNNCATAATCAGTCCCAGCAGTACC
AGCACACGTCTGAACTCTTTCCCTACACGACGCTCTTCCGATCTNNNCAGGTCACACGATCATATTGT
AGCACACGTCTGAACTCTTTCCCTACACGACGCTCTTCCGATCTNNNCAGGTCACACGATCATATTGT
AGCACACGTCTGAACTCTTTCCCTACACGACGCTCTTCCGATCTNNNCAGGTCACACGATCATATTGT
AGCACACGTCTGAACTCTTTCCCTACACGACGCTCTTCCGATCTNNNCAGGTCACACGATCATATTGT
AGCACACGTCTGAACTCTTTCCCTACACGACGCTCTTCCGATCTNNNCAGGTCACACGATCATATTGT

GAGCACACGTCTGAACTCTTTCCCTACACGACGCTCTTCCGATCTNNNGAGAATGGCTTGAGAGAACC

GAGCACACGTCTGAACTCTTTCCCTACACGACGCTCTTCCGATCTNNNGAGAATGGCTTGAGAGAACC

GAGCACACGTCTGAACTCTTTCCCTACACGACGCTCTTCCGATCTNNNGAGAATGGCTTGAGAGAACC

GAGCACACGTCTGAACTCTTTCCCTACACGACGCTCTTCCGATCTNNNGAGAATGGCTTGAGAGAACC

GAGCACACGTCTGAACTCTTTCCCTACACGACGCTCTTCCGATCTNNNGAGAATGGCTTGAGAGAACC

AGAGCACACGTCTGAACTCTTTCCCTACACGACGCTCTTCCGATCTNNNTCCATGTTGCTGCAAATTCAC

AGAGCACACGTCTGAACTCTTTCCCTACACGACGCTCTTCCGATCTNNNTCCATGTTGCTGCAAATTCAC

AGAGCACACGTCTGAACTCTTTCCCTACACGACGCTCTTCCGATCTNNNTCCATGTTGCTGCAAATTCAC

AGAGCACACGTCTGAACTCTTTCCCTACACGACGCTCTTCCGATCTNNNTCCATGTTGCTGCAAATTCAC

AGAGCACACGTCTGAACTCTTTCCCTACACGACGCTCTTCCGATCTNNNTCCATGTTGCTGCAAATTCAC

AGCACACGTCTGAACTCTTTCCCTACACGACGCTCTTCCGATCTNNNAAGCAGAAATAGATCCCGCCTT

AGCACACGTCTGAACTCTTTCCCTACACGACGCTCTTCCGATCTNNNAAGCAGAAATAGATCCCGCCTT

AGCACACGTCTGAACTCTTTCCCTACACGACGCTCTTCCGATCTNNNAAGCAGAAATAGATCCCGCCTT

AGCACACGTCTGAACTCTTTCCCTACACGACGCTCTTCCGATCTNNNAAGCAGAAATAGATCCCGCCTT

AGCACACGTCTGAACTCTTTCCCTACACGACGCTCTTCCGATCTNNNAAGCAGAAATAGATCCCGCCTT

GAGCACACGTCTGAACTCTTTCCCTACACGACGCTCTTCCGATCTNNNCTGAGGCAGGAGAACAGTGTA

GAGCACACGTCTGAACTCTTTCCCTACACGACGCTCTTCCGATCTNNNCTGAGGCAGGAGAACAGTGTA

GAGCACACGTCTGAACTCTTTCCCTACACGACGCTCTTCCGATCTNNNCTGAGGCAGGAGAACAGTGTA

GAGCACACGTCTGAACTCTTTCCCTACACGACGCTCTTCCGATCTNNNCTGAGGCAGGAGAACAGTGTA

GAGCACACGTCTGAACTCTTTCCCTACACGACGCTCTTCCGATCTNNNCTGAGGCAGGAGAACAGTGTA

AGAGCACACGTCTGAACTCTTTCCCTACACGACGCTCTTCCGATCTNNNGTTGTAATGAGCCAAGATCA

AGAGCACACGTCTGAACTCTTTCCCTACACGACGCTCTTCCGATCTNNNGTTGTAATGAGCCAAGATCA

AGAGCACACGTCTGAACTCTTTCCCTACACGACGCTCTTCCGATCTNNNGTTGTAATGAGCCAAGATCA

AGAGCACACGTCTGAACTCTTTCCCTACACGACGCTCTTCCGATCTNNNGTTGTAATGAGCCAAGATCA

AGAGCACACGTCTGAACTCTTTCCCTACACGACGCTCTTCCGATCTNNNGTTGTAATGAGCCAAGATCA

AGAGCACACGTCTGAACTCTTTCCCTACACGACGCTCTTCCGATCTNNNCCCGAAGACTCTCTTGTTCTA

AGAGCACACGTCTGAACTCTTTCCCTACACGACGCTCTTCCGATCTNNNCCCGAAGACTCTCTTGTTCTA

AGAGCACACGTCTGAACTCTTTCCCTACACGACGCTCTTCCGATCTNNNCCCGAAGACTCTCTTGTTCTA

AGAGCACACGTCTGAACTCTTTCCCTACACGACGCTCTTCCGATCTNNNCCCGAAGACTCTCTTGTTCTA

AGAGCACACGTCTGAACTCTTTCCCTACACGACGCTCTTCCGATCTNNNCCCGAAGACTCTCTTGTTCTA

AGAGCACACGTCTGAACTCTTTCCCTACACGACGCTCTTCCGATCTNNNACTCAGGGATGTCTTTAGGG

AGAGCACACGTCTGAACTCTTTCCCTACACGACGCTCTTCCGATCTNNNACTCAGGGATGTCTTTAGGG

AGAGCACACGTCTGAACTCTTTCCCTACACGACGCTCTTCCGATCTNNNACTCAGGGATGTCTTTAGGG

AGAGCACACGTCTGAACTCTTTCCCTACACGACGCTCTTCCGATCTNNNACTCAGGGATGTCTTTAGGG

AGAGCACACGTCTGAACTCTTTCCCTACACGACGCTCTTCCGATCTNNNACTCAGGGATGTCTTTAGGG

GAGCACACGTCTGAACTCTTTCCCTACACGACGCTCTTCCGATCTNNNGTAATCTCAGCTACTCGAGAG

GAGCACACGTCTGAACTCTTTCCCTACACGACGCTCTTCCGATCTNNNGTAATCTCAGCTACTCGAGAG

GAGCACACGTCTGAACTCTTTCCCTACACGACGCTCTTCCGATCTNNNGTAATCTCAGCTACTCGAGAG

GAGCACACGTCTGAACTCTTTCCCTACACGACGCTCTTCCGATCTNNNGTAATCTCAGCTACTCGAGAG

GAGCACACGTCTGAACTCTTTCCCTACACGACGCTCTTCCGATCTNNNGTAATCTCAGCTACTCGAGAG

AGCACACGTCTGAACTCTTTCCCTACACGACGCTCTTCCGATCTNNNTTGCACGTAGTTGCAGAATTCA

AGCACACGTCTGAACTCTTTCCCTACACGACGCTCTTCCGATCTNNNTTGCACGTAGTTGCAGAATTCA

AGCACACGTCTGAACTCTTTCCCTACACGACGCTCTTCCGATCTNNNTTGCACGTAGTTGCAGAATTCA

AGCACACGTCTGAACTCTTTCCCTACACGACGCTCTTCCGATCTNNNTTGCACGTAGTTGCAGAATTCA
AGCACACGTCTGAACTCTTTCCCTACACGACGCTCTTCCGATCTNNNTTGCACGTAGTTGCAGAATTCA
GAGCACACGTCTGAACTCTTTCCCTACACGACGCTCTTCCGATCTNNNGTGGGTTCAGCATTTTCTGTT
GAGCACACGTCTGAACTCTTTCCCTACACGACGCTCTTCCGATCTNNNGTGGGTTCAGCATTTTCTGTT
GAGCACACGTCTGAACTCTTTCCCTACACGACGCTCTTCCGATCTNNNGTGGGTTCAGCATTTTCTGTT
GAGCACACGTCTGAACTCTTTCCCTACACGACGCTCTTCCGATCTNNNGTGGGTTCAGCATTTTCTGTT
GAGCACACGTCTGAACTCTTTCCCTACACGACGCTCTTCCGATCTNNNGTGGGTTCAGCATTTTCTGTT
AGCACACGTCTGAACTCTTTCCCTACACGACGCTCTTCCGATCTNNNCCTCCCAGGCTGTTTTTCATTT
AGCACACGTCTGAACTCTTTCCCTACACGACGCTCTTCCGATCTNNNCCTCCCAGGCTGTTTTTCATTT
AGCACACGTCTGAACTCTTTCCCTACACGACGCTCTTCCGATCTNNNCCTCCCAGGCTGTTTTTCATTT
AGCACACGTCTGAACTCTTTCCCTACACGACGCTCTTCCGATCTNNNCCTCCCAGGCTGTTTTTCATTT
AGCACACGTCTGAACTCTTTCCCTACACGACGCTCTTCCGATCTNNNCCTCCCAGGCTGTTTTTCATTT
AGCACACGTCTGAACTCTTTCCCTACACGACGCTCTTCCGATCTNNNGCTCCTCTGTTGTGTATGCTTG
AGCACACGTCTGAACTCTTTCCCTACACGACGCTCTTCCGATCTNNNGCTCCTCTGTTGTGTATGCTTG
AGCACACGTCTGAACTCTTTCCCTACACGACGCTCTTCCGATCTNNNGCTCCTCTGTTGTGTATGCTTG
AGCACACGTCTGAACTCTTTCCCTACACGACGCTCTTCCGATCTNNNGCTCCTCTGTTGTGTATGCTTG
AGCACACGTCTGAACTCTTTCCCTACACGACGCTCTTCCGATCTNNNGCTCCTCTGTTGTGTATGCTTG
GAGCACACGTCTGAACTCTTTCCCTACACGACGCTCTTCCGATCTNNNAGCTACTCAGGAAGTTGAAG
GAGCACACGTCTGAACTCTTTCCCTACACGACGCTCTTCCGATCTNNNAGCTACTCAGGAAGTTGAAG
GAGCACACGTCTGAACTCTTTCCCTACACGACGCTCTTCCGATCTNNNAGCTACTCAGGAAGTTGAAG
GAGCACACGTCTGAACTCTTTCCCTACACGACGCTCTTCCGATCTNNNAGCTACTCAGGAAGTTGAAG
GAGCACACGTCTGAACTCTTTCCCTACACGACGCTCTTCCGATCTNNNAGCTACTCAGGAAGTTGAAG
GAGCACACGTCTGAACTCTTTCCCTACACGACGCTCTTCCGATCTNNNGTTCGGTTTTGTTTTCACGC
GAGCACACGTCTGAACTCTTTCCCTACACGACGCTCTTCCGATCTNNNGTTCGGTTTTGTTTTCACGC
GAGCACACGTCTGAACTCTTTCCCTACACGACGCTCTTCCGATCTNNNGTTCGGTTTTGTTTTCACGC
GAGCACACGTCTGAACTCTTTCCCTACACGACGCTCTTCCGATCTNNNGTTCGGTTTTGTTTTCACGC
GAGCACACGTCTGAACTCTTTCCCTACACGACGCTCTTCCGATCTNNNGTTCGGTTTTGTTTTCACGC
GCACACGTCTGAACTCTTTCCCTACACGACGCTCTTCCGATCTNNNATGCTGACAGTTTGTGGAGGTA
GCACACGTCTGAACTCTTTCCCTACACGACGCTCTTCCGATCTNNNATGCTGACAGTTTGTGGAGGTA
GCACACGTCTGAACTCTTTCCCTACACGACGCTCTTCCGATCTNNNATGCTGACAGTTTGTGGAGGTA
GCACACGTCTGAACTCTTTCCCTACACGACGCTCTTCCGATCTNNNATGCTGACAGTTTGTGGAGGTA
GCACACGTCTGAACTCTTTCCCTACACGACGCTCTTCCGATCTNNNATGCTGACAGTTTGTGGAGGTA
AGCACACGTCTGAACTCTTTCCCTACACGACGCTCTTCCGATCTNNNCCCAGCTAACTAAGTGCACAG
AGCACACGTCTGAACTCTTTCCCTACACGACGCTCTTCCGATCTNNNCCCAGCTAACTAAGTGCACAG
AGCACACGTCTGAACTCTTTCCCTACACGACGCTCTTCCGATCTNNNCCCAGCTAACTAAGTGCACAG
AGCACACGTCTGAACTCTTTCCCTACACGACGCTCTTCCGATCTNNNCCCAGCTAACTAAGTGCACAG
AGCACACGTCTGAACTCTTTCCCTACACGACGCTCTTCCGATCTNNNCCCAGCTAACTAAGTGCACAG
GAGCACACGTCTGAACTCTTTCCCTACACGACGCTCTTCCGATCTNNNGACTCCCTGTCAAACTGATG
GAGCACACGTCTGAACTCTTTCCCTACACGACGCTCTTCCGATCTNNNGACTCCCTGTCAAACTGATG
GAGCACACGTCTGAACTCTTTCCCTACACGACGCTCTTCCGATCTNNNGACTCCCTGTCAAACTGATG
GAGCACACGTCTGAACTCTTTCCCTACACGACGCTCTTCCGATCTNNNGACTCCCTGTCAAACTGATG
GAGCACACGTCTGAACTCTTTCCCTACACGACGCTCTTCCGATCTNNNGACTCCCTGTCAAACTGATG
AGCACACGTCTGAACTCTTTCCCTACACGACGCTCTTCCGATCTNNNGAAGTGAGTGAGTGACAGAC

AGCACACGTCTGAACTCTTTCCCTACACGACGCTCTTCCGATCTNNNGAAGTGAGTGAGTGACAGAC

AGCACACGTCTGAACTCTTTCCCTACACGACGCTCTTCCGATCTNNNGAAGTGAGTGAGTGACAGAC

AGCACACGTCTGAACTCTTTCCCTACACGACGCTCTTCCGATCTNNNGAAGTGAGTGAGTGACAGAC

AGCACACGTCTGAACTCTTTCCCTACACGACGCTCTTCCGATCTNNNGAAGTGAGTGAGTGACAGAC

GAGCACACGTCTGAACTCTTTCCCTACACGACGCTCTTCCGATCTNNNGACTTCTGGCTGCAGGTGT

GAGCACACGTCTGAACTCTTTCCCTACACGACGCTCTTCCGATCTNNNGACTTCTGGCTGCAGGTGT

GAGCACACGTCTGAACTCTTTCCCTACACGACGCTCTTCCGATCTNNNGACTTCTGGCTGCAGGTGT

GAGCACACGTCTGAACTCTTTCCCTACACGACGCTCTTCCGATCTNNNGACTTCTGGCTGCAGGTGT

GAGCACACGTCTGAACTCTTTCCCTACACGACGCTCTTCCGATCTNNNGACTTCTGGCTGCAGGTGT

AGCACACGTCTGAACTCTTTCCCTACACGACGCTCTTCCGATCTNNNCAGAGGTCTTGGGATGGTTTC

AGCACACGTCTGAACTCTTTCCCTACACGACGCTCTTCCGATCTNNNCAGAGGTCTTGGGATGGTTTC

AGCACACGTCTGAACTCTTTCCCTACACGACGCTCTTCCGATCTNNNCAGAGGTCTTGGGATGGTTTC

AGCACACGTCTGAACTCTTTCCCTACACGACGCTCTTCCGATCTNNNCAGAGGTCTTGGGATGGTTTC

AGCACACGTCTGAACTCTTTCCCTACACGACGCTCTTCCGATCTNNNCAGAGGTCTTGGGATGGTTTC

GCACACGTCTGAACTCTTTCCCTACACGACGCTCTTCCGATCTNNNACTTGCAATAGCTGAGTAGTAC

GCACACGTCTGAACTCTTTCCCTACACGACGCTCTTCCGATCTNNNACTTGCAATAGCTGAGTAGTAC

GCACACGTCTGAACTCTTTCCCTACACGACGCTCTTCCGATCTNNNACTTGCAATAGCTGAGTAGTAC

GCACACGTCTGAACTCTTTCCCTACACGACGCTCTTCCGATCTNNNACTTGCAATAGCTGAGTAGTAC

GCACACGTCTGAACTCTTTCCCTACACGACGCTCTTCCGATCTNNNACTTGCAATAGCTGAGTAGTAC

GCACACGTCTGAACTCTTTCCCTACACGACGCTCTTCCGATCTNNNTTTACCCTCTATTTTCACTTCCCA

GCACACGTCTGAACTCTTTCCCTACACGACGCTCTTCCGATCTNNNTTTACCCTCTATTTTCACTTCCCA

GCACACGTCTGAACTCTTTCCCTACACGACGCTCTTCCGATCTNNNTTTACCCTCTATTTTCACTTCCCA

GCACACGTCTGAACTCTTTCCCTACACGACGCTCTTCCGATCTNNNTTTACCCTCTATTTTCACTTCCCA

GCACACGTCTGAACTCTTTCCCTACACGACGCTCTTCCGATCTNNNTTTACCCTCTATTTTCACTTCCCA

AGCACACGTCTGAACTCTTTCCCTACACGACGCTCTTCCGATCTNNNTCTGACTCTTCTCTTTTCCCAGC

AGCACACGTCTGAACTCTTTCCCTACACGACGCTCTTCCGATCTNNNTCTGACTCTTCTCTTTTCCCAGC

AGCACACGTCTGAACTCTTTCCCTACACGACGCTCTTCCGATCTNNNTCTGACTCTTCTCTTTTCCCAGC

AGCACACGTCTGAACTCTTTCCCTACACGACGCTCTTCCGATCTNNNTCTGACTCTTCTCTTTTCCCAGC

AGCACACGTCTGAACTCTTTCCCTACACGACGCTCTTCCGATCTNNNTCTGACTCTTCTCTTTTCCCAGC

AGCACACGTCTGAACTCTTTCCCTACACGACGCTCTTCCGATCTNNNGCGACAGAGCAAGACTTCATC

AGCACACGTCTGAACTCTTTCCCTACACGACGCTCTTCCGATCTNNNGCGACAGAGCAAGACTTCATC

AGCACACGTCTGAACTCTTTCCCTACACGACGCTCTTCCGATCTNNNGCGACAGAGCAAGACTTCATC

AGCACACGTCTGAACTCTTTCCCTACACGACGCTCTTCCGATCTNNNGCGACAGAGCAAGACTTCATC

AGCACACGTCTGAACTCTTTCCCTACACGACGCTCTTCCGATCTNNNGCGACAGAGCAAGACTTCATC

AGCACACGTCTGAACTCTTTCCCTACACGACGCTCTTCCGATCTNNNGGACGTGAAATTTGCCAGTTG

AGCACACGTCTGAACTCTTTCCCTACACGACGCTCTTCCGATCTNNNGGACGTGAAATTTGCCAGTTG

AGCACACGTCTGAACTCTTTCCCTACACGACGCTCTTCCGATCTNNNGGACGTGAAATTTGCCAGTTG

AGCACACGTCTGAACTCTTTCCCTACACGACGCTCTTCCGATCTNNNGGACGTGAAATTTGCCAGTTG

AGCACACGTCTGAACTCTTTCCCTACACGACGCTCTTCCGATCTNNNGGACGTGAAATTTGCCAGTTG

AGCACACGTCTGAACTCTTTCCCTACACGACGCTCTTCCGATCTNNNAGGTTGCAGTGAGTTGAGAT

AGCACACGTCTGAACTCTTTCCCTACACGACGCTCTTCCGATCTNNNAGGTTGCAGTGAGTTGAGAT

AGCACACGTCTGAACTCTTTCCCTACACGACGCTCTTCCGATCTNNNAGGTTGCAGTGAGTTGAGAT

AGCACACGTCTGAACTCTTTCCCTACACGACGCTCTTCCGATCTNNNAGGTTGCAGTGAGTTGAGAT

AGCACACGTCTGAACTCTTTCCCTACACGACGCTCTTCCGATCTNNNAGGTTGCAGTGAGTTGAGAT

GCACACGTCTGAACTCTTTCCCTACACGACGCTCTTCCGATCTNNNCTGAGGCAGGAAAATTGCTTGA

GCACACGTCTGAACTCTTTCCCTACACGACGCTCTTCCGATCTNNNCTGAGGCAGGAAAATTGCTTGA

GCACACGTCTGAACTCTTTCCCTACACGACGCTCTTCCGATCTNNNCTGAGGCAGGAAAATTGCTTGA

GCACACGTCTGAACTCTTTCCCTACACGACGCTCTTCCGATCTNNNCTGAGGCAGGAAAATTGCTTGA

GCACACGTCTGAACTCTTTCCCTACACGACGCTCTTCCGATCTNNNCTGAGGCAGGAAAATTGCTTGA

CACACGTCTGAACTCTTTCCCTACACGACGCTCTTCCGATCTNNNGGGACTCACTGGGAATTGTAGT

CACACGTCTGAACTCTTTCCCTACACGACGCTCTTCCGATCTNNNGGGACTCACTGGGAATTGTAGT

CACACGTCTGAACTCTTTCCCTACACGACGCTCTTCCGATCTNNNGGGACTCACTGGGAATTGTAGT

CACACGTCTGAACTCTTTCCCTACACGACGCTCTTCCGATCTNNNGGGACTCACTGGGAATTGTAGT

CACACGTCTGAACTCTTTCCCTACACGACGCTCTTCCGATCTNNNGGGACTCACTGGGAATTGTAGT

AGCACACGTCTGAACTCTTTCCCTACACGACGCTCTTCCGATCTNNNCCAAAATAAAGATCAGGCTGG

AGCACACGTCTGAACTCTTTCCCTACACGACGCTCTTCCGATCTNNNCCAAAATAAAGATCAGGCTGG

AGCACACGTCTGAACTCTTTCCCTACACGACGCTCTTCCGATCTNNNCCAAAATAAAGATCAGGCTGG

AGCACACGTCTGAACTCTTTCCCTACACGACGCTCTTCCGATCTNNNCCAAAATAAAGATCAGGCTGG

AGCACACGTCTGAACTCTTTCCCTACACGACGCTCTTCCGATCTNNNCCAAAATAAAGATCAGGCTGG

AGCACACGTCTGAACTCTTTCCCTACACGACGCTCTTCCGATCTNNNTAGAAAGATATGGGGTGTGT

AGCACACGTCTGAACTCTTTCCCTACACGACGCTCTTCCGATCTNNNTAGAAAGATATGGGGTGTGT

AGCACACGTCTGAACTCTTTCCCTACACGACGCTCTTCCGATCTNNNTAGAAAGATATGGGGTGTGT

AGCACACGTCTGAACTCTTTCCCTACACGACGCTCTTCCGATCTNNNTAGAAAGATATGGGGTGTGT

AGCACACGTCTGAACTCTTTCCCTACACGACGCTCTTCCGATCTNNNTAGAAAGATATGGGGTGTGT

GCACACGTCTGAACTCTTTCCCTACACGACGCTCTTCCGATCTNNNTACAGCCAGCTACCTTGAGATT

GCACACGTCTGAACTCTTTCCCTACACGACGCTCTTCCGATCTNNNTACAGCCAGCTACCTTGAGATT

GCACACGTCTGAACTCTTTCCCTACACGACGCTCTTCCGATCTNNNTACAGCCAGCTACCTTGAGATT

GCACACGTCTGAACTCTTTCCCTACACGACGCTCTTCCGATCTNNNTACAGCCAGCTACCTTGAGATT

GCACACGTCTGAACTCTTTCCCTACACGACGCTCTTCCGATCTNNNTACAGCCAGCTACCTTGAGATT

GAGCACACGTCTGAACTCTTTCCCTACACGACGCTCTTCCGATCTNNNCAACTCAAGGACCGGACAAA

GAGCACACGTCTGAACTCTTTCCCTACACGACGCTCTTCCGATCTNNNCAACTCAAGGACCGGACAAA

GAGCACACGTCTGAACTCTTTCCCTACACGACGCTCTTCCGATCTNNNCAACTCAAGGACCGGACAAA

GAGCACACGTCTGAACTCTTTCCCTACACGACGCTCTTCCGATCTNNNCAACTCAAGGACCGGACAAA

GAGCACACGTCTGAACTCTTTCCCTACACGACGCTCTTCCGATCTNNNCAACTCAAGGACCGGACAAA

GCACACGTCTGAACTCTTTCCCTACACGACGCTCTTCCGATCTNNNATCCTTAAAGCTTCTGATTGTTT

GCACACGTCTGAACTCTTTCCCTACACGACGCTCTTCCGATCTNNNATCCTTAAAGCTTCTGATTGTTT

GCACACGTCTGAACTCTTTCCCTACACGACGCTCTTCCGATCTNNNATCCTTAAAGCTTCTGATTGTTT

GCACACGTCTGAACTCTTTCCCTACACGACGCTCTTCCGATCTNNNATCCTTAAAGCTTCTGATTGTTT

GCACACGTCTGAACTCTTTCCCTACACGACGCTCTTCCGATCTNNNATCCTTAAAGCTTCTGATTGTTT

GAGCACACGTCTGAACTCTTTCCCTACACGACGCTCTTCCGATCTNNNCTAGAGAGGGAGGCTACTG

GAGCACACGTCTGAACTCTTTCCCTACACGACGCTCTTCCGATCTNNNCTAGAGAGGGAGGCTACTG

GAGCACACGTCTGAACTCTTTCCCTACACGACGCTCTTCCGATCTNNNCTAGAGAGGGAGGCTACTG

GAGCACACGTCTGAACTCTTTCCCTACACGACGCTCTTCCGATCTNNNCTAGAGAGGGAGGCTACTG

GAGCACACGTCTGAACTCTTTCCCTACACGACGCTCTTCCGATCTNNNCTAGAGAGGGAGGCTACTG

AGCACACGTCTGAACTCTTTCCCTACACGACGCTCTTCCGATCTNNNGTATTCCCAGCTACTCAGGAA

AGCACACGTCTGAACTCTTTCCCTACACGACGCTCTTCCGATCTNNNGTATTCCCAGCTACTCAGGAA

AGCACACGTCTGAACTCTTTCCCTACACGACGCTCTTCCGATCTNNNGTATTCCCAGCTACTCAGGAA
AGCACACGTCTGAACTCTTTCCCTACACGACGCTCTTCCGATCTNNNGTATTCCCAGCTACTCAGGAA
AGCACACGTCTGAACTCTTTCCCTACACGACGCTCTTCCGATCTNNNGTATTCCCAGCTACTCAGGAA
GAGCACACGTCTGAACTCTTTCCCTACACGACGCTCTTCCGATCTNNNCTTCTCTCTAACCCCCAATGC
GAGCACACGTCTGAACTCTTTCCCTACACGACGCTCTTCCGATCTNNNCTTCTCTCTAACCCCCAATGC
GAGCACACGTCTGAACTCTTTCCCTACACGACGCTCTTCCGATCTNNNCTTCTCTCTAACCCCCAATGC
GAGCACACGTCTGAACTCTTTCCCTACACGACGCTCTTCCGATCTNNNCTTCTCTCTAACCCCCAATGC
GAGCACACGTCTGAACTCTTTCCCTACACGACGCTCTTCCGATCTNNNCTTCTCTCTAACCCCCAATGC
GAGCACACGTCTGAACTCTTTCCCTACACGACGCTCTTCCGATCTNNNCCCTGTGTTGGCAAGAGTAT
GAGCACACGTCTGAACTCTTTCCCTACACGACGCTCTTCCGATCTNNNCCCTGTGTTGGCAAGAGTAT
GAGCACACGTCTGAACTCTTTCCCTACACGACGCTCTTCCGATCTNNNCCCTGTGTTGGCAAGAGTAT
GAGCACACGTCTGAACTCTTTCCCTACACGACGCTCTTCCGATCTNNNCCCTGTGTTGGCAAGAGTAT
GAGCACACGTCTGAACTCTTTCCCTACACGACGCTCTTCCGATCTNNNCCCTGTGTTGGCAAGAGTAT
GAGCACACGTCTGAACTCTTTCCCTACACGACGCTCTTCCGATCTNNNGGCAGTGAAATGAATGAT
GAGCACACGTCTGAACTCTTTCCCTACACGACGCTCTTCCGATCTNNNGGCAGTGAAATGAATGAT
GAGCACACGTCTGAACTCTTTCCCTACACGACGCTCTTCCGATCTNNNGGCAGTGAAATGAATGAT
GAGCACACGTCTGAACTCTTTCCCTACACGACGCTCTTCCGATCTNNNGGCAGTGAAATGAATGAT
GAGCACACGTCTGAACTCTTTCCCTACACGACGCTCTTCCGATCTNNNGGCAGTGAAATGAATGAT
GCACACGTCTGAACTCTTTCCCTACACGACGCTCTTCCGATCTNNNGTTTTGAACTCCTGACCTCAAG
GCACACGTCTGAACTCTTTCCCTACACGACGCTCTTCCGATCTNNNGTTTTGAACTCCTGACCTCAAG
GCACACGTCTGAACTCTTTCCCTACACGACGCTCTTCCGATCTNNNGTTTTGAACTCCTGACCTCAAG
GCACACGTCTGAACTCTTTCCCTACACGACGCTCTTCCGATCTNNNGTTTTGAACTCCTGACCTCAAG
GCACACGTCTGAACTCTTTCCCTACACGACGCTCTTCCGATCTNNNGTTTTGAACTCCTGACCTCAAG
GAGCACACGTCTGAACTCTTTCCCTACACGACGCTCTTCCGATCTNNNCAAGTGAGCCTTCTTGGAAG
GAGCACACGTCTGAACTCTTTCCCTACACGACGCTCTTCCGATCTNNNCAAGTGAGCCTTCTTGGAAG
GAGCACACGTCTGAACTCTTTCCCTACACGACGCTCTTCCGATCTNNNCAAGTGAGCCTTCTTGGAAG
GAGCACACGTCTGAACTCTTTCCCTACACGACGCTCTTCCGATCTNNNCAAGTGAGCCTTCTTGGAAG
GAGCACACGTCTGAACTCTTTCCCTACACGACGCTCTTCCGATCTNNNCAAGTGAGCCTTCTTGGAAG
AGCACACGTCTGAACTCTTTCCCTACACGACGCTCTTCCGATCTNNNATCTGCCAGCACCTTGATTTTT
AGCACACGTCTGAACTCTTTCCCTACACGACGCTCTTCCGATCTNNNATCTGCCAGCACCTTGATTTTT
AGCACACGTCTGAACTCTTTCCCTACACGACGCTCTTCCGATCTNNNATCTGCCAGCACCTTGATTTTT
AGCACACGTCTGAACTCTTTCCCTACACGACGCTCTTCCGATCTNNNATCTGCCAGCACCTTGATTTTT
AGCACACGTCTGAACTCTTTCCCTACACGACGCTCTTCCGATCTNNNATCTGCCAGCACCTTGATTTTT
AGCACACGTCTGAACTCTTTCCCTACACGACGCTCTTCCGATCTNNNCTGTAATCCCAGCTACTCAGG
AGCACACGTCTGAACTCTTTCCCTACACGACGCTCTTCCGATCTNNNCTGTAATCCCAGCTACTCAGG
AGCACACGTCTGAACTCTTTCCCTACACGACGCTCTTCCGATCTNNNCTGTAATCCCAGCTACTCAGG
AGCACACGTCTGAACTCTTTCCCTACACGACGCTCTTCCGATCTNNNCTGTAATCCCAGCTACTCAGG
AGCACACGTCTGAACTCTTTCCCTACACGACGCTCTTCCGATCTNNNCTGTAATCCCAGCTACTCAGG
GAGCACACGTCTGAACTCTTTCCCTACACGACGCTCTTCCGATCTNNNTCCAAGCTTTCAATAGTAAAT
GAGCACACGTCTGAACTCTTTCCCTACACGACGCTCTTCCGATCTNNNTCCAAGCTTTCAATAGTAAAT
GAGCACACGTCTGAACTCTTTCCCTACACGACGCTCTTCCGATCTNNNTCCAAGCTTTCAATAGTAAAT
GAGCACACGTCTGAACTCTTTCCCTACACGACGCTCTTCCGATCTNNNTCCAAGCTTTCAATAGTAAAT
GAGCACACGTCTGAACTCTTTCCCTACACGACGCTCTTCCGATCTNNNTCCAAGCTTTCAATAGTAAAT

AGCACACGTCTGAACTCTTTCCCTACACGACGCTCTTCCGATCTNNNCCTATCTTGGACAATCCTCCAGG
AGCACACGTCTGAACTCTTTCCCTACACGACGCTCTTCCGATCTNNNCCTATCTTGGACAATCCTCCAGG
AGCACACGTCTGAACTCTTTCCCTACACGACGCTCTTCCGATCTNNNCCTATCTTGGACAATCCTCCAGG
AGCACACGTCTGAACTCTTTCCCTACACGACGCTCTTCCGATCTNNNCCTATCTTGGACAATCCTCCAGG
AGCACACGTCTGAACTCTTTCCCTACACGACGCTCTTCCGATCTNNNCCTATCTTGGACAATCCTCCAGG
GCACACGTCTGAACTCTTTCCCTACACGACGCTCTTCCGATCTNNNATCCCCACAGATTACAAATCCCCC
GCACACGTCTGAACTCTTTCCCTACACGACGCTCTTCCGATCTNNNATCCCCACAGATTACAAATCCCCC
GCACACGTCTGAACTCTTTCCCTACACGACGCTCTTCCGATCTNNNATCCCCACAGATTACAAATCCCCC
GCACACGTCTGAACTCTTTCCCTACACGACGCTCTTCCGATCTNNNATCCCCACAGATTACAAATCCCCC
GCACACGTCTGAACTCTTTCCCTACACGACGCTCTTCCGATCTNNNATCCCCACAGATTACAAATCCCCC
AGAGCACACGTCTGAACTCTTTCCCTACACGACGCTCTTCCGATCTNNNTGTTAGCTCTGTGCCTAACCA
AGAGCACACGTCTGAACTCTTTCCCTACACGACGCTCTTCCGATCTNNNTGTTAGCTCTGTGCCTAACCA
AGAGCACACGTCTGAACTCTTTCCCTACACGACGCTCTTCCGATCTNNNTGTTAGCTCTGTGCCTAACCA
AGAGCACACGTCTGAACTCTTTCCCTACACGACGCTCTTCCGATCTNNNTGTTAGCTCTGTGCCTAACCA
AGAGCACACGTCTGAACTCTTTCCCTACACGACGCTCTTCCGATCTNNNTGTTAGCTCTGTGCCTAACCA
AGCACACGTCTGAACTCTTTCCCTACACGACGCTCTTCCGATCTNNNGTATGGAATATAGCAGAGATG
AGCACACGTCTGAACTCTTTCCCTACACGACGCTCTTCCGATCTNNNGTATGGAATATAGCAGAGATG
AGCACACGTCTGAACTCTTTCCCTACACGACGCTCTTCCGATCTNNNGTATGGAATATAGCAGAGATG
AGCACACGTCTGAACTCTTTCCCTACACGACGCTCTTCCGATCTNNNGTATGGAATATAGCAGAGATG
AGCACACGTCTGAACTCTTTCCCTACACGACGCTCTTCCGATCTNNNGTATGGAATATAGCAGAGATG
AGAGCACACGTCTGAACTCTTTCCCTACACGACGCTCTTCCGATCTNNNGATCTTATTCCGGGTGGTAG
AGAGCACACGTCTGAACTCTTTCCCTACACGACGCTCTTCCGATCTNNNGATCTTATTCCGGGTGGTAG
AGAGCACACGTCTGAACTCTTTCCCTACACGACGCTCTTCCGATCTNNNGATCTTATTCCGGGTGGTAG
AGAGCACACGTCTGAACTCTTTCCCTACACGACGCTCTTCCGATCTNNNGATCTTATTCCGGGTGGTAG
AGAGCACACGTCTGAACTCTTTCCCTACACGACGCTCTTCCGATCTNNNGATCTTATTCCGGGTGGTAG
AGCACACGTCTGAACTCTTTCCCTACACGACGCTCTTCCGATCTNNNATGAACACACTCTTTAATTTCAT
AGCACACGTCTGAACTCTTTCCCTACACGACGCTCTTCCGATCTNNNATGAACACACTCTTTAATTTCAT
AGCACACGTCTGAACTCTTTCCCTACACGACGCTCTTCCGATCTNNNATGAACACACTCTTTAATTTCAT
AGCACACGTCTGAACTCTTTCCCTACACGACGCTCTTCCGATCTNNNATGAACACACTCTTTAATTTCAT
AGCACACGTCTGAACTCTTTCCCTACACGACGCTCTTCCGATCTNNNATGAACACACTCTTTAATTTCAT
AAGAGCACACGTCTGAACTCTTTCCCTACACGACGCTCTTCCGATCTNNNAAAAATTAGCCAGACGTGG
AAGAGCACACGTCTGAACTCTTTCCCTACACGACGCTCTTCCGATCTNNNAAAAATTAGCCAGACGTGG
AAGAGCACACGTCTGAACTCTTTCCCTACACGACGCTCTTCCGATCTNNNAAAAATTAGCCAGACGTGG
AAGAGCACACGTCTGAACTCTTTCCCTACACGACGCTCTTCCGATCTNNNAAAAATTAGCCAGACGTGG
AAGAGCACACGTCTGAACTCTTTCCCTACACGACGCTCTTCCGATCTNNNAAAAATTAGCCAGACGTGG
GAGCACACGTCTGAACTCTTTCCCTACACGACGCTCTTCCGATCTNNNCATAATATAAGTCAGCAGGTA
GAGCACACGTCTGAACTCTTTCCCTACACGACGCTCTTCCGATCTNNNCATAATATAAGTCAGCAGGTA
GAGCACACGTCTGAACTCTTTCCCTACACGACGCTCTTCCGATCTNNNCATAATATAAGTCAGCAGGTA
GAGCACACGTCTGAACTCTTTCCCTACACGACGCTCTTCCGATCTNNNCATAATATAAGTCAGCAGGTA
GAGCACACGTCTGAACTCTTTCCCTACACGACGCTCTTCCGATCTNNNCATAATATAAGTCAGCAGGTA
GCACACGTCTGAACTCTTTCCCTACACGACGCTCTTCCGATCTNNNCATGTGCATATTGTGGTTGGGT
GCACACGTCTGAACTCTTTCCCTACACGACGCTCTTCCGATCTNNNCATGTGCATATTGTGGTTGGGT
GCACACGTCTGAACTCTTTCCCTACACGACGCTCTTCCGATCTNNNCATGTGCATATTGTGGTTGGGT

GCACACGTCTGAACTCTTTCCCTACACGACGCTCTTCCGATCTNNNCATGTGCATATTGTGGTTGGGTT
GCACACGTCTGAACTCTTTCCCTACACGACGCTCTTCCGATCTNNNCATGTGCATATTGTGGTTGGGTT
GAGCACACGTCTGAACTCTTTCCCTACACGACGCTCTTCCGATCTNNNCTGTAGTCCCAGCTACTCAGGA
GAGCACACGTCTGAACTCTTTCCCTACACGACGCTCTTCCGATCTNNNCTGTAGTCCCAGCTACTCAGGA
GAGCACACGTCTGAACTCTTTCCCTACACGACGCTCTTCCGATCTNNNCTGTAGTCCCAGCTACTCAGGA
GAGCACACGTCTGAACTCTTTCCCTACACGACGCTCTTCCGATCTNNNCTGTAGTCCCAGCTACTCAGGA
GAGCACACGTCTGAACTCTTTCCCTACACGACGCTCTTCCGATCTNNNCTGTAGTCCCAGCTACTCAGGA
AGCACACGTCTGAACTCTTTCCCTACACGACGCTCTTCCGATCTNNNCCATCAGCAGAGAATGAAAGCA
AGCACACGTCTGAACTCTTTCCCTACACGACGCTCTTCCGATCTNNNCCATCAGCAGAGAATGAAAGCA
AGCACACGTCTGAACTCTTTCCCTACACGACGCTCTTCCGATCTNNNCCATCAGCAGAGAATGAAAGCA
AGCACACGTCTGAACTCTTTCCCTACACGACGCTCTTCCGATCTNNNCCATCAGCAGAGAATGAAAGCA
AGCACACGTCTGAACTCTTTCCCTACACGACGCTCTTCCGATCTNNNCCATCAGCAGAGAATGAAAGCA
AGCACACGTCTGAACTCTTTCCCTACACGACGCTCTTCCGATCTNNNGTCTTGAACTCCCGACCTCAGCA
AGCACACGTCTGAACTCTTTCCCTACACGACGCTCTTCCGATCTNNNGTCTTGAACTCCCGACCTCAGCA
AGCACACGTCTGAACTCTTTCCCTACACGACGCTCTTCCGATCTNNNGTCTTGAACTCCCGACCTCAGCA
AGCACACGTCTGAACTCTTTCCCTACACGACGCTCTTCCGATCTNNNGTCTTGAACTCCCGACCTCAGCA
AGCACACGTCTGAACTCTTTCCCTACACGACGCTCTTCCGATCTNNNGTCTTGAACTCCCGACCTCAGCA
AGCACACGTCTGAACTCTTTCCCTACACGACGCTCTTCCGATCTNNNTCTGGGAATGAAGTCACTGACT
AGCACACGTCTGAACTCTTTCCCTACACGACGCTCTTCCGATCTNNNTCTGGGAATGAAGTCACTGACT
AGCACACGTCTGAACTCTTTCCCTACACGACGCTCTTCCGATCTNNNTCTGGGAATGAAGTCACTGACT
AGCACACGTCTGAACTCTTTCCCTACACGACGCTCTTCCGATCTNNNTCTGGGAATGAAGTCACTGACT
AGCACACGTCTGAACTCTTTCCCTACACGACGCTCTTCCGATCTNNNTCTGGGAATGAAGTCACTGACT
GAGCACACGTCTGAACTCTTTCCCTACACGACGCTCTTCCGATCTNNNTAATTGTGTTTCTGTATAGGT
GAGCACACGTCTGAACTCTTTCCCTACACGACGCTCTTCCGATCTNNNTAATTGTGTTTCTGTATAGGT
GAGCACACGTCTGAACTCTTTCCCTACACGACGCTCTTCCGATCTNNNTAATTGTGTTTCTGTATAGGT
GAGCACACGTCTGAACTCTTTCCCTACACGACGCTCTTCCGATCTNNNTAATTGTGTTTCTGTATAGGT
GAGCACACGTCTGAACTCTTTCCCTACACGACGCTCTTCCGATCTNNNTAATTGTGTTTCTGTATAGGT
CACACGTCTGAACTCTTTCCCTACACGACGCTCTTCCGATCTNNNGGATGTCGTAACAACAGCTGTTTCA
CACACGTCTGAACTCTTTCCCTACACGACGCTCTTCCGATCTNNNGGATGTCGTAACAACAGCTGTTTCA
CACACGTCTGAACTCTTTCCCTACACGACGCTCTTCCGATCTNNNGGATGTCGTAACAACAGCTGTTTCA
CACACGTCTGAACTCTTTCCCTACACGACGCTCTTCCGATCTNNNGGATGTCGTAACAACAGCTGTTTCA
CACACGTCTGAACTCTTTCCCTACACGACGCTCTTCCGATCTNNNGGATGTCGTAACAACAGCTGTTTCA
AGAGCACACGTCTGAACTCTTTCCCTACACGACGCTCTTCCGATCTNNNTTTCTGACTGCTTTATGGGCT
AGAGCACACGTCTGAACTCTTTCCCTACACGACGCTCTTCCGATCTNNNTTTCTGACTGCTTTATGGGCT
AGAGCACACGTCTGAACTCTTTCCCTACACGACGCTCTTCCGATCTNNNTTTCTGACTGCTTTATGGGCT
AGAGCACACGTCTGAACTCTTTCCCTACACGACGCTCTTCCGATCTNNNTTTCTGACTGCTTTATGGGCT
AGAGCACACGTCTGAACTCTTTCCCTACACGACGCTCTTCCGATCTNNNTTTCTGACTGCTTTATGGGCT
GAGCACACGTCTGAACTCTTTCCCTACACGACGCTCTTCCGATCTNNNCAGCAAGAGGATTCTGCATCA
GAGCACACGTCTGAACTCTTTCCCTACACGACGCTCTTCCGATCTNNNCAGCAAGAGGATTCTGCATCA
GAGCACACGTCTGAACTCTTTCCCTACACGACGCTCTTCCGATCTNNNCAGCAAGAGGATTCTGCATCA
GAGCACACGTCTGAACTCTTTCCCTACACGACGCTCTTCCGATCTNNNCAGCAAGAGGATTCTGCATCA
GAGCACACGTCTGAACTCTTTCCCTACACGACGCTCTTCCGATCTNNNCAGCAAGAGGATTCTGCATCA
AGAGCACACGTCTGAACTCTTTCCCTACACGACGCTCTTCCGATCTNNNGGTTGCAGTGAGCCAAGAT

AGAGCACACGTCTGAACTCTTTCCCTACACGACGCTCTTCCGATCTNNNGGTTGCAGTGAGCCAAGAT
AGAGCACACGTCTGAACTCTTTCCCTACACGACGCTCTTCCGATCTNNNGGTTGCAGTGAGCCAAGAT
AGAGCACACGTCTGAACTCTTTCCCTACACGACGCTCTTCCGATCTNNNGGTTGCAGTGAGCCAAGAT
AGAGCACACGTCTGAACTCTTTCCCTACACGACGCTCTTCCGATCTNNNGGTTGCAGTGAGCCAAGAT
GAGCACACGTCTGAACTCTTTCCCTACACGACGCTCTTCCGATCTNNNCGTTTGTGGAAACTGCTCTACT
GAGCACACGTCTGAACTCTTTCCCTACACGACGCTCTTCCGATCTNNNCGTTTGTGGAAACTGCTCTACT
GAGCACACGTCTGAACTCTTTCCCTACACGACGCTCTTCCGATCTNNNCGTTTGTGGAAACTGCTCTACT
GAGCACACGTCTGAACTCTTTCCCTACACGACGCTCTTCCGATCTNNNCGTTTGTGGAAACTGCTCTACT
GAGCACACGTCTGAACTCTTTCCCTACACGACGCTCTTCCGATCTNNNCGTTTGTGGAAACTGCTCTACT
GCACACGTCTGAACTCTTTCCCTACACGACGCTCTTCCGATCTNNNCTGAAATCCCCACTACTCAGGAG
GCACACGTCTGAACTCTTTCCCTACACGACGCTCTTCCGATCTNNNCTGAAATCCCCACTACTCAGGAG
GCACACGTCTGAACTCTTTCCCTACACGACGCTCTTCCGATCTNNNCTGAAATCCCCACTACTCAGGAG
GCACACGTCTGAACTCTTTCCCTACACGACGCTCTTCCGATCTNNNCTGAAATCCCCACTACTCAGGAG
GCACACGTCTGAACTCTTTCCCTACACGACGCTCTTCCGATCTNNNCTGAAATCCCCACTACTCAGGAG
AGCACACGTCTGAACTCTTTCCCTACACGACGCTCTTCCGATCTNNNACACACAATGGAGGAAGGAAA
AGCACACGTCTGAACTCTTTCCCTACACGACGCTCTTCCGATCTNNNACACACAATGGAGGAAGGAAA
AGCACACGTCTGAACTCTTTCCCTACACGACGCTCTTCCGATCTNNNACACACAATGGAGGAAGGAAA
AGCACACGTCTGAACTCTTTCCCTACACGACGCTCTTCCGATCTNNNACACACAATGGAGGAAGGAAA
AGCACACGTCTGAACTCTTTCCCTACACGACGCTCTTCCGATCTNNNACACACAATGGAGGAAGGAAA
GAGCACACGTCTGAACTCTTTCCCTACACGACGCTCTTCCGATCTNNNAGCTCCGATGGGAAAGTACAA
GAGCACACGTCTGAACTCTTTCCCTACACGACGCTCTTCCGATCTNNNAGCTCCGATGGGAAAGTACAA
GAGCACACGTCTGAACTCTTTCCCTACACGACGCTCTTCCGATCTNNNAGCTCCGATGGGAAAGTACAA
GAGCACACGTCTGAACTCTTTCCCTACACGACGCTCTTCCGATCTNNNAGCTCCGATGGGAAAGTACAA
GAGCACACGTCTGAACTCTTTCCCTACACGACGCTCTTCCGATCTNNNAGCTCCGATGGGAAAGTACAA
AGAGCACACGTCTGAACTCTTTCCCTACACGACGCTCTTCCGATCTNNNTCTTTGAGTAAACAAATGAG
AGAGCACACGTCTGAACTCTTTCCCTACACGACGCTCTTCCGATCTNNNTCTTTGAGTAAACAAATGAG
AGAGCACACGTCTGAACTCTTTCCCTACACGACGCTCTTCCGATCTNNNTCTTTGAGTAAACAAATGAG
AGAGCACACGTCTGAACTCTTTCCCTACACGACGCTCTTCCGATCTNNNTCTTTGAGTAAACAAATGAG
AGAGCACACGTCTGAACTCTTTCCCTACACGACGCTCTTCCGATCTNNNTCTTTGAGTAAACAAATGAG
AGCACACGTCTGAACTCTTTCCCTACACGACGCTCTTCCGATCTNNNACAAACACACATCTCCACCTCTT
AGCACACGTCTGAACTCTTTCCCTACACGACGCTCTTCCGATCTNNNACAAACACACATCTCCACCTCTT
AGCACACGTCTGAACTCTTTCCCTACACGACGCTCTTCCGATCTNNNACAAACACACATCTCCACCTCTT
AGCACACGTCTGAACTCTTTCCCTACACGACGCTCTTCCGATCTNNNACAAACACACATCTCCACCTCTT
AGCACACGTCTGAACTCTTTCCCTACACGACGCTCTTCCGATCTNNNACAAACACACATCTCCACCTCTT
GAGCACACGTCTGAACTCTTTCCCTACACGACGCTCTTCCGATCTNNNTTATCTCCTCTTTTGCAACCCCA
GAGCACACGTCTGAACTCTTTCCCTACACGACGCTCTTCCGATCTNNNTTATCTCCTCTTTTGCAACCCCA
GAGCACACGTCTGAACTCTTTCCCTACACGACGCTCTTCCGATCTNNNTTATCTCCTCTTTTGCAACCCCA
GAGCACACGTCTGAACTCTTTCCCTACACGACGCTCTTCCGATCTNNNTTATCTCCTCTTTTGCAACCCCA
GAGCACACGTCTGAACTCTTTCCCTACACGACGCTCTTCCGATCTNNNTTATCTCCTCTTTTGCAACCCCA
AGAGCACACGTCTGAACTCTTTCCCTACACGACGCTCTTCCGATCTNNNGGATAGGATGGTATGGTAT
AGAGCACACGTCTGAACTCTTTCCCTACACGACGCTCTTCCGATCTNNNGGATAGGATGGTATGGTAT
AGAGCACACGTCTGAACTCTTTCCCTACACGACGCTCTTCCGATCTNNNGGATAGGATGGTATGGTAT
AGAGCACACGTCTGAACTCTTTCCCTACACGACGCTCTTCCGATCTNNNGGATAGGATGGTATGGTAT

AGAGCACACGTCTGAACTCTTTCCCTACACGACGCTCTTCCGATCTNNNGGATAGGATGGTATGGTAT
GAGCACACGTCTGAACTCTTTCCCTACACGACGCTCTTCCGATCTNNNGATGGACTGGAGGTAGAGA
GAGCACACGTCTGAACTCTTTCCCTACACGACGCTCTTCCGATCTNNNGATGGACTGGAGGTAGAGA
GAGCACACGTCTGAACTCTTTCCCTACACGACGCTCTTCCGATCTNNNGATGGACTGGAGGTAGAGA
GAGCACACGTCTGAACTCTTTCCCTACACGACGCTCTTCCGATCTNNNGATGGACTGGAGGTAGAGA
GAGCACACGTCTGAACTCTTTCCCTACACGACGCTCTTCCGATCTNNNGATGGACTGGAGGTAGAGA
GAGCACACGTCTGAACTCTTTCCCTACACGACGCTCTTCCGATCTNNNTGGCACAATTCACATGGAGTT
GAGCACACGTCTGAACTCTTTCCCTACACGACGCTCTTCCGATCTNNNTGGCACAATTCACATGGAGTT
GAGCACACGTCTGAACTCTTTCCCTACACGACGCTCTTCCGATCTNNNTGGCACAATTCACATGGAGTT
GAGCACACGTCTGAACTCTTTCCCTACACGACGCTCTTCCGATCTNNNTGGCACAATTCACATGGAGTT
GAGCACACGTCTGAACTCTTTCCCTACACGACGCTCTTCCGATCTNNNTGGCACAATTCACATGGAGTT
GAGCACACGTCTGAACTCTTTCCCTACACGACGCTCTTCCGATCTNNNCTGTAGTTCCAGCTACTTGGG
GAGCACACGTCTGAACTCTTTCCCTACACGACGCTCTTCCGATCTNNNCTGTAGTTCCAGCTACTTGGG
GAGCACACGTCTGAACTCTTTCCCTACACGACGCTCTTCCGATCTNNNCTGTAGTTCCAGCTACTTGGG
GAGCACACGTCTGAACTCTTTCCCTACACGACGCTCTTCCGATCTNNNCTGTAGTTCCAGCTACTTGGG
GAGCACACGTCTGAACTCTTTCCCTACACGACGCTCTTCCGATCTNNNCTGTAGTTCCAGCTACTTGGG
AGCACACGTCTGAACTCTTTCCCTACACGACGCTCTTCCGATCTNNNATTGACCTTCTGTTGTCTATGCA
AGCACACGTCTGAACTCTTTCCCTACACGACGCTCTTCCGATCTNNNATTGACCTTCTGTTGTCTATGCA
AGCACACGTCTGAACTCTTTCCCTACACGACGCTCTTCCGATCTNNNATTGACCTTCTGTTGTCTATGCA
AGCACACGTCTGAACTCTTTCCCTACACGACGCTCTTCCGATCTNNNATTGACCTTCTGTTGTCTATGCA
AGCACACGTCTGAACTCTTTCCCTACACGACGCTCTTCCGATCTNNNATTGACCTTCTGTTGTCTATGCA
CACACGTCTGAACTCTTTCCCTACACGACGCTCTTCCGATCTNNNACTGTTAGTATTCCTGGATGATGGA
CACACGTCTGAACTCTTTCCCTACACGACGCTCTTCCGATCTNNNACTGTTAGTATTCCTGGATGATGGA
CACACGTCTGAACTCTTTCCCTACACGACGCTCTTCCGATCTNNNACTGTTAGTATTCCTGGATGATGGA
CACACGTCTGAACTCTTTCCCTACACGACGCTCTTCCGATCTNNNACTGTTAGTATTCCTGGATGATGGA
CACACGTCTGAACTCTTTCCCTACACGACGCTCTTCCGATCTNNNACTGTTAGTATTCCTGGATGATGGA
GAGCACACGTCTGAACTCTTTCCCTACACGACGCTCTTCCGATCTNNNCTGCCCCGCCATTTTTTTTTTTT
GAGCACACGTCTGAACTCTTTCCCTACACGACGCTCTTCCGATCTNNNCTGCCCCGCCATTTTTTTTTTTT
GAGCACACGTCTGAACTCTTTCCCTACACGACGCTCTTCCGATCTNNNCTGCCCCGCCATTTTTTTTTTTT
GAGCACACGTCTGAACTCTTTCCCTACACGACGCTCTTCCGATCTNNNCTGCCCCGCCATTTTTTTTTTTT
GAGCACACGTCTGAACTCTTTCCCTACACGACGCTCTTCCGATCTNNNCTGCCCCGCCATTTTTTTTTTTT
AGCACACGTCTGAACTCTTTCCCTACACGACGCTCTTCCGATCTNNNCTGCTTTCCTTTCCTTCGTTTCCC
AGCACACGTCTGAACTCTTTCCCTACACGACGCTCTTCCGATCTNNNCTGCTTTCCTTTCCTTCGTTTCCC
AGCACACGTCTGAACTCTTTCCCTACACGACGCTCTTCCGATCTNNNCTGCTTTCCTTTCCTTCGTTTCCC
AGCACACGTCTGAACTCTTTCCCTACACGACGCTCTTCCGATCTNNNCTGCTTTCCTTTCCTTCGTTTCCC
AGCACACGTCTGAACTCTTTCCCTACACGACGCTCTTCCGATCTNNNCTGCTTTCCTTTCCTTCGTTTCCC
GAGCACACGTCTGAACTCTTTCCCTACACGACGCTCTTCCGATCTNNNGGGCAACAACAGCAAAACTCT
GAGCACACGTCTGAACTCTTTCCCTACACGACGCTCTTCCGATCTNNNGGGCAACAACAGCAAAACTCT
GAGCACACGTCTGAACTCTTTCCCTACACGACGCTCTTCCGATCTNNNGGGCAACAACAGCAAAACTCT
GAGCACACGTCTGAACTCTTTCCCTACACGACGCTCTTCCGATCTNNNGGGCAACAACAGCAAAACTCT
GAGCACACGTCTGAACTCTTTCCCTACACGACGCTCTTCCGATCTNNNGGGCAACAACAGCAAAACTCT
AGCACACGTCTGAACTCTTTCCCTACACGACGCTCTTCCGATCTNNNCTCCAACCCAACATCCTAAGTGA
AGCACACGTCTGAACTCTTTCCCTACACGACGCTCTTCCGATCTNNNCTCCAACCCAACATCCTAAGTGA

AGCACACGTCTGAACTCTTTCCCTACACGACGCTCTTCCGATCTNNNCTCCAACCCAACATCCTAAGTGA

AGCACACGTCTGAACTCTTTCCCTACACGACGCTCTTCCGATCTNNNCTCCAACCCAACATCCTAAGTGA

AGCACACGTCTGAACTCTTTCCCTACACGACGCTCTTCCGATCTNNNCTCCAACCCAACATCCTAAGTGA

GAGCACACGTCTGAACTCTTTCCCTACACGACGCTCTTCCGATCTNNNATGCTTGTAATCCCAGCGATT

GAGCACACGTCTGAACTCTTTCCCTACACGACGCTCTTCCGATCTNNNATGCTTGTAATCCCAGCGATT

GAGCACACGTCTGAACTCTTTCCCTACACGACGCTCTTCCGATCTNNNATGCTTGTAATCCCAGCGATT

GAGCACACGTCTGAACTCTTTCCCTACACGACGCTCTTCCGATCTNNNATGCTTGTAATCCCAGCGATT

GAGCACACGTCTGAACTCTTTCCCTACACGACGCTCTTCCGATCTNNNATGCTTGTAATCCCAGCGATT

GAGCACACGTCTGAACTCTTTCCCTACACGACGCTCTTCCGATCTNNNAGGTTACAGTGAGCTGAGATC

GAGCACACGTCTGAACTCTTTCCCTACACGACGCTCTTCCGATCTNNNAGGTTACAGTGAGCTGAGATC

GAGCACACGTCTGAACTCTTTCCCTACACGACGCTCTTCCGATCTNNNAGGTTACAGTGAGCTGAGATC

GAGCACACGTCTGAACTCTTTCCCTACACGACGCTCTTCCGATCTNNNAGGTTACAGTGAGCTGAGATC

GAGCACACGTCTGAACTCTTTCCCTACACGACGCTCTTCCGATCTNNNAGGTTACAGTGAGCTGAGATC

AGAGCACACGTCTGAACTCTTTCCCTACACGACGCTCTTCCGATCTNNNTAGACATAAGGGGGCACTA

AGAGCACACGTCTGAACTCTTTCCCTACACGACGCTCTTCCGATCTNNNTAGACATAAGGGGGCACTA

AGAGCACACGTCTGAACTCTTTCCCTACACGACGCTCTTCCGATCTNNNTAGACATAAGGGGGCACTA

AGAGCACACGTCTGAACTCTTTCCCTACACGACGCTCTTCCGATCTNNNTAGACATAAGGGGGCACTA

AGAGCACACGTCTGAACTCTTTCCCTACACGACGCTCTTCCGATCTNNNTAGACATAAGGGGGCACTA

GAGCACACGTCTGAACTCTTTCCCTACACGACGCTCTTCCGATCTNNNCACCTGTAATCCCAGCTACTCTC

GAGCACACGTCTGAACTCTTTCCCTACACGACGCTCTTCCGATCTNNNCACCTGTAATCCCAGCTACTCTC

GAGCACACGTCTGAACTCTTTCCCTACACGACGCTCTTCCGATCTNNNCACCTGTAATCCCAGCTACTCTC

GAGCACACGTCTGAACTCTTTCCCTACACGACGCTCTTCCGATCTNNNCACCTGTAATCCCAGCTACTCTC

GAGCACACGTCTGAACTCTTTCCCTACACGACGCTCTTCCGATCTNNNCACCTGTAATCCCAGCTACTCTC

GAGCACACGTCTGAACTCTTTCCCTACACGACGCTCTTCCGATCTNNNACATTCATTTTCTTAGCGGTGT

GAGCACACGTCTGAACTCTTTCCCTACACGACGCTCTTCCGATCTNNNACATTCATTTTCTTAGCGGTGT

GAGCACACGTCTGAACTCTTTCCCTACACGACGCTCTTCCGATCTNNNACATTCATTTTCTTAGCGGTGT

GAGCACACGTCTGAACTCTTTCCCTACACGACGCTCTTCCGATCTNNNACATTCATTTTCTTAGCGGTGT

GAGCACACGTCTGAACTCTTTCCCTACACGACGCTCTTCCGATCTNNNACATTCATTTTCTTAGCGGTGT

GAGCACACGTCTGAACTCTTTCCCTACACGACGCTCTTCCGATCTNNNTCAAGTGGAAGTAAAGGAGA

GAGCACACGTCTGAACTCTTTCCCTACACGACGCTCTTCCGATCTNNNTCAAGTGGAAGTAAAGGAGA

GAGCACACGTCTGAACTCTTTCCCTACACGACGCTCTTCCGATCTNNNTCAAGTGGAAGTAAAGGAGA

GAGCACACGTCTGAACTCTTTCCCTACACGACGCTCTTCCGATCTNNNTCAAGTGGAAGTAAAGGAGA

GAGCACACGTCTGAACTCTTTCCCTACACGACGCTCTTCCGATCTNNNTCAAGTGGAAGTAAAGGAGA

AGCACACGTCTGAACTCTTTCCCTACACGACGCTCTTCCGATCTNNNGCCAATATCACATCACTGCACTC

AGCACACGTCTGAACTCTTTCCCTACACGACGCTCTTCCGATCTNNNGCCAATATCACATCACTGCACTC

AGCACACGTCTGAACTCTTTCCCTACACGACGCTCTTCCGATCTNNNGCCAATATCACATCACTGCACTC

AGCACACGTCTGAACTCTTTCCCTACACGACGCTCTTCCGATCTNNNGCCAATATCACATCACTGCACTC

AGCACACGTCTGAACTCTTTCCCTACACGACGCTCTTCCGATCTNNNGCCAATATCACATCACTGCACTC

GAGCACACGTCTGAACTCTTTCCCTACACGACGCTCTTCCGATCTNNNCTACAGAGTGAGACTCCGTTT

GAGCACACGTCTGAACTCTTTCCCTACACGACGCTCTTCCGATCTNNNCTACAGAGTGAGACTCCGTTT

GAGCACACGTCTGAACTCTTTCCCTACACGACGCTCTTCCGATCTNNNCTACAGAGTGAGACTCCGTTT

GAGCACACGTCTGAACTCTTTCCCTACACGACGCTCTTCCGATCTNNNCTACAGAGTGAGACTCCGTTT

GAGCACACGTCTGAACTCTTTCCCTACACGACGCTCTTCCGATCTNNNCTACAGAGTGAGACTCCGTTT

GCACACGTCTGAACTCTTTCCCTACACGACGCTCTTCCGATCTNNNTCAGGCCTCAATGTACACCTAAC
GCACACGTCTGAACTCTTTCCCTACACGACGCTCTTCCGATCTNNNTCAGGCCTCAATGTACACCTAAC
GCACACGTCTGAACTCTTTCCCTACACGACGCTCTTCCGATCTNNNTCAGGCCTCAATGTACACCTAAC
GCACACGTCTGAACTCTTTCCCTACACGACGCTCTTCCGATCTNNNTCAGGCCTCAATGTACACCTAAC
GCACACGTCTGAACTCTTTCCCTACACGACGCTCTTCCGATCTNNNTCAGGCCTCAATGTACACCTAAC
GAGCACACGTCTGAACTCTTTCCCTACACGACGCTCTTCCGATCTNNNCACCTGAAGTCCCAGCTACTC
GAGCACACGTCTGAACTCTTTCCCTACACGACGCTCTTCCGATCTNNNCACCTGAAGTCCCAGCTACTC
GAGCACACGTCTGAACTCTTTCCCTACACGACGCTCTTCCGATCTNNNCACCTGAAGTCCCAGCTACTC
GAGCACACGTCTGAACTCTTTCCCTACACGACGCTCTTCCGATCTNNNCACCTGAAGTCCCAGCTACTC
GAGCACACGTCTGAACTCTTTCCCTACACGACGCTCTTCCGATCTNNNCACCTGAAGTCCCAGCTACTC
AGCACACGTCTGAACTCTTTCCCTACACGACGCTCTTCCGATCTNNNCTGAGGCAGGAGAATTGCTTG
AGCACACGTCTGAACTCTTTCCCTACACGACGCTCTTCCGATCTNNNCTGAGGCAGGAGAATTGCTTG
AGCACACGTCTGAACTCTTTCCCTACACGACGCTCTTCCGATCTNNNCTGAGGCAGGAGAATTGCTTG
AGCACACGTCTGAACTCTTTCCCTACACGACGCTCTTCCGATCTNNNCTGAGGCAGGAGAATTGCTTG
AGCACACGTCTGAACTCTTTCCCTACACGACGCTCTTCCGATCTNNNCTGAGGCAGGAGAATTGCTTG
AGAGCACACGTCTGAACTCTTTCCCTACACGACGCTCTTCCGATCTNNNTAAACCCTCCTCCATTTTTCA
AGAGCACACGTCTGAACTCTTTCCCTACACGACGCTCTTCCGATCTNNNTAAACCCTCCTCCATTTTTCA
AGAGCACACGTCTGAACTCTTTCCCTACACGACGCTCTTCCGATCTNNNTAAACCCTCCTCCATTTTTCA
AGAGCACACGTCTGAACTCTTTCCCTACACGACGCTCTTCCGATCTNNNTAAACCCTCCTCCATTTTTCA
AGAGCACACGTCTGAACTCTTTCCCTACACGACGCTCTTCCGATCTNNNTAAACCCTCCTCCATTTTTCA
GCACACGTCTGAACTCTTTCCCTACACGACGCTCTTCCGATCTNNNATTTGTGTGTGAGTGTGTGTGT
GCACACGTCTGAACTCTTTCCCTACACGACGCTCTTCCGATCTNNNATTTGTGTGTGAGTGTGTGTGT
GCACACGTCTGAACTCTTTCCCTACACGACGCTCTTCCGATCTNNNATTTGTGTGTGAGTGTGTGTGT
GCACACGTCTGAACTCTTTCCCTACACGACGCTCTTCCGATCTNNNATTTGTGTGTGAGTGTGTGTGT
GCACACGTCTGAACTCTTTCCCTACACGACGCTCTTCCGATCTNNNATTTGTGTGTGAGTGTGTGTGT
AGCACACGTCTGAACTCTTTCCCTACACGACGCTCTTCCGATCTNNNAACTCTGTGGGACAGATCTCA
AGCACACGTCTGAACTCTTTCCCTACACGACGCTCTTCCGATCTNNNAACTCTGTGGGACAGATCTCA
AGCACACGTCTGAACTCTTTCCCTACACGACGCTCTTCCGATCTNNNAACTCTGTGGGACAGATCTCA
AGCACACGTCTGAACTCTTTCCCTACACGACGCTCTTCCGATCTNNNAACTCTGTGGGACAGATCTCA
AGCACACGTCTGAACTCTTTCCCTACACGACGCTCTTCCGATCTNNNAACTCTGTGGGACAGATCTCA
AGCACACGTCTGAACTCTTTCCCTACACGACGCTCTTCCGATCTNNNCTGACGTAGGAGAATCGCATG
AGCACACGTCTGAACTCTTTCCCTACACGACGCTCTTCCGATCTNNNCTGACGTAGGAGAATCGCATG
AGCACACGTCTGAACTCTTTCCCTACACGACGCTCTTCCGATCTNNNCTGACGTAGGAGAATCGCATG
AGCACACGTCTGAACTCTTTCCCTACACGACGCTCTTCCGATCTNNNCTGACGTAGGAGAATCGCATG
AGCACACGTCTGAACTCTTTCCCTACACGACGCTCTTCCGATCTNNNCTGACGTAGGAGAATCGCATG
CACACGTCTGAACTCTTTCCCTACACGACGCTCTTCCGATCTNNNGAAGTGTCTGTTCATGTCCTTCGC
CACACGTCTGAACTCTTTCCCTACACGACGCTCTTCCGATCTNNNGAAGTGTCTGTTCATGTCCTTCGC
CACACGTCTGAACTCTTTCCCTACACGACGCTCTTCCGATCTNNNGAAGTGTCTGTTCATGTCCTTCGC
CACACGTCTGAACTCTTTCCCTACACGACGCTCTTCCGATCTNNNGAAGTGTCTGTTCATGTCCTTCGC
CACACGTCTGAACTCTTTCCCTACACGACGCTCTTCCGATCTNNNGAAGTGTCTGTTCATGTCCTTCGC
AGCACACGTCTGAACTCTTTCCCTACACGACGCTCTTCCGATCTNNNAGATTCCTGGCCTATTGGATTC
AGCACACGTCTGAACTCTTTCCCTACACGACGCTCTTCCGATCTNNNAGATTCCTGGCCTATTGGATTC
AGCACACGTCTGAACTCTTTCCCTACACGACGCTCTTCCGATCTNNNAGATTCCTGGCCTATTGGATTC

AGCACACGTCTGAACTCTTTCCCTACACGACGCTCTTCCGATCTNNNAGATTCCTGGCCTATTGGATTC

AGCACACGTCTGAACTCTTTCCCTACACGACGCTCTTCCGATCTNNNAGATTCCTGGCCTATTGGATTC

AGCACACGTCTGAACTCTTTCCCTACACGACGCTCTTCCGATCTNNNTCTCTCTCTCTCTGTGTGTGT

AGCACACGTCTGAACTCTTTCCCTACACGACGCTCTTCCGATCTNNNTCTCTCTCTCTCTGTGTGTGT

AGCACACGTCTGAACTCTTTCCCTACACGACGCTCTTCCGATCTNNNTCTCTCTCTCTCTGTGTGTGT

AGCACACGTCTGAACTCTTTCCCTACACGACGCTCTTCCGATCTNNNTCTCTCTCTCTCTGTGTGTGT

AGCACACGTCTGAACTCTTTCCCTACACGACGCTCTTCCGATCTNNNTCTCTCTCTCTCTGTGTGTGT

AGCACACGTCTGAACTCTTTCCCTACACGACGCTCTTCCGATCTNNNGAGGAAAGTGGAGGTGGAT

AGCACACGTCTGAACTCTTTCCCTACACGACGCTCTTCCGATCTNNNGAGGAAAGTGGAGGTGGAT

AGCACACGTCTGAACTCTTTCCCTACACGACGCTCTTCCGATCTNNNGAGGAAAGTGGAGGTGGAT

AGCACACGTCTGAACTCTTTCCCTACACGACGCTCTTCCGATCTNNNGAGGAAAGTGGAGGTGGAT

AGCACACGTCTGAACTCTTTCCCTACACGACGCTCTTCCGATCTNNNGAGGAAAGTGGAGGTGGAT

AGCACACGTCTGAACTCTTTCCCTACACGACGCTCTTCCGATCTNNNACAAATAGGTGTGCTAGGAGA

AGCACACGTCTGAACTCTTTCCCTACACGACGCTCTTCCGATCTNNNACAAATAGGTGTGCTAGGAGA

AGCACACGTCTGAACTCTTTCCCTACACGACGCTCTTCCGATCTNNNACAAATAGGTGTGCTAGGAGA

AGCACACGTCTGAACTCTTTCCCTACACGACGCTCTTCCGATCTNNNACAAATAGGTGTGCTAGGAGA

AGCACACGTCTGAACTCTTTCCCTACACGACGCTCTTCCGATCTNNNACAAATAGGTGTGCTAGGAGA

GAGCACACGTCTGAACTCTTTCCCTACACGACGCTCTTCCGATCTNNNGAGATCGAGACACAACACTC

GAGCACACGTCTGAACTCTTTCCCTACACGACGCTCTTCCGATCTNNNGAGATCGAGACACAACACTC

GAGCACACGTCTGAACTCTTTCCCTACACGACGCTCTTCCGATCTNNNGAGATCGAGACACAACACTC

GAGCACACGTCTGAACTCTTTCCCTACACGACGCTCTTCCGATCTNNNGAGATCGAGACACAACACTC

GAGCACACGTCTGAACTCTTTCCCTACACGACGCTCTTCCGATCTNNNGAGATCGAGACACAACACTC

AGCACACGTCTGAACTCTTTCCCTACACGACGCTCTTCCGATCTNNNGAAGCCTGTGTTTTTACCCACT

AGCACACGTCTGAACTCTTTCCCTACACGACGCTCTTCCGATCTNNNGAAGCCTGTGTTTTTACCCACT

AGCACACGTCTGAACTCTTTCCCTACACGACGCTCTTCCGATCTNNNGAAGCCTGTGTTTTTACCCACT

AGCACACGTCTGAACTCTTTCCCTACACGACGCTCTTCCGATCTNNNGAAGCCTGTGTTTTTACCCACT

AGCACACGTCTGAACTCTTTCCCTACACGACGCTCTTCCGATCTNNNGAAGCCTGTGTTTTTACCCACT

AGCACACGTCTGAACTCTTTCCCTACACGACGCTCTTCCGATCTNNNGAGACTGTAGCGAGACAAGA

AGCACACGTCTGAACTCTTTCCCTACACGACGCTCTTCCGATCTNNNGAGACTGTAGCGAGACAAGA

AGCACACGTCTGAACTCTTTCCCTACACGACGCTCTTCCGATCTNNNGAGACTGTAGCGAGACAAGA

AGCACACGTCTGAACTCTTTCCCTACACGACGCTCTTCCGATCTNNNGAGACTGTAGCGAGACAAGA

AGCACACGTCTGAACTCTTTCCCTACACGACGCTCTTCCGATCTNNNGAGACTGTAGCGAGACAAGA

CACACGTCTGAACTCTTTCCCTACACGACGCTCTTCCGATCTNNNAACTGTTGGTTATTTGTGTTATTC

CACACGTCTGAACTCTTTCCCTACACGACGCTCTTCCGATCTNNNAACTGTTGGTTATTTGTGTTATTC

CACACGTCTGAACTCTTTCCCTACACGACGCTCTTCCGATCTNNNAACTGTTGGTTATTTGTGTTATTC

CACACGTCTGAACTCTTTCCCTACACGACGCTCTTCCGATCTNNNAACTGTTGGTTATTTGTGTTATTC

CACACGTCTGAACTCTTTCCCTACACGACGCTCTTCCGATCTNNNAACTGTTGGTTATTTGTGTTATTC

AAGAGCACACGTCTGAACTCTTTCCCTACACGACGCTCTTCCGATCTNNNTCCCAAAGATGTGCATGTT

AAGAGCACACGTCTGAACTCTTTCCCTACACGACGCTCTTCCGATCTNNNTCCCAAAGATGTGCATGTT

AAGAGCACACGTCTGAACTCTTTCCCTACACGACGCTCTTCCGATCTNNNTCCCAAAGATGTGCATGTT

AAGAGCACACGTCTGAACTCTTTCCCTACACGACGCTCTTCCGATCTNNNTCCCAAAGATGTGCATGTT

AAGAGCACACGTCTGAACTCTTTCCCTACACGACGCTCTTCCGATCTNNNTCCCAAAGATGTGCATGTT

GAGCACACGTCTGAACTCTTTCCCTACACGACGCTCTTCCGATCTNNNACTCATCGCAGCTGTATCAAG

GAGCACACGTCTGAACTCTTTCCCTACACGACGCTCTTCCGATCTNNNACTCATCGCAGCTGTATCAAG
GAGCACACGTCTGAACTCTTTCCCTACACGACGCTCTTCCGATCTNNNACTCATCGCAGCTGTATCAAG
GAGCACACGTCTGAACTCTTTCCCTACACGACGCTCTTCCGATCTNNNACTCATCGCAGCTGTATCAAG
GAGCACACGTCTGAACTCTTTCCCTACACGACGCTCTTCCGATCTNNNACTCATCGCAGCTGTATCAAG
GAGCACACGTCTGAACTCTTTCCCTACACGACGCTCTTCCGATCTNNNAAGACCCATCAGTGTGCTGTA
GAGCACACGTCTGAACTCTTTCCCTACACGACGCTCTTCCGATCTNNNAAGACCCATCAGTGTGCTGTA
GAGCACACGTCTGAACTCTTTCCCTACACGACGCTCTTCCGATCTNNNAAGACCCATCAGTGTGCTGTA
GAGCACACGTCTGAACTCTTTCCCTACACGACGCTCTTCCGATCTNNNAAGACCCATCAGTGTGCTGTA
GAGCACACGTCTGAACTCTTTCCCTACACGACGCTCTTCCGATCTNNNAAGACCCATCAGTGTGCTGTA
AGCACACGTCTGAACTCTTTCCCTACACGACGCTCTTCCGATCTNNNCAGGGGGAAAACCAAATCACA
AGCACACGTCTGAACTCTTTCCCTACACGACGCTCTTCCGATCTNNNCAGGGGGAAAACCAAATCACA
AGCACACGTCTGAACTCTTTCCCTACACGACGCTCTTCCGATCTNNNCAGGGGGAAAACCAAATCACA
AGCACACGTCTGAACTCTTTCCCTACACGACGCTCTTCCGATCTNNNCAGGGGGAAAACCAAATCACA
AGCACACGTCTGAACTCTTTCCCTACACGACGCTCTTCCGATCTNNNCAGGGGGAAAACCAAATCACA
AAGAGCACACGTCTGAACTCTTTCCCTACACGACGCTCTTCCGATCTNNNACTCTTTGCAAGCCAGTGA
AAGAGCACACGTCTGAACTCTTTCCCTACACGACGCTCTTCCGATCTNNNACTCTTTGCAAGCCAGTGA
AAGAGCACACGTCTGAACTCTTTCCCTACACGACGCTCTTCCGATCTNNNACTCTTTGCAAGCCAGTGA
AAGAGCACACGTCTGAACTCTTTCCCTACACGACGCTCTTCCGATCTNNNACTCTTTGCAAGCCAGTGA
AAGAGCACACGTCTGAACTCTTTCCCTACACGACGCTCTTCCGATCTNNNACTCTTTGCAAGCCAGTGA
GAGCACACGTCTGAACTCTTTCCCTACACGACGCTCTTCCGATCTNNNGGCACCTGTAATCCCAGTTAC
GAGCACACGTCTGAACTCTTTCCCTACACGACGCTCTTCCGATCTNNNGGCACCTGTAATCCCAGTTAC
GAGCACACGTCTGAACTCTTTCCCTACACGACGCTCTTCCGATCTNNNGGCACCTGTAATCCCAGTTAC
GAGCACACGTCTGAACTCTTTCCCTACACGACGCTCTTCCGATCTNNNGGCACCTGTAATCCCAGTTAC
GAGCACACGTCTGAACTCTTTCCCTACACGACGCTCTTCCGATCTNNNGGCACCTGTAATCCCAGTTAC
CACACGTCTGAACTCTTTCCCTACACGACGCTCTTCCGATCTNNNTGTCACTCGTGGTAAATTTTATGTT
CACACGTCTGAACTCTTTCCCTACACGACGCTCTTCCGATCTNNNTGTCACTCGTGGTAAATTTTATGTT
CACACGTCTGAACTCTTTCCCTACACGACGCTCTTCCGATCTNNNTGTCACTCGTGGTAAATTTTATGTT
CACACGTCTGAACTCTTTCCCTACACGACGCTCTTCCGATCTNNNTGTCACTCGTGGTAAATTTTATGTT
CACACGTCTGAACTCTTTCCCTACACGACGCTCTTCCGATCTNNNTGTCACTCGTGGTAAATTTTATGTT
GAGCACACGTCTGAACTCTTTCCCTACACGACGCTCTTCCGATCTNNNTGAGTTGAGATTGTACCACTG
GAGCACACGTCTGAACTCTTTCCCTACACGACGCTCTTCCGATCTNNNTGAGTTGAGATTGTACCACTG
GAGCACACGTCTGAACTCTTTCCCTACACGACGCTCTTCCGATCTNNNTGAGTTGAGATTGTACCACTG
GAGCACACGTCTGAACTCTTTCCCTACACGACGCTCTTCCGATCTNNNTGAGTTGAGATTGTACCACTG
GAGCACACGTCTGAACTCTTTCCCTACACGACGCTCTTCCGATCTNNNTGAGTTGAGATTGTACCACTG
GAGCACACGTCTGAACTCTTTCCCTACACGACGCTCTTCCGATCTNNNGAGGCTGAGGCAGAAGAAT
GAGCACACGTCTGAACTCTTTCCCTACACGACGCTCTTCCGATCTNNNGAGGCTGAGGCAGAAGAAT
GAGCACACGTCTGAACTCTTTCCCTACACGACGCTCTTCCGATCTNNNGAGGCTGAGGCAGAAGAAT
GAGCACACGTCTGAACTCTTTCCCTACACGACGCTCTTCCGATCTNNNGAGGCTGAGGCAGAAGAAT
GAGCACACGTCTGAACTCTTTCCCTACACGACGCTCTTCCGATCTNNNGAGGCTGAGGCAGAAGAAT
AGCACACGTCTGAACTCTTTCCCTACACGACGCTCTTCCGATCTNNNATTTGTCCTCTGGGCTCTGAAA
AGCACACGTCTGAACTCTTTCCCTACACGACGCTCTTCCGATCTNNNATTTGTCCTCTGGGCTCTGAAA
AGCACACGTCTGAACTCTTTCCCTACACGACGCTCTTCCGATCTNNNATTTGTCCTCTGGGCTCTGAAA
AGCACACGTCTGAACTCTTTCCCTACACGACGCTCTTCCGATCTNNNATTTGTCCTCTGGGCTCTGAAA

AGCACACGTCTGAACTCTTTCCCTACACGACGCTCTTCCGATCTNNNATTTGTCCTCTGGGCTCTGAAA
GAGCACACGTCTGAACTCTTTCCCTACACGACGCTCTTCCGATCTNNNAAAAACTTTCCCTGACAGAAG
GAGCACACGTCTGAACTCTTTCCCTACACGACGCTCTTCCGATCTNNNAAAAACTTTCCCTGACAGAAG
GAGCACACGTCTGAACTCTTTCCCTACACGACGCTCTTCCGATCTNNNAAAAACTTTCCCTGACAGAAG
GAGCACACGTCTGAACTCTTTCCCTACACGACGCTCTTCCGATCTNNNAAAAACTTTCCCTGACAGAAG
GAGCACACGTCTGAACTCTTTCCCTACACGACGCTCTTCCGATCTNNNAAAAACTTTCCCTGACAGAAG
GAGCACACGTCTGAACTCTTTCCCTACACGACGCTCTTCCGATCTNNNAGAATTCAGGACCAACTTCA
GAGCACACGTCTGAACTCTTTCCCTACACGACGCTCTTCCGATCTNNNAGAATTCAGGACCAACTTCA
GAGCACACGTCTGAACTCTTTCCCTACACGACGCTCTTCCGATCTNNNAGAATTCAGGACCAACTTCA
GAGCACACGTCTGAACTCTTTCCCTACACGACGCTCTTCCGATCTNNNAGAATTCAGGACCAACTTCA
GAGCACACGTCTGAACTCTTTCCCTACACGACGCTCTTCCGATCTNNNAGAATTCAGGACCAACTTCA
AGCACACGTCTGAACTCTTTCCCTACACGACGCTCTTCCGATCTNNNTCGGTTTTGAAGGATAAGGGT
AGCACACGTCTGAACTCTTTCCCTACACGACGCTCTTCCGATCTNNNTCGGTTTTGAAGGATAAGGGT
AGCACACGTCTGAACTCTTTCCCTACACGACGCTCTTCCGATCTNNNTCGGTTTTGAAGGATAAGGGT
AGCACACGTCTGAACTCTTTCCCTACACGACGCTCTTCCGATCTNNNTCGGTTTTGAAGGATAAGGGT
AGCACACGTCTGAACTCTTTCCCTACACGACGCTCTTCCGATCTNNNTCGGTTTTGAAGGATAAGGGT
AAGAGCACACGTCTGAACTCTTTCCCTACACGACGCTCTTCCGATCTNNNGAACCTGCCTAACAGTGAG
AAGAGCACACGTCTGAACTCTTTCCCTACACGACGCTCTTCCGATCTNNNGAACCTGCCTAACAGTGAG
AAGAGCACACGTCTGAACTCTTTCCCTACACGACGCTCTTCCGATCTNNNGAACCTGCCTAACAGTGAG
AAGAGCACACGTCTGAACTCTTTCCCTACACGACGCTCTTCCGATCTNNNGAACCTGCCTAACAGTGAG
AAGAGCACACGTCTGAACTCTTTCCCTACACGACGCTCTTCCGATCTNNNGAACCTGCCTAACAGTGAG
GAGCACACGTCTGAACTCTTTCCCTACACGACGCTCTTCCGATCTNNNTCAAGATTCTTGCTCAATTTTA
GAGCACACGTCTGAACTCTTTCCCTACACGACGCTCTTCCGATCTNNNTCAAGATTCTTGCTCAATTTTA
GAGCACACGTCTGAACTCTTTCCCTACACGACGCTCTTCCGATCTNNNTCAAGATTCTTGCTCAATTTTA
GAGCACACGTCTGAACTCTTTCCCTACACGACGCTCTTCCGATCTNNNTCAAGATTCTTGCTCAATTTTA
GAGCACACGTCTGAACTCTTTCCCTACACGACGCTCTTCCGATCTNNNTCAAGATTCTTGCTCAATTTTA
AGCACACGTCTGAACTCTTTCCCTACACGACGCTCTTCCGATCTNNNAAAGTATTGCATTTCCCTCTGG
AGCACACGTCTGAACTCTTTCCCTACACGACGCTCTTCCGATCTNNNAAAGTATTGCATTTCCCTCTGG
AGCACACGTCTGAACTCTTTCCCTACACGACGCTCTTCCGATCTNNNAAAGTATTGCATTTCCCTCTGG
AGCACACGTCTGAACTCTTTCCCTACACGACGCTCTTCCGATCTNNNAAAGTATTGCATTTCCCTCTGG
AGCACACGTCTGAACTCTTTCCCTACACGACGCTCTTCCGATCTNNNAAAGTATTGCATTTCCCTCTGG
GAGCACACGTCTGAACTCTTTCCCTACACGACGCTCTTCCGATCTNNNCTCTGCAAAACTTGGGTTCAG
GAGCACACGTCTGAACTCTTTCCCTACACGACGCTCTTCCGATCTNNNCTCTGCAAAACTTGGGTTCAG
GAGCACACGTCTGAACTCTTTCCCTACACGACGCTCTTCCGATCTNNNCTCTGCAAAACTTGGGTTCAG
GAGCACACGTCTGAACTCTTTCCCTACACGACGCTCTTCCGATCTNNNCTCTGCAAAACTTGGGTTCAG
GAGCACACGTCTGAACTCTTTCCCTACACGACGCTCTTCCGATCTNNNCTCTGCAAAACTTGGGTTCAG
AGCACACGTCTGAACTCTTTCCCTACACGACGCTCTTCCGATCTNNNTCAATTTGAAAATACTCATCGA
AGCACACGTCTGAACTCTTTCCCTACACGACGCTCTTCCGATCTNNNTCAATTTGAAAATACTCATCGA
AGCACACGTCTGAACTCTTTCCCTACACGACGCTCTTCCGATCTNNNTCAATTTGAAAATACTCATCGA
AGCACACGTCTGAACTCTTTCCCTACACGACGCTCTTCCGATCTNNNTCAATTTGAAAATACTCATCGA
AGCACACGTCTGAACTCTTTCCCTACACGACGCTCTTCCGATCTNNNTCAATTTGAAAATACTCATCGA
GAGCACACGTCTGAACTCTTTCCCTACACGACGCTCTTCCGATCTNNNGACCTATTCACACCTGTCTCCT
GAGCACACGTCTGAACTCTTTCCCTACACGACGCTCTTCCGATCTNNNGACCTATTCACACCTGTCTCCT

AGCACACGTCTGAACTCTTTCCCTACACGACGCTCTTCCGATCTNNNGACCTATTCACACCTGTCTCCT
AGCACACGTCTGAACTCTTTCCCTACACGACGCTCTTCCGATCTNNNGACCTATTCACACCTGTCTCCT
AGCACACGTCTGAACTCTTTCCCTACACGACGCTCTTCCGATCTNNNGACCTATTCACACCTGTCTCCT
AGCACACGTCTGAACTCTTTCCCTACACGACGCTCTTCCGATCTNNNCAGGAGAGGTGTAGGTGTAC
AGCACACGTCTGAACTCTTTCCCTACACGACGCTCTTCCGATCTNNNCAGGAGAGGTGTAGGTGTAC
AGCACACGTCTGAACTCTTTCCCTACACGACGCTCTTCCGATCTNNNCAGGAGAGGTGTAGGTGTAC
AGCACACGTCTGAACTCTTTCCCTACACGACGCTCTTCCGATCTNNNCAGGAGAGGTGTAGGTGTAC
AGCACACGTCTGAACTCTTTCCCTACACGACGCTCTTCCGATCTNNNCAGGAGAGGTGTAGGTGTAC
GCACACGTCTGAACTCTTTCCCTACACGACGCTCTTCCGATCTNNNATGCCCAGCCACTTTTTTCTTT
GCACACGTCTGAACTCTTTCCCTACACGACGCTCTTCCGATCTNNNATGCCCAGCCACTTTTTTCTTT
GCACACGTCTGAACTCTTTCCCTACACGACGCTCTTCCGATCTNNNATGCCCAGCCACTTTTTTCTTT
GCACACGTCTGAACTCTTTCCCTACACGACGCTCTTCCGATCTNNNATGCCCAGCCACTTTTTTCTTT
GCACACGTCTGAACTCTTTCCCTACACGACGCTCTTCCGATCTNNNATGCCCAGCCACTTTTTTCTTT
GCACACGTCTGAACTCTTTCCCTACACGACGCTCTTCCGATCTNNNAGTGAGTCAAGGTAGTGCCATT
GCACACGTCTGAACTCTTTCCCTACACGACGCTCTTCCGATCTNNNAGTGAGTCAAGGTAGTGCCATT
GCACACGTCTGAACTCTTTCCCTACACGACGCTCTTCCGATCTNNNAGTGAGTCAAGGTAGTGCCATT
GCACACGTCTGAACTCTTTCCCTACACGACGCTCTTCCGATCTNNNAGTGAGTCAAGGTAGTGCCATT
GCACACGTCTGAACTCTTTCCCTACACGACGCTCTTCCGATCTNNNAGTGAGTCAAGGTAGTGCCATT
AGCACACGTCTGAACTCTTTCCCTACACGACGCTCTTCCGATCTNNNCTATGTTGCCTATGCTGGTCTC
AGCACACGTCTGAACTCTTTCCCTACACGACGCTCTTCCGATCTNNNCTATGTTGCCTATGCTGGTCTC
AGCACACGTCTGAACTCTTTCCCTACACGACGCTCTTCCGATCTNNNCTATGTTGCCTATGCTGGTCTC
AGCACACGTCTGAACTCTTTCCCTACACGACGCTCTTCCGATCTNNNCTATGTTGCCTATGCTGGTCTC
AGCACACGTCTGAACTCTTTCCCTACACGACGCTCTTCCGATCTNNNCTATGTTGCCTATGCTGGTCTC
AGCACACGTCTGAACTCTTTCCCTACACGACGCTCTTCCGATCTNNNCTGAGGCAGGAGAATCACTTG
AGCACACGTCTGAACTCTTTCCCTACACGACGCTCTTCCGATCTNNNCTGAGGCAGGAGAATCACTTG
AGCACACGTCTGAACTCTTTCCCTACACGACGCTCTTCCGATCTNNNCTGAGGCAGGAGAATCACTTG
AGCACACGTCTGAACTCTTTCCCTACACGACGCTCTTCCGATCTNNNCTGAGGCAGGAGAATCACTTG
AGCACACGTCTGAACTCTTTCCCTACACGACGCTCTTCCGATCTNNNCTGAGGCAGGAGAATCACTTG
GAGCACACGTCTGAACTCTTTCCCTACACGACGCTCTTCCGATCTNNNATGAAGGAGTGTTTTTGCCT
GAGCACACGTCTGAACTCTTTCCCTACACGACGCTCTTCCGATCTNNNATGAAGGAGTGTTTTTGCCT
GAGCACACGTCTGAACTCTTTCCCTACACGACGCTCTTCCGATCTNNNATGAAGGAGTGTTTTTGCCT
GAGCACACGTCTGAACTCTTTCCCTACACGACGCTCTTCCGATCTNNNATGAAGGAGTGTTTTTGCCT
GAGCACACGTCTGAACTCTTTCCCTACACGACGCTCTTCCGATCTNNNATGAAGGAGTGTTTTTGCCT
GCACACGTCTGAACTCTTTCCCTACACGACGCTCTTCCGATCTNNNATGAGAGATGAGAAATGCAGG
GCACACGTCTGAACTCTTTCCCTACACGACGCTCTTCCGATCTNNNATGAGAGATGAGAAATGCAGG
GCACACGTCTGAACTCTTTCCCTACACGACGCTCTTCCGATCTNNNATGAGAGATGAGAAATGCAGG
GCACACGTCTGAACTCTTTCCCTACACGACGCTCTTCCGATCTNNNATGAGAGATGAGAAATGCAGG
GCACACGTCTGAACTCTTTCCCTACACGACGCTCTTCCGATCTNNNATGAGAGATGAGAAATGCAGG
AGCACACGTCTGAACTCTTTCCCTACACGACGCTCTTCCGATCTNNNTGCCCAGTTTCATTTTGGTTGT
AGCACACGTCTGAACTCTTTCCCTACACGACGCTCTTCCGATCTNNNTGCCCAGTTTCATTTTGGTTGT
AGCACACGTCTGAACTCTTTCCCTACACGACGCTCTTCCGATCTNNNTGCCCAGTTTCATTTTGGTTGT
AGCACACGTCTGAACTCTTTCCCTACACGACGCTCTTCCGATCTNNNTGCCCAGTTTCATTTTGGTTGT
AGCACACGTCTGAACTCTTTCCCTACACGACGCTCTTCCGATCTNNNTGCCCAGTTTCATTTTGGTTGT

AGAGCACACGTCTGAACTCTTTCCCTACACGACGCTCTTCCGATCTNNNTCCCATCTGACCTCCTGTATT

AGAGCACACGTCTGAACTCTTTCCCTACACGACGCTCTTCCGATCTNNNTCCCATCTGACCTCCTGTATT

AGAGCACACGTCTGAACTCTTTCCCTACACGACGCTCTTCCGATCTNNNTCCCATCTGACCTCCTGTATT

AGAGCACACGTCTGAACTCTTTCCCTACACGACGCTCTTCCGATCTNNNTCCCATCTGACCTCCTGTATT

AGAGCACACGTCTGAACTCTTTCCCTACACGACGCTCTTCCGATCTNNNTCCCATCTGACCTCCTGTATT

GAGCACACGTCTGAACTCTTTCCCTACACGACGCTCTTCCGATCTNNNGGGTGGATCATTTGAGGTAA

GAGCACACGTCTGAACTCTTTCCCTACACGACGCTCTTCCGATCTNNNGGGTGGATCATTTGAGGTAA

GAGCACACGTCTGAACTCTTTCCCTACACGACGCTCTTCCGATCTNNNGGGTGGATCATTTGAGGTAA

GAGCACACGTCTGAACTCTTTCCCTACACGACGCTCTTCCGATCTNNNGGGTGGATCATTTGAGGTAA

GAGCACACGTCTGAACTCTTTCCCTACACGACGCTCTTCCGATCTNNNGGGTGGATCATTTGAGGTAA

AGCACACGTCTGAACTCTTTCCCTACACGACGCTCTTCCGATCTNNNACAAAAAGATGACTTGGCTGCT

AGCACACGTCTGAACTCTTTCCCTACACGACGCTCTTCCGATCTNNNACAAAAAGATGACTTGGCTGCT

AGCACACGTCTGAACTCTTTCCCTACACGACGCTCTTCCGATCTNNNACAAAAAGATGACTTGGCTGCT

AGCACACGTCTGAACTCTTTCCCTACACGACGCTCTTCCGATCTNNNACAAAAAGATGACTTGGCTGCT

AGCACACGTCTGAACTCTTTCCCTACACGACGCTCTTCCGATCTNNNACAAAAAGATGACTTGGCTGCT

AGAGCACACGTCTGAACTCTTTCCCTACACGACGCTCTTCCGATCTNNNCAAGATCCCACCACTGCACTC

AGAGCACACGTCTGAACTCTTTCCCTACACGACGCTCTTCCGATCTNNNCAAGATCCCACCACTGCACTC

AGAGCACACGTCTGAACTCTTTCCCTACACGACGCTCTTCCGATCTNNNCAAGATCCCACCACTGCACTC

AGAGCACACGTCTGAACTCTTTCCCTACACGACGCTCTTCCGATCTNNNCAAGATCCCACCACTGCACTC

AGAGCACACGTCTGAACTCTTTCCCTACACGACGCTCTTCCGATCTNNNCAAGATCCCACCACTGCACTC

AGCACACGTCTGAACTCTTTCCCTACACGACGCTCTTCCGATCTNNNGATTGTAGCTACTCAGGAGGCT

AGCACACGTCTGAACTCTTTCCCTACACGACGCTCTTCCGATCTNNNGATTGTAGCTACTCAGGAGGCT

AGCACACGTCTGAACTCTTTCCCTACACGACGCTCTTCCGATCTNNNGATTGTAGCTACTCAGGAGGCT

AGCACACGTCTGAACTCTTTCCCTACACGACGCTCTTCCGATCTNNNGATTGTAGCTACTCAGGAGGCT

AGCACACGTCTGAACTCTTTCCCTACACGACGCTCTTCCGATCTNNNGATTGTAGCTACTCAGGAGGCT

GAGCACACGTCTGAACTCTTTCCCTACACGACGCTCTTCCGATCTNNNGTGAGCTGAGATTGCACCATT

GAGCACACGTCTGAACTCTTTCCCTACACGACGCTCTTCCGATCTNNNGTGAGCTGAGATTGCACCATT

GAGCACACGTCTGAACTCTTTCCCTACACGACGCTCTTCCGATCTNNNGTGAGCTGAGATTGCACCATT

GAGCACACGTCTGAACTCTTTCCCTACACGACGCTCTTCCGATCTNNNGTGAGCTGAGATTGCACCATT

GAGCACACGTCTGAACTCTTTCCCTACACGACGCTCTTCCGATCTNNNGTGAGCTGAGATTGCACCATT

GAGCACACGTCTGAACTCTTTCCCTACACGACGCTCTTCCGATCTNNNGGGGGGCTCTCAAATATCATC

GAGCACACGTCTGAACTCTTTCCCTACACGACGCTCTTCCGATCTNNNGGGGGGCTCTCAAATATCATC

GAGCACACGTCTGAACTCTTTCCCTACACGACGCTCTTCCGATCTNNNGGGGGGCTCTCAAATATCATC

GAGCACACGTCTGAACTCTTTCCCTACACGACGCTCTTCCGATCTNNNGGGGGGCTCTCAAATATCATC

GAGCACACGTCTGAACTCTTTCCCTACACGACGCTCTTCCGATCTNNNGGGGGGCTCTCAAATATCATC

GAGCACACGTCTGAACTCTTTCCCTACACGACGCTCTTCCGATCTNNNCTTGCAGTGAGCTGAGATGG

GAGCACACGTCTGAACTCTTTCCCTACACGACGCTCTTCCGATCTNNNCTTGCAGTGAGCTGAGATGG

GAGCACACGTCTGAACTCTTTCCCTACACGACGCTCTTCCGATCTNNNCTTGCAGTGAGCTGAGATGG

GAGCACACGTCTGAACTCTTTCCCTACACGACGCTCTTCCGATCTNNNCTTGCAGTGAGCTGAGATGG

GAGCACACGTCTGAACTCTTTCCCTACACGACGCTCTTCCGATCTNNNCTTGCAGTGAGCTGAGATGG

AGCACACGTCTGAACTCTTTCCCTACACGACGCTCTTCCGATCTNNNTGAGATTACAGATGTGAGCCAC

AGCACACGTCTGAACTCTTTCCCTACACGACGCTCTTCCGATCTNNNTGAGATTACAGATGTGAGCCAC

AGCACACGTCTGAACTCTTTCCCTACACGACGCTCTTCCGATCTNNNTGAGATTACAGATGTGAGCCAC

AGCACACGTCTGAACTCTTTCCCTACACGACGCTCTTCCGATCTNNNTGAGATTACAGATGTGAGCCAC
AGCACACGTCTGAACTCTTTCCCTACACGACGCTCTTCCGATCTNNNTGAGATTACAGATGTGAGCCAC
GAGCACACGTCTGAACTCTTTCCCTACACGACGCTCTTCCGATCTNNNGAGGAAGGAAGGAAGGAAG
GAGCACACGTCTGAACTCTTTCCCTACACGACGCTCTTCCGATCTNNNGAGGAAGGAAGGAAGGAAG
GAGCACACGTCTGAACTCTTTCCCTACACGACGCTCTTCCGATCTNNNGAGGAAGGAAGGAAGGAAG
GAGCACACGTCTGAACTCTTTCCCTACACGACGCTCTTCCGATCTNNNGAGGAAGGAAGGAAGGAAG
GAGCACACGTCTGAACTCTTTCCCTACACGACGCTCTTCCGATCTNNNGAGGAAGGAAGGAAGGAAG
GCACACGTCTGAACTCTTTCCCTACACGACGCTCTTCCGATCTNNNAAGTCACCTGGGATGGCAAGCA
GCACACGTCTGAACTCTTTCCCTACACGACGCTCTTCCGATCTNNNAAGTCACCTGGGATGGCAAGCA
GCACACGTCTGAACTCTTTCCCTACACGACGCTCTTCCGATCTNNNAAGTCACCTGGGATGGCAAGCA
GCACACGTCTGAACTCTTTCCCTACACGACGCTCTTCCGATCTNNNAAGTCACCTGGGATGGCAAGCA
GCACACGTCTGAACTCTTTCCCTACACGACGCTCTTCCGATCTNNNAAGTCACCTGGGATGGCAAGCA
GCACACGTCTGAACTCTTTCCCTACACGACGCTCTTCCGATCTNNNCTCTGGAAATGAGCCAAGGACAA
GCACACGTCTGAACTCTTTCCCTACACGACGCTCTTCCGATCTNNNCTCTGGAAATGAGCCAAGGACAA
GCACACGTCTGAACTCTTTCCCTACACGACGCTCTTCCGATCTNNNCTCTGGAAATGAGCCAAGGACAA
GCACACGTCTGAACTCTTTCCCTACACGACGCTCTTCCGATCTNNNCTCTGGAAATGAGCCAAGGACAA
GCACACGTCTGAACTCTTTCCCTACACGACGCTCTTCCGATCTNNNCTCTGGAAATGAGCCAAGGACAA
GAGCACACGTCTGAACTCTTTCCCTACACGACGCTCTTCCGATCTNNNTGAGAATCACTTGAACCCAG
GAGCACACGTCTGAACTCTTTCCCTACACGACGCTCTTCCGATCTNNNTGAGAATCACTTGAACCCAG
GAGCACACGTCTGAACTCTTTCCCTACACGACGCTCTTCCGATCTNNNTGAGAATCACTTGAACCCAG
GAGCACACGTCTGAACTCTTTCCCTACACGACGCTCTTCCGATCTNNNTGAGAATCACTTGAACCCAG
GAGCACACGTCTGAACTCTTTCCCTACACGACGCTCTTCCGATCTNNNTGAGAATCACTTGAACCCAG
AGCACACGTCTGAACTCTTTCCCTACACGACGCTCTTCCGATCTNNNGGCCGTAGACCAAATATCAGG
AGCACACGTCTGAACTCTTTCCCTACACGACGCTCTTCCGATCTNNNGGCCGTAGACCAAATATCAGG
AGCACACGTCTGAACTCTTTCCCTACACGACGCTCTTCCGATCTNNNGGCCGTAGACCAAATATCAGG
AGCACACGTCTGAACTCTTTCCCTACACGACGCTCTTCCGATCTNNNGGCCGTAGACCAAATATCAGG
AGCACACGTCTGAACTCTTTCCCTACACGACGCTCTTCCGATCTNNNGGCCGTAGACCAAATATCAGG
AGCACACGTCTGAACTCTTTCCCTACACGACGCTCTTCCGATCTNNNTGTACCTTGCCCCATTCATACA
AGCACACGTCTGAACTCTTTCCCTACACGACGCTCTTCCGATCTNNNTGTACCTTGCCCCATTCATACA
AGCACACGTCTGAACTCTTTCCCTACACGACGCTCTTCCGATCTNNNTGTACCTTGCCCCATTCATACA
AGCACACGTCTGAACTCTTTCCCTACACGACGCTCTTCCGATCTNNNTGTACCTTGCCCCATTCATACA
AGCACACGTCTGAACTCTTTCCCTACACGACGCTCTTCCGATCTNNNTGTACCTTGCCCCATTCATACA
GCACACGTCTGAACTCTTTCCCTACACGACGCTCTTCCGATCTNNNAGATTCCAGATTGTGTTCTCCCCC
GCACACGTCTGAACTCTTTCCCTACACGACGCTCTTCCGATCTNNNAGATTCCAGATTGTGTTCTCCCCC
GCACACGTCTGAACTCTTTCCCTACACGACGCTCTTCCGATCTNNNAGATTCCAGATTGTGTTCTCCCCC
GCACACGTCTGAACTCTTTCCCTACACGACGCTCTTCCGATCTNNNAGATTCCAGATTGTGTTCTCCCCC
GCACACGTCTGAACTCTTTCCCTACACGACGCTCTTCCGATCTNNNAGATTCCAGATTGTGTTCTCCCCC
AGCACACGTCTGAACTCTTTCCCTACACGACGCTCTTCCGATCTNNNAAAATATGGAACCAGCCCAAAC
AGCACACGTCTGAACTCTTTCCCTACACGACGCTCTTCCGATCTNNNAAAATATGGAACCAGCCCAAAC
AGCACACGTCTGAACTCTTTCCCTACACGACGCTCTTCCGATCTNNNAAAATATGGAACCAGCCCAAAC
AGCACACGTCTGAACTCTTTCCCTACACGACGCTCTTCCGATCTNNNAAAATATGGAACCAGCCCAAAC
AGCACACGTCTGAACTCTTTCCCTACACGACGCTCTTCCGATCTNNNAAAATATGGAACCAGCCCAAAC
GAGCACACGTCTGAACTCTTTCCCTACACGACGCTCTTCCGATCTNNNAGCTCAGAACCTCTTTCTGCCT

GAGCACACGTCTGAACTCTTTCCCTACACGACGCTCTTCCGATCTNNNAGCTCAGAACCTCTTTCTGCCT
GAGCACACGTCTGAACTCTTTCCCTACACGACGCTCTTCCGATCTNNNAGCTCAGAACCTCTTTCTGCCT
GAGCACACGTCTGAACTCTTTCCCTACACGACGCTCTTCCGATCTNNNAGCTCAGAACCTCTTTCTGCCT
GAGCACACGTCTGAACTCTTTCCCTACACGACGCTCTTCCGATCTNNNAGCTCAGAACCTCTTTCTGCCT
GAGCACACGTCTGAACTCTTTCCCTACACGACGCTCTTCCGATCTNNNCCAAAGGTCCTTGAGTTTCTC
GAGCACACGTCTGAACTCTTTCCCTACACGACGCTCTTCCGATCTNNNCCAAAGGTCCTTGAGTTTCTC
GAGCACACGTCTGAACTCTTTCCCTACACGACGCTCTTCCGATCTNNNCCAAAGGTCCTTGAGTTTCTC
GAGCACACGTCTGAACTCTTTCCCTACACGACGCTCTTCCGATCTNNNCCAAAGGTCCTTGAGTTTCTC
GAGCACACGTCTGAACTCTTTCCCTACACGACGCTCTTCCGATCTNNNCCAAAGGTCCTTGAGTTTCTC
AGCACACGTCTGAACTCTTTCCCTACACGACGCTCTTCCGATCTNNNAGAGATTTGAACCCAGTAGTC
AGCACACGTCTGAACTCTTTCCCTACACGACGCTCTTCCGATCTNNNAGAGATTTGAACCCAGTAGTC
AGCACACGTCTGAACTCTTTCCCTACACGACGCTCTTCCGATCTNNNAGAGATTTGAACCCAGTAGTC
AGCACACGTCTGAACTCTTTCCCTACACGACGCTCTTCCGATCTNNNAGAGATTTGAACCCAGTAGTC
AGCACACGTCTGAACTCTTTCCCTACACGACGCTCTTCCGATCTNNNAGAGATTTGAACCCAGTAGTC
GAGCACACGTCTGAACTCTTTCCCTACACGACGCTCTTCCGATCTNNNACACTGGGGGTCAGAGAGTA
GAGCACACGTCTGAACTCTTTCCCTACACGACGCTCTTCCGATCTNNNACACTGGGGGTCAGAGAGTA
GAGCACACGTCTGAACTCTTTCCCTACACGACGCTCTTCCGATCTNNNACACTGGGGGTCAGAGAGTA
GAGCACACGTCTGAACTCTTTCCCTACACGACGCTCTTCCGATCTNNNACACTGGGGGTCAGAGAGTA
GAGCACACGTCTGAACTCTTTCCCTACACGACGCTCTTCCGATCTNNNACACTGGGGGTCAGAGAGTA
AGCACACGTCTGAACTCTTTCCCTACACGACGCTCTTCCGATCTNNNACAAGAATCCCACAAGTTCCAG
AGCACACGTCTGAACTCTTTCCCTACACGACGCTCTTCCGATCTNNNACAAGAATCCCACAAGTTCCAG
AGCACACGTCTGAACTCTTTCCCTACACGACGCTCTTCCGATCTNNNACAAGAATCCCACAAGTTCCAG
AGCACACGTCTGAACTCTTTCCCTACACGACGCTCTTCCGATCTNNNACAAGAATCCCACAAGTTCCAG
AGCACACGTCTGAACTCTTTCCCTACACGACGCTCTTCCGATCTNNNACAAGAATCCCACAAGTTCCAG
GAGCACACGTCTGAACTCTTTCCCTACACGACGCTCTTCCGATCTNNNCCCATATCTGAAGATAAATTT
GAGCACACGTCTGAACTCTTTCCCTACACGACGCTCTTCCGATCTNNNCCCATATCTGAAGATAAATTT
GAGCACACGTCTGAACTCTTTCCCTACACGACGCTCTTCCGATCTNNNCCCATATCTGAAGATAAATTT
GAGCACACGTCTGAACTCTTTCCCTACACGACGCTCTTCCGATCTNNNCCCATATCTGAAGATAAATTT
GAGCACACGTCTGAACTCTTTCCCTACACGACGCTCTTCCGATCTNNNCCCATATCTGAAGATAAATTT
AGCACACGTCTGAACTCTTTCCCTACACGACGCTCTTCCGATCTNNNGGGGATATGTTCACCTCTTTG
AGCACACGTCTGAACTCTTTCCCTACACGACGCTCTTCCGATCTNNNGGGGATATGTTCACCTCTTTG
AGCACACGTCTGAACTCTTTCCCTACACGACGCTCTTCCGATCTNNNGGGGATATGTTCACCTCTTTG
AGCACACGTCTGAACTCTTTCCCTACACGACGCTCTTCCGATCTNNNGGGGATATGTTCACCTCTTTG
AGCACACGTCTGAACTCTTTCCCTACACGACGCTCTTCCGATCTNNNGGGGATATGTTCACCTCTTTG
GCACACGTCTGAACTCTTTCCCTACACGACGCTCTTCCGATCTNNNAGAAAGTGAAAGGCAGAGGG
GCACACGTCTGAACTCTTTCCCTACACGACGCTCTTCCGATCTNNNAGAAAGTGAAAGGCAGAGGG
GCACACGTCTGAACTCTTTCCCTACACGACGCTCTTCCGATCTNNNAGAAAGTGAAAGGCAGAGGG
GCACACGTCTGAACTCTTTCCCTACACGACGCTCTTCCGATCTNNNAGAAAGTGAAAGGCAGAGGG
GCACACGTCTGAACTCTTTCCCTACACGACGCTCTTCCGATCTNNNAGAAAGTGAAAGGCAGAGGG
GCACACGTCTGAACTCTTTCCCTACACGACGCTCTTCCGATCTNNNGGAATCTGCCCTGTGTCGTCCAT
GCACACGTCTGAACTCTTTCCCTACACGACGCTCTTCCGATCTNNNGGAATCTGCCCTGTGTCGTCCAT
GCACACGTCTGAACTCTTTCCCTACACGACGCTCTTCCGATCTNNNGGAATCTGCCCTGTGTCGTCCAT
GCACACGTCTGAACTCTTTCCCTACACGACGCTCTTCCGATCTNNNGGAATCTGCCCTGTGTCGTCCAT

GCACACGTCTGAACTCTTTCCCTACACGACGCTCTTCCGATCTNNNGGAATCTGCCCTGTGTCGTCCAT

GCACACGTCTGAACTCTTTCCCTACACGACGCTCTTCCGATCTNNNTGCTTAGGGGACAAGAGTTTGAA

GCACACGTCTGAACTCTTTCCCTACACGACGCTCTTCCGATCTNNNTGCTTAGGGGACAAGAGTTTGAA

GCACACGTCTGAACTCTTTCCCTACACGACGCTCTTCCGATCTNNNTGCTTAGGGGACAAGAGTTTGAA

GCACACGTCTGAACTCTTTCCCTACACGACGCTCTTCCGATCTNNNTGCTTAGGGGACAAGAGTTTGAA

GCACACGTCTGAACTCTTTCCCTACACGACGCTCTTCCGATCTNNNTGCTTAGGGGACAAGAGTTTGAA

AGAGCACACGTCTGAACTCTTTCCCTACACGACGCTCTTCCGATCTNNNGAGGTTGCAGTGAGTTGAG

AGAGCACACGTCTGAACTCTTTCCCTACACGACGCTCTTCCGATCTNNNGAGGTTGCAGTGAGTTGAG

AGAGCACACGTCTGAACTCTTTCCCTACACGACGCTCTTCCGATCTNNNGAGGTTGCAGTGAGTTGAG

AGAGCACACGTCTGAACTCTTTCCCTACACGACGCTCTTCCGATCTNNNGAGGTTGCAGTGAGTTGAG

AGAGCACACGTCTGAACTCTTTCCCTACACGACGCTCTTCCGATCTNNNGAGGTTGCAGTGAGTTGAG

GAGCACACGTCTGAACTCTTTCCCTACACGACGCTCTTCCGATCTNNNGTTTCAGCTGACCTCAAGGACT

GAGCACACGTCTGAACTCTTTCCCTACACGACGCTCTTCCGATCTNNNGTTTCAGCTGACCTCAAGGACT

GAGCACACGTCTGAACTCTTTCCCTACACGACGCTCTTCCGATCTNNNGTTTCAGCTGACCTCAAGGACT

GAGCACACGTCTGAACTCTTTCCCTACACGACGCTCTTCCGATCTNNNGTTTCAGCTGACCTCAAGGACT

GAGCACACGTCTGAACTCTTTCCCTACACGACGCTCTTCCGATCTNNNGTTTCAGCTGACCTCAAGGACT

GAGCACACGTCTGAACTCTTTCCCTACACGACGCTCTTCCGATCTNNNGAATAAGCAGATCCATGGAG

GAGCACACGTCTGAACTCTTTCCCTACACGACGCTCTTCCGATCTNNNGAATAAGCAGATCCATGGAG

GAGCACACGTCTGAACTCTTTCCCTACACGACGCTCTTCCGATCTNNNGAATAAGCAGATCCATGGAG

GAGCACACGTCTGAACTCTTTCCCTACACGACGCTCTTCCGATCTNNNGAATAAGCAGATCCATGGAG

GAGCACACGTCTGAACTCTTTCCCTACACGACGCTCTTCCGATCTNNNGAATAAGCAGATCCATGGAG

GCACACGTCTGAACTCTTTCCCTACACGACGCTCTTCCGATCTNNNGGAGAAGTGGAATGCACACAAT

GCACACGTCTGAACTCTTTCCCTACACGACGCTCTTCCGATCTNNNGGAGAAGTGGAATGCACACAAT

GCACACGTCTGAACTCTTTCCCTACACGACGCTCTTCCGATCTNNNGGAGAAGTGGAATGCACACAAT

GCACACGTCTGAACTCTTTCCCTACACGACGCTCTTCCGATCTNNNGGAGAAGTGGAATGCACACAAT

GCACACGTCTGAACTCTTTCCCTACACGACGCTCTTCCGATCTNNNGGAGAAGTGGAATGCACACAAT

ACACGTCTGAACTCTTTCCCTACACGACGCTCTTCCGATCTNNNGTGTGTTGCCATAACAAAATACATC

ACACGTCTGAACTCTTTCCCTACACGACGCTCTTCCGATCTNNNGTGTGTTGCCATAACAAAATACATC

ACACGTCTGAACTCTTTCCCTACACGACGCTCTTCCGATCTNNNGTGTGTTGCCATAACAAAATACATC

ACACGTCTGAACTCTTTCCCTACACGACGCTCTTCCGATCTNNNGTGTGTTGCCATAACAAAATACATC

ACACGTCTGAACTCTTTCCCTACACGACGCTCTTCCGATCTNNNGTGTGTTGCCATAACAAAATACATC

AGAGCACACGTCTGAACTCTTTCCCTACACGACGCTCTTCCGATCTNNNGGTTGCAGTTAGCCAAGAT

AGAGCACACGTCTGAACTCTTTCCCTACACGACGCTCTTCCGATCTNNNGGTTGCAGTTAGCCAAGAT

AGAGCACACGTCTGAACTCTTTCCCTACACGACGCTCTTCCGATCTNNNGGTTGCAGTTAGCCAAGAT

AGAGCACACGTCTGAACTCTTTCCCTACACGACGCTCTTCCGATCTNNNGGTTGCAGTTAGCCAAGAT

AGAGCACACGTCTGAACTCTTTCCCTACACGACGCTCTTCCGATCTNNNGGTTGCAGTTAGCCAAGAT

AGAGCACACGTCTGAACTCTTTCCCTACACGACGCTCTTCCGATCTNNNAGGTGGAACAAACCTAAG

AGAGCACACGTCTGAACTCTTTCCCTACACGACGCTCTTCCGATCTNNNAGGTGGAACAAACCTAAG

AGAGCACACGTCTGAACTCTTTCCCTACACGACGCTCTTCCGATCTNNNAGGTGGAACAAACCTAAG

AGAGCACACGTCTGAACTCTTTCCCTACACGACGCTCTTCCGATCTNNNAGGTGGAACAAACCTAAG

AGAGCACACGTCTGAACTCTTTCCCTACACGACGCTCTTCCGATCTNNNAGGTGGAACAAACCTAAG

AGCACACGTCTGAACTCTTTCCCTACACGACGCTCTTCCGATCTNNNGAGGGGGAGAAGGGATGAAA

AGCACACGTCTGAACTCTTTCCCTACACGACGCTCTTCCGATCTNNNGAGGGGGAGAAGGGATGAAA

AGCACACGTCTGAACTCTTTCCCTACACGACGCTCTTCCGATCTNNNGAGGGGGAGAAGGGATGAAA
AGCACACGTCTGAACTCTTTCCCTACACGACGCTCTTCCGATCTNNNGAGGGGGAGAAGGGATGAAA
AGCACACGTCTGAACTCTTTCCCTACACGACGCTCTTCCGATCTNNNGAGGGGGAGAAGGGATGAAA
AGCACACGTCTGAACTCTTTCCCTACACGACGCTCTTCCGATCTNNNAGAGACATTTGGGGGTACTCA
AGCACACGTCTGAACTCTTTCCCTACACGACGCTCTTCCGATCTNNNAGAGACATTTGGGGGTACTCA
AGCACACGTCTGAACTCTTTCCCTACACGACGCTCTTCCGATCTNNNAGAGACATTTGGGGGTACTCA
AGCACACGTCTGAACTCTTTCCCTACACGACGCTCTTCCGATCTNNNAGAGACATTTGGGGGTACTCA
AGCACACGTCTGAACTCTTTCCCTACACGACGCTCTTCCGATCTNNNAGAGACATTTGGGGGTACTCA
AGCACACGTCTGAACTCTTTCCCTACACGACGCTCTTCCGATCTNNNACACCTAGAAGCAAGTCTGTAC
AGCACACGTCTGAACTCTTTCCCTACACGACGCTCTTCCGATCTNNNACACCTAGAAGCAAGTCTGTAC
AGCACACGTCTGAACTCTTTCCCTACACGACGCTCTTCCGATCTNNNACACCTAGAAGCAAGTCTGTAC
AGCACACGTCTGAACTCTTTCCCTACACGACGCTCTTCCGATCTNNNACACCTAGAAGCAAGTCTGTAC
AGCACACGTCTGAACTCTTTCCCTACACGACGCTCTTCCGATCTNNNACACCTAGAAGCAAGTCTGTAC
GAGCACACGTCTGAACTCTTTCCCTACACGACGCTCTTCCGATCTNNNCCTGAACTGTGATTTTGCCAC
GAGCACACGTCTGAACTCTTTCCCTACACGACGCTCTTCCGATCTNNNCCTGAACTGTGATTTTGCCAC
GAGCACACGTCTGAACTCTTTCCCTACACGACGCTCTTCCGATCTNNNCCTGAACTGTGATTTTGCCAC
GAGCACACGTCTGAACTCTTTCCCTACACGACGCTCTTCCGATCTNNNCCTGAACTGTGATTTTGCCAC
GAGCACACGTCTGAACTCTTTCCCTACACGACGCTCTTCCGATCTNNNCCTGAACTGTGATTTTGCCAC
GAGCACACGTCTGAACTCTTTCCCTACACGACGCTCTTCCGATCTNNNAGGAGGCTGAGGTAGGAGA
GAGCACACGTCTGAACTCTTTCCCTACACGACGCTCTTCCGATCTNNNAGGAGGCTGAGGTAGGAGA
GAGCACACGTCTGAACTCTTTCCCTACACGACGCTCTTCCGATCTNNNAGGAGGCTGAGGTAGGAGA
GAGCACACGTCTGAACTCTTTCCCTACACGACGCTCTTCCGATCTNNNAGGAGGCTGAGGTAGGAGA
GAGCACACGTCTGAACTCTTTCCCTACACGACGCTCTTCCGATCTNNNAGGAGGCTGAGGTAGGAGA
GAGCACACGTCTGAACTCTTTCCCTACACGACGCTCTTCCGATCTNNNTGAGGCAGCAAAATCACTTGA
GAGCACACGTCTGAACTCTTTCCCTACACGACGCTCTTCCGATCTNNNTGAGGCAGCAAAATCACTTGA
GAGCACACGTCTGAACTCTTTCCCTACACGACGCTCTTCCGATCTNNNTGAGGCAGCAAAATCACTTGA
GAGCACACGTCTGAACTCTTTCCCTACACGACGCTCTTCCGATCTNNNTGAGGCAGCAAAATCACTTGA
GAGCACACGTCTGAACTCTTTCCCTACACGACGCTCTTCCGATCTNNNTGAGGCAGCAAAATCACTTGA
AAGAGCACACGTCTGAACTCTTTCCCTACACGACGCTCTTCCGATCTNNNGAACCTGGAAGAACTGAG
AAGAGCACACGTCTGAACTCTTTCCCTACACGACGCTCTTCCGATCTNNNGAACCTGGAAGAACTGAG
AAGAGCACACGTCTGAACTCTTTCCCTACACGACGCTCTTCCGATCTNNNGAACCTGGAAGAACTGAG
AAGAGCACACGTCTGAACTCTTTCCCTACACGACGCTCTTCCGATCTNNNGAACCTGGAAGAACTGAG
AAGAGCACACGTCTGAACTCTTTCCCTACACGACGCTCTTCCGATCTNNNGAACCTGGAAGAACTGAG
AGCACACGTCTGAACTCTTTCCCTACACGACGCTCTTCCGATCTNNNAGAATCACTTGAACCCAGGAG
AGCACACGTCTGAACTCTTTCCCTACACGACGCTCTTCCGATCTNNNAGAATCACTTGAACCCAGGAG
AGCACACGTCTGAACTCTTTCCCTACACGACGCTCTTCCGATCTNNNAGAATCACTTGAACCCAGGAG
AGCACACGTCTGAACTCTTTCCCTACACGACGCTCTTCCGATCTNNNAGAATCACTTGAACCCAGGAG
AGCACACGTCTGAACTCTTTCCCTACACGACGCTCTTCCGATCTNNNAGAATCACTTGAACCCAGGAG
GAGCACACGTCTGAACTCTTTCCCTACACGACGCTCTTCCGATCTNNNTTTGGGCCTCCAGGAATACAA
GAGCACACGTCTGAACTCTTTCCCTACACGACGCTCTTCCGATCTNNNTTTGGGCCTCCAGGAATACAA
GAGCACACGTCTGAACTCTTTCCCTACACGACGCTCTTCCGATCTNNNTTTGGGCCTCCAGGAATACAA
GAGCACACGTCTGAACTCTTTCCCTACACGACGCTCTTCCGATCTNNNTTTGGGCCTCCAGGAATACAA
GAGCACACGTCTGAACTCTTTCCCTACACGACGCTCTTCCGATCTNNNTTTGGGCCTCCAGGAATACAA

AGCACACGTCTGAACTCTTTCCCTACACGACGCTCTTCCGATCTNNNGGCTATTTACTGTATTCAGGAA
AGCACACGTCTGAACTCTTTCCCTACACGACGCTCTTCCGATCTNNNGGCTATTTACTGTATTCAGGAA
AGCACACGTCTGAACTCTTTCCCTACACGACGCTCTTCCGATCTNNNGGCTATTTACTGTATTCAGGAA
AGCACACGTCTGAACTCTTTCCCTACACGACGCTCTTCCGATCTNNNGGCTATTTACTGTATTCAGGAA
AGCACACGTCTGAACTCTTTCCCTACACGACGCTCTTCCGATCTNNNGGCTATTTACTGTATTCAGGAA
AGCACACGTCTGAACTCTTTCCCTACACGACGCTCTTCCGATCTNNNATGGGAGTAGGATGGATTGAT
AGCACACGTCTGAACTCTTTCCCTACACGACGCTCTTCCGATCTNNNATGGGAGTAGGATGGATTGAT
AGCACACGTCTGAACTCTTTCCCTACACGACGCTCTTCCGATCTNNNATGGGAGTAGGATGGATTGAT
AGCACACGTCTGAACTCTTTCCCTACACGACGCTCTTCCGATCTNNNATGGGAGTAGGATGGATTGAT
AGCACACGTCTGAACTCTTTCCCTACACGACGCTCTTCCGATCTNNNATGGGAGTAGGATGGATTGAT
AGCACACGTCTGAACTCTTTCCCTACACGACGCTCTTCCGATCTNNNGTCTCCAGTGAGAATGCTTCCA
AGCACACGTCTGAACTCTTTCCCTACACGACGCTCTTCCGATCTNNNGTCTCCAGTGAGAATGCTTCCA
AGCACACGTCTGAACTCTTTCCCTACACGACGCTCTTCCGATCTNNNGTCTCCAGTGAGAATGCTTCCA
AGCACACGTCTGAACTCTTTCCCTACACGACGCTCTTCCGATCTNNNGTCTCCAGTGAGAATGCTTCCA
AGCACACGTCTGAACTCTTTCCCTACACGACGCTCTTCCGATCTNNNGTCTCCAGTGAGAATGCTTCCA
AGCACACGTCTGAACTCTTTCCCTACACGACGCTCTTCCGATCTNNNGGGAAAAGGTGTTTCAGGAAC
AGCACACGTCTGAACTCTTTCCCTACACGACGCTCTTCCGATCTNNNGGGAAAAGGTGTTTCAGGAAC
AGCACACGTCTGAACTCTTTCCCTACACGACGCTCTTCCGATCTNNNGGGAAAAGGTGTTTCAGGAAC
AGCACACGTCTGAACTCTTTCCCTACACGACGCTCTTCCGATCTNNNGGGAAAAGGTGTTTCAGGAAC
AGCACACGTCTGAACTCTTTCCCTACACGACGCTCTTCCGATCTNNNGGGAAAAGGTGTTTCAGGAAC
AGAGCACACGTCTGAACTCTTTCCCTACACGACGCTCTTCCGATCTNNNGGATTGCTTGAGCTCAGGAT
AGAGCACACGTCTGAACTCTTTCCCTACACGACGCTCTTCCGATCTNNNGGATTGCTTGAGCTCAGGAT
AGAGCACACGTCTGAACTCTTTCCCTACACGACGCTCTTCCGATCTNNNGGATTGCTTGAGCTCAGGAT
AGAGCACACGTCTGAACTCTTTCCCTACACGACGCTCTTCCGATCTNNNGGATTGCTTGAGCTCAGGAT
AGAGCACACGTCTGAACTCTTTCCCTACACGACGCTCTTCCGATCTNNNGGATTGCTTGAGCTCAGGAT
AGCACACGTCTGAACTCTTTCCCTACACGACGCTCTTCCGATCTNNNAACTGCCTTCCAACTCAAAACAC
AGCACACGTCTGAACTCTTTCCCTACACGACGCTCTTCCGATCTNNNAACTGCCTTCCAACTCAAAACAC
AGCACACGTCTGAACTCTTTCCCTACACGACGCTCTTCCGATCTNNNAACTGCCTTCCAACTCAAAACAC
AGCACACGTCTGAACTCTTTCCCTACACGACGCTCTTCCGATCTNNNAACTGCCTTCCAACTCAAAACAC
AGCACACGTCTGAACTCTTTCCCTACACGACGCTCTTCCGATCTNNNAACTGCCTTCCAACTCAAAACAC
GAGCACACGTCTGAACTCTTTCCCTACACGACGCTCTTCCGATCTNNNGAGGCTGAGGTGAGAGAATC
GAGCACACGTCTGAACTCTTTCCCTACACGACGCTCTTCCGATCTNNNGAGGCTGAGGTGAGAGAATC
GAGCACACGTCTGAACTCTTTCCCTACACGACGCTCTTCCGATCTNNNGAGGCTGAGGTGAGAGAATC
GAGCACACGTCTGAACTCTTTCCCTACACGACGCTCTTCCGATCTNNNGAGGCTGAGGTGAGAGAATC
GAGCACACGTCTGAACTCTTTCCCTACACGACGCTCTTCCGATCTNNNGAGGCTGAGGTGAGAGAATC
GAGCACACGTCTGAACTCTTTCCCTACACGACGCTCTTCCGATCTNNNGGAAGTAGTGTGTGCTTGGA
GAGCACACGTCTGAACTCTTTCCCTACACGACGCTCTTCCGATCTNNNGGAAGTAGTGTGTGCTTGGA
GAGCACACGTCTGAACTCTTTCCCTACACGACGCTCTTCCGATCTNNNGGAAGTAGTGTGTGCTTGGA
GAGCACACGTCTGAACTCTTTCCCTACACGACGCTCTTCCGATCTNNNGGAAGTAGTGTGTGCTTGGA
GAGCACACGTCTGAACTCTTTCCCTACACGACGCTCTTCCGATCTNNNGGAAGTAGTGTGTGCTTGGA
AGAGCACACGTCTGAACTCTTTCCCTACACGACGCTCTTCCGATCTNNNCATGCACACATTCACACAGA
AGAGCACACGTCTGAACTCTTTCCCTACACGACGCTCTTCCGATCTNNNCATGCACACATTCACACAGA
AGAGCACACGTCTGAACTCTTTCCCTACACGACGCTCTTCCGATCTNNNCATGCACACATTCACACAGA

AGAGCACACGTCTGAACTCTTTCCCTACACGACGCTCTTCCGATCTNNNCATGCACACATTCACACAGA

AGAGCACACGTCTGAACTCTTTCCCTACACGACGCTCTTCCGATCTNNNCATGCACACATTCACACAGA

AGAGCACACGTCTGAACTCTTTCCCTACACGACGCTCTTCCGATCTNNNCTGATCCCATCCTAGGCTAC

AGAGCACACGTCTGAACTCTTTCCCTACACGACGCTCTTCCGATCTNNNCTGATCCCATCCTAGGCTAC

AGAGCACACGTCTGAACTCTTTCCCTACACGACGCTCTTCCGATCTNNNCTGATCCCATCCTAGGCTAC

AGAGCACACGTCTGAACTCTTTCCCTACACGACGCTCTTCCGATCTNNNCTGATCCCATCCTAGGCTAC

AGAGCACACGTCTGAACTCTTTCCCTACACGACGCTCTTCCGATCTNNNCTGATCCCATCCTAGGCTAC

AGCACACGTCTGAACTCTTTCCCTACACGACGCTCTTCCGATCTNNNTGAGGGAGGAGAATCACTTG

AGCACACGTCTGAACTCTTTCCCTACACGACGCTCTTCCGATCTNNNTGAGGGAGGAGAATCACTTG

AGCACACGTCTGAACTCTTTCCCTACACGACGCTCTTCCGATCTNNNTGAGGGAGGAGAATCACTTG

AGCACACGTCTGAACTCTTTCCCTACACGACGCTCTTCCGATCTNNNTGAGGGAGGAGAATCACTTG

AGCACACGTCTGAACTCTTTCCCTACACGACGCTCTTCCGATCTNNNTGAGGGAGGAGAATCACTTG

GCACACGTCTGAACTCTTTCCCTACACGACGCTCTTCCGATCTNNNATCTAGGACAGCTTTGGCAAGA

GCACACGTCTGAACTCTTTCCCTACACGACGCTCTTCCGATCTNNNATCTAGGACAGCTTTGGCAAGA

GCACACGTCTGAACTCTTTCCCTACACGACGCTCTTCCGATCTNNNATCTAGGACAGCTTTGGCAAGA

GCACACGTCTGAACTCTTTCCCTACACGACGCTCTTCCGATCTNNNATCTAGGACAGCTTTGGCAAGA

GCACACGTCTGAACTCTTTCCCTACACGACGCTCTTCCGATCTNNNATCTAGGACAGCTTTGGCAAGA

GCACACGTCTGAACTCTTTCCCTACACGACGCTCTTCCGATCTNNNCTCCCTCTTTTTCTGTCTCCCAC

GCACACGTCTGAACTCTTTCCCTACACGACGCTCTTCCGATCTNNNCTCCCTCTTTTTCTGTCTCCCAC

GCACACGTCTGAACTCTTTCCCTACACGACGCTCTTCCGATCTNNNCTCCCTCTTTTTCTGTCTCCCAC

GCACACGTCTGAACTCTTTCCCTACACGACGCTCTTCCGATCTNNNCTCCCTCTTTTTCTGTCTCCCAC

GCACACGTCTGAACTCTTTCCCTACACGACGCTCTTCCGATCTNNNCTCCCTCTTTTTCTGTCTCCCAC

GAGCACACGTCTGAACTCTTTCCCTACACGACGCTCTTCCGATCTNNNCACTCAACAGGTTGCATTAG

GAGCACACGTCTGAACTCTTTCCCTACACGACGCTCTTCCGATCTNNNCACTCAACAGGTTGCATTAG

GAGCACACGTCTGAACTCTTTCCCTACACGACGCTCTTCCGATCTNNNCACTCAACAGGTTGCATTAG

GAGCACACGTCTGAACTCTTTCCCTACACGACGCTCTTCCGATCTNNNCACTCAACAGGTTGCATTAG

GAGCACACGTCTGAACTCTTTCCCTACACGACGCTCTTCCGATCTNNNCACTCAACAGGTTGCATTAG

GAGCACACGTCTGAACTCTTTCCCTACACGACGCTCTTCCGATCTNNNCCTATATATGTGGAGGGTGG

GAGCACACGTCTGAACTCTTTCCCTACACGACGCTCTTCCGATCTNNNCCTATATATGTGGAGGGTGG

GAGCACACGTCTGAACTCTTTCCCTACACGACGCTCTTCCGATCTNNNCCTATATATGTGGAGGGTGG

GAGCACACGTCTGAACTCTTTCCCTACACGACGCTCTTCCGATCTNNNCCTATATATGTGGAGGGTGG

GAGCACACGTCTGAACTCTTTCCCTACACGACGCTCTTCCGATCTNNNCCTATATATGTGGAGGGTGG

GCACACGTCTGAACTCTTTCCCTACACGACGCTCTTCCGATCTNNNTTTCTCCCAGTCTGTAGCTTGTC

GCACACGTCTGAACTCTTTCCCTACACGACGCTCTTCCGATCTNNNTTTCTCCCAGTCTGTAGCTTGTC

GCACACGTCTGAACTCTTTCCCTACACGACGCTCTTCCGATCTNNNTTTCTCCCAGTCTGTAGCTTGTC

GCACACGTCTGAACTCTTTCCCTACACGACGCTCTTCCGATCTNNNTTTCTCCCAGTCTGTAGCTTGTC

GCACACGTCTGAACTCTTTCCCTACACGACGCTCTTCCGATCTNNNTTTCTCCCAGTCTGTAGCTTGTC

GCACACGTCTGAACTCTTTCCCTACACGACGCTCTTCCGATCTNNNGGATCACACAGCGTGTAGTCTT

GCACACGTCTGAACTCTTTCCCTACACGACGCTCTTCCGATCTNNNGGATCACACAGCGTGTAGTCTT

GCACACGTCTGAACTCTTTCCCTACACGACGCTCTTCCGATCTNNNGGATCACACAGCGTGTAGTCTT

GCACACGTCTGAACTCTTTCCCTACACGACGCTCTTCCGATCTNNNGGATCACACAGCGTGTAGTCTT

GCACACGTCTGAACTCTTTCCCTACACGACGCTCTTCCGATCTNNNGGATCACACAGCGTGTAGTCTT

ACACGTCTGAACTCTTTCCCTACACGACGCTCTTCCGATCTNNNCTGCCTTTGTGAGATCAGAGGAG

CACACGTCTGAACTCTTTCCCTACACGACGCTCTTCCGATCTNNNCTGCCTTTGTGAGATCAGAGGAGC
CACACGTCTGAACTCTTTCCCTACACGACGCTCTTCCGATCTNNNCTGCCTTTGTGAGATCAGAGGAGC
CACACGTCTGAACTCTTTCCCTACACGACGCTCTTCCGATCTNNNCTGCCTTTGTGAGATCAGAGGAGC
CACACGTCTGAACTCTTTCCCTACACGACGCTCTTCCGATCTNNNCTGCCTTTGTGAGATCAGAGGAGC
GAGCACACGTCTGAACTCTTTCCCTACACGACGCTCTTCCGATCTNNNTGTCTCATAAACACACCAGCTG
GAGCACACGTCTGAACTCTTTCCCTACACGACGCTCTTCCGATCTNNNTGTCTCATAAACACACCAGCTG
GAGCACACGTCTGAACTCTTTCCCTACACGACGCTCTTCCGATCTNNNTGTCTCATAAACACACCAGCTG
GAGCACACGTCTGAACTCTTTCCCTACACGACGCTCTTCCGATCTNNNTGTCTCATAAACACACCAGCTG
GAGCACACGTCTGAACTCTTTCCCTACACGACGCTCTTCCGATCTNNNTGTCTCATAAACACACCAGCTG
AGCACACGTCTGAACTCTTTCCCTACACGACGCTCTTCCGATCTNNNAGGGTCTTACCATATTGCCCATG
AGCACACGTCTGAACTCTTTCCCTACACGACGCTCTTCCGATCTNNNAGGGTCTTACCATATTGCCCATG
AGCACACGTCTGAACTCTTTCCCTACACGACGCTCTTCCGATCTNNNAGGGTCTTACCATATTGCCCATG
AGCACACGTCTGAACTCTTTCCCTACACGACGCTCTTCCGATCTNNNAGGGTCTTACCATATTGCCCATG
AGCACACGTCTGAACTCTTTCCCTACACGACGCTCTTCCGATCTNNNAGGGTCTTACCATATTGCCCATG
GAGCACACGTCTGAACTCTTTCCCTACACGACGCTCTTCCGATCTNNNCTACTACACTCTAGCCTGGGTG
GAGCACACGTCTGAACTCTTTCCCTACACGACGCTCTTCCGATCTNNNCTACTACACTCTAGCCTGGGTG
GAGCACACGTCTGAACTCTTTCCCTACACGACGCTCTTCCGATCTNNNCTACTACACTCTAGCCTGGGTG
GAGCACACGTCTGAACTCTTTCCCTACACGACGCTCTTCCGATCTNNNCTACTACACTCTAGCCTGGGTG
GAGCACACGTCTGAACTCTTTCCCTACACGACGCTCTTCCGATCTNNNCTACTACACTCTAGCCTGGGTG
AGCACACGTCTGAACTCTTTCCCTACACGACGCTCTTCCGATCTNNNGAGAAGCTTTTCCCTTACCTCGA
AGCACACGTCTGAACTCTTTCCCTACACGACGCTCTTCCGATCTNNNGAGAAGCTTTTCCCTTACCTCGA
AGCACACGTCTGAACTCTTTCCCTACACGACGCTCTTCCGATCTNNNGAGAAGCTTTTCCCTTACCTCGA
AGCACACGTCTGAACTCTTTCCCTACACGACGCTCTTCCGATCTNNNGAGAAGCTTTTCCCTTACCTCGA
AGCACACGTCTGAACTCTTTCCCTACACGACGCTCTTCCGATCTNNNGAGAAGCTTTTCCCTTACCTCGA
GAGCACACGTCTGAACTCTTTCCCTACACGACGCTCTTCCGATCTNNNAGGAAAACGGGTATGAAGGA
GAGCACACGTCTGAACTCTTTCCCTACACGACGCTCTTCCGATCTNNNAGGAAAACGGGTATGAAGGA
GAGCACACGTCTGAACTCTTTCCCTACACGACGCTCTTCCGATCTNNNAGGAAAACGGGTATGAAGGA
GAGCACACGTCTGAACTCTTTCCCTACACGACGCTCTTCCGATCTNNNAGGAAAACGGGTATGAAGGA
GAGCACACGTCTGAACTCTTTCCCTACACGACGCTCTTCCGATCTNNNAGGAAAACGGGTATGAAGGA
AGAGCACACGTCTGAACTCTTTCCCTACACGACGCTCTTCCGATCTNNNAGTGCTAGGATTACAGGCAT
AGAGCACACGTCTGAACTCTTTCCCTACACGACGCTCTTCCGATCTNNNAGTGCTAGGATTACAGGCAT
AGAGCACACGTCTGAACTCTTTCCCTACACGACGCTCTTCCGATCTNNNAGTGCTAGGATTACAGGCAT
AGAGCACACGTCTGAACTCTTTCCCTACACGACGCTCTTCCGATCTNNNAGTGCTAGGATTACAGGCAT
AGAGCACACGTCTGAACTCTTTCCCTACACGACGCTCTTCCGATCTNNNAGTGCTAGGATTACAGGCAT
GAGCACACGTCTGAACTCTTTCCCTACACGACGCTCTTCCGATCTNNNGGAGTTTACATTCCATTTTATG
GAGCACACGTCTGAACTCTTTCCCTACACGACGCTCTTCCGATCTNNNGGAGTTTACATTCCATTTTATG
GAGCACACGTCTGAACTCTTTCCCTACACGACGCTCTTCCGATCTNNNGGAGTTTACATTCCATTTTATG
GAGCACACGTCTGAACTCTTTCCCTACACGACGCTCTTCCGATCTNNNGGAGTTTACATTCCATTTTATG
GAGCACACGTCTGAACTCTTTCCCTACACGACGCTCTTCCGATCTNNNGGAGTTTACATTCCATTTTATG
GCACACGTCTGAACTCTTTCCCTACACGACGCTCTTCCGATCTNNNATACAGCACCTCACTGCTCTTGCA
GCACACGTCTGAACTCTTTCCCTACACGACGCTCTTCCGATCTNNNATACAGCACCTCACTGCTCTTGCA
GCACACGTCTGAACTCTTTCCCTACACGACGCTCTTCCGATCTNNNATACAGCACCTCACTGCTCTTGCA
GCACACGTCTGAACTCTTTCCCTACACGACGCTCTTCCGATCTNNNATACAGCACCTCACTGCTCTTGCA

GCACACGTCTGAACTCTTTCCCTACACGACGCTCTTCCGATCTNNNATACAGCACCTCACTGCTCTTGCA
AGCACACGTCTGAACTCTTTCCCTACACGACGCTCTTCCGATCTNNNCATCATCCGCAATGACAATGAC
AGCACACGTCTGAACTCTTTCCCTACACGACGCTCTTCCGATCTNNNCATCATCCGCAATGACAATGAC
AGCACACGTCTGAACTCTTTCCCTACACGACGCTCTTCCGATCTNNNCATCATCCGCAATGACAATGAC
AGCACACGTCTGAACTCTTTCCCTACACGACGCTCTTCCGATCTNNNCATCATCCGCAATGACAATGAC
AGCACACGTCTGAACTCTTTCCCTACACGACGCTCTTCCGATCTNNNCATCATCCGCAATGACAATGAC
AGCACACGTCTGAACTCTTTCCCTACACGACGCTCTTCCGATCTNNNGCTCTGTCAAACTGTGTAGAGG
AGCACACGTCTGAACTCTTTCCCTACACGACGCTCTTCCGATCTNNNGCTCTGTCAAACTGTGTAGAGG
AGCACACGTCTGAACTCTTTCCCTACACGACGCTCTTCCGATCTNNNGCTCTGTCAAACTGTGTAGAGG
AGCACACGTCTGAACTCTTTCCCTACACGACGCTCTTCCGATCTNNNGCTCTGTCAAACTGTGTAGAGG
AGCACACGTCTGAACTCTTTCCCTACACGACGCTCTTCCGATCTNNNGCTCTGTCAAACTGTGTAGAGG
AGCACACGTCTGAACTCTTTCCCTACACGACGCTCTTCCGATCTNNNCCAAAGTGCTGGGATTACAGT
AGCACACGTCTGAACTCTTTCCCTACACGACGCTCTTCCGATCTNNNCCAAAGTGCTGGGATTACAGT
AGCACACGTCTGAACTCTTTCCCTACACGACGCTCTTCCGATCTNNNCCAAAGTGCTGGGATTACAGT
AGCACACGTCTGAACTCTTTCCCTACACGACGCTCTTCCGATCTNNNCCAAAGTGCTGGGATTACAGT
AGCACACGTCTGAACTCTTTCCCTACACGACGCTCTTCCGATCTNNNCCAAAGTGCTGGGATTACAGT
AGCACACGTCTGAACTCTTTCCCTACACGACGCTCTTCCGATCTNNNGGGCTCTTGTTTGGTTCCATAT
AGCACACGTCTGAACTCTTTCCCTACACGACGCTCTTCCGATCTNNNGGGCTCTTGTTTGGTTCCATAT
AGCACACGTCTGAACTCTTTCCCTACACGACGCTCTTCCGATCTNNNGGGCTCTTGTTTGGTTCCATAT
AGCACACGTCTGAACTCTTTCCCTACACGACGCTCTTCCGATCTNNNGGGCTCTTGTTTGGTTCCATAT
AGCACACGTCTGAACTCTTTCCCTACACGACGCTCTTCCGATCTNNNGGGCTCTTGTTTGGTTCCATAT
AGCACACGTCTGAACTCTTTCCCTACACGACGCTCTTCCGATCTNNNAAAGAGTGTCCATTCCCCACTTT
AGCACACGTCTGAACTCTTTCCCTACACGACGCTCTTCCGATCTNNNAAAGAGTGTCCATTCCCCACTTT
AGCACACGTCTGAACTCTTTCCCTACACGACGCTCTTCCGATCTNNNAAAGAGTGTCCATTCCCCACTTT
AGCACACGTCTGAACTCTTTCCCTACACGACGCTCTTCCGATCTNNNAAAGAGTGTCCATTCCCCACTTT
AGCACACGTCTGAACTCTTTCCCTACACGACGCTCTTCCGATCTNNNAAAGAGTGTCCATTCCCCACTTT
GAGCACACGTCTGAACTCTTTCCCTACACGACGCTCTTCCGATCTNNNGGATGACAGAGTGAGACTCC
GAGCACACGTCTGAACTCTTTCCCTACACGACGCTCTTCCGATCTNNNGGATGACAGAGTGAGACTCC
GAGCACACGTCTGAACTCTTTCCCTACACGACGCTCTTCCGATCTNNNGGATGACAGAGTGAGACTCC
GAGCACACGTCTGAACTCTTTCCCTACACGACGCTCTTCCGATCTNNNGGATGACAGAGTGAGACTCC
GAGCACACGTCTGAACTCTTTCCCTACACGACGCTCTTCCGATCTNNNGGATGACAGAGTGAGACTCC
GAGCACACGTCTGAACTCTTTCCCTACACGACGCTCTTCCGATCTNNNATTTCTGCAGCTGGCTTGAATT
GAGCACACGTCTGAACTCTTTCCCTACACGACGCTCTTCCGATCTNNNATTTCTGCAGCTGGCTTGAATT
GAGCACACGTCTGAACTCTTTCCCTACACGACGCTCTTCCGATCTNNNATTTCTGCAGCTGGCTTGAATT
GAGCACACGTCTGAACTCTTTCCCTACACGACGCTCTTCCGATCTNNNATTTCTGCAGCTGGCTTGAATT
GAGCACACGTCTGAACTCTTTCCCTACACGACGCTCTTCCGATCTNNNATTTCTGCAGCTGGCTTGAATT
AGCACACGTCTGAACTCTTTCCCTACACGACGCTCTTCCGATCTNNNCCCTTCTTCATACACTGCTCCATC
AGCACACGTCTGAACTCTTTCCCTACACGACGCTCTTCCGATCTNNNCCCTTCTTCATACACTGCTCCATC
AGCACACGTCTGAACTCTTTCCCTACACGACGCTCTTCCGATCTNNNCCCTTCTTCATACACTGCTCCATC
AGCACACGTCTGAACTCTTTCCCTACACGACGCTCTTCCGATCTNNNCCCTTCTTCATACACTGCTCCATC
AGCACACGTCTGAACTCTTTCCCTACACGACGCTCTTCCGATCTNNNCCCTTCTTCATACACTGCTCCATC
AGCACACGTCTGAACTCTTTCCCTACACGACGCTCTTCCGATCTNNNCTGCCTCTAGTTCCCATGACTTT
AGCACACGTCTGAACTCTTTCCCTACACGACGCTCTTCCGATCTNNNCTGCCTCTAGTTCCCATGACTTT

AGCACACGTCTGAACTCTTTCCCTACACGACGCTCTTCCGATCTNNNCTGCCTCTAGTTCCCATGACTTT
AGCACACGTCTGAACTCTTTCCCTACACGACGCTCTTCCGATCTNNNCTGCCTCTAGTTCCCATGACTTT
AGCACACGTCTGAACTCTTTCCCTACACGACGCTCTTCCGATCTNNNCTGCCTCTAGTTCCCATGACTTT
AGCACACGTCTGAACTCTTTCCCTACACGACGCTCTTCCGATCTNNNTCACTTTCCTTCTGGATGTTCC
AGCACACGTCTGAACTCTTTCCCTACACGACGCTCTTCCGATCTNNNTCACTTTCCTTCTGGATGTTCC
AGCACACGTCTGAACTCTTTCCCTACACGACGCTCTTCCGATCTNNNTCACTTTCCTTCTGGATGTTCC
AGCACACGTCTGAACTCTTTCCCTACACGACGCTCTTCCGATCTNNNTCACTTTCCTTCTGGATGTTCC
AGCACACGTCTGAACTCTTTCCCTACACGACGCTCTTCCGATCTNNNTCACTTTCCTTCTGGATGTTCC
GCACACGTCTGAACTCTTTCCCTACACGACGCTCTTCCGATCTNNNTGTCTGTCTCTCTCTGCTTGTTCC
GCACACGTCTGAACTCTTTCCCTACACGACGCTCTTCCGATCTNNNTGTCTGTCTCTCTCTGCTTGTTCC
GCACACGTCTGAACTCTTTCCCTACACGACGCTCTTCCGATCTNNNTGTCTGTCTCTCTCTGCTTGTTCC
GCACACGTCTGAACTCTTTCCCTACACGACGCTCTTCCGATCTNNNTGTCTGTCTCTCTCTGCTTGTTCC
GCACACGTCTGAACTCTTTCCCTACACGACGCTCTTCCGATCTNNNTGTCTGTCTCTCTCTGCTTGTTCC
GAGCACACGTCTGAACTCTTTCCCTACACGACGCTCTTCCGATCTNNNGAGGAGACGAGCAGGATCA
GAGCACACGTCTGAACTCTTTCCCTACACGACGCTCTTCCGATCTNNNGAGGAGACGAGCAGGATCA
GAGCACACGTCTGAACTCTTTCCCTACACGACGCTCTTCCGATCTNNNGAGGAGACGAGCAGGATCA
GAGCACACGTCTGAACTCTTTCCCTACACGACGCTCTTCCGATCTNNNGAGGAGACGAGCAGGATCA
GAGCACACGTCTGAACTCTTTCCCTACACGACGCTCTTCCGATCTNNNGAGGAGACGAGCAGGATCA
GAGCACACGTCTGAACTCTTTCCCTACACGACGCTCTTCCGATCTNNNATAACGCAAGGAAGGGAGA
GAGCACACGTCTGAACTCTTTCCCTACACGACGCTCTTCCGATCTNNNATAACGCAAGGAAGGGAGA
GAGCACACGTCTGAACTCTTTCCCTACACGACGCTCTTCCGATCTNNNATAACGCAAGGAAGGGAGA
GAGCACACGTCTGAACTCTTTCCCTACACGACGCTCTTCCGATCTNNNATAACGCAAGGAAGGGAGA
GAGCACACGTCTGAACTCTTTCCCTACACGACGCTCTTCCGATCTNNNATAACGCAAGGAAGGGAGA
AGCACACGTCTGAACTCTTTCCCTACACGACGCTCTTCCGATCTNNNACCCAGAGAGGAGTAGTACCTT
AGCACACGTCTGAACTCTTTCCCTACACGACGCTCTTCCGATCTNNNACCCAGAGAGGAGTAGTACCTT
AGCACACGTCTGAACTCTTTCCCTACACGACGCTCTTCCGATCTNNNACCCAGAGAGGAGTAGTACCTT
AGCACACGTCTGAACTCTTTCCCTACACGACGCTCTTCCGATCTNNNACCCAGAGAGGAGTAGTACCTT
AGCACACGTCTGAACTCTTTCCCTACACGACGCTCTTCCGATCTNNNACCCAGAGAGGAGTAGTACCTT
AGCACACGTCTGAACTCTTTCCCTACACGACGCTCTTCCGATCTNNNCTGAGACACGAGGATCACTTG
AGCACACGTCTGAACTCTTTCCCTACACGACGCTCTTCCGATCTNNNCTGAGACACGAGGATCACTTG
AGCACACGTCTGAACTCTTTCCCTACACGACGCTCTTCCGATCTNNNCTGAGACACGAGGATCACTTG
AGCACACGTCTGAACTCTTTCCCTACACGACGCTCTTCCGATCTNNNCTGAGACACGAGGATCACTTG
AGCACACGTCTGAACTCTTTCCCTACACGACGCTCTTCCGATCTNNNCTGAGACACGAGGATCACTTG
AGCACACGTCTGAACTCTTTCCCTACACGACGCTCTTCCGATCTNNNGGTGGGTCCAGTCATAAGGAT
AGCACACGTCTGAACTCTTTCCCTACACGACGCTCTTCCGATCTNNNGGTGGGTCCAGTCATAAGGAT
AGCACACGTCTGAACTCTTTCCCTACACGACGCTCTTCCGATCTNNNGGTGGGTCCAGTCATAAGGAT
AGCACACGTCTGAACTCTTTCCCTACACGACGCTCTTCCGATCTNNNGGTGGGTCCAGTCATAAGGAT
AGCACACGTCTGAACTCTTTCCCTACACGACGCTCTTCCGATCTNNNGGTGGGTCCAGTCATAAGGAT
GAGCACACGTCTGAACTCTTTCCCTACACGACGCTCTTCCGATCTNNNGGGCGACAGAGACTCTTGTC
GAGCACACGTCTGAACTCTTTCCCTACACGACGCTCTTCCGATCTNNNGGGCGACAGAGACTCTTGTC
GAGCACACGTCTGAACTCTTTCCCTACACGACGCTCTTCCGATCTNNNGGGCGACAGAGACTCTTGTC
GAGCACACGTCTGAACTCTTTCCCTACACGACGCTCTTCCGATCTNNNGGGCGACAGAGACTCTTGTC
GAGCACACGTCTGAACTCTTTCCCTACACGACGCTCTTCCGATCTNNNGGGCGACAGAGACTCTTGTC

AGAGCACACGTCTGAACTCTTTCCCTACACGACGCTCTTCCGATCTNNNTACGTATAGCTTGCCACCTT
AGAGCACACGTCTGAACTCTTTCCCTACACGACGCTCTTCCGATCTNNNTACGTATAGCTTGCCACCTT
AGAGCACACGTCTGAACTCTTTCCCTACACGACGCTCTTCCGATCTNNNTACGTATAGCTTGCCACCTT
AGAGCACACGTCTGAACTCTTTCCCTACACGACGCTCTTCCGATCTNNNTACGTATAGCTTGCCACCTT
AGAGCACACGTCTGAACTCTTTCCCTACACGACGCTCTTCCGATCTNNNTACGTATAGCTTGCCACCTT
AGCACACGTCTGAACTCTTTCCCTACACGACGCTCTTCCGATCTNNNTTGCAAACAGGGGAGATACA
AGCACACGTCTGAACTCTTTCCCTACACGACGCTCTTCCGATCTNNNTTGCAAACAGGGGAGATACA
AGCACACGTCTGAACTCTTTCCCTACACGACGCTCTTCCGATCTNNNTTGCAAACAGGGGAGATACA
AGCACACGTCTGAACTCTTTCCCTACACGACGCTCTTCCGATCTNNNTTGCAAACAGGGGAGATACA
AGCACACGTCTGAACTCTTTCCCTACACGACGCTCTTCCGATCTNNNTTGCAAACAGGGGAGATACA
CACACGTCTGAACTCTTTCCCTACACGACGCTCTTCCGATCTNNNTTGGATGGTAGCTGAGATGACA
CACACGTCTGAACTCTTTCCCTACACGACGCTCTTCCGATCTNNNTTGGATGGTAGCTGAGATGACA
CACACGTCTGAACTCTTTCCCTACACGACGCTCTTCCGATCTNNNTTGGATGGTAGCTGAGATGACA
CACACGTCTGAACTCTTTCCCTACACGACGCTCTTCCGATCTNNNTTGGATGGTAGCTGAGATGACA
CACACGTCTGAACTCTTTCCCTACACGACGCTCTTCCGATCTNNNTTGGATGGTAGCTGAGATGACA
GCACACGTCTGAACTCTTTCCCTACACGACGCTCTTCCGATCTNNNATTATATCAGAATAGCCTCTAGC
GCACACGTCTGAACTCTTTCCCTACACGACGCTCTTCCGATCTNNNATTATATCAGAATAGCCTCTAGC
GCACACGTCTGAACTCTTTCCCTACACGACGCTCTTCCGATCTNNNATTATATCAGAATAGCCTCTAGC
GCACACGTCTGAACTCTTTCCCTACACGACGCTCTTCCGATCTNNNATTATATCAGAATAGCCTCTAGC
GCACACGTCTGAACTCTTTCCCTACACGACGCTCTTCCGATCTNNNATTATATCAGAATAGCCTCTAGC
AGCACACGTCTGAACTCTTTCCCTACACGACGCTCTTCCGATCTNNNGCGCCTGTAATCCTAGTTACTC
AGCACACGTCTGAACTCTTTCCCTACACGACGCTCTTCCGATCTNNNGCGCCTGTAATCCTAGTTACTC
AGCACACGTCTGAACTCTTTCCCTACACGACGCTCTTCCGATCTNNNGCGCCTGTAATCCTAGTTACTC
AGCACACGTCTGAACTCTTTCCCTACACGACGCTCTTCCGATCTNNNGCGCCTGTAATCCTAGTTACTC
AGCACACGTCTGAACTCTTTCCCTACACGACGCTCTTCCGATCTNNNGCGCCTGTAATCCTAGTTACTC
AGAGCACACGTCTGAACTCTTTCCCTACACGACGCTCTTCCGATCTNNNTCCCCAATTATATGCTTATAT
AGAGCACACGTCTGAACTCTTTCCCTACACGACGCTCTTCCGATCTNNNTCCCCAATTATATGCTTATAT
AGAGCACACGTCTGAACTCTTTCCCTACACGACGCTCTTCCGATCTNNNTCCCCAATTATATGCTTATAT
AGAGCACACGTCTGAACTCTTTCCCTACACGACGCTCTTCCGATCTNNNTCCCCAATTATATGCTTATAT
AGAGCACACGTCTGAACTCTTTCCCTACACGACGCTCTTCCGATCTNNNTCCCCAATTATATGCTTATAT
GCACACGTCTGAACTCTTTCCCTACACGACGCTCTTCCGATCTNNNACTACTGACCTCTCCAGGAATCA
GCACACGTCTGAACTCTTTCCCTACACGACGCTCTTCCGATCTNNNACTACTGACCTCTCCAGGAATCA
GCACACGTCTGAACTCTTTCCCTACACGACGCTCTTCCGATCTNNNACTACTGACCTCTCCAGGAATCA
GCACACGTCTGAACTCTTTCCCTACACGACGCTCTTCCGATCTNNNACTACTGACCTCTCCAGGAATCA
GCACACGTCTGAACTCTTTCCCTACACGACGCTCTTCCGATCTNNNACTACTGACCTCTCCAGGAATCA
CACACGTCTGAACTCTTTCCCTACACGACGCTCTTCCGATCTNNNTTTAGTAGAGACGGGGTTTCACC
CACACGTCTGAACTCTTTCCCTACACGACGCTCTTCCGATCTNNNTTTAGTAGAGACGGGGTTTCACC
CACACGTCTGAACTCTTTCCCTACACGACGCTCTTCCGATCTNNNTTTAGTAGAGACGGGGTTTCACC
CACACGTCTGAACTCTTTCCCTACACGACGCTCTTCCGATCTNNNTTTAGTAGAGACGGGGTTTCACC
CACACGTCTGAACTCTTTCCCTACACGACGCTCTTCCGATCTNNNTTTAGTAGAGACGGGGTTTCACC
GAGCACACGTCTGAACTCTTTCCCTACACGACGCTCTTCCGATCTNNNAATGGCTTGATAGAGGAGTT
GAGCACACGTCTGAACTCTTTCCCTACACGACGCTCTTCCGATCTNNNAATGGCTTGATAGAGGAGTT
GAGCACACGTCTGAACTCTTTCCCTACACGACGCTCTTCCGATCTNNNAATGGCTTGATAGAGGAGTT

GAGCACACGTCTGAACTCTTTCCCTACACGACGCTCTTCCGATCTNNNAATGGCTTGATAGAGGAGTT
GAGCACACGTCTGAACTCTTTCCCTACACGACGCTCTTCCGATCTNNNAATGGCTTGATAGAGGAGTT
AGCACACGTCTGAACTCTTTCCCTACACGACGCTCTTCCGATCTNNNAGGCTGAGGCAGAATTACTTG
AGCACACGTCTGAACTCTTTCCCTACACGACGCTCTTCCGATCTNNNAGGCTGAGGCAGAATTACTTG
AGCACACGTCTGAACTCTTTCCCTACACGACGCTCTTCCGATCTNNNAGGCTGAGGCAGAATTACTTG
AGCACACGTCTGAACTCTTTCCCTACACGACGCTCTTCCGATCTNNNAGGCTGAGGCAGAATTACTTG
AGCACACGTCTGAACTCTTTCCCTACACGACGCTCTTCCGATCTNNNAGGCTGAGGCAGAATTACTTG
GCACACGTCTGAACTCTTTCCCTACACGACGCTCTTCCGATCTNNNACAAACAACCCCATCCAAAAGTG
GCACACGTCTGAACTCTTTCCCTACACGACGCTCTTCCGATCTNNNACAAACAACCCCATCCAAAAGTG
GCACACGTCTGAACTCTTTCCCTACACGACGCTCTTCCGATCTNNNACAAACAACCCCATCCAAAAGTG
GCACACGTCTGAACTCTTTCCCTACACGACGCTCTTCCGATCTNNNACAAACAACCCCATCCAAAAGTG
GCACACGTCTGAACTCTTTCCCTACACGACGCTCTTCCGATCTNNNACAAACAACCCCATCCAAAAGTG
GCACACGTCTGAACTCTTTCCCTACACGACGCTCTTCCGATCTNNNTGTGTGACTTCCTTTACTCAGAA
GCACACGTCTGAACTCTTTCCCTACACGACGCTCTTCCGATCTNNNTGTGTGACTTCCTTTACTCAGAA
GCACACGTCTGAACTCTTTCCCTACACGACGCTCTTCCGATCTNNNTGTGTGACTTCCTTTACTCAGAA
GCACACGTCTGAACTCTTTCCCTACACGACGCTCTTCCGATCTNNNTGTGTGACTTCCTTTACTCAGAA
GCACACGTCTGAACTCTTTCCCTACACGACGCTCTTCCGATCTNNNTGTGTGACTTCCTTTACTCAGAA
GCACACGTCTGAACTCTTTCCCTACACGACGCTCTTCCGATCTNNNGTGATGGAGCAAGACTCTGTCT
GCACACGTCTGAACTCTTTCCCTACACGACGCTCTTCCGATCTNNNGTGATGGAGCAAGACTCTGTCT
GCACACGTCTGAACTCTTTCCCTACACGACGCTCTTCCGATCTNNNGTGATGGAGCAAGACTCTGTCT
GCACACGTCTGAACTCTTTCCCTACACGACGCTCTTCCGATCTNNNGTGATGGAGCAAGACTCTGTCT
GCACACGTCTGAACTCTTTCCCTACACGACGCTCTTCCGATCTNNNGTGATGGAGCAAGACTCTGTCT
GAGCACACGTCTGAACTCTTTCCCTACACGACGCTCTTCCGATCTNNNCGCGTGGGTTACATTTTGAG
GAGCACACGTCTGAACTCTTTCCCTACACGACGCTCTTCCGATCTNNNCGCGTGGGTTACATTTTGAG
GAGCACACGTCTGAACTCTTTCCCTACACGACGCTCTTCCGATCTNNNCGCGTGGGTTACATTTTGAG
GAGCACACGTCTGAACTCTTTCCCTACACGACGCTCTTCCGATCTNNNCGCGTGGGTTACATTTTGAG
GAGCACACGTCTGAACTCTTTCCCTACACGACGCTCTTCCGATCTNNNCGCGTGGGTTACATTTTGAG
AGCACACGTCTGAACTCTTTCCCTACACGACGCTCTTCCGATCTNNNTTGTTATGGGGCACAACACTGT
AGCACACGTCTGAACTCTTTCCCTACACGACGCTCTTCCGATCTNNNTTGTTATGGGGCACAACACTGT
AGCACACGTCTGAACTCTTTCCCTACACGACGCTCTTCCGATCTNNNTTGTTATGGGGCACAACACTGT
AGCACACGTCTGAACTCTTTCCCTACACGACGCTCTTCCGATCTNNNTTGTTATGGGGCACAACACTGT
AGCACACGTCTGAACTCTTTCCCTACACGACGCTCTTCCGATCTNNNTTGTTATGGGGCACAACACTGT
AGCACACGTCTGAACTCTTTCCCTACACGACGCTCTTCCGATCTNNNGCCTGTAGTCCCAGCTACTTGC
AGCACACGTCTGAACTCTTTCCCTACACGACGCTCTTCCGATCTNNNGCCTGTAGTCCCAGCTACTTGC
AGCACACGTCTGAACTCTTTCCCTACACGACGCTCTTCCGATCTNNNGCCTGTAGTCCCAGCTACTTGC
AGCACACGTCTGAACTCTTTCCCTACACGACGCTCTTCCGATCTNNNGCCTGTAGTCCCAGCTACTTGC
AGCACACGTCTGAACTCTTTCCCTACACGACGCTCTTCCGATCTNNNGCCTGTAGTCCCAGCTACTTGC
ACACGTCTGAACTCTTTCCCTACACGACGCTCTTCCGATCTNNNGTGCTAGTCTCTGCTATGAGTCCCA
ACACGTCTGAACTCTTTCCCTACACGACGCTCTTCCGATCTNNNGTGCTAGTCTCTGCTATGAGTCCCA
ACACGTCTGAACTCTTTCCCTACACGACGCTCTTCCGATCTNNNGTGCTAGTCTCTGCTATGAGTCCCA
ACACGTCTGAACTCTTTCCCTACACGACGCTCTTCCGATCTNNNGTGCTAGTCTCTGCTATGAGTCCCA
ACACGTCTGAACTCTTTCCCTACACGACGCTCTTCCGATCTNNNGTGCTAGTCTCTGCTATGAGTCCCA
AGAGCACACGTCTGAACTCTTTCCCTACACGACGCTCTTCCGATCTNNNATCAAGCTGGAGGGAAAA

AGAGCACACGTCTGAACTCTTTCCCTACACGACGCTCTTCCGATCTNNNATCAAGCTGGAGGGAAAAA
AGAGCACACGTCTGAACTCTTTCCCTACACGACGCTCTTCCGATCTNNNATCAAGCTGGAGGGAAAAA
AGAGCACACGTCTGAACTCTTTCCCTACACGACGCTCTTCCGATCTNNNATCAAGCTGGAGGGAAAAA
AGAGCACACGTCTGAACTCTTTCCCTACACGACGCTCTTCCGATCTNNNATCAAGCTGGAGGGAAAAA
AGCACACGTCTGAACTCTTTCCCTACACGACGCTCTTCCGATCTNNNGTGTCTGCTTCTTAGGAGTCTCC
AGCACACGTCTGAACTCTTTCCCTACACGACGCTCTTCCGATCTNNNGTGTCTGCTTCTTAGGAGTCTCC
AGCACACGTCTGAACTCTTTCCCTACACGACGCTCTTCCGATCTNNNGTGTCTGCTTCTTAGGAGTCTCC
AGCACACGTCTGAACTCTTTCCCTACACGACGCTCTTCCGATCTNNNGTGTCTGCTTCTTAGGAGTCTCC
AGCACACGTCTGAACTCTTTCCCTACACGACGCTCTTCCGATCTNNNGTGTCTGCTTCTTAGGAGTCTCC
CACACGTCTGAACTCTTTCCCTACACGACGCTCTTCCGATCTNNNTACCTGGCTTCAAAGTGACAGAAC
CACACGTCTGAACTCTTTCCCTACACGACGCTCTTCCGATCTNNNTACCTGGCTTCAAAGTGACAGAAC
CACACGTCTGAACTCTTTCCCTACACGACGCTCTTCCGATCTNNNTACCTGGCTTCAAAGTGACAGAAC
CACACGTCTGAACTCTTTCCCTACACGACGCTCTTCCGATCTNNNTACCTGGCTTCAAAGTGACAGAAC
CACACGTCTGAACTCTTTCCCTACACGACGCTCTTCCGATCTNNNTACCTGGCTTCAAAGTGACAGAAC
GAGCACACGTCTGAACTCTTTCCCTACACGACGCTCTTCCGATCTNNNTTCCTTCCTTCCTTCCTTCCTTC
GAGCACACGTCTGAACTCTTTCCCTACACGACGCTCTTCCGATCTNNNTTCCTTCCTTCCTTCCTTCCTTC
GAGCACACGTCTGAACTCTTTCCCTACACGACGCTCTTCCGATCTNNNTTCCTTCCTTCCTTCCTTCCTTC
GAGCACACGTCTGAACTCTTTCCCTACACGACGCTCTTCCGATCTNNNTTCCTTCCTTCCTTCCTTCCTTC
GAGCACACGTCTGAACTCTTTCCCTACACGACGCTCTTCCGATCTNNNTTCCTTCCTTCCTTCCTTCCTTC
AGCACACGTCTGAACTCTTTCCCTACACGACGCTCTTCCGATCTNNNACCATGGTGTATTGATGTCCTGA
AGCACACGTCTGAACTCTTTCCCTACACGACGCTCTTCCGATCTNNNACCATGGTGTATTGATGTCCTGA
AGCACACGTCTGAACTCTTTCCCTACACGACGCTCTTCCGATCTNNNACCATGGTGTATTGATGTCCTGA
AGCACACGTCTGAACTCTTTCCCTACACGACGCTCTTCCGATCTNNNACCATGGTGTATTGATGTCCTGA
AGCACACGTCTGAACTCTTTCCCTACACGACGCTCTTCCGATCTNNNACCATGGTGTATTGATGTCCTGA
AGCACACGTCTGAACTCTTTCCCTACACGACGCTCTTCCGATCTNNNAGAGAGAGAGAGAGAGAAAG
AGCACACGTCTGAACTCTTTCCCTACACGACGCTCTTCCGATCTNNNAGAGAGAGAGAGAGAGAAAG
AGCACACGTCTGAACTCTTTCCCTACACGACGCTCTTCCGATCTNNNAGAGAGAGAGAGAGAGAAAG
AGCACACGTCTGAACTCTTTCCCTACACGACGCTCTTCCGATCTNNNAGAGAGAGAGAGAGAGAAAG
AGCACACGTCTGAACTCTTTCCCTACACGACGCTCTTCCGATCTNNNAGAGAGAGAGAGAGAGAAAG
GCACACGTCTGAACTCTTTCCCTACACGACGCTCTTCCGATCTNNNCCCATAAGCTCTACTTGGCTACACA
GCACACGTCTGAACTCTTTCCCTACACGACGCTCTTCCGATCTNNNCCCATAAGCTCTACTTGGCTACACA
GCACACGTCTGAACTCTTTCCCTACACGACGCTCTTCCGATCTNNNCCCATAAGCTCTACTTGGCTACACA
GCACACGTCTGAACTCTTTCCCTACACGACGCTCTTCCGATCTNNNCCCATAAGCTCTACTTGGCTACACA
GCACACGTCTGAACTCTTTCCCTACACGACGCTCTTCCGATCTNNNCCCATAAGCTCTACTTGGCTACACA
GAGCACACGTCTGAACTCTTTCCCTACACGACGCTCTTCCGATCTNNNAGATAATACCAGAGGCCAGCA
GAGCACACGTCTGAACTCTTTCCCTACACGACGCTCTTCCGATCTNNNAGATAATACCAGAGGCCAGCA
GAGCACACGTCTGAACTCTTTCCCTACACGACGCTCTTCCGATCTNNNAGATAATACCAGAGGCCAGCA
GAGCACACGTCTGAACTCTTTCCCTACACGACGCTCTTCCGATCTNNNAGATAATACCAGAGGCCAGCA
GAGCACACGTCTGAACTCTTTCCCTACACGACGCTCTTCCGATCTNNNAGATAATACCAGAGGCCAGCA
AGCACACGTCTGAACTCTTTCCCTACACGACGCTCTTCCGATCTNNNTGTCATTATTCAGTGCATGGCTC
AGCACACGTCTGAACTCTTTCCCTACACGACGCTCTTCCGATCTNNNTGTCATTATTCAGTGCATGGCTC
AGCACACGTCTGAACTCTTTCCCTACACGACGCTCTTCCGATCTNNNTGTCATTATTCAGTGCATGGCTC
AGCACACGTCTGAACTCTTTCCCTACACGACGCTCTTCCGATCTNNNTGTCATTATTCAGTGCATGGCTC

AGCACACGTCTGAACTCTTTCCCTACACGACGCTCTTCCGATCTNNNTGTCATTATTCAGTGCATGGCT
AGCACACGTCTGAACTCTTTCCCTACACGACGCTCTTCCGATCTNNNGCCTGTAGTCCCAGCTACTTGCA
AGCACACGTCTGAACTCTTTCCCTACACGACGCTCTTCCGATCTNNNGCCTGTAGTCCCAGCTACTTGCA
AGCACACGTCTGAACTCTTTCCCTACACGACGCTCTTCCGATCTNNNGCCTGTAGTCCCAGCTACTTGCA
AGCACACGTCTGAACTCTTTCCCTACACGACGCTCTTCCGATCTNNNGCCTGTAGTCCCAGCTACTTGCA
AGCACACGTCTGAACTCTTTCCCTACACGACGCTCTTCCGATCTNNNGCCTGTAGTCCCAGCTACTTGCA
AGCACACGTCTGAACTCTTTCCCTACACGACGCTCTTCCGATCTNNNCTGAGGCAGGAGAATTGCTTG
AGCACACGTCTGAACTCTTTCCCTACACGACGCTCTTCCGATCTNNNCTGAGGCAGGAGAATTGCTTG
AGCACACGTCTGAACTCTTTCCCTACACGACGCTCTTCCGATCTNNNCTGAGGCAGGAGAATTGCTTG
AGCACACGTCTGAACTCTTTCCCTACACGACGCTCTTCCGATCTNNNCTGAGGCAGGAGAATTGCTTG
AGCACACGTCTGAACTCTTTCCCTACACGACGCTCTTCCGATCTNNNCTGAGGCAGGAGAATTGCTTG
GAGCACACGTCTGAACTCTTTCCCTACACGACGCTCTTCCGATCTNNNAGAAGAGAGTGAGGATGACA
GAGCACACGTCTGAACTCTTTCCCTACACGACGCTCTTCCGATCTNNNAGAAGAGAGTGAGGATGACA
GAGCACACGTCTGAACTCTTTCCCTACACGACGCTCTTCCGATCTNNNAGAAGAGAGTGAGGATGACA
GAGCACACGTCTGAACTCTTTCCCTACACGACGCTCTTCCGATCTNNNAGAAGAGAGTGAGGATGACA
GAGCACACGTCTGAACTCTTTCCCTACACGACGCTCTTCCGATCTNNNAGAAGAGAGTGAGGATGACA
AGCACACGTCTGAACTCTTTCCCTACACGACGCTCTTCCGATCTNNNATAAGCAATACTGGGAACACG
AGCACACGTCTGAACTCTTTCCCTACACGACGCTCTTCCGATCTNNNATAAGCAATACTGGGAACACG
AGCACACGTCTGAACTCTTTCCCTACACGACGCTCTTCCGATCTNNNATAAGCAATACTGGGAACACG
AGCACACGTCTGAACTCTTTCCCTACACGACGCTCTTCCGATCTNNNATAAGCAATACTGGGAACACG
AGCACACGTCTGAACTCTTTCCCTACACGACGCTCTTCCGATCTNNNATAAGCAATACTGGGAACACG
GAGCACACGTCTGAACTCTTTCCCTACACGACGCTCTTCCGATCTNNNCATCTGTAGTCCCAGCTACTCA
GAGCACACGTCTGAACTCTTTCCCTACACGACGCTCTTCCGATCTNNNCATCTGTAGTCCCAGCTACTCA
GAGCACACGTCTGAACTCTTTCCCTACACGACGCTCTTCCGATCTNNNCATCTGTAGTCCCAGCTACTCA
GAGCACACGTCTGAACTCTTTCCCTACACGACGCTCTTCCGATCTNNNCATCTGTAGTCCCAGCTACTCA
GAGCACACGTCTGAACTCTTTCCCTACACGACGCTCTTCCGATCTNNNCATCTGTAGTCCCAGCTACTCA
GAGCACACGTCTGAACTCTTTCCCTACACGACGCTCTTCCGATCTNNNAATACAAAACTTAGCTGGGC
GAGCACACGTCTGAACTCTTTCCCTACACGACGCTCTTCCGATCTNNNAATACAAAACTTAGCTGGGC
GAGCACACGTCTGAACTCTTTCCCTACACGACGCTCTTCCGATCTNNNAATACAAAACTTAGCTGGGC
GAGCACACGTCTGAACTCTTTCCCTACACGACGCTCTTCCGATCTNNNAATACAAAACTTAGCTGGGC
GAGCACACGTCTGAACTCTTTCCCTACACGACGCTCTTCCGATCTNNNAATACAAAACTTAGCTGGGC
AGCACACGTCTGAACTCTTTCCCTACACGACGCTCTTCCGATCTNNNTCTGATGGCCTTTTCATACCACA
AGCACACGTCTGAACTCTTTCCCTACACGACGCTCTTCCGATCTNNNTCTGATGGCCTTTTCATACCACA
AGCACACGTCTGAACTCTTTCCCTACACGACGCTCTTCCGATCTNNNTCTGATGGCCTTTTCATACCACA
AGCACACGTCTGAACTCTTTCCCTACACGACGCTCTTCCGATCTNNNTCTGATGGCCTTTTCATACCACA
AGCACACGTCTGAACTCTTTCCCTACACGACGCTCTTCCGATCTNNNTCTGATGGCCTTTTCATACCACA
GCACACGTCTGAACTCTTTCCCTACACGACGCTCTTCCGATCTNNNTACCCATACACCATTGCAGTGTTC
GCACACGTCTGAACTCTTTCCCTACACGACGCTCTTCCGATCTNNNTACCCATACACCATTGCAGTGTTC
GCACACGTCTGAACTCTTTCCCTACACGACGCTCTTCCGATCTNNNTACCCATACACCATTGCAGTGTTC
GCACACGTCTGAACTCTTTCCCTACACGACGCTCTTCCGATCTNNNTACCCATACACCATTGCAGTGTTC
GCACACGTCTGAACTCTTTCCCTACACGACGCTCTTCCGATCTNNNTACCCATACACCATTGCAGTGTTC
GAGCACACGTCTGAACTCTTTCCCTACACGACGCTCTTCCGATCTNNNCTGTGATCATGCCACTACACTC
GAGCACACGTCTGAACTCTTTCCCTACACGACGCTCTTCCGATCTNNNCTGTGATCATGCCACTACACTC

AGCACACGTCTGAACTCTTTCCCTACACGACGCTCTTCCGATCTNNNCTGTGATCATGCCACTACACTC
AGCACACGTCTGAACTCTTTCCCTACACGACGCTCTTCCGATCTNNNCTGTGATCATGCCACTACACTC
AGCACACGTCTGAACTCTTTCCCTACACGACGCTCTTCCGATCTNNNCTGTGATCATGCCACTACACTC
AGCACACGTCTGAACTCTTTCCCTACACGACGCTCTTCCGATCTNNNGCAGAAGAATGGCATGAATCC
AGCACACGTCTGAACTCTTTCCCTACACGACGCTCTTCCGATCTNNNGCAGAAGAATGGCATGAATCC
AGCACACGTCTGAACTCTTTCCCTACACGACGCTCTTCCGATCTNNNGCAGAAGAATGGCATGAATCC
AGCACACGTCTGAACTCTTTCCCTACACGACGCTCTTCCGATCTNNNGCAGAAGAATGGCATGAATCC
AGCACACGTCTGAACTCTTTCCCTACACGACGCTCTTCCGATCTNNNGCAGAAGAATGGCATGAATCC
AGAGCACACGTCTGAACTCTTTCCCTACACGACGCTCTTCCGATCTNNNACCAAGGAATAGAGAATAA
AGCACACGTCTGAACTCTTTCCCTACACGACGCTCTTCCGATCTNNNTTTCTTGGTTAGTCTAATTAGT
GAGCACACGTCTGAACTCTTTCCCTACACGACGCTCTTCCGATCTNNNAGAAAGGATTGAGCATTCAT
GAGCACACGTCTGAACTCTTTCCCTACACGACGCTCTTCCGATCTNNNACTCTAGTTTTAGTCTAATCCC
AGCACACGTCTGAACTCTTTCCCTACACGACGCTCTTCCGATCTNNNTCCAAAAATATGATTTGAGTTT
GAGCACACGTCTGAACTCTTTCCCTACACGACGCTCTTCCGATCTNNNTCAAAGGAACAATCTAAGGTT
AAGAGCACACGTCTGAACTCTTTCCCTACACGACGCTCTTCCGATCTNNNACTTGACTGCATGTTTGA
GAGCACACGTCTGAACTCTTTCCCTACACGACGCTCTTCCGATCTNNNGGGTAAAATATTGACTCTCA
GAGCACACGTCTGAACTCTTTCCCTACACGACGCTCTTCCGATCTNNNGGGATATAATCACAAACTAAT
GAAGAGCACACGTCTGAACTCTTTCCCTACACGACGCTCTTCCGATCTNNNACTTAGATGTTCACGTA
AAGAGCACACGTCTGAACTCTTTCCCTACACGACGCTCTTCCGATCTNNNGGGCCTGAATCAACATAG
AGAGCACACGTCTGAACTCTTTCCCTACACGACGCTCTTCCGATCTNNNTCCTTTTCAAATGGTGAAC
AGCACACGTCTGAACTCTTTCCCTACACGACGCTCTTCCGATCTNNNATCAAGGAAATAAAAGAGGAC
GAAGAGCACACGTCTGAACTCTTTCCCTACACGACGCTCTTCCGATCTNNNCTGTCATACAGGAAGAG
GAGCACACGTCTGAACTCTTTCCCTACACGACGCTCTTCCGATCTNNNTGTATACACAAAACTGTATAC
GAGCACACGTCTGAACTCTTTCCCTACACGACGCTCTTCCGATCTNNNATCTCCCAGGTCCAAGCAATT
AGCACACGTCTGAACTCTTTCCCTACACGACGCTCTTCCGATCTNNNGAAGGATAAAATAAAAGTTGG
AAGAGCACACGTCTGAACTCTTTCCCTACACGACGCTCTTCCGATCTNNNTTCTCTCTTCTCCTGCTCTC
AGCACACGTCTGAACTCTTTCCCTACACGACGCTCTTCCGATCTNNNAGTCTCTTCTACTTCAAATAAA
GAGCACACGTCTGAACTCTTTCCCTACACGACGCTCTTCCGATCTNNNCATCATGTAGAACACAATGTA
GAAGAGCACACGTCTGAACTCTTTCCCTACACGACGCTCTTCCGATCTNNNACACTCAAGTCCAGAGG
GAAGAGCACACGTCTGAACTCTTTCCCTACACGACGCTCTTCCGATCTNNNATAAAGCCCTGTGCAGG
GAGCACACGTCTGAACTCTTTCCCTACACGACGCTCTTCCGATCTNNNAAACATGTACTTACTCTAAAG
GAAGAGCACACGTCTGAACTCTTTCCCTACACGACGCTCTTCCGATCTNNNGTTCCTCTCAAAAGTCTT
GAGCACACGTCTGAACTCTTTCCCTACACGACGCTCTTCCGATCTNNNCCAATTTGGCACTCTGTAGAT
GAAGAGCACACGTCTGAACTCTTTCCCTACACGACGCTCTTCCGATCTNNNCCTTGTTCTAACATTGGG
GAGCACACGTCTGAACTCTTTCCCTACACGACGCTCTTCCGATCTNNNCAAAAACGATATATCAGAGA
AAGAGCACACGTCTGAACTCTTTCCCTACACGACGCTCTTCCGATCTNNNTGGGATTCATTCATATTG
AGAGCACACGTCTGAACTCTTTCCCTACACGACGCTCTTCCGATCTNNNTGTGGTGTATTTGACACTAT
GAAGAGCACACGTCTGAACTCTTTCCCTACACGACGCTCTTCCGATCTNNNAACCCAGATCCAAATGCC
AGCACACGTCTGAACTCTTTCCCTACACGACGCTCTTCCGATCTNNNACTCTCAGGTACTAAGAAGAA
GAAGAGCACACGTCTGAACTCTTTCCCTACACGACGCTCTTCCGATCTNNNTATCTACCATTTCAGCAG
AGCACACGTCTGAACTCTTTCCCTACACGACGCTCTTCCGATCTNNNTCTTTTTCTTCTTTCTCTCAAAT
GAAGAGCACACGTCTGAACTCTTTCCCTACACGACGCTCTTCCGATCTNNNAGATACCGTCTGTTTGTT
AAGAGCACACGTCTGAACTCTTTCCCTACACGACGCTCTTCCGATCTNNNTCCTCCTACCTCTCTCTCCT

AAGAGCACACGTCTGAACTCTTTCCCTACACGACGCTCTTCCGATCTNNNTGAGGTTGTATAATGCAT
AGAGCACACGTCTGAACTCTTTCCCTACACGACGCTCTTCCGATCTNNNTGAGGAGAAGAAAGCTAT
AGCACACGTCTGAACTCTTTCCCTACACGACGCTCTTCCGATCTNNNACACTTACATAAGTTGTGTGG
GAGCACACGTCTGAACTCTTTCCCTACACGACGCTCTTCCGATCTNNNACTAGATGTCTAATAAGAGG
GAAGAGCACACGTCTGAACTCTTTCCCTACACGACGCTCTTCCGATCTNNNGGAGGGCCACCTTGAT
AAGAGCACACGTCTGAACTCTTTCCCTACACGACGCTCTTCCGATCTNNNGGTAACAAGAAACACTCC
GAAGAGCACACGTCTGAACTCTTTCCCTACACGACGCTCTTCCGATCTNNNTCAATTTCATCTACCTGCC
AGAGCACACGTCTGAACTCTTTCCCTACACGACGCTCTTCCGATCTNNNTGGGTTGATAGACCCTTAG
AGAGCACACGTCTGAACTCTTTCCCTACACGACGCTCTTCCGATCTNNNCCAAAGTCATCCTTATCTGT
AGAGCACACGTCTGAACTCTTTCCCTACACGACGCTCTTCCGATCTNNNCTTCTCCATAACCAGAGAGG
AGAGCACACGTCTGAACTCTTTCCCTACACGACGCTCTTCCGATCTNNNAGTGTTTCTCTCTCTCTCTCT
AGAGCACACGTCTGAACTCTTTCCCTACACGACGCTCTTCCGATCTNNNACAACAATTTTGTTTCCTCCT
AAGAGCACACGTCTGAACTCTTTCCCTACACGACGCTCTTCCGATCTNNNCGCAATTGTTCCTTAGTGT
AGAGCACACGTCTGAACTCTTTCCCTACACGACGCTCTTCCGATCTNNNTGGTGTACGAAAAACTTG
AAGAGCACACGTCTGAACTCTTTCCCTACACGACGCTCTTCCGATCTNNNTTTTCCCTTCCTTCCTTTG
GAAGAGCACACGTCTGAACTCTTTCCCTACACGACGCTCTTCCGATCTNNNGCCTGTACCTGTCAACTT
AGCACACGTCTGAACTCTTTCCCTACACGACGCTCTTCCGATCTNNNATCAATATATACATGCACACAC
AAGAGCACACGTCTGAACTCTTTCCCTACACGACGCTCTTCCGATCTNNNGGCCTCTGATTATCTTCTC
AAGAGCACACGTCTGAACTCTTTCCCTACACGACGCTCTTCCGATCTNNNGTTATTACTGGCTGTTGC
AAGAGCACACGTCTGAACTCTTTCCCTACACGACGCTCTTCCGATCTNNNCTTCGCACAGAGAGAAAG
GAGCACACGTCTGAACTCTTTCCCTACACGACGCTCTTCCGATCTNNNTGGTACATACCATGACCTATA
GAAGAGCACACGTCTGAACTCTTTCCCTACACGACGCTCTTCCGATCTNNNACAGGCTCAACCTAATTG
GAAGAGCACACGTCTGAACTCTTTCCCTACACGACGCTCTTCCGATCTNNNATACGCCTGTAGTCCCTA
AAGAGCACACGTCTGAACTCTTTCCCTACACGACGCTCTTCCGATCTNNNACTAAGTGCTAAGTCCCA
GAAGAGCACACGTCTGAACTCTTTCCCTACACGACGCTCTTCCGATCTNNNAAACGCCTTTCAGGAAC
AAGAGCACACGTCTGAACTCTTTCCCTACACGACGCTCTTCCGATCTNNNAAAACCCAAATGCTGTCG
GAAGAGCACACGTCTGAACTCTTTCCCTACACGACGCTCTTCCGATCTNNNACTACCCAATTGTTCCAA
GAGCACACGTCTGAACTCTTTCCCTACACGACGCTCTTCCGATCTNNNTCTGTTTTTGAGTTACAGCAG
GAAGAGCACACGTCTGAACTCTTTCCCTACACGACGCTCTTCCGATCTNNNCAACAAAAATGGACCCA
AAGAGCACACGTCTGAACTCTTTCCCTACACGACGCTCTTCCGATCTNNNACACTCTCCAATTCCATCC
AGCACACGTCTGAACTCTTTCCCTACACGACGCTCTTCCGATCTNNNTGTGTTAGGAGGAAAAAGAT
AAGAGCACACGTCTGAACTCTTTCCCTACACGACGCTCTTCCGATCTNNNCATGGTGTCATGGTGTTTT
GGAAGAGCACACGTCTGAACTCTTTCCCTACACGACGCTCTTCCGATCTNNNAAAGACATGTGGAAGC
AAGAGCACACGTCTGAACTCTTTCCCTACACGACGCTCTTCCGATCTNNNACACACAGCTTTTCCCAAA
AAGAGCACACGTCTGAACTCTTTCCCTACACGACGCTCTTCCGATCTNNNATGATCGTACACTACACT
AAGAGCACACGTCTGAACTCTTTCCCTACACGACGCTCTTCCGATCTNNNGGAGCTGCAGGGTAAAA
GAAGAGCACACGTCTGAACTCTTTCCCTACACGACGCTCTTCCGATCTNNNAGCTACTTCTTTGTGTAT
AGAGCACACGTCTGAACTCTTTCCCTACACGACGCTCTTCCGATCTNNNAGTGAGAACACTTATTAGT
GAAGAGCACACGTCTGAACTCTTTCCCTACACGACGCTCTTCCGATCTNNNGACAAGACCTATAGCGC
GAAGAGCACACGTCTGAACTCTTTCCCTACACGACGCTCTTCCGATCTNNNCCTTCACAGATTCCAGG
GAAGAGCACACGTCTGAACTCTTTCCCTACACGACGCTCTTCCGATCTNNNTGCACAATCCTTTCAAAT
AAGAGCACACGTCTGAACTCTTTCCCTACACGACGCTCTTCCGATCTNNNTCCTGATACTCTGCCAACT
AGAGCACACGTCTGAACTCTTTCCCTACACGACGCTCTTCCGATCTNNNAGGAAGGCCTTTGTCTTTCT

GAGCACACGTCTGAACTCTTTCCCTACACGACGCTCTTCCGATCTNNNTCTTCACATTGAGCATATGTAT
GGAAGAGCACACGTCTGAACTCTTTCCCTACACGACGCTCTTCCGATCTNNNAAGTTTCCTAGGAGGG
GAAGAGCACACGTCTGAACTCTTTCCCTACACGACGCTCTTCCGATCTNNNGGAGATCCTAGTCTTTA
GAAGAGCACACGTCTGAACTCTTTCCCTACACGACGCTCTTCCGATCTNNNCCATCCAGGTAATTGTGA
GGAAGAGCACACGTCTGAACTCTTTCCCTACACGACGCTCTTCCGATCTNNNATACATGATGTGGCCCT
GGAAGAGCACACGTCTGAACTCTTTCCCTACACGACGCTCTTCCGATCTNNNTGAGTGAGATTTTTCAC
AGCACACGTCTGAACTCTTTCCCTACACGACGCTCTTCCGATCTNNNACATTGTTCACAGTAGCCAAGA
AGCACACGTCTGAACTCTTTCCCTACACGACGCTCTTCCGATCTNNNAGTAAGGACCTACTCTTGTCAT
AAGAGCACACGTCTGAACTCTTTCCCTACACGACGCTCTTCCGATCTNNNGGCAGCTTTACTCTCCTTT
AGAGCACACGTCTGAACTCTTTCCCTACACGACGCTCTTCCGATCTNNNACACACACACACAGATCTAT
GAAGAGCACACGTCTGAACTCTTTCCCTACACGACGCTCTTCCGATCTNNNAATGCCAAAGTCTGGATG
GAGCACACGTCTGAACTCTTTCCCTACACGACGCTCTTCCGATCTNNNGCTATTATAGGTCCTTTTTACT
AGAGCACACGTCTGAACTCTTTCCCTACACGACGCTCTTCCGATCTNNNTCAGGAATATTGGCCTATA
AGAGCACACGTCTGAACTCTTTCCCTACACGACGCTCTTCCGATCTNNNTTGTATACAGGTTGCGCTCT
AGAGCACACGTCTGAACTCTTTCCCTACACGACGCTCTTCCGATCTNNNAGGTTTTCATGTTTCCAAGC
GAAGAGCACACGTCTGAACTCTTTCCCTACACGACGCTCTTCCGATCTNNNTGAACAACAACTGATCC
GAAGAGCACACGTCTGAACTCTTTCCCTACACGACGCTCTTCCGATCTNNNCAACGTTGACTAAAACAT
GAGCACACGTCTGAACTCTTTCCCTACACGACGCTCTTCCGATCTNNNCTCATGGTGGCTCACACTTAT
GAAGAGCACACGTCTGAACTCTTTCCCTACACGACGCTCTTCCGATCTNNNTTTAGTGGATCTGTAC
GAAGAGCACACGTCTGAACTCTTTCCCTACACGACGCTCTTCCGATCTNNNATGTATATCTCTGTGCGT
AAGAGCACACGTCTGAACTCTTTCCCTACACGACGCTCTTCCGATCTNNNCCTGGCCCACTAATGTACT
GAAGAGCACACGTCTGAACTCTTTCCCTACACGACGCTCTTCCGATCTNNNGCAGAGCTCTCACGGTT
GGAAGAGCACACGTCTGAACTCTTTCCCTACACGACGCTCTTCCGATCTNNNTCCAGGACTACTCTTGG
AGAGCACACGTCTGAACTCTTTCCCTACACGACGCTCTTCCGATCTNNNTCTCCATTGATTCTTGATTC
GAAGAGCACACGTCTGAACTCTTTCCCTACACGACGCTCTTCCGATCTNNNGTGGAAGAGGTACCCCA
GAAGAGCACACGTCTGAACTCTTTCCCTACACGACGCTCTTCCGATCTNNNATGTCATTGGCTTTCTG
GAAGAGCACACGTCTGAACTCTTTCCCTACACGACGCTCTTCCGATCTNNNATGCTGGTTTGCCTTCTG
GAGCACACGTCTGAACTCTTTCCCTACACGACGCTCTTCCGATCTNNNAGTCCAACAGGAAAATATCA
AGAGCACACGTCTGAACTCTTTCCCTACACGACGCTCTTCCGATCTNNNCTGGTCCATTTGCAATCAAT
AAGAGCACACGTCTGAACTCTTTCCCTACACGACGCTCTTCCGATCTNNNGACCATCCTCTTTAGCTCA
AAGAGCACACGTCTGAACTCTTTCCCTACACGACGCTCTTCCGATCTNNNCAGAGATAGTGGGGCAG
GGAAGAGCACACGTCTGAACTCTTTCCCTACACGACGCTCTTCCGATCTNNNACCCTGAAGAAGCATA
AAGAGCACACGTCTGAACTCTTTCCCTACACGACGCTCTTCCGATCTNNNGCTCTCAGAGTGGTGATT
AAGAGCACACGTCTGAACTCTTTCCCTACACGACGCTCTTCCGATCTNNNCTTGGGTCCCATGGAGGA
GAAGAGCACACGTCTGAACTCTTTCCCTACACGACGCTCTTCCGATCTNNNTTGTTGTTGAAACCATGA
GAAGAGCACACGTCTGAACTCTTTCCCTACACGACGCTCTTCCGATCTNNNTCATGTAGTCACACACAT
GAGCACACGTCTGAACTCTTTCCCTACACGACGCTCTTCCGATCTNNNATTCAATAGATATATATGCGT
AAGAGCACACGTCTGAACTCTTTCCCTACACGACGCTCTTCCGATCTNNNGCCTTCTCTGTCTTTCTGTC
GAAGAGCACACGTCTGAACTCTTTCCCTACACGACGCTCTTCCGATCTNNNATCATTCTGCGCTTACAGA
AGCACACGTCTGAACTCTTTCCCTACACGACGCTCTTCCGATCTNNNTGCAATAAATTCCAATCTATCG
AGCACACGTCTGAACTCTTTCCCTACACGACGCTCTTCCGATCTNNNTGGAGGAGAATTTTCCAAAAA
AGCACACGTCTGAACTCTTTCCCTACACGACGCTCTTCCGATCTNNNTGTATAAACATATAGTGTCTGT
GAAGAGCACACGTCTGAACTCTTTCCCTACACGACGCTCTTCCGATCTNNNTCGTATATGTGTGTTCAT

GAAGAGCACACGTCTGAACTCTTTCCCTACACGACGCTCTTCCGATCTNNNCTTCCTTTGGGAAGGG

AAGAGCACACGTCTGAACTCTTTCCCTACACGACGCTCTTCCGATCTNNNGGCTACTTGCCAGTCTTCA

GAAGAGCACACGTCTGAACTCTTTCCCTACACGACGCTCTTCCGATCTNNNCAGGGGAAATTTTCAA

GAGCACACGTCTGAACTCTTTCCCTACACGACGCTCTTCCGATCTNNNTGGTGTTTTAGCTTTCCTCTT

AAGAGCACACGTCTGAACTCTTTCCCTACACGACGCTCTTCCGATCTNNNAGAATTTCAAAGTCGGGA

GGAAGAGCACACGTCTGAACTCTTTCCCTACACGACGCTCTTCCGATCTNNNTGGGTCCCAGAAGAAA

GCACACGTCTGAACTCTTTCCCTACACGACGCTCTTCCGATCTNNNTCATCTTTTGGAAATGGAATAA

AGCACACGTCTGAACTCTTTCCCTACACGACGCTCTTCCGATCTNNNTTATACTTACATACTATTCCCTG

GAAGAGCACACGTCTGAACTCTTTCCCTACACGACGCTCTTCCGATCTNNNAGCAAGGAATAGGCAG

AAGAGCACACGTCTGAACTCTTTCCCTACACGACGCTCTTCCGATCTNNNTTTCTGTCTTTCTGCGTAT

AAGAGCACACGTCTGAACTCTTTCCCTACACGACGCTCTTCCGATCTNNNCGTATTTGCCACTTTCCAG

GAAGAGCACACGTCTGAACTCTTTCCCTACACGACGCTCTTCCGATCTNNNACAGCAGTCCAACTTCTG

GAAGAGCACACGTCTGAACTCTTTCCCTACACGACGCTCTTCCGATCTNNNATTTCCAAAAAGTATGG

GAAGAGCACACGTCTGAACTCTTTCCCTACACGACGCTCTTCCGATCTNNNTGAGGCTGTTCTTCCTG

AGCACACGTCTGAACTCTTTCCCTACACGACGCTCTTCCGATCTNNNAGCAGTAATTTTCACAAATTC

GAAGAGCACACGTCTGAACTCTTTCCCTACACGACGCTCTTCCGATCTNNNTCCTTGATAGCCTTGCCA

AGCACACGTCTGAACTCTTTCCCTACACGACGCTCTTCCGATCTNNNTGAAATTTCCTGATATTTTGG

GAAGAGCACACGTCTGAACTCTTTCCCTACACGACGCTCTTCCGATCTNNNCACTTTTTCCTCCCGCTTT

GAGCACACGTCTGAACTCTTTCCCTACACGACGCTCTTCCGATCTNNNTGTCACAAATTACCACAAAGT

AAGAGCACACGTCTGAACTCTTTCCCTACACGACGCTCTTCCGATCTNNNTCTCAAAAGCCCAACTGAG

GCACACGTCTGAACTCTTTCCCTACACGACGCTCTTCCGATCTNNNCGACATAGATGTAAAAATCCTC

GAAGAGCACACGTCTGAACTCTTTCCCTACACGACGCTCTTCCGATCTNNNGAAGTCTTCCCTGGGTG

GAAGAGCACACGTCTGAACTCTTTCCCTACACGACGCTCTTCCGATCTNNNAGAGTAAGTGTCATGG

GAAGAGCACACGTCTGAACTCTTTCCCTACACGACGCTCTTCCGATCTNNNGTGTGGGTCTGTGCAT

GAGCACACGTCTGAACTCTTTCCCTACACGACGCTCTTCCGATCTNNNTCTGTATACACGTCTCTCTCT

AAGAGCACACGTCTGAACTCTTTCCCTACACGACGCTCTTCCGATCTNNNGGAATGGAAGGGAAATG

GGAAGAGCACACGTCTGAACTCTTTCCCTACACGACGCTCTTCCGATCTNNNACAGACTGCATGGTAT

GCACACGTCTGAACTCTTTCCCTACACGACGCTCTTCCGATCTNNNGCAAATGATAATTTTGACCAGA

AAGAGCACACGTCTGAACTCTTTCCCTACACGACGCTCTTCCGATCTNNNTGCAGTCTTTGAAGAAC

GAGCACACGTCTGAACTCTTTCCCTACACGACGCTCTTCCGATCTNNNTGCTGAGTTACATTTACTCCA

GAAGAGCACACGTCTGAACTCTTTCCCTACACGACGCTCTTCCGATCTNNNTTGCAAACACATGACTT

AAGAGCACACGTCTGAACTCTTTCCCTACACGACGCTCTTCCGATCTNNNTATATGGTGCCCAACGAA

AAGAGCACACGTCTGAACTCTTTCCCTACACGACGCTCTTCCGATCTNNNTTTACCTGCCTTGGCCTAC

AAGAGCACACGTCTGAACTCTTTCCCTACACGACGCTCTTCCGATCTNNNAGCCACTAGGACCAGTA

GGAAGAGCACACGTCTGAACTCTTTCCCTACACGACGCTCTTCCGATCTNNNCATGACTGGCTTGCTTT

AAGAGCACACGTCTGAACTCTTTCCCTACACGACGCTCTTCCGATCTNNNGCATTAGATTCTCACTGG

AAGAGCACACGTCTGAACTCTTTCCCTACACGACGCTCTTCCGATCTNNNTGATTTCTCTGTCTCATC

AAGAGCACACGTCTGAACTCTTTCCCTACACGACGCTCTTCCGATCTNNNCTATACTCATGGCTCTGCA

AAGAGCACACGTCTGAACTCTTTCCCTACACGACGCTCTTCCGATCTNNNTGTCTCCCTTTTCTTAGCA

GAAGAGCACACGTCTGAACTCTTTCCCTACACGACGCTCTTCCGATCTNNNAGACCTAGCCCTCAGAT

AAGAGCACACGTCTGAACTCTTTCCCTACACGACGCTCTTCCGATCTNNNAGGCAAGGATGAAAACA

GAAGAGCACACGTCTGAACTCTTTCCCTACACGACGCTCTTCCGATCTNNNACTGAAAGACTGTCCTT

CACACGTCTGAACTCTTTCCCTACACGACGCTCTTCCGATCTNNNCCACCAACATTTTGATTCATTCTC

GAAGAGCACACGTCTGAACTCTTTCCCTACACGACGCTCTTCCGATCTNNNACACTCCCAGATTTGGG
AAGAGCACACGTCTGAACTCTTTCCCTACACGACGCTCTTCCGATCTNNNTATTTGTCTCGCCCATG
AGCACACGTCTGAACTCTTTCCCTACACGACGCTCTTCCGATCTNNNACACATACATACGTGTAGACA
GAAGAGCACACGTCTGAACTCTTTCCCTACACGACGCTCTTCCGATCTNNNCAGTTCTTAGAGCAAGT
AAGAGCACACGTCTGAACTCTTTCCCTACACGACGCTCTTCCGATCTNNNTATGTGTGGTGCAAGAA
AAGAGCACACGTCTGAACTCTTTCCCTACACGACGCTCTTCCGATCTNNNGTGCTCAGCAAAAGTCA
AGAGCACACGTCTGAACTCTTTCCCTACACGACGCTCTTCCGATCTNNNCCCTCTCTCTCTGGAGATAT
AAGAGCACACGTCTGAACTCTTTCCCTACACGACGCTCTTCCGATCTNNNTGTAGCACAATTCACAATT
AAGAGCACACGTCTGAACTCTTTCCCTACACGACGCTCTTCCGATCTNNNTTTCACTTTGCTTGGGAG
GCACACGTCTGAACTCTTTCCCTACACGACGCTCTTCCGATCTNNNTGACCATACAACTAATCTTGTTC
AGCACACGTCTGAACTCTTTCCCTACACGACGCTCTTCCGATCTNNNTGGATGGATAAACTCATACTA
AGCACACGTCTGAACTCTTTCCCTACACGACGCTCTTCCGATCTNNNTGAAAATCCAAATGTGTATCCC
AAGAGCACACGTCTGAACTCTTTCCCTACACGACGCTCTTCCGATCTNNNAGTACCCATGCTGTTCAA
GAGCACACGTCTGAACTCTTTCCCTACACGACGCTCTTCCGATCTNNNACATGTCACATTTGAGACTA
AAGAGCACACGTCTGAACTCTTTCCCTACACGACGCTCTTCCGATCTNNNTGTCATAGACACCACCACT
GAAGAGCACACGTCTGAACTCTTTCCCTACACGACGCTCTTCCGATCTNNNGGATTTGTGGGAGTTG
AAGAGCACACGTCTGAACTCTTTCCCTACACGACGCTCTTCCGATCTNNNGGTGATGAAGAGGTGCT
AGAGCACACGTCTGAACTCTTTCCCTACACGACGCTCTTCCGATCTNNNTTTGATCAAGTATTGAAATG
AGCACACGTCTGAACTCTTTCCCTACACGACGCTCTTCCGATCTNNNTTTTTCCGTAGGATAACGATT
AGCACACGTCTGAACTCTTTCCCTACACGACGCTCTTCCGATCTNNNATTTGTTATATATGGTGTGTT
GAGCACACGTCTGAACTCTTTCCCTACACGACGCTCTTCCGATCTNNNTGTGTGATTTAGAGATAACC
AGAGCACACGTCTGAACTCTTTCCCTACACGACGCTCTTCCGATCTNNNCCTAAAGGTGTCCTGTTTGA
AAGAGCACACGTCTGAACTCTTTCCCTACACGACGCTCTTCCGATCTNNNGAATCCACGGATGCAGAT
AGCACACGTCTGAACTCTTTCCCTACACGACGCTCTTCCGATCTNNNATAAAAATATGTGAAGCTTGAT
AAGAGCACACGTCTGAACTCTTTCCCTACACGACGCTCTTCCGATCTNNNTGATCCCCAATTCATGTCT
AAGAGCACACGTCTGAACTCTTTCCCTACACGACGCTCTTCCGATCTNNNAAAGCCATTTGCATTCCAT
AGCACACGTCTGAACTCTTTCCCTACACGACGCTCTTCCGATCTNNNAGCATATCTAGTCATAGAAAAG
AGCACACGTCTGAACTCTTTCCCTACACGACGCTCTTCCGATCTNNNAAAATAAGATGTGTATACGTT
AGCACACGTCTGAACTCTTTCCCTACACGACGCTCTTCCGATCTNNNACGCACATACATATATACCTGCT
AGCACACGTCTGAACTCTTTCCCTACACGACGCTCTTCCGATCTNNNGTCCAGAAAAGAAATCATAAA
GCACACGTCTGAACTCTTTCCCTACACGACGCTCTTCCGATCTNNNCCTTTATTCTCATAGGTTGCTAG
AGCACACGTCTGAACTCTTTCCCTACACGACGCTCTTCCGATCTNNNAGTTGAAATTTAGTTTTTCTTC
GAGCACACGTCTGAACTCTTTCCCTACACGACGCTCTTCCGATCTNNNCAGGGCTCAAAGAGAGTAA
AGCACACGTCTGAACTCTTTCCCTACACGACGCTCTTCCGATCTNNNAGAAGACCAATTATTGAAAAAC
AGAGCACACGTCTGAACTCTTTCCCTACACGACGCTCTTCCGATCTNNNACTTGAGCCACTGTGTCTG
AGAGCACACGTCTGAACTCTTTCCCTACACGACGCTCTTCCGATCTNNNTCATCTGGAGATGTTGGCAC
GAGCACACGTCTGAACTCTTTCCCTACACGACGCTCTTCCGATCTNNNTGCACATATTTAGTGTGCTAT
GAAGAGCACACGTCTGAACTCTTTCCCTACACGACGCTCTTCCGATCTNNNAGGTTACGTGAGCTAG
AAGAGCACACGTCTGAACTCTTTCCCTACACGACGCTCTTCCGATCTNNNATGGCCTGATTCAATTCT
AAGAGCACACGTCTGAACTCTTTCCCTACACGACGCTCTTCCGATCTNNNCAAGAGGTGTCTCCTCCT
AAGAGCACACGTCTGAACTCTTTCCCTACACGACGCTCTTCCGATCTNNNAATGCATTCTCTCCCTCTC
GCACACGTCTGAACTCTTTCCCTACACGACGCTCTTCCGATCTNNNACCAAGTGTTTGAATAGAAATA
AGAGCACACGTCTGAACTCTTTCCCTACACGACGCTCTTCCGATCTNNNACTTCCGAGTTTTCTCTCTCT

AGAGCACACGTCTGAACTCTTTCCCTACACGACGCTCTTCCGATCTNNNTAAGCTTACAATGCCAGGG
GAAGAGCACACGTCTGAACTCTTTCCCTACACGACGCTCTTCCGATCTNNNAATAAGAGGTAGCCCCT
AGAGCACACGTCTGAACTCTTTCCCTACACGACGCTCTTCCGATCTNNNTCTCATGGGAACAAGACCA
GAGCACACGTCTGAACTCTTTCCCTACACGACGCTCTTCCGATCTNNNACTTCCCTCTCTCTGTCTTTCT
AGAGCACACGTCTGAACTCTTTCCCTACACGACGCTCTTCCGATCTNNNAATTCAGGTTCTGTAGGTC
AAGAGCACACGTCTGAACTCTTTCCCTACACGACGCTCTTCCGATCTNNNATATCTCAGTTTCTGCGG
GGAAGAGCACACGTCTGAACTCTTTCCCTACACGACGCTCTTCCGATCTNNNAACAAGACAAGGATG
GCACACGTCTGAACTCTTTCCCTACACGACGCTCTTCCGATCTNNNAGGTCTCTTTTATACACACACAC
AGCACACGTCTGAACTCTTTCCCTACACGACGCTCTTCCGATCTNNNTGATGTAACAGTGAATAAAG
AGAGCACACGTCTGAACTCTTTCCCTACACGACGCTCTTCCGATCTNNNAGATCTTTTATCCTGCTGC
AGAGCACACGTCTGAACTCTTTCCCTACACGACGCTCTTCCGATCTNNNAAGATCCTAGAGGGTCAG
AGCACACGTCTGAACTCTTTCCCTACACGACGCTCTTCCGATCTNNNTGGTAAGAAAAGAGAGAAAA
AAGAGCACACGTCTGAACTCTTTCCCTACACGACGCTCTTCCGATCTNNNCTGGCTGACCTAGCCAAAG
AAGAGCACACGTCTGAACTCTTTCCCTACACGACGCTCTTCCGATCTNNNAAATGGGCTCTGTCTCTG
GGAAGAGCACACGTCTGAACTCTTTCCCTACACGACGCTCTTCCGATCTNNNTTGATTGGAGCCTCCT
GAAGAGCACACGTCTGAACTCTTTCCCTACACGACGCTCTTCCGATCTNNNAGAGGTAAAGGTGAAC
AGCACACGTCTGAACTCTTTCCCTACACGACGCTCTTCCGATCTNNNAGAAGAAAATCCTCCAAATGA
GAAGAGCACACGTCTGAACTCTTTCCCTACACGACGCTCTTCCGATCTNNNCTACCTCCAGGGAAGAG
GGAAGAGCACACGTCTGAACTCTTTCCCTACACGACGCTCTTCCGATCTNNNTTGAGCTTCCAAATCAG
GAAGAGCACACGTCTGAACTCTTTCCCTACACGACGCTCTTCCGATCTNNNTGCCACTTTCTAAAAAG
AAGAGCACACGTCTGAACTCTTTCCCTACACGACGCTCTTCCGATCTNNNTCCTGTAATTTGTATTACTC
AGAGCACACGTCTGAACTCTTTCCCTACACGACGCTCTTCCGATCTNNNTCTGTATCAATATTCATGCC
AGCACACGTCTGAACTCTTTCCCTACACGACGCTCTTCCGATCTNNNGAAAATGACTTTCAAACAGAA
AGAGCACACGTCTGAACTCTTTCCCTACACGACGCTCTTCCGATCTNNNTATAGCTCACACTCTCTCTC
AAGAGCACACGTCTGAACTCTTTCCCTACACGACGCTCTTCCGATCTNNNAAAGTGCATTGCTGGTAA
AAGAGCACACGTCTGAACTCTTTCCCTACACGACGCTCTTCCGATCTNNNAGCAAGACCTCATCTCAT
GGAAGAGCACACGTCTGAACTCTTTCCCTACACGACGCTCTTCCGATCTNNNTGCTCCCTACTTCTACCT
AAGAGCACACGTCTGAACTCTTTCCCTACACGACGCTCTTCCGATCTNNNGTGCCCTCATGATGGATT
AAGAGCACACGTCTGAACTCTTTCCCTACACGACGCTCTTCCGATCTNNNCTCTGGTTCCCAGGAGGC
AAGAGCACACGTCTGAACTCTTTCCCTACACGACGCTCTTCCGATCTNNNATGTCATTCTCTCTCTGTCC
AAGAGCACACGTCTGAACTCTTTCCCTACACGACGCTCTTCCGATCTNNNTTATTGGTCCTCTTTGTTC
AAGAGCACACGTCTGAACTCTTTCCCTACACGACGCTCTTCCGATCTNNNGAAGAAACACCATGTTGC
AAGAGCACACGTCTGAACTCTTTCCCTACACGACGCTCTTCCGATCTNNNGAAGGGCTGAGAAACCA
AAGAGCACACGTCTGAACTCTTTCCCTACACGACGCTCTTCCGATCTNNNTAGTCAAGCCTCATTTTTG
AAGAGCACACGTCTGAACTCTTTCCCTACACGACGCTCTTCCGATCTNNNTCACCTAATGCTGTGGCTT
AGCACACGTCTGAACTCTTTCCCTACACGACGCTCTTCCGATCTNNNACAGTTGTTATGAAAATACTC
GAAGAGCACACGTCTGAACTCTTTCCCTACACGACGCTCTTCCGATCTNNNACACTCCCAGATTTGGG
AGCACACGTCTGAACTCTTTCCCTACACGACGCTCTTCCGATCTNNNAGCACAATAACATATTACCTT
AGAGCACACGTCTGAACTCTTTCCCTACACGACGCTCTTCCGATCTNNNAAGATCCTAGAGGGTCAG
GGAAGAGCACACGTCTGAACTCTTTCCCTACACGACGCTCTTCCGATCTNNNGGCAGCTTTGTGGTGA
GCACACGTCTGAACTCTTTCCCTACACGACGCTCTTCCGATCTNNNAGATAGATTAGCTAGACAAACA
AGCACACGTCTGAACTCTTTCCCTACACGACGCTCTTCCGATCTNNNGCAAGTTACCATATTTATCAT
GAAGAGCACACGTCTGAACTCTTTCCCTACACGACGCTCTTCCGATCTNNNGTGGGGACACAGTAAC

GAGCACACGTCTGAACTCTTTCCCTACACGACGCTCTTCCGATCTNNNTGGGTAAACCTTTCTTTAGAAC
AAGAGCACACGTCTGAACTCTTTCCCTACACGACGCTCTTCCGATCTNNNTGTGTTCTTAGTTTTTGCCA
AAGAGCACACGTCTGAACTCTTTCCCTACACGACGCTCTTCCGATCTNNNCGCACATTTTATATGCGCA
AGCACACGTCTGAACTCTTTCCCTACACGACGCTCTTCCGATCTNNNTGTTTTATTTGGAGTAGAAAGT
GAGCACACGTCTGAACTCTTTCCCTACACGACGCTCTTCCGATCTNNNCACATGCCAATATGCTAATATA
AGCACACGTCTGAACTCTTTCCCTACACGACGCTCTTCCGATCTNNNTCCTCATAGTAGTTTTAGGTTAT
GGAAGAGCACACGTCTGAACTCTTTCCCTACACGACGCTCTTCCGATCTNNNTACCGGCCATTTCAAGCA
GAGCACACGTCTGAACTCTTTCCCTACACGACGCTCTTCCGATCTNNNTGCACTAGTTTTTGTAATTTTCT
GGAAGAGCACACGTCTGAACTCTTTCCCTACACGACGCTCTTCCGATCTNNNAGCTCCAGAAACTGATG
AAGAGCACACGTCTGAACTCTTTCCCTACACGACGCTCTTCCGATCTNNNTGCAAAACCTGTACTGTGTC
GAAGAGCACACGTCTGAACTCTTTCCCTACACGACGCTCTTCCGATCTNNNCCATAGTGCACAGTGATT
AGCACACGTCTGAACTCTTTCCCTACACGACGCTCTTCCGATCTNNNACTTCTGAACTCAATCAAATACA
GAGCACACGTCTGAACTCTTTCCCTACACGACGCTCTTCCGATCTNNNACTTTTGAGTTTTTGCTTAGC
GAAGAGCACACGTCTGAACTCTTTCCCTACACGACGCTCTTCCGATCTNNNAAATGACTGTTCCAGTTCC
GAAGAGCACACGTCTGAACTCTTTCCCTACACGACGCTCTTCCGATCTNNNTTGTATACAGGTTGCGCT
GAGCACACGTCTGAACTCTTTCCCTACACGACGCTCTTCCGATCTNNNTCTAGAATCACAGTACATGTTC
GAAGAGCACACGTCTGAACTCTTTCCCTACACGACGCTCTTCCGATCTNNNACTGGAAAATCTCCAAGC
GGAAGAGCACACGTCTGAACTCTTTCCCTACACGACGCTCTTCCGATCTNNNATGCAACTGAGAATAT
AAGAGCACACGTCTGAACTCTTTCCCTACACGACGCTCTTCCGATCTNNNGCCAGAAATGAGGGACAAA
AGAGCACACGTCTGAACTCTTTCCCTACACGACGCTCTTCCGATCTNNNAGACATTCTGATACCCTAAAA
GAGCACACGTCTGAACTCTTTCCCTACACGACGCTCTTCCGATCTNNNCCCATTTCCAACTAACATACTG
AGCACACGTCTGAACTCTTTCCCTACACGACGCTCTTCCGATCTNNNCACTAATATTTGGTTTACATGCT
GCACACGTCTGAACTCTTTCCCTACACGACGCTCTTCCGATCTNNNACAGGTAAAAAGAACATAACGAA
AGCACACGTCTGAACTCTTTCCCTACACGACGCTCTTCCGATCTNNNACTTTGGAAGTCAATATTCCCAA
AGCACACGTCTGAACTCTTTCCCTACACGACGCTCTTCCGATCTNNNGCATGTATACAGGGATGTATTT
AGAGCACACGTCTGAACTCTTTCCCTACACGACGCTCTTCCGATCTNNNTGAGTTTGGGGCATTTTTCT
AGAGCACACGTCTGAACTCTTTCCCTACACGACGCTCTTCCGATCTNNNTGTTAGCCCTCTATGATGTCT
AGCACACGTCTGAACTCTTTCCCTACACGACGCTCTTCCGATCTNNNTCTTGTATCCTACAACCTTATTGA
GAAGAGCACACGTCTGAACTCTTTCCCTACACGACGCTCTTCCGATCTNNNTAGAGCTGAACCAAAGG
AAGAGCACACGTCTGAACTCTTTCCCTACACGACGCTCTTCCGATCTNNNCGTTTGGTTAGTGAAGTTC
AGCACACGTCTGAACTCTTTCCCTACACGACGCTCTTCCGATCTNNNTCGATTCTGTGCTTTTTATATCT
GAGCACACGTCTGAACTCTTTCCCTACACGACGCTCTTCCGATCTNNNCAGGTCCTTTCTCAATATCATTT
AAGAGCACACGTCTGAACTCTTTCCCTACACGACGCTCTTCCGATCTNNNACCAAGTCATCCTATTCTTCT
AGCACACGTCTGAACTCTTTCCCTACACGACGCTCTTCCGATCTNNNACATGTTTTATCATCTAGAGCCT
AGCACACGTCTGAACTCTTTCCCTACACGACGCTCTTCCGATCTNNNTCTAGTAGATTTTCCTTTACATCT
AAGAGCACACGTCTGAACTCTTTCCCTACACGACGCTCTTCCGATCTNNNCCATGTGAAATTTGCTTCTG
GCACACGTCTGAACTCTTTCCCTACACGACGCTCTTCCGATCTNNNTGATAAATAGTAAAACCTCTCTGC
GAAGAGCACACGTCTGAACTCTTTCCCTACACGACGCTCTTCCGATCTNNNGCATGACTTACTGTGAGG
GAGCACACGTCTGAACTCTTTCCCTACACGACGCTCTTCCGATCTNNNACTGTTCTCATTTCACAGGCCT
GAAGAGCACACGTCTGAACTCTTTCCCTACACGACGCTCTTCCGATCTNNNAGGATGACTGTAATACCC
GAGCACACGTCTGAACTCTTTCCCTACACGACGCTCTTCCGATCTNNNACCAAATGTCACTGAATTGTA
AAGAGCACACGTCTGAACTCTTTCCCTACACGACGCTCTTCCGATCTNNNCAGTATGCTCTACAAGCAG
GAGCACACGTCTGAACTCTTTCCCTACACGACGCTCTTCCGATCTNNNGGGTGGGATAACAAGCAATA

GAGCACACGTCTGAACTCTTTCCCTACACGACGCTCTTCCGATCTNNNAAATATATATACATAGGTGT

GAAGAGCACACGTCTGAACTCTTTCCCTACACGACGCTCTTCCGATCTNNNCCCTCTAGTGTGAGTTG

AAGAGCACACGTCTGAACTCTTTCCCTACACGACGCTCTTCCGATCTNNNACATCTTCATTGCCCAAT

AGCACACGTCTGAACTCTTTCCCTACACGACGCTCTTCCGATCTNNNTGTTTACTATCTGTCCTTTTCA

AGAGCACACGTCTGAACTCTTTCCCTACACGACGCTCTTCCGATCTNNNAGCAAATGAAATCTACCTG

AAGAGCACACGTCTGAACTCTTTCCCTACACGACGCTCTTCCGATCTNNNTGACTTCCATTCAGGGA

GAAGAGCACACGTCTGAACTCTTTCCCTACACGACGCTCTTCCGATCTNNNTAGTCAAGCCTCATTTTT

GAAGAGCACACGTCTGAACTCTTTCCCTACACGACGCTCTTCCGATCTNNNGTACTCTGTGAGTGTGT

GAAGAGCACACGTCTGAACTCTTTCCCTACACGACGCTCTTCCGATCTNNNATGAGATGGCTTTTTAC

GAAGAGCACACGTCTGAACTCTTTCCCTACACGACGCTCTTCCGATCTNNNGGATGTATTCCCTCAGG

GAGCACACGTCTGAACTCTTTCCCTACACGACGCTCTTCCGATCTNNNTGCTAAGAGATTAGATTTA

AAGAGCACACGTCTGAACTCTTTCCCTACACGACGCTCTTCCGATCTNNNTTTCACCGTAGTGTGTGT

GAGCACACGTCTGAACTCTTTCCCTACACGACGCTCTTCCGATCTNNNCTTCAAAATGTTACTGTCAGG

GAGCACACGTCTGAACTCTTTCCCTACACGACGCTCTTCCGATCTNNNTTTTGATATAAGTCCATGGT

AGAGCACACGTCTGAACTCTTTCCCTACACGACGCTCTTCCGATCTNNNTCCTGTTTCTTCCTCTTCTCT

GAGCACACGTCTGAACTCTTTCCCTACACGACGCTCTTCCGATCTNNNAGTTCTAGTATCTCTCTCTCT

AGCACACGTCTGAACTCTTTCCCTACACGACGCTCTTCCGATCTNNNGCAAAATAGGTCTTATGACTT

AGAGCACACGTCTGAACTCTTTCCCTACACGACGCTCTTCCGATCTNNNTTGATGAGGAATCCTGAAT

AAGAGCACACGTCTGAACTCTTTCCCTACACGACGCTCTTCCGATCTNNNTCCCTCCTCTTTGTCCTCAT

GAGCACACGTCTGAACTCTTTCCCTACACGACGCTCTTCCGATCTNNNATAATATTCCAGGTGTGTGT

GAGCACACGTCTGAACTCTTTCCCTACACGACGCTCTTCCGATCTNNNTCTTCCTGTTCCTTGTATTGT

GAAGAGCACACGTCTGAACTCTTTCCCTACACGACGCTCTTCCGATCTNNNACAAGGTACATCATGCC

AAGAGCACACGTCTGAACTCTTTCCCTACACGACGCTCTTCCGATCTNNNTTATGTGGGTCCAAGCAG

AAGAGCACACGTCTGAACTCTTTCCCTACACGACGCTCTTCCGATCTNNNACAAAAACGATACTTGCA

AAGAGCACACGTCTGAACTCTTTCCCTACACGACGCTCTTCCGATCTNNNTGTACATTCTGTTTGAGG

AAGAGCACACGTCTGAACTCTTTCCCTACACGACGCTCTTCCGATCTNNNATGAATTGCCCTGAAGAC

GAAGAGCACACGTCTGAACTCTTTCCCTACACGACGCTCTTCCGATCTNNNCTGAGGCAGGAGAATC

GGAAGAGCACACGTCTGAACTCTTTCCCTACACGACGCTCTTCCGATCTNNNGGATAAATGTTGTGG

GCACACGTCTGAACTCTTTCCCTACACGACGCTCTTCCGATCTNNNTCCTTGATATGTCAATAACAGC

GCACACGTCTGAACTCTTTCCCTACACGACGCTCTTCCGATCTNNNTGTATAATGTGTTAGAATAAGT

GAAGAGCACACGTCTGAACTCTTTCCCTACACGACGCTCTTCCGATCTNNNATGCTTGTAATCCCAGC

GAAGAGCACACGTCTGAACTCTTTCCCTACACGACGCTCTTCCGATCTNNNGAGAATTGCTTGAACCT

AAGAGCACACGTCTGAACTCTTTCCCTACACGACGCTCTTCCGATCTNNNGAATGAGAATCGCTTGA

GAAGAGCACACGTCTGAACTCTTTCCCTACACGACGCTCTTCCGATCTNNNCTAAGGTAGGCCAGTC

AGCACACGTCTGAACTCTTTCCCTACACGACGCTCTTCCGATCTNNNACTTATTTGAACCTTCTCTCTT

GGAAGAGCACACGTCTGAACTCTTTCCCTACACGACGCTCTTCCGATCTNNNCAGGAGAATCACTTGA

AGCACACGTCTGAACTCTTTCCCTACACGACGCTCTTCCGATCTNNNAGCCAAAAAATAAAAGACAAT

AAGAGCACACGTCTGAACTCTTTCCCTACACGACGCTCTTCCGATCTNNNAGATTGCAGTGAGCTGA

AAGAGCACACGTCTGAACTCTTTCCCTACACGACGCTCTTCCGATCTNNNCTAATCTCAGCTACTCGA

AGCACACGTCTGAACTCTTTCCCTACACGACGCTCTTCCGATCTNNNTGCTCAGTAAATATTGTAGG

GCACACGTCTGAACTCTTTCCCTACACGACGCTCTTCCGATCTNNNACTATATATATGCATACACACAC

GCACACGTCTGAACTCTTTCCCTACACGACGCTCTTCCGATCTNNNACACATTATGTCTCATGATTAT

AAGAGCACACGTCTGAACTCTTTCCCTACACGACGCTCTTCCGATCTNNNCCAAGACCGAAAGGCCTA
GAGCACACGTCTGAACTCTTTCCCTACACGACGCTCTTCCGATCTNNNTGCATCTATTCTTCAAACTCA
AGAGCACACGTCTGAACTCTTTCCCTACACGACGCTCTTCCGATCTNNNTACTATGCCCAGCCGAAAA
AAGAGCACACGTCTGAACTCTTTCCCTACACGACGCTCTTCCGATCTNNNGTTACTTGAGCTTTGGAG
AAGAGCACACGTCTGAACTCTTTCCCTACACGACGCTCTTCCGATCTNNNATAAAACAGGCAAAGGTC
AAGAGCACACGTCTGAACTCTTTCCCTACACGACGCTCTTCCGATCTNNNGGCAGTTTGAAGACCAAT
AAGAGCACACGTCTGAACTCTTTCCCTACACGACGCTCTTCCGATCTNNNTCAAAATCAGGGCTCAAC
AGAGCACACGTCTGAACTCTTTCCCTACACGACGCTCTTCCGATCTNNNATTGAATGCACTGTGTTCTT
AAGAGCACACGTCTGAACTCTTTCCCTACACGACGCTCTTCCGATCTNNNTGATGAAGATTCCACTCA
AAGAGCACACGTCTGAACTCTTTCCCTACACGACGCTCTTCCGATCTNNNCCTTGGATAATTGAGACC
AGCACACGTCTGAACTCTTTCCCTACACGACGCTCTTCCGATCTNNNACAAGAATGTTTCAAAATGACT
AAGAGCACACGTCTGAACTCTTTCCCTACACGACGCTCTTCCGATCTNNNATGCCTGCTAAGGAATTT
GGAAGAGCACACGTCTGAACTCTTTCCCTACACGACGCTCTTCCGATCTNNNGTTACCTCCCAATAGAC
AGCACACGTCTGAACTCTTTCCCTACACGACGCTCTTCCGATCTNNNAGTTGAAAGAGATAGATATAC
AAGAGCACACGTCTGAACTCTTTCCCTACACGACGCTCTTCCGATCTNNNTTTGGAGTGGGGGTTGTT
AAGAGCACACGTCTGAACTCTTTCCCTACACGACGCTCTTCCGATCTNNNTAGGTACTCAGTGAACA
AGAGCACACGTCTGAACTCTTTCCCTACACGACGCTCTTCCGATCTNNNAAGAGGAGAGACGAAGA
GAGCACACGTCTGAACTCTTTCCCTACACGACGCTCTTCCGATCTNNNTGATTGATTCATTTGCTGTGC
AGAGCACACGTCTGAACTCTTTCCCTACACGACGCTCTTCCGATCTNNNCCCGGCCTCAAATCTTTATCT
GCACACGTCTGAACTCTTTCCCTACACGACGCTCTTCCGATCTNNNAGTGGCATTTATGTGTTTCTCTT
AGCACACGTCTGAACTCTTTCCCTACACGACGCTCTTCCGATCTNNNACTTGTGGATAAAAGTTACTA
AGCACACGTCTGAACTCTTTCCCTACACGACGCTCTTCCGATCTNNNTGTCCGAAGTAATTTGAAAGG
GAGCACACGTCTGAACTCTTTCCCTACACGACGCTCTTCCGATCTNNNACCAAAGGACACATGGTAAG
AGAGCACACGTCTGAACTCTTTCCCTACACGACGCTCTTCCGATCTNNNTTCTGTCCCGTTTTTGCCAT
GAGCACACGTCTGAACTCTTTCCCTACACGACGCTCTTCCGATCTNNNACACATACACAATTTTGCAAA
AGCACACGTCTGAACTCTTTCCCTACACGACGCTCTTCCGATCTNNNGTCATTGTTCTTGGTACTTTGT
AAGAGCACACGTCTGAACTCTTTCCCTACACGACGCTCTTCCGATCTNNNCAAAATGAAGAGCTCTGG
AAGAGCACACGTCTGAACTCTTTCCCTACACGACGCTCTTCCGATCTNNNCCATGTTCCTGTGAATGAC
GAAGAGCACACGTCTGAACTCTTTCCCTACACGACGCTCTTCCGATCTNNNAAAGTCACCTGGGATGA
AAGAGCACACGTCTGAACTCTTTCCCTACACGACGCTCTTCCGATCTNNNAACAAAAATCCCCAGGGA
AGCACACGTCTGAACTCTTTCCCTACACGACGCTCTTCCGATCTNNNAAAAAGTCCTGTTAGATGACA
AAGAGCACACGTCTGAACTCTTTCCCTACACGACGCTCTTCCGATCTNNNCATTGTAATACCCTTCTCCC
AGAGCACACGTCTGAACTCTTTCCCTACACGACGCTCTTCCGATCTNNNCACAACACATTTCTTCTTGCC
GAAGAGCACACGTCTGAACTCTTTCCCTACACGACGCTCTTCCGATCTNNNGATCATCCTGTCTCCGCC
AAGAGCACACGTCTGAACTCTTTCCCTACACGACGCTCTTCCGATCTNNNGAGGTTGCAGTGAGACG
AAGAGCACACGTCTGAACTCTTTCCCTACACGACGCTCTTCCGATCTNNNTGAATGGAAAAAGAAAAT
GCACACGTCTGAACTCTTTCCCTACACGACGCTCTTCCGATCTNNNTGTGGTTTATTTAGAATGACCA
AGCACACGTCTGAACTCTTTCCCTACACGACGCTCTTCCGATCTNNNGCCACCAATATATACACCTGGT
GAGCACACGTCTGAACTCTTTCCCTACACGACGCTCTTCCGATCTNNNAGTTACCTCTGAATGAAACA
AGAGCACACGTCTGAACTCTTTCCCTACACGACGCTCTTCCGATCTNNNGATCTGGCAGCAGGTTTTG
AAGAGCACACGTCTGAACTCTTTCCCTACACGACGCTCTTCCGATCTNNNCTCTTCAAGTGGTGGCCAG
GGAAGAGCACACGTCTGAACTCTTTCCCTACACGACGCTCTTCCGATCTNNNTGCCATTGATACACATT
AAGAGCACACGTCTGAACTCTTTCCCTACACGACGCTCTTCCGATCTNNNCTCTATGTGACCAGGCAG

AAGAGCACACGTCTGAACTCTTTCCCTACACGACGCTCTTCCGATCTNNNCCAAGACCGAAAGGCCTA
AGCACACGTCTGAACTCTTTCCCTACACGACGCTCTTCCGATCTNNNTGACTTACAAGGCAAAGAAA
AGAGCACACGTCTGAACTCTTTCCCTACACGACGCTCTTCCGATCTNNNCTGTGGCCTTGCCTATACTT
AAGAGCACACGTCTGAACTCTTTCCCTACACGACGCTCTTCCGATCTNNNGCTGTGAGCTCCAAGAAA
GAGCACACGTCTGAACTCTTTCCCTACACGACGCTCTTCCGATCTNNNTGAGTAGTTTTGTGTGTCTG
GAGCACACGTCTGAACTCTTTCCCTACACGACGCTCTTCCGATCTNNNTCAAATGTAGTGTATCCCTTT
AGCACACGTCTGAACTCTTTCCCTACACGACGCTCTTCCGATCTNNNAGGCAAATTCAGAAATACAGT
AAGAGCACACGTCTGAACTCTTTCCCTACACGACGCTCTTCCGATCTNNNACCTCCAAAAGCTTCCTCC
AGAGCACACGTCTGAACTCTTTCCCTACACGACGCTCTTCCGATCTNNNCACTTCAACCTCTGCCTTCA
AGAGCACACGTCTGAACTCTTTCCCTACACGACGCTCTTCCGATCTNNNGGCTTTGATCAACCACCAG
AAGAGCACACGTCTGAACTCTTTCCCTACACGACGCTCTTCCGATCTNNNTTCCTGATTCAGAACAGC
AAGAGCACACGTCTGAACTCTTTCCCTACACGACGCTCTTCCGATCTNNNGAGCCTGGACAAATTCCC
AGCACACGTCTGAACTCTTTCCCTACACGACGCTCTTCCGATCTNNNTGCTGTATCTGCTTATAATCTC
AAGAGCACACGTCTGAACTCTTTCCCTACACGACGCTCTTCCGATCTNNNGCATAGGAGATTGACAG
AGCACACGTCTGAACTCTTTCCCTACACGACGCTCTTCCGATCTNNNACATTTTCTCTCAACTCATTGTG
AGAGCACACGTCTGAACTCTTTCCCTACACGACGCTCTTCCGATCTNNNGGCATGGCTGTTGCTTCAT
AGAGCACACGTCTGAACTCTTTCCCTACACGACGCTCTTCCGATCTNNNTCAGTCCTTTTCCAAGTCCA
AGAGCACACGTCTGAACTCTTTCCCTACACGACGCTCTTCCGATCTNNNCCTTTGAAGTAGGTCCCAT
AAGAGCACACGTCTGAACTCTTTCCCTACACGACGCTCTTCCGATCTNNNAGCCCTACCTTGTGCATGT
AGAGCACACGTCTGAACTCTTTCCCTACACGACGCTCTTCCGATCTNNNTGGGTTTGGGTAAACATA
AAGAGCACACGTCTGAACTCTTTCCCTACACGACGCTCTTCCGATCTNNNGCAGTGATGAACAGGAA
AGAGCACACGTCTGAACTCTTTCCCTACACGACGCTCTTCCGATCTNNNTTGACGGCTCCAGAAGAC
AGAGCACACGTCTGAACTCTTTCCCTACACGACGCTCTTCCGATCTNNNAATTTTGCCGCATTTGCTTT
AGAGCACACGTCTGAACTCTTTCCCTACACGACGCTCTTCCGATCTNNNAGTCTTCTCTCCTGTCCCTT
GGAAGAGCACACGTCTGAACTCTTTCCCTACACGACGCTCTTCCGATCTNNNACCCTTTGCTTTCCCCAT
AAGAGCACACGTCTGAACTCTTTCCCTACACGACGCTCTTCCGATCTNNNTCCATCTTCTCACATAGCT
GAGCACACGTCTGAACTCTTTCCCTACACGACGCTCTTCCGATCTNNNTAGAACCTCCTATTCCACCAC
GAAGAGCACACGTCTGAACTCTTTCCCTACACGACGCTCTTCCGATCTNNNTTAGTGGGATCTCTCTC
AGAGCACACGTCTGAACTCTTTCCCTACACGACGCTCTTCCGATCTNNNAGCAGTATTCATAGTAGG
GAGCACACGTCTGAACTCTTTCCCTACACGACGCTCTTCCGATCTNNNAATCTGGCCCACATGCACAA
AAGAGCACACGTCTGAACTCTTTCCCTACACGACGCTCTTCCGATCTNNNGAAATCAGAACCCTAGTT
AGAGCACACGTCTGAACTCTTTCCCTACACGACGCTCTTCCGATCTNNNGGCCTAACCTGTGATAGCC
GAGCACACGTCTGAACTCTTTCCCTACACGACGCTCTTCCGATCTNNNACAATGCACTAGGAGATCAT
AGAGCACACGTCTGAACTCTTTCCCTACACGACGCTCTTCCGATCTNNNAACCTGTGTGACCATGGAA
AAGAGCACACGTCTGAACTCTTTCCCTACACGACGCTCTTCCGATCTNNNCTGAGTTTGCTTGGGGTT
AAGAGCACACGTCTGAACTCTTTCCCTACACGACGCTCTTCCGATCTNNNAATCAGACTGGGCGCAGA
AAGAGCACACGTCTGAACTCTTTCCCTACACGACGCTCTTCCGATCTNNNAACTTTTCCTCAAGAGCAC
GAGCACACGTCTGAACTCTTTCCCTACACGACGCTCTTCCGATCTNNNCCAAATGTCCCTACCTCTAAC
AGAGCACACGTCTGAACTCTTTCCCTACACGACGCTCTTCCGATCTNNNAGCAACTGATTTTGGTATG
AAGAGCACACGTCTGAACTCTTTCCCTACACGACGCTCTTCCGATCTNNNTCTCCCCAAGTAACTCACG
AGAGCACACGTCTGAACTCTTTCCCTACACGACGCTCTTCCGATCTNNNACCCTAGACAGACACCTTCC
AAGAGCACACGTCTGAACTCTTTCCCTACACGACGCTCTTCCGATCTNNNAGCTTTCTTCTGGTCCGA
GAGCACACGTCTGAACTCTTTCCCTACACGACGCTCTTCCGATCTNNNTCATGTTGTTGCAAATGACG

AGAGCACACGTCTGAACTCTTTCCCTACACGACGCTCTTCCGATCTNNNAGGAGGAAACTACCAACA
GAGCACACGTCTGAACTCTTTCCCTACACGACGCTCTTCCGATCTNNNACAGTTTCTCATGTTCTAGTC
AAGAGCACACGTCTGAACTCTTTCCCTACACGACGCTCTTCCGATCTNNNATCAGAAAGGACATGGA
AGAGCACACGTCTGAACTCTTTCCCTACACGACGCTCTTCCGATCTNNNAGTCCTTCCTCCTCCATTCTT
AGAGCACACGTCTGAACTCTTTCCCTACACGACGCTCTTCCGATCTNNNAACCAGTGTCACTTTCTGAA
AAGAGCACACGTCTGAACTCTTTCCCTACACGACGCTCTTCCGATCTNNNTCTCTAACAGCCATTCTG
AGCACACGTCTGAACTCTTTCCCTACACGACGCTCTTCCGATCTNNNTGTTTCTCTCCCTCCTCTTTCTT
AGAGCACACGTCTGAACTCTTTCCCTACACGACGCTCTTCCGATCTNNNTGTCACTATCACTGGTTTT
AGAGCACACGTCTGAACTCTTTCCCTACACGACGCTCTTCCGATCTNNNTATACACATGTGCACTTGG
GAGCACACGTCTGAACTCTTTCCCTACACGACGCTCTTCCGATCTNNNGGAAACACAGCTACTACTTC
AAGAGCACACGTCTGAACTCTTTCCCTACACGACGCTCTTCCGATCTNNNAGCTAAGACTACAGTGCC
AGAGCACACGTCTGAACTCTTTCCCTACACGACGCTCTTCCGATCTNNNTCCATGATTATGTTACCCCC
AAGAGCACACGTCTGAACTCTTTCCCTACACGACGCTCTTCCGATCTNNNTGGAATATGTGGAGTGT
AAGAGCACACGTCTGAACTCTTTCCCTACACGACGCTCTTCCGATCTNNNTGCCAATGTAGCCATGAT
AAGAGCACACGTCTGAACTCTTTCCCTACACGACGCTCTTCCGATCTNNNAGTCATGTCACCTTTGGC
GAAGAGCACACGTCTGAACTCTTTCCCTACACGACGCTCTTCCGATCTNNNAAGGAACAGCTGGGAT
AGAGCACACGTCTGAACTCTTTCCCTACACGACGCTCTTCCGATCTNNNCCTTTCCCCTATATGTGTGT
AAGAGCACACGTCTGAACTCTTTCCCTACACGACGCTCTTCCGATCTNNNGGATTTATACACATGCAC
GAAGAGCACACGTCTGAACTCTTTCCCTACACGACGCTCTTCCGATCTNNNCCAAAGCAAGAATGCAA
AGAGCACACGTCTGAACTCTTTCCCTACACGACGCTCTTCCGATCTNNNACATTCCATTCTGTGTGTGT
AAGAGCACACGTCTGAACTCTTTCCCTACACGACGCTCTTCCGATCTNNNTCAATTATGGTGCTCTCGA
AAGAGCACACGTCTGAACTCTTTCCCTACACGACGCTCTTCCGATCTNNNTGGGTGAAAGAATGAGA
AGAGCACACGTCTGAACTCTTTCCCTACACGACGCTCTTCCGATCTNNNCCATCAGGTGTTTAGCCTCA
AAGAGCACACGTCTGAACTCTTTCCCTACACGACGCTCTTCCGATCTNNNAGGGAGGGGTCCAAAAT
AGAGCACACGTCTGAACTCTTTCCCTACACGACGCTCTTCCGATCTNNNAGTCATCACCAAAGCCAAA
AGAGCACACGTCTGAACTCTTTCCCTACACGACGCTCTTCCGATCTNNNGGTGTTTTTCCAGCAGCAA
AGAGCACACGTCTGAACTCTTTCCCTACACGACGCTCTTCCGATCTNNNGCTGTTTTCTGGAGCTCTT
GAGCACACGTCTGAACTCTTTCCCTACACGACGCTCTTCCGATCTNNNTGACAGAGTAAGGTCCCTTC
GAAGAGCACACGTCTGAACTCTTTCCCTACACGACGCTCTTCCGATCTNNNGTCAAGATTTCAGGGAT
AAGAGCACACGTCTGAACTCTTTCCCTACACGACGCTCTTCCGATCTNNNTCCCAACTTTATCTAGAG
AGAGCACACGTCTGAACTCTTTCCCTACACGACGCTCTTCCGATCTNNNTGCCCTCTTTCCTGTACTCTG
AAGAGCACACGTCTGAACTCTTTCCCTACACGACGCTCTTCCGATCTNNNAGAGTTGGAAGCTTTGA
AGAGCACACGTCTGAACTCTTTCCCTACACGACGCTCTTCCGATCTNNNCAAAATCCAGAATCCCAGG
AGAGCACACGTCTGAACTCTTTCCCTACACGACGCTCTTCCGATCTNNNGGGATTAGGAGCCAGTAG
AAGAGCACACGTCTGAACTCTTTCCCTACACGACGCTCTTCCGATCTNNNTAGATTGGCATTCGTCAG
GAAGAGCACACGTCTGAACTCTTTCCCTACACGACGCTCTTCCGATCTNNNCCCAGCCAATGTAACTG
AAGAGCACACGTCTGAACTCTTTCCCTACACGACGCTCTTCCGATCTNNNATTGGGCCATGGTTTAGT
AGAGCACACGTCTGAACTCTTTCCCTACACGACGCTCTTCCGATCTNNNGGTGGAGAGAAATCCCAG
AAGAGCACACGTCTGAACTCTTTCCCTACACGACGCTCTTCCGATCTNNNAGATGTTTCAGCACCCTC
GGAAGAGCACACGTCTGAACTCTTTCCCTACACGACGCTCTTCCGATCTNNNTCCCAGTGAGTAAGT
AGAGCACACGTCTGAACTCTTTCCCTACACGACGCTCTTCCGATCTNNNACATTTTCCATTTCCCTCAC
GAAGAGCACACGTCTGAACTCTTTCCCTACACGACGCTCTTCCGATCTNNNATATGCCTTGGACTTTTT
AAGAGCACACGTCTGAACTCTTTCCCTACACGACGCTCTTCCGATCTNNNCTTTCACCTTATCCCCATC

AGCACACGTCTGAACTCTTTCCCTACACGACGCTCTTCCGATCTNNNCCAGTTGAATTTTGGAAATCTG
GAGCACACGTCTGAACTCTTTCCCTACACGACGCTCTTCCGATCTNNNACAGTAGTGAATGTTCTCTCT
GAGCACACGTCTGAACTCTTTCCCTACACGACGCTCTTCCGATCTNNNGGCAATTTACAAACAAAAC
AAGAGCACACGTCTGAACTCTTTCCCTACACGACGCTCTTCCGATCTNNNTGGTGGTCCCAACCACTAC
GAGCACACGTCTGAACTCTTTCCCTACACGACGCTCTTCCGATCTNNNACGGCAATAGTCCACTGAG
AGAGCACACGTCTGAACTCTTTCCCTACACGACGCTCTTCCGATCTNNNAGTCTCGACAGATAGGAT
AAGAGCACACGTCTGAACTCTTTCCCTACACGACGCTCTTCCGATCTNNNACTGTTGTGTTGGGGAAT
AAGAGCACACGTCTGAACTCTTTCCCTACACGACGCTCTTCCGATCTNNNTCTTGTTTGCCTCCTGTGT
AGAGCACACGTCTGAACTCTTTCCCTACACGACGCTCTTCCGATCTNNNACCTGGAATTTTCTCTCTCTG
AAGAGCACACGTCTGAACTCTTTCCCTACACGACGCTCTTCCGATCTNNNAAAACAGCTCTGGCGTAT
GAGCACACGTCTGAACTCTTTCCCTACACGACGCTCTTCCGATCTNNNCCGTGTAGGCTCACTGTAAG
AGAGCACACGTCTGAACTCTTTCCCTACACGACGCTCTTCCGATCTNNNGGGGTGAAACCACAGTTTT
AGAGCACACGTCTGAACTCTTTCCCTACACGACGCTCTTCCGATCTNNNAAGATTGAGGCTGCAGTG
GAAGAGCACACGTCTGAACTCTTTCCCTACACGACGCTCTTCCGATCTNNNGAGATGCCCCAGATAAA
AGAGCACACGTCTGAACTCTTTCCCTACACGACGCTCTTCCGATCTNNNATAGCATGTGGCTACTGCA
GCACACGTCTGAACTCTTTCCCTACACGACGCTCTTCCGATCTNNNTGCTTTGCTCATTATTTTGTCTC
GAAGAGCACACGTCTGAACTCTTTCCCTACACGACGCTCTTCCGATCTNNNTCCCCTAAACACCTTCCCA
AGAGCACACGTCTGAACTCTTTCCCTACACGACGCTCTTCCGATCTNNNGCTGATAGAGGATGAGCT
GAAGAGCACACGTCTGAACTCTTTCCCTACACGACGCTCTTCCGATCTNNNGCAAGGTACTGCCTGAT
AGAGCACACGTCTGAACTCTTTCCCTACACGACGCTCTTCCGATCTNNNCCCCGCATTCACCTCTTCTA
AGAGCACACGTCTGAACTCTTTCCCTACACGACGCTCTTCCGATCTNNNCTTTCAGTGGTCACATCCAG
GAGCACACGTCTGAACTCTTTCCCTACACGACGCTCTTCCGATCTNNNGGAAGAAACACCATAGTAAG
AAGAGCACACGTCTGAACTCTTTCCCTACACGACGCTCTTCCGATCTNNNACAAGTGGGTAAAAGAA
GAAGAGCACACGTCTGAACTCTTTCCCTACACGACGCTCTTCCGATCTNNNCCAAGCACAGAAGAGAG
GAGCACACGTCTGAACTCTTTCCCTACACGACGCTCTTCCGATCTNNNAGGAAAACTCATATGTCCCA
GAAGAGCACACGTCTGAACTCTTTCCCTACACGACGCTCTTCCGATCTNNNTGCTGCATAGAGTGGCT
AGAGCACACGTCTGAACTCTTTCCCTACACGACGCTCTTCCGATCTNNNCCTCCTGAACCTTGCTTTCT
AGAGCACACGTCTGAACTCTTTCCCTACACGACGCTCTTCCGATCTNNNGGCAGCAGCTAATGCTGA
AGAGCACACGTCTGAACTCTTTCCCTACACGACGCTCTTCCGATCTNNNACTGTCTTACTTTGCCTTGA
GAAGAGCACACGTCTGAACTCTTTCCCTACACGACGCTCTTCCGATCTNNNGTTGTGTGGTGTGCAAT
AAGAGCACACGTCTGAACTCTTTCCCTACACGACGCTCTTCCGATCTNNNGGAACACAGAACCAAACC
AAGAGCACACGTCTGAACTCTTTCCCTACACGACGCTCTTCCGATCTNNNTGAATTTCCCTTTGCCCCTA
AGCACACGTCTGAACTCTTTCCCTACACGACGCTCTTCCGATCTNNNTGTCTGCAAAACAAGGTAAAT
AAGAGCACACGTCTGAACTCTTTCCCTACACGACGCTCTTCCGATCTNNNTCCTTCCCCCACATGAAAG
AGAGCACACGTCTGAACTCTTTCCCTACACGACGCTCTTCCGATCTNNNCTGGTCCACATGTGTGAGC
AAGAGCACACGTCTGAACTCTTTCCCTACACGACGCTCTTCCGATCTNNNTATGCTGGAGGCTTTCGT
GCACACGTCTGAACTCTTTCCCTACACGACGCTCTTCCGATCTNNNAGGTCTCTTCTATTTTCCACCTC
AGCACACGTCTGAACTCTTTCCCTACACGACGCTCTTCCGATCTNNNACAACTACACATGTAAAAGA
AGAGCACACGTCTGAACTCTTTCCCTACACGACGCTCTTCCGATCTNNNTAAGACGCATGCATGTGA
AGAGCACACGTCTGAACTCTTTCCCTACACGACGCTCTTCCGATCTNNNTTTTGCCTTCCACTCTTCTTC
AAGAGCACACGTCTGAACTCTTTCCCTACACGACGCTCTTCCGATCTNNNAGAAAATGACCCATCTGC
GAAGAGCACACGTCTGAACTCTTTCCCTACACGACGCTCTTCCGATCTNNNTGTTGTATGGCTGAGCA
GAGCACACGTCTGAACTCTTTCCCTACACGACGCTCTTCCGATCTNNNGCCTCACCACTCCAATTTTCAA

GAAGAGCACACGTCTGAACTCTTTCCCTACACGACGCTCTTCCGATCTNNNACCATATTGTGCAGTACA
AAGAGCACACGTCTGAACTCTTTCCCTACACGACGCTCTTCCGATCTNNNTGTCTTGGGTGTTTGGTTT
AGAGCACACGTCTGAACTCTTTCCCTACACGACGCTCTTCCGATCTNNNAGAAAGAGTGAAACGCAG
AAGAGCACACGTCTGAACTCTTTCCCTACACGACGCTCTTCCGATCTNNNGCATCTAACATGGTACCGA
AGCACACGTCTGAACTCTTTCCCTACACGACGCTCTTCCGATCTNNNACCAGTGTATAATCAACGGAAC
AGAGCACACGTCTGAACTCTTTCCCTACACGACGCTCTTCCGATCTNNNCGAAGGATATATGTGTGTG
AGAGCACACGTCTGAACTCTTTCCCTACACGACGCTCTTCCGATCTNNNGGCCTCAAGAAATCCTCATT
AAGAGCACACGTCTGAACTCTTTCCCTACACGACGCTCTTCCGATCTNNNAGTTGCTCCTGAAGAAGAT
GAGCACACGTCTGAACTCTTTCCCTACACGACGCTCTTCCGATCTNNNAGCAACTGACCAAATTGCTAAA
AAGAGCACACGTCTGAACTCTTTCCCTACACGACGCTCTTCCGATCTNNNGGCCCATAGTCCAAGTGAA
AGAGCACACGTCTGAACTCTTTCCCTACACGACGCTCTTCCGATCTNNNGCAAATTTGTATGCAGAAG
AGAGCACACGTCTGAACTCTTTCCCTACACGACGCTCTTCCGATCTNNNCCTCTGCCTCTCTCAGGTCTC
GAGCACACGTCTGAACTCTTTCCCTACACGACGCTCTTCCGATCTNNNTGGATGCATCTTTGCTTTTCCC
AGAGCACACGTCTGAACTCTTTCCCTACACGACGCTCTTCCGATCTNNNGAGAAGCAGAGCCCAGCAT
AGCACACGTCTGAACTCTTTCCCTACACGACGCTCTTCCGATCTNNNATATATGTGCGTTCATGGATGT
GAAGAGCACACGTCTGAACTCTTTCCCTACACGACGCTCTTCCGATCTNNNACTAAATGCCAGGTCAGT
AAGAGCACACGTCTGAACTCTTTCCCTACACGACGCTCTTCCGATCTNNNTGGGTCCATTCATTTTGTGC
AGAGCACACGTCTGAACTCTTTCCCTACACGACGCTCTTCCGATCTNNNACAGCAGGAAGCCATTACTC
AGAGCACACGTCTGAACTCTTTCCCTACACGACGCTCTTCCGATCTNNNTGTGTAGCAAAGACAATTCT
AGAGCACACGTCTGAACTCTTTCCCTACACGACGCTCTTCCGATCTNNNTGTGTTCTACAAATTTTCCAA
AAGAGCACACGTCTGAACTCTTTCCCTACACGACGCTCTTCCGATCTNNNAGAAGTGAGGAAGCTGTC
AGCACACGTCTGAACTCTTTCCCTACACGACGCTCTTCCGATCTNNNTCAGTAGGAACAGTAATTTCATC
AAGAGCACACGTCTGAACTCTTTCCCTACACGACGCTCTTCCGATCTNNNAAGCCCTGTCAAGATTTCT
AAGAGCACACGTCTGAACTCTTTCCCTACACGACGCTCTTCCGATCTNNNACAAGAAAGACCAACAACT
AGCACACGTCTGAACTCTTTCCCTACACGACGCTCTTCCGATCTNNNTCTCTGTAATGCCTATGGAACTT
AAGAGCACACGTCTGAACTCTTTCCCTACACGACGCTCTTCCGATCTNNNTGTTCATTCCACAGATCAG
GAGCACACGTCTGAACTCTTTCCCTACACGACGCTCTTCCGATCTNNNTCCTACACATTTTCATGTTTTT
AGAGCACACGTCTGAACTCTTTCCCTACACGACGCTCTTCCGATCTNNNCAGTCCCAATGCTCACGAAC
AGCACACGTCTGAACTCTTTCCCTACACGACGCTCTTCCGATCTNNNAGAACAATGAACAGAAAAGAA
GAGCACACGTCTGAACTCTTTCCCTACACGACGCTCTTCCGATCTNNNTGAATACTCACCTCCAAAACAT
AGCACACGTCTGAACTCTTTCCCTACACGACGCTCTTCCGATCTNNNAGATAGATGATAAAGACCCATG
AAGAGCACACGTCTGAACTCTTTCCCTACACGACGCTCTTCCGATCTNNNGGGACCATCTTTTGCCTCAT
AGCACACGTCTGAACTCTTTCCCTACACGACGCTCTTCCGATCTNNNTCATTCTGTTATGATTCTCTACA
GAAGAGCACACGTCTGAACTCTTTCCCTACACGACGCTCTTCCGATCTNNNACCACGTATTTCTCACTCT
AGAGCACACGTCTGAACTCTTTCCCTACACGACGCTCTTCCGATCTNNNACACTGTCCTGTGATATTGT
GAGCACACGTCTGAACTCTTTCCCTACACGACGCTCTTCCGATCTNNNACATTGTGATTTTCCCTAAAAC
AGCACACGTCTGAACTCTTTCCCTACACGACGCTCTTCCGATCTNNNTCCAAAGTCATTCCAAAAGTAT
GCACACGTCTGAACTCTTTCCCTACACGACGCTCTTCCGATCTNNNACATGAAAAACCTGACAGATACA
AGAGCACACGTCTGAACTCTTTCCCTACACGACGCTCTTCCGATCTNNNGCTTGGGAGGCTGAGTTAT
AGCACACGTCTGAACTCTTTCCCTACACGACGCTCTTCCGATCTNNNGCTATGTAATTTTAGGCAACACA
GAGCACACGTCTGAACTCTTTCCCTACACGACGCTCTTCCGATCTNNNTGGATCATGGAATCACTTTCT
AGCACACGTCTGAACTCTTTCCCTACACGACGCTCTTCCGATCTNNNTGGTGCATGTCCTATATCAAAG
AGCACACGTCTGAACTCTTTCCCTACACGACGCTCTTCCGATCTNNNCAGAACCAACAAGTTTGATCAC

GAAGAGCACACGTCTGAACTCTTTCCCTACACGACGCTCTTCCGATCTNNNAAAGAAAGGCAGCAGG
AAGAGCACACGTCTGAACTCTTTCCCTACACGACGCTCTTCCGATCTNNNCTCATTCTAGCCCAGGTG
GCACACGTCTGAACTCTTTCCCTACACGACGCTCTTCCGATCTNNNAATATATGTGTTTGTATGTGTG
GCACACGTCTGAACTCTTTCCCTACACGACGCTCTTCCGATCTNNNTGTAACTAATACTGTACATCAC
GAAGAGCACACGTCTGAACTCTTTCCCTACACGACGCTCTTCCGATCTNNNAGCGAGTGGAATAAAA
GAGCACACGTCTGAACTCTTTCCCTACACGACGCTCTTCCGATCTNNNCCTTACATAAATTTTGCAGCT
AGAGCACACGTCTGAACTCTTTCCCTACACGACGCTCTTCCGATCTNNNTTTCTGGGTAACTTTGAGC
AGAGCACACGTCTGAACTCTTTCCCTACACGACGCTCTTCCGATCTNNNTCCCAGCCTTAGAGAGAAC
AGCACACGTCTGAACTCTTTCCCTACACGACGCTCTTCCGATCTNNNGCCAAGTAGAAGAGTACTAT
GAGCACACGTCTGAACTCTTTCCCTACACGACGCTCTTCCGATCTNNNTCAGCTCCACTTCATATGTTTT
AAGAGCACACGTCTGAACTCTTTCCCTACACGACGCTCTTCCGATCTNNNGGAGAAGGAAGGTGGG
GCACACGTCTGAACTCTTTCCCTACACGACGCTCTTCCGATCTNNNACTTTACTATTTGGAAGAAACGT
AGAGCACACGTCTGAACTCTTTCCCTACACGACGCTCTTCCGATCTNNNCACAAGATCTGCCCTCAAG
AGCACACGTCTGAACTCTTTCCCTACACGACGCTCTTCCGATCTNNNAGTTTTGCTATTTTCTTCTGTGT
AAGAGCACACGTCTGAACTCTTTCCCTACACGACGCTCTTCCGATCTNNNGCCTTTGGGCATAGTCAT
AGCACACGTCTGAACTCTTTCCCTACACGACGCTCTTCCGATCTNNNCAGTAAGCCATGATAGCACCT
GAGCACACGTCTGAACTCTTTCCCTACACGACGCTCTTCCGATCTNNNCTGTGACTTTAGGATGCTGT
AGAGCACACGTCTGAACTCTTTCCCTACACGACGCTCTTCCGATCTNNNAGGAACCCACCCAAGAAAA
AAGAGCACACGTCTGAACTCTTTCCCTACACGACGCTCTTCCGATCTNNNTTGTTTGTCTCTTCCACC
AGAGCACACGTCTGAACTCTTTCCCTACACGACGCTCTTCCGATCTNNNCACTGGAAAATAACCCAGC
AAGAGCACACGTCTGAACTCTTTCCCTACACGACGCTCTTCCGATCTNNNTCAAAAGGCTCTTCAGTG
AGCACACGTCTGAACTCTTTCCCTACACGACGCTCTTCCGATCTNNNTCACCAGTGCTCATGAATTCTT
AAGAGCACACGTCTGAACTCTTTCCCTACACGACGCTCTTCCGATCTNNNTGACAGAGCGAAACCCTA
AGAGCACACGTCTGAACTCTTTCCCTACACGACGCTCTTCCGATCTNNNTAAAGAGTTGCTTTGGAGC
GAAGAGCACACGTCTGAACTCTTTCCCTACACGACGCTCTTCCGATCTNNNTATACTCCTCAAAACACA
GCACACGTCTGAACTCTTTCCCTACACGACGCTCTTCCGATCTNNNTCTGTGGCTAGATAGTATTCCA
AGCACACGTCTGAACTCTTTCCCTACACGACGCTCTTCCGATCTNNNAGAACCAGATTGAGAATGTAA
AGAGCACACGTCTGAACTCTTTCCCTACACGACGCTCTTCCGATCTNNNATGCATGCTAAGATCATGC
AGAGCACACGTCTGAACTCTTTCCCTACACGACGCTCTTCCGATCTNNNTGAGCCCACCACTAAGAGA
AAGAGCACACGTCTGAACTCTTTCCCTACACGACGCTCTTCCGATCTNNNTGTATATATGGAACACCAT
GAGCACACGTCTGAACTCTTTCCCTACACGACGCTCTTCCGATCTNNNCCACATGGCTTAGTACTCAA
AAGAGCACACGTCTGAACTCTTTCCCTACACGACGCTCTTCCGATCTNNNTATATGTGAAATCGCCCT
AAGAGCACACGTCTGAACTCTTTCCCTACACGACGCTCTTCCGATCTNNNGCCCGGCCTTGAGTTTTA
AAGAGCACACGTCTGAACTCTTTCCCTACACGACGCTCTTCCGATCTNNNATGCATCCACTAAATGACC
AGCACACGTCTGAACTCTTTCCCTACACGACGCTCTTCCGATCTNNNTCCTAGGCAAAATATTAGGTG
GAGCACACGTCTGAACTCTTTCCCTACACGACGCTCTTCCGATCTNNNACACAGATAACCAGTCCTCC
GCACACGTCTGAACTCTTTCCCTACACGACGCTCTTCCGATCTNNNTAAAAGTGGCAGTTTCCCTTGA
AAGAGCACACGTCTGAACTCTTTCCCTACACGACGCTCTTCCGATCTNNNGAGGTCCAGGCAAAGCA
GAGCACACGTCTGAACTCTTTCCCTACACGACGCTCTTCCGATCTNNNAGTCCCACATTTCATTGGTAG
AGAGCACACGTCTGAACTCTTTCCCTACACGACGCTCTTCCGATCTNNNGAACAGCCTAAGCACATCCT
GAGCACACGTCTGAACTCTTTCCCTACACGACGCTCTTCCGATCTNNNTCCCTGAAATTAGTATTGGG
GCACACGTCTGAACTCTTTCCCTACACGACGCTCTTCCGATCTNNNACGTTGTTATAGGAGTATGGA
AAGAGCACACGTCTGAACTCTTTCCCTACACGACGCTCTTCCGATCTNNNATGTGTGTGTGCACTCTC

GAGCACACGTCTGAACTCTTTCCCTACACGACGCTCTTCCGATCTNNNAGGTCTTTAGCTGAAATGTAC
AGAGCACACGTCTGAACTCTTTCCCTACACGACGCTCTTCCGATCTNNNAGCCATGAACATTCTAAAGA
AGAGCACACGTCTGAACTCTTTCCCTACACGACGCTCTTCCGATCTNNNAGCAAACCATGAAAAGAAGA
GAAGAGCACACGTCTGAACTCTTTCCCTACACGACGCTCTTCCGATCTNNNAGTTCTCAGTTGTTGTCA
AGAGCACACGTCTGAACTCTTTCCCTACACGACGCTCTTCCGATCTNNNTCAAATTTAGGTCAGATTCA
GAGCACACGTCTGAACTCTTTCCCTACACGACGCTCTTCCGATCTNNNAGTCCAACCATCACTTGATTTA
AAGAGCACACGTCTGAACTCTTTCCCTACACGACGCTCTTCCGATCTNNNAGAACAGTCTCTGCATAACA
AGAGCACACGTCTGAACTCTTTCCCTACACGACGCTCTTCCGATCTNNNTGTTTACATAGAACCTTCCA
AAGAGCACACGTCTGAACTCTTTCCCTACACGACGCTCTTCCGATCTNNNTGCATTCTGAGATCAGATTA
AAGAGCACACGTCTGAACTCTTTCCCTACACGACGCTCTTCCGATCTNNNGAAGACAGAGCAAGCTG
AGAGCACACGTCTGAACTCTTTCCCTACACGACGCTCTTCCGATCTNNNGGAGGTATTGGTGTGAAGT
GAGCACACGTCTGAACTCTTTCCCTACACGACGCTCTTCCGATCTNNNTCAGTGGAAACATTCCCTTCAA
AGAGCACACGTCTGAACTCTTTCCCTACACGACGCTCTTCCGATCTNNNTCAAGACCCTACTTTGTATGA
AAGAGCACACGTCTGAACTCTTTCCCTACACGACGCTCTTCCGATCTNNNAACATAGGTTCATCGGCTG
GAGCACACGTCTGAACTCTTTCCCTACACGACGCTCTTCCGATCTNNNTCCTGAAATTCTTGTTTACTGT
GGAAGAGCACACGTCTGAACTCTTTCCCTACACGACGCTCTTCCGATCTNNNGCCAACTGGACAAATAG
GAGCACACGTCTGAACTCTTTCCCTACACGACGCTCTTCCGATCTNNNAGAAGTTTTACTTGGGTCAAT
GAGCACACGTCTGAACTCTTTCCCTACACGACGCTCTTCCGATCTNNNGGATAACAGGCCTTGATATT
AGCACACGTCTGAACTCTTTCCCTACACGACGCTCTTCCGATCTNNNTGGGATTTGTCATAATGAAGCA
AGCACACGTCTGAACTCTTTCCCTACACGACGCTCTTCCGATCTNNNAGGAGGTTTACAAACTCAGATT
AAGAGCACACGTCTGAACTCTTTCCCTACACGACGCTCTTCCGATCTNNNAAAATCCCAATGTCTCTGCA
AAGAGCACACGTCTGAACTCTTTCCCTACACGACGCTCTTCCGATCTNNNTATCCACCACTACCTCAGCT
AGAGCACACGTCTGAACTCTTTCCCTACACGACGCTCTTCCGATCTNNNACACAGGTAGGTTAGAAGG
AAGAGCACACGTCTGAACTCTTTCCCTACACGACGCTCTTCCGATCTNNNCCCTGTCCTGAACAGAGAAA
AAGAGCACACGTCTGAACTCTTTCCCTACACGACGCTCTTCCGATCTNNNCCCAGCCGATATCCTGTTC
AGAGCACACGTCTGAACTCTTTCCCTACACGACGCTCTTCCGATCTNNNTTAGCTCCTCAAATGCACCAA
AGAGCACACGTCTGAACTCTTTCCCTACACGACGCTCTTCCGATCTNNNACTATTTACATTCTTTGTACA
GAAGAGCACACGTCTGAACTCTTTCCCTACACGACGCTCTTCCGATCTNNNAAAGTAGCAAAGGAAAG
AGAGCACACGTCTGAACTCTTTCCCTACACGACGCTCTTCCGATCTNNNCCCCTCAAGATCCTTCCTCGA
AGCACACGTCTGAACTCTTTCCCTACACGACGCTCTTCCGATCTNNNACATTGGAAACACGATTGGATT
GGAAGAGCACACGTCTGAACTCTTTCCCTACACGACGCTCTTCCGATCTNNNCCCCCTCTAGCTTGCAAA
AGCACACGTCTGAACTCTTTCCCTACACGACGCTCTTCCGATCTNNNTCTTCAGAGAATTGTCTTTGTA
AAGAGCACACGTCTGAACTCTTTCCCTACACGACGCTCTTCCGATCTNNNTAGCTTTTGAAATCAGCCCA
AAGAGCACACGTCTGAACTCTTTCCCTACACGACGCTCTTCCGATCTNNNCAGAACTCCAGGCTCTCTG
GAGCACACGTCTGAACTCTTTCCCTACACGACGCTCTTCCGATCTNNNTGACATTTCCATAATGGTTAG
GAAGAGCACACGTCTGAACTCTTTCCCTACACGACGCTCTTCCGATCTNNNCCAGCAGTGTACAAAAGT
AGAGCACACGTCTGAACTCTTTCCCTACACGACGCTCTTCCGATCTNNNCAGAGAGAATATATATGTGT
AGCACACGTCTGAACTCTTTCCCTACACGACGCTCTTCCGATCTNNNACTCACTTGAACAGATTTGTAAG
GAAGAGCACACGTCTGAACTCTTTCCCTACACGACGCTCTTCCGATCTNNNCGTTTGTGGAGAGCTGG
AGCACACGTCTGAACTCTTTCCCTACACGACGCTCTTCCGATCTNNNTGGATTCTAATCTCTGGTGCAT
GAAGAGCACACGTCTGAACTCTTTCCCTACACGACGCTCTTCCGATCTNNNCCAGGAAGGAAGGCTTT
AGAGCACACGTCTGAACTCTTTCCCTACACGACGCTCTTCCGATCTNNNATGTCATTCTCTCTCTGTCCC
AGCACACGTCTGAACTCTTTCCCTACACGACGCTCTTCCGATCTNNNAGTAATTCAAAAAGGCCAATGT

GCACACGTCTGAACTCTTTCCCTACACGACGCTCTTCCGATCTNNNTGTATTACAGATTGTATTTTGCCA
AGCACACGTCTGAACTCTTTCCCTACACGACGCTCTTCCGATCTNNNTGTTGTTAGTGCATAGTTTGA
GCACACGTCTGAACTCTTTCCCTACACGACGCTCTTCCGATCTNNNACCTTGTTATTTATATTTGCGTCT
AGAGCACACGTCTGAACTCTTTCCCTACACGACGCTCTTCCGATCTNNNCTCCCCAACAGTGGTTGAAA
AAGAGCACACGTCTGAACTCTTTCCCTACACGACGCTCTTCCGATCTNNNTGCAACATTCTTTCATCTCA
GAAGAGCACACGTCTGAACTCTTTCCCTACACGACGCTCTTCCGATCTNNNTTTCCCCCTCCACATGCAT
GAGCACACGTCTGAACTCTTTCCCTACACGACGCTCTTCCGATCTNNNTCCATAGATGTGGATGGATA
GAAGAGCACACGTCTGAACTCTTTCCCTACACGACGCTCTTCCGATCTNNNTCTAGAGAGCCCAAGTC
AAGAGCACACGTCTGAACTCTTTCCCTACACGACGCTCTTCCGATCTNNNTGGCAATTTACATACACAC
GCACACGTCTGAACTCTTTCCCTACACGACGCTCTTCCGATCTNNNTCCTTAGAAATAAAGTGGAAAG
AGAGCACACGTCTGAACTCTTTCCCTACACGACGCTCTTCCGATCTNNNCTCCTTCAACAGAAGATGCT
AGAGCACACGTCTGAACTCTTTCCCTACACGACGCTCTTCCGATCTNNNTGGGCTAAGACTTCCTAGG
AGAGCACACGTCTGAACTCTTTCCCTACACGACGCTCTTCCGATCTNNNGGCACACAATTGTCCCTACA
AGCACACGTCTGAACTCTTTCCCTACACGACGCTCTTCCGATCTNNNGCACTCTTTACCTAGATTCACCT
GAAGAGCACACGTCTGAACTCTTTCCCTACACGACGCTCTTCCGATCTNNNTCTAATTCTTGGGGGCTT
AGAGCACACGTCTGAACTCTTTCCCTACACGACGCTCTTCCGATCTNNNGCTCCGTTCCACTTATTTCTC
AAGAGCACACGTCTGAACTCTTTCCCTACACGACGCTCTTCCGATCTNNNCCTGACAGTTGGACTGGG
AAGAGCACACGTCTGAACTCTTTCCCTACACGACGCTCTTCCGATCTNNNTTTTTCCCTTCCTTCCTTTGG
GCACACGTCTGAACTCTTTCCCTACACGACGCTCTTCCGATCTNNNAGCTGTAAATGTTTCTCTCTCTCT
AGAGCACACGTCTGAACTCTTTCCCTACACGACGCTCTTCCGATCTNNNTGCAAGATGAAAGTGTGTGT
AAGAGCACACGTCTGAACTCTTTCCCTACACGACGCTCTTCCGATCTNNNGCAGATCATGAGACTGCTA
GAAGAGCACACGTCTGAACTCTTTCCCTACACGACGCTCTTCCGATCTNNNCAGGTCTTTCTCAATAGCA
GAGCACACGTCTGAACTCTTTCCCTACACGACGCTCTTCCGATCTNNNACTTTGCTAAGCTAATAACA
GAAGAGCACACGTCTGAACTCTTTCCCTACACGACGCTCTTCCGATCTNNNCGCATAGCTCCTATCACCT
GAGCACACGTCTGAACTCTTTCCCTACACGACGCTCTTCCGATCTNNNAGAAGATTTTAGGTGCTGTTA
AAGAGCACACGTCTGAACTCTTTCCCTACACGACGCTCTTCCGATCTNNNACAGAACCAACAGAGACAG
GAGCACACGTCTGAACTCTTTCCCTACACGACGCTCTTCCGATCTNNNTTTTGGTTCTTTCCGAGTTGT
AGCACACGTCTGAACTCTTTCCCTACACGACGCTCTTCCGATCTNNNTCCTTTCAATAGTTTTTGACCCT
AAGAGCACACGTCTGAACTCTTTCCCTACACGACGCTCTTCCGATCTNNNAGGAAAACACATCAGGCT
GAAGAGCACACGTCTGAACTCTTTCCCTACACGACGCTCTTCCGATCTNNNATGAGTGATTCTGGGTGG
AGAGCACACGTCTGAACTCTTTCCCTACACGACGCTCTTCCGATCTNNNCCTGCAAGGAAAGACTATG
GAGCACACGTCTGAACTCTTTCCCTACACGACGCTCTTCCGATCTNNNGCTTGAAACTTGAAACAATGT
AAGAGCACACGTCTGAACTCTTTCCCTACACGACGCTCTTCCGATCTNNNAGCGAATGTTGAGTTGAC
AGAGCACACGTCTGAACTCTTTCCCTACACGACGCTCTTCCGATCTNNNAGGACATTTTGAGTACTGCT
GAGCACACGTCTGAACTCTTTCCCTACACGACGCTCTTCCGATCTNNNACAGGGCACTGTACTTGATT
AGAGCACACGTCTGAACTCTTTCCCTACACGACGCTCTTCCGATCTNNNTCAAAGCACTCTCCAAAGCCA
AGAGCACACGTCTGAACTCTTTCCCTACACGACGCTCTTCCGATCTNNNACAATGTAAGCTTCCAGGTA
AGAGCACACGTCTGAACTCTTTCCCTACACGACGCTCTTCCGATCTNNNTGGAAATCCTTAGATTGCCT
AGCACACGTCTGAACTCTTTCCCTACACGACGCTCTTCCGATCTNNNTGCTTTGTTTGTATGTTAGATT
AAGAGCACACGTCTGAACTCTTTCCCTACACGACGCTCTTCCGATCTNNNCTCAAGGAGAAAGGGCAT
AAGAGCACACGTCTGAACTCTTTCCCTACACGACGCTCTTCCGATCTNNNTCAGTTAGCTCTACATGGT
AGCACACGTCTGAACTCTTTCCCTACACGACGCTCTTCCGATCTNNNACATGAAGATCACTTCAAACTTA
AGCACACGTCTGAACTCTTTCCCTACACGACGCTCTTCCGATCTNNNAGCTTCTCTCATTACTTTTCACTG

AGCACACGTCTGAACTCTTTCCCTACACGACGCTCTTCCGATCTNNNAGATAACAAGTATGTGTATGT

AAGAGCACACGTCTGAACTCTTTCCCTACACGACGCTCTTCCGATCTNNNTGGAAATCCTTAGATTGCC

GAGCACACGTCTGAACTCTTTCCCTACACGACGCTCTTCCGATCTNNNAGAGAATCCTAATGAGAGAC

GAGCACACGTCTGAACTCTTTCCCTACACGACGCTCTTCCGATCTNNNTTCCAAAAGTTCAAACAGAAT

AAGAGCACACGTCTGAACTCTTTCCCTACACGACGCTCTTCCGATCTNNNTCTCAGAGAGCTTCTTCGG

AGAGCACACGTCTGAACTCTTTCCCTACACGACGCTCTTCCGATCTNNNTCTTTATACTCACCTGTGCCT

AAGAGCACACGTCTGAACTCTTTCCCTACACGACGCTCTTCCGATCTNNNGGCAACAGCTAATTCACA

GAGCACACGTCTGAACTCTTTCCCTACACGACGCTCTTCCGATCTNNNGCAAGAACTTTATCCATCCTT

AGCACACGTCTGAACTCTTTCCCTACACGACGCTCTTCCGATCTNNNTGCTTTGGAAATACATCAGGA

AGCACACGTCTGAACTCTTTCCCTACACGACGCTCTTCCGATCTNNNTCGATATTATTCAGATTTTCCG

AAGAGCACACGTCTGAACTCTTTCCCTACACGACGCTCTTCCGATCTNNNTGTAGTTGACAAGGAGA

AGCACACGTCTGAACTCTTTCCCTACACGACGCTCTTCCGATCTNNNACTCATTTTGGCATGTTGATTT

AGAGCACACGTCTGAACTCTTTCCCTACACGACGCTCTTCCGATCTNNNGGGCAAGGAAAAGTGTGT

AAGAGCACACGTCTGAACTCTTTCCCTACACGACGCTCTTCCGATCTNNNTGTTCTGCTTCCACTTTCA

AGAGCACACGTCTGAACTCTTTCCCTACACGACGCTCTTCCGATCTNNNTACTGTGCCTAAGCCATTTG

AGCACACGTCTGAACTCTTTCCCTACACGACGCTCTTCCGATCTNNNGCATTACTAAATAACTTGCAAA

AAGAGCACACGTCTGAACTCTTTCCCTACACGACGCTCTTCCGATCTNNNTTCCACAATCCCAGCTTTT

GAGCACACGTCTGAACTCTTTCCCTACACGACGCTCTTCCGATCTNNNTGCATTTTATCATGTTGGATG

AGCACACGTCTGAACTCTTTCCCTACACGACGCTCTTCCGATCTNNNGGAATGGATGGATACCTCCTT

AGCACACGTCTGAACTCTTTCCCTACACGACGCTCTTCCGATCTNNNAGTGCATTTTTATTTCCTCTCTC

AGAGCACACGTCTGAACTCTTTCCCTACACGACGCTCTTCCGATCTNNNGCTCTATGTGCTTGTTATTG

AGAGCACACGTCTGAACTCTTTCCCTACACGACGCTCTTCCGATCTNNNATTATGCCGTCCTGGGTAT

GAGCACACGTCTGAACTCTTTCCCTACACGACGCTCTTCCGATCTNNNAGGGAAAGAAAGAAAGAAA

GCACACGTCTGAACTCTTTCCCTACACGACGCTCTTCCGATCTNNNAGACTGCATCTCAAAAAAAAGAA

AGAGCACACGTCTGAACTCTTTCCCTACACGACGCTCTTCCGATCTNNNCTGAGGCAGGAGAATAGC

AGCACACGTCTGAACTCTTTCCCTACACGACGCTCTTCCGATCTNNNTGCCAATCACAAAAATGTTCAA

AAGAGCACACGTCTGAACTCTTTCCCTACACGACGCTCTTCCGATCTNNNACACCTGTAATCCCAGCTA

AAGAGCACACGTCTGAACTCTTTCCCTACACGACGCTCTTCCGATCTNNNTGTAGTCCCAGCTACTAG

AAGAGCACACGTCTGAACTCTTTCCCTACACGACGCTCTTCCGATCTNNNGTGATACAGGCTTCGGTG

GGAAGAGCACACGTCTGAACTCTTTCCCTACACGACGCTCTTCCGATCTNNNTGAGGCAAGAGAATG

GCACACGTCTGAACTCTTTCCCTACACGACGCTCTTCCGATCTNNNACGATGCATTTTGAGTTTAGTTT

AGCACACGTCTGAACTCTTTCCCTACACGACGCTCTTCCGATCTNNNCGTTGAAATGCAAGAAAATTTT

AGCACACGTCTGAACTCTTTCCCTACACGACGCTCTTCCGATCTNNNGCAATAAATATCCGTGTTTGT

AGCACACGTCTGAACTCTTTCCCTACACGACGCTCTTCCGATCTNNNAGTGCTTATGATATGTCAGACA

AAGAGCACACGTCTGAACTCTTTCCCTACACGACGCTCTTCCGATCTNNNAACTTGCAGTGAGCCAAG

GCACACGTCTGAACTCTTTCCCTACACGACGCTCTTCCGATCTNNNACTCCTGTCTTTTCTACATATTTGT

GCACACGTCTGAACTCTTTCCCTACACGACGCTCTTCCGATCTNNNTGCAGTACTATTTTGTTATACCAT

AAGAGCACACGTCTGAACTCTTTCCCTACACGACGCTCTTCCGATCTNNNGATTTCTTGAGCCTGAGA

AGCACACGTCTGAACTCTTTCCCTACACGACGCTCTTCCGATCTNNNACAAAAAAATTAGTGATTGCC

AGCACACGTCTGAACTCTTTCCCTACACGACGCTCTTCCGATCTNNNTGCAGAAATAGGTATCAACATT

AAGAGCACACGTCTGAACTCTTTCCCTACACGACGCTCTTCCGATCTNNNGAGAATCACTTGAACCCG

GAAGAGCACACGTCTGAACTCTTTCCCTACACGACGCTCTTCCGATCTNNNTCTGTAATCTCAGCTAG

AAGAGCACACGTCTGAACTCTTTCCCTACACGACGCTCTTCCGATCTNNNGTGCCTGTAGTCCCAGTT

AAGAGCACACGTCTGAACTCTTTCCCTACACGACGCTCTTCCGATCTNNNGGGAGGGATAGCATTGG
AAGAGCACACGTCTGAACTCTTTCCCTACACGACGCTCTTCCGATCTNNNATCCATTGAGCCTGGAAAG
AGCACACGTCTGAACTCTTTCCCTACACGACGCTCTTCCGATCTNNNAGATGCAAGATAAAAGTGATG
AGAGCACACGTCTGAACTCTTTCCCTACACGACGCTCTTCCGATCTNNNGAAATGTAGGTTGCAGTGA
GAGCACACGTCTGAACTCTTTCCCTACACGACGCTCTTCCGATCTNNNTCAAGTGTCAAAAACTATTGAA
GGAAGAGCACACGTCTGAACTCTTTCCCTACACGACGCTCTTCCGATCTNNNTTGCACAAGTGGGTGA
AGCACACGTCTGAACTCTTTCCCTACACGACGCTCTTCCGATCTNNNTCAGACAGAGATCTACATGAGA
AAGAGCACACGTCTGAACTCTTTCCCTACACGACGCTCTTCCGATCTNNNACCTAATGCTAGATGACAC
GAAGAGCACACGTCTGAACTCTTTCCCTACACGACGCTCTTCCGATCTNNNAGTCTGAAGCTGCAGTGT
AGCACACGTCTGAACTCTTTCCCTACACGACGCTCTTCCGATCTNNNACATCAAAAGTCATAGGCGTTT
AGCACACGTCTGAACTCTTTCCCTACACGACGCTCTTCCGATCTNNNCCACCTCTCTAATACTCAAAGTC
AAGAGCACACGTCTGAACTCTTTCCCTACACGACGCTCTTCCGATCTNNNGAGCTTACAGTGAGCAGAG
AAGAGCACACGTCTGAACTCTTTCCCTACACGACGCTCTTCCGATCTNNNATGATCATACCGCCACTACA
AAGAGCACACGTCTGAACTCTTTCCCTACACGACGCTCTTCCGATCTNNNGTCTCAGCTACTCCTAGGAG
GAAGAGCACACGTCTGAACTCTTTCCCTACACGACGCTCTTCCGATCTNNNGTTGCAGTGAGCTGAGA
CGGAAGAGCACACGTCTGAACTCTTTCCCTACACGACGCTCTTCCGATCTNNNCAGGAGAATCACCTAA
GAGCACACGTCTGAACTCTTTCCCTACACGACGCTCTTCCGATCTNNNTGGAGAAGTCAATATTTGAAT
GCACACGTCTGAACTCTTTCCCTACACGACGCTCTTCCGATCTNNNAGTGATAGAGGGAATTTATTCTT
AAGAGCACACGTCTGAACTCTTTCCCTACACGACGCTCTTCCGATCTNNNAAGGAGAATCATTTGAACG
AAGAGCACACGTCTGAACTCTTTCCCTACACGACGCTCTTCCGATCTNNNGAAGGAGAATGGTGAGAA
AGAGCACACGTCTGAACTCTTTCCCTACACGACGCTCTTCCGATCTNNNATACCCTTGTGTAGGTGTGT
AAGAGCACACGTCTGAACTCTTTCCCTACACGACGCTCTTCCGATCTNNNAAACATCCAGCCTGAATGT
GCACACGTCTGAACTCTTTCCCTACACGACGCTCTTCCGATCTNNNAGTTTATAAACTTCATGAAAGCCA
AGCACACGTCTGAACTCTTTCCCTACACGACGCTCTTCCGATCTNNNGCTTTAGCACAAATCATTCTACA
AGCACACGTCTGAACTCTTTCCCTACACGACGCTCTTCCGATCTNNNTGACAATGACAGTAGTCTTTCT
AGCACACGTCTGAACTCTTTCCCTACACGACGCTCTTCCGATCTNNNTTGGCATTTATGTAATACCATGA
GAAGAGCACACGTCTGAACTCTTTCCCTACACGACGCTCTTCCGATCTNNNTTACAGACACTGGTGAC
AAGAGCACACGTCTGAACTCTTTCCCTACACGACGCTCTTCCGATCTNNNAGACAGGAGAATCACTTG
AAGAGCACACGTCTGAACTCTTTCCCTACACGACGCTCTTCCGATCTNNNCTGAGGCAGGAAAATTGC
GAAGAGCACACGTCTGAACTCTTTCCCTACACGACGCTCTTCCGATCTNNNCAAGAGAATTGCTTGAA
AGAGCACACGTCTGAACTCTTTCCCTACACGACGCTCTTCCGATCTNNNTCCACAGAAAAAATGAGTG
GCACACGTCTGAACTCTTTCCCTACACGACGCTCTTCCGATCTNNNTGTGTGTATATATATGTGTGTGT
AAGAGCACACGTCTGAACTCTTTCCCTACACGACGCTCTTCCGATCTNNNGGTGGCAAGTATCTGTAA
AGAGCACACGTCTGAACTCTTTCCCTACACGACGCTCTTCCGATCTNNNGCTGCTCTAGTTCACTGCTC
AAGAGCACACGTCTGAACTCTTTCCCTACACGACGCTCTTCCGATCTNNNATTTTTGCTTTTGTTGCCTG
AGAGCACACGTCTGAACTCTTTCCCTACACGACGCTCTTCCGATCTNNNCCCTTCCTGATGATCCAGTCA
AGCACACGTCTGAACTCTTTCCCTACACGACGCTCTTCCGATCTNNNCCAGTATTCCTTCTTATGGAGTT
AGAGCACACGTCTGAACTCTTTCCCTACACGACGCTCTTCCGATCTNNNTCTGGTGATCTTCATTTCTTC
GAGCACACGTCTGAACTCTTTCCCTACACGACGCTCTTCCGATCTNNNTCCATTGACTATACATATCACT
GCACACGTCTGAACTCTTTCCCTACACGACGCTCTTCCGATCTNNNTGAAATGACTGAGAAGATACAT
AAGAGCACACGTCTGAACTCTTTCCCTACACGACGCTCTTCCGATCTNNNGTGAGTCTCCGTCTCAGGA
AGCACACGTCTGAACTCTTTCCCTACACGACGCTCTTCCGATCTNNNAGAAAACTCAAGAAGAAAATG
GAGCACACGTCTGAACTCTTTCCCTACACGACGCTCTTCCGATCTNNNAATGAACATTGCATTACTTTTT

AAGAGCACACGTCTGAACTCTTTCCCTACACGACGCTCTTCCGATCTNNNCTGTCAGGCTTGCTAAAG
AGAGCACACGTCTGAACTCTTTCCCTACACGACGCTCTTCCGATCTNNNCTGAGGCAGGAGAATTGT
AGCACACGTCTGAACTCTTTCCCTACACGACGCTCTTCCGATCTNNNTCGACCTGAATATTTGGAATA
AGAGCACACGTCTGAACTCTTTCCCTACACGACGCTCTTCCGATCTNNNTGAAATTGGCCACTGCACT
GCACACGTCTGAACTCTTTCCCTACACGACGCTCTTCCGATCTNNNGTCAAATGGAAATATTGTAATG
CACACGTCTGAACTCTTTCCCTACACGACGCTCTTCCGATCTNNNTCTTTCTTTCTCTTTCTTTCTCTTTC
AAGAGCACACGTCTGAACTCTTTCCCTACACGACGCTCTTCCGATCTNNNAATCGCTTGAACTCAGG
AAGAGCACACGTCTGAACTCTTTCCCTACACGACGCTCTTCCGATCTNNNCAAAACTGCCCTCCTGAAT
AGAGCACACGTCTGAACTCTTTCCCTACACGACGCTCTTCCGATCTNNNTGCTAAGAGCTGAAGAGA
AGAGCACACGTCTGAACTCTTTCCCTACACGACGCTCTTCCGATCTNNNGGAGGTTACAGTGAGCTG
AGAGCACACGTCTGAACTCTTTCCCTACACGACGCTCTTCCGATCTNNNTTCTGATTCAGTACTTCTGG
AAGAGCACACGTCTGAACTCTTTCCCTACACGACGCTCTTCCGATCTNNNGATCTCCTGACCTTGTGAT
GAAGAGCACACGTCTGAACTCTTTCCCTACACGACGCTCTTCCGATCTNNNATAGAAAGTGAGAGAC
GAGCACACGTCTGAACTCTTTCCCTACACGACGCTCTTCCGATCTNNNACATATCAAATGTGAGACAT
GAGCACACGTCTGAACTCTTTCCCTACACGACGCTCTTCCGATCTNNNCAGGAAAGAGCACAAGCAT
AAGAGCACACGTCTGAACTCTTTCCCTACACGACGCTCTTCCGATCTNNNCATTCCACACTCCAAAAGC
AGAGCACACGTCTGAACTCTTTCCCTACACGACGCTCTTCCGATCTNNNGCCTTCGTAATTGTGTGAA
AGAGCACACGTCTGAACTCTTTCCCTACACGACGCTCTTCCGATCTNNNAGGAAAGCTTTTCCTGTAAT
AGCACACGTCTGAACTCTTTCCCTACACGACGCTCTTCCGATCTNNNTCTGTATAGAAAGGCTTTGTT
GCACACGTCTGAACTCTTTCCCTACACGACGCTCTTCCGATCTNNNTGTTTTCTTCCCTGATCATATCCA
AGCACACGTCTGAACTCTTTCCCTACACGACGCTCTTCCGATCTNNNTGTGTGTATATGTGTGTGTGT
AAGAGCACACGTCTGAACTCTTTCCCTACACGACGCTCTTCCGATCTNNNAACTCAGGCAAAACACGT
AAGAGCACACGTCTGAACTCTTTCCCTACACGACGCTCTTCCGATCTNNNTAGGCCACAGTGACCTAT
GCACACGTCTGAACTCTTTCCCTACACGACGCTCTTCCGATCTNNNTGAAAAGAAAAAGAAATGCCCT
GAGCACACGTCTGAACTCTTTCCCTACACGACGCTCTTCCGATCTNNNTTCCATCCAGATTGCTGCAA
GAGCACACGTCTGAACTCTTTCCCTACACGACGCTCTTCCGATCTNNNTCAGAATGAGGAATCTGTAA
AGCACACGTCTGAACTCTTTCCCTACACGACGCTCTTCCGATCTNNNTTGAAAGAAAAGTCACACTAG
AGAGCACACGTCTGAACTCTTTCCCTACACGACGCTCTTCCGATCTNNNCTTGAAACAATTGCGCAAA
AGCACACGTCTGAACTCTTTCCCTACACGACGCTCTTCCGATCTNNNTCTGCCAGCTAAACTAATTTTG
AAGAGCACACGTCTGAACTCTTTCCCTACACGACGCTCTTCCGATCTNNNGCCAAGATCATGTTACTG
AGAGCACACGTCTGAACTCTTTCCCTACACGACGCTCTTCCGATCTNNNGGACAAGAGTGGGATTTG
GAGCACACGTCTGAACTCTTTCCCTACACGACGCTCTTCCGATCTNNNTGGACATAACATTTTCAACA
AGCACACGTCTGAACTCTTTCCCTACACGACGCTCTTCCGATCTNNNACAGGAGTTTTATGTGTATAT
AGAGCACACGTCTGAACTCTTTCCCTACACGACGCTCTTCCGATCTNNNAGATCTTCATGACCTAGGAT
AGAGCACACGTCTGAACTCTTTCCCTACACGACGCTCTTCCGATCTNNNCACACACACAATCCCAGTTC
AGCACACGTCTGAACTCTTTCCCTACACGACGCTCTTCCGATCTNNNTGTGTGTGTGTATATATGT
GAAGAGCACACGTCTGAACTCTTTCCCTACACGACGCTCTTCCGATCTNNNAGAGGGACAGAACTAA
AGCACACGTCTGAACTCTTTCCCTACACGACGCTCTTCCGATCTNNNAGGAAAGGAAGAGTAAAAAG
GAGCACACGTCTGAACTCTTTCCCTACACGACGCTCTTCCGATCTNNNACGAGAGTCCATTTCAAAAA
AGCACACGTCTGAACTCTTTCCCTACACGACGCTCTTCCGATCTNNNGCAAATATGTATTTCTTTGCTT
GCACACGTCTGAACTCTTTCCCTACACGACGCTCTTCCGATCTNNNTCTGACACACACATATATATACA
GCACACGTCTGAACTCTTTCCCTACACGACGCTCTTCCGATCTNNNCCAATGTTGGGAAAAAAGAAG
AGCACACGTCTGAACTCTTTCCCTACACGACGCTCTTCCGATCTNNNCAAACTATTATTCAACCAAATT

GAGCACACGTCTGAACTCTTTCCCTACACGACGCTCTTCCGATCTNNNACACAAAACATCATGAATGAA
GAGCACACGTCTGAACTCTTTCCCTACACGACGCTCTTCCGATCTNNNAGTTAGCTTTTTCTCACCATAT
GAAGAGCACACGTCTGAACTCTTTCCCTACACGACGCTCTTCCGATCTNNNTTTGCAATCCACCACAATC
GCACACGTCTGAACTCTTTCCCTACACGACGCTCTTCCGATCTNNNATAGATGCTGTAGTATCCATAGT
GAGCACACGTCTGAACTCTTTCCCTACACGACGCTCTTCCGATCTNNNTGTGTGACTGGTTAGTAAAAA
AAGAGCACACGTCTGAACTCTTTCCCTACACGACGCTCTTCCGATCTNNNAGATGGCAAATCATGGGA
AAGAGCACACGTCTGAACTCTTTCCCTACACGACGCTCTTCCGATCTNNNAGACATGTGTATACCATAC
GAGCACACGTCTGAACTCTTTCCCTACACGACGCTCTTCCGATCTNNNAATATTTGGGTGAGAAGGAA
AAGAGCACACGTCTGAACTCTTTCCCTACACGACGCTCTTCCGATCTNNNGATCTCAACTCACTGCAAC
GAAGAGCACACGTCTGAACTCTTTCCCTACACGACGCTCTTCCGATCTNNNTACAACACGCATCAGAAG
GAGCACACGTCTGAACTCTTTCCCTACACGACGCTCTTCCGATCTNNNTGAAACCATCTTTGAAGAATG
AGAGCACACGTCTGAACTCTTTCCCTACACGACGCTCTTCCGATCTNNNACTCAAATATCTCTGAGGTA
GAGCACACGTCTGAACTCTTTCCCTACACGACGCTCTTCCGATCTNNNAGGAATGAAATGTTTTCTGTC
GAAGAGCACACGTCTGAACTCTTTCCCTACACGACGCTCTTCCGATCTNNNACGCACGAAGGTATATTC
AGCACACGTCTGAACTCTTTCCCTACACGACGCTCTTCCGATCTNNNTCTTTCTTCTCTCTCTCTCTTTC
GAAGAGCACACGTCTGAACTCTTTCCCTACACGACGCTCTTCCGATCTNNNGAGATCACACCACTGTAC
AAGAGCACACGTCTGAACTCTTTCCCTACACGACGCTCTTCCGATCTNNNCTCCTTGACCTCCCAAAGTA
GCACACGTCTGAACTCTTTCCCTACACGACGCTCTTCCGATCTNNNTGGAAATTTCATTCTAGTGTAGTC
GAGCACACGTCTGAACTCTTTCCCTACACGACGCTCTTCCGATCTNNNTGTCTCAAACAAGAGAGAGAC
GAGCACACGTCTGAACTCTTTCCCTACACGACGCTCTTCCGATCTNNNACTCCATCTCAAAAAAGAAAGA
AGCACACGTCTGAACTCTTTCCCTACACGACGCTCTTCCGATCTNNNACTCTTGCTTTCTTTCTTTCTTTC
GAGCACACGTCTGAACTCTTTCCCTACACGACGCTCTTCCGATCTNNNATTCATGTATATATGTGTGTGT
AGCACACGTCTGAACTCTTTCCCTACACGACGCTCTTCCGATCTNNNGCCTACTCTAAGATAAGAAGTA
CGGAAGAGCACACGTCTGAACTCTTTCCCTACACGACGCTCTTCCGATCTNNNCACCATTGCACTCCAG
AAGAGCACACGTCTGAACTCTTTCCCTACACGACGCTCTTCCGATCTNNNTCAACAAAAATCATGCTGG
CGGAAGAGCACACGTCTGAACTCTTTCCCTACACGACGCTCTTCCGATCTNNNGGAGAATCACTTGAA
AGAGCACACGTCTGAACTCTTTCCCTACACGACGCTCTTCCGATCTNNNCAGTGTTTCTTACGCCAGTGT
CGGAAGAGCACACGTCTGAACTCTTTCCCTACACGACGCTCTTCCGATCTNNNGAGATAGGCCACTCCA
AGCACACGTCTGAACTCTTTCCCTACACGACGCTCTTCCGATCTNNNTCTTTCTCTCTCTCTCTTTCTTTCT
AGAGCACACGTCTGAACTCTTTCCCTACACGACGCTCTTCCGATCTNNNCCAGAGAAACTCTTCAGAG
GAAGAGCACACGTCTGAACTCTTTCCCTACACGACGCTCTTCCGATCTNNNTGGGCCCATAGAGTAGAC
AGCACACGTCTGAACTCTTTCCCTACACGACGCTCTTCCGATCTNNNTCTGCAGTTTGAATATGGTTTCC
AGAGCACACGTCTGAACTCTTTCCCTACACGACGCTCTTCCGATCTNNNAGAAATCATGTGAGGGGGT
AAGAGCACACGTCTGAACTCTTTCCCTACACGACGCTCTTCCGATCTNNNTAGCATGGTGACCACGGT
AGCACACGTCTGAACTCTTTCCCTACACGACGCTCTTCCGATCTNNNGGAGGATCCTTAGGTCAGGAG
AGAGCACACGTCTGAACTCTTTCCCTACACGACGCTCTTCCGATCTNNNAAGTCTTCAGGGTCTAGAGA
GCACACGTCTGAACTCTTTCCCTACACGACGCTCTTCCGATCTNNNTCAGATCCACATTTTCCCAGTTTC
AGAGCACACGTCTGAACTCTTTCCCTACACGACGCTCTTCCGATCTNNNACATGATCCAGGGACAGTGC
AGAGCACACGTCTGAACTCTTTCCCTACACGACGCTCTTCCGATCTNNNACCTTACTATGTTCCCAAAATC
AGAGCACACGTCTGAACTCTTTCCCTACACGACGCTCTTCCGATCTNNNGCTCACATGGCACCTCTCATA
GAGCACACGTCTGAACTCTTTCCCTACACGACGCTCTTCCGATCTNNNATAGAAGTAAGGGAGGAGG
AGAGCACACGTCTGAACTCTTTCCCTACACGACGCTCTTCCGATCTNNNCTGTACCTGTAATTGGACCC
AGAGCACACGTCTGAACTCTTTCCCTACACGACGCTCTTCCGATCTNNNACTGAAGGAGAGCATCATGA

GCACACGTCTGAACTCTTTCCCTACACGACGCTCTTCCGATCTNNNACCCACATATATACAGCATTGAA
AGAGCACACGTCTGAACTCTTTCCCTACACGACGCTCTTCCGATCTNNNGGAACTTTAGATCATGAAG
AGCACACGTCTGAACTCTTTCCCTACACGACGCTCTTCCGATCTNNNTGGAGGAGCAAAAAGGCAA
GAGCACACGTCTGAACTCTTTCCCTACACGACGCTCTTCCGATCTNNNGGGCTCTGGAACTTTGAATT
GAGCACACGTCTGAACTCTTTCCCTACACGACGCTCTTCCGATCTNNNGTAATTTCCTCTTGCTCCCCCT
AAGAGCACACGTCTGAACTCTTTCCCTACACGACGCTCTTCCGATCTNNNTGCGCAATTGATCTGTTTG
AGCACACGTCTGAACTCTTTCCCTACACGACGCTCTTCCGATCTNNNACCATGTGGACTTGATGTTCAT
AGCACACGTCTGAACTCTTTCCCTACACGACGCTCTTCCGATCTNNNGGAAGGCAAAGGAATATGAA
GAGCACACGTCTGAACTCTTTCCCTACACGACGCTCTTCCGATCTNNNTCACTGTAGCAGTCTTTCTTC
AGCACACGTCTGAACTCTTTCCCTACACGACGCTCTTCCGATCTNNNTCCTCAATGACAAGGCAATTT
AGAGCACACGTCTGAACTCTTTCCCTACACGACGCTCTTCCGATCTNNNCATTCCTCAGCCCTCTATCCT
AAGAGCACACGTCTGAACTCTTTCCCTACACGACGCTCTTCCGATCTNNNCTCCAGTCCTATTCTCCTTC
AAGAGCACACGTCTGAACTCTTTCCCTACACGACGCTCTTCCGATCTNNNACCTCCTCTGTGATATGGT
GAGCACACGTCTGAACTCTTTCCCTACACGACGCTCTTCCGATCTNNNACACGCCAGGACCTCTAAAAA
GAGCACACGTCTGAACTCTTTCCCTACACGACGCTCTTCCGATCTNNNGTGTTCCTAGACACCTTCTGCT
AGAGCACACGTCTGAACTCTTTCCCTACACGACGCTCTTCCGATCTNNNTGGCCTGGTTGTTTCCGTA
GAGCACACGTCTGAACTCTTTCCCTACACGACGCTCTTCCGATCTNNNTCATCTATACATGGGTTAGCA
AGAGCACACGTCTGAACTCTTTCCCTACACGACGCTCTTCCGATCTNNNTTGCAAGTCTCTGCATATTC
GAGCACACGTCTGAACTCTTTCCCTACACGACGCTCTTCCGATCTNNNCAGTAGGTCACTGCCTATCAA
GAGCACACGTCTGAACTCTTTCCCTACACGACGCTCTTCCGATCTNNNAACCAGGCATATTGTTGGAAG
AGAGCACACGTCTGAACTCTTTCCCTACACGACGCTCTTCCGATCTNNNGCTCTCAGAACTCAGCACAT
GAAGAGCACACGTCTGAACTCTTTCCCTACACGACGCTCTTCCGATCTNNNACTAAGTGCTAAGTCCCA
AGAGCACACGTCTGAACTCTTTCCCTACACGACGCTCTTCCGATCTNNNCCACTTTTGGGCTTTAGCTAC
AGAGCACACGTCTGAACTCTTTCCCTACACGACGCTCTTCCGATCTNNNGGAAATGATGTATGACAGC
AAGAGCACACGTCTGAACTCTTTCCCTACACGACGCTCTTCCGATCTNNNCCTTGGGTTAGGCATCTCA
AGAGCACACGTCTGAACTCTTTCCCTACACGACGCTCTTCCGATCTNNNAGGGAAAGGCAAAATTCAA
AAGAGCACACGTCTGAACTCTTTCCCTACACGACGCTCTTCCGATCTNNNTGCAGTATGATTCCAGGA
GAGCACACGTCTGAACTCTTTCCCTACACGACGCTCTTCCGATCTNNNGCTTACCTTGTGGGGTTGTT
GAAGAGCACACGTCTGAACTCTTTCCCTACACGACGCTCTTCCGATCTNNNCCCGGTCTCATTGAAGG
AGAGCACACGTCTGAACTCTTTCCCTACACGACGCTCTTCCGATCTNNNGCAAATATTGTCAACTGTGT
AGCACACGTCTGAACTCTTTCCCTACACGACGCTCTTCCGATCTNNNACTAATGTCGGTGTCTAGAACC
GAGCACACGTCTGAACTCTTTCCCTACACGACGCTCTTCCGATCTNNNGGAGTGTTTAGCTCAGTTAG
AGAGCACACGTCTGAACTCTTTCCCTACACGACGCTCTTCCGATCTNNNTTGAGGACTGCTGCCTTAT
AGAGCACACGTCTGAACTCTTTCCCTACACGACGCTCTTCCGATCTNNNAAGTGAAAAATCCATCCCCC
AGCACACGTCTGAACTCTTTCCCTACACGACGCTCTTCCGATCTNNNTCTACTTGCCTTTTTCTCTCTCT
AAGAGCACACGTCTGAACTCTTTCCCTACACGACGCTCTTCCGATCTNNNGAGCTGTTAGGCCTTGTG
AGAGCACACGTCTGAACTCTTTCCCTACACGACGCTCTTCCGATCTNNNGAACTCAAAGAAGTACTCCC
GAGCACACGTCTGAACTCTTTCCCTACACGACGCTCTTCCGATCTNNNCCCAGCTGTTTGATTGTTAT
AGAGCACACGTCTGAACTCTTTCCCTACACGACGCTCTTCCGATCTNNNAAAAGTCCCTCTTTGGAAT
GAGCACACGTCTGAACTCTTTCCCTACACGACGCTCTTCCGATCTNNNTTTACGTTCCGGGGATTATGA
AGAGCACACGTCTGAACTCTTTCCCTACACGACGCTCTTCCGATCTNNNACTTTTTGGACTACTGCTCA
AGAGCACACGTCTGAACTCTTTCCCTACACGACGCTCTTCCGATCTNNNATCACAAGCCTGGGGGCCT
AGCACACGTCTGAACTCTTTCCCTACACGACGCTCTTCCGATCTNNNTCTCAACTTATTCTTTCTCCTCG

AGCACACGTCTGAACTCTTTCCCTACACGACGCTCTTCCGATCTNNNACATATCCATAGTCAGGAGAG

GAGCACACGTCTGAACTCTTTCCCTACACGACGCTCTTCCGATCTNNNCACCGTATTGATAGGATAAG

GCACACGTCTGAACTCTTTCCCTACACGACGCTCTTCCGATCTNNNTGGGTAGTAAAACTGTGTCAAC

GCACACGTCTGAACTCTTTCCCTACACGACGCTCTTCCGATCTNNNTGCACTCTATTCTCTCTCTCTG

GAGCACACGTCTGAACTCTTTCCCTACACGACGCTCTTCCGATCTNNNAGTCCCCGTGTCAAGATCTA

GAGCACACGTCTGAACTCTTTCCCTACACGACGCTCTTCCGATCTNNNTCACCCTTGGCTTATTTTGC

AGCACACGTCTGAACTCTTTCCCTACACGACGCTCTTCCGATCTNNNACTGACACATCTATCACCACAC

AGCACACGTCTGAACTCTTTCCCTACACGACGCTCTTCCGATCTNNNTCAAGTGGACAAAATACCCAG

GAGCACACGTCTGAACTCTTTCCCTACACGACGCTCTTCCGATCTNNNAGCTGAATTTATCTATTGCCT

AGAGCACACGTCTGAACTCTTTCCCTACACGACGCTCTTCCGATCTNNNACATGCTATGTTTGGCTAT

AGAGCACACGTCTGAACTCTTTCCCTACACGACGCTCTTCCGATCTNNNAACTTGACTGGGCTAAGG

GAGCACACGTCTGAACTCTTTCCCTACACGACGCTCTTCCGATCTNNNTGTGCCCCTTCCCAATCAATA

AGAGCACACGTCTGAACTCTTTCCCTACACGACGCTCTTCCGATCTNNNAGTACACGTATGAACAGTG

AGCACACGTCTGAACTCTTTCCCTACACGACGCTCTTCCGATCTNNNCATTCTCTCACTTCAGCCTCCAA

GAGCACACGTCTGAACTCTTTCCCTACACGACGCTCTTCCGATCTNNNAAGTCCCTAAGGAGGGCTAG

AGAGCACACGTCTGAACTCTTTCCCTACACGACGCTCTTCCGATCTNNNAAAGACATGTGGAAGCGA

GAGCACACGTCTGAACTCTTTCCCTACACGACGCTCTTCCGATCTNNNTGCAATGGAAGTAGCATAAT

AGAGCACACGTCTGAACTCTTTCCCTACACGACGCTCTTCCGATCTNNNTTTGTACCAGGCATTGGGC

AGCACACGTCTGAACTCTTTCCCTACACGACGCTCTTCCGATCTNNNTGTCAAGAGAAAGACCACTTC

AGAGCACACGTCTGAACTCTTTCCCTACACGACGCTCTTCCGATCTNNNTTACTGTTGCCTTGTGTCC

GAGCACACGTCTGAACTCTTTCCCTACACGACGCTCTTCCGATCTNNNCCCTACAGCGTTTCCAGAAAG

GAGCACACGTCTGAACTCTTTCCCTACACGACGCTCTTCCGATCTNNNTGCAAGTTCGTCCCTGTAAC

AGAGCACACGTCTGAACTCTTTCCCTACACGACGCTCTTCCGATCTNNNGTCTCCATGCAATACAGTG

AGCACACGTCTGAACTCTTTCCCTACACGACGCTCTTCCGATCTNNNGCCCTATGCTAATCACAAAAA

AGCACACGTCTGAACTCTTTCCCTACACGACGCTCTTCCGATCTNNNAGGAGGCATATTGAAAGACTT

AGCACACGTCTGAACTCTTTCCCTACACGACGCTCTTCCGATCTNNNAGAGGTGGAAAATAATGTGG

GAGCACACGTCTGAACTCTTTCCCTACACGACGCTCTTCCGATCTNNNCTTGGGCTATAAACACTCCC

AAGAGCACACGTCTGAACTCTTTCCCTACACGACGCTCTTCCGATCTNNNTCTTCTCTTGTCAGTCAGT

GAGCACACGTCTGAACTCTTTCCCTACACGACGCTCTTCCGATCTNNNGGAGATGGTCAACAAATGC

AGAGCACACGTCTGAACTCTTTCCCTACACGACGCTCTTCCGATCTNNNTGTGGATAACACCCTGGAC

AGAGCACACGTCTGAACTCTTTCCCTACACGACGCTCTTCCGATCTNNNACCTCCCATACAACAGATGC

AGAGCACACGTCTGAACTCTTTCCCTACACGACGCTCTTCCGATCTNNNAAACATACACACACAGCCT

AGAGCACACGTCTGAACTCTTTCCCTACACGACGCTCTTCCGATCTNNNACAAATGTGTGTGTGAGAG

AGAGCACACGTCTGAACTCTTTCCCTACACGACGCTCTTCCGATCTNNNTGTCAGGTGATGTGTGAG

AGAGCACACGTCTGAACTCTTTCCCTACACGACGCTCTTCCGATCTNNNAGCAATTCAACGCTTCTCCT

GCACACGTCTGAACTCTTTCCCTACACGACGCTCTTCCGATCTNNNCCAACAAGATATTTTTGCAAGCC

AGAGCACACGTCTGAACTCTTTCCCTACACGACGCTCTTCCGATCTNNNTCCGGAATAGACTGGACTA

AAGAGCACACGTCTGAACTCTTTCCCTACACGACGCTCTTCCGATCTNNNACACAGCTCAGTACCTAG

AAGAGCACACGTCTGAACTCTTTCCCTACACGACGCTCTTCCGATCTNNNAAACCGGAAAGTGTGTT

AGCACACGTCTGAACTCTTTCCCTACACGACGCTCTTCCGATCTNNNCCTGAACAACTGAATAAAACCC

AGCACACGTCTGAACTCTTTCCCTACACGACGCTCTTCCGATCTNNNGTGTCTTGACAAGTGAAGTGT

AAGAGCACACGTCTGAACTCTTTCCCTACACGACGCTCTTCCGATCTNNNAACAAGACAAGGATGTCA

AGCACACGTCTGAACTCTTTCCCTACACGACGCTCTTCCGATCTNNNTGCTTTGAACACAGGTGGAT

AGAGCACACGTCTGAACTCTTTCCCTACACGACGCTCTTCCGATCTNNNTTCTTTGTGTACTTTCCGCAC
AGAGCACACGTCTGAACTCTTTCCCTACACGACGCTCTTCCGATCTNNNATGTGTTCACCAACTTCCTCT
AAGAGCACACGTCTGAACTCTTTCCCTACACGACGCTCTTCCGATCTNNNGCTAATGGTGAAATGGAG
AGAGCACACGTCTGAACTCTTTCCCTACACGACGCTCTTCCGATCTNNNGCATGCTCCTACTGTGCTGT
AAGAGCACACGTCTGAACTCTTTCCCTACACGACGCTCTTCCGATCTNNNTGGATGTTGAAATGGCCT
AGAGCACACGTCTGAACTCTTTCCCTACACGACGCTCTTCCGATCTNNNTAGCCTTGCCAAGTTGACA
GAGCACACGTCTGAACTCTTTCCCTACACGACGCTCTTCCGATCTNNNTGTTTTGTCTGGCTGCCTTC
AAGAGCACACGTCTGAACTCTTTCCCTACACGACGCTCTTCCGATCTNNNAAAAAGGCTTATTGGGGT
AAGAGCACACGTCTGAACTCTTTCCCTACACGACGCTCTTCCGATCTNNNCTCAGGGATGTTTCCCTC
GAGCACACGTCTGAACTCTTTCCCTACACGACGCTCTTCCGATCTNNNTTTGTAAAAAGCACACACACA
AAGAGCACACGTCTGAACTCTTTCCCTACACGACGCTCTTCCGATCTNNNAGCAATGCTTTGGATGAC
AAGAGCACACGTCTGAACTCTTTCCCTACACGACGCTCTTCCGATCTNNNTTTGTGGGTAAAGAGCC
AAGAGCACACGTCTGAACTCTTTCCCTACACGACGCTCTTCCGATCTNNNACACAAAGATTCCCAGCAG
AGAGCACACGTCTGAACTCTTTCCCTACACGACGCTCTTCCGATCTNNNCCATGTGCTTCAGGGAAGA
AGAGCACACGTCTGAACTCTTTCCCTACACGACGCTCTTCCGATCTNNNCCCGTCTCCTTCCCTCTTATC
AAGAGCACACGTCTGAACTCTTTCCCTACACGACGCTCTTCCGATCTNNNTCAGTATGCAGTCAGTGTG
AGAGCACACGTCTGAACTCTTTCCCTACACGACGCTCTTCCGATCTNNNGCACATGGGAGAAACAAA
AGAGCACACGTCTGAACTCTTTCCCTACACGACGCTCTTCCGATCTNNNGATACCAGGATAACAGTGC
AGAGCACACGTCTGAACTCTTTCCCTACACGACGCTCTTCCGATCTNNNACAGACTGGCCGTTTCTCAA
AGCACACGTCTGAACTCTTTCCCTACACGACGCTCTTCCGATCTNNNACCTCTGGTATAGTTCTTTTCTG
GAGCACACGTCTGAACTCTTTCCCTACACGACGCTCTTCCGATCTNNNAGCCTAGGAACTGGAGAAGT
GAGCACACGTCTGAACTCTTTCCCTACACGACGCTCTTCCGATCTNNNTGCTTGTATAATTCTGGGTGT
AGAGCACACGTCTGAACTCTTTCCCTACACGACGCTCTTCCGATCTNNNCAATTCAGTGCTCCTTTCCC
AAGAGCACACGTCTGAACTCTTTCCCTACACGACGCTCTTCCGATCTNNNATTCTGTCACTTCCCTTCTG
GAGCACACGTCTGAACTCTTTCCCTACACGACGCTCTTCCGATCTNNNACTTTGTTTTGTACTGGTTGTT
AAGAGCACACGTCTGAACTCTTTCCCTACACGACGCTCTTCCGATCTNNNTCTTTGTTCCTCATGTGTG
GAGCACACGTCTGAACTCTTTCCCTACACGACGCTCTTCCGATCTNNNGTCATGTTGGTGGGTCAGAA
AGCACACGTCTGAACTCTTTCCCTACACGACGCTCTTCCGATCTNNNTGAACTTTTGAGGACCCCACA
GAGCACACGTCTGAACTCTTTCCCTACACGACGCTCTTCCGATCTNNNTATTCATAAGCCAGCCCTGTG
AGCACACGTCTGAACTCTTTCCCTACACGACGCTCTTCCGATCTNNNAGCATACTAGCGTGTACATGT
AAGAGCACACGTCTGAACTCTTTCCCTACACGACGCTCTTCCGATCTNNNAGAGCCCTTTGTATTTTGG
AGAGCACACGTCTGAACTCTTTCCCTACACGACGCTCTTCCGATCTNNNTCAGCAGGAAATGAATAAG
AGCACACGTCTGAACTCTTTCCCTACACGACGCTCTTCCGATCTNNNAAAAGTAGAGATTATCAGAGT
AGAGCACACGTCTGAACTCTTTCCCTACACGACGCTCTTCCGATCTNNNGCACTCTGGCCTGGGTAAT
AAGAGCACACGTCTGAACTCTTTCCCTACACGACGCTCTTCCGATCTNNNCTTTGCTTTTATGCGTGTGT
GAGCACACGTCTGAACTCTTTCCCTACACGACGCTCTTCCGATCTNNNAGTGAAATAACCCATTTAGAA
AGCACACGTCTGAACTCTTTCCCTACACGACGCTCTTCCGATCTNNNAGGCCTAATACACATACACACA
GCACACGTCTGAACTCTTTCCCTACACGACGCTCTTCCGATCTNNNGTCAATGACACCAACAGAAACAT
GAAGAGCACACGTCTGAACTCTTTCCCTACACGACGCTCTTCCGATCTNNNATCTCTACAACCCTAGGG
AGAGCACACGTCTGAACTCTTTCCCTACACGACGCTCTTCCGATCTNNNTTTTCTGAAAGTCTTTGCACT
GCACACGTCTGAACTCTTTCCCTACACGACGCTCTTCCGATCTNNNAGATGTACCTTTGGATGTCTGTCT
GAAGAGCACACGTCTGAACTCTTTCCCTACACGACGCTCTTCCGATCTNNNTGACTTCAAGCCCATGTT
AGAGCACACGTCTGAACTCTTTCCCTACACGACGCTCTTCCGATCTNNNAGGGAAATGGAAGTAGTC

GAGCACACGTCTGAACTCTTTCCCTACACGACGCTCTTCCGATCTNNNTTCTGAAAGTGTTTTGATTTC
AGAGCACACGTCTGAACTCTTTCCCTACACGACGCTCTTCCGATCTNNNAAAATGCTAACACCACCACC
AGCACACGTCTGAACTCTTTCCCTACACGACGCTCTTCCGATCTNNNTCAAATCATCTTCCAGTTTCCCTG
AGCACACGTCTGAACTCTTTCCCTACACGACGCTCTTCCGATCTNNNCCAAAGAGTTTGAGAACTGAT
AGAGCACACGTCTGAACTCTTTCCCTACACGACGCTCTTCCGATCTNNNGAATTGGGATTCTCTCACTT
AGAGCACACGTCTGAACTCTTTCCCTACACGACGCTCTTCCGATCTNNNGCACCCCCTTTTAGCATGTA
AAGAGCACACGTCTGAACTCTTTCCCTACACGACGCTCTTCCGATCTNNNTGCTTTATTGAGGTGGAG
AGCACACGTCTGAACTCTTTCCCTACACGACGCTCTTCCGATCTNNNGGCTAAATCATACATACCAGGC
AGAGCACACGTCTGAACTCTTTCCCTACACGACGCTCTTCCGATCTNNNGGAACTGGGGGAACAGGA
GAAGAGCACACGTCTGAACTCTTTCCCTACACGACGCTCTTCCGATCTNNNAGAAGTAAAAGCCTAAG
AGAGCACACGTCTGAACTCTTTCCCTACACGACGCTCTTCCGATCTNNNTCCAAAACACCAGATGCAAA
GCACACGTCTGAACTCTTTCCCTACACGACGCTCTTCCGATCTNNNTGCTTAGAAGTCCTTGGTGTTCT
AGCACACGTCTGAACTCTTTCCCTACACGACGCTCTTCCGATCTNNNACTAAACTGATTTTTGCCATGG
GAGCACACGTCTGAACTCTTTCCCTACACGACGCTCTTCCGATCTNNNACAAAAACAAAAACAGGTGA
AGAGCACACGTCTGAACTCTTTCCCTACACGACGCTCTTCCGATCTNNNTGCCCACAATTACACTTCTAG
AGAGCACACGTCTGAACTCTTTCCCTACACGACGCTCTTCCGATCTNNNTTTCCCAGTCATGTAGACCC
AAGAGCACACGTCTGAACTCTTTCCCTACACGACGCTCTTCCGATCTNNNAGTTGAGAAAAGGGTGG
GAGCACACGTCTGAACTCTTTCCCTACACGACGCTCTTCCGATCTNNNACCATCTCTGTTTCTACTGAG
AAGAGCACACGTCTGAACTCTTTCCCTACACGACGCTCTTCCGATCTNNNGAAGTCTGCATGCTCTCCC
AGAGCACACGTCTGAACTCTTTCCCTACACGACGCTCTTCCGATCTNNNGAAACGCAAATACAGGTG
GAGCACACGTCTGAACTCTTTCCCTACACGACGCTCTTCCGATCTNNNTCAGTGTATGATCTAAAGACA
AGAGCACACGTCTGAACTCTTTCCCTACACGACGCTCTTCCGATCTNNNATATCAGGGTTTCAGAAGG
AGAGCACACGTCTGAACTCTTTCCCTACACGACGCTCTTCCGATCTNNNTCCGGTTCCCGAAAGTTTCT
AGAGCACACGTCTGAACTCTTTCCCTACACGACGCTCTTCCGATCTNNNCAGCTCTCAGGCATTCGAAA
GAGCACACGTCTGAACTCTTTCCCTACACGACGCTCTTCCGATCTNNNAGGCACACCTAAAATCCTAC
GAGCACACGTCTGAACTCTTTCCCTACACGACGCTCTTCCGATCTNNNACACCTACTAGAATTGCAACG
AAGAGCACACGTCTGAACTCTTTCCCTACACGACGCTCTTCCGATCTNNNACCCACACACATACATTTA
AGAGCACACGTCTGAACTCTTTCCCTACACGACGCTCTTCCGATCTNNNAAACCAAATGCCCAGTTATG
GAGCACACGTCTGAACTCTTTCCCTACACGACGCTCTTCCGATCTNNNTAGTCCTTGTGTTCTGCACTC
AAGAGCACACGTCTGAACTCTTTCCCTACACGACGCTCTTCCGATCTNNNATTGAGTGAAAAAGGCA
AAGAGCACACGTCTGAACTCTTTCCCTACACGACGCTCTTCCGATCTNNNACCAGTCCAGGAATAGAG
GAGCACACGTCTGAACTCTTTCCCTACACGACGCTCTTCCGATCTNNNACTCTATTAGGAGGAGTGA
GAAGAGCACACGTCTGAACTCTTTCCCTACACGACGCTCTTCCGATCTNNNAGGAAAGATGGGGATA
AAGAGCACACGTCTGAACTCTTTCCCTACACGACGCTCTTCCGATCTNNNGAACATCAGAAGTGCTGA
GAAGAGCACACGTCTGAACTCTTTCCCTACACGACGCTCTTCCGATCTNNNTGCGTTTGTCTCAGTAAG
AAGAGCACACGTCTGAACTCTTTCCCTACACGACGCTCTTCCGATCTNNNAAAGCATGGCTCTACCCAC
AGAGCACACGTCTGAACTCTTTCCCTACACGACGCTCTTCCGATCTNNNTCCTATTGTTCGATAGAAGT
AGAGCACACGTCTGAACTCTTTCCCTACACGACGCTCTTCCGATCTNNNATGGATTTCAGAGGCCCCA
GAGCACACGTCTGAACTCTTTCCCTACACGACGCTCTTCCGATCTNNNGTGGCTCCAATGACTTTGCT
GAGCACACGTCTGAACTCTTTCCCTACACGACGCTCTTCCGATCTNNNTCAAAGTCTTTTCCTGGGAAA
AAGAGCACACGTCTGAACTCTTTCCCTACACGACGCTCTTCCGATCTNNNGGGCAAGCTTGATCCCTAT
AGAGCACACGTCTGAACTCTTTCCCTACACGACGCTCTTCCGATCTNNNTTCTACCCTCATCACCCCCAT
AAGAGCACACGTCTGAACTCTTTCCCTACACGACGCTCTTCCGATCTNNNAGGTCACTGACAGGCAG

GAGCACACGTCTGAACTCTTTCCCTACACGACGCTCTTCCGATCTNNNTGCCTGTTGTACGTATATTCTT
GAGCACACGTCTGAACTCTTTCCCTACACGACGCTCTTCCGATCTNNNTCTCACTTCTTCTTGATATCCCA
AAGAGCACACGTCTGAACTCTTTCCCTACACGACGCTCTTCCGATCTNNNTGTAAGGAAGATCCAGGGG
AAGAGCACACGTCTGAACTCTTTCCCTACACGACGCTCTTCCGATCTNNNTGATGATTTGGGAGTAGG
AGAGCACACGTCTGAACTCTTTCCCTACACGACGCTCTTCCGATCTNNNATAATGGCAGGGAAAAAGG
AGAGCACACGTCTGAACTCTTTCCCTACACGACGCTCTTCCGATCTNNNTGAGGGTGGATCCCCTATAA
AGAGCACACGTCTGAACTCTTTCCCTACACGACGCTCTTCCGATCTNNNTAGTGTAGCGTGGGTGAATT
AGAGCACACGTCTGAACTCTTTCCCTACACGACGCTCTTCCGATCTNNNTGAAGTTGGCTCCTGTATCCC
AGAGCACACGTCTGAACTCTTTCCCTACACGACGCTCTTCCGATCTNNNATCTGCATTGTGCAACTTTCC
AAGAGCACACGTCTGAACTCTTTCCCTACACGACGCTCTTCCGATCTNNNTTCTCTGGGGACTGTGATG
AAGAGCACACGTCTGAACTCTTTCCCTACACGACGCTCTTCCGATCTNNNCAAGTCAGGTCATGAGGCA
AGAGCACACGTCTGAACTCTTTCCCTACACGACGCTCTTCCGATCTNNNCCTTTTTGGGGCTGAGTACT
AAGAGCACACGTCTGAACTCTTTCCCTACACGACGCTCTTCCGATCTNNNAAACATTTAGTGGTCCGTTC
AGAGCACACGTCTGAACTCTTTCCCTACACGACGCTCTTCCGATCTNNNGTCAACATGCAACCAAAAAG
AGAGCACACGTCTGAACTCTTTCCCTACACGACGCTCTTCCGATCTNNNGCTACCTTCTCCTTTTGTGAG
GAGCACACGTCTGAACTCTTTCCCTACACGACGCTCTTCCGATCTNNNGGTAGCCTGATGAAAGATGT
AGAGCACACGTCTGAACTCTTTCCCTACACGACGCTCTTCCGATCTNNNTGTGGGGAAGAGGAATCTC
AGAGCACACGTCTGAACTCTTTCCCTACACGACGCTCTTCCGATCTNNNGCTTGTGGGAGCACAGTTTT
AGCACACGTCTGAACTCTTTCCCTACACGACGCTCTTCCGATCTNNNACTTTGTCCTTGATATCCTTAGG
GAGCACACGTCTGAACTCTTTCCCTACACGACGCTCTTCCGATCTNNNAGAATCTGGGCATGAAATGG
AAGAGCACACGTCTGAACTCTTTCCCTACACGACGCTCTTCCGATCTNNNTCTCAGAGCCCGCCTAATTC
AGAGCACACGTCTGAACTCTTTCCCTACACGACGCTCTTCCGATCTNNNAAAGTCCTGCAACAGTAAGA
AGAGCACACGTCTGAACTCTTTCCCTACACGACGCTCTTCCGATCTNNNGCAGTTACTGGGCCTCATACA
AGAGCACACGTCTGAACTCTTTCCCTACACGACGCTCTTCCGATCTNNNAAGCAAAACTACCAGGTTCGG
AGAGCACACGTCTGAACTCTTTCCCTACACGACGCTCTTCCGATCTNNNACCAGTCAGGTTCATACCAGT
AGAGCACACGTCTGAACTCTTTCCCTACACGACGCTCTTCCGATCTNNNACCATGACTAGCCAGTTCAG
GAGCACACGTCTGAACTCTTTCCCTACACGACGCTCTTCCGATCTNNNCCATGTGACCAGTTACAGGTA
AAGAGCACACGTCTGAACTCTTTCCCTACACGACGCTCTTCCGATCTNNNAGCTGTGCCTATGGTATGCT
AAGAGCACACGTCTGAACTCTTTCCCTACACGACGCTCTTCCGATCTNNNGGAACAAGCCTAAATGCCC
AAGAGCACACGTCTGAACTCTTTCCCTACACGACGCTCTTCCGATCTNNNGCTATGACTGTGCACTTCTC
AAGAGCACACGTCTGAACTCTTTCCCTACACGACGCTCTTCCGATCTNNNAGTAACTCATTGTGAAGGT
GAGCACACGTCTGAACTCTTTCCCTACACGACGCTCTTCCGATCTNNNTCAACATACCTGTACACACACAC
GAAGAGCACACGTCTGAACTCTTTCCCTACACGACGCTCTTCCGATCTNNNGCCAGGACGGAACTTTTC
AGAGCACACGTCTGAACTCTTTCCCTACACGACGCTCTTCCGATCTNNNATGATGGAGGTCTGTCCAGT
AGAGCACACGTCTGAACTCTTTCCCTACACGACGCTCTTCCGATCTNNNATCACTACCCCACAAAAGCAC
GAGCACACGTCTGAACTCTTTCCCTACACGACGCTCTTCCGATCTNNNGAGGCTCTGTCTCAAAAGAAA
AAGAGCACACGTCTGAACTCTTTCCCTACACGACGCTCTTCCGATCTNNNTGCTGCTGGGTATGTACCA
AGAGCACACGTCTGAACTCTTTCCCTACACGACGCTCTTCCGATCTNNNAGTTTGAAGGACAAAGGGC
AAGAGCACACGTCTGAACTCTTTCCCTACACGACGCTCTTCCGATCTNNNAGCTGTGACACATTCCAAA
GAAGAGCACACGTCTGAACTCTTTCCCTACACGACGCTCTTCCGATCTNNNTTTTGGCTTCAAAAGGCTC
AGCACACGTCTGAACTCTTTCCCTACACGACGCTCTTCCGATCTNNNTGGAGTTCCAGGTAATAGTTTG
AAGAGCACACGTCTGAACTCTTTCCCTACACGACGCTCTTCCGATCTNNNTGAACTGTACAGGAAGGC
AAGAGCACACGTCTGAACTCTTTCCCTACACGACGCTCTTCCGATCTNNNTACAATCTACCGAGAGGG

AAGAGCACACGTCTGAACTCTTTCCCTACACGACGCTCTTCCGATCTNNNGTCAATACTACAGCGAG
AGAGCACACGTCTGAACTCTTTCCCTACACGACGCTCTTCCGATCTNNNGACATAATGGCTGTAGGG
AGCACACGTCTGAACTCTTTCCCTACACGACGCTCTTCCGATCTNNNGGAACTCCTCAGGGACAACTA
GCACACGTCTGAACTCTTTCCCTACACGACGCTCTTCCGATCTNNNTCCATCACCTCTAAAAGCTTCATT
GAGCACACGTCTGAACTCTTTCCCTACACGACGCTCTTCCGATCTNNNACCTAACAGTTCCCTTTCTTCA
GAGCACACGTCTGAACTCTTTCCCTACACGACGCTCTTCCGATCTNNNTGCCCTTGCCATTTATGTTTC
AGCACACGTCTGAACTCTTTCCCTACACGACGCTCTTCCGATCTNNNTCATAAAGAGGGAGAAGCCTT
AGAGCACACGTCTGAACTCTTTCCCTACACGACGCTCTTCCGATCTNNNACTTTTCAACCCATGAGGAA
AGAGCACACGTCTGAACTCTTTCCCTACACGACGCTCTTCCGATCTNNNCAGAGACATAGGAAGACC
GAGCACACGTCTGAACTCTTTCCCTACACGACGCTCTTCCGATCTNNNAAGGCACTTTCCTGTTTTCCC
AGAGCACACGTCTGAACTCTTTCCCTACACGACGCTCTTCCGATCTNNNACCCTTCTAACAACCTGGCA
AGAGCACACGTCTGAACTCTTTCCCTACACGACGCTCTTCCGATCTNNNACTATAGGTCTGTCTGGCC
GAGCACACGTCTGAACTCTTTCCCTACACGACGCTCTTCCGATCTNNNCAATCCCCTTGCATTGCTCAA
AAGAGCACACGTCTGAACTCTTTCCCTACACGACGCTCTTCCGATCTNNNAAGAACCCATCCCCCTACA
GAGCACACGTCTGAACTCTTTCCCTACACGACGCTCTTCCGATCTNNNAAAGCCTTCTCACTACTCCCA
GAGCACACGTCTGAACTCTTTCCCTACACGACGCTCTTCCGATCTNNNCAAACTAGCTCAGCTTGACT
GAGCACACGTCTGAACTCTTTCCCTACACGACGCTCTTCCGATCTNNNAGAAGCGCAGGGTGGAATT
GAAGAGCACACGTCTGAACTCTTTCCCTACACGACGCTCTTCCGATCTNNNGCCAAGGATGGCAAAG
AGAGCACACGTCTGAACTCTTTCCCTACACGACGCTCTTCCGATCTNNNTAAAAAGCACTGCACAAAC
AGAGCACACGTCTGAACTCTTTCCCTACACGACGCTCTTCCGATCTNNNTCCCTGTCTCACCACTGTTAC
AAGAGCACACGTCTGAACTCTTTCCCTACACGACGCTCTTCCGATCTNNNATAGTTTGTGAGCCACTGC
GAGCACACGTCTGAACTCTTTCCCTACACGACGCTCTTCCGATCTNNNTGTCTGGAATCTTTCACTTGG
AGCACACGTCTGAACTCTTTCCCTACACGACGCTCTTCCGATCTNNNGCACATGGCAGGTACTTGATA
AGAGCACACGTCTGAACTCTTTCCCTACACGACGCTCTTCCGATCTNNNCTGAGTTCACCCGTGTTGTC
AAGAGCACACGTCTGAACTCTTTCCCTACACGACGCTCTTCCGATCTNNNCCAGGTTGGCGTGTATGA
AAGAGCACACGTCTGAACTCTTTCCCTACACGACGCTCTTCCGATCTNNNGAGGCGTTTTGCCCATTA
AGCACACGTCTGAACTCTTTCCCTACACGACGCTCTTCCGATCTNNNGGCCAAGTTTAGAAACATTCC
AGAGCACACGTCTGAACTCTTTCCCTACACGACGCTCTTCCGATCTNNNACAAACAGAAGAGCACAGC
GAGCACACGTCTGAACTCTTTCCCTACACGACGCTCTTCCGATCTNNNCACCATAGCATGTATAGTAGG
AGAGCACACGTCTGAACTCTTTCCCTACACGACGCTCTTCCGATCTNNNAAGTACTGCATTGGCTTAG
AGCACACGTCTGAACTCTTTCCCTACACGACGCTCTTCCGATCTNNNCCTTATAGTCTTTAGGATTGCG
AAGAGCACACGTCTGAACTCTTTCCCTACACGACGCTCTTCCGATCTNNNCCATCTTTGACGGCAAACT
AGCACACGTCTGAACTCTTTCCCTACACGACGCTCTTCCGATCTNNNTCTTCCCAGGTTCAACAGATTTT
AGCACACGTCTGAACTCTTTCCCTACACGACGCTCTTCCGATCTNNNAGGAATCATCTGGAAAGAGA
GAGCACACGTCTGAACTCTTTCCCTACACGACGCTCTTCCGATCTNNNGTGCTATCAGGCCCAAAATC
AGAGCACACGTCTGAACTCTTTCCCTACACGACGCTCTTCCGATCTNNNGCCTTTCCACTGAGTTTTCCT
AGAGCACACGTCTGAACTCTTTCCCTACACGACGCTCTTCCGATCTNNNTAAACCAGGAGCACTTTGT
AGAGCACACGTCTGAACTCTTTCCCTACACGACGCTCTTCCGATCTNNNTGAACAGTCGACACATGTA
AGCACACGTCTGAACTCTTTCCCTACACGACGCTCTTCCGATCTNNNTTTGAAACCTTGCCTAGGGTC
AGAGCACACGTCTGAACTCTTTCCCTACACGACGCTCTTCCGATCTNNNACTGGGATTGGCTGATTTC
AGAGCACACGTCTGAACTCTTTCCCTACACGACGCTCTTCCGATCTNNNTCCAGTTGGCAGCTTCTGTG
AAGAGCACACGTCTGAACTCTTTCCCTACACGACGCTCTTCCGATCTNNNTAAGGTAATTGTGTGTGG
AGAGCACACGTCTGAACTCTTTCCCTACACGACGCTCTTCCGATCTNNNACAATGGTCACTACGAAGT

AGAGCACACGTCTGAACTCTTTCCCTACACGACGCTCTTCCGATCTNNNACAATGTCTATTCATGCGTGT
GAGCACACGTCTGAACTCTTTCCCTACACGACGCTCTTCCGATCTNNNTCTGATGCTACACATCCTTCCC
AGAGCACACGTCTGAACTCTTTCCCTACACGACGCTCTTCCGATCTNNNAGAAAGATGGCAAGACAAA
AGAGCACACGTCTGAACTCTTTCCCTACACGACGCTCTTCCGATCTNNNACCAGGAAACTTTTGAGATC
AAGAGCACACGTCTGAACTCTTTCCCTACACGACGCTCTTCCGATCTNNNATACATATCTCTGTGAGCT
AGAGCACACGTCTGAACTCTTTCCCTACACGACGCTCTTCCGATCTNNNGCTGTGCAGCAGGATTTTCA
AAGAGCACACGTCTGAACTCTTTCCCTACACGACGCTCTTCCGATCTNNNGTGAGTTTCCAAGCTCCAA
AGAGCACACGTCTGAACTCTTTCCCTACACGACGCTCTTCCGATCTNNNCACATCTGAGGATCTGGAG
AGAGCACACGTCTGAACTCTTTCCCTACACGACGCTCTTCCGATCTNNNTTATCTCTCAGACCCTGTGTC
AGAGCACACGTCTGAACTCTTTCCCTACACGACGCTCTTCCGATCTNNNTCATGCAGAAGAGATGCCAA
AGAGCACACGTCTGAACTCTTTCCCTACACGACGCTCTTCCGATCTNNNCATTATTTGGCAGGGGGAG
AAGAGCACACGTCTGAACTCTTTCCCTACACGACGCTCTTCCGATCTNNNTAATTATTGCCACCCCTTGG
AGAGCACACGTCTGAACTCTTTCCCTACACGACGCTCTTCCGATCTNNNTACGAATACCCTACTGGTTG
GAGCACACGTCTGAACTCTTTCCCTACACGACGCTCTTCCGATCTNNNAACCTGCTAGCCTCATCATTCC
AGAGCACACGTCTGAACTCTTTCCCTACACGACGCTCTTCCGATCTNNNTCTGAACTGTTTGAGTAGAT
AAGAGCACACGTCTGAACTCTTTCCCTACACGACGCTCTTCCGATCTNNNTGCTTTGTGTCAGATATGG
GAGCACACGTCTGAACTCTTTCCCTACACGACGCTCTTCCGATCTNNNAGTGAATAATGCATTGCTTCTC
AGAGCACACGTCTGAACTCTTTCCCTACACGACGCTCTTCCGATCTNNNCTATGGCACTCCCTTTCAAGT
AGAGCACACGTCTGAACTCTTTCCCTACACGACGCTCTTCCGATCTNNNGCCTTTACCCAGGTTCAGTCA
AGAGCACACGTCTGAACTCTTTCCCTACACGACGCTCTTCCGATCTNNNCCGTGAATACATTCGCAAAG
AGAGCACACGTCTGAACTCTTTCCCTACACGACGCTCTTCCGATCTNNNCTGCAAAGACCTAAGTCACA
GAGCACACGTCTGAACTCTTTCCCTACACGACGCTCTTCCGATCTNNNACATAGATCATCCACAAGCTGT
GAGCACACGTCTGAACTCTTTCCCTACACGACGCTCTTCCGATCTNNNTGGGCCAGAGTGATACTCTAA
AGCACACGTCTGAACTCTTTCCCTACACGACGCTCTTCCGATCTNNNTGCTAACAGAGTAGATCTTGTG
AAGAGCACACGTCTGAACTCTTTCCCTACACGACGCTCTTCCGATCTNNNCTCCTCTGCCTGTTTTCGAG
AGAGCACACGTCTGAACTCTTTCCCTACACGACGCTCTTCCGATCTNNNTGAATGAATGACGAACAGCA
AAGAGCACACGTCTGAACTCTTTCCCTACACGACGCTCTTCCGATCTNNNGTGCCATCTATGAACTTCAG
GAGCACACGTCTGAACTCTTTCCCTACACGACGCTCTTCCGATCTNNNCTATAGCTGCTGGAGCGACTT
AGAGCACACGTCTGAACTCTTTCCCTACACGACGCTCTTCCGATCTNNNTGGTTGAAAACAAGCCTGTA
AGCACACGTCTGAACTCTTTCCCTACACGACGCTCTTCCGATCTNNNTGAAAAATTAGTTATTGCTTGG
AGCACACGTCTGAACTCTTTCCCTACACGACGCTCTTCCGATCTNNNGCCACTGCAAACATAGTAATCTT
AAGAGCACACGTCTGAACTCTTTCCCTACACGACGCTCTTCCGATCTNNNCCAATCCAAGAGGAGCATC
AGAGCACACGTCTGAACTCTTTCCCTACACGACGCTCTTCCGATCTNNNGATTTCTGGTTCAGTGACCCT
AAGAGCACACGTCTGAACTCTTTCCCTACACGACGCTCTTCCGATCTNNNCACCCCGGAGAATTTCAAC
AAGAGCACACGTCTGAACTCTTTCCCTACACGACGCTCTTCCGATCTNNNACCATCAAATTCCTTGGGC
GAGCACACGTCTGAACTCTTTCCCTACACGACGCTCTTCCGATCTNNNGATCACGAGAGGCCATGAAGT
GAAGAGCACACGTCTGAACTCTTTCCCTACACGACGCTCTTCCGATCTNNNTCTGTACTGAAGCTACAT
AAGAGCACACGTCTGAACTCTTTCCCTACACGACGCTCTTCCGATCTNNNTAAAAATGTCACTGCTCCCC
AGAGCACACGTCTGAACTCTTTCCCTACACGACGCTCTTCCGATCTNNNACACAACTGATTTTGTGGGT
AAGAGCACACGTCTGAACTCTTTCCCTACACGACGCTCTTCCGATCTNNNGCTTATGCTACCCCTGTGT
AGAGCACACGTCTGAACTCTTTCCCTACACGACGCTCTTCCGATCTNNNAGTGTCTCCACTGTTGATTG
GAGCACACGTCTGAACTCTTTCCCTACACGACGCTCTTCCGATCTNNNAAATCATGGCTTCAGTTAGTG
AGAGCACACGTCTGAACTCTTTCCCTACACGACGCTCTTCCGATCTNNNGGCCAACATGTCACCTAACA

GAGCACACGTCTGAACTCTTTCCCTACACGACGCTCTTCCGATCTNNNAAAATACTTCAGGCTGGGTT
AGAGCACACGTCTGAACTCTTTCCCTACACGACGCTCTTCCGATCTNNNTCATGCTGTTTTCCCAGCTA
AGAGCACACGTCTGAACTCTTTCCCTACACGACGCTCTTCCGATCTNNNTTTGTACAATGAGGAGGTG
AAGAGCACACGTCTGAACTCTTTCCCTACACGACGCTCTTCCGATCTNNNTCCCCATGGGATATCTTC
AGAGCACACGTCTGAACTCTTTCCCTACACGACGCTCTTCCGATCTNNNTCACCAGGCACTCTGGAGA
GCACACGTCTGAACTCTTTCCCTACACGACGCTCTTCCGATCTNNNTGAGATATTCTCTGTGAGGACT
AGAGCACACGTCTGAACTCTTTCCCTACACGACGCTCTTCCGATCTNNNTAAGTCCCAAGCTGTCTGAG
AAGAGCACACGTCTGAACTCTTTCCCTACACGACGCTCTTCCGATCTNNNTGTGTCCTTATCCTTAGCTC
GCACACGTCTGAACTCTTTCCCTACACGACGCTCTTCCGATCTNNNTGAAAGGACCAAGTGAATAAA
AGCACACGTCTGAACTCTTTCCCTACACGACGCTCTTCCGATCTNNNATGGATTAGGTTTTGTTGTACA
AGAGCACACGTCTGAACTCTTTCCCTACACGACGCTCTTCCGATCTNNNCTTTTACCATTGGGGGTGTG
AAGAGCACACGTCTGAACTCTTTCCCTACACGACGCTCTTCCGATCTNNNCATGTGCCTGTCAGTGATG
AGAGCACACGTCTGAACTCTTTCCCTACACGACGCTCTTCCGATCTNNNCGCAGTCTTTTGAGTAGCTG
AGAGCACACGTCTGAACTCTTTCCCTACACGACGCTCTTCCGATCTNNNCAGATGCCAAGCTTTCTCTC
AGAGCACACGTCTGAACTCTTTCCCTACACGACGCTCTTCCGATCTNNNTGACCATGAAGTGCTTAGG
AGAGCACACGTCTGAACTCTTTCCCTACACGACGCTCTTCCGATCTNNNTTTCTGCCTGCTCAGTGATT
AGAGCACACGTCTGAACTCTTTCCCTACACGACGCTCTTCCGATCTNNNCTTGATTCTTGGGTCTGGGA
GAAGAGCACACGTCTGAACTCTTTCCCTACACGACGCTCTTCCGATCTNNNCCATGAATTAGATGCAG
AGAGCACACGTCTGAACTCTTTCCCTACACGACGCTCTTCCGATCTNNNAGCCACTAGGACCAGTAAT
GAGCACACGTCTGAACTCTTTCCCTACACGACGCTCTTCCGATCTNNNTTCTGATGTGAGGCTACCTTT
GCACACGTCTGAACTCTTTCCCTACACGACGCTCTTCCGATCTNNNAGATAATAGTCTCTGCAGCTGT
AGAGCACACGTCTGAACTCTTTCCCTACACGACGCTCTTCCGATCTNNNAGTTTCTTGTGAGAATCTAG
AGAGCACACGTCTGAACTCTTTCCCTACACGACGCTCTTCCGATCTNNNTCTATGGTCGTGTGAGCCA
AGAGCACACGTCTGAACTCTTTCCCTACACGACGCTCTTCCGATCTNNNAACCTCTGGCTTTCAATCTC
AGAGCACACGTCTGAACTCTTTCCCTACACGACGCTCTTCCGATCTNNNGCAACACAGAAGACAGTCT
AGAGCACACGTCTGAACTCTTTCCCTACACGACGCTCTTCCGATCTNNNCTACCAGTTCATCCATGTTGC
AGCACACGTCTGAACTCTTTCCCTACACGACGCTCTTCCGATCTNNNTCTGTCTGGTTTTATCCTTCAC
AGCACACGTCTGAACTCTTTCCCTACACGACGCTCTTCCGATCTNNNAACACAACTGCTATCAAAATA
AGCACACGTCTGAACTCTTTCCCTACACGACGCTCTTCCGATCTNNNAGTGACTGCAAATCAAATTTT
AGAGCACACGTCTGAACTCTTTCCCTACACGACGCTCTTCCGATCTNNNAGGCTCTACCACATAGACAA
AGCACACGTCTGAACTCTTTCCCTACACGACGCTCTTCCGATCTNNNCCATGTGACCAGTTACAGGTAA
AGCACACGTCTGAACTCTTTCCCTACACGACGCTCTTCCGATCTNNNCCTCTGAGGTTCCTTTTGGTAA
GAGCACACGTCTGAACTCTTTCCCTACACGACGCTCTTCCGATCTNNNGCATTTGGAATCAGGGACTA
GAAGAGCACACGTCTGAACTCTTTCCCTACACGACGCTCTTCCGATCTNNNGTTCTTTCAGAGAATGCC
AAGAGCACACGTCTGAACTCTTTCCCTACACGACGCTCTTCCGATCTNNNTGTCGAGCTTTGGATGCT
GAGCACACGTCTGAACTCTTTCCCTACACGACGCTCTTCCGATCTNNNTTCAATGGCTTCAGCTCCCATT
GAGCACACGTCTGAACTCTTTCCCTACACGACGCTCTTCCGATCTNNNAGTGAACTGTCTTTCTGGTT
GAGCACACGTCTGAACTCTTTCCCTACACGACGCTCTTCCGATCTNNNTGTGAAGCTACAGTTTTCTGG
AAGAGCACACGTCTGAACTCTTTCCCTACACGACGCTCTTCCGATCTNNNTCCTCCCTAGGGTCAGATA
AAGAGCACACGTCTGAACTCTTTCCCTACACGACGCTCTTCCGATCTNNNATAGTGTTTCCTCTGATGC
GAGCACACGTCTGAACTCTTTCCCTACACGACGCTCTTCCGATCTNNNCCAGATAGTGAACACGTACC
AGCACACGTCTGAACTCTTTCCCTACACGACGCTCTTCCGATCTNNNAGAGAGAGAGAAAGAGGAAG
AAGAGCACACGTCTGAACTCTTTCCCTACACGACGCTCTTCCGATCTNNNTCTTCTTATCTGTGGGGAG

GAGCACACGTCTGAACTCTTTCCCTACACGACGCTCTTCCGATCTNNNATCCAAGGGGAAGGAGTAGC
AGCACACGTCTGAACTCTTTCCCTACACGACGCTCTTCCGATCTNNNTGATGTAATATTTTACCCATCCCC
AGAGCACACGTCTGAACTCTTTCCCTACACGACGCTCTTCCGATCTNNNGGGTGTGTATGAGTGTGTG
GAGCACACGTCTGAACTCTTTCCCTACACGACGCTCTTCCGATCTNNNTCGTTCCTTTTTGTTTGTGACT
AAGAGCACACGTCTGAACTCTTTCCCTACACGACGCTCTTCCGATCTNNNTGAGAGAGAAGCTAGAGA
AAGAGCACACGTCTGAACTCTTTCCCTACACGACGCTCTTCCGATCTNNNGCCAAAATGCCCATCAAAC
AGAGCACACGTCTGAACTCTTTCCCTACACGACGCTCTTCCGATCTNNNAGTCTACTGTTCTTTACCCACC
AGAGCACACGTCTGAACTCTTTCCCTACACGACGCTCTTCCGATCTNNNACCTATTAGGAGTGCACAGA
AAGAGCACACGTCTGAACTCTTTCCCTACACGACGCTCTTCCGATCTNNNTACTTCCACGCAAAGGATC
GAGCACACGTCTGAACTCTTTCCCTACACGACGCTCTTCCGATCTNNNTATTTGGAGGAGGATGTGGC
AAGAGCACACGTCTGAACTCTTTCCCTACACGACGCTCTTCCGATCTNNNTGTGCGTCTGGAAAGTGA
AAGAGCACACGTCTGAACTCTTTCCCTACACGACGCTCTTCCGATCTNNNTGGGAATCCCTGTCTCTGC
AAGAGCACACGTCTGAACTCTTTCCCTACACGACGCTCTTCCGATCTNNNGAGGTTCTAAGTGGTGGC
GAGCACACGTCTGAACTCTTTCCCTACACGACGCTCTTCCGATCTNNNAGCAAGTGTCATAATCACGGA
AAGAGCACACGTCTGAACTCTTTCCCTACACGACGCTCTTCCGATCTNNNCTGAGGCTTCTGAGCTGAC
AGAGCACACGTCTGAACTCTTTCCCTACACGACGCTCTTCCGATCTNNNGTGATCAAACCAAAGCAGCA
AGAGCACACGTCTGAACTCTTTCCCTACACGACGCTCTTCCGATCTNNNCAGGGCTAACCAGAAAAGC
GAGCACACGTCTGAACTCTTTCCCTACACGACGCTCTTCCGATCTNNNACCTCCTTGGTAGGATGTATTC
AAGAGCACACGTCTGAACTCTTTCCCTACACGACGCTCTTCCGATCTNNNAGTCCTGTCTCCGATACTTC
AGAGCACACGTCTGAACTCTTTCCCTACACGACGCTCTTCCGATCTNNNCACTCCCAAATTGTGGTTCT
AAGAGCACACGTCTGAACTCTTTCCCTACACGACGCTCTTCCGATCTNNNACCTGTCAACAAACATGTG
AGAGCACACGTCTGAACTCTTTCCCTACACGACGCTCTTCCGATCTNNNTTTTATAACATCGGCGTGTG
GAAGAGCACACGTCTGAACTCTTTCCCTACACGACGCTCTTCCGATCTNNNTGGGAAGATGCCTTGCT
AGAGCACACGTCTGAACTCTTTCCCTACACGACGCTCTTCCGATCTNNNTAGTGTATTGGTCAGGATT
AGAGCACACGTCTGAACTCTTTCCCTACACGACGCTCTTCCGATCTNNNTCCTTTCTCTTCCTTGTTGGG
GAGCACACGTCTGAACTCTTTCCCTACACGACGCTCTTCCGATCTNNNGTGGATCTACACATACATGGA
AAGAGCACACGTCTGAACTCTTTCCCTACACGACGCTCTTCCGATCTNNNTCATCAATCAGGTGTGTGT
GAGCACACGTCTGAACTCTTTCCCTACACGACGCTCTTCCGATCTNNNAAAAGTTTGGGGAAGGGAA
AGAGCACACGTCTGAACTCTTTCCCTACACGACGCTCTTCCGATCTNNNCTGCGCTGATAAGATGCCAT
GAGCACACGTCTGAACTCTTTCCCTACACGACGCTCTTCCGATCTNNNAGGCAAAAATTAGTCCTAAGG
GAAGAGCACACGTCTGAACTCTTTCCCTACACGACGCTCTTCCGATCTNNNAGCCTCCTCTGGTAACTG
AGCACACGTCTGAACTCTTTCCCTACACGACGCTCTTCCGATCTNNNTGTTTTGACAAATGCTTGCTTTG
AGAGCACACGTCTGAACTCTTTCCCTACACGACGCTCTTCCGATCTNNNCCTGAGGGTCGATCTCATCA
AGCACACGTCTGAACTCTTTCCCTACACGACGCTCTTCCGATCTNNNTCCTAGTCGTGTTATTTTTCCAG
AGAGCACACGTCTGAACTCTTTCCCTACACGACGCTCTTCCGATCTNNNACAGCACATGCAAATGGATT
AGAGCACACGTCTGAACTCTTTCCCTACACGACGCTCTTCCGATCTNNNATCAGAGGTGAGATTTGGG
AGAGCACACGTCTGAACTCTTTCCCTACACGACGCTCTTCCGATCTNNNGCAGACTGCACATGTTCTGA
GAGCACACGTCTGAACTCTTTCCCTACACGACGCTCTTCCGATCTNNNATCCTAATTTCTCCCTCTTGTTC
AGCACACGTCTGAACTCTTTCCCTACACGACGCTCTTCCGATCTNNNTGGGTATGTTAGAACAAAACTC
AGCACACGTCTGAACTCTTTCCCTACACGACGCTCTTCCGATCTNNNAGGATGAGAACAATAGAATGA
AGCACACGTCTGAACTCTTTCCCTACACGACGCTCTTCCGATCTNNNTCTTGCTATAAATCACCAAACCAC
AGAGCACACGTCTGAACTCTTTCCCTACACGACGCTCTTCCGATCTNNNGCTGCAGCTTTCTTATCTGGA
AGAGCACACGTCTGAACTCTTTCCCTACACGACGCTCTTCCGATCTNNNTCCTGCAAATCCTATGTCTCC

GAGCACACGTCTGAACTCTTTCCCTACACGACGCTCTTCCGATCTNNNAGAAGACTGTAAGAAAAGCA
GAGCACACGTCTGAACTCTTTCCCTACACGACGCTCTTCCGATCTNNNGTGGGAATTCTCAGCCTCTGT
GAGCACACGTCTGAACTCTTTCCCTACACGACGCTCTTCCGATCTNNNGCACAAGAATTCCAAAAGGC
GAGCACACGTCTGAACTCTTTCCCTACACGACGCTCTTCCGATCTNNNTGGTATGAAAAGCTCATTGAG
AGCACACGTCTGAACTCTTTCCCTACACGACGCTCTTCCGATCTNNNTCAATTCTTGCATTGAATCTAT
AGAGCACACGTCTGAACTCTTTCCCTACACGACGCTCTTCCGATCTNNNTGTGTAGACTTCAGGGACA
AGAGCACACGTCTGAACTCTTTCCCTACACGACGCTCTTCCGATCTNNNTCCTCTCCCCTGATGTCCTAT
AGAGCACACGTCTGAACTCTTTCCCTACACGACGCTCTTCCGATCTNNNGGGACGGGAAAGGAAGA
AAGAGCACACGTCTGAACTCTTTCCCTACACGACGCTCTTCCGATCTNNNACAACGGACACCTTATTGA
AGAGCACACGTCTGAACTCTTTCCCTACACGACGCTCTTCCGATCTNNNGGTGGTTTTCCAGGTGGTA
AGAGCACACGTCTGAACTCTTTCCCTACACGACGCTCTTCCGATCTNNNGAGACTATGCCCAAAGCCA
AGCACACGTCTGAACTCTTTCCCTACACGACGCTCTTCCGATCTNNNTGTGTATGACATGATCCTTTTG
GCACACGTCTGAACTCTTTCCCTACACGACGCTCTTCCGATCTNNNTCTGGGATTATATCCTTCCATGC
GAGCACACGTCTGAACTCTTTCCCTACACGACGCTCTTCCGATCTNNNGCAAATGCCATTATTTCACGC
AGAGCACACGTCTGAACTCTTTCCCTACACGACGCTCTTCCGATCTNNNAAGATCATCGTCTGGTGAT
AGAGCACACGTCTGAACTCTTTCCCTACACGACGCTCTTCCGATCTNNNACATTGGAGGGGGAAAAG
GCACACGTCTGAACTCTTTCCCTACACGACGCTCTTCCGATCTNNNTGAGTTTCTGGTAGGAATTGGA
AGAGCACACGTCTGAACTCTTTCCCTACACGACGCTCTTCCGATCTNNNTGCTTCATCATTGGCACTCT
AGAGCACACGTCTGAACTCTTTCCCTACACGACGCTCTTCCGATCTNNNACCAGAGGACTACAACCCAC
AAGAGCACACGTCTGAACTCTTTCCCTACACGACGCTCTTCCGATCTNNNTAATGTCACAGAGCTCCAC
AGAGCACACGTCTGAACTCTTTCCCTACACGACGCTCTTCCGATCTNNNTTTGATACCCCTACCCCTCT
GAGCACACGTCTGAACTCTTTCCCTACACGACGCTCTTCCGATCTNNNTATACCAGCTCACAGAGGTCC
GCACACGTCTGAACTCTTTCCCTACACGACGCTCTTCCGATCTNNNCCTGAGGATTCTTTCTCTCTACC
AGCACACGTCTGAACTCTTTCCCTACACGACGCTCTTCCGATCTNNNTGTGGGATAGACACCAAAGTT
AAGAGCACACGTCTGAACTCTTTCCCTACACGACGCTCTTCCGATCTNNNTTGATCTTCCCAGCTTCAGT
AGCACACGTCTGAACTCTTTCCCTACACGACGCTCTTCCGATCTNNNTCTGTTACTCTAAGCAGGGTTA
GAGCACACGTCTGAACTCTTTCCCTACACGACGCTCTTCCGATCTNNNTGTGAGGAAGACATTACACAT
AGAGCACACGTCTGAACTCTTTCCCTACACGACGCTCTTCCGATCTNNNCTTCTCACATATGTCCAGCCT
AGCACACGTCTGAACTCTTTCCCTACACGACGCTCTTCCGATCTNNNTCTTGGTAAATGTGAGTAGAA
GAGCACACGTCTGAACTCTTTCCCTACACGACGCTCTTCCGATCTNNNTTTGTTTATGTTATGACGCATG
AGAGCACACGTCTGAACTCTTTCCCTACACGACGCTCTTCCGATCTNNNTCCCAGTGATCAAAAATACA
GAGCACACGTCTGAACTCTTTCCCTACACGACGCTCTTCCGATCTNNNCGTCTAATCCCATGGCTCAGA
GAGCACACGTCTGAACTCTTTCCCTACACGACGCTCTTCCGATCTNNNATACCATCCCACCACTCACTCA
AGAGCACACGTCTGAACTCTTTCCCTACACGACGCTCTTCCGATCTNNNAGAGAAGAAAATGTGGAAA
AGCACACGTCTGAACTCTTTCCCTACACGACGCTCTTCCGATCTNNNAGTTTATAGTCAGGGTACCTCA
AGAGCACACGTCTGAACTCTTTCCCTACACGACGCTCTTCCGATCTNNNTCTGGTAATGTAGGGAAG
GAAGAGCACACGTCTGAACTCTTTCCCTACACGACGCTCTTCCGATCTNNNTTTGACTGCAAGGCAGAA
AGAGCACACGTCTGAACTCTTTCCCTACACGACGCTCTTCCGATCTNNNACCAAAGGCAGAAAGAAGA
AGCACACGTCTGAACTCTTTCCCTACACGACGCTCTTCCGATCTNNNTTGAAACACGATGCTAAATGA
AGCACACGTCTGAACTCTTTCCCTACACGACGCTCTTCCGATCTNNNTCCCATCAATCTCAATTGCTGT
AGAGCACACGTCTGAACTCTTTCCCTACACGACGCTCTTCCGATCTNNNGGCTGAATTTGGGTCAGGT
AGCACACGTCTGAACTCTTTCCCTACACGACGCTCTTCCGATCTNNNACATATGTGATATGACCCCAAT
AGAGCACACGTCTGAACTCTTTCCCTACACGACGCTCTTCCGATCTNNNTGTGCTCTGAACCACCAGAT

GAGCACACGTCTGAACTCTTTCCCTACACGACGCTCTTCCGATCTNNNTCTTCTCCCTGGAAAAGCAA
AGCACACGTCTGAACTCTTTCCCTACACGACGCTCTTCCGATCTNNNTGGATTTCTCCTTTTAGTTCTG
GAGCACACGTCTGAACTCTTTCCCTACACGACGCTCTTCCGATCTNNNGCCCGGCCTATGTATCACAAT
GAGCACACGTCTGAACTCTTTCCCTACACGACGCTCTTCCGATCTNNNCATCCCTTCATTACCTGTGGT
AGCACACGTCTGAACTCTTTCCCTACACGACGCTCTTCCGATCTNNNCCAGAAACCTGCTCTCTCATTC
GAGCACACGTCTGAACTCTTTCCCTACACGACGCTCTTCCGATCTNNNTGAAGCTGTAAAATTACAAG
GAGCACACGTCTGAACTCTTTCCCTACACGACGCTCTTCCGATCTNNNATATTCTCCCTATCCTGCTGC
GAGCACACGTCTGAACTCTTTCCCTACACGACGCTCTTCCGATCTNNNGCCCCTGAACCTGTAGTCTCT
AGAGCACACGTCTGAACTCTTTCCCTACACGACGCTCTTCCGATCTNNNAGAGGGGCTATAATCCCAG
GAGCACACGTCTGAACTCTTTCCCTACACGACGCTCTTCCGATCTNNNGGCATTGCTTTTGTCCTGAC
GAGCACACGTCTGAACTCTTTCCCTACACGACGCTCTTCCGATCTNNNAGACTTTCTCACAGGGTAGA
GAGCACACGTCTGAACTCTTTCCCTACACGACGCTCTTCCGATCTNNNGTAATGTCCTCTGGGTTCATC
AGCACACGTCTGAACTCTTTCCCTACACGACGCTCTTCCGATCTNNNGGTAATGTCATTGAACCATCCT
AAGAGCACACGTCTGAACTCTTTCCCTACACGACGCTCTTCCGATCTNNNCAAAAAGCTTAGTGGCTT
AGAGCACACGTCTGAACTCTTTCCCTACACGACGCTCTTCCGATCTNNNGAACTTTGGTATGGTAGCC
AGCACACGTCTGAACTCTTTCCCTACACGACGCTCTTCCGATCTNNNAGTGTCCATAACTGTGGGAGC
AGAGCACACGTCTGAACTCTTTCCCTACACGACGCTCTTCCGATCTNNNCAACAAAGTGAGACCTCTG
GAGCACACGTCTGAACTCTTTCCCTACACGACGCTCTTCCGATCTNNNTCAGCTTCTGAGCAAGAGTT
GAGCACACGTCTGAACTCTTTCCCTACACGACGCTCTTCCGATCTNNNTTTGAGACAAGGTCGGTCTC
GAGCACACGTCTGAACTCTTTCCCTACACGACGCTCTTCCGATCTNNNTGTGGGTTTCATCACTAACC
AGAGCACACGTCTGAACTCTTTCCCTACACGACGCTCTTCCGATCTNNNGGTAGGGCCTTTGGAAGC
AGCACACGTCTGAACTCTTTCCCTACACGACGCTCTTCCGATCTNNNAGAACTGAAGCCTCAAATGG
GAGCACACGTCTGAACTCTTTCCCTACACGACGCTCTTCCGATCTNNNTTTTCTTCTTTCTTTCCCCCTTC
GAGCACACGTCTGAACTCTTTCCCTACACGACGCTCTTCCGATCTNNNATGAAAGTCTCCTTGCCTCTC
GAGCACACGTCTGAACTCTTTCCCTACACGACGCTCTTCCGATCTNNNTGCATGGTTTTCTCTCTCACA
GAGCACACGTCTGAACTCTTTCCCTACACGACGCTCTTCCGATCTNNNAGATTCTTACTTGAAATCCC
GAGCACACGTCTGAACTCTTTCCCTACACGACGCTCTTCCGATCTNNNACCTATCTGTCACCTTCCATA
GAGCACACGTCTGAACTCTTTCCCTACACGACGCTCTTCCGATCTNNNTTTCTCTTAGGTGACCTCTCC
GCACACGTCTGAACTCTTTCCCTACACGACGCTCTTCCGATCTNNNTCATGAGCCTGACATAATCAGG
AGAGCACACGTCTGAACTCTTTCCCTACACGACGCTCTTCCGATCTNNNATCCAGGTCCATTCGTTCCA
AGAGCACACGTCTGAACTCTTTCCCTACACGACGCTCTTCCGATCTNNNTTCTCTAGCTTCCCCTAAGC
AGCACACGTCTGAACTCTTTCCCTACACGACGCTCTTCCGATCTNNNAGTGGAAGAAAAGAAAACCTC
AGAGCACACGTCTGAACTCTTTCCCTACACGACGCTCTTCCGATCTNNNTTCTTATGAGCATGGGGCT
GAGCACACGTCTGAACTCTTTCCCTACACGACGCTCTTCCGATCTNNNAGAGTATTTTCCTTCTCACTCC
GCACACGTCTGAACTCTTTCCCTACACGACGCTCTTCCGATCTNNNTCTTTTCATCCTTTGGGTGGTTT
GCACACGTCTGAACTCTTTCCCTACACGACGCTCTTCCGATCTNNNACAAAAGTCTCAGAGGAAAGC
GAGCACACGTCTGAACTCTTTCCCTACACGACGCTCTTCCGATCTNNNCTGATCTCCTCAACTTTCCTG
AGAGCACACGTCTGAACTCTTTCCCTACACGACGCTCTTCCGATCTNNNTCACTCTGGAATGACTTTTT
AGCACACGTCTGAACTCTTTCCCTACACGACGCTCTTCCGATCTNNNTCCCAACAATGTTTTGAGATA
GAGCACACGTCTGAACTCTTTCCCTACACGACGCTCTTCCGATCTNNNTACCTGGCACTCATATTCACC
AGAGCACACGTCTGAACTCTTTCCCTACACGACGCTCTTCCGATCTNNNTGGCTCATTAGAAGGCCAA
AAGAGCACACGTCTGAACTCTTTCCCTACACGACGCTCTTCCGATCTNNNTGTTTCTTTGCTGAGCCCA
GAGCACACGTCTGAACTCTTTCCCTACACGACGCTCTTCCGATCTNNNACATGAGTTTTGGAAGGGAC

AGAGCACACGTCTGAACTCTTTCCCTACACGACGCTCTTCCGATCTNNNATCCAGTAAGGGCTCTGGA
GAGCACACGTCTGAACTCTTTCCCTACACGACGCTCTTCCGATCTNNNCTGCCTATGTTTCCTTCATGG
GAGCACACGTCTGAACTCTTTCCCTACACGACGCTCTTCCGATCTNNNCCAGAGACAGAGCAAAACAA
AGCACACGTCTGAACTCTTTCCCTACACGACGCTCTTCCGATCTNNNACATGAACAAGGACTTCAATAC
AGCACACGTCTGAACTCTTTCCCTACACGACGCTCTTCCGATCTNNNACACACAATTTCTGATGGACAA
AGCACACGTCTGAACTCTTTCCCTACACGACGCTCTTCCGATCTNNNCCTAACCAGAACTTCTCCCCTG
GAGCACACGTCTGAACTCTTTCCCTACACGACGCTCTTCCGATCTNNNGCAAAATCCCCAATACATGTC
AAGAGCACACGTCTGAACTCTTTCCCTACACGACGCTCTTCCGATCTNNNGGGTTTCCCCAGAGAAAG
AGAGCACACGTCTGAACTCTTTCCCTACACGACGCTCTTCCGATCTNNNACACAAAGTACTGAAAGAG
GAGCACACGTCTGAACTCTTTCCCTACACGACGCTCTTCCGATCTNNNTCCTGCAACTTTACTAAACCAA
AGCACACGTCTGAACTCTTTCCCTACACGACGCTCTTCCGATCTNNNACTATACTATGGTAGCATTTCC
AGAGCACACGTCTGAACTCTTTCCCTACACGACGCTCTTCCGATCTNNNTTGCATGTAGTCCCTTTCTCC
AGCACACGTCTGAACTCTTTCCCTACACGACGCTCTTCCGATCTNNNAGAGAACCACACTGACTATTTC
GAGCACACGTCTGAACTCTTTCCCTACACGACGCTCTTCCGATCTNNNTGCGTTATCTCTCTCTCTCT
AGAGCACACGTCTGAACTCTTTCCCTACACGACGCTCTTCCGATCTNNNAAAATCACTGCTCTAGCCAC
AGAGCACACGTCTGAACTCTTTCCCTACACGACGCTCTTCCGATCTNNNTGAAGTAAAACATTCAAGG
AGCACACGTCTGAACTCTTTCCCTACACGACGCTCTTCCGATCTNNNTCATCTTACAGCTTCAAGTTAC
AGAGCACACGTCTGAACTCTTTCCCTACACGACGCTCTTCCGATCTNNNTCTTCTGGCCTAATGTACTG
AGCACACGTCTGAACTCTTTCCCTACACGACGCTCTTCCGATCTNNNCCTTCTAGCTCTTCCTTTCTTCT
AAGAGCACACGTCTGAACTCTTTCCCTACACGACGCTCTTCCGATCTNNNTGGTTTTATTCCTCTGGCA
AGAGCACACGTCTGAACTCTTTCCCTACACGACGCTCTTCCGATCTNNNCCTGACAGGTCTAAGCACAT
GCACACGTCTGAACTCTTTCCCTACACGACGCTCTTCCGATCTNNNACTTCTGTCTTAGAATTTTTGCT
AGAGCACACGTCTGAACTCTTTCCCTACACGACGCTCTTCCGATCTNNNGAACTAAGGCACCCCAAAG
AGCACACGTCTGAACTCTTTCCCTACACGACGCTCTTCCGATCTNNNCAAGAATGCCAAGGTATTTCTT
AGAGCACACGTCTGAACTCTTTCCCTACACGACGCTCTTCCGATCTNNNCCTGTTGGCCATATACCTTT
AGAGCACACGTCTGAACTCTTTCCCTACACGACGCTCTTCCGATCTNNNCAAAACAAAACGAAATGCCC
AAGAGCACACGTCTGAACTCTTTCCCTACACGACGCTCTTCCGATCTNNNACTATGGTAGGGTGGTT
GAGCACACGTCTGAACTCTTTCCCTACACGACGCTCTTCCGATCTNNNTGGATAAAAGTGTCTCTCTCT
AGCACACGTCTGAACTCTTTCCCTACACGACGCTCTTCCGATCTNNNCCTTGGAGTTTGTAACTCACTT
GAGCACACGTCTGAACTCTTTCCCTACACGACGCTCTTCCGATCTNNNCCCTCAGTTCAGAAGATGTAT
GAGCACACGTCTGAACTCTTTCCCTACACGACGCTCTTCCGATCTNNNGATAACTGGCAAAACACTGT
GAGCACACGTCTGAACTCTTTCCCTACACGACGCTCTTCCGATCTNNNATGCCTGACAGGTTACCAAA
GAGCACACGTCTGAACTCTTTCCCTACACGACGCTCTTCCGATCTNNNCATGTCTGTCCCTTTAGCAGC
GAGCACACGTCTGAACTCTTTCCCTACACGACGCTCTTCCGATCTNNNATGAAATGACCATCAAAGGC
GAGCACACGTCTGAACTCTTTCCCTACACGACGCTCTTCCGATCTNNNACGAACTACAGAAAGGGTAA
GCACACGTCTGAACTCTTTCCCTACACGACGCTCTTCCGATCTNNNTCTGGGATTATATCCTTCCATGCC
AGAGCACACGTCTGAACTCTTTCCCTACACGACGCTCTTCCGATCTNNNATACCCATTCACTGCACATC
AGAGCACACGTCTGAACTCTTTCCCTACACGACGCTCTTCCGATCTNNNCCAGTGCACGAAAGGTCTAG
AGAGCACACGTCTGAACTCTTTCCCTACACGACGCTCTTCCGATCTNNNTGGCAAGACTGGATTCATA
AGAGCACACGTCTGAACTCTTTCCCTACACGACGCTCTTCCGATCTNNNAAAAAGTCCATTGTTGAGTA
AAGAGCACACGTCTGAACTCTTTCCCTACACGACGCTCTTCCGATCTNNNCTTGCGTACTTTGAGTTTC
GAGCACACGTCTGAACTCTTTCCCTACACGACGCTCTTCCGATCTNNNAGTTTTGAATTCCTGGTTCTT
GAGCACACGTCTGAACTCTTTCCCTACACGACGCTCTTCCGATCTNNNTGCACTCTAGGTAGTTGCTC

AAGAGCACACGTCTGAACTCTTTCCCTACACGACGCTCTTCCGATCTNNNTCTTCACCCTTGTAGTAGG
AAGAGCACACGTCTGAACTCTTTCCCTACACGACGCTCTTCCGATCTNNNTTTCATGTGAGGACAGT
AGCACACGTCTGAACTCTTTCCCTACACGACGCTCTTCCGATCTNNNCCTCTTTGTCTTTGTTTTTGGCA
GCACACGTCTGAACTCTTTCCCTACACGACGCTCTTCCGATCTNNNGGCCACAGAAGATTAGACTAAC
AGCACACGTCTGAACTCTTTCCCTACACGACGCTCTTCCGATCTNNNGCATCTAACGTATTTGCTACTT
GAGCACACGTCTGAACTCTTTCCCTACACGACGCTCTTCCGATCTNNNTGAATCTGCATGTGTGGTAT
GAGCACACGTCTGAACTCTTTCCCTACACGACGCTCTTCCGATCTNNNACAATCTGTTACAATGGTGG
AGAGCACACGTCTGAACTCTTTCCCTACACGACGCTCTTCCGATCTNNNTTTGACCTCAGATAATAGA
AGAGCACACGTCTGAACTCTTTCCCTACACGACGCTCTTCCGATCTNNNAGGTTTGCTATGTCCCCATT
AAGAGCACACGTCTGAACTCTTTCCCTACACGACGCTCTTCCGATCTNNNTAGTTTCTTGCTGAGGG
AGAGCACACGTCTGAACTCTTTCCCTACACGACGCTCTTCCGATCTNNNGACACCCATTCACACACCAA
GAGCACACGTCTGAACTCTTTCCCTACACGACGCTCTTCCGATCTNNNTCAGCTCCCTAGACAGAGTA
GAGCACACGTCTGAACTCTTTCCCTACACGACGCTCTTCCGATCTNNNTGATCTCATGGGGTTACATT
AAGAGCACACGTCTGAACTCTTTCCCTACACGACGCTCTTCCGATCTNNNTCTATTGAGATGGGGTAG
AAGAGCACACGTCTGAACTCTTTCCCTACACGACGCTCTTCCGATCTNNNTGCATTCCAAGTGTTCTCA
GAGCACACGTCTGAACTCTTTCCCTACACGACGCTCTTCCGATCTNNNGAGCGAGCAGCAGATAAGA
AGCACACGTCTGAACTCTTTCCCTACACGACGCTCTTCCGATCTNNNACTTGGTTCAAATGACAGCTGA
GAGCACACGTCTGAACTCTTTCCCTACACGACGCTCTTCCGATCTNNNTCAATCCTTAGATCAGTTCTT
AGCACACGTCTGAACTCTTTCCCTACACGACGCTCTTCCGATCTNNNCCCACAGATTGAACTGAAATCT
AGCACACGTCTGAACTCTTTCCCTACACGACGCTCTTCCGATCTNNNTCTGTATTGCTGCAGTCTTTGT
AGCACACGTCTGAACTCTTTCCCTACACGACGCTCTTCCGATCTNNNAAAACTTTGAATAAAGTGCTCA
GAGCACACGTCTGAACTCTTTCCCTACACGACGCTCTTCCGATCTNNNCAGGAAGACTGGTAATGGG
AGCACACGTCTGAACTCTTTCCCTACACGACGCTCTTCCGATCTNNNACTTCTACACAAGACCACATGA
AGAGCACACGTCTGAACTCTTTCCCTACACGACGCTCTTCCGATCTNNNTGTTAGAAGGGACAAGTTT
AAGAGCACACGTCTGAACTCTTTCCCTACACGACGCTCTTCCGATCTNNNCACAGTGGGAAAGAAAA
AGAGCACACGTCTGAACTCTTTCCCTACACGACGCTCTTCCGATCTNNNAGTGAAAAACAGGTGAGA
GAGCACACGTCTGAACTCTTTCCCTACACGACGCTCTTCCGATCTNNNACATGAGGCAATTCAGACAT
AGAGCACACGTCTGAACTCTTTCCCTACACGACGCTCTTCCGATCTNNNGTGGTTTTCCAGGGACAAA
GAGCACACGTCTGAACTCTTTCCCTACACGACGCTCTTCCGATCTNNNACACTCAGGAGAATAACAAGT
AGCACACGTCTGAACTCTTTCCCTACACGACGCTCTTCCGATCTNNNACTATCTTTCTAGAGGCATCCA
AGCACACGTCTGAACTCTTTCCCTACACGACGCTCTTCCGATCTNNNACCTCCCACATCAAAGATTTAC
AGCACACGTCTGAACTCTTTCCCTACACGACGCTCTTCCGATCTNNNTCTATACTAACCCAACCTCTAC
AGAGCACACGTCTGAACTCTTTCCCTACACGACGCTCTTCCGATCTNNNCTTCTGCCCCTACCCATAAA
AGAGCACACGTCTGAACTCTTTCCCTACACGACGCTCTTCCGATCTNNNTTGTGGTTTTGGGTGTTGG
AGCACACGTCTGAACTCTTTCCCTACACGACGCTCTTCCGATCTNNNAGTCAGACCCTAAAGACTATGT
AGAGCACACGTCTGAACTCTTTCCCTACACGACGCTCTTCCGATCTNNNTCCCAGAGAGGACTGCTAC
GAGCACACGTCTGAACTCTTTCCCTACACGACGCTCTTCCGATCTNNNCACAGTGGGAACAGTTCTTG
AGAGCACACGTCTGAACTCTTTCCCTACACGACGCTCTTCCGATCTNNNCAGGGGAAATTTTTCAAGA
AGAGCACACGTCTGAACTCTTTCCCTACACGACGCTCTTCCGATCTNNNTGCATACAACTTGTTCTCAG
AAGAGCACACGTCTGAACTCTTTCCCTACACGACGCTCTTCCGATCTNNNTCTAATTCTTGGGGGCTTA
AGAGCACACGTCTGAACTCTTTCCCTACACGACGCTCTTCCGATCTNNNATGAAGGACCCTTGTTCAT
GAGCACACGTCTGAACTCTTTCCCTACACGACGCTCTTCCGATCTNNNTCTATTTGGCCCTTTAGAATG
GAGCACACGTCTGAACTCTTTCCCTACACGACGCTCTTCCGATCTNNNAATTCATTCCAGGTTGCTGT

AGAGCACACGTCTGAACTCTTTCCCTACACGACGCTCTTCCGATCTNNNGGCTTTGTTTCTTACTCTTC
AGCACACGTCTGAACTCTTTCCCTACACGACGCTCTTCCGATCTNNNACTCATTCAGTACATATCACTG
AGAGCACACGTCTGAACTCTTTCCCTACACGACGCTCTTCCGATCTNNNAGGCCTTTACCTCTACAACT
GAGCACACGTCTGAACTCTTTCCCTACACGACGCTCTTCCGATCTNNNACATGATGTGTTCCAGGCTC
AGCACACGTCTGAACTCTTTCCCTACACGACGCTCTTCCGATCTNNNTCCATCTGTCTCAGAGTTTAG
AGAGCACACGTCTGAACTCTTTCCCTACACGACGCTCTTCCGATCTNNNTTATGCATGCACACAAAGCT
AGCACACGTCTGAACTCTTTCCCTACACGACGCTCTTCCGATCTNNNTGGGAAAATAATGGAATGGA
AGCACACGTCTGAACTCTTTCCCTACACGACGCTCTTCCGATCTNNNTGAGAGAAAAAGATGTCAAA
AGAGCACACGTCTGAACTCTTTCCCTACACGACGCTCTTCCGATCTNNNAGGCTCTGGCTGTACTCGA
AGCACACGTCTGAACTCTTTCCCTACACGACGCTCTTCCGATCTNNNTGAGTGTGTGTGTGTTACATA
GCACACGTCTGAACTCTTTCCCTACACGACGCTCTTCCGATCTNNNTGTTGTGGTAAAACCTGTGTCT
GAGCACACGTCTGAACTCTTTCCCTACACGACGCTCTTCCGATCTNNNTTTGGTGTCTCTCCTTCCAGTT
AGAGCACACGTCTGAACTCTTTCCCTACACGACGCTCTTCCGATCTNNNCAAGGCTTCTGTACACCTTCT
GAGCACACGTCTGAACTCTTTCCCTACACGACGCTCTTCCGATCTNNNCACTAAAGCCAAACTTCTGC
AGAGCACACGTCTGAACTCTTTCCCTACACGACGCTCTTCCGATCTNNNCAATGCCCAGGGAAATTGA
AAGAGCACACGTCTGAACTCTTTCCCTACACGACGCTCTTCCGATCTNNNGCAGCATGTCTCAGGTAT
AGCACACGTCTGAACTCTTTCCCTACACGACGCTCTTCCGATCTNNNACTTATCTTGTTCAAATTTCCC
AGCACACGTCTGAACTCTTTCCCTACACGACGCTCTTCCGATCTNNNTGATCTCGTATCTTGTGATTTTT
GAGCACACGTCTGAACTCTTTCCCTACACGACGCTCTTCCGATCTNNNCAATGTCTGGAAGAGAGAA
AGCACACGTCTGAACTCTTTCCCTACACGACGCTCTTCCGATCTNNNTACGTCTTCTAGAGAGCTTGTG
AGAGCACACGTCTGAACTCTTTCCCTACACGACGCTCTTCCGATCTNNNTGATTACCAGGGTCTGAGG
AGAGCACACGTCTGAACTCTTTCCCTACACGACGCTCTTCCGATCTNNNAAACTGTCCAGATGGTTCT
AAGAGCACACGTCTGAACTCTTTCCCTACACGACGCTCTTCCGATCTNNNTGTGCTCTGAACCACCAGA
GAGCACACGTCTGAACTCTTTCCCTACACGACGCTCTTCCGATCTNNNCCTAAAACAACGTGTGTGTGT
GCACACGTCTGAACTCTTTCCCTACACGACGCTCTTCCGATCTNNNAGTGTATTTATTGTTCCTGGGT
GAGCACACGTCTGAACTCTTTCCCTACACGACGCTCTTCCGATCTNNNACAACTCTACCATCCCCTAAC
AGCACACGTCTGAACTCTTTCCCTACACGACGCTCTTCCGATCTNNNTGGGTATGTCTTTATTTTGGGC
AGAGCACACGTCTGAACTCTTTCCCTACACGACGCTCTTCCGATCTNNNTGGCTAGGTCAGCATTTTTC
AGAGCACACGTCTGAACTCTTTCCCTACACGACGCTCTTCCGATCTNNNCCGAAGTAGTTGCAGCATTT
GAGCACACGTCTGAACTCTTTCCCTACACGACGCTCTTCCGATCTNNNCAGAAAATGCTGCCTTTCCCA
GCACACGTCTGAACTCTTTCCCTACACGACGCTCTTCCGATCTNNNTGGACTTGTGGGTTATTTTATG
AAGAGCACACGTCTGAACTCTTTCCCTACACGACGCTCTTCCGATCTNNNTTCTATCTGCAGTGCCAAT
GAGCACACGTCTGAACTCTTTCCCTACACGACGCTCTTCCGATCTNNNTCAATTTGGCCCGAATAGTA
GAGCACACGTCTGAACTCTTTCCCTACACGACGCTCTTCCGATCTNNNTCCGTCTTGACAACATTTCAA
AGAGCACACGTCTGAACTCTTTCCCTACACGACGCTCTTCCGATCTNNNCCCTGACTAGGTTGAAGTG
GCACACGTCTGAACTCTTTCCCTACACGACGCTCTTCCGATCTNNNACTAAGGTAACCAAATATCCCTC
GCACACGTCTGAACTCTTTCCCTACACGACGCTCTTCCGATCTNNNTCAGTGTATACTATCAGAAGGC
AGAGCACACGTCTGAACTCTTTCCCTACACGACGCTCTTCCGATCTNNNCCACCCTACCTGTTCTCACCT
AGAGCACACGTCTGAACTCTTTCCCTACACGACGCTCTTCCGATCTNNNCTGCGATCTCCTTGTCATAG
AGAGCACACGTCTGAACTCTTTCCCTACACGACGCTCTTCCGATCTNNNCAATGAAGAATTGCCCTCCA
AGCACACGTCTGAACTCTTTCCCTACACGACGCTCTTCCGATCTNNNTGCCTCATTTTGTCAGTAGTTTC
AGCACACGTCTGAACTCTTTCCCTACACGACGCTCTTCCGATCTNNNTCAATGGATGGTTGGATGAA
AGAGCACACGTCTGAACTCTTTCCCTACACGACGCTCTTCCGATCTNNNGCACTCCCATCAGGAGAGT

AGAGCACACGTCTGAACTCTTTCCCTACACGACGCTCTTCCGATCTNNNATGGCTGACCATTTCAGGAC
GAAGAGCACACGTCTGAACTCTTTCCCTACACGACGCTCTTCCGATCTNNNATCCCCTTTGGTGACAGG
AAGAGCACACGTCTGAACTCTTTCCCTACACGACGCTCTTCCGATCTNNNGCAAACACACTCACTCTCTG
AGAGCACACGTCTGAACTCTTTCCCTACACGACGCTCTTCCGATCTNNNACCCTCCTTGCATTCTCTTTAC
AGCACACGTCTGAACTCTTTCCCTACACGACGCTCTTCCGATCTNNNACAGATCAATGGAAGAGTATGC
AGCACACGTCTGAACTCTTTCCCTACACGACGCTCTTCCGATCTNNNACACACACAGATATGTACTCACA
AAGAGCACACGTCTGAACTCTTTCCCTACACGACGCTCTTCCGATCTNNNGGAACATAGGAAGTTCCAC
AGAGCACACGTCTGAACTCTTTCCCTACACGACGCTCTTCCGATCTNNNAGAAAGAGACAACTCTGTTCT
AGAGCACACGTCTGAACTCTTTCCCTACACGACGCTCTTCCGATCTNNNTGCTTGTGAAATGGAGAGTT
AGAGCACACGTCTGAACTCTTTCCCTACACGACGCTCTTCCGATCTNNNCACCCTGTCAATGTCTTCACT
AAGAGCACACGTCTGAACTCTTTCCCTACACGACGCTCTTCCGATCTNNNAAATAATTGTGTGCTTGGG
AAGAGCACACGTCTGAACTCTTTCCCTACACGACGCTCTTCCGATCTNNNATTGCAGATGGGCTCTTGT
AAGAGCACACGTCTGAACTCTTTCCCTACACGACGCTCTTCCGATCTNNNCCTCTCAACTACACAACCTAA
GAGCACACGTCTGAACTCTTTCCCTACACGACGCTCTTCCGATCTNNNTATTCAGACCACAGCATTCAG
AGAGCACACGTCTGAACTCTTTCCCTACACGACGCTCTTCCGATCTNNNTGTACAGACATGCACACAGA
AGAGCACACGTCTGAACTCTTTCCCTACACGACGCTCTTCCGATCTNNNCAACATGAAATTCGGTTGTGC
GAGCACACGTCTGAACTCTTTCCCTACACGACGCTCTTCCGATCTNNNTTTTGTGTATTCTTGCCCATG
GAGCACACGTCTGAACTCTTTCCCTACACGACGCTCTTCCGATCTNNNTCTCAGACTTTTGGTTTTTACC
AGAGCACACGTCTGAACTCTTTCCCTACACGACGCTCTTCCGATCTNNNCAAAAGGTAGGGGGACAAA
AAGAGCACACGTCTGAACTCTTTCCCTACACGACGCTCTTCCGATCTNNNCTCCACCATCAGATTTGGGA
AAGAGCACACGTCTGAACTCTTTCCCTACACGACGCTCTTCCGATCTNNNTTTGCACAGCACTCCTCTAA
AGAGCACACGTCTGAACTCTTTCCCTACACGACGCTCTTCCGATCTNNNCAGCTCCTGTAACACCTTGAG
AGCACACGTCTGAACTCTTTCCCTACACGACGCTCTTCCGATCTNNNTGTACAGATGTATACACACACAC
AAGAGCACACGTCTGAACTCTTTCCCTACACGACGCTCTTCCGATCTNNNAGCATTGTTACAGGCATTT
AAGAGCACACGTCTGAACTCTTTCCCTACACGACGCTCTTCCGATCTNNNGCCCAACTTTTCCTCCTTCCT
GAGCACACGTCTGAACTCTTTCCCTACACGACGCTCTTCCGATCTNNNTGGACTGCAATCTTTAGTTTCA
GAGCACACGTCTGAACTCTTTCCCTACACGACGCTCTTCCGATCTNNNGCCTTTCTCTTTCTTGCTGTCTT
AGAGCACACGTCTGAACTCTTTCCCTACACGACGCTCTTCCGATCTNNNAAGGAAATTACAGACCAGCA
AAGAGCACACGTCTGAACTCTTTCCCTACACGACGCTCTTCCGATCTNNNATAGCTGAGGTAGAGGTT
AAGAGCACACGTCTGAACTCTTTCCCTACACGACGCTCTTCCGATCTNNNGCTCTCCTAGGACACTGGG
GAAGAGCACACGTCTGAACTCTTTCCCTACACGACGCTCTTCCGATCTNNNTCAGCCTGTTGACTACAGT
AGAGCACACGTCTGAACTCTTTCCCTACACGACGCTCTTCCGATCTNNNTGGAGACTACCAGAGAAACA
AGCACACGTCTGAACTCTTTCCCTACACGACGCTCTTCCGATCTNNNTCCACATTTACATTCTCTCCCTCC
AGAGCACACGTCTGAACTCTTTCCCTACACGACGCTCTTCCGATCTNNNAGCGTTTCTTACATCAGCTCA
AAGAGCACACGTCTGAACTCTTTCCCTACACGACGCTCTTCCGATCTNNNCAGTGGGGAGAGACAACA
AAGAGCACACGTCTGAACTCTTTCCCTACACGACGCTCTTCCGATCTNNNTGGCAAGATGCAGACTAGA
AAGAGCACACGTCTGAACTCTTTCCCTACACGACGCTCTTCCGATCTNNNCCCAGATGCACCATTTCTCT
GAGCACACGTCTGAACTCTTTCCCTACACGACGCTCTTCCGATCTNNNGCACTAATGTTTAGTTCTGTGA
AAGAGCACACGTCTGAACTCTTTCCCTACACGACGCTCTTCCGATCTNNNGAAAGGGTAAAGAGCTGA
AGAGCACACGTCTGAACTCTTTCCCTACACGACGCTCTTCCGATCTNNNGACTGCTTACACCTTAGCTGC
AGAGCACACGTCTGAACTCTTTCCCTACACGACGCTCTTCCGATCTNNNCCTTCCTCCCTCCAATCTCTCT
GAGCACACGTCTGAACTCTTTCCCTACACGACGCTCTTCCGATCTNNNAGTCATAATGTGGTAAGGAA
AGCACACGTCTGAACTCTTTCCCTACACGACGCTCTTCCGATCTNNNTGGCTTGCGAATAAACAAATTA

AAGAGCACACGTCTGAACTCTTTCCCTACACGACGCTCTTCCGATCTNNNGAGAAAAGAAACAGCAG
GAGCACACGTCTGAACTCTTTCCCTACACGACGCTCTTCCGATCTNNNTTGCAAGACTGAGTTGTAGC
GAGCACACGTCTGAACTCTTTCCCTACACGACGCTCTTCCGATCTNNNACCTTGATCATGAAGAGTAA
GAGCACACGTCTGAACTCTTTCCCTACACGACGCTCTTCCGATCTNNNATTCTGAAAATGGCTGCACC
AGAGCACACGTCTGAACTCTTTCCCTACACGACGCTCTTCCGATCTNNNGTTGTGCGTTCCTAAACTC
AAGAGCACACGTCTGAACTCTTTCCCTACACGACGCTCTTCCGATCTNNNCTTTCTTGTCCTCTCCTCC
AGAGCACACGTCTGAACTCTTTCCCTACACGACGCTCTTCCGATCTNNNGGCATGCCAATACCTAGGA
AGCACACGTCTGAACTCTTTCCCTACACGACGCTCTTCCGATCTNNNTGTTTACTATCCAGCAGTCTT
AGAGCACACGTCTGAACTCTTTCCCTACACGACGCTCTTCCGATCTNNNCAACAAGAGTGAACCTCAG
AGCACACGTCTGAACTCTTTCCCTACACGACGCTCTTCCGATCTNNNAGAAACATTGAGAGATAGACA
AGAGCACACGTCTGAACTCTTTCCCTACACGACGCTCTTCCGATCTNNNGGAGAAAAATAATGCGAAG
AGCACACGTCTGAACTCTTTCCCTACACGACGCTCTTCCGATCTNNNCCTGCCCACTCATAAATCCATC
AGCACACGTCTGAACTCTTTCCCTACACGACGCTCTTCCGATCTNNNTCCTCACTGCCATTCCTCTATAT
AAGAGCACACGTCTGAACTCTTTCCCTACACGACGCTCTTCCGATCTNNNTGAAAGTAGTGGTTTCGG
AGAGCACACGTCTGAACTCTTTCCCTACACGACGCTCTTCCGATCTNNNCAGATCTGGTGAAAGCTAG
AGAGCACACGTCTGAACTCTTTCCCTACACGACGCTCTTCCGATCTNNNTTTTTGATCTGCAGTTGCGT
AGAGCACACGTCTGAACTCTTTCCCTACACGACGCTCTTCCGATCTNNNTAACAGAGTACCTTAGCCC
AAGAGCACACGTCTGAACTCTTTCCCTACACGACGCTCTTCCGATCTNNNAAATTTCTTGAGGCCAGAG
AGAGCACACGTCTGAACTCTTTCCCTACACGACGCTCTTCCGATCTNNNAAGTCTTCAGGGTCTAGAG
AGAGCACACGTCTGAACTCTTTCCCTACACGACGCTCTTCCGATCTNNNTTGGATTGTTGTTGCTCCTG
GCACACGTCTGAACTCTTTCCCTACACGACGCTCTTCCGATCTNNNAGTTTCCTTTCTTTTTCACTTGCC
GAGCACACGTCTGAACTCTTTCCCTACACGACGCTCTTCCGATCTNNNTGTGTACAATGAAATACTCAG
AAGAGCACACGTCTGAACTCTTTCCCTACACGACGCTCTTCCGATCTNNNTTGCTCTGCCCTGTAAGAA
AAGAGCACACGTCTGAACTCTTTCCCTACACGACGCTCTTCCGATCTNNNAATATGACTTACAGCTGTG
GAGCACACGTCTGAACTCTTTCCCTACACGACGCTCTTCCGATCTNNNCAGGACCTTACTGCCACTTTC
AAGAGCACACGTCTGAACTCTTTCCCTACACGACGCTCTTCCGATCTNNNGTGGAACACACTTCAAGC
AAGAGCACACGTCTGAACTCTTTCCCTACACGACGCTCTTCCGATCTNNNATTACCCATTTCAGAAGCG
GAGCACACGTCTGAACTCTTTCCCTACACGACGCTCTTCCGATCTNNNGTCATGCTCACCAAACATGC
AGAGCACACGTCTGAACTCTTTCCCTACACGACGCTCTTCCGATCTNNNTCTGCGCCTGAATTTCCTTA
GAGCACACGTCTGAACTCTTTCCCTACACGACGCTCTTCCGATCTNNNATTTACCAACTTGCATTTGTT
AAGAGCACACGTCTGAACTCTTTCCCTACACGACGCTCTTCCGATCTNNNTATGTGCTGATCAGTGTA
AAGAGCACACGTCTGAACTCTTTCCCTACACGACGCTCTTCCGATCTNNNATTGGTGCTTGCAAAGT
GAGCACACGTCTGAACTCTTTCCCTACACGACGCTCTTCCGATCTNNNTGCAACTCTGTGATGTTGTT
GAGCACACGTCTGAACTCTTTCCCTACACGACGCTCTTCCGATCTNNNCGTAACAGAGCGAGACTATG
AGAGCACACGTCTGAACTCTTTCCCTACACGACGCTCTTCCGATCTNNNTTCCCAACATTCCAGATCCC
GAGCACACGTCTGAACTCTTTCCCTACACGACGCTCTTCCGATCTNNNTCCTAGGGACAAGAGGAAG
GAGCACACGTCTGAACTCTTTCCCTACACGACGCTCTTCCGATCTNNNAAAAGAAGGTCTGCTCAGCT
AGAGCACACGTCTGAACTCTTTCCCTACACGACGCTCTTCCGATCTNNNAGAGGTATGGGGACAAAC
GAGCACACGTCTGAACTCTTTCCCTACACGACGCTCTTCCGATCTNNNTAGTGAAGACAGGACCCAG
GAGCACACGTCTGAACTCTTTCCCTACACGACGCTCTTCCGATCTNNNACAAATATCCTTGCACATCTC
AAGAGCACACGTCTGAACTCTTTCCCTACACGACGCTCTTCCGATCTNNNTAAGAGCCAAGCCAAGAA
AGAGCACACGTCTGAACTCTTTCCCTACACGACGCTCTTCCGATCTNNNTTTTTGCTTTGGCTGAGTTT
AGAGCACACGTCTGAACTCTTTCCCTACACGACGCTCTTCCGATCTNNNACATGTCCTCCTGTAGTCCC

AAGAGCACACGTCTGAACTCTTTCCCTACACGACGCTCTTCCGATCTNNNTGAATCCAGGAGGCAGAG
GAGCACACGTCTGAACTCTTTCCCTACACGACGCTCTTCCGATCTNNNTGTCACTTCTTTTGTTTTTTGG
AGAGCACACGTCTGAACTCTTTCCCTACACGACGCTCTTCCGATCTNNNCTCCTGTAATCCCAGCTACTT
AGAGCACACGTCTGAACTCTTTCCCTACACGACGCTCTTCCGATCTNNNTGAGGCAGGAGAATCACTT
AGAGCACACGTCTGAACTCTTTCCCTACACGACGCTCTTCCGATCTNNNGTGAAGGTTGTAGTGAGCT
AAGAGCACACGTCTGAACTCTTTCCCTACACGACGCTCTTCCGATCTNNNGAGAATGGTGTGAACTC
AGCACACGTCTGAACTCTTTCCCTACACGACGCTCTTCCGATCTNNNCTTCAAAAGCTGTGTAAAGTTA
GCACACGTCTGAACTCTTTCCCTACACGACGCTCTTCCGATCTNNNACAGTGAATAGAGAATTGGGA
GAGCACACGTCTGAACTCTTTCCCTACACGACGCTCTTCCGATCTNNNACAACCTAACATCATACCTAGT
AGCACACGTCTGAACTCTTTCCCTACACGACGCTCTTCCGATCTNNNTGGTAGGCATTTAGGTTCTCAA
AGCACACGTCTGAACTCTTTCCCTACACGACGCTCTTCCGATCTNNNCCCTAACTTTGAATAGAGAAG
AGAGCACACGTCTGAACTCTTTCCCTACACGACGCTCTTCCGATCTNNNAGCCAAATCCACAGAGTTA
GCACACGTCTGAACTCTTTCCCTACACGACGCTCTTCCGATCTNNNGCCATCTTTCTTAGTTTGACAAA
AGCACACGTCTGAACTCTTTCCCTACACGACGCTCTTCCGATCTNNNTCTTGAAGAGCTATTAGCACTC
GAGCACACGTCTGAACTCTTTCCCTACACGACGCTCTTCCGATCTNNNGAGGCAGGAAAATCACTTGA
AGAGCACACGTCTGAACTCTTTCCCTACACGACGCTCTTCCGATCTNNNTTGAGGCAGGAGAATCAC
GAGCACACGTCTGAACTCTTTCCCTACACGACGCTCTTCCGATCTNNNCTGAGGCAGGAGAATTGCTT
AAGAGCACACGTCTGAACTCTTTCCCTACACGACGCTCTTCCGATCTNNNATGCCTGTAATCCCAGTGA
GGAAGAGCACACGTCTGAACTCTTTCCCTACACGACGCTCTTCCGATCTNNNAGAATCACTTGAATCCA
GAGCACACGTCTGAACTCTTTCCCTACACGACGCTCTTCCGATCTNNNGAGGTTACAGTGAGCCGAGA
AAGAGCACACGTCTGAACTCTTTCCCTACACGACGCTCTTCCGATCTNNNAGGCTGAAGCAGGAGAAT
AAGAGCACACGTCTGAACTCTTTCCCTACACGACGCTCTTCCGATCTNNNCGAGGCAGAAGAATCACTT
AGCACACGTCTGAACTCTTTCCCTACACGACGCTCTTCCGATCTNNNTGAGATGCTTATTCCCAAATGT
AGCACACGTCTGAACTCTTTCCCTACACGACGCTCTTCCGATCTNNNTCTGTGGATTGTCTTTTCACTTT
AGAGCACACGTCTGAACTCTTTCCCTACACGACGCTCTTCCGATCTNNNAGTGAGCTGAGACCGTGATA
AGCACACGTCTGAACTCTTTCCCTACACGACGCTCTTCCGATCTNNNACTCAGAAAATACAAGCAGAGG
AAGAGCACACGTCTGAACTCTTTCCCTACACGACGCTCTTCCGATCTNNNGCAGGAGCATCACTTGAAT
GGAAGAGCACACGTCTGAACTCTTTCCCTACACGACGCTCTTCCGATCTNNNCTGTAGTCCCAGCTACT
GAGCACACGTCTGAACTCTTTCCCTACACGACGCTCTTCCGATCTNNNTCTAGAATGAGAAGAGTTCT
AAGAGCACACGTCTGAACTCTTTCCCTACACGACGCTCTTCCGATCTNNNATTTAGGCAATGAGGAGG
AGAGCACACGTCTGAACTCTTTCCCTACACGACGCTCTTCCGATCTNNNTTACAGTGAGCCAAAATCAT
AGAGCACACGTCTGAACTCTTTCCCTACACGACGCTCTTCCGATCTNNNCTGTATTCCCAGCTACTCAGG
AGCACACGTCTGAACTCTTTCCCTACACGACGCTCTTCCGATCTNNNGGCAAATTGAATCCAACAATAC
GAGCACACGTCTGAACTCTTTCCCTACACGACGCTCTTCCGATCTNNNAGGAGAATCGTTTGAACCCAT
GAGCACACGTCTGAACTCTTTCCCTACACGACGCTCTTCCGATCTNNNTTGAGTCCAGGAGTTCAAGGA
AAGAGCACACGTCTGAACTCTTTCCCTACACGACGCTCTTCCGATCTNNNCTGAGGCAGGAGAATCACT
GAGCACACGTCTGAACTCTTTCCCTACACGACGCTCTTCCGATCTNNNTTTTTTTTGAGATGGAGTTTC
GAGCACACGTCTGAACTCTTTCCCTACACGACGCTCTTCCGATCTNNNAGGTTCTTCCTACAGTACCAAA
GGAAGAGCACACGTCTGAACTCTTTCCCTACACGACGCTCTTCCGATCTNNNTTGCTTGAACATAGGA
AGAGCACACGTCTGAACTCTTTCCCTACACGACGCTCTTCCGATCTNNNTGTAGTCCCAGCTACTCAGTA
GAGCACACGTCTGAACTCTTTCCCTACACGACGCTCTTCCGATCTNNNATATCACAGCTACTTGGAGAG
AGAGCACACGTCTGAACTCTTTCCCTACACGACGCTCTTCCGATCTNNNGCGCATATCTGTAGTTCCAG
AGAGCACACGTCTGAACTCTTTCCCTACACGACGCTCTTCCGATCTNNNCTGTAGTCCCAGCTACTCAGG

AGAGCACACGTCTGAACTCTTTCCCTACACGACGCTCTTCCGATCTNNNAGGCAAGAGAATCCCTTGA
GAGCACACGTCTGAACTCTTTCCCTACACGACGCTCTTCCGATCTNNNACTAATTTGGAGTTTGGGTTT
GAAGAGCACACGTCTGAACTCTTTCCCTACACGACGCTCTTCCGATCTNNNGTAATCCCAGCTACTTAG
AAGAGCACACGTCTGAACTCTTTCCCTACACGACGCTCTTCCGATCTNNNCACACACACAGACACATAC
AGCACACGTCTGAACTCTTTCCCTACACGACGCTCTTCCGATCTNNNACAAGGAGACAAACTGAATAAT
GAAGAGCACACGTCTGAACTCTTTCCCTACACGACGCTCTTCCGATCTNNNGAGAATGGTGTGAACCT
GAGCACACGTCTGAACTCTTTCCCTACACGACGCTCTTCCGATCTNNNGAATCGCTTGAACCTAGGAG
GAGCACACGTCTGAACTCTTTCCCTACACGACGCTCTTCCGATCTNNNGTAATCCTAGCTACTTGGGA
AGAGCACACGTCTGAACTCTTTCCCTACACGACGCTCTTCCGATCTNNNCTGTAATCCCAGCTACTCAGG
AGAGCACACGTCTGAACTCTTTCCCTACACGACGCTCTTCCGATCTNNNCTGTAGTCCTAGCTACTCGG
GAGCACACGTCTGAACTCTTTCCCTACACGACGCTCTTCCGATCTNNNCTGTAATCTCAGCTACTCGGG
AGAGCACACGTCTGAACTCTTTCCCTACACGACGCTCTTCCGATCTNNNACAAAAATTAGCTGGGTGT
AAGAGCACACGTCTGAACTCTTTCCCTACACGACGCTCTTCCGATCTNNNTGTAGTCCCAGCTACTCAG
AAGAGCACACGTCTGAACTCTTTCCCTACACGACGCTCTTCCGATCTNNNCTTGCAGTGAGCTGAGAT
AGAGCACACGTCTGAACTCTTTCCCTACACGACGCTCTTCCGATCTNNNGTAATCTCAGCTACTCAGG
AGCACACGTCTGAACTCTTTCCCTACACGACGCTCTTCCGATCTNNNTGATGACATCAAATTCATACAC
GAGCACACGTCTGAACTCTTTCCCTACACGACGCTCTTCCGATCTNNNTGTGTGTGTGTACATTTAT
GAGCACACGTCTGAACTCTTTCCCTACACGACGCTCTTCCGATCTNNNAGCAATATCCCTTTGAGAAAG
AGAGCACACGTCTGAACTCTTTCCCTACACGACGCTCTTCCGATCTNNNAGAGAGAGAGAGAGAGAG
AGAGCACACGTCTGAACTCTTTCCCTACACGACGCTCTTCCGATCTNNNGCTGGTGCTCATGTATCCAA
AGAGCACACGTCTGAACTCTTTCCCTACACGACGCTCTTCCGATCTNNNTTGACACTGATACAATGCAC
GAGCACACGTCTGAACTCTTTCCCTACACGACGCTCTTCCGATCTNNNACTATATACTGTCTTCAGGGG
GAGCACACGTCTGAACTCTTTCCCTACACGACGCTCTTCCGATCTNNNTGGCACAATCAGTTTCTTCTTT
GCACACGTCTGAACTCTTTCCCTACACGACGCTCTTCCGATCTNNNTGAAAATTGAGTCTTTAGGGCT
AGAGCACACGTCTGAACTCTTTCCCTACACGACGCTCTTCCGATCTNNNCTGTAGTCCCAGCTACTCAG
AAGAGCACACGTCTGAACTCTTTCCCTACACGACGCTCTTCCGATCTNNNATCACTTGAACCCAGGAAG
AGAGCACACGTCTGAACTCTTTCCCTACACGACGCTCTTCCGATCTNNNTGTAGTCCTAGCTACTCCAG
AGAGCACACGTCTGAACTCTTTCCCTACACGACGCTCTTCCGATCTNNNAGAATGATGTCTCCCAGCTT
AGCACACGTCTGAACTCTTTCCCTACACGACGCTCTTCCGATCTNNNGGCTTCATCATTTTGTGATAGT
GAGCACACGTCTGAACTCTTTCCCTACACGACGCTCTTCCGATCTNNNCATGCACCTATAGTCCCAGCT
GAGCACACGTCTGAACTCTTTCCCTACACGACGCTCTTCCGATCTNNNAGGTGATAAATTTCTGTTGCT
AGCACACGTCTGAACTCTTTCCCTACACGACGCTCTTCCGATCTNNNAGCAATTTGAAAGGGTAAATG
AGCACACGTCTGAACTCTTTCCCTACACGACGCTCTTCCGATCTNNNGGACAAAACTCTAATTAGCTT
AGAGCACACGTCTGAACTCTTTCCCTACACGACGCTCTTCCGATCTNNNCTGAGATCATGCCACTGTA
AAGAGCACACGTCTGAACTCTTTCCCTACACGACGCTCTTCCGATCTNNNCTGAGACAGGAGAGTTG
AGCACACGTCTGAACTCTTTCCCTACACGACGCTCTTCCGATCTNNNTGCTTTTTCTTCAGAGAATCGT
AGCACACGTCTGAACTCTTTCCCTACACGACGCTCTTCCGATCTNNNTGATTGACAGAATAGTAGTGG
AGAGCACACGTCTGAACTCTTTCCCTACACGACGCTCTTCCGATCTNNNTTATTCTTCCTACCCTCCCTC
GAGCACACGTCTGAACTCTTTCCCTACACGACGCTCTTCCGATCTNNNGGCTGAGGCAGAAAAATCT
GCACACGTCTGAACTCTTTCCCTACACGACGCTCTTCCGATCTNNNTCAAGGTCAAATTGTACTTAGCC
AGCACACGTCTGAACTCTTTCCCTACACGACGCTCTTCCGATCTNNNACAAGTCATAAGGGTAACTTCA
AGAGCACACGTCTGAACTCTTTCCCTACACGACGCTCTTCCGATCTNNNATGAGGCAGGAGAATCGTT
AGAGCACACGTCTGAACTCTTTCCCTACACGACGCTCTTCCGATCTNNNCACCTGTAATCCCAGCTACT

AGCACACGTCTGAACTCTTTCCCTACACGACGCTCTTCCGATCTNNNTGACATTGTACTGAAGTTCTA
GAGCACACGTCTGAACTCTTTCCCTACACGACGCTCTTCCGATCTNNNCAGCTTCATCAGTATTGCTG
AGAGCACACGTCTGAACTCTTTCCCTACACGACGCTCTTCCGATCTNNNTTGACTATGAAGGGCCCCA
AGCACACGTCTGAACTCTTTCCCTACACGACGCTCTTCCGATCTNNNACAGATTCCTGGAACTTTTGT
GAGCACACGTCTGAACTCTTTCCCTACACGACGCTCTTCCGATCTNNNTTCTGTTCTCAGTTCAACTCC
AGAGCACACGTCTGAACTCTTTCCCTACACGACGCTCTTCCGATCTNNNAATAGAGCCAGACCATGTC
GAGCACACGTCTGAACTCTTTCCCTACACGACGCTCTTCCGATCTNNNTCACAGATAGAAGCCAATTT
AGAGCACACGTCTGAACTCTTTCCCTACACGACGCTCTTCCGATCTNNNTGTAATCACAGCTACTTGG
AGCACACGTCTGAACTCTTTCCCTACACGACGCTCTTCCGATCTNNNACTGGAAAAAAAAACAAAGA
AGAGCACACGTCTGAACTCTTTCCCTACACGACGCTCTTCCGATCTNNNTCCCAAGAGGAATGCATTT
AGAGCACACGTCTGAACTCTTTCCCTACACGACGCTCTTCCGATCTNNNTTTGCCAGTATGTAGACT
AGAGCACACGTCTGAACTCTTTCCCTACACGACGCTCTTCCGATCTNNNCGAGGCAAGAGAATCACT
GAGCACACGTCTGAACTCTTTCCCTACACGACGCTCTTCCGATCTNNNGTGAATCTTCTCTGACCCTCT
GCACACGTCTGAACTCTTTCCCTACACGACGCTCTTCCGATCTNNNTGACCTCTGGTGTTTTCATTATC
AGAGCACACGTCTGAACTCTTTCCCTACACGACGCTCTTCCGATCTNNNCAAAACAGCTGTTGACGAT
AGAGCACACGTCTGAACTCTTTCCCTACACGACGCTCTTCCGATCTNNNCATCTATTTGCTGCCCCGAA
GAGCACACGTCTGAACTCTTTCCCTACACGACGCTCTTCCGATCTNNNTAGAAGTCACTGCGGAGATG
GAGCACACGTCTGAACTCTTTCCCTACACGACGCTCTTCCGATCTNNNTGCCAAAATACCTTTCCCTTT
AGCACACGTCTGAACTCTTTCCCTACACGACGCTCTTCCGATCTNNNAGTCCATTAGCACACATAACAG
AGCACACGTCTGAACTCTTTCCCTACACGACGCTCTTCCGATCTNNNACAGCAGTGTACACATTAGT
GAGCACACGTCTGAACTCTTTCCCTACACGACGCTCTTCCGATCTNNNTCCATTTTAGGGATAGTCAA
GCACACGTCTGAACTCTTTCCCTACACGACGCTCTTCCGATCTNNNTGCTCTTACAGTAGGGAAACA
CACACGTCTGAACTCTTTCCCTACACGACGCTCTTCCGATCTNNNGAACAAGATCTGGTCTCAAAAA
AGCACACGTCTGAACTCTTTCCCTACACGACGCTCTTCCGATCTNNNAGGGTTCTCTAGAGGGATAG
AGAGCACACGTCTGAACTCTTTCCCTACACGACGCTCTTCCGATCTNNNTCCAATGTATTCTACTGTGC
GAGCACACGTCTGAACTCTTTCCCTACACGACGCTCTTCCGATCTNNNTCTTAGCTCACTGAAACCTCT
GCACACGTCTGAACTCTTTCCCTACACGACGCTCTTCCGATCTNNNACCTTTCTTCTCTCTCTCTCTC
AGAGCACACGTCTGAACTCTTTCCCTACACGACGCTCTTCCGATCTNNNATGATTTGTAACCCCTGAG
AGCACACGTCTGAACTCTTTCCCTACACGACGCTCTTCCGATCTNNNTCTTACATGTAGAAAACCTGAA
GAGCACACGTCTGAACTCTTTCCCTACACGACGCTCTTCCGATCTNNNTGTTATAGAATACCATATGC
AGCACACGTCTGAACTCTTTCCCTACACGACGCTCTTCCGATCTNNNACCCTGATATCAAAACTAGAC
AGCACACGTCTGAACTCTTTCCCTACACGACGCTCTTCCGATCTNNNTGTTCTTTTTTGGTTTTCATTG
GCACACGTCTGAACTCTTTCCCTACACGACGCTCTTCCGATCTNNNCACTCGCAACTACTTCTCTTCCT
AGCACACGTCTGAACTCTTTCCCTACACGACGCTCTTCCGATCTNNNTCTCTCTCTCTCTCTCTCTCT
AGCACACGTCTGAACTCTTTCCCTACACGACGCTCTTCCGATCTNNNGTGAGAGGGTTCACAATATCT
GCACACGTCTGAACTCTTTCCCTACACGACGCTCTTCCGATCTNNNATGCCTGTAATCCCAGCTACTTG
AAGAGCACACGTCTGAACTCTTTCCCTACACGACGCTCTTCCGATCTNNNAATGAGCCGAGATCATG
GCACACGTCTGAACTCTTTCCCTACACGACGCTCTTCCGATCTNNNACATATCCTATAAACCCAATGTC
GAGCACACGTCTGAACTCTTTCCCTACACGACGCTCTTCCGATCTNNNCCTGGTTATACCCCTCCTCAA
AGCACACGTCTGAACTCTTTCCCTACACGACGCTCTTCCGATCTNNNTCCTATAGTTGAAGCCACTGA
AAGAGCACACGTCTGAACTCTTTCCCTACACGACGCTCTTCCGATCTNNNCTCCAGTGTCTGTATGC
AGAGCACACGTCTGAACTCTTTCCCTACACGACGCTCTTCCGATCTNNNTATTTCCACCTTACCTCCAG
GAGCACACGTCTGAACTCTTTCCCTACACGACGCTCTTCCGATCTNNNTCCTCTATTCATGGCTTGATA

AGAGCACACGTCTGAACTCTTTCCCTACACGACGCTCTTCCGATCTNNNTGACTAGTGTTCCCCTATC
AGAGCACACGTCTGAACTCTTTCCCTACACGACGCTCTTCCGATCTNNNTCAAGGCTTCAGTGAACTG
GAGCACACGTCTGAACTCTTTCCCTACACGACGCTCTTCCGATCTNNNTCTGGATCTGTGGTCTTTGG
AGCACACGTCTGAACTCTTTCCCTACACGACGCTCTTCCGATCTNNNTGCCATTTCATCCTTTTGCTATT
AGAGCACACGTCTGAACTCTTTCCCTACACGACGCTCTTCCGATCTNNNGATTACTGCTTGAGTCTGG
GAGCACACGTCTGAACTCTTTCCCTACACGACGCTCTTCCGATCTNNNTGTTATTCTTGATAGGCCAAA
GAGCACACGTCTGAACTCTTTCCCTACACGACGCTCTTCCGATCTNNNTGATGGACACTTAGGTTGATT
AGCACACGTCTGAACTCTTTCCCTACACGACGCTCTTCCGATCTNNNTGAGAAGAGGTAAGGTCTAA
AGCACACGTCTGAACTCTTTCCCTACACGACGCTCTTCCGATCTNNNTCACTAAAAGGAAGACAGAAA
AGCACACGTCTGAACTCTTTCCCTACACGACGCTCTTCCGATCTNNNTCTTGTATTTTCTTCTCTCTCTCC
AGCACACGTCTGAACTCTTTCCCTACACGACGCTCTTCCGATCTNNNACAACAGAGAAAGACAGAGAG
AGAGCACACGTCTGAACTCTTTCCCTACACGACGCTCTTCCGATCTNNNTCTGGTCTCATGTTACATCC
GCACACGTCTGAACTCTTTCCCTACACGACGCTCTTCCGATCTNNNACTGACCATCATATCCCTATTGT
GAGCACACGTCTGAACTCTTTCCCTACACGACGCTCTTCCGATCTNNNGGTTGAGAGACAAATCAAGA
AGAGCACACGTCTGAACTCTTTCCCTACACGACGCTCTTCCGATCTNNNCCCAGCCCATGATTCCTTTTT
GCACACGTCTGAACTCTTTCCCTACACGACGCTCTTCCGATCTNNNTGTCCAGGTGAAAATATAGCTA
GGAAGAGCACACGTCTGAACTCTTTCCCTACACGACGCTCTTCCGATCTNNNAAAAGTTGCAGTGAGC
GAGCACACGTCTGAACTCTTTCCCTACACGACGCTCTTCCGATCTNNNTCTGACAAGGGTCTAGTATCC
GCACACGTCTGAACTCTTTCCCTACACGACGCTCTTCCGATCTNNNTCTGTAGGCTTGAGTTTGAGAC
GAGCACACGTCTGAACTCTTTCCCTACACGACGCTCTTCCGATCTNNNGCATCTGGCTTATTTCACTTA
AGCACACGTCTGAACTCTTTCCCTACACGACGCTCTTCCGATCTNNNTCTTTCTTCCTTCCAGTATGTTT
GAGCACACGTCTGAACTCTTTCCCTACACGACGCTCTTCCGATCTNNNACTCTGGAAGTTGTAAAAGC
GAGCACACGTCTGAACTCTTTCCCTACACGACGCTCTTCCGATCTNNNTGCAACTGATTTTTGTGGGT
AGCACACGTCTGAACTCTTTCCCTACACGACGCTCTTCCGATCTNNNAGGATTCATGTTCTTCTCATTA
AGCACACGTCTGAACTCTTTCCCTACACGACGCTCTTCCGATCTNNNTGCATTTGTGTGTGTATGTAT
AGAGCACACGTCTGAACTCTTTCCCTACACGACGCTCTTCCGATCTNNNTCTAAGAACTCCAAGGACAC
AAGAGCACACGTCTGAACTCTTTCCCTACACGACGCTCTTCCGATCTNNNCTGAGGCAGGAGAATCAC
AGAGCACACGTCTGAACTCTTTCCCTACACGACGCTCTTCCGATCTNNNGGGCAACAAGAGAGAAACT
GAGCACACGTCTGAACTCTTTCCCTACACGACGCTCTTCCGATCTNNNTGCCAAATACGTGATCTATG
AGAGCACACGTCTGAACTCTTTCCCTACACGACGCTCTTCCGATCTNNNCAGAGAAAGACCCTGTCTCA
GAGCACACGTCTGAACTCTTTCCCTACACGACGCTCTTCCGATCTNNNCCTCTCTTCCTATTTGTGTTT
GGAAGAGCACACGTCTGAACTCTTTCCCTACACGACGCTCTTCCGATCTNNNCAGTTCTAGAGGCAAG
AGCACACGTCTGAACTCTTTCCCTACACGACGCTCTTCCGATCTNNNAGAATGAATAAGACCTAGCA
AGCACACGTCTGAACTCTTTCCCTACACGACGCTCTTCCGATCTNNNAGATTCCATCTCACAAACAAA
GGAAGAGCACACGTCTGAACTCTTTCCCTACACGACGCTCTTCCGATCTNNNCTGAGGCACAAGAATC
AAGAGCACACGTCTGAACTCTTTCCCTACACGACGCTCTTCCGATCTNNNGTGAGCCAAGATCGCATC
GAGCACACGTCTGAACTCTTTCCCTACACGACGCTCTTCCGATCTNNNACAGATTTCAATTTGCTTCAT
AGAGCACACGTCTGAACTCTTTCCCTACACGACGCTCTTCCGATCTNNNAAATGCCAAGGGTTTTTGAA
GAGCACACGTCTGAACTCTTTCCCTACACGACGCTCTTCCGATCTNNNAAATAAGGTCAGCTAGGCTT
AAGAGCACACGTCTGAACTCTTTCCCTACACGACGCTCTTCCGATCTNNNAGACTCCGTCTCAAAAAAA
GGAAGAGCACACGTCTGAACTCTTTCCCTACACGACGCTCTTCCGATCTNNNGATCGTGTCACTCTAC
GAGCACACGTCTGAACTCTTTCCCTACACGACGCTCTTCCGATCTNNNCTGTAGTGCAGTGGTGTGAT
GCACACGTCTGAACTCTTTCCCTACACGACGCTCTTCCGATCTNNNAGCTCCATTTTCTCTCTTTCCTTC

GCACACGTCTGAACTCTTTCCCTACACGACGCTCTTCCGATCTNNNAGTCTCCCAAGTCTGATAAAGTG
AGAGCACACGTCTGAACTCTTTCCCTACACGACGCTCTTCCGATCTNNNAGTGTATGCTCTCACACTAG
AAGAGCACACGTCTGAACTCTTTCCCTACACGACGCTCTTCCGATCTNNNATCCCCTTTGGTGACAGGA
AGAGCACACGTCTGAACTCTTTCCCTACACGACGCTCTTCCGATCTNNNAGGGTGGCTAGCTGTAGCT
AGCACACGTCTGAACTCTTTCCCTACACGACGCTCTTCCGATCTNNNACACTCTGCTCAACTTTGCCTTT
GAGCACACGTCTGAACTCTTTCCCTACACGACGCTCTTCCGATCTNNNTACACCAAGCAACTCAATCCCA
GAGCACACGTCTGAACTCTTTCCCTACACGACGCTCTTCCGATCTNNNCTGGCTGACCTAGCCAAAGAT
AGCACACGTCTGAACTCTTTCCCTACACGACGCTCTTCCGATCTNNNTCAGCCAATCAATCAATCTGTC
AGAGCACACGTCTGAACTCTTTCCCTACACGACGCTCTTCCGATCTNNNACCTTGGAGTGGTTTTGGTA
AGCACACGTCTGAACTCTTTCCCTACACGACGCTCTTCCGATCTNNNAGAAATGTTACTGCTAAGCACC
AGAGCACACGTCTGAACTCTTTCCCTACACGACGCTCTTCCGATCTNNNGAAACAGTGGGCTTTGTTT
GAGCACACGTCTGAACTCTTTCCCTACACGACGCTCTTCCGATCTNNNCCAAATCTGGGCTTTACTTGTG
GAGCACACGTCTGAACTCTTTCCCTACACGACGCTCTTCCGATCTNNNATATTTGGGACACAGAACCTG
GAGCACACGTCTGAACTCTTTCCCTACACGACGCTCTTCCGATCTNNNGTGGTGTTCAGGGATTTAGG
GAGCACACGTCTGAACTCTTTCCCTACACGACGCTCTTCCGATCTNNNTGGAGCTAGTATCGATGCAG
GAGCACACGTCTGAACTCTTTCCCTACACGACGCTCTTCCGATCTNNNTCCCCTTCTCCAAGAAATAGC
AGCACACGTCTGAACTCTTTCCCTACACGACGCTCTTCCGATCTNNNACACATAGATCACATGGTACTC
GAGCACACGTCTGAACTCTTTCCCTACACGACGCTCTTCCGATCTNNNGAATCCAGGTGGGTAAACTG
GAGCACACGTCTGAACTCTTTCCCTACACGACGCTCTTCCGATCTNNNGTCACTTATTAGCCACAGGGC
GAGCACACGTCTGAACTCTTTCCCTACACGACGCTCTTCCGATCTNNNAGTTCTGTCCCTTTCTCTCTCCT
GAGCACACGTCTGAACTCTTTCCCTACACGACGCTCTTCCGATCTNNNTGCAGATTATGCCACTGCACTC
AGAGCACACGTCTGAACTCTTTCCCTACACGACGCTCTTCCGATCTNNNATTCACGGACCACTTTGATAC
GAGCACACGTCTGAACTCTTTCCCTACACGACGCTCTTCCGATCTNNNGAAACAACACGGCCACTAGG
AGCACACGTCTGAACTCTTTCCCTACACGACGCTCTTCCGATCTNNNTGGCATTGTTTGTTTGCCTAATC
AGCACACGTCTGAACTCTTTCCCTACACGACGCTCTTCCGATCTNNNGGCCAATGAACAAACTCTACAT
GAGCACACGTCTGAACTCTTTCCCTACACGACGCTCTTCCGATCTNNNGCATATGCTTCCTGGTTGAAA
GAGCACACGTCTGAACTCTTTCCCTACACGACGCTCTTCCGATCTNNNTCCTGATTGTTTACACACACAC
AGCACACGTCTGAACTCTTTCCCTACACGACGCTCTTCCGATCTNNNGCGAAGATCACATCACTGCATT
GAGCACACGTCTGAACTCTTTCCCTACACGACGCTCTTCCGATCTNNNGTGTCCCCTGGAGAAGATCA
GAGCACACGTCTGAACTCTTTCCCTACACGACGCTCTTCCGATCTNNNATCTCCCTGGGTGTAGAGAGA
AGAGCACACGTCTGAACTCTTTCCCTACACGACGCTCTTCCGATCTNNNTTGGGCATTTTTCCCCAGTAG
GAGCACACGTCTGAACTCTTTCCCTACACGACGCTCTTCCGATCTNNNGGAGGAGGCTGAGGAAAATT
GAGCACACGTCTGAACTCTTTCCCTACACGACGCTCTTCCGATCTNNNAGTGGAATCTAGGCAGGGA
AGAGCACACGTCTGAACTCTTTCCCTACACGACGCTCTTCCGATCTNNNCTTCCTTCAGGAACTGGTGC
GAGCACACGTCTGAACTCTTTCCCTACACGACGCTCTTCCGATCTNNNAGTGGTAGAGAGGGATCAAA
GAGCACACGTCTGAACTCTTTCCCTACACGACGCTCTTCCGATCTNNNTTCAGTACCCTTCAGTACCCCT
GAGCACACGTCTGAACTCTTTCCCTACACGACGCTCTTCCGATCTNNNGCACCCTTGATCTATGTCTCTC
AGCACACGTCTGAACTCTTTCCCTACACGACGCTCTTCCGATCTNNNTGCATTGTGGATCAATATTCTT
AGAGCACACGTCTGAACTCTTTCCCTACACGACGCTCTTCCGATCTNNNAAGCAAAGTGTTGTGGTAT
AAGAGCACACGTCTGAACTCTTTCCCTACACGACGCTCTTCCGATCTNNNGCCCAGGGGTTCAAGTTAT
AGAGCACACGTCTGAACTCTTTCCCTACACGACGCTCTTCCGATCTNNNTGAATGGCTTCAGAAGGTCT
AGAGCACACGTCTGAACTCTTTCCCTACACGACGCTCTTCCGATCTNNNACCCCAGTTTACATAAGTGTT
GAGCACACGTCTGAACTCTTTCCCTACACGACGCTCTTCCGATCTNNNCATGGCGTATTTTACAGCCAC

AAGAGCACACGTCTGAACTCTTTCCCTACACGACGCTCTTCCGATCTNNNAGGGATAGTGGGAAGTT
AGCACACGTCTGAACTCTTTCCCTACACGACGCTCTTCCGATCTNNNCAAGGAAGGAGGTGATGATG
GAGCACACGTCTGAACTCTTTCCCTACACGACGCTCTTCCGATCTNNNAGGAAGTAAAGGCCACAGT
AGAGCACACGTCTGAACTCTTTCCCTACACGACGCTCTTCCGATCTNNNCCAGGAAACAGGGGTTGA
AAGAGCACACGTCTGAACTCTTTCCCTACACGACGCTCTTCCGATCTNNNAGAGCAAAAGCCAAAATC
GAGCACACGTCTGAACTCTTTCCCTACACGACGCTCTTCCGATCTNNNGCAGCCAGTCTGTGTATTAG
AGCACACGTCTGAACTCTTTCCCTACACGACGCTCTTCCGATCTNNNTCATGTTGAACGATTGAACTG
GAGCACACGTCTGAACTCTTTCCCTACACGACGCTCTTCCGATCTNNNGTTGTGCTGCCAATATCTTCT
GAGCACACGTCTGAACTCTTTCCCTACACGACGCTCTTCCGATCTNNNCTACATCCCAACTGAGGGGT
GCACACGTCTGAACTCTTTCCCTACACGACGCTCTTCCGATCTNNNGGGCGTCTAGGTTGATTCTATC
GAGCACACGTCTGAACTCTTTCCCTACACGACGCTCTTCCGATCTNNNCCACCATCTTTCCCCAGTAAG
AGCACACGTCTGAACTCTTTCCCTACACGACGCTCTTCCGATCTNNNCCTGGCCCACTAATGTACTTTC
GAGCACACGTCTGAACTCTTTCCCTACACGACGCTCTTCCGATCTNNNAGTCCCTTTGGTAAGCAATC
GAGCACACGTCTGAACTCTTTCCCTACACGACGCTCTTCCGATCTNNNTTCCCCCACTCCTAACTTACA
GAGCACACGTCTGAACTCTTTCCCTACACGACGCTCTTCCGATCTNNNAGATCTGTACTCAATGGAAG
AGCACACGTCTGAACTCTTTCCCTACACGACGCTCTTCCGATCTNNNTACTTTCTTGCAGCCTGTGGTT
GAGCACACGTCTGAACTCTTTCCCTACACGACGCTCTTCCGATCTNNNATGGGGTGGGTTGATCTTTA
GAGCACACGTCTGAACTCTTTCCCTACACGACGCTCTTCCGATCTNNNTTGTTCCCAGGCCCTCTTAGA
GAGCACACGTCTGAACTCTTTCCCTACACGACGCTCTTCCGATCTNNNACAAATGGGCATGAGGGAA
AGCACACGTCTGAACTCTTTCCCTACACGACGCTCTTCCGATCTNNNTTTCAGTGATGTATGTGGAGA
GCACACGTCTGAACTCTTTCCCTACACGACGCTCTTCCGATCTNNNAGCCAACTAAGCCTTGTAGGTA
AGCACACGTCTGAACTCTTTCCCTACACGACGCTCTTCCGATCTNNNGCTCCTGTCAATTTCCACAGTC
AGAGCACACGTCTGAACTCTTTCCCTACACGACGCTCTTCCGATCTNNNCAGCAATGAGCTTGCTACCT
GAGCACACGTCTGAACTCTTTCCCTACACGACGCTCTTCCGATCTNNNCTTTCCTTCTTTCCTTCCACCC
AGAGCACACGTCTGAACTCTTTCCCTACACGACGCTCTTCCGATCTNNNTCTTTGCATGTTCACCACTG
AGCACACGTCTGAACTCTTTCCCTACACGACGCTCTTCCGATCTNNNAGAAAACAGTGAAGCCATGGT
AGCACACGTCTGAACTCTTTCCCTACACGACGCTCTTCCGATCTNNNCAGTTCCATGCTTCCATGTGAG
GAGCACACGTCTGAACTCTTTCCCTACACGACGCTCTTCCGATCTNNNTCCAAAGCAGGGCAAAACA
AGCACACGTCTGAACTCTTTCCCTACACGACGCTCTTCCGATCTNNNGAAGGTCAAAGTCCCTTTTCGG
AGAGCACACGTCTGAACTCTTTCCCTACACGACGCTCTTCCGATCTNNNATGAAGCCAGGGTCCTAAC
GAGCACACGTCTGAACTCTTTCCCTACACGACGCTCTTCCGATCTNNNACTTCTAGGAACAAGGAGTA
GAGCACACGTCTGAACTCTTTCCCTACACGACGCTCTTCCGATCTNNNTGTGACGGAGATGGGTATT
AGCACACGTCTGAACTCTTTCCCTACACGACGCTCTTCCGATCTNNNAATGTTTTTGTTTGTGGGCCT
GAGCACACGTCTGAACTCTTTCCCTACACGACGCTCTTCCGATCTNNNCAGTGATGCTTCCTTCAACCA
GAGCACACGTCTGAACTCTTTCCCTACACGACGCTCTTCCGATCTNNNTCCTTCTGCCTTTCAGGTTCAA
AGCACACGTCTGAACTCTTTCCCTACACGACGCTCTTCCGATCTNNNGTACTGCCAGGCAAAGATGAG
AGCACACGTCTGAACTCTTTCCCTACACGACGCTCTTCCGATCTNNNCCTCCAGCTCTATTCAAGTGGT
GCACACGTCTGAACTCTTTCCCTACACGACGCTCTTCCGATCTNNNAGCTGACAACAACTAACCTTTGG
AGCACACGTCTGAACTCTTTCCCTACACGACGCTCTTCCGATCTNNNTCACTGTGTCCTTATCTGGCCTT
GAGCACACGTCTGAACTCTTTCCCTACACGACGCTCTTCCGATCTNNNAATACATGACTTCAGGGTGG
AGCACACGTCTGAACTCTTTCCCTACACGACGCTCTTCCGATCTNNNACGCAATTCCTTTTCTGGGTAT
AGAGCACACGTCTGAACTCTTTCCCTACACGACGCTCTTCCGATCTNNNAACACAGAGAGCTGGTTTC
AGCACACGTCTGAACTCTTTCCCTACACGACGCTCTTCCGATCTNNNCCCTTCTATAGCAGCTACACA

GAGCACACGTCTGAACTCTTTCCCTACACGACGCTCTTCCGATCTNNNTGTTCAGTCTTCCATCTTGAA
GCACACGTCTGAACTCTTTCCCTACACGACGCTCTTCCGATCTNNNACTATCATGGTTCCCAGCAAAA
GAGCACACGTCTGAACTCTTTCCCTACACGACGCTCTTCCGATCTNNNTATCTCGACCCAATTAGCAT
AGCACACGTCTGAACTCTTTCCCTACACGACGCTCTTCCGATCTNNNGCCTGCGATCTTCCAGATTGT
AGCACACGTCTGAACTCTTTCCCTACACGACGCTCTTCCGATCTNNNACTCGAATGAAGCCTGTTTGG
GAGCACACGTCTGAACTCTTTCCCTACACGACGCTCTTCCGATCTNNNCCAGAGCTGGCAGTAGGATA
AGCACACGTCTGAACTCTTTCCCTACACGACGCTCTTCCGATCTNNNTTCTTACGCCCCTCTATAGCCA
GAGCACACGTCTGAACTCTTTCCCTACACGACGCTCTTCCGATCTNNNGGGGCCTTTTACAGGACAC
AGCACACGTCTGAACTCTTTCCCTACACGACGCTCTTCCGATCTNNNAGAGTTCTGTAGGCCAAGGTA
GAGCACACGTCTGAACTCTTTCCCTACACGACGCTCTTCCGATCTNNNTTTTTGTGTGAATACCCTGG
AGCACACGTCTGAACTCTTTCCCTACACGACGCTCTTCCGATCTNNNGCCAGGCCCTATTCTAAGTTCT
GAGCACACGTCTGAACTCTTTCCCTACACGACGCTCTTCCGATCTNNNAGATGTGCTTTGAAGGCTTT
AGCACACGTCTGAACTCTTTCCCTACACGACGCTCTTCCGATCTNNNATGTAGAGGCTGCTTTATGTG
AGAGCACACGTCTGAACTCTTTCCCTACACGACGCTCTTCCGATCTNNNTGCTCTTCACTGTGTAATGG
GCACACGTCTGAACTCTTTCCCTACACGACGCTCTTCCGATCTNNNGAAACAGAATATGGGTGGGG
AGAGCACACGTCTGAACTCTTTCCCTACACGACGCTCTTCCGATCTNNNCTTTACCTGATGCCCACCTA
AGCACACGTCTGAACTCTTTCCCTACACGACGCTCTTCCGATCTNNNTGTACCTGTACCCTCCTGTTTT
GCACACGTCTGAACTCTTTCCCTACACGACGCTCTTCCGATCTNNNAGTGTTCATGTTACTGGGAAG
GAGCACACGTCTGAACTCTTTCCCTACACGACGCTCTTCCGATCTNNNAAGGGGAGAGATTTATTTCC
GAGCACACGTCTGAACTCTTTCCCTACACGACGCTCTTCCGATCTNNNACCAAAATCCAGATGAGAGG
AGCACACGTCTGAACTCTTTCCCTACACGACGCTCTTCCGATCTNNNAACAGAGAGAAGGGAAGAG
AGCACACGTCTGAACTCTTTCCCTACACGACGCTCTTCCGATCTNNNTGCCAATTTGCCAATAACAACA
GCACACGTCTGAACTCTTTCCCTACACGACGCTCTTCCGATCTNNNCCACCCAAACTATCAATTGTTGC
GAGCACACGTCTGAACTCTTTCCCTACACGACGCTCTTCCGATCTNNNAGGGGCCGACTGTGTATGTA
AGCACACGTCTGAACTCTTTCCCTACACGACGCTCTTCCGATCTNNNTGTTTGACTATGTGCACTAGG
AGCACACGTCTGAACTCTTTCCCTACACGACGCTCTTCCGATCTNNNGGCATACTCTGGGATAGTAGG
GAGCACACGTCTGAACTCTTTCCCTACACGACGCTCTTCCGATCTNNNGGCTTTGGGCTAAGAAGGA
GAGCACACGTCTGAACTCTTTCCCTACACGACGCTCTTCCGATCTNNNGCAGGATCTTCTTGAGGTGT
GAGCACACGTCTGAACTCTTTCCCTACACGACGCTCTTCCGATCTNNNCCAAGCATTCTGGCAAGTTT
GAGCACACGTCTGAACTCTTTCCCTACACGACGCTCTTCCGATCTNNNTTCCTTGGAATTAGCCATTCC
GAGCACACGTCTGAACTCTTTCCCTACACGACGCTCTTCCGATCTNNNCTGGGCTAGTGCTGATCATCT
AGCACACGTCTGAACTCTTTCCCTACACGACGCTCTTCCGATCTNNNTGGATTGGAACTGCAATTTTG
GCACACGTCTGAACTCTTTCCCTACACGACGCTCTTCCGATCTNNNAGTAGTGCTGATTTCCAAATGCC
AGCACACGTCTGAACTCTTTCCCTACACGACGCTCTTCCGATCTNNNCTGCCTAACCATCAGACCATCA
GAGCACACGTCTGAACTCTTTCCCTACACGACGCTCTTCCGATCTNNNCCGTGAATACATTCGCAAAGT
GCACACGTCTGAACTCTTTCCCTACACGACGCTCTTCCGATCTNNNTGTTCTTGTCACCTAAGTTTCTG
GAGCACACGTCTGAACTCTTTCCCTACACGACGCTCTTCCGATCTNNNCGCCTGGTCAGAATCGTCATT
GAGCACACGTCTGAACTCTTTCCCTACACGACGCTCTTCCGATCTNNNCATGGATGTCTGCTTTAGG
AGAGCACACGTCTGAACTCTTTCCCTACACGACGCTCTTCCGATCTNNNCCAGTACACAGCCAACATA
GCACACGTCTGAACTCTTTCCCTACACGACGCTCTTCCGATCTNNNCAATCCTCATATGTGCATGAGTG
GAGCACACGTCTGAACTCTTTCCCTACACGACGCTCTTCCGATCTNNNTAACTTCTGGCCTCCCAGGTT
GCACACGTCTGAACTCTTTCCCTACACGACGCTCTTCCGATCTNNNGCTTTAGAGTTCACAGTGAGAC
GAGCACACGTCTGAACTCTTTCCCTACACGACGCTCTTCCGATCTNNNCCACCCCATTCAAAACAAATA

GAGCACACGTCTGAACTCTTTCCCTACACGACGCTCTTCCGATCTNNNTTGCAATTCCCTGTCCTTTGTAA
GAGCACACGTCTGAACTCTTTCCCTACACGACGCTCTTCCGATCTNNNGCGATTTTCAGAACTTACTGCG
AGAGCACACGTCTGAACTCTTTCCCTACACGACGCTCTTCCGATCTNNNCCTTTGATTGTCCCCACACAG
GAGCACACGTCTGAACTCTTTCCCTACACGACGCTCTTCCGATCTNNNAAAGTGCTGGGACTACACAAG
GAGCACACGTCTGAACTCTTTCCCTACACGACGCTCTTCCGATCTNNNTCCATGTTCCTGTGAATGACGA
AAGAGCACACGTCTGAACTCTTTCCCTACACGACGCTCTTCCGATCTNNNTAGGTGTTTGGTGGGAAAA
AGAGCACACGTCTGAACTCTTTCCCTACACGACGCTCTTCCGATCTNNNGAGGATCCCTTAGGCTGAG
AGCACACGTCTGAACTCTTTCCCTACACGACGCTCTTCCGATCTNNNGGAGTTTCCTTTCTTCTTCCACCT
AGCACACGTCTGAACTCTTTCCCTACACGACGCTCTTCCGATCTNNNCCTGTCACTGTGTGTGTCTCTGT
AGCACACGTCTGAACTCTTTCCCTACACGACGCTCTTCCGATCTNNNCCTCTAAATCTGAATGTAAGCC
AAGAGCACACGTCTGAACTCTTTCCCTACACGACGCTCTTCCGATCTNNNATGAAAGGCCCTGGAAATG
AGCACACGTCTGAACTCTTTCCCTACACGACGCTCTTCCGATCTNNNTCCTTTCTTGCTGGTAGACACA
GAGCACACGTCTGAACTCTTTCCCTACACGACGCTCTTCCGATCTNNNGGGTGAGTAGCATTCCATAGC
AGAGCACACGTCTGAACTCTTTCCCTACACGACGCTCTTCCGATCTNNNCCTACCACATGACGTTCCTTT
GAGCACACGTCTGAACTCTTTCCCTACACGACGCTCTTCCGATCTNNNATGAGAGGATCGCTTGTGTC
GAGCACACGTCTGAACTCTTTCCCTACACGACGCTCTTCCGATCTNNNCACAACACATTTCTTCTTGCCCC
GAGCACACGTCTGAACTCTTTCCCTACACGACGCTCTTCCGATCTNNNTGGTTTCCTGCCTTCAGAACAT
AGAGCACACGTCTGAACTCTTTCCCTACACGACGCTCTTCCGATCTNNNCTGTCACAGCAACTGTCCCAA
AGCACACGTCTGAACTCTTTCCCTACACGACGCTCTTCCGATCTNNNCTTTCTCTCTCTCTGTCTGTGGG
GAGCACACGTCTGAACTCTTTCCCTACACGACGCTCTTCCGATCTNNNTAGAGGAACTGTCAGGCTTCC
AGAGCACACGTCTGAACTCTTTCCCTACACGACGCTCTTCCGATCTNNNCTGCTCTCTTTTGGGACCATC
AGAGCACACGTCTGAACTCTTTCCCTACACGACGCTCTTCCGATCTNNNAGACCCTGGTACTTCCAGAA
AGAGCACACGTCTGAACTCTTTCCCTACACGACGCTCTTCCGATCTNNNCAGGAAACTGGCATTACGTC
AGCACACGTCTGAACTCTTTCCCTACACGACGCTCTTCCGATCTNNNTGACACTCCCTTTTGAACTTGTC
AGCACACGTCTGAACTCTTTCCCTACACGACGCTCTTCCGATCTNNNTAACAGATTTCACCTCACTGGG
GAGCACACGTCTGAACTCTTTCCCTACACGACGCTCTTCCGATCTNNNGTAAGAGGCAGGAAAGGTG
AGAGCACACGTCTGAACTCTTTCCCTACACGACGCTCTTCCGATCTNNNTCATCCCACAGATTCAAGAG
AGCACACGTCTGAACTCTTTCCCTACACGACGCTCTTCCGATCTNNNACAAAGGCATAAGAAATTGTG
GAGCACACGTCTGAACTCTTTCCCTACACGACGCTCTTCCGATCTNNNAGCAATATTTCTCCACGTGTGT
AGAGCACACGTCTGAACTCTTTCCCTACACGACGCTCTTCCGATCTNNNGGTTGATTTTGGTGTTACCA
AAGAGCACACGTCTGAACTCTTTCCCTACACGACGCTCTTCCGATCTNNNAATTCACTCTCCTGCCATTGT
AGCACACGTCTGAACTCTTTCCCTACACGACGCTCTTCCGATCTNNNTCTAAGAGGATGGAAGCAGTA
AGAGCACACGTCTGAACTCTTTCCCTACACGACGCTCTTCCGATCTNNNGAGTGAGGTTTGCTGCTTTC
AAGAGCACACGTCTGAACTCTTTCCCTACACGACGCTCTTCCGATCTNNNAGAGAAGAGGTGGTTGGT
AAGAGCACACGTCTGAACTCTTTCCCTACACGACGCTCTTCCGATCTNNNTCCAGTCTGGGAAAGACTT
AGCACACGTCTGAACTCTTTCCCTACACGACGCTCTTCCGATCTNNNTGGTGATTTTAGGGTTGTTGG
GAGCACACGTCTGAACTCTTTCCCTACACGACGCTCTTCCGATCTNNNTCACTCAACATTGTAGTCAGT
GAGCACACGTCTGAACTCTTTCCCTACACGACGCTCTTCCGATCTNNNACACAGAACATATCCACACACA
GAGCACACGTCTGAACTCTTTCCCTACACGACGCTCTTCCGATCTNNNCATCCATCTCCCCATCCCTCTAC
AGAGCACACGTCTGAACTCTTTCCCTACACGACGCTCTTCCGATCTNNNAGACACCTGCCTATCTCACCA
GAGCACACGTCTGAACTCTTTCCCTACACGACGCTCTTCCGATCTNNNGGGGGAGAACCTGATATAGT
AGAGCACACGTCTGAACTCTTTCCCTACACGACGCTCTTCCGATCTNNNGGCCTGTTTCCTCTAGCTTTT
AGAGCACACGTCTGAACTCTTTCCCTACACGACGCTCTTCCGATCTNNNGTGTCCTCCAGTTTCATCCACC

GAGCACACGTCTGAACTCTTTCCCTACACGACGCTCTTCCGATCTNNNTCTCACACGTCTCCTTCTCTCT
AGAGCACACGTCTGAACTCTTTCCCTACACGACGCTCTTCCGATCTNNNGAGAACTGCTCAGACATGG
GAGCACACGTCTGAACTCTTTCCCTACACGACGCTCTTCCGATCTNNNTCAAACACCCAGTCATTGCA
AGCACACGTCTGAACTCTTTCCCTACACGACGCTCTTCCGATCTNNNTGGAATGCCGCATATGTAAA
AGCACACGTCTGAACTCTTTCCCTACACGACGCTCTTCCGATCTNNNGAAGAGCAACAAGAGCAACA
AGAGCACACGTCTGAACTCTTTCCCTACACGACGCTCTTCCGATCTNNNAACCTAAATGCCGGTCAACA
AGAGCACACGTCTGAACTCTTTCCCTACACGACGCTCTTCCGATCTNNNGAGAACAACTGCTCTGGGG
AGCACACGTCTGAACTCTTTCCCTACACGACGCTCTTCCGATCTNNNGCTCACTACATATAGACACCCTG
GCACACGTCTGAACTCTTTCCCTACACGACGCTCTTCCGATCTNNNCACATTAGAATCCAAGGAGGG
AGCACACGTCTGAACTCTTTCCCTACACGACGCTCTTCCGATCTNNNTGAAGCATGCCTACAAGATTT
GAGCACACGTCTGAACTCTTTCCCTACACGACGCTCTTCCGATCTNNNGAGTTTGGAGTACAGCTGTG
AGCACACGTCTGAACTCTTTCCCTACACGACGCTCTTCCGATCTNNNTTGCTTGTTCTTTTTCTGTGCC
AGAGCACACGTCTGAACTCTTTCCCTACACGACGCTCTTCCGATCTNNNGTTTGTTCTAAGGTTGCGT
AGCACACGTCTGAACTCTTTCCCTACACGACGCTCTTCCGATCTNNNCCTATGGACTTGGGAAAGCAC
AGAGCACACGTCTGAACTCTTTCCCTACACGACGCTCTTCCGATCTNNNGCCACAAGTAGCTAGCAAAG
AGCACACGTCTGAACTCTTTCCCTACACGACGCTCTTCCGATCTNNNCGGATTCAGAATAGAGGGAG
GAGCACACGTCTGAACTCTTTCCCTACACGACGCTCTTCCGATCTNNNACGTGGATTTTATGCATTTGT
GAGCACACGTCTGAACTCTTTCCCTACACGACGCTCTTCCGATCTNNNTTTTCTCACACACCCATGTCA
GAGCACACGTCTGAACTCTTTCCCTACACGACGCTCTTCCGATCTNNNTACTTCAGCTTATCTTTGGCG
AGAGCACACGTCTGAACTCTTTCCCTACACGACGCTCTTCCGATCTNNNTACCCTGTGGAACTGTCCTT
AGAGCACACGTCTGAACTCTTTCCCTACACGACGCTCTTCCGATCTNNNTCTGTAATTCTTGCCCAGAC
AGCACACGTCTGAACTCTTTCCCTACACGACGCTCTTCCGATCTNNNACCTCAGAAAATTTGGCTTACA
AGCACACGTCTGAACTCTTTCCCTACACGACGCTCTTCCGATCTNNNACACTTTCAGTTTAGTGCACAC
GAGCACACGTCTGAACTCTTTCCCTACACGACGCTCTTCCGATCTNNNGAGGTGCTTCACCCTAACATC
GCACACGTCTGAACTCTTTCCCTACACGACGCTCTTCCGATCTNNNTGTGAGGAAGACATTACACATT
GAGCACACGTCTGAACTCTTTCCCTACACGACGCTCTTCCGATCTNNNTGTCTGTGCAAAACACAAAAC
AGCACACGTCTGAACTCTTTCCCTACACGACGCTCTTCCGATCTNNNACCTTGTGAAAGATCTGTGGC
GAGCACACGTCTGAACTCTTTCCCTACACGACGCTCTTCCGATCTNNNGAATTCCCCAAAGCAGCCTTA
GAGCACACGTCTGAACTCTTTCCCTACACGACGCTCTTCCGATCTNNNAGGCTGAGTGGCTTACTTAC
AGCACACGTCTGAACTCTTTCCCTACACGACGCTCTTCCGATCTNNNTGGGTCCCATAAAGGGGAAT
GCACACGTCTGAACTCTTTCCCTACACGACGCTCTTCCGATCTNNNAACCCAGGAATGCTTTTCCCAAG
GAGCACACGTCTGAACTCTTTCCCTACACGACGCTCTTCCGATCTNNNGTTGGCTGTGCTGCTTATTTT
GAGCACACGTCTGAACTCTTTCCCTACACGACGCTCTTCCGATCTNNNAACATGCTATTTGGGGGTTT
GCACACGTCTGAACTCTTTCCCTACACGACGCTCTTCCGATCTNNNAGCCAGGTTTCACCAGTTTTATG
GAGCACACGTCTGAACTCTTTCCCTACACGACGCTCTTCCGATCTNNNCTGCACCTGGAAAGTTCTAG
AGCACACGTCTGAACTCTTTCCCTACACGACGCTCTTCCGATCTNNNACAAAGTCTCTACCCTACTGGA
GAGCACACGTCTGAACTCTTTCCCTACACGACGCTCTTCCGATCTNNNTGGTCTCCTTGTGTATTGTCA
AGCACACGTCTGAACTCTTTCCCTACACGACGCTCTTCCGATCTNNNGCGTGGTGCCTTATTTGTTGA
GAGCACACGTCTGAACTCTTTCCCTACACGACGCTCTTCCGATCTNNNTGGAGTGAAATCCTGGTACT
GAGCACACGTCTGAACTCTTTCCCTACACGACGCTCTTCCGATCTNNNTCAGGACATTTCCTTCCTGTC
AGAGCACACGTCTGAACTCTTTCCCTACACGACGCTCTTCCGATCTNNNGACATTCTCTCTGAGGCCAG
AGAGCACACGTCTGAACTCTTTCCCTACACGACGCTCTTCCGATCTNNNGCCCTCAGGTCATTTTAGGA
GCACACGTCTGAACTCTTTCCCTACACGACGCTCTTCCGATCTNNNTGGTCTGAGATCTGTGTTCAACT

GCACACGTCTGAACTCTTTCCCTACACGACGCTCTTCCGATCTNNNACTGTGTATTTATGTGTTTGTG
AGCACACGTCTGAACTCTTTCCCTACACGACGCTCTTCCGATCTNNNCGTTTCAATGCCTCTCCTGAAT
AGCACACGTCTGAACTCTTTCCCTACACGACGCTCTTCCGATCTNNNAGCAGGATGCAGAAGCAAAT
GAGCACACGTCTGAACTCTTTCCCTACACGACGCTCTTCCGATCTNNNAAACTCAGAGGAAGGATGT
GAGCACACGTCTGAACTCTTTCCCTACACGACGCTCTTCCGATCTNNNTGCAGCCATGCCTAAAGATA
AGAGCACACGTCTGAACTCTTTCCCTACACGACGCTCTTCCGATCTNNNTTGAGTTCCCCTGCCAATCA
GAGCACACGTCTGAACTCTTTCCCTACACGACGCTCTTCCGATCTNNNTCCCTTTTCCTCCTGTTCCTCA
AGCACACGTCTGAACTCTTTCCCTACACGACGCTCTTCCGATCTNNNCCCTCTCTTCATCTTCACATGG
GAGCACACGTCTGAACTCTTTCCCTACACGACGCTCTTCCGATCTNNNTGAATGAAAAGGCAGGAAG
AGAGCACACGTCTGAACTCTTTCCCTACACGACGCTCTTCCGATCTNNNAGGTCACATTTGTGAGAAC
GAGCACACGTCTGAACTCTTTCCCTACACGACGCTCTTCCGATCTNNNTATCCAGCTAGGGACTTGGA
AGCACACGTCTGAACTCTTTCCCTACACGACGCTCTTCCGATCTNNNAAGGGCCTAACACTTCAGATT
AAGAGCACACGTCTGAACTCTTTCCCTACACGACGCTCTTCCGATCTNNNTTGACCTGGCTTCATAGG
AGCACACGTCTGAACTCTTTCCCTACACGACGCTCTTCCGATCTNNNGGTATCCAATTACATCCACGT
AGCACACGTCTGAACTCTTTCCCTACACGACGCTCTTCCGATCTNNNATGTGATTTTGTTGGGCACTG
AGCACACGTCTGAACTCTTTCCCTACACGACGCTCTTCCGATCTNNNTTGCATTCTTGTCTTCACAGGC
GAGCACACGTCTGAACTCTTTCCCTACACGACGCTCTTCCGATCTNNNCTACTTCCACGTCCAAACTTC
GCACACGTCTGAACTCTTTCCCTACACGACGCTCTTCCGATCTNNNTGTATCCTGAGACTGGGTTGAA
GCACACGTCTGAACTCTTTCCCTACACGACGCTCTTCCGATCTNNNCCTCCAACCCAACCAATATTGTC
AGCACACGTCTGAACTCTTTCCCTACACGACGCTCTTCCGATCTNNNCCCAACTAAGGGTACAAATG
AGCACACGTCTGAACTCTTTCCCTACACGACGCTCTTCCGATCTNNNACAGCAGATAATGTTCAAATG
AGCACACGTCTGAACTCTTTCCCTACACGACGCTCTTCCGATCTNNNGAAACGAGAGGATCGCTTGAC
GAGCACACGTCTGAACTCTTTCCCTACACGACGCTCTTCCGATCTNNNATGGATAAGTCTGCAAGTA
GAGCACACGTCTGAACTCTTTCCCTACACGACGCTCTTCCGATCTNNNGCAATTTGGTTCCTTCAAGC
GAGCACACGTCTGAACTCTTTCCCTACACGACGCTCTTCCGATCTNNNGTATACAGAGTTTGGGGCA
AGCACACGTCTGAACTCTTTCCCTACACGACGCTCTTCCGATCTNNNTTATTTCAGGTTTGGCCACTG
AGAGCACACGTCTGAACTCTTTCCCTACACGACGCTCTTCCGATCTNNNTTTTCTAACTTCAGCTGCCC
GCACACGTCTGAACTCTTTCCCTACACGACGCTCTTCCGATCTNNNTGTGAAGTTCCACAGTATCTGA
GCACACGTCTGAACTCTTTCCCTACACGACGCTCTTCCGATCTNNNGCTACAAAGTGCTTTACTGACA
AGAGCACACGTCTGAACTCTTTCCCTACACGACGCTCTTCCGATCTNNNAGCTGACCCATACTGCAGA
GAGCACACGTCTGAACTCTTTCCCTACACGACGCTCTTCCGATCTNNNACATACACAGAGTGAGCGA
AGCACACGTCTGAACTCTTTCCCTACACGACGCTCTTCCGATCTNNNTCAAAGACATGAAGATAGTC
AGCACACGTCTGAACTCTTTCCCTACACGACGCTCTTCCGATCTNNNACTGTCACTGTGGGGTTTTCT
AGCACACGTCTGAACTCTTTCCCTACACGACGCTCTTCCGATCTNNNGCCAAAGATCAGGGAACTCAT
AGCACACGTCTGAACTCTTTCCCTACACGACGCTCTTCCGATCTNNNTCCTGTGACATAAGAAATGCGT
AGCACACGTCTGAACTCTTTCCCTACACGACGCTCTTCCGATCTNNNAGGCCCCACTATCTAACCTTG
GAGCACACGTCTGAACTCTTTCCCTACACGACGCTCTTCCGATCTNNNGGGGCTCTAACATATAACAT
GAGCACACGTCTGAACTCTTTCCCTACACGACGCTCTTCCGATCTNNNACTAGAATCAGCCAATCTGG
AGAGCACACGTCTGAACTCTTTCCCTACACGACGCTCTTCCGATCTNNNTTCTCAGGAGATCTGGGGT
AGAGCACACGTCTGAACTCTTTCCCTACACGACGCTCTTCCGATCTNNNTACGTGAACTGATACACGTT
AGCACACGTCTGAACTCTTTCCCTACACGACGCTCTTCCGATCTNNNAGCCATGATTGCTTCCTCATTG
AGAGCACACGTCTGAACTCTTTCCCTACACGACGCTCTTCCGATCTNNNGGTCCTCATAAGCTTTCAC
GAGCACACGTCTGAACTCTTTCCCTACACGACGCTCTTCCGATCTNNNGTGTCCCTTCCTACTCACTTC

AGCACACGTCTGAACTCTTTCCCTACACGACGCTCTTCCGATCTNNNAGGCTAAACTGTGTAGTGTTTG
AGAGCACACGTCTGAACTCTTTCCCTACACGACGCTCTTCCGATCTNNNTCCACCACTATTTGGGGCTT
AGCACACGTCTGAACTCTTTCCCTACACGACGCTCTTCCGATCTNNNGCAACTCCAGGAAATTGATGTT
GAGCACACGTCTGAACTCTTTCCCTACACGACGCTCTTCCGATCTNNNAGTTCTCTAGGATGGAGCTG
AGCACACGTCTGAACTCTTTCCCTACACGACGCTCTTCCGATCTNNNAGGAGGAGATGTTAGTGACTC
GAGCACACGTCTGAACTCTTTCCCTACACGACGCTCTTCCGATCTNNNGATGGCAGGACAGATTTGG
AGCACACGTCTGAACTCTTTCCCTACACGACGCTCTTCCGATCTNNNGCTGTAACAGGAATCCCTACG
GAGCACACGTCTGAACTCTTTCCCTACACGACGCTCTTCCGATCTNNNGCAAGGTTCAAAAACTATGTG
GAGCACACGTCTGAACTCTTTCCCTACACGACGCTCTTCCGATCTNNNTTCTTCTCCCATAGGGCTCTGT
AGCACACGTCTGAACTCTTTCCCTACACGACGCTCTTCCGATCTNNNACCTACAGAGGAATCATACTGG
AGCACACGTCTGAACTCTTTCCCTACACGACGCTCTTCCGATCTNNNTTTTACATGTTTGGGTGGGTG
AGAGCACACGTCTGAACTCTTTCCCTACACGACGCTCTTCCGATCTNNNAGAGCTGGAAGAAGTGGA
AGAGCACACGTCTGAACTCTTTCCCTACACGACGCTCTTCCGATCTNNNATTGAAACGAGACGGCACA
GAGCACACGTCTGAACTCTTTCCCTACACGACGCTCTTCCGATCTNNNGGAAATGCTTACTGTGCCAAC
GAGCACACGTCTGAACTCTTTCCCTACACGACGCTCTTCCGATCTNNNGCAGTTCCAAGACCAATCAAT
AGAGCACACGTCTGAACTCTTTCCCTACACGACGCTCTTCCGATCTNNNTGTGTCCACTGATCACATTT
AGCACACGTCTGAACTCTTTCCCTACACGACGCTCTTCCGATCTNNNTCCTTTAGCCAAATCCTCAAAG
AGCACACGTCTGAACTCTTTCCCTACACGACGCTCTTCCGATCTNNNACCAATGTCAGCTCAATAGCAC
AAGAGCACACGTCTGAACTCTTTCCCTACACGACGCTCTTCCGATCTNNNTGTAAAGTTACGTGCGTGA
AGAGCACACGTCTGAACTCTTTCCCTACACGACGCTCTTCCGATCTNNNTAGAATCTGGCAGGGGAGG
AGAGCACACGTCTGAACTCTTTCCCTACACGACGCTCTTCCGATCTNNNTGTGAGATTGTGTTGGTGT
GAGCACACGTCTGAACTCTTTCCCTACACGACGCTCTTCCGATCTNNNACGATGGAGTGCCATGATCAA
AGAGCACACGTCTGAACTCTTTCCCTACACGACGCTCTTCCGATCTNNNTCATATTGTCACATTGGAAG
AGAGCACACGTCTGAACTCTTTCCCTACACGACGCTCTTCCGATCTNNNCCATCTCACAGCAGTTACTC
AGCACACGTCTGAACTCTTTCCCTACACGACGCTCTTCCGATCTNNNATCGGTGTTGATGGGGAAAGA
AGCACACGTCTGAACTCTTTCCCTACACGACGCTCTTCCGATCTNNNTCTTGTTTTGGTAGGGGTGTG
AGCACACGTCTGAACTCTTTCCCTACACGACGCTCTTCCGATCTNNNTGTAGTGGTCTCATCCCAGACC
AAGAGCACACGTCTGAACTCTTTCCCTACACGACGCTCTTCCGATCTNNNACATTTAGTCAGTGAGGGT
AGCACACGTCTGAACTCTTTCCCTACACGACGCTCTTCCGATCTNNNTCACTCTGCTTGAAAGATCAGC
GAGCACACGTCTGAACTCTTTCCCTACACGACGCTCTTCCGATCTNNNAGTAGAAAGGCAGGAGAGCA
AGAGCACACGTCTGAACTCTTTCCCTACACGACGCTCTTCCGATCTNNNGTGAGTGGAGATTGCATTC
AGAGCACACGTCTGAACTCTTTCCCTACACGACGCTCTTCCGATCTNNNAACTAACTTAGCTGCTCACCG
AAGAGCACACGTCTGAACTCTTTCCCTACACGACGCTCTTCCGATCTNNNAGTCCAGTGAGGCTGTGAT
GAGCACACGTCTGAACTCTTTCCCTACACGACGCTCTTCCGATCTNNNTGCTTGTTGTGTTATGACTTG
GAGCACACGTCTGAACTCTTTCCCTACACGACGCTCTTCCGATCTNNNAGTCTTGGCAGGTATGGACA
GAGCACACGTCTGAACTCTTTCCCTACACGACGCTCTTCCGATCTNNNTTCAATCTGAAGAGAAACCG
GAGCACACGTCTGAACTCTTTCCCTACACGACGCTCTTCCGATCTNNNACATAGAACAGGGAGGAGCT
AGCACACGTCTGAACTCTTTCCCTACACGACGCTCTTCCGATCTNNNTGATTTGGCCCTTGTTTTTCTTCA
AGCACACGTCTGAACTCTTTCCCTACACGACGCTCTTCCGATCTNNNGGGAGGGGGAGAGTTATCAT
AGAGCACACGTCTGAACTCTTTCCCTACACGACGCTCTTCCGATCTNNNATCCTATCATTTTCTTCTGGG
GAGCACACGTCTGAACTCTTTCCCTACACGACGCTCTTCCGATCTNNNTCTATACATAGACACACACACC
GAGCACACGTCTGAACTCTTTCCCTACACGACGCTCTTCCGATCTNNNTACAAACACAGAATGTCACCCC
AGCACACGTCTGAACTCTTTCCCTACACGACGCTCTTCCGATCTNNNAGAGTTTATCTGGAGGAACAA

GAGCACACGTCTGAACTCTTTCCCTACACGACGCTCTTCCGATCTNNNGCCCACATAAACCACTTACAC
GAGCACACGTCTGAACTCTTTCCCTACACGACGCTCTTCCGATCTNNNCACCTGTCTTTCTTGCTGAAG
GAGCACACGTCTGAACTCTTTCCCTACACGACGCTCTTCCGATCTNNNCTGACTACAGGTCAGGCAGA
AGCACACGTCTGAACTCTTTCCCTACACGACGCTCTTCCGATCTNNNACCTGATAGAACTAGTATGGCT
GAGCACACGTCTGAACTCTTTCCCTACACGACGCTCTTCCGATCTNNNAACGTTTCCGCCACTTTATCT
AGCACACGTCTGAACTCTTTCCCTACACGACGCTCTTCCGATCTNNNTGGATTTGATTGTGTTGGATC
AGCACACGTCTGAACTCTTTCCCTACACGACGCTCTTCCGATCTNNNAATGCACATAAACTGCATGGC
GAGCACACGTCTGAACTCTTTCCCTACACGACGCTCTTCCGATCTNNNTTTGTATCCTGCCCTTTACTG
GAGCACACGTCTGAACTCTTTCCCTACACGACGCTCTTCCGATCTNNNGTCAAAACAATGAGCACTCC
AGCACACGTCTGAACTCTTTCCCTACACGACGCTCTTCCGATCTNNNATCAGGGCAAGTAGAAAGCA
AGCACACGTCTGAACTCTTTCCCTACACGACGCTCTTCCGATCTNNNCACACGTTACCATGTCTGCCTA
AAGAGCACACGTCTGAACTCTTTCCCTACACGACGCTCTTCCGATCTNNNCACTCCCTTCACCCATCACA
AGAGCACACGTCTGAACTCTTTCCCTACACGACGCTCTTCCGATCTNNNGTTGGTAAAACAGTACACG
GAGCACACGTCTGAACTCTTTCCCTACACGACGCTCTTCCGATCTNNNAGGGACACTCTGTGTCTTTT
AAGAGCACACGTCTGAACTCTTTCCCTACACGACGCTCTTCCGATCTNNNCATTAGGGAGGAAGTCAC
GAGCACACGTCTGAACTCTTTCCCTACACGACGCTCTTCCGATCTNNNTCATCCAGGTTGTTGTGAAG
GAGCACACGTCTGAACTCTTTCCCTACACGACGCTCTTCCGATCTNNNGCCAGTGAGGATAGAGTGA
AGAGCACACGTCTGAACTCTTTCCCTACACGACGCTCTTCCGATCTNNNACCAGCTGAGATCCGTCTAT
AGCACACGTCTGAACTCTTTCCCTACACGACGCTCTTCCGATCTNNNATTGGCTGAAAAGACTACCCC
AGAGCACACGTCTGAACTCTTTCCCTACACGACGCTCTTCCGATCTNNNGCTCGGTTCCTTCAACCATT
AGCACACGTCTGAACTCTTTCCCTACACGACGCTCTTCCGATCTNNNTTTGACATATGAAGGGGTGT
AGCACACGTCTGAACTCTTTCCCTACACGACGCTCTTCCGATCTNNNTGTGTTTGGGAATAACCCAGT
GAGCACACGTCTGAACTCTTTCCCTACACGACGCTCTTCCGATCTNNNCCTGGAACCAATCCTCCTACA
AGAGCACACGTCTGAACTCTTTCCCTACACGACGCTCTTCCGATCTNNNGACCTAAGGTATGGCATGG
AGCACACGTCTGAACTCTTTCCCTACACGACGCTCTTCCGATCTNNNAGGGAGTAGAATGAGTTCTT
AGAGCACACGTCTGAACTCTTTCCCTACACGACGCTCTTCCGATCTNNNAGAATTATGGGTGTGTGTG
AGCACACGTCTGAACTCTTTCCCTACACGACGCTCTTCCGATCTNNNTGCAAAATTGGTATCCTGCTCT
AGCACACGTCTGAACTCTTTCCCTACACGACGCTCTTCCGATCTNNNAACAGGAAATTTCTCCCAAGG
AGCACACGTCTGAACTCTTTCCCTACACGACGCTCTTCCGATCTNNNTGAGGAGTTTTGCACTGACG
GAGCACACGTCTGAACTCTTTCCCTACACGACGCTCTTCCGATCTNNNCATTGCTGCATTGCTTTCCAA
AAGAGCACACGTCTGAACTCTTTCCCTACACGACGCTCTTCCGATCTNNNTCTGGTAAGGCCATGGG
GAGCACACGTCTGAACTCTTTCCCTACACGACGCTCTTCCGATCTNNNTGAAAGGTAATGGCTCTCAG
AGCACACGTCTGAACTCTTTCCCTACACGACGCTCTTCCGATCTNNNTGATTTTATCTGGGTCTCCTGG
AGCACACGTCTGAACTCTTTCCCTACACGACGCTCTTCCGATCTNNNGATCGTGAAAGCCCCAAAAAT
AGAGCACACGTCTGAACTCTTTCCCTACACGACGCTCTTCCGATCTNNNGGAGACGGGGTTTTGTGG
AGCACACGTCTGAACTCTTTCCCTACACGACGCTCTTCCGATCTNNNAAAGAAGAGAGGGGGAAGG
GCACACGTCTGAACTCTTTCCCTACACGACGCTCTTCCGATCTNNNACCTCCTTGGTAGGATGTATTG
AGCACACGTCTGAACTCTTTCCCTACACGACGCTCTTCCGATCTNNNAAAGTCGGCTCCACAAGAACT
AGAGCACACGTCTGAACTCTTTCCCTACACGACGCTCTTCCGATCTNNNTCAGGGAAAGAGCACTGG
GAGCACACGTCTGAACTCTTTCCCTACACGACGCTCTTCCGATCTNNNGCTTTGGGGAAGTTTGTTG
AGAGCACACGTCTGAACTCTTTCCCTACACGACGCTCTTCCGATCTNNNGCCATATTAGTCGGTGTTC
AAGAGCACACGTCTGAACTCTTTCCCTACACGACGCTCTTCCGATCTNNNGGCAGAGGCTAAAGAGG
GAGCACACGTCTGAACTCTTTCCCTACACGACGCTCTTCCGATCTNNNTTTCCCTGTTGTCTTCGAGCA

AGAGCACACGTCTGAACTCTTTCCCTACACGACGCTCTTCCGATCTNNNACAAGGTTGCAATTAGAGG
AGAGCACACGTCTGAACTCTTTCCCTACACGACGCTCTTCCGATCTNNNACAGAACAGGCCTTCCTTCT
GAGCACACGTCTGAACTCTTTCCCTACACGACGCTCTTCCGATCTNNNACTGTTCATGTCTTCACCCACTT
AAGAGCACACGTCTGAACTCTTTCCCTACACGACGCTCTTCCGATCTNNNTGGCACTTCCATTTTCATTG
AGAGCACACGTCTGAACTCTTTCCCTACACGACGCTCTTCCGATCTNNNCAAATCTGCCTTCAGACCTCA
AGAGCACACGTCTGAACTCTTTCCCTACACGACGCTCTTCCGATCTNNNTGTGGTGTAACACTCAAGAA
GAGCACACGTCTGAACTCTTTCCCTACACGACGCTCTTCCGATCTNNNGTCACTAAGCTGCCAAGTTCTC
AAGAGCACACGTCTGAACTCTTTCCCTACACGACGCTCTTCCGATCTNNNGCTAGTTTGAACCAGGCAT
AGAGCACACGTCTGAACTCTTTCCCTACACGACGCTCTTCCGATCTNNNTAGCAGGGTGAGACCTTGTC
AGAGCACACGTCTGAACTCTTTCCCTACACGACGCTCTTCCGATCTNNNGGTGTTGTCAACCTTTACTCC
GAGCACACGTCTGAACTCTTTCCCTACACGACGCTCTTCCGATCTNNNATGTGCAGTGTGGGATGAGA
GAGCACACGTCTGAACTCTTTCCCTACACGACGCTCTTCCGATCTNNNTTTGTCTATCAGTTCCAAGAGC
AAGAGCACACGTCTGAACTCTTTCCCTACACGACGCTCTTCCGATCTNNNTCTTTGTGGTAGTTGTTGG
GAGCACACGTCTGAACTCTTTCCCTACACGACGCTCTTCCGATCTNNNGCATCTCAGAACAAAAATCCCT
GAGCACACGTCTGAACTCTTTCCCTACACGACGCTCTTCCGATCTNNNGGAATTGTTCTTCTCTCAGAG
AGAGCACACGTCTGAACTCTTTCCCTACACGACGCTCTTCCGATCTNNNTCTTCGTCTTATGCTTTCACCT
AGCACACGTCTGAACTCTTTCCCTACACGACGCTCTTCCGATCTNNNAACGCTGCTAATTCATTCCTTTC
GAGCACACGTCTGAACTCTTTCCCTACACGACGCTCTTCCGATCTNNNTGCATTGCATGCTGATACCAA
AGAGCACACGTCTGAACTCTTTCCCTACACGACGCTCTTCCGATCTNNNTTGTTTGCTTCAGGTAATGG
AGAGCACACGTCTGAACTCTTTCCCTACACGACGCTCTTCCGATCTNNNACTCTGCCCATTGACTGATCT
AGAGCACACGTCTGAACTCTTTCCCTACACGACGCTCTTCCGATCTNNNCATGGAGACAGAGGCCAAG
AAGAGCACACGTCTGAACTCTTTCCCTACACGACGCTCTTCCGATCTNNNTGAAAGTAGTGGTTTCGGC
GAGCACACGTCTGAACTCTTTCCCTACACGACGCTCTTCCGATCTNNNAGAAGGGCTCACCATTTACTC
AGCACACGTCTGAACTCTTTCCCTACACGACGCTCTTCCGATCTNNNCAAATGGAAACCGTCAGAGTTC
GAGCACACGTCTGAACTCTTTCCCTACACGACGCTCTTCCGATCTNNNTCCGTGTCATGTCATTCCTGAA
AAGAGCACACGTCTGAACTCTTTCCCTACACGACGCTCTTCCGATCTNNNTGCAAGCATCTGTTTATGT
AGCACACGTCTGAACTCTTTCCCTACACGACGCTCTTCCGATCTNNNAACTTGTGGAGAAAAGCCAGA
AGAGCACACGTCTGAACTCTTTCCCTACACGACGCTCTTCCGATCTNNNTTTTGGGTTGATGGGTTCCA
GAGCACACGTCTGAACTCTTTCCCTACACGACGCTCTTCCGATCTNNNCTGGTGGTCTTTGACTTGTGG
AGAGCACACGTCTGAACTCTTTCCCTACACGACGCTCTTCCGATCTNNNGGGGAGACTGACTATCAAT
GAGCACACGTCTGAACTCTTTCCCTACACGACGCTCTTCCGATCTNNNACCATGAATAGCATGACCACA
AGAGCACACGTCTGAACTCTTTCCCTACACGACGCTCTTCCGATCTNNNACAGTAAACACAAACGAAC
GAGCACACGTCTGAACTCTTTCCCTACACGACGCTCTTCCGATCTNNNTCAACATACTCATCTTCCTCCTG
GAGCACACGTCTGAACTCTTTCCCTACACGACGCTCTTCCGATCTNNNGTACACAAGCTGGTATGTGTC
GAGCACACGTCTGAACTCTTTCCCTACACGACGCTCTTCCGATCTNNNCTGTCCTTGAGTAACCATCCTG
AGCACACGTCTGAACTCTTTCCCTACACGACGCTCTTCCGATCTNNNTTCACTGCAAATCAACTGGGTC
GAGCACACGTCTGAACTCTTTCCCTACACGACGCTCTTCCGATCTNNNAGGCCTTATCTGGAAAAAGCA
AGAGCACACGTCTGAACTCTTTCCCTACACGACGCTCTTCCGATCTNNNACCTGAACATTATAGCCCCAC
AGAGCACACGTCTGAACTCTTTCCCTACACGACGCTCTTCCGATCTNNNAGAGAAGTTGCTTTGGCCAT
AGCACACGTCTGAACTCTTTCCCTACACGACGCTCTTCCGATCTNNNTCAAGGAGAAGAAAGCAGTTCA
AGAGCACACGTCTGAACTCTTTCCCTACACGACGCTCTTCCGATCTNNNGAAGAAGAGCATTAGGGCC
AGAGCACACGTCTGAACTCTTTCCCTACACGACGCTCTTCCGATCTNNNGAGAGGAAAGGCCTGAAAC
GAGCACACGTCTGAACTCTTTCCCTACACGACGCTCTTCCGATCTNNNCCACTTTGGTGCTCTTGGAGAA

GAGCACACGTCTGAACTCTTTCCCTACACGACGCTCTTCCGATCTNNNAGGGTCTAAGTACTCCCAAGT
AGAGCACACGTCTGAACTCTTTCCCTACACGACGCTCTTCCGATCTNNNATTGTGGTGGCATGGAAAC
AGAGCACACGTCTGAACTCTTTCCCTACACGACGCTCTTCCGATCTNNNACCTCATGCATTTCCCTCTCT
AGCACACGTCTGAACTCTTTCCCTACACGACGCTCTTCCGATCTNNNCCTACAGTTTCCTACAAAGCGT
AGAGCACACGTCTGAACTCTTTCCCTACACGACGCTCTTCCGATCTNNNCTGTGCCACTATGGCAACAA
GAGCACACGTCTGAACTCTTTCCCTACACGACGCTCTTCCGATCTNNNAGTATGGCTTTTACAGACTG
AGCACACGTCTGAACTCTTTCCCTACACGACGCTCTTCCGATCTNNNGGAGAGCAAAACCCTGTCTAC
GAGCACACGTCTGAACTCTTTCCCTACACGACGCTCTTCCGATCTNNNTGTAAATGCGTGAGAAGAG
GAGCACACGTCTGAACTCTTTCCCTACACGACGCTCTTCCGATCTNNNGCAGGATTGACTGAAGGAGG
AGAGCACACGTCTGAACTCTTTCCCTACACGACGCTCTTCCGATCTNNNTTGACATGCACCTGTGGTTG
AGAGCACACGTCTGAACTCTTTCCCTACACGACGCTCTTCCGATCTNNNTGCCTTCACTCCTGAAGGAC
GAGCACACGTCTGAACTCTTTCCCTACACGACGCTCTTCCGATCTNNNGTTCGTGCAGGATGAATGG
GAGCACACGTCTGAACTCTTTCCCTACACGACGCTCTTCCGATCTNNNTCTGAAATACTAGGGTAGGG
GAGCACACGTCTGAACTCTTTCCCTACACGACGCTCTTCCGATCTNNNGTGGCAGTGCTATGATTTC
AGAGCACACGTCTGAACTCTTTCCCTACACGACGCTCTTCCGATCTNNNGGGAATGCCTTCATCAGAA
AGAGCACACGTCTGAACTCTTTCCCTACACGACGCTCTTCCGATCTNNNCAATAGTTTTGGGATGGCT
GAGCACACGTCTGAACTCTTTCCCTACACGACGCTCTTCCGATCTNNNATCAGGGTGATGTGTGTGTG
AGAGCACACGTCTGAACTCTTTCCCTACACGACGCTCTTCCGATCTNNNGCAGGCCCTTTCTTAGTACA
AGAGCACACGTCTGAACTCTTTCCCTACACGACGCTCTTCCGATCTNNNGAGAGAGAAGAACAGGCA
AGAGCACACGTCTGAACTCTTTCCCTACACGACGCTCTTCCGATCTNNNTCACTTGCACCGAGGCTAAG
AGCACACGTCTGAACTCTTTCCCTACACGACGCTCTTCCGATCTNNNCTTCAACTCCAGACCACCAACA
GAGCACACGTCTGAACTCTTTCCCTACACGACGCTCTTCCGATCTNNNGCTTGACAGGTTTCTAGAGT
AGAGCACACGTCTGAACTCTTTCCCTACACGACGCTCTTCCGATCTNNNGGTGCTCAGTTTGCCATTTT
AAGAGCACACGTCTGAACTCTTTCCCTACACGACGCTCTTCCGATCTNNNCTTACCAGCCAAGTTTCTG
AGCACACGTCTGAACTCTTTCCCTACACGACGCTCTTCCGATCTNNNTCGTTGCCAAATCATAGACACT
AGAGCACACGTCTGAACTCTTTCCCTACACGACGCTCTTCCGATCTNNNCACTGGAGCTTCCCAAATCA
AGCACACGTCTGAACTCTTTCCCTACACGACGCTCTTCCGATCTNNNCTCATTACAACTCTGCCTCTGGG
AGAGCACACGTCTGAACTCTTTCCCTACACGACGCTCTTCCGATCTNNNGAGGAACATGGATTTTCCG
AGAGCACACGTCTGAACTCTTTCCCTACACGACGCTCTTCCGATCTNNNTGTCCCCCTTATGGAGATGA
GAGCACACGTCTGAACTCTTTCCCTACACGACGCTCTTCCGATCTNNNCACATTCAGCTTCAGAAATGC
AGAGCACACGTCTGAACTCTTTCCCTACACGACGCTCTTCCGATCTNNNCTAAACAAAGGGCATGGT
GAGCACACGTCTGAACTCTTTCCCTACACGACGCTCTTCCGATCTNNNAATAACACTCCACTTTTCCCCG
AGCACACGTCTGAACTCTTTCCCTACACGACGCTCTTCCGATCTNNNCCTCTCAGTGTATGCTCACAAG
AGAGCACACGTCTGAACTCTTTCCCTACACGACGCTCTTCCGATCTNNNTCATTGAGAAGAAACCTTA
AGAGCACACGTCTGAACTCTTTCCCTACACGACGCTCTTCCGATCTNNNCAAAGGACCCCATGTGCTGA
AGAGCACACGTCTGAACTCTTTCCCTACACGACGCTCTTCCGATCTNNNTTCTGCCAATTTGGAGTGTG
AGCACACGTCTGAACTCTTTCCCTACACGACGCTCTTCCGATCTNNNCTTGTGTGTATCTGTATGTGCG
AGCACACGTCTGAACTCTTTCCCTACACGACGCTCTTCCGATCTNNNTCCTTCCTTGTGTGTGTGTGTTT
AAGAGCACACGTCTGAACTCTTTCCCTACACGACGCTCTTCCGATCTNNNACCCCACATCTACAAAGAA
AGAGCACACGTCTGAACTCTTTCCCTACACGACGCTCTTCCGATCTNNNCCCACCAAAGAGTGGGAAA
AGCACACGTCTGAACTCTTTCCCTACACGACGCTCTTCCGATCTNNNTGTAGCTCTCTGTGTATGTGTG
AGAGCACACGTCTGAACTCTTTCCCTACACGACGCTCTTCCGATCTNNNAGGCCAATGGAAAATAACA
GAGCACACGTCTGAACTCTTTCCCTACACGACGCTCTTCCGATCTNNNAACACCTCCATTTTGCTTTCCC

GAGCACACGTCTGAACTCTTTCCCTACACGACGCTCTTCCGATCTNNNTCCAAGGTTCTGTGAGGTAG
GAGCACACGTCTGAACTCTTTCCCTACACGACGCTCTTCCGATCTNNNTTTCCAAACTTTCAACTGGTT
AGAGCACACGTCTGAACTCTTTCCCTACACGACGCTCTTCCGATCTNNNATCTCCTAGGTTCATGCCTC
AGAGCACACGTCTGAACTCTTTCCCTACACGACGCTCTTCCGATCTNNNCTTGGGTCTTCTTGTACCTGT
AGAGCACACGTCTGAACTCTTTCCCTACACGACGCTCTTCCGATCTNNNGGAACACCTCCATTCCTCTT
GAGCACACGTCTGAACTCTTTCCCTACACGACGCTCTTCCGATCTNNNAATTCTGCATCTGTGTGTGTG
GAGCACACGTCTGAACTCTTTCCCTACACGACGCTCTTCCGATCTNNNAGAATTTCAAAGTCGGGAGC
AGCACACGTCTGAACTCTTTCCCTACACGACGCTCTTCCGATCTNNNACATGAAAGAACCAATTCAAG
GAGCACACGTCTGAACTCTTTCCCTACACGACGCTCTTCCGATCTNNNAGGAACAAGTTTTCTGGGTC
AGAGCACACGTCTGAACTCTTTCCCTACACGACGCTCTTCCGATCTNNNGACGCTAGTGAGCAAGGA
AGCACACGTCTGAACTCTTTCCCTACACGACGCTCTTCCGATCTNNNCTGTGCCCAGCCTACATAAGA
AGAGCACACGTCTGAACTCTTTCCCTACACGACGCTCTTCCGATCTNNNTGGGTTCAGTCTGATGATG
AGCACACGTCTGAACTCTTTCCCTACACGACGCTCTTCCGATCTNNNACAGTCTCTCTCTTTCTCACCCC
AGAGCACACGTCTGAACTCTTTCCCTACACGACGCTCTTCCGATCTNNNTGCAGGAGGTGTACTAGTT
AGCACACGTCTGAACTCTTTCCCTACACGACGCTCTTCCGATCTNNNACTGGCTCAGACAAGACTACA
CACACGTCTGAACTCTTTCCCTACACGACGCTCTTCCGATCTNNNGTGTGTGTGTCAATAAAAGGC
AGAGCACACGTCTGAACTCTTTCCCTACACGACGCTCTTCCGATCTNNNTGCATGCAAGGCCATCATTT
AGCACACGTCTGAACTCTTTCCCTACACGACGCTCTTCCGATCTNNNTGCATTCAAAAGGACGGGAA
AGCACACGTCTGAACTCTTTCCCTACACGACGCTCTTCCGATCTNNNTCTGAGAGGGCTGTTTTGTCC
AGAGCACACGTCTGAACTCTTTCCCTACACGACGCTCTTCCGATCTNNNCAAAATCAACCAGCTTGGCT
AGCACACGTCTGAACTCTTTCCCTACACGACGCTCTTCCGATCTNNNAATCTGGAATTCGCTGTGTCTC
GAGCACACGTCTGAACTCTTTCCCTACACGACGCTCTTCCGATCTNNNCTGCCACTGTCACAAGAAAGG
GAGCACACGTCTGAACTCTTTCCCTACACGACGCTCTTCCGATCTNNNTTACTGACAGTGGAGTTTGC
GAGCACACGTCTGAACTCTTTCCCTACACGACGCTCTTCCGATCTNNNATTTCTGCTGGGCAGGATAC
AAGAGCACACGTCTGAACTCTTTCCCTACACGACGCTCTTCCGATCTNNNGATCAAGCCAGGATTTGTT
AGCACACGTCTGAACTCTTTCCCTACACGACGCTCTTCCGATCTNNNATCTCAGCCAAGTTGTGCCTAT
AGCACACGTCTGAACTCTTTCCCTACACGACGCTCTTCCGATCTNNNAACAGTCTGATGTGTGTGTGT
AGCACACGTCTGAACTCTTTCCCTACACGACGCTCTTCCGATCTNNNAGATGAATCACTTTCATAACGG
AGCACACGTCTGAACTCTTTCCCTACACGACGCTCTTCCGATCTNNNCACAGGCTGCTTCATCTCTTTC
AGAGCACACGTCTGAACTCTTTCCCTACACGACGCTCTTCCGATCTNNNGGTTAGACTTACCCTCCTCCC
AGCACACGTCTGAACTCTTTCCCTACACGACGCTCTTCCGATCTNNNTGCAACAGTAAGGATCCATTT
GAGCACACGTCTGAACTCTTTCCCTACACGACGCTCTTCCGATCTNNNTTGAGAGCAGGGCCTCATATA
AGCACACGTCTGAACTCTTTCCCTACACGACGCTCTTCCGATCTNNNTAACCACAGAAAATGGTGCCCT
GAGCACACGTCTGAACTCTTTCCCTACACGACGCTCTTCCGATCTNNNACACGAAACAAATAGACATG
GAGCACACGTCTGAACTCTTTCCCTACACGACGCTCTTCCGATCTNNNGTGTGTTAGTGAAGGTTCTC
AGCACACGTCTGAACTCTTTCCCTACACGACGCTCTTCCGATCTNNNACGTCGCTCCAGTGTAAACAAA
AGAGCACACGTCTGAACTCTTTCCCTACACGACGCTCTTCCGATCTNNNATTGAAGGCTTCCCCTTCACT
GCACACGTCTGAACTCTTTCCCTACACGACGCTCTTCCGATCTNNNTCTGTGGCTAGATAGTATTCCAC
AGCACACGTCTGAACTCTTTCCCTACACGACGCTCTTCCGATCTNNNTGGCTTTCACTCGTTTTTACATG
AGAGCACACGTCTGAACTCTTTCCCTACACGACGCTCTTCCGATCTNNNGTTTGTCTTGGCTTCCCAGA
GAGCACACGTCTGAACTCTTTCCCTACACGACGCTCTTCCGATCTNNNAGAGGAAAGTCTGTGAATG
GAGCACACGTCTGAACTCTTTCCCTACACGACGCTCTTCCGATCTNNNAAGTGACATAAGAGTGAGG
GAGCACACGTCTGAACTCTTTCCCTACACGACGCTCTTCCGATCTNNNCAACAGTAAACTGTGGTAAG

GAGCACACGTCTGAACTCTTTCCCTACACGACGCTCTTCCGATCTNNNGCAGAAGCACAACTTCCACAA
GCACACGTCTGAACTCTTTCCCTACACGACGCTCTTCCGATCTNNNTGCAGTTCTTGTCCACCTAAAAGT
AGAGCACACGTCTGAACTCTTTCCCTACACGACGCTCTTCCGATCTNNNTCAGCGATGCATAACTCTGT
AGAGCACACGTCTGAACTCTTTCCCTACACGACGCTCTTCCGATCTNNNCACCAGTCCCACGTTAGAAC
AGCACACGTCTGAACTCTTTCCCTACACGACGCTCTTCCGATCTNNNAGATTGCCCTCACATCCATCCTA
AGCACACGTCTGAACTCTTTCCCTACACGACGCTCTTCCGATCTNNNTTTTCAGTGTCTCTCCCTCTCTC
GAGCACACGTCTGAACTCTTTCCCTACACGACGCTCTTCCGATCTNNNTTAGGCCTCTCTGATGAGGT
GAGCACACGTCTGAACTCTTTCCCTACACGACGCTCTTCCGATCTNNNAATCATCACTGCACTTTGAG
GAGCACACGTCTGAACTCTTTCCCTACACGACGCTCTTCCGATCTNNNAATGGTCACGTTCACTTCCTCA
GCACACGTCTGAACTCTTTCCCTACACGACGCTCTTCCGATCTNNNTGCAAAGTCATTCATATACTGCC
GAGCACACGTCTGAACTCTTTCCCTACACGACGCTCTTCCGATCTNNNCCCTCCCATTACTGTGGGGAA
AGAGCACACGTCTGAACTCTTTCCCTACACGACGCTCTTCCGATCTNNNGTGTGGGTTAGAATCCTGA
GAGCACACGTCTGAACTCTTTCCCTACACGACGCTCTTCCGATCTNNNATAGACCTACTGAGCCTCACC
GCACACGTCTGAACTCTTTCCCTACACGACGCTCTTCCGATCTNNNACGTAAAAACTAATTGCGAAAC
GAGCACACGTCTGAACTCTTTCCCTACACGACGCTCTTCCGATCTNNNCCTCACCTCCACTTCCCAAGTA
GAGCACACGTCTGAACTCTTTCCCTACACGACGCTCTTCCGATCTNNNGGCGACAATAGCAGAAACTC
AGAGCACACGTCTGAACTCTTTCCCTACACGACGCTCTTCCGATCTNNNACGAAACTGGACTGACGAA
GAGCACACGTCTGAACTCTTTCCCTACACGACGCTCTTCCGATCTNNNGTTAGTTCCAGCTTGGAGGTC
GCACACGTCTGAACTCTTTCCCTACACGACGCTCTTCCGATCTNNNTCATAAGTGTGTGCCTATGTGT
GAGCACACGTCTGAACTCTTTCCCTACACGACGCTCTTCCGATCTNNNTCCATAGGAGACCAGTGTATA
GAGCACACGTCTGAACTCTTTCCCTACACGACGCTCTTCCGATCTNNNTTTTGGTATTTTGGTGGCTGC
AGCACACGTCTGAACTCTTTCCCTACACGACGCTCTTCCGATCTNNNCGCGTGGCCCACTATTTTCTTAT
AGAGCACACGTCTGAACTCTTTCCCTACACGACGCTCTTCCGATCTNNNTTGCATGGTCTCTATGAGTC
AGAGCACACGTCTGAACTCTTTCCCTACACGACGCTCTTCCGATCTNNNACTTGAGCCTACCAAGCAG
GAGCACACGTCTGAACTCTTTCCCTACACGACGCTCTTCCGATCTNNNTCAAGGTATTAGATGCACCCA
GAGCACACGTCTGAACTCTTTCCCTACACGACGCTCTTCCGATCTNNNGCCTGGGCAATAGAAGTGA
GAGCACACGTCTGAACTCTTTCCCTACACGACGCTCTTCCGATCTNNNGTTTCCCCTGCAGATAAGGAA
GAGCACACGTCTGAACTCTTTCCCTACACGACGCTCTTCCGATCTNNNAGCTCTAGTACACCTTACACAT
GCACACGTCTGAACTCTTTCCCTACACGACGCTCTTCCGATCTNNNAGATCCTTTTGTAGTTGGTGCA
GAGCACACGTCTGAACTCTTTCCCTACACGACGCTCTTCCGATCTNNNCGTAGGCCGGTAAACGAAGT
GAGCACACGTCTGAACTCTTTCCCTACACGACGCTCTTCCGATCTNNNGAGTTCAAGCACGTCAAAAC
AGCACACGTCTGAACTCTTTCCCTACACGACGCTCTTCCGATCTNNNGGTCCTTCTTTCCTTCTCCTTTCC
AGAGCACACGTCTGAACTCTTTCCCTACACGACGCTCTTCCGATCTNNNCCCGTGTTATTCAAGGGTTA
GAGCACACGTCTGAACTCTTTCCCTACACGACGCTCTTCCGATCTNNNGCGATTGAGTGACTGCCTCA
AGAGCACACGTCTGAACTCTTTCCCTACACGACGCTCTTCCGATCTNNNCTGTGGGTGTGTTGTATTT
GAGCACACGTCTGAACTCTTTCCCTACACGACGCTCTTCCGATCTNNNCAGTTCTGCCATATCAGTGGC
GCACACGTCTGAACTCTTTCCCTACACGACGCTCTTCCGATCTNNNCCCCTGTTATAACCACCTACTCCA
GAGCACACGTCTGAACTCTTTCCCTACACGACGCTCTTCCGATCTNNNACATAACCTAACTGGCCCCTTG
GCACACGTCTGAACTCTTTCCCTACACGACGCTCTTCCGATCTNNNTGGAACTGCTCTTTGTCTTGTTC
GAGCACACGTCTGAACTCTTTCCCTACACGACGCTCTTCCGATCTNNNCCGTACTCCTAACACACATACA
GAGCACACGTCTGAACTCTTTCCCTACACGACGCTCTTCCGATCTNNNTCCCACAAACCATCCATAATG
AGCACACGTCTGAACTCTTTCCCTACACGACGCTCTTCCGATCTNNNTGATCTTCTTGGCTCATTGTCTC
GAGCACACGTCTGAACTCTTTCCCTACACGACGCTCTTCCGATCTNNNTACTCCAATGAAAGGCAAGA

GAGCACACGTCTGAACTCTTTCCCTACACGACGCTCTTCCGATCTNNNATACTGTGGGTTGGGCAGA
GAGCACACGTCTGAACTCTTTCCCTACACGACGCTCTTCCGATCTNNNTCATGTCCTCCTGATTCCATC
AGCACACGTCTGAACTCTTTCCCTACACGACGCTCTTCCGATCTNNNATGGATTCATGGGTGTGTG
GCACACGTCTGAACTCTTTCCCTACACGACGCTCTTCCGATCTNNNAACCTCAACAAATATGCCTTCA
AGCACACGTCTGAACTCTTTCCCTACACGACGCTCTTCCGATCTNNNCCGGAGGATCAGAATCAAGT
AGCACACGTCTGAACTCTTTCCCTACACGACGCTCTTCCGATCTNNNACAAATAGGACCAAATGCAAC
AGCACACGTCTGAACTCTTTCCCTACACGACGCTCTTCCGATCTNNNGGCTAGCATGTTTAGCATCA
AGAGCACACGTCTGAACTCTTTCCCTACACGACGCTCTTCCGATCTNNNTGAGGTGGAGGATCTAAG
GAGCACACGTCTGAACTCTTTCCCTACACGACGCTCTTCCGATCTNNNGATTTTTGTGTGCAGGGAG
GAGCACACGTCTGAACTCTTTCCCTACACGACGCTCTTCCGATCTNNNTCATTCTTTGAAAAATTGGG
GCACACGTCTGAACTCTTTCCCTACACGACGCTCTTCCGATCTNNNTTTATTGGAGACAGGCTCACG
GAGCACACGTCTGAACTCTTTCCCTACACGACGCTCTTCCGATCTNNNACGCTTCGTTTTGGCATGAA
AGAGCACACGTCTGAACTCTTTCCCTACACGACGCTCTTCCGATCTNNNGCAGCAGAACTAATATGGG
AGAGCACACGTCTGAACTCTTTCCCTACACGACGCTCTTCCGATCTNNNACCTACTAGCTCTGTGTGT
AGCACACGTCTGAACTCTTTCCCTACACGACGCTCTTCCGATCTNNNAGCCGTTGTATTCTGGAAACC
GAGCACACGTCTGAACTCTTTCCCTACACGACGCTCTTCCGATCTNNNGTTTCAGGCTGAGATGGGA
GAGCACACGTCTGAACTCTTTCCCTACACGACGCTCTTCCGATCTNNNGTTTGGAGCCACAGAGGAT
AGCACACGTCTGAACTCTTTCCCTACACGACGCTCTTCCGATCTNNNTGATTTGGATAATTGTTCCGG
AGCACACGTCTGAACTCTTTCCCTACACGACGCTCTTCCGATCTNNNCGACAAAGCAAAACACTGTCT
AAGAGCACACGTCTGAACTCTTTCCCTACACGACGCTCTTCCGATCTNNNTATGAGATTGAGCCAATG
AGCACACGTCTGAACTCTTTCCCTACACGACGCTCTTCCGATCTNNNAGTTGTGTGCGGATTTTGA
AGCACACGTCTGAACTCTTTCCCTACACGACGCTCTTCCGATCTNNNTCTTTGGATAGATTTGCCTCT
GCACACGTCTGAACTCTTTCCCTACACGACGCTCTTCCGATCTNNNACTGACCTGTGAAGTAGATGAT
GAGCACACGTCTGAACTCTTTCCCTACACGACGCTCTTCCGATCTNNNACCAGGTACCCCATTCAACAT
AGCACACGTCTGAACTCTTTCCCTACACGACGCTCTTCCGATCTNNNGCAGTGTTTTGTGCAAAGTCT
AGAGCACACGTCTGAACTCTTTCCCTACACGACGCTCTTCCGATCTNNNAATAGCAATTGCACATCCC
GAGCACACGTCTGAACTCTTTCCCTACACGACGCTCTTCCGATCTNNNCCTGGGTCTCCAGCTTACTGA
GCACACGTCTGAACTCTTTCCCTACACGACGCTCTTCCGATCTNNNCAGCTGCATCATCTGTTATTCC
GCACACGTCTGAACTCTTTCCCTACACGACGCTCTTCCGATCTNNNCAGAATAGCTCTCATGGGTTTG
AGCACACGTCTGAACTCTTTCCCTACACGACGCTCTTCCGATCTNNNAGTAGTAGGGCAAGCATACC
AGCACACGTCTGAACTCTTTCCCTACACGACGCTCTTCCGATCTNNNGGTGTTGGTCTACTCAGGGGT
AGAGCACACGTCTGAACTCTTTCCCTACACGACGCTCTTCCGATCTNNNAATCACATGGTTGCTGTGG
AGCACACGTCTGAACTCTTTCCCTACACGACGCTCTTCCGATCTNNNAGAAGCCACAAAAGCAAATGA
AGCACACGTCTGAACTCTTTCCCTACACGACGCTCTTCCGATCTNNNTGTACTTTGTATGCCTTGGGG
GAGCACACGTCTGAACTCTTTCCCTACACGACGCTCTTCCGATCTNNNTCCTGACCTTCTTCCTAATGG
AGAGCACACGTCTGAACTCTTTCCCTACACGACGCTCTTCCGATCTNNNGTGCTGAGTAAAACTGTGT
AGCACACGTCTGAACTCTTTCCCTACACGACGCTCTTCCGATCTNNNGGCTTTCTTCCCTTTGCTCTCAT
AGCACACGTCTGAACTCTTTCCCTACACGACGCTCTTCCGATCTNNNAGGTACAACCCAAGATCATCA
GAGCACACGTCTGAACTCTTTCCCTACACGACGCTCTTCCGATCTNNNCGCATGTGTCCTCTGGAATTG
AGCACACGTCTGAACTCTTTCCCTACACGACGCTCTTCCGATCTNNNAAGAAAAACCATCTCCCTTCCC
GAGCACACGTCTGAACTCTTTCCCTACACGACGCTCTTCCGATCTNNNACACATGGTCAAACCAGGAA
AGCACACGTCTGAACTCTTTCCCTACACGACGCTCTTCCGATCTNNNGGTGTGGAATGGTGTCCCATT
AGAGCACACGTCTGAACTCTTTCCCTACACGACGCTCTTCCGATCTNNNAACCATGTCTCAGCCTTCAC

AGCACACGTCTGAACTCTTTCCCTACACGACGCTCTTCCGATCTNNNCAATATGAACAAACGCACATG
AGCACACGTCTGAACTCTTTCCCTACACGACGCTCTTCCGATCTNNNACTGCACAGCATTGATACAGA
GAGCACACGTCTGAACTCTTTCCCTACACGACGCTCTTCCGATCTNNNATGAATTGCTCTTCACCGACC
GAGCACACGTCTGAACTCTTTCCCTACACGACGCTCTTCCGATCTNNNAATTAGTGTGTGTGTGTGCA
AGCACACGTCTGAACTCTTTCCCTACACGACGCTCTTCCGATCTNNNGCCTCTTAGCAGAGCCTTCAA
GAGCACACGTCTGAACTCTTTCCCTACACGACGCTCTTCCGATCTNNNGCTTAGTGATTCCCAACCTCC
GAGCACACGTCTGAACTCTTTCCCTACACGACGCTCTTCCGATCTNNNTGGATTAGCATGTCCAATGCC
AAGAGCACACGTCTGAACTCTTTCCCTACACGACGCTCTTCCGATCTNNNATTGGTGCAGCATGACAT
GAGCACACGTCTGAACTCTTTCCCTACACGACGCTCTTCCGATCTNNNATCCTCTTGCCTCGGCCTATA
AGAGCACACGTCTGAACTCTTTCCCTACACGACGCTCTTCCGATCTNNNTAGCTAACGAGGATGCAGT
AGAGCACACGTCTGAACTCTTTCCCTACACGACGCTCTTCCGATCTNNNAAAATCTCTGCCCTTAGGG
AGCACACGTCTGAACTCTTTCCCTACACGACGCTCTTCCGATCTNNNTGCAAGTACACTCTAATGTTC
GAGCACACGTCTGAACTCTTTCCCTACACGACGCTCTTCCGATCTNNNAGAGAACCACTGCTGCCAGT
AGCACACGTCTGAACTCTTTCCCTACACGACGCTCTTCCGATCTNNNAGGACAGCTTCCTCATTCTTTT
GAGCACACGTCTGAACTCTTTCCCTACACGACGCTCTTCCGATCTNNNTAAACACTAAAACCTCCGTG
GAGCACACGTCTGAACTCTTTCCCTACACGACGCTCTTCCGATCTNNNTGCCATTATCTCTTCACAGCA
AGCACACGTCTGAACTCTTTCCCTACACGACGCTCTTCCGATCTNNNACATTCTCAGGATAGTGGATTT
AGCACACGTCTGAACTCTTTCCCTACACGACGCTCTTCCGATCTNNNTTGGAGCTCAGAAGTTTTCTGG
GAGCACACGTCTGAACTCTTTCCCTACACGACGCTCTTCCGATCTNNNCTCACTTAGCAATGACACCCT
GAGCACACGTCTGAACTCTTTCCCTACACGACGCTCTTCCGATCTNNNTTTTCTCACACACCCATGTCA
AGAGCACACGTCTGAACTCTTTCCCTACACGACGCTCTTCCGATCTNNNGGGACTGTGTATGTGTGTC
GAGCACACGTCTGAACTCTTTCCCTACACGACGCTCTTCCGATCTNNNACCAATAACCCAAACTGTATC
GAGCACACGTCTGAACTCTTTCCCTACACGACGCTCTTCCGATCTNNNTGCTCCATCCTACTTGCTACT
AGAGCACACGTCTGAACTCTTTCCCTACACGACGCTCTTCCGATCTNNNGCAGAAGGAAGGCACGTAT
AGCACACGTCTGAACTCTTTCCCTACACGACGCTCTTCCGATCTNNNAGGTAAACCATTCTTATCAGAC
AGCACACGTCTGAACTCTTTCCCTACACGACGCTCTTCCGATCTNNNAGATTCTGCAGGCATCTTCACA
GAGCACACGTCTGAACTCTTTCCCTACACGACGCTCTTCCGATCTNNNTGAACAACAACTGATCCCTCC
AGAGCACACGTCTGAACTCTTTCCCTACACGACGCTCTTCCGATCTNNNTAGCTTCAGGCCAAAAGAC
AGCACACGTCTGAACTCTTTCCCTACACGACGCTCTTCCGATCTNNNGGGGTCATTGTAATGCCACCT
AGAGCACACGTCTGAACTCTTTCCCTACACGACGCTCTTCCGATCTNNNAGTTCAGTGTTTCATGGAC
AGAGCACACGTCTGAACTCTTTCCCTACACGACGCTCTTCCGATCTNNNTCCAAACACACAAAGTGGA
AGCACACGTCTGAACTCTTTCCCTACACGACGCTCTTCCGATCTNNNAAATTGAAAACCAGCATCTCTG
GAGCACACGTCTGAACTCTTTCCCTACACGACGCTCTTCCGATCTNNNCCCATCTGAGGGTATTTCTGT
GAGCACACGTCTGAACTCTTTCCCTACACGACGCTCTTCCGATCTNNNAATCCTGGATGACCTGGCAA
AGCACACGTCTGAACTCTTTCCCTACACGACGCTCTTCCGATCTNNNCCCTTTTTCACTCCTGATAGCG
AGCACACGTCTGAACTCTTTCCCTACACGACGCTCTTCCGATCTNNNTTTGCCCAGTGAACAGATGT
GAGCACACGTCTGAACTCTTTCCCTACACGACGCTCTTCCGATCTNNNTTGGTCAGTGCAACGTAACT
AGCACACGTCTGAACTCTTTCCCTACACGACGCTCTTCCGATCTNNNAGCTACCACCCAAAACATAGC
AGAGCACACGTCTGAACTCTTTCCCTACACGACGCTCTTCCGATCTNNNTGGAAAACTGCCTTATGACA
AGCACACGTCTGAACTCTTTCCCTACACGACGCTCTTCCGATCTNNNGAGGATGCCTAGATACGTCCA
AGCACACGTCTGAACTCTTTCCCTACACGACGCTCTTCCGATCTNNNAAAGCATAAAGGAGCCTTGCA
GAGCACACGTCTGAACTCTTTCCCTACACGACGCTCTTCCGATCTNNNTTTTCTGTTGAAGCTGTTGTT
GAGCACACGTCTGAACTCTTTCCCTACACGACGCTCTTCCGATCTNNNGCTGAAGCTTGTTTTTCCCC

AGCACACGTCTGAACTCTTTCCCTACACGACGCTCTTCCGATCTNNNTGTCTTTGGAGGGATTTGAGT
GAGCACACGTCTGAACTCTTTCCCTACACGACGCTCTTCCGATCTNNNTCAGAAGCTGCAGTTGTAA
GAGCACACGTCTGAACTCTTTCCCTACACGACGCTCTTCCGATCTNNNATCACTCATTAGCAATCCACC
GAGCACACGTCTGAACTCTTTCCCTACACGACGCTCTTCCGATCTNNNGTTTGCCAGTTGGCAGTCAA
GAGCACACGTCTGAACTCTTTCCCTACACGACGCTCTTCCGATCTNNNCTCTTTCCCTCTCCCCACAATC
AGAGCACACGTCTGAACTCTTTCCCTACACGACGCTCTTCCGATCTNNNTTTTATTGGGGCTACTCTCT
AGCACACGTCTGAACTCTTTCCCTACACGACGCTCTTCCGATCTNNNAACACATCCTCTGTCACCCATA
GAGCACACGTCTGAACTCTTTCCCTACACGACGCTCTTCCGATCTNNNCTCTCTCTCCAGCAACATCTG
GAGCACACGTCTGAACTCTTTCCCTACACGACGCTCTTCCGATCTNNNTAACTTGGCAGGAAACCTGT
GAGCACACGTCTGAACTCTTTCCCTACACGACGCTCTTCCGATCTNNNGCCAGGTTGCTAGAAAACTA
AGCACACGTCTGAACTCTTTCCCTACACGACGCTCTTCCGATCTNNNTGGCTTTCCATGGTAATTTGGT
GAGCACACGTCTGAACTCTTTCCCTACACGACGCTCTTCCGATCTNNNGAAGAGCAGAGGAGTAAC
GAGCACACGTCTGAACTCTTTCCCTACACGACGCTCTTCCGATCTNNNTGTTTGCTAGGTAGTTGTCT
GAGCACACGTCTGAACTCTTTCCCTACACGACGCTCTTCCGATCTNNNTCATGAGGGATGGTTACCCA
GAGCACACGTCTGAACTCTTTCCCTACACGACGCTCTTCCGATCTNNNTTCTCTCCACCCCTAGAGTGT
GAGCACACGTCTGAACTCTTTCCCTACACGACGCTCTTCCGATCTNNNGGTTGACATGGGCAGTAGG
GAGCACACGTCTGAACTCTTTCCCTACACGACGCTCTTCCGATCTNNNCTGGTTACACCCCCTTCCCTA
GAGCACACGTCTGAACTCTTTCCCTACACGACGCTCTTCCGATCTNNNTGTTCATATTTTGCCCCACTTG
AGAGCACACGTCTGAACTCTTTCCCTACACGACGCTCTTCCGATCTNNNACCTTGTGATAGAGCTGTC
AGCACACGTCTGAACTCTTTCCCTACACGACGCTCTTCCGATCTNNNACAGAGGTAGGGCAGCTAAC
AGAGCACACGTCTGAACTCTTTCCCTACACGACGCTCTTCCGATCTNNNAGGCTACCACACACTTTTTA
GAGCACACGTCTGAACTCTTTCCCTACACGACGCTCTTCCGATCTNNNTGCAATTCAAACTCCTACAAA
AGCACACGTCTGAACTCTTTCCCTACACGACGCTCTTCCGATCTNNNCCCAGCAATTCATCCCATCTCTC
AGCACACGTCTGAACTCTTTCCCTACACGACGCTCTTCCGATCTNNNAGAGAATCCTGTGTCATGATTG
CACACGTCTGAACTCTTTCCCTACACGACGCTCTTCCGATCTNNNGGATAGAGTAGCTAGGACTACA
AGAGCACACGTCTGAACTCTTTCCCTACACGACGCTCTTCCGATCTNNNGTGAGAATCTAACTGGTGG
AGCACACGTCTGAACTCTTTCCCTACACGACGCTCTTCCGATCTNNNTCCCTTCTCAATTCCCATACCAC
AAGAGCACACGTCTGAACTCTTTCCCTACACGACGCTCTTCCGATCTNNNGTGTCTCGCGCTTTCTCG
AGAGCACACGTCTGAACTCTTTCCCTACACGACGCTCTTCCGATCTNNNCCCTCAAGCGATTCACCCA
AGCACACGTCTGAACTCTTTCCCTACACGACGCTCTTCCGATCTNNNTGCCTTGATTCTGCATCTACTG
GCACACGTCTGAACTCTTTCCCTACACGACGCTCTTCCGATCTNNNTCTGCTCCTGGAGTAAGAAAGG
GAGCACACGTCTGAACTCTTTCCCTACACGACGCTCTTCCGATCTNNNCGCATGTAAGGTTAGTCAC
GAGCACACGTCTGAACTCTTTCCCTACACGACGCTCTTCCGATCTNNNAAGAAGAGCAGCTGGTGTT
GCACACGTCTGAACTCTTTCCCTACACGACGCTCTTCCGATCTNNNTCTGTTAGTTTTGAATTGGCCT
AGAGCACACGTCTGAACTCTTTCCCTACACGACGCTCTTCCGATCTNNNCAAGTGTGTATGTGGTGT
GAGCACACGTCTGAACTCTTTCCCTACACGACGCTCTTCCGATCTNNNTCCATCTTCTCACATAGCTAG
GCACACGTCTGAACTCTTTCCCTACACGACGCTCTTCCGATCTNNNTTCATCTCCTGGATTATGTGCTT
AGCACACGTCTGAACTCTTTCCCTACACGACGCTCTTCCGATCTNNNGTCTCCAACTTTACTGCAGAAG
GAGCACACGTCTGAACTCTTTCCCTACACGACGCTCTTCCGATCTNNNTGAGCCTGTTATTACCTTTCC
GAGCACACGTCTGAACTCTTTCCCTACACGACGCTCTTCCGATCTNNNGGCCTCCAAATACTTGGCTTT
GAGCACACGTCTGAACTCTTTCCCTACACGACGCTCTTCCGATCTNNNCAGTGGTGGTAGCAACAGA
AGCACACGTCTGAACTCTTTCCCTACACGACGCTCTTCCGATCTNNNATTACATGATTGGCGTTAGGC
GAGCACACGTCTGAACTCTTTCCCTACACGACGCTCTTCCGATCTNNNCTTCAGCCTGTTTGACAGAG

AGCACACGTCTGAACTCTTTCCCTACACGACGCTCTTCCGATCTNNNTTATCGCTTCTCAGCATTTGGC
AGCACACGTCTGAACTCTTTCCCTACACGACGCTCTTCCGATCTNNNTGGCACATATGAATACACACA
GAGCACACGTCTGAACTCTTTCCCTACACGACGCTCTTCCGATCTNNNGAGGGTATGATGGAAAAGC
AGCACACGTCTGAACTCTTTCCCTACACGACGCTCTTCCGATCTNNNGCTTTATTTGTCCTGGCCTCCT
GCACACGTCTGAACTCTTTCCCTACACGACGCTCTTCCGATCTNNNTCAGTCTGGGTGGCAGAATAAG
AGCACACGTCTGAACTCTTTCCCTACACGACGCTCTTCCGATCTNNNCATGCAGACATTGGATTCTTCT
GAGCACACGTCTGAACTCTTTCCCTACACGACGCTCTTCCGATCTNNNGGCAACATTGGCAAGATGAA
AGCACACGTCTGAACTCTTTCCCTACACGACGCTCTTCCGATCTNNNAGTCTAATGGTTTCTAAACCCC
GAGCACACGTCTGAACTCTTTCCCTACACGACGCTCTTCCGATCTNNNCCTAAATGTGCTTTTTGCATG
GCACACGTCTGAACTCTTTCCCTACACGACGCTCTTCCGATCTNNNACATTCAGGTTTTGCTGGTTTAG
AGCACACGTCTGAACTCTTTCCCTACACGACGCTCTTCCGATCTNNNAGTAGTATCAACCCCCACTTCAC
GAGCACACGTCTGAACTCTTTCCCTACACGACGCTCTTCCGATCTNNNAAAGAGAGAGAGTTCCTGAG
AGCACACGTCTGAACTCTTTCCCTACACGACGCTCTTCCGATCTNNNAACCAGCGAGTACTGGGTATG
AGCACACGTCTGAACTCTTTCCCTACACGACGCTCTTCCGATCTNNNACACATGCAGACAAATAAATG
GAGCACACGTCTGAACTCTTTCCCTACACGACGCTCTTCCGATCTNNNGTTCAGCCAGTAGCTAGAAGA
GAGCACACGTCTGAACTCTTTCCCTACACGACGCTCTTCCGATCTNNNCTGCTGCAACCTGGACTAGA
GAGCACACGTCTGAACTCTTTCCCTACACGACGCTCTTCCGATCTNNNCCTGTTATCCTTTTCAGGGCCT
GAGCACACGTCTGAACTCTTTCCCTACACGACGCTCTTCCGATCTNNNAAGGGATCTAGGGTCCTGT
GAGCACACGTCTGAACTCTTTCCCTACACGACGCTCTTCCGATCTNNNAGGCACTGATTCCAAGAAGA
AGCACACGTCTGAACTCTTTCCCTACACGACGCTCTTCCGATCTNNNAGAAGCTAGGAAGCCATCTGG
AGCACACGTCTGAACTCTTTCCCTACACGACGCTCTTCCGATCTNNNTGGCCAAAGCATTGAAAAGAT
GAGCACACGTCTGAACTCTTTCCCTACACGACGCTCTTCCGATCTNNNACTTCCAGTTGTTGGATTTTG
AGCACACGTCTGAACTCTTTCCCTACACGACGCTCTTCCGATCTNNNAGCTAGAATGGTTCTCAGAAAA
AGAGCACACGTCTGAACTCTTTCCCTACACGACGCTCTTCCGATCTNNNCTTCCTAAAGCCCCCACCAA
GAGCACACGTCTGAACTCTTTCCCTACACGACGCTCTTCCGATCTNNNTTTTGATGCCTGCTGGATCT
GAGCACACGTCTGAACTCTTTCCCTACACGACGCTCTTCCGATCTNNNTTCCAACATCTAGACCCTGCTG
GAGCACACGTCTGAACTCTTTCCCTACACGACGCTCTTCCGATCTNNNAAAGGCAATTTCTGTGTATG
GAGCACACGTCTGAACTCTTTCCCTACACGACGCTCTTCCGATCTNNNCCAGAGCCCCTGGAATTTATG
GAGCACACGTCTGAACTCTTTCCCTACACGACGCTCTTCCGATCTNNNTGTTCAGTCTCTGTCTGTCTCT
AGCACACGTCTGAACTCTTTCCCTACACGACGCTCTTCCGATCTNNNTCTGCTATAGTTTCCCTTCAGTC
AGAGCACACGTCTGAACTCTTTCCCTACACGACGCTCTTCCGATCTNNNAGGCATATTAGCAGATGAG
AGAGCACACGTCTGAACTCTTTCCCTACACGACGCTCTTCCGATCTNNNTGCTCAGATCTACAGGCCTA
GAGCACACGTCTGAACTCTTTCCCTACACGACGCTCTTCCGATCTNNNTCCTTGTATCCCTCAGTGTCTG
AGAGCACACGTCTGAACTCTTTCCCTACACGACGCTCTTCCGATCTNNNCTATGAATTGCTCCCCTACC
AGCACACGTCTGAACTCTTTCCCTACACGACGCTCTTCCGATCTNNNTCTTTCAATGATTTCTGGTGTG
GAGCACACGTCTGAACTCTTTCCCTACACGACGCTCTTCCGATCTNNNACTCTTTCTGATCTCAAGGGA
AGAGCACACGTCTGAACTCTTTCCCTACACGACGCTCTTCCGATCTNNNCCGGCCTTGCCAGTATTTTAT
GAGCACACGTCTGAACTCTTTCCCTACACGACGCTCTTCCGATCTNNNACTTTCCATTGCATATTGTTGC
GAGCACACGTCTGAACTCTTTCCCTACACGACGCTCTTCCGATCTNNNTGGGTATTGTTGAGTGAGAA
GAGCACACGTCTGAACTCTTTCCCTACACGACGCTCTTCCGATCTNNNGCTGTTGTAACTCCACTCAGG
AGCACACGTCTGAACTCTTTCCCTACACGACGCTCTTCCGATCTNNNCTGACCCAGGAGATTTACCAAG
GAGCACACGTCTGAACTCTTTCCCTACACGACGCTCTTCCGATCTNNNTCACATTTATACAGAGCCAAAC
AGCACACGTCTGAACTCTTTCCCTACACGACGCTCTTCCGATCTNNNACAGATTGCTGTATAGTAGGA

GAGCACACGTCTGAACTCTTTCCCTACACGACGCTCTTCCGATCTNNNACTACTAAAAGGAGGCTGAG
AGAGCACACGTCTGAACTCTTTCCCTACACGACGCTCTTCCGATCTNNNTGCAAATGCCATTACTCTGTG
AGCACACGTCTGAACTCTTTCCCTACACGACGCTCTTCCGATCTNNNGTGGTGTGATCTTAGGTCATTG
AGAGCACACGTCTGAACTCTTTCCCTACACGACGCTCTTCCGATCTNNNTCAGGCAGTGAAGTTTTTCC
AGAGCACACGTCTGAACTCTTTCCCTACACGACGCTCTTCCGATCTNNNTTCCAGACATGAGGTTGCTA
AGAGCACACGTCTGAACTCTTTCCCTACACGACGCTCTTCCGATCTNNNTCATCTCCAATCTACCTGCCCA
AGCACACGTCTGAACTCTTTCCCTACACGACGCTCTTCCGATCTNNNTGTCTGTGTACAACACTGATAG
AGAGCACACGTCTGAACTCTTTCCCTACACGACGCTCTTCCGATCTNNNAAGTTCAGTTCTTTTTCAATG
GAGCACACGTCTGAACTCTTTCCCTACACGACGCTCTTCCGATCTNNNAGACCTTACCCTGTCATGCAGA
AGCACACGTCTGAACTCTTTCCCTACACGACGCTCTTCCGATCTNNNTGGTCTCTCTCTCTCTTCTCTGTT
GAGCACACGTCTGAACTCTTTCCCTACACGACGCTCTTCCGATCTNNNGTCCCAACTACTACTCCCAAAG
AGCACACGTCTGAACTCTTTCCCTACACGACGCTCTTCCGATCTNNNTCAATCACTGAAACGAGTTACTG
AGAGCACACGTCTGAACTCTTTCCCTACACGACGCTCTTCCGATCTNNNTCCACTTTCACTGCTAATCACT
AGAGCACACGTCTGAACTCTTTCCCTACACGACGCTCTTCCGATCTNNNTTTGCTGGTTCTATCTTGGG
AGAGCACACGTCTGAACTCTTTCCCTACACGACGCTCTTCCGATCTNNNCTTGGCTACTGTGATAGCCCT
AGAGCACACGTCTGAACTCTTTCCCTACACGACGCTCTTCCGATCTNNNTGTCACTTTTGTCCTCTTCCAC
AGAGCACACGTCTGAACTCTTTCCCTACACGACGCTCTTCCGATCTNNNGGAAATGTTCTCATGGGCAA
GAGCACACGTCTGAACTCTTTCCCTACACGACGCTCTTCCGATCTNNNTGCCAGGGAGTGAAAATTAGA
GAGCACACGTCTGAACTCTTTCCCTACACGACGCTCTTCCGATCTNNNTTGATATTTTGCACTGCGGTG
AGAGCACACGTCTGAACTCTTTCCCTACACGACGCTCTTCCGATCTNNNGTATCAGCCTCTTGCTTCTAC
GAGCACACGTCTGAACTCTTTCCCTACACGACGCTCTTCCGATCTNNNGTTTCTTGTACTCTCCTGGAGG
AAGAGCACACGTCTGAACTCTTTCCCTACACGACGCTCTTCCGATCTNNNTCTGCCTTGTCTTTTTCAAG
AGAGCACACGTCTGAACTCTTTCCCTACACGACGCTCTTCCGATCTNNNAGTCTCAAATATAGACAAGC
AGAGCACACGTCTGAACTCTTTCCCTACACGACGCTCTTCCGATCTNNNGGGGAATTCCAGCTGATAGC
AGAGCACACGTCTGAACTCTTTCCCTACACGACGCTCTTCCGATCTNNNCCCCTTCTTCCAAAGTTCCTAC
AGCACACGTCTGAACTCTTTCCCTACACGACGCTCTTCCGATCTNNNACTGTTTGTTGGTCTGAATGTC
GCACACGTCTGAACTCTTTCCCTACACGACGCTCTTCCGATCTNNNCATGCCACCATGTTTTTGAACTGT
AAGAGCACACGTCTGAACTCTTTCCCTACACGACGCTCTTCCGATCTNNNGCTCTCTGCATCTCAATTCC
AAGAGCACACGTCTGAACTCTTTCCCTACACGACGCTCTTCCGATCTNNNTAGTAGGGATGCAGTAGC
AGCACACGTCTGAACTCTTTCCCTACACGACGCTCTTCCGATCTNNNCCAATGCTACTAGATAGGGGGC
GAGCACACGTCTGAACTCTTTCCCTACACGACGCTCTTCCGATCTNNNAGGCTCCAGGTACATAGAGTT
GAGCACACGTCTGAACTCTTTCCCTACACGACGCTCTTCCGATCTNNNTGTATATATTTGCCAGGGTTCT
AGCACACGTCTGAACTCTTTCCCTACACGACGCTCTTCCGATCTNNNTGCAATGTTCATTCCTGGTTGAC
AGCACACGTCTGAACTCTTTCCCTACACGACGCTCTTCCGATCTNNNTCAAGAAGACATTTTCCCCTTGC
GAGCACACGTCTGAACTCTTTCCCTACACGACGCTCTTCCGATCTNNNGGCAGCAGCTAATGCTGAATA
AAGAGCACACGTCTGAACTCTTTCCCTACACGACGCTCTTCCGATCTNNNGAGGTCAAAGCTCCCATCTT
AGCACACGTCTGAACTCTTTCCCTACACGACGCTCTTCCGATCTNNNTCATTTTGAAACCAAACAACACC
AGAGCACACGTCTGAACTCTTTCCCTACACGACGCTCTTCCGATCTNNNTGGTTGATGGGAAAAACAC
AGCACACGTCTGAACTCTTTCCCTACACGACGCTCTTCCGATCTNNNTTGTACCCAATTACAGAGCTTCA
AGCACACGTCTGAACTCTTTCCCTACACGACGCTCTTCCGATCTNNNTGCTGCAAAATGATTTGGTCTC
GAGCACACGTCTGAACTCTTTCCCTACACGACGCTCTTCCGATCTNNNACCCAGTGCCCCTTTGATATAC
AGCACACGTCTGAACTCTTTCCCTACACGACGCTCTTCCGATCTNNNGCGTTCATTGGTGAATGAATG
GAGCACACGTCTGAACTCTTTCCCTACACGACGCTCTTCCGATCTNNNACCCTGTAATTATTCGGTAAG

GAGCACACGTCTGAACTCTTTCCCTACACGACGCTCTTCCGATCTNNNATATAGGCATATGCTCACCTAT
GAGCACACGTCTGAACTCTTTCCCTACACGACGCTCTTCCGATCTNNNTAGATAGAGTTGTGTGTGTGT
AGCACACGTCTGAACTCTTTCCCTACACGACGCTCTTCCGATCTNNNTTAGGTTAGGTGCTCTTGGCTC
GAGCACACGTCTGAACTCTTTCCCTACACGACGCTCTTCCGATCTNNNTGTTTCCAAAGTCACAAAATGA
AGCACACGTCTGAACTCTTTCCCTACACGACGCTCTTCCGATCTNNNTCCACAATGCCTTACATTTTGCA
AGCACACGTCTGAACTCTTTCCCTACACGACGCTCTTCCGATCTNNNTCTTCCAAGCTGTGTTTCAAATG
GAGCACACGTCTGAACTCTTTCCCTACACGACGCTCTTCCGATCTNNNGATCCTGTCTCTCTCTCTCCT
AAGAGCACACGTCTGAACTCTTTCCCTACACGACGCTCTTCCGATCTNNNCACATAATTCCCAGAAGGTC
AGCACACGTCTGAACTCTTTCCCTACACGACGCTCTTCCGATCTNNNTGAAAGGTCTGTTGGACAAGA
AGCACACGTCTGAACTCTTTCCCTACACGACGCTCTTCCGATCTNNNGCTTGAAATCAGGTAGTGTTAG
GAGCACACGTCTGAACTCTTTCCCTACACGACGCTCTTCCGATCTNNNACTCCCTTTCTCATCTTCTCTTC
GAGCACACGTCTGAACTCTTTCCCTACACGACGCTCTTCCGATCTNNNCAGCTAGGACCACAGGTAAG
AAGAGCACACGTCTGAACTCTTTCCCTACACGACGCTCTTCCGATCTNNNAGGACAAAAGCTATAGGTG
AGAGCACACGTCTGAACTCTTTCCCTACACGACGCTCTTCCGATCTNNNACCACACACATTGGAGAGA
GAGCACACGTCTGAACTCTTTCCCTACACGACGCTCTTCCGATCTNNNAACAAAAACTGGACGGTAGT
AGAGCACACGTCTGAACTCTTTCCCTACACGACGCTCTTCCGATCTNNNGCGTCATACTTGCATTTTATT
AGAGCACACGTCTGAACTCTTTCCCTACACGACGCTCTTCCGATCTNNNCCTACTGCCACTACTACTACTG
AGCACACGTCTGAACTCTTTCCCTACACGACGCTCTTCCGATCTNNNCTTTAGCTTGCACAGCAGTCTCT
AGCACACGTCTGAACTCTTTCCCTACACGACGCTCTTCCGATCTNNNCAGCTTCACAACAAGGATGTTG
GCACACGTCTGAACTCTTTCCCTACACGACGCTCTTCCGATCTNNNCCCCTAGTGCTGTCCTTTGATAGA
GAGCACACGTCTGAACTCTTTCCCTACACGACGCTCTTCCGATCTNNNAGCCACTAAGTTTGCAGTAAT
AGAGCACACGTCTGAACTCTTTCCCTACACGACGCTCTTCCGATCTNNNGGAGATAGGTCATCTACTGT
GAGCACACGTCTGAACTCTTTCCCTACACGACGCTCTTCCGATCTNNNCCCTTCACAAACTAGGCCAAAC
GAGCACACGTCTGAACTCTTTCCCTACACGACGCTCTTCCGATCTNNNTTCTAAGTCAGAATCCCCAGGC
GAGCACACGTCTGAACTCTTTCCCTACACGACGCTCTTCCGATCTNNNCTACTGCCTTCCCTAGTTCTCC
GAGCACACGTCTGAACTCTTTCCCTACACGACGCTCTTCCGATCTNNNCTATTTGAAGGCATGGCACCT
GAGCACACGTCTGAACTCTTTCCCTACACGACGCTCTTCCGATCTNNNAATCCAATCACCTGGGGCAAA
GAGCACACGTCTGAACTCTTTCCCTACACGACGCTCTTCCGATCTNNNAGGGTTTAGGATGACAGGGT
AGCACACGTCTGAACTCTTTCCCTACACGACGCTCTTCCGATCTNNNGTGGGTAAATTCTCAGCCTTGC
AAGAGCACACGTCTGAACTCTTTCCCTACACGACGCTCTTCCGATCTNNNACCTTCCAAGGACATTCAAC
AGCACACGTCTGAACTCTTTCCCTACACGACGCTCTTCCGATCTNNNCTCCCTAAGCAAGACAGGAAAC
GAGCACACGTCTGAACTCTTTCCCTACACGACGCTCTTCCGATCTNNNTCCCATTGTGATTTCTCTCCTAG
GAGCACACGTCTGAACTCTTTCCCTACACGACGCTCTTCCGATCTNNNTCACATGAGGAAATTAGGCAA
AGAGCACACGTCTGAACTCTTTCCCTACACGACGCTCTTCCGATCTNNNCCTTGGCCCAATCAAGCCATT
AGCACACGTCTGAACTCTTTCCCTACACGACGCTCTTCCGATCTNNNTCTCCCTCACATTTCAGTACAACA
AGCACACGTCTGAACTCTTTCCCTACACGACGCTCTTCCGATCTNNNCTCCCCAACAGTGGTTGAAAAG
AGAGCACACGTCTGAACTCTTTCCCTACACGACGCTCTTCCGATCTNNNGCTAATCTAGCATGCCTGGCT
GAGCACACGTCTGAACTCTTTCCCTACACGACGCTCTTCCGATCTNNNTGTGACTAGTGCATGAGTTTCT
AAGAGCACACGTCTGAACTCTTTCCCTACACGACGCTCTTCCGATCTNNNATACATGTTATGCAGCCCCC
GAGCACACGTCTGAACTCTTTCCCTACACGACGCTCTTCCGATCTNNNGCTTCAGTTTTGGTAGGTTGG
GAGCACACGTCTGAACTCTTTCCCTACACGACGCTCTTCCGATCTNNNAGAGGGAAAAGCAGGAGTAC
AGCACACGTCTGAACTCTTTCCCTACACGACGCTCTTCCGATCTNNNTGGAAGAGTGGGTTGGAACAA
GAGCACACGTCTGAACTCTTTCCCTACACGACGCTCTTCCGATCTNNNAATAGACAACGAGTTGCCTGA

AGCACACGTCTGAACTCTTTCCCTACACGACGCTCTTCCGATCTNNNTCTTCAGAAAACAGGAATACA
GAGCACACGTCTGAACTCTTTCCCTACACGACGCTCTTCCGATCTNNNTAATGTCAGGTGAGGGAAG
AAGAGCACACGTCTGAACTCTTTCCCTACACGACGCTCTTCCGATCTNNNATGTGCAGATGTTCTGGG
GAGCACACGTCTGAACTCTTTCCCTACACGACGCTCTTCCGATCTNNNGAGAACATAAAGCAGATGCG
AGCACACGTCTGAACTCTTTCCCTACACGACGCTCTTCCGATCTNNNAGACAGACACCTCCTTATTGG
GAGCACACGTCTGAACTCTTTCCCTACACGACGCTCTTCCGATCTNNNACTGGAATGGTGGAAAAAG
GAGCACACGTCTGAACTCTTTCCCTACACGACGCTCTTCCGATCTNNNCAGAACCTCCTATGACAAGG
AGCACACGTCTGAACTCTTTCCCTACACGACGCTCTTCCGATCTNNNAACATAAAGGAAAACACAGGT
AGCACACGTCTGAACTCTTTCCCTACACGACGCTCTTCCGATCTNNNACAAAACTGTGTGTGCCTTCT
AGCACACGTCTGAACTCTTTCCCTACACGACGCTCTTCCGATCTNNNAGAAAATGCAGCATCCCTTGCT
AGCACACGTCTGAACTCTTTCCCTACACGACGCTCTTCCGATCTNNNTTCTCCAGCTCTCAGTCCTTCA
AAGAGCACACGTCTGAACTCTTTCCCTACACGACGCTCTTCCGATCTNNNCTGTGGATGCACCCCACA
GAGCACACGTCTGAACTCTTTCCCTACACGACGCTCTTCCGATCTNNNTCAGCATCTGCTTTAGTGAAA
GCACACGTCTGAACTCTTTCCCTACACGACGCTCTTCCGATCTNNNACCAGTGTTCTCCTTCCTTCCTTC
AGCACACGTCTGAACTCTTTCCCTACACGACGCTCTTCCGATCTNNNCTCCAGCAATTCTTCCCCATCT
GAGCACACGTCTGAACTCTTTCCCTACACGACGCTCTTCCGATCTNNNTAGTTACAGGGGTGGAGCA
AGCACACGTCTGAACTCTTTCCCTACACGACGCTCTTCCGATCTNNNCCAAATGTCCCTACCTCTAACA
GAGCACACGTCTGAACTCTTTCCCTACACGACGCTCTTCCGATCTNNNTGGAATGCATGTTTGTTTCAT
AGAGCACACGTCTGAACTCTTTCCCTACACGACGCTCTTCCGATCTNNNTAGAGGGTCTTCTCACCTTG
AGAGCACACGTCTGAACTCTTTCCCTACACGACGCTCTTCCGATCTNNNGATCGAGACCAGGGTTTTG
AAGAGCACACGTCTGAACTCTTTCCCTACACGACGCTCTTCCGATCTNNNCCAGGAAAGCAGCTCTTCA
AGAGCACACGTCTGAACTCTTTCCCTACACGACGCTCTTCCGATCTNNNTCAGTCACTCTCTCCCACTTG
AAGAGCACACGTCTGAACTCTTTCCCTACACGACGCTCTTCCGATCTNNNGTGAACACTCCCAGGCTAT
GAGCACACGTCTGAACTCTTTCCCTACACGACGCTCTTCCGATCTNNNTTGCATTCCACATGCTCCTCA
GAGCACACGTCTGAACTCTTTCCCTACACGACGCTCTTCCGATCTNNNTGAACCATCATAAGTCAGGTT
AGCACACGTCTGAACTCTTTCCCTACACGACGCTCTTCCGATCTNNNAGCATGGTCCTAGAATTTCCTC
GAGCACACGTCTGAACTCTTTCCCTACACGACGCTCTTCCGATCTNNNACGACCAGAGAAAAGAGTGA
GCACACGTCTGAACTCTTTCCCTACACGACGCTCTTCCGATCTNNNGTCACCCCTACAACACATAAACC
AGCACACGTCTGAACTCTTTCCCTACACGACGCTCTTCCGATCTNNNAGCCTGAGTGATGGAGTAAG
GAGCACACGTCTGAACTCTTTCCCTACACGACGCTCTTCCGATCTNNNAGCAAGATTCCAAACAGTAA
AGCACACGTCTGAACTCTTTCCCTACACGACGCTCTTCCGATCTNNNCCAATTGGCTGAAGACCTGGA
AGAGCACACGTCTGAACTCTTTCCCTACACGACGCTCTTCCGATCTNNNTCCTCTTTCTGTCTCTGCTGT
AAGAGCACACGTCTGAACTCTTTCCCTACACGACGCTCTTCCGATCTNNNCCTCGAGAGAAACAGACC
GGAAGAGCACACGTCTGAACTCTTTCCCTACACGACGCTCTTCCGATCTNNNTGCTGCAGGCAGGAT
GAGCACACGTCTGAACTCTTTCCCTACACGACGCTCTTCCGATCTNNNGTCCCCATGATAAGATTAGC
GCACACGTCTGAACTCTTTCCCTACACGACGCTCTTCCGATCTNNNGAAATCATCTACTCCATCCGCCC
AGCACACGTCTGAACTCTTTCCCTACACGACGCTCTTCCGATCTNNNGGAGAAAGGGAGGGAGAAG
AGAGCACACGTCTGAACTCTTTCCCTACACGACGCTCTTCCGATCTNNNCTAGAGTTGACAGTGAGGA
AGCACACGTCTGAACTCTTTCCCTACACGACGCTCTTCCGATCTNNNTGGGTTCCCACAGGTAAACCA
AGAGCACACGTCTGAACTCTTTCCCTACACGACGCTCTTCCGATCTNNNCGGGGAGACAAGAGAAAA
AGAGCACACGTCTGAACTCTTTCCCTACACGACGCTCTTCCGATCTNNNATCTTTTGCTCCCTGCATTG
GAGCACACGTCTGAACTCTTTCCCTACACGACGCTCTTCCGATCTNNNCTCCCTGGGTTGCTTTTTCAG
GAGCACACGTCTGAACTCTTTCCCTACACGACGCTCTTCCGATCTNNNTTTGAGACAAGGTCGGTCTC

GAGCACACGTCTGAACTCTTTCCCTACACGACGCTCTTCCGATCTNNNTGAAGTGTTCCAGGCAGAAA
GAGCACACGTCTGAACTCTTTCCCTACACGACGCTCTTCCGATCTNNNTGAGAAAATGACTTCTGATTG
GAGCACACGTCTGAACTCTTTCCCTACACGACGCTCTTCCGATCTNNNCTCTCCCTTTTTCTGGCTACA
AGAGCACACGTCTGAACTCTTTCCCTACACGACGCTCTTCCGATCTNNNGGAACCTGGCAAGACCTCAT
GAGCACACGTCTGAACTCTTTCCCTACACGACGCTCTTCCGATCTNNNTCTCAAATGGTTCAGGAGAAA
AGCACACGTCTGAACTCTTTCCCTACACGACGCTCTTCCGATCTNNNGGTACACATTCACATGTAGTTG
AGAGCACACGTCTGAACTCTTTCCCTACACGACGCTCTTCCGATCTNNNACTTCCCTGATGCACACTTT
GAGCACACGTCTGAACTCTTTCCCTACACGACGCTCTTCCGATCTNNNTCTTTCCATTGTGCCATCAACCA
GAGCACACGTCTGAACTCTTTCCCTACACGACGCTCTTCCGATCTNNNAAGTGCCAAACTCTCTCTCTCT
AAGAGCACACGTCTGAACTCTTTCCCTACACGACGCTCTTCCGATCTNNNTGGTGGCAACAGAAAGTA
GAGCACACGTCTGAACTCTTTCCCTACACGACGCTCTTCCGATCTNNNCTAGTCATGAGCTGCCATCTGT
AAGAGCACACGTCTGAACTCTTTCCCTACACGACGCTCTTCCGATCTNNNCACTCTGCAGACGGTGTTA
AGCACACGTCTGAACTCTTTCCCTACACGACGCTCTTCCGATCTNNNCCAGAAACTACTGGCAGGAACA
GAGCACACGTCTGAACTCTTTCCCTACACGACGCTCTTCCGATCTNNNCCCAAACTGTCTGCCAGTAATG
AGCACACGTCTGAACTCTTTCCCTACACGACGCTCTTCCGATCTNNNACTTCTTTCTCCTGGAACACAG
AGAGCACACGTCTGAACTCTTTCCCTACACGACGCTCTTCCGATCTNNNTCAGAGCCTCTCTCTCTCAACT
AGAGCACACGTCTGAACTCTTTCCCTACACGACGCTCTTCCGATCTNNNCACAGGGTGGGAACAAGAT
AGAGCACACGTCTGAACTCTTTCCCTACACGACGCTCTTCCGATCTNNNGGACCTATGGTGGGACTTCT
GAGCACACGTCTGAACTCTTTCCCTACACGACGCTCTTCCGATCTNNNATTTTCAGAGGAGATGGGGG
GAGCACACGTCTGAACTCTTTCCCTACACGACGCTCTTCCGATCTNNNATTTCAAACTGCCATTGGATG
GAGCACACGTCTGAACTCTTTCCCTACACGACGCTCTTCCGATCTNNNTCTCTTTACCCCCATTCCCCAA
GAGCACACGTCTGAACTCTTTCCCTACACGACGCTCTTCCGATCTNNNACAGAGCAAGACACCAGAAAA
AGAGCACACGTCTGAACTCTTTCCCTACACGACGCTCTTCCGATCTNNNGCTAGAAGACCTGGGGGCT
AGCACACGTCTGAACTCTTTCCCTACACGACGCTCTTCCGATCTNNNTGTCAGGCAAAAATAGCAAAAA
AAGAGCACACGTCTGAACTCTTTCCCTACACGACGCTCTTCCGATCTNNNGTGGGGGTGAGGAAATTG
AGAGCACACGTCTGAACTCTTTCCCTACACGACGCTCTTCCGATCTNNNACCCAGGTTTAGGATTTATG
AGAGCACACGTCTGAACTCTTTCCCTACACGACGCTCTTCCGATCTNNNACCTTGGCAGTGATTTCTTCT
AGCACACGTCTGAACTCTTTCCCTACACGACGCTCTTCCGATCTNNNTCTTGGTCAGTTGGATTTTGG
GAGCACACGTCTGAACTCTTTCCCTACACGACGCTCTTCCGATCTNNNAGTAGGTGACCCAGTGTTTT
GAGCACACGTCTGAACTCTTTCCCTACACGACGCTCTTCCGATCTNNNAGTCAAGACATTTGAGGAAA
AGAGCACACGTCTGAACTCTTTCCCTACACGACGCTCTTCCGATCTNNNTGTTGATGATCTAGTCAGCA
GAGCACACGTCTGAACTCTTTCCCTACACGACGCTCTTCCGATCTNNNTGACCTGTCCATTTTGATGCA
AGAGCACACGTCTGAACTCTTTCCCTACACGACGCTCTTCCGATCTNNNGGCCTGTAAGTGGGAATGAA
AAGAGCACACGTCTGAACTCTTTCCCTACACGACGCTCTTCCGATCTNNNCTCCATGTGCCAGATTCTGT
GAGCACACGTCTGAACTCTTTCCCTACACGACGCTCTTCCGATCTNNNTCCATCATGTAACCCTGTTGCA
GAGCACACGTCTGAACTCTTTCCCTACACGACGCTCTTCCGATCTNNNAGTGAGAAGTGAAGTGTTA
AGAGCACACGTCTGAACTCTTTCCCTACACGACGCTCTTCCGATCTNNNCTCTGAGACCTATGGGCAGG
GAGCACACGTCTGAACTCTTTCCCTACACGACGCTCTTCCGATCTNNNGTCCATGCATCAGAAACAACA
AGCACACGTCTGAACTCTTTCCCTACACGACGCTCTTCCGATCTNNNTGAGACTTCAAAAGGGATTGA
GAGCACACGTCTGAACTCTTTCCCTACACGACGCTCTTCCGATCTNNNTTTCTCCGTTTTTCAAACTGCC
AGCACACGTCTGAACTCTTTCCCTACACGACGCTCTTCCGATCTNNNGGATATTGGAAAGGCATCAGG
AGAGCACACGTCTGAACTCTTTCCCTACACGACGCTCTTCCGATCTNNNCTTTCCCTTTCCCCAACACTCT
AGAGCACACGTCTGAACTCTTTCCCTACACGACGCTCTTCCGATCTNNNGTGTCCATTATCAGTGGTCT

AGAGCACACGTCTGAACTCTTTCCCTACACGACGCTCTTCCGATCTNNNACAGACAATCCAGTCTTCTT
AGCACACGTCTGAACTCTTTCCCTACACGACGCTCTTCCGATCTNNNGCTGTGAATGCCTTTGCTTCAT
GAGCACACGTCTGAACTCTTTCCCTACACGACGCTCTTCCGATCTNNNAGAAAATACACGTCTGCTGC
GAGCACACGTCTGAACTCTTTCCCTACACGACGCTCTTCCGATCTNNNTGGCTATAAGAAGTGTGTG
AGCACACGTCTGAACTCTTTCCCTACACGACGCTCTTCCGATCTNNNTCTCAGTTTACTCACAGAAACG
GAGCACACGTCTGAACTCTTTCCCTACACGACGCTCTTCCGATCTNNNTGATCAGAGTACATCCCAGAT
AGCACACGTCTGAACTCTTTCCCTACACGACGCTCTTCCGATCTNNNGGAGGTACTGAGGTGAATTC
GAGCACACGTCTGAACTCTTTCCCTACACGACGCTCTTCCGATCTNNNAACATGCTACCAGCCAATAA
GAGCACACGTCTGAACTCTTTCCCTACACGACGCTCTTCCGATCTNNNATCTAATCTCCTGGCTGGAAG
AGCACACGTCTGAACTCTTTCCCTACACGACGCTCTTCCGATCTNNNAAGTGCTATGCAAATCAACTG
AGAGCACACGTCTGAACTCTTTCCCTACACGACGCTCTTCCGATCTNNNTGTCAAAGCAGACAGATGG
AGCACACGTCTGAACTCTTTCCCTACACGACGCTCTTCCGATCTNNNTGAATCTCCTGCTTTTTCCTGG
GAGCACACGTCTGAACTCTTTCCCTACACGACGCTCTTCCGATCTNNNCGTGAACCAAAGATTGGAAA
AGCACACGTCTGAACTCTTTCCCTACACGACGCTCTTCCGATCTNNNTTCCATTTGAAACCCTCTGGG
AGCACACGTCTGAACTCTTTCCCTACACGACGCTCTTCCGATCTNNNTGGAATGTCTTCTGACCCCAG
GAGCACACGTCTGAACTCTTTCCCTACACGACGCTCTTCCGATCTNNNTACCATCCTGGTGGGTGTGT
AGAGCACACGTCTGAACTCTTTCCCTACACGACGCTCTTCCGATCTNNNAAGAACAGTGATGGTGAA
AGCACACGTCTGAACTCTTTCCCTACACGACGCTCTTCCGATCTNNNATCATCCTTACCCTTGCTGAGA
AGCACACGTCTGAACTCTTTCCCTACACGACGCTCTTCCGATCTNNNAGTGCTTGACATCCTAGTTGG
AGCACACGTCTGAACTCTTTCCCTACACGACGCTCTTCCGATCTNNNTCTCTCTTTTGCTTTCTCTTGCA
AGCACACGTCTGAACTCTTTCCCTACACGACGCTCTTCCGATCTNNNTGACACAGTACTTTCTCTCACT
AGAGCACACGTCTGAACTCTTTCCCTACACGACGCTCTTCCGATCTNNNCCAGAAGAGGGGGAAGGG
AGCACACGTCTGAACTCTTTCCCTACACGACGCTCTTCCGATCTNNNTGGTGAAGGCATTAGAACAA
GAGCACACGTCTGAACTCTTTCCCTACACGACGCTCTTCCGATCTNNNAGTACTAGAAGAAACACACT
AGCACACGTCTGAACTCTTTCCCTACACGACGCTCTTCCGATCTNNNTCAAAGGAACATTGACCTGAG
AGCACACGTCTGAACTCTTTCCCTACACGACGCTCTTCCGATCTNNNTTATGTTCCTCCAAACAGCACC
AGCACACGTCTGAACTCTTTCCCTACACGACGCTCTTCCGATCTNNNTGACTTTCTTTTCATCTTGCAT
AGCACACGTCTGAACTCTTTCCCTACACGACGCTCTTCCGATCTNNNGCTGGTGTATAAGTCAGTAA
AGCACACGTCTGAACTCTTTCCCTACACGACGCTCTTCCGATCTNNNTGCCAAAACTTCTCACTGAAGA
GAGCACACGTCTGAACTCTTTCCCTACACGACGCTCTTCCGATCTNNNGGGTACAATTTGGGTACTGG
GAGCACACGTCTGAACTCTTTCCCTACACGACGCTCTTCCGATCTNNNGGGTACGTTGATCCTTGTTTT
GAGCACACGTCTGAACTCTTTCCCTACACGACGCTCTTCCGATCTNNNATACCGTGTTCCCATCTCTGA
AGCACACGTCTGAACTCTTTCCCTACACGACGCTCTTCCGATCTNNNGGATGGGGGCAGTGATAATG
GAGCACACGTCTGAACTCTTTCCCTACACGACGCTCTTCCGATCTNNNTCTGGTATTCTTCATACCACT
AGCACACGTCTGAACTCTTTCCCTACACGACGCTCTTCCGATCTNNNACTGCATCACAAACCTATGAC
GAGCACACGTCTGAACTCTTTCCCTACACGACGCTCTTCCGATCTNNNGGACAATGGATTTTTGGTCT
AGCACACGTCTGAACTCTTTCCCTACACGACGCTCTTCCGATCTNNNCAGCTTCACTCTGTCTCTGTCT
AGCACACGTCTGAACTCTTTCCCTACACGACGCTCTTCCGATCTNNNAGGTCTTAGCTTTGTTTATGTG
AGCACACGTCTGAACTCTTTCCCTACACGACGCTCTTCCGATCTNNNACACACTCACACCAAACACAAA
AGCACACGTCTGAACTCTTTCCCTACACGACGCTCTTCCGATCTNNNTTTTGAATGTTGACCTGGGTG
AAGAGCACACGTCTGAACTCTTTCCCTACACGACGCTCTTCCGATCTNNNCTTACACAGAGCCCAGACT
AGAGCACACGTCTGAACTCTTTCCCTACACGACGCTCTTCCGATCTNNNAACTCTGCATGAGGGGGA
GAGCACACGTCTGAACTCTTTCCCTACACGACGCTCTTCCGATCTNNNAAGCCTGGTCCATAAGGTTT

AGAGCACACGTCTGAACTCTTTCCCTACACGACGCTCTTCCGATCTNNNACCACCCTTATTTCTTCCTCT
GAGCACACGTCTGAACTCTTTCCCTACACGACGCTCTTCCGATCTNNNAGAAGACAACGCAAAGAGGT
GAGCACACGTCTGAACTCTTTCCCTACACGACGCTCTTCCGATCTNNNTTGCTCCCGTGAACATCACATA
AGAGCACACGTCTGAACTCTTTCCCTACACGACGCTCTTCCGATCTNNNCTCAAATCTGCCTTTCCTCCTC
AGCACACGTCTGAACTCTTTCCCTACACGACGCTCTTCCGATCTNNNTGGATAAGGAATACTTGCCCTG
AGAGCACACGTCTGAACTCTTTCCCTACACGACGCTCTTCCGATCTNNNCCAGCCTGGTCATAGAGAGA
AGCACACGTCTGAACTCTTTCCCTACACGACGCTCTTCCGATCTNNNTGAGTGAAAGATCCTAGACACA
GAGCACACGTCTGAACTCTTTCCCTACACGACGCTCTTCCGATCTNNNTTTCTACTTGGCTTTCCTAGGT
AGAGCACACGTCTGAACTCTTTCCCTACACGACGCTCTTCCGATCTNNNACTCAATGTTCAAGTTCTCCCT
GAGCACACGTCTGAACTCTTTCCCTACACGACGCTCTTCCGATCTNNNGCCTTATCCCAAATGTTGTCAC
GAGCACACGTCTGAACTCTTTCCCTACACGACGCTCTTCCGATCTNNNTACCCAACAGCTATGTGACTA
GAGCACACGTCTGAACTCTTTCCCTACACGACGCTCTTCCGATCTNNNCCTCAAATCATCCAGAAAGGCC
GAGCACACGTCTGAACTCTTTCCCTACACGACGCTCTTCCGATCTNNNAGGTTCTCCAGAAAAACCAAA
GAGCACACGTCTGAACTCTTTCCCTACACGACGCTCTTCCGATCTNNNTCAATACCTCATTGCCCACCTT
AGAGCACACGTCTGAACTCTTTCCCTACACGACGCTCTTCCGATCTNNNAGTGACTGTTTTCACCATTTCA
AGAGCACACGTCTGAACTCTTTCCCTACACGACGCTCTTCCGATCTNNNGACTTCAGAAAATTCTCTCTC
GCACACGTCTGAACTCTTTCCCTACACGACGCTCTTCCGATCTNNNTGCAATTGTCAGATGAATGAGAC
GAGCACACGTCTGAACTCTTTCCCTACACGACGCTCTTCCGATCTNNNGCACGAACAAGCAAAACCAGA
GAGCACACGTCTGAACTCTTTCCCTACACGACGCTCTTCCGATCTNNNAGAGTGAGAGAGAGATGTG
GAGCACACGTCTGAACTCTTTCCCTACACGACGCTCTTCCGATCTNNNATTCTCAGTTCTTTGCCTTCCAT
AGCACACGTCTGAACTCTTTCCCTACACGACGCTCTTCCGATCTNNNCCTCCCTCCCCCTTCTACATATCC
GAGCACACGTCTGAACTCTTTCCCTACACGACGCTCTTCCGATCTNNNAAGAATCAACTTTTCGCTTTG
GAGCACACGTCTGAACTCTTTCCCTACACGACGCTCTTCCGATCTNNNTCAAACAGTGCCACCAAGAAG
AAGAGCACACGTCTGAACTCTTTCCCTACACGACGCTCTTCCGATCTNNNACAAAGAGGAGCTGGTATC
GAGCACACGTCTGAACTCTTTCCCTACACGACGCTCTTCCGATCTNNNTCTATTCTGGAAAGCTGAGGC
GAGCACACGTCTGAACTCTTTCCCTACACGACGCTCTTCCGATCTNNNCAGGAAACACTCCTTTGGGCTA
AGCACACGTCTGAACTCTTTCCCTACACGACGCTCTTCCGATCTNNNTTGAACACTTTGTAAGTACCTGT
GCACACGTCTGAACTCTTTCCCTACACGACGCTCTTCCGATCTNNNAACTGTTCCTAATATGTTGTTTGC
GAGCACACGTCTGAACTCTTTCCCTACACGACGCTCTTCCGATCTNNNTGAGTGGTGATCATGCAGTAT
AGAGCACACGTCTGAACTCTTTCCCTACACGACGCTCTTCCGATCTNNNGCTAGTCTCAGCTCAGCAGA
AGAGCACACGTCTGAACTCTTTCCCTACACGACGCTCTTCCGATCTNNNACCTTGCTAAGAGGAGGTCT
GAGCACACGTCTGAACTCTTTCCCTACACGACGCTCTTCCGATCTNNNTGACACCACCTATGAAAGAGA
GAGCACACGTCTGAACTCTTTCCCTACACGACGCTCTTCCGATCTNNNAATGAAACTTGCCCTTTGGCA
GAGCACACGTCTGAACTCTTTCCCTACACGACGCTCTTCCGATCTNNNGGCACAAATTGCAGTGTACAT
GAGCACACGTCTGAACTCTTTCCCTACACGACGCTCTTCCGATCTNNNAAATGAGTGAGACCCTCCCTAC
AGAGCACACGTCTGAACTCTTTCCCTACACGACGCTCTTCCGATCTNNNCCTCCAGGTACTGAGAGATG
GAGCACACGTCTGAACTCTTTCCCTACACGACGCTCTTCCGATCTNNNTCTCAGGATGAGTTCAAGGTC
GAGCACACGTCTGAACTCTTTCCCTACACGACGCTCTTCCGATCTNNNAGCGCCAGGAAAAGTAAATCT
AGAGCACACGTCTGAACTCTTTCCCTACACGACGCTCTTCCGATCTNNNCTAGGACTTCTCGGCCCTCAT
AGAGCACACGTCTGAACTCTTTCCCTACACGACGCTCTTCCGATCTNNNCAAAATGAATGTCTCTCGCTC
AGAGCACACGTCTGAACTCTTTCCCTACACGACGCTCTTCCGATCTNNNGCTGTTGCCACTGAGTAAAA
AGCACACGTCTGAACTCTTTCCCTACACGACGCTCTTCCGATCTNNNAGGAGTAAATGGGAGTTTTCTG
AGAGCACACGTCTGAACTCTTTCCCTACACGACGCTCTTCCGATCTNNNTGGCATTTTTGTGGAGAGAG

GCACACGTCTGAACTCTTTCCCTACACGACGCTCTTCCGATCTNNNTCTCTGAATTTAGCTCAATGGTG
AGCACACGTCTGAACTCTTTCCCTACACGACGCTCTTCCGATCTNNNCACCTCCCAGAAAATGTTTGTT
AGAGCACACGTCTGAACTCTTTCCCTACACGACGCTCTTCCGATCTNNNCCCAGGTTCTGATGATTGC
GCACACGTCTGAACTCTTTCCCTACACGACGCTCTTCCGATCTNNNTGAGTGGTCTTGATAACCTTGT
GCACACGTCTGAACTCTTTCCCTACACGACGCTCTTCCGATCTNNNAGGGGTCTATACATACCATTTG
AGAGCACACGTCTGAACTCTTTCCCTACACGACGCTCTTCCGATCTNNNGCTCACTGAAATCGGAAGG
GAGCACACGTCTGAACTCTTTCCCTACACGACGCTCTTCCGATCTNNNTGTCCCATTCCTCTACCTACCA
GAGCACACGTCTGAACTCTTTCCCTACACGACGCTCTTCCGATCTNNNACCATGATTTTTGCTGTTTTCC
AGCACACGTCTGAACTCTTTCCCTACACGACGCTCTTCCGATCTNNNCTCCACACCTCACCTCCTCCAGC
GAGCACACGTCTGAACTCTTTCCCTACACGACGCTCTTCCGATCTNNNAGAAAAGTACTCCACGACAA
GCACACGTCTGAACTCTTTCCCTACACGACGCTCTTCCGATCTNNNGCACACTCTCATGCATTTTCTCT
GAGCACACGTCTGAACTCTTTCCCTACACGACGCTCTTCCGATCTNNNACAATCTTCACCAAACTCTAG
GCACACGTCTGAACTCTTTCCCTACACGACGCTCTTCCGATCTNNNTGCAATCATTTTCATGACAGGG
AGCACACGTCTGAACTCTTTCCCTACACGACGCTCTTCCGATCTNNNTGAATAAGGCTCCTGTGTTCTT
GCACACGTCTGAACTCTTTCCCTACACGACGCTCTTCCGATCTNNNAGGAAATGTACAAGTTGCTTTG
GAGCACACGTCTGAACTCTTTCCCTACACGACGCTCTTCCGATCTNNNTGCAAAGTATCATGGCTGCA
AGCACACGTCTGAACTCTTTCCCTACACGACGCTCTTCCGATCTNNNAGATTTCACCATCTTGAGCAGG
AGCACACGTCTGAACTCTTTCCCTACACGACGCTCTTCCGATCTNNNCATTCACACAAAGGTGACTGG
AGCACACGTCTGAACTCTTTCCCTACACGACGCTCTTCCGATCTNNNACAGTTACAAAGCACTAACGT
AGCACACGTCTGAACTCTTTCCCTACACGACGCTCTTCCGATCTNNNACCAAACACCTTCATGCCTAGA
AGCACACGTCTGAACTCTTTCCCTACACGACGCTCTTCCGATCTNNNAGTGCCAATCTTCTCTTATGCT
AGCACACGTCTGAACTCTTTCCCTACACGACGCTCTTCCGATCTNNNCAACGTTGACTAAAACATTGC
AGCACACGTCTGAACTCTTTCCCTACACGACGCTCTTCCGATCTNNNTGACTGTCATCTGGAATTTCTC
AGCACACGTCTGAACTCTTTCCCTACACGACGCTCTTCCGATCTNNNGCAATTGGAAAATGCTAAGG
AGAGCACACGTCTGAACTCTTTCCCTACACGACGCTCTTCCGATCTNNNACTTCCGCCTTTCCATTTCA
AGCACACGTCTGAACTCTTTCCCTACACGACGCTCTTCCGATCTNNNTCACCTCAAACTTGCTATACGAG
AGCACACGTCTGAACTCTTTCCCTACACGACGCTCTTCCGATCTNNNTTCTTTCGTTTTCAGCAGTTTGG
AGAGCACACGTCTGAACTCTTTCCCTACACGACGCTCTTCCGATCTNNNCTGGAATGCACTGTGAAGC
AGCACACGTCTGAACTCTTTCCCTACACGACGCTCTTCCGATCTNNNAGGAAGTGCTAACAAATGGTG
AGAGCACACGTCTGAACTCTTTCCCTACACGACGCTCTTCCGATCTNNNCCTCTCGATGTTGTAGTTGG
AAGAGCACACGTCTGAACTCTTTCCCTACACGACGCTCTTCCGATCTNNNTGCAGAAATTAGAGCAAC
AGAGCACACGTCTGAACTCTTTCCCTACACGACGCTCTTCCGATCTNNNGCCCTTATGAAAGAGGAGA
GAGCACACGTCTGAACTCTTTCCCTACACGACGCTCTTCCGATCTNNNGCAGCACTGAAGACTTTAGA
AGCACACGTCTGAACTCTTTCCCTACACGACGCTCTTCCGATCTNNNCTATATGTGGTTTGTGTGTGT
AGCACACGTCTGAACTCTTTCCCTACACGACGCTCTTCCGATCTNNNAGCCAAAAACCAGAATTAGCC
GAGCACACGTCTGAACTCTTTCCCTACACGACGCTCTTCCGATCTNNNACACAAAGTCCAAACAATCTG
GAGCACACGTCTGAACTCTTTCCCTACACGACGCTCTTCCGATCTNNNTGACAGAGCTGTATCAGTTGA
AGCACACGTCTGAACTCTTTCCCTACACGACGCTCTTCCGATCTNNNTGCTTGTCAGTTTTCTCTCTTCC
GCACACGTCTGAACTCTTTCCCTACACGACGCTCTTCCGATCTNNNTTGTTGGCAGCACATTCTAACCT
AGCACACGTCTGAACTCTTTCCCTACACGACGCTCTTCCGATCTNNNGGCAGGAAAAGGAGAGAAA
GAGCACACGTCTGAACTCTTTCCCTACACGACGCTCTTCCGATCTNNNACCAGGATGCTGGAATAGAG
GAGCACACGTCTGAACTCTTTCCCTACACGACGCTCTTCCGATCTNNNAAATGCATAGCTCCATGACCA
GAGCACACGTCTGAACTCTTTCCCTACACGACGCTCTTCCGATCTNNNAGTGCTCTTATTCAGGATCAG

GAGCACACGTCTGAACTCTTTCCCTACACGACGCTCTTCCGATCTNNNGTCCTCTTGGCATTTTGGGGT
AGCACACGTCTGAACTCTTTCCCTACACGACGCTCTTCCGATCTNNNTAGGCTGAACATCATTCCAGGC
AGCACACGTCTGAACTCTTTCCCTACACGACGCTCTTCCGATCTNNNTTATGTTGCCTGTCACAGTGTC
GAGCACACGTCTGAACTCTTTCCCTACACGACGCTCTTCCGATCTNNNGGGCCTTGTGATAGTGTGAGT
GAGCACACGTCTGAACTCTTTCCCTACACGACGCTCTTCCGATCTNNNTTGTGGTTTTGGGTGTTGGTT
AGCACACGTCTGAACTCTTTCCCTACACGACGCTCTTCCGATCTNNNGCACTTTTTGGCCTCATAAGCA
AGCACACGTCTGAACTCTTTCCCTACACGACGCTCTTCCGATCTNNNTGCCTAGTTTGTTTGGCTTCCAT
GAGCACACGTCTGAACTCTTTCCCTACACGACGCTCTTCCGATCTNNNTCAGGGCTTTATGTCTCAAATC
AGAGCACACGTCTGAACTCTTTCCCTACACGACGCTCTTCCGATCTNNNGCACCCCCTTTTAGCATGTAT
AGAGCACACGTCTGAACTCTTTCCCTACACGACGCTCTTCCGATCTNNNCCTCTCTTCTGAGCTGTTGTC
GCACACGTCTGAACTCTTTCCCTACACGACGCTCTTCCGATCTNNNACTGTGTTCTATGAGTTCTCTGCT
AGAGCACACGTCTGAACTCTTTCCCTACACGACGCTCTTCCGATCTNNNAACGAAGAACTTGTTTGTGG
GAGCACACGTCTGAACTCTTTCCCTACACGACGCTCTTCCGATCTNNNCCAAATTACCTTAGGTGAGGT
AGCACACGTCTGAACTCTTTCCCTACACGACGCTCTTCCGATCTNNNTGTAAATCCAGATTCCAAAACCA
AGCACACGTCTGAACTCTTTCCCTACACGACGCTCTTCCGATCTNNNTGGGTGTATATTCATCCCCAAA
AGAGCACACGTCTGAACTCTTTCCCTACACGACGCTCTTCCGATCTNNNTGGATAAAATCACTGCCAACA
GAGCACACGTCTGAACTCTTTCCCTACACGACGCTCTTCCGATCTNNNATCACATTACTCTCCACTCCCCC
GAGCACACGTCTGAACTCTTTCCCTACACGACGCTCTTCCGATCTNNNTGCTTGAAGAATCATGGTCCC
GAGCACACGTCTGAACTCTTTCCCTACACGACGCTCTTCCGATCTNNNGGAGGGGTCCATAGGCTTATC
GCACACGTCTGAACTCTTTCCCTACACGACGCTCTTCCGATCTNNNGGGAAAAAGCAAACTGTTTCCAC
GAGCACACGTCTGAACTCTTTCCCTACACGACGCTCTTCCGATCTNNNCCACACCAGCCACTAGTAAAGA
AGAGCACACGTCTGAACTCTTTCCCTACACGACGCTCTTCCGATCTNNNTGTGGCTGTATAGTACTCCAA
AGCACACGTCTGAACTCTTTCCCTACACGACGCTCTTCCGATCTNNNTTAGGGTCCTCCACAGTGTCAT
AAGAGCACACGTCTGAACTCTTTCCCTACACGACGCTCTTCCGATCTNNNACTGGACTTGAGGAAACA
AGCACACGTCTGAACTCTTTCCCTACACGACGCTCTTCCGATCTNNNTCTGCACTTATTCTTTGTCAGCC
AGAGCACACGTCTGAACTCTTTCCCTACACGACGCTCTTCCGATCTNNNTTCCCCCAAACTTTTGAAGAA
AGAGCACACGTCTGAACTCTTTCCCTACACGACGCTCTTCCGATCTNNNAGTGCTGAAAGACAAGCAA
AGCACACGTCTGAACTCTTTCCCTACACGACGCTCTTCCGATCTNNNAAAATGGAAAACAAATGTGCAT
AGCACACGTCTGAACTCTTTCCCTACACGACGCTCTTCCGATCTNNNTCCTAGTGGAAGATGATAAGCA
AGCACACGTCTGAACTCTTTCCCTACACGACGCTCTTCCGATCTNNNTGGAGGAAGATGCTAGTAGAG
GAGCACACGTCTGAACTCTTTCCCTACACGACGCTCTTCCGATCTNNNACCAACTCACAGCTTCTCTCTC
CACACGTCTGAACTCTTTCCCTACACGACGCTCTTCCGATCTNNNACAAAGAGAGGCCCTGTATCAAAA
AGCACACGTCTGAACTCTTTCCCTACACGACGCTCTTCCGATCTNNNAATTACTACCGTGACTGGGATG
AGAGCACACGTCTGAACTCTTTCCCTACACGACGCTCTTCCGATCTNNNACCATGACTAGCCAGTTCAGA
GAGCACACGTCTGAACTCTTTCCCTACACGACGCTCTTCCGATCTNNNAAAGACACAAAATGTGGCTGC
GAGCACACGTCTGAACTCTTTCCCTACACGACGCTCTTCCGATCTNNNTGTAGGTCTTTTACCTCCCTGC
AGCACACGTCTGAACTCTTTCCCTACACGACGCTCTTCCGATCTNNNCAGGATCAGGTCAGAAATGGAC
AGCACACGTCTGAACTCTTTCCCTACACGACGCTCTTCCGATCTNNNAAACAAGAGCAAGAAGAACTTC
AGCACACGTCTGAACTCTTTCCCTACACGACGCTCTTCCGATCTNNNCCATGTGGTATGTAGAGATGT
GAGCACACGTCTGAACTCTTTCCCTACACGACGCTCTTCCGATCTNNNTAAACCTCCCGAATCATGTGAC
AGAGCACACGTCTGAACTCTTTCCCTACACGACGCTCTTCCGATCTNNNGAACAACCCAAGACAATGTG
GAGCACACGTCTGAACTCTTTCCCTACACGACGCTCTTCCGATCTNNNCCAGCTTCAGCAGCTATAAAC
GAGCACACGTCTGAACTCTTTCCCTACACGACGCTCTTCCGATCTNNNTTCTCCCATAGCCTTCTACCAC

AGAGCACACGTCTGAACTCTTTCCCTACACGACGCTCTTCCGATCTNNNTCAAATTCCCCATTTGTTTCTC
AGCACACGTCTGAACTCTTTCCCTACACGACGCTCTTCCGATCTNNNACTCCTGTTCCTTCCTCAGTTAC
GAGCACACGTCTGAACTCTTTCCCTACACGACGCTCTTCCGATCTNNNCTGAGTACCCTGAGGAAGAG
AGCACACGTCTGAACTCTTTCCCTACACGACGCTCTTCCGATCTNNNTTATTGATGCCTAGCGTGTCTCA
AGAGCACACGTCTGAACTCTTTCCCTACACGACGCTCTTCCGATCTNNNTCATGGTATATGCACAGGAC
GAGCACACGTCTGAACTCTTTCCCTACACGACGCTCTTCCGATCTNNNTTGTAGATGGTGAGAGATGG
AGAGCACACGTCTGAACTCTTTCCCTACACGACGCTCTTCCGATCTNNNTGTCCAGTATGAACAGAACT
AGCACACGTCTGAACTCTTTCCCTACACGACGCTCTTCCGATCTNNNGCTACAGCCTTCTAAGGAAACA
AGAGCACACGTCTGAACTCTTTCCCTACACGACGCTCTTCCGATCTNNNTGGGAGAGGATCAATTTCTC
GAGCACACGTCTGAACTCTTTCCCTACACGACGCTCTTCCGATCTNNNGAATCTCCACCAACCATTCTGC
GAGCACACGTCTGAACTCTTTCCCTACACGACGCTCTTCCGATCTNNNAGAGAAGGTTTCCTTGCATTT
GAGCACACGTCTGAACTCTTTCCCTACACGACGCTCTTCCGATCTNNNTCAGTTGGATGTAGTGGTCCA
GAGCACACGTCTGAACTCTTTCCCTACACGACGCTCTTCCGATCTNNNGGCATCTCTGGAAGCCATATGA
GAGCACACGTCTGAACTCTTTCCCTACACGACGCTCTTCCGATCTNNNCACACACAGACATATAAACGTC
AGCACACGTCTGAACTCTTTCCCTACACGACGCTCTTCCGATCTNNNAGCCTGTTTCTCCTAGCATGTTC
AGAGCACACGTCTGAACTCTTTCCCTACACGACGCTCTTCCGATCTNNNCAGTGAGAGGTGCTTATTTC
AGCACACGTCTGAACTCTTTCCCTACACGACGCTCTTCCGATCTNNNACAAAAGCAATTCATTCGCACTG
AGCACACGTCTGAACTCTTTCCCTACACGACGCTCTTCCGATCTNNNTTTACCTTCTGCCAAATGGGGT
AGAGCACACGTCTGAACTCTTTCCCTACACGACGCTCTTCCGATCTNNNTGTTCATCTACCATTCCCTTCC
GAGCACACGTCTGAACTCTTTCCCTACACGACGCTCTTCCGATCTNNNTCTGTGTACATGGTGATGAAC
AGAGCACACGTCTGAACTCTTTCCCTACACGACGCTCTTCCGATCTNNNATGCCCAGTCCATCCAAGAGA
AGAGCACACGTCTGAACTCTTTCCCTACACGACGCTCTTCCGATCTNNNACACACATTGTCCTCTCTCTCT
AGCACACGTCTGAACTCTTTCCCTACACGACGCTCTTCCGATCTNNNCAGTAACTGTTCATTTGGCAATT
AGCACACGTCTGAACTCTTTCCCTACACGACGCTCTTCCGATCTNNNTCTCCTCTTTCTGGCTTGTGGAT
AGCACACGTCTGAACTCTTTCCCTACACGACGCTCTTCCGATCTNNNAGGTTTTGCCATAACACTCAGTC
AGAGCACACGTCTGAACTCTTTCCCTACACGACGCTCTTCCGATCTNNNTGTTGAATGAGGCAAATCAG
AGCACACGTCTGAACTCTTTCCCTACACGACGCTCTTCCGATCTNNNGGGTCATGCATCTTTCAACAAC
AGAGCACACGTCTGAACTCTTTCCCTACACGACGCTCTTCCGATCTNNNTTCATAATGAGGTGCAAGG
AGAGCACACGTCTGAACTCTTTCCCTACACGACGCTCTTCCGATCTNNNAAAAAGCACATCAACCACATT
GAGCACACGTCTGAACTCTTTCCCTACACGACGCTCTTCCGATCTNNNCGCTGTACCCAAAGTGCTATTC
AGCACACGTCTGAACTCTTTCCCTACACGACGCTCTTCCGATCTNNNCACTCAGACCCCATCTTACCCATC
AGCACACGTCTGAACTCTTTCCCTACACGACGCTCTTCCGATCTNNNAGCTTTATTTAGCACACACACAC
AGCACACGTCTGAACTCTTTCCCTACACGACGCTCTTCCGATCTNNNTGTGGGGAGAGTACCTCTTAG
GAGCACACGTCTGAACTCTTTCCCTACACGACGCTCTTCCGATCTNNNTTGAAGGTGAACCCAGAAGA
GAGCACACGTCTGAACTCTTTCCCTACACGACGCTCTTCCGATCTNNNTGGAATGGAACAGAGCAAAC
GAAGAGCACACGTCTGAACTCTTTCCCTACACGACGCTCTTCCGATCTNNNCGGAGGACAGCAGAGGT
GAGCACACGTCTGAACTCTTTCCCTACACGACGCTCTTCCGATCTNNNCAGGAGAATCTTGCAAAGCAG
AGAGCACACGTCTGAACTCTTTCCCTACACGACGCTCTTCCGATCTNNNCTTTGCACAATCCTGTGACCA
AGAGCACACGTCTGAACTCTTTCCCTACACGACGCTCTTCCGATCTNNNAGTTGCTCCTGAAGAAGATG
GAGCACACGTCTGAACTCTTTCCCTACACGACGCTCTTCCGATCTNNNGGGGAAATATGACAGGACTG
GAAGAGCACACGTCTGAACTCTTTCCCTACACGACGCTCTTCCGATCTNNNGCTCTGGGTAACACACTTC
AGCACACGTCTGAACTCTTTCCCTACACGACGCTCTTCCGATCTNNNCCAAGGGGAAGCCAAGAGATT
AGAGCACACGTCTGAACTCTTTCCCTACACGACGCTCTTCCGATCTNNNTGGAGTTAGATCCAAGATAC

GAGCACACGTCTGAACTCTTTCCCTACACGACGCTCTTCCGATCTNNNCCACCGTGACCAGCAGTTAT
AGCACACGTCTGAACTCTTTCCCTACACGACGCTCTTCCGATCTNNNCTCAATAAGCTCTCACTGCCTC
GAGCACACGTCTGAACTCTTTCCCTACACGACGCTCTTCCGATCTNNNAATGATGACCTACGGTTCCA
GCACACGTCTGAACTCTTTCCCTACACGACGCTCTTCCGATCTNNNAGTTGCATCATTTAGGGTACCA
AGCACACGTCTGAACTCTTTCCCTACACGACGCTCTTCCGATCTNNNCATTGCCACTATACCCTCAACT
AGCACACGTCTGAACTCTTTCCCTACACGACGCTCTTCCGATCTNNNCCAACTTCTCAAAAACCACACA
GAGCACACGTCTGAACTCTTTCCCTACACGACGCTCTTCCGATCTNNNTAGATACACATGCCCACATGC
AGAGCACACGTCTGAACTCTTTCCCTACACGACGCTCTTCCGATCTNNNGCAGAAAGTAAGGCAAGA
AGCACACGTCTGAACTCTTTCCCTACACGACGCTCTTCCGATCTNNNTGCAGCAATAACTGGAAATAG
AGCACACGTCTGAACTCTTTCCCTACACGACGCTCTTCCGATCTNNNACACTCACACTCAAACACTTGC
AGAGCACACGTCTGAACTCTTTCCCTACACGACGCTCTTCCGATCTNNNGATAAAGAACTGCTGTGCT
AGAGCACACGTCTGAACTCTTTCCCTACACGACGCTCTTCCGATCTNNNTGAAGCAGGCTCTTGTTTC
GAGCACACGTCTGAACTCTTTCCCTACACGACGCTCTTCCGATCTNNNTGTCCTCAGTGTCATTTTCCT
AGAGCACACGTCTGAACTCTTTCCCTACACGACGCTCTTCCGATCTNNNACCTAGTAAAGCAGTTGAA
AGCACACGTCTGAACTCTTTCCCTACACGACGCTCTTCCGATCTNNNTTTCGTATTTCACCCAGCCTTGT
AGAGCACACGTCTGAACTCTTTCCCTACACGACGCTCTTCCGATCTNNNGCTCATTTGTAATTAGGCCC
GCACACGTCTGAACTCTTTCCCTACACGACGCTCTTCCGATCTNNNTCCTCCCTCTCTATTCCTCTCTGT
GCACACGTCTGAACTCTTTCCCTACACGACGCTCTTCCGATCTNNNACCTTTTCTGAGCTGTAGTCCTC
GAGCACACGTCTGAACTCTTTCCCTACACGACGCTCTTCCGATCTNNNTGTGTTTTTGTTTCTTGCCAA
AGCACACGTCTGAACTCTTTCCCTACACGACGCTCTTCCGATCTNNNCAGGGACTTTGGAACACTTTG
AGAGCACACGTCTGAACTCTTTCCCTACACGACGCTCTTCCGATCTNNNCTTCAAGGCAGAAACCAGCT
AGAGCACACGTCTGAACTCTTTCCCTACACGACGCTCTTCCGATCTNNNTGGTTGTAATGTGTTTCCGG
AGCACACGTCTGAACTCTTTCCCTACACGACGCTCTTCCGATCTNNNATACATATCTCTGTGAGCTGC
AGCACACGTCTGAACTCTTTCCCTACACGACGCTCTTCCGATCTNNNTTGTGACTTGGGGAGTTTTCC
AGAGCACACGTCTGAACTCTTTCCCTACACGACGCTCTTCCGATCTNNNAGCTTGATGGGGGATGTA
AGCACACGTCTGAACTCTTTCCCTACACGACGCTCTTCCGATCTNNNCCCTGAAATACTCACATTCCCA
AGCACACGTCTGAACTCTTTCCCTACACGACGCTCTTCCGATCTNNNACCAACATTTCTGAAGAAGCA
GAGCACACGTCTGAACTCTTTCCCTACACGACGCTCTTCCGATCTNNNTTTTCACCTTCCTGAGACTTG
AGAGCACACGTCTGAACTCTTTCCCTACACGACGCTCTTCCGATCTNNNGTGGTTTTCAAATCTTCCC
GAGCACACGTCTGAACTCTTTCCCTACACGACGCTCTTCCGATCTNNNAGGGCTTGATATGTGCAAA
AGCACACGTCTGAACTCTTTCCCTACACGACGCTCTTCCGATCTNNNAGGTCACTCAGATGTGAACCA
GAGCACACGTCTGAACTCTTTCCCTACACGACGCTCTTCCGATCTNNNTGCCATGCCTTACACACACTA
AGCACACGTCTGAACTCTTTCCCTACACGACGCTCTTCCGATCTNNNGGAGGCAAACATGATTCACA
GAGCACACGTCTGAACTCTTTCCCTACACGACGCTCTTCCGATCTNNNACCCTTTTTGCCGTGTAACAG
AGAGCACACGTCTGAACTCTTTCCCTACACGACGCTCTTCCGATCTNNNTGGGGACCATTGTTCAACC
GAGCACACGTCTGAACTCTTTCCCTACACGACGCTCTTCCGATCTNNNAGATAACATTGACCATGTGCA
AAGAGCACACGTCTGAACTCTTTCCCTACACGACGCTCTTCCGATCTNNNGTGAAGTGAGGGAAACC
GAGCACACGTCTGAACTCTTTCCCTACACGACGCTCTTCCGATCTNNNAAGGATGAATCCCAGTTGCTT
AGCACACGTCTGAACTCTTTCCCTACACGACGCTCTTCCGATCTNNNCATGGAGCCACCTATTGCTTAC
GAGCACACGTCTGAACTCTTTCCCTACACGACGCTCTTCCGATCTNNNTCCCACTTCTTGGAATCACTC
AGAGCACACGTCTGAACTCTTTCCCTACACGACGCTCTTCCGATCTNNNAAGATAGACAGAAGGAGC
GAGCACACGTCTGAACTCTTTCCCTACACGACGCTCTTCCGATCTNNNAAGTTTGCCTCTAGGGACTGA
AGCACACGTCTGAACTCTTTCCCTACACGACGCTCTTCCGATCTNNNACATTACAAGGTTTGAAAGG

AGCACACGTCTGAACTCTTTCCCTACACGACGCTCTTCCGATCTNNNTGCTCTCTTTCACTCTTCCTCTCT
AGAGCACACGTCTGAACTCTTTCCCTACACGACGCTCTTCCGATCTNNNGGTGACCTGGGGATTCATTT
AGCACACGTCTGAACTCTTTCCCTACACGACGCTCTTCCGATCTNNNACCACCTAATAGTCAAAGGCAG
GAGCACACGTCTGAACTCTTTCCCTACACGACGCTCTTCCGATCTNNNTCTGGAGTTTTCATGGACTTT
GAGCACACGTCTGAACTCTTTCCCTACACGACGCTCTTCCGATCTNNNTTTTGCTAACCACAGTCCCA
AGAGCACACGTCTGAACTCTTTCCCTACACGACGCTCTTCCGATCTNNNCGAGCCAAATAATGGAAAC
AGAGCACACGTCTGAACTCTTTCCCTACACGACGCTCTTCCGATCTNNNTGGGAAGCTATAATTGGAG
AGAGCACACGTCTGAACTCTTTCCCTACACGACGCTCTTCCGATCTNNNAGTTGAGAAAAGGGTGGCG
AGAGCACACGTCTGAACTCTTTCCCTACACGACGCTCTTCCGATCTNNNGTGATTGTACCACCACAT
GAGCACACGTCTGAACTCTTTCCCTACACGACGCTCTTCCGATCTNNNGTGACTTGGATAACTCATCAG
GAGCACACGTCTGAACTCTTTCCCTACACGACGCTCTTCCGATCTNNNGGCCAGTTTTAGAAAGTTTG
AGCACACGTCTGAACTCTTTCCCTACACGACGCTCTTCCGATCTNNNCAACACAGTTCAAAATCTCTGC
AAGAGCACACGTCTGAACTCTTTCCCTACACGACGCTCTTCCGATCTNNNGTCCATGAGTGTGCATGG
GAGCACACGTCTGAACTCTTTCCCTACACGACGCTCTTCCGATCTNNNATCTTTCTCCCTCTCTCTCATCC
AGCACACGTCTGAACTCTTTCCCTACACGACGCTCTTCCGATCTNNNACGGGGACAAAGCTTCAAAG
AGCACACGTCTGAACTCTTTCCCTACACGACGCTCTTCCGATCTNNNGGGAATGGGGAGCTGAAAAG
GAGCACACGTCTGAACTCTTTCCCTACACGACGCTCTTCCGATCTNNNTTCAGATTTGCTAGCCCCTTCA
AGCACACGTCTGAACTCTTTCCCTACACGACGCTCTTCCGATCTNNNTGGAAGAGATCTTCGAGGGAA
AGAGCACACGTCTGAACTCTTTCCCTACACGACGCTCTTCCGATCTNNNATGCCCTTTATTAGGAGTTC
AGAGCACACGTCTGAACTCTTTCCCTACACGACGCTCTTCCGATCTNNNTACTAGGTCTGAGGTTCAG
GAGCACACGTCTGAACTCTTTCCCTACACGACGCTCTTCCGATCTNNNTTTGGACTTCAAGGATGCCA
AGAGCACACGTCTGAACTCTTTCCCTACACGACGCTCTTCCGATCTNNNCTGTGGGGAACCAAAGAGT
GAGCACACGTCTGAACTCTTTCCCTACACGACGCTCTTCCGATCTNNNTGCTAGACAAAGGCAATTAG
AGAGCACACGTCTGAACTCTTTCCCTACACGACGCTCTTCCGATCTNNNAGGCTGAAAATGCCTTGAGT
GAGCACACGTCTGAACTCTTTCCCTACACGACGCTCTTCCGATCTNNNATGGGATGCCTCTGCTGAGA
GAGCACACGTCTGAACTCTTTCCCTACACGACGCTCTTCCGATCTNNNTGGAGGAGATAGAGAGGG
AGCACACGTCTGAACTCTTTCCCTACACGACGCTCTTCCGATCTNNNGCTTTGCGAAACTTTTGTTTCAG
AGCACACGTCTGAACTCTTTCCCTACACGACGCTCTTCCGATCTNNNACTTCAAATCCAACTGGTGGG
AAGAGCACACGTCTGAACTCTTTCCCTACACGACGCTCTTCCGATCTNNNGTTATCAGCCAGCCTCATTG
AGAGCACACGTCTGAACTCTTTCCCTACACGACGCTCTTCCGATCTNNNAGAGAGGGCGACAGAGTA
AAGAGCACACGTCTGAACTCTTTCCCTACACGACGCTCTTCCGATCTNNNCACATGTCCTCTCCGAATC
GAGCACACGTCTGAACTCTTTCCCTACACGACGCTCTTCCGATCTNNNACTTTTCCTCTTCCCTTTAGG
AGCACACGTCTGAACTCTTTCCCTACACGACGCTCTTCCGATCTNNNCGGCATACCCTCCCTAAAGAAG
AGCACACGTCTGAACTCTTTCCCTACACGACGCTCTTCCGATCTNNNACATCAAATCTAGGGGATTCTT
AGAGCACACGTCTGAACTCTTTCCCTACACGACGCTCTTCCGATCTNNNCAAAGAAGGTTCTGTGCTTA
AGAGCACACGTCTGAACTCTTTCCCTACACGACGCTCTTCCGATCTNNNAGTTGATGGAGTTTGGCAT
AGAGCACACGTCTGAACTCTTTCCCTACACGACGCTCTTCCGATCTNNNTCAGAGCTGTGGTGCTCTTT
AGCACACGTCTGAACTCTTTCCCTACACGACGCTCTTCCGATCTNNNACCAAACCTTAGTGCCATATGA
AGAGCACACGTCTGAACTCTTTCCCTACACGACGCTCTTCCGATCTNNNACTGTAAACTCTAGAACAAC
AGCACACGTCTGAACTCTTTCCCTACACGACGCTCTTCCGATCTNNNACCATTATTGTTTTGTGTTCCCC
GAGCACACGTCTGAACTCTTTCCCTACACGACGCTCTTCCGATCTNNNGCCTCCTCTGACCTACACACAT
GAGCACACGTCTGAACTCTTTCCCTACACGACGCTCTTCCGATCTNNNACAACCCTATGGAAGAGTGA
AGAGCACACGTCTGAACTCTTTCCCTACACGACGCTCTTCCGATCTNNNCCGTGTAGAGATATTTCACC

GAGCACACGTCTGAACTCTTTCCCTACACGACGCTCTTCCGATCTNNNAATGCATCATGGCACATCTA
AGCACACGTCTGAACTCTTTCCCTACACGACGCTCTTCCGATCTNNNTAAAAGAATCACCCACAGCAC
AGAGCACACGTCTGAACTCTTTCCCTACACGACGCTCTTCCGATCTNNNAGCAGACAACTGATCACCC
GAGCACACGTCTGAACTCTTTCCCTACACGACGCTCTTCCGATCTNNNAACATATTGCAGACATCACC
AGCACACGTCTGAACTCTTTCCCTACACGACGCTCTTCCGATCTNNNTGCTTTCAGCTATCTCAGTCTG
GAGCACACGTCTGAACTCTTTCCCTACACGACGCTCTTCCGATCTNNNCAGTACAGCATTCTGTGCAT
AGCACACGTCTGAACTCTTTCCCTACACGACGCTCTTCCGATCTNNNTGAGCTTGGAGGTTACATGTT
AGCACACGTCTGAACTCTTTCCCTACACGACGCTCTTCCGATCTNNNTGCCTTCAGTCTTAGGTAATC
AGAGCACACGTCTGAACTCTTTCCCTACACGACGCTCTTCCGATCTNNNTCTTTGTGGCTGAATGGTA
GAGCACACGTCTGAACTCTTTCCCTACACGACGCTCTTCCGATCTNNNCCTCCAGGCAGCTTGCTTTTA
GAGCACACGTCTGAACTCTTTCCCTACACGACGCTCTTCCGATCTNNNAAGAAAGCAGGCACTCTTTT
GAGCACACGTCTGAACTCTTTCCCTACACGACGCTCTTCCGATCTNNNCCACAGTGAGAACTGAATG
AGCACACGTCTGAACTCTTTCCCTACACGACGCTCTTCCGATCTNNNAAGGGGAGGAACGGCTATAA
AGCACACGTCTGAACTCTTTCCCTACACGACGCTCTTCCGATCTNNNACTCTCTACATACCAGGCACAG
GAGCACACGTCTGAACTCTTTCCCTACACGACGCTCTTCCGATCTNNNACCCTTTGGAAGTTTCTTGAA
AGCACACGTCTGAACTCTTTCCCTACACGACGCTCTTCCGATCTNNNTCCTCCTCCTTCTTCTTCTCCTTC
GAGCACACGTCTGAACTCTTTCCCTACACGACGCTCTTCCGATCTNNNGCACACATACCGAGTCATATT
AGCACACGTCTGAACTCTTTCCCTACACGACGCTCTTCCGATCTNNNCTGAGGCAAAAGAACGACTT
AGCACACGTCTGAACTCTTTCCCTACACGACGCTCTTCCGATCTNNNTGCCTGTAATCCAAGCTACTCA
AGAGCACACGTCTGAACTCTTTCCCTACACGACGCTCTTCCGATCTNNNCCAGCTACTCTGAAGGCTA
GAGCACACGTCTGAACTCTTTCCCTACACGACGCTCTTCCGATCTNNNCTGTAATCCCAGCTACTCAGG
GAGCACACGTCTGAACTCTTTCCCTACACGACGCTCTTCCGATCTNNNAAATGCCCATAATCCCAGCT
GAGCACACGTCTGAACTCTTTCCCTACACGACGCTCTTCCGATCTNNNAGGTTGAGGCAGAAGAATC
GCACACGTCTGAACTCTTTCCCTACACGACGCTCTTCCGATCTNNNAGTGACTATAAACAATGTTGCA
AGCACACGTCTGAACTCTTTCCCTACACGACGCTCTTCCGATCTNNNTGGGGAGAAAGCTTTTCTTTT
AGCACACGTCTGAACTCTTTCCCTACACGACGCTCTTCCGATCTNNNACTCTAAAGATCCTGGACAGA
GAGCACACGTCTGAACTCTTTCCCTACACGACGCTCTTCCGATCTNNNGAGAATCACTTGAACCCAGA
GAGCACACGTCTGAACTCTTTCCCTACACGACGCTCTTCCGATCTNNNTGCCCATTTTCCATACTTAGTT
AAGAGCACACGTCTGAACTCTTTCCCTACACGACGCTCTTCCGATCTNNNCTGTAGTCCTAGCTACTCG
GCACACGTCTGAACTCTTTCCCTACACGACGCTCTTCCGATCTNNNACAAAATCCTCTTGGCTACATG
GCACACGTCTGAACTCTTTCCCTACACGACGCTCTTCCGATCTNNNGGGTTCATAAAGAAACACTGG
GAGCACACGTCTGAACTCTTTCCCTACACGACGCTCTTCCGATCTNNNAGAATCACTTGAAACTGGGA
AGCACACGTCTGAACTCTTTCCCTACACGACGCTCTTCCGATCTNNNCTGTAGTCCCAGCTACTCGAG
GCACACGTCTGAACTCTTTCCCTACACGACGCTCTTCCGATCTNNNACAAATCCAGGAGTTGGTTTT
AGCACACGTCTGAACTCTTTCCCTACACGACGCTCTTCCGATCTNNNTGGTCTATTCAGTTTCCTTGC
GAGCACACGTCTGAACTCTTTCCCTACACGACGCTCTTCCGATCTNNNTAAGGCAGGAGAATCACTG
AGAGCACACGTCTGAACTCTTTCCCTACACGACGCTCTTCCGATCTNNNTCAAGCAACCTACAGGCATA
GCACACGTCTGAACTCTTTCCCTACACGACGCTCTTCCGATCTNNNTGTGCCATAGACAAAATCTGTT
AGCACACGTCTGAACTCTTTCCCTACACGACGCTCTTCCGATCTNNNTGTATCAATAACCGACTTTGC
AGCACACGTCTGAACTCTTTCCCTACACGACGCTCTTCCGATCTNNNCACGATTGTGTCACTGTACTCC
AGAGCACACGTCTGAACTCTTTCCCTACACGACGCTCTTCCGATCTNNNCATGCCTGTAATCCCAGCT
GAGCACACGTCTGAACTCTTTCCCTACACGACGCTCTTCCGATCTNNNCACCTGTAGTCCCAGCTACT
AGAGCACACGTCTGAACTCTTTCCCTACACGACGCTCTTCCGATCTNNNCTCAGGAGGTTGAAGCAG

GCACACGTCTGAACTCTTTCCCTACACGACGCTCTTCCGATCTNNNGGTTGATTCCAATTTTTCAG

AGCACACGTCTGAACTCTTTCCCTACACGACGCTCTTCCGATCTNNNTGGATGAGATCTGGGAGAATT

GCACACGTCTGAACTCTTTCCCTACACGACGCTCTTCCGATCTNNNAGTGACTGAATCTGAAGATTTG

AGAGCACACGTCTGAACTCTTTCCCTACACGACGCTCTTCCGATCTNNNACGTTACTGGTATCTGAGG
AGCACACGTCTGAACTCTTTCCCTACACGACGCTCTTCCGATCTNNNGTTGGTTTTCATTTCTCTCCAG
AGCACACGTCTGAACTCTTTCCCTACACGACGCTCTTCCGATCTNNNCAGAGAAACAGAACCAATAGG
GAGCACACGTCTGAACTCTTTCCCTACACGACGCTCTTCCGATCTNNNTGTGTACACGTGTAGACATG
GAGCACACGTCTGAACTCTTTCCCTACACGACGCTCTTCCGATCTNNNACCATGAGAACACAGACACAT
GCACACGTCTGAACTCTTTCCCTACACGACGCTCTTCCGATCTNNNACCAATAGGAGAGAGAGAGAG
AGAGCACACGTCTGAACTCTTTCCCTACACGACGCTCTTCCGATCTNNNGTTGTCACAAAACCCATGG
AGCACACGTCTGAACTCTTTCCCTACACGACGCTCTTCCGATCTNNNTCTTACTTCTCAGGCCTTTGGA
AGCACACGTCTGAACTCTTTCCCTACACGACGCTCTTCCGATCTNNNTCCAGAGAAACAGAACCAACA
AGCACACGTCTGAACTCTTTCCCTACACGACGCTCTTCCGATCTNNNACTGAGATAGAACTGGGTTGA
GAGCACACGTCTGAACTCTTTCCCTACACGACGCTCTTCCGATCTNNNTCCTCCATTCCATTGGTTGTC
AGCACACGTCTGAACTCTTTCCCTACACGACGCTCTTCCGATCTNNNGTGCCTGGCTTATTTTACTTAG
AGCACACGTCTGAACTCTTTCCCTACACGACGCTCTTCCGATCTNNNGCAACTAAGACCCATAACTGAT
GAGCACACGTCTGAACTCTTTCCCTACACGACGCTCTTCCGATCTNNNACAGATACAGTGCAATCCCTA
GAGCACACGTCTGAACTCTTTCCCTACACGACGCTCTTCCGATCTNNNTTCCAGATTTGGGATCCTCA
AGCACACGTCTGAACTCTTTCCCTACACGACGCTCTTCCGATCTNNNAGGCATGTTACCTAAGTCACA
AGCACACGTCTGAACTCTTTCCCTACACGACGCTCTTCCGATCTNNNCCTCTAAGTCCTCCCAACTTTCC
AGCACACGTCTGAACTCTTTCCCTACACGACGCTCTTCCGATCTNNNACTTTAGGTCACTTTGATTCAG
AGAGCACACGTCTGAACTCTTTCCCTACACGACGCTCTTCCGATCTNNNCACACACACACACACACGAA
AGAGCACACGTCTGAACTCTTTCCCTACACGACGCTCTTCCGATCTNNNTGACAGAGCAAACTCCATC
AGCACACGTCTGAACTCTTTCCCTACACGACGCTCTTCCGATCTNNNCAAAGATTCCTGCACTTGTATG
GAGCACACGTCTGAACTCTTTCCCTACACGACGCTCTTCCGATCTNNNACAAGTTGTCAGGTTCCTAA
GAGCACACGTCTGAACTCTTTCCCTACACGACGCTCTTCCGATCTNNNCAAAACCAACCTGGAAGACT
AGCACACGTCTGAACTCTTTCCCTACACGACGCTCTTCCGATCTNNNTGAACAGAAAAAGTCAGTGTC
GAGCACACGTCTGAACTCTTTCCCTACACGACGCTCTTCCGATCTNNNATAGAAGCTCCTGAACACATG
GCACACGTCTGAACTCTTTCCCTACACGACGCTCTTCCGATCTNNNGGGGATGATACTCAGTACTACA
GAGCACACGTCTGAACTCTTTCCCTACACGACGCTCTTCCGATCTNNNACTCTTCCTCCAACCCAATTCT
AAGAGCACACGTCTGAACTCTTTCCCTACACGACGCTCTTCCGATCTNNNTCTTTCTGATGGGCATGA
GAAGAGCACACGTCTGAACTCTTTCCCTACACGACGCTCTTCCGATCTNNNCTCCAGTGCACCAAATG
GAGCACACGTCTGAACTCTTTCCCTACACGACGCTCTTCCGATCTNNNGGTGCCACCTATATGCTGATT
GAGCACACGTCTGAACTCTTTCCCTACACGACGCTCTTCCGATCTNNNCTCTATCGTCCTTCTCCCCAAT
GAGCACACGTCTGAACTCTTTCCCTACACGACGCTCTTCCGATCTNNNTGGTTTCTGCTGAAAGATTC
AGCACACGTCTGAACTCTTTCCCTACACGACGCTCTTCCGATCTNNNAGTTCCCACATATCTCTCTCTCC
AGCACACGTCTGAACTCTTTCCCTACACGACGCTCTTCCGATCTNNNTGGGTCAGATGATTCTCTCTCT
AGCACACGTCTGAACTCTTTCCCTACACGACGCTCTTCCGATCTNNNTGTTACTGACACAGAAAGGGA
GAGCACACGTCTGAACTCTTTCCCTACACGACGCTCTTCCGATCTNNNCCTAACATGGGGTCTAAGGA
GAGCACACGTCTGAACTCTTTCCCTACACGACGCTCTTCCGATCTNNNTCAGAGATGTTGACAAAGCA
AGCACACGTCTGAACTCTTTCCCTACACGACGCTCTTCCGATCTNNNTCCCAGTTAGACATACCTGAGA
GCACACGTCTGAACTCTTTCCCTACACGACGCTCTTCCGATCTNNNACAGTACACACATACACACACAG
GAGCACACGTCTGAACTCTTTCCCTACACGACGCTCTTCCGATCTNNNTCTTCAACAGAATTTCACTGG
AGCACACGTCTGAACTCTTTCCCTACACGACGCTCTTCCGATCTNNNTGAGCCATGATTATACTCCAGC
GAGCACACGTCTGAACTCTTTCCCTACACGACGCTCTTCCGATCTNNNTGACATCACATTGGTAGCTGG
GAGCACACGTCTGAACTCTTTCCCTACACGACGCTCTTCCGATCTNNNCAAACGTATCTGCATGTTTGT

AGAGCACACGTCTGAACTCTTTCCCTACACGACGCTCTTCCGATCTNNNTGTTCATCAGGGATATTGAC
AGCACACGTCTGAACTCTTTCCCTACACGACGCTCTTCCGATCTNNNACTTTTCCTAATACACACCCACAC
AGAGCACACGTCTGAACTCTTTCCCTACACGACGCTCTTCCGATCTNNNGGAGCCTAAGGAGACAGAA
GAGCACACGTCTGAACTCTTTCCCTACACGACGCTCTTCCGATCTNNNTCTCCAAATTTATGGCAACCTG
AGAGCACACGTCTGAACTCTTTCCCTACACGACGCTCTTCCGATCTNNNTGGATGGACCTGGAGGACA
GCACACGTCTGAACTCTTTCCCTACACGACGCTCTTCCGATCTNNNTGTTCCATTGGTCTATGTGTCTGT
AGAGCACACGTCTGAACTCTTTCCCTACACGACGCTCTTCCGATCTNNNGCAGAGAGATCAATCAAGA
AGCACACGTCTGAACTCTTTCCCTACACGACGCTCTTCCGATCTNNNTGAAGCCAGTATCACCAAATAC
AGAGCACACGTCTGAACTCTTTCCCTACACGACGCTCTTCCGATCTNNNAGCTACAGCAATCAGACAAG
AGAGCACACGTCTGAACTCTTTCCCTACACGACGCTCTTCCGATCTNNNTCCTGTCTCTCTCTGTCTCTCT
GAGCACACGTCTGAACTCTTTCCCTACACGACGCTCTTCCGATCTNNNGCAGACAGCCTATTGTGGGAT
GAGCACACGTCTGAACTCTTTCCCTACACGACGCTCTTCCGATCTNNNTGGAAGCATATCACATCCTGA
AGAGCACACGTCTGAACTCTTTCCCTACACGACGCTCTTCCGATCTNNNATACTGACACACACCACAACA
GAGCACACGTCTGAACTCTTTCCCTACACGACGCTCTTCCGATCTNNNATAGAAAAACAGGCAAATTGC
AGAGCACACGTCTGAACTCTTTCCCTACACGACGCTCTTCCGATCTNNNTATGTGCTGGGTTAGGAAGT
AGCACACGTCTGAACTCTTTCCCTACACGACGCTCTTCCGATCTNNNGGGTTTTCATCATGTTGACCAG
AGAGCACACGTCTGAACTCTTTCCCTACACGACGCTCTTCCGATCTNNNGAGAATCACTTGAACCTGGC
AGAGCACACGTCTGAACTCTTTCCCTACACGACGCTCTTCCGATCTNNNCCTCCAGGAGCTACTCTTTG
AGCACACGTCTGAACTCTTTCCCTACACGACGCTCTTCCGATCTNNNACACAGTAACAGTCTGATCTCT
GCACACGTCTGAACTCTTTCCCTACACGACGCTCTTCCGATCTNNNCCCAAATGAGATACATGCATAGC
GAGCACACGTCTGAACTCTTTCCCTACACGACGCTCTTCCGATCTNNNTCCTTGTCCTGAGAAATATGC
GAGCACACGTCTGAACTCTTTCCCTACACGACGCTCTTCCGATCTNNNGTGTGTGTGTATGTGTGTGT
GAGCACACGTCTGAACTCTTTCCCTACACGACGCTCTTCCGATCTNNNACCAATCCTAGGAAACCAGAA
AAGAGCACACGTCTGAACTCTTTCCCTACACGACGCTCTTCCGATCTNNNACAAGATAAGATGATCCTG
GAGCACACGTCTGAACTCTTTCCCTACACGACGCTCTTCCGATCTNNNACAGTCACAGATATCTGAGCT
GAGCACACGTCTGAACTCTTTCCCTACACGACGCTCTTCCGATCTNNNACTTGTTCATGGTGTTTTTCTC
GAGCACACGTCTGAACTCTTTCCCTACACGACGCTCTTCCGATCTNNNTGGCTTGGCTTACAGTTGAGT
GAGCACACGTCTGAACTCTTTCCCTACACGACGCTCTTCCGATCTNNNAGGTGATCTTACGTACAGATC
AAGAGCACACGTCTGAACTCTTTCCCTACACGACGCTCTTCCGATCTNNNACGCATGGATGTATGTACA
AGCACACGTCTGAACTCTTTCCCTACACGACGCTCTTCCGATCTNNNACCAGTCCTTGAAAGAATGAAA
GAGCACACGTCTGAACTCTTTCCCTACACGACGCTCTTCCGATCTNNNACTACTTTCCATCAACTGCCAC
AGAGCACACGTCTGAACTCTTTCCCTACACGACGCTCTTCCGATCTNNNAGTGTTACAGCAATCAATGG
GAGCACACGTCTGAACTCTTTCCCTACACGACGCTCTTCCGATCTNNNGCTATTTTGGTGGCATATGGT
GAGCACACGTCTGAACTCTTTCCCTACACGACGCTCTTCCGATCTNNNGGCTCACACCCAGAGGAATAT
AGCACACGTCTGAACTCTTTCCCTACACGACGCTCTTCCGATCTNNNGTGAAACTCTGTCTCAGGAAAA
GAGCACACGTCTGAACTCTTTCCCTACACGACGCTCTTCCGATCTNNNAGGAGAATTGCTTGATCCCAC
GAGCACACGTCTGAACTCTTTCCCTACACGACGCTCTTCCGATCTNNNCATTTTCTGGTAGCAGCAGTT
AGAGCACACGTCTGAACTCTTTCCCTACACGACGCTCTTCCGATCTNNNCAAAGGTTGCAGTGAACTGA
GAGCACACGTCTGAACTCTTTCCCTACACGACGCTCTTCCGATCTNNNATCGAGATCATACCACTGCACT
GAGCACACGTCTGAACTCTTTCCCTACACGACGCTCTTCCGATCTNNNGAGCAAGACTCTGTCTCAAAA
GCACACGTCTGAACTCTTTCCCTACACGACGCTCTTCCGATCTNNNTGTTGATTTTTGTATCCTGCAACTC
AGCACACGTCTGAACTCTTTCCCTACACGACGCTCTTCCGATCTNNNGCAACTGTGCCACATTACATTA
GAGCACACGTCTGAACTCTTTCCCTACACGACGCTCTTCCGATCTNNNTGTTCTCTCTCTCTCTCTCTGCA

GCACACGTCTGAACTCTTTCCCTACACGACGCTCTTCCGATCTNNNTGAAAGGCAGAAGAAAGCAAAA
GCACACGTCTGAACTCTTTCCCTACACGACGCTCTTCCGATCTNNNCAGAGGAACATCAGGGACCTTAC
CACACGTCTGAACTCTTTCCCTACACGACGCTCTTCCGATCTNNNAGAGAGAGAGAGAGAGAGAGAG
AGCACACGTCTGAACTCTTTCCCTACACGACGCTCTTCCGATCTNNNTCTCTCTCTGTCTCTCTGTCTCTG
AGCACACGTCTGAACTCTTTCCCTACACGACGCTCTTCCGATCTNNNCCAGAAGGGTAAAGATAGGAT
GAGCACACGTCTGAACTCTTTCCCTACACGACGCTCTTCCGATCTNNNAGTCCTAATCCCCATCCTCAGAA
GAGCACACGTCTGAACTCTTTCCCTACACGACGCTCTTCCGATCTNNNTGGAAAGTCTCAACTGCTACAC
GAGCACACGTCTGAACTCTTTCCCTACACGACGCTCTTCCGATCTNNNGGGTTAGCGCTGTCTCTTTTG
GAGCACACGTCTGAACTCTTTCCCTACACGACGCTCTTCCGATCTNNNGGGAATAACTGGCTTCCAGAC
GAGCACACGTCTGAACTCTTTCCCTACACGACGCTCTTCCGATCTNNNTCCTGCTAAAGTCTCCATCTTC
GAGCACACGTCTGAACTCTTTCCCTACACGACGCTCTTCCGATCTNNNCCAAAAGATAATCCCAGCCCTG
GAGCACACGTCTGAACTCTTTCCCTACACGACGCTCTTCCGATCTNNNCAATATGCAGCTTGCTTCACCA
AGCACACGTCTGAACTCTTTCCCTACACGACGCTCTTCCGATCTNNNATAGACCTCCTTTCCTCTCCTGCC
AGCACACGTCTGAACTCTTTCCCTACACGACGCTCTTCCGATCTNNNCAGCATGTGAGAGTGTGAGAG
GAGCACACGTCTGAACTCTTTCCCTACACGACGCTCTTCCGATCTNNNTCGATGGCACCAACTATCTGAA
AGCACACGTCTGAACTCTTTCCCTACACGACGCTCTTCCGATCTNNNTTGCTGCATTCCAGCCTAAAAGA
GAGCACACGTCTGAACTCTTTCCCTACACGACGCTCTTCCGATCTNNNTCTCCAGGGTTTTCCAAGAAA
GAGCACACGTCTGAACTCTTTCCCTACACGACGCTCTTCCGATCTNNNCCATGGTCGGCAGTCTTGATA
AGAGCACACGTCTGAACTCTTTCCCTACACGACGCTCTTCCGATCTNNNACAGATTCATACACACCTGCA
GAGCACACGTCTGAACTCTTTCCCTACACGACGCTCTTCCGATCTNNNACGGCGTCATTAGTACACAAC
GAGCACACGTCTGAACTCTTTCCCTACACGACGCTCTTCCGATCTNNNCTGAGCCCAAGAGTTCAAGTA
AGAGCACACGTCTGAACTCTTTCCCTACACGACGCTCTTCCGATCTNNNCTCAGATGCTTCCCTGTTCTT
AGCACACGTCTGAACTCTTTCCCTACACGACGCTCTTCCGATCTNNNCTCTAGCACAGTGGGCTTTCTTC
AGAGCACACGTCTGAACTCTTTCCCTACACGACGCTCTTCCGATCTNNNTTGAGCTTCCAAATCAGGTGC
GAGCACACGTCTGAACTCTTTCCCTACACGACGCTCTTCCGATCTNNNGCTACAGAGTTGTGATCATGC
AGCACACGTCTGAACTCTTTCCCTACACGACGCTCTTCCGATCTNNNGCTAATTCTGGGACTGTAGGCT
AGCACACGTCTGAACTCTTTCCCTACACGACGCTCTTCCGATCTNNNTATAAAAGAGGCTCCCCAAGGC
GAGCACACGTCTGAACTCTTTCCCTACACGACGCTCTTCCGATCTNNNCCAATAGGATGTGTGTGAGTG
AGCACACGTCTGAACTCTTTCCCTACACGACGCTCTTCCGATCTNNNCCACCTTCCTGTCAACTTGTCTCC
AGCACACGTCTGAACTCTTTCCCTACACGACGCTCTTCCGATCTNNNATATGTGACACACAGACACGTA
AGCACACGTCTGAACTCTTTCCCTACACGACGCTCTTCCGATCTNNNACGGATATTGGACCTTCAAACA
GAGCACACGTCTGAACTCTTTCCCTACACGACGCTCTTCCGATCTNNNGGCAAGGAGAAAGGACTAGA
GAGCACACGTCTGAACTCTTTCCCTACACGACGCTCTTCCGATCTNNNGTGTTAGCATGTCCACCTGAG
GAGCACACGTCTGAACTCTTTCCCTACACGACGCTCTTCCGATCTNNNAGGATGCACAGGGCTAAGTAC
GAGCACACGTCTGAACTCTTTCCCTACACGACGCTCTTCCGATCTNNNACGTTTTACCCCAGTTGCTTCA
GAGCACACGTCTGAACTCTTTCCCTACACGACGCTCTTCCGATCTNNNGTCTGTGAAGTGTGTGCCTGT
AGCACACGTCTGAACTCTTTCCCTACACGACGCTCTTCCGATCTNNNTTGAGTAGCCAAGTCAAGCAAC
GAGCACACGTCTGAACTCTTTCCCTACACGACGCTCTTCCGATCTNNNGGGTTTCCCCAGAGAAAGAGA
GCACACGTCTGAACTCTTTCCCTACACGACGCTCTTCCGATCTNNNCTCCTCCCTCATATCAGCCTCAACA
GAGCACACGTCTGAACTCTTTCCCTACACGACGCTCTTCCGATCTNNNGGTAGAGCTAGGACTGGTTCC
AGCACACGTCTGAACTCTTTCCCTACACGACGCTCTTCCGATCTNNNATACTGGTGGCCAAAGGTTTCTC
AGAGCACACGTCTGAACTCTTTCCCTACACGACGCTCTTCCGATCTNNNGCAAGAAGCCATGAGAAAGC
AGCACACGTCTGAACTCTTTCCCTACACGACGCTCTTCCGATCTNNNACGGAAACCCACACACATTGTT

AGCACACGTCTGAACTCTTTCCCTACACGACGCTCTTCCGATCTNNNAAAACAGGCCAGTGGACATTA
AGCACACGTCTGAACTCTTTCCCTACACGACGCTCTTCCGATCTNNNTTGTTGCACTGTCCTGAATGTG
AGCACACGTCTGAACTCTTTCCCTACACGACGCTCTTCCGATCTNNNCACTGGTAGATGTGCCATAGCT
GAGCACACGTCTGAACTCTTTCCCTACACGACGCTCTTCCGATCTNNNATCAGAAAGACACTCCTACCC
AGAGCACACGTCTGAACTCTTTCCCTACACGACGCTCTTCCGATCTNNNATTGACCATGTTGAGTTCGG
GAGCACACGTCTGAACTCTTTCCCTACACGACGCTCTTCCGATCTNNNTCGATGACCTTACATCCACCCA
AGAGCACACGTCTGAACTCTTTCCCTACACGACGCTCTTCCGATCTNNNTTATTGCCTCACCTAGTTAC
GAGCACACGTCTGAACTCTTTCCCTACACGACGCTCTTCCGATCTNNNCCCCTCACCCGATACATAGACA
GCACACGTCTGAACTCTTTCCCTACACGACGCTCTTCCGATCTNNNAAAAGAGAACCCAGAAGCAGCA
AGCACACGTCTGAACTCTTTCCCTACACGACGCTCTTCCGATCTNNNAGTCTGATATTGAAGTTGGCC
AGCACACGTCTGAACTCTTTCCCTACACGACGCTCTTCCGATCTNNNACTTCTCTCTGCCTGTGTCTTCT
GAGCACACGTCTGAACTCTTTCCCTACACGACGCTCTTCCGATCTNNNCGGGTGTTACAAACACTGCAA
AGCACACGTCTGAACTCTTTCCCTACACGACGCTCTTCCGATCTNNNTTTTCCAGAGCTGAGTACAGTG
AGCACACGTCTGAACTCTTTCCCTACACGACGCTCTTCCGATCTNNNTTTGATAGCACAGAAGGGTGA
AGCACACGTCTGAACTCTTTCCCTACACGACGCTCTTCCGATCTNNNAATCAGTCAGGGCTTTCCTTCA
AGCACACGTCTGAACTCTTTCCCTACACGACGCTCTTCCGATCTNNNTAAAACACTCTCCCAGCAAACCC
AGCACACGTCTGAACTCTTTCCCTACACGACGCTCTTCCGATCTNNNCAGACACAGACCCACTCACAAA
GCACACGTCTGAACTCTTTCCCTACACGACGCTCTTCCGATCTNNNTACAGTTGACCTGGGACCTTTGT
AGCACACGTCTGAACTCTTTCCCTACACGACGCTCTTCCGATCTNNNGGAAAATGGTTCTCTCCCTGAC
AGCACACGTCTGAACTCTTTCCCTACACGACGCTCTTCCGATCTNNNATAAACTTGAGGTAGGCCAGG
GAGCACACGTCTGAACTCTTTCCCTACACGACGCTCTTCCGATCTNNNACCTTCTCCAGAGAGACAGAA
GAGCACACGTCTGAACTCTTTCCCTACACGACGCTCTTCCGATCTNNNTGGCATCTGGGAGAGAACATT
GAGCACACGTCTGAACTCTTTCCCTACACGACGCTCTTCCGATCTNNNGTTCATGCGATTCTCCTGGTTC
GAGCACACGTCTGAACTCTTTCCCTACACGACGCTCTTCCGATCTNNNTAGTTACCAGGTATCAACCAT
AGAGCACACGTCTGAACTCTTTCCCTACACGACGCTCTTCCGATCTNNNTTGTCCTCTGCAAAGAACCT
AGCACACGTCTGAACTCTTTCCCTACACGACGCTCTTCCGATCTNNNTGAGCTTCATGCAAGAGAGTGA
GAGCACACGTCTGAACTCTTTCCCTACACGACGCTCTTCCGATCTNNNATCTGGAGGACAATTGGGAC
AGCACACGTCTGAACTCTTTCCCTACACGACGCTCTTCCGATCTNNNACTTCAGGGTGACCAAAACTTC
AGCACACGTCTGAACTCTTTCCCTACACGACGCTCTTCCGATCTNNNTGACAGGTTTTATTCCGCTCCTG
AGCACACGTCTGAACTCTTTCCCTACACGACGCTCTTCCGATCTNNNAGGTGGGTTACTGAAAGGAGA
AGCACACGTCTGAACTCTTTCCCTACACGACGCTCTTCCGATCTNNNCCCAACTCTTGTTTGGGGGTA
GAGCACACGTCTGAACTCTTTCCCTACACGACGCTCTTCCGATCTNNNTGGAAACGCTATTCTCAACCT
GAGCACACGTCTGAACTCTTTCCCTACACGACGCTCTTCCGATCTNNNGCACTGACAAACACACGCATA
GAGCACACGTCTGAACTCTTTCCCTACACGACGCTCTTCCGATCTNNNAGTTAGATGAGGTGGAGGA
AGCACACGTCTGAACTCTTTCCCTACACGACGCTCTTCCGATCTNNNGTGTCTCTCTCCTGCTTCCAGTA
AGCACACGTCTGAACTCTTTCCCTACACGACGCTCTTCCGATCTNNNTCAGTACACAGACATGCAACTG
GCACACGTCTGAACTCTTTCCCTACACGACGCTCTTCCGATCTNNNATGTGTCTTGGGGTTTCATTGCT
GAGCACACGTCTGAACTCTTTCCCTACACGACGCTCTTCCGATCTNNNCCTTCCACCTTTGCTGTTGAGA
AGCACACGTCTGAACTCTTTCCCTACACGACGCTCTTCCGATCTNNNATGTACCAGCTACCATCCATCCA
AGCACACGTCTGAACTCTTTCCCTACACGACGCTCTTCCGATCTNNNCCTGTGAAGGGACATTACTAAG
AGCACACGTCTGAACTCTTTCCCTACACGACGCTCTTCCGATCTNNNGAGAACATAAAGCAGATGCGT
GAGCACACGTCTGAACTCTTTCCCTACACGACGCTCTTCCGATCTNNNTCCTCCCTCTTTTTCCACACTCA
GCACACGTCTGAACTCTTTCCCTACACGACGCTCTTCCGATCTNNNTAATGTAGCGCATGACTGTGTG

GCACACGTCTGAACTCTTTCCCTACACGACGCTCTTCCGATCTNNNCACTGCTAAAGACAGAGCCAGA
AGCACACGTCTGAACTCTTTCCCTACACGACGCTCTTCCGATCTNNNGAACCCCCTGACATTCCACAAT
AGAGCACACGTCTGAACTCTTTCCCTACACGACGCTCTTCCGATCTNNNTACATGTAGCTGCATTGAAC
GCACACGTCTGAACTCTTTCCCTACACGACGCTCTTCCGATCTNNNCAGCATGATTCCTAAGGCTTCCA
AGCACACGTCTGAACTCTTTCCCTACACGACGCTCTTCCGATCTNNNTGCAATTGGCAATCCATCTGGT
AGCACACGTCTGAACTCTTTCCCTACACGACGCTCTTCCGATCTNNNTGGGACTAAATTCCTCCCCTCA
AGCACACGTCTGAACTCTTTCCCTACACGACGCTCTTCCGATCTNNNCGCAGCCTACACAATCAATAGC
AGCACACGTCTGAACTCTTTCCCTACACGACGCTCTTCCGATCTNNNTTGAGAATCACTGCGTTAGTGG
AGCACACGTCTGAACTCTTTCCCTACACGACGCTCTTCCGATCTNNNTAGGGCATGTTCTTTTGCTGAG
AGCACACGTCTGAACTCTTTCCCTACACGACGCTCTTCCGATCTNNNCCAAGAGCACTTTGAGCTCAGT
AGCACACGTCTGAACTCTTTCCCTACACGACGCTCTTCCGATCTNNNCAACTGGACCTCCTTGTGGAAA
GCACACGTCTGAACTCTTTCCCTACACGACGCTCTTCCGATCTNNNACAGTTTACACTGGCATCTGGA
AGCACACGTCTGAACTCTTTCCCTACACGACGCTCTTCCGATCTNNNAGCAACAGGCAGAATGGTTTG
AGCACACGTCTGAACTCTTTCCCTACACGACGCTCTTCCGATCTNNNATTCAACTCCAGGTAATGGTGG
AGCACACGTCTGAACTCTTTCCCTACACGACGCTCTTCCGATCTNNNCTCTTGTTCACCACCCGTTTTTC
GCACACGTCTGAACTCTTTCCCTACACGACGCTCTTCCGATCTNNNTTGTTTATTTGTGCAGACAGCTT
GCACACGTCTGAACTCTTTCCCTACACGACGCTCTTCCGATCTNNNGGGTCCCTCATATGTGCATAGT
GAGCACACGTCTGAACTCTTTCCCTACACGACGCTCTTCCGATCTNNNCAGGGGCGATATAGCATTCA
AGCACACGTCTGAACTCTTTCCCTACACGACGCTCTTCCGATCTNNNGAAATCCTTTACCTTCCCCCGAC
AGAGCACACGTCTGAACTCTTTCCCTACACGACGCTCTTCCGATCTNNNCAGATCTCTGGGATTCCTGT
GAGCACACGTCTGAACTCTTTCCCTACACGACGCTCTTCCGATCTNNNAAACTGCTCAGGGTCTCATT
AGCACACGTCTGAACTCTTTCCCTACACGACGCTCTTCCGATCTNNNTGAAGGAAGTTTTGCTTTGGG
GAGCACACGTCTGAACTCTTTCCCTACACGACGCTCTTCCGATCTNNNTTCTATGCACACCTCACATCAC
AGCACACGTCTGAACTCTTTCCCTACACGACGCTCTTCCGATCTNNNCAGTGTCCGAAATGAGGAACT
AGCACACGTCTGAACTCTTTCCCTACACGACGCTCTTCCGATCTNNNCTGTGGTAAATGGTACTGGGT
GAGCACACGTCTGAACTCTTTCCCTACACGACGCTCTTCCGATCTNNNTGTTCCAAAAGCCATAAGCCT
AGCACACGTCTGAACTCTTTCCCTACACGACGCTCTTCCGATCTNNNGTATTGGCTGGAAGAAACGC
AGCACACGTCTGAACTCTTTCCCTACACGACGCTCTTCCGATCTNNNGGAGAGATCCTTGTGTGTCTT
AGCACACGTCTGAACTCTTTCCCTACACGACGCTCTTCCGATCTNNNCCCCTTATGGCATCTTTGAGCAT
AGAGCACACGTCTGAACTCTTTCCCTACACGACGCTCTTCCGATCTNNNAGTTCCAACTCCTAGTCTCCT
AGCACACGTCTGAACTCTTTCCCTACACGACGCTCTTCCGATCTNNNGTAGAAAGCTGGGTGTTACAG
GAGCACACGTCTGAACTCTTTCCCTACACGACGCTCTTCCGATCTNNNACTGACACTGTACATCTGGTG
AGCACACGTCTGAACTCTTTCCCTACACGACGCTCTTCCGATCTNNNAGCTTCAGAGAAGATCTAGGC
AGCACACGTCTGAACTCTTTCCCTACACGACGCTCTTCCGATCTNNNCCTCCCTCCCTTTCCTAACTTCA
GCACACGTCTGAACTCTTTCCCTACACGACGCTCTTCCGATCTNNNTTTATTCTGGGCTCCTAGCACCT
GAGCACACGTCTGAACTCTTTCCCTACACGACGCTCTTCCGATCTNNNAGGTGGTGATAGGCTACCAT
AGCACACGTCTGAACTCTTTCCCTACACGACGCTCTTCCGATCTNNNGAGACCTCTCAGCATTCCTGAA
GAGCACACGTCTGAACTCTTTCCCTACACGACGCTCTTCCGATCTNNNTAAAGTTGCACTGTGACTGG
AGCACACGTCTGAACTCTTTCCCTACACGACGCTCTTCCGATCTNNNAAAGACATCCGAGACAGAACA
GCACACGTCTGAACTCTTTCCCTACACGACGCTCTTCCGATCTNNNAGGCAAGAACTAAAATCCACAG
GAGCACACGTCTGAACTCTTTCCCTACACGACGCTCTTCCGATCTNNNCTAAAGCCTTCATCCCACTGG
GCACACGTCTGAACTCTTTCCCTACACGACGCTCTTCCGATCTNNNGGTTAGCTAGTGGTTTGGCAGA
AGCACACGTCTGAACTCTTTCCCTACACGACGCTCTTCCGATCTNNNAGATCAATAGGGCCAGAAGT

AGCACACGTCTGAACTCTTTCCCTACACGACGCTCTTCCGATCTNNNAGTCACTTAGCCTGTGTGCTTA
GAGCACACGTCTGAACTCTTTCCCTACACGACGCTCTTCCGATCTNNNACTTCCGAAGGCCAAGGATA
AGCACACGTCTGAACTCTTTCCCTACACGACGCTCTTCCGATCTNNNGAGGTTGTTCTGTTGGTTGTT
AGCACACGTCTGAACTCTTTCCCTACACGACGCTCTTCCGATCTNNNTACCTACAAGATTTCCCCAGCA
AGCACACGTCTGAACTCTTTCCCTACACGACGCTCTTCCGATCTNNNAAACTGAAATGCTCATCAGGG
AGCACACGTCTGAACTCTTTCCCTACACGACGCTCTTCCGATCTNNNACACCTCTATGTGAGCAAACAC
AGCACACGTCTGAACTCTTTCCCTACACGACGCTCTTCCGATCTNNNGAACACCCTCGCTAGTTTCAAG
AGAGCACACGTCTGAACTCTTTCCCTACACGACGCTCTTCCGATCTNNNTCCATTTCTAGTGCTAGTCCC
AGCACACGTCTGAACTCTTTCCCTACACGACGCTCTTCCGATCTNNNGGAAGGTCAGTCTCTGACAAT
AGCACACGTCTGAACTCTTTCCCTACACGACGCTCTTCCGATCTNNNAGTTTCCTCTCTCTTTCCCTCCG
GCACACGTCTGAACTCTTTCCCTACACGACGCTCTTCCGATCTNNNTGGCTAACCCCTCACTTCCAATAC
GCACACGTCTGAACTCTTTCCCTACACGACGCTCTTCCGATCTNNNTCTAAAGGCATCACTCTCCCCATC
GAGCACACGTCTGAACTCTTTCCCTACACGACGCTCTTCCGATCTNNNTGTTCAGAAGTGGAACAGCA
AGCACACGTCTGAACTCTTTCCCTACACGACGCTCTTCCGATCTNNNCTGCTGTTTGCTCTCTCTCTCT
GAGCACACGTCTGAACTCTTTCCCTACACGACGCTCTTCCGATCTNNNCTCCTCTGATCTGGAACAGTG
GAGCACACGTCTGAACTCTTTCCCTACACGACGCTCTTCCGATCTNNNTAGTTTACCCCTAACTCCCCTG
AGCACACGTCTGAACTCTTTCCCTACACGACGCTCTTCCGATCTNNNCCTGTCGACGGAGTGAGACTTA
AGCACACGTCTGAACTCTTTCCCTACACGACGCTCTTCCGATCTNNNTCACTTCCACAAATCAGGCACT
AGCACACGTCTGAACTCTTTCCCTACACGACGCTCTTCCGATCTNNNCCAGTTATCTCTGTCTCTCTCGC
GAGCACACGTCTGAACTCTTTCCCTACACGACGCTCTTCCGATCTNNNTAGAAGCCTAGAGCTGTTGG
AGCACACGTCTGAACTCTTTCCCTACACGACGCTCTTCCGATCTNNNTTTGGAAGCACTGCAATGACA
AGCACACGTCTGAACTCTTTCCCTACACGACGCTCTTCCGATCTNNNATGTTTATGGTCTCTCGTCCCC
AGCACACGTCTGAACTCTTTCCCTACACGACGCTCTTCCGATCTNNNATCGGCTAATGAAGCCTGTGAT
AGCACACGTCTGAACTCTTTCCCTACACGACGCTCTTCCGATCTNNNAAACCTACTACGATCAGCAGCA
GAGCACACGTCTGAACTCTTTCCCTACACGACGCTCTTCCGATCTNNNTTTGAACAGAACAGGGCTTGT
AGCACACGTCTGAACTCTTTCCCTACACGACGCTCTTCCGATCTNNNTGTCTTTCCAACTCCAAAGCCA
AGAGCACACGTCTGAACTCTTTCCCTACACGACGCTCTTCCGATCTNNNAAATAAGCCAGCAAGGAGG
AGCACACGTCTGAACTCTTTCCCTACACGACGCTCTTCCGATCTNNNTGAGGGACAACAGTCTATAGG
AGAGCACACGTCTGAACTCTTTCCCTACACGACGCTCTTCCGATCTNNNTAGTAGTGTCATTGCACTCC
AGCACACGTCTGAACTCTTTCCCTACACGACGCTCTTCCGATCTNNNAAGTCCTAGGGTTCCCAAACCT
AGAGCACACGTCTGAACTCTTTCCCTACACGACGCTCTTCCGATCTNNNATGTGTGTAAAGGGGGCTT
GAGCACACGTCTGAACTCTTTCCCTACACGACGCTCTTCCGATCTNNNTCAGTCAGTTCTCAGAAGGG
AGAGCACACGTCTGAACTCTTTCCCTACACGACGCTCTTCCGATCTNNNCAGTTGGCTACTCTTGCTTC
AGCACACGTCTGAACTCTTTCCCTACACGACGCTCTTCCGATCTNNNCCTCAACTCACCTCACTCCCTATC
AGCACACGTCTGAACTCTTTCCCTACACGACGCTCTTCCGATCTNNNTGCTGTCTACCAAGGTGACTTT
GAGCACACGTCTGAACTCTTTCCCTACACGACGCTCTTCCGATCTNNNAGATTCGATGTGTACTCTTGC
AGCACACGTCTGAACTCTTTCCCTACACGACGCTCTTCCGATCTNNNCAAGCTCAGCGAATCCTTCTCTT
AGCACACGTCTGAACTCTTTCCCTACACGACGCTCTTCCGATCTNNNATACCTGAATGGCTGCTTATGC
GAGCACACGTCTGAACTCTTTCCCTACACGACGCTCTTCCGATCTNNNATTTGACCCATGGTCCCTTTAG
AGAGCACACGTCTGAACTCTTTCCCTACACGACGCTCTTCCGATCTNNNTGTCCTCACAAGGTAGGAG
GAGCACACGTCTGAACTCTTTCCCTACACGACGCTCTTCCGATCTNNNTCTGAGCACCAACTCATTCAAG
AGCACACGTCTGAACTCTTTCCCTACACGACGCTCTTCCGATCTNNNCAGTTATCTGTGAGCGTATGG
AGCACACGTCTGAACTCTTTCCCTACACGACGCTCTTCCGATCTNNNTGGCATGAAGATGGCACATTCT

AGAGCACACGTCTGAACTCTTTCCCTACACGACGCTCTTCCGATCTNNNCTATGTCTCACATGGCCTTT
AGCACACGTCTGAACTCTTTCCCTACACGACGCTCTTCCGATCTNNNCCTGAGTTGGCCTTTGACTGAA
GAGCACACGTCTGAACTCTTTCCCTACACGACGCTCTTCCGATCTNNNAAGAAATGTGCTCTACAGCT
GAGCACACGTCTGAACTCTTTCCCTACACGACGCTCTTCCGATCTNNNAGAAAGAAGAAGTCAAGGG
AGAGCACACGTCTGAACTCTTTCCCTACACGACGCTCTTCCGATCTNNNAGCAATCAGAGGGTTTTGG
AGCACACGTCTGAACTCTTTCCCTACACGACGCTCTTCCGATCTNNNTGTCATGGGGGTCTGTCTTGT
GAGCACACGTCTGAACTCTTTCCCTACACGACGCTCTTCCGATCTNNNTTGCCAAAACTAAATGCTGG
GAGCACACGTCTGAACTCTTTCCCTACACGACGCTCTTCCGATCTNNNTGAAAAGAACGTGAGAGGT
GAGCACACGTCTGAACTCTTTCCCTACACGACGCTCTTCCGATCTNNNACATCCTGGTCCTCTCTCTCTT
AGCACACGTCTGAACTCTTTCCCTACACGACGCTCTTCCGATCTNNNGTTGCCGATATACCCTATGACC
AGCACACGTCTGAACTCTTTCCCTACACGACGCTCTTCCGATCTNNNATGCTGTGCAAGAGAAGTGA
AGCACACGTCTGAACTCTTTCCCTACACGACGCTCTTCCGATCTNNNGGTTGTAAACTCAACTCGGAG
GAGCACACGTCTGAACTCTTTCCCTACACGACGCTCTTCCGATCTNNNAGAGCTGGAAGAAGTGGAG
AGCACACGTCTGAACTCTTTCCCTACACGACGCTCTTCCGATCTNNNATGTGTTTGCATGCCAGGTCTA
AGCACACGTCTGAACTCTTTCCCTACACGACGCTCTTCCGATCTNNNCTTACGCTTCCTTTTGGTGGAG
GAGCACACGTCTGAACTCTTTCCCTACACGACGCTCTTCCGATCTNNNGGGGAAGATATCTGCATCTG
AGCACACGTCTGAACTCTTTCCCTACACGACGCTCTTCCGATCTNNNTATCTGCTCAAATTGCGTGTGT
GAGCACACGTCTGAACTCTTTCCCTACACGACGCTCTTCCGATCTNNNAGACACCTGCCTATCTCACCAT
AGCACACGTCTGAACTCTTTCCCTACACGACGCTCTTCCGATCTNNNTCACACGCACACAGAAATACAC
GAGCACACGTCTGAACTCTTTCCCTACACGACGCTCTTCCGATCTNNNCTCATTCATCTCCCTCTCCCAGT
GAGCACACGTCTGAACTCTTTCCCTACACGACGCTCTTCCGATCTNNNGTGTGAATGGGCATGTGTT
GAGCACACGTCTGAACTCTTTCCCTACACGACGCTCTTCCGATCTNNNAGGCTGTTGCAGAGTAGAAT
GCACACGTCTGAACTCTTTCCCTACACGACGCTCTTCCGATCTNNNCCTAAGCAACGGAATGAAACCC
GAGCACACGTCTGAACTCTTTCCCTACACGACGCTCTTCCGATCTNNNCTCCAGGGAGAGATGATGTC
AGCACACGTCTGAACTCTTTCCCTACACGACGCTCTTCCGATCTNNNCAGGCACACACAAAGAAAGAT
AGCACACGTCTGAACTCTTTCCCTACACGACGCTCTTCCGATCTNNNGAGGGCAAAGAAGAGATCAC
AGCACACGTCTGAACTCTTTCCCTACACGACGCTCTTCCGATCTNNNAGGGCAATAGCCAGTTTTGTA
AGCACACGTCTGAACTCTTTCCCTACACGACGCTCTTCCGATCTNNNGTCACCTTCCAGAGAGAGAGA
GAGCACACGTCTGAACTCTTTCCCTACACGACGCTCTTCCGATCTNNNTCAGGAATGGGACCTCCTTC
GAGCACACGTCTGAACTCTTTCCCTACACGACGCTCTTCCGATCTNNNATCTTCCCCTGTGCTCTTTCTC
AGCACACGTCTGAACTCTTTCCCTACACGACGCTCTTCCGATCTNNNATTCAAGGGATCACTGTTCACC
GAGCACACGTCTGAACTCTTTCCCTACACGACGCTCTTCCGATCTNNNCACTTCCTCACCATCCTTTCCTG
GAGCACACGTCTGAACTCTTTCCCTACACGACGCTCTTCCGATCTNNNAGGATGCTCTAAGTACAGCC
AGCACACGTCTGAACTCTTTCCCTACACGACGCTCTTCCGATCTNNNACCACCTGCTATCTCATACCTCC
GAGCACACGTCTGAACTCTTTCCCTACACGACGCTCTTCCGATCTNNNAGTCTCCTAAAAAGAGCGTTG
GAGCACACGTCTGAACTCTTTCCCTACACGACGCTCTTCCGATCTNNNACTCAGAGTTTCCCTGATTCC
AGCACACGTCTGAACTCTTTCCCTACACGACGCTCTTCCGATCTNNNTTAGGGGATGCTCCTATCACT
AGCACACGTCTGAACTCTTTCCCTACACGACGCTCTTCCGATCTNNNTCTCATGCACACGAGTGTTTCA
AGAGCACACGTCTGAACTCTTTCCCTACACGACGCTCTTCCGATCTNNNTGACTTCCCAACAATCATGT
AGCACACGTCTGAACTCTTTCCCTACACGACGCTCTTCCGATCTNNNAATTAGAGGCTAGATGCCCCAC
GCACACGTCTGAACTCTTTCCCTACACGACGCTCTTCCGATCTNNNGTAGCTCTGTCTCTCTCCCTGTC
GAGCACACGTCTGAACTCTTTCCCTACACGACGCTCTTCCGATCTNNNGTCCCTCTGTCCCTCTATTCCT
AGCACACGTCTGAACTCTTTCCCTACACGACGCTCTTCCGATCTNNNTGCAACCTCAAGCCCTGTATAA

GAGCACACGTCTGAACTCTTTCCCTACACGACGCTCTTCCGATCTNNNTGAGAACCAGTCTGAACAA
AGCACACGTCTGAACTCTTTCCCTACACGACGCTCTTCCGATCTNNNGTCTCAATGAAGACGATGGG
GAGCACACGTCTGAACTCTTTCCCTACACGACGCTCTTCCGATCTNNNGATTTCTGCCTCAGCTCTTCT
AGCACACGTCTGAACTCTTTCCCTACACGACGCTCTTCCGATCTNNNTTCTGAAGCTCTTGGCTGAAG
GAGCACACGTCTGAACTCTTTCCCTACACGACGCTCTTCCGATCTNNNGGTTGCTACTGCTGCTTCAG
AGCACACGTCTGAACTCTTTCCCTACACGACGCTCTTCCGATCTNNNCTGCTCAGTGGATGGTTGTAT
AGCACACGTCTGAACTCTTTCCCTACACGACGCTCTTCCGATCTNNNCATCGGGTTGAGCTGAGAAG
GAGCACACGTCTGAACTCTTTCCCTACACGACGCTCTTCCGATCTNNNTTCAAGGTCTTTCCACTGTGC
AGCACACGTCTGAACTCTTTCCCTACACGACGCTCTTCCGATCTNNNATGGGTAAGTGTTGAAGAGG
GAGCACACGTCTGAACTCTTTCCCTACACGACGCTCTTCCGATCTNNNTGGGCTCTTTTAGTGTGTGT
AGCACACGTCTGAACTCTTTCCCTACACGACGCTCTTCCGATCTNNNAGGCAGAAGTCTTATAGCTGG
GCACACGTCTGAACTCTTTCCCTACACGACGCTCTTCCGATCTNNNAGTTTTTGTTGGGGCTTGAAGG
AGCACACGTCTGAACTCTTTCCCTACACGACGCTCTTCCGATCTNNNATCCCCACAGCAACCCTATAAG
GCACACGTCTGAACTCTTTCCCTACACGACGCTCTTCCGATCTNNNTGGTAGGTGTATTGCAAGTGG
AGCACACGTCTGAACTCTTTCCCTACACGACGCTCTTCCGATCTNNNGCAAGCAAGAAGTTAGTGCAG
AGCACACGTCTGAACTCTTTCCCTACACGACGCTCTTCCGATCTNNNGCTGTGAAAGCATGACACCAA
AGCACACGTCTGAACTCTTTCCCTACACGACGCTCTTCCGATCTNNNCACATTGCCACAAAGCCTGAAA
AGCACACGTCTGAACTCTTTCCCTACACGACGCTCTTCCGATCTNNNGTGGCCACACGAATCTAACAT
AGCACACGTCTGAACTCTTTCCCTACACGACGCTCTTCCGATCTNNNTCCTCTTCACATCAATAGGCTGA
AGCACACGTCTGAACTCTTTCCCTACACGACGCTCTTCCGATCTNNNAACCACAGCTGCTGATTTCTCA
AGCACACGTCTGAACTCTTTCCCTACACGACGCTCTTCCGATCTNNNGTTTGAGCTCTGGATGCTTCTG
AGCACACGTCTGAACTCTTTCCCTACACGACGCTCTTCCGATCTNNNATGTTAGGGCCCTCCAGAGAA
AGCACACGTCTGAACTCTTTCCCTACACGACGCTCTTCCGATCTNNNGGGCTCAGAAAGGTAACATAG
AGCACACGTCTGAACTCTTTCCCTACACGACGCTCTTCCGATCTNNNCAATGTGATGATCCAGGCACA
AGCACACGTCTGAACTCTTTCCCTACACGACGCTCTTCCGATCTNNNGAGGATCCCTTAGGCTGAGAA
AGCACACGTCTGAACTCTTTCCCTACACGACGCTCTTCCGATCTNNNAAAGTCATTCCCAGTCAAGGC
AGCACACGTCTGAACTCTTTCCCTACACGACGCTCTTCCGATCTNNNCCAAAATGAGATACGGCTGCT
AGCACACGTCTGAACTCTTTCCCTACACGACGCTCTTCCGATCTNNNATGACAAATGAGCTTCTGGGC
GCACACGTCTGAACTCTTTCCCTACACGACGCTCTTCCGATCTNNNAAATGAGCTCGCAGAACCTACA
GCACACGTCTGAACTCTTTCCCTACACGACGCTCTTCCGATCTNNNCCTATTCTCCAGAGTGGTGGGA
GAGCACACGTCTGAACTCTTTCCCTACACGACGCTCTTCCGATCTNNNGTGTGTTGTGTGCACATGCT
AGCACACGTCTGAACTCTTTCCCTACACGACGCTCTTCCGATCTNNNTTCAACTTGGCCCACTTGGAAA
AGCACACGTCTGAACTCTTTCCCTACACGACGCTCTTCCGATCTNNNTTGTTCATTGCCCTCTCCTTTCC
AGCACACGTCTGAACTCTTTCCCTACACGACGCTCTTCCGATCTNNNATGGACCCCAGGAATGGTTAC
GCACACGTCTGAACTCTTTCCCTACACGACGCTCTTCCGATCTNNNGCTGGAGAGGGAAAGAATGAC
AGCACACGTCTGAACTCTTTCCCTACACGACGCTCTTCCGATCTNNNGAAACCGGAATTCACCTTCCTG
GCACACGTCTGAACTCTTTCCCTACACGACGCTCTTCCGATCTNNNGTTTATAGCCTCTCCTCCCCTCCC
AGCACACGTCTGAACTCTTTCCCTACACGACGCTCTTCCGATCTNNNATCTGGTCTTCCCAAACTCATG
AGCACACGTCTGAACTCTTTCCCTACACGACGCTCTTCCGATCTNNNAGGGAGGGATGATCAAGTTG
GAGCACACGTCTGAACTCTTTCCCTACACGACGCTCTTCCGATCTNNNGCCATGGAAAACAGGTCATC
AGCACACGTCTGAACTCTTTCCCTACACGACGCTCTTCCGATCTNNNATCAAGGGAATCCTGGTTTCA
AGCACACGTCTGAACTCTTTCCCTACACGACGCTCTTCCGATCTNNNACAATTTCTTGGCATCTCTCCC
GCACACGTCTGAACTCTTTCCCTACACGACGCTCTTCCGATCTNNNGGTCAGACGTAGAATCCAGAC

AGCACACGTCTGAACTCTTTCCCTACACGACGCTCTTCCGATCTNNNACCTCCTGTATCAAACCCAGAGA
GAGCACACGTCTGAACTCTTTCCCTACACGACGCTCTTCCGATCTNNNGGTGGTGCTCTCCTGTAGTTTT
AGCACACGTCTGAACTCTTTCCCTACACGACGCTCTTCCGATCTNNNACACTGGTCCAAACCATTCTCAAG
GAGCACACGTCTGAACTCTTTCCCTACACGACGCTCTTCCGATCTNNNGCAAGGTACTGCCTGATGTAG
AGCACACGTCTGAACTCTTTCCCTACACGACGCTCTTCCGATCTNNNTCCCATCCGTCTAGAAGTGAAA
GAGCACACGTCTGAACTCTTTCCCTACACGACGCTCTTCCGATCTNNNCACCATCATCCCAAGGTATCCA
AGCACACGTCTGAACTCTTTCCCTACACGACGCTCTTCCGATCTNNNAGCTTTCATTGTGTTCTACTGGT
AGCACACGTCTGAACTCTTTCCCTACACGACGCTCTTCCGATCTNNNTATGGCTCTACAGAGAGAAGG
AGCACACGTCTGAACTCTTTCCCTACACGACGCTCTTCCGATCTNNNAATGAAAAGCATCACCTCCTCCA
AGCACACGTCTGAACTCTTTCCCTACACGACGCTCTTCCGATCTNNNACTGATTTCCCCAGCTTTCACTG
AGCACACGTCTGAACTCTTTCCCTACACGACGCTCTTCCGATCTNNNGTGAAGAAAGCTTGAGGTTTG
GAGCACACGTCTGAACTCTTTCCCTACACGACGCTCTTCCGATCTNNNTGGGGCATAGTAGAAACTGC
AGCACACGTCTGAACTCTTTCCCTACACGACGCTCTTCCGATCTNNNTAACTGTTCAGGATAGGGCGTC
AGCACACGTCTGAACTCTTTCCCTACACGACGCTCTTCCGATCTNNNGAGGGCCAATTCAGAAGCCAT
GAGCACACGTCTGAACTCTTTCCCTACACGACGCTCTTCCGATCTNNNGTGTGTGTGTTCACAAGCA
GAGCACACGTCTGAACTCTTTCCCTACACGACGCTCTTCCGATCTNNNGCAAAGGTTTTGAACATGGGC
AGCACACGTCTGAACTCTTTCCCTACACGACGCTCTTCCGATCTNNNGAAAAGGCCAACACCTCACAAT
GAGCACACGTCTGAACTCTTTCCCTACACGACGCTCTTCCGATCTNNNCTCTGCTCTCACTTTCATCACCC
AGCACACGTCTGAACTCTTTCCCTACACGACGCTCTTCCGATCTNNNGGATTCCGAAGGCAAAGAGCTA
GAGCACACGTCTGAACTCTTTCCCTACACGACGCTCTTCCGATCTNNNGATGATGTGGTGGGCATAAG
AGCACACGTCTGAACTCTTTCCCTACACGACGCTCTTCCGATCTNNNGATCTCCTTCAGCTGTCAACTCC
GAGCACACGTCTGAACTCTTTCCCTACACGACGCTCTTCCGATCTNNNAGACAGAAGAAAACTCTCCAA
AGCACACGTCTGAACTCTTTCCCTACACGACGCTCTTCCGATCTNNNAAGTTGGAGAGGAGGGAAACT
AGCACACGTCTGAACTCTTTCCCTACACGACGCTCTTCCGATCTNNNTCCAGGTAATGAGGCTTTAGGC
AGCACACGTCTGAACTCTTTCCCTACACGACGCTCTTCCGATCTNNNAGTCCCTTCCAAGGAATTAGCCA
AGCACACGTCTGAACTCTTTCCCTACACGACGCTCTTCCGATCTNNNTAGGCACAGAAGCCTCCTTTTTC
AGCACACGTCTGAACTCTTTCCCTACACGACGCTCTTCCGATCTNNNGAAACAGGGCTCTTTATCCAGG
GAGCACACGTCTGAACTCTTTCCCTACACGACGCTCTTCCGATCTNNNGCTTACCCTGCTGGAATATGCT
AGCACACGTCTGAACTCTTTCCCTACACGACGCTCTTCCGATCTNNNCCCCTCATTGAGAATCACTACGCC
AGCACACGTCTGAACTCTTTCCCTACACGACGCTCTTCCGATCTNNNTCTAACAAGCTCCTGGAAGGTG
GAGCACACGTCTGAACTCTTTCCCTACACGACGCTCTTCCGATCTNNNACGAGGTTCAGAAAGTGGCTT
AGCACACGTCTGAACTCTTTCCCTACACGACGCTCTTCCGATCTNNNTTTCAGCGGTAATACTTGCAGTT
AGCACACGTCTGAACTCTTTCCCTACACGACGCTCTTCCGATCTNNNGGTGCTTTCCTGTAATCCCACTT
AGCACACGTCTGAACTCTTTCCCTACACGACGCTCTTCCGATCTNNNTCTCAGAGAGCTTCTTCGGTCA
AGCACACGTCTGAACTCTTTCCCTACACGACGCTCTTCCGATCTNNNGGACGTAGGGTCCAACTTGAA
GAGCACACGTCTGAACTCTTTCCCTACACGACGCTCTTCCGATCTNNNCCCGTCACTCTACATATTCAGC
AGCACACGTCTGAACTCTTTCCCTACACGACGCTCTTCCGATCTNNNATTTTCCACCCTCATGATGCAGT
GAGCACACGTCTGAACTCTTTCCCTACACGACGCTCTTCCGATCTNNNGTGAACCTCATACAAAACCCCC
AGCACACGTCTGAACTCTTTCCCTACACGACGCTCTTCCGATCTNNNCATGTGAGACAGCACCTTGTCA
AGCACACGTCTGAACTCTTTCCCTACACGACGCTCTTCCGATCTNNNTACTTGCCCAATGAGGAGCTTT
AGCACACGTCTGAACTCTTTCCCTACACGACGCTCTTCCGATCTNNNCCTCTGTGACTGTAGGAACAAC
AGCACACGTCTGAACTCTTTCCCTACACGACGCTCTTCCGATCTNNNAAAGCTCTGCTTGGTCAGTTTGA
AGCACACGTCTGAACTCTTTCCCTACACGACGCTCTTCCGATCTNNNATATCTGAAGAACTCGCTGTGC

AGCACACGTCTGAACTCTTTCCCTACACGACGCTCTTCCGATCTNNNTTGTAGCCCCTACCAACAGAA
GAGCACACGTCTGAACTCTTTCCCTACACGACGCTCTTCCGATCTNNNCAGGGCTCTGCTACAAATGT
AGCACACGTCTGAACTCTTTCCCTACACGACGCTCTTCCGATCTNNNCTGGTGGTCACATAGGGAAT
AGCACACGTCTGAACTCTTTCCCTACACGACGCTCTTCCGATCTNNNCAACAGCAAGAGATGGTCTCC
AGCACACGTCTGAACTCTTTCCCTACACGACGCTCTTCCGATCTNNNACTCTTGCTCCTCTCACTACCAG
AGCACACGTCTGAACTCTTTCCCTACACGACGCTCTTCCGATCTNNNATGGAGGGGTATCACAACCAA
AGCACACGTCTGAACTCTTTCCCTACACGACGCTCTTCCGATCTNNNACTGTCAGCTCAACTGGGTTTT
AGCACACGTCTGAACTCTTTCCCTACACGACGCTCTTCCGATCTNNNCCTTTTTGTGACATCCTTGGTG
GAGCACACGTCTGAACTCTTTCCCTACACGACGCTCTTCCGATCTNNNTTATTCAACACACCTCTGGT
GAGCACACGTCTGAACTCTTTCCCTACACGACGCTCTTCCGATCTNNNGCCTGGGCAATAGAAGTGAG
GAGCACACGTCTGAACTCTTTCCCTACACGACGCTCTTCCGATCTNNNTTGCCATTCTCTTCCACAACA
AGCACACGTCTGAACTCTTTCCCTACACGACGCTCTTCCGATCTNNNTCAACATGGTAAGTTGGGGAC
GCACACGTCTGAACTCTTTCCCTACACGACGCTCTTCCGATCTNNNTTTCTTCGTCTTGACGGTACAGG
GAGCACACGTCTGAACTCTTTCCCTACACGACGCTCTTCCGATCTNNNTCTGACCTTAGGTAATTCGCC
AGCACACGTCTGAACTCTTTCCCTACACGACGCTCTTCCGATCTNNNGCAGAAGGTAGGCTTTTCCAT
GCACACGTCTGAACTCTTTCCCTACACGACGCTCTTCCGATCTNNNTCATCACTCAACCTCACTGTTGCC
AGCACACGTCTGAACTCTTTCCCTACACGACGCTCTTCCGATCTNNNTCAACTACCTCCATCCTAGGTC
GAGCACACGTCTGAACTCTTTCCCTACACGACGCTCTTCCGATCTNNNTAAATGGAGCAGTGAGTGA
AGCACACGTCTGAACTCTTTCCCTACACGACGCTCTTCCGATCTNNNACTTTGCATAAGCACACCACAA
AGCACACGTCTGAACTCTTTCCCTACACGACGCTCTTCCGATCTNNNCCTCTCAAGCTTCATCTCCAACC
AGCACACGTCTGAACTCTTTCCCTACACGACGCTCTTCCGATCTNNNTTCTGGGTGTGTTGTTTGTCT
GAGCACACGTCTGAACTCTTTCCCTACACGACGCTCTTCCGATCTNNNATCTAACCAACCCTTCTCCCTA
AGCACACGTCTGAACTCTTTCCCTACACGACGCTCTTCCGATCTNNNTGTCACTAAACGTCATGCATCT
AGCACACGTCTGAACTCTTTCCCTACACGACGCTCTTCCGATCTNNNTCTTTATACACAGCCCAGGCTC
AGCACACGTCTGAACTCTTTCCCTACACGACGCTCTTCCGATCTNNNCGGTCCTGGACACTAAACAAA
GAGCACACGTCTGAACTCTTTCCCTACACGACGCTCTTCCGATCTNNNGATGATCTGCCAGACACCAA
GAGCACACGTCTGAACTCTTTCCCTACACGACGCTCTTCCGATCTNNNACACAGGAGAAATCAGGAAC
AGCACACGTCTGAACTCTTTCCCTACACGACGCTCTTCCGATCTNNNACAGCTATAGGTACGGAAGT
AGCACACGTCTGAACTCTTTCCCTACACGACGCTCTTCCGATCTNNNAAACCTCTGTACAGGTATGTG
AGCACACGTCTGAACTCTTTCCCTACACGACGCTCTTCCGATCTNNNCATGCCTGATGTGCAACTACT
GAGCACACGTCTGAACTCTTTCCCTACACGACGCTCTTCCGATCTNNNACTCCTACATGTTTGTGTGTG
AGCACACGTCTGAACTCTTTCCCTACACGACGCTCTTCCGATCTNNNAGCCTGCTTTGTTTTGCATGTTT
AGAGCACACGTCTGAACTCTTTCCCTACACGACGCTCTTCCGATCTNNNGCTTCCAGGAATCTCAAAT
GAGCACACGTCTGAACTCTTTCCCTACACGACGCTCTTCCGATCTNNNTGCAGGCTAAGTATGTTCC
GAGCACACGTCTGAACTCTTTCCCTACACGACGCTCTTCCGATCTNNNCTGGGTCTGGAATCTGTGTC
AGCACACGTCTGAACTCTTTCCCTACACGACGCTCTTCCGATCTNNNCCTGTATTCAGGGAGCTTGCTT
AGCACACGTCTGAACTCTTTCCCTACACGACGCTCTTCCGATCTNNNACAGCAATTTCTGCTCTCTCAGG
AGCACACGTCTGAACTCTTTCCCTACACGACGCTCTTCCGATCTNNNAATGTCAGGGCACAGAATGT
AGCACACGTCTGAACTCTTTCCCTACACGACGCTCTTCCGATCTNNNAGCAAAAATGCAAGCCCTGAA
AGCACACGTCTGAACTCTTTCCCTACACGACGCTCTTCCGATCTNNNCCAGTATGAAGTGGTTGAACT
AGCACACGTCTGAACTCTTTCCCTACACGACGCTCTTCCGATCTNNNAGGCTTTTAGGCAGTCTAGTCC
AGCACACGTCTGAACTCTTTCCCTACACGACGCTCTTCCGATCTNNNAAAAAGCCAGACAGAGGAAA
GCACACGTCTGAACTCTTTCCCTACACGACGCTCTTCCGATCTNNNCAGTTTCCCTAAAGGGGAGAGG

GAGCACACGTCTGAACTCTTTCCCTACACGACGCTCTTCCGATCTNNNTACTGTCTCTCACCTTGATGGC
GCACACGTCTGAACTCTTTCCCTACACGACGCTCTTCCGATCTNNNCGTTGGTGGACATTTAGCTTCCT
GAGCACACGTCTGAACTCTTTCCCTACACGACGCTCTTCCGATCTNNNAATCAATGGCTCAGCAGGAAG
GAGCACACGTCTGAACTCTTTCCCTACACGACGCTCTTCCGATCTNNNAGGCAGAAGGAAGAGAAGA
GAGCACACGTCTGAACTCTTTCCCTACACGACGCTCTTCCGATCTNNNTGGCGACACCAATTTCCAACTT
AGCACACGTCTGAACTCTTTCCCTACACGACGCTCTTCCGATCTNNNACACCAATCCTCCCAAAGTAACA
GAGCACACGTCTGAACTCTTTCCCTACACGACGCTCTTCCGATCTNNNTATCTGGCCCACCTATTCCTAG
AGAGCACACGTCTGAACTCTTTCCCTACACGACGCTCTTCCGATCTNNNCTGGAACTTTTCTCCACAGG
GCACACGTCTGAACTCTTTCCCTACACGACGCTCTTCCGATCTNNNTCAATTTTGAAAGCCCTGGTGGT
AGCACACGTCTGAACTCTTTCCCTACACGACGCTCTTCCGATCTNNNGAGCTTTACCCTCATCTCCCTCT
AGCACACGTCTGAACTCTTTCCCTACACGACGCTCTTCCGATCTNNNTAAGTGAAGACGGGGTGCTGA
AGCACACGTCTGAACTCTTTCCCTACACGACGCTCTTCCGATCTNNNATCTGTGTCATCCTTTGGAAGCA
AGCACACGTCTGAACTCTTTCCCTACACGACGCTCTTCCGATCTNNNAAAAGCCTGTAAACCACCAACCA
CACACGTCTGAACTCTTTCCCTACACGACGCTCTTCCGATCTNNNCTCTAAGGATGTCCCCACCTTCCCA
GCACACGTCTGAACTCTTTCCCTACACGACGCTCTTCCGATCTNNNTTTCCTTCTCTTCCCCATGTGAGCA
GCACACGTCTGAACTCTTTCCCTACACGACGCTCTTCCGATCTNNNGTCTACGAAAATCACCGAGCACT
GCACACGTCTGAACTCTTTCCCTACACGACGCTCTTCCGATCTNNNGCTATTGCTGTGATGATGGTTG
AGCACACGTCTGAACTCTTTCCCTACACGACGCTCTTCCGATCTNNNACAGATGTTCAGGGAGGAATG
GCACACGTCTGAACTCTTTCCCTACACGACGCTCTTCCGATCTNNNCAAACTCACACAACCAGGAAGGA
AGCACACGTCTGAACTCTTTCCCTACACGACGCTCTTCCGATCTNNNGGCAATAGGAAAGTTTGAAGG
AGCACACGTCTGAACTCTTTCCCTACACGACGCTCTTCCGATCTNNNCACACATGCCTTGATAAAACGG
AGCACACGTCTGAACTCTTTCCCTACACGACGCTCTTCCGATCTNNNGGTCATTTCACACTTCTGCTTGG
GAGCACACGTCTGAACTCTTTCCCTACACGACGCTCTTCCGATCTNNNAGTTCCCTAGGTCCCTTTCACT
AGCACACGTCTGAACTCTTTCCCTACACGACGCTCTTCCGATCTNNNACCCAATTTCCTTCATGCTCACA
GAGCACACGTCTGAACTCTTTCCCTACACGACGCTCTTCCGATCTNNNTCATGCACATGTAAGTGTTTG
GAGCACACGTCTGAACTCTTTCCCTACACGACGCTCTTCCGATCTNNNATGGTCAGATTGATGGTCTGC
GAGCACACGTCTGAACTCTTTCCCTACACGACGCTCTTCCGATCTNNNCTCTACAGTGTGATGCTGGCT
AGCACACGTCTGAACTCTTTCCCTACACGACGCTCTTCCGATCTNNNCATCTCAGGTTCTGACAGTTGG
AGCACACGTCTGAACTCTTTCCCTACACGACGCTCTTCCGATCTNNNACTGGACATGGAAGAGTTGTC
AGAGCACACGTCTGAACTCTTTCCCTACACGACGCTCTTCCGATCTNNNGATGTGGCTCAGGAAGCTTA
AGCACACGTCTGAACTCTTTCCCTACACGACGCTCTTCCGATCTNNNGTTCAGGCACAGCTAGAATCCA
GAGCACACGTCTGAACTCTTTCCCTACACGACGCTCTTCCGATCTNNNAGGAATAGTGCCCATTGTGT
GAGCACACGTCTGAACTCTTTCCCTACACGACGCTCTTCCGATCTNNNAGGCTGCTTCTAGTCTTCTGC
GAGCACACGTCTGAACTCTTTCCCTACACGACGCTCTTCCGATCTNNNAGAATAAGCAGTATTGTGGC
GAGCACACGTCTGAACTCTTTCCCTACACGACGCTCTTCCGATCTNNNATCCATAATCAACAGAGCTGG
AGCACACGTCTGAACTCTTTCCCTACACGACGCTCTTCCGATCTNNNCATGCAAATGAGCATTTTGTGG
AGCACACGTCTGAACTCTTTCCCTACACGACGCTCTTCCGATCTNNNAGCAAAGTGAGAAAGAGCTGT
GAGCACACGTCTGAACTCTTTCCCTACACGACGCTCTTCCGATCTNNNAGTATGGCTTTTACAGACTGG
AGCACACGTCTGAACTCTTTCCCTACACGACGCTCTTCCGATCTNNNAATTCCCTTCACCCACTGCATTT
GAGCACACGTCTGAACTCTTTCCCTACACGACGCTCTTCCGATCTNNNCAGTGGTGTATTTGGCTACA
AGCACACGTCTGAACTCTTTCCCTACACGACGCTCTTCCGATCTNNNACCCTGTGTTGAATGATCATGG
GAGCACACGTCTGAACTCTTTCCCTACACGACGCTCTTCCGATCTNNNTTTAGCCTTCCTGTACACTCTGT
AGCACACGTCTGAACTCTTTCCCTACACGACGCTCTTCCGATCTNNNCTCCTTCTCCCACTTCCTCTCTTCA

GCACACGTCTGAACTCTTTCCCTACACGACGCTCTTCCGATCTNNNATAAGCAACCACAGATCCACACC

GCACACGTCTGAACTCTTTCCCTACACGACGCTCTTCCGATCTNNNTAGCAAAGCATTCCTTTGCCAC

AGCACACGTCTGAACTCTTTCCCTACACGACGCTCTTCCGATCTNNNTTACTACATTGCAGAAGCCCAG

GCACACGTCTGAACTCTTTCCCTACACGACGCTCTTCCGATCTNNNCAAGACTGAAACTCCGTGTCAA

GCACACGTCTGAACTCTTTCCCTACACGACGCTCTTCCGATCTNNNGTTTGCAAAGAGGTGGAGCAA

AGCACACGTCTGAACTCTTTCCCTACACGACGCTCTTCCGATCTNNNCTCTTGCCTGAACTATCCTACG

GCACACGTCTGAACTCTTTCCCTACACGACGCTCTTCCGATCTNNNCTCCCCACTCCCTAAAAAGCAAA

GCACACGTCTGAACTCTTTCCCTACACGACGCTCTTCCGATCTNNNTAAATGATGGCTGACAAGCCTC

AGCACACGTCTGAACTCTTTCCCTACACGACGCTCTTCCGATCTNNNTAGTGTCCTTGAAGGCTCTGG

GCACACGTCTGAACTCTTTCCCTACACGACGCTCTTCCGATCTNNNACTGTTGACTGCCTACCCAAAAT

GCACACGTCTGAACTCTTTCCCTACACGACGCTCTTCCGATCTNNNTTGCCTTAGTTTTTGGTTTGGA

GCACACGTCTGAACTCTTTCCCTACACGACGCTCTTCCGATCTNNNCAACCTCTACTGGTCATCTGCAC

AGCACACGTCTGAACTCTTTCCCTACACGACGCTCTTCCGATCTNNNGTTGCTTATTGGAAGGACATG

AGCACACGTCTGAACTCTTTCCCTACACGACGCTCTTCCGATCTNNNTAATGTGCATGCTCTCTCTCTCT

GAGCACACGTCTGAACTCTTTCCCTACACGACGCTCTTCCGATCTNNNACAAACTGAGGGAGAGGAA

GAGCACACGTCTGAACTCTTTCCCTACACGACGCTCTTCCGATCTNNNTGGGAGAAAATAATTGCAGT

AGCACACGTCTGAACTCTTTCCCTACACGACGCTCTTCCGATCTNNNAAATAAAGGAAGGGTGGTGG

AGAGCACACGTCTGAACTCTTTCCCTACACGACGCTCTTCCGATCTNNNCAAGGTCACAACAGATACT

AGCACACGTCTGAACTCTTTCCCTACACGACGCTCTTCCGATCTNNNACTTTTCTCAGGTGGCAAGTG

AGCACACGTCTGAACTCTTTCCCTACACGACGCTCTTCCGATCTNNNGGTGGGGTATTATGGCCACTT

GAGCACACGTCTGAACTCTTTCCCTACACGACGCTCTTCCGATCTNNNCCGCCCACCACTTTACCAAAT

AGCACACGTCTGAACTCTTTCCCTACACGACGCTCTTCCGATCTNNNAACCTACTCCAAGCATCCTTCCT

AGCACACGTCTGAACTCTTTCCCTACACGACGCTCTTCCGATCTNNNACTCCACACATGCACGATTCTT

GAGCACACGTCTGAACTCTTTCCCTACACGACGCTCTTCCGATCTNNNGCTTTTCAAAGAGGTGGTTT

AGCACACGTCTGAACTCTTTCCCTACACGACGCTCTTCCGATCTNNNATTCACAGCCTCTTCCTGAATG

AGCACACGTCTGAACTCTTTCCCTACACGACGCTCTTCCGATCTNNNGCCAGGTTCTTCCAAAGATGTC

GCACACGTCTGAACTCTTTCCCTACACGACGCTCTTCCGATCTNNNTGCATCTCCAGATTCACACCCTA

GCACACGTCTGAACTCTTTCCCTACACGACGCTCTTCCGATCTNNNATTCTAACCACTCCCTCTCCACCC

CACACGTCTGAACTCTTTCCCTACACGACGCTCTTCCGATCTNNNTCTTCCAAAATCAAAGCCCACTGC

GAGCACACGTCTGAACTCTTTCCCTACACGACGCTCTTCCGATCTNNNAAGGAAACCTCCAGCTTTGT

AGCACACGTCTGAACTCTTTCCCTACACGACGCTCTTCCGATCTNNNTTCTGAGGGGTCTACCTCCTA

AGCACACGTCTGAACTCTTTCCCTACACGACGCTCTTCCGATCTNNNTCAGCAACACAGCCAAATCTT

AGCACACGTCTGAACTCTTTCCCTACACGACGCTCTTCCGATCTNNNTCATTCTTTTTCCCCCATCCTGA

AGAGCACACGTCTGAACTCTTTCCCTACACGACGCTCTTCCGATCTNNNCACATATGGGAGGAACTCT

GAGCACACGTCTGAACTCTTTCCCTACACGACGCTCTTCCGATCTNNNACATGCAAAAGAGTGCTAGG

GCACACGTCTGAACTCTTTCCCTACACGACGCTCTTCCGATCTNNNAGAGCACACTATCCAGGTGATG

AGCACACGTCTGAACTCTTTCCCTACACGACGCTCTTCCGATCTNNNCTGTAGACCTTGGGATTTGGC

GAGCACACGTCTGAACTCTTTCCCTACACGACGCTCTTCCGATCTNNNTGGATGTGAGCAACTAGAG

AGCACACGTCTGAACTCTTTCCCTACACGACGCTCTTCCGATCTNNNTTGGTGTTTCCCCTCACTCCTT

AGCACACGTCTGAACTCTTTCCCTACACGACGCTCTTCCGATCTNNNCCACATATGCGCACACATCTCT

AGCACACGTCTGAACTCTTTCCCTACACGACGCTCTTCCGATCTNNNTCCAACTATTCTCTCCCCCAAC

GCACACGTCTGAACTCTTTCCCTACACGACGCTCTTCCGATCTNNNGCTCTCTATCTGGGCAGCAAAA

GCACACGTCTGAACTCTTTCCCTACACGACGCTCTTCCGATCTNNNCAAGGCATTACTTGCCAACTGT

AGCACACGTCTGAACTCTTTCCCTACACGACGCTCTTCCGATCTNNNATTGATTCTGAGGAGGCAGGG
AGCACACGTCTGAACTCTTTCCCTACACGACGCTCTTCCGATCTNNNTTCATTACCATCACTGCCACAG
GAGCACACGTCTGAACTCTTTCCCTACACGACGCTCTTCCGATCTNNNATTGTTGGTAGGAATCACAG
GAGCACACGTCTGAACTCTTTCCCTACACGACGCTCTTCCGATCTNNNGTATCAGCCTCTTGCTTCTAG
AGCACACGTCTGAACTCTTTCCCTACACGACGCTCTTCCGATCTNNNAGGAATCAGCTTTGAGGAGA
AGCACACGTCTGAACTCTTTCCCTACACGACGCTCTTCCGATCTNNNAGTTCTTTTGGGGGACCCTTT
AGCACACGTCTGAACTCTTTCCCTACACGACGCTCTTCCGATCTNNNGTGGGAGAAGATGGTACAAG
AGCACACGTCTGAACTCTTTCCCTACACGACGCTCTTCCGATCTNNNGGTCAGCAGGTTTCTGAGAAC
GAGCACACGTCTGAACTCTTTCCCTACACGACGCTCTTCCGATCTNNNCCTTTGTTCTCTCCTACTCCCC
AGCACACGTCTGAACTCTTTCCCTACACGACGCTCTTCCGATCTNNNTTCCCTGCCAACTACATCAACTC
GAGCACACGTCTGAACTCTTTCCCTACACGACGCTCTTCCGATCTNNNTGGCCACATACAGAACTGCTT
GAGCACACGTCTGAACTCTTTCCCTACACGACGCTCTTCCGATCTNNNTCTTTGTCTTTCTGGCTGGCTT
GCACACGTCTGAACTCTTTCCCTACACGACGCTCTTCCGATCTNNNGCACAAGAATTCCAAAAGGCAC
GAGCACACGTCTGAACTCTTTCCCTACACGACGCTCTTCCGATCTNNNCCTTATCTCCTACCCATCCCCAC
AGCACACGTCTGAACTCTTTCCCTACACGACGCTCTTCCGATCTNNNACAAGACTATCCCTGATGTCTC
GCACACGTCTGAACTCTTTCCCTACACGACGCTCTTCCGATCTNNNTCTCCAGCACTGTATCCACTTGA
GCACACGTCTGAACTCTTTCCCTACACGACGCTCTTCCGATCTNNNGCAGTGACCTTGTGTATTGTGT
AGCACACGTCTGAACTCTTTCCCTACACGACGCTCTTCCGATCTNNNCCTCAAAAGTGCAGTGAGTTC
GCACACGTCTGAACTCTTTCCCTACACGACGCTCTTCCGATCTNNNGTTAGTGGCCAGAATGTGGGA
GAGCACACGTCTGAACTCTTTCCCTACACGACGCTCTTCCGATCTNNNCCCCACAAAGATCCCAATAGA
GAGCACACGTCTGAACTCTTTCCCTACACGACGCTCTTCCGATCTNNNAGGAGATGCAGGCAAGGTA
GCACACGTCTGAACTCTTTCCCTACACGACGCTCTTCCGATCTNNNAGGTCCAAGCTGTTGAAGGAA
AGCACACGTCTGAACTCTTTCCCTACACGACGCTCTTCCGATCTNNNCACATTTGAGTTTTGCAGGGT
GAGCACACGTCTGAACTCTTTCCCTACACGACGCTCTTCCGATCTNNNACCTCGTCTGAAAAACAATGA
GCACACGTCTGAACTCTTTCCCTACACGACGCTCTTCCGATCTNNNCCTGTCTTTTTCCACCAGTGTCA
GAGCACACGTCTGAACTCTTTCCCTACACGACGCTCTTCCGATCTNNNCAGGTGCCAATTCCCCTTTAG
AGCACACGTCTGAACTCTTTCCCTACACGACGCTCTTCCGATCTNNNTATAATGTCCTCCGAGTTTGCC
AGCACACGTCTGAACTCTTTCCCTACACGACGCTCTTCCGATCTNNNTAGGGACTGAGACCAAAGGA
GAGCACACGTCTGAACTCTTTCCCTACACGACGCTCTTCCGATCTNNNGGGAGGGATGTAGGAGAG
GAGCACACGTCTGAACTCTTTCCCTACACGACGCTCTTCCGATCTNNNTAGCACATGCCCATAGTCCCA
AGCACACGTCTGAACTCTTTCCCTACACGACGCTCTTCCGATCTNNNCAGAATTTCCTCGTGGGTGGA
GAGCACACGTCTGAACTCTTTCCCTACACGACGCTCTTCCGATCTNNNGGAAGTAGAGGGCTAAGCG
GAGCACACGTCTGAACTCTTTCCCTACACGACGCTCTTCCGATCTNNNTGTACATTGACATTGGGGTT
AGCACACGTCTGAACTCTTTCCCTACACGACGCTCTTCCGATCTNNNGCTGGTTAGACAGCATGAGTG
AGCACACGTCTGAACTCTTTCCCTACACGACGCTCTTCCGATCTNNNGAACCCCAACAAGATACAGAG
GAGCACACGTCTGAACTCTTTCCCTACACGACGCTCTTCCGATCTNNNAATGCAACTCTCTTTTCTCCGG
GAGCACACGTCTGAACTCTTTCCCTACACGACGCTCTTCCGATCTNNNCTTGGCTCCATCTTCAGTCTA
AGCACACGTCTGAACTCTTTCCCTACACGACGCTCTTCCGATCTNNNGATTGGGACCTCAAAGTGAG
GCACACGTCTGAACTCTTTCCCTACACGACGCTCTTCCGATCTNNNGATTCCCTGAGCCACTTTCAGA
AGCACACGTCTGAACTCTTTCCCTACACGACGCTCTTCCGATCTNNNCTAACTGGGGGTTCTTTCTGC
GAGCACACGTCTGAACTCTTTCCCTACACGACGCTCTTCCGATCTNNNATGATAAGGGACCATGAAGG
AGCACACGTCTGAACTCTTTCCCTACACGACGCTCTTCCGATCTNNNTAAACACTAAAACCTCCGTGG
AGCACACGTCTGAACTCTTTCCCTACACGACGCTCTTCCGATCTNNNAGTCCGTGCATATCCAAGCAT

GAGCACACGTCTGAACTCTTTCCCTACACGACGCTCTTCCGATCTNNNCTGCATTGCCTTCTGTGGTAT
AGCACACGTCTGAACTCTTTCCCTACACGACGCTCTTCCGATCTNNNCCTGGTGATGAAATCTGTCCC
GCACACGTCTGAACTCTTTCCCTACACGACGCTCTTCCGATCTNNNTTGTTTTGGTTTGCCATTTCCCCC
AGCACACGTCTGAACTCTTTCCCTACACGACGCTCTTCCGATCTNNNAGACCCATCAGAGATTTGTCCC
AGCACACGTCTGAACTCTTTCCCTACACGACGCTCTTCCGATCTNNNGCTTACCGCAGAGATAAGACAC
AGCACACGTCTGAACTCTTTCCCTACACGACGCTCTTCCGATCTNNNAGGCAGTTGAGAGGAAAACC
GAGCACACGTCTGAACTCTTTCCCTACACGACGCTCTTCCGATCTNNNACCAGAACAGAAACACTTTGT
GCACACGTCTGAACTCTTTCCCTACACGACGCTCTTCCGATCTNNNCCATCTGGAAACAGGTTGAGGT
GCACACGTCTGAACTCTTTCCCTACACGACGCTCTTCCGATCTNNNTCTGACACATCATGGCCACAAAC
GCACACGTCTGAACTCTTTCCCTACACGACGCTCTTCCGATCTNNNATTTACCTACCACCTTGTAGTGG
AGCACACGTCTGAACTCTTTCCCTACACGACGCTCTTCCGATCTNNNAACTGACCATCTCAACCTACCC
GCACACGTCTGAACTCTTTCCCTACACGACGCTCTTCCGATCTNNNAGTTGGCAGAAGATAACCACTG
AGCACACGTCTGAACTCTTTCCCTACACGACGCTCTTCCGATCTNNNTCACACAACAGCAGGATCATC
AGCACACGTCTGAACTCTTTCCCTACACGACGCTCTTCCGATCTNNNATCTCTTTCTCTCTCGCTCTGGC
AGCACACGTCTGAACTCTTTCCCTACACGACGCTCTTCCGATCTNNNCCATGGCATGTAGGTAGGGT
GCACACGTCTGAACTCTTTCCCTACACGACGCTCTTCCGATCTNNNCCACTCCCTTCTCTCCCAAAATGC
AGCACACGTCTGAACTCTTTCCCTACACGACGCTCTTCCGATCTNNNCCAACATCCTTCCCTACTTCAGC
GCACACGTCTGAACTCTTTCCCTACACGACGCTCTTCCGATCTNNNGTTCAGGTCCTGTCGCCTAAAA
AGCACACGTCTGAACTCTTTCCCTACACGACGCTCTTCCGATCTNNNCAATTTGAATCTGGGCTTGGCT
GCACACGTCTGAACTCTTTCCCTACACGACGCTCTTCCGATCTNNNAGGGCTTACCAAGCTGAATGAT
AGCACACGTCTGAACTCTTTCCCTACACGACGCTCTTCCGATCTNNNTGATTGACAGTGGAGTAGCCA
GAGCACACGTCTGAACTCTTTCCCTACACGACGCTCTTCCGATCTNNNTCCACTTTCTCCTGAAACAGCT
GCACACGTCTGAACTCTTTCCCTACACGACGCTCTTCCGATCTNNNCCTGTCCCAAATACTTACCCAGCC
GCACACGTCTGAACTCTTTCCCTACACGACGCTCTTCCGATCTNNNAGACTGAGGCCCTACTTTGCTT
AGCACACGTCTGAACTCTTTCCCTACACGACGCTCTTCCGATCTNNNATTTCTCCCATGGTATGTGCGT
GAGCACACGTCTGAACTCTTTCCCTACACGACGCTCTTCCGATCTNNNAAGATCCAACTGGCATGATTG
GAGCACACGTCTGAACTCTTTCCCTACACGACGCTCTTCCGATCTNNNCAAAGAAAAGACCAGGCACA
GCACACGTCTGAACTCTTTCCCTACACGACGCTCTTCCGATCTNNNTCTCTACTTCTGCTTCTTGGGCT
AGCACACGTCTGAACTCTTTCCCTACACGACGCTCTTCCGATCTNNNAAGAGAAAAGAAGGGGCATG
AGCACACGTCTGAACTCTTTCCCTACACGACGCTCTTCCGATCTNNNTGATCCCAGAATTTCAAGGCC
GCACACGTCTGAACTCTTTCCCTACACGACGCTCTTCCGATCTNNNGCTGTTATGGCAGGAATGGGT
GAGCACACGTCTGAACTCTTTCCCTACACGACGCTCTTCCGATCTNNNGAGGAAAGACACAGATGCAC
AGCACACGTCTGAACTCTTTCCCTACACGACGCTCTTCCGATCTNNNATTGACAGGAGCCTGGGAAG
AGCACACGTCTGAACTCTTTCCCTACACGACGCTCTTCCGATCTNNNGCTGCAAAGAGGCTGATTTCA
GCACACGTCTGAACTCTTTCCCTACACGACGCTCTTCCGATCTNNNGTCATTTTCCTGGCTTGCAGATG
GCACACGTCTGAACTCTTTCCCTACACGACGCTCTTCCGATCTNNNGGAACCTTCTTGAGGCCATACA
GAGCACACGTCTGAACTCTTTCCCTACACGACGCTCTTCCGATCTNNNCTTGCAGCTACACATGGACTT
GCACACGTCTGAACTCTTTCCCTACACGACGCTCTTCCGATCTNNNGGAACACTCAGTCTCTTCAGCAG
GCACACGTCTGAACTCTTTCCCTACACGACGCTCTTCCGATCTNNNATGGTCAGGATGTGCTCTGGTT
AGCACACGTCTGAACTCTTTCCCTACACGACGCTCTTCCGATCTNNNCCTGGTTGGCTCAAAACTACCT
AGCACACGTCTGAACTCTTTCCCTACACGACGCTCTTCCGATCTNNNCTCTCTTCCATTGGTGCTCAGTC
AGCACACGTCTGAACTCTTTCCCTACACGACGCTCTTCCGATCTNNNATCCATAGTTGTGTACTGGGG
GCACACGTCTGAACTCTTTCCCTACACGACGCTCTTCCGATCTNNNTGGTTTCAATGGTCTAAGGCTG

AGCACACGTCTGAACTCTTTCCCTACACGACGCTCTTCCGATCTNNNACATAACCTAACTGGCCCCTTGA
AGCACACGTCTGAACTCTTTCCCTACACGACGCTCTTCCGATCTNNNATATGAATCACTTGGATGGCTG
GAGCACACGTCTGAACTCTTTCCCTACACGACGCTCTTCCGATCTNNNCTGTAGCTTCAGACCCATTGA
AGCACACGTCTGAACTCTTTCCCTACACGACGCTCTTCCGATCTNNNTTGTTCCAAAAAGCTGTGAAG
AGCACACGTCTGAACTCTTTCCCTACACGACGCTCTTCCGATCTNNNGAAATTTCTAGCCTCCCTGTGG
AGCACACGTCTGAACTCTTTCCCTACACGACGCTCTTCCGATCTNNNGGCTCTTCTCAAGCATTTTCCCT
AGCACACGTCTGAACTCTTTCCCTACACGACGCTCTTCCGATCTNNNGCCAGCAGGAAAGTGCAATAA
AGCACACGTCTGAACTCTTTCCCTACACGACGCTCTTCCGATCTNNNCCCTTCTACTGCAGCCAAATGAT
AGCACACGTCTGAACTCTTTCCCTACACGACGCTCTTCCGATCTNNNGACCAGAAGGGAGGAAAATGG
GCACACGTCTGAACTCTTTCCCTACACGACGCTCTTCCGATCTNNNTTTCAGCTTTATTTGCGTGGCAG
AGCACACGTCTGAACTCTTTCCCTACACGACGCTCTTCCGATCTNNNTTCAGTGGAGCTTCTGGACT
GCACACGTCTGAACTCTTTCCCTACACGACGCTCTTCCGATCTNNNGATGCAGATGGCCAATAGTGGA
AGCACACGTCTGAACTCTTTCCCTACACGACGCTCTTCCGATCTNNNCTTCCAGGGGATTCTCATGTTG
AGCACACGTCTGAACTCTTTCCCTACACGACGCTCTTCCGATCTNNNTCAAGTACCAACCTAGAGGCTT
GAGCACACGTCTGAACTCTTTCCCTACACGACGCTCTTCCGATCTNNNTCCTCTAAATTGCAATACGCCC
AGCACACGTCTGAACTCTTTCCCTACACGACGCTCTTCCGATCTNNNACTTCTGATGACCACTCCCTCAA
GCACACGTCTGAACTCTTTCCCTACACGACGCTCTTCCGATCTNNNGCAACCCTCTCTCTTTCTACAGGC
GCACACGTCTGAACTCTTTCCCTACACGACGCTCTTCCGATCTNNNTGGGTTGGAAGAGGTGTTTTTG
GAGCACACGTCTGAACTCTTTCCCTACACGACGCTCTTCCGATCTNNNGCATCCTTTTGTGTGACTGTG
GCACACGTCTGAACTCTTTCCCTACACGACGCTCTTCCGATCTNNNAGTCTTCACAAGCGCTGATCATT
AGCACACGTCTGAACTCTTTCCCTACACGACGCTCTTCCGATCTNNNTTTCCTCCCTATTATCGCCAGGT
GAGCACACGTCTGAACTCTTTCCCTACACGACGCTCTTCCGATCTNNNGGAGGGCACTAGTGAAGTCC
AGCACACGTCTGAACTCTTTCCCTACACGACGCTCTTCCGATCTNNNTACCCCATTGGCTTTGTTGTTG
AGCACACGTCTGAACTCTTTCCCTACACGACGCTCTTCCGATCTNNNTTTTCTCCAGCCTTTGCCTTCTTC
AGCACACGTCTGAACTCTTTCCCTACACGACGCTCTTCCGATCTNNNAAACCATGATACAGGTGCCAGA
GCACACGTCTGAACTCTTTCCCTACACGACGCTCTTCCGATCTNNNGTTCAGGTTGATGGGAGGGAA
GCACACGTCTGAACTCTTTCCCTACACGACGCTCTTCCGATCTNNNTACTTCTGTGAGCAAGAAGGAA
GCACACGTCTGAACTCTTTCCCTACACGACGCTCTTCCGATCTNNNGCAGACCATGACTTCCTTTGCTA
GCACACGTCTGAACTCTTTCCCTACACGACGCTCTTCCGATCTNNNTGCAAGTCACAAGCTGAATCCTT
GAGCACACGTCTGAACTCTTTCCCTACACGACGCTCTTCCGATCTNNNGGATGCCTCAGTTCTCTCTCT
AGCACACGTCTGAACTCTTTCCCTACACGACGCTCTTCCGATCTNNNGTAGAAAGAACTGACCAGCCA
GAGCACACGTCTGAACTCTTTCCCTACACGACGCTCTTCCGATCTNNNGGATTGGATGGGGATAAAT
AGCACACGTCTGAACTCTTTCCCTACACGACGCTCTTCCGATCTNNNGTCAGATCCCTCCCTCATTGTCT
AGCACACGTCTGAACTCTTTCCCTACACGACGCTCTTCCGATCTNNNCCTTGACATGTTAGTGTGGAGG
GAGCACACGTCTGAACTCTTTCCCTACACGACGCTCTTCCGATCTNNNCTCCCTCTGGGAGATAGTGTG
AGCACACGTCTGAACTCTTTCCCTACACGACGCTCTTCCGATCTNNNGTCCTCATGATGCATTGCCCTAA
AGCACACGTCTGAACTCTTTCCCTACACGACGCTCTTCCGATCTNNNCCAGCAGCCAAAGAGAAAACA
AGCACACGTCTGAACTCTTTCCCTACACGACGCTCTTCCGATCTNNNTATCTGCTCGGTCAATCAGGCTA
GAGCACACGTCTGAACTCTTTCCCTACACGACGCTCTTCCGATCTNNNATGGCAGTTGTTTCGTCTCTT
AGCACACGTCTGAACTCTTTCCCTACACGACGCTCTTCCGATCTNNNCCATGGCCTAGTTGTTGATTCCA
AGCACACGTCTGAACTCTTTCCCTACACGACGCTCTTCCGATCTNNNACCTTGGCAGTGATTTCTTCTCC
AGCACACGTCTGAACTCTTTCCCTACACGACGCTCTTCCGATCTNNNCGGATTCAGAATAGAGGGAGG
GAGCACACGTCTGAACTCTTTCCCTACACGACGCTCTTCCGATCTNNNACAGTTTGGGATGGCATTTA

GCACACGTCTGAACTCTTTCCCTACACGACGCTCTTCCGATCTNNNAGAATGATTATTCCCACACAGG
GCACACGTCTGAACTCTTTCCCTACACGACGCTCTTCCGATCTNNNCCATGATAACAAGGAGAAGCGC
AGCACACGTCTGAACTCTTTCCCTACACGACGCTCTTCCGATCTNNNTATCCCAGACAGGGAGAACAG
GCACACGTCTGAACTCTTTCCCTACACGACGCTCTTCCGATCTNNNTCTCTGGGTGGTCTTGCTCAAA
AGCACACGTCTGAACTCTTTCCCTACACGACGCTCTTCCGATCTNNNAAACTCCTAAGAGGGAGGCT
GCACACGTCTGAACTCTTTCCCTACACGACGCTCTTCCGATCTNNNGTTTCTGAGTTCATCCTGGCCTT
AGCACACGTCTGAACTCTTTCCCTACACGACGCTCTTCCGATCTNNNTCTTCTCTTTGCTCCATTGGCAT
AGCACACGTCTGAACTCTTTCCCTACACGACGCTCTTCCGATCTNNNCCAGAAGGAGAGAAGTGTGT
GAGCACACGTCTGAACTCTTTCCCTACACGACGCTCTTCCGATCTNNNGAGTAGCTTGGACTTTAGGG
AGCACACGTCTGAACTCTTTCCCTACACGACGCTCTTCCGATCTNNNGCCTTCATTGATGCGTTCAGAT
CACACGTCTGAACTCTTTCCCTACACGACGCTCTTCCGATCTNNNCTACCTTACTCACCTTTGCCTGCCA
AGCACACGTCTGAACTCTTTCCCTACACGACGCTCTTCCGATCTNNNTACATTTGTAGGCCCCTGTCAG
AGCACACGTCTGAACTCTTTCCCTACACGACGCTCTTCCGATCTNNNCTTTCTTCAGTGTGGAGCCTG
AGCACACGTCTGAACTCTTTCCCTACACGACGCTCTTCCGATCTNNNTTTCAGTTTCTGCTCTGGAGCT
AGCACACGTCTGAACTCTTTCCCTACACGACGCTCTTCCGATCTNNNTCTGCTGGTTCTCACTGATGGA
GCACACGTCTGAACTCTTTCCCTACACGACGCTCTTCCGATCTNNNCAGACTTTGAGCCCTGCTAACA
GCACACGTCTGAACTCTTTCCCTACACGACGCTCTTCCGATCTNNNATTTTGTGTCTTTCCCAAACGCC
GAGCACACGTCTGAACTCTTTCCCTACACGACGCTCTTCCGATCTNNNTCTTGGATGTGTATGTGCTTG
GAGCACACGTCTGAACTCTTTCCCTACACGACGCTCTTCCGATCTNNNGCTTGATTGTTGTAAGGCCTG
AGCACACGTCTGAACTCTTTCCCTACACGACGCTCTTCCGATCTNNNCTGAAGCAGTACCCTCACTTAG
GAGCACACGTCTGAACTCTTTCCCTACACGACGCTCTTCCGATCTNNNTAAAATCTGCCCCGATTGGG
GAGCACACGTCTGAACTCTTTCCCTACACGACGCTCTTCCGATCTNNNTAGGACCTTGCTCTTACCCAC
AGCACACGTCTGAACTCTTTCCCTACACGACGCTCTTCCGATCTNNNCTTGGCTAATCAGTACCACACC
AGCACACGTCTGAACTCTTTCCCTACACGACGCTCTTCCGATCTNNNGGAGACAAGATTGAGAGCCA
GCACACGTCTGAACTCTTTCCCTACACGACGCTCTTCCGATCTNNNGAGCCATTCACTTATTCCTGCCA
GAGCACACGTCTGAACTCTTTCCCTACACGACGCTCTTCCGATCTNNNGGACATCTTGTGTTGGCATTT
AGCACACGTCTGAACTCTTTCCCTACACGACGCTCTTCCGATCTNNNAAGGCATTTTGGCAATTCACTC
GCACACGTCTGAACTCTTTCCCTACACGACGCTCTTCCGATCTNNNGAAGCTCTTTCTGACTGGCCTA
AGCACACGTCTGAACTCTTTCCCTACACGACGCTCTTCCGATCTNNNCCAACGTACCACACACATACAC
AGCACACGTCTGAACTCTTTCCCTACACGACGCTCTTCCGATCTNNNATGAATCTCTGGCTGGACATA
AGCACACGTCTGAACTCTTTCCCTACACGACGCTCTTCCGATCTNNNCGGAAACTTCGGAATAGACCA
GCACACGTCTGAACTCTTTCCCTACACGACGCTCTTCCGATCTNNNGACTTGGCCTCTGGTTCATTTCA
GCACACGTCTGAACTCTTTCCCTACACGACGCTCTTCCGATCTNNNATTAGGCTTTGTTTGTCCCAGCA
AGCACACGTCTGAACTCTTTCCCTACACGACGCTCTTCCGATCTNNNGACGGGTGACATTTTCATCTC
GAGCACACGTCTGAACTCTTTCCCTACACGACGCTCTTCCGATCTNNNGACACAAACTCCTCCCACTCTC
GCACACGTCTGAACTCTTTCCCTACACGACGCTCTTCCGATCTNNNGCCTCCTTTACACAACCACCAAT
GAGCACACGTCTGAACTCTTTCCCTACACGACGCTCTTCCGATCTNNNAATGATATTCTCCCCACCTCCC
AGCACACGTCTGAACTCTTTCCCTACACGACGCTCTTCCGATCTNNNGCAAACAACATTGACATGGCT
AGCACACGTCTGAACTCTTTCCCTACACGACGCTCTTCCGATCTNNNATGCTAAGCTGAGGTTCCCAAT
AGCACACGTCTGAACTCTTTCCCTACACGACGCTCTTCCGATCTNNNAGTTTTCCTAAATGCCTGGAGC
GAGCACACGTCTGAACTCTTTCCCTACACGACGCTCTTCCGATCTNNNTGCAAAATCAAGAGGCATCC
AGCACACGTCTGAACTCTTTCCCTACACGACGCTCTTCCGATCTNNNTCTGCTGTAGTGGTTCCCAGA
GAGCACACGTCTGAACTCTTTCCCTACACGACGCTCTTCCGATCTNNNCCTGACAAGTTTGACACACAC

AGCACACGTCTGAACTCTTTCCCTACACGACGCTCTTCCGATCTNNNGCGGGCATACATATTCAGTCCA
GAGCACACGTCTGAACTCTTTCCCTACACGACGCTCTTCCGATCTNNNGCATGGGGTTTGCTTTCCTAAT
GAGCACACGTCTGAACTCTTTCCCTACACGACGCTCTTCCGATCTNNNAGCAATTCAACGCTTCTCCTCA
AGCACACGTCTGAACTCTTTCCCTACACGACGCTCTTCCGATCTNNNATATGCACAGGTTTTGCATCCCA
GAGCACACGTCTGAACTCTTTCCCTACACGACGCTCTTCCGATCTNNNTTCACCATCTGCTAAGTGATGC
AGAGCACACGTCTGAACTCTTTCCCTACACGACGCTCTTCCGATCTNNNCCTTCAGCTCTTTGGTCTTGTC
GAGCACACGTCTGAACTCTTTCCCTACACGACGCTCTTCCGATCTNNNAAACCTCCAACAAGGAGGGA
AGCACACGTCTGAACTCTTTCCCTACACGACGCTCTTCCGATCTNNNTCTGGAGCTCTGAAGTAAGTGT
GAGCACACGTCTGAACTCTTTCCCTACACGACGCTCTTCCGATCTNNNCAGCTGGTTACTCATGGTTCTC
AGCACACGTCTGAACTCTTTCCCTACACGACGCTCTTCCGATCTNNNCCTTTGTTCCATCGTGTGACTGA
AGCACACGTCTGAACTCTTTCCCTACACGACGCTCTTCCGATCTNNNGAACAGTTCTTGCCTGACAATG
GAGCACACGTCTGAACTCTTTCCCTACACGACGCTCTTCCGATCTNNNAATTTCATGATGCGGAAGGCT
GAGCACACGTCTGAACTCTTTCCCTACACGACGCTCTTCCGATCTNNNTGGATTGTGGAATGCTTAGCT
GAGCACACGTCTGAACTCTTTCCCTACACGACGCTCTTCCGATCTNNNGGTTTGGTTTGGTTGGTTTG
AGAGCACACGTCTGAACTCTTTCCCTACACGACGCTCTTCCGATCTNNNTGTGGGGGAAAGGTAGAAC
AGCACACGTCTGAACTCTTTCCCTACACGACGCTCTTCCGATCTNNNAATAAAGAGCACACACCAGAGC
GAGCACACGTCTGAACTCTTTCCCTACACGACGCTCTTCCGATCTNNNACCTAGGACCTCTCTAACTCCCT
AGCACACGTCTGAACTCTTTCCCTACACGACGCTCTTCCGATCTNNNATGGAAGGGAAGGGCAACTTT
AGCACACGTCTGAACTCTTTCCCTACACGACGCTCTTCCGATCTNNNTGATCATGGCAATCATATGGGA
GAGCACACGTCTGAACTCTTTCCCTACACGACGCTCTTCCGATCTNNNTCAGGACCCATGGTTTTCAGAA
GAGCACACGTCTGAACTCTTTCCCTACACGACGCTCTTCCGATCTNNNAGCACCTTCACCTCCTCAAGTA
GAGCACACGTCTGAACTCTTTCCCTACACGACGCTCTTCCGATCTNNNTCCAAGAACGGCTCAAAGATT
AGCACACGTCTGAACTCTTTCCCTACACGACGCTCTTCCGATCTNNNTGAGCAGACAGAGAAGTCTGG
GAGCACACGTCTGAACTCTTTCCCTACACGACGCTCTTCCGATCTNNNTGCAAAATAAAATCCCAGCACC
GAGCACACGTCTGAACTCTTTCCCTACACGACGCTCTTCCGATCTNNNTCATGGAAGACTTCTTCAAGCC
GCACACGTCTGAACTCTTTCCCTACACGACGCTCTTCCGATCTNNNCCAGGAACATTACTGCCTTGGAA
AGAGCACACGTCTGAACTCTTTCCCTACACGACGCTCTTCCGATCTNNNTTGTGGTGCACGTACTATTCC
GAGCACACGTCTGAACTCTTTCCCTACACGACGCTCTTCCGATCTNNNACAAATAATGCACCCACATCCC
GAGCACACGTCTGAACTCTTTCCCTACACGACGCTCTTCCGATCTNNNGTTTCCATCTTTCAGGCAACAC
AGCACACGTCTGAACTCTTTCCCTACACGACGCTCTTCCGATCTNNNAAAGGTGATTGTAAGCAAGCC
AGCACACGTCTGAACTCTTTCCCTACACGACGCTCTTCCGATCTNNNGAACACAGGTATGTGCCATCAC
GCACACGTCTGAACTCTTTCCCTACACGACGCTCTTCCGATCTNNNTTACTTACGATGCACCCCTCTCTCA
AGAGCACACGTCTGAACTCTTTCCCTACACGACGCTCTTCCGATCTNNNTTTAGGTACTGAGGGGCAAC
AGCACACGTCTGAACTCTTTCCCTACACGACGCTCTTCCGATCTNNNACAAGAGAAGACCTTCATGACC
AGCACACGTCTGAACTCTTTCCCTACACGACGCTCTTCCGATCTNNNGCATTTCCAAAGCATGTTGTGT
GCACACGTCTGAACTCTTTCCCTACACGACGCTCTTCCGATCTNNNGAGTACTAGGATTACAGGCAAG
GAGCACACGTCTGAACTCTTTCCCTACACGACGCTCTTCCGATCTNNNATTTGGGGTGGACACAAACAC
GAGCACACGTCTGAACTCTTTCCCTACACGACGCTCTTCCGATCTNNNGCCAATAAAGCGTGGTTCTCT
GAGCACACGTCTGAACTCTTTCCCTACACGACGCTCTTCCGATCTNNNAGTATACTACGCTGCTTCCCAG
GAGCACACGTCTGAACTCTTTCCCTACACGACGCTCTTCCGATCTNNNTGCTTGCATAGATGGACATTC
AGCACACGTCTGAACTCTTTCCCTACACGACGCTCTTCCGATCTNNNCATAAGCTGTTTCGGCCTTTTGG
AGCACACGTCTGAACTCTTTCCCTACACGACGCTCTTCCGATCTNNNGAACAGCAGTATTTTCCTTGCGT
GCACACGTCTGAACTCTTTCCCTACACGACGCTCTTCCGATCTNNNCTCACCTGAGCTCTGCCTTTATCC

GAGCACACGTCTGAACTCTTTCCCTACACGACGCTCTTCCGATCTNNNCCCTAGTCTTCTCAGTAGCCA
AGCACACGTCTGAACTCTTTCCCTACACGACGCTCTTCCGATCTNNNATAAGCTTGCGAAAGGGAAG
GAGCACACGTCTGAACTCTTTCCCTACACGACGCTCTTCCGATCTNNNAGTGCCCAGGAAAATACTTC
GAGCACACGTCTGAACTCTTTCCCTACACGACGCTCTTCCGATCTNNNACTCCTCTTCGGCATGAGACT
AGCACACGTCTGAACTCTTTCCCTACACGACGCTCTTCCGATCTNNNGTGTCTGCCTGGGTAGTAACT
GCACACGTCTGAACTCTTTCCCTACACGACGCTCTTCCGATCTNNNTCAGAAGCTGCAGTTGTAAGAA
AGCACACGTCTGAACTCTTTCCCTACACGACGCTCTTCCGATCTNNNCTCATCTCTTTTCAGCCCAGTC
AGCACACGTCTGAACTCTTTCCCTACACGACGCTCTTCCGATCTNNNCGTTGCTGATTCTGTCAAATGG
CACACGTCTGAACTCTTTCCCTACACGACGCTCTTCCGATCTNNNTCCTCTTTGTCCCTTGCAAAACAC
GCACACGTCTGAACTCTTTCCCTACACGACGCTCTTCCGATCTNNNTAGCTCTTTCTCTCTCTCGGTCCC
AGCACACGTCTGAACTCTTTCCCTACACGACGCTCTTCCGATCTNNNTTTGCTGTTGCTGAAAATGGT
AGCACACGTCTGAACTCTTTCCCTACACGACGCTCTTCCGATCTNNNTAACGAGGGCTCTGGGAGATA
GAGCACACGTCTGAACTCTTTCCCTACACGACGCTCTTCCGATCTNNNCAAGTGAGCATGCATTGACC
GCACACGTCTGAACTCTTTCCCTACACGACGCTCTTCCGATCTNNNCCCTCTGCACTTGGCTGAATTTT
AGCACACGTCTGAACTCTTTCCCTACACGACGCTCTTCCGATCTNNNCTACCACCACACCCAGCTCTATC
GAGCACACGTCTGAACTCTTTCCCTACACGACGCTCTTCCGATCTNNNGAAATGGACCCTTGGAGATG
GCACACGTCTGAACTCTTTCCCTACACGACGCTCTTCCGATCTNNNCTTCCCACCTTGGATATGCGTAG
GCACACGTCTGAACTCTTTCCCTACACGACGCTCTTCCGATCTNNNATTTGGACAAATGTCTCCCCCAC
GCACACGTCTGAACTCTTTCCCTACACGACGCTCTTCCGATCTNNNTAAAGTGGCCATTCTTCCTCAG
AGCACACGTCTGAACTCTTTCCCTACACGACGCTCTTCCGATCTNNNTAGGAAGAACAACCCTCCACTC
GCACACGTCTGAACTCTTTCCCTACACGACGCTCTTCCGATCTNNNTCTTGGAAACAATGCCCATGTG
GCACACGTCTGAACTCTTTCCCTACACGACGCTCTTCCGATCTNNNAGCTAGTACAAAGCACGCAGAA
AGCACACGTCTGAACTCTTTCCCTACACGACGCTCTTCCGATCTNNNTCAGTGACCGAAATGAACCAT
AGAGCACACGTCTGAACTCTTTCCCTACACGACGCTCTTCCGATCTNNNACATGTTGCCTGTACCAGTT
GAGCACACGTCTGAACTCTTTCCCTACACGACGCTCTTCCGATCTNNNAGAAATGGGACACAAAGA
GAGCACACGTCTGAACTCTTTCCCTACACGACGCTCTTCCGATCTNNNCCGGCCTGTAGTCTCTAGTTT
GCACACGTCTGAACTCTTTCCCTACACGACGCTCTTCCGATCTNNNGGAGGAGGCTGAGGAAAATTC
AGCACACGTCTGAACTCTTTCCCTACACGACGCTCTTCCGATCTNNNTGGACTTTTGCACTACAGGTG
AGCACACGTCTGAACTCTTTCCCTACACGACGCTCTTCCGATCTNNNATGCAGTGCCAAATAACATGG
AGCACACGTCTGAACTCTTTCCCTACACGACGCTCTTCCGATCTNNNAAAGGTCAGCTGAGTGATTTG
GAGCACACGTCTGAACTCTTTCCCTACACGACGCTCTTCCGATCTNNNACATGAACTTCTCAGAGCCAC
GAGCACACGTCTGAACTCTTTCCCTACACGACGCTCTTCCGATCTNNNCCTGACACAAAGACCTCACA
GCACACGTCTGAACTCTTTCCCTACACGACGCTCTTCCGATCTNNNCTCTCTCTGTTCCATGGTCAGGT
GCACACGTCTGAACTCTTTCCCTACACGACGCTCTTCCGATCTNNNCCATTTTCCTGTGGGTAGAGGT
AGCACACGTCTGAACTCTTTCCCTACACGACGCTCTTCCGATCTNNNTCCACCTTCTCTGTTTCTGACCT
CACACGTCTGAACTCTTTCCCTACACGACGCTCTTCCGATCTNNNTGGTGTCTTTTATTCCACAGAGC
GAGCACACGTCTGAACTCTTTCCCTACACGACGCTCTTCCGATCTNNNAAGCAGCTCAGTTTCCCATTC
GAGCACACGTCTGAACTCTTTCCCTACACGACGCTCTTCCGATCTNNNTGACAAGCCACACTAAGCTC
AGCACACGTCTGAACTCTTTCCCTACACGACGCTCTTCCGATCTNNNGCAGGATGGGAATCTCTTCTG
AGCACACGTCTGAACTCTTTCCCTACACGACGCTCTTCCGATCTNNNTTGAATCCAGCAAACCCATCAC
AGAGCACACGTCTGAACTCTTTCCCTACACGACGCTCTTCCGATCTNNNCCACTTGTTCAACTCCCGAA
AGCACACGTCTGAACTCTTTCCCTACACGACGCTCTTCCGATCTNNNACCTGTAAAACATGAGACTGC
AGCACACGTCTGAACTCTTTCCCTACACGACGCTCTTCCGATCTNNNCTTCTTTGGGGCTCTGAAAAT

AGCACACGTCTGAACTCTTTCCCTACACGACGCTCTTCCGATCTNNNTCCCAGCCTTAGAGAGAACTAG
GAGCACACGTCTGAACTCTTTCCCTACACGACGCTCTTCCGATCTNNNTCGTTTCTGTCTCCCTCTCTAC
GAGCACACGTCTGAACTCTTTCCCTACACGACGCTCTTCCGATCTNNNATACAGGCCTCTTGTGTGTA
AGCACACGTCTGAACTCTTTCCCTACACGACGCTCTTCCGATCTNNNTTCCATCCCGGGTGACAAACAT
AGCACACGTCTGAACTCTTTCCCTACACGACGCTCTTCCGATCTNNNGAGGCTTCAAAGAATTCAGAG
AGCACACGTCTGAACTCTTTCCCTACACGACGCTCTTCCGATCTNNNTCAAAGTGTGTGTGTGTATGC
GAGCACACGTCTGAACTCTTTCCCTACACGACGCTCTTCCGATCTNNNGCTTTGAATCCAGCTGCAACA
GAGCACACGTCTGAACTCTTTCCCTACACGACGCTCTTCCGATCTNNNGTGGACTGGAAAATTCAGCC
CACACGTCTGAACTCTTTCCCTACACGACGCTCTTCCGATCTNNNTGTGGTAATTGTGTACAGCAGG
GCACACGTCTGAACTCTTTCCCTACACGACGCTCTTCCGATCTNNNTTTCCTTCCCTCTTGTCCCTTCCC
AGCACACGTCTGAACTCTTTCCCTACACGACGCTCTTCCGATCTNNNTAAGAAGAGAGAGAGTGGGA
AGCACACGTCTGAACTCTTTCCCTACACGACGCTCTTCCGATCTNNNTGTACTCTGTTACCAGCTTTTCA
AGCACACGTCTGAACTCTTTCCCTACACGACGCTCTTCCGATCTNNNGCTGTTGCAGGTCCATCAACT
AGCACACGTCTGAACTCTTTCCCTACACGACGCTCTTCCGATCTNNNATTATGTTTGCAATGTGTGGG
AGCACACGTCTGAACTCTTTCCCTACACGACGCTCTTCCGATCTNNNCCTATCTCTGGGTGGATGCTG
AGCACACGTCTGAACTCTTTCCCTACACGACGCTCTTCCGATCTNNNAGAGATTCTCATACTCGGCCTC
AGCACACGTCTGAACTCTTTCCCTACACGACGCTCTTCCGATCTNNNCTTGGGTTCAGACTGTCTCCTTG
AGCACACGTCTGAACTCTTTCCCTACACGACGCTCTTCCGATCTNNNCCTGCTGTACTTCCCCTGTTTT
AGCACACGTCTGAACTCTTTCCCTACACGACGCTCTTCCGATCTNNNCAGCATTCCATCAGAGAAGCA
GAGCACACGTCTGAACTCTTTCCCTACACGACGCTCTTCCGATCTNNNCTATTCCTGGGGAAAACTCA
GAGCACACGTCTGAACTCTTTCCCTACACGACGCTCTTCCGATCTNNNAACTCACTCAAAGTAACACCG
AGCACACGTCTGAACTCTTTCCCTACACGACGCTCTTCCGATCTNNNTACAGGATGGCTGAGAAACTT
AGAGCACACGTCTGAACTCTTTCCCTACACGACGCTCTTCCGATCTNNNTTGGGTGAAAAATAAAGGG
AGCACACGTCTGAACTCTTTCCCTACACGACGCTCTTCCGATCTNNNCAATCCCACACCGATACCAAGA
AGAGCACACGTCTGAACTCTTTCCCTACACGACGCTCTTCCGATCTNNNAGCCATTGCGCCATCTATAT
GAGCACACGTCTGAACTCTTTCCCTACACGACGCTCTTCCGATCTNNNTTATAGTCCTGGCTTTGGCAC
GAGCACACGTCTGAACTCTTTCCCTACACGACGCTCTTCCGATCTNNNACCATCATCCTGGAGTTTTTC
GAGCACACGTCTGAACTCTTTCCCTACACGACGCTCTTCCGATCTNNNTGACCTACTCAGGTTTCACCAG
AGCACACGTCTGAACTCTTTCCCTACACGACGCTCTTCCGATCTNNNTTGATTCCTCACTGGCTTTGCTT
GAGCACACGTCTGAACTCTTTCCCTACACGACGCTCTTCCGATCTNNNTGCATAAAACATGGCACTGCT
GCACACGTCTGAACTCTTTCCCTACACGACGCTCTTCCGATCTNNNAATTTTATTCTGAGGGGGCTGT
AGAGCACACGTCTGAACTCTTTCCCTACACGACGCTCTTCCGATCTNNNCTGGACCCTAAACGAGTGT
GAGCACACGTCTGAACTCTTTCCCTACACGACGCTCTTCCGATCTNNNGGGGCCTAGTGATGGTGAT
GAGCACACGTCTGAACTCTTTCCCTACACGACGCTCTTCCGATCTNNNATCTAACCCCCTCTTCCTGTCAT
GAGCACACGTCTGAACTCTTTCCCTACACGACGCTCTTCCGATCTNNNTGTTCACAAGGTACTGCATAC
AGAGCACACGTCTGAACTCTTTCCCTACACGACGCTCTTCCGATCTNNNCGAGTTCATAATCCAGGGGA
GAGCACACGTCTGAACTCTTTCCCTACACGACGCTCTTCCGATCTNNNATACTCTAAGTCGTGCAGCCA
AGCACACGTCTGAACTCTTTCCCTACACGACGCTCTTCCGATCTNNNACTATGCTTTGCCTTCAACTCCG
GAGCACACGTCTGAACTCTTTCCCTACACGACGCTCTTCCGATCTNNNCCATCTAGAAACCACCAAGAG
AGCACACGTCTGAACTCTTTCCCTACACGACGCTCTTCCGATCTNNNTGAAGGCAGCTAAATGCTGAA
AGAGCACACGTCTGAACTCTTTCCCTACACGACGCTCTTCCGATCTNNNCAGGAAGGTGGCTTTGGTA
AGCACACGTCTGAACTCTTTCCCTACACGACGCTCTTCCGATCTNNNGCACATACACATGAGTTTTGG
AGCACACGTCTGAACTCTTTCCCTACACGACGCTCTTCCGATCTNNNTCTTAGGACTGGGCCAAGGAA

GCACACGTCTGAACTCTTTCCCTACACGACGCTCTTCCGATCTNNNACGTTTACCCCTGCACATTTTCTC
AGCACACGTCTGAACTCTTTCCCTACACGACGCTCTTCCGATCTNNNCAGCCATTACGCATCCTGTTTT
GAGCACACGTCTGAACTCTTTCCCTACACGACGCTCTTCCGATCTNNNTTCTCTCTCTTCTGCCTGTTGG
AGCACACGTCTGAACTCTTTCCCTACACGACGCTCTTCCGATCTNNNGCTACTTCATGCTACAGCCAGT
GCACACGTCTGAACTCTTTCCCTACACGACGCTCTTCCGATCTNNNGAGATTCCTTCCAGGCCAATGA
GAGCACACGTCTGAACTCTTTCCCTACACGACGCTCTTCCGATCTNNNCCAAATGTTGCCCTTTGGTTC
GAGCACACGTCTGAACTCTTTCCCTACACGACGCTCTTCCGATCTNNNTGTTTAGCCACAGTCACAGCT
AGCACACGTCTGAACTCTTTCCCTACACGACGCTCTTCCGATCTNNNTGGAGATGAACAGGCAGGAC
AGCACACGTCTGAACTCTTTCCCTACACGACGCTCTTCCGATCTNNNGCCATCAGGGTGATGTTCTTAC
GAGCACACGTCTGAACTCTTTCCCTACACGACGCTCTTCCGATCTNNNCATCACCTGAGCAGTTTCTAC
AGCACACGTCTGAACTCTTTCCCTACACGACGCTCTTCCGATCTNNNCTCACCCTCCTCAGTTTCAGAG
AGCACACGTCTGAACTCTTTCCCTACACGACGCTCTTCCGATCTNNNCCATGAAGATTCAGCACTGTG
GAGCACACGTCTGAACTCTTTCCCTACACGACGCTCTTCCGATCTNNNGGTGGCACCTACATTGCAAA
GCACACGTCTGAACTCTTTCCCTACACGACGCTCTTCCGATCTNNNTTGGATCCATCTGCAACTGAAG
GAGCACACGTCTGAACTCTTTCCCTACACGACGCTCTTCCGATCTNNNGCTAATGTCTTCCAGCTCCTTC
AGCACACGTCTGAACTCTTTCCCTACACGACGCTCTTCCGATCTNNNCCTTTGTGTGGCTCCTGATTTT
AGCACACGTCTGAACTCTTTCCCTACACGACGCTCTTCCGATCTNNNGGCTGTCCCTATAAGGGGGTA
AGCACACGTCTGAACTCTTTCCCTACACGACGCTCTTCCGATCTNNNCACACTTCTATTCTCCTTGCCCC
GCACACGTCTGAACTCTTTCCCTACACGACGCTCTTCCGATCTNNNTCTGTGCTTTGACGAAGTCATG
AGCACACGTCTGAACTCTTTCCCTACACGACGCTCTTCCGATCTNNNAAGCCACATTTGTCCCCATGA
AGCACACGTCTGAACTCTTTCCCTACACGACGCTCTTCCGATCTNNNGGAACTAACATTGCTGGAGA
GAGCACACGTCTGAACTCTTTCCCTACACGACGCTCTTCCGATCTNNNACTGGATCCATCATCCCTCAT
AGCACACGTCTGAACTCTTTCCCTACACGACGCTCTTCCGATCTNNNAGGAGTTCAAGGTCATGGGT
AGCACACGTCTGAACTCTTTCCCTACACGACGCTCTTCCGATCTNNNTGTAGTGTCTGTCCTTTTGGC
AGCACACGTCTGAACTCTTTCCCTACACGACGCTCTTCCGATCTNNNTAAGCCACTAGTGAGGGAACT
AGCACACGTCTGAACTCTTTCCCTACACGACGCTCTTCCGATCTNNNCACTGTAAATCCCTTGGCCTCTC
AGCACACGTCTGAACTCTTTCCCTACACGACGCTCTTCCGATCTNNNTTGGGCAATTCTGTTCTGCAT
AGCACACGTCTGAACTCTTTCCCTACACGACGCTCTTCCGATCTNNNCGAGGTGCTGCTGGATATGC
AGCACACGTCTGAACTCTTTCCCTACACGACGCTCTTCCGATCTNNNCTCTCGTCACTACCCTCCAGTT
AGCACACGTCTGAACTCTTTCCCTACACGACGCTCTTCCGATCTNNNTCTGGACAAAGTTGATCCTCTG
AGAGCACACGTCTGAACTCTTTCCCTACACGACGCTCTTCCGATCTNNNTCAGTGTAGTCAAGCCTTCC
GCACACGTCTGAACTCTTTCCCTACACGACGCTCTTCCGATCTNNNCCTAGATTGCACCACACTACTGC
AGCACACGTCTGAACTCTTTCCCTACACGACGCTCTTCCGATCTNNNAGAGCCAGGCAATGTGTTTGT
GAGCACACGTCTGAACTCTTTCCCTACACGACGCTCTTCCGATCTNNNTGGAATTGCTGGAGGCATGT
AGCACACGTCTGAACTCTTTCCCTACACGACGCTCTTCCGATCTNNNGCCTGGCCTTGTATGTCAATTC
GAGCACACGTCTGAACTCTTTCCCTACACGACGCTCTTCCGATCTNNNAACTGTAGGATACCCCTTGC
GAGCACACGTCTGAACTCTTTCCCTACACGACGCTCTTCCGATCTNNNTTCTTCAGAGGATGACCACC
AGCACACGTCTGAACTCTTTCCCTACACGACGCTCTTCCGATCTNNNACAGGTACCATCTGTAACCAC
GAGCACACGTCTGAACTCTTTCCCTACACGACGCTCTTCCGATCTNNNGATGGGAAATGCTTGCCTGT
GCACACGTCTGAACTCTTTCCCTACACGACGCTCTTCCGATCTNNNCTCACCCATTACCAGGTTCAAGG
AGAGCACACGTCTGAACTCTTTCCCTACACGACGCTCTTCCGATCTNNNAGAGACAAGGCTCACTGAG
AGCACACGTCTGAACTCTTTCCCTACACGACGCTCTTCCGATCTNNNCCCCTTCCCCTCTATTTGTTTG
GAGCACACGTCTGAACTCTTTCCCTACACGACGCTCTTCCGATCTNNNTGTGGAAGGAAGGGAGAAT

AGAGCACACGTCTGAACTCTTTCCCTACACGACGCTCTTCCGATCTNNNAGGTTTTAGGGAACTGGGT
GAGCACACGTCTGAACTCTTTCCCTACACGACGCTCTTCCGATCTNNNTGCAGACATGAAGGCAAGTC
AGCACACGTCTGAACTCTTTCCCTACACGACGCTCTTCCGATCTNNNGTTTGGATCTTCCTCCAACCTCC
AGAGCACACGTCTGAACTCTTTCCCTACACGACGCTCTTCCGATCTNNNCGTCTGGAAACCAGGATGT
GAGCACACGTCTGAACTCTTTCCCTACACGACGCTCTTCCGATCTNNNAAGATTTCCTCCTGCCCTGTAG
AGCACACGTCTGAACTCTTTCCCTACACGACGCTCTTCCGATCTNNNAACAAACGAATGTGACTTCTGC
AGAGCACACGTCTGAACTCTTTCCCTACACGACGCTCTTCCGATCTNNNAGCCTCAGAAACTCAATGGA
AGAGCACACGTCTGAACTCTTTCCCTACACGACGCTCTTCCGATCTNNNAGCAGAATACCAACAGTGCT
AGCACACGTCTGAACTCTTTCCCTACACGACGCTCTTCCGATCTNNNTCATTCCCTTTTGAATGGTTGGT
GAGCACACGTCTGAACTCTTTCCCTACACGACGCTCTTCCGATCTNNNTGGGTCATCACCAACATCATAA
GAGCACACGTCTGAACTCTTTCCCTACACGACGCTCTTCCGATCTNNNCCCTGCTCATTAGTTTGTGGGA
GAGCACACGTCTGAACTCTTTCCCTACACGACGCTCTTCCGATCTNNNATATCTGAGGTCTCCCAATCCC
AGCACACGTCTGAACTCTTTCCCTACACGACGCTCTTCCGATCTNNNTCATCTCCCGTTAGGTTAGCCAT
AGCACACGTCTGAACTCTTTCCCTACACGACGCTCTTCCGATCTNNNAACAACACAGAATTACCCACCCC
GAGCACACGTCTGAACTCTTTCCCTACACGACGCTCTTCCGATCTNNNACAACACAGTACAAACACACT
AGCACACGTCTGAACTCTTTCCCTACACGACGCTCTTCCGATCTNNNGGCAGAGCAGAAGAAAGGAA
GAGCACACGTCTGAACTCTTTCCCTACACGACGCTCTTCCGATCTNNNCACTGGGCTATTATCTACCGC
GAGCACACGTCTGAACTCTTTCCCTACACGACGCTCTTCCGATCTNNNACAGGAATTCACATGAGAGA
GAGCACACGTCTGAACTCTTTCCCTACACGACGCTCTTCCGATCTNNNCATACTGCGAGGATGACTTCT
AGCACACGTCTGAACTCTTTCCCTACACGACGCTCTTCCGATCTNNNTGCCCTTTTGGCATAACCTCATT
GAGCACACGTCTGAACTCTTTCCCTACACGACGCTCTTCCGATCTNNNTTCACCCGGATCTCAATTTTGCT
GAGCACACGTCTGAACTCTTTCCCTACACGACGCTCTTCCGATCTNNNAAACCACGTACACAAGAACCTG
AGCACACGTCTGAACTCTTTCCCTACACGACGCTCTTCCGATCTNNNACTCTCGTTGCTTCAGGATTCA
GAGCACACGTCTGAACTCTTTCCCTACACGACGCTCTTCCGATCTNNNATAATTTCCCTAGCCCCAGATG
GAGCACACGTCTGAACTCTTTCCCTACACGACGCTCTTCCGATCTNNNGGGAAAATGTCCATGATCCCA
GAGCACACGTCTGAACTCTTTCCCTACACGACGCTCTTCCGATCTNNNAGGACAGACTGAACGTCCTTA
GAGCACACGTCTGAACTCTTTCCCTACACGACGCTCTTCCGATCTNNNAGTGTTCCTCAGACCTGGTAAT
GAGCACACGTCTGAACTCTTTCCCTACACGACGCTCTTCCGATCTNNNTCTAAAATGAGCTGGCCAGG
AGCACACGTCTGAACTCTTTCCCTACACGACGCTCTTCCGATCTNNNGGTAAGCAGCCTGAAAGCATTT
GAGCACACGTCTGAACTCTTTCCCTACACGACGCTCTTCCGATCTNNNAAAAGTACATAACCCCTCTGC
AGAGCACACGTCTGAACTCTTTCCCTACACGACGCTCTTCCGATCTNNNATGGGGTGAGAGTGACTGT
AGCACACGTCTGAACTCTTTCCCTACACGACGCTCTTCCGATCTNNNTATCCTGAGGTTGCCTACAAGG
GAGCACACGTCTGAACTCTTTCCCTACACGACGCTCTTCCGATCTNNNATGTGTACCCAAGGGCAGAG
AGCACACGTCTGAACTCTTTCCCTACACGACGCTCTTCCGATCTNNNAAGTGATTGGATCTCTGAGGCTG
AGCACACGTCTGAACTCTTTCCCTACACGACGCTCTTCCGATCTNNNCAGCCCTTGAAGCTATCTGGTTT
GAGCACACGTCTGAACTCTTTCCCTACACGACGCTCTTCCGATCTNNNATATTCTTTCCCAGTCCCTGGCT
AGCACACGTCTGAACTCTTTCCCTACACGACGCTCTTCCGATCTNNNTGGAAGTAGAGCTGTCTCTCTG
GAGCACACGTCTGAACTCTTTCCCTACACGACGCTCTTCCGATCTNNNACTCTGTGAGCCAATTCCATG
AGCACACGTCTGAACTCTTTCCCTACACGACGCTCTTCCGATCTNNNCCTTATGAAACTCCAGCGAGGA
GAGCACACGTCTGAACTCTTTCCCTACACGACGCTCTTCCGATCTNNNCGTACACACACACATCCCTTAC
AGCACACGTCTGAACTCTTTCCCTACACGACGCTCTTCCGATCTNNNTGTGTCCTGCATCTTACATGTGC
GAGCACACGTCTGAACTCTTTCCCTACACGACGCTCTTCCGATCTNNNTTCATGGCAAAAGGCAGCAAA
AGCACACGTCTGAACTCTTTCCCTACACGACGCTCTTCCGATCTNNNTACCTAATGTTTGCCCCTAAGGC

GAGCACACGTCTGAACTCTTTCCCTACACGACGCTCTTCCGATCTNNNTGCATCTAATTCTGAGGGAC
AGCACACGTCTGAACTCTTTCCCTACACGACGCTCTTCCGATCTNNNTGCTAAGAATTGTGAACAGGC
GAGCACACGTCTGAACTCTTTCCCTACACGACGCTCTTCCGATCTNNNCCCCATTTCTCCACAACTTCT
GCACACGTCTGAACTCTTTCCCTACACGACGCTCTTCCGATCTNNNCTGTGGCCTTGCCTATACTTTGA
AGCACACGTCTGAACTCTTTCCCTACACGACGCTCTTCCGATCTNNNTAGCTTCTTCCTCCTTCCCCTTCC
AGCACACGTCTGAACTCTTTCCCTACACGACGCTCTTCCGATCTNNNCAACTCTGCTGGTTCTCTCCTT
AGCACACGTCTGAACTCTTTCCCTACACGACGCTCTTCCGATCTNNNCTCTGGAATTACACACAGGAGC
AGCACACGTCTGAACTCTTTCCCTACACGACGCTCTTCCGATCTNNNACTTATCCCTTGACTTCCGGTG
GAGCACACGTCTGAACTCTTTCCCTACACGACGCTCTTCCGATCTNNNTTATCTTGCATAGGCTATCCC
GAGCACACGTCTGAACTCTTTCCCTACACGACGCTCTTCCGATCTNNNGATGCTTGTGGGCTTAGAAA
AGCACACGTCTGAACTCTTTCCCTACACGACGCTCTTCCGATCTNNNATTTGAAGCCCTTGCTGGGTAA
AGCACACGTCTGAACTCTTTCCCTACACGACGCTCTTCCGATCTNNNCTGGAATGCACTGTGAAGCTA
AGCACACGTCTGAACTCTTTCCCTACACGACGCTCTTCCGATCTNNNTGCATCACAGGGTAGCATTTC
AGCACACGTCTGAACTCTTTCCCTACACGACGCTCTTCCGATCTNNNATCCAAGAGGTAGAGGGGAA
AGCACACGTCTGAACTCTTTCCCTACACGACGCTCTTCCGATCTNNNCGTCTAATCCCATGGCTCAGAG
GCACACGTCTGAACTCTTTCCCTACACGACGCTCTTCCGATCTNNNTAATGGTGATGCCATCTGTGTG
AGCACACGTCTGAACTCTTTCCCTACACGACGCTCTTCCGATCTNNNTGGCTGGCTTGTGTTTATGTT
GAGCACACGTCTGAACTCTTTCCCTACACGACGCTCTTCCGATCTNNNAGCTGGGGAGAGGGATATG
GCACACGTCTGAACTCTTTCCCTACACGACGCTCTTCCGATCTNNNAGAAAACAGGTCTGTGTGTGTG
AGCACACGTCTGAACTCTTTCCCTACACGACGCTCTTCCGATCTNNNGGGCTCTATAGGTGCTCTCTCT
GAGCACACGTCTGAACTCTTTCCCTACACGACGCTCTTCCGATCTNNNTACCCTCTCAGCTTACAGCCAT
GAGCACACGTCTGAACTCTTTCCCTACACGACGCTCTTCCGATCTNNNTAATGCTAGGAAGAGTGGG
GCACACGTCTGAACTCTTTCCCTACACGACGCTCTTCCGATCTNNNTTCCCCATCAAATGACAAGGTG
GAGCACACGTCTGAACTCTTTCCCTACACGACGCTCTTCCGATCTNNNCTCCCTGTCAAAGAGGTTCC
GAGCACACGTCTGAACTCTTTCCCTACACGACGCTCTTCCGATCTNNNAATAGCCTGCTACATGGCAAG
AGCACACGTCTGAACTCTTTCCCTACACGACGCTCTTCCGATCTNNNGTCAGTACCCCTAATCTCTGCC
AGCACACGTCTGAACTCTTTCCCTACACGACGCTCTTCCGATCTNNNATTTATCACTGCAAGCACCTCCA
GAGCACACGTCTGAACTCTTTCCCTACACGACGCTCTTCCGATCTNNNTTTCTCCACTACCAAAACTGG
AGCACACGTCTGAACTCTTTCCCTACACGACGCTCTTCCGATCTNNNTACTGGGTCGGTCATGGTTTCT
AGCACACGTCTGAACTCTTTCCCTACACGACGCTCTTCCGATCTNNNTCCAGTTCTATCCACGATGCAA
AGCACACGTCTGAACTCTTTCCCTACACGACGCTCTTCCGATCTNNNTCTCCATCACCCCTTTGTTTTGG
AGCACACGTCTGAACTCTTTCCCTACACGACGCTCTTCCGATCTNNNATGGCTTTGGTTCTTAGGTGG
AGCACACGTCTGAACTCTTTCCCTACACGACGCTCTTCCGATCTNNNGCCTTGCTACTCAGATTTCACCA
AGCACACGTCTGAACTCTTTCCCTACACGACGCTCTTCCGATCTNNNATAGGAGGGGGTTTCCCTTTCT
GAGCACACGTCTGAACTCTTTCCCTACACGACGCTCTTCCGATCTNNNAGAATAGAGAGCCAGAAGG
AGCACACGTCTGAACTCTTTCCCTACACGACGCTCTTCCGATCTNNNTACTGCTCCTTTCCAACACACA
GAGCACACGTCTGAACTCTTTCCCTACACGACGCTCTTCCGATCTNNNCACAACTGCCCAGATACCTA
AGCACACGTCTGAACTCTTTCCCTACACGACGCTCTTCCGATCTNNNATAAGAATCCATGTGGCCCAA
GAGCACACGTCTGAACTCTTTCCCTACACGACGCTCTTCCGATCTNNNAGTGCTGTGTCTTGACTTTTG
AGCACACGTCTGAACTCTTTCCCTACACGACGCTCTTCCGATCTNNNCTCTGCCTAAAAGATGGGAGC
AGCACACGTCTGAACTCTTTCCCTACACGACGCTCTTCCGATCTNNNCCCTCAAGGGGGAAATCAAA
GAGCACACGTCTGAACTCTTTCCCTACACGACGCTCTTCCGATCTNNNGGGGAGAGTTTTGTTGTCCC
AGAGCACACGTCTGAACTCTTTCCCTACACGACGCTCTTCCGATCTNNNCGTTGGCTTCTCAGATGAA

AGCACACGTCTGAACTCTTTCCCTACACGACGCTCTTCCGATCTNNNAATGTCCTCCAGCATTGTCTGTG
AGCACACGTCTGAACTCTTTCCCTACACGACGCTCTTCCGATCTNNNGGGCACCAGAGGCAATCTTATA
GAGCACACGTCTGAACTCTTTCCCTACACGACGCTCTTCCGATCTNNNGGGGACAATTACAGGCAGGA
AGAGCACACGTCTGAACTCTTTCCCTACACGACGCTCTTCCGATCTNNNTTTCTTAGGTGACCCTGGGG
AGCACACGTCTGAACTCTTTCCCTACACGACGCTCTTCCGATCTNNNTTCTAAGTCAGAATCCCCAGGC
GCACACGTCTGAACTCTTTCCCTACACGACGCTCTTCCGATCTNNNGTGGATACACTGTTGTTTGGTGC
GAGCACACGTCTGAACTCTTTCCCTACACGACGCTCTTCCGATCTNNNACTACACACCCCTTCTATCCCAC
AGCACACGTCTGAACTCTTTCCCTACACGACGCTCTTCCGATCTNNNACCTGCATATTTCACGAAGAGG
AGCACACGTCTGAACTCTTTCCCTACACGACGCTCTTCCGATCTNNNGCTGAAGGCTGTTCTATGGACAA
GCACACGTCTGAACTCTTTCCCTACACGACGCTCTTCCGATCTNNNACAGGATTTACCGGAAGAAGCAC
AGCACACGTCTGAACTCTTTCCCTACACGACGCTCTTCCGATCTNNNTGTGAAGAGTAGCACTTCTGTC
GAGCACACGTCTGAACTCTTTCCCTACACGACGCTCTTCCGATCTNNNTGACATACCCCCAACTCTCTCTC
GAGCACACGTCTGAACTCTTTCCCTACACGACGCTCTTCCGATCTNNNCTGTTTCCTGAGGCCCAACATT
GAGCACACGTCTGAACTCTTTCCCTACACGACGCTCTTCCGATCTNNNAGACCGATATAGCACTCACATC
AGCACACGTCTGAACTCTTTCCCTACACGACGCTCTTCCGATCTNNNTCTTTTGCTGTTCTGGTTCCTCC
GAGCACACGTCTGAACTCTTTCCCTACACGACGCTCTTCCGATCTNNNTTTTCAAATATTCCCCTCGCCCC
GAGCACACGTCTGAACTCTTTCCCTACACGACGCTCTTCCGATCTNNNACCCCATAAACCACACTCTCTCA
AGCACACGTCTGAACTCTTTCCCTACACGACGCTCTTCCGATCTNNNATTCTTGTCTGAGCAACCCACTCC
AGCACACGTCTGAACTCTTTCCCTACACGACGCTCTTCCGATCTNNNCCCCATGGAAGCCATGAAACATA
AGCACACGTCTGAACTCTTTCCCTACACGACGCTCTTCCGATCTNNNAACCTATACCCACCCTACAACCCC
AGAGCACACGTCTGAACTCTTTCCCTACACGACGCTCTTCCGATCTNNNGCAAGAGAGCTCCTTCAGGT
GCACACGTCTGAACTCTTTCCCTACACGACGCTCTTCCGATCTNNNTGGCTTGCAATCATGTGCAGAAT
AGCACACGTCTGAACTCTTTCCCTACACGACGCTCTTCCGATCTNNNGCTTTCCCCTGCAAATACCCATA
AGCACACGTCTGAACTCTTTCCCTACACGACGCTCTTCCGATCTNNNCCACACCTGTTACTGAGGGTAG
GAGCACACGTCTGAACTCTTTCCCTACACGACGCTCTTCCGATCTNNNTGGAACACTTTCCTTGACCGAT
AGCACACGTCTGAACTCTTTCCCTACACGACGCTCTTCCGATCTNNNATCTGGTAACTCTGGTGTGAGC
GAGCACACGTCTGAACTCTTTCCCTACACGACGCTCTTCCGATCTNNNCGCGTGCTGTAAGCTACCTAA
AGCACACGTCTGAACTCTTTCCCTACACGACGCTCTTCCGATCTNNNATGTCATACAAGTGCCAGAGCAC
GAGCACACGTCTGAACTCTTTCCCTACACGACGCTCTTCCGATCTNNNAGCCAACTCCTGACCTGTTTTT
GAGCACACGTCTGAACTCTTTCCCTACACGACGCTCTTCCGATCTNNNAGTTCAAAAGCCCTCAATCACC
AGCACACGTCTGAACTCTTTCCCTACACGACGCTCTTCCGATCTNNNGGCAATGCCAACCCTAAAGACTA
AGCACACGTCTGAACTCTTTCCCTACACGACGCTCTTCCGATCTNNNTGACCTCTCCTTCCATCCTCAAGC
GCACACGTCTGAACTCTTTCCCTACACGACGCTCTTCCGATCTNNNACCTCTCCTATCCTCTTTGCCACCA
AGAGCACACGTCTGAACTCTTTCCCTACACGACGCTCTTCCGATCTNNNAATAGCCCAAGGGGTCATCT
AGAGCACACGTCTGAACTCTTTCCCTACACGACGCTCTTCCGATCTNNNGGACTTGTGGTTTGCTCTTCC
GAGCACACGTCTGAACTCTTTCCCTACACGACGCTCTTCCGATCTNNNAGGAACTCTGGACGAAGACT
GAGCACACGTCTGAACTCTTTCCCTACACGACGCTCTTCCGATCTNNNAGCTAGAATTCAAGGGAGAGC
AGCACACGTCTGAACTCTTTCCCTACACGACGCTCTTCCGATCTNNNAGAAATGTTCTCTGCAGTTTTGT
GAGCACACGTCTGAACTCTTTCCCTACACGACGCTCTTCCGATCTNNNAGCCACTCAAAGGAACTTTCTG
GAGCACACGTCTGAACTCTTTCCCTACACGACGCTCTTCCGATCTNNNGTCACGTTTTCTGTTCCCTCCTT
AGAGCACACGTCTGAACTCTTTCCCTACACGACGCTCTTCCGATCTNNNCCATGGACAGTCTACCTCCTT
GAGCACACGTCTGAACTCTTTCCCTACACGACGCTCTTCCGATCTNNNTCTTTTGATCCCCCTCCTTGGAT
GAGCACACGTCTGAACTCTTTCCCTACACGACGCTCTTCCGATCTNNNAGTGGTCATTCTCACCTTTTCC

GCACACGTCTGAACTCTTTCCCTACACGACGCTCTTCCGATCTNNNAGCCTTACTTCCTCATCAGGGTTC
GAGCACACGTCTGAACTCTTTCCCTACACGACGCTCTTCCGATCTNNNACTTGGAAGAGGCCTTCAAAA
GAGCACACGTCTGAACTCTTTCCCTACACGACGCTCTTCCGATCTNNNTTCTACAACAGGCACTGGGGA
AGCACACGTCTGAACTCTTTCCCTACACGACGCTCTTCCGATCTNNNGGGTGGAATGTTCACACACACT
AGCACACGTCTGAACTCTTTCCCTACACGACGCTCTTCCGATCTNNNCAGTTATTGAGCTAATGGCCAG
AGCACACGTCTGAACTCTTTCCCTACACGACGCTCTTCCGATCTNNNAAGGAGAAGGAAGGACTGGGA
GAGCACACGTCTGAACTCTTTCCCTACACGACGCTCTTCCGATCTNNNTTCCAGGAGCTGAATTTTCCA
GAGCACACGTCTGAACTCTTTCCCTACACGACGCTCTTCCGATCTNNNCACTGAGATCAAAGCATAGCG
GAGCACACGTCTGAACTCTTTCCCTACACGACGCTCTTCCGATCTNNNAAGTGTTCCCAGTTTAGACCCT
GAGCACACGTCTGAACTCTTTCCCTACACGACGCTCTTCCGATCTNNNAGTGAAAGAGGAGGTGATG
AGAGCACACGTCTGAACTCTTTCCCTACACGACGCTCTTCCGATCTNNNGTGAAAATACCTCACCCCTGC
AGCACACGTCTGAACTCTTTCCCTACACGACGCTCTTCCGATCTNNNTGCGCAGCTTCATCACATACATA
GAGCACACGTCTGAACTCTTTCCCTACACGACGCTCTTCCGATCTNNNCGTGTCTGTTTGTGAGCATGA
GAGCACACGTCTGAACTCTTTCCCTACACGACGCTCTTCCGATCTNNNTAAGCTTGCCAAGTCACTGTCA
GAGCACACGTCTGAACTCTTTCCCTACACGACGCTCTTCCGATCTNNNGGTGTGATTGTGCTACTGTAC
AGCACACGTCTGAACTCTTTCCCTACACGACGCTCTTCCGATCTNNNTCCCTGGGCAATTGTACTCATTC
GCACACGTCTGAACTCTTTCCCTACACGACGCTCTTCCGATCTNNNTTGCCTTCACCAACACACATACACA
AGCACACGTCTGAACTCTTTCCCTACACGACGCTCTTCCGATCTNNNAAAGGGATGGAGAAACAGAGC
AGAGCACACGTCTGAACTCTTTCCCTACACGACGCTCTTCCGATCTNNNTCATAAATACACACAGCGGCA
AGAGCACACGTCTGAACTCTTTCCCTACACGACGCTCTTCCGATCTNNNGCACATGATCCAACGTTAGCA
AGCACACGTCTGAACTCTTTCCCTACACGACGCTCTTCCGATCTNNNTGCAGGGAATTTAGGAAAGAG
GAGCACACGTCTGAACTCTTTCCCTACACGACGCTCTTCCGATCTNNNTTGATCCCTCAAGTCCTGAATG
AGCACACGTCTGAACTCTTTCCCTACACGACGCTCTTCCGATCTNNNATTGCACCTTTAGGTAACCCAGC
GAGCACACGTCTGAACTCTTTCCCTACACGACGCTCTTCCGATCTNNNTCTTGAGCATGCACACATTTTC
AGCACACGTCTGAACTCTTTCCCTACACGACGCTCTTCCGATCTNNNGGGCATCAGGTTTTCTTTCATG
AGCACACGTCTGAACTCTTTCCCTACACGACGCTCTTCCGATCTNNNCTTTCCCCACTCCCCTCTTTCTACA
AGCACACGTCTGAACTCTTTCCCTACACGACGCTCTTCCGATCTNNNAATTCTCTCAAGGGTAGTGGGC
GAGCACACGTCTGAACTCTTTCCCTACACGACGCTCTTCCGATCTNNNAGGGTTCCCTCACAGATGAAT
AGAGCACACGTCTGAACTCTTTCCCTACACGACGCTCTTCCGATCTNNNCGGCCTCTTCTTGCACATTTCT
GCACACGTCTGAACTCTTTCCCTACACGACGCTCTTCCGATCTNNNTCACTGGCTCATTTCAACTGCATCA
AGCACACGTCTGAACTCTTTCCCTACACGACGCTCTTCCGATCTNNNCAGAGTTGTGTAAGAGCCTTGC
GAGCACACGTCTGAACTCTTTCCCTACACGACGCTCTTCCGATCTNNNGTTCCTTCCTCCATGGCATTCT
CACACGTCTGAACTCTTTCCCTACACGACGCTCTTCCGATCTNNNAAGTCATTATTCATACTCCCCTCTCCA
GAGCACACGTCTGAACTCTTTCCCTACACGACGCTCTTCCGATCTNNNAAAGCTCACCTATTGCCCTCAAC
GCACACGTCTGAACTCTTTCCCTACACGACGCTCTTCCGATCTNNNTGGTTTACTTTGTGGTTCTCGGGC
GAGCACACGTCTGAACTCTTTCCCTACACGACGCTCTTCCGATCTNNNGCAGGGTCCTAGAGAAAACCA
AGCACACGTCTGAACTCTTTCCCTACACGACGCTCTTCCGATCTNNNCACTCATCAACTCCAGGCAACTT
GCACACGTCTGAACTCTTTCCCTACACGACGCTCTTCCGATCTNNNTGCTGGGTCCCAAAATTATCGTTC
GAGCACACGTCTGAACTCTTTCCCTACACGACGCTCTTCCGATCTNNNCCCTTGCCATACAACAACCTAG
AGAGCACACGTCTGAACTCTTTCCCTACACGACGCTCTTCCGATCTNNNATCCAAGGGGAAGGAGTAG
AGCACACGTCTGAACTCTTTCCCTACACGACGCTCTTCCGATCTNNNGTACTGAGCGCAAATACCAAAC
AGCACACGTCTGAACTCTTTCCCTACACGACGCTCTTCCGATCTNNNCCTGACAGGTTTTACTACCCCTC
AGCACACGTCTGAACTCTTTCCCTACACGACGCTCTTCCGATCTNNNTTAGTGCAGAAGACTTGCTTGCT

GAGCACACGTCTGAACTCTTTCCCTACACGACGCTCTTCCGATCTNNNAACAGCAACAGAGCAATATCC
AGCACACGTCTGAACTCTTTCCCTACACGACGCTCTTCCGATCTNNNCGCCACCAATGGATCTGATTAC
AGCACACGTCTGAACTCTTTCCCTACACGACGCTCTTCCGATCTNNNTTTCTCATCACCACTCACCACACC
GAGCACACGTCTGAACTCTTTCCCTACACGACGCTCTTCCGATCTNNNGAATGGTTGTCATCAGCCCTA
AGCACACGTCTGAACTCTTTCCCTACACGACGCTCTTCCGATCTNNNTCGTTGGGTGACACTAGCATAT
GAGCACACGTCTGAACTCTTTCCCTACACGACGCTCTTCCGATCTNNNTGCCTTGTTTGGAAGACCTTGA
GAGCACACGTCTGAACTCTTTCCCTACACGACGCTCTTCCGATCTNNNAGCTGATTTTTACTGCGTCTCC
GAGCACACGTCTGAACTCTTTCCCTACACGACGCTCTTCCGATCTNNNAGGCACTGAGAAATCTTATAG
GAGCACACGTCTGAACTCTTTCCCTACACGACGCTCTTCCGATCTNNNCCAGGTTTATGTGCCTTTCTGG
AGAGCACACGTCTGAACTCTTTCCCTACACGACGCTCTTCCGATCTNNNGAAAGATGAATTGTGTGCG
AGCACACGTCTGAACTCTTTCCCTACACGACGCTCTTCCGATCTNNNCCACTCTGCCTGCTTATGTCATTC
GAGCACACGTCTGAACTCTTTCCCTACACGACGCTCTTCCGATCTNNNGAAGGCAGCTTCCTGTAAGTC
GAGCACACGTCTGAACTCTTTCCCTACACGACGCTCTTCCGATCTNNNACTTGAATCCTGCTTTTGCCTT
AGAGCACACGTCTGAACTCTTTCCCTACACGACGCTCTTCCGATCTNNNGCTAGACTGGGCCTCCTACAT
GAGCACACGTCTGAACTCTTTCCCTACACGACGCTCTTCCGATCTNNNTACCATGGAGTTAGGCACAAG
AGCACACGTCTGAACTCTTTCCCTACACGACGCTCTTCCGATCTNNNAAACAACCAACCAACCACCCAAT
AGCACACGTCTGAACTCTTTCCCTACACGACGCTCTTCCGATCTNNNGGAACGTGAAGTACCACTCTCA
AGCACACGTCTGAACTCTTTCCCTACACGACGCTCTTCCGATCTNNNTGGCATTGTCCTCAAAACATCCAG
AGCACACGTCTGAACTCTTTCCCTACACGACGCTCTTCCGATCTNNNTTCCCATCTTGCTGCTTCTTCTCC
AGCACACGTCTGAACTCTTTCCCTACACGACGCTCTTCCGATCTNNNGCTGATTCTATCTCTCCCCACCTCA
AGCACACGTCTGAACTCTTTCCCTACACGACGCTCTTCCGATCTNNNGTTATGGTGGGATGCGACTTCA
AGCACACGTCTGAACTCTTTCCCTACACGACGCTCTTCCGATCTNNNTTATTGCATCCCCATCAGTGTCC
AGCACACGTCTGAACTCTTTCCCTACACGACGCTCTTCCGATCTNNNGTGGTGCATTCCTCTCTTCCAAAG
AGCACACGTCTGAACTCTTTCCCTACACGACGCTCTTCCGATCTNNNAAGGTGAGAAGCCGTTATTGAC
GAGCACACGTCTGAACTCTTTCCCTACACGACGCTCTTCCGATCTNNNAAAGACATTTACAGGGCCAGG
GAGCACACGTCTGAACTCTTTCCCTACACGACGCTCTTCCGATCTNNNATTCAGCTGCTCAGTGTCATCA
GCACACGTCTGAACTCTTTCCCTACACGACGCTCTTCCGATCTNNNAGACCCATCAGAGATTTGTCCCAC
GAGCACACGTCTGAACTCTTTCCCTACACGACGCTCTTCCGATCTNNNGGGGCCAGAAAATGACAAGAT
GCACACGTCTGAACTCTTTCCCTACACGACGCTCTTCCGATCTNNNTGAACTTTTTGAGGACCCCACACCA
GCACACGTCTGAACTCTTTCCCTACACGACGCTCTTCCGATCTNNNACGTCGCTCCAGTGTAAACAAAA
AGCACACGTCTGAACTCTTTCCCTACACGACGCTCTTCCGATCTNNNTGTCCACATCATTTCCTTGCACA
AGCACACGTCTGAACTCTTTCCCTACACGACGCTCTTCCGATCTNNNAGTTGCACTGCTGTGAGAAAAG
AGCACACGTCTGAACTCTTTCCCTACACGACGCTCTTCCGATCTNNNCTGGGTAAAGAGTGAGGCTGTG
AGCACACGTCTGAACTCTTTCCCTACACGACGCTCTTCCGATCTNNNTTGCTTGTTCTTTTTCTGTGCCCC
AGCACACGTCTGAACTCTTTCCCTACACGACGCTCTTCCGATCTNNNTGAATGATCTGCATTTGGGTGG
AGCACACGTCTGAACTCTTTCCCTACACGACGCTCTTCCGATCTNNNGGGATCACCAGCACCTTATTGTT
GAGCACACGTCTGAACTCTTTCCCTACACGACGCTCTTCCGATCTNNNCATGGCAACTGGTTTCTCCCAT
GAGCACACGTCTGAACTCTTTCCCTACACGACGCTCTTCCGATCTNNNAGCTAACCTAAGAGAGCAGGC
AGCACACGTCTGAACTCTTTCCCTACACGACGCTCTTCCGATCTNNNTTTTATCCAGCTGAGAAGCAGG
GCACACGTCTGAACTCTTTCCCTACACGACGCTCTTCCGATCTNNNAGGTACTAGCCTCTCTCTCTCTCTC
GAGCACACGTCTGAACTCTTTCCCTACACGACGCTCTTCCGATCTNNNTCCTCTAACTCACTCGCTCTCTC
GAGCACACGTCTGAACTCTTTCCCTACACGACGCTCTTCCGATCTNNNTGTGTTGTTGCACCTACCCAAA
AGCACACGTCTGAACTCTTTCCCTACACGACGCTCTTCCGATCTNNNCCCAAGCTTGATAGGAGCTTTC

CACACGTCTGAACTCTTTCCCTACACGACGCTCTTCCGATCTNNNCCATCAGCTTGCTTTGGTCTGTAC
GCACACGTCTGAACTCTTTCCCTACACGACGCTCTTCCGATCTNNNCTGTTCCTAGAGCCTCGTTTTCC
GCACACGTCTGAACTCTTTCCCTACACGACGCTCTTCCGATCTNNNCACAGGAATGAGTGACTGTGG
AGCACACGTCTGAACTCTTTCCCTACACGACGCTCTTCCGATCTNNNGCCTGTTGTTCAAAGACACTTC
GCACACGTCTGAACTCTTTCCCTACACGACGCTCTTCCGATCTNNNAGGCAAGGATGAAAACAGCCA
AGCACACGTCTGAACTCTTTCCCTACACGACGCTCTTCCGATCTNNNGACAGCGATCAACGTGTAGT
AGCACACGTCTGAACTCTTTCCCTACACGACGCTCTTCCGATCTNNNGTGTTAGTGAAGGTTCTCC
GAGCACACGTCTGAACTCTTTCCCTACACGACGCTCTTCCGATCTNNNCCCTCAAGGAGTTTACACTC
AGCACACGTCTGAACTCTTTCCCTACACGACGCTCTTCCGATCTNNNTATTAGTAGAGACTGGCCTC
GCACACGTCTGAACTCTTTCCCTACACGACGCTCTTCCGATCTNNNGGATTCAACGTACCCTTCATGC
AGCACACGTCTGAACTCTTTCCCTACACGACGCTCTTCCGATCTNNNGCCTACCTGATCCAAGTAAAG
GCACACGTCTGAACTCTTTCCCTACACGACGCTCTTCCGATCTNNNGGTGGGGTAGTGGCTTATCCT
AGCACACGTCTGAACTCTTTCCCTACACGACGCTCTTCCGATCTNNNATGGTCACTGGAACATGGGTA
AGCACACGTCTGAACTCTTTCCCTACACGACGCTCTTCCGATCTNNNCACAAAGACTGCACCATGTTT
AGAGCACACGTCTGAACTCTTTCCCTACACGACGCTCTTCCGATCTNNNAATGCATTCACTCAGCAAGT
GCACACGTCTGAACTCTTTCCCTACACGACGCTCTTCCGATCTNNNTTGACTGTCAGAGTGTTGATGG
AGCACACGTCTGAACTCTTTCCCTACACGACGCTCTTCCGATCTNNNAGCAAATACTGAAGCTGTCTCC
AGCACACGTCTGAACTCTTTCCCTACACGACGCTCTTCCGATCTNNNTGAATTTGGAGAAGGCACCA
GCACACGTCTGAACTCTTTCCCTACACGACGCTCTTCCGATCTNNNGGCTCCAGCATCTTTCCTTCTTT
AGCACACGTCTGAACTCTTTCCCTACACGACGCTCTTCCGATCTNNNCAATTTGATGCAGTCTGATGC
AGCACACGTCTGAACTCTTTCCCTACACGACGCTCTTCCGATCTNNNATACAGCAGCATCTCTCTACGG
GCACACGTCTGAACTCTTTCCCTACACGACGCTCTTCCGATCTNNNAGAGGAGAGAATCAGCAAACA
AGCACACGTCTGAACTCTTTCCCTACACGACGCTCTTCCGATCTNNNCTATTGCCTTTGCAATCCCGAA
AGCACACGTCTGAACTCTTTCCCTACACGACGCTCTTCCGATCTNNNCCCTCCCATTACTGTGGGGAAT
AGCACACGTCTGAACTCTTTCCCTACACGACGCTCTTCCGATCTNNNCACTTCCTGGTTGGAGAGGAG
AGCACACGTCTGAACTCTTTCCCTACACGACGCTCTTCCGATCTNNNGCTCTTCTCTGCTAGTTCTGCT
AGCACACGTCTGAACTCTTTCCCTACACGACGCTCTTCCGATCTNNNCAAGGCTACCACTTTCTCTCCA
AGCACACGTCTGAACTCTTTCCCTACACGACGCTCTTCCGATCTNNNCAGCCAAGTTTGGAGATCCCAT
GAGCACACGTCTGAACTCTTTCCCTACACGACGCTCTTCCGATCTNNNAGAGTGTATAAGAGGGGG
GCACACGTCTGAACTCTTTCCCTACACGACGCTCTTCCGATCTNNNTTTCCCCAAGATGTCCTGGTTTG
AGCACACGTCTGAACTCTTTCCCTACACGACGCTCTTCCGATCTNNNTTGACTGGATATCTGGATTGC
AGCACACGTCTGAACTCTTTCCCTACACGACGCTCTTCCGATCTNNNACAGCATTCTGGTCCCATCTTT
AGCACACGTCTGAACTCTTTCCCTACACGACGCTCTTCCGATCTNNNCAGTATCTGCCTCCATCACCTT
AGCACACGTCTGAACTCTTTCCCTACACGACGCTCTTCCGATCTNNNTGTTTGTGCTGAGTTGTTTGC
GAGCACACGTCTGAACTCTTTCCCTACACGACGCTCTTCCGATCTNNNCTGTGGGGAACCAAAGAGT
AGCACACGTCTGAACTCTTTCCCTACACGACGCTCTTCCGATCTNNNAAACCATGACTGATCTATCCAG
GAGCACACGTCTGAACTCTTTCCCTACACGACGCTCTTCCGATCTNNNCCAGAGCAGTAGTTCCAAGA
GCACACGTCTGAACTCTTTCCCTACACGACGCTCTTCCGATCTNNNTATTTCTGGTGACAGCCTTTTGC
GCACACGTCTGAACTCTTTCCCTACACGACGCTCTTCCGATCTNNNACCCAGTCTCATTTCTGTCTCCA
AGCACACGTCTGAACTCTTTCCCTACACGACGCTCTTCCGATCTNNNCCCTAAGCCCCAACTAGAAAA
AGCACACGTCTGAACTCTTTCCCTACACGACGCTCTTCCGATCTNNNACAACACCGAATTCAAGCGATT
AGCACACGTCTGAACTCTTTCCCTACACGACGCTCTTCCGATCTNNNTTTCTACAGTCAAGCCAGTTTG
GCACACGTCTGAACTCTTTCCCTACACGACGCTCTTCCGATCTNNNGCCAGATGCTCCTTTATCTGGT

GAGCACACGTCTGAACTCTTTCCCTACACGACGCTCTTCCGATCTNNNTTCAATTATGAACAGGAGGG
AGCACACGTCTGAACTCTTTCCCTACACGACGCTCTTCCGATCTNNNGCCAGTGTACCCAGAGTGATT
AGCACACGTCTGAACTCTTTCCCTACACGACGCTCTTCCGATCTNNNTGGTAGTTGAGTGGCAAAATG
GAGCACACGTCTGAACTCTTTCCCTACACGACGCTCTTCCGATCTNNNATTCTTGTGCTTGAGGTGTGC
AGCACACGTCTGAACTCTTTCCCTACACGACGCTCTTCCGATCTNNNCAAAGTCATCATGCCTCCCTCAT
AGCACACGTCTGAACTCTTTCCCTACACGACGCTCTTCCGATCTNNNCTCTAGGGGAACCACAAAAAG
GCACACGTCTGAACTCTTTCCCTACACGACGCTCTTCCGATCTNNNGGTTCAGATTCTCGTAGCTCTGC
CACACGTCTGAACTCTTTCCCTACACGACGCTCTTCCGATCTNNNGATGGAGATGGGCAGAATCAAG
GCACACGTCTGAACTCTTTCCCTACACGACGCTCTTCCGATCTNNNCCCTGGTCTCAGATATGTCTCCC
GCACACGTCTGAACTCTTTCCCTACACGACGCTCTTCCGATCTNNNTCCTGTTCACTTGATCCCTTCCAC
GAGCACACGTCTGAACTCTTTCCCTACACGACGCTCTTCCGATCTNNNACAACAGTTGAGGGTTTTTGC
GAGCACACGTCTGAACTCTTTCCCTACACGACGCTCTTCCGATCTNNNACCGTGTGAGAAATTGTTTGG
AGCACACGTCTGAACTCTTTCCCTACACGACGCTCTTCCGATCTNNNCAAAGCTGGAAACAGCCCTAGA
AGCACACGTCTGAACTCTTTCCCTACACGACGCTCTTCCGATCTNNNTTGGCGTCAGTTACTGGAACTT
GAGCACACGTCTGAACTCTTTCCCTACACGACGCTCTTCCGATCTNNNTGCTCTGTGTAGACCCTGAT
GAGCACACGTCTGAACTCTTTCCCTACACGACGCTCTTCCGATCTNNNAGATCCAACATCCAACAACAA
AGCACACGTCTGAACTCTTTCCCTACACGACGCTCTTCCGATCTNNNTGGGGGCTCTATATTTAGTCGC
AGCACACGTCTGAACTCTTTCCCTACACGACGCTCTTCCGATCTNNNCTCTGCTCACAAGTCCTTCCTTC
GAGCACACGTCTGAACTCTTTCCCTACACGACGCTCTTCCGATCTNNNTCCCACTATGGAATGATGGG
GAGCACACGTCTGAACTCTTTCCCTACACGACGCTCTTCCGATCTNNNTAACTTGGTATGCGTTGCAAA
GCACACGTCTGAACTCTTTCCCTACACGACGCTCTTCCGATCTNNNAGCATGCTAACTTCTGACTTGGT
AGCACACGTCTGAACTCTTTCCCTACACGACGCTCTTCCGATCTNNNCATGCCGCTGGATAAGATTCCAT
GAGCACACGTCTGAACTCTTTCCCTACACGACGCTCTTCCGATCTNNNTTAGCACTCAACTGCTGTCTC
CACACGTCTGAACTCTTTCCCTACACGACGCTCTTCCGATCTNNNAAAGATCCAAAACACTTCTGCTTCC
GAGCACACGTCTGAACTCTTTCCCTACACGACGCTCTTCCGATCTNNNGGCAAGAAAATCACATTGCCC
AGCACACGTCTGAACTCTTTCCCTACACGACGCTCTTCCGATCTNNNGGCAGAGCTCCAAAGTATGTC
GCACACGTCTGAACTCTTTCCCTACACGACGCTCTTCCGATCTNNNTTTTCTGTTTCCTCAGTGAGCCCC
AGCACACGTCTGAACTCTTTCCCTACACGACGCTCTTCCGATCTNNNTGTTGGCTTGTAACCAATGACC
AGCACACGTCTGAACTCTTTCCCTACACGACGCTCTTCCGATCTNNNCTGCGATCTCCTTGTCATAGAG
AGCACACGTCTGAACTCTTTCCCTACACGACGCTCTTCCGATCTNNNTCATTGTCCTGGGCTACAGTTTT
GAGCACACGTCTGAACTCTTTCCCTACACGACGCTCTTCCGATCTNNNATATGTATCACACATGCCCCC
AGCACACGTCTGAACTCTTTCCCTACACGACGCTCTTCCGATCTNNNTGGTGTGTGGTGTGTATCTGT
GCACACGTCTGAACTCTTTCCCTACACGACGCTCTTCCGATCTNNNCAAAGAAAGGAAGATGGCGAG
GAGCACACGTCTGAACTCTTTCCCTACACGACGCTCTTCCGATCTNNNACACTTTGCTTACTTACACCCC
AGCACACGTCTGAACTCTTTCCCTACACGACGCTCTTCCGATCTNNNTTCCCAGGTCAATCTCTCTCTC
GAGCACACGTCTGAACTCTTTCCCTACACGACGCTCTTCCGATCTNNNAGTTCCCATTTCCTCCTTCCTC
GAGCACACGTCTGAACTCTTTCCCTACACGACGCTCTTCCGATCTNNNCATTCTGGTCATCCAGTCCTG
AGCACACGTCTGAACTCTTTCCCTACACGACGCTCTTCCGATCTNNNGAGCACTGGAGGCTGTCTATA
GAGCACACGTCTGAACTCTTTCCCTACACGACGCTCTTCCGATCTNNNCCCAGCAGCTTATGTGAATGT
AGCACACGTCTGAACTCTTTCCCTACACGACGCTCTTCCGATCTNNNGCAACACAAAATCACCACACAC
GCACACGTCTGAACTCTTTCCCTACACGACGCTCTTCCGATCTNNNCACTACAGTCATATCTGGCCCCT
GCACACGTCTGAACTCTTTCCCTACACGACGCTCTTCCGATCTNNNATCCATCACACCATTTCAGAGCCC
GCACACGTCTGAACTCTTTCCCTACACGACGCTCTTCCGATCTNNNCTTGTTCATCAAAAGCTGCACCA

AGCACACGTCTGAACTCTTTCCCTACACGACGCTCTTCCGATCTNNNTTTCTTCCCTCAATTTCTGCCCA
GAGCACACGTCTGAACTCTTTCCCTACACGACGCTCTTCCGATCTNNNAGGGAGACTGGGAATCAGG
GAGCACACGTCTGAACTCTTTCCCTACACGACGCTCTTCCGATCTNNNTGTGCTGTTTTCAGCTCTCTC
GAGCACACGTCTGAACTCTTTCCCTACACGACGCTCTTCCGATCTNNNTGGGAGGACTTAGGATTGT
AGCACACGTCTGAACTCTTTCCCTACACGACGCTCTTCCGATCTNNNACAGCTACACTCACTTCCTCCTG
AGCACACGTCTGAACTCTTTCCCTACACGACGCTCTTCCGATCTNNNCACCTGTAGAACTGGTAAACC
GCACACGTCTGAACTCTTTCCCTACACGACGCTCTTCCGATCTNNNATCTGAGAATCCCAAATGGCAC
GCACACGTCTGAACTCTTTCCCTACACGACGCTCTTCCGATCTNNNAACATGGTGTTCTCAGAGGCA
AGCACACGTCTGAACTCTTTCCCTACACGACGCTCTTCCGATCTNNNGAAGAGGCATCTGACAGCTT
AGCACACGTCTGAACTCTTTCCCTACACGACGCTCTTCCGATCTNNNCCTCCCCATCCCTTACTTATCCC
GCACACGTCTGAACTCTTTCCCTACACGACGCTCTTCCGATCTNNNAGATGTGGCAAAAACCAAACTG
GAGCACACGTCTGAACTCTTTCCCTACACGACGCTCTTCCGATCTNNNTGTTTCCAGGGGGTAAGGA
GAGCACACGTCTGAACTCTTTCCCTACACGACGCTCTTCCGATCTNNNACAACACCTAATCCTCACATG
GAGCACACGTCTGAACTCTTTCCCTACACGACGCTCTTCCGATCTNNNGAATTGAGACCACAGCCCTC
CACACGTCTGAACTCTTTCCCTACACGACGCTCTTCCGATCTNNNCTCTTTGAATATTGGCAGGCTGG
AGCACACGTCTGAACTCTTTCCCTACACGACGCTCTTCCGATCTNNNGGTCACTGGGTCTCTCTCTTTC
AGCACACGTCTGAACTCTTTCCCTACACGACGCTCTTCCGATCTNNNAACAGGGAAAAGGAGAGATG
AGCACACGTCTGAACTCTTTCCCTACACGACGCTCTTCCGATCTNNNTGCATTCAGAAGCTTGGCATT
AGCACACGTCTGAACTCTTTCCCTACACGACGCTCTTCCGATCTNNNTCCTGCTTCTTGTTTGACTTCCC
AGCACACGTCTGAACTCTTTCCCTACACGACGCTCTTCCGATCTNNNATTTCTTCCTCCATGTGTGGGT
AGCACACGTCTGAACTCTTTCCCTACACGACGCTCTTCCGATCTNNNTGGAATCCAGGAGCTGGTCA
AGCACACGTCTGAACTCTTTCCCTACACGACGCTCTTCCGATCTNNNTTCCTCCTTGACCCCTAAAGCAT
GAGCACACGTCTGAACTCTTTCCCTACACGACGCTCTTCCGATCTNNNCTTGAATTCACTGGGTTTGT
GCACACGTCTGAACTCTTTCCCTACACGACGCTCTTCCGATCTNNNAAGACCTCTCCTTCCCTTCTCTGC
GAGCACACGTCTGAACTCTTTCCCTACACGACGCTCTTCCGATCTNNNGTCCAGAGCAAGAATGGTAC
AGCACACGTCTGAACTCTTTCCCTACACGACGCTCTTCCGATCTNNNGAATCATCCCACTCAACACAGC
AGCACACGTCTGAACTCTTTCCCTACACGACGCTCTTCCGATCTNNNTTGCATTGAGGCCTGGGAAAT
AGCACACGTCTGAACTCTTTCCCTACACGACGCTCTTCCGATCTNNNTTGGTCTGGGTACATTGTGTG
GCACACGTCTGAACTCTTTCCCTACACGACGCTCTTCCGATCTNNNACAGGCTTGTGGTGTATATGT
AGCACACGTCTGAACTCTTTCCCTACACGACGCTCTTCCGATCTNNNAACATCTTAGGCTCTGCTTGA
AGCACACGTCTGAACTCTTTCCCTACACGACGCTCTTCCGATCTNNNAGCATTGCTGTGAACTTGTGA
GCACACGTCTGAACTCTTTCCCTACACGACGCTCTTCCGATCTNNNACTAAGAGCAGTCAGTCCCACTT
GCACACGTCTGAACTCTTTCCCTACACGACGCTCTTCCGATCTNNNACTGGAGATGGTCATACTGGCT
AGCACACGTCTGAACTCTTTCCCTACACGACGCTCTTCCGATCTNNNCAATTGCCATTCGTTCCAGTGT
GAGCACACGTCTGAACTCTTTCCCTACACGACGCTCTTCCGATCTNNNTGGTGCAACTTTGTGTCAGG
GAGCACACGTCTGAACTCTTTCCCTACACGACGCTCTTCCGATCTNNNTGCAATTCTTTTTAGCCTGG
AGCACACGTCTGAACTCTTTCCCTACACGACGCTCTTCCGATCTNNNTGAAGTCGAAAGAGGACCACA
AGCACACGTCTGAACTCTTTCCCTACACGACGCTCTTCCGATCTNNNAGGGATAGGGCCAGTGAGA
GCACACGTCTGAACTCTTTCCCTACACGACGCTCTTCCGATCTNNNTATCTCCTGCATCAGTTCCTCCCC
GAGCACACGTCTGAACTCTTTCCCTACACGACGCTCTTCCGATCTNNNCGGCCGTTCTCATGTATCTTG
GCACACGTCTGAACTCTTTCCCTACACGACGCTCTTCCGATCTNNNGTACAATCACCTTCCAGTGGCAT
GCACACGTCTGAACTCTTTCCCTACACGACGCTCTTCCGATCTNNNGTACTGAAGGCCTGCTATGAAC
AGCACACGTCTGAACTCTTTCCCTACACGACGCTCTTCCGATCTNNNTCTGGAGTGTGGTGGTGCAA

GAGCACACGTCTGAACTCTTTCCCTACACGACGCTCTTCCGATCTNNNCACTGCATTAGGGTTCTCCAG
AGCACACGTCTGAACTCTTTCCCTACACGACGCTCTTCCGATCTNNNTGAAAGTTGAACTGAACTCCCC
AGCACACGTCTGAACTCTTTCCCTACACGACGCTCTTCCGATCTNNNGTTTGTGTGCGTAAGTGCATG
AGCACACGTCTGAACTCTTTCCCTACACGACGCTCTTCCGATCTNNNCTGGGAAAGAGAAATGGTAGG
AGCACACGTCTGAACTCTTTCCCTACACGACGCTCTTCCGATCTNNNGTGGAACCCAATGCACTCTTGT
AGAGCACACGTCTGAACTCTTTCCCTACACGACGCTCTTCCGATCTNNNATAGTAACCCACAAACCCCTG
AGCACACGTCTGAACTCTTTCCCTACACGACGCTCTTCCGATCTNNNAAGGACACTCGGAGGAGAAGA
AGCACACGTCTGAACTCTTTCCCTACACGACGCTCTTCCGATCTNNNAGGTGATAAAGTCACTGAAGG
AGCACACGTCTGAACTCTTTCCCTACACGACGCTCTTCCGATCTNNNAGAGTGGATAGAAAAGGGCA
GAGCACACGTCTGAACTCTTTCCCTACACGACGCTCTTCCGATCTNNNATTCAGTAGGGTTGGGACAG
GAGCACACGTCTGAACTCTTTCCCTACACGACGCTCTTCCGATCTNNNGGTTGCTAATGTCTTCGGGAA
GAGCACACGTCTGAACTCTTTCCCTACACGACGCTCTTCCGATCTNNNGAGTTACACAGTTGGCAACAG
GAGCACACGTCTGAACTCTTTCCCTACACGACGCTCTTCCGATCTNNNGGCACTTTCAAAAGTCATGCTC
GAGCACACGTCTGAACTCTTTCCCTACACGACGCTCTTCCGATCTNNNAAGCATTCCTTGGTCTGGTTC
AGCACACGTCTGAACTCTTTCCCTACACGACGCTCTTCCGATCTNNNGCCAATATTGCGCCACTAAAGT
GAGCACACGTCTGAACTCTTTCCCTACACGACGCTCTTCCGATCTNNNTTTCTACCAAAGGGCTAAGTG
AGAGCACACGTCTGAACTCTTTCCCTACACGACGCTCTTCCGATCTNNNTCTGAGAGGGCTGTTTTGTC
GCACACGTCTGAACTCTTTCCCTACACGACGCTCTTCCGATCTNNNTGCATCCTGACTAATAAAGCCCA
AGCACACGTCTGAACTCTTTCCCTACACGACGCTCTTCCGATCTNNNTATGGGAGAGACCCAGTATGC
AGCACACGTCTGAACTCTTTCCCTACACGACGCTCTTCCGATCTNNNCCCTCCTCTGTCTCCCTTTTCATC
AGCACACGTCTGAACTCTTTCCCTACACGACGCTCTTCCGATCTNNNACCCTCTTCTTTCCTTAGCCACAC
AGCACACGTCTGAACTCTTTCCCTACACGACGCTCTTCCGATCTNNNAGATTCCAGAGCCAGAGGTGAA
GCACACGTCTGAACTCTTTCCCTACACGACGCTCTTCCGATCTNNNAAGTACCCATCCCAGTGTCTTGTC
AGCACACGTCTGAACTCTTTCCCTACACGACGCTCTTCCGATCTNNNGTGGGAGTAAGAAGAGGGAG
AGCACACGTCTGAACTCTTTCCCTACACGACGCTCTTCCGATCTNNNATGTGGATGGGAGAGTGTGAA
AGCACACGTCTGAACTCTTTCCCTACACGACGCTCTTCCGATCTNNNTAACTCCAGCCTCATCTCTGTCCC
AGCACACGTCTGAACTCTTTCCCTACACGACGCTCTTCCGATCTNNNGTTTACTATGTGCCCCTTCCCAG
AGCACACGTCTGAACTCTTTCCCTACACGACGCTCTTCCGATCTNNNCTCTGCCTTCGAATACCAGCTTT
AGAGCACACGTCTGAACTCTTTCCCTACACGACGCTCTTCCGATCTNNNTCCCCTATCTGTGTGGGTTTT
GAGCACACGTCTGAACTCTTTCCCTACACGACGCTCTTCCGATCTNNNTATGTTTCCCCTTCACCCTAGCA
AGCACACGTCTGAACTCTTTCCCTACACGACGCTCTTCCGATCTNNNTTGCTGACTTTGCCAACAATTCC
AGCACACGTCTGAACTCTTTCCCTACACGACGCTCTTCCGATCTNNNGAGGGTAAAGTCAAACCAGGC
AGCACACGTCTGAACTCTTTCCCTACACGACGCTCTTCCGATCTNNNCTAGGCTTTTCCAATGGGACCTC
GAGCACACGTCTGAACTCTTTCCCTACACGACGCTCTTCCGATCTNNNCCTCCGTGAGGAAGATTATGC
AGCACACGTCTGAACTCTTTCCCTACACGACGCTCTTCCGATCTNNNTCTCATAAGAACCTACCTGGCCC
GAGCACACGTCTGAACTCTTTCCCTACACGACGCTCTTCCGATCTNNNTGCAGTATTGGACACGACAGA
GAGCACACGTCTGAACTCTTTCCCTACACGACGCTCTTCCGATCTNNNAGTCTTTCCCTGTCTCTGTCTCT
GAGCACACGTCTGAACTCTTTCCCTACACGACGCTCTTCCGATCTNNNATTGTTTTCTGATGGGTGAGG
AGCACACGTCTGAACTCTTTCCCTACACGACGCTCTTCCGATCTNNNAGGATGGTAAGGATGAAAAAG
AGCACACGTCTGAACTCTTTCCCTACACGACGCTCTTCCGATCTNNNCCAGCCCTTTTCTTGTGCAATTTC
AGCACACGTCTGAACTCTTTCCCTACACGACGCTCTTCCGATCTNNNTTTCCTGTTGACTGAATGGTGGA
GAGCACACGTCTGAACTCTTTCCCTACACGACGCTCTTCCGATCTNNNTTTGGCTGTTTTTGGATGCTAC
AGCACACGTCTGAACTCTTTCCCTACACGACGCTCTTCCGATCTNNNAGCAAATCTTCCATGTTCCTCGC

AGCACACGTCTGAACTCTTTCCCTACACGACGCTCTTCCGATCTNNNTGTTCCTTCTCAACACCTTGTG
AGCACACGTCTGAACTCTTTCCCTACACGACGCTCTTCCGATCTNNNTCTCAAATGGTTCAGCAAAAA
AGCACACGTCTGAACTCTTTCCCTACACGACGCTCTTCCGATCTNNNAGGGAAGATAGTCGAGGAG
GAGCACACGTCTGAACTCTTTCCCTACACGACGCTCTTCCGATCTNNNCAGTGGTGTATTTGGCTACA
GCACACGTCTGAACTCTTTCCCTACACGACGCTCTTCCGATCTNNNTTTGGGTTTGTATGTCTTGCCCT
GCACACGTCTGAACTCTTTCCCTACACGACGCTCTTCCGATCTNNNCTGTACTCCTTGTTCTGCACCTC
GCACACGTCTGAACTCTTTCCCTACACGACGCTCTTCCGATCTNNNAAACCTAGAGTTGGATGTGCGT
GAGCACACGTCTGAACTCTTTCCCTACACGACGCTCTTCCGATCTNNNCACTTGTCTACAGCATCAGC
GCACACGTCTGAACTCTTTCCCTACACGACGCTCTTCCGATCTNNNATCCAGGTTGGTAGTTCTGTGG
GAGCACACGTCTGAACTCTTTCCCTACACGACGCTCTTCCGATCTNNNTTTGCTGTTGCTGAAAATGG
GCACACGTCTGAACTCTTTCCCTACACGACGCTCTTCCGATCTNNNAAGGTAAAGCCAAAGAAAGGC
GCACACGTCTGAACTCTTTCCCTACACGACGCTCTTCCGATCTNNNCTTCATCGGGCTTCAGTGGTTT
GCACACGTCTGAACTCTTTCCCTACACGACGCTCTTCCGATCTNNNACTCCTACTCTACTGAGACACGC
GCACACGTCTGAACTCTTTCCCTACACGACGCTCTTCCGATCTNNNTGCATGAAAACCTTCAGGTGGA
GCACACGTCTGAACTCTTTCCCTACACGACGCTCTTCCGATCTNNNGTCTGGAATGTTCCGCAATCTCT
AGCACACGTCTGAACTCTTTCCCTACACGACGCTCTTCCGATCTNNNAGCAAGGTCAATCCTGTCTCTC
AGCACACGTCTGAACTCTTTCCCTACACGACGCTCTTCCGATCTNNNGGGCCAAAGAAGGTGGTAAA
GAGCACACGTCTGAACTCTTTCCCTACACGACGCTCTTCCGATCTNNNTTGCCTGGGGTTTTTAGAGT
GAGCACACGTCTGAACTCTTTCCCTACACGACGCTCTTCCGATCTNNNTTTACCAAGGAACAGAGCCA
AGCACACGTCTGAACTCTTTCCCTACACGACGCTCTTCCGATCTNNNAACCACTGCCCTGGTAAAGAAA
AGCACACGTCTGAACTCTTTCCCTACACGACGCTCTTCCGATCTNNNATGTTTTCTTCCAACCTCTGGC
AGCACACGTCTGAACTCTTTCCCTACACGACGCTCTTCCGATCTNNNGATGCTTCCAGTCAGATGCTAC
GAGCACACGTCTGAACTCTTTCCCTACACGACGCTCTTCCGATCTNNNTTTTGAACCCACTAAGGAGC
AGCACACGTCTGAACTCTTTCCCTACACGACGCTCTTCCGATCTNNNTTGTTGTTCAACCCACGATGTC
AGCACACGTCTGAACTCTTTCCCTACACGACGCTCTTCCGATCTNNNAGAAAACCAAAGGATGCCTTC
GCACACGTCTGAACTCTTTCCCTACACGACGCTCTTCCGATCTNNNGGGTGGGAGACAGTAAAAGG
GCACACGTCTGAACTCTTTCCCTACACGACGCTCTTCCGATCTNNNCCGAGAAGCGACTGGTCATTAT
AGCACACGTCTGAACTCTTTCCCTACACGACGCTCTTCCGATCTNNNAGCCTAATCAAAGCCATTTGT
GAGCACACGTCTGAACTCTTTCCCTACACGACGCTCTTCCGATCTNNNAGAGAGGAACTATGCCTCA
GAGCACACGTCTGAACTCTTTCCCTACACGACGCTCTTCCGATCTNNNATTAGCCTTGGCGATCTTGT
AGCACACGTCTGAACTCTTTCCCTACACGACGCTCTTCCGATCTNNNAGGCAAAATCAGTGAAGCTG
GCACACGTCTGAACTCTTTCCCTACACGACGCTCTTCCGATCTNNNTGGATTCCAAATGGAGCCAAC
GAGCACACGTCTGAACTCTTTCCCTACACGACGCTCTTCCGATCTNNNAAACACTAGGGGAATTCAG
GAGCACACGTCTGAACTCTTTCCCTACACGACGCTCTTCCGATCTNNNTCCCCAGGTGAGAAAGAATG
AGCACACGTCTGAACTCTTTCCCTACACGACGCTCTTCCGATCTNNNGGGGTAGCGATCCTGTCATTT
AGCACACGTCTGAACTCTTTCCCTACACGACGCTCTTCCGATCTNNNCACACACAATGGAATGGCTTCT
AGCACACGTCTGAACTCTTTCCCTACACGACGCTCTTCCGATCTNNNAGTGAAGCTCACCTCACCCAT
GAGCACACGTCTGAACTCTTTCCCTACACGACGCTCTTCCGATCTNNNTGCTTGTTGTGTTATGACTTG
AGCACACGTCTGAACTCTTTCCCTACACGACGCTCTTCCGATCTNNNTTGCATGGTCTCTATGAGTGT
AAGAGCACACGTCTGAACTCTTTCCCTACACGACGCTCTTCCGATCTNNNGTGTGTAAACCAAGTCCCA
AGCACACGTCTGAACTCTTTCCCTACACGACGCTCTTCCGATCTNNNCCAGTGGCCTTCACACTTTTCT
AGCACACGTCTGAACTCTTTCCCTACACGACGCTCTTCCGATCTNNNCTATTAGTCCAAGGTTGGGCA
GAGCACACGTCTGAACTCTTTCCCTACACGACGCTCTTCCGATCTNNNACCAGACCTGAACCTACCAAT

AGCACACGTCTGAACTCTTTCCCTACACGACGCTCTTCCGATCTNNNATCAGGGTGATGTGTGTGTGT
GAGCACACGTCTGAACTCTTTCCCTACACGACGCTCTTCCGATCTNNNCAGTTCTGCCATATCAGTGGC
GAGCACACGTCTGAACTCTTTCCCTACACGACGCTCTTCCGATCTNNNTCCACTGGGATCTCAGAAATG
AGAGCACACGTCTGAACTCTTTCCCTACACGACGCTCTTCCGATCTNNNAGCAGAAACCACAATCTCTCT
GAGCACACGTCTGAACTCTTTCCCTACACGACGCTCTTCCGATCTNNNGGCCTGACGGGTAAAGCTATT
GAGCACACGTCTGAACTCTTTCCCTACACGACGCTCTTCCGATCTNNNACTGAATGCTTGCTCTCAAAG
GAGCACACGTCTGAACTCTTTCCCTACACGACGCTCTTCCGATCTNNNACTACCATGAACTAGCAGACA
AGCACACGTCTGAACTCTTTCCCTACACGACGCTCTTCCGATCTNNNTTGGTTTTCCATTCCATCCAGGT
GAGCACACGTCTGAACTCTTTCCCTACACGACGCTCTTCCGATCTNNNCCTCCTCACAGGTCTCCCTAAA
GAGCACACGTCTGAACTCTTTCCCTACACGACGCTCTTCCGATCTNNNTACATGTGTGTTTCTGGGTGC
GAGCACACGTCTGAACTCTTTCCCTACACGACGCTCTTCCGATCTNNNATGCAGGTATATGCAGGACT
GCACACGTCTGAACTCTTTCCCTACACGACGCTCTTCCGATCTNNNAGGAGCAATGGAGCAGATACTC
GAGCACACGTCTGAACTCTTTCCCTACACGACGCTCTTCCGATCTNNNTGGACAATAGGCAACTGGGT
AGAGCACACGTCTGAACTCTTTCCCTACACGACGCTCTTCCGATCTNNNATGGCATTGGTGAGTTGTT
GAGCACACGTCTGAACTCTTTCCCTACACGACGCTCTTCCGATCTNNNGTGTGATTGCCTAGAGATTG
AGCACACGTCTGAACTCTTTCCCTACACGACGCTCTTCCGATCTNNNATCCACTAGCCTCAGGGAACAT
GAGCACACGTCTGAACTCTTTCCCTACACGACGCTCTTCCGATCTNNNATTCTCATGGGTGCATTGCTT
AGCACACGTCTGAACTCTTTCCCTACACGACGCTCTTCCGATCTNNNCATGAGGTGGATGAAAGGAC
AGAGCACACGTCTGAACTCTTTCCCTACACGACGCTCTTCCGATCTNNNAATTGACTAGCTGGACTGTG
GAGCACACGTCTGAACTCTTTCCCTACACGACGCTCTTCCGATCTNNNAAATAGGTTTGGGAGGTTGA
GAGCACACGTCTGAACTCTTTCCCTACACGACGCTCTTCCGATCTNNNAGTTTGGAAAAGAGATCTGCT
AGAGCACACGTCTGAACTCTTTCCCTACACGACGCTCTTCCGATCTNNNACTTACACGTCTGGTACCTCT
AGCACACGTCTGAACTCTTTCCCTACACGACGCTCTTCCGATCTNNNTTGTCATAAATGCCTGGCAATG
AGCACACGTCTGAACTCTTTCCCTACACGACGCTCTTCCGATCTNNNTTGAAGAAGGGACTGAGCGAA
GAGCACACGTCTGAACTCTTTCCCTACACGACGCTCTTCCGATCTNNNGGTTCTCCAAGAGGTATCCAG
AGAGCACACGTCTGAACTCTTTCCCTACACGACGCTCTTCCGATCTNNNCAGGTGCTGCCTATGAGTAG
AGCACACGTCTGAACTCTTTCCCTACACGACGCTCTTCCGATCTNNNTCACCCACATTTGAAACAGGAC
AGCACACGTCTGAACTCTTTCCCTACACGACGCTCTTCCGATCTNNNTGAGCTAAGTACAGGACCCACA
GAGCACACGTCTGAACTCTTTCCCTACACGACGCTCTTCCGATCTNNNACAAGGGCCATGCTTTTGATT
GAGCACACGTCTGAACTCTTTCCCTACACGACGCTCTTCCGATCTNNNGCAGTCTTTTGTTTGCTGAGA
AGAGCACACGTCTGAACTCTTTCCCTACACGACGCTCTTCCGATCTNNNGACATCCTTCATTCCTAAGG
GAGCACACGTCTGAACTCTTTCCCTACACGACGCTCTTCCGATCTNNNGGAAAAAGCCCAGTTCTCAAT
AGAGCACACGTCTGAACTCTTTCCCTACACGACGCTCTTCCGATCTNNNAGGAGAAATTCAACAAGCC
AGAGCACACGTCTGAACTCTTTCCCTACACGACGCTCTTCCGATCTNNNCTGGACTACAGGAGGCCCTA
GAGCACACGTCTGAACTCTTTCCCTACACGACGCTCTTCCGATCTNNNCTCCTTCCCCAGGTTATCATTG
GCACACGTCTGAACTCTTTCCCTACACGACGCTCTTCCGATCTNNNTCCAAGTGTTTTCAGAAGTCCCC
GAGCACACGTCTGAACTCTTTCCCTACACGACGCTCTTCCGATCTNNNTTGTTCCAGGATGTGTCTGT
GAGCACACGTCTGAACTCTTTCCCTACACGACGCTCTTCCGATCTNNNATAGGGTTGACAGGTCTTCCC
AGAGCACACGTCTGAACTCTTTCCCTACACGACGCTCTTCCGATCTNNNAAACTGAGCACAGCTGATGA
GAGCACACGTCTGAACTCTTTCCCTACACGACGCTCTTCCGATCTNNNTCCTTCTGTGTTCCTCTTTGTG
GAGCACACGTCTGAACTCTTTCCCTACACGACGCTCTTCCGATCTNNNGGATACAAGGCATCTCAAGC
AGAGCACACGTCTGAACTCTTTCCCTACACGACGCTCTTCCGATCTNNNGGCCCTGAAGAGAAAAGA
AGCACACGTCTGAACTCTTTCCCTACACGACGCTCTTCCGATCTNNNCCTGCTGGTCTCCCAATTCTTTG

GAGCACACGTCTGAACTCTTTCCCTACACGACGCTCTTCCGATCTNNNTATGCCCATGCTCCAAAAACCT
GAGCACACGTCTGAACTCTTTCCCTACACGACGCTCTTCCGATCTNNNTCTCTCCTCATTGTGTACAACC
GCACACGTCTGAACTCTTTCCCTACACGACGCTCTTCCGATCTNNNAAACTCTTGCATCCTGGGAAACC
AGAGCACACGTCTGAACTCTTTCCCTACACGACGCTCTTCCGATCTNNNAGCATGTCTTTCTTTCCTCAT
GAGCACACGTCTGAACTCTTTCCCTACACGACGCTCTTCCGATCTNNNGCACCCCCTTTACTCTCTATCCT
AGAGCACACGTCTGAACTCTTTCCCTACACGACGCTCTTCCGATCTNNNTGTGTGTGTGTAGATTCCAG
GAGCACACGTCTGAACTCTTTCCCTACACGACGCTCTTCCGATCTNNNATGATGCTTTGGTTATTGCGG
GAGCACACGTCTGAACTCTTTCCCTACACGACGCTCTTCCGATCTNNNTAGCTCTTTGCTCCATTCAGTG
AGCACACGTCTGAACTCTTTCCCTACACGACGCTCTTCCGATCTNNNAAACATCTGGAGGACCTCTCTG
AGCACACGTCTGAACTCTTTCCCTACACGACGCTCTTCCGATCTNNNACTTCCCAGTATGAATGCACCA
AGCACACGTCTGAACTCTTTCCCTACACGACGCTCTTCCGATCTNNNTAGAACTGGCCCCTTGGGATAT
AGCACACGTCTGAACTCTTTCCCTACACGACGCTCTTCCGATCTNNNACATTCGAAATCAAGCTGTTGG
GAGCACACGTCTGAACTCTTTCCCTACACGACGCTCTTCCGATCTNNNTAAAATCGCTCAAGACACAGC
AGAGCACACGTCTGAACTCTTTCCCTACACGACGCTCTTCCGATCTNNNGCACTCAATTGCTTACTTCCA
AGCACACGTCTGAACTCTTTCCCTACACGACGCTCTTCCGATCTNNNAGTCCGGTAACTCATCTGTTTGC
GAGCACACGTCTGAACTCTTTCCCTACACGACGCTCTTCCGATCTNNNTCATTGAGAGACAAGGTCCTG
GAGCACACGTCTGAACTCTTTCCCTACACGACGCTCTTCCGATCTNNNCCCAAACTGTCTGCCAGTAAT
GAGCACACGTCTGAACTCTTTCCCTACACGACGCTCTTCCGATCTNNNCCGCTTTCTTTTCTCATGGTTG
AGCACACGTCTGAACTCTTTCCCTACACGACGCTCTTCCGATCTNNNTCTACGGAGGACTGGATGAAG
AGCACACGTCTGAACTCTTTCCCTACACGACGCTCTTCCGATCTNNNAGCGTTTTGACTTCTTGTTCCA
GAGCACACGTCTGAACTCTTTCCCTACACGACGCTCTTCCGATCTNNNACAAGAAGACAGGGATACG
GAGCACACGTCTGAACTCTTTCCCTACACGACGCTCTTCCGATCTNNNCTGGACAATAGAGCGAGTAA
AGAGCACACGTCTGAACTCTTTCCCTACACGACGCTCTTCCGATCTNNNTCCCTTCAACCACAGCTCTTA
AGAGCACACGTCTGAACTCTTTCCCTACACGACGCTCTTCCGATCTNNNAAAAGAGGGGGTCAGGGA
GAGCACACGTCTGAACTCTTTCCCTACACGACGCTCTTCCGATCTNNNGCTTGATGGGGGATGTAGAG
GAGCACACGTCTGAACTCTTTCCCTACACGACGCTCTTCCGATCTNNNTTCTTGCATTGTCTCCTTGCTG
AGAGCACACGTCTGAACTCTTTCCCTACACGACGCTCTTCCGATCTNNNAGGCCACATGGAGGAGTTA
AGCACACGTCTGAACTCTTTCCCTACACGACGCTCTTCCGATCTNNNCTTCCAGCCTTTGGACATGAAC
AGCACACGTCTGAACTCTTTCCCTACACGACGCTCTTCCGATCTNNNCCTCTGCATTAGTTTGGTCAGG
AGCACACGTCTGAACTCTTTCCCTACACGACGCTCTTCCGATCTNNNATCTGAATCTGAACAGTGGGT
AGCACACGTCTGAACTCTTTCCCTACACGACGCTCTTCCGATCTNNNACATGCTCCTCAGTCAACAACTC
AGAGCACACGTCTGAACTCTTTCCCTACACGACGCTCTTCCGATCTNNNTGACTGCTGCACTCTCTTTT
GAGCACACGTCTGAACTCTTTCCCTACACGACGCTCTTCCGATCTNNNAAAGCAGTTTGGTTGATGAG
GAGCACACGTCTGAACTCTTTCCCTACACGACGCTCTTCCGATCTNNNTCTGACTTTCCAAGTTGGCTG
GAGCACACGTCTGAACTCTTTCCCTACACGACGCTCTTCCGATCTNNNCCTAATGGTAGAAAGGCTTC
GAGCACACGTCTGAACTCTTTCCCTACACGACGCTCTTCCGATCTNNNATTTTCAGAGGAGATGGGGG
AGCACACGTCTGAACTCTTTCCCTACACGACGCTCTTCCGATCTNNNTTCCCATGTTAGAAGAGAGGG
AGCACACGTCTGAACTCTTTCCCTACACGACGCTCTTCCGATCTNNNGAAAGCAGCATCTGGGAAGGT
AGCACACGTCTGAACTCTTTCCCTACACGACGCTCTTCCGATCTNNNGCTCTGTGGTCAAGTCTAGGA
GAGCACACGTCTGAACTCTTTCCCTACACGACGCTCTTCCGATCTNNNACAAGACACAGACATCTTCCTG
GAGCACACGTCTGAACTCTTTCCCTACACGACGCTCTTCCGATCTNNNTACAGCACTCCAAGTCTCTGA
AGCACACGTCTGAACTCTTTCCCTACACGACGCTCTTCCGATCTNNNTGACTTACAGAGTTTCAGACTG
GAGCACACGTCTGAACTCTTTCCCTACACGACGCTCTTCCGATCTNNNTTAGACAAGCTGCAGGAAGA

AGCACACGTCTGAACTCTTTCCCTACACGACGCTCTTCCGATCTNNNAATGGGCAAAGTGGTGTTTTTC
AGAGCACACGTCTGAACTCTTTCCCTACACGACGCTCTTCCGATCTNNNTGCCATTGATACACATTGCCT
AGAGCACACGTCTGAACTCTTTCCCTACACGACGCTCTTCCGATCTNNNGCAGACTTGTTCAGAGAGC
GAGCACACGTCTGAACTCTTTCCCTACACGACGCTCTTCCGATCTNNNGCTTTGAATCCAGCTGCAACAT
AGCACACGTCTGAACTCTTTCCCTACACGACGCTCTTCCGATCTNNNTGTATGTGTGTGCGGTAGTGTT
AGAGCACACGTCTGAACTCTTTCCCTACACGACGCTCTTCCGATCTNNNGGGAACTCAGACAAATGGG
GAGCACACGTCTGAACTCTTTCCCTACACGACGCTCTTCCGATCTNNNAGTGGAGACAGACCCACAGAT
AGAGCACACGTCTGAACTCTTTCCCTACACGACGCTCTTCCGATCTNNNAATGCAAGGTGTATTAGCCA
AGAGCACACGTCTGAACTCTTTCCCTACACGACGCTCTTCCGATCTNNNCGGTGTCCTTACCACAGAAT
GAGCACACGTCTGAACTCTTTCCCTACACGACGCTCTTCCGATCTNNNGGCTGCAGTTGAGCTATGGTA
AGCACACGTCTGAACTCTTTCCCTACACGACGCTCTTCCGATCTNNNACCATAATTGAGCTTGAGCTGC
AAGAGCACACGTCTGAACTCTTTCCCTACACGACGCTCTTCCGATCTNNNCCTATACCTAGAGCCGGAG
AAGAGCACACGTCTGAACTCTTTCCCTACACGACGCTCTTCCGATCTNNNATGCCACTGCATGAAGCAA
AGAGCACACGTCTGAACTCTTTCCCTACACGACGCTCTTCCGATCTNNNGGCTGCAGGGTCATGTAAAA
GAGCACACGTCTGAACTCTTTCCCTACACGACGCTCTTCCGATCTNNNCCAAGTGCCTTGTTACAGTAG
AGAGCACACGTCTGAACTCTTTCCCTACACGACGCTCTTCCGATCTNNNCCAATTCCCTGCTTCCCTTCAA
AAGAGCACACGTCTGAACTCTTTCCCTACACGACGCTCTTCCGATCTNNNAGCAGTGAGCTCTAACAGA
GAGCACACGTCTGAACTCTTTCCCTACACGACGCTCTTCCGATCTNNNTAGGTCTACCTCTTTCCCAGCA
GAGCACACGTCTGAACTCTTTCCCTACACGACGCTCTTCCGATCTNNNACACAACACAGATCACAGACCA
AGCACACGTCTGAACTCTTTCCCTACACGACGCTCTTCCGATCTNNNTACCTGCAAAAACTCTCCACACC
AGAGCACACGTCTGAACTCTTTCCCTACACGACGCTCTTCCGATCTNNNGGGCTGCAAAACTGACCTAT
AGAGCACACGTCTGAACTCTTTCCCTACACGACGCTCTTCCGATCTNNNGATGTCCCTATTGCCATCACC
GAGCACACGTCTGAACTCTTTCCCTACACGACGCTCTTCCGATCTNNNATGAGAGCATGTGAGTGTGA
AGCACACGTCTGAACTCTTTCCCTACACGACGCTCTTCCGATCTNNNTAATTCCACAACAAAGGCACCC
AGCACACGTCTGAACTCTTTCCCTACACGACGCTCTTCCGATCTNNNAATGCCTGCTTGTGTCATTTTCC
AGCACACGTCTGAACTCTTTCCCTACACGACGCTCTTCCGATCTNNNAGTGCAGGGTGATTTGCAGTT
GAGCACACGTCTGAACTCTTTCCCTACACGACGCTCTTCCGATCTNNNTGCTTCCTTCCCAATCACTTTCA
AGCACACGTCTGAACTCTTTCCCTACACGACGCTCTTCCGATCTNNNGTTTCCCCTGCAGATAAGGAAG
AGCACACGTCTGAACTCTTTCCCTACACGACGCTCTTCCGATCTNNNTTCCCTCACTCTTTGGATGCTCTC
GAGCACACGTCTGAACTCTTTCCCTACACGACGCTCTTCCGATCTNNNCTCCTGCCATTCTCTTGTAAGG
AGCACACGTCTGAACTCTTTCCCTACACGACGCTCTTCCGATCTNNNTGCAAACCCTCTGAAAAACATG
GAGCACACGTCTGAACTCTTTCCCTACACGACGCTCTTCCGATCTNNNTAATGTGGCAGAAAGTGAAG
AGCACACGTCTGAACTCTTTCCCTACACGACGCTCTTCCGATCTNNNGTGTGTGTGTGTTCTGGTCTCT
AGCACACGTCTGAACTCTTTCCCTACACGACGCTCTTCCGATCTNNNAGAATCAGAAATGTTCCTCGGG
GAGCACACGTCTGAACTCTTTCCCTACACGACGCTCTTCCGATCTNNNCCCCTCTACTTGTCTGGAGATG
AGCACACGTCTGAACTCTTTCCCTACACGACGCTCTTCCGATCTNNNTGCAGAGAGGGAAAGAAACAG
GAGCACACGTCTGAACTCTTTCCCTACACGACGCTCTTCCGATCTNNNTGTCTCAAAAAGAGCAAAGGC
AGCACACGTCTGAACTCTTTCCCTACACGACGCTCTTCCGATCTNNNGCATTGCAAAAATATGGAACTG
AGAGCACACGTCTGAACTCTTTCCCTACACGACGCTCTTCCGATCTNNNTTACCACACATCTGGTTCCTCC
AGCACACGTCTGAACTCTTTCCCTACACGACGCTCTTCCGATCTNNNCTGGTAAGCAGCAGCATAGTTG
GAGCACACGTCTGAACTCTTTCCCTACACGACGCTCTTCCGATCTNNNTATAAATCAGATCCCCCGACCC
GAGCACACGTCTGAACTCTTTCCCTACACGACGCTCTTCCGATCTNNNCCTGGCTACATTTTGACACACC
AGCACACGTCTGAACTCTTTCCCTACACGACGCTCTTCCGATCTNNNTTGATGTTTCAGGCACACTGTCA

AGCACACGTCTGAACTCTTTCCCTACACGACGCTCTTCCGATCTNNNCCCTCTCCCCTACTCTTCCTTTCC
GCACACGTCTGAACTCTTTCCCTACACGACGCTCTTCCGATCTNNNTGGTAGGTGCACAGTGAATGT
AGCACACGTCTGAACTCTTTCCCTACACGACGCTCTTCCGATCTNNNAAATCCCCATGTGGCCAAAAA
AGCACACGTCTGAACTCTTTCCCTACACGACGCTCTTCCGATCTNNNCCGAGATTCCCTTAGGTGTGTT
GCACACGTCTGAACTCTTTCCCTACACGACGCTCTTCCGATCTNNNTCATTGCTTTGTTCTTGCTGCTGC
GCACACGTCTGAACTCTTTCCCTACACGACGCTCTTCCGATCTNNNAATTTTCCCGCCTTGTATGGTG
AGCACACGTCTGAACTCTTTCCCTACACGACGCTCTTCCGATCTNNNCCCTGAGGATACAGCCCTAAA
GAGCACACGTCTGAACTCTTTCCCTACACGACGCTCTTCCGATCTNNNCCAGATTCTGCAATAAAGCGG
GAGCACACGTCTGAACTCTTTCCCTACACGACGCTCTTCCGATCTNNNTAAAACCTGGATCCCGTTCA
AGCACACGTCTGAACTCTTTCCCTACACGACGCTCTTCCGATCTNNNTTGCTTATCCATTCAGCCTGTCC
GCACACGTCTGAACTCTTTCCCTACACGACGCTCTTCCGATCTNNNCGTGAGGCCTGAATGCAATATC
AGCACACGTCTGAACTCTTTCCCTACACGACGCTCTTCCGATCTNNNAACAGTACCACAGTACCATGT
AGCACACGTCTGAACTCTTTCCCTACACGACGCTCTTCCGATCTNNNCTCTAATCCAGGCAAAGAGGC
GAGCACACGTCTGAACTCTTTCCCTACACGACGCTCTTCCGATCTNNNCACTGCTGGTAGCCATCTAC
GAGCACACGTCTGAACTCTTTCCCTACACGACGCTCTTCCGATCTNNNACAGTGCCCATCCTTAGTTCT
AGCACACGTCTGAACTCTTTCCCTACACGACGCTCTTCCGATCTNNNATCTGTGTGTAGTGTGGGGG
GAGCACACGTCTGAACTCTTTCCCTACACGACGCTCTTCCGATCTNNNACGGACAGCTTCTAAGAGTG
AGCACACGTCTGAACTCTTTCCCTACACGACGCTCTTCCGATCTNNNCATATTTGCACACAGGCACATG
AGCACACGTCTGAACTCTTTCCCTACACGACGCTCTTCCGATCTNNNTGATTACCCTCAACACTCCCCA
CACACGTCTGAACTCTTTCCCTACACGACGCTCTTCCGATCTNNNGCTCAGAGCTGGAGTTGACTTAG
GCACACGTCTGAACTCTTTCCCTACACGACGCTCTTCCGATCTNNNTTGTCTTGTGTCTTGTTCCCTCCC
GAGCACACGTCTGAACTCTTTCCCTACACGACGCTCTTCCGATCTNNNTGTGTCTCAGAGAATTGGGA
AGCACACGTCTGAACTCTTTCCCTACACGACGCTCTTCCGATCTNNNGCTGTAGTCCTAAAACTCAACC
GAGCACACGTCTGAACTCTTTCCCTACACGACGCTCTTCCGATCTNNNTAAAGGGGCATGCAGTTTAC
GCACACGTCTGAACTCTTTCCCTACACGACGCTCTTCCGATCTNNNACCGCCTTTGTCTTTGTTTTTCCC
AGCACACGTCTGAACTCTTTCCCTACACGACGCTCTTCCGATCTNNNAGATCATGAGGGTAGGAGAG
AGCACACGTCTGAACTCTTTCCCTACACGACGCTCTTCCGATCTNNNCCACACATAACTCAGTCCGTGA
GCACACGTCTGAACTCTTTCCCTACACGACGCTCTTCCGATCTNNNAGGCCCCTTTACTTTGCTTGTTT
GCACACGTCTGAACTCTTTCCCTACACGACGCTCTTCCGATCTNNNGTCAGCAATGGCAAAATCAGCA
GAGCACACGTCTGAACTCTTTCCCTACACGACGCTCTTCCGATCTNNNACAGGCATACTTGTCCATTTT
AGAGCACACGTCTGAACTCTTTCCCTACACGACGCTCTTCCGATCTNNNAAGACAGGGTCCCATTCAT
GAGCACACGTCTGAACTCTTTCCCTACACGACGCTCTTCCGATCTNNNAGACCAAGTGTCTTGTGTTCC
AGAGCACACGTCTGAACTCTTTCCCTACACGACGCTCTTCCGATCTNNNGAATAAATGCCTGCTCCCA
AGCACACGTCTGAACTCTTTCCCTACACGACGCTCTTCCGATCTNNNACCAAGGAACTAGGGATTGG
GCACACGTCTGAACTCTTTCCCTACACGACGCTCTTCCGATCTNNNGGGGGTAAGAGGTGGTTAGAT
AGAGCACACGTCTGAACTCTTTCCCTACACGACGCTCTTCCGATCTNNNCTGATGCCCTTGATGGTTA
GAGCACACGTCTGAACTCTTTCCCTACACGACGCTCTTCCGATCTNNNTGATTACAGCACTGTGTGTGT
AGAGCACACGTCTGAACTCTTTCCCTACACGACGCTCTTCCGATCTNNNGTCAGTCCCCCATGGGATA
AGCACACGTCTGAACTCTTTCCCTACACGACGCTCTTCCGATCTNNNAGCAGTTGGATTTGTCTTCCCA
AGCACACGTCTGAACTCTTTCCCTACACGACGCTCTTCCGATCTNNNACAGTGGTGTGGACATTCGAA
GAGCACACGTCTGAACTCTTTCCCTACACGACGCTCTTCCGATCTNNNTAATTATTGCCACCCCTTGGCT
AGCACACGTCTGAACTCTTTCCCTACACGACGCTCTTCCGATCTNNNGTGATATCAGACTGGCCCTTGA
GAGCACACGTCTGAACTCTTTCCCTACACGACGCTCTTCCGATCTNNNGTCTTTCAGTTCCATCCTCTCC

AGCACACGTCTGAACTCTTTCCCTACACGACGCTCTTCCGATCTNNNTGCATCCTGAATGGATAAGGT
AGCACACGTCTGAACTCTTTCCCTACACGACGCTCTTCCGATCTNNNATTCTCCAGAGTGACAGAGCC
AGCACACGTCTGAACTCTTTCCCTACACGACGCTCTTCCGATCTNNNAATAGGAAATGTCAAAGCGG
AGCACACGTCTGAACTCTTTCCCTACACGACGCTCTTCCGATCTNNNCGTGGATGACCTTGGACATCA
GAGCACACGTCTGAACTCTTTCCCTACACGACGCTCTTCCGATCTNNNCCTGAGGTGGGAAGGAATT
AGAGCACACGTCTGAACTCTTTCCCTACACGACGCTCTTCCGATCTNNNAGATTCTACTGCTGTTGGCA
GAGCACACGTCTGAACTCTTTCCCTACACGACGCTCTTCCGATCTNNNTTAGGGAAGCAATGCAACTC
GAGCACACGTCTGAACTCTTTCCCTACACGACGCTCTTCCGATCTNNNATGACTGTTCCACTGGTATG
AGAGCACACGTCTGAACTCTTTCCCTACACGACGCTCTTCCGATCTNNNAAGTCCCCACCTTCATACAG
GAGCACACGTCTGAACTCTTTCCCTACACGACGCTCTTCCGATCTNNNTGCACGTCTGTGTATTTGTGT
AGAGCACACGTCTGAACTCTTTCCCTACACGACGCTCTTCCGATCTNNNATGTTGGGTGAAAGCTCCT
AGCACACGTCTGAACTCTTTCCCTACACGACGCTCTTCCGATCTNNNTCACGTTGCCACCTTACTTTCA
AAGAGCACACGTCTGAACTCTTTCCCTACACGACGCTCTTCCGATCTNNNACTGTACCTGCCCCCTCAC
AGAGCACACGTCTGAACTCTTTCCCTACACGACGCTCTTCCGATCTNNNGTGTTACCTCAGGCTATGT
AGAGCACACGTCTGAACTCTTTCCCTACACGACGCTCTTCCGATCTNNNTTTTCTCTTTCTCACGCACA
AGAGCACACGTCTGAACTCTTTCCCTACACGACGCTCTTCCGATCTNNNCATGAGGGAGACTTTGAGG
GAGCACACGTCTGAACTCTTTCCCTACACGACGCTCTTCCGATCTNNNGGCAAGAGAGCACACCAAG
AGCACACGTCTGAACTCTTTCCCTACACGACGCTCTTCCGATCTNNNTGGAGAGAGAAGAAGAGCT
AGCACACGTCTGAACTCTTTCCCTACACGACGCTCTTCCGATCTNNNCAGCTGTAGGAAAAGCCAGT
GAGCACACGTCTGAACTCTTTCCCTACACGACGCTCTTCCGATCTNNNGAGACTGTACACTACCAAGG
AGAGCACACGTCTGAACTCTTTCCCTACACGACGCTCTTCCGATCTNNNCTGCACAAGGGAGATGAA
AGAGCACACGTCTGAACTCTTTCCCTACACGACGCTCTTCCGATCTNNNAGAGACCCAGAGAGACAGT
GAGCACACGTCTGAACTCTTTCCCTACACGACGCTCTTCCGATCTNNNGTATGATTTTGGGGTGTGTG
GAGCACACGTCTGAACTCTTTCCCTACACGACGCTCTTCCGATCTNNNTGACATTGGTTGGAGAAACC
AGCACACGTCTGAACTCTTTCCCTACACGACGCTCTTCCGATCTNNNGTTGATGTCCTCCTGTGGCTAT
AGCACACGTCTGAACTCTTTCCCTACACGACGCTCTTCCGATCTNNNAAGAGGATCTTGTAGCCCAGG
AGCACACGTCTGAACTCTTTCCCTACACGACGCTCTTCCGATCTNNNTCAGGAGAGAGGGAGGAAT
AGCACACGTCTGAACTCTTTCCCTACACGACGCTCTTCCGATCTNNNAATGGCTACAGTCTTTGTGTTG
AGCACACGTCTGAACTCTTTCCCTACACGACGCTCTTCCGATCTNNNAGTGATGGTATGCACCTGTAT
AGCACACGTCTGAACTCTTTCCCTACACGACGCTCTTCCGATCTNNNTACACAAGCTAAAGCTGCACC
GCACACGTCTGAACTCTTTCCCTACACGACGCTCTTCCGATCTNNNAGAATTGGCAGGCATCACTTCT
AGCACACGTCTGAACTCTTTCCCTACACGACGCTCTTCCGATCTNNNACATTGCACAATGCACATAAG
AGCACACGTCTGAACTCTTTCCCTACACGACGCTCTTCCGATCTNNNGATTGTGGGGACCTATGTGT
GAGCACACGTCTGAACTCTTTCCCTACACGACGCTCTTCCGATCTNNNAACTTGCCTGACATAGGATT
GCACACGTCTGAACTCTTTCCCTACACGACGCTCTTCCGATCTNNNCAGCCCTTGAAGCTATCTGGTTT
AGAGCACACGTCTGAACTCTTTCCCTACACGACGCTCTTCCGATCTNNNCTATGTAGAACAGAGCTGC
AGAGCACACGTCTGAACTCTTTCCCTACACGACGCTCTTCCGATCTNNNATGATTAGTTGTGCCAGAT
GAGCACACGTCTGAACTCTTTCCCTACACGACGCTCTTCCGATCTNNNAAAATTATACCCCTGCCCCCA
GAGCACACGTCTGAACTCTTTCCCTACACGACGCTCTTCCGATCTNNNGAGCTCTAACTCTGAGCCAG
AGAGCACACGTCTGAACTCTTTCCCTACACGACGCTCTTCCGATCTNNNTGAACAAGCCCAAAATACT
AGCACACGTCTGAACTCTTTCCCTACACGACGCTCTTCCGATCTNNNGTGAGGCTTTGGGAGGAAAA
AGCACACGTCTGAACTCTTTCCCTACACGACGCTCTTCCGATCTNNNCCCTTTCTACTGCCTCTTCCTCT
GAGCACACGTCTGAACTCTTTCCCTACACGACGCTCTTCCGATCTNNNGGCTTTCTTCCCTTTGCTCTCA

GAGCACACGTCTGAACTCTTTCCCTACACGACGCTCTTCCGATCTNNNAGAAGCTGGGTTGGCAATAA
GAGCACACGTCTGAACTCTTTCCCTACACGACGCTCTTCCGATCTNNNGAGCTCAGGACAATAACATG
AGAGCACACGTCTGAACTCTTTCCCTACACGACGCTCTTCCGATCTNNNAAGGGCTTAGGTGTGATTT
AAGAGCACACGTCTGAACTCTTTCCCTACACGACGCTCTTCCGATCTNNNTTCTGAAAGCTCCCAATCT
AGAGCACACGTCTGAACTCTTTCCCTACACGACGCTCTTCCGATCTNNNTATCTGGGGAGGCTGATTA
AGAGCACACGTCTGAACTCTTTCCCTACACGACGCTCTTCCGATCTNNNAGGGGAAAGTGAGAGTTG
GAGCACACGTCTGAACTCTTTCCCTACACGACGCTCTTCCGATCTNNNTCATGTTGCCATGTGAATGTG
GAGCACACGTCTGAACTCTTTCCCTACACGACGCTCTTCCGATCTNNNAGAGCTTGGTGACATGCATT
AGCACACGTCTGAACTCTTTCCCTACACGACGCTCTTCCGATCTNNNCCAACTTGACCAAACTCACAGG
AGCACACGTCTGAACTCTTTCCCTACACGACGCTCTTCCGATCTNNNCAACATTGCAGCAAACAGCCAT
GAGCACACGTCTGAACTCTTTCCCTACACGACGCTCTTCCGATCTNNNCAGGCTTCATCCTCAAACCTTG
GAGCACACGTCTGAACTCTTTCCCTACACGACGCTCTTCCGATCTNNNCGGCACATATTTACCCAGAGG
GAGCACACGTCTGAACTCTTTCCCTACACGACGCTCTTCCGATCTNNNCCTCAGCTTTGCAATCTTAGG
GAGCACACGTCTGAACTCTTTCCCTACACGACGCTCTTCCGATCTNNNCCTTTGGTAAACTGTAAAGG
AGCACACGTCTGAACTCTTTCCCTACACGACGCTCTTCCGATCTNNNTCTTCTTATGTAACTGGGCACAA
AGCACACGTCTGAACTCTTTCCCTACACGACGCTCTTCCGATCTNNNAGTCCGAACAGATAAGCAGGG
GAGCACACGTCTGAACTCTTTCCCTACACGACGCTCTTCCGATCTNNNAGAATGTCTCCAAGGCTGTT
GAGCACACGTCTGAACTCTTTCCCTACACGACGCTCTTCCGATCTNNNCACATTCTTAGGGGCTTCCAC
AGAGCACACGTCTGAACTCTTTCCCTACACGACGCTCTTCCGATCTNNNATCTCTACAACCCTAGGGCT
AGAGCACACGTCTGAACTCTTTCCCTACACGACGCTCTTCCGATCTNNNCTATTGTCATCCCTTGGGCA
AGAGCACACGTCTGAACTCTTTCCCTACACGACGCTCTTCCGATCTNNNAAGCTCTGAATTGGACTGTC
GAGCACACGTCTGAACTCTTTCCCTACACGACGCTCTTCCGATCTNNNGTTATGCTCTTTGTCCACCCG
GAGCACACGTCTGAACTCTTTCCCTACACGACGCTCTTCCGATCTNNNTTTGTGTTGGAACTTGGGTC
AGCACACGTCTGAACTCTTTCCCTACACGACGCTCTTCCGATCTNNNTTAGGATGGAGTGAAGGCGA
AGCACACGTCTGAACTCTTTCCCTACACGACGCTCTTCCGATCTNNNTTGGAAGGAGGACCCTGAGA
GAGCACACGTCTGAACTCTTTCCCTACACGACGCTCTTCCGATCTNNNCCCAGCCAAGAACTTTCACTT
AGAGCACACGTCTGAACTCTTTCCCTACACGACGCTCTTCCGATCTNNNTTTGTCATAGAGACAGGGC
AGAGCACACGTCTGAACTCTTTCCCTACACGACGCTCTTCCGATCTNNNGCAGCCCTATATGTGTAGCT
GAGCACACGTCTGAACTCTTTCCCTACACGACGCTCTTCCGATCTNNNTGGGTATGAGCACTGTGACTT
GAGCACACGTCTGAACTCTTTCCCTACACGACGCTCTTCCGATCTNNNAGAGAATTTCAGACTTGAGC
GAGCACACGTCTGAACTCTTTCCCTACACGACGCTCTTCCGATCTNNNCTCAGGAGGCTGAACAAGGA
AGCACACGTCTGAACTCTTTCCCTACACGACGCTCTTCCGATCTNNNCCAGTAAAATGGCAGCAACAAG
AGCACACGTCTGAACTCTTTCCCTACACGACGCTCTTCCGATCTNNNGCACAATGAAAGCATTGCTCAG
AGAGCACACGTCTGAACTCTTTCCCTACACGACGCTCTTCCGATCTNNNAGATGATAGCCCAAGGACA
GAGCACACGTCTGAACTCTTTCCCTACACGACGCTCTTCCGATCTNNNGGATAACTCATGCTGCAGGG
GAGCACACGTCTGAACTCTTTCCCTACACGACGCTCTTCCGATCTNNNGCAACAATGGAGAGGGTTTC
GAGCACACGTCTGAACTCTTTCCCTACACGACGCTCTTCCGATCTNNNTTGGTCCCTGGAAGTGAACA
AGAGCACACGTCTGAACTCTTTCCCTACACGACGCTCTTCCGATCTNNNCCAGGAATTTGAGACCACCT
AGAGCACACGTCTGAACTCTTTCCCTACACGACGCTCTTCCGATCTNNNATTCTCTGCAGTGTCCTTTGG
GAGCACACGTCTGAACTCTTTCCCTACACGACGCTCTTCCGATCTNNNACTACTGGGTCCTGGGATGT
AGAGCACACGTCTGAACTCTTTCCCTACACGACGCTCTTCCGATCTNNNAGTGCCTAGTGCAGGATGT
AGCACACGTCTGAACTCTTTCCCTACACGACGCTCTTCCGATCTNNNTGACAACCTGGAAGTCATTGG
AGAGCACACGTCTGAACTCTTTCCCTACACGACGCTCTTCCGATCTNNNAACCAGCCAGCTTGCTCTAT

GAGCACACGTCTGAACTCTTTCCCTACACGACGCTCTTCCGATCTNNNCTGAATGGTTCTCTGATCCTG

AGCACACGTCTGAACTCTTTCCCTACACGACGCTCTTCCGATCTNNNCCATTCTATTGCAGGGCTCTCT

GAGCACACGTCTGAACTCTTTCCCTACACGACGCTCTTCCGATCTNNNGCATGTGAAACTGCAAAAGC

AGAGCACACGTCTGAACTCTTTCCCTACACGACGCTCTTCCGATCTNNNACATCAGGCCACTAAGGAAA

GAGCACACGTCTGAACTCTTTCCCTACACGACGCTCTTCCGATCTNNNGCCCAGTATGGGTCATAGGA

GAGCACACGTCTGAACTCTTTCCCTACACGACGCTCTTCCGATCTNNNGGATATTTTTGGCCCGTTGTT

GAGCACACGTCTGAACTCTTTCCCTACACGACGCTCTTCCGATCTNNNTCCTCACTCCTTCCTCTCTCTCT

AGCACACGTCTGAACTCTTTCCCTACACGACGCTCTTCCGATCTNNNGTATGCCCAAGTCAGCAAAATG

AGAGCACACGTCTGAACTCTTTCCCTACACGACGCTCTTCCGATCTNNNGTGGAGGGAAAGCAAATGC

GAGCACACGTCTGAACTCTTTCCCTACACGACGCTCTTCCGATCTNNNAGAGCCACTCCCTGCTTCTAAT

AGCACACGTCTGAACTCTTTCCCTACACGACGCTCTTCCGATCTNNNAATACATGCACAGCACACACAG

AGAGCACACGTCTGAACTCTTTCCCTACACGACGCTCTTCCGATCTNNNCACTAGGCAGAGGAAAAAG

AGAGCACACGTCTGAACTCTTTCCCTACACGACGCTCTTCCGATCTNNNCGGTGGAGAGAAACCTAAA

AGAGCACACGTCTGAACTCTTTCCCTACACGACGCTCTTCCGATCTNNNGAGGCTGGAAAACTGGCAA

AGAGCACACGTCTGAACTCTTTCCCTACACGACGCTCTTCCGATCTNNNGCCATCTCCAAGGGCCTTTTA

GAGCACACGTCTGAACTCTTTCCCTACACGACGCTCTTCCGATCTNNNTCCTTTCTCCCTTACCTGTCCTG

AGCACACGTCTGAACTCTTTCCCTACACGACGCTCTTCCGATCTNNNTTTCCTGTAAGGATGTGAGTGA

GAGCACACGTCTGAACTCTTTCCCTACACGACGCTCTTCCGATCTNNNCACAAGATGTAACAGCCATTG

AGCACACGTCTGAACTCTTTCCCTACACGACGCTCTTCCGATCTNNNACCTTGGAATTTGGACCAGTGA

GAGCACACGTCTGAACTCTTTCCCTACACGACGCTCTTCCGATCTNNNCCTCATGAGCAACATTCTTCCCA

GAGCACACGTCTGAACTCTTTCCCTACACGACGCTCTTCCGATCTNNNCCACTAATAGGATGAGGCAG

GAGCACACGTCTGAACTCTTTCCCTACACGACGCTCTTCCGATCTNNNGCTAGAAGAGCCCAGACTACA

AGAGCACACGTCTGAACTCTTTCCCTACACGACGCTCTTCCGATCTNNNTAGCAATACCCCCTCGGTACA

AGAGCACACGTCTGAACTCTTTCCCTACACGACGCTCTTCCGATCTNNNAGCTGAGTAGTCACACAGCA

AGCACACGTCTGAACTCTTTCCCTACACGACGCTCTTCCGATCTNNNTCCATCTATGTTGCCGCATGAAA

GAGCACACGTCTGAACTCTTTCCCTACACGACGCTCTTCCGATCTNNNAAATTTGGAGAAGCCAACAT

AGCACACGTCTGAACTCTTTCCCTACACGACGCTCTTCCGATCTNNNTTGGATCCATCTGCAACTGAAG

AGCACACGTCTGAACTCTTTCCCTACACGACGCTCTTCCGATCTNNNAAGTGGACTACTAAAGCAAGG

AGCACACGTCTGAACTCTTTCCCTACACGACGCTCTTCCGATCTNNNATCCCCCTGTATTCTGTAGGCAT

GAGCACACGTCTGAACTCTTTCCCTACACGACGCTCTTCCGATCTNNNCTACCTCCAGGGAAGAGACA

GAGCACACGTCTGAACTCTTTCCCTACACGACGCTCTTCCGATCTNNNCCATCCACAGCCTTTCCTGAAG

AGCACACGTCTGAACTCTTTCCCTACACGACGCTCTTCCGATCTNNNTATGAAGCCAGCCTATATGCCC

AGCACACGTCTGAACTCTTTCCCTACACGACGCTCTTCCGATCTNNNGCAGTATTTCTCACCAACTTGC

GAGCACACGTCTGAACTCTTTCCCTACACGACGCTCTTCCGATCTNNNACAGTCCCACACACTCTCTCTT

AGAGCACACGTCTGAACTCTTTCCCTACACGACGCTCTTCCGATCTNNNCTATGCATGGAGCAGAACA

AGCACACGTCTGAACTCTTTCCCTACACGACGCTCTTCCGATCTNNNTGCCCACAATCATAGTTCACTGC

AGCACACGTCTGAACTCTTTCCCTACACGACGCTCTTCCGATCTNNNCAAGGACTTGTCAGCCTCCATAA

GAGCACACGTCTGAACTCTTTCCCTACACGACGCTCTTCCGATCTNNNGTCAAAATAACAACTCCGCCCA

AGCACACGTCTGAACTCTTTCCCTACACGACGCTCTTCCGATCTNNNTTGCTGGTTCCTATGTCCCTTTG

GAGCACACGTCTGAACTCTTTCCCTACACGACGCTCTTCCGATCTNNNTGGTCTTCCCTGCTTTTCCTTT

AGCACACGTCTGAACTCTTTCCCTACACGACGCTCTTCCGATCTNNNAGTCTTTTGGGGTTGGCATAG

AGAGCACACGTCTGAACTCTTTCCCTACACGACGCTCTTCCGATCTNNNGTGTGGCCTGGTTCTCTGTAT

GAGCACACGTCTGAACTCTTTCCCTACACGACGCTCTTCCGATCTNNNAAAAGGAGGAGTGTGTGTGT

GAGCACACGTCTGAACTCTTTCCCTACACGACGCTCTTCCGATCTNNNTCCCATAGAGCCACAGAAGTT
GAGCACACGTCTGAACTCTTTCCCTACACGACGCTCTTCCGATCTNNNCCTTTCCATAGCCGAGAAAGC
AGCACACGTCTGAACTCTTTCCCTACACGACGCTCTTCCGATCTNNNTTTCAGGTTGCAGGTTTGGAA
AGCACACGTCTGAACTCTTTCCCTACACGACGCTCTTCCGATCTNNNTCCATTGTCTCCTTTCTTCCAGT
AGAGCACACGTCTGAACTCTTTCCCTACACGACGCTCTTCCGATCTNNNAGAGACCACACACACATGCA
AGAGCACACGTCTGAACTCTTTCCCTACACGACGCTCTTCCGATCTNNNAGTTGGCAAGATTTTGGGG
AGAGCACACGTCTGAACTCTTTCCCTACACGACGCTCTTCCGATCTNNNAAGGTGATGATCGACAAGT
GAGCACACGTCTGAACTCTTTCCCTACACGACGCTCTTCCGATCTNNNGGGCTCTCCCATTCTGTTTCA
GCACACGTCTGAACTCTTTCCCTACACGACGCTCTTCCGATCTNNNACGCTCTAATCCTAGCTCCACTTCA
AGAGCACACGTCTGAACTCTTTCCCTACACGACGCTCTTCCGATCTNNNAGATAGCCCCAGGAAAAGG
GAGCACACGTCTGAACTCTTTCCCTACACGACGCTCTTCCGATCTNNNGGGGACGCAGATGACATTCA
GAGCACACGTCTGAACTCTTTCCCTACACGACGCTCTTCCGATCTNNNCCCACCTACCTCACCCTTCTTA
GAGCACACGTCTGAACTCTTTCCCTACACGACGCTCTTCCGATCTNNNGTAATTTCCCAGGCAGTGAC
GAGCACACGTCTGAACTCTTTCCCTACACGACGCTCTTCCGATCTNNNGGGCCTTGGATGACATTATG
AGAGCACACGTCTGAACTCTTTCCCTACACGACGCTCTTCCGATCTNNNGGTTCTGTAGCCAAATGGGT
AGCACACGTCTGAACTCTTTCCCTACACGACGCTCTTCCGATCTNNNTCATCTAATTCCACAGTGCCACG
GAGCACACGTCTGAACTCTTTCCCTACACGACGCTCTTCCGATCTNNNCATTTCAGAGCTGAGTTCCTG
AGCACACGTCTGAACTCTTTCCCTACACGACGCTCTTCCGATCTNNNTTCCCAATAATGCTGACACTGG
GAGCACACGTCTGAACTCTTTCCCTACACGACGCTCTTCCGATCTNNNCCCGGATCTCTCTGTTTCTCTC
GAGCACACGTCTGAACTCTTTCCCTACACGACGCTCTTCCGATCTNNNGGCCCAAAATGGAAAACATA
GAGCACACGTCTGAACTCTTTCCCTACACGACGCTCTTCCGATCTNNNAATGCTTCCTTTCTCTTTTCCC
AGCACACGTCTGAACTCTTTCCCTACACGACGCTCTTCCGATCTNNNAAACCAGAACACGTAACTGCCT
GAGCACACGTCTGAACTCTTTCCCTACACGACGCTCTTCCGATCTNNNATCACACCAAACACCAGCAGAA
AGCACACGTCTGAACTCTTTCCCTACACGACGCTCTTCCGATCTNNNTAAAGCTCCTCTCACCAGATGG
AGCACACGTCTGAACTCTTTCCCTACACGACGCTCTTCCGATCTNNNCACACTCTTTCTACTGGCTGACC
GAGCACACGTCTGAACTCTTTCCCTACACGACGCTCTTCCGATCTNNNTTCGACCAAGACGTCATTACG
AGCACACGTCTGAACTCTTTCCCTACACGACGCTCTTCCGATCTNNNACCAACACTCCTCCCTCTATGTT
GAGCACACGTCTGAACTCTTTCCCTACACGACGCTCTTCCGATCTNNNGTTGCCTACTTAGGGTTCTGA
GAGCACACGTCTGAACTCTTTCCCTACACGACGCTCTTCCGATCTNNNTGCTGCTCTCTTAGACATAGC
GAGCACACGTCTGAACTCTTTCCCTACACGACGCTCTTCCGATCTNNNTGCTCTGACAGTGTCTAACCC
GAGCACACGTCTGAACTCTTTCCCTACACGACGCTCTTCCGATCTNNNATGGCATATGAGGTGTGTGT
GAGCACACGTCTGAACTCTTTCCCTACACGACGCTCTTCCGATCTNNNAGAGGAGAGAGTGGATTGG
GAGCACACGTCTGAACTCTTTCCCTACACGACGCTCTTCCGATCTNNNTGCTCATGATTCCCTTGGCTAT
AGCACACGTCTGAACTCTTTCCCTACACGACGCTCTTCCGATCTNNNTAGGCACATCTTACATGGCAACT
GAGCACACGTCTGAACTCTTTCCCTACACGACGCTCTTCCGATCTNNNCTTGGAATATCCGGGTTGCT
GAGCACACGTCTGAACTCTTTCCCTACACGACGCTCTTCCGATCTNNNCAGAATCCTGGTCTCTGAAG
GAGCACACGTCTGAACTCTTTCCCTACACGACGCTCTTCCGATCTNNNTACACCTCAACCCAGAACATAC
GAGCACACGTCTGAACTCTTTCCCTACACGACGCTCTTCCGATCTNNNTCATGGGCAACACATACACAC
AGCACACGTCTGAACTCTTTCCCTACACGACGCTCTTCCGATCTNNNCTTGCTCGGTCTCACTGTATTGT
AAGAGCACACGTCTGAACTCTTTCCCTACACGACGCTCTTCCGATCTNNNTCTAAGCATGTTACGTGAA
AGAGCACACGTCTGAACTCTTTCCCTACACGACGCTCTTCCGATCTNNNGCATGTTAGGTGGTTTTGT
GAGCACACGTCTGAACTCTTTCCCTACACGACGCTCTTCCGATCTNNNACTCTGTCTGTCCCCTAAAGGT
GAGCACACGTCTGAACTCTTTCCCTACACGACGCTCTTCCGATCTNNNTAAACTTTGCCAGAACTAGCG

AGCACACGTCTGAACTCTTTCCCTACACGACGCTCTTCCGATCTNNNAGGGTATGGCTTTGTTGTCAT

GCACACGTCTGAACTCTTTCCCTACACGACGCTCTTCCGATCTNNNGATGCAGATGGCCAATAGTGGA

AGCACACGTCTGAACTCTTTCCCTACACGACGCTCTTCCGATCTNNNGTTGTGTCAACTGCTGATATG

AGCACACGTCTGAACTCTTTCCCTACACGACGCTCTTCCGATCTNNNTTGAGACACATATCAGATCCC

AGCACACGTCTGAACTCTTTCCCTACACGACGCTCTTCCGATCTNNNCCTCCTCAACTTAGACAGTGG

GCACACGTCTGAACTCTTTCCCTACACGACGCTCTTCCGATCTNNNGATTTTGGCAGATGCCTTCCAC

AGCACACGTCTGAACTCTTTCCCTACACGACGCTCTTCCGATCTNNNAGGAGGTATTTGTACTGGGTG

GCACACGTCTGAACTCTTTCCCTACACGACGCTCTTCCGATCTNNNCAAGAACTACTTGTCAACCTCC

AGCACACGTCTGAACTCTTTCCCTACACGACGCTCTTCCGATCTNNNTCTCCCAAGTAACTCACGTCA

AGCACACGTCTGAACTCTTTCCCTACACGACGCTCTTCCGATCTNNNGTACACAAGCTGGTATGTGTG

GCACACGTCTGAACTCTTTCCCTACACGACGCTCTTCCGATCTNNNAACTATCCCAAGACACACTCTGC

AGCACACGTCTGAACTCTTTCCCTACACGACGCTCTTCCGATCTNNNGTTCCAAGGAACACATCCTCC

GAGCACACGTCTGAACTCTTTCCCTACACGACGCTCTTCCGATCTNNNAGAGAGAAGAAGAGCTCAG

GCACACGTCTGAACTCTTTCCCTACACGACGCTCTTCCGATCTNNNGATGCAGTCTACTCCCTGCTACT

AGCACACGTCTGAACTCTTTCCCTACACGACGCTCTTCCGATCTNNNTTTCTGGTGGGATATATCTGG

AGCACACGTCTGAACTCTTTCCCTACACGACGCTCTTCCGATCTNNNGGGTTTGGCTTGGAGCTAAT

AGAGCACACGTCTGAACTCTTTCCCTACACGACGCTCTTCCGATCTNNNGGCAAGTGTTTCAGATAGC

GCACACGTCTGAACTCTTTCCCTACACGACGCTCTTCCGATCTNNNTTCTGTGAGTGGGTCAAACTTG

AGCACACGTCTGAACTCTTTCCCTACACGACGCTCTTCCGATCTNNNGGGATGGGGTGAAAGGAAG

AGCACACGTCTGAACTCTTTCCCTACACGACGCTCTTCCGATCTNNNTTGCTGATGCCAGTGATTCAA

AGCACACGTCTGAACTCTTTCCCTACACGACGCTCTTCCGATCTNNNAACGATGGAAACTGTACCCTA

AGCACACGTCTGAACTCTTTCCCTACACGACGCTCTTCCGATCTNNNTCTCACTTTGGGCTTCAGCTTT

GAGCACACGTCTGAACTCTTTCCCTACACGACGCTCTTCCGATCTNNNTTTTCCTCGGAGATGTGTGT

GCACACGTCTGAACTCTTTCCCTACACGACGCTCTTCCGATCTNNNAAATAAGACAACATGGGCAGC

AGCACACGTCTGAACTCTTTCCCTACACGACGCTCTTCCGATCTNNNCTTGGGCAAAGGAGCAAGACT

AGCACACGTCTGAACTCTTTCCCTACACGACGCTCTTCCGATCTNNNATACCTGAAGTCCTGAAAGGT

AGCACACGTCTGAACTCTTTCCCTACACGACGCTCTTCCGATCTNNNAGGTCCCTCTTCCATTGTTCTC

GCACACGTCTGAACTCTTTCCCTACACGACGCTCTTCCGATCTNNNGAAGAAAGGAGAAGGAGAGC

GAGCACACGTCTGAACTCTTTCCCTACACGACGCTCTTCCGATCTNNNCATTGTGGCTTCCATCTTGGT

AGCACACGTCTGAACTCTTTCCCTACACGACGCTCTTCCGATCTNNNCCCTATGCATCTCTTTGGCCATT

AGCACACGTCTGAACTCTTTCCCTACACGACGCTCTTCCGATCTNNNCCTGCATAATTGTTCCGTGAAG

AGCACACGTCTGAACTCTTTCCCTACACGACGCTCTTCCGATCTNNNGAGGCAAGTGGAGAAGGAA

GCACACGTCTGAACTCTTTCCCTACACGACGCTCTTCCGATCTNNNAAAAGAAGTGAGCCCTCTTGTG

GAGCACACGTCTGAACTCTTTCCCTACACGACGCTCTTCCGATCTNNNGATCTGACAGTGGAAGGCAA

GCACACGTCTGAACTCTTTCCCTACACGACGCTCTTCCGATCTNNNTTCTCATTTTCCCACTCCCCTGAC

AGCACACGTCTGAACTCTTTCCCTACACGACGCTCTTCCGATCTNNNCGACAGACAGAGCAAAGCTCA

AGCACACGTCTGAACTCTTTCCCTACACGACGCTCTTCCGATCTNNNTATCAGTTTGGGGAGGGAAG

AGCACACGTCTGAACTCTTTCCCTACACGACGCTCTTCCGATCTNNNCAGCTGTGAAGTCTCTCTCACT

AGCACACGTCTGAACTCTTTCCCTACACGACGCTCTTCCGATCTNNNTCACTTCACTTCACCTCAACCCT

GCACACGTCTGAACTCTTTCCCTACACGACGCTCTTCCGATCTNNNATTGTGTGCATGTATTTGGTGTG

AGCACACGTCTGAACTCTTTCCCTACACGACGCTCTTCCGATCTNNNCTGCCTCTTGAATCCAGAACAG

AGCACACGTCTGAACTCTTTCCCTACACGACGCTCTTCCGATCTNNNGATGCCACTATACAACCACTGG

AGCACACGTCTGAACTCTTTCCCTACACGACGCTCTTCCGATCTNNNTATAAAGGGATAGCACCTGG

AGCACACGTCTGAACTCTTTCCCTACACGACGCTCTTCCGATCTNNNCACGCAGCCCTATGAGGTAAAA
GCACACGTCTGAACTCTTTCCCTACACGACGCTCTTCCGATCTNNNAAGAGTTACTGGGTCACCACTG
AGCACACGTCTGAACTCTTTCCCTACACGACGCTCTTCCGATCTNNNCACCAGAACACACTCACACATG
AGCACACGTCTGAACTCTTTCCCTACACGACGCTCTTCCGATCTNNNTCAATCTCCTCTTCCCCATCCTC
AGCACACGTCTGAACTCTTTCCCTACACGACGCTCTTCCGATCTNNNAATGTGCAAACTCCAGCATTTG
AGCACACGTCTGAACTCTTTCCCTACACGACGCTCTTCCGATCTNNNATTTCTCTACCTTCCCAGGCCAT
GAGCACACGTCTGAACTCTTTCCCTACACGACGCTCTTCCGATCTNNNTATGCCATCCACAGAAGTCTC
AGCACACGTCTGAACTCTTTCCCTACACGACGCTCTTCCGATCTNNNTGATCTCTGAGGGATGTGTGT
AGCACACGTCTGAACTCTTTCCCTACACGACGCTCTTCCGATCTNNNAGATATAACACTGCTGGCTGG
GAGCACACGTCTGAACTCTTTCCCTACACGACGCTCTTCCGATCTNNNGCTTCACTGGAATGAGAAGG
GAGCACACGTCTGAACTCTTTCCCTACACGACGCTCTTCCGATCTNNNGCAAGCACTCACTCTCCATTC
GAGCACACGTCTGAACTCTTTCCCTACACGACGCTCTTCCGATCTNNNGCTTGTCAAGAGATTCCTTCG
AGAGCACACGTCTGAACTCTTTCCCTACACGACGCTCTTCCGATCTNNNTGATGGCATTGAGCAGAAG
AGCACACGTCTGAACTCTTTCCCTACACGACGCTCTTCCGATCTNNNCCAAGTATCTGGCATGTACCAC
AGAGCACACGTCTGAACTCTTTCCCTACACGACGCTCTTCCGATCTNNNACATATTTGCTTAGCAGGG
GAGCACACGTCTGAACTCTTTCCCTACACGACGCTCTTCCGATCTNNNCAGGTACCTAACCTGTCTCTGG
AGCACACGTCTGAACTCTTTCCCTACACGACGCTCTTCCGATCTNNNGTTCCCAAACAAGTATGCTCCC
GAGCACACGTCTGAACTCTTTCCCTACACGACGCTCTTCCGATCTNNNCCAGACTGCTTGTAACAGTGC
AGCACACGTCTGAACTCTTTCCCTACACGACGCTCTTCCGATCTNNNTGACACTGAACAGGTCATTTG
AGCACACGTCTGAACTCTTTCCCTACACGACGCTCTTCCGATCTNNNAACCCAATTCAGCAAGGCCATT
AGCACACGTCTGAACTCTTTCCCTACACGACGCTCTTCCGATCTNNNTCCCAACCTGAGAAATTGTGT
AGCACACGTCTGAACTCTTTCCCTACACGACGCTCTTCCGATCTNNNATCCCTTTTCCTCCTGTTCCTCAC
GAGCACACGTCTGAACTCTTTCCCTACACGACGCTCTTCCGATCTNNNTAGGGTAGAAGAGGTGGAT
GAGCACACGTCTGAACTCTTTCCCTACACGACGCTCTTCCGATCTNNNGGTAGGAGGAGGAATGCAG
AGAGCACACGTCTGAACTCTTTCCCTACACGACGCTCTTCCGATCTNNNGGTGATTTCATACACGCACA
AGCACACGTCTGAACTCTTTCCCTACACGACGCTCTTCCGATCTNNNCATCATGCTCGGCTAAGAGTGT
GAGCACACGTCTGAACTCTTTCCCTACACGACGCTCTTCCGATCTNNNTGCCTGACTTAGCATAGTGTC
AAGAGCACACGTCTGAACTCTTTCCCTACACGACGCTCTTCCGATCTNNNAAGATTGCTTGTAGCCAAA
GAGCACACGTCTGAACTCTTTCCCTACACGACGCTCTTCCGATCTNNNCTTGCCTTCATTTGCATAGGG
GAGCACACGTCTGAACTCTTTCCCTACACGACGCTCTTCCGATCTNNNTGCCTTTCCTCAACAATTAGTG
AGAGCACACGTCTGAACTCTTTCCCTACACGACGCTCTTCCGATCTNNNCCACCATTTGTACAATGCTA
AGAGCACACGTCTGAACTCTTTCCCTACACGACGCTCTTCCGATCTNNNAAACACACATCTCTGTGCCT
GAGCACACGTCTGAACTCTTTCCCTACACGACGCTCTTCCGATCTNNNTTACAGGATGGTGGTTGCGA
AGAGCACACGTCTGAACTCTTTCCCTACACGACGCTCTTCCGATCTNNNAAACTCTTTCCTGCCCCTTCT
AGAGCACACGTCTGAACTCTTTCCCTACACGACGCTCTTCCGATCTNNNGGACTATAAAAGCCCTGGG
GAGCACACGTCTGAACTCTTTCCCTACACGACGCTCTTCCGATCTNNNCTCTCCTCTATCCCTGCCACTT
GAGCACACGTCTGAACTCTTTCCCTACACGACGCTCTTCCGATCTNNNATACCCATGGGGAAATAAGG
GAGCACACGTCTGAACTCTTTCCCTACACGACGCTCTTCCGATCTNNNCAATCCCACTCCTTAGCAGGA
GCACACGTCTGAACTCTTTCCCTACACGACGCTCTTCCGATCTNNNCTGGCCCTTTGGAGAAACTGAAA
AGAGCACACGTCTGAACTCTTTCCCTACACGACGCTCTTCCGATCTNNNCTGCAGAAGAGTTGGGAGA
GAGCACACGTCTGAACTCTTTCCCTACACGACGCTCTTCCGATCTNNNCCCACAGGACAAACACAAAAG
GCACACGTCTGAACTCTTTCCCTACACGACGCTCTTCCGATCTNNNGTGGTTTCTGTAGCGTGTGTAC
AGCACACGTCTGAACTCTTTCCCTACACGACGCTCTTCCGATCTNNNCCTTCTCCTCTCTCTCTGTCTGG

AGCACACGTCTGAACTCTTTCCCTACACGACGCTCTTCCGATCTNNNGGGTGTAGGGTATGTTCAGGG
AGCACACGTCTGAACTCTTTCCCTACACGACGCTCTTCCGATCTNNNGCATGGCTCAGGAGAAATGA
AGCACACGTCTGAACTCTTTCCCTACACGACGCTCTTCCGATCTNNNAGGAAGTCATCTCTCTCTCGCTT
AGCACACGTCTGAACTCTTTCCCTACACGACGCTCTTCCGATCTNNNTTCCCCAAACAACAACAGCAAT
GAGCACACGTCTGAACTCTTTCCCTACACGACGCTCTTCCGATCTNNNAACTTTTCCGTGAGCAAACAA
GAGCACACGTCTGAACTCTTTCCCTACACGACGCTCTTCCGATCTNNNTGCAGCCAGTCCAAGATTTCA
AGCACACGTCTGAACTCTTTCCCTACACGACGCTCTTCCGATCTNNNAAACCCAAGGTGAAACTGTAG
GAGCACACGTCTGAACTCTTTCCCTACACGACGCTCTTCCGATCTNNNAGGTGACTTGCTCCCTGTAGT
AGCACACGTCTGAACTCTTTCCCTACACGACGCTCTTCCGATCTNNNGGTCAGTAACTACGTCCCTTTT
AGAGCACACGTCTGAACTCTTTCCCTACACGACGCTCTTCCGATCTNNNAGGGTTCTTTCTACACTCCC
AGCACACGTCTGAACTCTTTCCCTACACGACGCTCTTCCGATCTNNNAACCTCACTTCCCAAAAATGCC
GCACACGTCTGAACTCTTTCCCTACACGACGCTCTTCCGATCTNNNTGCAGCACATGACTGAATTGGT
GAGCACACGTCTGAACTCTTTCCCTACACGACGCTCTTCCGATCTNNNAATTTGGGCAGGACATGGAC
AGCACACGTCTGAACTCTTTCCCTACACGACGCTCTTCCGATCTNNNGCAAAGAGCCACTCATACATCC
GCACACGTCTGAACTCTTTCCCTACACGACGCTCTTCCGATCTNNNGTGGCTTCATGCTGTTATCATGG
AGCACACGTCTGAACTCTTTCCCTACACGACGCTCTTCCGATCTNNNACAACTCTGTTGTGTGTGTGT
AGCACACGTCTGAACTCTTTCCCTACACGACGCTCTTCCGATCTNNNCTAGACAAGCCCAAAGTGTGTG
GCACACGTCTGAACTCTTTCCCTACACGACGCTCTTCCGATCTNNNGTGCACCAGGTAATTCAGACCAA
GAGCACACGTCTGAACTCTTTCCCTACACGACGCTCTTCCGATCTNNNGGGCCTCATTCAATCAGAGAA
GAGCACACGTCTGAACTCTTTCCCTACACGACGCTCTTCCGATCTNNNCACCACAAAAGAACACAAGG
GAGCACACGTCTGAACTCTTTCCCTACACGACGCTCTTCCGATCTNNNAACCTAAGCCTTCACCTCTCA
AGCACACGTCTGAACTCTTTCCCTACACGACGCTCTTCCGATCTNNNCTTTGGTGCTCCTGCTGTTTAG
AGCACACGTCTGAACTCTTTCCCTACACGACGCTCTTCCGATCTNNNTGACTTAGCCTGAGTTGGTCTC
AGCACACGTCTGAACTCTTTCCCTACACGACGCTCTTCCGATCTNNNAATCTCTTTTGTGCGCACTCTCT
AGAGCACACGTCTGAACTCTTTCCCTACACGACGCTCTTCCGATCTNNNTTTCTGTTGACAGCTCACAT
AGCACACGTCTGAACTCTTTCCCTACACGACGCTCTTCCGATCTNNNGAGGATCCCTTGGAATTCACCT
GAGCACACGTCTGAACTCTTTCCCTACACGACGCTCTTCCGATCTNNNCAGCAATTCTGTGAGAGGTA
AGAGCACACGTCTGAACTCTTTCCCTACACGACGCTCTTCCGATCTNNNTCACTACAGGGTGTGTGTG
AGCACACGTCTGAACTCTTTCCCTACACGACGCTCTTCCGATCTNNNAGGGAAGTTCCAACATTTTCCC
AGCACACGTCTGAACTCTTTCCCTACACGACGCTCTTCCGATCTNNNATCCCTGACAGCCATTAGTAGC
AGCACACGTCTGAACTCTTTCCCTACACGACGCTCTTCCGATCTNNNTGTGCAGGGTATAAGGTCTTTG
AGCACACGTCTGAACTCTTTCCCTACACGACGCTCTTCCGATCTNNNGTATTTCTTCCCTTCCCTCCCCTC
GAGCACACGTCTGAACTCTTTCCCTACACGACGCTCTTCCGATCTNNNCCTGACATGAAGGAGAAGTG
GCACACGTCTGAACTCTTTCCCTACACGACGCTCTTCCGATCTNNNGTTTCAGCAACACCAACTTGCTT
AGAGCACACGTCTGAACTCTTTCCCTACACGACGCTCTTCCGATCTNNNAGTGGAGATTTTGAAGGCA
AGCACACGTCTGAACTCTTTCCCTACACGACGCTCTTCCGATCTNNNTCCTTTCCTGACTGCTTGACCT
AGCACACGTCTGAACTCTTTCCCTACACGACGCTCTTCCGATCTNNNTCGCAGGGAGTAAAATGTGT
AGCACACGTCTGAACTCTTTCCCTACACGACGCTCTTCCGATCTNNNAACCTAAAATATGCCTGCCCCTG
AGCACACGTCTGAACTCTTTCCCTACACGACGCTCTTCCGATCTNNNAACTTCCCTAGCCCCTGAAAAT
GCACACGTCTGAACTCTTTCCCTACACGACGCTCTTCCGATCTNNNCCCAAATGTCATTATGCAGCACCC
AGAGCACACGTCTGAACTCTTTCCCTACACGACGCTCTTCCGATCTNNNTGTTTGACAGAGACAGAGA
GAGCACACGTCTGAACTCTTTCCCTACACGACGCTCTTCCGATCTNNNAATCTGTGTGGTCTGTCATGC
AGCACACGTCTGAACTCTTTCCCTACACGACGCTCTTCCGATCTNNNGCCATCTCTAGTGGACACACCT

AGCACACGTCTGAACTCTTTCCCTACACGACGCTCTTCCGATCTNNNTAATGTTGTGCAGAGTTGCTG
GAGCACACGTCTGAACTCTTTCCCTACACGACGCTCTTCCGATCTNNNTGTTATTTCCAGGCCAACCAC
AGCACACGTCTGAACTCTTTCCCTACACGACGCTCTTCCGATCTNNNACACTGCTTTTAGAGAATGCA
GAGCACACGTCTGAACTCTTTCCCTACACGACGCTCTTCCGATCTNNNTACAGTCAAGTGGAGGCAT
AGCACACGTCTGAACTCTTTCCCTACACGACGCTCTTCCGATCTNNNCCACAGCATTAGCTCTTGGCT
GCACACGTCTGAACTCTTTCCCTACACGACGCTCTTCCGATCTNNNAGGGCAAAAATCACAATCCCTGT
GAGCACACGTCTGAACTCTTTCCCTACACGACGCTCTTCCGATCTNNNGGTTTCTTCCAAGGCTAGTG
AGCACACGTCTGAACTCTTTCCCTACACGACGCTCTTCCGATCTNNNCCAGCAAATGTCCTATCCTGA
AGCACACGTCTGAACTCTTTCCCTACACGACGCTCTTCCGATCTNNNTCTATGGCCACAGCTCTAAGA
AGCACACGTCTGAACTCTTTCCCTACACGACGCTCTTCCGATCTNNNTGTGCATGAGCTGGACATCC
AGCACACGTCTGAACTCTTTCCCTACACGACGCTCTTCCGATCTNNNTTTCTAGTGGATGGGGCTTC
GAGCACACGTCTGAACTCTTTCCCTACACGACGCTCTTCCGATCTNNNCTTTTGCTAGTGGGCAGTGT
AGCACACGTCTGAACTCTTTCCCTACACGACGCTCTTCCGATCTNNNTTGTGAGAAAGAGCCAACTG
AGCACACGTCTGAACTCTTTCCCTACACGACGCTCTTCCGATCTNNNCTTTCCTCAAGTTCCACCAACC
GCACACGTCTGAACTCTTTCCCTACACGACGCTCTTCCGATCTNNNATTCTATGTGCACAGTGTCAGG
GCACACGTCTGAACTCTTTCCCTACACGACGCTCTTCCGATCTNNNGTTCCACCAATCCTGAGGGAAA
GCACACGTCTGAACTCTTTCCCTACACGACGCTCTTCCGATCTNNNTGGCTGAGGACTTGGATAGAAC
AGCACACGTCTGAACTCTTTCCCTACACGACGCTCTTCCGATCTNNNATGGGGCTCTATGATGGTGTC
GAGCACACGTCTGAACTCTTTCCCTACACGACGCTCTTCCGATCTNNNAAGTCATGGCACATAGATG
GAGCACACGTCTGAACTCTTTCCCTACACGACGCTCTTCCGATCTNNNAGGTTCAGCAGCTATTGTGA
AGCACACGTCTGAACTCTTTCCCTACACGACGCTCTTCCGATCTNNNACAGACATAGATACAGGCAGG
AGCACACGTCTGAACTCTTTCCCTACACGACGCTCTTCCGATCTNNNCAGGTCAAGTCCACAACCTTG
GCACACGTCTGAACTCTTTCCCTACACGACGCTCTTCCGATCTNNNTTGAAGTGCAACAGAACACAC
AGCACACGTCTGAACTCTTTCCCTACACGACGCTCTTCCGATCTNNNTGCAGTGTATTTTTCCCCTCA
GAGCACACGTCTGAACTCTTTCCCTACACGACGCTCTTCCGATCTNNNTTTTTACACCTGCCACCATTG
AGCACACGTCTGAACTCTTTCCCTACACGACGCTCTTCCGATCTNNNCCACACTACAACTGCCCATATC
GCACACGTCTGAACTCTTTCCCTACACGACGCTCTTCCGATCTNNNGTGGGTGACCTTGATTCTCTCT
GCACACGTCTGAACTCTTTCCCTACACGACGCTCTTCCGATCTNNNGCGGATGGAAAACACCACTTCT
GCACACGTCTGAACTCTTTCCCTACACGACGCTCTTCCGATCTNNNCAGCACTGTCACCTACATGATG
AGCACACGTCTGAACTCTTTCCCTACACGACGCTCTTCCGATCTNNNGAAGTAGCTTCTGCTAAGGG
AGCACACGTCTGAACTCTTTCCCTACACGACGCTCTTCCGATCTNNNGGGCAATTCTTCTGTTCCACA
GAGCACACGTCTGAACTCTTTCCCTACACGACGCTCTTCCGATCTNNNAGGTCCTTCAAACAGACACA
AGCACACGTCTGAACTCTTTCCCTACACGACGCTCTTCCGATCTNNNTTACACCCTGGACCTCATCTCT
AGCACACGTCTGAACTCTTTCCCTACACGACGCTCTTCCGATCTNNNGGACTGTTCAGTTTATGTCCCC
AGCACACGTCTGAACTCTTTCCCTACACGACGCTCTTCCGATCTNNNTCCCGAGAGGCAACCAGTATT
GAGCACACGTCTGAACTCTTTCCCTACACGACGCTCTTCCGATCTNNNAAGAGGGGCAGACAGAAAA
GAGCACACGTCTGAACTCTTTCCCTACACGACGCTCTTCCGATCTNNNGTGTGTCAGAATGAGTAGC
AGCACACGTCTGAACTCTTTCCCTACACGACGCTCTTCCGATCTNNNTGAAGCTTAGGAAACCAACCT
GAGCACACGTCTGAACTCTTTCCCTACACGACGCTCTTCCGATCTNNNACTGGCACTTAGGAAATTCA
AGCACACGTCTGAACTCTTTCCCTACACGACGCTCTTCCGATCTNNNAGTCTTGTTTCCCAAATGCTGG
GCACACGTCTGAACTCTTTCCCTACACGACGCTCTTCCGATCTNNNCCTGGCCCTGATGTGACTTTTA
AGCACACGTCTGAACTCTTTCCCTACACGACGCTCTTCCGATCTNNNTATGAAAAAGAAACCGGGGG
AGCACACGTCTGAACTCTTTCCCTACACGACGCTCTTCCGATCTNNNGTTTTAGTGGCACCTCCTCCA

GAGCACACGTCTGAACTCTTTCCCTACACGACGCTCTTCCGATCTNNNAATCAATGAAGGGAGGGAAA
GAGCACACGTCTGAACTCTTTCCCTACACGACGCTCTTCCGATCTNNNTCCAACTGTCAGAGTGTCAGG
GAGCACACGTCTGAACTCTTTCCCTACACGACGCTCTTCCGATCTNNNATGACAGAACGGAACTCTGA
GCACACGTCTGAACTCTTTCCCTACACGACGCTCTTCCGATCTNNNGAAGCCACAGGTAGAAGGAAGT
GAGCACACGTCTGAACTCTTTCCCTACACGACGCTCTTCCGATCTNNNCCATCCTACACACAGCAACACA
AGAGCACACGTCTGAACTCTTTCCCTACACGACGCTCTTCCGATCTNNNCCACAAGATTGCCGTGATGA
AGAGCACACGTCTGAACTCTTTCCCTACACGACGCTCTTCCGATCTNNNCTTTGTTCATAGCTTTGGGCC
GAGCACACGTCTGAACTCTTTCCCTACACGACGCTCTTCCGATCTNNNAGCCAGAAAATGAGATGAGG
GAGCACACGTCTGAACTCTTTCCCTACACGACGCTCTTCCGATCTNNNATCCTCTGTTTCATCTGGCAAC
AGAGCACACGTCTGAACTCTTTCCCTACACGACGCTCTTCCGATCTNNNCTCCACTCAAACCACTGGAG
GAGCACACGTCTGAACTCTTTCCCTACACGACGCTCTTCCGATCTNNNCAATCGAGATTGCCCTTGATG
AGAGCACACGTCTGAACTCTTTCCCTACACGACGCTCTTCCGATCTNNNTCTGGTTTGTAATGTCACAG
AGCACACGTCTGAACTCTTTCCCTACACGACGCTCTTCCGATCTNNNTCGTTTTCTTTCTCCCACCACAAC
AGCACACGTCTGAACTCTTTCCCTACACGACGCTCTTCCGATCTNNNTCAGCGGATTAGAGTGGGATA
AGCACACGTCTGAACTCTTTCCCTACACGACGCTCTTCCGATCTNNNTATGGCCTATCAACGTCTACTCG
AGCACACGTCTGAACTCTTTCCCTACACGACGCTCTTCCGATCTNNNGGCCATTACAGGAAGTGCTTTT
GAGCACACGTCTGAACTCTTTCCCTACACGACGCTCTTCCGATCTNNNCATCTGCATTATCTCAGGGTCC
GAGCACACGTCTGAACTCTTTCCCTACACGACGCTCTTCCGATCTNNNGCTTACCCACACATGTACACTCA
GCACACGTCTGAACTCTTTCCCTACACGACGCTCTTCCGATCTNNNGTACAAGGGGCTCAAGTCGTTTA
AGCACACGTCTGAACTCTTTCCCTACACGACGCTCTTCCGATCTNNNTAATTGCTTGATGGCCTTGAGC
GCACACGTCTGAACTCTTTCCCTACACGACGCTCTTCCGATCTNNNAGTCTAGGAATAAGGGCCTTGG
AGCACACGTCTGAACTCTTTCCCTACACGACGCTCTTCCGATCTNNNTACATTTGTAGGCCCCTGTCAG
GAGCACACGTCTGAACTCTTTCCCTACACGACGCTCTTCCGATCTNNNGTGGTGTCTCAACTCTCAAGT
GCACACGTCTGAACTCTTTCCCTACACGACGCTCTTCCGATCTNNNTAATCAAACCCTGGCTGAGGAGA
GAGCACACGTCTGAACTCTTTCCCTACACGACGCTCTTCCGATCTNNNCCATGACTGCTCATGCCATTTT
AGAGCACACGTCTGAACTCTTTCCCTACACGACGCTCTTCCGATCTNNNTTGGCTAGGAGACCAGGATT
GAGCACACGTCTGAACTCTTTCCCTACACGACGCTCTTCCGATCTNNNCCCCACTGCTTCATGTCTTACAC
AGCACACGTCTGAACTCTTTCCCTACACGACGCTCTTCCGATCTNNNACAAAACCTATGTGCTGTGCCTA
GAGCACACGTCTGAACTCTTTCCCTACACGACGCTCTTCCGATCTNNNTACTCAGGGCAGCTTACTTAG
GAGCACACGTCTGAACTCTTTCCCTACACGACGCTCTTCCGATCTNNNAGCTGTCAGATTGTTAGCTCT
GAGCACACGTCTGAACTCTTTCCCTACACGACGCTCTTCCGATCTNNNTTTCCAGTGTCTACCACCTCTGT
GAGCACACGTCTGAACTCTTTCCCTACACGACGCTCTTCCGATCTNNNCCACCTATTGAATTGTGCCAGC
GAGCACACGTCTGAACTCTTTCCCTACACGACGCTCTTCCGATCTNNNGGTAACTCACACTCATCCACCA
AGAGCACACGTCTGAACTCTTTCCCTACACGACGCTCTTCCGATCTNNNTCCTTGACAAGTGTTGATGG
GAGCACACGTCTGAACTCTTTCCCTACACGACGCTCTTCCGATCTNNNCAGAACGAGACCCTATCTCCCA
AGCACACGTCTGAACTCTTTCCCTACACGACGCTCTTCCGATCTNNNGCCAGTTCACGGTCTAGTTCAAG
AGCACACGTCTGAACTCTTTCCCTACACGACGCTCTTCCGATCTNNNGTTCTCTCCCCACCTCCAAAAAGC
AGCACACGTCTGAACTCTTTCCCTACACGACGCTCTTCCGATCTNNNGGATCACCAAAAGTCCTGAGAC
GAGCACACGTCTGAACTCTTTCCCTACACGACGCTCTTCCGATCTNNNTTAGGTTACCAAGGGATGGG
AGCACACGTCTGAACTCTTTCCCTACACGACGCTCTTCCGATCTNNNTCTCAAAGAGGCACACAAAGACA
AGCACACGTCTGAACTCTTTCCCTACACGACGCTCTTCCGATCTNNNCAAAATCCAGAATCCCAGGGTGC
GAGCACACGTCTGAACTCTTTCCCTACACGACGCTCTTCCGATCTNNNAAAAACATGAAGGGCCACACT
AGAGCACACGTCTGAACTCTTTCCCTACACGACGCTCTTCCGATCTNNNTCCCTAGTTCCTGGGTTGTGA

AGCACACGTCTGAACTCTTTCCCTACACGACGCTCTTCCGATCTNNNTTGTGCTAGGCTGTACTTCACAA
GCACACGTCTGAACTCTTTCCCTACACGACGCTCTTCCGATCTNNNTACCTATTAGCCAGTGTCTCCCCC
AGCACACGTCTGAACTCTTTCCCTACACGACGCTCTTCCGATCTNNNCTCCAGCCATTGCTAACTGCTTAG
AGCACACGTCTGAACTCTTTCCCTACACGACGCTCTTCCGATCTNNNGAAACAATTGCAGCCTGATTCCC
AGCACACGTCTGAACTCTTTCCCTACACGACGCTCTTCCGATCTNNNTGTTTTGTGTACACCCTGGTCAG
AGAGCACACGTCTGAACTCTTTCCCTACACGACGCTCTTCCGATCTNNNTGATCTTCACACCAGGAGGA
GAGCACACGTCTGAACTCTTTCCCTACACGACGCTCTTCCGATCTNNNTTCAGGTGCCATCACTTTTTCCA
AGCACACGTCTGAACTCTTTCCCTACACGACGCTCTTCCGATCTNNNTTGGTAACACTCATAGCACCTG
GCACACGTCTGAACTCTTTCCCTACACGACGCTCTTCCGATCTNNNCACCTGGGGTTTTGGAAATCAGT
AGCACACGTCTGAACTCTTTCCCTACACGACGCTCTTCCGATCTNNNTGGCTTTGTGATTTCTGCTTGAG
GAGCACACGTCTGAACTCTTTCCCTACACGACGCTCTTCCGATCTNNNAATGGGTGGCTAGGAGAATA
AGCACACGTCTGAACTCTTTCCCTACACGACGCTCTTCCGATCTNNNAGGGTGGCTAGCTGTAGCTATT
GCACACGTCTGAACTCTTTCCCTACACGACGCTCTTCCGATCTNNNAATCCTTACTGGATCATGTGCCCC
GAGCACACGTCTGAACTCTTTCCCTACACGACGCTCTTCCGATCTNNNCCTTATTCTTCCTCCAAGGTGG
AGCACACGTCTGAACTCTTTCCCTACACGACGCTCTTCCGATCTNNNAGCAACGACAAAAGCCAAAACA
GAGCACACGTCTGAACTCTTTCCCTACACGACGCTCTTCCGATCTNNNAGGAACAGAAAACTGGGGAG
GAGCACACGTCTGAACTCTTTCCCTACACGACGCTCTTCCGATCTNNNGAAGGGGGTTGTAAAGTGTG
AGCACACGTCTGAACTCTTTCCCTACACGACGCTCTTCCGATCTNNNAGTGAACACCACTGTTGACTAGG
GCACACGTCTGAACTCTTTCCCTACACGACGCTCTTCCGATCTNNNTCCCTCCTCACCAACACATACATCA
AGCACACGTCTGAACTCTTTCCCTACACGACGCTCTTCCGATCTNNNCTTTACTGGTGGTGATGGTGG
GAGCACACGTCTGAACTCTTTCCCTACACGACGCTCTTCCGATCTNNNGATTGCTAGTGAGGCTTGTGG
GAGCACACGTCTGAACTCTTTCCCTACACGACGCTCTTCCGATCTNNNGGACCAAAAGGAGCAATTCTG
AGCACACGTCTGAACTCTTTCCCTACACGACGCTCTTCCGATCTNNNGAGATGCTGGCCTGTGTAAGTA
AGCACACGTCTGAACTCTTTCCCTACACGACGCTCTTCCGATCTNNNGCCAAAGATGCTGTATCCTGCTA
AGCACACGTCTGAACTCTTTCCCTACACGACGCTCTTCCGATCTNNNAGCTCATCTCCAGTCTATACCCA
AGCACACGTCTGAACTCTTTCCCTACACGACGCTCTTCCGATCTNNNTCCAAGCACTCCAAAATCTCACC
AGCACACGTCTGAACTCTTTCCCTACACGACGCTCTTCCGATCTNNNCCTCACCCTCCTCCAAGTACATAC
AGCACACGTCTGAACTCTTTCCCTACACGACGCTCTTCCGATCTNNNCTCCTGGTGAGATAGATGCTTC
GAGCACACGTCTGAACTCTTTCCCTACACGACGCTCTTCCGATCTNNNGCACACTTACCTCTCACTACAGG
GAGCACACGTCTGAACTCTTTCCCTACACGACGCTCTTCCGATCTNNNGGAGCCTGATGAATTGTCTGT
GAGCACACGTCTGAACTCTTTCCCTACACGACGCTCTTCCGATCTNNNCTGAAACCGAGAGTTCTCCCAA
GCACACGTCTGAACTCTTTCCCTACACGACGCTCTTCCGATCTNNNTCCTCCACATTGTAAACCCTCCTCAT
AGCACACGTCTGAACTCTTTCCCTACACGACGCTCTTCCGATCTNNNGCAAGGAACAGGAACTAGATC
AGAGCACACGTCTGAACTCTTTCCCTACACGACGCTCTTCCGATCTNNNTGGCAATGGTGAGGAAACA
AGCACACGTCTGAACTCTTTCCCTACACGACGCTCTTCCGATCTNNNTCTGTAACTACACCACCAGCCTT
AGCACACGTCTGAACTCTTTCCCTACACGACGCTCTTCCGATCTNNNTGAAGAAGTCCTCTCTGCTCCAA
AGCACACGTCTGAACTCTTTCCCTACACGACGCTCTTCCGATCTNNNTTTCTGTCTTTGCTTGGCTCCATG
AGCACACGTCTGAACTCTTTCCCTACACGACGCTCTTCCGATCTNNNTGCAAACTGGCTTGTGATTTCT
AGCACACGTCTGAACTCTTTCCCTACACGACGCTCTTCCGATCTNNNTTGCCGGGTAAAGATCTAAGCA
AGCACACGTCTGAACTCTTTCCCTACACGACGCTCTTCCGATCTNNNATGCACCATGCAGGATTGCATA
GAGCACACGTCTGAACTCTTTCCCTACACGACGCTCTTCCGATCTNNNAACCTACTAGAAACCGTGGGTG
GCACACGTCTGAACTCTTTCCCTACACGACGCTCTTCCGATCTNNNTCCCTAAAATTGCACACCAGACCC
GAGCACACGTCTGAACTCTTTCCCTACACGACGCTCTTCCGATCTNNNTATCCTACAGGGAACAGGATG

GAGCACACGTCTGAACTCTTTCCCTACACGACGCTCTTCCGATCTNNNCAGGAAGGCTTCATCCTGCAT
GAGCACACGTCTGAACTCTTTCCCTACACGACGCTCTTCCGATCTNNNTATCCAGCTAGGGACTTGGAA
GAGCACACGTCTGAACTCTTTCCCTACACGACGCTCTTCCGATCTNNNACAGAATGAGCACAAAAGCCA
AGCACACGTCTGAACTCTTTCCCTACACGACGCTCTTCCGATCTNNNGCTTCCAGTTCTGTTCACATTGC
GAGCACACGTCTGAACTCTTTCCCTACACGACGCTCTTCCGATCTNNNCCATTTTGTGCAAGTCACAGAG
AGCACACGTCTGAACTCTTTCCCTACACGACGCTCTTCCGATCTNNNATCCTCAAAGCCAAATGTGGAC
GAGCACACGTCTGAACTCTTTCCCTACACGACGCTCTTCCGATCTNNNGAAAGAGTTCCAATGGCCAAA
AGCACACGTCTGAACTCTTTCCCTACACGACGCTCTTCCGATCTNNNATAAATGCCTCTTCTCATCCGCCC
GAGCACACGTCTGAACTCTTTCCCTACACGACGCTCTTCCGATCTNNNGATTTTCCCGAGACCCTACCA
GAGCACACGTCTGAACTCTTTCCCTACACGACGCTCTTCCGATCTNNNCACATTGTGTGGATCAAAGCCA
AGCACACGTCTGAACTCTTTCCCTACACGACGCTCTTCCGATCTNNNTAGCTACCTGTCCGTCCCTTATC
AGAGCACACGTCTGAACTCTTTCCCTACACGACGCTCTTCCGATCTNNNCGTGGTCACTCAGGGTCATA
GAGCACACGTCTGAACTCTTTCCCTACACGACGCTCTTCCGATCTNNNCCTGTGCCCTCTCATCATTCTCT
GAGCACACGTCTGAACTCTTTCCCTACACGACGCTCTTCCGATCTNNNTGTGGAATTCTTGTCCTAGCAC
AGCACACGTCTGAACTCTTTCCCTACACGACGCTCTTCCGATCTNNNCACCTGTGTCTCCTTGCTAGATC
GAGCACACGTCTGAACTCTTTCCCTACACGACGCTCTTCCGATCTNNNTGATGTACCAGGGAGACTGTG
GAGCACACGTCTGAACTCTTTCCCTACACGACGCTCTTCCGATCTNNNTCTGGGGATGGAGGTGGAA
GAGCACACGTCTGAACTCTTTCCCTACACGACGCTCTTCCGATCTNNNAGAGAGCCCCCAAGTTGATAC
AGCACACGTCTGAACTCTTTCCCTACACGACGCTCTTCCGATCTNNNAGAAACCAAGTCCAGTAGAGGC
AGCACACGTCTGAACTCTTTCCCTACACGACGCTCTTCCGATCTNNNGCTTCTCTGCCAAGTTCACAAAC
GAGCACACGTCTGAACTCTTTCCCTACACGACGCTCTTCCGATCTNNNGAAGACAGCCATAAGAAGGC
AGAGCACACGTCTGAACTCTTTCCCTACACGACGCTCTTCCGATCTNNNTCTGTGTTAGTCACGTTCTCCA
GAGCACACGTCTGAACTCTTTCCCTACACGACGCTCTTCCGATCTNNNAACCAGCTTAGAAGACAAAAG
AGCACACGTCTGAACTCTTTCCCTACACGACGCTCTTCCGATCTNNNTCAACCCTACCCAGAAATCTCCA
GAGCACACGTCTGAACTCTTTCCCTACACGACGCTCTTCCGATCTNNNCTGTGTACTCTGGTACTGTCCC
GAGCACACGTCTGAACTCTTTCCCTACACGACGCTCTTCCGATCTNNNACAGAGACACAACACAGGGA
AGCACACGTCTGAACTCTTTCCCTACACGACGCTCTTCCGATCTNNNACAATTACGAGAGAGGGGGAG
AGCACACGTCTGAACTCTTTCCCTACACGACGCTCTTCCGATCTNNNAGCAGTCATGGACCACAATTCTC
AGCACACGTCTGAACTCTTTCCCTACACGACGCTCTTCCGATCTNNNATGAAAAGCCAAAGAGGTGCTC
AGCACACGTCTGAACTCTTTCCCTACACGACGCTCTTCCGATCTNNNCAAATAGTCTCACAGGGAAGCC
GAGCACACGTCTGAACTCTTTCCCTACACGACGCTCTTCCGATCTNNNCTGGACAGGAGGAACAGTCTT
AGCACACGTCTGAACTCTTTCCCTACACGACGCTCTTCCGATCTNNNTATCATCATCTCCTCTGCCCACCC
GAGCACACGTCTGAACTCTTTCCCTACACGACGCTCTTCCGATCTNNNTAGGTGTGAAACAGCTTCTGC
AGCACACGTCTGAACTCTTTCCCTACACGACGCTCTTCCGATCTNNNGTGTATGCTAAGAGAGGGCATG
GAGCACACGTCTGAACTCTTTCCCTACACGACGCTCTTCCGATCTNNNCATTTCAGGCTTCCAGCAGAAG
GAGCACACGTCTGAACTCTTTCCCTACACGACGCTCTTCCGATCTNNNTGTTGAAAATCAGGCCACCTAC
GCACACGTCTGAACTCTTTCCCTACACGACGCTCTTCCGATCTNNNCACCTTGTCTGACATCTCCCTTGCA
GAGCACACGTCTGAACTCTTTCCCTACACGACGCTCTTCCGATCTNNNTTTCTCAATAGGGGTGTGTGT
AGCACACGTCTGAACTCTTTCCCTACACGACGCTCTTCCGATCTNNNTGAGGAAGTGTAGCTTCAAGG
GAGCACACGTCTGAACTCTTTCCCTACACGACGCTCTTCCGATCTNNNTTTCACTAGCACCTTCCAAAGC
GAGCACACGTCTGAACTCTTTCCCTACACGACGCTCTTCCGATCTNNNAATCACTGAATTCATGCTGGG
AGCACACGTCTGAACTCTTTCCCTACACGACGCTCTTCCGATCTNNNAACATGCTATTTGGGGGTTTCC
GAGCACACGTCTGAACTCTTTCCCTACACGACGCTCTTCCGATCTNNNCTTGACGTTTACCACATGTCCCT

AGCACACGTCTGAACTCTTTCCCTACACGACGCTCTTCCGATCTNNNAGATGCATGACAGCTGGTGAA
GAGCACACGTCTGAACTCTTTCCCTACACGACGCTCTTCCGATCTNNNTTACAAGCCACAGTACCACGT
GAGCACACGTCTGAACTCTTTCCCTACACGACGCTCTTCCGATCTNNNTTGTGCTGAAGTCTGCCTGTAT
AGAGCACACGTCTGAACTCTTTCCCTACACGACGCTCTTCCGATCTNNNGGTGAAGGTGTTCAGGTAG
GAGCACACGTCTGAACTCTTTCCCTACACGACGCTCTTCCGATCTNNNTGGAGGTCTTAGAGTACAGA
AGCACACGTCTGAACTCTTTCCCTACACGACGCTCTTCCGATCTNNNTTTCTGTCTTTTTCCTGCTGCCT
GAGCACACGTCTGAACTCTTTCCCTACACGACGCTCTTCCGATCTNNNACACAACCAATCATGAGACAG
GAGCACACGTCTGAACTCTTTCCCTACACGACGCTCTTCCGATCTNNNTTTCTTCATCCCGCCATCCTTA
GAGCACACGTCTGAACTCTTTCCCTACACGACGCTCTTCCGATCTNNNGAGCATTGTAGGTATTGCCC
GAGCACACGTCTGAACTCTTTCCCTACACGACGCTCTTCCGATCTNNNTGCTGGATGTGGATTGAGCTT
GAGCACACGTCTGAACTCTTTCCCTACACGACGCTCTTCCGATCTNNNAGTCCTCCCTTCTCAGCCTCAT
GAGCACACGTCTGAACTCTTTCCCTACACGACGCTCTTCCGATCTNNNACATTGGGACTAAGTGTGGCA
AGCACACGTCTGAACTCTTTCCCTACACGACGCTCTTCCGATCTNNNCTGCTTCACCTCAGTTCTCAACA
AGCACACGTCTGAACTCTTTCCCTACACGACGCTCTTCCGATCTNNNAGTTTCTTTGGGAATGCAGGA
AGCACACGTCTGAACTCTTTCCCTACACGACGCTCTTCCGATCTNNNAGCGGAAACATTGCAAACACAA
GAGCACACGTCTGAACTCTTTCCCTACACGACGCTCTTCCGATCTNNNAGACCCTGCAGCTATCATAAG
AGAGCACACGTCTGAACTCTTTCCCTACACGACGCTCTTCCGATCTNNNTGACATGGTCAATACTTCTG
GAGCACACGTCTGAACTCTTTCCCTACACGACGCTCTTCCGATCTNNNTGCTTGTGGCTGGTACAAGTT
AGCACACGTCTGAACTCTTTCCCTACACGACGCTCTTCCGATCTNNNTACCTGTGTGCTACTCATTGTC
CACACGTCTGAACTCTTTCCCTACACGACGCTCTTCCGATCTNNNTGCAGGTTTTCTTTCAATGGGTGC
GAGCACACGTCTGAACTCTTTCCCTACACGACGCTCTTCCGATCTNNNTCCCAGCCACTAATGCACTTA
AGCACACGTCTGAACTCTTTCCCTACACGACGCTCTTCCGATCTNNNAGTGAAGCAACCCTTCTCTCTCT
AGAGCACACGTCTGAACTCTTTCCCTACACGACGCTCTTCCGATCTNNNCAGGCCACTCTCAACTAAGTG
GAGCACACGTCTGAACTCTTTCCCTACACGACGCTCTTCCGATCTNNNAAGCTGCTGTTTGGAGAGCTA
AGCACACGTCTGAACTCTTTCCCTACACGACGCTCTTCCGATCTNNNAGGGAAGTAAGGAAGTAGCCA
GAGCACACGTCTGAACTCTTTCCCTACACGACGCTCTTCCGATCTNNNGAATCCATCACCACAGCAGAC
AGCACACGTCTGAACTCTTTCCCTACACGACGCTCTTCCGATCTNNNACATGGGGCTTTGGTAAAGAA
AGAGCACACGTCTGAACTCTTTCCCTACACGACGCTCTTCCGATCTNNNATAGTGGTTTCCTGGAACTG
AGAGCACACGTCTGAACTCTTTCCCTACACGACGCTCTTCCGATCTNNNGATGCCAGAAAAATCCCCTG
GAGCACACGTCTGAACTCTTTCCCTACACGACGCTCTTCCGATCTNNNTGACTGACTACAACTATGGCA
GAGCACACGTCTGAACTCTTTCCCTACACGACGCTCTTCCGATCTNNNATCTGTTTCAGCAATGACACC
GCACACGTCTGAACTCTTTCCCTACACGACGCTCTTCCGATCTNNNCCTGGCTACCACCTCTCTTTCTACA
GAGCACACGTCTGAACTCTTTCCCTACACGACGCTCTTCCGATCTNNNGGCAGAGCAGAAGAAAGGAA
AGCACACGTCTGAACTCTTTCCCTACACGACGCTCTTCCGATCTNNNACTCTCACATGAAGAGAACCCCA
AGAGCACACGTCTGAACTCTTTCCCTACACGACGCTCTTCCGATCTNNNTGTTTACTGCATCTGTATGG
AGCACACGTCTGAACTCTTTCCCTACACGACGCTCTTCCGATCTNNNTGCAAAATAAAATCCCAGCACCA
AGAGCACACGTCTGAACTCTTTCCCTACACGACGCTCTTCCGATCTNNNTTTCCTTCTTGCCTTCTTCCCA
AGCACACGTCTGAACTCTTTCCCTACACGACGCTCTTCCGATCTNNNCTTAGACAGGAGACCTACACCC
GAGCACACGTCTGAACTCTTTCCCTACACGACGCTCTTCCGATCTNNNTAAGGGGAGGAGAGAGAGA
GAGCACACGTCTGAACTCTTTCCCTACACGACGCTCTTCCGATCTNNNCAGAATTTGCACTCCAGGCTA
GAGCACACGTCTGAACTCTTTCCCTACACGACGCTCTTCCGATCTNNNAGAGATGAGACTTTGGAGTC
GAGCACACGTCTGAACTCTTTCCCTACACGACGCTCTTCCGATCTNNNTATTCAGTGCCAGCAGCTATCT
AGAGCACACGTCTGAACTCTTTCCCTACACGACGCTCTTCCGATCTNNNTGATTACGTTTTTCCAAGGCT

AGCACACGTCTGAACTCTTTCCCTACACGACGCTCTTCCGATCTNNNCTCATGATAGGATTGGGGGTC
GCACACGTCTGAACTCTTTCCCTACACGACGCTCTTCCGATCTNNNAAAAAGCCCTCCATCCATGTCTCC
AGCACACGTCTGAACTCTTTCCCTACACGACGCTCTTCCGATCTNNNTTTGAGGCTGGAGTAAGCAG
AGCACACGTCTGAACTCTTTCCCTACACGACGCTCTTCCGATCTNNNTTGTGACAAATGCCTCAGTGT
AGCACACGTCTGAACTCTTTCCCTACACGACGCTCTTCCGATCTNNNCCATTTGCACCACTTTCCAAGT
AGCACACGTCTGAACTCTTTCCCTACACGACGCTCTTCCGATCTNNNAATCCCTCCCCACATTCTCTCTC
GAGCACACGTCTGAACTCTTTCCCTACACGACGCTCTTCCGATCTNNNGTCACCATCATGTGGCAATTT
GAGCACACGTCTGAACTCTTTCCCTACACGACGCTCTTCCGATCTNNNGTCAGGATTGAAACTGGCT
GAGCACACGTCTGAACTCTTTCCCTACACGACGCTCTTCCGATCTNNNGGATGTGACTAGGAGGGAG
GAGCACACGTCTGAACTCTTTCCCTACACGACGCTCTTCCGATCTNNNGGAAGCAGAGATTGCAGAG
AGCACACGTCTGAACTCTTTCCCTACACGACGCTCTTCCGATCTNNNAGAAGACATACACGCACACACT
AGCACACGTCTGAACTCTTTCCCTACACGACGCTCTTCCGATCTNNNATTTGGCATTTGGGATGTGTG
GAGCACACGTCTGAACTCTTTCCCTACACGACGCTCTTCCGATCTNNNTCTACTGGGAACAGAATCCT
GAGCACACGTCTGAACTCTTTCCCTACACGACGCTCTTCCGATCTNNNGCCAATGAGCTGTGAATGAA
AGCACACGTCTGAACTCTTTCCCTACACGACGCTCTTCCGATCTNNNATGTTTGGTCCCAGAGATGGG
AGCACACGTCTGAACTCTTTCCCTACACGACGCTCTTCCGATCTNNNCCATATCACTCCCCAAATGCGA
AGCACACGTCTGAACTCTTTCCCTACACGACGCTCTTCCGATCTNNNTGGCCTACATGGAGGTAACAG
AGCACACGTCTGAACTCTTTCCCTACACGACGCTCTTCCGATCTNNNATAGTCACACAAACACCCAACC
GAGCACACGTCTGAACTCTTTCCCTACACGACGCTCTTCCGATCTNNNGCAGTTCTGTTCTCAACATCT
AGCACACGTCTGAACTCTTTCCCTACACGACGCTCTTCCGATCTNNNAGACCATGTTCTCAACAGGACA
GCACACGTCTGAACTCTTTCCCTACACGACGCTCTTCCGATCTNNNTGGTTTCCAGTACTCCTGGTTCT
GCACACGTCTGAACTCTTTCCCTACACGACGCTCTTCCGATCTNNNTGGTCCCACAGTTCTAGCTCATA
AGAGCACACGTCTGAACTCTTTCCCTACACGACGCTCTTCCGATCTNNNGCTCAACATTGGCCTGCTTT
AGCACACGTCTGAACTCTTTCCCTACACGACGCTCTTCCGATCTNNNACCAAAACATGGACATTTTCCC
AGCACACGTCTGAACTCTTTCCCTACACGACGCTCTTCCGATCTNNNGTCAATATCTTGTGTTGGCGCT
AGCACACGTCTGAACTCTTTCCCTACACGACGCTCTTCCGATCTNNNTATGGAGCTCAGAGAGGTGA
GAGCACACGTCTGAACTCTTTCCCTACACGACGCTCTTCCGATCTNNNACCTTCCAAAGTCGTACAGCT
GAGCACACGTCTGAACTCTTTCCCTACACGACGCTCTTCCGATCTNNNTCTCCATGAAGACAGGGAAG
AGCACACGTCTGAACTCTTTCCCTACACGACGCTCTTCCGATCTNNNAACTCTCCATTTTGGAAGCAG
GCACACGTCTGAACTCTTTCCCTACACGACGCTCTTCCGATCTNNNAATCTCACAGCTCTAAGTGGCAG
GCACACGTCTGAACTCTTTCCCTACACGACGCTCTTCCGATCTNNNAGCCCTTCTACAGTGGCATAAG
AGCACACGTCTGAACTCTTTCCCTACACGACGCTCTTCCGATCTNNNGTCAAGCACCAAATGGAAACC
AGCACACGTCTGAACTCTTTCCCTACACGACGCTCTTCCGATCTNNNCGAAAAGCACAAGATTCAACCC
GAGCACACGTCTGAACTCTTTCCCTACACGACGCTCTTCCGATCTNNNCACAAATGTGCTTGCCTCTCTT
AGAGCACACGTCTGAACTCTTTCCCTACACGACGCTCTTCCGATCTNNNTAGATTGTGGCTTTCCAGA
AGAGCACACGTCTGAACTCTTTCCCTACACGACGCTCTTCCGATCTNNNGAGGCCTGGTACTGTGACT
AGCACACGTCTGAACTCTTTCCCTACACGACGCTCTTCCGATCTNNNGGCTCAGGTATGACTGGTGAA
GAGCACACGTCTGAACTCTTTCCCTACACGACGCTCTTCCGATCTNNNCAACCCATCTAAGGGGATTCA
AGCACACGTCTGAACTCTTTCCCTACACGACGCTCTTCCGATCTNNNTAAGTTGGCCACGTTTAGACAG
AGCACACGTCTGAACTCTTTCCCTACACGACGCTCTTCCGATCTNNNGACAGAGTGGCTTGAACACTA
AGCACACGTCTGAACTCTTTCCCTACACGACGCTCTTCCGATCTNNNTGGAGTAGGGATAATCACACA
GCACACGTCTGAACTCTTTCCCTACACGACGCTCTTCCGATCTNNNTCCTGTGTATCTGGCTCTGGTTT
GCACACGTCTGAACTCTTTCCCTACACGACGCTCTTCCGATCTNNNTCCAGCAGATCAGAACTCTTTGC

AGCACACGTCTGAACTCTTTCCCTACACGACGCTCTTCCGATCTNNNTCCTCTGGAACCTAACGTTTTCAC
AGCACACGTCTGAACTCTTTCCCTACACGACGCTCTTCCGATCTNNNAAGCACTGTCTGTGAGTAATGG
GCACACGTCTGAACTCTTTCCCTACACGACGCTCTTCCGATCTNNNACTTTTTCTCCCTCTCTCCTGCTCA
AGCACACGTCTGAACTCTTTCCCTACACGACGCTCTTCCGATCTNNNTAAAATCCAGTGCCCTCCCACTA
AGCACACGTCTGAACTCTTTCCCTACACGACGCTCTTCCGATCTNNNGTCGAAGTTGCAGAGAGCAAG
AGCACACGTCTGAACTCTTTCCCTACACGACGCTCTTCCGATCTNNNGCCCTCATCTCTTTCTCTCCTCTCA
AGCACACGTCTGAACTCTTTCCCTACACGACGCTCTTCCGATCTNNNGAGCCTTCTCAGTTCTTTCCCAG
GAGCACACGTCTGAACTCTTTCCCTACACGACGCTCTTCCGATCTNNNGAGAGGACACACTCTAGTCCA
GAGCACACGTCTGAACTCTTTCCCTACACGACGCTCTTCCGATCTNNNACCTCCGGAGACTTGATTCAC
AGCACACGTCTGAACTCTTTCCCTACACGACGCTCTTCCGATCTNNNTAAACAGTACTGGTCAGTGGCC
GAGCACACGTCTGAACTCTTTCCCTACACGACGCTCTTCCGATCTNNNAGCCTGGGGATCACTGTATTT
AGAGCACACGTCTGAACTCTTTCCCTACACGACGCTCTTCCGATCTNNNATGGTATATCTGAGCCCTCCA
GAGCACACGTCTGAACTCTTTCCCTACACGACGCTCTTCCGATCTNNNAAGGAAAAACTGTGTGTGTG
AGCACACGTCTGAACTCTTTCCCTACACGACGCTCTTCCGATCTNNNGGGAGGCTGAAGGAACAAAG
AGCACACGTCTGAACTCTTTCCCTACACGACGCTCTTCCGATCTNNNTAAAACACCAACCTTTCACCCCAC
GAGCACACGTCTGAACTCTTTCCCTACACGACGCTCTTCCGATCTNNNGCCTAAAAGGTGAAACCCCCAT
AGCACACGTCTGAACTCTTTCCCTACACGACGCTCTTCCGATCTNNNTGATGACATTTTGGTTGTGCGT
AGCACACGTCTGAACTCTTTCCCTACACGACGCTCTTCCGATCTNNNAGCTGAACAAACCTGAGAGGG
GAGCACACGTCTGAACTCTTTCCCTACACGACGCTCTTCCGATCTNNNTTCTCCCTCTGGAATTTGCAAC
AGCACACGTCTGAACTCTTTCCCTACACGACGCTCTTCCGATCTNNNTGGGAAAACACAGTGCAGTTCA
AGCACACGTCTGAACTCTTTCCCTACACGACGCTCTTCCGATCTNNNGATCTAGGAGGCACGCATGAT
GAGCACACGTCTGAACTCTTTCCCTACACGACGCTCTTCCGATCTNNNAGAAATGGGCACCTTAGTACT
GAGCACACGTCTGAACTCTTTCCCTACACGACGCTCTTCCGATCTNNNTTTTCTTTCCCTTGCCTCTCACC
AGAGCACACGTCTGAACTCTTTCCCTACACGACGCTCTTCCGATCTNNNCAACCATACAGGGGCCAAAT
GAGCACACGTCTGAACTCTTTCCCTACACGACGCTCTTCCGATCTNNNGGGGATTGGAACAATGCCTG
GAGCACACGTCTGAACTCTTTCCCTACACGACGCTCTTCCGATCTNNNCAGAGCCCACAAACAATCTTCC
AGAGCACACGTCTGAACTCTTTCCCTACACGACGCTCTTCCGATCTNNNCATTCACTGCTCCACATCATCC
AGCACACGTCTGAACTCTTTCCCTACACGACGCTCTTCCGATCTNNNACACTGTTTGTGGTAGGAAAG
AGAGCACACGTCTGAACTCTTTCCCTACACGACGCTCTTCCGATCTNNNCATGAGCAGTGAATCTCACCA
AGCACACGTCTGAACTCTTTCCCTACACGACGCTCTTCCGATCTNNNGTATACCAAGCCATCCACCTTCCC
GAGCACACGTCTGAACTCTTTCCCTACACGACGCTCTTCCGATCTNNNAGTCACTTAGCTGGTCTTCCTG
AGCACACGTCTGAACTCTTTCCCTACACGACGCTCTTCCGATCTNNNAGTTGTCATGCTGGTGGATGTT
AGCACACGTCTGAACTCTTTCCCTACACGACGCTCTTCCGATCTNNNTGAGATTTACAAGAGGGCCTG
GAGCACACGTCTGAACTCTTTCCCTACACGACGCTCTTCCGATCTNNNAGTGCAGACATTGCATTTGAG
GAGCACACGTCTGAACTCTTTCCCTACACGACGCTCTTCCGATCTNNNGCTGGGGGTAGAAGAAAAAC
AGCACACGTCTGAACTCTTTCCCTACACGACGCTCTTCCGATCTNNNAATAGTCAGCCTGGTCCACAGAA
GAGCACACGTCTGAACTCTTTCCCTACACGACGCTCTTCCGATCTNNNAAAGGGAGAAGAAGAGGTG
GAGCACACGTCTGAACTCTTTCCCTACACGACGCTCTTCCGATCTNNNCTGTGTTCCAGTGTTACAACCC
AGAGCACACGTCTGAACTCTTTCCCTACACGACGCTCTTCCGATCTNNNTGTGCTGATAGAAACACCTG
GAGCACACGTCTGAACTCTTTCCCTACACGACGCTCTTCCGATCTNNNGAGGAAGGAATGCTGGAGA
GAGCACACGTCTGAACTCTTTCCCTACACGACGCTCTTCCGATCTNNNTATAAATGGGAAGTGGCTGG
AGAGCACACGTCTGAACTCTTTCCCTACACGACGCTCTTCCGATCTNNNCAAGGAAGAAGGGACAGGC
AGCACACGTCTGAACTCTTTCCCTACACGACGCTCTTCCGATCTNNNACCTGATCTGCAAACAATCCTG

AGCACACGTCTGAACTCTTTCCCTACACGACGCTCTTCCGATCTNNNGGCAATCGGACAATGTAGGAC
AGCACACGTCTGAACTCTTTCCCTACACGACGCTCTTCCGATCTNNNGTGCTTAGGCTTACTCAGTGCT
GCACACGTCTGAACTCTTTCCCTACACGACGCTCTTCCGATCTNNNGACTTTGACAGCAAGTGTGTCT
GAGCACACGTCTGAACTCTTTCCCTACACGACGCTCTTCCGATCTNNNTTTGTTCCACTGGGAAAGC
AGCACACGTCTGAACTCTTTCCCTACACGACGCTCTTCCGATCTNNNTGCACCCACATACACACACATT
GCACACGTCTGAACTCTTTCCCTACACGACGCTCTTCCGATCTNNNATAGCCTTTACTTACGTGCCAGC
AGCACACGTCTGAACTCTTTCCCTACACGACGCTCTTCCGATCTNNNTACTTACAGATCCTGGAGGGA
AGCACACGTCTGAACTCTTTCCCTACACGACGCTCTTCCGATCTNNNATTTGATGCAGTGACATGGGT
GAGCACACGTCTGAACTCTTTCCCTACACGACGCTCTTCCGATCTNNNGGACCATGGAATGTGCTTTG
AGCACACGTCTGAACTCTTTCCCTACACGACGCTCTTCCGATCTNNNGTCCGAAGACCTCTTTTGTCTC
GCACACGTCTGAACTCTTTCCCTACACGACGCTCTTCCGATCTNNNATGTCAAGAATTTTTGCTGGTGT
AGCACACGTCTGAACTCTTTCCCTACACGACGCTCTTCCGATCTNNNCCTCTACAAGCTTCTGATGCGT
GAGCACACGTCTGAACTCTTTCCCTACACGACGCTCTTCCGATCTNNNGGTACACACCTTGGCTAAGA
AGCACACGTCTGAACTCTTTCCCTACACGACGCTCTTCCGATCTNNNGTGCAGCAAGCTTGTGTAATA
GAGCACACGTCTGAACTCTTTCCCTACACGACGCTCTTCCGATCTNNNGCTCTAGACTTGCTTCTGCTT
AGCACACGTCTGAACTCTTTCCCTACACGACGCTCTTCCGATCTNNNCCCAGATAACCCCTATTGCCA
GCACACGTCTGAACTCTTTCCCTACACGACGCTCTTCCGATCTNNNAGATGAAGGAGGGCAGAGAA
AGCACACGTCTGAACTCTTTCCCTACACGACGCTCTTCCGATCTNNNTCGCAGGTACTTGGTGGTAGT
GAGCACACGTCTGAACTCTTTCCCTACACGACGCTCTTCCGATCTNNNTGAATCTCACCATTGATGGG
GAGCACACGTCTGAACTCTTTCCCTACACGACGCTCTTCCGATCTNNNATGCAACCATGCTATACTTGG
CACACGTCTGAACTCTTTCCCTACACGACGCTCTTCCGATCTNNNAAGAATCCGGAAGACCAGGAGA
GCACACGTCTGAACTCTTTCCCTACACGACGCTCTTCCGATCTNNNATCTGATTGCCAGGTTGATCTG
GCACACGTCTGAACTCTTTCCCTACACGACGCTCTTCCGATCTNNNCAGGGGAGCTCTCTGTAAAGA
GCACACGTCTGAACTCTTTCCCTACACGACGCTCTTCCGATCTNNNCCAGGTGGAGAAGGGAGAATG
GAGCACACGTCTGAACTCTTTCCCTACACGACGCTCTTCCGATCTNNNTGTAAATGGGATTGGAGGC
GCACACGTCTGAACTCTTTCCCTACACGACGCTCTTCCGATCTNNNTCTCAAGTCAGCAGACCCCAAA
AGCACACGTCTGAACTCTTTCCCTACACGACGCTCTTCCGATCTNNNAGGAAGGAATTTGGGCTGAG
GCACACGTCTGAACTCTTTCCCTACACGACGCTCTTCCGATCTNNNCCCAAACTGACACCTTTGTGTGT
GAGCACACGTCTGAACTCTTTCCCTACACGACGCTCTTCCGATCTNNNATACACCAATGTGATGGTGC
GAGCACACGTCTGAACTCTTTCCCTACACGACGCTCTTCCGATCTNNNACAGTGGGTACAGTTACTCC
AGAGCACACGTCTGAACTCTTTCCCTACACGACGCTCTTCCGATCTNNNAATTTCCAGTTGGGAAAGG
GCACACGTCTGAACTCTTTCCCTACACGACGCTCTTCCGATCTNNNACTTTCCTTGTCTCTAGCCCTGT
GCACACGTCTGAACTCTTTCCCTACACGACGCTCTTCCGATCTNNNTCCAATAAAGAACATGCATCCAG
GAGCACACGTCTGAACTCTTTCCCTACACGACGCTCTTCCGATCTNNNAGTAGGCCTCATTGGTCATTC
AGCACACGTCTGAACTCTTTCCCTACACGACGCTCTTCCGATCTNNNAGAAAGTAAAGCTGCCTTGCCT
AGCACACGTCTGAACTCTTTCCCTACACGACGCTCTTCCGATCTNNNAAATCAGGTTCCTTCTCCCCAG
GAGCACACGTCTGAACTCTTTCCCTACACGACGCTCTTCCGATCTNNNTTCAAGCAGCACACCAAGTAA
AGCACACGTCTGAACTCTTTCCCTACACGACGCTCTTCCGATCTNNNTGCCCTATCCTGGTCTCTACCTA
AGCACACGTCTGAACTCTTTCCCTACACGACGCTCTTCCGATCTNNNAGAGACAGAGCACCATGACTT
GAGCACACGTCTGAACTCTTTCCCTACACGACGCTCTTCCGATCTNNNCGTTTCCAACAGTTCCATGTGT
AGCACACGTCTGAACTCTTTCCCTACACGACGCTCTTCCGATCTNNNAAGATCAGGCTTCCCTCAGAA
AGCACACGTCTGAACTCTTTCCCTACACGACGCTCTTCCGATCTNNNCCACTGAGAGTAAATAAGGGC
GAGCACACGTCTGAACTCTTTCCCTACACGACGCTCTTCCGATCTNNNTCCCAGGACATTGGACCTAG

GAGCACACGTCTGAACTCTTTCCCTACACGACGCTCTTCCGATCTNNNAAGCCTCCCGGAAATGATTCT
AGCACACGTCTGAACTCTTTCCCTACACGACGCTCTTCCGATCTNNNATCATATTCCAGTGTGCCTACCT
AGCACACGTCTGAACTCTTTCCCTACACGACGCTCTTCCGATCTNNNACCTACTTTTCTGCTTTCCCCACA
GAGCACACGTCTGAACTCTTTCCCTACACGACGCTCTTCCGATCTNNNGGAATTTGTGTGTGTGTACCT
GAGCACACGTCTGAACTCTTTCCCTACACGACGCTCTTCCGATCTNNNGGGTCAAAGGAGAAAAGGCA
GAGCACACGTCTGAACTCTTTCCCTACACGACGCTCTTCCGATCTNNNGCATGAATGAAGTGGTAGCC
AGCACACGTCTGAACTCTTTCCCTACACGACGCTCTTCCGATCTNNNAGGGTGTCTAAATGGTCCCAGT
GAGCACACGTCTGAACTCTTTCCCTACACGACGCTCTTCCGATCTNNNACCCAGCAAAATGTACAAACA
AGCACACGTCTGAACTCTTTCCCTACACGACGCTCTTCCGATCTNNNTCTGACATTCCTTGCCACAAACA
GAGCACACGTCTGAACTCTTTCCCTACACGACGCTCTTCCGATCTNNNGAATGGTGTCCTGAGCTTCTG
GAGCACACGTCTGAACTCTTTCCCTACACGACGCTCTTCCGATCTNNNTGAGGGATTGGAACTGAGGT
GAGCACACGTCTGAACTCTTTCCCTACACGACGCTCTTCCGATCTNNNTCCATAGCTTCCCCACTGTAGT
GAGCACACGTCTGAACTCTTTCCCTACACGACGCTCTTCCGATCTNNNTTTGTTCCACTGGGAAAGCT
GAGCACACGTCTGAACTCTTTCCCTACACGACGCTCTTCCGATCTNNNTGATCCTCTACCGAGAGAGCA
GAGCACACGTCTGAACTCTTTCCCTACACGACGCTCTTCCGATCTNNNAATCATCACTGCACTTTGAGG
AGCACACGTCTGAACTCTTTCCCTACACGACGCTCTTCCGATCTNNNAACCCCATTTCCTACACACATGC
AGCACACGTCTGAACTCTTTCCCTACACGACGCTCTTCCGATCTNNNCCCATTTGCTTCCTCATTTGCTCCA
GAGCACACGTCTGAACTCTTTCCCTACACGACGCTCTTCCGATCTNNNAAGAGGGCAAACCAAACAGGA
GCACACGTCTGAACTCTTTCCCTACACGACGCTCTTCCGATCTNNNTACCATTTGGGTTTGTTGGGTGT
AGAGCACACGTCTGAACTCTTTCCCTACACGACGCTCTTCCGATCTNNNGCCTGAATAGCCCATTTTTCT
AGCACACGTCTGAACTCTTTCCCTACACGACGCTCTTCCGATCTNNNTGGGACTTAGCTTCCACAAACA
AGAGCACACGTCTGAACTCTTTCCCTACACGACGCTCTTCCGATCTNNNCCAGAATTCAGCCTTGGAGT
AGCACACGTCTGAACTCTTTCCCTACACGACGCTCTTCCGATCTNNNGAAAGGCAGTTGGAACTTGTGT
AGCACACGTCTGAACTCTTTCCCTACACGACGCTCTTCCGATCTNNNAAGCCCACCAATTGATGAGTGA
GAGCACACGTCTGAACTCTTTCCCTACACGACGCTCTTCCGATCTNNNTCCTGGCTTCTACATGGTCAAG
GAGCACACGTCTGAACTCTTTCCCTACACGACGCTCTTCCGATCTNNNCACTGGCTTGCACTCACTATGA
GAGCACACGTCTGAACTCTTTCCCTACACGACGCTCTTCCGATCTNNNATAACCACATCACACGCCATAC
GAGCACACGTCTGAACTCTTTCCCTACACGACGCTCTTCCGATCTNNNGCCGGAGATCAGTAAGTTGG
GAGCACACGTCTGAACTCTTTCCCTACACGACGCTCTTCCGATCTNNNTTCTGATTGATGGCCTGCCTTA
GAGCACACGTCTGAACTCTTTCCCTACACGACGCTCTTCCGATCTNNNGAGACAAGTGAAAGCCGAAG
GAGCACACGTCTGAACTCTTTCCCTACACGACGCTCTTCCGATCTNNNTTTCCCTCTACAAATCAGCGTG
AGCACACGTCTGAACTCTTTCCCTACACGACGCTCTTCCGATCTNNNCCACAGAACAGAGCGTCAAAAA
AGCACACGTCTGAACTCTTTCCCTACACGACGCTCTTCCGATCTNNNCTGGATTTAGCCTCAGACCAAC
GAGCACACGTCTGAACTCTTTCCCTACACGACGCTCTTCCGATCTNNNAGATGTCTCTGTCTTCTGCCAG
GAGCACACGTCTGAACTCTTTCCCTACACGACGCTCTTCCGATCTNNNGGGTGTGTTTTGATGGACCTG
AGCACACGTCTGAACTCTTTCCCTACACGACGCTCTTCCGATCTNNNCTGTACAATGGGGAAAGTGAG
AGCACACGTCTGAACTCTTTCCCTACACGACGCTCTTCCGATCTNNNGCTTCAACCTCCTTCTAGCTGTC
AGCACACGTCTGAACTCTTTCCCTACACGACGCTCTTCCGATCTNNNTGTTGGAGACAACCACAAGTCA
AGCACACGTCTGAACTCTTTCCCTACACGACGCTCTTCCGATCTNNNCTACCTGAGTTGTCACCGGAAA
AGAGCACACGTCTGAACTCTTTCCCTACACGACGCTCTTCCGATCTNNNTGCATGTTAGCACCCAAAAG
GCACACGTCTGAACTCTTTCCCTACACGACGCTCTTCCGATCTNNNTGGGAACACATAAAAGTCTGTCA
GAGCACACGTCTGAACTCTTTCCCTACACGACGCTCTTCCGATCTNNNTTTGGTAGGTTGGCATGTTTC
AGCACACGTCTGAACTCTTTCCCTACACGACGCTCTTCCGATCTNNNTGTGTTTTGGCCAAGAGGTCAA

GAGCACACGTCTGAACTCTTTCCCTACACGACGCTCTTCCGATCTNNNATCTCCAACCCCATTCAGATT
AGCACACGTCTGAACTCTTTCCCTACACGACGCTCTTCCGATCTNNNACATTCAAACATGCTCCCTCTCT
AGCACACGTCTGAACTCTTTCCCTACACGACGCTCTTCCGATCTNNNAAACTGCTATCTCACCTCCCACT
GAGCACACGTCTGAACTCTTTCCCTACACGACGCTCTTCCGATCTNNNCATGACCCATTCTCTGTTCCCT
GCACACGTCTGAACTCTTTCCCTACACGACGCTCTTCCGATCTNNNACAAATGGTACACAGGCCAAAC
AGAGCACACGTCTGAACTCTTTCCCTACACGACGCTCTTCCGATCTNNNATGAAGGACCCTTGTTCATG
GAGCACACGTCTGAACTCTTTCCCTACACGACGCTCTTCCGATCTNNNCCATTCTAAGCTCACATTCGCC
AGCACACGTCTGAACTCTTTCCCTACACGACGCTCTTCCGATCTNNNTTGGTAAACACTACCAACCAG
AGCACACGTCTGAACTCTTTCCCTACACGACGCTCTTCCGATCTNNNTAACAGGTTGAATGCAGAGGC
GAGCACACGTCTGAACTCTTTCCCTACACGACGCTCTTCCGATCTNNNCTCCCAGGGGAACTTATTCA
AGCACACGTCTGAACTCTTTCCCTACACGACGCTCTTCCGATCTNNNGCCCATCAATCAACAAGAGTG
AGCACACGTCTGAACTCTTTCCCTACACGACGCTCTTCCGATCTNNNCTGGAGAAGTGGGTACCTTTGT
GAGCACACGTCTGAACTCTTTCCCTACACGACGCTCTTCCGATCTNNNTCACTCCTCCCTTTTCTTCGTCT
GCACACGTCTGAACTCTTTCCCTACACGACGCTCTTCCGATCTNNNACAGTTACATTCCACCTTGCCTG
AGCACACGTCTGAACTCTTTCCCTACACGACGCTCTTCCGATCTNNNCTGTGGCTGGAGTTGAAATAG
AGCACACGTCTGAACTCTTTCCCTACACGACGCTCTTCCGATCTNNNTACCCTCAACCTACAGAAGCAC
AGCACACGTCTGAACTCTTTCCCTACACGACGCTCTTCCGATCTNNNTACACACATAACACCCAGACAC
AGCACACGTCTGAACTCTTTCCCTACACGACGCTCTTCCGATCTNNNTGTAAGGAAGATCCAGGGCTT
GAGCACACGTCTGAACTCTTTCCCTACACGACGCTCTTCCGATCTNNNTATGGGCTTTCAGGGGATAG
AGAGCACACGTCTGAACTCTTTCCCTACACGACGCTCTTCCGATCTNNNGTCATAAGCCTCTGTCCACA
AGCACACGTCTGAACTCTTTCCCTACACGACGCTCTTCCGATCTNNNGTGGTTTATCCATGGAAGGGC
AGAGCACACGTCTGAACTCTTTCCCTACACGACGCTCTTCCGATCTNNNTTATCTGTTGGATGCTCCCT
AGCACACGTCTGAACTCTTTCCCTACACGACGCTCTTCCGATCTNNNAGGATTTAGGGATGGAGGTT
GAGCACACGTCTGAACTCTTTCCCTACACGACGCTCTTCCGATCTNNNAAGCCCAGAGAGTATTGTTC
GAGCACACGTCTGAACTCTTTCCCTACACGACGCTCTTCCGATCTNNNATGCTTCCATGTTATAGCCCA
AGCACACGTCTGAACTCTTTCCCTACACGACGCTCTTCCGATCTNNNCATAGGAAAACGCTGGAAGCA
AGCACACGTCTGAACTCTTTCCCTACACGACGCTCTTCCGATCTNNNTCCTCAGTGGAAACTGTGTTAG
GAGCACACGTCTGAACTCTTTCCCTACACGACGCTCTTCCGATCTNNNTGGATGGTTCAGGGATAGTT
GAGCACACGTCTGAACTCTTTCCCTACACGACGCTCTTCCGATCTNNNCCCTGCAGAGACAGACTAGTG
GAGCACACGTCTGAACTCTTTCCCTACACGACGCTCTTCCGATCTNNNTGGTTTCAGGTGGTTTTGGC
AGAGCACACGTCTGAACTCTTTCCCTACACGACGCTCTTCCGATCTNNNTTCAAACCATGTCTGGACAA
GAGCACACGTCTGAACTCTTTCCCTACACGACGCTCTTCCGATCTNNNCTTGTGTGTATCTTGGTCGT
GCACACGTCTGAACTCTTTCCCTACACGACGCTCTTCCGATCTNNNGGAAGATTGAGAAGCTACTGG
GAGCACACGTCTGAACTCTTTCCCTACACGACGCTCTTCCGATCTNNNCCAAATAGTTCCTGCATGTGT
GAGCACACGTCTGAACTCTTTCCCTACACGACGCTCTTCCGATCTNNNTACCAGGAATACGGCAGCAA
GAGCACACGTCTGAACTCTTTCCCTACACGACGCTCTTCCGATCTNNNGTTGGGCATTTGTGTGTGTTT
AGCACACGTCTGAACTCTTTCCCTACACGACGCTCTTCCGATCTNNNTTCATGCTAGTCTGACCACAGG
AGCACACGTCTGAACTCTTTCCCTACACGACGCTCTTCCGATCTNNNAGCACTTCTGACTGATCCTCCTT
AGCACACGTCTGAACTCTTTCCCTACACGACGCTCTTCCGATCTNNNCCCAGCCTCCTTCTTCTTTTTCTC
GCACACGTCTGAACTCTTTCCCTACACGACGCTCTTCCGATCTNNNTCTGAGTTCCTAACCACTGAGCA
GCACACGTCTGAACTCTTTCCCTACACGACGCTCTTCCGATCTNNNTTTAGCTCTGGGGCTGGTGTTAT
GAGCACACGTCTGAACTCTTTCCCTACACGACGCTCTTCCGATCTNNNACTTCAGCCCTGAACTATGG
GCACACGTCTGAACTCTTTCCCTACACGACGCTCTTCCGATCTNNNGCAGGGTCTTGTCACATCTTCAA

AGCACACGTCTGAACTCTTTCCCTACACGACGCTCTTCCGATCTNNNTAATAATGCATGGTGCCCACAG
GCACACGTCTGAACTCTTTCCCTACACGACGCTCTTCCGATCTNNNTTTCTTCCTCCTGCTGCTTTGAACA
AGCACACGTCTGAACTCTTTCCCTACACGACGCTCTTCCGATCTNNNTGTTATCTTCTCCCCCTCATTGC
GAGCACACGTCTGAACTCTTTCCCTACACGACGCTCTTCCGATCTNNNGGCTAAAACCCAGCTGATTTC
AGCACACGTCTGAACTCTTTCCCTACACGACGCTCTTCCGATCTNNNGTTTCCTAATGCCTGGGAGATG
AGCACACGTCTGAACTCTTTCCCTACACGACGCTCTTCCGATCTNNNACCCAACTCAGACAACCACTGA
AGAGCACACGTCTGAACTCTTTCCCTACACGACGCTCTTCCGATCTNNNAGTTGGATTGGAAGGTGG
GAGCACACGTCTGAACTCTTTCCCTACACGACGCTCTTCCGATCTNNNCACATCCTACGCTCACATACCT
GAGCACACGTCTGAACTCTTTCCCTACACGACGCTCTTCCGATCTNNNTTGTCTTGTTTCCCAGAGAA
AGCACACGTCTGAACTCTTTCCCTACACGACGCTCTTCCGATCTNNNTGCCCACAAATATAGAGGGGA
GAGCACACGTCTGAACTCTTTCCCTACACGACGCTCTTCCGATCTNNNCTGAGCAGCTGGAAAGTACA
AGCACACGTCTGAACTCTTTCCCTACACGACGCTCTTCCGATCTNNNAGAACAGACATGTCCATAGCC
AGCACACGTCTGAACTCTTTCCCTACACGACGCTCTTCCGATCTNNNGCCTATATCTCCAGGATGGCTC
GAGCACACGTCTGAACTCTTTCCCTACACGACGCTCTTCCGATCTNNNAAGAATTGGCTCCTTCACCTTC
AGCACACGTCTGAACTCTTTCCCTACACGACGCTCTTCCGATCTNNNTATCAGCACAGGTCCTGGAACA
GAGCACACGTCTGAACTCTTTCCCTACACGACGCTCTTCCGATCTNNNAAGGAAAAACTGTGTGTGTG
AGCACACGTCTGAACTCTTTCCCTACACGACGCTCTTCCGATCTNNNTTGTGCCCATTGTTTCTTGTCCT
GAGCACACGTCTGAACTCTTTCCCTACACGACGCTCTTCCGATCTNNNCTATTCTCTGGGTTTGTGCTG
GCACACGTCTGAACTCTTTCCCTACACGACGCTCTTCCGATCTNNNTGAGAGAGGCGAAGAATGAAG
GCACACGTCTGAACTCTTTCCCTACACGACGCTCTTCCGATCTNNNAGCTTATGGAGGATCATTTTGTT
GAGCACACGTCTGAACTCTTTCCCTACACGACGCTCTTCCGATCTNNNATGGAACAGAAAATGCACAA
AGCACACGTCTGAACTCTTTCCCTACACGACGCTCTTCCGATCTNNNCGAAGAAACCTTTTTCCCTGCAC
GAGCACACGTCTGAACTCTTTCCCTACACGACGCTCTTCCGATCTNNNAAAACTCCACAAGCCCCAGAA
GAGCACACGTCTGAACTCTTTCCCTACACGACGCTCTTCCGATCTNNNGACACACCAGTACACACACAC
GAGCACACGTCTGAACTCTTTCCCTACACGACGCTCTTCCGATCTNNNACAAGACTGTGTGACTTCTTG
GAGCACACGTCTGAACTCTTTCCCTACACGACGCTCTTCCGATCTNNNCTGTCTCTCTGGCGTCATTTGT
AGCACACGTCTGAACTCTTTCCCTACACGACGCTCTTCCGATCTNNNGTACACTGTGATCCCACAGTCA
AGCACACGTCTGAACTCTTTCCCTACACGACGCTCTTCCGATCTNNNTGCGGATACAAAACCTTCTAGC
GAGCACACGTCTGAACTCTTTCCCTACACGACGCTCTTCCGATCTNNNACAGATGACGGACTGTAGGA
AGCACACGTCTGAACTCTTTCCCTACACGACGCTCTTCCGATCTNNNTTTCAGATGCAGGCACATGTTT
GAGCACACGTCTGAACTCTTTCCCTACACGACGCTCTTCCGATCTNNNGTCTGTAAGGGGTCACCAAT
AGCACACGTCTGAACTCTTTCCCTACACGACGCTCTTCCGATCTNNNCGCAGTGGTTACACACATCTA
GAGCACACGTCTGAACTCTTTCCCTACACGACGCTCTTCCGATCTNNNCAGAACATAGCATCAGCACG
AAGAGCACACGTCTGAACTCTTTCCCTACACGACGCTCTTCCGATCTNNNCCAGCCCCTGAAAAGATTT
GAGCACACGTCTGAACTCTTTCCCTACACGACGCTCTTCCGATCTNNNAACCATTCATGGCAACAGGT
GAGCACACGTCTGAACTCTTTCCCTACACGACGCTCTTCCGATCTNNNTCCAGCAAGGCTGCTAAAAGT
AGCACACGTCTGAACTCTTTCCCTACACGACGCTCTTCCGATCTNNNGCACTTGGCAGGATTTGTATG
AGCACACGTCTGAACTCTTTCCCTACACGACGCTCTTCCGATCTNNNACCTGCATGGCACGAATAAGA
AGAGCACACGTCTGAACTCTTTCCCTACACGACGCTCTTCCGATCTNNNGCAAATTGCCCAAGGTAACA
GCACACGTCTGAACTCTTTCCCTACACGACGCTCTTCCGATCTNNNATAGGTGGGTTTTGAACATGG
AGCACACGTCTGAACTCTTTCCCTACACGACGCTCTTCCGATCTNNNATGCAACCAACACCTCTTGTCT
GAGCACACGTCTGAACTCTTTCCCTACACGACGCTCTTCCGATCTNNNTATTGCCCTGAGCAGAGATGA
AGCACACGTCTGAACTCTTTCCCTACACGACGCTCTTCCGATCTNNNTTATGCAACCTAAGAAATGCC

AGCACACGTCTGAACTCTTTCCCTACACGACGCTCTTCCGATCTNNNGGCCTAAACTGCCATCACCATT
GCACACGTCTGAACTCTTTCCCTACACGACGCTCTTCCGATCTNNNAGATCCTTCCTCATCTGCCACAA
GCACACGTCTGAACTCTTTCCCTACACGACGCTCTTCCGATCTNNNGAGGAGACAAGGTGAGAGGAA
GAGCACACGTCTGAACTCTTTCCCTACACGACGCTCTTCCGATCTNNNGGATTGATGTTCACTGTCAG
GAGCACACGTCTGAACTCTTTCCCTACACGACGCTCTTCCGATCTNNNTCAAGTGTGTCCCAAAGAAC
AGCACACGTCTGAACTCTTTCCCTACACGACGCTCTTCCGATCTNNNTACACAGAGACAGAAGGTCAG
AGCACACGTCTGAACTCTTTCCCTACACGACGCTCTTCCGATCTNNNCATGTGGCTCACCGTTTTCTTA
AGCACACGTCTGAACTCTTTCCCTACACGACGCTCTTCCGATCTNNNGTTCTTTTGGTTGTTGGTGGTG
AGCACACGTCTGAACTCTTTCCCTACACGACGCTCTTCCGATCTNNNTCCAGGCCAAAAAGAAGTTGA
AGCACACGTCTGAACTCTTTCCCTACACGACGCTCTTCCGATCTNNNAATCCTTTCAGGTGGCCCTTTTT
GCACACGTCTGAACTCTTTCCCTACACGACGCTCTTCCGATCTNNNGGCTAGGCACAGATGTGAGAAA
AGCACACGTCTGAACTCTTTCCCTACACGACGCTCTTCCGATCTNNNCTCATGAGAATGCTCTTGCATG
AGCACACGTCTGAACTCTTTCCCTACACGACGCTCTTCCGATCTNNNTCAATTACCAGATGCACTCCTG
AGCACACGTCTGAACTCTTTCCCTACACGACGCTCTTCCGATCTNNNGAATGCACCAGAACAGGTCTC
AGCACACGTCTGAACTCTTTCCCTACACGACGCTCTTCCGATCTNNNGCCACTCAAGCAGAATCTCTA
AGCACACGTCTGAACTCTTTCCCTACACGACGCTCTTCCGATCTNNNAGATTTCTGCCTTGGGGGAA
AGCACACGTCTGAACTCTTTCCCTACACGACGCTCTTCCGATCTNNNATTATACCTTTCCCCGTCCCAGT
GAGCACACGTCTGAACTCTTTCCCTACACGACGCTCTTCCGATCTNNNGAAGCACTTCTCCTTGCTTTC
GAGCACACGTCTGAACTCTTTCCCTACACGACGCTCTTCCGATCTNNNAGGGAATCTCCTCATGGAAC
AGCACACGTCTGAACTCTTTCCCTACACGACGCTCTTCCGATCTNNNCCACCTCTGAAATGCCAGTGAT
AGCACACGTCTGAACTCTTTCCCTACACGACGCTCTTCCGATCTNNNCTCTTTCCTTGTCGGAGCTTGTA
GCACACGTCTGAACTCTTTCCCTACACGACGCTCTTCCGATCTNNNTCAGTCCAGGGATTGAGAGAAC
AGCACACGTCTGAACTCTTTCCCTACACGACGCTCTTCCGATCTNNNACACACGCGGTTTAGCAATAT
AGCACACGTCTGAACTCTTTCCCTACACGACGCTCTTCCGATCTNNNCTTCACTATGTCCAATGCAGG
GAGCACACGTCTGAACTCTTTCCCTACACGACGCTCTTCCGATCTNNNCAGGTTTGAGCTGTCACTTCC
AGCACACGTCTGAACTCTTTCCCTACACGACGCTCTTCCGATCTNNNAACATCTTCCAGATTAGTGCG
GCACACGTCTGAACTCTTTCCCTACACGACGCTCTTCCGATCTNNNGGCCCATGAATGCGTATTTTCA
AGCACACGTCTGAACTCTTTCCCTACACGACGCTCTTCCGATCTNNNGCTGGATTTATAGGACCATG
AGCACACGTCTGAACTCTTTCCCTACACGACGCTCTTCCGATCTNNNTAAGTGGAAAAACCCCAGCT
AGCACACGTCTGAACTCTTTCCCTACACGACGCTCTTCCGATCTNNNGAACCCTCCAGGAGACCATAA
GAGCACACGTCTGAACTCTTTCCCTACACGACGCTCTTCCGATCTNNNCAGACTAGAGTGGAGGAAG
AGCACACGTCTGAACTCTTTCCCTACACGACGCTCTTCCGATCTNNNGTTCTGTAATTCGTGCGTGTG
AGCACACGTCTGAACTCTTTCCCTACACGACGCTCTTCCGATCTNNNCTGCCTCACACTTACTGTGGAA
AGAGCACACGTCTGAACTCTTTCCCTACACGACGCTCTTCCGATCTNNNTCACCCGTCTTCTTCTAAGG
AGCACACGTCTGAACTCTTTCCCTACACGACGCTCTTCCGATCTNNNGGTCACAATCACCATGATTCG
AGCACACGTCTGAACTCTTTCCCTACACGACGCTCTTCCGATCTNNNTGGGGCTGGACTTCTACACTT
GCACACGTCTGAACTCTTTCCCTACACGACGCTCTTCCGATCTNNNCCCATTGTCTGAATGGGATAAG
AGCACACGTCTGAACTCTTTCCCTACACGACGCTCTTCCGATCTNNNAGTTCAGTTGGCACTGTTGTTG
GAGCACACGTCTGAACTCTTTCCCTACACGACGCTCTTCCGATCTNNNGCTGCCCTAGCAAAGGTCTA
GCACACGTCTGAACTCTTTCCCTACACGACGCTCTTCCGATCTNNNCCCTCTACCCTCCTTCTGGAAAAC
AGCACACGTCTGAACTCTTTCCCTACACGACGCTCTTCCGATCTNNNCTCCATCTACCTTTCTGGTCCCT
GAGCACACGTCTGAACTCTTTCCCTACACGACGCTCTTCCGATCTNNNCCTGGGACAAAGCAGAACCT
AGCACACGTCTGAACTCTTTCCCTACACGACGCTCTTCCGATCTNNNCATTTTGGTCATGCACATCCCT

AGCACACGTCTGAACTCTTTCCCTACACGACGCTCTTCCGATCTNNNTCTCAAGCCTATCCTCTCCATC
GAGCACACGTCTGAACTCTTTCCCTACACGACGCTCTTCCGATCTNNNGCCAGGATAGGATGCACATT
GCACACGTCTGAACTCTTTCCCTACACGACGCTCTTCCGATCTNNNAGTGGGTGTTGCTTTTCACTCA
AGAGCACACGTCTGAACTCTTTCCCTACACGACGCTCTTCCGATCTNNNTCATGCACTGCAAGGCATA
AGAGCACACGTCTGAACTCTTTCCCTACACGACGCTCTTCCGATCTNNNGAATTGGGACTTTGGGTG
AGCACACGTCTGAACTCTTTCCCTACACGACGCTCTTCCGATCTNNNAGCTGAACCAGGAAACAACTT
GAGCACACGTCTGAACTCTTTCCCTACACGACGCTCTTCCGATCTNNNTTGTGACAGTTTGGAGTAGA
AGCACACGTCTGAACTCTTTCCCTACACGACGCTCTTCCGATCTNNNACTATACACACACTCCCCTTCCC
CACACGTCTGAACTCTTTCCCTACACGACGCTCTTCCGATCTNNNGTACCTCAGAGGACACATCTCCA
GAGCACACGTCTGAACTCTTTCCCTACACGACGCTCTTCCGATCTNNNTGAGGCAGAATTGCCAGAAC
AGCACACGTCTGAACTCTTTCCCTACACGACGCTCTTCCGATCTNNNGAACAGTTGTAGAGGGCTTGT
AGCACACGTCTGAACTCTTTCCCTACACGACGCTCTTCCGATCTNNNCCTCTGCAAAAATCACAGCCTT
AGCACACGTCTGAACTCTTTCCCTACACGACGCTCTTCCGATCTNNNGAAACTCAGGCTGGGTAACA
GAGCACACGTCTGAACTCTTTCCCTACACGACGCTCTTCCGATCTNNNAGACCTTCCAAAACAGGATT
GCACACGTCTGAACTCTTTCCCTACACGACGCTCTTCCGATCTNNNTGCCCTTGCCATTTATGTTTCCT
AGCACACGTCTGAACTCTTTCCCTACACGACGCTCTTCCGATCTNNNACATTTTCTGACCCCCACAACTC
GAGCACACGTCTGAACTCTTTCCCTACACGACGCTCTTCCGATCTNNNCAAAACACTCTGCAACACCA
AGCACACGTCTGAACTCTTTCCCTACACGACGCTCTTCCGATCTNNNGGCTGGTGTCACCTCTTACATT
AGCACACGTCTGAACTCTTTCCCTACACGACGCTCTTCCGATCTNNNCAATTCAGTGCTCCTTTCCCTC
AGCACACGTCTGAACTCTTTCCCTACACGACGCTCTTCCGATCTNNNAGGCAGGTAACAGTAGTCCTC
AGCACACGTCTGAACTCTTTCCCTACACGACGCTCTTCCGATCTNNNGTATAGCTGGAGTGGAGGCT
AGCACACGTCTGAACTCTTTCCCTACACGACGCTCTTCCGATCTNNNTTTGTAGGGAAAACCAGCATC
GAGCACACGTCTGAACTCTTTCCCTACACGACGCTCTTCCGATCTNNNTCATCATTGGCTTTGTTTTG
GCACACGTCTGAACTCTTTCCCTACACGACGCTCTTCCGATCTNNNTGCATGCTCATTCACTTTAGGCA
GCACACGTCTGAACTCTTTCCCTACACGACGCTCTTCCGATCTNNNGCCTTTGTGGTCTGTTTGCTTTT
GCACACGTCTGAACTCTTTCCCTACACGACGCTCTTCCGATCTNNNGGGGTCTCACTCCGAATCATTAC
GAGCACACGTCTGAACTCTTTCCCTACACGACGCTCTTCCGATCTNNNCAGTTCCTGGCACAAACTAAT
CACACGTCTGAACTCTTTCCCTACACGACGCTCTTCCGATCTNNNCTTTGCAAGAACCTCTGTCTGCC
GAGCACACGTCTGAACTCTTTCCCTACACGACGCTCTTCCGATCTNNNTAAACTTTGCCAGAACTAGC
GAGCACACGTCTGAACTCTTTCCCTACACGACGCTCTTCCGATCTNNNTGAGCTCTGCAAAGTGAGTA
AGCACACGTCTGAACTCTTTCCCTACACGACGCTCTTCCGATCTNNNGGTAAAAGTAGGGCTCTTGG
AGCACACGTCTGAACTCTTTCCCTACACGACGCTCTTCCGATCTNNNTTAGCCGCCTTATTTTGGCAG
GCACACGTCTGAACTCTTTCCCTACACGACGCTCTTCCGATCTNNNAAAGAGGGTTAGGAGGTCTTG
AGCACACGTCTGAACTCTTTCCCTACACGACGCTCTTCCGATCTNNNACAGGCAATCATTTAGCCTCCT
AGCACACGTCTGAACTCTTTCCCTACACGACGCTCTTCCGATCTNNNTGTTGTTATGCATATGTGGCT
AGCACACGTCTGAACTCTTTCCCTACACGACGCTCTTCCGATCTNNNCCCTCTGTTCTCTCATTCCCCAT
AGCACACGTCTGAACTCTTTCCCTACACGACGCTCTTCCGATCTNNNTCTCCTCTACTGTCCCAGAAA
GCACACGTCTGAACTCTTTCCCTACACGACGCTCTTCCGATCTNNNGTGCTTCATGTATCCTCTGGCTT
GAGCACACGTCTGAACTCTTTCCCTACACGACGCTCTTCCGATCTNNNACTAAGCTGCATTGTAAGCG
AGCACACGTCTGAACTCTTTCCCTACACGACGCTCTTCCGATCTNNNTGTCCTCGATATAGCTGGGTG
AGCACACGTCTGAACTCTTTCCCTACACGACGCTCTTCCGATCTNNNGCGACTTTTCTCTGCTTTCCATT
GAGCACACGTCTGAACTCTTTCCCTACACGACGCTCTTCCGATCTNNNTGTCTGGCCTTGTTCTCATCT
AGCACACGTCTGAACTCTTTCCCTACACGACGCTCTTCCGATCTNNNTTGCTAACTTACCTGCAATGCC

AGCACACGTCTGAACTCTTTCCCTACACGACGCTCTTCCGATCTNNNAGTGTTGGCTTATTGCTTTGC
AGCACACGTCTGAACTCTTTCCCTACACGACGCTCTTCCGATCTNNNACAAGAAGGCAGAAAGAGGA
AGCACACGTCTGAACTCTTTCCCTACACGACGCTCTTCCGATCTNNNATGCCACCTTTGAAATCACTTG
AGAGCACACGTCTGAACTCTTTCCCTACACGACGCTCTTCCGATCTNNNAAATTTGGGGGTGTAGCT
AGCACACGTCTGAACTCTTTCCCTACACGACGCTCTTCCGATCTNNNATCAGTCACTTCTCTGGCCTCT
GCACACGTCTGAACTCTTTCCCTACACGACGCTCTTCCGATCTNNNGGCTCGAGCATCTGTATCATCTG
AGCACACGTCTGAACTCTTTCCCTACACGACGCTCTTCCGATCTNNNCAGCATTCCATCAGAGAAGCA
AGCACACGTCTGAACTCTTTCCCTACACGACGCTCTTCCGATCTNNNTCAGGCATGTGCTATGTGAAT
AGCACACGTCTGAACTCTTTCCCTACACGACGCTCTTCCGATCTNNNATAAACCAAACCCCAGTGACG
AGCACACGTCTGAACTCTTTCCCTACACGACGCTCTTCCGATCTNNNGTCTGCAGAGACATAAAGTG
AGCACACGTCTGAACTCTTTCCCTACACGACGCTCTTCCGATCTNNNAAAGGGAGATAGTTGCATCCA
AGCACACGTCTGAACTCTTTCCCTACACGACGCTCTTCCGATCTNNNTTCTGGGCAGGGACTGTATTGT
AGCACACGTCTGAACTCTTTCCCTACACGACGCTCTTCCGATCTNNNTAAAATCGCTCAAGACACAGC
AGCACACGTCTGAACTCTTTCCCTACACGACGCTCTTCCGATCTNNNCACTGGCTGACACACCCTATATT
AGCACACGTCTGAACTCTTTCCCTACACGACGCTCTTCCGATCTNNNTCAAAGTTGGTGTCAAGGTCA
AGCACACGTCTGAACTCTTTCCCTACACGACGCTCTTCCGATCTNNNTTTGAGGAGTTCCAGAGGTAG
AGCACACGTCTGAACTCTTTCCCTACACGACGCTCTTCCGATCTNNNTTAGCCAACCCCTCTTTATCCCA
AGCACACGTCTGAACTCTTTCCCTACACGACGCTCTTCCGATCTNNNCTGGAAGCATCTGCATCCTTCTT
GAGCACACGTCTGAACTCTTTCCCTACACGACGCTCTTCCGATCTNNNGCTCTGGCTAATGGAAGTGA
AGCACACGTCTGAACTCTTTCCCTACACGACGCTCTTCCGATCTNNNGTAAATTGCCCAGCAGAATG
GCACACGTCTGAACTCTTTCCCTACACGACGCTCTTCCGATCTNNNTGGGAAGTAACAGTAGTTGGT
AGCACACGTCTGAACTCTTTCCCTACACGACGCTCTTCCGATCTNNNTGAACCTTACTAGGAAACACA
AGCACACGTCTGAACTCTTTCCCTACACGACGCTCTTCCGATCTNNNCAGGGCAGATGCAAGGAATA
AGCACACGTCTGAACTCTTTCCCTACACGACGCTCTTCCGATCTNNNATATTTGGGACACAGAACCTG
AGCACACGTCTGAACTCTTTCCCTACACGACGCTCTTCCGATCTNNNGAGGCTCTCAACAGTGTAATG
AGCACACGTCTGAACTCTTTCCCTACACGACGCTCTTCCGATCTNNNTGGACACATAGGAAGTGTGT
AGCACACGTCTGAACTCTTTCCCTACACGACGCTCTTCCGATCTNNNGGCACTGTCTGTGTTTCTCTCT
GCACACGTCTGAACTCTTTCCCTACACGACGCTCTTCCGATCTNNNGCAGGAAGCACATCTACTAGCT
AGCACACGTCTGAACTCTTTCCCTACACGACGCTCTTCCGATCTNNNAAGGCAGTTATCAGCAGGCAA
AGCACACGTCTGAACTCTTTCCCTACACGACGCTCTTCCGATCTNNNTGTGGGTACTAGGAGTTGGTT
AGCACACGTCTGAACTCTTTCCCTACACGACGCTCTTCCGATCTNNNGCCTGAGTGATAGAGAGCATC
AGCACACGTCTGAACTCTTTCCCTACACGACGCTCTTCCGATCTNNNATATACTCTTCATGCCCCCACCA
AGCACACGTCTGAACTCTTTCCCTACACGACGCTCTTCCGATCTNNNATTGGGGAAAAATGCCCATCT
GCACACGTCTGAACTCTTTCCCTACACGACGCTCTTCCGATCTNNNTTCTTATCTCGGGTCACCACCTT
AGCACACGTCTGAACTCTTTCCCTACACGACGCTCTTCCGATCTNNNTTCCAAACACTTGAGGTCAGA
AGCACACGTCTGAACTCTTTCCCTACACGACGCTCTTCCGATCTNNNGACTTGAAATCCTGCCTGGAA
AGCACACGTCTGAACTCTTTCCCTACACGACGCTCTTCCGATCTNNNCCCCGACCCTACATAGTACAC
AGCACACGTCTGAACTCTTTCCCTACACGACGCTCTTCCGATCTNNNTCCTAGAATCAGTCCTCCCAAC
GAGCACACGTCTGAACTCTTTCCCTACACGACGCTCTTCCGATCTNNNGTACACGTATGAACAGTGCT
AGCACACGTCTGAACTCTTTCCCTACACGACGCTCTTCCGATCTNNNGGCTAGACACACTGAAGGAAA
AGAGCACACGTCTGAACTCTTTCCCTACACGACGCTCTTCCGATCTNNNTGACTATCGGCCATTGTAA
AGCACACGTCTGAACTCTTTCCCTACACGACGCTCTTCCGATCTNNNTTTCTAGAGTGTTTGCCTCCTG
AGAGCACACGTCTGAACTCTTTCCCTACACGACGCTCTTCCGATCTNNNTAGGAAGGGTTGAGTGAC

GAGCACACGTCTGAACTCTTTCCCTACACGACGCTCTTCCGATCTNNNCCCGTGTTATTCAAGGGTTA
AGCACACGTCTGAACTCTTTCCCTACACGACGCTCTTCCGATCTNNNTCGTACCTGTCACCTCCAAGATT
AGCACACGTCTGAACTCTTTCCCTACACGACGCTCTTCCGATCTNNNCTTGCAGATGGTTCACATGACC
AGCACACGTCTGAACTCTTTCCCTACACGACGCTCTTCCGATCTNNNTACTCATCCTGCAGACTTTGGG
AGCACACGTCTGAACTCTTTCCCTACACGACGCTCTTCCGATCTNNNCAAGAACTTCCAGGAATCAGTG
AGCACACGTCTGAACTCTTTCCCTACACGACGCTCTTCCGATCTNNNTATATTCCATACCCCTCCCCACC
AGAGCACACGTCTGAACTCTTTCCCTACACGACGCTCTTCCGATCTNNNCTTGAGTGACCTGTACCCAA
AGCACACGTCTGAACTCTTTCCCTACACGACGCTCTTCCGATCTNNNTGGGTTCACTGCCTTGTTTACT
AGCACACGTCTGAACTCTTTCCCTACACGACGCTCTTCCGATCTNNNTGAAGCTGCCCCTGTTTTGTATT
AGCACACGTCTGAACTCTTTCCCTACACGACGCTCTTCCGATCTNNNTCAAAAGGGAGGAAGGGAGA
AGCACACGTCTGAACTCTTTCCCTACACGACGCTCTTCCGATCTNNNCCTCCCCATTGTGGTTTAGGAT
GCACACGTCTGAACTCTTTCCCTACACGACGCTCTTCCGATCTNNNGTTTTGGAGATTTCTTGGCTGG
AGCACACGTCTGAACTCTTTCCCTACACGACGCTCTTCCGATCTNNNTACTCTGGAACATTCTGGCATC
AGCACACGTCTGAACTCTTTCCCTACACGACGCTCTTCCGATCTNNNGGAGAAGAGAGCTGTTGTGTG
AGCACACGTCTGAACTCTTTCCCTACACGACGCTCTTCCGATCTNNNGTTAGACAGCATTAGCTCCCAG
GAGCACACGTCTGAACTCTTTCCCTACACGACGCTCTTCCGATCTNNNATGAAACCAGAGAGCTTGTG
AGCACACGTCTGAACTCTTTCCCTACACGACGCTCTTCCGATCTNNNCACTGCTTCCCCTTGCCTTATAG
AGCACACGTCTGAACTCTTTCCCTACACGACGCTCTTCCGATCTNNNTACTCAGGGCAGCTTACTTAG
AGCACACGTCTGAACTCTTTCCCTACACGACGCTCTTCCGATCTNNNCAAGTACATGGGTCTCACGAA
AGCACACGTCTGAACTCTTTCCCTACACGACGCTCTTCCGATCTNNNTAAATGAAGCGAGACAACACC
AGCACACGTCTGAACTCTTTCCCTACACGACGCTCTTCCGATCTNNNGGAAACTGCTAGTGCGTGATA
GCACACGTCTGAACTCTTTCCCTACACGACGCTCTTCCGATCTNNNAACTCCCTTACAGGGGAGAAAA
AGCACACGTCTGAACTCTTTCCCTACACGACGCTCTTCCGATCTNNNCCATCGTGAGGGTTTTGAGGT
AGCACACGTCTGAACTCTTTCCCTACACGACGCTCTTCCGATCTNNNAGGAGACACGAGAAAGCTTAG
AGCACACGTCTGAACTCTTTCCCTACACGACGCTCTTCCGATCTNNNGGCTGTTGAGGCTGAAACATC
GAGCACACGTCTGAACTCTTTCCCTACACGACGCTCTTCCGATCTNNNAGAGCAGAGATGGCAAATTC
GCACACGTCTGAACTCTTTCCCTACACGACGCTCTTCCGATCTNNNAGCTTTTCATTAGCTCAGCCTCG
AGCACACGTCTGAACTCTTTCCCTACACGACGCTCTTCCGATCTNNNTCTATGGTCGTGTGAGCCAAA
GCACACGTCTGAACTCTTTCCCTACACGACGCTCTTCCGATCTNNNAGCCCTTACTACCCACATACTCCC
AGCACACGTCTGAACTCTTTCCCTACACGACGCTCTTCCGATCTNNNGGTCTGTTGTTGACTGGTACTC
AGCACACGTCTGAACTCTTTCCCTACACGACGCTCTTCCGATCTNNNATCCAGGATGTGATGTATGGG
AGAGCACACGTCTGAACTCTTTCCCTACACGACGCTCTTCCGATCTNNNAGCTAGCACCATGGGAAAA
AGCACACGTCTGAACTCTTTCCCTACACGACGCTCTTCCGATCTNNNGCCATTGTGCAACATACACAA
AGCACACGTCTGAACTCTTTCCCTACACGACGCTCTTCCGATCTNNNAAAACACCATGAGTGCTTCTGC
AGCACACGTCTGAACTCTTTCCCTACACGACGCTCTTCCGATCTNNNGTGGCATGTCTGTGTGTGTGA
AGCACACGTCTGAACTCTTTCCCTACACGACGCTCTTCCGATCTNNNCAAGCACATATCCCACAATGCA
GAGCACACGTCTGAACTCTTTCCCTACACGACGCTCTTCCGATCTNNNCCACAGTGAATTTTGGTGGT
AGCACACGTCTGAACTCTTTCCCTACACGACGCTCTTCCGATCTNNNGTTGGGATATGTGGGAAAGT
GAGCACACGTCTGAACTCTTTCCCTACACGACGCTCTTCCGATCTNNNGTGTCTGGTACAAAGGGTAA
AGCACACGTCTGAACTCTTTCCCTACACGACGCTCTTCCGATCTNNNCCTGATACCCTCCTCTTATGGCA
AGCACACGTCTGAACTCTTTCCCTACACGACGCTCTTCCGATCTNNNGCATAGCAATGGGGGTGGAA
AGCACACGTCTGAACTCTTTCCCTACACGACGCTCTTCCGATCTNNNCCCTTTCTCTAGCCTCTTCACCT
AGCACACGTCTGAACTCTTTCCCTACACGACGCTCTTCCGATCTNNNTGTTTCCTCTGCCTTCATTGCAT

AGCACACGTCTGAACTCTTTCCCTACACGACGCTCTTCCGATCTNNNACATTCTTCCCTGACTGTAAAGC
AGCACACGTCTGAACTCTTTCCCTACACGACGCTCTTCCGATCTNNNGTTGAGTGGCTCTGATCCCAAA
AGCACACGTCTGAACTCTTTCCCTACACGACGCTCTTCCGATCTNNNTGAGAGACCTAGCTACAAACA
AGCACACGTCTGAACTCTTTCCCTACACGACGCTCTTCCGATCTNNNCAGAGTGTGATGGTTCGTCTG
AGCACACGTCTGAACTCTTTCCCTACACGACGCTCTTCCGATCTNNNCTGTTCTCTCTCTCTTCCCCCTTCA
GAGCACACGTCTGAACTCTTTCCCTACACGACGCTCTTCCGATCTNNNGAAACTCAGCCTCCTCTGTAA
GCACACGTCTGAACTCTTTCCCTACACGACGCTCTTCCGATCTNNNCGCTTTCCCTCTAGCTTTCTCTCCA
AGCACACGTCTGAACTCTTTCCCTACACGACGCTCTTCCGATCTNNNCTTGAGTGCAGTGTCCCTATCA
AGCACACGTCTGAACTCTTTCCCTACACGACGCTCTTCCGATCTNNNCCGTACACTTACAAGTGAGGCAA
AGCACACGTCTGAACTCTTTCCCTACACGACGCTCTTCCGATCTNNNCTAGGCTTTTCCAATGGGACCTC
GCACACGTCTGAACTCTTTCCCTACACGACGCTCTTCCGATCTNNNCATAGCAGAGGGTAAACCACCA
AGCACACGTCTGAACTCTTTCCCTACACGACGCTCTTCCGATCTNNNACTCTGCTGTATGTATCTGTGGT
AGCACACGTCTGAACTCTTTCCCTACACGACGCTCTTCCGATCTNNNGGACTGTTAGTATTGCTTGGG
AGCACACGTCTGAACTCTTTCCCTACACGACGCTCTTCCGATCTNNNTGTTGAACGTTCCTCCATACTGT
GAGCACACGTCTGAACTCTTTCCCTACACGACGCTCTTCCGATCTNNNGGACCTACTCGGTACTTGGAT
GAGCACACGTCTGAACTCTTTCCCTACACGACGCTCTTCCGATCTNNNTGTGCCCGGTATGGTTTTCAAT
AGCACACGTCTGAACTCTTTCCCTACACGACGCTCTTCCGATCTNNNTACAGGGGCTATGGCTATGAA
AGCACACGTCTGAACTCTTTCCCTACACGACGCTCTTCCGATCTNNNACACAGCTTGGGCTAAAGTGAA
GCACACGTCTGAACTCTTTCCCTACACGACGCTCTTCCGATCTNNNACACACAGCTTTTCCCAAAGTTAC
GCACACGTCTGAACTCTTTCCCTACACGACGCTCTTCCGATCTNNNTCCCTTTACAAGTGGTGGAGCTA
GCACACGTCTGAACTCTTTCCCTACACGACGCTCTTCCGATCTNNNACAGTCAACAGAGCACGTAACAT
AGAGCACACGTCTGAACTCTTTCCCTACACGACGCTCTTCCGATCTNNNGTCTCCAGACCTGCCCAGATA
AGCACACGTCTGAACTCTTTCCCTACACGACGCTCTTCCGATCTNNNGAGATCTCTGCTTGGAATGACC
GCACACGTCTGAACTCTTTCCCTACACGACGCTCTTCCGATCTNNNACTTAGTGGTGGTAATGGTGGG
AGCACACGTCTGAACTCTTTCCCTACACGACGCTCTTCCGATCTNNNCTTCATCTGGGGAATAAGGTGC
AGCACACGTCTGAACTCTTTCCCTACACGACGCTCTTCCGATCTNNNGGTTTCTCTTGTCCCCCAACATA
GAGCACACGTCTGAACTCTTTCCCTACACGACGCTCTTCCGATCTNNNCATATCCGTGCCTGTGTATGT
AGCACACGTCTGAACTCTTTCCCTACACGACGCTCTTCCGATCTNNNTATACCTGTCAGGGGTGGTGT
AGCACACGTCTGAACTCTTTCCCTACACGACGCTCTTCCGATCTNNNACCCACATCATTGTGTCTTCCAC
AGAGCACACGTCTGAACTCTTTCCCTACACGACGCTCTTCCGATCTNNNCTAAATGGGTTGTGGGGAG
GAGCACACGTCTGAACTCTTTCCCTACACGACGCTCTTCCGATCTNNNGAATCCAGCAACCAACCAGTA
AGCACACGTCTGAACTCTTTCCCTACACGACGCTCTTCCGATCTNNNAATGAGAAGGTTGCTTCTGCCT
AGCACACGTCTGAACTCTTTCCCTACACGACGCTCTTCCGATCTNNNTATCCAGGATCCCCAAGGACAA
AGAGCACACGTCTGAACTCTTTCCCTACACGACGCTCTTCCGATCTNNNTGATCCACTGATACAGAACCT
AGCACACGTCTGAACTCTTTCCCTACACGACGCTCTTCCGATCTNNNGCCAAGTTCTCATGTGCTCTGA
GAGCACACGTCTGAACTCTTTCCCTACACGACGCTCTTCCGATCTNNNCCAAGAGGCTCTAAGACGAA
GAGCACACGTCTGAACTCTTTCCCTACACGACGCTCTTCCGATCTNNNTATACACACTGCACTCCAAGCA
AGCACACGTCTGAACTCTTTCCCTACACGACGCTCTTCCGATCTNNNTAAAGGCAACTATTGCCCACTCC
GAGCACACGTCTGAACTCTTTCCCTACACGACGCTCTTCCGATCTNNNACAATCAAGCAAATCTGGAG
GAGCACACGTCTGAACTCTTTCCCTACACGACGCTCTTCCGATCTNNNCACCTAGCAAATAACAGGGGT
AGAGCACACGTCTGAACTCTTTCCCTACACGACGCTCTTCCGATCTNNNACACAGATCCAGGCACAAAG
GAGCACACGTCTGAACTCTTTCCCTACACGACGCTCTTCCGATCTNNNAAATTTCTTGAGGCCAGAGTT
AGCACACGTCTGAACTCTTTCCCTACACGACGCTCTTCCGATCTNNNATCTGGGCTCAAGAAATCCACA

AGAGCACACGTCTGAACTCTTTCCCTACACGACGCTCTTCCGATCTNNNGTTGGGCAGAGGGATTAGT
GAGCACACGTCTGAACTCTTTCCCTACACGACGCTCTTCCGATCTNNNGGTAGATTCTGCAGTTTGGG
AGAGCACACGTCTGAACTCTTTCCCTACACGACGCTCTTCCGATCTNNNCGAACTCATACTGCAGATCCT
AGCACACGTCTGAACTCTTTCCCTACACGACGCTCTTCCGATCTNNNTGTAGCAACTCTGGATGCCAAT
AGCACACGTCTGAACTCTTTCCCTACACGACGCTCTTCCGATCTNNNTGTTCGCAACAGAAAAACACGA
AGCACACGTCTGAACTCTTTCCCTACACGACGCTCTTCCGATCTNNNGTTTGATTGTGTCTTGCTCTGCC
AGCACACGTCTGAACTCTTTCCCTACACGACGCTCTTCCGATCTNNNTGTGTGTATAGTTCACAGTGCC
GCACACGTCTGAACTCTTTCCCTACACGACGCTCTTCCGATCTNNNTCACTCACACTAGCAAATGAAAG
GCACACGTCTGAACTCTTTCCCTACACGACGCTCTTCCGATCTNNNTGTGTGACTGCAAGTAAGCATGT
AGCACACGTCTGAACTCTTTCCCTACACGACGCTCTTCCGATCTNNNCTCTCTGGAAAGGCAGGGTTTT
GAGCACACGTCTGAACTCTTTCCCTACACGACGCTCTTCCGATCTNNNACCCGGCACGATTTATTTGGT
AGCACACGTCTGAACTCTTTCCCTACACGACGCTCTTCCGATCTNNNTTCCTTCCCTCATGGAGGTCTTT
AGCACACGTCTGAACTCTTTCCCTACACGACGCTCTTCCGATCTNNNTGCTTCTGGGGACAGGAATTAC
AGCACACGTCTGAACTCTTTCCCTACACGACGCTCTTCCGATCTNNNAAGGGCCAAGACAGCTATTTCC
AGCACACGTCTGAACTCTTTCCCTACACGACGCTCTTCCGATCTNNNAGGTGAACAGAGAAGAGCTGC
AGCACACGTCTGAACTCTTTCCCTACACGACGCTCTTCCGATCTNNNGTTTGGGGATCACTGAAGAGG
AGCACACGTCTGAACTCTTTCCCTACACGACGCTCTTCCGATCTNNNTGCAACCAAGTTCAAGGTATGC
GAGCACACGTCTGAACTCTTTCCCTACACGACGCTCTTCCGATCTNNNTCCTGATGGAACCAGCCTAAAT
GCACACGTCTGAACTCTTTCCCTACACGACGCTCTTCCGATCTNNNAGCAACAAATCCAATCTGTCCGTC
GAGCACACGTCTGAACTCTTTCCCTACACGACGCTCTTCCGATCTNNNAGCAGCTGCACATGAACATTTT
AGCACACGTCTGAACTCTTTCCCTACACGACGCTCTTCCGATCTNNNTTGTGCCATTTCATGTCCTTCCAC
AGCACACGTCTGAACTCTTTCCCTACACGACGCTCTTCCGATCTNNNAACATGCTCATGCTTCCATTACT
GAGCACACGTCTGAACTCTTTCCCTACACGACGCTCTTCCGATCTNNNGTGGGCAGTGCTATGATTTCT
GAGCACACGTCTGAACTCTTTCCCTACACGACGCTCTTCCGATCTNNNTATTCCTAACAGCGTACAAGG
GCACACGTCTGAACTCTTTCCCTACACGACGCTCTTCCGATCTNNNCTGCAGGTGTATGTTACTGGCAT
GCACACGTCTGAACTCTTTCCCTACACGACGCTCTTCCGATCTNNNTGTATGGGAAAGTGTCCACAACG
AGCACACGTCTGAACTCTTTCCCTACACGACGCTCTTCCGATCTNNNTGTGCCCTTAGGCAGTTTCTTTT
AGCACACGTCTGAACTCTTTCCCTACACGACGCTCTTCCGATCTNNNTCTGGGGAGTGTGTGTCTGTAA
GCACACGTCTGAACTCTTTCCCTACACGACGCTCTTCCGATCTNNNTCTTTGGGTACCTGACCTCTTCCCA
GAGCACACGTCTGAACTCTTTCCCTACACGACGCTCTTCCGATCTNNNATGTACCCGTTTTGTGTGTGT
GAGCACACGTCTGAACTCTTTCCCTACACGACGCTCTTCCGATCTNNNAGTAAGCAGCCACTGACATCA
GAGCACACGTCTGAACTCTTTCCCTACACGACGCTCTTCCGATCTNNNAAACCCCAAAACAAACCAACC
AGCACACGTCTGAACTCTTTCCCTACACGACGCTCTTCCGATCTNNNAAATATCCACTGCCCTTGGACAC
GAGCACACGTCTGAACTCTTTCCCTACACGACGCTCTTCCGATCTNNNTTCTTGCCTAGAAAGCTTAGCC
AGCACACGTCTGAACTCTTTCCCTACACGACGCTCTTCCGATCTNNNCCCCAAGTTTTCAACAGACATG
GAGCACACGTCTGAACTCTTTCCCTACACGACGCTCTTCCGATCTNNNTGCAGATGCTAGAAAGTGCT
GCACACGTCTGAACTCTTTCCCTACACGACGCTCTTCCGATCTNNNTCTAGCTTTGAGGACAGAGGGT
AGAGCACACGTCTGAACTCTTTCCCTACACGACGCTCTTCCGATCTNNNAAGCAGGTTGCAGAATACCT
GAGCACACGTCTGAACTCTTTCCCTACACGACGCTCTTCCGATCTNNNACCATGTGCAAGAATAAGCTC
AGCACACGTCTGAACTCTTTCCCTACACGACGCTCTTCCGATCTNNNGGGAGCCAAACTCGAAAAATA
GAGCACACGTCTGAACTCTTTCCCTACACGACGCTCTTCCGATCTNNNCTGCCCTCTGTCACTTCCAATT
AGCACACGTCTGAACTCTTTCCCTACACGACGCTCTTCCGATCTNNNTGATTTGTCACTGGGGTATTGC
GAGCACACGTCTGAACTCTTTCCCTACACGACGCTCTTCCGATCTNNNTCCAGCAAGGCTGCTAAAAGT

AGCACACGTCTGAACTCTTTCCCTACACGACGCTCTTCCGATCTNNNGAGCTGTGTTAGTCTGGGTTC
GCACACGTCTGAACTCTTTCCCTACACGACGCTCTTCCGATCTNNNACCTGGACCAATTGGAAAGGAT
GCACACGTCTGAACTCTTTCCCTACACGACGCTCTTCCGATCTNNNCAAAACCAAATTCCCGACCTTCG
AGCACACGTCTGAACTCTTTCCCTACACGACGCTCTTCCGATCTNNNGATGAACCACGGATCCTGCTA
AGCACACGTCTGAACTCTTTCCCTACACGACGCTCTTCCGATCTNNNTGGCATCAAAGACTACTGGAA
GCACACGTCTGAACTCTTTCCCTACACGACGCTCTTCCGATCTNNNGCTGAAGGCTGTTCTATGGACA
AGCACACGTCTGAACTCTTTCCCTACACGACGCTCTTCCGATCTNNNCCATTCTGCTCATTCAGTCCTGT
GAGCACACGTCTGAACTCTTTCCCTACACGACGCTCTTCCGATCTNNNTTTCTGTCCCCTAGAATCCTGC
GCACACGTCTGAACTCTTTCCCTACACGACGCTCTTCCGATCTNNNCTTCCTGGTCACTTTCTTGCTGT
AGCACACGTCTGAACTCTTTCCCTACACGACGCTCTTCCGATCTNNNGTCCTTGTTTCCAGCTTTTCTGG
AGCACACGTCTGAACTCTTTCCCTACACGACGCTCTTCCGATCTNNNGAACTCAAGTGATCCTGCCACT
AGCACACGTCTGAACTCTTTCCCTACACGACGCTCTTCCGATCTNNNTTGTATATATCCCACCCCTGCCC
GCACACGTCTGAACTCTTTCCCTACACGACGCTCTTCCGATCTNNNGGTGAGAAGTCTACAGGGCTTT
AGCACACGTCTGAACTCTTTCCCTACACGACGCTCTTCCGATCTNNNTGAATGACCTCAATAGAGCCCC
AGCACACGTCTGAACTCTTTCCCTACACGACGCTCTTCCGATCTNNNTCTTTTCACCTGGCTCCTGAAAC
AGCACACGTCTGAACTCTTTCCCTACACGACGCTCTTCCGATCTNNNGAAAGCTCATGAAGGGTGGGA
AGCACACGTCTGAACTCTTTCCCTACACGACGCTCTTCCGATCTNNNTAAAGCCTGGTGAGAACTGGG
GAGCACACGTCTGAACTCTTTCCCTACACGACGCTCTTCCGATCTNNNAGCTCAGCTCACTCCAATCTC
GAGCACACGTCTGAACTCTTTCCCTACACGACGCTCTTCCGATCTNNNCTCACTCCAGAGGCTTCATTT
AGCACACGTCTGAACTCTTTCCCTACACGACGCTCTTCCGATCTNNNCCCTGAATATTTTTGCCCACACG
GCACACGTCTGAACTCTTTCCCTACACGACGCTCTTCCGATCTNNNCTGGGAGAGGCATGGCAAAAAT
AGCACACGTCTGAACTCTTTCCCTACACGACGCTCTTCCGATCTNNNAAGCAAACCACAGAAAACCAAG
AGCACACGTCTGAACTCTTTCCCTACACGACGCTCTTCCGATCTNNNGCACACGCTGTCCTGTTGTTTA
AGCACACGTCTGAACTCTTTCCCTACACGACGCTCTTCCGATCTNNNTGGTATACATGGCTCTATGCG
AGCACACGTCTGAACTCTTTCCCTACACGACGCTCTTCCGATCTNNNTATCCTAGCACTCTGCCTTCCAA
GAGCACACGTCTGAACTCTTTCCCTACACGACGCTCTTCCGATCTNNNATGGAACGTGGCAGAAGTGA
AGCACACGTCTGAACTCTTTCCCTACACGACGCTCTTCCGATCTNNNCCCAGGGAAATGAGGCAAGTT
GAGCACACGTCTGAACTCTTTCCCTACACGACGCTCTTCCGATCTNNNTAATGGAAGCTGGTCATCAG
AGCACACGTCTGAACTCTTTCCCTACACGACGCTCTTCCGATCTNNNGAGCCACTGAATAGAGGAGCT
AGCACACGTCTGAACTCTTTCCCTACACGACGCTCTTCCGATCTNNNCAATCTGTCTATCTGGCCCCCT
GAGCACACGTCTGAACTCTTTCCCTACACGACGCTCTTCCGATCTNNNAATCTTCACACACAGGATGAG
AGCACACGTCTGAACTCTTTCCCTACACGACGCTCTTCCGATCTNNNGGAAGCTTGCCACTGAAAGAC
GAGCACACGTCTGAACTCTTTCCCTACACGACGCTCTTCCGATCTNNNAAATTTCAGATGCTGGAAAC
AGCACACGTCTGAACTCTTTCCCTACACGACGCTCTTCCGATCTNNNAGTTGGGAAGGAGAGGTAAG
AGCACACGTCTGAACTCTTTCCCTACACGACGCTCTTCCGATCTNNNATGTCTGCAAACACCTTCCATCA
GAGCACACGTCTGAACTCTTTCCCTACACGACGCTCTTCCGATCTNNNCCAAGCAATACTGTCTCCCTTC
AGCACACGTCTGAACTCTTTCCCTACACGACGCTCTTCCGATCTNNNGGGAAACCAGTGTTCTGGGTT
AGCACACGTCTGAACTCTTTCCCTACACGACGCTCTTCCGATCTNNNTTAGTCTCAAGGCTTCGGTCTCT
AGCACACGTCTGAACTCTTTCCCTACACGACGCTCTTCCGATCTNNNATGGCTTGCTATTGCAGGGTTT
AGCACACGTCTGAACTCTTTCCCTACACGACGCTCTTCCGATCTNNNTCAGATGCCCCAGCTGAATGTA
AGCACACGTCTGAACTCTTTCCCTACACGACGCTCTTCCGATCTNNNTTCATTTCCTAATGTCCTGGGG
AGCACACGTCTGAACTCTTTCCCTACACGACGCTCTTCCGATCTNNNTGGCCTGCAGATTTTGAGACTT
GAGCACACGTCTGAACTCTTTCCCTACACGACGCTCTTCCGATCTNNNTCATTTCTCCCCTCTACTTCCCC

AGAGCACACGTCTGAACTCTTTCCCTACACGACGCTCTTCCGATCTNNNCATTCCTTTCCTGTCCCTTTG
AGCACACGTCTGAACTCTTTCCCTACACGACGCTCTTCCGATCTNNNCTGGACTAGTTGAGGCCATTCA
AGCACACGTCTGAACTCTTTCCCTACACGACGCTCTTCCGATCTNNNGCACATGCAATGCAGAGATCC
GAGCACACGTCTGAACTCTTTCCCTACACGACGCTCTTCCGATCTNNNCCACTGAACCACCCAAGATTG
GAGCACACGTCTGAACTCTTTCCCTACACGACGCTCTTCCGATCTNNNGACTTAGCATGTCCTCAGCCA
GCACACGTCTGAACTCTTTCCCTACACGACGCTCTTCCGATCTNNNTTCCCTCCAATATCTGTCTGCTCA
AGCACACGTCTGAACTCTTTCCCTACACGACGCTCTTCCGATCTNNNTCACCTATTTGGTAGTTCCCGGT
AGCACACGTCTGAACTCTTTCCCTACACGACGCTCTTCCGATCTNNNAAAAGGTGGGGGACCACTACAT
AGCACACGTCTGAACTCTTTCCCTACACGACGCTCTTCCGATCTNNNGCCTTCAGAACCTCCTTCATGTT
GCACACGTCTGAACTCTTTCCCTACACGACGCTCTTCCGATCTNNNGAGGGCGCTAGTTCTAAAGATG
AGCACACGTCTGAACTCTTTCCCTACACGACGCTCTTCCGATCTNNNGCAGCATTAGAAAGAGGCAGA
GAGCACACGTCTGAACTCTTTCCCTACACGACGCTCTTCCGATCTNNNTAAATGCACCATGGAGGTGA
AGCACACGTCTGAACTCTTTCCCTACACGACGCTCTTCCGATCTNNNCGACTACAGGAGCTCGTCACTAT
GCACACGTCTGAACTCTTTCCCTACACGACGCTCTTCCGATCTNNNTCGCCCTCCTCTACCCATATAGAC
GCACACGTCTGAACTCTTTCCCTACACGACGCTCTTCCGATCTNNNAAAAACTCTGCAGGAGGTAGGG
AGCACACGTCTGAACTCTTTCCCTACACGACGCTCTTCCGATCTNNNGCAACAGTGACCAAGACACAAA
GCACACGTCTGAACTCTTTCCCTACACGACGCTCTTCCGATCTNNNTCTGAACCCATCCCACTCCTAAGC
AGCACACGTCTGAACTCTTTCCCTACACGACGCTCTTCCGATCTNNNGTAAACTCAGTTGCACCCATTGC
AGCACACGTCTGAACTCTTTCCCTACACGACGCTCTTCCGATCTNNNAAGGCCTTCCCTCACTACATCTG
AGCACACGTCTGAACTCTTTCCCTACACGACGCTCTTCCGATCTNNNTCTGGGAGCTATTTTGTGGCTT
AGCACACGTCTGAACTCTTTCCCTACACGACGCTCTTCCGATCTNNNCTCCCTAATGTAGGTCAGCCTC
GAGCACACGTCTGAACTCTTTCCCTACACGACGCTCTTCCGATCTNNNAATGAGGCAAACAGATCAAT
GCACACGTCTGAACTCTTTCCCTACACGACGCTCTTCCGATCTNNNTGGCTCACATTAGTACCTTGCAT
AGCACACGTCTGAACTCTTTCCCTACACGACGCTCTTCCGATCTNNNTTCTTTCCTGCTTCAGTTGTCCC
GAGCACACGTCTGAACTCTTTCCCTACACGACGCTCTTCCGATCTNNNGGTTGGTCAGGCTTTGTCTCT
GAGCACACGTCTGAACTCTTTCCCTACACGACGCTCTTCCGATCTNNNGGCTGTCTGCTGTCCATATTT
AGCACACGTCTGAACTCTTTCCCTACACGACGCTCTTCCGATCTNNNACCTCATGCAAGGCCAAATGTA
GCACACGTCTGAACTCTTTCCCTACACGACGCTCTTCCGATCTNNNCACATGACCATGCCAGCTAAGAT
AGAGCACACGTCTGAACTCTTTCCCTACACGACGCTCTTCCGATCTNNNATTAGAAGGTAGCTGGAGG
GAGCACACGTCTGAACTCTTTCCCTACACGACGCTCTTCCGATCTNNNCCCTCAGTACGCAGTAGCTTT
GAGCACACGTCTGAACTCTTTCCCTACACGACGCTCTTCCGATCTNNNCCACAGGACGCACCTGTAATA
AGCACACGTCTGAACTCTTTCCCTACACGACGCTCTTCCGATCTNNNTGGAATTCAACCAGCCCTAACT
AGCACACGTCTGAACTCTTTCCCTACACGACGCTCTTCCGATCTNNNTACTGTTGCTACCTTTTGCATCT
AGCACACGTCTGAACTCTTTCCCTACACGACGCTCTTCCGATCTNNNCTTCCTGGGAGAGGACTGAAG
GCACACGTCTGAACTCTTTCCCTACACGACGCTCTTCCGATCTNNNTTCTGGCACCTCTCACTCCATTAC
AGCACACGTCTGAACTCTTTCCCTACACGACGCTCTTCCGATCTNNNAACCTGGGGTCCAAGGATAGA
AGCACACGTCTGAACTCTTTCCCTACACGACGCTCTTCCGATCTNNNACAAGAACACTAAATGCTGTGT
AGCACACGTCTGAACTCTTTCCCTACACGACGCTCTTCCGATCTNNNTTAGCTTTCCAACATCCCCCTTG
GAGCACACGTCTGAACTCTTTCCCTACACGACGCTCTTCCGATCTNNNTGATGGTCTGTTTTCTTTTGCC
GAGCACACGTCTGAACTCTTTCCCTACACGACGCTCTTCCGATCTNNNATGGTCCATGAGAGACAGAA
AGCACACGTCTGAACTCTTTCCCTACACGACGCTCTTCCGATCTNNNTACACAAACACAAAAACAGGCCC
GAGCACACGTCTGAACTCTTTCCCTACACGACGCTCTTCCGATCTNNNATACACAACCCAAGGAAGTCT
AGCACACGTCTGAACTCTTTCCCTACACGACGCTCTTCCGATCTNNNAGTTGGGCACCCATATTTCACT

AGCACACGTCTGAACTCTTTCCCTACACGACGCTCTTCCGATCTNNNAGAAAAGGTGGCTTTCAGACT
AGCACACGTCTGAACTCTTTCCCTACACGACGCTCTTCCGATCTNNNAAGAGTGTATCAGTCAGGGT
GAGCACACGTCTGAACTCTTTCCCTACACGACGCTCTTCCGATCTNNNTTGTTTTCCTCTGAGCTGCC
GCACACGTCTGAACTCTTTCCCTACACGACGCTCTTCCGATCTNNNAAATATGGATGCAGGGTGTGTG
GCACACGTCTGAACTCTTTCCCTACACGACGCTCTTCCGATCTNNNGTCAAACAGCTGAAGCCAAAGA
GAGCACACGTCTGAACTCTTTCCCTACACGACGCTCTTCCGATCTNNNTACACAAACATGCATGCCTTT
AGCACACGTCTGAACTCTTTCCCTACACGACGCTCTTCCGATCTNNNGAAAATGAAATTTGCCAGCGG
AGCACACGTCTGAACTCTTTCCCTACACGACGCTCTTCCGATCTNNNTCCCAATAATGAGCCTTTCTG
AGCACACGTCTGAACTCTTTCCCTACACGACGCTCTTCCGATCTNNNCAAGGGTCAGTACAGAAAAG
GCACACGTCTGAACTCTTTCCCTACACGACGCTCTTCCGATCTNNNCCACAGAAGACAGTGTCCACAA
GCACACGTCTGAACTCTTTCCCTACACGACGCTCTTCCGATCTNNNAGACAGTTTGGAACAACCCAA
AGCACACGTCTGAACTCTTTCCCTACACGACGCTCTTCCGATCTNNNAACTTGTACACGGAAACACCT
AGCACACGTCTGAACTCTTTCCCTACACGACGCTCTTCCGATCTNNNGTCTACAGAAGCCCCCTGCTA
AGCACACGTCTGAACTCTTTCCCTACACGACGCTCTTCCGATCTNNNAGGAAGCTTCTGGTTCCTACC
AGCACACGTCTGAACTCTTTCCCTACACGACGCTCTTCCGATCTNNNCCTCTTTTGGCGCTTCTCTTTT
GCACACGTCTGAACTCTTTCCCTACACGACGCTCTTCCGATCTNNNTAAATGGACTTGAAGGGGCAT
AGCACACGTCTGAACTCTTTCCCTACACGACGCTCTTCCGATCTNNNTGGCTTCCTCCTGGTTATGTA
GAGCACACGTCTGAACTCTTTCCCTACACGACGCTCTTCCGATCTNNNGATTGAGCAACATTGCCTTA
AGCACACGTCTGAACTCTTTCCCTACACGACGCTCTTCCGATCTNNNAATTCCCAGGTTCTTTCCAGTG
AGCACACGTCTGAACTCTTTCCCTACACGACGCTCTTCCGATCTNNNCTCCGCAAGTATAAGGAGGAA
AGCACACGTCTGAACTCTTTCCCTACACGACGCTCTTCCGATCTNNNTGCTTTTACATGTCCCAAGCTC
GCACACGTCTGAACTCTTTCCCTACACGACGCTCTTCCGATCTNNNTGTATTTCAACTTGCACCCACCT
GAGCACACGTCTGAACTCTTTCCCTACACGACGCTCTTCCGATCTNNNGAAACTGGCCTTACCGGAGT
GAGCACACGTCTGAACTCTTTCCCTACACGACGCTCTTCCGATCTNNNTCAGCTTTACCTTACAGGCA
AGCACACGTCTGAACTCTTTCCCTACACGACGCTCTTCCGATCTNNNTTTGTGTTCTTGTCTGGCCTTG
GAGCACACGTCTGAACTCTTTCCCTACACGACGCTCTTCCGATCTNNNTACCAGAGTGACTTAGAGCT
GCACACGTCTGAACTCTTTCCCTACACGACGCTCTTCCGATCTNNNTTCTCTCTTGCCTATCCCTACCG
GCACACGTCTGAACTCTTTCCCTACACGACGCTCTTCCGATCTNNNCCCACCTGACTTGAAAACAATG
AGCACACGTCTGAACTCTTTCCCTACACGACGCTCTTCCGATCTNNNTGAGTAAATGCATGCACAAG
GCACACGTCTGAACTCTTTCCCTACACGACGCTCTTCCGATCTNNNGTGGGTGTCATAGCAACTAGA
AGCACACGTCTGAACTCTTTCCCTACACGACGCTCTTCCGATCTNNNAAACCAAAACCGAGACACACA
AGCACACGTCTGAACTCTTTCCCTACACGACGCTCTTCCGATCTNNNAATATCCTCTGTGTCCCTCTCCC
AGCACACGTCTGAACTCTTTCCCTACACGACGCTCTTCCGATCTNNNAGCCCCTGCTTTTTCCTTTCTAC
AGCACACGTCTGAACTCTTTCCCTACACGACGCTCTTCCGATCTNNNTCGTTTATGGGACTGGACAGC
GCACACGTCTGAACTCTTTCCCTACACGACGCTCTTCCGATCTNNNGGGTTGCAGGTAAAAACACCT
GAGCACACGTCTGAACTCTTTCCCTACACGACGCTCTTCCGATCTNNNGGGTGGGGGTATGAATCTA
AGCACACGTCTGAACTCTTTCCCTACACGACGCTCTTCCGATCTNNNGAACTGTGGCCTTCTTCCCTAA
AGCACACGTCTGAACTCTTTCCCTACACGACGCTCTTCCGATCTNNNGCTGAGCTTATTGTGAGGAAG
GCACACGTCTGAACTCTTTCCCTACACGACGCTCTTCCGATCTNNNCATCCTCCCAAGTTCATGGACAC
GCACACGTCTGAACTCTTTCCCTACACGACGCTCTTCCGATCTNNNCCTGGTAGCAAGCAAGAAGAC
AGCACACGTCTGAACTCTTTCCCTACACGACGCTCTTCCGATCTNNNCAAGCAAACAAGTTTCGCCTC
AGCACACGTCTGAACTCTTTCCCTACACGACGCTCTTCCGATCTNNNTTTGGAATCCAGATCACCCTCC
AGCACACGTCTGAACTCTTTCCCTACACGACGCTCTTCCGATCTNNNTGAAATACGGTAAGCTCCCCTC

GAGCACACGTCTGAACTCTTTCCCTACACGACGCTCTTCCGATCTNNNAGAGCTTAGATCTGTCTTCCC
AGCACACGTCTGAACTCTTTCCCTACACGACGCTCTTCCGATCTNNNCATGTGGACCAAGTCTAGAA
AGCACACGTCTGAACTCTTTCCCTACACGACGCTCTTCCGATCTNNNGAGGATTTGCGCTTGTGTATG
GAGCACACGTCTGAACTCTTTCCCTACACGACGCTCTTCCGATCTNNNAGAAACTGGGGAGGTGAGG
GAGCACACGTCTGAACTCTTTCCCTACACGACGCTCTTCCGATCTNNNAGTCTCCCAGTCTTGGGTATG
GAGCACACGTCTGAACTCTTTCCCTACACGACGCTCTTCCGATCTNNNCACTGTTCCAATGCCCATTTCA
GAGCACACGTCTGAACTCTTTCCCTACACGACGCTCTTCCGATCTNNNAAATTCTTTGCACTGTCCACCC
GAGCACACGTCTGAACTCTTTCCCTACACGACGCTCTTCCGATCTNNNGGTAGTTAGATGTCCCATCCT
AGCACACGTCTGAACTCTTTCCCTACACGACGCTCTTCCGATCTNNNTTCATTCTTGCTGAGCGAGTAC
AGCACACGTCTGAACTCTTTCCCTACACGACGCTCTTCCGATCTNNNAGTGAAAGGTGCTATCTTGCT
GCACACGTCTGAACTCTTTCCCTACACGACGCTCTTCCGATCTNNNAATCCAGTGTTACCTGCCACTTC
AGCACACGTCTGAACTCTTTCCCTACACGACGCTCTTCCGATCTNNNCAGCGTCTCACTCTGGTTTCTA
AGCACACGTCTGAACTCTTTCCCTACACGACGCTCTTCCGATCTNNNTGAGGTCTGTCGTTGACCAAA
AGCACACGTCTGAACTCTTTCCCTACACGACGCTCTTCCGATCTNNNGCCTAGAAACCTGTCACATGAG
GAGCACACGTCTGAACTCTTTCCCTACACGACGCTCTTCCGATCTNNNTTCCATTGTGAGGAAACTGG
GAGCACACGTCTGAACTCTTTCCCTACACGACGCTCTTCCGATCTNNNTTGCCAGCCATTCTGTTTGTAG
AGCACACGTCTGAACTCTTTCCCTACACGACGCTCTTCCGATCTNNNATTTGGTTCCTTGCAAAGTGCT
AGCACACGTCTGAACTCTTTCCCTACACGACGCTCTTCCGATCTNNNAGTGTTGGGGTAGTTTAGCAA
AGCACACGTCTGAACTCTTTCCCTACACGACGCTCTTCCGATCTNNNAGTTCAACTGATCATGGCTGG
GAGCACACGTCTGAACTCTTTCCCTACACGACGCTCTTCCGATCTNNNGATCAGGGAGGAAGCAACA
AGCACACGTCTGAACTCTTTCCCTACACGACGCTCTTCCGATCTNNNAAATGGGCTTCAAGTGATCAG
GAGCACACGTCTGAACTCTTTCCCTACACGACGCTCTTCCGATCTNNNTAAGGATGGTAGCTAAAGGG
GAGCACACGTCTGAACTCTTTCCCTACACGACGCTCTTCCGATCTNNNGAGTTCTCTCTGTGTGTGCAT
AGCACACGTCTGAACTCTTTCCCTACACGACGCTCTTCCGATCTNNNTTTCAAATGCTTGCTGCTCTTGC
GCACACGTCTGAACTCTTTCCCTACACGACGCTCTTCCGATCTNNNAAATCTACCTGTGAGCTGGAAC
AGAGCACACGTCTGAACTCTTTCCCTACACGACGCTCTTCCGATCTNNNCTTGGGGCTTGCATGTTGTT
GAGCACACGTCTGAACTCTTTCCCTACACGACGCTCTTCCGATCTNNNGGTGCTGAGGTGTTTCTTCT
GAGCACACGTCTGAACTCTTTCCCTACACGACGCTCTTCCGATCTNNNACTGCTGATTACAGGTTGTGT
GAGCACACGTCTGAACTCTTTCCCTACACGACGCTCTTCCGATCTNNNTTGCTGAGAGAGTAGGTGTC
GAGCACACGTCTGAACTCTTTCCCTACACGACGCTCTTCCGATCTNNNGCCCTTATGTGACATCATTCG
GAGCACACGTCTGAACTCTTTCCCTACACGACGCTCTTCCGATCTNNNTAACCCTAAGGAGGGCTTAC
AGCACACGTCTGAACTCTTTCCCTACACGACGCTCTTCCGATCTNNNATTCTTTGGATTCTGGGGTCCC
GAGCACACGTCTGAACTCTTTCCCTACACGACGCTCTTCCGATCTNNNGTGACACCCCATCACTTTAGTC
AGCACACGTCTGAACTCTTTCCCTACACGACGCTCTTCCGATCTNNNCAGGGCAAGGGGTGTCTTTTAT
GAGCACACGTCTGAACTCTTTCCCTACACGACGCTCTTCCGATCTNNNGGACAAACCTTACATCCCCTTC
GAGCACACGTCTGAACTCTTTCCCTACACGACGCTCTTCCGATCTNNNTAACGCTGGAGATGGAGGTT
GCACACGTCTGAACTCTTTCCCTACACGACGCTCTTCCGATCTNNNCGCACTATAACTGCCCTACTTCCC
AGCACACGTCTGAACTCTTTCCCTACACGACGCTCTTCCGATCTNNNGGTATTCACCCTGGAGAAGCAA
GCACACGTCTGAACTCTTTCCCTACACGACGCTCTTCCGATCTNNNTTTGGCTCCTGCTTCATCTTCTCC
AGCACACGTCTGAACTCTTTCCCTACACGACGCTCTTCCGATCTNNNGAAAAGAGTGTGTGTGTGTCCC
GAGCACACGTCTGAACTCTTTCCCTACACGACGCTCTTCCGATCTNNNAGCAGTGTTTCCCAGGAATG
AGCACACGTCTGAACTCTTTCCCTACACGACGCTCTTCCGATCTNNNTGGATGGGGGCAAATAATCAC
AGCACACGTCTGAACTCTTTCCCTACACGACGCTCTTCCGATCTNNNTTGCTCAACTACAGCAATCCCAT

AGCACACGTCTGAACTCTTTCCCTACACGACGCTCTTCCGATCTNNNAAGTCAAAACAGGTCCAAGAG
AGCACACGTCTGAACTCTTTCCCTACACGACGCTCTTCCGATCTNNNTAACCATGGGAATTTCCTGGAC
AGCACACGTCTGAACTCTTTCCCTACACGACGCTCTTCCGATCTNNNGCCCATTGCAGTATTTTGTTGC
AGCACACGTCTGAACTCTTTCCCTACACGACGCTCTTCCGATCTNNNCAATCCCACACCGATACCAAGAC
AGCACACGTCTGAACTCTTTCCCTACACGACGCTCTTCCGATCTNNNAACCCTCTGGTGCTGTCAAAAA
GAGCACACGTCTGAACTCTTTCCCTACACGACGCTCTTCCGATCTNNNGGAGAATGGGGGAAGAAGC
GCACACGTCTGAACTCTTTCCCTACACGACGCTCTTCCGATCTNNNTGCCTGCCAGACATTTTTCTTTG
AGCACACGTCTGAACTCTTTCCCTACACGACGCTCTTCCGATCTNNNTGCAAACTAACCTCACACCCTTC
AGCACACGTCTGAACTCTTTCCCTACACGACGCTCTTCCGATCTNNNTAGTGAGCTTCCTACGTTGGCT
AGCACACGTCTGAACTCTTTCCCTACACGACGCTCTTCCGATCTNNNAGGGTATTGCTTTCCTAGCCTG
AGCACACGTCTGAACTCTTTCCCTACACGACGCTCTTCCGATCTNNNTCCTTGGCCAATTCCACAACTTT
AGCACACGTCTGAACTCTTTCCCTACACGACGCTCTTCCGATCTNNNTTGCTGGAAGAACTCATGGCT
GCACACGTCTGAACTCTTTCCCTACACGACGCTCTTCCGATCTNNNTTTGGTTGGCTTTCACTGTCTGA
GCACACGTCTGAACTCTTTCCCTACACGACGCTCTTCCGATCTNNNCTTGTGTGGTTTGGAGTGAGT
AGCACACGTCTGAACTCTTTCCCTACACGACGCTCTTCCGATCTNNNCAGGCCAGGGACAAATAACT
AGCACACGTCTGAACTCTTTCCCTACACGACGCTCTTCCGATCTNNNTGGACAACACTGCTCTACATTC
GAGCACACGTCTGAACTCTTTCCCTACACGACGCTCTTCCGATCTNNNAAGGCCAGTAACACTGAGAG
AGCACACGTCTGAACTCTTTCCCTACACGACGCTCTTCCGATCTNNNAACCTCACAAACCAGCTTCACT
AGCACACGTCTGAACTCTTTCCCTACACGACGCTCTTCCGATCTNNNCACCATGTAATCTCCCAAGAGC
AGCACACGTCTGAACTCTTTCCCTACACGACGCTCTTCCGATCTNNNAGCCTGAAATTGGCTGAACTT
AGCACACGTCTGAACTCTTTCCCTACACGACGCTCTTCCGATCTNNNTGCATAGGGCCACCTTCTTACT
GAGCACACGTCTGAACTCTTTCCCTACACGACGCTCTTCCGATCTNNNAGTCCACAGAGCATCAGAACA
GCACACGTCTGAACTCTTTCCCTACACGACGCTCTTCCGATCTNNNTGCCAAGTGTGTTTCTTGCTACT
AGCACACGTCTGAACTCTTTCCCTACACGACGCTCTTCCGATCTNNNGCATGAGTGCTTAGCTGTCTCT
AGCACACGTCTGAACTCTTTCCCTACACGACGCTCTTCCGATCTNNNTCAGCTCTCTCTTACCTAGTGC
AGCACACGTCTGAACTCTTTCCCTACACGACGCTCTTCCGATCTNNNTACCCTTGACTACCTGGGAAG
AGCACACGTCTGAACTCTTTCCCTACACGACGCTCTTCCGATCTNNNGCAGAAGCACAACTTCCACAAT
GCACACGTCTGAACTCTTTCCCTACACGACGCTCTTCCGATCTNNNAACAGTAAGTTACCTTGCACCCG
AGCACACGTCTGAACTCTTTCCCTACACGACGCTCTTCCGATCTNNNCAGAGTATCCACTCCCAACAAG
AGCACACGTCTGAACTCTTTCCCTACACGACGCTCTTCCGATCTNNNAACTTCAACTGCTACCCAGACAC
AGCACACGTCTGAACTCTTTCCCTACACGACGCTCTTCCGATCTNNNCCCAATATTTCTGCAGTGCAAC
AGCACACGTCTGAACTCTTTCCCTACACGACGCTCTTCCGATCTNNNTAAGTTCATTCAGTGTACGTG
AGCACACGTCTGAACTCTTTCCCTACACGACGCTCTTCCGATCTNNNATATCCAGCTTGTGTGTGTGT
AGCACACGTCTGAACTCTTTCCCTACACGACGCTCTTCCGATCTNNNCCTTGGAAATATGTCCCCCTGT
GCACACGTCTGAACTCTTTCCCTACACGACGCTCTTCCGATCTNNNAGCTGTTAGCTCAGTGATCCTCT
GCACACGTCTGAACTCTTTCCCTACACGACGCTCTTCCGATCTNNNTACATGGCCCAGTGTTGATAAG
AGCACACGTCTGAACTCTTTCCCTACACGACGCTCTTCCGATCTNNNTGTCCCAGCTTTCAATGTCATCA
GAGCACACGTCTGAACTCTTTCCCTACACGACGCTCTTCCGATCTNNNGCTGCAAAGTCTTAGAACCCA
GCACACGTCTGAACTCTTTCCCTACACGACGCTCTTCCGATCTNNNTCTCCTTCCTCATCTCCTACCCTCA
GCACACGTCTGAACTCTTTCCCTACACGACGCTCTTCCGATCTNNNCCATCTCTCCCAACCTCCATACACA
AGCACACGTCTGAACTCTTTCCCTACACGACGCTCTTCCGATCTNNNTGACCGAGTTCTGAGCTCTATG
AGCACACGTCTGAACTCTTTCCCTACACGACGCTCTTCCGATCTNNNCCAAAAGTCCCCCAAAGCATAG
GCACACGTCTGAACTCTTTCCCTACACGACGCTCTTCCGATCTNNNGACCCAGAGGATCTGGTTAGAG

GCACACGTCTGAACTCTTTCCCTACACGACGCTCTTCCGATCTNNNGGCCTGGGTAATCTCTCTCTCTC
AGCACACGTCTGAACTCTTTCCCTACACGACGCTCTTCCGATCTNNNCTGAAGAATATTTCCCTGCCCC
AGCACACGTCTGAACTCTTTCCCTACACGACGCTCTTCCGATCTNNNCTACAGTGAAGGAGTGAAGG
GCACACGTCTGAACTCTTTCCCTACACGACGCTCTTCCGATCTNNNTCCGGATCCAGTATCCTTACAGTC
AGCACACGTCTGAACTCTTTCCCTACACGACGCTCTTCCGATCTNNNAAGCCTGTATCAGTCTGGTGG
AGCACACGTCTGAACTCTTTCCCTACACGACGCTCTTCCGATCTNNNAGTTTACTGCGTAGTGCTTTGG
GCACACGTCTGAACTCTTTCCCTACACGACGCTCTTCCGATCTNNNGACACTGAGGCTTTGTGTGATG
AGCACACGTCTGAACTCTTTCCCTACACGACGCTCTTCCGATCTNNNAGGCATCATCCAAGGGATTA
AGCACACGTCTGAACTCTTTCCCTACACGACGCTCTTCCGATCTNNNCCCGGCCAGTTCTGATTCTTAT
AGCACACGTCTGAACTCTTTCCCTACACGACGCTCTTCCGATCTNNNGCTATTTTGGCCACACAACCA
AGCACACGTCTGAACTCTTTCCCTACACGACGCTCTTCCGATCTNNNAAAAGAGGAGGTGATAGCTCA
GCACACGTCTGAACTCTTTCCCTACACGACGCTCTTCCGATCTNNNCCTGGGTGTCCAGATTATGGAT
AGCACACGTCTGAACTCTTTCCCTACACGACGCTCTTCCGATCTNNNTTGATCTTGGGTCATCTGTGT
GAGCACACGTCTGAACTCTTTCCCTACACGACGCTCTTCCGATCTNNNTCAGGTCTGACCCAGTCATA
GAGCACACGTCTGAACTCTTTCCCTACACGACGCTCTTCCGATCTNNNAGCTCACATGAAAACATCCCC
GCACACGTCTGAACTCTTTCCCTACACGACGCTCTTCCGATCTNNNCTTCTGTTCTCCTTCAAGCCACAC
GCACACGTCTGAACTCTTTCCCTACACGACGCTCTTCCGATCTNNNACCTCCTTTATCCTGTCAGCCTTC
GCACACGTCTGAACTCTTTCCCTACACGACGCTCTTCCGATCTNNNTGCCTATTTGGGCTTCTCTCTCTC
CACACGTCTGAACTCTTTCCCTACACGACGCTCTTCCGATCTNNNGAAAGCACCACATCAAAATGGCA
GCACACGTCTGAACTCTTTCCCTACACGACGCTCTTCCGATCTNNNGGGCAAGAATATGGCTGTGCTA
GCACACGTCTGAACTCTTTCCCTACACGACGCTCTTCCGATCTNNNGACAATGTATTCCAGAGGCAGG
GAGCACACGTCTGAACTCTTTCCCTACACGACGCTCTTCCGATCTNNNTGTGATACAGCGTCTACTGG
AGCACACGTCTGAACTCTTTCCCTACACGACGCTCTTCCGATCTNNNCTTTTCTTGCTCTCGGCATCCTT
AGCACACGTCTGAACTCTTTCCCTACACGACGCTCTTCCGATCTNNNGGGACTTAGAGGGTCTACTG
AGCACACGTCTGAACTCTTTCCCTACACGACGCTCTTCCGATCTNNNATTGGTGGCTAGAGCTTCAGA
AGCACACGTCTGAACTCTTTCCCTACACGACGCTCTTCCGATCTNNNATACTGCCATGAAAGCCAGTC
AGCACACGTCTGAACTCTTTCCCTACACGACGCTCTTCCGATCTNNNCATGGTTCATCAGGGAAGCTGT
GCACACGTCTGAACTCTTTCCCTACACGACGCTCTTCCGATCTNNNGAACATGTCTTCCCTTGTAGGGC
AGCACACGTCTGAACTCTTTCCCTACACGACGCTCTTCCGATCTNNNACTCTTTCCCACAGTCAAGCTC
AGCACACGTCTGAACTCTTTCCCTACACGACGCTCTTCCGATCTNNNAAGCTTGCACTTCTGAGCCTTTT
GCACACGTCTGAACTCTTTCCCTACACGACGCTCTTCCGATCTNNNGCAAAGGACAGTTCTGCCTAAG
AGCACACGTCTGAACTCTTTCCCTACACGACGCTCTTCCGATCTNNNCACTCTGTACTCACATGGGCTTT
GCACACGTCTGAACTCTTTCCCTACACGACGCTCTTCCGATCTNNNCCTACATAATCCATCACTGGGCG
GCACACGTCTGAACTCTTTCCCTACACGACGCTCTTCCGATCTNNNGGGGGTGGTTACACAGATGTA
GCACACGTCTGAACTCTTTCCCTACACGACGCTCTTCCGATCTNNNGGAACTATCCCCTCTGACTGGAG
AGCACACGTCTGAACTCTTTCCCTACACGACGCTCTTCCGATCTNNNAAGTGATGTGGGGCAAGTTA
AGCACACGTCTGAACTCTTTCCCTACACGACGCTCTTCCGATCTNNNAATGTGAGGGTGCTTGATTCTG
AGCACACGTCTGAACTCTTTCCCTACACGACGCTCTTCCGATCTNNNATGTTGTACCTTGGTGGCTTCA
AGCACACGTCTGAACTCTTTCCCTACACGACGCTCTTCCGATCTNNNAAAAGGGTGCAGTCTGTAGGA
GAGCACACGTCTGAACTCTTTCCCTACACGACGCTCTTCCGATCTNNNGACCTCCATAAAGTTCACCGC
AGCACACGTCTGAACTCTTTCCCTACACGACGCTCTTCCGATCTNNNTGCTTCCCATGTAGTGTGTTTC
GCACACGTCTGAACTCTTTCCCTACACGACGCTCTTCCGATCTNNNAGCACAGGCAGAATTTCTTCAAA
GCACACGTCTGAACTCTTTCCCTACACGACGCTCTTCCGATCTNNNCCACTTGTTCAACTCCCGAAGAA

GAGCACACGTCTGAACTCTTTCCCTACACGACGCTCTTCCGATCTNNNTAAGGTAAGGGTTGTGTGTG
AGCACACGTCTGAACTCTTTCCCTACACGACGCTCTTCCGATCTNNNACCTTCTTCTGACCGCATTTGTTC
GAGCACACGTCTGAACTCTTTCCCTACACGACGCTCTTCCGATCTNNNAATCTCATTGTTTCCCCAATGC
AGCACACGTCTGAACTCTTTCCCTACACGACGCTCTTCCGATCTNNNAGTGAAGGAAAAACAGCAGGG
AGCACACGTCTGAACTCTTTCCCTACACGACGCTCTTCCGATCTNNNGTCCTGTAGGATCATGCACACTG
AGCACACGTCTGAACTCTTTCCCTACACGACGCTCTTCCGATCTNNNTTGGATTAGTCAGGGTTCAGG
AGCACACGTCTGAACTCTTTCCCTACACGACGCTCTTCCGATCTNNNTGCATTAGGGAACAAATGGGA
GAGCACACGTCTGAACTCTTTCCCTACACGACGCTCTTCCGATCTNNNTTGCAGATAGACCCCTTCTCAC
AGCACACGTCTGAACTCTTTCCCTACACGACGCTCTTCCGATCTNNNGATGAATAAGGACAGTGACG
GAGCACACGTCTGAACTCTTTCCCTACACGACGCTCTTCCGATCTNNNTGCCTGGCCCACAGATAGATA
AGCACACGTCTGAACTCTTTCCCTACACGACGCTCTTCCGATCTNNNTACACATGCCTGAGTGTAAGCA
AGCACACGTCTGAACTCTTTCCCTACACGACGCTCTTCCGATCTNNNTGAACTAACACAAGCAGCCAA
AGCACACGTCTGAACTCTTTCCCTACACGACGCTCTTCCGATCTNNNGACCTGTTTGATTTTCCCACCCT
AGCACACGTCTGAACTCTTTCCCTACACGACGCTCTTCCGATCTNNNGCAAGGACAAAAGTAACACAG
AGCACACGTCTGAACTCTTTCCCTACACGACGCTCTTCCGATCTNNNAAATTGCTGGTGGAAACACTTG
GAGCACACGTCTGAACTCTTTCCCTACACGACGCTCTTCCGATCTNNNGATAGGATGTCCAGGATGG
AGCACACGTCTGAACTCTTTCCCTACACGACGCTCTTCCGATCTNNNGCAAAGATAGAAGCAGAGAG
GAGCACACGTCTGAACTCTTTCCCTACACGACGCTCTTCCGATCTNNNTTTGTAGAATTGCCTGACCCA
AGCACACGTCTGAACTCTTTCCCTACACGACGCTCTTCCGATCTNNNATCTGCTCATGTGTTCCTTCTGC
GAGCACACGTCTGAACTCTTTCCCTACACGACGCTCTTCCGATCTNNNTTCTCAGATGTGTGGGAAAA
AGCACACGTCTGAACTCTTTCCCTACACGACGCTCTTCCGATCTNNNCGACAGACACCCTGTCATACAAA
GCACACGTCTGAACTCTTTCCCTACACGACGCTCTTCCGATCTNNNTTTTCCTTCTTCGCTCCTCTAGCC
AGCACACGTCTGAACTCTTTCCCTACACGACGCTCTTCCGATCTNNNTATAATGCTTACGGACCCAGTG
AGCACACGTCTGAACTCTTTCCCTACACGACGCTCTTCCGATCTNNNAAACGTAGTTCAGACCCAGAT
AGAGCACACGTCTGAACTCTTTCCCTACACGACGCTCTTCCGATCTNNNTCAAACATGGAAGGTAAGC
AGCACACGTCTGAACTCTTTCCCTACACGACGCTCTTCCGATCTNNNACATGCAACTCACACACACAGA
AGCACACGTCTGAACTCTTTCCCTACACGACGCTCTTCCGATCTNNNGCACTTATGTCAAGCAGGGAT
GAGCACACGTCTGAACTCTTTCCCTACACGACGCTCTTCCGATCTNNNTGCCAGTTTATCTTACAGGCA
GCACACGTCTGAACTCTTTCCCTACACGACGCTCTTCCGATCTNNNGCTGTAGTGGGGTGGTTTTTAC
GAGCACACGTCTGAACTCTTTCCCTACACGACGCTCTTCCGATCTNNNTGCCTGACCATTCAAATCCATG
GAGCACACGTCTGAACTCTTTCCCTACACGACGCTCTTCCGATCTNNNCTTCCAGTAACTCCAGACCTTG
GAGCACACGTCTGAACTCTTTCCCTACACGACGCTCTTCCGATCTNNNCCTGCATTCTTACCTTTGCAAG
AGCACACGTCTGAACTCTTTCCCTACACGACGCTCTTCCGATCTNNNTGGGAATGGTGCTTCAGAGAA
GAGCACACGTCTGAACTCTTTCCCTACACGACGCTCTTCCGATCTNNNAGCATGCACATAACCCAGAGA
AGAGCACACGTCTGAACTCTTTCCCTACACGACGCTCTTCCGATCTNNNGCTTGGAAGAAAGTTCCGT
AGAGCACACGTCTGAACTCTTTCCCTACACGACGCTCTTCCGATCTNNNCCACGGATCATACTGCCTCAA
GAGCACACGTCTGAACTCTTTCCCTACACGACGCTCTTCCGATCTNNNGTGCCCTAACCTTTTCTCTCCTG
GAGCACACGTCTGAACTCTTTCCCTACACGACGCTCTTCCGATCTNNNGGGAATCCCAGCTGCCTAAA
GAGCACACGTCTGAACTCTTTCCCTACACGACGCTCTTCCGATCTNNNAACTTTACCTGCCTCTGCTTTC
GAGCACACGTCTGAACTCTTTCCCTACACGACGCTCTTCCGATCTNNNTAACAGGGGTGACTTAGAGG
GAGCACACGTCTGAACTCTTTCCCTACACGACGCTCTTCCGATCTNNNGGAAGCTCCAATGAGGACTG
GAGCACACGTCTGAACTCTTTCCCTACACGACGCTCTTCCGATCTNNNATCTCTCATAGGCTCTACAGG
AGCACACGTCTGAACTCTTTCCCTACACGACGCTCTTCCGATCTNNNAGTTAGCATTCAGCAGTCTCCA

GAGCACACGTCTGAACTCTTTCCCTACACGACGCTCTTCCGATCTNNNAGAAGGGTATTCTGATCTGG
AGCACACGTCTGAACTCTTTCCCTACACGACGCTCTTCCGATCTNNNGAACCCCCATTCCCATAGGTTT
GCACACGTCTGAACTCTTTCCCTACACGACGCTCTTCCGATCTNNNCTTGATGCATGGGCTGAAATTG
GCACACGTCTGAACTCTTTCCCTACACGACGCTCTTCCGATCTNNNAGGATGTCCTAACAGGACCTAG
AGCACACGTCTGAACTCTTTCCCTACACGACGCTCTTCCGATCTNNNAACTCTCCAGGTAAGTTGGCT
AGCACACGTCTGAACTCTTTCCCTACACGACGCTCTTCCGATCTNNNCAGGTGCAATGATGTTACCTGG
AGCACACGTCTGAACTCTTTCCCTACACGACGCTCTTCCGATCTNNNAAAAGGGACAGCAGTTTGGC
GAGCACACGTCTGAACTCTTTCCCTACACGACGCTCTTCCGATCTNNNGGCGCACAGACAAAGAAAGT
GCACACGTCTGAACTCTTTCCCTACACGACGCTCTTCCGATCTNNNCTATCTGGTTGGTGCTACTGCA
AGAGCACACGTCTGAACTCTTTCCCTACACGACGCTCTTCCGATCTNNNCCTTTGTATACGGGGTGCT
AGCACACGTCTGAACTCTTTCCCTACACGACGCTCTTCCGATCTNNNACTGCCCTCATACCAACTCTCAA
GCACACGTCTGAACTCTTTCCCTACACGACGCTCTTCCGATCTNNNTGCCCTCATTTGTATCCAGTCTC
AGAGCACACGTCTGAACTCTTTCCCTACACGACGCTCTTCCGATCTNNNTGCTGAGTGAAACCAAATC
GAGCACACGTCTGAACTCTTTCCCTACACGACGCTCTTCCGATCTNNNGCTCTCTTTACGGTGCTTACC
AGCACACGTCTGAACTCTTTCCCTACACGACGCTCTTCCGATCTNNNATGTGAAGGAGACCCTGACAT
GAGCACACGTCTGAACTCTTTCCCTACACGACGCTCTTCCGATCTNNNTGCATCCTGCACTTTTTCTCA
AGAGCACACGTCTGAACTCTTTCCCTACACGACGCTCTTCCGATCTNNNAGACTGTCAACTAAAAGAGT
AGCACACGTCTGAACTCTTTCCCTACACGACGCTCTTCCGATCTNNNATCTTTTGCTCCCTGCATTGTTT
AGCACACGTCTGAACTCTTTCCCTACACGACGCTCTTCCGATCTNNNAAATCCAAAAGCCCCATCCTCT
GAGCACACGTCTGAACTCTTTCCCTACACGACGCTCTTCCGATCTNNNAGGAAGGGGCTCCGTTAGA
AGCACACGTCTGAACTCTTTCCCTACACGACGCTCTTCCGATCTNNNACCACCAAACGACAGTTGAGA
AGCACACGTCTGAACTCTTTCCCTACACGACGCTCTTCCGATCTNNNCTGGAAAGCTTGGAAGATTG
AGAGCACACGTCTGAACTCTTTCCCTACACGACGCTCTTCCGATCTNNNGCTTTGAAAGGCAAGCAA
AGAGCACACGTCTGAACTCTTTCCCTACACGACGCTCTTCCGATCTNNNATGGCCTCCTAAAAGCTTCC
GAGCACACGTCTGAACTCTTTCCCTACACGACGCTCTTCCGATCTNNNAGACACCAACCCTGTATGAA
AGCACACGTCTGAACTCTTTCCCTACACGACGCTCTTCCGATCTNNNAGTTCTCTAGGATGGAGCTGT
AGCACACGTCTGAACTCTTTCCCTACACGACGCTCTTCCGATCTNNNGTCTCTGCCATACTTTTCCTGG
AGCACACGTCTGAACTCTTTCCCTACACGACGCTCTTCCGATCTNNNTGAAGCAAGCATTGACAGACT
AGCACACGTCTGAACTCTTTCCCTACACGACGCTCTTCCGATCTNNNTCTCTCTGTGTTGTCTTATGCCT
AGCACACGTCTGAACTCTTTCCCTACACGACGCTCTTCCGATCTNNNTTCTGCTCATATCCTGTGCTGT
GCACACGTCTGAACTCTTTCCCTACACGACGCTCTTCCGATCTNNNTCCTGCAAATCTGAACTCTTCCG
GCACACGTCTGAACTCTTTCCCTACACGACGCTCTTCCGATCTNNNAAGAGCCTTGTGGGAAGAGAGA
AGCACACGTCTGAACTCTTTCCCTACACGACGCTCTTCCGATCTNNNTTATCCGGGCTGAGAGATCCT
AGCACACGTCTGAACTCTTTCCCTACACGACGCTCTTCCGATCTNNNTGGTTACTGTAGTTTGCTGAGC
GCACACGTCTGAACTCTTTCCCTACACGACGCTCTTCCGATCTNNNTTTCTAAGTGCTTACGTGCTCAA
GCACACGTCTGAACTCTTTCCCTACACGACGCTCTTCCGATCTNNNAGAGCCTGCAATGTTGATGTTG
GAGCACACGTCTGAACTCTTTCCCTACACGACGCTCTTCCGATCTNNNGTTGGCATGCAAGGATGTAT
AGCACACGTCTGAACTCTTTCCCTACACGACGCTCTTCCGATCTNNNTCAGTGATTCCATCCTCCCACT
AGCACACGTCTGAACTCTTTCCCTACACGACGCTCTTCCGATCTNNNATCACCAGCGAGGAGACAGAA
GCACACGTCTGAACTCTTTCCCTACACGACGCTCTTCCGATCTNNNCATGACGCCTGTCATAACTGCTA
GCACACGTCTGAACTCTTTCCCTACACGACGCTCTTCCGATCTNNNCACTATGCAGTTATCTGAATGGC
AGCACACGTCTGAACTCTTTCCCTACACGACGCTCTTCCGATCTNNNCAGCGCCTGTAACTACATAAGC
AGAGCACACGTCTGAACTCTTTCCCTACACGACGCTCTTCCGATCTNNNCCAAACCAAACAGTACTTCCC

GAGCACACGTCTGAACTCTTTCCCTACACGACGCTCTTCCGATCTNNNACAGCACCACACACACACATA
GAGCACACGTCTGAACTCTTTCCCTACACGACGCTCTTCCGATCTNNNCCACCGGATGAACTCCAAAAT
GAGCACACGTCTGAACTCTTTCCCTACACGACGCTCTTCCGATCTNNNAAAGTCTGAAGCCAAAGAGG
GCACACGTCTGAACTCTTTCCCTACACGACGCTCTTCCGATCTNNNGGCAAGTAGAATGCTGATGGGA
AGCACACGTCTGAACTCTTTCCCTACACGACGCTCTTCCGATCTNNNAGACAGAGCCACTGGAGAAGA
AGCACACGTCTGAACTCTTTCCCTACACGACGCTCTTCCGATCTNNNTTATGGAGCCGTTCAGACACAA
AGCACACGTCTGAACTCTTTCCCTACACGACGCTCTTCCGATCTNNNAGCAATAGATGCTGTGGCTTGT
AGCACACGTCTGAACTCTTTCCCTACACGACGCTCTTCCGATCTNNNATTCCATTGTCACCAAACTGCCT
GAGCACACGTCTGAACTCTTTCCCTACACGACGCTCTTCCGATCTNNNAACCTTTCCCCAGCAAAGAGA
GAGCACACGTCTGAACTCTTTCCCTACACGACGCTCTTCCGATCTNNNAAAAGGGAAATAGGGGGAC
AGCACACGTCTGAACTCTTTCCCTACACGACGCTCTTCCGATCTNNNCTCTTCGAGGCCCATCTACAATG
GAGCACACGTCTGAACTCTTTCCCTACACGACGCTCTTCCGATCTNNNATTTGCCCTCTCATGTTCAATG
GAGCACACGTCTGAACTCTTTCCCTACACGACGCTCTTCCGATCTNNNTGTTTGTCATGGACTTGGTAC
GCACACGTCTGAACTCTTTCCCTACACGACGCTCTTCCGATCTNNNTCTAGAGATGGAGGCACAACCA
GCACACGTCTGAACTCTTTCCCTACACGACGCTCTTCCGATCTNNNGTTTCCCTCTCACCAGCCAATAGC
GAGCACACGTCTGAACTCTTTCCCTACACGACGCTCTTCCGATCTNNNGAAGACCGCAGTAAACACTGA
GCACACGTCTGAACTCTTTCCCTACACGACGCTCTTCCGATCTNNNTCTGAGTTTCCATCACTGCTTGG
AGAGCACACGTCTGAACTCTTTCCCTACACGACGCTCTTCCGATCTNNNTGTGATGGGGAGGTTATTC
GAGCACACGTCTGAACTCTTTCCCTACACGACGCTCTTCCGATCTNNNCAAGAGCGACACTCTGTCGA
GAGCACACGTCTGAACTCTTTCCCTACACGACGCTCTTCCGATCTNNNACAGATCACAACCCTGCCTCTA
AGCACACGTCTGAACTCTTTCCCTACACGACGCTCTTCCGATCTNNNTGGCTGCTCAATTTACCACCTTA
GAGCACACGTCTGAACTCTTTCCCTACACGACGCTCTTCCGATCTNNNGAGGCAGACAAAATCGCACT
AGCACACGTCTGAACTCTTTCCCTACACGACGCTCTTCCGATCTNNNAGAGATGGCACATTCTGCAAA
GAGCACACGTCTGAACTCTTTCCCTACACGACGCTCTTCCGATCTNNNACATGCCCCTAGCTTACTAGC
AGCACACGTCTGAACTCTTTCCCTACACGACGCTCTTCCGATCTNNNTGGAGAGAGCTCACCTATTGCT
GAGCACACGTCTGAACTCTTTCCCTACACGACGCTCTTCCGATCTNNNCCTTCCTCCTCTGAAACTGCAT
AGCACACGTCTGAACTCTTTCCCTACACGACGCTCTTCCGATCTNNNATTCATTCCTTCCCCTCCACCTCC
GAGCACACGTCTGAACTCTTTCCCTACACGACGCTCTTCCGATCTNNNGACAAAGTCCACTGGAAGGA
AGAGCACACGTCTGAACTCTTTCCCTACACGACGCTCTTCCGATCTNNNTTATCTGGGCTGGAGCTAC
AGCACACGTCTGAACTCTTTCCCTACACGACGCTCTTCCGATCTNNNACACATAAGGCTGAAGGAAGT
AGAGCACACGTCTGAACTCTTTCCCTACACGACGCTCTTCCGATCTNNNGACTGGGATTGGTTCCACTC
GAGCACACGTCTGAACTCTTTCCCTACACGACGCTCTTCCGATCTNNNCAGGGACAGTGTAATCTTGG
GAGCACACGTCTGAACTCTTTCCCTACACGACGCTCTTCCGATCTNNNAGTGGACAAAACACGGGAAC
GAGCACACGTCTGAACTCTTTCCCTACACGACGCTCTTCCGATCTNNNTCATGTTCAATTGGTGCTTCC
AGAGCACACGTCTGAACTCTTTCCCTACACGACGCTCTTCCGATCTNNNCACTTAGCTGAAGGGATGG
GAGCACACGTCTGAACTCTTTCCCTACACGACGCTCTTCCGATCTNNNCTACCAGTTCATCCATGTTGCC
GAGCACACGTCTGAACTCTTTCCCTACACGACGCTCTTCCGATCTNNNTTCTTTTGGTTTAGGGGCTTG
GAGCACACGTCTGAACTCTTTCCCTACACGACGCTCTTCCGATCTNNNTGTGTGCATGTGTGAGCATA
GAGCACACGTCTGAACTCTTTCCCTACACGACGCTCTTCCGATCTNNNCAGACTGGGTGTGTAATCAA
GAGCACACGTCTGAACTCTTTCCCTACACGACGCTCTTCCGATCTNNNAAACAAAGGATGCTTCCCTCC
GAGCACACGTCTGAACTCTTTCCCTACACGACGCTCTTCCGATCTNNNATATTTACTGGCTTGCCCCAAG
AGAGCACACGTCTGAACTCTTTCCCTACACGACGCTCTTCCGATCTNNNGCAGAAACATCCTCCTTCAG
AGCACACGTCTGAACTCTTTCCCTACACGACGCTCTTCCGATCTNNNTATTGCTAACAGGTCAGCATGG

GAGCACACGTCTGAACTCTTTCCCTACACGACGCTCTTCCGATCTNNNAGGAGACACGAGAAAGCTTAC
AGCACACGTCTGAACTCTTTCCCTACACGACGCTCTTCCGATCTNNNAGCAACTGTATCAATCAGCTGTC
GAGCACACGTCTGAACTCTTTCCCTACACGACGCTCTTCCGATCTNNNATTCCCCATACACACACACAAG
GAGCACACGTCTGAACTCTTTCCCTACACGACGCTCTTCCGATCTNNNACATTCTGAGAGAGAGGGACA
AGAGCACACGTCTGAACTCTTTCCCTACACGACGCTCTTCCGATCTNNNTTCTCACCACTCTCTCTCTTCCT
AGCACACGTCTGAACTCTTTCCCTACACGACGCTCTTCCGATCTNNNTTCCATGCATTTCTCCAAGTGCT
GAGCACACGTCTGAACTCTTTCCCTACACGACGCTCTTCCGATCTNNNAGGCAGAGCAGAAACGTAGA
GCACACGTCTGAACTCTTTCCCTACACGACGCTCTTCCGATCTNNNCTCCTCTGCCTGTTTTCGAGTTTCA
GAGCACACGTCTGAACTCTTTCCCTACACGACGCTCTTCCGATCTNNNAAATCCTGTGATCCTGTGAATC
GAGCACACGTCTGAACTCTTTCCCTACACGACGCTCTTCCGATCTNNNTTGATGCAGTTGATGTAAATC
AGCACACGTCTGAACTCTTTCCCTACACGACGCTCTTCCGATCTNNNGAACAGAAGCAGACCACACAGA
AGCACACGTCTGAACTCTTTCCCTACACGACGCTCTTCCGATCTNNNCAGCAGAGGTGATGTTGTACCA
AGCACACGTCTGAACTCTTTCCCTACACGACGCTCTTCCGATCTNNNGCTATAAGGCTGATGTGTGGG
AGCACACGTCTGAACTCTTTCCCTACACGACGCTCTTCCGATCTNNNGGTAGGAAACAGCAACAGCAA
AGCACACGTCTGAACTCTTTCCCTACACGACGCTCTTCCGATCTNNNCCTCTCTTTTTCTTTCACGCCCTCA
GAGCACACGTCTGAACTCTTTCCCTACACGACGCTCTTCCGATCTNNNCCATGCCATGTGAAGGACATT
GAGCACACGTCTGAACTCTTTCCCTACACGACGCTCTTCCGATCTNNNCTGGAGAGAGCTACATACCCAC
AGCACACGTCTGAACTCTTTCCCTACACGACGCTCTTCCGATCTNNNTTCACTGCAAATCAACTGGGTCA
GAGCACACGTCTGAACTCTTTCCCTACACGACGCTCTTCCGATCTNNNGTGGATTTGTAACAGTGTCCC
GAGCACACGTCTGAACTCTTTCCCTACACGACGCTCTTCCGATCTNNNTCTCTGACCACACAGAAAGTTG
AGCACACGTCTGAACTCTTTCCCTACACGACGCTCTTCCGATCTNNNTCCTTACAGCAAAGCAAAACGGA
GAGCACACGTCTGAACTCTTTCCCTACACGACGCTCTTCCGATCTNNNCCGTCAAACCCCTTCCTTACTTC
AGAGCACACGTCTGAACTCTTTCCCTACACGACGCTCTTCCGATCTNNNCCCATAAATGTGCATGGGGA
AGCACACGTCTGAACTCTTTCCCTACACGACGCTCTTCCGATCTNNNTCTTCTCAAGTTTAGCGGCCATC
AGCACACGTCTGAACTCTTTCCCTACACGACGCTCTTCCGATCTNNNTGTGCCATGCATTCATGTTCTCTC
GAGCACACGTCTGAACTCTTTCCCTACACGACGCTCTTCCGATCTNNNAGTAGAAAGATGAAGTGGGT
AGCACACGTCTGAACTCTTTCCCTACACGACGCTCTTCCGATCTNNNACATTGTACGTGCTTTGGGACAT
AGCACACGTCTGAACTCTTTCCCTACACGACGCTCTTCCGATCTNNNTATGTCAGGAAACACACACACGC
GAGCACACGTCTGAACTCTTTCCCTACACGACGCTCTTCCGATCTNNNCCCTTCTCTGTCTGCATACTTCC
AGCACACGTCTGAACTCTTTCCCTACACGACGCTCTTCCGATCTNNNAAGGGAAGGAGGAATCTAAGC
AGCACACGTCTGAACTCTTTCCCTACACGACGCTCTTCCGATCTNNNCATTTGCTGCAACACAGACACTT
AGCACACGTCTGAACTCTTTCCCTACACGACGCTCTTCCGATCTNNNTGCATTCTGATGCTCTGCTGAAA
GAGCACACGTCTGAACTCTTTCCCTACACGACGCTCTTCCGATCTNNNTTGCTGTCACAACTACCACCAT
AGCACACGTCTGAACTCTTTCCCTACACGACGCTCTTCCGATCTNNNACACAGCAAGGAAAAGAACTG
GAGCACACGTCTGAACTCTTTCCCTACACGACGCTCTTCCGATCTNNNTAACCCCATTTGAGCATGACC
AGAGCACACGTCTGAACTCTTTCCCTACACGACGCTCTTCCGATCTNNNTGTGACTTACTGGCCCAACTT
AAGAGCACACGTCTGAACTCTTTCCCTACACGACGCTCTTCCGATCTNNNTACCTCTCTCCATGTCCCCTC
GAGCACACGTCTGAACTCTTTCCCTACACGACGCTCTTCCGATCTNNNGTGGTCGTGTCCAGTTATTCA
GAGCACACGTCTGAACTCTTTCCCTACACGACGCTCTTCCGATCTNNNGTCCCCTAGTCTCTAGTTCCCT
AGCACACGTCTGAACTCTTTCCCTACACGACGCTCTTCCGATCTNNNATCAAAGAAAGCTTTGGCCCTCT
GAGCACACGTCTGAACTCTTTCCCTACACGACGCTCTTCCGATCTNNNATTCCATTTAGCATCTCCCACC
GAGCACACGTCTGAACTCTTTCCCTACACGACGCTCTTCCGATCTNNNATGGAGAAAATAACTGGGCT
AGAGCACACGTCTGAACTCTTTCCCTACACGACGCTCTTCCGATCTNNNAGAAGTCGAGCCAAAACCAA

AGCACACGTCTGAACTCTTTCCCTACACGACGCTCTTCCGATCTNNNTCGTTGGGTGACACTAGCATA
AGCACACGTCTGAACTCTTTCCCTACACGACGCTCTTCCGATCTNNNCACCCCAAAAACTCACACCAGT
AGCACACGTCTGAACTCTTTCCCTACACGACGCTCTTCCGATCTNNNAGTCTCTCATGTGTGTGAGGG
AGCACACGTCTGAACTCTTTCCCTACACGACGCTCTTCCGATCTNNNTGAGGAGAAACAAGTGCAAG
AGCACACGTCTGAACTCTTTCCCTACACGACGCTCTTCCGATCTNNNTGCCGTCAATATGGGTTTTCCT
AGCACACGTCTGAACTCTTTCCCTACACGACGCTCTTCCGATCTNNNATCCTAATGCACACACTTGCAC
AGCACACGTCTGAACTCTTTCCCTACACGACGCTCTTCCGATCTNNNTTCTAGCAGTGCAATGTTCCAG
GAGCACACGTCTGAACTCTTTCCCTACACGACGCTCTTCCGATCTNNNAGGTGTCAGAGGTTTTGAT
GAGCACACGTCTGAACTCTTTCCCTACACGACGCTCTTCCGATCTNNNTGCAACAAGCAAAGAGATTG
GAGCACACGTCTGAACTCTTTCCCTACACGACGCTCTTCCGATCTNNNGGGACTGATCATAAATGCAG
GAGCACACGTCTGAACTCTTTCCCTACACGACGCTCTTCCGATCTNNNAGTGTCCATCAACCAATGAAG
GAGCACACGTCTGAACTCTTTCCCTACACGACGCTCTTCCGATCTNNNTTATGGACTGGGAGGAAGA
GAGCACACGTCTGAACTCTTTCCCTACACGACGCTCTTCCGATCTNNNTGGTAGATAGGATCTGCTGG
AGCACACGTCTGAACTCTTTCCCTACACGACGCTCTTCCGATCTNNNCTTCCCAGGGTTGTTGTGAAG
AGAGCACACGTCTGAACTCTTTCCCTACACGACGCTCTTCCGATCTNNNTGCCAAACCCTGATAGGTAG
AGCACACGTCTGAACTCTTTCCCTACACGACGCTCTTCCGATCTNNNTGTTGAGCTGGTATGCAGAAA
AGAGCACACGTCTGAACTCTTTCCCTACACGACGCTCTTCCGATCTNNNATTGATGTGGACAAGACTG
AGAGCACACGTCTGAACTCTTTCCCTACACGACGCTCTTCCGATCTNNNACCTACTTTTGGATGGTTT
AGCACACGTCTGAACTCTTTCCCTACACGACGCTCTTCCGATCTNNNTGTTGTGAGATGGCCACAAGT
GAGCACACGTCTGAACTCTTTCCCTACACGACGCTCTTCCGATCTNNNCCCATATGACTAGTTTCCCAG
GAGCACACGTCTGAACTCTTTCCCTACACGACGCTCTTCCGATCTNNNAGCATTTCTCACTGACAGCAA
GAGCACACGTCTGAACTCTTTCCCTACACGACGCTCTTCCGATCTNNNTTTTCTTGCGTGGGAGTGTT
AGCACACGTCTGAACTCTTTCCCTACACGACGCTCTTCCGATCTNNNTAACCCTAAGGAGGGCTTACG
AGCACACGTCTGAACTCTTTCCCTACACGACGCTCTTCCGATCTNNNCGGAGAGGGACCATGTCTCTA
GAGCACACGTCTGAACTCTTTCCCTACACGACGCTCTTCCGATCTNNNGGGTTAGTGGTGGTGGAG
AGAGCACACGTCTGAACTCTTTCCCTACACGACGCTCTTCCGATCTNNNCAAATCAAGCAGAGGACAT
GAGCACACGTCTGAACTCTTTCCCTACACGACGCTCTTCCGATCTNNNATGTCTTGGAGAAAGCAAGC
GAGCACACGTCTGAACTCTTTCCCTACACGACGCTCTTCCGATCTNNNCTCCCTAAAGTGTCTGCCACA
GAGCACACGTCTGAACTCTTTCCCTACACGACGCTCTTCCGATCTNNNTAACAGTCAGGGTACCACCA
GAGCACACGTCTGAACTCTTTCCCTACACGACGCTCTTCCGATCTNNNTTCTTGTGCATCCTCTATGTG
AGCACACGTCTGAACTCTTTCCCTACACGACGCTCTTCCGATCTNNNCCACTAAGAGAATGCCCAATGG
AGAGCACACGTCTGAACTCTTTCCCTACACGACGCTCTTCCGATCTNNNACAGACAGTCGTTTCTCTCCT
AGCACACGTCTGAACTCTTTCCCTACACGACGCTCTTCCGATCTNNNATGGGCTCCCAACAGATTGAA
AGCACACGTCTGAACTCTTTCCCTACACGACGCTCTTCCGATCTNNNCAAAGGCTGCAAAATGAGTGT
AGCACACGTCTGAACTCTTTCCCTACACGACGCTCTTCCGATCTNNNGCAGGAGAATGCTTGATCCCA
AGAGCACACGTCTGAACTCTTTCCCTACACGACGCTCTTCCGATCTNNNAGGAATTCAATCCCACTCCC
AGCACACGTCTGAACTCTTTCCCTACACGACGCTCTTCCGATCTNNNTTTCAGGTTGGTTGCTATGTCC
GAGCACACGTCTGAACTCTTTCCCTACACGACGCTCTTCCGATCTNNNTCACAAGTCTCTGACACAACA
GAGCACACGTCTGAACTCTTTCCCTACACGACGCTCTTCCGATCTNNNACTGGATTCAGTCACGCCTTT
AGCACACGTCTGAACTCTTTCCCTACACGACGCTCTTCCGATCTNNNTGGAGTGATTCTTTTCAGCAG
GAGCACACGTCTGAACTCTTTCCCTACACGACGCTCTTCCGATCTNNNAAAAATGAAGCTTTGCAGCA
AGCACACGTCTGAACTCTTTCCCTACACGACGCTCTTCCGATCTNNNTCTGCTTCTGTGCAACTTCTGTT
AGCACACGTCTGAACTCTTTCCCTACACGACGCTCTTCCGATCTNNNGATCACCTACCTTCTGGTGACC

GAGCACACGTCTGAACTCTTTCCCTACACGACGCTCTTCCGATCTNNNAGTTGTGGTGCTTAGGGAA
GAGCACACGTCTGAACTCTTTCCCTACACGACGCTCTTCCGATCTNNNGCCACAACGCTAGTCATAGT
GCACACGTCTGAACTCTTTCCCTACACGACGCTCTTCCGATCTNNNAGAAGGATTGGGTCAGGTGAT
AGCACACGTCTGAACTCTTTCCCTACACGACGCTCTTCCGATCTNNNACAGGCTTTGCATATTTCACCG
AGCACACGTCTGAACTCTTTCCCTACACGACGCTCTTCCGATCTNNNACTACGCTAAGCACTTTACCT
GAGCACACGTCTGAACTCTTTCCCTACACGACGCTCTTCCGATCTNNNGAGAGCTCTGATTGGCCAA
AGCACACGTCTGAACTCTTTCCCTACACGACGCTCTTCCGATCTNNNTTACAAGCAGTCCAAGAGCAT
AGCACACGTCTGAACTCTTTCCCTACACGACGCTCTTCCGATCTNNNTTTGTCTTTTGCAGGCACTCT
AGCACACGTCTGAACTCTTTCCCTACACGACGCTCTTCCGATCTNNNACACAGGCTCGTTCATTCACA
AGCACACGTCTGAACTCTTTCCCTACACGACGCTCTTCCGATCTNNNAAACTTTCTCTCTCACCCCGGTA
AGCACACGTCTGAACTCTTTCCCTACACGACGCTCTTCCGATCTNNNCAGCTCTTAGAAAAACACAG
AGCACACGTCTGAACTCTTTCCCTACACGACGCTCTTCCGATCTNNNTAAAGCTCACACTTGGAAGCA
GCACACGTCTGAACTCTTTCCCTACACGACGCTCTTCCGATCTNNNAATGATGGGGGACAGACATTG
AGAGCACACGTCTGAACTCTTTCCCTACACGACGCTCTTCCGATCTNNNTGAACTTTGGCCATCTACAG
AGCACACGTCTGAACTCTTTCCCTACACGACGCTCTTCCGATCTNNNGAGGCAGGTACAGCACAGAA
GCACACGTCTGAACTCTTTCCCTACACGACGCTCTTCCGATCTNNNTGTTCCCCTCCTTCTTTTTCTGCC
AGCACACGTCTGAACTCTTTCCCTACACGACGCTCTTCCGATCTNNNTGATAACACCAGTAAGCTGTG
GAGCACACGTCTGAACTCTTTCCCTACACGACGCTCTTCCGATCTNNNGGAAAAAGCAGCTATCTGAT
GAGCACACGTCTGAACTCTTTCCCTACACGACGCTCTTCCGATCTNNNTACGCCAACTTCTTATGCTGT
AGCACACGTCTGAACTCTTTCCCTACACGACGCTCTTCCGATCTNNNTACACAAGCATGTATGCACAG
GAGCACACGTCTGAACTCTTTCCCTACACGACGCTCTTCCGATCTNNNTTCTTGACCTTGAGATCCGAC
AGCACACGTCTGAACTCTTTCCCTACACGACGCTCTTCCGATCTNNNTTGATATTTGTTTGGCAGGCAG
AGCACACGTCTGAACTCTTTCCCTACACGACGCTCTTCCGATCTNNNATGGCTTGCTATTGCAGGGTT
AGCACACGTCTGAACTCTTTCCCTACACGACGCTCTTCCGATCTNNNCTTTAGGAAGAGGACTGTGG
AGCACACGTCTGAACTCTTTCCCTACACGACGCTCTTCCGATCTNNNTGAAGAAGTCCTCTCTGCTCCA
AGCACACGTCTGAACTCTTTCCCTACACGACGCTCTTCCGATCTNNNATAAGGAGTGCTGAGGAAG
AGCACACGTCTGAACTCTTTCCCTACACGACGCTCTTCCGATCTNNNGTATTGCTAATCTCTGGGGTG
AGCACACGTCTGAACTCTTTCCCTACACGACGCTCTTCCGATCTNNNAAGGAAGGTCCAAGTCTTTGCT
GCACACGTCTGAACTCTTTCCCTACACGACGCTCTTCCGATCTNNNTTGTGTTTTGTTTTGAGGGCCA
AGCACACGTCTGAACTCTTTCCCTACACGACGCTCTTCCGATCTNNNCCGGCCTATCAGTGGGTTTTT
AGCACACGTCTGAACTCTTTCCCTACACGACGCTCTTCCGATCTNNNTTGGGTCGTAAGGGTGGATTT
GAGCACACGTCTGAACTCTTTCCCTACACGACGCTCTTCCGATCTNNNCAGGTATCAGATAAGTGTGG
AGCACACGTCTGAACTCTTTCCCTACACGACGCTCTTCCGATCTNNNTGGTTTGAAATTGCAAGTGTC
AGCACACGTCTGAACTCTTTCCCTACACGACGCTCTTCCGATCTNNNCTTGCCTCATCCCCCATTACTT
AGCACACGTCTGAACTCTTTCCCTACACGACGCTCTTCCGATCTNNNCCTTTCGGGATAGTCGCAAGA
AGCACACGTCTGAACTCTTTCCCTACACGACGCTCTTCCGATCTNNNTCAATGAGTGACCTTTTTGCAG
AGCACACGTCTGAACTCTTTCCCTACACGACGCTCTTCCGATCTNNNTGTGAAGGAAAAACCTGGTGA
AGCACACGTCTGAACTCTTTCCCTACACGACGCTCTTCCGATCTNNNATGGGACTTTCAGTGGTAAAC
AGCACACGTCTGAACTCTTTCCCTACACGACGCTCTTCCGATCTNNNCATGCACCATCACCCTGGATAA
GAGCACACGTCTGAACTCTTTCCCTACACGACGCTCTTCCGATCTNNNCTAAGGATGCTTGTGTTGCC
GAGCACACGTCTGAACTCTTTCCCTACACGACGCTCTTCCGATCTNNNGAAATGCTGCTCTGATAGGG
AGCACACGTCTGAACTCTTTCCCTACACGACGCTCTTCCGATCTNNNGTAAGCTCCAGTCCATGTCCAA
AGCACACGTCTGAACTCTTTCCCTACACGACGCTCTTCCGATCTNNNCTGAGCACAGTCTCTCTCTCTC

GAGCACACGTCTGAACTCTTTCCCTACACGACGCTCTTCCGATCTNNNTTCCTTCCAAGACAAAGCCCA
GAGCACACGTCTGAACTCTTTCCCTACACGACGCTCTTCCGATCTNNNGCCCAGCCTAAGATCCTATGT
AGAGCACACGTCTGAACTCTTTCCCTACACGACGCTCTTCCGATCTNNNGGAGACAGAAGGAGAAAG
AGCACACGTCTGAACTCTTTCCCTACACGACGCTCTTCCGATCTNNNTCCTAGAAGCAGCTTTCCCAAT
AGCACACGTCTGAACTCTTTCCCTACACGACGCTCTTCCGATCTNNNTTGATTTTTCTCTCCCCCACC
GAGCACACGTCTGAACTCTTTCCCTACACGACGCTCTTCCGATCTNNNTACGGGCTAAGAAGAGATG
GAGCACACGTCTGAACTCTTTCCCTACACGACGCTCTTCCGATCTNNNCGTTTCCCACCCAGAGAAGAT
GAGCACACGTCTGAACTCTTTCCCTACACGACGCTCTTCCGATCTNNNTGAACCTGAGCTCTGATCCT
AGCACACGTCTGAACTCTTTCCCTACACGACGCTCTTCCGATCTNNNCTGCCTTCAAAGGCCTACAATC
AGCACACGTCTGAACTCTTTCCCTACACGACGCTCTTCCGATCTNNNCCTTGGGGGTTTTGCTAGGAT
AGCACACGTCTGAACTCTTTCCCTACACGACGCTCTTCCGATCTNNNTGCCAGAACAAATGCACTAGG
GAGCACACGTCTGAACTCTTTCCCTACACGACGCTCTTCCGATCTNNNCATTCCTCTCTGCCTCTACCAG
GAGCACACGTCTGAACTCTTTCCCTACACGACGCTCTTCCGATCTNNNAATTTGGGTTGACTACCACCA
AGCACACGTCTGAACTCTTTCCCTACACGACGCTCTTCCGATCTNNNGGGCTCCAACAGAGAAACAG
GAGCACACGTCTGAACTCTTTCCCTACACGACGCTCTTCCGATCTNNNGCTGAATTTCAACCATCCCCA
AGCACACGTCTGAACTCTTTCCCTACACGACGCTCTTCCGATCTNNNAACAACTGCTCAGTTCCTTCCT
GCACACGTCTGAACTCTTTCCCTACACGACGCTCTTCCGATCTNNNCTATGGTGCCAGATTTAGGCAG
AGCACACGTCTGAACTCTTTCCCTACACGACGCTCTTCCGATCTNNNGCAACGATGCCATTGGAAAGA
AGCACACGTCTGAACTCTTTCCCTACACGACGCTCTTCCGATCTNNNACCTCCAGTCCTTCTTTACCCT
AGCACACGTCTGAACTCTTTCCCTACACGACGCTCTTCCGATCTNNNATTCTTCACGGTGCTGTATCCT
GAGCACACGTCTGAACTCTTTCCCTACACGACGCTCTTCCGATCTNNNGTAGACCTGAGGGACCATTC
AGCACACGTCTGAACTCTTTCCCTACACGACGCTCTTCCGATCTNNNGGAAAATCTGGCAACATTGAG
GAGCACACGTCTGAACTCTTTCCCTACACGACGCTCTTCCGATCTNNNGCCACCTTTTAGCATCCACCT
AGCACACGTCTGAACTCTTTCCCTACACGACGCTCTTCCGATCTNNNAGGCCTTAGCCAAACAAATCCA
AGCACACGTCTGAACTCTTTCCCTACACGACGCTCTTCCGATCTNNNTGGCCCTGAAGATCCAGCTAT
AGCACACGTCTGAACTCTTTCCCTACACGACGCTCTTCCGATCTNNNTGTGGATGAAGGTTGTCTGGG
AGCACACGTCTGAACTCTTTCCCTACACGACGCTCTTCCGATCTNNNCCCACTAGATGCTCTTGGAGAG
AGCACACGTCTGAACTCTTTCCCTACACGACGCTCTTCCGATCTNNNCCAGGGAAAGGACATGATCT
GAGCACACGTCTGAACTCTTTCCCTACACGACGCTCTTCCGATCTNNNATATAGGAGGGACTGAGTG
AGCACACGTCTGAACTCTTTCCCTACACGACGCTCTTCCGATCTNNNGCTGACTACCAAGCCATCATT
AGCACACGTCTGAACTCTTTCCCTACACGACGCTCTTCCGATCTNNNATCCCAAAGGCCTCTCTTGACT
AGCACACGTCTGAACTCTTTCCCTACACGACGCTCTTCCGATCTNNNATAGTCACATAGGGGCTTAGG
AGCACACGTCTGAACTCTTTCCCTACACGACGCTCTTCCGATCTNNNCCAAATGTCCACCGACAGATA
GAGCACACGTCTGAACTCTTTCCCTACACGACGCTCTTCCGATCTNNNGGTCCTATTTTGGCTCCTTCT
AGCACACGTCTGAACTCTTTCCCTACACGACGCTCTTCCGATCTNNNCAAAGTCTTGGCTGCTCTGTCT
GAGCACACGTCTGAACTCTTTCCCTACACGACGCTCTTCCGATCTNNNGACACATACATGCGGGTCTG
GCACACGTCTGAACTCTTTCCCTACACGACGCTCTTCCGATCTNNNCCCCTCCCTTGTTTCATTAGCTCC
GAGCACACGTCTGAACTCTTTCCCTACACGACGCTCTTCCGATCTNNNTTGTAGGAGACCACCTTCTC
GCACACGTCTGAACTCTTTCCCTACACGACGCTCTTCCGATCTNNNTTATACCCCGTTACATCCCTCCCC
AGCACACGTCTGAACTCTTTCCCTACACGACGCTCTTCCGATCTNNNTAATGATTGGGCTCTGGCTGA
GAGCACACGTCTGAACTCTTTCCCTACACGACGCTCTTCCGATCTNNNAGCATGCTGGCATATATCCT
AGCACACGTCTGAACTCTTTCCCTACACGACGCTCTTCCGATCTNNNAAATAGCCCATAGTGTGTGTG
AGCACACGTCTGAACTCTTTCCCTACACGACGCTCTTCCGATCTNNNATAGGGCCCTGCTTTGTCATCT

AGCACACGTCTGAACTCTTTCCCTACACGACGCTCTTCCGATCTNNNGGGGATGGAGGCAGATGAAT
AGCACACGTCTGAACTCTTTCCCTACACGACGCTCTTCCGATCTNNNAAAATGATAGCCGTCCTGTTG
GAGCACACGTCTGAACTCTTTCCCTACACGACGCTCTTCCGATCTNNNATGGGCTAATCCTTATCTGG
AGCACACGTCTGAACTCTTTCCCTACACGACGCTCTTCCGATCTNNNGAGGGAGGTGGACAAATGG
GAGCACACGTCTGAACTCTTTCCCTACACGACGCTCTTCCGATCTNNNCTCCCTGAAGTTGCTTGGGA
AGCACACGTCTGAACTCTTTCCCTACACGACGCTCTTCCGATCTNNNAATCCTTCTTTCTGTGCTCTGCC
GCACACGTCTGAACTCTTTCCCTACACGACGCTCTTCCGATCTNNNCCAGCTGGTCCTAAACAGAAAC
AGCACACGTCTGAACTCTTTCCCTACACGACGCTCTTCCGATCTNNNCCTAGCTTGGGGTGCAGAAAT
GAGCACACGTCTGAACTCTTTCCCTACACGACGCTCTTCCGATCTNNNAGTCAGGGTTTCTCAGACAG
GAGCACACGTCTGAACTCTTTCCCTACACGACGCTCTTCCGATCTNNNTGACTTCTGTGTGAGTGTGTT
AGCACACGTCTGAACTCTTTCCCTACACGACGCTCTTCCGATCTNNNTGCCCTCACATCCATCCTATCTT
AGCACACGTCTGAACTCTTTCCCTACACGACGCTCTTCCGATCTNNNCTTGAGGTTTGGGGATGAGT
GAGCACACGTCTGAACTCTTTCCCTACACGACGCTCTTCCGATCTNNNGCCTGTATCCCTTTCCATAAG
GCACACGTCTGAACTCTTTCCCTACACGACGCTCTTCCGATCTNNNTGATGCCATAATCTCCAAAGCC
AGCACACGTCTGAACTCTTTCCCTACACGACGCTCTTCCGATCTNNNTCAAGATCCAGCCTGTGGTTTA
AGCACACGTCTGAACTCTTTCCCTACACGACGCTCTTCCGATCTNNNTATAGGGCTGTGTGGGAACA
AGCACACGTCTGAACTCTTTCCCTACACGACGCTCTTCCGATCTNNNTCCTGTCACTTACAGAAGCTCC
GAGCACACGTCTGAACTCTTTCCCTACACGACGCTCTTCCGATCTNNNTCCCACAGATCCTCCATGAAG
AGCACACGTCTGAACTCTTTCCCTACACGACGCTCTTCCGATCTNNNAAGCAAGGTTGTGATTTGACT
GCACACGTCTGAACTCTTTCCCTACACGACGCTCTTCCGATCTNNNCAAACTTTAGCCTGTTTGGGCAC
GCACACGTCTGAACTCTTTCCCTACACGACGCTCTTCCGATCTNNNTGGAGAACTGCAAGACAGTAGG
GCACACGTCTGAACTCTTTCCCTACACGACGCTCTTCCGATCTNNNAGACCTTTGTACACAGCCTACCT
GCACACGTCTGAACTCTTTCCCTACACGACGCTCTTCCGATCTNNNCCACCCAATCTGCCTCTATCACT
GAGCACACGTCTGAACTCTTTCCCTACACGACGCTCTTCCGATCTNNNGTCCTCTGCTTCTTCGTGTTCA
GCACACGTCTGAACTCTTTCCCTACACGACGCTCTTCCGATCTNNNAGGCAGGCCTGTAAATTATCTGC
AGCACACGTCTGAACTCTTTCCCTACACGACGCTCTTCCGATCTNNNTTCTGCCCACCATACAGCTACT
AGCACACGTCTGAACTCTTTCCCTACACGACGCTCTTCCGATCTNNNTTTGCCAGTTCCCTAGTTTCCAC
AGCACACGTCTGAACTCTTTCCCTACACGACGCTCTTCCGATCTNNNTACAACCCACTGACATCCCTGA
GCACACGTCTGAACTCTTTCCCTACACGACGCTCTTCCGATCTNNNCAAGGGCATTCTTTTGCCTTCAG
GCACACGTCTGAACTCTTTCCCTACACGACGCTCTTCCGATCTNNNTTATCCATACACACCATGCCAGC
GAGCACACGTCTGAACTCTTTCCCTACACGACGCTCTTCCGATCTNNNAGGGGATGAATGTGTGCTTT
GCACACGTCTGAACTCTTTCCCTACACGACGCTCTTCCGATCTNNNAAATGCCTCCTAAGGGTGTTCT
GAGCACACGTCTGAACTCTTTCCCTACACGACGCTCTTCCGATCTNNNTTGATTACGCTCTCCTGCCTTT
AGCACACGTCTGAACTCTTTCCCTACACGACGCTCTTCCGATCTNNNCTGGCACATGTCTACTAGAGG
GCACACGTCTGAACTCTTTCCCTACACGACGCTCTTCCGATCTNNNCTGCCCTTTTCTCACTTCCTCTCC
GAGCACACGTCTGAACTCTTTCCCTACACGACGCTCTTCCGATCTNNNACATGAAGCTCATGCAGTACC
AGCACACGTCTGAACTCTTTCCCTACACGACGCTCTTCCGATCTNNNCACAGTCATCCCTCCCTAAGAC
GAGCACACGTCTGAACTCTTTCCCTACACGACGCTCTTCCGATCTNNNGTAATACAGGCATGTGCCACA
GAGCACACGTCTGAACTCTTTCCCTACACGACGCTCTTCCGATCTNNNCACTGAACACAGGGAAGTCA
AGCACACGTCTGAACTCTTTCCCTACACGACGCTCTTCCGATCTNNNTCGTCTCTTGTATCCTACACGCA
GAGCACACGTCTGAACTCTTTCCCTACACGACGCTCTTCCGATCTNNNCTCCATACAGCTTTCCATGG
AGCACACGTCTGAACTCTTTCCCTACACGACGCTCTTCCGATCTNNNCTATGCTCATTGGCGTAGCACA
AGCACACGTCTGAACTCTTTCCCTACACGACGCTCTTCCGATCTNNNGATCACCTGTGTGTGATCTGCT

AGCACACGTCTGAACTCTTTCCCTACACGACGCTCTTCCGATCTNNNTTTCCCAATGCCTACACACAGA
AGCACACGTCTGAACTCTTTCCCTACACGACGCTCTTCCGATCTNNNCCACACCTTGGAAATGGATTTG
AGAGCACACGTCTGAACTCTTTCCCTACACGACGCTCTTCCGATCTNNNAAGACTGAATGTGGTGTGT
AGCACACGTCTGAACTCTTTCCCTACACGACGCTCTTCCGATCTNNNCAGAATCCAGAGAGGAAGGG
AGCACACGTCTGAACTCTTTCCCTACACGACGCTCTTCCGATCTNNNTGGATTAGCATGTCCAATGCCT
GAGCACACGTCTGAACTCTTTCCCTACACGACGCTCTTCCGATCTNNNTGGGCAATAGCAAGTCCCTAT
AGCACACGTCTGAACTCTTTCCCTACACGACGCTCTTCCGATCTNNNAGGAAATCACCTCACCCCAAGTA
GAGCACACGTCTGAACTCTTTCCCTACACGACGCTCTTCCGATCTNNNCACCCACCGTACTCCATATACCT
GAGCACACGTCTGAACTCTTTCCCTACACGACGCTCTTCCGATCTNNNGTCTCTGGTTTTGTGCAGTTTG
GAGCACACGTCTGAACTCTTTCCCTACACGACGCTCTTCCGATCTNNNGGAGACACCCTAAATGTGTCC
AGCACACGTCTGAACTCTTTCCCTACACGACGCTCTTCCGATCTNNNGTAAGAGGTAGCTCTGCCCTAC
GAGCACACGTCTGAACTCTTTCCCTACACGACGCTCTTCCGATCTNNNGTCATTTTGTACACGGATGTG
GAGCACACGTCTGAACTCTTTCCCTACACGACGCTCTTCCGATCTNNNATCTAGAATCCCAGCCTAGTG
GAGCACACGTCTGAACTCTTTCCCTACACGACGCTCTTCCGATCTNNNGCACCATGCATACTCTCAAGTC
AGCACACGTCTGAACTCTTTCCCTACACGACGCTCTTCCGATCTNNNTTGAGAGCAGGGCCTCATATAC
AGCACACGTCTGAACTCTTTCCCTACACGACGCTCTTCCGATCTNNNGGATTCGGATCAAAGGAGCTG
AGCACACGTCTGAACTCTTTCCCTACACGACGCTCTTCCGATCTNNNCGTCTACTCAGTGTCACGGTTTT
GAGCACACGTCTGAACTCTTTCCCTACACGACGCTCTTCCGATCTNNNCAGAAAAAGGACACTCTGAG
AGCACACGTCTGAACTCTTTCCCTACACGACGCTCTTCCGATCTNNNGTCCTTGCCTAATGACCACAGA
AGCACACGTCTGAACTCTTTCCCTACACGACGCTCTTCCGATCTNNNAGGAAAGCTTTTAGCCCTCATG
GAGCACACGTCTGAACTCTTTCCCTACACGACGCTCTTCCGATCTNNNACACACAGCTTCTAACAACCAA
AGCACACGTCTGAACTCTTTCCCTACACGACGCTCTTCCGATCTNNNATGATTTGCTTCGCATAGACAG
GAGCACACGTCTGAACTCTTTCCCTACACGACGCTCTTCCGATCTNNNTATGTCTCCGTGCTTACCAGT
AGCACACGTCTGAACTCTTTCCCTACACGACGCTCTTCCGATCTNNNTGACTATTTGCACATCTGGCTCC
AGAGCACACGTCTGAACTCTTTCCCTACACGACGCTCTTCCGATCTNNNCCCCAAGAGTGTCAATTTC
GCACACGTCTGAACTCTTTCCCTACACGACGCTCTTCCGATCTNNNCTCTGCTCCAGTTCCTCAACATCC
GAGCACACGTCTGAACTCTTTCCCTACACGACGCTCTTCCGATCTNNNCCCAAAAGTATGACCATGTG
AGAGCACACGTCTGAACTCTTTCCCTACACGACGCTCTTCCGATCTNNNTTCCAGCCACTGACAGAGTA
AGCACACGTCTGAACTCTTTCCCTACACGACGCTCTTCCGATCTNNNAGTTTTCACCGTCTGCTTCAGT
GAGCACACGTCTGAACTCTTTCCCTACACGACGCTCTTCCGATCTNNNCACATCCACGTTCTAGGCAGA
GAGCACACGTCTGAACTCTTTCCCTACACGACGCTCTTCCGATCTNNNTGGGTTTCAGCCTAAAGCTAC
AGAGCACACGTCTGAACTCTTTCCCTACACGACGCTCTTCCGATCTNNNAATGGAATTTGCCACAGAG
GAGCACACGTCTGAACTCTTTCCCTACACGACGCTCTTCCGATCTNNNGCATGGTGGTAGGTATGATT
GAGCACACGTCTGAACTCTTTCCCTACACGACGCTCTTCCGATCTNNNTGGAGACGAGTTCATGAGTT
AGCACACGTCTGAACTCTTTCCCTACACGACGCTCTTCCGATCTNNNTTAGCCGTCTTTCCCCTATTTGC
GAGCACACGTCTGAACTCTTTCCCTACACGACGCTCTTCCGATCTNNNAGCTGTGAGAGGGTGGTAA
GAGCACACGTCTGAACTCTTTCCCTACACGACGCTCTTCCGATCTNNNAGTGCACTAATAGGGGACAC
AGCACACGTCTGAACTCTTTCCCTACACGACGCTCTTCCGATCTNNNGGCAGCCAAAGGGTGAATCTA
AGCACACGTCTGAACTCTTTCCCTACACGACGCTCTTCCGATCTNNNAGCAAAACAAAACCCTGCAAGA
GAGCACACGTCTGAACTCTTTCCCTACACGACGCTCTTCCGATCTNNNGTACAGAGGACATCACATGC
AGCACACGTCTGAACTCTTTCCCTACACGACGCTCTTCCGATCTNNNTAACAGCAAGATATAGGCCCT
AGCACACGTCTGAACTCTTTCCCTACACGACGCTCTTCCGATCTNNNCACCTTTTCCTCTGTTCCAGTCA
AGCACACGTCTGAACTCTTTCCCTACACGACGCTCTTCCGATCTNNNGGATATGCACTCAAAACACTG

GAGCACACGTCTGAACTCTTTCCCTACACGACGCTCTTCCGATCTNNNGTCCCTAAAGGAATGCTGTCA
AGCACACGTCTGAACTCTTTCCCTACACGACGCTCTTCCGATCTNNNCTATGCCAAGCACTCATTCAGGT
AGCACACGTCTGAACTCTTTCCCTACACGACGCTCTTCCGATCTNNNCTTAGCAAGTGGCTTCCAACTCT
AGCACACGTCTGAACTCTTTCCCTACACGACGCTCTTCCGATCTNNNTTGCTTCCAAGATTTCCAGCAGT
AGCACACGTCTGAACTCTTTCCCTACACGACGCTCTTCCGATCTNNNACACATTCATATGCAGTTGCCCA
GAGCACACGTCTGAACTCTTTCCCTACACGACGCTCTTCCGATCTNNNTACGCTGCTTCTACCAAGTGAT
GAGCACACGTCTGAACTCTTTCCCTACACGACGCTCTTCCGATCTNNNTGTGTAAGAGCATGGGGGA
GAGCACACGTCTGAACTCTTTCCCTACACGACGCTCTTCCGATCTNNNAGAGATTGCATGGGTAGAGC
GAGCACACGTCTGAACTCTTTCCCTACACGACGCTCTTCCGATCTNNNACTCAGTGATTGCCAAGGAAG
GAGCACACGTCTGAACTCTTTCCCTACACGACGCTCTTCCGATCTNNNTCCAGTGACACATATGCACTC
GCACACGTCTGAACTCTTTCCCTACACGACGCTCTTCCGATCTNNNCCTAGTTTGATATGGCTGGAGG
GAGCACACGTCTGAACTCTTTCCCTACACGACGCTCTTCCGATCTNNNATGATGTGGTATGTATGTGTG
AGCACACGTCTGAACTCTTTCCCTACACGACGCTCTTCCGATCTNNNGCCAGCTGAGCCTACTCTTTTA
AGCACACGTCTGAACTCTTTCCCTACACGACGCTCTTCCGATCTNNNTGCAGGAAGAAGGAACAAAGG
GAGCACACGTCTGAACTCTTTCCCTACACGACGCTCTTCCGATCTNNNGCCTTGCTGTACTTTAGAAGG
GAGCACACGTCTGAACTCTTTCCCTACACGACGCTCTTCCGATCTNNNTCTTACTGGAGCTACCATAGG
AGAGCACACGTCTGAACTCTTTCCCTACACGACGCTCTTCCGATCTNNNAAGGGGTAACTTAGCAGGG
GAGCACACGTCTGAACTCTTTCCCTACACGACGCTCTTCCGATCTNNNGCAACACAATCCTCTTCATCCC
AGCACACGTCTGAACTCTTTCCCTACACGACGCTCTTCCGATCTNNNAGCATGCTTTTGTATGAGGCA
AGCACACGTCTGAACTCTTTCCCTACACGACGCTCTTCCGATCTNNNGGAAAAGGGAGGTGAAGGAA
GAGCACACGTCTGAACTCTTTCCCTACACGACGCTCTTCCGATCTNNNTCATCTTGTGTTTGGACAGTG
GAGCACACGTCTGAACTCTTTCCCTACACGACGCTCTTCCGATCTNNNAGCAGATAGTGTCCTGGTAG
AGCACACGTCTGAACTCTTTCCCTACACGACGCTCTTCCGATCTNNNTAGGAGTTGAAAACACACGCAC
AGCACACGTCTGAACTCTTTCCCTACACGACGCTCTTCCGATCTNNNTGTAGCTGAGGGAGAATGCTA
AGCACACGTCTGAACTCTTTCCCTACACGACGCTCTTCCGATCTNNNTTCTTTTCTGCCTTCCCCAACTCCA
AGCACACGTCTGAACTCTTTCCCTACACGACGCTCTTCCGATCTNNNGAGGAAGATCAGGAAATCCCAG
GAGCACACGTCTGAACTCTTTCCCTACACGACGCTCTTCCGATCTNNNAGGGAGAGGAGATTGCTCAA
AGCACACGTCTGAACTCTTTCCCTACACGACGCTCTTCCGATCTNNNAGGCCCAGGTCGTCATACTTAT
GAGCACACGTCTGAACTCTTTCCCTACACGACGCTCTTCCGATCTNNNTGCCTCACTACTCCTGGAATTG
GAGCACACGTCTGAACTCTTTCCCTACACGACGCTCTTCCGATCTNNNCACACATTTTCCCAGCACATTCA
GAGCACACGTCTGAACTCTTTCCCTACACGACGCTCTTCCGATCTNNNAAGCTTACTAGGGAATGGAG
AGCACACGTCTGAACTCTTTCCCTACACGACGCTCTTCCGATCTNNNACCACTAATCCACTTCCTTGACCC
AGCACACGTCTGAACTCTTTCCCTACACGACGCTCTTCCGATCTNNNAGACAACCACAATTCTCTGAGC
AGCACACGTCTGAACTCTTTCCCTACACGACGCTCTTCCGATCTNNNAGCTTGGGCTGAATAATGAGCA
GAGCACACGTCTGAACTCTTTCCCTACACGACGCTCTTCCGATCTNNNAGAGGTAATGCATACATGGAA
AGCACACGTCTGAACTCTTTCCCTACACGACGCTCTTCCGATCTNNNAACTCAGTCTTTGGAGGGGCAT
GAGCACACGTCTGAACTCTTTCCCTACACGACGCTCTTCCGATCTNNNGAAAAACAACGGGGCCTGTAT
GCACACGTCTGAACTCTTTCCCTACACGACGCTCTTCCGATCTNNNGCAGGACCCTACATTATTTGCCAC
GCACACGTCTGAACTCTTTCCCTACACGACGCTCTTCCGATCTNNNCCTTGTTCACTCAAAGAGCACCTC
GAGCACACGTCTGAACTCTTTCCCTACACGACGCTCTTCCGATCTNNNGCTTTGCCTTCTGTGCTCTATG
GAGCACACGTCTGAACTCTTTCCCTACACGACGCTCTTCCGATCTNNNAGTCAAGCATCCAAAGACAGC
AGCACACGTCTGAACTCTTTCCCTACACGACGCTCTTCCGATCTNNNGAAAGGCACATGTGACAAAGAC
AGCACACGTCTGAACTCTTTCCCTACACGACGCTCTTCCGATCTNNNCCAGGTATCATTTTCTGAGCCC

GCACACGTCTGAACTCTTTCCCTACACGACGCTCTTCCGATCTNNNACATGGGGCTTTGGTAAAGAA

CACACGTCTGAACTCTTTCCCTACACGACGCTCTTCCGATCTNNNTCCTGTCTTATCTCCTTGGCTCCCA

GCACACGTCTGAACTCTTTCCCTACACGACGCTCTTCCGATCTNNNCCCTCTCTTCATCTTCACATGGCC

AGCACACGTCTGAACTCTTTCCCTACACGACGCTCTTCCGATCTNNNTTGGTTGTGTAGCAATCCCTC

AGCACACGTCTGAACTCTTTCCCTACACGACGCTCTTCCGATCTNNNGCAGGCCTTCTTACTGAATCCT

GAGCACACGTCTGAACTCTTTCCCTACACGACGCTCTTCCGATCTNNNTGGGGAGGGACATAAACTTC

AGCACACGTCTGAACTCTTTCCCTACACGACGCTCTTCCGATCTNNNGAGAAGTTTCCCCAAGGTTGG

AGCACACGTCTGAACTCTTTCCCTACACGACGCTCTTCCGATCTNNNGTCCAATGCCACTGGTGTTCT

AGCACACGTCTGAACTCTTTCCCTACACGACGCTCTTCCGATCTNNNTGAGGCATGTGAATGTCTTG

GCACACGTCTGAACTCTTTCCCTACACGACGCTCTTCCGATCTNNNTGAGACACACAAAGCACTGTG

GAGCACACGTCTGAACTCTTTCCCTACACGACGCTCTTCCGATCTNNNGTTTACGTTAGTCCTGTGGG

AGCACACGTCTGAACTCTTTCCCTACACGACGCTCTTCCGATCTNNNGGAAAGTACGTGGCATGGAA

GAGCACACGTCTGAACTCTTTCCCTACACGACGCTCTTCCGATCTNNNTGTGACGGAGGAAAATAAG

AGCACACGTCTGAACTCTTTCCCTACACGACGCTCTTCCGATCTNNNTCAACATGATGCACTAGAGGG

CACACGTCTGAACTCTTTCCCTACACGACGCTCTTCCGATCTNNNAGGACAAATATGTGCCTTGAGC

AGCACACGTCTGAACTCTTTCCCTACACGACGCTCTTCCGATCTNNNATGGGTTGGAATTTGTCTGC

AGCACACGTCTGAACTCTTTCCCTACACGACGCTCTTCCGATCTNNNCCTCCCAGTACCTACCTTTCTCC

AGCACACGTCTGAACTCTTTCCCTACACGACGCTCTTCCGATCTNNNTGAGACTGACAGAGAAGTCCC

AGCACACGTCTGAACTCTTTCCCTACACGACGCTCTTCCGATCTNNNCTGAACCTGAAAACAGGTCTG

AGCACACGTCTGAACTCTTTCCCTACACGACGCTCTTCCGATCTNNNATTCCACTTCTCTGCCTCCAGTA

AGCACACGTCTGAACTCTTTCCCTACACGACGCTCTTCCGATCTNNNAGTGAGAGCCACAAATTCCTC

AGCACACGTCTGAACTCTTTCCCTACACGACGCTCTTCCGATCTNNNAACAAGCCTTTTAGAGCAGGT

AGCACACGTCTGAACTCTTTCCCTACACGACGCTCTTCCGATCTNNNATAATCTCCCCCACCACACTTCT

AGCACACGTCTGAACTCTTTCCCTACACGACGCTCTTCCGATCTNNNATGTGAAAGAGATGCTGTGAC

AGCACACGTCTGAACTCTTTCCCTACACGACGCTCTTCCGATCTNNNACACCCTGCTGCATCATTTGTA

GCACACGTCTGAACTCTTTCCCTACACGACGCTCTTCCGATCTNNNTGCTTCCTTTACGTGGCTTTCTTC

AGCACACGTCTGAACTCTTTCCCTACACGACGCTCTTCCGATCTNNNTTTCCTAGTCCTGGTACTTGC

GCACACGTCTGAACTCTTTCCCTACACGACGCTCTTCCGATCTNNNTACCTAAGGTCCATCGTCAGGT

GCACACGTCTGAACTCTTTCCCTACACGACGCTCTTCCGATCTNNNATCTCTCCCTCACACACAGAAGT

GCACACGTCTGAACTCTTTCCCTACACGACGCTCTTCCGATCTNNNGCTGAAGGCTGTCTCTCTCTTTC

GCACACGTCTGAACTCTTTCCCTACACGACGCTCTTCCGATCTNNNTCCCATAGGAATGAAACCTCAA

CACACGTCTGAACTCTTTCCCTACACGACGCTCTTCCGATCTNNNTATTCTGCAGTTCAAAGGCCTGCC

AGCACACGTCTGAACTCTTTCCCTACACGACGCTCTTCCGATCTNNNTAGAAACCCAGGTCAGACACA

GAGCACACGTCTGAACTCTTTCCCTACACGACGCTCTTCCGATCTNNNGCAAAGAAACACTCGGAGTT

AGCACACGTCTGAACTCTTTCCCTACACGACGCTCTTCCGATCTNNNACCCTGAGTGAATTGCCTAAAG

AGCACACGTCTGAACTCTTTCCCTACACGACGCTCTTCCGATCTNNNTATTCCTGTGTGCATGCCTACC

AGCACACGTCTGAACTCTTTCCCTACACGACGCTCTTCCGATCTNNNAGCAGGAAATCCTGTGCATTC

GAGCACACGTCTGAACTCTTTCCCTACACGACGCTCTTCCGATCTNNNGTCAACACTGCTGAAGTTG

AGCACACGTCTGAACTCTTTCCCTACACGACGCTCTTCCGATCTNNNGTCCCTGGCAAGCATGAATTT

AGCACACGTCTGAACTCTTTCCCTACACGACGCTCTTCCGATCTNNNACGTGCACATATGGGAAAAT

AGCACACGTCTGAACTCTTTCCCTACACGACGCTCTTCCGATCTNNNTACCTGAAGTGGCATTCGATG

AGCACACGTCTGAACTCTTTCCCTACACGACGCTCTTCCGATCTNNNCGTTGGAATTCACACATCTCAG

GCACACGTCTGAACTCTTTCCCTACACGACGCTCTTCCGATCTNNNCACAGCCACTCTAGCTGATTCTC

GCACACGTCTGAACTCTTTCCCTACACGACGCTCTTCCGATCTNNNCAGCTACCACCCTTCATCAATCCC
GCACACGTCTGAACTCTTTCCCTACACGACGCTCTTCCGATCTNNNAGCACTATTTGGATTCTGCTTGT
AGCACACGTCTGAACTCTTTCCCTACACGACGCTCTTCCGATCTNNNTCAACGCCTTTCTCTCTCTCTCT
AGCACACGTCTGAACTCTTTCCCTACACGACGCTCTTCCGATCTNNNTCACTCTATGATCCTGGGCAAC
AGCACACGTCTGAACTCTTTCCCTACACGACGCTCTTCCGATCTNNNGCCTGTACATCAAAACAAAGC
AGCACACGTCTGAACTCTTTCCCTACACGACGCTCTTCCGATCTNNNTGTGTGTAAGTCATTGCCAACA
AGAGCACACGTCTGAACTCTTTCCCTACACGACGCTCTTCCGATCTNNNATGTCTGCAGGCTACATTCA
AGCACACGTCTGAACTCTTTCCCTACACGACGCTCTTCCGATCTNNNGCACACAGGTCCACTTCTTAG
GAGCACACGTCTGAACTCTTTCCCTACACGACGCTCTTCCGATCTNNNTCCTTGGTTGCTAACCTCCTG
GAGCACACGTCTGAACTCTTTCCCTACACGACGCTCTTCCGATCTNNNGAAACAGAGTAGTAGCAGC
AGAGCACACGTCTGAACTCTTTCCCTACACGACGCTCTTCCGATCTNNNATGGGAAGTGCACTGTACT
AGCACACGTCTGAACTCTTTCCCTACACGACGCTCTTCCGATCTNNNCCATCTCTCCTGGAGCCTAGTT
GAGCACACGTCTGAACTCTTTCCCTACACGACGCTCTTCCGATCTNNNTGTGCCTTCAGAGCTTCTCTT
AGCACACGTCTGAACTCTTTCCCTACACGACGCTCTTCCGATCTNNNGAGCTCTGCCCATGAGAGACT
AGCACACGTCTGAACTCTTTCCCTACACGACGCTCTTCCGATCTNNNCTGGAAAGATGTTGGTAGGG
AGCACACGTCTGAACTCTTTCCCTACACGACGCTCTTCCGATCTNNNTTGCAAGTTTATTGCACGCAT
AGCACACGTCTGAACTCTTTCCCTACACGACGCTCTTCCGATCTNNNATATATGCCAGTCACACTCCCC
AGCACACGTCTGAACTCTTTCCCTACACGACGCTCTTCCGATCTNNNTCGTACAAACCAGCCACTAAAG
GCACACGTCTGAACTCTTTCCCTACACGACGCTCTTCCGATCTNNNATTGAAGGGGATAGGGTGCTT
GCACACGTCTGAACTCTTTCCCTACACGACGCTCTTCCGATCTNNNATTTCTAGCCCAGTCAACCTGGA
GCACACGTCTGAACTCTTTCCCTACACGACGCTCTTCCGATCTNNNCTTTGGTCTCTGTTCCTCCTCCCC
AGCACACGTCTGAACTCTTTCCCTACACGACGCTCTTCCGATCTNNNAGAAGTGGCTTCCAAACTGTGT
GCACACGTCTGAACTCTTTCCCTACACGACGCTCTTCCGATCTNNNAGACTGACCCCTAAGTGCAAGAT
AGCACACGTCTGAACTCTTTCCCTACACGACGCTCTTCCGATCTNNNAGGATTTCAATGCCAAAGTCCC
GAGCACACGTCTGAACTCTTTCCCTACACGACGCTCTTCCGATCTNNNCACAGAGAATCACCTGCACA
AGCACACGTCTGAACTCTTTCCCTACACGACGCTCTTCCGATCTNNNTCTCCTTCTTGAGCACCTGATG
AGCACACGTCTGAACTCTTTCCCTACACGACGCTCTTCCGATCTNNNGGCCTATGGGAAAGAGTTGT
GCACACGTCTGAACTCTTTCCCTACACGACGCTCTTCCGATCTNNNGGTTTTCCTAATCTCCATGCCCCC
AGCACACGTCTGAACTCTTTCCCTACACGACGCTCTTCCGATCTNNNTCTCTCTCAGAAATCAACCCCG
AGCACACGTCTGAACTCTTTCCCTACACGACGCTCTTCCGATCTNNNGGATCTTCTCTGAGCATGCCTA
GAGCACACGTCTGAACTCTTTCCCTACACGACGCTCTTCCGATCTNNNGTGTCCCTTCCTACTCACTTCC
GAGCACACGTCTGAACTCTTTCCCTACACGACGCTCTTCCGATCTNNNCACTGCTGTCAGTAAGCATA
AGCACACGTCTGAACTCTTTCCCTACACGACGCTCTTCCGATCTNNNAGAAAAGTAGGCCAGTGTGT
GAGCACACGTCTGAACTCTTTCCCTACACGACGCTCTTCCGATCTNNNGTGTTCCCTATCTCTTTCCTCG
AGCACACGTCTGAACTCTTTCCCTACACGACGCTCTTCCGATCTNNNATTCTTTCCTACACCAGGGGTG
AGCACACGTCTGAACTCTTTCCCTACACGACGCTCTTCCGATCTNNNTGACCCACAGATCATAGCCTTC
AGCACACGTCTGAACTCTTTCCCTACACGACGCTCTTCCGATCTNNNGCAATTTTTGTTGTCCCATGG
AGCACACGTCTGAACTCTTTCCCTACACGACGCTCTTCCGATCTNNNTTTCATGGATGGAAAAAGCCA
GCACACGTCTGAACTCTTTCCCTACACGACGCTCTTCCGATCTNNNCCATGAAGATGATGTCCCACAG
AGCACACGTCTGAACTCTTTCCCTACACGACGCTCTTCCGATCTNNNATGCCTCTGTCTCATCCTCTCT
AGCACACGTCTGAACTCTTTCCCTACACGACGCTCTTCCGATCTNNNTCCCTATGTCTGCATTGCTCCT
AGCACACGTCTGAACTCTTTCCCTACACGACGCTCTTCCGATCTNNNAAGTGGGTCATTCCCACAGAC
AGCACACGTCTGAACTCTTTCCCTACACGACGCTCTTCCGATCTNNNATGTGTGAGAGACAGCGAGT

GAGCACACGTCTGAACTCTTTCCCTACACGACGCTCTTCCGATCTNNNGGTGTGGGATATGGTACTTG
AGCACACGTCTGAACTCTTTCCCTACACGACGCTCTTCCGATCTNNNGGGTACTTGGGTATCTTCACCC
AGCACACGTCTGAACTCTTTCCCTACACGACGCTCTTCCGATCTNNNTATACTCAAGAGAGACCCAGG
AGCACACGTCTGAACTCTTTCCCTACACGACGCTCTTCCGATCTNNNTGAAACAATGGCCTTTGGAGCA
GAGCACACGTCTGAACTCTTTCCCTACACGACGCTCTTCCGATCTNNNATCAATCCACTGCCTATGGGGT
GAGCACACGTCTGAACTCTTTCCCTACACGACGCTCTTCCGATCTNNNCAGAGAGTGGCACCCTTATAG
GAGCACACGTCTGAACTCTTTCCCTACACGACGCTCTTCCGATCTNNNAGGCAAATGGATTTCTTGAG
AGCACACGTCTGAACTCTTTCCCTACACGACGCTCTTCCGATCTNNNCATTCCTTGGCTTGCAAAGCAT
GAGCACACGTCTGAACTCTTTCCCTACACGACGCTCTTCCGATCTNNNTTAGTAGGGTCAGAGGCAGT
GCACACGTCTGAACTCTTTCCCTACACGACGCTCTTCCGATCTNNNATGGCTGTCTCATTCCAACCATCC
AGCACACGTCTGAACTCTTTCCCTACACGACGCTCTTCCGATCTNNNTGTAACATTAGGCCTGTTCCAG
GCACACGTCTGAACTCTTTCCCTACACGACGCTCTTCCGATCTNNNAAGCAGTGGTGCCTAAAGCTATC
AGCACACGTCTGAACTCTTTCCCTACACGACGCTCTTCCGATCTNNNGATGAGTGTGAGTAGTGCTGCA
AGCACACGTCTGAACTCTTTCCCTACACGACGCTCTTCCGATCTNNNAGACATGATTGCTAAGGTCCAA
GAGCACACGTCTGAACTCTTTCCCTACACGACGCTCTTCCGATCTNNNAAAAACAAGCCTGCTTGAGCA
AGCACACGTCTGAACTCTTTCCCTACACGACGCTCTTCCGATCTNNNAACATCCTGGGTACATGCACTT
AGCACACGTCTGAACTCTTTCCCTACACGACGCTCTTCCGATCTNNNAAAGCTACAAGTCCCAGTGCAA
AGCACACGTCTGAACTCTTTCCCTACACGACGCTCTTCCGATCTNNNGCAGTTCCAAGACCAATCAATG
AGCACACGTCTGAACTCTTTCCCTACACGACGCTCTTCCGATCTNNNATCTCATTGCATTACGGCCAGT
GAGCACACGTCTGAACTCTTTCCCTACACGACGCTCTTCCGATCTNNNCCTATCCCACTTCAGGGAAAG
GAGCACACGTCTGAACTCTTTCCCTACACGACGCTCTTCCGATCTNNNTTTCTCTCTCTCTCCCCCAGATG
AGCACACGTCTGAACTCTTTCCCTACACGACGCTCTTCCGATCTNNNCACCAGGCATCAGCAATAAGCTA
AGCACACGTCTGAACTCTTTCCCTACACGACGCTCTTCCGATCTNNNTCACTTCCTTCTGACCCTCCTTTC
GAGCACACGTCTGAACTCTTTCCCTACACGACGCTCTTCCGATCTNNNACCACCTTACCCCAGTAACGTA
GCACACGTCTGAACTCTTTCCCTACACGACGCTCTTCCGATCTNNNCTTGGGCTGAACTATCCTCAGTG
GAGCACACGTCTGAACTCTTTCCCTACACGACGCTCTTCCGATCTNNNTCAGATTCTCCTGAATCACCCC
AGCACACGTCTGAACTCTTTCCCTACACGACGCTCTTCCGATCTNNNATTCTTGGCAAAGAGGTGCTCT
AAGAGCACACGTCTGAACTCTTTCCCTACACGACGCTCTTCCGATCTNNNAGTCAAAGTCCACACCTATT
AGCACACGTCTGAACTCTTTCCCTACACGACGCTCTTCCGATCTNNNATCTTTGGGCTGTGATTGTTCCA
AGCACACGTCTGAACTCTTTCCCTACACGACGCTCTTCCGATCTNNNAACACGCAACCTTCTTCTTGTCT
GCACACGTCTGAACTCTTTCCCTACACGACGCTCTTCCGATCTNNNGCAGTAGAGGTTTCCTGGTTTC
AGCACACGTCTGAACTCTTTCCCTACACGACGCTCTTCCGATCTNNNAGGGAAGGCATCAGTTCCTTA
AGCACACGTCTGAACTCTTTCCCTACACGACGCTCTTCCGATCTNNNCTGTCTGTTTTCCGTGACTGTCT
GCACACGTCTGAACTCTTTCCCTACACGACGCTCTTCCGATCTNNNAACCAGGCTGCTTCTTTTGAGACC
AGCACACGTCTGAACTCTTTCCCTACACGACGCTCTTCCGATCTNNNTCTCTCTGATCACACCCCACTTCCA
AGCACACGTCTGAACTCTTTCCCTACACGACGCTCTTCCGATCTNNNCACAGATGATCCACGTGTTGCTA
GAGCACACGTCTGAACTCTTTCCCTACACGACGCTCTTCCGATCTNNNAATGTACTGATGCCACTGAAC
AGCACACGTCTGAACTCTTTCCCTACACGACGCTCTTCCGATCTNNNTGCAGAACTGTGAAGTCCTCCT
AGCACACGTCTGAACTCTTTCCCTACACGACGCTCTTCCGATCTNNNGAGTGAGATCCACTTTGGAGAC
AGCACACGTCTGAACTCTTTCCCTACACGACGCTCTTCCGATCTNNNATTTACCACACTACAGGGGTG
GCACACGTCTGAACTCTTTCCCTACACGACGCTCTTCCGATCTNNNAGTGAAATGTGAAGCAGCAAGG
GCACACGTCTGAACTCTTTCCCTACACGACGCTCTTCCGATCTNNNACTGTCATCTCCTTGGTATGGGG
GAGCACACGTCTGAACTCTTTCCCTACACGACGCTCTTCCGATCTNNNCCCTCAAGCGATTCACCCACTT

GAGCACACGTCTGAACTCTTTCCCTACACGACGCTCTTCCGATCTNNNACACAGCCAGATTTGGAACCA
GCACACGTCTGAACTCTTTCCCTACACGACGCTCTTCCGATCTNNNTTTGTTCTTTCCCAGCCCTACACCA
AGCACACGTCTGAACTCTTTCCCTACACGACGCTCTTCCGATCTNNNATACTGTGTGTGGATGTAGGG
GAGCACACGTCTGAACTCTTTCCCTACACGACGCTCTTCCGATCTNNNTTCTCCCAGAGGCAGTGACTTT
GAGCACACGTCTGAACTCTTTCCCTACACGACGCTCTTCCGATCTNNNCACCCAGGCCATAAACCTCAAA
GAGCACACGTCTGAACTCTTTCCCTACACGACGCTCTTCCGATCTNNNTAAGCTCCACCCCCTAAAGATC
AGCACACGTCTGAACTCTTTCCCTACACGACGCTCTTCCGATCTNNNATTACCAACATTGCCTTTTGCCCC
AGAGCACACGTCTGAACTCTTTCCCTACACGACGCTCTTCCGATCTNNNAAGCAAGGACAATATGTGG
AGCACACGTCTGAACTCTTTCCCTACACGACGCTCTTCCGATCTNNNCTGAGCCTTGCCAGAAGAGTAA
AGCACACGTCTGAACTCTTTCCCTACACGACGCTCTTCCGATCTNNNACACTGTGTGCTTTCTTGCTTTCC
GAGCACACGTCTGAACTCTTTCCCTACACGACGCTCTTCCGATCTNNNCCAGTTTTGCAGTCTGACACCT
AGCACACGTCTGAACTCTTTCCCTACACGACGCTCTTCCGATCTNNNAAAGTCAGAGACAGGAAGCGG
AGAGCACACGTCTGAACTCTTTCCCTACACGACGCTCTTCCGATCTNNNATGTTTGCAGTTGACAGACTC
GAGCACACGTCTGAACTCTTTCCCTACACGACGCTCTTCCGATCTNNNGAGGATGTGAGGTTAGGAGC
AGCACACGTCTGAACTCTTTCCCTACACGACGCTCTTCCGATCTNNNCCTAGGTTTTGCGTGAGAGTC
GAGCACACGTCTGAACTCTTTCCCTACACGACGCTCTTCCGATCTNNNTGTGGTAGGAGGATCCTTGAC
GAGCACACGTCTGAACTCTTTCCCTACACGACGCTCTTCCGATCTNNNGGGAAAAGGGGACGAAATGC
AGCACACGTCTGAACTCTTTCCCTACACGACGCTCTTCCGATCTNNNCCAGGTAGGAAGTGATTTCTGC
GAGCACACGTCTGAACTCTTTCCCTACACGACGCTCTTCCGATCTNNNTACATTTCAGGAGACCAAGCA
AGCACACGTCTGAACTCTTTCCCTACACGACGCTCTTCCGATCTNNNCATCCCTTCATTACCTGTGGTGT
AGCACACGTCTGAACTCTTTCCCTACACGACGCTCTTCCGATCTNNNTTCTGGAAAGTTCCTCCTCCCC
GCACACGTCTGAACTCTTTCCCTACACGACGCTCTTCCGATCTNNNGTTCAGCTGTCATTGCTCTCTGTC
GAGCACACGTCTGAACTCTTTCCCTACACGACGCTCTTCCGATCTNNNCCATGACTTGACCAGCCTAACA
AGCACACGTCTGAACTCTTTCCCTACACGACGCTCTTCCGATCTNNNTTTGTGGAACTGACAAAGGCTG
GCACACGTCTGAACTCTTTCCCTACACGACGCTCTTCCGATCTNNNTATGTTGACGCTCAAACCACACTC
GAGCACACGTCTGAACTCTTTCCCTACACGACGCTCTTCCGATCTNNNAGCTCAAGAAGTGCAGAAGTA
GAGCACACGTCTGAACTCTTTCCCTACACGACGCTCTTCCGATCTNNNTGTTTCCCATAGGTTGGTCTCC
GAGCACACGTCTGAACTCTTTCCCTACACGACGCTCTTCCGATCTNNNATACCTAGTGGCTCTCTCCCTCT
GAGCACACGTCTGAACTCTTTCCCTACACGACGCTCTTCCGATCTNNNGGCCATGTGAGCCAATTCCTTA
GAGCACACGTCTGAACTCTTTCCCTACACGACGCTCTTCCGATCTNNNCCACCGTGACCAGCAGTTATTT
AGCACACGTCTGAACTCTTTCCCTACACGACGCTCTTCCGATCTNNNTGAGGGTCAACTCTCTACAGAC
AGCACACGTCTGAACTCTTTCCCTACACGACGCTCTTCCGATCTNNNTCTTGGTCCCAACACAGAGGTT
AGCACACGTCTGAACTCTTTCCCTACACGACGCTCTTCCGATCTNNNCATGGTGATGTCCAGATTCCCA
GAGCACACGTCTGAACTCTTTCCCTACACGACGCTCTTCCGATCTNNNCTGCTGCCTCATAACCCTACAT
AGCACACGTCTGAACTCTTTCCCTACACGACGCTCTTCCGATCTNNNTCCATACTGAGACCCTTCACGTT
GCACACGTCTGAACTCTTTCCCTACACGACGCTCTTCCGATCTNNNGATGTGACTTCACAGGCACAGAT
AGCACACGTCTGAACTCTTTCCCTACACGACGCTCTTCCGATCTNNNATCTTATGCACACACACACGGAC
GAGCACACGTCTGAACTCTTTCCCTACACGACGCTCTTCCGATCTNNNAGGCTGGGTAAGTATGAAGA
GAGCACACGTCTGAACTCTTTCCCTACACGACGCTCTTCCGATCTNNNAAACAGATCAGACGAGAAGCC
AGCACACGTCTGAACTCTTTCCCTACACGACGCTCTTCCGATCTNNNCCATCCACGACCTCTCTCTCTAAC
GAGCACACGTCTGAACTCTTTCCCTACACGACGCTCTTCCGATCTNNNCACCTGAAAAATCCCTTTTGCCC
AGCACACGTCTGAACTCTTTCCCTACACGACGCTCTTCCGATCTNNNAGCTTTGAGTTTGTGTATTGCTC
AGCACACGTCTGAACTCTTTCCCTACACGACGCTCTTCCGATCTNNNGCTTGGTGTATGTCTGTGGTCT

CACACGTCTGAACTCTTTCCCTACACGACGCTCTTCCGATCTNNNTGGTGGGAGTAGTTATGTGGGA
GAGCACACGTCTGAACTCTTTCCCTACACGACGCTCTTCCGATCTNNNATGTTCTGGATACGGGAAGT
AGCACACGTCTGAACTCTTTCCCTACACGACGCTCTTCCGATCTNNNAGAATCAGTTCTAGGGTGCCT
AGCACACGTCTGAACTCTTTCCCTACACGACGCTCTTCCGATCTNNNTGAAGGAGGCCACATTTTACT
GCACACGTCTGAACTCTTTCCCTACACGACGCTCTTCCGATCTNNNCTATGCTTCCTCTGAAGGCACTG
GCACACGTCTGAACTCTTTCCCTACACGACGCTCTTCCGATCTNNNTCCTATTCCTACAGTGCGTCCAT
GCACACGTCTGAACTCTTTCCCTACACGACGCTCTTCCGATCTNNNGCTTCCTTTCAGGGTTTGTGGA
GCACACGTCTGAACTCTTTCCCTACACGACGCTCTTCCGATCTNNNGGCCTCCTCCTTCAAGACTTCTAC
GCACACGTCTGAACTCTTTCCCTACACGACGCTCTTCCGATCTNNNAGTAAGTACTCAAACGCCTGCTG
CACACGTCTGAACTCTTTCCCTACACGACGCTCTTCCGATCTNNNTTCCATGGTAAACACTGTCAGCC
CACACGTCTGAACTCTTTCCCTACACGACGCTCTTCCGATCTNNNAGCTATTCATGCACACTCACACC
GCACACGTCTGAACTCTTTCCCTACACGACGCTCTTCCGATCTNNNCCTTTCCCCTCAACGTGTACAATC
GCACACGTCTGAACTCTTTCCCTACACGACGCTCTTCCGATCTNNNCCGTGTTTCACATTTGGCTTTGT
GCACACGTCTGAACTCTTTCCCTACACGACGCTCTTCCGATCTNNNGGGAAGAGTTTTTCTCCTCTGG
AGCACACGTCTGAACTCTTTCCCTACACGACGCTCTTCCGATCTNNNTCCGATACAGCACAGTGATTAC
GCACACGTCTGAACTCTTTCCCTACACGACGCTCTTCCGATCTNNNAACTCTAGGACTGGGGCAAATG
AGCACACGTCTGAACTCTTTCCCTACACGACGCTCTTCCGATCTNNNATCTCCATGCCTATCCTCAGCTC
AGCACACGTCTGAACTCTTTCCCTACACGACGCTCTTCCGATCTNNNCATAAGGCTTTGGGATCCTGT
GCACACGTCTGAACTCTTTCCCTACACGACGCTCTTCCGATCTNNNTCTGCTGCATCTCTGTGTTTCTC
GCACACGTCTGAACTCTTTCCCTACACGACGCTCTTCCGATCTNNNCTGTGAAATGCCCAAAAGCACA
GCACACGTCTGAACTCTTTCCCTACACGACGCTCTTCCGATCTNNNAAACAGGCAGAAATCCCCTGAA
GCACACGTCTGAACTCTTTCCCTACACGACGCTCTTCCGATCTNNNTTCCCATCACTGCTTCCCTCTAAC
CACACGTCTGAACTCTTTCCCTACACGACGCTCTTCCGATCTNNNGTCTTTGATCTGCCATTCTGGGAC
AGCACACGTCTGAACTCTTTCCCTACACGACGCTCTTCCGATCTNNNCTTCTTGGCCATGACATCTTCC
GAGCACACGTCTGAACTCTTTCCCTACACGACGCTCTTCCGATCTNNNTGCTTGGTTTCTCCTTGATGC
GAGCACACGTCTGAACTCTTTCCCTACACGACGCTCTTCCGATCTNNNTCCTTGAAAAGTTCAGCAAC
AGCACACGTCTGAACTCTTTCCCTACACGACGCTCTTCCGATCTNNNTTACACTACATGCACACACATG
GCACACGTCTGAACTCTTTCCCTACACGACGCTCTTCCGATCTNNNCCAGACCCTTGTAAGTGCTCAAA
AGCACACGTCTGAACTCTTTCCCTACACGACGCTCTTCCGATCTNNNCCTTTGCACTTGGACGGGAAT
AGCACACGTCTGAACTCTTTCCCTACACGACGCTCTTCCGATCTNNNCCTTGGACCAGTATTGAAATG
AGCACACGTCTGAACTCTTTCCCTACACGACGCTCTTCCGATCTNNNGCCAAGCTCTTTAGAGCCTAC
AGCACACGTCTGAACTCTTTCCCTACACGACGCTCTTCCGATCTNNNGTAGGGAAAGGAGACAGAA
AGCACACGTCTGAACTCTTTCCCTACACGACGCTCTTCCGATCTNNNACAAGATCACTGGAAAGAGCC
GCACACGTCTGAACTCTTTCCCTACACGACGCTCTTCCGATCTNNNAAGAAGTCTCCCACCCCTAGGT
AGCACACGTCTGAACTCTTTCCCTACACGACGCTCTTCCGATCTNNNGGGGGATATAAAGGCTTGGT
GAGCACACGTCTGAACTCTTTCCCTACACGACGCTCTTCCGATCTNNNTAATTCTTGGCACCCAGAAC
AGCACACGTCTGAACTCTTTCCCTACACGACGCTCTTCCGATCTNNNATGTCCATCCATCCCGTCCTAAT
AGCACACGTCTGAACTCTTTCCCTACACGACGCTCTTCCGATCTNNNGGAATCACATCACACACATGC
GCACACGTCTGAACTCTTTCCCTACACGACGCTCTTCCGATCTNNNAAGAAGGGAAATTCCTACGCAC
GCACACGTCTGAACTCTTTCCCTACACGACGCTCTTCCGATCTNNNCTACAGCACAGTAGCAAGACCA
AGCACACGTCTGAACTCTTTCCCTACACGACGCTCTTCCGATCTNNNATCTCTGTCCCTCTTCCCCATT
AGCACACGTCTGAACTCTTTCCCTACACGACGCTCTTCCGATCTNNNAAATCATGCGTCTACACTCCAG
AGCACACGTCTGAACTCTTTCCCTACACGACGCTCTTCCGATCTNNNGACATTCTCTCTGAGGCCAGTT

GAGCACACGTCTGAACTCTTTCCCTACACGACGCTCTTCCGATCTNNNCAAACATGAGGCATTTGTTC
AGCACACGTCTGAACTCTTTCCCTACACGACGCTCTTCCGATCTNNNTAACTCCCACTCCCATTCCTGAG
GAGCACACGTCTGAACTCTTTCCCTACACGACGCTCTTCCGATCTNNNCCCTCACTCATACTTCTCTCCG
AGCACACGTCTGAACTCTTTCCCTACACGACGCTCTTCCGATCTNNNTGGGAGCAAAACACCATTACAC
AGCACACGTCTGAACTCTTTCCCTACACGACGCTCTTCCGATCTNNNCCTGAGCCTCACTACCTTCCTA
GAGCACACGTCTGAACTCTTTCCCTACACGACGCTCTTCCGATCTNNNGCAGTCACGGGATTTGCTAC
GAGCACACGTCTGAACTCTTTCCCTACACGACGCTCTTCCGATCTNNNAGGGTTGGTAAATGCTAGG
GAGCACACGTCTGAACTCTTTCCCTACACGACGCTCTTCCGATCTNNNACTGAGCTCTAATCTGGGCT
GAGCACACGTCTGAACTCTTTCCCTACACGACGCTCTTCCGATCTNNNTTCTTCCCTCCCACACTCTGTA
GAGCACACGTCTGAACTCTTTCCCTACACGACGCTCTTCCGATCTNNNGTACACATGCACACAAGGAG
AGCACACGTCTGAACTCTTTCCCTACACGACGCTCTTCCGATCTNNNATCTTGGGGATCAGCAAGGTT
AGCACACGTCTGAACTCTTTCCCTACACGACGCTCTTCCGATCTNNNCCATGACTGCTCATGCCATTTT
AGCACACGTCTGAACTCTTTCCCTACACGACGCTCTTCCGATCTNNNTGTATGGGGTTGCAGTCAGAG
AGCACACGTCTGAACTCTTTCCCTACACGACGCTCTTCCGATCTNNNATGCTAGCTCAGCCTCTAGACA
AGCACACGTCTGAACTCTTTCCCTACACGACGCTCTTCCGATCTNNNCCTCCTCCTTCCATTGCTTTCTCC
AGCACACGTCTGAACTCTTTCCCTACACGACGCTCTTCCGATCTNNNCCTGGCCTTGTGTCCTTATCTT
AGCACACGTCTGAACTCTTTCCCTACACGACGCTCTTCCGATCTNNNGCAAAACTGAGGTTGACAAAG
AGCACACGTCTGAACTCTTTCCCTACACGACGCTCTTCCGATCTNNNGCCTGCTTAGAGTAGCAGTAAC
AGCACACGTCTGAACTCTTTCCCTACACGACGCTCTTCCGATCTNNNTTTTCCCTACCTTTTCCCCAGC
AGCACACGTCTGAACTCTTTCCCTACACGACGCTCTTCCGATCTNNNCCACAGGGATGAAGTCAGTGA
GAGCACACGTCTGAACTCTTTCCCTACACGACGCTCTTCCGATCTNNNTTGAGTTCCCCTGCCAATCAA
AGCACACGTCTGAACTCTTTCCCTACACGACGCTCTTCCGATCTNNNAGGCATCCAAATGTTTCCTCCT
AGCACACGTCTGAACTCTTTCCCTACACGACGCTCTTCCGATCTNNNCCAGCAAGGGTTTTAGAGCCT
AGCACACGTCTGAACTCTTTCCCTACACGACGCTCTTCCGATCTNNNGAAAGTCACAAGTGCCTCCTCA
AGCACACGTCTGAACTCTTTCCCTACACGACGCTCTTCCGATCTNNNGACAAACTCTCACTCTTGGGCT
AGCACACGTCTGAACTCTTTCCCTACACGACGCTCTTCCGATCTNNNACCAAACACCAAGTCCCTCTTTC
GCACACGTCTGAACTCTTTCCCTACACGACGCTCTTCCGATCTNNNCCAGCTTTGTCAGCAACATTCAG
AGCACACGTCTGAACTCTTTCCCTACACGACGCTCTTCCGATCTNNNGTGGTAGTAAAGGAAGCCCC
AGCACACGTCTGAACTCTTTCCCTACACGACGCTCTTCCGATCTNNNAACAATATTCTCAGGGCATCGG
GAGCACACGTCTGAACTCTTTCCCTACACGACGCTCTTCCGATCTNNNAAGAAAGATGCCATGACGTG
GAGCACACGTCTGAACTCTTTCCCTACACGACGCTCTTCCGATCTNNNAGTGAAGCCTTCTTACTCTTC
GCACACGTCTGAACTCTTTCCCTACACGACGCTCTTCCGATCTNNNTGTCTGTGTGTGTGTTGAGTGA
GAGCACACGTCTGAACTCTTTCCCTACACGACGCTCTTCCGATCTNNNTGACTAGTGGGTGGATAGT
AGCACACGTCTGAACTCTTTCCCTACACGACGCTCTTCCGATCTNNNGCAAAATTTGTTTCAGCAGAGC
GAGCACACGTCTGAACTCTTTCCCTACACGACGCTCTTCCGATCTNNNAAATTGCCCTCTCTCTCCCTCT
GAGCACACGTCTGAACTCTTTCCCTACACGACGCTCTTCCGATCTNNNGAAGCAAAGGGTCATCAGG
GCACACGTCTGAACTCTTTCCCTACACGACGCTCTTCCGATCTNNNGCTGGGGCAAATGGCTACTAAA
GAGCACACGTCTGAACTCTTTCCCTACACGACGCTCTTCCGATCTNNNAAGCCACATACATTCCAAAGC
AGCACACGTCTGAACTCTTTCCCTACACGACGCTCTTCCGATCTNNNTAGGGTCTTGTCAGCCTTCATC
AGCACACGTCTGAACTCTTTCCCTACACGACGCTCTTCCGATCTNNNACACAGGAAGGCATTTCTTAG
AGCACACGTCTGAACTCTTTCCCTACACGACGCTCTTCCGATCTNNNCTGCAACAAATTGCCATCGACT
AGCACACGTCTGAACTCTTTCCCTACACGACGCTCTTCCGATCTNNNGATTGCAAGCATTTGCCAGAA
AGCACACGTCTGAACTCTTTCCCTACACGACGCTCTTCCGATCTNNNGCTGGCCTTGGTATTAGTAGT

GAGCACACGTCTGAACTCTTTCCCTACACGACGCTCTTCCGATCTNNNCGCAGCATTGTGTCATATGG
AGCACACGTCTGAACTCTTTCCCTACACGACGCTCTTCCGATCTNNNCCGACTTAGGAGCAAACAATCC
AGCACACGTCTGAACTCTTTCCCTACACGACGCTCTTCCGATCTNNNATTGCGGGCTCTTATCTCTCTC
AGCACACGTCTGAACTCTTTCCCTACACGACGCTCTTCCGATCTNNNACTGGAAGACGAACTTGGAG
GAGCACACGTCTGAACTCTTTCCCTACACGACGCTCTTCCGATCTNNNAGCTCCTAAAGAGTTGTGAG
AGCACACGTCTGAACTCTTTCCCTACACGACGCTCTTCCGATCTNNNCCCTCTCCCCTGATTTCACATCT
GAGCACACGTCTGAACTCTTTCCCTACACGACGCTCTTCCGATCTNNNCACTGCACACAGACATATCCA
AGCACACGTCTGAACTCTTTCCCTACACGACGCTCTTCCGATCTNNNTCCCAATGTCTCTGGGGTTTAG
GAGCACACGTCTGAACTCTTTCCCTACACGACGCTCTTCCGATCTNNNGCACGAGTTTTGCTTTGTAG
AGCACACGTCTGAACTCTTTCCCTACACGACGCTCTTCCGATCTNNNACCAACAAACAAGTGGCTCGTA
AGCACACGTCTGAACTCTTTCCCTACACGACGCTCTTCCGATCTNNNAAACTAAAGTCAGTCACTGGCC
GCACACGTCTGAACTCTTTCCCTACACGACGCTCTTCCGATCTNNNTTTGAATGTACACGGTGTCAGG
AGCACACGTCTGAACTCTTTCCCTACACGACGCTCTTCCGATCTNNNAAAACACTAGTAACCCCACTGC
AGCACACGTCTGAACTCTTTCCCTACACGACGCTCTTCCGATCTNNNACCATAAGTGTTGGTTTGGGT
AGCACACGTCTGAACTCTTTCCCTACACGACGCTCTTCCGATCTNNNCCTGTGAGATAGATGGCCAGA
AGCACACGTCTGAACTCTTTCCCTACACGACGCTCTTCCGATCTNNNACCTCAGAGCTCCTGTCATTAG
AGCACACGTCTGAACTCTTTCCCTACACGACGCTCTTCCGATCTNNNAAAAGGGAGACTGGGGAGA
AGCACACGTCTGAACTCTTTCCCTACACGACGCTCTTCCGATCTNNNCAAGAGCAGAGTTGTAAACT
AGCACACGTCTGAACTCTTTCCCTACACGACGCTCTTCCGATCTNNNGTAGAGAGTGGTGGAAATGG
AGCACACGTCTGAACTCTTTCCCTACACGACGCTCTTCCGATCTNNNTCTTCAACCTGCTGCTGTCATAC
AGCACACGTCTGAACTCTTTCCCTACACGACGCTCTTCCGATCTNNNGAGCATTGTAGGTATTGCCCC
GAGCACACGTCTGAACTCTTTCCCTACACGACGCTCTTCCGATCTNNNAAACGCCTTTCAGGAACTAG
AGCACACGTCTGAACTCTTTCCCTACACGACGCTCTTCCGATCTNNNCAGAGGTTGCAGGAGAGATT
AGCACACGTCTGAACTCTTTCCCTACACGACGCTCTTCCGATCTNNNTTCTCCTATCAGGTAGTGTGCG
AGCACACGTCTGAACTCTTTCCCTACACGACGCTCTTCCGATCTNNNATTCCTCCCACAGCAAAGAACA
GAGCACACGTCTGAACTCTTTCCCTACACGACGCTCTTCCGATCTNNNCAACAATCTGACTTGCCTCAG
GCACACGTCTGAACTCTTTCCCTACACGACGCTCTTCCGATCTNNNAATCAAGCTGGAATTAGGCCTG
AGCACACGTCTGAACTCTTTCCCTACACGACGCTCTTCCGATCTNNNTTGTTCCAATGGTTCCAGCAAC
AGCACACGTCTGAACTCTTTCCCTACACGACGCTCTTCCGATCTNNNTTCCACCGATGGTAAACCAGA
GCACACGTCTGAACTCTTTCCCTACACGACGCTCTTCCGATCTNNNAAAGAGAGACCACATGGAAAG
GCACACGTCTGAACTCTTTCCCTACACGACGCTCTTCCGATCTNNNTCTTCCTCTTCTCTGCTGAAGTTC
AGCACACGTCTGAACTCTTTCCCTACACGACGCTCTTCCGATCTNNNGACATAGTCAGGGAAGCCTGT
GCACACGTCTGAACTCTTTCCCTACACGACGCTCTTCCGATCTNNNCCTTTGCCCTCAGGACTATCTCTC
AGCACACGTCTGAACTCTTTCCCTACACGACGCTCTTCCGATCTNNNTCAGATGGTTGGGGTACTGTC
GCACACGTCTGAACTCTTTCCCTACACGACGCTCTTCCGATCTNNNCTCCTCTTTACTGGGGCTTTCACC
AGCACACGTCTGAACTCTTTCCCTACACGACGCTCTTCCGATCTNNNCCCTACTAGATCAGCACCAAGG
AGCACACGTCTGAACTCTTTCCCTACACGACGCTCTTCCGATCTNNNTCAAATGCCTCCTCCTCCACTAA
AGCACACGTCTGAACTCTTTCCCTACACGACGCTCTTCCGATCTNNNCCCACATACTTATCCATGGCAGG
GCACACGTCTGAACTCTTTCCCTACACGACGCTCTTCCGATCTNNNGATGCCTTCTTGTGGCTTTTTCCC
AGCACACGTCTGAACTCTTTCCCTACACGACGCTCTTCCGATCTNNNACTGCTCTAATGGACAGAAACC
AGAGCACACGTCTGAACTCTTTCCCTACACGACGCTCTTCCGATCTNNNGAGACATGGAGGAGTTTGG
AGCACACGTCTGAACTCTTTCCCTACACGACGCTCTTCCGATCTNNNCTTCAAACCAATCAGGGTGGCT
GCACACGTCTGAACTCTTTCCCTACACGACGCTCTTCCGATCTNNNGGCCATCCAAGACACATACACAT

GCACACGTCTGAACTCTTTCCCTACACGACGCTCTTCCGATCTNNNATTAGCCTCAGGCCGTTAGTTTG
AGCACACGTCTGAACTCTTTCCCTACACGACGCTCTTCCGATCTNNNCGCTATTGGCTCTTGACCTGAAT
GAGCACACGTCTGAACTCTTTCCCTACACGACGCTCTTCCGATCTNNNATGTACTTACAAGCTCCATGT
GAGCACACGTCTGAACTCTTTCCCTACACGACGCTCTTCCGATCTNNNAAACCAGAAGGAGTTGAGCA
AGCACACGTCTGAACTCTTTCCCTACACGACGCTCTTCCGATCTNNNAAGTCCTCTTTGGCAGATACAG
AGAGCACACGTCTGAACTCTTTCCCTACACGACGCTCTTCCGATCTNNNACCCCCAAATTAGGAGACTT
AGCACACGTCTGAACTCTTTCCCTACACGACGCTCTTCCGATCTNNNCTCAGTATGGGTGTATGAGCA
AGAGCACACGTCTGAACTCTTTCCCTACACGACGCTCTTCCGATCTNNNCCCCAATCCAATTCCCCTTGAA
GAGCACACGTCTGAACTCTTTCCCTACACGACGCTCTTCCGATCTNNNGCTAGCTCCTGTGTTTCCATCA
AGCACACGTCTGAACTCTTTCCCTACACGACGCTCTTCCGATCTNNNAGACAATGAGGGCTTTGGATT
AGAGCACACGTCTGAACTCTTTCCCTACACGACGCTCTTCCGATCTNNNTGTGAGGAAGCTACTCCAA
AGCACACGTCTGAACTCTTTCCCTACACGACGCTCTTCCGATCTNNNATGTGGTGTTTTCTGCCTGGAA
AGCACACGTCTGAACTCTTTCCCTACACGACGCTCTTCCGATCTNNNTATGTGACCTTCACCTGACATC
GAGCACACGTCTGAACTCTTTCCCTACACGACGCTCTTCCGATCTNNNGTAGGGATGGGGGATTCTGA
AGCACACGTCTGAACTCTTTCCCTACACGACGCTCTTCCGATCTNNNTAGCCTGGAGGTGTGTCATTTT
AGCACACGTCTGAACTCTTTCCCTACACGACGCTCTTCCGATCTNNNTCACAGCTTTTGGAGCTTCCATT
AGCACACGTCTGAACTCTTTCCCTACACGACGCTCTTCCGATCTNNNATGCCACACACATATCTATGCA
AGCACACGTCTGAACTCTTTCCCTACACGACGCTCTTCCGATCTNNNCTAGGGTTGTGTGGGTAAAGC
AGCACACGTCTGAACTCTTTCCCTACACGACGCTCTTCCGATCTNNNCTCATGTGGTCCTCCTTAGGTG
AGCACACGTCTGAACTCTTTCCCTACACGACGCTCTTCCGATCTNNNAAGGCCTCTGGAGAGATCAGT
AGCACACGTCTGAACTCTTTCCCTACACGACGCTCTTCCGATCTNNNGAACACACCTCAGAGTTACCAC
GAGCACACGTCTGAACTCTTTCCCTACACGACGCTCTTCCGATCTNNNTGGCTGAGATGCATATGAGA
GCACACGTCTGAACTCTTTCCCTACACGACGCTCTTCCGATCTNNNCTGTGATCCTCCTTTTCCAACCACA
GAGCACACGTCTGAACTCTTTCCCTACACGACGCTCTTCCGATCTNNNACACAGAGGTTGTGTTTCTCT
GAGCACACGTCTGAACTCTTTCCCTACACGACGCTCTTCCGATCTNNNAATGTGACACAGTTCCAGTG
GAGCACACGTCTGAACTCTTTCCCTACACGACGCTCTTCCGATCTNNNCCTTGTAGTCAAGTGGAACC
AGCACACGTCTGAACTCTTTCCCTACACGACGCTCTTCCGATCTNNNCTAGTGGTCCTACGTCCTCATCT
AGCACACGTCTGAACTCTTTCCCTACACGACGCTCTTCCGATCTNNNAACTGAAATGTACACCGGTGA
AGCACACGTCTGAACTCTTTCCCTACACGACGCTCTTCCGATCTNNNCCAGTGGTGAGAAACATGGTGA
AGCACACGTCTGAACTCTTTCCCTACACGACGCTCTTCCGATCTNNNATTATTCTGCACAAGCAGAGGCA
AGCACACGTCTGAACTCTTTCCCTACACGACGCTCTTCCGATCTNNNCCCAGAAGCCACTTCATACAGA
GAGCACACGTCTGAACTCTTTCCCTACACGACGCTCTTCCGATCTNNNCTAGCTGGGTGGGTTACACTT
AGCACACGTCTGAACTCTTTCCCTACACGACGCTCTTCCGATCTNNNCTGAAATCCCACCACCCAAAGAT
GCACACGTCTGAACTCTTTCCCTACACGACGCTCTTCCGATCTNNNGGTTGAGATGATTACCTCGGAC
GCACACGTCTGAACTCTTTCCCTACACGACGCTCTTCCGATCTNNNTACAGAGTATAGCCACTTGCGTG
GAGCACACGTCTGAACTCTTTCCCTACACGACGCTCTTCCGATCTNNNCCTACCACATGACGTTCCTTTG
GCACACGTCTGAACTCTTTCCCTACACGACGCTCTTCCGATCTNNNTAGAAACCCTGCCTTCAGCTAGT
GAGCACACGTCTGAACTCTTTCCCTACACGACGCTCTTCCGATCTNNNCAAGCTGTAACAAGGTGCATT
GCACACGTCTGAACTCTTTCCCTACACGACGCTCTTCCGATCTNNNTTATCCCCTTGATCTTGCCTCCACA
AGCACACGTCTGAACTCTTTCCCTACACGACGCTCTTCCGATCTNNNACAGTGGCAGAGCAAAACTAT
AGCACACGTCTGAACTCTTTCCCTACACGACGCTCTTCCGATCTNNNTAGTGGGAAGGAAGGAGGAA
AGCACACGTCTGAACTCTTTCCCTACACGACGCTCTTCCGATCTNNNATGGTTGTGGGTCTTTTCTGCA
AGCACACGTCTGAACTCTTTCCCTACACGACGCTCTTCCGATCTNNNGTCTATGAAGGTCTGCGGATT

AGCACACGTCTGAACTCTTTCCCTACACGACGCTCTTCCGATCTNNNGTCCTTACTGTCAGCATGAACCA
GCACACGTCTGAACTCTTTCCCTACACGACGCTCTTCCGATCTNNNTTTGCAGCTGTCTTCTCCTTACGCA
GCACACGTCTGAACTCTTTCCCTACACGACGCTCTTCCGATCTNNNTAAGACAGGGAAAGGGGACTGA
GAGCACACGTCTGAACTCTTTCCCTACACGACGCTCTTCCGATCTNNNGGTGGCAGGTGGTGATAAT
AGCACACGTCTGAACTCTTTCCCTACACGACGCTCTTCCGATCTNNNACCGTAATACCGTCTCCCAATG
GAGCACACGTCTGAACTCTTTCCCTACACGACGCTCTTCCGATCTNNNGCACAAAAGCACCTTCTCTCTC
AGCACACGTCTGAACTCTTTCCCTACACGACGCTCTTCCGATCTNNNTGAACCCAGATTGTAGCACAGA
GCACACGTCTGAACTCTTTCCCTACACGACGCTCTTCCGATCTNNNATTTGGACAAATGTCTCCCCCACC
GCACACGTCTGAACTCTTTCCCTACACGACGCTCTTCCGATCTNNNAACCCTGCTTCTTGGCATTTTTGC
GCACACGTCTGAACTCTTTCCCTACACGACGCTCTTCCGATCTNNNGGTGTTTAGGTGAGACAACCTC
GCACACGTCTGAACTCTTTCCCTACACGACGCTCTTCCGATCTNNNTATCTCTCATGCTCACCTCTGGGC
GAGCACACGTCTGAACTCTTTCCCTACACGACGCTCTTCCGATCTNNNCTTCTCCAGCCTAGCAGTGTT
AGCACACGTCTGAACTCTTTCCCTACACGACGCTCTTCCGATCTNNNTGGCTTCTCTTCTCTCAATCTCTC
AGCACACGTCTGAACTCTTTCCCTACACGACGCTCTTCCGATCTNNNTACCTGTTGGATTATGAAGGG
AGCACACGTCTGAACTCTTTCCCTACACGACGCTCTTCCGATCTNNNTTGGAATCAATTGCTCTGGCACT
GCACACGTCTGAACTCTTTCCCTACACGACGCTCTTCCGATCTNNNCAGCTCAGAGTGTAGGTTAGGC
GAGCACACGTCTGAACTCTTTCCCTACACGACGCTCTTCCGATCTNNNTGTATTACCTCGAGTAGGGC
AGCACACGTCTGAACTCTTTCCCTACACGACGCTCTTCCGATCTNNNCAAGCTCCCATTTTCCCTCTCTCC
GCACACGTCTGAACTCTTTCCCTACACGACGCTCTTCCGATCTNNNCATTCTCCAAGAGGGAGTGCAAA
GCACACGTCTGAACTCTTTCCCTACACGACGCTCTTCCGATCTNNNTTGTGCTTGTAGAACCTGCTGTTC
AGCACACGTCTGAACTCTTTCCCTACACGACGCTCTTCCGATCTNNNACCACATAACTCAGAGTAGGG
GCACACGTCTGAACTCTTTCCCTACACGACGCTCTTCCGATCTNNNTAGAGGCTATCACATCCGAAGCA
GAGCACACGTCTGAACTCTTTCCCTACACGACGCTCTTCCGATCTNNNACACTCTTACAACCACACACAC
AGAGCACACGTCTGAACTCTTTCCCTACACGACGCTCTTCCGATCTNNNAGGGACTGCTTGGATACTG
GAGCACACGTCTGAACTCTTTCCCTACACGACGCTCTTCCGATCTNNNTCTCAATGCATGGTCACTGGA
AGAGCACACGTCTGAACTCTTTCCCTACACGACGCTCTTCCGATCTNNNTCTCAGGTTCTCTTTCCCTCT
GAGCACACGTCTGAACTCTTTCCCTACACGACGCTCTTCCGATCTNNNTTGCCTCCTCTCCTTCCCATAAA
GAGCACACGTCTGAACTCTTTCCCTACACGACGCTCTTCCGATCTNNNGCCTGCCTGTCTCCATATCATT
GCACACGTCTGAACTCTTTCCCTACACGACGCTCTTCCGATCTNNNTGCCAAAGGTGTTCAATGAGTG
GAGCACACGTCTGAACTCTTTCCCTACACGACGCTCTTCCGATCTNNNTTTGTGTTCCACACAGCTTGGT
GAGCACACGTCTGAACTCTTTCCCTACACGACGCTCTTCCGATCTNNNACCAAGAGAGTAGATGAGG
GAGCACACGTCTGAACTCTTTCCCTACACGACGCTCTTCCGATCTNNNATGGAGAATGGGAAATCGTG
AGAGCACACGTCTGAACTCTTTCCCTACACGACGCTCTTCCGATCTNNNTTGATTGATGCACCTTCTCAC
AGCACACGTCTGAACTCTTTCCCTACACGACGCTCTTCCGATCTNNNCCACCTACCTAAACCAGTGTGTC
AGAGCACACGTCTGAACTCTTTCCCTACACGACGCTCTTCCGATCTNNNAAAAAGCTCCACTTTTTGAG
GAGCACACGTCTGAACTCTTTCCCTACACGACGCTCTTCCGATCTNNNGGTCTCCCTCTCTCTTCACACAT
GAGCACACGTCTGAACTCTTTCCCTACACGACGCTCTTCCGATCTNNNTGTCCACTGCAGCATCATATTC
GAGCACACGTCTGAACTCTTTCCCTACACGACGCTCTTCCGATCTNNNCCACGAGCTTCCTGAAGTCTAA
AGCACACGTCTGAACTCTTTCCCTACACGACGCTCTTCCGATCTNNNAAGGTAGAGAAGTGTTCCTCCC
GAGCACACGTCTGAACTCTTTCCCTACACGACGCTCTTCCGATCTNNNCCCTTTTGGTCTCTCTTGTGTC
AGCACACGTCTGAACTCTTTCCCTACACGACGCTCTTCCGATCTNNNACTTTTACTCAGTGCAGTTTGGC
AGCACACGTCTGAACTCTTTCCCTACACGACGCTCTTCCGATCTNNNGAAGGCCTGAGAACCAAGAAC
GCACACGTCTGAACTCTTTCCCTACACGACGCTCTTCCGATCTNNNGCTGATAGAGGATGAGCTGCAA

AGCACACGTCTGAACTCTTTCCCTACACGACGCTCTTCCGATCTNNNCAATATGTCATGCTTCACGCAC
AGAGCACACGTCTGAACTCTTTCCCTACACGACGCTCTTCCGATCTNNNGATTTTTGGCAGATGCCTT
AGAGCACACGTCTGAACTCTTTCCCTACACGACGCTCTTCCGATCTNNNGCATCACGGTAACCATAAAG
AGAGCACACGTCTGAACTCTTTCCCTACACGACGCTCTTCCGATCTNNNAACTACTCTATGCGTGCCTCT
AGCACACGTCTGAACTCTTTCCCTACACGACGCTCTTCCGATCTNNNAGGGGAAAGAGGGCAAGATA
AGAGCACACGTCTGAACTCTTTCCCTACACGACGCTCTTCCGATCTNNNGCTCTTTTGGAAAGCCCTCT
GAGCACACGTCTGAACTCTTTCCCTACACGACGCTCTTCCGATCTNNNCAAACTCCATCCCATCCTTCCT
AGAGCACACGTCTGAACTCTTTCCCTACACGACGCTCTTCCGATCTNNNAGGCTGTTGTGTGAGGAA
GAGCACACGTCTGAACTCTTTCCCTACACGACGCTCTTCCGATCTNNNCTGTTCATGTACACACACCCA
AGCACACGTCTGAACTCTTTCCCTACACGACGCTCTTCCGATCTNNNGCACAGAACCAATCAGAGGGT
AGCACACGTCTGAACTCTTTCCCTACACGACGCTCTTCCGATCTNNNTGTGAGTTGTGCATGTAGTCTG
GAGCACACGTCTGAACTCTTTCCCTACACGACGCTCTTCCGATCTNNNATAATCCCAAAATGCAGTGGC
AGAGCACACGTCTGAACTCTTTCCCTACACGACGCTCTTCCGATCTNNNGACCTAAAGGCCAAGCAAG
GAGCACACGTCTGAACTCTTTCCCTACACGACGCTCTTCCGATCTNNNGAGAGAACCTGAAAGGAAG
AGCACACGTCTGAACTCTTTCCCTACACGACGCTCTTCCGATCTNNNTTGCACATACACACACACAAAG
AGCACACGTCTGAACTCTTTCCCTACACGACGCTCTTCCGATCTNNNTTGCACACAGCCAGGATTATGT
GAGCACACGTCTGAACTCTTTCCCTACACGACGCTCTTCCGATCTNNNAGAAGCTTCAGATCCTGGAA
GAGCACACGTCTGAACTCTTTCCCTACACGACGCTCTTCCGATCTNNNCAATATGCTTGCCTCAAGGGT
GAGCACACGTCTGAACTCTTTCCCTACACGACGCTCTTCCGATCTNNNTGTCTTCCTTCCAGTGCTGTCT
AGAGCACACGTCTGAACTCTTTCCCTACACGACGCTCTTCCGATCTNNNAGAAGCCCTCCCAGAGATG
GAGCACACGTCTGAACTCTTTCCCTACACGACGCTCTTCCGATCTNNNCGGGTTGGATCAAAAATACG
GAGCACACGTCTGAACTCTTTCCCTACACGACGCTCTTCCGATCTNNNGAAAGGCTTATCAGTGAGG
AGCACACGTCTGAACTCTTTCCCTACACGACGCTCTTCCGATCTNNNACCCATAGCTCAGAACACATGA
GAGCACACGTCTGAACTCTTTCCCTACACGACGCTCTTCCGATCTNNNCCATCAGGCCTACGAAGAGTA
AGCACACGTCTGAACTCTTTCCCTACACGACGCTCTTCCGATCTNNNGCTTTGCTTGGTGCTTCATTAG
GAGCACACGTCTGAACTCTTTCCCTACACGACGCTCTTCCGATCTNNNGAAGGATCCCTTCCCTCATCCT
GAGCACACGTCTGAACTCTTTCCCTACACGACGCTCTTCCGATCTNNNTGGAATATGTGGAGTGTGTG
AGCACACGTCTGAACTCTTTCCCTACACGACGCTCTTCCGATCTNNNTGATATGTCACTGCTCCCCAGA
AGCACACGTCTGAACTCTTTCCCTACACGACGCTCTTCCGATCTNNNAGCCTCGAATGATTTCTCTCCT
GCACACGTCTGAACTCTTTCCCTACACGACGCTCTTCCGATCTNNNCGTATGCACAAGCTCACACAATC
GAGCACACGTCTGAACTCTTTCCCTACACGACGCTCTTCCGATCTNNNCTCAAGTAAGAAGTTCCGGAG
AGCACACGTCTGAACTCTTTCCCTACACGACGCTCTTCCGATCTNNNCTTTTTCTCCTACGTGCCCTTCA
GAGCACACGTCTGAACTCTTTCCCTACACGACGCTCTTCCGATCTNNNGAGCCTCTCCCCTTTACAATGT
GAGCACACGTCTGAACTCTTTCCCTACACGACGCTCTTCCGATCTNNNTCTTCCCAGACCTAGACTGTG
GAGCACACGTCTGAACTCTTTCCCTACACGACGCTCTTCCGATCTNNNTTTCACATTGACCACTCCACCA
GAGCACACGTCTGAACTCTTTCCCTACACGACGCTCTTCCGATCTNNNCATGGCAGGCTTGTATGCAA
AGCACACGTCTGAACTCTTTCCCTACACGACGCTCTTCCGATCTNNNGAGGGCCTGAGAGAGAAAAA
AGCACACGTCTGAACTCTTTCCCTACACGACGCTCTTCCGATCTNNNTCTGTGGTTGATATTGCTGCCT
AGCACACGTCTGAACTCTTTCCCTACACGACGCTCTTCCGATCTNNNCCCATGCAACATAGGGGATGA
GAGCACACGTCTGAACTCTTTCCCTACACGACGCTCTTCCGATCTNNNAGTCCCTGTGATAGCTCCTGA
AGCACACGTCTGAACTCTTTCCCTACACGACGCTCTTCCGATCTNNNCTGACAATTCCAGATCCTCAGG
GAGCACACGTCTGAACTCTTTCCCTACACGACGCTCTTCCGATCTNNNATCCGTATTAGTTGGGGTCCT
GAGCACACGTCTGAACTCTTTCCCTACACGACGCTCTTCCGATCTNNNCCAATGGAGGGTCAGAGAAA

GAGCACACGTCTGAACTCTTTCCCTACACGACGCTCTTCCGATCTNNNTTGTTGCCATGTGGTAATGC
GCACACGTCTGAACTCTTTCCCTACACGACGCTCTTCCGATCTNNNTCCTCCCCTCTTTTTCTGCTACAC
GAGCACACGTCTGAACTCTTTCCCTACACGACGCTCTTCCGATCTNNNGAGTTGGAGCAGGACTAAG
AGCACACGTCTGAACTCTTTCCCTACACGACGCTCTTCCGATCTNNNGAAGCTCAAGTGGCAAGGAAC
AGCACACGTCTGAACTCTTTCCCTACACGACGCTCTTCCGATCTNNNATGAATGGTCTTGGGGAACAT
GCACACGTCTGAACTCTTTCCCTACACGACGCTCTTCCGATCTNNNTCCAAGATGGATTCGTTAGCAG
AGCACACGTCTGAACTCTTTCCCTACACGACGCTCTTCCGATCTNNNTGCAAAGTCCTCCATCTTGAGT
CACACGTCTGAACTCTTTCCCTACACGACGCTCTTCCGATCTNNNTTCCCAGGAAGGTAGGAAAGAG
AGCACACGTCTGAACTCTTTCCCTACACGACGCTCTTCCGATCTNNNCCTCCATTACATTGTGACCAGG
AGCACACGTCTGAACTCTTTCCCTACACGACGCTCTTCCGATCTNNNTCAGCCTCTGAAAGCCCTGATT
AGCACACGTCTGAACTCTTTCCCTACACGACGCTCTTCCGATCTNNNTCATGCCAGTCCCTCTATTGGA
GAGCACACGTCTGAACTCTTTCCCTACACGACGCTCTTCCGATCTNNNAGCTCTTGATTTTGGCACAGT
GAGCACACGTCTGAACTCTTTCCCTACACGACGCTCTTCCGATCTNNNGCAAGAGTAAAGCACTGTTC
GAGCACACGTCTGAACTCTTTCCCTACACGACGCTCTTCCGATCTNNNATTTGGAAGGCCACCATGTG
AGAGCACACGTCTGAACTCTTTCCCTACACGACGCTCTTCCGATCTNNNTGCCACTGGTGGAAATTGT
AGAGCACACGTCTGAACTCTTTCCCTACACGACGCTCTTCCGATCTNNNGATTCAATAGCACACCAGC
GAGCACACGTCTGAACTCTTTCCCTACACGACGCTCTTCCGATCTNNNATTCTGACAGGGCCAATTCAT
AGCACACGTCTGAACTCTTTCCCTACACGACGCTCTTCCGATCTNNNTCACCTTGTCTACACACACATG
GAGCACACGTCTGAACTCTTTCCCTACACGACGCTCTTCCGATCTNNNTTGAGTGACACAAAAGAGT
GAGCACACGTCTGAACTCTTTCCCTACACGACGCTCTTCCGATCTNNNGGAGTTGATCTGGTTTCAAT
AGCACACGTCTGAACTCTTTCCCTACACGACGCTCTTCCGATCTNNNAAAAGATGTGTGTGGGGAGT
AGAGCACACGTCTGAACTCTTTCCCTACACGACGCTCTTCCGATCTNNNTCCCTATTTGTAAGTGCAGC
GAGCACACGTCTGAACTCTTTCCCTACACGACGCTCTTCCGATCTNNNGGGAAGAAGGACAAGGAG
AGCACACGTCTGAACTCTTTCCCTACACGACGCTCTTCCGATCTNNNGGGTATGATAGCGGGTTCCTG
AGCACACGTCTGAACTCTTTCCCTACACGACGCTCTTCCGATCTNNNTTCGGGTTTGGGTTTCACAAG
GCACACGTCTGAACTCTTTCCCTACACGACGCTCTTCCGATCTNNNATGCCCAAAATCTCTGCTTTGGA
GAGCACACGTCTGAACTCTTTCCCTACACGACGCTCTTCCGATCTNNNTCCACCCTTTTGCTTTTGCAT
AGCACACGTCTGAACTCTTTCCCTACACGACGCTCTTCCGATCTNNNAAACCAACAACCCATAATCCCC
GAGCACACGTCTGAACTCTTTCCCTACACGACGCTCTTCCGATCTNNNCTAAGCTCCACTGATGTGCTC
GAGCACACGTCTGAACTCTTTCCCTACACGACGCTCTTCCGATCTNNNTACTCAGCACACACAAACACA
GAGCACACGTCTGAACTCTTTCCCTACACGACGCTCTTCCGATCTNNNCCTCCCTCCCTTTCCTAACTTCA
GCACACGTCTGAACTCTTTCCCTACACGACGCTCTTCCGATCTNNNGCTGTTCCCCTTACCTAGAATGC
AAGAGCACACGTCTGAACTCTTTCCCTACACGACGCTCTTCCGATCTNNNGCAGCATCCTTTGTCCTCTG
AGAGCACACGTCTGAACTCTTTCCCTACACGACGCTCTTCCGATCTNNNGCAAGGCAGAATGAGCAAA
AGAGCACACGTCTGAACTCTTTCCCTACACGACGCTCTTCCGATCTNNNCATGAAAAGGTGGAGCAGA
GAGCACACGTCTGAACTCTTTCCCTACACGACGCTCTTCCGATCTNNNACATGCAGGATGGGGTAGAA
AGAGCACACGTCTGAACTCTTTCCCTACACGACGCTCTTCCGATCTNNNACTGTTGCAGTCCGGAAAC
AGCACACGTCTGAACTCTTTCCCTACACGACGCTCTTCCGATCTNNNGACAAACCAGCAAATGAGCAA
GAGCACACGTCTGAACTCTTTCCCTACACGACGCTCTTCCGATCTNNNACGCTATAGAGCCTGGAGTTT
GAGCACACGTCTGAACTCTTTCCCTACACGACGCTCTTCCGATCTNNNTGTCCCATCAATCAGTGTAGT
GAGCACACGTCTGAACTCTTTCCCTACACGACGCTCTTCCGATCTNNNATGGCAGCCTCATTAGAGGA
AGCACACGTCTGAACTCTTTCCCTACACGACGCTCTTCCGATCTNNNTGCACCACTGTCATAAACACAC
AGCACACGTCTGAACTCTTTCCCTACACGACGCTCTTCCGATCTNNNGTCTGATGGATTTTTGCTCTG

AGCACACGTCTGAACTCTTTCCCTACACGACGCTCTTCCGATCTNNNGGAGGTGGGTATTGGTCAAGT
GAGCACACGTCTGAACTCTTTCCCTACACGACGCTCTTCCGATCTNNNAGCTGGTTTGTACATCAAAGG
AGCACACGTCTGAACTCTTTCCCTACACGACGCTCTTCCGATCTNNNATGCATCTGGAAAGCAAAAAG
GAGCACACGTCTGAACTCTTTCCCTACACGACGCTCTTCCGATCTNNNAGTTTACAGCCTAAGCCCCTGT
GAGCACACGTCTGAACTCTTTCCCTACACGACGCTCTTCCGATCTNNNTCTCTGTCCAGCAGGAAGAAG
AGAGCACACGTCTGAACTCTTTCCCTACACGACGCTCTTCCGATCTNNNAGGGAGGAATGTGTCAGCTA
AGAGCACACGTCTGAACTCTTTCCCTACACGACGCTCTTCCGATCTNNNACGATCAGGAATGCAGAGG
AGAGCACACGTCTGAACTCTTTCCCTACACGACGCTCTTCCGATCTNNNACACTTTCCTCCATCTCTCCTC
AGAGCACACGTCTGAACTCTTTCCCTACACGACGCTCTTCCGATCTNNNAGCCAGCAGCTCATCTTTTTC
GAGCACACGTCTGAACTCTTTCCCTACACGACGCTCTTCCGATCTNNNGGAGTGCAGAATAAGGGTGA
GAGCACACGTCTGAACTCTTTCCCTACACGACGCTCTTCCGATCTNNNTCCTAGCATCTCAGGGCATAA
AGAGCACACGTCTGAACTCTTTCCCTACACGACGCTCTTCCGATCTNNNTGGTGGGAAAGTCAGAAAA
GAGCACACGTCTGAACTCTTTCCCTACACGACGCTCTTCCGATCTNNNACCAACATCAAGGTGAGGAA
AGAGCACACGTCTGAACTCTTTCCCTACACGACGCTCTTCCGATCTNNNAGATTGCTTCTGACAGCACA
AGAGCACACGTCTGAACTCTTTCCCTACACGACGCTCTTCCGATCTNNNTGGTTAGTCAATGTGTGCTCT
AGAGCACACGTCTGAACTCTTTCCCTACACGACGCTCTTCCGATCTNNNAGTAGGGGATGAGGGAATA
GAGCACACGTCTGAACTCTTTCCCTACACGACGCTCTTCCGATCTNNNATTATCATACCACCCAGAGGC
AGAGCACACGTCTGAACTCTTTCCCTACACGACGCTCTTCCGATCTNNNACAGGTGCATTAGGATCTCTC
AGCACACGTCTGAACTCTTTCCCTACACGACGCTCTTCCGATCTNNNGGTGGGGCTCTATACTACAGGA
AGCACACGTCTGAACTCTTTCCCTACACGACGCTCTTCCGATCTNNNACTTCTAGGCAGTTTTGTGGAG
GAGCACACGTCTGAACTCTTTCCCTACACGACGCTCTTCCGATCTNNNGGTATGTGTGTGAAGTGGGA
AGCACACGTCTGAACTCTTTCCCTACACGACGCTCTTCCGATCTNNNAGTACCCTTACAGGAAGAGCCA
AGAGCACACGTCTGAACTCTTTCCCTACACGACGCTCTTCCGATCTNNNCATTGAGGAAGTGGGGGAA
AGCACACGTCTGAACTCTTTCCCTACACGACGCTCTTCCGATCTNNNAGGAACTGTGAACACTCCAAAC
GAGCACACGTCTGAACTCTTTCCCTACACGACGCTCTTCCGATCTNNNACACAGCAAAACCTTGTCTGG
AGCACACGTCTGAACTCTTTCCCTACACGACGCTCTTCCGATCTNNNGTACATGTCTCTAGGCAGAGGC
AGCACACGTCTGAACTCTTTCCCTACACGACGCTCTTCCGATCTNNNCCTTCATGCTGATGACTTCCAGT
AGCACACGTCTGAACTCTTTCCCTACACGACGCTCTTCCGATCTNNNACAGTCCTTTGTTGCCTGATGTT
GAGCACACGTCTGAACTCTTTCCCTACACGACGCTCTTCCGATCTNNNTTCCTCATGATCTCTATTCCCC
GAGCACACGTCTGAACTCTTTCCCTACACGACGCTCTTCCGATCTNNNCAACTCTGTCCCTTTGAGGAAC
GAGCACACGTCTGAACTCTTTCCCTACACGACGCTCTTCCGATCTNNNTGCTGCCTACTCAGTCTCTTTC
GAGCACACGTCTGAACTCTTTCCCTACACGACGCTCTTCCGATCTNNNTCTTGCAGGGAGAGTAAGGT
AGCACACGTCTGAACTCTTTCCCTACACGACGCTCTTCCGATCTNNNCTGAGAGAGGGAGGGATTGAG
GAGCACACGTCTGAACTCTTTCCCTACACGACGCTCTTCCGATCTNNNTGCTCGGACCCTGATTACTATC
AGCACACGTCTGAACTCTTTCCCTACACGACGCTCTTCCGATCTNNNGCCTGGCTTCATATCCAAAAGTG
GAGCACACGTCTGAACTCTTTCCCTACACGACGCTCTTCCGATCTNNNGCTTTGCTTTTGTGAGGCTTTC
AGCACACGTCTGAACTCTTTCCCTACACGACGCTCTTCCGATCTNNNCCAGGGTGTTGATGTCTCCCTTA
GCACACGTCTGAACTCTTTCCCTACACGACGCTCTTCCGATCTNNNGAGGCTCTCCTAGACGAAGAGTT
GAGCACACGTCTGAACTCTTTCCCTACACGACGCTCTTCCGATCTNNNAGGAAAGTGTGCTTACAGAA
AGCACACGTCTGAACTCTTTCCCTACACGACGCTCTTCCGATCTNNNCCCCATATACCCCATATCCCAGC
GAGCACACGTCTGAACTCTTTCCCTACACGACGCTCTTCCGATCTNNNTCACTGCATGGTGACCACAAT
GAGCACACGTCTGAACTCTTTCCCTACACGACGCTCTTCCGATCTNNNATGGAAAATGGGATACCCCAG
GAGCACACGTCTGAACTCTTTCCCTACACGACGCTCTTCCGATCTNNNGTGTGGCATGTGTAAGAGAG

AGCACACGTCTGAACTCTTTCCCTACACGACGCTCTTCCGATCTNNNGGATTATTTGGTCCCTCTGCC
GAGCACACGTCTGAACTCTTTCCCTACACGACGCTCTTCCGATCTNNNCCCTTTTGCTGGGCATAAAGT
AGCACACGTCTGAACTCTTTCCCTACACGACGCTCTTCCGATCTNNNACAGGCTTGAAAGTCTCTGTCT
GAGCACACGTCTGAACTCTTTCCCTACACGACGCTCTTCCGATCTNNNTCCTTTCTCATTGTGCCAGTTC
GCACACGTCTGAACTCTTTCCCTACACGACGCTCTTCCGATCTNNNTACAGTGCCTTGATTCACAACCCC
GCACACGTCTGAACTCTTTCCCTACACGACGCTCTTCCGATCTNNNCATGCTGTTCTCATGGTGGAGTT
AGCACACGTCTGAACTCTTTCCCTACACGACGCTCTTCCGATCTNNNTACTCAGCTCTTCTGGGTTTGG
AGCACACGTCTGAACTCTTTCCCTACACGACGCTCTTCCGATCTNNNTTTTCCCCCAAATTCCTTCCCTG
GAGCACACGTCTGAACTCTTTCCCTACACGACGCTCTTCCGATCTNNNTTAGGAGCTGGTTGAAAAGG
GCACACGTCTGAACTCTTTCCCTACACGACGCTCTTCCGATCTNNNCAGTGGGTGGGTACTCAAATAG
GCACACGTCTGAACTCTTTCCCTACACGACGCTCTTCCGATCTNNNATGCGGCTATTTCCAAGAGTGT
GAGCACACGTCTGAACTCTTTCCCTACACGACGCTCTTCCGATCTNNNGTGGGGGTGATACTCTCCAC
GCACACGTCTGAACTCTTTCCCTACACGACGCTCTTCCGATCTNNNGCAGATTTTAGGCAGCTCTCCTT
GCACACGTCTGAACTCTTTCCCTACACGACGCTCTTCCGATCTNNNAGTTCCAGCCAACAGAACATGA
AGAGCACACGTCTGAACTCTTTCCCTACACGACGCTCTTCCGATCTNNNGCCTAATCCAATCCGACTGT
AGCACACGTCTGAACTCTTTCCCTACACGACGCTCTTCCGATCTNNNTCTTGGCACCTCAATACCAATC
GAGCACACGTCTGAACTCTTTCCCTACACGACGCTCTTCCGATCTNNNTTTCACTGGAGCAAAGAGTT
GAGCACACGTCTGAACTCTTTCCCTACACGACGCTCTTCCGATCTNNNGGGAGAAATCATGGGAACC
GAGCACACGTCTGAACTCTTTCCCTACACGACGCTCTTCCGATCTNNNTCTAGTTGCTAGTTACACAGC
AGCACACGTCTGAACTCTTTCCCTACACGACGCTCTTCCGATCTNNNAAGTTGGTATGCCAGCAAAAA
GAGCACACGTCTGAACTCTTTCCCTACACGACGCTCTTCCGATCTNNNTCGCTTTATAACAGCCAGAGT
GAGCACACGTCTGAACTCTTTCCCTACACGACGCTCTTCCGATCTNNNACCAAGCAAGGGTGAAAGC
GAGCACACGTCTGAACTCTTTCCCTACACGACGCTCTTCCGATCTNNNGGGACGGGAAAGGAAGAAA
GAGCACACGTCTGAACTCTTTCCCTACACGACGCTCTTCCGATCTNNNCCTCTATCTGCAAGAAGGACA
GAGCACACGTCTGAACTCTTTCCCTACACGACGCTCTTCCGATCTNNNTGCTCTGCATCTTTGTCACCAA
GAGCACACGTCTGAACTCTTTCCCTACACGACGCTCTTCCGATCTNNNTCACTTTCTGTATCCCACAGG
AGCACACGTCTGAACTCTTTCCCTACACGACGCTCTTCCGATCTNNNTGGTAGGCTATTTGGCTCTACA
GAGCACACGTCTGAACTCTTTCCCTACACGACGCTCTTCCGATCTNNNACTGCAGTCTTCATTGCTGCT
GAGCACACGTCTGAACTCTTTCCCTACACGACGCTCTTCCGATCTNNNAAGGTTGGCATTCCATTTACC
AGCACACGTCTGAACTCTTTCCCTACACGACGCTCTTCCGATCTNNNATCATGTGTGTGGGTAGGAAG
GAGCACACGTCTGAACTCTTTCCCTACACGACGCTCTTCCGATCTNNNTTGAGTGACTGAATTGCTAGG
AGCACACGTCTGAACTCTTTCCCTACACGACGCTCTTCCGATCTNNNGTGTGTGTAAGTGATGCCAGT
GCACACGTCTGAACTCTTTCCCTACACGACGCTCTTCCGATCTNNNCAATCATTGACCAGCCATGGAGA
GAGCACACGTCTGAACTCTTTCCCTACACGACGCTCTTCCGATCTNNNAGCAATGGAGGCTGTCAAA
AGCACACGTCTGAACTCTTTCCCTACACGACGCTCTTCCGATCTNNNATGGATAACATGTCCTCACTCC
GAGCACACGTCTGAACTCTTTCCCTACACGACGCTCTTCCGATCTNNNCTTCAGTCTTTGCCTCCCATA
GAGCACACGTCTGAACTCTTTCCCTACACGACGCTCTTCCGATCTNNNTCTGGAAAAGGCAAACAATAC
AGCACACGTCTGAACTCTTTCCCTACACGACGCTCTTCCGATCTNNNATGCTGTGTTCTCCATCTTTGC
GAGCACACGTCTGAACTCTTTCCCTACACGACGCTCTTCCGATCTNNNGGGAATCCGATCTGAAAGAC
AGCACACGTCTGAACTCTTTCCCTACACGACGCTCTTCCGATCTNNNGTGTCAAAGAGAGCCAAGGCT
GAGCACACGTCTGAACTCTTTCCCTACACGACGCTCTTCCGATCTNNNGATCCATGCAGATCCAGTCTC
GCACACGTCTGAACTCTTTCCCTACACGACGCTCTTCCGATCTNNNATTCCAAGCTGGGGAAGGAAG
GAGCACACGTCTGAACTCTTTCCCTACACGACGCTCTTCCGATCTNNNACTCTTTCTCTCTCCTGTGGCT

AGCACACGTCTGAACTCTTTCCCTACACGACGCTCTTCCGATCTNNNATCTGGAGCTAAAATGAGCCT
GCACACGTCTGAACTCTTTCCCTACACGACGCTCTTCCGATCTNNNCTTGGATCTCTCTCCCTCTCTCCC
AGCACACGTCTGAACTCTTTCCCTACACGACGCTCTTCCGATCTNNNCTTCTAGTCTGGTGGTTGACAC
AGCACACGTCTGAACTCTTTCCCTACACGACGCTCTTCCGATCTNNNGGAGAGAAAAAGGGAGTAGG
CACACGTCTGAACTCTTTCCCTACACGACGCTCTTCCGATCTNNNTCCTTTCTGCTCAATTACAGGGCC
AGCACACGTCTGAACTCTTTCCCTACACGACGCTCTTCCGATCTNNNTATGATCCCAAAGCATTCCAG
GCACACGTCTGAACTCTTTCCCTACACGACGCTCTTCCGATCTNNNCTCTATGTGAGCATCTTGGGGT
CACACGTCTGAACTCTTTCCCTACACGACGCTCTTCCGATCTNNNCTTTGTTCCTTTCCTGGTCTGGGC
AGCACACGTCTGAACTCTTTCCCTACACGACGCTCTTCCGATCTNNNCTTGCTCGGTCTCACTGTATTG
GAGCACACGTCTGAACTCTTTCCCTACACGACGCTCTTCCGATCTNNNCTTTGTGTGAAAAGGACAGG
AGCACACGTCTGAACTCTTTCCCTACACGACGCTCTTCCGATCTNNNTATGGTGGAATATGCTGGCCC
AGCACACGTCTGAACTCTTTCCCTACACGACGCTCTTCCGATCTNNNTGATCTGAGCTGTAAGGGAGG
GAGCACACGTCTGAACTCTTTCCCTACACGACGCTCTTCCGATCTNNNAGTGGCAAAACCACAATTAC
AGCACACGTCTGAACTCTTTCCCTACACGACGCTCTTCCGATCTNNNGAGTGTACCTGAAACCAGTCTG
AGCACACGTCTGAACTCTTTCCCTACACGACGCTCTTCCGATCTNNNTTTGTAGCCTTGCCTTTATGCC
AGCACACGTCTGAACTCTTTCCCTACACGACGCTCTTCCGATCTNNNGGGTCCATGAAGGCAGAGAT
AGCACACGTCTGAACTCTTTCCCTACACGACGCTCTTCCGATCTNNNGTCCCACAGAGGTTTCTCTCTC
GAGCACACGTCTGAACTCTTTCCCTACACGACGCTCTTCCGATCTNNNGGTGATGCCTTGAATGAGG
AGCACACGTCTGAACTCTTTCCCTACACGACGCTCTTCCGATCTNNNTGGTGTCACTCAAATTTTGCCC
GAGCACACGTCTGAACTCTTTCCCTACACGACGCTCTTCCGATCTNNNATCACTTCCTCCAGGTTGAAA
GAGCACACGTCTGAACTCTTTCCCTACACGACGCTCTTCCGATCTNNNCTACGTTCTGGGAAGGGAAG
AGCACACGTCTGAACTCTTTCCCTACACGACGCTCTTCCGATCTNNNTAGGTACTCACACCTCCCTTTC
GAGCACACGTCTGAACTCTTTCCCTACACGACGCTCTTCCGATCTNNNAGAGAAGCAAACGCAAATCC
GAGCACACGTCTGAACTCTTTCCCTACACGACGCTCTTCCGATCTNNNTCAACCCTAACACTTTTCCCC
AGCACACGTCTGAACTCTTTCCCTACACGACGCTCTTCCGATCTNNNGGAAGTCTACCCTGCTCTCTCA
GAGCACACGTCTGAACTCTTTCCCTACACGACGCTCTTCCGATCTNNNTTGGGTGATGTAGCCAGAAC
GAGCACACGTCTGAACTCTTTCCCTACACGACGCTCTTCCGATCTNNNTAGAATACATGCACCCACTGC
AGCACACGTCTGAACTCTTTCCCTACACGACGCTCTTCCGATCTNNNATGAGGGGCTTTCCTATCCTGT
AGAGCACACGTCTGAACTCTTTCCCTACACGACGCTCTTCCGATCTNNNGACAGTCTTTCTCTCTCCTTG
AGAGCACACGTCTGAACTCTTTCCCTACACGACGCTCTTCCGATCTNNNATTAGGTCATAAGGGTGGG
AGAGCACACGTCTGAACTCTTTCCCTACACGACGCTCTTCCGATCTNNNATGGGGAGAGTTGGTTATG
AGAGCACACGTCTGAACTCTTTCCCTACACGACGCTCTTCCGATCTNNNCTGGATGGGAATCCTGCTT
AGCACACGTCTGAACTCTTTCCCTACACGACGCTCTTCCGATCTNNNGGTTCAACCAAACCAGAGACA
GAGCACACGTCTGAACTCTTTCCCTACACGACGCTCTTCCGATCTNNNTTTGTAGGGAAAACCAGCAT
GAGCACACGTCTGAACTCTTTCCCTACACGACGCTCTTCCGATCTNNNTGATGATTTGGGAGTAGGG
GAGCACACGTCTGAACTCTTTCCCTACACGACGCTCTTCCGATCTNNNTGTGCCCTGTGTGAAAACAG
AGCACACGTCTGAACTCTTTCCCTACACGACGCTCTTCCGATCTNNNGCAGTGAGACCTCTCTCTCTCT
GCACACGTCTGAACTCTTTCCCTACACGACGCTCTTCCGATCTNNNTCAAAGCAGGCGACTAGTAGAG
AGCACACGTCTGAACTCTTTCCCTACACGACGCTCTTCCGATCTNNNCCTGCTTACACACTATCACCAC
GAGCACACGTCTGAACTCTTTCCCTACACGACGCTCTTCCGATCTNNNTCCACCCCAAAGTAGATAGA
GAGCACACGTCTGAACTCTTTCCCTACACGACGCTCTTCCGATCTNNNCCCCTTGGACCCAAGATTACA
AGCACACGTCTGAACTCTTTCCCTACACGACGCTCTTCCGATCTNNNCTCAGGTAGGTGTCCTTATCCC
GCACACGTCTGAACTCTTTCCCTACACGACGCTCTTCCGATCTNNNCTCTCCATCCCATACAGCCAAACC

AGCACACGTCTGAACTCTTTCCCTACACGACGCTCTTCCGATCTNNNTGTCCTAACTGACTGGCAGAA
GAGCACACGTCTGAACTCTTTCCCTACACGACGCTCTTCCGATCTNNNCACCCAATACTGAGGAGGCT
GAGCACACGTCTGAACTCTTTCCCTACACGACGCTCTTCCGATCTNNNGTGTTAGTCAAGTCTGACCC
GAGCACACGTCTGAACTCTTTCCCTACACGACGCTCTTCCGATCTNNNTGTACACTGTGCAGAGGTAA
AGCACACGTCTGAACTCTTTCCCTACACGACGCTCTTCCGATCTNNNCTTCCATGAGCTCAGCTTGACT
GAGCACACGTCTGAACTCTTTCCCTACACGACGCTCTTCCGATCTNNNACAGGTCAGTAGGAATCGTT
GAGCACACGTCTGAACTCTTTCCCTACACGACGCTCTTCCGATCTNNNTCCACATGACAAAACACTGG
AGCACACGTCTGAACTCTTTCCCTACACGACGCTCTTCCGATCTNNNTAACCAAATGCCCACCAGTCA
AGCACACGTCTGAACTCTTTCCCTACACGACGCTCTTCCGATCTNNNGGCCAATAACTACCTGGGGA
GCACACGTCTGAACTCTTTCCCTACACGACGCTCTTCCGATCTNNNTTATTGTCAAGAAGCAGAGCA
GAGCACACGTCTGAACTCTTTCCCTACACGACGCTCTTCCGATCTNNNAAAGAAACTCTCTCCTCCCTC
GCACACGTCTGAACTCTTTCCCTACACGACGCTCTTCCGATCTNNNCTAACTCTCTGAGCTCTCCCTCC
AGCACACGTCTGAACTCTTTCCCTACACGACGCTCTTCCGATCTNNNGTGAAAGGCACTGGAGGTT
GAGCACACGTCTGAACTCTTTCCCTACACGACGCTCTTCCGATCTNNNATTCCTTCTGGGCCCTTTCAA
AGCACACGTCTGAACTCTTTCCCTACACGACGCTCTTCCGATCTNNNATTCCACATCTGTTTTCCCCACC
AGCACACGTCTGAACTCTTTCCCTACACGACGCTCTTCCGATCTNNNTGCCTCTGCTGTCCTGTATCTA
GCACACGTCTGAACTCTTTCCCTACACGACGCTCTTCCGATCTNNNCCCTCTCAAACCATTCCCTTCTG
AGCACACGTCTGAACTCTTTCCCTACACGACGCTCTTCCGATCTNNNGAACTTAGAGTGGTAAGCTG
AGCACACGTCTGAACTCTTTCCCTACACGACGCTCTTCCGATCTNNNTAAGCTTGGTTTTCTGACGCC
GAGCACACGTCTGAACTCTTTCCCTACACGACGCTCTTCCGATCTNNNGCTGAATCCTGAGTAACAGA
AGCACACGTCTGAACTCTTTCCCTACACGACGCTCTTCCGATCTNNNTGCAAACCCTTCATTCACAAGG
AGCACACGTCTGAACTCTTTCCCTACACGACGCTCTTCCGATCTNNNCCAAAGCCTGGGGTTTTCATC
GCACACGTCTGAACTCTTTCCCTACACGACGCTCTTCCGATCTNNNTCTTCTCTCTCTCCCTGTCCTC
GCACACGTCTGAACTCTTTCCCTACACGACGCTCTTCCGATCTNNNACACAAAGAATGCACAAGCATG
AGCACACGTCTGAACTCTTTCCCTACACGACGCTCTTCCGATCTNNNAGGGTGGCTAAAAATGTGG
AGCACACGTCTGAACTCTTTCCCTACACGACGCTCTTCCGATCTNNNAAGTGCTTCTCTGAGGAGGG
AGCACACGTCTGAACTCTTTCCCTACACGACGCTCTTCCGATCTNNNGACATTCCGAGTATGCGTGTG
AGCACACGTCTGAACTCTTTCCCTACACGACGCTCTTCCGATCTNNNAATTTTAGGCTTTGGGGCTG
AGCACACGTCTGAACTCTTTCCCTACACGACGCTCTTCCGATCTNNNTGTAAGAAGAGGAAAGGAG
AGCACACGTCTGAACTCTTTCCCTACACGACGCTCTTCCGATCTNNNTAGCTGTATCAGGTACTGCAG
GCACACGTCTGAACTCTTTCCCTACACGACGCTCTTCCGATCTNNNGCAAGGGAAATTCACACACACA
AGCACACGTCTGAACTCTTTCCCTACACGACGCTCTTCCGATCTNNNAATCCGTTGGGTTCACTCCTTT
AGCACACGTCTGAACTCTTTCCCTACACGACGCTCTTCCGATCTNNNCGACAATGGGTGATAGTGGG
GCACACGTCTGAACTCTTTCCCTACACGACGCTCTTCCGATCTNNNGCATAGTACATGGGTGTCCTCA
AGCACACGTCTGAACTCTTTCCCTACACGACGCTCTTCCGATCTNNNAGCCCCAACAAAAGTTTGTCT
GCACACGTCTGAACTCTTTCCCTACACGACGCTCTTCCGATCTNNNATGAGCATGTACACACACACAGT
GCACACGTCTGAACTCTTTCCCTACACGACGCTCTTCCGATCTNNNTCCCCCACTCAAGCATTTGAATC
AGCACACGTCTGAACTCTTTCCCTACACGACGCTCTTCCGATCTNNNTAGGGGTGAAGGACATCTGA
GCACACGTCTGAACTCTTTCCCTACACGACGCTCTTCCGATCTNNNTAACACAGCTCTCACCACCATTC
AGCACACGTCTGAACTCTTTCCCTACACGACGCTCTTCCGATCTNNNTTCCTGTAGGAACTCTCCAGG
GCACACGTCTGAACTCTTTCCCTACACGACGCTCTTCCGATCTNNNGGCATCTTGGTCTCCCATAGGTA
AGAGCACACGTCTGAACTCTTTCCCTACACGACGCTCTTCCGATCTNNNCTTCCCCTTCACTTCTTCAGT
GAGCACACGTCTGAACTCTTTCCCTACACGACGCTCTTCCGATCTNNNTGCAAGCATCTGTTTATGTG

AGAGCACACGTCTGAACTCTTTCCCTACACGACGCTCTTCCGATCTNNNAAAGGAACCAGCACATGAG
AGAGCACACGTCTGAACTCTTTCCCTACACGACGCTCTTCCGATCTNNNAGTACCCTATTCAGCAGAG
GAGCACACGTCTGAACTCTTTCCCTACACGACGCTCTTCCGATCTNNNCATGCTCCTCAACTTGTGAC
AGAGCACACGTCTGAACTCTTTCCCTACACGACGCTCTTCCGATCTNNNGAATTCTCTTTTCCTGGCTT
AAGAGCACACGTCTGAACTCTTTCCCTACACGACGCTCTTCCGATCTNNNCCAAACAGATAGAAGGGC
GAGCACACGTCTGAACTCTTTCCCTACACGACGCTCTTCCGATCTNNNCTCTGGGAAAGAATCTGGC
AGCACACGTCTGAACTCTTTCCCTACACGACGCTCTTCCGATCTNNNTAGCTGTATCAGGTACTGCAG
AGCACACGTCTGAACTCTTTCCCTACACGACGCTCTTCCGATCTNNNTCTGATGCAAAACTCGACCTT
AGCACACGTCTGAACTCTTTCCCTACACGACGCTCTTCCGATCTNNNTACAGTTGAAAATATGGGCCC
AGCACACGTCTGAACTCTTTCCCTACACGACGCTCTTCCGATCTNNNTTCCCATGTCCTCCTCTTCTCTC
AGAGCACACGTCTGAACTCTTTCCCTACACGACGCTCTTCCGATCTNNNTGATGTGAGGAATGTCCC
AGAGCACACGTCTGAACTCTTTCCCTACACGACGCTCTTCCGATCTNNNTTTGAGTAAACCAGGCCC
GAGCACACGTCTGAACTCTTTCCCTACACGACGCTCTTCCGATCTNNNACCCACTACAGGATATCTGGC
GCACACGTCTGAACTCTTTCCCTACACGACGCTCTTCCGATCTNNNTTTCATCCTTCCCTCCTCCTTCTCA
AGCACACGTCTGAACTCTTTCCCTACACGACGCTCTTCCGATCTNNNCAGTCTGACACCTGCTCTTTTCA
AGCACACGTCTGAACTCTTTCCCTACACGACGCTCTTCCGATCTNNNTGGAGCAGAGGTGCACTATC
GAGCACACGTCTGAACTCTTTCCCTACACGACGCTCTTCCGATCTNNNACTAGATGATTCCCAAGGGG
AGCACACGTCTGAACTCTTTCCCTACACGACGCTCTTCCGATCTNNNTTTAGACACCACCTCCCACAGAT
AGCACACGTCTGAACTCTTTCCCTACACGACGCTCTTCCGATCTNNNAGCTGGTTTGTACATCAAAGG
GAGCACACGTCTGAACTCTTTCCCTACACGACGCTCTTCCGATCTNNNCTTGCATGATACAGGGAGGC
AGCACACGTCTGAACTCTTTCCCTACACGACGCTCTTCCGATCTNNNCCCTGGACTGGAACACAAATT
AGCACACGTCTGAACTCTTTCCCTACACGACGCTCTTCCGATCTNNNATAGATGGGGGCCAAAATGTT
GAGCACACGTCTGAACTCTTTCCCTACACGACGCTCTTCCGATCTNNNTGAGCTGTGCATGGAGAAAA
GAGCACACGTCTGAACTCTTTCCCTACACGACGCTCTTCCGATCTNNNATGTTGGTCATGTTCTGCTTC
GAGCACACGTCTGAACTCTTTCCCTACACGACGCTCTTCCGATCTNNNATCCAGGAACTCAGATCTACC
AGCACACGTCTGAACTCTTTCCCTACACGACGCTCTTCCGATCTNNNCTTCTTTGTCTCTGTCCCTCCCT
CACACGTCTGAACTCTTTCCCTACACGACGCTCTTCCGATCTNNNGTGTATATATGTATGTGTGAGAC
GAGCACACGTCTGAACTCTTTCCCTACACGACGCTCTTCCGATCTNNNAGAATAGGGAAGGGGAAGG
GAGCACACGTCTGAACTCTTTCCCTACACGACGCTCTTCCGATCTNNNGCTACACTCAGCACCTGTTAC
GCACACGTCTGAACTCTTTCCCTACACGACGCTCTTCCGATCTNNNTCAGCAAGAGGGGAAAAGAGA
AGCACACGTCTGAACTCTTTCCCTACACGACGCTCTTCCGATCTNNNACCAGAGGGATGAAAAGGAA
AGAGCACACGTCTGAACTCTTTCCCTACACGACGCTCTTCCGATCTNNNATTCTGAGCTCCTTCTTTGCA
AGAGCACACGTCTGAACTCTTTCCCTACACGACGCTCTTCCGATCTNNNAGATTGAGATCTCTGCGGA
GCACACGTCTGAACTCTTTCCCTACACGACGCTCTTCCGATCTNNNAGAAGCTAGGAAGCCATCTGGA
GCACACGTCTGAACTCTTTCCCTACACGACGCTCTTCCGATCTNNNAAAGAGCAGATGGAGCAGAAT
AAGAGCACACGTCTGAACTCTTTCCCTACACGACGCTCTTCCGATCTNNNAAGCATTTTATTGAACGGG
AGAGCACACGTCTGAACTCTTTCCCTACACGACGCTCTTCCGATCTNNNGTTTGGCCTGAGTTTCTGG
AGAGCACACGTCTGAACTCTTTCCCTACACGACGCTCTTCCGATCTNNNACATACATGTTGAGATCCTG
GAGCACACGTCTGAACTCTTTCCCTACACGACGCTCTTCCGATCTNNNTCCCCAGAACTCAAGGGTAAT
GAGCACACGTCTGAACTCTTTCCCTACACGACGCTCTTCCGATCTNNNTTTTGGGTTGATGGGTTCCA
GAGCACACGTCTGAACTCTTTCCCTACACGACGCTCTTCCGATCTNNNGTGAGGAGTGAGAAAGAGG
GAGCACACGTCTGAACTCTTTCCCTACACGACGCTCTTCCGATCTNNNAAGAAAGGGTGGTGAGACA
GAGCACACGTCTGAACTCTTTCCCTACACGACGCTCTTCCGATCTNNNTTACTGTTGTGAGCCAGTCTT

GAGCACACGTCTGAACTCTTTCCCTACACGACGCTCTTCCGATCTNNNTCAAGGATGTGTAGGTGCAT
AGCACACGTCTGAACTCTTTCCCTACACGACGCTCTTCCGATCTNNNTTCTTCTAAAGCCCAAGGTAGG
AGAGCACACGTCTGAACTCTTTCCCTACACGACGCTCTTCCGATCTNNNTGCTGCTTCATGAATGACCC
GAGCACACGTCTGAACTCTTTCCCTACACGACGCTCTTCCGATCTNNNGATAGCGTCAGATCCAGACAC
AGCACACGTCTGAACTCTTTCCCTACACGACGCTCTTCCGATCTNNNTCAGCCCATACAAAACTGACCTG
GAGCACACGTCTGAACTCTTTCCCTACACGACGCTCTTCCGATCTNNNCCTTCCTTTCACAACCCCTTCT
AGAGCACACGTCTGAACTCTTTCCCTACACGACGCTCTTCCGATCTNNNAGATGTTTCTGTGCATGTG
GAGCACACGTCTGAACTCTTTCCCTACACGACGCTCTTCCGATCTNNNGGTCTGCGAAAGATGAGTTA
AGCACACGTCTGAACTCTTTCCCTACACGACGCTCTTCCGATCTNNNAGAGGCATGTGTACTTACGGC
AGAGCACACGTCTGAACTCTTTCCCTACACGACGCTCTTCCGATCTNNNAAACCACTAACTCTCACGCA
AGAGCACACGTCTGAACTCTTTCCCTACACGACGCTCTTCCGATCTNNNCTTGTGTTTTCAGATGGGT
AGAGCACACGTCTGAACTCTTTCCCTACACGACGCTCTTCCGATCTNNNTTGCATTTGCAACAAAGGA
AGCACACGTCTGAACTCTTTCCCTACACGACGCTCTTCCGATCTNNNTGGCCTGCCAGGATTTAGAGA
GCACACGTCTGAACTCTTTCCCTACACGACGCTCTTCCGATCTNNNGGGAGTGTATACGTGGGAGAG
GAGCACACGTCTGAACTCTTTCCCTACACGACGCTCTTCCGATCTNNNCATGGAGGGCATCTCTTTCTC
AGAGCACACGTCTGAACTCTTTCCCTACACGACGCTCTTCCGATCTNNNATTGCGGTAACTCTGAGTG
AGCACACGTCTGAACTCTTTCCCTACACGACGCTCTTCCGATCTNNNTACATACACACAAAAGGGTGC
AGCACACGTCTGAACTCTTTCCCTACACGACGCTCTTCCGATCTNNNAGAAAACACTATGCCCACTCTC
AGAGCACACGTCTGAACTCTTTCCCTACACGACGCTCTTCCGATCTNNNAGCACAGTACAGCAAACTTG
GAGCACACGTCTGAACTCTTTCCCTACACGACGCTCTTCCGATCTNNNCTGGACAACATCGTAAAGCT
GAGCACACGTCTGAACTCTTTCCCTACACGACGCTCTTCCGATCTNNNAAAATGCAGAGTAATGGTGG
AGAGCACACGTCTGAACTCTTTCCCTACACGACGCTCTTCCGATCTNNNCATGCCTGACATGCAGCAA
GAGCACACGTCTGAACTCTTTCCCTACACGACGCTCTTCCGATCTNNNCTTTCTCTCCCTCCATCCCCTT
GAGCACACGTCTGAACTCTTTCCCTACACGACGCTCTTCCGATCTNNNACCTACCATAAGCTGTACTCAG
AGAGCACACGTCTGAACTCTTTCCCTACACGACGCTCTTCCGATCTNNNGACGGCAGAATGAAACCTT
GAGCACACGTCTGAACTCTTTCCCTACACGACGCTCTTCCGATCTNNNTATCCAAGCTAAGAAGCACG
AGAGCACACGTCTGAACTCTTTCCCTACACGACGCTCTTCCGATCTNNNTGCTTATAATTTTGTCAGCG
AGAGCACACGTCTGAACTCTTTCCCTACACGACGCTCTTCCGATCTNNNTCTGTCAAGCATCATCTGCTT
GAGCACACGTCTGAACTCTTTCCCTACACGACGCTCTTCCGATCTNNNCGCACATTCCCTCTCTCTCTTT
AGCACACGTCTGAACTCTTTCCCTACACGACGCTCTTCCGATCTNNNGAATGCTACATATGACCCTGCC
AGCACACGTCTGAACTCTTTCCCTACACGACGCTCTTCCGATCTNNNCTGTGCGTACAGCTACTCTCAT
AGAGCACACGTCTGAACTCTTTCCCTACACGACGCTCTTCCGATCTNNNTCTGTAAGTACTTTGCCTGT
GAGCACACGTCTGAACTCTTTCCCTACACGACGCTCTTCCGATCTNNNGTGCTGCCCAAAGACAGAAT
GCACACGTCTGAACTCTTTCCCTACACGACGCTCTTCCGATCTNNNCTAGCACCTCCATCCTACCTCTCC
GAGCACACGTCTGAACTCTTTCCCTACACGACGCTCTTCCGATCTNNNGGCAATGTTTGCCATGAAAG
GAGCACACGTCTGAACTCTTTCCCTACACGACGCTCTTCCGATCTNNNTCCTATCTGAATGGCTACGCT
AGCACACGTCTGAACTCTTTCCCTACACGACGCTCTTCCGATCTNNNAACGGTATGTTATATGGGTGC
GAGCACACGTCTGAACTCTTTCCCTACACGACGCTCTTCCGATCTNNNATGTTGGGAAACAGCTGGGT
AGCACACGTCTGAACTCTTTCCCTACACGACGCTCTTCCGATCTNNNCCTTGCCCATTTTGCCCTACTAT
GAGCACACGTCTGAACTCTTTCCCTACACGACGCTCTTCCGATCTNNNCCTTGGGACACTTTTGTTTG
GAGCACACGTCTGAACTCTTTCCCTACACGACGCTCTTCCGATCTNNNGTTTACAAGGGTGGAGACAG
GAGCACACGTCTGAACTCTTTCCCTACACGACGCTCTTCCGATCTNNNGGGCAAAGCTTATCTACTGGC
GAGCACACGTCTGAACTCTTTCCCTACACGACGCTCTTCCGATCTNNNGAAGGTCCAGGAACAGAGG

AGAGCACACGTCTGAACTCTTTCCCTACACGACGCTCTTCCGATCTNNNAGGCACCATCAAAAATGACC
GAGCACACGTCTGAACTCTTTCCCTACACGACGCTCTTCCGATCTNNNGGAGCACTCTGGAATTCTCTCT
AGAGCACACGTCTGAACTCTTTCCCTACACGACGCTCTTCCGATCTNNNTCAATCGTGACGGTTTTATG
AGCACACGTCTGAACTCTTTCCCTACACGACGCTCTTCCGATCTNNNAAGCTAAGTCAGTAAGCCACAG
AGCACACGTCTGAACTCTTTCCCTACACGACGCTCTTCCGATCTNNNTTATTCTGTCACCAAGGTGTGC
AGCACACGTCTGAACTCTTTCCCTACACGACGCTCTTCCGATCTNNNAGAAAATATGAGTGCACATGC
AGCACACGTCTGAACTCTTTCCCTACACGACGCTCTTCCGATCTNNNTATGACCTTCACTCTGGGACAG
AGCACACGTCTGAACTCTTTCCCTACACGACGCTCTTCCGATCTNNNAGCTGTTGCCACAAATTCTGAA
AGAGCACACGTCTGAACTCTTTCCCTACACGACGCTCTTCCGATCTNNNTAAAAAGAGCCTGTGGTTGC
AGCACACGTCTGAACTCTTTCCCTACACGACGCTCTTCCGATCTNNNTTTTATCCGCACACATTCACCCA
GAGCACACGTCTGAACTCTTTCCCTACACGACGCTCTTCCGATCTNNNGCCTCCAGGTTTCCTTGTAGAT
GAGCACACGTCTGAACTCTTTCCCTACACGACGCTCTTCCGATCTNNNACACAGTGGCTAAAGAGAGA
AGAGCACACGTCTGAACTCTTTCCCTACACGACGCTCTTCCGATCTNNNTAAGAGGGCCCAGTACTCAC
AGAGCACACGTCTGAACTCTTTCCCTACACGACGCTCTTCCGATCTNNNGGTTGGCTATCACAAAACTG
AGAGCACACGTCTGAACTCTTTCCCTACACGACGCTCTTCCGATCTNNNCCACTGCTAGCTCTCTCCTCAT
GAGCACACGTCTGAACTCTTTCCCTACACGACGCTCTTCCGATCTNNNGAAGCACATGGGCAAGGTAT
GAGCACACGTCTGAACTCTTTCCCTACACGACGCTCTTCCGATCTNNNCTCCTCCAGGAATCTTTGGCTT
AGCACACGTCTGAACTCTTTCCCTACACGACGCTCTTCCGATCTNNNAAATCCCCTCTTCTCACTCTCCCCA
AGCACACGTCTGAACTCTTTCCCTACACGACGCTCTTCCGATCTNNNCTGATGCTGAAGGGTAGAGGA
AGAGCACACGTCTGAACTCTTTCCCTACACGACGCTCTTCCGATCTNNNGTGCATGAAGATAGGGCCTT
GAGCACACGTCTGAACTCTTTCCCTACACGACGCTCTTCCGATCTNNNTAAAAGAGTGTGGCTCCTCCT
AGAGCACACGTCTGAACTCTTTCCCTACACGACGCTCTTCCGATCTNNNCTGGAGAGTAGGAAGTGAG
GAGCACACGTCTGAACTCTTTCCCTACACGACGCTCTTCCGATCTNNNTGAAGTGTGTGTGTGTGTGT
AGCACACGTCTGAACTCTTTCCCTACACGACGCTCTTCCGATCTNNNCCCAACATCATCAGAAGAGCCA
AGAGCACACGTCTGAACTCTTTCCCTACACGACGCTCTTCCGATCTNNNATTGTTACTGTTTTCGCAGG
AGCACACGTCTGAACTCTTTCCCTACACGACGCTCTTCCGATCTNNNGGGACCATGAGAAGCATGAAT
AGCACACGTCTGAACTCTTTCCCTACACGACGCTCTTCCGATCTNNNCCTTACACGCTCTGTGAACATCA
AGCACACGTCTGAACTCTTTCCCTACACGACGCTCTTCCGATCTNNNTGCTCTGCTTACTTACCTACCCC
GAGCACACGTCTGAACTCTTTCCCTACACGACGCTCTTCCGATCTNNNGTAAGAGCAAGCAGAGACCCA
AGCACACGTCTGAACTCTTTCCCTACACGACGCTCTTCCGATCTNNNTCCTGGGTTTGTGTTGTTGCAT
GAGCACACGTCTGAACTCTTTCCCTACACGACGCTCTTCCGATCTNNNCTGTTGCAGCCAATTCTTTCA
GAGCACACGTCTGAACTCTTTCCCTACACGACGCTCTTCCGATCTNNNATCCCTGGCTCCTAGTTCAAT
GAGCACACGTCTGAACTCTTTCCCTACACGACGCTCTTCCGATCTNNNGCCACAGAGTGGAGATTTTCA
AGCACACGTCTGAACTCTTTCCCTACACGACGCTCTTCCGATCTNNNTACCTACAGTCAACCCTGGACAC
AGAGCACACGTCTGAACTCTTTCCCTACACGACGCTCTTCCGATCTNNNGGAGACCGAAGCTACTTTTT
GAGCACACGTCTGAACTCTTTCCCTACACGACGCTCTTCCGATCTNNNCCTTAGCCATCTTAGGGAAG
AGCACACGTCTGAACTCTTTCCCTACACGACGCTCTTCCGATCTNNNTATTTGAATGAACGGGCCCTCA
AGCACACGTCTGAACTCTTTCCCTACACGACGCTCTTCCGATCTNNNGTCTGCCTCCCCATCATACATTC
GAGCACACGTCTGAACTCTTTCCCTACACGACGCTCTTCCGATCTNNNGCTTACCATGCATGTGGACCTA
GAGCACACGTCTGAACTCTTTCCCTACACGACGCTCTTCCGATCTNNNAAAGCCATCTGTTACAGCACT
AGCACACGTCTGAACTCTTTCCCTACACGACGCTCTTCCGATCTNNNGGGTTGTAAGGGTGTGTTGAT
AGAGCACACGTCTGAACTCTTTCCCTACACGACGCTCTTCCGATCTNNNGGGAGATTCTGTATAGCCAT
GAGCACACGTCTGAACTCTTTCCCTACACGACGCTCTTCCGATCTNNNAGTATTCACGGACATTGCATC

AGCACACGTCTGAACTCTTTCCCTACACGACGCTCTTCCGATCTNNNCTCACTTGTTCTGTTCTCCCCTG

AGAGCACACGTCTGAACTCTTTCCCTACACGACGCTCTTCCGATCTNNNCCAGGAGGTAGACTTAGCA

AGCACACGTCTGAACTCTTTCCCTACACGACGCTCTTCCGATCTNNNAAATGTATGCGTTCTACCCCCA

GAGCACACGTCTGAACTCTTTCCCTACACGACGCTCTTCCGATCTNNNTCAGGTGCCCATCAGTGGAT

AGCACACGTCTGAACTCTTTCCCTACACGACGCTCTTCCGATCTNNNAATCTGAATCCCTTGGCTGTGA

AGCACACGTCTGAACTCTTTCCCTACACGACGCTCTTCCGATCTNNNCACACCACCACACCCTGATTTAT

GCACACGTCTGAACTCTTTCCCTACACGACGCTCTTCCGATCTNNNAAAGTCAGTCTCTCTCTCTCCCTT

AGCACACGTCTGAACTCTTTCCCTACACGACGCTCTTCCGATCTNNNAGACAGGGTCTTCCAATAGGC

GCACACGTCTGAACTCTTTCCCTACACGACGCTCTTCCGATCTNNNAGCATTGAAAAGCTGTGTGGGT

GCACACGTCTGAACTCTTTCCCTACACGACGCTCTTCCGATCTNNNTAAAGCACCCAAGAAAAACCAC

AGCACACGTCTGAACTCTTTCCCTACACGACGCTCTTCCGATCTNNNCAGAAATCCTACCTTCCTTGCCA

AGAGCACACGTCTGAACTCTTTCCCTACACGACGCTCTTCCGATCTNNNGGTTGCAAAGTTTCTCGAG

GCACACGTCTGAACTCTTTCCCTACACGACGCTCTTCCGATCTNNNTATCTCATCTCCTTTGCCCTCCCCA

AGCACACGTCTGAACTCTTTCCCTACACGACGCTCTTCCGATCTNNNGGTGGTCTCACAGGTCTTTGTT

AGCACACGTCTGAACTCTTTCCCTACACGACGCTCTTCCGATCTNNNTGAGGAAGAGGTAGCTGGAA

GAGCACACGTCTGAACTCTTTCCCTACACGACGCTCTTCCGATCTNNNACAAAGATCCATTCCCTCCCA

AGCACACGTCTGAACTCTTTCCCTACACGACGCTCTTCCGATCTNNNACAGAACATGTGTAGCACTTC

AGCACACGTCTGAACTCTTTCCCTACACGACGCTCTTCCGATCTNNNCTCTTTAGGTCAGCGGTGATTC

GAGCACACGTCTGAACTCTTTCCCTACACGACGCTCTTCCGATCTNNNAGTATACCCATGATTCCAGCC

AGCACACGTCTGAACTCTTTCCCTACACGACGCTCTTCCGATCTNNNAGCACAAAATTGACAAAAAGA

AGCACACGTCTGAACTCTTTCCCTACACGACGCTCTTCCGATCTNNNCAGCAAGCTGAGCTATGTCCTT

GAGCACACGTCTGAACTCTTTCCCTACACGACGCTCTTCCGATCTNNNCCAAGAGCCTTTTGTTCAGG

AGCACACGTCTGAACTCTTTCCCTACACGACGCTCTTCCGATCTNNNATTGGGCATGAGTGGTTACAC

GAGCACACGTCTGAACTCTTTCCCTACACGACGCTCTTCCGATCTNNNCTGTGGCTCTCTGTTTGCCTT

AGCACACGTCTGAACTCTTTCCCTACACGACGCTCTTCCGATCTNNNCATGGTACGCAACCAGTGTGT

AGCACACGTCTGAACTCTTTCCCTACACGACGCTCTTCCGATCTNNNAGTGAAGCAACCCTTCTCTCTCT

GAGCACACGTCTGAACTCTTTCCCTACACGACGCTCTTCCGATCTNNNAAGCTAAGCCAAATAAGAG

AGCACACGTCTGAACTCTTTCCCTACACGACGCTCTTCCGATCTNNNTAAGGTAATACCAGTGTGCAG

AGCACACGTCTGAACTCTTTCCCTACACGACGCTCTTCCGATCTNNNTTCACACACCTGCATGTAAGCA

GAGCACACGTCTGAACTCTTTCCCTACACGACGCTCTTCCGATCTNNNCCAATCAAGGACTTCAGAGT

AGCACACGTCTGAACTCTTTCCCTACACGACGCTCTTCCGATCTNNNTCCTGTTCTAAGGCAAGGCAA

AGCACACGTCTGAACTCTTTCCCTACACGACGCTCTTCCGATCTNNNAGGGATCTGCTTCTGAACTGG

AGCACACGTCTGAACTCTTTCCCTACACGACGCTCTTCCGATCTNNNTCCAGCATGTTTACACACACAC

GAGCACACGTCTGAACTCTTTCCCTACACGACGCTCTTCCGATCTNNNCAAGATCCACTCCAGGCTTTCT

GAGCACACGTCTGAACTCTTTCCCTACACGACGCTCTTCCGATCTNNNGGCATCTGAGTGTTCCATGC

AGCACACGTCTGAACTCTTTCCCTACACGACGCTCTTCCGATCTNNNGCGCCTTATCTCTGTCTCTCTCT

GAGCACACGTCTGAACTCTTTCCCTACACGACGCTCTTCCGATCTNNNTTTTTCCAGCTCCCATTTTGC

GAGCACACGTCTGAACTCTTTCCCTACACGACGCTCTTCCGATCTNNNTGTGCTTGTTTTTGCAGATGT

GAGCACACGTCTGAACTCTTTCCCTACACGACGCTCTTCCGATCTNNNACAAGACCCTAGCAACTCTCT

AGCACACGTCTGAACTCTTTCCCTACACGACGCTCTTCCGATCTNNNGCTCAGTGTGGACATGGAAAT

GAGCACACGTCTGAACTCTTTCCCTACACGACGCTCTTCCGATCTNNNGAAGGAGCTTGCAGAGCATA

AGCACACGTCTGAACTCTTTCCCTACACGACGCTCTTCCGATCTNNNAACTTTGGTGGCTGTCCTAAGT

GAGCACACGTCTGAACTCTTTCCCTACACGACGCTCTTCCGATCTNNNATGCCCCAGAGATCTGGTTTT

GAGCACACGTCTGAACTCTTTCCCTACACGACGCTCTTCCGATCTNNNGAAGAAGTCCTGATGTCACT
AGCACACGTCTGAACTCTTTCCCTACACGACGCTCTTCCGATCTNNNACATGAGACAGAGGGGGTAG
GAGCACACGTCTGAACTCTTTCCCTACACGACGCTCTTCCGATCTNNNGCCCAGGCAACAATGGTAAA
GAGCACACGTCTGAACTCTTTCCCTACACGACGCTCTTCCGATCTNNNCTCCACAGAGTAGCATTTGA
AGCACACGTCTGAACTCTTTCCCTACACGACGCTCTTCCGATCTNNNTAGGATTGCAAGTGAGTGGT
AGCACACGTCTGAACTCTTTCCCTACACGACGCTCTTCCGATCTNNNGTTTCTGGTTCTTGCTGAAGTG
GAGCACACGTCTGAACTCTTTCCCTACACGACGCTCTTCCGATCTNNNGTTGGGCGCTCTTTCTTACA
AGCACACGTCTGAACTCTTTCCCTACACGACGCTCTTCCGATCTNNNCCCCACATGGTTTTGAAAGAG
GCACACGTCTGAACTCTTTCCCTACACGACGCTCTTCCGATCTNNNGCTAAGGGACAGACTTGTTACG
GAGCACACGTCTGAACTCTTTCCCTACACGACGCTCTTCCGATCTNNNGTAAAATGAACCTGTTGGCA
GAGCACACGTCTGAACTCTTTCCCTACACGACGCTCTTCCGATCTNNNGCAAGATCCATGTTGCTGAG
GCACACGTCTGAACTCTTTCCCTACACGACGCTCTTCCGATCTNNNTTTGTGACCTGTGAAGGTGGT
GCACACGTCTGAACTCTTTCCCTACACGACGCTCTTCCGATCTNNNCTGAGACCCTGTCTCTAAACTGC
AGCACACGTCTGAACTCTTTCCCTACACGACGCTCTTCCGATCTNNNAGAAGGGCCTTATCCACATCC
GCACACGTCTGAACTCTTTCCCTACACGACGCTCTTCCGATCTNNNTGGTGTTGAAAGTGCTTCTTGA
GCACACGTCTGAACTCTTTCCCTACACGACGCTCTTCCGATCTNNNAATAGCTCAGAATGTGCCAGAC
GAGCACACGTCTGAACTCTTTCCCTACACGACGCTCTTCCGATCTNNNCTAACCTCCCCTCAGGCTATGT
AGCACACGTCTGAACTCTTTCCCTACACGACGCTCTTCCGATCTNNNACTTCTATCTCAGGCTCGTCCT
GAGCACACGTCTGAACTCTTTCCCTACACGACGCTCTTCCGATCTNNNGGGGTTCTCTGGGTTTTCTT
GCACACGTCTGAACTCTTTCCCTACACGACGCTCTTCCGATCTNNNATGTTTTCTCCTTGCAAACGCTG
AGAGCACACGTCTGAACTCTTTCCCTACACGACGCTCTTCCGATCTNNNAGCCCATGCTAGTAGACCTC
AGCACACGTCTGAACTCTTTCCCTACACGACGCTCTTCCGATCTNNNGAGCAAATTCTGCAAGCTGAA
AGCACACGTCTGAACTCTTTCCCTACACGACGCTCTTCCGATCTNNNCCCAGCTGCCTATTCTGAGTTT
AGCACACGTCTGAACTCTTTCCCTACACGACGCTCTTCCGATCTNNNCTCCAGCTCCCACTGTAGACTA
AGCACACGTCTGAACTCTTTCCCTACACGACGCTCTTCCGATCTNNNCTGTTACAACACCCCAGCTTCAT
GAGCACACGTCTGAACTCTTTCCCTACACGACGCTCTTCCGATCTNNNCCCTCTGGGTCAGACAATCA
GAGCACACGTCTGAACTCTTTCCCTACACGACGCTCTTCCGATCTNNNTCGAAGGTGTACTTTTCCCT
AGCACACGTCTGAACTCTTTCCCTACACGACGCTCTTCCGATCTNNNCTGGAATTCCCAGCTATGCAT
AGCACACGTCTGAACTCTTTCCCTACACGACGCTCTTCCGATCTNNNTCTGCTTCCTTCCCACTCTTCTT
GAGCACACGTCTGAACTCTTTCCCTACACGACGCTCTTCCGATCTNNNAAGGTTTATCAAAAGCACTG
AGCACACGTCTGAACTCTTTCCCTACACGACGCTCTTCCGATCTNNNAACCAAGAACCAACAGGAGAC
AGCACACGTCTGAACTCTTTCCCTACACGACGCTCTTCCGATCTNNNCCTCCCTGGGACTATGTCTCAA
AGCACACGTCTGAACTCTTTCCCTACACGACGCTCTTCCGATCTNNNCAAACAAGCCAACAAACCCAA
GAGCACACGTCTGAACTCTTTCCCTACACGACGCTCTTCCGATCTNNNCCGTGATTGTACACAACGCTC
AGCACACGTCTGAACTCTTTCCCTACACGACGCTCTTCCGATCTNNNAGATCAAGCTGGGAAGCAAA
GAGCACACGTCTGAACTCTTTCCCTACACGACGCTCTTCCGATCTNNNGCTGCTGAATGCCTTCCAACT
AGCACACGTCTGAACTCTTTCCCTACACGACGCTCTTCCGATCTNNNGCTTGTGTCACCCTACTCTCAAC
AGCACACGTCTGAACTCTTTCCCTACACGACGCTCTTCCGATCTNNNCTCACCACCTAGGACATGGAG
AGCACACGTCTGAACTCTTTCCCTACACGACGCTCTTCCGATCTNNNTATTTCCAGTCTGCCAGGCATT
GCACACGTCTGAACTCTTTCCCTACACGACGCTCTTCCGATCTNNNCTTTATGCAAATTGGCCTCGCTG
AGCACACGTCTGAACTCTTTCCCTACACGACGCTCTTCCGATCTNNNTTCCTCCCTGCATGGATTCTGT
GAGCACACGTCTGAACTCTTTCCCTACACGACGCTCTTCCGATCTNNNCTAGGTTGGCCTCTCTCTCTC
AGCACACGTCTGAACTCTTTCCCTACACGACGCTCTTCCGATCTNNNAAGGGTCTACGGACTTCAACA

AGCACACGTCTGAACTCTTTCCCTACACGACGCTCTTCCGATCTNNNGTCTTGGGCTAAACCGATGAA
AGCACACGTCTGAACTCTTTCCCTACACGACGCTCTTCCGATCTNNNATAGCACACATGCCTCCAAACT
AGCACACGTCTGAACTCTTTCCCTACACGACGCTCTTCCGATCTNNNCACATGGCTTTCAGGAGGTCA
GCACACGTCTGAACTCTTTCCCTACACGACGCTCTTCCGATCTNNNATCTAAGTACCCCTTATACACGC
GCACACGTCTGAACTCTTTCCCTACACGACGCTCTTCCGATCTNNNTGGTGCTTTGGTTATCCTTGGTC
AGCACACGTCTGAACTCTTTCCCTACACGACGCTCTTCCGATCTNNNAGAAGTCCTCTTCCCTATCAGC
AGCACACGTCTGAACTCTTTCCCTACACGACGCTCTTCCGATCTNNNTGCTGAGTGTATCCAAATCCTG
GCACACGTCTGAACTCTTTCCCTACACGACGCTCTTCCGATCTNNNAGCTTGCAGGGAATATCTGTTC
AGCACACGTCTGAACTCTTTCCCTACACGACGCTCTTCCGATCTNNNGTGCTTGCACCATTTACAATCC
AGCACACGTCTGAACTCTTTCCCTACACGACGCTCTTCCGATCTNNNGTTCCCATTCTGCTTTTGAGC
AGCACACGTCTGAACTCTTTCCCTACACGACGCTCTTCCGATCTNNNGGAAGTGAACACGGCTGATTA
GCACACGTCTGAACTCTTTCCCTACACGACGCTCTTCCGATCTNNNTGTTCAAGAGACCAGGCTACTC
AGCACACGTCTGAACTCTTTCCCTACACGACGCTCTTCCGATCTNNNCACGAACTCACACACACAACTT
AGCACACGTCTGAACTCTTTCCCTACACGACGCTCTTCCGATCTNNNAAGGGTATCACATGAAACCTG
AGCACACGTCTGAACTCTTTCCCTACACGACGCTCTTCCGATCTNNNTGTCTTGGTTCCTGAATGGTA
GAGCACACGTCTGAACTCTTTCCCTACACGACGCTCTTCCGATCTNNNAGATGTCTAGAACCTGCTCT
GCACACGTCTGAACTCTTTCCCTACACGACGCTCTTCCGATCTNNNAAACCATGAAACCTCCATGTCC
GAGCACACGTCTGAACTCTTTCCCTACACGACGCTCTTCCGATCTNNNCCCCTAAGGCCTTCAGTCTA
GCACACGTCTGAACTCTTTCCCTACACGACGCTCTTCCGATCTNNNCCCACTCCTGTAATCTCAGCTTCC
AGCACACGTCTGAACTCTTTCCCTACACGACGCTCTTCCGATCTNNNGAATGGACAGCTTCTGTATGG
AGCACACGTCTGAACTCTTTCCCTACACGACGCTCTTCCGATCTNNNCTGTGCTTCACTCTCTCATCACA
AGCACACGTCTGAACTCTTTCCCTACACGACGCTCTTCCGATCTNNNTGCAAGCTGGTCAGAAACTCA
AGCACACGTCTGAACTCTTTCCCTACACGACGCTCTTCCGATCTNNNCAACCCAGGTTGCCAAAGTTC
GCACACGTCTGAACTCTTTCCCTACACGACGCTCTTCCGATCTNNNCACACAATGAGCAGCATGGAAA
AGCACACGTCTGAACTCTTTCCCTACACGACGCTCTTCCGATCTNNNTAGGGTGGCCTAAAAGACACT
AGCACACGTCTGAACTCTTTCCCTACACGACGCTCTTCCGATCTNNNTTAGGCTACCTGAGGCTGAAG
GCACACGTCTGAACTCTTTCCCTACACGACGCTCTTCCGATCTNNNGAAGGTAAGCGATCCTTCTGAC
GCACACGTCTGAACTCTTTCCCTACACGACGCTCTTCCGATCTNNNACGTCTACCAGAGTTCATTTGG
AGCACACGTCTGAACTCTTTCCCTACACGACGCTCTTCCGATCTNNNTTCCCAGAGGTAGCTGATTTG
AGCACACGTCTGAACTCTTTCCCTACACGACGCTCTTCCGATCTNNNGCACAGCACAGGAATGAATAG
AGCACACGTCTGAACTCTTTCCCTACACGACGCTCTTCCGATCTNNNTAGCTAGAAATAGGCCTGGA
AGCACACGTCTGAACTCTTTCCCTACACGACGCTCTTCCGATCTNNNTGTAAGGTTGCTGGGGAGAG
AGCACACGTCTGAACTCTTTCCCTACACGACGCTCTTCCGATCTNNNAGCATGCTATCATTTGTGTGG
AGCACACGTCTGAACTCTTTCCCTACACGACGCTCTTCCGATCTNNNTTGTTTAGCCAGTCACAAGAG
AGCACACGTCTGAACTCTTTCCCTACACGACGCTCTTCCGATCTNNNAATGAGAGAGAGAGAGCAG
GAGCACACGTCTGAACTCTTTCCCTACACGACGCTCTTCCGATCTNNNAACTTCTCCAAGAGCAAGCA
GCACACGTCTGAACTCTTTCCCTACACGACGCTCTTCCGATCTNNNTTTATTGAGTGTTGCTGTTGGC
GCACACGTCTGAACTCTTTCCCTACACGACGCTCTTCCGATCTNNNAACACACAGACTTGTCCTACTG
AGCACACGTCTGAACTCTTTCCCTACACGACGCTCTTCCGATCTNNNCCCACCCCCAATCAGAAAGAA
GAGCACACGTCTGAACTCTTTCCCTACACGACGCTCTTCCGATCTNNNCCAGGAAAGCAGCTCTTCAT
AGCACACGTCTGAACTCTTTCCCTACACGACGCTCTTCCGATCTNNNCTTTTGGACACACACTCTCCAG
GAGCACACGTCTGAACTCTTTCCCTACACGACGCTCTTCCGATCTNNNAGCATTGAAAAATTGGTGC
AGCACACGTCTGAACTCTTTCCCTACACGACGCTCTTCCGATCTNNNACTTCCTCCTCCCTCCAGAAAA

AGCACACGTCTGAACTCTTTCCCTACACGACGCTCTTCCGATCTNNNAGGGTAGCAGGTAGCTGTATGT
GAGCACACGTCTGAACTCTTTCCCTACACGACGCTCTTCCGATCTNNNCTGTTCTGATCCTCCAAGCTGT
AGCACACGTCTGAACTCTTTCCCTACACGACGCTCTTCCGATCTNNNTTGCTATTCCACTCTGCAGTCAC
GAGCACACGTCTGAACTCTTTCCCTACACGACGCTCTTCCGATCTNNNTTGACAGAAAAGGGACAAAG
GAGCACACGTCTGAACTCTTTCCCTACACGACGCTCTTCCGATCTNNNTCCTCCTCTACCTTCCTGTAAG
GAGCACACGTCTGAACTCTTTCCCTACACGACGCTCTTCCGATCTNNNGGGTGGGTGGAAATGTGTG
AGAGCACACGTCTGAACTCTTTCCCTACACGACGCTCTTCCGATCTNNNAGGTAGGTGGACAATCCAA
GAGCACACGTCTGAACTCTTTCCCTACACGACGCTCTTCCGATCTNNNCACACTCCAGCAAAACTCTGTC
GAGCACACGTCTGAACTCTTTCCCTACACGACGCTCTTCCGATCTNNNTCAGGCCAGATTTGACCAACA
AGCACACGTCTGAACTCTTTCCCTACACGACGCTCTTCCGATCTNNNAAGCACAGGTTACACAGACAG
GAGCACACGTCTGAACTCTTTCCCTACACGACGCTCTTCCGATCTNNNATCCATGAAACCTTCCTCAGTG
AGAGCACACGTCTGAACTCTTTCCCTACACGACGCTCTTCCGATCTNNNCACACAATCTGTCCTCACCTCA
AGCACACGTCTGAACTCTTTCCCTACACGACGCTCTTCCGATCTNNNTGTCCATCCTCTCTCTGTCTCTG
GAGCACACGTCTGAACTCTTTCCCTACACGACGCTCTTCCGATCTNNNAAGGGACAAGGACTGGGAAC
AGAGCACACGTCTGAACTCTTTCCCTACACGACGCTCTTCCGATCTNNNGAAGTTGAGGCAAACAGCAC
GAGCACACGTCTGAACTCTTTCCCTACACGACGCTCTTCCGATCTNNNTGCACTGTGTTGAATTTTGGTC
GAGCACACGTCTGAACTCTTTCCCTACACGACGCTCTTCCGATCTNNNATGTTCCTCTCCTCTCAGCATG
GAGCACACGTCTGAACTCTTTCCCTACACGACGCTCTTCCGATCTNNNTCTCTCCCACTTTGGAGTTAGCA
AGCACACGTCTGAACTCTTTCCCTACACGACGCTCTTCCGATCTNNNCTTGTGGTAAGGAGTGGTTTGC
AGCACACGTCTGAACTCTTTCCCTACACGACGCTCTTCCGATCTNNNCAAGGTCTTTGCCAGAGTGAGA
AGCACACGTCTGAACTCTTTCCCTACACGACGCTCTTCCGATCTNNNAAGCTTGGAACACAAGAGTGGA
GCACACGTCTGAACTCTTTCCCTACACGACGCTCTTCCGATCTNNNGCTTCAAGGATTGTTGTGCCTAC
GAGCACACGTCTGAACTCTTTCCCTACACGACGCTCTTCCGATCTNNNAATGGCATAAAACACACACGC
GAGCACACGTCTGAACTCTTTCCCTACACGACGCTCTTCCGATCTNNNACCTTGGAATTTGGACCAGTG
AGCACACGTCTGAACTCTTTCCCTACACGACGCTCTTCCGATCTNNNTAACTACCATGGTCCATTGCCTC
AGCACACGTCTGAACTCTTTCCCTACACGACGCTCTTCCGATCTNNNCATATCTGTAGGCATGAGCCACC
GAGCACACGTCTGAACTCTTTCCCTACACGACGCTCTTCCGATCTNNNACATCCTCAAACCAGGCCTTCTA
AGCACACGTCTGAACTCTTTCCCTACACGACGCTCTTCCGATCTNNNGAAGACAGAGAGGCAGGAGA
AGCACACGTCTGAACTCTTTCCCTACACGACGCTCTTCCGATCTNNNGCTCTATGAGGGGGAGTGTTTT
AGCACACGTCTGAACTCTTTCCCTACACGACGCTCTTCCGATCTNNNGAGGACAGACAGAAAGCAAGC
GAGCACACGTCTGAACTCTTTCCCTACACGACGCTCTTCCGATCTNNNGACTTGATACAAGCCACAGGG
GCACACGTCTGAACTCTTTCCCTACACGACGCTCTTCCGATCTNNNCCCAAGAACCGGGAACTCTAATG
AGAGCACACGTCTGAACTCTTTCCCTACACGACGCTCTTCCGATCTNNNGGAAAAGGCTGAGAAGGA
AGAGCACACGTCTGAACTCTTTCCCTACACGACGCTCTTCCGATCTNNNGTCATTTTGGCTTCTTTCCCTC
AGCACACGTCTGAACTCTTTCCCTACACGACGCTCTTCCGATCTNNNTGATGATGGCAACAACCTGGAA
AGAGCACACGTCTGAACTCTTTCCCTACACGACGCTCTTCCGATCTNNNATGCATCTGTCTTTCGTGGA
GAGCACACGTCTGAACTCTTTCCCTACACGACGCTCTTCCGATCTNNNATGTCCAGGCTACTCTCTCATC
GAGCACACGTCTGAACTCTTTCCCTACACGACGCTCTTCCGATCTNNNAGCAAGCTATGAACCAGGAG
AGCACACGTCTGAACTCTTTCCCTACACGACGCTCTTCCGATCTNNNAGGGATTCCCCAAAGGAATCAC
AGCACACGTCTGAACTCTTTCCCTACACGACGCTCTTCCGATCTNNNAGGAATGACAAGAGTGGGAAT
AGCACACGTCTGAACTCTTTCCCTACACGACGCTCTTCCGATCTNNNGAGAATGAGAGTGGGGCAAAC
GAGCACACGTCTGAACTCTTTCCCTACACGACGCTCTTCCGATCTNNNGAACAGAAACTTGGGTGACAA
AGAGCACACGTCTGAACTCTTTCCCTACACGACGCTCTTCCGATCTNNNGGCTGCTGTTTTGTGTACGT

GAGCACACGTCTGAACTCTTTCCCTACACGACGCTCTTCCGATCTNNNGAGGGAGCACCAAGAATGAT
AGCACACGTCTGAACTCTTTCCCTACACGACGCTCTTCCGATCTNNNTGTTCATTATCTCTGGTGTGTGC
GAGCACACGTCTGAACTCTTTCCCTACACGACGCTCTTCCGATCTNNNTCCAGGCATGTCCATGTAGGT
GAGCACACGTCTGAACTCTTTCCCTACACGACGCTCTTCCGATCTNNNTGAGAGGGGACTTACAGGGA
GAGCACACGTCTGAACTCTTTCCCTACACGACGCTCTTCCGATCTNNNGGTCCCAGGGAGAATGCAATT
AGCACACGTCTGAACTCTTTCCCTACACGACGCTCTTCCGATCTNNNGAAGGGAAAACGGAGGAAGAA
GAGCACACGTCTGAACTCTTTCCCTACACGACGCTCTTCCGATCTNNNGGGATTAGGAGCCAGTAGGC
AGAGCACACGTCTGAACTCTTTCCCTACACGACGCTCTTCCGATCTNNNGAACTTGTAGCTGTCCACACC
AGCACACGTCTGAACTCTTTCCCTACACGACGCTCTTCCGATCTNNNTTCCCGGTGTACAGAAACACAA
GAGCACACGTCTGAACTCTTTCCCTACACGACGCTCTTCCGATCTNNNCCCCCTCTTCTGTGATCAGTCT
AGCACACGTCTGAACTCTTTCCCTACACGACGCTCTTCCGATCTNNNTGACATCTAGCAAGAAGTTGGG
GAGCACACGTCTGAACTCTTTCCCTACACGACGCTCTTCCGATCTNNNATCCCTCTTCCCATCAAGAGCA
GAGCACACGTCTGAACTCTTTCCCTACACGACGCTCTTCCGATCTNNNAAATTTCTAGGATGCTTGCGG
GAGCACACGTCTGAACTCTTTCCCTACACGACGCTCTTCCGATCTNNNATGTAAAGTGCTTACCCTGTG
GAGCACACGTCTGAACTCTTTCCCTACACGACGCTCTTCCGATCTNNNGCTCTGGGTAACACACTTCTT
AGAGCACACGTCTGAACTCTTTCCCTACACGACGCTCTTCCGATCTNNNTACATCGTTCCAAAACAGGTC
AGAGCACACGTCTGAACTCTTTCCCTACACGACGCTCTTCCGATCTNNNCAATGGAGGAAGGTGACAC
GAGCACACGTCTGAACTCTTTCCCTACACGACGCTCTTCCGATCTNNNAGCTTCACTCAGGAAGTGGCA
GAGCACACGTCTGAACTCTTTCCCTACACGACGCTCTTCCGATCTNNNGCCAAAGAATCAAAACCGACA
AGAGCACACGTCTGAACTCTTTCCCTACACGACGCTCTTCCGATCTNNNGCTGTCAACATGTATCCCTTC
AGCACACGTCTGAACTCTTTCCCTACACGACGCTCTTCCGATCTNNNCGGACACACACACATACATCCAT
AGAGCACACGTCTGAACTCTTTCCCTACACGACGCTCTTCCGATCTNNNAGTCTAGCCAAGAGAGCTCA
AGAGCACACGTCTGAACTCTTTCCCTACACGACGCTCTTCCGATCTNNNTACGAATACCCTACTGGTTGC
GAGCACACGTCTGAACTCTTTCCCTACACGACGCTCTTCCGATCTNNNCCAGAAGAGCTCTGTGCTGTTA
GAGCACACGTCTGAACTCTTTCCCTACACGACGCTCTTCCGATCTNNNGCCTGATTGGTGAATCACTCT
AGAGCACACGTCTGAACTCTTTCCCTACACGACGCTCTTCCGATCTNNNATGCCAGTCCTACATACATGC
GAGCACACGTCTGAACTCTTTCCCTACACGACGCTCTTCCGATCTNNNAGTACCCTTACAGGAAGAGCC
GAGCACACGTCTGAACTCTTTCCCTACACGACGCTCTTCCGATCTNNNTGTCCTTGTTGTCCTTGTTCTG
GAGCACACGTCTGAACTCTTTCCCTACACGACGCTCTTCCGATCTNNNCCCGTGAGATGAAAGACACA
AGAGCACACGTCTGAACTCTTTCCCTACACGACGCTCTTCCGATCTNNNTAACCACCTGGTCTTGCTCTT
AGCACACGTCTGAACTCTTTCCCTACACGACGCTCTTCCGATCTNNNAATTCCCAACCACCACTTTGGAA
AGAGCACACGTCTGAACTCTTTCCCTACACGACGCTCTTCCGATCTNNNATGCACATGTGTGTATGTGT
AGAGCACACGTCTGAACTCTTTCCCTACACGACGCTCTTCCGATCTNNNACCCCTAAGAAGTCCATAGAT
AGAGCACACGTCTGAACTCTTTCCCTACACGACGCTCTTCCGATCTNNNTATGGCATGATTTGAGTGCTC
AGAGCACACGTCTGAACTCTTTCCCTACACGACGCTCTTCCGATCTNNNTGATAAGAAATCGGTGAGC
AGCACACGTCTGAACTCTTTCCCTACACGACGCTCTTCCGATCTNNNCTAACACAACCCTTTCCCAGGAC
GAGCACACGTCTGAACTCTTTCCCTACACGACGCTCTTCCGATCTNNNAATCATCATGAAACCCTCTGGC
AGCACACGTCTGAACTCTTTCCCTACACGACGCTCTTCCGATCTNNNACTGCCTTTGCCTAGTCACAAAA
GAGCACACGTCTGAACTCTTTCCCTACACGACGCTCTTCCGATCTNNNGCCACCGCACATAAATGTTCAA
GAGCACACGTCTGAACTCTTTCCCTACACGACGCTCTTCCGATCTNNNTTGAGGAGGAAAGGCATGTTA
GAGCACACGTCTGAACTCTTTCCCTACACGACGCTCTTCCGATCTNNNTTCTTACTGCTCTCCTCTACCCG
AGAGCACACGTCTGAACTCTTTCCCTACACGACGCTCTTCCGATCTNNNAAGAAAGGAGTGTTCACAG
AGCACACGTCTGAACTCTTTCCCTACACGACGCTCTTCCGATCTNNNGCTCTGTATCAGTCACGGTTCT

AGCACACGTCTGAACTCTTTCCCTACACGACGCTCTTCCGATCTNNNTTCAAGTGGCTACCTGAGAGTT
AGCACACGTCTGAACTCTTTCCCTACACGACGCTCTTCCGATCTNNNATGAACAGTCTGGTGCATGTCT
GAGCACACGTCTGAACTCTTTCCCTACACGACGCTCTTCCGATCTNNNGAAAGTGGAAACAGCCTCCTG
GAGCACACGTCTGAACTCTTTCCCTACACGACGCTCTTCCGATCTNNNGAGCTCCAAGTGCATTTTCAG
GAGCACACGTCTGAACTCTTTCCCTACACGACGCTCTTCCGATCTNNNTGTGACCAAGAAGCCAACGAT
AGAGCACACGTCTGAACTCTTTCCCTACACGACGCTCTTCCGATCTNNNAACCAACCCCTAAATGATGG
GAGCACACGTCTGAACTCTTTCCCTACACGACGCTCTTCCGATCTNNNTATTACTGGCCTCCCGTCCTAA
GAGCACACGTCTGAACTCTTTCCCTACACGACGCTCTTCCGATCTNNNATTTTGTCTGCCATACATGCTG
GAGCACACGTCTGAACTCTTTCCCTACACGACGCTCTTCCGATCTNNNTTGGCTTAGTGGGGAAGCATA
GAGCACACGTCTGAACTCTTTCCCTACACGACGCTCTTCCGATCTNNNACCTTGACAATCTGCCTCCAAA
GAGCACACGTCTGAACTCTTTCCCTACACGACGCTCTTCCGATCTNNNGGAGCTGTTTCTTTGGAGAG
AGAGCACACGTCTGAACTCTTTCCCTACACGACGCTCTTCCGATCTNNNTGAAACACATTACAGAGGG
GAGCACACGTCTGAACTCTTTCCCTACACGACGCTCTTCCGATCTNNNTGATCTGTAGTGGTGTGTGTG
GAGCACACGTCTGAACTCTTTCCCTACACGACGCTCTTCCGATCTNNNAACAGTCTACTTTGGAAGGCA
GAGCACACGTCTGAACTCTTTCCCTACACGACGCTCTTCCGATCTNNNTTTTGGGACTTTACCCAACTCC
AGCACACGTCTGAACTCTTTCCCTACACGACGCTCTTCCGATCTNNNGCAGGAGCTGGGTCTTTTACAT
AGCACACGTCTGAACTCTTTCCCTACACGACGCTCTTCCGATCTNNNTTCAGGTGCCATCACTTTTTCCAG
GAGCACACGTCTGAACTCTTTCCCTACACGACGCTCTTCCGATCTNNNGTCAAACAGAGAGCAATTGAG
AGCACACGTCTGAACTCTTTCCCTACACGACGCTCTTCCGATCTNNNTGTTGGTATTTCAGGTGGTGCA
GAGCACACGTCTGAACTCTTTCCCTACACGACGCTCTTCCGATCTNNNAAAAATGCACTTGTGCCCTTAG
GAGCACACGTCTGAACTCTTTCCCTACACGACGCTCTTCCGATCTNNNGGTCCATGAAGGCCAAGTAAG
AGCACACGTCTGAACTCTTTCCCTACACGACGCTCTTCCGATCTNNNAGGTGATTCTCCACCTCTCAGTG
AGCACACGTCTGAACTCTTTCCCTACACGACGCTCTTCCGATCTNNNAACATGTTCGAAGCAGAAACCC
GAGCACACGTCTGAACTCTTTCCCTACACGACGCTCTTCCGATCTNNNTGTGTCTGTAATTCCCACCCCTA
GAGCACACGTCTGAACTCTTTCCCTACACGACGCTCTTCCGATCTNNNTCCAGGCAAATGTGTAGGTGA
AGAGCACACGTCTGAACTCTTTCCCTACACGACGCTCTTCCGATCTNNNTTAGCACTCAACTGCTGTCTC
AGAGCACACGTCTGAACTCTTTCCCTACACGACGCTCTTCCGATCTNNNTACTGATTTGCCCTGTGTGTG
AGAGCACACGTCTGAACTCTTTCCCTACACGACGCTCTTCCGATCTNNNAAAAAGGCCAAACTGCTAAG
GAGCACACGTCTGAACTCTTTCCCTACACGACGCTCTTCCGATCTNNNAGCATCAAGACACAGAGATGC
GAGCACACGTCTGAACTCTTTCCCTACACGACGCTCTTCCGATCTNNNCTGGGCCAGTAAAGAGATGC
AAGAGCACACGTCTGAACTCTTTCCCTACACGACGCTCTTCCGATCTNNNCCCATTGGCATTTCCCAAAT
GAGCACACGTCTGAACTCTTTCCCTACACGACGCTCTTCCGATCTNNNCATCTCTCTTCCATCCCCTAGCA
AGAGCACACGTCTGAACTCTTTCCCTACACGACGCTCTTCCGATCTNNNGCTGTAGTGAGGAAAGGAG
GAGCACACGTCTGAACTCTTTCCCTACACGACGCTCTTCCGATCTNNNGTGTATTTCCTTGACACCCTCCC
AGAGCACACGTCTGAACTCTTTCCCTACACGACGCTCTTCCGATCTNNNGTGCCTTGTAGAAGGAAGGT
GAGCACACGTCTGAACTCTTTCCCTACACGACGCTCTTCCGATCTNNNACTGTTTCTCAAACAAGAGGGG
GAGCACACGTCTGAACTCTTTCCCTACACGACGCTCTTCCGATCTNNNTTCGGTTTCTGATTTGGAGGT
GAGCACACGTCTGAACTCTTTCCCTACACGACGCTCTTCCGATCTNNNTTTCCTTGGCCCATACAGTGTT
AGCACACGTCTGAACTCTTTCCCTACACGACGCTCTTCCGATCTNNNTAGGGTCTTGTGTGTGCTCTCAT
AGAGCACACGTCTGAACTCTTTCCCTACACGACGCTCTTCCGATCTNNNAGCAAGAACAGCAGAGGAA
AGCACACGTCTGAACTCTTTCCCTACACGACGCTCTTCCGATCTNNNCCCAAGGGAGGGAAGGTGTAA
AGAGCACACGTCTGAACTCTTTCCCTACACGACGCTCTTCCGATCTNNNAGTCATGCTGCAAAGTTATG
GAGCACACGTCTGAACTCTTTCCCTACACGACGCTCTTCCGATCTNNNGAACCAAGAATGCATCATCGG

AGAGCACACGTCTGAACTCTTTCCCTACACGACGCTCTTCCGATCTNNNATACCAAGTGGACCGAAGG
GAGCACACGTCTGAACTCTTTCCCTACACGACGCTCTTCCGATCTNNNATGTACAAGGCAAGCAGCATC
AGAGCACACGTCTGAACTCTTTCCCTACACGACGCTCTTCCGATCTNNNCATGGCATCCTGAGATGGA
AGAGCACACGTCTGAACTCTTTCCCTACACGACGCTCTTCCGATCTNNNTCAGAGCTGACATTTGCTCC
GAGCACACGTCTGAACTCTTTCCCTACACGACGCTCTTCCGATCTNNNACCTCTTCCAAATCACACACAC
AGAGCACACGTCTGAACTCTTTCCCTACACGACGCTCTTCCGATCTNNNTGTTTAGAGAACAGTGTCCCT
GAGCACACGTCTGAACTCTTTCCCTACACGACGCTCTTCCGATCTNNNCAGTCCTACCAAGACACAACCA
GAGCACACGTCTGAACTCTTTCCCTACACGACGCTCTTCCGATCTNNNTCCCTGCAGTACTGTTTACCAC
AGCACACGTCTGAACTCTTTCCCTACACGACGCTCTTCCGATCTNNNTGCCTCTCACCCTTTACTAAGGT
GAGCACACGTCTGAACTCTTTCCCTACACGACGCTCTTCCGATCTNNNCTTCTACCCAATGTGCCAAGGT
AGAGCACACGTCTGAACTCTTTCCCTACACGACGCTCTTCCGATCTNNNCCAGAGTTGGAAAGTAAACC
GAGCACACGTCTGAACTCTTTCCCTACACGACGCTCTTCCGATCTNNNGCCATACCAAAGAGCCATTCG
AAGAGCACACGTCTGAACTCTTTCCCTACACGACGCTCTTCCGATCTNNNCCATGCCTCCCAAGGGATAT
GAGCACACGTCTGAACTCTTTCCCTACACGACGCTCTTCCGATCTNNNTCCTCCTGGAAGCTCATTGAGA
AGCACACGTCTGAACTCTTTCCCTACACGACGCTCTTCCGATCTNNNCAGACACTGAGAATGGCCATGA
AGCACACGTCTGAACTCTTTCCCTACACGACGCTCTTCCGATCTNNNTGCAGAGGGCTCATCTCTAATC
GAGCACACGTCTGAACTCTTTCCCTACACGACGCTCTTCCGATCTNNNGTCACGTTTTCTGTTCCCTCCTT
GAGCACACGTCTGAACTCTTTCCCTACACGACGCTCTTCCGATCTNNNAGAGGGAATGTGCTATGTGT
AGCACACGTCTGAACTCTTTCCCTACACGACGCTCTTCCGATCTNNNAAGTCAGGTGGTGGTTTTGTTC
AGAGCACACGTCTGAACTCTTTCCCTACACGACGCTCTTCCGATCTNNNAGCCTGATTGGGGTAAGTT
GAGCACACGTCTGAACTCTTTCCCTACACGACGCTCTTCCGATCTNNNAAAGACAGGGTGAGAGGAGA
AGAGCACACGTCTGAACTCTTTCCCTACACGACGCTCTTCCGATCTNNNGCCACGTTTGTTCAGAGGAC
AGAGCACACGTCTGAACTCTTTCCCTACACGACGCTCTTCCGATCTNNNCTCTGAGTACAGTCGTGTGTC
AGAGCACACGTCTGAACTCTTTCCCTACACGACGCTCTTCCGATCTNNNGGCATTTTGGGGGTGAAGA
GAGCACACGTCTGAACTCTTTCCCTACACGACGCTCTTCCGATCTNNNTTAGAACGCCCTACTCTCTTCCC
AGCACACGTCTGAACTCTTTCCCTACACGACGCTCTTCCGATCTNNNGCATTCCTGCAGCTCTAGTTTCT
GAGCACACGTCTGAACTCTTTCCCTACACGACGCTCTTCCGATCTNNNATCTGGACAGGTTAGACCAG
AGCACACGTCTGAACTCTTTCCCTACACGACGCTCTTCCGATCTNNNCTGGCACGAGCATTTTGAGTAT
AGAGCACACGTCTGAACTCTTTCCCTACACGACGCTCTTCCGATCTNNNTTCTGGTCCAAAATGCAACCA
AGAGCACACGTCTGAACTCTTTCCCTACACGACGCTCTTCCGATCTNNNACGGCAATGAAACCAGGTAG
GAGCACACGTCTGAACTCTTTCCCTACACGACGCTCTTCCGATCTNNNGTGAGTTTCCAAGCTCCAAAG
GAGCACACGTCTGAACTCTTTCCCTACACGACGCTCTTCCGATCTNNNCCTGCTGATCTACAGCAGTGTT
AGAGCACACGTCTGAACTCTTTCCCTACACGACGCTCTTCCGATCTNNNACAAAATAGTGGGGGTTCTC
GAGCACACGTCTGAACTCTTTCCCTACACGACGCTCTTCCGATCTNNNCTGACGTGTCTCATGGAAAGG
GAGCACACGTCTGAACTCTTTCCCTACACGACGCTCTTCCGATCTNNNGGTTGATTTTGGTGTTACCAG
AGAGCACACGTCTGAACTCTTTCCCTACACGACGCTCTTCCGATCTNNNGTATTGGCTGCTGTGGAAGA
AGCACACGTCTGAACTCTTTCCCTACACGACGCTCTTCCGATCTNNNTAAGGCCTTGAATGGGAGAGT
AGCACACGTCTGAACTCTTTCCCTACACGACGCTCTTCCGATCTNNNCTAGTAAGCAGTGGACTGGCAT
AGAGCACACGTCTGAACTCTTTCCCTACACGACGCTCTTCCGATCTNNNCAATCCTGAGATTCCCAACCCT
GAGCACACGTCTGAACTCTTTCCCTACACGACGCTCTTCCGATCTNNNTTCCAGCCTCTCAGCAAAATAC
GAGCACACGTCTGAACTCTTTCCCTACACGACGCTCTTCCGATCTNNNTAATCATGTTGTGCTCACCCCAC
AGCACACGTCTGAACTCTTTCCCTACACGACGCTCTTCCGATCTNNNTGCACCAATTTCTTGCTAACCCTC
AGCACACGTCTGAACTCTTTCCCTACACGACGCTCTTCCGATCTNNNGTTGGCTCTGTTCGTTCACTAGA

GAGCACACGTCTGAACTCTTTCCCTACACGACGCTCTTCCGATCTNNNCTATTGAGGTCCCAGAGAGG
AGCACACGTCTGAACTCTTTCCCTACACGACGCTCTTCCGATCTNNNGTTTGCTGCTAAGTGTACTCCC
GAGCACACGTCTGAACTCTTTCCCTACACGACGCTCTTCCGATCTNNNGGAAAGTCCCTGTCCCAACAA
GAGCACACGTCTGAACTCTTTCCCTACACGACGCTCTTCCGATCTNNNATAGCCACCACGTCTCTCTGT
AGCACACGTCTGAACTCTTTCCCTACACGACGCTCTTCCGATCTNNNAGTCACTTAGCCTGTGTGCTTA
GAGCACACGTCTGAACTCTTTCCCTACACGACGCTCTTCCGATCTNNNGGAGTGTTTGCCTAACTGTCC
AGCACACGTCTGAACTCTTTCCCTACACGACGCTCTTCCGATCTNNNCTGCATCATCCCCAGCACATAA
GAGCACACGTCTGAACTCTTTCCCTACACGACGCTCTTCCGATCTNNNAGCCTTTGTGTTTGGATGGA
AGCACACGTCTGAACTCTTTCCCTACACGACGCTCTTCCGATCTNNNATCAACAGTGAGAAATGCCCTC
AGAGCACACGTCTGAACTCTTTCCCTACACGACGCTCTTCCGATCTNNNACGGGAGGAGAGTGTTTCT
AGCACACGTCTGAACTCTTTCCCTACACGACGCTCTTCCGATCTNNNTACATACCAAATCAGGGTGGGC
GAGCACACGTCTGAACTCTTTCCCTACACGACGCTCTTCCGATCTNNNCAAATTTGAGGCTGCAGTGA
AGCACACGTCTGAACTCTTTCCCTACACGACGCTCTTCCGATCTNNNGCTACAGGGGAAAAGGTGTGT
GAGCACACGTCTGAACTCTTTCCCTACACGACGCTCTTCCGATCTNNNCCCAAGGACATTCATCACTGG
AGAGCACACGTCTGAACTCTTTCCCTACACGACGCTCTTCCGATCTNNNCTGCACAAGGGAGATGAAC
AGCACACGTCTGAACTCTTTCCCTACACGACGCTCTTCCGATCTNNNTGCTAAGGAGGTTGGGGGAA
AGCACACGTCTGAACTCTTTCCCTACACGACGCTCTTCCGATCTNNNAGGCTTTTACATTCCCTGTGCT
GAGCACACGTCTGAACTCTTTCCCTACACGACGCTCTTCCGATCTNNNGCTCCTTTGAACACACTCCAGT
AGCACACGTCTGAACTCTTTCCCTACACGACGCTCTTCCGATCTNNNACGTGTGTGTAAGTAGCAATCC
AGCACACGTCTGAACTCTTTCCCTACACGACGCTCTTCCGATCTNNNAGCTGTACCAGTGGAGATGGA
AGCACACGTCTGAACTCTTTCCCTACACGACGCTCTTCCGATCTNNNTATTTGCCTGGCACGAACAAAA
GAGCACACGTCTGAACTCTTTCCCTACACGACGCTCTTCCGATCTNNNTGCTGACATACCTGGCAACAA
AGCACACGTCTGAACTCTTTCCCTACACGACGCTCTTCCGATCTNNNCGGAAGCTGTGTTTCACTGTAG
AGCACACGTCTGAACTCTTTCCCTACACGACGCTCTTCCGATCTNNNACTCTAAACCATAGCAGACTGC
GAGCACACGTCTGAACTCTTTCCCTACACGACGCTCTTCCGATCTNNNTTGCTCAGGGTCATGTGTCTG
AGCACACGTCTGAACTCTTTCCCTACACGACGCTCTTCCGATCTNNNATTCCCTGGCAACCACATTCATA
GAGCACACGTCTGAACTCTTTCCCTACACGACGCTCTTCCGATCTNNNAGCCTCGTCTCAGTAACTACC
GAGCACACGTCTGAACTCTTTCCCTACACGACGCTCTTCCGATCTNNNAATATACACTGAGGGCCATGA
AGCACACGTCTGAACTCTTTCCCTACACGACGCTCTTCCGATCTNNNAAGGTGAGCCAGCTTCTCATTT
AGCACACGTCTGAACTCTTTCCCTACACGACGCTCTTCCGATCTNNNCTGCCTCCATGATCCTCTTCAAC
AGCACACGTCTGAACTCTTTCCCTACACGACGCTCTTCCGATCTNNNCATGTCCTGGATAGCTGCATGT
GAGCACACGTCTGAACTCTTTCCCTACACGACGCTCTTCCGATCTNNNGTATGTGGACAGCATGAGAG
GAGCACACGTCTGAACTCTTTCCCTACACGACGCTCTTCCGATCTNNNCCCTCCCTGTCATGTTTCTCTC
AGAGCACACGTCTGAACTCTTTCCCTACACGACGCTCTTCCGATCTNNNCAAGAGTTTCCTGAGGCTAC
AGAGCACACGTCTGAACTCTTTCCCTACACGACGCTCTTCCGATCTNNNTGTAAATTTCCAGGCCCTTCA
GAGCACACGTCTGAACTCTTTCCCTACACGACGCTCTTCCGATCTNNNTCTCCTTAGTCCACTGATGCCA
AGAGCACACGTCTGAACTCTTTCCCTACACGACGCTCTTCCGATCTNNNAAAATCTCTGCCCTTAGGGA
AGCACACGTCTGAACTCTTTCCCTACACGACGCTCTTCCGATCTNNNCTCCTACCATATTCCCCTCCCTG
AGCACACGTCTGAACTCTTTCCCTACACGACGCTCTTCCGATCTNNNACGCTCTGAGTTTATCATCGCTC
AGAGCACACGTCTGAACTCTTTCCCTACACGACGCTCTTCCGATCTNNNCCTAAGTCAGTCCCATTGAA
AGCACACGTCTGAACTCTTTCCCTACACGACGCTCTTCCGATCTNNNAGTGGGTGACAGAACCTAGAC
AGCACACGTCTGAACTCTTTCCCTACACGACGCTCTTCCGATCTNNNTATTCCAGTTTTGGGTTGTGGC
GAGCACACGTCTGAACTCTTTCCCTACACGACGCTCTTCCGATCTNNNGAGATGTCTGGGGTGAATG

GAGCACACGTCTGAACTCTTTCCCTACACGACGCTCTTCCGATCTNNNTAGTCAAGTGGTGTTGAGA
AGCACACGTCTGAACTCTTTCCCTACACGACGCTCTTCCGATCTNNNCTATGGTTCAGTTACAGCAGG
AGCACACGTCTGAACTCTTTCCCTACACGACGCTCTTCCGATCTNNNAGTGGCTGTCTTCTTCATGTG
GAGCACACGTCTGAACTCTTTCCCTACACGACGCTCTTCCGATCTNNNGAGAGTGATTCGTGCCAAGA
AGCACACGTCTGAACTCTTTCCCTACACGACGCTCTTCCGATCTNNNAGCATGAGAAAGTGAGAACC
AGCACACGTCTGAACTCTTTCCCTACACGACGCTCTTCCGATCTNNNTTCTGATGTTCCACCCTTCTGG
GAGCACACGTCTGAACTCTTTCCCTACACGACGCTCTTCCGATCTNNNCAAAACTCCCATGGCAGCAT
GCACACGTCTGAACTCTTTCCCTACACGACGCTCTTCCGATCTNNNTTCTGAAGCAAGTGATTGTCCC
GAGCACACGTCTGAACTCTTTCCCTACACGACGCTCTTCCGATCTNNNGGCTATGTCTCAAACACCCCT
GCACACGTCTGAACTCTTTCCCTACACGACGCTCTTCCGATCTNNNTGCTTCTCTGGTACAGGTTTCCT
GCACACGTCTGAACTCTTTCCCTACACGACGCTCTTCCGATCTNNNCCTCCTTTGGTTTCTAGGAGCT
GCACACGTCTGAACTCTTTCCCTACACGACGCTCTTCCGATCTNNNGCATGCCTACTAAATGCCAGCT
GAGCACACGTCTGAACTCTTTCCCTACACGACGCTCTTCCGATCTNNNTTCTGTGGTGCTCTCTTGCA
AGCACACGTCTGAACTCTTTCCCTACACGACGCTCTTCCGATCTNNNTATTCGCCCATCCTTCAACACA
GAGCACACGTCTGAACTCTTTCCCTACACGACGCTCTTCCGATCTNNNAGTGACTGTGCTAAGGAGG
GAGCACACGTCTGAACTCTTTCCCTACACGACGCTCTTCCGATCTNNNAATTTGCAAGCCCTTTACTGT
GAGCACACGTCTGAACTCTTTCCCTACACGACGCTCTTCCGATCTNNNAATTCAGGCTCAGATGCTCT
GAGCACACGTCTGAACTCTTTCCCTACACGACGCTCTTCCGATCTNNNTGCTGGGTAATCTTGTAGAA
GAGCACACGTCTGAACTCTTTCCCTACACGACGCTCTTCCGATCTNNNTTTTCCTCCTCCCCAAACATG
GAGCACACGTCTGAACTCTTTCCCTACACGACGCTCTTCCGATCTNNNAGGATATGAACAGGAATTG
AGCACACGTCTGAACTCTTTCCCTACACGACGCTCTTCCGATCTNNNCCATACAAACATCCAAAGCCTG
AGCACACGTCTGAACTCTTTCCCTACACGACGCTCTTCCGATCTNNNACGTATTTTGTGTGTGGCTGTT
GAGCACACGTCTGAACTCTTTCCCTACACGACGCTCTTCCGATCTNNNCATAAGCACAGGGTTACCG
GAGCACACGTCTGAACTCTTTCCCTACACGACGCTCTTCCGATCTNNNGAGCCCCACCACAAAAACAAA
GAGCACACGTCTGAACTCTTTCCCTACACGACGCTCTTCCGATCTNNNTGAGGCTGGATTCAATATC
AGCACACGTCTGAACTCTTTCCCTACACGACGCTCTTCCGATCTNNNCTTGAGGGTGCTCTTGTCTTG
AGCACACGTCTGAACTCTTTCCCTACACGACGCTCTTCCGATCTNNNTGGCACAGAGACAGAGAGAA
AGCACACGTCTGAACTCTTTCCCTACACGACGCTCTTCCGATCTNNNGCCTCAGCCAACCAAGTAAAT
GAGCACACGTCTGAACTCTTTCCCTACACGACGCTCTTCCGATCTNNNGGTGAAGTAGAGGAAGTGA
GAGCACACGTCTGAACTCTTTCCCTACACGACGCTCTTCCGATCTNNNATGGTACCCGTTGAGCCATTT
AGCACACGTCTGAACTCTTTCCCTACACGACGCTCTTCCGATCTNNNGAAGTTCCTCTCTGGGCTAAT
AGCACACGTCTGAACTCTTTCCCTACACGACGCTCTTCCGATCTNNNATATCCTGACAACACACACCCT
AGCACACGTCTGAACTCTTTCCCTACACGACGCTCTTCCGATCTNNNTGCTTCCAGACCTTTCAGTTG
GAGCACACGTCTGAACTCTTTCCCTACACGACGCTCTTCCGATCTNNNATGAAATCCAGGCCTGTGGA
GAGCACACGTCTGAACTCTTTCCCTACACGACGCTCTTCCGATCTNNNACTGCTGCTATAGCTATCCTT
AGAGCACACGTCTGAACTCTTTCCCTACACGACGCTCTTCCGATCTNNNTCCAGGCCTCAGAGACATC
AGCACACGTCTGAACTCTTTCCCTACACGACGCTCTTCCGATCTNNNTCTGTAATTCTTGCCCAGACA
GCACACGTCTGAACTCTTTCCCTACACGACGCTCTTCCGATCTNNNGTAGCTGGGCAACGTCTAGTTT
GAGCACACGTCTGAACTCTTTCCCTACACGACGCTCTTCCGATCTNNNTGAGGCACCTTCCAGAGAAG
AGCACACGTCTGAACTCTTTCCCTACACGACGCTCTTCCGATCTNNNAGCAAAGACTTGGAACCAAA
GAGCACACGTCTGAACTCTTTCCCTACACGACGCTCTTCCGATCTNNNGGAAACCAGAGACACAAAA
AGCACACGTCTGAACTCTTTCCCTACACGACGCTCTTCCGATCTNNNTTACTTCCCATCACCTGTCCCA
AGCACACGTCTGAACTCTTTCCCTACACGACGCTCTTCCGATCTNNNCACACGTGACAAACTCCACCT

GAGCACACGTCTGAACTCTTTCCCTACACGACGCTCTTCCGATCTNNNGAAGAGGCTAAATGAGGCA
GAGCACACGTCTGAACTCTTTCCCTACACGACGCTCTTCCGATCTNNNCAAAGCAGCATGAGATCTCC
GAGCACACGTCTGAACTCTTTCCCTACACGACGCTCTTCCGATCTNNNCCAGAGTGAGACCCTGAAA
AGCACACGTCTGAACTCTTTCCCTACACGACGCTCTTCCGATCTNNNGTCCTTCCTGTGGTCTCTACCT
AGCACACGTCTGAACTCTTTCCCTACACGACGCTCTTCCGATCTNNNCCTTCTGTGCACTTCCTACTCT
GAGCACACGTCTGAACTCTTTCCCTACACGACGCTCTTCCGATCTNNNATTCAGCTGACAGCCAATTCC
GCACACGTCTGAACTCTTTCCCTACACGACGCTCTTCCGATCTNNNAGCTCTAAGAGGTGGACATGA
GAGCACACGTCTGAACTCTTTCCCTACACGACGCTCTTCCGATCTNNNCCAGAGAAACCAAACCTGTG
GCACACGTCTGAACTCTTTCCCTACACGACGCTCTTCCGATCTNNNTTTCTCTGCCTCCTCACCATACAC
AGCACACGTCTGAACTCTTTCCCTACACGACGCTCTTCCGATCTNNNTAGATGGGCGAGTTTGTGGA
GAGCACACGTCTGAACTCTTTCCCTACACGACGCTCTTCCGATCTNNNAGGCCAGACTGGAAGAAGA
AGCACACGTCTGAACTCTTTCCCTACACGACGCTCTTCCGATCTNNNCCTCTTTTGCGCTCTCTCTGTA
GAGCACACGTCTGAACTCTTTCCCTACACGACGCTCTTCCGATCTNNNATAACTGAGAAACAGTGGC
GAGCACACGTCTGAACTCTTTCCCTACACGACGCTCTTCCGATCTNNNTGGCATAGTTTTGGCTCTAG
AGCACACGTCTGAACTCTTTCCCTACACGACGCTCTTCCGATCTNNNCCAGGAATGATAGACTTTGGG
AGCACACGTCTGAACTCTTTCCCTACACGACGCTCTTCCGATCTNNNAAGCTATGGAGGTTGGAGTG
GAGCACACGTCTGAACTCTTTCCCTACACGACGCTCTTCCGATCTNNNCAGGGCTGAAACCTTCTCTCA
GCACACGTCTGAACTCTTTCCCTACACGACGCTCTTCCGATCTNNNCATCTGCAGCACCTCTATTTCCAC
GAGCACACGTCTGAACTCTTTCCCTACACGACGCTCTTCCGATCTNNNATCTAACCCCCTCTTCCTGTCA
GAGCACACGTCTGAACTCTTTCCCTACACGACGCTCTTCCGATCTNNNATCCACCCATCATCCTCAGCA
AGCACACGTCTGAACTCTTTCCCTACACGACGCTCTTCCGATCTNNNCAAGCTCAGGTTGGACACTAC
AGCACACGTCTGAACTCTTTCCCTACACGACGCTCTTCCGATCTNNNTAGATCTCTTCCTCCAACCCCTG
GAGCACACGTCTGAACTCTTTCCCTACACGACGCTCTTCCGATCTNNNATGATTGCACTATGTGCTCCA
AGCACACGTCTGAACTCTTTCCCTACACGACGCTCTTCCGATCTNNNCTATGCCAGACATCATGCTAGG
AGCACACGTCTGAACTCTTTCCCTACACGACGCTCTTCCGATCTNNNTTCCTCTCTCTTCTCCTCCTGGT
AGCACACGTCTGAACTCTTTCCCTACACGACGCTCTTCCGATCTNNNCAAAGATTTCCGTCCTTCTGCC
AGAGCACACGTCTGAACTCTTTCCCTACACGACGCTCTTCCGATCTNNNTTTGTGTAAAGCCTCAGAG
AGCACACGTCTGAACTCTTTCCCTACACGACGCTCTTCCGATCTNNNCGTCTTCATTTTGGTGGTGG
AGCACACGTCTGAACTCTTTCCCTACACGACGCTCTTCCGATCTNNNATACACAGCAAGGCAATGGG
AGCACACGTCTGAACTCTTTCCCTACACGACGCTCTTCCGATCTNNNACAAGGTGGAGGTGGATTAG
GAGCACACGTCTGAACTCTTTCCCTACACGACGCTCTTCCGATCTNNNGTGTGGGTTAGAATCCTGAG
AGAGCACACGTCTGAACTCTTTCCCTACACGACGCTCTTCCGATCTNNNAAAGAAGAGCTGGGGCAA
GAGCACACGTCTGAACTCTTTCCCTACACGACGCTCTTCCGATCTNNNGCAGAGTGAGGCTTTGTGTC
AGAGCACACGTCTGAACTCTTTCCCTACACGACGCTCTTCCGATCTNNNGACCTGAAGAACCAGGTAG
AGCACACGTCTGAACTCTTTCCCTACACGACGCTCTTCCGATCTNNNGCAACAATCCCATTCCCAATAC
AGCACACGTCTGAACTCTTTCCCTACACGACGCTCTTCCGATCTNNNGGGATGTTCCTCCCTCTGTAT
GAGCACACGTCTGAACTCTTTCCCTACACGACGCTCTTCCGATCTNNNAGTGGAGATTTTGAAGGCA
AGCACACGTCTGAACTCTTTCCCTACACGACGCTCTTCCGATCTNNNAGAGACATCCTGAGGCTAAG
GAGCACACGTCTGAACTCTTTCCCTACACGACGCTCTTCCGATCTNNNTTCAAACCTCCTGAACACACC
GCACACGTCTGAACTCTTTCCCTACACGACGCTCTTCCGATCTNNNCCCCTCCCATATTGTGGTAAG
AGCACACGTCTGAACTCTTTCCCTACACGACGCTCTTCCGATCTNNNCTGGGCTGAACAACCACAGA
GAGCACACGTCTGAACTCTTTCCCTACACGACGCTCTTCCGATCTNNNGGAATTTTAGCCTGCTTCCC
GAGCACACGTCTGAACTCTTTCCCTACACGACGCTCTTCCGATCTNNNCACACACAGACCTTCTCTTCA

AGAGCACACGTCTGAACTCTTTCCCTACACGACGCTCTTCCGATCTNNNCTGCACCTGGACAACTTTCT
GAGCACACGTCTGAACTCTTTCCCTACACGACGCTCTTCCGATCTNNNTCAGGTAGATCACACACACA
GAGCACACGTCTGAACTCTTTCCCTACACGACGCTCTTCCGATCTNNNGCTTGGCTATTTCTAGGCAGA
AGAGCACACGTCTGAACTCTTTCCCTACACGACGCTCTTCCGATCTNNNATGGATGGATAGTTGGGA
AGCACACGTCTGAACTCTTTCCCTACACGACGCTCTTCCGATCTNNNGTGATGATCAGGGATGCAGT
AGAGCACACGTCTGAACTCTTTCCCTACACGACGCTCTTCCGATCTNNNGGTCAGCAAAGGAGAACCT
GAGCACACGTCTGAACTCTTTCCCTACACGACGCTCTTCCGATCTNNNGAAAGCGGCTGACCATTTGT
GAGCACACGTCTGAACTCTTTCCCTACACGACGCTCTTCCGATCTNNNTAAACATGTACACAACCCTG
AGCACACGTCTGAACTCTTTCCCTACACGACGCTCTTCCGATCTNNNCTTGGTCTAAGGGTCACCAT
GAGCACACGTCTGAACTCTTTCCCTACACGACGCTCTTCCGATCTNNNGTAGTCCCACATGTACACCAA
AGCACACGTCTGAACTCTTTCCCTACACGACGCTCTTCCGATCTNNNAAAGGGCCTACTCAGGAAAA
GAGCACACGTCTGAACTCTTTCCCTACACGACGCTCTTCCGATCTNNNTTACGTGTTTTGGTGATGGTG
AGCACACGTCTGAACTCTTTCCCTACACGACGCTCTTCCGATCTNNNCCTGAAGTGTGTGAAGAAACC
GAGCACACGTCTGAACTCTTTCCCTACACGACGCTCTTCCGATCTNNNTTTCATGCAAGGTTTGTGTCA
GAGCACACGTCTGAACTCTTTCCCTACACGACGCTCTTCCGATCTNNNACACCCATGCACACACTTTGTA
GAGCACACGTCTGAACTCTTTCCCTACACGACGCTCTTCCGATCTNNNTCCATTACATCTATCCCCCACC
GCACACGTCTGAACTCTTTCCCTACACGACGCTCTTCCGATCTNNNTTCCTGTGAAAAACTGTTTGGGT
AGCACACGTCTGAACTCTTTCCCTACACGACGCTCTTCCGATCTNNNGGCGTGTAGTTCTGGAATTCTG
AGCACACGTCTGAACTCTTTCCCTACACGACGCTCTTCCGATCTNNNTCTGGGATCCTCTTTTAGGGCA
AGCACACGTCTGAACTCTTTCCCTACACGACGCTCTTCCGATCTNNNATTCTGCCAAAGAGAAAGTGG
AGCACACGTCTGAACTCTTTCCCTACACGACGCTCTTCCGATCTNNNAATGTTTTGCAGAGAGGGACT
AGCACACGTCTGAACTCTTTCCCTACACGACGCTCTTCCGATCTNNNATGGTACTGCACACAGTAGTT
GAGCACACGTCTGAACTCTTTCCCTACACGACGCTCTTCCGATCTNNNGGCTGACCTCAACTCAGAACT
AGCACACGTCTGAACTCTTTCCCTACACGACGCTCTTCCGATCTNNNATTTATGGCTGACTGGCATAC
GAGCACACGTCTGAACTCTTTCCCTACACGACGCTCTTCCGATCTNNNTGAGTGGTTTTGCTCCCTTTT
AGCACACGTCTGAACTCTTTCCCTACACGACGCTCTTCCGATCTNNNACCTAGAAAGAGGGACCATCA
AGCACACGTCTGAACTCTTTCCCTACACGACGCTCTTCCGATCTNNNATCTCCACTGCTCCCATACACTT
AGCACACGTCTGAACTCTTTCCCTACACGACGCTCTTCCGATCTNNNATTTCCCTGTGCCCTTTGTTAGC
GAGCACACGTCTGAACTCTTTCCCTACACGACGCTCTTCCGATCTNNNCAGAAAGCCTGGAAGCTGAA
AGAGCACACGTCTGAACTCTTTCCCTACACGACGCTCTTCCGATCTNNNAGAACCCACGAACACAATTC
GAGCACACGTCTGAACTCTTTCCCTACACGACGCTCTTCCGATCTNNNTTATTGATGGAGGGGTGGA
GAGCACACGTCTGAACTCTTTCCCTACACGACGCTCTTCCGATCTNNNAGGGACCAGTATTGTCACTC
GAGCACACGTCTGAACTCTTTCCCTACACGACGCTCTTCCGATCTNNNTGTCCAATGTTCCATAAGGGC
AGCACACGTCTGAACTCTTTCCCTACACGACGCTCTTCCGATCTNNNATGATGGCACTGGAGTTCTTT
AGCACACGTCTGAACTCTTTCCCTACACGACGCTCTTCCGATCTNNNCACAGAGCAACAGCATCATAG
AGCACACGTCTGAACTCTTTCCCTACACGACGCTCTTCCGATCTNNNACACAGAATAAACAGCCAGGC
AGCACACGTCTGAACTCTTTCCCTACACGACGCTCTTCCGATCTNNNCAGGAACCAGTGCATACATTC
GAGCACACGTCTGAACTCTTTCCCTACACGACGCTCTTCCGATCTNNNCAACTCCAGCAGAGAAAGGG
GAGCACACGTCTGAACTCTTTCCCTACACGACGCTCTTCCGATCTNNNATCTCACAGGTAGGGAGCAA
AGCACACGTCTGAACTCTTTCCCTACACGACGCTCTTCCGATCTNNNTCCCATACTCACTGATGCTCCAT
AGCACACGTCTGAACTCTTTCCCTACACGACGCTCTTCCGATCTNNNGGTCTGAGTGCAATTCCCAAT
AGCACACGTCTGAACTCTTTCCCTACACGACGCTCTTCCGATCTNNNGCAGTTACTGGGCCTCATACAA
AGCACACGTCTGAACTCTTTCCCTACACGACGCTCTTCCGATCTNNNATGTCTGGCACTTTTGGGTATG

AGAGCACACGTCTGAACTCTTTCCCTACACGACGCTCTTCCGATCTNNNGCTGCTTCTCTCTAAGCTTGA
GAGCACACGTCTGAACTCTTTCCCTACACGACGCTCTTCCGATCTNNNAAGCTGAGAATGAAGTGCAG
AGCACACGTCTGAACTCTTTCCCTACACGACGCTCTTCCGATCTNNNCCAGCTGTATCCCTGTTGCTAGA
AGCACACGTCTGAACTCTTTCCCTACACGACGCTCTTCCGATCTNNNAGGCATGACAGTTCTAGCCACT
AGAGCACACGTCTGAACTCTTTCCCTACACGACGCTCTTCCGATCTNNNAAGATATCCTGAGAGAGGTC
GAGCACACGTCTGAACTCTTTCCCTACACGACGCTCTTCCGATCTNNNGCCTATATTCACAGACGCCTGA
AGCACACGTCTGAACTCTTTCCCTACACGACGCTCTTCCGATCTNNNTGTAAGGGACAGAGAGGGAG
AGCACACGTCTGAACTCTTTCCCTACACGACGCTCTTCCGATCTNNNAATACGTGCCCATTGATGGATA
AGCACACGTCTGAACTCTTTCCCTACACGACGCTCTTCCGATCTNNNTCTGTAAAACATACCCACCCAGA
AGCACACGTCTGAACTCTTTCCCTACACGACGCTCTTCCGATCTNNNTTTTCTCTCCTCAACCCATCGTCCA
GAGCACACGTCTGAACTCTTTCCCTACACGACGCTCTTCCGATCTNNNGCACGTTCCCTTATAGACTGA
GAGCACACGTCTGAACTCTTTCCCTACACGACGCTCTTCCGATCTNNNTGCTTCCAAACAAGAATGTGA
GAGCACACGTCTGAACTCTTTCCCTACACGACGCTCTTCCGATCTNNNCTTCCTGAACACACACAGGATC
AGCACACGTCTGAACTCTTTCCCTACACGACGCTCTTCCGATCTNNNTGTATGACAGGATGCAGCTAGG
AGCACACGTCTGAACTCTTTCCCTACACGACGCTCTTCCGATCTNNNCCTACTCAACCAGAGGGGAAGA
AGCACACGTCTGAACTCTTTCCCTACACGACGCTCTTCCGATCTNNNTCCCCTTTATGTGTACGTCCCTT
AGCACACGTCTGAACTCTTTCCCTACACGACGCTCTTCCGATCTNNNGCCCCAGGACAGTTGTGTATTA
AGCACACGTCTGAACTCTTTCCCTACACGACGCTCTTCCGATCTNNNCCCTCTTTCCTCTCTTATGCTGCC
AGCACACGTCTGAACTCTTTCCCTACACGACGCTCTTCCGATCTNNNACAATGATGGTAGGGCTTCAA
AGCACACGTCTGAACTCTTTCCCTACACGACGCTCTTCCGATCTNNNTGGTTTTGGGAGAGGAGGTAC
AGCACACGTCTGAACTCTTTCCCTACACGACGCTCTTCCGATCTNNNTTTCATGACGCATCACTCTACCT
GAGCACACGTCTGAACTCTTTCCCTACACGACGCTCTTCCGATCTNNNCAGGTGCACACGTTATGACTA
GAGCACACGTCTGAACTCTTTCCCTACACGACGCTCTTCCGATCTNNNTCTGCTTGGATGGCAGATGTA
GAGCACACGTCTGAACTCTTTCCCTACACGACGCTCTTCCGATCTNNNAGCCATGGCATAATGGGAAAC
GAGCACACGTCTGAACTCTTTCCCTACACGACGCTCTTCCGATCTNNNTGAAGACTGTGTGTGTGTGTC
GAGCACACGTCTGAACTCTTTCCCTACACGACGCTCTTCCGATCTNNNTGCTTACCTGTCCCACTTCACTA
GAGCACACGTCTGAACTCTTTCCCTACACGACGCTCTTCCGATCTNNNAAAAGTGCCCTGAAGGAAACT
GAGCACACGTCTGAACTCTTTCCCTACACGACGCTCTTCCGATCTNNNATAAAAGTCCAAGGGAGGTGG
AGAGCACACGTCTGAACTCTTTCCCTACACGACGCTCTTCCGATCTNNNGAAGGCTTTGCTGTCTGGAA
GAGCACACGTCTGAACTCTTTCCCTACACGACGCTCTTCCGATCTNNNGACCTTTCTGTCATGGCTCACA
GAGCACACGTCTGAACTCTTTCCCTACACGACGCTCTTCCGATCTNNNACCTGAACATTATAGCCCCACC
AGAGCACACGTCTGAACTCTTTCCCTACACGACGCTCTTCCGATCTNNNCTGGCCTTTGGAAGTAGGTT
GAGCACACGTCTGAACTCTTTCCCTACACGACGCTCTTCCGATCTNNNATACACACTCTCTTGAGCCAAA
GAGCACACGTCTGAACTCTTTCCCTACACGACGCTCTTCCGATCTNNNAGACCCAAGCAGACATAGACA
AGCACACGTCTGAACTCTTTCCCTACACGACGCTCTTCCGATCTNNNGAACGAGAGGCGTGTCAGAAA
GAGCACACGTCTGAACTCTTTCCCTACACGACGCTCTTCCGATCTNNNAAGAGACCTGCTAACACACAC
AGCACACGTCTGAACTCTTTCCCTACACGACGCTCTTCCGATCTNNNCAACTGTTCCTAGGCTGAATCCC
AGCACACGTCTGAACTCTTTCCCTACACGACGCTCTTCCGATCTNNNTAGGGAAGTAGGGCGTTGAAT
GAGCACACGTCTGAACTCTTTCCCTACACGACGCTCTTCCGATCTNNNAGCTTCAAGTCCTTCAAGACA
GAGCACACGTCTGAACTCTTTCCCTACACGACGCTCTTCCGATCTNNNTGAATATGCCAGTTTGGATGCA
AGCACACGTCTGAACTCTTTCCCTACACGACGCTCTTCCGATCTNNNCCCATCCTATCCCATCTTGTCCTC
GAGCACACGTCTGAACTCTTTCCCTACACGACGCTCTTCCGATCTNNNCCCAGGATTGAGTTCTCTGCTT
AGCACACGTCTGAACTCTTTCCCTACACGACGCTCTTCCGATCTNNNAACTTCTGTGCTTGATACCCTGC

GAGCACACGTCTGAACTCTTTCCCTACACGACGCTCTTCCGATCTNNNCAGTTGAGTGAGGAACAGAG
GAGCACACGTCTGAACTCTTTCCCTACACGACGCTCTTCCGATCTNNNAGTAAATGAGGAGGGCATCCA
GAGCACACGTCTGAACTCTTTCCCTACACGACGCTCTTCCGATCTNNNTTTTGTCTGCCTTGTGCTCTCAT
GAGCACACGTCTGAACTCTTTCCCTACACGACGCTCTTCCGATCTNNNCTCTGCTCTCCAACCTATGAGG
AGAGCACACGTCTGAACTCTTTCCCTACACGACGCTCTTCCGATCTNNNAAGAATAGAGATGGGCTGT
GAGCACACGTCTGAACTCTTTCCCTACACGACGCTCTTCCGATCTNNNCTCTCCTCTTGTAGATGGCAGG
AGCACACGTCTGAACTCTTTCCCTACACGACGCTCTTCCGATCTNNNCCTGCATCTTCTGCTACCATCTC
AGCACACGTCTGAACTCTTTCCCTACACGACGCTCTTCCGATCTNNNTTTGACAAGGCCTCTTACTGTC
GCACACGTCTGAACTCTTTCCCTACACGACGCTCTTCCGATCTNNNACACCAACAAGACCACAGGAGTA
AGCACACGTCTGAACTCTTTCCCTACACGACGCTCTTCCGATCTNNNACATACATGCACATACACCCTGG
AGCACACGTCTGAACTCTTTCCCTACACGACGCTCTTCCGATCTNNNTCCAGACCATCCTTCACCTTTCA
AGCACACGTCTGAACTCTTTCCCTACACGACGCTCTTCCGATCTNNNGGCAGCAGGATGATAGGTAG
AGCACACGTCTGAACTCTTTCCCTACACGACGCTCTTCCGATCTNNNCTTCTCTTGCTCATCCCATCCTGC
GAGCACACGTCTGAACTCTTTCCCTACACGACGCTCTTCCGATCTNNNCATACCAAGTGTCTCGATGCT
AGCACACGTCTGAACTCTTTCCCTACACGACGCTCTTCCGATCTNNNGTAGGAACCTATAGACCGCAC
GAGCACACGTCTGAACTCTTTCCCTACACGACGCTCTTCCGATCTNNNATTTGTACTGTGTATGCTCCA
GAGCACACGTCTGAACTCTTTCCCTACACGACGCTCTTCCGATCTNNNCATTACTGCGATGAGATCCTG
AGCACACGTCTGAACTCTTTCCCTACACGACGCTCTTCCGATCTNNNAGTTGGCACAGAAGGTGAAGA
GAGCACACGTCTGAACTCTTTCCCTACACGACGCTCTTCCGATCTNNNGCAAGGTTGTAGAGATCCCAA
GAGCACACGTCTGAACTCTTTCCCTACACGACGCTCTTCCGATCTNNNTGCAACCGTTGGGTTGAGATA
GAGCACACGTCTGAACTCTTTCCCTACACGACGCTCTTCCGATCTNNNAGCTCTGTGCTGGAAATTCTG
GCACACGTCTGAACTCTTTCCCTACACGACGCTCTTCCGATCTNNNAGCCACAACCATGTGATTTCGAT
AGCACACGTCTGAACTCTTTCCCTACACGACGCTCTTCCGATCTNNNCAGATTTTGCCACTTACACCTGC
GAGCACACGTCTGAACTCTTTCCCTACACGACGCTCTTCCGATCTNNNTAAGGACAATCGAAACCCTCC
GAGCACACGTCTGAACTCTTTCCCTACACGACGCTCTTCCGATCTNNNGATGGCAGGACAGATTTGGA
AGAGCACACGTCTGAACTCTTTCCCTACACGACGCTCTTCCGATCTNNNCTAGCCATGAGAAAGTCAAG
AGAGCACACGTCTGAACTCTTTCCCTACACGACGCTCTTCCGATCTNNNATCTCTCTCTTTCTCCCATCCG
AGAGCACACGTCTGAACTCTTTCCCTACACGACGCTCTTCCGATCTNNNTTTGGGCCATGTCTGTTTCAG
AGAGCACACGTCTGAACTCTTTCCCTACACGACGCTCTTCCGATCTNNNCTCTCTGCTTTGCATCTGGAC
GAGCACACGTCTGAACTCTTTCCCTACACGACGCTCTTCCGATCTNNNAACCTGGCAACATTTGGTAAA
GAGCACACGTCTGAACTCTTTCCCTACACGACGCTCTTCCGATCTNNNAAGTCATCCATTTGCAGGGAA
AGCACACGTCTGAACTCTTTCCCTACACGACGCTCTTCCGATCTNNNTCAGATAGGCTTCCCCTACCTTC
AGAGCACACGTCTGAACTCTTTCCCTACACGACGCTCTTCCGATCTNNNAACCCCCTAACGTGGATGAA
AGAGCACACGTCTGAACTCTTTCCCTACACGACGCTCTTCCGATCTNNNGCTGGAGCAAAAAGCAATG
GAGCACACGTCTGAACTCTTTCCCTACACGACGCTCTTCCGATCTNNNTGTGAACATAACATCAGGGA
GCACACGTCTGAACTCTTTCCCTACACGACGCTCTTCCGATCTNNNACCTAACCTGCTACTAGCTTGACA
GAGCACACGTCTGAACTCTTTCCCTACACGACGCTCTTCCGATCTNNNTACTCCAATGAAAGGCAAGAG
AAGAGCACACGTCTGAACTCTTTCCCTACACGACGCTCTTCCGATCTNNNCCCTTTCTTGTTTTAGGGTG
AAGAGCACACGTCTGAACTCTTTCCCTACACGACGCTCTTCCGATCTNNNGACAAATGAAACTGGCAG
GAGCACACGTCTGAACTCTTTCCCTACACGACGCTCTTCCGATCTNNNTAGGTAAGGTTCCCCCAGACA
AGCACACGTCTGAACTCTTTCCCTACACGACGCTCTTCCGATCTNNNAGAAATGGGAAGTGTTTGGGT
AGAGCACACGTCTGAACTCTTTCCCTACACGACGCTCTTCCGATCTNNNGACTGGTACCACAGGAGTT
GAGCACACGTCTGAACTCTTTCCCTACACGACGCTCTTCCGATCTNNNTCTCTCCCCTTTCTCATTCAACA

AGCACACGTCTGAACTCTTTCCCTACACGACGCTCTTCCGATCTNNNATATGCACAGGTTTTGCATCCC
GAGCACACGTCTGAACTCTTTCCCTACACGACGCTCTTCCGATCTNNNTAGCTAACGAGGATGCAGTG
AGCACACGTCTGAACTCTTTCCCTACACGACGCTCTTCCGATCTNNNGGTGAGAATGCTGTCTGTCTG
GAGCACACGTCTGAACTCTTTCCCTACACGACGCTCTTCCGATCTNNNGACCCTACCTCTCAGTCTCTCA
AGCACACGTCTGAACTCTTTCCCTACACGACGCTCTTCCGATCTNNNTCCTTTCCTTCTGTCTTCCATGCC
GAGCACACGTCTGAACTCTTTCCCTACACGACGCTCTTCCGATCTNNNGATGCCTGGATACCCAGAGT
GAGCACACGTCTGAACTCTTTCCCTACACGACGCTCTTCCGATCTNNNGCTACCCTAGTTTGGCTTTCT
AGCACACGTCTGAACTCTTTCCCTACACGACGCTCTTCCGATCTNNNGGGCACCAGAGGCAATCTTAT
GAGCACACGTCTGAACTCTTTCCCTACACGACGCTCTTCCGATCTNNNCACCCACACACATACTCATGC
AGCACACGTCTGAACTCTTTCCCTACACGACGCTCTTCCGATCTNNNCCCTTTCTCTAGCCTCTTCACCT
AGCACACGTCTGAACTCTTTCCCTACACGACGCTCTTCCGATCTNNNGTAGCCAAGGCTGATAGAGAG
GAGCACACGTCTGAACTCTTTCCCTACACGACGCTCTTCCGATCTNNNACATTCTCACACAGCTTCTGT
AGCACACGTCTGAACTCTTTCCCTACACGACGCTCTTCCGATCTNNNGCCGAGGCATCTGTCTCAAAAT
AGCACACGTCTGAACTCTTTCCCTACACGACGCTCTTCCGATCTNNNAGCATCAGGCCTATAGCAGACT
AGCACACGTCTGAACTCTTTCCCTACACGACGCTCTTCCGATCTNNNTGTGTCCGTGCATATGTGTGT
AGCACACGTCTGAACTCTTTCCCTACACGACGCTCTTCCGATCTNNNTTGCCTATAGGTGTGTGAGGT
GAGCACACGTCTGAACTCTTTCCCTACACGACGCTCTTCCGATCTNNNTTCAGCATAAGGCAGTTTGGG
AGCACACGTCTGAACTCTTTCCCTACACGACGCTCTTCCGATCTNNNAAAGAGCAGATGGAGCAGAA
GAGCACACGTCTGAACTCTTTCCCTACACGACGCTCTTCCGATCTNNNCAGTTTATGGACAGAAGCTG
GCACACGTCTGAACTCTTTCCCTACACGACGCTCTTCCGATCTNNNTCCGGAATAGACTGGACTACAG
CACACGTCTGAACTCTTTCCCTACACGACGCTCTTCCGATCTNNNAGCCAAATGGCCAGTTACAGAAA
AGAGCACACGTCTGAACTCTTTCCCTACACGACGCTCTTCCGATCTNNNGAGGGACAGTTCATATTGG
AGCACACGTCTGAACTCTTTCCCTACACGACGCTCTTCCGATCTNNNAGTGTGCCCATTTCTCTCTCTCT
GAGCACACGTCTGAACTCTTTCCCTACACGACGCTCTTCCGATCTNNNAGACTGAGATTTGCCTGTGT
GAGCACACGTCTGAACTCTTTCCCTACACGACGCTCTTCCGATCTNNNATGCAAGTATGCTGTCTTGA
GAGCACACGTCTGAACTCTTTCCCTACACGACGCTCTTCCGATCTNNNCTGTTGGCCTGTCTATTGGTC
AGAGCACACGTCTGAACTCTTTCCCTACACGACGCTCTTCCGATCTNNNGTGTGAATATTGGTCTGCG
AGCACACGTCTGAACTCTTTCCCTACACGACGCTCTTCCGATCTNNNTGGGCACACATTTGTGTTACAG
GAGCACACGTCTGAACTCTTTCCCTACACGACGCTCTTCCGATCTNNNAGCAATGGCTGGAAACACTT
GAGCACACGTCTGAACTCTTTCCCTACACGACGCTCTTCCGATCTNNNTTAGGGAGTTTATTCATTCCC
AGCACACGTCTGAACTCTTTCCCTACACGACGCTCTTCCGATCTNNNGCTTTTACATGGGTGTTCCCAC
AGCACACGTCTGAACTCTTTCCCTACACGACGCTCTTCCGATCTNNNCTGAATCAGCCATCAGGGAAG
AGCACACGTCTGAACTCTTTCCCTACACGACGCTCTTCCGATCTNNNTCAATGAGTTCACCAGACAAC
GAGCACACGTCTGAACTCTTTCCCTACACGACGCTCTTCCGATCTNNNGACACAGAGAGCTGAAAGAC
AGCACACGTCTGAACTCTTTCCCTACACGACGCTCTTCCGATCTNNNCTAGTGTGCAGATCCAAGAGA
GAGCACACGTCTGAACTCTTTCCCTACACGACGCTCTTCCGATCTNNNACAAAACACCACTATGCCCAC
AGCACACGTCTGAACTCTTTCCCTACACGACGCTCTTCCGATCTNNNTTCAGTAGGACACACCACCTGA
GAGCACACGTCTGAACTCTTTCCCTACACGACGCTCTTCCGATCTNNNAGGAACCAGAGGCTGTGTAG
AGCACACGTCTGAACTCTTTCCCTACACGACGCTCTTCCGATCTNNNACCCTCACAACTTCCTCACACTC
AGAGCACACGTCTGAACTCTTTCCCTACACGACGCTCTTCCGATCTNNNCCCACTATCCCCAATCAGCTT
AGCACACGTCTGAACTCTTTCCCTACACGACGCTCTTCCGATCTNNNAGAGCCAGCATTATTTGTGCC
AGCACACGTCTGAACTCTTTCCCTACACGACGCTCTTCCGATCTNNNAGGCCTTTTCCATACTCACACA
AGCACACGTCTGAACTCTTTCCCTACACGACGCTCTTCCGATCTNNNGAGACTAAGCTGTCTGGCTTG

AGCACACGTCTGAACTCTTTCCCTACACGACGCTCTTCCGATCTNNNTGAGATCACTTTGGGCAACAA
GCACACGTCTGAACTCTTTCCCTACACGACGCTCTTCCGATCTNNNAATGTTCTTGTGGGGACGGTAT
AGCACACGTCTGAACTCTTTCCCTACACGACGCTCTTCCGATCTNNNATGTGTGAGCCTGATGGCTTA
GAGCACACGTCTGAACTCTTTCCCTACACGACGCTCTTCCGATCTNNNTGACCCTAAGATTTTCGCCAG
GCACACGTCTGAACTCTTTCCCTACACGACGCTCTTCCGATCTNNNGGAGGATGCAATATTACCAGGG
GAGCACACGTCTGAACTCTTTCCCTACACGACGCTCTTCCGATCTNNNAGCCCACATAGACATGAACAC
AGCACACGTCTGAACTCTTTCCCTACACGACGCTCTTCCGATCTNNNTTGGTATTCTTCATCACATTGC
AGCACACGTCTGAACTCTTTCCCTACACGACGCTCTTCCGATCTNNNATATATACAGGGTGCGTGCGT
AGCACACGTCTGAACTCTTTCCCTACACGACGCTCTTCCGATCTNNNACAGCAGGACATGTGTGCTA
GAGCACACGTCTGAACTCTTTCCCTACACGACGCTCTTCCGATCTNNNCTCCACTGCAGAAAGGAGAC
GAGCACACGTCTGAACTCTTTCCCTACACGACGCTCTTCCGATCTNNNCAATCTCAGACCCTTCACAGC
AGCACACGTCTGAACTCTTTCCCTACACGACGCTCTTCCGATCTNNNCAAGCCAAGGCTGCAATAAGC
GAGCACACGTCTGAACTCTTTCCCTACACGACGCTCTTCCGATCTNNNGGCAGGGACCAGATCATACA
GAGCACACGTCTGAACTCTTTCCCTACACGACGCTCTTCCGATCTNNNAGCTTTCTCCAAGGCTGTGT
GAGCACACGTCTGAACTCTTTCCCTACACGACGCTCTTCCGATCTNNNTCACATAGCGGCAAGGTGT
GAGCACACGTCTGAACTCTTTCCCTACACGACGCTCTTCCGATCTNNNTCCCAACTTTTCCTTTGTCCC
AGCACACGTCTGAACTCTTTCCCTACACGACGCTCTTCCGATCTNNNCTTGAAGGGTTGAGTGAAGG
GAGCACACGTCTGAACTCTTTCCCTACACGACGCTCTTCCGATCTNNNGGGATGGGTAGGATCTAAC
AGCACACGTCTGAACTCTTTCCCTACACGACGCTCTTCCGATCTNNNGCATTTCTGGAGTAGGATGG
AGCACACGTCTGAACTCTTTCCCTACACGACGCTCTTCCGATCTNNNTGGGGATGACAGAATGATGA
GAGCACACGTCTGAACTCTTTCCCTACACGACGCTCTTCCGATCTNNNTATCTTCTCGGCATGGCTTT
AGCACACGTCTGAACTCTTTCCCTACACGACGCTCTTCCGATCTNNNTCATAGCTCTACAGCTTACTGG
GAGCACACGTCTGAACTCTTTCCCTACACGACGCTCTTCCGATCTNNNTATAAACGTGGCCAGTTCTC
AGCACACGTCTGAACTCTTTCCCTACACGACGCTCTTCCGATCTNNNATTGCAATGTATGAGGCATCG
AGCACACGTCTGAACTCTTTCCCTACACGACGCTCTTCCGATCTNNNATGTACCTTACCAGTGGCACT
AGCACACGTCTGAACTCTTTCCCTACACGACGCTCTTCCGATCTNNNCCACACATACACATTCCCAACC
AGCACACGTCTGAACTCTTTCCCTACACGACGCTCTTCCGATCTNNNGAACAGGATTCAGGCTCTGT
AGCACACGTCTGAACTCTTTCCCTACACGACGCTCTTCCGATCTNNNATGCGTGCTCTCTCTCCATGTA
AGCACACGTCTGAACTCTTTCCCTACACGACGCTCTTCCGATCTNNNACCAAGGAGAGAAGTGTCCTT
AGCACACGTCTGAACTCTTTCCCTACACGACGCTCTTCCGATCTNNNTGGACAGCGGTAAGATGAAG
GCACACGTCTGAACTCTTTCCCTACACGACGCTCTTCCGATCTNNNCTGTTCCCACCAGATCACAAGA
AGCACACGTCTGAACTCTTTCCCTACACGACGCTCTTCCGATCTNNNCTGGTGAGGACACGTTACAC
AGCACACGTCTGAACTCTTTCCCTACACGACGCTCTTCCGATCTNNNTTGTTCTTTGAGTTAGCCAGG
AGCACACGTCTGAACTCTTTCCCTACACGACGCTCTTCCGATCTNNNACACACTCAATTTGAAACCGCA
AGCACACGTCTGAACTCTTTCCCTACACGACGCTCTTCCGATCTNNNGCTAGTAGGAGTGTTTGCCTG
GCACACGTCTGAACTCTTTCCCTACACGACGCTCTTCCGATCTNNNCTGCCTTGAATTCCATCAAGCCT
AGAGCACACGTCTGAACTCTTTCCCTACACGACGCTCTTCCGATCTNNNACCAGCCCAATTCTACTCTC
GAGCACACGTCTGAACTCTTTCCCTACACGACGCTCTTCCGATCTNNNGGAATCAGTTCAGCAAGAG
GCACACGTCTGAACTCTTTCCCTACACGACGCTCTTCCGATCTNNNCCCTCCTGCATCACTCATTTTCA
AGCACACGTCTGAACTCTTTCCCTACACGACGCTCTTCCGATCTNNNTTCTCATATGCCTTTCCCCTCCA
AGCACACGTCTGAACTCTTTCCCTACACGACGCTCTTCCGATCTNNNGCCCAAAGAGCAGGTGAGAT
GAGCACACGTCTGAACTCTTTCCCTACACGACGCTCTTCCGATCTNNNGGCAAAGTCTGACCTCATAG
AGCACACGTCTGAACTCTTTCCCTACACGACGCTCTTCCGATCTNNNATTCCCAGGCTTTTCTCTTGG

GCACACGTCTGAACTCTTTCCCTACACGACGCTCTTCCGATCTNNNTTCAGAGGTGTTGAAGGAATGG
AGCACACGTCTGAACTCTTTCCCTACACGACGCTCTTCCGATCTNNNCTACACACCACACACACTTACCC
GAGCACACGTCTGAACTCTTTCCCTACACGACGCTCTTCCGATCTNNNGACAATGAGAACACCTCTGGT
AGCACACGTCTGAACTCTTTCCCTACACGACGCTCTTCCGATCTNNNTGAATGTTTTTCAGGACCTGTG
AGCACACGTCTGAACTCTTTCCCTACACGACGCTCTTCCGATCTNNNAACAGCCAGATACTCCTCACGT
GAGCACACGTCTGAACTCTTTCCCTACACGACGCTCTTCCGATCTNNNCTAGGAGTTCTGGGCTTAGTC
GAGCACACGTCTGAACTCTTTCCCTACACGACGCTCTTCCGATCTNNNAAACAGGAGGCTCTCTTTGTT
AGAGCACACGTCTGAACTCTTTCCCTACACGACGCTCTTCCGATCTNNNTATTGGTACTTACCTGCCCCC
GAGCACACGTCTGAACTCTTTCCCTACACGACGCTCTTCCGATCTNNNGTAGTGGCTTGGTCCTAAGG
AGAGCACACGTCTGAACTCTTTCCCTACACGACGCTCTTCCGATCTNNNTTCGAGATCATCCTGGGTCTA
AGCACACGTCTGAACTCTTTCCCTACACGACGCTCTTCCGATCTNNNCCATGGATACCAAGGAATGAGC
GCACACGTCTGAACTCTTTCCCTACACGACGCTCTTCCGATCTNNNAAAGATAAAGCCCCAAAGGCCA
GAGCACACGTCTGAACTCTTTCCCTACACGACGCTCTTCCGATCTNNNACTCCTTCCTTACCAAAGTGGC
AGCACACGTCTGAACTCTTTCCCTACACGACGCTCTTCCGATCTNNNGAAGTAGTGCAGGTGATCAAG
GAGCACACGTCTGAACTCTTTCCCTACACGACGCTCTTCCGATCTNNNGTCAGGCGGGATATTTGAGT
AGCACACGTCTGAACTCTTTCCCTACACGACGCTCTTCCGATCTNNNGCGGGTATATGCAACTTTGGG
AGAGCACACGTCTGAACTCTTTCCCTACACGACGCTCTTCCGATCTNNNCCTAACTTGCAGAGGGACAC
GAGCACACGTCTGAACTCTTTCCCTACACGACGCTCTTCCGATCTNNNGTGACAGGTATGGCAGACAG
GAGCACACGTCTGAACTCTTTCCCTACACGACGCTCTTCCGATCTNNNAATATCACAGGGCACCCACAT
AGCACACGTCTGAACTCTTTCCCTACACGACGCTCTTCCGATCTNNNACCGAAGAACATGAACACACAC
AGAGCACACGTCTGAACTCTTTCCCTACACGACGCTCTTCCGATCTNNNGCTGGTGGAGGTAGCATAG
GAGCACACGTCTGAACTCTTTCCCTACACGACGCTCTTCCGATCTNNNCTGCTAATCCAGAAGACTTGC
GAGCACACGTCTGAACTCTTTCCCTACACGACGCTCTTCCGATCTNNNTTTGTTCTGTGAAGCAAGGT
AGCACACGTCTGAACTCTTTCCCTACACGACGCTCTTCCGATCTNNNATTGAGTGTGGAGAGGCAAAC
AGAGCACACGTCTGAACTCTTTCCCTACACGACGCTCTTCCGATCTNNNTGAGACACAAACTTGGAAAC
AGAGCACACGTCTGAACTCTTTCCCTACACGACGCTCTTCCGATCTNNNTGCAAGGAGAACCTTGGGA
GAGCACACGTCTGAACTCTTTCCCTACACGACGCTCTTCCGATCTNNNGTCTCTCTTGTGCGTTCTCTCT
GAGCACACGTCTGAACTCTTTCCCTACACGACGCTCTTCCGATCTNNNACATATGTGTCCTCTTTCCCCA
GAGCACACGTCTGAACTCTTTCCCTACACGACGCTCTTCCGATCTNNNACTCTTGTTACAGGTACCCTCC
AGAGCACACGTCTGAACTCTTTCCCTACACGACGCTCTTCCGATCTNNNCCAATGCTGTTCCCTTCTCTTC
GAGCACACGTCTGAACTCTTTCCCTACACGACGCTCTTCCGATCTNNNGAGGTGATTATCCTCACTGCT
GCACACGTCTGAACTCTTTCCCTACACGACGCTCTTCCGATCTNNNCCTCAGGTTACTCAGGTTTCTGCC
AGCACACGTCTGAACTCTTTCCCTACACGACGCTCTTCCGATCTNNNACTCCTTTGGACAGCCATAGTCT
GAGCACACGTCTGAACTCTTTCCCTACACGACGCTCTTCCGATCTNNNTAAATGCTTTTGGAGCATCAG
AGAGCACACGTCTGAACTCTTTCCCTACACGACGCTCTTCCGATCTNNNGGTTATGGGTCTTTGCTTTG
GAGCACACGTCTGAACTCTTTCCCTACACGACGCTCTTCCGATCTNNNTGGCCAAAAACAGGGTCAAA
AGCACACGTCTGAACTCTTTCCCTACACGACGCTCTTCCGATCTNNNAATTTTGCTATGGGTTGCTGGC
AGAGCACACGTCTGAACTCTTTCCCTACACGACGCTCTTCCGATCTNNNCCTGAGAATGGAGGGTAAA
AGCACACGTCTGAACTCTTTCCCTACACGACGCTCTTCCGATCTNNNTTCCTTCTCAACAGGGAGAGGT
GCACACGTCTGAACTCTTTCCCTACACGACGCTCTTCCGATCTNNNGCCTTCATCTTCAAATGGAGTGTC
GAGCACACGTCTGAACTCTTTCCCTACACGACGCTCTTCCGATCTNNNGCAAAGCCATCTATTTCCATGC
GAGCACACGTCTGAACTCTTTCCCTACACGACGCTCTTCCGATCTNNNGGCTATTGCCACACGTTTTTCA
AGCACACGTCTGAACTCTTTCCCTACACGACGCTCTTCCGATCTNNNCAGAGAACAGGACCCTAGGAC

AGCACACGTCTGAACTCTTTCCCTACACGACGCTCTTCCGATCTNNNCGGGTCTATCTGTGCTTGTTG
GAGCACACGTCTGAACTCTTTCCCTACACGACGCTCTTCCGATCTNNNAGGCTTTCTTCTGTTCTCTGT
GAGCACACGTCTGAACTCTTTCCCTACACGACGCTCTTCCGATCTNNNTCTTAGGACTGGGCCAAGGA
GAGCACACGTCTGAACTCTTTCCCTACACGACGCTCTTCCGATCTNNNTGGAGTACTGCAAAACCCAC
AGCACACGTCTGAACTCTTTCCCTACACGACGCTCTTCCGATCTNNNATAGCACACCTTGGCTGGATTT
AGCACACGTCTGAACTCTTTCCCTACACGACGCTCTTCCGATCTNNNTGCTCTGCTTTTCTGCTTTACCT
GAGCACACGTCTGAACTCTTTCCCTACACGACGCTCTTCCGATCTNNNTCCCTGCAGCTAGGGTTTATC
AGCACACGTCTGAACTCTTTCCCTACACGACGCTCTTCCGATCTNNNCGACTTTCTGGAAGAGGAAGC
GAGCACACGTCTGAACTCTTTCCCTACACGACGCTCTTCCGATCTNNNTGGAGGAGATCCTAATGGGC
GCACACGTCTGAACTCTTTCCCTACACGACGCTCTTCCGATCTNNNATTGCACATTCTTGTGCTATGCC
AGCACACGTCTGAACTCTTTCCCTACACGACGCTCTTCCGATCTNNNGGGATGGAGGTGAGTAGGTC
GAGCACACGTCTGAACTCTTTCCCTACACGACGCTCTTCCGATCTNNNTCACAAGAATGATTCCAGCC
AGCACACGTCTGAACTCTTTCCCTACACGACGCTCTTCCGATCTNNNCAGTGCCAAGGTTCTGAGACT
GCACACGTCTGAACTCTTTCCCTACACGACGCTCTTCCGATCTNNNGGACTTTTTGACCTCCAGCCTAC
GAGCACACGTCTGAACTCTTTCCCTACACGACGCTCTTCCGATCTNNNATCTTGGGTTCCTTGGTTTCA
AGCACACGTCTGAACTCTTTCCCTACACGACGCTCTTCCGATCTNNNATGTGGGGTGGTATAAAGGG
AGCACACGTCTGAACTCTTTCCCTACACGACGCTCTTCCGATCTNNNACCAGTCAAATTAGAAGCGCT
GAGCACACGTCTGAACTCTTTCCCTACACGACGCTCTTCCGATCTNNNACAATCAAGAGTGCCATTAG
AGCACACGTCTGAACTCTTTCCCTACACGACGCTCTTCCGATCTNNNAGATACCTAGCTTTGTGCGTG
AGCACACGTCTGAACTCTTTCCCTACACGACGCTCTTCCGATCTNNNGATTTTGGTGCAACCTTGGGA
AGCACACGTCTGAACTCTTTCCCTACACGACGCTCTTCCGATCTNNNTCATGGCTCTGCAGATTTTACC
GAGCACACGTCTGAACTCTTTCCCTACACGACGCTCTTCCGATCTNNNCTCTTGCTGTCTCCTTACACAC
GAGCACACGTCTGAACTCTTTCCCTACACGACGCTCTTCCGATCTNNNTGCAGAATCAACTTGAAGCA
GAGCACACGTCTGAACTCTTTCCCTACACGACGCTCTTCCGATCTNNNAGCATTGCCATGGAAGTTGG
AGCACACGTCTGAACTCTTTCCCTACACGACGCTCTTCCGATCTNNNGTGTGTGTACCTGGCTAAGAA
AGCACACGTCTGAACTCTTTCCCTACACGACGCTCTTCCGATCTNNNCCTAGAACATGGGTCTGGGTT
GAGCACACGTCTGAACTCTTTCCCTACACGACGCTCTTCCGATCTNNNAGCAGTGTTTCGTTTCTGGT
AGCACACGTCTGAACTCTTTCCCTACACGACGCTCTTCCGATCTNNNAACTCAATTCTTCAAGACGCCA
AGCACACGTCTGAACTCTTTCCCTACACGACGCTCTTCCGATCTNNNCGTCTGCTTCCAATTATCCTGG
AGCACACGTCTGAACTCTTTCCCTACACGACGCTCTTCCGATCTNNNAAGTGGGAGGTAGAGGGAA
GCACACGTCTGAACTCTTTCCCTACACGACGCTCTTCCGATCTNNNTGGATTCAAATCCCCTTGGATCA
GCACACGTCTGAACTCTTTCCCTACACGACGCTCTTCCGATCTNNNGGGCTTTGAACAGTGTCTACTG
GAGCACACGTCTGAACTCTTTCCCTACACGACGCTCTTCCGATCTNNNTTTACTATGCTGGGCGACAA
AGCACACGTCTGAACTCTTTCCCTACACGACGCTCTTCCGATCTNNNAGCTCCTACACTCCCTTCTTACA
GAGCACACGTCTGAACTCTTTCCCTACACGACGCTCTTCCGATCTNNNTTACAATGTGCTGCTAGGCAG
AGCACACGTCTGAACTCTTTCCCTACACGACGCTCTTCCGATCTNNNTAATGCCTTGGAGGGATTTGG
GAGCACACGTCTGAACTCTTTCCCTACACGACGCTCTTCCGATCTNNNCCCACTACATACCCACCGTCTT
AGCACACGTCTGAACTCTTTCCCTACACGACGCTCTTCCGATCTNNNGTGGCTTACCAGGAAGAACTC
AGCACACGTCTGAACTCTTTCCCTACACGACGCTCTTCCGATCTNNNCAGACCCACAGGGAAATTCTG
GAGCACACGTCTGAACTCTTTCCCTACACGACGCTCTTCCGATCTNNNCTGAGTGGGTGAGAGTTGA
GAGCACACGTCTGAACTCTTTCCCTACACGACGCTCTTCCGATCTNNNAGAGGAAACGTGTACACACA
AGCACACGTCTGAACTCTTTCCCTACACGACGCTCTTCCGATCTNNNCCTGCCATCTAAACCCATAACCC
GAGCACACGTCTGAACTCTTTCCCTACACGACGCTCTTCCGATCTNNNATTCCTTGGTAGAAGTGCAA

GCACACGTCTGAACTCTTTCCCTACACGACGCTCTTCCGATCTNNNAAAGCCCTGAGGATCTGTAGGA
GAGCACACGTCTGAACTCTTTCCCTACACGACGCTCTTCCGATCTNNNAAAACGATGTCTGTCCACTCA
AGCACACGTCTGAACTCTTTCCCTACACGACGCTCTTCCGATCTNNNAAGCACTCAGTGACCATTAGCA
AGCACACGTCTGAACTCTTTCCCTACACGACGCTCTTCCGATCTNNNTATTCTCTCTTGTGCACTCGTGG
AGCACACGTCTGAACTCTTTCCCTACACGACGCTCTTCCGATCTNNNAGACTTCTGCAATGTCAAGTGG
GAGCACACGTCTGAACTCTTTCCCTACACGACGCTCTTCCGATCTNNNAGTAGCTGGGATTCTGGCATA
AGCACACGTCTGAACTCTTTCCCTACACGACGCTCTTCCGATCTNNNGGAGGACCCTTGTTGTTCAACT
GAGCACACGTCTGAACTCTTTCCCTACACGACGCTCTTCCGATCTNNNAGTCCATAAATGCAGGGAGA
AGCACACGTCTGAACTCTTTCCCTACACGACGCTCTTCCGATCTNNNAGATCCAGAAGGTAAGGAAG
AGCACACGTCTGAACTCTTTCCCTACACGACGCTCTTCCGATCTNNNTACCATGGAGTTCTATCAGGG
AGCACACGTCTGAACTCTTTCCCTACACGACGCTCTTCCGATCTNNNTTGGGATGAAGCTCTGTGTTG
AGCACACGTCTGAACTCTTTCCCTACACGACGCTCTTCCGATCTNNNGGCGGGTCACCTAATCTCTCTA
AGCACACGTCTGAACTCTTTCCCTACACGACGCTCTTCCGATCTNNNATACAGATGCAACCAAAGCAC
AGCACACGTCTGAACTCTTTCCCTACACGACGCTCTTCCGATCTNNNACAACTGACCCTTCACGTAGGT
AGCACACGTCTGAACTCTTTCCCTACACGACGCTCTTCCGATCTNNNCTTGTGTATGTGTGTGTCCAGG
AGCACACGTCTGAACTCTTTCCCTACACGACGCTCTTCCGATCTNNNCATGTTCAGTACGCACGCAATT
GAGCACACGTCTGAACTCTTTCCCTACACGACGCTCTTCCGATCTNNNGGTCAATTTCTTGGGGGATT
AGCACACGTCTGAACTCTTTCCCTACACGACGCTCTTCCGATCTNNNGGGAGACCTAGCTTCCTTTTC
GAGCACACGTCTGAACTCTTTCCCTACACGACGCTCTTCCGATCTNNNAGGGAGCAGCAGATGTGTAT
AGCACACGTCTGAACTCTTTCCCTACACGACGCTCTTCCGATCTNNNACTTTGCCTAAAGGAGCATTC
GAGCACACGTCTGAACTCTTTCCCTACACGACGCTCTTCCGATCTNNNGCAAGAAACCAGATTGGAT
GAGCACACGTCTGAACTCTTTCCCTACACGACGCTCTTCCGATCTNNNATCTGCAGTGTGAGGCAATG
GAGCACACGTCTGAACTCTTTCCCTACACGACGCTCTTCCGATCTNNNGTCCGTTGCTCATTCTCTCTCT
GAGCACACGTCTGAACTCTTTCCCTACACGACGCTCTTCCGATCTNNNGACCTATGTTGCCAGCTTCTC
GAGCACACGTCTGAACTCTTTCCCTACACGACGCTCTTCCGATCTNNNAGATTTCCTTGGATGCCAGAA
AGCACACGTCTGAACTCTTTCCCTACACGACGCTCTTCCGATCTNNNTATGGCCTTTAGGCAAATGGCT
AGCACACGTCTGAACTCTTTCCCTACACGACGCTCTTCCGATCTNNNCCTGGCCCCCTAATTTCACATA
AGCACACGTCTGAACTCTTTCCCTACACGACGCTCTTCCGATCTNNNTATTATGGCTGCTGCACTGGT
GAGCACACGTCTGAACTCTTTCCCTACACGACGCTCTTCCGATCTNNNCGAAACCCTGTCTCTGCAATG
GAGCACACGTCTGAACTCTTTCCCTACACGACGCTCTTCCGATCTNNNCTTTGTGACTGTGAGAACCCT
AGCACACGTCTGAACTCTTTCCCTACACGACGCTCTTCCGATCTNNNTTAGGCCAGTTTCACTCCACCTT
AGCACACGTCTGAACTCTTTCCCTACACGACGCTCTTCCGATCTNNNTGCCCTAATCTTGGATTTCTGG
GAGCACACGTCTGAACTCTTTCCCTACACGACGCTCTTCCGATCTNNNAATGTGCTTGACAGTCACACT
GAGCACACGTCTGAACTCTTTCCCTACACGACGCTCTTCCGATCTNNNGTCCAGCTGCACAAGAAGTAT
AGCACACGTCTGAACTCTTTCCCTACACGACGCTCTTCCGATCTNNNTGTAAGTGGAAGACAGGCAGA
AGCACACGTCTGAACTCTTTCCCTACACGACGCTCTTCCGATCTNNNGAACATGTACCTGCACTGTCCT
AGCACACGTCTGAACTCTTTCCCTACACGACGCTCTTCCGATCTNNNGCTGAGACCCTACACGTTTTTCT
AGCACACGTCTGAACTCTTTCCCTACACGACGCTCTTCCGATCTNNNGAAACAATTGCAGCCTGATTCC
AGCACACGTCTGAACTCTTTCCCTACACGACGCTCTTCCGATCTNNNAGATGAGGGTTTGGAATCTTG
GCACACGTCTGAACTCTTTCCCTACACGACGCTCTTCCGATCTNNNGGAGATGTACTGTGGAAAGCA
GAGCACACGTCTGAACTCTTTCCCTACACGACGCTCTTCCGATCTNNNTGAGATGTACCAAATCCCTGT
AGCACACGTCTGAACTCTTTCCCTACACGACGCTCTTCCGATCTNNNTTTTTCATTGCCTCGATCGTCCT
GCACACGTCTGAACTCTTTCCCTACACGACGCTCTTCCGATCTNNNGATTGATGAGATCCTGCAAGG

AGCACACGTCTGAACTCTTTCCCTACACGACGCTCTTCCGATCTNNNCAAGGGGGTGGGCCAATAAA
AGCACACGTCTGAACTCTTTCCCTACACGACGCTCTTCCGATCTNNNTAGTTGTCCAAGGATGTTTTCC
GCACACGTCTGAACTCTTTCCCTACACGACGCTCTTCCGATCTNNNTAGTGGTAAGAGGGAAGGAG
AGCACACGTCTGAACTCTTTCCCTACACGACGCTCTTCCGATCTNNNGAGGCAGGTTTCTCTCTCTCTC
AGCACACGTCTGAACTCTTTCCCTACACGACGCTCTTCCGATCTNNNCTAACAGTAGCTTGGGCAAG
AGCACACGTCTGAACTCTTTCCCTACACGACGCTCTTCCGATCTNNNAAGCATCCCACAGTGACAGA
AGCACACGTCTGAACTCTTTCCCTACACGACGCTCTTCCGATCTNNNCATCTGGCCCTTTATTCTTCCC
AGCACACGTCTGAACTCTTTCCCTACACGACGCTCTTCCGATCTNNNGGCTTCATCACCAGACCAGAAA
AGCACACGTCTGAACTCTTTCCCTACACGACGCTCTTCCGATCTNNNTTCAGTCCTGTCTCCCAAGGTA
GCACACGTCTGAACTCTTTCCCTACACGACGCTCTTCCGATCTNNNTGTCTTACTGGTGTGGTTGTAG
AGCACACGTCTGAACTCTTTCCCTACACGACGCTCTTCCGATCTNNNTCCATCTACTCTCTTGCTCCCA
GCACACGTCTGAACTCTTTCCCTACACGACGCTCTTCCGATCTNNNGGTTCCTAAACACGCTCTCTCTC
GCACACGTCTGAACTCTTTCCCTACACGACGCTCTTCCGATCTNNNTAACAGCAGCTGGATGGATGA
AGCACACGTCTGAACTCTTTCCCTACACGACGCTCTTCCGATCTNNNCACATTGAGAACACGTTCGGTT
GCACACGTCTGAACTCTTTCCCTACACGACGCTCTTCCGATCTNNNGTTGCCAAGCTTTTGGTGGAAA
AGCACACGTCTGAACTCTTTCCCTACACGACGCTCTTCCGATCTNNNTTTGACAGAACAGGTCAGCAC
AGCACACGTCTGAACTCTTTCCCTACACGACGCTCTTCCGATCTNNNTGAGATAGAGTGCCATGGAA
AGCACACGTCTGAACTCTTTCCCTACACGACGCTCTTCCGATCTNNNTTCTTTTATCTCCCACTGCACCC
AGCACACGTCTGAACTCTTTCCCTACACGACGCTCTTCCGATCTNNNCAGCTCACACTTCAGTCACTCT
GCACACGTCTGAACTCTTTCCCTACACGACGCTCTTCCGATCTNNNCTGAAGCCTCAGACCTTGGAAAG
GAGCACACGTCTGAACTCTTTCCCTACACGACGCTCTTCCGATCTNNNCCTGTGCGAAGACTTTAGGT
AGCACACGTCTGAACTCTTTCCCTACACGACGCTCTTCCGATCTNNNTAATGTCAGGTGAGGGAAGA
GCACACGTCTGAACTCTTTCCCTACACGACGCTCTTCCGATCTNNNATTGACATTCCAGGAAGATGGG
AGCACACGTCTGAACTCTTTCCCTACACGACGCTCTTCCGATCTNNNAAGCCACTCTCCGAAAACAAGA
AGCACACGTCTGAACTCTTTCCCTACACGACGCTCTTCCGATCTNNNGACTGGAGCACGTAGGATGT
GAGCACACGTCTGAACTCTTTCCCTACACGACGCTCTTCCGATCTNNNACAGCATGTGGTCATGTTGG
GCACACGTCTGAACTCTTTCCCTACACGACGCTCTTCCGATCTNNNTTGCAAAACAAACCCCTGGACTA
GCACACGTCTGAACTCTTTCCCTACACGACGCTCTTCCGATCTNNNCCTCTCTCCCCTCTGTCTCTCTACA
AGCACACGTCTGAACTCTTTCCCTACACGACGCTCTTCCGATCTNNNCAACAAGGAATCCAGTCTGAC
AGCACACGTCTGAACTCTTTCCCTACACGACGCTCTTCCGATCTNNNTCCCTTCAGAGGGAGAGAATG
AGCACACGTCTGAACTCTTTCCCTACACGACGCTCTTCCGATCTNNNGCCAGGGTTCTATGGAGAAAC
AGAGCACACGTCTGAACTCTTTCCCTACACGACGCTCTTCCGATCTNNNAAGTCTTCCTGGAGCAGAG
AGAGCACACGTCTGAACTCTTTCCCTACACGACGCTCTTCCGATCTNNNAAGGGGTGAAAACTGCTT
GAGCACACGTCTGAACTCTTTCCCTACACGACGCTCTTCCGATCTNNNAAGGAGGTGATGTGAGAGT
AGCACACGTCTGAACTCTTTCCCTACACGACGCTCTTCCGATCTNNNGCAGGGGCAGAGGAAAATAG
AGCACACGTCTGAACTCTTTCCCTACACGACGCTCTTCCGATCTNNNCTCCTGGTTCCCATTACTAGCCT
AGCACACGTCTGAACTCTTTCCCTACACGACGCTCTTCCGATCTNNNTGGGTACAACAGGAGGAGAG
GCACACGTCTGAACTCTTTCCCTACACGACGCTCTTCCGATCTNNNTTTGTCCACTGTTATGCCTTGCT
AGAGCACACGTCTGAACTCTTTCCCTACACGACGCTCTTCCGATCTNNNCCCTCTGGACAAAAAGCTG
GAGCACACGTCTGAACTCTTTCCCTACACGACGCTCTTCCGATCTNNNCACAGGGAAAACCTAAACTG
GAGCACACGTCTGAACTCTTTCCCTACACGACGCTCTTCCGATCTNNNTCTCTTACGCACACACACA
AGCACACGTCTGAACTCTTTCCCTACACGACGCTCTTCCGATCTNNNCCCCAACTGACATTCTAAGGA
GCACACGTCTGAACTCTTTCCCTACACGACGCTCTTCCGATCTNNNCAAACTAGCGTCTTCTCTGGGT

AGCACACGTCTGAACTCTTTCCCTACACGACGCTCTTCCGATCTNNNCCCTACAATACACAGGACACCC
GAGCACACGTCTGAACTCTTTCCCTACACGACGCTCTTCCGATCTNNNGTGCTCCAGGACTTATCTTT
GAGCACACGTCTGAACTCTTTCCCTACACGACGCTCTTCCGATCTNNNAGCAGTCAGGGCAAGATAGA
GAGCACACGTCTGAACTCTTTCCCTACACGACGCTCTTCCGATCTNNNTTGATTCAAGGCAGGATTC
AGCACACGTCTGAACTCTTTCCCTACACGACGCTCTTCCGATCTNNNCATTTATATTGGCTGCCTGGT
AGCACACGTCTGAACTCTTTCCCTACACGACGCTCTTCCGATCTNNNTGTTGGGAGCTACACCAATAG
GAGCACACGTCTGAACTCTTTCCCTACACGACGCTCTTCCGATCTNNNTGAAGAGGCTGTTCTGAGGT
AGCACACGTCTGAACTCTTTCCCTACACGACGCTCTTCCGATCTNNNAAAAGGAGGAGTGTGTGTGT
GAGCACACGTCTGAACTCTTTCCCTACACGACGCTCTTCCGATCTNNNAAAGCAGACAAACCATTTGC
GAGCACACGTCTGAACTCTTTCCCTACACGACGCTCTTCCGATCTNNNTGGGAATCCCTGTCTCTGCA
GAGCACACGTCTGAACTCTTTCCCTACACGACGCTCTTCCGATCTNNNTGCACATAGACCAGTAATCCC
GAGCACACGTCTGAACTCTTTCCCTACACGACGCTCTTCCGATCTNNNCACTCTCTCTAGCCTGAGTGT
AGCACACGTCTGAACTCTTTCCCTACACGACGCTCTTCCGATCTNNNGGGGGAGCTTAGAGTACATG
GAGCACACGTCTGAACTCTTTCCCTACACGACGCTCTTCCGATCTNNNCTTCAGTTTCAGGCACAGTGC
AGCACACGTCTGAACTCTTTCCCTACACGACGCTCTTCCGATCTNNNTGAAACTTGAGACGGTGGTTC
AGCACACGTCTGAACTCTTTCCCTACACGACGCTCTTCCGATCTNNNTTGGTCAGTAGCTACACAAAT
GAGCACACGTCTGAACTCTTTCCCTACACGACGCTCTTCCGATCTNNNCTGATTACAGAAACGACGGT
GAGCACACGTCTGAACTCTTTCCCTACACGACGCTCTTCCGATCTNNNGGCTGCTGTTCTTACACAAT
AGAGCACACGTCTGAACTCTTTCCCTACACGACGCTCTTCCGATCTNNNTACCCTCACAACATGGCAGT
AGCACACGTCTGAACTCTTTCCCTACACGACGCTCTTCCGATCTNNNTCACAAGAAAGATTCTGGTCTC
AGCACACGTCTGAACTCTTTCCCTACACGACGCTCTTCCGATCTNNNACAGGCTTCCACTTTTTCCACAT
AGCACACGTCTGAACTCTTTCCCTACACGACGCTCTTCCGATCTNNNTGTATTCCTAACCCTACCCCAGC
GAGCACACGTCTGAACTCTTTCCCTACACGACGCTCTTCCGATCTNNNAACTATTGTGTGTGTGCGTGT
AGCACACGTCTGAACTCTTTCCCTACACGACGCTCTTCCGATCTNNNCTAGTGGCTCTAAAGCATGAC
GCACACGTCTGAACTCTTTCCCTACACGACGCTCTTCCGATCTNNNGGCAGACATACTCAGACCTCTCT
GAGCACACGTCTGAACTCTTTCCCTACACGACGCTCTTCCGATCTNNNGGTGCAGTTCTGGTCTTCTCA
AGCACACGTCTGAACTCTTTCCCTACACGACGCTCTTCCGATCTNNNTGAGGTAGGAACCTGCAAGT
GAGCACACGTCTGAACTCTTTCCCTACACGACGCTCTTCCGATCTNNNAATGCTAAATGCTAAAGGCG
GAGCACACGTCTGAACTCTTTCCCTACACGACGCTCTTCCGATCTNNNGTACACTTCTGGGGTCAGTC
AGCACACGTCTGAACTCTTTCCCTACACGACGCTCTTCCGATCTNNNAATCACTTAGCACAACACCTGG
GAGCACACGTCTGAACTCTTTCCCTACACGACGCTCTTCCGATCTNNNTTTCAGCCTCTCTACTCTGCTG
GAGCACACGTCTGAACTCTTTCCCTACACGACGCTCTTCCGATCTNNNGATTTCTGTCAGACTGGTGA
GAGCACACGTCTGAACTCTTTCCCTACACGACGCTCTTCCGATCTNNNTACCCAGCCATCCTTGCCTAT
AGCACACGTCTGAACTCTTTCCCTACACGACGCTCTTCCGATCTNNNATCTTTGCAGATTGGCTGTCTG
GAGCACACGTCTGAACTCTTTCCCTACACGACGCTCTTCCGATCTNNNGTGTCGAGTTCTCCAGCTGT
AGAGCACACGTCTGAACTCTTTCCCTACACGACGCTCTTCCGATCTNNNTGAGGAATTTCTGGATGTG
GAGCACACGTCTGAACTCTTTCCCTACACGACGCTCTTCCGATCTNNNCTGGCTGGTAGAACTGTGAA
GAGCACACGTCTGAACTCTTTCCCTACACGACGCTCTTCCGATCTNNNCATGTGGTGTTTGAACATGG
GAGCACACGTCTGAACTCTTTCCCTACACGACGCTCTTCCGATCTNNNCTCTATTCTGCCACCCTCCTA
GCACACGTCTGAACTCTTTCCCTACACGACGCTCTTCCGATCTNNNGGCCATCTCTCCCTGGATAATCT
GAGCACACGTCTGAACTCTTTCCCTACACGACGCTCTTCCGATCTNNNGTGAGTGCACACGAGTATG
GAGCACACGTCTGAACTCTTTCCCTACACGACGCTCTTCCGATCTNNNACCAACTCCAAAGCATGTGTT
GAGCACACGTCTGAACTCTTTCCCTACACGACGCTCTTCCGATCTNNNTATGGCCAGGTGGTTCAATT

AGCACACGTCTGAACTCTTTCCCTACACGACGCTCTTCCGATCTNNNTGACTTCAGTGATCCTCCAAGG
AGCACACGTCTGAACTCTTTCCCTACACGACGCTCTTCCGATCTNNNGATTTCTGCCCAGAAGAGGAG
AGCACACGTCTGAACTCTTTCCCTACACGACGCTCTTCCGATCTNNNTGCTGAGAGGGAAGAAAACAA
GAGCACACGTCTGAACTCTTTCCCTACACGACGCTCTTCCGATCTNNNTGCCTGGCTGAAATTTCTCCTT
AGAGCACACGTCTGAACTCTTTCCCTACACGACGCTCTTCCGATCTNNNGCTTGGCTCAAACTCTCTCT
GAGCACACGTCTGAACTCTTTCCCTACACGACGCTCTTCCGATCTNNNAGGAAAGGAAATGCTGTCTCT
AGAGCACACGTCTGAACTCTTTCCCTACACGACGCTCTTCCGATCTNNNGTCATCTCATTGGTGGTCAC
GAGCACACGTCTGAACTCTTTCCCTACACGACGCTCTTCCGATCTNNNGTCACCAGGTCACCATTCAAA
GAGCACACGTCTGAACTCTTTCCCTACACGACGCTCTTCCGATCTNNNTGGTTTTCCAGAGGCTATGA
AGAGCACACGTCTGAACTCTTTCCCTACACGACGCTCTTCCGATCTNNNATGACACACACAAGCAGACA
AGAGCACACGTCTGAACTCTTTCCCTACACGACGCTCTTCCGATCTNNNCCTCATCTTCGTCACTTTCAG
AGCACACGTCTGAACTCTTTCCCTACACGACGCTCTTCCGATCTNNNGGGTCATGAAGGAGAAATGT
AGCACACGTCTGAACTCTTTCCCTACACGACGCTCTTCCGATCTNNNGGTGTGCAACGAGGTGAGAA
AGCACACGTCTGAACTCTTTCCCTACACGACGCTCTTCCGATCTNNNACGTTCTTCTCACTCCCTCATCA
GAGCACACGTCTGAACTCTTTCCCTACACGACGCTCTTCCGATCTNNNATGGGGACTTCTCTCTCTCTCC
AGCACACGTCTGAACTCTTTCCCTACACGACGCTCTTCCGATCTNNNCCTTACTTTCTCAGCTCAGTGG
AGCACACGTCTGAACTCTTTCCCTACACGACGCTCTTCCGATCTNNNGTTTTCCCCTTGTCTCTCCTTGG
AGCACACGTCTGAACTCTTTCCCTACACGACGCTCTTCCGATCTNNNCAACCTGGCTATGTCAGAGCAT
GAGCACACGTCTGAACTCTTTCCCTACACGACGCTCTTCCGATCTNNNAGTGTTCAGGGAAAACAGTC
AGAGCACACGTCTGAACTCTTTCCCTACACGACGCTCTTCCGATCTNNNAAAAAGCATGGGGTGTGTG
GAGCACACGTCTGAACTCTTTCCCTACACGACGCTCTTCCGATCTNNNTCAAAAGCCATATACGCCCAC
AGCACACGTCTGAACTCTTTCCCTACACGACGCTCTTCCGATCTNNNGAACAAATGGGAGGAACTCCT
AGAGCACACGTCTGAACTCTTTCCCTACACGACGCTCTTCCGATCTNNNTACAGCCTAGTGAGGATCCA
GAGCACACGTCTGAACTCTTTCCCTACACGACGCTCTTCCGATCTNNNCATATTGAGCCAAGGGAAGG
GAGCACACGTCTGAACTCTTTCCCTACACGACGCTCTTCCGATCTNNNGGGTAGATGCTTTGAAATGG
GAGCACACGTCTGAACTCTTTCCCTACACGACGCTCTTCCGATCTNNNTCGTTCAGGGTGTGTTTTGT
GAGCACACGTCTGAACTCTTTCCCTACACGACGCTCTTCCGATCTNNNTAGATCATGTGGTAGGGTTT
AGAGCACACGTCTGAACTCTTTCCCTACACGACGCTCTTCCGATCTNNNTCAGCGATGCATAACTCTGT
AGAGCACACGTCTGAACTCTTTCCCTACACGACGCTCTTCCGATCTNNNCTTACCTCATTGCAGCTGTCT
GAGCACACGTCTGAACTCTTTCCCTACACGACGCTCTTCCGATCTNNNCTGAGAAGTGAGGCAGGAG
AGCACACGTCTGAACTCTTTCCCTACACGACGCTCTTCCGATCTNNNACGGGTTTTCTTTTTGCATTGG
GAGCACACGTCTGAACTCTTTCCCTACACGACGCTCTTCCGATCTNNNTAAATCTGGGCCACAGAGCAA
GAGCACACGTCTGAACTCTTTCCCTACACGACGCTCTTCCGATCTNNNTTCTCAGGCCCACTTCCTGTAT
GAGCACACGTCTGAACTCTTTCCCTACACGACGCTCTTCCGATCTNNNCCAGTGTGTTCATGGGCATCA
GAGCACACGTCTGAACTCTTTCCCTACACGACGCTCTTCCGATCTNNNAAAAAGTCCCCATGCTACACA
GAGCACACGTCTGAACTCTTTCCCTACACGACGCTCTTCCGATCTNNNGCCCTCCCTTGATGGTATGTAT
GAGCACACGTCTGAACTCTTTCCCTACACGACGCTCTTCCGATCTNNNCTGTGGGTGTGTTGTATTTGT
GAGCACACGTCTGAACTCTTTCCCTACACGACGCTCTTCCGATCTNNNGCACTGTGAATCTCCAAGAA
GAGCACACGTCTGAACTCTTTCCCTACACGACGCTCTTCCGATCTNNNTTTCAGTCTTCATGTGTGTGG
GCACACGTCTGAACTCTTTCCCTACACGACGCTCTTCCGATCTNNNTGTTTTTGTGTACACCCTGGTCA
AGCACACGTCTGAACTCTTTCCCTACACGACGCTCTTCCGATCTNNNATGAAGATTTGGCCATCTTCCCC
AGAGCACACGTCTGAACTCTTTCCCTACACGACGCTCTTCCGATCTNNNCTAAGTCAAGGGAGTGATG
GAGCACACGTCTGAACTCTTTCCCTACACGACGCTCTTCCGATCTNNNCAGACACTGTGACAGATCAT

AGCACACGTCTGAACTCTTTCCCTACACGACGCTCTTCCGATCTNNNCTGGACCACATCTGGAATGAA
AGCACACGTCTGAACTCTTTCCCTACACGACGCTCTTCCGATCTNNNGCACTCTGTTTCAGTCTGGAG
GAGCACACGTCTGAACTCTTTCCCTACACGACGCTCTTCCGATCTNNNAACATAGGTTCATCGGCTGT
AGCACACGTCTGAACTCTTTCCCTACACGACGCTCTTCCGATCTNNNGGCTCATAGTACATTTCCCCA
GAGCACACGTCTGAACTCTTTCCCTACACGACGCTCTTCCGATCTNNNGGTGTCCGTGGCTTATCTTT
GAGCACACGTCTGAACTCTTTCCCTACACGACGCTCTTCCGATCTNNNGGGCACTGGTTTGTGATACT
AGCACACGTCTGAACTCTTTCCCTACACGACGCTCTTCCGATCTNNNTGCAGACAGAATGAAGCCAAG
GCACACGTCTGAACTCTTTCCCTACACGACGCTCTTCCGATCTNNNATGTACAGAAAGATGACGCTG
AGCACACGTCTGAACTCTTTCCCTACACGACGCTCTTCCGATCTNNNAGAGAAGTCTCCAATAAGGG
GCACACGTCTGAACTCTTTCCCTACACGACGCTCTTCCGATCTNNNGCACAGTCTCCGATTTCTGGATT
AGCACACGTCTGAACTCTTTCCCTACACGACGCTCTTCCGATCTNNNTAAAGGTTCCCATGTGTCTGG
AGCACACGTCTGAACTCTTTCCCTACACGACGCTCTTCCGATCTNNNGTTACATGTGAAACAGAGCAC
AGCACACGTCTGAACTCTTTCCCTACACGACGCTCTTCCGATCTNNNCCCTTTGAGGATGGCCTGAAA
AGCACACGTCTGAACTCTTTCCCTACACGACGCTCTTCCGATCTNNNATGAGAATAGGTCTCATGGG
GAGCACACGTCTGAACTCTTTCCCTACACGACGCTCTTCCGATCTNNNTGAGGGACATACAGATTGCT
AGCACACGTCTGAACTCTTTCCCTACACGACGCTCTTCCGATCTNNNTCGTAATGGGGATACAGAAAG
AGCACACGTCTGAACTCTTTCCCTACACGACGCTCTTCCGATCTNNNCCAGGAATGGGGATTGCCAAT
GAGCACACGTCTGAACTCTTTCCCTACACGACGCTCTTCCGATCTNNNCTCAGCACAATGCTACTGGG
GAGCACACGTCTGAACTCTTTCCCTACACGACGCTCTTCCGATCTNNNACAGGTATTGCATACCAAGC
GAGCACACGTCTGAACTCTTTCCCTACACGACGCTCTTCCGATCTNNNTAATGGATCTACACTCCGGCG
AGCACACGTCTGAACTCTTTCCCTACACGACGCTCTTCCGATCTNNNACAGCTGTAGGAAAACCAAAC
GAGCACACGTCTGAACTCTTTCCCTACACGACGCTCTTCCGATCTNNNAATCAGTGAACAGACACACA
GAGCACACGTCTGAACTCTTTCCCTACACGACGCTCTTCCGATCTNNNGGAAGGCTTTCCTGGAGAA
AGCACACGTCTGAACTCTTTCCCTACACGACGCTCTTCCGATCTNNNCAAAGAAAGGCTGGAGCACA
GAGCACACGTCTGAACTCTTTCCCTACACGACGCTCTTCCGATCTNNNTTATCCCATGGTGGAAGGAG
GCACACGTCTGAACTCTTTCCCTACACGACGCTCTTCCGATCTNNNAGCTGGGACTACATGTGAATGT
GAGCACACGTCTGAACTCTTTCCCTACACGACGCTCTTCCGATCTNNNTCTAATCTATCTGCCTGAGGG
AGCACACGTCTGAACTCTTTCCCTACACGACGCTCTTCCGATCTNNNTAATCCAAGGCGATCCACAAA
AGCACACGTCTGAACTCTTTCCCTACACGACGCTCTTCCGATCTNNNAGGACGGAGAGAATGTGAA
AGCACACGTCTGAACTCTTTCCCTACACGACGCTCTTCCGATCTNNNAGGGTTTGGTTTTGTTGTTTG
AGCACACGTCTGAACTCTTTCCCTACACGACGCTCTTCCGATCTNNNTTTCCTGAATGATGGCAACCCT
GCACACGTCTGAACTCTTTCCCTACACGACGCTCTTCCGATCTNNNAGCTAATCACAGGCAGGACCTA
GAGCACACGTCTGAACTCTTTCCCTACACGACGCTCTTCCGATCTNNNACCGAGCTGATCAAAAGACA
GCACACGTCTGAACTCTTTCCCTACACGACGCTCTTCCGATCTNNNGCTTCAGGTCTCACATGTACCA
AGCACACGTCTGAACTCTTTCCCTACACGACGCTCTTCCGATCTNNNACATGGGAATTTTCTGGGGAC
GCACACGTCTGAACTCTTTCCCTACACGACGCTCTTCCGATCTNNNTCTTTGAAGGAAGCCTCAGCAT
AGCACACGTCTGAACTCTTTCCCTACACGACGCTCTTCCGATCTNNNCCGGGTATTTACTGTGTAGAG
GCACACGTCTGAACTCTTTCCCTACACGACGCTCTTCCGATCTNNNAAGGATTGGGACCTCCAAAGCT
AGCACACGTCTGAACTCTTTCCCTACACGACGCTCTTCCGATCTNNNTGGAGGAAGCTCTGATCCAA
AGCACACGTCTGAACTCTTTCCCTACACGACGCTCTTCCGATCTNNNCTGCATATCATCCTCCCACTCCT
AGCACACGTCTGAACTCTTTCCCTACACGACGCTCTTCCGATCTNNNCTACTGCCTCGCAATACCAAA
AGCACACGTCTGAACTCTTTCCCTACACGACGCTCTTCCGATCTNNNTCTCTTCCACCCCCAAAAGAA
GCACACGTCTGAACTCTTTCCCTACACGACGCTCTTCCGATCTNNNATTCACTACACACACATGTGGA

GAGCACACGTCTGAACTCTTTCCCTACACGACGCTCTTCCGATCTNNNCCAAGCAAAAGTTGAAGAGCC
GAGCACACGTCTGAACTCTTTCCCTACACGACGCTCTTCCGATCTNNNTGGGAAGGCAAATGTGTTTT
AGAGCACACGTCTGAACTCTTTCCCTACACGACGCTCTTCCGATCTNNNACACAGTCCTTACTTCCTGAC
GAGCACACGTCTGAACTCTTTCCCTACACGACGCTCTTCCGATCTNNNACCATTCTTCACTTTCCTCAGG
AGCACACGTCTGAACTCTTTCCCTACACGACGCTCTTCCGATCTNNNTGCTCAAGCATGAAAGAGAGC
GCACACGTCTGAACTCTTTCCCTACACGACGCTCTTCCGATCTNNNCTTTGACCCTGGGTGGTTCTTTC
GAGCACACGTCTGAACTCTTTCCCTACACGACGCTCTTCCGATCTNNNAGGGCAAGTGACTGCATTCT
GAGCACACGTCTGAACTCTTTCCCTACACGACGCTCTTCCGATCTNNNGTGACAACAAACCACACACAC
AGCACACGTCTGAACTCTTTCCCTACACGACGCTCTTCCGATCTNNNTAAAAATGTCCCTTTTGGCACG
AGCACACGTCTGAACTCTTTCCCTACACGACGCTCTTCCGATCTNNNTTATGGCCTTCCTCAGCCTAACT
GAGCACACGTCTGAACTCTTTCCCTACACGACGCTCTTCCGATCTNNNAAGTAGCAGAGCTGGAATTT
AGCACACGTCTGAACTCTTTCCCTACACGACGCTCTTCCGATCTNNNAGTTGGATCCCAAAATGGATC
AGCACACGTCTGAACTCTTTCCCTACACGACGCTCTTCCGATCTNNNACAAGAGCTGCCCTACTTACAG
AGCACACGTCTGAACTCTTTCCCTACACGACGCTCTTCCGATCTNNNGCTTCTCCTGCCTTTCTCAAAGT
GAGCACACGTCTGAACTCTTTCCCTACACGACGCTCTTCCGATCTNNNGGAGAGATCAGCCACCTTTTC
GAGCACACGTCTGAACTCTTTCCCTACACGACGCTCTTCCGATCTNNNACAACTGTTAGAAGTGCCCTG
AGAGCACACGTCTGAACTCTTTCCCTACACGACGCTCTTCCGATCTNNNAGACCCATAGCAAACTTGG
GCACACGTCTGAACTCTTTCCCTACACGACGCTCTTCCGATCTNNNACTGTTCCAGGGCATAGTCTCAT
GAGCACACGTCTGAACTCTTTCCCTACACGACGCTCTTCCGATCTNNNTTGTGCTTTTTCCCCAGTTTGT
GAGCACACGTCTGAACTCTTTCCCTACACGACGCTCTTCCGATCTNNNTTCTGAGAATACACCACTGGG
AGCACACGTCTGAACTCTTTCCCTACACGACGCTCTTCCGATCTNNNAGAGACCCATTTCCTCCAATCC
GAGCACACGTCTGAACTCTTTCCCTACACGACGCTCTTCCGATCTNNNGTCGACAAAAAGCTACACTGC
AGAGCACACGTCTGAACTCTTTCCCTACACGACGCTCTTCCGATCTNNNTAGCTTCCATAAAAGCACCA
AGCACACGTCTGAACTCTTTCCCTACACGACGCTCTTCCGATCTNNNTCATGAGTGTTCCTGCTGTTTC
GAGCACACGTCTGAACTCTTTCCCTACACGACGCTCTTCCGATCTNNNAGCAAGGTCAATCCTGTCTCT
AGCACACGTCTGAACTCTTTCCCTACACGACGCTCTTCCGATCTNNNTGGACTTCCACAAGTTTCAGCT
AGAGCACACGTCTGAACTCTTTCCCTACACGACGCTCTTCCGATCTNNNTCAAAGTCAACGCAAAGAGA
GAGCACACGTCTGAACTCTTTCCCTACACGACGCTCTTCCGATCTNNNCATGCCCCTTGGAATGTAGAC
AGCACACGTCTGAACTCTTTCCCTACACGACGCTCTTCCGATCTNNNATGGAAGGGAATTCAACTGTG
AGCACACGTCTGAACTCTTTCCCTACACGACGCTCTTCCGATCTNNNCCAATCTTATTCAGAGCCCTCCA
AGAGCACACGTCTGAACTCTTTCCCTACACGACGCTCTTCCGATCTNNNAGCAATTGTACATGGGTGT
GAGCACACGTCTGAACTCTTTCCCTACACGACGCTCTTCCGATCTNNNCCCATTCAGCTTGTCACTTGT
AGCACACGTCTGAACTCTTTCCCTACACGACGCTCTTCCGATCTNNNAGAGAAGTTGCTTTGGCCATTC
GAGCACACGTCTGAACTCTTTCCCTACACGACGCTCTTCCGATCTNNNCTAGCTTGGTGTCTCCTTCAAC
AGAGCACACGTCTGAACTCTTTCCCTACACGACGCTCTTCCGATCTNNNTCATTTCACTACCGTATGCCG
AGCACACGTCTGAACTCTTTCCCTACACGACGCTCTTCCGATCTNNNATGCACCCATAGTGTGCTAGAG
AGCACACGTCTGAACTCTTTCCCTACACGACGCTCTTCCGATCTNNNTTCAGGTGAAAGTGGCTAAGT
AGCACACGTCTGAACTCTTTCCCTACACGACGCTCTTCCGATCTNNNAGGCTGTAGAGTTCTTTGCAT
AGCACACGTCTGAACTCTTTCCCTACACGACGCTCTTCCGATCTNNNAGATGGAGGAAGAGCCAATCC
GAGCACACGTCTGAACTCTTTCCCTACACGACGCTCTTCCGATCTNNNTTTACGAGTGCATGTGTCTGT
AGAGCACACGTCTGAACTCTTTCCCTACACGACGCTCTTCCGATCTNNNTTCAGTAAACTTCATGGGGC
AGAGCACACGTCTGAACTCTTTCCCTACACGACGCTCTTCCGATCTNNNAATGTGAGAGGAATCACGC
GAGCACACGTCTGAACTCTTTCCCTACACGACGCTCTTCCGATCTNNNTGGTAGCAGCATGAGTATCT

GAGCACACGTCTGAACTCTTTCCCTACACGACGCTCTTCCGATCTNNNCCAAATGGGTACCCCTCAATC
AGCACACGTCTGAACTCTTTCCCTACACGACGCTCTTCCGATCTNNNCCTGGCCCTCTACCTTCATTTTT
AGCACACGTCTGAACTCTTTCCCTACACGACGCTCTTCCGATCTNNNCCTGCAAACACACAAACACAGA
AGCACACGTCTGAACTCTTTCCCTACACGACGCTCTTCCGATCTNNNTGTTCGGTCTACTCCATGAACA
AGCACACGTCTGAACTCTTTCCCTACACGACGCTCTTCCGATCTNNNTTAGCCTACAGGCTCTGTCTCA
AGCACACGTCTGAACTCTTTCCCTACACGACGCTCTTCCGATCTNNNCCAGGGTGAGCAATCCAAAAG
AGCACACGTCTGAACTCTTTCCCTACACGACGCTCTTCCGATCTNNNGTGGCCAGGAATTGAAAGAG
GCACACGTCTGAACTCTTTCCCTACACGACGCTCTTCCGATCTNNNTAGCTGCTTCTCTATCCACTCCC
GCACACGTCTGAACTCTTTCCCTACACGACGCTCTTCCGATCTNNNTCTGCTGCATAAATTGCACCTCTC
AGCACACGTCTGAACTCTTTCCCTACACGACGCTCTTCCGATCTNNNTCCCTAGCTGCCACATTCTACTT
GCACACGTCTGAACTCTTTCCCTACACGACGCTCTTCCGATCTNNNTATCCTGGAACTGGAAGATGG
AGCACACGTCTGAACTCTTTCCCTACACGACGCTCTTCCGATCTNNNCCCTGAAGGAAGCCAAACTAC
GCACACGTCTGAACTCTTTCCCTACACGACGCTCTTCCGATCTNNNACAACAAAAGCCCAGTGAGACA
AGCACACGTCTGAACTCTTTCCCTACACGACGCTCTTCCGATCTNNNGAACACGCTAGGATCTGTACG
AGCACACGTCTGAACTCTTTCCCTACACGACGCTCTTCCGATCTNNNTGCTTCCTAGGTATGTAAGGC
GAGCACACGTCTGAACTCTTTCCCTACACGACGCTCTTCCGATCTNNNCTGTGACACTTGGCATGGTTT
AGAGCACACGTCTGAACTCTTTCCCTACACGACGCTCTTCCGATCTNNNGTTGGTTTCCGAAGCTCCAA
GAGCACACGTCTGAACTCTTTCCCTACACGACGCTCTTCCGATCTNNNATCTCAAGTGGCTTGTCTCTC
AGCACACGTCTGAACTCTTTCCCTACACGACGCTCTTCCGATCTNNNCCTGAGAATGCCCTTGACTGTTT
AGCACACGTCTGAACTCTTTCCCTACACGACGCTCTTCCGATCTNNNGGTCTATGGAGAGAAGGAGC
AGCACACGTCTGAACTCTTTCCCTACACGACGCTCTTCCGATCTNNNTTTAGGGTCTCTCATGCTGCCAT
AGCACACGTCTGAACTCTTTCCCTACACGACGCTCTTCCGATCTNNNTTCGACCTTTGATGACTGGACA
GAGCACACGTCTGAACTCTTTCCCTACACGACGCTCTTCCGATCTNNNTATACTCCTCAAAACACACACG
GAGCACACGTCTGAACTCTTTCCCTACACGACGCTCTTCCGATCTNNNGCTCGGGTTTCTAAGGGAGA
AGCACACGTCTGAACTCTTTCCCTACACGACGCTCTTCCGATCTNNNCTTACCCTCAGTGTTCTCCATG
AGCACACGTCTGAACTCTTTCCCTACACGACGCTCTTCCGATCTNNNCAGAAAGCAAGGAAGTGCTTG
GAGCACACGTCTGAACTCTTTCCCTACACGACGCTCTTCCGATCTNNNAAGCTCATCACCAGTGAGTAG
AGCACACGTCTGAACTCTTTCCCTACACGACGCTCTTCCGATCTNNNCACGGCCCAAGAGTGATGAAT
GCACACGTCTGAACTCTTTCCCTACACGACGCTCTTCCGATCTNNNGTGCCTGCCTATGGGTGAAAAT
GCACACGTCTGAACTCTTTCCCTACACGACGCTCTTCCGATCTNNNTTCACTCCATTCCCCACACACATC
AGCACACGTCTGAACTCTTTCCCTACACGACGCTCTTCCGATCTNNNTTTCTCCTCCACAACTCACCTCT
AGCACACGTCTGAACTCTTTCCCTACACGACGCTCTTCCGATCTNNNGGCTTGAAATGAACAGGGAAT
AGCACACGTCTGAACTCTTTCCCTACACGACGCTCTTCCGATCTNNNAAATTCCCCTGAACTCACTGTG
AGAGCACACGTCTGAACTCTTTCCCTACACGACGCTCTTCCGATCTNNNACACAAATGGGATCAGGAC
GCACACGTCTGAACTCTTTCCCTACACGACGCTCTTCCGATCTNNNCCAAAGTCTCACAGCCAGTTAG
AGCACACGTCTGAACTCTTTCCCTACACGACGCTCTTCCGATCTNNNTGCCGGCATTCTTCTTTCCTATT
AGAGCACACGTCTGAACTCTTTCCCTACACGACGCTCTTCCGATCTNNNGAGTTGATCTCCTGGAGGT
GAGCACACGTCTGAACTCTTTCCCTACACGACGCTCTTCCGATCTNNNCCCAGACCACCTCAGTATGAA
AGCACACGTCTGAACTCTTTCCCTACACGACGCTCTTCCGATCTNNNTACCTGGAACATGGGTTAGGA
GCACACGTCTGAACTCTTTCCCTACACGACGCTCTTCCGATCTNNNAAGAGTGGGGACAAAACACTGT
GAGCACACGTCTGAACTCTTTCCCTACACGACGCTCTTCCGATCTNNNACTAAGAATACTCCACCAAGG
GAGCACACGTCTGAACTCTTTCCCTACACGACGCTCTTCCGATCTNNNTCTGATGCCACTTCCCAAAACA
AGAGCACACGTCTGAACTCTTTCCCTACACGACGCTCTTCCGATCTNNNACGAGGCAAGAATGAAAGT

AGCACACGTCTGAACTCTTTCCCTACACGACGCTCTTCCGATCTNNNTGCTAACACTTGGCATCAAGTCA
GAGCACACGTCTGAACTCTTTCCCTACACGACGCTCTTCCGATCTNNNCCTGAGGCTGGGAGAACATTT
GAGCACACGTCTGAACTCTTTCCCTACACGACGCTCTTCCGATCTNNNGATGGTGACGTGGGATGTCAT
GAGCACACGTCTGAACTCTTTCCCTACACGACGCTCTTCCGATCTNNNTCTTGAACTACTGGACTGTGG
AGAGCACACGTCTGAACTCTTTCCCTACACGACGCTCTTCCGATCTNNNTGGTCAACCATGTCATCATCT
GAGCACACGTCTGAACTCTTTCCCTACACGACGCTCTTCCGATCTNNNTGGATTAGAGTAAGGGGAG
GAGCACACGTCTGAACTCTTTCCCTACACGACGCTCTTCCGATCTNNNCAATGGGTACAGTCAGACAG
AGCACACGTCTGAACTCTTTCCCTACACGACGCTCTTCCGATCTNNNACAGACAGCAGATGGTAGGAC
GAGCACACGTCTGAACTCTTTCCCTACACGACGCTCTTCCGATCTNNNTGGAGTTAGAGAGCTCCTTGT
GAGCACACGTCTGAACTCTTTCCCTACACGACGCTCTTCCGATCTNNNCTACATGAAGGAAACAAACGC
AGCACACGTCTGAACTCTTTCCCTACACGACGCTCTTCCGATCTNNNTGACGTCTGTAGCACACACAAA
AGAGCACACGTCTGAACTCTTTCCCTACACGACGCTCTTCCGATCTNNNTTCACTGAAGTGATGTAAAG
GAGCACACGTCTGAACTCTTTCCCTACACGACGCTCTTCCGATCTNNNCAGCTCTTTCCTGTTACCCTTCC
AGCACACGTCTGAACTCTTTCCCTACACGACGCTCTTCCGATCTNNNACACATCTTACATGCAGAGGCA
AGCACACGTCTGAACTCTTTCCCTACACGACGCTCTTCCGATCTNNNTACAGTGAGCTGATACTGCGCT
GAGCACACGTCTGAACTCTTTCCCTACACGACGCTCTTCCGATCTNNNTTTTCACCAGAGCTAAACCCCA
AGCACACGTCTGAACTCTTTCCCTACACGACGCTCTTCCGATCTNNNACTGACCCTTCCTTCTCTACCTGC
AGCACACGTCTGAACTCTTTCCCTACACGACGCTCTTCCGATCTNNNACTGTTCATGTCTTCACCCACTT
GAGCACACGTCTGAACTCTTTCCCTACACGACGCTCTTCCGATCTNNNTAACAGGCGACAGACGACGA
AGCACACGTCTGAACTCTTTCCCTACACGACGCTCTTCCGATCTNNNAGGGGTTTGGAGAAGTCACAG
GAGCACACGTCTGAACTCTTTCCCTACACGACGCTCTTCCGATCTNNNGCAAGCATGTCCTCTTATGGTG
GAGCACACGTCTGAACTCTTTCCCTACACGACGCTCTTCCGATCTNNNTTATCCGGGCTGAGAGATCCT
AGCACACGTCTGAACTCTTTCCCTACACGACGCTCTTCCGATCTNNNTTCTTCTCCCATAGGGCTCTGTC
GAGCACACGTCTGAACTCTTTCCCTACACGACGCTCTTCCGATCTNNNGCTCATACATCGTGTGTTCTT
AGCACACGTCTGAACTCTTTCCCTACACGACGCTCTTCCGATCTNNNTCCAGGACATAGGTCAAAGAC
AGCACACGTCTGAACTCTTTCCCTACACGACGCTCTTCCGATCTNNNGGACAATCTGTGTCGATTGAG
GAGCACACGTCTGAACTCTTTCCCTACACGACGCTCTTCCGATCTNNNTGGGTTCCCACAGGTAAACCA
GAGCACACGTCTGAACTCTTTCCCTACACGACGCTCTTCCGATCTNNNCTTTTGCTCCCTCTCTCCAAGAC
AGAGCACACGTCTGAACTCTTTCCCTACACGACGCTCTTCCGATCTNNNATTTGGTTGGCTCTGGAAT
AGCACACGTCTGAACTCTTTCCCTACACGACGCTCTTCCGATCTNNNACGTGAATATTGACAGGGGCTT
AGAGCACACGTCTGAACTCTTTCCCTACACGACGCTCTTCCGATCTNNNCAGCCCAGCCCTATATCTTGG
GAGCACACGTCTGAACTCTTTCCCTACACGACGCTCTTCCGATCTNNNGGGAAAGAATGAACAAAGCA
GAGCACACGTCTGAACTCTTTCCCTACACGACGCTCTTCCGATCTNNNGCATGTTAGGTGGTTTTGTGT
GAGCACACGTCTGAACTCTTTCCCTACACGACGCTCTTCCGATCTNNNAGAAACATGTAGGAGACGAA
GAGCACACGTCTGAACTCTTTCCCTACACGACGCTCTTCCGATCTNNNCCACTTCCACACTCCTAACTCAC
GCACACGTCTGAACTCTTTCCCTACACGACGCTCTTCCGATCTNNNCATCTGACTGTGTAGCACTTCCCC
GAGCACACGTCTGAACTCTTTCCCTACACGACGCTCTTCCGATCTNNNAACCCATCAAAAGACAGTTCG
GAGCACACGTCTGAACTCTTTCCCTACACGACGCTCTTCCGATCTNNNACAACCCTGATTCCATTGCTCT
GAGCACACGTCTGAACTCTTTCCCTACACGACGCTCTTCCGATCTNNNCCAAGTCGAAGTTGCCAAAGA
AGCACACGTCTGAACTCTTTCCCTACACGACGCTCTTCCGATCTNNNACTTTTGTGATGTGGAAAGTG
GAGCACACGTCTGAACTCTTTCCCTACACGACGCTCTTCCGATCTNNNACTGGTCTGACTACACCCCTTA
AGCACACGTCTGAACTCTTTCCCTACACGACGCTCTTCCGATCTNNNAAAGGTAACCAAGTGCTGCTA
AGCACACGTCTGAACTCTTTCCCTACACGACGCTCTTCCGATCTNNNCACAATCTCCTGAGCCAATTCCA

AGAGCACACGTCTGAACTCTTTCCCTACACGACGCTCTTCCGATCTNNNGCATAGGACAAGGACCCAAT
AGCACACGTCTGAACTCTTTCCCTACACGACGCTCTTCCGATCTNNNTTTGCCATGTGGGAGTTGGTAA
GAGCACACGTCTGAACTCTTTCCCTACACGACGCTCTTCCGATCTNNNTGAACAGAACCTCTCCTCTTCG
AGCACACGTCTGAACTCTTTCCCTACACGACGCTCTTCCGATCTNNNTACTGAACTGTGTCCTCCAACCA
AGAGCACACGTCTGAACTCTTTCCCTACACGACGCTCTTCCGATCTNNNCAGTCACCTATTCCTGCTGCT
AGAGCACACGTCTGAACTCTTTCCCTACACGACGCTCTTCCGATCTNNNGTCCAGGGGTCCATATACAC
AGAGCACACGTCTGAACTCTTTCCCTACACGACGCTCTTCCGATCTNNNTGCAAGTTGATCAGCCTAGT
GAGCACACGTCTGAACTCTTTCCCTACACGACGCTCTTCCGATCTNNNACCTTCCCAAGGTGTCAAAAAG
GAGCACACGTCTGAACTCTTTCCCTACACGACGCTCTTCCGATCTNNNATAGCCTGTCCAAATTGTCCCA
GAGCACACGTCTGAACTCTTTCCCTACACGACGCTCTTCCGATCTNNNTGATGATCCTCTCCTGTTGCTG
AGAGCACACGTCTGAACTCTTTCCCTACACGACGCTCTTCCGATCTNNNTCCATGGGATGTATCAGTGA
GAGCACACGTCTGAACTCTTTCCCTACACGACGCTCTTCCGATCTNNNCACCAGTCCCACGTTAGAACA
GAGCACACGTCTGAACTCTTTCCCTACACGACGCTCTTCCGATCTNNNTGGGGGATTGTTTTTGTGCT
GAGCACACGTCTGAACTCTTTCCCTACACGACGCTCTTCCGATCTNNNTGTACATGGACTTCCCTCTCCT
GAGCACACGTCTGAACTCTTTCCCTACACGACGCTCTTCCGATCTNNNAGTACAACCGTAGGAGTAGTG
GCACACGTCTGAACTCTTTCCCTACACGACGCTCTTCCGATCTNNNTCAAAGTTTGGTGGGGAAAACC
AGAGCACACGTCTGAACTCTTTCCCTACACGACGCTCTTCCGATCTNNNAGAATGCAGGGGATATGCA
GAGCACACGTCTGAACTCTTTCCCTACACGACGCTCTTCCGATCTNNNGCACTGGCTCACTGAATGGAA
AGAGCACACGTCTGAACTCTTTCCCTACACGACGCTCTTCCGATCTNNNTGTGGTACTATTCTGGGAGG
AGCACACGTCTGAACTCTTTCCCTACACGACGCTCTTCCGATCTNNNATCACCAGAAGTCTGTCTTCCCC
GAGCACACGTCTGAACTCTTTCCCTACACGACGCTCTTCCGATCTNNNGCTGCAGTAACCCATGATTCAG
AGAGCACACGTCTGAACTCTTTCCCTACACGACGCTCTTCCGATCTNNNGCCCACAACGACAGTCTTTTT
GAGCACACGTCTGAACTCTTTCCCTACACGACGCTCTTCCGATCTNNNATACTCCACCAATAGTCCCCCAG
GAGCACACGTCTGAACTCTTTCCCTACACGACGCTCTTCCGATCTNNNAGGCACGCAGAAATATACCTT
AGCACACGTCTGAACTCTTTCCCTACACGACGCTCTTCCGATCTNNNTTTCTCTTTGCCTCTGCTTTCGTG
GCACACGTCTGAACTCTTTCCCTACACGACGCTCTTCCGATCTNNNTCCCCTACAGAATAGAAAGCCCCC
GAGCACACGTCTGAACTCTTTCCCTACACGACGCTCTTCCGATCTNNNCCTCCTTTGAGCAACCTTTGAG
GAGCACACGTCTGAACTCTTTCCCTACACGACGCTCTTCCGATCTNNNGGAAGAAGGTGACCTGGACC
AGCACACGTCTGAACTCTTTCCCTACACGACGCTCTTCCGATCTNNNAGAACCCAGAAGCCTCTTCTCTT
GAGCACACGTCTGAACTCTTTCCCTACACGACGCTCTTCCGATCTNNNGATTGAGCAATGCCAGGCTAA
GAGCACACGTCTGAACTCTTTCCCTACACGACGCTCTTCCGATCTNNNTTCTATACAGTTGCACAATGC
AGCACACGTCTGAACTCTTTCCCTACACGACGCTCTTCCGATCTNNNGCACACAGCACTCTCTGTTTCTA
AGCACACGTCTGAACTCTTTCCCTACACGACGCTCTTCCGATCTNNNTCTTTGGCCATAAGGGGAGACA
AGAGCACACGTCTGAACTCTTTCCCTACACGACGCTCTTCCGATCTNNNCTCAAAACTTTGGGCCAGTCA
GAGCACACGTCTGAACTCTTTCCCTACACGACGCTCTTCCGATCTNNNTCTGGGTTACCAGGTAGAGA
GAGCACACGTCTGAACTCTTTCCCTACACGACGCTCTTCCGATCTNNNTCTGGGATGCTATCAAGAGTG
GAGCACACGTCTGAACTCTTTCCCTACACGACGCTCTTCCGATCTNNNTGTTTGAGTTGTAGCTTACTG
GCACACGTCTGAACTCTTTCCCTACACGACGCTCTTCCGATCTNNNCTTTCCCATCAGCCCTTCAGTTTCA
GCACACGTCTGAACTCTTTCCCTACACGACGCTCTTCCGATCTNNNTAGGCTTCCTGTCCCACCTAAAAC
AGCACACGTCTGAACTCTTTCCCTACACGACGCTCTTCCGATCTNNNACGAAAGCAAAATGGTCAAAGG
GAGCACACGTCTGAACTCTTTCCCTACACGACGCTCTTCCGATCTNNNCCAAAGCCTGACAGATAGACA
AGCACACGTCTGAACTCTTTCCCTACACGACGCTCTTCCGATCTNNNTAGCCACTTCCTGGCTTGTAGAT
AGCACACGTCTGAACTCTTTCCCTACACGACGCTCTTCCGATCTNNNTTCCTCTCCTCACTCTCCTTGACC

AGAGCACACGTCTGAACTCTTTCCCTACACGACGCTCTTCCGATCTNNNCCCAGAGAAAGAGCCATTAG
AGCACACGTCTGAACTCTTTCCCTACACGACGCTCTTCCGATCTNNNTAAAATACCTAGTACGGGCCGG
GAGCACACGTCTGAACTCTTTCCCTACACGACGCTCTTCCGATCTNNNAATCGGGAGAAAGAGACAGC
AGAGCACACGTCTGAACTCTTTCCCTACACGACGCTCTTCCGATCTNNNGAAAAGATAAGCTCCCTGCC
AGCACACGTCTGAACTCTTTCCCTACACGACGCTCTTCCGATCTNNNACAGACCACAAGCTGAATTACGA
GAGCACACGTCTGAACTCTTTCCCTACACGACGCTCTTCCGATCTNNNAGAAGATGCAAACAGCTGGGA
AGAGCACACGTCTGAACTCTTTCCCTACACGACGCTCTTCCGATCTNNNGGGGACAGAGTGAGATCCA
AGAGCACACGTCTGAACTCTTTCCCTACACGACGCTCTTCCGATCTNNNACTTGAACTCACTTCTGTCCA
GAGCACACGTCTGAACTCTTTCCCTACACGACGCTCTTCCGATCTNNNTACAGGTGTGTTGTGGAGTTT
GAGCACACGTCTGAACTCTTTCCCTACACGACGCTCTTCCGATCTNNNGCAAACTCAGGGTGATCAAGC
AGCACACGTCTGAACTCTTTCCCTACACGACGCTCTTCCGATCTNNNTTGTTCACACAGAAAAACGAAC
AGCACACGTCTGAACTCTTTCCCTACACGACGCTCTTCCGATCTNNNAAACAAAGATTGGCATGCAGG
AGCACACGTCTGAACTCTTTCCCTACACGACGCTCTTCCGATCTNNNCTGGGAAGCCATTATTCTGTCCA
AGCACACGTCTGAACTCTTTCCCTACACGACGCTCTTCCGATCTNNNGAGAACTTCTACATTCCCCAGC
AGAGCACACGTCTGAACTCTTTCCCTACACGACGCTCTTCCGATCTNNNGCTCAAACCCAGATCTTCCAA
GAGCACACGTCTGAACTCTTTCCCTACACGACGCTCTTCCGATCTNNNAAATGGCATGGTGTTTGTGTC
AGCACACGTCTGAACTCTTTCCCTACACGACGCTCTTCCGATCTNNNTGAGTTCCACCCATGAAAAAGCA
AGAGCACACGTCTGAACTCTTTCCCTACACGACGCTCTTCCGATCTNNNGATGGGGTAGTTTGAGCATC
AGCACACGTCTGAACTCTTTCCCTACACGACGCTCTTCCGATCTNNNATGAACATTTCGATCTGAGGGC
GAGCACACGTCTGAACTCTTTCCCTACACGACGCTCTTCCGATCTNNNTGCCCAATCCCAACACAAATAC
AGCACACGTCTGAACTCTTTCCCTACACGACGCTCTTCCGATCTNNNCTCCAGCAATTCTTCCCCATCTCCA
GCACACGTCTGAACTCTTTCCCTACACGACGCTCTTCCGATCTNNNTGTTTTCCACAAGCTCAAGTTCCC
AGCACACGTCTGAACTCTTTCCCTACACGACGCTCTTCCGATCTNNNTGGGACCTACAACTCAAGATGG
GCACACGTCTGAACTCTTTCCCTACACGACGCTCTTCCGATCTNNNTCTCCCTTCCTTTTCTCTTCCCACAT
AGCACACGTCTGAACTCTTTCCCTACACGACGCTCTTCCGATCTNNNTGAGCCTAAGGGAAAAATCTGC
GCACACGTCTGAACTCTTTCCCTACACGACGCTCTTCCGATCTNNNGACACACACTCTTGGTTCCAGTTC
GAGCACACGTCTGAACTCTTTCCCTACACGACGCTCTTCCGATCTNNNCAGCCCAGATTCAAGGAGGTA
AGCACACGTCTGAACTCTTTCCCTACACGACGCTCTTCCGATCTNNNTGGCATGTGCCAATCACCTATTT
GCACACGTCTGAACTCTTTCCCTACACGACGCTCTTCCGATCTNNNACCCAGTGCCCCTTTGATATACTC
GAGCACACGTCTGAACTCTTTCCCTACACGACGCTCTTCCGATCTNNNATGACAGTGGAGATGGCATT
GAGCACACGTCTGAACTCTTTCCCTACACGACGCTCTTCCGATCTNNNCCCGCCTGTATTCCATAAACAG
GAGCACACGTCTGAACTCTTTCCCTACACGACGCTCTTCCGATCTNNNCAGAGCCGTAATGAAGACACA
GAGCACACGTCTGAACTCTTTCCCTACACGACGCTCTTCCGATCTNNNGGAATATGGAGACCCAAGTCC
AGCACACGTCTGAACTCTTTCCCTACACGACGCTCTTCCGATCTNNNACCCTGGTTCTCTCAACTGGTTA
GAGCACACGTCTGAACTCTTTCCCTACACGACGCTCTTCCGATCTNNNACACATAGGCAACATAGGCAC
GAGCACACGTCTGAACTCTTTCCCTACACGACGCTCTTCCGATCTNNNTCCGTGTCATGTCATTCCTGAA
AGCACACGTCTGAACTCTTTCCCTACACGACGCTCTTCCGATCTNNNTGGTTTCCACATGTTCCCCAAAA
AGCACACGTCTGAACTCTTTCCCTACACGACGCTCTTCCGATCTNNNTCAATCTTTTCTCCTTGCCCCACCA
AGCACACGTCTGAACTCTTTCCCTACACGACGCTCTTCCGATCTNNNGTCCCAGAATGCTTTGCAATGTT
GCACACGTCTGAACTCTTTCCCTACACGACGCTCTTCCGATCTNNNGCCCAGAGACAACTTGATTGTGA
GCACACGTCTGAACTCTTTCCCTACACGACGCTCTTCCGATCTNNNCTGACAATAGCCTCCTCTGTGCTC
AGCACACGTCTGAACTCTTTCCCTACACGACGCTCTTCCGATCTNNNTTGGTCATCCTTCCTCACCATCTC
AGCACACGTCTGAACTCTTTCCCTACACGACGCTCTTCCGATCTNNNTTTCAAGTTGGCTGTTCTCCACAC

AGCACACGTCTGAACTCTTTCCCTACACGACGCTCTTCCGATCTNNNTCCCTCTCTCTCTGAATCTCCCTCA
AGCACACGTCTGAACTCTTTCCCTACACGACGCTCTTCCGATCTNNNTTGACACTGTATTCCCTCTGCCTC
GAGCACACGTCTGAACTCTTTCCCTACACGACGCTCTTCCGATCTNNNATTTAGGGTGTTCTTCCACCCC
AGCACACGTCTGAACTCTTTCCCTACACGACGCTCTTCCGATCTNNNAAGAGCTTCAGCAAAGGGTCAA
AGCACACGTCTGAACTCTTTCCCTACACGACGCTCTTCCGATCTNNNAGAAGCTGAATGCTTCTGAATG
AGCACACGTCTGAACTCTTTCCCTACACGACGCTCTTCCGATCTNNNTGATGCCTTTTGCATCTCTCTCTC
AGCACACGTCTGAACTCTTTCCCTACACGACGCTCTTCCGATCTNNNCGCTTCATGTCTCAGATTTGTCC
AGCACACGTCTGAACTCTTTCCCTACACGACGCTCTTCCGATCTNNNAAATGCCAGGGTTGCAATATCA
AGCACACGTCTGAACTCTTTCCCTACACGACGCTCTTCCGATCTNNNTGCTTAGATTCAGTGGACAATC
GAGCACACGTCTGAACTCTTTCCCTACACGACGCTCTTCCGATCTNNNTTTCCAGTGCTAAGGCATTCCA
GAGCACACGTCTGAACTCTTTCCCTACACGACGCTCTTCCGATCTNNNACCATGCTAACGTGTCTACTTC
AGCACACGTCTGAACTCTTTCCCTACACGACGCTCTTCCGATCTNNNTGGGGTCATGAGAAGACAAATC
GCACACGTCTGAACTCTTTCCCTACACGACGCTCTTCCGATCTNNNAAACTGTTTTTATGGACCGTGCCC
AGAGCACACGTCTGAACTCTTTCCCTACACGACGCTCTTCCGATCTNNNAACCTCTAATTTGGTTTGCCA
AGAGCACACGTCTGAACTCTTTCCCTACACGACGCTCTTCCGATCTNNNCACAACCTCTGACATGGCAA
GAGCACACGTCTGAACTCTTTCCCTACACGACGCTCTTCCGATCTNNNTCCATATTGGGGCTGTGAACTC
GAGCACACGTCTGAACTCTTTCCCTACACGACGCTCTTCCGATCTNNNTCGACCTGGTCATTTGTACATG
AGAGCACACGTCTGAACTCTTTCCCTACACGACGCTCTTCCGATCTNNNTTGATAGCACAGTCTCCTGT
GAGCACACGTCTGAACTCTTTCCCTACACGACGCTCTTCCGATCTNNNGCTTAGGCCAAAGAAGTAAGC
GAGCACACGTCTGAACTCTTTCCCTACACGACGCTCTTCCGATCTNNNACAAAACCACAAAACGCAGAT
AGCACACGTCTGAACTCTTTCCCTACACGACGCTCTTCCGATCTNNNTTGTGCACAGCCTCTGTAAATCA
GAGCACACGTCTGAACTCTTTCCCTACACGACGCTCTTCCGATCTNNNAATGTCAGGAAAAGGACATGC
GAGCACACGTCTGAACTCTTTCCCTACACGACGCTCTTCCGATCTNNNTCATGAATACCTAGCATGTGCC
GCACACGTCTGAACTCTTTCCCTACACGACGCTCTTCCGATCTNNNCCCATGGAAATGACTGAACAGGT
AGCACACGTCTGAACTCTTTCCCTACACGACGCTCTTCCGATCTNNNGGACACTTCTGCTTCCTGATTG
AGCACACGTCTGAACTCTTTCCCTACACGACGCTCTTCCGATCTNNNCTGCCTTTTTCCTCCCTCTCTTCCA
GAGCACACGTCTGAACTCTTTCCCTACACGACGCTCTTCCGATCTNNNTGATGTATGTTGGCTATGGTC
AGCACACGTCTGAACTCTTTCCCTACACGACGCTCTTCCGATCTNNNAGCCTGGATCAAAAGTCAGGA
AGCACACGTCTGAACTCTTTCCCTACACGACGCTCTTCCGATCTNNNTTCTGCCCACCCAATATCCTTCTC
AGAGCACACGTCTGAACTCTTTCCCTACACGACGCTCTTCCGATCTNNNTTCTACTGGATTAGGGTGGG
AGCACACGTCTGAACTCTTTCCCTACACGACGCTCTTCCGATCTNNNGCAGATGCAAAAAGGGTATGG
GAGCACACGTCTGAACTCTTTCCCTACACGACGCTCTTCCGATCTNNNAGCTGTGCCTATGGTATGCTA
GAGCACACGTCTGAACTCTTTCCCTACACGACGCTCTTCCGATCTNNNCCTGCCCTCCTTATGATATGCAC
GCACACGTCTGAACTCTTTCCCTACACGACGCTCTTCCGATCTNNNAGAAGATATGGCAGTTCCAGTGC
AGCACACGTCTGAACTCTTTCCCTACACGACGCTCTTCCGATCTNNNCCTCATATCGGGTTTCAGCAGA
GAGCACACGTCTGAACTCTTTCCCTACACGACGCTCTTCCGATCTNNNACCCCTCACACTGGTATGTTTT
AGCACACGTCTGAACTCTTTCCCTACACGACGCTCTTCCGATCTNNNTCTGGATCTGGGTGGTTGATAC
AGCACACGTCTGAACTCTTTCCCTACACGACGCTCTTCCGATCTNNNCATGCCAATCCCAGAAAGGTGTA
AGCACACGTCTGAACTCTTTCCCTACACGACGCTCTTCCGATCTNNNAGTGTTTTGCTTTCGTTCTTCCT
AGCACACGTCTGAACTCTTTCCCTACACGACGCTCTTCCGATCTNNNGGAAGATGGCATTTCATAGCCCT
AGCACACGTCTGAACTCTTTCCCTACACGACGCTCTTCCGATCTNNNCAGTTCTCCCACCTCAACCTATGC
AGCACACGTCTGAACTCTTTCCCTACACGACGCTCTTCCGATCTNNNTCTCGCTGTCTTCCTATGTGA
AGCACACGTCTGAACTCTTTCCCTACACGACGCTCTTCCGATCTNNNTTCTGTTGCATCTGACAGGCTTT

AGCACACGTCTGAACTCTTTCCCTACACGACGCTCTTCCGATCTNNNTCCATTGTTTGTCTGGTGCTGT
GCACACGTCTGAACTCTTTCCCTACACGACGCTCTTCCGATCTNNNTTTCCACCCAAAAGGAATCCTCC
AGCACACGTCTGAACTCTTTCCCTACACGACGCTCTTCCGATCTNNNAAACGTTTTCTGGAGGTCCTT
AGCACACGTCTGAACTCTTTCCCTACACGACGCTCTTCCGATCTNNNGCTGAGCAGAAAATTCAGGCA
GCACACGTCTGAACTCTTTCCCTACACGACGCTCTTCCGATCTNNNCCCATTTCCTGCCTTCTACAACAC
GCACACGTCTGAACTCTTTCCCTACACGACGCTCTTCCGATCTNNNACATGTGTTTCACTAGTCTGCCA
GCACACGTCTGAACTCTTTCCCTACACGACGCTCTTCCGATCTNNNTGTGTAGTACTCCATTGGCTGGA
GCACACGTCTGAACTCTTTCCCTACACGACGCTCTTCCGATCTNNNCTTTCTGGTGTGCTACCTTCTCC
AGCACACGTCTGAACTCTTTCCCTACACGACGCTCTTCCGATCTNNNCTCCTGTGTCTCTTTGACAACCC
GCACACGTCTGAACTCTTTCCCTACACGACGCTCTTCCGATCTNNNCCTGCTCTAGTAAAGGAACCCCT
GAGCACACGTCTGAACTCTTTCCCTACACGACGCTCTTCCGATCTNNNGGATACCGGGAAGGTTTGT
GCACACGTCTGAACTCTTTCCCTACACGACGCTCTTCCGATCTNNNAGGCACTATCCTAGTTGTTGGG
GAGCACACGTCTGAACTCTTTCCCTACACGACGCTCTTCCGATCTNNNACATAGTTGCCAAGGCTTTGT
GCACACGTCTGAACTCTTTCCCTACACGACGCTCTTCCGATCTNNNGTGAATGAAATGCTTGGAGCAG
GCACACGTCTGAACTCTTTCCCTACACGACGCTCTTCCGATCTNNNATTGTCCCTTTCCGCTTTCCTTTC
AGCACACGTCTGAACTCTTTCCCTACACGACGCTCTTCCGATCTNNNACAGTACCATTTCACAGCCCAA
GAGCACACGTCTGAACTCTTTCCCTACACGACGCTCTTCCGATCTNNNTAAAGGAAAGGTTCCAGCCC
GCACACGTCTGAACTCTTTCCCTACACGACGCTCTTCCGATCTNNNAGGATTGAAGCCTCTCTCTCTG
AGCACACGTCTGAACTCTTTCCCTACACGACGCTCTTCCGATCTNNNTTTGGGGTAAAAATGCCACAT
GCACACGTCTGAACTCTTTCCCTACACGACGCTCTTCCGATCTNNNCAGCTCTCCTCAGCAAATTCAGT
AGCACACGTCTGAACTCTTTCCCTACACGACGCTCTTCCGATCTNNNAATGTGGAAATTAGTTCCTCTG
GAGCACACGTCTGAACTCTTTCCCTACACGACGCTCTTCCGATCTNNNGTCTCCACAGTCATTCTTGGC
GCACACGTCTGAACTCTTTCCCTACACGACGCTCTTCCGATCTNNNGAACTTGGTACCTTTCCTGCCTC
GCACACGTCTGAACTCTTTCCCTACACGACGCTCTTCCGATCTNNNCCTGTGGTTCTGTGATGTCTGTT
GCACACGTCTGAACTCTTTCCCTACACGACGCTCTTCCGATCTNNNTCTCTGCTCTCTCTCCTCTCTCAC
AGCACACGTCTGAACTCTTTCCCTACACGACGCTCTTCCGATCTNNNCCTTCTGGGGCTGATTTCAGTT
GCACACGTCTGAACTCTTTCCCTACACGACGCTCTTCCGATCTNNNTGGATTGTGTGCGTATGCAGTT
GCACACGTCTGAACTCTTTCCCTACACGACGCTCTTCCGATCTNNNCTCTTAGAGACCATCCACTGCAC
AGCACACGTCTGAACTCTTTCCCTACACGACGCTCTTCCGATCTNNNATTCATCATCTTCCACACGCAC
GAGCACACGTCTGAACTCTTTCCCTACACGACGCTCTTCCGATCTNNNTCGCTTTATAACAGCCAGAGT
GAGCACACGTCTGAACTCTTTCCCTACACGACGCTCTTCCGATCTNNNAAATAGCTCAGCCCCTTTTG
GAGCACACGTCTGAACTCTTTCCCTACACGACGCTCTTCCGATCTNNNGAGGGTTAGCTGTCCTGAAA
AGCACACGTCTGAACTCTTTCCCTACACGACGCTCTTCCGATCTNNNGTTGGGCCTTAGGAGTAAAT
AGCACACGTCTGAACTCTTTCCCTACACGACGCTCTTCCGATCTNNNTGAGGGTCCCTGAAAGAAAT
AGCACACGTCTGAACTCTTTCCCTACACGACGCTCTTCCGATCTNNNCTGCCTGCCTCTCTTCCTACATA
GCACACGTCTGAACTCTTTCCCTACACGACGCTCTTCCGATCTNNNTTCTGGCTGATGAAGGTGTTCT
GAGCACACGTCTGAACTCTTTCCCTACACGACGCTCTTCCGATCTNNNAAGTGATTTGGAAAACATGC
AGCACACGTCTGAACTCTTTCCCTACACGACGCTCTTCCGATCTNNNGCACTCTGTGTGTGTGAGAGA
GAGCACACGTCTGAACTCTTTCCCTACACGACGCTCTTCCGATCTNNNTTCCTCTCTCCTCTTCTCTGAC
GAGCACACGTCTGAACTCTTTCCCTACACGACGCTCTTCCGATCTNNNGAGAAGTTTCCCCAAGGTTG
AGCACACGTCTGAACTCTTTCCCTACACGACGCTCTTCCGATCTNNNCGAAGAGCCTGACATAAACA
AGCACACGTCTGAACTCTTTCCCTACACGACGCTCTTCCGATCTNNNACAAAGGATCATACTGCAGCC
AGCACACGTCTGAACTCTTTCCCTACACGACGCTCTTCCGATCTNNNGTCACTGAATCACATGGGTAG

GAGCACACGTCTGAACTCTTTCCCTACACGACGCTCTTCCGATCTNNNCAATAAGCCACAAACACACAC
GAGCACACGTCTGAACTCTTTCCCTACACGACGCTCTTCCGATCTNNNAGCATTTCCCTTTGTGTTTAGC
GAGCACACGTCTGAACTCTTTCCCTACACGACGCTCTTCCGATCTNNNAGAGTGTCCAGAAATGGCTTT
AGCACACGTCTGAACTCTTTCCCTACACGACGCTCTTCCGATCTNNNCTGCTCAAAATGACTGATGGG
AGCACACGTCTGAACTCTTTCCCTACACGACGCTCTTCCGATCTNNNTTAGGAGGCCCCAAACTTCAAA
GAGCACACGTCTGAACTCTTTCCCTACACGACGCTCTTCCGATCTNNNCTAATGCCAGCATGATCCCTGT
AAGAGCACACGTCTGAACTCTTTCCCTACACGACGCTCTTCCGATCTNNNACACAAAAAGCAAAGCTTG
AGCACACGTCTGAACTCTTTCCCTACACGACGCTCTTCCGATCTNNNTGTACACTTCATGTCCCAGACCTC
AGCACACGTCTGAACTCTTTCCCTACACGACGCTCTTCCGATCTNNNACATCTTGCATGGATAAGGTG
GAGCACACGTCTGAACTCTTTCCCTACACGACGCTCTTCCGATCTNNNACAAAAGGGAAACTAAGGTCA
AGAGCACACGTCTGAACTCTTTCCCTACACGACGCTCTTCCGATCTNNNCGGTTCTCCAGAGAAACAAC
AGAGCACACGTCTGAACTCTTTCCCTACACGACGCTCTTCCGATCTNNNGACATGGGGTGAAGATGTC
GAGCACACGTCTGAACTCTTTCCCTACACGACGCTCTTCCGATCTNNNTCCCTCTCTGCCCTTTCAACTAT
AGAGCACACGTCTGAACTCTTTCCCTACACGACGCTCTTCCGATCTNNNTATGCCTTTTACCTCACCATG
GAGCACACGTCTGAACTCTTTCCCTACACGACGCTCTTCCGATCTNNNTTCTTATATCTGCGTGTGTGTG
GAGCACACGTCTGAACTCTTTCCCTACACGACGCTCTTCCGATCTNNNAGCATGAGAGCTGCTAAAACC
AGCACACGTCTGAACTCTTTCCCTACACGACGCTCTTCCGATCTNNNATGGAATCTCCCTGTCCCATCTC
GAGCACACGTCTGAACTCTTTCCCTACACGACGCTCTTCCGATCTNNNGAAAGTCATGTAGTCCCCACA
AGCACACGTCTGAACTCTTTCCCTACACGACGCTCTTCCGATCTNNNCAACAGCATCCTCACATAGGCTT
GCACACGTCTGAACTCTTTCCCTACACGACGCTCTTCCGATCTNNNATTGCACATTCTTGTGCTATGCCC
AGAGCACACGTCTGAACTCTTTCCCTACACGACGCTCTTCCGATCTNNNAGCCAAGCATGGTTTCTCCTA
AGCACACGTCTGAACTCTTTCCCTACACGACGCTCTTCCGATCTNNNCTGCCACAGAAGCAAATTCACTG
AGAGCACACGTCTGAACTCTTTCCCTACACGACGCTCTTCCGATCTNNNACATGATAACATCTGGGTG
AGAGCACACGTCTGAACTCTTTCCCTACACGACGCTCTTCCGATCTNNNGCATAAGTAGGTCACAGGGA
GAGCACACGTCTGAACTCTTTCCCTACACGACGCTCTTCCGATCTNNNATGCTCTTGAGTTGCTGATGG
AGAGCACACGTCTGAACTCTTTCCCTACACGACGCTCTTCCGATCTNNNTCCTTCAGGACATGTTTGTG
GAGCACACGTCTGAACTCTTTCCCTACACGACGCTCTTCCGATCTNNNATGTTTGCGTGTCTGGCTTAT
GAGCACACGTCTGAACTCTTTCCCTACACGACGCTCTTCCGATCTNNNTATGAGAACTAGGGCACACTC
AGCACACGTCTGAACTCTTTCCCTACACGACGCTCTTCCGATCTNNNAACTCTTAGGAGGATGCCCAACA
AGAGCACACGTCTGAACTCTTTCCCTACACGACGCTCTTCCGATCTNNNCAAGTGCCTGTGTCAAGTAA
AGAGCACACGTCTGAACTCTTTCCCTACACGACGCTCTTCCGATCTNNNTCTGTTGGCTTGATTTTCATC
AAGAGCACACGTCTGAACTCTTTCCCTACACGACGCTCTTCCGATCTNNNAAGGACCCTCCTTTGTTTCT
GAGCACACGTCTGAACTCTTTCCCTACACGACGCTCTTCCGATCTNNNGGCTATGCTCTGGAGAGAGA
AGAGCACACGTCTGAACTCTTTCCCTACACGACGCTCTTCCGATCTNNNCATCTACCCCTTTGTGAAGGA
AGAGCACACGTCTGAACTCTTTCCCTACACGACGCTCTTCCGATCTNNNAATCTTCTGTCCATCGCCCTTT
GAGCACACGTCTGAACTCTTTCCCTACACGACGCTCTTCCGATCTNNNGTTTTGCCTGGTTTGGCTCTTT
GAGCACACGTCTGAACTCTTTCCCTACACGACGCTCTTCCGATCTNNNTGGCTATTTTAGGATCTGTGC
AGCACACGTCTGAACTCTTTCCCTACACGACGCTCTTCCGATCTNNNCCCTTCTTTTGCCCTCTTCAACTCA
GAGCACACGTCTGAACTCTTTCCCTACACGACGCTCTTCCGATCTNNNAATCTTAGGTGGTGCAGAGG
AGCACACGTCTGAACTCTTTCCCTACACGACGCTCTTCCGATCTNNNTGGTTTCTGGGAGAGGATGTT
AGCACACGTCTGAACTCTTTCCCTACACGACGCTCTTCCGATCTNNNTAGTTAGAGCTGCCTGGCTAAG
GAGCACACGTCTGAACTCTTTCCCTACACGACGCTCTTCCGATCTNNNCACTTTTATTGTACCAGCCACG
GAGCACACGTCTGAACTCTTTCCCTACACGACGCTCTTCCGATCTNNNTGTGGCATTATGAAGAGTGTC

GAGCACACGTCTGAACTCTTTCCCTACACGACGCTCTTCCGATCTNNNTCTGTCCAGCCTCAAAGACCAT
AGAGCACACGTCTGAACTCTTTCCCTACACGACGCTCTTCCGATCTNNNCCTGTCTCAGAAAACATGGCT
AGAGCACACGTCTGAACTCTTTCCCTACACGACGCTCTTCCGATCTNNNTGGGAAAACTGAGGGATTCA
AGCACACGTCTGAACTCTTTCCCTACACGACGCTCTTCCGATCTNNNACCTGTACTTCTGCCTGGAATCT
GCACACGTCTGAACTCTTTCCCTACACGACGCTCTTCCGATCTNNNGTGGCAGATATTTGAAGGCTG
AGAGCACACGTCTGAACTCTTTCCCTACACGACGCTCTTCCGATCTNNNGAATCCATTCAGGTCCGACTC
GAGCACACGTCTGAACTCTTTCCCTACACGACGCTCTTCCGATCTNNNGACCCTCACAATGTCGTTGTTC
GAGCACACGTCTGAACTCTTTCCCTACACGACGCTCTTCCGATCTNNNGAATACTTACAGGGCTCCAGT
AGCACACGTCTGAACTCTTTCCCTACACGACGCTCTTCCGATCTNNNTCTAAAGGGTGTTGGAGCCCAT
GAGCACACGTCTGAACTCTTTCCCTACACGACGCTCTTCCGATCTNNNATTGTGACAACTTACCCCACCA
GCACACGTCTGAACTCTTTCCCTACACGACGCTCTTCCGATCTNNNTGTTGGCTTCTCTCTATGCTCCTCA
AGCACACGTCTGAACTCTTTCCCTACACGACGCTCTTCCGATCTNNNTCTTTGAGCACTATGGTCTTGC
GAGCACACGTCTGAACTCTTTCCCTACACGACGCTCTTCCGATCTNNNAGTGGTGGAATAGATGCCTT
AGCACACGTCTGAACTCTTTCCCTACACGACGCTCTTCCGATCTNNNCAGCATGGATGGAAAGACAGA
AGCACACGTCTGAACTCTTTCCCTACACGACGCTCTTCCGATCTNNNGACTTGCAGAACAGTAGGTGTC
GAGCACACGTCTGAACTCTTTCCCTACACGACGCTCTTCCGATCTNNNTCGAGATAGAGCTCGTCTAGT
AGCACACGTCTGAACTCTTTCCCTACACGACGCTCTTCCGATCTNNNAGGCAAATGACCCTCATACCTCT
GAGCACACGTCTGAACTCTTTCCCTACACGACGCTCTTCCGATCTNNNTGTAGATCCCCCTACACTTCCT
GAGCACACGTCTGAACTCTTTCCCTACACGACGCTCTTCCGATCTNNNCCGCCTCACTTTCTTTCTCTCTC
GAGCACACGTCTGAACTCTTTCCCTACACGACGCTCTTCCGATCTNNNCTGAGAATTTGGCAAAGGGCA
GAGCACACGTCTGAACTCTTTCCCTACACGACGCTCTTCCGATCTNNNCCTTCGCTTCTGTTCAATGCTTC
GAGCACACGTCTGAACTCTTTCCCTACACGACGCTCTTCCGATCTNNNTGTGCGTCTGGAAAGTGAAT
AGAGCACACGTCTGAACTCTTTCCCTACACGACGCTCTTCCGATCTNNNAAAGTCACTGATCATGCCCCT
GAGCACACGTCTGAACTCTTTCCCTACACGACGCTCTTCCGATCTNNNAATCAAAATGCCCCTGGAAGT
AGCACACGTCTGAACTCTTTCCCTACACGACGCTCTTCCGATCTNNNATTTTCCAGAACCCTCTCACCC
GAGCACACGTCTGAACTCTTTCCCTACACGACGCTCTTCCGATCTNNNGGTTGAAGATACTGGGTGGT
AGCACACGTCTGAACTCTTTCCCTACACGACGCTCTTCCGATCTNNNGGAAAGAGATGCAGAAAAGCC
AGCACACGTCTGAACTCTTTCCCTACACGACGCTCTTCCGATCTNNNCTTGGCTATTCTTGGAGCTGTG
AGAGCACACGTCTGAACTCTTTCCCTACACGACGCTCTTCCGATCTNNNGTTCCCTTTGCAATGGATGTG
AGCACACGTCTGAACTCTTTCCCTACACGACGCTCTTCCGATCTNNNGTGTTCACTCAGAACCCTTGTCC
GAGCACACGTCTGAACTCTTTCCCTACACGACGCTCTTCCGATCTNNNCAACTTGCAACAAGAGGAATC
AGCACACGTCTGAACTCTTTCCCTACACGACGCTCTTCCGATCTNNNACCCGTTCTTCCAGAATAGGTCA
AGAGCACACGTCTGAACTCTTTCCCTACACGACGCTCTTCCGATCTNNNCTTACACAGAGCCCAGACTTT
AGCACACGTCTGAACTCTTTCCCTACACGACGCTCTTCCGATCTNNNAAACAAAGTTCAAAACAGGCAG
AGCACACGTCTGAACTCTTTCCCTACACGACGCTCTTCCGATCTNNNTAACTAGGGGAGGAAGCCTTA
AGCACACGTCTGAACTCTTTCCCTACACGACGCTCTTCCGATCTNNNCATAAACTGCCCAGATGGACAC
GAGCACACGTCTGAACTCTTTCCCTACACGACGCTCTTCCGATCTNNNTACATTTCAAGTGCTGTCCCTC
AGAGCACACGTCTGAACTCTTTCCCTACACGACGCTCTTCCGATCTNNNAGATAAAGAGCATTATGTCC
AGCACACGTCTGAACTCTTTCCCTACACGACGCTCTTCCGATCTNNNGAACCATCATAAGTTGGGACCC
AGCACACGTCTGAACTCTTTCCCTACACGACGCTCTTCCGATCTNNNCTTCATTCAGTGTCTGGGGTCT
AGCACACGTCTGAACTCTTTCCCTACACGACGCTCTTCCGATCTNNNTCCTCTATCCCCACAACCAACTTC
AGCACACGTCTGAACTCTTTCCCTACACGACGCTCTTCCGATCTNNNTATAAACCGTGCACTCTCCCTCT
AGCACACGTCTGAACTCTTTCCCTACACGACGCTCTTCCGATCTNNNTCCACCACTATTTGGGGCTTTTT

AGCACACGTCTGAACTCTTTCCCTACACGACGCTCTTCCGATCTNNNCTAACTCCTCACCTCCCTCCAT
GCACACGTCTGAACTCTTTCCCTACACGACGCTCTTCCGATCTNNNAACTTACTGTGTTTCAGGGCCA
AGCACACGTCTGAACTCTTTCCCTACACGACGCTCTTCCGATCTNNNAAGGTGCTGGACATCTTTTCA
GCACACGTCTGAACTCTTTCCCTACACGACGCTCTTCCGATCTNNNGCTCTGGCTTCACTCTAAACCT
GCACACGTCTGAACTCTTTCCCTACACGACGCTCTTCCGATCTNNNTGCCAAAGCTCAAGTCTTCAAC
GAGCACACGTCTGAACTCTTTCCCTACACGACGCTCTTCCGATCTNNNGCGCAGAACCATCTATTTG
GAGCACACGTCTGAACTCTTTCCCTACACGACGCTCTTCCGATCTNNNCATTAGGAGCTGTAGACTG
GCACACGTCTGAACTCTTTCCCTACACGACGCTCTTCCGATCTNNNTGCTGAGGTGCTTTGTAAACTC
GAGCACACGTCTGAACTCTTTCCCTACACGACGCTCTTCCGATCTNNNTATTTCAGCATCTGCCTCTTC
GCACACGTCTGAACTCTTTCCCTACACGACGCTCTTCCGATCTNNNTCATCAACTCATGGCCAATCCTG
CACACGTCTGAACTCTTTCCCTACACGACGCTCTTCCGATCTNNNCAGGTGCACATATTAGCCAGAG
GCACACGTCTGAACTCTTTCCCTACACGACGCTCTTCCGATCTNNNATCCCTAGTGGACGACTGAAAC
GCACACGTCTGAACTCTTTCCCTACACGACGCTCTTCCGATCTNNNCCACTAATCTAAGGGCAACCCA
AGCACACGTCTGAACTCTTTCCCTACACGACGCTCTTCCGATCTNNNGGGATGCTCCCATCCTACTATG
GCACACGTCTGAACTCTTTCCCTACACGACGCTCTTCCGATCTNNNACATCCCACACACACATCTCTCTC
GCACACGTCTGAACTCTTTCCCTACACGACGCTCTTCCGATCTNNNCCTCTGGGGTCTTTGAGATTGT
AGCACACGTCTGAACTCTTTCCCTACACGACGCTCTTCCGATCTNNNGCATTAGGGTGGAAAAATGG
AGCACACGTCTGAACTCTTTCCCTACACGACGCTCTTCCGATCTNNNTCCACATACAGACAAACGTGCA
GAGCACACGTCTGAACTCTTTCCCTACACGACGCTCTTCCGATCTNNNCATCCAACCTCAATCCTCAGC
AGAGCACACGTCTGAACTCTTTCCCTACACGACGCTCTTCCGATCTNNNTCTCTGGCTCATCTTCTCCTA
AGCACACGTCTGAACTCTTTCCCTACACGACGCTCTTCCGATCTNNNCTGGGCAACACTCCAGTGAAA
AGCACACGTCTGAACTCTTTCCCTACACGACGCTCTTCCGATCTNNNCACACAACACTTACATGCCACA
GAGCACACGTCTGAACTCTTTCCCTACACGACGCTCTTCCGATCTNNNAAGCTAACCCCCAGAACTTC
GAGCACACGTCTGAACTCTTTCCCTACACGACGCTCTTCCGATCTNNNAGGGGAAGAAGGCATATGG
CACACGTCTGAACTCTTTCCCTACACGACGCTCTTCCGATCTNNNACCTGATATCCAGGGCCATTTCTC
AGCACACGTCTGAACTCTTTCCCTACACGACGCTCTTCCGATCTNNNAATCACACACAAAGGGAAAGC
GAGCACACGTCTGAACTCTTTCCCTACACGACGCTCTTCCGATCTNNNAATACCCTGTGTGTGTGGAT
AGCACACGTCTGAACTCTTTCCCTACACGACGCTCTTCCGATCTNNNGTGCAATAGTATGGCTGGGTC
GCACACGTCTGAACTCTTTCCCTACACGACGCTCTTCCGATCTNNNTAATCCCTGAGCTGCATCCAAA
GCACACGTCTGAACTCTTTCCCTACACGACGCTCTTCCGATCTNNNTACATACCTCAAAACCCAGAGG
AGCACACGTCTGAACTCTTTCCCTACACGACGCTCTTCCGATCTNNNGGGTAAGCTAGTGGTAGTTCT
GCACACGTCTGAACTCTTTCCCTACACGACGCTCTTCCGATCTNNNGGGGCGGAAGCTAGTTTTACTT
GAGCACACGTCTGAACTCTTTCCCTACACGACGCTCTTCCGATCTNNNATGTGAGCCTTCCTCTGATTC
GAGCACACGTCTGAACTCTTTCCCTACACGACGCTCTTCCGATCTNNNCCTGTGCGAAGACTTTAGGT
GCACACGTCTGAACTCTTTCCCTACACGACGCTCTTCCGATCTNNNAGAAACTGTCCTTGAGAGGGT
GAGCACACGTCTGAACTCTTTCCCTACACGACGCTCTTCCGATCTNNNAAGAACCCATCCCCCTACAAC
AGCACACGTCTGAACTCTTTCCCTACACGACGCTCTTCCGATCTNNNATTCCCTTCCATGCCTCCTTAGT
GCACACGTCTGAACTCTTTCCCTACACGACGCTCTTCCGATCTNNNGACAATTACATACACGCACGCA
AGCACACGTCTGAACTCTTTCCCTACACGACGCTCTTCCGATCTNNNTGCCTTTAGTGTCCAAGGCAT
GAGCACACGTCTGAACTCTTTCCCTACACGACGCTCTTCCGATCTNNNCGTCTTGCCTATCCTGAAACC
AGCACACGTCTGAACTCTTTCCCTACACGACGCTCTTCCGATCTNNNTGTAGATGAAGACAGCACTG
AGCACACGTCTGAACTCTTTCCCTACACGACGCTCTTCCGATCTNNNAGCTTCAAAGACACAGAAGG
AGCACACGTCTGAACTCTTTCCCTACACGACGCTCTTCCGATCTNNNGGCTAGATCAATGGGCTCTA

GAGCACACGTCTGAACTCTTTCCCTACACGACGCTCTTCCGATCTNNNGCTCTCCCGGTTTTTGTTTTTG
AGCACACGTCTGAACTCTTTCCCTACACGACGCTCTTCCGATCTNNNTGGGGCCAAAGGAAAATAACC
GAGCACACGTCTGAACTCTTTCCCTACACGACGCTCTTCCGATCTNNNAGAGCTCTATGTGGGTCCATG
AGCACACGTCTGAACTCTTTCCCTACACGACGCTCTTCCGATCTNNNTCACGTACATCCTCCTTCATCCAC
GCACACGTCTGAACTCTTTCCCTACACGACGCTCTTCCGATCTNNNTGCAAGCCCTTTATTCTGGCATTC
GAGCACACGTCTGAACTCTTTCCCTACACGACGCTCTTCCGATCTNNNTGGTGTCTCCATGACCTACCAT
AGCACACGTCTGAACTCTTTCCCTACACGACGCTCTTCCGATCTNNNTCTTATTTCTGCCCACCCACTTCCA
GAGCACACGTCTGAACTCTTTCCCTACACGACGCTCTTCCGATCTNNNCATACATCAAACCCCACAACACC
GAGCACACGTCTGAACTCTTTCCCTACACGACGCTCTTCCGATCTNNNCTGTGTCGCAGCACATGAAGT
AGAGCACACGTCTGAACTCTTTCCCTACACGACGCTCTTCCGATCTNNNTAGATCCCTCGTTTGTTTTGG
AGCACACGTCTGAACTCTTTCCCTACACGACGCTCTTCCGATCTNNNGTTGTCTTTGCAAGTTCTTGGCT
AGCACACGTCTGAACTCTTTCCCTACACGACGCTCTTCCGATCTNNNCAAAGTGTGGCCACTCCTCAATA
AGCACACGTCTGAACTCTTTCCCTACACGACGCTCTTCCGATCTNNNGGGTCACATCCTGCACTATACCA
GAGCACACGTCTGAACTCTTTCCCTACACGACGCTCTTCCGATCTNNNAAGGTTTGTGTGAGAATCAA
AGAGCACACGTCTGAACTCTTTCCCTACACGACGCTCTTCCGATCTNNNCCGCTTCTGGTACTCTGATGA
GAGCACACGTCTGAACTCTTTCCCTACACGACGCTCTTCCGATCTNNNAAAGCCTGATACACACCAAGC
GAGCACACGTCTGAACTCTTTCCCTACACGACGCTCTTCCGATCTNNNTCTTCTTTCCTGACTTCCCAGG
GAGCACACGTCTGAACTCTTTCCCTACACGACGCTCTTCCGATCTNNNCCTGTTGCTGCAGTTGATTTCT
GAGCACACGTCTGAACTCTTTCCCTACACGACGCTCTTCCGATCTNNNCAGAGTGAGCTAGAAATACGC
GAGCACACGTCTGAACTCTTTCCCTACACGACGCTCTTCCGATCTNNNGACTTCACCCATGATGTACCA
AGCACACGTCTGAACTCTTTCCCTACACGACGCTCTTCCGATCTNNNTCAAGCCTCACACCTATTTTGGG
GAGCACACGTCTGAACTCTTTCCCTACACGACGCTCTTCCGATCTNNNTTAGTGAGGCTTGAGACCCCT
GAGCACACGTCTGAACTCTTTCCCTACACGACGCTCTTCCGATCTNNNGCTGATCTTGAAGCCCTTGAG
AGCACACGTCTGAACTCTTTCCCTACACGACGCTCTTCCGATCTNNNCACTCAGTTGAGGGCCAATTCT
GAGCACACGTCTGAACTCTTTCCCTACACGACGCTCTTCCGATCTNNNATGAGGAAGAAAAGGGGGAC
AGCACACGTCTGAACTCTTTCCCTACACGACGCTCTTCCGATCTNNNGGGTGACATCTTTCACAATCGGA
AGCACACGTCTGAACTCTTTCCCTACACGACGCTCTTCCGATCTNNNCATCTAGGGACTGCAGAGGAGA
GAGCACACGTCTGAACTCTTTCCCTACACGACGCTCTTCCGATCTNNNGTCATCACCCTTCCAATCTCCTC
GAGCACACGTCTGAACTCTTTCCCTACACGACGCTCTTCCGATCTNNNGCCAGCACAAAAATGAGTTACC
AGAGCACACGTCTGAACTCTTTCCCTACACGACGCTCTTCCGATCTNNNTTGTTAGAAACCGACAGGTC
GAGCACACGTCTGAACTCTTTCCCTACACGACGCTCTTCCGATCTNNNAAAGGTTCTTGGTGGTACTGC
GCACACGTCTGAACTCTTTCCCTACACGACGCTCTTCCGATCTNNNCCTATCTTTATGCCCATGCCTCCCA
GAGCACACGTCTGAACTCTTTCCCTACACGACGCTCTTCCGATCTNNNAGGCCACTCAACCATTTCATTG
AGCACACGTCTGAACTCTTTCCCTACACGACGCTCTTCCGATCTNNNTGTGGATTTGCAGGCATTCCTTA
AGCACACGTCTGAACTCTTTCCCTACACGACGCTCTTCCGATCTNNNTAGACAAGCAAGACAGCTTCCAC
AGCACACGTCTGAACTCTTTCCCTACACGACGCTCTTCCGATCTNNNCCTTCTGCCTCAGGTGGAATTTT
GAGCACACGTCTGAACTCTTTCCCTACACGACGCTCTTCCGATCTNNNGGTAGTTGTTGTGCTGTGGGT
AGCACACGTCTGAACTCTTTCCCTACACGACGCTCTTCCGATCTNNNTCAGTTGAGGCATCTTCTGCTTC
GAGCACACGTCTGAACTCTTTCCCTACACGACGCTCTTCCGATCTNNNAGTCAGTCACAAACAAGTCAC
GAGCACACGTCTGAACTCTTTCCCTACACGACGCTCTTCCGATCTNNNGAATGAAGGATGCCCTGGAA
AGCACACGTCTGAACTCTTTCCCTACACGACGCTCTTCCGATCTNNNTGTCTTTGCATCCATCATCCAAT
GAGCACACGTCTGAACTCTTTCCCTACACGACGCTCTTCCGATCTNNNCCTATCACAATGCAAGCTCCAG
AGAGCACACGTCTGAACTCTTTCCCTACACGACGCTCTTCCGATCTNNNCAGAAAGCAAAGGACCTCA

AGCACACGTCTGAACTCTTTCCCTACACGACGCTCTTCCGATCTNNNATCCACTAAGCCCAGCAGAAA
GAGCACACGTCTGAACTCTTTCCCTACACGACGCTCTTCCGATCTNNNCCATGATAGCTGTGACATAG
GAGCACACGTCTGAACTCTTTCCCTACACGACGCTCTTCCGATCTNNNCAAGAAACAGAAATGCGCTC
AGCACACGTCTGAACTCTTTCCCTACACGACGCTCTTCCGATCTNNNGTAGCAAAAGACCATGCAGG
GCACACGTCTGAACTCTTTCCCTACACGACGCTCTTCCGATCTNNNTACATACATATACGGGACGGG
GAGCACACGTCTGAACTCTTTCCCTACACGACGCTCTTCCGATCTNNNACAGCTGGAGGAGACTGAA
GAGCACACGTCTGAACTCTTTCCCTACACGACGCTCTTCCGATCTNNNATAGCACAGCTAGCATCAAG
AGCACACGTCTGAACTCTTTCCCTACACGACGCTCTTCCGATCTNNNGAAGACACTTCCCACCTTCAT
AGCACACGTCTGAACTCTTTCCCTACACGACGCTCTTCCGATCTNNNTCCCTGCCCTCAAGAAACAAA
GAGCACACGTCTGAACTCTTTCCCTACACGACGCTCTTCCGATCTNNNCACTTGTCTACAGCATCAGC
AGCACACGTCTGAACTCTTTCCCTACACGACGCTCTTCCGATCTNNNCCTAGCTCTTCTGTCTAGCCCT
AGCACACGTCTGAACTCTTTCCCTACACGACGCTCTTCCGATCTNNNGGTAAACCATCATCACCATTGC
AGCACACGTCTGAACTCTTTCCCTACACGACGCTCTTCCGATCTNNNCAAAGTGCCACCAAAAGCTCA
AGCACACGTCTGAACTCTTTCCCTACACGACGCTCTTCCGATCTNNNGAAGCCAAAACTGTGAACTGC
AGCACACGTCTGAACTCTTTCCCTACACGACGCTCTTCCGATCTNNNTACTGTGTCTCACAGAGAAGGC
GCACACGTCTGAACTCTTTCCCTACACGACGCTCTTCCGATCTNNNGAGGAGAGCAGCAGTCTAGTT
AGCACACGTCTGAACTCTTTCCCTACACGACGCTCTTCCGATCTNNNGACAAATACCCCAGTTTCCCTC
AGCACACGTCTGAACTCTTTCCCTACACGACGCTCTTCCGATCTNNNATTGGGGACTAACTTGGATTG
AGCACACGTCTGAACTCTTTCCCTACACGACGCTCTTCCGATCTNNNCTTCAGCTTCCCTCACTTTACC
AGCACACGTCTGAACTCTTTCCCTACACGACGCTCTTCCGATCTNNNTTCCACCACAGTACACTGTTTCC
GCACACGTCTGAACTCTTTCCCTACACGACGCTCTTCCGATCTNNNGCTTGTGTTGAGAGAGAGTGA
AGCACACGTCTGAACTCTTTCCCTACACGACGCTCTTCCGATCTNNNGGAAGACTGAAGGGGAGAG
GAGCACACGTCTGAACTCTTTCCCTACACGACGCTCTTCCGATCTNNNTCATCTCCTCCTGAGAGATCC
CACACGTCTGAACTCTTTCCCTACACGACGCTCTTCCGATCTNNNATTAGGTAGTCCCAGCACTTTGG
CACACGTCTGAACTCTTTCCCTACACGACGCTCTTCCGATCTNNNTTTGCTGTTGTGTTGGAGAGGT
GCACACGTCTGAACTCTTTCCCTACACGACGCTCTTCCGATCTNNNATCGCCAGTGTTTCTCTCTTTGCC
GCACACGTCTGAACTCTTTCCCTACACGACGCTCTTCCGATCTNNNAGGCAGATGGAACAGAAAGGT
AGCACACGTCTGAACTCTTTCCCTACACGACGCTCTTCCGATCTNNNCACCACTGTACAATGAGACCC
GCACACGTCTGAACTCTTTCCCTACACGACGCTCTTCCGATCTNNNAAGGGATTTGGTCACTGGGATT
AGCACACGTCTGAACTCTTTCCCTACACGACGCTCTTCCGATCTNNNTCATAGGAAGTGTTGCAGGG
GAGCACACGTCTGAACTCTTTCCCTACACGACGCTCTTCCGATCTNNNTGCAGACCTATTCACAATTGC
AGCACACGTCTGAACTCTTTCCCTACACGACGCTCTTCCGATCTNNNATAACAAGACAGGGGATGGCT
AGCACACGTCTGAACTCTTTCCCTACACGACGCTCTTCCGATCTNNNAGATGGAATGTAAACCATGCC
AGCACACGTCTGAACTCTTTCCCTACACGACGCTCTTCCGATCTNNNGTTCAGAAGTCTCAGAAACCCC
AGCACACGTCTGAACTCTTTCCCTACACGACGCTCTTCCGATCTNNNCTTCCCTCCCTTCTCCTTTTCTGC
GCACACGTCTGAACTCTTTCCCTACACGACGCTCTTCCGATCTNNNATGCATCCAGAAAAGTGCCTTG
GCACACGTCTGAACTCTTTCCCTACACGACGCTCTTCCGATCTNNNGAGGGGGTACTCATTCAGTCCA
AGCACACGTCTGAACTCTTTCCCTACACGACGCTCTTCCGATCTNNNGTATGTGGGAGGGAAGTAGG
GAGCACACGTCTGAACTCTTTCCCTACACGACGCTCTTCCGATCTNNNATGAGAATAGGTCTCATGGG
AGCACACGTCTGAACTCTTTCCCTACACGACGCTCTTCCGATCTNNNTCCTTTTGAGGAAAAAGACCC
AGCACACGTCTGAACTCTTTCCCTACACGACGCTCTTCCGATCTNNNTGGCAACAAAATCACTCCTGA
GCACACGTCTGAACTCTTTCCCTACACGACGCTCTTCCGATCTNNNTTGTGTGCCCTTCCATGAATGTT
AGCACACGTCTGAACTCTTTCCCTACACGACGCTCTTCCGATCTNNNATGCCATCTCCCCCAAATTCTT

AGCACACGTCTGAACTCTTTCCCTACACGACGCTCTTCCGATCTNNNACAAAGTTGCTCCTCTCGTTCTG
GAGCACACGTCTGAACTCTTTCCCTACACGACGCTCTTCCGATCTNNNCCTCCCTATCGCTCTACTCCATT
AGCACACGTCTGAACTCTTTCCCTACACGACGCTCTTCCGATCTNNNGCCCATAAAGAGAATTTGCTGG
AGCACACGTCTGAACTCTTTCCCTACACGACGCTCTTCCGATCTNNNGGCAGCACTAACCTCATTCTAG
GAGCACACGTCTGAACTCTTTCCCTACACGACGCTCTTCCGATCTNNNAGCATGGAATGAGCGGTTTT
AGCACACGTCTGAACTCTTTCCCTACACGACGCTCTTCCGATCTNNNATATCATCAAATGACTGTGGG
AGCACACGTCTGAACTCTTTCCCTACACGACGCTCTTCCGATCTNNNAAAACCTGTCTCCAAACAGACCA
GAGCACACGTCTGAACTCTTTCCCTACACGACGCTCTTCCGATCTNNNTGGTTCTGTCTGTTACATCCCC
GAGCACACGTCTGAACTCTTTCCCTACACGACGCTCTTCCGATCTNNNCAAGGCAGGACCTTCTGACTG
GAGCACACGTCTGAACTCTTTCCCTACACGACGCTCTTCCGATCTNNNTTGCCTGATTGAGAACCAACC
AGAGCACACGTCTGAACTCTTTCCCTACACGACGCTCTTCCGATCTNNNCCAATGCAAGATACACACACA
AGAGCACACGTCTGAACTCTTTCCCTACACGACGCTCTTCCGATCTNNNTGATTGAATCCAACAATGCGT
GAGCACACGTCTGAACTCTTTCCCTACACGACGCTCTTCCGATCTNNNTTGGCCCCAAAAAGTACACTTG
AGCACACGTCTGAACTCTTTCCCTACACGACGCTCTTCCGATCTNNNATTTACACACATGCACGCTCTCTG
GAGCACACGTCTGAACTCTTTCCCTACACGACGCTCTTCCGATCTNNNACAGGGATGTAGGACGTAGG
AGCACACGTCTGAACTCTTTCCCTACACGACGCTCTTCCGATCTNNNCTGGTTTACAGATGGCAGTCTC
AGCACACGTCTGAACTCTTTCCCTACACGACGCTCTTCCGATCTNNNAGTTTCAAGTAGCCAACTGAGA
AGCACACGTCTGAACTCTTTCCCTACACGACGCTCTTCCGATCTNNNTTTGATAGTCTGCTGTTGAGCCG
AGCACACGTCTGAACTCTTTCCCTACACGACGCTCTTCCGATCTNNNGTCTAGCATTCTTTCACTGCTGG
GCACACGTCTGAACTCTTTCCCTACACGACGCTCTTCCGATCTNNNGGCCAGGATTCATAGGCTCTCTA
GAGCACACGTCTGAACTCTTTCCCTACACGACGCTCTTCCGATCTNNNAACCTTCCCTAGCTGCTCACAT
AGCACACGTCTGAACTCTTTCCCTACACGACGCTCTTCCGATCTNNNTCCACTCATTCATGCTTCCTGTGC
GAGCACACGTCTGAACTCTTTCCCTACACGACGCTCTTCCGATCTNNNATCTTTGGTCTGTTCAGCGCT
AGCACACGTCTGAACTCTTTCCCTACACGACGCTCTTCCGATCTNNNCCTGAGCCAGTGTACCATTTGAA
GAGCACACGTCTGAACTCTTTCCCTACACGACGCTCTTCCGATCTNNNCTCCACAGCAGTGAGACCTTTA
AGCACACGTCTGAACTCTTTCCCTACACGACGCTCTTCCGATCTNNNTATATGTACACTGCTGACCCCCA
AGCACACGTCTGAACTCTTTCCCTACACGACGCTCTTCCGATCTNNNTAGCCTGTGGGCCTTTTCTCTAT
AGAGCACACGTCTGAACTCTTTCCCTACACGACGCTCTTCCGATCTNNNACTATGCTTTGCCTTCAACTCC
AGCACACGTCTGAACTCTTTCCCTACACGACGCTCTTCCGATCTNNNATGCTCTTTTGGCCATGCTTTTCC
AGCACACGTCTGAACTCTTTCCCTACACGACGCTCTTCCGATCTNNNCAATTTCACTCCGTGGTCTTCCT
GAGCACACGTCTGAACTCTTTCCCTACACGACGCTCTTCCGATCTNNNGGGAGAGACAGGGAGAAAC
AGCACACGTCTGAACTCTTTCCCTACACGACGCTCTTCCGATCTNNNTGTAAACATGCGGGAATTTGGG
GAGCACACGTCTGAACTCTTTCCCTACACGACGCTCTTCCGATCTNNNAGCTAATGAAGCCTGACCATG
AGCACACGTCTGAACTCTTTCCCTACACGACGCTCTTCCGATCTNNNCATGCTTCCATTGCACCAAACATG
GAGCACACGTCTGAACTCTTTCCCTACACGACGCTCTTCCGATCTNNNAACCACAAGAGGACTGCAGTA
GAGCACACGTCTGAACTCTTTCCCTACACGACGCTCTTCCGATCTNNNCCCAGAGATGGAACCCTCTACA
GAGCACACGTCTGAACTCTTTCCCTACACGACGCTCTTCCGATCTNNNGTTGATGTCCTCCTGTGGCTAT
AGCACACGTCTGAACTCTTTCCCTACACGACGCTCTTCCGATCTNNNGCTGCATTGAGAACCAAGATG
AGAGCACACGTCTGAACTCTTTCCCTACACGACGCTCTTCCGATCTNNNATGGCCAGAAGCAATGACTA
AGAGCACACGTCTGAACTCTTTCCCTACACGACGCTCTTCCGATCTNNNGAAGTTGTGGATCTTCTCCC
GAGCACACGTCTGAACTCTTTCCCTACACGACGCTCTTCCGATCTNNNTGGGAAAACACAGTGCAGTTG
GAGCACACGTCTGAACTCTTTCCCTACACGACGCTCTTCCGATCTNNNTCAGAGCTTACTTTGGTGGTTG
AGCACACGTCTGAACTCTTTCCCTACACGACGCTCTTCCGATCTNNNCCAAGCAGCCTTCCCTAATGAAT

AGAGCACACGTCTGAACTCTTTCCCTACACGACGCTCTTCCGATCTNNNCCACACATGTGATAAACCTG
AGCACACGTCTGAACTCTTTCCCTACACGACGCTCTTCCGATCTNNNTTACAAATTGGGGGAGAAGGG
GAGCACACGTCTGAACTCTTTCCCTACACGACGCTCTTCCGATCTNNNCCTCTGGAGGAACTCTGACCT
AGCACACGTCTGAACTCTTTCCCTACACGACGCTCTTCCGATCTNNNTTTACCATCTTCTGAGCAGCCAC
AGAGCACACGTCTGAACTCTTTCCCTACACGACGCTCTTCCGATCTNNNGCATGCAGCAATTCATCTAG
GAGCACACGTCTGAACTCTTTCCCTACACGACGCTCTTCCGATCTNNNCGTCCAGCTGAAACGTGTGTA
AGCACACGTCTGAACTCTTTCCCTACACGACGCTCTTCCGATCTNNNTCATTGGTTCATCAGTGGGAA
GAGCACACGTCTGAACTCTTTCCCTACACGACGCTCTTCCGATCTNNNGTCATTCTGAAGGGGAGTAG
AGCACACGTCTGAACTCTTTCCCTACACGACGCTCTTCCGATCTNNNTCCACTTCTCAAGCTGGAACAA
GAGCACACGTCTGAACTCTTTCCCTACACGACGCTCTTCCGATCTNNNTAGGATGAAGGGGCATCATG
AGCACACGTCTGAACTCTTTCCCTACACGACGCTCTTCCGATCTNNNTAGAGTAGTGGTCTCCAACTCC
GAGCACACGTCTGAACTCTTTCCCTACACGACGCTCTTCCGATCTNNNTACCTTGAGGTATGACTGTGC
AGCACACGTCTGAACTCTTTCCCTACACGACGCTCTTCCGATCTNNNCCAGAGAAATTGGCCTCTTTGG
AGCACACGTCTGAACTCTTTCCCTACACGACGCTCTTCCGATCTNNNGTGGTTGGCTATCTGCCTTTCTT
AGCACACGTCTGAACTCTTTCCCTACACGACGCTCTTCCGATCTNNNAGTCTTTCCCTGTCTCTGTCTCTC
AGCACACGTCTGAACTCTTTCCCTACACGACGCTCTTCCGATCTNNNTCCATTTATCCAGGGAGCAAAG
AGCACACGTCTGAACTCTTTCCCTACACGACGCTCTTCCGATCTNNNGTTCCCTTTGCAATGGATGTGA
AGCACACGTCTGAACTCTTTCCCTACACGACGCTCTTCCGATCTNNNTTTGGGTGGTGTCACTTTTGG
GAGCACACGTCTGAACTCTTTCCCTACACGACGCTCTTCCGATCTNNNGGCAACTGTCATCATCAACTG
AGCACACGTCTGAACTCTTTCCCTACACGACGCTCTTCCGATCTNNNTCTGGAAAACTGGAATTTGGG
GCACACGTCTGAACTCTTTCCCTACACGACGCTCTTCCGATCTNNNCCTTCCTTACCATTGCTCCCAAGCA
AGCACACGTCTGAACTCTTTCCCTACACGACGCTCTTCCGATCTNNNTGTTGTTTTTCATCCTTGGCAG
AGCACACGTCTGAACTCTTTCCCTACACGACGCTCTTCCGATCTNNNGTGTGAGGCCCAACAAATGAA
GAGCACACGTCTGAACTCTTTCCCTACACGACGCTCTTCCGATCTNNNAGAAGTCTTGTGATGGTTGG
AGCACACGTCTGAACTCTTTCCCTACACGACGCTCTTCCGATCTNNNTAGAGGAACTGTCAGGCTTCCA
AGCACACGTCTGAACTCTTTCCCTACACGACGCTCTTCCGATCTNNNGAAAGAAAGCCGATCTTTGTGC
GAGCACACGTCTGAACTCTTTCCCTACACGACGCTCTTCCGATCTNNNCCTGTGGTGATGTCAGGCTTT
AGCACACGTCTGAACTCTTTCCCTACACGACGCTCTTCCGATCTNNNTGTGTTGACTGTTGAATGAGG
GAGCACACGTCTGAACTCTTTCCCTACACGACGCTCTTCCGATCTNNNAGCAGTTTCTACTGCACCCTC
AGCACACGTCTGAACTCTTTCCCTACACGACGCTCTTCCGATCTNNNTGAGAGAGATCTTCCACCGAAC
GAGCACACGTCTGAACTCTTTCCCTACACGACGCTCTTCCGATCTNNNAGGGGTTTTTAGACCAATGCA
GAGCACACGTCTGAACTCTTTCCCTACACGACGCTCTTCCGATCTNNNTACCTGAAGATAGTGCCAGTC
GAGCACACGTCTGAACTCTTTCCCTACACGACGCTCTTCCGATCTNNNTGCTCATCACTCAAACACAACC
GAGCACACGTCTGAACTCTTTCCCTACACGACGCTCTTCCGATCTNNNACCAGCACTTCCAAGAACACAA
AGCACACGTCTGAACTCTTTCCCTACACGACGCTCTTCCGATCTNNNCCCTTCCAATGCAGAAACTCAGA
GAGCACACGTCTGAACTCTTTCCCTACACGACGCTCTTCCGATCTNNNTCAGGGACATTGATGGGCTTG
AGCACACGTCTGAACTCTTTCCCTACACGACGCTCTTCCGATCTNNNAATTAGCCAGATGTTGTGACCC
GAGCACACGTCTGAACTCTTTCCCTACACGACGCTCTTCCGATCTNNNTTCTAGCTGGCAAACACACCAT
AGCACACGTCTGAACTCTTTCCCTACACGACGCTCTTCCGATCTNNNTCACTCTCCCAAATGCAGTAAAA
AGCACACGTCTGAACTCTTTCCCTACACGACGCTCTTCCGATCTNNNCTTTGGCCAACTCAGGGACTTAG
GAGCACACGTCTGAACTCTTTCCCTACACGACGCTCTTCCGATCTNNNTAGGAGAGTGAGCTGGAAG
AGCACACGTCTGAACTCTTTCCCTACACGACGCTCTTCCGATCTNNNCCGAGTGAAAACTTCCTGCCATA
AGCACACGTCTGAACTCTTTCCCTACACGACGCTCTTCCGATCTNNNGATACTTGTTCCAAAGCACACG

AGCACACGTCTGAACTCTTTCCCTACACGACGCTCTTCCGATCTNNNCTGGAACCAGATCAATACCAGG
AGCACACGTCTGAACTCTTTCCCTACACGACGCTCTTCCGATCTNNNTCTTGCTTCCTTGCTACACAACA
AGAGCACACGTCTGAACTCTTTCCCTACACGACGCTCTTCCGATCTNNNAGTTTGTTGTCTGGCTCTTCA
GAGCACACGTCTGAACTCTTTCCCTACACGACGCTCTTCCGATCTNNNAAATGCAGTCTGTGCAGAGTT
AGCACACGTCTGAACTCTTTCCCTACACGACGCTCTTCCGATCTNNNATATCACCTTCCAGAGTTACGTT
GCACACGTCTGAACTCTTTCCCTACACGACGCTCTTCCGATCTNNNCCAAGCACCAAGGATGGGTAAT
AGCACACGTCTGAACTCTTTCCCTACACGACGCTCTTCCGATCTNNNACAGGTATGCACAACATCTCTC
GCACACGTCTGAACTCTTTCCCTACACGACGCTCTTCCGATCTNNNATGCTTTCAGGTCTTCAAGCCAA
AGCACACGTCTGAACTCTTTCCCTACACGACGCTCTTCCGATCTNNNAAAAATTACTGCCTGGGGTGA
GCACACGTCTGAACTCTTTCCCTACACGACGCTCTTCCGATCTNNNGGTGCTTGCTGAAGATAAGGGA
AGCACACGTCTGAACTCTTTCCCTACACGACGCTCTTCCGATCTNNNCTAAAAATCTCCAGGCACCCTGC
GAGCACACGTCTGAACTCTTTCCCTACACGACGCTCTTCCGATCTNNNCAGCAACAGCTTCCTTCCTGTA
AGCACACGTCTGAACTCTTTCCCTACACGACGCTCTTCCGATCTNNNCTCCTCCTCGTTCACAAAGTCAG
AGCACACGTCTGAACTCTTTCCCTACACGACGCTCTTCCGATCTNNNCCTTGCTACTGGAACTCAGCTT
AGCACACGTCTGAACTCTTTCCCTACACGACGCTCTTCCGATCTNNNGAAAGGCCACTATATCACCAGG
AGCACACGTCTGAACTCTTTCCCTACACGACGCTCTTCCGATCTNNNATGTGAGTTTCATACCCAGGTG
AGCACACGTCTGAACTCTTTCCCTACACGACGCTCTTCCGATCTNNNACACACAACTTAGGCTTCCTTCC
AGCACACGTCTGAACTCTTTCCCTACACGACGCTCTTCCGATCTNNNGCACTCCCTTTTCAGATTCTCCA
GCACACGTCTGAACTCTTTCCCTACACGACGCTCTTCCGATCTNNNAGCTGAGGAGCTCACAGACTAA
GCACACGTCTGAACTCTTTCCCTACACGACGCTCTTCCGATCTNNNCTTTCCTCCATGGCATTCACAGACA
GCACACGTCTGAACTCTTTCCCTACACGACGCTCTTCCGATCTNNNCACCATTGATTCTACCCCCGATGCA
GCACACGTCTGAACTCTTTCCCTACACGACGCTCTTCCGATCTNNNTATCCCTACACTCCTACACCCACCA
AGCACACGTCTGAACTCTTTCCCTACACGACGCTCTTCCGATCTNNNCTAGACCCCAATCAAAAGCCTCC
GCACACGTCTGAACTCTTTCCCTACACGACGCTCTTCCGATCTNNNTGCTACTGCCATTACTGCATTGTC
AGCACACGTCTGAACTCTTTCCCTACACGACGCTCTTCCGATCTNNNCTAAGCTCTGGCAGGGAGAAAT
GAGCACACGTCTGAACTCTTTCCCTACACGACGCTCTTCCGATCTNNNCATCCCTTTTCCTTCCTCCTCC
AGCACACGTCTGAACTCTTTCCCTACACGACGCTCTTCCGATCTNNNGCAAAGCTCCTCTCTACTCTTCC
AGCACACGTCTGAACTCTTTCCCTACACGACGCTCTTCCGATCTNNNTACACTCAAACCACTCGGCAAA
GAGCACACGTCTGAACTCTTTCCCTACACGACGCTCTTCCGATCTNNNGAGAGGGAAGTGAATTGCAG
AGAGCACACGTCTGAACTCTTTCCCTACACGACGCTCTTCCGATCTNNNGGACTTCCACACTCGCCTATT
GAGCACACGTCTGAACTCTTTCCCTACACGACGCTCTTCCGATCTNNNGGAAACCAAAGGGGGTCAAA
GAGCACACGTCTGAACTCTTTCCCTACACGACGCTCTTCCGATCTNNNTGGGTTTGACCACTTTCTCCTT
GAGCACACGTCTGAACTCTTTCCCTACACGACGCTCTTCCGATCTNNNACCCAAATCCACGCATGTGATA
AGCACACGTCTGAACTCTTTCCCTACACGACGCTCTTCCGATCTNNNACTCTCCCTCTTTTTCCTTCCCCCA
GAGCACACGTCTGAACTCTTTCCCTACACGACGCTCTTCCGATCTNNNCCGCAACCTCCTTTATTACTGG
GAGCACACGTCTGAACTCTTTCCCTACACGACGCTCTTCCGATCTNNNAGGCCTGAAGTATTGCTGACC
AGCACACGTCTGAACTCTTTCCCTACACGACGCTCTTCCGATCTNNNCCTCCCTCCCTCAATCTTACCTTC
GAGCACACGTCTGAACTCTTTCCCTACACGACGCTCTTCCGATCTNNNAGCAAATTGCAGAGCCGTAT
AGCACACGTCTGAACTCTTTCCCTACACGACGCTCTTCCGATCTNNNTTTGTTTTGGTGTCTGTCTCCCC
AGCACACGTCTGAACTCTTTCCCTACACGACGCTCTTCCGATCTNNNCAGTCTCCCTTCCTTAGTCCCTTC
AGCACACGTCTGAACTCTTTCCCTACACGACGCTCTTCCGATCTNNNTATTCCCCTGCACAGAGATGTC
AGCACACGTCTGAACTCTTTCCCTACACGACGCTCTTCCGATCTNNNTGACAACACTCAAGCTGAGAAC
GCACACGTCTGAACTCTTTCCCTACACGACGCTCTTCCGATCTNNNTCAAAACCAGAAACGGAACTACA

AGCACACGTCTGAACTCTTTCCCTACACGACGCTCTTCCGATCTNNNTATGAACAGGGCCTACCAGAA
AGCACACGTCTGAACTCTTTCCCTACACGACGCTCTTCCGATCTNNNACGCTTCAAAGGGACAAAGAA
GAGCACACGTCTGAACTCTTTCCCTACACGACGCTCTTCCGATCTNNNTTTTACGTGGCAAAATGTAC
AGAGCACACGTCTGAACTCTTTCCCTACACGACGCTCTTCCGATCTNNNGAAGGCCATGCAGACAAA
AGCACACGTCTGAACTCTTTCCCTACACGACGCTCTTCCGATCTNNNCCAGAGCCCCTGGAATTTATG
GAGCACACGTCTGAACTCTTTCCCTACACGACGCTCTTCCGATCTNNNTACTAGGTCTGAGGTTCAGG
GAGCACACGTCTGAACTCTTTCCCTACACGACGCTCTTCCGATCTNNNGGACAACAGAAAGCACCCAC
AGCACACGTCTGAACTCTTTCCCTACACGACGCTCTTCCGATCTNNNGACTTGGCTGTTAGGATGGG
GCACACGTCTGAACTCTTTCCCTACACGACGCTCTTCCGATCTNNNATCCTGCGCTCACATTTCTTCTT
GCACACGTCTGAACTCTTTCCCTACACGACGCTCTTCCGATCTNNNTTAGTATCCACGGAAGGACCTG
GAGCACACGTCTGAACTCTTTCCCTACACGACGCTCTTCCGATCTNNNTGTACCAGTCTCTCTCTGCAA
AGCACACGTCTGAACTCTTTCCCTACACGACGCTCTTCCGATCTNNNAACCTGCCTCTGGGTTTTTGTT
AGCACACGTCTGAACTCTTTCCCTACACGACGCTCTTCCGATCTNNNGCCCTTTAGAGGCAAACACAT
GCACACGTCTGAACTCTTTCCCTACACGACGCTCTTCCGATCTNNNACGTTTTAGCAGATGATGTGGT
GCACACGTCTGAACTCTTTCCCTACACGACGCTCTTCCGATCTNNNGTGACAGGGCCTATATGTGCAT
GCACACGTCTGAACTCTTTCCCTACACGACGCTCTTCCGATCTNNNCGTTAGCATGGTGGAGTATGT
AGCACACGTCTGAACTCTTTCCCTACACGACGCTCTTCCGATCTNNNGCTGAGTGGATCAGCTCAAAG
AGCACACGTCTGAACTCTTTCCCTACACGACGCTCTTCCGATCTNNNCTGAGACTGACCCATGTCATGT
AGCACACGTCTGAACTCTTTCCCTACACGACGCTCTTCCGATCTNNNATCAAATCCAGTAGAGCCAAG
GCACACGTCTGAACTCTTTCCCTACACGACGCTCTTCCGATCTNNNCTGCTCAGTTAGCTTAGTCGTC
AGCACACGTCTGAACTCTTTCCCTACACGACGCTCTTCCGATCTNNNGAAATCAGAAAGGGCTGGTA
GCACACGTCTGAACTCTTTCCCTACACGACGCTCTTCCGATCTNNNTTCAAAAGACCCCAGACCCAAT
GCACACGTCTGAACTCTTTCCCTACACGACGCTCTTCCGATCTNNNAGGCTGGGATGGATACTACTG
GCACACGTCTGAACTCTTTCCCTACACGACGCTCTTCCGATCTNNNATTGCTACTCTTGTGGGAGAGG
AGCACACGTCTGAACTCTTTCCCTACACGACGCTCTTCCGATCTNNNTGCCTGGCAGTAATGATTGGA
GCACACGTCTGAACTCTTTCCCTACACGACGCTCTTCCGATCTNNNAAAATAGACACACATATGCACG
AGCACACGTCTGAACTCTTTCCCTACACGACGCTCTTCCGATCTNNNCTGTTCTGTGAGGTTGTTCAG
GAGCACACGTCTGAACTCTTTCCCTACACGACGCTCTTCCGATCTNNNTGCTGATACGCTGATGGAAT
GCACACGTCTGAACTCTTTCCCTACACGACGCTCTTCCGATCTNNNCACTGCCACCACAATTTCATCTG
AGCACACGTCTGAACTCTTTCCCTACACGACGCTCTTCCGATCTNNNACTCCCCACTCTCATTTACTCCC
AGCACACGTCTGAACTCTTTCCCTACACGACGCTCTTCCGATCTNNNTTTGGATGCCTTTTCTGAGGT
GCACACGTCTGAACTCTTTCCCTACACGACGCTCTTCCGATCTNNNCCACTACCAGTACCTGTCTCCTCC
AGCACACGTCTGAACTCTTTCCCTACACGACGCTCTTCCGATCTNNNTTACCAGGTACGGTGATAAG
AGCACACGTCTGAACTCTTTCCCTACACGACGCTCTTCCGATCTNNNCCCCAGAATCCAGGACTTCCTA
AGCACACGTCTGAACTCTTTCCCTACACGACGCTCTTCCGATCTNNNGCATTTATGTGACCAGCCAAA
GCACACGTCTGAACTCTTTCCCTACACGACGCTCTTCCGATCTNNNGGTGGGCTAAGTTTCTGTCTCT
CACACGTCTGAACTCTTTCCCTACACGACGCTCTTCCGATCTNNNAAAGTCTTCCCCTGAATGCCTTGC
AGCACACGTCTGAACTCTTTCCCTACACGACGCTCTTCCGATCTNNNGTGGTCCTCAAATCCCTGCATA
AGCACACGTCTGAACTCTTTCCCTACACGACGCTCTTCCGATCTNNNTTCCCATCTCTGTGTTCCCATCT
GCACACGTCTGAACTCTTTCCCTACACGACGCTCTTCCGATCTNNNGAAGAGCAGATGGTTGGCTAC
GCACACGTCTGAACTCTTTCCCTACACGACGCTCTTCCGATCTNNNCTTGGTCATACCTGCAATCTGCT
GCACACGTCTGAACTCTTTCCCTACACGACGCTCTTCCGATCTNNNGCAGTTTCGTCAGAGGTAAAGG
GCACACGTCTGAACTCTTTCCCTACACGACGCTCTTCCGATCTNNNGTTTCCCAGGTAACCTTCAGCTA

GCACACGTCTGAACTCTTTCCCTACACGACGCTCTTCCGATCTNNNCTTTGGCTCTGTTTCTTTCTGCGCA
AGCACACGTCTGAACTCTTTCCCTACACGACGCTCTTCCGATCTNNNGCCTAGCACATTGGAGAATGACA
GAGCACACGTCTGAACTCTTTCCCTACACGACGCTCTTCCGATCTNNNGGATATGCACATGCACACAA
GCACACGTCTGAACTCTTTCCCTACACGACGCTCTTCCGATCTNNNTGGGTTCTTACCTCTCCTCTGTGC
AGCACACGTCTGAACTCTTTCCCTACACGACGCTCTTCCGATCTNNNGCTGTTTCTAGCAGGCACCTATT
AGCACACGTCTGAACTCTTTCCCTACACGACGCTCTTCCGATCTNNNATGAAATTGCAGGCAGTGTTT
GAGCACACGTCTGAACTCTTTCCCTACACGACGCTCTTCCGATCTNNNGATCCTTGATTTGGTCCCTGT
AGCACACGTCTGAACTCTTTCCCTACACGACGCTCTTCCGATCTNNNGGCAGCACAGACACTTAGGATT
AGAGCACACGTCTGAACTCTTTCCCTACACGACGCTCTTCCGATCTNNNAAGGAAGAACTCCATCTTCC
AGCACACGTCTGAACTCTTTCCCTACACGACGCTCTTCCGATCTNNNGCATGAGAGTTTGCTCTCTGTG
AGAGCACACGTCTGAACTCTTTCCCTACACGACGCTCTTCCGATCTNNNTTGAGGGATGCCAGGATAA
AGCACACGTCTGAACTCTTTCCCTACACGACGCTCTTCCGATCTNNNAAAATGAGTATACGCGCACCTG
AGCACACGTCTGAACTCTTTCCCTACACGACGCTCTTCCGATCTNNNAGAGCCAATCTCACATGCTTTGT
GCACACGTCTGAACTCTTTCCCTACACGACGCTCTTCCGATCTNNNCCTGGGAATAACTCCCTCCTCTTC
GAGCACACGTCTGAACTCTTTCCCTACACGACGCTCTTCCGATCTNNNAGTGGGACCTCATGTGTACCAA
GCACACGTCTGAACTCTTTCCCTACACGACGCTCTTCCGATCTNNNTCCAAATCTCTACTAAGCCGGGG
GCACACGTCTGAACTCTTTCCCTACACGACGCTCTTCCGATCTNNNTGGAACCATGGTCTTTTCCCTTGC
GAGCACACGTCTGAACTCTTTCCCTACACGACGCTCTTCCGATCTNNNCATCTGGGAAACCAAGCTGT
GAGCACACGTCTGAACTCTTTCCCTACACGACGCTCTTCCGATCTNNNGGGGCATGGAAAGGAATAGA
GCACACGTCTGAACTCTTTCCCTACACGACGCTCTTCCGATCTNNNCTGACCCAGGAGATTTACCAAGC
AGCACACGTCTGAACTCTTTCCCTACACGACGCTCTTCCGATCTNNNTCCTAGAAGGCAATGCACCATT
AGCACACGTCTGAACTCTTTCCCTACACGACGCTCTTCCGATCTNNNCTTTTCCTCCAGGGCTTTGTGAT
AGCACACGTCTGAACTCTTTCCCTACACGACGCTCTTCCGATCTNNNCCCAACCAGGAAACTAGAGTGA
AGCACACGTCTGAACTCTTTCCCTACACGACGCTCTTCCGATCTNNNCAATGTCCCGGAAACAAAGTCCT
AGCACACGTCTGAACTCTTTCCCTACACGACGCTCTTCCGATCTNNNTGATACAATATGGCTTGTGCCC
AGAGCACACGTCTGAACTCTTTCCCTACACGACGCTCTTCCGATCTNNNTTTCTCTGATCAACAGCCACA
AGCACACGTCTGAACTCTTTCCCTACACGACGCTCTTCCGATCTNNNATAGACCTACTGAGCCTCACCT
AGCACACGTCTGAACTCTTTCCCTACACGACGCTCTTCCGATCTNNNTGCTGTAAGTGCCTCTCTAAGT
AGCACACGTCTGAACTCTTTCCCTACACGACGCTCTTCCGATCTNNNTGGTATGACAGAGGGAAACAG
GAGCACACGTCTGAACTCTTTCCCTACACGACGCTCTTCCGATCTNNNCTCATGGAGCAGATTCCAGGA
AGCACACGTCTGAACTCTTTCCCTACACGACGCTCTTCCGATCTNNNCCTCAGTACCTTTACCCATTGCCC
AGCACACGTCTGAACTCTTTCCCTACACGACGCTCTTCCGATCTNNNGCAAAGAAAAGAGAAGGGCAA
GAGCACACGTCTGAACTCTTTCCCTACACGACGCTCTTCCGATCTNNNTGTCTCAAAGAGCCTTTCTCTG
GAGCACACGTCTGAACTCTTTCCCTACACGACGCTCTTCCGATCTNNNAACCGATGGTTGTTTCCTGA
AGCACACGTCTGAACTCTTTCCCTACACGACGCTCTTCCGATCTNNNGCACACACACACACCTTGAGATA
AGCACACGTCTGAACTCTTTCCCTACACGACGCTCTTCCGATCTNNNTGGCTGGGGCAATTAGAAAGA
GAGCACACGTCTGAACTCTTTCCCTACACGACGCTCTTCCGATCTNNNCCCAGGAAGTATGTTTGGTA
AGCACACGTCTGAACTCTTTCCCTACACGACGCTCTTCCGATCTNNNCAGCTCTCAGGCATTCGAAAAA
AGCACACGTCTGAACTCTTTCCCTACACGACGCTCTTCCGATCTNNNTATTTCTGGTGACAGCCTTTTG
GAGCACACGTCTGAACTCTTTCCCTACACGACGCTCTTCCGATCTNNNAATTTTTGCTCCCTCCCCATCT
AGAGCACACGTCTGAACTCTTTCCCTACACGACGCTCTTCCGATCTNNNCCACAGAAGCCAACAATCAA
AGCACACGTCTGAACTCTTTCCCTACACGACGCTCTTCCGATCTNNNTTTGCGCATGTGTCCACTTTTA
GAGCACACGTCTGAACTCTTTCCCTACACGACGCTCTTCCGATCTNNNTACATTCTGCTTTCCTGGTGGT

AGAGCACACGTCTGAACTCTTTCCCTACACGACGCTCTTCCGATCTNNNACCTTCCAAGGACATTCAACT
AGCACACGTCTGAACTCTTTCCCTACACGACGCTCTTCCGATCTNNNCCAACCCCTGTATCACACAAACA
GAGCACACGTCTGAACTCTTTCCCTACACGACGCTCTTCCGATCTNNNACCCCAGAACCTAAAGGACTG
GAGCACACGTCTGAACTCTTTCCCTACACGACGCTCTTCCGATCTNNNACACATGGCCTTCTGTTAGTGT
GAGCACACGTCTGAACTCTTTCCCTACACGACGCTCTTCCGATCTNNNGGTAATTGGCCCAGTGATTTT
GAGCACACGTCTGAACTCTTTCCCTACACGACGCTCTTCCGATCTNNNAAGTCTCTTCTCCCAGGCTGAT
AGCACACGTCTGAACTCTTTCCCTACACGACGCTCTTCCGATCTNNNAGTCAGATGTGTTTTTGTGCAG
AGCACACGTCTGAACTCTTTCCCTACACGACGCTCTTCCGATCTNNNTTTACTCCTTAGACACCACCCCC
AGCACACGTCTGAACTCTTTCCCTACACGACGCTCTTCCGATCTNNNATGAGCTAGGGGTTCATGAGA
GAGCACACGTCTGAACTCTTTCCCTACACGACGCTCTTCCGATCTNNNGTGCGAGGTATGGATAGAAG
AGAGCACACGTCTGAACTCTTTCCCTACACGACGCTCTTCCGATCTNNNTTTTGGCATGGTCTGCCTTT
GAGCACACGTCTGAACTCTTTCCCTACACGACGCTCTTCCGATCTNNNTACCAGAAGTTTAGGCCTAGG
GCACACGTCTGAACTCTTTCCCTACACGACGCTCTTCCGATCTNNNTGACAAGTCATCCAAATGCACCAC
AGCACACGTCTGAACTCTTTCCCTACACGACGCTCTTCCGATCTNNNTTTTTCCTCAGGTGAGGCTGGAC
GAGCACACGTCTGAACTCTTTCCCTACACGACGCTCTTCCGATCTNNNCAGCCCGGCTCTAACAATAAGT
AGCACACGTCTGAACTCTTTCCCTACACGACGCTCTTCCGATCTNNNTGCTTTGTGTCAGATATGGCTC
GAGCACACGTCTGAACTCTTTCCCTACACGACGCTCTTCCGATCTNNNACAGTAGTTCCCAGTTCCATCC
GAGCACACGTCTGAACTCTTTCCCTACACGACGCTCTTCCGATCTNNNTCTATTGGAGAAGCCTGCCTA
AGCACACGTCTGAACTCTTTCCCTACACGACGCTCTTCCGATCTNNNTATAGACGCTTGAGGCCAGGAT
GAGCACACGTCTGAACTCTTTCCCTACACGACGCTCTTCCGATCTNNNGGCCTCAGAATAGATCTCCCTC
AGCACACGTCTGAACTCTTTCCCTACACGACGCTCTTCCGATCTNNNCAGCCTTCTGCTTTCAGTCTACA
AGCACACGTCTGAACTCTTTCCCTACACGACGCTCTTCCGATCTNNNTTCTGCTGTTCAAGTTTGGCTCT
AGAGCACACGTCTGAACTCTTTCCCTACACGACGCTCTTCCGATCTNNNGGGGTAAAGGGCCATGATTT
GCACACGTCTGAACTCTTTCCCTACACGACGCTCTTCCGATCTNNNATGGCGATGAAGTGAAGACAGA
GAGCACACGTCTGAACTCTTTCCCTACACGACGCTCTTCCGATCTNNNACCCACAGAAACTGAAAGACA
AGCACACGTCTGAACTCTTTCCCTACACGACGCTCTTCCGATCTNNNTGTAGGCAGTCTCTCTCTCTCAG
AGCACACGTCTGAACTCTTTCCCTACACGACGCTCTTCCGATCTNNNATTCTCTCCCTGTTGAATGCTCC
GAGCACACGTCTGAACTCTTTCCCTACACGACGCTCTTCCGATCTNNNGTGACATTGACAGAAGCCACT
AGCACACGTCTGAACTCTTTCCCTACACGACGCTCTTCCGATCTNNNCTCAGATTGTTCAGCGGTTCCAT
GAGCACACGTCTGAACTCTTTCCCTACACGACGCTCTTCCGATCTNNNGCCTGGGGTCCTAGAGATTTA
AGAGCACACGTCTGAACTCTTTCCCTACACGACGCTCTTCCGATCTNNNAGCAGAAAACCTATCCCCAAC
GAGCACACGTCTGAACTCTTTCCCTACACGACGCTCTTCCGATCTNNNTAGACTTACCCCAGGAAAGGC
AAGAGCACACGTCTGAACTCTTTCCCTACACGACGCTCTTCCGATCTNNNATGAGGAACCTGCAGATAC
GAGCACACGTCTGAACTCTTTCCCTACACGACGCTCTTCCGATCTNNNAGAAATTACTCCAGGAGCACC
AGCACACGTCTGAACTCTTTCCCTACACGACGCTCTTCCGATCTNNNCTGTCTTTCTCTGGGGCTGATTC
GAGCACACGTCTGAACTCTTTCCCTACACGACGCTCTTCCGATCTNNNTTCATTTTGCTAGAGGCTCAGC
GAGCACACGTCTGAACTCTTTCCCTACACGACGCTCTTCCGATCTNNNGCAGTGCTTGGCAATTGAACTA
GAGCACACGTCTGAACTCTTTCCCTACACGACGCTCTTCCGATCTNNNCTGAATGGAAGCTATCCTGGC
AGCACACGTCTGAACTCTTTCCCTACACGACGCTCTTCCGATCTNNNAGGCAATAGACCAGAGCCCATT
AGCACACGTCTGAACTCTTTCCCTACACGACGCTCTTCCGATCTNNNCTTACAGAGGGACTGGTGTGTT
GAGCACACGTCTGAACTCTTTCCCTACACGACGCTCTTCCGATCTNNNAGAGTGATGGAATGTAGCCCA
AGCACACGTCTGAACTCTTTCCCTACACGACGCTCTTCCGATCTNNNTCATTCATGTTCCCGAGCAATCC
AGCACACGTCTGAACTCTTTCCCTACACGACGCTCTTCCGATCTNNNATAAATCACAGCCAGCACTGTG

AGCACACGTCTGAACTCTTTCCCTACACGACGCTCTTCCGATCTNNNGTCTCTTGGAAGGGGTGTAGG
AGAGCACACGTCTGAACTCTTTCCCTACACGACGCTCTTCCGATCTNNNGCTAAACCAGGGTTCTTTG
GAGCACACGTCTGAACTCTTTCCCTACACGACGCTCTTCCGATCTNNNAGTGCTCTGGCTCTGAAGACT
AGCACACGTCTGAACTCTTTCCCTACACGACGCTCTTCCGATCTNNNAGAGTGTGGGAGAGGAAGGA
AGCACACGTCTGAACTCTTTCCCTACACGACGCTCTTCCGATCTNNNTGCATGTTCACCTCCTTTACACA
GAGCACACGTCTGAACTCTTTCCCTACACGACGCTCTTCCGATCTNNNATGTTGCTGGCATGTAAACT
GAGCACACGTCTGAACTCTTTCCCTACACGACGCTCTTCCGATCTNNNAGAGCAGACAGGTTGCTTCTT
AGCACACGTCTGAACTCTTTCCCTACACGACGCTCTTCCGATCTNNNTGCCCTGCTGTCACTCAATAAAA
GAGCACACGTCTGAACTCTTTCCCTACACGACGCTCTTCCGATCTNNNAAGGTGGTGTGTTAGGGTTC
AGCACACGTCTGAACTCTTTCCCTACACGACGCTCTTCCGATCTNNNGGAGAAGAGAGCTGTTGTGTG
GAGCACACGTCTGAACTCTTTCCCTACACGACGCTCTTCCGATCTNNNATGGCAAACCCAGAAATCAG
AGCACACGTCTGAACTCTTTCCCTACACGACGCTCTTCCGATCTNNNCTACTGCACTTCAGCAAGACACT
AGAGCACACGTCTGAACTCTTTCCCTACACGACGCTCTTCCGATCTNNNTTTGCTGCAAGACAAGGTTT
AGCACACGTCTGAACTCTTTCCCTACACGACGCTCTTCCGATCTNNNGGGGTCCACATGCCAATAATTT
AGCACACGTCTGAACTCTTTCCCTACACGACGCTCTTCCGATCTNNNCATTCACATCCTTGTCCCCAGAG
AGCACACGTCTGAACTCTTTCCCTACACGACGCTCTTCCGATCTNNNGTTTGGGGAGGCTAAAGCAGT
AGCACACGTCTGAACTCTTTCCCTACACGACGCTCTTCCGATCTNNNCTTCCCTCATTTACAGACCCTGC
AGCACACGTCTGAACTCTTTCCCTACACGACGCTCTTCCGATCTNNNGAATGGTTCTGCTCTGTTTGCT
AGCACACGTCTGAACTCTTTCCCTACACGACGCTCTTCCGATCTNNNCCTTGAACTTCTGAGGAGGTGA
AGCACACGTCTGAACTCTTTCCCTACACGACGCTCTTCCGATCTNNNCGTAATTTCCCCATGCCCATGTA
AGCACACGTCTGAACTCTTTCCCTACACGACGCTCTTCCGATCTNNNAAAAACAGTCAGTGGCTCATTG
GAGCACACGTCTGAACTCTTTCCCTACACGACGCTCTTCCGATCTNNNCACCTCTCTGCCCTCATGTTTTT
GAGCACACGTCTGAACTCTTTCCCTACACGACGCTCTTCCGATCTNNNTCGCTATCAAAATCCCAGGAG
AGCACACGTCTGAACTCTTTCCCTACACGACGCTCTTCCGATCTNNNAGACTGCTGGATCAGGTGAAAT
GAGCACACGTCTGAACTCTTTCCCTACACGACGCTCTTCCGATCTNNNTGGGTGTCCATGGTAGGGAA
GAGCACACGTCTGAACTCTTTCCCTACACGACGCTCTTCCGATCTNNNGGGATGTTCCTCCCTCTGTATG
GAGCACACGTCTGAACTCTTTCCCTACACGACGCTCTTCCGATCTNNNATCAGTCATTGCTTGGCTAGT
AGCACACGTCTGAACTCTTTCCCTACACGACGCTCTTCCGATCTNNNTCAATGAACCTGAGGTAGGGTA
GAGCACACGTCTGAACTCTTTCCCTACACGACGCTCTTCCGATCTNNNTGGGGTGTGTGTGTGTAGAG
GAGCACACGTCTGAACTCTTTCCCTACACGACGCTCTTCCGATCTNNNACTGAAAAACTACAAGCCCAC
AGAGCACACGTCTGAACTCTTTCCCTACACGACGCTCTTCCGATCTNNNCTCAGGAAGTCTCCCTCCTTCT
AGAGCACACGTCTGAACTCTTTCCCTACACGACGCTCTTCCGATCTNNNAGCTTCCTCAGATTGAGGCT
AGAGCACACGTCTGAACTCTTTCCCTACACGACGCTCTTCCGATCTNNNTGGCAGCCTCTTAGGAAAAT
GAGCACACGTCTGAACTCTTTCCCTACACGACGCTCTTCCGATCTNNNGAGGGCTTTGCTCTATACATG
AGCACACGTCTGAACTCTTTCCCTACACGACGCTCTTCCGATCTNNNCAACTTCTCCACTCAACCCTCAG
AGCACACGTCTGAACTCTTTCCCTACACGACGCTCTTCCGATCTNNNATTTATCTTTCCCAGAGCCCAGC
GCACACGTCTGAACTCTTTCCCTACACGACGCTCTTCCGATCTNNNACCAACTTGTCTTTGTCATCGGTCA
AGCACACGTCTGAACTCTTTCCCTACACGACGCTCTTCCGATCTNNNGCATGGGGTCTGTGAAACTGTA
AGCACACGTCTGAACTCTTTCCCTACACGACGCTCTTCCGATCTNNNTGTGCCTCCTCCTTCTCATTTTCCA
GAGCACACGTCTGAACTCTTTCCCTACACGACGCTCTTCCGATCTNNNAACTTGCCACCTCTCACAAACA
GAGCACACGTCTGAACTCTTTCCCTACACGACGCTCTTCCGATCTNNNTTCCAGGGAACAGACACTAGA
AGCACACGTCTGAACTCTTTCCCTACACGACGCTCTTCCGATCTNNNGAGTGTGTGCGCATGTTCTTTTA
AGCACACGTCTGAACTCTTTCCCTACACGACGCTCTTCCGATCTNNNATGGGAAAGACTGGTGTCTGG

GAGCACACGTCTGAACTCTTTCCCTACACGACGCTCTTCCGATCTNNNAAGGGCAAGGACTGATATAC
AGCACACGTCTGAACTCTTTCCCTACACGACGCTCTTCCGATCTNNNTTGTGGAAAGGAGCGTATCTG
AGCACACGTCTGAACTCTTTCCCTACACGACGCTCTTCCGATCTNNNATTGCTCTTTAGCCACACTGCTT
GAGCACACGTCTGAACTCTTTCCCTACACGACGCTCTTCCGATCTNNNTCGGTTTTATAGGCCAACAAG
GAGCACACGTCTGAACTCTTTCCCTACACGACGCTCTTCCGATCTNNNCGTGTGCATGTTTCAAATGTTC
GAGCACACGTCTGAACTCTTTCCCTACACGACGCTCTTCCGATCTNNNATATGCTCATCTGGCCTTTCCC
GAGCACACGTCTGAACTCTTTCCCTACACGACGCTCTTCCGATCTNNNGATCACCAGTGGCCAATAACG
AGCACACGTCTGAACTCTTTCCCTACACGACGCTCTTCCGATCTNNNATTTTAGGAAGAGCTGGCTGC
GAGCACACGTCTGAACTCTTTCCCTACACGACGCTCTTCCGATCTNNNATATGCTAGTGTGTCCTCCCCA
GAGCACACGTCTGAACTCTTTCCCTACACGACGCTCTTCCGATCTNNNTCATGAGAAGAGTGTTGCAGA
AGCACACGTCTGAACTCTTTCCCTACACGACGCTCTTCCGATCTNNNTGTAAAGCATTTGGAGACCAGC
GAGCACACGTCTGAACTCTTTCCCTACACGACGCTCTTCCGATCTNNNAGGTTGATGCTACCTGGGGTA
GAGCACACGTCTGAACTCTTTCCCTACACGACGCTCTTCCGATCTNNNGCAGAGGTGGGGTAGAATTC
GCACACGTCTGAACTCTTTCCCTACACGACGCTCTTCCGATCTNNNTCTGCCTACTTTCTCCAGCTTCTCA
AGCACACGTCTGAACTCTTTCCCTACACGACGCTCTTCCGATCTNNNCTTGTCTCATGGGGCAGACATT
AGCACACGTCTGAACTCTTTCCCTACACGACGCTCTTCCGATCTNNNAGCTGAACCCTTTGGTTCTCAAG
GAGCACACGTCTGAACTCTTTCCCTACACGACGCTCTTCCGATCTNNNCAAATGGTCCACTTTTTCCCGT
AGAGCACACGTCTGAACTCTTTCCCTACACGACGCTCTTCCGATCTNNNGGATTGCAGTCAGCTTGTTT
AGCACACGTCTGAACTCTTTCCCTACACGACGCTCTTCCGATCTNNNCAGAACACAGGAAGGGTTTTG
GCACACGTCTGAACTCTTTCCCTACACGACGCTCTTCCGATCTNNNGCCCGAGAGCTGAAGTTTCATTA
GAGCACACGTCTGAACTCTTTCCCTACACGACGCTCTTCCGATCTNNNTCTCAGTGAAGTGCCCGATTTT
GCACACGTCTGAACTCTTTCCCTACACGACGCTCTTCCGATCTNNNCTATCCACAGGACACAAACCAGG
GAGCACACGTCTGAACTCTTTCCCTACACGACGCTCTTCCGATCTNNNCTGGATCTACATACAGCCTTCCC
AGCACACGTCTGAACTCTTTCCCTACACGACGCTCTTCCGATCTNNNCTGCCCTGAGGCTTTTCACTAGC
AGCACACGTCTGAACTCTTTCCCTACACGACGCTCTTCCGATCTNNNTCCAGCATTGGTCATCAAGAACT
GAGCACACGTCTGAACTCTTTCCCTACACGACGCTCTTCCGATCTNNNCTACCATGCCTAAGCTGCTTTCA
AGCACACGTCTGAACTCTTTCCCTACACGACGCTCTTCCGATCTNNNCTGGGCAACATAGAGACAAGG
AGAGCACACGTCTGAACTCTTTCCCTACACGACGCTCTTCCGATCTNNNAACATGACATAGGAGACACA
GAGCACACGTCTGAACTCTTTCCCTACACGACGCTCTTCCGATCTNNNCACAGGCTCAGAGAGCAATGA
GAGCACACGTCTGAACTCTTTCCCTACACGACGCTCTTCCGATCTNNNACTCACAGGTCTTACTGGTCCA
GAGCACACGTCTGAACTCTTTCCCTACACGACGCTCTTCCGATCTNNNTAATAATGGGAGAGAGCCTG
AGAGCACACGTCTGAACTCTTTCCCTACACGACGCTCTTCCGATCTNNNCCCAGAATACCCCAAGATGG
GAGCACACGTCTGAACTCTTTCCCTACACGACGCTCTTCCGATCTNNNTAGGCTCAGAAAGACTTCCCA
AGCACACGTCTGAACTCTTTCCCTACACGACGCTCTTCCGATCTNNNCCTGTAGTAAGAACCTAGCGCCT
GAGCACACGTCTGAACTCTTTCCCTACACGACGCTCTTCCGATCTNNNTGGCTTCCTCCTGGTTATGTAC
AGCACACGTCTGAACTCTTTCCCTACACGACGCTCTTCCGATCTNNNTGATTGGGCCAGATTTTGTGTG
GAGCACACGTCTGAACTCTTTCCCTACACGACGCTCTTCCGATCTNNNAAGTGTAATGCACAGACACCT
AGCACACGTCTGAACTCTTTCCCTACACGACGCTCTTCCGATCTNNNGCCAGAGCCCCATCTAAAAATC
GAGCACACGTCTGAACTCTTTCCCTACACGACGCTCTTCCGATCTNNNTTGAGACAGCCTTTAGCACAGT
AGCACACGTCTGAACTCTTTCCCTACACGACGCTCTTCCGATCTNNNAGCTCTTGCTGGAATGGATAGG
AGCACACGTCTGAACTCTTTCCCTACACGACGCTCTTCCGATCTNNNTCTCATGCCTGGCCAGATAAAA
AGCACACGTCTGAACTCTTTCCCTACACGACGCTCTTCCGATCTNNNCTTGTGCCTTCTGGGAAAGAAC
AGCACACGTCTGAACTCTTTCCCTACACGACGCTCTTCCGATCTNNNTCATCTTGTTGTCTAACGCCTGC

AGCACACGTCTGAACTCTTTCCCTACACGACGCTCTTCCGATCTNNNGCTCTTCTACGTTTCACTGTCGTC
AGAGCACACGTCTGAACTCTTTCCCTACACGACGCTCTTCCGATCTNNNCTGTGCTCCTGGAAGTTGTTT
GAGCACACGTCTGAACTCTTTCCCTACACGACGCTCTTCCGATCTNNNGATTACAGACAACCTTGGTGCC
GCACACGTCTGAACTCTTTCCCTACACGACGCTCTTCCGATCTNNNCAAGTCTTGACCTACAGCTGACCC
GCACACGTCTGAACTCTTTCCCTACACGACGCTCTTCCGATCTNNNGGCAGGGTGATCGTGTCATATT
AGCACACGTCTGAACTCTTTCCCTACACGACGCTCTTCCGATCTNNNTGCACTAGGATGTCACTGCTTTG
AGAGCACACGTCTGAACTCTTTCCCTACACGACGCTCTTCCGATCTNNNATGCCCAAAGTACCATACCAC
AGCACACGTCTGAACTCTTTCCCTACACGACGCTCTTCCGATCTNNNAAATCCTCTTGGTGCAAGCTGAA
AGAGCACACGTCTGAACTCTTTCCCTACACGACGCTCTTCCGATCTNNNTGCAGTTTCAGTGGGACATA
GAGCACACGTCTGAACTCTTTCCCTACACGACGCTCTTCCGATCTNNNAGCCAGTTATCTTGGGTCTTGC
GAGCACACGTCTGAACTCTTTCCCTACACGACGCTCTTCCGATCTNNNAACAGCAGCTCATTCCTTGTCT
GAGCACACGTCTGAACTCTTTCCCTACACGACGCTCTTCCGATCTNNNGGGATTCCAACAGGAGCAATC
GAGCACACGTCTGAACTCTTTCCCTACACGACGCTCTTCCGATCTNNNCTGAGTTTCTGGATTGCAGGG
AGAGCACACGTCTGAACTCTTTCCCTACACGACGCTCTTCCGATCTNNNTTACAGTTCCGGCCTCTCAGT
AGCACACGTCTGAACTCTTTCCCTACACGACGCTCTTCCGATCTNNNGCTGTGCTTCTCTCAGTCTTCAAC
AGCACACGTCTGAACTCTTTCCCTACACGACGCTCTTCCGATCTNNNGTCTTAGAGGGGCATGATGGT
AGCACACGTCTGAACTCTTTCCCTACACGACGCTCTTCCGATCTNNNAACATCGTGCCATTTTCCAATGC
GAGCACACGTCTGAACTCTTTCCCTACACGACGCTCTTCCGATCTNNNGCAGTGATACTTTGGACGTCA
AGCACACGTCTGAACTCTTTCCCTACACGACGCTCTTCCGATCTNNNCCTAGTTTGGGAAGCAGAGACA
AGCACACGTCTGAACTCTTTCCCTACACGACGCTCTTCCGATCTNNNCCATTCTGGAAAGCAATTGGCAC
AGCACACGTCTGAACTCTTTCCCTACACGACGCTCTTCCGATCTNNNTCAGTTCTTTGTGAAGCCACGTT
AGCACACGTCTGAACTCTTTCCCTACACGACGCTCTTCCGATCTNNNTTGCATAGCACATGTCCTCTCTG
GAGCACACGTCTGAACTCTTTCCCTACACGACGCTCTTCCGATCTNNNAACCTTTGAGTGCCATAGCAA
AGCACACGTCTGAACTCTTTCCCTACACGACGCTCTTCCGATCTNNNGGCTTTTCCTCTCCATACCTCCTC
AGCACACGTCTGAACTCTTTCCCTACACGACGCTCTTCCGATCTNNNGTATACCTGGTGAAAGCAGCAC
AGAGCACACGTCTGAACTCTTTCCCTACACGACGCTCTTCCGATCTNNNCTAGATCCTGTCTTGGAGCCT
AGCACACGTCTGAACTCTTTCCCTACACGACGCTCTTCCGATCTNNNATTGCCAGTCAGTGTCTGTCTT
AGCACACGTCTGAACTCTTTCCCTACACGACGCTCTTCCGATCTNNNTGAGGGACCACAGGGGAATTTT
AGCACACGTCTGAACTCTTTCCCTACACGACGCTCTTCCGATCTNNNCCTGTTAGTTTCAGAGGCTGTC
GAGCACACGTCTGAACTCTTTCCCTACACGACGCTCTTCCGATCTNNNCCCAAAAACAGTCATGCAAGTC
AGCACACGTCTGAACTCTTTCCCTACACGACGCTCTTCCGATCTNNNAGTGTCTTGTACCCAGCCTATG
AGCACACGTCTGAACTCTTTCCCTACACGACGCTCTTCCGATCTNNNCAAAACAGGAAATGGACAGTGG
GAGCACACGTCTGAACTCTTTCCCTACACGACGCTCTTCCGATCTNNNGATCTGCCGGGAGAGGAGAA
AGCACACGTCTGAACTCTTTCCCTACACGACGCTCTTCCGATCTNNNCTACCACATACAAACGCCACTGT
AGCACACGTCTGAACTCTTTCCCTACACGACGCTCTTCCGATCTNNNGGTAGGTTGCAGAGCATAGGA
AGCACACGTCTGAACTCTTTCCCTACACGACGCTCTTCCGATCTNNNACGGAAGCAAGGAAGGAATGT
GAGCACACGTCTGAACTCTTTCCCTACACGACGCTCTTCCGATCTNNNAGAGCCAACCCCTCACTTCTTA
AGAGCACACGTCTGAACTCTTTCCCTACACGACGCTCTTCCGATCTNNNACAGAGCAGCAGCAATTACA
AGCACACGTCTGAACTCTTTCCCTACACGACGCTCTTCCGATCTNNNATCGAAGAGTTCAACGTAGCCT
AGAGCACACGTCTGAACTCTTTCCCTACACGACGCTCTTCCGATCTNNNCTTTGACAAATGTAGGGGCC
GAGCACACGTCTGAACTCTTTCCCTACACGACGCTCTTCCGATCTNNNAATACAGGTCTGAAAGTCGTG
AGCACACGTCTGAACTCTTTCCCTACACGACGCTCTTCCGATCTNNNAGTCCTATGTCAACATTCAGCGT
AGAGCACACGTCTGAACTCTTTCCCTACACGACGCTCTTCCGATCTNNNAGGTTTGTCAGCAACTTGTG

AGCACACGTCTGAACTCTTTCCCTACACGACGCTCTTCCGATCTNNNAGGGCATAAGGATGGATGG
GCACACGTCTGAACTCTTTCCCTACACGACGCTCTTCCGATCTNNNAAGTTTGGCATTTCCTTGCCTCT
GAGCACACGTCTGAACTCTTTCCCTACACGACGCTCTTCCGATCTNNNCTGCACTGAACCTGCATAGAT
AGCACACGTCTGAACTCTTTCCCTACACGACGCTCTTCCGATCTNNNACCAGGAGATCCAAAGACTGA
AGCACACGTCTGAACTCTTTCCCTACACGACGCTCTTCCGATCTNNNAACAAAACCCCGTCTCAAAATG
GCACACGTCTGAACTCTTTCCCTACACGACGCTCTTCCGATCTNNNTCACCTCTTCTCAGACAACCACTC
AGCACACGTCTGAACTCTTTCCCTACACGACGCTCTTCCGATCTNNNTCAAAGTACACGGTAGGTAGG
CACACGTCTGAACTCTTTCCCTACACGACGCTCTTCCGATCTNNNCACAGTGCCTGGCTTATAGACAA
AGCACACGTCTGAACTCTTTCCCTACACGACGCTCTTCCGATCTNNNTAGGAGCCAGACCTGTCTCAA
GCACACGTCTGAACTCTTTCCCTACACGACGCTCTTCCGATCTNNNTCCTGGCAGTCCTAAAACAAACC
GCACACGTCTGAACTCTTTCCCTACACGACGCTCTTCCGATCTNNNGACTAAAACTCAGCCAGCAGGA
AGCACACGTCTGAACTCTTTCCCTACACGACGCTCTTCCGATCTNNNAGAGCTCAATAAACCCAAGGC
GCACACGTCTGAACTCTTTCCCTACACGACGCTCTTCCGATCTNNNATATCTCAGTTTCTGCGGGTCCA
AGCACACGTCTGAACTCTTTCCCTACACGACGCTCTTCCGATCTNNNTGTCCAACAGAGGAAACACAT
AGCACACGTCTGAACTCTTTCCCTACACGACGCTCTTCCGATCTNNNTACCAGCGCTAGGATCACTATG
GAGCACACGTCTGAACTCTTTCCCTACACGACGCTCTTCCGATCTNNNCCCCAACTCTGTGCCAATAAA
AGCACACGTCTGAACTCTTTCCCTACACGACGCTCTTCCGATCTNNNTCAGCAGCTCTGGAGTATTCCT
GAGCACACGTCTGAACTCTTTCCCTACACGACGCTCTTCCGATCTNNNGTCGAGAGAGTGGTGACTAC
GCACACGTCTGAACTCTTTCCCTACACGACGCTCTTCCGATCTNNNCGTGGTCATAACAGAGCACTGA
GCACACGTCTGAACTCTTTCCCTACACGACGCTCTTCCGATCTNNNTCAAACGATACCTTGAACACTCA
AGCACACGTCTGAACTCTTTCCCTACACGACGCTCTTCCGATCTNNNAACTTTGGGGAATACCAACCC
AGCACACGTCTGAACTCTTTCCCTACACGACGCTCTTCCGATCTNNNATGTGCAGTGTGGGATGAGAT
GCACACGTCTGAACTCTTTCCCTACACGACGCTCTTCCGATCTNNNCAACCTTCCTACCTCCTGCAAAAC
GCACACGTCTGAACTCTTTCCCTACACGACGCTCTTCCGATCTNNNTCCCCTTGACTGTCTCTCAAAGTC
AGCACACGTCTGAACTCTTTCCCTACACGACGCTCTTCCGATCTNNNAGGGAACTTCTTTTGCTCCCT
AGCACACGTCTGAACTCTTTCCCTACACGACGCTCTTCCGATCTNNNATGTGCCAGGAACTACCTTAT
AGAGCACACGTCTGAACTCTTTCCCTACACGACGCTCTTCCGATCTNNNTGCCATATCTCAATGAAAG
GCACACGTCTGAACTCTTTCCCTACACGACGCTCTTCCGATCTNNNACAGATGGATGTAGGGGCTCAA
AGCACACGTCTGAACTCTTTCCCTACACGACGCTCTTCCGATCTNNNTTGCAGAGATAGAGCCATCGG
GAGCACACGTCTGAACTCTTTCCCTACACGACGCTCTTCCGATCTNNNTGACAAAGGTAAGCTCAGG
GAGCACACGTCTGAACTCTTTCCCTACACGACGCTCTTCCGATCTNNNTCCTTGCCTTTCTTACCCCTGT
AGCACACGTCTGAACTCTTTCCCTACACGACGCTCTTCCGATCTNNNAATGCGTTCACCATGTAACAGG
GAGCACACGTCTGAACTCTTTCCCTACACGACGCTCTTCCGATCTNNNTTTCTCCTGCCTCTCAGCCTA
GAGCACACGTCTGAACTCTTTCCCTACACGACGCTCTTCCGATCTNNNAGCTTTCTTCTGGTCCGACTT
GAGCACACGTCTGAACTCTTTCCCTACACGACGCTCTTCCGATCTNNNAACTGTCTCATGCCAAGCTG
GAGCACACGTCTGAACTCTTTCCCTACACGACGCTCTTCCGATCTNNNACAATGGAGTGGGTCATGAA
GAGCACACGTCTGAACTCTTTCCCTACACGACGCTCTTCCGATCTNNNCAGTCAACATGACAGTCACC
AGCACACGTCTGAACTCTTTCCCTACACGACGCTCTTCCGATCTNNNCTGGCTGAATGTAATGTGTCC
AGCACACGTCTGAACTCTTTCCCTACACGACGCTCTTCCGATCTNNNAGCCATTACAAGAAGGGAAGC
AGCACACGTCTGAACTCTTTCCCTACACGACGCTCTTCCGATCTNNNAACAACTTTACCTGAAGCACCG
GCACACGTCTGAACTCTTTCCCTACACGACGCTCTTCCGATCTNNNGTAAGCCAGGATTCACCAGGAA
AGCACACGTCTGAACTCTTTCCCTACACGACGCTCTTCCGATCTNNNATCAAAGGGCAAAGCATTTCA
AGCACACGTCTGAACTCTTTCCCTACACGACGCTCTTCCGATCTNNNGGGTTTTCTGTCACTCTGCTGT

AGCACACGTCTGAACTCTTTCCCTACACGACGCTCTTCCGATCTNNNTCTTCCCCTACACAGACACACAT
AGAGCACACGTCTGAACTCTTTCCCTACACGACGCTCTTCCGATCTNNNTTGAAGTAATCCTCCTGCCT
GCACACGTCTGAACTCTTTCCCTACACGACGCTCTTCCGATCTNNNCTGAGACACAGGCATATGCAAC
GCACACGTCTGAACTCTTTCCCTACACGACGCTCTTCCGATCTNNNGGGTCCTCACTTTCCTTGAACTC
GAGCACACGTCTGAACTCTTTCCCTACACGACGCTCTTCCGATCTNNNAAAATACATGCGTTGGAGGT
AGCACACGTCTGAACTCTTTCCCTACACGACGCTCTTCCGATCTNNNTGAGGAGGGAGGTAACTTCCA
AGCACACGTCTGAACTCTTTCCCTACACGACGCTCTTCCGATCTNNNCACAACAGGATGCAAGAAAAC
GAGCACACGTCTGAACTCTTTCCCTACACGACGCTCTTCCGATCTNNNGAGAAAGTTGTCTGGGCTTG
GCACACGTCTGAACTCTTTCCCTACACGACGCTCTTCCGATCTNNNAAGCCCACAGAAGAACAGGTTT
GCACACGTCTGAACTCTTTCCCTACACGACGCTCTTCCGATCTNNNAGAAAAGAATCCAGCCCTCATG
AGCACACGTCTGAACTCTTTCCCTACACGACGCTCTTCCGATCTNNNATCAGAGGTGAGATTTGGGT
GAGCACACGTCTGAACTCTTTCCCTACACGACGCTCTTCCGATCTNNNAGAGGTCTACTCATTGCTCAC
AGCACACGTCTGAACTCTTTCCCTACACGACGCTCTTCCGATCTNNNTATCCATCACCTCACACAGGGA
AGCACACGTCTGAACTCTTTCCCTACACGACGCTCTTCCGATCTNNNCACACAGAGACCCATTCAGAC
AGCACACGTCTGAACTCTTTCCCTACACGACGCTCTTCCGATCTNNNTCCTCTAGTCACCAAAGAAACC
GCACACGTCTGAACTCTTTCCCTACACGACGCTCTTCCGATCTNNNGTCTCCATGGTTTGCTCACAGAT
CACACGTCTGAACTCTTTCCCTACACGACGCTCTTCCGATCTNNNTGAATGAGGCTGTACCTTGCTTC
AGCACACGTCTGAACTCTTTCCCTACACGACGCTCTTCCGATCTNNNATCCTGACACCAAAACTGTGC
CACACGTCTGAACTCTTTCCCTACACGACGCTCTTCCGATCTNNNATATGTGCTCTCCCCCTCTCCTACA
AGCACACGTCTGAACTCTTTCCCTACACGACGCTCTTCCGATCTNNNTTCAACTCTGGAGGCAAACCA
GCACACGTCTGAACTCTTTCCCTACACGACGCTCTTCCGATCTNNNCCTTATCCCTCTGTTTCTCTGCGC
GCACACGTCTGAACTCTTTCCCTACACGACGCTCTTCCGATCTNNNAACCTGGACTGCAAATCACACAT
GAGCACACGTCTGAACTCTTTCCCTACACGACGCTCTTCCGATCTNNNATCCAGACCATGACCCACTTT
AGCACACGTCTGAACTCTTTCCCTACACGACGCTCTTCCGATCTNNNAGTAGAATTTCAGGCCTGTCT
AGCACACGTCTGAACTCTTTCCCTACACGACGCTCTTCCGATCTNNNTAACTAATAGCTCCCTCGTGG
AGCACACGTCTGAACTCTTTCCCTACACGACGCTCTTCCGATCTNNNGTCCTTCAAGCTCCGTTTTCTC
AGCACACGTCTGAACTCTTTCCCTACACGACGCTCTTCCGATCTNNNACACACACAGATAAGTGGAAG
GAGCACACGTCTGAACTCTTTCCCTACACGACGCTCTTCCGATCTNNNGTCTAGTCTCCCTGTTCCCAA
AGCACACGTCTGAACTCTTTCCCTACACGACGCTCTTCCGATCTNNNGGTGTCACAGCTCTCCAGAA
AGCACACGTCTGAACTCTTTCCCTACACGACGCTCTTCCGATCTNNNTTTCTGCCATTTCTGACTGAG
AGCACACGTCTGAACTCTTTCCCTACACGACGCTCTTCCGATCTNNNAGCCACTGTGTACTTAGCATGT
GCACACGTCTGAACTCTTTCCCTACACGACGCTCTTCCGATCTNNNGGTGGCAATTGGTGGTCTTCTA
AGCACACGTCTGAACTCTTTCCCTACACGACGCTCTTCCGATCTNNNGCAGTCCATGGTTCACCAAAA
AGCACACGTCTGAACTCTTTCCCTACACGACGCTCTTCCGATCTNNNCTGACTTGGCCTCTTGTTCTC
GCACACGTCTGAACTCTTTCCCTACACGACGCTCTTCCGATCTNNNACACTAAAAGGAAATGGTGGG
AGCACACGTCTGAACTCTTTCCCTACACGACGCTCTTCCGATCTNNNTAGTCATAGGGAATGTGTTGT
AGCACACGTCTGAACTCTTTCCCTACACGACGCTCTTCCGATCTNNNCAACTATATGTGTCACCTCCCG
GCACACGTCTGAACTCTTTCCCTACACGACGCTCTTCCGATCTNNNGTTTCTTCCTCCAGGAGTGCTTC
AGCACACGTCTGAACTCTTTCCCTACACGACGCTCTTCCGATCTNNNCCTCAGGTTCTGGAGGAATAC
GCACACGTCTGAACTCTTTCCCTACACGACGCTCTTCCGATCTNNNCGCACCTGGCCATCTTTTATCTT
GCACACGTCTGAACTCTTTCCCTACACGACGCTCTTCCGATCTNNNCTGAAAACTTGGTGGGAACTT
GCACACGTCTGAACTCTTTCCCTACACGACGCTCTTCCGATCTNNNGGTCTCCTTCTTGCTGACTGTAG
AGCACACGTCTGAACTCTTTCCCTACACGACGCTCTTCCGATCTNNNTCCCCGTTCTTATGAGCCTTAC

GCACACGTCTGAACTCTTTCCCTACACGACGCTCTTCCGATCTNNNCAAGGGATTTCTCCCCTTGTAGCC
GCACACGTCTGAACTCTTTCCCTACACGACGCTCTTCCGATCTNNNACACAGCACTTATGTGTTTGCACC
AGCACACGTCTGAACTCTTTCCCTACACGACGCTCTTCCGATCTNNNATCCACCTGGGTTTTGATTTGCT
AGCACACGTCTGAACTCTTTCCCTACACGACGCTCTTCCGATCTNNNTGGAGCCCTGATTTCACTCAATC
GCACACGTCTGAACTCTTTCCCTACACGACGCTCTTCCGATCTNNNGGTAAGTAGGAGTTGCATGGGT
GAGCACACGTCTGAACTCTTTCCCTACACGACGCTCTTCCGATCTNNNTGCTGGAGCAACTTGATATCT
GAGCACACGTCTGAACTCTTTCCCTACACGACGCTCTTCCGATCTNNNACCCTCAATAACCAGGACCATC
GAGCACACGTCTGAACTCTTTCCCTACACGACGCTCTTCCGATCTNNNAGTTTTGGGGGTTGTTTGACC
AGAGCACACGTCTGAACTCTTTCCCTACACGACGCTCTTCCGATCTNNNAACTTCCCCGTGTAACTACCT
AGCACACGTCTGAACTCTTTCCCTACACGACGCTCTTCCGATCTNNNATCTTTCTTGCACAGGTCCACTCC
AGAGCACACGTCTGAACTCTTTCCCTACACGACGCTCTTCCGATCTNNNCCTATTCCAGGGAGGGAAA
AGCACACGTCTGAACTCTTTCCCTACACGACGCTCTTCCGATCTNNNTGTCCCAATCCCACACAGATACT
GAGCACACGTCTGAACTCTTTCCCTACACGACGCTCTTCCGATCTNNNCAGTCCTACCAAGACACAACC
AGCACACGTCTGAACTCTTTCCCTACACGACGCTCTTCCGATCTNNNTCATCTTGGTGTGAACCAATCC
AGCACACGTCTGAACTCTTTCCCTACACGACGCTCTTCCGATCTNNNACCTCCAAGTCTCTCAGCTTCAT
GAGCACACGTCTGAACTCTTTCCCTACACGACGCTCTTCCGATCTNNNATCCTCACCAAAAATGGTGCTC
AGCACACGTCTGAACTCTTTCCCTACACGACGCTCTTCCGATCTNNNGAGGTTGTGAGCCGAAATCGT
GCACACGTCTGAACTCTTTCCCTACACGACGCTCTTCCGATCTNNNTATGCAGTCTGAGGTTCAAAGCC
AGCACACGTCTGAACTCTTTCCCTACACGACGCTCTTCCGATCTNNNTGGGAGAGACTAGCATCTCTGT
AGAGCACACGTCTGAACTCTTTCCCTACACGACGCTCTTCCGATCTNNNACTCCAAGACTTTCCCCAGAA
GAGCACACGTCTGAACTCTTTCCCTACACGACGCTCTTCCGATCTNNNAACTCAGAAGTGAAGATGCCT
GAGCACACGTCTGAACTCTTTCCCTACACGACGCTCTTCCGATCTNNNTCAAGAGAAGACCCCTCTCACA
AGCACACGTCTGAACTCTTTCCCTACACGACGCTCTTCCGATCTNNNGTACTTTTGAAACAGGGCTTGC
GAGCACACGTCTGAACTCTTTCCCTACACGACGCTCTTCCGATCTNNNTATTCAGGAGGCTTAGGCGGA
GAGCACACGTCTGAACTCTTTCCCTACACGACGCTCTTCCGATCTNNNTTCTGTATCTGTCATCGGCACA
GAGCACACGTCTGAACTCTTTCCCTACACGACGCTCTTCCGATCTNNNATGTGCTCGGCTAACAGTTTT
GAGCACACGTCTGAACTCTTTCCCTACACGACGCTCTTCCGATCTNNNCGCCTATTTGCACTGGGAATAC
AGCACACGTCTGAACTCTTTCCCTACACGACGCTCTTCCGATCTNNNATACCCATTCACTGCACATCAGC
AGCACACGTCTGAACTCTTTCCCTACACGACGCTCTTCCGATCTNNNACATATCCCCATAACAGCTCCCTC
AGCACACGTCTGAACTCTTTCCCTACACGACGCTCTTCCGATCTNNNAGCCTGGGCAATAGCATAAGA
AGCACACGTCTGAACTCTTTCCCTACACGACGCTCTTCCGATCTNNNATGTGTTTAGTCTCACCCACTGC
AGCACACGTCTGAACTCTTTCCCTACACGACGCTCTTCCGATCTNNNTACAAAGAGCAGCACGAGATA
AGCACACGTCTGAACTCTTTCCCTACACGACGCTCTTCCGATCTNNNTTTCTGCTCCAACCAGCAAGAAT
GAGCACACGTCTGAACTCTTTCCCTACACGACGCTCTTCCGATCTNNNGGGCTACCCCAAACAATATCAC
GAGCACACGTCTGAACTCTTTCCCTACACGACGCTCTTCCGATCTNNNGACTACTGGCTTTGTATGTGG
AGAGCACACGTCTGAACTCTTTCCCTACACGACGCTCTTCCGATCTNNNTCCCAGATCAGCAAACATTTT
GAGCACACGTCTGAACTCTTTCCCTACACGACGCTCTTCCGATCTNNNTCCTGGTGTCCAAGGAAAGAA
GAGCACACGTCTGAACTCTTTCCCTACACGACGCTCTTCCGATCTNNNGCTGACATTGAGACAGACAGA
AGCACACGTCTGAACTCTTTCCCTACACGACGCTCTTCCGATCTNNNTCCCATGAGAAGACACTTACCT
AGCACACGTCTGAACTCTTTCCCTACACGACGCTCTTCCGATCTNNNCCTTCCAGGACTGTGAGAAGAC
AGAGCACACGTCTGAACTCTTTCCCTACACGACGCTCTTCCGATCTNNNAGTGGGGAGAACTAAGACAC
GAGCACACGTCTGAACTCTTTCCCTACACGACGCTCTTCCGATCTNNNGTTTCTGTGGCTCCATCTATCC
AGAGCACACGTCTGAACTCTTTCCCTACACGACGCTCTTCCGATCTNNNTGCAGTGGTAGCAAAAGCAA

AGCACACGTCTGAACTCTTTCCCTACACGACGCTCTTCCGATCTNNNAGGCTGGGGGTGTTCTAATAT
AGCACACGTCTGAACTCTTTCCCTACACGACGCTCTTCCGATCTNNNTTTTCGGAGCAAGTTGAGGTTG
AGCACACGTCTGAACTCTTTCCCTACACGACGCTCTTCCGATCTNNNCTTTCAGAAGGAGGATGGTTC
AGAGCACACGTCTGAACTCTTTCCCTACACGACGCTCTTCCGATCTNNNTGCAGAAGAAATTCTACCCTG
AGAGCACACGTCTGAACTCTTTCCCTACACGACGCTCTTCCGATCTNNNCTGCCAGAACTACCAGTACTA
AGAGCACACGTCTGAACTCTTTCCCTACACGACGCTCTTCCGATCTNNNTGCTGAACTAGTTGCTTGCA
AGAGCACACGTCTGAACTCTTTCCCTACACGACGCTCTTCCGATCTNNNTCCTGAAGACAAAGCAGTG
AGAGCACACGTCTGAACTCTTTCCCTACACGACGCTCTTCCGATCTNNNTTACCACCTTACATGCTGCCT
GCACACGTCTGAACTCTTTCCCTACACGACGCTCTTCCGATCTNNNACTCTCCTGTCCTCATCTCACTGCA
GAGCACACGTCTGAACTCTTTCCCTACACGACGCTCTTCCGATCTNNNTACACCACACCTGAAGGAGA
GAGCACACGTCTGAACTCTTTCCCTACACGACGCTCTTCCGATCTNNNGGAGGATTTGTTTGTGAAG
AGAGCACACGTCTGAACTCTTTCCCTACACGACGCTCTTCCGATCTNNNTTGATTGGAGCCTCCTGATT
AGCACACGTCTGAACTCTTTCCCTACACGACGCTCTTCCGATCTNNNTTCTGCAATACAACTTCCTGCCA
AGCACACGTCTGAACTCTTTCCCTACACGACGCTCTTCCGATCTNNNCTAATTACTACCCCCACTCCCCA
AGAGCACACGTCTGAACTCTTTCCCTACACGACGCTCTTCCGATCTNNNTGAAGAGCTGACCCATAGTG
GAGCACACGTCTGAACTCTTTCCCTACACGACGCTCTTCCGATCTNNNCTGTGCCTGTTCCTCTCTTTCTC
AGCACACGTCTGAACTCTTTCCCTACACGACGCTCTTCCGATCTNNNCTGCTGGTCAACCACTATGTGT
GAGCACACGTCTGAACTCTTTCCCTACACGACGCTCTTCCGATCTNNNGGTTTCTCTGTCTCTCCACACA
GAGCACACGTCTGAACTCTTTCCCTACACGACGCTCTTCCGATCTNNNCCTGAGGATTCATCCACCTGA
AGCACACGTCTGAACTCTTTCCCTACACGACGCTCTTCCGATCTNNNCATGCATGGACACACACAATAC
AGCACACGTCTGAACTCTTTCCCTACACGACGCTCTTCCGATCTNNNTCCCATTCAGTCCATTGTAGGG
GAGCACACGTCTGAACTCTTTCCCTACACGACGCTCTTCCGATCTNNNTGTTTTCTGTGTGGAAAGGG
GCACACGTCTGAACTCTTTCCCTACACGACGCTCTTCCGATCTNNNGAGAAAACTATGGCTGGAGACC
AGCACACGTCTGAACTCTTTCCCTACACGACGCTCTTCCGATCTNNNTCTCACCCAGATGTTCTGCTGTA
GAGCACACGTCTGAACTCTTTCCCTACACGACGCTCTTCCGATCTNNNCGTGCACAGACACGCAGAAAA
GAGCACACGTCTGAACTCTTTCCCTACACGACGCTCTTCCGATCTNNNTAGAAATGAGCCCCCAAGTGA
AGCACACGTCTGAACTCTTTCCCTACACGACGCTCTTCCGATCTNNNTTTCCAGATGCAGAAGCCAAAGT
GAGCACACGTCTGAACTCTTTCCCTACACGACGCTCTTCCGATCTNNNGACAAATCCCATGAGATGGCA
AGCACACGTCTGAACTCTTTCCCTACACGACGCTCTTCCGATCTNNNGCCTAAAATTCACCCCCTTTGGA
GAGCACACGTCTGAACTCTTTCCCTACACGACGCTCTTCCGATCTNNNCAATCCTGAGAAACTTGGAG
AGAGCACACGTCTGAACTCTTTCCCTACACGACGCTCTTCCGATCTNNNGCATCATGCTGTGTCTTGAA
AGCACACGTCTGAACTCTTTCCCTACACGACGCTCTTCCGATCTNNNCATGTAGGAGTGGAAGCTTTCG
AGAGCACACGTCTGAACTCTTTCCCTACACGACGCTCTTCCGATCTNNNCCTGGATGAAAAGGCAGAGT
AGCACACGTCTGAACTCTTTCCCTACACGACGCTCTTCCGATCTNNNCTGTTGTGTTCCTGACATTGTG
GCACACGTCTGAACTCTTTCCCTACACGACGCTCTTCCGATCTNNNAATTTCATTGGAAGGAGGGGTC
AGCACACGTCTGAACTCTTTCCCTACACGACGCTCTTCCGATCTNNNTATATCCTAATGGGAGGCGAA
GCACACGTCTGAACTCTTTCCCTACACGACGCTCTTCCGATCTNNNTCAGGGTACTAGCACTGTTGGTT
GAGCACACGTCTGAACTCTTTCCCTACACGACGCTCTTCCGATCTNNNAGGGTATCCACAAAGCAATGA
GAGCACACGTCTGAACTCTTTCCCTACACGACGCTCTTCCGATCTNNNGGTGCTGTCTTGTGTGGTAGA
AGAGCACACGTCTGAACTCTTTCCCTACACGACGCTCTTCCGATCTNNNGAAGGAAGATTCCAGGCAGA
GAGCACACGTCTGAACTCTTTCCCTACACGACGCTCTTCCGATCTNNNTTCCGTCTTTCCTCTAATGTGG
GAGCACACGTCTGAACTCTTTCCCTACACGACGCTCTTCCGATCTNNNCTCACTCTTTTCCCACTGCTGAA
GAGCACACGTCTGAACTCTTTCCCTACACGACGCTCTTCCGATCTNNNATCGGAAATGTTGGCCTGTA

GAGCACACGTCTGAACTCTTTCCCTACACGACGCTCTTCCGATCTNNNCCAGTACCTTTGCTGTGAATG

GAGCACACGTCTGAACTCTTTCCCTACACGACGCTCTTCCGATCTNNNGATCTTCCTTGCGTTTCCTTCC

GAGCACACGTCTGAACTCTTTCCCTACACGACGCTCTTCCGATCTNNNATAGCAACAACCAAAGTCCCCT

GCACACGTCTGAACTCTTTCCCTACACGACGCTCTTCCGATCTNNNAGAGGAAAACGTAAGCAGAACC

AGCACACGTCTGAACTCTTTCCCTACACGACGCTCTTCCGATCTNNNCTCTCCCTCCCACTTCTTTCAC

GAGCACACGTCTGAACTCTTTCCCTACACGACGCTCTTCCGATCTNNNACAGCAGCTTCTCTCTGTCTCT

AGCACACGTCTGAACTCTTTCCCTACACGACGCTCTTCCGATCTNNNGCCTAAGTCCAGAAGCAGGTTA

GCACACGTCTGAACTCTTTCCCTACACGACGCTCTTCCGATCTNNNATCTTGCTTCTCATTCTGTGGGCCA

GCACACGTCTGAACTCTTTCCCTACACGACGCTCTTCCGATCTNNNAAAAAGTCTGAGGGCCTGGTTA

GAGCACACGTCTGAACTCTTTCCCTACACGACGCTCTTCCGATCTNNNAAATGGTCGAATTATGGGCGT

AGCACACGTCTGAACTCTTTCCCTACACGACGCTCTTCCGATCTNNNGCCTGGAGCATCTTATAGTGCA

GAGCACACGTCTGAACTCTTTCCCTACACGACGCTCTTCCGATCTNNNCCCTGTAACTGCACCAAACTTC

GAGCACACGTCTGAACTCTTTCCCTACACGACGCTCTTCCGATCTNNNTGCTAAATCTCATGGGGCCATT

AGCACACGTCTGAACTCTTTCCCTACACGACGCTCTTCCGATCTNNNGGGCCCTGGCTTTTCATTAGTT

AGAGCACACGTCTGAACTCTTTCCCTACACGACGCTCTTCCGATCTNNNGTGGGTTTTACGACAGCACA

AGCACACGTCTGAACTCTTTCCCTACACGACGCTCTTCCGATCTNNNTCCTCCTATTCCATTACCACCGCC

GAGCACACGTCTGAACTCTTTCCCTACACGACGCTCTTCCGATCTNNNTAGAACTGGCCCCTTGGGATA

GAGCACACGTCTGAACTCTTTCCCTACACGACGCTCTTCCGATCTNNNGAGGGAAGACATTCTCTCTCC

GAGCACACGTCTGAACTCTTTCCCTACACGACGCTCTTCCGATCTNNNCCGGGCTTCCTGTTTAGTCTCT

AGCACACGTCTGAACTCTTTCCCTACACGACGCTCTTCCGATCTNNNTATTTGCATGGGCTCCTTAGCTG

GAGCACACGTCTGAACTCTTTCCCTACACGACGCTCTTCCGATCTNNNATCCTTCTTTCCCAACCATCAGC

AGCACACGTCTGAACTCTTTCCCTACACGACGCTCTTCCGATCTNNNCATGTTCAGTACGCACGCAATT

AGCACACGTCTGAACTCTTTCCCTACACGACGCTCTTCCGATCTNNNGCCTTGTGAGATCTGTTCATTG

AGCACACGTCTGAACTCTTTCCCTACACGACGCTCTTCCGATCTNNNACTTTTTCGGTTCTGGCTGTCTTC

AGAGCACACGTCTGAACTCTTTCCCTACACGACGCTCTTCCGATCTNNNAACCTCATTGATATGCATCAC

AAGAGCACACGTCTGAACTCTTTCCCTACACGACGCTCTTCCGATCTNNNCATACATCTGTCCCCAACAA

GAGCACACGTCTGAACTCTTTCCCTACACGACGCTCTTCCGATCTNNNTAGCTCTGAGAACATAGTGGC

GAGCACACGTCTGAACTCTTTCCCTACACGACGCTCTTCCGATCTNNNGCCTTCTTGCTCATGAAACTGA

GAGCACACGTCTGAACTCTTTCCCTACACGACGCTCTTCCGATCTNNNTAGAATCATCCATTCCCATCGC

GAGCACACGTCTGAACTCTTTCCCTACACGACGCTCTTCCGATCTNNNGAGATCTCGTGCTCTCTCTCTC

GAGCACACGTCTGAACTCTTTCCCTACACGACGCTCTTCCGATCTNNNGATTAGGCATACGCCATCCTGA

GAGCACACGTCTGAACTCTTTCCCTACACGACGCTCTTCCGATCTNNNTTGTCTGACAGTTTCCTCAGCA

AGCACACGTCTGAACTCTTTCCCTACACGACGCTCTTCCGATCTNNNATGAGAAAGGGGTGAGGATGT

GAGCACACGTCTGAACTCTTTCCCTACACGACGCTCTTCCGATCTNNNGCGTGTCTATGTCGATTTGCA

AGAGCACACGTCTGAACTCTTTCCCTACACGACGCTCTTCCGATCTNNNTGGATGTGGTAGGATGAAG

GCACACGTCTGAACTCTTTCCCTACACGACGCTCTTCCGATCTNNNTACACCTAATGGGCAAACAAGCA

AGAGCACACGTCTGAACTCTTTCCCTACACGACGCTCTTCCGATCTNNNCACAGATACTCTGGGCACGTA

AGCACACGTCTGAACTCTTTCCCTACACGACGCTCTTCCGATCTNNNGTAAGGTTACTGTGTTAGGCCC

AGAGCACACGTCTGAACTCTTTCCCTACACGACGCTCTTCCGATCTNNNCCCCCAAGAGTGTCAATTTCT

GCACACGTCTGAACTCTTTCCCTACACGACGCTCTTCCGATCTNNNTCTACTGTTGTGTCAGGGTCACTC

AGCACACGTCTGAACTCTTTCCCTACACGACGCTCTTCCGATCTNNNCTGCATCTAGAATGGGAGTGTC

AGAGCACACGTCTGAACTCTTTCCCTACACGACGCTCTTCCGATCTNNNTAAGAAAACGACGACGACA

GAGCACACGTCTGAACTCTTTCCCTACACGACGCTCTTCCGATCTNNNACCATCACACCCTTTAGGACAC

GAGCACACGTCTGAACTCTTTCCCTACACGACGCTCTTCCGATCTNNNAGCTTTTGACTTTTGGCCACAT
AGCACACGTCTGAACTCTTTCCCTACACGACGCTCTTCCGATCTNNNTGTCCAACTCTCTCTCTCACACAC
AGCACACGTCTGAACTCTTTCCCTACACGACGCTCTTCCGATCTNNNAGCAATAAGCACTAGAAGGAG
GAGCACACGTCTGAACTCTTTCCCTACACGACGCTCTTCCGATCTNNNTCTAATGATGTGTGTGTGTGC
GAGCACACGTCTGAACTCTTTCCCTACACGACGCTCTTCCGATCTNNNGCAGAAAGAGCAAGGAAGAC
AGCACACGTCTGAACTCTTTCCCTACACGACGCTCTTCCGATCTNNNCCTGCTATCTTGTTGCAGGACTT
AGCACACGTCTGAACTCTTTCCCTACACGACGCTCTTCCGATCTNNNTTAGCGCATCGATTTCTTCCTCAC
AGCACACGTCTGAACTCTTTCCCTACACGACGCTCTTCCGATCTNNNTATCTGGTCTCTGGGGTCATTGC
GAGCACACGTCTGAACTCTTTCCCTACACGACGCTCTTCCGATCTNNNTACCTGTTCACTTCTGTCACCTG
GAGCACACGTCTGAACTCTTTCCCTACACGACGCTCTTCCGATCTNNNAGAGCCTTGTGTGTTATGGCA
AGCACACGTCTGAACTCTTTCCCTACACGACGCTCTTCCGATCTNNNTCTATTCATGACCTACCCCCGACC
GAGCACACGTCTGAACTCTTTCCCTACACGACGCTCTTCCGATCTNNNTTGAGCTAGTAGGAGAGCCAC
AGAGCACACGTCTGAACTCTTTCCCTACACGACGCTCTTCCGATCTNNNTGTAGCTTTGAGTTTGGTGA
GAGCACACGTCTGAACTCTTTCCCTACACGACGCTCTTCCGATCTNNNTGGATGCTACACTGCATCTTCA
GAGCACACGTCTGAACTCTTTCCCTACACGACGCTCTTCCGATCTNNNACAAGGACAGACTGGAGTTTG
GAGCACACGTCTGAACTCTTTCCCTACACGACGCTCTTCCGATCTNNNTTTGTATGCCCCTCTTTGCTTC
AGCACACGTCTGAACTCTTTCCCTACACGACGCTCTTCCGATCTNNNAGGACAGTAGGACACACACAGA
AGCACACGTCTGAACTCTTTCCCTACACGACGCTCTTCCGATCTNNNAACAGTCCCAGAAAGGAAAAG
GAGCACACGTCTGAACTCTTTCCCTACACGACGCTCTTCCGATCTNNNGAGCAAGCTTACCTGATGTCCT
GAGCACACGTCTGAACTCTTTCCCTACACGACGCTCTTCCGATCTNNNGCCAAAAAGGCAAGTCATGGA
GAGCACACGTCTGAACTCTTTCCCTACACGACGCTCTTCCGATCTNNNATTGTGTTCGAGTATCACCCA
GAGCACACGTCTGAACTCTTTCCCTACACGACGCTCTTCCGATCTNNNAAAACACCCAAACCCCACAAAA
GAGCACACGTCTGAACTCTTTCCCTACACGACGCTCTTCCGATCTNNNAGAGAAAGTCAGGAGAATGT
GAGCACACGTCTGAACTCTTTCCCTACACGACGCTCTTCCGATCTNNNCATGGACTTTCCCCTGACTTAG
GAGCACACGTCTGAACTCTTTCCCTACACGACGCTCTTCCGATCTNNNGTGCAGCACAGAACATCCTCA
AGCACACGTCTGAACTCTTTCCCTACACGACGCTCTTCCGATCTNNNGGGCATAGAAAGACCCCATCTC
AGCACACGTCTGAACTCTTTCCCTACACGACGCTCTTCCGATCTNNNTCTGGCTAACAGCTTTCTGAACT
GAGCACACGTCTGAACTCTTTCCCTACACGACGCTCTTCCGATCTNNNTCTACAACGGCAACCACTTTCA
AGCACACGTCTGAACTCTTTCCCTACACGACGCTCTTCCGATCTNNNACGCATCTCCCAAAATACACACAC
GAGCACACGTCTGAACTCTTTCCCTACACGACGCTCTTCCGATCTNNNCAGGAGGCATCGAGGTAGAG
AGAGCACACGTCTGAACTCTTTCCCTACACGACGCTCTTCCGATCTNNNTAAGTGTGAGGTAAGGTGG
GCACACGTCTGAACTCTTTCCCTACACGACGCTCTTCCGATCTNNNTGTGGCAAGGCAGTATGATTGTT
AGAGCACACGTCTGAACTCTTTCCCTACACGACGCTCTTCCGATCTNNNGAGCTCAAGTAGAGTGGTTG
GAGCACACGTCTGAACTCTTTCCCTACACGACGCTCTTCCGATCTNNNATGTATATCTCTGTGCGTGTGC
GAGCACACGTCTGAACTCTTTCCCTACACGACGCTCTTCCGATCTNNNACAAAATGGTCCATGCCTCTG
GAGCACACGTCTGAACTCTTTCCCTACACGACGCTCTTCCGATCTNNNATCCATGGTGGATGATGCTAG
GAGCACACGTCTGAACTCTTTCCCTACACGACGCTCTTCCGATCTNNNTAACCATCATGCAAGGGCAAA
AGCACACGTCTGAACTCTTTCCCTACACGACGCTCTTCCGATCTNNNTTAGTAATACCACCACACCACCA
AGCACACGTCTGAACTCTTTCCCTACACGACGCTCTTCCGATCTNNNTAGGAATGCTACCTCCCACTCAC
GAGCACACGTCTGAACTCTTTCCCTACACGACGCTCTTCCGATCTNNNACACAATGAGGGGATTGGAGA
GAGCACACGTCTGAACTCTTTCCCTACACGACGCTCTTCCGATCTNNNACCTGAGATGTCCTCCTTTCAC
AGCACACGTCTGAACTCTTTCCCTACACGACGCTCTTCCGATCTNNNTGAAGGAAAAAGAAGTGGGCC
GAGCACACGTCTGAACTCTTTCCCTACACGACGCTCTTCCGATCTNNNCACCAACACACAATTCCCCTGTA

AGCACACGTCTGAACTCTTTCCCTACACGACGCTCTTCCGATCTNNNAGGAGATCAGAGCTGTCACT
GAGCACACGTCTGAACTCTTTCCCTACACGACGCTCTTCCGATCTNNNCACACTATGACAGCCAAGAAG
GCACACGTCTGAACTCTTTCCCTACACGACGCTCTTCCGATCTNNNAAGGCCCAGTTCTCCTCTCATTA
AGCACACGTCTGAACTCTTTCCCTACACGACGCTCTTCCGATCTNNNGAGTAGTCTGCACTCCCATGTT
GAGCACACGTCTGAACTCTTTCCCTACACGACGCTCTTCCGATCTNNNACCGAAGTCAGACACCAAGG
GCACACGTCTGAACTCTTTCCCTACACGACGCTCTTCCGATCTNNNCTTGCTCACCCTTCTCTTAGCAG
GAGCACACGTCTGAACTCTTTCCCTACACGACGCTCTTCCGATCTNNNGACCTACCAGTGGGACTTTC
AGCACACGTCTGAACTCTTTCCCTACACGACGCTCTTCCGATCTNNNTCAGAGTTGCTGGGCTATTTCA
AGCACACGTCTGAACTCTTTCCCTACACGACGCTCTTCCGATCTNNNCCTGGCCTGATTTGGATTTTC
GCACACGTCTGAACTCTTTCCCTACACGACGCTCTTCCGATCTNNNGGGAGTGTATACGTGGGAGA
GCACACGTCTGAACTCTTTCCCTACACGACGCTCTTCCGATCTNNNTGGTAAGTGGTCTCTGGGTTA
AGCACACGTCTGAACTCTTTCCCTACACGACGCTCTTCCGATCTNNNTAAGCACAGAGCCAAACACCTT
GAGCACACGTCTGAACTCTTTCCCTACACGACGCTCTTCCGATCTNNNTAGCTTCAAACAGAAGTCGG
GCACACGTCTGAACTCTTTCCCTACACGACGCTCTTCCGATCTNNNTATTTGCTTCCCATGTCCCCAG
AGCACACGTCTGAACTCTTTCCCTACACGACGCTCTTCCGATCTNNNTCATATTGCAGTTGTCCCTCGA
GAGCACACGTCTGAACTCTTTCCCTACACGACGCTCTTCCGATCTNNNTACATTGGAGGAGCAGACTG
AGCACACGTCTGAACTCTTTCCCTACACGACGCTCTTCCGATCTNNNTCCAACCATTCCTTCTTCTCTGC
GCACACGTCTGAACTCTTTCCCTACACGACGCTCTTCCGATCTNNNTTGGGAGAAAGGGGTCAGTGT
AGAGCACACGTCTGAACTCTTTCCCTACACGACGCTCTTCCGATCTNNNATTGGTGCAGCATGACAT
GAGCACACGTCTGAACTCTTTCCCTACACGACGCTCTTCCGATCTNNNGGGGGATCAAACTGCCAACT
AGCACACGTCTGAACTCTTTCCCTACACGACGCTCTTCCGATCTNNNGTGGATGATGTGTCCTTTCAC
GAGCACACGTCTGAACTCTTTCCCTACACGACGCTCTTCCGATCTNNNCAAAAAGGGACAGCCAAAGT
GCACACGTCTGAACTCTTTCCCTACACGACGCTCTTCCGATCTNNNTTCCAACCTTGCAGAGAGGGTA
GAGCACACGTCTGAACTCTTTCCCTACACGACGCTCTTCCGATCTNNNCCTTCAATAGGCTTCTTGCCC
AGCACACGTCTGAACTCTTTCCCTACACGACGCTCTTCCGATCTNNNGGGGGTCCTGTCTTCTTCTGT
AGCACACGTCTGAACTCTTTCCCTACACGACGCTCTTCCGATCTNNNCAGTGGGGAGAGACAACAAT
AGCACACGTCTGAACTCTTTCCCTACACGACGCTCTTCCGATCTNNNGGCTATAGTGAGCCTCAAAGG
AGCACACGTCTGAACTCTTTCCCTACACGACGCTCTTCCGATCTNNNCTTCCACTTCCAATAGCTGGC
AGCACACGTCTGAACTCTTTCCCTACACGACGCTCTTCCGATCTNNNATATAACTAGAGGCCGAGCCA
GAGCACACGTCTGAACTCTTTCCCTACACGACGCTCTTCCGATCTNNNCAAACATGTCAGGAATTTGG
AGCACACGTCTGAACTCTTTCCCTACACGACGCTCTTCCGATCTNNNAAGAATGGTTGGAACGGTTG
AGCACACGTCTGAACTCTTTCCCTACACGACGCTCTTCCGATCTNNNATGAAGCTGTACTGGGATTTG
AGCACACGTCTGAACTCTTTCCCTACACGACGCTCTTCCGATCTNNNTTGCATTCTGAAGACAAGCCAG
AGCACACGTCTGAACTCTTTCCCTACACGACGCTCTTCCGATCTNNNTTTTTCTCGCCGGGTAGTAGTT
AGCACACGTCTGAACTCTTTCCCTACACGACGCTCTTCCGATCTNNNTGCACCACACAAATACACTCAC
AGCACACGTCTGAACTCTTTCCCTACACGACGCTCTTCCGATCTNNNAAGAGTGGGGCTGTGATCTTA
AGCACACGTCTGAACTCTTTCCCTACACGACGCTCTTCCGATCTNNNATAGAGGAGAGGTTGAGGA
AGCACACGTCTGAACTCTTTCCCTACACGACGCTCTTCCGATCTNNNTCCTCTGAGGGCCTGAATACAA
GAGCACACGTCTGAACTCTTTCCCTACACGACGCTCTTCCGATCTNNNCTGGCTTCCTCCAAAGTTAGC
AGCACACGTCTGAACTCTTTCCCTACACGACGCTCTTCCGATCTNNNATACTGTGCTGTTTCTGGAGC
GAGCACACGTCTGAACTCTTTCCCTACACGACGCTCTTCCGATCTNNNCGGTGAGGAATTCAAATGT
AGCACACGTCTGAACTCTTTCCCTACACGACGCTCTTCCGATCTNNNTCACATCCACAGTACACATGCA
GAGCACACGTCTGAACTCTTTCCCTACACGACGCTCTTCCGATCTNNNGCTCTGCTTGTTTCATTTGGT

GCACACGTCTGAACTCTTTCCCTACACGACGCTCTTCCGATCTNNNCAGGGGAATGCCCAATACTTCT
CACACGTCTGAACTCTTTCCCTACACGACGCTCTTCCGATCTNNNTGGGGAATTCTAATGCACCCAAC
GAGCACACGTCTGAACTCTTTCCCTACACGACGCTCTTCCGATCTNNNATCTATGCGAGTGTGTTTGT
AGCACACGTCTGAACTCTTTCCCTACACGACGCTCTTCCGATCTNNNAAAGCTAACCGGAACATCACA
GCACACGTCTGAACTCTTTCCCTACACGACGCTCTTCCGATCTNNNTAGATCAATGCGCAGTCTGGTT
AGCACACGTCTGAACTCTTTCCCTACACGACGCTCTTCCGATCTNNNGGACTGGTGTTGACTTTGGAT
GCACACGTCTGAACTCTTTCCCTACACGACGCTCTTCCGATCTNNNACACCTGGGAATTAGAAGGGG
AGCACACGTCTGAACTCTTTCCCTACACGACGCTCTTCCGATCTNNNACACAGTGAGACCCCCTTTAC
GCACACGTCTGAACTCTTTCCCTACACGACGCTCTTCCGATCTNNNAAGGACTTTGTCTGCAAGCATC
GAGCACACGTCTGAACTCTTTCCCTACACGACGCTCTTCCGATCTNNNTGGGATCTGTAAAGTTAGA
GAGCACACGTCTGAACTCTTTCCCTACACGACGCTCTTCCGATCTNNNAGCAAAGTAATGGCTCCCTC
AGCACACGTCTGAACTCTTTCCCTACACGACGCTCTTCCGATCTNNNGCCAGACTACGTCTCTCTCTCT
AGAGCACACGTCTGAACTCTTTCCCTACACGACGCTCTTCCGATCTNNNCCGGCCTCGTGTATTGATTT
GCACACGTCTGAACTCTTTCCCTACACGACGCTCTTCCGATCTNNNTTACTGAAAACAGCCCAGAGGA
AGCACACGTCTGAACTCTTTCCCTACACGACGCTCTTCCGATCTNNNGGTCAGTAAGAAGGGAGACT
GCACACGTCTGAACTCTTTCCCTACACGACGCTCTTCCGATCTNNNGCTTCCACCAAACACCACATTT
GCACACGTCTGAACTCTTTCCCTACACGACGCTCTTCCGATCTNNNACTCAGACAGGAACACACACAA
CACACGTCTGAACTCTTTCCCTACACGACGCTCTTCCGATCTNNNAGGTCACAGTCATAAGCCTCCAA
AGCACACGTCTGAACTCTTTCCCTACACGACGCTCTTCCGATCTNNNATCTGTGACGAAGCTGTGATG
AGCACACGTCTGAACTCTTTCCCTACACGACGCTCTTCCGATCTNNNGTTTCTACACTTAGGCCTGGAC
AGCACACGTCTGAACTCTTTCCCTACACGACGCTCTTCCGATCTNNNAATCCAGGCTTTCCCAAATGTG
GCACACGTCTGAACTCTTTCCCTACACGACGCTCTTCCGATCTNNNCTCCACAAAATTAGGCCATGCTG
GCACACGTCTGAACTCTTTCCCTACACGACGCTCTTCCGATCTNNNTCCTGGAATAGACTGCATGTCA
AGCACACGTCTGAACTCTTTCCCTACACGACGCTCTTCCGATCTNNNTTTCTAACTGGCATGTGGAG
GCACACGTCTGAACTCTTTCCCTACACGACGCTCTTCCGATCTNNNACATGAGGGTTTTAGTGTCCTG
GAGCACACGTCTGAACTCTTTCCCTACACGACGCTCTTCCGATCTNNNCTTGTGTCTTCCTGATCCCAC
GAGCACACGTCTGAACTCTTTCCCTACACGACGCTCTTCCGATCTNNNTCATTCACAGGGAATCTATG
AGCACACGTCTGAACTCTTTCCCTACACGACGCTCTTCCGATCTNNNCATTTATAGGCCCCATACCCC
AGCACACGTCTGAACTCTTTCCCTACACGACGCTCTTCCGATCTNNNTACATGCACACATAAATGCAGG
AGCACACGTCTGAACTCTTTCCCTACACGACGCTCTTCCGATCTNNNCCACATGCTCAAAGGGTACATC
AGCACACGTCTGAACTCTTTCCCTACACGACGCTCTTCCGATCTNNNTGGCCTCAGTTACTGCAAACAA
AGCACACGTCTGAACTCTTTCCCTACACGACGCTCTTCCGATCTNNNAGAACATAGATCTCTCCCCCTG
GAGCACACGTCTGAACTCTTTCCCTACACGACGCTCTTCCGATCTNNNAATCTAAGGAGAAGCCAGG
AGAGCACACGTCTGAACTCTTTCCCTACACGACGCTCTTCCGATCTNNNAAGAAAGCACAACACTAAG
GCACACGTCTGAACTCTTTCCCTACACGACGCTCTTCCGATCTNNNTTTCAGGCTCTGATATTTGGCCC
AGCACACGTCTGAACTCTTTCCCTACACGACGCTCTTCCGATCTNNNCATTCATCATTGGTGGAACGC
GCACACGTCTGAACTCTTTCCCTACACGACGCTCTTCCGATCTNNNTTTATTTGAGGGAAAGCCGCCA
GCACACGTCTGAACTCTTTCCCTACACGACGCTCTTCCGATCTNNNTTGCCAAAGCCTTAGGACTTCT
GAGCACACGTCTGAACTCTTTCCCTACACGACGCTCTTCCGATCTNNNTCAAGTTTGGTTCAGCACTAG
AGCACACGTCTGAACTCTTTCCCTACACGACGCTCTTCCGATCTNNNGACCAAAAGCTAGTCAGCACT
GCACACGTCTGAACTCTTTCCCTACACGACGCTCTTCCGATCTNNNCTCCGCCCAAGTTCTTCTAACTCC
GAGCACACGTCTGAACTCTTTCCCTACACGACGCTCTTCCGATCTNNNCCCAATAACCTGATGCACATG
AGCACACGTCTGAACTCTTTCCCTACACGACGCTCTTCCGATCTNNNAATCTAACTGTCCATCAAGGG

AGCACACGTCTGAACTCTTTCCCTACACGACGCTCTTCCGATCTNNNGCTTGCATTTTGATGGAGCATG
GAGCACACGTCTGAACTCTTTCCCTACACGACGCTCTTCCGATCTNNNACCACTTCCCCAATCTCTCTCT
AGCACACGTCTGAACTCTTTCCCTACACGACGCTCTTCCGATCTNNNTGCCAATAAAGACGCTTGTCAA
GAGCACACGTCTGAACTCTTTCCCTACACGACGCTCTTCCGATCTNNNGAAGGTAGCTGACATTGGC
AGCACACGTCTGAACTCTTTCCCTACACGACGCTCTTCCGATCTNNNCATGAGTGAAGACCACTCACG
AGCACACGTCTGAACTCTTTCCCTACACGACGCTCTTCCGATCTNNNTAATCAGAACATCCCACAGTG
GAGCACACGTCTGAACTCTTTCCCTACACGACGCTCTTCCGATCTNNNGCTAGCTGAACCAGGAAACA
AGCACACGTCTGAACTCTTTCCCTACACGACGCTCTTCCGATCTNNNTCATTGAGTGGTGGGATTTCAC
GAGCACACGTCTGAACTCTTTCCCTACACGACGCTCTTCCGATCTNNNGTGGGCAGTAAAACAGGAC
AGCACACGTCTGAACTCTTTCCCTACACGACGCTCTTCCGATCTNNNTGGAGCAGGAATTTGAACTG
GCACACGTCTGAACTCTTTCCCTACACGACGCTCTTCCGATCTNNNAGAACCCAGAAGCCTCTTCTCTT
AGCACACGTCTGAACTCTTTCCCTACACGACGCTCTTCCGATCTNNNAGGCTCCTCATCAGATTCCCAT
GAGCACACGTCTGAACTCTTTCCCTACACGACGCTCTTCCGATCTNNNGGAGGAGTTGAGTGGAAA
AGCACACGTCTGAACTCTTTCCCTACACGACGCTCTTCCGATCTNNNTCGATGACATTCTCACCTCTGT
AGCACACGTCTGAACTCTTTCCCTACACGACGCTCTTCCGATCTNNNGTGTGTGTGTATGTGAACCCT
GAGCACACGTCTGAACTCTTTCCCTACACGACGCTCTTCCGATCTNNNGCACAGAGGCAGACACTGA
AGCACACGTCTGAACTCTTTCCCTACACGACGCTCTTCCGATCTNNNACATGAGGATACAGAGGTGG
AGCACACGTCTGAACTCTTTCCCTACACGACGCTCTTCCGATCTNNNAGCAAGTAGGGGAGAAGGAT
AGCACACGTCTGAACTCTTTCCCTACACGACGCTCTTCCGATCTNNNGCTCCTACTGAATCCAGAAAC
AGCACACGTCTGAACTCTTTCCCTACACGACGCTCTTCCGATCTNNNTATGGACAGTAAGCAAGCCCA
AGCACACGTCTGAACTCTTTCCCTACACGACGCTCTTCCGATCTNNNTTGCCACCTGTGGCTTTTTCTAT
GAGCACACGTCTGAACTCTTTCCCTACACGACGCTCTTCCGATCTNNNAGGTTCTCTTGAGAGACAGA
AGCACACGTCTGAACTCTTTCCCTACACGACGCTCTTCCGATCTNNNGACAATGCCCCCTCTAGACAAT
AGCACACGTCTGAACTCTTTCCCTACACGACGCTCTTCCGATCTNNNTGGATGTGGACAGACTAGAGC
AGCACACGTCTGAACTCTTTCCCTACACGACGCTCTTCCGATCTNNNCCAGAATTTTCTTCACTGTGGG
AGCACACGTCTGAACTCTTTCCCTACACGACGCTCTTCCGATCTNNNGATGGCTTGGATGGAAGGAA
GCACACGTCTGAACTCTTTCCCTACACGACGCTCTTCCGATCTNNNCCTTAGCTGTTGTTGGCATCTTC
GAGCACACGTCTGAACTCTTTCCCTACACGACGCTCTTCCGATCTNNNAGGAAACAAGGCTTACCTAG
AGCACACGTCTGAACTCTTTCCCTACACGACGCTCTTCCGATCTNNNAGCCTTTTTCATCACAGGGAAG
GCACACGTCTGAACTCTTTCCCTACACGACGCTCTTCCGATCTNNNGGACCTCAGAGAGACCTGAAAG
GAGCACACGTCTGAACTCTTTCCCTACACGACGCTCTTCCGATCTNNNCATGGGTTATCTGTGACGA
AGCACACGTCTGAACTCTTTCCCTACACGACGCTCTTCCGATCTNNNTTGCTTTTCCCTTTCTTTTGCCC
AGCACACGTCTGAACTCTTTCCCTACACGACGCTCTTCCGATCTNNNATGGACCCTCTTGAGCAACTGA
AGCACACGTCTGAACTCTTTCCCTACACGACGCTCTTCCGATCTNNNGCCAGACAGGAGGTTACTTAG
GAGCACACGTCTGAACTCTTTCCCTACACGACGCTCTTCCGATCTNNNAACATCTCCTTCCACTCCTCCT
AGCACACGTCTGAACTCTTTCCCTACACGACGCTCTTCCGATCTNNNAGGTGGCTGGAATTCTCTCTCT
AGCACACGTCTGAACTCTTTCCCTACACGACGCTCTTCCGATCTNNNCTGAGTGTGTGTGTGTCTGGA
GAGCACACGTCTGAACTCTTTCCCTACACGACGCTCTTCCGATCTNNNTTGTTCCCCATCCACCACATA
GAGCACACGTCTGAACTCTTTCCCTACACGACGCTCTTCCGATCTNNNAAATGATTTGGGGCAGGTC
GAGCACACGTCTGAACTCTTTCCCTACACGACGCTCTTCCGATCTNNNTTCAAAGAGATGCCTTCTTCG
GAGCACACGTCTGAACTCTTTCCCTACACGACGCTCTTCCGATCTNNNAAGCTCTCCTAGCGTCAGAGT
AGCACACGTCTGAACTCTTTCCCTACACGACGCTCTTCCGATCTNNNCTAGACGGAAGCTTGGAGCA
AGCACACGTCTGAACTCTTTCCCTACACGACGCTCTTCCGATCTNNNATTCTGAGGCATGCTACCTTTG

AGCACACGTCTGAACTCTTTCCCTACACGACGCTCTTCCGATCTNNNTGAGCAACCACTCAAGAGCAAA
AGCACACGTCTGAACTCTTTCCCTACACGACGCTCTTCCGATCTNNNAACATTACCCCTCTCATACCTGA
GCACACGTCTGAACTCTTTCCCTACACGACGCTCTTCCGATCTNNNGCATTGCATTCAGGAGGTTAGG
AGCACACGTCTGAACTCTTTCCCTACACGACGCTCTTCCGATCTNNNCTTGCCTTCAAGGAATTATGGC
GAGCACACGTCTGAACTCTTTCCCTACACGACGCTCTTCCGATCTNNNAAATGAGGCTGCCTTCTAGG
AGAGCACACGTCTGAACTCTTTCCCTACACGACGCTCTTCCGATCTNNNTAGTGACTTCTTCAGGGGTA
AGAGCACACGTCTGAACTCTTTCCCTACACGACGCTCTTCCGATCTNNNTGGGAAGGTGCATCTCAGT
GAGCACACGTCTGAACTCTTTCCCTACACGACGCTCTTCCGATCTNNNCTGGCCAACAGAGACAGTTAC
GAGCACACGTCTGAACTCTTTCCCTACACGACGCTCTTCCGATCTNNNCCCTACCACAGGCTCATCATT
AGAGCACACGTCTGAACTCTTTCCCTACACGACGCTCTTCCGATCTNNNAAAACAACTCAGCATTCTGG
GAGCACACGTCTGAACTCTTTCCCTACACGACGCTCTTCCGATCTNNNAATGACCTTTCAGGTGGTGCA
AGCACACGTCTGAACTCTTTCCCTACACGACGCTCTTCCGATCTNNNCCCACTTTCAGAACCTGCTCTTT
GCACACGTCTGAACTCTTTCCCTACACGACGCTCTTCCGATCTNNNAGAGAAGAAAACCCTGAACCAG
AGAGCACACGTCTGAACTCTTTCCCTACACGACGCTCTTCCGATCTNNNTAGCCATATTTGGAGGATG
AGCACACGTCTGAACTCTTTCCCTACACGACGCTCTTCCGATCTNNNTCTAAGGTTGAGCAAATGCATA
AGCACACGTCTGAACTCTTTCCCTACACGACGCTCTTCCGATCTNNNCCACTAGGCCTCTTTTGAATGCT
AGCACACGTCTGAACTCTTTCCCTACACGACGCTCTTCCGATCTNNNATCATTTGCCCATTTTGAATTGC
AGCACACGTCTGAACTCTTTCCCTACACGACGCTCTTCCGATCTNNNTCATGCAGGGTAGGAAAGACA
GAGCACACGTCTGAACTCTTTCCCTACACGACGCTCTTCCGATCTNNNTAATTTGCACCTAGTCCTGTG
GAGCACACGTCTGAACTCTTTCCCTACACGACGCTCTTCCGATCTNNNTAAGCACTGTGTACAGCAAG
GAGCACACGTCTGAACTCTTTCCCTACACGACGCTCTTCCGATCTNNNACAGCACCTGGTCCAATTTTCT
AGCACACGTCTGAACTCTTTCCCTACACGACGCTCTTCCGATCTNNNCTGGTCACATATGCAGAAGCCA
GAGCACACGTCTGAACTCTTTCCCTACACGACGCTCTTCCGATCTNNNCGTCTACAACAACACCCTCAAC
AGAGCACACGTCTGAACTCTTTCCCTACACGACGCTCTTCCGATCTNNNGCATCCTGATCCAGGTACAA
AGCACACGTCTGAACTCTTTCCCTACACGACGCTCTTCCGATCTNNNCACCTCTACCTGTCTGAACTTGC
AGAGCACACGTCTGAACTCTTTCCCTACACGACGCTCTTCCGATCTNNNGCATCAGATAAGCCAGAGGT
GAGCACACGTCTGAACTCTTTCCCTACACGACGCTCTTCCGATCTNNNCCTTGGCAGTGACTTTGAAAC
GAGCACACGTCTGAACTCTTTCCCTACACGACGCTCTTCCGATCTNNNAGTTCACAAGTGTGCTCTGGT
GAGCACACGTCTGAACTCTTTCCCTACACGACGCTCTTCCGATCTNNNTCTCAACACCATGGAAAAGTT
AGCACACGTCTGAACTCTTTCCCTACACGACGCTCTTCCGATCTNNNTGTCCAGTTCCCAACAATTGCAT
GAGCACACGTCTGAACTCTTTCCCTACACGACGCTCTTCCGATCTNNNTAGCCATGGTTTCCTTCATCTG
GAGCACACGTCTGAACTCTTTCCCTACACGACGCTCTTCCGATCTNNNACCTAGACTGACTTCAGGGAA
AGAGCACACGTCTGAACTCTTTCCCTACACGACGCTCTTCCGATCTNNNCACTCGTGTGTCCTTCATCAT
GAGCACACGTCTGAACTCTTTCCCTACACGACGCTCTTCCGATCTNNNTTAGCCTAGCACAACAGTACA
GAGCACACGTCTGAACTCTTTCCCTACACGACGCTCTTCCGATCTNNNCTATCAAACAAATGCCGAGGT
AGAGCACACGTCTGAACTCTTTCCCTACACGACGCTCTTCCGATCTNNNAGAGAAAAGAAACAGCAGA
GAGCACACGTCTGAACTCTTTCCCTACACGACGCTCTTCCGATCTNNNATGTTTGGGTATGGAGCTTGT
GAGCACACGTCTGAACTCTTTCCCTACACGACGCTCTTCCGATCTNNNAAGTTCAAAGCCTGGTGAGA
AGAGCACACGTCTGAACTCTTTCCCTACACGACGCTCTTCCGATCTNNNGGGGATGGAGTACACAGA
AGCACACGTCTGAACTCTTTCCCTACACGACGCTCTTCCGATCTNNNAACTCCTCAGCAAAGTCGATAG
GAGCACACGTCTGAACTCTTTCCCTACACGACGCTCTTCCGATCTNNNGACGTTATTTGGAGGTAACG
GAGCACACGTCTGAACTCTTTCCCTACACGACGCTCTTCCGATCTNNNCACAGAATCACTCAGGATGCC
GAGCACACGTCTGAACTCTTTCCCTACACGACGCTCTTCCGATCTNNNCTCCCCAAGGTTTTGGAAAAG

GAGCACACGTCTGAACTCTTTCCCTACACGACGCTCTTCCGATCTNNNGTTCAGTAGGAATTCGCTCA
GAGCACACGTCTGAACTCTTTCCCTACACGACGCTCTTCCGATCTNNNATCCCTACTGCCTTCTGGACTT
GAGCACACGTCTGAACTCTTTCCCTACACGACGCTCTTCCGATCTNNNGAATCCTTTTCACAACAGGCA
AGAGCACACGTCTGAACTCTTTCCCTACACGACGCTCTTCCGATCTNNNTAATACAAATGGGCTGACTG
AGCACACGTCTGAACTCTTTCCCTACACGACGCTCTTCCGATCTNNNGCCAAAGTTGGAACAAAGTGT
AGAGCACACGTCTGAACTCTTTCCCTACACGACGCTCTTCCGATCTNNNGATTCTGACTGTAGTGCTTC
AGAGCACACGTCTGAACTCTTTCCCTACACGACGCTCTTCCGATCTNNNTGAAGGGAAAACACAAAAA
AGAGCACACGTCTGAACTCTTTCCCTACACGACGCTCTTCCGATCTNNNCCCCAGCCTTTCATGCTCTATT
AGCACACGTCTGAACTCTTTCCCTACACGACGCTCTTCCGATCTNNNGAAGAGGGGGAGAAAGGAGA
AGCACACGTCTGAACTCTTTCCCTACACGACGCTCTTCCGATCTNNNCGACTTGAACTTCTCAAATGCC
AGCACACGTCTGAACTCTTTCCCTACACGACGCTCTTCCGATCTNNNACTGCCCCTGGATCATCTGTATT
GAGCACACGTCTGAACTCTTTCCCTACACGACGCTCTTCCGATCTNNNACCAAGTTAGTGAGCACAGCA
GAGCACACGTCTGAACTCTTTCCCTACACGACGCTCTTCCGATCTNNNCCTGGCATCCTTCCTGTATCCTA
AGAGCACACGTCTGAACTCTTTCCCTACACGACGCTCTTCCGATCTNNNAATGGAGACTACCATGACGT
GAGCACACGTCTGAACTCTTTCCCTACACGACGCTCTTCCGATCTNNNTGATGTTGTGGTGACGATGA
GAGCACACGTCTGAACTCTTTCCCTACACGACGCTCTTCCGATCTNNNGTAAGCGTTTCTGTCTCTCCCT
GAGCACACGTCTGAACTCTTTCCCTACACGACGCTCTTCCGATCTNNNGCCAATGCCTGTTTGGATTAC
GAGCACACGTCTGAACTCTTTCCCTACACGACGCTCTTCCGATCTNNNAAACAGAAGTCTACCACCCCT
AGCACACGTCTGAACTCTTTCCCTACACGACGCTCTTCCGATCTNNNGTTTGCCAGTTGGCAGTCAAAA
GAGCACACGTCTGAACTCTTTCCCTACACGACGCTCTTCCGATCTNNNGTAGACCCCAAAGCACTATGT
GAGCACACGTCTGAACTCTTTCCCTACACGACGCTCTTCCGATCTNNNAAACTCACACACACTCTCTCCCT
GAGCACACGTCTGAACTCTTTCCCTACACGACGCTCTTCCGATCTNNNAACAAGGCCTGCAAATAGAA
AGCACACGTCTGAACTCTTTCCCTACACGACGCTCTTCCGATCTNNNGATGTGGGAGGAAAACATGA
AGCACACGTCTGAACTCTTTCCCTACACGACGCTCTTCCGATCTNNNCAACCTTATTTGTCCTGACAGCC
GAGCACACGTCTGAACTCTTTCCCTACACGACGCTCTTCCGATCTNNNACCAGGAAACTTTTGAGATGG
GAGCACACGTCTGAACTCTTTCCCTACACGACGCTCTTCCGATCTNNNAGCCTGCAATAAACTTGGTGT
GAGCACACGTCTGAACTCTTTCCCTACACGACGCTCTTCCGATCTNNNGACAGTGACAGCTGTAGCAGA
GCACACGTCTGAACTCTTTCCCTACACGACGCTCTTCCGATCTNNNCAGAGCATGTCTGTCTGTCTGTC
GAGCACACGTCTGAACTCTTTCCCTACACGACGCTCTTCCGATCTNNNGTGTGGTAGTCTGGGTAGTCT
AGAGCACACGTCTGAACTCTTTCCCTACACGACGCTCTTCCGATCTNNNTCATGAGGCTGCTTTCTCAGT
AGAGCACACGTCTGAACTCTTTCCCTACACGACGCTCTTCCGATCTNNNTGATCTTTCTGCTGTCTAGC
GAGCACACGTCTGAACTCTTTCCCTACACGACGCTCTTCCGATCTNNNTGTTTGAGGGGAAGCAAGTA
AGCACACGTCTGAACTCTTTCCCTACACGACGCTCTTCCGATCTNNNGGAAGGCAACAGATATTGCAG
AGCACACGTCTGAACTCTTTCCCTACACGACGCTCTTCCGATCTNNNATCAATTGCCAAATGTACCCAC
AGCACACGTCTGAACTCTTTCCCTACACGACGCTCTTCCGATCTNNNGCATGCAATTTTCTGCAAAGCT
AGCACACGTCTGAACTCTTTCCCTACACGACGCTCTTCCGATCTNNNTATCTATAAGTGCGTGTGTGC
AGCACACGTCTGAACTCTTTCCCTACACGACGCTCTTCCGATCTNNNCAGAGGCAACTTTAGCCTCCAA
GAGCACACGTCTGAACTCTTTCCCTACACGACGCTCTTCCGATCTNNNTGACATGGGCCCCTTTGATAT
AGCACACGTCTGAACTCTTTCCCTACACGACGCTCTTCCGATCTNNNACTTACTGATGTGTTTCCCCTGT
AGCACACGTCTGAACTCTTTCCCTACACGACGCTCTTCCGATCTNNNCTTATAAGCATCCAGCCTCTGCC
GAGCACACGTCTGAACTCTTTCCCTACACGACGCTCTTCCGATCTNNNCTGTCTGCAGCTTCTATCTTCC
GAGCACACGTCTGAACTCTTTCCCTACACGACGCTCTTCCGATCTNNNATTCCTTCCTAGGAGCCCATCA
GAGCACACGTCTGAACTCTTTCCCTACACGACGCTCTTCCGATCTNNNGCTGCTGTCCTGATGTCTTGT

GAGCACACGTCTGAACTCTTTCCCTACACGACGCTCTTCCGATCTNNNGACCTTCTGAGTTTGAGGCAG
AGCACACGTCTGAACTCTTTCCCTACACGACGCTCTTCCGATCTNNNCCATGAATCCCAGCAATTAGCC
AGCACACGTCTGAACTCTTTCCCTACACGACGCTCTTCCGATCTNNNTACAGTGGCCAGTCAGATTTTG
GAGCACACGTCTGAACTCTTTCCCTACACGACGCTCTTCCGATCTNNNTCCCTTGATAGCCACTTTGAG
AGCACACGTCTGAACTCTTTCCCTACACGACGCTCTTCCGATCTNNNGCAGGATTGACTGAAGGAGGT
AGAGCACACGTCTGAACTCTTTCCCTACACGACGCTCTTCCGATCTNNNTAGAAAGGCTGGTGACTTA
GAGCACACGTCTGAACTCTTTCCCTACACGACGCTCTTCCGATCTNNNGGTGAGGATTTGGGGACAA
GAGCACACGTCTGAACTCTTTCCCTACACGACGCTCTTCCGATCTNNNTCACTCTGGTGTCTGCTATCT
AGAGCACACGTCTGAACTCTTTCCCTACACGACGCTCTTCCGATCTNNNGAGTTGAATGAGGTCATTG
AAGAGCACACGTCTGAACTCTTTCCCTACACGACGCTCTTCCGATCTNNNTTACTGCCCTCTCCTCACTT
GAGCACACGTCTGAACTCTTTCCCTACACGACGCTCTTCCGATCTNNNGGGGATATGTTCTCAGGCTT
AGAGCACACGTCTGAACTCTTTCCCTACACGACGCTCTTCCGATCTNNNTCCCGCCCCCAAATCAAATAT
GAGCACACGTCTGAACTCTTTCCCTACACGACGCTCTTCCGATCTNNNCCTACTGCTTACATCCCAAGCT
AGCACACGTCTGAACTCTTTCCCTACACGACGCTCTTCCGATCTNNNGCTGACTGCCAGGTTAGAATGA
AGAGCACACGTCTGAACTCTTTCCCTACACGACGCTCTTCCGATCTNNNGCCCATCTTCCATTGGTGTAT
AGAGCACACGTCTGAACTCTTTCCCTACACGACGCTCTTCCGATCTNNNGGCAGCATTGTGAAAACCCA
AGCACACGTCTGAACTCTTTCCCTACACGACGCTCTTCCGATCTNNNAAAGTGTCCCACATCCTTTTCCC
GAGCACACGTCTGAACTCTTTCCCTACACGACGCTCTTCCGATCTNNNTAATGGAGGAGGGTGGTTT
GAGCACACGTCTGAACTCTTTCCCTACACGACGCTCTTCCGATCTNNNACTGACATGCCCTTGACAAGT
GAGCACACGTCTGAACTCTTTCCCTACACGACGCTCTTCCGATCTNNNTGCAAATGTTTTCCTCAGTTG
AGCACACGTCTGAACTCTTTCCCTACACGACGCTCTTCCGATCTNNNCTGAGGCACATTCAGATCTTCC
GAGCACACGTCTGAACTCTTTCCCTACACGACGCTCTTCCGATCTNNNGCTGCTGACTTGGAAAAACTC
GAGCACACGTCTGAACTCTTTCCCTACACGACGCTCTTCCGATCTNNNACTTTTTCTTGGAGGCTGACA
AGAGCACACGTCTGAACTCTTTCCCTACACGACGCTCTTCCGATCTNNNTGTATCCCAAAGAACAGGTT
AGCACACGTCTGAACTCTTTCCCTACACGACGCTCTTCCGATCTNNNAGAGCTTGCAGTGAGTAGACA
AGAGCACACGTCTGAACTCTTTCCCTACACGACGCTCTTCCGATCTNNNTTAGGGAGGCAGAAGCAA
AGAGCACACGTCTGAACTCTTTCCCTACACGACGCTCTTCCGATCTNNNCTGGGCTTTGGTTTGGAGTA
AGCACACGTCTGAACTCTTTCCCTACACGACGCTCTTCCGATCTNNNACGAACAAACAACCAACCAACA
AGCACACGTCTGAACTCTTTCCCTACACGACGCTCTTCCGATCTNNNCTTACAGACACCACACGCACAT
AGAGCACACGTCTGAACTCTTTCCCTACACGACGCTCTTCCGATCTNNNTGTTCAAGAGGAGTTTTGG
AGAGCACACGTCTGAACTCTTTCCCTACACGACGCTCTTCCGATCTNNNGTACAGCTTTACTGAAGCAC
GAGCACACGTCTGAACTCTTTCCCTACACGACGCTCTTCCGATCTNNNGGTGACAGAGCTGGACACTG
GAGCACACGTCTGAACTCTTTCCCTACACGACGCTCTTCCGATCTNNNGCAAGGTTGCTGGATATTGA
GAGCACACGTCTGAACTCTTTCCCTACACGACGCTCTTCCGATCTNNNATGGAAGGGGAAGACGGAC
GAGCACACGTCTGAACTCTTTCCCTACACGACGCTCTTCCGATCTNNNTGAGTTGGAGAGACAGCCAA
AGAGCACACGTCTGAACTCTTTCCCTACACGACGCTCTTCCGATCTNNNTACTGACATGGCAACATTTA
GCACACGTCTGAACTCTTTCCCTACACGACGCTCTTCCGATCTNNNCCATCAGCTTGCTTTGGTCTGTAC
GCACACGTCTGAACTCTTTCCCTACACGACGCTCTTCCGATCTNNNCAGAGCCAGAGTTAGTAGGGAG
AGAGCACACGTCTGAACTCTTTCCCTACACGACGCTCTTCCGATCTNNNAGATGTGTCCCGTCTCCTCTT
AGCACACGTCTGAACTCTTTCCCTACACGACGCTCTTCCGATCTNNNAACCTCAGAGGAAGAATGGGT
AAGAGCACACGTCTGAACTCTTTCCCTACACGACGCTCTTCCGATCTNNNATGAAATCAGTGTTGCCTT
GAGCACACGTCTGAACTCTTTCCCTACACGACGCTCTTCCGATCTNNNATGAGGCTCACCAAATGTGTG
GAGCACACGTCTGAACTCTTTCCCTACACGACGCTCTTCCGATCTNNNACAACCTGTGTTCGTCAATGG

GAGCACACGTCTGAACTCTTTCCCTACACGACGCTCTTCCGATCTNNNGTGAGCGTTCCTCCTCTTTTC
AGCACACGTCTGAACTCTTTCCCTACACGACGCTCTTCCGATCTNNNGGAGGAGGAGGAGTATGAG
AGCACACGTCTGAACTCTTTCCCTACACGACGCTCTTCCGATCTNNNCCTTTCCCTATTTTGGGCACAC
AGCACACGTCTGAACTCTTTCCCTACACGACGCTCTTCCGATCTNNNTGACGGAACGAAGAGGGATT
AGAGCACACGTCTGAACTCTTTCCCTACACGACGCTCTTCCGATCTNNNTTTGAAAGGTTCTCAGGAC
GAGCACACGTCTGAACTCTTTCCCTACACGACGCTCTTCCGATCTNNNGAAGTCAGGTGGGAAAGG
AGCACACGTCTGAACTCTTTCCCTACACGACGCTCTTCCGATCTNNNCTTTTCCTTTGCTTTGAGGGC
AGCACACGTCTGAACTCTTTCCCTACACGACGCTCTTCCGATCTNNNTTCTGCACAATCAGCATTACC
GAGCACACGTCTGAACTCTTTCCCTACACGACGCTCTTCCGATCTNNNGGGTAAACAGGCCTGAAAA
GAGCACACGTCTGAACTCTTTCCCTACACGACGCTCTTCCGATCTNNNAGTGGTAATGCAAGAGGAG
GCACACGTCTGAACTCTTTCCCTACACGACGCTCTTCCGATCTNNNTCGCGAATGGATCTCCTTCTTTC
GCACACGTCTGAACTCTTTCCCTACACGACGCTCTTCCGATCTNNNCGTGGGGACTATGGTGACTAC
GAGCACACGTCTGAACTCTTTCCCTACACGACGCTCTTCCGATCTNNNCATATTCATACACACGCACGC
AGAGCACACGTCTGAACTCTTTCCCTACACGACGCTCTTCCGATCTNNNTCTTGTTTGCTTCCCGGCTTT
GAGCACACGTCTGAACTCTTTCCCTACACGACGCTCTTCCGATCTNNNATGGGAGGATGTGACAGAA
AGAGCACACGTCTGAACTCTTTCCCTACACGACGCTCTTCCGATCTNNNAGCTCATAAGTGCTTGGGC
GAGCACACGTCTGAACTCTTTCCCTACACGACGCTCTTCCGATCTNNNCTACACACCCAATCACATGGG
GCACACGTCTGAACTCTTTCCCTACACGACGCTCTTCCGATCTNNNCTTTCACCTTATCCCCCATCTGCC
AGCACACGTCTGAACTCTTTCCCTACACGACGCTCTTCCGATCTNNNCAACATTCACAGTCACGTTCAC
AGCACACGTCTGAACTCTTTCCCTACACGACGCTCTTCCGATCTNNNCTTAGCCAGCCTGATGATGTC
GAGCACACGTCTGAACTCTTTCCCTACACGACGCTCTTCCGATCTNNNTAGCAATACCCCCTCGGTACA
AGCACACGTCTGAACTCTTTCCCTACACGACGCTCTTCCGATCTNNNCACCCAGGCCTTGTGCATATAAT
AAGAGCACACGTCTGAACTCTTTCCCTACACGACGCTCTTCCGATCTNNNTGGAGTTGGAAAATGAAA
AGAGCACACGTCTGAACTCTTTCCCTACACGACGCTCTTCCGATCTNNNTTCAACAGCTGGAGCATGA
AGCACACGTCTGAACTCTTTCCCTACACGACGCTCTTCCGATCTNNNTGAGGGGAAATGGATGGGGT
GAGCACACGTCTGAACTCTTTCCCTACACGACGCTCTTCCGATCTNNNGAATTGCCTGTGAATGCCCA
AGCACACGTCTGAACTCTTTCCCTACACGACGCTCTTCCGATCTNNNTCATCTCATAACCCTCTGACCCC
GAGCACACGTCTGAACTCTTTCCCTACACGACGCTCTTCCGATCTNNNCCCCATCTCCCAAAACAAGAA
AGCACACGTCTGAACTCTTTCCCTACACGACGCTCTTCCGATCTNNNACCTCCCTCTCTCTTCTCCCTTA
GAGCACACGTCTGAACTCTTTCCCTACACGACGCTCTTCCGATCTNNNGGCAACTTCCTCATGCTTTTG
GAGCACACGTCTGAACTCTTTCCCTACACGACGCTCTTCCGATCTNNNATGTATGAAGGGGGAAAAA
AGCACACGTCTGAACTCTTTCCCTACACGACGCTCTTCCGATCTNNNAGGATCCAAGGGTGACAGGT
AGCACACGTCTGAACTCTTTCCCTACACGACGCTCTTCCGATCTNNNCCAGCTTGCATTCTAAGTGCTT
GAGCACACGTCTGAACTCTTTCCCTACACGACGCTCTTCCGATCTNNNTGATAGCCTGCCTCTCTTCTCT
GAGCACACGTCTGAACTCTTTCCCTACACGACGCTCTTCCGATCTNNNATCAGAATCCCAACAGAGGG
GAGCACACGTCTGAACTCTTTCCCTACACGACGCTCTTCCGATCTNNNAATGTGAGTGGGCTGTCAAT
AGCACACGTCTGAACTCTTTCCCTACACGACGCTCTTCCGATCTNNNGCCAAATCAGGAATGCAATAG
GCACACGTCTGAACTCTTTCCCTACACGACGCTCTTCCGATCTNNNGATGAGTTCTGGGAACACTTGG
GAGCACACGTCTGAACTCTTTCCCTACACGACGCTCTTCCGATCTNNNAAAAACGCCTGGCTAGAGTT
AGCACACGTCTGAACTCTTTCCCTACACGACGCTCTTCCGATCTNNNCACAAGATCTGCCCTCAAGATC
AGCACACGTCTGAACTCTTTCCCTACACGACGCTCTTCCGATCTNNNGGGGAGAACTTGCATTCTGGA
GCACACGTCTGAACTCTTTCCCTACACGACGCTCTTCCGATCTNNNGGGGCATTATCCTTCACCACAA
GAGCACACGTCTGAACTCTTTCCCTACACGACGCTCTTCCGATCTNNNTCTGAAGAGCTCAATACTGT

iAGCACACGTCTGAACTCTTTCCCTACACGACGCTCTTCCGATCTNNNGACAGGTTTGGAATCAGGCTT
iAGCACACGTCTGAACTCTTTCCCTACACGACGCTCTTCCGATCTNNNTCTCTAACAGCTTGCAGGTCAA
AGAGCACACGTCTGAACTCTTTCCCTACACGACGCTCTTCCGATCTNNNACCTTGTTGGAGTACAGAAG
iAGCACACGTCTGAACTCTTTCCCTACACGACGCTCTTCCGATCTNNNTTTTAGCCTGATTCCACAGCCA
AGCACACGTCTGAACTCTTTCCCTACACGACGCTCTTCCGATCTNNNAGTGTCTGAGCATAGAGTTGG
AGCACACGTCTGAACTCTTTCCCTACACGACGCTCTTCCGATCTNNNTGGTGAAAGGCACCTAGAACA
AGCACACGTCTGAACTCTTTCCCTACACGACGCTCTTCCGATCTNNNTAGAATCCCAAGCCAAGTCAGG
AGCACACGTCTGAACTCTTTCCCTACACGACGCTCTTCCGATCTNNNCAGAGGAGGCAAACTGTGATG
GAGCACACGTCTGAACTCTTTCCCTACACGACGCTCTTCCGATCTNNNTGAAGCTGTGTTGTTCAGAG
GAGCACACGTCTGAACTCTTTCCCTACACGACGCTCTTCCGATCTNNNGGGTGAAAGTTCTGTGTACG
AGAGCACACGTCTGAACTCTTTCCCTACACGACGCTCTTCCGATCTNNNTGGCTGTCATGTAGCAATCC
iAGCACACGTCTGAACTCTTTCCCTACACGACGCTCTTCCGATCTNNNGGCCATCTGCTCAGTAGGTCT
AGCACACGTCTGAACTCTTTCCCTACACGACGCTCTTCCGATCTNNNGGGAGATAGGATGGAGACACC
AGAGCACACGTCTGAACTCTTTCCCTACACGACGCTCTTCCGATCTNNNTGCGTTTGTCTCAGTAAGCA
AGAGCACACGTCTGAACTCTTTCCCTACACGACGCTCTTCCGATCTNNNTAACTTTCAGGACAGGTAG
iAGCACACGTCTGAACTCTTTCCCTACACGACGCTCTTCCGATCTNNNGTGAACACCCCAGTGGTCAAT
GAGCACACGTCTGAACTCTTTCCCTACACGACGCTCTTCCGATCTNNNGGTCTATCTTTCACCCCTCCCTT
iAGCACACGTCTGAACTCTTTCCCTACACGACGCTCTTCCGATCTNNNTGACATTATCGTTTGCCAGTCC
iAGCACACGTCTGAACTCTTTCCCTACACGACGCTCTTCCGATCTNNNTGACATACCCCCAACTCTCTCTC
AGCACACGTCTGAACTCTTTCCCTACACGACGCTCTTCCGATCTNNNCTGTTTCTCCTAACATCCTGGGC
iAGCACACGTCTGAACTCTTTCCCTACACGACGCTCTTCCGATCTNNNCAGTAGTGTTGAGCAGAGCTG
GAGCACACGTCTGAACTCTTTCCCTACACGACGCTCTTCCGATCTNNNGTTTACTTCTCCCTCTGCCTGT
GAGCACACGTCTGAACTCTTTCCCTACACGACGCTCTTCCGATCTNNNGAGGGAAAGGAGGTGAGAG
GAGCACACGTCTGAACTCTTTCCCTACACGACGCTCTTCCGATCTNNNTTGTGTTGATGGTAACCCAAA
GAGCACACGTCTGAACTCTTTCCCTACACGACGCTCTTCCGATCTNNNGCCTCTGCAGTCCAGATTTGA
GAGCACACGTCTGAACTCTTTCCCTACACGACGCTCTTCCGATCTNNNAACTTGCCTAGTCACCCCTACA
GAGCACACGTCTGAACTCTTTCCCTACACGACGCTCTTCCGATCTNNNGTCCAGGTGACTTGTGTATG
AGCACACGTCTGAACTCTTTCCCTACACGACGCTCTTCCGATCTNNNTATTGATCCGCCCCAATTGTCTC
GAGCACACGTCTGAACTCTTTCCCTACACGACGCTCTTCCGATCTNNNCCAGCCAGAAGATGACCACAT
GAGCACACGTCTGAACTCTTTCCCTACACGACGCTCTTCCGATCTNNNGGGAGCCGTACACAAATCTCA
AGAGCACACGTCTGAACTCTTTCCCTACACGACGCTCTTCCGATCTNNNACTACTGAGCATACCTAGGT
GAGCACACGTCTGAACTCTTTCCCTACACGACGCTCTTCCGATCTNNNGGATGAGGACACACATGCTA
iAGCACACGTCTGAACTCTTTCCCTACACGACGCTCTTCCGATCTNNNACCAAAGCACTGTACACTTCAA
GAGCACACGTCTGAACTCTTTCCCTACACGACGCTCTTCCGATCTNNNTTGGAAGTTTGGTGAGGTGT
iAGCACACGTCTGAACTCTTTCCCTACACGACGCTCTTCCGATCTNNNAGCCTAGTCTAACATCACCACA
AGCACACGTCTGAACTCTTTCCCTACACGACGCTCTTCCGATCTNNNAGAGGTGTATTCCCCTTTGAGG
AGCACACGTCTGAACTCTTTCCCTACACGACGCTCTTCCGATCTNNNCTGGATCTTGGACACTGTCTGG
AGAGCACACGTCTGAACTCTTTCCCTACACGACGCTCTTCCGATCTNNNGATGCAGTTTGATCCCTGGC
AAGAGCACACGTCTGAACTCTTTCCCTACACGACGCTCTTCCGATCTNNNCTCCATAACCCCTCCCCCATT
AGAGCACACGTCTGAACTCTTTCCCTACACGACGCTCTTCCGATCTNNNGGTGTTGTAGGATTCGGTTT
AGAGCACACGTCTGAACTCTTTCCCTACACGACGCTCTTCCGATCTNNNAACTGACCCCTGTTTGCACT
GAGCACACGTCTGAACTCTTTCCCTACACGACGCTCTTCCGATCTNNNTGGATTTCAGGTGATTGCGAT
AGCACACGTCTGAACTCTTTCCCTACACGACGCTCTTCCGATCTNNNTTAGGGTCAGACCAGTGCCTTT

GAGCACACGTCTGAACTCTTTCCCTACACGACGCTCTTCCGATCTNNNCTAGAACCCGATCCACGTCTT
AGAGCACACGTCTGAACTCTTTCCCTACACGACGCTCTTCCGATCTNNNACCCTTGCGATGAAAGGTG
AGAGCACACGTCTGAACTCTTTCCCTACACGACGCTCTTCCGATCTNNNATGATTAGTTGTGCCAGAT
GCACACGTCTGAACTCTTTCCCTACACGACGCTCTTCCGATCTNNNCATTCCAGTGCATCATCTTGTGC
AGCACACGTCTGAACTCTTTCCCTACACGACGCTCTTCCGATCTNNNCAGCTGGTGGGATTCTAACCT
AGCACACGTCTGAACTCTTTCCCTACACGACGCTCTTCCGATCTNNNGGCTCTTGCTTCCATACCTTCTC
AGCACACGTCTGAACTCTTTCCCTACACGACGCTCTTCCGATCTNNNATAGACAGCCTACAGCCAGACA
GAGCACACGTCTGAACTCTTTCCCTACACGACGCTCTTCCGATCTNNNCCTGGACCAGACAGCAAAGA
AGCACACGTCTGAACTCTTTCCCTACACGACGCTCTTCCGATCTNNNTCCAGAGGGGAAGAGGATATT
AGCACACGTCTGAACTCTTTCCCTACACGACGCTCTTCCGATCTNNNCGTCAGCACAAGAAATTGCTC
GAGCACACGTCTGAACTCTTTCCCTACACGACGCTCTTCCGATCTNNNAGCTCGTGAGAATACTGCTT
GAGCACACGTCTGAACTCTTTCCCTACACGACGCTCTTCCGATCTNNNTTCTCTCCACCCCTAGAGTGTT
GAGCACACGTCTGAACTCTTTCCCTACACGACGCTCTTCCGATCTNNNATGTGGGTTTATGGATTGCT
AGCACACGTCTGAACTCTTTCCCTACACGACGCTCTTCCGATCTNNNTCTCTTTGTAGGGGCAGCATTT
AGCACACGTCTGAACTCTTTCCCTACACGACGCTCTTCCGATCTNNNACAGGATCTGGTCCTACAGGA
AGCACACGTCTGAACTCTTTCCCTACACGACGCTCTTCCGATCTNNNTCAGCAAACCCCAAAAAGAATG
GAGCACACGTCTGAACTCTTTCCCTACACGACGCTCTTCCGATCTNNNTGTTGGTGGTCTCAAGTCTGT
AGCACACGTCTGAACTCTTTCCCTACACGACGCTCTTCCGATCTNNNATCATGTTCTGGACTTCCTACG
AGCACACGTCTGAACTCTTTCCCTACACGACGCTCTTCCGATCTNNNGGAGGAAGGGTTGCATTTTAG
GCACACGTCTGAACTCTTTCCCTACACGACGCTCTTCCGATCTNNNACGTTCAGCCATGGAGATGATT
AGCACACGTCTGAACTCTTTCCCTACACGACGCTCTTCCGATCTNNNCTCTCCCCTTTTTCTGGCTACAC
AGCACACGTCTGAACTCTTTCCCTACACGACGCTCTTCCGATCTNNNAGAGACAACATGGAAAGGAAG
AGCACACGTCTGAACTCTTTCCCTACACGACGCTCTTCCGATCTNNNTTTCTTTTGGTGGCAGGGAGT
GAGCACACGTCTGAACTCTTTCCCTACACGACGCTCTTCCGATCTNNNGTGGGGAGATGGTTTCTCTC
AGCACACGTCTGAACTCTTTCCCTACACGACGCTCTTCCGATCTNNNACACATATCTACTGGATCGTGG
GAGCACACGTCTGAACTCTTTCCCTACACGACGCTCTTCCGATCTNNNAGGACAGTTCAGAGATGCTG
GAGCACACGTCTGAACTCTTTCCCTACACGACGCTCTTCCGATCTNNNGAGCTCTGTGATTAGCAGGA
AGAGCACACGTCTGAACTCTTTCCCTACACGACGCTCTTCCGATCTNNNTGAAGCCCTACGTATCCAAA
GAGCACACGTCTGAACTCTTTCCCTACACGACGCTCTTCCGATCTNNNGGGCTCTCCCATTCTGTTTCA
AGCACACGTCTGAACTCTTTCCCTACACGACGCTCTTCCGATCTNNNTTCCCATTCCAGCCTCACAAAA
AGCACACGTCTGAACTCTTTCCCTACACGACGCTCTTCCGATCTNNNAGGATGACTGTAATACCCAGC
GAGCACACGTCTGAACTCTTTCCCTACACGACGCTCTTCCGATCTNNNTCTGCTTTATTCCTCTCTGGCC
GAGCACACGTCTGAACTCTTTCCCTACACGACGCTCTTCCGATCTNNNACCACACCCTACCACTTCACTA
AGCACACGTCTGAACTCTTTCCCTACACGACGCTCTTCCGATCTNNNGCCTACAACATAGACCATGTCCC
GAGCACACGTCTGAACTCTTTCCCTACACGACGCTCTTCCGATCTNNNAAACCACCTAGTGTTACAGG
AGAGCACACGTCTGAACTCTTTCCCTACACGACGCTCTTCCGATCTNNNAGATGGCCTAGTAAAGCAT
AGAGCACACGTCTGAACTCTTTCCCTACACGACGCTCTTCCGATCTNNNAGTGATTGCTGACTGTTCAG
AGCACACGTCTGAACTCTTTCCCTACACGACGCTCTTCCGATCTNNNCCATGTGCACTGTAGGTAGCA
AGAGCACACGTCTGAACTCTTTCCCTACACGACGCTCTTCCGATCTNNNATTTTCCCCAATGCCCACAAA
AGAGCACACGTCTGAACTCTTTCCCTACACGACGCTCTTCCGATCTNNNTGGCTTCTTATGCCTAACAC
GAGCACACGTCTGAACTCTTTCCCTACACGACGCTCTTCCGATCTNNNACTGGTGCTCTTGAAAGACA
GAGCACACGTCTGAACTCTTTCCCTACACGACGCTCTTCCGATCTNNNTTGTCTAGGCAGAAGTGGT
GAGCACACGTCTGAACTCTTTCCCTACACGACGCTCTTCCGATCTNNNATATTCTTGAGAGCTTGGCCA

GAGCACACGTCTGAACTCTTTCCCTACACGACGCTCTTCCGATCTNNNAAGGTCACTTTTCTCTCACAG
GAGCACACGTCTGAACTCTTTCCCTACACGACGCTCTTCCGATCTNNNCCTATTTCCCAACTCCACTGCTC
GAGCACACGTCTGAACTCTTTCCCTACACGACGCTCTTCCGATCTNNNGCCTATATACATGCGCACACA
GAGCACACGTCTGAACTCTTTCCCTACACGACGCTCTTCCGATCTNNNTTAGGGGATTCTGGGGAGAA
GAGCACACGTCTGAACTCTTTCCCTACACGACGCTCTTCCGATCTNNNCCATCATTAGCCACAATGCCA
AGAGCACACGTCTGAACTCTTTCCCTACACGACGCTCTTCCGATCTNNNGCAGGACAACTTTGTGCTTTT
AGCACACGTCTGAACTCTTTCCCTACACGACGCTCTTCCGATCTNNNACAGACACCAAGCAATATGCTC
GAGCACACGTCTGAACTCTTTCCCTACACGACGCTCTTCCGATCTNNNATGAGGGTCTCTGCTGTTCTT
AGCACACGTCTGAACTCTTTCCCTACACGACGCTCTTCCGATCTNNNCAGTGTCCTGAGAGCTACTACCA
AGAGCACACGTCTGAACTCTTTCCCTACACGACGCTCTTCCGATCTNNNCATGGAGGCCATACTGAAGT
GAGCACACGTCTGAACTCTTTCCCTACACGACGCTCTTCCGATCTNNNGCTTCATAAGCATTGGATGA
GAGCACACGTCTGAACTCTTTCCCTACACGACGCTCTTCCGATCTNNNTACCTGTAATGGCATGTGTGC
GAGCACACGTCTGAACTCTTTCCCTACACGACGCTCTTCCGATCTNNNGATCTGGCAGCAGGTTTTGTT
GAGCACACGTCTGAACTCTTTCCCTACACGACGCTCTTCCGATCTNNNAATGGCAGATGGGGACTTGT
AGCACACGTCTGAACTCTTTCCCTACACGACGCTCTTCCGATCTNNNTGTCCCAGAATGTAGGTCAATG
GAGCACACGTCTGAACTCTTTCCCTACACGACGCTCTTCCGATCTNNNCAGGCATGGTTTTTGAATGGT
AGCACACGTCTGAACTCTTTCCCTACACGACGCTCTTCCGATCTNNNCACATACACACTCTCGCTCTCAA
AGCACACGTCTGAACTCTTTCCCTACACGACGCTCTTCCGATCTNNNCACAGGTCCACATGTACACACA
GAGCACACGTCTGAACTCTTTCCCTACACGACGCTCTTCCGATCTNNNTTTCTCCTTCTGTCTCCTCCCTG
AGAGCACACGTCTGAACTCTTTCCCTACACGACGCTCTTCCGATCTNNNCTTCAGTTCAGGCACCACTTCT
AGCACACGTCTGAACTCTTTCCCTACACGACGCTCTTCCGATCTNNNCTCCCATTTTTGGCGAAGACAA
GAGCACACGTCTGAACTCTTTCCCTACACGACGCTCTTCCGATCTNNNAACAAGCCTCCATCCTGAAAA
AGCACACGTCTGAACTCTTTCCCTACACGACGCTCTTCCGATCTNNNCCATGGTGAGTGGAATGTTGGA
AGCACACGTCTGAACTCTTTCCCTACACGACGCTCTTCCGATCTNNNAGAATGTTCAATGGTTGGCTGA
AGCACACGTCTGAACTCTTTCCCTACACGACGCTCTTCCGATCTNNNGTCACCTGCACACATACAGACA
GAGCACACGTCTGAACTCTTTCCCTACACGACGCTCTTCCGATCTNNNGCTAATGTCCTGACAAGAAGG
GAGCACACGTCTGAACTCTTTCCCTACACGACGCTCTTCCGATCTNNNCTGAAGAGGCTGCAAGAAGA
GAGCACACGTCTGAACTCTTTCCCTACACGACGCTCTTCCGATCTNNNCTCTGCCTCCTCCACTATGAA
GAGCACACGTCTGAACTCTTTCCCTACACGACGCTCTTCCGATCTNNNAATAGGATGGATGTGTGCGT
AGAGCACACGTCTGAACTCTTTCCCTACACGACGCTCTTCCGATCTNNNTACCCCAATGAATGTGCCAT
GAGCACACGTCTGAACTCTTTCCCTACACGACGCTCTTCCGATCTNNNTGTTGGATGTCTCCCTGCTCTA
AGCACACGTCTGAACTCTTTCCCTACACGACGCTCTTCCGATCTNNNATCTTGTGGACAGCTTGGGAAT
GAGCACACGTCTGAACTCTTTCCCTACACGACGCTCTTCCGATCTNNNTCACATCTCATTGTATCCAGG
GAGCACACGTCTGAACTCTTTCCCTACACGACGCTCTTCCGATCTNNNGGCATCTGGGTGTTTCCTAA
GAGCACACGTCTGAACTCTTTCCCTACACGACGCTCTTCCGATCTNNNGAACAAAGGCCATGGGGAAA
AGCACACGTCTGAACTCTTTCCCTACACGACGCTCTTCCGATCTNNNATGTTAGACACACATGCACTG
AGCACACGTCTGAACTCTTTCCCTACACGACGCTCTTCCGATCTNNNTCCCTAGCCGATATCCACAGAA
AGCACACGTCTGAACTCTTTCCCTACACGACGCTCTTCCGATCTNNNTCTTTCTTTCCAAGCAGGTTGACA
AGAGCACACGTCTGAACTCTTTCCCTACACGACGCTCTTCCGATCTNNNCTGCAGAGAAACCAGTAGGA
GAGCACACGTCTGAACTCTTTCCCTACACGACGCTCTTCCGATCTNNNGAGAATTTGCTCAGTGGGAG
AGAGCACACGTCTGAACTCTTTCCCTACACGACGCTCTTCCGATCTNNNTCTCATTTCCCAGAGCTTTCA
AGAGCACACGTCTGAACTCTTTCCCTACACGACGCTCTTCCGATCTNNNGGTTTTCACCACGGTGTTTCA
AGCACACGTCTGAACTCTTTCCCTACACGACGCTCTTCCGATCTNNNGCTTTGCTCCCAATCTCTCTCTC

GAGCACACGTCTGAACTCTTTCCCTACACGACGCTCTTCCGATCTNNNTCAGTTTGATGTGTTTCTGGG
AGCACACGTCTGAACTCTTTCCCTACACGACGCTCTTCCGATCTNNNTAGCTAGCAGCAGGATCAACTC
GAGCACACGTCTGAACTCTTTCCCTACACGACGCTCTTCCGATCTNNNCCGGAGAAGGTCCTCAAAGTA
GCACACGTCTGAACTCTTTCCCTACACGACGCTCTTCCGATCTNNNGAAAGTCACCCTCTCCCCACTATC
GAGCACACGTCTGAACTCTTTCCCTACACGACGCTCTTCCGATCTNNNGGACCATGATTATCCCATGCCT
AGAGCACACGTCTGAACTCTTTCCCTACACGACGCTCTTCCGATCTNNNAGTAACCGCTATTTGTTGAG
GAGCACACGTCTGAACTCTTTCCCTACACGACGCTCTTCCGATCTNNNTAGGTGTTTGGTGGGAAAAC
AGCACACGTCTGAACTCTTTCCCTACACGACGCTCTTCCGATCTNNNTGTATGTGTGTGCGGTAGTGT
GAGCACACGTCTGAACTCTTTCCCTACACGACGCTCTTCCGATCTNNNTGGATGGCAAAGAACACATC
AGAGCACACGTCTGAACTCTTTCCCTACACGACGCTCTTCCGATCTNNNATTCTGCTGCTGTCCCCTAG
GAGCACACGTCTGAACTCTTTCCCTACACGACGCTCTTCCGATCTNNNACCTGGGACAGTAACCACATT
AGCACACGTCTGAACTCTTTCCCTACACGACGCTCTTCCGATCTNNNTTCTCCTGTAGGTACAGAATTG
AGCACACGTCTGAACTCTTTCCCTACACGACGCTCTTCCGATCTNNNAAGTGACATAAGAGTGAGGGC
GAGCACACGTCTGAACTCTTTCCCTACACGACGCTCTTCCGATCTNNNGCTAACTGGCTGCAAGAAACT
GAGCACACGTCTGAACTCTTTCCCTACACGACGCTCTTCCGATCTNNNATATGGCATTGTTGGACCCTG
AGAGCACACGTCTGAACTCTTTCCCTACACGACGCTCTTCCGATCTNNNATCTGTTCAGCAGGGAGAGA
GAGCACACGTCTGAACTCTTTCCCTACACGACGCTCTTCCGATCTNNNTGGCTCCCCTGATAAACCATCT
GAGCACACGTCTGAACTCTTTCCCTACACGACGCTCTTCCGATCTNNNCTTTTTCTAGAACCTGCCCCCA
AGAGCACACGTCTGAACTCTTTCCCTACACGACGCTCTTCCGATCTNNNCCATTCCAGGCAATCTGAAG
AGCACACGTCTGAACTCTTTCCCTACACGACGCTCTTCCGATCTNNNACTCCACAGATGTACTGTAGCG
AGAGCACACGTCTGAACTCTTTCCCTACACGACGCTCTTCCGATCTNNNTACTCTGAAATGGTTCCCAAA
AGAGCACACGTCTGAACTCTTTCCCTACACGACGCTCTTCCGATCTNNNTGCTCAGCATATCAAAGCAC
AGCACACGTCTGAACTCTTTCCCTACACGACGCTCTTCCGATCTNNNCATCTGCCAGGAATCCCTTTCTC
GAGCACACGTCTGAACTCTTTCCCTACACGACGCTCTTCCGATCTNNNCAAGTGCTGTGGTTTGGATG
GAGCACACGTCTGAACTCTTTCCCTACACGACGCTCTTCCGATCTNNNAAGCCCTGTCAAGATTTCTGA
AGCACACGTCTGAACTCTTTCCCTACACGACGCTCTTCCGATCTNNNTAACATATTCCCCAAGTTGCCCC
AGAGCACACGTCTGAACTCTTTCCCTACACGACGCTCTTCCGATCTNNNATCCTGTGTAAAGACAGGCA
AGCACACGTCTGAACTCTTTCCCTACACGACGCTCTTCCGATCTNNNTGTGTCACTTCTCTCAGGTCCTC
GAGCACACGTCTGAACTCTTTCCCTACACGACGCTCTTCCGATCTNNNGATGCTCTCCTTTAGGTCTGT
AGCACACGTCTGAACTCTTTCCCTACACGACGCTCTTCCGATCTNNNAAACTCAGCCTCAAGGGAAAG
AGCACACGTCTGAACTCTTTCCCTACACGACGCTCTTCCGATCTNNNGCAACTTCTGCACTTGAACTGA
GAGCACACGTCTGAACTCTTTCCCTACACGACGCTCTTCCGATCTNNNACTTCTGTGCCATCATGTGTAC
GAGCACACGTCTGAACTCTTTCCCTACACGACGCTCTTCCGATCTNNNCTGGGCCACATAGGAAGATT
AGAGCACACGTCTGAACTCTTTCCCTACACGACGCTCTTCCGATCTNNNGCCAGGATTTTTATCTGTGT
AGCACACGTCTGAACTCTTTCCCTACACGACGCTCTTCCGATCTNNNCCAAATTCTCCTCTCCTGCATCC
AGAGCACACGTCTGAACTCTTTCCCTACACGACGCTCTTCCGATCTNNNAAGATGCCATGAGAGCAAAG
GAGCACACGTCTGAACTCTTTCCCTACACGACGCTCTTCCGATCTNNNCTCCAGGAGGAAAACCAGAG
GAGCACACGTCTGAACTCTTTCCCTACACGACGCTCTTCCGATCTNNNTTATCAGAAGGAGGGATGGG
GCACACGTCTGAACTCTTTCCCTACACGACGCTCTTCCGATCTNNNTCTCCTCATCTGTGAAATGCACCC
GAGCACACGTCTGAACTCTTTCCCTACACGACGCTCTTCCGATCTNNNGTGCCCTCAATAAGGCACATA
GAGCACACGTCTGAACTCTTTCCCTACACGACGCTCTTCCGATCTNNNATTCATGCTAGCAAGCTCTCC
GAGCACACGTCTGAACTCTTTCCCTACACGACGCTCTTCCGATCTNNNGGTCATCTCTCAGTCTGATT
GAGCACACGTCTGAACTCTTTCCCTACACGACGCTCTTCCGATCTNNNAACTAATGGTGGCCTCTCTGT

AGCACACGTCTGAACTCTTTCCCTACACGACGCTCTTCCGATCTNNNCTTCCCCGAACAGCAGTTCATAA
GAGCACACGTCTGAACTCTTTCCCTACACGACGCTCTTCCGATCTNNNCCCAATATAGCAACAGGGGAA
GAGCACACGTCTGAACTCTTTCCCTACACGACGCTCTTCCGATCTNNNAGCCAACTTGTCTTCTAGTACC
AGCACACGTCTGAACTCTTTCCCTACACGACGCTCTTCCGATCTNNNCCTTAGTAACTTTCTGGTGGCC
AGCACACGTCTGAACTCTTTCCCTACACGACGCTCTTCCGATCTNNNGATTTCTGCAGGAGGAACCAGA
AGCACACGTCTGAACTCTTTCCCTACACGACGCTCTTCCGATCTNNNCAATCCCTGCCTCAGTAAACTCC
GAGCACACGTCTGAACTCTTTCCCTACACGACGCTCTTCCGATCTNNNCACTGAGTTTAGTTGGGGAT
GCACACGTCTGAACTCTTTCCCTACACGACGCTCTTCCGATCTNNNGAGGTGCAGGAATGTAGGACAA
AGAGCACACGTCTGAACTCTTTCCCTACACGACGCTCTTCCGATCTNNNCAGGTCCCCGAAAGAATGA
GCACACGTCTGAACTCTTTCCCTACACGACGCTCTTCCGATCTNNNCTGTCCCCCAAAAATGCATACCTC
GAGCACACGTCTGAACTCTTTCCCTACACGACGCTCTTCCGATCTNNNCCAAGCCAGAGCATTGATACCA
AGCACACGTCTGAACTCTTTCCCTACACGACGCTCTTCCGATCTNNNGATGTGGACATCCCAAACCAAG
GAGCACACGTCTGAACTCTTTCCCTACACGACGCTCTTCCGATCTNNNGGATGGAGTTGGGAGGATGC
AGCACACGTCTGAACTCTTTCCCTACACGACGCTCTTCCGATCTNNNTGTAAGTGTAGGCCTCTCTCTG
GAGCACACGTCTGAACTCTTTCCCTACACGACGCTCTTCCGATCTNNNACAATTAGGACCTTGCACCCAT
AGCACACGTCTGAACTCTTTCCCTACACGACGCTCTTCCGATCTNNNCCAGTGGCATAATGGCAAATG
AGAGCACACGTCTGAACTCTTTCCCTACACGACGCTCTTCCGATCTNNNAGAGATGATCCTTAGGCAGA
GAGCACACGTCTGAACTCTTTCCCTACACGACGCTCTTCCGATCTNNNCCCAACAGACAGAGGTTGGAA
GAGCACACGTCTGAACTCTTTCCCTACACGACGCTCTTCCGATCTNNNTGGGGACAAGAAGAAACCAT
GAGCACACGTCTGAACTCTTTCCCTACACGACGCTCTTCCGATCTNNNACATTGCAGTTCTCTCACTGGT
AGAGCACACGTCTGAACTCTTTCCCTACACGACGCTCTTCCGATCTNNNTTATCATGGGTGACTTTACCC
AGAGCACACGTCTGAACTCTTTCCCTACACGACGCTCTTCCGATCTNNNTGGTGTATCTATTGTCTGGCC
AGAGCACACGTCTGAACTCTTTCCCTACACGACGCTCTTCCGATCTNNNTTTTGACATGACACTCCCACA
GCACACGTCTGAACTCTTTCCCTACACGACGCTCTTCCGATCTNNNATTGCATGTCTGTGTGTATGGGT
GAGCACACGTCTGAACTCTTTCCCTACACGACGCTCTTCCGATCTNNNTCAAGTACTAACCATGGGCTTC
GAGCACACGTCTGAACTCTTTCCCTACACGACGCTCTTCCGATCTNNNTGGCCATAAGCCAAATCCAAC
AGAGCACACGTCTGAACTCTTTCCCTACACGACGCTCTTCCGATCTNNNTAGCCCCAGCTCTTTATTTGC
GAGCACACGTCTGAACTCTTTCCCTACACGACGCTCTTCCGATCTNNNAAGATGTTTCAGCACCCTCCAA
AGAGCACACGTCTGAACTCTTTCCCTACACGACGCTCTTCCGATCTNNNACTCCACAACATAGGATGGG
AGCACACGTCTGAACTCTTTCCCTACACGACGCTCTTCCGATCTNNNAGGGTCAGCAAAGTCAGGAAA
GAGCACACGTCTGAACTCTTTCCCTACACGACGCTCTTCCGATCTNNNTGCATATACCATCGCCTCCAAA
GCACACGTCTGAACTCTTTCCCTACACGACGCTCTTCCGATCTNNNTAAAACACAATCCCAACCTCCCCA
AGCACACGTCTGAACTCTTTCCCTACACGACGCTCTTCCGATCTNNNATTTCCTGTACCTCATGGAGCCT
GAGCACACGTCTGAACTCTTTCCCTACACGACGCTCTTCCGATCTNNNGTAGGTGTTTTGCCATCGCATT
GAGCACACGTCTGAACTCTTTCCCTACACGACGCTCTTCCGATCTNNNGGCAGACCCCTACTCCAATATC
GAGCACACGTCTGAACTCTTTCCCTACACGACGCTCTTCCGATCTNNNAACCCAATATGTTCTGCATCCA
AGAGCACACGTCTGAACTCTTTCCCTACACGACGCTCTTCCGATCTNNNTGTGCAACCCAGTTTGTGGT
AGCACACGTCTGAACTCTTTCCCTACACGACGCTCTTCCGATCTNNNACATCCAAATCAGGAAATCGGC
AGCACACGTCTGAACTCTTTCCCTACACGACGCTCTTCCGATCTNNNAGCCCTTCTACAGTGGCATAAG
AGAGCACACGTCTGAACTCTTTCCCTACACGACGCTCTTCCGATCTNNNTTCCAAAAGGAATGATGGGA
AGCACACGTCTGAACTCTTTCCCTACACGACGCTCTTCCGATCTNNNGCATGACCAGACCCCAGTAAAA
GCACACGTCTGAACTCTTTCCCTACACGACGCTCTTCCGATCTNNNGGTGAAGATACACACTCATGCCTC
AGCACACGTCTGAACTCTTTCCCTACACGACGCTCTTCCGATCTNNNACACTTAGGAGAGACCAGGAG

AGCACACGTCTGAACTCTTTCCCTACACGACGCTCTTCCGATCTNNNCGCTCTCTCTCTCTCTCTCTCC
GAGCACACGTCTGAACTCTTTCCCTACACGACGCTCTTCCGATCTNNNAACCTTTTGAAAACCCAGCACT
AGCACACGTCTGAACTCTTTCCCTACACGACGCTCTTCCGATCTNNNAGAGAGAGAGAGAGAGAGAG
AGAGCACACGTCTGAACTCTTTCCCTACACGACGCTCTTCCGATCTNNNGGGCTGAGTGAAGAGAATC
CACACGTCTGAACTCTTTCCCTACACGACGCTCTTCCGATCTNNNGCATTTACCAGAAGCCTCTCTCTCA
AGCACACGTCTGAACTCTTTCCCTACACGACGCTCTTCCGATCTNNNGCAACAGGATGAGACTTTGTCA
AGAGCACACGTCTGAACTCTTTCCCTACACGACGCTCTTCCGATCTNNNGCTTGTGTTGAGGATAAGG
AGCACACGTCTGAACTCTTTCCCTACACGACGCTCTTCCGATCTNNNGTCAGCACAAAAGAAGTCAGGA
GCACACGTCTGAACTCTTTCCCTACACGACGCTCTTCCGATCTNNNGGTCTTTCTCCCTTTCTCCCTTTCAT
AGCACACGTCTGAACTCTTTCCCTACACGACGCTCTTCCGATCTNNNCAGAGGTTGACATCTGTGGAGT
AGCACACGTCTGAACTCTTTCCCTACACGACGCTCTTCCGATCTNNNGTGCTCTTGGGTTGAGTACCTA
GAGCACACGTCTGAACTCTTTCCCTACACGACGCTCTTCCGATCTNNNCTTTCTTTCTCTGCCTCCTCCATG
AGAGCACACGTCTGAACTCTTTCCCTACACGACGCTCTTCCGATCTNNNAAGACCCCTTGAGTCTAGGA
AGCACACGTCTGAACTCTTTCCCTACACGACGCTCTTCCGATCTNNNGATCACTTGAGCTCAGGAGTTC
CACACGTCTGAACTCTTTCCCTACACGACGCTCTTCCGATCTNNNTGTCTGTCTGTCTCTGTCTCTCTCAT
GAGCACACGTCTGAACTCTTTCCCTACACGACGCTCTTCCGATCTNNNGAGAGACAGAGACGAGAGAC
AGCACACGTCTGAACTCTTTCCCTACACGACGCTCTTCCGATCTNNNTTAGCATCTCCCTTTCCATCCTGC
GAGCACACGTCTGAACTCTTTCCCTACACGACGCTCTTCCGATCTNNNAGACAGGAGAATTGCTTGAAG
GAGCACACGTCTGAACTCTTTCCCTACACGACGCTCTTCCGATCTNNNGAATCATTCCCAAAACCCCCTT
GAGCACACGTCTGAACTCTTTCCCTACACGACGCTCTTCCGATCTNNNGAATTGGAGAGAGGAGAGA
GAGCACACGTCTGAACTCTTTCCCTACACGACGCTCTTCCGATCTNNNGAGCAAGACCTTGTCTCAAAC
GAGCACACGTCTGAACTCTTTCCCTACACGACGCTCTTCCGATCTNNNCACTTGAGCTCAGGAGTTTGA
AGCACACGTCTGAACTCTTTCCCTACACGACGCTCTTCCGATCTNNNAGAGAGAGAGGGAGAGAGAG
AGCACACGTCTGAACTCTTTCCCTACACGACGCTCTTCCGATCTNNNGGGGAGAAGAGAGAGAGAGA
GAGCACACGTCTGAACTCTTTCCCTACACGACGCTCTTCCGATCTNNNGAGCGAGACTCCGTCTATAAA
GAGCACACGTCTGAACTCTTTCCCTACACGACGCTCTTCCGATCTNNNCTCTCTCTCTCACACACACAC
GAGCACACGTCTGAACTCTTTCCCTACACGACGCTCTTCCGATCTNNNACACCTGGCCAGATACCTTTTT
GAGCACACGTCTGAACTCTTTCCCTACACGACGCTCTTCCGATCTNNNCAGCCCTTTGAGAGAGAGAG
AGAGCACACGTCTGAACTCTTTCCCTACACGACGCTCTTCCGATCTNNNCATGTCTGAATCCCAGCACTT
AGCACACGTCTGAACTCTTTCCCTACACGACGCTCTTCCGATCTNNNAAACAAACAAGAAGCTTACCGC
GAGCACACGTCTGAACTCTTTCCCTACACGACGCTCTTCCGATCTNNNATTGTGCCATTGTACTCCATCC
AGCACACGTCTGAACTCTTTCCCTACACGACGCTCTTCCGATCTNNNGGGACCAGGGTGTGTGAATAT
GAGCACACGTCTGAACTCTTTCCCTACACGACGCTCTTCCGATCTNNNCAGGCAACATAGTGAGACCTG
GAGCACACGTCTGAACTCTTTCCCTACACGACGCTCTTCCGATCTNNNCAACAAGAGCGAAACTGTCTC
AGCACACGTCTGAACTCTTTCCCTACACGACGCTCTTCCGATCTNNNGGATCACTTGAGGTCAGGAGTT
GAGCACACGTCTGAACTCTTTCCCTACACGACGCTCTTCCGATCTNNNTTCCAAAGTGCTGGGATTAGA
CACACGTCTGAACTCTTTCCCTACACGACGCTCTTCCGATCTNNNAGCGAGTTTATAGACTTTGGACTC
GAGCACACGTCTGAACTCTTTCCCTACACGACGCTCTTCCGATCTNNNGAGAGAGAGAGAGACGAG
CACACGTCTGAACTCTTTCCCTACACGACGCTCTTCCGATCTNNNGCAAACATACTGCAGTAACAGGAC
GAGCACACGTCTGAACTCTTTCCCTACACGACGCTCTTCCGATCTNNNCCAACCTGATCACAGAAGTTTC
GAGCACACGTCTGAACTCTTTCCCTACACGACGCTCTTCCGATCTNNNAGTCATCCAGGACTAGCAGTA
GAGCACACGTCTGAACTCTTTCCCTACACGACGCTCTTCCGATCTNNNATCAACATGCATACACAAGCAC
GAGCACACGTCTGAACTCTTTCCCTACACGACGCTCTTCCGATCTNNNAGCCTGAGCAACATAGTAAGA

AGCACACGTCTGAACTCTTTCCCTACACGACGCTCTTCCGATCTNNNGAACCTGGGAGAATTGCTTGA

GAGCACACGTCTGAACTCTTTCCCTACACGACGCTCTTCCGATCTNNNGCGAGGCCCTGTCTCAATAAA

AGCACACGTCTGAACTCTTTCCCTACACGACGCTCTTCCGATCTNNNCCCGCAGTGAGTTTTTAGTTGT

AGCACACGTCTGAACTCTTTCCCTACACGACGCTCTTCCGATCTNNNCACCAGCAAACACAGTTATTCGT

AGCACACGTCTGAACTCTTTCCCTACACGACGCTCTTCCGATCTNNNACCTCATCCTTCAAGTGCTCTTT

AGCACACGTCTGAACTCTTTCCCTACACGACGCTCTTCCGATCTNNNACCTCCTTTCTGCTTTCTGGAAT

GAGCACACGTCTGAACTCTTTCCCTACACGACGCTCTTCCGATCTNNNGTGAAAAGGAGGCAGGTTTT

AGCACACGTCTGAACTCTTTCCCTACACGACGCTCTTCCGATCTNNNACTTGAACAGGCTATGGTTCCA

AGCACACGTCTGAACTCTTTCCCTACACGACGCTCTTCCGATCTNNNGGAGAAAAGTTCAGACAACGT

AGCACACGTCTGAACTCTTTCCCTACACGACGCTCTTCCGATCTNNNATGGCCTAATATGTAGTGGCTG

GAGCACACGTCTGAACTCTTTCCCTACACGACGCTCTTCCGATCTNNNTGGCATGCCTTTGAACTTTGA

GAGCACACGTCTGAACTCTTTCCCTACACGACGCTCTTCCGATCTNNNATTCTGCATTGGTTGGACAGA

GAGCACACGTCTGAACTCTTTCCCTACACGACGCTCTTCCGATCTNNNCACCACTGTACTCTTAGCCTGA
AGAGCACACGTCTGAACTCTTTCCCTACACGACGCTCTTCCGATCTNNNTGAGGTTGGAGGATTCTTTG
GAGCACACGTCTGAACTCTTTCCCTACACGACGCTCTTCCGATCTNNNAGAATCACTTGAACTGGAGAC
AGCACACGTCTGAACTCTTTCCCTACACGACGCTCTTCCGATCTNNNCAAAATCCTGCCACTGTACTCCA
GCACACGTCTGAACTCTTTCCCTACACGACGCTCTTCCGATCTNNNCCCAGCTACTACAGAGGCTAAAG
GAGCACACGTCTGAACTCTTTCCCTACACGACGCTCTTCCGATCTNNNAGGCAGGAGAATCATTTGAAC
AGCACACGTCTGAACTCTTTCCCTACACGACGCTCTTCCGATCTNNNCTGTAGTCCCAGCTACTCAGGA
AGCACACGTCTGAACTCTTTCCCTACACGACGCTCTTCCGATCTNNNGATTACCCTCTCAATCGCATTGC
AGCACACGTCTGAACTCTTTCCCTACACGACGCTCTTCCGATCTNNNGAGGCAAGAGAATCACTTGAA
AGCACACGTCTGAACTCTTTCCCTACACGACGCTCTTCCGATCTNNNGGGAGACTGAGGCAAGATAAT
GAGCACACGTCTGAACTCTTTCCCTACACGACGCTCTTCCGATCTNNNGCATGCCTATAATCCCAGCTAC
AGAGCACACGTCTGAACTCTTTCCCTACACGACGCTCTTCCGATCTNNNTACAAAATATAGCCAGGCGT
AGCACACGTCTGAACTCTTTCCCTACACGACGCTCTTCCGATCTNNNGATTGTTTGAGCCTGGGAGAT
GAGCACACGTCTGAACTCTTTCCCTACACGACGCTCTTCCGATCTNNNATGAGCACCTATAGTCCCAGC
GAGCACACGTCTGAACTCTTTCCCTACACGACGCTCTTCCGATCTNNNGAGAACCGTGATTATGCCATTC
AGCACACGTCTGAACTCTTTCCCTACACGACGCTCTTCCGATCTNNNTGTAATCCTAGCACTTTGGGAG
AGAGCACACGTCTGAACTCTTTCCCTACACGACGCTCTTCCGATCTNNNGGAGAATGGCATGAACCTG
AGCACACGTCTGAACTCTTTCCCTACACGACGCTCTTCCGATCTNNNCTGAGGCAGGAGAATAGCTTG
GAGCACACGTCTGAACTCTTTCCCTACACGACGCTCTTCCGATCTNNNCACCTGTAATCCCAGATAGTCC
GAGCACACGTCTGAACTCTTTCCCTACACGACGCTCTTCCGATCTNNNACCTTCCCTCCACTATTGTCCTA
GAGCACACGTCTGAACTCTTTCCCTACACGACGCTCTTCCGATCTNNNATGATCACACCACTGCACTCTA
AGAGCACACGTCTGAACTCTTTCCCTACACGACGCTCTTCCGATCTNNNGAGGTTGGAGTGAGCTGAG
AGCACACGTCTGAACTCTTTCCCTACACGACGCTCTTCCGATCTNNNCTGAGGCAGGAGAATTGCTTGA
GAGCACACGTCTGAACTCTTTCCCTACACGACGCTCTTCCGATCTNNNGTCAGGAGATCGAGACCATTC
AGCACACGTCTGAACTCTTTCCCTACACGACGCTCTTCCGATCTNNNCTGAGGTAGGAGGATCACTTGA
GAGCACACGTCTGAACTCTTTCCCTACACGACGCTCTTCCGATCTNNNGTAGAGTTTGCAGTGAGCTG
GAGCACACGTCTGAACTCTTTCCCTACACGACGCTCTTCCGATCTNNNTGAGGCAGAAGAATTGCTTGA
GAGCACACGTCTGAACTCTTTCCCTACACGACGCTCTTCCGATCTNNNAGAAGTTGCAGTCAGTGGAG
AGCACACGTCTGAACTCTTTCCCTACACGACGCTCTTCCGATCTNNNGCTGAGACAGGAGAATCACTTG
AGAGCACACGTCTGAACTCTTTCCCTACACGACGCTCTTCCGATCTNNNGATCACTTTAGCCCAGGAGT
AGAGCACACGTCTGAACTCTTTCCCTACACGACGCTCTTCCGATCTNNNCAGGAGAATTGTTTTGAGCC
GAGCACACGTCTGAACTCTTTCCCTACACGACGCTCTTCCGATCTNNNCCTGGGTGACAAGAGTAAGA
GAGCACACGTCTGAACTCTTTCCCTACACGACGCTCTTCCGATCTNNNGGGCAACAGAGTGAAATTCCA
AGCACACGTCTGAACTCTTTCCCTACACGACGCTCTTCCGATCTNNNTCAAGTAATTCTCCCACCTCAGC
AGCACACGTCTGAACTCTTTCCCTACACGACGCTCTTCCGATCTNNNAGAGAAAGAAAAAACAGCCTGC
AGCACACGTCTGAACTCTTTCCCTACACGACGCTCTTCCGATCTNNNAATGAAGTCTGTTTCCAAGCAG
AGCACACGTCTGAACTCTTTCCCTACACGACGCTCTTCCGATCTNNNTCCTGCACATTCATACCATGGAAC
GAGCACACGTCTGAACTCTTTCCCTACACGACGCTCTTCCGATCTNNNTCACCTGACAGGCTTTAGAATG
AGCACACGTCTGAACTCTTTCCCTACACGACGCTCTTCCGATCTNNNAACACCACCAACAACATTGACTT
AGCACACGTCTGAACTCTTTCCCTACACGACGCTCTTCCGATCTNNNGGTTCAAGCGATTCTCATGTCTC
AGCACACGTCTGAACTCTTTCCCTACACGACGCTCTTCCGATCTNNNGGGCTGGATGATGACCTACAAT
GAGCACACGTCTGAACTCTTTCCCTACACGACGCTCTTCCGATCTNNNAATGAAATGGGAAACAGCAG
GAGCACACGTCTGAACTCTTTCCCTACACGACGCTCTTCCGATCTNNNTCTGATTGAGCACTGGGTTCT

AGAGCACACGTCTGAACTCTTTCCCTACACGACGCTCTTCCGATCTNNNTGGTAGGCAGAGGTTTCAAT
AGCACACGTCTGAACTCTTTCCCTACACGACGCTCTTCCGATCTNNNTGAGACACGAGAATCACTTGAG
AGCACACGTCTGAACTCTTTCCCTACACGACGCTCTTCCGATCTNNNGAGGCCCATCTCTACAAGAAA
GAGCACACGTCTGAACTCTTTCCCTACACGACGCTCTTCCGATCTNNNCAGTAAGCTGAGATTGCACCA
AGAGCACACGTCTGAACTCTTTCCCTACACGACGCTCTTCCGATCTNNNAGACAGGAGAATCACATGA
GAGCACACGTCTGAACTCTTTCCCTACACGACGCTCTTCCGATCTNNNCTGTAGTCTGAGCTTCTCTGG
GAGCACACGTCTGAACTCTTTCCCTACACGACGCTCTTCCGATCTNNNGAAACAGGAGAATCGCTTGA
AGCACACGTCTGAACTCTTTCCCTACACGACGCTCTTCCGATCTNNNACGACTGTAATTGCAGCACTTT
AGAGCACACGTCTGAACTCTTTCCCTACACGACGCTCTTCCGATCTNNNCTGTAGTCCCAGCTACTCAG
GAGCACACGTCTGAACTCTTTCCCTACACGACGCTCTTCCGATCTNNNTGTGCCTGTAATTCCAGCTAC
GAGCACACGTCTGAACTCTTTCCCTACACGACGCTCTTCCGATCTNNNGAGGTTGAGTGAGCCGATAT
AGCACACGTCTGAACTCTTTCCCTACACGACGCTCTTCCGATCTNNNAGGCTGAGACATAAGAATCGC
GAGCACACGTCTGAACTCTTTCCCTACACGACGCTCTTCCGATCTNNNGTAATCCCAGCTATTCAGGAG
GAGCACACGTCTGAACTCTTTCCCTACACGACGCTCTTCCGATCTNNNGCATCCCAGTGTTTATTACCA
AGCACACGTCTGAACTCTTTCCCTACACGACGCTCTTCCGATCTNNNATTGGTATCCTTACACACAGGG
GAGCACACGTCTGAACTCTTTCCCTACACGACGCTCTTCCGATCTNNNTGGACGCATGACTTTACAGAG
AGCACACGTCTGAACTCTTTCCCTACACGACGCTCTTCCGATCTNNNAGTCACACACAACCTTCTTCTCC
AGCACACGTCTGAACTCTTTCCCTACACGACGCTCTTCCGATCTNNNCCAGCATGCCTGACTCCATATAT
GAGCACACGTCTGAACTCTTTCCCTACACGACGCTCTTCCGATCTNNNTTCCAACTAGCTGGGATTACA
GCACACGTCTGAACTCTTTCCCTACACGACGCTCTTCCGATCTNNNTGTTCCTGGAAAAAGAGGACCA
GAGCACACGTCTGAACTCTTTCCCTACACGACGCTCTTCCGATCTNNNATGCCCACAAGTAGAATTGT
GAGCACACGTCTGAACTCTTTCCCTACACGACGCTCTTCCGATCTNNNTGCTGTTCACAGACAGTACAT
GAGCACACGTCTGAACTCTTTCCCTACACGACGCTCTTCCGATCTNNNGCCAGTATTACAGACATAGGC
GAGCACACGTCTGAACTCTTTCCCTACACGACGCTCTTCCGATCTNNNCCCACAGAAGACAGCTGTAAG
AGAGCACACGTCTGAACTCTTTCCCTACACGACGCTCTTCCGATCTNNNTTGGTTTTTTTTGCAGCTTCG
AGCACACGTCTGAACTCTTTCCCTACACGACGCTCTTCCGATCTNNNGCAAGGGAGAAAGTCAGAGA
AGCACACGTCTGAACTCTTTCCCTACACGACGCTCTTCCGATCTNNNAGAGAGTTTGGTTGGGAGGAA
GAGCACACGTCTGAACTCTTTCCCTACACGACGCTCTTCCGATCTNNNTCAAGTCGTGATATTCCCAGT
AGAGCACACGTCTGAACTCTTTCCCTACACGACGCTCTTCCGATCTNNNAAATAGGCACAAGGGTAGT
AGCACACGTCTGAACTCTTTCCCTACACGACGCTCTTCCGATCTNNNAGTCCTGTTCTCCAATCCACTTA
GAGCACACGTCTGAACTCTTTCCCTACACGACGCTCTTCCGATCTNNNGGGCTCTTGCTGCTTTATTTCT
AGCACACGTCTGAACTCTTTCCCTACACGACGCTCTTCCGATCTNNNAGTAGGTAGGACTACAGGCAT
AGCACACGTCTGAACTCTTTCCCTACACGACGCTCTTCCGATCTNNNTACTGGTACCACAAGTGTTTCT
AGCACACGTCTGAACTCTTTCCCTACACGACGCTCTTCCGATCTNNNTTCAGTTCTGAGCCACATGATC
AGCACACGTCTGAACTCTTTCCCTACACGACGCTCTTCCGATCTNNNAATCATCCAGCTAAGCTTCTCC
GAGCACACGTCTGAACTCTTTCCCTACACGACGCTCTTCCGATCTNNNTGTAGGGACAGGAAAGGAA
GCACACGTCTGAACTCTTTCCCTACACGACGCTCTTCCGATCTNNNGGGATGCAAAATCAAAGAAGG
AGCACACGTCTGAACTCTTTCCCTACACGACGCTCTTCCGATCTNNNTTCCCTCTGACTTCCCTGTCATA
GAGCACACGTCTGAACTCTTTCCCTACACGACGCTCTTCCGATCTNNNTCCTGTTCCTTCTTAGCCTGTT
AGCACACGTCTGAACTCTTTCCCTACACGACGCTCTTCCGATCTNNNTGAGCACACTTCCTAAGGTCAA
GAGCACACGTCTGAACTCTTTCCCTACACGACGCTCTTCCGATCTNNNCTTCCAAGTGGCAAGATTCCAT
AGCACACGTCTGAACTCTTTCCCTACACGACGCTCTTCCGATCTNNNGCCCATGGATGAACTTAGTGA
GCACACGTCTGAACTCTTTCCCTACACGACGCTCTTCCGATCTNNNTGTTTTGGGCCATTTCAAATCCA

GAGCACACGTCTGAACTCTTTCCCTACACGACGCTCTTCCGATCTNNNAGGAACAAGAATCACCTGAAC
AGCACACGTCTGAACTCTTTCCCTACACGACGCTCTTCCGATCTNNNCTCTGGCTCACTCCTGTAATACC
AGCACACGTCTGAACTCTTTCCCTACACGACGCTCTTCCGATCTNNNTGTAGAGCTAATGCCCCTTTAC
AGCACACGTCTGAACTCTTTCCCTACACGACGCTCTTCCGATCTNNNAAATGCTGAGACTCCAAGTCCA
GAGCACACGTCTGAACTCTTTCCCTACACGACGCTCTTCCGATCTNNNCAGCCATATAAGGAAACACTG
AGCACACGTCTGAACTCTTTCCCTACACGACGCTCTTCCGATCTNNNCACCTCAGGTACAACATGGAAGA
AGCACACGTCTGAACTCTTTCCCTACACGACGCTCTTCCGATCTNNNTTCAAGTAGTTCTCCGTCTCAGC
GAGCACACGTCTGAACTCTTTCCCTACACGACGCTCTTCCGATCTNNNAGAATCCTGGGGCCATTTTCTA
GCACACGTCTGAACTCTTTCCCTACACGACGCTCTTCCGATCTNNNTGTTTTCACAATGTTGTCAGTTCA
AGCACACGTCTGAACTCTTTCCCTACACGACGCTCTTCCGATCTNNNGTAACTGGAAGAAACAGGGGCT
AGCACACGTCTGAACTCTTTCCCTACACGACGCTCTTCCGATCTNNNTGTCACTAGGACCAGAACAGAT
AGCACACGTCTGAACTCTTTCCCTACACGACGCTCTTCCGATCTNNNACAGAGCTATGCCTTTGGATCA
GAGCACACGTCTGAACTCTTTCCCTACACGACGCTCTTCCGATCTNNNTGACACTCAGCCCAATCATTAC
AGCACACGTCTGAACTCTTTCCCTACACGACGCTCTTCCGATCTNNNCTTAGTCAAAGTCAGGTGGGT
AGCACACGTCTGAACTCTTTCCCTACACGACGCTCTTCCGATCTNNNCAACCATCCTCTCAACTCTAGGG
AGCACACGTCTGAACTCTTTCCCTACACGACGCTCTTCCGATCTNNNCTTCCTGGAGCTTCGACAGTATT
AGCACACGTCTGAACTCTTTCCCTACACGACGCTCTTCCGATCTNNNGCCTCCTGAATAGCTGGGATTA
GAGCACACGTCTGAACTCTTTCCCTACACGACGCTCTTCCGATCTNNNGTTCCTTTACCTTCCACAAGTC
AGCACACGTCTGAACTCTTTCCCTACACGACGCTCTTCCGATCTNNNCATTGCTGATGTGCTTGTCACT
AGCACACGTCTGAACTCTTTCCCTACACGACGCTCTTCCGATCTNNNATCCTCCATCACACATACTACGCC
GAGCACACGTCTGAACTCTTTCCCTACACGACGCTCTTCCGATCTNNNGCAAGAGAATCACTTGAACCC
GAGCACACGTCTGAACTCTTTCCCTACACGACGCTCTTCCGATCTNNNTAAGTGCTTGAACTGAAGAG
CACACGTCTGAACTCTTTCCCTACACGACGCTCTTCCGATCTNNNTGGCAGAAGTAATGGTATGTCCCC
AGCACACGTCTGAACTCTTTCCCTACACGACGCTCTTCCGATCTNNNCATGCCTGTAATCCCAGCTACTC
GAGCACACGTCTGAACTCTTTCCCTACACGACGCTCTTCCGATCTNNNTTTACTCCATCCCACTTTGCTGT
GAGCACACGTCTGAACTCTTTCCCTACACGACGCTCTTCCGATCTNNNGCAACATGGTGAAATCCTGTC
AAGAGCACACGTCTGAACTCTTTCCCTACACGACGCTCTTCCGATCTNNNTAGTCCCAGCTACTCATGAG
AGAGCACACGTCTGAACTCTTTCCCTACACGACGCTCTTCCGATCTNNNAGAATGGCATGAACCTGGG
GAGCACACGTCTGAACTCTTTCCCTACACGACGCTCTTCCGATCTNNNCAATGCAGTGAGCCATGATCAT
GAGCACACGTCTGAACTCTTTCCCTACACGACGCTCTTCCGATCTNNNCTGAGGCAGGAGAATCACTTG
GAGCACACGTCTGAACTCTTTCCCTACACGACGCTCTTCCGATCTNNNCAAGAGAATTGCTTGAACCCAG
AGAGCACACGTCTGAACTCTTTCCCTACACGACGCTCTTCCGATCTNNNGAAATCGAGACCATCCTGGCT
AGCACACGTCTGAACTCTTTCCCTACACGACGCTCTTCCGATCTNNNCTGAGGCAGGAGAATCACTTG
GAGCACACGTCTGAACTCTTTCCCTACACGACGCTCTTCCGATCTNNNGCATGCCTGTAATCCCAGATAC
GCACACGTCTGAACTCTTTCCCTACACGACGCTCTTCCGATCTNNNGTGAACTACTGGGGTCAAGTGA
GAGCACACGTCTGAACTCTTTCCCTACACGACGCTCTTCCGATCTNNNCACAGCTGACTAAAACATTGCC
AGCACACGTCTGAACTCTTTCCCTACACGACGCTCTTCCGATCTNNNCATTCTGTGTCCTTTGGCATCTG
AGCACACGTCTGAACTCTTTCCCTACACGACGCTCTTCCGATCTNNNTTGTGGGTCTATAGGGATAGG
AGCACACGTCTGAACTCTTTCCCTACACGACGCTCTTCCGATCTNNNTAAGCACAGATGTCTGAAGAGC
AGCACACGTCTGAACTCTTTCCCTACACGACGCTCTTCCGATCTNNNAGCTCCTGGAATCAGAAGACAA
AGCACACGTCTGAACTCTTTCCCTACACGACGCTCTTCCGATCTNNNTCAAGCAATTCTCCTACCTCAGC
AGCACACGTCTGAACTCTTTCCCTACACGACGCTCTTCCGATCTNNNCAGTCTGGTCTCAAACTCCTGAC
AGCACACGTCTGAACTCTTTCCCTACACGACGCTCTTCCGATCTNNNAAGTAGCTGGTATTACAGGCAC

AGCACACGTCTGAACTCTTTCCCTACACGACGCTCTTCCGATCTNNNAAGACCCATCAGTGTGCTGTAT
GAGCACACGTCTGAACTCTTTCCCTACACGACGCTCTTCCGATCTNNNGACGACTGTAATCCCAGCTA
AGCACACGTCTGAACTCTTTCCCTACACGACGCTCTTCCGATCTNNNTGCTCATAACTACCTCCCAAA
AGCACACGTCTGAACTCTTTCCCTACACGACGCTCTTCCGATCTNNNAGCCGAGATTATGCCACTGTA
GAGCACACGTCTGAACTCTTTCCCTACACGACGCTCTTCCGATCTNNNGATGTGGTGGCTCATACCTG
AGCACACGTCTGAACTCTTTCCCTACACGACGCTCTTCCGATCTNNNGTAATCCTGGCTACTCAGAAG
GCACACGTCTGAACTCTTTCCCTACACGACGCTCTTCCGATCTNNNATGCCTTGTAATCCCAGCTACTC
GAGCACACGTCTGAACTCTTTCCCTACACGACGCTCTTCCGATCTNNNTTTGTCTTCTGGGACTAGTGT
GCACACGTCTGAACTCTTTCCCTACACGACGCTCTTCCGATCTNNNCAGAAGAATTGCTTGAACCCAG
AGCACACGTCTGAACTCTTTCCCTACACGACGCTCTTCCGATCTNNNTTGAACCCAGGAGACAGAGA
AGCACACGTCTGAACTCTTTCCCTACACGACGCTCTTCCGATCTNNNCTGTAGTTCCAGCTACTTGGG
AGCACACGTCTGAACTCTTTCCCTACACGACGCTCTTCCGATCTNNNTGGAGGCTGCAATAAGACAT
AGCACACGTCTGAACTCTTTCCCTACACGACGCTCTTCCGATCTNNNACATGCCTGTAATCCCAGCTAT
GAGCACACGTCTGAACTCTTTCCCTACACGACGCTCTTCCGATCTNNNGTCAGGAGAATAGCTTGAAC
AGCACACGTCTGAACTCTTTCCCTACACGACGCTCTTCCGATCTNNNGGTAAAGGTATCACTTGAGCC
AGCACACGTCTGAACTCTTTCCCTACACGACGCTCTTCCGATCTNNNGAGAATGACGTGAACCTAGG
AGCACACGTCTGAACTCTTTCCCTACACGACGCTCTTCCGATCTNNNCTGAACTCCAGCCTAGGCAAT
AGCACACGTCTGAACTCTTTCCCTACACGACGCTCTTCCGATCTNNNGAGATCACGCCATTTCATTCC
AGCACACGTCTGAACTCTTTCCCTACACGACGCTCTTCCGATCTNNNGGAGTTTGCAGTGAGTGGAG
GAGCACACGTCTGAACTCTTTCCCTACACGACGCTCTTCCGATCTNNNCACGGATCACAAGGTCACTA
AGCACACGTCTGAACTCTTTCCCTACACGACGCTCTTCCGATCTNNNGGGATCACTAGGTCAGGAGT
GAGCACACGTCTGAACTCTTTCCCTACACGACGCTCTTCCGATCTNNNAGGCAGGTGGATTATGAAGT
AGCACACGTCTGAACTCTTTCCCTACACGACGCTCTTCCGATCTNNNAGAAAAGTGATGAGATGGCT
GAGCACACGTCTGAACTCTTTCCCTACACGACGCTCTTCCGATCTNNNGAGAATGGAGTGAACTCAG
AGCACACGTCTGAACTCTTTCCCTACACGACGCTCTTCCGATCTNNNGCAACAAAAGCAGAACTCCATC
AGCACACGTCTGAACTCTTTCCCTACACGACGCTCTTCCGATCTNNNGTTGCTGTGAGTCGAGATTA
AAGAGCACACGTCTGAACTCTTTCCCTACACGACGCTCTTCCGATCTNNNAGTTTGAGGCTGCAGTAA
AGCACACGTCTGAACTCTTTCCCTACACGACGCTCTTCCGATCTNNNGCCTGTATTCCCAGCTACTCTA
GAGCACACGTCTGAACTCTTTCCCTACACGACGCTCTTCCGATCTNNNATTGCTTGAGCCTAGGAGTT
AGCACACGTCTGAACTCTTTCCCTACACGACGCTCTTCCGATCTNNNTTGGTGAGTTAGGGCTGAATCT
AGCACACGTCTGAACTCTTTCCCTACACGACGCTCTTCCGATCTNNNGGTTCCCATGTCATGCAAAACT
AGCACACGTCTGAACTCTTTCCCTACACGACGCTCTTCCGATCTNNNTTCAAGCAATTGTTCTGCCTCAG
AGCACACGTCTGAACTCTTTCCCTACACGACGCTCTTCCGATCTNNNTGTTTCTGGGAATGTCCATGG
AGCACACGTCTGAACTCTTTCCCTACACGACGCTCTTCCGATCTNNNGTTCATACCATTCTCCTGCCTCA
GCACACGTCTGAACTCTTTCCCTACACGACGCTCTTCCGATCTNNNGTGCTGGTTTGCAGTTTCTTTTG
GCACACGTCTGAACTCTTTCCCTACACGACGCTCTTCCGATCTNNNGCTCAGATCATCTTCCCACTTCAC
AGCACACGTCTGAACTCTTTCCCTACACGACGCTCTTCCGATCTNNNACATTCAACTCTTGGGTGGAAG
AGCACACGTCTGAACTCTTTCCCTACACGACGCTCTTCCGATCTNNNCCAATTGGCCTGAACTCTTCAA
AGCACACGTCTGAACTCTTTCCCTACACGACGCTCTTCCGATCTNNNCAAAGCAGGACAAGAGACCAT
AGCACACGTCTGAACTCTTTCCCTACACGACGCTCTTCCGATCTNNNCACTCTGTCCTTTTTCAAGCAGG
AGCACACGTCTGAACTCTTTCCCTACACGACGCTCTTCCGATCTNNNTTCAAGCAATTTTCCTGCCTCA
AGCACACGTCTGAACTCTTTCCCTACACGACGCTCTTCCGATCTNNNTAGAAGCACCTTCGAGCTGAT
GCACACGTCTGAACTCTTTCCCTACACGACGCTCTTCCGATCTNNNCAAACTCCTGAGCTCAAGCAAT

AGCACACGTCTGAACTCTTTCCCTACACGACGCTCTTCCGATCTNNNACAAAAAGTAGGTTGCCAAG
AGCACACGTCTGAACTCTTTCCCTACACGACGCTCTTCCGATCTNNNTCAGCCAAAATAGGATTCAAG
GAGCACACGTCTGAACTCTTTCCCTACACGACGCTCTTCCGATCTNNNAGACAGGAGAATCACTTGAA
AGCACACGTCTGAACTCTTTCCCTACACGACGCTCTTCCGATCTNNNATAAGGGCCAGCCTTCAAATC
AGCACACGTCTGAACTCTTTCCCTACACGACGCTCTTCCGATCTNNNGTAATCCCAGCTACTCAGGAAG
AGCACACGTCTGAACTCTTTCCCTACACGACGCTCTTCCGATCTNNNCTCAGTTTCTGCAACCCCAATT
AGCACACGTCTGAACTCTTTCCCTACACGACGCTCTTCCGATCTNNNCACCACTCCCAGCTAATTTTG
AGCACACGTCTGAACTCTTTCCCTACACGACGCTCTTCCGATCTNNNGATTCAAGCAATTCTCCTGCG
GAGCACACGTCTGAACTCTTTCCCTACACGACGCTCTTCCGATCTNNNAGATGAGCACATTTGGCCTC
GAGCACACGTCTGAACTCTTTCCCTACACGACGCTCTTCCGATCTNNNTTGGTTTTTGTTGAGTTTTG
GAGCACACGTCTGAACTCTTTCCCTACACGACGCTCTTCCGATCTNNNAAAGCTTGCCCAGATAAGAG
AGCACACGTCTGAACTCTTTCCCTACACGACGCTCTTCCGATCTNNNATGGAGTCATTTGGTTGCCT
AGAGCACACGTCTGAACTCTTTCCCTACACGACGCTCTTCCGATCTNNNTTTCCTAATGTGCCCAATGA
AGCACACGTCTGAACTCTTTCCCTACACGACGCTCTTCCGATCTNNNCTTGGTGAACTGCAAAGCCTA
GCACACGTCTGAACTCTTTCCCTACACGACGCTCTTCCGATCTNNNCAGACACCAGACACCACTAAAC
AGCACACGTCTGAACTCTTTCCCTACACGACGCTCTTCCGATCTNNNTGGGATTTGGATGATTGGTG
AGCACACGTCTGAACTCTTTCCCTACACGACGCTCTTCCGATCTNNNAGAGAGAAATGAACTGCTCC
AGCACACGTCTGAACTCTTTCCCTACACGACGCTCTTCCGATCTNNNGTTCTCATCATTCATCCCCCAC
AGCACACGTCTGAACTCTTTCCCTACACGACGCTCTTCCGATCTNNNTGGTTTACCGAAAGACCAGG
AGCACACGTCTGAACTCTTTCCCTACACGACGCTCTTCCGATCTNNNAATCTGCTATCAAACCTCTCTG
GCACACGTCTGAACTCTTTCCCTACACGACGCTCTTCCGATCTNNNTTCCCGTATCACCGTCTCTTTTC
GAGCACACGTCTGAACTCTTTCCCTACACGACGCTCTTCCGATCTNNNAATGTCTTTCGGACCTTCCAG
AGCACACGTCTGAACTCTTTCCCTACACGACGCTCTTCCGATCTNNNTTTGCATTGTGCTGAACCTAG
AGCACACGTCTGAACTCTTTCCCTACACGACGCTCTTCCGATCTNNNTTTGTGTCCAGTGAGCTACAA
GCACACGTCTGAACTCTTTCCCTACACGACGCTCTTCCGATCTNNNAGATGTTTGGTTTGAACAGTG
GCACACGTCTGAACTCTTTCCCTACACGACGCTCTTCCGATCTNNNACATGACAGATAGCATACCACC
GCACACGTCTGAACTCTTTCCCTACACGACGCTCTTCCGATCTNNNTTCCCAGGTTCAAGTGATTCTC
AGCACACGTCTGAACTCTTTCCCTACACGACGCTCTTCCGATCTNNNACTCTCTTGTTAGGGGCTAAT
GCACACGTCTGAACTCTTTCCCTACACGACGCTCTTCCGATCTNNNGCACTGTAGCCTAGGTCTCAAA
GCACACGTCTGAACTCTTTCCCTACACGACGCTCTTCCGATCTNNNAGTGGATGGATGAATGGATG
AGCACACGTCTGAACTCTTTCCCTACACGACGCTCTTCCGATCTNNNGGTGGAAGTCAGTGGTTACA
GCACACGTCTGAACTCTTTCCCTACACGACGCTCTTCCGATCTNNNGCCTGAACTTTTTCTGCATCTCC
GAGCACACGTCTGAACTCTTTCCCTACACGACGCTCTTCCGATCTNNNTTGCATTTTTTTCGCCAATCG
GAGCACACGTCTGAACTCTTTCCCTACACGACGCTCTTCCGATCTNNNATGTCTATCACCACAAGGAG
GCACACGTCTGAACTCTTTCCCTACACGACGCTCTTCCGATCTNNNGGATTATGCTGTCCTTGTAGGC
GCACACGTCTGAACTCTTTCCCTACACGACGCTCTTCCGATCTNNNCTGCTTACCTCCATTGGCTAAG
GCACACGTCTGAACTCTTTCCCTACACGACGCTCTTCCGATCTNNNTCCTTCTCACATTTCCTCAGCAA
GAGCACACGTCTGAACTCTTTCCCTACACGACGCTCTTCCGATCTNNNATTTACAGCTACTTCATCGCC
GAGCACACGTCTGAACTCTTTCCCTACACGACGCTCTTCCGATCTNNNTCTCTCTCTCTCTCTCTCTCT
GAGCACACGTCTGAACTCTTTCCCTACACGACGCTCTTCCGATCTNNNTTTTGGAACATGCTTGCCCTA
AGCACACGTCTGAACTCTTTCCCTACACGACGCTCTTCCGATCTNNNGTGATAGAGAGCAAGACCCTG
AGCACACGTCTGAACTCTTTCCCTACACGACGCTCTTCCGATCTNNNGTGAGCCAAGATCATGCTATT
AGCACACGTCTGAACTCTTTCCCTACACGACGCTCTTCCGATCTNNNGAGGTTACAGTGAGCCAAGAT

GAGCACACGTCTGAACTCTTTCCCTACACGACGCTCTTCCGATCTNNNACCACACCCCGCTAACTTTATA
AGCACACGTCTGAACTCTTTCCCTACACGACGCTCTTCCGATCTNNNTTTGCTTTCAACCTAAATGGGCC
AGCACACGTCTGAACTCTTTCCCTACACGACGCTCTTCCGATCTNNNTAGGGAGGGAGTTTCAGCATTT
GAGCACACGTCTGAACTCTTTCCCTACACGACGCTCTTCCGATCTNNNTGCTGATGTACTTTGGGATGC
AGCACACGTCTGAACTCTTTCCCTACACGACGCTCTTCCGATCTNNNTAGAACAGCAGCTATCAAGGAG
AGCACACGTCTGAACTCTTTCCCTACACGACGCTCTTCCGATCTNNNAGGGTGTCAGTCTATTGCTCAT
AGCACACGTCTGAACTCTTTCCCTACACGACGCTCTTCCGATCTNNNCGTATTTCTGTTGCCCAATCTGC
GAGCACACGTCTGAACTCTTTCCCTACACGACGCTCTTCCGATCTNNNTTCTGTGTCCTTTTCTTTGCCAC
GAGCACACGTCTGAACTCTTTCCCTACACGACGCTCTTCCGATCTNNNGTTTCTATGACTGGTAGTGGG
AGCACACGTCTGAACTCTTTCCCTACACGACGCTCTTCCGATCTNNNAGTAGAGATGGGCCACCTTATT
AGCACACGTCTGAACTCTTTCCCTACACGACGCTCTTCCGATCTNNNCCTCCTCAAGTAGCTGGGATTA
AGCACACGTCTGAACTCTTTCCCTACACGACGCTCTTCCGATCTNNNATTTTTTCCAGCTGTCACCACTGC
GAGCACACGTCTGAACTCTTTCCCTACACGACGCTCTTCCGATCTNNNCTTGAACTCCTGACCTCAAGTC
AGCACACGTCTGAACTCTTTCCCTACACGACGCTCTTCCGATCTNNNGGCCCACATGCATTTTAGAAACA
AGCACACGTCTGAACTCTTTCCCTACACGACGCTCTTCCGATCTNNNGAGGAGTTGCTAGTACTGGCA
GAGCACACGTCTGAACTCTTTCCCTACACGACGCTCTTCCGATCTNNNGATTACAGGAGTGAGCCACTC
GAGCACACGTCTGAACTCTTTCCCTACACGACGCTCTTCCGATCTNNNCATGCACATTACAGTCTGAGC
AGCACACGTCTGAACTCTTTCCCTACACGACGCTCTTCCGATCTNNNTCACTTCCAATCACTGGCACTATC
AGCACACGTCTGAACTCTTTCCCTACACGACGCTCTTCCGATCTNNNTCACTTCCAATCACTGGCACTATC
GAGCACACGTCTGAACTCTTTCCCTACACGACGCTCTTCCGATCTNNNACACAATGCTGAATGCTGTGA
AGCACACGTCTGAACTCTTTCCCTACACGACGCTCTTCCGATCTNNNTAGTGGAGAGAAAGCCCAGAAA
GAGCACACGTCTGAACTCTTTCCCTACACGACGCTCTTCCGATCTNNNTGAGATGATAGTGCCTTTGGT
GAGCACACGTCTGAACTCTTTCCCTACACGACGCTCTTCCGATCTNNNCGACACTCCGTCTCCAAAAAAA
AGCACACGTCTGAACTCTTTCCCTACACGACGCTCTTCCGATCTNNNAGTCATGTGCTGTTTCTTTAGCC
GAGCACACGTCTGAACTCTTTCCCTACACGACGCTCTTCCGATCTNNNGATGGGGTTTCATCACATTGG
AGCACACGTCTGAACTCTTTCCCTACACGACGCTCTTCCGATCTNNNAGAAAACCCTTGTGGCAGTTAG
GAGCACACGTCTGAACTCTTTCCCTACACGACGCTCTTCCGATCTNNNTGCAGTTTGGCTAGCATCTCTT
AGCACACGTCTGAACTCTTTCCCTACACGACGCTCTTCCGATCTNNNAGGGAGCTTGAAAATGCATTTC
AGCACACGTCTGAACTCTTTCCCTACACGACGCTCTTCCGATCTNNNCCTACACATTATCAGGCCTTTGG
AGCACACGTCTGAACTCTTTCCCTACACGACGCTCTTCCGATCTNNNTTCACTGTGATAAGTTCCAGGC
AGCACACGTCTGAACTCTTTCCCTACACGACGCTCTTCCGATCTNNNTGGCTCCAAAATTGGTTGGTTA
GAGCACACGTCTGAACTCTTTCCCTACACGACGCTCTTCCGATCTNNNCACTTGAGGTCAGAGGTTCAA
AGCACACGTCTGAACTCTTTCCCTACACGACGCTCTTCCGATCTNNNGGTGGCATATGCTCATAGTCTC
GAGCACACGTCTGAACTCTTTCCCTACACGACGCTCTTCCGATCTNNNGGTGGAAGTTTCAATGAGCCA
GAGCACACGTCTGAACTCTTTCCCTACACGACGCTCTTCCGATCTNNNTGCCTGTAATTCCAGCTACTCA
AGCACACGTCTGAACTCTTTCCCTACACGACGCTCTTCCGATCTNNNGGGAGACAGAGTGAGACTTCA
GAGCACACGTCTGAACTCTTTCCCTACACGACGCTCTTCCGATCTNNNGAATGACTGAATGAGCTGCAC
AGCACACGTCTGAACTCTTTCCCTACACGACGCTCTTCCGATCTNNNAGAATCACTTGAATCCAGGAGC
AGCACACGTCTGAACTCTTTCCCTACACGACGCTCTTCCGATCTNNNAAGATCGCTCCATTACACTCCAG
AGCACACGTCTGAACTCTTTCCCTACACGACGCTCTTCCGATCTNNNGTCTGTAGTCCTAGCTACCAGG
GAGCACACGTCTGAACTCTTTCCCTACACGACGCTCTTCCGATCTNNNCGAAACCCCGTCTGTACTAAA
GAGCACACGTCTGAACTCTTTCCCTACACGACGCTCTTCCGATCTNNNCCTGGGGGAACTACAATTCAA
GAGCACACGTCTGAACTCTTTCCCTACACGACGCTCTTCCGATCTNNNACAAGACAACCTATCCCTTGGT

AGCACACGTCTGAACTCTTTCCCTACACGACGCTCTTCCGATCTNNNAGCCAAGTACTTGATTGCCTACT
GAGCACACGTCTGAACTCTTTCCCTACACGACGCTCTTCCGATCTNNNCAGAAATGTGCACCCGTAATC
AGCACACGTCTGAACTCTTTCCCTACACGACGCTCTTCCGATCTNNNCAACAGTCCTAAGTCCACATCCA
AGCACACGTCTGAACTCTTTCCCTACACGACGCTCTTCCGATCTNNNATGCCCAGCTAAAGAACTCTTCT
CACGTCTGAACTCTTTCCCTACACGACGCTCTTCCGATCTNNNAGACTCTACCAAAAAACTCTTACACAC
GAGCACACGTCTGAACTCTTTCCCTACACGACGCTCTTCCGATCTNNNAAATCTCACCCACTGATCTAGC
AGCACACGTCTGAACTCTTTCCCTACACGACGCTCTTCCGATCTNNNACTCCATCTTTTGTCAGAGGCT
AGCACACGTCTGAACTCTTTCCCTACACGACGCTCTTCCGATCTNNNCTCCTGAGTAGCTGGGACTACA
AGCACACGTCTGAACTCTTTCCCTACACGACGCTCTTCCGATCTNNNCTCTCCTTTGTGCCCTATTTCCTC
AGCACACGTCTGAACTCTTTCCCTACACGACGCTCTTCCGATCTNNNCCACCACGTCCAGCTAATTTTTT
GAGCACACGTCTGAACTCTTTCCCTACACGACGCTCTTCCGATCTNNNCAATGATTGACTGGCATCAG
AGCACACGTCTGAACTCTTTCCCTACACGACGCTCTTCCGATCTNNNTCCTCTTACCACTCCCTCATTCTC
GCACACGTCTGAACTCTTTCCCTACACGACGCTCTTCCGATCTNNNATGGTAGGTGTTTTCTCTCAGGC
GAGCACACGTCTGAACTCTTTCCCTACACGACGCTCTTCCGATCTNNNCATTTCACTAAGTGCTGCCCAA
GCACACGTCTGAACTCTTTCCCTACACGACGCTCTTCCGATCTNNNTGAATAGGTGGAACTACAGGTG
AGAGCACACGTCTGAACTCTTTCCCTACACGACGCTCTTCCGATCTNNNCCTTTCATGTGTGTCTGAGT
GAGCACACGTCTGAACTCTTTCCCTACACGACGCTCTTCCGATCTNNNGAGAAGGGTCTCTTGGCTGAA
GAGCACACGTCTGAACTCTTTCCCTACACGACGCTCTTCCGATCTNNNGATATTTTCCTGCTGGACAAG
GAGCACACGTCTGAACTCTTTCCCTACACGACGCTCTTCCGATCTNNNCAATGGGTGTGTTCAAAGCTC
GAGCACACGTCTGAACTCTTTCCCTACACGACGCTCTTCCGATCTNNNGATTATAGCCATGTGCCACCA
AGCACACGTCTGAACTCTTTCCCTACACGACGCTCTTCCGATCTNNNTGATCTCCTGACCTTGTATGTG
GCACACGTCTGAACTCTTTCCCTACACGACGCTCTTCCGATCTNNNCACTGGCTCCTTACTCTTGAACTC
AGCACACGTCTGAACTCTTTCCCTACACGACGCTCTTCCGATCTNNNGATCTCTCCATGCTCAGGTGAT
AGCACACGTCTGAACTCTTTCCCTACACGACGCTCTTCCGATCTNNNCGATGAGCCTCTGATATTGGAC
AGCACACGTCTGAACTCTTTCCCTACACGACGCTCTTCCGATCTNNNCCACTGTGCATGGATTCCTTTTT
AGCACACGTCTGAACTCTTTCCCTACACGACGCTCTTCCGATCTNNNTTGCATATGTTGAACCAGCCTT
AGCACACGTCTGAACTCTTTCCCTACACGACGCTCTTCCGATCTNNNACCCACAAAACTGTAGCTACAT
AGCACACGTCTGAACTCTTTCCCTACACGACGCTCTTCCGATCTNNNGGTGTGGGTAAGAAATGCATT
GCACACGTCTGAACTCTTTCCCTACACGACGCTCTTCCGATCTNNNATATCCTTTTCTAGTCCTGGGCCC
GAGCACACGTCTGAACTCTTTCCCTACACGACGCTCTTCCGATCTNNNTGCAACTCCTAAGCTCAAGTGA
GAGCACACGTCTGAACTCTTTCCCTACACGACGCTCTTCCGATCTNNNCCATTTGAGTCTCCTGTGAGGA
GCACACGTCTGAACTCTTTCCCTACACGACGCTCTTCCGATCTNNNCAAAAAGGCTTTTGTGGCTACTA
GAGCACACGTCTGAACTCTTTCCCTACACGACGCTCTTCCGATCTNNNGGCAGCAAATCACACTCTGAAT
AGCACACGTCTGAACTCTTTCCCTACACGACGCTCTTCCGATCTNNNCTTCATCAGTGAGAAAGGAGC
AGCACACGTCTGAACTCTTTCCCTACACGACGCTCTTCCGATCTNNNGTCGTGATGTGCTCTACTCTAGC
GAGCACACGTCTGAACTCTTTCCCTACACGACGCTCTTCCGATCTNNNGGATTACAAGTGTGTACCA
GAGCACACGTCTGAACTCTTTCCCTACACGACGCTCTTCCGATCTNNNGGGTTTGACAAAAGGTACAA
GCACACGTCTGAACTCTTTCCCTACACGACGCTCTTCCGATCTNNNTGCACTCAGTGAAATCAGGAGA
AGCACACGTCTGAACTCTTTCCCTACACGACGCTCTTCCGATCTNNNTGGATTATGCCACTACACTCCAG
AGAGCACACGTCTGAACTCTTTCCCTACACGACGCTCTTCCGATCTNNNGGAAGTTGTAGTGAGCCGAG
AGAGCACACGTCTGAACTCTTTCCCTACACGACGCTCTTCCGATCTNNNTAGGGCAGGAGAATCACTTG
GAGCACACGTCTGAACTCTTTCCCTACACGACGCTCTTCCGATCTNNNTTGAGACGGGAGAATTGCTT
AGCACACGTCTGAACTCTTTCCCTACACGACGCTCTTCCGATCTNNNCTGAGGCAGGAGAATCACTTGA

AGCACACGTCTGAACTCTTTCCCTACACGACGCTCTTCCGATCTNNNCTGGGTGAGAAGAAACCAACTT
AGAGCACACGTCTGAACTCTTTCCCTACACGACGCTCTTCCGATCTNNNACCGGTACATGATACTTGAC
AGCACACGTCTGAACTCTTTCCCTACACGACGCTCTTCCGATCTNNNGTGCCCCAACATCCATTCTTTT
GAGCACACGTCTGAACTCTTTCCCTACACGACGCTCTTCCGATCTNNNCACAGGGGAGTGTCTCAAAT
AGCACACGTCTGAACTCTTTCCCTACACGACGCTCTTCCGATCTNNNGCCACATCTTCAGTACAAAGAG
AGAGCACACGTCTGAACTCTTTCCCTACACGACGCTCTTCCGATCTNNNGGTGAGCCAAGATTGTGCT
AGCACACGTCTGAACTCTTTCCCTACACGACGCTCTTCCGATCTNNNGATTGAGACCATCCTGGCTAA
AGCACACGTCTGAACTCTTTCCCTACACGACGCTCTTCCGATCTNNNCTAGAGTAGCTGGGATTACAG
GAGCACACGTCTGAACTCTTTCCCTACACGACGCTCTTCCGATCTNNNGCCACAGAGTAAGACTCCAT
AGAGCACACGTCTGAACTCTTTCCCTACACGACGCTCTTCCGATCTNNNAAGCCTTGGGTGATATATG
AGCACACGTCTGAACTCTTTCCCTACACGACGCTCTTCCGATCTNNNAGGTGGGTTCGAAGATTTCCTT
GAGCACACGTCTGAACTCTTTCCCTACACGACGCTCTTCCGATCTNNNCTGGCTTGGTGGAATAAAAG
GCACACGTCTGAACTCTTTCCCTACACGACGCTCTTCCGATCTNNNCCTGGGTGATACTCTCTCTCTC
GAGCACACGTCTGAACTCTTTCCCTACACGACGCTCTTCCGATCTNNNCAGGCTGGCTAACATGGTAA
GCACACGTCTGAACTCTTTCCCTACACGACGCTCTTCCGATCTNNNTGTCTCTCTGTCTCTCTGTCTGTC
GCACACGTCTGAACTCTTTCCCTACACGACGCTCTTCCGATCTNNNTGAGATTTACACTGCATGCTGG
AGCACACGTCTGAACTCTTTCCCTACACGACGCTCTTCCGATCTNNNTTCCCATCTGAAACCATGGAGA
GAGCACACGTCTGAACTCTTTCCCTACACGACGCTCTTCCGATCTNNNAGGGAGAGAGAGAGAGAGAG
AGAGCACACGTCTGAACTCTTTCCCTACACGACGCTCTTCCGATCTNNNAGATTGCTAACACAGTTTTG
AGCACACGTCTGAACTCTTTCCCTACACGACGCTCTTCCGATCTNNNAAAGGGAGAAAGGGAGAGA
AGCACACGTCTGAACTCTTTCCCTACACGACGCTCTTCCGATCTNNNCCACGGCCACCTAAGTATTTGT
AGCACACGTCTGAACTCTTTCCCTACACGACGCTCTTCCGATCTNNNTGATTGGAATTTCCTGCAGGAG
AGCACACGTCTGAACTCTTTCCCTACACGACGCTCTTCCGATCTNNNTCTTCCCTTACTTTCTCCCAAGC
AGCACACGTCTGAACTCTTTCCCTACACGACGCTCTTCCGATCTNNNAGAGAGAGAGAGAGAGAGG
AGCACACGTCTGAACTCTTTCCCTACACGACGCTCTTCCGATCTNNNAGGGAGAGAGAGAGAGAGAG
GAGCACACGTCTGAACTCTTTCCCTACACGACGCTCTTCCGATCTNNNAGTAGAAGAGGAAGCAGGTA
GAGCACACGTCTGAACTCTTTCCCTACACGACGCTCTTCCGATCTNNNTTGAGGCTGCAGTAAACTCA
AGAGCACACGTCTGAACTCTTTCCCTACACGACGCTCTTCCGATCTNNNCATTCTGAGTAGCCTGGGGT
GAGCACACGTCTGAACTCTTTCCCTACACGACGCTCTTCCGATCTNNNAAATCAAGCCATCGCACTCCAT
AGCACACGTCTGAACTCTTTCCCTACACGACGCTCTTCCGATCTNNNTAAACCTCTTTTGGGGGAGCTT
AGCACACGTCTGAACTCTTTCCCTACACGACGCTCTTCCGATCTNNNGAACATGGCCACACTCATTCAT
GCACACGTCTGAACTCTTTCCCTACACGACGCTCTTCCGATCTNNNCCAGAATCTTGGTTTCCAATCCCC
GAGCACACGTCTGAACTCTTTCCCTACACGACGCTCTTCCGATCTNNNAGAGAATGTTTTTGCGCATGT
AGCACACGTCTGAACTCTTTCCCTACACGACGCTCTTCCGATCTNNNCAGAGTAAGACCCTGTCCCAAA
GAGCACACGTCTGAACTCTTTCCCTACACGACGCTCTTCCGATCTNNNCCTACTGGTCTGTTGACTACC
AGCACACGTCTGAACTCTTTCCCTACACGACGCTCTTCCGATCTNNNCATGGGTGTGAAGTGGTATCT
AGCACACGTCTGAACTCTTTCCCTACACGACGCTCTTCCGATCTNNNCTGGGAAATGGTAGAGCTGGA
GAGCACACGTCTGAACTCTTTCCCTACACGACGCTCTTCCGATCTNNNTTTTTTGAGAGACAGGGTTT
AGCACACGTCTGAACTCTTTCCCTACACGACGCTCTTCCGATCTNNNTGTTCTTCCTGTCTTTCCCCTTTC
AGCACACGTCTGAACTCTTTCCCTACACGACGCTCTTCCGATCTNNNGGAGCAGAAAAAGCAAGACTG
AGCACACGTCTGAACTCTTTCCCTACACGACGCTCTTCCGATCTNNNCATCCTCATTTTGGTTGAGCACA
GCACACGTCTGAACTCTTTCCCTACACGACGCTCTTCCGATCTNNNTTTTGGAATGGCAGCATTCAGA
AGCACACGTCTGAACTCTTTCCCTACACGACGCTCTTCCGATCTNNNTTTTGTTCCTCAAGTGCACCAA

AGCACACGTCTGAACTCTTTCCCTACACGACGCTCTTCCGATCTNNNGAGCTTCCAGGTCATGTTTAG
AGCACACGTCTGAACTCTTTCCCTACACGACGCTCTTCCGATCTNNNTAGATGCTGGTGTTCAACATG
GCACACGTCTGAACTCTTTCCCTACACGACGCTCTTCCGATCTNNNGGTCCTTTCACTCCTTTCCTCAA
GCACACGTCTGAACTCTTTCCCTACACGACGCTCTTCCGATCTNNNTCAGGTCTGAGTTTGCACAAGA
AGCACACGTCTGAACTCTTTCCCTACACGACGCTCTTCCGATCTNNNGAGGTTGCAGTGAACCAAGAT
GAGCACACGTCTGAACTCTTTCCCTACACGACGCTCTTCCGATCTNNNAGAGAACTGTTTGAACCTGG
GAGCACACGTCTGAACTCTTTCCCTACACGACGCTCTTCCGATCTNNNGAAGATCTTTCCACTGCACTC
GCACACGTCTGAACTCTTTCCCTACACGACGCTCTTCCGATCTNNNCCAAGATTGCACCACTGTACTT
GAGCACACGTCTGAACTCTTTCCCTACACGACGCTCTTCCGATCTNNNCTGAGGCAGGAAAATTGCTT
AGCACACGTCTGAACTCTTTCCCTACACGACGCTCTTCCGATCTNNNCCTAGTGAGACTGCAGAGAGA
AGCACACGTCTGAACTCTTTCCCTACACGACGCTCTTCCGATCTNNNACTGTTGGCTGAGAAATCAT
GCACACGTCTGAACTCTTTCCCTACACGACGCTCTTCCGATCTNNNGCCAAGATTATGTCACTGCACTC
AGCACACGTCTGAACTCTTTCCCTACACGACGCTCTTCCGATCTNNNGGAGATTGCAGTGAGCCAAG
AGCACACGTCTGAACTCTTTCCCTACACGACGCTCTTCCGATCTNNNTTTTCACCTAGCAAAGCTCACA
GCACACGTCTGAACTCTTTCCCTACACGACGCTCTTCCGATCTNNNCTGTAGTCCCAGCTACTCAGGA
AGCACACGTCTGAACTCTTTCCCTACACGACGCTCTTCCGATCTNNNAGCATAACTGACAGAGCAAGA
AGCACACGTCTGAACTCTTTCCCTACACGACGCTCTTCCGATCTNNNGGGATGAGAATTGCTTGAAC
AGCACACGTCTGAACTCTTTCCCTACACGACGCTCTTCCGATCTNNNGCACATGCTTGTAATACCAGC
AGCACACGTCTGAACTCTTTCCCTACACGACGCTCTTCCGATCTNNNCTCAGGAGGCTGAAGCATAAG
AGCACACGTCTGAACTCTTTCCCTACACGACGCTCTTCCGATCTNNNAGGTTTCAGTGAGCCAAGAT
GAGCACACGTCTGAACTCTTTCCCTACACGACGCTCTTCCGATCTNNNTCGGAATAAGAATGGCAAA
GAGCACACGTCTGAACTCTTTCCCTACACGACGCTCTTCCGATCTNNNAGAGGTTGCAGTAAGCTGA
AGCACACGTCTGAACTCTTTCCCTACACGACGCTCTTCCGATCTNNNCTATGGTCCCAGCTACTCAGG
AGCACACGTCTGAACTCTTTCCCTACACGACGCTCTTCCGATCTNNNCAAGCTGGAGCACAATGAAAT
AGCACACGTCTGAACTCTTTCCCTACACGACGCTCTTCCGATCTNNNTTTGGGTGCAACTGTGTAAA
AGAGCACACGTCTGAACTCTTTCCCTACACGACGCTCTTCCGATCTNNNTAGTTGCCAAGCCTTCACTG
AGCACACGTCTGAACTCTTTCCCTACACGACGCTCTTCCGATCTNNNAAGAAATTGTCTCTGGAGGG
AGCACACGTCTGAACTCTTTCCCTACACGACGCTCTTCCGATCTNNNCAGCTTTGAATCCTTCCATCAG
GCACACGTCTGAACTCTTTCCCTACACGACGCTCTTCCGATCTNNNTTAGCTGTGGGTTTGTTGTATG
AGCACACGTCTGAACTCTTTCCCTACACGACGCTCTTCCGATCTNNNTGTGAATAGTGATGATGTGC
AGCACACGTCTGAACTCTTTCCCTACACGACGCTCTTCCGATCTNNNTCCTGAGTAGCTGGGATTAC
AGCACACGTCTGAACTCTTTCCCTACACGACGCTCTTCCGATCTNNNAAAGTGCTGGGATTACAAGCA
AGCACACGTCTGAACTCTTTCCCTACACGACGCTCTTCCGATCTNNNTATCACCTATTCAACACCCTCCC
AGCACACGTCTGAACTCTTTCCCTACACGACGCTCTTCCGATCTNNNGCTCCGTAGTTCTTAGGATGG
GCACACGTCTGAACTCTTTCCCTACACGACGCTCTTCCGATCTNNNGAGATAGCAGAGTGACTAAGG
GCACACGTCTGAACTCTTTCCCTACACGACGCTCTTCCGATCTNNNGGGGTTTTTGTGGTTCCATAG
GAGCACACGTCTGAACTCTTTCCCTACACGACGCTCTTCCGATCTNNNAGGAGGCATATAGATACAG
GCACACGTCTGAACTCTTTCCCTACACGACGCTCTTCCGATCTNNNTGAGATGATTGTGTTGGAGCA
GAGCACACGTCTGAACTCTTTCCCTACACGACGCTCTTCCGATCTNNNCATCTCTGCTGCTTTCCTTATG
GCACACGTCTGAACTCTTTCCCTACACGACGCTCTTCCGATCTNNNTATGGCATTCTTGGGAGTGTGA
GCACACGTCTGAACTCTTTCCCTACACGACGCTCTTCCGATCTNNNACAGGCAGGACAATGAGTAACA
GCACACGTCTGAACTCTTTCCCTACACGACGCTCTTCCGATCTNNNTTGGCATTTTGATAGGGATTGC
AGCACACGTCTGAACTCTTTCCCTACACGACGCTCTTCCGATCTNNNTAACATAACCAAGAGGCATCC

AGCACACGTCTGAACTCTTTCCCTACACGACGCTCTTCCGATCTNNNGCTTGGAGAATGAGTGCAAA
AGCACACGTCTGAACTCTTTCCCTACACGACGCTCTTCCGATCTNNNGTTCAAGAGATTCTCCTGCCTC
AGCACACGTCTGAACTCTTTCCCTACACGACGCTCTTCCGATCTNNNCTGAGGCACAAGAATCACCTG
AGCACACGTCTGAACTCTTTCCCTACACGACGCTCTTCCGATCTNNNATCCAGGAATGAATACACCGTC
GAGCACACGTCTGAACTCTTTCCCTACACGACGCTCTTCCGATCTNNNGCAAGACCCAGACTTTTCTTT
AGCACACGTCTGAACTCTTTCCCTACACGACGCTCTTCCGATCTNNNAACCACCAAAAGCACTCACTTT
GCACACGTCTGAACTCTTTCCCTACACGACGCTCTTCCGATCTNNNTGGCTACAGTAGACTAGCTTTG
GAGCACACGTCTGAACTCTTTCCCTACACGACGCTCTTCCGATCTNNNTCTCATTGCCCAGAATTGAGT
AGCACACGTCTGAACTCTTTCCCTACACGACGCTCTTCCGATCTNNNTCAGCTGTGGAATGCAAATGTT
AGCACACGTCTGAACTCTTTCCCTACACGACGCTCTTCCGATCTNNNCTTCCTAGGCTCAAGTGATCCTC
AGCACACGTCTGAACTCTTTCCCTACACGACGCTCTTCCGATCTNNNAAGAAGACAGAGTGCCCTGTA
GAGCACACGTCTGAACTCTTTCCCTACACGACGCTCTTCCGATCTNNNCAGGACATGCTACCCTAAAAC
GCACACGTCTGAACTCTTTCCCTACACGACGCTCTTCCGATCTNNNCACTTCTCCCCACAGAATAACCAC
GCACACGTCTGAACTCTTTCCCTACACGACGCTCTTCCGATCTNNNGCCCAAGCTAAGACAGTAACAA
AGCACACGTCTGAACTCTTTCCCTACACGACGCTCTTCCGATCTNNNAGCAGGAGAATCTCTTGAACC
AGCACACGTCTGAACTCTTTCCCTACACGACGCTCTTCCGATCTNNNGGAGGTTACAGTGAGCCAAG
GAGCACACGTCTGAACTCTTTCCCTACACGACGCTCTTCCGATCTNNNACGCTGAGGAAGAAGAATCA
AGCACACGTCTGAACTCTTTCCCTACACGACGCTCTTCCGATCTNNNTGTGCCTGTAATCCCAGCTATT
AGCACACGTCTGAACTCTTTCCCTACACGACGCTCTTCCGATCTNNNGAGCCAAGATTGCATCACTGTA
GAGCACACGTCTGAACTCTTTCCCTACACGACGCTCTTCCGATCTNNNTGTAGTCCCAGCTACTTAGGA
AGCACACGTCTGAACTCTTTCCCTACACGACGCTCTTCCGATCTNNNCTCCTTTCCCCAACCAGCTATAT
AGCACACGTCTGAACTCTTTCCCTACACGACGCTCTTCCGATCTNNNGCTCCAAGTCAGGTAAGACAG
AGCACACGTCTGAACTCTTTCCCTACACGACGCTCTTCCGATCTNNNGCAGGAGAATCATTTGAACCT
GAGCACACGTCTGAACTCTTTCCCTACACGACGCTCTTCCGATCTNNNCAAAATTAGTCAGGCGTGGT
GCACACGTCTGAACTCTTTCCCTACACGACGCTCTTCCGATCTNNNGTAATCCCAGCTACTTAGGAGG
AGCACACGTCTGAACTCTTTCCCTACACGACGCTCTTCCGATCTNNNTAGATGATGATCAGGCACAGT
AGCACACGTCTGAACTCTTTCCCTACACGACGCTCTTCCGATCTNNNTGAGCTCACGAGTTTGAGACT
AGCACACGTCTGAACTCTTTCCCTACACGACGCTCTTCCGATCTNNNACCAGAGACCTTTGTTCACTTG
GAGCACACGTCTGAACTCTTTCCCTACACGACGCTCTTCCGATCTNNNCGGTAAACTGAGCTGGGAG
GAGCACACGTCTGAACTCTTTCCCTACACGACGCTCTTCCGATCTNNNGGGCAACAAGAATGAAACTC
AGCACACGTCTGAACTCTTTCCCTACACGACGCTCTTCCGATCTNNNCTGTAGTCCCAGCTACTTTGGA
GAGCACACGTCTGAACTCTTTCCCTACACGACGCTCTTCCGATCTNNNCCTGTCCTACACGGTGTTCTA
AGCACACGTCTGAACTCTTTCCCTACACGACGCTCTTCCGATCTNNNCTGAGGCAGGAGAATCACTTG
GCACACGTCTGAACTCTTTCCCTACACGACGCTCTTCCGATCTNNNGGAGAATAGCGTGAACCTGGA
GAGCACACGTCTGAACTCTTTCCCTACACGACGCTCTTCCGATCTNNNGTAGTGAGCCGAGATCATGC
AGAGCACACGTCTGAACTCTTTCCCTACACGACGCTCTTCCGATCTNNNAGAATGGCATGAACCCAGG
GAGCACACGTCTGAACTCTTTCCCTACACGACGCTCTTCCGATCTNNNTGCCTGTAATTCCAGCTACTC
AGCACACGTCTGAACTCTTTCCCTACACGACGCTCTTCCGATCTNNNGAATCGTTTGAACCTGGGAAG
AGCACACGTCTGAACTCTTTCCCTACACGACGCTCTTCCGATCTNNNCTGTAGTCCCAGCTACTCGAG
GAGCACACGTCTGAACTCTTTCCCTACACGACGCTCTTCCGATCTNNNAGGAGAATCAGTTCCTTGAAC
AGCACACGTCTGAACTCTTTCCCTACACGACGCTCTTCCGATCTNNNCTTAGGCAACAAGAGCGAAAC
AGCACACGTCTGAACTCTTTCCCTACACGACGCTCTTCCGATCTNNNCAGGAGGCTGAGACAGGATA
GAGCACACGTCTGAACTCTTTCCCTACACGACGCTCTTCCGATCTNNNTCTCGAACTCCTAGCCTCAAG

AGCACACGTCTGAACTCTTTCCCTACACGACGCTCTTCCGATCTNNNGCACCTGTAGTCTCAGCTACTTG
GAGCACACGTCTGAACTCTTTCCCTACACGACGCTCTTCCGATCTNNNCTGTGGTCCTAGCTACTCAGG
AGCACACGTCTGAACTCTTTCCCTACACGACGCTCTTCCGATCTNNNTGTGATGGAAAGATTGCCTGA
GAGCACACGTCTGAACTCTTTCCCTACACGACGCTCTTCCGATCTNNNAGTGAGTAGAGATTGTGCCAC
AGCACACGTCTGAACTCTTTCCCTACACGACGCTCTTCCGATCTNNNTGCCTGTAATCTCAGCTACTTGC
GAGCACACGTCTGAACTCTTTCCCTACACGACGCTCTTCCGATCTNNNCTGTAATTCCAGCTACTCGGGA
AGCACACGTCTGAACTCTTTCCCTACACGACGCTCTTCCGATCTNNNCCACTGCACCCAGCTATTGTATTC
AGCACACGTCTGAACTCTTTCCCTACACGACGCTCTTCCGATCTNNNTTCCCTCGTTTTGTGTTTGTCTG
GAGCACACGTCTGAACTCTTTCCCTACACGACGCTCTTCCGATCTNNNAAGGGTACAGAGAGGTGGAA
AGCACACGTCTGAACTCTTTCCCTACACGACGCTCTTCCGATCTNNNGATGGTCTCAATCTCCTGACCTC
GAGCACACGTCTGAACTCTTTCCCTACACGACGCTCTTCCGATCTNNNTCTCCCATGTGCATCATTCAAG
AGCACACGTCTGAACTCTTTCCCTACACGACGCTCTTCCGATCTNNNGCTTTTTCTGAAATTGAGCAGG
AGCACACGTCTGAACTCTTTCCCTACACGACGCTCTTCCGATCTNNNTTCCGAAACAGCTGGGATTACA
AGCACACGTCTGAACTCTTTCCCTACACGACGCTCTTCCGATCTNNNCAGGCTGGTCTCTATCTCTTGAC
AGCACACGTCTGAACTCTTTCCCTACACGACGCTCTTCCGATCTNNNCTGGTGAAGATGCTGTGAACAT
AGCACACGTCTGAACTCTTTCCCTACACGACGCTCTTCCGATCTNNNTGCTTTGCTATCCCACATTCTCTC
AGCACACGTCTGAACTCTTTCCCTACACGACGCTCTTCCGATCTNNNTGTCTCTTTCACCATAGCCAGAG
AGCACACGTCTGAACTCTTTCCCTACACGACGCTCTTCCGATCTNNNCTGCCTCCTAAAGTTCTGGGAT
AGCACACGTCTGAACTCTTTCCCTACACGACGCTCTTCCGATCTNNNTCAAGTCTGCTGCTGAACTTTTC
GAGCACACGTCTGAACTCTTTCCCTACACGACGCTCTTCCGATCTNNNTCTCCCTCCCTGAATAGTTCCTT
GAGCACACGTCTGAACTCTTTCCCTACACGACGCTCTTCCGATCTNNNGGAAATGTTTTTTGGGGGGA
AGCACACGTCTGAACTCTTTCCCTACACGACGCTCTTCCGATCTNNNCTCATTGCAGCCTAGACTTCCTG
AGCACACGTCTGAACTCTTTCCCTACACGACGCTCTTCCGATCTNNNCAAACTACCTGCCTCAGCTTTTC
AGCACACGTCTGAACTCTTTCCCTACACGACGCTCTTCCGATCTNNNACGACTGGCTGATGGTCTTATT
AGCACACGTCTGAACTCTTTCCCTACACGACGCTCTTCCGATCTNNNGCTGGTCTCAAATTTCTGACGTC
GAGCACACGTCTGAACTCTTTCCCTACACGACGCTCTTCCGATCTNNNTATTACCAGTGGAGAAAGGGT
AGCACACGTCTGAACTCTTTCCCTACACGACGCTCTTCCGATCTNNNGATTGCCTAAGGACAGGAGTT
AGCACACGTCTGAACTCTTTCCCTACACGACGCTCTTCCGATCTNNNTTGGCTCTGTATTCACTCTTCCCC
AGCACACGTCTGAACTCTTTCCCTACACGACGCTCTTCCGATCTNNNGGAAAGCAATGCTATGATCACC
AGCACACGTCTGAACTCTTTCCCTACACGACGCTCTTCCGATCTNNNAGTGAGCCAAGATTGCATCATT
AGAGCACACGTCTGAACTCTTTCCCTACACGACGCTCTTCCGATCTNNNGAGCTTGTAGTGAGCCGAG
AGCACACGTCTGAACTCTTTCCCTACACGACGCTCTTCCGATCTNNNTTGAACTCAGGAGGTGGAGGT
GAGCACACGTCTGAACTCTTTCCCTACACGACGCTCTTCCGATCTNNNGAAGGATCACTCGAGCTTAG
AGCACACGTCTGAACTCTTTCCCTACACGACGCTCTTCCGATCTNNNTTCATCTGATGCCAGTTCCATCA
AGCACACGTCTGAACTCTTTCCCTACACGACGCTCTTCCGATCTNNNTGAGGTAGGAGAATGGTGTGA
GAGCACACGTCTGAACTCTTTCCCTACACGACGCTCTTCCGATCTNNNGCAGGAGAATCACTTGAACCT
AGCACACGTCTGAACTCTTTCCCTACACGACGCTCTTCCGATCTNNNCTGAGGCAGAGAATTGCTTGA
AGCACACGTCTGAACTCTTTCCCTACACGACGCTCTTCCGATCTNNNATCACTTGAATCCAGGAGGCAA
GAGCACACGTCTGAACTCTTTCCCTACACGACGCTCTTCCGATCTNNNAGTGAGCCAAGATTGCATGAT
AGAGCACACGTCTGAACTCTTTCCCTACACGACGCTCTTCCGATCTNNNGGAGGTTGTAGTGAGCCAAC
GAGCACACGTCTGAACTCTTTCCCTACACGACGCTCTTCCGATCTNNNGAATCGCTTGAACATGGGATG
GAGCACACGTCTGAACTCTTTCCCTACACGACGCTCTTCCGATCTNNNCTGTAATCCCAGCTACTCAGGA
GCACACGTCTGAACTCTTTCCCTACACGACGCTCTTCCGATCTNNNGACAGTGTTAGGTGGCACAAAA

AGCACACGTCTGAACTCTTTCCCTACACGACGCTCTTCCGATCTNNNTATCTGTTTAGGTCCCAGGTTG
GAGCACACGTCTGAACTCTTTCCCTACACGACGCTCTTCCGATCTNNNCACCTGGCCTATTGTTACTCT
AGCACACGTCTGAACTCTTTCCCTACACGACGCTCTTCCGATCTNNNAAACCAGACCACCTAAAGCAAA
GAGCACACGTCTGAACTCTTTCCCTACACGACGCTCTTCCGATCTNNNTCTTGTTCTGTGTTTTCTCCAG
AGCACACGTCTGAACTCTTTCCCTACACGACGCTCTTCCGATCTNNNATCACCTTCCACCACGAGTAAA
GAGCACACGTCTGAACTCTTTCCCTACACGACGCTCTTCCGATCTNNNGGTGGGATCCTTATCAGGGTT
AGCACACGTCTGAACTCTTTCCCTACACGACGCTCTTCCGATCTNNNAGTTTATGGAGCCAATGCCCTA
AGCACACGTCTGAACTCTTTCCCTACACGACGCTCTTCCGATCTNNNGATGGTCTCAATCTCCTGACCT
GAGCACACGTCTGAACTCTTTCCCTACACGACGCTCTTCCGATCTNNNAAGTGTAGAAGCTTCCATCCC
AGCACACGTCTGAACTCTTTCCCTACACGACGCTCTTCCGATCTNNNGGTCTCGAACTCCTAACCTCAA
AGCACACGTCTGAACTCTTTCCCTACACGACGCTCTTCCGATCTNNNACTTGGGAATAAGGGGAATG
GAGCACACGTCTGAACTCTTTCCCTACACGACGCTCTTCCGATCTNNNTTGAGGCAGGAGAATCACTT
GAGCACACGTCTGAACTCTTTCCCTACACGACGCTCTTCCGATCTNNNCTGGAACTCCTGACTTCAAGT
AGCACACGTCTGAACTCTTTCCCTACACGACGCTCTTCCGATCTNNNATACAGTAGTGCACTCCTGTGA
GAGCACACGTCTGAACTCTTTCCCTACACGACGCTCTTCCGATCTNNNTTACAGCATTTTGAGCCACTG
AGCACACGTCTGAACTCTTTCCCTACACGACGCTCTTCCGATCTNNNAATACCACCATATCATGCCTGCA
GAGCACACGTCTGAACTCTTTCCCTACACGACGCTCTTCCGATCTNNNAGGTTCAAGTGATTCTTCTGC
AGCACACGTCTGAACTCTTTCCCTACACGACGCTCTTCCGATCTNNNGGTGATCCGTGATGTGAAGA
GAGCACACGTCTGAACTCTTTCCCTACACGACGCTCTTCCGATCTNNNACCCTGTAACACTCCAATACC
GCACACGTCTGAACTCTTTCCCTACACGACGCTCTTCCGATCTNNNAGAAGGACAGTGCATGAATGA
GAGCACACGTCTGAACTCTTTCCCTACACGACGCTCTTCCGATCTNNNCCCATGGAAGGTCATCGTAT
AGCACACGTCTGAACTCTTTCCCTACACGACGCTCTTCCGATCTNNNCACCACCATGCCTGGATAATTT
GAGCACACGTCTGAACTCTTTCCCTACACGACGCTCTTCCGATCTNNNTGTTCCTGATCTTTGGGGGA
AGAGCACACGTCTGAACTCTTTCCCTACACGACGCTCTTCCGATCTNNNCTGTAATCCCAGCTACTCAG
AGCACACGTCTGAACTCTTTCCCTACACGACGCTCTTCCGATCTNNNGAGGCAGGATAATTGCTTGAA
AGCACACGTCTGAACTCTTTCCCTACACGACGCTCTTCCGATCTNNNAGGGAGGAGAATTTCTTGAA
AGCACACGTCTGAACTCTTTCCCTACACGACGCTCTTCCGATCTNNNACTAAAAGAAGGTGGGTTTG
AGCACACGTCTGAACTCTTTCCCTACACGACGCTCTTCCGATCTNNNAGAATCGTTTGAACCTAGGAGG
GAGCACACGTCTGAACTCTTTCCCTACACGACGCTCTTCCGATCTNNNGTTGCAGTGAGCCAAGATCAT
AGCACACGTCTGAACTCTTTCCCTACACGACGCTCTTCCGATCTNNNGGCATCTGTAATCCCAGCTACT
GAGCACACGTCTGAACTCTTTCCCTACACGACGCTCTTCCGATCTNNNGACCCCCTTCTCCAAAAACAA
GAGCACACGTCTGAACTCTTTCCCTACACGACGCTCTTCCGATCTNNNGACCCCCTTCTCCAAAAACAA
AGAGCACACGTCTGAACTCTTTCCCTACACGACGCTCTTCCGATCTNNNAGGTTGCAGTTAGTTGAGA
GAGCACACGTCTGAACTCTTTCCCTACACGACGCTCTTCCGATCTNNNCAGTCCCAACAATGTGGCTTA
AGCACACGTCTGAACTCTTTCCCTACACGACGCTCTTCCGATCTNNNGAATCTCTGTGTGTTAGGCAGG
GAGCACACGTCTGAACTCTTTCCCTACACGACGCTCTTCCGATCTNNNAAGGCAGGATAATTGCTTGA
GCACACGTCTGAACTCTTTCCCTACACGACGCTCTTCCGATCTNNNAGAAGTTGCAGTGAGCCAAGAT
GAGCACACGTCTGAACTCTTTCCCTACACGACGCTCTTCCGATCTNNNGCAACAAGAGCGAAACTTCAT
AGCACACGTCTGAACTCTTTCCCTACACGACGCTCTTCCGATCTNNNCACCTGAGGTCAAGAGTTTGA
AGAGCACACGTCTGAACTCTTTCCCTACACGACGCTCTTCCGATCTNNNCACCTGTAATCCCAGCTACTC
GAGCACACGTCTGAACTCTTTCCCTACACGACGCTCTTCCGATCTNNNTTGCTTGAGCTCAGGAGTTCT
GAGCACACGTCTGAACTCTTTCCCTACACGACGCTCTTCCGATCTNNNACCAGCCTGGAAAAGATGGT
GAGCACACGTCTGAACTCTTTCCCTACACGACGCTCTTCCGATCTNNNACTTGGGAGGATAAGACAG

GAGCACACGTCTGAACTCTTTCCCTACACGACGCTCTTCCGATCTNNNGAGAATGGTGTGAAGCTGG
AGAGCACACGTCTGAACTCTTTCCCTACACGACGCTCTTCCGATCTNNNTAGGTGACAGAGTAAGACC
AGCACACGTCTGAACTCTTTCCCTACACGACGCTCTTCCGATCTNNNCTGTAATCCCAGCTACTCAGGA
AGCACACGTCTGAACTCTTTCCCTACACGACGCTCTTCCGATCTNNNGATTGCAGTGAGCTGAGATTG
GCACACGTCTGAACTCTTTCCCTACACGACGCTCTTCCGATCTNNNCATTGTCCAGTTGACTGTGCATT
AGAGCACACGTCTGAACTCTTTCCCTACACGACGCTCTTCCGATCTNNNTTCCTTTCTTCCTCCCTCTCT
AGCACACGTCTGAACTCTTTCCCTACACGACGCTCTTCCGATCTNNNTGCCAAACTAATCCAAAGCAACT
GAGCACACGTCTGAACTCTTTCCCTACACGACGCTCTTCCGATCTNNNCCTCCTGGATTCAAGCGATTCT
GCACACGTCTGAACTCTTTCCCTACACGACGCTCTTCCGATCTNNNTCTGCCTGGCATGTTTTGAAATT
GAGCACACGTCTGAACTCTTTCCCTACACGACGCTCTTCCGATCTNNNGGGCCGGTAGATTCGTTACTA
AGCACACGTCTGAACTCTTTCCCTACACGACGCTCTTCCGATCTNNNCCTGTTCCCCAAAAACCTATGG
AGCACACGTCTGAACTCTTTCCCTACACGACGCTCTTCCGATCTNNNCCTTCAGTTTCCTCTAGCCTTGT
GCACACGTCTGAACTCTTTCCCTACACGACGCTCTTCCGATCTNNNCACATGTCTCTCATCCTCCAACAC
AGCACACGTCTGAACTCTTTCCCTACACGACGCTCTTCCGATCTNNNATTCAATTGATCTGCCTGCCTTG
GAGCACACGTCTGAACTCTTTCCCTACACGACGCTCTTCCGATCTNNNGATCCTCTCACCTCAGCCTCTA
GAGCACACGTCTGAACTCTTTCCCTACACGACGCTCTTCCGATCTNNNGCCCCTCTATCTTTCTATTCCC
GAGCACACGTCTGAACTCTTTCCCTACACGACGCTCTTCCGATCTNNNTCCCACCTCAGTCTCCTAAGTA
AGCACACGTCTGAACTCTTTCCCTACACGACGCTCTTCCGATCTNNNTTCAGCCCCAGCAGTAAAAATTG
AGCACACGTCTGAACTCTTTCCCTACACGACGCTCTTCCGATCTNNNACCCTTTGTCCACATTCTTTCCAC
AGCACACGTCTGAACTCTTTCCCTACACGACGCTCTTCCGATCTNNNTGCCCTCTCAGCTACTTTTCTTTC
GAGCACACGTCTGAACTCTTTCCCTACACGACGCTCTTCCGATCTNNNTTCAAGTGATTCTCCTGCCTCA
AGCACACGTCTGAACTCTTTCCCTACACGACGCTCTTCCGATCTNNNCTCAGCCTCCTAAAGTGTTTGG
GAGCACACGTCTGAACTCTTTCCCTACACGACGCTCTTCCGATCTNNNTCTCCTGTGCTAACTCATTAGG
AGAGCACACGTCTGAACTCTTTCCCTACACGACGCTCTTCCGATCTNNNACTGCTTCCTAGTCGCTTTCT
GAGCACACGTCTGAACTCTTTCCCTACACGACGCTCTTCCGATCTNNNTTCTCTGGGCCTTCATTTTCTC
AGCACACGTCTGAACTCTTTCCCTACACGACGCTCTTCCGATCTNNNTGCTTGTCCTCTTACTCTGCTTT
AGCACACGTCTGAACTCTTTCCCTACACGACGCTCTTCCGATCTNNNAAAGGAATGCACTTTGAGGTC
AGCACACGTCTGAACTCTTTCCCTACACGACGCTCTTCCGATCTNNNCCACACTGACTTCCACAATGATT
GCACACGTCTGAACTCTTTCCCTACACGACGCTCTTCCGATCTNNNCCCACTGGTTTTTTGAACTGACA
GAGCACACGTCTGAACTCTTTCCCTACACGACGCTCTTCCGATCTNNNCTTCTCCCATGGTTTCTGCAAT
AGCACACGTCTGAACTCTTTCCCTACACGACGCTCTTCCGATCTNNNAGCTAGAACTACAGGCGCATAT
AGCACACGTCTGAACTCTTTCCCTACACGACGCTCTTCCGATCTNNNAAGTGATTCTCCTGTTTCAGCC
AGCACACGTCTGAACTCTTTCCCTACACGACGCTCTTCCGATCTNNNTGGTCTTGAACTCCTGACTTCAG
AGCACACGTCTGAACTCTTTCCCTACACGACGCTCTTCCGATCTNNNGGAGTAAGAAGGTGGGAGAA
AGCACACGTCTGAACTCTTTCCCTACACGACGCTCTTCCGATCTNNNGCCTTTTCTTTCATGCAGCTTCT
AGCACACGTCTGAACTCTTTCCCTACACGACGCTCTTCCGATCTNNNACTTCCTGTACCTTGCTTTCACT
GAGCACACGTCTGAACTCTTTCCCTACACGACGCTCTTCCGATCTNNNGTGCATTCACTGACCTCCAAAT
GAGCACACGTCTGAACTCTTTCCCTACACGACGCTCTTCCGATCTNNNGTGGGGTTCTAAAGAGGATC
AGCACACGTCTGAACTCTTTCCCTACACGACGCTCTTCCGATCTNNNGGTATCATTCCTTCAGAGGCTG
AGCACACGTCTGAACTCTTTCCCTACACGACGCTCTTCCGATCTNNNTTGAACTCCTGGCTTCAAGTGA
GAGCACACGTCTGAACTCTTTCCCTACACGACGCTCTTCCGATCTNNNGTATCCAGCATGCTCTCATCTC
AGCACACGTCTGAACTCTTTCCCTACACGACGCTCTTCCGATCTNNNGCAGGGAAGAGGAAGCAAAAT
GCACACGTCTGAACTCTTTCCCTACACGACGCTCTTCCGATCTNNNCACAGCACCCTCTAACATACCAA

GAGCACACGTCTGAACTCTTTCCCTACACGACGCTCTTCCGATCTNNNTCCTCAAAGTGACTTGTTTAG
AGCACACGTCTGAACTCTTTCCCTACACGACGCTCTTCCGATCTNNNTTTAGTAGAGACGGGGTTTC
AGCACACGTCTGAACTCTTTCCCTACACGACGCTCTTCCGATCTNNNCAAATGTGGAGCTGACTTTTC
GAGCACACGTCTGAACTCTTTCCCTACACGACGCTCTTCCGATCTNNNGTAATCTCAGCTACTCAGGA
AGCACACGTCTGAACTCTTTCCCTACACGACGCTCTTCCGATCTNNNAGACAGGAGAATCACTTGAAC
GAGCACACGTCTGAACTCTTTCCCTACACGACGCTCTTCCGATCTNNNCTGAGGCAGGAGAATCACTT
GAGCACACGTCTGAACTCTTTCCCTACACGACGCTCTTCCGATCTNNNGCAACAGAGCAAGATTCCATC
AGCACACGTCTGAACTCTTTCCCTACACGACGCTCTTCCGATCTNNNCCTGGGTGAAAAAGTGAGACT
AGCACACGTCTGAACTCTTTCCCTACACGACGCTCTTCCGATCTNNNGGTGCCTGTAATCCTGCTACTA
GAGCACACGTCTGAACTCTTTCCCTACACGACGCTCTTCCGATCTNNNAAAGTTGCAGTGAGCTAGGA
GCACACGTCTGAACTCTTTCCCTACACGACGCTCTTCCGATCTNNNCTGTAGTCCCAGCTACTCAGGAG
GAGCACACGTCTGAACTCTTTCCCTACACGACGCTCTTCCGATCTNNNGAGACTGTACCACTGTACTC
AGCACACGTCTGAACTCTTTCCCTACACGACGCTCTTCCGATCTNNNGGCATTTTGATTGGCTAGCCT
GAGCACACGTCTGAACTCTTTCCCTACACGACGCTCTTCCGATCTNNNTTGAGGCAGGAGAATCACTT
AGCACACGTCTGAACTCTTTCCCTACACGACGCTCTTCCGATCTNNNTCCAGCCTAGATGACAGAGAA
AGCACACGTCTGAACTCTTTCCCTACACGACGCTCTTCCGATCTNNNCCCTGTGCCTAGAATATCTGCT
GAGCACACGTCTGAACTCTTTCCCTACACGACGCTCTTCCGATCTNNNGGCTGAGGCACAAGAATTAC
GAGCACACGTCTGAACTCTTTCCCTACACGACGCTCTTCCGATCTNNNGAAGAATTGCTTGAACCCAG
AGCACACGTCTGAACTCTTTCCCTACACGACGCTCTTCCGATCTNNNCTGAGGCAGGAGAATCACTTG
AGCACACGTCTGAACTCTTTCCCTACACGACGCTCTTCCGATCTNNNTGTAGGTATGACAGAGCCCAA
AGAGCACACGTCTGAACTCTTTCCCTACACGACGCTCTTCCGATCTNNNCCTGTAATCCCAGCTACTCAA
GAGCACACGTCTGAACTCTTTCCCTACACGACGCTCTTCCGATCTNNNAGAATCATTTGAACCCCGGA
GAGCACACGTCTGAACTCTTTCCCTACACGACGCTCTTCCGATCTNNNAGGAGAATTGTTTGAACCTG
GAGCACACGTCTGAACTCTTTCCCTACACGACGCTCTTCCGATCTNNNACTTGTAATCCCTGCTACTTG
AGCACACGTCTGAACTCTTTCCCTACACGACGCTCTTCCGATCTNNNCCAGGCAACAAAAGCAAAACTT
AGAGCACACGTCTGAACTCTTTCCCTACACGACGCTCTTCCGATCTNNNGCAGGAGAATTGCTTGAAC
AGCACACGTCTGAACTCTTTCCCTACACGACGCTCTTCCGATCTNNNCTGTAAACCCAGCTACTCAGGA
AGAGCACACGTCTGAACTCTTTCCCTACACGACGCTCTTCCGATCTNNNGAGGTTGCAGTGAACCAAG
GAGCACACGTCTGAACTCTTTCCCTACACGACGCTCTTCCGATCTNNNGATTGTGCCACTATACTCCAG
AGCACACGTCTGAACTCTTTCCCTACACGACGCTCTTCCGATCTNNNTGTAATCCCAGCACTTTAGGAG
GAGCACACGTCTGAACTCTTTCCCTACACGACGCTCTTCCGATCTNNNATAGTGAAACCCCGTCCCTACT
GAGCACACGTCTGAACTCTTTCCCTACACGACGCTCTTCCGATCTNNNCTGTAATCCCAGCTACTCAGG
AGCACACGTCTGAACTCTTTCCCTACACGACGCTCTTCCGATCTNNNAGAGCTTGCAGTAAGTCGAGA
AGCACACGTCTGAACTCTTTCCCTACACGACGCTCTTCCGATCTNNNGGCAACAGAGTGAGACTTTGT
GAGCACACGTCTGAACTCTTTCCCTACACGACGCTCTTCCGATCTNNNGATCACAAGGTCAGGAGATC
AGCACACGTCTGAACTCTTTCCCTACACGACGCTCTTCCGATCTNNNGAGATTGCTGTGAGCCAAGA
AGCACACGTCTGAACTCTTTCCCTACACGACGCTCTTCCGATCTNNNTCCCAACACTTTAGGAGGTCA
GCACACGTCTGAACTCTTTCCCTACACGACGCTCTTCCGATCTNNNGGCAAGAAAATCACTTGAACCC
GAGCACACGTCTGAACTCTTTCCCTACACGACGCTCTTCCGATCTNNNCAACATAGCAAGACCCTGTC
GAGCACACGTCTGAACTCTTTCCCTACACGACGCTCTTCCGATCTNNNTGAGGCAGGAGAATCTCTTG
AGCACACGTCTGAACTCTTTCCCTACACGACGCTCTTCCGATCTNNNCCTGTAATCCCACCTACTCAGG
AGCACACGTCTGAACTCTTTCCCTACACGACGCTCTTCCGATCTNNNCTGAGACAGGAGAATGGCATG
GCACACGTCTGAACTCTTTCCCTACACGACGCTCTTCCGATCTNNNCAGAGGTTGTAGTGAGCCAAGA

GAGCACACGTCTGAACTCTTTCCCTACACGACGCTCTTCCGATCTNNNAAAAAGAATTTGGATGAGGC
AGCACACGTCTGAACTCTTTCCCTACACGACGCTCTTCCGATCTNNNTCTGACCATACCACTGTACTCCA
AGAGCACACGTCTGAACTCTTTCCCTACACGACGCTCTTCCGATCTNNNACACTTGTAAGCCCAGCTATT
AGCACACGTCTGAACTCTTTCCCTACACGACGCTCTTCCGATCTNNNACCTTCTGGGCTCAATTCATCTT
AGCACACGTCTGAACTCTTTCCCTACACGACGCTCTTCCGATCTNNNGATGGTCTTGATCTCCTGACCTC
AGCACACGTCTGAACTCTTTCCCTACACGACGCTCTTCCGATCTNNNCAACTGAGCTAACTGGCTATTG
GCACACGTCTGAACTCTTTCCCTACACGACGCTCTTCCGATCTNNNCATGAACTGTACATGGTCAGTGC
AGCACACGTCTGAACTCTTTCCCTACACGACGCTCTTCCGATCTNNNACATCCTCTGTGCTTTACCCATT
GCACACGTCTGAACTCTTTCCCTACACGACGCTCTTCCGATCTNNNTTCTGTCTTTGGAAAGTCACGGA
AGCACACGTCTGAACTCTTTCCCTACACGACGCTCTTCCGATCTNNNTCTGCTCTTTTCTGCTTCTCCATC
GCACACGTCTGAACTCTTTCCCTACACGACGCTCTTCCGATCTNNNGTTTCCCCCATCTTGTTCTTGTGC
GAGCACACGTCTGAACTCTTTCCCTACACGACGCTCTTCCGATCTNNNGACCTTCCAAAGTGCTGGGAT
GAGCACACGTCTGAACTCTTTCCCTACACGACGCTCTTCCGATCTNNNAGAGCAACTTCCAGAAGAAAC
GAGCACACGTCTGAACTCTTTCCCTACACGACGCTCTTCCGATCTNNNAATGGCGTAATCTTGGCTCACT
AGCACACGTCTGAACTCTTTCCCTACACGACGCTCTTCCGATCTNNNTTCAGGTCCAGGTTCTAATGTGC
GAGCACACGTCTGAACTCTTTCCCTACACGACGCTCTTCCGATCTNNNATGTGTCCAGTACAGTTAGGA
GAGCACACGTCTGAACTCTTTCCCTACACGACGCTCTTCCGATCTNNNGAGTAGCTGGGATTACAAGG
AGCACACGTCTGAACTCTTTCCCTACACGACGCTCTTCCGATCTNNNAGCAAACACTGATTCTCCACTCA
AGCACACGTCTGAACTCTTTCCCTACACGACGCTCTTCCGATCTNNNCACACAGCGTTCAAATTTCCATC
AGCACACGTCTGAACTCTTTCCCTACACGACGCTCTTCCGATCTNNNCTCAGAGAATCCACCCAACTTGG
GCACACGTCTGAACTCTTTCCCTACACGACGCTCTTCCGATCTNNNCCTCTAAAGGCACATACCCAGTTC
AGCACACGTCTGAACTCTTTCCCTACACGACGCTCTTCCGATCTNNNAAAGTGCTGGAGTTACAGGTGT
AGCACACGTCTGAACTCTTTCCCTACACGACGCTCTTCCGATCTNNNGATGAGATCTCCCTATGTTGCCC
AGCACACGTCTGAACTCTTTCCCTACACGACGCTCTTCCGATCTNNNCCACGCCCTGCTAATTTTTGTAT
GAGCACACGTCTGAACTCTTTCCCTACACGACGCTCTTCCGATCTNNNGAGTGCTGGGATTACAAGCA
GCACACGTCTGAACTCTTTCCCTACACGACGCTCTTCCGATCTNNNTTTTGAATTTTGTGCCATGTGGG
AGCACACGTCTGAACTCTTTCCCTACACGACGCTCTTCCGATCTNNNAAAATGGCCAAAATCCTTGCTC
GAGCACACGTCTGAACTCTTTCCCTACACGACGCTCTTCCGATCTNNNTGGTAGAGACAGGGTTTCTCC
GCACACGTCTGAACTCTTTCCCTACACGACGCTCTTCCGATCTNNNCTGCATGTCAGTGAGGAGTATG
GAGCACACGTCTGAACTCTTTCCCTACACGACGCTCTTCCGATCTNNNGCACTTCCTTCCCTTTCTCCTTA
AGCACACGTCTGAACTCTTTCCCTACACGACGCTCTTCCGATCTNNNTCTGAAACGGTGAGCCTTTTTC
AGCACACGTCTGAACTCTTTCCCTACACGACGCTCTTCCGATCTNNNCAGGATAGGAGCTCTGACCATT
AGCACACGTCTGAACTCTTTCCCTACACGACGCTCTTCCGATCTNNNCCCTAGTAGTTGAGATTACGCGA
AGCACACGTCTGAACTCTTTCCCTACACGACGCTCTTCCGATCTNNNGTTATCTTTTATCCCTTGCGCCCC
GAGCACACGTCTGAACTCTTTCCCTACACGACGCTCTTCCGATCTNNNGTCTTTTCAGGACAAGGAAGC
AGCACACGTCTGAACTCTTTCCCTACACGACGCTCTTCCGATCTNNNAGTCCTCCTGCTGTAGAGAGAA
GAGCACACGTCTGAACTCTTTCCCTACACGACGCTCTTCCGATCTNNNCCACTGCACTGGCTAACTTTTT
AGCACACGTCTGAACTCTTTCCCTACACGACGCTCTTCCGATCTNNNACTTTGTGCTGATCCCTACGATT
AGCACACGTCTGAACTCTTTCCCTACACGACGCTCTTCCGATCTNNNCTGTGTACTAGGACTACAGGCA
AGCACACGTCTGAACTCTTTCCCTACACGACGCTCTTCCGATCTNNNTCTTGGATCACTACAACCTCTGC
AGCACACGTCTGAACTCTTTCCCTACACGACGCTCTTCCGATCTNNNAACAGCCAAGTGTTATTCTGCAC
AGCACACGTCTGAACTCTTTCCCTACACGACGCTCTTCCGATCTNNNGAACATGAGGTTGCTCAGGAAT
GCACACGTCTGAACTCTTTCCCTACACGACGCTCTTCCGATCTNNNTGTTCTGTACTACCCAAGATGCC

GAGCACACGTCTGAACTCTTTCCCTACACGACGCTCTTCCGATCTNNNCTTCTACCTGATGGCCAGTGA
AGAGCACACGTCTGAACTCTTTCCCTACACGACGCTCTTCCGATCTNNNCCCCAAAACTTTCACTAGCT
GAGCACACGTCTGAACTCTTTCCCTACACGACGCTCTTCCGATCTNNNAAGTGTTGGGATTACAGGCA
AGCACACGTCTGAACTCTTTCCCTACACGACGCTCTTCCGATCTNNNAATAGAAGGAAGCAGCAGTGA
AGCACACGTCTGAACTCTTTCCCTACACGACGCTCTTCCGATCTNNNAAGCACAGACCCAAAAACATT
AGCACACGTCTGAACTCTTTCCCTACACGACGCTCTTCCGATCTNNNGATCACAGCTCACTATAGCCA
AGCACACGTCTGAACTCTTTCCCTACACGACGCTCTTCCGATCTNNNTGAGGTAGGAGAATCGCTTGA
AGCACACGTCTGAACTCTTTCCCTACACGACGCTCTTCCGATCTNNNAGAATTGCTTGAATTTGGGA
GAGCACACGTCTGAACTCTTTCCCTACACGACGCTCTTCCGATCTNNNGGAGAATCACTTGAACCTGG
AGCACACGTCTGAACTCTTTCCCTACACGACGCTCTTCCGATCTNNNTGTGATGAAACTCCGTATCCA
GAGCACACGTCTGAACTCTTTCCCTACACGACGCTCTTCCGATCTNNNCTGAGGCAGGAGAATCACTT
GCACACGTCTGAACTCTTTCCCTACACGACGCTCTTCCGATCTNNNGGCAGCAAAGTGAGACTCTAT
GAGCACACGTCTGAACTCTTTCCCTACACGACGCTCTTCCGATCTNNNCCAAGATCATGTCATTGCAC
AGCACACGTCTGAACTCTTTCCCTACACGACGCTCTTCCGATCTNNNTGCCTGTAATCCCAGTTACTT
AGCACACGTCTGAACTCTTTCCCTACACGACGCTCTTCCGATCTNNNGAGGTTGCAGTGAGCTGAGA
GAGCACACGTCTGAACTCTTTCCCTACACGACGCTCTTCCGATCTNNNGGCAACAAAGTGAGACTCTG
GCACACGTCTGAACTCTTTCCCTACACGACGCTCTTCCGATCTNNNCTGAGGCAGAGAATTGCTTGAA
AGCACACGTCTGAACTCTTTCCCTACACGACGCTCTTCCGATCTNNNGTTTTACAAGAAACTCCCAGG
GCACACGTCTGAACTCTTTCCCTACACGACGCTCTTCCGATCTNNNGAGGATTGCTTGAACCTAGGA
AGCACACGTCTGAACTCTTTCCCTACACGACGCTCTTCCGATCTNNNCTGTAGTCACAGCTACTCGGG
GCACACGTCTGAACTCTTTCCCTACACGACGCTCTTCCGATCTNNNCTTCGCTCCCATGCTAAGAAGTA
GAGCACACGTCTGAACTCTTTCCCTACACGACGCTCTTCCGATCTNNNGGTAGAGGTTACAGTGAGC
GAGCACACGTCTGAACTCTTTCCCTACACGACGCTCTTCCGATCTNNNCAAGATCGTGCCATTACACT
GAGCACACGTCTGAACTCTTTCCCTACACGACGCTCTTCCGATCTNNNGGTGACAGAGAAAGACCTTG
GAGCACACGTCTGAACTCTTTCCCTACACGACGCTCTTCCGATCTNNNCTGAGACAGGTGAATTGCTT
GCACACGTCTGAACTCTTTCCCTACACGACGCTCTTCCGATCTNNNAGAACTGTTTGAACTCAGGAG
AGCACACGTCTGAACTCTTTCCCTACACGACGCTCTTCCGATCTNNNATGGTGGCTCATACCTGTAAT
GCACACGTCTGAACTCTTTCCCTACACGACGCTCTTCCGATCTNNNAGGCAAGAGAATCACTTGAACC
GCACACGTCTGAACTCTTTCCCTACACGACGCTCTTCCGATCTNNNCGTAATCCAAGCTACTCAGGAG
AGCACACGTCTGAACTCTTTCCCTACACGACGCTCTTCCGATCTNNNGAGGATCACTTGAACACAGGA
AGCACACGTCTGAACTCTTTCCCTACACGACGCTCTTCCGATCTNNNGTGCCTGTAGTCCCAGTTACTT
GCACACGTCTGAACTCTTTCCCTACACGACGCTCTTCCGATCTNNNCAGGAGGTGGAGGTTATATGA
GAGCACACGTCTGAACTCTTTCCCTACACGACGCTCTTCCGATCTNNNAGGAGAATCACTTGAACCCA
AGAGCACACGTCTGAACTCTTTCCCTACACGACGCTCTTCCGATCTNNNTTGGGATGTTGAGGTGGAT
AGCACACGTCTGAACTCTTTCCCTACACGACGCTCTTCCGATCTNNNGGTATGGTAGCTCATGCCTGTA
AGCACACGTCTGAACTCTTTCCCTACACGACGCTCTTCCGATCTNNNTCCGAGAAACACCCAAGAATG
GCACACGTCTGAACTCTTTCCCTACACGACGCTCTTCCGATCTNNNCAGAAGAATTGCTTGAACCCAG
AGCACACGTCTGAACTCTTTCCCTACACGACGCTCTTCCGATCTNNNCTGAGGCAGGAGAATCACTTG
AGCACACGTCTGAACTCTTTCCCTACACGACGCTCTTCCGATCTNNNCCGTAATCCCAGCTACTCAGAA
AGCACACGTCTGAACTCTTTCCCTACACGACGCTCTTCCGATCTNNNCTTGAACCTAGGAGGCAGATG
GCACACGTCTGAACTCTTTCCCTACACGACGCTCTTCCGATCTNNNTGGTGACAGAATGAGACTCCAT
AGCACACGTCTGAACTCTTTCCCTACACGACGCTCTTCCGATCTNNNAGAATTGCTTGAATCTCGGAG
AGAGCACACGTCTGAACTCTTTCCCTACACGACGCTCTTCCGATCTNNNGGAGGCTGAGACAGAAGA

AGAGCACACGTCTGAACTCTTTCCCTACACGACGCTCTTCCGATCTNNNAAGCAGGAGAATCACTTGAA
GAGCACACGTCTGAACTCTTTCCCTACACGACGCTCTTCCGATCTNNNTGAGATCAGGAGTTTGAGAC
GAGCACACGTCTGAACTCTTTCCCTACACGACGCTCTTCCGATCTNNNGGATCACCTGAGGTAAGGAG
GAGCACACGTCTGAACTCTTTCCCTACACGACGCTCTTCCGATCTNNNAAGCCCTTGTTGAAAAATGCTG
GAGCACACGTCTGAACTCTTTCCCTACACGACGCTCTTCCGATCTNNNAAGATCACTTGAAGCCAGGAG
AGCACACGTCTGAACTCTTTCCCTACACGACGCTCTTCCGATCTNNNCTGTAATCCCAGCTACTCAGGA
AGAGCACACGTCTGAACTCTTTCCCTACACGACGCTCTTCCGATCTNNNCTGTAGTCCCAGCTACTCCACT
GCACACGTCTGAACTCTTTCCCTACACGACGCTCTTCCGATCTNNNCTGAGGCAGGAGAATCACTTGAA
AGCACACGTCTGAACTCTTTCCCTACACGACGCTCTTCCGATCTNNNGGTGGGAGAATTGCTTGAGTC
GCACACGTCTGAACTCTTTCCCTACACGACGCTCTTCCGATCTNNNCCCCTAGGAGCAATGGTTCAATA
AGCACACGTCTGAACTCTTTCCCTACACGACGCTCTTCCGATCTNNNGAAAATGGCATGAACCTGGGA
GAGCACACGTCTGAACTCTTTCCCTACACGACGCTCTTCCGATCTNNNGTAATCCCAGCTACTTAGGAG
AGAGCACACGTCTGAACTCTTTCCCTACACGACGCTCTTCCGATCTNNNCTAGGAGTTTGAGACCATCCT
AGCACACGTCTGAACTCTTTCCCTACACGACGCTCTTCCGATCTNNNCTGTAGTCCCAGCTACTCAGGA
AGAGCACACGTCTGAACTCTTTCCCTACACGACGCTCTTCCGATCTNNNGAGGCAGGATAATTGCTTGA
AGCACACGTCTGAACTCTTTCCCTACACGACGCTCTTCCGATCTNNNGCAGCCTTCAAACTAAGTCAGT
AGCACACGTCTGAACTCTTTCCCTACACGACGCTCTTCCGATCTNNNTGCTCCTAACTGAACACACAAG
AGCACACGTCTGAACTCTTTCCCTACACGACGCTCTTCCGATCTNNNGAGAACCACGCTATGCTCTCTAT
AGCACACGTCTGAACTCTTTCCCTACACGACGCTCTTCCGATCTNNNCCATGTAAGACGTGATTTGCTCC
GAGCACACGTCTGAACTCTTTCCCTACACGACGCTCTTCCGATCTNNNGACACCAAAACCACAAGTGAC
GAGCACACGTCTGAACTCTTTCCCTACACGACGCTCTTCCGATCTNNNACTGGCTTCTTTCACGTAGCAT
AGCACACGTCTGAACTCTTTCCCTACACGACGCTCTTCCGATCTNNNAGTGCTGGGATTAGAAGTGTG
AGCACACGTCTGAACTCTTTCCCTACACGACGCTCTTCCGATCTNNNGGGATTCCCTGCAGAAGAGTAA
AGCACACGTCTGAACTCTTTCCCTACACGACGCTCTTCCGATCTNNNTGCAGACATTACATTTTCCCTGC
AGCACACGTCTGAACTCTTTCCCTACACGACGCTCTTCCGATCTNNNTAGTTTCCTGTGTCATCATGGG
AGCACACGTCTGAACTCTTTCCCTACACGACGCTCTTCCGATCTNNNAGTAAGCCGAGATCATACCACT
AGCACACGTCTGAACTCTTTCCCTACACGACGCTCTTCCGATCTNNNCTGAAGCATGAAACTCGCTTGA
AGCACACGTCTGAACTCTTTCCCTACACGACGCTCTTCCGATCTNNNCTGAGGCAGGAGAATCACTTG
GAGCACACGTCTGAACTCTTTCCCTACACGACGCTCTTCCGATCTNNNATGATCACACCACTGCACTCTA
GAGCACACGTCTGAACTCTTTCCCTACACGACGCTCTTCCGATCTNNNGAGAATCACTTGAACCCAGGA
AGCACACGTCTGAACTCTTTCCCTACACGACGCTCTTCCGATCTNNNGAGACAGGGTTTCACCATTTTG
AGCACACGTCTGAACTCTTTCCCTACACGACGCTCTTCCGATCTNNNTAGGCCAGTAAGCGTGTCTATT
AGCACACGTCTGAACTCTTTCCCTACACGACGCTCTTCCGATCTNNNTCCTACCTATCAGCTAGCTACTCC
GAGCACACGTCTGAACTCTTTCCCTACACGACGCTCTTCCGATCTNNNGCAGTGAGGTACAATCAGAG
GAGCACACGTCTGAACTCTTTCCCTACACGACGCTCTTCCGATCTNNNATCAGAGACTAGGATTGCAAC
AGCACACGTCTGAACTCTTTCCCTACACGACGCTCTTCCGATCTNNNGCATGGCATCCACTTTTCAAGAT
GAGCACACGTCTGAACTCTTTCCCTACACGACGCTCTTCCGATCTNNNTGGCTGTTCTTTAGGCTTCTCT
GAGCACACGTCTGAACTCTTTCCCTACACGACGCTCTTCCGATCTNNNGGCACACTGTAAGCACTCAGTA
GAGCACACGTCTGAACTCTTTCCCTACACGACGCTCTTCCGATCTNNNCTTGAGGAAGACATGAAGCTT
AGAGCACACGTCTGAACTCTTTCCCTACACGACGCTCTTCCGATCTNNNGTAACCCAACCTAATCCCACC
GAGCACACGTCTGAACTCTTTCCCTACACGACGCTCTTCCGATCTNNNTGAGCATCTATCAATCCAAGCC
AGCACACGTCTGAACTCTTTCCCTACACGACGCTCTTCCGATCTNNNAGGGCAAATTAGACCCAAATCCT
AGCACACGTCTGAACTCTTTCCCTACACGACGCTCTTCCGATCTNNNTCTGATTTCTGTGTTCTCTGCCC

AGCACACGTCTGAACTCTTTCCCTACACGACGCTCTTCCGATCTNNNGCAGGTATGACAGAGCCATTT
AGCACACGTCTGAACTCTTTCCCTACACGACGCTCTTCCGATCTNNNTGGCTGTACCATAGTGAAGTC
GCACACGTCTGAACTCTTTCCCTACACGACGCTCTTCCGATCTNNNCCAAGGTTCAAGCAATTCTCCTG
AGCACACGTCTGAACTCTTTCCCTACACGACGCTCTTCCGATCTNNNTCTCCCAGAGACTAGTGGACTT
GCACACGTCTGAACTCTTTCCCTACACGACGCTCTTCCGATCTNNNATCCACCATGTCCCAATTTCAGA
GCACACGTCTGAACTCTTTCCCTACACGACGCTCTTCCGATCTNNNGGAGGGTCCAGCTCTAGTTAG
GCACACGTCTGAACTCTTTCCCTACACGACGCTCTTCCGATCTNNNACTTCTTGAATGTTCTGCAGTG
GCACACGTCTGAACTCTTTCCCTACACGACGCTCTTCCGATCTNNNTGACCTGGAATACTGTAAGCAC
AGCACACGTCTGAACTCTTTCCCTACACGACGCTCTTCCGATCTNNNGAGTTTCATCTGTGTTGATGG
AGCACACGTCTGAACTCTTTCCCTACACGACGCTCTTCCGATCTNNNGTCCAAGATCACTGTGACAGT
GAGCACACGTCTGAACTCTTTCCCTACACGACGCTCTTCCGATCTNNNTATCTGGAACTACAGGCATA
GCACACGTCTGAACTCTTTCCCTACACGACGCTCTTCCGATCTNNNCAGTATTACCCTGGGTTAGCCA
AGCACACGTCTGAACTCTTTCCCTACACGACGCTCTTCCGATCTNNNTGTAGTGGTGCAATCTTGACT
AGCACACGTCTGAACTCTTTCCCTACACGACGCTCTTCCGATCTNNNAATGGGGTTGCTGGATCATAT
GAGCACACGTCTGAACTCTTTCCCTACACGACGCTCTTCCGATCTNNNATAGTGTCCTTGGAAAACTG
GCACACGTCTGAACTCTTTCCCTACACGACGCTCTTCCGATCTNNNAGACAAAACCAACTAACCAACCA
GAGCACACGTCTGAACTCTTTCCCTACACGACGCTCTTCCGATCTNNNGTGTGATCTCAGCTCATTGCA
GCACACGTCTGAACTCTTTCCCTACACGACGCTCTTCCGATCTNNNGTGGGATAGAGTGACTCATTC
GAGCACACGTCTGAACTCTTTCCCTACACGACGCTCTTCCGATCTNNNTGAATGCCAAACTAACAGTC
GAGCACACGTCTGAACTCTTTCCCTACACGACGCTCTTCCGATCTNNNATTCTAATGTGCAGCCAAGGT
GCACACGTCTGAACTCTTTCCCTACACGACGCTCTTCCGATCTNNNAAATCACCTGTCTTCTGCATTGC
GCACACGTCTGAACTCTTTCCCTACACGACGCTCTTCCGATCTNNNTGCCCAGACTAAGAGTGCTTTTC
GCACACGTCTGAACTCTTTCCCTACACGACGCTCTTCCGATCTNNNGCTGGGCCTGTAATGTTTACAT
AGCACACGTCTGAACTCTTTCCCTACACGACGCTCTTCCGATCTNNNACCTTTTGAGCACCAACATGAT
GCACACGTCTGAACTCTTTCCCTACACGACGCTCTTCCGATCTNNNATTGGCCGTGAGTGTATCTAGT
AGCACACGTCTGAACTCTTTCCCTACACGACGCTCTTCCGATCTNNNCAACAGAAGCAGCACTAAGAT
GCACACGTCTGAACTCTTTCCCTACACGACGCTCTTCCGATCTNNNTCCATCACAATGTTTGTTGCACT
AGCACACGTCTGAACTCTTTCCCTACACGACGCTCTTCCGATCTNNNAGCCATCCCAGTTTTCTTTCTTT
AGCACACGTCTGAACTCTTTCCCTACACGACGCTCTTCCGATCTNNNCATACACACACACACACACACA
GCACACGTCTGAACTCTTTCCCTACACGACGCTCTTCCGATCTNNNATTTTCCTTGTCCTCAGTGGTGA
GCACACGTCTGAACTCTTTCCCTACACGACGCTCTTCCGATCTNNNTAGTCCTTGCTGCTAAGATGTG
AGCACACGTCTGAACTCTTTCCCTACACGACGCTCTTCCGATCTNNNCTTGAACTCCTGACCTCGTGAT
GCACACGTCTGAACTCTTTCCCTACACGACGCTCTTCCGATCTNNNGCTGTAAAAATGAATGAGTGCA
GCACACGTCTGAACTCTTTCCCTACACGACGCTCTTCCGATCTNNNCTGTCTGACACTGTCTTGGATCA
GCACACGTCTGAACTCTTTCCCTACACGACGCTCTTCCGATCTNNNTCTCTTCACTTCTTCCCTTCGTGC
GAGCACACGTCTGAACTCTTTCCCTACACGACGCTCTTCCGATCTNNNGGTGTGATCGTAATTCACTG
GAGCACACGTCTGAACTCTTTCCCTACACGACGCTCTTCCGATCTNNNGGTTCAAGTGATTCTTCTGCC
GCACACGTCTGAACTCTTTCCCTACACGACGCTCTTCCGATCTNNNCAAACATCCCCGCTTCTTTTTCTC
GCACACGTCTGAACTCTTTCCCTACACGACGCTCTTCCGATCTNNNCATTTCAGTGGCACTCTCTCTCT
GCACACGTCTGAACTCTTTCCCTACACGACGCTCTTCCGATCTNNNCGGCTTGTTCATAAGTTGCATG
AGAGCACACGTCTGAACTCTTTCCCTACACGACGCTCTTCCGATCTNNNCGTCTCCAGTGAGCTATGA
GAGCACACGTCTGAACTCTTTCCCTACACGACGCTCTTCCGATCTNNNATGACAGTGCAAGAGCCTAT
GCACACGTCTGAACTCTTTCCCTACACGACGCTCTTCCGATCTNNNCTGAGGCAGGAGAATCACTTGA

GCACACGTCTGAACTCTTTCCCTACACGACGCTCTTCCGATCTNNNTGGGCAAGAGCTTTTAGGAGAT
AGCACACGTCTGAACTCTTTCCCTACACGACGCTCTTCCGATCTNNNTGAGATTGTGTCACTGCACTG
GAGCACACGTCTGAACTCTTTCCCTACACGACGCTCTTCCGATCTNNNCTGAGGCAGGAGAATCACTT
AGCACACGTCTGAACTCTTTCCCTACACGACGCTCTTCCGATCTNNNAGAGTATTGAGGCAGGAGAA
GAGCACACGTCTGAACTCTTTCCCTACACGACGCTCTTCCGATCTNNNGAGGCAGGAGAATTGCTTG
AGCACACGTCTGAACTCTTTCCCTACACGACGCTCTTCCGATCTNNNTTGCTTGAATCTGGGAGAATT
GAGCACACGTCTGAACTCTTTCCCTACACGACGCTCTTCCGATCTNNNCTGGGAAACATAGAGAAAC
AGCACACGTCTGAACTCTTTCCCTACACGACGCTCTTCCGATCTNNNGAGGCTAAGACAGGAGAATTG
AGCACACGTCTGAACTCTTTCCCTACACGACGCTCTTCCGATCTNNNTATAGTCCCAGCTACTCGAGAG
GCACACGTCTGAACTCTTTCCCTACACGACGCTCTTCCGATCTNNNTGAGGCTGTAGTGAGTTGTGAT
GAGCACACGTCTGAACTCTTTCCCTACACGACGCTCTTCCGATCTNNNGAGCTTGCAGTGAGCCAAG
AGCACACGTCTGAACTCTTTCCCTACACGACGCTCTTCCGATCTNNNGAAGTTGCAGTGAGCTGAGAT
GCACACGTCTGAACTCTTTCCCTACACGACGCTCTTCCGATCTNNNTAAGGCAGGAGAATCGCTTGA
GAGCACACGTCTGAACTCTTTCCCTACACGACGCTCTTCCGATCTNNNTGTAATCACAGCTACTCGGG
GCACACGTCTGAACTCTTTCCCTACACGACGCTCTTCCGATCTNNNCACCAGCAGTTAGGCTCTTCTAT
GAGCACACGTCTGAACTCTTTCCCTACACGACGCTCTTCCGATCTNNNTGTAATCCTAGCACTTTGGGA
GAGCACACGTCTGAACTCTTTCCCTACACGACGCTCTTCCGATCTNNNTGTAGTCCTAGCTACTCGGA
AGCACACGTCTGAACTCTTTCCCTACACGACGCTCTTCCGATCTNNNCTGAGGCAGGAGAATTGCTAG
AGCACACGTCTGAACTCTTTCCCTACACGACGCTCTTCCGATCTNNNGAGGTTGCAGTGAGACAAGA
GAGCACACGTCTGAACTCTTTCCCTACACGACGCTCTTCCGATCTNNNAAGTTGAGGCAGGAGAATT
GAGCACACGTCTGAACTCTTTCCCTACACGACGCTCTTCCGATCTNNNGGATCACAAGGTCAGGAGA
GCACACGTCTGAACTCTTTCCCTACACGACGCTCTTCCGATCTNNNGGATCACTTGAGGTCAGGAGT
GCACACGTCTGAACTCTTTCCCTACACGACGCTCTTCCGATCTNNNATACAAAAATTAGCTGGGCGTG
GAGCACACGTCTGAACTCTTTCCCTACACGACGCTCTTCCGATCTNNNGATTGCTTGAAGCCAGGAGT
GCACACGTCTGAACTCTTTCCCTACACGACGCTCTTCCGATCTNNNGAATAATTGCTTGAACCCGGGA
AGCACACGTCTGAACTCTTTCCCTACACGACGCTCTTCCGATCTNNNTGTAGTCCTATCTACTCGGGAG
GAGCACACGTCTGAACTCTTTCCCTACACGACGCTCTTCCGATCTNNNCTGTAATCCCAGCTACTCAGG
AGCACACGTCTGAACTCTTTCCCTACACGACGCTCTTCCGATCTNNNTCATACAAACAGGGTCTGGAG
GAGCACACGTCTGAACTCTTTCCCTACACGACGCTCTTCCGATCTNNNGAGGTTGCAGTGAGCTGAG
GCACACGTCTGAACTCTTTCCCTACACGACGCTCTTCCGATCTNNNCTGGGATGACTCATCTCTGTTCC
GAGCACACGTCTGAACTCTTTCCCTACACGACGCTCTTCCGATCTNNNAGGTAGGAGAACCACCTAAG
GAGCACACGTCTGAACTCTTTCCCTACACGACGCTCTTCCGATCTNNNATCTGCAGAGTGGATGTCAT
AGAGCACACGTCTGAACTCTTTCCCTACACGACGCTCTTCCGATCTNNNGCCACAAACTGTGCCCATAT
GAGCACACGTCTGAACTCTTTCCCTACACGACGCTCTTCCGATCTNNNAGGAGGCTGAGGTAGGAG
GCACACGTCTGAACTCTTTCCCTACACGACGCTCTTCCGATCTNNNTGAGGCATGAGAATTGCTTGA
AGCACACGTCTGAACTCTTTCCCTACACGACGCTCTTCCGATCTNNNGGAGGCTGAGGTAGGAAGA
GCACACGTCTGAACTCTTTCCCTACACGACGCTCTTCCGATCTNNNCTGAGGCAGGAGAATGACATG
AGCACACGTCTGAACTCTTTCCCTACACGACGCTCTTCCGATCTNNNGAGAATTGCTTGAACCCAGG
AGCACACGTCTGAACTCTTTCCCTACACGACGCTCTTCCGATCTNNNCTGAGGCAAAAGAATCGCTT
GAGCACACGTCTGAACTCTTTCCCTACACGACGCTCTTCCGATCTNNNGGCTGAGACAGAAGAATTG
AGCACACGTCTGAACTCTTTCCCTACACGACGCTCTTCCGATCTNNNATACAAATCTTGAGCTGCCAGG
GCACACGTCTGAACTCTTTCCCTACACGACGCTCTTCCGATCTNNNGTGGTGAGTGCCTATAATCCCA
GAGCACACGTCTGAACTCTTTCCCTACACGACGCTCTTCCGATCTNNNCAGGAGAATGCCAAAAGAA

AGCACACGTCTGAACTCTTTCCCTACACGACGCTCTTCCGATCTNNNAAATTATTGCCAGCCACCTGAA
AGCACACGTCTGAACTCTTTCCCTACACGACGCTCTTCCGATCTNNNGTAATCCTAGCTACTTGGGAG
AGCACACGTCTGAACTCTTTCCCTACACGACGCTCTTCCGATCTNNNCTGTAGTCCCAGCTACTCAGG
AGCACACGTCTGAACTCTTTCCCTACACGACGCTCTTCCGATCTNNNATTCCCACCTACTCAGAAGGTT
GAGCACACGTCTGAACTCTTTCCCTACACGACGCTCTTCCGATCTNNNAATGCAGGAGAATTGCTTG
AGCACACGTCTGAACTCTTTCCCTACACGACGCTCTTCCGATCTNNNCAGGAGAATTGCTTGAACTTG
AGCACACGTCTGAACTCTTTCCCTACACGACGCTCTTCCGATCTNNNAGGAGAATTGCTTGAACCTGG
AGCACACGTCTGAACTCTTTCCCTACACGACGCTCTTCCGATCTNNNGTAATCTCAGCTACTCACGAGG
GAGCACACGTCTGAACTCTTTCCCTACACGACGCTCTTCCGATCTNNNGTGGATCACAAGGTCAGGA
GAGCACACGTCTGAACTCTTTCCCTACACGACGCTCTTCCGATCTNNNTTTGCGACATTACACTCTAGC
GCACACGTCTGAACTCTTTCCCTACACGACGCTCTTCCGATCTNNNCTTGGAAGGCAGAGAAGGAAG
AGCACACGTCTGAACTCTTTCCCTACACGACGCTCTTCCGATCTNNNTAATCCCATCTACTCAGGAGGC
GAGCACACGTCTGAACTCTTTCCCTACACGACGCTCTTCCGATCTNNNATGGTAGTGCACGCCTATAA
AGCACACGTCTGAACTCTTTCCCTACACGACGCTCTTCCGATCTNNNTGTGATCCTAGCACTTTGGGAA
GAGCACACGTCTGAACTCTTTCCCTACACGACGCTCTTCCGATCTNNNTGAGAATCACTTGAACCCAG
GCACACGTCTGAACTCTTTCCCTACACGACGCTCTTCCGATCTNNNCTGAGGTGGGAGAATTGCTTG
AGCACACGTCTGAACTCTTTCCCTACACGACGCTCTTCCGATCTNNNGGCAAGAATGGAGTCTAATCC
AGCACACGTCTGAACTCTTTCCCTACACGACGCTCTTCCGATCTNNNTACATGCCTGTAATCCCAGCTA
AGCACACGTCTGAACTCTTTCCCTACACGACGCTCTTCCGATCTNNNCGCACCTATAATCCCAGCTACA
AGCACACGTCTGAACTCTTTCCCTACACGACGCTCTTCCGATCTNNNGCTGAAAGAGGAGGATCACTA
GCACACGTCTGAACTCTTTCCCTACACGACGCTCTTCCGATCTNNNCCACTGGGAAGGAGTATTCACT
AGCACACGTCTGAACTCTTTCCCTACACGACGCTCTTCCGATCTNNNAGGTCTCAAATGTGTCTCTGAG
AGCACACGTCTGAACTCTTTCCCTACACGACGCTCTTCCGATCTNNNGAGTCTGGGACATAGAAGATG
AGCACACGTCTGAACTCTTTCCCTACACGACGCTCTTCCGATCTNNNTGAGGGGCTCTCTGTTTTCTA
AGCACACGTCTGAACTCTTTCCCTACACGACGCTCTTCCGATCTNNNGTACTCCACTTGAGCTCTCTGA
AGCACACGTCTGAACTCTTTCCCTACACGACGCTCTTCCGATCTNNNGAGTTTCACTCTTCTTGCCCAG
GCACACGTCTGAACTCTTTCCCTACACGACGCTCTTCCGATCTNNNAGGCTGCTGAGACTGTATTTCTT
AGCACACGTCTGAACTCTTTCCCTACACGACGCTCTTCCGATCTNNNCCCAAAGGGCTAGGATTATAG
GAGCACACGTCTGAACTCTTTCCCTACACGACGCTCTTCCGATCTNNNTTCTTTCCTTCCTTCCCTCCTTC
GAGCACACGTCTGAACTCTTTCCCTACACGACGCTCTTCCGATCTNNNGCAACATAGTGAGACCCCAT
AGCACACGTCTGAACTCTTTCCCTACACGACGCTCTTCCGATCTNNNCACCATGCCCAGCTAATTTCTT
AGCACACGTCTGAACTCTTTCCCTACACGACGCTCTTCCGATCTNNNACCCTGCATTTTTCCTTCCTCAT
AGCACACGTCTGAACTCTTTCCCTACACGACGCTCTTCCGATCTNNNCTATGTGTGTGTGTGTGTGTG
AGCACACGTCTGAACTCTTTCCCTACACGACGCTCTTCCGATCTNNNAGACAGTGACAAGAACATGGT
AGCACACGTCTGAACTCTTTCCCTACACGACGCTCTTCCGATCTNNNCAGCCATCTAATCACACAACTG
AGCACACGTCTGAACTCTTTCCCTACACGACGCTCTTCCGATCTNNNTAGTGAGTGCAGGAGTTTGAG
AGCACACGTCTGAACTCTTTCCCTACACGACGCTCTTCCGATCTNNNGAACTCCTGACCTCATGATCCA
AGCACACGTCTGAACTCTTTCCCTACACGACGCTCTTCCGATCTNNNCCCAGCCTTTTTTGTTTTGTGTT
AGCACACGTCTGAACTCTTTCCCTACACGACGCTCTTCCGATCTNNNATATGCTGCTGGATTTGGTTTG
GAGCACACGTCTGAACTCTTTCCCTACACGACGCTCTTCCGATCTNNNGTAATTGTGGCCTGGAGTTT
GAGCACACGTCTGAACTCTTTCCCTACACGACGCTCTTCCGATCTNNNGTCCACCTTGCATCTTCCAATT
AGCACACGTCTGAACTCTTTCCCTACACGACGCTCTTCCGATCTNNNCAGATCTGTGGACCATTCAAA
AGCACACGTCTGAACTCTTTCCCTACACGACGCTCTTCCGATCTNNNACTGCCTCCACCCTTGTAAATA

AGCACACGTCTGAACTCTTTCCCTACACGACGCTCTTCCGATCTNNNATCCAGTCCTCAAGCAACCTAA

AGCACACGTCTGAACTCTTTCCCTACACGACGCTCTTCCGATCTNNNGAAAGATGGGTTGGATTGGCA

GAGCACACGTCTGAACTCTTTCCCTACACGACGCTCTTCCGATCTNNNGCTGTCCCTGATCAAGGCTAT

AGCACACGTCTGAACTCTTTCCCTACACGACGCTCTTCCGATCTNNNCTGTAATCCTAGCTACCCGGAA

GAGCACACGTCTGAACTCTTTCCCTACACGACGCTCTTCCGATCTNNNAGGATCATGCCATTGTACTCCA

GAGCACACGTCTGAACTCTTTCCCTACACGACGCTCTTCCGATCTNNNCACCTGTAATCCCAGCTACTCA

GAGCACACGTCTGAACTCTTTCCCTACACGACGCTCTTCCGATCTNNNCTGTAATCCCAGCTACTCAGGA

AGCACACGTCTGAACTCTTTCCCTACACGACGCTCTTCCGATCTNNNAACAAAAAAAAACCAGGGGG

GAGCACACGTCTGAACTCTTTCCCTACACGACGCTCTTCCGATCTNNNGGAGAATCACTTGAACCCAGG

AGAGCACACGTCTGAACTCTTTCCCTACACGACGCTCTTCCGATCTNNNGATCACGCCACTGTACTCTA

GAGCACACGTCTGAACTCTTTCCCTACACGACGCTCTTCCGATCTNNNGAGGATTACTTGAACCCAGGA

AGCACACGTCTGAACTCTTTCCCTACACGACGCTCTTCCGATCTNNNCATTGCACTTCAGCCTGGATTA

AGCACACGTCTGAACTCTTTCCCTACACGACGCTCTTCCGATCTNNNGGAGAATTGCTTGAACTCTGCA

GAGCACACGTCTGAACTCTTTCCCTACACGACGCTCTTCCGATCTNNNATGACAAAAAATTAGCTGGG

GAGCACACGTCTGAACTCTTTCCCTACACGACGCTCTTCCGATCTNNNCTTCCAAAGTGCTGGGATTA

AGCACACGTCTGAACTCTTTCCCTACACGACGCTCTTCCGATCTNNNCAGGAGTTAGAGGCAGAAGG

GAGCACACGTCTGAACTCTTTCCCTACACGACGCTCTTCCGATCTNNNGGAAGCAGAGGTTGTAGTAA

AGAGCACACGTCTGAACTCTTTCCCTACACGACGCTCTTCCGATCTNNNACCTGTAAATCCCAGCTACTT

GAGCACACGTCTGAACTCTTTCCCTACACGACGCTCTTCCGATCTNNNGGTGACAGAGTGAGACTCCA

AGAGCACACGTCTGAACTCTTTCCCTACACGACGCTCTTCCGATCTNNNCAGGAGAATTGCTTGAACTT

AGCACACGTCTGAACTCTTTCCCTACACGACGCTCTTCCGATCTNNNGAGAATCGCTTGAACCTGAGA

GAGCACACGTCTGAACTCTTTCCCTACACGACGCTCTTCCGATCTNNNAGTGAGCTGAGATTGCGAGA

GAGCACACGTCTGAACTCTTTCCCTACACGACGCTCTTCCGATCTNNNAGGAGAATCGCTTGAACTGA

GAGCACACGTCTGAACTCTTTCCCTACACGACGCTCTTCCGATCTNNNCGGAGAATGGTGTGAAACTA

GAGCACACGTCTGAACTCTTTCCCTACACGACGCTCTTCCGATCTNNNCCTGAGTGACACAATGAGACT

AGAGCACACGTCTGAACTCTTTCCCTACACGACGCTCTTCCGATCTNNNGAGATCACACCACTACACTC

GAGCACACGTCTGAACTCTTTCCCTACACGACGCTCTTCCGATCTNNNGAGACAGGAGACTCACTTGA

AGCACACGTCTGAACTCTTTCCCTACACGACGCTCTTCCGATCTNNNGGAGGCAAAAGTTTTGATGAG

GAGCACACGTCTGAACTCTTTCCCTACACGACGCTCTTCCGATCTNNNCACTTGAGGTCAGGAGTTTG

AGCACACGTCTGAACTCTTTCCCTACACGACGCTCTTCCGATCTNNNTGTAATAAGCTGAGATCGTGC

AGCACACGTCTGAACTCTTTCCCTACACGACGCTCTTCCGATCTNNNGGTGTTGAAGTGAGACTCTGT

GAGCACACGTCTGAACTCTTTCCCTACACGACGCTCTTCCGATCTNNNCTCGAACTGCTGACCTCATGA

AGAGCACACGTCTGAACTCTTTCCCTACACGACGCTCTTCCGATCTNNNGGAATTCAATGTTGCAGTG

AGCACACGTCTGAACTCTTTCCCTACACGACGCTCTTCCGATCTNNNCTGAGGCAGGAGAATAGCTTGA

AGCACACGTCTGAACTCTTTCCCTACACGACGCTCTTCCGATCTNNNCTGTAATCCCAGCTACTTGAGG

GAGCACACGTCTGAACTCTTTCCCTACACGACGCTCTTCCGATCTNNNACAAAAAATTAGACCGGCGTT

AGAGCACACGTCTGAACTCTTTCCCTACACGACGCTCTTCCGATCTNNNCACCTGTAATCTCAGCTACC

GAGCACACGTCTGAACTCTTTCCCTACACGACGCTCTTCCGATCTNNNAGAGGATGAGTTGAACCCTA

GAGCACACGTCTGAACTCTTTCCCTACACGACGCTCTTCCGATCTNNNGAGAATCTCTTGAACCCAGGA

AGCACACGTCTGAACTCTTTCCCTACACGACGCTCTTCCGATCTNNNGGAGATTGCAGTGAGCCAAAA

GAGCACACGTCTGAACTCTTTCCCTACACGACGCTCTTCCGATCTNNNGTGACACAGCAAGACTCTGAT

AGCACACGTCTGAACTCTTTCCCTACACGACGCTCTTCCGATCTNNNGGTGACAGAGTGAGACTCCAT

GAGCACACGTCTGAACTCTTTCCCTACACGACGCTCTTCCGATCTNNNAAGAGAATCGTTTGAACTCG

GAGCACACGTCTGAACTCTTTCCCTACACGACGCTCTTCCGATCTNNNTTGTTGTTCCAGCTACTCAGGA
AGCACACGTCTGAACTCTTTCCCTACACGACGCTCTTCCGATCTNNNCCTGCCTATAATCCCAGCTACTC
GCACACGTCTGAACTCTTTCCCTACACGACGCTCTTCCGATCTNNNGAAGTTGCAGTGAACCGATAGT
AGAGCACACGTCTGAACTCTTTCCCTACACGACGCTCTTCCGATCTNNNGTGGTGTATGCCTATAGTCC
AGAGCACACGTCTGAACTCTTTCCCTACACGACGCTCTTCCGATCTNNNCTGAGGCAGGAGAATTGCT
GAGCACACGTCTGAACTCTTTCCCTACACGACGCTCTTCCGATCTNNNGCACCTGTAGTTCCAGCTACTA
AGCACACGTCTGAACTCTTTCCCTACACGACGCTCTTCCGATCTNNNGTCTTTGGTCCATCTGGCTACT
GAGCACACGTCTGAACTCTTTCCCTACACGACGCTCTTCCGATCTNNNGACTACAAGTGCAAACCACCA
AGCACACGTCTGAACTCTTTCCCTACACGACGCTCTTCCGATCTNNNGCTTCAGACACCCCTACTTTCTA
AGCACACGTCTGAACTCTTTCCCTACACGACGCTCTTCCGATCTNNNGCCTCCCAGGTTCAATCTATTCT
AGCACACGTCTGAACTCTTTCCCTACACGACGCTCTTCCGATCTNNNGGCACACTGCTTACGTATCTTG
AGCACACGTCTGAACTCTTTCCCTACACGACGCTCTTCCGATCTNNNGCTACCTTCTCATTGTGTCCTCA
AGCACACGTCTGAACTCTTTCCCTACACGACGCTCTTCCGATCTNNNGAGGTTGCAGTAGCTGAGATCA
AGCACACGTCTGAACTCTTTCCCTACACGACGCTCTTCCGATCTNNNCCAGAGTGGCTTCTTTCTTACCT
AGCACACGTCTGAACTCTTTCCCTACACGACGCTCTTCCGATCTNNNACAGCAGGCTAGGGAATGATTA
AGCACACGTCTGAACTCTTTCCCTACACGACGCTCTTCCGATCTNNNCCAAAACCATGTGTAAGCCACTA
AGCACACGTCTGAACTCTTTCCCTACACGACGCTCTTCCGATCTNNNAATTTTTGGGATCACAGGCATC
GAGCACACGTCTGAACTCTTTCCCTACACGACGCTCTTCCGATCTNNNTAGGCAGGATTTCACTCTGTC
AGCACACGTCTGAACTCTTTCCCTACACGACGCTCTTCCGATCTNNNAGAGTGATGCCCTGTCTCAAAA
AGCACACGTCTGAACTCTTTCCCTACACGACGCTCTTCCGATCTNNNCATGACCCCATGCCTGATTTATA
GAGCACACGTCTGAACTCTTTCCCTACACGACGCTCTTCCGATCTNNNTACCAGGTGTCTCCATTTTGA
AGCACACGTCTGAACTCTTTCCCTACACGACGCTCTTCCGATCTNNNGTTTTGGAGCCAGATTCAAACCA
GAGCACACGTCTGAACTCTTTCCCTACACGACGCTCTTCCGATCTNNNGAGGTTGAAGGGGAACATGA
GAGCACACGTCTGAACTCTTTCCCTACACGACGCTCTTCCGATCTNNNTCCTAAAGTGCTGGGATTACA
GCACACGTCTGAACTCTTTCCCTACACGACGCTCTTCCGATCTNNNGCCTCCTGAGTAGCTGAGATTAC
CACGTCTGAACTCTTTCCCTACACGACGCTCTTCCGATCTNNNAGTTTAGCCACCATAGAATAGTCAGT
GAGCACACGTCTGAACTCTTTCCCTACACGACGCTCTTCCGATCTNNNCTCAAGTGATTCTCCTGCCTCA
GAGCACACGTCTGAACTCTTTCCCTACACGACGCTCTTCCGATCTNNNGAGTCTGGTTTCACGGAAGAC
AGCACACGTCTGAACTCTTTCCCTACACGACGCTCTTCCGATCTNNNTTGGAGCTCAGGGTGATAAAAC
AGCACACGTCTGAACTCTTTCCCTACACGACGCTCTTCCGATCTNNNCGGCCTTCCAAAGTACTGGTATT
AGCACACGTCTGAACTCTTTCCCTACACGACGCTCTTCCGATCTNNNTGGAATATTCTTCCTGCGTCTG
AGCACACGTCTGAACTCTTTCCCTACACGACGCTCTTCCGATCTNNNGACCGAGGTTCTACTGATTGTC
AGCACACGTCTGAACTCTTTCCCTACACGACGCTCTTCCGATCTNNNTCCCAGGTTCAAGAATTCTCCTG
AGCACACGTCTGAACTCTTTCCCTACACGACGCTCTTCCGATCTNNNAATGGGGAAAAGGTAGCTGGA
GAGCACACGTCTGAACTCTTTCCCTACACGACGCTCTTCCGATCTNNNATTCTTCCAGTCCACGAGTATG
AGCACACGTCTGAACTCTTTCCCTACACGACGCTCTTCCGATCTNNNCCAGTGCTAGCTCAGTTATGCTA
AGCACACGTCTGAACTCTTTCCCTACACGACGCTCTTCCGATCTNNNAGGTTCAAGTGATTCTCCTGCT
GAGCACACGTCTGAACTCTTTCCCTACACGACGCTCTTCCGATCTNNNAGTGGTACGATCTCAAGTTCA
GAGCACACGTCTGAACTCTTTCCCTACACGACGCTCTTCCGATCTNNNACAAGAGTATCCAGCTGATCTC
GAGCACACGTCTGAACTCTTTCCCTACACGACGCTCTTCCGATCTNNNTGCCACCACACCTAGCTAATTT
GCACACGTCTGAACTCTTTCCCTACACGACGCTCTTCCGATCTNNNAGCCTGGGAACTACTAAAATGG
GAGCACACGTCTGAACTCTTTCCCTACACGACGCTCTTCCGATCTNNNAGACAAAGCAAGACTCCAACT
AGCACACGTCTGAACTCTTTCCCTACACGACGCTCTTCCGATCTNNNTCAGCTTGACATCATCCTGTTGA

AGCACACGTCTGAACTCTTTCCCTACACGACGCTCTTCCGATCTNNNTGCCTTCACCAGCATAGTGATT

AGCACACGTCTGAACTCTTTCCCTACACGACGCTCTTCCGATCTNNNGTAGTAGAGATGGGTGTTTG

GCACACGTCTGAACTCTTTCCCTACACGACGCTCTTCCGATCTNNNGTTTCACCATGTTAGCCAGGATG

AGCACACGTCTGAACTCTTTCCCTACACGACGCTCTTCCGATCTNNNATAAAAGGGGAGTGGAGGT

AGCACACGTCTGAACTCTTTCCCTACACGACGCTCTTCCGATCTNNNGTTAGGCAGAATGGTCTCGAT

GCACACGTCTGAACTCTTTCCCTACACGACGCTCTTCCGATCTNNNCGGAAGAAGCTGAGTATCACA

AGCACACGTCTGAACTCTTTCCCTACACGACGCTCTTCCGATCTNNNGATGAGATTCTCACCCTTTTG

AGCACACGTCTGAACTCTTTCCCTACACGACGCTCTTCCGATCTNNNGCATACGTAGTGACAATGCCT

AGCACACGTCTGAACTCTTTCCCTACACGACGCTCTTCCGATCTNNNATAGAACTACAGGTGTGAGC

GAGCACACGTCTGAACTCTTTCCCTACACGACGCTCTTCCGATCTNNNCTGAGATCACATCACTGCACT

GCACACGTCTGAACTCTTTCCCTACACGACGCTCTTCCGATCTNNNCTGTAGTCCTAGCTACTCGGGA

GCACACGTCTGAACTCTTTCCCTACACGACGCTCTTCCGATCTNNNAGCCTGGGTAATAGAGTGAGA

GCACACGTCTGAACTCTTTCCCTACACGACGCTCTTCCGATCTNNNGTCAAGGTCATAGTGAGCCATG

AGCACACGTCTGAACTCTTTCCCTACACGACGCTCTTCCGATCTNNNACCATGGAATACTACACAGCTG

GAGCACACGTCTGAACTCTTTCCCTACACGACGCTCTTCCGATCTNNNAAGGATACAGCAACCAAAAC

GAGCACACGTCTGAACTCTTTCCCTACACGACGCTCTTCCGATCTNNNTGCTTGTAATCCCAGCTATTC

GAGCACACGTCTGAACTCTTTCCCTACACGACGCTCTTCCGATCTNNNCTGAGGCAGGAGAATAGCAT

AGCACACGTCTGAACTCTTTCCCTACACGACGCTCTTCCGATCTNNNGAGGCAGGAGAATCACTTGA

AGCACACGTCTGAACTCTTTCCCTACACGACGCTCTTCCGATCTNNNCGCTGCTACTCAAAGAAGTTC

AGCACACGTCTGAACTCTTTCCCTACACGACGCTCTTCCGATCTNNNTCATGTCACATGCAAAGACACA

AGCACACGTCTGAACTCTTTCCCTACACGACGCTCTTCCGATCTNNNCTGAGACAGGAGAATGGCATG

CACACGTCTGAACTCTTTCCCTACACGACGCTCTTCCGATCTNNNCTGTAGTCCCAGCTACTCAGGAG

GCACACGTCTGAACTCTTTCCCTACACGACGCTCTTCCGATCTNNNCTTGAACCCAAAAGGCAGAGGT

AGCACACGTCTGAACTCTTTCCCTACACGACGCTCTTCCGATCTNNNCTGTAGTACCAGCTACTCGGGA

AGCACACGTCTGAACTCTTTCCCTACACGACGCTCTTCCGATCTNNNGGTGACAAGAGTGAGACCTC

GAGCACACGTCTGAACTCTTTCCCTACACGACGCTCTTCCGATCTNNNCTGAGGCAGGAGAATTGCTT

GAGCACACGTCTGAACTCTTTCCCTACACGACGCTCTTCCGATCTNNNTAGGAGAATTGCTTGAACCT

AGCACACGTCTGAACTCTTTCCCTACACGACGCTCTTCCGATCTNNNCTGAGGCAGGATAATCGCTTG

AGCACACGTCTGAACTCTTTCCCTACACGACGCTCTTCCGATCTNNNGGCAACAGAATGAGACTCCAT

GAGCACACGTCTGAACTCTTTCCCTACACGACGCTCTTCCGATCTNNNACCAAATAGCCAAGGCAATC

GAGCACACGTCTGAACTCTTTCCCTACACGACGCTCTTCCGATCTNNNAGAATCACTTGAACCTGGGA

AGCACACGTCTGAACTCTTTCCCTACACGACGCTCTTCCGATCTNNNGAGAATTGCTTGAACCCGGAA

GAGCACACGTCTGAACTCTTTCCCTACACGACGCTCTTCCGATCTNNNTTTCAGTTACTTGGGAGGCT

AGCACACGTCTGAACTCTTTCCCTACACGACGCTCTTCCGATCTNNNCTGGGTTACTAAGCAAGACT

AGCACACGTCTGAACTCTTTCCCTACACGACGCTCTTCCGATCTNNNGCAACAAGAGCAAAACTGCAT

AGCACACGTCTGAACTCTTTCCCTACACGACGCTCTTCCGATCTNNNTGAGGCAGGAGAATCACTTGT

GCACACGTCTGAACTCTTTCCCTACACGACGCTCTTCCGATCTNNNTGTGGCAGGAGAATCACTTGA

AGCACACGTCTGAACTCTTTCCCTACACGACGCTCTTCCGATCTNNNAGTAGCTGGGATTATAGGCAT

AGCACACGTCTGAACTCTTTCCCTACACGACGCTCTTCCGATCTNNNGTGAGTAGAGATTGTGCCATT

AGCACACGTCTGAACTCTTTCCCTACACGACGCTCTTCCGATCTNNNCTATAATCCCAGCTACTGAGG

AGCACACGTCTGAACTCTTTCCCTACACGACGCTCTTCCGATCTNNNGAGAATCACTTGAACCCAGGA

GAGCACACGTCTGAACTCTTTCCCTACACGACGCTCTTCCGATCTNNNGAGCTTGCAGTGAGCCAAT

AGCACACGTCTGAACTCTTTCCCTACACGACGCTCTTCCGATCTNNNGGCAACAAGAGCAAAAACTCT

GAGCACACGTCTGAACTCTTTCCCTACACGACGCTCTTCCGATCTNNNCGTGTAATCCCAGCTATTCAG
GAGCACACGTCTGAACTCTTTCCCTACACGACGCTCTTCCGATCTNNNGGTTGAAGTAAACTGAGAT
GAGCACACGTCTGAACTCTTTCCCTACACGACGCTCTTCCGATCTNNNTGAGGTGGAAGGATCGTTT
AGCACACGTCTGAACTCTTTCCCTACACGACGCTCTTCCGATCTNNNTAGAAATCAGGATACGTGGG
GAGCACACGTCTGAACTCTTTCCCTACACGACGCTCTTCCGATCTNNNTGTAATTCCAGCTACTTGGG
AGCACACGTCTGAACTCTTTCCCTACACGACGCTCTTCCGATCTNNNCTCACAGCTATAATCCCAGCACT
AGCACACGTCTGAACTCTTTCCCTACACGACGCTCTTCCGATCTNNNGGAGAATTCCTTGAACCCAG
GAGCACACGTCTGAACTCTTTCCCTACACGACGCTCTTCCGATCTNNNTTCAGGACATAGGCATGGAC
GAGCACACGTCTGAACTCTTTCCCTACACGACGCTCTTCCGATCTNNNTGCCTATAATCCCAGCACTTT
AGCACACGTCTGAACTCTTTCCCTACACGACGCTCTTCCGATCTNNNTCAGGAGAATCACTTGAACCC
AGCACACGTCTGAACTCTTTCCCTACACGACGCTCTTCCGATCTNNNACAAAAATTAGCTGGGTGTGG
GAGCACACGTCTGAACTCTTTCCCTACACGACGCTCTTCCGATCTNNNAGTGAGTCCTAATCATGCCAC
AGCACACGTCTGAACTCTTTCCCTACACGACGCTCTTCCGATCTNNNGTTCAAGACCAGTCTAGGCAA
GAGCACACGTCTGAACTCTTTCCCTACACGACGCTCTTCCGATCTNNNGTGACAGAGTGAGACTCCGT
GAGCACACGTCTGAACTCTTTCCCTACACGACGCTCTTCCGATCTNNNCAGGTAGAGGTTGTAGTGA
AGCACACGTCTGAACTCTTTCCCTACACGACGCTCTTCCGATCTNNNGAGGTTGTAGTGAGCCGAGA
GAGCACACGTCTGAACTCTTTCCCTACACGACGCTCTTCCGATCTNNNTGAGGCAGGAGAATCACTT
GAGCACACGTCTGAACTCTTTCCCTACACGACGCTCTTCCGATCTNNNAGAATTGCTTGAATCTGGG
GAGCACACGTCTGAACTCTTTCCCTACACGACGCTCTTCCGATCTNNNGCTCACACCTGTAATCCCAGT
AGCACACGTCTGAACTCTTTCCCTACACGACGCTCTTCCGATCTNNNCAGCCTAAGAGTCCCCAGAAT
AGCACACGTCTGAACTCTTTCCCTACACGACGCTCTTCCGATCTNNNAAGATCTCACTCTGTTGTCCAG
AGCACACGTCTGAACTCTTTCCCTACACGACGCTCTTCCGATCTNNNCCCTGTCACCTTCCTGAATTGTA
AGCACACGTCTGAACTCTTTCCCTACACGACGCTCTTCCGATCTNNNTTTCTGGGAGTTAGTTGGCAC
AGCACACGTCTGAACTCTTTCCCTACACGACGCTCTTCCGATCTNNNTTCAGCCTTTCTTTTTTGTGGG
AGCACACGTCTGAACTCTTTCCCTACACGACGCTCTTCCGATCTNNNACCGGCTGTAGCTATTCTTTTC
AGCACACGTCTGAACTCTTTCCCTACACGACGCTCTTCCGATCTNNNTGTTTCCTAACTTTGTGTCTGG
AGCACACGTCTGAACTCTTTCCCTACACGACGCTCTTCCGATCTNNNGGTTCAAGCGATTCTCTTGTCT
AGCACACGTCTGAACTCTTTCCCTACACGACGCTCTTCCGATCTNNNGAAGGAGGAGATGGGAGGAA
GCACACGTCTGAACTCTTTCCCTACACGACGCTCTTCCGATCTNNNCTGTTTTCTTCCTCTTCCTTCGCC
AGCACACGTCTGAACTCTTTCCCTACACGACGCTCTTCCGATCTNNNAAGCCTCTGTTGCTCATTTTCA
AGCACACGTCTGAACTCTTTCCCTACACGACGCTCTTCCGATCTNNNAGCAGACCCTTACAAGCTTCTT
AGCACACGTCTGAACTCTTTCCCTACACGACGCTCTTCCGATCTNNNTAGCTACTGTATCCTGAGCCCTT
AGCACACGTCTGAACTCTTTCCCTACACGACGCTCTTCCGATCTNNNAAGCATTTCTGGCACCCTGTAT
AGCACACGTCTGAACTCTTTCCCTACACGACGCTCTTCCGATCTNNNCTCAAGTGATCTTCCCACCTCA
AGCACACGTCTGAACTCTTTCCCTACACGACGCTCTTCCGATCTNNNATCTCAGCTCACTTCAACCTCTG
AGCACACGTCTGAACTCTTTCCCTACACGACGCTCTTCCGATCTNNNCCATCCCTGTTTCCACCTATTCA
AGCACACGTCTGAACTCTTTCCCTACACGACGCTCTTCCGATCTNNNCAGGCTTGGATCTTCTTTCTCT
AGCACACGTCTGAACTCTTTCCCTACACGACGCTCTTCCGATCTNNNAAACCAACTCTCCCTGAGGTT
GCACACGTCTGAACTCTTTCCCTACACGACGCTCTTCCGATCTNNNGCTCAAGTAATCCTCCCACTTCA
AGCACACGTCTGAACTCTTTCCCTACACGACGCTCTTCCGATCTNNNTCTCAAAGTGCTGGGATTACA
AGCACACGTCTGAACTCTTTCCCTACACGACGCTCTTCCGATCTNNNGGAGGTCTTAGCCATATGCCA
AGCACACGTCTGAACTCTTTCCCTACACGACGCTCTTCCGATCTNNNAAAACCCCCAGATTAGCACTGA
GCACACGTCTGAACTCTTTCCCTACACGACGCTCTTCCGATCTNNNTCCACTGCAAGCATTACTCTCTT

AGCACACGTCTGAACTCTTTCCCTACACGACGCTCTTCCGATCTNNNGTGGGAACAGACACTATCTCCT
AGCACACGTCTGAACTCTTTCCCTACACGACGCTCTTCCGATCTNNNGGTCAGCAGAGAGAGAGGAA
AGCACACGTCTGAACTCTTTCCCTACACGACGCTCTTCCGATCTNNNTCATTTTCCCCTTGATCTGAGCA
AGCACACGTCTGAACTCTTTCCCTACACGACGCTCTTCCGATCTNNNCTGTAGGGTGTAGCCAAAGAA
AGCACACGTCTGAACTCTTTCCCTACACGACGCTCTTCCGATCTNNNACATCCTTTTCCCAACCTGTAGG
AGCACACGTCTGAACTCTTTCCCTACACGACGCTCTTCCGATCTNNNTTCCAGGTGTTCAGACTACCAAT
GAGCACACGTCTGAACTCTTTCCCTACACGACGCTCTTCCGATCTNNNAAACAATTCTCCTGCCTCATCCT
AGCACACGTCTGAACTCTTTCCCTACACGACGCTCTTCCGATCTNNNAGGAGTGGTGAGAGTGGATA
GAGCACACGTCTGAACTCTTTCCCTACACGACGCTCTTCCGATCTNNNTCAAATCCACACCTCAAGTCTC
GAGCACACGTCTGAACTCTTTCCCTACACGACGCTCTTCCGATCTNNNTGTCTGGGTGATTCAAGAGA
AGCACACGTCTGAACTCTTTCCCTACACGACGCTCTTCCGATCTNNNGCTGATGGCATTGACATAGGT
AGCACACGTCTGAACTCTTTCCCTACACGACGCTCTTCCGATCTNNNAAATCACTCTGCAACTAGTGGG
AGCACACGTCTGAACTCTTTCCCTACACGACGCTCTTCCGATCTNNNCCAAAATGCTGGGATTACAAG
GAGCACACGTCTGAACTCTTTCCCTACACGACGCTCTTCCGATCTNNNACACTGCCTTTACACATGACAA
AGCACACGTCTGAACTCTTTCCCTACACGACGCTCTTCCGATCTNNNTTATTGGGAGGCTGTCTCTGTT
AGCACACGTCTGAACTCTTTCCCTACACGACGCTCTTCCGATCTNNNTTACAGGTATGAGCCACCGTAT
AGCACACGTCTGAACTCTTTCCCTACACGACGCTCTTCCGATCTNNNGTACTTTTTCAACCCATGCCTCCC
AGCACACGTCTGAACTCTTTCCCTACACGACGCTCTTCCGATCTNNNAGTGGTGTGATCATGGCTTACT
GAGCACACGTCTGAACTCTTTCCCTACACGACGCTCTTCCGATCTNNNCCAGAAGCAATTGGGAGTCAA
AGCACACGTCTGAACTCTTTCCCTACACGACGCTCTTCCGATCTNNNGTCCTAATTCACCATCCCTGTGT
GAGCACACGTCTGAACTCTTTCCCTACACGACGCTCTTCCGATCTNNNCCACCACCATACCCAGCTAATTA
GAGCACACGTCTGAACTCTTTCCCTACACGACGCTCTTCCGATCTNNNAGTGATCATCTTGCCTCAACCT
AGCACACGTCTGAACTCTTTCCCTACACGACGCTCTTCCGATCTNNNTGCAACTTGTAACTTGTAGCTG
AGCACACGTCTGAACTCTTTCCCTACACGACGCTCTTCCGATCTNNNCCTCAGTCTGCCAAGTAACTAG
AGCACACGTCTGAACTCTTTCCCTACACGACGCTCTTCCGATCTNNNGAAGTCTGGGCAGGTAGAATG
GAGCACACGTCTGAACTCTTTCCCTACACGACGCTCTTCCGATCTNNNGAGAACTCACTATGTGTCAGG
GAGCACACGTCTGAACTCTTTCCCTACACGACGCTCTTCCGATCTNNNCCCAGATTTGCTTCCTTTCAGT
GAGCACACGTCTGAACTCTTTCCCTACACGACGCTCTTCCGATCTNNNGGTGGAATAGAAACGCATCA
GCACACGTCTGAACTCTTTCCCTACACGACGCTCTTCCGATCTNNNTCTCACCAACCTGCTCTTTCTTTCA
GAGCACACGTCTGAACTCTTTCCCTACACGACGCTCTTCCGATCTNNNAGCAGCTGGAATTTCAAGTGT
AGCACACGTCTGAACTCTTTCCCTACACGACGCTCTTCCGATCTNNNTTCCTTGAGCAGAAGAAAATG
AGCACACGTCTGAACTCTTTCCCTACACGACGCTCTTCCGATCTNNNGCCTGTAGAATCACAAGCCAAT
GAGCACACGTCTGAACTCTTTCCCTACACGACGCTCTTCCGATCTNNNGGAATGAGTAGATCGCATGT
AGCACACGTCTGAACTCTTTCCCTACACGACGCTCTTCCGATCTNNNCAAATGCCCATTTTGATTCCCCT
GAGCACACGTCTGAACTCTTTCCCTACACGACGCTCTTCCGATCTNNNCTTCTGTAAACCCTTCAGTCCCA
GAGCACACGTCTGAACTCTTTCCCTACACGACGCTCTTCCGATCTNNNTAGATTTACTGCAGCCTCCACA
AGCACACGTCTGAACTCTTTCCCTACACGACGCTCTTCCGATCTNNNGGGGAAAGCAAAGTTTAGAAC
GAGCACACGTCTGAACTCTTTCCCTACACGACGCTCTTCCGATCTNNNAATCTCGAAACTCTTCACCCCT
AGCACACGTCTGAACTCTTTCCCTACACGACGCTCTTCCGATCTNNNCTCCTGAGTAGCTGGGATTACA
GAGCACACGTCTGAACTCTTTCCCTACACGACGCTCTTCCGATCTNNNAAAACTGGGTGCACACATTTA
AGCACACGTCTGAACTCTTTCCCTACACGACGCTCTTCCGATCTNNNAACAGCTGCATTGGAGTTTTCT
GAGCACACGTCTGAACTCTTTCCCTACACGACGCTCTTCCGATCTNNNGTTGAGCTGTGAAGGGCTTTA
AGCACACGTCTGAACTCTTTCCCTACACGACGCTCTTCCGATCTNNNATCACACCTTACTCTCCTCCTCA

AGCACACGTCTGAACTCTTTCCCTACACGACGCTCTTCCGATCTNNNGAACAATGTAATAAGGGCCAG
GAGCACACGTCTGAACTCTTTCCCTACACGACGCTCTTCCGATCTNNNGGCAGTTTCGTGGTATGAA
GAGCACACGTCTGAACTCTTTCCCTACACGACGCTCTTCCGATCTNNNAATGTTGGCATTACAGGTGTG
AGCACACGTCTGAACTCTTTCCCTACACGACGCTCTTCCGATCTNNNCTTCCAAAGTGCTGGGATTACA
GCACACGTCTGAACTCTTTCCCTACACGACGCTCTTCCGATCTNNNTGGTTGGGGTCAAGGACATTTAT
GAGCACACGTCTGAACTCTTTCCCTACACGACGCTCTTCCGATCTNNNTTGCTCTATTGGTCCATGCATCT
AGCACACGTCTGAACTCTTTCCCTACACGACGCTCTTCCGATCTNNNTGTCCCTTTGATGGTAGCAGAT
GAGCACACGTCTGAACTCTTTCCCTACACGACGCTCTTCCGATCTNNNAATCAGCCTTCTGGAGAGACT
AGCACACGTCTGAACTCTTTCCCTACACGACGCTCTTCCGATCTNNNTACCTGACTCTTGTTGCTAGCTC
AGCACACGTCTGAACTCTTTCCCTACACGACGCTCTTCCGATCTNNNCGTCCACCTGTAATCCTAGCTA
AGCACACGTCTGAACTCTTTCCCTACACGACGCTCTTCCGATCTNNNCGTCCACCTGTAATCCTAGCTA
AGAGCACACGTCTGAACTCTTTCCCTACACGACGCTCTTCCGATCTNNNCTGCCCTAGAAACATGCTTC
AGCACACGTCTGAACTCTTTCCCTACACGACGCTCTTCCGATCTNNNCTGAGGCAGGAGAATCACTTGA
AGCACACGTCTGAACTCTTTCCCTACACGACGCTCTTCCGATCTNNNGGAACTTGTAGTGAGCCCAGAT
GAGCACACGTCTGAACTCTTTCCCTACACGACGCTCTTCCGATCTNNNCATCTGTAACCCCAGCTACTCA
GAGCACACGTCTGAACTCTTTCCCTACACGACGCTCTTCCGATCTNNNCTGAGGCAGGAGAATCACTTG
AGCACACGTCTGAACTCTTTCCCTACACGACGCTCTTCCGATCTNNNGAGGCAGAAGAATCACTTGAA
GAGCACACGTCTGAACTCTTTCCCTACACGACGCTCTTCCGATCTNNNTGCCTGTAATCTCAGCTACTTG
AGCACACGTCTGAACTCTTTCCCTACACGACGCTCTTCCGATCTNNNGGCAGGAGGATTACTTGAACT
AGCACACGTCTGAACTCTTTCCCTACACGACGCTCTTCCGATCTNNNCTGAGGCAGGAGAATCACTTG
GAGCACACGTCTGAACTCTTTCCCTACACGACGCTCTTCCGATCTNNNGCATGCCTGTAATCCTAGCTAC
GCACACGTCTGAACTCTTTCCCTACACGACGCTCTTCCGATCTNNNCAGGAGAATCACTTGAACCTGGA
GAGCACACGTCTGAACTCTTTCCCTACACGACGCTCTTCCGATCTNNNGGAGTTTGAGGTTCGGTAA
GAGCACACGTCTGAACTCTTTCCCTACACGACGCTCTTCCGATCTNNNGGTGCCTATAATCCCAGCTACT
GAGCACACGTCTGAACTCTTTCCCTACACGACGCTCTTCCGATCTNNNTAGAGGTTGCAGTGAGCTGA
GAGCACACGTCTGAACTCTTTCCCTACACGACGCTCTTCCGATCTNNNCTGAGGCAGGAGAATGGTTT
GAGCACACGTCTGAACTCTTTCCCTACACGACGCTCTTCCGATCTNNNCAAGAAGTCTGGGCTTTGTT
AGCACACGTCTGAACTCTTTCCCTACACGACGCTCTTCCGATCTNNNTGGAGGTTGCAGTAAACTGAG
GAGCACACGTCTGAACTCTTTCCCTACACGACGCTCTTCCGATCTNNNAATCCCAGTTACTCAGGAAGCT
GAGCACACGTCTGAACTCTTTCCCTACACGACGCTCTTCCGATCTNNNATGACTTGAGCCCAGAGATT
AGCACACGTCTGAACTCTTTCCCTACACGACGCTCTTCCGATCTNNNAACATTGTACCACTGCACTCCA
AGAGCACACGTCTGAACTCTTTCCCTACACGACGCTCTTCCGATCTNNNAGTTTGCACTGAGTCAAGAT
GAGCACACGTCTGAACTCTTTCCCTACACGACGCTCTTCCGATCTNNNCCTAGGCAACAGAGTGAGAC
GAGCACACGTCTGAACTCTTTCCCTACACGACGCTCTTCCGATCTNNNGTAGTGAGCCATGATTGCATC
GAGCACACGTCTGAACTCTTTCCCTACACGACGCTCTTCCGATCTNNNCACATGAGGTCAGGAGTTTGA
GCACACGTCTGAACTCTTTCCCTACACGACGCTCTTCCGATCTNNNGGAGAATCACTTGAACCCGAGA
AGCACACGTCTGAACTCTTTCCCTACACGACGCTCTTCCGATCTNNNTTCCAGCTATTTGGATGGCTAC
GAGCACACGTCTGAACTCTTTCCCTACACGACGCTCTTCCGATCTNNNGGCATCCATAATCCCAGCTAC
AGCACACGTCTGAACTCTTTCCCTACACGACGCTCTTCCGATCTNNNCTGTAATCCCAGCTACTCAGGA
GAGCACACGTCTGAACTCTTTCCCTACACGACGCTCTTCCGATCTNNNTGAAGTCAGGAGTTTGAGAC
GCACACGTCTGAACTCTTTCCCTACACGACGCTCTTCCGATCTNNNTGACAATCGCTTGAACCCAAAAG
GAGCACACGTCTGAACTCTTTCCCTACACGACGCTCTTCCGATCTNNNGATGTGCCTATAGTCCCAGCTA
GAGCACACGTCTGAACTCTTTCCCTACACGACGCTCTTCCGATCTNNNAATTTGCTGGTACAAAGGCA

AGCACACGTCTGAACTCTTTCCCTACACGACGCTCTTCCGATCTNNNCTAAGTAGACTGAGGTGGGA
AGCACACGTCTGAACTCTTTCCCTACACGACGCTCTTCCGATCTNNNCTGAGGTAGGAGGATTGCTT
GAGCACACGTCTGAACTCTTTCCCTACACGACGCTCTTCCGATCTNNNTGAGTCCAAGAGTTTGAGA
AGCACACGTCTGAACTCTTTCCCTACACGACGCTCTTCCGATCTNNNCTCCTGTAATCCCAACTACTCG
GAGCACACGTCTGAACTCTTTCCCTACACGACGCTCTTCCGATCTNNNTGTAATTCCAGCTACTTGGG
GCACACGTCTGAACTCTTTCCCTACACGACGCTCTTCCGATCTNNNCTATGGTCACAGCTACTTGGGA
GAGCACACGTCTGAACTCTTTCCCTACACGACGCTCTTCCGATCTNNNGAGAATCGCTTGAACTCAGG
GCACACGTCTGAACTCTTTCCCTACACGACGCTCTTCCGATCTNNNGAGTTTGCAGTGAGCCAAGATT
AGCACACGTCTGAACTCTTTCCCTACACGACGCTCTTCCGATCTNNNCACTCCTGTGATCTCAGCTACT
GAGCACACGTCTGAACTCTTTCCCTACACGACGCTCTTCCGATCTNNNTGAGATCACATCATTGCACTC
AGCACACGTCTGAACTCTTTCCCTACACGACGCTCTTCCGATCTNNNCAGAGGTTACAGTGAGCTGA
AGCACACGTCTGAACTCTTTCCCTACACGACGCTCTTCCGATCTNNNGAAATCGTACCATTGCACTCC
GCACACGTCTGAACTCTTTCCCTACACGACGCTCTTCCGATCTNNNGTGATCCTAGCTACTCAGGAAG
GAGCACACGTCTGAACTCTTTCCCTACACGACGCTCTTCCGATCTNNNACATGTCTGTAATCCCAGCAG
AGCACACGTCTGAACTCTTTCCCTACACGACGCTCTTCCGATCTNNNCTGAGGCAGGAGAATGGTGT
AGCACACGTCTGAACTCTTTCCCTACACGACGCTCTTCCGATCTNNNCTGTAGTTCCAGCTACTCGGG
GAGCACACGTCTGAACTCTTTCCCTACACGACGCTCTTCCGATCTNNNTGCACTGAGCCAAGATCATAT
AGCACACGTCTGAACTCTTTCCCTACACGACGCTCTTCCGATCTNNNAAGAATCACTTGAACCCAGGA
AGCACACGTCTGAACTCTTTCCCTACACGACGCTCTTCCGATCTNNNCTGGGGAAGGAGAATCACTT
GCACACGTCTGAACTCTTTCCCTACACGACGCTCTTCCGATCTNNNGTTGTAGTGAGCCAAGATCATG
AGCACACGTCTGAACTCTTTCCCTACACGACGCTCTTCCGATCTNNNCTGTAGTCCCAGCTACTCCAGA
AGAGCACACGTCTGAACTCTTTCCCTACACGACGCTCTTCCGATCTNNNCTGTAGTCCCAGCTACTCAG
GCACACGTCTGAACTCTTTCCCTACACGACGCTCTTCCGATCTNNNGAGCTTGCAGTGAGTCGAGAT
AGCACACGTCTGAACTCTTTCCCTACACGACGCTCTTCCGATCTNNNTACGGTCATTTGGCATTTGAA
AGCACACGTCTGAACTCTTTCCCTACACGACGCTCTTCCGATCTNNNATCATGCCACTGTACTTTAGCC
GAGCACACGTCTGAACTCTTTCCCTACACGACGCTCTTCCGATCTNNNTGTAGTCTCAGCTACTCAGGA
AAGAGCACACGTCTGAACTCTTTCCCTACACGACGCTCTTCCGATCTNNNGCCTGTAGTCCCAGCTACT
AGCACACGTCTGAACTCTTTCCCTACACGACGCTCTTCCGATCTNNNCTGAGGCAGGAGAATCACTT
GAGCACACGTCTGAACTCTTTCCCTACACGACGCTCTTCCGATCTNNNCTGTTGTACCCCAGACTGTTG
GCACACGTCTGAACTCTTTCCCTACACGACGCTCTTCCGATCTNNNCTGAGATCACATCACTGCACTC
AGCACACGTCTGAACTCTTTCCCTACACGACGCTCTTCCGATCTNNNGTTGAGGCATGAGAATTGCT
AGCACACGTCTGAACTCTTTCCCTACACGACGCTCTTCCGATCTNNNCAACAGCCATACTCCCTATCCT
AGCACACGTCTGAACTCTTTCCCTACACGACGCTCTTCCGATCTNNNAGAATTGCTTGAACCCAGGAG
AGCACACGTCTGAACTCTTTCCCTACACGACGCTCTTCCGATCTNNNTAACTACTGAGGAGGCTGAG
AGCACACGTCTGAACTCTTTCCCTACACGACGCTCTTCCGATCTNNNATGCTTGTAATCCCAGCTACTC
GCACACGTCTGAACTCTTTCCCTACACGACGCTCTTCCGATCTNNNGTGACAGAGTGAGACTCCATCA
AGCACACGTCTGAACTCTTTCCCTACACGACGCTCTTCCGATCTNNNGCTATATCCGTCTCCTCATGCT
AGCACACGTCTGAACTCTTTCCCTACACGACGCTCTTCCGATCTNNNCAAGAGAATTGCTTGAACCTG
GCACACGTCTGAACTCTTTCCCTACACGACGCTCTTCCGATCTNNNCAGTGAGCTGAGATCAAGTCA
AGCACACGTCTGAACTCTTTCCCTACACGACGCTCTTCCGATCTNNNGCAAGACCCGGTCTCTACTAA
GAGCACACGTCTGAACTCTTTCCCTACACGACGCTCTTCCGATCTNNNGCAACAGACCAAAACTCCATC
AGCACACGTCTGAACTCTTTCCCTACACGACGCTCTTCCGATCTNNNCCCTGGGTGATAGAGTGAGA
GCACACGTCTGAACTCTTTCCCTACACGACGCTCTTCCGATCTNNNTCAGGTAGGAGAATCGCTTGA

AGCACACGTCTGAACTCTTTCCCTACACGACGCTCTTCCGATCTNNNGAGGCAGGACAATTTCTTGAA
GAGCACACGTCTGAACTCTTTCCCTACACGACGCTCTTCCGATCTNNNACTCAAAGGTAAGCAGAGGA
GAGCACACGTCTGAACTCTTTCCCTACACGACGCTCTTCCGATCTNNNGCCTATAATCCCAGCTTCTCAG
GAGCACACGTCTGAACTCTTTCCCTACACGACGCTCTTCCGATCTNNNCAGAAGAATCGCTTGAACTTG
AGCACACGTCTGAACTCTTTCCCTACACGACGCTCTTCCGATCTNNNATAGCCACCTACAGCACTTACTG
GAGCACACGTCTGAACTCTTTCCCTACACGACGCTCTTCCGATCTNNNCTGGTGCTAGGACTGTGAGTA
AGCACACGTCTGAACTCTTTCCCTACACGACGCTCTTCCGATCTNNNCAAACACACTTTCGGGTGCTATT
GAGCACACGTCTGAACTCTTTCCCTACACGACGCTCTTCCGATCTNNNCTCCTGCCTTAGCTTTCCAAAT
AGCACACGTCTGAACTCTTTCCCTACACGACGCTCTTCCGATCTNNNCCACTTACAGATGAGGATGCTG
AGCACACGTCTGAACTCTTTCCCTACACGACGCTCTTCCGATCTNNNGCAGATTGTCCTGGGATTTTCT
GAGCACACGTCTGAACTCTTTCCCTACACGACGCTCTTCCGATCTNNNCTTCCTGGGTTCAAGGGATTC
AGCACACGTCTGAACTCTTTCCCTACACGACGCTCTTCCGATCTNNNCTCTTGTGGCCTTTCCTCTATGG
GAGCACACGTCTGAACTCTTTCCCTACACGACGCTCTTCCGATCTNNNAAAGCAGTTTCCCCAGTGATA
GCACACGTCTGAACTCTTTCCCTACACGACGCTCTTCCGATCTNNNAGTTACCCCTGTCCTGAATCCTA
AGCACACGTCTGAACTCTTTCCCTACACGACGCTCTTCCGATCTNNNATTCTCCTATCTCAGCCTCCTGA
AGCACACGTCTGAACTCTTTCCCTACACGACGCTCTTCCGATCTNNNCCCAGCCCCCATAATTACTTTTTC
AGCACACGTCTGAACTCTTTCCCTACACGACGCTCTTCCGATCTNNNCTGAGCCATCTGTCTGTTTTGT
AGCACACGTCTGAACTCTTTCCCTACACGACGCTCTTCCGATCTNNNGGTCTTCTGTAAGTTTTCTGGC
GAGCACACGTCTGAACTCTTTCCCTACACGACGCTCTTCCGATCTNNNCACCTCCATCTCCATCTTCCAAA
GAGCACACGTCTGAACTCTTTCCCTACACGACGCTCTTCCGATCTNNNAGATAAGTGAATGCCTGGCT
GAGCACACGTCTGAACTCTTTCCCTACACGACGCTCTTCCGATCTNNNATGTTGTCCAGGCTGTACTCA
AGCACACGTCTGAACTCTTTCCCTACACGACGCTCTTCCGATCTNNNTGGTACTAACTTTTCCCTCAACA
GAGCACACGTCTGAACTCTTTCCCTACACGACGCTCTTCCGATCTNNNCACCATGCCCAGCTAATTTTT
GAGCACACGTCTGAACTCTTTCCCTACACGACGCTCTTCCGATCTNNNTCCCTTAGTTTCACCCCTATCA
AGCACACGTCTGAACTCTTTCCCTACACGACGCTCTTCCGATCTNNNACCCTGTGTCAAAAAAATACCC
AGCACACGTCTGAACTCTTTCCCTACACGACGCTCTTCCGATCTNNNCCCTTTACTCAGGTTACTGCTGT
AGCACACGTCTGAACTCTTTCCCTACACGACGCTCTTCCGATCTNNNTAGCTCATTGTAGCCCTGAACT
GAGCACACGTCTGAACTCTTTCCCTACACGACGCTCTTCCGATCTNNNCTAACACCACCCACAGTCAACTA
AGCACACGTCTGAACTCTTTCCCTACACGACGCTCTTCCGATCTNNNTCTACGCACAATGCACAAATGTT
AGCACACGTCTGAACTCTTTCCCTACACGACGCTCTTCCGATCTNNNAAATGTTGCATCACAGTTTCCGT
GAGCACACGTCTGAACTCTTTCCCTACACGACGCTCTTCCGATCTNNNCCCAGTCATCCAGCTATGTTTTT
GAGCACACGTCTGAACTCTTTCCCTACACGACGCTCTTCCGATCTNNNTATTTCTCCACATCCTCTCCAG
GAGCACACGTCTGAACTCTTTCCCTACACGACGCTCTTCCGATCTNNNTTTAGACAGTCTGGCTCTATC
GAGCACACGTCTGAACTCTTTCCCTACACGACGCTCTTCCGATCTNNNCCTCCTTCCTATGAGCAGAGA
GAGCACACGTCTGAACTCTTTCCCTACACGACGCTCTTCCGATCTNNNAGCTAATTCTCTCCCCTACCAC
AGCACACGTCTGAACTCTTTCCCTACACGACGCTCTTCCGATCTNNNGCACTAGGCTCTGGAGAGATA
GAGCACACGTCTGAACTCTTTCCCTACACGACGCTCTTCCGATCTNNNTGAACTGAGTGAAAGTACGT
AGCACACGTCTGAACTCTTTCCCTACACGACGCTCTTCCGATCTNNNGTACTAAACACTGACTCAGGCC
AGCACACGTCTGAACTCTTTCCCTACACGACGCTCTTCCGATCTNNNCTCCCAAGTAGCTGAGACTACA
AGCACACGTCTGAACTCTTTCCCTACACGACGCTCTTCCGATCTNNNTGATCCATGATGGTCAAGGAC
AGCACACGTCTGAACTCTTTCCCTACACGACGCTCTTCCGATCTNNNCTGTCATGGGATTGAGACTTG
AGAGCACACGTCTGAACTCTTTCCCTACACGACGCTCTTCCGATCTNNNAGGAGAAACTGCTGGGAAT
AGAGCACACGTCTGAACTCTTTCCCTACACGACGCTCTTCCGATCTNNNCCAGGCATTTGAGACTACAG

AGCACACGTCTGAACTCTTTCCCTACACGACGCTCTTCCGATCTNNNGAGGTTGCAGTAAGCCAAGA
AGCACACGTCTGAACTCTTTCCCTACACGACGCTCTTCCGATCTNNNAGAATCGCTTGAACTAGGGAG
AGCACACGTCTGAACTCTTTCCCTACACGACGCTCTTCCGATCTNNNATAATCCCAGCACTTTAGGAG
AGCACACGTCTGAACTCTTTCCCTACACGACGCTCTTCCGATCTNNNCTGAGGCAGGAGAATCACTTG
GAGCACACGTCTGAACTCTTTCCCTACACGACGCTCTTCCGATCTNNNTGTAATCCCAGCACTTTGAGA
AGCACACGTCTGAACTCTTTCCCTACACGACGCTCTTCCGATCTNNNTATTTGGAAGCTAAGGCCAGA
GAGCACACGTCTGAACTCTTTCCCTACACGACGCTCTTCCGATCTNNNAGAATCACTTGAACCCAGCAG
AGCACACGTCTGAACTCTTTCCCTACACGACGCTCTTCCGATCTNNNAGTAAGCTGTGATCATGGCAC
AGAGCACACGTCTGAACTCTTTCCCTACACGACGCTCTTCCGATCTNNNGATTGCAGTGAGCTGAGAT
AGCACACGTCTGAACTCTTTCCCTACACGACGCTCTTCCGATCTNNNATCACATCATTGCACTCTAGCCT
AGCACACGTCTGAACTCTTTCCCTACACGACGCTCTTCCGATCTNNNTGAGCAGAGATCATACCATTG
GAGCACACGTCTGAACTCTTTCCCTACACGACGCTCTTCCGATCTNNNGAGAATCGCTTGAAACCAGA
AGAGCACACGTCTGAACTCTTTCCCTACACGACGCTCTTCCGATCTNNNAGGAGGTGGAAGTTTCAGT
AGCACACGTCTGAACTCTTTCCCTACACGACGCTCTTCCGATCTNNNGCTGAGGTGGTAGAATAGCT
AGCACACGTCTGAACTCTTTCCCTACACGACGCTCTTCCGATCTNNNGAGAATTGCCTGAACCTAGGA
GCACACGTCTGAACTCTTTCCCTACACGACGCTCTTCCGATCTNNNCTGTAATCCCAGCTACTCAGGA
AGCACACGTCTGAACTCTTTCCCTACACGACGCTCTTCCGATCTNNNAGGAGAATTGCTTGAACTGGG
GAGCACACGTCTGAACTCTTTCCCTACACGACGCTCTTCCGATCTNNNGGCTGAGGCAAGAGAATCAT
AGCACACGTCTGAACTCTTTCCCTACACGACGCTCTTCCGATCTNNNCACCTGTAATCCCAGCTACTCAA
GAGCACACGTCTGAACTCTTTCCCTACACGACGCTCTTCCGATCTNNNGAGCAACACTCCATCTCAAAA
GAGCACACGTCTGAACTCTTTCCCTACACGACGCTCTTCCGATCTNNNGGCAACAGAGTAAGACCTTG
GCACACGTCTGAACTCTTTCCCTACACGACGCTCTTCCGATCTNNNCCTGTAATTCCAGCTACTTGGGA
AGCACACGTCTGAACTCTTTCCCTACACGACGCTCTTCCGATCTNNNGAGACAGGAGACTAGCATGA
GAGCACACGTCTGAACTCTTTCCCTACACGACGCTCTTCCGATCTNNNGCAACATGATGAAACCCTGTC
GAGCACACGTCTGAACTCTTTCCCTACACGACGCTCTTCCGATCTNNNGGATCACTTGAGGCCATGAG
AGCACACGTCTGAACTCTTTCCCTACACGACGCTCTTCCGATCTNNNCAGGAGAATCACTTGAACCCA
GAGCACACGTCTGAACTCTTTCCCTACACGACGCTCTTCCGATCTNNNGTGATGCACGTCTGTAATCC
GAGCACACGTCTGAACTCTTTCCCTACACGACGCTCTTCCGATCTNNNGGAAGTTGCAATGAGCCAA
GAGCACACGTCTGAACTCTTTCCCTACACGACGCTCTTCCGATCTNNNCTGAGGCAGGAGAATCACTT
GCACACGTCTGAACTCTTTCCCTACACGACGCTCTTCCGATCTNNNCTGAGGCAGGAGAATTGCTTGA
AGAGCACACGTCTGAACTCTTTCCCTACACGACGCTCTTCCGATCTNNNAATGAGCCAAATTACACCAC
GCACACGTCTGAACTCTTTCCCTACACGACGCTCTTCCGATCTNNNCTGTAATCCCAGCTACTGAGGA
GAGCACACGTCTGAACTCTTTCCCTACACGACGCTCTTCCGATCTNNNTGAGGCAGAAGAATTGCTTG
GAGCACACGTCTGAACTCTTTCCCTACACGACGCTCTTCCGATCTNNNCTCTGAGCCTCAGTTTCCTCAT
GCACACGTCTGAACTCTTTCCCTACACGACGCTCTTCCGATCTNNNATGCTGTGATTTTTGCCATCAGG
AGCACACGTCTGAACTCTTTCCCTACACGACGCTCTTCCGATCTNNNGCCCCAAGTAACATAGGAGAG
AGCACACGTCTGAACTCTTTCCCTACACGACGCTCTTCCGATCTNNNTCACACCACTACATTCCAGTCTG
AGCACACGTCTGAACTCTTTCCCTACACGACGCTCTTCCGATCTNNNATGAACCCAGAGGAAAGAGC
AAGAGCACACGTCTGAACTCTTTCCCTACACGACGCTCTTCCGATCTNNNGAGAATCGCTTGAACCTA
AGCACACGTCTGAACTCTTTCCCTACACGACGCTCTTCCGATCTNNNAGGTTATGGCCCTATATCCTG
AGCACACGTCTGAACTCTTTCCCTACACGACGCTCTTCCGATCTNNNGTAATCCCAGCTACTCAGAAG
AGCACACGTCTGAACTCTTTCCCTACACGACGCTCTTCCGATCTNNNGAGTTTCCATTGAGCTGAGGT
AGCACACGTCTGAACTCTTTCCCTACACGACGCTCTTCCGATCTNNNTGTAATCTCAGCAACTCAGGA

```
GCACACGTCTGAACTCTTTCCCTACACGACGCTCTTCCGATCTNNNCATCTTGCCTCTAGTGGACAAGG
GAGCACACGTCTGAACTCTTTCCCTACACGACGCTCTTCCGATCTNNNGGGAGAAAGAGCACAGAGG
AGCACACGTCTGAACTCTTTCCCTACACGACGCTCTTCCGATCTNNNGCTGGGATTACAGACATAAG
GAGCACACGTCTGAACTCTTTCCCTACACGACGCTCTTCCGATCTNNNTGTTGAAGAGCCTCATTCAGT
AGCACACGTCTGAACTCTTTCCCTACACGACGCTCTTCCGATCTNNNCACATAAGAGAGCACGGACCT
AGCACACGTCTGAACTCTTTCCCTACACGACGCTCTTCCGATCTNNNGGATTTCCCCCTATTTTCCCACT
AGCACACGTCTGAACTCTTTCCCTACACGACGCTCTTCCGATCTNNNTCCAACCCATCCCCTTTATCAA
GCACACGTCTGAACTCTTTCCCTACACGACGCTCTTCCGATCTNNNAGCCAGATGATTTTCCAAAGCA
AGCACACGTCTGAACTCTTTCCCTACACGACGCTCTTCCGATCTNNNACCTCGCCTGGTCTTTTTTTTT
AGCACACGTCTGAACTCTTTCCCTACACGACGCTCTTCCGATCTNNNATATGCCCAGAAAGTGTCAGG
GAGCACACGTCTGAACTCTTTCCCTACACGACGCTCTTCCGATCTNNNTTACACTGCAGACTCTTCAGA
AGCACACGTCTGAACTCTTTCCCTACACGACGCTCTTCCGATCTNNNAAACTCTAACTCCTGGGCTCAA
AGCACACGTCTGAACTCTTTCCCTACACGACGCTCTTCCGATCTNNNCATCAGACTATTGCCACAGGG
AGCACACGTCTGAACTCTTTCCCTACACGACGCTCTTCCGATCTNNNAGACTTTGTTCAGGGGTCTG
AGCACACGTCTGAACTCTTTCCCTACACGACGCTCTTCCGATCTNNNTAAGTGATCCAGTGATACATG
AGCACACGTCTGAACTCTTTCCCTACACGACGCTCTTCCGATCTNNNCCCTCCTGTCCCTTCTTTTTCTT
AGCACACGTCTGAACTCTTTCCCTACACGACGCTCTTCCGATCTNNNACGGTTTTTGCCATTGAGCATA
AGCACACGTCTGAACTCTTTCCCTACACGACGCTCTTCCGATCTNNNTCTAGCACACACTGATCTTGCAT
GCACACGTCTGAACTCTTTCCCTACACGACGCTCTTCCGATCTNNNAAGCAGACCCACATAGACTTGA
AGCACACGTCTGAACTCTTTCCCTACACGACGCTCTTCCGATCTNNNACAAGAGTAATACCAGGGCTC
AGCACACGTCTGAACTCTTTCCCTACACGACGCTCTTCCGATCTNNNTCAAGTGATACTACTGCCTCAG
AGCACACGTCTGAACTCTTTCCCTACACGACGCTCTTCCGATCTNNNCACCACACCCAGCTAATTTTG
GAGCACACGTCTGAACTCTTTCCCTACACGACGCTCTTCCGATCTNNNAGTGCTGGGATTACAGGAAT
GAGCACACGTCTGAACTCTTTCCCTACACGACGCTCTTCCGATCTNNNCTCTCGAGTAGCTGGGACTA
AGCACACGTCTGAACTCTTTCCCTACACGACGCTCTTCCGATCTNNNGATGGTCTCAATCTCCTGACCT
AGCACACGTCTGAACTCTTTCCCTACACGACGCTCTTCCGATCTNNNCGTCTCTGTATTGCCCAGTCTA
AGCACACGTCTGAACTCTTTCCCTACACGACGCTCTTCCGATCTNNNCTGAGGCAGGAGAATCACTTG
GAGCACACGTCTGAACTCTTTCCCTACACGACGCTCTTCCGATCTNNNGCAGGAGAATTGCTTGAAC
AGCACACGTCTGAACTCTTTCCCTACACGACGCTCTTCCGATCTNNNCCCAACCAATTGAAACCACTGT
AGCACACGTCTGAACTCTTTCCCTACACGACGCTCTTCCGATCTNNNGTGCCTATAGTCCCAGCTACTT
GAGCACACGTCTGAACTCTTTCCCTACACGACGCTCTTCCGATCTNNNCTATCATTTTGGGAGGCTGA
AGCACACGTCTGAACTCTTTCCCTACACGACGCTCTTCCGATCTNNNGGCACACATGTAATTCCAGCTA
GAGCACACGTCTGAACTCTTTCCCTACACGACGCTCTTCCGATCTNNNTGAAGTAGGAGAGCCACTT
AGCACACGTCTGAACTCTTTCCCTACACGACGCTCTTCCGATCTNNNGATTGCTTATGCCCAGGAGTT
AGCACACGTCTGAACTCTTTCCCTACACGACGCTCTTCCGATCTNNNTGTAATCCTAGCACTTTGGGA
AGCACACGTCTGAACTCTTTCCCTACACGACGCTCTTCCGATCTNNNGAGGCAGGAGAATAGCTTGA
AGCACACGTCTGAACTCTTTCCCTACACGACGCTCTTCCGATCTNNNGAGGCAGAAGAATCGCTTGA
AGAGCACACGTCTGAACTCTTTCCCTACACGACGCTCTTCCGATCTNNNGGAGGCTGAGAGAGAAGA
AGCACACGTCTGAACTCTTTCCCTACACGACGCTCTTCCGATCTNNNCTGTAATCCCAGCTACTCAGG
GAGCACACGTCTGAACTCTTTCCCTACACGACGCTCTTCCGATCTNNNCCTTGTAATACCAGCACCTTG
GCACACGTCTGAACTCTTTCCCTACACGACGCTCTTCCGATCTNNNAGACAGGAGAATCACTTGAACC
AGCACACGTCTGAACTCTTTCCCTACACGACGCTCTTCCGATCTNNNGGAGGCTGAAGCAGAATAAT
AGCACACGTCTGAACTCTTTCCCTACACGACGCTCTTCCGATCTNNNGAGGCAGGAGAATTGTTTGA
```

GAGCACACGTCTGAACTCTTTCCCTACACGACGCTCTTCCGATCTNNNTCTCAACACTTTGGGAGGCT
AGCACACGTCTGAACTCTTTCCCTACACGACGCTCTTCCGATCTNNNGGTCATGATCACCTCATTGGA
AGCACACGTCTGAACTCTTTCCCTACACGACGCTCTTCCGATCTNNNGAAGTTGCAGTGAGCCAAGA
AGCACACGTCTGAACTCTTTCCCTACACGACGCTCTTCCGATCTNNNCAGCTACTCAGAAGCAGGAG
AGCACACGTCTGAACTCTTTCCCTACACGACGCTCTTCCGATCTNNNCAAGAAGATTGCTTGAACCTG
AGCACACGTCTGAACTCTTTCCCTACACGACGCTCTTCCGATCTNNNCTGTAGTCCCAGCTACTCAGA
AGCACACGTCTGAACTCTTTCCCTACACGACGCTCTTCCGATCTNNNCCTAGCCAACATAGTGAAACC
AGCACACGTCTGAACTCTTTCCCTACACGACGCTCTTCCGATCTNNNTTGGGAGGCTAAAGCATGAG
GAGCACACGTCTGAACTCTTTCCCTACACGACGCTCTTCCGATCTNNNGCATGTGCCTGTAATCCTAG
AGCACACGTCTGAACTCTTTCCCTACACGACGCTCTTCCGATCTNNNACCTTCCCTCCACTATTGTCCTA
GCACACGTCTGAACTCTTTCCCTACACGACGCTCTTCCGATCTNNNGCAGTGACCTATGATTGCATCA
AGCACACGTCTGAACTCTTTCCCTACACGACGCTCTTCCGATCTNNNGAGTTCAAGACCAGCCTAGA
AGCACACGTCTGAACTCTTTCCCTACACGACGCTCTTCCGATCTNNNGCAAGAGAATAGCTTGAACC
AGCACACGTCTGAACTCTTTCCCTACACGACGCTCTTCCGATCTNNNTGAGCTGAGACACCACTGTATT
GCACACGTCTGAACTCTTTCCCTACACGACGCTCTTCCGATCTNNNGTGGATCACGAGGTCAGTAG
AGCACACGTCTGAACTCTTTCCCTACACGACGCTCTTCCGATCTNNNGGGAGACTGAGTCAGGAGA
GCACACGTCTGAACTCTTTCCCTACACGACGCTCTTCCGATCTNNNATCTTGGCTTACTGCAACCTCTG
GCACACGTCTGAACTCTTTCCCTACACGACGCTCTTCCGATCTNNNGAGTGCAATAGTGTGATCATG
GAGCACACGTCTGAACTCTTTCCCTACACGACGCTCTTCCGATCTNNNTCTGACTACCCAAGACATCC
GCACACGTCTGAACTCTTTCCCTACACGACGCTCTTCCGATCTNNNTTAGGATTACAGCTGTGAGCC
GAGCACACGTCTGAACTCTTTCCCTACACGACGCTCTTCCGATCTNNNTAGATGCGTAGGTAGATTC
AGCACACGTCTGAACTCTTTCCCTACACGACGCTCTTCCGATCTNNNGACAGCATACAGTTGGATCAT
AGCACACGTCTGAACTCTTTCCCTACACGACGCTCTTCCGATCTNNNTTGGAATCAAAGGAAGGTCA
AGCACACGTCTGAACTCTTTCCCTACACGACGCTCTTCCGATCTNNNTACAGACGGGAATTCACCATG
GCACACGTCTGAACTCTTTCCCTACACGACGCTCTTCCGATCTNNNTAGCTAGCTGGAATGAAGTGG
GCACACGTCTGAACTCTTTCCCTACACGACGCTCTTCCGATCTNNNGAAGGAAGTGGGGAAAGAGA
GAGCACACGTCTGAACTCTTTCCCTACACGACGCTCTTCCGATCTNNNAGAAGTCAGGTCCCTGGTTC
GCACACGTCTGAACTCTTTCCCTACACGACGCTCTTCCGATCTNNNGAGGACGATCAAGAAAACACA
GCACACGTCTGAACTCTTTCCCTACACGACGCTCTTCCGATCTNNNAAGTCCAATTTGTGTGTTTCGG
GCACACGTCTGAACTCTTTCCCTACACGACGCTCTTCCGATCTNNNTGTAATAAGGCCCTGGTTTCC
AGCACACGTCTGAACTCTTTCCCTACACGACGCTCTTCCGATCTNNNAAACCCATCACTTCACTGCTTT
GCACACGTCTGAACTCTTTCCCTACACGACGCTCTTCCGATCTNNNGTAGCTGGGACTTTAGGCATGA
AGCACACGTCTGAACTCTTTCCCTACACGACGCTCTTCCGATCTNNNCTTCCAGGCTCATTCATGTTGT
AGCACACGTCTGAACTCTTTCCCTACACGACGCTCTTCCGATCTNNNGTATGGGTCAAGGATAGGAG
GCACACGTCTGAACTCTTTCCCTACACGACGCTCTTCCGATCTNNNTCTCCAATAGCAACAGTGTCA
AGCACACGTCTGAACTCTTTCCCTACACGACGCTCTTCCGATCTNNNCCAGCCTCTCTGTTGTTTCTTT
AGCACACGTCTGAACTCTTTCCCTACACGACGCTCTTCCGATCTNNNGCTCCACACAAAGCAAGTAAGT
AGCACACGTCTGAACTCTTTCCCTACACGACGCTCTTCCGATCTNNNGTCTGCTTTGCCTTTTTGCATT
GCACACGTCTGAACTCTTTCCCTACACGACGCTCTTCCGATCTNNNTCAAAGAGAGTCAAGCCAGATG
GAGCACACGTCTGAACTCTTTCCCTACACGACGCTCTTCCGATCTNNNATTCTCCAAGACCGGCTATCT
AGCACACGTCTGAACTCTTTCCCTACACGACGCTCTTCCGATCTNNNATAGTGAGCTGAGATTGCACC
AGCACACGTCTGAACTCTTTCCCTACACGACGCTCTTCCGATCTNNNAAAATTGCGCCATTACACTCCA
GAGCACACGTCTGAACTCTTTCCCTACACGACGCTCTTCCGATCTNNNAGACAGGAGAATTGCTTGA

GAGCACACGTCTGAACTCTTTCCCTACACGACGCTCTTCCGATCTNNNCTGTAGTCCCAGCTACTCAGGA
GAGCACACGTCTGAACTCTTTCCCTACACGACGCTCTTCCGATCTNNNGGCACAAGAATCATTTGAACC
GAGCACACGTCTGAACTCTTTCCCTACACGACGCTCTTCCGATCTNNNGTACCTGTAGTCCCAGCTACTT
AGCACACGTCTGAACTCTTTCCCTACACGACGCTCTTCCGATCTNNNGAGCTTGCAATGAGCCGAGATA
GAGCACACGTCTGAACTCTTTCCCTACACGACGCTCTTCCGATCTNNNCTGAGGCAGGAGAATCACTTC
AGCACACGTCTGAACTCTTTCCCTACACGACGCTCTTCCGATCTNNNTGTGTCTGTCTCCAGTGTAAAGC
AGCACACGTCTGAACTCTTTCCCTACACGACGCTCTTCCGATCTNNNGCAGCATGGCAAAATCCTATCTC
AGCACACGTCTGAACTCTTTCCCTACACGACGCTCTTCCGATCTNNNGACAAAAGAATCGCTTGAACCCA
GAGCACACGTCTGAACTCTTTCCCTACACGACGCTCTTCCGATCTNNNTCAAGACAAGCCTGGGAAACA
AGCACACGTCTGAACTCTTTCCCTACACGACGCTCTTCCGATCTNNNTCAGAAGGCATCATGTTACCCA
AGCACACGTCTGAACTCTTTCCCTACACGACGCTCTTCCGATCTNNNATGCCTCTAATCCCAGCTACTTG
GCACACGTCTGAACTCTTTCCCTACACGACGCTCTTCCGATCTNNNAGTGAGCCAAGATCATGACACTC
GAGCACACGTCTGAACTCTTTCCCTACACGACGCTCTTCCGATCTNNNAGTGAGTGGAGATCATGCCA
GAGCACACGTCTGAACTCTTTCCCTACACGACGCTCTTCCGATCTNNNCTGCTTGATATTGAGGCATTG
GAGCACACGTCTGAACTCTTTCCCTACACGACGCTCTTCCGATCTNNNATCTCAGTGGCTGTTTCCAGA
GAGCACACGTCTGAACTCTTTCCCTACACGACGCTCTTCCGATCTNNNCACCTGTAGTACCAGCTACTTG
AGCACACGTCTGAACTCTTTCCCTACACGACGCTCTTCCGATCTNNNTGTAGTCCCAGCTACTCAGGAG
AGAGCACACGTCTGAACTCTTTCCCTACACGACGCTCTTCCGATCTNNNTGAGGCTGGAGAATCACTTC
GAGCACACGTCTGAACTCTTTCCCTACACGACGCTCTTCCGATCTNNNAAGACATCACAGAATCTCCAG
AGAGCACACGTCTGAACTCTTTCCCTACACGACGCTCTTCCGATCTNNNGAGCGAGACTCCGTTTCAAA
AGCACACGTCTGAACTCTTTCCCTACACGACGCTCTTCCGATCTNNNCTGAGACAGGAGAATCGCTTG
AGAGCACACGTCTGAACTCTTTCCCTACACGACGCTCTTCCGATCTNNNTGGGAGACAGAGTAAGACTC
AGCACACGTCTGAACTCTTTCCCTACACGACGCTCTTCCGATCTNNNCTTGAATCGCTTGAACCTGGAA
AGAGCACACGTCTGAACTCTTTCCCTACACGACGCTCTTCCGATCTNNNTGGGAGGAAGATGTTGCAG
AGCACACGTCTGAACTCTTTCCCTACACGACGCTCTTCCGATCTNNNTCTGTTATGGTCCTTCCCACTTGC
GAGCACACGTCTGAACTCTTTCCCTACACGACGCTCTTCCGATCTNNNAAGGTTGCAGTAAGCCAAGAT
GAGCACACGTCTGAACTCTTTCCCTACACGACGCTCTTCCGATCTNNNTGAGGGAGTAGAAGTGCTTG
GAGCACACGTCTGAACTCTTTCCCTACACGACGCTCTTCCGATCTNNNGGCTGAGACATGAGAAATGCT
GAGCACACGTCTGAACTCTTTCCCTACACGACGCTCTTCCGATCTNNNCTTGGGTTCCTGAGTCTCTTGT
AGCACACGTCTGAACTCTTTCCCTACACGACGCTCTTCCGATCTNNNGCCTGAATCACCTTCTACCTTCA
GAGCACACGTCTGAACTCTTTCCCTACACGACGCTCTTCCGATCTNNNGAGAGTGTAGGGATGAAGAG
AGCACACGTCTGAACTCTTTCCCTACACGACGCTCTTCCGATCTNNNCACTCACTCACAGCAACTTTCAG
GAGCACACGTCTGAACTCTTTCCCTACACGACGCTCTTCCGATCTNNNAATTCCATCAGGACCACTAAC
AGCACACGTCTGAACTCTTTCCCTACACGACGCTCTTCCGATCTNNNGGTGGTGCTTTCTTTCTTGACTC
AGCACACGTCTGAACTCTTTCCCTACACGACGCTCTTCCGATCTNNNGCCCAGGTACACAGATTCACTA
AGCACACGTCTGAACTCTTTCCCTACACGACGCTCTTCCGATCTNNNTTGTAGTAGAGACGGGTTTCAC
AGCACACGTCTGAACTCTTTCCCTACACGACGCTCTTCCGATCTNNNGTAGCTAGGACTACAGGCACAT
AGCACACGTCTGAACTCTTTCCCTACACGACGCTCTTCCGATCTNNNCATGGAAGCAGACTAGAGCATC
AGCACACGTCTGAACTCTTTCCCTACACGACGCTCTTCCGATCTNNNGGCAACTACCATTCTGCTCTTTCC
AGCACACGTCTGAACTCTTTCCCTACACGACGCTCTTCCGATCTNNNATCGCAACTCTATGTTAGCTCCA
GAGCACACGTCTGAACTCTTTCCCTACACGACGCTCTTCCGATCTNNNCATATGGCCACAGAAGTCCTG
AGCACACGTCTGAACTCTTTCCCTACACGACGCTCTTCCGATCTNNNTTTTTCCATATGCTTGTTGGCCAC
AGCACACGTCTGAACTCTTTCCCTACACGACGCTCTTCCGATCTNNNCTCACTCTTGCCCTTTCACATTG

GCACACGTCTGAACTCTTTCCCTACACGACGCTCTTCCGATCTNNNTAGATAGGACTATGGCATGTG
AGCACACGTCTGAACTCTTTCCCTACACGACGCTCTTCCGATCTNNNCAGGGGAAGCTTCTTAGTCCTT
GAGCACACGTCTGAACTCTTTCCCTACACGACGCTCTTCCGATCTNNNGGATTACAGGCTTGAGCTACC
AGCACACGTCTGAACTCTTTCCCTACACGACGCTCTTCCGATCTNNNTTGATACTAAAGCTGTCCACGCA
AGCACACGTCTGAACTCTTTCCCTACACGACGCTCTTCCGATCTNNNAGGTTGTTCAGTTTCCGTGTAG
GAGCACACGTCTGAACTCTTTCCCTACACGACGCTCTTCCGATCTNNNAGGCTACATTGTTGTGTTTTG
GCACACGTCTGAACTCTTTCCCTACACGACGCTCTTCCGATCTNNNCTGACAACCCACATTGCAGATTT
GCACACGTCTGAACTCTTTCCCTACACGACGCTCTTCCGATCTNNNGCCTCGCAAAATACTGGGATTAC
AGCACACGTCTGAACTCTTTCCCTACACGACGCTCTTCCGATCTNNNGACAGCAATGGCAGTAATAGC
AGCACACGTCTGAACTCTTTCCCTACACGACGCTCTTCCGATCTNNNTCTGCTAGGGGCCAGATATTTT
GAGCACACGTCTGAACTCTTTCCCTACACGACGCTCTTCCGATCTNNNGACAGAGTCTCACTCCATCACT
GCACACGTCTGAACTCTTTCCCTACACGACGCTCTTCCGATCTNNNTTGATGGTAGAACTGTAGTCACC
GCACACGTCTGAACTCTTTCCCTACACGACGCTCTTCCGATCTNNNCTGCTACAGCTGTCATGGAAAAC
AGCACACGTCTGAACTCTTTCCCTACACGACGCTCTTCCGATCTNNNGGTGTGGACGAGGTAATGTAA
AGCACACGTCTGAACTCTTTCCCTACACGACGCTCTTCCGATCTNNNCAAGTCTGCTGTTACGGCTTTTT
AGAGCACACGTCTGAACTCTTTCCCTACACGACGCTCTTCCGATCTNNNCCTAGTAGCTGGGACTATAG
AGCACACGTCTGAACTCTTTCCCTACACGACGCTCTTCCGATCTNNNTAGTCTTGCTTTGGCTATGGAG
GCACACGTCTGAACTCTTTCCCTACACGACGCTCTTCCGATCTNNNTCAGTTATGCCTGAAATCTGCAA
AGCACACGTCTGAACTCTTTCCCTACACGACGCTCTTCCGATCTNNNTATTGGAATCAGCCCACATTGTC
GAGCACACGTCTGAACTCTTTCCCTACACGACGCTCTTCCGATCTNNNAAAGGATGCTTCCAGGGTGT
GAGCACACGTCTGAACTCTTTCCCTACACGACGCTCTTCCGATCTNNNTAAGGCTGGAGTCATGACATA
GAGCACACGTCTGAACTCTTTCCCTACACGACGCTCTTCCGATCTNNNTGGGTCAAATGGTGGTTCTAC
GAGCACACGTCTGAACTCTTTCCCTACACGACGCTCTTCCGATCTNNNACAATATCCTGGTCCATTGTCA
GAGCACACGTCTGAACTCTTTCCCTACACGACGCTCTTCCGATCTNNNCCCTGTCTTGAGTACTGTAGCT
GAGCACACGTCTGAACTCTTTCCCTACACGACGCTCTTCCGATCTNNNAGCTCATCATCAGGTGGAAAC
GAGCACACGTCTGAACTCTTTCCCTACACGACGCTCTTCCGATCTNNNGAAGAGGAAGAGAGTGGGA
GCACACGTCTGAACTCTTTCCCTACACGACGCTCTTCCGATCTNNNGGGGGCTATAAGTCCTTTTCACA
AGCACACGTCTGAACTCTTTCCCTACACGACGCTCTTCCGATCTNNNCCATTACTGCACCACTACACTCT
GAGCACACGTCTGAACTCTTTCCCTACACGACGCTCTTCCGATCTNNNTTCACTTATCTGCTGACCTTCC
GAGCACACGTCTGAACTCTTTCCCTACACGACGCTCTTCCGATCTNNNGCTTGCACACAGGAATTCAAG
GCACACGTCTGAACTCTTTCCCTACACGACGCTCTTCCGATCTNNNGGGCAACTCCATCTCAAAAACAAC
GAGCACACGTCTGAACTCTTTCCCTACACGACGCTCTTCCGATCTNNNGGTGGATCACAAAGTCAGGA
AGAGCACACGTCTGAACTCTTTCCCTACACGACGCTCTTCCGATCTNNNGCCAAAATCACACCACTACAC
GCACACGTCTGAACTCTTTCCCTACACGACGCTCTTCCGATCTNNNCTGAGGCAGAAGAATGGTGTGA
GAGCACACGTCTGAACTCTTTCCCTACACGACGCTCTTCCGATCTNNNTCACTTCCAGCCAAAACAATGT
GAGCACACGTCTGAACTCTTTCCCTACACGACGCTCTTCCGATCTNNNCTGAGGCAGAAGAATCCCTTG
AGCACACGTCTGAACTCTTTCCCTACACGACGCTCTTCCGATCTNNNAGGACTGCTTGACGTGATACAA
GAGCACACGTCTGAACTCTTTCCCTACACGACGCTCTTCCGATCTNNNGTCTGTAGTCCCAGCTACTAGC
GCACACGTCTGAACTCTTTCCCTACACGACGCTCTTCCGATCTNNNACAAAAATTAGCTGGCCATGGTC
AGCACACGTCTGAACTCTTTCCCTACACGACGCTCTTCCGATCTNNNCAACAAGAGCAAAAACTCCGTC
AGCACACGTCTGAACTCTTTCCCTACACGACGCTCTTCCGATCTNNNTAAACTACGGGCAGACAAATG
GAGCACACGTCTGAACTCTTTCCCTACACGACGCTCTTCCGATCTNNNCTGGGCAACAAGAATGAATC
GAGCACACGTCTGAACTCTTTCCCTACACGACGCTCTTCCGATCTNNNAAAAAAACATCCAGATTGCGC

GAGCACACGTCTGAACTCTTTCCCTACACGACGCTCTTCCGATCTNNNGAGGCAGGAGAATTGCTTG
GAGCACACGTCTGAACTCTTTCCCTACACGACGCTCTTCCGATCTNNNACCTGTAGTCCTAGCTACTCT
AGAGCACACGTCTGAACTCTTTCCCTACACGACGCTCTTCCGATCTNNNCCTGGGTGATAAGAGCAA
AGCACACGTCTGAACTCTTTCCCTACACGACGCTCTTCCGATCTNNNCTGGAGCAGTCTTGTTGTTTG
AGCACACGTCTGAACTCTTTCCCTACACGACGCTCTTCCGATCTNNNTGTCTGTAATCCCAGCTATTCG
AGCACACGTCTGAACTCTTTCCCTACACGACGCTCTTCCGATCTNNNGGCAGAAGAATTGCTTGAACC
AGCACACGTCTGAACTCTTTCCCTACACGACGCTCTTCCGATCTNNNTATAGTCTCAGCTACTCGGGA
AGCACACGTCTGAACTCTTTCCCTACACGACGCTCTTCCGATCTNNNGTAATCACAGCTACTAGGGA
GCACACGTCTGAACTCTTTCCCTACACGACGCTCTTCCGATCTNNNATACTGTCACTCTGGGGGTTAG
GAGCACACGTCTGAACTCTTTCCCTACACGACGCTCTTCCGATCTNNNCTGAGGCATGAAAATCGCTT
AGCACACGTCTGAACTCTTTCCCTACACGACGCTCTTCCGATCTNNNCGATACAAAAGTAACAGGCTG
AGCACACGTCTGAACTCTTTCCCTACACGACGCTCTTCCGATCTNNNCACCTGTCATCTCAGCTACTCA
GCACACGTCTGAACTCTTTCCCTACACGACGCTCTTCCGATCTNNNGTACAACAAACTCCCATCACAC
GCACACGTCTGAACTCTTTCCCTACACGACGCTCTTCCGATCTNNNGCCTATTTTACCAGGTCAGGGA
AGCACACGTCTGAACTCTTTCCCTACACGACGCTCTTCCGATCTNNNGTACATTGCAGTGAGCCAAGA
GCACACGTCTGAACTCTTTCCCTACACGACGCTCTTCCGATCTNNNCTGAGGCAGGAAAATTGCTTG
GAGCACACGTCTGAACTCTTTCCCTACACGACGCTCTTCCGATCTNNNACACGAGAATTGCTTCAACT
GAGCACACGTCTGAACTCTTTCCCTACACGACGCTCTTCCGATCTNNNTCATGAGGTCAGGAGTTTGA
GAGCACACGTCTGAACTCTTTCCCTACACGACGCTCTTCCGATCTNNNGGTGCTAGTAATCCCAGCTA
CACACGTCTGAACTCTTTCCCTACACGACGCTCTTCCGATCTNNNCATGCACCTGTGATCTCACCTACC
AGCACACGTCTGAACTCTTTCCCTACACGACGCTCTTCCGATCTNNNCTGAGGCAGGAGAATCACTTG
GAGCACACGTCTGAACTCTTTCCCTACACGACGCTCTTCCGATCTNNNAGAATCACTTGAACCCAGGA
GAGCACACGTCTGAACTCTTTCCCTACACGACGCTCTTCCGATCTNNNCTGAGGCAGGAGAATCACTT
GCACACGTCTGAACTCTTTCCCTACACGACGCTCTTCCGATCTNNNTGAGGCAGGAGAATTGCTTGT
AGCACACGTCTGAACTCTTTCCCTACACGACGCTCTTCCGATCTNNNACGCCAGTAATTCCAGCATTTT
GAGCACACGTCTGAACTCTTTCCCTACACGACGCTCTTCCGATCTNNNCTGAGGTAGGAGGATTGCT
AGCACACGTCTGAACTCTTTCCCTACACGACGCTCTTCCGATCTNNNAAGAGAATCGTTTGAACCCA
AGAGCACACGTCTGAACTCTTTCCCTACACGACGCTCTTCCGATCTNNNAGAGATTGCAGTGAGCCAA
GCACACGTCTGAACTCTTTCCCTACACGACGCTCTTCCGATCTNNNCTAGGCAGGAGAACCTCTTGAA
GAGCACACGTCTGAACTCTTTCCCTACACGACGCTCTTCCGATCTNNNTTACTTGAACCTGGGAGTT
AGCACACGTCTGAACTCTTTCCCTACACGACGCTCTTCCGATCTNNNAATTCCAGCTACTTGGGAGAC
AGCACACGTCTGAACTCTTTCCCTACACGACGCTCTTCCGATCTNNNCACAATGTGCACATGTACCCT
GCACACGTCTGAACTCTTTCCCTACACGACGCTCTTCCGATCTNNNGCTGTGATATAAGTCCCCCTCT
GCACACGTCTGAACTCTTTCCCTACACGACGCTCTTCCGATCTNNNAGTTTGGTGCTTCTCGACTGTA
GCACACGTCTGAACTCTTTCCCTACACGACGCTCTTCCGATCTNNNAGCTTACTCTTGACCAGAAGCTT
GCACACGTCTGAACTCTTTCCCTACACGACGCTCTTCCGATCTNNNTGGTTCCTTGGAGCAATTCAGT
AGCACACGTCTGAACTCTTTCCCTACACGACGCTCTTCCGATCTNNNGTGTATCTGTCTGCATGCCTCT
AGCACACGTCTGAACTCTTTCCCTACACGACGCTCTTCCGATCTNNNGCCACCACAAAAATATGAACC
AGCACACGTCTGAACTCTTTCCCTACACGACGCTCTTCCGATCTNNNTGTAACTCTGGCAGAAATCAG
AGCACACGTCTGAACTCTTTCCCTACACGACGCTCTTCCGATCTNNNCAGTGAAGACAGAGAGTGGA
GAGCACACGTCTGAACTCTTTCCCTACACGACGCTCTTCCGATCTNNNTATGCTTTGCAACAAAATGGT
AGCACACGTCTGAACTCTTTCCCTACACGACGCTCTTCCGATCTNNNCCTCCCTACCTCTCTCTCTCTTT
AGCACACGTCTGAACTCTTTCCCTACACGACGCTCTTCCGATCTNNNTCCTGAAGGTCCTGTCATGGA

AGCACACGTCTGAACTCTTTCCCTACACGACGCTCTTCCGATCTNNNCAATCAGCAGCCAGGTATAGA
AGCACACGTCTGAACTCTTTCCCTACACGACGCTCTTCCGATCTNNNGCAACCCCTGGTTTTTTTGTTT
AGCACACGTCTGAACTCTTTCCCTACACGACGCTCTTCCGATCTNNNGCCCTCGTTTGTGTTTTTCATTC
AGCACACGTCTGAACTCTTTCCCTACACGACGCTCTTCCGATCTNNNAACACAAACCTCTCCAACAGTCT
AGCACACGTCTGAACTCTTTCCCTACACGACGCTCTTCCGATCTNNNGTGTGTGTGTGTGTGTGT
GCACACGTCTGAACTCTTTCCCTACACGACGCTCTTCCGATCTNNNTGGGTCAGACATTTCCATTCAA
AGCACACGTCTGAACTCTTTCCCTACACGACGCTCTTCCGATCTNNNGCCGGTTACTGGAGAAAACAA
AGCACACGTCTGAACTCTTTCCCTACACGACGCTCTTCCGATCTNNNGGCTGGTCTCAAACTTCTCAAA
AGCACACGTCTGAACTCTTTCCCTACACGACGCTCTTCCGATCTNNNCAAGCTGGTCTCAAACTCATGA
AGCACACGTCTGAACTCTTTCCCTACACGACGCTCTTCCGATCTNNNATGTTAGAAAGGGCCTTCCAT
AGCACACGTCTGAACTCTTTCCCTACACGACGCTCTTCCGATCTNNNGTGACTGTTCAGGTCTTTTGCT
GAGCACACGTCTGAACTCTTTCCCTACACGACGCTCTTCCGATCTNNNCACCACCACACAGATTTTTCTC
AGCACACGTCTGAACTCTTTCCCTACACGACGCTCTTCCGATCTNNNTCCAAAGTGCTAGGATTACAGG
AGCACACGTCTGAACTCTTTCCCTACACGACGCTCTTCCGATCTNNNTCAAATCATTTGCTCCTCCCTGA
GCACACGTCTGAACTCTTTCCCTACACGACGCTCTTCCGATCTNNNAAAAACATCACCAAAACCCCCTG
GCACACGTCTGAACTCTTTCCCTACACGACGCTCTTCCGATCTNNNTTCACCACAGGGGAAAATTCTCT
AGCACACGTCTGAACTCTTTCCCTACACGACGCTCTTCCGATCTNNNGTACTCAGAAGGGTCTTGCTTC
AGCACACGTCTGAACTCTTTCCCTACACGACGCTCTTCCGATCTNNNTCACCAAGCTCTTCAGTTCCTAC
GCACACGTCTGAACTCTTTCCCTACACGACGCTCTTCCGATCTNNNACAGAGACACGTCTAGATTGCT
AGCACACGTCTGAACTCTTTCCCTACACGACGCTCTTCCGATCTNNNAGCTCCTTGAGTGGTATAATG
AGCACACGTCTGAACTCTTTCCCTACACGACGCTCTTCCGATCTNNNGCTTGCTACAGGAACTAGACAG
AGCACACGTCTGAACTCTTTCCCTACACGACGCTCTTCCGATCTNNNAGAGACACAGAGCAAGTAAGT
AGCACACGTCTGAACTCTTTCCCTACACGACGCTCTTCCGATCTNNNCTGATCTCCTCAAAGAACCAGCT
AGCACACGTCTGAACTCTTTCCCTACACGACGCTCTTCCGATCTNNNGACCCTAAGAAACAACGTTTG
AGCACACGTCTGAACTCTTTCCCTACACGACGCTCTTCCGATCTNNNTACAAAAAATTAGCCGGGCAT
AGCACACGTCTGAACTCTTTCCCTACACGACGCTCTTCCGATCTNNNCATGCCTGTAGTCCTAGCTACT
AGCACACGTCTGAACTCTTTCCCTACACGACGCTCTTCCGATCTNNNCTCTGGAAAGCAGAATGGAG
GAGCACACGTCTGAACTCTTTCCCTACACGACGCTCTTCCGATCTNNNCTGAGGCAGGAGAATCACTT
GAGCACACGTCTGAACTCTTTCCCTACACGACGCTCTTCCGATCTNNNGGCATGTTCTCACTCATTCTT
AGCACACGTCTGAACTCTTTCCCTACACGACGCTCTTCCGATCTNNNAGACAGGAGAATTGCTTGAA
GAGCACACGTCTGAACTCTTTCCCTACACGACGCTCTTCCGATCTNNNCCTATAGTCCCAGCTACTCGG
GAGCACACGTCTGAACTCTTTCCCTACACGACGCTCTTCCGATCTNNNGAGCTTGTAGAGAGCCGAGA
GAGCACACGTCTGAACTCTTTCCCTACACGACGCTCTTCCGATCTNNNCCCAAAGTGCTAGGGTTACA
GAGCACACGTCTGAACTCTTTCCCTACACGACGCTCTTCCGATCTNNNTTCTTGAAAACTGGAGGTG
GAGCACACGTCTGAACTCTTTCCCTACACGACGCTCTTCCGATCTNNNAGGCAGAAGAATCACTTGAA
GAGCACACGTCTGAACTCTTTCCCTACACGACGCTCTTCCGATCTNNNCTAAGGCAGGAGAATCGCTT
AGCACACGTCTGAACTCTTTCCCTACACGACGCTCTTCCGATCTNNNCTTGAACCAGAGTGTCAGAGGT
AGCACACGTCTGAACTCTTTCCCTACACGACGCTCTTCCGATCTNNNGCAGAGGTTGTAGTGAACCAA
AGCACACGTCTGAACTCTTTCCCTACACGACGCTCTTCCGATCTNNNCTAGCCTGGACGAAAGAAACT
GAGCACACGTCTGAACTCTTTCCCTACACGACGCTCTTCCGATCTNNNTCACGAGGTCAGGAATTCAA
GAGCACACGTCTGAACTCTTTCCCTACACGACGCTCTTCCGATCTNNNGACCAATAGTTCAAGACCAG
GCACACGTCTGAACTCTTTCCCTACACGACGCTCTTCCGATCTNNNTGAGGCAGGAGAATCTCTTGAA
AGCACACGTCTGAACTCTTTCCCTACACGACGCTCTTCCGATCTNNNATCCTATCAAATCGCCAGTGTC

AGCACACGTCTGAACTCTTTCCCTACACGACGCTCTTCCGATCTNNNCTGTAGTCCCAGCTACTCAGGA
GCACACGTCTGAACTCTTTCCCTACACGACGCTCTTCCGATCTNNNGAAAAATCACTTGAACCCGGGA
AGCACACGTCTGAACTCTTTCCCTACACGACGCTCTTCCGATCTNNNGAAACTGCAGTGAGCCATGA
AGCACACGTCTGAACTCTTTCCCTACACGACGCTCTTCCGATCTNNNTGTAATCACAGTCACTCAGGA
GAGCACACGTCTGAACTCTTTCCCTACACGACGCTCTTCCGATCTNNNTTGAACTGGGGAGACAGAG
GCACACGTCTGAACTCTTTCCCTACACGACGCTCTTCCGATCTNNNCTGAGGCAGGAGAATTGCTTGA
GCACACGTCTGAACTCTTTCCCTACACGACGCTCTTCCGATCTNNNAATGGTGTGAACCTGAACCTAG
AGAGCACACGTCTGAACTCTTTCCCTACACGACGCTCTTCCGATCTNNNGAGGCAGGAGAATGGTGT
GCACACGTCTGAACTCTTTCCCTACACGACGCTCTTCCGATCTNNNTGGGCAACATAGTGAAACCCTA
GCACACGTCTGAACTCTTTCCCTACACGACGCTCTTCCGATCTNNNCTGAGACAGGAGAATGGCATGA
AGCACACGTCTGAACTCTTTCCCTACACGACGCTCTTCCGATCTNNNCCAAAATCAAGCCACTGTACTC
AGCACACGTCTGAACTCTTTCCCTACACGACGCTCTTCCGATCTNNNGGCAACAGAGTAAGACTCCA
AGCACACGTCTGAACTCTTTCCCTACACGACGCTCTTCCGATCTNNNATGCCTTGAGGTCAAAAAGA
AGCACACGTCTGAACTCTTTCCCTACACGACGCTCTTCCGATCTNNNGTAATACCAGCATTTTGGGAG
GAGCACACGTCTGAACTCTTTCCCTACACGACGCTCTTCCGATCTNNNGCTGATAACCCACTTCATGTG
AGCACACGTCTGAACTCTTTCCCTACACGACGCTCTTCCGATCTNNNAGAGCAATCAGGCAAGAGA
GAGCACACGTCTGAACTCTTTCCCTACACGACGCTCTTCCGATCTNNNCTGAGACAGGAGAATGGTG
AGCACACGTCTGAACTCTTTCCCTACACGACGCTCTTCCGATCTNNNAGCAGGCCTTTATTCCCAACT
AGCACACGTCTGAACTCTTTCCCTACACGACGCTCTTCCGATCTNNNATAGGCCGCACAACTATAATCC
AGCACACGTCTGAACTCTTTCCCTACACGACGCTCTTCCGATCTNNNCTGTAGTCCCAGCTACTCACA
AGCACACGTCTGAACTCTTTCCCTACACGACGCTCTTCCGATCTNNNAAATACAAAAATTAGCCGGG
GAGCACACGTCTGAACTCTTTCCCTACACGACGCTCTTCCGATCTNNNCTGTAATTGCAGCACTTTGG
GAGCACACGTCTGAACTCTTTCCCTACACGACGCTCTTCCGATCTNNNGAATGGCATGAATATTGGA
AGCACACGTCTGAACTCTTTCCCTACACGACGCTCTTCCGATCTNNNGCTGAGATGGGAGAATCACT
AGCACACGTCTGAACTCTTTCCCTACACGACGCTCTTCCGATCTNNNGGAGAATCGCTTGAACCAGA
AGCACACGTCTGAACTCTTTCCCTACACGACGCTCTTCCGATCTNNNCTGTTTCTGGCACCAGCTATTT
AGCACACGTCTGAACTCTTTCCCTACACGACGCTCTTCCGATCTNNNAGTGAGTTAGAGGTTGCAGT
AGCACACGTCTGAACTCTTTCCCTACACGACGCTCTTCCGATCTNNNTTGCTTGAATCCAGAAGGCAA
AGCACACGTCTGAACTCTTTCCCTACACGACGCTCTTCCGATCTNNNCCTGTAGTCCCAGCTACTAGG
AGCACACGTCTGAACTCTTTCCCTACACGACGCTCTTCCGATCTNNNAAGAGATATTCAAGCCAGGG
GCACACGTCTGAACTCTTTCCCTACACGACGCTCTTCCGATCTNNNAACATAGCAAGACCCCGTCTCTA
AGCACACGTCTGAACTCTTTCCCTACACGACGCTCTTCCGATCTNNNACCACCACACCCAGCTAATTT
GCACACGTCTGAACTCTTTCCCTACACGACGCTCTTCCGATCTNNNTGAGGCAAGAAAATCGCTTGA
GCACACGTCTGAACTCTTTCCCTACACGACGCTCTTCCGATCTNNNCACCTGTAATCCCAGCTACTCAG
GAGCACACGTCTGAACTCTTTCCCTACACGACGCTCTTCCGATCTNNNAGAAACACAAGTCTTGTCCA
AGCACACGTCTGAACTCTTTCCCTACACGACGCTCTTCCGATCTNNNGAGATCGCATCACTACACTCCA
GAGCACACGTCTGAACTCTTTCCCTACACGACGCTCTTCCGATCTNNNTGAGAATCACTTGAGCCAGG
GAGCACACGTCTGAACTCTTTCCCTACACGACGCTCTTCCGATCTNNNTGTAGTACCAGCTACTCAGG
GCACACGTCTGAACTCTTTCCCTACACGACGCTCTTCCGATCTNNNGGCTCTCTTGGTCCCTAAAGAA
AGCACACGTCTGAACTCTTTCCCTACACGACGCTCTTCCGATCTNNNCACAAATGTTGACAAGCAAAG
GCACACGTCTGAACTCTTTCCCTACACGACGCTCTTCCGATCTNNNGATCATAGCTCCCTGCATCCTTG
GCACACGTCTGAACTCTTTCCCTACACGACGCTCTTCCGATCTNNNCTCTCTCTTCCCTGTCCAAACTCC
AGCACACGTCTGAACTCTTTCCCTACACGACGCTCTTCCGATCTNNNCACCAGCACACACATAATGTCT

GAGCACACGTCTGAACTCTTTCCCTACACGACGCTCTTCCGATCTNNNTAAACTGCATGGGAGGAAAA
GCACACGTCTGAACTCTTTCCCTACACGACGCTCTTCCGATCTNNNCTGTTTCCCTTTTCATGCCTTCCC
AGCACACGTCTGAACTCTTTCCCTACACGACGCTCTTCCGATCTNNNGCTCCTTGCATTTTGTACCACTT
AGCACACGTCTGAACTCTTTCCCTACACGACGCTCTTCCGATCTNNNGTCTTTTCCTAGCTTTGCTGGT
AGCACACGTCTGAACTCTTTCCCTACACGACGCTCTTCCGATCTNNNGAACTTGGGAGCAGCTTAGCT
GCACACGTCTGAACTCTTTCCCTACACGACGCTCTTCCGATCTNNNCCCACTCAAAGGAAAACAGACA
AGCACACGTCTGAACTCTTTCCCTACACGACGCTCTTCCGATCTNNNGCCTGACTGTATATGGGAGGA
GAGCACACGTCTGAACTCTTTCCCTACACGACGCTCTTCCGATCTNNNCATGACAGGTGCTGTGAACA
GCACACGTCTGAACTCTTTCCCTACACGACGCTCTTCCGATCTNNNCCCTCCACAGATACACAGATAGC
GCACACGTCTGAACTCTTTCCCTACACGACGCTCTTCCGATCTNNNGATTCAGTAGGTCTGGAATGG
AGCACACGTCTGAACTCTTTCCCTACACGACGCTCTTCCGATCTNNNTTTCCTCAGTTGGTTAGTCCCA
AGCACACGTCTGAACTCTTTCCCTACACGACGCTCTTCCGATCTNNNCCGCCTTTCTCACTAACTCTTCT
AGCACACGTCTGAACTCTTTCCCTACACGACGCTCTTCCGATCTNNNAAAGTGCTGGGATTACAGGTG
AGCACACGTCTGAACTCTTTCCCTACACGACGCTCTTCCGATCTNNNAGGAAGACATAGACCACACCCT
AGCACACGTCTGAACTCTTTCCCTACACGACGCTCTTCCGATCTNNNAGCTGTCACAGAGTATGGTCT
AGCACACGTCTGAACTCTTTCCCTACACGACGCTCTTCCGATCTNNNAAACTTCTCCCTGGACATCCAA
AGCACACGTCTGAACTCTTTCCCTACACGACGCTCTTCCGATCTNNNAAGTTGGATTTCAAAGGAGCA
AGCACACGTCTGAACTCTTTCCCTACACGACGCTCTTCCGATCTNNNCTAGAATTCTGCTCTCCTGGG
AGCACACGTCTGAACTCTTTCCCTACACGACGCTCTTCCGATCTNNNCATTGTTGGGATTTGTCTGACC
AGCACACGTCTGAACTCTTTCCCTACACGACGCTCTTCCGATCTNNNATTGGTAGAGATGGGGTTTC
AGCACACGTCTGAACTCTTTCCCTACACGACGCTCTTCCGATCTNNNCTGCTGACCCTCCTGTATCAAAT
GAGCACACGTCTGAACTCTTTCCCTACACGACGCTCTTCCGATCTNNNATTTTCCTGTGTGTGTGTGTG
AGCACACGTCTGAACTCTTTCCCTACACGACGCTCTTCCGATCTNNNCCTGGCTGCCCTTGTTCTTTATA
AGCACACGTCTGAACTCTTTCCCTACACGACGCTCTTCCGATCTNNNCTCACTGCAACCTCAAATTCCTG
AGCACACGTCTGAACTCTTTCCCTACACGACGCTCTTCCGATCTNNNATCAACCCGTCGTCTACATTAG
GAGCACACGTCTGAACTCTTTCCCTACACGACGCTCTTCCGATCTNNNCTCCTGAGTAGCTGGGATTA
GAGCACACGTCTGAACTCTTTCCCTACACGACGCTCTTCCGATCTNNNGAGAATGGTGTGAACCTGG
GAGCACACGTCTGAACTCTTTCCCTACACGACGCTCTTCCGATCTNNNGAGAATGGTGTGAACCTGG
GAGCACACGTCTGAACTCTTTCCCTACACGACGCTCTTCCGATCTNNNTCTGGAAACGTACAACCTTTC
AGCACACGTCTGAACTCTTTCCCTACACGACGCTCTTCCGATCTNNNTTCTGGATCCAAGCATTTCCGT
AGCACACGTCTGAACTCTTTCCCTACACGACGCTCTTCCGATCTNNNCTGAGGCAGGAGAATCACTTG
GCACACGTCTGAACTCTTTCCCTACACGACGCTCTTCCGATCTNNNCTGTAGTCCCAGCTACTCAGGAG
AGCACACGTCTGAACTCTTTCCCTACACGACGCTCTTCCGATCTNNNTGCACACCTATAGTCCAAGCTA
GCACACGTCTGAACTCTTTCCCTACACGACGCTCTTCCGATCTNNNAATGTGCTGGCAGATTTGATGT
AGCACACGTCTGAACTCTTTCCCTACACGACGCTCTTCCGATCTNNNAGAGTTTGTAGTGAGCCGAG
AGCACACGTCTGAACTCTTTCCCTACACGACGCTCTTCCGATCTNNNGCAATAAGAGTGAGTCTCCGTC
AGCACACGTCTGAACTCTTTCCCTACACGACGCTCTTCCGATCTNNNGCTGTAGAAAACCTGAGCAACA
GAGCACACGTCTGAACTCTTTCCCTACACGACGCTCTTCCGATCTNNNTCACAAGGTCAGGAGTTTGA
GAGCACACGTCTGAACTCTTTCCCTACACGACGCTCTTCCGATCTNNNAGGAGGCTGAGACAAGAGAA
AGCACACGTCTGAACTCTTTCCCTACACGACGCTCTTCCGATCTNNNATGTTGCAGTGAGTTGAGATC
AGCACACGTCTGAACTCTTTCCCTACACGACGCTCTTCCGATCTNNNCTCATGCCTATAATCCCAGCAC
AGCACACGTCTGAACTCTTTCCCTACACGACGCTCTTCCGATCTNNNAAAAGAGTAAAAACCGGGCAT
AGCACACGTCTGAACTCTTTCCCTACACGACGCTCTTCCGATCTNNNAATTCCAGCTACTTGGGAGAC

AGCACACGTCTGAACTCTTTCCCTACACGACGCTCTTCCGATCTNNNGAGGCAGAAGAATCGCTTGA
AGCACACGTCTGAACTCTTTCCCTACACGACGCTCTTCCGATCTNNNGATCAAGACCATCCTGGCTAAC
AGCACACGTCTGAACTCTTTCCCTACACGACGCTCTTCCGATCTNNNGTAGGAGAATCTCTTGAGCCC
GAGCACACGTCTGAACTCTTTCCCTACACGACGCTCTTCCGATCTNNNCGCCTGTAATCCTAGCTACTC
AGAGCACACGTCTGAACTCTTTCCCTACACGACGCTCTTCCGATCTNNNGTAATCCCAGCTACTTAGGA
AGCACACGTCTGAACTCTTTCCCTACACGACGCTCTTCCGATCTNNNTAGGCAGGAGAATCACTTGAA
GAGCACACGTCTGAACTCTTTCCCTACACGACGCTCTTCCGATCTNNNTAATCCCAGCTACTTGCAAGG
AGCACACGTCTGAACTCTTTCCCTACACGACGCTCTTCCGATCTNNNGACCTTCTGACCTCGTGATCTA
GAGCACACGTCTGAACTCTTTCCCTACACGACGCTCTTCCGATCTNNNCTGTAATCCCAGCTACCAAGG
GAGCACACGTCTGAACTCTTTCCCTACACGACGCTCTTCCGATCTNNNGTCAAGACTACAGTGAGCCA
CACACGTCTGAACTCTTTCCCTACACGACGCTCTTCCGATCTNNNGAGGTTGCAGTGAGCTGAGATA
GAGCACACGTCTGAACTCTTTCCCTACACGACGCTCTTCCGATCTNNNTGAGGTCAGGAGTTCAAGAC
AGCACACGTCTGAACTCTTTCCCTACACGACGCTCTTCCGATCTNNNTGAGGCAGGAGTATCACTTGA
AGCACACGTCTGAACTCTTTCCCTACACGACGCTCTTCCGATCTNNNGAGGCTAAGGCAAGAGAATCA
GAGCACACGTCTGAACTCTTTCCCTACACGACGCTCTTCCGATCTNNNTTGAGCCCAGAAGTTCATGA
GAGCACACGTCTGAACTCTTTCCCTACACGACGCTCTTCCGATCTNNNCTTGTAGTCTCAGCTACTTGG
GAGCACACGTCTGAACTCTTTCCCTACACGACGCTCTTCCGATCTNNNGCTGAAACCCCGACTCTACTA
AGCACACGTCTGAACTCTTTCCCTACACGACGCTCTTCCGATCTNNNGAGGTTGCAGTGAGCTAAGAT
GAGCACACGTCTGAACTCTTTCCCTACACGACGCTCTTCCGATCTNNNCTTGAGGTTAGGAGTTCGAG
AGCACACGTCTGAACTCTTTCCCTACACGACGCTCTTCCGATCTNNNGAGGTTGTAGTGAGCTGAGTA
GAGCACACGTCTGAACTCTTTCCCTACACGACGCTCTTCCGATCTNNNTGCCTGTAATCCCAACTACTT
GAGCACACGTCTGAACTCTTTCCCTACACGACGCTCTTCCGATCTNNNCTAATCCCAGTTACTCAGGAG
GAGCACACGTCTGAACTCTTTCCCTACACGACGCTCTTCCGATCTNNNGGTTGCAGTGAGCTAAGATT
AGCACACGTCTGAACTCTTTCCCTACACGACGCTCTTCCGATCTNNNCACCTGTAATTCCAGCACTTTG
AGCACACGTCTGAACTCTTTCCCTACACGACGCTCTTCCGATCTNNNGATAACATCACTGCATTCCAGC
GAGCACACGTCTGAACTCTTTCCCTACACGACGCTCTTCCGATCTNNNGAGTTGCTGGTGACATTTCA
AGCACACGTCTGAACTCTTTCCCTACACGACGCTCTTCCGATCTNNNCTGTTGCTTTGACTCAGAGCCT
AGCACACGTCTGAACTCTTTCCCTACACGACGCTCTTCCGATCTNNNTTTTTTTTCAGGGGGAGACAG
GAGCACACGTCTGAACTCTTTCCCTACACGACGCTCTTCCGATCTNNNATAAGTGTTTGGGAGGCAAG
AGCACACGTCTGAACTCTTTCCCTACACGACGCTCTTCCGATCTNNNCGTCGACTGGACTCTTATTTTG
AGCACACGTCTGAACTCTTTCCCTACACGACGCTCTTCCGATCTNNNCTATACTTCAGCCTGGGACCTT
GCACACGTCTGAACTCTTTCCCTACACGACGCTCTTCCGATCTNNNACTGTCTACTCTGTGTCTTCCCT
GCACACGTCTGAACTCTTTCCCTACACGACGCTCTTCCGATCTNNNCTGGATGTTGCTGACAGTGTTT
GCACACGTCTGAACTCTTTCCCTACACGACGCTCTTCCGATCTNNNCCACTGTACCCGGCTAATTTTG
GAGCACACGTCTGAACTCTTTCCCTACACGACGCTCTTCCGATCTNNNACCACCACACCTGGCTAATTA
AGCACACGTCTGAACTCTTTCCCTACACGACGCTCTTCCGATCTNNNTGTTGTTTGTCAGCTTGCCTAT
GCACACGTCTGAACTCTTTCCCTACACGACGCTCTTCCGATCTNNNTGTGCATGTTCTTTCAGACTGA
AGCACACGTCTGAACTCTTTCCCTACACGACGCTCTTCCGATCTNNNCCCTCGTTTTGGGGAAGAACAT
AGCACACGTCTGAACTCTTTCCCTACACGACGCTCTTCCGATCTNNNATCACAGCTCATTACAGCCTTGA
GAGCACACGTCTGAACTCTTTCCCTACACGACGCTCTTCCGATCTNNNCGTCAGACACCACCTAATGAA
GAGCACACGTCTGAACTCTTTCCCTACACGACGCTCTTCCGATCTNNNATTGCTTGAGGTCAGGAGTT
GAGCACACGTCTGAACTCTTTCCCTACACGACGCTCTTCCGATCTNNNTGGACAAACCTGAGTGTGA
GAGCACACGTCTGAACTCTTTCCCTACACGACGCTCTTCCGATCTNNNAGAAAAAGGGAACAGGGTC

AGCACACGTCTGAACTCTTTCCCTACACGACGCTCTTCCGATCTNNNCCAGGGCTATATAGGAACTGTG
AGCACACGTCTGAACTCTTTCCCTACACGACGCTCTTCCGATCTNNNTTTTGAACGCAGCTTCCCAAAT
GAGCACACGTCTGAACTCTTTCCCTACACGACGCTCTTCCGATCTNNNAGCAAACATCTCTCACCTGGAT
AGCACACGTCTGAACTCTTTCCCTACACGACGCTCTTCCGATCTNNNAGCAGGAGTGTCCCTTTAGTTTT
GAGCACACGTCTGAACTCTTTCCCTACACGACGCTCTTCCGATCTNNNGACACTAATCTTGGTGGACCCA
AGCACACGTCTGAACTCTTTCCCTACACGACGCTCTTCCGATCTNNNCCAAAGTGCTGGGATTACAAGT
GAGCACACGTCTGAACTCTTTCCCTACACGACGCTCTTCCGATCTNNNGTTCAAGTAGTTCTCCTGCCTC
AGCACACGTCTGAACTCTTTCCCTACACGACGCTCTTCCGATCTNNNGCTAATTGAATGGCCCTGAAAG
AGCACACGTCTGAACTCTTTCCCTACACGACGCTCTTCCGATCTNNNATTACCATTCAATCTCCCCCACCA
GAGCACACGTCTGAACTCTTTCCCTACACGACGCTCTTCCGATCTNNNACAGACACAAATCTCCTGATCC
AGAGCACACGTCTGAACTCTTTCCCTACACGACGCTCTTCCGATCTNNNATCCACAGGGTTCTATCCAAT
GAGCACACGTCTGAACTCTTTCCCTACACGACGCTCTTCCGATCTNNNTGGCTAGTACAATGAATCTGG
GAGCACACGTCTGAACTCTTTCCCTACACGACGCTCTTCCGATCTNNNACAAGTGCTCACCATAACACTC
AGCACACGTCTGAACTCTTTCCCTACACGACGCTCTTCCGATCTNNNCCCACACACTAACTCCCTCAAAT
AGCACACGTCTGAACTCTTTCCCTACACGACGCTCTTCCGATCTNNNGATTGTATCTGCTCTGCCTCACT
AGCACACGTCTGAACTCTTTCCCTACACGACGCTCTTCCGATCTNNNGATGGGGTGAAGGACATCTGA
AGAGCACACGTCTGAACTCTTTCCCTACACGACGCTCTTCCGATCTNNNTCCAGAGTAGCTAGGACTACA
AGCACACGTCTGAACTCTTTCCCTACACGACGCTCTTCCGATCTNNNTTTTCATGTGTTTTTTGGCTGCA
GAGCACACGTCTGAACTCTTTCCCTACACGACGCTCTTCCGATCTNNNATTCTCCTGCCTTAGCCTCCTAA
GAGCACACGTCTGAACTCTTTCCCTACACGACGCTCTTCCGATCTNNNGGAAAAAAGAAGGAAGCTCC
GAGCACACGTCTGAACTCTTTCCCTACACGACGCTCTTCCGATCTNNNCGCAGAGTGTAAGAGAGTGA
GAGCACACGTCTGAACTCTTTCCCTACACGACGCTCTTCCGATCTNNNAAGACATCGTTGAGCAGAGAG
GAGCACACGTCTGAACTCTTTCCCTACACGACGCTCTTCCGATCTNNNGTTTCACCATGTTAGTCAGGCT
GAGCACACGTCTGAACTCTTTCCCTACACGACGCTCTTCCGATCTNNNTGAGCTTTGTACGAAATGCTG
AGCACACGTCTGAACTCTTTCCCTACACGACGCTCTTCCGATCTNNNACAATATTGCAATCTGGGCTGT
GAGCACACGTCTGAACTCTTTCCCTACACGACGCTCTTCCGATCTNNNTCGTTGTCCCTGGTCCAATTATT
AGCACACGTCTGAACTCTTTCCCTACACGACGCTCTTCCGATCTNNNCTGTAGGTTGCATGTTCACACTG
AGCACACGTCTGAACTCTTTCCCTACACGACGCTCTTCCGATCTNNNGTTCAATACAGAGAGGGTCTCCC
AGCACACGTCTGAACTCTTTCCCTACACGACGCTCTTCCGATCTNNNTATCAGGAACAACAGCACCAAG
GCACACGTCTGAACTCTTTCCCTACACGACGCTCTTCCGATCTNNNCTAAGGCCGGAGAATCACTTGAA
AGAGCACACGTCTGAACTCTTTCCCTACACGACGCTCTTCCGATCTNNNGGTGCCTGTAATACCAGCTA
GAGCACACGTCTGAACTCTTTCCCTACACGACGCTCTTCCGATCTNNNAGCTGAGATCATACCCCTACA
GAGCACACGTCTGAACTCTTTCCCTACACGACGCTCTTCCGATCTNNNCCCTTGAAGACCCATTTCATCCT
AGCACACGTCTGAACTCTTTCCCTACACGACGCTCTTCCGATCTNNNCTGAGACTGGAGAATCGCTTG
GAGCACACGTCTGAACTCTTTCCCTACACGACGCTCTTCCGATCTNNNCAGCAACAACAGTGAGACTTT
AGCACACGTCTGAACTCTTTCCCTACACGACGCTCTTCCGATCTNNNCTTGCAGTGAGCTAAGATTGTG
AGCACACGTCTGAACTCTTTCCCTACACGACGCTCTTCCGATCTNNNCTTGAACCTAGGAGGCAGATGT
GAGCACACGTCTGAACTCTTTCCCTACACGACGCTCTTCCGATCTNNNCACTTGAGCCTAAGTAGGTGG
GAGCACACGTCTGAACTCTTTCCCTACACGACGCTCTTCCGATCTNNNGAGACCAGCCTAGGTAACATA
GAGCACACGTCTGAACTCTTTCCCTACACGACGCTCTTCCGATCTNNNCCCACTCTTGCCTTTACTTTGTG
AGCACACGTCTGAACTCTTTCCCTACACGACGCTCTTCCGATCTNNNCTGTAGTCCCAGCTACTCAGGA
AGAGCACACGTCTGAACTCTTTCCCTACACGACGCTCTTCCGATCTNNNAAGGCTGTAGTCAGTGGTG
AGCACACGTCTGAACTCTTTCCCTACACGACGCTCTTCCGATCTNNNCTGTAGTCCCAGCTATTCAGGAC

AGCACACGTCTGAACTCTTTCCCTACACGACGCTCTTCCGATCTNNNCTGAGATTGTACCACTGCACTCC
AGAGCACACGTCTGAACTCTTTCCCTACACGACGCTCTTCCGATCTNNNCTAAGGCAGGAGAATCGCT
AGCACACGTCTGAACTCTTTCCCTACACGACGCTCTTCCGATCTNNNCTGAGGCAGGAGAATCACTTGA
GAGCACACGTCTGAACTCTTTCCCTACACGACGCTCTTCCGATCTNNNGCAGAAGAATCACTTGAACCCA
AGCACACGTCTGAACTCTTTCCCTACACGACGCTCTTCCGATCTNNNAGTGAGCTGAGATTGTACCACT
AGAGCACACGTCTGAACTCTTTCCCTACACGACGCTCTTCCGATCTNNNAATCCAGCCTGGGTAACACA
GAGCACACGTCTGAACTCTTTCCCTACACGACGCTCTTCCGATCTNNNTTACACAGGACAATCGCTTGA
AGCACACGTCTGAACTCTTTCCCTACACGACGCTCTTCCGATCTNNNCAGAGTGAGTAAGCCTCCATCT
GAGCACACGTCTGAACTCTTTCCCTACACGACGCTCTTCCGATCTNNNATCACTTCACTTCGGGAGTTT
AGAGCACACGTCTGAACTCTTTCCCTACACGACGCTCTTCCGATCTNNNTGAGGCAGGAGAATCACTTG
AGCACACGTCTGAACTCTTTCCCTACACGACGCTCTTCCGATCTNNNAGGCAGGAGAATCATTTGAACC
GAGCACACGTCTGAACTCTTTCCCTACACGACGCTCTTCCGATCTNNNCAAGACAGCGAAACCCTATCT
AGCACACGTCTGAACTCTTTCCCTACACGACGCTCTTCCGATCTNNNCTGTAATCCCAGCTACTCAGGA
AGCACACGTCTGAACTCTTTCCCTACACGACGCTCTTCCGATCTNNNATCCTAGTACTTTGGGAGGCCA
GAGCACACGTCTGAACTCTTTCCCTACACGACGCTCTTCCGATCTNNNAATCCCAACTATTCAGGAGGCT
GAGCACACGTCTGAACTCTTTCCCTACACGACGCTCTTCCGATCTNNNCTGGGAGTTCAAGCTGTAGT
GCACACGTCTGAACTCTTTCCCTACACGACGCTCTTCCGATCTNNNTAAGAGGCTGAAGCAGGAGAAT
GAGCACACGTCTGAACTCTTTCCCTACACGACGCTCTTCCGATCTNNNTATGCCTGTAATCCCAGCACTT
GCACACGTCTGAACTCTTTCCCTACACGACGCTCTTCCGATCTNNNGAGATAGCACCACTACACTCCAG
AGCACACGTCTGAACTCTTTCCCTACACGACGCTCTTCCGATCTNNNAGTACTTTGGGAAGCCAAGGT
AGCACACGTCTGAACTCTTTCCCTACACGACGCTCTTCCGATCTNNNTGAGGGAGGAGAATCACTTGA
GAGCACACGTCTGAACTCTTTCCCTACACGACGCTCTTCCGATCTNNNGATCACTTGAGCTCAGGAGTT
AGCACACGTCTGAACTCTTTCCCTACACGACGCTCTTCCGATCTNNNCACGAGAATTGCTTGAACTCAG
AGCACACGTCTGAACTCTTTCCCTACACGACGCTCTTCCGATCTNNNCAGCCCAGACAACAGAGTAAGA
AGCACACGTCTGAACTCTTTCCCTACACGACGCTCTTCCGATCTNNNTGAGGTGAAGAGTTTGAGACCA
AGCACACGTCTGAACTCTTTCCCTACACGACGCTCTTCCGATCTNNNCTGAGGCAGGAGAATTGACTG
GAGCACACGTCTGAACTCTTTCCCTACACGACGCTCTTCCGATCTNNNGGCAAAACCCCGTCTCTACTAA
AGCACACGTCTGAACTCTTTCCCTACACGACGCTCTTCCGATCTNNNGCTTTGAGGGCATTTTACCTGA
GAGCACACGTCTGAACTCTTTCCCTACACGACGCTCTTCCGATCTNNNGAGGCTGAATCAGGAGAATCA
GAGCACACGTCTGAACTCTTTCCCTACACGACGCTCTTCCGATCTNNNTGATGCTGAAGAACTGCTTGA
GAGCACACGTCTGAACTCTTTCCCTACACGACGCTCTTCCGATCTNNNTCTGTTTGTGAGACCAGTTAC
GAGCACACGTCTGAACTCTTTCCCTACACGACGCTCTTCCGATCTNNNCACTTGAGGTCAGGAGTTCAA
AGCACACGTCTGAACTCTTTCCCTACACGACGCTCTTCCGATCTNNNTTCTGTGAAAGTTTGAAAGCCCT
AGCACACGTCTGAACTCTTTCCCTACACGACGCTCTTCCGATCTNNNGGATGTTGCAGTGAGCTGAGAT
GCACACGTCTGAACTCTTTCCCTACACGACGCTCTTCCGATCTNNNCACCAATAATCCCAGCTACTCCACA
GAGCACACGTCTGAACTCTTTCCCTACACGACGCTCTTCCGATCTNNNGGGAATCACTTGAAACCAGGA
GAGCACACGTCTGAACTCTTTCCCTACACGACGCTCTTCCGATCTNNNGAATTGCTTGAACCTAGGAGT
GAGCACACGTCTGAACTCTTTCCCTACACGACGCTCTTCCGATCTNNNCAGGAGAATTGCTTGAACATC
AGAGCACACGTCTGAACTCTTTCCCTACACGACGCTCTTCCGATCTNNNCTAGGCAACAAGAGCAAAAC
GAGCACACGTCTGAACTCTTTCCCTACACGACGCTCTTCCGATCTNNNAATCCAACTTGCCACTTGTTTCC
AGCACACGTCTGAACTCTTTCCCTACACGACGCTCTTCCGATCTNNNCATGAAAACCCACTCATCATGCT
GAGCACACGTCTGAACTCTTTCCCTACACGACGCTCTTCCGATCTNNNAGACCTTTGCAGTGAGTGTTA
GAGCACACGTCTGAACTCTTTCCCTACACGACGCTCTTCCGATCTNNNCTCAGGTGATCCTCCCATCTTA

GCACACGTCTGAACTCTTTCCCTACACGACGCTCTTCCGATCTNNNAGTAGCCAGGACTACAGGTATG
GCACACGTCTGAACTCTTTCCCTACACGACGCTCTTCCGATCTNNNTGGGATTGCTGAGTCAAATGG
AGCACACGTCTGAACTCTTTCCCTACACGACGCTCTTCCGATCTNNNGGCTGAAGCTAATTCCCTCTAC
GCACACGTCTGAACTCTTTCCCTACACGACGCTCTTCCGATCTNNNCCAGTGAGAAGAAGCCTTGCAT
AGCACACGTCTGAACTCTTTCCCTACACGACGCTCTTCCGATCTNNNAGCATCACTTTGGGGTTAGGAT
GCACACGTCTGAACTCTTTCCCTACACGACGCTCTTCCGATCTNNNTGTTGGATTACAGTAGTGAGCC
AGCACACGTCTGAACTCTTTCCCTACACGACGCTCTTCCGATCTNNNCCAGGATCAGTCAATTGGTTTG
AGCACACGTCTGAACTCTTTCCCTACACGACGCTCTTCCGATCTNNNTTTGGTCCTTGGGCTATGTTCT
GCACACGTCTGAACTCTTTCCCTACACGACGCTCTTCCGATCTNNNGTCACAAAGACTTCTACACAGG
AGCACACGTCTGAACTCTTTCCCTACACGACGCTCTTCCGATCTNNNGGCAGGAAACTTAGGTAGGG
AGCACACGTCTGAACTCTTTCCCTACACGACGCTCTTCCGATCTNNNGATGGTCTTGATCTCCTGACC
GCACACGTCTGAACTCTTTCCCTACACGACGCTCTTCCGATCTNNNTAAGGAGAAACTAAGGTGGAG
AGCACACGTCTGAACTCTTTCCCTACACGACGCTCTTCCGATCTNNNCTGAGTAGCTGGGATTACAGG
CACACGTCTGAACTCTTTCCCTACACGACGCTCTTCCGATCTNNNAGCAGTCCTTCCCTAAAGAAAGG
GAGCACACGTCTGAACTCTTTCCCTACACGACGCTCTTCCGATCTNNNCATACCATCATGCCCAGCAAA
AGCACACGTCTGAACTCTTTCCCTACACGACGCTCTTCCGATCTNNNTCCGATCACTTTGTGTAAGTC
AGCACACGTCTGAACTCTTTCCCTACACGACGCTCTTCCGATCTNNNGGTAGCTGGAACTACAGTCAT
GCACACGTCTGAACTCTTTCCCTACACGACGCTCTTCCGATCTNNNGGTGGGTGCTAATGTAAATGG
AGCACACGTCTGAACTCTTTCCCTACACGACGCTCTTCCGATCTNNNGCTCAAGCTATCTTCTGCACTC
GAGCACACGTCTGAACTCTTTCCCTACACGACGCTCTTCCGATCTNNNAGAACCAAATGACTGGGAAG
AGCACACGTCTGAACTCTTTCCCTACACGACGCTCTTCCGATCTNNNTGCAGCCTGCTATTATACTCTC
AGCACACGTCTGAACTCTTTCCCTACACGACGCTCTTCCGATCTNNNATGTGTACCTCACTTTCCTTCCC
GAGCACACGTCTGAACTCTTTCCCTACACGACGCTCTTCCGATCTNNNCTCAAGTGATCCTCCTGTCTC
AGCACACGTCTGAACTCTTTCCCTACACGACGCTCTTCCGATCTNNNGGCAACAATAAGCATGGAACT
GAGCACACGTCTGAACTCTTTCCCTACACGACGCTCTTCCGATCTNNNCTTTAGCCAAACCACAGACAC
AGCACACGTCTGAACTCTTTCCCTACACGACGCTCTTCCGATCTNNNGCTTCCTCACCAAAGTCAGAAA
AGCACACGTCTGAACTCTTTCCCTACACGACGCTCTTCCGATCTNNNGGTTAGAACTGCTCCATTCTTG
GAGCACACGTCTGAACTCTTTCCCTACACGACGCTCTTCCGATCTNNNTGTAGAAATGGTTGGGATCC
AGCACACGTCTGAACTCTTTCCCTACACGACGCTCTTCCGATCTNNNCACACTTGCTCTACCAAAAAGC
AGAGCACACGTCTGAACTCTTTCCCTACACGACGCTCTTCCGATCTNNNACACCTACTATGTACCCACTA
AGCACACGTCTGAACTCTTTCCCTACACGACGCTCTTCCGATCTNNNAACGTTATTGCTGCTGGTGTA
AGCACACGTCTGAACTCTTTCCCTACACGACGCTCTTCCGATCTNNNCTCATTGTAGCCTCAGACTCAG
AGCACACGTCTGAACTCTTTCCCTACACGACGCTCTTCCGATCTNNNCCCAGAGATTACGTCCTATTGG
GCACACGTCTGAACTCTTTCCCTACACGACGCTCTTCCGATCTNNNGGGTGTGTGGGTTGAATGAAT
AGCACACGTCTGAACTCTTTCCCTACACGACGCTCTTCCGATCTNNNATCCCACTTTACCTTCCTTGCAT
AGCACACGTCTGAACTCTTTCCCTACACGACGCTCTTCCGATCTNNNCCCAGGTTCAAGCAATTCTTCTG
GCACACGTCTGAACTCTTTCCCTACACGACGCTCTTCCGATCTNNNTGTATTTGGGGAAGTGGGAGA
GCACACGTCTGAACTCTTTCCCTACACGACGCTCTTCCGATCTNNNTGATCTCTTGACCTCATGATCCG
AGCACACGTCTGAACTCTTTCCCTACACGACGCTCTTCCGATCTNNNCCACCAAGCCCAGCTAATTTTTT
AGCACACGTCTGAACTCTTTCCCTACACGACGCTCTTCCGATCTNNNCACGGTTTGGAAAATGACTAG
AGCACACGTCTGAACTCTTTCCCTACACGACGCTCTTCCGATCTNNNAATGAGGACAGCTGTGTTCTG
AGCACACGTCTGAACTCTTTCCCTACACGACGCTCTTCCGATCTNNNGCAAGCAAGCACCTTCTTCATA
GCACACGTCTGAACTCTTTCCCTACACGACGCTCTTCCGATCTNNNAAGGTCTAACAGATTGGTCCCA

AGCACACGTCTGAACTCTTTCCCTACACGACGCTCTTCCGATCTNNNACCTTCCCAATGACCACTTGAA
GCACACGTCTGAACTCTTTCCCTACACGACGCTCTTCCGATCTNNNCATGGTCTTTTCCTGAAACTGCA
AGCACACGTCTGAACTCTTTCCCTACACGACGCTCTTCCGATCTNNNTTTCTTTCTTTTGTGTGTGGTG
AGCACACGTCTGAACTCTTTCCCTACACGACGCTCTTCCGATCTNNNTGTTTCTGTTGGTTAGCTCACC
GCACACGTCTGAACTCTTTCCCTACACGACGCTCTTCCGATCTNNNTATCCAAGACACCCCACCAAAAA
GCACACGTCTGAACTCTTTCCCTACACGACGCTCTTCCGATCTNNNGCTGTAGGCCAGAGTAAAAAG
GAGCACACGTCTGAACTCTTTCCCTACACGACGCTCTTCCGATCTNNNATCCTCTTCTGCACTGATGGT
AGCACACGTCTGAACTCTTTCCCTACACGACGCTCTTCCGATCTNNNTGCGATCATAGCTCACTGTAT
AGCACACGTCTGAACTCTTTCCCTACACGACGCTCTTCCGATCTNNNAAGGTGTAAGCCCAAAAGAGT
AGCACACGTCTGAACTCTTTCCCTACACGACGCTCTTCCGATCTNNNATGAGTAGTTGGCAATGCAT
GAGCACACGTCTGAACTCTTTCCCTACACGACGCTCTTCCGATCTNNNCCTAGGTGACAGAACAAGA
GAGCACACGTCTGAACTCTTTCCCTACACGACGCTCTTCCGATCTNNNCAGAGCTTGTAGTGAGCAG
AGCACACGTCTGAACTCTTTCCCTACACGACGCTCTTCCGATCTNNNCAAGAGGTAGAGGCTATGGTG
AGCACACGTCTGAACTCTTTCCCTACACGACGCTCTTCCGATCTNNNGAGAATTGTTTGAACCTGGGA
AGCACACGTCTGAACTCTTTCCCTACACGACGCTCTTCCGATCTNNNCTGAGGCAGGAGAATTGCTTG
GCACACGTCTGAACTCTTTCCCTACACGACGCTCTTCCGATCTNNNCTGTAATCCCAGCTACTTGGAGG
AGCACACGTCTGAACTCTTTCCCTACACGACGCTCTTCCGATCTNNNCTGAGGCAGGAGAATAGCTT
GAGCACACGTCTGAACTCTTTCCCTACACGACGCTCTTCCGATCTNNNCTGGGTCAGAACGAGATTCC
AGAGCACACGTCTGAACTCTTTCCCTACACGACGCTCTTCCGATCTNNNGAGACAGGAGAATTGCTTG
AGCACACGTCTGAACTCTTTCCCTACACGACGCTCTTCCGATCTNNNCACCTGTAATCCCAGCTACTCAG
AGCACACGTCTGAACTCTTTCCCTACACGACGCTCTTCCGATCTNNNCAGAAGAATTGCTTGAACCTGG
GCACACGTCTGAACTCTTTCCCTACACGACGCTCTTCCGATCTNNNAGGGAATTGCTTGAATCCAGGA
AGCACACGTCTGAACTCTTTCCCTACACGACGCTCTTCCGATCTNNNACATGTAATCCCAGCTACTCAG
AGCACACGTCTGAACTCTTTCCCTACACGACGCTCTTCCGATCTNNNAGAGGTTGCAGTGAACTGAG
GAGCACACGTCTGAACTCTTTCCCTACACGACGCTCTTCCGATCTNNNAAGAAAAGCAAAAAGAGGC
GAGCACACGTCTGAACTCTTTCCCTACACGACGCTCTTCCGATCTNNNGATGCCTGTTATCCCAGCTAC
GAGCACACGTCTGAACTCTTTCCCTACACGACGCTCTTCCGATCTNNNGAGTATCACTTGAACCTGGG
GAGCACACGTCTGAACTCTTTCCCTACACGACGCTCTTCCGATCTNNNCTGAGGCAGGAGAATAGCTT
GAGCACACGTCTGAACTCTTTCCCTACACGACGCTCTTCCGATCTNNNCTCATGCCTGTAATCCTAGCA
GAGCACACGTCTGAACTCTTTCCCTACACGACGCTCTTCCGATCTNNNCTCACACCTGTAATCTCAGCA
AGCACACGTCTGAACTCTTTCCCTACACGACGCTCTTCCGATCTNNNCTTGAACCCAGGACACAGAGA
AGCACACGTCTGAACTCTTTCCCTACACGACGCTCTTCCGATCTNNNCAAGATCACGCCACTTCATTCA
AGCACACGTCTGAACTCTTTCCCTACACGACGCTCTTCCGATCTNNNAAGGTTTGATTTCATGCCTGGT
AGCACACGTCTGAACTCTTTCCCTACACGACGCTCTTCCGATCTNNNCAGAAGAATTGCTCAAACCTGG
GAGCACACGTCTGAACTCTTTCCCTACACGACGCTCTTCCGATCTNNNGAGAGTTGCTTGAACCTAGG
AGCACACGTCTGAACTCTTTCCCTACACGACGCTCTTCCGATCTNNNTGTAATCCCAGGTACTCAGGA
AGCACACGTCTGAACTCTTTCCCTACACGACGCTCTTCCGATCTNNNGAGTTTGCAGTAGGCTGAGA
AGCACACGTCTGAACTCTTTCCCTACACGACGCTCTTCCGATCTNNNCTGAGACAGGAGAATCGCTTG
AGCACACGTCTGAACTCTTTCCCTACACGACGCTCTTCCGATCTNNNGGTAGAAGGAAGAAGCACA
GCACACGTCTGAACTCTTTCCCTACACGACGCTCTTCCGATCTNNNCTGTAGTCCTAGCTACTCGGGAG
AGCACACGTCTGAACTCTTTCCCTACACGACGCTCTTCCGATCTNNNGTGACACAGCAAGACTCTTTC
AGCACACGTCTGAACTCTTTCCCTACACGACGCTCTTCCGATCTNNNTGAGGCAGGAGAATCTCTTG
AGCACACGTCTGAACTCTTTCCCTACACGACGCTCTTCCGATCTNNNAAGAAGAGCAAAACCCTGTCTC

AGAGCACACGTCTGAACTCTTTCCCTACACGACGCTCTTCCGATCTNNNCTGTAGTCCCAGCTACTCAGG
GAGCACACGTCTGAACTCTTTCCCTACACGACGCTCTTCCGATCTNNNAGCTTACAGTGAGCCAAGATCA
AGCACACGTCTGAACTCTTTCCCTACACGACGCTCTTCCGATCTNNNGAGGCAGAAGAATCGTTTGAA
AGAGCACACGTCTGAACTCTTTCCCTACACGACGCTCTTCCGATCTNNNCTGAGGCAGGAGAATCACT
GAGCACACGTCTGAACTCTTTCCCTACACGACGCTCTTCCGATCTNNNGTACCTGTAATCCCAGCTACTG
AGCACACGTCTGAACTCTTTCCCTACACGACGCTCTTCCGATCTNNNCAAGGCAGGAGAATTGCTTGAA
GAGCACACGTCTGAACTCTTTCCCTACACGACGCTCTTCCGATCTNNNCACCTGTAATCCCAGCTACTCA
GAGCACACGTCTGAACTCTTTCCCTACACGACGCTCTTCCGATCTNNNGTTGCAGTAAGCTGAGATTGC
AAGAGCACACGTCTGAACTCTTTCCCTACACGACGCTCTTCCGATCTNNNGAAGGTTGCAGTGAGTCG
AGCACACGTCTGAACTCTTTCCCTACACGACGCTCTTCCGATCTNNNCTCAGCTACTTGAGAGGCTAAG
AGCACACGTCTGAACTCTTTCCCTACACGACGCTCTTCCGATCTNNNTGGAGATTGCAGTGAACTGAG
GCACACGTCTGAACTCTTTCCCTACACGACGCTCTTCCGATCTNNNTGAGCTACTGCCCTTGACATATCC
AGAGCACACGTCTGAACTCTTTCCCTACACGACGCTCTTCCGATCTNNNATCTGTAATCTCAGCTGCTCA
GCACACGTCTGAACTCTTTCCCTACACGACGCTCTTCCGATCTNNNACACTTGTAATCCCAGCTACTCGG
GAGCACACGTCTGAACTCTTTCCCTACACGACGCTCTTCCGATCTNNNGAGGTTGCAGTGAGTTGATTG
AGCACACGTCTGAACTCTTTCCCTACACGACGCTCTTCCGATCTNNNAGAATTTCTTGAGGCCAGGAG
AGCACACGTCTGAACTCTTTCCCTACACGACGCTCTTCCGATCTNNNGCATGCCTATAATCCCAGCTACT
AGCACACGTCTGAACTCTTTCCCTACACGACGCTCTTCCGATCTNNNTTCCCATCTACTCGGTAGGTAGG
AGCACACGTCTGAACTCTTTCCCTACACGACGCTCTTCCGATCTNNNCTATAATCCCAGCTACTCGGCAG
AGCACACGTCTGAACTCTTTCCCTACACGACGCTCTTCCGATCTNNNCTGAGGCAGGAGAATTGCTTGA
AGCACACGTCTGAACTCTTTCCCTACACGACGCTCTTCCGATCTNNNGCCTGACTTTCTGACTCCAAACTG
AGCACACGTCTGAACTCTTTCCCTACACGACGCTCTTCCGATCTNNNGGCCCAGCCTAAAAATTCCAAA
GAGCACACGTCTGAACTCTTTCCCTACACGACGCTCTTCCGATCTNNNCCACCATGCCCTGCTAAATTTT
GAGCACACGTCTGAACTCTTTCCCTACACGACGCTCTTCCGATCTNNNTAATGGCATGATCTCAGCTCAG
AGCACACGTCTGAACTCTTTCCCTACACGACGCTCTTCCGATCTNNNCCAGATCCTACAGATGCACATCA
AGCACACGTCTGAACTCTTTCCCTACACGACGCTCTTCCGATCTNNNTTACAATAACGGCCTCTCACTCCG
AGCACACGTCTGAACTCTTTCCCTACACGACGCTCTTCCGATCTNNNCCAAACAGTCCTCAGGTACAAGA
AGCACACGTCTGAACTCTTTCCCTACACGACGCTCTTCCGATCTNNNAGTATTGCACTGAGTGAGCTGA
GAGCACACGTCTGAACTCTTTCCCTACACGACGCTCTTCCGATCTNNNTCACCACGCCCAACTCATTTATA
AGCACACGTCTGAACTCTTTCCCTACACGACGCTCTTCCGATCTNNNAATCTCCGCCTTCCTAGTTCAAG
AGCACACGTCTGAACTCTTTCCCTACACGACGCTCTTCCGATCTNNNAAAAGAGACCATGTGTGACCCTT
GAGCACACGTCTGAACTCTTTCCCTACACGACGCTCTTCCGATCTNNNGGGACTACAGGCATGCATATG
AGCACACGTCTGAACTCTTTCCCTACACGACGCTCTTCCGATCTNNNCTCCCATGAAGCGTATCTCAGAG
AGCACACGTCTGAACTCTTTCCCTACACGACGCTCTTCCGATCTNNNATAAGGAAGGAGTGTCTGGGA
AGCACACGTCTGAACTCTTTCCCTACACGACGCTCTTCCGATCTNNNTTCAAGCAATTCTCCTGCCTCAG
AGCACACGTCTGAACTCTTTCCCTACACGACGCTCTTCCGATCTNNNGGTGACAGCCTAACAGAGAAGA
AGCACACGTCTGAACTCTTTCCCTACACGACGCTCTTCCGATCTNNNGAGGAGTTGCTAGTACTGGCA
AGCACACGTCTGAACTCTTTCCCTACACGACGCTCTTCCGATCTNNNAAACTTCGGCTTGCATTAGGTT
AGCACACGTCTGAACTCTTTCCCTACACGACGCTCTTCCGATCTNNNAAAATTGTTCTTCCTGCAGGGTG
GAGCACACGTCTGAACTCTTTCCCTACACGACGCTCTTCCGATCTNNNCTCAAGCAATCCTCCCACTTCAG
GAGCACACGTCTGAACTCTTTCCCTACACGACGCTCTTCCGATCTNNNTGACAACGAAAGCCATCTCCT
GAGCACACGTCTGAACTCTTTCCCTACACGACGCTCTTCCGATCTNNNAAATGTAAAATGGCGTGACTG
AGCACACGTCTGAACTCTTTCCCTACACGACGCTCTTCCGATCTNNNAGATGTCCCTCAAGTGAATTGG

AGCACACGTCTGAACTCTTTCCCTACACGACGCTCTTCCGATCTNNNGTCTAATTGGCTCACAGTTCTG
GAGCACACGTCTGAACTCTTTCCCTACACGACGCTCTTCCGATCTNNNTGATGACCCTATCTCATACGC
AGCACACGTCTGAACTCTTTCCCTACACGACGCTCTTCCGATCTNNNTAGCAAACCATCATGCCCTCTT
GAGCACACGTCTGAACTCTTTCCCTACACGACGCTCTTCCGATCTNNNGACTACATGCATGTGCCACTA
AGCACACGTCTGAACTCTTTCCCTACACGACGCTCTTCCGATCTNNNCTTTCCTGCTATTCCGCAAGTT
AGCACACGTCTGAACTCTTTCCCTACACGACGCTCTTCCGATCTNNNAGGGAGTTGGACTCATGGTTT
AGCACACGTCTGAACTCTTTCCCTACACGACGCTCTTCCGATCTNNNCTAATGCCACATGTTCCATGCT
AGCACACGTCTGAACTCTTTCCCTACACGACGCTCTTCCGATCTNNNTCTGCCGAGGAAGTCAGTTTTT
AGCACACGTCTGAACTCTTTCCCTACACGACGCTCTTCCGATCTNNNAAGCACTCAGTAACCAGCTGT
AGCACACGTCTGAACTCTTTCCCTACACGACGCTCTTCCGATCTNNNCCTAGCACTTTGAAAAGCCAA
GAGCACACGTCTGAACTCTTTCCCTACACGACGCTCTTCCGATCTNNNCAGCATGAGTGACAGAGTG
GAGCACACGTCTGAACTCTTTCCCTACACGACGCTCTTCCGATCTNNNCAGGAGAATCACTTGAACCC
AGCACACGTCTGAACTCTTTCCCTACACGACGCTCTTCCGATCTNNNATCACTTGAATTCAGGAGGCT
AGCACACGTCTGAACTCTTTCCCTACACGACGCTCTTCCGATCTNNNAGTGAGCTATGATAGTGCCACT
AGCACACGTCTGAACTCTTTCCCTACACGACGCTCTTCCGATCTNNNCACCTGTAATCCCAGCTACTCA
GCACACGTCTGAACTCTTTCCCTACACGACGCTCTTCCGATCTNNNCGAGATCATATCACTGCACTCCA
AGCACACGTCTGAACTCTTTCCCTACACGACGCTCTTCCGATCTNNNACATAGTGAGACTGAGACCCT
GAGCACACGTCTGAACTCTTTCCCTACACGACGCTCTTCCGATCTNNNGGGAGAATCACTTGAACCCA
AGCACACGTCTGAACTCTTTCCCTACACGACGCTCTTCCGATCTNNNTGGCAACAGAGTAAGACTCCA
GCACACGTCTGAACTCTTTCCCTACACGACGCTCTTCCGATCTNNNGAGAATGGCTTGAACCCAGAAA
GAGCACACGTCTGAACTCTTTCCCTACACGACGCTCTTCCGATCTNNNGAGATTGCAGTGAGCCAAGA
GAGCACACGTCTGAACTCTTTCCCTACACGACGCTCTTCCGATCTNNNACCTGGGCAACAAAATTGAA
GAGCACACGTCTGAACTCTTTCCCTACACGACGCTCTTCCGATCTNNNATAAGAATCGCTTGAACCTGG
AGAGCACACGTCTGAACTCTTTCCCTACACGACGCTCTTCCGATCTNNNGAGATTGCACTGAGCTGAG
AGCACACGTCTGAACTCTTTCCCTACACGACGCTCTTCCGATCTNNNCCCAGGAGTTCAAGGCTATAG
GAGCACACGTCTGAACTCTTTCCCTACACGACGCTCTTCCGATCTNNNGAGACAGGAGAATTGCTTG
GCACACGTCTGAACTCTTTCCCTACACGACGCTCTTCCGATCTNNNTGTAGTCCCAGTTACTCAGGAG
GAGCACACGTCTGAACTCTTTCCCTACACGACGCTCTTCCGATCTNNNCCTGGGAGTTTGAGAATGCAT
AGCACACGTCTGAACTCTTTCCCTACACGACGCTCTTCCGATCTNNNCAGATACTTGGTAGGCTGAGG
GAGCACACGTCTGAACTCTTTCCCTACACGACGCTCTTCCGATCTNNNCGTGCCTATAGTTCCAGCTA
GAGCACACGTCTGAACTCTTTCCCTACACGACGCTCTTCCGATCTNNNCCAAAGCAGGAGAATCGTTT
GAGCACACGTCTGAACTCTTTCCCTACACGACGCTCTTCCGATCTNNNTTGAACAGGGAAGAGAGAG
AGCACACGTCTGAACTCTTTCCCTACACGACGCTCTTCCGATCTNNNATGACAGACCAATTGCTTGAAC
GAGCACACGTCTGAACTCTTTCCCTACACGACGCTCTTCCGATCTNNNATCCTAGCTACTCAGAAGCCT
AGCACACGTCTGAACTCTTTCCCTACACGACGCTCTTCCGATCTNNNGGGAACAGAGTGAGATTCC
GCACACGTCTGAACTCTTTCCCTACACGACGCTCTTCCGATCTNNNTGCCTGTAATACCAGCTACTTGG
GAGCACACGTCTGAACTCTTTCCCTACACGACGCTCTTCCGATCTNNNGAGGCATGAGAATTGCTTGT
AGAGCACACGTCTGAACTCTTTCCCTACACGACGCTCTTCCGATCTNNNCGAGGTCAGGAGATCAAGA
AGAGCACACGTCTGAACTCTTTCCCTACACGACGCTCTTCCGATCTNNNTCATGCTACTACTGCACTCC
AGCACACGTCTGAACTCTTTCCCTACACGACGCTCTTCCGATCTNNNTCATGCCTGTAATCCCAACACTT
AGCACACGTCTGAACTCTTTCCCTACACGACGCTCTTCCGATCTNNNCATGAGAATTGCTTGAACTGG
GAGCACACGTCTGAACTCTTTCCCTACACGACGCTCTTCCGATCTNNNGAGAATGGCATGAATCCAGG
AGCACACGTCTGAACTCTTTCCCTACACGACGCTCTTCCGATCTNNNTTCACTTGTAATCCCAGCTACCC

AGCACACGTCTGAACTCTTTCCCTACACGACGCTCTTCCGATCTNNNGCTTGGATGACAGAGTGAGA
GCACACGTCTGAACTCTTTCCCTACACGACGCTCTTCCGATCTNNNAGACAGGAGAATCACTTGAAC
GAGCACACGTCTGAACTCTTTCCCTACACGACGCTCTTCCGATCTNNNAATTGTGCCAAGCTGTGAG
AGCACACGTCTGAACTCTTTCCCTACACGACGCTCTTCCGATCTNNNGGTGCCTGTAATCCAGCTACT
GAGCACACGTCTGAACTCTTTCCCTACACGACGCTCTTCCGATCTNNNCTGAGGCAGGAGAATTGCTT
AGCACACGTCTGAACTCTTTCCCTACACGACGCTCTTCCGATCTNNNTGACTGTAATCCCAGCTGCTTA
GAGCACACGTCTGAACTCTTTCCCTACACGACGCTCTTCCGATCTNNNGAGCTTGCAGTGAGCCAAG
AGCACACGTCTGAACTCTTTCCCTACACGACGCTCTTCCGATCTNNNGAGACAGGAAAATCGGTTGA
AGAGCACACGTCTGAACTCTTTCCCTACACGACGCTCTTCCGATCTNNNCTGAGGCAGGAGAATCACT
GAGCACACGTCTGAACTCTTTCCCTACACGACGCTCTTCCGATCTNNNACCTGGGCAACAAAATTGAA
AGCACACGTCTGAACTCTTTCCCTACACGACGCTCTTCCGATCTNNNGTGGTGTATGCCTGTAGTCCT
GCACACGTCTGAACTCTTTCCCTACACGACGCTCTTCCGATCTNNNCTGTAATCCCAGCTACTCAGGAG
AGCACACGTCTGAACTCTTTCCCTACACGACGCTCTTCCGATCTNNNTAGGAGAATTGCTTGAACCTG
GAGCACACGTCTGAACTCTTTCCCTACACGACGCTCTTCCGATCTNNNCAAGAGTCACCAAGTACCTG
AGCACACGTCTGAACTCTTTCCCTACACGACGCTCTTCCGATCTNNNGTAATCCCAGTTACTCAGGAG
AGCACACGTCTGAACTCTTTCCCTACACGACGCTCTTCCGATCTNNNGAGGTTGCAGTGAATCAAGA
GAGCACACGTCTGAACTCTTTCCCTACACGACGCTCTTCCGATCTNNNCAGGAGATTCACTTGAACCT
GAGCACACGTCTGAACTCTTTCCCTACACGACGCTCTTCCGATCTNNNTGAACCTAAGAGGAGGAGT
AGCACACGTCTGAACTCTTTCCCTACACGACGCTCTTCCGATCTNNNTGGCTCATACCTGCAATTCTA
GAGCACACGTCTGAACTCTTTCCCTACACGACGCTCTTCCGATCTNNNCCTGGGCAACAAGAGAGAA
GCACACGTCTGAACTCTTTCCCTACACGACGCTCTTCCGATCTNNNACATCGTTCCCTGATTGCCTAAT
AGCACACGTCTGAACTCTTTCCCTACACGACGCTCTTCCGATCTNNNTTGGTTACAAGAGTTAGCTCC
AGCACACGTCTGAACTCTTTCCCTACACGACGCTCTTCCGATCTNNNTGTTTGCAGTCCTGATGTTACC
GCACACGTCTGAACTCTTTCCCTACACGACGCTCTTCCGATCTNNNACTTGCTTTATGACCACAATGG
GAGCACACGTCTGAACTCTTTCCCTACACGACGCTCTTCCGATCTNNNGAGAGAGGGGCAAAGTCTC
AGCACACGTCTGAACTCTTTCCCTACACGACGCTCTTCCGATCTNNNTGCTTCTCTTTTAGGATGTGTG
AGCACACGTCTGAACTCTTTCCCTACACGACGCTCTTCCGATCTNNNCCATTCTAACAGGTGTGAGGT
GCACACGTCTGAACTCTTTCCCTACACGACGCTCTTCCGATCTNNNCTGGGTTCAAGTGATTCTCTTG
GCACACGTCTGAACTCTTTCCCTACACGACGCTCTTCCGATCTNNNCAACTTCCCTGGCTCAAATGAT
AGCACACGTCTGAACTCTTTCCCTACACGACGCTCTTCCGATCTNNNGCTGATTGTTTCTTGCCTACCA
GAGCACACGTCTGAACTCTTTCCCTACACGACGCTCTTCCGATCTNNNCAGGAAGGACTTCACTAGGC
GCACACGTCTGAACTCTTTCCCTACACGACGCTCTTCCGATCTNNNCAGAAACAAGTTGGGAAGAGC
GAGCACACGTCTGAACTCTTTCCCTACACGACGCTCTTCCGATCTNNNGCATGGCTACCTCCCAACTAT
AGCACACGTCTGAACTCTTTCCCTACACGACGCTCTTCCGATCTNNNTATTTTTTTGAGATGGGCGG
GCACACGTCTGAACTCTTTCCCTACACGACGCTCTTCCGATCTNNNTAGAGCCAGTCTAGTGGTGATG
AGCACACGTCTGAACTCTTTCCCTACACGACGCTCTTCCGATCTNNNACTGATGGAAATGAAGACCA
AGCACACGTCTGAACTCTTTCCCTACACGACGCTCTTCCGATCTNNNATTGGGCTTGTCGATTGCTTTT
AGCACACGTCTGAACTCTTTCCCTACACGACGCTCTTCCGATCTNNNGCCTATTGTCCCTGAAGCATTT
GCACACGTCTGAACTCTTTCCCTACACGACGCTCTTCCGATCTNNNACAGATGCACACCACCATATCTT
AGCACACGTCTGAACTCTTTCCCTACACGACGCTCTTCCGATCTNNNTGTTTTGACGGGAGAATTCAA
AGCACACGTCTGAACTCTTTCCCTACACGACGCTCTTCCGATCTNNNGGCAACCTTTGTATCAACAGC
GAGCACACGTCTGAACTCTTTCCCTACACGACGCTCTTCCGATCTNNNTGGGCCCGAATCAATGGTAA
AGCACACGTCTGAACTCTTTCCCTACACGACGCTCTTCCGATCTNNNGGCTAGGACTATGAGCACATA

AGCACACGTCTGAACTCTTTCCCTACACGACGCTCTTCCGATCTNNNTGTGTTTCGAAAAAGCCACACT
AGCACACGTCTGAACTCTTTCCCTACACGACGCTCTTCCGATCTNNNGCTGTCTGATGAGGACTAGCTT
GAGCACACGTCTGAACTCTTTCCCTACACGACGCTCTTCCGATCTNNNGCGTGACTACATTCAGGTTTGA
GAGCACACGTCTGAACTCTTTCCCTACACGACGCTCTTCCGATCTNNNATGGTCTTAGCTCACTGCAACT
AGCACACGTCTGAACTCTTTCCCTACACGACGCTCTTCCGATCTNNNTACCCATTACAGGTTGACCATCC
GAGCACACGTCTGAACTCTTTCCCTACACGACGCTCTTCCGATCTNNNTTTCTTCCTGGGCTCTTAGTTC
GAGCACACGTCTGAACTCTTTCCCTACACGACGCTCTTCCGATCTNNNTTACAAGAGTTGCACATGGTTC
GAGCACACGTCTGAACTCTTTCCCTACACGACGCTCTTCCGATCTNNNTTGTCATCCTGGTTATGGGAG
GAGCACACGTCTGAACTCTTTCCCTACACGACGCTCTTCCGATCTNNNGCAGGAGTGGAGCCAAAAAA
AGCACACGTCTGAACTCTTTCCCTACACGACGCTCTTCCGATCTNNNAAGCTCGGAAGTGGTAAAAGC
AGCACACGTCTGAACTCTTTCCCTACACGACGCTCTTCCGATCTNNNCAAGACAGCCTTTGTGTACCAT
AGCACACGTCTGAACTCTTTCCCTACACGACGCTCTTCCGATCTNNNATTGACGAGTTGACAAGGAACT
AGCACACGTCTGAACTCTTTCCCTACACGACGCTCTTCCGATCTNNNTCAAGAGGTTCTTCTGTCTCAG
AGCACACGTCTGAACTCTTTCCCTACACGACGCTCTTCCGATCTNNNTACATCATTGGTTGTCTGAGGCA
AGCACACGTCTGAACTCTTTCCCTACACGACGCTCTTCCGATCTNNNGCACAGGCAACACAACAAAAAA
GAGCACACGTCTGAACTCTTTCCCTACACGACGCTCTTCCGATCTNNNATGCCCTACCACATTCCTCTTTT
GAGCACACGTCTGAACTCTTTCCCTACACGACGCTCTTCCGATCTNNNTGTGACTCATTTCAAACTCAGA
GAGCACACGTCTGAACTCTTTCCCTACACGACGCTCTTCCGATCTNNNGCAGTGTCTCCTCTTTCTCTCTC
AGCACACGTCTGAACTCTTTCCCTACACGACGCTCTTCCGATCTNNNCCTTAGCCTCAGGAACATCTGA
GAGCACACGTCTGAACTCTTTCCCTACACGACGCTCTTCCGATCTNNNCTTCTTCCATCCCTTTGTCCTCA
GAGCACACGTCTGAACTCTTTCCCTACACGACGCTCTTCCGATCTNNNTTCATGTATTCACTGGAGCAG
GAGCACACGTCTGAACTCTTTCCCTACACGACGCTCTTCCGATCTNNNTCCCAGGTATCAAGCCTAGTA
GAGCACACGTCTGAACTCTTTCCCTACACGACGCTCTTCCGATCTNNNGTTCTGGTGAGTACCCTCTTCA
GAGCACACGTCTGAACTCTTTCCCTACACGACGCTCTTCCGATCTNNNCCTGCGACCTCAATTCTCTGAT
AGCACACGTCTGAACTCTTTCCCTACACGACGCTCTTCCGATCTNNNCCTCGCCCACCTCCAAATAATATC
GAGCACACGTCTGAACTCTTTCCCTACACGACGCTCTTCCGATCTNNNCTTCCCTTCCCTTTCTCTCTCTC
AGCACACGTCTGAACTCTTTCCCTACACGACGCTCTTCCGATCTNNNTCACACTGAATCTTGTCTGTTGT
AGCACACGTCTGAACTCTTTCCCTACACGACGCTCTTCCGATCTNNNATGACACAACCACTTTGGAACAC
AGCACACGTCTGAACTCTTTCCCTACACGACGCTCTTCCGATCTNNNAGTGCTGGGATTACAGATGTGA
AGCACACGTCTGAACTCTTTCCCTACACGACGCTCTTCCGATCTNNNGAGTGCTGGGATTATAGGCATC
AGCACACGTCTGAACTCTTTCCCTACACGACGCTCTTCCGATCTNNNCGAAGCCCAGCCAATTTTTTGTA
GAGCACACGTCTGAACTCTTTCCCTACACGACGCTCTTCCGATCTNNNCTCAACTCCCTCTGACTTCAGT
AGCACACGTCTGAACTCTTTCCCTACACGACGCTCTTCCGATCTNNNCCCTAACCCCAGTAACCTCAAAAC
GAGCACACGTCTGAACTCTTTCCCTACACGACGCTCTTCCGATCTNNNTAAATGGAGTAGCCCAGTACT
GCACACGTCTGAACTCTTTCCCTACACGACGCTCTTCCGATCTNNNTGCCTTTGCTTTCTTTTCAATTCCC
AGCACACGTCTGAACTCTTTCCCTACACGACGCTCTTCCGATCTNNNTCTCCCTACCCCGTCATTCATATC
AGCACACGTCTGAACTCTTTCCCTACACGACGCTCTTCCGATCTNNNCACTGCATCCTTGAATTCCAGTG
GAGCACACGTCTGAACTCTTTCCCTACACGACGCTCTTCCGATCTNNNGGTGGTAAGTGGCACCTTATC
GAGCACACGTCTGAACTCTTTCCCTACACGACGCTCTTCCGATCTNNNTCCTACTTACCAAGGAAGGGT
AGCACACGTCTGAACTCTTTCCCTACACGACGCTCTTCCGATCTNNNCTGTAATCCCAGCTACTTGGCAC
AGCACACGTCTGAACTCTTTCCCTACACGACGCTCTTCCGATCTNNNCAGGAGACTTGCTTGAACCTGA
GAGCACACGTCTGAACTCTTTCCCTACACGACGCTCTTCCGATCTNNNGGAGAATCACTTGAACCCAGT
AGAGCACACGTCTGAACTCTTTCCCTACACGACGCTCTTCCGATCTNNNAATCACTTGAACTCAGGAGG

GAGCACACGTCTGAACTCTTTCCCTACACGACGCTCTTCCGATCTNNNTGAGGCAGGAGAATCACTTG
GAGCACACGTCTGAACTCTTTCCCTACACGACGCTCTTCCGATCTNNNAGATAGTTTGAGACCAGCCCTT
GAGCACACGTCTGAACTCTTTCCCTACACGACGCTCTTCCGATCTNNNCATGAGAATTGCTTGAACCTG
AGCACACGTCTGAACTCTTTCCCTACACGACGCTCTTCCGATCTNNNAGGCAGAAGAATCACTTGAAC
GAGCACACGTCTGAACTCTTTCCCTACACGACGCTCTTCCGATCTNNNGAGAATCACTTGAACTCGGGA
GAGCACACGTCTGAACTCTTTCCCTACACGACGCTCTTCCGATCTNNNTGTAGTCCTAGCTACTCTGGAG
AGAGCACACGTCTGAACTCTTTCCCTACACGACGCTCTTCCGATCTNNNCAGGAGAATGTCATGAACCC
GAGCACACGTCTGAACTCTTTCCCTACACGACGCTCTTCCGATCTNNNGTGGATCATGAGGTCAGGAC
GAGCACACGTCTGAACTCTTTCCCTACACGACGCTCTTCCGATCTNNNGCACATGCCTATAATCCCAGCT
AGAGCACACGTCTGAACTCTTTCCCTACACGACGCTCTTCCGATCTNNNGGGTGATAGAATGAGACTCC
AGAGCACACGTCTGAACTCTTTCCCTACACGACGCTCTTCCGATCTNNNGTAAAAGTTCTTCCTGGCTGC
AGCACACGTCTGAACTCTTTCCCTACACGACGCTCTTCCGATCTNNNCTCCAGCCTTGTCAAAAAGAGT
AAGAGCACACGTCTGAACTCTTTCCCTACACGACGCTCTTCCGATCTNNNGCACCTGTAGTCCTAGCTA
GCACACGTCTGAACTCTTTCCCTACACGACGCTCTTCCGATCTNNNTTGCAGTAAGCCGAGATAGTACC
GAGCACACGTCTGAACTCTTTCCCTACACGACGCTCTTCCGATCTNNNAGCACAGTAGCATGTACGTGT
GAGCACACGTCTGAACTCTTTCCCTACACGACGCTCTTCCGATCTNNNGGAGGCTGAGACAGGAAAAT
GCACACGTCTGAACTCTTTCCCTACACGACGCTCTTCCGATCTNNNGGTGACAAAGTGAGACTCCATCTC
AGCACACGTCTGAACTCTTTCCCTACACGACGCTCTTCCGATCTNNNTGAGGCAGGAGAATCACTTGA
GAGCACACGTCTGAACTCTTTCCCTACACGACGCTCTTCCGATCTNNNTGAGAGACAAGCCTGGAATAT
AGCACACGTCTGAACTCTTTCCCTACACGACGCTCTTCCGATCTNNNTGTAGTCCCAGCTACTTAGGAG
AGCACACGTCTGAACTCTTTCCCTACACGACGCTCTTCCGATCTNNNTGAGGCAAGAGAATTGCTTGA
AGAGCACACGTCTGAACTCTTTCCCTACACGACGCTCTTCCGATCTNNNTGTAATCCTAGCACTTTGGGA
AGCACACGTCTGAACTCTTTCCCTACACGACGCTCTTCCGATCTNNNGCCAAGATCATGCCATTGTACTC
AGCACACGTCTGAACTCTTTCCCTACACGACGCTCTTCCGATCTNNNTGGGAGGAAGAGAGAGAACCT
AGCACACGTCTGAACTCTTTCCCTACACGACGCTCTTCCGATCTNNNAGAATCATATGAACCCAGGAGG
AGCACACGTCTGAACTCTTTCCCTACACGACGCTCTTCCGATCTNNNGAGAATGGTGTAAACCTGGGA
AGCACACGTCTGAACTCTTTCCCTACACGACGCTCTTCCGATCTNNNAGCAAAGGGAATCGCTAAGAT
AGCACACGTCTGAACTCTTTCCCTACACGACGCTCTTCCGATCTNNNAGATGTTGTGGTGAGCTGAGA
GAGCACACGTCTGAACTCTTTCCCTACACGACGCTCTTCCGATCTNNNAAGAGAATTGCTTGAACCCAG
GAGCACACGTCTGAACTCTTTCCCTACACGACGCTCTTCCGATCTNNNCTGTAATCCCAGCTACCAAGGA
GAGCACACGTCTGAACTCTTTCCCTACACGACGCTCTTCCGATCTNNNAAGATATCAAGACCATCCTGG
AGCACACGTCTGAACTCTTTCCCTACACGACGCTCTTCCGATCTNNNGAGACAGGACAATTGCTTGAAC
GAGCACACGTCTGAACTCTTTCCCTACACGACGCTCTTCCGATCTNNNTACTTGGGAGGTTGAGACAGC
AGCACACGTCTGAACTCTTTCCCTACACGACGCTCTTCCGATCTNNNAAACCCATATTCAGGAGGCTGAG
AGCACACGTCTGAACTCTTTCCCTACACGACGCTCTTCCGATCTNNNAGAATTGCTTGAACCCCAGAGAC
AGAGCACACGTCTGAACTCTTTCCCTACACGACGCTCTTCCGATCTNNNCTGAGGCAGGAGAATCACTT
AGCACACGTCTGAACTCTTTCCCTACACGACGCTCTTCCGATCTNNNCAAGGTAGTCTCTGCCTATGCA
GAGCACACGTCTGAACTCTTTCCCTACACGACGCTCTTCCGATCTNNNCTAGGCAACAGAGTGAGACTT
GAGCACACGTCTGAACTCTTTCCCTACACGACGCTCTTCCGATCTNNNATTACCCGAGCTCAGGAATTT
AGCACACGTCTGAACTCTTTCCCTACACGACGCTCTTCCGATCTNNNGTAATCTCAGCTACTCAGGAGGC
GAGCACACGTCTGAACTCTTTCCCTACACGACGCTCTTCCGATCTNNNGAGGTTGCAGTAAGCCAAGA
AGCACACGTCTGAACTCTTTCCCTACACGACGCTCTTCCGATCTNNNGCTGAGACAGGAGAATAGCTTC
AGCACACGTCTGAACTCTTTCCCTACACGACGCTCTTCCGATCTNNNCAGAAGATCAAAGCTGCAGTGA

AGCACACGTCTGAACTCTTTCCCTACACGACGCTCTTCCGATCTNNNTTCTACAATTGGGATGCCAAGG
GAGCACACGTCTGAACTCTTTCCCTACACGACGCTCTTCCGATCTNNNATCAAGAGCATAAGGACAGA
AGCACACGTCTGAACTCTTTCCCTACACGACGCTCTTCCGATCTNNNCCAACATGGTGAAATCCTGTCT
AAGAGCACACGTCTGAACTCTTTCCCTACACGACGCTCTTCCGATCTNNNGAGAATCATTTGAACCCGG
AGCACACGTCTGAACTCTTTCCCTACACGACGCTCTTCCGATCTNNNGCACCTGTAAACCCAGTTACTT
GCACACGTCTGAACTCTTTCCCTACACGACGCTCTTCCGATCTNNNAATCATCTGAATCCTGAGGGTGG
AGCACACGTCTGAACTCTTTCCCTACACGACGCTCTTCCGATCTNNNATCCTAGTACTTTGGGAGTCCG
AGAGCACACGTCTGAACTCTTTCCCTACACGACGCTCTTCCGATCTNNNCAGTGATCTGAGATGGTGA
GAGCACACGTCTGAACTCTTTCCCTACACGACGCTCTTCCGATCTNNNCTATAATCCCAGCTGCTCAGG
AGAGCACACGTCTGAACTCTTTCCCTACACGACGCTCTTCCGATCTNNNGCACATGCCTATAATCCCAG
GAGCACACGTCTGAACTCTTTCCCTACACGACGCTCTTCCGATCTNNNCTGTAATCCCAGCTACTCAGG
GAGCACACGTCTGAACTCTTTCCCTACACGACGCTCTTCCGATCTNNNGCAAGAGAATCACTTGAACT
AGCACACGTCTGAACTCTTTCCCTACACGACGCTCTTCCGATCTNNNGTAATCCCAGCTACTAAGGAGG
AGCACACGTCTGAACTCTTTCCCTACACGACGCTCTTCCGATCTNNNGTCCAAACACTTAGGGAGGCT
GAGCACACGTCTGAACTCTTTCCCTACACGACGCTCTTCCGATCTNNNCAACAAAGACTGAAACTGCGT
GAGCACACGTCTGAACTCTTTCCCTACACGACGCTCTTCCGATCTNNNCTCACATCTGTAATCCCAGCAC
AGCACACGTCTGAACTCTTTCCCTACACGACGCTCTTCCGATCTNNNAGGCAGGAGAATCACTTGAAT
GAGCACACGTCTGAACTCTTTCCCTACACGACGCTCTTCCGATCTNNNCAAAAATTAGCCGGGCATTAT
AGAGCACACGTCTGAACTCTTTCCCTACACGACGCTCTTCCGATCTNNNAGAATTGCTTGAACCCAGGA
AGCACACGTCTGAACTCTTTCCCTACACGACGCTCTTCCGATCTNNNCAGATTTCCCCTTCTAACCTGCT
GAGCACACGTCTGAACTCTTTCCCTACACGACGCTCTTCCGATCTNNNGAGGCAAGGGAATTTCTTGA
GAGCACACGTCTGAACTCTTTCCCTACACGACGCTCTTCCGATCTNNNAAGTTGTAGTGCACCAAGATT
AGAGCACACGTCTGAACTCTTTCCCTACACGACGCTCTTCCGATCTNNNTACGCCTGTAATTCCAGCTAC
GAGCACACGTCTGAACTCTTTCCCTACACGACGCTCTTCCGATCTNNNCTGTAGTCCCAGCTACTCAGA
AGCACACGTCTGAACTCTTTCCCTACACGACGCTCTTCCGATCTNNNGAGTAGCTGGGATTATAGGTG
AGCACACGTCTGAACTCTTTCCCTACACGACGCTCTTCCGATCTNNNTCTCTCTCTCTCTCTCTCCCTC
AGCACACGTCTGAACTCTTTCCCTACACGACGCTCTTCCGATCTNNNAGATTACAGGTGTGAGCCATTG
AGCACACGTCTGAACTCTTTCCCTACACGACGCTCTTCCGATCTNNNTGCAGAAACTTGCTCAAAACAC
AGCACACGTCTGAACTCTTTCCCTACACGACGCTCTTCCGATCTNNNCTCATACACCTGTTGGCTGTTTG
AGCACACGTCTGAACTCTTTCCCTACACGACGCTCTTCCGATCTNNNCACCCTCCCTACATACAGATGTG
AGCACACGTCTGAACTCTTTCCCTACACGACGCTCTTCCGATCTNNNGGTTTGCAAGGATGTTCTTTTC
AGCACACGTCTGAACTCTTTCCCTACACGACGCTCTTCCGATCTNNNTGAGATTACAGGTGCCCACTAA
AGCACACGTCTGAACTCTTTCCCTACACGACGCTCTTCCGATCTNNNGCCCAGCTAAATTCTGAGGTTT
AGCACACGTCTGAACTCTTTCCCTACACGACGCTCTTCCGATCTNNNAGGTTGCGGTTTACATAGAGT
GAGCACACGTCTGAACTCTTTCCCTACACGACGCTCTTCCGATCTNNNGTGACACAGCTGAGTTCCAT
GAGCACACGTCTGAACTCTTTCCCTACACGACGCTCTTCCGATCTNNNTGGTCTCGATTTCCTGACTTTG
AGCACACGTCTGAACTCTTTCCCTACACGACGCTCTTCCGATCTNNNGTGCCCAGCCAACAATATAACT
GCACACGTCTGAACTCTTTCCCTACACGACGCTCTTCCGATCTNNNCCAGGCTTGCCACTAAATTGAAA
AGCACACGTCTGAACTCTTTCCCTACACGACGCTCTTCCGATCTNNNCAGGAAAAGAAGAGTCTGGGG
AGCACACGTCTGAACTCTTTCCCTACACGACGCTCTTCCGATCTNNNTGGGACCAGAAAAAAGAGTCA
GAGCACACGTCTGAACTCTTTCCCTACACGACGCTCTTCCGATCTNNNACCTTCCTAGAACACAGCAAA
AGCACACGTCTGAACTCTTTCCCTACACGACGCTCTTCCGATCTNNNTCATGGCTGTTAGCTCTTGAAC
AGCACACGTCTGAACTCTTTCCCTACACGACGCTCTTCCGATCTNNNTACAGAGATTGCCCATATGCCT

AGCACACGTCTGAACTCTTTCCCTACACGACGCTCTTCCGATCTNNNGCTGCCTGCAATTAGATGAATG
AGCACACGTCTGAACTCTTTCCCTACACGACGCTCTTCCGATCTNNNAAACATGCTTACCTACAGGGGA
AGCACACGTCTGAACTCTTTCCCTACACGACGCTCTTCCGATCTNNNTGATTAGGCTGGTCTTGAACT
AGCACACGTCTGAACTCTTTCCCTACACGACGCTCTTCCGATCTNNNGAGAGGAAGCGAGAGAGAGA
AGCACACGTCTGAACTCTTTCCCTACACGACGCTCTTCCGATCTNNNCTCAAATGATCCTCTTGCCTCAG
GAGCACACGTCTGAACTCTTTCCCTACACGACGCTCTTCCGATCTNNNTTTATGATCCTTTGCTCCTTGC
AGCACACGTCTGAACTCTTTCCCTACACGACGCTCTTCCGATCTNNNGCCCACATAGTAGCCTCACAATA
AGCACACGTCTGAACTCTTTCCCTACACGACGCTCTTCCGATCTNNNGCTCACTGGAAAGAGAGCAAGA
GAGCACACGTCTGAACTCTTTCCCTACACGACGCTCTTCCGATCTNNNTTTGCCTTGGGTAGCCTTTAC
GAGCACACGTCTGAACTCTTTCCCTACACGACGCTCTTCCGATCTNNNACGGGTCTGTTTATGGTCATT
AGCACACGTCTGAACTCTTTCCCTACACGACGCTCTTCCGATCTNNNACTTCTGGGTTCAAGTCATCCT
GAGCACACGTCTGAACTCTTTCCCTACACGACGCTCTTCCGATCTNNNGGGTTTATCAAACATCTGGAG
GAGCACACGTCTGAACTCTTTCCCTACACGACGCTCTTCCGATCTNNNCTATAGCCTCCTCCTGAGTAGC
GAGCACACGTCTGAACTCTTTCCCTACACGACGCTCTTCCGATCTNNNCAGAGCTAAAGGAGAGGACA
AGCACACGTCTGAACTCTTTCCCTACACGACGCTCTTCCGATCTNNNGCCTCCCAAGTAGCTGAGATTA
GAGCACACGTCTGAACTCTTTCCCTACACGACGCTCTTCCGATCTNNNTCTTGCTCAGTGGCATGATCA
AGCACACGTCTGAACTCTTTCCCTACACGACGCTCTTCCGATCTNNNTGTTCCCCCTTCACTATCTCTCTC
GCACACGTCTGAACTCTTTCCCTACACGACGCTCTTCCGATCTNNNCTACAGGCGTGAACATCACTCTT
GAGCACACGTCTGAACTCTTTCCCTACACGACGCTCTTCCGATCTNNNTGCACTCTCATCATCACCAGAA
GAGCACACGTCTGAACTCTTTCCCTACACGACGCTCTTCCGATCTNNNCGGCTTCTACTGGGAATGATA
GAGCACACGTCTGAACTCTTTCCCTACACGACGCTCTTCCGATCTNNNAGTCACCTTGGGTACACTTTCT
AGCACACGTCTGAACTCTTTCCCTACACGACGCTCTTCCGATCTNNNCAAGTAGCTGGGACTATAGGCA
AGCACACGTCTGAACTCTTTCCCTACACGACGCTCTTCCGATCTNNNGGTATCCCCATTCCACTTTAGG
AGCACACGTCTGAACTCTTTCCCTACACGACGCTCTTCCGATCTNNNGAGACTTCAGCTGATCCAGTAG
AGCACACGTCTGAACTCTTTCCCTACACGACGCTCTTCCGATCTNNNCAGAAGGCTGTGTACAAGGTA
GCACACGTCTGAACTCTTTCCCTACACGACGCTCTTCCGATCTNNNTTCAGATACCTTTGGGTTTGGGT
GAGCACACGTCTGAACTCTTTCCCTACACGACGCTCTTCCGATCTNNNGAAGTACACTTGTGCTGAAGC
GAGCACACGTCTGAACTCTTTCCCTACACGACGCTCTTCCGATCTNNNAGGAGAATTGCTTGAATACG
AGCACACGTCTGAACTCTTTCCCTACACGACGCTCTTCCGATCTNNNGCACTACAGACTGGCACATAGT
AGCACACGTCTGAACTCTTTCCCTACACGACGCTCTTCCGATCTNNNATCTGGGTTCTTGTCTCATGAC
AGCACACGTCTGAACTCTTTCCCTACACGACGCTCTTCCGATCTNNNCTGTAGTTCCAGCTACTCGAGA
AGCACACGTCTGAACTCTTTCCCTACACGACGCTCTTCCGATCTNNNAAGTAGGGTTTCCAAGGAGTC
AGCACACGTCTGAACTCTTTCCCTACACGACGCTCTTCCGATCTNNNTTGTTTGAACTCGGGAGGTAG
AGAGCACACGTCTGAACTCTTTCCCTACACGACGCTCTTCCGATCTNNNCTTACAGTGAGCTGAGATCG
GAGCACACGTCTGAACTCTTTCCCTACACGACGCTCTTCCGATCTNNNGCATGCCTGTAATCCTAGCTA
GAGCACACGTCTGAACTCTTTCCCTACACGACGCTCTTCCGATCTNNNGAGAATGGCATGAACCTGGGA
GAGCACACGTCTGAACTCTTTCCCTACACGACGCTCTTCCGATCTNNNTGAGGCAGGAGAATCTCTTGA
GAGCACACGTCTGAACTCTTTCCCTACACGACGCTCTTCCGATCTNNNGGTTGTAGTGATCTGAGATC
GAGCACACGTCTGAACTCTTTCCCTACACGACGCTCTTCCGATCTNNNGAGACTGAGGATGCAATGA
GAGCACACGTCTGAACTCTTTCCCTACACGACGCTCTTCCGATCTNNNCATGGTGAGGCATGCTATCTA
GAGCACACGTCTGAACTCTTTCCCTACACGACGCTCTTCCGATCTNNNTGGGTAACATAGTGAGACCC
AGCACACGTCTGAACTCTTTCCCTACACGACGCTCTTCCGATCTNNNACCTTCCCTCCACTATTGTCCTA
GAGCACACGTCTGAACTCTTTCCCTACACGACGCTCTTCCGATCTNNNGGCTAAGGCAGGAAAATCAC

GAGCACACGTCTGAACTCTTTCCCTACACGACGCTCTTCCGATCTNNNGTAATCTCAGCTACTCAGGAG
GAGCACACGTCTGAACTCTTTCCCTACACGACGCTCTTCCGATCTNNNTGAGATGGGAGAATCACTGG
AGCACACGTCTGAACTCTTTCCCTACACGACGCTCTTCCGATCTNNNTCTGTAATCCTAGCTACTCGGG
GAGCACACGTCTGAACTCTTTCCCTACACGACGCTCTTCCGATCTNNNGAGATCGCACCAGTTCATTC
GCACACGTCTGAACTCTTTCCCTACACGACGCTCTTCCGATCTNNNGGAGTGAGCTGAAATTGTGTC
GAGCACACGTCTGAACTCTTTCCCTACACGACGCTCTTCCGATCTNNNCCTAAGTGACAGCAAGACTC
GAGCACACGTCTGAACTCTTTCCCTACACGACGCTCTTCCGATCTNNNGGCAACAGAGCGAGATTATG
GAGCACACGTCTGAACTCTTTCCCTACACGACGCTCTTCCGATCTNNNTGAGCTGAGATCATGCCATT
AGAGCACACGTCTGAACTCTTTCCCTACACGACGCTCTTCCGATCTNNNGCCTGTAATCCCAGCTACTC
AGAGCACACGTCTGAACTCTTTCCCTACACGACGCTCTTCCGATCTNNNCTGAGGCAGGAGAATCGTT
AGCACACGTCTGAACTCTTTCCCTACACGACGCTCTTCCGATCTNNNGAGACAGGAGAATTGCTTGAA
GAGCACACGTCTGAACTCTTTCCCTACACGACGCTCTTCCGATCTNNNACCCTAGGCAGTGTTTCTCAA
AAGAGCACACGTCTGAACTCTTTCCCTACACGACGCTCTTCCGATCTNNNGAGCTGAGATTGTGCCAC
GAGCACACGTCTGAACTCTTTCCCTACACGACGCTCTTCCGATCTNNNCCAAGATCATGTCATTGCACT
AGCACACGTCTGAACTCTTTCCCTACACGACGCTCTTCCGATCTNNNGAGGCTACAGTGAGCTGAGAT
AGAGCACACGTCTGAACTCTTTCCCTACACGACGCTCTTCCGATCTNNNTGCACACCTGTAATCCTAGC
AGCACACGTCTGAACTCTTTCCCTACACGACGCTCTTCCGATCTNNNGTGACAGAGTGAGACTCCTTC
GAGCACACGTCTGAACTCTTTCCCTACACGACGCTCTTCCGATCTNNNCACTTGAGGTCAGGAGTTCA
AGCACACGTCTGAACTCTTTCCCTACACGACGCTCTTCCGATCTNNNCGCCTATAATCCCAGCTACTCAG
GAGCACACGTCTGAACTCTTTCCCTACACGACGCTCTTCCGATCTNNNAGACATGAGAATCGCTTGAAC
AGCACACGTCTGAACTCTTTCCCTACACGACGCTCTTCCGATCTNNNAAGAATTGCTTGATCTGGGAG
AGAGCACACGTCTGAACTCTTTCCCTACACGACGCTCTTCCGATCTNNNCATGCCTGTAATCCCAGCTA
GAGCACACGTCTGAACTCTTTCCCTACACGACGCTCTTCCGATCTNNNCTGTAGTCCCAGCTACTCAGG
AGCACACGTCTGAACTCTTTCCCTACACGACGCTCTTCCGATCTNNNGTGGGAGGATCACTTGAGTTC
GAGCACACGTCTGAACTCTTTCCCTACACGACGCTCTTCCGATCTNNNGAATCGCTTGAACCTGGAAA
GAGCACACGTCTGAACTCTTTCCCTACACGACGCTCTTCCGATCTNNNGAGAAAAAGAGACTGGAGC
AGCACACGTCTGAACTCTTTCCCTACACGACGCTCTTCCGATCTNNNTCTGTTCTCTGAAGATCTCCCC
AGCACACGTCTGAACTCTTTCCCTACACGACGCTCTTCCGATCTNNNCTGGGTAACACAGTGACTCTC
AGCACACGTCTGAACTCTTTCCCTACACGACGCTCTTCCGATCTNNNCACTGTGAGACTTGAGAGCTC
GAGCACACGTCTGAACTCTTTCCCTACACGACGCTCTTCCGATCTNNNGGAACATCAGGAACAATGCC
GCACACGTCTGAACTCTTTCCCTACACGACGCTCTTCCGATCTNNNACACCATGAATGTTTGCTGGAAG
GAGCACACGTCTGAACTCTTTCCCTACACGACGCTCTTCCGATCTNNNTCAACCTAGAAGCCTTTCCTG
AGCACACGTCTGAACTCTTTCCCTACACGACGCTCTTCCGATCTNNNCTAAACGGTTTACTCTGTCCCC
GAGCACACGTCTGAACTCTTTCCCTACACGACGCTCTTCCGATCTNNNCTGGCCTTGTTCCTTACATTTG
GAGCACACGTCTGAACTCTTTCCCTACACGACGCTCTTCCGATCTNNNAAAAGACTTTAGGGAGCTGG
GAGCACACGTCTGAACTCTTTCCCTACACGACGCTCTTCCGATCTNNNTGGATGTTCAACAAGTGAGA
GAGCACACGTCTGAACTCTTTCCCTACACGACGCTCTTCCGATCTNNNTGTAATCCCAGCAGTTTAGGA
GAGCACACGTCTGAACTCTTTCCCTACACGACGCTCTTCCGATCTNNNTATGTAGTACCGAACTGTGTG
GAGCACACGTCTGAACTCTTTCCCTACACGACGCTCTTCCGATCTNNNGGTGACAGAGTGAGACTCCA
GAGCACACGTCTGAACTCTTTCCCTACACGACGCTCTTCCGATCTNNNTGACAGACACATGTGATCCTA
GAGCACACGTCTGAACTCTTTCCCTACACGACGCTCTTCCGATCTNNNTGGGTGACAGAATGAAACTC
GAGCACACGTCTGAACTCTTTCCCTACACGACGCTCTTCCGATCTNNNGGCAACATGGTGAAACATT
GAGCACACGTCTGAACTCTTTCCCTACACGACGCTCTTCCGATCTNNNTGTCCATTCATTCAAACCACGT

AGCACACGTCTGAACTCTTTCCCTACACGACGCTCTTCCGATCTNNNTGAGAGACTACTGGGAACTCAG
GAGCACACGTCTGAACTCTTTCCCTACACGACGCTCTTCCGATCTNNNGCAACGTAGCAACAAAGGTTT
GAGCACACGTCTGAACTCTTTCCCTACACGACGCTCTTCCGATCTNNNGACCAAGGCTGTCCATTTTGA
GAGCACACGTCTGAACTCTTTCCCTACACGACGCTCTTCCGATCTNNNCAGAAGAATTGCTTGAACCCAG
AGAGCACACGTCTGAACTCTTTCCCTACACGACGCTCTTCCGATCTNNNATAAATGAACCCAACCCCTCC
GAGCACACGTCTGAACTCTTTCCCTACACGACGCTCTTCCGATCTNNNCTGTAGTCCCAGCTACTCAGGA
AGCACACGTCTGAACTCTTTCCCTACACGACGCTCTTCCGATCTNNNGGCCTGGTAAGCTGCAAATATTT
AGCACACGTCTGAACTCTTTCCCTACACGACGCTCTTCCGATCTNNNTCTTCCTTGCAACCTAGAATGCTG
AGCACACGTCTGAACTCTTTCCCTACACGACGCTCTTCCGATCTNNNGTGGTAGCTAAGGGGTTTACTG
AGCACACGTCTGAACTCTTTCCCTACACGACGCTCTTCCGATCTNNNGAGTGCACAACTGTTTCCTTCTC
AGCACACGTCTGAACTCTTTCCCTACACGACGCTCTTCCGATCTNNNATCTCATGTGCCCTAAGTACAGG
AGCACACGTCTGAACTCTTTCCCTACACGACGCTCTTCCGATCTNNNATGAACCTGGGAGATGGAAGT
AGCACACGTCTGAACTCTTTCCCTACACGACGCTCTTCCGATCTNNNCTGTAGTTCCAGCTACTTGGGAG
GAGCACACGTCTGAACTCTTTCCCTACACGACGCTCTTCCGATCTNNNTGTGAGCATACCACTGCATTCT
AGCACACGTCTGAACTCTTTCCCTACACGACGCTCTTCCGATCTNNNAGCCCTTCTTGCTGTTTTGATTT
AGAGCACACGTCTGAACTCTTTCCCTACACGACGCTCTTCCGATCTNNNGCCTAGTTGACAGAGCAAGA
AAGAGCACACGTCTGAACTCTTTCCCTACACGACGCTCTTCCGATCTNNNGAGGCAGGAGAATGGTG
GAGCACACGTCTGAACTCTTTCCCTACACGACGCTCTTCCGATCTNNNCTGAGGCAGGAGAATGTCATG
AGCACACGTCTGAACTCTTTCCCTACACGACGCTCTTCCGATCTNNNAGTGGCACAATCACAACTCATT
GCACACGTCTGAACTCTTTCCCTACACGACGCTCTTCCGATCTNNNGAGAATCACTCGAACCTGGGAAG
AGCACACGTCTGAACTCTTTCCCTACACGACGCTCTTCCGATCTNNNAATACACACTAAGAACGGGCTG
GAGCACACGTCTGAACTCTTTCCCTACACGACGCTCTTCCGATCTNNNTGCTGGCCTATAATCCCAGTTA
AGCACACGTCTGAACTCTTTCCCTACACGACGCTCTTCCGATCTNNNAGAGCTTGCATTGAGCCAAAAT
GCACACGTCTGAACTCTTTCCCTACACGACGCTCTTCCGATCTNNNGATCAATTAGAGCCTGGGAGGT
AAGAGCACACGTCTGAACTCTTTCCCTACACGACGCTCTTCCGATCTNNNCTGTAATCCCAGCTACTCAG
AGCACACGTCTGAACTCTTTCCCTACACGACGCTCTTCCGATCTNNNCTGAGGCAGGAGAATCACTTGA
AGCACACGTCTGAACTCTTTCCCTACACGACGCTCTTCCGATCTNNNGAGAATGGTGTGAAACTGGGA
GAGCACACGTCTGAACTCTTTCCCTACACGACGCTCTTCCGATCTNNNGAGGCAGGAAAATGCTGTGAA
GAGCACACGTCTGAACTCTTTCCCTACACGACGCTCTTCCGATCTNNNTGACAAGTCTGGGAAGAGATT
AGCACACGTCTGAACTCTTTCCCTACACGACGCTCTTCCGATCTNNNCACAAATGTTGACAAGCAAAGG
AGCACACGTCTGAACTCTTTCCCTACACGACGCTCTTCCGATCTNNNGATGGTCTCAATCTCCTGACCTC
GAGCACACGTCTGAACTCTTTCCCTACACGACGCTCTTCCGATCTNNNATGGTTCCAATAGAGGACCAT
AGCACACGTCTGAACTCTTTCCCTACACGACGCTCTTCCGATCTNNNACTGTTCTTGTGGGGAAGCTAA
AGCACACGTCTGAACTCTTTCCCTACACGACGCTCTTCCGATCTNNNTCGGAGTAGCTAGGACTATAGG
GAGCACACGTCTGAACTCTTTCCCTACACGACGCTCTTCCGATCTNNNTTCAGAACAGTTCCATCACCTC
GAGCACACGTCTGAACTCTTTCCCTACACGACGCTCTTCCGATCTNNNCATGTTCTAGGAGTTCCAGACC
GCACACGTCTGAACTCTTTCCCTACACGACGCTCTTCCGATCTNNNTCTTTGTGTTCTTTCAAATGGGTG
AGCACACGTCTGAACTCTTTCCCTACACGACGCTCTTCCGATCTNNNCGCACCTGGCCTAGAGTAATATT
AGCACACGTCTGAACTCTTTCCCTACACGACGCTCTTCCGATCTNNNATTTTAGAGACAGCAGACTCGGG
GAGCACACGTCTGAACTCTTTCCCTACACGACGCTCTTCCGATCTNNNAGCCAGCTCTCTAAAGAAGAA
AGAGCACACGTCTGAACTCTTTCCCTACACGACGCTCTTCCGATCTNNNTCAAGGTTATGCTGGCCTCAT
AGCACACGTCTGAACTCTTTCCCTACACGACGCTCTTCCGATCTNNNAAGTGTTGGGATTACAGGTGTG
GAGCACACGTCTGAACTCTTTCCCTACACGACGCTCTTCCGATCTNNNTGGTTACTGCTCTGTTATTGG

AGCACACGTCTGAACTCTTTCCCTACACGACGCTCTTCCGATCTNNNCTCGAATTCCTGACCTCGTGAT
AGCACACGTCTGAACTCTTTCCCTACACGACGCTCTTCCGATCTNNNAGATGAGATTTGGGTGAGGA
AGCACACGTCTGAACTCTTTCCCTACACGACGCTCTTCCGATCTNNNACTTCAGCCTCTTGAGTAACTG
GCACACGTCTGAACTCTTTCCCTACACGACGCTCTTCCGATCTNNNTCCCAAAGTGACAGGATTACAG
AGCACACGTCTGAACTCTTTCCCTACACGACGCTCTTCCGATCTNNNAAGGGCAGAACAAACATGAA
AGCACACGTCTGAACTCTTTCCCTACACGACGCTCTTCCGATCTNNNCCTCTGCTGTCTCACTAGTTAG
AGCACACGTCTGAACTCTTTCCCTACACGACGCTCTTCCGATCTNNNGCTTGGTGAGGACTGTGTAA
GCACACGTCTGAACTCTTTCCCTACACGACGCTCTTCCGATCTNNNTCCTGAGTAGCTGGGATTACA
GCACACGTCTGAACTCTTTCCCTACACGACGCTCTTCCGATCTNNNAATCACCTGGTCATTGTCTTTG
GCACACGTCTGAACTCTTTCCCTACACGACGCTCTTCCGATCTNNNCCTGGAAAACAAGAGCGAAAC
AGCACACGTCTGAACTCTTTCCCTACACGACGCTCTTCCGATCTNNNTGGGCAACAGAGTGATAACTT
AGCACACGTCTGAACTCTTTCCCTACACGACGCTCTTCCGATCTNNNCGCACACCTATAGTTCCAGCTA
AGCACACGTCTGAACTCTTTCCCTACACGACGCTCTTCCGATCTNNNAATCACTTGAACTCAGGAGGT
AGCACACGTCTGAACTCTTTCCCTACACGACGCTCTTCCGATCTNNNCTGTAATCCCAGCTACTCAGG
GAGCACACGTCTGAACTCTTTCCCTACACGACGCTCTTCCGATCTNNNCATTTGAGCCCAAGAAACAG
AGCACACGTCTGAACTCTTTCCCTACACGACGCTCTTCCGATCTNNNGCTGTGATTGCACTACTGTGT
GCACACGTCTGAACTCTTTCCCTACACGACGCTCTTCCGATCTNNNACTCCCACTAACAGCGTAAAG
AGCACACGTCTGAACTCTTTCCCTACACGACGCTCTTCCGATCTNNNCTCCTGAGTAGCTGGGACTAC
AGCACACGTCTGAACTCTTTCCCTACACGACGCTCTTCCGATCTNNNCTGGAGTTCAGTGGCATGATT
AGCACACGTCTGAACTCTTTCCCTACACGACGCTCTTCCGATCTNNNCTCACTGTATCCTTGACCTCC
AGCACACGTCTGAACTCTTTCCCTACACGACGCTCTTCCGATCTNNNCAGGATTTGAGAACAGCTTG
AGCACACGTCTGAACTCTTTCCCTACACGACGCTCTTCCGATCTNNNTTGTGTCTACGTTCACCAGTG
AGCACACGTCTGAACTCTTTCCCTACACGACGCTCTTCCGATCTNNNTGGCTAACACAGTGAAATCCT
GAGCACACGTCTGAACTCTTTCCCTACACGACGCTCTTCCGATCTNNNCTTGAGCCCAAGGAGTATGA
AGCACACGTCTGAACTCTTTCCCTACACGACGCTCTTCCGATCTNNNACACACACACACACACACTA
AGCACACGTCTGAACTCTTTCCCTACACGACGCTCTTCCGATCTNNNCCAGCATGTGGTATTGTCAAA
GCACACGTCTGAACTCTTTCCCTACACGACGCTCTTCCGATCTNNNGGAGTCCTAAGTAACCATGGTG
GAGCACACGTCTGAACTCTTTCCCTACACGACGCTCTTCCGATCTNNNATACACCAACAGCAAACAAG
AGCACACGTCTGAACTCTTTCCCTACACGACGCTCTTCCGATCTNNNCTCCAGTTCCATCCAACTTGTT
AGCACACGTCTGAACTCTTTCCCTACACGACGCTCTTCCGATCTNNNTGAGGCAGGAGAATAGCTTGA
AGCACACGTCTGAACTCTTTCCCTACACGACGCTCTTCCGATCTNNNTGAGACAGGAGAATCGCTTG
GAGCACACGTCTGAACTCTTTCCCTACACGACGCTCTTCCGATCTNNNGTCTGTCTTTCACACACACACA
GCACACGTCTGAACTCTTTCCCTACACGACGCTCTTCCGATCTNNNACTCCTGTATTCACTGCAGCATT
GCACACGTCTGAACTCTTTCCCTACACGACGCTCTTCCGATCTNNNTGATGGGTCTATGGGTCTGTA
AGCACACGTCTGAACTCTTTCCCTACACGACGCTCTTCCGATCTNNNAGGAATGAGGTTGGTAGGAG
AGCACACGTCTGAACTCTTTCCCTACACGACGCTCTTCCGATCTNNNCTATCATGGCTCACTGCAACCT
GCACACGTCTGAACTCTTTCCCTACACGACGCTCTTCCGATCTNNNAACCCCTTCTCTACTAACAACCC
AGCACACGTCTGAACTCTTTCCCTACACGACGCTCTTCCGATCTNNNGATTCCCTTCAACAAATGGTC
GAGCACACGTCTGAACTCTTTCCCTACACGACGCTCTTCCGATCTNNNTGGCAAAACCTCATCTCTACC
GCACACGTCTGAACTCTTTCCCTACACGACGCTCTTCCGATCTNNNACAGTGGGAGGAGAAGAATG
GAGCACACGTCTGAACTCTTTCCCTACACGACGCTCTTCCGATCTNNNGAAGAAAAGCTGCCCAGTTC
AGCACACGTCTGAACTCTTTCCCTACACGACGCTCTTCCGATCTNNNGGTAACATAGCAAGACCCCTTC
AGCACACGTCTGAACTCTTTCCCTACACGACGCTCTTCCGATCTNNNGAGGACCTGCTGTAATTCCCTA

GAGCACACGTCTGAACTCTTTCCCTACACGACGCTCTTCCGATCTNNNGTGTCCAATGATGAAACTGG
CACACGTCTGAACTCTTTCCCTACACGACGCTCTTCCGATCTNNNACACATACCCCAAATGCTTACCTCA
AGCACACGTCTGAACTCTTTCCCTACACGACGCTCTTCCGATCTNNNCCTGAACACTCACTACTTCCCAT
AGCACACGTCTGAACTCTTTCCCTACACGACGCTCTTCCGATCTNNNACAACAACTAGGATGGGTGGA
AGCACACGTCTGAACTCTTTCCCTACACGACGCTCTTCCGATCTNNNCAATCTTGGCTTACTGCAACCTC
GCACACGTCTGAACTCTTTCCCTACACGACGCTCTTCCGATCTNNNCCCCTACAAGCTTTACTGCACTA
GAGCACACGTCTGAACTCTTTCCCTACACGACGCTCTTCCGATCTNNNTTCTGACTGTGTAGGTCTGT
AGCACACGTCTGAACTCTTTCCCTACACGACGCTCTTCCGATCTNNNCAGAGGTTGCAGAGAACTGA
GAGCACACGTCTGAACTCTTTCCCTACACGACGCTCTTCCGATCTNNNGTCAACAGTCCCTGAAAAACA
GCACACGTCTGAACTCTTTCCCTACACGACGCTCTTCCGATCTNNNCTTAGGGTGTGTGTTTCAATGC
GAGCACACGTCTGAACTCTTTCCCTACACGACGCTCTTCCGATCTNNNATACCATCACAAAACCCCCAAG
GAGCACACGTCTGAACTCTTTCCCTACACGACGCTCTTCCGATCTNNNTGGATTACTTGAGCTCAGGC
GCACACGTCTGAACTCTTTCCCTACACGACGCTCTTCCGATCTNNNACAGAGAGAGAGAGAGAGAGA
GCACACGTCTGAACTCTTTCCCTACACGACGCTCTTCCGATCTNNNAGAGAGAGAGGGAGAGAGAG
GCACACGTCTGAACTCTTTCCCTACACGACGCTCTTCCGATCTNNNTGGTACTTAGCCTGCATAGTCAC
AGAGCACACGTCTGAACTCTTTCCCTACACGACGCTCTTCCGATCTNNNAGAGATCGCGTCACTGTAT
AGAGCACACGTCTGAACTCTTTCCCTACACGACGCTCTTCCGATCTNNNGTGAGTGTGTGGAAGAGA
AGCACACGTCTGAACTCTTTCCCTACACGACGCTCTTCCGATCTNNNTCATTGGTTGGGTTCTGGTGA
GCACACGTCTGAACTCTTTCCCTACACGACGCTCTTCCGATCTNNNTCCAGGGTTCACATACTATCCCT
GAGCACACGTCTGAACTCTTTCCCTACACGACGCTCTTCCGATCTNNNATATGGAGAGAGAGAGAG
AGAGCACACGTCTGAACTCTTTCCCTACACGACGCTCTTCCGATCTNNNAGGAAGCAGATGAAAAGT
CACACGTCTGAACTCTTTCCCTACACGACGCTCTTCCGATCTNNNCTCTCTCTCTCTCTCGCTCTCCAT
AGCACACGTCTGAACTCTTTCCCTACACGACGCTCTTCCGATCTNNNTGCTCTTCTGTTCTCAGTTTGGA
GAGCACACGTCTGAACTCTTTCCCTACACGACGCTCTTCCGATCTNNNCTAAGGTGGGAGGATCACTT
AGAGCACACGTCTGAACTCTTTCCCTACACGACGCTCTTCCGATCTNNNCAATGCAACAGAGGCCAAA
GAGCACACGTCTGAACTCTTTCCCTACACGACGCTCTTCCGATCTNNNCTGTACTCCAGCCTGAGTAAC
AGCACACGTCTGAACTCTTTCCCTACACGACGCTCTTCCGATCTNNNTTCCCCTCCATTTCCATACTGTG
AGCACACGTCTGAACTCTTTCCCTACACGACGCTCTTCCGATCTNNNCCAGGCCAGAACTTTCCTTTTT
AGCACACGTCTGAACTCTTTCCCTACACGACGCTCTTCCGATCTNNNAAAGTGGGAATTTCTTGCCCT
AGCACACGTCTGAACTCTTTCCCTACACGACGCTCTTCCGATCTNNNGTATCGAACTCTTGACCTCGTG
AGCACACGTCTGAACTCTTTCCCTACACGACGCTCTTCCGATCTNNNAACGTGTTCTCAAAATGGCTC
AGCACACGTCTGAACTCTTTCCCTACACGACGCTCTTCCGATCTNNNAGAAAGATTTCTGCCCAAACC
AGCACACGTCTGAACTCTTTCCCTACACGACGCTCTTCCGATCTNNNCAGCTACACTTTGTACCTCTGG
GAGCACACGTCTGAACTCTTTCCCTACACGACGCTCTTCCGATCTNNNCTTAGGCAGGGTCTTACTGT
GAGCACACGTCTGAACTCTTTCCCTACACGACGCTCTTCCGATCTNNNTTGAAGTTCCTGCCTCTGTCT
AGCACACGTCTGAACTCTTTCCCTACACGACGCTCTTCCGATCTNNNCTGTCTCTTGCTGCTTTCAAGA
AGCACACGTCTGAACTCTTTCCCTACACGACGCTCTTCCGATCTNNNGTGTGAGTACAAGAGTCCAGA
AGCACACGTCTGAACTCTTTCCCTACACGACGCTCTTCCGATCTNNNAAATACTACTGGCCTTGTCTGC
GCACACGTCTGAACTCTTTCCCTACACGACGCTCTTCCGATCTNNNCTGAAGCATGCATGGATCTCAA
AGCACACGTCTGAACTCTTTCCCTACACGACGCTCTTCCGATCTNNNAGTGCTAGTATTACAGGCGTGA
AGCACACGTCTGAACTCTTTCCCTACACGACGCTCTTCCGATCTNNNTTTCTGTCTCAACAAGTCCCTCC
GAGCACACGTCTGAACTCTTTCCCTACACGACGCTCTTCCGATCTNNNGCCTTTTTGACTGATGGTTTG
AGCACACGTCTGAACTCTTTCCCTACACGACGCTCTTCCGATCTNNNTGTGTATTAGAGGGCTCACTG

AGCACACGTCTGAACTCTTTCCCTACACGACGCTCTTCCGATCTNNNATGAAGAACCTGTGGTGTTG
AGCACACGTCTGAACTCTTTCCCTACACGACGCTCTTCCGATCTNNNTTTTTTGAGTGGTGCAATCTCG
ACACGTCTGAACTCTTTCCCTACACGACGCTCTTCCGATCTNNNTTGAAGTTATGTTTCTAGGTGCTTC
AGCACACGTCTGAACTCTTTCCCTACACGACGCTCTTCCGATCTNNNGACCTCCCCTTTGGTACTCTTTT
AGCACACGTCTGAACTCTTTCCCTACACGACGCTCTTCCGATCTNNNTGTTTCCAGCTGAACTTAGCAG
AGCACACGTCTGAACTCTTTCCCTACACGACGCTCTTCCGATCTNNNCACCTGTAATCCCAGCTACTTG
AGCACACGTCTGAACTCTTTCCCTACACGACGCTCTTCCGATCTNNNCTTGCAGTAAGCTGAGATCAG
AGAGCACACGTCTGAACTCTTTCCCTACACGACGCTCTTCCGATCTNNNAGAATTTGCAGTGAGCCGA
GCACACGTCTGAACTCTTTCCCTACACGACGCTCTTCCGATCTNNNCTGAAGTGGGAAGATCACTGG
AGCACACGTCTGAACTCTTTCCCTACACGACGCTCTTCCGATCTNNNAGGTTGCAGTGAGCTGATATC
AGCACACGTCTGAACTCTTTCCCTACACGACGCTCTTCCGATCTNNNACACCTGTAATCCCAGCACTTTG
GAGCACACGTCTGAACTCTTTCCCTACACGACGCTCTTCCGATCTNNNGGAAATTGCAGTGAGCCAAG
GAGCACACGTCTGAACTCTTTCCCTACACGACGCTCTTCCGATCTNNNAAAAATTAGCTACTGGGGAG
GCACACGTCTGAACTCTTTCCCTACACGACGCTCTTCCGATCTNNNCAAGAGGCTGAGGTAGGAGAA
AGCACACGTCTGAACTCTTTCCCTACACGACGCTCTTCCGATCTNNNGAGCTTTCAGTGAGCCGAGAT
GAGCACACGTCTGAACTCTTTCCCTACACGACGCTCTTCCGATCTNNNCTGTAATCCCAGCTACTCAGG
AGCACACGTCTGAACTCTTTCCCTACACGACGCTCTTCCGATCTNNNAGATTGTGGCACTGTACTCTA
AGCACACGTCTGAACTCTTTCCCTACACGACGCTCTTCCGATCTNNNGGCACCTGTAATCCTAGCTACT
AGCACACGTCTGAACTCTTTCCCTACACGACGCTCTTCCGATCTNNNCAGATTGAGACCATCCTGGCT
AGCACACGTCTGAACTCTTTCCCTACACGACGCTCTTCCGATCTNNNCTGGGCAACATATGAGACTCC
GAGCACACGTCTGAACTCTTTCCCTACACGACGCTCTTCCGATCTNNNTCACAAGGTCAGGAGTTCAA
AGCACACGTCTGAACTCTTTCCCTACACGACGCTCTTCCGATCTNNNTGTTTGGGAGGTGAATGTTCT
AGCACACGTCTGAACTCTTTCCCTACACGACGCTCTTCCGATCTNNNGGGTAACAAGCTGTGATTGCA
GAGCACACGTCTGAACTCTTTCCCTACACGACGCTCTTCCGATCTNNNGGCAAGACCCAGAATAAGAC
AGCACACGTCTGAACTCTTTCCCTACACGACGCTCTTCCGATCTNNNACCAACAGTAACCAAGCTGAG
AGCACACGTCTGAACTCTTTCCCTACACGACGCTCTTCCGATCTNNNGGGCAACAAGAACAAAACTC
AGCACACGTCTGAACTCTTTCCCTACACGACGCTCTTCCGATCTNNNCTTCGGGTGTTCATTCAAAAGC
AGCACACGTCTGAACTCTTTCCCTACACGACGCTCTTCCGATCTNNNCGCCTATAATCCCAGCTACTCAG
AGCACACGTCTGAACTCTTTCCCTACACGACGCTCTTCCGATCTNNNACTGGGAAGTGTGAGTTATA
AGCACACGTCTGAACTCTTTCCCTACACGACGCTCTTCCGATCTNNNTCCTGTCCTTTGATCCTTTCCTG
AGCACACGTCTGAACTCTTTCCCTACACGACGCTCTTCCGATCTNNNGATGCTCGAAGTCTGAATTCTG
GCACACGTCTGAACTCTTTCCCTACACGACGCTCTTCCGATCTNNNCAAGGGCTCAATCCAATCTTTC
AGCACACGTCTGAACTCTTTCCCTACACGACGCTCTTCCGATCTNNNCTTCACACATCAGTCAGTGGAG
AGCACACGTCTGAACTCTTTCCCTACACGACGCTCTTCCGATCTNNNGCACAGTTTCAGAGAAGGAC
GCACACGTCTGAACTCTTTCCCTACACGACGCTCTTCCGATCTNNNCTAACAGGTGTGAGTTGTTCCC
AGCACACGTCTGAACTCTTTCCCTACACGACGCTCTTCCGATCTNNNTGATGTAGGTACTTGAGGCTC
AGCACACGTCTGAACTCTTTCCCTACACGACGCTCTTCCGATCTNNNAGTAGGGCTTAGGACACAAAT
AGCACACGTCTGAACTCTTTCCCTACACGACGCTCTTCCGATCTNNNGATATCCTGACTCGTGATCCAC
AGCACACGTCTGAACTCTTTCCCTACACGACGCTCTTCCGATCTNNNATCAACTTCTTCCCAACTCCTGT
GAGCACACGTCTGAACTCTTTCCCTACACGACGCTCTTCCGATCTNNNCTTCTTGTAGCAGCTACGAGT
AGCACACGTCTGAACTCTTTCCCTACACGACGCTCTTCCGATCTNNNTCTTCACTGTCACCTCACTAAC
GAGCACACGTCTGAACTCTTTCCCTACACGACGCTCTTCCGATCTNNNCAATCAAAAGAGTGAGGCAG
AGCACACGTCTGAACTCTTTCCCTACACGACGCTCTTCCGATCTNNNCCCGAAATGGAAAACCTCTGA

AGCACACGTCTGAACTCTTTCCCTACACGACGCTCTTCCGATCTNNNCTGCAGGTTTCTGTGCCTTTTA
AGCACACGTCTGAACTCTTTCCCTACACGACGCTCTTCCGATCTNNNTGGTCTTTGTCCCTATTTCCTGG
GAGCACACGTCTGAACTCTTTCCCTACACGACGCTCTTCCGATCTNNNCTCACGAGTAGCTGGGACTA
AGCACACGTCTGAACTCTTTCCCTACACGACGCTCTTCCGATCTNNNTGGAAAAGTTAGTGGATGGC
GAGCACACGTCTGAACTCTTTCCCTACACGACGCTCTTCCGATCTNNNGCTGACCACAAAGAAACTCTG
AGCACACGTCTGAACTCTTTCCCTACACGACGCTCTTCCGATCTNNNGTAGAGACAGAGTTTCACCGT
GAGCACACGTCTGAACTCTTTCCCTACACGACGCTCTTCCGATCTNNNCTGATTCAGCAGGACTAAGT
AGCACACGTCTGAACTCTTTCCCTACACGACGCTCTTCCGATCTNNNTGTGTGTGTGTGTGTGTAT
AGCACACGTCTGAACTCTTTCCCTACACGACGCTCTTCCGATCTNNNGACTACAGAACTACAGCCACC
GCACACGTCTGAACTCTTTCCCTACACGACGCTCTTCCGATCTNNNTCTGTGAGCATACCCTTTCGATT
GAGCACACGTCTGAACTCTTTCCCTACACGACGCTCTTCCGATCTNNNGCAGTGATGCGATCATAGTT
GCACACGTCTGAACTCTTTCCCTACACGACGCTCTTCCGATCTNNNCTGGGTTTCAAGCACAAAACTG
GCACACGTCTGAACTCTTTCCCTACACGACGCTCTTCCGATCTNNNTGCCTAAAGAGAGACGTAGAA
GCACACGTCTGAACTCTTTCCCTACACGACGCTCTTCCGATCTNNNCAGGAGGAGGAAGAAAAGTCC
AGCACACGTCTGAACTCTTTCCCTACACGACGCTCTTCCGATCTNNNCTTTGTCTGCAACTGCTCCTAG
GAGCACACGTCTGAACTCTTTCCCTACACGACGCTCTTCCGATCTNNNCATTGGAGTCCCTATTGGTCT
AGCACACGTCTGAACTCTTTCCCTACACGACGCTCTTCCGATCTNNNCTTATGTGCTCATTGGCCATCT
AGCACACGTCTGAACTCTTTCCCTACACGACGCTCTTCCGATCTNNNTACATCCCCTGCTGAGACTTAG
AGCACACGTCTGAACTCTTTCCCTACACGACGCTCTTCCGATCTNNNGCTTCTTTACACTCTGGGGCTA
AGCACACGTCTGAACTCTTTCCCTACACGACGCTCTTCCGATCTNNNAAATGGGGCTAGTGTGACTA
GCACACGTCTGAACTCTTTCCCTACACGACGCTCTTCCGATCTNNNATTGCCATCTGGATTTTTGTGG
GCACACGTCTGAACTCTTTCCCTACACGACGCTCTTCCGATCTNNNCTTGACTCACTACACAACCTCCA
AGCACACGTCTGAACTCTTTCCCTACACGACGCTCTTCCGATCTNNNTCAGATGATCCTCCCAATTCAG
AGCACACGTCTGAACTCTTTCCCTACACGACGCTCTTCCGATCTNNNGAGACATGAGCAGAGACCTGA
AGCACACGTCTGAACTCTTTCCCTACACGACGCTCTTCCGATCTNNNATGTAAGACATGCCTTGCTTCC
AGCACACGTCTGAACTCTTTCCCTACACGACGCTCTTCCGATCTNNNAGATGCAAATAACTCCATGGC
GCACACGTCTGAACTCTTTCCCTACACGACGCTCTTCCGATCTNNNAGCCACTTCTGATGGGTCTAAA
AGCACACGTCTGAACTCTTTCCCTACACGACGCTCTTCCGATCTNNNGCAGAGAAGGAGAAGAGAG
GAGCACACGTCTGAACTCTTTCCCTACACGACGCTCTTCCGATCTNNNCATATGCCATCATGCCCAGTT
AGCACACGTCTGAACTCTTTCCCTACACGACGCTCTTCCGATCTNNNAACCCCAGTTCCTTCATTTCAG
GCACACGTCTGAACTCTTTCCCTACACGACGCTCTTCCGATCTNNNCAGAATAGACCATGCCACTTCCT
AGCACACGTCTGAACTCTTTCCCTACACGACGCTCTTCCGATCTNNNAGCATTCAAGGGAGTCTTCCT
GCACACGTCTGAACTCTTTCCCTACACGACGCTCTTCCGATCTNNNCTCAAACAAACTTCAGAGCAGG
GCACACGTCTGAACTCTTTCCCTACACGACGCTCTTCCGATCTNNNCTCGATCTCCTGACCTCATGAT
AGCACACGTCTGAACTCTTTCCCTACACGACGCTCTTCCGATCTNNNCTGGAACAAAGGCTATAGTTG
AGCACACGTCTGAACTCTTTCCCTACACGACGCTCTTCCGATCTNNNCTGTAGTCCCAGCTACTCAGG
GAGCACACGTCTGAACTCTTTCCCTACACGACGCTCTTCCGATCTNNNCTGTAGTCCCAGCTACTCGAG
GCACACGTCTGAACTCTTTCCCTACACGACGCTCTTCCGATCTNNNCAGAAGGTTGAAGCTGCAGTA
GAGCACACGTCTGAACTCTTTCCCTACACGACGCTCTTCCGATCTNNNTGGCTATCTGTGATGTCACC
GAGCACACGTCTGAACTCTTTCCCTACACGACGCTCTTCCGATCTNNNCTGTGAGGTATTTCAAGCAG
GAGCACACGTCTGAACTCTTTCCCTACACGACGCTCTTCCGATCTNNNTTGGGTATAGTGGCATGTAC
AGCACACGTCTGAACTCTTTCCCTACACGACGCTCTTCCGATCTNNNAATGAAAATCGCTTGAACCAG
GCACACGTCTGAACTCTTTCCCTACACGACGCTCTTCCGATCTNNNGTGAAACCCTGGCTCTACTAAA

AGCACACGTCTGAACTCTTTCCCTACACGACGCTCTTCCGATCTNNNCTGTAGTCCCAGCTACTCAGG
AGCACACGTCTGAACTCTTTCCCTACACGACGCTCTTCCGATCTNNNTGGCTGAGGCAAGAGAATTA
AGCACACGTCTGAACTCTTTCCCTACACGACGCTCTTCCGATCTNNNATTGCTTGAGCCTAGGAGTTC
GCACACGTCTGAACTCTTTCCCTACACGACGCTCTTCCGATCTNNNAAGAATCACTTGAACCCAGGAG
GAGCACACGTCTGAACTCTTTCCCTACACGACGCTCTTCCGATCTNNNGGTAGGAGAATGGCTTGAA
AGCACACGTCTGAACTCTTTCCCTACACGACGCTCTTCCGATCTNNNGGCAACAGATTGAGACTCCAT
GAGCACACGTCTGAACTCTTTCCCTACACGACGCTCTTCCGATCTNNNCTGTAGTCCCAGCTACTCAGG
GAGCACACGTCTGAACTCTTTCCCTACACGACGCTCTTCCGATCTNNNTCTGCATCAAATCCAACCTCT
GAGCACACGTCTGAACTCTTTCCCTACACGACGCTCTTCCGATCTNNNCTGTAGTCCCAGCTACTCAGG
AGCACACGTCTGAACTCTTTCCCTACACGACGCTCTTCCGATCTNNNAGGCTGAGGTAGGAGTATCA
GAGCACACGTCTGAACTCTTTCCCTACACGACGCTCTTCCGATCTNNNGCAGAAGAATCACTTGAACC
GCACACGTCTGAACTCTTTCCCTACACGACGCTCTTCCGATCTNNNGCACATGCCTATAATCCCAGCTA
AGCACACGTCTGAACTCTTTCCCTACACGACGCTCTTCCGATCTNNNGGGACAAGAGTGAGACTTCA
GAGCACACGTCTGAACTCTTTCCCTACACGACGCTCTTCCGATCTNNNCACCTGTAATCCCAGCTACTC
GCACACGTCTGAACTCTTTCCCTACACGACGCTCTTCCGATCTNNNCAGAAGTTGCAATGAGCCAAGA
AGCACACGTCTGAACTCTTTCCCTACACGACGCTCTTCCGATCTNNNGCCACATCCAGCCACGATATTA
GAGCACACGTCTGAACTCTTTCCCTACACGACGCTCTTCCGATCTNNNGCTCATGCCTGTAATCTCATTG
AGCACACGTCTGAACTCTTTCCCTACACGACGCTCTTCCGATCTNNNAATACAAAAAATTAGCCGGGC
GAGCACACGTCTGAACTCTTTCCCTACACGACGCTCTTCCGATCTNNNACACAAGCCTGTAATTCCAGC
AGCACACGTCTGAACTCTTTCCCTACACGACGCTCTTCCGATCTNNNAAACCGGGACAAACTGATCTC
AGCACACGTCTGAACTCTTTCCCTACACGACGCTCTTCCGATCTNNNTGCATGTAATCCCAGCTACTCA
GAGCACACGTCTGAACTCTTTCCCTACACGACGCTCTTCCGATCTNNNCATGCCTGTAGACCCAGCTAT
GAGCACACGTCTGAACTCTTTCCCTACACGACGCTCTTCCGATCTNNNCTGAGGCAGGAGAATCACTT
GAGCACACGTCTGAACTCTTTCCCTACACGACGCTCTTCCGATCTNNNAGATTGTGTCACTGTACTCCA
AGCACACGTCTGAACTCTTTCCCTACACGACGCTCTTCCGATCTNNNTTTGAGACTAGTGTGGGCAA
AGCACACGTCTGAACTCTTTCCCTACACGACGCTCTTCCGATCTNNNCTTGAACCTGGGATGCAGAAG
AAGAGCACACGTCTGAACTCTTTCCCTACACGACGCTCTTCCGATCTNNNGCAGGAGAATTGCTTGAA
AGCACACGTCTGAACTCTTTCCCTACACGACGCTCTTCCGATCTNNNGAATTGCTTGAAATGGGAGGT
AGCACACGTCTGAACTCTTTCCCTACACGACGCTCTTCCGATCTNNNCAGCCTGGCTAACATGGTAAA
AGCACACGTCTGAACTCTTTCCCTACACGACGCTCTTCCGATCTNNNAGAATCACTTGAACCTGGGAG
GAGCACACGTCTGAACTCTTTCCCTACACGACGCTCTTCCGATCTNNNGTAGTTCCAGCTACTCAGAA
GAGCACACGTCTGAACTCTTTCCCTACACGACGCTCTTCCGATCTNNNCAAGAGAATTGCTTGAACCC
GAGCACACGTCTGAACTCTTTCCCTACACGACGCTCTTCCGATCTNNNCTGTAGTCCCAGCTACTCAGG
GAGCACACGTCTGAACTCTTTCCCTACACGACGCTCTTCCGATCTNNNTGGAAAAAGATATTCCTGAG
AGAGCACACGTCTGAACTCTTTCCCTACACGACGCTCTTCCGATCTNNNCATCCTGGCTAACATGGTGA
GAGCACACGTCTGAACTCTTTCCCTACACGACGCTCTTCCGATCTNNNCTGAGATGGGAGGATCACTT
AGCACACGTCTGAACTCTTTCCCTACACGACGCTCTTCCGATCTNNNCTTGGGACCACTGTACTTTCTG
AGCACACGTCTGAACTCTTTCCCTACACGACGCTCTTCCGATCTNNNGTCAAGATACAAAACAGGCCA
AGAGCACACGTCTGAACTCTTTCCCTACACGACGCTCTTCCGATCTNNNGGGTAGCTGAGGTAGGAG
AGCACACGTCTGAACTCTTTCCCTACACGACGCTCTTCCGATCTNNNAGTCCTAGATACTCAGGAGGC
GCACACGTCTGAACTCTTTCCCTACACGACGCTCTTCCGATCTNNNAAATCTCTTGAACAGGGGAGGT
AGCACACGTCTGAACTCTTTCCCTACACGACGCTCTTCCGATCTNNNCTGAGGTGAGTGGTTTACCTG
GCACACGTCTGAACTCTTTCCCTACACGACGCTCTTCCGATCTNNNTAAGGCAGGAGAATCGCTTGAA

GAGCACACGTCTGAACTCTTTCCCTACACGACGCTCTTCCGATCTNNNAGGTTGTGATGAGCTGAGA
GAGCACACGTCTGAACTCTTTCCCTACACGACGCTCTTCCGATCTNNNGGCAACAAGAGCAAAACTTT
GAGCACACGTCTGAACTCTTTCCCTACACGACGCTCTTCCGATCTNNNTATAATCCCAGCTACACGGA
AGCACACGTCTGAACTCTTTCCCTACACGACGCTCTTCCGATCTNNNAAAAATTAGCCAGGCGTGGT
AGCACACGTCTGAACTCTTTCCCTACACGACGCTCTTCCGATCTNNNGAGAATCACTTGAACCAGGGA
AGCACACGTCTGAACTCTTTCCCTACACGACGCTCTTCCGATCTNNNCACACCTGTAATCCCAGCTACT
GAGCACACGTCTGAACTCTTTCCCTACACGACGCTCTTCCGATCTNNNAGGAAGGAGAATTGCTTGAA
GAGCACACGTCTGAACTCTTTCCCTACACGACGCTCTTCCGATCTNNNTGGTCTTGAACTCTTGACCTC
GCACACGTCTGAACTCTTTCCCTACACGACGCTCTTCCGATCTNNNGTCTTTTATCCCTCATCCCCCTCC
GAGCACACGTCTGAACTCTTTCCCTACACGACGCTCTTCCGATCTNNNCCTAGGAGTTCAAGGCTGTA
AGCACACGTCTGAACTCTTTCCCTACACGACGCTCTTCCGATCTNNNTGGAAGAAGCCCATGAGATCA
GCACACGTCTGAACTCTTTCCCTACACGACGCTCTTCCGATCTNNNCGCCCGGTCGGTGATATATATA
AGCACACGTCTGAACTCTTTCCCTACACGACGCTCTTCCGATCTNNNATGAACCCTGACCCAAACATG
GCACACGTCTGAACTCTTTCCCTACACGACGCTCTTCCGATCTNNNAGCAAGAACCTACTGAAGCTGA
AGCACACGTCTGAACTCTTTCCCTACACGACGCTCTTCCGATCTNNNAATCCCCTGGAAATGTGTCTC
GCACACGTCTGAACTCTTTCCCTACACGACGCTCTTCCGATCTNNNCTGGGGTGGAGCCTTAGATAAT
GCACACGTCTGAACTCTTTCCCTACACGACGCTCTTCCGATCTNNNCTTTATCTGCCCTTCCAGAGTGG
GAGCACACGTCTGAACTCTTTCCCTACACGACGCTCTTCCGATCTNNNATCTCCTGACATTGTGATCCA
AGCACACGTCTGAACTCTTTCCCTACACGACGCTCTTCCGATCTNNNGGGTCACAGAAATAGACACTC
AGCACACGTCTGAACTCTTTCCCTACACGACGCTCTTCCGATCTNNNCCCATTCCTGCTTTCATTCACA
AGCACACGTCTGAACTCTTTCCCTACACGACGCTCTTCCGATCTNNNCACTTGCACATTGTTGATGGG
GCACACGTCTGAACTCTTTCCCTACACGACGCTCTTCCGATCTNNNCTCCAACAGTGCTGGGATTACAG
GCACACGTCTGAACTCTTTCCCTACACGACGCTCTTCCGATCTNNNACAACTTCCTAAAGCGCTAGTCA
AGCACACGTCTGAACTCTTTCCCTACACGACGCTCTTCCGATCTNNNAGAGATGGGGTTTCACTCTGT
AGCACACGTCTGAACTCTTTCCCTACACGACGCTCTTCCGATCTNNNGCAAACGAACACAAAAGAACA
GCACACGTCTGAACTCTTTCCCTACACGACGCTCTTCCGATCTNNNTCAGAAGACTGAGGAAAGTGCT
AGCACACGTCTGAACTCTTTCCCTACACGACGCTCTTCCGATCTNNNAACCATGTCTTCACCTTCCTTC
GCACACGTCTGAACTCTTTCCCTACACGACGCTCTTCCGATCTNNNCACAGCTTCGCAAATGGAGTAA
AGCACACGTCTGAACTCTTTCCCTACACGACGCTCTTCCGATCTNNNATAAACATTATTCCTGGGCCCG
AGCACACGTCTGAACTCTTTCCCTACACGACGCTCTTCCGATCTNNNCCTTTCCTGTCCTGTTACTGTCA
AGCACACGTCTGAACTCTTTCCCTACACGACGCTCTTCCGATCTNNNGCAGGAGAATTATTTGAACCC
GAGCACACGTCTGAACTCTTTCCCTACACGACGCTCTTCCGATCTNNNTGAGGCAGGAGAATCACTTG
AGCACACGTCTGAACTCTTTCCCTACACGACGCTCTTCCGATCTNNNGAGCTTGCAGTGAATTGAGAT
GAGCACACGTCTGAACTCTTTCCCTACACGACGCTCTTCCGATCTNNNGAAGCAGGAAAATAGCTCG
GAGCACACGTCTGAACTCTTTCCCTACACGACGCTCTTCCGATCTNNNGAATCACTTGAACTCAGAAG
GAGCACACGTCTGAACTCTTTCCCTACACGACGCTCTTCCGATCTNNNGTGAGCTGAGATTGTGCTAT
GAGCACACGTCTGAACTCTTTCCCTACACGACGCTCTTCCGATCTNNNATACAAAAATTAGCCGAGCG
GCACACGTCTGAACTCTTTCCCTACACGACGCTCTTCCGATCTNNNCAGGAGTTCAAAGTTGCAGTGA
AGCACACGTCTGAACTCTTTCCCTACACGACGCTCTTCCGATCTNNNTAGGTGACTGAGCGAGACTC
AGCACACGTCTGAACTCTTTCCCTACACGACGCTCTTCCGATCTNNNGGCAACAGAGTAAGACTGTCT
GAGCACACGTCTGAACTCTTTCCCTACACGACGCTCTTCCGATCTNNNTGAGCTCAGAAGTTTGAGAC
AGCACACGTCTGAACTCTTTCCCTACACGACGCTCTTCCGATCTNNNTAGCTCATGTCTGTAGTCCCAG
GAGCACACGTCTGAACTCTTTCCCTACACGACGCTCTTCCGATCTNNNCTGAGGCAGGAGAATAGCA

AGCACACGTCTGAACTCTTTCCCTACACGACGCTCTTCCGATCTNNNAGTGAGCTGAGATTGTGAGGT

GAGCACACGTCTGAACTCTTTCCCTACACGACGCTCTTCCGATCTNNNGAGGCAGGAGAATTGCTTG

AGCACACGTCTGAACTCTTTCCCTACACGACGCTCTTCCGATCTNNNCTGTAGTCCCAGCTACTTGAG

GCACACGTCTGAACTCTTTCCCTACACGACGCTCTTCCGATCTNNNTGAGCTGAGAACACACCATTAC

AGCACACGTCTGAACTCTTTCCCTACACGACGCTCTTCCGATCTNNNCTACACTCCAGTCTGGGCAATA

AGCACACGTCTGAACTCTTTCCCTACACGACGCTCTTCCGATCTNNNTGTTTGATGCTGCAGTGAACT

GAGCACACGTCTGAACTCTTTCCCTACACGACGCTCTTCCGATCTNNNACCTGTAATCCCAGCACTTTA

AGCACACGTCTGAACTCTTTCCCTACACGACGCTCTTCCGATCTNNNCTGCTCTCAATGTACTCTGGAG

AGCACACGTCTGAACTCTTTCCCTACACGACGCTCTTCCGATCTNNNCTGAGGCAGGAGAATCACTTG

AGCACACGTCTGAACTCTTTCCCTACACGACGCTCTTCCGATCTNNNTTTAGTAGAGACAGGGTTTCG

AGCACACGTCTGAACTCTTTCCCTACACGACGCTCTTCCGATCTNNNTCTGTTTCAAGAGGTTTAGGCC

AGCACACGTCTGAACTCTTTCCCTACACGACGCTCTTCCGATCTNNNCTCTGCATGTCTAACACACTCC

GAGCACACGTCTGAACTCTTTCCCTACACGACGCTCTTCCGATCTNNNCAACTTGGATGTTGCTGGTGT

GAGCACACGTCTGAACTCTTTCCCTACACGACGCTCTTCCGATCTNNNGTTTCTGATTCTCTGGGTCTG

GAGCACACGTCTGAACTCTTTCCCTACACGACGCTCTTCCGATCTNNNCTGTGATAGGGACTTTGAGG

GCACACGTCTGAACTCTTTCCCTACACGACGCTCTTCCGATCTNNNCATGAGAAGTCTGAGCTCTCCA

GAGCACACGTCTGAACTCTTTCCCTACACGACGCTCTTCCGATCTNNNCCTTGCAGCTGGTTTTCTTTTC

GAGCACACGTCTGAACTCTTTCCCTACACGACGCTCTTCCGATCTNNNCTCCCAAGTAGCTGGAACTAT

GAGCACACGTCTGAACTCTTTCCCTACACGACGCTCTTCCGATCTNNNCTGTCACAAATGGATCACGTG

GCACACGTCTGAACTCTTTCCCTACACGACGCTCTTCCGATCTNNNGGCTAAACAGCAATGAGAAGCA

AGCACACGTCTGAACTCTTTCCCTACACGACGCTCTTCCGATCTNNNCTATCATAGCTCACTGCAGCCT

AGCACACGTCTGAACTCTTTCCCTACACGACGCTCTTCCGATCTNNNCTCCTCACCTCACTATACTGTG

AGCACACGTCTGAACTCTTTCCCTACACGACGCTCTTCCGATCTNNNAGTCAGACGCATCTCAATCTCT

GCACACGTCTGAACTCTTTCCCTACACGACGCTCTTCCGATCTNNNAGGTTCAAGTGATTCTCATGCC

AGCACACGTCTGAACTCTTTCCCTACACGACGCTCTTCCGATCTNNNCTCAATCATGGGGAATCTGGT

GCACACGTCTGAACTCTTTCCCTACACGACGCTCTTCCGATCTNNNGGTGGGATTATAGACATGAGC

GCACACGTCTGAACTCTTTCCCTACACGACGCTCTTCCGATCTNNNCGCCTGAGGTCAAACCCATATAT

AGCACACGTCTGAACTCTTTCCCTACACGACGCTCTTCCGATCTNNNAAATTGCTTTGGACTGTCAGA

GAGCACACGTCTGAACTCTTTCCCTACACGACGCTCTTCCGATCTNNNCCGAGTAGCTGAGATTACAG

GCACACGTCTGAACTCTTTCCCTACACGACGCTCTTCCGATCTNNNAAAGTGCTGGGATTACAAGCTT

AGCACACGTCTGAACTCTTTCCCTACACGACGCTCTTCCGATCTNNNGCTCCAAGTCTGTCTGAATCCT

AGCACACGTCTGAACTCTTTCCCTACACGACGCTCTTCCGATCTNNNGCTCCAAGTCTGTCTGAATCCT

AGCACACGTCTGAACTCTTTCCCTACACGACGCTCTTCCGATCTNNNCCCCCAATTCAAAGCAAAGAGT

AGCACACGTCTGAACTCTTTCCCTACACGACGCTCTTCCGATCTNNNCACCAGTGTAGAGAAAGCTAC

AGCACACGTCTGAACTCTTTCCCTACACGACGCTCTTCCGATCTNNNTGCAGTCTTTCTCCACTAGCTT

AGCACACGTCTGAACTCTTTCCCTACACGACGCTCTTCCGATCTNNNGAAAGCCTTAGTTTGATGCCA

GAGCACACGTCTGAACTCTTTCCCTACACGACGCTCTTCCGATCTNNNCTCACACCTGTAATCTCAGC

GAGCACACGTCTGAACTCTTTCCCTACACGACGCTCTTCCGATCTNNNTGGGCAATAGAATGAGACT

GAGCACACGTCTGAACTCTTTCCCTACACGACGCTCTTCCGATCTNNNCTATGAGGTGGAGATTGCAG

GAGCACACGTCTGAACTCTTTCCCTACACGACGCTCTTCCGATCTNNNGTAATCCCAGCTACTCAGTAG

AGCACACGTCTGAACTCTTTCCCTACACGACGCTCTTCCGATCTNNNATGATCATGCCACTTCACTCCAT

GAGCACACGTCTGAACTCTTTCCCTACACGACGCTCTTCCGATCTNNNATGCATGTAATCCCAGCACTTT

GAGCACACGTCTGAACTCTTTCCCTACACGACGCTCTTCCGATCTNNNTGCAGTGAGATAAGATCATG

AGAGCACACGTCTGAACTCTTTCCCTACACGACGCTCTTCCGATCTNNNGGAAGCTGAGACAGAATTG
AGCACACGTCTGAACTCTTTCCCTACACGACGCTCTTCCGATCTNNNACACCTGTAATCCCAGCTACTT
GAGCACACGTCTGAACTCTTTCCCTACACGACGCTCTTCCGATCTNNNCTGAGGCAGGAGAATCACTT
AGCACACGTCTGAACTCTTTCCCTACACGACGCTCTTCCGATCTNNNAGAATCACTTGAACCAGGGAG
GAGCACACGTCTGAACTCTTTCCCTACACGACGCTCTTCCGATCTNNNTAGGCACCTGTAATCCAAGCT
AGCACACGTCTGAACTCTTTCCCTACACGACGCTCTTCCGATCTNNNCTAGGCAACATAGGGAGACA
GAGCACACGTCTGAACTCTTTCCCTACACGACGCTCTTCCGATCTNNNGGCACCTATAATCCCAGCTA
AGAGCACACGTCTGAACTCTTTCCCTACACGACGCTCTTCCGATCTNNNTGTAGTCTCAGCTACTCAGG
AGCACACGTCTGAACTCTTTCCCTACACGACGCTCTTCCGATCTNNNATCTTGAACTGGGCTCAAGTG
GCACACGTCTGAACTCTTTCCCTACACGACGCTCTTCCGATCTNNNGAGGCAGGAGAATTGTTTGAAC
AGCACACGTCTGAACTCTTTCCCTACACGACGCTCTTCCGATCTNNNGAGGTTGTAGTGAGCGGAGAT
AGCACACGTCTGAACTCTTTCCCTACACGACGCTCTTCCGATCTNNNGACCCTCTCCTTGAATTCCTCT
AGCACACGTCTGAACTCTTTCCCTACACGACGCTCTTCCGATCTNNNGTAGTGAACCAAGATTGTGC
AGCACACGTCTGAACTCTTTCCCTACACGACGCTCTTCCGATCTNNNCGAGATCACACCATTGTACTCC
GAGCACACGTCTGAACTCTTTCCCTACACGACGCTCTTCCGATCTNNNGCACCTATAGTCCCAGTTACT
AGCACACGTCTGAACTCTTTCCCTACACGACGCTCTTCCGATCTNNNGATGCAGTGAGTTGAGATCA
AGAGCACACGTCTGAACTCTTTCCCTACACGACGCTCTTCCGATCTNNNGAGGCAGGAGAATCGCTT
AGCACACGTCTGAACTCTTTCCCTACACGACGCTCTTCCGATCTNNNAGAAGAATCGCCTGAACCCAA
AGCACACGTCTGAACTCTTTCCCTACACGACGCTCTTCCGATCTNNNCAGAGGTTGTCGTTAGCTGA
AGAGCACACGTCTGAACTCTTTCCCTACACGACGCTCTTCCGATCTNNNTGAGAATCACTTGAACCCAG
GAGCACACGTCTGAACTCTTTCCCTACACGACGCTCTTCCGATCTNNNAAATATTAGCTGGGTGTGGT
AGCACACGTCTGAACTCTTTCCCTACACGACGCTCTTCCGATCTNNNACCTAGGCTCAAAACCAGGATA
AGCACACGTCTGAACTCTTTCCCTACACGACGCTCTTCCGATCTNNNTTGCAAGGTAGGTTTCTCCTCT
AGCACACGTCTGAACTCTTTCCCTACACGACGCTCTTCCGATCTNNNACTAACTCAGCTTCCCCAATAG
AGCACACGTCTGAACTCTTTCCCTACACGACGCTCTTCCGATCTNNNGTCCCTCACATTCCATGCATCT
AGCACACGTCTGAACTCTTTCCCTACACGACGCTCTTCCGATCTNNNGATACACTCCATGTCGAGCTCA
GCACACGTCTGAACTCTTTCCCTACACGACGCTCTTCCGATCTNNNTGAGTAGCTGAGATTACAGGCA
AGCACACGTCTGAACTCTTTCCCTACACGACGCTCTTCCGATCTNNNCTGTCACTCCCTATCAGTGTTCC
AGCACACGTCTGAACTCTTTCCCTACACGACGCTCTTCCGATCTNNNTAGCATTCCATTTGTGCCACATC
AGCACACGTCTGAACTCTTTCCCTACACGACGCTCTTCCGATCTNNNGAGTAGTCTCTGGCTTTAGGC
AGCACACGTCTGAACTCTTTCCCTACACGACGCTCTTCCGATCTNNNCTCTAATACCTGGGCTCAAGCA
AGCACACGTCTGAACTCTTTCCCTACACGACGCTCTTCCGATCTNNNACCATGTTGCATGTTGTTGAT
AGCACACGTCTGAACTCTTTCCCTACACGACGCTCTTCCGATCTNNNGGCCAGAACAACTGTAATCAA
AGCACACGTCTGAACTCTTTCCCTACACGACGCTCTTCCGATCTNNNCAGGGTAATGTTGGCCTCAGA
GAGCACACGTCTGAACTCTTTCCCTACACGACGCTCTTCCGATCTNNNTGCTATTCCTTCTGCCTGAAAT
AGCACACGTCTGAACTCTTTCCCTACACGACGCTCTTCCGATCTNNNCTAGATTGACAGTGGCTTTCCC
AGCACACGTCTGAACTCTTTCCCTACACGACGCTCTTCCGATCTNNNATAAAGCTCTACCTCATGGGC
AGCACACGTCTGAACTCTTTCCCTACACGACGCTCTTCCGATCTNNNACTTTAGCTAGTCTCCCACCTTT
AGCACACGTCTGAACTCTTTCCCTACACGACGCTCTTCCGATCTNNNAATTGGGCCATCAAATGTGTCT
CACACGTCTGAACTCTTTCCCTACACGACGCTCTTCCGATCTNNNTGAGTGCCAGCTTAGAATTCCAT
AGCACACGTCTGAACTCTTTCCCTACACGACGCTCTTCCGATCTNNNAACAGGTGATATGAATGGCAG
AGCACACGTCTGAACTCTTTCCCTACACGACGCTCTTCCGATCTNNNTTCTGGAAAAGTCTGGTGTCAC
AGCACACGTCTGAACTCTTTCCCTACACGACGCTCTTCCGATCTNNNTCAGTTCTTGTTTCCCATGAGCT

AGCACACGTCTGAACTCTTTCCCTACACGACGCTCTTCCGATCTNNNTAAACAGCCAAGTTGTGAACA
GCACACGTCTGAACTCTTTCCCTACACGACGCTCTTCCGATCTNNNTTAGCTGAAGGGAAATGACGT
AGCACACGTCTGAACTCTTTCCCTACACGACGCTCTTCCGATCTNNNTTATTCTCATTCCCTACCCCGT
GAGCACACGTCTGAACTCTTTCCCTACACGACGCTCTTCCGATCTNNNCCCTTTCCTTCTCTTTTGTGTG
AGCACACGTCTGAACTCTTTCCCTACACGACGCTCTTCCGATCTNNNCATGCCTCCTTCTACAGTACCAG
AGCACACGTCTGAACTCTTTCCCTACACGACGCTCTTCCGATCTNNNCCAAAGTGCTGGGATAACAGG
AGCACACGTCTGAACTCTTTCCCTACACGACGCTCTTCCGATCTNNNGTCTACTCTGGGCCATCTTGAT
GCACACGTCTGAACTCTTTCCCTACACGACGCTCTTCCGATCTNNNAACCTCCCAAGTAGCTGAGACTA
AGCACACGTCTGAACTCTTTCCCTACACGACGCTCTTCCGATCTNNNACACTTTCATCCCCTCAAACAG
AGCACACGTCTGAACTCTTTCCCTACACGACGCTCTTCCGATCTNNNAATCTCCTGACCTCATGATCTG
GAGCACACGTCTGAACTCTTTCCCTACACGACGCTCTTCCGATCTNNNAAGTCCAAAGTGCTGGGATT
AGCACACGTCTGAACTCTTTCCCTACACGACGCTCTTCCGATCTNNNAGTGTTGAGATTACAGGCCTG
GCACACGTCTGAACTCTTTCCCTACACGACGCTCTTCCGATCTNNNGATCCTACAGATGCACTTCACCA
AGCACACGTCTGAACTCTTTCCCTACACGACGCTCTTCCGATCTNNNTGTTTCAGTACTCAGCTCAGG
AGCACACGTCTGAACTCTTTCCCTACACGACGCTCTTCCGATCTNNNGTCTGAGGGGTAATTGTACT
AGCACACGTCTGAACTCTTTCCCTACACGACGCTCTTCCGATCTNNNTATAAGAGGCCAAGCTGGGT
GAGCACACGTCTGAACTCTTTCCCTACACGACGCTCTTCCGATCTNNNCTCCTGAGTAGCTGGGATTA
AGCACACGTCTGAACTCTTTCCCTACACGACGCTCTTCCGATCTNNNATTGAAGCTACTTTTGGCCCTT
AGCACACGTCTGAACTCTTTCCCTACACGACGCTCTTCCGATCTNNNGGAGTCCCTCCTTTCCAATTGAT
CACACGTCTGAACTCTTTCCCTACACGACGCTCTTCCGATCTNNNTGCTGTGATTTTGATTGGTATTAC
AGCACACGTCTGAACTCTTTCCCTACACGACGCTCTTCCGATCTNNNAGTTGAGACTACAGGTATGCA
AGCACACGTCTGAACTCTTTCCCTACACGACGCTCTTCCGATCTNNNGCAGAGCCAAACCATATCATT
AGCACACGTCTGAACTCTTTCCCTACACGACGCTCTTCCGATCTNNNGAACTCCTAACCTCAGGTGATC
AGCACACGTCTGAACTCTTTCCCTACACGACGCTCTTCCGATCTNNNTAGATTGGCAACTGCAGTAA
AGCACACGTCTGAACTCTTTCCCTACACGACGCTCTTCCGATCTNNNCAAAGGTCTCCACAAGTGAACT
AGCACACGTCTGAACTCTTTCCCTACACGACGCTCTTCCGATCTNNNAATTTGTGAGGTGGGAAAAC
AGCACACGTCTGAACTCTTTCCCTACACGACGCTCTTCCGATCTNNNATACAATACCCCCTAACAACGC
GAGCACACGTCTGAACTCTTTCCCTACACGACGCTCTTCCGATCTNNNATCTCCTGTAAATCTGGGACG
AGCACACGTCTGAACTCTTTCCCTACACGACGCTCTTCCGATCTNNNCCCTTCATTTTTCTACACTGGG
AGCACACGTCTGAACTCTTTCCCTACACGACGCTCTTCCGATCTNNNGCCACCATGCTAGGCTAATTTT
GCACACGTCTGAACTCTTTCCCTACACGACGCTCTTCCGATCTNNNAGGGAAAGAGGCTTCCGTTTTT
AGAGCACACGTCTGAACTCTTTCCCTACACGACGCTCTTCCGATCTNNNATACAAGCAGGGTGAGCAT
AGCACACGTCTGAACTCTTTCCCTACACGACGCTCTTCCGATCTNNNCAAGACACTCCAAAGGCAAGAA
GCACACGTCTGAACTCTTTCCCTACACGACGCTCTTCCGATCTNNNTTGCTGTATCCCAGAGGTTTTGA
AGCACACGTCTGAACTCTTTCCCTACACGACGCTCTTCCGATCTNNNGATTCAAGCCGTTCTTATGCCT
AGCACACGTCTGAACTCTTTCCCTACACGACGCTCTTCCGATCTNNNGGAGAGCACCGTTCAAAATTT
AGCACACGTCTGAACTCTTTCCCTACACGACGCTCTTCCGATCTNNNATACTTGGGTTGTTTCCACCTC
AGCACACGTCTGAACTCTTTCCCTACACGACGCTCTTCCGATCTNNNGCATAATCATGGCTCACTGAA
AGCACACGTCTGAACTCTTTCCCTACACGACGCTCTTCCGATCTNNNCACCTGTAATCCCAGCTACTCA
GCACACGTCTGAACTCTTTCCCTACACGACGCTCTTCCGATCTNNNCAGCTTGGGTGACAGAGTAAG
AGCACACGTCTGAACTCTTTCCCTACACGACGCTCTTCCGATCTNNNAGGATCACTTGAGGTCAGCAA
GAGCACACGTCTGAACTCTTTCCCTACACGACGCTCTTCCGATCTNNNCGAGACAGGAGAATTGCTCA
AGCACACGTCTGAACTCTTTCCCTACACGACGCTCTTCCGATCTNNNCTGAGGTGGGAGAATCACTT

AGCACACGTCTGAACTCTTTCCCTACACGACGCTCTTCCGATCTNNNTACATGCCTGTAATCCCAGCTA
GAGCACACGTCTGAACTCTTTCCCTACACGACGCTCTTCCGATCTNNNTTTTCCCCAGGAATTCAAGAC
AGCACACGTCTGAACTCTTTCCCTACACGACGCTCTTCCGATCTNNNCTGTAATCCCAGCTACTCAGG
AGCACACGTCTGAACTCTTTCCCTACACGACGCTCTTCCGATCTNNNGTAGTCTCAGCTACTCAGGAAG
AGCACACGTCTGAACTCTTTCCCTACACGACGCTCTTCCGATCTNNNTGTAGTCTCAGCTACTCAGGA
GAGCACACGTCTGAACTCTTTCCCTACACGACGCTCTTCCGATCTNNNGAGGTTGCAGTAAGCCAAG
GCACACGTCTGAACTCTTTCCCTACACGACGCTCTTCCGATCTNNNTGAGGCAGGAGAATCTCTTGAA
GAGCACACGTCTGAACTCTTTCCCTACACGACGCTCTTCCGATCTNNNATTTGCCCATAGCTGATCTGG
GAGCACACGTCTGAACTCTTTCCCTACACGACGCTCTTCCGATCTNNNGAGGCAGAAGAATCACTTGA
AGCACACGTCTGAACTCTTTCCCTACACGACGCTCTTCCGATCTNNNGGAGAATCACTTGAACCTGGG
GAGCACACGTCTGAACTCTTTCCCTACACGACGCTCTTCCGATCTNNNGTGACACAGCGAGACTCTGT
GAGCACACGTCTGAACTCTTTCCCTACACGACGCTCTTCCGATCTNNNATGCTTGTAATCCCAGCTACT
GAGCACACGTCTGAACTCTTTCCCTACACGACGCTCTTCCGATCTNNNGGGAGAATCGCTTGAAACCA
AGCACACGTCTGAACTCTTTCCCTACACGACGCTCTTCCGATCTNNNAAGTTGCAGTGAACTGAGATT
AGCACACGTCTGAACTCTTTCCCTACACGACGCTCTTCCGATCTNNNGGCTAACACCTGTAATCCCATC
AGCACACGTCTGAACTCTTTCCCTACACGACGCTCTTCCGATCTNNNACTCCTGTAATCCCAGCACTTT
GAGCACACGTCTGAACTCTTTCCCTACACGACGCTCTTCCGATCTNNNCTGAGGCAGAAGAATGGTGT
GAGCACACGTCTGAACTCTTTCCCTACACGACGCTCTTCCGATCTNNNACCATCTCACACCAGTCAGAA
GAGCACACGTCTGAACTCTTTCCCTACACGACGCTCTTCCGATCTNNNTGTGATCACAGCTAAATGGG
GCACACGTCTGAACTCTTTCCCTACACGACGCTCTTCCGATCTNNNGGGAGAACTGCTTGAATTCAGG
AGCACACGTCTGAACTCTTTCCCTACACGACGCTCTTCCGATCTNNNAGAGGTTGCAGTGAACTCAG
AGCACACGTCTGAACTCTTTCCCTACACGACGCTCTTCCGATCTNNNATACAGGAAGCATGGTGGTTT
GAGCACACGTCTGAACTCTTTCCCTACACGACGCTCTTCCGATCTNNNTGCAATCAAAACCACAGTGA
GAGCACACGTCTGAACTCTTTCCCTACACGACGCTCTTCCGATCTNNNGCAGGAGAATCATTTGAAC
GAGCACACGTCTGAACTCTTTCCCTACACGACGCTCTTCCGATCTNNNAGGTTACAATGAGCCAAGAC
AGCACACGTCTGAACTCTTTCCCTACACGACGCTCTTCCGATCTNNNCTGAGGCAGGAGAATCACTTG
GCACACGTCTGAACTCTTTCCCTACACGACGCTCTTCCGATCTNNNCGTGAAGTTGAAGCTGCAATGA
AGCACACGTCTGAACTCTTTCCCTACACGACGCTCTTCCGATCTNNNAGGCAGGAGAATAACTTGAAC
GAGCACACGTCTGAACTCTTTCCCTACACGACGCTCTTCCGATCTNNNTGAGGAAGGAGAATCGCTT
AGCACACGTCTGAACTCTTTCCCTACACGACGCTCTTCCGATCTNNNGCCCCTTCCAATATGATTGCAT
AGCACACGTCTGAACTCTTTCCCTACACGACGCTCTTCCGATCTNNNCTGTAGTCCCAGCTACTCGGA
AGCACACGTCTGAACTCTTTCCCTACACGACGCTCTTCCGATCTNNNCACCTGTAATCCCAGCTACTTG
AGCACACGTCTGAACTCTTTCCCTACACGACGCTCTTCCGATCTNNNAGAGGTTGCAGTGAGCTAAG
GAGCACACGTCTGAACTCTTTCCCTACACGACGCTCTTCCGATCTNNNTACAAAAATTAGCCAGGCAG
AGCACACGTCTGAACTCTTTCCCTACACGACGCTCTTCCGATCTNNNAGGTAGGAGAATTGCTTGAA
GAGCACACGTCTGAACTCTTTCCCTACACGACGCTCTTCCGATCTNNNAGCAACAGAACAAGACTCCAT
GAGCACACGTCTGAACTCTTTCCCTACACGACGCTCTTCCGATCTNNNTGCACCTGTAATCCCAGTTA
GCACACGTCTGAACTCTTTCCCTACACGACGCTCTTCCGATCTNNNATGGACTCTCTAAGTGTGCATCC
GCACACGTCTGAACTCTTTCCCTACACGACGCTCTTCCGATCTNNNGAGACAGGAGAATTGCTTGAA
AGCACACGTCTGAACTCTTTCCCTACACGACGCTCTTCCGATCTNNNAGTTATGGCTATTTCGAGCTGC
AGCACACGTCTGAACTCTTTCCCTACACGACGCTCTTCCGATCTNNNTTGTCATATGAATCAGCACCTC
AGCACACGTCTGAACTCTTTCCCTACACGACGCTCTTCCGATCTNNNCCAGCCTTCAGAACTGTGAGAA
AGCACACGTCTGAACTCTTTCCCTACACGACGCTCTTCCGATCTNNNGTACAGTGGCACGATCTCTAC

AGCACACGTCTGAACTCTTTCCCTACACGACGCTCTTCCGATCTNNNGTTGGTCTCAAACTCCTGACCT
AGCACACGTCTGAACTCTTTCCCTACACGACGCTCTTCCGATCTNNNTTCTGGGATTATAGGTGCAAG
GAGCACACGTCTGAACTCTTTCCCTACACGACGCTCTTCCGATCTNNNAGGTTCAAGCAATTCTCCTGT
GAGCACACGTCTGAACTCTTTCCCTACACGACGCTCTTCCGATCTNNNGCCTCGGTGTGATGTTTTGATA
AGAGCACACGTCTGAACTCTTTCCCTACACGACGCTCTTCCGATCTNNNACTCATGCAATTTGCCATGT
GAGCACACGTCTGAACTCTTTCCCTACACGACGCTCTTCCGATCTNNNACAGAGTAGAGTGTTGCATA
AGCACACGTCTGAACTCTTTCCCTACACGACGCTCTTCCGATCTNNNTGTCTTAGCTACTCAGGAGGTG
AGCACACGTCTGAACTCTTTCCCTACACGACGCTCTTCCGATCTNNNCACCTCGCCTGGCTAGTTTATAT
GAGCACACGTCTGAACTCTTTCCCTACACGACGCTCTTCCGATCTNNNGCACACACAGAGGACATTCTTT
GAGCACACGTCTGAACTCTTTCCCTACACGACGCTCTTCCGATCTNNNTAGACAGAGTCTCACTCCATT
AGCACACGTCTGAACTCTTTCCCTACACGACGCTCTTCCGATCTNNNGGCTCTTGATTCTGCTTCCAAAA
AGCACACGTCTGAACTCTTTCCCTACACGACGCTCTTCCGATCTNNNTGGGTTTTCAGAATGGATTTG
AGCACACGTCTGAACTCTTTCCCTACACGACGCTCTTCCGATCTNNNAGCTGAACTGCTTCTTCTTCCTT
AGCACACGTCTGAACTCTTTCCCTACACGACGCTCTTCCGATCTNNNTCCTGAGTAGGTGGGATTACAG
AGCACACGTCTGAACTCTTTCCCTACACGACGCTCTTCCGATCTNNNCAGTTATTTCTCCAAGGAGCCCT
GAGCACACGTCTGAACTCTTTCCCTACACGACGCTCTTCCGATCTNNNGGACATGTGAGCTAGCATCCT
AGCACACGTCTGAACTCTTTCCCTACACGACGCTCTTCCGATCTNNNCTCAAACTCCCAATCTCAGGTGA
GCACACGTCTGAACTCTTTCCCTACACGACGCTCTTCCGATCTNNNTAACACAAGGTTGGGGAAGATG
AGCACACGTCTGAACTCTTTCCCTACACGACGCTCTTCCGATCTNNNAGAAAAGAGAGCCAAGCATCT
AGCACACGTCTGAACTCTTTCCCTACACGACGCTCTTCCGATCTNNNTTGTACCAGATGACACCATTGG
GAGCACACGTCTGAACTCTTTCCCTACACGACGCTCTTCCGATCTNNNAAAGTGTTGGGATTACAAGC
AGCACACGTCTGAACTCTTTCCCTACACGACGCTCTTCCGATCTNNNAAAGTGCTGGGATTACAAGCAT
AGCACACGTCTGAACTCTTTCCCTACACGACGCTCTTCCGATCTNNNCTGGCTCTAGTCTGTTTCTCCAG
GAGCACACGTCTGAACTCTTTCCCTACACGACGCTCTTCCGATCTNNNTGGGATTACAGGTGTGAGTC
AGCACACGTCTGAACTCTTTCCCTACACGACGCTCTTCCGATCTNNNCTGGCCTTGAACTATCTGGCTAT
AGCACACGTCTGAACTCTTTCCCTACACGACGCTCTTCCGATCTNNNCAATCACGGCTCACTGAAACTTC
AGCACACGTCTGAACTCTTTCCCTACACGACGCTCTTCCGATCTNNNCTGGAACTCCTGACCTCAAATGA
GAGCACACGTCTGAACTCTTTCCCTACACGACGCTCTTCCGATCTNNNGGGTATGAAGAGGGACTTGA
AGCACACGTCTGAACTCTTTCCCTACACGACGCTCTTCCGATCTNNNGGATTACAGGTGTGAGTCACT
GAGCACACGTCTGAACTCTTTCCCTACACGACGCTCTTCCGATCTNNNTCTTGAATGCTGGCCTCAAGTA
AGCACACGTCTGAACTCTTTCCCTACACGACGCTCTTCCGATCTNNNCGTCCCTCTAATTCGGAGCTAAA
AGCACACGTCTGAACTCTTTCCCTACACGACGCTCTTCCGATCTNNNAGGCCCTCCACAATGTTATTCTT
GAGCACACGTCTGAACTCTTTCCCTACACGACGCTCTTCCGATCTNNNTACATGTTGGCCATTTGTGTGT
AGCACACGTCTGAACTCTTTCCCTACACGACGCTCTTCCGATCTNNNTTCAGTTGGCTCAAGATTCCGTA
AGCACACGTCTGAACTCTTTCCCTACACGACGCTCTTCCGATCTNNNCTCCCGGGTTCAAACAATTCTT
AGCACACGTCTGAACTCTTTCCCTACACGACGCTCTTCCGATCTNNNAGCCATCGGTGCTCTTAGAATA
AGCACACGTCTGAACTCTTTCCCTACACGACGCTCTTCCGATCTNNNTCCCAAAGTGCTGAGATTACAG
GAGCACACGTCTGAACTCTTTCCCTACACGACGCTCTTCCGATCTNNNACTACGAAGTAAGAGAGACA
AGCACACGTCTGAACTCTTTCCCTACACGACGCTCTTCCGATCTNNNTCTTACCCGCTCAGAGTTTCATG
AGCACACGTCTGAACTCTTTCCCTACACGACGCTCTTCCGATCTNNNAGAAGAACCACCTAGGAAAAG
GAGCACACGTCTGAACTCTTTCCCTACACGACGCTCTTCCGATCTNNNCTGACCTCCTGTCTCTCTCTTT
AGCACACGTCTGAACTCTTTCCCTACACGACGCTCTTCCGATCTNNNAATTGCAGGATACCAAGAGGGA
AGCACACGTCTGAACTCTTTCCCTACACGACGCTCTTCCGATCTNNNCAGTCAGGGCTGTGAGAGAAT

GAGCACACGTCTGAACTCTTTCCCTACACGACGCTCTTCCGATCTNNNCTTTCTTCCTTCTGTCTCCCTCC
AGCACACGTCTGAACTCTTTCCCTACACGACGCTCTTCCGATCTNNNTCAATCTCCTGACCTTGTGATC
GAGCACACGTCTGAACTCTTTCCCTACACGACGCTCTTCCGATCTNNNAGACAGGAGAATTGCTTGAA
GAGCACACGTCTGAACTCTTTCCCTACACGACGCTCTTCCGATCTNNNTTGTAGTGAGCCTAGATCAT
AGCACACGTCTGAACTCTTTCCCTACACGACGCTCTTCCGATCTNNNCTGTAGTCCCAGCTTCTCAGGA
AGCACACGTCTGAACTCTTTCCCTACACGACGCTCTTCCGATCTNNNAGATGGGAGAATCACTTGAA
AGCACACGTCTGAACTCTTTCCCTACACGACGCTCTTCCGATCTNNNTGAGCTGAGATTATTCCACTGC
AGCACACGTCTGAACTCTTTCCCTACACGACGCTCTTCCGATCTNNNAATCCCAGCTTCTTGAGAGAC
GAGCACACGTCTGAACTCTTTCCCTACACGACGCTCTTCCGATCTNNNGAGAATCACCTGAACCCAGT
AGCACACGTCTGAACTCTTTCCCTACACGACGCTCTTCCGATCTNNNGCTCATGCCTATTATCTCAGCA
GAGCACACGTCTGAACTCTTTCCCTACACGACGCTCTTCCGATCTNNNCTGTAATCACAGCACTTTGGG
GCACACGTCTGAACTCTTTCCCTACACGACGCTCTTCCGATCTNNNGGTGACAGAGTGAGACTCCATC
GCACACGTCTGAACTCTTTCCCTACACGACGCTCTTCCGATCTNNNAGAAGAATTTCAAAGCCTTCGC
AGCACACGTCTGAACTCTTTCCCTACACGACGCTCTTCCGATCTNNNTTGTGCGGACATACGTTTTCAA
AGCACACGTCTGAACTCTTTCCCTACACGACGCTCTTCCGATCTNNNATATCATGCCACTGTACTCCAG
GCACACGTCTGAACTCTTTCCCTACACGACGCTCTTCCGATCTNNNCTGAGGCAGAGAATTGCTTGA
GCACACGTCTGAACTCTTTCCCTACACGACGCTCTTCCGATCTNNNGAGCTTGCAGTGAGCCAAGAT
AGCACACGTCTGAACTCTTTCCCTACACGACGCTCTTCCGATCTNNNTGAGCTGAGATCACACCATTAC
AGCACACGTCTGAACTCTTTCCCTACACGACGCTCTTCCGATCTNNNGGGAGGTTGAGACAGGAGA
AGCACACGTCTGAACTCTTTCCCTACACGACGCTCTTCCGATCTNNNGAGAATACCTTGAACCTGGG
AGCACACGTCTGAACTCTTTCCCTACACGACGCTCTTCCGATCTNNNGAGGATTTCTTGAGCTCAGGA
AGCACACGTCTGAACTCTTTCCCTACACGACGCTCTTCCGATCTNNNGAGGTTGCAGTGAGTTGAGA
GAGCACACGTCTGAACTCTTTCCCTACACGACGCTCTTCCGATCTNNNTAGTCTCAACTACTCAGGAG
GCACACGTCTGAACTCTTTCCCTACACGACGCTCTTCCGATCTNNNGGCTGAAGCAGAAGAATCACTT
GCACACGTCTGAACTCTTTCCCTACACGACGCTCTTCCGATCTNNNGTAGTCCCAGCTACTCATTAGGC
AGCACACGTCTGAACTCTTTCCCTACACGACGCTCTTCCGATCTNNNCTGAGGCAGGAGAATCACTT
AGCACACGTCTGAACTCTTTCCCTACACGACGCTCTTCCGATCTNNNGGGCAATAGAGTGAGACTCCA
GCACACGTCTGAACTCTTTCCCTACACGACGCTCTTCCGATCTNNNGAGGCAGGAGATTCTCTTGATC
AGCACACGTCTGAACTCTTTCCCTACACGACGCTCTTCCGATCTNNNAACCACAGAGAGCCTTTGATG
AGCACACGTCTGAACTCTTTCCCTACACGACGCTCTTCCGATCTNNNAATGCCTGTAATCCCAGCTAC
GCACACGTCTGAACTCTTTCCCTACACGACGCTCTTCCGATCTNNNTGCACGACTGAATGACACTTCT
GAGCACACGTCTGAACTCTTTCCCTACACGACGCTCTTCCGATCTNNNCACACCTGTAGTTCCAGCTAC
AGCACACGTCTGAACTCTTTCCCTACACGACGCTCTTCCGATCTNNNAGGAGAATTGCTTGAACTCG
GAGCACACGTCTGAACTCTTTCCCTACACGACGCTCTTCCGATCTNNNTGGGAGGATTGATTGAACC
GCACACGTCTGAACTCTTTCCCTACACGACGCTCTTCCGATCTNNNCTGTAGTCCCAGCTACTCAGAA
AGCACACGTCTGAACTCTTTCCCTACACGACGCTCTTCCGATCTNNNTGAGTTGGAAGGATCACTTG
AGCACACGTCTGAACTCTTTCCCTACACGACGCTCTTCCGATCTNNNCTTAGAAGGCTGAGGCAAGA
AGCACACGTCTGAACTCTTTCCCTACACGACGCTCTTCCGATCTNNNGAGGCAGGAAAATCACTTGA
GCACACGTCTGAACTCTTTCCCTACACGACGCTCTTCCGATCTNNNAATCACTTGAGGTCAGGAGTT
AGCACACGTCTGAACTCTTTCCCTACACGACGCTCTTCCGATCTNNNGTAACAGAGTGGGACTCCAT
GCACACGTCTGAACTCTTTCCCTACACGACGCTCTTCCGATCTNNNCACCTGTAATCCCAGCTACTCAG
GAGCACACGTCTGAACTCTTTCCCTACACGACGCTCTTCCGATCTNNNCTGAGGCAGGAGAATCACTT
GCACACGTCTGAACTCTTTCCCTACACGACGCTCTTCCGATCTNNNCTGTAGTCTCAGCTACGAAGGA

AGCACACGTCTGAACTCTTTCCCTACACGACGCTCTTCCGATCTNNNGAAGTTGCAGTGAACCGAGAT
GAGCACACGTCTGAACTCTTTCCCTACACGACGCTCTTCCGATCTNNNGAGGTTGCAGCAATCTGAGA
GAGCACACGTCTGAACTCTTTCCCTACACGACGCTCTTCCGATCTNNNAGGAGAATTGCTTGAACCCAG
GCACACGTCTGAACTCTTTCCCTACACGACGCTCTTCCGATCTNNNAGTTGCCTATGGTAAAGCTTGGA
GAGCACACGTCTGAACTCTTTCCCTACACGACGCTCTTCCGATCTNNNGAGTTTGCAGTGAGCTGAGAT
AGCACACGTCTGAACTCTTTCCCTACACGACGCTCTTCCGATCTNNNGTAGTCCCAGCTACTTAGGAA
GAGCACACGTCTGAACTCTTTCCCTACACGACGCTCTTCCGATCTNNNGCCTGTAGTCTTAGTTACTCG
AGCACACGTCTGAACTCTTTCCCTACACGACGCTCTTCCGATCTNNNTCTGTAATCCCAGCTACTCGGTA
GAGCACACGTCTGAACTCTTTCCCTACACGACGCTCTTCCGATCTNNNTCCCAACATGCTAGGATTACAG
GAGCACACGTCTGAACTCTTTCCCTACACGACGCTCTTCCGATCTNNNGAATCATTTGAATCCAGGAGG
GAGCACACGTCTGAACTCTTTCCCTACACGACGCTCTTCCGATCTNNNTGAGCAAAGATCACACCACTA
AGCACACGTCTGAACTCTTTCCCTACACGACGCTCTTCCGATCTNNNTATAGTCCCAGCTACTCAGGAG
GAGCACACGTCTGAACTCTTTCCCTACACGACGCTCTTCCGATCTNNNAGCTACTTGGAAGGCTAAGAT
AGCACACGTCTGAACTCTTTCCCTACACGACGCTCTTCCGATCTNNNTATAGTCCCAGCCACTTGGAAA
GCACACGTCTGAACTCTTTCCCTACACGACGCTCTTCCGATCTNNNCAACGGGAGAAGGGGAATCTAA
AGCACACGTCTGAACTCTTTCCCTACACGACGCTCTTCCGATCTNNNCCCACTCCTTGTCTGTACATTCA
AGCACACGTCTGAACTCTTTCCCTACACGACGCTCTTCCGATCTNNNTATATCCAGTCTCAGTCTCGGG
AGCACACGTCTGAACTCTTTCCCTACACGACGCTCTTCCGATCTNNNAAGCTGAGGTATTCTGGCAAT
GAGCACACGTCTGAACTCTTTCCCTACACGACGCTCTTCCGATCTNNNTGTAAATGTTCCTGTGTCTGCA
AGCACACGTCTGAACTCTTTCCCTACACGACGCTCTTCCGATCTNNNCTCTCTCTGTACCCTTCCTTCCTC
AGCACACGTCTGAACTCTTTCCCTACACGACGCTCTTCCGATCTNNNTCCCTCGTTGTATACCAGACAGA
GAGCACACGTCTGAACTCTTTCCCTACACGACGCTCTTCCGATCTNNNTTTGACAGGGTCTTCCTCTGT
GAGCACACGTCTGAACTCTTTCCCTACACGACGCTCTTCCGATCTNNNTCCTCTATAGAGCAGTGTCTC
GCACACGTCTGAACTCTTTCCCTACACGACGCTCTTCCGATCTNNNAGCCTCCCAAAGTGCTAGAATTA
AGCACACGTCTGAACTCTTTCCCTACACGACGCTCTTCCGATCTNNNCCCCTTGCAGATCTAATGTCAGA
AGCACACGTCTGAACTCTTTCCCTACACGACGCTCTTCCGATCTNNNCTGCTCCATACATTCGTCCTGAC
AGCACACGTCTGAACTCTTTCCCTACACGACGCTCTTCCGATCTNNNGAAGCAATAAACAGAACTGCC
GAGCACACGTCTGAACTCTTTCCCTACACGACGCTCTTCCGATCTNNNCATGGTGTCACATGCCTGTAAT
AGAGCACACGTCTGAACTCTTTCCCTACACGACGCTCTTCCGATCTNNNGCAAGAGAATCACTTGAACC
GAGCACACGTCTGAACTCTTTCCCTACACGACGCTCTTCCGATCTNNNTGTAGTCCTAGCTACTTGGGA
GCACACGTCTGAACTCTTTCCCTACACGACGCTCTTCCGATCTNNNCCACCCTGGGAGAAAAGAATGA
GAGCACACGTCTGAACTCTTTCCCTACACGACGCTCTTCCGATCTNNNGGATCACCTGAGGTCAAGAG
AGAGCACACGTCTGAACTCTTTCCCTACACGACGCTCTTCCGATCTNNNGGTAGTAGTGGTGCACCTGT
GAGCACACGTCTGAACTCTTTCCCTACACGACGCTCTTCCGATCTNNNGAGACTGAGATGGGAGAATT
GAGCACACGTCTGAACTCTTTCCCTACACGACGCTCTTCCGATCTNNNCACATGCCTGTAATCCCAGTA
GAGCACACGTCTGAACTCTTTCCCTACACGACGCTCTTCCGATCTNNNTGTGCCTGTAATCCCAACTACT
AGCACACGTCTGAACTCTTTCCCTACACGACGCTCTTCCGATCTNNNGTGCACCTGTAATCCCAGTTACT
GAGCACACGTCTGAACTCTTTCCCTACACGACGCTCTTCCGATCTNNNCTGTAATCCCAGCTACTCAGGA
AGAGCACACGTCTGAACTCTTTCCCTACACGACGCTCTTCCGATCTNNNTGTGTGTCTGTAATCCCAGC
AGCACACGTCTGAACTCTTTCCCTACACGACGCTCTTCCGATCTNNNACACCTGTAATCCCAGCACTTTG
AGCACACGTCTGAACTCTTTCCCTACACGACGCTCTTCCGATCTNNNGATTATGGCTTGAACCCAGGAG
GAGCACACGTCTGAACTCTTTCCCTACACGACGCTCTTCCGATCTNNNAATCTCAAGGGACCTCGAATA
AGCACACGTCTGAACTCTTTCCCTACACGACGCTCTTCCGATCTNNNCATGCCTGTAATCCCAGCTACTC

AGCACACGTCTGAACTCTTTCCCTACACGACGCTCTTCCGATCTNNNACCATTTTATTTTGCCCCTCCTT
AGCACACGTCTGAACTCTTTCCCTACACGACGCTCTTCCGATCTNNNTGGAGTGCAGTAGTACCATCA
GCACACGTCTGAACTCTTTCCCTACACGACGCTCTTCCGATCTNNNATTCAAGTGATTCTCCTGCCTCA
GAGCACACGTCTGAACTCTTTCCCTACACGACGCTCTTCCGATCTNNNAACTTGAAAGCATTCTCTGGT
AGCACACGTCTGAACTCTTTCCCTACACGACGCTCTTCCGATCTNNNCCTTTCATCTAAAGCAAGGTG
AGCACACGTCTGAACTCTTTCCCTACACGACGCTCTTCCGATCTNNNTCATGTAATCTTCCCACCTCAG
GCACACGTCTGAACTCTTTCCCTACACGACGCTCTTCCGATCTNNNGGACATCCTTGTTTTGTTCCTG
AGCACACGTCTGAACTCTTTCCCTACACGACGCTCTTCCGATCTNNNAGACTGAAACCTTGACATCCCT
GCACACGTCTGAACTCTTTCCCTACACGACGCTCTTCCGATCTNNNAGAGCTTTCCTTCATCCCTCAAG
AGCACACGTCTGAACTCTTTCCCTACACGACGCTCTTCCGATCTNNNGTGCAATCTTGACTCACTGCTA
AGCACACGTCTGAACTCTTTCCCTACACGACGCTCTTCCGATCTNNNGCTCTTACCATTGCTTTCCACT
AGCACACGTCTGAACTCTTTCCCTACACGACGCTCTTCCGATCTNNNATCCCATGACTTTACCCTACAC
GCACACGTCTGAACTCTTTCCCTACACGACGCTCTTCCGATCTNNNAGAAGTGGGGTTGCTGAATCA
AGCACACGTCTGAACTCTTTCCCTACACGACGCTCTTCCGATCTNNNTGAGACTGGGTTTTGCTCTAAC
GCACACGTCTGAACTCTTTCCCTACACGACGCTCTTCCGATCTNNNAGGTTGACACATGGAAAGTCTC
AGCACACGTCTGAACTCTTTCCCTACACGACGCTCTTCCGATCTNNNCTGAGTAGCTGGGACTACAG
GCACACGTCTGAACTCTTTCCCTACACGACGCTCTTCCGATCTNNNGGAAGTCATAGGAGAATGGAG
GCACACGTCTGAACTCTTTCCCTACACGACGCTCTTCCGATCTNNNCTTGAACCCGAGTGATTCAGAG
AGCACACGTCTGAACTCTTTCCCTACACGACGCTCTTCCGATCTNNNTTTAGTAGAGACGGGGTTTCA
AGCACACGTCTGAACTCTTTCCCTACACGACGCTCTTCCGATCTNNNAAAAGATCTTCCCATCTTGGCC
GCACACGTCTGAACTCTTTCCCTACACGACGCTCTTCCGATCTNNNATCTTAGCAAAGGAGAGGGCAA
GCACACGTCTGAACTCTTTCCCTACACGACGCTCTTCCGATCTNNNGATCTCAGCTCATTGCAAACTCC
GCACACGTCTGAACTCTTTCCCTACACGACGCTCTTCCGATCTNNNGGTAAAAGGTGAGGGAGGAG
AGCACACGTCTGAACTCTTTCCCTACACGACGCTCTTCCGATCTNNNCTCCAGGTGCAGGTTGTCATT
AGCACACGTCTGAACTCTTTCCCTACACGACGCTCTTCCGATCTNNNTCTTGCTTTTCTCCCTCTTCTCC
GCACACGTCTGAACTCTTTCCCTACACGACGCTCTTCCGATCTNNNCTCCCAAAGGGCTAGGATTACA
AGCACACGTCTGAACTCTTTCCCTACACGACGCTCTTCCGATCTNNNTTTATGCATTCTGCCATTCTG
AGAGCACACGTCTGAACTCTTTCCCTACACGACGCTCTTCCGATCTNNNAATGCTTTTTGTTGTTTGCCC
AGCACACGTCTGAACTCTTTCCCTACACGACGCTCTTCCGATCTNNNTCCAAATGCAAAACCACTTCCTT
GCACACGTCTGAACTCTTTCCCTACACGACGCTCTTCCGATCTNNNGGCTATGGAGAGCATGCTTTTT
AGCACACGTCTGAACTCTTTCCCTACACGACGCTCTTCCGATCTNNNGTCAAGCTATCCTCCTGACTCA
AGCACACGTCTGAACTCTTTCCCTACACGACGCTCTTCCGATCTNNNTTTCATGTTTGTTGGCCACTT
GCACACGTCTGAACTCTTTCCCTACACGACGCTCTTCCGATCTNNNGTACTTCCTGCTTCGTCTCTCAG
ACACGTCTGAACTCTTTCCCTACACGACGCTCTTCCGATCTNNNTCACGGAAATTTTATGTTACTAGTG
AGCACACGTCTGAACTCTTTCCCTACACGACGCTCTTCCGATCTNNNTGTCTCCTAGAATTCAACACAC
GCACACGTCTGAACTCTTTCCCTACACGACGCTCTTCCGATCTNNNCAACCTCTGTCCTCTGAGTTCAA
AGCACACGTCTGAACTCTTTCCCTACACGACGCTCTTCCGATCTNNNTCTGGCCTTGGTTGTCTTCAAT
GCACACGTCTGAACTCTTTCCCTACACGACGCTCTTCCGATCTNNNACTAACAGTGAACCTCTCAGCT
AGCACACGTCTGAACTCTTTCCCTACACGACGCTCTTCCGATCTNNNTATTTGCTGCCATACTGGAGTG
AGCACACGTCTGAACTCTTTCCCTACACGACGCTCTTCCGATCTNNNGCTTGCAGTTGTTCTAGCAGT
GCACACGTCTGAACTCTTTCCCTACACGACGCTCTTCCGATCTNNNGTAGAGTGCTTTGGCTTTGTTC
AGCACACGTCTGAACTCTTTCCCTACACGACGCTCTTCCGATCTNNNATCCATCTTCAACCTCCTGAGT
AGCACACGTCTGAACTCTTTCCCTACACGACGCTCTTCCGATCTNNNTTCACTCCTTGACGGTTTCTTT

GAGCACACGTCTGAACTCTTTCCCTACACGACGCTCTTCCGATCTNNNGTCTTATGCCCTATTGTCACCC

GAGCACACGTCTGAACTCTTTCCCTACACGACGCTCTTCCGATCTNNNTTTCTTTGATTGAGTGCCTTGG

GAGCACACGTCTGAACTCTTTCCCTACACGACGCTCTTCCGATCTNNNACTGAGGGTAAGCCACATCAA

AGCACACGTCTGAACTCTTTCCCTACACGACGCTCTTCCGATCTNNNAAAAGTGAGTTTGGCCCTCTCT

GAGCACACGTCTGAACTCTTTCCCTACACGACGCTCTTCCGATCTNNNGGCCAAAGTAAAAGGAATGC

GAGCACACGTCTGAACTCTTTCCCTACACGACGCTCTTCCGATCTNNNTAAAGTGTTCCTCTCCACTGCA

AGCACACGTCTGAACTCTTTCCCTACACGACGCTCTTCCGATCTNNNTGCATCTTGTCCCCACAAATCTA

AGCACACGTCTGAACTCTTTCCCTACACGACGCTCTTCCGATCTNNNGCCAAAACAGACTCTCAAGCAT

AGCACACGTCTGAACTCTTTCCCTACACGACGCTCTTCCGATCTNNNCATGAAGGTTCCAATGTCTCCAC

AGCACACGTCTGAACTCTTTCCCTACACGACGCTCTTCCGATCTNNNTTTGAAGCTTGTTTCCATATGCC

AGCACACGTCTGAACTCTTTCCCTACACGACGCTCTTCCGATCTNNNCTGTGCCTCTGGGGTATTATTCA

GAGCACACGTCTGAACTCTTTCCCTACACGACGCTCTTCCGATCTNNNACCCCAATCAATCTAGAACACG

AGCACACGTCTGAACTCTTTCCCTACACGACGCTCTTCCGATCTNNNTAGATACTATTGTGTGCGTCCGC

AGCACACGTCTGAACTCTTTCCCTACACGACGCTCTTCCGATCTNNNGGATTTCTCAGGACAGTGAGTG

AGCACACGTCTGAACTCTTTCCCTACACGACGCTCTTCCGATCTNNNAAGCAGACAATAGGACACACTC

AGCACACGTCTGAACTCTTTCCCTACACGACGCTCTTCCGATCTNNNCTCTTGTAAGGTAGGTCTGGTG

AGCACACGTCTGAACTCTTTCCCTACACGACGCTCTTCCGATCTNNNCTCGTGGAAGAGCATAGTCTAC

GAGCACACGTCTGAACTCTTTCCCTACACGACGCTCTTCCGATCTNNNGTCTGACTTTTGACTCCCAAGG

AGCACACGTCTGAACTCTTTCCCTACACGACGCTCTTCCGATCTNNNTCAGTGTTAGCCCTGTTGAAGA

GAGCACACGTCTGAACTCTTTCCCTACACGACGCTCTTCCGATCTNNNTCCTCCTCTACCTTCTAAAAGCT

AGAGCACACGTCTGAACTCTTTCCCTACACGACGCTCTTCCGATCTNNNTCAACAAATTTTGCCCCGTAC

GAGCACACGTCTGAACTCTTTCCCTACACGACGCTCTTCCGATCTNNNGGCAACTTTGTGTTTTTCTTCG

GAGCACACGTCTGAACTCTTTCCCTACACGACGCTCTTCCGATCTNNNATCACTTGAATCCAGGAGGCA

GAGCACACGTCTGAACTCTTTCCCTACACGACGCTCTTCCGATCTNNNGCAACAGAGTAAGACTCCCTC

AGCACACGTCTGAACTCTTTCCCTACACGACGCTCTTCCGATCTNNNTAATCCCAGTTACTCAGGAGGC

AGAGCACACGTCTGAACTCTTTCCCTACACGACGCTCTTCCGATCTNNNGCTGCTCTGTTTTGTTTTGCT

AGAGCACACGTCTGAACTCTTTCCCTACACGACGCTCTTCCGATCTNNNAGGCAGAAGAATCACTTGAA

GCACACGTCTGAACTCTTTCCCTACACGACGCTCTTCCGATCTNNNGGGAGGGGTTCTCATAGAGTAG

GCACACGTCTGAACTCTTTCCCTACACGACGCTCTTCCGATCTNNNACCAGGATTATGCCACTGTACTC

AGCACACGTCTGAACTCTTTCCCTACACGACGCTCTTCCGATCTNNNCCAACATGGTGAAACACTGTCTC

AGCACACGTCTGAACTCTTTCCCTACACGACGCTCTTCCGATCTNNNATGAGAATCGCTTGAATCCAGG

GAGCACACGTCTGAACTCTTTCCCTACACGACGCTCTTCCGATCTNNNGAAGTTGCAGTGAACTGAGAT

GAGCACACGTCTGAACTCTTTCCCTACACGACGCTCTTCCGATCTNNNCTGTAGTCCCAGCTACTCAGG

AGCACACGTCTGAACTCTTTCCCTACACGACGCTCTTCCGATCTNNNCTGAGACAGGAGAATCGCTTGA

AGCACACGTCTGAACTCTTTCCCTACACGACGCTCTTCCGATCTNNNTGTAATTCCAGCTACTTGGGAG

GAGCACACGTCTGAACTCTTTCCCTACACGACGCTCTTCCGATCTNNNGGAAAGTACTACAAGGGCACA

AGCACACGTCTGAACTCTTTCCCTACACGACGCTCTTCCGATCTNNNGTGACAGAGCAAGACTACGTCT

GCACACGTCTGAACTCTTTCCCTACACGACGCTCTTCCGATCTNNNACGTGTTTATCTGCTGACCTTCTC

GAGCACACGTCTGAACTCTTTCCCTACACGACGCTCTTCCGATCTNNNCAACATGGAGAAACCCCATCT

GAGCACACGTCTGAACTCTTTCCCTACACGACGCTCTTCCGATCTNNNGAGAATGGCATGAATCCAGG

GAGCACACGTCTGAACTCTTTCCCTACACGACGCTCTTCCGATCTNNNAATTGCTTGAACCTGGAAAGT

AGAGCACACGTCTGAACTCTTTCCCTACACGACGCTCTTCCGATCTNNNAGAGGTTGCATGAGCTAAGA

AGCACACGTCTGAACTCTTTCCCTACACGACGCTCTTCCGATCTNNNGAGGTTGCATTGAGCTGAGATC

AGCACACGTCTGAACTCTTTCCCTACACGACGCTCTTCCGATCTNNNCTGTAATCCCAGCTACTCAGG
AGAGCACACGTCTGAACTCTTTCCCTACACGACGCTCTTCCGATCTNNNGAGAATCGCTTGAACTCAG
AGCACACGTCTGAACTCTTTCCCTACACGACGCTCTTCCGATCTNNNCCAGGAGTTTGAGGTTGGAGT
AGCACACGTCTGAACTCTTTCCCTACACGACGCTCTTCCGATCTNNNCTTGAGCCTAGGAGTTCCAGA
AGCACACGTCTGAACTCTTTCCCTACACGACGCTCTTCCGATCTNNNTCGAGGCTGCAATGAACTATG
AGCACACGTCTGAACTCTTTCCCTACACGACGCTCTTCCGATCTNNNTGAGGCAGGAGAATCACTTG
GAGCACACGTCTGAACTCTTTCCCTACACGACGCTCTTCCGATCTNNNGTGAAACCCGCTCTCTACTAA
AGCACACGTCTGAACTCTTTCCCTACACGACGCTCTTCCGATCTNNNGCCTGTAATCCTAGCATTTTGG
AGCACACGTCTGAACTCTTTCCCTACACGACGCTCTTCCGATCTNNNGAGTCAGTAGCTACTCAGAAG
AGCACACGTCTGAACTCTTTCCCTACACGACGCTCTTCCGATCTNNNCAGAGCTTGTAGTGAGCAGAG
AGCACACGTCTGAACTCTTTCCCTACACGACGCTCTTCCGATCTNNNCAACATAGCAAACCCCCTCTCT
AGCACACGTCTGAACTCTTTCCCTACACGACGCTCTTCCGATCTNNNAACAAAGTGAAGTGAGACCC
AGCACACGTCTGAACTCTTTCCCTACACGACGCTCTTCCGATCTNNNAGTGCTGGGATTATAGGCGT
GCACACGTCTGAACTCTTTCCCTACACGACGCTCTTCCGATCTNNNGAGGCAAGAGAATCACATGAA
GCACACGTCTGAACTCTTTCCCTACACGACGCTCTTCCGATCTNNNACTGTAATCCCAGCTACTCAGG
AGCACACGTCTGAACTCTTTCCCTACACGACGCTCTTCCGATCTNNNGTTATGGTAAGCAGTGACTGT
GAGCACACGTCTGAACTCTTTCCCTACACGACGCTCTTCCGATCTNNNAGATCGAGTCACTGCATTCTA
AGCACACGTCTGAACTCTTTCCCTACACGACGCTCTTCCGATCTNNNCAGAAGTTGCAATGAGCCAA
AGCACACGTCTGAACTCTTTCCCTACACGACGCTCTTCCGATCTNNNCAAGATCACACCACTGAACTCC
GCACACGTCTGAACTCTTTCCCTACACGACGCTCTTCCGATCTNNNCTGAGGCAGGATAATGGTGTG
AGCACACGTCTGAACTCTTTCCCTACACGACGCTCTTCCGATCTNNNCTGGGCAAAAGAACAAGACT
AGCACACGTCTGAACTCTTTCCCTACACGACGCTCTTCCGATCTNNNCTGAGGCAGGAGAATCACTTG
AGCACACGTCTGAACTCTTTCCCTACACGACGCTCTTCCGATCTNNNGAATTGCTTGAGCCCAGGAAT
AGCACACGTCTGAACTCTTTCCCTACACGACGCTCTTCCGATCTNNNAGGATTGCTTGAGTCCAGGAA
AGCACACGTCTGAACTCTTTCCCTACACGACGCTCTTCCGATCTNNNTCACTTGATCCCAGGAGTTCAA
AGCACACGTCTGAACTCTTTCCCTACACGACGCTCTTCCGATCTNNNCTGTAATCCCAGCTACTAGGCA
AGCACACGTCTGAACTCTTTCCCTACACGACGCTCTTCCGATCTNNNAAGACCCATCAGTGTGCTGTA
AGCACACGTCTGAACTCTTTCCCTACACGACGCTCTTCCGATCTNNNCGGAGATCACACCATTGTACT
GCACACGTCTGAACTCTTTCCCTACACGACGCTCTTCCGATCTNNNATTTTGAGGCTGCAGTGAGCTA
AGCACACGTCTGAACTCTTTCCCTACACGACGCTCTTCCGATCTNNNAACTAAGGAGGTTGAGGATG
AGCACACGTCTGAACTCTTTCCCTACACGACGCTCTTCCGATCTNNNAGGTTGCAGTGAGTCAAGAT
GAGCACACGTCTGAACTCTTTCCCTACACGACGCTCTTCCGATCTNNNATACAAAAATTAGCTGGGC
AGCACACGTCTGAACTCTTTCCCTACACGACGCTCTTCCGATCTNNNGGTACATGCCTGTCATCCCATA
GCACACGTCTGAACTCTTTCCCTACACGACGCTCTTCCGATCTNNNGCAACAGAGAAAGACTGTCTCC
GAGCACACGTCTGAACTCTTTCCCTACACGACGCTCTTCCGATCTNNNAGGAGAATTGCTTGAACCTG
GAGCACACGTCTGAACTCTTTCCCTACACGACGCTCTTCCGATCTNNNAGGAAGGAGAATCACTTGAA
AGCACACGTCTGAACTCTTTCCCTACACGACGCTCTTCCGATCTNNNGAGAATTGCTTGAGCCTAGGA
GAGCACACGTCTGAACTCTTTCCCTACACGACGCTCTTCCGATCTNNNGCCTGTAATCCCAGATACTC
GAGCACACGTCTGAACTCTTTCCCTACACGACGCTCTTCCGATCTNNNTAGTCTCAGCTACTCAGAAG
AGAGCACACGTCTGAACTCTTTCCCTACACGACGCTCTTCCGATCTNNNCTGTAGTCCCAGCTACTCGA
AGCACACGTCTGAACTCTTTCCCTACACGACGCTCTTCCGATCTNNNGAGAATGGTGTAAACCTGGG
GAGCACACGTCTGAACTCTTTCCCTACACGACGCTCTTCCGATCTNNNAATAAGCTGTGATTGTGCC
GCACACGTCTGAACTCTTTCCCTACACGACGCTCTTCCGATCTNNNCATGAGAATTGCTTGAACCTGG

GAGCACACGTCTGAACTCTTTCCCTACACGACGCTCTTCCGATCTNNNACCTTGCTGAACTCACTGTAGT
AGCACACGTCTGAACTCTTTCCCTACACGACGCTCTTCCGATCTNNNGTAGCTGATTTTGGTGAGGCTT
AGCACACGTCTGAACTCTTTCCCTACACGACGCTCTTCCGATCTNNNCATGATCGTGCTACTACACTCCA
AGCACACGTCTGAACTCTTTCCCTACACGACGCTCTTCCGATCTNNNTTTTAGTAGAGGCGGGGTTT
GAGCACACGTCTGAACTCTTTCCCTACACGACGCTCTTCCGATCTNNNTGAAAGCCAGTATCAGATGGC
GAGCACACGTCTGAACTCTTTCCCTACACGACGCTCTTCCGATCTNNNATGGGAATAGGCAAGACACC
AGCACACGTCTGAACTCTTTCCCTACACGACGCTCTTCCGATCTNNNTGTTTCATATGAGAATGCGTCCT
GAGCACACGTCTGAACTCTTTCCCTACACGACGCTCTTCCGATCTNNNAGTAGAGATGGGGTTTTGCAT
AGCACACGTCTGAACTCTTTCCCTACACGACGCTCTTCCGATCTNNNCTCCCGGGTTCAAGTGATTCTTA
AGCACACGTCTGAACTCTTTCCCTACACGACGCTCTTCCGATCTNNNCTCCCTTGTCTCACTTTCCTCTCCA
AGCACACGTCTGAACTCTTTCCCTACACGACGCTCTTCCGATCTNNNTTTAGAGTGATGTTTCCCGCTGA
AGCACACGTCTGAACTCTTTCCCTACACGACGCTCTTCCGATCTNNNTGATCTCTTGACCTCGTGATCTC
AGCACACGTCTGAACTCTTTCCCTACACGACGCTCTTCCGATCTNNNCAGTTACTGTGCACTCCTCCTAC
AGCACACGTCTGAACTCTTTCCCTACACGACGCTCTTCCGATCTNNNCTCCACCTCATTTTTTGGCTCTC
GAGCACACGTCTGAACTCTTTCCCTACACGACGCTCTTCCGATCTNNNATCCAAAATTCCTGATCCCTGC
AGCACACGTCTGAACTCTTTCCCTACACGACGCTCTTCCGATCTNNNGTGCTGGAATTATAGGCATCAC
AGCACACGTCTGAACTCTTTCCCTACACGACGCTCTTCCGATCTNNNACAAAGTGCTAGGATTACAGG
AGCACACGTCTGAACTCTTTCCCTACACGACGCTCTTCCGATCTNNNTAAAGTGCTGGGATTACAGGCA
GAGCACACGTCTGAACTCTTTCCCTACACGACGCTCTTCCGATCTNNNTGTGCCAGGCTAACATTTTGTT
AGCACACGTCTGAACTCTTTCCCTACACGACGCTCTTCCGATCTNNNCCTGGCTAACTGGTGTCTTTTCT
AGCACACGTCTGAACTCTTTCCCTACACGACGCTCTTCCGATCTNNNAGTGCTGGGATTACAGACATGA
AGCACACGTCTGAACTCTTTCCCTACACGACGCTCTTCCGATCTNNNATTTTCCTTGCAGCAATCTCCTG
AGCACACGTCTGAACTCTTTCCCTACACGACGCTCTTCCGATCTNNNTGACAAACCATAGCTGGGTACT
GAGCACACGTCTGAACTCTTTCCCTACACGACGCTCTTCCGATCTNNNCCATCGTGGGCATAAATCAAC
AGCACACGTCTGAACTCTTTCCCTACACGACGCTCTTCCGATCTNNNCTCTCTGCATTGGCTGTTCTCTAC
GAGCACACGTCTGAACTCTTTCCCTACACGACGCTCTTCCGATCTNNNCACCCGGCTGTTTTGTTTTCATA
AGCACACGTCTGAACTCTTTCCCTACACGACGCTCTTCCGATCTNNNCAGTGACTCTGTAAGCTGACAG
GAGCACACGTCTGAACTCTTTCCCTACACGACGCTCTTCCGATCTNNNACTCCTGACCTCAAATGATCCA
AGCACACGTCTGAACTCTTTCCCTACACGACGCTCTTCCGATCTNNNCTAAGGGCCTATGATCCTAGGG
AGCACACGTCTGAACTCTTTCCCTACACGACGCTCTTCCGATCTNNNAGTGCTGGGATTACAGGCATAA
GAGCACACGTCTGAACTCTTTCCCTACACGACGCTCTTCCGATCTNNNTTTGAGATTGAGTTTCGCCCTT
AGCACACGTCTGAACTCTTTCCCTACACGACGCTCTTCCGATCTNNNCTCAAAGTGTTGGGATTACAGC
AGCACACGTCTGAACTCTTTCCCTACACGACGCTCTTCCGATCTNNNTCTGCCTTTCTTCCTTCTTCCTCC
AGCACACGTCTGAACTCTTTCCCTACACGACGCTCTTCCGATCTNNNAACTCTGCATGGTAGGTCATTG
AGCACACGTCTGAACTCTTTCCCTACACGACGCTCTTCCGATCTNNNATATCTCACACCCTCAACGTAGC
AGCACACGTCTGAACTCTTTCCCTACACGACGCTCTTCCGATCTNNNACCATGTTCTTCATACACAGCCA
AGCACACGTCTGAACTCTTTCCCTACACGACGCTCTTCCGATCTNNNCCAGCTGACTTACTGCCATCTTA
GAGCACACGTCTGAACTCTTTCCCTACACGACGCTCTTCCGATCTNNNACAGTGGACAATCACAGAACA
AGAGCACACGTCTGAACTCTTTCCCTACACGACGCTCTTCCGATCTNNNACCTAAAGCCATTTCCTAGGT
GAGCACACGTCTGAACTCTTTCCCTACACGACGCTCTTCCGATCTNNNGGGGTTCTCAGTTTTCCAGATC
AGCACACGTCTGAACTCTTTCCCTACACGACGCTCTTCCGATCTNNNTCCATTCTGTTTTTTCCCCAACTCA
AGAGCACACGTCTGAACTCTTTCCCTACACGACGCTCTTCCGATCTNNNAGTGATCCTTCCACTTTAGCCT
AGCACACGTCTGAACTCTTTCCCTACACGACGCTCTTCCGATCTNNNGCAGGAGTCTCTTTCTAACCTG

AGCACACGTCTGAACTCTTTCCCTACACGACGCTCTTCCGATCTNNNCTCCAAAAGTGCTGGGATTACA
AGCACACGTCTGAACTCTTTCCCTACACGACGCTCTTCCGATCTNNNACTGTTGGACTATTCTTGGCCAT
GCACACGTCTGAACTCTTTCCCTACACGACGCTCTTCCGATCTNNNAGACCCCAGCATTTCCCATAAAAC
GAGCACACGTCTGAACTCTTTCCCTACACGACGCTCTTCCGATCTNNNTGGAGTGACCATAGTTTTCTG
AGCACACGTCTGAACTCTTTCCCTACACGACGCTCTTCCGATCTNNNGCATGACTCCTTAGACCCCTTAG
GAGCACACGTCTGAACTCTTTCCCTACACGACGCTCTTCCGATCTNNNCCGTAATTCCAGCTACTCAGGA
GAGCACACGTCTGAACTCTTTCCCTACACGACGCTCTTCCGATCTNNNAGATTGTGGTGAGCTGAGATT
AGCACACGTCTGAACTCTTTCCCTACACGACGCTCTTCCGATCTNNNTGGACGAAACAGTGAGATTCCA
GAGCACACGTCTGAACTCTTTCCCTACACGACGCTCTTCCGATCTNNNAGACATGAGACTTGCATGAAC
GAGCACACGTCTGAACTCTTTCCCTACACGACGCTCTTCCGATCTNNNTGTAGTCCCAGCTACTAGGAA
GCACACGTCTGAACTCTTTCCCTACACGACGCTCTTCCGATCTNNNGAGACTCACTTGAATCCAGGAGG
AGCACACGTCTGAACTCTTTCCCTACACGACGCTCTTCCGATCTNNNTGGGTGATAGAGTGAGACTGT
AGAGCACACGTCTGAACTCTTTCCCTACACGACGCTCTTCCGATCTNNNCTGAGGCAGGAGAATAGCT
GAGCACACGTCTGAACTCTTTCCCTACACGACGCTCTTCCGATCTNNNGGTGACAGAAAGCTAGACTC
AGCACACGTCTGAACTCTTTCCCTACACGACGCTCTTCCGATCTNNNGTGACAGAGTGAGACTCCGTAT
AGCACACGTCTGAACTCTTTCCCTACACGACGCTCTTCCGATCTNNNTGCAGTGGCTCATTCATACCTAT
AGCACACGTCTGAACTCTTTCCCTACACGACGCTCTTCCGATCTNNNCTGAGGCAGGAGAATCACTTGA
AGAGCACACGTCTGAACTCTTTCCCTACACGACGCTCTTCCGATCTNNNAAAAAAAATTAGCCAGGCGT
AGAGCACACGTCTGAACTCTTTCCCTACACGACGCTCTTCCGATCTNNNGGTGATGGAGTGAGACTCAC
AGAGCACACGTCTGAACTCTTTCCCTACACGACGCTCTTCCGATCTNNNTGTGAGACTTGTAATGCCAG
AGCACACGTCTGAACTCTTTCCCTACACGACGCTCTTCCGATCTNNNTGGTAATAAAACCCACCTGGAG
GAGCACACGTCTGAACTCTTTCCCTACACGACGCTCTTCCGATCTNNNGCAGATTACCTGAGCTCAAGA
GAGCACACGTCTGAACTCTTTCCCTACACGACGCTCTTCCGATCTNNNTGAGCCAGGATCTCAATATTG
GAGCACACGTCTGAACTCTTTCCCTACACGACGCTCTTCCGATCTNNNGTTGCAGTGAGCTGTGATCAT
AGCACACGTCTGAACTCTTTCCCTACACGACGCTCTTCCGATCTNNNGAGAGGCTGAGACAGGATAAT
GAGCACACGTCTGAACTCTTTCCCTACACGACGCTCTTCCGATCTNNNCTGAGACACGAAAATCGCTTG
GAGCACACGTCTGAACTCTTTCCCTACACGACGCTCTTCCGATCTNNNAATTATAGGCTGGTTGGGGG
AGCACACGTCTGAACTCTTTCCCTACACGACGCTCTTCCGATCTNNNAAGAATTGCTTCAACCCAGGAG
GAGCACACGTCTGAACTCTTTCCCTACACGACGCTCTTCCGATCTNNNCAGAGAATTCCTTGAACCTGGC
GAGCACACGTCTGAACTCTTTCCCTACACGACGCTCTTCCGATCTNNNTGAAGTCAGGAGTTCAAGAC
GAGCACACGTCTGAACTCTTTCCCTACACGACGCTCTTCCGATCTNNNAAGGATAAGCTCACTGTCAGT
GAGCACACGTCTGAACTCTTTCCCTACACGACGCTCTTCCGATCTNNNAGATACCTGTAGTCCCAGCTAC
GAGCACACGTCTGAACTCTTTCCCTACACGACGCTCTTCCGATCTNNNAATGAGCTGAGATCGCACTAC
GAGCACACGTCTGAACTCTTTCCCTACACGACGCTCTTCCGATCTNNNGCTATGATTGATCACACCACTG
GAGCACACGTCTGAACTCTTTCCCTACACGACGCTCTTCCGATCTNNNGCTTGAGGTTGGAAGTTTGA
GCACACGTCTGAACTCTTTCCCTACACGACGCTCTTCCGATCTNNNGTGCCTGTAGTCCCAGCTATTAG
GCACACGTCTGAACTCTTTCCCTACACGACGCTCTTCCGATCTNNNTATACGCCTGTAATCCCAGCTACC
AGAGCACACGTCTGAACTCTTTCCCTACACGACGCTCTTCCGATCTNNNGTAATCTCAGCTACTCAGGA
GAGCACACGTCTGAACTCTTTCCCTACACGACGCTCTTCCGATCTNNNACACCTGTAGTCCCAGATACTC
AGAGCACACGTCTGAACTCTTTCCCTACACGACGCTCTTCCGATCTNNNCTGAGGCAGGAGAATCACTT
GAGCACACGTCTGAACTCTTTCCCTACACGACGCTCTTCCGATCTNNNCTGTAATTCCAGCTGCTCAGG
GAGCACACGTCTGAACTCTTTCCCTACACGACGCTCTTCCGATCTNNNGAACTTGCAGTGAGCTGAGAT
GAGCACACGTCTGAACTCTTTCCCTACACGACGCTCTTCCGATCTNNNTAAGGCAGGAGAATCGCTTGA

GCACACGTCTGAACTCTTTCCCTACACGACGCTCTTCCGATCTNNNCTGTAGTCCCAGCTACTCAGGAG
GAGCACACGTCTGAACTCTTTCCCTACACGACGCTCTTCCGATCTNNNTTTGAATCCAGGAGGCAGAG
AGCACACGTCTGAACTCTTTCCCTACACGACGCTCTTCCGATCTNNNGGGCAACATAGAGAGACCTT
AGCACACGTCTGAACTCTTTCCCTACACGACGCTCTTCCGATCTNNNGTTCACACCTGTAATGCCAACT
AGCACACGTCTGAACTCTTTCCCTACACGACGCTCTTCCGATCTNNNGCAAGACCCTGTCTCAAAACAA
GAGCACACGTCTGAACTCTTTCCCTACACGACGCTCTTCCGATCTNNNACACACCTATAGTCCCAGCT
GAGCACACGTCTGAACTCTTTCCCTACACGACGCTCTTCCGATCTNNNAAAAATTGGCAGGAGAATC
AGCACACGTCTGAACTCTTTCCCTACACGACGCTCTTCCGATCTNNNCTGGAGGTTGAGGTTACAGTG
AGCACACGTCTGAACTCTTTCCCTACACGACGCTCTTCCGATCTNNNGGAGACAGAGCAAGAACTTG
AGCACACGTCTGAACTCTTTCCCTACACGACGCTCTTCCGATCTNNNCAGCCTGAGCAACAAAAGTGA
AGCACACGTCTGAACTCTTTCCCTACACGACGCTCTTCCGATCTNNNGCTGAGGCAGGATAATCACT
GAGCACACGTCTGAACTCTTTCCCTACACGACGCTCTTCCGATCTNNNCACAGGAAGACTTGATTCAG
GCACACGTCTGAACTCTTTCCCTACACGACGCTCTTCCGATCTNNNAAGTACCCAGTTACCCAGTCTCA
AGCACACGTCTGAACTCTTTCCCTACACGACGCTCTTCCGATCTNNNGGAATTCTCCTAACCTGGAAA
AGCACACGTCTGAACTCTTTCCCTACACGACGCTCTTCCGATCTNNNTATCTGCCCACCTCAGTTTCTT
GAGCACACGTCTGAACTCTTTCCCTACACGACGCTCTTCCGATCTNNNCCTGTTTCCTCACACATTTGC
AGCACACGTCTGAACTCTTTCCCTACACGACGCTCTTCCGATCTNNNCTGGCTTGTCATTGGAACAG
AGCACACGTCTGAACTCTTTCCCTACACGACGCTCTTCCGATCTNNNGCCAGATTGCAGAGGATCATA
AGCACACGTCTGAACTCTTTCCCTACACGACGCTCTTCCGATCTNNNAGTATAGTGGCCTCCAGCTCTA
AGCACACGTCTGAACTCTTTCCCTACACGACGCTCTTCCGATCTNNNTGTGTACATAAGAAAGCGCCT
AGCACACGTCTGAACTCTTTCCCTACACGACGCTCTTCCGATCTNNNAACTTCCAGGCTGAACCAGAT
AGCACACGTCTGAACTCTTTCCCTACACGACGCTCTTCCGATCTNNNAAATGGAAGTACATTCGGTCA
GCACACGTCTGAACTCTTTCCCTACACGACGCTCTTCCGATCTNNNCTGGGACTATATGTATGGGCCA
AGCACACGTCTGAACTCTTTCCCTACACGACGCTCTTCCGATCTNNNCCCTCTATCTCCTATGTCACAG
AGCACACGTCTGAACTCTTTCCCTACACGACGCTCTTCCGATCTNNNGAGTGGATTGCATGCTTTCTCT
GAGCACACGTCTGAACTCTTTCCCTACACGACGCTCTTCCGATCTNNNGAGGACTTGAGTATACACGG
GAGCACACGTCTGAACTCTTTCCCTACACGACGCTCTTCCGATCTNNNAGTGCTGGGATTACAGACAT
AGCACACGTCTGAACTCTTTCCCTACACGACGCTCTTCCGATCTNNNTTCAATCTATCTCAGCTCCAGC
AGCACACGTCTGAACTCTTTCCCTACACGACGCTCTTCCGATCTNNNCCAGCAAGTTTTGTTTCTCCTGA
GAGCACACGTCTGAACTCTTTCCCTACACGACGCTCTTCCGATCTNNNCCCATGACACCATCTCACTAG
AGCACACGTCTGAACTCTTTCCCTACACGACGCTCTTCCGATCTNNNCACCACGCCTTGCTAATTTCATA
AGCACACGTCTGAACTCTTTCCCTACACGACGCTCTTCCGATCTNNNCAGGGACTTACTGTGAGTCAG
AGCACACGTCTGAACTCTTTCCCTACACGACGCTCTTCCGATCTNNNTCGATCTCCTGACCTCATGATCC
AGCACACGTCTGAACTCTTTCCCTACACGACGCTCTTCCGATCTNNNGGAATGGTTCCTGTACCTAGA
AGCACACGTCTGAACTCTTTCCCTACACGACGCTCTTCCGATCTNNNAAAAGTCATGATTTCCCCAGG
AGCACACGTCTGAACTCTTTCCCTACACGACGCTCTTCCGATCTNNNCAGTTCCGCAGTCATGTTTAC
AGCACACGTCTGAACTCTTTCCCTACACGACGCTCTTCCGATCTNNNCTGTGTTTGGCTTCTTTCACTGA
AGCACACGTCTGAACTCTTTCCCTACACGACGCTCTTCCGATCTNNNTACAAAAATTAGCCGGGCAGA
AGCACACGTCTGAACTCTTTCCCTACACGACGCTCTTCCGATCTNNNCTCAAGTGATCTTCCTGCCCTA
AGCACACGTCTGAACTCTTTCCCTACACGACGCTCTTCCGATCTNNNGGTTGAAGTAGTTCTCTGCCTC
AGCACACGTCTGAACTCTTTCCCTACACGACGCTCTTCCGATCTNNNGAAAGTCCAAGGAACCTCTGT
AGCACACGTCTGAACTCTTTCCCTACACGACGCTCTTCCGATCTNNNAGCAGAGTAACACCTTGTACA
AGCACACGTCTGAACTCTTTCCCTACACGACGCTCTTCCGATCTNNNGAGCCCCTATCAGAGCAACATA

AGCACACGTCTGAACTCTTTCCCTACACGACGCTCTTCCGATCTNNNAACATTTTCATGAAGGAGGCA
GCACACGTCTGAACTCTTTCCCTACACGACGCTCTTCCGATCTNNNTCACTATGTGATATGCCTGCTCC
AGCACACGTCTGAACTCTTTCCCTACACGACGCTCTTCCGATCTNNNTCAGAATTGTTTGCTGAGACA
GAGCACACGTCTGAACTCTTTCCCTACACGACGCTCTTCCGATCTNNNAAATTTGTGCTCCCTCTTGCTT
AGCACACGTCTGAACTCTTTCCCTACACGACGCTCTTCCGATCTNNNGGGATTATAGGAGTGAGCCA
AGCACACGTCTGAACTCTTTCCCTACACGACGCTCTTCCGATCTNNNTGGCCTGCTATATGTCCAAAA
AGCACACGTCTGAACTCTTTCCCTACACGACGCTCTTCCGATCTNNNACCAAGCCATTCATGAGGAATC
GCACACGTCTGAACTCTTTCCCTACACGACGCTCTTCCGATCTNNNTGGTACAGCCTCATTACCATTCT
GCACACGTCTGAACTCTTTCCCTACACGACGCTCTTCCGATCTNNNAGCATTGCTGTTTCTTTGACATT
GAGCACACGTCTGAACTCTTTCCCTACACGACGCTCTTCCGATCTNNNTGAGGCTAAGCAAAGGAGA
AGCACACGTCTGAACTCTTTCCCTACACGACGCTCTTCCGATCTNNNTTGGGAGCTCTATGAATTCAG
GAGCACACGTCTGAACTCTTTCCCTACACGACGCTCTTCCGATCTNNNAGAAGATCTCTGTGGAACTG
AGCACACGTCTGAACTCTTTCCCTACACGACGCTCTTCCGATCTNNNTCAGCCCTTACAAAAACAAAGC
AGCACACGTCTGAACTCTTTCCCTACACGACGCTCTTCCGATCTNNNAAAAGTCCCATTTCTCCCATGG
AGCACACGTCTGAACTCTTTCCCTACACGACGCTCTTCCGATCTNNNGGGAATTTGTGTTGGCAGTAT
AGCACACGTCTGAACTCTTTCCCTACACGACGCTCTTCCGATCTNNNCACTACTTGTCTTCTCAGCCTGT
AGCACACGTCTGAACTCTTTCCCTACACGACGCTCTTCCGATCTNNNTTTGAGAGAAGAGCAGGAGT
GCACACGTCTGAACTCTTTCCCTACACGACGCTCTTCCGATCTNNNCCCCTGAACCTCAATCTCCTTACC
AGCACACGTCTGAACTCTTTCCCTACACGACGCTCTTCCGATCTNNNGTCTCGAATTTCTGACCTCAGG
AGCACACGTCTGAACTCTTTCCCTACACGACGCTCTTCCGATCTNNNAGTTCCGAGTCTGTGGAGTTT
AGCACACGTCTGAACTCTTTCCCTACACGACGCTCTTCCGATCTNNNGCTAGAGTACAGTGGCACAA
AGCACACGTCTGAACTCTTTCCCTACACGACGCTCTTCCGATCTNNNAGTGCTAGGATTACAGGCAT
GCACACGTCTGAACTCTTTCCCTACACGACGCTCTTCCGATCTNNNATGGTCTATGTTAGGGTTGGCA
AGCACACGTCTGAACTCTTTCCCTACACGACGCTCTTCCGATCTNNNCGCCCGCCTTTTTTTTTCTTTTT
AGCACACGTCTGAACTCTTTCCCTACACGACGCTCTTCCGATCTNNNGAGAAAGCAAATCTTCCTGTG
AGCACACGTCTGAACTCTTTCCCTACACGACGCTCTTCCGATCTNNNTGAGTAGCTAGGACTACAGGT
AGCACACGTCTGAACTCTTTCCCTACACGACGCTCTTCCGATCTNNNTAGGTTCAAGTGATTCTCCTG
AGCACACGTCTGAACTCTTTCCCTACACGACGCTCTTCCGATCTNNNGCCAAAGTCATTAGCAGGCAT
AGCACACGTCTGAACTCTTTCCCTACACGACGCTCTTCCGATCTNNNGTTTCACCATGTTAGCCAGGAT
AGCACACGTCTGAACTCTTTCCCTACACGACGCTCTTCCGATCTNNNACATTACACACATATGGCTTGC
GAGCACACGTCTGAACTCTTTCCCTACACGACGCTCTTCCGATCTNNNCTAAGGCAGGAGAATCGCTT
AGCACACGTCTGAACTCTTTCCCTACACGACGCTCTTCCGATCTNNNGTTGCAGTAAGCCAATATCACC
AGCACACGTCTGAACTCTTTCCCTACACGACGCTCTTCCGATCTNNNCTGTAGTTCCAGCTACTCAGG
GCACACGTCTGAACTCTTTCCCTACACGACGCTCTTCCGATCTNNNATGAACCTGGGAGACGGAAAT
GCACACGTCTGAACTCTTTCCCTACACGACGCTCTTCCGATCTNNNGGAGACTAAGCAAGACTACCTCG
GAGCACACGTCTGAACTCTTTCCCTACACGACGCTCTTCCGATCTNNNCAGGAGGATAAGGCAGGAG
GAGCACACGTCTGAACTCTTTCCCTACACGACGCTCTTCCGATCTNNNCTGAGGTGGGAGGATCATTT
AGCACACGTCTGAACTCTTTCCCTACACGACGCTCTTCCGATCTNNNAGAGCTTGCAGTGAGCCAATAT
AGCACACGTCTGAACTCTTTCCCTACACGACGCTCTTCCGATCTNNNCGAAACACAGCAAAACCTCAT
AGCACACGTCTGAACTCTTTCCCTACACGACGCTCTTCCGATCTNNNCTGTAGACCCAGCTACTCCAGA
GCACACGTCTGAACTCTTTCCCTACACGACGCTCTTCCGATCTNNNTGTAGTACCAGCTACTCAGGAG
GCACACGTCTGAACTCTTTCCCTACACGACGCTCTTCCGATCTNNNCAGAGGTTACAGTGAGCCAAGA
GCACACGTCTGAACTCTTTCCCTACACGACGCTCTTCCGATCTNNNCTGTAGTCCCAGCTACTCAGGAG

AGCACACGTCTGAACTCTTTCCCTACACGACGCTCTTCCGATCTNNNCAGAAGAATTGCTTGAACCTG
AGAGCACACGTCTGAACTCTTTCCCTACACGACGCTCTTCCGATCTNNNATGGCAAGAGAATTGCTTG
AGAGCACACGTCTGAACTCTTTCCCTACACGACGCTCTTCCGATCTNNNGAGAATCGCTTGAACCAGA
GCACACGTCTGAACTCTTTCCCTACACGACGCTCTTCCGATCTNNNCTGAGGCAGAAGAATGGTGTG
GCACACGTCTGAACTCTTTCCCTACACGACGCTCTTCCGATCTNNNGACTGGGAGAGAGAGTGAGA
GAGCACACGTCTGAACTCTTTCCCTACACGACGCTCTTCCGATCTNNNGAATTGCTTGAACCCTAGAA
GCACACGTCTGAACTCTTTCCCTACACGACGCTCTTCCGATCTNNNTGAGGTAGAGAGTTGCTTGAA
AGCACACGTCTGAACTCTTTCCCTACACGACGCTCTTCCGATCTNNNCAGAAGAATTGCTTGAACCTG
GAGCACACGTCTGAACTCTTTCCCTACACGACGCTCTTCCGATCTNNNGAGATTGTGCCATTGTACTC
AGCACACGTCTGAACTCTTTCCCTACACGACGCTCTTCCGATCTNNNGTGCCTAAAATCCCAGCTACT
AGCACACGTCTGAACTCTTTCCCTACACGACGCTCTTCCGATCTNNNGAGAATCGCTTGAATCCTGGA
GCACACGTCTGAACTCTTTCCCTACACGACGCTCTTCCGATCTNNNAAGGAGAATCTATTGAACCCGG
AGAGCACACGTCTGAACTCTTTCCCTACACGACGCTCTTCCGATCTNNNCTAAGGCAGGAGAATGACG
AGCACACGTCTGAACTCTTTCCCTACACGACGCTCTTCCGATCTNNNCACTACATTCCAGTCTGGTGA
AGCACACGTCTGAACTCTTTCCCTACACGACGCTCTTCCGATCTNNNGTTACAGTGAGCCGTGATCAA
GCACACGTCTGAACTCTTTCCCTACACGACGCTCTTCCGATCTNNNGAGAATCACTTGAACTCGGGA
GAGCACACGTCTGAACTCTTTCCCTACACGACGCTCTTCCGATCTNNNCCAAGATCATGCCATTGAACT
GAGCACACGTCTGAACTCTTTCCCTACACGACGCTCTTCCGATCTNNNTGTAGTCCCAGCTACTAAGG
GAGCACACGTCTGAACTCTTTCCCTACACGACGCTCTTCCGATCTNNNTACTTGAAAGGCTGAGATGG
AGCACACGTCTGAACTCTTTCCCTACACGACGCTCTTCCGATCTNNNGTAAGCTATGATCATGCCACTG
AGCACACGTCTGAACTCTTTCCCTACACGACGCTCTTCCGATCTNNNCCAAGATTGTGCCATTGTACT
AGCACACGTCTGAACTCTTTCCCTACACGACGCTCTTCCGATCTNNNAGAAGTTGCAGTGAACCGAG
AGCACACGTCTGAACTCTTTCCCTACACGACGCTCTTCCGATCTNNNGGTGCCTGTAATCCTAGCTACT
GAGCACACGTCTGAACTCTTTCCCTACACGACGCTCTTCCGATCTNNNTGAGGCAGAGAATTGCTTGA
GCACACGTCTGAACTCTTTCCCTACACGACGCTCTTCCGATCTNNNGGAGCTTACAGTGAGCCAAGAT
AGCACACGTCTGAACTCTTTCCCTACACGACGCTCTTCCGATCTNNNTCATGCCACTATACTCTGAAGC
GCACACGTCTGAACTCTTTCCCTACACGACGCTCTTCCGATCTNNNTGAGGCAGGAGAATCTCTTGA
AGCACACGTCTGAACTCTTTCCCTACACGACGCTCTTCCGATCTNNNCAACATAGTGAAGCCCTGTCT
AGCACACGTCTGAACTCTTTCCCTACACGACGCTCTTCCGATCTNNNGAGCTTGCAGTGAACCGAGAT
GCACACGTCTGAACTCTTTCCCTACACGACGCTCTTCCGATCTNNNGGCCACAAGAATGAAACTCCAT
GAGCACACGTCTGAACTCTTTCCCTACACGACGCTCTTCCGATCTNNNTTGAATCCAGGAGGTGAAG
GCACACGTCTGAACTCTTTCCCTACACGACGCTCTTCCGATCTNNNGGCGACAGAGTGAGATCCTAT
AGCACACGTCTGAACTCTTTCCCTACACGACGCTCTTCCGATCTNNNACCTTCCCTCCACTATTGTCCT
GAGCACACGTCTGAACTCTTTCCCTACACGACGCTCTTCCGATCTNNNGGGTGATGGAATGAGACTCT
GAGCACACGTCTGAACTCTTTCCCTACACGACGCTCTTCCGATCTNNNCAGCCTGGGCAATATAGCAA
AGCACACGTCTGAACTCTTTCCCTACACGACGCTCTTCCGATCTNNNAAGGCAACATAGTGAGACCT
GAGCACACGTCTGAACTCTTTCCCTACACGACGCTCTTCCGATCTNNNTAAGGCAGAAGAATCGCTTG
GCACACGTCTGAACTCTTTCCCTACACGACGCTCTTCCGATCTNNNCTGAGGCAGGAAAATTGCATGA
AGAGCACACGTCTGAACTCTTTCCCTACACGACGCTCTTCCGATCTNNNGAGGTGAGAGGATGGCTT
GCACACGTCTGAACTCTTTCCCTACACGACGCTCTTCCGATCTNNNTAGCAACAGAGTGAGACTCCAT
AGCACACGTCTGAACTCTTTCCCTACACGACGCTCTTCCGATCTNNNGGCTAAGGTGGGAGGATAA
AGCACACGTCTGAACTCTTTCCCTACACGACGCTCTTCCGATCTNNNACTTCTTTGTTCAGGTGTCCTC
GCACACGTCTGAACTCTTTCCCTACACGACGCTCTTCCGATCTNNNGTTTGAGACCAGTTGAGAACA

GAGCACACGTCTGAACTCTTTCCCTACACGACGCTCTTCCGATCTNNNTCAGACCTAGCCACAATTTTG
GAGCACACGTCTGAACTCTTTCCCTACACGACGCTCTTCCGATCTNNNCTGAGGCAGGAGAATCACTT
AGCACACGTCTGAACTCTTTCCCTACACGACGCTCTTCCGATCTNNNAGAGGTTGCAGTGAGCTAAGA
GAGCACACGTCTGAACTCTTTCCCTACACGACGCTCTTCCGATCTNNNTCTGGGTGATGGAGTGATAC
AGAGCACACGTCTGAACTCTTTCCCTACACGACGCTCTTCCGATCTNNNAGCCTAGGTTACAGGGTAA
CACACGTCTGAACTCTTTCCCTACACGACGCTCTTCCGATCTNNNTTAGGACCAAATGTGCTACCTCCC
GAGCACACGTCTGAACTCTTTCCCTACACGACGCTCTTCCGATCTNNNCATGAGAATTGCTTGAACCCA
AGCACACGTCTGAACTCTTTCCCTACACGACGCTCTTCCGATCTNNNCGGAATTCCCTGTCTATTCTGG
AGCACACGTCTGAACTCTTTCCCTACACGACGCTCTTCCGATCTNNNGAGGAATCACTTCAACCTGGG
AGCACACGTCTGAACTCTTTCCCTACACGACGCTCTTCCGATCTNNNCTGTAGTCCCAGCTACTTGAGA
AGCACACGTCTGAACTCTTTCCCTACACGACGCTCTTCCGATCTNNNATAGCTCATGCCTGTAATCCCA
GAGCACACGTCTGAACTCTTTCCCTACACGACGCTCTTCCGATCTNNNATACAAAAATTGGCCGGACA
GCACACGTCTGAACTCTTTCCCTACACGACGCTCTTCCGATCTNNNAGAATCACTTGAACCCAGGAGA
GAGCACACGTCTGAACTCTTTCCCTACACGACGCTCTTCCGATCTNNNGCGTAACAGAGCAAGATTC
AGCACACGTCTGAACTCTTTCCCTACACGACGCTCTTCCGATCTNNNCTGTAGTCCCAGCTACTCAGGA
GAGCACACGTCTGAACTCTTTCCCTACACGACGCTCTTCCGATCTNNNCACATGAGGTCAGGAGTTTG
AGCACACGTCTGAACTCTTTCCCTACACGACGCTCTTCCGATCTNNNCATGCCTGTAGTACCAGCTACT
AGCACACGTCTGAACTCTTTCCCTACACGACGCTCTTCCGATCTNNNTAATTCCATCCAGGTTGCTGTG
GCACACGTCTGAACTCTTTCCCTACACGACGCTCTTCCGATCTNNNAAACCTGAGTCTGAATCCTGGT
AGCACACGTCTGAACTCTTTCCCTACACGACGCTCTTCCGATCTNNNATTTGGTCAGGCTAGTCTCGA
GCACACGTCTGAACTCTTTCCCTACACGACGCTCTTCCGATCTNNNACAGGCCTTTGATCAGAGGTAA
GCACACGTCTGAACTCTTTCCCTACACGACGCTCTTCCGATCTNNNACGCCCGGCCTCTATTTTTATAA
AGCACACGTCTGAACTCTTTCCCTACACGACGCTCTTCCGATCTNNNTCCTGAATAGCTGAGAATAGG
AGCACACGTCTGAACTCTTTCCCTACACGACGCTCTTCCGATCTNNNCACAGAAATACCAGAGGTGTG
AGCACACGTCTGAACTCTTTCCCTACACGACGCTCTTCCGATCTNNNCTTCCTGTCTTCCCCACTCATTT
GAGCACACGTCTGAACTCTTTCCCTACACGACGCTCTTCCGATCTNNNTAGTTGCAGGATGGTACTTG
GCACACGTCTGAACTCTTTCCCTACACGACGCTCTTCCGATCTNNNCATCAGCTGTTGGGTTACATGT
GCACACGTCTGAACTCTTTCCCTACACGACGCTCTTCCGATCTNNNATCGCTGCCAGCATTTATAAAC
GCACACGTCTGAACTCTTTCCCTACACGACGCTCTTCCGATCTNNNAGAAAATGTTTGGAACCCACGT
AGCACACGTCTGAACTCTTTCCCTACACGACGCTCTTCCGATCTNNNGGATGTTCTGCTCTCACTGATG
AGCACACGTCTGAACTCTTTCCCTACACGACGCTCTTCCGATCTNNNTTTTAGTAGAGGCGGGGTTT
AGCACACGTCTGAACTCTTTCCCTACACGACGCTCTTCCGATCTNNNTGTGACTTGAATTAGGAGGCA
AGCACACGTCTGAACTCTTTCCCTACACGACGCTCTTCCGATCTNNNGGCCATCTGTATAGCTTCTTTG
AGCACACGTCTGAACTCTTTCCCTACACGACGCTCTTCCGATCTNNNCTGGAAACCCGTGGACATATAC
GCACACGTCTGAACTCTTTCCCTACACGACGCTCTTCCGATCTNNNAGTGCTAGGATTACAGGCATG
GAGCACACGTCTGAACTCTTTCCCTACACGACGCTCTTCCGATCTNNNGTGTACTCTGGTCTCTTGGGA
GCACACGTCTGAACTCTTTCCCTACACGACGCTCTTCCGATCTNNNTAGGCAGTTTTTCAACCCTCACT
GCACACGTCTGAACTCTTTCCCTACACGACGCTCTTCCGATCTNNNGGTAAGCAAAGCCAAGATGGTT
AGCACACGTCTGAACTCTTTCCCTACACGACGCTCTTCCGATCTNNNCTCAAACTCCTGACCTCATGTGA
GAGCACACGTCTGAACTCTTTCCCTACACGACGCTCTTCCGATCTNNNAAGTCTGGGATTACAGGCAT
GCACACGTCTGAACTCTTTCCCTACACGACGCTCTTCCGATCTNNNACAGTGCACCTATAATCAGGGA
GCACACGTCTGAACTCTTTCCCTACACGACGCTCTTCCGATCTNNNTGGCCCAACACTGACTTCTAAAT
GCACACGTCTGAACTCTTTCCCTACACGACGCTCTTCCGATCTNNNAGACAGGAGACAAAGTGGCTA

GAGCACACGTCTGAACTCTTTCCCTACACGACGCTCTTCCGATCTNNNCCTGTGTTACTTTGACATGCTG
AGCACACGTCTGAACTCTTTCCCTACACGACGCTCTTCCGATCTNNNAAGGGGATGTAGTATGCTGTTG
GAGCACACGTCTGAACTCTTTCCCTACACGACGCTCTTCCGATCTNNNTCAATCATCCTGACCTCGTGAT
AGCACACGTCTGAACTCTTTCCCTACACGACGCTCTTCCGATCTNNNGGATGACAGCACACCTCTTTTTT
AGCACACGTCTGAACTCTTTCCCTACACGACGCTCTTCCGATCTNNNTAGCTCACTGCAGCCTAGATTT
AGCACACGTCTGAACTCTTTCCCTACACGACGCTCTTCCGATCTNNNCCAATCCAGCCCTCTACAATACA
AGCACACGTCTGAACTCTTTCCCTACACGACGCTCTTCCGATCTNNNTTGAACTCCTGACCTCATGATC
AGCACACGTCTGAACTCTTTCCCTACACGACGCTCTTCCGATCTNNNTAAAGCAATCCTCCCATCTCAG
AGCACACGTCTGAACTCTTTCCCTACACGACGCTCTTCCGATCTNNNGTCCCTAAAGGCTCAAAAGTCCT
GAGCACACGTCTGAACTCTTTCCCTACACGACGCTCTTCCGATCTNNNATGCTCCAAGTGCTACTCTTCA
GCACACGTCTGAACTCTTTCCCTACACGACGCTCTTCCGATCTNNNAGGAGAAAAGCCAGAATCCTCAA
AGCACACGTCTGAACTCTTTCCCTACACGACGCTCTTCCGATCTNNNCAGTAAAGAGAGCCTCACAAGG
GAGCACACGTCTGAACTCTTTCCCTACACGACGCTCTTCCGATCTNNNGTTCATGATCATGGCTCATTG
AGCACACGTCTGAACTCTTTCCCTACACGACGCTCTTCCGATCTNNNAGTGGCACAATTATGGCTCATT
AGCACACGTCTGAACTCTTTCCCTACACGACGCTCTTCCGATCTNNNCAAACTCCTGGGATCTAGTGAC
AGCACACGTCTGAACTCTTTCCCTACACGACGCTCTTCCGATCTNNNAGACTTCAGAGCACTTTCACAG
AGCACACGTCTGAACTCTTTCCCTACACGACGCTCTTCCGATCTNNNAGTGTGGCACATAGTAAGAGCT
GAGCACACGTCTGAACTCTTTCCCTACACGACGCTCTTCCGATCTNNNTCTTGCCTGCCACTCTTAGAAA
GAGCACACGTCTGAACTCTTTCCCTACACGACGCTCTTCCGATCTNNNCACCACCATGCCTGACTAATTT
AGCACACGTCTGAACTCTTTCCCTACACGACGCTCTTCCGATCTNNNAATGTCTGTTCATGTCCTTTGCC
AGCACACGTCTGAACTCTTTCCCTACACGACGCTCTTCCGATCTNNNGAAAGGTGGGTTGGTGTAAAT
AGCACACGTCTGAACTCTTTCCCTACACGACGCTCTTCCGATCTNNNAACTAGTAAAAACTTGCTGCAG
AGCACACGTCTGAACTCTTTCCCTACACGACGCTCTTCCGATCTNNNCCAGTGCTGAGACTATAGGTG
AGCACACGTCTGAACTCTTTCCCTACACGACGCTCTTCCGATCTNNNGCCAGAAGACAATGGAGTGGT
AGCACACGTCTGAACTCTTTCCCTACACGACGCTCTTCCGATCTNNNGAGTATCTGGGACTAAGGCACC
GAGCACACGTCTGAACTCTTTCCCTACACGACGCTCTTCCGATCTNNNCTTTGTGGAAGAAGTTTGGCT
GAGCACACGTCTGAACTCTTTCCCTACACGACGCTCTTCCGATCTNNNCTGTGCTGGGTTTCTGTCTTCT
AGCACACGTCTGAACTCTTTCCCTACACGACGCTCTTCCGATCTNNNAGTATAAAACCCAGCTAGCAG
GAGCACACGTCTGAACTCTTTCCCTACACGACGCTCTTCCGATCTNNNAAAAATCAGTGGATCCAGGA
AGCACACGTCTGAACTCTTTCCCTACACGACGCTCTTCCGATCTNNNCCCTCCTCAAGAAGTGCCTTATG
GAGCACACGTCTGAACTCTTTCCCTACACGACGCTCTTCCGATCTNNNTTTGGGGGTTTACTAAGCTTG
CACACGTCTGAACTCTTTCCCTACACGACGCTCTTCCGATCTNNNTGTGAGTTTTTCAGCTCTATCAGA
AGCACACGTCTGAACTCTTTCCCTACACGACGCTCTTCCGATCTNNNGTAGGTGCTCAATAAATGCAGG
GAGCACACGTCTGAACTCTTTCCCTACACGACGCTCTTCCGATCTNNNCTCCCAAGTAGCTGGAACTAC
AGCACACGTCTGAACTCTTTCCCTACACGACGCTCTTCCGATCTNNNAGCCTGTTCCTGTGTTCTTTAGA
AGCACACGTCTGAACTCTTTCCCTACACGACGCTCTTCCGATCTNNNCATCTTACCTCAGTGCTCCTCTCC
GAGCACACGTCTGAACTCTTTCCCTACACGACGCTCTTCCGATCTNNNCCAGAAGCAATTGGGAGTCAA
AGCACACGTCTGAACTCTTTCCCTACACGACGCTCTTCCGATCTNNNGATTACAGGAGCCTACCACCAT
AGCACACGTCTGAACTCTTTCCCTACACGACGCTCTTCCGATCTNNNTATGCTTGTTCTATTCCCCCCTCC
AGCACACGTCTGAACTCTTTCCCTACACGACGCTCTTCCGATCTNNNAGCCTACTGCCTTCTTCTTTTGTC
AGCACACGTCTGAACTCTTTCCCTACACGACGCTCTTCCGATCTNNNTCCTACTGACTCCCTGTAGTGAA
GCACACGTCTGAACTCTTTCCCTACACGACGCTCTTCCGATCTNNNGATTTCAGATTGAGAAGCCTGG
AGCACACGTCTGAACTCTTTCCCTACACGACGCTCTTCCGATCTNNNCACATCTGTCATTCACTCTGTGC

AGCACACGTCTGAACTCTTTCCCTACACGACGCTCTTCCGATCTNNNTGTTTAGTAGAGACAGGCTGG
AGCACACGTCTGAACTCTTTCCCTACACGACGCTCTTCCGATCTNNNCTCGAAGTCCTGACCTCATGATC
GAGCACACGTCTGAACTCTTTCCCTACACGACGCTCTTCCGATCTNNNCCCTGCTGGTTTTGAATTTCTG
GAGCACACGTCTGAACTCTTTCCCTACACGACGCTCTTCCGATCTNNNTATTTGTGGCTCTCTCCTTCCTC
AGCACACGTCTGAACTCTTTCCCTACACGACGCTCTTCCGATCTNNNTCTGGGTTCAAGTGATTCTCCT
GAGCACACGTCTGAACTCTTTCCCTACACGACGCTCTTCCGATCTNNNTGTAGTGTCTGAGTTCTCTGT
GAGCACACGTCTGAACTCTTTCCCTACACGACGCTCTTCCGATCTNNNTGAAAGCTCACTGTCAGAGTT
AGCACACGTCTGAACTCTTTCCCTACACGACGCTCTTCCGATCTNNNACCTCAACCCATTGTCATATTGAC
AGCACACGTCTGAACTCTTTCCCTACACGACGCTCTTCCGATCTNNNCTGAGTTCACACAATTCTCCTGCC
GAGCACACGTCTGAACTCTTTCCCTACACGACGCTCTTCCGATCTNNNTTCTGCTCCCACCTCCTCTATAA
AGCACACGTCTGAACTCTTTCCCTACACGACGCTCTTCCGATCTNNNTGTCAAATAAAGCAGGGAATGC
GAGCACACGTCTGAACTCTTTCCCTACACGACGCTCTTCCGATCTNNNAATAGCCACTGTACTCCAACCTC
AGCACACGTCTGAACTCTTTCCCTACACGACGCTCTTCCGATCTNNNGGAGAGCTGGGAGTCTTTTTCA
AGCACACGTCTGAACTCTTTCCCTACACGACGCTCTTCCGATCTNNNTGGGTAGCTGAAATTACAGGCA
AGCACACGTCTGAACTCTTTCCCTACACGACGCTCTTCCGATCTNNNCCTCTGACTTTCCTCCATCCCTAC
GAGCACACGTCTGAACTCTTTCCCTACACGACGCTCTTCCGATCTNNNTCAATTCCATCTGCTCCCAAAG
GAGCACACGTCTGAACTCTTTCCCTACACGACGCTCTTCCGATCTNNNGCTGTGTATTTGGCACCTTTCT
GAGCACACGTCTGAACTCTTTCCCTACACGACGCTCTTCCGATCTNNNACAGATTGTTGGGGAGAAAT
AGCACACGTCTGAACTCTTTCCCTACACGACGCTCTTCCGATCTNNNCCACTGTGCCAAGCCTAAAAAAT
AGCACACGTCTGAACTCTTTCCCTACACGACGCTCTTCCGATCTNNNGAGTGGCTGGGACTATAGGAAA
GAGCACACGTCTGAACTCTTTCCCTACACGACGCTCTTCCGATCTNNNGTCGAGGGTGACAACTGATGT
AGCACACGTCTGAACTCTTTCCCTACACGACGCTCTTCCGATCTNNNTAAATACCTGAGACTTGGCGCA
AGCACACGTCTGAACTCTTTCCCTACACGACGCTCTTCCGATCTNNNAGTAGCCTTGAATGACCTTTTGT
AGCACACGTCTGAACTCTTTCCCTACACGACGCTCTTCCGATCTNNNCAGCAGCCATAAAACCAGTAACC
GAGCACACGTCTGAACTCTTTCCCTACACGACGCTCTTCCGATCTNNNCAGGGGCTCTGCATTTTTACTT
AGCACACGTCTGAACTCTTTCCCTACACGACGCTCTTCCGATCTNNNTGTGAGGGTCCTATTGCAATTC
AGCACACGTCTGAACTCTTTCCCTACACGACGCTCTTCCGATCTNNNGCCACTGCTCAAGAATGACTTAT
GCACACGTCTGAACTCTTTCCCTACACGACGCTCTTCCGATCTNNNGGCACATGTCTTCAGGACTTACTC
GAGCACACGTCTGAACTCTTTCCCTACACGACGCTCTTCCGATCTNNNAACACATGGCTTTTGGACTTGT
AGCACACGTCTGAACTCTTTCCCTACACGACGCTCTTCCGATCTNNNGATGCAATCATAGCTCACTGCAG
GAGCACACGTCTGAACTCTTTCCCTACACGACGCTCTTCCGATCTNNNATCACTTGAAGGCCTATGCTT
GCACACGTCTGAACTCTTTCCCTACACGACGCTCTTCCGATCTNNNAAAACATTGACATTTCCCTGCGGC
AGCACACGTCTGAACTCTTTCCCTACACGACGCTCTTCCGATCTNNNTTATCTGCACTGAACTGTCCCTC
AGCACACGTCTGAACTCTTTCCCTACACGACGCTCTTCCGATCTNNNTGTCCTCAGACTTTCTGGTTCAG
AGCACACGTCTGAACTCTTTCCCTACACGACGCTCTTCCGATCTNNNCATGCCCGGCTAGTTTTTTGTAT
AGCACACGTCTGAACTCTTTCCCTACACGACGCTCTTCCGATCTNNNTGGGAGTTAGAGTGTTCAAGAC
AGCACACGTCTGAACTCTTTCCCTACACGACGCTCTTCCGATCTNNNGGTCAAGAGAGGCCAAAGTACT
AGCACACGTCTGAACTCTTTCCCTACACGACGCTCTTCCGATCTNNNTATGTTCCTCCTCCTCCTTCCTTCA
AGCACACGTCTGAACTCTTTCCCTACACGACGCTCTTCCGATCTNNNGATCTTGGCTCACTGCAAACTC
GAGCACACGTCTGAACTCTTTCCCTACACGACGCTCTTCCGATCTNNNATAGTGAGCCAAGATCATGCCA
AGCACACGTCTGAACTCTTTCCCTACACGACGCTCTTCCGATCTNNNCCTGGAACAACAAGAGCAAAAC
AGCACACGTCTGAACTCTTTCCCTACACGACGCTCTTCCGATCTNNNTATAGTCCCAGCTACTCAGGAG
AGCACACGTCTGAACTCTTTCCCTACACGACGCTCTTCCGATCTNNNTCCAGTAAGCCATATTTGTGCCT

AGCACACGTCTGAACTCTTTCCCTACACGACGCTCTTCCGATCTNNNCTGTAGTCCCAGCTACTCAGG
AGCACACGTCTGAACTCTTTCCCTACACGACGCTCTTCCGATCTNNNAGAATTGCTTGAATCCAGGA
AGCACACGTCTGAACTCTTTCCCTACACGACGCTCTTCCGATCTNNNAGAATTGCTTGAATCCAGGA
AGCACACGTCTGAACTCTTTCCCTACACGACGCTCTTCCGATCTNNNCTGAGGCAGGAGAATCACTTG
AGCACACGTCTGAACTCTTTCCCTACACGACGCTCTTCCGATCTNNNCATGCCTGTAATCCCAGCTACT
GAGCACACGTCTGAACTCTTTCCCTACACGACGCTCTTCCGATCTNNNAAAAATTAGCCGGGTGAGAT
AGAGCACACGTCTGAACTCTTTCCCTACACGACGCTCTTCCGATCTNNNATGTACCTGTAGTCCCAGCT
AGCACACGTCTGAACTCTTTCCCTACACGACGCTCTTCCGATCTNNNGACACCTGTAGTCCCAGCTATT
GAGCACACGTCTGAACTCTTTCCCTACACGACGCTCTTCCGATCTNNNTGATGCAGGAGAATTGCTT
GAGCACACGTCTGAACTCTTTCCCTACACGACGCTCTTCCGATCTNNNGACAGATCACTTGAGCCTAG
AGCACACGTCTGAACTCTTTCCCTACACGACGCTCTTCCGATCTNNNGCTTCTGGTTCAAGGAACATG
AGCACACGTCTGAACTCTTTCCCTACACGACGCTCTTCCGATCTNNNGGTGACAAAGTGAGACTCTG
AGCACACGTCTGAACTCTTTCCCTACACGACGCTCTTCCGATCTNNNTTATGCTTCTGTGCTCCTACCT
AGCACACGTCTGAACTCTTTCCCTACACGACGCTCTTCCGATCTNNNAGAATCACTTGAACTCAGGAG
AGCACACGTCTGAACTCTTTCCCTACACGACGCTCTTCCGATCTNNNGTAGTTCCAGCTACTTGAGAG
AGCACACGTCTGAACTCTTTCCCTACACGACGCTCTTCCGATCTNNNGCATGGCTTTGAAGATGGGA
GCACACGTCTGAACTCTTTCCCTACACGACGCTCTTCCGATCTNNNCTAACTACTCAGGAGGCAGAG
GAGCACACGTCTGAACTCTTTCCCTACACGACGCTCTTCCGATCTNNNCCTAGGAGGCAGAACTTGTA
GAGCACACGTCTGAACTCTTTCCCTACACGACGCTCTTCCGATCTNNNTGTGGGTTCTCTCTTGGAAT
AGAGCACACGTCTGAACTCTTTCCCTACACGACGCTCTTCCGATCTNNNAGCTACATTACAGGCTGAG
AGCACACGTCTGAACTCTTTCCCTACACGACGCTCTTCCGATCTNNNGTATTCCCAGCTACTCGAGAA
AGCACACGTCTGAACTCTTTCCCTACACGACGCTCTTCCGATCTNNNCTGAGCCAGGAGAATCACTTG
GAGCACACGTCTGAACTCTTTCCCTACACGACGCTCTTCCGATCTNNNTAGGTGGATTGCTTGAGCTT
AGCACACGTCTGAACTCTTTCCCTACACGACGCTCTTCCGATCTNNNGAGGTTACAGTGAGCCGAGA
GAGCACACGTCTGAACTCTTTCCCTACACGACGCTCTTCCGATCTNNNCTTCAGTGAGTCATGATTGT
AGCACACGTCTGAACTCTTTCCCTACACGACGCTCTTCCGATCTNNNATACCATTACACCATTGCCCTC
AGCACACGTCTGAACTCTTTCCCTACACGACGCTCTTCCGATCTNNNTACAGTGAGCTGAGATGTAC
AGCACACGTCTGAACTCTTTCCCTACACGACGCTCTTCCGATCTNNNCATTTTGGCAGTTCCTTTTCG
AGCACACGTCTGAACTCTTTCCCTACACGACGCTCTTCCGATCTNNNAATTCCAGCTACTCAGGAGG
AGCACACGTCTGAACTCTTTCCCTACACGACGCTCTTCCGATCTNNNAGTGAGCTGAAATTGCACCA
GCACACGTCTGAACTCTTTCCCTACACGACGCTCTTCCGATCTNNNGCAACACAGCAAGATTCTGTCT
AGCACACGTCTGAACTCTTTCCCTACACGACGCTCTTCCGATCTNNNGGGAGGATCTCTTGAACCTAG
AGCACACGTCTGAACTCTTTCCCTACACGACGCTCTTCCGATCTNNNCCCTGGCCGAATTATCAATTG
GCACACGTCTGAACTCTTTCCCTACACGACGCTCTTCCGATCTNNNACACACACCTATAATCCCAGAG
GAGCACACGTCTGAACTCTTTCCCTACACGACGCTCTTCCGATCTNNNAACAAGAGCGAAAACTCCGT
AGCACACGTCTGAACTCTTTCCCTACACGACGCTCTTCCGATCTNNNCAGTAAGCCAAGATTGCACTA
AGCACACGTCTGAACTCTTTCCCTACACGACGCTCTTCCGATCTNNNTATAGTCCCAGCTACTCAGGA
AGCACACGTCTGAACTCTTTCCCTACACGACGCTCTTCCGATCTNNNCAGAGCTTGTAGTGAGCCAA
GAGCACACGTCTGAACTCTTTCCCTACACGACGCTCTTCCGATCTNNNAGAGGTTGCAGTGAGTCAA
AGCACACGTCTGAACTCTTTCCCTACACGACGCTCTTCCGATCTNNNGCCTCCTCTCTTTCTCTCGAATT
AGCACACGTCTGAACTCTTTCCCTACACGACGCTCTTCCGATCTNNNGTAATCTCAGCTACTCAGGAG
AGCACACGTCTGAACTCTTTCCCTACACGACGCTCTTCCGATCTNNNCACTTGAGACCAGGAGTTCA
AGCACACGTCTGAACTCTTTCCCTACACGACGCTCTTCCGATCTNNNCACCTGTAGTCCCAGCTATTCA

AGCACACGTCTGAACTCTTTCCCTACACGACGCTCTTCCGATCTNNNGGCAACAGAATGATACCCTGT
AGCACACGTCTGAACTCTTTCCCTACACGACGCTCTTCCGATCTNNNCTGTAGTCCCAGCTATTCAGGA
GAGCACACGTCTGAACTCTTTCCCTACACGACGCTCTTCCGATCTNNNAGAATTGCTTGAACCTGGGAG
GAGCACACGTCTGAACTCTTTCCCTACACGACGCTCTTCCGATCTNNNACACCTGTAATCCCAGCTACTT
GCACACGTCTGAACTCTTTCCCTACACGACGCTCTTCCGATCTNNNTACCTGTAATCCCACCACTTTGGCA
GAGCACACGTCTGAACTCTTTCCCTACACGACGCTCTTCCGATCTNNNTTAGGGATGGGAAAAGAGGA
AGAGCACACGTCTGAACTCTTTCCCTACACGACGCTCTTCCGATCTNNNTACACGCCTGTAATCTCAGCT
GAGCACACGTCTGAACTCTTTCCCTACACGACGCTCTTCCGATCTNNNTATAGTCCCAGCTACTCGGAA
AGCACACGTCTGAACTCTTTCCCTACACGACGCTCTTCCGATCTNNNAATGCCTATCAACGGTAGACTG
AGCACACGTCTGAACTCTTTCCCTACACGACGCTCTTCCGATCTNNNAGATTGCAGTGAGCCAAAATCA
AGCACACGTCTGAACTCTTTCCCTACACGACGCTCTTCCGATCTNNNGAGATCACACCACTACACTCCAG
AGCACACGTCTGAACTCTTTCCCTACACGACGCTCTTCCGATCTNNNTTGAACCCAGGAAGTAGAGGT
GAGCACACGTCTGAACTCTTTCCCTACACGACGCTCTTCCGATCTNNNCTGAGGCAGGAGAATCACTT
GCACACGTCTGAACTCTTTCCCTACACGACGCTCTTCCGATCTNNNGGTGACAGTGTGAGATTCAGTC
GAGCACACGTCTGAACTCTTTCCCTACACGACGCTCTTCCGATCTNNNCTGTAGTCCCAGCTACTCAGGA
AGCACACGTCTGAACTCTTTCCCTACACGACGCTCTTCCGATCTNNNTGGCAACATAATGAGACCTCCT
GAGCACACGTCTGAACTCTTTCCCTACACGACGCTCTTCCGATCTNNNCGAGATCTCAGCATTGTACTC
AGCACACGTCTGAACTCTTTCCCTACACGACGCTCTTCCGATCTNNNGGGTGTGTGATAGGCAATTGTT
AGAGCACACGTCTGAACTCTTTCCCTACACGACGCTCTTCCGATCTNNNAAGAATCGCTTGAACCCTAG
GAGCACACGTCTGAACTCTTTCCCTACACGACGCTCTTCCGATCTNNNACACCTGTAATCCCAGCTATTC
GAGCACACGTCTGAACTCTTTCCCTACACGACGCTCTTCCGATCTNNNTGGGTGACAAGAACAAAACT
AGCACACGTCTGAACTCTTTCCCTACACGACGCTCTTCCGATCTNNNCTGAGGCAGGAGAATCACTTGA
AGCACACGTCTGAACTCTTTCCCTACACGACGCTCTTCCGATCTNNNCTTGGGAGGGCTATGAAGAAG
AGCACACGTCTGAACTCTTTCCCTACACGACGCTCTTCCGATCTNNNCGAGGCAGGAGAATCATTTGA
AGAGCACACGTCTGAACTCTTTCCCTACACGACGCTCTTCCGATCTNNNGAGAATCGCTTGTACCTAGG
AGCACACGTCTGAACTCTTTCCCTACACGACGCTCTTCCGATCTNNNCAACTACTAAGGAAGCTGAGG
GAGCACACGTCTGAACTCTTTCCCTACACGACGCTCTTCCGATCTNNNAGGTAGGACAATTGCTTGAA
AGAGCACACGTCTGAACTCTTTCCCTACACGACGCTCTTCCGATCTNNNCTGAGACAGGAGAATCGCT
AGCACACGTCTGAACTCTTTCCCTACACGACGCTCTTCCGATCTNNNAGTGAGCCGAGATCATGCTATT
AGCACACGTCTGAACTCTTTCCCTACACGACGCTCTTCCGATCTNNNCTTAGCAGGCTGAGTCAAGAG
GAGCACACGTCTGAACTCTTTCCCTACACGACGCTCTTCCGATCTNNNACACCTGTAATCCCAGCTACTT
AGCACACGTCTGAACTCTTTCCCTACACGACGCTCTTCCGATCTNNNCAGGAGAATCACTTGAACCCAG
AGCACACGTCTGAACTCTTTCCCTACACGACGCTCTTCCGATCTNNNTGTAATCCCAGCACTTTGAGAG
AGCACACGTCTGAACTCTTTCCCTACACGACGCTCTTCCGATCTNNNTGAGAACCACTTGAACCTGAGA
AGCACACGTCTGAACTCTTTCCCTACACGACGCTCTTCCGATCTNNNTTACAGTGAGCTGAGATTCCAC
GAGCACACGTCTGAACTCTTTCCCTACACGACGCTCTTCCGATCTNNNTGGCTCAGTCTCTTCTGTTCTT
AGCACACGTCTGAACTCTTTCCCTACACGACGCTCTTCCGATCTNNNGCAGGAGAATTGCTTGAATCCA
GAGCACACGTCTGAACTCTTTCCCTACACGACGCTCTTCCGATCTNNNACTTAGCTGGAAGTATCGCTT
GAGCACACGTCTGAACTCTTTCCCTACACGACGCTCTTCCGATCTNNNGTAGTACACGCCTGTAGTCTCA
GAGCACACGTCTGAACTCTTTCCCTACACGACGCTCTTCCGATCTNNNCACCTGTAATCCCAGCTACTCA
AGCACACGTCTGAACTCTTTCCCTACACGACGCTCTTCCGATCTNNNCTGAGGCAGGAGAATAGCTTGA
GAGCACACGTCTGAACTCTTTCCCTACACGACGCTCTTCCGATCTNNNGAGGCAGGAGAATTGCTTGA
AGCACACGTCTGAACTCTTTCCCTACACGACGCTCTTCCGATCTNNNGAGAATTTTGAACCCAGGAG

AGCACACGTCTGAACTCTTTCCCTACACGACGCTCTTCCGATCTNNNTGGCGTAATCATAGCTCAGTG
AGCACACGTCTGAACTCTTTCCCTACACGACGCTCTTCCGATCTNNNGCATGATCTCAGTTCACCACAA
GCACACGTCTGAACTCTTTCCCTACACGACGCTCTTCCGATCTNNNTGCAATAAGTCAGGGTACATG
AGCACACGTCTGAACTCTTTCCCTACACGACGCTCTTCCGATCTNNNAAACTAGTTTCAGAGACCACG
AGCACACGTCTGAACTCTTTCCCTACACGACGCTCTTCCGATCTNNNGTAAACCCATGAAACAACGAG
AGCACACGTCTGAACTCTTTCCCTACACGACGCTCTTCCGATCTNNNCAGAAAATTGCAGTCACAGGG
AGCACACGTCTGAACTCTTTCCCTACACGACGCTCTTCCGATCTNNNTATTGAGGCAGGAGAATTGGG
GAGCACACGTCTGAACTCTTTCCCTACACGACGCTCTTCCGATCTNNNGGACAACTGGCTGTAACTAG
AGCACACGTCTGAACTCTTTCCCTACACGACGCTCTTCCGATCTNNNAAACTAGCAGGTATTTGCCCA
AGCACACGTCTGAACTCTTTCCCTACACGACGCTCTTCCGATCTNNNGTAAATAGCGCTGCAATGACC
AGCACACGTCTGAACTCTTTCCCTACACGACGCTCTTCCGATCTNNNCGGCTTCCTTTCCAGTTTGTTA
GCACACGTCTGAACTCTTTCCCTACACGACGCTCTTCCGATCTNNNATCATAACTCACTGCAGCCTCAA
GCACACGTCTGAACTCTTTCCCTACACGACGCTCTTCCGATCTNNNCAGAGTTGTATCTGCTGGAAG
GCACACGTCTGAACTCTTTCCCTACACGACGCTCTTCCGATCTNNNTGACTTCCAGTTTCCTCCATCTG
GAGCACACGTCTGAACTCTTTCCCTACACGACGCTCTTCCGATCTNNNCCTTCCTCCTCTCTCTCTCTC
GAGCACACGTCTGAACTCTTTCCCTACACGACGCTCTTCCGATCTNNNCTGGGGCACTGTTTCTTTTCT
AGCACACGTCTGAACTCTTTCCCTACACGACGCTCTTCCGATCTNNNGTCGGGCTTCAAGGGAAATTT
GCACACGTCTGAACTCTTTCCCTACACGACGCTCTTCCGATCTNNNAGTTCTCCTGAGAGCGTTTCTTT
AGCACACGTCTGAACTCTTTCCCTACACGACGCTCTTCCGATCTNNNCAAGGTGAGTTCTACAATGTG
AGCACACGTCTGAACTCTTTCCCTACACGACGCTCTTCCGATCTNNNACCCATACCCCCAGGAGTTATT
AGCACACGTCTGAACTCTTTCCCTACACGACGCTCTTCCGATCTNNNACAGAAAGCAGCCAAAAACTA
GCACACGTCTGAACTCTTTCCCTACACGACGCTCTTCCGATCTNNNTCCAGAAGGCCTAAGAGGTCAT
GCACACGTCTGAACTCTTTCCCTACACGACGCTCTTCCGATCTNNNCATGAAGTCATGCACTTTGGGA
AGCACACGTCTGAACTCTTTCCCTACACGACGCTCTTCCGATCTNNNGCCAAAATGTGTGTATAGGAG
GCACACGTCTGAACTCTTTCCCTACACGACGCTCTTCCGATCTNNNGTTTGAATTACCACGGCCTATG
AGCACACGTCTGAACTCTTTCCCTACACGACGCTCTTCCGATCTNNNTAGTGAGTAGCTGGGACTAC
AGCACACGTCTGAACTCTTTCCCTACACGACGCTCTTCCGATCTNNNATGGATGCATGAGTCATTCTTT
AGCACACGTCTGAACTCTTTCCCTACACGACGCTCTTCCGATCTNNNTATGCTTGCAGCTTGCTTTTTC
AGCACACGTCTGAACTCTTTCCCTACACGACGCTCTTCCGATCTNNNTTAGGACTCCCCAGTACCTTG
AGCACACGTCTGAACTCTTTCCCTACACGACGCTCTTCCGATCTNNNGCTCAAGGCACTGAATGAATG
AGCACACGTCTGAACTCTTTCCCTACACGACGCTCTTCCGATCTNNNAAAGGTGGCATACTGTATGTG
AGCACACGTCTGAACTCTTTCCCTACACGACGCTCTTCCGATCTNNNATTGTTGTGCAGCCAATCTTC
GAGCACACGTCTGAACTCTTTCCCTACACGACGCTCTTCCGATCTNNNGGAACTCCTGACCTCGTGATT
GCACACGTCTGAACTCTTTCCCTACACGACGCTCTTCCGATCTNNNAGCTGGAAGTACAGGTGTATA
AGCACACGTCTGAACTCTTTCCCTACACGACGCTCTTCCGATCTNNNGGAGTGTGAGGAGCATAAACT
AGCACACGTCTGAACTCTTTCCCTACACGACGCTCTTCCGATCTNNNGAATGAGGCAGCTCAAACTC
GAGCACACGTCTGAACTCTTTCCCTACACGACGCTCTTCCGATCTNNNCTTCTTAGGAGAAACCCCTTC
GAGCACACGTCTGAACTCTTTCCCTACACGACGCTCTTCCGATCTNNNAGAATGTGAGCAACAACTTG
AGCACACGTCTGAACTCTTTCCCTACACGACGCTCTTCCGATCTNNNGTCTTGTTTTCCTGCTCATGTC
GAGCACACGTCTGAACTCTTTCCCTACACGACGCTCTTCCGATCTNNNGAATGCAGCTGGAAATGAC
AGAGCACACGTCTGAACTCTTTCCCTACACGACGCTCTTCCGATCTNNNGCTATGACAGATCGATGGG
GAGCACACGTCTGAACTCTTTCCCTACACGACGCTCTTCCGATCTNNNCAAAACAACCAGCTTCCTGA
GAGCACACGTCTGAACTCTTTCCCTACACGACGCTCTTCCGATCTNNNCCCCATGAAGTCTCTGAATCC

AGAGCACACGTCTGAACTCTTTCCCTACACGACGCTCTTCCGATCTNNNGTTTCGGAAGTGCCTACCAA
GAGCACACGTCTGAACTCTTTCCCTACACGACGCTCTTCCGATCTNNNGGTTACCTGCTCTCTCTCTCT
AGCACACGTCTGAACTCTTTCCCTACACGACGCTCTTCCGATCTNNNGCCTTCAAGACCCTTGATCATCT
GAGCACACGTCTGAACTCTTTCCCTACACGACGCTCTTCCGATCTNNNTCCTGTAATAGCCTAACCTTC
GAGCACACGTCTGAACTCTTTCCCTACACGACGCTCTTCCGATCTNNNCCTGCTCTTTCTCCTAGTGGAA
AGCACACGTCTGAACTCTTTCCCTACACGACGCTCTTCCGATCTNNNTAAGGACTCTTTACCCCTTCCCT
AGCACACGTCTGAACTCTTTCCCTACACGACGCTCTTCCGATCTNNNATTATGTGTGGGTATTGACGG
GAGCACACGTCTGAACTCTTTCCCTACACGACGCTCTTCCGATCTNNNGAGCACTTTGGACCTCTTTTGT
AGCACACGTCTGAACTCTTTCCCTACACGACGCTCTTCCGATCTNNNTGGAGTCACTCTTGATTGCTGA
AGCACACGTCTGAACTCTTTCCCTACACGACGCTCTTCCGATCTNNNTTCATCCAGAGACACACACAGA
AGAGCACACGTCTGAACTCTTTCCCTACACGACGCTCTTCCGATCTNNNGGACTCTAGAAAGCCCAGGT
AGCACACGTCTGAACTCTTTCCCTACACGACGCTCTTCCGATCTNNNTCATCCAGCATTGTCCATCTCAA
GAGCACACGTCTGAACTCTTTCCCTACACGACGCTCTTCCGATCTNNNACCTTTGAGAATTTCCCACTG
AGAGCACACGTCTGAACTCTTTCCCTACACGACGCTCTTCCGATCTNNNACCAAGAACACCAAGGAGA
AGAGCACACGTCTGAACTCTTTCCCTACACGACGCTCTTCCGATCTNNNATCCTCACCCCTTCTCTGTAG
GAGCACACGTCTGAACTCTTTCCCTACACGACGCTCTTCCGATCTNNNATGTAAACATTGCTGGCTCAG
GAGCACACGTCTGAACTCTTTCCCTACACGACGCTCTTCCGATCTNNNGGAAGATTCCTGGGTCACACA
AGCACACGTCTGAACTCTTTCCCTACACGACGCTCTTCCGATCTNNNTGTGGAAGCATTGAGTTACTT
GAGCACACGTCTGAACTCTTTCCCTACACGACGCTCTTCCGATCTNNNCAAGTAGTACCTGACACTCCA
GAGCACACGTCTGAACTCTTTCCCTACACGACGCTCTTCCGATCTNNNCCTTTTCTCCTCATGCCAGTCTA
GAGCACACGTCTGAACTCTTTCCCTACACGACGCTCTTCCGATCTNNNGATCCCTCTGACCAGTGACTGA
GAGCACACGTCTGAACTCTTTCCCTACACGACGCTCTTCCGATCTNNNTGCCTGACAGTTCAAGATGAG
AGCACACGTCTGAACTCTTTCCCTACACGACGCTCTTCCGATCTNNNTCCAGTGTGAATGAATGCATGT
GAGCACACGTCTGAACTCTTTCCCTACACGACGCTCTTCCGATCTNNNAGGCCGAATCAATTAGAGTCC
AGCACACGTCTGAACTCTTTCCCTACACGACGCTCTTCCGATCTNNNCTGTTGAGCACTCCCTATCCTTT
AGCACACGTCTGAACTCTTTCCCTACACGACGCTCTTCCGATCTNNNAAAAACAGTCACCACCTCCTCAT
GAGCACACGTCTGAACTCTTTCCCTACACGACGCTCTTCCGATCTNNNCTAAGGATTTCACCATGCCCTC
GAGCACACGTCTGAACTCTTTCCCTACACGACGCTCTTCCGATCTNNNAGGACTCCTATTTCCAAAGGCT
AGAGCACACGTCTGAACTCTTTCCCTACACGACGCTCTTCCGATCTNNNGAGAAGGAAGTAGGCCAGA
GAGCACACGTCTGAACTCTTTCCCTACACGACGCTCTTCCGATCTNNNTAATCCCAATCAAACTCCCAGC
AGCACACGTCTGAACTCTTTCCCTACACGACGCTCTTCCGATCTNNNTGTGTCTTTTACAGAGACGGGT
GAGCACACGTCTGAACTCTTTCCCTACACGACGCTCTTCCGATCTNNNACCCAGAGATAGGACCAGTAT
GAGCACACGTCTGAACTCTTTCCCTACACGACGCTCTTCCGATCTNNNCGCGTTCCAGAATTCTCAAACA
AGAGCACACGTCTGAACTCTTTCCCTACACGACGCTCTTCCGATCTNNNCTCCCAAGTAGCTGGAACTA
AGCACACGTCTGAACTCTTTCCCTACACGACGCTCTTCCGATCTNNNCAAGGCAGGAGAATTGCTTGA
AGAGCACACGTCTGAACTCTTTCCCTACACGACGCTCTTCCGATCTNNNTAACCAAGATCAGAGCAGAA
GAGCACACGTCTGAACTCTTTCCCTACACGACGCTCTTCCGATCTNNNAATCTCTTGAACTCAGGAGGCA
AGCACACGTCTGAACTCTTTCCCTACACGACGCTCTTCCGATCTNNNGAAGTAGGAGAATCACCTGAG
GAGCACACGTCTGAACTCTTTCCCTACACGACGCTCTTCCGATCTNNNACAGGGCCAAATGGAAAAGA
AGCACACGTCTGAACTCTTTCCCTACACGACGCTCTTCCGATCTNNNGCCAAGTCCACTCTCTTTTTGTT
AGCACACGTCTGAACTCTTTCCCTACACGACGCTCTTCCGATCTNNNCAGCTATAATCCCAGCACGTTTG
GAGCACACGTCTGAACTCTTTCCCTACACGACGCTCTTCCGATCTNNNCTTTCCAGGTTCACGTCATTCT
AGCACACGTCTGAACTCTTTCCCTACACGACGCTCTTCCGATCTNNNAGGTTCCCAGTCTCTCCTCATTC

AGCACACGTCTGAACTCTTTCCCTACACGACGCTCTTCCGATCTNNNTCCATCACCTCACCTACTTACCAC
AGCACACGTCTGAACTCTTTCCCTACACGACGCTCTTCCGATCTNNNCTTCAGGAGGCCAACTAGAACA
AGCACACGTCTGAACTCTTTCCCTACACGACGCTCTTCCGATCTNNNCCGAGGTGGAAGGATTACTTGA
AGCACACGTCTGAACTCTTTCCCTACACGACGCTCTTCCGATCTNNNCCCAAAATGTTCCAGTGAGCATT
AGCACACGTCTGAACTCTTTCCCTACACGACGCTCTTCCGATCTNNNTGTGCTCTGGGAAATCCTGATA
AGAGCACACGTCTGAACTCTTTCCCTACACGACGCTCTTCCGATCTNNNACACCTGTAATCCCAGCTACTT
AGCACACGTCTGAACTCTTTCCCTACACGACGCTCTTCCGATCTNNNCTGTGTGTACTCGGGTAGGTTA
AGCACACGTCTGAACTCTTTCCCTACACGACGCTCTTCCGATCTNNNATCTCAGCTTATTGCAACCTCCAC
AGCACACGTCTGAACTCTTTCCCTACACGACGCTCTTCCGATCTNNNGGTCTCGAACTTCTGACCTTAG
AGCACACGTCTGAACTCTTTCCCTACACGACGCTCTTCCGATCTNNNCTTGAGGGTAGGAGTTTGAGA
AGAGCACACGTCTGAACTCTTTCCCTACACGACGCTCTTCCGATCTNNNTGTAATCAATCCCAGCTACTC
AGCACACGTCTGAACTCTTTCCCTACACGACGCTCTTCCGATCTNNNTTAGTTCTGGTTCCCTCTCCCTT
AGCACACGTCTGAACTCTTTCCCTACACGACGCTCTTCCGATCTNNNTCCCTCTCTCTCTCTCTCTCTCA
AGCACACGTCTGAACTCTTTCCCTACACGACGCTCTTCCGATCTNNNAGGTGGAGGATCTCTACATGG
GAGCACACGTCTGAACTCTTTCCCTACACGACGCTCTTCCGATCTNNNAAGCAGGAGAATCACTTGAAC
AGAGCACACGTCTGAACTCTTTCCCTACACGACGCTCTTCCGATCTNNNGCCCTGCTGCTAATTTTCCA
AGAGCACACGTCTGAACTCTTTCCCTACACGACGCTCTTCCGATCTNNNAGAACTCCCCTTGTTGACTTT
GAGCACACGTCTGAACTCTTTCCCTACACGACGCTCTTCCGATCTNNNCGACGCTATTCACAGTAGCAAA
AGCACACGTCTGAACTCTTTCCCTACACGACGCTCTTCCGATCTNNNAAGAAGAAAGGCAGAGATGGC
GAGCACACGTCTGAACTCTTTCCCTACACGACGCTCTTCCGATCTNNNGAGTTCAAGACCAGCCTGGAT
GAGCACACGTCTGAACTCTTTCCCTACACGACGCTCTTCCGATCTNNNCTAGTAGCTGGGATTACAGGC
GAGCACACGTCTGAACTCTTTCCCTACACGACGCTCTTCCGATCTNNNGAGTCCAGGAGTTTGAGACCA
GAGCACACGTCTGAACTCTTTCCCTACACGACGCTCTTCCGATCTNNNTTGACACAGCCTGGGCAATATA
GAGCACACGTCTGAACTCTTTCCCTACACGACGCTCTTCCGATCTNNNAGCTGCATGTCTGTGTATCTG
GAGCACACGTCTGAACTCTTTCCCTACACGACGCTCTTCCGATCTNNNCAGAGTGAGACCCTGTCTGAA
GAGCACACGTCTGAACTCTTTCCCTACACGACGCTCTTCCGATCTNNNCCAGAGTCCCCCTTCTCTACTAA
AGAGCACACGTCTGAACTCTTTCCCTACACGACGCTCTTCCGATCTNNNCTGAGGCAGGAGAATCACTT
GAGCACACGTCTGAACTCTTTCCCTACACGACGCTCTTCCGATCTNNNAGGGTAGAGGCTATCTCACTG
GAGCACACGTCTGAACTCTTTCCCTACACGACGCTCTTCCGATCTNNNGATTCCCTTCAACAAATGGTCG
GAGCACACGTCTGAACTCTTTCCCTACACGACGCTCTTCCGATCTNNNCCACATTAGGCTCCTGAATAGC
AGAGCACACGTCTGAACTCTTTCCCTACACGACGCTCTTCCGATCTNNNCTGAGGCAGGAGAATTGCTT
GAGCACACGTCTGAACTCTTTCCCTACACGACGCTCTTCCGATCTNNNGAGCAAGACCCTGTCTCAAAA
AGCACACGTCTGAACTCTTTCCCTACACGACGCTCTTCCGATCTNNNTTACTATCTGGCCCTGCACATAC
GAGCACACGTCTGAACTCTTTCCCTACACGACGCTCTTCCGATCTNNNGGCAACAAAATGAGACCATG
GAGCACACGTCTGAACTCTTTCCCTACACGACGCTCTTCCGATCTNNNAGTGTAGGGACGCAACTATGA
AGCACACGTCTGAACTCTTTCCCTACACGACGCTCTTCCGATCTNNNATCAGTTGGGCAGGAGTGTATT
AGCACACGTCTGAACTCTTTCCCTACACGACGCTCTTCCGATCTNNNAAGTAGCTGGGATTACAGGCA
GAGCACACGTCTGAACTCTTTCCCTACACGACGCTCTTCCGATCTNNNTGAGGAGAGACAGAGAGAGA
AGCACACGTCTGAACTCTTTCCCTACACGACGCTCTTCCGATCTNNNAGACCTCCATAATCACATGAGCC
GAGCACACGTCTGAACTCTTTCCCTACACGACGCTCTTCCGATCTNNNCAGAGCAGGTTCTTTTTGGGT
GAGCACACGTCTGAACTCTTTCCCTACACGACGCTCTTCCGATCTNNNCCAGCTACTTTTGAAGGTGCCT
AGCACACGTCTGAACTCTTTCCCTACACGACGCTCTTCCGATCTNNNCACGAGAATTGCTTGAATCTGGC
AGCACACGTCTGAACTCTTTCCCTACACGACGCTCTTCCGATCTNNNAGAGAACCATTTGAAGCTGGGA

GAGCACACGTCTGAACTCTTTCCCTACACGACGCTCTTCCGATCTNNNATCGCTTGAAGCTAGGAGTT
AGCACACGTCTGAACTCTTTCCCTACACGACGCTCTTCCGATCTNNNAATTGCTTGAACTCGGAATGT
AGCACACGTCTGAACTCTTTCCCTACACGACGCTCTTCCGATCTNNNCCCAATCCTTTGTGCGGTTATTA
AGCACACGTCTGAACTCTTTCCCTACACGACGCTCTTCCGATCTNNNTCTCTTGAACCAGGGAAATGG
AGCACACGTCTGAACTCTTTCCCTACACGACGCTCTTCCGATCTNNNTAAGGTAGTTTTCACTGGGAG
GCACACGTCTGAACTCTTTCCCTACACGACGCTCTTCCGATCTNNNAAGTTCCAGTGAGCCAAGATTG
GAGCACACGTCTGAACTCTTTCCCTACACGACGCTCTTCCGATCTNNNTTCTCCCCTTCTCATTACCACA
AGCACACGTCTGAACTCTTTCCCTACACGACGCTCTTCCGATCTNNNAGAGAGAGAGAGAGAGAGA
GAGCACACGTCTGAACTCTTTCCCTACACGACGCTCTTCCGATCTNNNAGACCTAGACACCTGAGAGA
AGCACACGTCTGAACTCTTTCCCTACACGACGCTCTTCCGATCTNNNGGCTAAGGCACAAGAATCACTT
AGCACACGTCTGAACTCTTTCCCTACACGACGCTCTTCCGATCTNNNGTGTGTGTGTGTGTGTGTA
GCACACGTCTGAACTCTTTCCCTACACGACGCTCTTCCGATCTNNNTCCCTCTTTCTCTCTGTCTCTGTC
GCACACGTCTGAACTCTTTCCCTACACGACGCTCTTCCGATCTNNNACCATGAGAGAGAGAGATCGTG
AGCACACGTCTGAACTCTTTCCCTACACGACGCTCTTCCGATCTNNNAGTCCCTTCCCAGACTCAGAAT
GAGCACACGTCTGAACTCTTTCCCTACACGACGCTCTTCCGATCTNNNACACACAGAGACACACATGT
CACACGTCTGAACTCTTTCCCTACACGACGCTCTTCCGATCTNNNCTCCTTCTGATTGGCTCTTTCCTCA
GCACACGTCTGAACTCTTTCCCTACACGACGCTCTTCCGATCTNNNCAACAGAGCAAGACTGTCTCAA
AGCACACGTCTGAACTCTTTCCCTACACGACGCTCTTCCGATCTNNNTCTCTTTTCCCTCCTTCCTTCCC
AAGAGCACACGTCTGAACTCTTTCCCTACACGACGCTCTTCCGATCTNNNGTATGATCCCACCACTGTA
GAGCACACGTCTGAACTCTTTCCCTACACGACGCTCTTCCGATCTNNNACAAAGCAGTTCCAGAGAGA
AGCACACGTCTGAACTCTTTCCCTACACGACGCTCTTCCGATCTNNNAGTTTACTGCTAATGGGGTGG
GCACACGTCTGAACTCTTTCCCTACACGACGCTCTTCCGATCTNNNGGCTCACAAGATAAACACTCTGC
AGCACACGTCTGAACTCTTTCCCTACACGACGCTCTTCCGATCTNNNAGAATGTGGCTGTGAATCCAT
AGCACACGTCTGAACTCTTTCCCTACACGACGCTCTTCCGATCTNNNAGGGAGAGAGAGAGAGAGA
AGCACACGTCTGAACTCTTTCCCTACACGACGCTCTTCCGATCTNNNTAAAACACACACACACACACAC
GAGCACACGTCTGAACTCTTTCCCTACACGACGCTCTTCCGATCTNNNTTGCAAACTTCTCAAACACAG
AGCACACGTCTGAACTCTTTCCCTACACGACGCTCTTCCGATCTNNNGGAGGAAGAGATCAAGAAGC
AGCACACGTCTGAACTCTTTCCCTACACGACGCTCTTCCGATCTNNNAGTTACCTTTCTTCCCAGTTGC
AGCACACGTCTGAACTCTTTCCCTACACGACGCTCTTCCGATCTNNNGTTCCCACCTAATCTCTTCCTCC
AGAGCACACGTCTGAACTCTTTCCCTACACGACGCTCTTCCGATCTNNNAAAGAGCTGAAGAGTGTA
AAGAGCACACGTCTGAACTCTTTCCCTACACGACGCTCTTCCGATCTNNNGGGAAGGAGAGAGAGAG
GCACACGTCTGAACTCTTTCCCTACACGACGCTCTTCCGATCTNNNAGGGAACTGAGATTGTGCTACT
GAGCACACGTCTGAACTCTTTCCCTACACGACGCTCTTCCGATCTNNNACACCTGTAATCCCAGCACTT
AGCACACGTCTGAACTCTTTCCCTACACGACGCTCTTCCGATCTNNNATAGGAGTTTGAGACAAGCC
AGCACACGTCTGAACTCTTTCCCTACACGACGCTCTTCCGATCTNNNGGATACAGACTGGACAGAGC
CACACGTCTGAACTCTTTCCCTACACGACGCTCTTCCGATCTNNNTCCCCAAATCAAGTCAAAGTGGA
AGCACACGTCTGAACTCTTTCCCTACACGACGCTCTTCCGATCTNNNGCAGGAGAATCACTTGAACCA
AGCACACGTCTGAACTCTTTCCCTACACGACGCTCTTCCGATCTNNNATTCTTTTATGACCCTGTGGC
AGCACACGTCTGAACTCTTTCCCTACACGACGCTCTTCCGATCTNNNGACTCAGCCTCTATTGTCCAG
AGCACACGTCTGAACTCTTTCCCTACACGACGCTCTTCCGATCTNNNGGAGAATGGCATTCGAGTTC
GAGCACACGTCTGAACTCTTTCCCTACACGACGCTCTTCCGATCTNNNCCGACCTCTTTTGAAAGCTG
AGCACACGTCTGAACTCTTTCCCTACACGACGCTCTTCCGATCTNNNATCCCCTTAGTTTGTGAACACC
GAGCACACGTCTGAACTCTTTCCCTACACGACGCTCTTCCGATCTNNNAGGAAAGGGACTGACGTTA

GCACACGTCTGAACTCTTTCCCTACACGACGCTCTTCCGATCTNNNGTCCAGACCGGTACTTTACTACC
AGCACACGTCTGAACTCTTTCCCTACACGACGCTCTTCCGATCTNNNTGCTCATTTATGCACTGACAT
AGCACACGTCTGAACTCTTTCCCTACACGACGCTCTTCCGATCTNNNCCCAACCAGATACCAAGTTCC
AGCACACGTCTGAACTCTTTCCCTACACGACGCTCTTCCGATCTNNNCTGAAGAATAAGGTGCCAGG
GCACACGTCTGAACTCTTTCCCTACACGACGCTCTTCCGATCTNNNCCACAGTACTGTATCTGGAGCT
GAGCACACGTCTGAACTCTTTCCCTACACGACGCTCTTCCGATCTNNNGGTTCTCACTTTTCAACCAAG
AGCACACGTCTGAACTCTTTCCCTACACGACGCTCTTCCGATCTNNNGGAGAAGAATGGCCAGTACA
GAGCACACGTCTGAACTCTTTCCCTACACGACGCTCTTCCGATCTNNNATTCCTTGGGTCTTTGCTTTT
AGCACACGTCTGAACTCTTTCCCTACACGACGCTCTTCCGATCTNNNTTCCATCCATTCATCTGTCCCAG
GCACACGTCTGAACTCTTTCCCTACACGACGCTCTTCCGATCTNNNGGTGATAGAGCAAAACCCTGT
AGCACACGTCTGAACTCTTTCCCTACACGACGCTCTTCCGATCTNNNTGGCTGGTTGTAAGATGAAT
AGCACACGTCTGAACTCTTTCCCTACACGACGCTCTTCCGATCTNNNTGGGTGATAGATCGAGACTC
GCACACGTCTGAACTCTTTCCCTACACGACGCTCTTCCGATCTNNNATGTTCAAATCACGTAGGGGT
GCACACGTCTGAACTCTTTCCCTACACGACGCTCTTCCGATCTNNNGGGTCTGTATGAGTGCAAGAG
AGCACACGTCTGAACTCTTTCCCTACACGACGCTCTTCCGATCTNNNCTCTATCCCTCAGCTCTCCTTC
AGCACACGTCTGAACTCTTTCCCTACACGACGCTCTTCCGATCTNNNCACATCTCATATGGTGGCAGA
AGCACACGTCTGAACTCTTTCCCTACACGACGCTCTTCCGATCTNNNGGACCCTGCTATTGCTGATGAT
CACACGTCTGAACTCTTTCCCTACACGACGCTCTTCCGATCTNNNAGGAGATGGTACCTCATTGTCCT
GAGCACACGTCTGAACTCTTTCCCTACACGACGCTCTTCCGATCTNNNTTGAGTTGGGTGGTCTCTA
AGCACACGTCTGAACTCTTTCCCTACACGACGCTCTTCCGATCTNNNATTGCTCACCCTGACCTAAAAC
AGCACACGTCTGAACTCTTTCCCTACACGACGCTCTTCCGATCTNNNTAAGAATGAGGTTAGGGCAA
GAGCACACGTCTGAACTCTTTCCCTACACGACGCTCTTCCGATCTNNNGATGTATAGGCCACTGTGG
GAGCACACGTCTGAACTCTTTCCCTACACGACGCTCTTCCGATCTNNNAGCCATAGCAGTACTGTGTC
GCACACGTCTGAACTCTTTCCCTACACGACGCTCTTCCGATCTNNNAGATGCCATAGGAAGGAGAGA
AGCACACGTCTGAACTCTTTCCCTACACGACGCTCTTCCGATCTNNNGGGTAACAGAGCAAGACCCTA
GCACACGTCTGAACTCTTTCCCTACACGACGCTCTTCCGATCTNNNTCCTTCTGAGACCTGGATCTTCT
AGCACACGTCTGAACTCTTTCCCTACACGACGCTCTTCCGATCTNNNCAAACCACCTCATCTCACTACC
AGAGCACACGTCTGAACTCTTTCCCTACACGACGCTCTTCCGATCTNNNACCAAAAATCCATTGCACA
GAGCACACGTCTGAACTCTTTCCCTACACGACGCTCTTCCGATCTNNNGTACTTCTGTAAGGCCTGTT
GCACACGTCTGAACTCTTTCCCTACACGACGCTCTTCCGATCTNNNTGATCTCCTGACCTTGTGATCTG
CACACGTCTGAACTCTTTCCCTACACGACGCTCTTCCGATCTNNNCATGACCCCTGTCCTAGAAACAA
AGCACACGTCTGAACTCTTTCCCTACACGACGCTCTTCCGATCTNNNAGAAACAGGACCAATACAGCC
AGCACACGTCTGAACTCTTTCCCTACACGACGCTCTTCCGATCTNNNGCTGATAACTCTCTCGGTCTCT
AGCACACGTCTGAACTCTTTCCCTACACGACGCTCTTCCGATCTNNNCCTCCAGGAAACTAGACCCAAT
GAGCACACGTCTGAACTCTTTCCCTACACGACGCTCTTCCGATCTNNNGCCCCTTCTACTCACACATAC
GCACACGTCTGAACTCTTTCCCTACACGACGCTCTTCCGATCTNNNTCTTCCTTCCAGATCTTCAGAGC
GCACACGTCTGAACTCTTTCCCTACACGACGCTCTTCCGATCTNNNCCCTGTCCCTCATTTACCCAAATC
AGCACACGTCTGAACTCTTTCCCTACACGACGCTCTTCCGATCTNNNCTCACGAGTAGCTAAGACCAC
CACACGTCTGAACTCTTTCCCTACACGACGCTCTTCCGATCTNNNTTTTCAGTGGACAGAGCTACGAA
AGCACACGTCTGAACTCTTTCCCTACACGACGCTCTTCCGATCTNNNTTTTCAAGTGGCTATTGCTGCT
AGCACACGTCTGAACTCTTTCCCTACACGACGCTCTTCCGATCTNNNTGGCAAAGTCTAATAAAGCCC
AGCACACGTCTGAACTCTTTCCCTACACGACGCTCTTCCGATCTNNNATAGTGACTCCAAGAAACTGG
GAGCACACGTCTGAACTCTTTCCCTACACGACGCTCTTCCGATCTNNNGATATGCTGTGAAGGCTGAG

AGCACACGTCTGAACTCTTTCCCTACACGACGCTCTTCCGATCTNNNTGACTCTCATGCCTGCCTATCTA
GAGCACACGTCTGAACTCTTTCCCTACACGACGCTCTTCCGATCTNNNCCCCCAGTCTTAGGAAACATG
AGCACACGTCTGAACTCTTTCCCTACACGACGCTCTTCCGATCTNNNAAGCCAGGTAACACATCCTGTT
AGAGCACACGTCTGAACTCTTTCCCTACACGACGCTCTTCCGATCTNNNGAGACAGACAGACAGACA
AGCACACGTCTGAACTCTTTCCCTACACGACGCTCTTCCGATCTNNNAGCTTCTCTGCAAGTTCCCTTT
AGCACACGTCTGAACTCTTTCCCTACACGACGCTCTTCCGATCTNNNCTGGGCTTCTCTGGAAATGTA
AGCACACGTCTGAACTCTTTCCCTACACGACGCTCTTCCGATCTNNNGAAGATTGAGGCAGGAGGTT
GAGCACACGTCTGAACTCTTTCCCTACACGACGCTCTTCCGATCTNNNAAAGCACACAACCAGAAAGA
AGCACACGTCTGAACTCTTTCCCTACACGACGCTCTTCCGATCTNNNTGGTGTTGGAGATGGAAAGA
GAGCACACGTCTGAACTCTTTCCCTACACGACGCTCTTCCGATCTNNNTGCCAAAGAGCTGGTCATTA
AGCACACGTCTGAACTCTTTCCCTACACGACGCTCTTCCGATCTNNNGTGGTTCATTCCAAACATGAG
AGCACACGTCTGAACTCTTTCCCTACACGACGCTCTTCCGATCTNNNCACTCATACATGTGGTTGCACA
AGCACACGTCTGAACTCTTTCCCTACACGACGCTCTTCCGATCTNNNGCCTACACTCTACACCTCTATGC
AGCACACGTCTGAACTCTTTCCCTACACGACGCTCTTCCGATCTNNNAAGAAGAGATGAGGGAGGTA
AGCACACGTCTGAACTCTTTCCCTACACGACGCTCTTCCGATCTNNNGCCTACCATGTGAGGACTTGA
GAGCACACGTCTGAACTCTTTCCCTACACGACGCTCTTCCGATCTNNNTGTTTTTCCCTGCACAAAGGT
AGCACACGTCTGAACTCTTTCCCTACACGACGCTCTTCCGATCTNNNTCTCTCAAATTTACAGGGTGGG
AGCACACGTCTGAACTCTTTCCCTACACGACGCTCTTCCGATCTNNNACAGGGTGCTCAGATATTTGG
AGAGCACACGTCTGAACTCTTTCCCTACACGACGCTCTTCCGATCTNNNGATAAAGACAAGGGAGGT
AGCACACGTCTGAACTCTTTCCCTACACGACGCTCTTCCGATCTNNNGGTTGCAATGAGCCAAGATCA
AGCACACGTCTGAACTCTTTCCCTACACGACGCTCTTCCGATCTNNNGGTAATCCAAGCTACTCAGGT
AGAGCACACGTCTGAACTCTTTCCCTACACGACGCTCTTCCGATCTNNNGAGAATTGCTTGAGCCTGT
AGAGCACACGTCTGAACTCTTTCCCTACACGACGCTCTTCCGATCTNNNGACACAGTGAGACTCTGCA
AGAGCACACGTCTGAACTCTTTCCCTACACGACGCTCTTCCGATCTNNNCACCTGTAGTCCCAGCTACT
AGCACACGTCTGAACTCTTTCCCTACACGACGCTCTTCCGATCTNNNAGGCATGAGTCTCTTGAATCC
AGCACACGTCTGAACTCTTTCCCTACACGACGCTCTTCCGATCTNNNCTGTAATCCCAGCTACACAGG
GAGCACACGTCTGAACTCTTTCCCTACACGACGCTCTTCCGATCTNNNTGGCAAAACTGCATCTCTACA
AGCACACGTCTGAACTCTTTCCCTACACGACGCTCTTCCGATCTNNNTATTGGAGGGAATCAGTGTCC
AGCACACGTCTGAACTCTTTCCCTACACGACGCTCTTCCGATCTNNNACCTTCAAGAATGAGTTGTCG
GCACACGTCTGAACTCTTTCCCTACACGACGCTCTTCCGATCTNNNTGCAGGTAACACATTTTAGCCTG
AGAGCACACGTCTGAACTCTTTCCCTACACGACGCTCTTCCGATCTNNNCCAATGCTGTCAGTAATTTG
GAGCACACGTCTGAACTCTTTCCCTACACGACGCTCTTCCGATCTNNNAGCACAATTCACCGTTGCAA
AGCACACGTCTGAACTCTTTCCCTACACGACGCTCTTCCGATCTNNNGGACGGAGAGAAAGAGAGAG
AGCACACGTCTGAACTCTTTCCCTACACGACGCTCTTCCGATCTNNNGGGGTCACTGTGACTCTCATA
AGCACACGTCTGAACTCTTTCCCTACACGACGCTCTTCCGATCTNNNTGTGTCCTGAGAGCACAAATAC
GCACACGTCTGAACTCTTTCCCTACACGACGCTCTTCCGATCTNNNAGAACTCAAATTCTTGCCACTGC
AGCACACGTCTGAACTCTTTCCCTACACGACGCTCTTCCGATCTNNNTCCCTCTCTCTCTCTCTCTCTC
AGCACACGTCTGAACTCTTTCCCTACACGACGCTCTTCCGATCTNNNAGGATTGCTTGAACTCAGGA
AGCACACGTCTGAACTCTTTCCCTACACGACGCTCTTCCGATCTNNNTGAAACCCCCCTGTCTCTACAAA
AGCACACGTCTGAACTCTTTCCCTACACGACGCTCTTCCGATCTNNNAATGAGTTGTGTTCATGCCAC
GAGCACACGTCTGAACTCTTTCCCTACACGACGCTCTTCCGATCTNNNACAGGGCCAAATGGAAAAGA
GAGCACACGTCTGAACTCTTTCCCTACACGACGCTCTTCCGATCTNNNAATCCTATCCTGGGCAACATG
AGCACACGTCTGAACTCTTTCCCTACACGACGCTCTTCCGATCTNNNCCCTGTCTCTCTCTCTCTCTCC

AGAGCACACGTCTGAACTCTTTCCCTACACGACGCTCTTCCGATCTNNNAAGTTGCAGTAAGCCGAGAT
AGCACACGTCTGAACTCTTTCCCTACACGACGCTCTTCCGATCTNNNCTGGGATTATAGACACCTGCCA
GAGCACACGTCTGAACTCTTTCCCTACACGACGCTCTTCCGATCTNNNCAGCAAGAAACTAGAGCTCCA
GAGCACACGTCTGAACTCTTTCCCTACACGACGCTCTTCCGATCTNNNTGATCTCCAATTCCATCCAGGT
GAGCACACGTCTGAACTCTTTCCCTACACGACGCTCTTCCGATCTNNNAGGTTGCAATGAGCCTAAAT
GAGCACACGTCTGAACTCTTTCCCTACACGACGCTCTTCCGATCTNNNTCACAAGGTCAGGAGTTTGAG
GCACACGTCTGAACTCTTTCCCTACACGACGCTCTTCCGATCTNNNGAAGCATTCGTGGTTTTTCAGGA
AGAGCACACGTCTGAACTCTTTCCCTACACGACGCTCTTCCGATCTNNNTCCCAGAAAGAGAGAACAA
AGCACACGTCTGAACTCTTTCCCTACACGACGCTCTTCCGATCTNNNCTCCTATCTCGTCCTTGTTCAGG
GCACACGTCTGAACTCTTTCCCTACACGACGCTCTTCCGATCTNNNACTACAGGTATGTGTCACCATG
GAGCACACGTCTGAACTCTTTCCCTACACGACGCTCTTCCGATCTNNNGCTTACTAAAACCTCTGCCTCC
AGCACACGTCTGAACTCTTTCCCTACACGACGCTCTTCCGATCTNNNGGCTAATGGCTCTTCAGACATCT
GAGCACACGTCTGAACTCTTTCCCTACACGACGCTCTTCCGATCTNNNAGGGACATGATCTCAGTTCAC
AGCACACGTCTGAACTCTTTCCCTACACGACGCTCTTCCGATCTNNNGGCTGATGATGCTACAACGAAT
GAGCACACGTCTGAACTCTTTCCCTACACGACGCTCTTCCGATCTNNNTGATTAGGCGGCTTATCCAAT
GAGCACACGTCTGAACTCTTTCCCTACACGACGCTCTTCCGATCTNNNAGTCATGATTACCCCATTGCAC
GAGCACACGTCTGAACTCTTTCCCTACACGACGCTCTTCCGATCTNNNAGAATCACTTGAACCCAGGAG
GAGCACACGTCTGAACTCTTTCCCTACACGACGCTCTTCCGATCTNNNCCGTCTCTACCAAAAACACACA
GAGCACACGTCTGAACTCTTTCCCTACACGACGCTCTTCCGATCTNNNGGCACCCCGTCTCTACTAAAAA
AGCACACGTCTGAACTCTTTCCCTACACGACGCTCTTCCGATCTNNNTGCTGCGAATGCCATTATTTCAT
AGCACACGTCTGAACTCTTTCCCTACACGACGCTCTTCCGATCTNNNTAGTTGTTGTTAGTTCCCCACCC
AGCACACGTCTGAACTCTTTCCCTACACGACGCTCTTCCGATCTNNNGTCCAGTGTACTGTAGGACGTT
AGCACACGTCTGAACTCTTTCCCTACACGACGCTCTTCCGATCTNNNCGATAACAAATGCTGGCCAAGA
GAGCACACGTCTGAACTCTTTCCCTACACGACGCTCTTCCGATCTNNNCTGTAATCCCAGCACTTTGGA
GAGCACACGTCTGAACTCTTTCCCTACACGACGCTCTTCCGATCTNNNTTGCTCTCTCTCTCTCTCTGTCT
GAGCACACGTCTGAACTCTTTCCCTACACGACGCTCTTCCGATCTNNNTGGGTCAGTGGAATCAACTTC
AGCACACGTCTGAACTCTTTCCCTACACGACGCTCTTCCGATCTNNNTTTTGGGGGGTGCTTACTTATG
AGCACACGTCTGAACTCTTTCCCTACACGACGCTCTTCCGATCTNNNGCCCGGCCTCTTTTTGTTTTTAT
AGCACACGTCTGAACTCTTTCCCTACACGACGCTCTTCCGATCTNNNTTATTGCTACCTCTTCCTCCCAG
GAGCACACGTCTGAACTCTTTCCCTACACGACGCTCTTCCGATCTNNNTGTGGTCATTGATCAGGTGTT
GAGCACACGTCTGAACTCTTTCCCTACACGACGCTCTTCCGATCTNNNCCTGAGGCATAGGAATAGCTT
AGCACACGTCTGAACTCTTTCCCTACACGACGCTCTTCCGATCTNNNGAGAATCTCTTGAACCAGGGAG
GAGCACACGTCTGAACTCTTTCCCTACACGACGCTCTTCCGATCTNNNACTTCATTCCTTTTTGAGGCTG
AGCACACGTCTGAACTCTTTCCCTACACGACGCTCTTCCGATCTNNNTTCTTCCTTCCTCTTCAGCTTCCC
GAGCACACGTCTGAACTCTTTCCCTACACGACGCTCTTCCGATCTNNNCTGAGGCAAAAGAATCGCTTG
GAGCACACGTCTGAACTCTTTCCCTACACGACGCTCTTCCGATCTNNNGAGTAAAAGAAAGCAAGGGG
AGCACACGTCTGAACTCTTTCCCTACACGACGCTCTTCCGATCTNNNTTTTTTTTTCTCCTGTGCCCGAG
AGCACACGTCTGAACTCTTTCCCTACACGACGCTCTTCCGATCTNNNATTTCAGCCCCCACAATTTCATG
AGCACACGTCTGAACTCTTTCCCTACACGACGCTCTTCCGATCTNNNGAGATTGCACCATTGTACTCCA
AGCACACGTCTGAACTCTTTCCCTACACGACGCTCTTCCGATCTNNNTATGTCCTGTTCTGATGCCAACA
GAGCACACGTCTGAACTCTTTCCCTACACGACGCTCTTCCGATCTNNNCAGTTGGCTCCTGTGTTACTTT
GAGCACACGTCTGAACTCTTTCCCTACACGACGCTCTTCCGATCTNNNGTAGCTCACTGTAGCCTCAAAC
GAGCACACGTCTGAACTCTTTCCCTACACGACGCTCTTCCGATCTNNNGGGTTCAAGTGATTCTCCTGT

AGCACACGTCTGAACTCTTTCCCTACACGACGCTCTTCCGATCTNNNGCAGGAGAATTGCTTGAACTCA
GAGCACACGTCTGAACTCTTTCCCTACACGACGCTCTTCCGATCTNNNCTGGGAAGCCAAAAGACAAAA
GAGCACACGTCTGAACTCTTTCCCTACACGACGCTCTTCCGATCTNNNTCTCTCTCTCACACACACTCATG
AGCACACGTCTGAACTCTTTCCCTACACGACGCTCTTCCGATCTNNNAATGAGCCTTGATCACACCATT
GCACACGTCTGAACTCTTTCCCTACACGACGCTCTTCCGATCTNNNATGATCTGCCACATACTGTCACC
AGCACACGTCTGAACTCTTTCCCTACACGACGCTCTTCCGATCTNNNTCTCTCTCTCTCTCTCGCAAACA
AGCACACGTCTGAACTCTTTCCCTACACGACGCTCTTCCGATCTNNNAGGGTTCTTACTTTGTCACCCA
GAGCACACGTCTGAACTCTTTCCCTACACGACGCTCTTCCGATCTNNNAAAGGATCAATTCAGCAAGG
AGCACACGTCTGAACTCTTTCCCTACACGACGCTCTTCCGATCTNNNTGATTCTCCTATCTCAGCCTCCC
GCACACGTCTGAACTCTTTCCCTACACGACGCTCTTCCGATCTNNNTGTGATCATACAACTGCACTCCA
GAGCACACGTCTGAACTCTTTCCCTACACGACGCTCTTCCGATCTNNNCTGTAATCCTACCACTTCGGGA
GAGCACACGTCTGAACTCTTTCCCTACACGACGCTCTTCCGATCTNNNGAGGATTGCTTGAGACTAGG
GAGCACACGTCTGAACTCTTTCCCTACACGACGCTCTTCCGATCTNNNAGTGAGCTGAGATTGTGCTAC
GAGCACACGTCTGAACTCTTTCCCTACACGACGCTCTTCCGATCTNNNTCTCAGCTACTTGGGAGACTGA
AGCACACGTCTGAACTCTTTCCCTACACGACGCTCTTCCGATCTNNNTGATGGGCTCTCCTAGATTGTT
GAGCACACGTCTGAACTCTTTCCCTACACGACGCTCTTCCGATCTNNNCTGTAGTCCCAGCTACTCGAGA
GAGCACACGTCTGAACTCTTTCCCTACACGACGCTCTTCCGATCTNNNAACAGTTTAGGAGTGCCGAG
GCACACGTCTGAACTCTTTCCCTACACGACGCTCTTCCGATCTNNNCAGTAGGCAGATCATGGGACTAC
GAGCACACGTCTGAACTCTTTCCCTACACGACGCTCTTCCGATCTNNNAATACACACACACACACACACA
GAGCACACGTCTGAACTCTTTCCCTACACGACGCTCTTCCGATCTNNNCAATCCTCAACTTGGGGCTTT
GAGCACACGTCTGAACTCTTTCCCTACACGACGCTCTTCCGATCTNNNCACACCTGGGCAATATAGTGA
GAGCACACGTCTGAACTCTTTCCCTACACGACGCTCTTCCGATCTNNNAGCAAGGCTCCGTCTAAAAAA
GAGCACACGTCTGAACTCTTTCCCTACACGACGCTCTTCCGATCTNNNCTATAGTTCCAGCTGCTCCAGA
GAGCACACGTCTGAACTCTTTCCCTACACGACGCTCTTCCGATCTNNNTGGCTGGAGATGAAGTTCAAC
AGAGCACACGTCTGAACTCTTTCCCTACACGACGCTCTTCCGATCTNNNCCGCCATGAATCTTCCAAAAA
GAGCACACGTCTGAACTCTTTCCCTACACGACGCTCTTCCGATCTNNNACTCCCCAAATTCTCTTTCTCCC
GAGCACACGTCTGAACTCTTTCCCTACACGACGCTCTTCCGATCTNNNTATTAGCCAGGGTTCTCCAGA
AGCACACGTCTGAACTCTTTCCCTACACGACGCTCTTCCGATCTNNNGATCACTTCAGCCCAGGAATTT
GAGCACACGTCTGAACTCTTTCCCTACACGACGCTCTTCCGATCTNNNCTGAGGCAGGAGAATTGCTT
GAGCACACGTCTGAACTCTTTCCCTACACGACGCTCTTCCGATCTNNNCACAAGGGCCAAAGTAACTAA
AGCACACGTCTGAACTCTTTCCCTACACGACGCTCTTCCGATCTNNNTTCCTTTGGGCAACTTGAAAAC
GAGCACACGTCTGAACTCTTTCCCTACACGACGCTCTTCCGATCTNNNACCTGTCTGTCTCTCTCTCTC
GAGCACACGTCTGAACTCTTTCCCTACACGACGCTCTTCCGATCTNNNCTGACGAAGGAGAATCGCTTG
AGCACACGTCTGAACTCTTTCCCTACACGACGCTCTTCCGATCTNNNATGGCAAAACCCCATCTCTACAT
AGCACACGTCTGAACTCTTTCCCTACACGACGCTCTTCCGATCTNNNGAGAATCACTTGAACCTGGGA
AGCACACGTCTGAACTCTTTCCCTACACGACGCTCTTCCGATCTNNNCTAAGGTGGGAAGATCACTTG
GAGCACACGTCTGAACTCTTTCCCTACACGACGCTCTTCCGATCTNNNGTGAGCTGAGATCACACCACT
GAGCACACGTCTGAACTCTTTCCCTACACGACGCTCTTCCGATCTNNNTCTCTAGTAGCTGAGACTGCA
GAGCACACGTCTGAACTCTTTCCCTACACGACGCTCTTCCGATCTNNNATTCCTGAACTGGCAAGAAGA
GAGCACACGTCTGAACTCTTTCCCTACACGACGCTCTTCCGATCTNNNTGACTATCAGAAACACCTGCCA
AGCACACGTCTGAACTCTTTCCCTACACGACGCTCTTCCGATCTNNNCAACTACATTTGCACCAACCTAA
GAGCACACGTCTGAACTCTTTCCCTACACGACGCTCTTCCGATCTNNNAAAGGGATGGAGATACTGGA
GAGCACACGTCTGAACTCTTTCCCTACACGACGCTCTTCCGATCTNNNAAAGGGATGGAGATACTGGA

AGCACACGTCTGAACTCTTTCCCTACACGACGCTCTTCCGATCTNNNATGGCGAAAACTGTCTCTACC
AGCACACGTCTGAACTCTTTCCCTACACGACGCTCTTCCGATCTNNNTGCTGTTCAATAAATGTGGGG
GAGCACACGTCTGAACTCTTTCCCTACACGACGCTCTTCCGATCTNNNCAAGAGAATTGCTTGAACTC
AGCACACGTCTGAACTCTTTCCCTACACGACGCTCTTCCGATCTNNNAGTGTCCATCATTCCCATGAAT
GCACACGTCTGAACTCTTTCCCTACACGACGCTCTTCCGATCTNNNAGAGAACCATTTGAAGCTGGG
AGCACACGTCTGAACTCTTTCCCTACACGACGCTCTTCCGATCTNNNTCTCTGAAACCTGCTAAACAGC
AGCACACGTCTGAACTCTTTCCCTACACGACGCTCTTCCGATCTNNNCTCTCCTTTTCCCCAATACACAC
AGCACACGTCTGAACTCTTTCCCTACACGACGCTCTTCCGATCTNNNGGACTACAGGTGTATGCCATC
AGCACACGTCTGAACTCTTTCCCTACACGACGCTCTTCCGATCTNNNAAACAATCCTCATGCCTCAGTCT
AAGAGCACACGTCTGAACTCTTTCCCTACACGACGCTCTTCCGATCTNNNAATGCTGGGATTACAGGC
GAGCACACGTCTGAACTCTTTCCCTACACGACGCTCTTCCGATCTNNNGAGACCAGGAGTTCAAAACA
AGCACACGTCTGAACTCTTTCCCTACACGACGCTCTTCCGATCTNNNCTCCCAGGTTCAAGTGATTCTC
AGCACACGTCTGAACTCTTTCCCTACACGACGCTCTTCCGATCTNNNTAGGGCAGGAGAATCACTTG
GAGCACACGTCTGAACTCTTTCCCTACACGACGCTCTTCCGATCTNNNTGATGCTCTACTGGCTTTTCC
GAGCACACGTCTGAACTCTTTCCCTACACGACGCTCTTCCGATCTNNNGCAAGACTCCCTGTCCAAAA
AGCACACGTCTGAACTCTTTCCCTACACGACGCTCTTCCGATCTNNNTCTCCAAAACCCACAATGAGTC
AGCACACGTCTGAACTCTTTCCCTACACGACGCTCTTCCGATCTNNNCCCTGAGAGAGAGAGAGAGAG
AGCACACGTCTGAACTCTTTCCCTACACGACGCTCTTCCGATCTNNNCTGTAAGCCTAGACTGTGCCAT
AGCACACGTCTGAACTCTTTCCCTACACGACGCTCTTCCGATCTNNNCTGAGGCAGGAGAATCACTTG
AGCACACGTCTGAACTCTTTCCCTACACGACGCTCTTCCGATCTNNNGGTATCAAAACCAGAGAAAGG
AGCACACGTCTGAACTCTTTCCCTACACGACGCTCTTCCGATCTNNNATTCTCCAGCTCCTTTTCTTCCCC
AGCACACGTCTGAACTCTTTCCCTACACGACGCTCTTCCGATCTNNNCTCTTTTGTGTGTGTGTGTGTG
GAGCACACGTCTGAACTCTTTCCCTACACGACGCTCTTCCGATCTNNNAGAGAGAGAGAGAGAGAGG
GCACACGTCTGAACTCTTTCCCTACACGACGCTCTTCCGATCTNNNCTATGGTGAGTAGATTTTGCGC
AGCACACGTCTGAACTCTTTCCCTACACGACGCTCTTCCGATCTNNNACCTGTCTGTCTGCATGTTCAT
CACACGTCTGAACTCTTTCCCTACACGACGCTCTTCCGATCTNNNGTTGAGGTAGGAGGATTGCTTG
AGCACACGTCTGAACTCTTTCCCTACACGACGCTCTTCCGATCTNNNTGTTTTCACATGTGCTTTGTAG
AGAGCACACGTCTGAACTCTTTCCCTACACGACGCTCTTCCGATCTNNNCACCAAAATAAACTCACGGG
GCACACGTCTGAACTCTTTCCCTACACGACGCTCTTCCGATCTNNNCTGAAGTGTCTCCTACCTTTCCCC
AGAGCACACGTCTGAACTCTTTCCCTACACGACGCTCTTCCGATCTNNNTATCTTATCCCCATCTCCCAG
CACACGTCTGAACTCTTTCCCTACACGACGCTCTTCCGATCTNNNTCAAACCAGGACTCCAACAATCAC
GAGCACACGTCTGAACTCTTTCCCTACACGACGCTCTTCCGATCTNNNAAGGAGAGAGAGAAAGGG
AAGAGCACACGTCTGAACTCTTTCCCTACACGACGCTCTTCCGATCTNNNCATCCTGGCTAACAAGGT
AGCACACGTCTGAACTCTTTCCCTACACGACGCTCTTCCGATCTNNNCCCATTGCAATGCTTATTCCTG
CACACGTCTGAACTCTTTCCCTACACGACGCTCTTCCGATCTNNNATAATGGGGGAAGTCAAGTGCA
AGCACACGTCTGAACTCTTTCCCTACACGACGCTCTTCCGATCTNNNCTGGCCATTTCCTTTCTCTCTCTC
GCACACGTCTGAACTCTTTCCCTACACGACGCTCTTCCGATCTNNNTGAAAACTGGGATTTCCGTTTCC
AGCACACGTCTGAACTCTTTCCCTACACGACGCTCTTCCGATCTNNNTCCACCCTGGACATCATTTTCAT
AGAGCACACGTCTGAACTCTTTCCCTACACGACGCTCTTCCGATCTNNNACTGGATATCTTGGAGTGG
AGCACACGTCTGAACTCTTTCCCTACACGACGCTCTTCCGATCTNNNCTCGTGATAGGGTTCTCATGAC
AGCACACGTCTGAACTCTTTCCCTACACGACGCTCTTCCGATCTNNNTTACGTGTGTACATGCATGTG
AGCACACGTCTGAACTCTTTCCCTACACGACGCTCTTCCGATCTNNNAGGGAGGCAAATCACTTGAAT
CACACGTCTGAACTCTTTCCCTACACGACGCTCTTCCGATCTNNNAGCCTCTCTAATTGTGTGAACCAC

AGCACACGTCTGAACTCTTTCCCTACACGACGCTCTTCCGATCTNNNAATCCAGTTTCCCTTCCTCCATCCA
GAGCACACGTCTGAACTCTTTCCCTACACGACGCTCTTCCGATCTNNNCAGACTGCCTGGTTTCAATCTT
AGAGCACACGTCTGAACTCTTTCCCTACACGACGCTCTTCCGATCTNNNAGAGAGAGAGAGAGAGGG
GAGCACACGTCTGAACTCTTTCCCTACACGACGCTCTTCCGATCTNNNCTTTTCCAAACTCGATGCCACT
AGCACACGTCTGAACTCTTTCCCTACACGACGCTCTTCCGATCTNNNAGATGAGATTGGTCCACTGCAT
CACACGTCTGAACTCTTTCCCTACACGACGCTCTTCCGATCTNNNCATCACACCATTGCTACTGGTTGCA
AGCACACGTCTGAACTCTTTCCCTACACGACGCTCTTCCGATCTNNNAGCAGAGACAGAAATCAGACAG
GCACACGTCTGAACTCTTTCCCTACACGACGCTCTTCCGATCTNNNGTTTTGAGTTCCCTAACCCCTCTC
AGCACACGTCTGAACTCTTTCCCTACACGACGCTCTTCCGATCTNNNGAGGCAGAGAGAGAGAGAGA
GAGCACACGTCTGAACTCTTTCCCTACACGACGCTCTTCCGATCTNNNTATGCCTGGGGAAATATTGCA
GAGCACACGTCTGAACTCTTTCCCTACACGACGCTCTTCCGATCTNNNGCAACAGAGCAAGAATCCATCT
AGCACACGTCTGAACTCTTTCCCTACACGACGCTCTTCCGATCTNNNAGTTATGCAGTTCCTGTCCTTCC
GAGCACACGTCTGAACTCTTTCCCTACACGACGCTCTTCCGATCTNNNCCCAAAGCCTTCTCAGAGACTA
GAGCACACGTCTGAACTCTTTCCCTACACGACGCTCTTCCGATCTNNNAGTTTCAGCTATTCGGGAGACT
GAGCACACGTCTGAACTCTTTCCCTACACGACGCTCTTCCGATCTNNNAGTGATCCTCCTACCTCAGCAT
GAGCACACGTCTGAACTCTTTCCCTACACGACGCTCTTCCGATCTNNNATCACTTGAACCTGGGAGATG
GCACACGTCTGAACTCTTTCCCTACACGACGCTCTTCCGATCTNNNCAGGCTTGTAATCCCAGCTACTCC
GAGCACACGTCTGAACTCTTTCCCTACACGACGCTCTTCCGATCTNNNTCGGCAATAAGAGCAAAATTCC
AGCACACGTCTGAACTCTTTCCCTACACGACGCTCTTCCGATCTNNNCAGGTGCCATCTCTTATTGACCT
AGCACACGTCTGAACTCTTTCCCTACACGACGCTCTTCCGATCTNNNATTATGCTGGCTTGGCTTCTTTC
AGCACACGTCTGAACTCTTTCCCTACACGACGCTCTTCCGATCTNNNCAGCACAGCTAGGAAGGAGTTT
AGCACACGTCTGAACTCTTTCCCTACACGACGCTCTTCCGATCTNNNCAGAACATTTCATCCAAGGGCTC
AGCACACGTCTGAACTCTTTCCCTACACGACGCTCTTCCGATCTNNNGGTAAGCCAAGATCACACCATTG
AGCACACGTCTGAACTCTTTCCCTACACGACGCTCTTCCGATCTNNNTATCCTTCTCTACCAGGTCCCTC
GAGCACACGTCTGAACTCTTTCCCTACACGACGCTCTTCCGATCTNNNGCCTGTAGTCCTAGCTACTCCT
AGCACACGTCTGAACTCTTTCCCTACACGACGCTCTTCCGATCTNNNTATTCCCAGCTACTCAGGAGACT
AAGAGCACACGTCTGAACTCTTTCCCTACACGACGCTCTTCCGATCTNNNTGTAGTACCACCTACTCTGC
AGCACACGTCTGAACTCTTTCCCTACACGACGCTCTTCCGATCTNNNAGGTAGGAGAATCGCTTGAAT
AGCACACGTCTGAACTCTTTCCCTACACGACGCTCTTCCGATCTNNNCTGAGGCACAACAATCACTTGAC
GAGCACACGTCTGAACTCTTTCCCTACACGACGCTCTTCCGATCTNNNTATGACCTGGAAGCCAATTCTG
AGCACACGTCTGAACTCTTTCCCTACACGACGCTCTTCCGATCTNNNCCAAAGTGCTGGGATTACAGGT
AGAGCACACGTCTGAACTCTTTCCCTACACGACGCTCTTCCGATCTNNNGAAGTTGCAGTGAGCTGAG
GAGCACACGTCTGAACTCTTTCCCTACACGACGCTCTTCCGATCTNNNTGGAGGTTGAAGTGAGCAAA
AGAGCACACGTCTGAACTCTTTCCCTACACGACGCTCTTCCGATCTNNNGGCATATGTAGTCCCAGCTA
AGAGCACACGTCTGAACTCTTTCCCTACACGACGCTCTTCCGATCTNNNAGAGGTTGCAGTGAATGGAC
AGCACACGTCTGAACTCTTTCCCTACACGACGCTCTTCCGATCTNNNGGAGGCTGTGGTGAGTTATGA
GAGCACACGTCTGAACTCTTTCCCTACACGACGCTCTTCCGATCTNNNATTCCCAAGCGCAGAGATTCTA
AGCACACGTCTGAACTCTTTCCCTACACGACGCTCTTCCGATCTNNNGGATCATAGACCCTCCTACTCCA
GAGCACACGTCTGAACTCTTTCCCTACACGACGCTCTTCCGATCTNNNGAAAGTGGGCAGTGGAAAAA
AGCACACGTCTGAACTCTTTCCCTACACGACGCTCTTCCGATCTNNNGAATGAGAGGGGACATCTGAG
AGCACACGTCTGAACTCTTTCCCTACACGACGCTCTTCCGATCTNNNCTCGATCTCCTGACCTTGTGATC
GAGCACACGTCTGAACTCTTTCCCTACACGACGCTCTTCCGATCTNNNAATTGCCGGGATTGTAGGAAT
AGCACACGTCTGAACTCTTTCCCTACACGACGCTCTTCCGATCTNNNCAGTGGTACGATCACAGCTTAC

AGCACACGTCTGAACTCTTTCCCTACACGACGCTCTTCCGATCTNNNAGTGCTGGGATTATAGGTGTG
AGCACACGTCTGAACTCTTTCCCTACACGACGCTCTTCCGATCTNNNCAGCTGGGGAAGAGGAACTAT
AGCACACGTCTGAACTCTTTCCCTACACGACGCTCTTCCGATCTNNNTGGTCTTGAACTCCTGACATCAC
AGCACACGTCTGAACTCTTTCCCTACACGACGCTCTTCCGATCTNNNTGCACTTTAGCTCACAGATTGA
AGCACACGTCTGAACTCTTTCCCTACACGACGCTCTTCCGATCTNNNACCATCCTTATATTCCTGCCTGG
GAGCACACGTCTGAACTCTTTCCCTACACGACGCTCTTCCGATCTNNNAGCAAGGTTTCTATTTCCTCCT
AGCACACGTCTGAACTCTTTCCCTACACGACGCTCTTCCGATCTNNNAACAATCTTTGGGCTTACAGAC
AGCACACGTCTGAACTCTTTCCCTACACGACGCTCTTCCGATCTNNNGCTAGCTCCTGGAGTATGGTA
GAGCACACGTCTGAACTCTTTCCCTACACGACGCTCTTCCGATCTNNNGGTCTCCCTTTCTCTCTCTCTCT
AGCACACGTCTGAACTCTTTCCCTACACGACGCTCTTCCGATCTNNNAAAACCTACTTTGAATGCCACCC
AGCACACGTCTGAACTCTTTCCCTACACGACGCTCTTCCGATCTNNNTCCTTTCCCTTTCCACTCACATTCA
GCACACGTCTGAACTCTTTCCCTACACGACGCTCTTCCGATCTNNNCTTCTGGCTTCAAGCAATCCTTCC
GAGCACACGTCTGAACTCTTTCCCTACACGACGCTCTTCCGATCTNNNTCCTGAAATTTCCAGCGTATTG
GAGCACACGTCTGAACTCTTTCCCTACACGACGCTCTTCCGATCTNNNCTCAAACAATTCTCCTGCCTCA
AGCACACGTCTGAACTCTTTCCCTACACGACGCTCTTCCGATCTNNNGTAGCATGATCTTGGTTCACTG
GAGCACACGTCTGAACTCTTTCCCTACACGACGCTCTTCCGATCTNNNGTGCTGGATGAATCTAACTC
AGCACACGTCTGAACTCTTTCCCTACACGACGCTCTTCCGATCTNNNTGATCAGTGGTGGCATTGAGAT
AGCACACGTCTGAACTCTTTCCCTACACGACGCTCTTCCGATCTNNNTTCCTGGGTTCAAGCTATTCTCC
GAGCACACGTCTGAACTCTTTCCCTACACGACGCTCTTCCGATCTNNNTTCACGCCATTCTCCTTTCTCAG
AGCACACGTCTGAACTCTTTCCCTACACGACGCTCTTCCGATCTNNNAATGCAAACTATTTGGGGTGT
AGAGCACACGTCTGAACTCTTTCCCTACACGACGCTCTTCCGATCTNNNCGGATCCCAAATCAATGCAC
GAGCACACGTCTGAACTCTTTCCCTACACGACGCTCTTCCGATCTNNNTGGAGATGGGGTTTCATCATG
AGCACACGTCTGAACTCTTTCCCTACACGACGCTCTTCCGATCTNNNAGTGCTGAGATTACAGAGGTG
AGCACACGTCTGAACTCTTTCCCTACACGACGCTCTTCCGATCTNNNAAAGTGCTGGAATTACAGGCA
AGCACACGTCTGAACTCTTTCCCTACACGACGCTCTTCCGATCTNNNTTTAGTAGAGACCGGGTTTCAC
GAGCACACGTCTGAACTCTTTCCCTACACGACGCTCTTCCGATCTNNNGGCTTCTTGGGCTCCTAAAGA
GAGCACACGTCTGAACTCTTTCCCTACACGACGCTCTTCCGATCTNNNCTGGAGTACAGTGGCACAATC
AGCACACGTCTGAACTCTTTCCCTACACGACGCTCTTCCGATCTNNNTGTCAGCCTGAAGTCTGAATGA
GAGCACACGTCTGAACTCTTTCCCTACACGACGCTCTTCCGATCTNNNTCCGGTAGGCAAAAGATTACT
AGCACACGTCTGAACTCTTTCCCTACACGACGCTCTTCCGATCTNNNCCAGGCCTCAGTTTTCATGTTTT
AGCACACGTCTGAACTCTTTCCCTACACGACGCTCTTCCGATCTNNNTTTGCAGGTGAAGGTAGACAA
GAGCACACGTCTGAACTCTTTCCCTACACGACGCTCTTCCGATCTNNNCAAGCAACAGTTTATTTCCCG
GCACACGTCTGAACTCTTTCCCTACACGACGCTCTTCCGATCTNNNAAGGTCACAGAGCAGGTAAGAT
AGCACACGTCTGAACTCTTTCCCTACACGACGCTCTTCCGATCTNNNAGAATTGGTCTGCAATTCCACTC
GAGCACACGTCTGAACTCTTTCCCTACACGACGCTCTTCCGATCTNNNCTGGGATTATAGGTGTGAGCC
GAGCACACGTCTGAACTCTTTCCCTACACGACGCTCTTCCGATCTNNNGCCGCTGTGCTACATCTTTTTT
AGCACACGTCTGAACTCTTTCCCTACACGACGCTCTTCCGATCTNNNGAAGAGTAGAACCCACCTTGGA
AGCACACGTCTGAACTCTTTCCCTACACGACGCTCTTCCGATCTNNNCCCTCAGCAAATCCCTTCTAGAA
AGCACACGTCTGAACTCTTTCCCTACACGACGCTCTTCCGATCTNNNTCTACTAGGCGGGGAAACTCAT
AGCACACGTCTGAACTCTTTCCCTACACGACGCTCTTCCGATCTNNNGCCCGGCCTTGATACAATATTTT
AGCACACGTCTGAACTCTTTCCCTACACGACGCTCTTCCGATCTNNNGTTTCACCATATTGGCCAGGATC
GAGCACACGTCTGAACTCTTTCCCTACACGACGCTCTTCCGATCTNNNTAGGGTTCATCCATGTCAAGCA
GAGCACACGTCTGAACTCTTTCCCTACACGACGCTCTTCCGATCTNNNCTGGACTAAAGCGATTCTCCTG

AGCACACGTCTGAACTCTTTCCCTACACGACGCTCTTCCGATCTNNNCCTAAAGTGCTGGGATTACAGC
GAGCACACGTCTGAACTCTTTCCCTACACGACGCTCTTCCGATCTNNNGGAAAATGGGGAGAGGTGAT
GAGCACACGTCTGAACTCTTTCCCTACACGACGCTCTTCCGATCTNNNCAAGTGTACCTGGGCTCACTA
AGCACACGTCTGAACTCTTTCCCTACACGACGCTCTTCCGATCTNNNTGATCACTAAAGGGCATGGGAT
GCACACGTCTGAACTCTTTCCCTACACGACGCTCTTCCGATCTNNNCAGACTCTGTTCTGGGAAGGAAC
GAGCACACGTCTGAACTCTTTCCCTACACGACGCTCTTCCGATCTNNNTGAAATGACCCTTCAAAGTCG
AGCACACGTCTGAACTCTTTCCCTACACGACGCTCTTCCGATCTNNNCACAAATGATTCTCCCACCTCAG
AGCACACGTCTGAACTCTTTCCCTACACGACGCTCTTCCGATCTNNNCATAGTCGGGGTAAGTGCTAG
GAGCACACGTCTGAACTCTTTCCCTACACGACGCTCTTCCGATCTNNNTCCGTAACAGCTTCCCTTTTCTC
AGCACACGTCTGAACTCTTTCCCTACACGACGCTCTTCCGATCTNNNTTCTCCACCCTCTTTGTTCTCTGC
GAGCACACGTCTGAACTCTTTCCCTACACGACGCTCTTCCGATCTNNNCCTGGGCAATAAGAGTGAAAC
GAGCACACGTCTGAACTCTTTCCCTACACGACGCTCTTCCGATCTNNNTGCAACCCCTAATCTCATTGTCA
GAGCACACGTCTGAACTCTTTCCCTACACGACGCTCTTCCGATCTNNNGTGGATCATGAGGTCAGGAG
AGAGCACACGTCTGAACTCTTTCCCTACACGACGCTCTTCCGATCTNNNCACCTGTAATCCCAGCTACTC
GAGCACACGTCTGAACTCTTTCCCTACACGACGCTCTTCCGATCTNNNAGGCATGAGAATCTCTGAACCT
GAGCACACGTCTGAACTCTTTCCCTACACGACGCTCTTCCGATCTNNNCTCTCTGCTTCTAAGATGGCGA
AGCACACGTCTGAACTCTTTCCCTACACGACGCTCTTCCGATCTNNNCATAGAAGGCAGAGGCAGATG
GAGCACACGTCTGAACTCTTTCCCTACACGACGCTCTTCCGATCTNNNAGGGAGGAGAATCATTTGAA
AGCACACGTCTGAACTCTTTCCCTACACGACGCTCTTCCGATCTNNNTGCTAAAATGTTTGGGGCCAA
AGCACACGTCTGAACTCTTTCCCTACACGACGCTCTTCCGATCTNNNAGTTATGGATGGGGATAACCAG
AGCACACGTCTGAACTCTTTCCCTACACGACGCTCTTCCGATCTNNNCTGAGATCATGCCACTGAACTC
AGCACACGTCTGAACTCTTTCCCTACACGACGCTCTTCCGATCTNNNGAGGCAGGAGAATTGCTTGAA
AGCACACGTCTGAACTCTTTCCCTACACGACGCTCTTCCGATCTNNNGGTGCCTATAATCCCAGCTACTC
AGAGCACACGTCTGAACTCTTTCCCTACACGACGCTCTTCCGATCTNNNCTAAGGCTGGAGAATCGCTT
AGCACACGTCTGAACTCTTTCCCTACACGACGCTCTTCCGATCTNNNTGATGAAGAAAGCAGCCATGTT
GAGCACACGTCTGAACTCTTTCCCTACACGACGCTCTTCCGATCTNNNTGGGCAACATGGTGAAATCCT
GAGCACACGTCTGAACTCTTTCCCTACACGACGCTCTTCCGATCTNNNTATTCCAGAGGCTGAGGATTT
AGAGCACACGTCTGAACTCTTTCCCTACACGACGCTCTTCCGATCTNNNGGAGAATTGCTTGAACCCGA
GAGCACACGTCTGAACTCTTTCCCTACACGACGCTCTTCCGATCTNNNTGAGACAGAATACCTTGAACC
GAGCACACGTCTGAACTCTTTCCCTACACGACGCTCTTCCGATCTNNNGTGGGTGCTTTATGTTGGCTA
AGCACACGTCTGAACTCTTTCCCTACACGACGCTCTTCCGATCTNNNGAGATGGGCTTTCATCATGTTC
AGCACACGTCTGAACTCTTTCCCTACACGACGCTCTTCCGATCTNNNACCTGTAATCCCAGCTAATCAG
GCACACGTCTGAACTCTTTCCCTACACGACGCTCTTCCGATCTNNNGATGAATAGCTACTGCCTCTGCT
AGCACACGTCTGAACTCTTTCCCTACACGACGCTCTTCCGATCTNNNGTGAGTACAGGGTTGCAAAGTT
AGCACACGTCTGAACTCTTTCCCTACACGACGCTCTTCCGATCTNNNTCTCCTGCTCATTTTCTTTGTCTC
GAGCACACGTCTGAACTCTTTCCCTACACGACGCTCTTCCGATCTNNNAGAGCTAGTTACAAAGGCTTG
GAGCACACGTCTGAACTCTTTCCCTACACGACGCTCTTCCGATCTNNNCTCTCAGACTAGACCAGGAAC
AGCACACGTCTGAACTCTTTCCCTACACGACGCTCTTCCGATCTNNNCAGGAGAATCACTTGAACCCAG
AGAGCACACGTCTGAACTCTTTCCCTACACGACGCTCTTCCGATCTNNNTGGGTGACAAATTGAGACTC
GAGCACACGTCTGAACTCTTTCCCTACACGACGCTCTTCCGATCTNNNTGCCTGTAATTCCAGCATTTTG
AGCACACGTCTGAACTCTTTCCCTACACGACGCTCTTCCGATCTNNNCTATAGAGACCCATCCTCCAGCT
AGCACACGTCTGAACTCTTTCCCTACACGACGCTCTTCCGATCTNNNAATACTGAGAGCCTATGTGCAGA
AGCACACGTCTGAACTCTTTCCCTACACGACGCTCTTCCGATCTNNNTGAGTTGGGTCTTGGAGGAAA

GCACACGTCTGAACTCTTTCCCTACACGACGCTCTTCCGATCTNNNGTTTGCATGTGTTGAAGGACAG
AGCACACGTCTGAACTCTTTCCCTACACGACGCTCTTCCGATCTNNNCTGGAACAACAGGTGCATTCT
AGCACACGTCTGAACTCTTTCCCTACACGACGCTCTTCCGATCTNNNTGAAACCTTGCTGAAAGAAGC
AGCACACGTCTGAACTCTTTCCCTACACGACGCTCTTCCGATCTNNNAGTAAGCTGAGATAGTGCCAC
AGCACACGTCTGAACTCTTTCCCTACACGACGCTCTTCCGATCTNNNTTCCTATCCCAGCACTTTGGAA
GAGCACACGTCTGAACTCTTTCCCTACACGACGCTCTTCCGATCTNNNGGCGACAGAGCAAGAATCTA
AGCACACGTCTGAACTCTTTCCCTACACGACGCTCTTCCGATCTNNNTAACTACCATTCAGCCCAGCAA
AGAGCACACGTCTGAACTCTTTCCCTACACGACGCTCTTCCGATCTNNNGAGAATGGTGTGAACCTA
AGCACACGTCTGAACTCTTTCCCTACACGACGCTCTTCCGATCTNNNGAGATGGCACCACTATACTCC
AGCACACGTCTGAACTCTTTCCCTACACGACGCTCTTCCGATCTNNNCCCAGCCTGAATTCAGTATTGG
GCACACGTCTGAACTCTTTCCCTACACGACGCTCTTCCGATCTNNNTGGTTTGAACTCCTGACCTCAT
GCACACGTCTGAACTCTTTCCCTACACGACGCTCTTCCGATCTNNNCTAGTGGATACGAAGTGGTGT
AGCACACGTCTGAACTCTTTCCCTACACGACGCTCTTCCGATCTNNNGGCAGTAGAATCACTTGATCCC
AGCACACGTCTGAACTCTTTCCCTACACGACGCTCTTCCGATCTNNNAATCTCAACTACCCAGGAAGCT
AGCACACGTCTGAACTCTTTCCCTACACGACGCTCTTCCGATCTNNNGTAGTCCCAGCTATTCAGGAA
GCACACGTCTGAACTCTTTCCCTACACGACGCTCTTCCGATCTNNNCAGGAGAATTGCTTGAACTTGG
AGCACACGTCTGAACTCTTTCCCTACACGACGCTCTTCCGATCTNNNGTCAGGAGAATTGCTTGAACTC
GCACACGTCTGAACTCTTTCCCTACACGACGCTCTTCCGATCTNNNCTCAGGTACTTGGGAGGATTGA
AGCACACGTCTGAACTCTTTCCCTACACGACGCTCTTCCGATCTNNNAGATCGAGACCATCCTGGCTA
AGCACACGTCTGAACTCTTTCCCTACACGACGCTCTTCCGATCTNNNCTGTAATCCCAGCTACTCAGGA
GAGCACACGTCTGAACTCTTTCCCTACACGACGCTCTTCCGATCTNNNATGCAGGAGAATCACTTGA
GCACACGTCTGAACTCTTTCCCTACACGACGCTCTTCCGATCTNNNACAGACAAAGCCCTACAGTCATT
GAGCACACGTCTGAACTCTTTCCCTACACGACGCTCTTCCGATCTNNNATACCACCATGCCCAGCTAAT
AGCACACGTCTGAACTCTTTCCCTACACGACGCTCTTCCGATCTNNNAAGTGCTAGAATTACAGGCGTG
GAGCACACGTCTGAACTCTTTCCCTACACGACGCTCTTCCGATCTNNNAGTAGCTGGGACTACATACA
GAGCACACGTCTGAACTCTTTCCCTACACGACGCTCTTCCGATCTNNNAAAGATTACACCACTGCACTC
AGCACACGTCTGAACTCTTTCCCTACACGACGCTCTTCCGATCTNNNAGTGAGCCAAGATCATACCAC
GAGCACACGTCTGAACTCTTTCCCTACACGACGCTCTTCCGATCTNNNGCAGGTGGATAACTTGAGGT
GAGCACACGTCTGAACTCTTTCCCTACACGACGCTCTTCCGATCTNNNGAACACTGGTCTTGCAGTCTA
AGCACACGTCTGAACTCTTTCCCTACACGACGCTCTTCCGATCTNNNGCACCTGTAATCTCAGCTACTC
GCACACGTCTGAACTCTTTCCCTACACGACGCTCTTCCGATCTNNNCAGAGGTTGTAGTGAGCCAAGA
GCACACGTCTGAACTCTTTCCCTACACGACGCTCTTCCGATCTNNNGAACTACTGGGCTCAAGCAATT
GAGCACACGTCTGAACTCTTTCCCTACACGACGCTCTTCCGATCTNNNGAGACAGAGTGAGACTCCTC
GAGCACACGTCTGAACTCTTTCCCTACACGACGCTCTTCCGATCTNNNATGCCTGTAATCCCAGCTACTT
AGCACACGTCTGAACTCTTTCCCTACACGACGCTCTTCCGATCTNNNTGATCATGTCACTACACTCCAG
AGCACACGTCTGAACTCTTTCCCTACACGACGCTCTTCCGATCTNNNGATTGCAGTGACCCAAGATC
GAGCACACGTCTGAACTCTTTCCCTACACGACGCTCTTCCGATCTNNNGTAGTCCATCACTTCCCAGAA
GCACACGTCTGAACTCTTTCCCTACACGACGCTCTTCCGATCTNNNATCAAGTCATCATGCCCAGCTAA
AGCACACGTCTGAACTCTTTCCCTACACGACGCTCTTCCGATCTNNNTGGTCAGTCCACTTTGATGTG
AGCACACGTCTGAACTCTTTCCCTACACGACGCTCTTCCGATCTNNNCACCTGTAATCCCAGCTACTTG
AGCACACGTCTGAACTCTTTCCCTACACGACGCTCTTCCGATCTNNNCTCTAGTCCCAGCTACTCAGGA
AGCACACGTCTGAACTCTTTCCCTACACGACGCTCTTCCGATCTNNNGCAACAAGAGCAAAACTGCAT
GCACACGTCTGAACTCTTTCCCTACACGACGCTCTTCCGATCTNNNTGAGGCAGGAGAATCTCTTGA

GAGCACACGTCTGAACTCTTTCCCTACACGACGCTCTTCCGATCTNNNGAGATTGCAGTGAGCTGAGA
GAGCACACGTCTGAACTCTTTCCCTACACGACGCTCTTCCGATCTNNNGAAGGATAAAGATGTTCGGG
AGAGCACACGTCTGAACTCTTTCCCTACACGACGCTCTTCCGATCTNNNACGTGTTTATCTGCTGACCTT
GAGCACACGTCTGAACTCTTTCCCTACACGACGCTCTTCCGATCTNNNTTCCTGGAAACACACATCATCC
GAAGAGCACACGTCTGAACTCTTTCCCTACACGACGCTCTTCCGATCTNNNGAGGCTGAGGCAGGAG
GAGCACACGTCTGAACTCTTTCCCTACACGACGCTCTTCCGATCTNNNTGTAGTCCTAGCTATTCGGGA
GAGCACACGTCTGAACTCTTTCCCTACACGACGCTCTTCCGATCTNNNCTGAGGCAGGAGAATCACTTC
GAGCACACGTCTGAACTCTTTCCCTACACGACGCTCTTCCGATCTNNNGAACCAAGATCGAGCTATTGC
GAGCACACGTCTGAACTCTTTCCCTACACGACGCTCTTCCGATCTNNNTTACAGTGAGCTACAGTCACAC
AGCACACGTCTGAACTCTTTCCCTACACGACGCTCTTCCGATCTNNNGGTGCCTATAGTCCCAGCTATT
GAGCACACGTCTGAACTCTTTCCCTACACGACGCTCTTCCGATCTNNNTGAGGAAGGCAGATCACTTTC
GAGCACACGTCTGAACTCTTTCCCTACACGACGCTCTTCCGATCTNNNACCATCCTCAGTTTCAGAGATG
GAGCACACGTCTGAACTCTTTCCCTACACGACGCTCTTCCGATCTNNNATGATTTGCTTGAGGTCAGGA
GAGCACACGTCTGAACTCTTTCCCTACACGACGCTCTTCCGATCTNNNATTCCCAAAGTAATCCCAGCAC
GAGCACACGTCTGAACTCTTTCCCTACACGACGCTCTTCCGATCTNNNCTGTAGTCCCAGCTACTCAGGA
GAGCACACGTCTGAACTCTTTCCCTACACGACGCTCTTCCGATCTNNNGAAGCTGTGGTGACTCAAGAT
GAGCACACGTCTGAACTCTTTCCCTACACGACGCTCTTCCGATCTNNNTATGCCTGTAATCCCAGCACTT
GAGCACACGTCTGAACTCTTTCCCTACACGACGCTCTTCCGATCTNNNTTCAGCCTAAGCAACAGAGTGA
GCACACGTCTGAACTCTTTCCCTACACGACGCTCTTCCGATCTNNNGGATCACTTGAGCTTGGAAGGTA
AGCACACGTCTGAACTCTTTCCCTACACGACGCTCTTCCGATCTNNNATTGTTTGAGTCCAGGATGTCCA
AGCACACGTCTGAACTCTTTCCCTACACGACGCTCTTCCGATCTNNNCTGAGGCAGGAGAATCACTTGA
GAGCACACGTCTGAACTCTTTCCCTACACGACGCTCTTCCGATCTNNNCTGAGGCAGGAGAATTGCTT
AGCACACGTCTGAACTCTTTCCCTACACGACGCTCTTCCGATCTNNNCTGAGGCAGGAGAATCACTTGA
AGCACACGTCTGAACTCTTTCCCTACACGACGCTCTTCCGATCTNNNGACATTAGAATCGCTTGAACCC
AGAGCACACGTCTGAACTCTTTCCCTACACGACGCTCTTCCGATCTNNNGAGGTAGGAGAACAGTGTG
AGCACACGTCTGAACTCTTTCCCTACACGACGCTCTTCCGATCTNNNCTGTAGTCCCAGCTACTTGGAA
AGCACACGTCTGAACTCTTTCCCTACACGACGCTCTTCCGATCTNNNCTGTAATCCCAGCTACACAGGA
AGCACACGTCTGAACTCTTTCCCTACACGACGCTCTTCCGATCTNNNTCCTAGCTACTTGGGAGACTGA
GAGCACACGTCTGAACTCTTTCCCTACACGACGCTCTTCCGATCTNNNTAGTCCCAGCTACTCGGTACTT
GAGCACACGTCTGAACTCTTTCCCTACACGACGCTCTTCCGATCTNNNATAATCCCAGCTACTTTGGAG
GAGCACACGTCTGAACTCTTTCCCTACACGACGCTCTTCCGATCTNNNGTAATCTCAGCTACTCAGGAG
AGAGCACACGTCTGAACTCTTTCCCTACACGACGCTCTTCCGATCTNNNCAGTCTGGCCAACATAGTGA
AGCACACGTCTGAACTCTTTCCCTACACGACGCTCTTCCGATCTNNNTGAGGCAGGAGGAGAATTACT
GAGCACACGTCTGAACTCTTTCCCTACACGACGCTCTTCCGATCTNNNACAGAGCCAAGATGACATGAC
AGCACACGTCTGAACTCTTTCCCTACACGACGCTCTTCCGATCTNNNAGTGAGCCAAGATCATACCACT
AGAGCACACGTCTGAACTCTTTCCCTACACGACGCTCTTCCGATCTNNNCCTGGAGAACAAGAGCAAAA
AGCACACGTCTGAACTCTTTCCCTACACGACGCTCTTCCGATCTNNNGAGGCTGAGGAAGAAGAATCA
GAGCACACGTCTGAACTCTTTCCCTACACGACGCTCTTCCGATCTNNNCAAAATGCTGGGATTACAGG
AGCACACGTCTGAACTCTTTCCCTACACGACGCTCTTCCGATCTNNNAGAGGTCTAGGTTGTAGTGAG
AGCACACGTCTGAACTCTTTCCCTACACGACGCTCTTCCGATCTNNNTGCTTGTAATCCCAGCTACTCAG
GAGCACACGTCTGAACTCTTTCCCTACACGACGCTCTTCCGATCTNNNTGTAGTCTCAGCTACTCGGGA
AGCACACGTCTGAACTCTTTCCCTACACGACGCTCTTCCGATCTNNNCTCCCAAGACTGAACAAGGAAC
AGCACACGTCTGAACTCTTTCCCTACACGACGCTCTTCCGATCTNNNCTGAGGCAGGAGAATCACTTG

AGCACACGTCTGAACTCTTTCCCTACACGACGCTCTTCCGATCTNNNCAAGGTGGGAAGATCATTTGA
GAGCACACGTCTGAACTCTTTCCCTACACGACGCTCTTCCGATCTNNNGAGATAGCTCCATTGCACTC
AGAGCACACGTCTGAACTCTTTCCCTACACGACGCTCTTCCGATCTNNNTGGGAGGCTGAGATAAGA
AGAGCACACGTCTGAACTCTTTCCCTACACGACGCTCTTCCGATCTNNNGGAGGTTACAGTGAGCCAA
GAGCACACGTCTGAACTCTTTCCCTACACGACGCTCTTCCGATCTNNNGAGGTGGGAGGATCACTTGA
GCACACGTCTGAACTCTTTCCCTACACGACGCTCTTCCGATCTNNNCGTGAATAGCTACAGAGGTCAC
GAGCACACGTCTGAACTCTTTCCCTACACGACGCTCTTCCGATCTNNNAGGCAAGAGAATCACTTGAA
GCACACGTCTGAACTCTTTCCCTACACGACGCTCTTCCGATCTNNNGTTGCAATAAGCTGAGATCGCAT
AGCACACGTCTGAACTCTTTCCCTACACGACGCTCTTCCGATCTNNNGAAGTTGCAGTGAGCTGAGAT
GCACACGTCTGAACTCTTTCCCTACACGACGCTCTTCCGATCTNNNATGGAAGAGGGAGAACAATGT
GAGCACACGTCTGAACTCTTTCCCTACACGACGCTCTTCCGATCTNNNAGGAAATCAACGAATGCGAA
GAGCACACGTCTGAACTCTTTCCCTACACGACGCTCTTCCGATCTNNNCTGAGGCAGGAGAATCACTT
GAGCACACGTCTGAACTCTTTCCCTACACGACGCTCTTCCGATCTNNNGAGGTTGCATTGAGCTGAGA
GAGCACACGTCTGAACTCTTTCCCTACACGACGCTCTTCCGATCTNNNCTGAGGCTGGAGAATTGCT
GAGCACACGTCTGAACTCTTTCCCTACACGACGCTCTTCCGATCTNNNCTGAGGCAGGAGAATCACTT
GCACACGTCTGAACTCTTTCCCTACACGACGCTCTTCCGATCTNNNGGACAACAAAGTGAGACACTG
AGCACACGTCTGAACTCTTTCCCTACACGACGCTCTTCCGATCTNNNCACTTGTAATCCCAGCACTTTG
GAGCACACGTCTGAACTCTTTCCCTACACGACGCTCTTCCGATCTNNNGGTGACAAATCAAGACTCCGT
GAGCACACGTCTGAACTCTTTCCCTACACGACGCTCTTCCGATCTNNNCGCCTATAATCCCAGCTACTCA
GAGCACACGTCTGAACTCTTTCCCTACACGACGCTCTTCCGATCTNNNCTGAGGCAGAGAATTGCTTG
AGCACACGTCTGAACTCTTTCCCTACACGACGCTCTTCCGATCTNNNAAAACCGCACCACTGCATTCTA
GAGCACACGTCTGAACTCTTTCCCTACACGACGCTCTTCCGATCTNNNCTGAGGCTGGAGAATCACT
AGCACACGTCTGAACTCTTTCCCTACACGACGCTCTTCCGATCTNNNTGAGGTCAGAAGTGTGAGAA
GAGCACACGTCTGAACTCTTTCCCTACACGACGCTCTTCCGATCTNNNCTGAGGCAGGAGAATAGCTT
AGCACACGTCTGAACTCTTTCCCTACACGACGCTCTTCCGATCTNNNGAGAACTTGGCCTGGCAAAAT
GAGCACACGTCTGAACTCTTTCCCTACACGACGCTCTTCCGATCTNNNTGAGGCAGGAGAATCTCTTG
AGCACACGTCTGAACTCTTTCCCTACACGACGCTCTTCCGATCTNNNGTGTCCACAGACTGGTCTTAGT
AGCACACGTCTGAACTCTTTCCCTACACGACGCTCTTCCGATCTNNNCACGGTGTGGAAAGACTTAGG
GCACACGTCTGAACTCTTTCCCTACACGACGCTCTTCCGATCTNNNATTACTACTGGTGTGAGCCACT
GAGCACACGTCTGAACTCTTTCCCTACACGACGCTCTTCCGATCTNNNTGGTCAGAGTATCACGAAAT
AGCACACGTCTGAACTCTTTCCCTACACGACGCTCTTCCGATCTNNNAAGGCTGACTTAGATGAACTG
GAGCACACGTCTGAACTCTTTCCCTACACGACGCTCTTCCGATCTNNNTTGAGTGGGCCTTAGTTACTT
AGCACACGTCTGAACTCTTTCCCTACACGACGCTCTTCCGATCTNNNCCAAGTATCTGGGACTACAGG
GCACACGTCTGAACTCTTTCCCTACACGACGCTCTTCCGATCTNNNCTGCCGGTTAGGACAATCAAAA
AGCACACGTCTGAACTCTTTCCCTACACGACGCTCTTCCGATCTNNNCTTGAACTCCTGACCTCGTGAT
GAGCACACGTCTGAACTCTTTCCCTACACGACGCTCTTCCGATCTNNNGTAGTAAGTAGCTGGGACCA
GCACACGTCTGAACTCTTTCCCTACACGACGCTCTTCCGATCTNNNAAGTAATCCTCCTACCTCAGCCT
GAGCACACGTCTGAACTCTTTCCCTACACGACGCTCTTCCGATCTNNNTAATGGAAATGGGCACTGTA
GCACACGTCTGAACTCTTTCCCTACACGACGCTCTTCCGATCTNNNACTCCAAAGAAGGGATTGTGTG
GCACACGTCTGAACTCTTTCCCTACACGACGCTCTTCCGATCTNNNAAGCCAGAGTGTCTTCTTACCT
GCACACGTCTGAACTCTTTCCCTACACGACGCTCTTCCGATCTNNNTTTGGGGGAAAGAAAATGAAG
GCACACGTCTGAACTCTTTCCCTACACGACGCTCTTCCGATCTNNNTGGCTTTCTCCATAAGCTTCACT
GAGCACACGTCTGAACTCTTTCCCTACACGACGCTCTTCCGATCTNNNTTGAGATGGAGTCTTGCTCTG

AGCACACGTCTGAACTCTTTCCCTACACGACGCTCTTCCGATCTNNNTTCCCAAGTAGCTGCGATTACA
GAGCACACGTCTGAACTCTTTCCCTACACGACGCTCTTCCGATCTNNNCATGTGACCAGAACTGAGCT
AGCACACGTCTGAACTCTTTCCCTACACGACGCTCTTCCGATCTNNNCTTTCCTAGCCTTGGTGTCCTA
AGCACACGTCTGAACTCTTTCCCTACACGACGCTCTTCCGATCTNNNGTGATCTCTCCTCACTGTAACCC
AGCACACGTCTGAACTCTTTCCCTACACGACGCTCTTCCGATCTNNNGTTCAAGCGATTCTCCTACCTC
AGCACACGTCTGAACTCTTTCCCTACACGACGCTCTTCCGATCTNNNGGGGAACAGGTGGTATTTGGT
AGCACACGTCTGAACTCTTTCCCTACACGACGCTCTTCCGATCTNNNAAAACCAGGTTCATTAGAGGC
AGCACACGTCTGAACTCTTTCCCTACACGACGCTCTTCCGATCTNNNGTTCATACCATTCTCCTGCCTCA
AGCACACGTCTGAACTCTTTCCCTACACGACGCTCTTCCGATCTNNNTATGTGAGTTGAGAGGATGCT
AGCACACGTCTGAACTCTTTCCCTACACGACGCTCTTCCGATCTNNNGACCTCCCAGTCTCTAGGATCAT
AGCACACGTCTGAACTCTTTCCCTACACGACGCTCTTCCGATCTNNNTGGCTCTTGAATTGAATTCCTG
AGCACACGTCTGAACTCTTTCCCTACACGACGCTCTTCCGATCTNNNATATGCCACACTGACAACACTGT
GAGCACACGTCTGAACTCTTTCCCTACACGACGCTCTTCCGATCTNNNAGATCCGTTTTTCTTGCTCTC
AGAGCACACGTCTGAACTCTTTCCCTACACGACGCTCTTCCGATCTNNNCCGCTTCCTAACACATTAGCT
GAGCACACGTCTGAACTCTTTCCCTACACGACGCTCTTCCGATCTNNNCCTCCTGAGTAGCTGGGACT
AGCACACGTCTGAACTCTTTCCCTACACGACGCTCTTCCGATCTNNNTCTGATTATAGTTTGCCAGGCC
GCACACGTCTGAACTCTTTCCCTACACGACGCTCTTCCGATCTNNNCAATTCCACTTTCCCTTTGCCTTC
AGCACACGTCTGAACTCTTTCCCTACACGACGCTCTTCCGATCTNNNTCCTAACCTGAAGTGATCCTCCT
AGCACACGTCTGAACTCTTTCCCTACACGACGCTCTTCCGATCTNNNGAAGGTAAGAGGAGGCAAAA
GAGCACACGTCTGAACTCTTTCCCTACACGACGCTCTTCCGATCTNNNCTACTCTAGGCTAGGAAGTG
GAGCACACGTCTGAACTCTTTCCCTACACGACGCTCTTCCGATCTNNNTGTTGCTGCAAATGGAGGAA
GAGCACACGTCTGAACTCTTTCCCTACACGACGCTCTTCCGATCTNNNCTGCTCTTAGTGACTCTTGCTG
GAGCACACGTCTGAACTCTTTCCCTACACGACGCTCTTCCGATCTNNNTTCCTTTGTCATCCAGGACACT
AGCACACGTCTGAACTCTTTCCCTACACGACGCTCTTCCGATCTNNNGGTTCAAGTGATTCTCCTGCTTC
AGCACACGTCTGAACTCTTTCCCTACACGACGCTCTTCCGATCTNNNCAGCTGCCTTCTACTTTCCTTTC
GAGCACACGTCTGAACTCTTTCCCTACACGACGCTCTTCCGATCTNNNAGTGAGGTCCTAGGGAGAAA
AGCACACGTCTGAACTCTTTCCCTACACGACGCTCTTCCGATCTNNNCCTTCCAGTTGAGAGTGACTGT
AGCACACGTCTGAACTCTTTCCCTACACGACGCTCTTCCGATCTNNNGAGTAGCTGGGACTACAGTCA
GAGCACACGTCTGAACTCTTTCCCTACACGACGCTCTTCCGATCTNNNATATTCCTGTCAGCAGTGCAT
GAGCACACGTCTGAACTCTTTCCCTACACGACGCTCTTCCGATCTNNNACTTAGTGTTTGTCCCAGGAG
GAGCACACGTCTGAACTCTTTCCCTACACGACGCTCTTCCGATCTNNNGCTACCACACCTGGCTAATTTT
AGCACACGTCTGAACTCTTTCCCTACACGACGCTCTTCCGATCTNNNCAGCTCCTGGATTCCTTGATTTT
AGCACACGTCTGAACTCTTTCCCTACACGACGCTCTTCCGATCTNNNGCTGGGATTACATACATGAGCC
AGCACACGTCTGAACTCTTTCCCTACACGACGCTCTTCCGATCTNNNACAGTGATGTGAGTGAACTTG
AGCACACGTCTGAACTCTTTCCCTACACGACGCTCTTCCGATCTNNNTCAGTGCATGACATACCTCGAA
GAGCACACGTCTGAACTCTTTCCCTACACGACGCTCTTCCGATCTNNNTCCTGAGTAGCTAGGACTACA
AGCACACGTCTGAACTCTTTCCCTACACGACGCTCTTCCGATCTNNNCAAGTAGCTGCTGGGACTATA
AGCACACGTCTGAACTCTTTCCCTACACGACGCTCTTCCGATCTNNNCCAGGCTTATCTTGAACTCCAG
GAGCACACGTCTGAACTCTTTCCCTACACGACGCTCTTCCGATCTNNNTTCTGCATTTCCCTAGGTTCTC
GAGCACACGTCTGAACTCTTTCCCTACACGACGCTCTTCCGATCTNNNATGAGGTGGCAATTAGGACT
AGCACACGTCTGAACTCTTTCCCTACACGACGCTCTTCCGATCTNNNCCCTCCTGCATTTCCTGTCTTAT
AGCACACGTCTGAACTCTTTCCCTACACGACGCTCTTCCGATCTNNNGGGGATGTAGAACCTGCATAT
AGCACACGTCTGAACTCTTTCCCTACACGACGCTCTTCCGATCTNNNCTTAGACTGCATGTCCAGAGTG

GAGCACACGTCTGAACTCTTTCCCTACACGACGCTCTTCCGATCTNNNTATTTGGAATGCTGGCCTGAG
GAGCACACGTCTGAACTCTTTCCCTACACGACGCTCTTCCGATCTNNNAATACGAAGTCCAAGAGAGC
AGCACACGTCTGAACTCTTTCCCTACACGACGCTCTTCCGATCTNNNTCCTCCATTGTTCTTCAGCCTAC
GCACACGTCTGAACTCTTTCCCTACACGACGCTCTTCCGATCTNNNACCTAAAGTGCTGGGATTACAG
AGCACACGTCTGAACTCTTTCCCTACACGACGCTCTTCCGATCTNNNGCAGAGCAGAGTAGAAATTAC
AGCACACGTCTGAACTCTTTCCCTACACGACGCTCTTCCGATCTNNNTACCTCATCTCCCTCCCATATCCA
AGCACACGTCTGAACTCTTTCCCTACACGACGCTCTTCCGATCTNNNTGAAGTGTGATAGTTGTTGGG
GCACACGTCTGAACTCTTTCCCTACACGACGCTCTTCCGATCTNNNGGACATTCTCCATAACTGCCTCTCA
AGCACACGTCTGAACTCTTTCCCTACACGACGCTCTTCCGATCTNNNTTGGCCCAAATGAAAAAGATGC
GAGCACACGTCTGAACTCTTTCCCTACACGACGCTCTTCCGATCTNNNCAGATTAGCTGGGACTACATG
GAGCACACGTCTGAACTCTTTCCCTACACGACGCTCTTCCGATCTNNNCACTAGTCTAGGCCTGGAAGTT
AGCACACGTCTGAACTCTTTCCCTACACGACGCTCTTCCGATCTNNNCACTGGAGAAGGCAGTAGAGA
GAGCACACGTCTGAACTCTTTCCCTACACGACGCTCTTCCGATCTNNNGGGTTTAGTGTAATCACAACG
GAGCACACGTCTGAACTCTTTCCCTACACGACGCTCTTCCGATCTNNNTCCTTGTGCCACCTAAAAATTG
GAGCACACGTCTGAACTCTTTCCCTACACGACGCTCTTCCGATCTNNNAGCAACCCAAATCTCTCTTGAC
AGCACACGTCTGAACTCTTTCCCTACACGACGCTCTTCCGATCTNNNATTCGTTGCATCTTCTGGCTTTCC
AGCACACGTCTGAACTCTTTCCCTACACGACGCTCTTCCGATCTNNNAACAGCTATGCAGAATAAAAGC
AGCACACGTCTGAACTCTTTCCCTACACGACGCTCTTCCGATCTNNNAGTCAGAAGATCTAGGTGTCAC
AGCACACGTCTGAACTCTTTCCCTACACGACGCTCTTCCGATCTNNNTGTGTTACTCTAAGCTGCCAAGT
GAGCACACGTCTGAACTCTTTCCCTACACGACGCTCTTCCGATCTNNNGCATGATCTCAGATCACTGCAA
AGCACACGTCTGAACTCTTTCCCTACACGACGCTCTTCCGATCTNNNTTTCCCTATGCTGTCCTTGTGA
AGCACACGTCTGAACTCTTTCCCTACACGACGCTCTTCCGATCTNNNATGTACTAGCAGCACACTGTTG
AGCACACGTCTGAACTCTTTCCCTACACGACGCTCTTCCGATCTNNNAGGAATCACTATCTATGGCAGC
AGCACACGTCTGAACTCTTTCCCTACACGACGCTCTTCCGATCTNNNCAGGGTCTGCAGTTTCATTTGT
AGCACACGTCTGAACTCTTTCCCTACACGACGCTCTTCCGATCTNNNTCTTCCTTTCTGATGCTCCAAGTC
AGCACACGTCTGAACTCTTTCCCTACACGACGCTCTTCCGATCTNNNTTGCAAAGGGAGAAGAAAACA
GAGCACACGTCTGAACTCTTTCCCTACACGACGCTCTTCCGATCTNNNGCTTTCCAGACAGCTTAGACA
GAGCACACGTCTGAACTCTTTCCCTACACGACGCTCTTCCGATCTNNNCTAAGCCAAATGAGCAGCCAAA
GAGCACACGTCTGAACTCTTTCCCTACACGACGCTCTTCCGATCTNNNCATGGGATGTGTTTCCATTTG
AGCACACGTCTGAACTCTTTCCCTACACGACGCTCTTCCGATCTNNNAATGTCGCAATCTAGGCTCACTC
AGCACACGTCTGAACTCTTTCCCTACACGACGCTCTTCCGATCTNNNGTCCCAGCTGTTATTATCTGGCT
GAGCACACGTCTGAACTCTTTCCCTACACGACGCTCTTCCGATCTNNNTGCTCTGCTCTCAGGTCTTTTT
GAGCACACGTCTGAACTCTTTCCCTACACGACGCTCTTCCGATCTNNNTGTGTTGAGTTGAACTTTGCAC
AGCACACGTCTGAACTCTTTCCCTACACGACGCTCTTCCGATCTNNNCTCCTCAGTAGCTAAGACTGGG
GAGCACACGTCTGAACTCTTTCCCTACACGACGCTCTTCCGATCTNNNCATGTAGCTTCGGTGAAATGT
AGCACACGTCTGAACTCTTTCCCTACACGACGCTCTTCCGATCTNNNCAAAAAATCTCTGCCCAGTCCAA
AGCACACGTCTGAACTCTTTCCCTACACGACGCTCTTCCGATCTNNNTTTGCCCTGAAAACAATTCCTCC
AGCACACGTCTGAACTCTTTCCCTACACGACGCTCTTCCGATCTNNNCTGGAATTACAGGCATGAACCA
GAGCACACGTCTGAACTCTTTCCCTACACGACGCTCTTCCGATCTNNNTGGTGTGACATGTCTGTTCAG
GAGCACACGTCTGAACTCTTTCCCTACACGACGCTCTTCCGATCTNNNATTGAACACCAAGGAAGTAGC
GAGCACACGTCTGAACTCTTTCCCTACACGACGCTCTTCCGATCTNNNAGAGAACAAAGAGAGGGTTC
AGCACACGTCTGAACTCTTTCCCTACACGACGCTCTTCCGATCTNNNTCTCATTCTGCACGTTGTCTTTT
AGCACACGTCTGAACTCTTTCCCTACACGACGCTCTTCCGATCTNNNCTGCCAGTGTCATTTCCTGTTTT

GAGCACACGTCTGAACTCTTTCCCTACACGACGCTCTTCCGATCTNNNTTGCTCTCTCATTCACTCATGCT
AGCACACGTCTGAACTCTTTCCCTACACGACGCTCTTCCGATCTNNNCTCAAGTGATTCCCCTGCCTTAG
AGCACACGTCTGAACTCTTTCCCTACACGACGCTCTTCCGATCTNNNCTGTTTACAGAGGATTGCAACCT
AGCACACGTCTGAACTCTTTCCCTACACGACGCTCTTCCGATCTNNNTTTCCCATGGGGCTTTGTACTAA
AGCACACGTCTGAACTCTTTCCCTACACGACGCTCTTCCGATCTNNNTGACATGATCACATCTCACCACA
AGCACACGTCTGAACTCTTTCCCTACACGACGCTCTTCCGATCTNNNGGTGGAGAGAGAGAGAGAGAG
AGAGCACACGTCTGAACTCTTTCCCTACACGACGCTCTTCCGATCTNNNTGTCCTTGACTTCTCAACAGC
AGCACACGTCTGAACTCTTTCCCTACACGACGCTCTTCCGATCTNNNTCTCTACTTCATGCTGCTGAGTC
AGCACACGTCTGAACTCTTTCCCTACACGACGCTCTTCCGATCTNNNTTAGAGAAATCCTGCCTGCCTTT
AGCACACGTCTGAACTCTTTCCCTACACGACGCTCTTCCGATCTNNNGCCCAAGGTTACCACGTATAGT
AGCACACGTCTGAACTCTTTCCCTACACGACGCTCTTCCGATCTNNNTTCCACAGGTTCTCTCTCTCTCTC
AGCACACGTCTGAACTCTTTCCCTACACGACGCTCTTCCGATCTNNNAGCTGTATCCTGCAAATTTTGAT
AGCACACGTCTGAACTCTTTCCCTACACGACGCTCTTCCGATCTNNNGAACTTGCAATGTGAGGTGAGA
GAGCACACGTCTGAACTCTTTCCCTACACGACGCTCTTCCGATCTNNNTTGCGTTTACTTCCAGAGATC
GAGCACACGTCTGAACTCTTTCCCTACACGACGCTCTTCCGATCTNNNTCAAGTTTCACCAGTTGTTCCA
AGCACACGTCTGAACTCTTTCCCTACACGACGCTCTTCCGATCTNNNCTCTCAAGTAGCTGGGATTACG
AGCACACGTCTGAACTCTTTCCCTACACGACGCTCTTCCGATCTNNNGTCTTGAACTCCTGACCTGAGA
GAGCACACGTCTGAACTCTTTCCCTACACGACGCTCTTCCGATCTNNNGCACTTTCCCACTCAAGTACCT
AGCACACGTCTGAACTCTTTCCCTACACGACGCTCTTCCGATCTNNNGCAAGTCTGCTGGTAATGACTTC
AGCACACGTCTGAACTCTTTCCCTACACGACGCTCTTCCGATCTNNNCTCTGTGGGAGGTTTTTTTCGT
GCACACGTCTGAACTCTTTCCCTACACGACGCTCTTCCGATCTNNNAGTGCTGGGATAACAGATGTGA
GAGCACACGTCTGAACTCTTTCCCTACACGACGCTCTTCCGATCTNNNTCCATTCTCTCCTTCATTTGCCA
AGCACACGTCTGAACTCTTTCCCTACACGACGCTCTTCCGATCTNNNTTTTCAGTAGAGACGGGGGTTT
AGCACACGTCTGAACTCTTTCCCTACACGACGCTCTTCCGATCTNNNCTACTCGGGAGGCAGTAGAATT
GAGCACACGTCTGAACTCTTTCCCTACACGACGCTCTTCCGATCTNNNTACAAAAAAATCAGCTGGGCG
AGCACACGTCTGAACTCTTTCCCTACACGACGCTCTTCCGATCTNNNCTGAGATAGCACCACTGTACTCC
GAGCACACGTCTGAACTCTTTCCCTACACGACGCTCTTCCGATCTNNNCGCACCTATAGTCTCAGCTACT
GAGCACACGTCTGAACTCTTTCCCTACACGACGCTCTTCCGATCTNNNGGTGTATGCCTGTAATCTCAGC
GAGCACACGTCTGAACTCTTTCCCTACACGACGCTCTTCCGATCTNNNCAACATGGTGAAACTCCCTCTC
GAGCACACGTCTGAACTCTTTCCCTACACGACGCTCTTCCGATCTNNNCTGTAGTCTCAGCTACTTGGG
GCACACGTCTGAACTCTTTCCCTACACGACGCTCTTCCGATCTNNNAGCTGTGTAGAAGGTCTGGAATC
GAGCACACGTCTGAACTCTTTCCCTACACGACGCTCTTCCGATCTNNNGAGAACTGCTTGAACACAGG
GAGCACACGTCTGAACTCTTTCCCTACACGACGCTCTTCCGATCTNNNGCACCTATAGTCCCAGCTACT
AGAGCACACGTCTGAACTCTTTCCCTACACGACGCTCTTCCGATCTNNNCTGAGACAGGAGAATGGCA
GAGCACACGTCTGAACTCTTTCCCTACACGACGCTCTTCCGATCTNNNCTGTAATGCCAGCTACTCAGG
AGAGCACACGTCTGAACTCTTTCCCTACACGACGCTCTTCCGATCTNNNTGTCCATTTCATAGGCATTCA
AGCACACGTCTGAACTCTTTCCCTACACGACGCTCTTCCGATCTNNNAGTGAGTCATGATCATGCCACTC
GAGCACACGTCTGAACTCTTTCCCTACACGACGCTCTTCCGATCTNNNTGTGTGTGTATCCCTTGCTTCA
GAGCACACGTCTGAACTCTTTCCCTACACGACGCTCTTCCGATCTNNNTGTAATCCCAGCACTTTGTGA
GAGCACACGTCTGAACTCTTTCCCTACACGACGCTCTTCCGATCTNNNCACATGCCTATAGTCCCAGGTA
AGCACACGTCTGAACTCTTTCCCTACACGACGCTCTTCCGATCTNNNCGTATCATGAGGTCAGGAGATC
GAGCACACGTCTGAACTCTTTCCCTACACGACGCTCTTCCGATCTNNNTTACCATTGATACCTCCGTGAC
GAGCACACGTCTGAACTCTTTCCCTACACGACGCTCTTCCGATCTNNNCAGAAGAATTGCTTGAACCCA

GCACACGTCTGAACTCTTTCCCTACACGACGCTCTTCCGATCTNNNGTTCCAGTAAGCTGAGATTGTG
GCACACGTCTGAACTCTTTCCCTACACGACGCTCTTCCGATCTNNNCTGTAATCCCAGCTACACAGGA
AGCACACGTCTGAACTCTTTCCCTACACGACGCTCTTCCGATCTNNNGTCCTTGTAGTCCCAGCTACTT
GCACACGTCTGAACTCTTTCCCTACACGACGCTCTTCCGATCTNNNAGGAGAATTGCTTGAACCCAG
AGCACACGTCTGAACTCTTTCCCTACACGACGCTCTTCCGATCTNNNAATACAAAAATTAGCTGGGC
GAGCACACGTCTGAACTCTTTCCCTACACGACGCTCTTCCGATCTNNNCTTGCAGTGAGATGAGATTG
GAGCACACGTCTGAACTCTTTCCCTACACGACGCTCTTCCGATCTNNNGGGAGAATTGCTTGAACTCT
GAGCACACGTCTGAACTCTTTCCCTACACGACGCTCTTCCGATCTNNNAGAATCACTTGAACCCAGGA
GAGCACACGTCTGAACTCTTTCCCTACACGACGCTCTTCCGATCTNNNGAGGTTAGGGGATTGAGA
GAGCACACGTCTGAACTCTTTCCCTACACGACGCTCTTCCGATCTNNNCAACAAGGGCAAAACTCTGT
AGCACACGTCTGAACTCTTTCCCTACACGACGCTCTTCCGATCTNNNGGGCAACAGAATGAGACTCA
GCACACGTCTGAACTCTTTCCCTACACGACGCTCTTCCGATCTNNNGGGAGGGATAGCATTGGGAG
AGCACACGTCTGAACTCTTTCCCTACACGACGCTCTTCCGATCTNNNTGTAGTCACAGCTACTTGGGA
AGCACACGTCTGAACTCTTTCCCTACACGACGCTCTTCCGATCTNNNGCAGGAGAATCACTTGAACAC
AGCACACGTCTGAACTCTTTCCCTACACGACGCTCTTCCGATCTNNNAAAGTGAAAATGCAGACAAGG
AGCACACGTCTGAACTCTTTCCCTACACGACGCTCTTCCGATCTNNNAGGAGAATTGCTTGAACCCA
AGCACACGTCTGAACTCTTTCCCTACACGACGCTCTTCCGATCTNNNAGTAGGTTGAGGCAGGAGAA
GCACACGTCTGAACTCTTTCCCTACACGACGCTCTTCCGATCTNNNACATGCACTTGTAATCCCAGCTA
GCACACGTCTGAACTCTTTCCCTACACGACGCTCTTCCGATCTNNNTGAGAGAGGAGAATTGCTTGA
AGCACACGTCTGAACTCTTTCCCTACACGACGCTCTTCCGATCTNNNCACCTGTAATCTCAGCTACTCG
AGCACACGTCTGAACTCTTTCCCTACACGACGCTCTTCCGATCTNNNTGGAAAATCACTTGTACCCGG
GAGCACACGTCTGAACTCTTTCCCTACACGACGCTCTTCCGATCTNNNGAAGATCACTTGAGTCCAGG
AGCACACGTCTGAACTCTTTCCCTACACGACGCTCTTCCGATCTNNNGATTATTTGAGCCTGGGAGG
AGCACACGTCTGAACTCTTTCCCTACACGACGCTCTTCCGATCTNNNGAATCATTTGAACCTGAGAGG
GAGCACACGTCTGAACTCTTTCCCTACACGACGCTCTTCCGATCTNNNTAGTCCTAGCTACTCACAAG
GAGCACACGTCTGAACTCTTTCCCTACACGACGCTCTTCCGATCTNNNTGCTTGTAATCCCAGCTACTC
GCACACGTCTGAACTCTTTCCCTACACGACGCTCTTCCGATCTNNNGACTACTCTATGCCTGGAGAAG
AGCACACGTCTGAACTCTTTCCCTACACGACGCTCTTCCGATCTNNNGTCTTGAAATCCTGACCTCGTG
AGCACACGTCTGAACTCTTTCCCTACACGACGCTCTTCCGATCTNNNTTGCAATGAACCAAGATCATG
AGCACACGTCTGAACTCTTTCCCTACACGACGCTCTTCCGATCTNNNCTGTAGTCCCAGCTACTCAGG
AGCACACGTCTGAACTCTTTCCCTACACGACGCTCTTCCGATCTNNNAGAAACATTTTGGAGCCAACT
AGCACACGTCTGAACTCTTTCCCTACACGACGCTCTTCCGATCTNNNCACCTGTAATCCCTGATGCTCG
GCACACGTCTGAACTCTTTCCCTACACGACGCTCTTCCGATCTNNNGTGACAGTGCAAGACTTTGTCT
AGCACACGTCTGAACTCTTTCCCTACACGACGCTCTTCCGATCTNNNCTGGAGGCTGAGACATGAGA
AGCACACGTCTGAACTCTTTCCCTACACGACGCTCTTCCGATCTNNNTCTTGAACCTGGGAGATAGAG
AGCACACGTCTGAACTCTTTCCCTACACGACGCTCTTCCGATCTNNNGGATCACAGGGTCAAGAGAT
AGCACACGTCTGAACTCTTTCCCTACACGACGCTCTTCCGATCTNNNGAGAATCACTTGAACCCAGGA
GAGCACACGTCTGAACTCTTTCCCTACACGACGCTCTTCCGATCTNNNGAGCGAAAATCATGGCAATA
AGCACACGTCTGAACTCTTTCCCTACACGACGCTCTTCCGATCTNNNGCATATGCCTGTAATCCCAGCT
AGCACACGTCTGAACTCTTTCCCTACACGACGCTCTTCCGATCTNNNTTTAGCAATTTGTGGCTAGGC
AGCACACGTCTGAACTCTTTCCCTACACGACGCTCTTCCGATCTNNNATGATCAACATGGAGAAACC
AGCACACGTCTGAACTCTTTCCCTACACGACGCTCTTCCGATCTNNNAGCCAAAATAGACAGGAGCA
AGCACACGTCTGAACTCTTTCCCTACACGACGCTCTTCCGATCTNNNCTGTAGTCCTAGCTACTCGGG

GAGCACACGTCTGAACTCTTTCCCTACACGACGCTCTTCCGATCTNNNTGCCTGTAGTCCCAGCTACTA
GAGCACACGTCTGAACTCTTTCCCTACACGACGCTCTTCCGATCTNNNAGACATGGCGAAACTTCATCT
GAGCACACGTCTGAACTCTTTCCCTACACGACGCTCTTCCGATCTNNNGCAGTGAGCTGAGGTTATA
GAGCACACGTCTGAACTCTTTCCCTACACGACGCTCTTCCGATCTNNNGTAGTGAGCTGAGATGATGC
AGCACACGTCTGAACTCTTTCCCTACACGACGCTCTTCCGATCTNNNAAAAAACCCTGAAAAACCCAC
AGCACACGTCTGAACTCTTTCCCTACACGACGCTCTTCCGATCTNNNGGCACCTATAATCCCAGCTACT
AGCACACGTCTGAACTCTTTCCCTACACGACGCTCTTCCGATCTNNNGCTGTAATGACGTAATCGCAC
AGCACACGTCTGAACTCTTTCCCTACACGACGCTCTTCCGATCTNNNCTCACACCTGTAATCTCAGCACT
GAGCACACGTCTGAACTCTTTCCCTACACGACGCTCTTCCGATCTNNNTCTGTAATCTCAGCTACTCGG
GAGCACACGTCTGAACTCTTTCCCTACACGACGCTCTTCCGATCTNNNTGGTGTGCTTCTGTAATC
GAGCACACGTCTGAACTCTTTCCCTACACGACGCTCTTCCGATCTNNNGAGGTTGCAGTGAGCCAAG
AGCACACGTCTGAACTCTTTCCCTACACGACGCTCTTCCGATCTNNNGAGAATCGCTTGAACACATGA
GAGCACACGTCTGAACTCTTTCCCTACACGACGCTCTTCCGATCTNNNCTGTAGTCCCAGCTACTCAGG
AGCACACGTCTGAACTCTTTCCCTACACGACGCTCTTCCGATCTNNNCTGTAGTCCCAGCTACTGTGG
GAGCACACGTCTGAACTCTTTCCCTACACGACGCTCTTCCGATCTNNNAAAGAACAATGGGAAAGAG
AGCACACGTCTGAACTCTTTCCCTACACGACGCTCTTCCGATCTNNNAGAATCAGATACCTGGCCAGTT
GAGCACACGTCTGAACTCTTTCCCTACACGACGCTCTTCCGATCTNNNGAGAATCACTTGAACCTGGG
AGCACACGTCTGAACTCTTTCCCTACACGACGCTCTTCCGATCTNNNAGAATAACTTGAACCCAGGAG
GCACACGTCTGAACTCTTTCCCTACACGACGCTCTTCCGATCTNNNTGAGCTGAGATCATACCATTGC
GAGCACACGTCTGAACTCTTTCCCTACACGACGCTCTTCCGATCTNNNGGGGCATGAGAATTGCTTGA
GAGCACACGTCTGAACTCTTTCCCTACACGACGCTCTTCCGATCTNNNAGCCTGAAAGTAAAGGGATG
GAGCACACGTCTGAACTCTTTCCCTACACGACGCTCTTCCGATCTNNNGAGGCAGGAGAATTGTTTG
GCACACGTCTGAACTCTTTCCCTACACGACGCTCTTCCGATCTNNNTTGTGTGTGAATGCATGCACAT
GAGCACACGTCTGAACTCTTTCCCTACACGACGCTCTTCCGATCTNNNGCACCTGTAATCCCAACTACT
AGCACACGTCTGAACTCTTTCCCTACACGACGCTCTTCCGATCTNNNGCCTGTAATCCCAGCTATTAGG
GAGCACACGTCTGAACTCTTTCCCTACACGACGCTCTTCCGATCTNNNTGATCTGTTCTGAAGCAGG
GCACACGTCTGAACTCTTTCCCTACACGACGCTCTTCCGATCTNNNTATAGTCCCAGCTACTCGAGAG
GAGCACACGTCTGAACTCTTTCCCTACACGACGCTCTTCCGATCTNNNAGGGAGGAGAATTGATTGA
AGCACACGTCTGAACTCTTTCCCTACACGACGCTCTTCCGATCTNNNCTGTGATTCCAGCTACTTGGGA
AGCACACGTCTGAACTCTTTCCCTACACGACGCTCTTCCGATCTNNNGATCTTGCAGTGAGCCGAGAT
GAGCACACGTCTGAACTCTTTCCCTACACGACGCTCTTCCGATCTNNNTGAACCCTGGAGTTGGAGTT
GCACACGTCTGAACTCTTTCCCTACACGACGCTCTTCCGATCTNNNACCTTACTTCCTGGGCTTGAATT
GAGCACACGTCTGAACTCTTTCCCTACACGACGCTCTTCCGATCTNNNGCCTGTAATCTCAGCTACTCA
GCACACGTCTGAACTCTTTCCCTACACGACGCTCTTCCGATCTNNNGGACAAAGAAGTTATGCCATGG
AGAGCACACGTCTGAACTCTTTCCCTACACGACGCTCTTCCGATCTNNNTGTAGTCCCAGCTACTCAGG
GAGCACACGTCTGAACTCTTTCCCTACACGACGCTCTTCCGATCTNNNCTGAGGCAGGAGAATCACTT
GAGCACACGTCTGAACTCTTTCCCTACACGACGCTCTTCCGATCTNNNCTGTAGTCCCAGGTATTCCGG
GAGCACACGTCTGAACTCTTTCCCTACACGACGCTCTTCCGATCTNNNAGGAGAATGGCATGAATTGG
GAGCACACGTCTGAACTCTTTCCCTACACGACGCTCTTCCGATCTNNNAGGAATTGCTTGAACCTAGG
GAGCACACGTCTGAACTCTTTCCCTACACGACGCTCTTCCGATCTNNNTGGACTCACACAGCAGTAAT
GAGCACACGTCTGAACTCTTTCCCTACACGACGCTCTTCCGATCTNNNTGAGAGACAGAGTGAGACTC
AGAGCACACGTCTGAACTCTTTCCCTACACGACGCTCTTCCGATCTNNNAGAATCACTTGAACCCAGG
GAGCACACGTCTGAACTCTTTCCCTACACGACGCTCTTCCGATCTNNNGTGATAGAGGAAGACACCGT

AGCACACGTCTGAACTCTTTCCCTACACGACGCTCTTCCGATCTNNNGATCACTTGAGGCTAGGAGTC
AGCACACGTCTGAACTCTTTCCCTACACGACGCTCTTCCGATCTNNNCTAGGAGACAGAGCAAGACTGT
AGCACACGTCTGAACTCTTTCCCTACACGACGCTCTTCCGATCTNNNCTGAGTCAGGAGAATGGTGTG
AGCACACGTCTGAACTCTTTCCCTACACGACGCTCTTCCGATCTNNNTCCAAAAATTAGCCAGTGGGAA
AGCACACGTCTGAACTCTTTCCCTACACGACGCTCTTCCGATCTNNNCCATTCCATTCACTGAGCCATTG
GAGCACACGTCTGAACTCTTTCCCTACACGACGCTCTTCCGATCTNNNGTGTATAGGCTCCATGGTCCA
GAGCACACGTCTGAACTCTTTCCCTACACGACGCTCTTCCGATCTNNNGGTTGTGGTGAGTCAAGATT
GAGCACACGTCTGAACTCTTTCCCTACACGACGCTCTTCCGATCTNNNGGTGACAGAGTGAGACTCCAT
GAGCACACGTCTGAACTCTTTCCCTACACGACGCTCTTCCGATCTNNNTGTAGTCCCAGCTACTAGAGA
AGCACACGTCTGAACTCTTTCCCTACACGACGCTCTTCCGATCTNNNAAAAAAAGAAGCAGGCACTCCA
GAGCACACGTCTGAACTCTTTCCCTACACGACGCTCTTCCGATCTNNNCTTGCAATGAGCTGAGATCAT
GAGCACACGTCTGAACTCTTTCCCTACACGACGCTCTTCCGATCTNNNGAGAATTGCTTGAACCAGGGA
AGCACACGTCTGAACTCTTTCCCTACACGACGCTCTTCCGATCTNNNCACCTGTAATCCCAGCTACTCAG
AGCACACGTCTGAACTCTTTCCCTACACGACGCTCTTCCGATCTNNNAGGATAATCGCTTGAACCCAGA
AGAGCACACGTCTGAACTCTTTCCCTACACGACGCTCTTCCGATCTNNNCTGTAGTCCCAGCTACTTGGA
AGCACACGTCTGAACTCTTTCCCTACACGACGCTCTTCCGATCTNNNGAGACATGAGACTTGCTTGAAC
AGCACACGTCTGAACTCTTTCCCTACACGACGCTCTTCCGATCTNNNGAGAATGATTTGAACCTGCGA
AGCACACGTCTGAACTCTTTCCCTACACGACGCTCTTCCGATCTNNNGAGCAAGACTCCGTCTCAAAAAA
AGCACACGTCTGAACTCTTTCCCTACACGACGCTCTTCCGATCTNNNAGTAGGAGAATCGCTTGAACCT
AGCACACGTCTGAACTCTTTCCCTACACGACGCTCTTCCGATCTNNNCTCAGGGCAATACATTCTGAGC
GAGCACACGTCTGAACTCTTTCCCTACACGACGCTCTTCCGATCTNNNCCCCTCTATTGAAGTCCCAATCA
AGCACACGTCTGAACTCTTTCCCTACACGACGCTCTTCCGATCTNNNGACCAACTCTGTCCCCACTTTTAC
AGCACACGTCTGAACTCTTTCCCTACACGACGCTCTTCCGATCTNNNGATGCAAATCCCAGCTCATCTTC
GAGCACACGTCTGAACTCTTTCCCTACACGACGCTCTTCCGATCTNNNCAGGATGAGGGCTTAGTGTG
GAGCACACGTCTGAACTCTTTCCCTACACGACGCTCTTCCGATCTNNNATTGGCCCTTTGTACCTCTTCTT
GAGCACACGTCTGAACTCTTTCCCTACACGACGCTCTTCCGATCTNNNCCATGTTTGTGCAAAGGACAT
AGCACACGTCTGAACTCTTTCCCTACACGACGCTCTTCCGATCTNNNATTGTTGTTTCAGCTTCGATGTC
AGCACACGTCTGAACTCTTTCCCTACACGACGCTCTTCCGATCTNNNCAGCGTTTTGATACAAGCCAGAA
GCACACGTCTGAACTCTTTCCCTACACGACGCTCTTCCGATCTNNNACAATCATGGTGGAAAGCAAAG
AGAGCACACGTCTGAACTCTTTCCCTACACGACGCTCTTCCGATCTNNNTGTAGTCCCAGCTACTTGGA
GCACACGTCTGAACTCTTTCCCTACACGACGCTCTTCCGATCTNNNGAGAGTGCAGTGAGTCATGATC
AGCACACGTCTGAACTCTTTCCCTACACGACGCTCTTCCGATCTNNNGGGCAAGAAGAGCAAAACTTC
GAGCACACGTCTGAACTCTTTCCCTACACGACGCTCTTCCGATCTNNNCACTTGAGCCCAGGAGTTTGA
GAGCACACGTCTGAACTCTTTCCCTACACGACGCTCTTCCGATCTNNNCATATGCCTGTAGTCCCAGCTA
AGCACACGTCTGAACTCTTTCCCTACACGACGCTCTTCCGATCTNNNCTTTGCTGACCCCTAAAATGACC
AGAGCACACGTCTGAACTCTTTCCCTACACGACGCTCTTCCGATCTNNNGAAAATCGCTTGAATCTGGG
GAGCACACGTCTGAACTCTTTCCCTACACGACGCTCTTCCGATCTNNNAAATTAGCTGGGTGTGGTAG
AGCACACGTCTGAACTCTTTCCCTACACGACGCTCTTCCGATCTNNNGCCTCCTAAATAGCTGGGACTA
GAGCACACGTCTGAACTCTTTCCCTACACGACGCTCTTCCGATCTNNNCTGAGGCAAGAGAATGGTGT
GAGCACACGTCTGAACTCTTTCCCTACACGACGCTCTTCCGATCTNNNGCAAGGGACAAAAGAATGCT
GAGCACACGTCTGAACTCTTTCCCTACACGACGCTCTTCCGATCTNNNGTCTTGGGCAGTTCTTTACAAC
GAGCACACGTCTGAACTCTTTCCCTACACGACGCTCTTCCGATCTNNNTTGCATTGAACCAAGATCACA
AGCACACGTCTGAACTCTTTCCCTACACGACGCTCTTCCGATCTNNNTTTGAAACCATGTTGTGTGAGG

GAGCACACGTCTGAACTCTTTCCCTACACGACGCTCTTCCGATCTNNNTGTAATCCCAGCTACTCAGGA
AGCACACGTCTGAACTCTTTCCCTACACGACGCTCTTCCGATCTNNNAAGGTGGGAGAATCTCTTGA
GAGCACACGTCTGAACTCTTTCCCTACACGACGCTCTTCCGATCTNNNGGTCTTCTCCTTGGAGTTCTG
AGCACACGTCTGAACTCTTTCCCTACACGACGCTCTTCCGATCTNNNATAAGCAGAGATCATGCCTTTG
AGCACACGTCTGAACTCTTTCCCTACACGACGCTCTTCCGATCTNNNCATTGGCAGCAAGAAAGAAGT
GAGCACACGTCTGAACTCTTTCCCTACACGACGCTCTTCCGATCTNNNGCTGAGGCATGAGAATCTCT
AGCACACGTCTGAACTCTTTCCCTACACGACGCTCTTCCGATCTNNNCTGTAGTACCAGCTACTCGGG
AGCACACGTCTGAACTCTTTCCCTACACGACGCTCTTCCGATCTNNNACCATTGCCCATCCTAAAGTTA
AGCACACGTCTGAACTCTTTCCCTACACGACGCTCTTCCGATCTNNNGCGACAGAATGACAGAATGAG
AGCACACGTCTGAACTCTTTCCCTACACGACGCTCTTCCGATCTNNNCTAGCTGCAGTATAGAGGATG
GAGCACACGTCTGAACTCTTTCCCTACACGACGCTCTTCCGATCTNNNGCTACAATGAGCTGTGATCA
AGCACACGTCTGAACTCTTTCCCTACACGACGCTCTTCCGATCTNNNCTGTAGTCCCAGCTACTCAGG
AGCACACGTCTGAACTCTTTCCCTACACGACGCTCTTCCGATCTNNNTTGTTAGTAGAGACGGGGTTT
AGCACACGTCTGAACTCTTTCCCTACACGACGCTCTTCCGATCTNNNGAGGCAAGAGAATCACTTGA
AGCACACGTCTGAACTCTTTCCCTACACGACGCTCTTCCGATCTNNNGAGCTACTGCACCTGGTCTAT
GAGCACACGTCTGAACTCTTTCCCTACACGACGCTCTTCCGATCTNNNAGCGAGACTCTGCCAAAAAA
AGCACACGTCTGAACTCTTTCCCTACACGACGCTCTTCCGATCTNNNATTAGGGTTCACTCCTGGTGTT
AGCACACGTCTGAACTCTTTCCCTACACGACGCTCTTCCGATCTNNNGAGTTTGAGACTAGCCTGAC
GAGCACACGTCTGAACTCTTTCCCTACACGACGCTCTTCCGATCTNNNCTCCTGAGTAGCTGGGACTA
AGCACACGTCTGAACTCTTTCCCTACACGACGCTCTTCCGATCTNNNAGATCCTCCCACTTACATCCTCA
AGCACACGTCTGAACTCTTTCCCTACACGACGCTCTTCCGATCTNNNTGCAGTGAGCCAAGATTATGC
GAGCACACGTCTGAACTCTTTCCCTACACGACGCTCTTCCGATCTNNNGACAATCACTTGAGAACCTG
AGCACACGTCTGAACTCTTTCCCTACACGACGCTCTTCCGATCTNNNGGAGACACAACAAGACTCCAT
GCACACGTCTGAACTCTTTCCCTACACGACGCTCTTCCGATCTNNNGCTGTTTCCCCCACTAATCTGTAC
AGCACACGTCTGAACTCTTTCCCTACACGACGCTCTTCCGATCTNNNTAGGCCCACATTGTTTCTTCAGT
GAGCACACGTCTGAACTCTTTCCCTACACGACGCTCTTCCGATCTNNNGAGAATCACTTGAACCCAGG
AGCACACGTCTGAACTCTTTCCCTACACGACGCTCTTCCGATCTNNNCTGTAGTTCCAGCTACTTGGG
AGCACACGTCTGAACTCTTTCCCTACACGACGCTCTTCCGATCTNNNCTTGCAGTTCCTCTACTTGGG
GAGCACACGTCTGAACTCTTTCCCTACACGACGCTCTTCCGATCTNNNAGTGAGCTGAGATTGTACCA
GCACACGTCTGAACTCTTTCCCTACACGACGCTCTTCCGATCTNNNCTGAGATAGAGCACTGCACATC
GCACACGTCTGAACTCTTTCCCTACACGACGCTCTTCCGATCTNNNGGAGAATGACGTGAATCCGTG
AGCACACGTCTGAACTCTTTCCCTACACGACGCTCTTCCGATCTNNNAGAATCACTTGAACTCAGGAG
AGCACACGTCTGAACTCTTTCCCTACACGACGCTCTTCCGATCTNNNTACAGTCTCTGCCTGACCATTTC
AGCACACGTCTGAACTCTTTCCCTACACGACGCTCTTCCGATCTNNNACACCTGTATTCCCAGCTACTT
GAGCACACGTCTGAACTCTTTCCCTACACGACGCTCTTCCGATCTNNNAATCACTTGAACCTGGGAAG
GAGCACACGTCTGAACTCTTTCCCTACACGACGCTCTTCCGATCTNNNTCCTAGCACTTTGGGAAGTCA
AGCACACGTCTGAACTCTTTCCCTACACGACGCTCTTCCGATCTNNNAGAATCACTTGTACCCAGTAGG
AGCACACGTCTGAACTCTTTCCCTACACGACGCTCTTCCGATCTNNNCAAGATTCCACCATTGCATTCC
AGCACACGTCTGAACTCTTTCCCTACACGACGCTCTTCCGATCTNNNATTTGTGCTGCTTCCTGCTTTT
AGCACACGTCTGAACTCTTTCCCTACACGACGCTCTTCCGATCTNNNTCTTCACCCAACATATTCGCTCA
GCACACGTCTGAACTCTTTCCCTACACGACGCTCTTCCGATCTNNNCTCCAACACCTAATACAGTGCCT
GCACACGTCTGAACTCTTTCCCTACACGACGCTCTTCCGATCTNNNGATATGCAGCTTCCTTTCCACAC
AGCACACGTCTGAACTCTTTCCCTACACGACGCTCTTCCGATCTNNNGTGGCATTCTGGGCTTTGTAA

AGCACACGTCTGAACTCTTTCCCTACACGACGCTCTTCCGATCTNNNTTGAACTCCTGGCTTCAAGCAAT
GAGCACACGTCTGAACTCTTTCCCTACACGACGCTCTTCCGATCTNNNTATGTTGTGCAGCAATCACTTC
GCACACGTCTGAACTCTTTCCCTACACGACGCTCTTCCGATCTNNNGCCAAGATAGCACCATTGAACTC
AGAGCACACGTCTGAACTCTTTCCCTACACGACGCTCTTCCGATCTNNNGCTGAAGTAGGAGAATCGC
GAGCACACGTCTGAACTCTTTCCCTACACGACGCTCTTCCGATCTNNNTACATGCCTGTAATCCCAGCTA
AGCACACGTCTGAACTCTTTCCCTACACGACGCTCTTCCGATCTNNNCACCTGTAATCCCAGCTACTTGG
AGCACACGTCTGAACTCTTTCCCTACACGACGCTCTTCCGATCTNNNAAAGGTTCCAGTGAGCTGAGA
AGCACACGTCTGAACTCTTTCCCTACACGACGCTCTTCCGATCTNNNGGCTCCTCCTACTTCAGTCAATC
AGCACACGTCTGAACTCTTTCCCTACACGACGCTCTTCCGATCTNNNCCCAAAAGTTCACATTGTGGACA
GAGCACACGTCTGAACTCTTTCCCTACACGACGCTCTTCCGATCTNNNACGGAGTTTCACTACATTGTCC
GAGCACACGTCTGAACTCTTTCCCTACACGACGCTCTTCCGATCTNNNGGCCCTTGTTCATTCATTCTCTC
GAGCACACGTCTGAACTCTTTCCCTACACGACGCTCTTCCGATCTNNNGAGTCTCTCTCTCTGTCACCCA
AGCACACGTCTGAACTCTTTCCCTACACGACGCTCTTCCGATCTNNNGTATCTTGGGTGTCCTTCCACAT
AGCACACGTCTGAACTCTTTCCCTACACGACGCTCTTCCGATCTNNNAGGAGAGATGCTGATTGGACA
GAGCACACGTCTGAACTCTTTCCCTACACGACGCTCTTCCGATCTNNNTTGTGGTCTCAACTACTTGGG
AGAGCACACGTCTGAACTCTTTCCCTACACGACGCTCTTCCGATCTNNNTGTGGTGGTGAATGCCTGTA
GAGCACACGTCTGAACTCTTTCCCTACACGACGCTCTTCCGATCTNNNGAGGCAAGACAATCACTTGAA
AGCACACGTCTGAACTCTTTCCCTACACGACGCTCTTCCGATCTNNNTAGAGTTGTTTCATCCCTGGCTG
GAGCACACGTCTGAACTCTTTCCCTACACGACGCTCTTCCGATCTNNNGTAATCCCAGCTACTTAGGAGG
AGCACACGTCTGAACTCTTTCCCTACACGACGCTCTTCCGATCTNNNGAAATATATCTGGCTGGACCCCA
GAGCACACGTCTGAACTCTTTCCCTACACGACGCTCTTCCGATCTNNNATCATACATCACTGCACCTTCGA
GCACACGTCTGAACTCTTTCCCTACACGACGCTCTTCCGATCTNNNTGCCCTTCTTTGCCTTTTTTGATCA
GAGCACACGTCTGAACTCTTTCCCTACACGACGCTCTTCCGATCTNNNAGATCCACCACAACGTACATGA
AGAGCACACGTCTGAACTCTTTCCCTACACGACGCTCTTCCGATCTNNNAGTGCTGGGATTATAGGCAT
AGCACACGTCTGAACTCTTTCCCTACACGACGCTCTTCCGATCTNNNTTTTAGGGAAACTTGGGGTCAC
AGAGCACACGTCTGAACTCTTTCCCTACACGACGCTCTTCCGATCTNNNGAGAATCTGGGATTACAGGC
AGCACACGTCTGAACTCTTTCCCTACACGACGCTCTTCCGATCTNNNATTTTAGTCCAGTGAGGCCCATT
AGCACACGTCTGAACTCTTTCCCTACACGACGCTCTTCCGATCTNNNTGTAATCCCAGGTACTCAGGAG
AGCACACGTCTGAACTCTTTCCCTACACGACGCTCTTCCGATCTNNNGAACACCCATACCATCACTGCTA
GAGCACACGTCTGAACTCTTTCCCTACACGACGCTCTTCCGATCTNNNTGACTGTCTTCCCAGTGTTATC
AGCACACGTCTGAACTCTTTCCCTACACGACGCTCTTCCGATCTNNNTGGTGACTAGCCTTGAGACTTT
AGCACACGTCTGAACTCTTTCCCTACACGACGCTCTTCCGATCTNNNCTTTCCCATTACATCTCCCACCAC
GAGCACACGTCTGAACTCTTTCCCTACACGACGCTCTTCCGATCTNNNGGTTGTAGTAAATGCTGCCTCA
GAGCACACGTCTGAACTCTTTCCCTACACGACGCTCTTCCGATCTNNNTGCCTTGGATATGACACAGAC
AGAGCACACGTCTGAACTCTTTCCCTACACGACGCTCTTCCGATCTNNNCATTTGTAAGTTCAGCTGGT
AGAGCACACGTCTGAACTCTTTCCCTACACGACGCTCTTCCGATCTNNNGCACCTTCTACAGAAACCTC
AGCACACGTCTGAACTCTTTCCCTACACGACGCTCTTCCGATCTNNNCTGAGGCAGGAGAATCACTTGA
GAGCACACGTCTGAACTCTTTCCCTACACGACGCTCTTCCGATCTNNNAATTCGAGGTTGCAGTGAGG
GAGCACACGTCTGAACTCTTTCCCTACACGACGCTCTTCCGATCTNNNTGTCTCTGTCTCTGTCTGTCTCT
GAGCACACGTCTGAACTCTTTCCCTACACGACGCTCTTCCGATCTNNNGCAACATGGTGAAACTCTTGT
GAGCACACGTCTGAACTCTTTCCCTACACGACGCTCTTCCGATCTNNNCCAAGATTGTGCCATTGTACTC
GAGCACACGTCTGAACTCTTTCCCTACACGACGCTCTTCCGATCTNNNCAGCCTGGACAACAAAAGAAA
GAGCACACGTCTGAACTCTTTCCCTACACGACGCTCTTCCGATCTNNNGACACCAGAGACCACTTTGAGT

GAGCACACGTCTGAACTCTTTCCCTACACGACGCTCTTCCGATCTNNNGAGGCAAGAGAATCGCATGA
AGCACACGTCTGAACTCTTTCCCTACACGACGCTCTTCCGATCTNNNAGGATTGCTTGAGTCCAGGAAT
AGCACACGTCTGAACTCTTTCCCTACACGACGCTCTTCCGATCTNNNCTGAGGCAGGAGAATCACTTGA
GAGCACACGTCTGAACTCTTTCCCTACACGACGCTCTTCCGATCTNNNACAGAAGAAGCTCTCGAATCTC
GAGCACACGTCTGAACTCTTTCCCTACACGACGCTCTTCCGATCTNNNATATAGCTCTACTGCAGGTGG
GAGCACACGTCTGAACTCTTTCCCTACACGACGCTCTTCCGATCTNNNCTGCAGTGAGCTGATATATCCC
AGCACACGTCTGAACTCTTTCCCTACACGACGCTCTTCCGATCTNNNATCTCCCTGAAACCTGTTTCCTCC
GAGCACACGTCTGAACTCTTTCCCTACACGACGCTCTTCCGATCTNNNAGAGGTTGCAGTAAGCCAAG
AGCACACGTCTGAACTCTTTCCCTACACGACGCTCTTCCGATCTNNNCCTGCATCTCTCTCTCTCTCTCA
AGCACACGTCTGAACTCTTTCCCTACACGACGCTCTTCCGATCTNNNTGGGTGACATAGAGAGACTCC
GAGCACACGTCTGAACTCTTTCCCTACACGACGCTCTTCCGATCTNNNCAGGAAAATCACTTGAACGGA
GCACACGTCTGAACTCTTTCCCTACACGACGCTCTTCCGATCTNNNGAAGGTTGCAGTGAGCCAAAAT
AGCACACGTCTGAACTCTTTCCCTACACGACGCTCTTCCGATCTNNNGTGTCAAGGGTGGAAAAACTCA
GAGCACACGTCTGAACTCTTTCCCTACACGACGCTCTTCCGATCTNNNGCCTATGTGCACTCTCTCCTAA
GCACACGTCTGAACTCTTTCCCTACACGACGCTCTTCCGATCTNNNGTTGGGAAGCGTCAATGAAATG
AGCACACGTCTGAACTCTTTCCCTACACGACGCTCTTCCGATCTNNNCCGAAAATGCCGAGTACTTTAG
GAGCACACGTCTGAACTCTTTCCCTACACGACGCTCTTCCGATCTNNNCTGCTGTCATATTGCATTCTCCC
AGCACACGTCTGAACTCTTTCCCTACACGACGCTCTTCCGATCTNNNAGCCTATCCAGTTTACAGTGACT
AGCACACGTCTGAACTCTTTCCCTACACGACGCTCTTCCGATCTNNNACTCTTGGCACTCTGATTCTCTG
AGAGCACACGTCTGAACTCTTTCCCTACACGACGCTCTTCCGATCTNNNAATGAAGTACAGGGTGCTA
GAGCACACGTCTGAACTCTTTCCCTACACGACGCTCTTCCGATCTNNNAGGCAGGAACATCACTTGAAT
AGAGCACACGTCTGAACTCTTTCCCTACACGACGCTCTTCCGATCTNNNCACATCCGTAGAGAGGCTCTA
AGCACACGTCTGAACTCTTTCCCTACACGACGCTCTTCCGATCTNNNTCAGAACTCTTGGACTCCAAAG
GCACACGTCTGAACTCTTTCCCTACACGACGCTCTTCCGATCTNNNTGAGCTATCACACCATTGCACTCC
AGCACACGTCTGAACTCTTTCCCTACACGACGCTCTTCCGATCTNNNGGGTGATCTTGACTGTAGGGA
AGCACACGTCTGAACTCTTTCCCTACACGACGCTCTTCCGATCTNNNCCTATGAGGCTCTCACAGTTTGT
AGCACACGTCTGAACTCTTTCCCTACACGACGCTCTTCCGATCTNNNTTTAGAACTCCCAACCTCAGGTC
GAGCACACGTCTGAACTCTTTCCCTACACGACGCTCTTCCGATCTNNNCTGGAGTGTAGTGGTGCAATT
GAGCACACGTCTGAACTCTTTCCCTACACGACGCTCTTCCGATCTNNNCAGCTATACACGGAGACAACA
AGCACACGTCTGAACTCTTTCCCTACACGACGCTCTTCCGATCTNNNACAGAGCGAGATTCCATCTAAC
AGCACACGTCTGAACTCTTTCCCTACACGACGCTCTTCCGATCTNNNGGTGTTTTGGTTCTGCTGGTATC
AGCACACGTCTGAACTCTTTCCCTACACGACGCTCTTCCGATCTNNNATTTCCACGTTCCTTGTACCAGA
AGCACACGTCTGAACTCTTTCCCTACACGACGCTCTTCCGATCTNNNAGACGAATCTTGGGATATGCCA
GAGCACACGTCTGAACTCTTTCCCTACACGACGCTCTTCCGATCTNNNGGAGTACAGTGACACGATCTT
GAGCACACGTCTGAACTCTTTCCCTACACGACGCTCTTCCGATCTNNNGCACCTTGATCTTGGACTTCCT
AGCACACGTCTGAACTCTTTCCCTACACGACGCTCTTCCGATCTNNNAAATACCGACTCCAGAGGTTTC
GAGCACACGTCTGAACTCTTTCCCTACACGACGCTCTTCCGATCTNNNAGCCCAAAGTTGAAGTCAAAT
AGCACACGTCTGAACTCTTTCCCTACACGACGCTCTTCCGATCTNNNTCTCCAGCATCTCTCTTCTGAC
GCACACGTCTGAACTCTTTCCCTACACGACGCTCTTCCGATCTNNNTCTGTCCCTATGTCTCTGTGTCTCA
GAGCACACGTCTGAACTCTTTCCCTACACGACGCTCTTCCGATCTNNNATTTCCTTTCTTCTGCTTTGGG
AGCACACGTCTGAACTCTTTCCCTACACGACGCTCTTCCGATCTNNNCCGAGCAAGTTGGATGTAAACA
GAGCACACGTCTGAACTCTTTCCCTACACGACGCTCTTCCGATCTNNNAGAGTTTGCAGTGAGCCAAG
AGCACACGTCTGAACTCTTTCCCTACACGACGCTCTTCCGATCTNNNCAAGTACTCTGGGCAAAAAGTC

GAGCACACGTCTGAACTCTTTCCCTACACGACGCTCTTCCGATCTNNNTATAGTCCCAGCTACTCAGGA
GAGCACACGTCTGAACTCTTTCCCTACACGACGCTCTTCCGATCTNNNTATCTTTCTGTCTGTCCACCTGC
GAGCACACGTCTGAACTCTTTCCCTACACGACGCTCTTCCGATCTNNNAGGATGGGTAGAAATGAGA
AGCACACGTCTGAACTCTTTCCCTACACGACGCTCTTCCGATCTNNNGTTCCATCTACTCATGAGGCTG
GAGCACACGTCTGAACTCTTTCCCTACACGACGCTCTTCCGATCTNNNGAGAAAGACACACACACACA
AGCACACGTCTGAACTCTTTCCCTACACGACGCTCTTCCGATCTNNNCTGAGGTGAGAGAACTGCTTG
GAGCACACGTCTGAACTCTTTCCCTACACGACGCTCTTCCGATCTNNNACAAAACCTCCGTGATAATGG
AGCACACGTCTGAACTCTTTCCCTACACGACGCTCTTCCGATCTNNNTTGCAATGAGTCAAGATCATGC
AGAGCACACGTCTGAACTCTTTCCCTACACGACGCTCTTCCGATCTNNNTATCATCTGCCTGAACTCAGC
AGAGCACACGTCTGAACTCTTTCCCTACACGACGCTCTTCCGATCTNNNGACACCTGTAATCCCAGCTAC
GAGCACACGTCTGAACTCTTTCCCTACACGACGCTCTTCCGATCTNNNAGCTGCAACCCTTTATAACTGG
AGAGCACACGTCTGAACTCTTTCCCTACACGACGCTCTTCCGATCTNNNGGCCTCTCAAATCCCATCCATA
GAGCACACGTCTGAACTCTTTCCCTACACGACGCTCTTCCGATCTNNNCTGTAATCCCAGCACTTTGGA
AGCACACGTCTGAACTCTTTCCCTACACGACGCTCTTCCGATCTNNNCTGGTCTCAAACTCCTGACTTCA
AGCACACGTCTGAACTCTTTCCCTACACGACGCTCTTCCGATCTNNNCAGACAAAAGCCACTCAATACC
AGAGCACACGTCTGAACTCTTTCCCTACACGACGCTCTTCCGATCTNNNGGCACCTATAATCCCAGCTAC
GCACACGTCTGAACTCTTTCCCTACACGACGCTCTTCCGATCTNNNCAGACCTTAGAACTTGTCAGCCT
AGCACACGTCTGAACTCTTTCCCTACACGACGCTCTTCCGATCTNNNAGTGAGCTGAGATTGTGTCACT
AGCACACGTCTGAACTCTTTCCCTACACGACGCTCTTCCGATCTNNNGAGGCAGGAGAATGGTGTGA
GAGCACACGTCTGAACTCTTTCCCTACACGACGCTCTTCCGATCTNNNGCAAGGTTAGCTCTCTCTCT
GAGCACACGTCTGAACTCTTTCCCTACACGACGCTCTTCCGATCTNNNCCCTGTCCATCACTCCTATCCTA
GAGCACACGTCTGAACTCTTTCCCTACACGACGCTCTTCCGATCTNNNCCGAAAGTGCTGTTTAGGGTT
AGCACACGTCTGAACTCTTTCCCTACACGACGCTCTTCCGATCTNNNATTTGTGTAACCACCACCCAGAT
AGCACACGTCTGAACTCTTTCCCTACACGACGCTCTTCCGATCTNNNCACACATCACTCAAGTCACCCTAC
AGAGCACACGTCTGAACTCTTTCCCTACACGACGCTCTTCCGATCTNNNGAGGCACAGAAAGAGACAG
AGCACACGTCTGAACTCTTTCCCTACACGACGCTCTTCCGATCTNNNATGGAGAAACCCCGTCTCTACT
GAGCACACGTCTGAACTCTTTCCCTACACGACGCTCTTCCGATCTNNNGGGCAAACAGGCTTTATATGC
AAGAGCACACGTCTGAACTCTTTCCCTACACGACGCTCTTCCGATCTNNNGAGGCAGGGAGAATAGGT
GAGCACACGTCTGAACTCTTTCCCTACACGACGCTCTTCCGATCTNNNGTGGGATTCCTGGATTGTATC
GAGCACACGTCTGAACTCTTTCCCTACACGACGCTCTTCCGATCTNNNCTGTAGTCCCAGCTACTCAGG
AGCACACGTCTGAACTCTTTCCCTACACGACGCTCTTCCGATCTNNNTAATGGGAATTCGTAGTGTGCC
GAGCACACGTCTGAACTCTTTCCCTACACGACGCTCTTCCGATCTNNNCCCTGCTTTTTTTGGTCCAGA
AGAGCACACGTCTGAACTCTTTCCCTACACGACGCTCTTCCGATCTNNNAGGGGAAAGACTTACAGAG
AGAGCACACGTCTGAACTCTTTCCCTACACGACGCTCTTCCGATCTNNNCTACCATGCTTCATAGAGACC
AGCACACGTCTGAACTCTTTCCCTACACGACGCTCTTCCGATCTNNNAAAGGTCTACACAAAGGCAGTC
AGCACACGTCTGAACTCTTTCCCTACACGACGCTCTTCCGATCTNNNTGACAGCTTTTCACCTACTCCTCC
AGCACACGTCTGAACTCTTTCCCTACACGACGCTCTTCCGATCTNNNGTTTCAACAAACTGTCTGTGGCT
GAGCACACGTCTGAACTCTTTCCCTACACGACGCTCTTCCGATCTNNNTGAATGAGAATTTACAGTGGC
AGCACACGTCTGAACTCTTTCCCTACACGACGCTCTTCCGATCTNNNTCTCACAGTGGATGGTCACAATC
AAGAGCACACGTCTGAACTCTTTCCCTACACGACGCTCTTCCGATCTNNNAATGAGTCATCACACTCCTC
AGCACACGTCTGAACTCTTTCCCTACACGACGCTCTTCCGATCTNNNATGCTCAGATTTTCTTGCCAGTG
AGCACACGTCTGAACTCTTTCCCTACACGACGCTCTTCCGATCTNNNGGAGGTTGCTGTTAGCCTAGAA
GAGCACACGTCTGAACTCTTTCCCTACACGACGCTCTTCCGATCTNNNAAAGGTCTGTCTGTCTGTCTG

AGAGCACACGTCTGAACTCTTTCCCTACACGACGCTCTTCCGATCTNNNGGCTGCAATCTGAAAGGAA
AGCACACGTCTGAACTCTTTCCCTACACGACGCTCTTCCGATCTNNNGCTAGTGCTGCAATCTCTTGAT
GAGCACACGTCTGAACTCTTTCCCTACACGACGCTCTTCCGATCTNNNTTTGGAGTTGAGCCCTTCAG
GAGCACACGTCTGAACTCTTTCCCTACACGACGCTCTTCCGATCTNNNTCAGGTTGAGTAGACTGTCC
AGCACACGTCTGAACTCTTTCCCTACACGACGCTCTTCCGATCTNNNCCATTGCTATGCACTGAATGAG
AGCACACGTCTGAACTCTTTCCCTACACGACGCTCTTCCGATCTNNNGCTTCCAAAGTAGCTGCAACTA
GAGCACACGTCTGAACTCTTTCCCTACACGACGCTCTTCCGATCTNNNTCACTTGAACCTGGAGACAA
GAGCACACGTCTGAACTCTTTCCCTACACGACGCTCTTCCGATCTNNNTCTGCCTTTCCCATTTAGATGC
AGCACACGTCTGAACTCTTTCCCTACACGACGCTCTTCCGATCTNNNTCTTGCTCCTTTGCCTCCTAAAT
AGAGCACACGTCTGAACTCTTTCCCTACACGACGCTCTTCCGATCTNNNCAGCATGGCCATTTTCACGA
GAGCACACGTCTGAACTCTTTCCCTACACGACGCTCTTCCGATCTNNNAGAGATCCCCATACTCTACCG
GAGCACACGTCTGAACTCTTTCCCTACACGACGCTCTTCCGATCTNNNTGCATCCTCCCTGTAACATTCT
GAGCACACGTCTGAACTCTTTCCCTACACGACGCTCTTCCGATCTNNNACCAGCAGAGTTTTGCCTTTT
GCACACGTCTGAACTCTTTCCCTACACGACGCTCTTCCGATCTNNNGCTGAAAACCACCTTGAATGTG
GCACACGTCTGAACTCTTTCCCTACACGACGCTCTTCCGATCTNNNGATTCGGTAGGCCTAGGATGAT
GCACACGTCTGAACTCTTTCCCTACACGACGCTCTTCCGATCTNNNCCAGAGTTCCAAGCAATACTCCT
AGCACACGTCTGAACTCTTTCCCTACACGACGCTCTTCCGATCTNNNTTACTTCCTTCTGCACTGCATCT
GAGCACACGTCTGAACTCTTTCCCTACACGACGCTCTTCCGATCTNNNGCTGTTTTTGGTGTTTTGGTG
AGCACACGTCTGAACTCTTTCCCTACACGACGCTCTTCCGATCTNNNGTGCCATCACAACATACTGCAT
CACACGTCTGAACTCTTTCCCTACACGACGCTCTTCCGATCTNNNGATCACAGCAGGAGACACTCAAG
AGCACACGTCTGAACTCTTTCCCTACACGACGCTCTTCCGATCTNNNGCCGATAGCCCAACTACTTGAT
AGCACACGTCTGAACTCTTTCCCTACACGACGCTCTTCCGATCTNNNGCTTCTCACCTCTCAGTGACTA
GCACACGTCTGAACTCTTTCCCTACACGACGCTCTTCCGATCTNNNAAAGGCTTGCCGGGGATTTTAT
GAGCACACGTCTGAACTCTTTCCCTACACGACGCTCTTCCGATCTNNNCTTGAACATTTGTGGCAGAG
GCACACGTCTGAACTCTTTCCCTACACGACGCTCTTCCGATCTNNNACCTGAGAAGTAGCCAAAACCA
GAGCACACGTCTGAACTCTTTCCCTACACGACGCTCTTCCGATCTNNNGAGGCAGGAGAATTGTTTG
AGCACACGTCTGAACTCTTTCCCTACACGACGCTCTTCCGATCTNNNTGGGAGTCATGTTCAGCTTAC
AGCACACGTCTGAACTCTTTCCCTACACGACGCTCTTCCGATCTNNNCAACAAGAGCGAAACTGTGTC
AGCACACGTCTGAACTCTTTCCCTACACGACGCTCTTCCGATCTNNNCCCATCAGCCTGATCAGATGTAT
AGCACACGTCTGAACTCTTTCCCTACACGACGCTCTTCCGATCTNNNTCAAACCATTGCACTTTAGCCTG
GAGCACACGTCTGAACTCTTTCCCTACACGACGCTCTTCCGATCTNNNGGTTGTACCACTGTACTCTAG
AGCACACGTCTGAACTCTTTCCCTACACGACGCTCTTCCGATCTNNNACCATTGCTATCTACTTCCCTG
GCACACGTCTGAACTCTTTCCCTACACGACGCTCTTCCGATCTNNNCTGAGGCAGAAAAATCGCTTGA
GAGCACACGTCTGAACTCTTTCCCTACACGACGCTCTTCCGATCTNNNGTAATCCCAGCTACTTGAGAG
AGCACACGTCTGAACTCTTTCCCTACACGACGCTCTTCCGATCTNNNTGCAGCATTCTCTTTGGTTTAG
AGAGCACACGTCTGAACTCTTTCCCTACACGACGCTCTTCCGATCTNNNACCCCATTCCCCTACATAAAA
AGAGCACACGTCTGAACTCTTTCCCTACACGACGCTCTTCCGATCTNNNCTGTAATTCCAGCTACTCGG
AGAGCACACGTCTGAACTCTTTCCCTACACGACGCTCTTCCGATCTNNNGTAATCCCAGCTACTCTGTA
GCACACGTCTGAACTCTTTCCCTACACGACGCTCTTCCGATCTNNNGTGAAAGTCCCTTCTGCAACATC
GAGCACACGTCTGAACTCTTTCCCTACACGACGCTCTTCCGATCTNNNGCATATTGCGCAGAACTATGT
GAGCACACGTCTGAACTCTTTCCCTACACGACGCTCTTCCGATCTNNNGGAGGATACAGTGAGAACC
AGCACACGTCTGAACTCTTTCCCTACACGACGCTCTTCCGATCTNNNCACCTGTAATCCCAGCTACTCA
AGCACACGTCTGAACTCTTTCCCTACACGACGCTCTTCCGATCTNNNAGTGAGCCAAGATCATGTCAC

GAGCACACGTCTGAACTCTTTCCCTACACGACGCTCTTCCGATCTNNNGCAGAAGAATCACTTGAACC
GCACACGTCTGAACTCTTTCCCTACACGACGCTCTTCCGATCTNNNCCTCCGTAATAGTGTGAGCCAAT
AGCACACGTCTGAACTCTTTCCCTACACGACGCTCTTCCGATCTNNNTGTGTAAAGGAAAGGTTTGGA
AGCACACGTCTGAACTCTTTCCCTACACGACGCTCTTCCGATCTNNNTGTAGTCTGTAGTCCCAGCTAC
GAGCACACGTCTGAACTCTTTCCCTACACGACGCTCTTCCGATCTNNNAAGAACTGCATCATTTGGCTG
GAGCACACGTCTGAACTCTTTCCCTACACGACGCTCTTCCGATCTNNNACCCTGTGTCAAACAAACAAA
AGCACACGTCTGAACTCTTTCCCTACACGACGCTCTTCCGATCTNNNAAAATCTGAGACCCCGAAATG
AGCACACGTCTGAACTCTTTCCCTACACGACGCTCTTCCGATCTNNNACTCTTCTCTCATGTTTTGGCCT
AGCACACGTCTGAACTCTTTCCCTACACGACGCTCTTCCGATCTNNNTCATGGACTCCCTAACTCTCCTT
AGCACACGTCTGAACTCTTTCCCTACACGACGCTCTTCCGATCTNNNCTACAAAGCAAGACTCCATCTC
GAGCACACGTCTGAACTCTTTCCCTACACGACGCTCTTCCGATCTNNNTCCCTCGCTCTTTTCCTTTTCTC
GAGCACACGTCTGAACTCTTTCCCTACACGACGCTCTTCCGATCTNNNCTGTAATCCCAGCACTTTGGA
AGAGCACACGTCTGAACTCTTTCCCTACACGACGCTCTTCCGATCTNNNGCCTGTAATCCCAACTACTA
AGCACACGTCTGAACTCTTTCCCTACACGACGCTCTTCCGATCTNNNAGAATTGCTTGAACCTGGGAG
AGCACACGTCTGAACTCTTTCCCTACACGACGCTCTTCCGATCTNNNATGTCCTTTTCAAATGCTCCTGC
AGCACACGTCTGAACTCTTTCCCTACACGACGCTCTTCCGATCTNNNGTAATCCTAGCTACTTGGGAG
GAGCACACGTCTGAACTCTTTCCCTACACGACGCTCTTCCGATCTNNNCCTGATATCTGGGATTACAGG
AGCACACGTCTGAACTCTTTCCCTACACGACGCTCTTCCGATCTNNNATGCCTGTAATTCCAGCACTTTG
GCACACGTCTGAACTCTTTCCCTACACGACGCTCTTCCGATCTNNNATGCCTGTAATCCCAGCTACTTG
AGAGCACACGTCTGAACTCTTTCCCTACACGACGCTCTTCCGATCTNNNGAGAACCACTTGAACCCAG
GAGCACACGTCTGAACTCTTTCCCTACACGACGCTCTTCCGATCTNNNACTCAAGCCTGGGTGATAGA
GCACACGTCTGAACTCTTTCCCTACACGACGCTCTTCCGATCTNNNCTCAAGCAATCCTCCCACCTTAG
AGCACACGTCTGAACTCTTTCCCTACACGACGCTCTTCCGATCTNNNGGCTCAAATGATCCTCCTACCT
GCACACGTCTGAACTCTTTCCCTACACGACGCTCTTCCGATCTNNNGCTCAATTTTACAGGCTGCTCTT
GCACACGTCTGAACTCTTTCCCTACACGACGCTCTTCCGATCTNNNGCCATTCTTTTATGCCTTGCCTTC
GAGCACACGTCTGAACTCTTTCCCTACACGACGCTCTTCCGATCTNNNGCTGATTGTTTCTTTTGCTGT
AGCACACGTCTGAACTCTTTCCCTACACGACGCTCTTCCGATCTNNNTGGGGTTCATAAGCACAAGAA
AGCACACGTCTGAACTCTTTCCCTACACGACGCTCTTCCGATCTNNNTCAGTTTTTACCAGGGCTCTCTC
AGCACACGTCTGAACTCTTTCCCTACACGACGCTCTTCCGATCTNNNGAGATGGCAGCACTGTACTAC
AGCACACGTCTGAACTCTTTCCCTACACGACGCTCTTCCGATCTNNNGTTGAATTCTAGCTTTGCCACCT
AGCACACGTCTGAACTCTTTCCCTACACGACGCTCTTCCGATCTNNNCTGAGTAGCTGGGACTAAAGG
GAGCACACGTCTGAACTCTTTCCCTACACGACGCTCTTCCGATCTNNNCCTGTTTTCTCCAGTACAGTCC
AGAGCACACGTCTGAACTCTTTCCCTACACGACGCTCTTCCGATCTNNNGTGTTGGGATTACAAGTGT
AGCACACGTCTGAACTCTTTCCCTACACGACGCTCTTCCGATCTNNNGAGAGAGACACACAGTGGAGT
AGCACACGTCTGAACTCTTTCCCTACACGACGCTCTTCCGATCTNNNGATCTCATGATCCACCCACCTTG
AGCACACGTCTGAACTCTTTCCCTACACGACGCTCTTCCGATCTNNNTGTAATGAGCTGTGATCACACC
GCACACGTCTGAACTCTTTCCCTACACGACGCTCTTCCGATCTNNNGCCTGCTCAATTCCAAAGTACAA
AGCACACGTCTGAACTCTTTCCCTACACGACGCTCTTCCGATCTNNNGCGACAGAGAGAGACTCCATAT
GAGCACACGTCTGAACTCTTTCCCTACACGACGCTCTTCCGATCTNNNTGCTCTGATAGGGACTCTCTT
GCACACGTCTGAACTCTTTCCCTACACGACGCTCTTCCGATCTNNNGGCACCTATAATCCCAGCTACTT
GAGCACACGTCTGAACTCTTTCCCTACACGACGCTCTTCCGATCTNNNAGCATATGAGAGGTTCCATTC
GAGCACACGTCTGAACTCTTTCCCTACACGACGCTCTTCCGATCTNNNTGGCCAACATGGTGAAACTCT
GAGCACACGTCTGAACTCTTTCCCTACACGACGCTCTTCCGATCTNNNGGATCGCTTGAGGTCAAAAG

GCACACGTCTGAACTCTTTCCCTACACGACGCTCTTCCGATCTNNNCTCCTGAGTAGCTGGGATTACA
GAGCACACGTCTGAACTCTTTCCCTACACGACGCTCTTCCGATCTNNNAATCCGCACAACAAACCTTA
GAGCACACGTCTGAACTCTTTCCCTACACGACGCTCTTCCGATCTNNNTAGCTGTACTCACGCAGAATT
AGCACACGTCTGAACTCTTTCCCTACACGACGCTCTTCCGATCTNNNAACATAGTGAGACCTCGTCCC
GAGCACACGTCTGAACTCTTTCCCTACACGACGCTCTTCCGATCTNNNGTAATTCCAGCTACTCAGGA
AGCACACGTCTGAACTCTTTCCCTACACGACGCTCTTCCGATCTNNNTCTCTGGGTGACTATCCACTC
AGCACACGTCTGAACTCTTTCCCTACACGACGCTCTTCCGATCTNNNTGTAATCCCACCACTTTGAGAG
AGCACACGTCTGAACTCTTTCCCTACACGACGCTCTTCCGATCTNNNCTACACAGCGAAGACTCTGTC
GCACACGTCTGAACTCTTTCCCTACACGACGCTCTTCCGATCTNNNGAAGATACTCAGCACCAGGTTC
GCACACGTCTGAACTCTTTCCCTACACGACGCTCTTCCGATCTNNNATGTCTGTCCTCAAGGTTCATCT
GCACACGTCTGAACTCTTTCCCTACACGACGCTCTTCCGATCTNNNCTCGATCTCCTGACCTTGTGATC
AGCACACGTCTGAACTCTTTCCCTACACGACGCTCTTCCGATCTNNNAATGTTTCACTTTGGCCAAGG
GCACACGTCTGAACTCTTTCCCTACACGACGCTCTTCCGATCTNNNGTGACCATTTTCTCACTCAAGG
AGCACACGTCTGAACTCTTTCCCTACACGACGCTCTTCCGATCTNNNAGGCAGGATAATCACATGAAC
AGCACACGTCTGAACTCTTTCCCTACACGACGCTCTTCCGATCTNNNGCTGAGGCAGGAGAATTACTT
GCACACGTCTGAACTCTTTCCCTACACGACGCTCTTCCGATCTNNNGTGACAGAGCAAGACCTTGTTT
GAGCACACGTCTGAACTCTTTCCCTACACGACGCTCTTCCGATCTNNNAGTAACTCCACAGGCAAAGC
AGCACACGTCTGAACTCTTTCCCTACACGACGCTCTTCCGATCTNNNTTCACCAGCACAATCATAGCTC
GCACACGTCTGAACTCTTTCCCTACACGACGCTCTTCCGATCTNNNACCCCTTTCTCTACATCTTTGCA
GCACACGTCTGAACTCTTTCCCTACACGACGCTCTTCCGATCTNNNTCACCAAATACCTTCCTTGCATTC
GCACACGTCTGAACTCTTTCCCTACACGACGCTCTTCCGATCTNNNCTAGGATTACAAGCATGCACCA
AGCACACGTCTGAACTCTTTCCCTACACGACGCTCTTCCGATCTNNNGGGCAACAGAGTGAGATCTTG
AGCACACGTCTGAACTCTTTCCCTACACGACGCTCTTCCGATCTNNNGTGTGTGTGTGTGTGTGT
GAGCACACGTCTGAACTCTTTCCCTACACGACGCTCTTCCGATCTNNNTGGTGCCAAACTATGAAGA
GCACACGTCTGAACTCTTTCCCTACACGACGCTCTTCCGATCTNNNTGTGTCCTGAGAGCACAAATAC
AGCACACGTCTGAACTCTTTCCCTACACGACGCTCTTCCGATCTNNNCAGCCTCGAACTCATGGGTAAT
GCACACGTCTGAACTCTTTCCCTACACGACGCTCTTCCGATCTNNNGAAGAACTCAGTTACTGGAGC
AGCACACGTCTGAACTCTTTCCCTACACGACGCTCTTCCGATCTNNNCTTAGCAACAGAGTGAGACTC
AGCACACGTCTGAACTCTTTCCCTACACGACGCTCTTCCGATCTNNNCAATAAGCTGAGATTGTGGCA
GCACACGTCTGAACTCTTTCCCTACACGACGCTCTTCCGATCTNNNGTATTGTTGTGCACCAATCCCTC
GCACACGTCTGAACTCTTTCCCTACACGACGCTCTTCCGATCTNNNCAGGAGGACACATTTTCTTGAGC
AGCACACGTCTGAACTCTTTCCCTACACGACGCTCTTCCGATCTNNNGGACTATACACACACACACA
AGCACACGTCTGAACTCTTTCCCTACACGACGCTCTTCCGATCTNNNGGGTTCAAGTGATTCTCCTGT
AGCACACGTCTGAACTCTTTCCCTACACGACGCTCTTCCGATCTNNNTTATGTTGGGATTGACTAGGG
GAGCACACGTCTGAACTCTTTCCCTACACGACGCTCTTCCGATCTNNNACAGGGCCAAATGGAAAAGA
AGCACACGTCTGAACTCTTTCCCTACACGACGCTCTTCCGATCTNNNGGGTGACAGAGAAAGACTCC
AGCACACGTCTGAACTCTTTCCCTACACGACGCTCTTCCGATCTNNNGAGGTTGTGATGAGCCGAGA
AGCACACGTCTGAACTCTTTCCCTACACGACGCTCTTCCGATCTNNNGGAACTGGAAGGAAAGAAAA
GAGCACACGTCTGAACTCTTTCCCTACACGACGCTCTTCCGATCTNNNGTTGGTTGAAATCAGGCATG
AGCACACGTCTGAACTCTTTCCCTACACGACGCTCTTCCGATCTNNNGATTGGCAAACTGATGGATGC
AGCACACGTCTGAACTCTTTCCCTACACGACGCTCTTCCGATCTNNNTCCCTAGTAGCTGGGACTATA
GAGCACACGTCTGAACTCTTTCCCTACACGACGCTCTTCCGATCTNNNGTCTCCCCTAAAGAGATGCTC
AGCACACGTCTGAACTCTTTCCCTACACGACGCTCTTCCGATCTNNNTGTAGTGACCTTCTCTGTCTG

GCACACGTCTGAACTCTTTCCCTACACGACGCTCTTCCGATCTNNNCTGTAGTCCCAGCTACTCAGGAG
AGCACACGTCTGAACTCTTTCCCTACACGACGCTCTTCCGATCTNNNGTCCCTTCATCTCAGCATTAGGT
AAGAGCACACGTCTGAACTCTTTCCCTACACGACGCTCTTCCGATCTNNNAGGCAGAAGAATCACTTG
GCACACGTCTGAACTCTTTCCCTACACGACGCTCTTCCGATCTNNNCACAGTGCTGGGATTATAGGA
AGCACACGTCTGAACTCTTTCCCTACACGACGCTCTTCCGATCTNNNACATCATTGCACTCTAGCCTGA
AGCACACGTCTGAACTCTTTCCCTACACGACGCTCTTCCGATCTNNNCATGTGGTTTCTGCATCCTCATT
AGCACACGTCTGAACTCTTTCCCTACACGACGCTCTTCCGATCTNNNACTGCACACATAGTCTTTGTTG
AGCACACGTCTGAACTCTTTCCCTACACGACGCTCTTCCGATCTNNNCTGAGAGATGGCATTTTCTGT
AGCACACGTCTGAACTCTTTCCCTACACGACGCTCTTCCGATCTNNNATGTCACCACACCTGGCTAATT
GAGCACACGTCTGAACTCTTTCCCTACACGACGCTCTTCCGATCTNNNGCTTTGTTCGTTTTGCTCAGG
AGCACACGTCTGAACTCTTTCCCTACACGACGCTCTTCCGATCTNNNCTTCGCAGACCTCATGTTTTCA
AGCACACGTCTGAACTCTTTCCCTACACGACGCTCTTCCGATCTNNNAGCGAGACAAGATCACACTAC
GAGCACACGTCTGAACTCTTTCCCTACACGACGCTCTTCCGATCTNNNCAACTTGCTCAGAAATGGGT
AGCACACGTCTGAACTCTTTCCCTACACGACGCTCTTCCGATCTNNNCCTTTCCCACTCTGATATGCTCA
AGCACACGTCTGAACTCTTTCCCTACACGACGCTCTTCCGATCTNNNCAGACTAGGCAACATAGGGA
GAGCACACGTCTGAACTCTTTCCCTACACGACGCTCTTCCGATCTNNNACAACTTTCCAGTTTCCTGAG
GCACACGTCTGAACTCTTTCCCTACACGACGCTCTTCCGATCTNNNAGCACTTTGAGAGAATGAGAG
GAGCACACGTCTGAACTCTTTCCCTACACGACGCTCTTCCGATCTNNNAGGCCATGAAGATTCAGCTG
AGCACACGTCTGAACTCTTTCCCTACACGACGCTCTTCCGATCTNNNAGCATCCTTGTCTTGTTCCAGAT
GCACACGTCTGAACTCTTTCCCTACACGACGCTCTTCCGATCTNNNTCCCTCTGAAGATCCTGGTACTT
AGCACACGTCTGAACTCTTTCCCTACACGACGCTCTTCCGATCTNNNGCAAGACCCTGACTCAAAAAC
GAGCACACGTCTGAACTCTTTCCCTACACGACGCTCTTCCGATCTNNNCTCTCTCTCTCACACACACA
GCACACGTCTGAACTCTTTCCCTACACGACGCTCTTCCGATCTNNNTGCTTGGTAGAGTCTGCATTGA
GCACACGTCTGAACTCTTTCCCTACACGACGCTCTTCCGATCTNNNACCTGGGGATGTATTCTTTTGG
AGCACACGTCTGAACTCTTTCCCTACACGACGCTCTTCCGATCTNNNGAGAATCACTGGAACTGAGG
GAGCACACGTCTGAACTCTTTCCCTACACGACGCTCTTCCGATCTNNNGAGGGAGGAAAATTGCTTG
GCACACGTCTGAACTCTTTCCCTACACGACGCTCTTCCGATCTNNNGCCACACCCTGCTAATTTCTTTCC
GAGCACACGTCTGAACTCTTTCCCTACACGACGCTCTTCCGATCTNNNGATGCAGGAGAATCGTTTGA
AGCACACGTCTGAACTCTTTCCCTACACGACGCTCTTCCGATCTNNNAACAAAAACAGGACCCAGAGA
AGCACACGTCTGAACTCTTTCCCTACACGACGCTCTTCCGATCTNNNTGCACATCACCATGATAGGCT
AGCACACGTCTGAACTCTTTCCCTACACGACGCTCTTCCGATCTNNNGAATCGTTTGAATCTGGAGGC
AGAGCACACGTCTGAACTCTTTCCCTACACGACGCTCTTCCGATCTNNNCAGTTACCCAATCCTAGGCC
GCACACGTCTGAACTCTTTCCCTACACGACGCTCTTCCGATCTNNNAGTCTTTCTCTCTCTCTCCCCCA
GAGCACACGTCTGAACTCTTTCCCTACACGACGCTCTTCCGATCTNNNGACACCTGTAATCCCAGCACT
AGCACACGTCTGAACTCTTTCCCTACACGACGCTCTTCCGATCTNNNGAGGGAGGAAAATTGCTTGA
GAGCACACGTCTGAACTCTTTCCCTACACGACGCTCTTCCGATCTNNNGGATGGTCTTGAACTTCTGAC
GAGCACACGTCTGAACTCTTTCCCTACACGACGCTCTTCCGATCTNNNGTGAGAGAGAAAATGAGAG
GAGCACACGTCTGAACTCTTTCCCTACACGACGCTCTTCCGATCTNNNATTCAAATCCATCACTTTCGG
GAGCACACGTCTGAACTCTTTCCCTACACGACGCTCTTCCGATCTNNNCATCTCTGACTCTGCATAGTC
AGCACACGTCTGAACTCTTTCCCTACACGACGCTCTTCCGATCTNNNAGTTAGGTCATGAGAGTGAAG
AGCACACGTCTGAACTCTTTCCCTACACGACGCTCTTCCGATCTNNNGAAGAGTCAGGTCCCACTATA
GAGCACACGTCTGAACTCTTTCCCTACACGACGCTCTTCCGATCTNNNTTCACCTTCCACCATGATTGT
AGCACACGTCTGAACTCTTTCCCTACACGACGCTCTTCCGATCTNNNGGCTGCAGTGAGCTAAGATCA

AGCACACGTCTGAACTCTTTCCCTACACGACGCTCTTCCGATCTNNNGCATGTGTTTTTGCTGGGTTTT
AGCACACGTCTGAACTCTTTCCCTACACGACGCTCTTCCGATCTNNNCGGAGCAAGACTCTGTCTCAAA
AGCACACGTCTGAACTCTTTCCCTACACGACGCTCTTCCGATCTNNNTGACCAAAGCATCCTTACATGG
GCACACGTCTGAACTCTTTCCCTACACGACGCTCTTCCGATCTNNNCGTGTGTGTTTGTTTGTGTGTT
AGCACACGTCTGAACTCTTTCCCTACACGACGCTCTTCCGATCTNNNACACATACGCGTTTTTTGTGTTT
GAGCACACGTCTGAACTCTTTCCCTACACGACGCTCTTCCGATCTNNNTCCTCTCTTCTCTCTCTCTCT
GAGCACACGTCTGAACTCTTTCCCTACACGACGCTCTTCCGATCTNNNTCCATTCAGGACATCGGGAT
GCACACGTCTGAACTCTTTCCCTACACGACGCTCTTCCGATCTNNNGGCAACATAGTGAGATCCTGAG
AGCACACGTCTGAACTCTTTCCCTACACGACGCTCTTCCGATCTNNNCTGAGGCAGGAGAATCACTTG
AGCACACGTCTGAACTCTTTCCCTACACGACGCTCTTCCGATCTNNNAGACAGGAAAATCGCTTGATC
GCACACGTCTGAACTCTTTCCCTACACGACGCTCTTCCGATCTNNNTCACATGGTTACAAGGTCAAGC
AGCACACGTCTGAACTCTTTCCCTACACGACGCTCTTCCGATCTNNNGTTGCATTGAGCTGAGATTGT
GAGCACACGTCTGAACTCTTTCCCTACACGACGCTCTTCCGATCTNNNATTATGTTCCTTTGGAGGTGC
GAGCACACGTCTGAACTCTTTCCCTACACGACGCTCTTCCGATCTNNNTATGCTGTTAGGTTTGGTTT
AGCACACGTCTGAACTCTTTCCCTACACGACGCTCTTCCGATCTNNNGAGTAGCTGGGATTACAGGC
GAGCACACGTCTGAACTCTTTCCCTACACGACGCTCTTCCGATCTNNNTAGCTGAGGCACAAGAATCA
AGCACACGTCTGAACTCTTTCCCTACACGACGCTCTTCCGATCTNNNGTGCCTATAGTCCCAGCTACTT
AGCACACGTCTGAACTCTTTCCCTACACGACGCTCTTCCGATCTNNNGATTGCTTGAACTGAGGCATT
AGAGCACACGTCTGAACTCTTTCCCTACACGACGCTCTTCCGATCTNNNCTTGCAGTGAGCCGAGATC
GCACACGTCTGAACTCTTTCCCTACACGACGCTCTTCCGATCTNNNAGAGGCTGACCAATGAAATGG
GCACACGTCTGAACTCTTTCCCTACACGACGCTCTTCCGATCTNNNAGCTTACAGATGGTAGGTTGTG
GCACACGTCTGAACTCTTTCCCTACACGACGCTCTTCCGATCTNNNATCCTAGCATTTTTGGAGGCTG
AGCACACGTCTGAACTCTTTCCCTACACGACGCTCTTCCGATCTNNNCTGTAATCCTGGCTACTTGGGA
AGCACACGTCTGAACTCTTTCCCTACACGACGCTCTTCCGATCTNNNATTGCAATTAGGACCAACCAAG
AGCACACGTCTGAACTCTTTCCCTACACGACGCTCTTCCGATCTNNNAAAGAGGAGGTTAGGACACA
GCACACGTCTGAACTCTTTCCCTACACGACGCTCTTCCGATCTNNNAGTGGCTGTTTTTTCTGATGAC
AGCACACGTCTGAACTCTTTCCCTACACGACGCTCTTCCGATCTNNNCTAATGACCCAGGGAACACAG
GAGCACACGTCTGAACTCTTTCCCTACACGACGCTCTTCCGATCTNNNGTTGTTTTCGCTTCAGGTCTG
GAGCACACGTCTGAACTCTTTCCCTACACGACGCTCTTCCGATCTNNNACCCTGACACTAACACCAGAA
AGCACACGTCTGAACTCTTTCCCTACACGACGCTCTTCCGATCTNNNTTCTTGGGATCCAGATTGCAGA
GAGCACACGTCTGAACTCTTTCCCTACACGACGCTCTTCCGATCTNNNGATCATGAGGTCAGGAGTTC
AGCACACGTCTGAACTCTTTCCCTACACGACGCTCTTCCGATCTNNNTTTTTTTTTGTCCCCCCCAACAA
AGCACACGTCTGAACTCTTTCCCTACACGACGCTCTTCCGATCTNNNACAAGAAACTGTTACCTTTGG
AGCACACGTCTGAACTCTTTCCCTACACGACGCTCTTCCGATCTNNNTCTCTTGCTTCCCATGTTGAAGA
GCACACGTCTGAACTCTTTCCCTACACGACGCTCTTCCGATCTNNNGAATCGCTTGAACCTGGAAAGA
GAGCACACGTCTGAACTCTTTCCCTACACGACGCTCTTCCGATCTNNNGCACTTAGAGTAACGTGTTCA
GAGCACACGTCTGAACTCTTTCCCTACACGACGCTCTTCCGATCTNNNCCACAAATTTGCTGTGTGAGT
GAGCACACGTCTGAACTCTTTCCCTACACGACGCTCTTCCGATCTNNNGAGCTACCATGCCCAGCTAAT
GAGCACACGTCTGAACTCTTTCCCTACACGACGCTCTTCCGATCTNNNGGTCCCAAGCATTTCTGATA
GAGCACACGTCTGAACTCTTTCCCTACACGACGCTCTTCCGATCTNNNGGAGCAATGGCATTTGAAAG
AGCACACGTCTGAACTCTTTCCCTACACGACGCTCTTCCGATCTNNNGCTCATGCCTGTAATCTCAACA
AGAGCACACGTCTGAACTCTTTCCCTACACGACGCTCTTCCGATCTNNNAACTCCAAAACTCTTTCCCC
AGCACACGTCTGAACTCTTTCCCTACACGACGCTCTTCCGATCTNNNAAATGCCCATCAATCAACGAGT

GAGCACACGTCTGAACTCTTTCCCTACACGACGCTCTTCCGATCTNNNAGTGAGCCAAGATCATGACAC
GAGCACACGTCTGAACTCTTTCCCTACACGACGCTCTTCCGATCTNNNAGCTATGATCACATCACTGCAG
GAGCACACGTCTGAACTCTTTCCCTACACGACGCTCTTCCGATCTNNNGGGTAACAAAGCAAGAACCTG
AGCACACGTCTGAACTCTTTCCCTACACGACGCTCTTCCGATCTNNNATCGGTTTACTCAAGCCACTTCTG
GAGCACACGTCTGAACTCTTTCCCTACACGACGCTCTTCCGATCTNNNCTGCTTTTACAGTTGTGCCAGA
GAGCACACGTCTGAACTCTTTCCCTACACGACGCTCTTCCGATCTNNNGATGAGGCAGGAGAATCACTT
AGCACACGTCTGAACTCTTTCCCTACACGACGCTCTTCCGATCTNNNCTGTAGTCCCAGCTACTCAGGA
GAGCACACGTCTGAACTCTTTCCCTACACGACGCTCTTCCGATCTNNNGGATAGCTTGTGCAGGGAAA
GAGCACACGTCTGAACTCTTTCCCTACACGACGCTCTTCCGATCTNNNATAAACCCCCACCTATACACAC
GAGCACACGTCTGAACTCTTTCCCTACACGACGCTCTTCCGATCTNNNCATGATCCTCAGGGGCATCAA
GAGCACACGTCTGAACTCTTTCCCTACACGACGCTCTTCCGATCTNNNAATGGCTGATTCTACTAGCGG
AGCACACGTCTGAACTCTTTCCCTACACGACGCTCTTCCGATCTNNNTGGATGGGAAATGAGTCTGTG
AGCACACGTCTGAACTCTTTCCCTACACGACGCTCTTCCGATCTNNNGTAATCCCAGCTACTCAGGAAG
GAGCACACGTCTGAACTCTTTCCCTACACGACGCTCTTCCGATCTNNNGGAGGATTGCTTGAGTCTAGG
GAGCACACGTCTGAACTCTTTCCCTACACGACGCTCTTCCGATCTNNNGATTGAGACCATCCTGGCTAAG
GAGCACACGTCTGAACTCTTTCCCTACACGACGCTCTTCCGATCTNNNTTTCTTTTGCTCTCCCCTGATG
AGCACACGTCTGAACTCTTTCCCTACACGACGCTCTTCCGATCTNNNCACCTGTAGTTCCAGCTACTCAG
AGCACACGTCTGAACTCTTTCCCTACACGACGCTCTTCCGATCTNNNTTAGAGTGATCCATTAGCTCCG
AGCACACGTCTGAACTCTTTCCCTACACGACGCTCTTCCGATCTNNNTGGATTTTCTAGGGCAGGTCTT
GCACACGTCTGAACTCTTTCCCTACACGACGCTCTTCCGATCTNNNAACCTCTGAAATTCTCCCCATGGC
AGCACACGTCTGAACTCTTTCCCTACACGACGCTCTTCCGATCTNNNGGATACAATCCTAGCTCACTGC
GAGCACACGTCTGAACTCTTTCCCTACACGACGCTCTTCCGATCTNNNAAGCTCTGTCTGCATGAGATTG
GAGCACACGTCTGAACTCTTTCCCTACACGACGCTCTTCCGATCTNNNGAGGCAGGATAATTGCTTGA
GAGCACACGTCTGAACTCTTTCCCTACACGACGCTCTTCCGATCTNNNGCAAGATTATAGCTCACTGCAG
AGCACACGTCTGAACTCTTTCCCTACACGACGCTCTTCCGATCTNNNTCAGGAGACCCTTCCTTAGTCTT
GAGCACACGTCTGAACTCTTTCCCTACACGACGCTCTTCCGATCTNNNCAGAGGTTGAAGTGAGCTGA
AGCACACGTCTGAACTCTTTCCCTACACGACGCTCTTCCGATCTNNNCAGATCACTTGAGCTCAGGAGT
AGCACACGTCTGAACTCTTTCCCTACACGACGCTCTTCCGATCTNNNCTGCACTTGTATGTTTGACAGC
GCACACGTCTGAACTCTTTCCCTACACGACGCTCTTCCGATCTNNNTGGCTCCCCATTTCTTTCTAAGTCA
GAGCACACGTCTGAACTCTTTCCCTACACGACGCTCTTCCGATCTNNNGGTGACAAAGGAAGACTCTGT
AGAGCACACGTCTGAACTCTTTCCCTACACGACGCTCTTCCGATCTNNNGGCTAAGGCAGGAAAATCA
AGCACACGTCTGAACTCTTTCCCTACACGACGCTCTTCCGATCTNNNGTTGAGCTGAGATCACACCATT
GAGCACACGTCTGAACTCTTTCCCTACACGACGCTCTTCCGATCTNNNTGAGGCAGGAGAATCACTTGA
GAGCACACGTCTGAACTCTTTCCCTACACGACGCTCTTCCGATCTNNNGAATCACTTGAACCTGGGAG
CACACGTCTGAACTCTTTCCCTACACGACGCTCTTCCGATCTNNNCCATGTTGTTGCAAATGTCAGGAC
AGCACACGTCTGAACTCTTTCCCTACACGACGCTCTTCCGATCTNNNTGGAGTGTGGTGGTATGATCTT
AGCACACGTCTGAACTCTTTCCCTACACGACGCTCTTCCGATCTNNNTGCTGCTGTGTGTTAGTGACTA
AAGAGCACACGTCTGAACTCTTTCCCTACACGACGCTCTTCCGATCTNNNGGTCTTGTTCTATGGTAGG
AGCACACGTCTGAACTCTTTCCCTACACGACGCTCTTCCGATCTNNNCAACTACAGCCTGAACTTGAAGG
GAGCACACGTCTGAACTCTTTCCCTACACGACGCTCTTCCGATCTNNNAGTAGCTAGGACTAAAGGCAT
AGCACACGTCTGAACTCTTTCCCTACACGACGCTCTTCCGATCTNNNTCTCTCTCTGTCTCTCTGTCTCCC
GAGCACACGTCTGAACTCTTTCCCTACACGACGCTCTTCCGATCTNNNCCTGAAGGGCAAATGCTATAT
GAGCACACGTCTGAACTCTTTCCCTACACGACGCTCTTCCGATCTNNNCTGAGGTGGGAGAATGGTGT

AGCACACGTCTGAACTCTTTCCCTACACGACGCTCTTCCGATCTNNNTGAGGTAGGAGAATCGCTTG
AGCACACGTCTGAACTCTTTCCCTACACGACGCTCTTCCGATCTNNNATTCATGTGATTCTCCACCTCCC
AGCACACGTCTGAACTCTTTCCCTACACGACGCTCTTCCGATCTNNNAGAATTGCTTGAATCCAGGAG
AGCACACGTCTGAACTCTTTCCCTACACGACGCTCTTCCGATCTNNNGCTTACCTAACCTCCACAAGTC
AGCACACGTCTGAACTCTTTCCCTACACGACGCTCTTCCGATCTNNNTTAGCACAGTTCCTGGCACAT
AGCACACGTCTGAACTCTTTCCCTACACGACGCTCTTCCGATCTNNNACTTCCCAGGTTCAAGCAATTT
AGCACACGTCTGAACTCTTTCCCTACACGACGCTCTTCCGATCTNNNACCTTCTTGCCTTGGTATCTCT
GCACACGTCTGAACTCTTTCCCTACACGACGCTCTTCCGATCTNNNGTAGCTGGGACTACAGGCATAT
AGCACACGTCTGAACTCTTTCCCTACACGACGCTCTTCCGATCTNNNAAAATTAGCTGGGCGTGATG
AGCACACGTCTGAACTCTTTCCCTACACGACGCTCTTCCGATCTNNNACTCATACCTCCTCCTCCTTCTC
GCACACGTCTGAACTCTTTCCCTACACGACGCTCTTCCGATCTNNNTTGAGGGGAAAGAGAGACAG
GAGCACACGTCTGAACTCTTTCCCTACACGACGCTCTTCCGATCTNNNTATACCCACCCATGATTCCTTG
AGCACACGTCTGAACTCTTTCCCTACACGACGCTCTTCCGATCTNNNACCAGAGACCTTTGTTCACTTG
AGCACACGTCTGAACTCTTTCCCTACACGACGCTCTTCCGATCTNNNGCCAGAGAATTGCTTGAACTG
GCACACGTCTGAACTCTTTCCCTACACGACGCTCTTCCGATCTNNNCACGGTGAAAACCCTTCTCTACT
GAGCACACGTCTGAACTCTTTCCCTACACGACGCTCTTCCGATCTNNNGGCAACATAGAGAGATCCTG
GAGCACACGTCTGAACTCTTTCCCTACACGACGCTCTTCCGATCTNNNTGTAATTCCAGCTACTTGGG
GCACACGTCTGAACTCTTTCCCTACACGACGCTCTTCCGATCTNNNGAGGGAGAGGCAGAGAAAAT
AGCACACGTCTGAACTCTTTCCCTACACGACGCTCTTCCGATCTNNNACTAAAGCAGGAGAATCGCTT
GCACACGTCTGAACTCTTTCCCTACACGACGCTCTTCCGATCTNNNAGATATTTAGCAGCACGTCTGA
AGCACACGTCTGAACTCTTTCCCTACACGACGCTCTTCCGATCTNNNAGAACTGCATATATTGGGTCCC
AGCACACGTCTGAACTCTTTCCCTACACGACGCTCTTCCGATCTNNNGCAGATCACTTGAGGTAAGG
AGCACACGTCTGAACTCTTTCCCTACACGACGCTCTTCCGATCTNNNAGTAATTCTAGCACGTTGGGA
GAGCACACGTCTGAACTCTTTCCCTACACGACGCTCTTCCGATCTNNNCAGCAATCTCACTACTGGGTG
GCACACGTCTGAACTCTTTCCCTACACGACGCTCTTCCGATCTNNNGAGAATGGCTTGAACCCAGTA
GCACACGTCTGAACTCTTTCCCTACACGACGCTCTTCCGATCTNNNACCACTGCTAATATCTTGGCAC
CACACGTCTGAACTCTTTCCCTACACGACGCTCTTCCGATCTNNNATTGATTGTGTCCAGGAGGTTGA
GAGCACACGTCTGAACTCTTTCCCTACACGACGCTCTTCCGATCTNNNTGGAAACAACAACAACGAC
AGCACACGTCTGAACTCTTTCCCTACACGACGCTCTTCCGATCTNNNGCAACAAGAGCAACACTTTGT
GCACACGTCTGAACTCTTTCCCTACACGACGCTCTTCCGATCTNNNTTGGTTTACAATTGCAACCTTGC
GAGCACACGTCTGAACTCTTTCCCTACACGACGCTCTTCCGATCTNNNAGGTTGAGATGGGAGAATT
GCACACGTCTGAACTCTTTCCCTACACGACGCTCTTCCGATCTNNNAGTCCCAACTTCAGATGATCCAC
AGCACACGTCTGAACTCTTTCCCTACACGACGCTCTTCCGATCTNNNTGTGTGTGTGTGTGTGTAGA
AGCACACGTCTGAACTCTTTCCCTACACGACGCTCTTCCGATCTNNNGGAAACCCCGTCTCTCCTAAAA
CACACGTCTGAACTCTTTCCCTACACGACGCTCTTCCGATCTNNNATCTTTCTAACATGGGCCCACATC
AGCACACGTCTGAACTCTTTCCCTACACGACGCTCTTCCGATCTNNNTTGGCTCTCTCCAACAAGTTCT
AGCACACGTCTGAACTCTTTCCCTACACGACGCTCTTCCGATCTNNNTCTGGGCAACAAGAGTGATA
AGCACACGTCTGAACTCTTTCCCTACACGACGCTCTTCCGATCTNNNTCAAGTCTGGAGTTCCTTAGCT
GAGCACACGTCTGAACTCTTTCCCTACACGACGCTCTTCCGATCTNNNGAGAGTTGTGGGGCCTAGA
AGCACACGTCTGAACTCTTTCCCTACACGACGCTCTTCCGATCTNNNCTCCTCATCTTTCTGTGACCACA
AGCACACGTCTGAACTCTTTCCCTACACGACGCTCTTCCGATCTNNNATTGACTTTTGCTGTTGTTGCT
AGCACACGTCTGAACTCTTTCCCTACACGACGCTCTTCCGATCTNNNCATTCCCACCAACAGTGCATAA
GAGCACACGTCTGAACTCTTTCCCTACACGACGCTCTTCCGATCTNNNCCAAGCAGAGACTCTTGTTCT

AGCACACGTCTGAACTCTTTCCCTACACGACGCTCTTCCGATCTNNNTCCCTCTCTCTCTCTCTCTCTC
AGCACACGTCTGAACTCTTTCCCTACACGACGCTCTTCCGATCTNNNGGCCATTATCCATAGAAGGG
AGCACACGTCTGAACTCTTTCCCTACACGACGCTCTTCCGATCTNNNCTGAGGCAGGAGAATCACTTG
AGCACACGTCTGAACTCTTTCCCTACACGACGCTCTTCCGATCTNNNTGTTTGAACCCAGGAGTTTGA
GAGCACACGTCTGAACTCTTTCCCTACACGACGCTCTTCCGATCTNNNGCCCATCAATCAATGAGTGG
GCACACGTCTGAACTCTTTCCCTACACGACGCTCTTCCGATCTNNNGCTTCTGAAAATATGGCGATGC
GCACACGTCTGAACTCTTTCCCTACACGACGCTCTTCCGATCTNNNTTTCGCATCAATGTTCATCAGG
AGCACACGTCTGAACTCTTTCCCTACACGACGCTCTTCCGATCTNNNAATTAGCTGGATGTGGTAGT
AGCACACGTCTGAACTCTTTCCCTACACGACGCTCTTCCGATCTNNNCAGGGTGATACTCCGTCTGAA
AGCACACGTCTGAACTCTTTCCCTACACGACGCTCTTCCGATCTNNNCAGGCCCTGTCTCAAAAAACAA
AGCACACGTCTGAACTCTTTCCCTACACGACGCTCTTCCGATCTNNNCCAGCTTGCTTTCCCTTTTTTTC
AGCACACGTCTGAACTCTTTCCCTACACGACGCTCTTCCGATCTNNNCTCAAGGGATCCTCTTGCTTCA
AGCACACGTCTGAACTCTTTCCCTACACGACGCTCTTCCGATCTNNNGAGGCATAAGCACACAATTTG
AGCACACGTCTGAACTCTTTCCCTACACGACGCTCTTCCGATCTNNNAACATGGTGAAATCCCGTCTG
AGCACACGTCTGAACTCTTTCCCTACACGACGCTCTTCCGATCTNNNCAGAATTGCTTGAACCCAGGA
GCACACGTCTGAACTCTTTCCCTACACGACGCTCTTCCGATCTNNNCTGAGGCAGAGAACCATTTGAA
AGCACACGTCTGAACTCTTTCCCTACACGACGCTCTTCCGATCTNNNCAACTCTTTGGGAGGCTGAAA
GAGCACACGTCTGAACTCTTTCCCTACACGACGCTCTTCCGATCTNNNACAGCTCTCTCTCTTTACCTC
GAGCACACGTCTGAACTCTTTCCCTACACGACGCTCTTCCGATCTNNNATACAAAAATTAGCCGGGCA
AGCACACGTCTGAACTCTTTCCCTACACGACGCTCTTCCGATCTNNNCCATGGAGTTGGAGTCTACA
AGCACACGTCTGAACTCTTTCCCTACACGACGCTCTTCCGATCTNNNTAGCTCTTTTTCCCTCCCTTTCC
GCACACGTCTGAACTCTTTCCCTACACGACGCTCTTCCGATCTNNNCCAGGGATGGTTTGTTTTTGGT
AGCACACGTCTGAACTCTTTCCCTACACGACGCTCTTCCGATCTNNNCCCACCTTGGACCATTTTCTTTT
AGCACACGTCTGAACTCTTTCCCTACACGACGCTCTTCCGATCTNNNTGTAATCCCAGCTACTCAGGAG
AGCACACGTCTGAACTCTTTCCCTACACGACGCTCTTCCGATCTNNNCTGCATGTGAGGAAAAACTGA
AGCACACGTCTGAACTCTTTCCCTACACGACGCTCTTCCGATCTNNNGAGGGTACAGCAACAAACAC
AGCACACGTCTGAACTCTTTCCCTACACGACGCTCTTCCGATCTNNNACTTGAGGTCAGGAGTTCAA
AGCACACGTCTGAACTCTTTCCCTACACGACGCTCTTCCGATCTNNNGGCAACAACAGCGAAACTCTA
AGCACACGTCTGAACTCTTTCCCTACACGACGCTCTTCCGATCTNNNCTGGTTGCCTCTACACCTAGTA
GCACACGTCTGAACTCTTTCCCTACACGACGCTCTTCCGATCTNNNCAGCTTTTCTCCATGTTGGTGA
GCACACGTCTGAACTCTTTCCCTACACGACGCTCTTCCGATCTNNNAGGAGTCTTGCCTCACTATAGGT
GCACACGTCTGAACTCTTTCCCTACACGACGCTCTTCCGATCTNNNAGGAAGCTGAGGCAGGAAAA
GCACACGTCTGAACTCTTTCCCTACACGACGCTCTTCCGATCTNNNGGCTCCTTCAAAACCTCATTCAG
GAGCACACGTCTGAACTCTTTCCCTACACGACGCTCTTCCGATCTNNNCTGGGCAATAGAGGGAGAT
AGCACACGTCTGAACTCTTTCCCTACACGACGCTCTTCCGATCTNNNTGGAAATCAGACCGTGAACAT
GCACACGTCTGAACTCTTTCCCTACACGACGCTCTTCCGATCTNNNCCCTTATTCACTCTCCTGCCTTTC
AGCACACGTCTGAACTCTTTCCCTACACGACGCTCTTCCGATCTNNNCCAACATGGTGAAATCCCATCT
GCACACGTCTGAACTCTTTCCCTACACGACGCTCTTCCGATCTNNNAACATTTTGGGAGGCTGAGGTA
GAGCACACGTCTGAACTCTTTCCCTACACGACGCTCTTCCGATCTNNNCTTGAGCCCAGGAGTTCAAG
GCACACGTCTGAACTCTTTCCCTACACGACGCTCTTCCGATCTNNNCTTCCCTGTTCCTGGGTAGTAAC
GCACACGTCTGAACTCTTTCCCTACACGACGCTCTTCCGATCTNNNGCAGATCACTTGAAGTCAGGA
AGCACACGTCTGAACTCTTTCCCTACACGACGCTCTTCCGATCTNNNTTTGCCTTCATCATGTGTGTGA
CACACGTCTGAACTCTTTCCCTACACGACGCTCTTCCGATCTNNNATCTTTCTAACATGGGCCCACATCA

GAGCACACGTCTGAACTCTTTCCCTACACGACGCTCTTCCGATCTNNNAGACAGGAGAATCACTTGAA
AGAGCACACGTCTGAACTCTTTCCCTACACGACGCTCTTCCGATCTNNNGAATGTCGTGAACCTGGGA
AGCACACGTCTGAACTCTTTCCCTACACGACGCTCTTCCGATCTNNNTGTAATTCCAGCACTTTGGGAG
GAGCACACGTCTGAACTCTTTCCCTACACGACGCTCTTCCGATCTNNNGAAGCATATAGAACCTGGCA
AGCACACGTCTGAACTCTTTCCCTACACGACGCTCTTCCGATCTNNNCTGTAATCCCAGCTACTGAGGA
GCACACGTCTGAACTCTTTCCCTACACGACGCTCTTCCGATCTNNNAACAGAGAAAGCAACATCAGCA
GAGCACACGTCTGAACTCTTTCCCTACACGACGCTCTTCCGATCTNNNAGATTGCAATGAGCTGAGTT
AGCACACGTCTGAACTCTTTCCCTACACGACGCTCTTCCGATCTNNNTGCTTGAAAACGGAAAATGGA
GCACACGTCTGAACTCTTTCCCTACACGACGCTCTTCCGATCTNNNCTACACAGGTCTGAGTCACACTC
GCACACGTCTGAACTCTTTCCCTACACGACGCTCTTCCGATCTNNNGGAGGTTACAGTGAGCCAAGAT
GAGCACACGTCTGAACTCTTTCCCTACACGACGCTCTTCCGATCTNNNCTTGCAGTAAGCCAAGATTG
GAGCACACGTCTGAACTCTTTCCCTACACGACGCTCTTCCGATCTNNNAATTAGCTGGGCATGGTGGA
AGCACACGTCTGAACTCTTTCCCTACACGACGCTCTTCCGATCTNNNGAGTTTGCAGTGAGCTGAGAT
AGCACACGTCTGAACTCTTTCCCTACACGACGCTCTTCCGATCTNNNAGAACCCACAGACAGAAGAAA
AGCACACGTCTGAACTCTTTCCCTACACGACGCTCTTCCGATCTNNNAAAGAATTTCTTCCCTCCCAGCT
GAGCACACGTCTGAACTCTTTCCCTACACGACGCTCTTCCGATCTNNNGGATAGAGAGGGAGGGAGA
GAGCACACGTCTGAACTCTTTCCCTACACGACGCTCTTCCGATCTNNNAATGTGCCTTACAATGAGCCA
AGAGCACACGTCTGAACTCTTTCCCTACACGACGCTCTTCCGATCTNNNATTACTCTCCCTTAGTTGGC
AGCACACGTCTGAACTCTTTCCCTACACGACGCTCTTCCGATCTNNNAAGGCATGAGAATTGCTTGAA
GAGCACACGTCTGAACTCTTTCCCTACACGACGCTCTTCCGATCTNNNCAGGAGTTCAAAGTTGCAGT
AGCACACGTCTGAACTCTTTCCCTACACGACGCTCTTCCGATCTNNNGCAACAGATCGAGACTCTGAT
GAGCACACGTCTGAACTCTTTCCCTACACGACGCTCTTCCGATCTNNNGAGAGAGAGACAGAGAGAG
CACACGTCTGAACTCTTTCCCTACACGACGCTCTTCCGATCTNNNCCTGCTTAGGGAAGGACATTTCTC
GAGCACACGTCTGAACTCTTTCCCTACACGACGCTCTTCCGATCTNNNGAGATTGCCCCACTGCACTAT
AGAGCACACGTCTGAACTCTTTCCCTACACGACGCTCTTCCGATCTNNNAAAGGGAAAGAGAGGAGA
AGCACACGTCTGAACTCTTTCCCTACACGACGCTCTTCCGATCTNNNTTTACAAGAAAACCGGGCATGT
GCACACGTCTGAACTCTTTCCCTACACGACGCTCTTCCGATCTNNNTGACTGTAATCACCTTGTTGTGC
GAGCACACGTCTGAACTCTTTCCCTACACGACGCTCTTCCGATCTNNNACCTGTATCTTCCCCTTACCTG
AGCACACGTCTGAACTCTTTCCCTACACGACGCTCTTCCGATCTNNNGAAGATGCCAAAGAGTTGAGT
AAGAGCACACGTCTGAACTCTTTCCCTACACGACGCTCTTCCGATCTNNNGGTTTAGACAATGGGAGG
GAGCACACGTCTGAACTCTTTCCCTACACGACGCTCTTCCGATCTNNNGATGTTTAGGTGTGTTCTGTG
GCACACGTCTGAACTCTTTCCCTACACGACGCTCTTCCGATCTNNNCTTCCTTCCTTCCTTCCTTCCTCCAT
GAGCACACGTCTGAACTCTTTCCCTACACGACGCTCTTCCGATCTNNNGACAGAAGAATTGCTTGAGC
AGCACACGTCTGAACTCTTTCCCTACACGACGCTCTTCCGATCTNNNGGTGACAAAGTGAGACTCCGTA
GAGCACACGTCTGAACTCTTTCCCTACACGACGCTCTTCCGATCTNNNGAGCTGAAATATCATGCCACT
AGCACACGTCTGAACTCTTTCCCTACACGACGCTCTTCCGATCTNNNTCTTTCCCTGTGTTCCCTGAAAA
AGCACACGTCTGAACTCTTTCCCTACACGACGCTCTTCCGATCTNNNTGAGGTAGGAGAATGGTGTGA
GCACACGTCTGAACTCTTTCCCTACACGACGCTCTTCCGATCTNNNGGTGTCTCTCTGTCTCTCTCTCTCA
GAAGAGCACACGTCTGAACTCTTTCCCTACACGACGCTCTTCCGATCTNNNCTACATTCCAGCCTGGG
AGCACACGTCTGAACTCTTTCCCTACACGACGCTCTTCCGATCTNNNGTCTGAGATAGGAGTTTGCCA
AGCACACGTCTGAACTCTTTCCCTACACGACGCTCTTCCGATCTNNNAGAGAGAGAGAGAGAGAGA
GAGCACACGTCTGAACTCTTTCCCTACACGACGCTCTTCCGATCTNNNGGCTCAATTGATGTCCAGGA
AGCACACGTCTGAACTCTTTCCCTACACGACGCTCTTCCGATCTNNNTACAACCTAACCCAAGAACCTG

AGAGCACACGTCTGAACTCTTTCCCTACACGACGCTCTTCCGATCTNNNGAAGGAAGGAAGGAAGG
GAGCACACGTCTGAACTCTTTCCCTACACGACGCTCTTCCGATCTNNNTAGGAAACAGTACCATGGTC
GCACACGTCTGAACTCTTTCCCTACACGACGCTCTTCCGATCTNNNGTGCAGAGAAAGAGAGAAAGG
CACACGTCTGAACTCTTTCCCTACACGACGCTCTTCCGATCTNNNCTTGCCATACATCTGGATTTACC
AGCACACGTCTGAACTCTTTCCCTACACGACGCTCTTCCGATCTNNNCCTAGCCCATTCTCATGCTAAC
AGCACACGTCTGAACTCTTTCCCTACACGACGCTCTTCCGATCTNNNTGGAATTTCCATGTGTAGGTTC
AGCACACGTCTGAACTCTTTCCCTACACGACGCTCTTCCGATCTNNNCAGAGAATTTCTTCACAGCCAG
AGCACACGTCTGAACTCTTTCCCTACACGACGCTCTTCCGATCTNNNCACATGCTGTAGGAAAAACGG
AGCACACGTCTGAACTCTTTCCCTACACGACGCTCTTCCGATCTNNNATCAGTTGGCTGCAACTATTTG
GAGCACACGTCTGAACTCTTTCCCTACACGACGCTCTTCCGATCTNNNCAAAGATTGCAGTGAGCTGA
AGCACACGTCTGAACTCTTTCCCTACACGACGCTCTTCCGATCTNNNCTTTCTCCCCTTCTCCTCCTTTCC
GCACACGTCTGAACTCTTTCCCTACACGACGCTCTTCCGATCTNNNTTATGGGTGTGTGTGTGTGTGT
GCACACGTCTGAACTCTTTCCCTACACGACGCTCTTCCGATCTNNNTTATGGGTGTGTGTGTGTGTGT
AGAGCACACGTCTGAACTCTTTCCCTACACGACGCTCTTCCGATCTNNNTCAAGAAGAGAGCCTAGG
GAGCACACGTCTGAACTCTTTCCCTACACGACGCTCTTCCGATCTNNNCTGTGTCATCCTCCAGAACAT
AGCACACGTCTGAACTCTTTCCCTACACGACGCTCTTCCGATCTNNNGGAGGGCATACTTCCAAACTC
AGCACACGTCTGAACTCTTTCCCTACACGACGCTCTTCCGATCTNNNGGCAACAGAGTGAGACTTCAT
AGCACACGTCTGAACTCTTTCCCTACACGACGCTCTTCCGATCTNNNGGGTGACAGAATGAGACTCC
GCACACGTCTGAACTCTTTCCCTACACGACGCTCTTCCGATCTNNNTTGAGGCAGGAGAATCACTTG
GAGCACACGTCTGAACTCTTTCCCTACACGACGCTCTTCCGATCTNNNCGGTAAGCTGAGATTGTACC
AGCACACGTCTGAACTCTTTCCCTACACGACGCTCTTCCGATCTNNNGGTGACAGAGTGAGACTCCAT
GAGCACACGTCTGAACTCTTTCCCTACACGACGCTCTTCCGATCTNNNTTGACCAAGCAGTCAGAAAA
GCACACGTCTGAACTCTTTCCCTACACGACGCTCTTCCGATCTNNNGACACATGTGATTGCACTTAGG
GCACACGTCTGAACTCTTTCCCTACACGACGCTCTTCCGATCTNNNTGAGAATCACTTGAACCCAGGA
AGCACACGTCTGAACTCTTTCCCTACACGACGCTCTTCCGATCTNNNGTAATACCAGCAATTTGGGAG
AGCACACGTCTGAACTCTTTCCCTACACGACGCTCTTCCGATCTNNNCAACAAGAGCCAAACTGTGTC
GCACACGTCTGAACTCTTTCCCTACACGACGCTCTTCCGATCTNNNCCCCTTCTTCATATCTGGTGTT
AGCACACGTCTGAACTCTTTCCCTACACGACGCTCTTCCGATCTNNNGTGCCTGTAGTCCTAGCTACTT
AGCACACGTCTGAACTCTTTCCCTACACGACGCTCTTCCGATCTNNNGCAAAGTATCCAGTTTCTCTCG
GAGCACACGTCTGAACTCTTTCCCTACACGACGCTCTTCCGATCTNNNTTCACTACATAACACTGGGCC
AGCACACGTCTGAACTCTTTCCCTACACGACGCTCTTCCGATCTNNNCTGAGGCAGGAGAATTGCTT
AGCACACGTCTGAACTCTTTCCCTACACGACGCTCTTCCGATCTNNNGAGACACCAGACTCTGTTCTC
AGCACACGTCTGAACTCTTTCCCTACACGACGCTCTTCCGATCTNNNTCCTGAGTAGATGGGACCTAC
GCACACGTCTGAACTCTTTCCCTACACGACGCTCTTCCGATCTNNNCTTTCCACCGGACATTTGTGATC
AGCACACGTCTGAACTCTTTCCCTACACGACGCTCTTCCGATCTNNNTATAGCTAGGTGTGGGATTGC
GAGCACACGTCTGAACTCTTTCCCTACACGACGCTCTTCCGATCTNNNCCAACATGGTGAAACTCTGTC
AGCACACGTCTGAACTCTTTCCCTACACGACGCTCTTCCGATCTNNNGTAGTTGGGATTACAGTTGCC
AGCACACGTCTGAACTCTTTCCCTACACGACGCTCTTCCGATCTNNNGGGTGACTGGTTGCATCAAA
GAGCACACGTCTGAACTCTTTCCCTACACGACGCTCTTCCGATCTNNNTAAGAGGGCATGTGTTCCAG
GCACACGTCTGAACTCTTTCCCTACACGACGCTCTTCCGATCTNNNAAAATGGATCATAGGCGCTTCA
CACACGTCTGAACTCTTTCCCTACACGACGCTCTTCCGATCTNNNTCACCACAGTGTCATCTCTTCTGC
AGCACACGTCTGAACTCTTTCCCTACACGACGCTCTTCCGATCTNNNGGCTTTTTGTTGTTTGAAGCTC
GAGCACACGTCTGAACTCTTTCCCTACACGACGCTCTTCCGATCTNNNTGAGGCAGGAGAATCTCTTG

GAGCACACGTCTGAACTCTTTCCCTACACGACGCTCTTCCGATCTNNNGCCGCAGCTAATTTTCTAACTC
GAGCACACGTCTGAACTCTTTCCCTACACGACGCTCTTCCGATCTNNNCACCAGTAGTCCCAGCTACTTG
GAGCACACGTCTGAACTCTTTCCCTACACGACGCTCTTCCGATCTNNNTGACTCCAGACCTCTTGTTCAAC
GAGCACACGTCTGAACTCTTTCCCTACACGACGCTCTTCCGATCTNNNGAGGCAGCAGAATCACTTGAA
AGCACACGTCTGAACTCTTTCCCTACACGACGCTCTTCCGATCTNNNGAAAGTGCTGGGATTACAGGT
GCACACGTCTGAACTCTTTCCCTACACGACGCTCTTCCGATCTNNNTAATCCTAGCTACTCAGGAGGCTG
AGAGCACACGTCTGAACTCTTTCCCTACACGACGCTCTTCCGATCTNNNGCAAGGCAGGCAGGATTTAT
AGCACACGTCTGAACTCTTTCCCTACACGACGCTCTTCCGATCTNNNCTGTAGTCCCAGCTACTCAGGA
GAGCACACGTCTGAACTCTTTCCCTACACGACGCTCTTCCGATCTNNNATATGCACCTGTAGTCCCAGCT
GAGCACACGTCTGAACTCTTTCCCTACACGACGCTCTTCCGATCTNNNTAAGTACCACTGAGACAGGAG
AGCACACGTCTGAACTCTTTCCCTACACGACGCTCTTCCGATCTNNNAAATACAAAAGTTAGTCCGGG
GAGCACACGTCTGAACTCTTTCCCTACACGACGCTCTTCCGATCTNNNGCGATACAGCAAGACTCAGTC
GAGCACACGTCTGAACTCTTTCCCTACACGACGCTCTTCCGATCTNNNTTGAGGCAGGAGAATCACTT
GAGCACACGTCTGAACTCTTTCCCTACACGACGCTCTTCCGATCTNNNCACCATGGAGTACTACTCAACC
GAGCACACGTCTGAACTCTTTCCCTACACGACGCTCTTCCGATCTNNNGCAGAGTTCCAATGACCATAA
GCACACGTCTGAACTCTTTCCCTACACGACGCTCTTCCGATCTNNNTGAAGGTTGCAGTGAACTGAGA
AGCACACGTCTGAACTCTTTCCCTACACGACGCTCTTCCGATCTNNNGACTTGAACACAGCAGACAACA
GAGCACACGTCTGAACTCTTTCCCTACACGACGCTCTTCCGATCTNNNATGTTGGTCAGATTGGTCTCG
GAGCACACGTCTGAACTCTTTCCCTACACGACGCTCTTCCGATCTNNNCTCTTGTCTGTAATCCCAGCAG
AGAGCACACGTCTGAACTCTTTCCCTACACGACGCTCTTCCGATCTNNNCTACTCAGGAGAATGGGGTG
GAGCACACGTCTGAACTCTTTCCCTACACGACGCTCTTCCGATCTNNNATGCCTGTAATCCCAGCACTTT
AGAGCACACGTCTGAACTCTTTCCCTACACGACGCTCTTCCGATCTNNNTCTGTAATCCTAGCTACTCGG
AGCACACGTCTGAACTCTTTCCCTACACGACGCTCTTCCGATCTNNNCTGAGGCAGAGAATTGCTTGA
GAGCACACGTCTGAACTCTTTCCCTACACGACGCTCTTCCGATCTNNNGATCACGAGGTCAGGAGATTG
GCACACGTCTGAACTCTTTCCCTACACGACGCTCTTCCGATCTNNNAGGCAGAGGTCTCAAAAATCAG
AGAGCACACGTCTGAACTCTTTCCCTACACGACGCTCTTCCGATCTNNNTGGAAGAGCAGAGTGGATT
AGCACACGTCTGAACTCTTTCCCTACACGACGCTCTTCCGATCTNNNAAGACCCATCAGTGTGCTGTAT
GAGCACACGTCTGAACTCTTTCCCTACACGACGCTCTTCCGATCTNNNGAGGCAGGAGAATTGCTTGA
GAGCACACGTCTGAACTCTTTCCCTACACGACGCTCTTCCGATCTNNNTGATGAAGAACAGCAGCACAA
AGCACACGTCTGAACTCTTTCCCTACACGACGCTCTTCCGATCTNNNGCACATCTGTAATCCCAGCTACTC
GAGCACACGTCTGAACTCTTTCCCTACACGACGCTCTTCCGATCTNNNTTACTTGAACCTGGAAGTGGA
AGCACACGTCTGAACTCTTTCCCTACACGACGCTCTTCCGATCTNNNCTGTAGTCCCAGCTACTCAGGAC
GAGCACACGTCTGAACTCTTTCCCTACACGACGCTCTTCCGATCTNNNAGAGGTTGCAGTGAACTGAG
GAGCACACGTCTGAACTCTTTCCCTACACGACGCTCTTCCGATCTNNNGTGTTGCAGCAAGTCTCTTTCT
GAGCACACGTCTGAACTCTTTCCCTACACGACGCTCTTCCGATCTNNNTAAAATGGCTGCTTTGGAACAC
AGCACACGTCTGAACTCTTTCCCTACACGACGCTCTTCCGATCTNNNTCAGGAATCAGAATCGCCATAC
AGCACACGTCTGAACTCTTTCCCTACACGACGCTCTTCCGATCTNNNGTGACAGAATCTCGCTATGTTG
AGCACACGTCTGAACTCTTTCCCTACACGACGCTCTTCCGATCTNNNGCCTAACACCCAAGTCTTGAATT
GAGCACACGTCTGAACTCTTTCCCTACACGACGCTCTTCCGATCTNNNTGTGAGTAGGTGTATGTGTG
AGCACACGTCTGAACTCTTTCCCTACACGACGCTCTTCCGATCTNNNTGGTTGTTATACTCTGGAGAGC
AGCACACGTCTGAACTCTTTCCCTACACGACGCTCTTCCGATCTNNNCCTGAGTAGCTGGGATTACAGT
AGCACACGTCTGAACTCTTTCCCTACACGACGCTCTTCCGATCTNNNTTTTTGACAGGCTGCTTTTCACA
AGCACACGTCTGAACTCTTTCCCTACACGACGCTCTTCCGATCTNNNGCTCTTTGAAGGGAGGGATAC

AGCACACGTCTGAACTCTTTCCCTACACGACGCTCTTCCGATCTNNNTCCTCTTGGGCTCATAGTACTT
GCACACGTCTGAACTCTTTCCCTACACGACGCTCTTCCGATCTNNNTAGGGGCTCTATTCTCATGACC
AGCACACGTCTGAACTCTTTCCCTACACGACGCTCTTCCGATCTNNNCATACATGCAGCATACACCATG
AGCACACGTCTGAACTCTTTCCCTACACGACGCTCTTCCGATCTNNNGTAATCCCAGCTACTCAGGAA
AGCACACGTCTGAACTCTTTCCCTACACGACGCTCTTCCGATCTNNNTAGACTGGAGCTGTTCCTATT
AGCACACGTCTGAACTCTTTCCCTACACGACGCTCTTCCGATCTNNNCGTGTTAGTCAGGATGGTCTT
AGCACACGTCTGAACTCTTTCCCTACACGACGCTCTTCCGATCTNNNTGTGGTGTAATGTGTCTAGCT
AGCACACGTCTGAACTCTTTCCCTACACGACGCTCTTCCGATCTNNNTGTCCGTCCTATCTCACTAACC
AGCACACGTCTGAACTCTTTCCCTACACGACGCTCTTCCGATCTNNNCCTCCCATTCTGCAACTACTTG
AGCACACGTCTGAACTCTTTCCCTACACGACGCTCTTCCGATCTNNNGAAATTCACACTTCCTGCTGA
GCACACGTCTGAACTCTTTCCCTACACGACGCTCTTCCGATCTNNNTGCTCGAACCATGAACCTAAGA
AGCACACGTCTGAACTCTTTCCCTACACGACGCTCTTCCGATCTNNNAGTGCTGGGATTACAGATGTG
GCACACGTCTGAACTCTTTCCCTACACGACGCTCTTCCGATCTNNNCTTCACCTCCTTCCAGTTCAAGA
GCACACGTCTGAACTCTTTCCCTACACGACGCTCTTCCGATCTNNNAAACTAGGTCTTCAAGGGGCAA
GCACACGTCTGAACTCTTTCCCTACACGACGCTCTTCCGATCTNNNTCGATTGTTCTCACCTTCCTCTC
AGCACACGTCTGAACTCTTTCCCTACACGACGCTCTTCCGATCTNNNCTGAGGCAGGAGAATCACTT
AGCACACGTCTGAACTCTTTCCCTACACGACGCTCTTCCGATCTNNNCAAGGTCACAGATCATGCCAT
AGCACACGTCTGAACTCTTTCCCTACACGACGCTCTTCCGATCTNNNCCTCTCCTGTGCATACTCATGT
GAGCACACGTCTGAACTCTTTCCCTACACGACGCTCTTCCGATCTNNNTTCTTGACCATATGGGGGTT
AGCACACGTCTGAACTCTTTCCCTACACGACGCTCTTCCGATCTNNNGACTGTCCTGGTGTTACATAG
GCACACGTCTGAACTCTTTCCCTACACGACGCTCTTCCGATCTNNNAGGGTCTAGCATGTGAGAATG
GCACACGTCTGAACTCTTTCCCTACACGACGCTCTTCCGATCTNNNACTGTGACAGGCGATAATGGT
AGCACACGTCTGAACTCTTTCCCTACACGACGCTCTTCCGATCTNNNTTTGCCATGTGAACAAGAGCA
AGCACACGTCTGAACTCTTTCCCTACACGACGCTCTTCCGATCTNNNCTCCACTAATGGCAATAACCAG
AGCACACGTCTGAACTCTTTCCCTACACGACGCTCTTCCGATCTNNNTGAGGTAGCACTTTGAAAGAC
AGCACACGTCTGAACTCTTTCCCTACACGACGCTCTTCCGATCTNNNACCAATTCATTCTTCCACCAAC
AGCACACGTCTGAACTCTTTCCCTACACGACGCTCTTCCGATCTNNNCCACTGAATTCCTGAGTCTCC
GCACACGTCTGAACTCTTTCCCTACACGACGCTCTTCCGATCTNNNCTCAGAACCTGGTCACATGTAC
GAGCACACGTCTGAACTCTTTCCCTACACGACGCTCTTCCGATCTNNNAGACTCTGTTCTGATTCCTGC
AGCACACGTCTGAACTCTTTCCCTACACGACGCTCTTCCGATCTNNNGCGGAAGTGACACAGTACAA
GCACACGTCTGAACTCTTTCCCTACACGACGCTCTTCCGATCTNNNGCACATACCTGCACACAAATGT
AGCACACGTCTGAACTCTTTCCCTACACGACGCTCTTCCGATCTNNNAGGTAGGGCCTCTCAGAAGT
AGCACACGTCTGAACTCTTTCCCTACACGACGCTCTTCCGATCTNNNTGGGGAGAGTCTGTTTAGGA
AGCACACGTCTGAACTCTTTCCCTACACGACGCTCTTCCGATCTNNNTGGCAATGATGGAAAGGGT
AGCACACGTCTGAACTCTTTCCCTACACGACGCTCTTCCGATCTNNNGCATACCAGGAAAACATCACC
AGCACACGTCTGAACTCTTTCCCTACACGACGCTCTTCCGATCTNNNGCAGCCTCCTCCATTGTATTAG
GCACACGTCTGAACTCTTTCCCTACACGACGCTCTTCCGATCTNNNCCATGATTATACCACTGCGTTCC
AGCACACGTCTGAACTCTTTCCCTACACGACGCTCTTCCGATCTNNNACAACATTGGGAAAAACAAGG
GCACACGTCTGAACTCTTTCCCTACACGACGCTCTTCCGATCTNNNGCAAATTCAGTTTCAGGACTGC
GAGCACACGTCTGAACTCTTTCCCTACACGACGCTCTTCCGATCTNNNTTGATAACCCTCTTTGGACTC
AGCACACGTCTGAACTCTTTCCCTACACGACGCTCTTCCGATCTNNNGGTCAGAAAAAGGCAATGAG
AGCACACGTCTGAACTCTTTCCCTACACGACGCTCTTCCGATCTNNNTTCCAAGTCAGGCAAGGCATAT
AGCACACGTCTGAACTCTTTCCCTACACGACGCTCTTCCGATCTNNNGATCTCTCTATGTGCGTTCTCG

GAGCACACGTCTGAACTCTTTCCCTACACGACGCTCTTCCGATCTNNNAATTCCCCCATGTTTTCCTCAA
AGCACACGTCTGAACTCTTTCCCTACACGACGCTCTTCCGATCTNNNGACTTGAAGCAGCTATGAACCA
GAGCACACGTCTGAACTCTTTCCCTACACGACGCTCTTCCGATCTNNNGCAGTACTGTTTCCCTAACACC
AGCACACGTCTGAACTCTTTCCCTACACGACGCTCTTCCGATCTNNNCTTTGCCTGTTCCTGTTCTGATGC
AGCACACGTCTGAACTCTTTCCCTACACGACGCTCTTCCGATCTNNNCCCACAAGTTACTTTGCCACTTC
AGCACACGTCTGAACTCTTTCCCTACACGACGCTCTTCCGATCTNNNTGCCCAAATGTACTTCCATCTGA
AGAGCACACGTCTGAACTCTTTCCCTACACGACGCTCTTCCGATCTNNNCAGCCTTTGGAAGCTAAGA
GAGCACACGTCTGAACTCTTTCCCTACACGACGCTCTTCCGATCTNNNGTCTGCCTCCTTTCCAAAAAAC
GAGCACACGTCTGAACTCTTTCCCTACACGACGCTCTTCCGATCTNNNAATGCTTTCAAAGTACGGGTT
AGCACACGTCTGAACTCTTTCCCTACACGACGCTCTTCCGATCTNNNGCCCTGTAACCTGACTTCTCTTA
AGCACACGTCTGAACTCTTTCCCTACACGACGCTCTTCCGATCTNNNGAGTTTCTCAGTCTGGTCTTTG
GAGCACACGTCTGAACTCTTTCCCTACACGACGCTCTTCCGATCTNNNTGCACTTTCTCTCTCTGTCTCTC
AGCACACGTCTGAACTCTTTCCCTACACGACGCTCTTCCGATCTNNNATGGACATTTCCTCCAAGATGCT
GAGCACACGTCTGAACTCTTTCCCTACACGACGCTCTTCCGATCTNNNTCAAACTGCAAGATGCTTTGT
AGCACACGTCTGAACTCTTTCCCTACACGACGCTCTTCCGATCTNNNTTGCTGGCAAGGTAGAGAGAAA
GAGCACACGTCTGAACTCTTTCCCTACACGACGCTCTTCCGATCTNNNTTTCCCCATAAGGTGACCAGAA
AGCACACGTCTGAACTCTTTCCCTACACGACGCTCTTCCGATCTNNNGAGCATCAGAGAAGGAAGGTG
AGCACACGTCTGAACTCTTTCCCTACACGACGCTCTTCCGATCTNNNGAAGGAAAGCCAATGCAAATCC
GAGCACACGTCTGAACTCTTTCCCTACACGACGCTCTTCCGATCTNNNTTTGCCATTTGACCAGACTCTT
AGCACACGTCTGAACTCTTTCCCTACACGACGCTCTTCCGATCTNNNTTGCCTGTCTCTCTCTTTCTGTC
AGCACACGTCTGAACTCTTTCCCTACACGACGCTCTTCCGATCTNNNTTGACTTCCATTGCTGCTTTCAG
GAGCACACGTCTGAACTCTTTCCCTACACGACGCTCTTCCGATCTNNNAAACTGAGGAAACGGAAACA
GAGCACACGTCTGAACTCTTTCCCTACACGACGCTCTTCCGATCTNNNCCAGCAAAAGAAACCTTCACCA
AGCACACGTCTGAACTCTTTCCCTACACGACGCTCTTCCGATCTNNNTCACAAACTTGACAGGAAGGAC
GAGCACACGTCTGAACTCTTTCCCTACACGACGCTCTTCCGATCTNNNGCTCCCCATGTTACTAGAACAC
GAGCACACGTCTGAACTCTTTCCCTACACGACGCTCTTCCGATCTNNNACCTGGAGTCATGGATAGTCA
AGCACACGTCTGAACTCTTTCCCTACACGACGCTCTTCCGATCTNNNAACGTCACACTCAGGAACTCAAT
GAGCACACGTCTGAACTCTTTCCCTACACGACGCTCTTCCGATCTNNNGAGAAGCACAGAGGACTCTAC
GAGCACACGTCTGAACTCTTTCCCTACACGACGCTCTTCCGATCTNNNACAGACGTCACCTGGAGGATA
AGCACACGTCTGAACTCTTTCCCTACACGACGCTCTTCCGATCTNNNGCTCTCACATTTCTTCAGAAGCA
AGCACACGTCTGAACTCTTTCCCTACACGACGCTCTTCCGATCTNNNTCCAGGAAGTAAAATCTGGGTA
AGAGCACACGTCTGAACTCTTTCCCTACACGACGCTCTTCCGATCTNNNCAGGTGAGGACAAAAGGAA
GAGCACACGTCTGAACTCTTTCCCTACACGACGCTCTTCCGATCTNNNAAATAGTCTTGCAGAGGAGTC
AGCACACGTCTGAACTCTTTCCCTACACGACGCTCTTCCGATCTNNNTATGTCCGCTGTCACCAAGGATC
AGCACACGTCTGAACTCTTTCCCTACACGACGCTCTTCCGATCTNNNATTGCTCAAAGATGATGCTGTG
GAGCACACGTCTGAACTCTTTCCCTACACGACGCTCTTCCGATCTNNNTTTCCTGGAAAGAGTACAGCTC
GAGCACACGTCTGAACTCTTTCCCTACACGACGCTCTTCCGATCTNNNTGGTCAGAAGACACCCTTTTAC
GAGCACACGTCTGAACTCTTTCCCTACACGACGCTCTTCCGATCTNNNGCTTTACATGTCCAAAAGGCA
GAGCACACGTCTGAACTCTTTCCCTACACGACGCTCTTCCGATCTNNNATCTAAGCCCCAGTAACAAGTC
GAGCACACGTCTGAACTCTTTCCCTACACGACGCTCTTCCGATCTNNNCTCAATAAAACCTCAGCCTGTG
AGCACACGTCTGAACTCTTTCCCTACACGACGCTCTTCCGATCTNNNGCTCCATCATTCACGTTCTAAGC
GAGCACACGTCTGAACTCTTTCCCTACACGACGCTCTTCCGATCTNNNGGCCTCACCTTAGACCAACTA
GAGCACACGTCTGAACTCTTTCCCTACACGACGCTCTTCCGATCTNNNGTGGGACTTTCCGGCTTCTATT

GAGCACACGTCTGAACTCTTTCCCTACACGACGCTCTTCCGATCTNNNAGACTGAACTGGCATAGCAG
AGCACACGTCTGAACTCTTTCCCTACACGACGCTCTTCCGATCTNNNGCTTTGTTCAGAGGCTCCTTTTC
AGCACACGTCTGAACTCTTTCCCTACACGACGCTCTTCCGATCTNNNAAAATTGGGCAGAAAAGAGCA
AGCACACGTCTGAACTCTTTCCCTACACGACGCTCTTCCGATCTNNNACCATTAGGAAAGTCAGTGCAG
GCACACGTCTGAACTCTTTCCCTACACGACGCTCTTCCGATCTNNNAAGAACAGAAGTCCCTGCAGTAG
AGAGCACACGTCTGAACTCTTTCCCTACACGACGCTCTTCCGATCTNNNTGTGATGCAACCCGGAATAG
AGCACACGTCTGAACTCTTTCCCTACACGACGCTCTTCCGATCTNNNAGGTTGACGAGGAAAAGAGAC
AGCACACGTCTGAACTCTTTCCCTACACGACGCTCTTCCGATCTNNNTTTCCTTTTTCTTCTCCCCTCCCCA
GCACACGTCTGAACTCTTTCCCTACACGACGCTCTTCCGATCTNNNCTCCAAACCAGCCTCCTAGTAAGC
GAGCACACGTCTGAACTCTTTCCCTACACGACGCTCTTCCGATCTNNNACTAGAAAGTGGGGATAAGG
GAGCACACGTCTGAACTCTTTCCCTACACGACGCTCTTCCGATCTNNNAAACAAGTGGATAGTAGAGG
AGAGCACACGTCTGAACTCTTTCCCTACACGACGCTCTTCCGATCTNNNGCAGAGGAAGTTGCAGTGA
GAGCACACGTCTGAACTCTTTCCCTACACGACGCTCTTCCGATCTNNNGCTGGCAGTTGTAATAGTCAG
GCACACGTCTGAACTCTTTCCCTACACGACGCTCTTCCGATCTNNNGAATAGGGGGTTCCTTGGAGAA
AGAGCACACGTCTGAACTCTTTCCCTACACGACGCTCTTCCGATCTNNNCTGATCTGGGCTGCTCTATGA
GAGCACACGTCTGAACTCTTTCCCTACACGACGCTCTTCCGATCTNNNATGCTTGACTATTAGCTCCAGC
GAGCACACGTCTGAACTCTTTCCCTACACGACGCTCTTCCGATCTNNNGAGAGAGAGCTTGTGTGTGTG
GAGCACACGTCTGAACTCTTTCCCTACACGACGCTCTTCCGATCTNNNGGGGGACAAAAGCTAAGAG
AGCACACGTCTGAACTCTTTCCCTACACGACGCTCTTCCGATCTNNNAAACATCCACAGAGCTCTTCAC
AGCACACGTCTGAACTCTTTCCCTACACGACGCTCTTCCGATCTNNNTCACTTACACCAGAGTTCAGCAA
AGCACACGTCTGAACTCTTTCCCTACACGACGCTCTTCCGATCTNNNAAAGTAAGGGGAGCCACATTC
AGCACACGTCTGAACTCTTTCCCTACACGACGCTCTTCCGATCTNNNATGGGGTGTCTACTTACCAGG
AGCACACGTCTGAACTCTTTCCCTACACGACGCTCTTCCGATCTNNNCCTGGAATTTGGTATGAGAGG
GAGCACACGTCTGAACTCTTTCCCTACACGACGCTCTTCCGATCTNNNGTGTGGAGGAGCATACAGAT
GAGCACACGTCTGAACTCTTTCCCTACACGACGCTCTTCCGATCTNNNATAATTGCATCCCTTTCCCTGC
AGCACACGTCTGAACTCTTTCCCTACACGACGCTCTTCCGATCTNNNGGCCTCTGCTGGGATAATTAGA
AGCACACGTCTGAACTCTTTCCCTACACGACGCTCTTCCGATCTNNNAAACTATCGTGTACATGCATGC
GCACACGTCTGAACTCTTTCCCTACACGACGCTCTTCCGATCTNNNGGGAGAAGAATATCACCTTCCCC
AGCACACGTCTGAACTCTTTCCCTACACGACGCTCTTCCGATCTNNNGAGGGTAGAATCAACTCCATGC
GAGCACACGTCTGAACTCTTTCCCTACACGACGCTCTTCCGATCTNNNCATTTGGTGAGCCCTGAAGAA
GAGCACACGTCTGAACTCTTTCCCTACACGACGCTCTTCCGATCTNNNTAACCTCAAGTTCTTCTCCTGG
AGAGCACACGTCTGAACTCTTTCCCTACACGACGCTCTTCCGATCTNNNGGGAGGGGGATATTCAAA
AGCACACGTCTGAACTCTTTCCCTACACGACGCTCTTCCGATCTNNNGTTCAACTTTCCACTTACACGCAC
AGCACACGTCTGAACTCTTTCCCTACACGACGCTCTTCCGATCTNNNTGCAGTCAGGACTCAGGAATTA
AGCACACGTCTGAACTCTTTCCCTACACGACGCTCTTCCGATCTNNNTAAACGGGAAGAATGACTCCCA
GAGCACACGTCTGAACTCTTTCCCTACACGACGCTCTTCCGATCTNNNTCTAGTTTTTCTGTCTCCACCC
AAGAGCACACGTCTGAACTCTTTCCCTACACGACGCTCTTCCGATCTNNNTGATAAGATTGAGGGCAAC
GAGCACACGTCTGAACTCTTTCCCTACACGACGCTCTTCCGATCTNNNATCTCCCAAAATACAGGTGGCA
AGAGCACACGTCTGAACTCTTTCCCTACACGACGCTCTTCCGATCTNNNTGGCAAGACATCCTAGAAGA
AGCACACGTCTGAACTCTTTCCCTACACGACGCTCTTCCGATCTNNNATGCCTATCACATCAGCATCCTT
AGCACACGTCTGAACTCTTTCCCTACACGACGCTCTTCCGATCTNNNGCTGACACTGCAAAGAGAAAGA
AGCACACGTCTGAACTCTTTCCCTACACGACGCTCTTCCGATCTNNNTGTTTGCTTGTTTGTTTTCCCCTC
AGCACACGTCTGAACTCTTTCCCTACACGACGCTCTTCCGATCTNNNCAGACTTGTACTGGGACTACAC

GAGCACACGTCTGAACTCTTTCCCTACACGACGCTCTTCCGATCTNNNTGCGTGTGATGGACATATCTG
AGCACACGTCTGAACTCTTTCCCTACACGACGCTCTTCCGATCTNNNCCATATGACTAGCTCCAGATGCA
AGCACACGTCTGAACTCTTTCCCTACACGACGCTCTTCCGATCTNNNGCTTATTGGGGGTGTTACTGGT
GCACACGTCTGAACTCTTTCCCTACACGACGCTCTTCCGATCTNNNCTTCCCCTCTAGGCTACTCTTCTCA
AGAGCACACGTCTGAACTCTTTCCCTACACGACGCTCTTCCGATCTNNNGTGACTTGAGCATTCACAG
GAGCACACGTCTGAACTCTTTCCCTACACGACGCTCTTCCGATCTNNNTTCAACCCCTTGACAAACACTT
GCACACGTCTGAACTCTTTCCCTACACGACGCTCTTCCGATCTNNNAAGCTTTCTTCCCTGTGTTGTTGC
AGCACACGTCTGAACTCTTTCCCTACACGACGCTCTTCCGATCTNNNACAACACCATCTTAGTCCAAGGT
GAGCACACGTCTGAACTCTTTCCCTACACGACGCTCTTCCGATCTNNNATTTGTAAAACCAGGTGAAGC
GAGCACACGTCTGAACTCTTTCCCTACACGACGCTCTTCCGATCTNNNTTTTCTCCCTCTCTCTCCCTCTTC
GAGCACACGTCTGAACTCTTTCCCTACACGACGCTCTTCCGATCTNNNTGACTGAAGGCATCAAACAGA
AGCACACGTCTGAACTCTTTCCCTACACGACGCTCTTCCGATCTNNNGACTCCCCACCATTTTCTTCTCTCA
AGCACACGTCTGAACTCTTTCCCTACACGACGCTCTTCCGATCTNNNAGGGACTTTGAGACAATACAG
GAGCACACGTCTGAACTCTTTCCCTACACGACGCTCTTCCGATCTNNNTGGCAAGAAAAAGTGGTATC
GCACACGTCTGAACTCTTTCCCTACACGACGCTCTTCCGATCTNNNCTGGTCTTCTGTGTTCTCCTCTCCA
GAGCACACGTCTGAACTCTTTCCCTACACGACGCTCTTCCGATCTNNNGGACTTTGCTTCTCTTTTGCCTT
AGCACACGTCTGAACTCTTTCCCTACACGACGCTCTTCCGATCTNNNAGCTTTCCTTTTCTCATGTCCCTC
GAGCACACGTCTGAACTCTTTCCCTACACGACGCTCTTCCGATCTNNNAGAAATGCAAATTACTGGGCC
AGCACACGTCTGAACTCTTTCCCTACACGACGCTCTTCCGATCTNNNACACGGACAAATGCACATGTATC
AGCACACGTCTGAACTCTTTCCCTACACGACGCTCTTCCGATCTNNNCTCATTGGCCCCATATGTCCTTCC
GAGCACACGTCTGAACTCTTTCCCTACACGACGCTCTTCCGATCTNNNTGCTCCTTTTCTGGGATCTTTG
AGCACACGTCTGAACTCTTTCCCTACACGACGCTCTTCCGATCTNNNGGTAATTCTTCAGTTGGCTTCC
AGCACACGTCTGAACTCTTTCCCTACACGACGCTCTTCCGATCTNNNTTTGATGTTCCAAGAGGTACCCC
GAGCACACGTCTGAACTCTTTCCCTACACGACGCTCTTCCGATCTNNNAGAGGGTTTCAGTTCTTCAGT
GAGCACACGTCTGAACTCTTTCCCTACACGACGCTCTTCCGATCTNNNCCCTCTCATGCTCTAATCCATC
AGCACACGTCTGAACTCTTTCCCTACACGACGCTCTTCCGATCTNNNAAAGAAGTCCCCTTTCCAGACTC
AGAGCACACGTCTGAACTCTTTCCCTACACGACGCTCTTCCGATCTNNNCACTGGGGGATCTTAGGGA
AGCACACGTCTGAACTCTTTCCCTACACGACGCTCTTCCGATCTNNNCATCACAAAAAGCCCTGGTGTA
AGCACACGTCTGAACTCTTTCCCTACACGACGCTCTTCCGATCTNNNTGGGAAGAGACTGGCATTTGAA
GAGCACACGTCTGAACTCTTTCCCTACACGACGCTCTTCCGATCTNNNCAGTGTCAATGGCCAAGCTTT
AGCACACGTCTGAACTCTTTCCCTACACGACGCTCTTCCGATCTNNNGTTTTCAGTAAACTCCAGCCCAC
GAGCACACGTCTGAACTCTTTCCCTACACGACGCTCTTCCGATCTNNNTTCACTCCTGGTGTTCGTGTTA
AGCACACGTCTGAACTCTTTCCCTACACGACGCTCTTCCGATCTNNNGTTCAAGCAATTCTTCTGCCTCA
GAGCACACGTCTGAACTCTTTCCCTACACGACGCTCTTCCGATCTNNNATAAACCATTGCCAAACCTGAC
AGAGCACACGTCTGAACTCTTTCCCTACACGACGCTCTTCCGATCTNNNGGAGATAGCGTGAACATTTC
AGCACACGTCTGAACTCTTTCCCTACACGACGCTCTTCCGATCTNNNTGATTAGTCAAGGCTGGACAGA
AGCACACGTCTGAACTCTTTCCCTACACGACGCTCTTCCGATCTNNNCTGCCTCCATTCTTCTTCTTCCTCA
AGCACACGTCTGAACTCTTTCCCTACACGACGCTCTTCCGATCTNNNGGATTACTGGCATGAGTCACTC
GAGCACACGTCTGAACTCTTTCCCTACACGACGCTCTTCCGATCTNNNCCCTTCTCCAGCTCTCTCTTCTA
AGCACACGTCTGAACTCTTTCCCTACACGACGCTCTTCCGATCTNNNCCTTGTCACATGGCCTCATAGTA
GAGCACACGTCTGAACTCTTTCCCTACACGACGCTCTTCCGATCTNNNAAATGGAAATGCTACCAGCTTC
AGAGCACACGTCTGAACTCTTTCCCTACACGACGCTCTTCCGATCTNNNACAGGGCCAAATGGAAAAGA
AGAGCACACGTCTGAACTCTTTCCCTACACGACGCTCTTCCGATCTNNNACAGGGCCAAATGGAAAAGA

AGCACACGTCTGAACTCTTTCCCTACACGACGCTCTTCCGATCTNNNGATCACGAGGTCAGGAGTTCA
GCACACGTCTGAACTCTTTCCCTACACGACGCTCTTCCGATCTNNNTTGGCTCCTAAATACCAGGAAC
GAGCACACGTCTGAACTCTTTCCCTACACGACGCTCTTCCGATCTNNNGCAGGAGAATTGCTTGAAC
AGCACACGTCTGAACTCTTTCCCTACACGACGCTCTTCCGATCTNNNTCCCTTAGTTCCTTGACACATG
GCACACGTCTGAACTCTTTCCCTACACGACGCTCTTCCGATCTNNNCCCTCTTTCATACCTAGACACGTC
AGCACACGTCTGAACTCTTTCCCTACACGACGCTCTTCCGATCTNNNCATGACAGCAGTGAAAACAG
GCACACGTCTGAACTCTTTCCCTACACGACGCTCTTCCGATCTNNNTTTGTAAATGGCAGAGCAGCT
GCACACGTCTGAACTCTTTCCCTACACGACGCTCTTCCGATCTNNNTTCCCTCTCTCCAGCAAATCATC
AGCACACGTCTGAACTCTTTCCCTACACGACGCTCTTCCGATCTNNNTGGGTAAAGAAAACATGGTG
AGCACACGTCTGAACTCTTTCCCTACACGACGCTCTTCCGATCTNNNAATGCTCATTGTAGTTTGGCC
AGCACACGTCTGAACTCTTTCCCTACACGACGCTCTTCCGATCTNNNGCACAAGAAAGTGAGAGGAC
AGCACACGTCTGAACTCTTTCCCTACACGACGCTCTTCCGATCTNNNATCCTATATGCTTCTCTCCCCCA
AGCACACGTCTGAACTCTTTCCCTACACGACGCTCTTCCGATCTNNNGGAGTCCAGTCCTGAAAATCA
AGCACACGTCTGAACTCTTTCCCTACACGACGCTCTTCCGATCTNNNACTTTGCGGAAGATAATGGG
GCACACGTCTGAACTCTTTCCCTACACGACGCTCTTCCGATCTNNNCTTTCCAGCCAGTTGAAGGAATG
GCACACGTCTGAACTCTTTCCCTACACGACGCTCTTCCGATCTNNNCTGCTAAGAAACGCAAGGAGA
AGCACACGTCTGAACTCTTTCCCTACACGACGCTCTTCCGATCTNNNGGCTCATCTTGGTCCACTGTT
AGCACACGTCTGAACTCTTTCCCTACACGACGCTCTTCCGATCTNNNACTGTCAGTCTGCCAGCATAAA
AGAGCACACGTCTGAACTCTTTCCCTACACGACGCTCTTCCGATCTNNNGTGTCTCACAATGGGTCTA
AGCACACGTCTGAACTCTTTCCCTACACGACGCTCTTCCGATCTNNNCACCATCATTGCTAGAGGTCA
GAGCACACGTCTGAACTCTTTCCCTACACGACGCTCTTCCGATCTNNNTATGGTACCAGCTACTCGAG
GCACACGTCTGAACTCTTTCCCTACACGACGCTCTTCCGATCTNNNCCTCTAGAGAACCCTGACTGACA
AGCACACGTCTGAACTCTTTCCCTACACGACGCTCTTCCGATCTNNNGAGGATTTTTGCGTCGATGTT
AGCACACGTCTGAACTCTTTCCCTACACGACGCTCTTCCGATCTNNNGTGTGTGTGTGTGTGTGTGT
GCACACGTCTGAACTCTTTCCCTACACGACGCTCTTCCGATCTNNNTGTGATTCTCAAGCATGGAGAG
AGCACACGTCTGAACTCTTTCCCTACACGACGCTCTTCCGATCTNNNGATAATGGCTTCCAACTCCATG
AGCACACGTCTGAACTCTTTCCCTACACGACGCTCTTCCGATCTNNNCTCAGGTGATCCTCCCACTTCA
AGCACACGTCTGAACTCTTTCCCTACACGACGCTCTTCCGATCTNNNAGTAAGCTGAGATCATGCCA
GCACACGTCTGAACTCTTTCCCTACACGACGCTCTTCCGATCTNNNTTACTGCCTCCTACAACACCAAA
GAGCACACGTCTGAACTCTTTCCCTACACGACGCTCTTCCGATCTNNNGTGGGAGAATTGTTTGAAC
AGCACACGTCTGAACTCTTTCCCTACACGACGCTCTTCCGATCTNNNCACCTGTAATCCCAGCTACTTG
AGCACACGTCTGAACTCTTTCCCTACACGACGCTCTTCCGATCTNNNCACCCTACTGAGTCAGAATCA
AGCACACGTCTGAACTCTTTCCCTACACGACGCTCTTCCGATCTNNNAAATGAAGTCTCTCCGTGTGT
AGCACACGTCTGAACTCTTTCCCTACACGACGCTCTTCCGATCTNNNAAGGTTCAATCCAACAAGCAG
AGCACACGTCTGAACTCTTTCCCTACACGACGCTCTTCCGATCTNNNTCAGGTAGCTGAGACTATAGG
GCACACGTCTGAACTCTTTCCCTACACGACGCTCTTCCGATCTNNNAGCCAGACTCCATCTCAAAACA
AGCACACGTCTGAACTCTTTCCCTACACGACGCTCTTCCGATCTNNNGCATCACTTGAAGCCAGAAGT
GCACACGTCTGAACTCTTTCCCTACACGACGCTCTTCCGATCTNNNCCTCGATTCCTCTCTCACTTTGTC
AGCACACGTCTGAACTCTTTCCCTACACGACGCTCTTCCGATCTNNNTATTATTGGCTGAAGGTGAG
AGCACACGTCTGAACTCTTTCCCTACACGACGCTCTTCCGATCTNNNTGAGGTAGGAGAATCGCTTGA
AGCACACGTCTGAACTCTTTCCCTACACGACGCTCTTCCGATCTNNNTGGATCTTGTCATAGCGACTG
AGCACACGTCTGAACTCTTTCCCTACACGACGCTCTTCCGATCTNNNTTCTCTACAACTTCCAGGTCTG
AGCACACGTCTGAACTCTTTCCCTACACGACGCTCTTCCGATCTNNNAGACAAGAAACCAAAAAAGC

AGCACACGTCTGAACTCTTTCCCTACACGACGCTCTTCCGATCTNNNAATCCTAGTAGTTTGCTGGTG
GAGCACACGTCTGAACTCTTTCCCTACACGACGCTCTTCCGATCTNNNTACTCTGTGTGTGTGTGTGTG
GAGCACACGTCTGAACTCTTTCCCTACACGACGCTCTTCCGATCTNNNCCCCAAACCTAGCAGAAAACA
AGCACACGTCTGAACTCTTTCCCTACACGACGCTCTTCCGATCTNNNTTGGTTGATACTGGCTGCAAA
GAGCACACGTCTGAACTCTTTCCCTACACGACGCTCTTCCGATCTNNNTTCAGTCTCTCCCTCTCTTTCC
AGAGCACACGTCTGAACTCTTTCCCTACACGACGCTCTTCCGATCTNNNTGGATATTGCAGTGAACTG
GAGCACACGTCTGAACTCTTTCCCTACACGACGCTCTTCCGATCTNNNCCAACATAGCAAAACCCCATC
AGCACACGTCTGAACTCTTTCCCTACACGACGCTCTTCCGATCTNNNTCTCAGCCTCCATAAATGCGTA
GCACACGTCTGAACTCTTTCCCTACACGACGCTCTTCCGATCTNNNGCCCACAAAATCAACCTGAAACT
AGCACACGTCTGAACTCTTTCCCTACACGACGCTCTTCCGATCTNNNCGCTTTCCCTGAATCCTAGACT
GAGCACACGTCTGAACTCTTTCCCTACACGACGCTCTTCCGATCTNNNATGTTCAAGCCACTGCACTCT
AGCACACGTCTGAACTCTTTCCCTACACGACGCTCTTCCGATCTNNNCTCCCAAGTAGCTAGGAACAC
AGCACACGTCTGAACTCTTTCCCTACACGACGCTCTTCCGATCTNNNCTCCTGAGCAGCTAGGATTAC
AGCACACGTCTGAACTCTTTCCCTACACGACGCTCTTCCGATCTNNNTAATTGTGAGGGCTGTGTGT
GAGCACACGTCTGAACTCTTTCCCTACACGACGCTCTTCCGATCTNNNCACAGGACAGGACCAAAGT
AGCACACGTCTGAACTCTTTCCCTACACGACGCTCTTCCGATCTNNNACTCTCTAGCAGCTGAGACTAC
GAGCACACGTCTGAACTCTTTCCCTACACGACGCTCTTCCGATCTNNNCACTGCACAAACTACCTTTGG
AGAGCACACGTCTGAACTCTTTCCCTACACGACGCTCTTCCGATCTNNNCCCTGTTGTGATAACAGCG
GAGCACACGTCTGAACTCTTTCCCTACACGACGCTCTTCCGATCTNNNTCTAGGGTTCTAAGGCACAA
GAGCACACGTCTGAACTCTTTCCCTACACGACGCTCTTCCGATCTNNNTAAAATGCACACACACACACA
GCACACGTCTGAACTCTTTCCCTACACGACGCTCTTCCGATCTNNNAAGGCACTAGGTGGTCTATTTG
AGCACACGTCTGAACTCTTTCCCTACACGACGCTCTTCCGATCTNNNAATGCCCATGAATCAACAACTG
GCACACGTCTGAACTCTTTCCCTACACGACGCTCTTCCGATCTNNNTTTTGGTGGCAGGGTTTGTTTT
AGCACACGTCTGAACTCTTTCCCTACACGACGCTCTTCCGATCTNNNATCTGGAACTCTTGGCCTCAAA
AGCACACGTCTGAACTCTTTCCCTACACGACGCTCTTCCGATCTNNNTCCAATTCCATCTAGGTTGCTG
AGCACACGTCTGAACTCTTTCCCTACACGACGCTCTTCCGATCTNNNGGGTAGTGCCAGATGGGATA
AGCACACGTCTGAACTCTTTCCCTACACGACGCTCTTCCGATCTNNNTGGCTAACGGCTCTTCAGATAT
AGAGCACACGTCTGAACTCTTTCCCTACACGACGCTCTTCCGATCTNNNCCTATCCTCAGCTCCAGACT
GAGCACACGTCTGAACTCTTTCCCTACACGACGCTCTTCCGATCTNNNCTGCTTTTACAAATGGGGTG
GAGCACACGTCTGAACTCTTTCCCTACACGACGCTCTTCCGATCTNNNTATTCCTGGCCGGGAGTGTAT
GAGCACACGTCTGAACTCTTTCCCTACACGACGCTCTTCCGATCTNNNCATAGCAAGACTCTGTCTCTG
AGCACACGTCTGAACTCTTTCCCTACACGACGCTCTTCCGATCTNNNAGACAAGAGAATCGCTTGAAC
GAGCACACGTCTGAACTCTTTCCCTACACGACGCTCTTCCGATCTNNNGTAATCCAAGCTACTCTGGA
AGCACACGTCTGAACTCTTTCCCTACACGACGCTCTTCCGATCTNNNGAGGCAGGAAGATTGCTTGA
AGCACACGTCTGAACTCTTTCCCTACACGACGCTCTTCCGATCTNNNCCGTGTCCTTCCATACCCTAAAA
CACACGTCTGAACTCTTTCCCTACACGACGCTCTTCCGATCTNNNTCCGTTAGTTGGGCTTTTGTTTCC
AGCACACGTCTGAACTCTTTCCCTACACGACGCTCTTCCGATCTNNNTCTCAACACCATGTCACATCCAT
AGCACACGTCTGAACTCTTTCCCTACACGACGCTCTTCCGATCTNNNAAATACTGGGGACTGCTGCAA
AGCACACGTCTGAACTCTTTCCCTACACGACGCTCTTCCGATCTNNNACTCCCGAGTAGCTAGGATTAC
AAGAGCACACGTCTGAACTCTTTCCCTACACGACGCTCTTCCGATCTNNNAGCATAACAGGGGTTTAG
GAGCACACGTCTGAACTCTTTCCCTACACGACGCTCTTCCGATCTNNNACCCATGTCTGTGTGTCTGTA
AGCACACGTCTGAACTCTTTCCCTACACGACGCTCTTCCGATCTNNNGTCAATCACTTGAGGTCAGGA
GCACACGTCTGAACTCTTTCCCTACACGACGCTCTTCCGATCTNNNATTGAGCTGAGATTGCGTCAAT

GAGCACACGTCTGAACTCTTTCCCTACACGACGCTCTTCCGATCTNNNCTGAAGTTCCATCTCTACCCC
GCACACGTCTGAACTCTTTCCCTACACGACGCTCTTCCGATCTNNNAGTCATTCCCTCTTCCTCTCTCTC
AGCACACGTCTGAACTCTTTCCCTACACGACGCTCTTCCGATCTNNNATGTTGGCTTACTGCAACCTCT
AGCACACGTCTGAACTCTTTCCCTACACGACGCTCTTCCGATCTNNNGTGTTTAGACTTGGGTGCCAT
AGCACACGTCTGAACTCTTTCCCTACACGACGCTCTTCCGATCTNNNGAACTCCTGGCCTGAAGTGAT
AGCACACGTCTGAACTCTTTCCCTACACGACGCTCTTCCGATCTNNNACGTCAGACACTCTACAAAGCA
AAGAGCACACGTCTGAACTCTTTCCCTACACGACGCTCTTCCGATCTNNNAAAATGTTCAGCACTTTAG
GCACACGTCTGAACTCTTTCCCTACACGACGCTCTTCCGATCTNNNCTTGAACCTAGGAAGTTGAGG
GCACACGTCTGAACTCTTTCCCTACACGACGCTCTTCCGATCTNNNTGAGATCTCACCACTGTACTCCA
AGCACACGTCTGAACTCTTTCCCTACACGACGCTCTTCCGATCTNNNATAACAATCAGCGAGATCAG
GAGCACACGTCTGAACTCTTTCCCTACACGACGCTCTTCCGATCTNNNACACAGTGAAACCCCATCTCT
GCACACGTCTGAACTCTTTCCCTACACGACGCTCTTCCGATCTNNNTCCTACCTGTGTTCCAAAGTCAC
AGCACACGTCTGAACTCTTTCCCTACACGACGCTCTTCCGATCTNNNTGAAGCTGAGAGCCAAATCA
AGCACACGTCTGAACTCTTTCCCTACACGACGCTCTTCCGATCTNNNAATTCTCTAGGGCTCGATGGT
GAGCACACGTCTGAACTCTTTCCCTACACGACGCTCTTCCGATCTNNNTGAGGTAGGAGAATCGCTT
GAGCACACGTCTGAACTCTTTCCCTACACGACGCTCTTCCGATCTNNNCCTCATGTGGGTTAGGACTT
AGCACACGTCTGAACTCTTTCCCTACACGACGCTCTTCCGATCTNNNGAGAAAATTCCAGGCAGCAG
AGCACACGTCTGAACTCTTTCCCTACACGACGCTCTTCCGATCTNNNAGTTTGGGGGAAAAACAATG
GCACACGTCTGAACTCTTTCCCTACACGACGCTCTTCCGATCTNNNTAGTTCGGTACTTGGAGGCTG
AGCACACGTCTGAACTCTTTCCCTACACGACGCTCTTCCGATCTNNNAACCTTTTACCCCAGATTATG
AGCACACGTCTGAACTCTTTCCCTACACGACGCTCTTCCGATCTNNNCTCTTGCTTTTTGGGACCAGAA
GCACACGTCTGAACTCTTTCCCTACACGACGCTCTTCCGATCTNNNCATGCTTCTGTAGAGGACTTGG
GCACACGTCTGAACTCTTTCCCTACACGACGCTCTTCCGATCTNNNATACCAGATGTGAATGTGGTG
GAGCACACGTCTGAACTCTTTCCCTACACGACGCTCTTCCGATCTNNNGAGAATCACTTGAACCTGGG
GAGCACACGTCTGAACTCTTTCCCTACACGACGCTCTTCCGATCTNNNCAACATAGTGAGACTCTGGC
GAGCACACGTCTGAACTCTTTCCCTACACGACGCTCTTCCGATCTNNNGCCTGAGCAACATATCGAA
AGCACACGTCTGAACTCTTTCCCTACACGACGCTCTTCCGATCTNNNGCAGAACTGAGATGACAGAG
GCACACGTCTGAACTCTTTCCCTACACGACGCTCTTCCGATCTNNNCTGTCGAAATTGGCTAGCATGA
GAGCACACGTCTGAACTCTTTCCCTACACGACGCTCTTCCGATCTNNNACCTAGATGCTAACACACAC
AGCACACGTCTGAACTCTTTCCCTACACGACGCTCTTCCGATCTNNNTTACTGTCAAGGAGAAGGGCT
CACACGTCTGAACTCTTTCCCTACACGACGCTCTTCCGATCTNNNGGCCTTGAACATCTAACACACAC
AGCACACGTCTGAACTCTTTCCCTACACGACGCTCTTCCGATCTNNNCCAGATTCAAGCAATTCTCCTG
AGCACACGTCTGAACTCTTTCCCTACACGACGCTCTTCCGATCTNNNTTCCAAGCAAACATCTTTGGTG
AGCACACGTCTGAACTCTTTCCCTACACGACGCTCTTCCGATCTNNNCTGCCTCAGCTTCTCAAGTAGA
GAGCACACGTCTGAACTCTTTCCCTACACGACGCTCTTCCGATCTNNNCTGAGGTGGGAGAATTGCT
GCACACGTCTGAACTCTTTCCCTACACGACGCTCTTCCGATCTNNNTTCAAGTTGTCTGGAGTGTCTG
GCACACGTCTGAACTCTTTCCCTACACGACGCTCTTCCGATCTNNNAGAGAAAAAGGTGGACACAG
GAGCACACGTCTGAACTCTTTCCCTACACGACGCTCTTCCGATCTNNNACAATACAGCTAGAAGGCC
AGCACACGTCTGAACTCTTTCCCTACACGACGCTCTTCCGATCTNNNGTTAGCAACCGTACATCACAA
GAGCACACGTCTGAACTCTTTCCCTACACGACGCTCTTCCGATCTNNNTCAATTAGGGGCTAGGGCAA
GAGCACACGTCTGAACTCTTTCCCTACACGACGCTCTTCCGATCTNNNGGGAAAGAGAGGGATGAG
AGCACACGTCTGAACTCTTTCCCTACACGACGCTCTTCCGATCTNNNTTAGCCAAGATCATGCCATCGT
AGCACACGTCTGAACTCTTTCCCTACACGACGCTCTTCCGATCTNNNAAGATTGTGCATTGAACTCCA

GAGCACACGTCTGAACTCTTTCCCTACACGACGCTCTTCCGATCTNNNTAATCCCAGTTACTCAGGAGG
AGCACACGTCTGAACTCTTTCCCTACACGACGCTCTTCCGATCTNNNGCAGAATCTATGGACACAAAGG
GAGCACACGTCTGAACTCTTTCCCTACACGACGCTCTTCCGATCTNNNCACACACACACACACACATTA
AGCACACGTCTGAACTCTTTCCCTACACGACGCTCTTCCGATCTNNNGGTTGAATTACGACCGGTTTTC
GAGCACACGTCTGAACTCTTTCCCTACACGACGCTCTTCCGATCTNNNCCCCAATCTGGTGTGTGTGTA
AAGAGCACACGTCTGAACTCTTTCCCTACACGACGCTCTTCCGATCTNNNATCTGTCCCATCAGCCAACA
GAGCACACGTCTGAACTCTTTCCCTACACGACGCTCTTCCGATCTNNNCCTGTTCTTCCATCCCTGTATCC
AGCACACGTCTGAACTCTTTCCCTACACGACGCTCTTCCGATCTNNNGAGGTTGCAGTAAGCCAAGAT
AGCACACGTCTGAACTCTTTCCCTACACGACGCTCTTCCGATCTNNNGTGTGATGAAAGCCAAAGGAC
GAGCACACGTCTGAACTCTTTCCCTACACGACGCTCTTCCGATCTNNNGAGGCCGAGACAGGAATATCA
AGCACACGTCTGAACTCTTTCCCTACACGACGCTCTTCCGATCTNNNACAGAGCCAGATTCCATCTCAAA
GAGCACACGTCTGAACTCTTTCCCTACACGACGCTCTTCCGATCTNNNTTGATGTCAAAACGAAGCACA
GAGCACACGTCTGAACTCTTTCCCTACACGACGCTCTTCCGATCTNNNTCTCTCCCTCTCTCTCTCTCTC
AGCACACGTCTGAACTCTTTCCCTACACGACGCTCTTCCGATCTNNNCAGTCTTGCATTCCCAGGAAAA
AGCACACGTCTGAACTCTTTCCCTACACGACGCTCTTCCGATCTNNNCAGGCATATCACTTGAGGTCAG
GAGCACACGTCTGAACTCTTTCCCTACACGACGCTCTTCCGATCTNNNTATAGTCCCAGCTACTCAGGAG
GAGCACACGTCTGAACTCTTTCCCTACACGACGCTCTTCCGATCTNNNCTTTTGACCTTGTGATCTGCCT
GAGCACACGTCTGAACTCTTTCCCTACACGACGCTCTTCCGATCTNNNTGGGTAAGCTTTTTGATGTGC
AGCACACGTCTGAACTCTTTCCCTACACGACGCTCTTCCGATCTNNNGGTAGAGGTTGCAGTGAGTCT
GAGCACACGTCTGAACTCTTTCCCTACACGACGCTCTTCCGATCTNNNAGACGCCGTCTCAAAAAAAAAA
AGCACACGTCTGAACTCTTTCCCTACACGACGCTCTTCCGATCTNNNGGGAGCCAGTCACATTTTACTGT
GAGCACACGTCTGAACTCTTTCCCTACACGACGCTCTTCCGATCTNNNCAGTTCCCTCCCATTACTGTGAG
AGAGCACACGTCTGAACTCTTTCCCTACACGACGCTCTTCCGATCTNNNCGCCTGGCCACTATTTTTTTT
GCACACGTCTGAACTCTTTCCCTACACGACGCTCTTCCGATCTNNNGTTTGGTCTCGTACAACAAAGGG
AGCACACGTCTGAACTCTTTCCCTACACGACGCTCTTCCGATCTNNNAGGTTATTTTCACAACCCCTGGT
AGAGCACACGTCTGAACTCTTTCCCTACACGACGCTCTTCCGATCTNNNGCTATGATCACAACACTGCAC
AGCACACGTCTGAACTCTTTCCCTACACGACGCTCTTCCGATCTNNNTACACATGTGAGCCGATTTCTCA
AGCACACGTCTGAACTCTTTCCCTACACGACGCTCTTCCGATCTNNNACACTTTCCAGAAACTGCACATG
AGCACACGTCTGAACTCTTTCCCTACACGACGCTCTTCCGATCTNNNGAGGTTGCAATGGACCGAGAT
AGCACACGTCTGAACTCTTTCCCTACACGACGCTCTTCCGATCTNNNTAAACACGCACATACACACACAC
GAGCACACGTCTGAACTCTTTCCCTACACGACGCTCTTCCGATCTNNNTAAACACACACACACACACACA
GAGCACACGTCTGAACTCTTTCCCTACACGACGCTCTTCCGATCTNNNCTGCTTCTTGGTACGCTGATTT
GAGCACACGTCTGAACTCTTTCCCTACACGACGCTCTTCCGATCTNNNTGAAATTCCTACCCAACACAGT
AGCACACGTCTGAACTCTTTCCCTACACGACGCTCTTCCGATCTNNNAGATACCCCTTTGCCTCAATGTT
AGCACACGTCTGAACTCTTTCCCTACACGACGCTCTTCCGATCTNNNTGCTCAGTTTGCAGAAAGATTG
GCACACGTCTGAACTCTTTCCCTACACGACGCTCTTCCGATCTNNNCCCCAAAGCAATAACCTAGGGATG
GAGCACACGTCTGAACTCTTTCCCTACACGACGCTCTTCCGATCTNNNAAATATGGAACCAGCCCAAAT
GCACACGTCTGAACTCTTTCCCTACACGACGCTCTTCCGATCTNNNGGTAGGAGAATCACTTGAACCCA
GAGCACACGTCTGAACTCTTTCCCTACACGACGCTCTTCCGATCTNNNATGTGTTCTTGCCAAAACTCGA
AGCACACGTCTGAACTCTTTCCCTACACGACGCTCTTCCGATCTNNNTTGACTCTCTTCCCCACTTCCTACA
GAGCACACGTCTGAACTCTTTCCCTACACGACGCTCTTCCGATCTNNNTCACTTTCCAGAGTCTCTGTTC
AGCACACGTCTGAACTCTTTCCCTACACGACGCTCTTCCGATCTNNNAAAGTGCTGGGATTACAGGAG
GAGCACACGTCTGAACTCTTTCCCTACACGACGCTCTTCCGATCTNNNCCAACACACACAGACACATACA

AGCACACGTCTGAACTCTTTCCCTACACGACGCTCTTCCGATCTNNNGATGCTGATGCCTTCCCTATGTA
GAGCACACGTCTGAACTCTTTCCCTACACGACGCTCTTCCGATCTNNNTTGAGGCAGGAGAATCACTTG
AGCACACGTCTGAACTCTTTCCCTACACGACGCTCTTCCGATCTNNNACAAGCTATTATCAGCAGTGCA
AGCACACGTCTGAACTCTTTCCCTACACGACGCTCTTCCGATCTNNNAAAGTGCTGGGATTACAGGTG
GAGCACACGTCTGAACTCTTTCCCTACACGACGCTCTTCCGATCTNNNTACAGTGGTGCAATCAGAACT
AGCACACGTCTGAACTCTTTCCCTACACGACGCTCTTCCGATCTNNNTGCAGGTGAGTCAAGTACCTTA
GAGCACACGTCTGAACTCTTTCCCTACACGACGCTCTTCCGATCTNNNGGGTAACAGAGTGAGATCCT
GAGCACACGTCTGAACTCTTTCCCTACACGACGCTCTTCCGATCTNNNATAGTCCTAGCTACTCAGGAG
GAGCACACGTCTGAACTCTTTCCCTACACGACGCTCTTCCGATCTNNNCTGAGGCAGGACAATCACTTG
GAGCACACGTCTGAACTCTTTCCCTACACGACGCTCTTCCGATCTNNNGGTAACAAGAGGGAGACTCCA
AGCACACGTCTGAACTCTTTCCCTACACGACGCTCTTCCGATCTNNNCACTTGAGGTCAGGAGTTCAAG
GAGCACACGTCTGAACTCTTTCCCTACACGACGCTCTTCCGATCTNNNCTCGCCTCTTCCTCATTAGAGT
AAGAGCACACGTCTGAACTCTTTCCCTACACGACGCTCTTCCGATCTNNNCCTCTCCAAGAAACACCCAA
AGAGCACACGTCTGAACTCTTTCCCTACACGACGCTCTTCCGATCTNNNCATCTGTGGAAGCTCTCTTCT
AGCACACGTCTGAACTCTTTCCCTACACGACGCTCTTCCGATCTNNNCTCCAACTCCATCCAAATTGCTG
GCACACGTCTGAACTCTTTCCCTACACGACGCTCTTCCGATCTNNNTAAGAGAGGAATCCCAGGGGTT
AGCACACGTCTGAACTCTTTCCCTACACGACGCTCTTCCGATCTNNNGTGTGTGTGTGTGTGTACAT
AGCACACGTCTGAACTCTTTCCCTACACGACGCTCTTCCGATCTNNNGGTGACAGAGAGATACCCTGTT
GAGCACACGTCTGAACTCTTTCCCTACACGACGCTCTTCCGATCTNNNAGTGCCATAAAGGAAGTGCTA
AGCACACGTCTGAACTCTTTCCCTACACGACGCTCTTCCGATCTNNNTATGTCTGTCCTGGGGTCTGATT
GAGCACACGTCTGAACTCTTTCCCTACACGACGCTCTTCCGATCTNNNGGAAACAGTTGGTCCAGGATT
GCACACGTCTGAACTCTTTCCCTACACGACGCTCTTCCGATCTNNNCATACAAACACACATGCCCCTTTCA
GAGCACACGTCTGAACTCTTTCCCTACACGACGCTCTTCCGATCTNNNATGGGGTTTCACTATGTTGGC
GAGCACACGTCTGAACTCTTTCCCTACACGACGCTCTTCCGATCTNNNGGAAGATTGCATGAGCTTGG
GAGCACACGTCTGAACTCTTTCCCTACACGACGCTCTTCCGATCTNNNGAGCAAGACTCTGCCTCAAAA
AGCACACGTCTGAACTCTTTCCCTACACGACGCTCTTCCGATCTNNNAGAGTTTGCAGTGAGCCAAGA
GAGCACACGTCTGAACTCTTTCCCTACACGACGCTCTTCCGATCTNNNGGAGGTTGCAGTGAACAGAG
GAGCACACGTCTGAACTCTTTCCCTACACGACGCTCTTCCGATCTNNNGCAACATAGCAAGACCATCTCT
GAGCACACGTCTGAACTCTTTCCCTACACGACGCTCTTCCGATCTNNNACTTGAATCTGAGAAGCGGAG
AGCACACGTCTGAACTCTTTCCCTACACGACGCTCTTCCGATCTNNNATCTACAGTTTTGGGGTGGAGT
GAGCACACGTCTGAACTCTTTCCCTACACGACGCTCTTCCGATCTNNNCATTGTCACTGTTCAAAGGTCC
GCACACGTCTGAACTCTTTCCCTACACGACGCTCTTCCGATCTNNNTGAGAATCCATTTCAGCCTCTGGC
AGCACACGTCTGAACTCTTTCCCTACACGACGCTCTTCCGATCTNNNGAGGCATTAGAATCGCTTGAAC
GAGCACACGTCTGAACTCTTTCCCTACACGACGCTCTTCCGATCTNNNGCCACTACTGCACTATAGCCTA
GAGCACACGTCTGAACTCTTTCCCTACACGACGCTCTTCCGATCTNNNAGGCTTAGGCACGAGAATTG
GCACACGTCTGAACTCTTTCCCTACACGACGCTCTTCCGATCTNNNGCCTCTCCAATTCAGCAGAAATCC
GAGCACACGTCTGAACTCTTTCCCTACACGACGCTCTTCCGATCTNNNTACTCCCAATACACACACACAG
AGCACACGTCTGAACTCTTTCCCTACACGACGCTCTTCCGATCTNNNAGTACAGAGAGTGAGCAGTGA
AGCACACGTCTGAACTCTTTCCCTACACGACGCTCTTCCGATCTNNNGTGTGCTTGGTGTAGGTAGTG
GAGCACACGTCTGAACTCTTTCCCTACACGACGCTCTTCCGATCTNNNAGTGCTGGGATTACAGACTTG
GAGCACACGTCTGAACTCTTTCCCTACACGACGCTCTTCCGATCTNNNCGCCTGGCCACTATTTTTTCTTT
AGAGCACACGTCTGAACTCTTTCCCTACACGACGCTCTTCCGATCTNNNAAGATTCCTGCTCTCCTGTTG
GAGCACACGTCTGAACTCTTTCCCTACACGACGCTCTTCCGATCTNNNCACCTGTAATCCCAGCATTTTG

CACACGTCTGAACTCTTTCCCTACACGACGCTCTTCCGATCTNNNATGGTCTCTCTCTCTCTCTCCCAT
AGCACACGTCTGAACTCTTTCCCTACACGACGCTCTTCCGATCTNNNCCTTCTTGCTGGCAATTTCAGA
GCACACGTCTGAACTCTTTCCCTACACGACGCTCTTCCGATCTNNNTATTTCTGCAGCTATGTGCCTCT
GAGCACACGTCTGAACTCTTTCCCTACACGACGCTCTTCCGATCTNNNGCTTACACCTATAACCCAGCA
AGCACACGTCTGAACTCTTTCCCTACACGACGCTCTTCCGATCTNNNCAAGAGAGGGAGTAAGAGAG
GCACACGTCTGAACTCTTTCCCTACACGACGCTCTTCCGATCTNNNACTCTCTATGAGGGTCTCTCCAC
AGCACACGTCTGAACTCTTTCCCTACACGACGCTCTTCCGATCTNNNCGAACTCCAATCTCTCTCTCTCC
AGCACACGTCTGAACTCTTTCCCTACACGACGCTCTTCCGATCTNNNTAAAGAGAGAAACAGAGGG
GCACACGTCTGAACTCTTTCCCTACACGACGCTCTTCCGATCTNNNGCCTTTGAACTCAAGACTGCAA
GAGCACACGTCTGAACTCTTTCCCTACACGACGCTCTTCCGATCTNNNACTGGATATCTTGGAGAGG
AGCACACGTCTGAACTCTTTCCCTACACGACGCTCTTCCGATCTNNNTATGAGAGAGAGAGAGGCA
GAGCACACGTCTGAACTCTTTCCCTACACGACGCTCTTCCGATCTNNNACTTGAGGTCAGGAGTTCAA
GCACACGTCTGAACTCTTTCCCTACACGACGCTCTTCCGATCTNNNAACCATGAGCGAGAGAAAGAG
GCACACGTCTGAACTCTTTCCCTACACGACGCTCTTCCGATCTNNNCAATCGTGTTGGTTTCTGGTGA
AGCACACGTCTGAACTCTTTCCCTACACGACGCTCTTCCGATCTNNNAATAAGACGAACTGGAGCCTT
GAGCACACGTCTGAACTCTTTCCCTACACGACGCTCTTCCGATCTNNNGTGTGGAGCAGCTAACTCT
AGCACACGTCTGAACTCTTTCCCTACACGACGCTCTTCCGATCTNNNCTCTCTTTCTCGCTGTCTCTCTC
GCACACGTCTGAACTCTTTCCCTACACGACGCTCTTCCGATCTNNNCTGGCCATTTCCTTTCTCTCTCTC
GCACACGTCTGAACTCTTTCCCTACACGACGCTCTTCCGATCTNNNCAGGGGGAAAAATAGTTGAAG
GAGCACACGTCTGAACTCTTTCCCTACACGACGCTCTTCCGATCTNNNAGATATTTGGGAAGCAGAG
AGCACACGTCTGAACTCTTTCCCTACACGACGCTCTTCCGATCTNNNTGTTTGTGTGTGTGTGCATAT
GAGCACACGTCTGAACTCTTTCCCTACACGACGCTCTTCCGATCTNNNGGTTCCCAAGAGAAACAGAG
AGCACACGTCTGAACTCTTTCCCTACACGACGCTCTTCCGATCTNNNGTGCTGTCTCGTGATAGAGTT
CACACGTCTGAACTCTTTCCCTACACGACGCTCTTCCGATCTNNNGGTAGTAACCCTTGCAAAGCAAA
AGCACACGTCTGAACTCTTTCCCTACACGACGCTCTTCCGATCTNNNAAATTTTTCTTGGAGGGGGA
GCACACGTCTGAACTCTTTCCCTACACGACGCTCTTCCGATCTNNNAGCATGAGCAATAGAGTGAGA
GCACACGTCTGAACTCTTTCCCTACACGACGCTCTTCCGATCTNNNTCTCTCTCTCTCTCTCTCTCCC
AGCACACGTCTGAACTCTTTCCCTACACGACGCTCTTCCGATCTNNNTGGATGCTTTGATTGCTCCAAT
AGCACACGTCTGAACTCTTTCCCTACACGACGCTCTTCCGATCTNNNTAGAAGTGTATGGCATTTCCCC
AGAGCACACGTCTGAACTCTTTCCCTACACGACGCTCTTCCGATCTNNNTTGATGACGGGAGAGACTT
AGCACACGTCTGAACTCTTTCCCTACACGACGCTCTTCCGATCTNNNTAGAGATGGGTTTTCGCCATG
AAGAGCACACGTCTGAACTCTTTCCCTACACGACGCTCTTCCGATCTNNNGGGAAGGAAGGGAGAAA
GAGCACACGTCTGAACTCTTTCCCTACACGACGCTCTTCCGATCTNNNGGGAAAAGGCATGTGTTGA
GCACACGTCTGAACTCTTTCCCTACACGACGCTCTTCCGATCTNNNTTTCTCTCTCTCTCTCTCCCCCA
AGCACACGTCTGAACTCTTTCCCTACACGACGCTCTTCCGATCTNNNGCCAAGTGGTCCTGGTCAATA
GCACACGTCTGAACTCTTTCCCTACACGACGCTCTTCCGATCTNNNCTCTGCAAAGACCCTCCTTGATA
GCACACGTCTGAACTCTTTCCCTACACGACGCTCTTCCGATCTNNNCATTAGGAGGGTCAGTCTGTCT
CACACGTCTGAACTCTTTCCCTACACGACGCTCTTCCGATCTNNNTGAGGAAGGAGAAAGTGTTTG
AGCACACGTCTGAACTCTTTCCCTACACGACGCTCTTCCGATCTNNNAAGGCTCAACAAAACAAAAAG
AGCACACGTCTGAACTCTTTCCCTACACGACGCTCTTCCGATCTNNNATGCTCTCACAGTCTGTTGACA
AGCACACGTCTGAACTCTTTCCCTACACGACGCTCTTCCGATCTNNNTGAACTTGGTTCCTACAGGAC
GAGCACACGTCTGAACTCTTTCCCTACACGACGCTCTTCCGATCTNNNTCAGAGTTCTCTCTCTCAGAG
GAGCACACGTCTGAACTCTTTCCCTACACGACGCTCTTCCGATCTNNNAGGAAGGGCGGAGACAAA

AGCACACGTCTGAACTCTTTCCCTACACGACGCTCTTCCGATCTNNNTTTAGGAGAATGGCTGTTCAG
GAGCACACGTCTGAACTCTTTCCCTACACGACGCTCTTCCGATCTNNNGAGAGGTTTTTCCCAGCATAA
GAGCACACGTCTGAACTCTTTCCCTACACGACGCTCTTCCGATCTNNNTATCCCCAACAATGCCAGGAAA
GAGCACACGTCTGAACTCTTTCCCTACACGACGCTCTTCCGATCTNNNGAGTGGCTTCCATAGAAAAG
GAGCACACGTCTGAACTCTTTCCCTACACGACGCTCTTCCGATCTNNNTGCATTTTCCGTAGACATTCCT
AGCACACGTCTGAACTCTTTCCCTACACGACGCTCTTCCGATCTNNNTTTCTGCCACTGAAATGTGAAC
AAGAGCACACGTCTGAACTCTTTCCCTACACGACGCTCTTCCGATCTNNNTGGATGCAAGACAAGTTTT
GAGCACACGTCTGAACTCTTTCCCTACACGACGCTCTTCCGATCTNNNAGGAAGCAAATAAGCCAGTCT
AGCACACGTCTGAACTCTTTCCCTACACGACGCTCTTCCGATCTNNNGATTTTGGAGCCTTTGGGATGT
GAGCACACGTCTGAACTCTTTCCCTACACGACGCTCTTCCGATCTNNNGGGGTATGGTGGAGTTTTTT
AGCACACGTCTGAACTCTTTCCCTACACGACGCTCTTCCGATCTNNNTAGTTGGGCAACAAGAGTGAAA
GAGCACACGTCTGAACTCTTTCCCTACACGACGCTCTTCCGATCTNNNGAGGGAGAGGAAGTATGCTG
GAGCACACGTCTGAACTCTTTCCCTACACGACGCTCTTCCGATCTNNNTTATGGCTTCTAGGCTGGCTAT
GCACACGTCTGAACTCTTTCCCTACACGACGCTCTTCCGATCTNNNGAGCAGGTAGAAAGGAGAGGA
GAGCACACGTCTGAACTCTTTCCCTACACGACGCTCTTCCGATCTNNNCTACAGAAGTCCCAAGAGCAC
AGCACACGTCTGAACTCTTTCCCTACACGACGCTCTTCCGATCTNNNAGTCTAAAGGAGCAATGTGGT
AGAGCACACGTCTGAACTCTTTCCCTACACGACGCTCTTCCGATCTNNNTTGTGGCTAAAAGCAGCAGT
AGCACACGTCTGAACTCTTTCCCTACACGACGCTCTTCCGATCTNNNGGACTCAGAAGAAAGAAGGTG
AGCACACGTCTGAACTCTTTCCCTACACGACGCTCTTCCGATCTNNNCCTGGAAATGTGCATGTCAAGT
AGAGCACACGTCTGAACTCTTTCCCTACACGACGCTCTTCCGATCTNNNGTGACAAAACTAGGGGTAGA
AGCACACGTCTGAACTCTTTCCCTACACGACGCTCTTCCGATCTNNNAGAATGCCCGTAGACTCACAAAT
GAGCACACGTCTGAACTCTTTCCCTACACGACGCTCTTCCGATCTNNNGCTTTCAGGTGAGTTTCTGCA
AGAGCACACGTCTGAACTCTTTCCCTACACGACGCTCTTCCGATCTNNNCCTCGGATTGCCTGTGAAATA
AGCACACGTCTGAACTCTTTCCCTACACGACGCTCTTCCGATCTNNNTTCACCAGGCAGATAAGGATA
GAGCACACGTCTGAACTCTTTCCCTACACGACGCTCTTCCGATCTNNNAAACTGGTGAAGAGCTACACA
GAGCACACGTCTGAACTCTTTCCCTACACGACGCTCTTCCGATCTNNNTGCCCCTTTTCTGTTATACAGG
AGCACACGTCTGAACTCTTTCCCTACACGACGCTCTTCCGATCTNNNGCCTGCTTTCAGTAGACAGATG
GAGCACACGTCTGAACTCTTTCCCTACACGACGCTCTTCCGATCTNNNTGCCTGTAGCTTAGGAATCAT
AGCACACGTCTGAACTCTTTCCCTACACGACGCTCTTCCGATCTNNNGTTTGAAAGGTGGGGATCTCT
AGAGCACACGTCTGAACTCTTTCCCTACACGACGCTCTTCCGATCTNNNCAGAGATGATGGGGAAAGA
AGAGCACACGTCTGAACTCTTTCCCTACACGACGCTCTTCCGATCTNNNCCAAAGAACTTGAAGACAG
AGAGCACACGTCTGAACTCTTTCCCTACACGACGCTCTTCCGATCTNNNGAAGAGATGGAACCCAAAG
AGAGCACACGTCTGAACTCTTTCCCTACACGACGCTCTTCCGATCTNNNCCAGTTCCAGATAGACAGAC
AGCACACGTCTGAACTCTTTCCCTACACGACGCTCTTCCGATCTNNNCAATGTGGTAAGTGCTGCAAAG
AGAGCACACGTCTGAACTCTTTCCCTACACGACGCTCTTCCGATCTNNNTCCCTGGACACAATTATTGGA
GAGCACACGTCTGAACTCTTTCCCTACACGACGCTCTTCCGATCTNNNTATCTATCTCTCTCTCTCCCCC
AGAGCACACGTCTGAACTCTTTCCCTACACGACGCTCTTCCGATCTNNNGCTTACCTGAAGAATGTGCAT
GAGCACACGTCTGAACTCTTTCCCTACACGACGCTCTTCCGATCTNNNTCCTTCCATTTCTGATGACACCT
AGCACACGTCTGAACTCTTTCCCTACACGACGCTCTTCCGATCTNNNCCACTCCTAGGCATTTACTGACA
GAGCACACGTCTGAACTCTTTCCCTACACGACGCTCTTCCGATCTNNNCAGGTCACCAGATTGCATGTT
GAGCACACGTCTGAACTCTTTCCCTACACGACGCTCTTCCGATCTNNNATAAGCTTCTTGCCTGTTCCAG
AGCACACGTCTGAACTCTTTCCCTACACGACGCTCTTCCGATCTNNNGTGCAGCTGCTATTTCATACCAG
GAGCACACGTCTGAACTCTTTCCCTACACGACGCTCTTCCGATCTNNNAAGGAGCAAGGATAGGGA

AGCACACGTCTGAACTCTTTCCCTACACGACGCTCTTCCGATCTNNNGTCAGGGATCTCAGAAATCCAC
AGCACACGTCTGAACTCTTTCCCTACACGACGCTCTTCCGATCTNNNCTGAGCTTCTCATTTCCCAGAG
AGCACACGTCTGAACTCTTTCCCTACACGACGCTCTTCCGATCTNNNAATCACTTGAGCCTAGGAGTT
GAGCACACGTCTGAACTCTTTCCCTACACGACGCTCTTCCGATCTNNNTGACTCCCTCTGATATGTAGC
AGCACACGTCTGAACTCTTTCCCTACACGACGCTCTTCCGATCTNNNCTTATACTCTGCTCACTCTGCCC
AGCACACGTCTGAACTCTTTCCCTACACGACGCTCTTCCGATCTNNNGTAGAGAGTGCTAGTTGCCT
AGCACACGTCTGAACTCTTTCCCTACACGACGCTCTTCCGATCTNNNTCCCCCATCTCTTATCTAGCTGA
AGCACACGTCTGAACTCTTTCCCTACACGACGCTCTTCCGATCTNNNTGAAGCACATTCTCTATGTCCC
AGCACACGTCTGAACTCTTTCCCTACACGACGCTCTTCCGATCTNNNACTAAGAACAGAACTGCAGCT
AGCACACGTCTGAACTCTTTCCCTACACGACGCTCTTCCGATCTNNNTGAAACAGTCACCCTCAGAAA
GAGCACACGTCTGAACTCTTTCCCTACACGACGCTCTTCCGATCTNNNACAGAGCCCAGTCTCTAAAAC
GAGCACACGTCTGAACTCTTTCCCTACACGACGCTCTTCCGATCTNNNATCTACGCAAAGATGGGAGA
AGCACACGTCTGAACTCTTTCCCTACACGACGCTCTTCCGATCTNNNACAACCCAAGACCTTGTCTCAA
AGCACACGTCTGAACTCTTTCCCTACACGACGCTCTTCCGATCTNNNTCAGATTCTTTCTCTTGTGGG
GAGCACACGTCTGAACTCTTTCCCTACACGACGCTCTTCCGATCTNNNATCTATAAGCCCATGTTTGCC
GAGCACACGTCTGAACTCTTTCCCTACACGACGCTCTTCCGATCTNNNAGGTCTAGGACTCCAAATCC
AGCACACGTCTGAACTCTTTCCCTACACGACGCTCTTCCGATCTNNNATACCACGCAGTTCTACCTTCA
GCACACGTCTGAACTCTTTCCCTACACGACGCTCTTCCGATCTNNNTCTCTTGTCTTCCTCTCCCTCTCC
GCACACGTCTGAACTCTTTCCCTACACGACGCTCTTCCGATCTNNNACCTCCCTCCTCTGAGTTCTTTAC
AGCACACGTCTGAACTCTTTCCCTACACGACGCTCTTCCGATCTNNNTAGCAGCTGGGATTATGGTAT
AGCACACGTCTGAACTCTTTCCCTACACGACGCTCTTCCGATCTNNNTTTTTCAGGAGCCACAGAGAG
AGCACACGTCTGAACTCTTTCCCTACACGACGCTCTTCCGATCTNNNCTTTCATGTGTTCAGTACTGGC
AGCACACGTCTGAACTCTTTCCCTACACGACGCTCTTCCGATCTNNNAGGTATACTCTACAAGTCAGG
AGCACACGTCTGAACTCTTTCCCTACACGACGCTCTTCCGATCTNNNACACACACACACCCCTCTTTATT
GAGCACACGTCTGAACTCTTTCCCTACACGACGCTCTTCCGATCTNNNGCTGTATGTGGGAAGGGAG
AGCACACGTCTGAACTCTTTCCCTACACGACGCTCTTCCGATCTNNNAGTGTGTATCTGTCTTCCTCC
AGCACACGTCTGAACTCTTTCCCTACACGACGCTCTTCCGATCTNNNAGGGCCACAGCAACATATCTA
GCACACGTCTGAACTCTTTCCCTACACGACGCTCTTCCGATCTNNNGCTGGATGTAACCCATTTTTGCT
GCACACGTCTGAACTCTTTCCCTACACGACGCTCTTCCGATCTNNNCGTTGCTCTACTCTCGTTCTCTCC
GAGCACACGTCTGAACTCTTTCCCTACACGACGCTCTTCCGATCTNNNGCCCTCTCGTTGATGTATACT
AGCACACGTCTGAACTCTTTCCCTACACGACGCTCTTCCGATCTNNNAGGAGCTGATTCTTCCTAAAG
AGCACACGTCTGAACTCTTTCCCTACACGACGCTCTTCCGATCTNNNGAAGCCTATGTTGGTAGGGA
GCACACGTCTGAACTCTTTCCCTACACGACGCTCTTCCGATCTNNNTAGTGAGCCATAATTGTGCCACT
AGCACACGTCTGAACTCTTTCCCTACACGACGCTCTTCCGATCTNNNCTTTTTTACTGCAACCTCGTCCA
GCACACGTCTGAACTCTTTCCCTACACGACGCTCTTCCGATCTNNNAACTTGACCCAGATATACCTGCC
AGCACACGTCTGAACTCTTTCCCTACACGACGCTCTTCCGATCTNNNATGAAGTTGGCCCTTTCGTAGA
AGCACACGTCTGAACTCTTTCCCTACACGACGCTCTTCCGATCTNNNCATCAGGGGATCCAGCTAATAC
AGCACACGTCTGAACTCTTTCCCTACACGACGCTCTTCCGATCTNNNCTCTATCCATGTGAGTCGGTG
GAGCACACGTCTGAACTCTTTCCCTACACGACGCTCTTCCGATCTNNNTCTGTCAGAGGGCTTATTTCA
GAGCACACGTCTGAACTCTTTCCCTACACGACGCTCTTCCGATCTNNNCTTCTCTCTTTTGCTGGCTGT
AGCACACGTCTGAACTCTTTCCCTACACGACGCTCTTCCGATCTNNNCACCCATAGAAGAACCTCACAC
GCACACGTCTGAACTCTTTCCCTACACGACGCTCTTCCGATCTNNNATAGCATGGACCACGTCATATCC
GCACACGTCTGAACTCTTTCCCTACACGACGCTCTTCCGATCTNNNAAATGCAGTCTCCATGTGGTCTA

GAGCACACGTCTGAACTCTTTCCCTACACGACGCTCTTCCGATCTNNNCTAATTAGCTGCAGGAAAAG

AGCACACGTCTGAACTCTTTCCCTACACGACGCTCTTCCGATCTNNNCTTGATTAGCCTAGTCAGTCCC

AGCACACGTCTGAACTCTTTCCCTACACGACGCTCTTCCGATCTNNNCCATGAAAATTCCACTTGTGG

AGCACACGTCTGAACTCTTTCCCTACACGACGCTCTTCCGATCTNNNCCAGCAAGATACAGAACTGGG

GAGCACACGTCTGAACTCTTTCCCTACACGACGCTCTTCCGATCTNNNATGCAACCTCAGAATCACACT

AGCACACGTCTGAACTCTTTCCCTACACGACGCTCTTCCGATCTNNNCAAGAGTCCTATCCTGACGTCA

AGCACACGTCTGAACTCTTTCCCTACACGACGCTCTTCCGATCTNNNGGTTTTTTGGAGACACAGA

AGCACACGTCTGAACTCTTTCCCTACACGACGCTCTTCCGATCTNNNGGTATCAATATAGGGCTTGCC

AGCACACGTCTGAACTCTTTCCCTACACGACGCTCTTCCGATCTNNNGGGCTCTTGTCTTTGTTCTTCA

GAGCACACGTCTGAACTCTTTCCCTACACGACGCTCTTCCGATCTNNNATTCCTCTGTTCTGGCCTTTT

AGCACACGTCTGAACTCTTTCCCTACACGACGCTCTTCCGATCTNNNTCTCTTTCCAGATGAGACATGC

AGCACACGTCTGAACTCTTTCCCTACACGACGCTCTTCCGATCTNNNCCAATGGGTTGAGGGTTCAGA

AGCACACGTCTGAACTCTTTCCCTACACGACGCTCTTCCGATCTNNNGGATGAAGAGAACTGAACAG

AGCACACGTCTGAACTCTTTCCCTACACGACGCTCTTCCGATCTNNNAAGTAGGCACTGTTGTTGAG

AGCACACGTCTGAACTCTTTCCCTACACGACGCTCTTCCGATCTNNNATGACAAAACCAGTGCTTGTCT

AGCACACGTCTGAACTCTTTCCCTACACGACGCTCTTCCGATCTNNNACGATGCTGTTTCTCTCTCTCC

GAGCACACGTCTGAACTCTTTCCCTACACGACGCTCTTCCGATCTNNNCACGATAATTGTTGGAAGCT

GCACACGTCTGAACTCTTTCCCTACACGACGCTCTTCCGATCTNNNAAAACGGTTTGCATTTCCCTTCA

AGCACACGTCTGAACTCTTTCCCTACACGACGCTCTTCCGATCTNNNGATCACGATGTCTCCATTCCTGA

GAGCACACGTCTGAACTCTTTCCCTACACGACGCTCTTCCGATCTNNNTTTCAGAGAGCCTTGAATGA

GAGCACACGTCTGAACTCTTTCCCTACACGACGCTCTTCCGATCTNNNTGGACCTCTTGGACATGCATA

AGCACACGTCTGAACTCTTTCCCTACACGACGCTCTTCCGATCTNNNGGAATAGAATCTGGGGGCCAT

AGCACACGTCTGAACTCTTTCCCTACACGACGCTCTTCCGATCTNNNACAGGTATCCTAGAGAGCACT

AGCACACGTCTGAACTCTTTCCCTACACGACGCTCTTCCGATCTNNNGAACTGGGATCTGATCTGGG

AGCACACGTCTGAACTCTTTCCCTACACGACGCTCTTCCGATCTNNNTCTTTAGCCAAAGGGGAGGAA

AGCACACGTCTGAACTCTTTCCCTACACGACGCTCTTCCGATCTNNNTTAGTAGGCACCCTTTCTGGA

GAGCACACGTCTGAACTCTTTCCCTACACGACGCTCTTCCGATCTNNNTACTCATCTCCCGAGTGTACT

GCACACGTCTGAACTCTTTCCCTACACGACGCTCTTCCGATCTNNNACGAAAAACTGTGTATGTGTGC

GCACACGTCTGAACTCTTTCCCTACACGACGCTCTTCCGATCTNNNGAATTGCTTGAGCCCAGGATTT

AGCACACGTCTGAACTCTTTCCCTACACGACGCTCTTCCGATCTNNNTCTCTTTTCCTCTCTTTGGGGTG

AGCACACGTCTGAACTCTTTCCCTACACGACGCTCTTCCGATCTNNNAAGAGAGTTGACAGCAAAAT

AGCACACGTCTGAACTCTTTCCCTACACGACGCTCTTCCGATCTNNNGGGTTCTCCTTTGTCCTTTCTCT

GAGCACACGTCTGAACTCTTTCCCTACACGACGCTCTTCCGATCTNNNAGGAAAGAACAAACAACGG

GCACACGTCTGAACTCTTTCCCTACACGACGCTCTTCCGATCTNNNGTGCCTTTGTGTGTCTTACTGTC

GAGCACACGTCTGAACTCTTTCCCTACACGACGCTCTTCCGATCTNNNTTCCACTGCAGTTTGAAACAT

AGCACACGTCTGAACTCTTTCCCTACACGACGCTCTTCCGATCTNNNCTCATACCACTTCAGAGCCTTGT

GCACACGTCTGAACTCTTTCCCTACACGACGCTCTTCCGATCTNNNACAGTGGAAGAAGAAGTGTCG

AGCACACGTCTGAACTCTTTCCCTACACGACGCTCTTCCGATCTNNNTCCTAGGGCTACAACTTTGCAT

AGCACACGTCTGAACTCTTTCCCTACACGACGCTCTTCCGATCTNNNCATGGAAGATTTGAGTCCTGG

GCACACGTCTGAACTCTTTCCCTACACGACGCTCTTCCGATCTNNNTTGCCCTTCTGGATGAAATCTCAC

AGCACACGTCTGAACTCTTTCCCTACACGACGCTCTTCCGATCTNNNTTGCTTAGATCCACACTCACAC

AGCACACGTCTGAACTCTTTCCCTACACGACGCTCTTCCGATCTNNNCAATTGAATCAACAGCACCCA

AGCACACGTCTGAACTCTTTCCCTACACGACGCTCTTCCGATCTNNNTCACCTCCGAGTTTTCAGACAA

GCACACGTCTGAACTCTTTCCCTACACGACGCTCTTCCGATCTNNNGAGCTTCTAAACCCACCTCAGA
GCACACGTCTGAACTCTTTCCCTACACGACGCTCTTCCGATCTNNNAAACAAGCAAGTCAGAGGATCC
AGCACACGTCTGAACTCTTTCCCTACACGACGCTCTTCCGATCTNNNTGAGCCACATAGCAAGATCTCA
AGCACACGTCTGAACTCTTTCCCTACACGACGCTCTTCCGATCTNNNAGGTGCTGGGAAAGAAAGAA
GCACACGTCTGAACTCTTTCCCTACACGACGCTCTTCCGATCTNNNGGGGAGGGGCAATATTTTCAA
AGCACACGTCTGAACTCTTTCCCTACACGACGCTCTTCCGATCTNNNGGGCCTGAGTTATTTGGGCTA
GCACACGTCTGAACTCTTTCCCTACACGACGCTCTTCCGATCTNNNAGGGCTTCTACTACTGGGAGAT
GCACACGTCTGAACTCTTTCCCTACACGACGCTCTTCCGATCTNNNTGTTGAGCCATGTGCAATAGAG
GAGCACACGTCTGAACTCTTTCCCTACACGACGCTCTTCCGATCTNNNTACCTCCTTCTGAACTTCTGT
AGCACACGTCTGAACTCTTTCCCTACACGACGCTCTTCCGATCTNNNCAGACTGAGAGCTATTGGGT
GCACACGTCTGAACTCTTTCCCTACACGACGCTCTTCCGATCTNNNTGTAGGTTCAGATGAGACTGCA
AGCACACGTCTGAACTCTTTCCCTACACGACGCTCTTCCGATCTNNNGTCAGCATGGCAGATAAGTCA
GCACACGTCTGAACTCTTTCCCTACACGACGCTCTTCCGATCTNNNAGACCTGAGGTAGAGGACAAA
AGCACACGTCTGAACTCTTTCCCTACACGACGCTCTTCCGATCTNNNGATCCATGGATGGGTTTGGA
AGCACACGTCTGAACTCTTTCCCTACACGACGCTCTTCCGATCTNNNGTTTCCAGAGGAGAGTTGGA
AGCACACGTCTGAACTCTTTCCCTACACGACGCTCTTCCGATCTNNNGCTATAGCCTGGAGTTGTAGA
AGCACACGTCTGAACTCTTTCCCTACACGACGCTCTTCCGATCTNNNTCAGTAATAGGTTTCCCTTGG
AGCACACGTCTGAACTCTTTCCCTACACGACGCTCTTCCGATCTNNNATTCAATTCACCTGGCTGTCTT
GCACACGTCTGAACTCTTTCCCTACACGACGCTCTTCCGATCTNNNAAACCCACATGACAAATCAGCTG
AGCACACGTCTGAACTCTTTCCCTACACGACGCTCTTCCGATCTNNNGGCATGAGAATGAAAATGGC
AGCACACGTCTGAACTCTTTCCCTACACGACGCTCTTCCGATCTNNNTTTCCTGTCTCTTGCTGATGTG
AGCACACGTCTGAACTCTTTCCCTACACGACGCTCTTCCGATCTNNNTATTGGAGTGTTGCTTCTCCCA
AGCACACGTCTGAACTCTTTCCCTACACGACGCTCTTCCGATCTNNNTGGAAAAACAAGCCAAGGAGG
AGCACACGTCTGAACTCTTTCCCTACACGACGCTCTTCCGATCTNNNTTAGAATGAACTGATGGGAGC
AGCACACGTCTGAACTCTTTCCCTACACGACGCTCTTCCGATCTNNNTCTTTGGTGCTTACAGAACTT
AGCACACGTCTGAACTCTTTCCCTACACGACGCTCTTCCGATCTNNNTAGAAAGTGGTAGCAGCAGA
GAGCACACGTCTGAACTCTTTCCCTACACGACGCTCTTCCGATCTNNNTCATCTGCCCAGACATGTACA
AGCACACGTCTGAACTCTTTCCCTACACGACGCTCTTCCGATCTNNNGGGAACACAATGCCAACACAT
AGCACACGTCTGAACTCTTTCCCTACACGACGCTCTTCCGATCTNNNATTTTGCCTCCTACAGTCTTGG
AGCACACGTCTGAACTCTTTCCCTACACGACGCTCTTCCGATCTNNNCATCCAGGTCAGGGAGGATAT
GCACACGTCTGAACTCTTTCCCTACACGACGCTCTTCCGATCTNNNTTCAAGCCCCGAAAAAATGAGA
AGCACACGTCTGAACTCTTTCCCTACACGACGCTCTTCCGATCTNNNGCAGAACAGTCAAGACACTGA
AGCACACGTCTGAACTCTTTCCCTACACGACGCTCTTCCGATCTNNNATGTCCAGAAAAGTGAAAGCA
AGAGCACACGTCTGAACTCTTTCCCTACACGACGCTCTTCCGATCTNNNGCCTCAAATCTCTGCTCTCT
GCACACGTCTGAACTCTTTCCCTACACGACGCTCTTCCGATCTNNNTATGAGATCTGCCATTGCACTCC
AGCACACGTCTGAACTCTTTCCCTACACGACGCTCTTCCGATCTNNNTCTCAGTCCCTATCTGTCCTCTG
AGCACACGTCTGAACTCTTTCCCTACACGACGCTCTTCCGATCTNNNACGTTGATGCTGGAGAGAGAA
AGCACACGTCTGAACTCTTTCCCTACACGACGCTCTTCCGATCTNNNGAATGAACACAAAAAGCAGC
AGCACACGTCTGAACTCTTTCCCTACACGACGCTCTTCCGATCTNNNCCCACCCTGTTTTCAAAACTGTT
AGCACACGTCTGAACTCTTTCCCTACACGACGCTCTTCCGATCTNNNTGCCTGGCATATCAGTAAGTG
AGCACACGTCTGAACTCTTTCCCTACACGACGCTCTTCCGATCTNNNAAACACATGCCATCACTTCCTT
AGCACACGTCTGAACTCTTTCCCTACACGACGCTCTTCCGATCTNNNGGTCTCCACATCATAAATGCTG
AGCACACGTCTGAACTCTTTCCCTACACGACGCTCTTCCGATCTNNNAAGGGAGAGAGGGGACTTCT

AGCACACGTCTGAACTCTTTCCCTACACGACGCTCTTCCGATCTNNNGTCCTCACTGATAAATGGGCT
GCACACGTCTGAACTCTTTCCCTACACGACGCTCTTCCGATCTNNNACAGAGGGAGAGAAAGAGAA
GCACACGTCTGAACTCTTTCCCTACACGACGCTCTTCCGATCTNNNTGGATGGATGGAAATAAATGC
AGCACACGTCTGAACTCTTTCCCTACACGACGCTCTTCCGATCTNNNTTGGAATTGATGTGGGGTTTT
GCACACGTCTGAACTCTTTCCCTACACGACGCTCTTCCGATCTNNNTGTCTTTTGGACAGAACTGGG
AGCACACGTCTGAACTCTTTCCCTACACGACGCTCTTCCGATCTNNNACAACCATTCATAGTTCATGAG
GAGCACACGTCTGAACTCTTTCCCTACACGACGCTCTTCCGATCTNNNGGAATGAAGTGCCCATTTTG
GAGCACACGTCTGAACTCTTTCCCTACACGACGCTCTTCCGATCTNNNGACAGGAGCAAAAAGGACA
AGCACACGTCTGAACTCTTTCCCTACACGACGCTCTTCCGATCTNNNCCACCTCCTAATGGGAATTCAC
AGCACACGTCTGAACTCTTTCCCTACACGACGCTCTTCCGATCTNNNGAGAGTACAAACCCACCCTTCA
AGCACACGTCTGAACTCTTTCCCTACACGACGCTCTTCCGATCTNNNGTTCTCAACCAGACTGTAGGG
AGCACACGTCTGAACTCTTTCCCTACACGACGCTCTTCCGATCTNNNTGCTTTGGAATCTGTGCTTTT
AGCACACGTCTGAACTCTTTCCCTACACGACGCTCTTCCGATCTNNNCTGATTCTTCCTGAGTCCTCTG
AGCACACGTCTGAACTCTTTCCCTACACGACGCTCTTCCGATCTNNNATAAATCAACTGTCCAGTGGC
AGCACACGTCTGAACTCTTTCCCTACACGACGCTCTTCCGATCTNNNCCCCTTGGTCCAGCTAATAATG
GAGCACACGTCTGAACTCTTTCCCTACACGACGCTCTTCCGATCTNNNGGGCATGCACAATAGGAAAA
GCACACGTCTGAACTCTTTCCCTACACGACGCTCTTCCGATCTNNNCCCTGGTGAATCCTATGGTCTAG
GCACACGTCTGAACTCTTTCCCTACACGACGCTCTTCCGATCTNNNAGTGCCTGATTTCCCTCATGAAT
GAGCACACGTCTGAACTCTTTCCCTACACGACGCTCTTCCGATCTNNNGACAGATAAACTCTCCTGGC
CACACGTCTGAACTCTTTCCCTACACGACGCTCTTCCGATCTNNNAAAGTGCTCATCTCGTACCACTTC
GCACACGTCTGAACTCTTTCCCTACACGACGCTCTTCCGATCTNNNCTGTGTGTCTGTGTGTCCATTT
AGCACACGTCTGAACTCTTTCCCTACACGACGCTCTTCCGATCTNNNTCTTGCCTAGCATTCTGTCCTTT
AGAGCACACGTCTGAACTCTTTCCCTACACGACGCTCTTCCGATCTNNNGAGATCTCAATTCCAGCCCT
AGCACACGTCTGAACTCTTTCCCTACACGACGCTCTTCCGATCTNNNTACCACAACTTCCCAGAACAG
GCACACGTCTGAACTCTTTCCCTACACGACGCTCTTCCGATCTNNNGGGTACTTTGCAATCTGAGCTTC
AGCACACGTCTGAACTCTTTCCCTACACGACGCTCTTCCGATCTNNNAACTGCAGCACAAACCCATAA
AGCACACGTCTGAACTCTTTCCCTACACGACGCTCTTCCGATCTNNNTGTCCATTCATGTACCACCACA
AGCACACGTCTGAACTCTTTCCCTACACGACGCTCTTCCGATCTNNNCAGGTTTACGTCATTCCCCAAA
AGCACACGTCTGAACTCTTTCCCTACACGACGCTCTTCCGATCTNNNGCTTTTGGGCCATTGTTTTGTT
AGCACACGTCTGAACTCTTTCCCTACACGACGCTCTTCCGATCTNNNAGGGAACAGCTGTACTTCCTC
AGCACACGTCTGAACTCTTTCCCTACACGACGCTCTTCCGATCTNNNCCTTCTTATTGCTGTTGCCACA
AGAGCACACGTCTGAACTCTTTCCCTACACGACGCTCTTCCGATCTNNNTTTAGCACTCAAATTCTGCC
AGCACACGTCTGAACTCTTTCCCTACACGACGCTCTTCCGATCTNNNAGAAGAGGCAAAGCTGAGATT
GCACACGTCTGAACTCTTTCCCTACACGACGCTCTTCCGATCTNNNCTGCAGTAAGCTATGATTGCAC
GCACACGTCTGAACTCTTTCCCTACACGACGCTCTTCCGATCTNNNACAAAAATTAGCTGGGTGTGGT
GCACACGTCTGAACTCTTTCCCTACACGACGCTCTTCCGATCTNNNGCAATGTTAGAGGCTTCACTGA
AGCACACGTCTGAACTCTTTCCCTACACGACGCTCTTCCGATCTNNNATTTCCTCGTCCTAATAGGCCT
AGCACACGTCTGAACTCTTTCCCTACACGACGCTCTTCCGATCTNNNACCTGGTCTGTCAATCACATTG
AGCACACGTCTGAACTCTTTCCCTACACGACGCTCTTCCGATCTNNNGAACAGTGACCAATACAGGA
GCACACGTCTGAACTCTTTCCCTACACGACGCTCTTCCGATCTNNNTGTGTGTCCTGAGAGCACAAAT
AGCACACGTCTGAACTCTTTCCCTACACGACGCTCTTCCGATCTNNNCTCTTGAGTAGCTGGGACAAC
AGCACACGTCTGAACTCTTTCCCTACACGACGCTCTTCCGATCTNNNAGCCAGTGAAAGGTATTGCTG
AGCACACGTCTGAACTCTTTCCCTACACGACGCTCTTCCGATCTNNNTTGGTTCCCTTTCCTTTCCTCTT

AGCACACGTCTGAACTCTTTCCCTACACGACGCTCTTCCGATCTNNNGCCTCTCAAGTACCTGGGATTA
AGCACACGTCTGAACTCTTTCCCTACACGACGCTCTTCCGATCTNNNACGGTATTAGGGTTCTCCAGA
GCACACGTCTGAACTCTTTCCCTACACGACGCTCTTCCGATCTNNNCTGGCAGACCTGAACATCATAG
GCACACGTCTGAACTCTTTCCCTACACGACGCTCTTCCGATCTNNNTCCCAAAGTGCTAGACTTACAG
GAGCACACGTCTGAACTCTTTCCCTACACGACGCTCTTCCGATCTNNNAAGTAGCTGGGATTACAGGC
GCACACGTCTGAACTCTTTCCCTACACGACGCTCTTCCGATCTNNNCACAATCTATGTTCCTCTGCTGC
GAGCACACGTCTGAACTCTTTCCCTACACGACGCTCTTCCGATCTNNNCCAATAGGGTGCTTTTCCTCT
GCACACGTCTGAACTCTTTCCCTACACGACGCTCTTCCGATCTNNNACAATTCACTGAGCTCGTTCTCT
GCACACGTCTGAACTCTTTCCCTACACGACGCTCTTCCGATCTNNNCTGAGGGAGGAGGATCACTTG
AGCACACGTCTGAACTCTTTCCCTACACGACGCTCTTCCGATCTNNNCCTCCAGTTCTGTGATTTTG
AGCACACGTCTGAACTCTTTCCCTACACGACGCTCTTCCGATCTNNNCCTGTATTAGTCAGGGTTCTC
AGCACACGTCTGAACTCTTTCCCTACACGACGCTCTTCCGATCTNNNTAGTCAGGGTTCTCTAGAGGG
AGCACACGTCTGAACTCTTTCCCTACACGACGCTCTTCCGATCTNNNTGAATTCAAGTTTCCTCCATGC
GCACACGTCTGAACTCTTTCCCTACACGACGCTCTTCCGATCTNNNAGCCTCTGTTGATTTCTCCTCAGC
GAGCACACGTCTGAACTCTTTCCCTACACGACGCTCTTCCGATCTNNNAAAGAAGTCACTGAGAACCT
AGCACACGTCTGAACTCTTTCCCTACACGACGCTCTTCCGATCTNNNCCTGGGTGACAAAGTGAGAAT
GAGCACACGTCTGAACTCTTTCCCTACACGACGCTCTTCCGATCTNNNTATAAGGTGGAGGTTGCAG
AGCACACGTCTGAACTCTTTCCCTACACGACGCTCTTCCGATCTNNNGGCAACTAAGTGAGATCCTGT
AGCACACGTCTGAACTCTTTCCCTACACGACGCTCTTCCGATCTNNNACATAGTGGTGTGTGCATGT
AGCACACGTCTGAACTCTTTCCCTACACGACGCTCTTCCGATCTNNNACACCTGTAATCTCAGCACTCT
GCACACGTCTGAACTCTTTCCCTACACGACGCTCTTCCGATCTNNNATTACCATGATTTGTCCCTGGCC
GCACACGTCTGAACTCTTTCCCTACACGACGCTCTTCCGATCTNNNTGAGGTGGGAGAATTGCTTGA
AGCACACGTCTGAACTCTTTCCCTACACGACGCTCTTCCGATCTNNNAGTCTAGCTACTTGGGAGGC
AGCACACGTCTGAACTCTTTCCCTACACGACGCTCTTCCGATCTNNNGCTTGGACAAGAGAGTGAG
GAGCACACGTCTGAACTCTTTCCCTACACGACGCTCTTCCGATCTNNNACAGGGCCAAATGGAAAAG
GAGCACACGTCTGAACTCTTTCCCTACACGACGCTCTTCCGATCTNNNTAATCCTAGTACTTTGGGAG
AGCACACGTCTGAACTCTTTCCCTACACGACGCTCTTCCGATCTNNNGGCAACATAGCAAAATCCCAT
AGCACACGTCTGAACTCTTTCCCTACACGACGCTCTTCCGATCTNNNGGCAGGAGGATTTGCAAGAAT
AGCACACGTCTGAACTCTTTCCCTACACGACGCTCTTCCGATCTNNNTCAAGGTTTCTGCAAATGCCAT
AGCACACGTCTGAACTCTTTCCCTACACGACGCTCTTCCGATCTNNNGTTCCTTATGATTTGCCTTGCG
AGCACACGTCTGAACTCTTTCCCTACACGACGCTCTTCCGATCTNNNTCTCAAGGGGAATGCTTTATG
AGCACACGTCTGAACTCTTTCCCTACACGACGCTCTTCCGATCTNNNCTGATGCCTGTAAGAAGTTGC
GCACACGTCTGAACTCTTTCCCTACACGACGCTCTTCCGATCTNNNTTCACATCATTCTCCTGCCTCAGC
AGCACACGTCTGAACTCTTTCCCTACACGACGCTCTTCCGATCTNNNATTGTCTGTCTCCTTTCCTTGG
AGCACACGTCTGAACTCTTTCCCTACACGACGCTCTTCCGATCTNNNGCCACAGATATCTCCAGTGAC
AGCACACGTCTGAACTCTTTCCCTACACGACGCTCTTCCGATCTNNNTATGGTGTGTGTGTGTGAGA
GAGCACACGTCTGAACTCTTTCCCTACACGACGCTCTTCCGATCTNNNTCAGCCTAAATGCTCATCAAC
GCACACGTCTGAACTCTTTCCCTACACGACGCTCTTCCGATCTNNNGAGAAAGACTCAGGTAACCTCC
AGCACACGTCTGAACTCTTTCCCTACACGACGCTCTTCCGATCTNNNAAGCTTTTTGATGTGCTGTTG
AGAGCACACGTCTGAACTCTTTCCCTACACGACGCTCTTCCGATCTNNNAGTGATGTCCTTACAGTCT
AGCACACGTCTGAACTCTTTCCCTACACGACGCTCTTCCGATCTNNNATCTCAGAGCTTTGGGAGTTG
GAGCACACGTCTGAACTCTTTCCCTACACGACGCTCTTCCGATCTNNNTATCATTGCACCACTGAGTTC
GCACACGTCTGAACTCTTTCCCTACACGACGCTCTTCCGATCTNNNGCTGAAGGAGCTTCTGGATTTA

AGCACACGTCTGAACTCTTTCCCTACACGACGCTCTTCCGATCTNNNTATATGTCTCTTGGTTGGCACC
AGCACACGTCTGAACTCTTTCCCTACACGACGCTCTTCCGATCTNNNTGTTACCTACCTTGGCAGCTTA
AGCACACGTCTGAACTCTTTCCCTACACGACGCTCTTCCGATCTNNNTGCATTAGTCAGAGTTCTCCT
GAGCACACGTCTGAACTCTTTCCCTACACGACGCTCTTCCGATCTNNNTGCAAGTCTCATGGTAAACA
GAGCACACGTCTGAACTCTTTCCCTACACGACGCTCTTCCGATCTNNNGGCTGAGGTAAGAGAATTG
GCACACGTCTGAACTCTTTCCCTACACGACGCTCTTCCGATCTNNNGAGGCAGAAGAATCACTTGAAC
AGCACACGTCTGAACTCTTTCCCTACACGACGCTCTTCCGATCTNNNTGGAAAGCAGTTTGGCATGAT
AGCACACGTCTGAACTCTTTCCCTACACGACGCTCTTCCGATCTNNNGACCAACATAGTGAGACCCCAT
GCACACGTCTGAACTCTTTCCCTACACGACGCTCTTCCGATCTNNNCCTCTTTTCATCCCCTAGCAACA
GAGCACACGTCTGAACTCTTTCCCTACACGACGCTCTTCCGATCTNNNATTTATATGCCTCCCCACAGA
AGCACACGTCTGAACTCTTTCCCTACACGACGCTCTTCCGATCTNNNAGTCTTCCTTTCAGAGATGAC
AGCACACGTCTGAACTCTTTCCCTACACGACGCTCTTCCGATCTNNNCTAGTCTCGAACTCCTGACCTC
GAGCACACGTCTGAACTCTTTCCCTACACGACGCTCTTCCGATCTNNNTGTCTCTCTCTCTCTGTCTC
AGCACACGTCTGAACTCTTTCCCTACACGACGCTCTTCCGATCTNNNAGATCACCTGATGTTGGGAGT
GAGCACACGTCTGAACTCTTTCCCTACACGACGCTCTTCCGATCTNNNCTAAGGCAGGAGGATCACTT
GAGCACACGTCTGAACTCTTTCCCTACACGACGCTCTTCCGATCTNNNCCTACACACAAACACACAGA
GCACACGTCTGAACTCTTTCCCTACACGACGCTCTTCCGATCTNNNCTCCATCTCTACCACACACACAT
GAGCACACGTCTGAACTCTTTCCCTACACGACGCTCTTCCGATCTNNNAGAGCTGAACTGAAGGAGA
AGCACACGTCTGAACTCTTTCCCTACACGACGCTCTTCCGATCTNNNCACACATGTTTATGGTGGCAC
GCACACGTCTGAACTCTTTCCCTACACGACGCTCTTCCGATCTNNNAGGCTGAACTCCTTTTGTAGGT
AGCACACGTCTGAACTCTTTCCCTACACGACGCTCTTCCGATCTNNNACTCTTTCAGTTGGTTCCTGTG
AGCACACGTCTGAACTCTTTCCCTACACGACGCTCTTCCGATCTNNNTGTGTGTGTGTGTGTGTAG
GCACACGTCTGAACTCTTTCCCTACACGACGCTCTTCCGATCTNNNCAAACAATTCTCCTGCCTTAGCC
AGCACACGTCTGAACTCTTTCCCTACACGACGCTCTTCCGATCTNNNCCCACAAAGCTAAAGGCATCTT
AGCACACGTCTGAACTCTTTCCCTACACGACGCTCTTCCGATCTNNNCCTGACCTATCTGACCATACGT
AGCACACGTCTGAACTCTTTCCCTACACGACGCTCTTCCGATCTNNNCAGGATAAGCCTCAGGAATTG
AGCACACGTCTGAACTCTTTCCCTACACGACGCTCTTCCGATCTNNNGCAGTGGTATCTACCCCATCAT
AGCACACGTCTGAACTCTTTCCCTACACGACGCTCTTCCGATCTNNNCTGTTGACATTTGGACAGTGC
GAGCACACGTCTGAACTCTTTCCCTACACGACGCTCTTCCGATCTNNNCGTTCCTAAGGAAGCAGGTT
AGCACACGTCTGAACTCTTTCCCTACACGACGCTCTTCCGATCTNNNGCCCAGGAGTTCAAGGTTATA
GCACACGTCTGAACTCTTTCCCTACACGACGCTCTTCCGATCTNNNCAAGGATATTGGCCAGCAGTTT
GCACACGTCTGAACTCTTTCCCTACACGACGCTCTTCCGATCTNNNCACCGGAGCCGTATCAATACTA
AGCACACGTCTGAACTCTTTCCCTACACGACGCTCTTCCGATCTNNNGGGGAAAGCATGTTTATGTG
AGCACACGTCTGAACTCTTTCCCTACACGACGCTCTTCCGATCTNNNCAGCATAGTGAAACCCTGTCT
AGCACACGTCTGAACTCTTTCCCTACACGACGCTCTTCCGATCTNNNAGAAGAATCACTTGAACCCAG
GAGCACACGTCTGAACTCTTTCCCTACACGACGCTCTTCCGATCTNNNCCAGGAGTTCAAGATAAGCC
AGCACACGTCTGAACTCTTTCCCTACACGACGCTCTTCCGATCTNNNTGCCTTTTTCCTCACATGACAA
AGCACACGTCTGAACTCTTTCCCTACACGACGCTCTTCCGATCTNNNAAAATTAGCCAGGCATAGTGG
GCACACGTCTGAACTCTTTCCCTACACGACGCTCTTCCGATCTNNNAAGAGGGACTAGGACATTGCTT
AGCACACGTCTGAACTCTTTCCCTACACGACGCTCTTCCGATCTNNNTTTGTTTTGCAGGAACCACACA
GAGCACACGTCTGAACTCTTTCCCTACACGACGCTCTTCCGATCTNNNAGGGAATTCCAAATCTCATG
AGCACACGTCTGAACTCTTTCCCTACACGACGCTCTTCCGATCTNNNTGAGGCCAGCATTATCCTGAT
AGCACACGTCTGAACTCTTTCCCTACACGACGCTCTTCCGATCTNNNCTGAGACAGGCGAATCACTTG

GAGCACACGTCTGAACTCTTTCCCTACACGACGCTCTTCCGATCTNNNAAATATGGAACCAGCCCAAAT
AGCACACGTCTGAACTCTTTCCCTACACGACGCTCTTCCGATCTNNNTCTAGCTCCATCCATGTTGCTAC
GAGCACACGTCTGAACTCTTTCCCTACACGACGCTCTTCCGATCTNNNGTGTGTGTGTGTGTGTGTAT
AGCACACGTCTGAACTCTTTCCCTACACGACGCTCTTCCGATCTNNNCAGGAGGAGGAAAAAGGCAT
AGCACACGTCTGAACTCTTTCCCTACACGACGCTCTTCCGATCTNNNCTTTGGGAGTTCAAGGTGAGA
AGCACACGTCTGAACTCTTTCCCTACACGACGCTCTTCCGATCTNNNAAGACAAGAAAAGGCAGAGTG
AGCACACGTCTGAACTCTTTCCCTACACGACGCTCTTCCGATCTNNNAGCAAAACCCTGTCTCTACCAA
GAGCACACGTCTGAACTCTTTCCCTACACGACGCTCTTCCGATCTNNNCTATGATCACACCACTGCACT
AGCACACGTCTGAACTCTTTCCCTACACGACGCTCTTCCGATCTNNNCAGAGATAGAAAGAGAGGG
GAGCACACGTCTGAACTCTTTCCCTACACGACGCTCTTCCGATCTNNNCGAGACTCCCTCTCAAACAAA
GAGCACACGTCTGAACTCTTTCCCTACACGACGCTCTTCCGATCTNNNGGCACTGAATCACTGGCTAAT
AGCACACGTCTGAACTCTTTCCCTACACGACGCTCTTCCGATCTNNNGGAGATTGCAGTGAGCCAAGA
AGAGCACACGTCTGAACTCTTTCCCTACACGACGCTCTTCCGATCTNNNAGAGATGGACTATTAGGCA
GAGCACACGTCTGAACTCTTTCCCTACACGACGCTCTTCCGATCTNNNGATTACAGTCTAGTGGCAG
AGCACACGTCTGAACTCTTTCCCTACACGACGCTCTTCCGATCTNNNACATGCACGCATATGTTCATTG
GAGCACACGTCTGAACTCTTTCCCTACACGACGCTCTTCCGATCTNNNCTGAGATCCAAGGACTGAGA
AGAGCACACGTCTGAACTCTTTCCCTACACGACGCTCTTCCGATCTNNNACACACAGTGAGATCCTGAG
GAGCACACGTCTGAACTCTTTCCCTACACGACGCTCTTCCGATCTNNNTGAGGCAGGAGAATCTCTTG
AGCACACGTCTGAACTCTTTCCCTACACGACGCTCTTCCGATCTNNNGAAGGTTGAGGTGGGAGAAT
AGCACACGTCTGAACTCTTTCCCTACACGACGCTCTTCCGATCTNNNATACTCCTGCTGTGGTTGCTTTA
AGCACACGTCTGAACTCTTTCCCTACACGACGCTCTTCCGATCTNNNCAGCAGCATGAAGAAATCCAA
GAGCACACGTCTGAACTCTTTCCCTACACGACGCTCTTCCGATCTNNNCACCACACCTGGCCAGAAATA
GAGCACACGTCTGAACTCTTTCCCTACACGACGCTCTTCCGATCTNNNTCTTTGTGTGTGTGTGTGT
GCACACGTCTGAACTCTTTCCCTACACGACGCTCTTCCGATCTNNNTCCCATGAGAACTTCCACATAGC
AGAGCACACGTCTGAACTCTTTCCCTACACGACGCTCTTCCGATCTNNNACTGGCTCTAACTTGGAGAT
AGCACACGTCTGAACTCTTTCCCTACACGACGCTCTTCCGATCTNNNTCCCAAAAACATTGCCAATCAG
AGCACACGTCTGAACTCTTTCCCTACACGACGCTCTTCCGATCTNNNCCACTGATTCAAGGTCTCTCAC
AGCACACGTCTGAACTCTTTCCCTACACGACGCTCTTCCGATCTNNNAATCCTAGTAGTTTGCTGGTG
GAGCACACGTCTGAACTCTTTCCCTACACGACGCTCTTCCGATCTNNNCTCCCATCGCTAAGACACTAA
GAGCACACGTCTGAACTCTTTCCCTACACGACGCTCTTCCGATCTNNNCATCTCTGCCTCCATGGTTAC
GAGCACACGTCTGAACTCTTTCCCTACACGACGCTCTTCCGATCTNNNAGGAGAACCATTTGAACCTG
AGCACACGTCTGAACTCTTTCCCTACACGACGCTCTTCCGATCTNNNGGTTTCTGAAAAGTGGCAA
GCACACGTCTGAACTCTTTCCCTACACGACGCTCTTCCGATCTNNNGCCTGTGGCTTTTTCAAAAGGT
GAGCACACGTCTGAACTCTTTCCCTACACGACGCTCTTCCGATCTNNNCTGTAATCCCAGCTACTCAGG
GAGCACACGTCTGAACTCTTTCCCTACACGACGCTCTTCCGATCTNNNAATGCTCATCAAACAACGAG
GAGCACACGTCTGAACTCTTTCCCTACACGACGCTCTTCCGATCTNNNGATTGCTTTGAGCTCAGGAG
GAGCACACGTCTGAACTCTTTCCCTACACGACGCTCTTCCGATCTNNNGTTCGCTTGAAGCCAGAAGT
AGCACACGTCTGAACTCTTTCCCTACACGACGCTCTTCCGATCTNNNCTTAGGATAATGGCCTCCAGCT
AGCACACGTCTGAACTCTTTCCCTACACGACGCTCTTCCGATCTNNNCACTGCAGGTATTCCAACCACT
AGCACACGTCTGAACTCTTTCCCTACACGACGCTCTTCCGATCTNNNCTCTAGGAGACAGAGGGAGG
GCACACGTCTGAACTCTTTCCCTACACGACGCTCTTCCGATCTNNNATGATCCCTTCCCCTTCATTCACC
AGCACACGTCTGAACTCTTTCCCTACACGACGCTCTTCCGATCTNNNCTCCCCGACTATATTGCATCCA
GAGCACACGTCTGAACTCTTTCCCTACACGACGCTCTTCCGATCTNNNAGGTTGAGTCAGGAGAATTG

GAGCACACGTCTGAACTCTTTCCCTACACGACGCTCTTCCGATCTNNNAGTGAGCCTAGATCATGCCATT
GAGCACACGTCTGAACTCTTTCCCTACACGACGCTCTTCCGATCTNNNAACTCAGTCCCACCCTAAGGA
GAGCACACGTCTGAACTCTTTCCCTACACGACGCTCTTCCGATCTNNNCCTATGAAGATTCAGGGGCCT
AGAGCACACGTCTGAACTCTTTCCCTACACGACGCTCTTCCGATCTNNNTGAATTCACGGGGTCACTTAT
GAGCACACGTCTGAACTCTTTCCCTACACGACGCTCTTCCGATCTNNNTCATCTTTTCCTGATCCAACCC
GCACACGTCTGAACTCTTTCCCTACACGACGCTCTTCCGATCTNNNGTTGAGGATCCCATCTTACCCAAC
AGCACACGTCTGAACTCTTTCCCTACACGACGCTCTTCCGATCTNNNCTTTGTGTCTGTGTGTGTGTG
AGAGCACACGTCTGAACTCTTTCCCTACACGACGCTCTTCCGATCTNNNACAAGAGCGAACTCCATCTCA
GAGCACACGTCTGAACTCTTTCCCTACACGACGCTCTTCCGATCTNNNGTCCCCCTAGAGAACCTTGACT
GAGCACACGTCTGAACTCTTTCCCTACACGACGCTCTTCCGATCTNNNTCCTATGTTTGTTGCAGGACTG
AGCACACGTCTGAACTCTTTCCCTACACGACGCTCTTCCGATCTNNNCATCCTGGCTAACACAGTGAAA
GAGCACACGTCTGAACTCTTTCCCTACACGACGCTCTTCCGATCTNNNCCCCACCCAGAAATCTGATCAT
GAGCACACGTCTGAACTCTTTCCCTACACGACGCTCTTCCGATCTNNNAGTTGCAGTGAGCCAAGATCT
AGCACACGTCTGAACTCTTTCCCTACACGACGCTCTTCCGATCTNNNTGAAACTTGGGCAATGAGTCCT
AGCACACGTCTGAACTCTTTCCCTACACGACGCTCTTCCGATCTNNNCCCCTTTTCTTTGCCCTCTCTATCA
AGCACACGTCTGAACTCTTTCCCTACACGACGCTCTTCCGATCTNNNTAGGTGACAGAGTGATACCCTC
GAGCACACGTCTGAACTCTTTCCCTACACGACGCTCTTCCGATCTNNNAACTCCTGAGCTCAAGTGATCT
AGCACACGTCTGAACTCTTTCCCTACACGACGCTCTTCCGATCTNNNCAAGTGTTTTGGAGGGGGATA
AGAGCACACGTCTGAACTCTTTCCCTACACGACGCTCTTCCGATCTNNNGAGCAAGACTCCGTCTCAAAA
AGCACACGTCTGAACTCTTTCCCTACACGACGCTCTTCCGATCTNNNTCCCTACTTGGCCTTCTCTGATA
AGCACACGTCTGAACTCTTTCCCTACACGACGCTCTTCCGATCTNNNGGCTCACATTGTATTTCCACTGC
GAGCACACGTCTGAACTCTTTCCCTACACGACGCTCTTCCGATCTNNNCAGGAGAATTGCTTCAACCCAT
GAGCACACGTCTGAACTCTTTCCCTACACGACGCTCTTCCGATCTNNNTATACTTGGCCCCTTCCCTTTC
GAGCACACGTCTGAACTCTTTCCCTACACGACGCTCTTCCGATCTNNNGGCAACATAGCAAGACTCCAT
AGCACACGTCTGAACTCTTTCCCTACACGACGCTCTTCCGATCTNNNAGGTTGCAGTGAGTCAAGATCA
AGAGCACACGTCTGAACTCTTTCCCTACACGACGCTCTTCCGATCTNNNATGGAGAGAGACAGACAGA
AGAGCACACGTCTGAACTCTTTCCCTACACGACGCTCTTCCGATCTNNNCAGAGAATCACTTGAACCTC
GAGCACACGTCTGAACTCTTTCCCTACACGACGCTCTTCCGATCTNNNTGAGCCAAGATGGTGCCATTA
AGCACACGTCTGAACTCTTTCCCTACACGACGCTCTTCCGATCTNNNGTCAAACCCAGGAACTCAAATGC
GAGCACACGTCTGAACTCTTTCCCTACACGACGCTCTTCCGATCTNNNCACCACACCTGGCTATTTTTCTG
GAGCACACGTCTGAACTCTTTCCCTACACGACGCTCTTCCGATCTNNNTGTGGATAGTGTGATGGCATT
GAGCACACGTCTGAACTCTTTCCCTACACGACGCTCTTCCGATCTNNNATCCAGGTTGGCCCAGTATAA
GAGCACACGTCTGAACTCTTTCCCTACACGACGCTCTTCCGATCTNNNCCAGAGGAAAACCTCATGTAC
AGCACACGTCTGAACTCTTTCCCTACACGACGCTCTTCCGATCTNNNAGCTCTCAATGAAAACAAATGC
AGCACACGTCTGAACTCTTTCCCTACACGACGCTCTTCCGATCTNNNAAGTAGCAGAGACTAAAGGCA
AGCACACGTCTGAACTCTTTCCCTACACGACGCTCTTCCGATCTNNNCTCTCAGGCATAACAGAGACCA
AGCACACGTCTGAACTCTTTCCCTACACGACGCTCTTCCGATCTNNNCTCTCTCTCTCTGTGTGTGTGTC
AGAGCACACGTCTGAACTCTTTCCCTACACGACGCTCTTCCGATCTNNNGACCCTGATTGAAGAGTGTC
AGCACACGTCTGAACTCTTTCCCTACACGACGCTCTTCCGATCTNNNTCCCAGATCCAAAATCCCCAAATC
GAGCACACGTCTGAACTCTTTCCCTACACGACGCTCTTCCGATCTNNNCCAGGATCCCTTTCCTTTTGCTA
GAGCACACGTCTGAACTCTTTCCCTACACGACGCTCTTCCGATCTNNNGTGGATCATGAGGTCAGGAG
AGCACACGTCTGAACTCTTTCCCTACACGACGCTCTTCCGATCTNNNCCACATCTCCTGTCATGGATCAT
GAGCACACGTCTGAACTCTTTCCCTACACGACGCTCTTCCGATCTNNNGAGTTTGAGACCAGCCTGACT

GCACACGTCTGAACTCTTTCCCTACACGACGCTCTTCCGATCTNNNCTGCAATGGCTTTACAGTGAGA
AGCACACGTCTGAACTCTTTCCCTACACGACGCTCTTCCGATCTNNNCAAGTAATTCAAGACCAGCCT
GCACACGTCTGAACTCTTTCCCTACACGACGCTCTTCCGATCTNNNTCTCAAACTTGTGCAACCATCAC
GAGCACACGTCTGAACTCTTTCCCTACACGACGCTCTTCCGATCTNNNGTGTGTGTGCGTGTGTGTAT
AGCACACGTCTGAACTCTTTCCCTACACGACGCTCTTCCGATCTNNNATGCGTCTAAGTTTCACCTACG
AGAGCACACGTCTGAACTCTTTCCCTACACGACGCTCTTCCGATCTNNNTTTGTCCAGTCCCCCAAACA
AGAGCACACGTCTGAACTCTTTCCCTACACGACGCTCTTCCGATCTNNNAGAAGCAGAGAGAGACA
AGCACACGTCTGAACTCTTTCCCTACACGACGCTCTTCCGATCTNNNGCTCCTCCCTAGCTCATTCTAT
AGCACACGTCTGAACTCTTTCCCTACACGACGCTCTTCCGATCTNNNGCTGACTCCTGCAATTCCAATA
GAGCACACGTCTGAACTCTTTCCCTACACGACGCTCTTCCGATCTNNNTGTCCCATTGATGGCAGTAAT
AGCACACGTCTGAACTCTTTCCCTACACGACGCTCTTCCGATCTNNNCGGTGGTTCACGTCTGTAATT
GAGCACACGTCTGAACTCTTTCCCTACACGACGCTCTTCCGATCTNNNTGTGTGTATATGTGTGTGTG
GAGCACACGTCTGAACTCTTTCCCTACACGACGCTCTTCCGATCTNNNAGTAAGCCAAGACTACACCA
AGCACACGTCTGAACTCTTTCCCTACACGACGCTCTTCCGATCTNNNATCTCGAATTGCCTCTGACCTTT
AGCACACGTCTGAACTCTTTCCCTACACGACGCTCTTCCGATCTNNNGTTCCTTATGATTTGCCTTGCG
GCACACGTCTGAACTCTTTCCCTACACGACGCTCTTCCGATCTNNNGTGTGTATGTCTGTGTGTGTGT
GCACACGTCTGAACTCTTTCCCTACACGACGCTCTTCCGATCTNNNAGATCCTGTTAGTCACTACGGG
GCACACGTCTGAACTCTTTCCCTACACGACGCTCTTCCGATCTNNNTTGTTTTCCTGACCCTGTGTGT
GCACACGTCTGAACTCTTTCCCTACACGACGCTCTTCCGATCTNNNGACAAAGCAAGACCCTACCTCT
AGCACACGTCTGAACTCTTTCCCTACACGACGCTCTTCCGATCTNNNCGAGAGAATCACCCTTTGCTTT
AGCACACGTCTGAACTCTTTCCCTACACGACGCTCTTCCGATCTNNNATGTCCAATTGTTCCAGTACCG
AGCACACGTCTGAACTCTTTCCCTACACGACGCTCTTCCGATCTNNNACACTGCTTTTACCACCTAACC
AGCACACGTCTGAACTCTTTCCCTACACGACGCTCTTCCGATCTNNNCCATGATTGGACTAGGATGGG
GCACACGTCTGAACTCTTTCCCTACACGACGCTCTTCCGATCTNNNATAGCCTACTGTTGACTGGAAG
AGCACACGTCTGAACTCTTTCCCTACACGACGCTCTTCCGATCTNNNCTTGTAAGTGGACCTGTGGA
AGCACACGTCTGAACTCTTTCCCTACACGACGCTCTTCCGATCTNNNCGTACATACACACACGATCTCC
AGCACACGTCTGAACTCTTTCCCTACACGACGCTCTTCCGATCTNNNCGAGATCGTGTCATTGTACTC
GCACACGTCTGAACTCTTTCCCTACACGACGCTCTTCCGATCTNNNATGCTTTAGGCTGTAGACCACA
GAGCACACGTCTGAACTCTTTCCCTACACGACGCTCTTCCGATCTNNNCTCTTCTCTGTTTGCCCCACT
CACACGTCTGAACTCTTTCCCTACACGACGCTCTTCCGATCTNNNTGCCAGAACAAGAAGTCTCTCTCC
AGCACACGTCTGAACTCTTTCCCTACACGACGCTCTTCCGATCTNNNCTCCTACCTTGGTCTCCTGAGT
GAGCACACGTCTGAACTCTTTCCCTACACGACGCTCTTCCGATCTNNNGGTGGAATCTAAATGTGAC
GAGCACACGTCTGAACTCTTTCCCTACACGACGCTCTTCCGATCTNNNAGACATTGCAGTGAGCTGA
AGCACACGTCTGAACTCTTTCCCTACACGACGCTCTTCCGATCTNNNAATGCAGCCTACCTACATTTCC
AGCACACGTCTGAACTCTTTCCCTACACGACGCTCTTCCGATCTNNNGGTGACAAGAGCGAAATTCT
GAGCACACGTCTGAACTCTTTCCCTACACGACGCTCTTCCGATCTNNNCAGAAGAATTGCTTGAACCT
GCACACGTCTGAACTCTTTCCCTACACGACGCTCTTCCGATCTNNNGGGTAAGAGAGGAACAGACTT
GAGCACACGTCTGAACTCTTTCCCTACACGACGCTCTTCCGATCTNNNGTTTTAGCCCAGGGTTTCAA
GAGCACACGTCTGAACTCTTTCCCTACACGACGCTCTTCCGATCTNNNGAGCAAGACTCCGTCTCGAA
AGCACACGTCTGAACTCTTTCCCTACACGACGCTCTTCCGATCTNNNGCTTTGACTCTGCTGTCCATTT
GAGCACACGTCTGAACTCTTTCCCTACACGACGCTCTTCCGATCTNNNTTCCCATCAAATGCACAGCTT
GCACACGTCTGAACTCTTTCCCTACACGACGCTCTTCCGATCTNNNAATTTTCCTCTGCCACACCTGTA
GCACACGTCTGAACTCTTTCCCTACACGACGCTCTTCCGATCTNNNACAGTCACACAATAAGAAGGT

AGCACACGTCTGAACTCTTTCCCTACACGACGCTCTTCCGATCTNNNTATAAGGCTAGCATGGAGTGT
AGCACACGTCTGAACTCTTTCCCTACACGACGCTCTTCCGATCTNNNTGGGGACAGTTCTATAGACAG
GAGCACACGTCTGAACTCTTTCCCTACACGACGCTCTTCCGATCTNNNAGAGTAGGACCCAAAAATCC
GAGCACACGTCTGAACTCTTTCCCTACACGACGCTCTTCCGATCTNNNATACCAGCCTGGACAATATAG
GCACACGTCTGAACTCTTTCCCTACACGACGCTCTTCCGATCTNNNCTTCCAAACTCAACTTTCCACCCC
GAGCACACGTCTGAACTCTTTCCCTACACGACGCTCTTCCGATCTNNNGATCAAGACCATCCTGGCTAA
AGCACACGTCTGAACTCTTTCCCTACACGACGCTCTTCCGATCTNNNGAAACAGAAACCCGTGAAGAC
AGCACACGTCTGAACTCTTTCCCTACACGACGCTCTTCCGATCTNNNATCCTGGTGTTAGCTTTTAGGC
GCACACGTCTGAACTCTTTCCCTACACGACGCTCTTCCGATCTNNNTATGATGACACCATGCACTCCAG
AGCACACGTCTGAACTCTTTCCCTACACGACGCTCTTCCGATCTNNNAACCCCCTATGGATACTGAAG
GAGCACACGTCTGAACTCTTTCCCTACACGACGCTCTTCCGATCTNNNTGGCTGTCCTATTTACACACAC
AGCACACGTCTGAACTCTTTCCCTACACGACGCTCTTCCGATCTNNNTGGGCATAGGAAGCTTTTTCAT
AGCACACGTCTGAACTCTTTCCCTACACGACGCTCTTCCGATCTNNNCCTGGACAGATTTTTTCAGCCA
GAGCACACGTCTGAACTCTTTCCCTACACGACGCTCTTCCGATCTNNNTGTTTGTGTGTGTATGTGTG
GAGCACACGTCTGAACTCTTTCCCTACACGACGCTCTTCCGATCTNNNACTCCTTAGCCTTCTGAGTAC
GAGCACACGTCTGAACTCTTTCCCTACACGACGCTCTTCCGATCTNNNGGGTGGGTGTTATGAATTGG
AGCACACGTCTGAACTCTTTCCCTACACGACGCTCTTCCGATCTNNNTCAAGGGATTCTTCTGTCTCAG
AGCACACGTCTGAACTCTTTCCCTACACGACGCTCTTCCGATCTNNNGCAGATGACATGCATGATCTTG
AGCACACGTCTGAACTCTTTCCCTACACGACGCTCTTCCGATCTNNNGTTCAGAATGTTCACAAGGGG
GAGCACACGTCTGAACTCTTTCCCTACACGACGCTCTTCCGATCTNNNCTCTTCTTCTACCCCTTTGGCAT
GAGCACACGTCTGAACTCTTTCCCTACACGACGCTCTTCCGATCTNNNCAACAGGATTTGGAAGAAGG
GAGCACACGTCTGAACTCTTTCCCTACACGACGCTCTTCCGATCTNNNACACACATGGACTCTTTCTCA
GAGCACACGTCTGAACTCTTTCCCTACACGACGCTCTTCCGATCTNNNAGATGGGGTTTCACTATGTTG
AGAGCACACGTCTGAACTCTTTCCCTACACGACGCTCTTCCGATCTNNNACCCATAGTGAGAGAGAGA
AGCACACGTCTGAACTCTTTCCCTACACGACGCTCTTCCGATCTNNNGAGTCAGGTGTATGGCCTTTT
GAGCACACGTCTGAACTCTTTCCCTACACGACGCTCTTCCGATCTNNNGATCAAGACCATCCTGGCTA
AGCACACGTCTGAACTCTTTCCCTACACGACGCTCTTCCGATCTNNNTTGCTTGAAGCTAGGAGTTGG
AGCACACGTCTGAACTCTTTCCCTACACGACGCTCTTCCGATCTNNNAGCCTGGATAATAGAGCAAGA
GAGCACACGTCTGAACTCTTTCCCTACACGACGCTCTTCCGATCTNNNCAAAGCGTTGGAATTACAGG
AGCACACGTCTGAACTCTTTCCCTACACGACGCTCTTCCGATCTNNNATCTTTTGAGCCCAGGAGTTTG
AGCACACGTCTGAACTCTTTCCCTACACGACGCTCTTCCGATCTNNNAACTCAGTTTGTTCTAGGAGGG
AGAGCACACGTCTGAACTCTTTCCCTACACGACGCTCTTCCGATCTNNNTTCAGGTTGGCTTTCAGAAG
AGCACACGTCTGAACTCTTTCCCTACACGACGCTCTTCCGATCTNNNCTCTGGGTTAGCTACTTGGTTG
AGCACACGTCTGAACTCTTTCCCTACACGACGCTCTTCCGATCTNNNCTCTGGGTTAGCTACTTGGTTG
GAGCACACGTCTGAACTCTTTCCCTACACGACGCTCTTCCGATCTNNNCAGAGAGGCCGATAACAGG
GAGCACACGTCTGAACTCTTTCCCTACACGACGCTCTTCCGATCTNNNCTCTACCTTGACCTGAATGCAC
AGAGCACACGTCTGAACTCTTTCCCTACACGACGCTCTTCCGATCTNNNCTGTGTGTGTCTGCATATGT
AGCACACGTCTGAACTCTTTCCCTACACGACGCTCTTCCGATCTNNNCGTCCCAAAACAACAACAACAA
AGAGCACACGTCTGAACTCTTTCCCTACACGACGCTCTTCCGATCTNNNAGGCAGACCCCATCTCAAAA
AGCACACGTCTGAACTCTTTCCCTACACGACGCTCTTCCGATCTNNNGAGTACTTGAGCTCAGGAGTT
AGCACACGTCTGAACTCTTTCCCTACACGACGCTCTTCCGATCTNNNTGCTTGAGGTGATGATGGATA
AGCACACGTCTGAACTCTTTCCCTACACGACGCTCTTCCGATCTNNNGACAACCAGAGTGAGACTGTG
GAGCACACGTCTGAACTCTTTCCCTACACGACGCTCTTCCGATCTNNNCCTGGGCAATAAGAGCAAAA

AGCACACGTCTGAACTCTTTCCCTACACGACGCTCTTCCGATCTNNNTGGATAATGAAGTGCAGGGT
AGCACACGTCTGAACTCTTTCCCTACACGACGCTCTTCCGATCTNNNCATAGCTTACTGCAACGTTGAC
GAGCACACGTCTGAACTCTTTCCCTACACGACGCTCTTCCGATCTNNNTTTGAAGTTGGGTAGCATGA
AGCACACGTCTGAACTCTTTCCCTACACGACGCTCTTCCGATCTNNNTCTGCCACCAAGGATAATTTCA
AGCACACGTCTGAACTCTTTCCCTACACGACGCTCTTCCGATCTNNNATATCCCACCACCTGATCCTAC
AGCACACGTCTGAACTCTTTCCCTACACGACGCTCTTCCGATCTNNNCTGCCAAGTAGCTGGGACTAT
AGCACACGTCTGAACTCTTTCCCTACACGACGCTCTTCCGATCTNNNCTCAAGCAATCCTCCAACCTCA
AGCACACGTCTGAACTCTTTCCCTACACGACGCTCTTCCGATCTNNNAAATACTGACATACCGCTAGG
AGCACACGTCTGAACTCTTTCCCTACACGACGCTCTTCCGATCTNNNGATGCTGATGCCTTCCCTATGT
GCACACGTCTGAACTCTTTCCCTACACGACGCTCTTCCGATCTNNNAGCAAACCTGTCTCTAAACCTGT
GCACACGTCTGAACTCTTTCCCTACACGACGCTCTTCCGATCTNNNCAAGCCTGTCAACATCAAGAAG
AGCACACGTCTGAACTCTTTCCCTACACGACGCTCTTCCGATCTNNNAAGCACTATACTAGGCAATGG
GAGCACACGTCTGAACTCTTTCCCTACACGACGCTCTTCCGATCTNNNATAATTTATGGGCTTTGGCT
GAGCACACGTCTGAACTCTTTCCCTACACGACGCTCTTCCGATCTNNNATTTCACTGTGTTGACCAAG
GAGCACACGTCTGAACTCTTTCCCTACACGACGCTCTTCCGATCTNNNGCTAAAGCATTTGGAGTGTG
GCACACGTCTGAACTCTTTCCCTACACGACGCTCTTCCGATCTNNNTGCTTTGTTTTGCTTTGTTCTGC
AGCACACGTCTGAACTCTTTCCCTACACGACGCTCTTCCGATCTNNNGGTCTTGAACTCCTAACCTCAG
AGAGCACACGTCTGAACTCTTTCCCTACACGACGCTCTTCCGATCTNNNATAAAGATGGGTTCCCAGG
AGCACACGTCTGAACTCTTTCCCTACACGACGCTCTTCCGATCTNNNGTGGGTTGCCTTTTCACTTTCTT
GAGCACACGTCTGAACTCTTTCCCTACACGACGCTCTTCCGATCTNNNTCTCACCTCACCCTCTTGAGTA
GCACACGTCTGAACTCTTTCCCTACACGACGCTCTTCCGATCTNNNAAAGGAGTGAACAAATGCATCG
AGCACACGTCTGAACTCTTTCCCTACACGACGCTCTTCCGATCTNNNTGTAATCTCAGCACTTGTGGAG
AGCACACGTCTGAACTCTTTCCCTACACGACGCTCTTCCGATCTNNNCAGCTCTGCAGTACAATGACAT
AGAGCACACGTCTGAACTCTTTCCCTACACGACGCTCTTCCGATCTNNNAAGACCGGTCCACATAAAA
AGCACACGTCTGAACTCTTTCCCTACACGACGCTCTTCCGATCTNNNGATCTCTGGAGCCTAGGAGTTT
GAGCACACGTCTGAACTCTTTCCCTACACGACGCTCTTCCGATCTNNNGAAGTTGCAGTGAGCTAAGA
GCACACGTCTGAACTCTTTCCCTACACGACGCTCTTCCGATCTNNNACCCAGACAATAAGCCTCATTCT
AGCACACGTCTGAACTCTTTCCCTACACGACGCTCTTCCGATCTNNNTGGTGAAAGGGCATTCTAAGT
AGCACACGTCTGAACTCTTTCCCTACACGACGCTCTTCCGATCTNNNTCTCCATAATTGCATGAGCCAGT
AGCACACGTCTGAACTCTTTCCCTACACGACGCTCTTCCGATCTNNNTGTGTTTTCATATACGCTCCCC
GAGCACACGTCTGAACTCTTTCCCTACACGACGCTCTTCCGATCTNNNGTGTGTGTGTTGACAATA
GCACACGTCTGAACTCTTTCCCTACACGACGCTCTTCCGATCTNNNGAATGGGTAGCATGCTCATTCA
AGCACACGTCTGAACTCTTTCCCTACACGACGCTCTTCCGATCTNNNCTTCCTCTTCTGCATTCCAAAC
GAGCACACGTCTGAACTCTTTCCCTACACGACGCTCTTCCGATCTNNNGGATATAGCCCAGGTCTAA
AGCACACGTCTGAACTCTTTCCCTACACGACGCTCTTCCGATCTNNNCAGATCATGAGACTTCTTGGG
GCACACGTCTGAACTCTTTCCCTACACGACGCTCTTCCGATCTNNNATAGCAAGACCCCACCTCTACAA
GCACACGTCTGAACTCTTTCCCTACACGACGCTCTTCCGATCTNNNCCAAGCTTACCACAAGGCATAA
AGCACACGTCTGAACTCTTTCCCTACACGACGCTCTTCCGATCTNNNGCGAAACTCCATCTCTCTCTCT
GCACACGTCTGAACTCTTTCCCTACACGACGCTCTTCCGATCTNNNCCAATGACCCAATTCCAAAGCAT
AGCACACGTCTGAACTCTTTCCCTACACGACGCTCTTCCGATCTNNNTCAGCCTTTCTCTCTCTCTCTC
GAGCACACGTCTGAACTCTTTCCCTACACGACGCTCTTCCGATCTNNNCTGAGCAACATGGTGAAACT
GCACACGTCTGAACTCTTTCCCTACACGACGCTCTTCCGATCTNNNATCAGAAATTCTAGGGGCAGGA
GCACACGTCTGAACTCTTTCCCTACACGACGCTCTTCCGATCTNNNGCCCATGATGTGGCATGAATTA

AGCACACGTCTGAACTCTTTCCCTACACGACGCTCTTCCGATCTNNNAACCCCACTCTTCTTCTCCTCTTC
AGCACACGTCTGAACTCTTTCCCTACACGACGCTCTTCCGATCTNNNTCAATGTTGCCGATGCTGTATT
GAGCACACGTCTGAACTCTTTCCCTACACGACGCTCTTCCGATCTNNNTGGTTCCCCCACATACATAAC
AGCACACGTCTGAACTCTTTCCCTACACGACGCTCTTCCGATCTNNNTAAGGTAGTTTTCACTGGGAG
GAGCACACGTCTGAACTCTTTCCCTACACGACGCTCTTCCGATCTNNNAGAGCCAGACCTTGTCTCAAA
GCACACGTCTGAACTCTTTCCCTACACGACGCTCTTCCGATCTNNNAATCACTTGAACTCAGGAGGTG
GAGCACACGTCTGAACTCTTTCCCTACACGACGCTCTTCCGATCTNNNCTGAGATCACACCACTGCACT
GAGCACACGTCTGAACTCTTTCCCTACACGACGCTCTTCCGATCTNNNTGTCTGTCTGTCTGTCTCATT
CACACGTCTGAACTCTTTCCCTACACGACGCTCTTCCGATCTNNNAGAATTGCTTGAACCCAGGAGTC
GAGCACACGTCTGAACTCTTTCCCTACACGACGCTCTTCCGATCTNNNCAGTAGCTGGGTAAAGAAGT
AGCACACGTCTGAACTCTTTCCCTACACGACGCTCTTCCGATCTNNNGACTACAGGCATTCACCACCAT
GAGCACACGTCTGAACTCTTTCCCTACACGACGCTCTTCCGATCTNNNCCAGTTGCAATTTCCCCTTTTC
GCACACGTCTGAACTCTTTCCCTACACGACGCTCTTCCGATCTNNNAGGACACAAGGAAGAAGAAAGT
GAGCACACGTCTGAACTCTTTCCCTACACGACGCTCTTCCGATCTNNNACACCATCTATAGCTCCTTCT
AGAGCACACGTCTGAACTCTTTCCCTACACGACGCTCTTCCGATCTNNNTGTGGTTTCCTCCCTAATTC
AAGAGCACACGTCTGAACTCTTTCCCTACACGACGCTCTTCCGATCTNNNGAAGGAAGAAGAAACCA
AGCACACGTCTGAACTCTTTCCCTACACGACGCTCTTCCGATCTNNNACTCCAGCCTAGGTGACAGAAT
GCACACGTCTGAACTCTTTCCCTACACGACGCTCTTCCGATCTNNNGGTGTAGTGGTTCACGCCTATA
AGCACACGTCTGAACTCTTTCCCTACACGACGCTCTTCCGATCTNNNTGAAGTTCAAGTTGCGTTTAGG
AGCACACGTCTGAACTCTTTCCCTACACGACGCTCTTCCGATCTNNNCCTCACTCTTACTCTCTCCCAC
GAGCACACGTCTGAACTCTTTCCCTACACGACGCTCTTCCGATCTNNNTACCACCATGTCCGGCTAATTT
AGAGCACACGTCTGAACTCTTTCCCTACACGACGCTCTTCCGATCTNNNCATTCTCTCCTGGGACAAGT
GCACACGTCTGAACTCTTTCCCTACACGACGCTCTTCCGATCTNNNTGAGTGGACACAGACTTTTGGA
GAGCACACGTCTGAACTCTTTCCCTACACGACGCTCTTCCGATCTNNNGAGGTTGCAGTAAGCCAAGA
AGCACACGTCTGAACTCTTTCCCTACACGACGCTCTTCCGATCTNNNAATTCAGTACTAGGTGCCAAG
AGCACACGTCTGAACTCTTTCCCTACACGACGCTCTTCCGATCTNNNACATTGCCTCTTAGTTCACACCT
AGCACACGTCTGAACTCTTTCCCTACACGACGCTCTTCCGATCTNNNAGTGAGCAGAGATCAGAGATT
AGCACACGTCTGAACTCTTTCCCTACACGACGCTCTTCCGATCTNNNTAGGTGCTGCAGATGGATAAG
AGCACACGTCTGAACTCTTTCCCTACACGACGCTCTTCCGATCTNNNACCTCTACTTGTGGGGATGGA
CACACGTCTGAACTCTTTCCCTACACGACGCTCTTCCGATCTNNNGTGGAGGTTGTAGTAAGCCAAGA
AGCACACGTCTGAACTCTTTCCCTACACGACGCTCTTCCGATCTNNNCATTCTTCGGAAGGTGCAATC
AGAGCACACGTCTGAACTCTTTCCCTACACGACGCTCTTCCGATCTNNNTCCAAGATCAGTCAGCACAT
AGCACACGTCTGAACTCTTTCCCTACACGACGCTCTTCCGATCTNNNTCACACAGAGAATAGCAGACC
GCACACGTCTGAACTCTTTCCCTACACGACGCTCTTCCGATCTNNNTGTTTACAGAGCCTTTCCTTGGT
GAGCACACGTCTGAACTCTTTCCCTACACGACGCTCTTCCGATCTNNNGCAGAATCATGAGCCCCATAA
GAGCACACGTCTGAACTCTTTCCCTACACGACGCTCTTCCGATCTNNNTGAGAAGCAATTACAAGGGT
GAGCACACGTCTGAACTCTTTCCCTACACGACGCTCTTCCGATCTNNNGGGTCTCTCTCTCTCTCTCTC
AGCACACGTCTGAACTCTTTCCCTACACGACGCTCTTCCGATCTNNNGAAGACCAGGTCACACAGATT
AGAGCACACGTCTGAACTCTTTCCCTACACGACGCTCTTCCGATCTNNNGGATAGTGGGTCAAAAGA
AGAGCACACGTCTGAACTCTTTCCCTACACGACGCTCTTCCGATCTNNNGGAGTCTCCAGAGAAACAG
AGCACACGTCTGAACTCTTTCCCTACACGACGCTCTTCCGATCTNNNAAAGTGCTGGGATTACAGGAG
GAGCACACGTCTGAACTCTTTCCCTACACGACGCTCTTCCGATCTNNNAGTGAGCCAAGATCGTATGA
AGCACACGTCTGAACTCTTTCCCTACACGACGCTCTTCCGATCTNNNAATGGAGGAGACGGAAATCC

AGCACACGTCTGAACTCTTTCCCTACACGACGCTCTTCCGATCTNNNTCTCTTTCTGGCGTCCTCATATG
AGCACACGTCTGAACTCTTTCCCTACACGACGCTCTTCCGATCTNNNCTGGGCTGAACATAGTGAGAT
AGCACACGTCTGAACTCTTTCCCTACACGACGCTCTTCCGATCTNNNATAGCTAGCCCCTCTGTTTTCTG
GCACACGTCTGAACTCTTTCCCTACACGACGCTCTTCCGATCTNNNCTGATGACTGATTTGGTTGCCTG
AGCACACGTCTGAACTCTTTCCCTACACGACGCTCTTCCGATCTNNNCAAGAGAACAGAGTACCAGCTC
AGCACACGTCTGAACTCTTTCCCTACACGACGCTCTTCCGATCTNNNATTTGTGTCTCCTGGGTCTCTC
AGCACACGTCTGAACTCTTTCCCTACACGACGCTCTTCCGATCTNNNTGCCTAGAATTACTTTCGCCACA
AGAGCACACGTCTGAACTCTTTCCCTACACGACGCTCTTCCGATCTNNNCAGAGCCCCAAGTAGATATT
GAGCACACGTCTGAACTCTTTCCCTACACGACGCTCTTCCGATCTNNNCAGCACAGTCATAGTGGTAGT
GAGCACACGTCTGAACTCTTTCCCTACACGACGCTCTTCCGATCTNNNAAAGGAAGGGAAAGGAAGG
AGCACACGTCTGAACTCTTTCCCTACACGACGCTCTTCCGATCTNNNGGATTACCAGGCAGAGACTCT
AGAGCACACGTCTGAACTCTTTCCCTACACGACGCTCTTCCGATCTNNNGAACTAGGGTAAGAACTCAG
CACACGTCTGAACTCTTTCCCTACACGACGCTCTTCCGATCTNNNTGATGGTGAGTGAGTTTTCAGGA
AGCACACGTCTGAACTCTTTCCCTACACGACGCTCTTCCGATCTNNNAGAATATGTGTGTGTGTGTGT
AGCACACGTCTGAACTCTTTCCCTACACGACGCTCTTCCGATCTNNNGAAAAGACACACACACACACACA
AGCACACGTCTGAACTCTTTCCCTACACGACGCTCTTCCGATCTNNNTAGAGAGAGGGAGAGAGAAG
AAGAGCACACGTCTGAACTCTTTCCCTACACGACGCTCTTCCGATCTNNNGCGGACGTCAAGACAGAC
AGCACACGTCTGAACTCTTTCCCTACACGACGCTCTTCCGATCTNNNCCCTTTGATCTAAGTACCACCCA
AGCACACGTCTGAACTCTTTCCCTACACGACGCTCTTCCGATCTNNNATCCTCCGGTTCTTTCAAACACC
AGCACACGTCTGAACTCTTTCCCTACACGACGCTCTTCCGATCTNNNATCGAGTTCCCTGATGTACACAC
AGAGCACACGTCTGAACTCTTTCCCTACACGACGCTCTTCCGATCTNNNAGTAAGGAGGATGAGAGAC
AGCACACGTCTGAACTCTTTCCCTACACGACGCTCTTCCGATCTNNNATCCCAGTATTGGTGCTCAGATC
AGCACACGTCTGAACTCTTTCCCTACACGACGCTCTTCCGATCTNNNCAGGTAGGCAGGTGTTTTTCTT
AGCACACGTCTGAACTCTTTCCCTACACGACGCTCTTCCGATCTNNNCTGAAGGAGAGATGCAAAGGA
GAGCACACGTCTGAACTCTTTCCCTACACGACGCTCTTCCGATCTNNNGCTTCCCCTAGAGACAAGTAA
GAGCACACGTCTGAACTCTTTCCCTACACGACGCTCTTCCGATCTNNNAAAGAAAGAAAGAGGAGGT
GCACACGTCTGAACTCTTTCCCTACACGACGCTCTTCCGATCTNNNTCTCCTGGTAGTGATACCCTTGTC
AGCACACGTCTGAACTCTTTCCCTACACGACGCTCTTCCGATCTNNNGAGCTCTCTTTCATGCACTCTCTC
AGCACACGTCTGAACTCTTTCCCTACACGACGCTCTTCCGATCTNNNTATCTCCTTCCAACCAAACCCAGC
AGCACACGTCTGAACTCTTTCCCTACACGACGCTCTTCCGATCTNNNAGAGAGAGAGAGAGAGAGGC
GAGCACACGTCTGAACTCTTTCCCTACACGACGCTCTTCCGATCTNNNAAATGCCCATCAATCAACGAG
AGCACACGTCTGAACTCTTTCCCTACACGACGCTCTTCCGATCTNNNCTCTTCCCTCACTTTCTCCCAAAC
AAGAGCACACGTCTGAACTCTTTCCCTACACGACGCTCTTCCGATCTNNNCCTAGGCAACAAGAGCAAA
GCACACGTCTGAACTCTTTCCCTACACGACGCTCTTCCGATCTNNNAACACCAGCCAAGATTCCAAATG
AGAGCACACGTCTGAACTCTTTCCCTACACGACGCTCTTCCGATCTNNNGGATGTGAAGCAGGTTTGT
GAGCACACGTCTGAACTCTTTCCCTACACGACGCTCTTCCGATCTNNNGTATGGTGGTTATCAGGGGC
GAGCACACGTCTGAACTCTTTCCCTACACGACGCTCTTCCGATCTNNNAAAACAAGAGGTGTGAGAGC
AGCACACGTCTGAACTCTTTCCCTACACGACGCTCTTCCGATCTNNNGTCAATTCCCATCCTTTGCCAAA
GCACACGTCTGAACTCTTTCCCTACACGACGCTCTTCCGATCTNNNGACAGCTCATGGTGCTTTTTTCTC
AGCACACGTCTGAACTCTTTCCCTACACGACGCTCTTCCGATCTNNNTTCACCCTTAGTCACTGACAAGG
AGCACACGTCTGAACTCTTTCCCTACACGACGCTCTTCCGATCTNNNGCCTTCTCCCTGTCTAGCTTATG
AGCACACGTCTGAACTCTTTCCCTACACGACGCTCTTCCGATCTNNNTATTTCGTGTGTGTGTGTGTGT
GCACACGTCTGAACTCTTTCCCTACACGACGCTCTTCCGATCTNNNTCTGTTTTCCTAGGGCTAAGTGC

CACACGTCTGAACTCTTTCCCTACACGACGCTCTTCCGATCTNNNAGTAGTCTGCTCTTGTGATGTGT
GAGCACACGTCTGAACTCTTTCCCTACACGACGCTCTTCCGATCTNNNTTTTACCTCTTTCTTTGGGGCA
GAGCACACGTCTGAACTCTTTCCCTACACGACGCTCTTCCGATCTNNNAGAGAGAGAGAGAGAGAGA
AGAGCACACGTCTGAACTCTTTCCCTACACGACGCTCTTCCGATCTNNNTCAGGGTGTAAACCTTAGC
GCACACGTCTGAACTCTTTCCCTACACGACGCTCTTCCGATCTNNNATAATGTGCTTAGGAAGGTGCC
AGAGCACACGTCTGAACTCTTTCCCTACACGACGCTCTTCCGATCTNNNGGAGACAATTACCCCACAG
AGCACACGTCTGAACTCTTTCCCTACACGACGCTCTTCCGATCTNNNGAGTTACTCCTGCTGAAGTTCC
AGCACACGTCTGAACTCTTTCCCTACACGACGCTCTTCCGATCTNNNAGTGCGTATATACAGTGTGTGT
GAGCACACGTCTGAACTCTTTCCCTACACGACGCTCTTCCGATCTNNNATTAGATCCCCTTCTTCAGCTG
AGCACACGTCTGAACTCTTTCCCTACACGACGCTCTTCCGATCTNNNTGTTCTGGAATAGGAATGGAC
AGCACACGTCTGAACTCTTTCCCTACACGACGCTCTTCCGATCTNNNCCCACTCCCCTTCAAAAGTATCA
AGCACACGTCTGAACTCTTTCCCTACACGACGCTCTTCCGATCTNNNTGGAGTGCTTTAGTGTACATG
GAGCACACGTCTGAACTCTTTCCCTACACGACGCTCTTCCGATCTNNNCCTTTCACTTCTGCCCTCATACT
AGCACACGTCTGAACTCTTTCCCTACACGACGCTCTTCCGATCTNNNGCCACCCCAGTCTAAAAATTCAC
AGCACACGTCTGAACTCTTTCCCTACACGACGCTCTTCCGATCTNNNCAGCTGCCTCTGCTACAGATAA
GAGCACACGTCTGAACTCTTTCCCTACACGACGCTCTTCCGATCTNNNAATGGAATGTGATGGCAGTG
GAGCACACGTCTGAACTCTTTCCCTACACGACGCTCTTCCGATCTNNNCTTGGAAAGGGATCTAGAAG
AGAGCACACGTCTGAACTCTTTCCCTACACGACGCTCTTCCGATCTNNNGACTTCCTCATCTTACCACTG
AGCACACGTCTGAACTCTTTCCCTACACGACGCTCTTCCGATCTNNNTTCTTCTATTGGAGGCAACCACA
AGCACACGTCTGAACTCTTTCCCTACACGACGCTCTTCCGATCTNNNAATTTCTTGAGAAGGCCAGGG
AGAGCACACGTCTGAACTCTTTCCCTACACGACGCTCTTCCGATCTNNNAGGAGGTGAGATATTCGAG
AGCACACGTCTGAACTCTTTCCCTACACGACGCTCTTCCGATCTNNNTGCCTATCTGTGATGCCTTGAT
AGCACACGTCTGAACTCTTTCCCTACACGACGCTCTTCCGATCTNNNTGGGAAACTTCCTGACCATCTT
GAGCACACGTCTGAACTCTTTCCCTACACGACGCTCTTCCGATCTNNNCTAGATTTTAGTGCCATGGC
AGCACACGTCTGAACTCTTTCCCTACACGACGCTCTTCCGATCTNNNTCTTGTGGGGTTTTGTGTATAC
GAGCACACGTCTGAACTCTTTCCCTACACGACGCTCTTCCGATCTNNNTGAAGAGTTCATAACCAAGC
GCACACGTCTGAACTCTTTCCCTACACGACGCTCTTCCGATCTNNNTAACTACCCTTTGCTTCCTGTCCC
GAGCACACGTCTGAACTCTTTCCCTACACGACGCTCTTCCGATCTNNNCACACTTCCCAATGGTGTAAA
AGCACACGTCTGAACTCTTTCCCTACACGACGCTCTTCCGATCTNNNGCTATCTTCTCAGCCACATGGA
GAGCACACGTCTGAACTCTTTCCCTACACGACGCTCTTCCGATCTNNNAGGGTTCTCCAGAGATACAGA
GAGCACACGTCTGAACTCTTTCCCTACACGACGCTCTTCCGATCTNNNCTTATAATAGATACCCCCCGC
AGCACACGTCTGAACTCTTTCCCTACACGACGCTCTTCCGATCTNNNAGTCAAGTTGTCCGCTGAAAT
GAGCACACGTCTGAACTCTTTCCCTACACGACGCTCTTCCGATCTNNNGCTTCGGATCTGAACTAGAG
AGCACACGTCTGAACTCTTTCCCTACACGACGCTCTTCCGATCTNNNAATCAGAAGCCCAAGACCAATC
GAGCACACGTCTGAACTCTTTCCCTACACGACGCTCTTCCGATCTNNNTGTGTGTGTGTGTGTGTGTG
AGCACACGTCTGAACTCTTTCCCTACACGACGCTCTTCCGATCTNNNTGGGGATAGAGAATGGAGAG
GCACACGTCTGAACTCTTTCCCTACACGACGCTCTTCCGATCTNNNGAAAATTTGCTGCTAGGAGAG
AGCACACGTCTGAACTCTTTCCCTACACGACGCTCTTCCGATCTNNNCCACAATTGGGCACAGGAATT
GAGCACACGTCTGAACTCTTTCCCTACACGACGCTCTTCCGATCTNNNGTTCTGTGGGCCAAACTTTTA
AGCACACGTCTGAACTCTTTCCCTACACGACGCTCTTCCGATCTNNNGGAGATGGAGTTTCAAGGCAG
AGCACACGTCTGAACTCTTTCCCTACACGACGCTCTTCCGATCTNNNGAAAGAAAGAGAGAGAGTG
AGCACACGTCTGAACTCTTTCCCTACACGACGCTCTTCCGATCTNNNTCAAGGTTCCCCAGAAGAACA
AGCACACGTCTGAACTCTTTCCCTACACGACGCTCTTCCGATCTNNNTAACTATGTCTGTCTTGGTGCC

GAGCACACGTCTGAACTCTTTCCCTACACGACGCTCTTCCGATCTNNNCGTACAATATCCCTCCCAGTA
AGCACACGTCTGAACTCTTTCCCTACACGACGCTCTTCCGATCTNNNTGGCCTGAAACGTCACTTTTTTT
GAGCACACGTCTGAACTCTTTCCCTACACGACGCTCTTCCGATCTNNNACTGATCTCTGGGCTTCAACT
AGAGCACACGTCTGAACTCTTTCCCTACACGACGCTCTTCCGATCTNNNCTCATCTCTGACTGACCATG
GCACACGTCTGAACTCTTTCCCTACACGACGCTCTTCCGATCTNNNTCTCCAGGCAGTGTAATGTTTGT
GAGCACACGTCTGAACTCTTTCCCTACACGACGCTCTTCCGATCTNNNGCCGGAACTGTCCAATATCTT
GCACACGTCTGAACTCTTTCCCTACACGACGCTCTTCCGATCTNNNTGGCACAAACATGTCATGGTAA
AGCACACGTCTGAACTCTTTCCCTACACGACGCTCTTCCGATCTNNNTAGATGAAGGTCCTGCCCAGAA
AGCACACGTCTGAACTCTTTCCCTACACGACGCTCTTCCGATCTNNNAACTCTACCCCTCCTTCTATCC
AGCACACGTCTGAACTCTTTCCCTACACGACGCTCTTCCGATCTNNNATAAAAGGTGGTCCCCTTCCTT
GAGCACACGTCTGAACTCTTTCCCTACACGACGCTCTTCCGATCTNNNGGAGGCTGAGGTAAGAGAA
AGCACACGTCTGAACTCTTTCCCTACACGACGCTCTTCCGATCTNNNAGGGGCTCAGAGAAAATAAC
AGAGCACACGTCTGAACTCTTTCCCTACACGACGCTCTTCCGATCTNNNTCAACAAACACTCACTGTTCT
GAGCACACGTCTGAACTCTTTCCCTACACGACGCTCTTCCGATCTNNNAGGAATCGTAAAACAGTGCT
AAGAGCACACGTCTGAACTCTTTCCCTACACGACGCTCTTCCGATCTNNNAGCGTTACTGACAGACTTC
AAGAGCACACGTCTGAACTCTTTCCCTACACGACGCTCTTCCGATCTNNNACTCCTGCAGAAGAAACCT
GAGCACACGTCTGAACTCTTTCCCTACACGACGCTCTTCCGATCTNNNCTATAATATGCACCCCCTTCCC
GAGCACACGTCTGAACTCTTTCCCTACACGACGCTCTTCCGATCTNNNGCAGAATACTGGAGAGAAG
AGAGCACACGTCTGAACTCTTTCCCTACACGACGCTCTTCCGATCTNNNATTGTCCAGTGCTTCAACCT
AGAGCACACGTCTGAACTCTTTCCCTACACGACGCTCTTCCGATCTNNNCTATTGAAGGAACAGTGAG
GAGCACACGTCTGAACTCTTTCCCTACACGACGCTCTTCCGATCTNNNACATAGCTATGGAGTGAAGG
AGCACACGTCTGAACTCTTTCCCTACACGACGCTCTTCCGATCTNNNCAAGAGTGTGAGGTAGCACTG
GAGCACACGTCTGAACTCTTTCCCTACACGACGCTCTTCCGATCTNNNTCTTCTGGCTAGGGTTGTAG
AGAGCACACGTCTGAACTCTTTCCCTACACGACGCTCTTCCGATCTNNNTGTGCCTGCTTACTCACCTAA
GAGCACACGTCTGAACTCTTTCCCTACACGACGCTCTTCCGATCTNNNAGGAATCCTCACCAAACAGTC
GAGCACACGTCTGAACTCTTTCCCTACACGACGCTCTTCCGATCTNNNGCCGCCATGTATTTCTTTAGG
AGCACACGTCTGAACTCTTTCCCTACACGACGCTCTTCCGATCTNNNGACATTACCCTCCTCCAACTCTCC
GAGCACACGTCTGAACTCTTTCCCTACACGACGCTCTTCCGATCTNNNCCAGAGAACAGCACAGTCATA
AGAGCACACGTCTGAACTCTTTCCCTACACGACGCTCTTCCGATCTNNNTATGCTGTTGGACCAAAGCT
AGCACACGTCTGAACTCTTTCCCTACACGACGCTCTTCCGATCTNNNAGCATGAGTCCCATTGTTCTCT
AGCACACGTCTGAACTCTTTCCCTACACGACGCTCTTCCGATCTNNNGGCCTCACTTGATGTGTTTCTT
AGCACACGTCTGAACTCTTTCCCTACACGACGCTCTTCCGATCTNNNAGACTGGATAATGGCCTGTGTT
GAGCACACGTCTGAACTCTTTCCCTACACGACGCTCTTCCGATCTNNNGCCATTTCCAACACTGTCTCTT
AGCACACGTCTGAACTCTTTCCCTACACGACGCTCTTCCGATCTNNNGAACAGAGAGAGAAAGGCAG
GAGCACACGTCTGAACTCTTTCCCTACACGACGCTCTTCCGATCTNNNCACATCTGCCTCTTCCAAGTG
AGCACACGTCTGAACTCTTTCCCTACACGACGCTCTTCCGATCTNNNTATTTCCTCAGGAAGTGCCACA
AGCACACGTCTGAACTCTTTCCCTACACGACGCTCTTCCGATCTNNNTTTTCCCCATTGCTGACAAGTTG
GAGCACACGTCTGAACTCTTTCCCTACACGACGCTCTTCCGATCTNNNCCCCTTCTTAGAAGGCTTGCT
AGCACACGTCTGAACTCTTTCCCTACACGACGCTCTTCCGATCTNNNCTCCTGCCATGAACATTCTCTCT
GAGCACACGTCTGAACTCTTTCCCTACACGACGCTCTTCCGATCTNNNCAGCTGGAACAATCTCTCTCT
AGCACACGTCTGAACTCTTTCCCTACACGACGCTCTTCCGATCTNNNTGTCTCTGTCTCTGTCTCTGTCT
AGCACACGTCTGAACTCTTTCCCTACACGACGCTCTTCCGATCTNNNCACGTGCATGTGTGGATATAA
AGCACACGTCTGAACTCTTTCCCTACACGACGCTCTTCCGATCTNNNTGGTGACATACGAAAGAGCCAT

AGCACACGTCTGAACTCTTTCCCTACACGACGCTCTTCCGATCTNNNACAATGGGAGATGGGATTCTC
GCACACGTCTGAACTCTTTCCCTACACGACGCTCTTCCGATCTNNNATGACAAGGTCAGAGGCACTAA
AGCACACGTCTGAACTCTTTCCCTACACGACGCTCTTCCGATCTNNNTGTGCATGTTTATGTGTGTGCA
AGCACACGTCTGAACTCTTTCCCTACACGACGCTCTTCCGATCTNNNGCAGACAAAAAGAAAGAGGG
GAGCACACGTCTGAACTCTTTCCCTACACGACGCTCTTCCGATCTNNNTGAACTGAGAGGCAAAGAGA
AGCACACGTCTGAACTCTTTCCCTACACGACGCTCTTCCGATCTNNNCCTCTCTCTCCCTTACTCACTCT
AAGAGCACACGTCTGAACTCTTTCCCTACACGACGCTCTTCCGATCTNNNTCTCTCCAGCATCTCTCTTC
GAGCACACGTCTGAACTCTTTCCCTACACGACGCTCTTCCGATCTNNNAAAAGCAACTGCCTTTCACCA
AGCACACGTCTGAACTCTTTCCCTACACGACGCTCTTCCGATCTNNNACCACACCCATATCTCTCTCTCT
AGCACACGTCTGAACTCTTTCCCTACACGACGCTCTTCCGATCTNNNAGCACAGGCAAATTTACCAGTT
AGCACACGTCTGAACTCTTTCCCTACACGACGCTCTTCCGATCTNNNGCACAGAGCAAACACTTGAAA
GAGCACACGTCTGAACTCTTTCCCTACACGACGCTCTTCCGATCTNNNGTTCCCATTGATCCCTAGCTC
AGCACACGTCTGAACTCTTTCCCTACACGACGCTCTTCCGATCTNNNATGTGCATTGCTTTTGTGGAGT
AGCACACGTCTGAACTCTTTCCCTACACGACGCTCTTCCGATCTNNNTGAAATTAGCAGACGCACATG
GCACACGTCTGAACTCTTTCCCTACACGACGCTCTTCCGATCTNNNGCAGAATTGCTTGAACCTGAGA
GCACACGTCTGAACTCTTTCCCTACACGACGCTCTTCCGATCTNNNTCTATCCCCCTTGTCAAGAGGTA
AGCACACGTCTGAACTCTTTCCCTACACGACGCTCTTCCGATCTNNNAACTGTCCTGGTCACCTCTGTA
AGCACACGTCTGAACTCTTTCCCTACACGACGCTCTTCCGATCTNNNAGAGAGAGGGGGAGAGAGA
AGCACACGTCTGAACTCTTTCCCTACACGACGCTCTTCCGATCTNNNATAGCAAGGATGTCAGTGTGG
AGCACACGTCTGAACTCTTTCCCTACACGACGCTCTTCCGATCTNNNACAGAAAGAATGGGCTTAGG
AGCACACGTCTGAACTCTTTCCCTACACGACGCTCTTCCGATCTNNNTCAGATTTTCCTAGTCCCTCTG
AGCACACGTCTGAACTCTTTCCCTACACGACGCTCTTCCGATCTNNNTCCCACTCAATCTTCTCTCTCTC
AGAGCACACGTCTGAACTCTTTCCCTACACGACGCTCTTCCGATCTNNNTGGGTGAGTGGGAAGATA
GAGCACACGTCTGAACTCTTTCCCTACACGACGCTCTTCCGATCTNNNACATGAAATCAAACCGAAGC
GAGCACACGTCTGAACTCTTTCCCTACACGACGCTCTTCCGATCTNNNATCGTGTTCCCTTCTGTGTAC
GCACACGTCTGAACTCTTTCCCTACACGACGCTCTTCCGATCTNNNCGTAGAAGGAGGATCAGAGGA
GCACACGTCTGAACTCTTTCCCTACACGACGCTCTTCCGATCTNNNAGCAATAGCAAACATCAGGCAT
AGCACACGTCTGAACTCTTTCCCTACACGACGCTCTTCCGATCTNNNGTCCTAGACCCCTGAAGAAAC
GCACACGTCTGAACTCTTTCCCTACACGACGCTCTTCCGATCTNNNAAATAAGCAAAGAGGACATGCC
GAGCACACGTCTGAACTCTTTCCCTACACGACGCTCTTCCGATCTNNNATGCTCATTAGTCCAGAAAG
GAGCACACGTCTGAACTCTTTCCCTACACGACGCTCTTCCGATCTNNNCAGAGTGGAGCGTGTTTGAA
AGAGCACACGTCTGAACTCTTTCCCTACACGACGCTCTTCCGATCTNNNCAGTTTCTCCTTGACTCGAG
AGCACACGTCTGAACTCTTTCCCTACACGACGCTCTTCCGATCTNNNATATAATACACACACATGCGC
AGCACACGTCTGAACTCTTTCCCTACACGACGCTCTTCCGATCTNNNTGAGAAACACTCCCTTAGGCTT
AGCACACGTCTGAACTCTTTCCCTACACGACGCTCTTCCGATCTNNNAGCAGTATCAATGGCAGTGG
AGAGCACACGTCTGAACTCTTTCCCTACACGACGCTCTTCCGATCTNNNTGCTCCCTAGACCTTGTCTG
AGCACACGTCTGAACTCTTTCCCTACACGACGCTCTTCCGATCTNNNTTCTGCAAGCCCAGTCTAAATC
AAGAGCACACGTCTGAACTCTTTCCCTACACGACGCTCTTCCGATCTNNNGTTTGTGATTACTTGGGG
AGCACACGTCTGAACTCTTTCCCTACACGACGCTCTTCCGATCTNNNAAAGAATGAATGTGCCCGTCT
AGCACACGTCTGAACTCTTTCCCTACACGACGCTCTTCCGATCTNNNAAGTGATTGTAGGAGCTGTGG
AGCACACGTCTGAACTCTTTCCCTACACGACGCTCTTCCGATCTNNNCATGGAACTTCTTAGCTTCCAC
AGCACACGTCTGAACTCTTTCCCTACACGACGCTCTTCCGATCTNNNAGAGAGGACCATTTTGAGTCA
AGCACACGTCTGAACTCTTTCCCTACACGACGCTCTTCCGATCTNNNCTATGTGCCTGTTGTGTGCATT

GAGCACACGTCTGAACTCTTTCCCTACACGACGCTCTTCCGATCTNNNTCCAAAACTCAGGGTTGTAG
GAGCACACGTCTGAACTCTTTCCCTACACGACGCTCTTCCGATCTNNNAGCAGTTGAAGCATACTAGG
GCACACGTCTGAACTCTTTCCCTACACGACGCTCTTCCGATCTNNNTTCCAGGTGAGAGAGAGAGAGA
AGCACACGTCTGAACTCTTTCCCTACACGACGCTCTTCCGATCTNNNGTGTCATATAAGGCCAGGGAG
AGCACACGTCTGAACTCTTTCCCTACACGACGCTCTTCCGATCTNNNTAATGATTGAGGCTGGGTGAT
AGCACACGTCTGAACTCTTTCCCTACACGACGCTCTTCCGATCTNNNAGCTCTGGACCCTAGAACTGTA
AGCACACGTCTGAACTCTTTCCCTACACGACGCTCTTCCGATCTNNNGTTTCAGAAGGAGAGCTTGC
GAGCACACGTCTGAACTCTTTCCCTACACGACGCTCTTCCGATCTNNNCAGACCTAAGGGCTGAATGA
GCACACGTCTGAACTCTTTCCCTACACGACGCTCTTCCGATCTNNNTTAGCAACCAGTAGAGCCAGAG
AGCACACGTCTGAACTCTTTCCCTACACGACGCTCTTCCGATCTNNNCAACAGCAAATGCCTGTGTTC
GCACACGTCTGAACTCTTTCCCTACACGACGCTCTTCCGATCTNNNGTTGGTGACTGTTTCTCAAAGG
AGAGCACACGTCTGAACTCTTTCCCTACACGACGCTCTTCCGATCTNNNGGAGCACTGATCAATGCGA
GAGCACACGTCTGAACTCTTTCCCTACACGACGCTCTTCCGATCTNNNCGTGTTTTCTGCACATTTTCTG
GAGCACACGTCTGAACTCTTTCCCTACACGACGCTCTTCCGATCTNNNCTCTGTTAGGTGTTATGTGG
GCACACGTCTGAACTCTTTCCCTACACGACGCTCTTCCGATCTNNNTTTCTAGGGCCAAGCAAAGAGA
AGCACACGTCTGAACTCTTTCCCTACACGACGCTCTTCCGATCTNNNTCCTTCTTCCCCCTCTCTAGAAG
AGCACACGTCTGAACTCTTTCCCTACACGACGCTCTTCCGATCTNNNGTTTCTTCTGGGTGTGAATGCT
GCACACGTCTGAACTCTTTCCCTACACGACGCTCTTCCGATCTNNNGATGAAATGCTGCATAGGTGG
GAGCACACGTCTGAACTCTTTCCCTACACGACGCTCTTCCGATCTNNNAAGCCATACCACTCTATTCAG
GAGCACACGTCTGAACTCTTTCCCTACACGACGCTCTTCCGATCTNNNTGCTGATTGAGAGTACACAC
AGCACACGTCTGAACTCTTTCCCTACACGACGCTCTTCCGATCTNNNTGAAATGGAGAGGTAAGGCT
GCACACGTCTGAACTCTTTCCCTACACGACGCTCTTCCGATCTNNNCTGGATGTGGGCCTTCTCTTTTA
AGCACACGTCTGAACTCTTTCCCTACACGACGCTCTTCCGATCTNNNTTTGAAACACCAGCTTGCAGAA
GCACACGTCTGAACTCTTTCCCTACACGACGCTCTTCCGATCTNNNTGGGGGTAATGGTAAAAGCAA
AGCACACGTCTGAACTCTTTCCCTACACGACGCTCTTCCGATCTNNNTAGGAGCTCAGATTCCATTGT
AGCACACGTCTGAACTCTTTCCCTACACGACGCTCTTCCGATCTNNNTCCCAAAAACAGGAGGGAAGA
GCACACGTCTGAACTCTTTCCCTACACGACGCTCTTCCGATCTNNNTGTCTGTCTGTCTCTCTCTGTCT
AGCACACGTCTGAACTCTTTCCCTACACGACGCTCTTCCGATCTNNNCCAATGTATGCACTCAACATCC
GAGCACACGTCTGAACTCTTTCCCTACACGACGCTCTTCCGATCTNNNTCTTCCTTGCCCCTCACAATTTA
GAGCACACGTCTGAACTCTTTCCCTACACGACGCTCTTCCGATCTNNNGAGGTGCAAAGCTCAGTTTGA
AGCACACGTCTGAACTCTTTCCCTACACGACGCTCTTCCGATCTNNNTCAAATGGGCTGAGCTAGTAA
AGCACACGTCTGAACTCTTTCCCTACACGACGCTCTTCCGATCTNNNTCTATCCCCCATTTCTGGTTGT
GAGCACACGTCTGAACTCTTTCCCTACACGACGCTCTTCCGATCTNNNATATGTTCCATGGCAGGTGT
GAGCACACGTCTGAACTCTTTCCCTACACGACGCTCTTCCGATCTNNNCCGCTTCAGATGTTCATGTTT
GAGCACACGTCTGAACTCTTTCCCTACACGACGCTCTTCCGATCTNNNAGTTTAGCAAGCAGATTTGG
AGCACACGTCTGAACTCTTTCCCTACACGACGCTCTTCCGATCTNNNTTAGGTGGGATTAGATGTGTG
GCACACGTCTGAACTCTTTCCCTACACGACGCTCTTCCGATCTNNNACACAAATGCAGGCAGAATGTT
AGCACACGTCTGAACTCTTTCCCTACACGACGCTCTTCCGATCTNNNACTTACCGGAAGAAAAGCGAG
AGCACACGTCTGAACTCTTTCCCTACACGACGCTCTTCCGATCTNNNGGTTCCACCTCACTTATGTGTC
AGCACACGTCTGAACTCTTTCCCTACACGACGCTCTTCCGATCTNNNTCTCTTTCTGTCCCTCATGCATG
GCACACGTCTGAACTCTTTCCCTACACGACGCTCTTCCGATCTNNNGGTGACTAGACAAGGACCCAAA
AGAGCACACGTCTGAACTCTTTCCCTACACGACGCTCTTCCGATCTNNNAGGCTGGAGTGTTCAAATA
GAGCACACGTCTGAACTCTTTCCCTACACGACGCTCTTCCGATCTNNNGTGTAGGGTAACACTCCACA

GAGCACACGTCTGAACTCTTTCCCTACACGACGCTCTTCCGATCTNNNCTGTCAACACATACACACACA
CACACGTCTGAACTCTTTCCCTACACGACGCTCTTCCGATCTNNNTCCAGCACTCTGAGCTTTTAGTTC
AGAGCACACGTCTGAACTCTTTCCCTACACGACGCTCTTCCGATCTNNNAACTGTCTGGTCGGTCATTG
AGCACACGTCTGAACTCTTTCCCTACACGACGCTCTTCCGATCTNNNAGGGTTCTCTAGAGGGACAG
AGCACACGTCTGAACTCTTTCCCTACACGACGCTCTTCCGATCTNNNGACAGCAGGTTGTAGGACTTC
AGCACACGTCTGAACTCTTTCCCTACACGACGCTCTTCCGATCTNNNCAGTTTGTTCTGGTTCCATGTC
AGCACACGTCTGAACTCTTTCCCTACACGACGCTCTTCCGATCTNNNTGTTTTCGCATGCTTCATTCCTT
AGCACACGTCTGAACTCTTTCCCTACACGACGCTCTTCCGATCTNNNCCTTGGAGATCAGAGAAGCTC
AGCACACGTCTGAACTCTTTCCCTACACGACGCTCTTCCGATCTNNNTTTGTGTGGTAGACTGTCACT
GAGCACACGTCTGAACTCTTTCCCTACACGACGCTCTTCCGATCTNNNTACCCTCTAGTGTTGCATAGG
AGCACACGTCTGAACTCTTTCCCTACACGACGCTCTTCCGATCTNNNCCTTTGATGTCATGCCAGAGAG
AGCACACGTCTGAACTCTTTCCCTACACGACGCTCTTCCGATCTNNNAAGGCTTCTGAGAAATCCCAC
AGCACACGTCTGAACTCTTTCCCTACACGACGCTCTTCCGATCTNNNGTTTAGTCTGCACTGGAAGGA
GAGCACACGTCTGAACTCTTTCCCTACACGACGCTCTTCCGATCTNNNGCTAAGAGGAAGTGAGTCT
AGCACACGTCTGAACTCTTTCCCTACACGACGCTCTTCCGATCTNNNCTCTACAACCACCCCTCAGTTAC
AGCACACGTCTGAACTCTTTCCCTACACGACGCTCTTCCGATCTNNNAGATCTGGCAGGAAGTTGATG
GAGCACACGTCTGAACTCTTTCCCTACACGACGCTCTTCCGATCTNNNAACTCATTTAGTGGCAGAGCT
GCACACGTCTGAACTCTTTCCCTACACGACGCTCTTCCGATCTNNNTTGGTAATAGCACCTCCCACAAA
AGCACACGTCTGAACTCTTTCCCTACACGACGCTCTTCCGATCTNNNAGCTTGTAGCCATCATCTTGGA
AGCACACGTCTGAACTCTTTCCCTACACGACGCTCTTCCGATCTNNNCTCCTCCCTATTTTCCAGTGAG
AGCACACGTCTGAACTCTTTCCCTACACGACGCTCTTCCGATCTNNNGCTGACTTTCGGAAATGAATC
AGCACACGTCTGAACTCTTTCCCTACACGACGCTCTTCCGATCTNNNAGAAGTCAGAACATGCTGCCTT
AGAGCACACGTCTGAACTCTTTCCCTACACGACGCTCTTCCGATCTNNNGTGTGGGCTTGTACGAAAT
GAGCACACGTCTGAACTCTTTCCCTACACGACGCTCTTCCGATCTNNNCCTCCCTCTGTTTGCCTGAAAT
GCACACGTCTGAACTCTTTCCCTACACGACGCTCTTCCGATCTNNNAGGAATAAAATGGGAGGCAGG
AGCACACGTCTGAACTCTTTCCCTACACGACGCTCTTCCGATCTNNNGCTCCAGAGACAGAGACAGA
AGCACACGTCTGAACTCTTTCCCTACACGACGCTCTTCCGATCTNNNGAAAGGGGGTAGACAGGTAA
GAGCACACGTCTGAACTCTTTCCCTACACGACGCTCTTCCGATCTNNNAAGAGGAGGGTTCAGAGAG
GAGCACACGTCTGAACTCTTTCCCTACACGACGCTCTTCCGATCTNNNAGCTGCAAGGAACAGGAAC
AGCACACGTCTGAACTCTTTCCCTACACGACGCTCTTCCGATCTNNNCCTGCATTTTGCTCCTGCTATA
AGAGCACACGTCTGAACTCTTTCCCTACACGACGCTCTTCCGATCTNNNCCAAAGGGCAGTTCTCGGA
AGCACACGTCTGAACTCTTTCCCTACACGACGCTCTTCCGATCTNNNCTACTGGACAATGGTTTCCTC
AGCACACGTCTGAACTCTTTCCCTACACGACGCTCTTCCGATCTNNNTTGATAGGCAAATGCATCCCA
AGCACACGTCTGAACTCTTTCCCTACACGACGCTCTTCCGATCTNNNCCGATCACTGAGCATTGAAAC
GAGCACACGTCTGAACTCTTTCCCTACACGACGCTCTTCCGATCTNNNATGTGAGGTGTGTGTGTGTG
AGCACACGTCTGAACTCTTTCCCTACACGACGCTCTTCCGATCTNNNGCTGCTCTTGTTGTAAGTTTGC
GCACACGTCTGAACTCTTTCCCTACACGACGCTCTTCCGATCTNNNAACTCCTCCTTCTGTCTGTCTCTC
GAGCACACGTCTGAACTCTTTCCCTACACGACGCTCTTCCGATCTNNNGGGAAAGAGAGAGAGAG
GAGCACACGTCTGAACTCTTTCCCTACACGACGCTCTTCCGATCTNNNCATTGGTCGTATGATTGGCA
AGCACACGTCTGAACTCTTTCCCTACACGACGCTCTTCCGATCTNNNAAGAGGTGGGAAAAAAGCAA
AGCACACGTCTGAACTCTTTCCCTACACGACGCTCTTCCGATCTNNNCTTCCTCTTTCTCCTGACTCACCC
AGCACACGTCTGAACTCTTTCCCTACACGACGCTCTTCCGATCTNNNAGGATTGTTGTGGCTTTTTGG
AGCACACGTCTGAACTCTTTCCCTACACGACGCTCTTCCGATCTNNNAACTCAACAGAGAGGAGGAG

AGCACACGTCTGAACTCTTTCCCTACACGACGCTCTTCCGATCTNNNAGACGTGGCATTCATCTTGTTT
GCACACGTCTGAACTCTTTCCCTACACGACGCTCTTCCGATCTNNNTTTCAAATATGAAGACCTCCGCA
AGCACACGTCTGAACTCTTTCCCTACACGACGCTCTTCCGATCTNNNTACACACACAGACCTACACACA
GAGCACACGTCTGAACTCTTTCCCTACACGACGCTCTTCCGATCTNNNTTTGAGCCCAGTTTCGTTTTG
AGAGCACACGTCTGAACTCTTTCCCTACACGACGCTCTTCCGATCTNNNCTGCATCATAGTGAGCAAT
AGCACACGTCTGAACTCTTTCCCTACACGACGCTCTTCCGATCTNNNACATTTACTCAGGCTCCTCCA
AGCACACGTCTGAACTCTTTCCCTACACGACGCTCTTCCGATCTNNNTGCATGTGTTCAACTCTGGTTT
AGCACACGTCTGAACTCTTTCCCTACACGACGCTCTTCCGATCTNNNGCATGGAATCAGGAGTCAGA
AGCACACGTCTGAACTCTTTCCCTACACGACGCTCTTCCGATCTNNNAGCCTCTCTGAAATTTCTGCCAT
AGCACACGTCTGAACTCTTTCCCTACACGACGCTCTTCCGATCTNNNTCTTGTTACCCAGGAGATCAAT
AGCACACGTCTGAACTCTTTCCCTACACGACGCTCTTCCGATCTNNNGGTATGTAGTAGCTGCTTGGT
AGCACACGTCTGAACTCTTTCCCTACACGACGCTCTTCCGATCTNNNTGATCCCTGCAGCATATTCTCT
GCACACGTCTGAACTCTTTCCCTACACGACGCTCTTCCGATCTNNNTACAGATCCTGGGACTTTTCAG
GCACACGTCTGAACTCTTTCCCTACACGACGCTCTTCCGATCTNNNGCAAAGTGCAAAGTAAAGGTGT
GCACACGTCTGAACTCTTTCCCTACACGACGCTCTTCCGATCTNNNACTTCTCTCATTGAGCAGTTGGT
GCACACGTCTGAACTCTTTCCCTACACGACGCTCTTCCGATCTNNNGAACCTTTTAGGGCAGTTTTCCC
GAGCACACGTCTGAACTCTTTCCCTACACGACGCTCTTCCGATCTNNNTCTGCCAACAAAAGAGCAAG
AGCACACGTCTGAACTCTTTCCCTACACGACGCTCTTCCGATCTNNNACTGGATAAATGCCTGTTTCC
GAGCACACGTCTGAACTCTTTCCCTACACGACGCTCTTCCGATCTNNNGGCTAAATGAAAGGAGGAG
AGCACACGTCTGAACTCTTTCCCTACACGACGCTCTTCCGATCTNNNGCAGTTCTCAGGGGTCTAAAT
AGCACACGTCTGAACTCTTTCCCTACACGACGCTCTTCCGATCTNNNAGGGACTTGAGCATCCATGAA
AGCACACGTCTGAACTCTTTCCCTACACGACGCTCTTCCGATCTNNNTGGTGCCATTTACTGGGTAAA
AGCACACGTCTGAACTCTTTCCCTACACGACGCTCTTCCGATCTNNNGGACTTTGTGGAGGTGTTTTT
AGCACACGTCTGAACTCTTTCCCTACACGACGCTCTTCCGATCTNNNGTGTGTGTGTGCATTATGTGT
GCACACGTCTGAACTCTTTCCCTACACGACGCTCTTCCGATCTNNNTCTCCAAGCCATAGTTGGTTGA
AGCACACGTCTGAACTCTTTCCCTACACGACGCTCTTCCGATCTNNNTTCAAAGATGGAATTACGCGC
AGCACACGTCTGAACTCTTTCCCTACACGACGCTCTTCCGATCTNNNAATCTACTCCGGTGATGTTTGC
AGCACACGTCTGAACTCTTTCCCTACACGACGCTCTTCCGATCTNNNCACACACACACTCTCTCTCTCTC
AGCACACGTCTGAACTCTTTCCCTACACGACGCTCTTCCGATCTNNNGGCTTCACAGAGTTTTGGGAA
AGCACACGTCTGAACTCTTTCCCTACACGACGCTCTTCCGATCTNNNCTATGTGTGTGTGTGTGTGTG
AGCACACGTCTGAACTCTTTCCCTACACGACGCTCTTCCGATCTNNNGCTGATGATCGAGGTATTGCA
AGCACACGTCTGAACTCTTTCCCTACACGACGCTCTTCCGATCTNNNTGGGGCTTTGCACAAAAATAA
AGCACACGTCTGAACTCTTTCCCTACACGACGCTCTTCCGATCTNNNTGAGTACCACAGTCCTAAGGA
AGCACACGTCTGAACTCTTTCCCTACACGACGCTCTTCCGATCTNNNTTGGCACATATTTGCGCTAAG
AGCACACGTCTGAACTCTTTCCCTACACGACGCTCTTCCGATCTNNNCATATCTTTCAGGGGAGCAGG
GCACACGTCTGAACTCTTTCCCTACACGACGCTCTTCCGATCTNNNGTGAATTTATGCCAGATCACCC
AGCACACGTCTGAACTCTTTCCCTACACGACGCTCTTCCGATCTNNNAACTGTATGTGTGTGTGTGTG
AGCACACGTCTGAACTCTTTCCCTACACGACGCTCTTCCGATCTNNNAAGAAGCACTGTCACAGGAT
AGCACACGTCTGAACTCTTTCCCTACACGACGCTCTTCCGATCTNNNTTAGAAGCCACGTTCCTAACT
GCACACGTCTGAACTCTTTCCCTACACGACGCTCTTCCGATCTNNNCTTCTGTCTCCATGTCTCCAGAGC
AGCACACGTCTGAACTCTTTCCCTACACGACGCTCTTCCGATCTNNNCACATGCCACACATCACATACA
GCACACGTCTGAACTCTTTCCCTACACGACGCTCTTCCGATCTNNNTGTACTTTTGCAGGGCAGTTCTA
AGCACACGTCTGAACTCTTTCCCTACACGACGCTCTTCCGATCTNNNTGCAGGAAAGACAGGTAAAG

GCACACGTCTGAACTCTTTCCCTACACGACGCTCTTCCGATCTNNNGGCTTTCTTCTTGGTTTGCAGA
GCACACGTCTGAACTCTTTCCCTACACGACGCTCTTCCGATCTNNNGCTTCATGGTCCCTTTTCTTTCCC
GCACACGTCTGAACTCTTTCCCTACACGACGCTCTTCCGATCTNNNTACTGACACCAAAACTGGGTCA
GAGCACACGTCTGAACTCTTTCCCTACACGACGCTCTTCCGATCTNNNACTCCTGATCTCAAGTGATCT
GAGCACACGTCTGAACTCTTTCCCTACACGACGCTCTTCCGATCTNNNCACTTATCGAGATAGGAGG
GCACACGTCTGAACTCTTTCCCTACACGACGCTCTTCCGATCTNNNTCTCCAATTCTCCACTCTCCTCTC
GCACACGTCTGAACTCTTTCCCTACACGACGCTCTTCCGATCTNNNTCTCTATCTATCCAAGCCCCCTC
AGCACACGTCTGAACTCTTTCCCTACACGACGCTCTTCCGATCTNNNCTGGGATTGGGTAACTCAGAG
GCACACGTCTGAACTCTTTCCCTACACGACGCTCTTCCGATCTNNNACCCTTTTTAGCCCTGACCTATG
CACACGTCTGAACTCTTTCCCTACACGACGCTCTTCCGATCTNNNAGCAGTTACTCCAGGGAGAAAT
GCACACGTCTGAACTCTTTCCCTACACGACGCTCTTCCGATCTNNNCAACTCCTTTACCTCCCATTCCAC
AGCACACGTCTGAACTCTTTCCCTACACGACGCTCTTCCGATCTNNNCTTATAGGATCTGGGGTGTA
GAGCACACGTCTGAACTCTTTCCCTACACGACGCTCTTCCGATCTNNNCCATAGTCTTACCCCAGTGCT
GCACACGTCTGAACTCTTTCCCTACACGACGCTCTTCCGATCTNNNCTGAGAGTAGATTTTAGGGGC
AGCACACGTCTGAACTCTTTCCCTACACGACGCTCTTCCGATCTNNNCTACACTTGTCCTTGTGGGAG
AGCACACGTCTGAACTCTTTCCCTACACGACGCTCTTCCGATCTNNNACCAATCAGCACTCTGTGTCTA
AGCACACGTCTGAACTCTTTCCCTACACGACGCTCTTCCGATCTNNNATGGTAGCTCTGAGGCTGAA
GCACACGTCTGAACTCTTTCCCTACACGACGCTCTTCCGATCTNNNAAAACAACATTGGCGTAGACCA
GAGCACACGTCTGAACTCTTTCCCTACACGACGCTCTTCCGATCTNNNTGTGAAATCCAAGCATAATG
GCACACGTCTGAACTCTTTCCCTACACGACGCTCTTCCGATCTNNNAATAAAGCTTCCTCTCTCCACCCC
AGCACACGTCTGAACTCTTTCCCTACACGACGCTCTTCCGATCTNNNTGTAGTCCCAGCTACTGAGGA
AGCACACGTCTGAACTCTTTCCCTACACGACGCTCTTCCGATCTNNNAGTAGCTAGGATTACAGGCAT
AGAGCACACGTCTGAACTCTTTCCCTACACGACGCTCTTCCGATCTNNNCTCTGAGTCCTGAGATAAA
GCACACGTCTGAACTCTTTCCCTACACGACGCTCTTCCGATCTNNNTCAGCTTCAAGCAATTCTTCTGC
AGCACACGTCTGAACTCTTTCCCTACACGACGCTCTTCCGATCTNNNATGGCCTCATTAGAGACATTG
AGCACACGTCTGAACTCTTTCCCTACACGACGCTCTTCCGATCTNNNAGTATAGAGATGCACCACCAT
AGCACACGTCTGAACTCTTTCCCTACACGACGCTCTTCCGATCTNNNTTCACTTACAGTAACAAGCCC
AGCACACGTCTGAACTCTTTCCCTACACGACGCTCTTCCGATCTNNNAAGCATAGAGGAGTGCATCTT
GCACACGTCTGAACTCTTTCCCTACACGACGCTCTTCCGATCTNNNGTGTTTTGTTCTTAGGCCTTCCC
AGCACACGTCTGAACTCTTTCCCTACACGACGCTCTTCCGATCTNNNCCATCATCCCTCATGTGTCTTCA
AGCACACGTCTGAACTCTTTCCCTACACGACGCTCTTCCGATCTNNNGTTCTCTAGAGAAGCAGAGC
AGCACACGTCTGAACTCTTTCCCTACACGACGCTCTTCCGATCTNNNTGATGTCCGGTCTTTGTGTGTA
GCACACGTCTGAACTCTTTCCCTACACGACGCTCTTCCGATCTNNNTCTTTGACGTTTGATGCCGTATG
GCACACGTCTGAACTCTTTCCCTACACGACGCTCTTCCGATCTNNNCTGAGGCAGGAGAATCACTTG
GCACACGTCTGAACTCTTTCCCTACACGACGCTCTTCCGATCTNNNAAGGAATCCATATTCTCAGCCC
GAGCACACGTCTGAACTCTTTCCCTACACGACGCTCTTCCGATCTNNNCTTGAGCTGAGGAGTTCTAG
GCACACGTCTGAACTCTTTCCCTACACGACGCTCTTCCGATCTNNNTCTGATCTCTCTTGCTTTTCCCC
GCACACGTCTGAACTCTTTCCCTACACGACGCTCTTCCGATCTNNNTGTGTGTGTGTGTATACTACG
AGCACACGTCTGAACTCTTTCCCTACACGACGCTCTTCCGATCTNNNGGTTCATGCCTATAATCCCAC
AGCACACGTCTGAACTCTTTCCCTACACGACGCTCTTCCGATCTNNNAGCACTCCTATACACCAACAAC
GAGCACACGTCTGAACTCTTTCCCTACACGACGCTCTTCCGATCTNNNCAGTTACTAAGGAGGCTGA
GAGCACACGTCTGAACTCTTTCCCTACACGACGCTCTTCCGATCTNNNCTGAGACAGGAGAATCGCTT
GAGCACACGTCTGAACTCTTTCCCTACACGACGCTCTTCCGATCTNNNGCAACAAAGTGAGACTCTGT

AGCACACGTCTGAACTCTTTCCCTACACGACGCTCTTCCGATCTNNNTATACACACACACACACCCAAC
AGCACACGTCTGAACTCTTTCCCTACACGACGCTCTTCCGATCTNNNGGCTTGGAAACTGCACATTTC
AGCACACGTCTGAACTCTTTCCCTACACGACGCTCTTCCGATCTNNNGAAATCATGCCATGTACTCCAG
AGCACACGTCTGAACTCTTTCCCTACACGACGCTCTTCCGATCTNNNACAAAGCAGGACTCCATCTCA
AGCACACGTCTGAACTCTTTCCCTACACGACGCTCTTCCGATCTNNNCTCCATGCGGCAGGTATGTAT
AGCACACGTCTGAACTCTTTCCCTACACGACGCTCTTCCGATCTNNNATGAGAATTTCTTGAACCGGG
GAGCACACGTCTGAACTCTTTCCCTACACGACGCTCTTCCGATCTNNNATGAACAAACGAACACACAC
AGCACACGTCTGAACTCTTTCCCTACACGACGCTCTTCCGATCTNNNCATCTTCTTCTCCTAGCCATCCC
AGCACACGTCTGAACTCTTTCCCTACACGACGCTCTTCCGATCTNNNCAGCTCAAATGCCCATCAATCA
AGCACACGTCTGAACTCTTTCCCTACACGACGCTCTTCCGATCTNNNGCCTCCAAAGTAGCTGGTACT
GAGCACACGTCTGAACTCTTTCCCTACACGACGCTCTTCCGATCTNNNCTTCCTGCATGGCCAAAAG
GCACACGTCTGAACTCTTTCCCTACACGACGCTCTTCCGATCTNNNCAGCTATGGGTTTTTGGTAAGG
GAGCACACGTCTGAACTCTTTCCCTACACGACGCTCTTCCGATCTNNNCAGCTCCTGGGTTTATCAATT
AGCACACGTCTGAACTCTTTCCCTACACGACGCTCTTCCGATCTNNNTCAAAAGGATCAACTGTCCGA
AGCACACGTCTGAACTCTTTCCCTACACGACGCTCTTCCGATCTNNNACAGCTCTGCTTTCACATCTACA
AGCACACGTCTGAACTCTTTCCCTACACGACGCTCTTCCGATCTNNNTGGTGGGATCTCAGCTCACTA
AGCACACGTCTGAACTCTTTCCCTACACGACGCTCTTCCGATCTNNNTTAGGTCTGTGAGGGAAGGA
GAGCACACGTCTGAACTCTTTCCCTACACGACGCTCTTCCGATCTNNNTGTGTATGTCTGTGTCTGT
AGAGCACACGTCTGAACTCTTTCCCTACACGACGCTCTTCCGATCTNNNATGCCAGGTAAGCAAGGA
GCACACGTCTGAACTCTTTCCCTACACGACGCTCTTCCGATCTNNNCTCTTGGGTTGAGTACCTGAGA
GAGCACACGTCTGAACTCTTTCCCTACACGACGCTCTTCCGATCTNNNTGGTACCATGTCCAAATCCA
AGCACACGTCTGAACTCTTTCCCTACACGACGCTCTTCCGATCTNNNACCCAGCCTGGAGTACTTTTT
GAGCACACGTCTGAACTCTTTCCCTACACGACGCTCTTCCGATCTNNNTATGATTGTGTCACTGCACTC
GAGCACACGTCTGAACTCTTTCCCTACACGACGCTCTTCCGATCTNNNTCCCAAAGTGCTAGGATTAC
AGCACACGTCTGAACTCTTTCCCTACACGACGCTCTTCCGATCTNNNTTCTTGAACACACACACACACA
AGCACACGTCTGAACTCTTTCCCTACACGACGCTCTTCCGATCTNNNATGCCTGTAATCCCAATGCTTT
AGCACACGTCTGAACTCTTTCCCTACACGACGCTCTTCCGATCTNNNGTTGCAGTGAGCCAAGATCAT
GCACACGTCTGAACTCTTTCCCTACACGACGCTCTTCCGATCTNNNGGTTTCACCATGTCAAACTCCTG
AGCACACGTCTGAACTCTTTCCCTACACGACGCTCTTCCGATCTNNNGGTCACACTTGTAATCCCAAC
GAGCACACGTCTGAACTCTTTCCCTACACGACGCTCTTCCGATCTNNNAGGTGCTGGAAATACAACAC
AGCACACGTCTGAACTCTTTCCCTACACGACGCTCTTCCGATCTNNNCATCCTGGCTAACACAGTGAAA
AGCACACGTCTGAACTCTTTCCCTACACGACGCTCTTCCGATCTNNNTGACAGCAAGACTCCATCTCA
AGCACACGTCTGAACTCTTTCCCTACACGACGCTCTTCCGATCTNNNCATCAAGGAGCAGGAACTCTT
AGCACACGTCTGAACTCTTTCCCTACACGACGCTCTTCCGATCTNNNCATATACATGCACACACGACCA
AGCACACGTCTGAACTCTTTCCCTACACGACGCTCTTCCGATCTNNNTATTTGCCACCTGATCACTGTC
GCACACGTCTGAACTCTTTCCCTACACGACGCTCTTCCGATCTNNNTATCATGACAAGGCCTAGGGTT
GAGCACACGTCTGAACTCTTTCCCTACACGACGCTCTTCCGATCTNNNGGGCAGAGAAAAGAAAAGG
AGCACACGTCTGAACTCTTTCCCTACACGACGCTCTTCCGATCTNNNGACCTAGAAAGTCAGTGAGCA
AGCACACGTCTGAACTCTTTCCCTACACGACGCTCTTCCGATCTNNNGCTCAAAAACTCTAAGCAGCA
GAGCACACGTCTGAACTCTTTCCCTACACGACGCTCTTCCGATCTNNNTCTCTCTCTCTCTCTCAGAC
AGCACACGTCTGAACTCTTTCCCTACACGACGCTCTTCCGATCTNNNCCTCCCTCCCTTTTCCTCTATCT
CACACGTCTGAACTCTTTCCCTACACGACGCTCTTCCGATCTNNNGGCTGCAGAGAGCTATGATCATA
AGCACACGTCTGAACTCTTTCCCTACACGACGCTCTTCCGATCTNNNTGGGTGACAGAACAACATCGT

GAGCACACGTCTGAACTCTTTCCCTACACGACGCTCTTCCGATCTNNNGCGCCCGGCAGTATGTATATA
AGCACACGTCTGAACTCTTTCCCTACACGACGCTCTTCCGATCTNNNCAGTTCCATCCATGTTGATGCA
AGAGCACACGTCTGAACTCTTTCCCTACACGACGCTCTTCCGATCTNNNAGGAAAAGGCATACATTGAC
GAGCACACGTCTGAACTCTTTCCCTACACGACGCTCTTCCGATCTNNNATGATCATAGCTCATGGCAAC
GAGCACACGTCTGAACTCTTTCCCTACACGACGCTCTTCCGATCTNNNGTGAGTACCTTATTTGTGCTGC
GAGCACACGTCTGAACTCTTTCCCTACACGACGCTCTTCCGATCTNNNACTTCCTATGCTTGTCCTTCCTG
AGCACACGTCTGAACTCTTTCCCTACACGACGCTCTTCCGATCTNNNCTGGAGCAGATTGAACAGAAA
GAGCACACGTCTGAACTCTTTCCCTACACGACGCTCTTCCGATCTNNNTGAAACACCCGTCTCTACCAAA
AGCACACGTCTGAACTCTTTCCCTACACGACGCTCTTCCGATCTNNNGTGTGTGTGTGTGCATATGA
AGCACACGTCTGAACTCTTTCCCTACACGACGCTCTTCCGATCTNNNTTGTAATGCACAGGGTAGTCTC
GAGCACACGTCTGAACTCTTTCCCTACACGACGCTCTTCCGATCTNNNAGGTTGCTGTGAATGCCATTTT
GAGCACACGTCTGAACTCTTTCCCTACACGACGCTCTTCCGATCTNNNGAGAACCCTCTCCTACCAAAA
GAGCACACGTCTGAACTCTTTCCCTACACGACGCTCTTCCGATCTNNNAAATCTGTGCTTTCGTTGCTTCA
GAGCACACGTCTGAACTCTTTCCCTACACGACGCTCTTCCGATCTNNNCAAGCCAAGAGCCAAATCAAG
AGAGCACACGTCTGAACTCTTTCCCTACACGACGCTCTTCCGATCTNNNCCATATCAGCAGGGAAGCAA
GAGCACACGTCTGAACTCTTTCCCTACACGACGCTCTTCCGATCTNNNACATGTTACAAGCACAGGGAT
GAGCACACGTCTGAACTCTTTCCCTACACGACGCTCTTCCGATCTNNNTGCAACAGAGATGGCTTTGAA
AGCACACGTCTGAACTCTTTCCCTACACGACGCTCTTCCGATCTNNNTGTTGTGTTCAATTTTTTCCCCTC
GAGCACACGTCTGAACTCTTTCCCTACACGACGCTCTTCCGATCTNNNGCTTGCTTTCTAGTGGGGAGA
AGAGCACACGTCTGAACTCTTTCCCTACACGACGCTCTTCCGATCTNNNTTGAGCATGGGAGTTCAAGC
GAGCACACGTCTGAACTCTTTCCCTACACGACGCTCTTCCGATCTNNNCTGAGATCATGCCACTGAACT
AAGAGCACACGTCTGAACTCTTTCCCTACACGACGCTCTTCCGATCTNNNTTTCCAAAACAGACATCGA
GAGCACACGTCTGAACTCTTTCCCTACACGACGCTCTTCCGATCTNNNATTGGGCAACATAATGAGACC
AGAGCACACGTCTGAACTCTTTCCCTACACGACGCTCTTCCGATCTNNNATAAAGGAGTAGCAGAAAG
GAGCACACGTCTGAACTCTTTCCCTACACGACGCTCTTCCGATCTNNNCGACAGAGCAAGATTCCGTCT
GCACACGTCTGAACTCTTTCCCTACACGACGCTCTTCCGATCTNNNTGCACATTTATGCCAGAACTTCAC
GAGCACACGTCTGAACTCTTTCCCTACACGACGCTCTTCCGATCTNNNATTGGCACTTGTAATCCCAGC
AGCACACGTCTGAACTCTTTCCCTACACGACGCTCTTCCGATCTNNNTCTTGAAACTGGAAGGCAGAG
AGCACACGTCTGAACTCTTTCCCTACACGACGCTCTTCCGATCTNNNCTCCCGGGCTTGAATTTTTTTCA
AGCACACGTCTGAACTCTTTCCCTACACGACGCTCTTCCGATCTNNNAGACCCTGCCTCAAAAAAAGAG
GCACACGTCTGAACTCTTTCCCTACACGACGCTCTTCCGATCTNNNGTTTGTACCCATTCAGCAACCTCC
AGCACACGTCTGAACTCTTTCCCTACACGACGCTCTTCCGATCTNNNAATTGTTGGTAAGCATGTGCAC
AGCACACGTCTGAACTCTTTCCCTACACGACGCTCTTCCGATCTNNNGGAACTGTGTGAGGTAGGTAA
GCACACGTCTGAACTCTTTCCCTACACGACGCTCTTCCGATCTNNNTGGAGTTTCTGTTGGGTCCTTAG
GAGCACACGTCTGAACTCTTTCCCTACACGACGCTCTTCCGATCTNNNTTTCTGAGGACCCCAGAGATA
GAGCACACGTCTGAACTCTTTCCCTACACGACGCTCTTCCGATCTNNNGCCTGTCACTTCCACTACACATA
GAGCACACGTCTGAACTCTTTCCCTACACGACGCTCTTCCGATCTNNNTGAGCCATGATTATACCACTGC
AGCACACGTCTGAACTCTTTCCCTACACGACGCTCTTCCGATCTNNNACCATTTGTTTGTGTCCTCTCTT
GAGCACACGTCTGAACTCTTTCCCTACACGACGCTCTTCCGATCTNNNATCCCAGAAACTCTGTGTGTGT
AGCACACGTCTGAACTCTTTCCCTACACGACGCTCTTCCGATCTNNNGTACAACCACCATGGAGAAGAG
AGCACACGTCTGAACTCTTTCCCTACACGACGCTCTTCCGATCTNNNTGGGAAGTCTTGAAATAGAAAT
AGCACACGTCTGAACTCTTTCCCTACACGACGCTCTTCCGATCTNNNTGTGTGTAGAGAGACAGTCTTC
AGAGCACACGTCTGAACTCTTTCCCTACACGACGCTCTTCCGATCTNNNGTGTTCTCCAGGTCTTAGCTC

AGCACACGTCTGAACTCTTTCCCTACACGACGCTCTTCCGATCTNNNCAACTGCAGGGCTTGTTGAAT
AGCACACGTCTGAACTCTTTCCCTACACGACGCTCTTCCGATCTNNNACCCCCAAATCTAGTCTGACTC
AGCACACGTCTGAACTCTTTCCCTACACGACGCTCTTCCGATCTNNNGGGAAAGTAAACTTGTGCAC
GAGCACACGTCTGAACTCTTTCCCTACACGACGCTCTTCCGATCTNNNTGTGGAATCAACCTGAGTGT
AGCACACGTCTGAACTCTTTCCCTACACGACGCTCTTCCGATCTNNNTGATCACATATGCCAACTCCCAT
AGCACACGTCTGAACTCTTTCCCTACACGACGCTCTTCCGATCTNNNCTGCTTCTTCCGATCTCTCTCTC
AGCACACGTCTGAACTCTTTCCCTACACGACGCTCTTCCGATCTNNNTAAAACCAGGACTCCATCACTG
AGCACACGTCTGAACTCTTTCCCTACACGACGCTCTTCCGATCTNNNACCAATGACATTCTTCACAGGA
AGCACACGTCTGAACTCTTTCCCTACACGACGCTCTTCCGATCTNNNCCAAGACCTAGGCACTAAACT
GCACACGTCTGAACTCTTTCCCTACACGACGCTCTTCCGATCTNNNCCTCCATCCCTATAAACATCCCCCA
GAGCACACGTCTGAACTCTTTCCCTACACGACGCTCTTCCGATCTNNNGGAAGGCTAAGGTGAGACA
GAGCACACGTCTGAACTCTTTCCCTACACGACGCTCTTCCGATCTNNNCACTCTCTTCCACCATTCCCTA
AGAGCACACGTCTGAACTCTTTCCCTACACGACGCTCTTCCGATCTNNNAAAGCAAACTGAGGAGAG
AGCACACGTCTGAACTCTTTCCCTACACGACGCTCTTCCGATCTNNNCACATAGCACTCACAGCATAC
AGCACACGTCTGAACTCTTTCCCTACACGACGCTCTTCCGATCTNNNCTGGAGGCTGAAATAGGAGGA
AGCACACGTCTGAACTCTTTCCCTACACGACGCTCTTCCGATCTNNNCCTCCAGCTCCATCCAAATTACA
AGCACACGTCTGAACTCTTTCCCTACACGACGCTCTTCCGATCTNNNTGAAAAAAAGGTGGGAGGGT
AGCACACGTCTGAACTCTTTCCCTACACGACGCTCTTCCGATCTNNNCCAACTGGTGAGAAGAACAG
AGCACACGTCTGAACTCTTTCCCTACACGACGCTCTTCCGATCTNNNCCAGGTTGTGTGAATGCCATT
GAGCACACGTCTGAACTCTTTCCCTACACGACGCTCTTCCGATCTNNNCATGCTTGAAGTTTGAGACA
AGCACACGTCTGAACTCTTTCCCTACACGACGCTCTTCCGATCTNNNAGAATCAGCCCCAGATTTATT
GCACACGTCTGAACTCTTTCCCTACACGACGCTCTTCCGATCTNNNGTGAGGGATGAGTGTAGCAGT
AGCACACGTCTGAACTCTTTCCCTACACGACGCTCTTCCGATCTNNNTTACCAGGTTGTAAGGCTCAA
AGCACACGTCTGAACTCTTTCCCTACACGACGCTCTTCCGATCTNNNTGTTTTTTCTGTCATGGGTTG
AGCACACGTCTGAACTCTTTCCCTACACGACGCTCTTCCGATCTNNNAAAGGGTGGAAAGGAGAAGT
AGCACACGTCTGAACTCTTTCCCTACACGACGCTCTTCCGATCTNNNTCTTTTATCTTAGCCAGGGAGC
AGCACACGTCTGAACTCTTTCCCTACACGACGCTCTTCCGATCTNNNGAGAGTGACAGGCAGTAAGG
AGCACACGTCTGAACTCTTTCCCTACACGACGCTCTTCCGATCTNNNCCAGATCATGGTACGTCTCAGA
AGCACACGTCTGAACTCTTTCCCTACACGACGCTCTTCCGATCTNNNACTACTTGCCTCTTCTCCATGTG
AGCACACGTCTGAACTCTTTCCCTACACGACGCTCTTCCGATCTNNNGTTATAAGCCTCACCAACACA
AGCACACGTCTGAACTCTTTCCCTACACGACGCTCTTCCGATCTNNNACACACTTAGTCAAGAGGAG
AGCACACGTCTGAACTCTTTCCCTACACGACGCTCTTCCGATCTNNNGAGGTTGCAGTAAGCCAAGA
GAGCACACGTCTGAACTCTTTCCCTACACGACGCTCTTCCGATCTNNNCTCAGCAACATAGTATGACCC
GAGCACACGTCTGAACTCTTTCCCTACACGACGCTCTTCCGATCTNNNGTTGTTTGAACATGGGTAG
AGCACACGTCTGAACTCTTTCCCTACACGACGCTCTTCCGATCTNNNTGAGGTAGGAGAATGGTGTG
AGCACACGTCTGAACTCTTTCCCTACACGACGCTCTTCCGATCTNNNCAGCAAAGCAATACTCCCTCT
AGCACACGTCTGAACTCTTTCCCTACACGACGCTCTTCCGATCTNNNTCCTTATACTTGGGATCTGTG
AGCACACGTCTGAACTCTTTCCCTACACGACGCTCTTCCGATCTNNNGGAAATACACCACTGGTTCTC
AGCACACGTCTGAACTCTTTCCCTACACGACGCTCTTCCGATCTNNNTCCAGAGCAAGATTTCTGTCT
GAGCACACGTCTGAACTCTTTCCCTACACGACGCTCTTCCGATCTNNNCAGTAATGTTGGAAGGACC
AGCACACGTCTGAACTCTTTCCCTACACGACGCTCTTCCGATCTNNNAATTGGACTGAATTGCACCACT
GCACACGTCTGAACTCTTTCCCTACACGACGCTCTTCCGATCTNNNGCCTACTCTGTGTGTGTGTGTA
AGCACACGTCTGAACTCTTTCCCTACACGACGCTCTTCCGATCTNNNCTCAAGCGATCCTCTCACTTCAG

AGAGCACACGTCTGAACTCTTTCCCTACACGACGCTCTTCCGATCTNNNGGCACTAGTAGTTTCACCCT
AGCACACGTCTGAACTCTTTCCCTACACGACGCTCTTCCGATCTNNNGTGTGTGTGTGTGTGTGTGTT
AGCACACGTCTGAACTCTTTCCCTACACGACGCTCTTCCGATCTNNNAACTGACCCTATGATCCAAGCA
GCACACGTCTGAACTCTTTCCCTACACGACGCTCTTCCGATCTNNNTCTTGCACAATGCCTGGGATATA
AGCACACGTCTGAACTCTTTCCCTACACGACGCTCTTCCGATCTNNNCCACGTTGCTACAAAAGACAT
GCACACGTCTGAACTCTTTCCCTACACGACGCTCTTCCGATCTNNNGGTCCTGGTGTGAATAAGAGG
AGCACACGTCTGAACTCTTTCCCTACACGACGCTCTTCCGATCTNNNTGTTGTGATCAGTGGGAATG
AGCACACGTCTGAACTCTTTCCCTACACGACGCTCTTCCGATCTNNNTACTGTCTAGGTTCCCCACTAG
GCACACGTCTGAACTCTTTCCCTACACGACGCTCTTCCGATCTNNNTGTAGGTGGACATTCACATGTG
CACACGTCTGAACTCTTTCCCTACACGACGCTCTTCCGATCTNNNTTAGCATTTCATTGTCACTGGGCC
GCACACGTCTGAACTCTTTCCCTACACGACGCTCTTCCGATCTNNNGGATCATGAGGTCAGGAATTC
GCACACGTCTGAACTCTTTCCCTACACGACGCTCTTCCGATCTNNNCCTTTCTCCAGGCTCATACACAT
GAGCACACGTCTGAACTCTTTCCCTACACGACGCTCTTCCGATCTNNNAAGAACGTGCCACACAATA
GCACACGTCTGAACTCTTTCCCTACACGACGCTCTTCCGATCTNNNAATCAACCTAAGTGCTCATCGG
AGCACACGTCTGAACTCTTTCCCTACACGACGCTCTTCCGATCTNNNCATCCTGGCTAACATGGTGAA
AGCACACGTCTGAACTCTTTCCCTACACGACGCTCTTCCGATCTNNNTTAGTGTCACTGCAAATGACCC
AGCACACGTCTGAACTCTTTCCCTACACGACGCTCTTCCGATCTNNNATCTGCCACTGTAAGGAGAGT
GAGCACACGTCTGAACTCTTTCCCTACACGACGCTCTTCCGATCTNNNATTCTTTGGGAAGCAGTGGT
GAGCACACGTCTGAACTCTTTCCCTACACGACGCTCTTCCGATCTNNNAAGAGAGAAGGACAGAGA
AGCACACGTCTGAACTCTTTCCCTACACGACGCTCTTCCGATCTNNNCCCCAACTAGTTTTGGCAAA
AAGAGCACACGTCTGAACTCTTTCCCTACACGACGCTCTTCCGATCTNNNAGAGAGTCCCCCTATAAA
GAGCACACGTCTGAACTCTTTCCCTACACGACGCTCTTCCGATCTNNNTACACACACACACACACAAAC
AGCACACGTCTGAACTCTTTCCCTACACGACGCTCTTCCGATCTNNNAAATGCCCATCAATCAACGAGT
AGCACACGTCTGAACTCTTTCCCTACACGACGCTCTTCCGATCTNNNAAGTACTAGTCTAGGCCTGGG
GAGCACACGTCTGAACTCTTTCCCTACACGACGCTCTTCCGATCTNNNGAGGACCTGAATTTATTTGG
GAGCACACGTCTGAACTCTTTCCCTACACGACGCTCTTCCGATCTNNNTGTACCTTGACTGTGGTAGT
AGCACACGTCTGAACTCTTTCCCTACACGACGCTCTTCCGATCTNNNCTCCTGGAATCACAGCCTCATA
GAGCACACGTCTGAACTCTTTCCCTACACGACGCTCTTCCGATCTNNNGGCACTGGGTTTCTACTGTA
GAGCACACGTCTGAACTCTTTCCCTACACGACGCTCTTCCGATCTNNNTCCCAAGTGACTGAGTTGAC
GAGCACACGTCTGAACTCTTTCCCTACACGACGCTCTTCCGATCTNNNTGGAGCAATACGAGAAGAGA
GCACACGTCTGAACTCTTTCCCTACACGACGCTCTTCCGATCTNNNTAAAATCTGTGGGAGTGTCCTG
AGCACACGTCTGAACTCTTTCCCTACACGACGCTCTTCCGATCTNNNCCTGGCCCAGAAGTTTTTGTT
AGCACACGTCTGAACTCTTTCCCTACACGACGCTCTTCCGATCTNNNTTGAAGACCAAAGGAGAGAG
GCACACGTCTGAACTCTTTCCCTACACGACGCTCTTCCGATCTNNNACAAATTTTTCCACCTTTGGCCA
AGCACACGTCTGAACTCTTTCCCTACACGACGCTCTTCCGATCTNNNGAATTACAGGCATGAACCACC
AGCACACGTCTGAACTCTTTCCCTACACGACGCTCTTCCGATCTNNNGACTTATCCTTGGATGAGCAC
GCACACGTCTGAACTCTTTCCCTACACGACGCTCTTCCGATCTNNNCTGTCTCTCTCTCCCTCTCTCTCC
AGCACACGTCTGAACTCTTTCCCTACACGACGCTCTTCCGATCTNNNCCTATAGGGAGGCTGATGTTG
GCACACGTCTGAACTCTTTCCCTACACGACGCTCTTCCGATCTNNNGAAGCGGTTTCACTATGTTGAC
AGCACACGTCTGAACTCTTTCCCTACACGACGCTCTTCCGATCTNNNATTCTCAGGTCAGAAAGTGCA
AGCACACGTCTGAACTCTTTCCCTACACGACGCTCTTCCGATCTNNNCAGAAGTTCAAGACCACCCTAG
GCACACGTCTGAACTCTTTCCCTACACGACGCTCTTCCGATCTNNNGGCAGATTCACTTGAGGTCAGT
AGCACACGTCTGAACTCTTTCCCTACACGACGCTCTTCCGATCTNNNCTAGATGCTTTCCAACCTTCCCT

GCACACGTCTGAACTCTTTCCCTACACGACGCTCTTCCGATCTNNNATAAGATTTGCTTGCTAGGCAG
GAGCACACGTCTGAACTCTTTCCCTACACGACGCTCTTCCGATCTNNNTGAGGGAGTAGATGGGCTA
AGCACACGTCTGAACTCTTTCCCTACACGACGCTCTTCCGATCTNNNACTAAATCATCCCAAGGCCACT
AGCACACGTCTGAACTCTTTCCCTACACGACGCTCTTCCGATCTNNNACTAAATCATCCCAAGGCCACT
AGCACACGTCTGAACTCTTTCCCTACACGACGCTCTTCCGATCTNNNGCTTCTGAGTAGCTGGAACTA
GAGCACACGTCTGAACTCTTTCCCTACACGACGCTCTTCCGATCTNNNTTCAAACAACATTGCCAGGA
AGCACACGTCTGAACTCTTTCCCTACACGACGCTCTTCCGATCTNNNCAGCAGCAGAAAGGGACTAA
GAGCACACGTCTGAACTCTTTCCCTACACGACGCTCTTCCGATCTNNNCTCAGGAGTTTGAGACTAG
GAGCACACGTCTGAACTCTTTCCCTACACGACGCTCTTCCGATCTNNNGAGATGGCTCTACTGCTTTCC
GAGCACACGTCTGAACTCTTTCCCTACACGACGCTCTTCCGATCTNNNCTCCATCTCACCTGTAATCCCA
GCACACGTCTGAACTCTTTCCCTACACGACGCTCTTCCGATCTNNNGCAAGACCCCATCTCCAAAAAAT
AGAGCACACGTCTGAACTCTTTCCCTACACGACGCTCTTCCGATCTNNNAAGGAAGAGAAGCAAGG
GCACACGTCTGAACTCTTTCCCTACACGACGCTCTTCCGATCTNNNGGCAATAGAGTGAGACTCCAT
AGCACACGTCTGAACTCTTTCCCTACACGACGCTCTTCCGATCTNNNTGAACTAAAGGCGTATCTCCA
AGCACACGTCTGAACTCTTTCCCTACACGACGCTCTTCCGATCTNNNTCCCACATTCATACCACACAT
GCACACGTCTGAACTCTTTCCCTACACGACGCTCTTCCGATCTNNNCTTTCCCATTACATCTCCCACCAC
AGCACACGTCTGAACTCTTTCCCTACACGACGCTCTTCCGATCTNNNTTCAAGTGATTCTCCCGCCTTA
AGCACACGTCTGAACTCTTTCCCTACACGACGCTCTTCCGATCTNNNCCAAAGTACATGAACATGCACG
AGCACACGTCTGAACTCTTTCCCTACACGACGCTCTTCCGATCTNNNGATGTTTTCTGCTCCCCTCTCTG
GAGCACACGTCTGAACTCTTTCCCTACACGACGCTCTTCCGATCTNNNTGCTAGGCTTTGTATTTCCTG
GCACACGTCTGAACTCTTTCCCTACACGACGCTCTTCCGATCTNNNAGATCCTGTTAGTCACTACGGG
AGCACACGTCTGAACTCTTTCCCTACACGACGCTCTTCCGATCTNNNGCTAGAGTTGGCAATCTGAGG
GCACACGTCTGAACTCTTTCCCTACACGACGCTCTTCCGATCTNNNGAATAGCCTCTTCCTGCTTCACT
AGCACACGTCTGAACTCTTTCCCTACACGACGCTCTTCCGATCTNNNACAGGAAGAGACTGTCTCAAA
GAGCACACGTCTGAACTCTTTCCCTACACGACGCTCTTCCGATCTNNNTATATCTCATGCAGCTTGTGT
GAGCACACGTCTGAACTCTTTCCCTACACGACGCTCTTCCGATCTNNNCACAGTTCTGAAGGCTAGGA
AGCACACGTCTGAACTCTTTCCCTACACGACGCTCTTCCGATCTNNNCAGGTTTTTGGTGAGAGGAC
GCACACGTCTGAACTCTTTCCCTACACGACGCTCTTCCGATCTNNNAGTTTACTGCTAATGGGGTGGA
GAGCACACGTCTGAACTCTTTCCCTACACGACGCTCTTCCGATCTNNNGCGTGTAACAAACCACTCCA
AGCACACGTCTGAACTCTTTCCCTACACGACGCTCTTCCGATCTNNNGGAAGAGGAAGTCCATTAGGC
GCACACGTCTGAACTCTTTCCCTACACGACGCTCTTCCGATCTNNNACCACTATCAGCCATTTCTACCTC
GAGCACACGTCTGAACTCTTTCCCTACACGACGCTCTTCCGATCTNNNAGGTAGGAGAATCGTTTGAA
GAGCACACGTCTGAACTCTTTCCCTACACGACGCTCTTCCGATCTNNNAATGGACTGAGGCCAACTTT
AGCACACGTCTGAACTCTTTCCCTACACGACGCTCTTCCGATCTNNNAGCAAGCAGTATTCTTTGTGCA
AGCACACGTCTGAACTCTTTCCCTACACGACGCTCTTCCGATCTNNNTCTCTCTCGTCTCTCTGTCTCTC
GAGCACACGTCTGAACTCTTTCCCTACACGACGCTCTTCCGATCTNNNGGAGACACAGGATCTCTCTCT
GAGCACACGTCTGAACTCTTTCCCTACACGACGCTCTTCCGATCTNNNATGCCTGATGATCTGAAGTG
AGCACACGTCTGAACTCTTTCCCTACACGACGCTCTTCCGATCTNNNCTGTTCTCTGACTGGTTCCTCTC
AGCACACGTCTGAACTCTTTCCCTACACGACGCTCTTCCGATCTNNNAATGGCTGATTCTAGGATTGG
GCACACGTCTGAACTCTTTCCCTACACGACGCTCTTCCGATCTNNNGCAGGATAATCGTTTGAACCAG
GAGCACACGTCTGAACTCTTTCCCTACACGACGCTCTTCCGATCTNNNTGCCTGACACACAGTGGATT
GCACACGTCTGAACTCTTTCCCTACACGACGCTCTTCCGATCTNNNCACTCATGCTCCATTCTTTGTCAC
GCACACGTCTGAACTCTTTCCCTACACGACGCTCTTCCGATCTNNNCACTCATCTCCATCCGCATTTTCC

GAGCACACGTCTGAACTCTTTCCCTACACGACGCTCTTCCGATCTNNNCACAAGGTTTGGAGTTTGAG
AGCACACGTCTGAACTCTTTCCCTACACGACGCTCTTCCGATCTNNNCTGGCCTGGACTTTTGATTAC
AGCACACGTCTGAACTCTTTCCCTACACGACGCTCTTCCGATCTNNNTTACAACAAATTTCCAGCCCCA
AGCACACGTCTGAACTCTTTCCCTACACGACGCTCTTCCGATCTNNNGATTTCATGTCCTCCACAGGT
AGCACACGTCTGAACTCTTTCCCTACACGACGCTCTTCCGATCTNNNGGCAAGAGAATTGCTTGAAT
GCACACGTCTGAACTCTTTCCCTACACGACGCTCTTCCGATCTNNNCCCTCATCCTATCCAATCAACCA
GCACACGTCTGAACTCTTTCCCTACACGACGCTCTTCCGATCTNNNCCCGGCAAATTCCTTCATTTTTG
AGCACACGTCTGAACTCTTTCCCTACACGACGCTCTTCCGATCTNNNTTCCTGACCTCAGTTGATCCAT
AGCACACGTCTGAACTCTTTCCCTACACGACGCTCTTCCGATCTNNNTTCCTTACGCTTTATCTTGTGCA
GAGCACACGTCTGAACTCTTTCCCTACACGACGCTCTTCCGATCTNNNAGTTGGAGGAGAAAGCTAC
AGCACACGTCTGAACTCTTTCCCTACACGACGCTCTTCCGATCTNNNTCCTGTGTCCTCTCCTCTTCTT
AGCACACGTCTGAACTCTTTCCCTACACGACGCTCTTCCGATCTNNNCAAGAGAGTGACACTCCATCT
GAGCACACGTCTGAACTCTTTCCCTACACGACGCTCTTCCGATCTNNNTGAGGTCAGGAGTTCGTGA
AGCACACGTCTGAACTCTTTCCCTACACGACGCTCTTCCGATCTNNNTCCTGTCTATGTTTCCACCACT
GCACACGTCTGAACTCTTTCCCTACACGACGCTCTTCCGATCTNNNGGGGTGTGTCCTTAGTCTTCAA
AGCACACGTCTGAACTCTTTCCCTACACGACGCTCTTCCGATCTNNNGTTTACTGTGGCACTGTTCACA
AGCACACGTCTGAACTCTTTCCCTACACGACGCTCTTCCGATCTNNNTGCTGCAAAGGACATGATTTT
AGCACACGTCTGAACTCTTTCCCTACACGACGCTCTTCCGATCTNNNGGTGTTTGATCATTTGCAACA
AGCACACGTCTGAACTCTTTCCCTACACGACGCTCTTCCGATCTNNNTAGATCGTATGTGAGAGGTC
GAGCACACGTCTGAACTCTTTCCCTACACGACGCTCTTCCGATCTNNNAAAGAATAGAGACAGGGG
AGCACACGTCTGAACTCTTTCCCTACACGACGCTCTTCCGATCTNNNACTCTGAGGATTTGACACTGG
AGCACACGTCTGAACTCTTTCCCTACACGACGCTCTTCCGATCTNNNAGTCAGGAACCAAACCCACATT
AGCACACGTCTGAACTCTTTCCCTACACGACGCTCTTCCGATCTNNNCGTCAAACCAGGCTAATTTGT
AGCACACGTCTGAACTCTTTCCCTACACGACGCTCTTCCGATCTNNNATTTTAGAAGCTACTGGGGG
AGCACACGTCTGAACTCTTTCCCTACACGACGCTCTTCCGATCTNNNACCACATCCAGCTAATTGTGT
AGCACACGTCTGAACTCTTTCCCTACACGACGCTCTTCCGATCTNNNTGAGAAATAATGTGGCATGG
AGCACACGTCTGAACTCTTTCCCTACACGACGCTCTTCCGATCTNNNGAGAATTGCTAGAACCTGGGA
AGCACACGTCTGAACTCTTTCCCTACACGACGCTCTTCCGATCTNNNTGGGTGACAGAGTGAGATTT
AGCACACGTCTGAACTCTTTCCCTACACGACGCTCTTCCGATCTNNNGCATCCAGCCACAGAGAGATA
AGCACACGTCTGAACTCTTTCCCTACACGACGCTCTTCCGATCTNNNGTAACAGAGTGAGATCCTGT
AGCACACGTCTGAACTCTTTCCCTACACGACGCTCTTCCGATCTNNNCCTGAAACCCAGCATTCAAGTT
AGCACACGTCTGAACTCTTTCCCTACACGACGCTCTTCCGATCTNNNTTTGCAGTGAGCTAAGATTGT
AGCACACGTCTGAACTCTTTCCCTACACGACGCTCTTCCGATCTNNNCCAATAATGTTGGAGCAGAG
GAGCACACGTCTGAACTCTTTCCCTACACGACGCTCTTCCGATCTNNNGGAAGGAGAATCACTTGAA
AGCACACGTCTGAACTCTTTCCCTACACGACGCTCTTCCGATCTNNNCCTGAGGTTAGGAGTTTGAG
GCACACGTCTGAACTCTTTCCCTACACGACGCTCTTCCGATCTNNNCCCAGCCCCAAAAAGCTTTTTA
GCACACGTCTGAACTCTTTCCCTACACGACGCTCTTCCGATCTNNNATTCATTCTGCCAATCCACGAAC
AGCACACGTCTGAACTCTTTCCCTACACGACGCTCTTCCGATCTNNNGGAGGATACGGTGAGCTAAG
AGCACACGTCTGAACTCTTTCCCTACACGACGCTCTTCCGATCTNNNGACAAGAGGACCAAGTGCTAA
GCACACGTCTGAACTCTTTCCCTACACGACGCTCTTCCGATCTNNNACAGCCTCATTTTTCATGTGGT
AGCACACGTCTGAACTCTTTCCCTACACGACGCTCTTCCGATCTNNNCCCTCACAAACACAACTCCTCT
GCACACGTCTGAACTCTTTCCCTACACGACGCTCTTCCGATCTNNNCACAGAGGTAAACCATGGTCTG
GCACACGTCTGAACTCTTTCCCTACACGACGCTCTTCCGATCTNNNTGCTTGGGGTTTTGAAATGGT

GAGCACACGTCTGAACTCTTTCCCTACACGACGCTCTTCCGATCTNNNTTCAAGGTCTCAGGTGATTGG
AGCACACGTCTGAACTCTTTCCCTACACGACGCTCTTCCGATCTNNNACAGTGCAAGACTCCATCTCAAA
GCACACGTCTGAACTCTTTCCCTACACGACGCTCTTCCGATCTNNNTACACACACACATATGCATGCACC
AGCACACGTCTGAACTCTTTCCCTACACGACGCTCTTCCGATCTNNNCCAGCTGGGTAATGAGAGGAAA
GAGCACACGTCTGAACTCTTTCCCTACACGACGCTCTTCCGATCTNNNTACACACTCCACACACCATACA
GCACACGTCTGAACTCTTTCCCTACACGACGCTCTTCCGATCTNNNGAGGCATGAGAATAGCTTGAAC
GAGCACACGTCTGAACTCTTTCCCTACACGACGCTCTTCCGATCTNNNGAGATCAGGCCACTGCATTCTA
GAGCACACGTCTGAACTCTTTCCCTACACGACGCTCTTCCGATCTNNNAGACCACCTGTATTAGTCTGGC
GAGCACACGTCTGAACTCTTTCCCTACACGACGCTCTTCCGATCTNNNCTGGTCTTGAACTCTTGGGATC
AGCACACGTCTGAACTCTTTCCCTACACGACGCTCTTCCGATCTNNNGGGACTCTTTCTTAGGTGTCAA
AGCACACGTCTGAACTCTTTCCCTACACGACGCTCTTCCGATCTNNNGAGCCTGTCTCAAAAACAAGCA
AGCACACGTCTGAACTCTTTCCCTACACGACGCTCTTCCGATCTNNNCACCGGCTGCAACCAATTATTAT
AGCACACGTCTGAACTCTTTCCCTACACGACGCTCTTCCGATCTNNNTCATCACTGTTGATCTGGTCTG
AGCACACGTCTGAACTCTTTCCCTACACGACGCTCTTCCGATCTNNNGGTGCTTTTGGACATCGACTAC
GAGCACACGTCTGAACTCTTTCCCTACACGACGCTCTTCCGATCTNNNAGGTGAAGAAGCTGTGTTCTC
AGCACACGTCTGAACTCTTTCCCTACACGACGCTCTTCCGATCTNNNCCCTCAGCATCCTGATATTCGAG
GAGCACACGTCTGAACTCTTTCCCTACACGACGCTCTTCCGATCTNNNAAAGGGATGGAGATACTGGA
AGCACACGTCTGAACTCTTTCCCTACACGACGCTCTTCCGATCTNNNCTTTGGGGAATGCTTGATCTGT
AGCACACGTCTGAACTCTTTCCCTACACGACGCTCTTCCGATCTNNNAGTGAGACTCTGCCTAAAAAAC
GAGCACACGTCTGAACTCTTTCCCTACACGACGCTCTTCCGATCTNNNTCCACCCGTCCTAATTTGAATG
AGCACACGTCTGAACTCTTTCCCTACACGACGCTCTTCCGATCTNNNTGAAGCCAGCATCATCCTGATAA
GAGCACACGTCTGAACTCTTTCCCTACACGACGCTCTTCCGATCTNNNAGGTAGAAATCATCATCCCCCT
GAGCACACGTCTGAACTCTTTCCCTACACGACGCTCTTCCGATCTNNNCCAGCCTGTCCTTATCTTCTGAC
AGCACACGTCTGAACTCTTTCCCTACACGACGCTCTTCCGATCTNNNTGCCCCTTACTGAAAAGTGACAT
GAGCACACGTCTGAACTCTTTCCCTACACGACGCTCTTCCGATCTNNNGTGTGTGTGTGTGTATGTC
GAGCACACGTCTGAACTCTTTCCCTACACGACGCTCTTCCGATCTNNNGTGGCAAAATCACGTCTCTGA
AGCACACGTCTGAACTCTTTCCCTACACGACGCTCTTCCGATCTNNNATAGTCCCACTTTGCTCAGGAAC
AGCACACGTCTGAACTCTTTCCCTACACGACGCTCTTCCGATCTNNNAGAGTTGCCAGATCTTCAGGTT
GAGCACACGTCTGAACTCTTTCCCTACACGACGCTCTTCCGATCTNNNGAGTTGTGACCCTAACATTGCA
AGAGCACACGTCTGAACTCTTTCCCTACACGACGCTCTTCCGATCTNNNTCATGAACAGTGCTATGCTG
GAGCACACGTCTGAACTCTTTCCCTACACGACGCTCTTCCGATCTNNNCCCAGAGGTTAGTTTCTCCAGA
GAGCACACGTCTGAACTCTTTCCCTACACGACGCTCTTCCGATCTNNNGGAAACAGACCAAGACTCCATC
GAGCACACGTCTGAACTCTTTCCCTACACGACGCTCTTCCGATCTNNNGTGATCAGGCTACTACACTCCA
GAGCACACGTCTGAACTCTTTCCCTACACGACGCTCTTCCGATCTNNNCCAGAGAAAAAGAACACACGC
GAGCACACGTCTGAACTCTTTCCCTACACGACGCTCTTCCGATCTNNNAGACCTCGGTTTTACACACAC
GAGCACACGTCTGAACTCTTTCCCTACACGACGCTCTTCCGATCTNNNAATGAGACTCTGACACACACA
AGAGCACACGTCTGAACTCTTTCCCTACACGACGCTCTTCCGATCTNNNGAATTTGGGAAGCATGGTG
AGCACACGTCTGAACTCTTTCCCTACACGACGCTCTTCCGATCTNNNGCAGCCTACTTGTATCATGCTTC
GAGCACACGTCTGAACTCTTTCCCTACACGACGCTCTTCCGATCTNNNATATATTTGCGTGTGTGTGTG
AGAGCACACGTCTGAACTCTTTCCCTACACGACGCTCTTCCGATCTNNNAGAGTCTGGCACATACATGA
GAGCACACGTCTGAACTCTTTCCCTACACGACGCTCTTCCGATCTNNNACAGGACTTATCTGCTTGTGCT
AGCACACGTCTGAACTCTTTCCCTACACGACGCTCTTCCGATCTNNNTTTTTGCTGGGATTACAGGTGT
GCACACGTCTGAACTCTTTCCCTACACGACGCTCTTCCGATCTNNNCAAAAACTAGGTCCCTCCCATCAC

GAGCACACGTCTGAACTCTTTCCCTACACGACGCTCTTCCGATCTNNNAGCCTAGGCAACAAGAGTGA
GAGCACACGTCTGAACTCTTTCCCTACACGACGCTCTTCCGATCTNNNGACCCGTTTCATGGAAGACAAT
AGCACACGTCTGAACTCTTTCCCTACACGACGCTCTTCCGATCTNNNAGGGATTTGAAAGTTCCTGAG
GAGCACACGTCTGAACTCTTTCCCTACACGACGCTCTTCCGATCTNNNCTCAGGAGTTCGAGAACAGT
GAGCACACGTCTGAACTCTTTCCCTACACGACGCTCTTCCGATCTNNNTGGATAGGACAGAAGTCATCA
GAGCACACGTCTGAACTCTTTCCCTACACGACGCTCTTCCGATCTNNNTTGCACAGCAGCTTTACTAAT
GAGCACACGTCTGAACTCTTTCCCTACACGACGCTCTTCCGATCTNNNGCTGGATTACTTGAGGTCAGG
GAGCACACGTCTGAACTCTTTCCCTACACGACGCTCTTCCGATCTNNNGGCACAGTGGCTTACATCTGT
GAGCACACGTCTGAACTCTTTCCCTACACGACGCTCTTCCGATCTNNNGCAATAAGCCAAGATGGTACC
GAGCACACGTCTGAACTCTTTCCCTACACGACGCTCTTCCGATCTNNNGAGGGTACAGCAACAAACACA
AGCACACGTCTGAACTCTTTCCCTACACGACGCTCTTCCGATCTNNNGCAATACTTTGTGACCCTGTCTC
AGCACACGTCTGAACTCTTTCCCTACACGACGCTCTTCCGATCTNNNTCTCTACACACACACACACACAAC
AGCACACGTCTGAACTCTTTCCCTACACGACGCTCTTCCGATCTNNNAGGAGAATTGCTTGTACCTGTG
AGCACACGTCTGAACTCTTTCCCTACACGACGCTCTTCCGATCTNNNGCTTGGCAAAAGAGACACTTCA
AGCACACGTCTGAACTCTTTCCCTACACGACGCTCTTCCGATCTNNNGCTCTGCATCCTCACAAGAAAT
GCACACGTCTGAACTCTTTCCCTACACGACGCTCTTCCGATCTNNNAGGTCTCTATTCTGTTCCATTGGG
AGAGCACACGTCTGAACTCTTTCCCTACACGACGCTCTTCCGATCTNNNAACTCTGCCACGCTTTATCAT
AGCACACGTCTGAACTCTTTCCCTACACGACGCTCTTCCGATCTNNNCCAGTCCTCCTCAGTCGTTATTA
GAGCACACGTCTGAACTCTTTCCCTACACGACGCTCTTCCGATCTNNNCTCCTCTCTCTCACTCTGTCTC
GAGCACACGTCTGAACTCTTTCCCTACACGACGCTCTTCCGATCTNNNGACATTGAAAAGTGAGGGCT
AGCACACGTCTGAACTCTTTCCCTACACGACGCTCTTCCGATCTNNNGAGTAGCTGGGATTATAGGTG
AGCACACGTCTGAACTCTTTCCCTACACGACGCTCTTCCGATCTNNNTCAATGTTGCCGATGCTGTATT
AGCACACGTCTGAACTCTTTCCCTACACGACGCTCTTCCGATCTNNNAAAGAGAGCGAGAGAGAGAG
AGCACACGTCTGAACTCTTTCCCTACACGACGCTCTTCCGATCTNNNGTCTCTGTATGACTTGGAGCGA
AGCACACGTCTGAACTCTTTCCCTACACGACGCTCTTCCGATCTNNNTCTCCAAAACCCACAATGAGTCA
GAGCACACGTCTGAACTCTTTCCCTACACGACGCTCTTCCGATCTNNNCAGCCTGGATAACATAGGGA
GAGCACACGTCTGAACTCTTTCCCTACACGACGCTCTTCCGATCTNNNAGATTGTACCACTGCACTCTA
AGCACACGTCTGAACTCTTTCCCTACACGACGCTCTTCCGATCTNNNACGCCCGGCTAATTTTTTGTATT
AGCACACGTCTGAACTCTTTCCCTACACGACGCTCTTCCGATCTNNNTTGTAGTCCCAGCACTTTGGAA
AGCACACGTCTGAACTCTTTCCCTACACGACGCTCTTCCGATCTNNNTCTCCAAAACCCACAATGAGTCA
AGCACACGTCTGAACTCTTTCCCTACACGACGCTCTTCCGATCTNNNTGTAACAAAGTCCTCTCTCAGCC
AGCACACGTCTGAACTCTTTCCCTACACGACGCTCTTCCGATCTNNNGTCACCCTTGTTCCATTTGTGTC
GAGCACACGTCTGAACTCTTTCCCTACACGACGCTCTTCCGATCTNNNGAGAATCACTTGAACCCAGGA
AGCACACGTCTGAACTCTTTCCCTACACGACGCTCTTCCGATCTNNNAAAGTATTTGGGTGTGCTGACA
AGAGCACACGTCTGAACTCTTTCCCTACACGACGCTCTTCCGATCTNNNCACTGTGCCTGACTTTAGGTT
AGCACACGTCTGAACTCTTTCCCTACACGACGCTCTTCCGATCTNNNACAGACACAATGCTTCTGAAAC
AGCACACGTCTGAACTCTTTCCCTACACGACGCTCTTCCGATCTNNNTTCTGGTGGGGCATGGTTATAA
GAGCACACGTCTGAACTCTTTCCCTACACGACGCTCTTCCGATCTNNNTCAGAGGTCAACCAATCAAGA
AGCACACGTCTGAACTCTTTCCCTACACGACGCTCTTCCGATCTNNNCTAGTTTGGGCCCAAATCCCTAT
AGCACACGTCTGAACTCTTTCCCTACACGACGCTCTTCCGATCTNNNTGCCAGAAATGACTAGAAGCAC
AGCACACGTCTGAACTCTTTCCCTACACGACGCTCTTCCGATCTNNNTATGCCTGTAATCCCAGCACTTT
GAGCACACGTCTGAACTCTTTCCCTACACGACGCTCTTCCGATCTNNNATATGTGTGTGTGTGTGTGTG
AGCACACGTCTGAACTCTTTCCCTACACGACGCTCTTCCGATCTNNNAACCTCCCAGACTCAAGTGATTC

AGCACACGTCTGAACTCTTTCCCTACACGACGCTCTTCCGATCTNNNACCATATTGGACAGTGTAGCTC
GAGCACACGTCTGAACTCTTTCCCTACACGACGCTCTTCCGATCTNNNCAACAGTGCGAAACTCCATCA
AGCACACGTCTGAACTCTTTCCCTACACGACGCTCTTCCGATCTNNNGAGGGTCTCTCTCCACTTTCTT
AGCACACGTCTGAACTCTTTCCCTACACGACGCTCTTCCGATCTNNNTAGGCCCCTATTTGACAATGTG
AGCACACGTCTGAACTCTTTCCCTACACGACGCTCTTCCGATCTNNNCAACAGAGTGAGACCCTGTCTA
AGCACACGTCTGAACTCTTTCCCTACACGACGCTCTTCCGATCTNNNGCAAGACTCCGTCTCAAAACAA
AGCACACGTCTGAACTCTTTCCCTACACGACGCTCTTCCGATCTNNNCTGGAACACTTCTCTTCCTGTCT
GCACACGTCTGAACTCTTTCCCTACACGACGCTCTTCCGATCTNNNTCCTGGTAACCTTGTGCATTACA
GAGCACACGTCTGAACTCTTTCCCTACACGACGCTCTTCCGATCTNNNAGCTTGGTGGCTGTATTCTAA
GCACACGTCTGAACTCTTTCCCTACACGACGCTCTTCCGATCTNNNTTATAGCTGGCTTTGGGGAAAG
GAGCACACGTCTGAACTCTTTCCCTACACGACGCTCTTCCGATCTNNNCTATTGCACTCCAGCCTAAGT
GAGCACACGTCTGAACTCTTTCCCTACACGACGCTCTTCCGATCTNNNGGTTTGCAGAACACCAGAATC
AGCACACGTCTGAACTCTTTCCCTACACGACGCTCTTCCGATCTNNNGGAGCCAGAAGAGAGATGCTT
GAGCACACGTCTGAACTCTTTCCCTACACGACGCTCTTCCGATCTNNNACAACAGCAAAACTCGAACTCA
AAGAGCACACGTCTGAACTCTTTCCCTACACGACGCTCTTCCGATCTNNNAAGTAATGGGGCCTTCAGA
AGCACACGTCTGAACTCTTTCCCTACACGACGCTCTTCCGATCTNNNATGCTGATACATTGTTTGGGG
AGCACACGTCTGAACTCTTTCCCTACACGACGCTCTTCCGATCTNNNTTTCTCTTCCTCCTCCTCTTCCTCA
GCACACGTCTGAACTCTTTCCCTACACGACGCTCTTCCGATCTNNNATGATCACTTGAAGCCAGGAGTT
GCACACGTCTGAACTCTTTCCCTACACGACGCTCTTCCGATCTNNNTCCTGTCTCTCTTTCCTCTGTCTCA
AGCACACGTCTGAACTCTTTCCCTACACGACGCTCTTCCGATCTNNNAGAGAATTCCCCAGCAAGTCAA
AGCACACGTCTGAACTCTTTCCCTACACGACGCTCTTCCGATCTNNNTGGCATCTAGAAGTGATTTCCCA
CACACGTCTGAACTCTTTCCCTACACGACGCTCTTCCGATCTNNNCCACCAGAGTCTCTTGTCTCTTCCAT
GAGCACACGTCTGAACTCTTTCCCTACACGACGCTCTTCCGATCTNNNCCTTTCTCCTTCCTTCCTTCCTC
GCACACGTCTGAACTCTTTCCCTACACGACGCTCTTCCGATCTNNNGACTGAGTCCCTGGGAATATTCG
AAGAGCACACGTCTGAACTCTTTCCCTACACGACGCTCTTCCGATCTNNNAAAAACTGAGAGAGGCCA
AGAGCACACGTCTGAACTCTTTCCCTACACGACGCTCTTCCGATCTNNNAAGCCATCCACAAAACAGAT
GCACACGTCTGAACTCTTTCCCTACACGACGCTCTTCCGATCTNNNAAGAAAGACTTTGATGTCCCCGT
GAGCACACGTCTGAACTCTTTCCCTACACGACGCTCTTCCGATCTNNNTCTCCCAAACAAATGGAGTCT
GAGCACACGTCTGAACTCTTTCCCTACACGACGCTCTTCCGATCTNNNCAAGCTTTTCTGCTGGACTCA
AGCACACGTCTGAACTCTTTCCCTACACGACGCTCTTCCGATCTNNNAAAGTTTTGTGATCATGGGCA
GAAGAGCACACGTCTGAACTCTTTCCCTACACGACGCTCTTCCGATCTNNNGTGACAGAGTGAGACCC
GAGCACACGTCTGAACTCTTTCCCTACACGACGCTCTTCCGATCTNNNAAGACCATGTCTGTCTGTCTG
AGAGCACACGTCTGAACTCTTTCCCTACACGACGCTCTTCCGATCTNNNATATGGAGAGAGAGAGAG
GAGCACACGTCTGAACTCTTTCCCTACACGACGCTCTTCCGATCTNNNCTCCTGGGTATGCATCTACTG
AGCACACGTCTGAACTCTTTCCCTACACGACGCTCTTCCGATCTNNNTCCATCTAGCACGGCTCTAAATC
AGCACACGTCTGAACTCTTTCCCTACACGACGCTCTTCCGATCTNNNAATCTTTGAGGGCTTGTGATTG
AGCACACGTCTGAACTCTTTCCCTACACGACGCTCTTCCGATCTNNNCAAGGTTGCAGGACATAAAGC
AGCACACGTCTGAACTCTTTCCCTACACGACGCTCTTCCGATCTNNNCATAATGCAGTGCACACAGAGA
AGCACACGTCTGAACTCTTTCCCTACACGACGCTCTTCCGATCTNNNGCAAAGTGCTCTTACTGATGT
AGAGCACACGTCTGAACTCTTTCCCTACACGACGCTCTTCCGATCTNNNTCTGGCCAAACTCTCAGTAA
GCACACGTCTGAACTCTTTCCCTACACGACGCTCTTCCGATCTNNNTGGCATAGGGAAAATCCAAGTG
AGCACACGTCTGAACTCTTTCCCTACACGACGCTCTTCCGATCTNNNACGTGCTCTCATGACTACTTTGT
AGCACACGTCTGAACTCTTTCCCTACACGACGCTCTTCCGATCTNNNTCTCATCTCAGTCTCCAGGTA

AGCACACGTCTGAACTCTTTCCCTACACGACGCTCTTCCGATCTNNNCATCTCACTTTGGACCTCCTTGA
AGCACACGTCTGAACTCTTTCCCTACACGACGCTCTTCCGATCTNNNAATATGCCCACACCTTTGAATG
AGCACACGTCTGAACTCTTTCCCTACACGACGCTCTTCCGATCTNNNGCCAAAGTGCTGATATTACAG
GAGCACACGTCTGAACTCTTTCCCTACACGACGCTCTTCCGATCTNNNAGAAAGAGAGAGAGAGAGG
AGCACACGTCTGAACTCTTTCCCTACACGACGCTCTTCCGATCTNNNCCAACCCACGCTGATTTTTCTTA
AAGAGCACACGTCTGAACTCTTTCCCTACACGACGCTCTTCCGATCTNNNAGGAAGGAGACAGAGAG
AGCACACGTCTGAACTCTTTCCCTACACGACGCTCTTCCGATCTNNNCCTAGGCAACAAGAGCAAAACT
GAGCACACGTCTGAACTCTTTCCCTACACGACGCTCTTCCGATCTNNNAGTTGTGGATTTTCTCTCTCTC
AAGAGCACACGTCTGAACTCTTTCCCTACACGACGCTCTTCCGATCTNNNGAGGTTGCAATGAGCCGA
AGCACACGTCTGAACTCTTTCCCTACACGACGCTCTTCCGATCTNNNGAGAATTGCTTGAACCCAGGA
AGCACACGTCTGAACTCTTTCCCTACACGACGCTCTTCCGATCTNNNTTCTTCCTTCCTTCCTTCCTTCCC
AGAGCACACGTCTGAACTCTTTCCCTACACGACGCTCTTCCGATCTNNNCCAGAAGAACCTCAGGATCA
AGCACACGTCTGAACTCTTTCCCTACACGACGCTCTTCCGATCTNNNTGATGGCTGTGGCTGGTAAAT
GAGCACACGTCTGAACTCTTTCCCTACACGACGCTCTTCCGATCTNNNACCAAGTCCAAATTCATCCTCC
GAGCACACGTCTGAACTCTTTCCCTACACGACGCTCTTCCGATCTNNNGAGAGAGAGAGAGTGTGTG
AGCACACGTCTGAACTCTTTCCCTACACGACGCTCTTCCGATCTNNNACACAGTACCAAACCATAAGCCT
GAGCACACGTCTGAACTCTTTCCCTACACGACGCTCTTCCGATCTNNNGAGCAAGACTCCGTCTCAAAA
AGCACACGTCTGAACTCTTTCCCTACACGACGCTCTTCCGATCTNNNGTCAAGATTGTCCCAATGCACT
GAGCACACGTCTGAACTCTTTCCCTACACGACGCTCTTCCGATCTNNNAATGAACTGTGTTTGCACCACT
AGCACACGTCTGAACTCTTTCCCTACACGACGCTCTTCCGATCTNNNGATGAGAGGACGTGTCTCTTC
GCACACGTCTGAACTCTTTCCCTACACGACGCTCTTCCGATCTNNNGCAATGGCTCCATTCATTCTCAAC
AGCACACGTCTGAACTCTTTCCCTACACGACGCTCTTCCGATCTNNNAACGTGCTGAGAGATAAACTGC
AGCACACGTCTGAACTCTTTCCCTACACGACGCTCTTCCGATCTNNNAGTGGCATCTTATCTTCCAGCA
AGCACACGTCTGAACTCTTTCCCTACACGACGCTCTTCCGATCTNNNACTGAATAGCCTCTGTGTGTGT
AGAGCACACGTCTGAACTCTTTCCCTACACGACGCTCTTCCGATCTNNNATGGAGAGAGAGATGGAG
AGCACACGTCTGAACTCTTTCCCTACACGACGCTCTTCCGATCTNNNCTTCTCACTGTAGCCTCACATG
AAGAGCACACGTCTGAACTCTTTCCCTACACGACGCTCTTCCGATCTNNNTGTAATTCCAGCTACTTGG
AGCACACGTCTGAACTCTTTCCCTACACGACGCTCTTCCGATCTNNNATGTGTGTGTGTGTGTGTATG
AGCACACGTCTGAACTCTTTCCCTACACGACGCTCTTCCGATCTNNNATGAAATCTGGTCTGGTCTGCT
AGCACACGTCTGAACTCTTTCCCTACACGACGCTCTTCCGATCTNNNATGTTTAGATTTGGGTCCCCCTC
GCACACGTCTGAACTCTTTCCCTACACGACGCTCTTCCGATCTNNNCTAAGAAGGCAGCCAGGTCATTA
AAGAGCACACGTCTGAACTCTTTCCCTACACGACGCTCTTCCGATCTNNNTAATGAAGATGTGCCCAGG
CACACGTCTGAACTCTTTCCCTACACGACGCTCTTCCGATCTNNNGCATTTTATTCTCAAGGCCCCTGCA
AGCACACGTCTGAACTCTTTCCCTACACGACGCTCTTCCGATCTNNNGCATTCTCATGGCCACCATATTC
GAGCACACGTCTGAACTCTTTCCCTACACGACGCTCTTCCGATCTNNNTGGGTGACTTGCTTACCTGATT
AGCACACGTCTGAACTCTTTCCCTACACGACGCTCTTCCGATCTNNNGAGGAGTGAGGACCTTCTAGG
AGAGCACACGTCTGAACTCTTTCCCTACACGACGCTCTTCCGATCTNNNATGCATCTATGACTTTCCTCC
AGCACACGTCTGAACTCTTTCCCTACACGACGCTCTTCCGATCTNNNACAAACAGGCACAACAGTGTAA
GAGCACACGTCTGAACTCTTTCCCTACACGACGCTCTTCCGATCTNNNAAGGGAGATTTGAGAATGGT
GAGCACACGTCTGAACTCTTTCCCTACACGACGCTCTTCCGATCTNNNCTATCTCTGTTTTCCCACCCGAA
AGCACACGTCTGAACTCTTTCCCTACACGACGCTCTTCCGATCTNNNGAAAGTGGGGGAAAACAGGA
GAGCACACGTCTGAACTCTTTCCCTACACGACGCTCTTCCGATCTNNNGCAACTACACAGGAAACAGTT
AGCACACGTCTGAACTCTTTCCCTACACGACGCTCTTCCGATCTNNNAGGGAGAGAGAGAGAGAGAGA

AGCACACGTCTGAACTCTTTCCCTACACGACGCTCTTCCGATCTNNNTGCCTCTTTCCATCTCTCCTCTA
GCACACGTCTGAACTCTTTCCCTACACGACGCTCTTCCGATCTNNNTCACCATCTTCCTCTACCTCACTCA
GCACACGTCTGAACTCTTTCCCTACACGACGCTCTTCCGATCTNNNTATAAAAGAGTCCTGAGGCTGG
AGCACACGTCTGAACTCTTTCCCTACACGACGCTCTTCCGATCTNNNTCAGCACAGATCTAACAGTCCA
GAGCACACGTCTGAACTCTTTCCCTACACGACGCTCTTCCGATCTNNNTCAGCAGACTTACACTACTAG
AGCACACGTCTGAACTCTTTCCCTACACGACGCTCTTCCGATCTNNNCCCAATGTTAGGTCCTCTTGT
AGCACACGTCTGAACTCTTTCCCTACACGACGCTCTTCCGATCTNNNGCCTGAGTGGGGAATTGAACT
GAGCACACGTCTGAACTCTTTCCCTACACGACGCTCTTCCGATCTNNNCCTGCTCCCAAGACTGGAAAT
AGCACACGTCTGAACTCTTTCCCTACACGACGCTCTTCCGATCTNNNAGAGTCTCAGGGTTGGCAAAT
AGCACACGTCTGAACTCTTTCCCTACACGACGCTCTTCCGATCTNNNAATGAACAGGTGGCTACTTGG
AGCACACGTCTGAACTCTTTCCCTACACGACGCTCTTCCGATCTNNNGATCCAGCGACCTTTTTTGTG
AGAGCACACGTCTGAACTCTTTCCCTACACGACGCTCTTCCGATCTNNNGCTGAGAAAGTCACAGATC
AGCACACGTCTGAACTCTTTCCCTACACGACGCTCTTCCGATCTNNNCAGCCACTCACTGTAATACCTG
AGCACACGTCTGAACTCTTTCCCTACACGACGCTCTTCCGATCTNNNGCAGTAGCTAATCAGGACCAG
GCACACGTCTGAACTCTTTCCCTACACGACGCTCTTCCGATCTNNNCTATGACTGTTTCCACGTTGCTG
AGCACACGTCTGAACTCTTTCCCTACACGACGCTCTTCCGATCTNNNGCATTTGACAGGAGCTCTGAA
GAGCACACGTCTGAACTCTTTCCCTACACGACGCTCTTCCGATCTNNNTGACACACCCACATCATAACA
AGCACACGTCTGAACTCTTTCCCTACACGACGCTCTTCCGATCTNNNGAGCTTGCAGATATGGATTTG
GCACACGTCTGAACTCTTTCCCTACACGACGCTCTTCCGATCTNNNTTTCTCCACAATAGGCACACAGT
CACACGTCTGAACTCTTTCCCTACACGACGCTCTTCCGATCTNNNCTATAGACCCCTGGATCGTTTCCC
GAGCACACGTCTGAACTCTTTCCCTACACGACGCTCTTCCGATCTNNNCCCATACTCCTTACCAAAGCC
GCACACGTCTGAACTCTTTCCCTACACGACGCTCTTCCGATCTNNNCAACTCTCGAAAATTCTCAGCCA
AGCACACGTCTGAACTCTTTCCCTACACGACGCTCTTCCGATCTNNNTCTTGCTCGAATGTTCTCACTC
AGCACACGTCTGAACTCTTTCCCTACACGACGCTCTTCCGATCTNNNGAGATAATCACATGAACCCGG
GAGCACACGTCTGAACTCTTTCCCTACACGACGCTCTTCCGATCTNNNACCTTCAACATCAGACTCAGA
AGCACACGTCTGAACTCTTTCCCTACACGACGCTCTTCCGATCTNNNTCCGAGTCCTACCTTAGTACCA
AGCACACGTCTGAACTCTTTCCCTACACGACGCTCTTCCGATCTNNNCAGCCAGCCTCCATAATCACAT
AGCACACGTCTGAACTCTTTCCCTACACGACGCTCTTCCGATCTNNNCTTACAAACAAACCCAGTCCTC
GAGCACACGTCTGAACTCTTTCCCTACACGACGCTCTTCCGATCTNNNTGCTGAGAGAGAGAGAGAG
GAGCACACGTCTGAACTCTTTCCCTACACGACGCTCTTCCGATCTNNNTAATCATCTCCCAAAGGCCTG
GAGCACACGTCTGAACTCTTTCCCTACACGACGCTCTTCCGATCTNNNGATGAGAAAGCTGAGCCACA
GAGCACACGTCTGAACTCTTTCCCTACACGACGCTCTTCCGATCTNNNATCAATAAGGAGGACTCTTG
AGCACACGTCTGAACTCTTTCCCTACACGACGCTCTTCCGATCTNNNGCAGGAAATAACATGGGAGG
AGCACACGTCTGAACTCTTTCCCTACACGACGCTCTTCCGATCTNNNACAGACTCATCTCACGGTTTT
GAGCACACGTCTGAACTCTTTCCCTACACGACGCTCTTCCGATCTNNNTCTGTAGTTCCAGCTACTCAG
AGCACACGTCTGAACTCTTTCCCTACACGACGCTCTTCCGATCTNNNTTTCCACACTCTAGAATGCCC
AGCACACGTCTGAACTCTTTCCCTACACGACGCTCTTCCGATCTNNNACAACTGGCTCTACATGAGTC
GAGCACACGTCTGAACTCTTTCCCTACACGACGCTCTTCCGATCTNNNGTAAGGATGGACATGAGAC
AGCACACGTCTGAACTCTTTCCCTACACGACGCTCTTCCGATCTNNNAGATCACATGAACCCAGCAGA
AGCACACGTCTGAACTCTTTCCCTACACGACGCTCTTCCGATCTNNNACCTCATGTCTCTCTTCTTTGCC
GAGCACACGTCTGAACTCTTTCCCTACACGACGCTCTTCCGATCTNNNATGACAGATTTGAAGAGTG
AGCACACGTCTGAACTCTTTCCCTACACGACGCTCTTCCGATCTNNNCAGAATTGCCATCTGATTGAG
AGCACACGTCTGAACTCTTTCCCTACACGACGCTCTTCCGATCTNNNGCAGAGCATCACAGGTATAAA

GAGCACACGTCTGAACTCTTTCCCTACACGACGCTCTTCCGATCTNNNCTAGCCTGGTTCTTAGGGTAT
GAGCACACGTCTGAACTCTTTCCCTACACGACGCTCTTCCGATCTNNNGAGTGGCACTAACTAGAGTCT
GAGCACACGTCTGAACTCTTTCCCTACACGACGCTCTTCCGATCTNNNAGAGATGAACACACAGCATC
AGAGCACACGTCTGAACTCTTTCCCTACACGACGCTCTTCCGATCTNNNGTTCCTACTTCCTATGCTCCCA
GAGCACACGTCTGAACTCTTTCCCTACACGACGCTCTTCCGATCTNNNAGTGAGAGACTGCATCACCA
AGCACACGTCTGAACTCTTTCCCTACACGACGCTCTTCCGATCTNNNCTGGTTACAAATCTGTCCCCTCA
GAGCACACGTCTGAACTCTTTCCCTACACGACGCTCTTCCGATCTNNNTAGCTGGCATCTTTTCCTTCCA
AGCACACGTCTGAACTCTTTCCCTACACGACGCTCTTCCGATCTNNNACTAGGAGATGAGTCGAAGGG
AGCACACGTCTGAACTCTTTCCCTACACGACGCTCTTCCGATCTNNNATTCTTCTCCATCACCATCGCTTC
GAGCACACGTCTGAACTCTTTCCCTACACGACGCTCTTCCGATCTNNNATGAGCTATGATGGCACTACT
AGCACACGTCTGAACTCTTTCCCTACACGACGCTCTTCCGATCTNNNCCAAGTTTCCTCAGAAAAGCAG
GCACACGTCTGAACTCTTTCCCTACACGACGCTCTTCCGATCTNNNGGATGAAAGTTAGGCTGGACAG
AGAGCACACGTCTGAACTCTTTCCCTACACGACGCTCTTCCGATCTNNNTGGTTCTTGTTTTTGTGCTC
GCACACGTCTGAACTCTTTCCCTACACGACGCTCTTCCGATCTNNNACATGTATGTAGAGGGGGGAAC
GAGCACACGTCTGAACTCTTTCCCTACACGACGCTCTTCCGATCTNNNGAGGACACATGTTTGCCAAGA
AGCACACGTCTGAACTCTTTCCCTACACGACGCTCTTCCGATCTNNNCTGCATGTGTAGGAATGACCAT
AGCACACGTCTGAACTCTTTCCCTACACGACGCTCTTCCGATCTNNNATACTCACTGCGTTAGCTAGAG
AGCACACGTCTGAACTCTTTCCCTACACGACGCTCTTCCGATCTNNNCCTGGCCTGGATTTCTTTTTGAC
GAGCACACGTCTGAACTCTTTCCCTACACGACGCTCTTCCGATCTNNNCCCTAAGGTCAGAGCAACAGA
GAGCACACGTCTGAACTCTTTCCCTACACGACGCTCTTCCGATCTNNNTATGTGCTCAAAACCACACACA
GAGCACACGTCTGAACTCTTTCCCTACACGACGCTCTTCCGATCTNNNCCCTTCAAAGGTTTCTGACTCCT
AGCACACGTCTGAACTCTTTCCCTACACGACGCTCTTCCGATCTNNNGGCTACTCCATTCACTCTTTCCT
AGCACACGTCTGAACTCTTTCCCTACACGACGCTCTTCCGATCTNNNTGTTTATTTGCATGGCCCACAAT
GCACACGTCTGAACTCTTTCCCTACACGACGCTCTTCCGATCTNNNTCTTTGGGACCAGCTAGGAGATA
GAGCACACGTCTGAACTCTTTCCCTACACGACGCTCTTCCGATCTNNNTTTTGAGTAGGGGTCCTGTGA
AGCACACGTCTGAACTCTTTCCCTACACGACGCTCTTCCGATCTNNNCCAATCCCGTGTGTGTGTACAT
AGCACACGTCTGAACTCTTTCCCTACACGACGCTCTTCCGATCTNNNTTCTCATCTGGGCAAGTAACAC
AGCACACGTCTGAACTCTTTCCCTACACGACGCTCTTCCGATCTNNNAATAAGCTGTTTTGCAGGCACA
GAGCACACGTCTGAACTCTTTCCCTACACGACGCTCTTCCGATCTNNNCCTCTTTCTCCTCTTGACTCCTG
GAGCACACGTCTGAACTCTTTCCCTACACGACGCTCTTCCGATCTNNNCAACATCCATGTGCTCTTTTCC
GAGCACACGTCTGAACTCTTTCCCTACACGACGCTCTTCCGATCTNNNATTTTGACACAGGGACACACA
AGCACACGTCTGAACTCTTTCCCTACACGACGCTCTTCCGATCTNNNGTTGGAAATGATGACAGCACAC
AGCACACGTCTGAACTCTTTCCCTACACGACGCTCTTCCGATCTNNNTCTCTGTCACAAAGAAACCTGGA
AGCACACGTCTGAACTCTTTCCCTACACGACGCTCTTCCGATCTNNNACCCAAATGTTTATGCATTGCCA
AGCACACGTCTGAACTCTTTCCCTACACGACGCTCTTCCGATCTNNNATTGTGTTTTAGGAGCTGTGTG
AGCACACGTCTGAACTCTTTCCCTACACGACGCTCTTCCGATCTNNNTCACCAGGAATGTATGAGTGTC
AGAGCACACGTCTGAACTCTTTCCCTACACGACGCTCTTCCGATCTNNNCACTGATGGACTTGGGAAAG
AGAGCACACGTCTGAACTCTTTCCCTACACGACGCTCTTCCGATCTNNNTCCTATGATTTCCTCCCCTTCC
AGCACACGTCTGAACTCTTTCCCTACACGACGCTCTTCCGATCTNNNGCCAGAACTTCACACTTCTAAGC
AGCACACGTCTGAACTCTTTCCCTACACGACGCTCTTCCGATCTNNNTTCCCTTGGTAATGCATCTGTGT
AGCACACGTCTGAACTCTTTCCCTACACGACGCTCTTCCGATCTNNNCTTATCCCACCTTCTGTTCTGGTC
GAGCACACGTCTGAACTCTTTCCCTACACGACGCTCTTCCGATCTNNNTGGGAAAAGCAGATTAGACCT
GAGCACACGTCTGAACTCTTTCCCTACACGACGCTCTTCCGATCTNNNGGAAGTTTGAGCTGACTTTCC

AGCACACGTCTGAACTCTTTCCCTACACGACGCTCTTCCGATCTNNNAGTAGCCCTAATGCACTTTCCA
AGCACACGTCTGAACTCTTTCCCTACACGACGCTCTTCCGATCTNNNCTTGTGGGATTGTGGGATTT
GCACACGTCTGAACTCTTTCCCTACACGACGCTCTTCCGATCTNNNTATACCCTAAACCTCAGCATCGC
AGCACACGTCTGAACTCTTTCCCTACACGACGCTCTTCCGATCTNNNGTGACAATTTAGAGGGGGGA
GCACACGTCTGAACTCTTTCCCTACACGACGCTCTTCCGATCTNNNGAAAGTACCTGTCTCAAGGGGT
AGCACACGTCTGAACTCTTTCCCTACACGACGCTCTTCCGATCTNNNCCCTAACCCCTGTGTGATTCAA
GAGCACACGTCTGAACTCTTTCCCTACACGACGCTCTTCCGATCTNNNCTAACTCAACCTCGGGACAG
GAGCACACGTCTGAACTCTTTCCCTACACGACGCTCTTCCGATCTNNNATTTGTACCAGCAAACCAAG
AGCACACGTCTGAACTCTTTCCCTACACGACGCTCTTCCGATCTNNNCTCTCTGGTGGTGGACATCAT
GCACACGTCTGAACTCTTTCCCTACACGACGCTCTTCCGATCTNNNTTCTCCTCATTGCTACTTTTGCTC
GAGCACACGTCTGAACTCTTTCCCTACACGACGCTCTTCCGATCTNNNTGAGTGTGATGAGTCTGGA
AGCACACGTCTGAACTCTTTCCCTACACGACGCTCTTCCGATCTNNNGGAAGGGCTATGTGGAGAAG
AGCACACGTCTGAACTCTTTCCCTACACGACGCTCTTCCGATCTNNNTCAGACAGGTCTTAGAGTCCA
AGCACACGTCTGAACTCTTTCCCTACACGACGCTCTTCCGATCTNNNATGCTGGACCCAAAATGAATC
AGCACACGTCTGAACTCTTTCCCTACACGACGCTCTTCCGATCTNNNGCAAGCCCAGAAATAGTGTTT
AAGAGCACACGTCTGAACTCTTTCCCTACACGACGCTCTTCCGATCTNNNCCCCTGTCTCTTTGAATCCT
AGCACACGTCTGAACTCTTTCCCTACACGACGCTCTTCCGATCTNNNGAGAAAAAGGGGAAGGTGC
GCACACGTCTGAACTCTTTCCCTACACGACGCTCTTCCGATCTNNNATCTGACAATGCCTTCCCATCTT
AGCACACGTCTGAACTCTTTCCCTACACGACGCTCTTCCGATCTNNNTAGCCTCTTCTTTTCCTGGAAG
AGCACACGTCTGAACTCTTTCCCTACACGACGCTCTTCCGATCTNNNTCTTTTACTGACTAGCACGTGG
AGCACACGTCTGAACTCTTTCCCTACACGACGCTCTTCCGATCTNNNGCACAAGTCAAGCTGTTTCTT
GAGCACACGTCTGAACTCTTTCCCTACACGACGCTCTTCCGATCTNNNCAGAGCCTTCTTGGGACTGT
CACACGTCTGAACTCTTTCCCTACACGACGCTCTTCCGATCTNNNTAAGAATGAGGTTAGGGCAAGG
AGCACACGTCTGAACTCTTTCCCTACACGACGCTCTTCCGATCTNNNTGGACTAAGACACACACTCACA
AGCACACGTCTGAACTCTTTCCCTACACGACGCTCTTCCGATCTNNNTTTTCAATACGGGAGAGCCAG
GCACACGTCTGAACTCTTTCCCTACACGACGCTCTTCCGATCTNNNCTTGTGGTTGTGTGTGTGTGTA
GAGCACACGTCTGAACTCTTTCCCTACACGACGCTCTTCCGATCTNNNATGCACTGTGTCTGGTGTAG
GAGCACACGTCTGAACTCTTTCCCTACACGACGCTCTTCCGATCTNNNGAAACAGGACTGCATGGAT
AGCACACGTCTGAACTCTTTCCCTACACGACGCTCTTCCGATCTNNNACAATGTGAACACTCCTATGG
GCACACGTCTGAACTCTTTCCCTACACGACGCTCTTCCGATCTNNNCTATCTACCATTCAGGACCTGGC
AGCACACGTCTGAACTCTTTCCCTACACGACGCTCTTCCGATCTNNNGCTCAAAGCACTTGTGGGAAA
GCACACGTCTGAACTCTTTCCCTACACGACGCTCTTCCGATCTNNNCATGCAAGACTGTGTGAAAGGA
AGCACACGTCTGAACTCTTTCCCTACACGACGCTCTTCCGATCTNNNTCCCAAACTTTCCACAGTATCGT
AGCACACGTCTGAACTCTTTCCCTACACGACGCTCTTCCGATCTNNNACCCAAAAGCACTATCTTCAAC
GAGCACACGTCTGAACTCTTTCCCTACACGACGCTCTTCCGATCTNNNAAGCTGCTATGCTGTGAATC
AGCACACGTCTGAACTCTTTCCCTACACGACGCTCTTCCGATCTNNNTGTGCTGGGTTTAGAAGTAGC
GCACACGTCTGAACTCTTTCCCTACACGACGCTCTTCCGATCTNNNAACTGGAGAAAGTCAGTGCAT
GAGCACACGTCTGAACTCTTTCCCTACACGACGCTCTTCCGATCTNNNAATGAAGGGAATGCAGAGT
GAGCACACGTCTGAACTCTTTCCCTACACGACGCTCTTCCGATCTNNNGCCACATTCATTCAGAGCTTG
AGCACACGTCTGAACTCTTTCCCTACACGACGCTCTTCCGATCTNNNAAAATGACAATTTGGGGTTG
AGCACACGTCTGAACTCTTTCCCTACACGACGCTCTTCCGATCTNNNAAAGTCCGTACCAGTGGGATA
GAGCACACGTCTGAACTCTTTCCCTACACGACGCTCTTCCGATCTNNNGTGTTTGCCTTTCTCCTCTTGG
GAGCACACGTCTGAACTCTTTCCCTACACGACGCTCTTCCGATCTNNNATGTAGCTGAAGGGGAACA

AGCACACGTCTGAACTCTTTCCCTACACGACGCTCTTCCGATCTNNNATTCTGGAGCTACACTTTGGG
AGAGCACACGTCTGAACTCTTTCCCTACACGACGCTCTTCCGATCTNNNATAGAGGTTTAGTGCACAC
AGCACACGTCTGAACTCTTTCCCTACACGACGCTCTTCCGATCTNNNACATGCATGTCTCTCTCTCTGT
GCACACGTCTGAACTCTTTCCCTACACGACGCTCTTCCGATCTNNNGGGGATGCTCTTTCTAGAGGTT
GCACACGTCTGAACTCTTTCCCTACACGACGCTCTTCCGATCTNNNGGCAGGCACCTCAAATTCTAATC
GAGCACACGTCTGAACTCTTTCCCTACACGACGCTCTTCCGATCTNNNGTTTTCCTCTGTCAATCATGG
AGCACACGTCTGAACTCTTTCCCTACACGACGCTCTTCCGATCTNNNACCCAAAGGTCTGTTATACCCT
GAGCACACGTCTGAACTCTTTCCCTACACGACGCTCTTCCGATCTNNNTAGAGGGGAGGCTTCTGAAT
GCACACGTCTGAACTCTTTCCCTACACGACGCTCTTCCGATCTNNNGAGGTACACATGTCCAAGAGCA
AGCACACGTCTGAACTCTTTCCCTACACGACGCTCTTCCGATCTNNNAAAAACTGCAGAACCCTCCAC
GAGCACACGTCTGAACTCTTTCCCTACACGACGCTCTTCCGATCTNNNAGAAGCCAAACAGGAGAAG
AGCACACGTCTGAACTCTTTCCCTACACGACGCTCTTCCGATCTNNNGTAGAGCCATCACCTCAGAA
AGCACACGTCTGAACTCTTTCCCTACACGACGCTCTTCCGATCTNNNGTTGTCTTCTCTCCCTAACCCTC
GCACACGTCTGAACTCTTTCCCTACACGACGCTCTTCCGATCTNNNAGAGATAGAGTTGGGGGTTG
GAGCACACGTCTGAACTCTTTCCCTACACGACGCTCTTCCGATCTNNNTGCGTAAGATAAGCTCTGTC
GAGCACACGTCTGAACTCTTTCCCTACACGACGCTCTTCCGATCTNNNGGAAGGAAGAGGCTACGTA
AGAGCACACGTCTGAACTCTTTCCCTACACGACGCTCTTCCGATCTNNNTAGGGAACACGTGCTGAAC
AGCACACGTCTGAACTCTTTCCCTACACGACGCTCTTCCGATCTNNNCTTGGGTATTAGACTAGACG
AGCACACGTCTGAACTCTTTCCCTACACGACGCTCTTCCGATCTNNNAAGACAATTCTCTGCCTCTCCT
AGCACACGTCTGAACTCTTTCCCTACACGACGCTCTTCCGATCTNNNTGCACCCCTAAATCCTTCACTTT
AGCACACGTCTGAACTCTTTCCCTACACGACGCTCTTCCGATCTNNNCGGGCTTTTACTTATGGCAGA
GCACACGTCTGAACTCTTTCCCTACACGACGCTCTTCCGATCTNNNTGATACTGCATGATCCCTTCTG
AGCACACGTCTGAACTCTTTCCCTACACGACGCTCTTCCGATCTNNNCAGAGAACATGGAAACCTGG
AGCACACGTCTGAACTCTTTCCCTACACGACGCTCTTCCGATCTNNNATTGGGAAGTCAAATGTTGC
AGCACACGTCTGAACTCTTTCCCTACACGACGCTCTTCCGATCTNNNTTGCTTGTCTCATTTGGTCTTC
GAGCACACGTCTGAACTCTTTCCCTACACGACGCTCTTCCGATCTNNNGACTCTGGCTCTTCTACTGAC
GCACACGTCTGAACTCTTTCCCTACACGACGCTCTTCCGATCTNNNCCAGCGGGTGAATATCTAGGTA
AGCACACGTCTGAACTCTTTCCCTACACGACGCTCTTCCGATCTNNNCTGAAGGAAATGGCTTTCTGC
AGCACACGTCTGAACTCTTTCCCTACACGACGCTCTTCCGATCTNNNGATCATGGCAGAATTGTGCAT
GCACACGTCTGAACTCTTTCCCTACACGACGCTCTTCCGATCTNNNCACACATACCAGACAAACCACA
AGCACACGTCTGAACTCTTTCCCTACACGACGCTCTTCCGATCTNNNAGAAAGTAAGAATGGAGCCCC
GCACACGTCTGAACTCTTTCCCTACACGACGCTCTTCCGATCTNNNAAATCTCAGTCTCTCTCTCCCCA
AGCACACGTCTGAACTCTTTCCCTACACGACGCTCTTCCGATCTNNNCTTTTCAACACCACCACCAGAT
GAGCACACGTCTGAACTCTTTCCCTACACGACGCTCTTCCGATCTNNNATCTGAGGGTGAGTAGTGC
AGAGCACACGTCTGAACTCTTTCCCTACACGACGCTCTTCCGATCTNNNAGAGATCTGACATCACATG
AGCACACGTCTGAACTCTTTCCCTACACGACGCTCTTCCGATCTNNNTCTGCTTTTTGAGTTATGTGCG
GAGCACACGTCTGAACTCTTTCCCTACACGACGCTCTTCCGATCTNNNTCCAGATTTCCGTGTGTGTAT
GAGCACACGTCTGAACTCTTTCCCTACACGACGCTCTTCCGATCTNNNCCCACACACAATGGTTACAC
GCACACGTCTGAACTCTTTCCCTACACGACGCTCTTCCGATCTNNNCAAAGTGTGGTCACCTCAATGT
AGCACACGTCTGAACTCTTTCCCTACACGACGCTCTTCCGATCTNNNCAACAAGAAAGTGGTGGTGG
GCACACGTCTGAACTCTTTCCCTACACGACGCTCTTCCGATCTNNNCCCCGTGGAAGAAAGGCTATAT
AGAGCACACGTCTGAACTCTTTCCCTACACGACGCTCTTCCGATCTNNNAAGCCAATAGAAGCTCCAA
AGCACACGTCTGAACTCTTTCCCTACACGACGCTCTTCCGATCTNNNTGCAGCTGATGTTTCTTTCCATT

GAGCACACGTCTGAACTCTTTCCCTACACGACGCTCTTCCGATCTNNNGACATGGATGATGGGAATG
CACACGTCTGAACTCTTTCCCTACACGACGCTCTTCCGATCTNNNTACTGACCCCAGCTTCATTCTTCC
AGCACACGTCTGAACTCTTTCCCTACACGACGCTCTTCCGATCTNNNCTGACTCAGGAGATTGCTCTG
GCACACGTCTGAACTCTTTCCCTACACGACGCTCTTCCGATCTNNNTTGTTCTCTTCAGTGCATTGGTG
GAGCACACGTCTGAACTCTTTCCCTACACGACGCTCTTCCGATCTNNNGTGTGTGTGTAAGGGCTTCA
AGCACACGTCTGAACTCTTTCCCTACACGACGCTCTTCCGATCTNNNGCTTGGCTGCTCTGTCTATAGA
AGCACACGTCTGAACTCTTTCCCTACACGACGCTCTTCCGATCTNNNATTCCTCAGTCAGATTACTCG
AGCACACGTCTGAACTCTTTCCCTACACGACGCTCTTCCGATCTNNNTGTGTTAGTCTTCTCCAGGGA
GCACACGTCTGAACTCTTTCCCTACACGACGCTCTTCCGATCTNNNAGCCTCTATAACCATGTGAACC
AGCACACGTCTGAACTCTTTCCCTACACGACGCTCTTCCGATCTNNNAGGAGGTCCAGGTAAGAAGT
AGCACACGTCTGAACTCTTTCCCTACACGACGCTCTTCCGATCTNNNAAATTTTCAAACAGTTGCCCA
AGCACACGTCTGAACTCTTTCCCTACACGACGCTCTTCCGATCTNNNAGGTGTGACTTAGGGAGGA
GAGCACACGTCTGAACTCTTTCCCTACACGACGCTCTTCCGATCTNNNACTGGCCCTCTCTAACTACCT
AGCACACGTCTGAACTCTTTCCCTACACGACGCTCTTCCGATCTNNNATTTTGCTCCCTCTCCATGAAGT
GAGCACACGTCTGAACTCTTTCCCTACACGACGCTCTTCCGATCTNNNTGGCACGTCCCAAACATTATC
AGCACACGTCTGAACTCTTTCCCTACACGACGCTCTTCCGATCTNNNGAGAGAGGCTGGCAGATGTA
GCACACGTCTGAACTCTTTCCCTACACGACGCTCTTCCGATCTNNNAGCAAAACAGAATGCCTCACAA
AGCACACGTCTGAACTCTTTCCCTACACGACGCTCTTCCGATCTNNNCAAGGGAGTGCCATTTTTGTG
GAGCACACGTCTGAACTCTTTCCCTACACGACGCTCTTCCGATCTNNNAGGTATGAATTTTGCAAGGT
GAGCACACGTCTGAACTCTTTCCCTACACGACGCTCTTCCGATCTNNNGAAGTCAAAAAAATCCCCCA
AGCACACGTCTGAACTCTTTCCCTACACGACGCTCTTCCGATCTNNNGCTGCGGTATTGATTCTACAC
GCACACGTCTGAACTCTTTCCCTACACGACGCTCTTCCGATCTNNNTGTCTTCCAGGAAAGGATACAG
AGCACACGTCTGAACTCTTTCCCTACACGACGCTCTTCCGATCTNNNACTGCAGATCTCAGAACTTGT
GCACACGTCTGAACTCTTTCCCTACACGACGCTCTTCCGATCTNNNCTGTTGTCTTCCAAACATGCAC
AGCACACGTCTGAACTCTTTCCCTACACGACGCTCTTCCGATCTNNNGAAGGAAAATGGCAGAGACA
AGCACACGTCTGAACTCTTTCCCTACACGACGCTCTTCCGATCTNNNGACCAATGCCCCTGTAGAAGA
AGCACACGTCTGAACTCTTTCCCTACACGACGCTCTTCCGATCTNNNTCTATCCTAGAGGGGCTGGAA
GCACACGTCTGAACTCTTTCCCTACACGACGCTCTTCCGATCTNNNAGCAGCTGTGTTAGGAGTTCTT
AGCACACGTCTGAACTCTTTCCCTACACGACGCTCTTCCGATCTNNNCTCCAGCATTGACCAAAATGT
GCACACGTCTGAACTCTTTCCCTACACGACGCTCTTCCGATCTNNNTATGATCAGCTGTGTGTACGCAT
AGAGCACACGTCTGAACTCTTTCCCTACACGACGCTCTTCCGATCTNNNTGGGGAATACTTTATGCA
GAGCACACGTCTGAACTCTTTCCCTACACGACGCTCTTCCGATCTNNNTTGCAGATCATGGGACTTCT
AGCACACGTCTGAACTCTTTCCCTACACGACGCTCTTCCGATCTNNNGGAAAATCTGGGCTTTTCTTC
AGCACACGTCTGAACTCTTTCCCTACACGACGCTCTTCCGATCTNNNAACTCTGGCATGTAAACTGG
GAGCACACGTCTGAACTCTTTCCCTACACGACGCTCTTCCGATCTNNNGAGAACCCCCATCTTAGATTC
AGCACACGTCTGAACTCTTTCCCTACACGACGCTCTTCCGATCTNNNAGATTATGGCAGAAGGGATG
GCACACGTCTGAACTCTTTCCCTACACGACGCTCTTCCGATCTNNNGGAACTGGGACATCTACATTCC
GAGCACACGTCTGAACTCTTTCCCTACACGACGCTCTTCCGATCTNNNCGGTGTCATCTCAAGGGTTC
AGCACACGTCTGAACTCTTTCCCTACACGACGCTCTTCCGATCTNNNCTTTGCTCATGTTCTCAGAAGC
AGCACACGTCTGAACTCTTTCCCTACACGACGCTCTTCCGATCTNNNTGACTACTTCCGTGGGATATTC
AGCACACGTCTGAACTCTTTCCCTACACGACGCTCTTCCGATCTNNNTCTAAGGTACCAGCTTGGGAG
AGCACACGTCTGAACTCTTTCCCTACACGACGCTCTTCCGATCTNNNCCTGGGAGGTTGAGGGTATA
GCACACGTCTGAACTCTTTCCCTACACGACGCTCTTCCGATCTNNNTTCCAAGGAGAAAGGAAAGCA

AGCACACGTCTGAACTCTTTCCCTACACGACGCTCTTCCGATCTNNNGTAAGTTCTCCTCGATCTCCAGC
AGCACACGTCTGAACTCTTTCCCTACACGACGCTCTTCCGATCTNNNACTGATGGGTATTCACAGGTT
GAGCACACGTCTGAACTCTTTCCCTACACGACGCTCTTCCGATCTNNNCCTGTATCTGCAGTTCAGGAC
GAGCACACGTCTGAACTCTTTCCCTACACGACGCTCTTCCGATCTNNNACTGCACCTCAAAACAACAAC
GAGCACACGTCTGAACTCTTTCCCTACACGACGCTCTTCCGATCTNNNCAAATCCTTCCCATTCCCAAAC
AGCACACGTCTGAACTCTTTCCCTACACGACGCTCTTCCGATCTNNNATGGGACTGGGTAATAGTCAAC
GAGCACACGTCTGAACTCTTTCCCTACACGACGCTCTTCCGATCTNNNAGGTGTTCCATCATTCTGGG
AGCACACGTCTGAACTCTTTCCCTACACGACGCTCTTCCGATCTNNNCGCACCTGGTCAACAATATCTT
GAGCACACGTCTGAACTCTTTCCCTACACGACGCTCTTCCGATCTNNNAGCCACATGCAGGTAGTAGT
GAGCACACGTCTGAACTCTTTCCCTACACGACGCTCTTCCGATCTNNNAGCCATCTGGAAACCTCTTTA
GAGCACACGTCTGAACTCTTTCCCTACACGACGCTCTTCCGATCTNNNACTCTCCCTTCTCAAATGGTTG
GCACACGTCTGAACTCTTTCCCTACACGACGCTCTTCCGATCTNNNGCCCGAGACTCATGTAGTTTCTT
GAGCACACGTCTGAACTCTTTCCCTACACGACGCTCTTCCGATCTNNNACTCTCTTCAGTGTGGGATGA
GCACACGTCTGAACTCTTTCCCTACACGACGCTCTTCCGATCTNNNGAGAAGACCCCAAAAACTTCTGC
GAGCACACGTCTGAACTCTTTCCCTACACGACGCTCTTCCGATCTNNNGATAATGAGAGACTGCAGCC
AGCACACGTCTGAACTCTTTCCCTACACGACGCTCTTCCGATCTNNNCTGAATCTCAACTACACAGGCC
GCACACGTCTGAACTCTTTCCCTACACGACGCTCTTCCGATCTNNNACATGGACTGAGAGCTGTTTGA
GAGCACACGTCTGAACTCTTTCCCTACACGACGCTCTTCCGATCTNNNTGCCTTTTCTCTTCTGTCTGTGT
AGCACACGTCTGAACTCTTTCCCTACACGACGCTCTTCCGATCTNNNAGGTGAGGACTTACTTTACGTG
GCACACGTCTGAACTCTTTCCCTACACGACGCTCTTCCGATCTNNNCCTAGAACTTGATTTTCGTGGCA
GAGCACACGTCTGAACTCTTTCCCTACACGACGCTCTTCCGATCTNNNATTCCAACACATGTTCCCTCTG
GAGCACACGTCTGAACTCTTTCCCTACACGACGCTCTTCCGATCTNNNAGGTTGGGATGATCTGGTGT
AGCACACGTCTGAACTCTTTCCCTACACGACGCTCTTCCGATCTNNNGCAGGAATGCCTAGGTCAGAA
AGCACACGTCTGAACTCTTTCCCTACACGACGCTCTTCCGATCTNNNTTCATACCATGAGGATGCAAGG
GAGCACACGTCTGAACTCTTTCCCTACACGACGCTCTTCCGATCTNNNGCCACTTTGGGTTCTGTTTAT
GAGCACACGTCTGAACTCTTTCCCTACACGACGCTCTTCCGATCTNNNAGCCTGTCATAGCAAAGAGA
AGCACACGTCTGAACTCTTTCCCTACACGACGCTCTTCCGATCTNNNGCTTGACACATATTGAAGAGGC
AGCACACGTCTGAACTCTTTCCCTACACGACGCTCTTCCGATCTNNNTGTGCACTCTCTAAATCTTGCCT
GAGCACACGTCTGAACTCTTTCCCTACACGACGCTCTTCCGATCTNNNTTACTTGAGCCTTCGAGGAAG
AGAGCACACGTCTGAACTCTTTCCCTACACGACGCTCTTCCGATCTNNNAAGTGATCAACAAGCAGGT
AGCACACGTCTGAACTCTTTCCCTACACGACGCTCTTCCGATCTNNNACACAGCAGCTGAAACTAATTT
AGCACACGTCTGAACTCTTTCCCTACACGACGCTCTTCCGATCTNNNCATTGCCCAGAAAGAACAGACA
GAGCACACGTCTGAACTCTTTCCCTACACGACGCTCTTCCGATCTNNNAACACAAATCTGACTTCAGCT
GAGCACACGTCTGAACTCTTTCCCTACACGACGCTCTTCCGATCTNNNGGACACTGAGTAACAAAACTG
GAGCACACGTCTGAACTCTTTCCCTACACGACGCTCTTCCGATCTNNNTGCAGCCCTTCTGTGAATATG
GCACACGTCTGAACTCTTTCCCTACACGACGCTCTTCCGATCTNNNTGGAGAAGGGACACCACTAATT
GCACACGTCTGAACTCTTTCCCTACACGACGCTCTTCCGATCTNNNTCCAAGTTTGTCCATTTGCAGTC
GAGCACACGTCTGAACTCTTTCCCTACACGACGCTCTTCCGATCTNNNAATGAGCACATCTAGGCAATT
GAGCACACGTCTGAACTCTTTCCCTACACGACGCTCTTCCGATCTNNNATGGAGGTTTCAAAAGCTTGT
GAGCACACGTCTGAACTCTTTCCCTACACGACGCTCTTCCGATCTNNNTCTGCCCTCCACAGGTAGTATA
GAGCACACGTCTGAACTCTTTCCCTACACGACGCTCTTCCGATCTNNNTGGTGACATAAGATACAGGT
AGCACACGTCTGAACTCTTTCCCTACACGACGCTCTTCCGATCTNNNTAAAGCCAACTACACCTCCACA
GAGCACACGTCTGAACTCTTTCCCTACACGACGCTCTTCCGATCTNNNTGAATGAGGACTTGGACTTG

AGCACACGTCTGAACTCTTTCCCTACACGACGCTCTTCCGATCTNNNGGGGAAAAGTGTTCATGTTG
AGCACACGTCTGAACTCTTTCCCTACACGACGCTCTTCCGATCTNNNCAATGTGACTGTGCTTCATGA
GAGCACACGTCTGAACTCTTTCCCTACACGACGCTCTTCCGATCTNNNGGTCTGTCTTGATCACGTAG
AGCACACGTCTGAACTCTTTCCCTACACGACGCTCTTCCGATCTNNNACCTGTGACAAAGTTACGTGA
GAGCACACGTCTGAACTCTTTCCCTACACGACGCTCTTCCGATCTNNNCTGCCTCTTTGTTTACCTGTG
AGCACACGTCTGAACTCTTTCCCTACACGACGCTCTTCCGATCTNNNTTGTCCTGCCAAGTTTTGTCTT
AGCACACGTCTGAACTCTTTCCCTACACGACGCTCTTCCGATCTNNNATTCTGTCTTGAGGTGCTCATG
AGCACACGTCTGAACTCTTTCCCTACACGACGCTCTTCCGATCTNNNCGCCCAGCCTAGTTTTTTCTTTT
AGCACACGTCTGAACTCTTTCCCTACACGACGCTCTTCCGATCTNNNTAGATGGACCCCACAGCTAAAA
AGCACACGTCTGAACTCTTTCCCTACACGACGCTCTTCCGATCTNNNTTGTCAGCCTGTAATTAGGTG
AGCACACGTCTGAACTCTTTCCCTACACGACGCTCTTCCGATCTNNNCTCCTCCATACACACATGCTCTT
CACACGTCTGAACTCTTTCCCTACACGACGCTCTTCCGATCTNNNGTTTGTCATCTCAGGTCTAGGCA
GAGCACACGTCTGAACTCTTTCCCTACACGACGCTCTTCCGATCTNNNGAAAGAACAGGCATGGGGA
GAGCACACGTCTGAACTCTTTCCCTACACGACGCTCTTCCGATCTNNNATTATTTGTGTAGGCTCCTGG
AGCACACGTCTGAACTCTTTCCCTACACGACGCTCTTCCGATCTNNNATATCTGGTTTCTTAGGCCCCA
AGCACACGTCTGAACTCTTTCCCTACACGACGCTCTTCCGATCTNNNAGGCAGCCTCCTGTTTATAAA
GAGCACACGTCTGAACTCTTTCCCTACACGACGCTCTTCCGATCTNNNATTCATGATGAAGGTCTGTG
GAGCACACGTCTGAACTCTTTCCCTACACGACGCTCTTCCGATCTNNNCTAGGGACATGAGCTTGAG
AGCACACGTCTGAACTCTTTCCCTACACGACGCTCTTCCGATCTNNNACAGTCAAAGTCCTCAACTCCA
AGCACACGTCTGAACTCTTTCCCTACACGACGCTCTTCCGATCTNNNTTTGAATTGGAGGGGACTTTC
AGCACACGTCTGAACTCTTTCCCTACACGACGCTCTTCCGATCTNNNACCCTAGACTTCTCAGATGTA
GAGCACACGTCTGAACTCTTTCCCTACACGACGCTCTTCCGATCTNNNCTATTGGCTGTATCGTTCATG
GAGCACACGTCTGAACTCTTTCCCTACACGACGCTCTTCCGATCTNNNTAGGATGCCAGTAATACCTC
GAGCACACGTCTGAACTCTTTCCCTACACGACGCTCTTCCGATCTNNNGCTATTTATTGAACACGCCTG
AGCACACGTCTGAACTCTTTCCCTACACGACGCTCTTCCGATCTNNNCTACTCAAGCTGCCAGGAAAC
AGCACACGTCTGAACTCTTTCCCTACACGACGCTCTTCCGATCTNNNTCAAATTGCTCCTGGGGATAA
AGCACACGTCTGAACTCTTTCCCTACACGACGCTCTTCCGATCTNNNTGTGTCTTTTACAGAGACGGG
GAGCACACGTCTGAACTCTTTCCCTACACGACGCTCTTCCGATCTNNNTTTTTTGAAGAGAGGGGGC
AGCACACGTCTGAACTCTTTCCCTACACGACGCTCTTCCGATCTNNNCTCCCAACCCTTTCAAATCCAAC
AGCACACGTCTGAACTCTTTCCCTACACGACGCTCTTCCGATCTNNNGGGGAAGTTCTCTGAACTCAA
GCACACGTCTGAACTCTTTCCCTACACGACGCTCTTCCGATCTNNNATTAGTCAAGGCTGGACAGAGT
AGCACACGTCTGAACTCTTTCCCTACACGACGCTCTTCCGATCTNNNGCAAAGAGGTTGTAACTTGAG
AGCACACGTCTGAACTCTTTCCCTACACGACGCTCTTCCGATCTNNNTGTGTGTGTACGTACCATGAA
GAGCACACGTCTGAACTCTTTCCCTACACGACGCTCTTCCGATCTNNNATTCTCCTACCTCACCCTCTTG
GCACACGTCTGAACTCTTTCCCTACACGACGCTCTTCCGATCTNNNTTCAAGCAATTTTCCTGCCTCAG
GCACACGTCTGAACTCTTTCCCTACACGACGCTCTTCCGATCTNNNGGAATCTCTCTCTCTCCCTCTC
AGCACACGTCTGAACTCTTTCCCTACACGACGCTCTTCCGATCTNNNGGCAACTCTTCATCCTTTCAAG
AGCACACGTCTGAACTCTTTCCCTACACGACGCTCTTCCGATCTNNNAGGCATTACCAAGGAATACG
AGCACACGTCTGAACTCTTTCCCTACACGACGCTCTTCCGATCTNNNATAGGTATATCCCACCACACC
GCACACGTCTGAACTCTTTCCCTACACGACGCTCTTCCGATCTNNNCTTCTTTACAGCTGTTTGGGAG
AGCACACGTCTGAACTCTTTCCCTACACGACGCTCTTCCGATCTNNNCTCCATCTCCCAAAGTGTTGAG
AGCACACGTCTGAACTCTTTCCCTACACGACGCTCTTCCGATCTNNNAATGGTGCTAAGGGTGCTAT
AGCACACGTCTGAACTCTTTCCCTACACGACGCTCTTCCGATCTNNNCACTTTGCAGACTGGGTCTAAT

GAGCACACGTCTGAACTCTTTCCCTACACGACGCTCTTCCGATCTNNNATATTCTTGCTGAGGGAAAG
GAGCACACGTCTGAACTCTTTCCCTACACGACGCTCTTCCGATCTNNNCAAGTATCACACAAACACACC
AGCACACGTCTGAACTCTTTCCCTACACGACGCTCTTCCGATCTNNNCAGGCAGAGACTCTTGTTCTCT
AGCACACGTCTGAACTCTTTCCCTACACGACGCTCTTCCGATCTNNNGCATGCCACTATGCCAGCTAAT
AGCACACGTCTGAACTCTTTCCCTACACGACGCTCTTCCGATCTNNNGGATCCAGCTCCTCCTCTACATA
AGCACACGTCTGAACTCTTTCCCTACACGACGCTCTTCCGATCTNNNTTCCTGCTGCAGTCTCTGTAAA
AGCACACGTCTGAACTCTTTCCCTACACGACGCTCTTCCGATCTNNNTCCCTGGCTTATTTCACTTACCA
GAGCACACGTCTGAACTCTTTCCCTACACGACGCTCTTCCGATCTNNNTCCTCTGTGTGTTCTCTTACTG
GAGCACACGTCTGAACTCTTTCCCTACACGACGCTCTTCCGATCTNNNTTCCCTTACTACCAGTACCCAG
GCACACGTCTGAACTCTTTCCCTACACGACGCTCTTCCGATCTNNNTGATTAGTCAAGGCTGGACAGA
GAGCACACGTCTGAACTCTTTCCCTACACGACGCTCTTCCGATCTNNNCCATTGTAGAATCAGGCAACA
GAGCACACGTCTGAACTCTTTCCCTACACGACGCTCTTCCGATCTNNNCTTTGGTGATAAAGTGAAGCC
AGCACACGTCTGAACTCTTTCCCTACACGACGCTCTTCCGATCTNNNTCTGTCTATAGTCAGACCAGGG
GCACACGTCTGAACTCTTTCCCTACACGACGCTCTTCCGATCTNNNAAGAACAACTTCAGTGGCAAAC
GAGCACACGTCTGAACTCTTTCCCTACACGACGCTCTTCCGATCTNNNCTGAGGTGGGTGGATTGATT
AGCACACGTCTGAACTCTTTCCCTACACGACGCTCTTCCGATCTNNNCAAATAGAGAGCAGATGGAG
GAGCACACGTCTGAACTCTTTCCCTACACGACGCTCTTCCGATCTNNNTAAGCTGTCAGGTCTGCTGTA
GAGCACACGTCTGAACTCTTTCCCTACACGACGCTCTTCCGATCTNNNCTCTCTCTCACACACACACAT
AGCACACGTCTGAACTCTTTCCCTACACGACGCTCTTCCGATCTNNNGTCTTTGCCTCTGTCTCTCTCTCC
AGCACACGTCTGAACTCTTTCCCTACACGACGCTCTTCCGATCTNNNTTGTGATAGACTAGGGCCAGG
GAGCACACGTCTGAACTCTTTCCCTACACGACGCTCTTCCGATCTNNNGTTGGCCACTCATAGAGGTTC
AGCACACGTCTGAACTCTTTCCCTACACGACGCTCTTCCGATCTNNNACATTCACTCTACCAAGGGTTC
AGCACACGTCTGAACTCTTTCCCTACACGACGCTCTTCCGATCTNNNAATTCAGGAGGTAGAGGTTGC
GAGCACACGTCTGAACTCTTTCCCTACACGACGCTCTTCCGATCTNNNTCTCAAAAAAAACCACACCCCT
AGCACACGTCTGAACTCTTTCCCTACACGACGCTCTTCCGATCTNNNGATCACTTGAGCCCATGAGTTT
AGCACACGTCTGAACTCTTTCCCTACACGACGCTCTTCCGATCTNNNCATGCAGTTTTGGTTCAGTAGC
AGCACACGTCTGAACTCTTTCCCTACACGACGCTCTTCCGATCTNNNCGCTGTTCCTCCATGAAACTGA
AGCACACGTCTGAACTCTTTCCCTACACGACGCTCTTCCGATCTNNNAGTATCGAAGACACACACACAC
AGCACACGTCTGAACTCTTTCCCTACACGACGCTCTTCCGATCTNNNAATGGAGTCTACTCTGGATCC
AGCACACGTCTGAACTCTTTCCCTACACGACGCTCTTCCGATCTNNNCACATACACACACAGACACACA
AGAGCACACGTCTGAACTCTTTCCCTACACGACGCTCTTCCGATCTNNNTGTAGATTTTGAACTTGCCA
GAGCACACGTCTGAACTCTTTCCCTACACGACGCTCTTCCGATCTNNNCCCATGTTGATAGCAGCACAAT
GAGCACACGTCTGAACTCTTTCCCTACACGACGCTCTTCCGATCTNNNTCCAGCCCTGCATATCCTAGA
GAGCACACGTCTGAACTCTTTCCCTACACGACGCTCTTCCGATCTNNNGGAATTCTTGGCAGTTACAC
AGCACACGTCTGAACTCTTTCCCTACACGACGCTCTTCCGATCTNNNAACCACAAAGCCATCATAACAG
GAGCACACGTCTGAACTCTTTCCCTACACGACGCTCTTCCGATCTNNNTGCAACATCAACTCCTACCAG
AGCACACGTCTGAACTCTTTCCCTACACGACGCTCTTCCGATCTNNNCTCCATAAAGCTGACAGGCAAG
AGAGCACACGTCTGAACTCTTTCCCTACACGACGCTCTTCCGATCTNNNTAAAAGCAATCACTCTGGGG
GAGCACACGTCTGAACTCTTTCCCTACACGACGCTCTTCCGATCTNNNTCCAAGAAGCTATGCCTGAAA
GAGCACACGTCTGAACTCTTTCCCTACACGACGCTCTTCCGATCTNNNGCATGGTCCTGGATGATTTT
GAGCACACGTCTGAACTCTTTCCCTACACGACGCTCTTCCGATCTNNNGCCAGCCAAGTTTCTTTACAC
GAGCACACGTCTGAACTCTTTCCCTACACGACGCTCTTCCGATCTNNNCTAGCTTACCTATCTCCCCACC
GCACACGTCTGAACTCTTTCCCTACACGACGCTCTTCCGATCTNNNCACCTCCCACCCTAATCACTTTTC

GCACACGTCTGAACTCTTTCCCTACACGACGCTCTTCCGATCTNNNCCTGCATTGTATGTGTCTGTGTC
AGCACACGTCTGAACTCTTTCCCTACACGACGCTCTTCCGATCTNNNAGAGATGCACCACCATATCCAA
AGCACACGTCTGAACTCTTTCCCTACACGACGCTCTTCCGATCTNNNACAGAGTGAGACTCCACTGAG
GAGCACACGTCTGAACTCTTTCCCTACACGACGCTCTTCCGATCTNNNCACCAACAGTGTCCAAAGATT
AGCACACGTCTGAACTCTTTCCCTACACGACGCTCTTCCGATCTNNNTTCCATCCATGTTGCTGCAAAA
GCACACGTCTGAACTCTTTCCCTACACGACGCTCTTCCGATCTNNNCTCTGTATTCCAGCCTAGGTGAC
AGCACACGTCTGAACTCTTTCCCTACACGACGCTCTTCCGATCTNNNGACTACAGGCACATGTCACCAT
GCACACGTCTGAACTCTTTCCCTACACGACGCTCTTCCGATCTNNNAATGTATCAGCTGTGTGCCCTTA
AGCACACGTCTGAACTCTTTCCCTACACGACGCTCTTCCGATCTNNNAAACATACAACCAAAAGCCCCA
AGCACACGTCTGAACTCTTTCCCTACACGACGCTCTTCCGATCTNNNGGAGTGTTTTGTAGCTCTGCT
GAGCACACGTCTGAACTCTTTCCCTACACGACGCTCTTCCGATCTNNNCTGCCCAATCTAGGTAGCCTT
GAGCACACGTCTGAACTCTTTCCCTACACGACGCTCTTCCGATCTNNNGGCTGTGGTCTCTTTTTTACC
GAGCACACGTCTGAACTCTTTCCCTACACGACGCTCTTCCGATCTNNNGTCCAAGAGATTCTCATGCCT
AGCACACGTCTGAACTCTTTCCCTACACGACGCTCTTCCGATCTNNNCCGGTTGCTGTGAATGTTATCA
GCACACGTCTGAACTCTTTCCCTACACGACGCTCTTCCGATCTNNNGTCTTCCCCCAAATCCATTCTCTCA
AGCACACGTCTGAACTCTTTCCCTACACGACGCTCTTCCGATCTNNNAATTCCATTGAGGTTGCTGCAA
AGAGCACACGTCTGAACTCTTTCCCTACACGACGCTCTTCCGATCTNNNGAGTTTGAGGCTGTAGTGAC
GAGCACACGTCTGAACTCTTTCCCTACACGACGCTCTTCCGATCTNNNAGTGGTGTGATCATAGCTCA
AGAGCACACGTCTGAACTCTTTCCCTACACGACGCTCTTCCGATCTNNNAGAGAGGGAGTGTTAGCTC
AGCACACGTCTGAACTCTTTCCCTACACGACGCTCTTCCGATCTNNNGTAAGGGTGTTGGATTTTGCCA
GAGCACACGTCTGAACTCTTTCCCTACACGACGCTCTTCCGATCTNNNTCTCTGGTGTCACAGACAAGA
GAGCACACGTCTGAACTCTTTCCCTACACGACGCTCTTCCGATCTNNNGCAAGACTCCATCTCGAAAAA
GCACACGTCTGAACTCTTTCCCTACACGACGCTCTTCCGATCTNNNACCCTTCTGATGTACAACTGTTT
AGCACACGTCTGAACTCTTTCCCTACACGACGCTCTTCCGATCTNNNGGTGGATCACTTGAGATCAGG
GAGCACACGTCTGAACTCTTTCCCTACACGACGCTCTTCCGATCTNNNGAACTGGAGAACCAAGAGCA
AGCACACGTCTGAACTCTTTCCCTACACGACGCTCTTCCGATCTNNNAGGCCAACATCATCTTGATACC
GAGCACACGTCTGAACTCTTTCCCTACACGACGCTCTTCCGATCTNNNTGATGATCTCTATCTCCTGGC
GAGCACACGTCTGAACTCTTTCCCTACACGACGCTCTTCCGATCTNNNCGAGATCACACCATTGAACTC
AGCACACGTCTGAACTCTTTCCCTACACGACGCTCTTCCGATCTNNNAGCCTGGGTGATAAAGTGAGA
AGAGCACACGTCTGAACTCTTTCCCTACACGACGCTCTTCCGATCTNNNGTATCCAGCAACTCCTTCCTT
GAGCACACGTCTGAACTCTTTCCCTACACGACGCTCTTCCGATCTNNNTCCGTAGCATTTCTGTACACC
AGCACACGTCTGAACTCTTTCCCTACACGACGCTCTTCCGATCTNNNAACATGCTCACTTGGAAGTCTC
GAGCACACGTCTGAACTCTTTCCCTACACGACGCTCTTCCGATCTNNNAAGGCCCTGTCTCAAAAACAA
GAGCACACGTCTGAACTCTTTCCCTACACGACGCTCTTCCGATCTNNNTTCTCAGTGCAAAATTGGCAG
GAGCACACGTCTGAACTCTTTCCCTACACGACGCTCTTCCGATCTNNNAAAATCCCTCTCACACACTTAC
AGCACACGTCTGAACTCTTTCCCTACACGACGCTCTTCCGATCTNNNAAGCTAGAGTAACCAACACAG
GAGCACACGTCTGAACTCTTTCCCTACACGACGCTCTTCCGATCTNNNCAACAACATAGTTGGGCAAT
GAGCACACGTCTGAACTCTTTCCCTACACGACGCTCTTCCGATCTNNNATGGCAGAGATTTTGCAGAT
GCACACGTCTGAACTCTTTCCCTACACGACGCTCTTCCGATCTNNNTTCCTCTTTCTACTGGCCACTTCC
GCACACGTCTGAACTCTTTCCCTACACGACGCTCTTCCGATCTNNNGCAGTGATACAGGGTAGACTCA
GAGCACACGTCTGAACTCTTTCCCTACACGACGCTCTTCCGATCTNNNAAAGAACAAGGTTGTAGCCT
GAGCACACGTCTGAACTCTTTCCCTACACGACGCTCTTCCGATCTNNNAGCCCAAATGTGCATCAATCA
AGCACACGTCTGAACTCTTTCCCTACACGACGCTCTTCCGATCTNNNGAAGAGGAGATGCCATGTCTC

AGAGCACACGTCTGAACTCTTTCCCTACACGACGCTCTTCCGATCTNNNTCCCTGTCAGAAGCTGTTTCT
AGCACACGTCTGAACTCTTTCCCTACACGACGCTCTTCCGATCTNNNTCTCTCTCTCTCTCTCCCTCTCA
GAGCACACGTCTGAACTCTTTCCCTACACGACGCTCTTCCGATCTNNNCTTCATCCCTACGCACACACATA
AGCACACGTCTGAACTCTTTCCCTACACGACGCTCTTCCGATCTNNNCCAGTCAATACACACACACACAC
AGCACACGTCTGAACTCTTTCCCTACACGACGCTCTTCCGATCTNNNTAGTCTCCAATGTCATCCAGGTC
AGCACACGTCTGAACTCTTTCCCTACACGACGCTCTTCCGATCTNNNGTGCGCATGTGTATTATGTA
AGCACACGTCTGAACTCTTTCCCTACACGACGCTCTTCCGATCTNNNAAAGAACAGCGTGTGTATGTG
GAGCACACGTCTGAACTCTTTCCCTACACGACGCTCTTCCGATCTNNNACTTCCCTTCCAGAACCACTTTT
AGCACACGTCTGAACTCTTTCCCTACACGACGCTCTTCCGATCTNNNTCCATGCCCTTCTCTTAGCTTTT
GAGCACACGTCTGAACTCTTTCCCTACACGACGCTCTTCCGATCTNNNTGGGTTCTCCAAGGAAACAGA
AGCACACGTCTGAACTCTTTCCCTACACGACGCTCTTCCGATCTNNNGCCTCTTGAGTAGCAGGGATTA
AGCACACGTCTGAACTCTTTCCCTACACGACGCTCTTCCGATCTNNNTTGTGGTTTACTGATGTGAGCA
AGCACACGTCTGAACTCTTTCCCTACACGACGCTCTTCCGATCTNNNGGCCAAAACCACATAAGTGCATA
AGCACACGTCTGAACTCTTTCCCTACACGACGCTCTTCCGATCTNNNCACATATTCCCTGTCTCTTCCCCA
GAGCACACGTCTGAACTCTTTCCCTACACGACGCTCTTCCGATCTNNNGATTCAGCAGGGTGAGTGTAT
GAGCACACGTCTGAACTCTTTCCCTACACGACGCTCTTCCGATCTNNNTGGGTATGTCTTTATTAGCAG
AGCACACGTCTGAACTCTTTCCCTACACGACGCTCTTCCGATCTNNNGCATGGGACTTTTTGTCTTCCAT
GAGCACACGTCTGAACTCTTTCCCTACACGACGCTCTTCCGATCTNNNAGGGTTCTCCAGACAGACAAA
GAGCACACGTCTGAACTCTTTCCCTACACGACGCTCTTCCGATCTNNNTTCTGTGTGTACATCGCTCTGA
AGCACACGTCTGAACTCTTTCCCTACACGACGCTCTTCCGATCTNNNAAACTCACTGGGGATTTGACAG
AGCACACGTCTGAACTCTTTCCCTACACGACGCTCTTCCGATCTNNNAAACTTGGTCTGTGTTTGTGTG
GAGCACACGTCTGAACTCTTTCCCTACACGACGCTCTTCCGATCTNNNATCCAGGTCCTAGAAGTCTTG
AGCACACGTCTGAACTCTTTCCCTACACGACGCTCTTCCGATCTNNNGTGTGTGTGTGGTTATGTA
AGCACACGTCTGAACTCTTTCCCTACACGACGCTCTTCCGATCTNNNGTTCCCCAAAGAAACAGAACCAC
AGCACACGTCTGAACTCTTTCCCTACACGACGCTCTTCCGATCTNNNAAAGACCGTTTCCAAAACTGAC
AGAGCACACGTCTGAACTCTTTCCCTACACGACGCTCTTCCGATCTNNNAGATATTCAAGAGCAGCCTA
AGAGCACACGTCTGAACTCTTTCCCTACACGACGCTCTTCCGATCTNNNAGAGGTTTAGGAGTAAAAC
GAGCACACGTCTGAACTCTTTCCCTACACGACGCTCTTCCGATCTNNNTCCTGTGCATTGTAGGACGTT
AGCACACGTCTGAACTCTTTCCCTACACGACGCTCTTCCGATCTNNNTCGCTTGAGGTCAGTAGTTCAA
GAGCACACGTCTGAACTCTTTCCCTACACGACGCTCTTCCGATCTNNNATGTGTCAGGCTTGTGAATTT
GAGCACACGTCTGAACTCTTTCCCTACACGACGCTCTTCCGATCTNNNAAGAGGTTGCAGTGAGGTGA
AGAGCACACGTCTGAACTCTTTCCCTACACGACGCTCTTCCGATCTNNNTCCTCTCTCTCTCTTCATCTGCA
AGCACACGTCTGAACTCTTTCCCTACACGACGCTCTTCCGATCTNNNCAAGGTTGGAGGATCACTTGA
GAGCACACGTCTGAACTCTTTCCCTACACGACGCTCTTCCGATCTNNNTGTGGTGATGGAACTACACA
GAGCACACGTCTGAACTCTTTCCCTACACGACGCTCTTCCGATCTNNNAGTGAGCCAAGATTGAGCTAC
GAGCACACGTCTGAACTCTTTCCCTACACGACGCTCTTCCGATCTNNNAGAGTCTGGCTTGGTCTTTTCA
GAGCACACGTCTGAACTCTTTCCCTACACGACGCTCTTCCGATCTNNNAGAACCAATGAGTATGGACAC
GCACACGTCTGAACTCTTTCCCTACACGACGCTCTTCCGATCTNNNCGGATCTCTTCAGCGATATGGAA
AGAGCACACGTCTGAACTCTTTCCCTACACGACGCTCTTCCGATCTNNNTTCCCCACAAAAGTCCTTCAG
AGCACACGTCTGAACTCTTTCCCTACACGACGCTCTTCCGATCTNNNGAGACTCCGTTCTGTGTTGTGT
AGCACACGTCTGAACTCTTTCCCTACACGACGCTCTTCCGATCTNNNTGGTTGATTTGTCTTCATAGGC
AGCACACGTCTGAACTCTTTCCCTACACGACGCTCTTCCGATCTNNNTCAGTCACCACAATCAGTACAGT
GAGCACACGTCTGAACTCTTTCCCTACACGACGCTCTTCCGATCTNNNGCAATGAGCCTATATTACGCC

AGCACACGTCTGAACTCTTTCCCTACACGACGCTCTTCCGATCTNNNCTCTGTGCAGGTGTACCATTTTT
AGCACACGTCTGAACTCTTTCCCTACACGACGCTCTTCCGATCTNNNCAGCATGGTTAGGCTCAAAGAT
AGCACACGTCTGAACTCTTTCCCTACACGACGCTCTTCCGATCTNNNGCTCAGTTTTCCCTGCACTATTC
AGCACACGTCTGAACTCTTTCCCTACACGACGCTCTTCCGATCTNNNACTTTCCATAGTGGTCACATCCA
AGCACACGTCTGAACTCTTTCCCTACACGACGCTCTTCCGATCTNNNTGCTGCCATTTCTTTAGGAAAG
GAGCACACGTCTGAACTCTTTCCCTACACGACGCTCTTCCGATCTNNNGTGAAGATGTGATTGTGTGT
AGAGCACACGTCTGAACTCTTTCCCTACACGACGCTCTTCCGATCTNNNAGAAGCAAGGAAGGCAAG
GAGCACACGTCTGAACTCTTTCCCTACACGACGCTCTTCCGATCTNNNACAAACAGTTATTGAGCACCTC
AGCACACGTCTGAACTCTTTCCCTACACGACGCTCTTCCGATCTNNNTGCTCATGGGTAGGAAGAATCA
AGCACACGTCTGAACTCTTTCCCTACACGACGCTCTTCCGATCTNNNTGGTATGTACCCTTGATGTGGT
GAGCACACGTCTGAACTCTTTCCCTACACGACGCTCTTCCGATCTNNNGGCTTACTCACCTTGTCACCTA
GAGCACACGTCTGAACTCTTTCCCTACACGACGCTCTTCCGATCTNNNACAGAGTTAGACCCTGTCTCA
GAGCACACGTCTGAACTCTTTCCCTACACGACGCTCTTCCGATCTNNNCCAACCTACATGTCCATCAGTC
GCACACGTCTGAACTCTTTCCCTACACGACGCTCTTCCGATCTNNNCTGAAGCACAAACACACCAGATTC
GCACACGTCTGAACTCTTTCCCTACACGACGCTCTTCCGATCTNNNGGAAGGTGCTCTGTTGTTACCTA
GAGCACACGTCTGAACTCTTTCCCTACACGACGCTCTTCCGATCTNNNTGAAATACAGCAAAAGCAGCA
GAGCACACGTCTGAACTCTTTCCCTACACGACGCTCTTCCGATCTNNNATATCTGGTTTTACTGGCCTGC
GAGCACACGTCTGAACTCTTTCCCTACACGACGCTCTTCCGATCTNNNATAGCCCAAGGGAAAATCACA
AGCACACGTCTGAACTCTTTCCCTACACGACGCTCTTCCGATCTNNNTATGGGGAGGATTCTCATGCAT
AGAGCACACGTCTGAACTCTTTCCCTACACGACGCTCTTCCGATCTNNNTGAGCTCAAGAGTTCAAGAC
GAGCACACGTCTGAACTCTTTCCCTACACGACGCTCTTCCGATCTNNNCAGAGATAGAAAGAGAGGGC
AGCACACGTCTGAACTCTTTCCCTACACGACGCTCTTCCGATCTNNNTTCCTATCTGGGGGCTAAGAACT
AGAGCACACGTCTGAACTCTTTCCCTACACGACGCTCTTCCGATCTNNNACAACAATAGGCAAGCCAAG
GAGCACACGTCTGAACTCTTTCCCTACACGACGCTCTTCCGATCTNNNCAACAGTGACCAAGCTGAGAA
AGCACACGTCTGAACTCTTTCCCTACACGACGCTCTTCCGATCTNNNTCAAGGGAGATTTTGGCCTTAC
GCACACGTCTGAACTCTTTCCCTACACGACGCTCTTCCGATCTNNNTGGACTTGAACTAGTTCTCCAGC
AGCACACGTCTGAACTCTTTCCCTACACGACGCTCTTCCGATCTNNNCTCAGCTACTTGAGAGGCTTAG
AGCACACGTCTGAACTCTTTCCCTACACGACGCTCTTCCGATCTNNNGTCAGGTGGTTTGTAGATCTCC
GAGCACACGTCTGAACTCTTTCCCTACACGACGCTCTTCCGATCTNNNTGTGTGTGTGTGTAGAGAGA
GAGCACACGTCTGAACTCTTTCCCTACACGACGCTCTTCCGATCTNNNGTGACAAGTGAGACTCTGACT
GAGCACACGTCTGAACTCTTTCCCTACACGACGCTCTTCCGATCTNNNTGGCTAATTTGGTGTCTTCTT
GAGCACACGTCTGAACTCTTTCCCTACACGACGCTCTTCCGATCTNNNATGAAAGACCAAGACAGCAAT
GAGCACACGTCTGAACTCTTTCCCTACACGACGCTCTTCCGATCTNNNCCAGAGAAACAGAACCAGAAC
AGCACACGTCTGAACTCTTTCCCTACACGACGCTCTTCCGATCTNNNTAAAAATCATCCCAGGTTGCTG
AGCACACGTCTGAACTCTTTCCCTACACGACGCTCTTCCGATCTNNNAAACAACACAGGTTTGAACTGC
AGCACACGTCTGAACTCTTTCCCTACACGACGCTCTTCCGATCTNNNTCCAAATGATCCTCCAACCTCAGC
GAGCACACGTCTGAACTCTTTCCCTACACGACGCTCTTCCGATCTNNNATAGTGTGTGTGTGTGTGTGT
AGCACACGTCTGAACTCTTTCCCTACACGACGCTCTTCCGATCTNNNAATCCTAGTAGTTTGCTGGTGC
GAGCACACGTCTGAACTCTTTCCCTACACGACGCTCTTCCGATCTNNNGATATAAGCAGCCGGGTACTC
AGCACACGTCTGAACTCTTTCCCTACACGACGCTCTTCCGATCTNNNGGTACCCCTGAAACTAAAAGCC
AGAGCACACGTCTGAACTCTTTCCCTACACGACGCTCTTCCGATCTNNNGACTGCAGTAATGGCCTGAA
GAGCACACGTCTGAACTCTTTCCCTACACGACGCTCTTCCGATCTNNNACAATAGTACCTTGCCCATGCT
GAGCACACGTCTGAACTCTTTCCCTACACGACGCTCTTCCGATCTNNNGATCAAGACCATCCTGGCTAA

GCACACGTCTGAACTCTTTCCCTACACGACGCTCTTCCGATCTNNNAGCCCCAATTGAGACAAGAATGT
AGCACACGTCTGAACTCTTTCCCTACACGACGCTCTTCCGATCTNNNTAACACATCTTGGCCTGTGGTTA
AGCACACGTCTGAACTCTTTCCCTACACGACGCTCTTCCGATCTNNNTTCTGTAAGGCCAGCATCATTCT
AGCACACGTCTGAACTCTTTCCCTACACGACGCTCTTCCGATCTNNNTCTCTGGAACTCCACTAACCTC
AGCACACGTCTGAACTCTTTCCCTACACGACGCTCTTCCGATCTNNNAGGTTGCAGTTAGCCAAGATCT
GAGCACACGTCTGAACTCTTTCCCTACACGACGCTCTTCCGATCTNNNCTGCCAATGAACTCAAATGCTC
GAGCACACGTCTGAACTCTTTCCCTACACGACGCTCTTCCGATCTNNNCAGGCATGTTTATAGCAGCAC
AGCACACGTCTGAACTCTTTCCCTACACGACGCTCTTCCGATCTNNNACACAAGACCAACAAGGCTAAG
GAGCACACGTCTGAACTCTTTCCCTACACGACGCTCTTCCGATCTNNNTGAGTGGGAGAATCACTTGAA
GAGCACACGTCTGAACTCTTTCCCTACACGACGCTCTTCCGATCTNNNGAGAATCACTTGAACCCAGGA
AGCACACGTCTGAACTCTTTCCCTACACGACGCTCTTCCGATCTNNNAGCAGACTACAGGTATGTGTGT
AGCACACGTCTGAACTCTTTCCCTACACGACGCTCTTCCGATCTNNNTCAATGCAAAAGTGGTGATGTC
AGCACACGTCTGAACTCTTTCCCTACACGACGCTCTTCCGATCTNNNGGGTAACAGATCGAGACTCCA
AGCACACGTCTGAACTCTTTCCCTACACGACGCTCTTCCGATCTNNNGCTACTGTCCCACTATCTTCTGGC
AGCACACGTCTGAACTCTTTCCCTACACGACGCTCTTCCGATCTNNNCTCAGGTGATTCTCCAGCCTAA
AAGAGCACACGTCTGAACTCTTTCCCTACACGACGCTCTTCCGATCTNNNGACACCTACTGCAGACACAA
AGCACACGTCTGAACTCTTTCCCTACACGACGCTCTTCCGATCTNNNAGAACTCTGCTTCTCTTGAGCAA
GAGCACACGTCTGAACTCTTTCCCTACACGACGCTCTTCCGATCTNNNTTTTGCCAACTGTACATGCAC
GCACACGTCTGAACTCTTTCCCTACACGACGCTCTTCCGATCTNNNTTATCTGCCCAGCATCCTTTCTCA
GAGCACACGTCTGAACTCTTTCCCTACACGACGCTCTTCCGATCTNNNGAGGCATGAGAATCGCTTGAA
AGCACACGTCTGAACTCTTTCCCTACACGACGCTCTTCCGATCTNNNCCCTTTCCTTCCACTAAATCCCAC
GCACACGTCTGAACTCTTTCCCTACACGACGCTCTTCCGATCTNNNATTTCATTTGTTCCCACCAGCTGCA
AGAGCACACGTCTGAACTCTTTCCCTACACGACGCTCTTCCGATCTNNNGAGAATCACGTGAACCCAGC
GAGCACACGTCTGAACTCTTTCCCTACACGACGCTCTTCCGATCTNNNGGTACAGTGGCTCACATCTGTA
AGCACACGTCTGAACTCTTTCCCTACACGACGCTCTTCCGATCTNNNAACATGACAAAACCCCGTCTCTA
AGCACACGTCTGAACTCTTTCCCTACACGACGCTCTTCCGATCTNNNGAGGTTTCGGCAATGAGTTTAC
AGCACACGTCTGAACTCTTTCCCTACACGACGCTCTTCCGATCTNNNATGTGCTTCTCTGTCCATAGTGC
GAGCACACGTCTGAACTCTTTCCCTACACGACGCTCTTCCGATCTNNNTGGTAGGCAAAATAGCAAGA
AGCACACGTCTGAACTCTTTCCCTACACGACGCTCTTCCGATCTNNNTTATGTCCAAGAAACCCTACGC
GCACACGTCTGAACTCTTTCCCTACACGACGCTCTTCCGATCTNNNCTACTCCTCATAGTCTGTGTGGCC
AGCACACGTCTGAACTCTTTCCCTACACGACGCTCTTCCGATCTNNNTGTGCTTTCAGACACACATGTAT
AGCACACGTCTGAACTCTTTCCCTACACGACGCTCTTCCGATCTNNNATCCTTTCCTTTCACAGTAGCCCC
AGCACACGTCTGAACTCTTTCCCTACACGACGCTCTTCCGATCTNNNGCACACACATGTTTATAGCAGC
AGCACACGTCTGAACTCTTTCCCTACACGACGCTCTTCCGATCTNNNACAAGTCATGTTCAGGGACTCAA
GAGCACACGTCTGAACTCTTTCCCTACACGACGCTCTTCCGATCTNNNTGCAATTTGCCCTTGAGAGAT
GAGCACACGTCTGAACTCTTTCCCTACACGACGCTCTTCCGATCTNNNGCGAAACTCCGTCTCAAAGAAA
AGCACACGTCTGAACTCTTTCCCTACACGACGCTCTTCCGATCTNNNCATGCAATAGCTTGGGTTTTCTC
AGCACACGTCTGAACTCTTTCCCTACACGACGCTCTTCCGATCTNNNCTGCATTCACACCAACATCTGTTC
GAGCACACGTCTGAACTCTTTCCCTACACGACGCTCTTCCGATCTNNNCAGAAAGGTTCGTCAGAGGC
AGCACACGTCTGAACTCTTTCCCTACACGACGCTCTTCCGATCTNNNGGTTTGCAATGATTCCAACAGGA
GAGCACACGTCTGAACTCTTTCCCTACACGACGCTCTTCCGATCTNNNGAGGAGGAAAGACAATGCCA
AGCACACGTCTGAACTCTTTCCCTACACGACGCTCTTCCGATCTNNNGGCTCCGTCTTGAAAGAACAAA
AGCACACGTCTGAACTCTTTCCCTACACGACGCTCTTCCGATCTNNNTGTCCTGTAAAAGCATTACCCCAC

AGCACACGTCTGAACTCTTTCCCTACACGACGCTCTTCCGATCTNNNGGGCGACAGAGTGAGACATA
AGCACACGTCTGAACTCTTTCCCTACACGACGCTCTTCCGATCTNNNGAATTAGCAAAGCCTGGGAAG
AGCACACGTCTGAACTCTTTCCCTACACGACGCTCTTCCGATCTNNNGTACTTAGCTCTCAGTAGCACC
GAGCACACGTCTGAACTCTTTCCCTACACGACGCTCTTCCGATCTNNNCTGTGACCCAGTATTTCCACT
GAGCACACGTCTGAACTCTTTCCCTACACGACGCTCTTCCGATCTNNNTGGGCAACATAATGAGACAC
AGCACACGTCTGAACTCTTTCCCTACACGACGCTCTTCCGATCTNNNTGTGAATGAGTGGAGATAGG
AGCACACGTCTGAACTCTTTCCCTACACGACGCTCTTCCGATCTNNNTATCAGTCAGGCTTCTCCAGA
AGCACACGTCTGAACTCTTTCCCTACACGACGCTCTTCCGATCTNNNTTGTTATCAGGCACACAAGCAT
AGCACACGTCTGAACTCTTTCCCTACACGACGCTCTTCCGATCTNNNTCTCTCTGTTTCTCTCTCTCCCCC
GCACACGTCTGAACTCTTTCCCTACACGACGCTCTTCCGATCTNNNCACACATACAGACACACACACAC
AGCACACGTCTGAACTCTTTCCCTACACGACGCTCTTCCGATCTNNNGAGGAAAGTGGAAGCTGCAA
AGCACACGTCTGAACTCTTTCCCTACACGACGCTCTTCCGATCTNNNTCCTTAGGTCCCAAAAGACACA
AGCACACGTCTGAACTCTTTCCCTACACGACGCTCTTCCGATCTNNNGGAAAACTTGGACACAGACAC
GAGCACACGTCTGAACTCTTTCCCTACACGACGCTCTTCCGATCTNNNGCCAGTGCAGGAAAAATACC
AGCACACGTCTGAACTCTTTCCCTACACGACGCTCTTCCGATCTNNNGTAACCCTCTCCATCACCTCAA
AGAGCACACGTCTGAACTCTTTCCCTACACGACGCTCTTCCGATCTNNNGAGTTGCTGCTTTGAGAGA
GAGCACACGTCTGAACTCTTTCCCTACACGACGCTCTTCCGATCTNNNAATATTCCAATCGCATGCTCG
CACACGTCTGAACTCTTTCCCTACACGACGCTCTTCCGATCTNNNTGTAGTCATTGTCAGTACACATAC
GAGCACACGTCTGAACTCTTTCCCTACACGACGCTCTTCCGATCTNNNCTACGACAGGTGTTCCAGAG
AGCACACGTCTGAACTCTTTCCCTACACGACGCTCTTCCGATCTNNNCTTTCACATCCTTGGGTAGAGG
AGCACACGTCTGAACTCTTTCCCTACACGACGCTCTTCCGATCTNNNAGTGAGTGAGTGAGTGAGAG
AGCACACGTCTGAACTCTTTCCCTACACGACGCTCTTCCGATCTNNNCCAAAGTACTGGGATTATGGG
AGCACACGTCTGAACTCTTTCCCTACACGACGCTCTTCCGATCTNNNCAGAGAAGCAAATTCGCCATG
AGCACACGTCTGAACTCTTTCCCTACACGACGCTCTTCCGATCTNNNTTGATTGATAATGTGGCCAGC
AGCACACGTCTGAACTCTTTCCCTACACGACGCTCTTCCGATCTNNNTGATCTTTTGAGAGTGTGTGT
AGCACACGTCTGAACTCTTTCCCTACACGACGCTCTTCCGATCTNNNCACACTGGGAGAAGAATGGA
AGCACACGTCTGAACTCTTTCCCTACACGACGCTCTTCCGATCTNNNCTAGCCAGTTATCCCAGCATCA
AGCACACGTCTGAACTCTTTCCCTACACGACGCTCTTCCGATCTNNNCAGTAGCTGAGACTACAGGTG
AGCACACGTCTGAACTCTTTCCCTACACGACGCTCTTCCGATCTNNNGCTGACTTCAGTCTCCATGAG
AGCACACGTCTGAACTCTTTCCCTACACGACGCTCTTCCGATCTNNNCCCTTCTACACATCTTAGCTGG
AGCACACGTCTGAACTCTTTCCCTACACGACGCTCTTCCGATCTNNNTCCAGCTCTGTGTGATTGGTT
AGCACACGTCTGAACTCTTTCCCTACACGACGCTCTTCCGATCTNNNATGAGAGTGAGTGTGTGTGT
GAGCACACGTCTGAACTCTTTCCCTACACGACGCTCTTCCGATCTNNNTTGTCTGATGGGAGAAAGAG
AGCACACGTCTGAACTCTTTCCCTACACGACGCTCTTCCGATCTNNNAAAGACGGAGACTCTGTTTCA
AGCACACGTCTGAACTCTTTCCCTACACGACGCTCTTCCGATCTNNNTCAAGCTCTCTCTCTCTCAGACA
GAGCACACGTCTGAACTCTTTCCCTACACGACGCTCTTCCGATCTNNNATTTGGCAGGAGATACACAC
AGCACACGTCTGAACTCTTTCCCTACACGACGCTCTTCCGATCTNNNATTGCAGCACTATTCACAATGG
AGCACACGTCTGAACTCTTTCCCTACACGACGCTCTTCCGATCTNNNACCAAAACAGCATGGTACTGGT
AGCACACGTCTGAACTCTTTCCCTACACGACGCTCTTCCGATCTNNNGGAGGTTACAGTGAGCTGAG
AGCACACGTCTGAACTCTTTCCCTACACGACGCTCTTCCGATCTNNNAAAGAGCAAGACCCTGTCTCA
AGCACACGTCTGAACTCTTTCCCTACACGACGCTCTTCCGATCTNNNCTGGGTGACAAAGCAAGATTC
AGCACACGTCTGAACTCTTTCCCTACACGACGCTCTTCCGATCTNNNATATTTGGAAGTACATCCCGG
GAGCACACGTCTGAACTCTTTCCCTACACGACGCTCTTCCGATCTNNNGCACGAACATTTGCAGAAAG

GCACACGTCTGAACTCTTTCCCTACACGACGCTCTTCCGATCTNNNGCTGAGGAAGGAGGATCATTTGT
GAGCACACGTCTGAACTCTTTCCCTACACGACGCTCTTCCGATCTNNNATATGGAACCAGACCAAATGC
AGCACACGTCTGAACTCTTTCCCTACACGACGCTCTTCCGATCTNNNTGCTACCAACTATCACCACTTCCC
AGCACACGTCTGAACTCTTTCCCTACACGACGCTCTTCCGATCTNNNAAATTGCCCAGTCTCAGGTATG
GCACACGTCTGAACTCTTTCCCTACACGACGCTCTTCCGATCTNNNTCTCTTTCCCTACCTTCCTGACA
GAGCACACGTCTGAACTCTTTCCCTACACGACGCTCTTCCGATCTNNNCTCCTGAGTAGCTGGGACTAC
GAGCACACGTCTGAACTCTTTCCCTACACGACGCTCTTCCGATCTNNNGCACATAGCAGGGGTTGAAT
AGAGCACACGTCTGAACTCTTTCCCTACACGACGCTCTTCCGATCTNNNCCGTGGTGGACCTCATATTGA
GAGCACACGTCTGAACTCTTTCCCTACACGACGCTCTTCCGATCTNNNGCATTGACTCATTTCCACCTTG
AGCACACGTCTGAACTCTTTCCCTACACGACGCTCTTCCGATCTNNNGTGTGTTGTGTATGTGTGTGT
AGCACACGTCTGAACTCTTTCCCTACACGACGCTCTTCCGATCTNNNGTAAGCCAAGATTGTGCCACTAC
GAGCACACGTCTGAACTCTTTCCCTACACGACGCTCTTCCGATCTNNNTTCTCTGTTGTTGGAAGGAAG
GAGCACACGTCTGAACTCTTTCCCTACACGACGCTCTTCCGATCTNNNTGAGAGTCACTGTTCTATGTG
GAGCACACGTCTGAACTCTTTCCCTACACGACGCTCTTCCGATCTNNNCAATGGCCACCGTCTATTTCTT
AGAGCACACGTCTGAACTCTTTCCCTACACGACGCTCTTCCGATCTNNNTGTCAAGGAAGAGTTCACAA
AGCACACGTCTGAACTCTTTCCCTACACGACGCTCTTCCGATCTNNNTGAGGCAAGAAGATCACTTGA
GAGCACACGTCTGAACTCTTTCCCTACACGACGCTCTTCCGATCTNNNCTGGATGAATGAGAGTGAGC
GAGCACACGTCTGAACTCTTTCCCTACACGACGCTCTTCCGATCTNNNGTCTGTCCTTAGACTCTGCTCTC
GAGCACACGTCTGAACTCTTTCCCTACACGACGCTCTTCCGATCTNNNATCCATCTTGTTTCTGCCATTGC
GAGCACACGTCTGAACTCTTTCCCTACACGACGCTCTTCCGATCTNNNCACTTGAGGTCAGGAGTTCAA
AGCACACGTCTGAACTCTTTCCCTACACGACGCTCTTCCGATCTNNNCTGCCCAGTTGATTACCCTTTTGC
AGCACACGTCTGAACTCTTTCCCTACACGACGCTCTTCCGATCTNNNTATGGAACCAGCTCAAATTCCC
GAGCACACGTCTGAACTCTTTCCCTACACGACGCTCTTCCGATCTNNNTAGGTACCAGGTCACTGTTCTC
GAGCACACGTCTGAACTCTTTCCCTACACGACGCTCTTCCGATCTNNNAATATGGAATCAGCCCAAATG
AGCACACGTCTGAACTCTTTCCCTACACGACGCTCTTCCGATCTNNNATTTGGGGCTGGACAATTCTTT
AGCACACGTCTGAACTCTTTCCCTACACGACGCTCTTCCGATCTNNNATACGTACAATCCCAGCACTTCA
AGCACACGTCTGAACTCTTTCCCTACACGACGCTCTTCCGATCTNNNTGATGCAGGAGAATTGCTTGA
AGCACACGTCTGAACTCTTTCCCTACACGACGCTCTTCCGATCTNNNGGCTCTACATCCCTTCAGCTTAG
AGCACACGTCTGAACTCTTTCCCTACACGACGCTCTTCCGATCTNNNGTGTGTATGTGTGTGTGTGTGA
GCACACGTCTGAACTCTTTCCCTACACGACGCTCTTCCGATCTNNNAAAAGACCCATCACTGGTACCTCC
AGCACACGTCTGAACTCTTTCCCTACACGACGCTCTTCCGATCTNNNACCTCAAGTGGCATTGTGAAAT
AGCACACGTCTGAACTCTTTCCCTACACGACGCTCTTCCGATCTNNNAGATCCTGTTAGTCACTACGGA
GCACACGTCTGAACTCTTTCCCTACACGACGCTCTTCCGATCTNNNGCTGCTCCTGGACTCATTTTTTTCA
GAGCACACGTCTGAACTCTTTCCCTACACGACGCTCTTCCGATCTNNNGATGGCTTCCAGGTTCATTCA
AGCACACGTCTGAACTCTTTCCCTACACGACGCTCTTCCGATCTNNNAAAGAACGAGGAGATGGGAA
AGCACACGTCTGAACTCTTTCCCTACACGACGCTCTTCCGATCTNNNAACTGTTCTAGCACCGTTTGTTC
GCACACGTCTGAACTCTTTCCCTACACGACGCTCTTCCGATCTNNNAGAGTGGGACCCTGAATCAAAAA
AGCACACGTCTGAACTCTTTCCCTACACGACGCTCTTCCGATCTNNNACACCTGTAATCTCAGCACTCTG
GAGCACACGTCTGAACTCTTTCCCTACACGACGCTCTTCCGATCTNNNCATGGCAGGAGGATAACTTGA
AGCACACGTCTGAACTCTTTCCCTACACGACGCTCTTCCGATCTNNNAGTGAGCTATGATCATGCCACTG
AGCACACGTCTGAACTCTTTCCCTACACGACGCTCTTCCGATCTNNNCATCTGTGAATAACCACTGCACC
AGCACACGTCTGAACTCTTTCCCTACACGACGCTCTTCCGATCTNNNAAAATATTGGACATCCCTGCCC
GAGCACACGTCTGAACTCTTTCCCTACACGACGCTCTTCCGATCTNNNGAGCGAAACTCTGTCTCAAAAA

GAGCACACGTCTGAACTCTTTCCCTACACGACGCTCTTCCGATCTNNNCAGCTCAAATGCCCATCAATCA
GAGCACACGTCTGAACTCTTTCCCTACACGACGCTCTTCCGATCTNNNGACTGTAATCCCAGCTACTCA
AGCACACGTCTGAACTCTTTCCCTACACGACGCTCTTCCGATCTNNNGGAGGATCACTTGACCAAGAGT
AGCACACGTCTGAACTCTTTCCCTACACGACGCTCTTCCGATCTNNNGAGTTCAAGACTAGCCTGGTCA
GAGCACACGTCTGAACTCTTTCCCTACACGACGCTCTTCCGATCTNNNGCCCCTCTATGGATGGTTGAA
AGCACACGTCTGAACTCTTTCCCTACACGACGCTCTTCCGATCTNNNCTGAGACAGGAGAATCGCTTGA
GCACACGTCTGAACTCTTTCCCTACACGACGCTCTTCCGATCTNNNCGTAGGGCTGGACTGTACTATAT
GAGCACACGTCTGAACTCTTTCCCTACACGACGCTCTTCCGATCTNNNATAACACACACACACACACACA
GAGCACACGTCTGAACTCTTTCCCTACACGACGCTCTTCCGATCTNNNGGATGATGGAGTGAGACTCT
GAGCACACGTCTGAACTCTTTCCCTACACGACGCTCTTCCGATCTNNNATCTCTCATCCCATTGTGCCTA
GAGCACACGTCTGAACTCTTTCCCTACACGACGCTCTTCCGATCTNNNTACAGGCCCTGATCTCAAGAA
GCACACGTCTGAACTCTTTCCCTACACGACGCTCTTCCGATCTNNNAGCCTCCTAAGTAGCTGGAACTA
GAGCACACGTCTGAACTCTTTCCCTACACGACGCTCTTCCGATCTNNNTGGGTGACAAATTGAGACCCT
AGAGCACACGTCTGAACTCTTTCCCTACACGACGCTCTTCCGATCTNNNTGTGTCATTATGGGATGCCT
GAGCACACGTCTGAACTCTTTCCCTACACGACGCTCTTCCGATCTNNNCTGGAGTGCAAAGGTGTGAT
GAGCACACGTCTGAACTCTTTCCCTACACGACGCTCTTCCGATCTNNNAGGTTCTCCTCCTCAAATGAA
AGCACACGTCTGAACTCTTTCCCTACACGACGCTCTTCCGATCTNNNATAGGGAGGTTGTTTGTGTGT
GAGCACACGTCTGAACTCTTTCCCTACACGACGCTCTTCCGATCTNNNTGTGGACCATGAGTGTGTATA
AGCACACGTCTGAACTCTTTCCCTACACGACGCTCTTCCGATCTNNNGATCTCAGCTTACTGCAACCTCT
AGCACACGTCTGAACTCTTTCCCTACACGACGCTCTTCCGATCTNNNCTAACGGCTCTTCAGACATCTGT
AGCACACGTCTGAACTCTTTCCCTACACGACGCTCTTCCGATCTNNNAGCAAGACCCTGTCTCAAAAGA
GAGCACACGTCTGAACTCTTTCCCTACACGACGCTCTTCCGATCTNNNGGATATTAGCGGGAAGAGCT
AGCACACGTCTGAACTCTTTCCCTACACGACGCTCTTCCGATCTNNNTACATCATCCCGAAGCATGCATA
GAGCACACGTCTGAACTCTTTCCCTACACGACGCTCTTCCGATCTNNNAGGTAGGTAGGTAGGGAGG
GAGCACACGTCTGAACTCTTTCCCTACACGACGCTCTTCCGATCTNNNACTGGGTCCAAAAAGTCTTAG
GAGCACACGTCTGAACTCTTTCCCTACACGACGCTCTTCCGATCTNNNTGTGACACGAGGCATTTCTTTT
GAGCACACGTCTGAACTCTTTCCCTACACGACGCTCTTCCGATCTNNNCGCCCAGCCAGCCATATATATA
AGCACACGTCTGAACTCTTTCCCTACACGACGCTCTTCCGATCTNNNCGTCTGTAATCCCAGCATTTTGG
GAGCACACGTCTGAACTCTTTCCCTACACGACGCTCTTCCGATCTNNNGCAACAGAGCAAGACACTATC
GAGCACACGTCTGAACTCTTTCCCTACACGACGCTCTTCCGATCTNNNGCTTAGACGACAGAGTGAGA
GAGCACACGTCTGAACTCTTTCCCTACACGACGCTCTTCCGATCTNNNAGGGTTGTCCAGAGAAACAG
AGAGCACACGTCTGAACTCTTTCCCTACACGACGCTCTTCCGATCTNNNCTTATTAGCTCCCCCAAACAC
GAGCACACGTCTGAACTCTTTCCCTACACGACGCTCTTCCGATCTNNNAAAAAGGAAGACCAGAGAAG
GAGCACACGTCTGAACTCTTTCCCTACACGACGCTCTTCCGATCTNNNAGAAACTGCTGTATTGCTCAG
AGCACACGTCTGAACTCTTTCCCTACACGACGCTCTTCCGATCTNNNTAGACTTCTTTGTGGTTCCCCAG
AGCACACGTCTGAACTCTTTCCCTACACGACGCTCTTCCGATCTNNNTCATGTGAATGCATGCAGACAT
GAGCACACGTCTGAACTCTTTCCCTACACGACGCTCTTCCGATCTNNNCTGAGGCAGGAGAATCACTT
AGCACACGTCTGAACTCTTTCCCTACACGACGCTCTTCCGATCTNNNCAGGAGGTTGATGCTACAGTGA
GAGCACACGTCTGAACTCTTTCCCTACACGACGCTCTTCCGATCTNNNAAACTGTGAACCAGCCTAAAT
AGAGCACACGTCTGAACTCTTTCCCTACACGACGCTCTTCCGATCTNNNGCTTACAATGCAGTTGGAGT
AGCACACGTCTGAACTCTTTCCCTACACGACGCTCTTCCGATCTNNNGGTTGGTAATTGTGTGCACCTA
GAGCACACGTCTGAACTCTTTCCCTACACGACGCTCTTCCGATCTNNNCTCTTGGGATGTCTGTGAGGA
AGCACACGTCTGAACTCTTTCCCTACACGACGCTCTTCCGATCTNNNGAACTCCGTATTTTGGCATGCA

AGCACACGTCTGAACTCTTTCCCTACACGACGCTCTTCCGATCTNNNTGTGTGGTTGAATTTGGTTTG
AGCACACGTCTGAACTCTTTCCCTACACGACGCTCTTCCGATCTNNNACAGAGCAAGACCTTGTCTCAA
AGCACACGTCTGAACTCTTTCCCTACACGACGCTCTTCCGATCTNNNGCACACACACCCAGCTAATTTT
GAGCACACGTCTGAACTCTTTCCCTACACGACGCTCTTCCGATCTNNNTTAGTAGCTGAGACTACAGG
GAGCACACGTCTGAACTCTTTCCCTACACGACGCTCTTCCGATCTNNNGTTGGCATAGCAACCTGATTT
AGCACACGTCTGAACTCTTTCCCTACACGACGCTCTTCCGATCTNNNGTTCAATTCCATCTTCCTGGCAG
AGCACACGTCTGAACTCTTTCCCTACACGACGCTCTTCCGATCTNNNTTGCTCTGCTTTACTGGACTCA
GCACACGTCTGAACTCTTTCCCTACACGACGCTCTTCCGATCTNNNGAGGTTTCTCTCCCCTTGTACTG
GAGCACACGTCTGAACTCTTTCCCTACACGACGCTCTTCCGATCTNNNTCTTTGCACATATGTGGGGAA
GAGCACACGTCTGAACTCTTTCCCTACACGACGCTCTTCCGATCTNNNCACAGTCAGTTCACCAATCAT
AGCACACGTCTGAACTCTTTCCCTACACGACGCTCTTCCGATCTNNNGAAGGTTGCATTGAGCCAAGA
GAGCACACGTCTGAACTCTTTCCCTACACGACGCTCTTCCGATCTNNNGAGCGATACTCCTTCTCAAC
GAGCACACGTCTGAACTCTTTCCCTACACGACGCTCTTCCGATCTNNNTGAGATGGGAAGATCACTTG
AGCACACGTCTGAACTCTTTCCCTACACGACGCTCTTCCGATCTNNNCAAGAGGTCAGGAGTTCAAG
AGCACACGTCTGAACTCTTTCCCTACACGACGCTCTTCCGATCTNNNTCAGAAGCTGGTCCTTCAATGA
AGCACACGTCTGAACTCTTTCCCTACACGACGCTCTTCCGATCTNNNACTCTGAAAAAACCACAACCCT
CACACGTCTGAACTCTTTCCCTACACGACGCTCTTCCGATCTNNNGCTTCTCAGGACTGACTCATCTCC
AGCACACGTCTGAACTCTTTCCCTACACGACGCTCTTCCGATCTNNNCCTCTTTCACCTCATAACCACCA
GAGCACACGTCTGAACTCTTTCCCTACACGACGCTCTTCCGATCTNNNCCCAAATGCCCATCAATCAAT
GAGCACACGTCTGAACTCTTTCCCTACACGACGCTCTTCCGATCTNNNTCCTGGAATCAACCTAAGTG
AGCACACGTCTGAACTCTTTCCCTACACGACGCTCTTCCGATCTNNNGTGAAATTTGGGTCACTGATG
AGAGCACACGTCTGAACTCTTTCCCTACACGACGCTCTTCCGATCTNNNAGAAGAACAGAAATCAGGC
AGCACACGTCTGAACTCTTTCCCTACACGACGCTCTTCCGATCTNNNTGATGTCTCCTTTCCTTTGCTG
GAGCACACGTCTGAACTCTTTCCCTACACGACGCTCTTCCGATCTNNNATCTTTTTTGTGGCTGGAGG
GAGCACACGTCTGAACTCTTTCCCTACACGACGCTCTTCCGATCTNNNTTCAAGCTATTCTCCTGCCTCA
AGCACACGTCTGAACTCTTTCCCTACACGACGCTCTTCCGATCTNNNGTAGAGATCTTTCACCTCCCTG
GAGCACACGTCTGAACTCTTTCCCTACACGACGCTCTTCCGATCTNNNAACGGAATGCAAAAGATGA
AGCACACGTCTGAACTCTTTCCCTACACGACGCTCTTCCGATCTNNNTCATGTGTATGTCTGTGTGTG
AGCACACGTCTGAACTCTTTCCCTACACGACGCTCTTCCGATCTNNNGAGAATCCCTGAGAAGCTTGTG
AGCACACGTCTGAACTCTTTCCCTACACGACGCTCTTCCGATCTNNNATGGACACACATTCATCCAAGG
AGCACACGTCTGAACTCTTTCCCTACACGACGCTCTTCCGATCTNNNTCTGTTATTGTGGAGGGAACA
GAGCACACGTCTGAACTCTTTCCCTACACGACGCTCTTCCGATCTNNNCTTGAGCATCCACAGACTTA
GAGCACACGTCTGAACTCTTTCCCTACACGACGCTCTTCCGATCTNNNTTTCCCTTTCTCTCTGCCAAGT
GAGCACACGTCTGAACTCTTTCCCTACACGACGCTCTTCCGATCTNNNTAAGGATGTCTTTGCCACTAC
AGCACACGTCTGAACTCTTTCCCTACACGACGCTCTTCCGATCTNNNTTGCCCATCTCAAACACATTCAC
AAGAGCACACGTCTGAACTCTTTCCCTACACGACGCTCTTCCGATCTNNNAAGACACAGCTTCTAGTCG
AGCACACGTCTGAACTCTTTCCCTACACGACGCTCTTCCGATCTNNNGAAGCAGTGAATTGTTGACAC
GAGCACACGTCTGAACTCTTTCCCTACACGACGCTCTTCCGATCTNNNCTAACGGATCTCAGACAGTG
GAGCACACGTCTGAACTCTTTCCCTACACGACGCTCTTCCGATCTNNNCATAGCTCATTGCAGCCTTG
AGCACACGTCTGAACTCTTTCCCTACACGACGCTCTTCCGATCTNNNCTCTGTCACTGACCCAAGACTAA
GAGCACACGTCTGAACTCTTTCCCTACACGACGCTCTTCCGATCTNNNGAAGGAGAAAAAGAAAGCC
GAGCACACGTCTGAACTCTTTCCCTACACGACGCTCTTCCGATCTNNNCCCTGCAAAGGACATAATCTT
AGCACACGTCTGAACTCTTTCCCTACACGACGCTCTTCCGATCTNNNCTCCTAGGCTCATGTGATCTTC

GAGCACACGTCTGAACTCTTTCCCTACACGACGCTCTTCCGATCTNNNGTTTTCTCTTCCGTGAAATGCC
AGCACACGTCTGAACTCTTTCCCTACACGACGCTCTTCCGATCTNNNGAGGGTCTCTCTCCACTTTCTTC
GAGCACACGTCTGAACTCTTTCCCTACACGACGCTCTTCCGATCTNNNTAGGATGACAGAACAACAGAC
AAGAGCACACGTCTGAACTCTTTCCCTACACGACGCTCTTCCGATCTNNNCAAAGATTCCTGGCCCTGTA
AGCACACGTCTGAACTCTTTCCCTACACGACGCTCTTCCGATCTNNNACAATCCAATGCCCATCAGTAGA
GAGCACACGTCTGAACTCTTTCCCTACACGACGCTCTTCCGATCTNNNAGTGGCTCACAGTACTTTGAG
AGCACACGTCTGAACTCTTTCCCTACACGACGCTCTTCCGATCTNNNTCCTGGCTAACATGGTGAAATC
AGCACACGTCTGAACTCTTTCCCTACACGACGCTCTTCCGATCTNNNGTGTCCTGTTACAAGTCCTAGG
AGCACACGTCTGAACTCTTTCCCTACACGACGCTCTTCCGATCTNNNAAGAATCTTTCTAAGGGGGCTC
GAGCACACGTCTGAACTCTTTCCCTACACGACGCTCTTCCGATCTNNNCTGAGGCAGGAGAATTGCTTC
AGCACACGTCTGAACTCTTTCCCTACACGACGCTCTTCCGATCTNNNTGAATCCAGGAGTTCAAGACCA
AGCACACGTCTGAACTCTTTCCCTACACGACGCTCTTCCGATCTNNNGGAAACACACCTGGCCAAAAAAT
GAGCACACGTCTGAACTCTTTCCCTACACGACGCTCTTCCGATCTNNNAGCCTAGAAGAGCTGACATTT
AGAGCACACGTCTGAACTCTTTCCCTACACGACGCTCTTCCGATCTNNNGTGAGGAAATCGCAGTCAAC
AGCACACGTCTGAACTCTTTCCCTACACGACGCTCTTCCGATCTNNNAACAGAGGGTAAAGTGGAAAG
AGCACACGTCTGAACTCTTTCCCTACACGACGCTCTTCCGATCTNNNGCCACAGAGTGAGACTATGTCT
GAGCACACGTCTGAACTCTTTCCCTACACGACGCTCTTCCGATCTNNNCTGAGGCAGGAGAATAGCTTC
AGCACACGTCTGAACTCTTTCCCTACACGACGCTCTTCCGATCTNNNAAGTGTTTGCTGTTGTTCTGCT
GAGCACACGTCTGAACTCTTTCCCTACACGACGCTCTTCCGATCTNNNTCAATCTAAGTGCCCATCAAC
GAGCACACGTCTGAACTCTTTCCCTACACGACGCTCTTCCGATCTNNNTGTCAGGGGTTAGCTTTCTCT
GAGCACACGTCTGAACTCTTTCCCTACACGACGCTCTTCCGATCTNNNCTGCTCACCTTGACTTTCCAAA
AGCACACGTCTGAACTCTTTCCCTACACGACGCTCTTCCGATCTNNNCCATAATCACATGAGCCAGTTC
AGCACACGTCTGAACTCTTTCCCTACACGACGCTCTTCCGATCTNNNAGAGCAGAAGGGCAGGAATA
GCACACGTCTGAACTCTTTCCCTACACGACGCTCTTCCGATCTNNNCTAGCCAGGGTGAGAGAACAAA
GAGCACACGTCTGAACTCTTTCCCTACACGACGCTCTTCCGATCTNNNAAGTAACGGTACATCAGCCTCT
AAGAGCACACGTCTGAACTCTTTCCCTACACGACGCTCTTCCGATCTNNNACAAACAAAGGCCAGTTGC
GAGCACACGTCTGAACTCTTTCCCTACACGACGCTCTTCCGATCTNNNAGGCTATTCATGTCTGTGTGT
AGCACACGTCTGAACTCTTTCCCTACACGACGCTCTTCCGATCTNNNCCCTGTCTCTCTCTCTCTCTCTCCA
GAGCACACGTCTGAACTCTTTCCCTACACGACGCTCTTCCGATCTNNNAAGCCAGTGGTGAAAATAGAC
AGCACACGTCTGAACTCTTTCCCTACACGACGCTCTTCCGATCTNNNCTCTCTCTCTCTCACACACACACC
AGCACACGTCTGAACTCTTTCCCTACACGACGCTCTTCCGATCTNNNCACTTGAGGTCAGAAGTTCGAG
GAGCACACGTCTGAACTCTTTCCCTACACGACGCTCTTCCGATCTNNNAACATGGCAAAACCCTGTCTCT
AGCACACGTCTGAACTCTTTCCCTACACGACGCTCTTCCGATCTNNNAATCAAACCCCCTACTCCCAAAG
AGAGCACACGTCTGAACTCTTTCCCTACACGACGCTCTTCCGATCTNNNACCCTGTCTCTCTTTCTCTGTC
GAGCACACGTCTGAACTCTTTCCCTACACGACGCTCTTCCGATCTNNNTCCCTCTCTCTCTCTCTCTCTCTC
AGCACACGTCTGAACTCTTTCCCTACACGACGCTCTTCCGATCTNNNGCAAGACTCCATCTCAAACACAC
GCACACGTCTGAACTCTTTCCCTACACGACGCTCTTCCGATCTNNNAAGTGCTGGGATTACAGGCATAA
AGAGCACACGTCTGAACTCTTTCCCTACACGACGCTCTTCCGATCTNNNATGAGCTGATAGCCTGAACT
GCACACGTCTGAACTCTTTCCCTACACGACGCTCTTCCGATCTNNNTAGGAGAATCACTTGAACCTGGC
GAGCACACGTCTGAACTCTTTCCCTACACGACGCTCTTCCGATCTNNNTGTGTGTGTCTGCCTGTGTATA
AGCACACGTCTGAACTCTTTCCCTACACGACGCTCTTCCGATCTNNNCCTACTCCAGGGGTTTCTATCAA
AGAGCACACGTCTGAACTCTTTCCCTACACGACGCTCTTCCGATCTNNNAAGGCTACTTTCTTCGTACAG
AGAGCACACGTCTGAACTCTTTCCCTACACGACGCTCTTCCGATCTNNNGTCTGGGATAGAAGGACTG

GCACACGTCTGAACTCTTTCCCTACACGACGCTCTTCCGATCTNNNCCCCAACCCCTTTTGTTTTGATTC
AGCACACGTCTGAACTCTTTCCCTACACGACGCTCTTCCGATCTNNNCCTGAGTAGCTGGGATTATAG
AGCACACGTCTGAACTCTTTCCCTACACGACGCTCTTCCGATCTNNNCGAATGCCGTTGATTCATTCC
AGCACACGTCTGAACTCTTTCCCTACACGACGCTCTTCCGATCTNNNGCTTTTGAGGTTAGGACTCTC
AGCACACGTCTGAACTCTTTCCCTACACGACGCTCTTCCGATCTNNNATGTGTGTGTATGTGTGTGTG
AGCACACGTCTGAACTCTTTCCCTACACGACGCTCTTCCGATCTNNNCTTAGACAGAGTCCCAGGCATT
AGAGCACACGTCTGAACTCTTTCCCTACACGACGCTCTTCCGATCTNNNTCATGAACAGTGCTATGCTG
AGAGCACACGTCTGAACTCTTTCCCTACACGACGCTCTTCCGATCTNNNTTACCCTGAACGAAAGACCT
AGCACACGTCTGAACTCTTTCCCTACACGACGCTCTTCCGATCTNNNTGCCAAACAAATCATTCACCTC
GAGCACACGTCTGAACTCTTTCCCTACACGACGCTCTTCCGATCTNNNGGTGGAGGTTATAGTGAGCT
AGCACACGTCTGAACTCTTTCCCTACACGACGCTCTTCCGATCTNNNGGAACACACTCCTAACTGCCTA
GCACACGTCTGAACTCTTTCCCTACACGACGCTCTTCCGATCTNNNCTGAGGCAGAGAACCATTTGAA
AGCACACGTCTGAACTCTTTCCCTACACGACGCTCTTCCGATCTNNNTTCTGGAGATGCAAGGAAATG
AGCACACGTCTGAACTCTTTCCCTACACGACGCTCTTCCGATCTNNNCAGAAAAAGACAGTGCCCAAG
AGCACACGTCTGAACTCTTTCCCTACACGACGCTCTTCCGATCTNNNTGGGTTCCACATCCGTAAATTC
AGCACACGTCTGAACTCTTTCCCTACACGACGCTCTTCCGATCTNNNTATTCTGTTCCCACCTTTCCCACC
AGCACACGTCTGAACTCTTTCCCTACACGACGCTCTTCCGATCTNNNTCCCTCCTCAGTTTTCAAGTAGC
GCACACGTCTGAACTCTTTCCCTACACGACGCTCTTCCGATCTNNNCCAGTCTCTAACACCAGGAGTTC
GAGCACACGTCTGAACTCTTTCCCTACACGACGCTCTTCCGATCTNNNCTCCCTCCCTAACACATTCTACC
GCACACGTCTGAACTCTTTCCCTACACGACGCTCTTCCGATCTNNNTGACAAAGGAAACCATTTTGGG
AGCACACGTCTGAACTCTTTCCCTACACGACGCTCTTCCGATCTNNNAAGGAGCTTGGACTTCATTCTC
GAGCACACGTCTGAACTCTTTCCCTACACGACGCTCTTCCGATCTNNNCCTCCCGGGTTCAAGATATTCT
AGCACACGTCTGAACTCTTTCCCTACACGACGCTCTTCCGATCTNNNGCCTCCCAAGTAGCTAGGATTA
GAGCACACGTCTGAACTCTTTCCCTACACGACGCTCTTCCGATCTNNNGCAATGGAGTGATCATGGTT
AGCACACGTCTGAACTCTTTCCCTACACGACGCTCTTCCGATCTNNNAATCTCTCTCTCTCTCTCTCCCCC
AGCACACGTCTGAACTCTTTCCCTACACGACGCTCTTCCGATCTNNNTCTACAGATGCAATGCAATCCC
GAGCACACGTCTGAACTCTTTCCCTACACGACGCTCTTCCGATCTNNNGATACGCTTGAGGAACACGA
GAGCACACGTCTGAACTCTTTCCCTACACGACGCTCTTCCGATCTNNNGAAAGTTGCAGTGAGCTGA
AGCACACGTCTGAACTCTTTCCCTACACGACGCTCTTCCGATCTNNNTTTCAAATGTCCTCAGTGCCAA
GAGCACACGTCTGAACTCTTTCCCTACACGACGCTCTTCCGATCTNNNTCACTAATCATCAGGGAAAG
AGCACACGTCTGAACTCTTTCCCTACACGACGCTCTTCCGATCTNNNTAGGATAAGGGTCTCCAGCTT
GAGCACACGTCTGAACTCTTTCCCTACACGACGCTCTTCCGATCTNNNCTGTGGATCAATGAGTCCCTT
AGCACACGTCTGAACTCTTTCCCTACACGACGCTCTTCCGATCTNNNCTTTTGCTTCTGGGACTGTCTT
AGCACACGTCTGAACTCTTTCCCTACACGACGCTCTTCCGATCTNNNGCAGCAGCTGTTGTAGAAAGT
AGCACACGTCTGAACTCTTTCCCTACACGACGCTCTTCCGATCTNNNCTACAGTGAGCTGAGATCATG
GAGCACACGTCTGAACTCTTTCCCTACACGACGCTCTTCCGATCTNNNGCGACAGAGCAAGATTGTCT
GAGCACACGTCTGAACTCTTTCCCTACACGACGCTCTTCCGATCTNNNCACTTGAGACCAGGAGTTTG
AGCACACGTCTGAACTCTTTCCCTACACGACGCTCTTCCGATCTNNNCATCTCGGATCACTGTAAGCTC
GCACACGTCTGAACTCTTTCCCTACACGACGCTCTTCCGATCTNNNGGTCTTGAATTGCTGACCTCAA
AGCACACGTCTGAACTCTTTCCCTACACGACGCTCTTCCGATCTNNNACAGAAGTTGCAGTGAACTGA
AGCACACGTCTGAACTCTTTCCCTACACGACGCTCTTCCGATCTNNNAAGTAGCTGGGATTACAGGCA
AGCACACGTCTGAACTCTTTCCCTACACGACGCTCTTCCGATCTNNNTTCAGTTCTCTCCCTCTCTGTCTC
GAGCACACGTCTGAACTCTTTCCCTACACGACGCTCTTCCGATCTNNNTACTTTTGCTGTGTCCCACAA

GAGCACACGTCTGAACTCTTTCCCTACACGACGCTCTTCCGATCTNNNCATGGGCTGTATTTTGCTGAG
AGAGCACACGTCTGAACTCTTTCCCTACACGACGCTCTTCCGATCTNNNCTGACCTGTGTGTGTGTCTG
AGCACACGTCTGAACTCTTTCCCTACACGACGCTCTTCCGATCTNNNCTACCACGCCCAGCTAATTTTT
GAGCACACGTCTGAACTCTTTCCCTACACGACGCTCTTCCGATCTNNNGTGACCAGCTGCAATAAGTGT
GAGCACACGTCTGAACTCTTTCCCTACACGACGCTCTTCCGATCTNNNAAAAACAAAGGGGTGTGTGT
AGAGCACACGTCTGAACTCTTTCCCTACACGACGCTCTTCCGATCTNNNAGCAGATGTTGGTGTGTCTA
AGCACACGTCTGAACTCTTTCCCTACACGACGCTCTTCCGATCTNNNGGCTTCCCTATTTTTCCTGATGC
GAGCACACGTCTGAACTCTTTCCCTACACGACGCTCTTCCGATCTNNNCTAAAGAGCTGGGATTACAG
GAGCACACGTCTGAACTCTTTCCCTACACGACGCTCTTCCGATCTNNNGATTGTGTGTGGGGAGATT
GAGCACACGTCTGAACTCTTTCCCTACACGACGCTCTTCCGATCTNNNCTCATACTGTGACCAGACCCAA
AGCACACGTCTGAACTCTTTCCCTACACGACGCTCTTCCGATCTNNNTTCAGTTCTCATGTCAGGGCTTT
AGCACACGTCTGAACTCTTTCCCTACACGACGCTCTTCCGATCTNNNATTAGCCCATCTCTCCCATGAAT
GAGCACACGTCTGAACTCTTTCCCTACACGACGCTCTTCCGATCTNNNAAACTAGGTAGCTGCAGGAA
GAGCACACGTCTGAACTCTTTCCCTACACGACGCTCTTCCGATCTNNNCGAGACGTTTTCCATCCCATTT
AGCACACGTCTGAACTCTTTCCCTACACGACGCTCTTCCGATCTNNNTGTAATCCCAGCTACTTGGAAG
GAGCACACGTCTGAACTCTTTCCCTACACGACGCTCTTCCGATCTNNNCAAGTAGCTGGGATTACATGT
GCACACGTCTGAACTCTTTCCCTACACGACGCTCTTCCGATCTNNNGTGTTTCAATGCCTCTGCTCAAGC
GAGCACACGTCTGAACTCTTTCCCTACACGACGCTCTTCCGATCTNNNACTGTGGTGACTGTCTTTGTA
AGAGCACACGTCTGAACTCTTTCCCTACACGACGCTCTTCCGATCTNNNGACAGGAGTTCCATACCAGT
AGCACACGTCTGAACTCTTTCCCTACACGACGCTCTTCCGATCTNNNGCATGAGTCACTCTCCAACCTTA
AGCACACGTCTGAACTCTTTCCCTACACGACGCTCTTCCGATCTNNNATGGAGAACAAGGCCTCTGAA
GAGCACACGTCTGAACTCTTTCCCTACACGACGCTCTTCCGATCTNNNTGCTGCAGCTTATAGTCATCA
AGCACACGTCTGAACTCTTTCCCTACACGACGCTCTTCCGATCTNNNCTATCTGGGAATAATGAGGCGC
AGCACACGTCTGAACTCTTTCCCTACACGACGCTCTTCCGATCTNNNGCAGGATTATGTAGGGTTGGA
AGCACACGTCTGAACTCTTTCCCTACACGACGCTCTTCCGATCTNNNTCCTTATGTCTTCCTCCCTCCTTCA
AGCACACGTCTGAACTCTTTCCCTACACGACGCTCTTCCGATCTNNNGAAAGACCTGAGCTGAACTGA
AGAGCACACGTCTGAACTCTTTCCCTACACGACGCTCTTCCGATCTNNNACATCCTAAGTGCTTGCCATA
GAGCACACGTCTGAACTCTTTCCCTACACGACGCTCTTCCGATCTNNNGCACTGTGGAAAACTTTGAG
GAGCACACGTCTGAACTCTTTCCCTACACGACGCTCTTCCGATCTNNNCGCCTGGCCACTAATTTTTCTT
AGCACACGTCTGAACTCTTTCCCTACACGACGCTCTTCCGATCTNNNTTGTACCTGGTAGATGGAGGTT
GAGCACACGTCTGAACTCTTTCCCTACACGACGCTCTTCCGATCTNNNCACACACACACAGAGAGAGA
AGCACACGTCTGAACTCTTTCCCTACACGACGCTCTTCCGATCTNNNAACATAGCAAGATCAGCCCCTA
AGCACACGTCTGAACTCTTTCCCTACACGACGCTCTTCCGATCTNNNGGTTCATACCTGTAATCCCAGCA
GAGCACACGTCTGAACTCTTTCCCTACACGACGCTCTTCCGATCTNNNGCTAAAATCACTCCTCGCATGT
AGCACACGTCTGAACTCTTTCCCTACACGACGCTCTTCCGATCTNNNTGGGAGGTGATTGAATCATGGA
GAGCACACGTCTGAACTCTTTCCCTACACGACGCTCTTCCGATCTNNNCTCCACCCTTCTCTTTTGACTCA
AGCACACGTCTGAACTCTTTCCCTACACGACGCTCTTCCGATCTNNNTATCAGCCTTTTCCCCTGATTCC
AGCACACGTCTGAACTCTTTCCCTACACGACGCTCTTCCGATCTNNNGGCTCATAGCTCCCTGTATTAG
AGCACACGTCTGAACTCTTTCCCTACACGACGCTCTTCCGATCTNNNAAAAAGAGGAGACAGAGACTG
AAGAGCACACGTCTGAACTCTTTCCCTACACGACGCTCTTCCGATCTNNNTCTTACTGAGTGTGTGTGT
GAGCACACGTCTGAACTCTTTCCCTACACGACGCTCTTCCGATCTNNNAGGCATCTCTCTCTACTCTGTGT
AGCACACGTCTGAACTCTTTCCCTACACGACGCTCTTCCGATCTNNNATCACAGTTTCAGCACTACCACA
GCACACGTCTGAACTCTTTCCCTACACGACGCTCTTCCGATCTNNNCACTCACCTTAGGTCTAAGCCAG

GAGCACACGTCTGAACTCTTTCCCTACACGACGCTCTTCCGATCTNNNATAGTAGGAGTGGAGAGGA
GAGCACACGTCTGAACTCTTTCCCTACACGACGCTCTTCCGATCTNNNGATCACAGGTAATGTCTCCAG
GCACACGTCTGAACTCTTTCCCTACACGACGCTCTTCCGATCTNNNCCTGGCTTACTTATCCAGTAGGG
GAGCACACGTCTGAACTCTTTCCCTACACGACGCTCTTCCGATCTNNNGACCCAGAGTAGTCCTAGTG
GAGCACACGTCTGAACTCTTTCCCTACACGACGCTCTTCCGATCTNNNAGAGAGAGGGAGAGAGAG
GAGCACACGTCTGAACTCTTTCCCTACACGACGCTCTTCCGATCTNNNCAATCATGGCAGAAGGTGAA
GAGCACACGTCTGAACTCTTTCCCTACACGACGCTCTTCCGATCTNNNGAGCAAGACTCCGTCTCAAAA
GAGCACACGTCTGAACTCTTTCCCTACACGACGCTCTTCCGATCTNNNTCTCTTACACACACACACACAC
AGCACACGTCTGAACTCTTTCCCTACACGACGCTCTTCCGATCTNNNTTTCTTCCTTTCTTCCCTCCCTCC
AGCACACGTCTGAACTCTTTCCCTACACGACGCTCTTCCGATCTNNNCTAGCCCCTTTTCAAGATGGAG
AGCACACGTCTGAACTCTTTCCCTACACGACGCTCTTCCGATCTNNNATTCATTCAACCAGTTTGCCCAG
AGCACACGTCTGAACTCTTTCCCTACACGACGCTCTTCCGATCTNNNCACACAGGAATGTACAGTCGA
AGCACACGTCTGAACTCTTTCCCTACACGACGCTCTTCCGATCTNNNCTCTTCCTGACTTGGAGACTG
AGCACACGTCTGAACTCTTTCCCTACACGACGCTCTTCCGATCTNNNAGCTCTCACCCCTATCTCTTTCTC
GAGCACACGTCTGAACTCTTTCCCTACACGACGCTCTTCCGATCTNNNAGGCTACAGTAACCAAAACAG
GAGCACACGTCTGAACTCTTTCCCTACACGACGCTCTTCCGATCTNNNTGAAGGAGAAGCAAGCATGT
GAGCACACGTCTGAACTCTTTCCCTACACGACGCTCTTCCGATCTNNNGTCCTTCAGGTTTCAGGATCC
AGCACACGTCTGAACTCTTTCCCTACACGACGCTCTTCCGATCTNNNACCCACCTGAAGAAACTGATGT
GCACACGTCTGAACTCTTTCCCTACACGACGCTCTTCCGATCTNNNAGCCTGAGCAACATAGTAAGACC
GCACACGTCTGAACTCTTTCCCTACACGACGCTCTTCCGATCTNNNATGAATGGGATTAGCGCCCTTAT
GCACACGTCTGAACTCTTTCCCTACACGACGCTCTTCCGATCTNNNTCCTTTTTCCTTCCTTCCTTCCTCAT
AGCACACGTCTGAACTCTTTCCCTACACGACGCTCTTCCGATCTNNNGCCAAGTTTCATACACACACAC
GAGCACACGTCTGAACTCTTTCCCTACACGACGCTCTTCCGATCTNNNAGAGGGAGAGAGAGAGAGA
AGCACACGTCTGAACTCTTTCCCTACACGACGCTCTTCCGATCTNNNGGCCAACATGGTGAAAATACA
AGCACACGTCTGAACTCTTTCCCTACACGACGCTCTTCCGATCTNNNTTGGAAGTTCTGTGTACGTGGA
GCACACGTCTGAACTCTTTCCCTACACGACGCTCTTCCGATCTNNNAGATTTTTCACAGACTGCCTCCTC
AGCACACGTCTGAACTCTTTCCCTACACGACGCTCTTCCGATCTNNNCCACGTGGTTTTTGGTAGGATT
AGCACACGTCTGAACTCTTTCCCTACACGACGCTCTTCCGATCTNNNCTGGAGAACATACAACCAGGAG
AGCACACGTCTGAACTCTTTCCCTACACGACGCTCTTCCGATCTNNNGTCCTTTTTATATGGCGTCTGG
GAGCACACGTCTGAACTCTTTCCCTACACGACGCTCTTCCGATCTNNNTAGTGAACACTATTCCTGCTG
GAGCACACGTCTGAACTCTTTCCCTACACGACGCTCTTCCGATCTNNNGTTGCAATGAGCTGAGATCA
CACACGTCTGAACTCTTTCCCTACACGACGCTCTTCCGATCTNNNTAACCAAGAACTATTCCAGGGGG
AGCACACGTCTGAACTCTTTCCCTACACGACGCTCTTCCGATCTNNNAGAGAAGGTGTGTTACGGTTC
AGCACACGTCTGAACTCTTTCCCTACACGACGCTCTTCCGATCTNNNTCCTTTCCCCTTTCTTTTCTGCTC
AGCACACGTCTGAACTCTTTCCCTACACGACGCTCTTCCGATCTNNNGGTTGCAATGAGCTGATATCG
GAGCACACGTCTGAACTCTTTCCCTACACGACGCTCTTCCGATCTNNNACCAACTGGATGCAGTATCAT
GAGCACACGTCTGAACTCTTTCCCTACACGACGCTCTTCCGATCTNNNTTTCTTTCTCTCTCTCCCTCCCTC
AGCACACGTCTGAACTCTTTCCCTACACGACGCTCTTCCGATCTNNNTGTCTCTGTCTCTCTCTCTCTCTC
AGCACACGTCTGAACTCTTTCCCTACACGACGCTCTTCCGATCTNNNCCCAACAACTAGTAGCTGCAGA
AGCACACGTCTGAACTCTTTCCCTACACGACGCTCTTCCGATCTNNNCACACACACACACACACACATAT
AGCACACGTCTGAACTCTTTCCCTACACGACGCTCTTCCGATCTNNNCTGTGTTCCTAGTCCCCAGAAAG
AGCACACGTCTGAACTCTTTCCCTACACGACGCTCTTCCGATCTNNNTGACCTATGATCATGCACTCCAG
AGCACACGTCTGAACTCTTTCCCTACACGACGCTCTTCCGATCTNNNCTGAGGCAGGAGAATCACTTG

AGCACACGTCTGAACTCTTTCCCTACACGACGCTCTTCCGATCTNNNCTTCCTCAAAAGCAAGACCACT
GAGCACACGTCTGAACTCTTTCCCTACACGACGCTCTTCCGATCTNNNACATTGCATGTATGTGTGTG
AGAGCACACGTCTGAACTCTTTCCCTACACGACGCTCTTCCGATCTNNNCAAGAGCGAAATTCCGTCT
CACACGTCTGAACTCTTTCCCTACACGACGCTCTTCCGATCTNNNCATTCTATCAGCCCCTCTCTGTCCA
GAGCACACGTCTGAACTCTTTCCCTACACGACGCTCTTCCGATCTNNNTCACAGCCAAACCATATCAGT
AGCACACGTCTGAACTCTTTCCCTACACGACGCTCTTCCGATCTNNNATGGAGAGACTAGGGTTTTG
GCACACGTCTGAACTCTTTCCCTACACGACGCTCTTCCGATCTNNNCATTCCTCCAAACACACACACACC
GAGCACACGTCTGAACTCTTTCCCTACACGACGCTCTTCCGATCTNNNACTTTGATCTCTCTCTCGCTCT
AGCACACGTCTGAACTCTTTCCCTACACGACGCTCTTCCGATCTNNNGCTCTCTCTCTCTCTCTGTCTCTC
GCACACGTCTGAACTCTTTCCCTACACGACGCTCTTCCGATCTNNNCCTTCTCTCTCTCCCTCTCTCATC
AGAGCACACGTCTGAACTCTTTCCCTACACGACGCTCTTCCGATCTNNNGTGTATGTGTGTGTGTGTG
AGCACACGTCTGAACTCTTTCCCTACACGACGCTCTTCCGATCTNNNAACCATGTGCTACCATAACCAC
AGCACACGTCTGAACTCTTTCCCTACACGACGCTCTTCCGATCTNNNGGCAGGAGAAGGTCAGAGAA
GCACACGTCTGAACTCTTTCCCTACACGACGCTCTTCCGATCTNNNTTATGGGTGTGTGTGTGTGTGT
AGCACACGTCTGAACTCTTTCCCTACACGACGCTCTTCCGATCTNNNTCTCTCTCACACACACACACAAA
GCACACGTCTGAACTCTTTCCCTACACGACGCTCTTCCGATCTNNNATTTGCCTCTCCTCTTTCCTCTCCA
AGCACACGTCTGAACTCTTTCCCTACACGACGCTCTTCCGATCTNNNCCACAGTATACATCTTGGGGCT
GCACACGTCTGAACTCTTTCCCTACACGACGCTCTTCCGATCTNNNGAATTCGGCCAGTTTATGTCCTG
AGCACACGTCTGAACTCTTTCCCTACACGACGCTCTTCCGATCTNNNCTACGGAGACAGCACAGAAAG
GCACACGTCTGAACTCTTTCCCTACACGACGCTCTTCCGATCTNNNATATGTGTGTGTGTGTGTGTGT
GAGCACACGTCTGAACTCTTTCCCTACACGACGCTCTTCCGATCTNNNAGAATTGCTTGAGTCCAGGA
AGCACACGTCTGAACTCTTTCCCTACACGACGCTCTTCCGATCTNNNTGTTCAACCTAGTGGCTACTCA
AGCACACGTCTGAACTCTTTCCCTACACGACGCTCTTCCGATCTNNNTCCCCAGGTGTATTAGGGTTCT
AGCACACGTCTGAACTCTTTCCCTACACGACGCTCTTCCGATCTNNNTTTTTAGTAGAGGCCGGGTTT
AGCACACGTCTGAACTCTTTCCCTACACGACGCTCTTCCGATCTNNNTTTGACCCCCCAGAGTTTTACA
GAGCACACGTCTGAACTCTTTCCCTACACGACGCTCTTCCGATCTNNNTATCTCACACACACACACACAC
GAGCACACGTCTGAACTCTTTCCCTACACGACGCTCTTCCGATCTNNNGGTGACTCAAAAGTTGAAGT
GAGCACACGTCTGAACTCTTTCCCTACACGACGCTCTTCCGATCTNNNTGCAGACTTTCAATTCTCAGT
GCACACGTCTGAACTCTTTCCCTACACGACGCTCTTCCGATCTNNNCTCGAGTCTGACAACAAGAAGG
AGCACACGTCTGAACTCTTTCCCTACACGACGCTCTTCCGATCTNNNGATGATCCACGCTCTAGATGT
AGCACACGTCTGAACTCTTTCCCTACACGACGCTCTTCCGATCTNNNTCAGACACTTTTCAAACCCTTCC
CACACGTCTGAACTCTTTCCCTACACGACGCTCTTCCGATCTNNNTCTACCTAAAGCCAGAGTGATGC
AGAGCACACGTCTGAACTCTTTCCCTACACGACGCTCTTCCGATCTNNNCTGGAAGAAGAGGCCATTA
AGCACACGTCTGAACTCTTTCCCTACACGACGCTCTTCCGATCTNNNACCCTTACAACTCCATGAGTGT
AGCACACGTCTGAACTCTTTCCCTACACGACGCTCTTCCGATCTNNNTTGCAATGACCTAGACTCAGAC
AGCACACGTCTGAACTCTTTCCCTACACGACGCTCTTCCGATCTNNNCCTTGTCCTGCAGAAATTCACT
GAGCACACGTCTGAACTCTTTCCCTACACGACGCTCTTCCGATCTNNNGGGAAGGAGACAGAAGGCA
GAGCACACGTCTGAACTCTTTCCCTACACGACGCTCTTCCGATCTNNNACACATACACAGACACTCCCAT
GAGCACACGTCTGAACTCTTTCCCTACACGACGCTCTTCCGATCTNNNTGTCAGTGCAATGGAGGTAA
GAGCACACGTCTGAACTCTTTCCCTACACGACGCTCTTCCGATCTNNNTATAACTAACCCTCCCCCGAT
GAGCACACGTCTGAACTCTTTCCCTACACGACGCTCTTCCGATCTNNNCTAGCCAGTCCCTGTCAAACT
GAGCACACGTCTGAACTCTTTCCCTACACGACGCTCTTCCGATCTNNNTGATGGCCCATTTGGTTACAG
GAGCACACGTCTGAACTCTTTCCCTACACGACGCTCTTCCGATCTNNNACATCGTAGTGCTGTTTACCA

AGCACACGTCTGAACTCTTTCCCTACACGACGCTCTTCCGATCTNNNGGTGATGATGTGCGGATATCT
AGCACACGTCTGAACTCTTTCCCTACACGACGCTCTTCCGATCTNNNCTGTCCATCCATGCATCCATCTA
AGCACACGTCTGAACTCTTTCCCTACACGACGCTCTTCCGATCTNNNTTTTCTACAAAATTGACCACGCA
AGCACACGTCTGAACTCTTTCCCTACACGACGCTCTTCCGATCTNNNTGACAAGTGGTTGAAGTGAGT
GAGCACACGTCTGAACTCTTTCCCTACACGACGCTCTTCCGATCTNNNTGGGTGAAGACGCATGATTG
GAGCACACGTCTGAACTCTTTCCCTACACGACGCTCTTCCGATCTNNNTGGTTCGTCTGTGAAATGAG
GAGCACACGTCTGAACTCTTTCCCTACACGACGCTCTTCCGATCTNNNCAATCCTCCCCCAGATTCTGAA
AGCACACGTCTGAACTCTTTCCCTACACGACGCTCTTCCGATCTNNNTTTCTCTCTCTCCCTCTCTCTCCC
AGCACACGTCTGAACTCTTTCCCTACACGACGCTCTTCCGATCTNNNGGAAAGAGTTCTGCTACCACAG
GAGCACACGTCTGAACTCTTTCCCTACACGACGCTCTTCCGATCTNNNGACACTAGCAATACAGAGGT
AGCACACGTCTGAACTCTTTCCCTACACGACGCTCTTCCGATCTNNNACTCCTGGCATTGTACTAGCAA
AGCACACGTCTGAACTCTTTCCCTACACGACGCTCTTCCGATCTNNNAAATGAGCAGGGTCTACTTGT
GAGCACACGTCTGAACTCTTTCCCTACACGACGCTCTTCCGATCTNNNCTTATTCTAGAGTCGAGGGT
AGCACACGTCTGAACTCTTTCCCTACACGACGCTCTTCCGATCTNNNATGTCTGAACCCAAAAATCTGC
GAGCACACGTCTGAACTCTTTCCCTACACGACGCTCTTCCGATCTNNNGGGACAAACATGAGCTGTGA
GAGCACACGTCTGAACTCTTTCCCTACACGACGCTCTTCCGATCTNNNGCAGATGGTGTTGAGAATTA
GAGCACACGTCTGAACTCTTTCCCTACACGACGCTCTTCCGATCTNNNAGAATGTGACAGCTTCTCCTT
GCACACGTCTGAACTCTTTCCCTACACGACGCTCTTCCGATCTNNNGTCTTAGCCCTGTGGTTTTGTCT
GAGCACACGTCTGAACTCTTTCCCTACACGACGCTCTTCCGATCTNNNGGCCTTCTCTCTCTCATTCTCTG
AGCACACGTCTGAACTCTTTCCCTACACGACGCTCTTCCGATCTNNNGGGGGGAAGAAGCAAAAAA
GAGCACACGTCTGAACTCTTTCCCTACACGACGCTCTTCCGATCTNNNACTAGGAGACAGACAGACAG
GCACACGTCTGAACTCTTTCCCTACACGACGCTCTTCCGATCTNNNTGGAGGTAGGGAAGATGGTTT
GAGCACACGTCTGAACTCTTTCCCTACACGACGCTCTTCCGATCTNNNAGAAGATGTGAATGGGAAA
AGAGCACACGTCTGAACTCTTTCCCTACACGACGCTCTTCCGATCTNNNATGAAGAGCCATACTACTCC
GAGCACACGTCTGAACTCTTTCCCTACACGACGCTCTTCCGATCTNNNAGTTACTCATCCTACCACCCAG
AGCACACGTCTGAACTCTTTCCCTACACGACGCTCTTCCGATCTNNNGATAATTGTACTCCAGCCTGG
GAGCACACGTCTGAACTCTTTCCCTACACGACGCTCTTCCGATCTNNNTGATTGTTTGCCCAAGGAAA
AGCACACGTCTGAACTCTTTCCCTACACGACGCTCTTCCGATCTNNNTACTAAAAGCCCTACAACACCC
AGCACACGTCTGAACTCTTTCCCTACACGACGCTCTTCCGATCTNNNGACTTCCTGTCCTTTCACATCCT
GAGCACACGTCTGAACTCTTTCCCTACACGACGCTCTTCCGATCTNNNGCAACAGAGCAATATCCCCAA
GCACACGTCTGAACTCTTTCCCTACACGACGCTCTTCCGATCTNNNACCACACAAGAATGGTCTGAAC
AGCACACGTCTGAACTCTTTCCCTACACGACGCTCTTCCGATCTNNNCCTCAGTATTTCTTGGGGAAG
GAGCACACGTCTGAACTCTTTCCCTACACGACGCTCTTCCGATCTNNNAAGCACTCCACACAACATTGA
GAGCACACGTCTGAACTCTTTCCCTACACGACGCTCTTCCGATCTNNNGCATTGATTGAGCTGAGATTG
AGCACACGTCTGAACTCTTTCCCTACACGACGCTCTTCCGATCTNNNTACTTTTGTTTGCTGTTTGCCCA
AGCACACGTCTGAACTCTTTCCCTACACGACGCTCTTCCGATCTNNNTAGAACTGATTGTCTTGGGGCA
AGCACACGTCTGAACTCTTTCCCTACACGACGCTCTTCCGATCTNNNAGACTGCTCCAATCTTTTGAGG
AGCACACGTCTGAACTCTTTCCCTACACGACGCTCTTCCGATCTNNNTGTCATGGGCAAGAACTGGAT
AAGAGCACACGTCTGAACTCTTTCCCTACACGACGCTCTTCCGATCTNNNACTCTTCACATGCACGTTT
AGCACACGTCTGAACTCTTTCCCTACACGACGCTCTTCCGATCTNNNTGTGAGGACCTGAAGTTCCAA
GAGCACACGTCTGAACTCTTTCCCTACACGACGCTCTTCCGATCTNNNCAGCTCACACATCTCCCTCTAT
GAGCACACGTCTGAACTCTTTCCCTACACGACGCTCTTCCGATCTNNNCCTTTGAAAACCAACTGACTC
AGCACACGTCTGAACTCTTTCCCTACACGACGCTCTTCCGATCTNNNTATGGTAACCCCTAGGCCCTTA

GAGCACACGTCTGAACTCTTTCCCTACACGACGCTCTTCCGATCTNNNATACAAAAAGCAACTCCCTGT
AGAGCACACGTCTGAACTCTTTCCCTACACGACGCTCTTCCGATCTNNNTTTTCGGATTGTTTGCAGGA
AGAGCACACGTCTGAACTCTTTCCCTACACGACGCTCTTCCGATCTNNNAGCAGACAAAATCCCTGACT
AGAGCACACGTCTGAACTCTTTCCCTACACGACGCTCTTCCGATCTNNNATCCACAGAGCTTTGTTTCA
AAGAGCACACGTCTGAACTCTTTCCCTACACGACGCTCTTCCGATCTNNNTCAGTGTGGTGGAAGGAG
GAGCACACGTCTGAACTCTTTCCCTACACGACGCTCTTCCGATCTNNNATGGAAACCACACAAATGTCC
GAGCACACGTCTGAACTCTTTCCCTACACGACGCTCTTCCGATCTNNNCCTGCTGCTCAATTACTTGCAT
GAGCACACGTCTGAACTCTTTCCCTACACGACGCTCTTCCGATCTNNNCGAGGAGTTTGAGGTTACAG
GAGCACACGTCTGAACTCTTTCCCTACACGACGCTCTTCCGATCTNNNATGCTGTGCTTCTTCAGTTTT
GAGCACACGTCTGAACTCTTTCCCTACACGACGCTCTTCCGATCTNNNTAGAAGGTAGGACAGAAGCC
AGAGCACACGTCTGAACTCTTTCCCTACACGACGCTCTTCCGATCTNNNCTGCCCTAGAGTTCTGTAGCA
AGCACACGTCTGAACTCTTTCCCTACACGACGCTCTTCCGATCTNNNGTACCTGGCACATAGTAGTTG
AGCACACGTCTGAACTCTTTCCCTACACGACGCTCTTCCGATCTNNNTCTTGGTCCAAAGACAGGTCAA
AGCACACGTCTGAACTCTTTCCCTACACGACGCTCTTCCGATCTNNNTCCACTTTCTTGAGTGCCACTAG
AGCACACGTCTGAACTCTTTCCCTACACGACGCTCTTCCGATCTNNNGTTCCAGGAGTCTTCCCTCATTT
GAGCACACGTCTGAACTCTTTCCCTACACGACGCTCTTCCGATCTNNNGAAAATGTCAGGTCTACAGC
GCACACGTCTGAACTCTTTCCCTACACGACGCTCTTCCGATCTNNNTGTCGTGCAACCTATGGATGTTA
AGAGCACACGTCTGAACTCTTTCCCTACACGACGCTCTTCCGATCTNNNAGTAGGACTTGTCAAGGGA
AGCACACGTCTGAACTCTTTCCCTACACGACGCTCTTCCGATCTNNNGTCCGTCCCCATTCTATATGGTC
AGCACACGTCTGAACTCTTTCCCTACACGACGCTCTTCCGATCTNNNATGAGAAGCTCTGCTCATCTCT
AGCACACGTCTGAACTCTTTCCCTACACGACGCTCTTCCGATCTNNNCTCCTGAGCAGCTGGGATTATA
AGCACACGTCTGAACTCTTTCCCTACACGACGCTCTTCCGATCTNNNAGTGTGGGATGGCAAGATAAA
AGCACACGTCTGAACTCTTTCCCTACACGACGCTCTTCCGATCTNNNATATGCATGTGTGTGTGTGTGT
GAGCACACGTCTGAACTCTTTCCCTACACGACGCTCTTCCGATCTNNNCAGATGTCAGTGGATGTCAG
GAGCACACGTCTGAACTCTTTCCCTACACGACGCTCTTCCGATCTNNNTTTCAACTTGCTTTTCCTGGTG
GAGCACACGTCTGAACTCTTTCCCTACACGACGCTCTTCCGATCTNNNTGTGAGGTAAGACGAAAGAG
GAGCACACGTCTGAACTCTTTCCCTACACGACGCTCTTCCGATCTNNNCAGATCAGACTTTGCTGCCAA
AGCACACGTCTGAACTCTTTCCCTACACGACGCTCTTCCGATCTNNNAATATTTGCCTGCTCCCCACTA
GAGCACACGTCTGAACTCTTTCCCTACACGACGCTCTTCCGATCTNNNGGGCTCAATGTCCAGCTAAGA
GAGCACACGTCTGAACTCTTTCCCTACACGACGCTCTTCCGATCTNNNCACTAGAAGCTCGGTGATGCT
AGCACACGTCTGAACTCTTTCCCTACACGACGCTCTTCCGATCTNNNTCTCTTATGACCCAGTTCTCCCC
GAGCACACGTCTGAACTCTTTCCCTACACGACGCTCTTCCGATCTNNNGCAGCCATCCAAGAAAGATCA
AGCACACGTCTGAACTCTTTCCCTACACGACGCTCTTCCGATCTNNNGTGGTGGGCAGAATTCTAACAT
GAGCACACGTCTGAACTCTTTCCCTACACGACGCTCTTCCGATCTNNNGAATTACACCATGGGCTTTCTG
GAGCACACGTCTGAACTCTTTCCCTACACGACGCTCTTCCGATCTNNNGCCTTCTCAGCTGACAGAGAA
AGCACACGTCTGAACTCTTTCCCTACACGACGCTCTTCCGATCTNNNTTTGCTGATACTGTCTCCTGGTC
AGAGCACACGTCTGAACTCTTTCCCTACACGACGCTCTTCCGATCTNNNGGGTCTCTCTCTCTCTCTCTCT
GAGCACACGTCTGAACTCTTTCCCTACACGACGCTCTTCCGATCTNNNTGGGCAACATAGTGAGTCTCA
AGCACACGTCTGAACTCTTTCCCTACACGACGCTCTTCCGATCTNNNATATTCTGAAGGTGGAGCCAA
AGAGCACACGTCTGAACTCTTTCCCTACACGACGCTCTTCCGATCTNNNGCTAGAGAGGAGGAAAGA
AGAGCACACGTCTGAACTCTTTCCCTACACGACGCTCTTCCGATCTNNNTTCACTTGGATGACATGCCCT
GAGCACACGTCTGAACTCTTTCCCTACACGACGCTCTTCCGATCTNNNTTGACCTCCTGAATCCACACAAT
AGCACACGTCTGAACTCTTTCCCTACACGACGCTCTTCCGATCTNNNGAAATGCAAAGATCTGAGGCCA

AGCACACGTCTGAACTCTTTCCCTACACGACGCTCTTCCGATCTNNNAATGCCCATAATTAGCTTTGCC

GCACACGTCTGAACTCTTTCCCTACACGACGCTCTTCCGATCTNNNTACACACAGACACACAAGCATCT

AGCACACGTCTGAACTCTTTCCCTACACGACGCTCTTCCGATCTNNNTTCAGAGCCTAGAGACTCCATC

AGCACACGTCTGAACTCTTTCCCTACACGACGCTCTTCCGATCTNNNCAGTCAAGCTTGGGGATTTGT

AGCACACGTCTGAACTCTTTCCCTACACGACGCTCTTCCGATCTNNNTGTTCTTCCGATGACACTTCCTT

AGAGCACACGTCTGAACTCTTTCCCTACACGACGCTCTTCCGATCTNNNGTTTTCTGCCCACATTACCAG

GCACACGTCTGAACTCTTTCCCTACACGACGCTCTTCCGATCTNNNGGGAAAACAACTTCCTCTTCAG

AGCACACGTCTGAACTCTTTCCCTACACGACGCTCTTCCGATCTNNNAGAGGACCTGGGCATTTATTC

GCACACGTCTGAACTCTTTCCCTACACGACGCTCTTCCGATCTNNNTCCTGCCCTTGTACTCAAATTGA

AGAGCACACGTCTGAACTCTTTCCCTACACGACGCTCTTCCGATCTNNNTTCTAAAAAAGACAACCCAC

AGCACACGTCTGAACTCTTTCCCTACACGACGCTCTTCCGATCTNNNGGAAAACAAATTCTTGGGGTG

AGCACACGTCTGAACTCTTTCCCTACACGACGCTCTTCCGATCTNNNAGCATGTGTTGAAGACTGGTA

GCACACGTCTGAACTCTTTCCCTACACGACGCTCTTCCGATCTNNNGTAGACCTCTCTCTAAGTGCACG

GAGCACACGTCTGAACTCTTTCCCTACACGACGCTCTTCCGATCTNNNTAGGCTAAATGTCCGGTGCTT

GAGCACACGTCTGAACTCTTTCCCTACACGACGCTCTTCCGATCTNNNTCTCTCTCTCTCTCTCTCTCA

AGCACACGTCTGAACTCTTTCCCTACACGACGCTCTTCCGATCTNNNTAGCATACTGCCTAACACACTG

GAGCACACGTCTGAACTCTTTCCCTACACGACGCTCTTCCGATCTNNNAGGGCTGAGACATTGTCTTT

GAGCACACGTCTGAACTCTTTCCCTACACGACGCTCTTCCGATCTNNNTTCCCCAGAATTGATAGTGCA

AGCACACGTCTGAACTCTTTCCCTACACGACGCTCTTCCGATCTNNNCTCTTCCTTTTTTCATGGGGCT

AGCACACGTCTGAACTCTTTCCCTACACGACGCTCTTCCGATCTNNNATCCTACGGCTTTAGCTCTTCAG

GCACACGTCTGAACTCTTTCCCTACACGACGCTCTTCCGATCTNNNTGGTAACATACATTGGTGAGCC

GAGCACACGTCTGAACTCTTTCCCTACACGACGCTCTTCCGATCTNNNCCAGGGGCAGAATAGTCAT

GAGCACACGTCTGAACTCTTTCCCTACACGACGCTCTTCCGATCTNNNAAAGAAAACACACTCTGGCA

GAGCACACGTCTGAACTCTTTCCCTACACGACGCTCTTCCGATCTNNNGGTGCTCCTGTACTGACTACA

AGCACACGTCTGAACTCTTTCCCTACACGACGCTCTTCCGATCTNNNGTGGATCTCAACTGGGAGTGA

GCACACGTCTGAACTCTTTCCCTACACGACGCTCTTCCGATCTNNNCCCTTGCTCTTCTCACTCTCTTTC

GCACACGTCTGAACTCTTTCCCTACACGACGCTCTTCCGATCTNNNACTCTTAGCTTCTTTGGTGCAGT

AGCACACGTCTGAACTCTTTCCCTACACGACGCTCTTCCGATCTNNNCATCAAACAGCTCACTTGACCA

GCACACGTCTGAACTCTTTCCCTACACGACGCTCTTCCGATCTNNNCACACACACACACACACACATAT

GCACACGTCTGAACTCTTTCCCTACACGACGCTCTTCCGATCTNNNACTCAGGGACCAACAAAATTCC

AGCACACGTCTGAACTCTTTCCCTACACGACGCTCTTCCGATCTNNNTTTCTCTCCCGGTCTACATCTCT

AGCACACGTCTGAACTCTTTCCCTACACGACGCTCTTCCGATCTNNNTGGCAATCAAGGAATCACTTG

AGCACACGTCTGAACTCTTTCCCTACACGACGCTCTTCCGATCTNNNTGAACTGTCTTTCCCACTCTCTG

GAGCACACGTCTGAACTCTTTCCCTACACGACGCTCTTCCGATCTNNNAAGCCATTTTGGACCTACTAG

GCACACGTCTGAACTCTTTCCCTACACGACGCTCTTCCGATCTNNNCCCTCGACTCTGAAGGTAAAGA

GCACACGTCTGAACTCTTTCCCTACACGACGCTCTTCCGATCTNNNCACGTTCACCTGGAAACCTTATT

GAGCACACGTCTGAACTCTTTCCCTACACGACGCTCTTCCGATCTNNNCCGTGGAGAAAGGAATGTG

GAGCACACGTCTGAACTCTTTCCCTACACGACGCTCTTCCGATCTNNNATTGTTGATTACCAAGAGGC

AGCACACGTCTGAACTCTTTCCCTACACGACGCTCTTCCGATCTNNNCAGAGCAAGACCATGTCTCAGA

GAGCACACGTCTGAACTCTTTCCCTACACGACGCTCTTCCGATCTNNNCCTCACAGAAAGAAGTGGCTA

GCACACGTCTGAACTCTTTCCCTACACGACGCTCTTCCGATCTNNNCCCTCCTGTTGGTTTGAAATTGA

AGAGCACACGTCTGAACTCTTTCCCTACACGACGCTCTTCCGATCTNNNGACACCTACTGCAGACACA

AGCACACGTCTGAACTCTTTCCCTACACGACGCTCTTCCGATCTNNNGAGGTTCCTGGATCTAGCAAC

GAGCACACGTCTGAACTCTTTCCCTACACGACGCTCTTCCGATCTNNNTCACTGAATCCCTTTAGCCAG

GAGCACACGTCTGAACTCTTTCCCTACACGACGCTCTTCCGATCTNNNTGGGGACAGCTTCAATGATG

AGCACACGTCTGAACTCTTTCCCTACACGACGCTCTTCCGATCTNNNGTAAGAAAAGGAGGAGGCAG

AGCACACGTCTGAACTCTTTCCCTACACGACGCTCTTCCGATCTNNNCCAGGTACAAGCAGGCAATAA

AGCACACGTCTGAACTCTTTCCCTACACGACGCTCTTCCGATCTNNNCAGTGTGAGTTGCAGATAGGT

AGCACACGTCTGAACTCTTTCCCTACACGACGCTCTTCCGATCTNNNTCTAGACAAAGAACCAACAG

AGCACACGTCTGAACTCTTTCCCTACACGACGCTCTTCCGATCTNNNGCAGTTCTTGTTTTCCCTCCTTG

GAGCACACGTCTGAACTCTTTCCCTACACGACGCTCTTCCGATCTNNNTCATCATGTTGACTTTCCCTC

AGCACACGTCTGAACTCTTTCCCTACACGACGCTCTTCCGATCTNNNCCTTAGTGTTAGCTGAGGACC

AGCACACGTCTGAACTCTTTCCCTACACGACGCTCTTCCGATCTNNNCCAGTTTCTCACGTATCCTTCCA

GAGCACACGTCTGAACTCTTTCCCTACACGACGCTCTTCCGATCTNNNTACTGCTCAAACCATCCTGGA

GCACACGTCTGAACTCTTTCCCTACACGACGCTCTTCCGATCTNNNGGCATTGATAAGTCTTCCTTGG

GAGCACACGTCTGAACTCTTTCCCTACACGACGCTCTTCCGATCTNNNATGTACTTGTTTGGTGTGGC

GAGCACACGTCTGAACTCTTTCCCTACACGACGCTCTTCCGATCTNNNGATGGTTTCCCAAAAGAAGG

GCACACGTCTGAACTCTTTCCCTACACGACGCTCTTCCGATCTNNNTCCCCCATCTGTCATCAAAATGT

AGCACACGTCTGAACTCTTTCCCTACACGACGCTCTTCCGATCTNNNTATACAGGGTTCCCTTCTTGTG

AGCACACGTCTGAACTCTTTCCCTACACGACGCTCTTCCGATCTNNNCTAGGAAAAGGGTCCGCCATA

AGCACACGTCTGAACTCTTTCCCTACACGACGCTCTTCCGATCTNNNCCCTTCGGCTGTCTTATTTTTGG

GCACACGTCTGAACTCTTTCCCTACACGACGCTCTTCCGATCTNNNTTTCTTCTGAGGCCACAGACTAG

AGCACACGTCTGAACTCTTTCCCTACACGACGCTCTTCCGATCTNNNGGAGACTTCTGTTTGGGGATT

AGCACACGTCTGAACTCTTTCCCTACACGACGCTCTTCCGATCTNNNTGCCATTTATTGGGTGCTGAA

AGCACACGTCTGAACTCTTTCCCTACACGACGCTCTTCCGATCTNNNTCTGTGGGAATTTGAGAGTGC

GAGCACACGTCTGAACTCTTTCCCTACACGACGCTCTTCCGATCTNNNTTTCTCATCTTCCCAGCTGAC

GAGCACACGTCTGAACTCTTTCCCTACACGACGCTCTTCCGATCTNNNATTCTCGTCGACTCAGTCATC

AGCACACGTCTGAACTCTTTCCCTACACGACGCTCTTCCGATCTNNNTTGATGAAACCTGTGAAATGG

AGCACACGTCTGAACTCTTTCCCTACACGACGCTCTTCCGATCTNNNGTTCTTCAGATTGCGGAACAC

GCACACGTCTGAACTCTTTCCCTACACGACGCTCTTCCGATCTNNNGACCACTTGTGATGTTGAGCAT

GCACACGTCTGAACTCTTTCCCTACACGACGCTCTTCCGATCTNNNAAATCTCTTGGGCCTCTTTTCCT

AGCACACGTCTGAACTCTTTCCCTACACGACGCTCTTCCGATCTNNNATGTTTACCCATGCTATGCCTCT

AGCACACGTCTGAACTCTTTCCCTACACGACGCTCTTCCGATCTNNNGAAAGAGAGGCATTCACAGC

AGCACACGTCTGAACTCTTTCCCTACACGACGCTCTTCCGATCTNNNGATGGAGCAAAAACCCTCAC

GCACACGTCTGAACTCTTTCCCTACACGACGCTCTTCCGATCTNNNTCCTGATGTTCCTTGGTTTGTG

GAGCACACGTCTGAACTCTTTCCCTACACGACGCTCTTCCGATCTNNNGCCAAGGGAAATGGAAGCA

AGCACACGTCTGAACTCTTTCCCTACACGACGCTCTTCCGATCTNNNTCCCCACTGAACTACTTCCTA

AGCACACGTCTGAACTCTTTCCCTACACGACGCTCTTCCGATCTNNNGGTGACAGCTCTCTCTCTTTC

AGCACACGTCTGAACTCTTTCCCTACACGACGCTCTTCCGATCTNNNATCCTCTACCCAGTCTCAGGT

GAGCACACGTCTGAACTCTTTCCCTACACGACGCTCTTCCGATCTNNNGCTCCTTGGGGCTTACATAC

AGCACACGTCTGAACTCTTTCCCTACACGACGCTCTTCCGATCTNNNTGTGGAACCCTAGGATATGGA

AGCACACGTCTGAACTCTTTCCCTACACGACGCTCTTCCGATCTNNNGCACTATTCCCAACAGCCAAAA

AGCACACGTCTGAACTCTTTCCCTACACGACGCTCTTCCGATCTNNNTCTCACATAACAAGAAGCCTGG

AGCACACGTCTGAACTCTTTCCCTACACGACGCTCTTCCGATCTNNNATGGGAATGACTTTGAAGCT

GAGCACACGTCTGAACTCTTTCCCTACACGACGCTCTTCCGATCTNNNGACAGAGACCAGGAGAGAA

GCACACGTCTGAACTCTTTCCCTACACGACGCTCTTCCGATCTNNNTGGCAGGGCTGAGAATAAATCA

GAGCACACGTCTGAACTCTTTCCCTACACGACGCTCTTCCGATCTNNNGAGCAAGTGTTGGAAGTAA

GAGCACACGTCTGAACTCTTTCCCTACACGACGCTCTTCCGATCTNNNCTGATGCCCCAAACATACAA

GAGCACACGTCTGAACTCTTTCCCTACACGACGCTCTTCCGATCTNNNATGAAAGGGGAGTGACAGT

AGCACACGTCTGAACTCTTTCCCTACACGACGCTCTTCCGATCTNNNTTGAGCTTGCTCTTGGGACAT

AGCACACGTCTGAACTCTTTCCCTACACGACGCTCTTCCGATCTNNNGTGCTCTGTGTACGTTTTCAC

AGCACACGTCTGAACTCTTTCCCTACACGACGCTCTTCCGATCTNNNAGCTGACTACCTTTCAAGCAAC

AGCACACGTCTGAACTCTTTCCCTACACGACGCTCTTCCGATCTNNNAATGTTTGTGTTTGGTGTGAG

GAGCACACGTCTGAACTCTTTCCCTACACGACGCTCTTCCGATCTNNNTGTGGAGGTGGCTTCTAAAG

AGCACACGTCTGAACTCTTTCCCTACACGACGCTCTTCCGATCTNNNTTATACCTCCTGCTACCCCAAGA

AGCACACGTCTGAACTCTTTCCCTACACGACGCTCTTCCGATCTNNNGGAATCAAGGGTGAGAGACA

AGCACACGTCTGAACTCTTTCCCTACACGACGCTCTTCCGATCTNNNTCTTCTCTTTTTCCACGCACAC

AGCACACGTCTGAACTCTTTCCCTACACGACGCTCTTCCGATCTNNNCAAACTCTGCACCACCACATTAG

AGCACACGTCTGAACTCTTTCCCTACACGACGCTCTTCCGATCTNNNGATCCCCTGAACACCTCTTCATT

AGCACACGTCTGAACTCTTTCCCTACACGACGCTCTTCCGATCTNNNCTGGGTGACACTTTCCCTAATC

AGCACACGTCTGAACTCTTTCCCTACACGACGCTCTTCCGATCTNNNTTGGCTTGGATGTTGGGACT

GCACACGTCTGAACTCTTTCCCTACACGACGCTCTTCCGATCTNNNGTCTTGTCTCTCTGTCTCTGCTCC

CACACGTCTGAACTCTTTCCCTACACGACGCTCTTCCGATCTNNNCATTCCAGCTTCATCTTTTGCCAC

AGCACACGTCTGAACTCTTTCCCTACACGACGCTCTTCCGATCTNNNTGAAAATGGAAACCACAGCT

GAGCACACGTCTGAACTCTTTCCCTACACGACGCTCTTCCGATCTNNNAAGCCTGTGTTCATCCTGTTT

GAGCACACGTCTGAACTCTTTCCCTACACGACGCTCTTCCGATCTNNNGCTTATTGTAGCCAGGAACG

GAGCACACGTCTGAACTCTTTCCCTACACGACGCTCTTCCGATCTNNNAAAGGGGTATTGTTGTCCTC

CACACGTCTGAACTCTTTCCCTACACGACGCTCTTCCGATCTNNNAAGACTTCACTGAGGGGTCAAAG

AGCACACGTCTGAACTCTTTCCCTACACGACGCTCTTCCGATCTNNNCAGGCCTTCAGACAAACAGAA

GAGCACACGTCTGAACTCTTTCCCTACACGACGCTCTTCCGATCTNNNCAGCATCTATTCCCTCCTTCCT

CACACGTCTGAACTCTTTCCCTACACGACGCTCTTCCGATCTNNNTGCTTTCTTCCTTTCCTCCTACCCA

GCACACGTCTGAACTCTTTCCCTACACGACGCTCTTCCGATCTNNNGGCTCAATCAAATACACAAACC

GAGCACACGTCTGAACTCTTTCCCTACACGACGCTCTTCCGATCTNNNCAACCCCATTTGTGAAGACTT

AGCACACGTCTGAACTCTTTCCCTACACGACGCTCTTCCGATCTNNNATCTATGGCAGCTCCTCTTCTTC

AGCACACGTCTGAACTCTTTCCCTACACGACGCTCTTCCGATCTNNNATGATTGCCAGTCAGTGACTC

AGCACACGTCTGAACTCTTTCCCTACACGACGCTCTTCCGATCTNNNAGGTCGAATGCACACTTCTAC

GCACACGTCTGAACTCTTTCCCTACACGACGCTCTTCCGATCTNNNTACACTATCAGTTTTCCTGGGCC

AGCACACGTCTGAACTCTTTCCCTACACGACGCTCTTCCGATCTNNNCCTCAGGTTGGGTCTATCCTTT

GAGCACACGTCTGAACTCTTTCCCTACACGACGCTCTTCCGATCTNNNGATTCTTCGGTGTCACAAACA

AGCACACGTCTGAACTCTTTCCCTACACGACGCTCTTCCGATCTNNNCCAGGTCAAAAGACCCTTGAA

GCACACGTCTGAACTCTTTCCCTACACGACGCTCTTCCGATCTNNNCCAGTGTTTGAGAACCAGTAGG

AGCACACGTCTGAACTCTTTCCCTACACGACGCTCTTCCGATCTNNNGATTTCAGTGATGCCAACGTCT

GAGCACACGTCTGAACTCTTTCCCTACACGACGCTCTTCCGATCTNNNCCAGTATTCTTCCCTTCTCCAC

AGCACACGTCTGAACTCTTTCCCTACACGACGCTCTTCCGATCTNNNCGACTCTTCAGACTGTGGACTT

AGCACACGTCTGAACTCTTTCCCTACACGACGCTCTTCCGATCTNNNCTCTGACCACCATTCTTCCTTCA

AGCACACGTCTGAACTCTTTCCCTACACGACGCTCTTCCGATCTNNNAGGGCTTGATTTCAGGAGTTC

GCACACGTCTGAACTCTTTCCCTACACGACGCTCTTCCGATCTNNNCCTTTTTGCGAATTGCTAAAGCC

GAGCACACGTCTGAACTCTTTCCCTACACGACGCTCTTCCGATCTNNNTTTACGGGGAAAAAAATGG

GAGCACACGTCTGAACTCTTTCCCTACACGACGCTCTTCCGATCTNNNTTTTCTGCCTCACAAACACCA

GAGCACACGTCTGAACTCTTTCCCTACACGACGCTCTTCCGATCTNNNAAGAGGGAGAAGGTGCAAT
GAGCACACGTCTGAACTCTTTCCCTACACGACGCTCTTCCGATCTNNNGGAAGATGATTTCCAGGCAC
AGAGCACACGTCTGAACTCTTTCCCTACACGACGCTCTTCCGATCTNNNCCATGGAGGAGAAAAGAG
AGCACACGTCTGAACTCTTTCCCTACACGACGCTCTTCCGATCTNNNTTTCATGCTGACTGGGGAAGA
AGCACACGTCTGAACTCTTTCCCTACACGACGCTCTTCCGATCTNNNTCATGACCTGTCCCGTTTTATGT
AGCACACGTCTGAACTCTTTCCCTACACGACGCTCTTCCGATCTNNNCCTGGCTACAAACTTGCATTTC
AGCACACGTCTGAACTCTTTCCCTACACGACGCTCTTCCGATCTNNNATTCATTAGTGGGGTTGGACT
AGCACACGTCTGAACTCTTTCCCTACACGACGCTCTTCCGATCTNNNAAAAGCTGGTAGTATCACACC
AGCACACGTCTGAACTCTTTCCCTACACGACGCTCTTCCGATCTNNNGCTGAAGTGACTCACCAAAAT
GAGCACACGTCTGAACTCTTTCCCTACACGACGCTCTTCCGATCTNNNGCTATGGAAAATCCTTGTCTC
GAGCACACGTCTGAACTCTTTCCCTACACGACGCTCTTCCGATCTNNNGGTCTCTCTCTCTCTGTCTCT
AGCACACGTCTGAACTCTTTCCCTACACGACGCTCTTCCGATCTNNNGAGGGGCATGAAGAGTGAAG
AGCACACGTCTGAACTCTTTCCCTACACGACGCTCTTCCGATCTNNNACTGTTGTGGCCAGAGACTAA
GAGCACACGTCTGAACTCTTTCCCTACACGACGCTCTTCCGATCTNNNCTCTTGTTCCCTAGACGACTAG
AGAGCACACGTCTGAACTCTTTCCCTACACGACGCTCTTCCGATCTNNNTACTGGATCATCTCTAGGC
AGCACACGTCTGAACTCTTTCCCTACACGACGCTCTTCCGATCTNNNCAAGAGTCAGATTGCCCCATTT
GCACACGTCTGAACTCTTTCCCTACACGACGCTCTTCCGATCTNNNTCTTCTGGCCACCTTCTAACTTCC
GCACACGTCTGAACTCTTTCCCTACACGACGCTCTTCCGATCTNNNAGAACTGAGCTGGCCTTTATGA
AGCACACGTCTGAACTCTTTCCCTACACGACGCTCTTCCGATCTNNNTCCAGACTGTCTTTGGACTTGA
GAGCACACGTCTGAACTCTTTCCCTACACGACGCTCTTCCGATCTNNNATGGTGAATGTACAAGACCA
GAGCACACGTCTGAACTCTTTCCCTACACGACGCTCTTCCGATCTNNNGAGTAATGGGGAGTGTTAG
AGCACACGTCTGAACTCTTTCCCTACACGACGCTCTTCCGATCTNNNCTCCTAATTGAACCCAGACTGCT
GAGCACACGTCTGAACTCTTTCCCTACACGACGCTCTTCCGATCTNNNAGCAACTGATTATTGGCAGT
AGAGCACACGTCTGAACTCTTTCCCTACACGACGCTCTTCCGATCTNNNGGGCTCTTACTACACAGAAA
AGCACACGTCTGAACTCTTTCCCTACACGACGCTCTTCCGATCTNNNCCACAAAAGACTGCAAAAAAGG
AGCACACGTCTGAACTCTTTCCCTACACGACGCTCTTCCGATCTNNNTTCATTTGCATGCTAAAGCCCT
GAGCACACGTCTGAACTCTTTCCCTACACGACGCTCTTCCGATCTNNNGTACCTTGCTACCTTGTCCATC
GAGCACACGTCTGAACTCTTTCCCTACACGACGCTCTTCCGATCTNNNTCTACTTTGAACAAATCCCCC
GCACACGTCTGAACTCTTTCCCTACACGACGCTCTTCCGATCTNNNTAATCACACACACACACACACAC
AGCACACGTCTGAACTCTTTCCCTACACGACGCTCTTCCGATCTNNNACTTGGTAGCCTCAAACAGCTA
AGCACACGTCTGAACTCTTTCCCTACACGACGCTCTTCCGATCTNNNAGGCTTATATGGGTTTGGGAG
GAGCACACGTCTGAACTCTTTCCCTACACGACGCTCTTCCGATCTNNNGCCACCTTGATAGTCTTGGAC
GAGCACACGTCTGAACTCTTTCCCTACACGACGCTCTTCCGATCTNNNGATATGCTGTGAAGGCTGA
GCACACGTCTGAACTCTTTCCCTACACGACGCTCTTCCGATCTNNNAGCTGCTGACAAGATGTATTTG
AGCACACGTCTGAACTCTTTCCCTACACGACGCTCTTCCGATCTNNNAGAGATCCAATGAGCAACAGG
GAGCACACGTCTGAACTCTTTCCCTACACGACGCTCTTCCGATCTNNNGATACCACATGGAGGAATGG
AGCACACGTCTGAACTCTTTCCCTACACGACGCTCTTCCGATCTNNNGACCATCAGGAAATTGCCTCTT
GAGCACACGTCTGAACTCTTTCCCTACACGACGCTCTTCCGATCTNNNGGAGGATGTCTAGGCTTACAT
AGCACACGTCTGAACTCTTTCCCTACACGACGCTCTTCCGATCTNNNTTCCAGAAGGTTAGAAGTGAG
GAGCACACGTCTGAACTCTTTCCCTACACGACGCTCTTCCGATCTNNNCTTTGCATTGGATTCAGCAGG
GCACACGTCTGAACTCTTTCCCTACACGACGCTCTTCCGATCTNNNAGTGTGAACAGAGGCATAAACC
AGCACACGTCTGAACTCTTTCCCTACACGACGCTCTTCCGATCTNNNTTCAAGCTCTAAGACCAGAAG
AGCACACGTCTGAACTCTTTCCCTACACGACGCTCTTCCGATCTNNNCCACACAAAAACACAGGTTCAC

GAGCACACGTCTGAACTCTTTCCCTACACGACGCTCTTCCGATCTNNNAACCAAAAATGCCCGTGTTT
AGCACACGTCTGAACTCTTTCCCTACACGACGCTCTTCCGATCTNNNCTCTCTCTCTCTCTCCACACAC
AGCACACGTCTGAACTCTTTCCCTACACGACGCTCTTCCGATCTNNNCTTGGCTGTGATGGTCCTTTTT
AGCACACGTCTGAACTCTTTCCCTACACGACGCTCTTCCGATCTNNNAGACTTGTCAGCCTCTGCAATT
AGCACACGTCTGAACTCTTTCCCTACACGACGCTCTTCCGATCTNNNAGGTCAATTACACTGAAGATG
GAGCACACGTCTGAACTCTTTCCCTACACGACGCTCTTCCGATCTNNNTCCCTTCTTCTCCCTTTCTCTC
GCACACGTCTGAACTCTTTCCCTACACGACGCTCTTCCGATCTNNNGGGAAGTTTGAGGGTAGGTTG
AGCACACGTCTGAACTCTTTCCCTACACGACGCTCTTCCGATCTNNNCTCATATTTGGACCTTCCCAGC
AGCACACGTCTGAACTCTTTCCCTACACGACGCTCTTCCGATCTNNNACCAAACCAAGAGCTCACATTT
AGCACACGTCTGAACTCTTTCCCTACACGACGCTCTTCCGATCTNNNAGCCTCCTGAGTAGCTAGGAT
GCACACGTCTGAACTCTTTCCCTACACGACGCTCTTCCGATCTNNNTGCTGAGAATAGAAGGCTTCCT
GCACACGTCTGAACTCTTTCCCTACACGACGCTCTTCCGATCTNNNGGTTATGTGGGAGTATGGGTG
CACACGTCTGAACTCTTTCCCTACACGACGCTCTTCCGATCTNNNCTTACAGAACTGTACCTCTGCCCC
CACACGTCTGAACTCTTTCCCTACACGACGCTCTTCCGATCTNNNGCTCACTCCAAAACAATGACCTCC
CACACGTCTGAACTCTTTCCCTACACGACGCTCTTCCGATCTNNNCACTCTTCCCTTGGGTTTTTACGC
GAGCACACGTCTGAACTCTTTCCCTACACGACGCTCTTCCGATCTNNNGTTAGAACAACTTGTGTCAG
AGCACACGTCTGAACTCTTTCCCTACACGACGCTCTTCCGATCTNNNGCTTCAGAGTATGTGGGGTA
AGCACACGTCTGAACTCTTTCCCTACACGACGCTCTTCCGATCTNNNAGCTATCACTTGCAAGCTTAG
GCACACGTCTGAACTCTTTCCCTACACGACGCTCTTCCGATCTNNNCAAGTGAGCACCTGAGTACAGT
AGCACACGTCTGAACTCTTTCCCTACACGACGCTCTTCCGATCTNNNGCTGAGGCAAGATGAAAAAG
GCACACGTCTGAACTCTTTCCCTACACGACGCTCTTCCGATCTNNNTCCATACTGCCCCTACTCATACAC
GAGCACACGTCTGAACTCTTTCCCTACACGACGCTCTTCCGATCTNNNGATTCCTTTTTGTGTTGGCT
AGCACACGTCTGAACTCTTTCCCTACACGACGCTCTTCCGATCTNNNGAATACCAAAGCCAGTTACAC
GCACACGTCTGAACTCTTTCCCTACACGACGCTCTTCCGATCTNNNGGCATCACATTTTCCAGAAAGC
AGCACACGTCTGAACTCTTTCCCTACACGACGCTCTTCCGATCTNNNTGTGTCCAGTCTAGGAAAAGG
AGCACACGTCTGAACTCTTTCCCTACACGACGCTCTTCCGATCTNNNTTCTGACCTCAAAGGAAGGG
GAGCACACGTCTGAACTCTTTCCCTACACGACGCTCTTCCGATCTNNNTCCCTTTGTGTTTGTGTTTGT
AGCACACGTCTGAACTCTTTCCCTACACGACGCTCTTCCGATCTNNNCACACATGTGGTTACCAGCTAC
GAGCACACGTCTGAACTCTTTCCCTACACGACGCTCTTCCGATCTNNNGGCCAACTCTTACTCAGATCA
GAGCACACGTCTGAACTCTTTCCCTACACGACGCTCTTCCGATCTNNNAGGTTCACTGTGAATGTCAG
AGCACACGTCTGAACTCTTTCCCTACACGACGCTCTTCCGATCTNNNAGAAGTTCTCTACGGTTCATG
CACACGTCTGAACTCTTTCCCTACACGACGCTCTTCCGATCTNNNCTGAAGCCACACACCAAAGGATA
AGCACACGTCTGAACTCTTTCCCTACACGACGCTCTTCCGATCTNNNCTAGTATGGCTGAAGCAGACT
AGCACACGTCTGAACTCTTTCCCTACACGACGCTCTTCCGATCTNNNGCTTCTAGGGCGGATAAGTTG
GCACACGTCTGAACTCTTTCCCTACACGACGCTCTTCCGATCTNNNGGATGGGGTGTCTCTCTTACAA
AGCACACGTCTGAACTCTTTCCCTACACGACGCTCTTCCGATCTNNNGGGGTTGAAAAGATGGGGTA
AGCACACGTCTGAACTCTTTCCCTACACGACGCTCTTCCGATCTNNNATAGAGACTCTGTCCTCCCCAA
GCACACGTCTGAACTCTTTCCCTACACGACGCTCTTCCGATCTNNNCTGACTGTAAAAGAAATGGGG
GCACACGTCTGAACTCTTTCCCTACACGACGCTCTTCCGATCTNNNAGAAATCTGTGGGATGGTAGT
AGCACACGTCTGAACTCTTTCCCTACACGACGCTCTTCCGATCTNNNGTATGATGCAAACTAAGGCAG
AGCACACGTCTGAACTCTTTCCCTACACGACGCTCTTCCGATCTNNNACAGCAGTCATCTCCTTTTGAG
AGCACACGTCTGAACTCTTTCCCTACACGACGCTCTTCCGATCTNNNTAGCCCCCAAGAAAAACCCTA
GCACACGTCTGAACTCTTTCCCTACACGACGCTCTTCCGATCTNNNACATTCAGTCCAGCCTCAAGAA

AGCACACGTCTGAACTCTTTCCCTACACGACGCTCTTCCGATCTNNNCTTCCTGGAAAGAGCTACTGACC
GAGCACACGTCTGAACTCTTTCCCTACACGACGCTCTTCCGATCTNNNGAGTGAAGACTGTCCCTGCTA
GAGCACACGTCTGAACTCTTTCCCTACACGACGCTCTTCCGATCTNNNAGTCTGTAATGCCAGCATTTTC
GCACACGTCTGAACTCTTTCCCTACACGACGCTCTTCCGATCTNNNGTCAGTGCAGTTTTTCAGTTGGT
AGCACACGTCTGAACTCTTTCCCTACACGACGCTCTTCCGATCTNNNTTACCCTAACAAGATGACACTGT
AGAGCACACGTCTGAACTCTTTCCCTACACGACGCTCTTCCGATCTNNNCTCAGATTGCTTTCTTCTGTG
AGAGCACACGTCTGAACTCTTTCCCTACACGACGCTCTTCCGATCTNNNTCCTCATGCTACCTTCATAACC
AGCACACGTCTGAACTCTTTCCCTACACGACGCTCTTCCGATCTNNNCAAAAGTCGTGAACTCCAGAACC
AGCACACGTCTGAACTCTTTCCCTACACGACGCTCTTCCGATCTNNNTGGTCCAGTTTAGGGAACAGTT
GCACACGTCTGAACTCTTTCCCTACACGACGCTCTTCCGATCTNNNATTTCTGTTCCTAACTGGCACCTC
GAGCACACGTCTGAACTCTTTCCCTACACGACGCTCTTCCGATCTNNNGGAACTGGATGGGGAGACAA
GAGCACACGTCTGAACTCTTTCCCTACACGACGCTCTTCCGATCTNNNGGGAGTTACAATGGGACTGG
AGCACACGTCTGAACTCTTTCCCTACACGACGCTCTTCCGATCTNNNAGCACATCTACGTTCTTTGCAAA
AGCACACGTCTGAACTCTTTCCCTACACGACGCTCTTCCGATCTNNNTTCCTCAGCTGAAGGGCTAAAAT
AGCACACGTCTGAACTCTTTCCCTACACGACGCTCTTCCGATCTNNNGAAGGCTACAGTGTTACCTAGC
AGCACACGTCTGAACTCTTTCCCTACACGACGCTCTTCCGATCTNNNCACATACCCATAGTCCCAGCTACC
GAGCACACGTCTGAACTCTTTCCCTACACGACGCTCTTCCGATCTNNNACAGTGATCACCCCTATTCTTG
AGAGCACACGTCTGAACTCTTTCCCTACACGACGCTCTTCCGATCTNNNTGGCTTTGAACTTTGTCCATC
GAGCACACGTCTGAACTCTTTCCCTACACGACGCTCTTCCGATCTNNNACTAAGGCAGGAGATTTGCTT
AGCACACGTCTGAACTCTTTCCCTACACGACGCTCTTCCGATCTNNNCAGTGTTGCCAGAAACCTTCAAT
GAGCACACGTCTGAACTCTTTCCCTACACGACGCTCTTCCGATCTNNNCAAGAGCAGAGAAGGTCTACA
AGCACACGTCTGAACTCTTTCCCTACACGACGCTCTTCCGATCTNNNCGTTGCCACTTTTTTCTAGGGAA
AGCACACGTCTGAACTCTTTCCCTACACGACGCTCTTCCGATCTNNNGCTATATTCCTGTGTGTGTGTGC
AGCACACGTCTGAACTCTTTCCCTACACGACGCTCTTCCGATCTNNNAGACACGTGTGTACACAATTGT
GAGCACACGTCTGAACTCTTTCCCTACACGACGCTCTTCCGATCTNNNATAAAAATGGAACCATGGCCA
GAGCACACGTCTGAACTCTTTCCCTACACGACGCTCTTCCGATCTNNNATTCCCTTGGTGTCAGTATCTG
GCACACGTCTGAACTCTTTCCCTACACGACGCTCTTCCGATCTNNNGCTATGTATCCACCCCTCTCAAGC
AGCACACGTCTGAACTCTTTCCCTACACGACGCTCTTCCGATCTNNNCAACAGACGTCAGGGACATTTT
AGCACACGTCTGAACTCTTTCCCTACACGACGCTCTTCCGATCTNNNTTCCAGATGTTTCTTCCCAAGGT
AGCACACGTCTGAACTCTTTCCCTACACGACGCTCTTCCGATCTNNNATCTGTTTGTGTGTGTGTGTGTG
AGCACACGTCTGAACTCTTTCCCTACACGACGCTCTTCCGATCTNNNGGGGTTGTAAATGCACCAATCA
GAGCACACGTCTGAACTCTTTCCCTACACGACGCTCTTCCGATCTNNNCTTCTCTGAGCACCTATGGACT
GAGCACACGTCTGAACTCTTTCCCTACACGACGCTCTTCCGATCTNNNTCTTTTTGGGTTGGATCCATTG
AGCACACGTCTGAACTCTTTCCCTACACGACGCTCTTCCGATCTNNNTGTGTCTTTTACAGAGACGGGT
AGAGCACACGTCTGAACTCTTTCCCTACACGACGCTCTTCCGATCTNNNAAAAGATGCAACTGCTCAGG
AGCACACGTCTGAACTCTTTCCCTACACGACGCTCTTCCGATCTNNNGCACAGCGATCTTCTCAAACAAT
GAGCACACGTCTGAACTCTTTCCCTACACGACGCTCTTCCGATCTNNNCACTGTAGAAGGAGACCCAAA
AGCACACGTCTGAACTCTTTCCCTACACGACGCTCTTCCGATCTNNNCCCTGGACTCTCTACCCATTAGA
GAGCACACGTCTGAACTCTTTCCCTACACGACGCTCTTCCGATCTNNNTGACACAGCAAGAATCTGTCTC
GAGCACACGTCTGAACTCTTTCCCTACACGACGCTCTTCCGATCTNNNCCCAAAGTGTGTGAGAGAGAG
AAGAGCACACGTCTGAACTCTTTCCCTACACGACGCTCTTCCGATCTNNNTTCATCCTCCCTCTTCCTACC
AGCACACGTCTGAACTCTTTCCCTACACGACGCTCTTCCGATCTNNNGAACTTGTCCAAAGTCACACAGC
GAGCACACGTCTGAACTCTTTCCCTACACGACGCTCTTCCGATCTNNNACCAGTCATTTCACTCTAGGA

AGCACACGTCTGAACTCTTTCCCTACACGACGCTCTTCCGATCTNNNACCCTGAGTTGTAGTATTTGC
AGCACACGTCTGAACTCTTTCCCTACACGACGCTCTTCCGATCTNNNAGGTAATGATAGTGGACGCA
AGCACACGTCTGAACTCTTTCCCTACACGACGCTCTTCCGATCTNNNTTGCTTGTTGTGCTGTTATTCT
GCACACGTCTGAACTCTTTCCCTACACGACGCTCTTCCGATCTNNNGGCCCCAATTATCAATGCTCATC
AGCACACGTCTGAACTCTTTCCCTACACGACGCTCTTCCGATCTNNNGGTCTCCAACTGTTCAGATGT
AGCACACGTCTGAACTCTTTCCCTACACGACGCTCTTCCGATCTNNNGCACAAAATTGAAGGAGCCAG
AGCACACGTCTGAACTCTTTCCCTACACGACGCTCTTCCGATCTNNNATGGATTTGTTTCTGGGAGCT
AGAGCACACGTCTGAACTCTTTCCCTACACGACGCTCTTCCGATCTNNNACCAACTCTCCCACTGAAA
AGCACACGTCTGAACTCTTTCCCTACACGACGCTCTTCCGATCTNNNACAATTCACTGAGCTCGTTCTC
GCACACGTCTGAACTCTTTCCCTACACGACGCTCTTCCGATCTNNNACAATTCACTGAGCTCGTTCTC
AGCACACGTCTGAACTCTTTCCCTACACGACGCTCTTCCGATCTNNNACAATTCACTGAGCTCGTTCTC
AGCACACGTCTGAACTCTTTCCCTACACGACGCTCTTCCGATCTNNNCAACAGAATCTTGAGTCCCCA
AGCACACGTCTGAACTCTTTCCCTACACGACGCTCTTCCGATCTNNNGCTCTCTCTCTCTCTCTCTCT
AGCACACGTCTGAACTCTTTCCCTACACGACGCTCTTCCGATCTNNNTTGCCGTGATCGCAATTATTT
AGCACACGTCTGAACTCTTTCCCTACACGACGCTCTTCCGATCTNNNCCACTAGCGTAAGAGCAATTC
AGCACACGTCTGAACTCTTTCCCTACACGACGCTCTTCCGATCTNNNCTTCTGTGATCCCCACTCCTAC
AGCACACGTCTGAACTCTTTCCCTACACGACGCTCTTCCGATCTNNNTCTGACTTGGGCTACACAATTG
AGCACACGTCTGAACTCTTTCCCTACACGACGCTCTTCCGATCTNNNCCTCAGGCCTTCTAAGACTGAA
GAGCACACGTCTGAACTCTTTCCCTACACGACGCTCTTCCGATCTNNNGCACTGTGGAAAACTTTGAG
GCACACGTCTGAACTCTTTCCCTACACGACGCTCTTCCGATCTNNNAGTGTGGTAACTTAGGTAGCAG
AGCACACGTCTGAACTCTTTCCCTACACGACGCTCTTCCGATCTNNNGAGATCACACCACTACACTCCA
CACACGTCTGAACTCTTTCCCTACACGACGCTCTTCCGATCTNNNGCCAAATCATGCAGAGCTTTACTC
GAGCACACGTCTGAACTCTTTCCCTACACGACGCTCTTCCGATCTNNNATGTATATGCACCCAACACT
GAGCACACGTCTGAACTCTTTCCCTACACGACGCTCTTCCGATCTNNNCCATAGGGCCATGTGTATTCA
GCACACGTCTGAACTCTTTCCCTACACGACGCTCTTCCGATCTNNNAGACTGCTTAGGGTTTCCTGAA
GAGCACACGTCTGAACTCTTTCCCTACACGACGCTCTTCCGATCTNNNGAGATCGAGAACATCCTGGC
GCACACGTCTGAACTCTTTCCCTACACGACGCTCTTCCGATCTNNNTCTTCTTTTCCAGGACAGCATCA
AGCACACGTCTGAACTCTTTCCCTACACGACGCTCTTCCGATCTNNNTGTACTCTAGCCAAACACCACA
GCACACGTCTGAACTCTTTCCCTACACGACGCTCTTCCGATCTNNNTTTGAGACCAGCCTGACCAACA
GAGCACACGTCTGAACTCTTTCCCTACACGACGCTCTTCCGATCTNNNTGCCTAGCACTTACTAGTTCA
AGCACACGTCTGAACTCTTTCCCTACACGACGCTCTTCCGATCTNNNAGATTGCAGGAAAACCCAAGA
AGCACACGTCTGAACTCTTTCCCTACACGACGCTCTTCCGATCTNNNGGTGACAGAGTGAGACTCCAT
AGCACACGTCTGAACTCTTTCCCTACACGACGCTCTTCCGATCTNNNCTGACTTGGTGACTTTCTCTGC
AGAGCACACGTCTGAACTCTTTCCCTACACGACGCTCTTCCGATCTNNNGGTGGGAGGATGGCTTGA
GAGCACACGTCTGAACTCTTTCCCTACACGACGCTCTTCCGATCTNNNCAGATCACTTGAGCTCAGGAG
GAGCACACGTCTGAACTCTTTCCCTACACGACGCTCTTCCGATCTNNNGAGAATCTCTTGAACCTGGGA
GCACACGTCTGAACTCTTTCCCTACACGACGCTCTTCCGATCTNNNACTGTCCTGGTCATTGCAGTATT
GAGCACACGTCTGAACTCTTTCCCTACACGACGCTCTTCCGATCTNNNATATGTGTGTGTGTGTGT
GAGCACACGTCTGAACTCTTTCCCTACACGACGCTCTTCCGATCTNNNACTGACCTAAGCACTGAACAC
GCACACGTCTGAACTCTTTCCCTACACGACGCTCTTCCGATCTNNNCCTCCATGTTAGGCACATATGGA
GAGCACACGTCTGAACTCTTTCCCTACACGACGCTCTTCCGATCTNNNATTTTCTTCCTAGGGCCAAGT
AGCACACGTCTGAACTCTTTCCCTACACGACGCTCTTCCGATCTNNNCTGTAATCCCAGCAGGTTGTGA
AGCACACGTCTGAACTCTTTCCCTACACGACGCTCTTCCGATCTNNNAAACTCCTGGACTCAACTGATC

GAGCACACGTCTGAACTCTTTCCCTACACGACGCTCTTCCGATCTNNNGATCACTTGAGCTCAGGAGT
AGCACACGTCTGAACTCTTTCCCTACACGACGCTCTTCCGATCTNNNCTGGGAACTCCGTCTCAAAAAA
GAGCACACGTCTGAACTCTTTCCCTACACGACGCTCTTCCGATCTNNNAAGGAAAGGCAGGAATCTA
AGCACACGTCTGAACTCTTTCCCTACACGACGCTCTTCCGATCTNNNCAAGATCACACCACTGTACTCCA
AGCACACGTCTGAACTCTTTCCCTACACGACGCTCTTCCGATCTNNNATAATCCTAGCACTTTGGGAG
GCACACGTCTGAACTCTTTCCCTACACGACGCTCTTCCGATCTNNNCTACAGTGGGACCTTAGCTACA
AGCACACGTCTGAACTCTTTCCCTACACGACGCTCTTCCGATCTNNNCTGAGGCAGGAGAATCACTTG
GCACACGTCTGAACTCTTTCCCTACACGACGCTCTTCCGATCTNNNAGGTCCTCCCTACTTTTCAGGTA
AGCACACGTCTGAACTCTTTCCCTACACGACGCTCTTCCGATCTNNNTGTCTAGAAAGGACTGCTCCAT
GAGCACACGTCTGAACTCTTTCCCTACACGACGCTCTTCCGATCTNNNTGCCCTTCCTCTTCCTTCTTTA
AGCACACGTCTGAACTCTTTCCCTACACGACGCTCTTCCGATCTNNNCTGATGCCTGTAAGAAGTTGC
AGCACACGTCTGAACTCTTTCCCTACACGACGCTCTTCCGATCTNNNCCATCATACACACACACACACAC
GCACACGTCTGAACTCTTTCCCTACACGACGCTCTTCCGATCTNNNACATGGCTAGACCCTCATCTCTA
GCACACGTCTGAACTCTTTCCCTACACGACGCTCTTCCGATCTNNNTTCAAGTGATTCTCCTGCCTCAG
GAGCACACGTCTGAACTCTTTCCCTACACGACGCTCTTCCGATCTNNNCTTAGCATTTCTCTCATTCCCC
AGCACACGTCTGAACTCTTTCCCTACACGACGCTCTTCCGATCTNNNTTCCTGCAAAGGACATGATCTG
AGCACACGTCTGAACTCTTTCCCTACACGACGCTCTTCCGATCTNNNAAAGGGTACTAAGCAGGTCA
AGCACACGTCTGAACTCTTTCCCTACACGACGCTCTTCCGATCTNNNCAAATACCTTTGTGCGATCGTC
AGCACACGTCTGAACTCTTTCCCTACACGACGCTCTTCCGATCTNNNATTCCATTGGTCTATGTGCCTG
AGCACACGTCTGAACTCTTTCCCTACACGACGCTCTTCCGATCTNNNACATGAAGAGCTTCCCAAAAGG
AGCACACGTCTGAACTCTTTCCCTACACGACGCTCTTCCGATCTNNNTCTGTGTGTGCATCTATGTGTG
AGCACACGTCTGAACTCTTTCCCTACACGACGCTCTTCCGATCTNNNGGTGCTAAGAAATTACCTTGGC
AGCACACGTCTGAACTCTTTCCCTACACGACGCTCTTCCGATCTNNNCTGGGTTTCTTCAGAGAGTGTG
GAGCACACGTCTGAACTCTTTCCCTACACGACGCTCTTCCGATCTNNNAATAAAGTGTGTTGCCTCCAG
GAGCACACGTCTGAACTCTTTCCCTACACGACGCTCTTCCGATCTNNNTCCCCAGATTTGGTTTACACG
AGCACACGTCTGAACTCTTTCCCTACACGACGCTCTTCCGATCTNNNCTATTCTAAGGCCCAATTGCAG
GAGCACACGTCTGAACTCTTTCCCTACACGACGCTCTTCCGATCTNNNCCACCACCCCAGCTATTGTATT
GAGCACACGTCTGAACTCTTTCCCTACACGACGCTCTTCCGATCTNNNGTGACAAAGCAATACCCTGTC
AGCACACGTCTGAACTCTTTCCCTACACGACGCTCTTCCGATCTNNNTCGAGATCATCCTGGCTAACAT
AGAGCACACGTCTGAACTCTTTCCCTACACGACGCTCTTCCGATCTNNNGAGCAAGACTCTGTCTCCAA
GCACACGTCTGAACTCTTTCCCTACACGACGCTCTTCCGATCTNNNCATGAACACACACTCAGACACAC
AGCACACGTCTGAACTCTTTCCCTACACGACGCTCTTCCGATCTNNNATTGCAAAACTATGGAACCAG
GAGCACACGTCTGAACTCTTTCCCTACACGACGCTCTTCCGATCTNNNTACACCAACAACATCCAAGCT
AGCACACGTCTGAACTCTTTCCCTACACGACGCTCTTCCGATCTNNNCGCCAAGACAATAGCCCATTTA
AGCACACGTCTGAACTCTTTCCCTACACGACGCTCTTCCGATCTNNNTCATGAGGTCAGGAGATGGA
AGAGCACACGTCTGAACTCTTTCCCTACACGACGCTCTTCCGATCTNNNAAACATGCTCAGTGAAAGA
GAGCACACGTCTGAACTCTTTCCCTACACGACGCTCTTCCGATCTNNNGGATAGGTGAGCTCAAACAC
GAGCACACGTCTGAACTCTTTCCCTACACGACGCTCTTCCGATCTNNNTATATCTTCTTTCTCGTGCACG
AGCACACGTCTGAACTCTTTCCCTACACGACGCTCTTCCGATCTNNNTTTGGCCATTGAGAGCTCTTTT
AGCACACGTCTGAACTCTTTCCCTACACGACGCTCTTCCGATCTNNNATGACAGGGAGGAAACGATCA
AGCACACGTCTGAACTCTTTCCCTACACGACGCTCTTCCGATCTNNNGGCTGAGGTGAAAGTATTGCT
GAGCACACGTCTGAACTCTTTCCCTACACGACGCTCTTCCGATCTNNNGAGGCAAGAGGATCACTTGA
GCACACGTCTGAACTCTTTCCCTACACGACGCTCTTCCGATCTNNNTATGACTGGGGGAACACACTAA

AGAGCACACGTCTGAACTCTTTCCCTACACGACGCTCTTCCGATCTNNNAGTTGAAGACCCGAATACAC
AGCACACGTCTGAACTCTTTCCCTACACGACGCTCTTCCGATCTNNNTCATGCCTGAAATCCTAGCACTT
AGAGCACACGTCTGAACTCTTTCCCTACACGACGCTCTTCCGATCTNNNGGCAAAACCCCGTCTCTACTA
AGAGCACACGTCTGAACTCTTTCCCTACACGACGCTCTTCCGATCTNNNGCCTCCTTTTCTTACAGCACTT
GAGCACACGTCTGAACTCTTTCCCTACACGACGCTCTTCCGATCTNNNCATTCTCTCTCAGGACACACACA
AGCACACGTCTGAACTCTTTCCCTACACGACGCTCTTCCGATCTNNNCTGCAGCTGTTTGAAGGATCTTC
GAGCACACGTCTGAACTCTTTCCCTACACGACGCTCTTCCGATCTNNNAGGTAGGTGGATCACTTGAG
GAGCACACGTCTGAACTCTTTCCCTACACGACGCTCTTCCGATCTNNNGAAAATCTCTTCCCCCTGGAGA
AGCACACGTCTGAACTCTTTCCCTACACGACGCTCTTCCGATCTNNNGCAAATGCTACACCATTTCCTGT
AGCACACGTCTGAACTCTTTCCCTACACGACGCTCTTCCGATCTNNNCCAACAGTGAATTCTATCCCCCA
AGCACACGTCTGAACTCTTTCCCTACACGACGCTCTTCCGATCTNNNGTACAGCACTCACCAACATTCAG
AGCACACGTCTGAACTCTTTCCCTACACGACGCTCTTCCGATCTNNNATTTACGATTGGGAGTCTGAGG
GAGCACACGTCTGAACTCTTTCCCTACACGACGCTCTTCCGATCTNNNCTCTCTCTCTGTCTCTCTCAG
GAGCACACGTCTGAACTCTTTCCCTACACGACGCTCTTCCGATCTNNNCTGGTCTCGAATTTCTGACCTC
GAGCACACGTCTGAACTCTTTCCCTACACGACGCTCTTCCGATCTNNNGATTGCTTGAGCCTGGGAATT
GAGCACACGTCTGAACTCTTTCCCTACACGACGCTCTTCCGATCTNNNATTGGCAATGCAACATTTTAG
GAGCACACGTCTGAACTCTTTCCCTACACGACGCTCTTCCGATCTNNNCTTAGCTACTTGCACCAAAGCT
AGCACACGTCTGAACTCTTTCCCTACACGACGCTCTTCCGATCTNNNACACGGTATATATACGTGTGTG
AGCACACGTCTGAACTCTTTCCCTACACGACGCTCTTCCGATCTNNNTTTATACCAGGGGCTTGCTCA
GAGCACACGTCTGAACTCTTTCCCTACACGACGCTCTTCCGATCTNNNCTTGTGCTTTGTCTAGTAGGG
AGCACACGTCTGAACTCTTTCCCTACACGACGCTCTTCCGATCTNNNGTAGTGTGATGTGGCTAGTGT
AGCACACGTCTGAACTCTTTCCCTACACGACGCTCTTCCGATCTNNNGCTTTTCACAAAAAACGGAAGCC
AAGAGCACACGTCTGAACTCTTTCCCTACACGACGCTCTTCCGATCTNNNCCCAAGATAGTCATGTGGA
AGCACACGTCTGAACTCTTTCCCTACACGACGCTCTTCCGATCTNNNGCAGAGTAAGACCCTGTCTCAA
GAGCACACGTCTGAACTCTTTCCCTACACGACGCTCTTCCGATCTNNNTGTGTACCTATGTGTGTGTGTG
AGAGCACACGTCTGAACTCTTTCCCTACACGACGCTCTTCCGATCTNNNGCACAAGAGAGAGACTGTCA
AGCACACGTCTGAACTCTTTCCCTACACGACGCTCTTCCGATCTNNNATGAGCAACTCCAAAAAGTGCT
GAGCACACGTCTGAACTCTTTCCCTACACGACGCTCTTCCGATCTNNNCTCAAGTGATCCTCCTCCATGA
AGAGCACACGTCTGAACTCTTTCCCTACACGACGCTCTTCCGATCTNNNCAGGGTAAGGTTTGGGTAA
GAGCACACGTCTGAACTCTTTCCCTACACGACGCTCTTCCGATCTNNNCCAAGTAGCTGGGACTACATG
GAGCACACGTCTGAACTCTTTCCCTACACGACGCTCTTCCGATCTNNNGATTACAGTCATGTGCCACCA
GAGCACACGTCTGAACTCTTTCCCTACACGACGCTCTTCCGATCTNNNGAGAATCACTTGAACCTGGGA
AGCACACGTCTGAACTCTTTCCCTACACGACGCTCTTCCGATCTNNNCGGTGTCCTGACTGACAAAAAT
GAGCACACGTCTGAACTCTTTCCCTACACGACGCTCTTCCGATCTNNNACGATTGCAAAATCATGGAAG
AGCACACGTCTGAACTCTTTCCCTACACGACGCTCTTCCGATCTNNNGAGTTAGCTCTAGTGGATAGGG
AGCACACGTCTGAACTCTTTCCCTACACGACGCTCTTCCGATCTNNNTGGTTTTGCCAGAGTTCCTAAAG
AGCACACGTCTGAACTCTTTCCCTACACGACGCTCTTCCGATCTNNNACCCATCAGATAATCCATCGTGT
AGCACACGTCTGAACTCTTTCCCTACACGACGCTCTTCCGATCTNNNGGATTACAGGCATGAGTCACTG
GAGCACACGTCTGAACTCTTTCCCTACACGACGCTCTTCCGATCTNNNAAGAAGCCATATGTCCCACAC
AGCACACGTCTGAACTCTTTCCCTACACGACGCTCTTCCGATCTNNNTCACTGTGGCCTCAAAAATCCTA
AGCACACGTCTGAACTCTTTCCCTACACGACGCTCTTCCGATCTNNNCCCCTACTTATAGGCACCTCTCT
AGCACACGTCTGAACTCTTTCCCTACACGACGCTCTTCCGATCTNNNTATTCTCAGGCCAGGACTTTGAG
GAGCACACGTCTGAACTCTTTCCCTACACGACGCTCTTCCGATCTNNNTCATCCCCTTACACTCACACATG

AGCACACGTCTGAACTCTTTCCCTACACGACGCTCTTCCGATCTNNNGGACTGACTTTGGAACTGGG
AGCACACGTCTGAACTCTTTCCCTACACGACGCTCTTCCGATCTNNNAAGAATCTCTTGAACCCAGGA
GAGCACACGTCTGAACTCTTTCCCTACACGACGCTCTTCCGATCTNNNAGGAGAATCATTTGAACCCG
GAGCACACGTCTGAACTCTTTCCCTACACGACGCTCTTCCGATCTNNNCTAGGCAACATAGTGAGACC
AGCACACGTCTGAACTCTTTCCCTACACGACGCTCTTCCGATCTNNNTGTAGTCCCAGCTACTTAGGA
AGCACACGTCTGAACTCTTTCCCTACACGACGCTCTTCCGATCTNNNATGAGAATCGCTTGAACTCAG
AGCACACGTCTGAACTCTTTCCCTACACGACGCTCTTCCGATCTNNNGGCAGTTTGTTCTTCTACTGAG
AGCACACGTCTGAACTCTTTCCCTACACGACGCTCTTCCGATCTNNNACTCTACCATGTTACCTGCACT
GAGCACACGTCTGAACTCTTTCCCTACACGACGCTCTTCCGATCTNNNATAGATGCAAATTGGGAGG
GAGCACACGTCTGAACTCTTTCCCTACACGACGCTCTTCCGATCTNNNTTCTTTCTGGACTTCCTGGGA
AGCACACGTCTGAACTCTTTCCCTACACGACGCTCTTCCGATCTNNNACAGGAGAAAAGAAAAGGCC
GCACACGTCTGAACTCTTTCCCTACACGACGCTCTTCCGATCTNNNTGTGGTGGCTCACATCTGTATA
AGCACACGTCTGAACTCTTTCCCTACACGACGCTCTTCCGATCTNNNGCAGGAGAGAGTAAAAAGGA
AGCACACGTCTGAACTCTTTCCCTACACGACGCTCTTCCGATCTNNNCTATCACACACACACACACAC
AGCACACGTCTGAACTCTTTCCCTACACGACGCTCTTCCGATCTNNNCGAGACATGTAGGGATTATGG
AGCACACGTCTGAACTCTTTCCCTACACGACGCTCTTCCGATCTNNNTGACAGTCAGCTTGTATCACT
AGCACACGTCTGAACTCTTTCCCTACACGACGCTCTTCCGATCTNNNCCTAGGATAATGGCCTCCAGTT
AGCACACGTCTGAACTCTTTCCCTACACGACGCTCTTCCGATCTNNNAGTTCTGTGCTAATTTGGGAG
GAGCACACGTCTGAACTCTTTCCCTACACGACGCTCTTCCGATCTNNNCCAGGGAATTAGGTGTCTGT
GCACACGTCTGAACTCTTTCCCTACACGACGCTCTTCCGATCTNNNATCTTACTGGGGTGTGGTTTGT
GCACACGTCTGAACTCTTTCCCTACACGACGCTCTTCCGATCTNNNTTTGGCCCCATTATCTTCCTACCC
AGCACACGTCTGAACTCTTTCCCTACACGACGCTCTTCCGATCTNNNTCCTGGTTCAATCGATTCTCCT
AGAGCACACGTCTGAACTCTTTCCCTACACGACGCTCTTCCGATCTNNNGGAGGTACTTTCTCAGCGA
GAGCACACGTCTGAACTCTTTCCCTACACGACGCTCTTCCGATCTNNNATAAGGGATGCCAAAACCTC
GAGCACACGTCTGAACTCTTTCCCTACACGACGCTCTTCCGATCTNNNCTGGGCGAAAGAGCTATAT
GCACACGTCTGAACTCTTTCCCTACACGACGCTCTTCCGATCTNNNTGTTGTGCTCTCTCTCTCTCTC
AGCACACGTCTGAACTCTTTCCCTACACGACGCTCTTCCGATCTNNNCTCCTGGGTAGCTGGGATTACA
GAGCACACGTCTGAACTCTTTCCCTACACGACGCTCTTCCGATCTNNNCCCTACTGAGTCAGAATCACC
AAGAGCACACGTCTGAACTCTTTCCCTACACGACGCTCTTCCGATCTNNNGAACTGATGATGTGTGCT
AGCACACGTCTGAACTCTTTCCCTACACGACGCTCTTCCGATCTNNNGCTTCCTGATAAGCCTCTAGC
AGCACACGTCTGAACTCTTTCCCTACACGACGCTCTTCCGATCTNNNAGGCCATTGTGTTTCAAAACC
GAGCACACGTCTGAACTCTTTCCCTACACGACGCTCTTCCGATCTNNNAGACACATGCTTTCTCTGTG
AGAGCACACGTCTGAACTCTTTCCCTACACGACGCTCTTCCGATCTNNNACGTTTCCATTTTCAAAGGA
GAGCACACGTCTGAACTCTTTCCCTACACGACGCTCTTCCGATCTNNNGGATCACTTGAGGTCAGGA
AGCACACGTCTGAACTCTTTCCCTACACGACGCTCTTCCGATCTNNNGCTGTCTAAATGTGGCCAAGA
AGCACACGTCTGAACTCTTTCCCTACACGACGCTCTTCCGATCTNNNGGTGAAGCCCCATCTCTACTAA
GCACACGTCTGAACTCTTTCCCTACACGACGCTCTTCCGATCTNNNGTGACAGCTCTTAGTGTACAGC
GAGCACACGTCTGAACTCTTTCCCTACACGACGCTCTTCCGATCTNNNACCTATCACCTGAAAAACTG
GAGCACACGTCTGAACTCTTTCCCTACACGACGCTCTTCCGATCTNNNCACTGACTCTCTTTTCGGACT
GAGCACACGTCTGAACTCTTTCCCTACACGACGCTCTTCCGATCTNNNTGTTTGTTGCAGCACTATTC
AGCACACGTCTGAACTCTTTCCCTACACGACGCTCTTCCGATCTNNNTGGGATGAGATCTCTCCTCTC
AGCACACGTCTGAACTCTTTCCCTACACGACGCTCTTCCGATCTNNNCAGAGATAGAAAGAGAGGG
AGCACACGTCTGAACTCTTTCCCTACACGACGCTCTTCCGATCTNNNCAGAGATAGAAAGAGAGGG

AGCACACGTCTGAACTCTTTCCCTACACGACGCTCTTCCGATCTNNNCTCATGCCTATAATCCCAGCACT
AGCACACGTCTGAACTCTTTCCCTACACGACGCTCTTCCGATCTNNNATAGTAAAACCCCTGCATCCCTG
AGCACACGTCTGAACTCTTTCCCTACACGACGCTCTTCCGATCTNNNTGTGGTTTCTGGGGTAACATGA
GAGCACACGTCTGAACTCTTTCCCTACACGACGCTCTTCCGATCTNNNCAGGAGAATCACTTGAACCCA
GAGCACACGTCTGAACTCTTTCCCTACACGACGCTCTTCCGATCTNNNTCACGGATGCCCATTTTGTTTA
GAGCACACGTCTGAACTCTTTCCCTACACGACGCTCTTCCGATCTNNNTTCCACCAAAAAGACACACACA
AGAGCACACGTCTGAACTCTTTCCCTACACGACGCTCTTCCGATCTNNNTGGTGACAAGGATACCAAAG
AGCACACGTCTGAACTCTTTCCCTACACGACGCTCTTCCGATCTNNNACGGATTTTATCACTTCCCAGCT
GAGCACACGTCTGAACTCTTTCCCTACACGACGCTCTTCCGATCTNNNAGTGCTGGGATTATAGGCATG
AGAGCACACGTCTGAACTCTTTCCCTACACGACGCTCTTCCGATCTNNNTGGCTGCTGCATTTTACAATG
AGCACACGTCTGAACTCTTTCCCTACACGACGCTCTTCCGATCTNNNCCAGTACAAATGCCCATCAATCA
AGAGCACACGTCTGAACTCTTTCCCTACACGACGCTCTTCCGATCTNNNCAAGTCTCTCTTGGCCTCAG
GAGCACACGTCTGAACTCTTTCCCTACACGACGCTCTTCCGATCTNNNAAATGTGGAACCAGCCTAAATG
GCACACGTCTGAACTCTTTCCCTACACGACGCTCTTCCGATCTNNNCACCACGCTCAGCTATTTTTTGTCA
AGCACACGTCTGAACTCTTTCCCTACACGACGCTCTTCCGATCTNNNAGTGTTTTCAAGAGGCCCTACA
AGCACACGTCTGAACTCTTTCCCTACACGACGCTCTTCCGATCTNNNCCAGAACTCCCACGTATGTCTAC
AGCACACGTCTGAACTCTTTCCCTACACGACGCTCTTCCGATCTNNNTGATTTCCCATTGCTCAGCTACT
AGCACACGTCTGAACTCTTTCCCTACACGACGCTCTTCCGATCTNNNTGTAAAACTGTGTGTGTGTGTG
GAGCACACGTCTGAACTCTTTCCCTACACGACGCTCTTCCGATCTNNNGGGAGCATGTAACCCTAAGAG
AGAGCACACGTCTGAACTCTTTCCCTACACGACGCTCTTCCGATCTNNNGCCATACCCAGCTTCTAGGAA
GAGCACACGTCTGAACTCTTTCCCTACACGACGCTCTTCCGATCTNNNAGATAGCATCCTGAAACCTTCC
AGCACACGTCTGAACTCTTTCCCTACACGACGCTCTTCCGATCTNNNGGAAATCCTACCGACCACATTCT
AGCACACGTCTGAACTCTTTCCCTACACGACGCTCTTCCGATCTNNNTGTCTGACTCCCATGTGACTAA
GAGCACACGTCTGAACTCTTTCCCTACACGACGCTCTTCCGATCTNNNATCCAAGGTACAACATGAGGA
AGCACACGTCTGAACTCTTTCCCTACACGACGCTCTTCCGATCTNNNTGGTGAGTATGTGGAGAAAAG
AGAGCACACGTCTGAACTCTTTCCCTACACGACGCTCTTCCGATCTNNNACTTCAACCATTGATGACTCC
AGCACACGTCTGAACTCTTTCCCTACACGACGCTCTTCCGATCTNNNCCTCAGTGGTGGTTTGATCTACA
GAGCACACGTCTGAACTCTTTCCCTACACGACGCTCTTCCGATCTNNNACGCACACACATATACACACA
GAGCACACGTCTGAACTCTTTCCCTACACGACGCTCTTCCGATCTNNNGAGCGAGACTCCACCTACTAA
GCACACGTCTGAACTCTTTCCCTACACGACGCTCTTCCGATCTNNNCCCAGAGACAGAGGTACTAGGAA
GAGCACACGTCTGAACTCTTTCCCTACACGACGCTCTTCCGATCTNNNCCAACATGATGAAACCCTGTCT
GAGCACACGTCTGAACTCTTTCCCTACACGACGCTCTTCCGATCTNNNCTGTAGTCCCAGCTACTCAGG
AGCACACGTCTGAACTCTTTCCCTACACGACGCTCTTCCGATCTNNNGCTCCATTGTCTGCAAGTTTTGA
AGCACACGTCTGAACTCTTTCCCTACACGACGCTCTTCCGATCTNNNCAACAGAGTGAGACCTTGTGTG
AGCACACGTCTGAACTCTTTCCCTACACGACGCTCTTCCGATCTNNNAGCCTCTTTCTCTCTGTCTCTCTCA
GAGCACACGTCTGAACTCTTTCCCTACACGACGCTCTTCCGATCTNNNTGCACAATGAAGTGGATCATT
AGAGCACACGTCTGAACTCTTTCCCTACACGACGCTCTTCCGATCTNNNAGGCCCAGTGTAATCAGAT
AGCACACGTCTGAACTCTTTCCCTACACGACGCTCTTCCGATCTNNNTATGCTTTCCAGAGAGGGAGAG
GCACACGTCTGAACTCTTTCCCTACACGACGCTCTTCCGATCTNNNACACACACACACACACAACTTTTCA
GAGCACACGTCTGAACTCTTTCCCTACACGACGCTCTTCCGATCTNNNATTTTGTGGGCTCTGACTTACC
GAGCACACGTCTGAACTCTTTCCCTACACGACGCTCTTCCGATCTNNNATACAAAGCATGAGCATGGCA
AGCACACGTCTGAACTCTTTCCCTACACGACGCTCTTCCGATCTNNNCTGAAGTGTTCTTGCCTGTCTG
AGCACACGTCTGAACTCTTTCCCTACACGACGCTCTTCCGATCTNNNGCCGATACAGCACTGTTTTCATT

AGCACACGTCTGAACTCTTTCCCTACACGACGCTCTTCCGATCTNNNTTTTGCCTTCTACCCCATGAGAT
GAGCACACGTCTGAACTCTTTCCCTACACGACGCTCTTCCGATCTNNNAAAGTGGCCATTTGAATACGA
AGCACACGTCTGAACTCTTTCCCTACACGACGCTCTTCCGATCTNNNTCACTGTAACCTCAAACTCCTGG
GAGCACACGTCTGAACTCTTTCCCTACACGACGCTCTTCCGATCTNNNTATCACAGCAGAGCCAAGACA
GCACACGTCTGAACTCTTTCCCTACACGACGCTCTTCCGATCTNNNAGAGCAACGAGAATGAAGGGAA
AGCACACGTCTGAACTCTTTCCCTACACGACGCTCTTCCGATCTNNNTGGCTAAGTGTGCGTGTATATG
AGAGCACACGTCTGAACTCTTTCCCTACACGACGCTCTTCCGATCTNNNTAAACTGAAGGGCCTAGGT
AGAGCACACGTCTGAACTCTTTCCCTACACGACGCTCTTCCGATCTNNNTGTGTGTGTGTGAGAGA
AGAGCACACGTCTGAACTCTTTCCCTACACGACGCTCTTCCGATCTNNNCTCTGCACAACTCAAGCTTGA
AGCACACGTCTGAACTCTTTCCCTACACGACGCTCTTCCGATCTNNNCTAGACTAGCCTGGGCAACATAC
GAGCACACGTCTGAACTCTTTCCCTACACGACGCTCTTCCGATCTNNNATCATACATAAAGGGGGGACT
AGCACACGTCTGAACTCTTTCCCTACACGACGCTCTTCCGATCTNNNGTGAGGCCCTGTCTCTACAAATT
AGAGCACACGTCTGAACTCTTTCCCTACACGACGCTCTTCCGATCTNNNCTAGGCGTCTCCTCTTTCTGTA
GAGCACACGTCTGAACTCTTTCCCTACACGACGCTCTTCCGATCTNNNGGGTTGCTAGAGGCTAAGAGT
GAGCACACGTCTGAACTCTTTCCCTACACGACGCTCTTCCGATCTNNNGGTGACAGAATGATACTCTGC
AGCACACGTCTGAACTCTTTCCCTACACGACGCTCTTCCGATCTNNNAGCTGAAATCAAGAAGGCAATC
AGCACACGTCTGAACTCTTTCCCTACACGACGCTCTTCCGATCTNNNCAGTCTGGCCAATATGGTGAAA
AGCACACGTCTGAACTCTTTCCCTACACGACGCTCTTCCGATCTNNNGCTATGATTGTGCCACTCATTCC
GAGCACACGTCTGAACTCTTTCCCTACACGACGCTCTTCCGATCTNNNGGTTGTTTTCACATGCTCAGT
GAGCACACGTCTGAACTCTTTCCCTACACGACGCTCTTCCGATCTNNNTTTCAAACACACACACACACAC
AGCACACGTCTGAACTCTTTCCCTACACGACGCTCTTCCGATCTNNNTTTGAAATATATCGGCTGGGCA
GAGCACACGTCTGAACTCTTTCCCTACACGACGCTCTTCCGATCTNNNGTGAGAAAGCAAGACCCTGTT
GAGCACACGTCTGAACTCTTTCCCTACACGACGCTCTTCCGATCTNNNAGGCTGAAGCAGGAGGAGAT
AGCACACGTCTGAACTCTTTCCCTACACGACGCTCTTCCGATCTNNNCGTCCCCTTCTCTTGTCAAACTA
GAGCACACGTCTGAACTCTTTCCCTACACGACGCTCTTCCGATCTNNNATCTGGGCATGTATACACCACA
AGCACACGTCTGAACTCTTTCCCTACACGACGCTCTTCCGATCTNNNTCTGAGTCCCCAGAGTCCATTAT
GAGCACACGTCTGAACTCTTTCCCTACACGACGCTCTTCCGATCTNNNTATTCTCTGCATCCACACCAAC
AGCACACGTCTGAACTCTTTCCCTACACGACGCTCTTCCGATCTNNNGCCCTTTCTATTTGCAGTTGGTT
GAGCACACGTCTGAACTCTTTCCCTACACGACGCTCTTCCGATCTNNNAGACTGCAGTGAAAAAACTGC
GAGCACACGTCTGAACTCTTTCCCTACACGACGCTCTTCCGATCTNNNTGGGTCAGATTTTCATCCCAGT
GAGCACACGTCTGAACTCTTTCCCTACACGACGCTCTTCCGATCTNNNCCTCTGCAGGGACTAAAGAAA
GAGCACACGTCTGAACTCTTTCCCTACACGACGCTCTTCCGATCTNNNCCTAGCACGTTCTTTGGCCTAT
GAGCACACGTCTGAACTCTTTCCCTACACGACGCTCTTCCGATCTNNNCCACTGTGCCTGGCCTATTTAT
GCACACGTCTGAACTCTTTCCCTACACGACGCTCTTCCGATCTNNNCCCCCCTCACTTTCTTTCTCTTTCAT
GAGCACACGTCTGAACTCTTTCCCTACACGACGCTCTTCCGATCTNNNATGTTATGCTCAGGGGTTTGT
AGCACACGTCTGAACTCTTTCCCTACACGACGCTCTTCCGATCTNNNCCTTTTCTCTGGCTTTGAATGCA
AGCACACGTCTGAACTCTTTCCCTACACGACGCTCTTCCGATCTNNNTCTCTAGTAGCTGGGACTACAG
AGCACACGTCTGAACTCTTTCCCTACACGACGCTCTTCCGATCTNNNGGGACTATAAGTGTGTGCCAT
AGCACACGTCTGAACTCTTTCCCTACACGACGCTCTTCCGATCTNNNTCCCATTTATTCTGTGGCAGTG
GAGCACACGTCTGAACTCTTTCCCTACACGACGCTCTTCCGATCTNNNCCAAATTTGAGGCCTTCGAAAC
GAGCACACGTCTGAACTCTTTCCCTACACGACGCTCTTCCGATCTNNNTTTGCTCACATTATGCCAGCTT
GAGCACACGTCTGAACTCTTTCCCTACACGACGCTCTTCCGATCTNNNTCAAAAAAACCTTGGTCTCCAC
AGCACACGTCTGAACTCTTTCCCTACACGACGCTCTTCCGATCTNNNTCCACTTGTTCTAGTCTACCCAAC

GAGCACACGTCTGAACTCTTTCCCTACACGACGCTCTTCCGATCTNNNATCCTGGAGAATCTTGTAGTG
AGCACACGTCTGAACTCTTTCCCTACACGACGCTCTTCCGATCTNNNTCACCTGTGTCTTCCTATGTGTG
AGCACACGTCTGAACTCTTTCCCTACACGACGCTCTTCCGATCTNNNGGGAGGCTGAGACATGAGAAT
AGAGCACACGTCTGAACTCTTTCCCTACACGACGCTCTTCCGATCTNNNCGATTGAGCCTAGGAGTTTG
AGCACACGTCTGAACTCTTTCCCTACACGACGCTCTTCCGATCTNNNTGAGATTGCACCATTGTACTCCA
AGCACACGTCTGAACTCTTTCCCTACACGACGCTCTTCCGATCTNNNGCCATGTTATTTCCTGGGGAGA
GAGCACACGTCTGAACTCTTTCCCTACACGACGCTCTTCCGATCTNNNTGAAATGAAACAAGAGGCAG
GAGCACACGTCTGAACTCTTTCCCTACACGACGCTCTTCCGATCTNNNGCAACAGAGTGACACCCTATT
GAGCACACGTCTGAACTCTTTCCCTACACGACGCTCTTCCGATCTNNNGCAATCTGCATGGTGCTTGTA
AGCACACGTCTGAACTCTTTCCCTACACGACGCTCTTCCGATCTNNNAGTGATCATGTGGAATCTCACC
GAGCACACGTCTGAACTCTTTCCCTACACGACGCTCTTCCGATCTNNNCTTCATGGCATGGCATGTTAG
GAGCACACGTCTGAACTCTTTCCCTACACGACGCTCTTCCGATCTNNNTGGGCAACATAGCACGATTCT
AGAGCACACGTCTGAACTCTTTCCCTACACGACGCTCTTCCGATCTNNNCAGCCTGGCCAACATAGTAAA
AGCACACGTCTGAACTCTTTCCCTACACGACGCTCTTCCGATCTNNNCCACCTCCCAGATTCAAGTGATT
GAGCACACGTCTGAACTCTTTCCCTACACGACGCTCTTCCGATCTNNNAAGGAGACCCTCTTACCTTTTG
GAGCACACGTCTGAACTCTTTCCCTACACGACGCTCTTCCGATCTNNNTAATATGGTAACGGGCTATGG
AGCACACGTCTGAACTCTTTCCCTACACGACGCTCTTCCGATCTNNNCTGCACCTTGATGATGGATTTCC
GAGCACACGTCTGAACTCTTTCCCTACACGACGCTCTTCCGATCTNNNTAGGTCCTGCTCCTGTCTAGTA
AGCACACGTCTGAACTCTTTCCCTACACGACGCTCTTCCGATCTNNNTTTCCTTCCAATGACAACACGTG
AGCACACGTCTGAACTCTTTCCCTACACGACGCTCTTCCGATCTNNNCCTCCTTGGTTAGATGTATTCCT
AGAGCACACGTCTGAACTCTTTCCCTACACGACGCTCTTCCGATCTNNNCCCACATGCTCGTATTTGAGA
GAGCACACGTCTGAACTCTTTCCCTACACGACGCTCTTCCGATCTNNNAAAACATCTTCCCTGGACCATCA
AGCACACGTCTGAACTCTTTCCCTACACGACGCTCTTCCGATCTNNNACGCTACTTTTCTTCTGTCGAGT
AGCACACGTCTGAACTCTTTCCCTACACGACGCTCTTCCGATCTNNNTTGAGAGTTGTGGCTGTGGAA
AGAGCACACGTCTGAACTCTTTCCCTACACGACGCTCTTCCGATCTNNNGCAGAGACTCTTGTTCTCTTC
AGCACACGTCTGAACTCTTTCCCTACACGACGCTCTTCCGATCTNNNGAGCGAAACTCCATCTCTCTCTC
GAGCACACGTCTGAACTCTTTCCCTACACGACGCTCTTCCGATCTNNNCTGGCCAAGATGGAGAAAAC
AGCACACGTCTGAACTCTTTCCCTACACGACGCTCTTCCGATCTNNNATTCTCAGGTCAGAAAGTGCAC
GAGCACACGTCTGAACTCTTTCCCTACACGACGCTCTTCCGATCTNNNGAGAGGATGATGATAGCTGC
AGCACACGTCTGAACTCTTTCCCTACACGACGCTCTTCCGATCTNNNAGATCTAGAAACCAGCTGTGCT
AGAGCACACGTCTGAACTCTTTCCCTACACGACGCTCTTCCGATCTNNNGCAAGATAGAGCCCGTCTCA
AGCACACGTCTGAACTCTTTCCCTACACGACGCTCTTCCGATCTNNNGGGCTTTCCAGAGAAAAAGAA
GAGCACACGTCTGAACTCTTTCCCTACACGACGCTCTTCCGATCTNNNAGAATGACTTGAACCTGGTAG
GAGCACACGTCTGAACTCTTTCCCTACACGACGCTCTTCCGATCTNNNCATGGGAATCGTTTGAACTTG
AGCACACGTCTGAACTCTTTCCCTACACGACGCTCTTCCGATCTNNNCACCACCATGGCCGCTATATATA
GAGCACACGTCTGAACTCTTTCCCTACACGACGCTCTTCCGATCTNNNAGACCCCGTCTCAGGAAAAAA
AGCACACGTCTGAACTCTTTCCCTACACGACGCTCTTCCGATCTNNNAAGCAGGGTCTCAAATACACAC
AGCACACGTCTGAACTCTTTCCCTACACGACGCTCTTCCGATCTNNNCACACACTCTCTCTCACTCTCACC
GAGCACACGTCTGAACTCTTTCCCTACACGACGCTCTTCCGATCTNNNCCTCTTCTCACTACTGCACTTGT
GAGCACACGTCTGAACTCTTTCCCTACACGACGCTCTTCCGATCTNNNCATTTGAGCCCAGAAGTTCAA
AGCACACGTCTGAACTCTTTCCCTACACGACGCTCTTCCGATCTNNNAGCCCACTGAAAGCCATATTAG
AGCACACGTCTGAACTCTTTCCCTACACGACGCTCTTCCGATCTNNNACCCCATTGTAAGTTGAGGAACA
GAGCACACGTCTGAACTCTTTCCCTACACGACGCTCTTCCGATCTNNNCGGGCTGTCTTTAGAGAAGTT

AGCACACGTCTGAACTCTTTCCCTACACGACGCTCTTCCGATCTNNNTGCTTTTAGCTGATGTGTGTG
GAGCACACGTCTGAACTCTTTCCCTACACGACGCTCTTCCGATCTNNNCAGCCTGGGTAACAAGAGTG
AGCACACGTCTGAACTCTTTCCCTACACGACGCTCTTCCGATCTNNNTTCCAACCAGCTTACTGCAAAT
GCACACGTCTGAACTCTTTCCCTACACGACGCTCTTCCGATCTNNNCCTCATATTGAAACTCCCTCCGT
GAGCACACGTCTGAACTCTTTCCCTACACGACGCTCTTCCGATCTNNNCTCTCTCCTACCCCATCAA
AGCACACGTCTGAACTCTTTCCCTACACGACGCTCTTCCGATCTNNNGAGTCTGAGAGTTAGTTACG
GCACACGTCTGAACTCTTTCCCTACACGACGCTCTTCCGATCTNNNTTTCTTAGGTATTACCGGGGCTG
GCACACGTCTGAACTCTTTCCCTACACGACGCTCTTCCGATCTNNNTAAACCTCAGTTTCCTCCTCTGCC
AGCACACGTCTGAACTCTTTCCCTACACGACGCTCTTCCGATCTNNNTTTTCCCATGGGTAAAAGGGG
GAGCACACGTCTGAACTCTTTCCCTACACGACGCTCTTCCGATCTNNNGATTGGCCCAGGTAACCATA
AGCACACGTCTGAACTCTTTCCCTACACGACGCTCTTCCGATCTNNNTGCGTATGTAGCATGAAATC
GCACACGTCTGAACTCTTTCCCTACACGACGCTCTTCCGATCTNNNGGTCTGCAGATTTCAAGTAACG
AGCACACGTCTGAACTCTTTCCCTACACGACGCTCTTCCGATCTNNNTTGGTTCAGAAGTGGGTAAG
AGCACACGTCTGAACTCTTTCCCTACACGACGCTCTTCCGATCTNNNCCCAGCCAAAATTCATCACTTC
GCACACGTCTGAACTCTTTCCCTACACGACGCTCTTCCGATCTNNNAGGTGTTCTCAGTACATGCACA
AGCACACGTCTGAACTCTTTCCCTACACGACGCTCTTCCGATCTNNNCCAACGTGGTAAAACCTCATC
GAGCACACGTCTGAACTCTTTCCCTACACGACGCTCTTCCGATCTNNNTCTTGTCCTCACTTGGATGGA
AGCACACGTCTGAACTCTTTCCCTACACGACGCTCTTCCGATCTNNNCATAGCGAAACCCTGACTCTAC
AGCACACGTCTGAACTCTTTCCCTACACGACGCTCTTCCGATCTNNNAGCCAGACTAGATCGCAGTATT
GCACACGTCTGAACTCTTTCCCTACACGACGCTCTTCCGATCTNNNCAGGATCTTCATGCCTCTCTCTC
AGCACACGTCTGAACTCTTTCCCTACACGACGCTCTTCCGATCTNNNTCCAAAAAATTGAGGAGGAG
GCACACGTCTGAACTCTTTCCCTACACGACGCTCTTCCGATCTNNNCCTCAGGAGGGAAGAAAAGTC
AGCACACGTCTGAACTCTTTCCCTACACGACGCTCTTCCGATCTNNNGGGCTCATCTTTGTTCAGCAAT
AGCACACGTCTGAACTCTTTCCCTACACGACGCTCTTCCGATCTNNNGTTTCTCCACTTCACTCCGTTT
GCACACGTCTGAACTCTTTCCCTACACGACGCTCTTCCGATCTNNNATTGCCTGTCTACGTTTCTCCAT
AGCACACGTCTGAACTCTTTCCCTACACGACGCTCTTCCGATCTNNNTATGGCCAAGTGGATTCCTAC
AGAGCACACGTCTGAACTCTTTCCCTACACGACGCTCTTCCGATCTNNNACTCATATCAGCAACAAAT
GCACACGTCTGAACTCTTTCCCTACACGACGCTCTTCCGATCTNNNCTCACTACAATCTCCACCTCCTG
AGCACACGTCTGAACTCTTTCCCTACACGACGCTCTTCCGATCTNNNGCAACTGAGGTTTGAAGAGA
AGCACACGTCTGAACTCTTTCCCTACACGACGCTCTTCCGATCTNNNAAGGACAGCACAGTGACTGAT
GAGCACACGTCTGAACTCTTTCCCTACACGACGCTCTTCCGATCTNNNAGGCAGGTAAATAAACATG
GCACACGTCTGAACTCTTTCCCTACACGACGCTCTTCCGATCTNNNAGATCCTGTTAGTCACTACGGG
AGCACACGTCTGAACTCTTTCCCTACACGACGCTCTTCCGATCTNNNAACCTTTTACCCCAGATTATGC
AGCACACGTCTGAACTCTTTCCCTACACGACGCTCTTCCGATCTNNNCTCCTGAGTAGCTGGGATTAC
AGCACACGTCTGAACTCTTTCCCTACACGACGCTCTTCCGATCTNNNCTGATCCACTACTGGCTTCTCT
AGCACACGTCTGAACTCTTTCCCTACACGACGCTCTTCCGATCTNNNGAGTGGTTGACTTGAGGTGA
AGCACACGTCTGAACTCTTTCCCTACACGACGCTCTTCCGATCTNNNCAAAGGCTACCTGAAAGTTTG
AGCACACGTCTGAACTCTTTCCCTACACGACGCTCTTCCGATCTNNNGGGCTCAAGTGATCCTCCTAA
AGCACACGTCTGAACTCTTTCCCTACACGACGCTCTTCCGATCTNNNAGGTGTGCGGTCTAAGAATG
GCACACGTCTGAACTCTTTCCCTACACGACGCTCTTCCGATCTNNNGATTACAGGTGTGAGCCATTGT
AGCACACGTCTGAACTCTTTCCCTACACGACGCTCTTCCGATCTNNNCATCTGAAAAATTCCCGTGGA
AGCACACGTCTGAACTCTTTCCCTACACGACGCTCTTCCGATCTNNNAATGAGGAATCTGGAGCCCA
AGCACACGTCTGAACTCTTTCCCTACACGACGCTCTTCCGATCTNNNGCCTCACCCATTTTACCCTTCT

AGCACACGTCTGAACTCTTTCCCTACACGACGCTCTTCCGATCTNNNCGTGTGCATGTGTGTATGTAT
AGCACACGTCTGAACTCTTTCCCTACACGACGCTCTTCCGATCTNNNCCTCTGGAGGCTGAAATATGA
AGCACACGTCTGAACTCTTTCCCTACACGACGCTCTTCCGATCTNNNACTGGGAGAGCTGGAAAATTGA
AGCACACGTCTGAACTCTTTCCCTACACGACGCTCTTCCGATCTNNNATAAAACAATTCTGCAGCCTGG
GAGCACACGTCTGAACTCTTTCCCTACACGACGCTCTTCCGATCTNNNCAACATACCAAGACCTCGTCT
AGCACACGTCTGAACTCTTTCCCTACACGACGCTCTTCCGATCTNNNGCAACATAGCAAGACCCTGTTT
AGCACACGTCTGAACTCTTTCCCTACACGACGCTCTTCCGATCTNNNTGAGGTAGGAGAACTGCTTGA
AGCACACGTCTGAACTCTTTCCCTACACGACGCTCTTCCGATCTNNNTATCACCCTGATACCAAAGCCA
GAGCACACGTCTGAACTCTTTCCCTACACGACGCTCTTCCGATCTNNNGCAAGTGGCCATGACAAACTT
GAGCACACGTCTGAACTCTTTCCCTACACGACGCTCTTCCGATCTNNNCTCTTGTGGTCAGGAGTTCAA
AGCACACGTCTGAACTCTTTCCCTACACGACGCTCTTCCGATCTNNNTGTGCTTGTAATCCCAGCTACTT
AGAGCACACGTCTGAACTCTTTCCCTACACGACGCTCTTCCGATCTNNNACTCTGTCTCTCTCTCTCTC
AGCACACGTCTGAACTCTTTCCCTACACGACGCTCTTCCGATCTNNNGAGTACAATGGAAGTGAGAG
AGCACACGTCTGAACTCTTTCCCTACACGACGCTCTTCCGATCTNNNAAACACACATGAACACATGCACA
AGAGCACACGTCTGAACTCTTTCCCTACACGACGCTCTTCCGATCTNNNTTCATCCATCCATCCATCCATC
GAGCACACGTCTGAACTCTTTCCCTACACGACGCTCTTCCGATCTNNNCAGATCACTTGAGCTCAGGA
GAGCACACGTCTGAACTCTTTCCCTACACGACGCTCTTCCGATCTNNNCCTCTGCCTCTACCCTAGACATA
AGCACACGTCTGAACTCTTTCCCTACACGACGCTCTTCCGATCTNNNACACACACACAAAATACACACCA
AGCACACGTCTGAACTCTTTCCCTACACGACGCTCTTCCGATCTNNNCAAAGTCATCCTGAAGACTGGGA
AGCACACGTCTGAACTCTTTCCCTACACGACGCTCTTCCGATCTNNNACCAACCTGGGCAATATAGTGA
GAGCACACGTCTGAACTCTTTCCCTACACGACGCTCTTCCGATCTNNNAAGACTAACTTCCCTGGAGCAA
AGCACACGTCTGAACTCTTTCCCTACACGACGCTCTTCCGATCTNNNCCTGTCTTAGCCAATTCCCCTAA
GAGCACACGTCTGAACTCTTTCCCTACACGACGCTCTTCCGATCTNNNTAGTTCATGAAACCCACCACA
GAGCACACGTCTGAACTCTTTCCCTACACGACGCTCTTCCGATCTNNNTTTGGGCTAAGTTGGTAGGT
AGCACACGTCTGAACTCTTTCCCTACACGACGCTCTTCCGATCTNNNATTAGTCAAGGCTGGACAGAGT
GAGCACACGTCTGAACTCTTTCCCTACACGACGCTCTTCCGATCTNNNATGGAACCAGCCTAAATGTCCA
AGCACACGTCTGAACTCTTTCCCTACACGACGCTCTTCCGATCTNNNGTATTTGGCTTTCTGTTCCTGCA
AGCACACGTCTGAACTCTTTCCCTACACGACGCTCTTCCGATCTNNNAATGGAAGTGATTTTTGCATGC
AGAGCACACGTCTGAACTCTTTCCCTACACGACGCTCTTCCGATCTNNNGTGGCACTTCCTTGATTCTGA
AGAGCACACGTCTGAACTCTTTCCCTACACGACGCTCTTCCGATCTNNNAAACAGCCCCTAAAGGTCAA
AGCACACGTCTGAACTCTTTCCCTACACGACGCTCTTCCGATCTNNNGGTGACAGAGTGAGACTCCAT
AGCACACGTCTGAACTCTTTCCCTACACGACGCTCTTCCGATCTNNNTGAGACCATGTGCATACCTCAT
GAGCACACGTCTGAACTCTTTCCCTACACGACGCTCTTCCGATCTNNNGGGAGAGAGAGAGAGAGAG
AGAGCACACGTCTGAACTCTTTCCCTACACGACGCTCTTCCGATCTNNNTGCCTGGCTAATGTTTTCTGT
AGCACACGTCTGAACTCTTTCCCTACACGACGCTCTTCCGATCTNNNATTCAGCTCCCTTCTCTTTCCTGC
AGCACACGTCTGAACTCTTTCCCTACACGACGCTCTTCCGATCTNNNGCATGCTGGGATCTGTAGTCTA
AGCACACGTCTGAACTCTTTCCCTACACGACGCTCTTCCGATCTNNNGGGATGAGGAAAGTGGCTTA
AGCACACGTCTGAACTCTTTCCCTACACGACGCTCTTCCGATCTNNNGTATGTGTGTGTGTGTTTCTGG
GCACACGTCTGAACTCTTTCCCTACACGACGCTCTTCCGATCTNNNATGTGCCTTTCAGACTTCAAAGC
AGCACACGTCTGAACTCTTTCCCTACACGACGCTCTTCCGATCTNNNAACTTCCTAGGCTCCAAGAACAG
GAGCACACGTCTGAACTCTTTCCCTACACGACGCTCTTCCGATCTNNNATCCTCTGGAACAAGAACACA
GAGCACACGTCTGAACTCTTTCCCTACACGACGCTCTTCCGATCTNNNCTGCCTTCTTCCTTCTTCTCTGT
AGCACACGTCTGAACTCTTTCCCTACACGACGCTCTTCCGATCTNNNTTATGAAGCCAGCATCATCCTGA

GAGCACACGTCTGAACTCTTTCCCTACACGACGCTCTTCCGATCTNNNACAGAGGGAAAATGCTGCAA
GAGCACACGTCTGAACTCTTTCCCTACACGACGCTCTTCCGATCTNNNAAAAAGACAACAACAACCACC
AGCACACGTCTGAACTCTTTCCCTACACGACGCTCTTCCGATCTNNNTAAGGTGGGTGGATCATTTGA
AGAGCACACGTCTGAACTCTTTCCCTACACGACGCTCTTCCGATCTNNNTGGCACATGACAGGAGTTTC
AGCACACGTCTGAACTCTTTCCCTACACGACGCTCTTCCGATCTNNNGAGCAAGACTCTACCTCAAAAC
AGCACACGTCTGAACTCTTTCCCTACACGACGCTCTTCCGATCTNNNTAGGATGACTCTGACCCCATCAT
AGAGCACACGTCTGAACTCTTTCCCTACACGACGCTCTTCCGATCTNNNGAAACATGGCGAAACTCCAT
AGCACACGTCTGAACTCTTTCCCTACACGACGCTCTTCCGATCTNNNCAAACACACACAAACACACGAA
AGCACACGTCTGAACTCTTTCCCTACACGACGCTCTTCCGATCTNNNGAAGAGTGGAACCCTGTCTCAA
AGCACACGTCTGAACTCTTTCCCTACACGACGCTCTTCCGATCTNNNCTCCACTTCCCCAAAAAGTAGA
AGCACACGTCTGAACTCTTTCCCTACACGACGCTCTTCCGATCTNNNGCAAGTTCATGTGCTTAGACCT
AGAGCACACGTCTGAACTCTTTCCCTACACGACGCTCTTCCGATCTNNNTGTAGTCCCAACAGTTTAGG
GAGCACACGTCTGAACTCTTTCCCTACACGACGCTCTTCCGATCTNNNGGGGAGGGGATACATGAGA
AGCACACGTCTGAACTCTTTCCCTACACGACGCTCTTCCGATCTNNNACACTCTCTGACATCCCTGATTG
AGCACACGTCTGAACTCTTTCCCTACACGACGCTCTTCCGATCTNNNGCTTTTCTTGGCTTCTAGTGACC
GAGCACACGTCTGAACTCTTTCCCTACACGACGCTCTTCCGATCTNNNAGATTGCTATCCACTGTGTGT
AGCACACGTCTGAACTCTTTCCCTACACGACGCTCTTCCGATCTNNNTGCTCTCAAAGTTCTGCTCAGAT
GAGCACACGTCTGAACTCTTTCCCTACACGACGCTCTTCCGATCTNNNTGCTCATGTTCTCTCTCTCTCT
AGCACACGTCTGAACTCTTTCCCTACACGACGCTCTTCCGATCTNNNTGAAGGTGGCTGTTTTAGGAA
GAGCACACGTCTGAACTCTTTCCCTACACGACGCTCTTCCGATCTNNNCCAATTCTACCTCTCTCCAGCT
GAGCACACGTCTGAACTCTTTCCCTACACGACGCTCTTCCGATCTNNNACACATACACACAAATGGCAC
GAGCACACGTCTGAACTCTTTCCCTACACGACGCTCTTCCGATCTNNNAAAATGTGTGTGTGTGTGTG
GAGCACACGTCTGAACTCTTTCCCTACACGACGCTCTTCCGATCTNNNTTTCAAACAGCAGCACAGAGT
AGCACACGTCTGAACTCTTTCCCTACACGACGCTCTTCCGATCTNNNGTGTGTGTGAAATGTAGTCTCG
AGCACACGTCTGAACTCTTTCCCTACACGACGCTCTTCCGATCTNNNAAGCTCTTAGAACTCACACCCAG
AGCACACGTCTGAACTCTTTCCCTACACGACGCTCTTCCGATCTNNNGAGACTGAGGTGAGCAGAGAT
AGCACACGTCTGAACTCTTTCCCTACACGACGCTCTTCCGATCTNNNCATGTATGTGGGCAGAGGATA
AGAGCACACGTCTGAACTCTTTCCCTACACGACGCTCTTCCGATCTNNNAGACTGCTATGGCTTTGACA
GAGCACACGTCTGAACTCTTTCCCTACACGACGCTCTTCCGATCTNNNCCAGCGGGATATATACACACA
AGAGCACACGTCTGAACTCTTTCCCTACACGACGCTCTTCCGATCTNNNTTCTTTCATCCATCCACCCTGT
GAGCACACGTCTGAACTCTTTCCCTACACGACGCTCTTCCGATCTNNNCTAAGCAACAAAGAGAGACCC
AGCACACGTCTGAACTCTTTCCCTACACGACGCTCTTCCGATCTNNNGATTTTGACCACTTCTCTCTGCCC
GAGCACACGTCTGAACTCTTTCCCTACACGACGCTCTTCCGATCTNNNTTGGACGTAACAAGGACTAAG
GAGCACACGTCTGAACTCTTTCCCTACACGACGCTCTTCCGATCTNNNCTCTAAACAGTGCTTGGGAAT
GAGCACACGTCTGAACTCTTTCCCTACACGACGCTCTTCCGATCTNNNTCTCCATAATTGCTGAGCCAGT
AGCACACGTCTGAACTCTTTCCCTACACGACGCTCTTCCGATCTNNNACACACAGACATACACACACCAT
AGAGCACACGTCTGAACTCTTTCCCTACACGACGCTCTTCCGATCTNNNGTAATCCCAGCTACTTCGGA
GCACACGTCTGAACTCTTTCCCTACACGACGCTCTTCCGATCTNNNGGCTTGTTTCTTCCCAAAATCCAC
AGCACACGTCTGAACTCTTTCCCTACACGACGCTCTTCCGATCTNNNGAGTCACACTACACTCTAGCCT
AGCACACGTCTGAACTCTTTCCCTACACGACGCTCTTCCGATCTNNNTTCATTTGCTGTTGCTGCTGTA
AGCACACGTCTGAACTCTTTCCCTACACGACGCTCTTCCGATCTNNNCACAGTGTTTTCCCCACTAGAG
AGCACACGTCTGAACTCTTTCCCTACACGACGCTCTTCCGATCTNNNGAAGAAAGTCATGCAGAACGT
GAGCACACGTCTGAACTCTTTCCCTACACGACGCTCTTCCGATCTNNNACATTTAGGTGTGACCATTGA

GAGCACACGTCTGAACTCTTTCCCTACACGACGCTCTTCCGATCTNNNCTTCAGGAAAGTTTCAGGAC
GAGCACACGTCTGAACTCTTTCCCTACACGACGCTCTTCCGATCTNNNCTAGACATCTAGCAGGCTCA
AGCACACGTCTGAACTCTTTCCCTACACGACGCTCTTCCGATCTNNNCCCCTACTCCAATATGACTGGT
AGCACACGTCTGAACTCTTTCCCTACACGACGCTCTTCCGATCTNNNTATAAATTGGAAAGCAGTGGG
CACACGTCTGAACTCTTTCCCTACACGACGCTCTTCCGATCTNNNTAAATTTTTTGGGGGGACAGGG
GAGCACACGTCTGAACTCTTTCCCTACACGACGCTCTTCCGATCTNNNACTGGCTGAACTGACTGAAA
GAGCACACGTCTGAACTCTTTCCCTACACGACGCTCTTCCGATCTNNNGGGTTGTCTCTCTCTCTCT
AGCACACGTCTGAACTCTTTCCCTACACGACGCTCTTCCGATCTNNNTCCTGGCAATTTAGTTCAAACA
GAGCACACGTCTGAACTCTTTCCCTACACGACGCTCTTCCGATCTNNNTTCCTTGCTTCACTTTCCTCAG
GAGCACACGTCTGAACTCTTTCCCTACACGACGCTCTTCCGATCTNNNGATCTGTTGTTTCTCCCGTAC
GAGCACACGTCTGAACTCTTTCCCTACACGACGCTCTTCCGATCTNNNTGTGGACGACCTGATAACTA
AGCACACGTCTGAACTCTTTCCCTACACGACGCTCTTCCGATCTNNNTATTGTCAGTGGGGTGTTGTC
AGCACACGTCTGAACTCTTTCCCTACACGACGCTCTTCCGATCTNNNTCTGGCATTTTGTAGTTGAAG
AGCACACGTCTGAACTCTTTCCCTACACGACGCTCTTCCGATCTNNNAAGGCGTATGGATGGGAAA
GCACACGTCTGAACTCTTTCCCTACACGACGCTCTTCCGATCTNNNGAATCTCAGGCACTTGTGCATTT
AGCACACGTCTGAACTCTTTCCCTACACGACGCTCTTCCGATCTNNNGCAGTCCCATTTACAATAGCC
AGCACACGTCTGAACTCTTTCCCTACACGACGCTCTTCCGATCTNNNATAAAAATTAGCCGAGCGTG
GCACACGTCTGAACTCTTTCCCTACACGACGCTCTTCCGATCTNNNACACAGACACAGACACACATACA
AGCACACGTCTGAACTCTTTCCCTACACGACGCTCTTCCGATCTNNNGGAAAATCACTTGAAACCGGG
AGCACACGTCTGAACTCTTTCCCTACACGACGCTCTTCCGATCTNNNCCTGCCTGGGAAATAAAGTG
AGCACACGTCTGAACTCTTTCCCTACACGACGCTCTTCCGATCTNNNAGGTACATTGTTGCTGGAGAG
AGCACACGTCTGAACTCTTTCCCTACACGACGCTCTTCCGATCTNNNAAATGCTTTGTGTGTGTGTGTG
GCACACGTCTGAACTCTTTCCCTACACGACGCTCTTCCGATCTNNNATTACACACACACACACACACC
AGCACACGTCTGAACTCTTTCCCTACACGACGCTCTTCCGATCTNNNTACATGACATCCTGGAAAAGGC
AGCACACGTCTGAACTCTTTCCCTACACGACGCTCTTCCGATCTNNNTGTAGATCATGCAGCACCTTT
AGCACACGTCTGAACTCTTTCCCTACACGACGCTCTTCCGATCTNNNTACAAGACAAAGCAGCAATGG
AGCACACGTCTGAACTCTTTCCCTACACGACGCTCTTCCGATCTNNNACACTTAGGGACTCTCTCTCCTC
AGCACACGTCTGAACTCTTTCCCTACACGACGCTCTTCCGATCTNNNGTGCTAAAAGCTTAGACCAG
AGCACACGTCTGAACTCTTTCCCTACACGACGCTCTTCCGATCTNNNGAACCTGAGATGTGAAGTTGC
AGCACACGTCTGAACTCTTTCCCTACACGACGCTCTTCCGATCTNNNCCTGTCTCTCTCTCTCTCACACA
AGCACACGTCTGAACTCTTTCCCTACACGACGCTCTTCCGATCTNNNTCATCGAATCAACCTGTGTCCA
AGCACACGTCTGAACTCTTTCCCTACACGACGCTCTTCCGATCTNNNAGATTGCTGCAAATGCCATGA
AGCACACGTCTGAACTCTTTCCCTACACGACGCTCTTCCGATCTNNNGCTTTTGTACAAATCGTCCAGC
AGCACACGTCTGAACTCTTTCCCTACACGACGCTCTTCCGATCTNNNCTAGAAGCAGGAGAATCGCTT
GAGCACACGTCTGAACTCTTTCCCTACACGACGCTCTTCCGATCTNNNAGTGAGCTAAGACTGTGCCA
GAGCACACGTCTGAACTCTTTCCCTACACGACGCTCTTCCGATCTNNNTCCATTCCCTATCACCTCTCTG
AGCACACGTCTGAACTCTTTCCCTACACGACGCTCTTCCGATCTNNNGATCTGGGACATCTCAACCTCA
GAGCACACGTCTGAACTCTTTCCCTACACGACGCTCTTCCGATCTNNNTGCCAGTCTCCATAATCATGT
AGCACACGTCTGAACTCTTTCCCTACACGACGCTCTTCCGATCTNNNCTCACAGCAGCCTCAAATTTCA
GAGCACACGTCTGAACTCTTTCCCTACACGACGCTCTTCCGATCTNNNAATGGGGGAAAAGACCATG
GAGCACACGTCTGAACTCTTTCCCTACACGACGCTCTTCCGATCTNNNTGTGTAGAGAGAGGGAACA
GAGCACACGTCTGAACTCTTTCCCTACACGACGCTCTTCCGATCTNNNGATCTGGTCCCTTCCTCACAC
AGCACACGTCTGAACTCTTTCCCTACACGACGCTCTTCCGATCTNNNTCTATATGCGCATACATGTGT

AGCACACGTCTGAACTCTTTCCCTACACGACGCTCTTCCGATCTNNNGAGCAAGACTCCATCTCAAAA
AGCACACGTCTGAACTCTTTCCCTACACGACGCTCTTCCGATCTNNNTATAGCTGTGTGTGTCTGTGT
AGCACACGTCTGAACTCTTTCCCTACACGACGCTCTTCCGATCTNNNCCTCCCAAGTAGCTGAAACTAC
AGCACACGTCTGAACTCTTTCCCTACACGACGCTCTTCCGATCTNNNGAGTAGCTGAGACTACAGGC
AGCACACGTCTGAACTCTTTCCCTACACGACGCTCTTCCGATCTNNNCTCAATCTCCTGACCTCGTGAT
AGCACACGTCTGAACTCTTTCCCTACACGACGCTCTTCCGATCTNNNGTTCGAGTAATTCTCCTGCCTC
GAGCACACGTCTGAACTCTTTCCCTACACGACGCTCTTCCGATCTNNNTAAGACCAGAGGCGGACACT
GCACACGTCTGAACTCTTTCCCTACACGACGCTCTTCCGATCTNNNCCAGAAGAGGAGGACAGAAAG
GAGCACACGTCTGAACTCTTTCCCTACACGACGCTCTTCCGATCTNNNAGACTCTGTCTCACACACACA
AGCACACGTCTGAACTCTTTCCCTACACGACGCTCTTCCGATCTNNNCCCTGTCTCTCTCTCTCTCTCTCC
AGAGCACACGTCTGAACTCTTTCCCTACACGACGCTCTTCCGATCTNNNTTATCTCCTGACCTCCTGATC
GAGCACACGTCTGAACTCTTTCCCTACACGACGCTCTTCCGATCTNNNGGCAACAAGAACGAAACTGT
GAGCACACGTCTGAACTCTTTCCCTACACGACGCTCTTCCGATCTNNNAGCAAGACTCCATCTCAAACA
AGCACACGTCTGAACTCTTTCCCTACACGACGCTCTTCCGATCTNNNAGTCTATGGCAGTTTGACCACA
AGCACACGTCTGAACTCTTTCCCTACACGACGCTCTTCCGATCTNNNCTGAAGTGGGAAGATCACCTG
AGCACACGTCTGAACTCTTTCCCTACACGACGCTCTTCCGATCTNNNCACACACACACACACACATACA
AGCACACGTCTGAACTCTTTCCCTACACGACGCTCTTCCGATCTNNNCAGCACTTTTGGAGGTTGAGA
AGCACACGTCTGAACTCTTTCCCTACACGACGCTCTTCCGATCTNNNTCCAAACTGGAAGGGAAGAA
AGCACACGTCTGAACTCTTTCCCTACACGACGCTCTTCCGATCTNNNCCCCTGCCCTGTTCTAGAAATA
GCACACGTCTGAACTCTTTCCCTACACGACGCTCTTCCGATCTNNNCTCTCTATAGTCCTAGGCCTGCT
AGCACACGTCTGAACTCTTTCCCTACACGACGCTCTTCCGATCTNNNTGAGTTCAGGAGTTTGAGACC
AGCACACGTCTGAACTCTTTCCCTACACGACGCTCTTCCGATCTNNNACACACACACACACACTCTTGAG
AGCACACGTCTGAACTCTTTCCCTACACGACGCTCTTCCGATCTNNNGGTGGCAACCTCATTTCTAAC
GAGCACACGTCTGAACTCTTTCCCTACACGACGCTCTTCCGATCTNNNGCCCGGCCCATAGAATTATTT
GCACACGTCTGAACTCTTTCCCTACACGACGCTCTTCCGATCTNNNGCCTCTCTCTGCAAACATTGTTT
AGCACACGTCTGAACTCTTTCCCTACACGACGCTCTTCCGATCTNNNCTGAATTGAGAGGGACTAGC
AGCACACGTCTGAACTCTTTCCCTACACGACGCTCTTCCGATCTNNNACTGTGATCTTGAATGAGGC
CACACGTCTGAACTCTTTCCCTACACGACGCTCTTCCGATCTNNNACCCATCATAACTTGCCAACTCTCA
AGCACACGTCTGAACTCTTTCCCTACACGACGCTCTTCCGATCTNNNCTGGAGTGCAGTGGTATGATC
AGAGCACACGTCTGAACTCTTTCCCTACACGACGCTCTTCCGATCTNNNACTTGAATCACGATGATGG
AGCACACGTCTGAACTCTTTCCCTACACGACGCTCTTCCGATCTNNNGATAAAGCTGTGAAGGTGG
GAGCACACGTCTGAACTCTTTCCCTACACGACGCTCTTCCGATCTNNNCTGAGGCAGAGGAATTGCTT
GAGCACACGTCTGAACTCTTTCCCTACACGACGCTCTTCCGATCTNNNACCCGAATATTTGTGTGTGT
GCACACGTCTGAACTCTTTCCCTACACGACGCTCTTCCGATCTNNNTTTCTAATTGGTGGGATGAGG
AGCACACGTCTGAACTCTTTCCCTACACGACGCTCTTCCGATCTNNNGTCTGACACCTGGAGTTGTAT
GCACACGTCTGAACTCTTTCCCTACACGACGCTCTTCCGATCTNNNGTACTGGGATTACAGGAGTGA
AGCACACGTCTGAACTCTTTCCCTACACGACGCTCTTCCGATCTNNNCTCGCACTCTCTTTCTCTCTCTCC
GAGCACACGTCTGAACTCTTTCCCTACACGACGCTCTTCCGATCTNNNTATGGTGCTTTATCCTGCCAT
AGCACACGTCTGAACTCTTTCCCTACACGACGCTCTTCCGATCTNNNCTGCGTGTGTGTATCTCTTTGT
CACACGTCTGAACTCTTTCCCTACACGACGCTCTTCCGATCTNNNGACTCCGTCTCAAACAAACAACA
GCACACGTCTGAACTCTTTCCCTACACGACGCTCTTCCGATCTNNNTCTCCACACATGTGTTGTTGTTG
AGCACACGTCTGAACTCTTTCCCTACACGACGCTCTTCCGATCTNNNAAATACTGACATACCGCTAGG
AGCACACGTCTGAACTCTTTCCCTACACGACGCTCTTCCGATCTNNNAAAGCCACAGTAACCAAAACA

GAGCACACGTCTGAACTCTTTCCCTACACGACGCTCTTCCGATCTNNNGATACCATTCCCCCATTCAGG
GAGCACACGTCTGAACTCTTTCCCTACACGACGCTCTTCCGATCTNNNTTCGTGTTTGTTGCAGCACTA
GAGCACACGTCTGAACTCTTTCCCTACACGACGCTCTTCCGATCTNNNTCCTCCCTTCCTCTCTTTCTCAA
GAGCACACGTCTGAACTCTTTCCCTACACGACGCTCTTCCGATCTNNNGCGAGATTCTGCCTCCAAAA
AGCACACGTCTGAACTCTTTCCCTACACGACGCTCTTCCGATCTNNNGTGGCAAAACCCCATCTCTACT
GAGCACACGTCTGAACTCTTTCCCTACACGACGCTCTTCCGATCTNNNCTGAGGCAGAGAACCGTTTG
GAGCACACGTCTGAACTCTTTCCCTACACGACGCTCTTCCGATCTNNNAAACCCAGGAACTCACAGGTA
GAGCACACGTCTGAACTCTTTCCCTACACGACGCTCTTCCGATCTNNNCCTGGTTGTTAGAGTGAGAC
AGCACACGTCTGAACTCTTTCCCTACACGACGCTCTTCCGATCTNNNCATTCCAGCTTAGGTGACAGAG
GAGCACACGTCTGAACTCTTTCCCTACACGACGCTCTTCCGATCTNNNCCTGGGCAATAATAGCGAAA
AGCACACGTCTGAACTCTTTCCCTACACGACGCTCTTCCGATCTNNNCAGAGGTTGTAGTGAGCTGAG
AGCACACGTCTGAACTCTTTCCCTACACGACGCTCTTCCGATCTNNNGGTTCAGGAGATGACAAGTG
AGCACACGTCTGAACTCTTTCCCTACACGACGCTCTTCCGATCTNNNCATGGTCAGGAGCAGAAGTTTT
GAGCACACGTCTGAACTCTTTCCCTACACGACGCTCTTCCGATCTNNNTTGACCATGGTGAGGAGTTC
GAGCACACGTCTGAACTCTTTCCCTACACGACGCTCTTCCGATCTNNNGCACTGTGGAAAACTTTGAG
GAGCACACGTCTGAACTCTTTCCCTACACGACGCTCTTCCGATCTNNNCGGTGCCACTATTTTCAAAAG
GAGCACACGTCTGAACTCTTTCCCTACACGACGCTCTTCCGATCTNNNAAAGATTTCTTCTGGGTCGTG
GAGCACACGTCTGAACTCTTTCCCTACACGACGCTCTTCCGATCTNNNACACATGGCTTGTTTGGGAAT
GAGCACACGTCTGAACTCTTTCCCTACACGACGCTCTTCCGATCTNNNGCCACTACACTCAGCCAGATA
GAGCACACGTCTGAACTCTTTCCCTACACGACGCTCTTCCGATCTNNNAGAGGTTTCAGTGAGCCAAGA
AGCACACGTCTGAACTCTTTCCCTACACGACGCTCTTCCGATCTNNNCACCCTTTTCCTTTGGACCTTTCC
AGCACACGTCTGAACTCTTTCCCTACACGACGCTCTTCCGATCTNNNCCTGGGCAAGAAGAGTAAAAC
AGAGCACACGTCTGAACTCTTTCCCTACACGACGCTCTTCCGATCTNNNGCCAAGATAGTGAAACCCTG
AGCACACGTCTGAACTCTTTCCCTACACGACGCTCTTCCGATCTNNNCGGTAAGTATGATGGAGGTA
GCACACGTCTGAACTCTTTCCCTACACGACGCTCTTCCGATCTNNNACCTACAGAGAGGAACAGGCTA
GAGCACACGTCTGAACTCTTTCCCTACACGACGCTCTTCCGATCTNNNGAGATCACACTGCTGTACTCC
GCACACGTCTGAACTCTTTCCCTACACGACGCTCTTCCGATCTNNNCCTGAGCAATAGAGTGAAACCC
AGCACACGTCTGAACTCTTTCCCTACACGACGCTCTTCCGATCTNNNGTATGTGTCTGTGTGTGTGTG
AGCACACGTCTGAACTCTTTCCCTACACGACGCTCTTCCGATCTNNNTGGGAAGTACATTGATGGACC
AGAGCACACGTCTGAACTCTTTCCCTACACGACGCTCTTCCGATCTNNNATGTGTGTGTGTGTGAATG
AGCACACGTCTGAACTCTTTCCCTACACGACGCTCTTCCGATCTNNNTTCAAGTGATTCTCCTGCCTCAG
AGCACACGTCTGAACTCTTTCCCTACACGACGCTCTTCCGATCTNNNCTACAAGGATCAATAGCAAGG
AGCACACGTCTGAACTCTTTCCCTACACGACGCTCTTCCGATCTNNNCAGGAGAATTGCTTTGAACCTG
GAGCACACGTCTGAACTCTTTCCCTACACGACGCTCTTCCGATCTNNNGACTCCAGCTTATGCAGTACA
AGAGCACACGTCTGAACTCTTTCCCTACACGACGCTCTTCCGATCTNNNCAAGACCCTTTCTCCAACAAC
AGCACACGTCTGAACTCTTTCCCTACACGACGCTCTTCCGATCTNNNATCTCATCCAGGTCACTGCAAA
AGCACACGTCTGAACTCTTTCCCTACACGACGCTCTTCCGATCTNNNCAGGCACATTTCATACACACAC
AGAGCACACGTCTGAACTCTTTCCCTACACGACGCTCTTCCGATCTNNNTGACAGCTTTGTCTAGTCAC
AGCACACGTCTGAACTCTTTCCCTACACGACGCTCTTCCGATCTNNNTGACCCTGGTAGAGTAGAGAC
AGCACACGTCTGAACTCTTTCCCTACACGACGCTCTTCCGATCTNNNTAAGTAGCTGGGACTACAGGC
AGCACACGTCTGAACTCTTTCCCTACACGACGCTCTTCCGATCTNNNCTCAAGTGATCCTCTTGCCTCA
GAGCACACGTCTGAACTCTTTCCCTACACGACGCTCTTCCGATCTNNNTCCCATTTATGTTCCCACCAAC
GAGCACACGTCTGAACTCTTTCCCTACACGACGCTCTTCCGATCTNNNATCTGGGCCAAGAAAAGCTT

GAGCACACGTCTGAACTCTTTCCCTACACGACGCTCTTCCGATCTNNNGTATGCCACCATACCCTGCTAA
GAGCACACGTCTGAACTCTTTCCCTACACGACGCTCTTCCGATCTNNNGCGACAGAGTGAGACTTCATT
AGCACACGTCTGAACTCTTTCCCTACACGACGCTCTTCCGATCTNNNGACAATCGCTTGACTCCCATAGA
GAGCACACGTCTGAACTCTTTCCCTACACGACGCTCTTCCGATCTNNNCCACGCCCAGCTAATTTTTGT
GAGCACACGTCTGAACTCTTTCCCTACACGACGCTCTTCCGATCTNNNCCCTGTCTCTCTCTCTCTCTCC
GAGCACACGTCTGAACTCTTTCCCTACACGACGCTCTTCCGATCTNNNGAGATCACACCACTACACTCCA
AGCACACGTCTGAACTCTTTCCCTACACGACGCTCTTCCGATCTNNNGGTGGGCAGGATTCTAATAAG
AGCACACGTCTGAACTCTTTCCCTACACGACGCTCTTCCGATCTNNNCTGGCCCAACTGTAACCTCTAAA
GAGCACACGTCTGAACTCTTTCCCTACACGACGCTCTTCCGATCTNNNCATTGCATATACACACACACCC
GAGCACACGTCTGAACTCTTTCCCTACACGACGCTCTTCCGATCTNNNGATTCTCACAATCCAGTCCCTG
GAGCACACGTCTGAACTCTTTCCCTACACGACGCTCTTCCGATCTNNNGCAACTAACGGACTCATGCAAT
AGCACACGTCTGAACTCTTTCCCTACACGACGCTCTTCCGATCTNNNCAACTTGTCTTGCAGTCATCTGG
AGCACACGTCTGAACTCTTTCCCTACACGACGCTCTTCCGATCTNNNTGTTTTGACTGACCCACTTCTCT
GAGCACACGTCTGAACTCTTTCCCTACACGACGCTCTTCCGATCTNNNGTGTGTGTGTGTTTTGAGA
AGCACACGTCTGAACTCTTTCCCTACACGACGCTCTTCCGATCTNNNCAACGTTGAAAGGGAGAGTCCT
AGCACACGTCTGAACTCTTTCCCTACACGACGCTCTTCCGATCTNNNGAAGGTTGCAGTGAGCTGAGA
AGCACACGTCTGAACTCTTTCCCTACACGACGCTCTTCCGATCTNNNGGGCGATAAGAGAGAAACTCC
GAGCACACGTCTGAACTCTTTCCCTACACGACGCTCTTCCGATCTNNNTCTGTCAACCATGCTACAAGT
AGCACACGTCTGAACTCTTTCCCTACACGACGCTCTTCCGATCTNNNGTGACAGAGCAAGACTAGGTCT
AGAGCACACGTCTGAACTCTTTCCCTACACGACGCTCTTCCGATCTNNNAAAACAATGTGTGTGTGTCC
GAGCACACGTCTGAACTCTTTCCCTACACGACGCTCTTCCGATCTNNNTTAGTAACACGAAGTCCAGTCC
AGCACACGTCTGAACTCTTTCCCTACACGACGCTCTTCCGATCTNNNTTCCAAGAAAGGACAGATGGC
AGCACACGTCTGAACTCTTTCCCTACACGACGCTCTTCCGATCTNNNCATACATATGCGTGTGTGTGTG
GAGCACACGTCTGAACTCTTTCCCTACACGACGCTCTTCCGATCTNNNTTCAGCTCAGTATGTGTGTGTG
GAGCACACGTCTGAACTCTTTCCCTACACGACGCTCTTCCGATCTNNNTGAGCCACTGCAGCTATAAAA
AGCACACGTCTGAACTCTTTCCCTACACGACGCTCTTCCGATCTNNNTCAAGTGATCCTTACACCTCAGG
AGCACACGTCTGAACTCTTTCCCTACACGACGCTCTTCCGATCTNNNGACAGAGCAAGACCTTGTCTCT
AGCACACGTCTGAACTCTTTCCCTACACGACGCTCTTCCGATCTNNNGTGACACAGCTAGACTCTGTCTC
GCACACGTCTGAACTCTTTCCCTACACGACGCTCTTCCGATCTNNNCATTACACTCTAGCCTGGGTGACC
GCACACGTCTGAACTCTTTCCCTACACGACGCTCTTCCGATCTNNNTGATAGTGAATGAGCTCTTGCGA
AGCACACGTCTGAACTCTTTCCCTACACGACGCTCTTCCGATCTNNNGTGCCTATAATCCCAGCTACTCG
GAGCACACGTCTGAACTCTTTCCCTACACGACGCTCTTCCGATCTNNNAAAGAGAGAGAGAGAGAGAG
GAGCACACGTCTGAACTCTTTCCCTACACGACGCTCTTCCGATCTNNNATGCAGACTGATTGAAGCTCT
GAGCACACGTCTGAACTCTTTCCCTACACGACGCTCTTCCGATCTNNNATATGTGTGTGTGTGTGTGT
GAGCACACGTCTGAACTCTTTCCCTACACGACGCTCTTCCGATCTNNNGGCTGTATTAGGGTTCTCCAG
AGAGCACACGTCTGAACTCTTTCCCTACACGACGCTCTTCCGATCTNNNGAGCAAGACTCCGTCTCAAAA
GAGCACACGTCTGAACTCTTTCCCTACACGACGCTCTTCCGATCTNNNCAGAATGCGACCATGTCTCAA
GAGCACACGTCTGAACTCTTTCCCTACACGACGCTCTTCCGATCTNNNGAGGCAGGAAAATCACTTGA
GAGCACACGTCTGAACTCTTTCCCTACACGACGCTCTTCCGATCTNNNACAACAGGAGTCCATGTGTCAT
GAGCACACGTCTGAACTCTTTCCCTACACGACGCTCTTCCGATCTNNNCACTTGAGACCAGGAGTTCAA
GAGCACACGTCTGAACTCTTTCCCTACACGACGCTCTTCCGATCTNNNCCCATCTCTCTCTCTCTCTCTGT
AGCACACGTCTGAACTCTTTCCCTACACGACGCTCTTCCGATCTNNNTCAAGACTGCAGTGAGCCATAA
GAGCACACGTCTGAACTCTTTCCCTACACGACGCTCTTCCGATCTNNNCAGGAGAATTGCTTGAAACTC

CACACGTCTGAACTCTTTCCCTACACGACGCTCTTCCGATCTNNNCAAGCTGGGTCAATGATTACACA
AGCACACGTCTGAACTCTTTCCCTACACGACGCTCTTCCGATCTNNNAGTTGTATGCTGAGTGGAAG
GCACACGTCTGAACTCTTTCCCTACACGACGCTCTTCCGATCTNNNTCACCTTTGGGTCAGAAAAGCA
AGCACACGTCTGAACTCTTTCCCTACACGACGCTCTTCCGATCTNNNGGCTCTTGAATGTCTGCTCTTT
GCACACGTCTGAACTCTTTCCCTACACGACGCTCTTCCGATCTNNNAATGTTGTGGTTCTTGCACACTG
AGCACACGTCTGAACTCTTTCCCTACACGACGCTCTTCCGATCTNNNATAGGAGAGACAGGAGAGA
AGCACACGTCTGAACTCTTTCCCTACACGACGCTCTTCCGATCTNNNGCTAGTCAGGGAGTTCTCATG
AGCACACGTCTGAACTCTTTCCCTACACGACGCTCTTCCGATCTNNNGCACAAACACACACACATACAG
GCACACGTCTGAACTCTTTCCCTACACGACGCTCTTCCGATCTNNNAAGATTCCATTCCTAGTGAGGGC
AAGAGCACACGTCTGAACTCTTTCCCTACACGACGCTCTTCCGATCTNNNAGCAAGACTCCTCATCAAA
GCACACGTCTGAACTCTTTCCCTACACGACGCTCTTCCGATCTNNNGGCAGAAACTCTTGTTGTCTTCC
AGCACACGTCTGAACTCTTTCCCTACACGACGCTCTTCCGATCTNNNAAGTTTGTTTTCTACCCCTCCCC
AGAGCACACGTCTGAACTCTTTCCCTACACGACGCTCTTCCGATCTNNNTATAGAGAGAGAGAGCACC
GAGCACACGTCTGAACTCTTTCCCTACACGACGCTCTTCCGATCTNNNTGGAGCTGTTGTGGGATTAT
AGCACACGTCTGAACTCTTTCCCTACACGACGCTCTTCCGATCTNNNGCTCATCTGACTCTACTTTGCCA
GCACACGTCTGAACTCTTTCCCTACACGACGCTCTTCCGATCTNNNTTACCATTTACACACGCACACACC
GAGCACACGTCTGAACTCTTTCCCTACACGACGCTCTTCCGATCTNNNACTCATGGTCATTTCAGGCTC
AGCACACGTCTGAACTCTTTCCCTACACGACGCTCTTCCGATCTNNNTGGCCCAGAGTAACAGAAAA
AGAGCACACGTCTGAACTCTTTCCCTACACGACGCTCTTCCGATCTNNNGCTCACAATGTCAAACCCTT
AGCACACGTCTGAACTCTTTCCCTACACGACGCTCTTCCGATCTNNNCAACATCATTTGCCCCAAACTCT
GCACACGTCTGAACTCTTTCCCTACACGACGCTCTTCCGATCTNNNTTTGGGGGTATAGTGTGCAAT
AGCACACGTCTGAACTCTTTCCCTACACGACGCTCTTCCGATCTNNNGGCAAAGAGGAAAAGAAAGA
AGCACACGTCTGAACTCTTTCCCTACACGACGCTCTTCCGATCTNNNAAGAATCACTTGAACCTGGGA
AGAGCACACGTCTGAACTCTTTCCCTACACGACGCTCTTCCGATCTNNNAAAATTAGCTGGACGTGGT
CACACGTCTGAACTCTTTCCCTACACGACGCTCTTCCGATCTNNNCTCTCTCTGTCTCTCGCTCTTCAT
AGAGCACACGTCTGAACTCTTTCCCTACACGACGCTCTTCCGATCTNNNAATTTGCCTTCTGTCCTAAG
GAGCACACGTCTGAACTCTTTCCCTACACGACGCTCTTCCGATCTNNNCTCTTCTCCTGCGTCTTCCATA
AGCACACGTCTGAACTCTTTCCCTACACGACGCTCTTCCGATCTNNNGGATCACTTCAGTCCAGGAGTT
GCACACGTCTGAACTCTTTCCCTACACGACGCTCTTCCGATCTNNNTGAAGCCCCTACTATTGTTGTGT
GCACACGTCTGAACTCTTTCCCTACACGACGCTCTTCCGATCTNNNACATGCGTAGTCTTCTTGGAATG
AGAGCACACGTCTGAACTCTTTCCCTACACGACGCTCTTCCGATCTNNNATGGTCAATGCATGTGTGTT
AGCACACGTCTGAACTCTTTCCCTACACGACGCTCTTCCGATCTNNNAAAGAAGCCTGAAAGAGATGG
GAGCACACGTCTGAACTCTTTCCCTACACGACGCTCTTCCGATCTNNNAAGATCACAGTGGGCCAGTT
AGCACACGTCTGAACTCTTTCCCTACACGACGCTCTTCCGATCTNNNCAGGGACACCATTACTCAGGAT
AGCACACGTCTGAACTCTTTCCCTACACGACGCTCTTCCGATCTNNNGGGAAGGCTCCAGGAAAATCA
GAGCACACGTCTGAACTCTTTCCCTACACGACGCTCTTCCGATCTNNNTAGATGGAGTCATGAGAGTG
GCACACGTCTGAACTCTTTCCCTACACGACGCTCTTCCGATCTNNNAGATTGAGAGAGAGCGAGAGA
AAGAGCACACGTCTGAACTCTTTCCCTACACGACGCTCTTCCGATCTNNNCTTGTAGACAGCCCCTTCT
GAGCACACGTCTGAACTCTTTCCCTACACGACGCTCTTCCGATCTNNNGCTGCATAATTCGCAATTGCA
AGCACACGTCTGAACTCTTTCCCTACACGACGCTCTTCCGATCTNNNGAGTTCTTCTCTCCAACTTCCGT
AGCACACGTCTGAACTCTTTCCCTACACGACGCTCTTCCGATCTNNNCTTTGTCAAGCACTTGGTTCCA
AGCACACGTCTGAACTCTTTCCCTACACGACGCTCTTCCGATCTNNNTCTCTCTTTGATCTGCTCGCTTT
GAGCACACGTCTGAACTCTTTCCCTACACGACGCTCTTCCGATCTNNNACAGTGTGTGTGTTTGTAAG

GAGCACACGTCTGAACTCTTTCCCTACACGACGCTCTTCCGATCTNNNTGGGCAACATAGTGAGACTT
AGCACACGTCTGAACTCTTTCCCTACACGACGCTCTTCCGATCTNNNGCCTGATGCAGAGCAGTCTAA
AGCACACGTCTGAACTCTTTCCCTACACGACGCTCTTCCGATCTNNNAATCACCATCACTCCAGCATGA
GCACACGTCTGAACTCTTTCCCTACACGACGCTCTTCCGATCTNNNCCAGAAGATTCAGTTCCTGGTG
AGCACACGTCTGAACTCTTTCCCTACACGACGCTCTTCCGATCTNNNCTTGCTGTCTTCTTCCTGTGACC
GCACACGTCTGAACTCTTTCCCTACACGACGCTCTTCCGATCTNNNTCTCACTCTCTGTCTCTCTCCTCC
GCACACGTCTGAACTCTTTCCCTACACGACGCTCTTCCGATCTNNNTAGGATAGCTGAATTTGGGGT
AGCACACGTCTGAACTCTTTCCCTACACGACGCTCTTCCGATCTNNNGAATGGTTCTGTGATGCTATG
GAGCACACGTCTGAACTCTTTCCCTACACGACGCTCTTCCGATCTNNNGATTGCACCAATCTCCAAATG
CACACGTCTGAACTCTTTCCCTACACGACGCTCTTCCGATCTNNNGCTTGGTGTCTCTCTCTCTCTCTCA
AGCACACGTCTGAACTCTTTCCCTACACGACGCTCTTCCGATCTNNNTCAAACACACACACACAGAGA
AGCACACGTCTGAACTCTTTCCCTACACGACGCTCTTCCGATCTNNNCCAAGAGTTTGAGGCTGTAGT
GAGCACACGTCTGAACTCTTTCCCTACACGACGCTCTTCCGATCTNNNGGGCATCTCAACGATTTTT
AGCACACGTCTGAACTCTTTCCCTACACGACGCTCTTCCGATCTNNNCTCAAAAAGGAAGGAAGGG
AGCACACGTCTGAACTCTTTCCCTACACGACGCTCTTCCGATCTNNNTTAGAGGTTTCCAGGTGTTTG
CACACGTCTGAACTCTTTCCCTACACGACGCTCTTCCGATCTNNNGTCTGTGTGTTTGTCTGTGTGTG
AGCACACGTCTGAACTCTTTCCCTACACGACGCTCTTCCGATCTNNNCAGATTGCTCGATCTTTCCCCT
GCACACGTCTGAACTCTTTCCCTACACGACGCTCTTCCGATCTNNNATATGTGTGTGTGTGTGTGTG
AGAGCACACGTCTGAACTCTTTCCCTACACGACGCTCTTCCGATCTNNNAGAAAGGAAAGAGAGAG
AGCACACGTCTGAACTCTTTCCCTACACGACGCTCTTCCGATCTNNNGGGAGAGAGAGAGAGAGA
GAGCACACGTCTGAACTCTTTCCCTACACGACGCTCTTCCGATCTNNNATCACTTGAATCCAGAAGGC
AGAGCACACGTCTGAACTCTTTCCCTACACGACGCTCTTCCGATCTNNNGGGAGAGAGAGAGAGAG
CACACGTCTGAACTCTTTCCCTACACGACGCTCTTCCGATCTNNNACAAACCTATGCACATACCCACT
AGCACACGTCTGAACTCTTTCCCTACACGACGCTCTTCCGATCTNNNGAGTGGAACGTGCTTCAGAAA
AGCACACGTCTGAACTCTTTCCCTACACGACGCTCTTCCGATCTNNNCTAACTACTGGTAGGCCTTTG
AGCACACGTCTGAACTCTTTCCCTACACGACGCTCTTCCGATCTNNNGGTCATGAGCATAGAACCCTC
AGCACACGTCTGAACTCTTTCCCTACACGACGCTCTTCCGATCTNNNTCTGTCTTCTTCATTTGGGACC
AGCACACGTCTGAACTCTTTCCCTACACGACGCTCTTCCGATCTNNNACTTCTCATAACTGTGGCCAAG
GCACACGTCTGAACTCTTTCCCTACACGACGCTCTTCCGATCTNNNGCTCTCTGTCTCTCTCCAACAAT
GCACACGTCTGAACTCTTTCCCTACACGACGCTCTTCCGATCTNNNGGTAGTCTCCCATAATCCACTGC
GCACACGTCTGAACTCTTTCCCTACACGACGCTCTTCCGATCTNNNTGGATTCTGGAATGCAAATCAG
GCACACGTCTGAACTCTTTCCCTACACGACGCTCTTCCGATCTNNNTCACCCAGACTTGCTGTTATGTA
GCACACGTCTGAACTCTTTCCCTACACGACGCTCTTCCGATCTNNNTGTCTCTACTGTCCTTGATAGCC
GCACACGTCTGAACTCTTTCCCTACACGACGCTCTTCCGATCTNNNTCGACTCAGTCAACAAAAGGTC
AGCACACGTCTGAACTCTTTCCCTACACGACGCTCTTCCGATCTNNNGGGTTTGTAGCAAATGTCTG
AGAGCACACGTCTGAACTCTTTCCCTACACGACGCTCTTCCGATCTNNNCCGAAAGTGTTTTGCCTTA
AGCACACGTCTGAACTCTTTCCCTACACGACGCTCTTCCGATCTNNNTGTGACTCTTGAGGAGAAACC
AGCACACGTCTGAACTCTTTCCCTACACGACGCTCTTCCGATCTNNNAGGCACTTTCCTGAATTCACTG
GCACACGTCTGAACTCTTTCCCTACACGACGCTCTTCCGATCTNNNTTCTTCATGGCTTGTAGATGGCT
AGCACACGTCTGAACTCTTTCCCTACACGACGCTCTTCCGATCTNNNTCGCATTGCTAGAATCTCCCAT
AGCACACGTCTGAACTCTTTCCCTACACGACGCTCTTCCGATCTNNNAGACCCTAGAGAAACAAAGCT
GCACACGTCTGAACTCTTTCCCTACACGACGCTCTTCCGATCTNNNCAGGAGGAGCTGTGTATTTCCT
AGCACACGTCTGAACTCTTTCCCTACACGACGCTCTTCCGATCTNNNATGTGTCTGTCTGTCTGTCTG

AGCACACGTCTGAACTCTTTCCCTACACGACGCTCTTCCGATCTNNNTTACTGCCGGTCTAGTGATCC
GCACACGTCTGAACTCTTTCCCTACACGACGCTCTTCCGATCTNNNAACTGGATTCACAACCATGCTTG
AGCACACGTCTGAACTCTTTCCCTACACGACGCTCTTCCGATCTNNNGTATCCAGATTTCAGCAGGC
GAGCACACGTCTGAACTCTTTCCCTACACGACGCTCTTCCGATCTNNNTCCCTCTCTCCTCTGAATAAG
AGAGCACACGTCTGAACTCTTTCCCTACACGACGCTCTTCCGATCTNNNAACTGAAGCAGAGAAGGG
AGCACACGTCTGAACTCTTTCCCTACACGACGCTCTTCCGATCTNNNAGAAGGTCTGCAATCTCACAC
GAGCACACGTCTGAACTCTTTCCCTACACGACGCTCTTCCGATCTNNNCTGATACACACACACACACAC
AGCACACGTCTGAACTCTTTCCCTACACGACGCTCTTCCGATCTNNNTAAGCACCTTGTAGAATGTG
GCACACGTCTGAACTCTTTCCCTACACGACGCTCTTCCGATCTNNNGACAAGGACCAGGAGAGATTC
AGCACACGTCTGAACTCTTTCCCTACACGACGCTCTTCCGATCTNNNAAGGTGCAGTCAGATTTAGAG
AGCACACGTCTGAACTCTTTCCCTACACGACGCTCTTCCGATCTNNNTCCTCCACATGGATGAAGGTTT
AGCACACGTCTGAACTCTTTCCCTACACGACGCTCTTCCGATCTNNNAGCTATCACTTGCAAGCTTAGG
GCACACGTCTGAACTCTTTCCCTACACGACGCTCTTCCGATCTNNNCCTAACCTCCTTCCTCCATTTCCCA
AGCACACGTCTGAACTCTTTCCCTACACGACGCTCTTCCGATCTNNNGAGATATGCATGTGTGTGTG
AGCACACGTCTGAACTCTTTCCCTACACGACGCTCTTCCGATCTNNNAAACATCCTAGCCAGGTCAAC
AGCACACGTCTGAACTCTTTCCCTACACGACGCTCTTCCGATCTNNNTGCCCAAACTAAGAGTGTGAA
AGCACACGTCTGAACTCTTTCCCTACACGACGCTCTTCCGATCTNNNGCAGGAGGGTGAGTTCCATTA
GAGCACACGTCTGAACTCTTTCCCTACACGACGCTCTTCCGATCTNNNCAGTTTTGTGCGTGTGTGTG
AGCACACGTCTGAACTCTTTCCCTACACGACGCTCTTCCGATCTNNNACTCAGGTCTAGATCTCTGGGA
AGCACACGTCTGAACTCTTTCCCTACACGACGCTCTTCCGATCTNNNTTGAGATGGAGTCTTGCTCTG
AGCACACGTCTGAACTCTTTCCCTACACGACGCTCTTCCGATCTNNNTCAATGATTACAACGTTTGGCC
AGCACACGTCTGAACTCTTTCCCTACACGACGCTCTTCCGATCTNNNCAATTTCTCTTTAGGCCTGGCA
GCACACGTCTGAACTCTTTCCCTACACGACGCTCTTCCGATCTNNNATTGTATCTCCCAGAATTCCCGG
AGCACACGTCTGAACTCTTTCCCTACACGACGCTCTTCCGATCTNNNAAGAACAAAGCTTCCACAGTGT
GAGCACACGTCTGAACTCTTTCCCTACACGACGCTCTTCCGATCTNNNCCACCAAAACTGAAGACCCAT
GCACACGTCTGAACTCTTTCCCTACACGACGCTCTTCCGATCTNNNCTGGAAACTGACCCCAAATAGG
AGCACACGTCTGAACTCTTTCCCTACACGACGCTCTTCCGATCTNNNCTGTTGAGTAGCTGGGACTA
AGCACACGTCTGAACTCTTTCCCTACACGACGCTCTTCCGATCTNNNCAGAGAATCACTTGAACCTGG
AGCACACGTCTGAACTCTTTCCCTACACGACGCTCTTCCGATCTNNNACCTTCCTCAACACAACAGAT
AGCACACGTCTGAACTCTTTCCCTACACGACGCTCTTCCGATCTNNNGATTGAGAAATCGGATGGTTG
AGCACACGTCTGAACTCTTTCCCTACACGACGCTCTTCCGATCTNNNGAGCTCAGACCTGTCTAATCCT
AGCACACGTCTGAACTCTTTCCCTACACGACGCTCTTCCGATCTNNNCTATCCACAGGACAGCAGTTT
AGCACACGTCTGAACTCTTTCCCTACACGACGCTCTTCCGATCTNNNGCCAGAAATTAGGAGGCAAA
AGCACACGTCTGAACTCTTTCCCTACACGACGCTCTTCCGATCTNNNAACAATAGCAACAATTCGCTGG
AGAGCACACGTCTGAACTCTTTCCCTACACGACGCTCTTCCGATCTNNNGCCTTGTTCCTTCTCTAGG
AGCACACGTCTGAACTCTTTCCCTACACGACGCTCTTCCGATCTNNNGATCATCTCTCCAAGTCCTCAC
GAGCACACGTCTGAACTCTTTCCCTACACGACGCTCTTCCGATCTNNNGGCAAGGTGAGCTTGTTTTG
AGCACACGTCTGAACTCTTTCCCTACACGACGCTCTTCCGATCTNNNACAAGATTGTAAGAAAGCCCC
AGCACACGTCTGAACTCTTTCCCTACACGACGCTCTTCCGATCTNNNCTCTTCCTTCCTTCCTTCCTTCCC
GCACACGTCTGAACTCTTTCCCTACACGACGCTCTTCCGATCTNNNGTTTCACTGTGTTAGCCAGGATG
GAGCACACGTCTGAACTCTTTCCCTACACGACGCTCTTCCGATCTNNNCCTCATTTATCTACTCCCAGG
GCACACGTCTGAACTCTTTCCCTACACGACGCTCTTCCGATCTNNNTTTGGTTTCCTTTGTGTCACGAG
GCACACGTCTGAACTCTTTCCCTACACGACGCTCTTCCGATCTNNNAGAAACAGGAACAAAGGCAGA

AGCACACGTCTGAACTCTTTCCCTACACGACGCTCTTCCGATCTNNNTCATGGGTAACACAGAAAGCAC
AGCACACGTCTGAACTCTTTCCCTACACGACGCTCTTCCGATCTNNNGTGCCTATAATCCCAGCTACTCG
AGCACACGTCTGAACTCTTTCCCTACACGACGCTCTTCCGATCTNNNGGTAATGTCTACCAAGGGTCCT
AAGAGCACACGTCTGAACTCTTTCCCTACACGACGCTCTTCCGATCTNNNCAATCATGGCAGAAGGTG
AGCACACGTCTGAACTCTTTCCCTACACGACGCTCTTCCGATCTNNNGAGCCGTGATCAAAACACTACA
AGCACACGTCTGAACTCTTTCCCTACACGACGCTCTTCCGATCTNNNAGGTTGCAGTGAGTGAAGATC
AGCACACGTCTGAACTCTTTCCCTACACGACGCTCTTCCGATCTNNNTTGACTGATACAGTGTCCACCA
AGCACACGTCTGAACTCTTTCCCTACACGACGCTCTTCCGATCTNNNTTGTCTGGCTTCTTTGTTTGCAT
GAGCACACGTCTGAACTCTTTCCCTACACGACGCTCTTCCGATCTNNNGTTGCAATGAGCTGAGATCT
AGCACACGTCTGAACTCTTTCCCTACACGACGCTCTTCCGATCTNNNGCCCTAGGGAGCTAATATAAG
GAGCACACGTCTGAACTCTTTCCCTACACGACGCTCTTCCGATCTNNNGCTTCTGTAATTGCATGAGCCA
AGAGCACACGTCTGAACTCTTTCCCTACACGACGCTCTTCCGATCTNNNGAGAAAGAGAGAGAAGGCA
GAGCACACGTCTGAACTCTTTCCCTACACGACGCTCTTCCGATCTNNNCTCCCCTCAAAGCAAGTGAAAA
AGAGCACACGTCTGAACTCTTTCCCTACACGACGCTCTTCCGATCTNNNGGTGACAGAGTGAGACGAA
GAGCACACGTCTGAACTCTTTCCCTACACGACGCTCTTCCGATCTNNNCCACTCCTGGTACCAAAATCTG
GAGCACACGTCTGAACTCTTTCCCTACACGACGCTCTTCCGATCTNNNAGTGGAAAGTGGAAACTCAC
AGCACACGTCTGAACTCTTTCCCTACACGACGCTCTTCCGATCTNNNTCTCTCTCTCTCCCTCTCTCTCAT
GAGCACACGTCTGAACTCTTTCCCTACACGACGCTCTTCCGATCTNNNTCAAGCATCAAGACCATTCAGG
GAGCACACGTCTGAACTCTTTCCCTACACGACGCTCTTCCGATCTNNNGCCCATAATAGCAAGCAATGG
GAGCACACGTCTGAACTCTTTCCCTACACGACGCTCTTCCGATCTNNNATGTGTGTGTGTGTGTGTGT
AGCACACGTCTGAACTCTTTCCCTACACGACGCTCTTCCGATCTNNNACACAGGTACACACACACACATA
AGCACACGTCTGAACTCTTTCCCTACACGACGCTCTTCCGATCTNNNAAACATAGTAAGTAGCTGGCCC
AGCACACGTCTGAACTCTTTCCCTACACGACGCTCTTCCGATCTNNNCTAGCCTAGGTGACAGAGTGA
AGAGCACACGTCTGAACTCTTTCCCTACACGACGCTCTTCCGATCTNNNCTGCCTCAGTGCTCTTTTCAA
AAGAGCACACGTCTGAACTCTTTCCCTACACGACGCTCTTCCGATCTNNNGAGGTTGTAGTGAGCCAG
AAGAGCACACGTCTGAACTCTTTCCCTACACGACGCTCTTCCGATCTNNNGAAAGGGAGAGAGAGAG
AGCACACGTCTGAACTCTTTCCCTACACGACGCTCTTCCGATCTNNNCTGTGTCTCTCTCTCTGTGTGTC
GCACACGTCTGAACTCTTTCCCTACACGACGCTCTTCCGATCTNNNAGAAGCAGCTAGGGTTAGTGTT
AGCACACGTCTGAACTCTTTCCCTACACGACGCTCTTCCGATCTNNNCTCTCTCTCTCTCTTCCCTCCCA
GCACACGTCTGAACTCTTTCCCTACACGACGCTCTTCCGATCTNNNCTGATGCTCACATTCGTGGTCTA
GAGCACACGTCTGAACTCTTTCCCTACACGACGCTCTTCCGATCTNNNATACCATTTGTTCTGCTGGGT
CACACGTCTGAACTCTTTCCCTACACGACGCTCTTCCGATCTNNNTGCAGGCAAGTTATCTCCTCTTTCA
AGCACACGTCTGAACTCTTTCCCTACACGACGCTCTTCCGATCTNNNCTTCTCTCTGCCTCAGTTTCTGT
GCACACGTCTGAACTCTTTCCCTACACGACGCTCTTCCGATCTNNNCTTTCTGCTGCTGTTTACCTTCCCA
AAGAGCACACGTCTGAACTCTTTCCCTACACGACGCTCTTCCGATCTNNNCTGAGGTGGGAGGATTAC
AAGAGCACACGTCTGAACTCTTTCCCTACACGACGCTCTTCCGATCTNNNCCCTGAGACCTTGTCAGAA
AGCACACGTCTGAACTCTTTCCCTACACGACGCTCTTCCGATCTNNNGAGAGAGGGAGAGTGAGAGA
AGAGCACACGTCTGAACTCTTTCCCTACACGACGCTCTTCCGATCTNNNCATAGGGGTGCTTACGTCAT
AGCACACGTCTGAACTCTTTCCCTACACGACGCTCTTCCGATCTNNNGGAAGCAAGCACCTTCTTCAAA
GAGCACACGTCTGAACTCTTTCCCTACACGACGCTCTTCCGATCTNNNGAAATACTGGGGGGGAAAGA
GAGCACACGTCTGAACTCTTTCCCTACACGACGCTCTTCCGATCTNNNTTCTCACTGTGTCTCCATATGG
AGCACACGTCTGAACTCTTTCCCTACACGACGCTCTTCCGATCTNNNTCCTGGTTTTGGTATTAGGGTG
GAGCACACGTCTGAACTCTTTCCCTACACGACGCTCTTCCGATCTNNNCTCTCTCTCTCTCCCTCTCTCC

GAGCACACGTCTGAACTCTTTCCCTACACGACGCTCTTCCGATCTNNNGGAGTATTTACTGGTGCCTG
GCACACGTCTGAACTCTTTCCCTACACGACGCTCTTCCGATCTNNNTCCCTCTCTCTCTCTCTCTCTCAT
AGAGCACACGTCTGAACTCTTTCCCTACACGACGCTCTTCCGATCTNNNCCAGGAATTTCAGGCAGTAG
GAGCACACGTCTGAACTCTTTCCCTACACGACGCTCTTCCGATCTNNNTCCACCAGAGGAAAAAGAACA
AGCACACGTCTGAACTCTTTCCCTACACGACGCTCTTCCGATCTNNNCATGCCATCAGTAGGAAAGGCT
AAGAGCACACGTCTGAACTCTTTCCCTACACGACGCTCTTCCGATCTNNNAGCCTGGATAAAAGAGCAA
GAGCACACGTCTGAACTCTTTCCCTACACGACGCTCTTCCGATCTNNNCTGAGGTCTTGGGTAAGTCAC
AGCACACGTCTGAACTCTTTCCCTACACGACGCTCTTCCGATCTNNNCTCTCTAAAGCTCTTGGTCCCC
GAGCACACGTCTGAACTCTTTCCCTACACGACGCTCTTCCGATCTNNNCATGCCATCATACCTGCTGAGA
AGCACACGTCTGAACTCTTTCCCTACACGACGCTCTTCCGATCTNNNGCAGTGATGGTTGCTCAATATC
AGCACACGTCTGAACTCTTTCCCTACACGACGCTCTTCCGATCTNNNTTCTTTCTGAGTCCTCACATGGG
AAGAGCACACGTCTGAACTCTTTCCCTACACGACGCTCTTCCGATCTNNNAGAGAAAGGAGAGAAAG
AGCACACGTCTGAACTCTTTCCCTACACGACGCTCTTCCGATCTNNNTGGGAGGTAAGGAGATCATGA
AGAGCACACGTCTGAACTCTTTCCCTACACGACGCTCTTCCGATCTNNNAGCCAGTTCAGAAGTATGCT
AGCACACGTCTGAACTCTTTCCCTACACGACGCTCTTCCGATCTNNNAAACCTGTGAGAAGGCATATGC
AGCACACGTCTGAACTCTTTCCCTACACGACGCTCTTCCGATCTNNNAGCAGAAGGCCTTGATAGAACA
AGCACACGTCTGAACTCTTTCCCTACACGACGCTCTTCCGATCTNNNCCCTCCCTCCCTTTTCTTTCTTTCA
AGCACACGTCTGAACTCTTTCCCTACACGACGCTCTTCCGATCTNNNTGCACACCTGTAGTCTCAGTTAC
AGCACACGTCTGAACTCTTTCCCTACACGACGCTCTTCCGATCTNNNGTGTGTGTGTGTGTGTGTA
GAGCACACGTCTGAACTCTTTCCCTACACGACGCTCTTCCGATCTNNNAAATGCCACCTTCTCCCTGTAT
AAGAGCACACGTCTGAACTCTTTCCCTACACGACGCTCTTCCGATCTNNNCTGTAATTCCAGCTACTCGG
AGCACACGTCTGAACTCTTTCCCTACACGACGCTCTTCCGATCTNNNTTCCTACTGTTAGCCTTGGTGTC
GAGCACACGTCTGAACTCTTTCCCTACACGACGCTCTTCCGATCTNNNTGCCTCTGTCTCTTTCTCTCTCT
AAGAGCACACGTCTGAACTCTTTCCCTACACGACGCTCTTCCGATCTNNNGTGTAGTTCAAAGGCCTGA
AGCACACGTCTGAACTCTTTCCCTACACGACGCTCTTCCGATCTNNNTCCCAAACAAATGGAGTCTCTCT
AGCACACGTCTGAACTCTTTCCCTACACGACGCTCTTCCGATCTNNNTCAACTTGACCCTTCTACTCCACC
AGCACACGTCTGAACTCTTTCCCTACACGACGCTCTTCCGATCTNNNTTGTGGATAAGCTGTCTCTGCAC
GAGCACACGTCTGAACTCTTTCCCTACACGACGCTCTTCCGATCTNNNTTTTCCAGCAAAGACATTGAGC
GAGCACACGTCTGAACTCTTTCCCTACACGACGCTCTTCCGATCTNNNGGGGGAAAAAAAACAGCAGA
GCACACGTCTGAACTCTTTCCCTACACGACGCTCTTCCGATCTNNNGCTTGACCGCTGTGAAAGTATTT
AGCACACGTCTGAACTCTTTCCCTACACGACGCTCTTCCGATCTNNNTAATGTGTGTGTGTGTGTGTG
GAGCACACGTCTGAACTCTTTCCCTACACGACGCTCTTCCGATCTNNNAAAAAGAGAGAGGGAGAGAC
AGCACACGTCTGAACTCTTTCCCTACACGACGCTCTTCCGATCTNNNTGATGTCCTCATGAGAATGTGC
GAGCACACGTCTGAACTCTTTCCCTACACGACGCTCTTCCGATCTNNNTCATCACTGTTGATCTGGTCTG
AGCACACGTCTGAACTCTTTCCCTACACGACGCTCTTCCGATCTNNNCAGAGGTTGCAGTGAGATGAG
GAGCACACGTCTGAACTCTTTCCCTACACGACGCTCTTCCGATCTNNNTACATTAGGGCCAGGTAGAG
CACACGTCTGAACTCTTTCCCTACACGACGCTCTTCCGATCTNNNTGAGCCACTATATGCCTCTCTCTCA
GAGCACACGTCTGAACTCTTTCCCTACACGACGCTCTTCCGATCTNNNAGAGAGAGAGAGAGAGAGA
AGCACACGTCTGAACTCTTTCCCTACACGACGCTCTTCCGATCTNNNACAGAACGAGACTGTCTCTCTC
GAGCACACGTCTGAACTCTTTCCCTACACGACGCTCTTCCGATCTNNNTAAAACCACCGGGATAACAGA
AGCACACGTCTGAACTCTTTCCCTACACGACGCTCTTCCGATCTNNNTGGGGACTTTTGCTGAAGTAA
AGCACACGTCTGAACTCTTTCCCTACACGACGCTCTTCCGATCTNNNGTCTCACTGGACAAGACCTGATT
AGCACACGTCTGAACTCTTTCCCTACACGACGCTCTTCCGATCTNNNGAGATGGAGAATAAGGCTGAC

AGAGCACACGTCTGAACTCTTTCCCTACACGACGCTCTTCCGATCTNNNCATTGATGGCCTGAATGTCT
GAGCACACGTCTGAACTCTTTCCCTACACGACGCTCTTCCGATCTNNNACTTTCCAGGAGCTAAAGGAA
GAGCACACGTCTGAACTCTTTCCCTACACGACGCTCTTCCGATCTNNNTGATCATGCCACTATACTCCA
GCACACGTCTGAACTCTTTCCCTACACGACGCTCTTCCGATCTNNNATTACACCACCAGCTTTCCTAGG
GAGCACACGTCTGAACTCTTTCCCTACACGACGCTCTTCCGATCTNNNAGAGCGAGATCCTGTCTGTA
GAGCACACGTCTGAACTCTTTCCCTACACGACGCTCTTCCGATCTNNNAAGAGGTCAAAGGCTGCTAT
GAGCACACGTCTGAACTCTTTCCCTACACGACGCTCTTCCGATCTNNNGAAATGATGAGAGGGAAGG
GAGCACACGTCTGAACTCTTTCCCTACACGACGCTCTTCCGATCTNNNAGTGCTGGGATTATAGGCAT
AGCACACGTCTGAACTCTTTCCCTACACGACGCTCTTCCGATCTNNNACCAAAGGAAGCAAGAGAGA
AAGAGCACACGTCTGAACTCTTTCCCTACACGACGCTCTTCCGATCTNNNGGGACACCTCAGGAAACT
GAGCACACGTCTGAACTCTTTCCCTACACGACGCTCTTCCGATCTNNNAGAGAGACACAGAGAGTCAC
AGCACACGTCTGAACTCTTTCCCTACACGACGCTCTTCCGATCTNNNGGGTGACATAGTGAGACTCT
GCACACGTCTGAACTCTTTCCCTACACGACGCTCTTCCGATCTNNNTTATGGGTGTGTGTGTGTGTGTT
AGCACACGTCTGAACTCTTTCCCTACACGACGCTCTTCCGATCTNNNCCATACATCCAGTTCCAGTTCCA
AGAGCACACGTCTGAACTCTTTCCCTACACGACGCTCTTCCGATCTNNNAGAGAGAGAGAGAGGAA
GAGCACACGTCTGAACTCTTTCCCTACACGACGCTCTTCCGATCTNNNAGAGAGAGAGAGAGAGGGA
GAGCACACGTCTGAACTCTTTCCCTACACGACGCTCTTCCGATCTNNNTGGGATGTCTCCACTTCAGTT
GCACACGTCTGAACTCTTTCCCTACACGACGCTCTTCCGATCTNNNGTCTCTCTCTCTTCCCTGTCTCTC
AGAGCACACGTCTGAACTCTTTCCCTACACGACGCTCTTCCGATCTNNNCAGCCTTAGTGACAGAGGA
GCACACGTCTGAACTCTTTCCCTACACGACGCTCTTCCGATCTNNNTTGCATACTTCTGGAGGTTGGA
AGCACACGTCTGAACTCTTTCCCTACACGACGCTCTTCCGATCTNNNATAGAAGTTCAAGACCAGCCTG
GAGCACACGTCTGAACTCTTTCCCTACACGACGCTCTTCCGATCTNNNTTGATTGAAATGGCTGCTTTC
GAGCACACGTCTGAACTCTTTCCCTACACGACGCTCTTCCGATCTNNNCGACCTGATCTCAGCAAAAAC
AGCACACGTCTGAACTCTTTCCCTACACGACGCTCTTCCGATCTNNNTGTAAAGAATGGGGGCTGAGA
AGCACACGTCTGAACTCTTTCCCTACACGACGCTCTTCCGATCTNNNTTTGAGGCTGCAGTGAGCTATT
AGCACACGTCTGAACTCTTTCCCTACACGACGCTCTTCCGATCTNNNGATCTCTCACCCATGTCAGAGA
GAGCACACGTCTGAACTCTTTCCCTACACGACGCTCTTCCGATCTNNNAAGACTGTGGCTTTTGTCTTG
AGCACACGTCTGAACTCTTTCCCTACACGACGCTCTTCCGATCTNNNGACCTATCAGACTCCAGCATCT
AGCACACGTCTGAACTCTTTCCCTACACGACGCTCTTCCGATCTNNNAGAGACACCAGAAATCACTTGC
GAGCACACGTCTGAACTCTTTCCCTACACGACGCTCTTCCGATCTNNNTTGGGGCACATGTCATAGCTA
AGCACACGTCTGAACTCTTTCCCTACACGACGCTCTTCCGATCTNNNCTGGATATCACCCAACTGGAG
GCACACGTCTGAACTCTTTCCCTACACGACGCTCTTCCGATCTNNNTACCCTGAAAGCTTGTCCTTTGT
GAGCACACGTCTGAACTCTTTCCCTACACGACGCTCTTCCGATCTNNNGAGGAGACCAGAAGACTACA
AGCACACGTCTGAACTCTTTCCCTACACGACGCTCTTCCGATCTNNNCCACTTTTGTTTCCTGTCTGGT
GAGCACACGTCTGAACTCTTTCCCTACACGACGCTCTTCCGATCTNNNCTCTGGGATAGAACACAACCA
GCACACGTCTGAACTCTTTCCCTACACGACGCTCTTCCGATCTNNNTCTTCCTAGCTTGTAGGAGACCA
GAGCACACGTCTGAACTCTTTCCCTACACGACGCTCTTCCGATCTNNNTGTGTGTGGAAATGTGTATG
AGCACACGTCTGAACTCTTTCCCTACACGACGCTCTTCCGATCTNNNATGTGTATGTGTGTGTGTGTG
AGAGCACACGTCTGAACTCTTTCCCTACACGACGCTCTTCCGATCTNNNTGGGAGTCTTCCTTGATTCT
AGCACACGTCTGAACTCTTTCCCTACACGACGCTCTTCCGATCTNNNAGTTGAGTTAGGCTGAACTCC
AGCACACGTCTGAACTCTTTCCCTACACGACGCTCTTCCGATCTNNNGGCTTCATTTCAAGCTCTGAT
GAGCACACGTCTGAACTCTTTCCCTACACGACGCTCTTCCGATCTNNNCCTCCCTTACATCCTTCCAAAC
AGCACACGTCTGAACTCTTTCCCTACACGACGCTCTTCCGATCTNNNTACCTGTAGTCCCAGCTACTTG

GAGCACACGTCTGAACTCTTTCCCTACACGACGCTCTTCCGATCTNNNTTTGCAGACTCATAGAGATG
AGCACACGTCTGAACTCTTTCCCTACACGACGCTCTTCCGATCTNNNAGACTCAGGCAAGAAATGTCC
GCACACGTCTGAACTCTTTCCCTACACGACGCTCTTCCGATCTNNNCTAGCAGCTTCTCAGATCCCAAT
GCACACGTCTGAACTCTTTCCCTACACGACGCTCTTCCGATCTNNNCTGCAGACTTCAGACTTACCAG
GAGCACACGTCTGAACTCTTTCCCTACACGACGCTCTTCCGATCTNNNTGAGCTCAATGAAGAGGAT
GAGCACACGTCTGAACTCTTTCCCTACACGACGCTCTTCCGATCTNNNTCAGATCCTGGAGATATTGCT
AGCACACGTCTGAACTCTTTCCCTACACGACGCTCTTCCGATCTNNNGTGCTTGTGTCATCTCATCCCT
GCACACGTCTGAACTCTTTCCCTACACGACGCTCTTCCGATCTNNNACAGTTTGAGCTTCTGGAAATG
GAGCACACGTCTGAACTCTTTCCCTACACGACGCTCTTCCGATCTNNNCGTGGCAGTGAAGGATCTAA
GAGCACACGTCTGAACTCTTTCCCTACACGACGCTCTTCCGATCTNNNGGCAGGTGATTGCTATTTCC
GAGCACACGTCTGAACTCTTTCCCTACACGACGCTCTTCCGATCTNNNGCAAATCACAAGGTCAGGAG
GAGCACACGTCTGAACTCTTTCCCTACACGACGCTCTTCCGATCTNNNCTATTCCTGCAGAGAGAACC
GCACACGTCTGAACTCTTTCCCTACACGACGCTCTTCCGATCTNNNCTGGAGTCTGGTCTCATTGGAT
GCACACGTCTGAACTCTTTCCCTACACGACGCTCTTCCGATCTNNNTCACACCATTATACTCCAGCCTG
GCACACGTCTGAACTCTTTCCCTACACGACGCTCTTCCGATCTNNNTTTGACAGCATCTCTTTGTGCAG
GAGCACACGTCTGAACTCTTTCCCTACACGACGCTCTTCCGATCTNNNTGGAAGTTCTTCGATGAGTT
GCACACGTCTGAACTCTTTCCCTACACGACGCTCTTCCGATCTNNNTTTGGTATGTCCTTTTGTGGG
GAGCACACGTCTGAACTCTTTCCCTACACGACGCTCTTCCGATCTNNNAGGCCACTTCACTGTTGTCAT
GAGCACACGTCTGAACTCTTTCCCTACACGACGCTCTTCCGATCTNNNCTCTGTGTCTAGCTCAGGGAT
AGCACACGTCTGAACTCTTTCCCTACACGACGCTCTTCCGATCTNNNAGGCTGGAGTACAATGACACA
AGCACACGTCTGAACTCTTTCCCTACACGACGCTCTTCCGATCTNNNGAGATCACGCCACTGAACTCT
CACACGTCTGAACTCTTTCCCTACACGACGCTCTTCCGATCTNNNAGCTCATCATGGATCTGGAACTC
GCACACGTCTGAACTCTTTCCCTACACGACGCTCTTCCGATCTNNNCTGACTTCCAGGTGAATCCTCT
AGCACACGTCTGAACTCTTTCCCTACACGACGCTCTTCCGATCTNNNACTCTCTCTCTCTCTCTCTCT
GAGCACACGTCTGAACTCTTTCCCTACACGACGCTCTTCCGATCTNNNCACCACACCCAGCTAATTTTG
AGCACACGTCTGAACTCTTTCCCTACACGACGCTCTTCCGATCTNNNCTGGTCTCAAGATATCCTCCAG
GAGCACACGTCTGAACTCTTTCCCTACACGACGCTCTTCCGATCTNNNGTAGAAAGTGGGAAAACTG
AGCACACGTCTGAACTCTTTCCCTACACGACGCTCTTCCGATCTNNNGGACTTCCGATACAAGAGATG
GCACACGTCTGAACTCTTTCCCTACACGACGCTCTTCCGATCTNNNCTGAGGCAGTTGAATTGCTTG
AGCACACGTCTGAACTCTTTCCCTACACGACGCTCTTCCGATCTNNNTGACTTCCCATTTCACTCAGTGT
AGCACACGTCTGAACTCTTTCCCTACACGACGCTCTTCCGATCTNNNCTTCGAGTCCCCTCTGTATTCA
AGCACACGTCTGAACTCTTTCCCTACACGACGCTCTTCCGATCTNNNCTGTTTCTTCTTCCGGTAGTGG
GAGCACACGTCTGAACTCTTTCCCTACACGACGCTCTTCCGATCTNNNTAAAAGAAAAACACAGGCTG
GAGCACACGTCTGAACTCTTTCCCTACACGACGCTCTTCCGATCTNNNTTAGCATTACCCTTTCTGCCCA
AGCACACGTCTGAACTCTTTCCCTACACGACGCTCTTCCGATCTNNNCTCTACCAAAAATTAGCTGGGC
GAGCACACGTCTGAACTCTTTCCCTACACGACGCTCTTCCGATCTNNNTCTTTGGCTTTACCCACTGA
GCACACGTCTGAACTCTTTCCCTACACGACGCTCTTCCGATCTNNNCTCCATCTCTCTCTCTCTCCCC
GCACACGTCTGAACTCTTTCCCTACACGACGCTCTTCCGATCTNNNCCCCTGAATGGTACTTCTAAGG
AGCACACGTCTGAACTCTTTCCCTACACGACGCTCTTCCGATCTNNNTGTTATTTGGAAATGCGGCC
GAGCACACGTCTGAACTCTTTCCCTACACGACGCTCTTCCGATCTNNNACCTGCAGTAAAACCTTGAA
AGCACACGTCTGAACTCTTTCCCTACACGACGCTCTTCCGATCTNNNGCTCTGGTTGTTTGAAAGTGT
GCACACGTCTGAACTCTTTCCCTACACGACGCTCTTCCGATCTNNNGTTAGCCGGGATGATCTCAATCT
GCACACGTCTGAACTCTTTCCCTACACGACGCTCTTCCGATCTNNNACGGCCAAGCATTTATTCAGTT

GAGCACACGTCTGAACTCTTTCCCTACACGACGCTCTTCCGATCTNNNTTTCAGTCTGATCCTGAGGG
GAGCACACGTCTGAACTCTTTCCCTACACGACGCTCTTCCGATCTNNNCCTCAAAACCAAAAACCCACA
AGCACACGTCTGAACTCTTTCCCTACACGACGCTCTTCCGATCTNNNACTGCGAGAGAGAAAGAGA
GCACACGTCTGAACTCTTTCCCTACACGACGCTCTTCCGATCTNNNGGTAGCAAGAGAGGGATTGA
GCACACGTCTGAACTCTTTCCCTACACGACGCTCTTCCGATCTNNNCATCTCTGATGAGGCCTCTCTTCC
GCACACGTCTGAACTCTTTCCCTACACGACGCTCTTCCGATCTNNNGGCAGAGACAGAGAGAGAAA
GAGCACACGTCTGAACTCTTTCCCTACACGACGCTCTTCCGATCTNNNGCAGTCCTCTCCATAGAACT
AGCACACGTCTGAACTCTTTCCCTACACGACGCTCTTCCGATCTNNNGGCCATGTCAACAGTGGAAAT
GAGCACACGTCTGAACTCTTTCCCTACACGACGCTCTTCCGATCTNNNAGAGAACGAGAGAAAGAG
AGAGCACACGTCTGAACTCTTTCCCTACACGACGCTCTTCCGATCTNNNCAACCATGGCATCATTTTCT
AGAGCACACGTCTGAACTCTTTCCCTACACGACGCTCTTCCGATCTNNNCAACCATGGCATCATTTTCT
AGAGCACACGTCTGAACTCTTTCCCTACACGACGCTCTTCCGATCTNNNAGGAATGCCAATCCAGTCT
AGAGCACACGTCTGAACTCTTTCCCTACACGACGCTCTTCCGATCTNNNTCTCTGTGTATGTGTGTGT
AGCACACGTCTGAACTCTTTCCCTACACGACGCTCTTCCGATCTNNNTGCTTTCAACATTTTCTTGCCC
GAGCACACGTCTGAACTCTTTCCCTACACGACGCTCTTCCGATCTNNNCAGCCTGGTCAATATAGCG
GCACACGTCTGAACTCTTTCCCTACACGACGCTCTTCCGATCTNNNAGAGGCATCTCATCGACAAAAG
AGCACACGTCTGAACTCTTTCCCTACACGACGCTCTTCCGATCTNNNCAGAAAACTGTCCCCCAACTTT
AGCACACGTCTGAACTCTTTCCCTACACGACGCTCTTCCGATCTNNNAAAAGAGAATTTGCATGCAG
AGCACACGTCTGAACTCTTTCCCTACACGACGCTCTTCCGATCTNNNAGGGAGAGAGAGAGAGAGA
GCACACGTCTGAACTCTTTCCCTACACGACGCTCTTCCGATCTNNNGCTGGTCCAAGTTATAGAACCC
AGCACACGTCTGAACTCTTTCCCTACACGACGCTCTTCCGATCTNNNAGAGAGAGAGAGAGAGAGG
AGCACACGTCTGAACTCTTTCCCTACACGACGCTCTTCCGATCTNNNCGAGAGAGAGAGAGAGAGA
GCACACGTCTGAACTCTTTCCCTACACGACGCTCTTCCGATCTNNNGTTCCAGCCTGTTTGTGATCTTC
AGCACACGTCTGAACTCTTTCCCTACACGACGCTCTTCCGATCTNNNAAGGACAGGAGAAGATGCGT
AGCACACGTCTGAACTCTTTCCCTACACGACGCTCTTCCGATCTNNNAGAGTGTGTGTGTGTGTGTGT
AGCACACGTCTGAACTCTTTCCCTACACGACGCTCTTCCGATCTNNNTGTGCAAGACATAGTACAGCT
GCACACGTCTGAACTCTTTCCCTACACGACGCTCTTCCGATCTNNNCACAGAACCTGCCAATAAACTG
GCACACGTCTGAACTCTTTCCCTACACGACGCTCTTCCGATCTNNNAGAGCAAACAAGCTGAAAACC
GCACACGTCTGAACTCTTTCCCTACACGACGCTCTTCCGATCTNNNGAGAACTAGCTCTTGTCTGCTC
GCACACGTCTGAACTCTTTCCCTACACGACGCTCTTCCGATCTNNNGTATGAACTGGAGAGCAGGG
GCACACGTCTGAACTCTTTCCCTACACGACGCTCTTCCGATCTNNNAGGAGAGGAGAGAGAGAGAG
AGCACACGTCTGAACTCTTTCCCTACACGACGCTCTTCCGATCTNNNACAATGGCTAGACCTCCATGA
AGAGCACACGTCTGAACTCTTTCCCTACACGACGCTCTTCCGATCTNNNGTAATCTCAGCTACTCTGG
GCACACGTCTGAACTCTTTCCCTACACGACGCTCTTCCGATCTNNNGGTGACAGAGTGAGACTCCAT
AGCACACGTCTGAACTCTTTCCCTACACGACGCTCTTCCGATCTNNNGTGTGCTCAGTGTAAGAGGA
GCACACGTCTGAACTCTTTCCCTACACGACGCTCTTCCGATCTNNNGATAGGGTTCTCAAGAGCTCTG
GAGCACACGTCTGAACTCTTTCCCTACACGACGCTCTTCCGATCTNNNTTTCTCTCTCTTCCTCCCTCCTT
GAGCACACGTCTGAACTCTTTCCCTACACGACGCTCTTCCGATCTNNNTTCAGGATGCAAACACAACC
GCACACGTCTGAACTCTTTCCCTACACGACGCTCTTCCGATCTNNNGTGCTTTAGGCCTCAGAAAACA
GCACACGTCTGAACTCTTTCCCTACACGACGCTCTTCCGATCTNNNTTCCTTCCTTCCTTCCTTCCTTCCA
CACACGTCTGAACTCTTTCCCTACACGACGCTCTTCCGATCTNNNATACTACTCTCCTCTGTCCAGGG
CACACGTCTGAACTCTTTCCCTACACGACGCTCTTCCGATCTNNNAGCAGTGATGGTCTTTCTTTCTTC
GAGCACACGTCTGAACTCTTTCCCTACACGACGCTCTTCCGATCTNNNATTGCAAAGCTGGGTGTTTA

GCACACGTCTGAACTCTTTCCCTACACGACGCTCTTCCGATCTNNNACTGATCAAGAGACGGCAACAT
GAGCACACGTCTGAACTCTTTCCCTACACGACGCTCTTCCGATCTNNNATGAGAGAAGGCTTTCCCA
AGCACACGTCTGAACTCTTTCCCTACACGACGCTCTTCCGATCTNNNATGGAAAGAGAGATGGAGAG
GCACACGTCTGAACTCTTTCCCTACACGACGCTCTTCCGATCTNNNAGAGAGAGAGAGAGAGAGGG
AGCACACGTCTGAACTCTTTCCCTACACGACGCTCTTCCGATCTNNNGAGGTTGCAGTGAGCTGAG
AGCACACGTCTGAACTCTTTCCCTACACGACGCTCTTCCGATCTNNNCCTACCTCAACCTCTCAAGTAG
AGCACACGTCTGAACTCTTTCCCTACACGACGCTCTTCCGATCTNNNTCTCTCTCTCTCTCTCTCTCC
GCACACGTCTGAACTCTTTCCCTACACGACGCTCTTCCGATCTNNNTGGGAGAAGCTGAATAAGATG
CACACGTCTGAACTCTTTCCCTACACGACGCTCTTCCGATCTNNNACTCACTTACTTAGGGCCAATCCC
AGCACACGTCTGAACTCTTTCCCTACACGACGCTCTTCCGATCTNNNAGGTAGCTGAAAGAACACTC
AGCACACGTCTGAACTCTTTCCCTACACGACGCTCTTCCGATCTNNNGGGCTGGAATAGAACAAAAA
GCACACGTCTGAACTCTTTCCCTACACGACGCTCTTCCGATCTNNNTTGTTTCTGTGTGTGTGTGTGT
GAGCACACGTCTGAACTCTTTCCCTACACGACGCTCTTCCGATCTNNNACAAATGTCAAAGAGGTGG
GCACACGTCTGAACTCTTTCCCTACACGACGCTCTTCCGATCTNNNGGATCACGAGGTCAGGAGTTA
AGCACACGTCTGAACTCTTTCCCTACACGACGCTCTTCCGATCTNNNACCTGTTGTTGAACTTCACGTA
GAGCACACGTCTGAACTCTTTCCCTACACGACGCTCTTCCGATCTNNNTCAATACCTGGTGAGGAGT
GAGCACACGTCTGAACTCTTTCCCTACACGACGCTCTTCCGATCTNNNAGGCCAAGACCAGAAAGAA
GAGCACACGTCTGAACTCTTTCCCTACACGACGCTCTTCCGATCTNNNCCAAAGAAGGAAGGAAGG
GCACACGTCTGAACTCTTTCCCTACACGACGCTCTTCCGATCTNNNAGGCAGAGACTCTTGTTCTCTT
AGCACACGTCTGAACTCTTTCCCTACACGACGCTCTTCCGATCTNNNGACCCTCAGCTTCAAGGTATCT
GCACACGTCTGAACTCTTTCCCTACACGACGCTCTTCCGATCTNNNCCTGCTTCAGAGAGATGATGAC
AGCACACGTCTGAACTCTTTCCCTACACGACGCTCTTCCGATCTNNNAAAGGGGAGGGAGAGAGAAA
AGCACACGTCTGAACTCTTTCCCTACACGACGCTCTTCCGATCTNNNATGTGTTTGTGTGTGTGACTG
GCACACGTCTGAACTCTTTCCCTACACGACGCTCTTCCGATCTNNNGAGCAGGACTCTGTCAGAAAG
GCACACGTCTGAACTCTTTCCCTACACGACGCTCTTCCGATCTNNNCTTTCCTTCCCTTCCTTCTCCTCCA
AGCACACGTCTGAACTCTTTCCCTACACGACGCTCTTCCGATCTNNNCCCTAATCTGGTAACACTGACG
AAGAGCACACGTCTGAACTCTTTCCCTACACGACGCTCTTCCGATCTNNNAGTGTTTGCTCAGAATGC
GCACACGTCTGAACTCTTTCCCTACACGACGCTCTTCCGATCTNNNCTCCAGAGAGAGAGAGAGAGA
GCACACGTCTGAACTCTTTCCCTACACGACGCTCTTCCGATCTNNNATGGAGAAGGGTTCGTCTAGA
GCACACGTCTGAACTCTTTCCCTACACGACGCTCTTCCGATCTNNNTGGAAGGTCTGTGTGTCTGAT
AGCACACGTCTGAACTCTTTCCCTACACGACGCTCTTCCGATCTNNNGGTGACAATGGTGAAACTCT
AGCACACGTCTGAACTCTTTCCCTACACGACGCTCTTCCGATCTNNNAGAGGTCTCAGAGGTACCAT
AGCACACGTCTGAACTCTTTCCCTACACGACGCTCTTCCGATCTNNNTACCTCTCAGCCCATAAGACAG
AGAGCACACGTCTGAACTCTTTCCCTACACGACGCTCTTCCGATCTNNNATTTCATGACCAACAACCTT
GAGCACACGTCTGAACTCTTTCCCTACACGACGCTCTTCCGATCTNNNAGACCCAAATACTGTGATGC
GCACACGTCTGAACTCTTTCCCTACACGACGCTCTTCCGATCTNNNATGCGGAAAAAAATCAGTCCCA
AGCACACGTCTGAACTCTTTCCCTACACGACGCTCTTCCGATCTNNNGGATTCAAGACTAGCCTGGAC
AGCACACGTCTGAACTCTTTCCCTACACGACGCTCTTCCGATCTNNNGGGAGGAAGAGGTAAGTCCA
AGCACACGTCTGAACTCTTTCCCTACACGACGCTCTTCCGATCTNNNCTCAGATCTTCCTGCTGGACTT
CACACGTCTGAACTCTTTCCCTACACGACGCTCTTCCGATCTNNNTGTGGATGAATATTTGCTTCCCT
AGCACACGTCTGAACTCTTTCCCTACACGACGCTCTTCCGATCTNNNAATGAGAGAGAGATGACGGG
AGCACACGTCTGAACTCTTTCCCTACACGACGCTCTTCCGATCTNNNCCTGGAGAGTGAACAACAAG
AGCACACGTCTGAACTCTTTCCCTACACGACGCTCTTCCGATCTNNNCATCAGCCCCAGCATTTACATT

GCACACGTCTGAACTCTTTCCCTACACGACGCTCTTCCGATCTNNNGGGACTAGAAACAGGACTCACT
GAGCACACGTCTGAACTCTTTCCCTACACGACGCTCTTCCGATCTNNNTTCTTCTCTGTCTCTGTCTCCG
GAGCACACGTCTGAACTCTTTCCCTACACGACGCTCTTCCGATCTNNNCAGAAAGTGGGGAGAAGGA
AGCACACGTCTGAACTCTTTCCCTACACGACGCTCTTCCGATCTNNNAATGGACAAATTGGGAGGGAG
AGAGCACACGTCTGAACTCTTTCCCTACACGACGCTCTTCCGATCTNNNGGGAAGAGAGAGAGAGAGA
AGAGCACACGTCTGAACTCTTTCCCTACACGACGCTCTTCCGATCTNNNGTGCAGAGTTTTACAAAGG
GCACACGTCTGAACTCTTTCCCTACACGACGCTCTTCCGATCTNNNTCTCTCTCTCCCTCTCTCTCTCAT
AGCACACGTCTGAACTCTTTCCCTACACGACGCTCTTCCGATCTNNNTAGTTATTGCAGGAGTGGGTT
AGCACACGTCTGAACTCTTTCCCTACACGACGCTCTTCCGATCTNNNCACACACACACACACACACATAT
GAGCACACGTCTGAACTCTTTCCCTACACGACGCTCTTCCGATCTNNNGGGATGGAGTGAGAGCTTTT
AGCACACGTCTGAACTCTTTCCCTACACGACGCTCTTCCGATCTNNNTCCAGTGCCTCTAGCTATGATCT
GAGCACACGTCTGAACTCTTTCCCTACACGACGCTCTTCCGATCTNNNGAAGGAAGGAAGGAAGGAA
AGCACACGTCTGAACTCTTTCCCTACACGACGCTCTTCCGATCTNNNCTGAGCAAGACTCTGTCTCAAA
AGCACACGTCTGAACTCTTTCCCTACACGACGCTCTTCCGATCTNNNCATTACTTGGGACCAGGAGTT
GAGCACACGTCTGAACTCTTTCCCTACACGACGCTCTTCCGATCTNNNTGAGATTGCAGTCAGCTGAG
GAGCACACGTCTGAACTCTTTCCCTACACGACGCTCTTCCGATCTNNNAGGATAGGAGTGTGTGTGA
AAGAGCACACGTCTGAACTCTTTCCCTACACGACGCTCTTCCGATCTNNNTCTAACACTCAGTCACTGCT
GAGCACACGTCTGAACTCTTTCCCTACACGACGCTCTTCCGATCTNNNTGGGAAGAGAGAGAGAAGG
AAGAGCACACGTCTGAACTCTTTCCCTACACGACGCTCTTCCGATCTNNNGCGAAACTCCGTCAAGAAA
GAGCACACGTCTGAACTCTTTCCCTACACGACGCTCTTCCGATCTNNNGCAAGGGAGGAGCAAGTCAT
AGCACACGTCTGAACTCTTTCCCTACACGACGCTCTTCCGATCTNNNTACAGTCTATCTCAGGGAAGGC
GCACACGTCTGAACTCTTTCCCTACACGACGCTCTTCCGATCTNNNCTCACACACACACACATACACC
AGCACACGTCTGAACTCTTTCCCTACACGACGCTCTTCCGATCTNNNCACCGTGGATTTTTGTCCTGAA
GAGCACACGTCTGAACTCTTTCCCTACACGACGCTCTTCCGATCTNNNCCACACATGAGAAAAATGCCC
AGCACACGTCTGAACTCTTTCCCTACACGACGCTCTTCCGATCTNNNGTATCACCTTTTCAGTGAGGGCT
AGAGCACACGTCTGAACTCTTTCCCTACACGACGCTCTTCCGATCTNNNAGTACCTAGGGCTACAGAT
AGCACACGTCTGAACTCTTTCCCTACACGACGCTCTTCCGATCTNNNAATTCACTGTTTATGGCATTGC
GCACACGTCTGAACTCTTTCCCTACACGACGCTCTTCCGATCTNNNATGCCTCTTCAGTCCTTCATGTTC
GAGCACACGTCTGAACTCTTTCCCTACACGACGCTCTTCCGATCTNNNTGGTTACAAGCTACTGCAAAT
AGCACACGTCTGAACTCTTTCCCTACACGACGCTCTTCCGATCTNNNTACAATGTGTGTGTGTGTGTGT
AGCACACGTCTGAACTCTTTCCCTACACGACGCTCTTCCGATCTNNNTATTCTGGAGGCTGAAGCACAA
AGCACACGTCTGAACTCTTTCCCTACACGACGCTCTTCCGATCTNNNAATAAAATCTGCCAGTGCCATG
GAGCACACGTCTGAACTCTTTCCCTACACGACGCTCTTCCGATCTNNNAAACAATGGACTACTACACCC
AGCACACGTCTGAACTCTTTCCCTACACGACGCTCTTCCGATCTNNNGCAACTATGATCCCCAGATTCCT
AGCACACGTCTGAACTCTTTCCCTACACGACGCTCTTCCGATCTNNNGAGTCTGTTTATCTGCACTGGG
GAGCACACGTCTGAACTCTTTCCCTACACGACGCTCTTCCGATCTNNNTGTACTGCTTTGATGGTCTTG
GAGCACACGTCTGAACTCTTTCCCTACACGACGCTCTTCCGATCTNNNCCCCACAAAATAAGAAGCAAG
GCACACGTCTGAACTCTTTCCCTACACGACGCTCTTCCGATCTNNNAGTTTTAGCTACCTTGCTCTGCT
GAGCACACGTCTGAACTCTTTCCCTACACGACGCTCTTCCGATCTNNNGAGATCACACCACTGCACTCT
AGCACACGTCTGAACTCTTTCCCTACACGACGCTCTTCCGATCTNNNCTGATCTGGTCTGCTCTTTCCAT
GAGCACACGTCTGAACTCTTTCCCTACACGACGCTCTTCCGATCTNNNCTTCGATGTAGAACACACACA
GAGCACACGTCTGAACTCTTTCCCTACACGACGCTCTTCCGATCTNNNTGTAGGGGAGGTAGTCAGTC
AGCACACGTCTGAACTCTTTCCCTACACGACGCTCTTCCGATCTNNNTCTTCTTTCCTCCCCCTCCTATTC

AGCACACGTCTGAACTCTTTCCCTACACGACGCTCTTCCGATCTNNNTCCTTCCTTCCTTCCTTCCTTTCCA
GAGCACACGTCTGAACTCTTTCCCTACACGACGCTCTTCCGATCTNNNCTTGAGAGTGAATGTCTGGCT
GAGCACACGTCTGAACTCTTTCCCTACACGACGCTCTTCCGATCTNNNGGGATTGGAACAGGGATCTTA
GAGCACACGTCTGAACTCTTTCCCTACACGACGCTCTTCCGATCTNNNAAGCTGACCCTGATGGATTCA
GAGCACACGTCTGAACTCTTTCCCTACACGACGCTCTTCCGATCTNNNAGCAAGCTCTGAGGTGCTAAA
AGCACACGTCTGAACTCTTTCCCTACACGACGCTCTTCCGATCTNNNCTGAGAAAAGACACCAGAGAG
AGCACACGTCTGAACTCTTTCCCTACACGACGCTCTTCCGATCTNNNCATTGTGAATGTGGGTAATGT
GAGCACACGTCTGAACTCTTTCCCTACACGACGCTCTTCCGATCTNNNTCTCTTTCACACACACACACA
GAGCACACGTCTGAACTCTTTCCCTACACGACGCTCTTCCGATCTNNNACTGCAAAAACCACAACTTTGC
GAGCACACGTCTGAACTCTTTCCCTACACGACGCTCTTCCGATCTNNNCTCCACGCTGTCTTTCTAAACAC
GAGCACACGTCTGAACTCTTTCCCTACACGACGCTCTTCCGATCTNNNCAGACCACTGGACCAAAACTAC
GAGCACACGTCTGAACTCTTTCCCTACACGACGCTCTTCCGATCTNNNTGTTCACAGAATTGATGCTTG
AGAGCACACGTCTGAACTCTTTCCCTACACGACGCTCTTCCGATCTNNNCCTAGGTGAGTCTGCTCTTCT
GAGCACACGTCTGAACTCTTTCCCTACACGACGCTCTTCCGATCTNNNTGGGATGATAGTTTCACCATGC
AGAGCACACGTCTGAACTCTTTCCCTACACGACGCTCTTCCGATCTNNNGTGATGGAGTGAGACTCTCT
AGCACACGTCTGAACTCTTTCCCTACACGACGCTCTTCCGATCTNNNATTTGCAGTTGGTCACCTTCTTG
AGCACACGTCTGAACTCTTTCCCTACACGACGCTCTTCCGATCTNNNCTTTTGCTGGATCTGCTCTTCCC
AGCACACGTCTGAACTCTTTCCCTACACGACGCTCTTCCGATCTNNNGTGGGTTTCCCTTTTTCCTTGTTC
AGCACACGTCTGAACTCTTTCCCTACACGACGCTCTTCCGATCTNNNCTCTTTGTGTGTGTGTGTGTG
AGCACACGTCTGAACTCTTTCCCTACACGACGCTCTTCCGATCTNNNCTGTAATACCAGCTACTCGGGA
GCACACGTCTGAACTCTTTCCCTACACGACGCTCTTCCGATCTNNNCCTGGTACTGACAACTAAGCCAAC
GAGCACACGTCTGAACTCTTTCCCTACACGACGCTCTTCCGATCTNNNAAAAACACTTGTTCCCTTGCCT
GAGCACACGTCTGAACTCTTTCCCTACACGACGCTCTTCCGATCTNNNAGAGAGTGAGATGGTCTTAG
GAGCACACGTCTGAACTCTTTCCCTACACGACGCTCTTCCGATCTNNNTAGAGCCTTGCTTTTGTTCTCC
AGCACACGTCTGAACTCTTTCCCTACACGACGCTCTTCCGATCTNNNGGGAGGCAGAAGTAGAATAGA
AGCACACGTCTGAACTCTTTCCCTACACGACGCTCTTCCGATCTNNNTACCTGAGAAACACACACACACA
AGCACACGTCTGAACTCTTTCCCTACACGACGCTCTTCCGATCTNNNTCTCAAGTACACACACACACACA
AGCACACGTCTGAACTCTTTCCCTACACGACGCTCTTCCGATCTNNNGAGAAAGAGAGATGGGAGAG
AGCACACGTCTGAACTCTTTCCCTACACGACGCTCTTCCGATCTNNNTTCATAGTTCATGAGCCACACAC
AGCACACGTCTGAACTCTTTCCCTACACGACGCTCTTCCGATCTNNNTTCCAAGTAGCTGGGATTACAG
GAGCACACGTCTGAACTCTTTCCCTACACGACGCTCTTCCGATCTNNNCTGTGTGTGTATGTGTGTGTG
AGCACACGTCTGAACTCTTTCCCTACACGACGCTCTTCCGATCTNNNTTCTGTACCTCTCAGAACTCCCTC
AGAGCACACGTCTGAACTCTTTCCCTACACGACGCTCTTCCGATCTNNNAGACAAGGCTTCATTGGAGC
AGCACACGTCTGAACTCTTTCCCTACACGACGCTCTTCCGATCTNNNGTGTGTGTGTGTGTGTATG
AGAGCACACGTCTGAACTCTTTCCCTACACGACGCTCTTCCGATCTNNNTTGAGTGTATTGTGAGAGGC
AGCACACGTCTGAACTCTTTCCCTACACGACGCTCTTCCGATCTNNNGAATCCTGGATTCCTTTTGCTGC
GAGCACACGTCTGAACTCTTTCCCTACACGACGCTCTTCCGATCTNNNTTTTGCTTTTCTCACAAGCTCC
GAGCACACGTCTGAACTCTTTCCCTACACGACGCTCTTCCGATCTNNNTTTTCATGGTGGCTTTCCAAA
GAGCACACGTCTGAACTCTTTCCCTACACGACGCTCTTCCGATCTNNNGTTCACCCCTTGCTAGTAGGAA
AGCACACGTCTGAACTCTTTCCCTACACGACGCTCTTCCGATCTNNNTCTTTTTCGCACTTCTTTGCACTCA
GCACACGTCTGAACTCTTTCCCTACACGACGCTCTTCCGATCTNNNTTCAAGGTGAAGAAGGGAGACA
GAGCACACGTCTGAACTCTTTCCCTACACGACGCTCTTCCGATCTNNNGTGCTTCCAAATCGAGAATGC
AGCACACGTCTGAACTCTTTCCCTACACGACGCTCTTCCGATCTNNNCATCCTGAAATGCCCTTTGGAA

AGCACACGTCTGAACTCTTTCCCTACACGACGCTCTTCCGATCTNNNGCCACCATGCCTAGCTAATTTT
AGCACACGTCTGAACTCTTTCCCTACACGACGCTCTTCCGATCTNNNCGGGTACTATTACGCCCCATAT
GCACACGTCTGAACTCTTTCCCTACACGACGCTCTTCCGATCTNNNTGCCAAGAGGAAAAGAAAAAA
AGCACACGTCTGAACTCTTTCCCTACACGACGCTCTTCCGATCTNNNTCACCCCTTTTCTCCCTTCAAAA
GAGCACACGTCTGAACTCTTTCCCTACACGACGCTCTTCCGATCTNNNGGCCTTTTATACTGAAGATG
AGCACACGTCTGAACTCTTTCCCTACACGACGCTCTTCCGATCTNNNTGGTTACACCCTCTGAAGGAT
AGCACACGTCTGAACTCTTTCCCTACACGACGCTCTTCCGATCTNNNCGGCCGTTACGTTATTTTCTGTT
AGCACACGTCTGAACTCTTTCCCTACACGACGCTCTTCCGATCTNNNGAACTTGAGTTCCTGCAAACCT
AGCACACGTCTGAACTCTTTCCCTACACGACGCTCTTCCGATCTNNNCACCACGCTGGCTATTGTATTT
AGCACACGTCTGAACTCTTTCCCTACACGACGCTCTTCCGATCTNNNCTGCATCATTCAGCTCTGTGAA
GCACACGTCTGAACTCTTTCCCTACACGACGCTCTTCCGATCTNNNGCTCCCAACTAAAAGGTCTGAG
AGCACACGTCTGAACTCTTTCCCTACACGACGCTCTTCCGATCTNNNATGTGGTGGCTCATGTCTGTA
AGCACACGTCTGAACTCTTTCCCTACACGACGCTCTTCCGATCTNNNCATTGCGTCTCTCTCTCTCTCC
CACACGTCTGAACTCTTTCCCTACACGACGCTCTTCCGATCTNNNGCTCTCTGTGTCAGTCTAGAACCC
AGCACACGTCTGAACTCTTTCCCTACACGACGCTCTTCCGATCTNNNCCAATTTACGACGTCACTGAA
AGCACACGTCTGAACTCTTTCCCTACACGACGCTCTTCCGATCTNNNCCAGCCATATAGAGATCACCT
GCACACGTCTGAACTCTTTCCCTACACGACGCTCTTCCGATCTNNNTCTCTCTCTTTTGTGTGTTCGGA
AGCACACGTCTGAACTCTTTCCCTACACGACGCTCTTCCGATCTNNNAACGTGTCTGATTTGCGAAAC
AGCACACGTCTGAACTCTTTCCCTACACGACGCTCTTCCGATCTNNNGAGAAGGAAGGAAGTCACG
GCACACGTCTGAACTCTTTCCCTACACGACGCTCTTCCGATCTNNNGACTCCTCAAAAACACACACACA
AGCACACGTCTGAACTCTTTCCCTACACGACGCTCTTCCGATCTNNNAGGAGATGCTGTAATGGACT
AGCACACGTCTGAACTCTTTCCCTACACGACGCTCTTCCGATCTNNNATAAACTTTCCATGCTGGGTT
AGCACACGTCTGAACTCTTTCCCTACACGACGCTCTTCCGATCTNNNAGCATTCCATCACCACCTAGAT
GAGCACACGTCTGAACTCTTTCCCTACACGACGCTCTTCCGATCTNNNGAACCACAGACGTTCCCATCT
AGCACACGTCTGAACTCTTTCCCTACACGACGCTCTTCCGATCTNNNCACCCCTCCTCACCTAACCTATA
AGCACACGTCTGAACTCTTTCCCTACACGACGCTCTTCCGATCTNNNTTTGGTGCCTAGTTGATTTGC
AGCACACGTCTGAACTCTTTCCCTACACGACGCTCTTCCGATCTNNNTTTTTTTCTTCACTACGCGACCC
GAGCACACGTCTGAACTCTTTCCCTACACGACGCTCTTCCGATCTNNNGCCCTAAAGAATCCTAGCTTG
GAGCACACGTCTGAACTCTTTCCCTACACGACGCTCTTCCGATCTNNNCAACTTCGGACTACACTTCCC
AGCACACGTCTGAACTCTTTCCCTACACGACGCTCTTCCGATCTNNNTCCCTCATTCTCTCCCATTCTCT
AGCACACGTCTGAACTCTTTCCCTACACGACGCTCTTCCGATCTNNNGGAGCTCTGTTGTTGTAGTGG
AGCACACGTCTGAACTCTTTCCCTACACGACGCTCTTCCGATCTNNNGAACTGAGATCACACCATTGC
AGCACACGTCTGAACTCTTTCCCTACACGACGCTCTTCCGATCTNNNATACATGACGTCCAGAGTTGC
AGCACACGTCTGAACTCTTTCCCTACACGACGCTCTTCCGATCTNNNCTTCCTTCCTTCCTTCCTTCCTC
GAGCACACGTCTGAACTCTTTCCCTACACGACGCTCTTCCGATCTNNNCCTCAAAACCAAAAACCCACA
GAGCACACGTCTGAACTCTTTCCCTACACGACGCTCTTCCGATCTNNNCAAGCTAGAGAACCACAAAA
GCACACGTCTGAACTCTTTCCCTACACGACGCTCTTCCGATCTNNNAGCCACTCATGATCTACAACACA
AGCACACGTCTGAACTCTTTCCCTACACGACGCTCTTCCGATCTNNNTAGAGAGAGAGAGAGAGAGAC
GAGCACACGTCTGAACTCTTTCCCTACACGACGCTCTTCCGATCTNNNAGAGAGAGAGAGAGAGAGAG
GCACACGTCTGAACTCTTTCCCTACACGACGCTCTTCCGATCTNNNGGGAGTGTGTGTGTGTGTTTAT
AGCACACGTCTGAACTCTTTCCCTACACGACGCTCTTCCGATCTNNNACAAGGAGCAGACTGGAGAA
CACACGTCTGAACTCTTTCCCTACACGACGCTCTTCCGATCTNNNGCAGAGAAAGCACAGTCAATGA
CACACGTCTGAACTCTTTCCCTACACGACGCTCTTCCGATCTNNNGCAATATGGATAAGTGCTGGCA

AGCACACGTCTGAACTCTTTCCCTACACGACGCTCTTCCGATCTNNNACTTTTGTGTGGGAGCTATAGT

GCACACGTCTGAACTCTTTCCCTACACGACGCTCTTCCGATCTNNNAAATCAATATCTGGGTGCTCGG

AGAGCACACGTCTGAACTCTTTCCCTACACGACGCTCTTCCGATCTNNNGAGAGTGAACAATTGGGT

AGCACACGTCTGAACTCTTTCCCTACACGACGCTCTTCCGATCTNNNGCTCAAAAAAGTCAGTCCTGC

AGCACACGTCTGAACTCTTTCCCTACACGACGCTCTTCCGATCTNNNTTCAACACACACAAACACACAC

AGAGCACACGTCTGAACTCTTTCCCTACACGACGCTCTTCCGATCTNNNAGCGAGACCCTGCCTAAAAA

AGCACACGTCTGAACTCTTTCCCTACACGACGCTCTTCCGATCTNNNTACACACAAACACACACACACA

GCACACGTCTGAACTCTTTCCCTACACGACGCTCTTCCGATCTNNNTCCTCCTGTGTGAGAGAGAAGA

GAGCACACGTCTGAACTCTTTCCCTACACGACGCTCTTCCGATCTNNNACTCTGACACAGTGACTGATG

GCACACGTCTGAACTCTTTCCCTACACGACGCTCTTCCGATCTNNNTCTCTCTCTCTCCCTCTCTCTCAT

AGCACACGTCTGAACTCTTTCCCTACACGACGCTCTTCCGATCTNNNTCTCTCTCTCTCACTCACTCAC

AGCACACGTCTGAACTCTTTCCCTACACGACGCTCTTCCGATCTNNNGTCTCTCTCTCCCTCTCTCTCTC

GCACACGTCTGAACTCTTTCCCTACACGACGCTCTTCCGATCTNNNACGAAGGCCTTTCCTCCAATAAA

AGCACACGTCTGAACTCTTTCCCTACACGACGCTCTTCCGATCTNNNTTTCTCATCAGTCTCCACACTGG

AGCACACGTCTGAACTCTTTCCCTACACGACGCTCTTCCGATCTNNNGAGCAAGACTCCACCTCAAAAA

AGCACACGTCTGAACTCTTTCCCTACACGACGCTCTTCCGATCTNNNATCATGGGAGTAGGTTCCTGGT

AGCACACGTCTGAACTCTTTCCCTACACGACGCTCTTCCGATCTNNNTACCTGAGCTTATTCACTTCCCG

GAGCACACGTCTGAACTCTTTCCCTACACGACGCTCTTCCGATCTNNNAAAGAAAACTGTACTCCCCGG

GCACACGTCTGAACTCTTTCCCTACACGACGCTCTTCCGATCTNNNCACACCTGTAATCCCAGCTACTCC

AGCACACGTCTGAACTCTTTCCCTACACGACGCTCTTCCGATCTNNNAGCTATTCAGGAAGATGAGGC

AGCACACGTCTGAACTCTTTCCCTACACGACGCTCTTCCGATCTNNNTTTATCTGGAAGTTGGAGTGG

AGCACACGTCTGAACTCTTTCCCTACACGACGCTCTTCCGATCTNNNACTTTAGTTTGGGGAGAGAGA

AGAGCACACGTCTGAACTCTTTCCCTACACGACGCTCTTCCGATCTNNNTACTCTTTGTCCTCCATCTG

GAGCACACGTCTGAACTCTTTCCCTACACGACGCTCTTCCGATCTNNNAGTGAGCTGAGATTGTACCA

AGCACACGTCTGAACTCTTTCCCTACACGACGCTCTTCCGATCTNNNCTTGTAGCCACGTCTACCTGAA

GAGCACACGTCTGAACTCTTTCCCTACACGACGCTCTTCCGATCTNNNAGATGACACAGCAAGACTCTG

GAGCACACGTCTGAACTCTTTCCCTACACGACGCTCTTCCGATCTNNNTGACACAGCAAGACTCTATCA

GAGCACACGTCTGAACTCTTTCCCTACACGACGCTCTTCCGATCTNNNGATCATACCACTACACTCCAG

AGCACACGTCTGAACTCTTTCCCTACACGACGCTCTTCCGATCTNNNACCCTTTACTTACATTCCCCAAA

GCACACGTCTGAACTCTTTCCCTACACGACGCTCTTCCGATCTNNNTGAGGCAAAAGGAACACTCAAA

AGAGCACACGTCTGAACTCTTTCCCTACACGACGCTCTTCCGATCTNNNAGTGAGCCGAGATCTTACC

AGCACACGTCTGAACTCTTTCCCTACACGACGCTCTTCCGATCTNNNGGTGGCTCAGAAGAGAAAGAG

GAGCACACGTCTGAACTCTTTCCCTACACGACGCTCTTCCGATCTNNNTAGCTGATGAAGTTGTAGGA

AAGAGCACACGTCTGAACTCTTTCCCTACACGACGCTCTTCCGATCTNNNCTGAGGTGGGAGGATCA

GAGCACACGTCTGAACTCTTTCCCTACACGACGCTCTTCCGATCTNNNTTGGGGGAGGAAGAAGAAT

AGCACACGTCTGAACTCTTTCCCTACACGACGCTCTTCCGATCTNNNAGCCTTGGACTAAAACTATGCC

AGCACACGTCTGAACTCTTTCCCTACACGACGCTCTTCCGATCTNNNGTTTGTTTCATTGCTTCCTTGCC

AGCACACGTCTGAACTCTTTCCCTACACGACGCTCTTCCGATCTNNNGGTTTCAGTGTGCCTAGATCTC

AGCACACGTCTGAACTCTTTCCCTACACGACGCTCTTCCGATCTNNNTACTGCACCATTGTACTCCAGT

AGCACACGTCTGAACTCTTTCCCTACACGACGCTCTTCCGATCTNNNAGAGAGAGAGAGAGAGAGAGA

AGAGCACACGTCTGAACTCTTTCCCTACACGACGCTCTTCCGATCTNNNCTTGTTCCTGAGCAAAGGTG

AGCACACGTCTGAACTCTTTCCCTACACGACGCTCTTCCGATCTNNNTGATCTCTAAGGCATCTTCCAG

GAGCACACGTCTGAACTCTTTCCCTACACGACGCTCTTCCGATCTNNNTGGGTGACAGAATGAGATCC

GCACACGTCTGAACTCTTTCCCTACACGACGCTCTTCCGATCTNNNGTTTGCTACCATGTTGGCTTTGA
AGCACACGTCTGAACTCTTTCCCTACACGACGCTCTTCCGATCTNNNCTCGGTGATTCTTGCAGACTAG
CACACGTCTGAACTCTTTCCCTACACGACGCTCTTCCGATCTNNNTGAGGCTGTCATAATGGGATCAG
GCACACGTCTGAACTCTTTCCCTACACGACGCTCTTCCGATCTNNNCTCTCTCTCTCACACACACACC
AGAGCACACGTCTGAACTCTTTCCCTACACGACGCTCTTCCGATCTNNNTCTCAAATTCCTGGACTCAAG
GAGCACACGTCTGAACTCTTTCCCTACACGACGCTCTTCCGATCTNNNTGGGTCAGATTTTCATCCCAG
GCACACGTCTGAACTCTTTCCCTACACGACGCTCTTCCGATCTNNNTGGATTAGCCATTTCCAAGGTGA
GAGCACACGTCTGAACTCTTTCCCTACACGACGCTCTTCCGATCTNNNAAATCGTTGTGTGTGTGTGT
AGCACACGTCTGAACTCTTTCCCTACACGACGCTCTTCCGATCTNNNAGAGGTTGCAGTGAGTTGAGA
AGCACACGTCTGAACTCTTTCCCTACACGACGCTCTTCCGATCTNNNGTCTGCTCCCTCTGTCTCTTTTAC
GCACACGTCTGAACTCTTTCCCTACACGACGCTCTTCCGATCTNNNGGGTTTATGGGAGGAAGTCAG
GCACACGTCTGAACTCTTTCCCTACACGACGCTCTTCCGATCTNNNAATGCTATTCTCAACACCACCCTC
GCACACGTCTGAACTCTTTCCCTACACGACGCTCTTCCGATCTNNNATATCTCTCTCTCTCCCCATGCCC
AGCACACGTCTGAACTCTTTCCCTACACGACGCTCTTCCGATCTNNNGTGGAACGTGAAGGAGAAAC
AGCACACGTCTGAACTCTTTCCCTACACGACGCTCTTCCGATCTNNNATGATCACACACCACTGTACTC
AAGAGCACACGTCTGAACTCTTTCCCTACACGACGCTCTTCCGATCTNNNATTGCTTGAGCCTAGGAGT
GAGCACACGTCTGAACTCTTTCCCTACACGACGCTCTTCCGATCTNNNACCCTCATCTCTGTCTCTATGG
GCACACGTCTGAACTCTTTCCCTACACGACGCTCTTCCGATCTNNNAGTACACCTTCTCCTTCTGTCCTC
AGCACACGTCTGAACTCTTTCCCTACACGACGCTCTTCCGATCTNNNTATGGTGGTGCTACAAGAAGT
AGCACACGTCTGAACTCTTTCCCTACACGACGCTCTTCCGATCTNNNCCAGGAGATTCCAACTGCTACA
AGAGCACACGTCTGAACTCTTTCCCTACACGACGCTCTTCCGATCTNNNACCCTTGTCCTGCAATGAGT
AGAGCACACGTCTGAACTCTTTCCCTACACGACGCTCTTCCGATCTNNNGCGATTCCTCAGCTCTCAAT
AGCACACGTCTGAACTCTTTCCCTACACGACGCTCTTCCGATCTNNNGAGAGAGAGAGAGAGATGTG
AGAGCACACGTCTGAACTCTTTCCCTACACGACGCTCTTCCGATCTNNNGGAGAAGGGAGTGAGAAA
AGCACACGTCTGAACTCTTTCCCTACACGACGCTCTTCCGATCTNNNACCAAAATTGCCTAAAGCTGGA
AGCACACGTCTGAACTCTTTCCCTACACGACGCTCTTCCGATCTNNNACCCAGTGACAGAGATAGACA
GCACACGTCTGAACTCTTTCCCTACACGACGCTCTTCCGATCTNNNGATACGTGTGTTTGTGTGTGTG
GCACACGTCTGAACTCTTTCCCTACACGACGCTCTTCCGATCTNNNTTCCTCCAATGCTGTTCTCATGAC
AGCACACGTCTGAACTCTTTCCCTACACGACGCTCTTCCGATCTNNNGCGAAAAAGCAAGTCTCCATC
AGCACACGTCTGAACTCTTTCCCTACACGACGCTCTTCCGATCTNNNTGACAGGAAAAGTCCCAGATCA
AAGAGCACACGTCTGAACTCTTTCCCTACACGACGCTCTTCCGATCTNNNTATTGCAGCATAACCTAGC
AGCACACGTCTGAACTCTTTCCCTACACGACGCTCTTCCGATCTNNNTCCTCCTTCCCTCTTTCTTTCTTC
AGCACACGTCTGAACTCTTTCCCTACACGACGCTCTTCCGATCTNNNATAATCTGGGTCTTTGGGGCT
AGAGCACACGTCTGAACTCTTTCCCTACACGACGCTCTTCCGATCTNNNTCCCTCTCTCTCTCTCTCTC
GAGCACACGTCTGAACTCTTTCCCTACACGACGCTCTTCCGATCTNNNCTGTGTTAGGGTTCTTCAGG
GAGCACACGTCTGAACTCTTTCCCTACACGACGCTCTTCCGATCTNNNCCTTTCTGTGTAAGAGTGGG
AGCACACGTCTGAACTCTTTCCCTACACGACGCTCTTCCGATCTNNNTCCTTCCTTCCTTCTTTCCTTCCC
GAGCACACGTCTGAACTCTTTCCCTACACGACGCTCTTCCGATCTNNNTCTCAGCCTTCAAAACTGTGA
AGAGCACACGTCTGAACTCTTTCCCTACACGACGCTCTTCCGATCTNNNTAGTGGGAAGTGAAAATTG
GAGCACACGTCTGAACTCTTTCCCTACACGACGCTCTTCCGATCTNNNGATGCCTCAATCTGCCTTCTTT
AGCACACGTCTGAACTCTTTCCCTACACGACGCTCTTCCGATCTNNNGAGAGAGAGAGAGAGAGAGA
AGAGCACACGTCTGAACTCTTTCCCTACACGACGCTCTTCCGATCTNNNGAAGAGACAGAGCAAACAG
AAGAGCACACGTCTGAACTCTTTCCCTACACGACGCTCTTCCGATCTNNNGAGAATTGCTTGAATCCGG

AGCACACGTCTGAACTCTTTCCCTACACGACGCTCTTCCGATCTNNNGCCGATAATCTGTGCAGAAACA
GAGCACACGTCTGAACTCTTTCCCTACACGACGCTCTTCCGATCTNNNATCTCAGAGAGATAGGGAGC
AGCACACGTCTGAACTCTTTCCCTACACGACGCTCTTCCGATCTNNNTATATCGGACACACACACACACA
AGCACACGTCTGAACTCTTTCCCTACACGACGCTCTTCCGATCTNNNTTTCTTACAGTTCTGGAGGCTGC
GAGCACACGTCTGAACTCTTTCCCTACACGACGCTCTTCCGATCTNNNAACCTTGACTTGCCTCCTTCTT
GAGCACACGTCTGAACTCTTTCCCTACACGACGCTCTTCCGATCTNNNTCTGTCTGTGCAGGAACTAGA
AGCACACGTCTGAACTCTTTCCCTACACGACGCTCTTCCGATCTNNNGAACCTGAGATGTGAAGTTGCA
GAGCACACGTCTGAACTCTTTCCCTACACGACGCTCTTCCGATCTNNNACCCTAACACAGATCTACCAGC
AGCACACGTCTGAACTCTTTCCCTACACGACGCTCTTCCGATCTNNNGGTGGCTCATAACAGAGAGAG
GAGCACACGTCTGAACTCTTTCCCTACACGACGCTCTTCCGATCTNNNCAGAGAGGGAGAGAGAAAGC
GAGCACACGTCTGAACTCTTTCCCTACACGACGCTCTTCCGATCTNNNCTACAGGGTTGCTTGTGTCAC
AGCACACGTCTGAACTCTTTCCCTACACGACGCTCTTCCGATCTNNNCAGGTCATCGTCAAACATCTCCT
GAGCACACGTCTGAACTCTTTCCCTACACGACGCTCTTCCGATCTNNNACAGGCTGCTATAATTGCCCAT
AGAGCACACGTCTGAACTCTTTCCCTACACGACGCTCTTCCGATCTNNNCTGAGAGGGAGAGAGAGAG
AGAGCACACGTCTGAACTCTTTCCCTACACGACGCTCTTCCGATCTNNNAGAGGCTTGTTTCTTCCTAGC
AGCACACGTCTGAACTCTTTCCCTACACGACGCTCTTCCGATCTNNNGATGGAGTGCTGGAGATAGCT
AGCACACGTCTGAACTCTTTCCCTACACGACGCTCTTCCGATCTNNNCCTCCCTCCCTCTTTTCTTTTCTCA
GAGCACACGTCTGAACTCTTTCCCTACACGACGCTCTTCCGATCTNNNGAGAAACAGGGCTGGGTATA
AGCACACGTCTGAACTCTTTCCCTACACGACGCTCTTCCGATCTNNNCATTTGACTCTGCATTCCCACTGC
CACACGTCTGAACTCTTTCCCTACACGACGCTCTTCCGATCTNNNTGTGCATTTGATGTTTCTCTGGGCA
AGCACACGTCTGAACTCTTTCCCTACACGACGCTCTTCCGATCTNNNACACACTCTCTCTCTCTCTCTCCCA
GAGCACACGTCTGAACTCTTTCCCTACACGACGCTCTTCCGATCTNNNCCTGGGCAATAAGAGGGAAAG
AGCACACGTCTGAACTCTTTCCCTACACGACGCTCTTCCGATCTNNNACAGTGCTGGGATTACAGTTGT
AGCACACGTCTGAACTCTTTCCCTACACGACGCTCTTCCGATCTNNNGTCTCGAATTCCTGGACTCAAG
AGCACACGTCTGAACTCTTTCCCTACACGACGCTCTTCCGATCTNNNAGGGGCTTACCATCTCAAAGAA
GAGCACACGTCTGAACTCTTTCCCTACACGACGCTCTTCCGATCTNNNGAGGTTGCAGTGAGCCAATAT
AGAGCACACGTCTGAACTCTTTCCCTACACGACGCTCTTCCGATCTNNNATATTGGCCAGGTTGGTCTT
AGCACACGTCTGAACTCTTTCCCTACACGACGCTCTTCCGATCTNNNCATATGTGTGTGTGTGTGTGTG
GAGCACACGTCTGAACTCTTTCCCTACACGACGCTCTTCCGATCTNNNACCCATTCATCCTCATTCTTCCC
AGCACACGTCTGAACTCTTTCCCTACACGACGCTCTTCCGATCTNNNACTGCACCTTTTTTCCTCTAGGA
AAGAGCACACGTCTGAACTCTTTCCCTACACGACGCTCTTCCGATCTNNNCAGCCTGGGAAAGAAGAG
GCACACGTCTGAACTCTTTCCCTACACGACGCTCTTCCGATCTNNNGTGGCTGTTTTTCCCCTTTAGTCCA
AGCACACGTCTGAACTCTTTCCCTACACGACGCTCTTCCGATCTNNNACAAGCAAGCTTTTCAAGTCGTT
GAGCACACGTCTGAACTCTTTCCCTACACGACGCTCTTCCGATCTNNNGTCCGACATTTGGTGTATGCA
AGCACACGTCTGAACTCTTTCCCTACACGACGCTCTTCCGATCTNNNTCTTAGAAATCCCCAACACAGCA
GAGCACACGTCTGAACTCTTTCCCTACACGACGCTCTTCCGATCTNNNGCAACAGAATGAAGCAGAGG
GCACACGTCTGAACTCTTTCCCTACACGACGCTCTTCCGATCTNNNAAAGAACCTGGAGTCTGATGTCC
GAGCACACGTCTGAACTCTTTCCCTACACGACGCTCTTCCGATCTNNNGTGACCTATTGGGAAGCCCTTT
GAGCACACGTCTGAACTCTTTCCCTACACGACGCTCTTCCGATCTNNNAAGTTATGCTGGTGCTTCAAAG
AGCACACGTCTGAACTCTTTCCCTACACGACGCTCTTCCGATCTNNNAAATCACCTAAGCCTGGGAAGT
GAGCACACGTCTGAACTCTTTCCCTACACGACGCTCTTCCGATCTNNNAAGGTTGAGTAGAGGTGCTT
GAGCACACGTCTGAACTCTTTCCCTACACGACGCTCTTCCGATCTNNNCCCCTGATACTGCACCTCTAAA
AGCACACGTCTGAACTCTTTCCCTACACGACGCTCTTCCGATCTNNNCCATGTCCTAAATCCCGGAATCT

AGCACACGTCTGAACTCTTTCCCTACACGACGCTCTTCCGATCTNNNTGATGTGAGGCTTCTGTCAACA
AGCACACGTCTGAACTCTTTCCCTACACGACGCTCTTCCGATCTNNNGGCATCTGAGGAGAAGAGCA
AGCACACGTCTGAACTCTTTCCCTACACGACGCTCTTCCGATCTNNNAAGCAGATGCAGTAAGAAGGT
GAGCACACGTCTGAACTCTTTCCCTACACGACGCTCTTCCGATCTNNNGAGACTAGAGGTGTGATTTT
AGCACACGTCTGAACTCTTTCCCTACACGACGCTCTTCCGATCTNNNTAGGGCTGGGGTCCTTATAAG
AGCACACGTCTGAACTCTTTCCCTACACGACGCTCTTCCGATCTNNNCGGGAGTGGTTGTTAGAAGAA
AGCACACGTCTGAACTCTTTCCCTACACGACGCTCTTCCGATCTNNNGTACCTGGAACTCTGAGAAAG
AGCACACGTCTGAACTCTTTCCCTACACGACGCTCTTCCGATCTNNNTGGTTGGTGAGAGTCTCCTAT
AGCACACGTCTGAACTCTTTCCCTACACGACGCTCTTCCGATCTNNNGGATGCCAGCAATCACTTTCTTT
AGCACACGTCTGAACTCTTTCCCTACACGACGCTCTTCCGATCTNNNAGTTGGAAAGTAGACAGGGCT
GCACACGTCTGAACTCTTTCCCTACACGACGCTCTTCCGATCTNNNCCCACTCCCCATGTAAATACCAA
GCACACGTCTGAACTCTTTCCCTACACGACGCTCTTCCGATCTNNNAGTCCAATTCCCTTCCCTTTGAA
AGCACACGTCTGAACTCTTTCCCTACACGACGCTCTTCCGATCTNNNAGGCTTTTTCTCAAAGTGCCAT
AGCACACGTCTGAACTCTTTCCCTACACGACGCTCTTCCGATCTNNNTACTTTGAGAGAGAGACGGCA
AGCACACGTCTGAACTCTTTCCCTACACGACGCTCTTCCGATCTNNNAGACATGTTACACGTCATGGA
GCACACGTCTGAACTCTTTCCCTACACGACGCTCTTCCGATCTNNNTCACCTGGGTTATGCTTGCAATA
CACACGTCTGAACTCTTTCCCTACACGACGCTCTTCCGATCTNNNAGCAAGTGACTTCATGAGAAATA
GAGCACACGTCTGAACTCTTTCCCTACACGACGCTCTTCCGATCTNNNAAGACAAACACACACACAC
GCACACGTCTGAACTCTTTCCCTACACGACGCTCTTCCGATCTNNNCCTCTATCTTGTTTCCACCACCAC
GAGCACACGTCTGAACTCTTTCCCTACACGACGCTCTTCCGATCTNNNTTCATCCCGGTTTGCTCTCTTT
GCACACGTCTGAACTCTTTCCCTACACGACGCTCTTCCGATCTNNNTTGAAGTCCTGGCCAATAAAGG
GAGCACACGTCTGAACTCTTTCCCTACACGACGCTCTTCCGATCTNNNGTATTTCAGATCCAGTCCTCG
AGCACACGTCTGAACTCTTTCCCTACACGACGCTCTTCCGATCTNNNCCATGGAGAGAGAGATGAAT
GAGCACACGTCTGAACTCTTTCCCTACACGACGCTCTTCCGATCTNNNAGCATGGTGTTCATTCTCTCT
AGCACACGTCTGAACTCTTTCCCTACACGACGCTCTTCCGATCTNNNGTCTTGGTGCGGTCTTCACTAT
GAGCACACGTCTGAACTCTTTCCCTACACGACGCTCTTCCGATCTNNNTGATCTGAGGATGTGGGATT
AGAGCACACGTCTGAACTCTTTCCCTACACGACGCTCTTCCGATCTNNNGAGGTTGTAGCAAGCCAAG
GCACACGTCTGAACTCTTTCCCTACACGACGCTCTTCCGATCTNNNAAAAGGGAAAGTAGGAGCCAG
GAGCACACGTCTGAACTCTTTCCCTACACGACGCTCTTCCGATCTNNNATGGGAAAAAGCCTGGATG
GCACACGTCTGAACTCTTTCCCTACACGACGCTCTTCCGATCTNNNCAAACTGCTGACCTCATGTGAT
GAGCACACGTCTGAACTCTTTCCCTACACGACGCTCTTCCGATCTNNNTTTTCTAACAGGACTCTGAG
AGAGCACACGTCTGAACTCTTTCCCTACACGACGCTCTTCCGATCTNNNGCTGCCACCTATTGTTTTCA
AGCACACGTCTGAACTCTTTCCCTACACGACGCTCTTCCGATCTNNNCCTCGTTTCTTCCTAAGGCTTCT
GAGCACACGTCTGAACTCTTTCCCTACACGACGCTCTTCCGATCTNNNGCCTCTTGGTTCAAACGATTC
AGCACACGTCTGAACTCTTTCCCTACACGACGCTCTTCCGATCTNNNAACAAAGGTTTGATGCCACTCA
GAGCACACGTCTGAACTCTTTCCCTACACGACGCTCTTCCGATCTNNNTGAGAGTTTCTATTTAGGCCG
GCACACGTCTGAACTCTTTCCCTACACGACGCTCTTCCGATCTNNNCTCCTGAGTAGCTGGGATTACA
GCACACGTCTGAACTCTTTCCCTACACGACGCTCTTCCGATCTNNNGCCTCCTGAGTAGCTGAGATTA
GAGCACACGTCTGAACTCTTTCCCTACACGACGCTCTTCCGATCTNNNGTAATCCCAGCTACTCAGAAG
GCACACGTCTGAACTCTTTCCCTACACGACGCTCTTCCGATCTNNNGTTGGGGATTCACTTTCCATTG
AGCACACGTCTGAACTCTTTCCCTACACGACGCTCTTCCGATCTNNNACGCCCGGCTAATTTTTGTATT
AGCACACGTCTGAACTCTTTCCCTACACGACGCTCTTCCGATCTNNNAAACACACACACACCACATATG
GCACACGTCTGAACTCTTTCCCTACACGACGCTCTTCCGATCTNNNGCAAAGGCCAGATATGGAGTA

AGCACACGTCTGAACTCTTTCCCTACACGACGCTCTTCCGATCTNNNTGGGGATTTCTCACCTCTTTCTG
AGCACACGTCTGAACTCTTTCCCTACACGACGCTCTTCCGATCTNNNCAGGGCAGAGGGTAACAATAT
GAGCACACGTCTGAACTCTTTCCCTACACGACGCTCTTCCGATCTNNNCAATCAGGGCTTTCGAGGAA
AGCACACGTCTGAACTCTTTCCCTACACGACGCTCTTCCGATCTNNNCTGCCTCCTAGGTTCAAGTGATT
AGCACACGTCTGAACTCTTTCCCTACACGACGCTCTTCCGATCTNNNCTACCTTCTTCCAAACCAAGGGA
GAGCACACGTCTGAACTCTTTCCCTACACGACGCTCTTCCGATCTNNNTGTCCATAATCCCAGCTACTTG
GAGCACACGTCTGAACTCTTTCCCTACACGACGCTCTTCCGATCTNNNCTTCTGGTAGGTCACAACTCC
GAGCACACGTCTGAACTCTTTCCCTACACGACGCTCTTCCGATCTNNNATACAAAAATTAGCCGAGCG
GAGCACACGTCTGAACTCTTTCCCTACACGACGCTCTTCCGATCTNNNATGAAGAAGAGGGAGAAGG
GAGCACACGTCTGAACTCTTTCCCTACACGACGCTCTTCCGATCTNNNATCTAAAGAAACAACAGCAGC
AGCACACGTCTGAACTCTTTCCCTACACGACGCTCTTCCGATCTNNNGATATGCTTCTTCACCTTTCCCGC
AGCACACGTCTGAACTCTTTCCCTACACGACGCTCTTCCGATCTNNNTAAAATGGGCCAATCAGCTCTCT
AGCACACGTCTGAACTCTTTCCCTACACGACGCTCTTCCGATCTNNNGGAACATGAAGGCAGACTAGC
GCACACGTCTGAACTCTTTCCCTACACGACGCTCTTCCGATCTNNNCAAAAGTTGTCTCTCCCCCAGATC
GAGCACACGTCTGAACTCTTTCCCTACACGACGCTCTTCCGATCTNNNACTAAGGAAGGGGCATGAAA
GAGCACACGTCTGAACTCTTTCCCTACACGACGCTCTTCCGATCTNNNGCCATACACTCACATATGCAAC
AGCACACGTCTGAACTCTTTCCCTACACGACGCTCTTCCGATCTNNNCCTGGCCCAACCTTCATATGTAA
GAGCACACGTCTGAACTCTTTCCCTACACGACGCTCTTCCGATCTNNNCAGTTGAGGTCAGGAGTTTGA
GAGCACACGTCTGAACTCTTTCCCTACACGACGCTCTTCCGATCTNNNATCCAAAAGACGGATACAGAC
AGCACACGTCTGAACTCTTTCCCTACACGACGCTCTTCCGATCTNNNAAGTACGTTAGTGTGTGGGAGT
AGCACACGTCTGAACTCTTTCCCTACACGACGCTCTTCCGATCTNNNTGTGGTAGTGCACACCTGTAAT
GAGCACACGTCTGAACTCTTTCCCTACACGACGCTCTTCCGATCTNNNTCAACTAAAGCCAGGAATGAG
GAGCACACGTCTGAACTCTTTCCCTACACGACGCTCTTCCGATCTNNNTCACTTGAGTCTGGGACATTG
AGCACACGTCTGAACTCTTTCCCTACACGACGCTCTTCCGATCTNNNACTACTGTGAGAATGGCACCAA
AGCACACGTCTGAACTCTTTCCCTACACGACGCTCTTCCGATCTNNNGACATTGAGCCACTACACTCCAG
AGCACACGTCTGAACTCTTTCCCTACACGACGCTCTTCCGATCTNNNTGTGTCTTTTACAGAGACGGGT
AGCACACGTCTGAACTCTTTCCCTACACGACGCTCTTCCGATCTNNNTGTGTCTTTTACAGAGACGGGT
AGCACACGTCTGAACTCTTTCCCTACACGACGCTCTTCCGATCTNNNCAAGTACTCACCTGTGCTGTAG
AGCACACGTCTGAACTCTTTCCCTACACGACGCTCTTCCGATCTNNNATCATTTCAGTCTTAGGGGACG
AGCACACGTCTGAACTCTTTCCCTACACGACGCTCTTCCGATCTNNNGGGTCCTGCTACTCTTTCTTTGA
AGCACACGTCTGAACTCTTTCCCTACACGACGCTCTTCCGATCTNNNGATCACTTGAGCCTAGGAGTTC
AGAGCACACGTCTGAACTCTTTCCCTACACGACGCTCTTCCGATCTNNNAGTCTCTGCTCTCATCTCAGA
AGCACACGTCTGAACTCTTTCCCTACACGACGCTCTTCCGATCTNNNCTGTAATATCCCAGCTACTCGGG
AGCACACGTCTGAACTCTTTCCCTACACGACGCTCTTCCGATCTNNNGGATTGGGTTTTCAGGGACTC
AGCACACGTCTGAACTCTTTCCCTACACGACGCTCTTCCGATCTNNNAACAGAATCCGCCTGACTACAAT
AGAGCACACGTCTGAACTCTTTCCCTACACGACGCTCTTCCGATCTNNNGTCTTTTCTGCTACTCCACCTC
GAGCACACGTCTGAACTCTTTCCCTACACGACGCTCTTCCGATCTNNNATGTGGGATTTGTACCTAGCA
AGCACACGTCTGAACTCTTTCCCTACACGACGCTCTTCCGATCTNNNTCAAGCTATTCTCCTGCCTCATC
AGCACACGTCTGAACTCTTTCCCTACACGACGCTCTTCCGATCTNNNAAGACCTTGTGGCATTTTTGCTT
AGAGCACACGTCTGAACTCTTTCCCTACACGACGCTCTTCCGATCTNNNTGGAAGAGACAGGGAACGT
GAGCACACGTCTGAACTCTTTCCCTACACGACGCTCTTCCGATCTNNNACAAAACTCTGCAAAAGAACC
GAGCACACGTCTGAACTCTTTCCCTACACGACGCTCTTCCGATCTNNNACCTGATCCCAAATCAAAAAGC
AGAGCACACGTCTGAACTCTTTCCCTACACGACGCTCTTCCGATCTNNNTTTAGTAGAGACGGGGTTTC

AGCACACGTCTGAACTCTTTCCCTACACGACGCTCTTCCGATCTNNNTCTTGAACTCGTGACCTCATGTT
GCACACGTCTGAACTCTTTCCCTACACGACGCTCTTCCGATCTNNNTTTATTTTTGTAGAGCTGGGGGC
AGCACACGTCTGAACTCTTTCCCTACACGACGCTCTTCCGATCTNNNGTTTCTTTCCAGTGTCCTCATGCC
AGCACACGTCTGAACTCTTTCCCTACACGACGCTCTTCCGATCTNNNAAACAGACCAATCAGCTCTCTGT
GAGCACACGTCTGAACTCTTTCCCTACACGACGCTCTTCCGATCTNNNTGAATGAAGACACAGGAGTC
GAGCACACGTCTGAACTCTTTCCCTACACGACGCTCTTCCGATCTNNNTCCCTTCTGGACAGTTTGCTT
GAGCACACGTCTGAACTCTTTCCCTACACGACGCTCTTCCGATCTNNNAGGAGCTACTGGAAATCTGA
GAGCACACGTCTGAACTCTTTCCCTACACGACGCTCTTCCGATCTNNNAATTAGTGCAGCTGTAACCCA
GAGCACACGTCTGAACTCTTTCCCTACACGACGCTCTTCCGATCTNNNTAGGAAAAGAGGTCAGACCC
AGCACACGTCTGAACTCTTTCCCTACACGACGCTCTTCCGATCTNNNCTTTCTCTCCCTTCCTTCCTTCCCA
CACACGTCTGAACTCTTTCCCTACACGACGCTCTTCCGATCTNNNCGGATGAATGACTGGGGTTTAGA
GAGCACACGTCTGAACTCTTTCCCTACACGACGCTCTTCCGATCTNNNATCACTGGCTGATCTCTCTCTC
GAGCACACGTCTGAACTCTTTCCCTACACGACGCTCTTCCGATCTNNNATTTTACAACCCTTGCTTGCCT
GAGCACACGTCTGAACTCTTTCCCTACACGACGCTCTTCCGATCTNNNCTCTCAGGCAATGTTCTTCTCC
AAGAGCACACGTCTGAACTCTTTCCCTACACGACGCTCTTCCGATCTNNNCTATCCCATTCCCCTCTTCCT
AGAGCACACGTCTGAACTCTTTCCCTACACGACGCTCTTCCGATCTNNNACTCATAAAACACTCCCAGCA
AGCACACGTCTGAACTCTTTCCCTACACGACGCTCTTCCGATCTNNNGGGAGATTGAGATAAAGAGG
AGCACACGTCTGAACTCTTTCCCTACACGACGCTCTTCCGATCTNNNCTGACTGCCAACTTCTCATTGTG
GAGCACACGTCTGAACTCTTTCCCTACACGACGCTCTTCCGATCTNNNAAGAGAGAGACAGAGAGTG
AAGAGCACACGTCTGAACTCTTTCCCTACACGACGCTCTTCCGATCTNNNGAGGCAGGAGAATTGTTT
GAGCACACGTCTGAACTCTTTCCCTACACGACGCTCTTCCGATCTNNNTACCCCTAAGACCAAGCAACTT
AAGAGCACACGTCTGAACTCTTTCCCTACACGACGCTCTTCCGATCTNNNGATGATAGCTTGAGCATGC
AGCACACGTCTGAACTCTTTCCCTACACGACGCTCTTCCGATCTNNNACCTTGTCTCTCTCTCTCTGTCTC
GAGCACACGTCTGAACTCTTTCCCTACACGACGCTCTTCCGATCTNNNGCCCGATTTTTTCAGTGCTTTT
AGCACACGTCTGAACTCTTTCCCTACACGACGCTCTTCCGATCTNNNGGGCCCTAATAGAACAAAAAGC
GCACACGTCTGAACTCTTTCCCTACACGACGCTCTTCCGATCTNNNGTGGGGCCCTCATTATAGATCTGC
GAGCACACGTCTGAACTCTTTCCCTACACGACGCTCTTCCGATCTNNNGGAGACAATGAAAGGAGGTT
AGCACACGTCTGAACTCTTTCCCTACACGACGCTCTTCCGATCTNNNTTTGATCTGCCAGTGAACAGAC
AAGAGCACACGTCTGAACTCTTTCCCTACACGACGCTCTTCCGATCTNNNAGCTGGTAATTGAAGCTGT
AGAGCACACGTCTGAACTCTTTCCCTACACGACGCTCTTCCGATCTNNNACAAGGTTATAAGGGTTGGC
AGCACACGTCTGAACTCTTTCCCTACACGACGCTCTTCCGATCTNNNGGACTTCTTGGCCTCCATAATCA
GAGCACACGTCTGAACTCTTTCCCTACACGACGCTCTTCCGATCTNNNATACAATGGGAGGGAGACAG
AGCACACGTCTGAACTCTTTCCCTACACGACGCTCTTCCGATCTNNNATTGTGCTGCGTCTGTTTCATT
GAGCACACGTCTGAACTCTTTCCCTACACGACGCTCTTCCGATCTNNNATATGGTCCCACCTTCTTCCCTA
GAGCACACGTCTGAACTCTTTCCCTACACGACGCTCTTCCGATCTNNNAGGTGAATGCAGTACAATCGG
AGCACACGTCTGAACTCTTTCCCTACACGACGCTCTTCCGATCTNNNGTGTGTGTATGTGTGTGTGTGA
AAGAGCACACGTCTGAACTCTTTCCCTACACGACGCTCTTCCGATCTNNNTAGGAGGCTGAGAATTGCT
GCACACGTCTGAACTCTTTCCCTACACGACGCTCTTCCGATCTNNNGGTGGGCCTTTGAGAGATATTCA
GAGCACACGTCTGAACTCTTTCCCTACACGACGCTCTTCCGATCTNNNCGCATTTCATGAATAGTGAGC
GAGCACACGTCTGAACTCTTTCCCTACACGACGCTCTTCCGATCTNNNCATGAAAGGTGATGTGGCTTC
AGAGCACACGTCTGAACTCTTTCCCTACACGACGCTCTTCCGATCTNNNGGGCAATCCTAATGAGAGA
GCACACGTCTGAACTCTTTCCCTACACGACGCTCTTCCGATCTNNNATTCTTGTGTGTGTGTGTGTGTG
AAGAGCACACGTCTGAACTCTTTCCCTACACGACGCTCTTCCGATCTNNNTTCTAGTGAGTGGAGATTC

AGCACACGTCTGAACTCTTTCCCTACACGACGCTCTTCCGATCTNNNTGCCATTTTACACACACACACA
AGCACACGTCTGAACTCTTTCCCTACACGACGCTCTTCCGATCTNNNTCCCTCTCTCTCTTTCTCTCCA
AGCACACGTCTGAACTCTTTCCCTACACGACGCTCTTCCGATCTNNNCCATTCTGTGCTCACTCACTAGT
AGCACACGTCTGAACTCTTTCCCTACACGACGCTCTTCCGATCTNNNGGCCATAACTTCAGACTTCTCCA
AGCACACGTCTGAACTCTTTCCCTACACGACGCTCTTCCGATCTNNNCTGGAGGCTGAAATAGGAGGA
GCACACGTCTGAACTCTTTCCCTACACGACGCTCTTCCGATCTNNNTGTGGACTTCACCTTGTGATAG
AGAGCACACGTCTGAACTCTTTCCCTACACGACGCTCTTCCGATCTNNNGCTTCCAGAGCCCAACTATT
AGCACACGTCTGAACTCTTTCCCTACACGACGCTCTTCCGATCTNNNTGTCTTACGGTGCAGGTGAATA
AGCACACGTCTGAACTCTTTCCCTACACGACGCTCTTCCGATCTNNNAGAAGGAAGAAGGGAGAGAG
GCACACGTCTGAACTCTTTCCCTACACGACGCTCTTCCGATCTNNNAAGCACATCTTACCGTGGTAGAG
GAGCACACGTCTGAACTCTTTCCCTACACGACGCTCTTCCGATCTNNNGGAGGAGTAGGGAGAGAAT
AGCACACGTCTGAACTCTTTCCCTACACGACGCTCTTCCGATCTNNNAACATGTCTTCCCATGACCAGAA
GAGCACACGTCTGAACTCTTTCCCTACACGACGCTCTTCCGATCTNNNAAGGCTGAATAGGGTGAAG
AGAGCACACGTCTGAACTCTTTCCCTACACGACGCTCTTCCGATCTNNNATAGGAAGTGTGTGTGTGT
GAGCACACGTCTGAACTCTTTCCCTACACGACGCTCTTCCGATCTNNNTGTCTCTGTCTGTCTCTGTCTC
AGCACACGTCTGAACTCTTTCCCTACACGACGCTCTTCCGATCTNNNTCTCTCCTTCTGTGTCTCACAC
GAGCACACGTCTGAACTCTTTCCCTACACGACGCTCTTCCGATCTNNNCTGAGGCAGGAGAATGACAT
AGCACACGTCTGAACTCTTTCCCTACACGACGCTCTTCCGATCTNNNTCTCTCTCTCCCTCTCTCTCTCA
GAGCACACGTCTGAACTCTTTCCCTACACGACGCTCTTCCGATCTNNNAGATTGTGACTCTGTACTCCA
GAGCACACGTCTGAACTCTTTCCCTACACGACGCTCTTCCGATCTNNNTTATCAGGCAGGGACTCTTGT
GAGCACACGTCTGAACTCTTTCCCTACACGACGCTCTTCCGATCTNNNCACATACACACACACACACACA
AGCACACGTCTGAACTCTTTCCCTACACGACGCTCTTCCGATCTNNNGGAGTGCATGGGATATAGAA
GAGCACACGTCTGAACTCTTTCCCTACACGACGCTCTTCCGATCTNNNGGGGAGGCCTTAGGAAACAT
GAGCACACGTCTGAACTCTTTCCCTACACGACGCTCTTCCGATCTNNNTGTCTGTCTCTCTCTCTCTCT
GAGCACACGTCTGAACTCTTTCCCTACACGACGCTCTTCCGATCTNNNTCCATAAAGGGCCAGAAAGT
AAGAGCACACGTCTGAACTCTTTCCCTACACGACGCTCTTCCGATCTNNNAGGGAATGTTCCTTTGTCT
GAGCACACGTCTGAACTCTTTCCCTACACGACGCTCTTCCGATCTNNNTCAATCTCAATCCCTCTCCCTCT
AGCACACGTCTGAACTCTTTCCCTACACGACGCTCTTCCGATCTNNNTCCTCTCTCTCTGTCTCTCTGTC
GCACACGTCTGAACTCTTTCCCTACACGACGCTCTTCCGATCTNNNATACTTGCAAACTCGATTCCCCTC
AAGAGCACACGTCTGAACTCTTTCCCTACACGACGCTCTTCCGATCTNNNTGCCAGCCATTATTCTAGG
AGCACACGTCTGAACTCTTTCCCTACACGACGCTCTTCCGATCTNNNGTGAAGTCTCTAGGCACTCTCT
GAGCACACGTCTGAACTCTTTCCCTACACGACGCTCTTCCGATCTNNNGTGCCTGCTAGTTTTTCCCTTT
GAGCACACGTCTGAACTCTTTCCCTACACGACGCTCTTCCGATCTNNNCACACACACACTCTCTCTCTCT
GAGCACACGTCTGAACTCTTTCCCTACACGACGCTCTTCCGATCTNNNGGAGGTTGTAGTGAGCTGA
GAGCACACGTCTGAACTCTTTCCCTACACGACGCTCTTCCGATCTNNNCTGGTCTTGAACTCCTGACAT
GCACACGTCTGAACTCTTTCCCTACACGACGCTCTTCCGATCTNNNGCATATTTCATGCTGCCTACTGG
AGCACACGTCTGAACTCTTTCCCTACACGACGCTCTTCCGATCTNNNGCTTTGGTGGTACTTTGTGGAT
AGCACACGTCTGAACTCTTTCCCTACACGACGCTCTTCCGATCTNNNTCTCTGTCTCTGTCTCTCTGTC
AGCACACGTCTGAACTCTTTCCCTACACGACGCTCTTCCGATCTNNNATCTCAACCACACCCCAGATTAC
GAGCACACGTCTGAACTCTTTCCCTACACGACGCTCTTCCGATCTNNNAGGACTGTCTTTACATGGCAG
GAGCACACGTCTGAACTCTTTCCCTACACGACGCTCTTCCGATCTNNNGAAAGATCCCGTCTTCCCCTAA
AGAGCACACGTCTGAACTCTTTCCCTACACGACGCTCTTCCGATCTNNNCTGTAGTTCCAGCTATTCGG
AGAGCACACGTCTGAACTCTTTCCCTACACGACGCTCTTCCGATCTNNNTTCCTTCCTTCCTTCCTTCCTT

GCACACGTCTGAACTCTTTCCCTACACGACGCTCTTCCGATCTNNNGGGAGTCAGTGAACTAAGACCT
AGAGCACACGTCTGAACTCTTTCCCTACACGACGCTCTTCCGATCTNNNGATGTGTGTGTATGTGTGT
GAGCACACGTCTGAACTCTTTCCCTACACGACGCTCTTCCGATCTNNNAGGTCTGTAGAGTGAGAGCA
AGCACACGTCTGAACTCTTTCCCTACACGACGCTCTTCCGATCTNNNCACACACACACACACACACATAT
AGCACACGTCTGAACTCTTTCCCTACACGACGCTCTTCCGATCTNNNATCCCTTGCTTCTTTCCATCCTT
AGCACACGTCTGAACTCTTTCCCTACACGACGCTCTTCCGATCTNNNGGCCAAAATGAGACCAGAACTT
GCACACGTCTGAACTCTTTCCCTACACGACGCTCTTCCGATCTNNNCCTTCCCCTTCTCCTTTCTTCTCCA
AGCACACGTCTGAACTCTTTCCCTACACGACGCTCTTCCGATCTNNNGAAGCAACTTGATGATGTCAC
AGCACACGTCTGAACTCTTTCCCTACACGACGCTCTTCCGATCTNNNTCAGTACCTAGAAATGACCAG
AGCACACGTCTGAACTCTTTCCCTACACGACGCTCTTCCGATCTNNNACATTGAGAAAAGGAGGAAG
AGCACACGTCTGAACTCTTTCCCTACACGACGCTCTTCCGATCTNNNTAAAGGCTTTGGCTAACAGGTC
CACACGTCTGAACTCTTTCCCTACACGACGCTCTTCCGATCTNNNAATCCATTGTCAGAGTCTGGCATC
GCACACGTCTGAACTCTTTCCCTACACGACGCTCTTCCGATCTNNNCCTTGGTTGGCAGATGACTATCT
GAGCACACGTCTGAACTCTTTCCCTACACGACGCTCTTCCGATCTNNNTTCTTAGGGAAGTTGGCCAC
AGCACACGTCTGAACTCTTTCCCTACACGACGCTCTTCCGATCTNNNAGATCAAGACTGCAGTGAACCA
AAGAGCACACGTCTGAACTCTTTCCCTACACGACGCTCTTCCGATCTNNNAAAGAAAGAGTAGCGAG
AGCACACGTCTGAACTCTTTCCCTACACGACGCTCTTCCGATCTNNNCCAGTTCCACTAAGAAATCCCC
AGCACACGTCTGAACTCTTTCCCTACACGACGCTCTTCCGATCTNNNCAGATGTAACGCGCGCAATATA
GAGCACACGTCTGAACTCTTTCCCTACACGACGCTCTTCCGATCTNNNCAGGATAACAAGGCCTTCTCC
GAGCACACGTCTGAACTCTTTCCCTACACGACGCTCTTCCGATCTNNNTCAGCTCAGTCTTCTCTCTCT
CACACGTCTGAACTCTTTCCCTACACGACGCTCTTCCGATCTNNNGTCTCTGACCTTCAGTTTCCTCACA
GAGCACACGTCTGAACTCTTTCCCTACACGACGCTCTTCCGATCTNNNCCCCCTAACTCAGAGACAGTT
AGCACACGTCTGAACTCTTTCCCTACACGACGCTCTTCCGATCTNNNGAAATGTTTGCATCCAGGAGG
AGCACACGTCTGAACTCTTTCCCTACACGACGCTCTTCCGATCTNNNGGAAGTCTGAGATCTAGGTGT
AGAGCACACGTCTGAACTCTTTCCCTACACGACGCTCTTCCGATCTNNNCACACAGAGGAACCAGTAAG
AGCACACGTCTGAACTCTTTCCCTACACGACGCTCTTCCGATCTNNNACTCATAAGGCTGAAGCACAA
AGCACACGTCTGAACTCTTTCCCTACACGACGCTCTTCCGATCTNNNCAGCTGAAGACTGAGATGAAC
AGAGCACACGTCTGAACTCTTTCCCTACACGACGCTCTTCCGATCTNNNCTCACGAGTTTGAGACTAGC
CACACGTCTGAACTCTTTCCCTACACGACGCTCTTCCGATCTNNNCCCAGATAGAACGAAAAGGTGGA
AGCACACGTCTGAACTCTTTCCCTACACGACGCTCTTCCGATCTNNNCTATAGTTGGGGTCACGTGAG
AGCACACGTCTGAACTCTTTCCCTACACGACGCTCTTCCGATCTNNNAGCTTTAGGTGGCTTAGAACA
GAGCACACGTCTGAACTCTTTCCCTACACGACGCTCTTCCGATCTNNNAGCGAGAGTAAGAGAGAGA
AAGAGCACACGTCTGAACTCTTTCCCTACACGACGCTCTTCCGATCTNNNGTGTGTGTGAGAGAGA
AGCACACGTCTGAACTCTTTCCCTACACGACGCTCTTCCGATCTNNNCCGGGTGAAGACAGAGGAAA
GAGCACACGTCTGAACTCTTTCCCTACACGACGCTCTTCCGATCTNNNGTGTGTGTGTCTGGAAGGCT
GCACACGTCTGAACTCTTTCCCTACACGACGCTCTTCCGATCTNNNCACACACACACACACACACATAT
AAGAGCACACGTCTGAACTCTTTCCCTACACGACGCTCTTCCGATCTNNNTGCCTGTAATCTCAGCTACT
GAGCACACGTCTGAACTCTTTCCCTACACGACGCTCTTCCGATCTNNNAGGGTACATGTTTTATCCCCT
AGCACACGTCTGAACTCTTTCCCTACACGACGCTCTTCCGATCTNNNTCCCAATTGTGTGTATGTGTGG
AGAGCACACGTCTGAACTCTTTCCCTACACGACGCTCTTCCGATCTNNNGCAGAGAGAGAGAGAGAGA
GAGCACACGTCTGAACTCTTTCCCTACACGACGCTCTTCCGATCTNNNTGTCTCTTTTGCTCACCACTCT
AGCACACGTCTGAACTCTTTCCCTACACGACGCTCTTCCGATCTNNNCAGAAAGGCGGGAGATGATTA
GAGCACACGTCTGAACTCTTTCCCTACACGACGCTCTTCCGATCTNNNACATCTGTGAATAGCCACTGT

AGCACACGTCTGAACTCTTTCCCTACACGACGCTCTTCCGATCTNNNCTGTAGGACAAAGTGAGACCCT
GAGCACACGTCTGAACTCTTTCCCTACACGACGCTCTTCCGATCTNNNCTAGCTTTAGGTCTGACCCAG
GAGCACACGTCTGAACTCTTTCCCTACACGACGCTCTTCCGATCTNNNTGCTTTTCTCACTTCAAGCTTG
GCACACGTCTGAACTCTTTCCCTACACGACGCTCTTCCGATCTNNNAAAGCCTGCTTTTCTCCTCTCTTCA
GAGCACACGTCTGAACTCTTTCCCTACACGACGCTCTTCCGATCTNNNTTCTTGCCTCCAGGTATTCATCC
CACACGTCTGAACTCTTTCCCTACACGACGCTCTTCCGATCTNNNCCGAGGTCAACAAATCAACAACAC
AGAGCACACGTCTGAACTCTTTCCCTACACGACGCTCTTCCGATCTNNNACCACTTGAATTCAGTTGCAC
AGCACACGTCTGAACTCTTTCCCTACACGACGCTCTTCCGATCTNNNTTGTTGGCTTACCCCTGTAATCC
GAGCACACGTCTGAACTCTTTCCCTACACGACGCTCTTCCGATCTNNNATGATCCATGATTGCACAACTC
AGCACACGTCTGAACTCTTTCCCTACACGACGCTCTTCCGATCTNNNCCCATGGTGAAAGTAGAAGGA
GAGCACACGTCTGAACTCTTTCCCTACACGACGCTCTTCCGATCTNNNCTGAAGGCTGTAGTGAGCTA
AGAGCACACGTCTGAACTCTTTCCCTACACGACGCTCTTCCGATCTNNNAAGTCAGCGTATGAAGTGGA
AAGAGCACACGTCTGAACTCTTTCCCTACACGACGCTCTTCCGATCTNNNCAGGAGAATCACTTGAACC
GAGCACACGTCTGAACTCTTTCCCTACACGACGCTCTTCCGATCTNNNATACCCTGAGAATGGTAGAGG
AGCACACGTCTGAACTCTTTCCCTACACGACGCTCTTCCGATCTNNNATGTGTCTTAGGACTGGAAAG
GAGCACACGTCTGAACTCTTTCCCTACACGACGCTCTTCCGATCTNNNAAGAGGTGGACTTGGGATTC
AGCACACGTCTGAACTCTTTCCCTACACGACGCTCTTCCGATCTNNNGGTAAAGGGGTCAATTCAGCA
AGAGCACACGTCTGAACTCTTTCCCTACACGACGCTCTTCCGATCTNNNGAGGGCAGATCTTACTAGTC
AGCACACGTCTGAACTCTTTCCCTACACGACGCTCTTCCGATCTNNNTCATCCATTCAACCAGTTTGCACC
AGAGCACACGTCTGAACTCTTTCCCTACACGACGCTCTTCCGATCTNNNAAATCAGCTTACCACAGTGCT
AGCACACGTCTGAACTCTTTCCCTACACGACGCTCTTCCGATCTNNNTGTACCACTGTCTTCTGGAGTTC
AGCACACGTCTGAACTCTTTCCCTACACGACGCTCTTCCGATCTNNNCAGGTTTCTCCAGAGGAACAGA
GAGCACACGTCTGAACTCTTTCCCTACACGACGCTCTTCCGATCTNNNTCTGTCTGTCTGTCTCTGTCTCT
GAGCACACGTCTGAACTCTTTCCCTACACGACGCTCTTCCGATCTNNNTACATTGTGCTCAACCTTCCTCA
GAGCACACGTCTGAACTCTTTCCCTACACGACGCTCTTCCGATCTNNNGAGAGACGGAGAGAGAGAGA
AGCACACGTCTGAACTCTTTCCCTACACGACGCTCTTCCGATCTNNNGCTTATAGATGGCTGCCTTCTCA
AGCACACGTCTGAACTCTTTCCCTACACGACGCTCTTCCGATCTNNNAGAGGGAGAGAGAGAGAGAG
AGCACACGTCTGAACTCTTTCCCTACACGACGCTCTTCCGATCTNNNGTGGAACTTCTTAGCCTCCCTAAC
GAGCACACGTCTGAACTCTTTCCCTACACGACGCTCTTCCGATCTNNNTTCTTCCTCTTCCTCTTCCTCCTC
AAGAGCACACGTCTGAACTCTTTCCCTACACGACGCTCTTCCGATCTNNNGAGTGAGACTCCGTCAGAA
AGAGCACACGTCTGAACTCTTTCCCTACACGACGCTCTTCCGATCTNNNTGAGAAAGAGAGAGCGAGA
AGCACACGTCTGAACTCTTTCCCTACACGACGCTCTTCCGATCTNNNAGGTTGCAGTGAGTGAAGATCA
AAGAGCACACGTCTGAACTCTTTCCCTACACGACGCTCTTCCGATCTNNNTTCAGCTACTTGAGAGACT
AGAGCACACGTCTGAACTCTTTCCCTACACGACGCTCTTCCGATCTNNNGCAATCCCAGTGATAGTAGG
AGCACACGTCTGAACTCTTTCCCTACACGACGCTCTTCCGATCTNNNAGATTACAAGTGTGAGCCACCA
GCACACGTCTGAACTCTTTCCCTACACGACGCTCTTCCGATCTNNNTAACATCTAAGAAGGCAGCCAGG
AGCACACGTCTGAACTCTTTCCCTACACGACGCTCTTCCGATCTNNNACTGAAGAATGCATCAAAGCCA
AGCACACGTCTGAACTCTTTCCCTACACGACGCTCTTCCGATCTNNNTAGATGAAACTCTGTGCCACTGC
GAGCACACGTCTGAACTCTTTCCCTACACGACGCTCTTCCGATCTNNNTGCTCAGTCATATTGGGTGAC
AGCACACGTCTGAACTCTTTCCCTACACGACGCTCTTCCGATCTNNNACTGATTTATTGGCCAGGGTTCT
AAGAGCACACGTCTGAACTCTTTCCCTACACGACGCTCTTCCGATCTNNNCTACCTGGAGTGCTTACCCT
GAGCACACGTCTGAACTCTTTCCCTACACGACGCTCTTCCGATCTNNNCCTCTCTCTCTTTGTCCCTCTCT
GCACACGTCTGAACTCTTTCCCTACACGACGCTCTTCCGATCTNNNTCCAGGGGTAGAAGAAGATGGA

GAGCACACGTCTGAACTCTTTCCCTACACGACGCTCTTCCGATCTNNNAGGCTCTGTTTCTCCATTTGT
AAGAGCACACGTCTGAACTCTTTCCCTACACGACGCTCTTCCGATCTNNNAGAGAGAGAGAGAGAGG
GAGCACACGTCTGAACTCTTTCCCTACACGACGCTCTTCCGATCTNNNATGACAGAGAGCTAGGTACG
GAGCACACGTCTGAACTCTTTCCCTACACGACGCTCTTCCGATCTNNNCAGATTCAAACCACATCTGGC
GCACACGTCTGAACTCTTTCCCTACACGACGCTCTTCCGATCTNNNATAGTAACATTCCCAGCTCGTCC
GCACACGTCTGAACTCTTTCCCTACACGACGCTCTTCCGATCTNNNACGGTCATGTGCTTTCCTTTTTC
AGCACACGTCTGAACTCTTTCCCTACACGACGCTCTTCCGATCTNNNATTGTACCATCCATCCCCCTTTC
AGCACACGTCTGAACTCTTTCCCTACACGACGCTCTTCCGATCTNNNAACTCCAGAGGTTCAAGTCAT
GCACACGTCTGAACTCTTTCCCTACACGACGCTCTTCCGATCTNNNCCTTTCAATCTTCCCACCCTGTAC
AGCACACGTCTGAACTCTTTCCCTACACGACGCTCTTCCGATCTNNNGAGTTGACTTTCTCCCAAAAG
GAGCACACGTCTGAACTCTTTCCCTACACGACGCTCTTCCGATCTNNNACACTTGGTATAGGAACATG
AGCACACGTCTGAACTCTTTCCCTACACGACGCTCTTCCGATCTNNNAGACTTCTCAGCCTCCAGAAT
AGCACACGTCTGAACTCTTTCCCTACACGACGCTCTTCCGATCTNNNCTGGGAAAGCCAAGCAAGATA
GAGCACACGTCTGAACTCTTTCCCTACACGACGCTCTTCCGATCTNNNAGGGTTCCCTAGGGAGATA
AAGAGCACACGTCTGAACTCTTTCCCTACACGACGCTCTTCCGATCTNNNAATACACACACACACACAC
GAGCACACGTCTGAACTCTTTCCCTACACGACGCTCTTCCGATCTNNNAAAAGCTACCTCCAGGGAAG
GCACACGTCTGAACTCTTTCCCTACACGACGCTCTTCCGATCTNNNAACAGCTAGTTGGCAAAAAGCA
AGCACACGTCTGAACTCTTTCCCTACACGACGCTCTTCCGATCTNNNACTCCAGAACAAGTGGTCATAC
AGCACACGTCTGAACTCTTTCCCTACACGACGCTCTTCCGATCTNNNTCTTACAATAGACTGCTGCTG
AGCACACGTCTGAACTCTTTCCCTACACGACGCTCTTCCGATCTNNNATTGTTTCTTTCCTGCCTGGTT
GCACACGTCTGAACTCTTTCCCTACACGACGCTCTTCCGATCTNNNACTTCTAGGCTTCTCCTCACTCT
AGCACACGTCTGAACTCTTTCCCTACACGACGCTCTTCCGATCTNNNCTGTCACATACAAATAGGCAC
AGCACACGTCTGAACTCTTTCCCTACACGACGCTCTTCCGATCTNNNATCTGCAGGAAGACATGAAA
GCACACGTCTGAACTCTTTCCCTACACGACGCTCTTCCGATCTNNNATGTCTGTCTTCTCACTGTGTCC
AGCACACGTCTGAACTCTTTCCCTACACGACGCTCTTCCGATCTNNNGAATGAAGACAGGTGTGTGTG
GAGCACACGTCTGAACTCTTTCCCTACACGACGCTCTTCCGATCTNNNGCCAGCTTTTACAGTGACAC
AGCACACGTCTGAACTCTTTCCCTACACGACGCTCTTCCGATCTNNNGCTCTCTTTCTGGTTTGCAGAT
GCACACGTCTGAACTCTTTCCCTACACGACGCTCTTCCGATCTNNNGTTTTTCATCTGCCTGGAATGCT
GCACACGTCTGAACTCTTTCCCTACACGACGCTCTTCCGATCTNNNTAGGAGCCATGTAAGGTCGAC
GCACACGTCTGAACTCTTTCCCTACACGACGCTCTTCCGATCTNNNCCCACACCACAACCTTCTATCTTC
AGAGCACACGTCTGAACTCTTTCCCTACACGACGCTCTTCCGATCTNNNATCGCTGTGGTCCTAAAGGT
GCACACGTCTGAACTCTTTCCCTACACGACGCTCTTCCGATCTNNNTAGGCTGTGATTTTTCCCCCTAG
AGCACACGTCTGAACTCTTTCCCTACACGACGCTCTTCCGATCTNNNAGGATATGGCCAAGTGACCAT
AGCACACGTCTGAACTCTTTCCCTACACGACGCTCTTCCGATCTNNNGATGAGGAATGTGTGTGTGTG
AGCACACGTCTGAACTCTTTCCCTACACGACGCTCTTCCGATCTNNNTGACACCTTCAGAGCTAACATC
AGCACACGTCTGAACTCTTTCCCTACACGACGCTCTTCCGATCTNNNGGAATGGGATTAGTGCCCTTG
GAGCACACGTCTGAACTCTTTCCCTACACGACGCTCTTCCGATCTNNNCACACACCGTAAAACCAATTG
AGAGCACACGTCTGAACTCTTTCCCTACACGACGCTCTTCCGATCTNNNTGTCAATATGGGGTTCCTCT
AGCACACGTCTGAACTCTTTCCCTACACGACGCTCTTCCGATCTNNNTGACCCTGGTCTTCTAGTGTA
AGCACACGTCTGAACTCTTTCCCTACACGACGCTCTTCCGATCTNNNATGACTGTGCCTCTACATTCCA
GAGCACACGTCTGAACTCTTTCCCTACACGACGCTCTTCCGATCTNNNAGTTGCCTATTCATGTAGCCA
GAGCACACGTCTGAACTCTTTCCCTACACGACGCTCTTCCGATCTNNNATGCCATCATGTCAGTAATCC
GAGCACACGTCTGAACTCTTTCCCTACACGACGCTCTTCCGATCTNNNACTCCCAAATTCCATTTCATCC

GAGCACACGTCTGAACTCTTTCCCTACACGACGCTCTTCCGATCTNNNAGGTTTTTCCATCTGCTCTC
AGCACACGTCTGAACTCTTTCCCTACACGACGCTCTTCCGATCTNNNCCGGGCTGATCTGTTCTCTTATT
AGCACACGTCTGAACTCTTTCCCTACACGACGCTCTTCCGATCTNNNTAGAAGTGTGTGTGTGTGTGT
AGCACACGTCTGAACTCTTTCCCTACACGACGCTCTTCCGATCTNNNGTGTGTGTGTGTGTGTGTGT
AGCACACGTCTGAACTCTTTCCCTACACGACGCTCTTCCGATCTNNNCGACCTGTCTGCAAAAATCTTG
AGCACACGTCTGAACTCTTTCCCTACACGACGCTCTTCCGATCTNNNGGGGAGGTGGAAGGATAGA
AGCACACGTCTGAACTCTTTCCCTACACGACGCTCTTCCGATCTNNNTCTGGAGAACACAGTCACAAC
AGAGCACACGTCTGAACTCTTTCCCTACACGACGCTCTTCCGATCTNNNGTATGCTTAGAGCCTGTGTT
GAGCACACGTCTGAACTCTTTCCCTACACGACGCTCTTCCGATCTNNNTATTCCAAAAGCGTGTGTGT
AGCACACGTCTGAACTCTTTCCCTACACGACGCTCTTCCGATCTNNNGACCAGGTCTTGCAAGTATCT
GAGCACACGTCTGAACTCTTTCCCTACACGACGCTCTTCCGATCTNNNAGAAGTGAATTACGGATTG
AGCACACGTCTGAACTCTTTCCCTACACGACGCTCTTCCGATCTNNNGAGAACAAAGGGGAGAGTA
AGCACACGTCTGAACTCTTTCCCTACACGACGCTCTTCCGATCTNNNGATGTGCGTGAATGTGTATG
GAGCACACGTCTGAACTCTTTCCCTACACGACGCTCTTCCGATCTNNNGAGAATCACACACACACACA
AGCACACGTCTGAACTCTTTCCCTACACGACGCTCTTCCGATCTNNNCGACTTCTGAACTGGTGGCTAT
AGCACACGTCTGAACTCTTTCCCTACACGACGCTCTTCCGATCTNNNGGCAGATTACACACACTCTAG
AGCACACGTCTGAACTCTTTCCCTACACGACGCTCTTCCGATCTNNNGACCAACACCATATGGAAGTTG
GAGCACACGTCTGAACTCTTTCCCTACACGACGCTCTTCCGATCTNNNTGTAATCCCAGCATTTTGGAA
AGCACACGTCTGAACTCTTTCCCTACACGACGCTCTTCCGATCTNNNATTTTTTCTTTGCTCTCACGCCA
AGCACACGTCTGAACTCTTTCCCTACACGACGCTCTTCCGATCTNNNTCTGCCAAGACACTTAGGAACT
AGCACACGTCTGAACTCTTTCCCTACACGACGCTCTTCCGATCTNNNTGAGTTTACAGAGTGGGCAAT
AGCACACGTCTGAACTCTTTCCCTACACGACGCTCTTCCGATCTNNNCTAGCTTTGGAAACCTGCTCA
GAGCACACGTCTGAACTCTTTCCCTACACGACGCTCTTCCGATCTNNNGCTGGGTTACAAAGTTGATT
AGCACACGTCTGAACTCTTTCCCTACACGACGCTCTTCCGATCTNNNCTCATACACACACACACACACA
GCACACGTCTGAACTCTTTCCCTACACGACGCTCTTCCGATCTNNNTCTCCATTCTGATGACAACAGCA
GAGCACACGTCTGAACTCTTTCCCTACACGACGCTCTTCCGATCTNNNGACCGAGAAGAAAAGAAAA
GCACACGTCTGAACTCTTTCCCTACACGACGCTCTTCCGATCTNNNAAATCTGCTTGAACAACCTGTG
GAGCACACGTCTGAACTCTTTCCCTACACGACGCTCTTCCGATCTNNNGAGAAGAGCATAGGGTCAT
AGCACACGTCTGAACTCTTTCCCTACACGACGCTCTTCCGATCTNNNATACAAAAATTAGCCGGGCATG
AGCACACGTCTGAACTCTTTCCCTACACGACGCTCTTCCGATCTNNNAAAACAGACTAGTCAGCCCTCT
AGCACACGTCTGAACTCTTTCCCTACACGACGCTCTTCCGATCTNNNGCAGATGCTGTAACAAATTCT
AGCACACGTCTGAACTCTTTCCCTACACGACGCTCTTCCGATCTNNNCCCTCACCTTCCTCTTCCCTATA
AGCACACGTCTGAACTCTTTCCCTACACGACGCTCTTCCGATCTNNNGAAACTCAGTGTGTGATGTTG
AGAGCACACGTCTGAACTCTTTCCCTACACGACGCTCTTCCGATCTNNNATCACAAAACATAGCCACCC
AGCACACGTCTGAACTCTTTCCCTACACGACGCTCTTCCGATCTNNNCACCTTCATCCTCTTTTTCTGG
AGCACACGTCTGAACTCTTTCCCTACACGACGCTCTTCCGATCTNNNCTAACCAGAAACCTTGCAACCA
AGCACACGTCTGAACTCTTTCCCTACACGACGCTCTTCCGATCTNNNGTGAAAGACATTTCTTAGGG
AGCACACGTCTGAACTCTTTCCCTACACGACGCTCTTCCGATCTNNNAACTTGCAACTGGTGTGTGTAT
AGCACACGTCTGAACTCTTTCCCTACACGACGCTCTTCCGATCTNNNATGGACATTTCCTGGATGAA
AGCACACGTCTGAACTCTTTCCCTACACGACGCTCTTCCGATCTNNNGTTGTTTTCTTTGTTCCCTCCCC
AGCACACGTCTGAACTCTTTCCCTACACGACGCTCTTCCGATCTNNNCCTGCTCTCACTCCAAACTACAT
AGCACACGTCTGAACTCTTTCCCTACACGACGCTCTTCCGATCTNNNTCCAGGTTTCCTACTTGAATGC
GAGCACACGTCTGAACTCTTTCCCTACACGACGCTCTTCCGATCTNNNGTGGTGAAAGAGTGGCTTC

AGCACACGTCTGAACTCTTTCCCTACACGACGCTCTTCCGATCTNNNTTTGGAAATTTTGCAGCCTGA
GAGCACACGTCTGAACTCTTTCCCTACACGACGCTCTTCCGATCTNNNGCAGACTCCAGGAGAGATT
GAGCACACGTCTGAACTCTTTCCCTACACGACGCTCTTCCGATCTNNNGCCTGAGGAATCTTTAGACC
AGCACACGTCTGAACTCTTTCCCTACACGACGCTCTTCCGATCTNNNGTGGAGTGAAAAGCCTGATG
GCACACGTCTGAACTCTTTCCCTACACGACGCTCTTCCGATCTNNNTGGTGGAAGTATGGCATCATCA
GAGCACACGTCTGAACTCTTTCCCTACACGACGCTCTTCCGATCTNNNCAGGGAAAGCTTTTGTCCAG
GCACACGTCTGAACTCTTTCCCTACACGACGCTCTTCCGATCTNNNTGATGCTTGGCAAAGTTTATGG
GAGCACACGTCTGAACTCTTTCCCTACACGACGCTCTTCCGATCTNNNGTGTCACACATTTCCAAGATG
AGCACACGTCTGAACTCTTTCCCTACACGACGCTCTTCCGATCTNNNGCAGCATGTCAGCATTACCTTA
GCACACGTCTGAACTCTTTCCCTACACGACGCTCTTCCGATCTNNNTGTGTGTGTAGCTGGGATTAC
AGCACACGTCTGAACTCTTTCCCTACACGACGCTCTTCCGATCTNNNACTCCCCCAGAATGTAAAGCA
AGCACACGTCTGAACTCTTTCCCTACACGACGCTCTTCCGATCTNNNGCAACAGAGCAAGAATCCATC
GAGCACACGTCTGAACTCTTTCCCTACACGACGCTCTTCCGATCTNNNTTTTTGTGTGTGTGTGACAG
GCACACGTCTGAACTCTTTCCCTACACGACGCTCTTCCGATCTNNNTCCTCTTCCTAATAAGCCTGGGA
GCACACGTCTGAACTCTTTCCCTACACGACGCTCTTCCGATCTNNNTTATTCTCTTATCTGGCCCCACCC
AGCACACGTCTGAACTCTTTCCCTACACGACGCTCTTCCGATCTNNNGCTCAGGTCTGAATGAGTTC
GAGCACACGTCTGAACTCTTTCCCTACACGACGCTCTTCCGATCTNNNAGCTACATGATTGAACAGGA
GCACACGTCTGAACTCTTTCCCTACACGACGCTCTTCCGATCTNNNCCGGAAAGGAAATAAACAGCC
AGCACACGTCTGAACTCTTTCCCTACACGACGCTCTTCCGATCTNNNCAACTGCAGGTGGATCATTTT
GAGCACACGTCTGAACTCTTTCCCTACACGACGCTCTTCCGATCTNNNTGCAGTCAAGAACCCATCACT
AGCACACGTCTGAACTCTTTCCCTACACGACGCTCTTCCGATCTNNNGAGAATCGCTTGAACTCGAG
AGCACACGTCTGAACTCTTTCCCTACACGACGCTCTTCCGATCTNNNGGCTCACCAGGAAATCTCACTT
GCACACGTCTGAACTCTTTCCCTACACGACGCTCTTCCGATCTNNNTGCCATCATGCCTAGCAATTTTT
AGCACACGTCTGAACTCTTTCCCTACACGACGCTCTTCCGATCTNNNGATTTCTCATCCACTACCCTCC
GCACACGTCTGAACTCTTTCCCTACACGACGCTCTTCCGATCTNNNGTAGAGATGGGGTTTCACGAT
AGCACACGTCTGAACTCTTTCCCTACACGACGCTCTTCCGATCTNNNCGTCTTGCGAACTCTGATTCA
AGCACACGTCTGAACTCTTTCCCTACACGACGCTCTTCCGATCTNNNGTTCAGTGGCCACCTGTTTATT
GAGCACACGTCTGAACTCTTTCCCTACACGACGCTCTTCCGATCTNNNCGACAACAACAACAAAAACA
GCACACGTCTGAACTCTTTCCCTACACGACGCTCTTCCGATCTNNNCCGCCCATAGCATGTGTTTTATT
GCACACGTCTGAACTCTTTCCCTACACGACGCTCTTCCGATCTNNNCAGTGGTGGTCTTCCAGTATCA
AGAGCACACGTCTGAACTCTTTCCCTACACGACGCTCTTCCGATCTNNNGGATAATGTGAGTTGATG
AGCACACGTCTGAACTCTTTCCCTACACGACGCTCTTCCGATCTNNNACGTGATAGAAATGTGCAGG
CACACGTCTGAACTCTTTCCCTACACGACGCTCTTCCGATCTNNNAGAATCAAAGCCAGACTCCTCTG
GCACACGTCTGAACTCTTTCCCTACACGACGCTCTTCCGATCTNNNTGTAATCCCAACACTTCAGGAG
AGCACACGTCTGAACTCTTTCCCTACACGACGCTCTTCCGATCTNNNTTGTCAAAAGAATTCCGAGTG
AGCACACGTCTGAACTCTTTCCCTACACGACGCTCTTCCGATCTNNNCCAAGTATCTGGGACTACAG
AGCACACGTCTGAACTCTTTCCCTACACGACGCTCTTCCGATCTNNNTGCTCTGTAAAAACGCACCAAT
AGCACACGTCTGAACTCTTTCCCTACACGACGCTCTTCCGATCTNNNCCGCACCTGGCTAATTTTTGTA
GCACACGTCTGAACTCTTTCCCTACACGACGCTCTTCCGATCTNNNCACCACACCCAGCTAATTTTGTT
AGCACACGTCTGAACTCTTTCCCTACACGACGCTCTTCCGATCTNNNCAGGCATCTCCTTCCGATAAT
GCACACGTCTGAACTCTTTCCCTACACGACGCTCTTCCGATCTNNNAAAAAAAAAGCCTCAAGACGCC
GAGCACACGTCTGAACTCTTTCCCTACACGACGCTCTTCCGATCTNNNGGATGATCTCTTGGTTCCCCT
AGCACACGTCTGAACTCTTTCCCTACACGACGCTCTTCCGATCTNNNCCAAGATTGGTCACAGGAGAT

GCACACGTCTGAACTCTTTCCCTACACGACGCTCTTCCGATCTNNNTCTGTCTCTATTGGCTGACAACA
AGCACACGTCTGAACTCTTTCCCTACACGACGCTCTTCCGATCTNNNGGCGAAACCTCGTCTCTACTAA
GAGCACACGTCTGAACTCTTTCCCTACACGACGCTCTTCCGATCTNNNTGCTTCTTTGGAGGAAAGCA
GCACACGTCTGAACTCTTTCCCTACACGACGCTCTTCCGATCTNNNAAGAAGAACGATTGGAAAGAC
AGCACACGTCTGAACTCTTTCCCTACACGACGCTCTTCCGATCTNNNGCCTCAGAGATTACAGCCTAA
AGAGCACACGTCTGAACTCTTTCCCTACACGACGCTCTTCCGATCTNNNGGCCCATGCTGTCTTGTAGA
AGCACACGTCTGAACTCTTTCCCTACACGACGCTCTTCCGATCTNNNACCCCTTCGGCCTTTTTTTTTT
AGCACACGTCTGAACTCTTTCCCTACACGACGCTCTTCCGATCTNNNCCTTTCTCAAATAGAGCACTG
AGCACACGTCTGAACTCTTTCCCTACACGACGCTCTTCCGATCTNNNAATAAGGGTTTGCAGGGGAG
AGCACACGTCTGAACTCTTTCCCTACACGACGCTCTTCCGATCTNNNACTCTGTCTCCTTTGTTTGGTG
AGCACACGTCTGAACTCTTTCCCTACACGACGCTCTTCCGATCTNNNTGGAGGAAGCCTGAGTTACT
AGCACACGTCTGAACTCTTTCCCTACACGACGCTCTTCCGATCTNNNTTTTCGGCAGGGAAATTGTG
AGCACACGTCTGAACTCTTTCCCTACACGACGCTCTTCCGATCTNNNTTTCTCTCTCTCTCTCTCCCCC
GCACACGTCTGAACTCTTTCCCTACACGACGCTCTTCCGATCTNNNGTCCTGTTACTCCCTCATCAGTG
CACACGTCTGAACTCTTTCCCTACACGACGCTCTTCCGATCTNNNCAAGACAGTGCAAGTGATTCTGG
GAGCACACGTCTGAACTCTTTCCCTACACGACGCTCTTCCGATCTNNNTCCTCTCTCTCTCTCTCTCCT
AGCACACGTCTGAACTCTTTCCCTACACGACGCTCTTCCGATCTNNNCAGGTTGGTTGGTCTTGAAAT
GAGCACACGTCTGAACTCTTTCCCTACACGACGCTCTTCCGATCTNNNGTTCTGTAGTTCGTCCTCTCT
GCACACGTCTGAACTCTTTCCCTACACGACGCTCTTCCGATCTNNNATGTGTTCACTCGACTTGTTCCT
AGCACACGTCTGAACTCTTTCCCTACACGACGCTCTTCCGATCTNNNTTGTCACTTCACTCCAGTCTGA
GCACACGTCTGAACTCTTTCCCTACACGACGCTCTTCCGATCTNNNGAGAGAGGGTGAGAGAGAGA
AGCACACGTCTGAACTCTTTCCCTACACGACGCTCTTCCGATCTNNNGTGAGGGTTTGTTACTTGGG
GAGCACACGTCTGAACTCTTTCCCTACACGACGCTCTTCCGATCTNNNCTTGAGCCCAAAAGTTTGAG
AGCACACGTCTGAACTCTTTCCCTACACGACGCTCTTCCGATCTNNNCATGGATGGATGGATGGATG
GAGCACACGTCTGAACTCTTTCCCTACACGACGCTCTTCCGATCTNNNACAGGGCCAAATGGAAAAG
GCACACGTCTGAACTCTTTCCCTACACGACGCTCTTCCGATCTNNNCAATTCCAGCTACTTGAGAGGC
AGCACACGTCTGAACTCTTTCCCTACACGACGCTCTTCCGATCTNNNGGACCATCATTGTCTTCAGGA
GAGCACACGTCTGAACTCTTTCCCTACACGACGCTCTTCCGATCTNNNAATTTCATAAGGGGTTGGCT
GCACACGTCTGAACTCTTTCCCTACACGACGCTCTTCCGATCTNNNGAAGAGAGAGAAAGAGTCAG
AGCACACGTCTGAACTCTTTCCCTACACGACGCTCTTCCGATCTNNNTCCTGTTCCATTTCTCCTTGGA
GCACACGTCTGAACTCTTTCCCTACACGACGCTCTTCCGATCTNNNTAGATCCCTTGAGCCTAGGAGT
AGCACACGTCTGAACTCTTTCCCTACACGACGCTCTTCCGATCTNNNCATCACCCCAATATGCCACATT
GCACACGTCTGAACTCTTTCCCTACACGACGCTCTTCCGATCTNNNTCTCTGACTTGGGTAAACTGCT
GCACACGTCTGAACTCTTTCCCTACACGACGCTCTTCCGATCTNNNCTCTTCCCACCTTCATAACCCAA
GAGCACACGTCTGAACTCTTTCCCTACACGACGCTCTTCCGATCTNNNTGGCATGTTACTTCTGAGGT
AGCACACGTCTGAACTCTTTCCCTACACGACGCTCTTCCGATCTNNNATGCCATTATGCTGTAGCCTA
AGCACACGTCTGAACTCTTTCCCTACACGACGCTCTTCCGATCTNNNACCAAGCTGCCTGGAAATAGA
AGCACACGTCTGAACTCTTTCCCTACACGACGCTCTTCCGATCTNNNTGGCATTGAGGTAAAGGAAG
GAGCACACGTCTGAACTCTTTCCCTACACGACGCTCTTCCGATCTNNNCGTACACACACACACAGAAA
AGCACACGTCTGAACTCTTTCCCTACACGACGCTCTTCCGATCTNNNTTTTCTGTGTGTGTGTGTGTG
AGCACACGTCTGAACTCTTTCCCTACACGACGCTCTTCCGATCTNNNATGGCTCAGAGAGGTAAATG
GAGCACACGTCTGAACTCTTTCCCTACACGACGCTCTTCCGATCTNNNTATACACACACACACACACAC
GAGCACACGTCTGAACTCTTTCCCTACACGACGCTCTTCCGATCTNNNATGAGGAGCAGCTGAAATG

AGCACACGTCTGAACTCTTTCCCTACACGACGCTCTTCCGATCTNNNGGGGAGAGAGAGAAAGAGA

AGCACACGTCTGAACTCTTTCCCTACACGACGCTCTTCCGATCTNNNGTTTCACCATGTTAGCCAGGAT

CACACGTCTGAACTCTTTCCCTACACGACGCTCTTCCGATCTNNNCATTCTTGCCTCATCCTCTCTGTCA

AGCACACGTCTGAACTCTTTCCCTACACGACGCTCTTCCGATCTNNNACACACACATATTCTCAGCCTGT

GCACACGTCTGAACTCTTTCCCTACACGACGCTCTTCCGATCTNNNGCTTTGGACTAAGCCCTATACAT

GCACACGTCTGAACTCTTTCCCTACACGACGCTCTTCCGATCTNNNAAGAATGTCCCTCACTCATGCAT

GAGCACACGTCTGAACTCTTTCCCTACACGACGCTCTTCCGATCTNNNACTCCTGTAATCCCACCACTTT

AGCACACGTCTGAACTCTTTCCCTACACGACGCTCTTCCGATCTNNNGTCTGTCTGTCTGTCTGTCTGT

AGCACACGTCTGAACTCTTTCCCTACACGACGCTCTTCCGATCTNNNTGGATCAGGACCCCTTTTCTTTT

GAGCACACGTCTGAACTCTTTCCCTACACGACGCTCTTCCGATCTNNNCACTTGTAGTCCCAGCTACTC

GCACACGTCTGAACTCTTTCCCTACACGACGCTCTTCCGATCTNNNAGAGTTTCACTATCCCTAGGCC

AGCACACGTCTGAACTCTTTCCCTACACGACGCTCTTCCGATCTNNNATATAGGAGGCATGGCTGTTG

AGCACACGTCTGAACTCTTTCCCTACACGACGCTCTTCCGATCTNNNCTGTAGATGTGCTCTCACCAAG

GCACACGTCTGAACTCTTTCCCTACACGACGCTCTTCCGATCTNNNGGGCCTGACTAGGGTGAGATAT

AGAGCACACGTCTGAACTCTTTCCCTACACGACGCTCTTCCGATCTNNNGTGTCTGTGTGTGAGAGAG

GAGCACACGTCTGAACTCTTTCCCTACACGACGCTCTTCCGATCTNNNCATACACACACACACACACAC

GAGCACACGTCTGAACTCTTTCCCTACACGACGCTCTTCCGATCTNNNTAGAGTTTGTGTGTGTGTGC

GAGCACACGTCTGAACTCTTTCCCTACACGACGCTCTTCCGATCTNNNGCTGTGGACTCCGAAGATGA

AGCACACGTCTGAACTCTTTCCCTACACGACGCTCTTCCGATCTNNNTTTCTATCTTTCCCTCCCTCCCT

GAGCACACGTCTGAACTCTTTCCCTACACGACGCTCTTCCGATCTNNNGATAGAGTTCTCACGAGAGC

GAGCACACGTCTGAACTCTTTCCCTACACGACGCTCTTCCGATCTNNNGAAGTAAATTGGCCACTTCA

AGCACACGTCTGAACTCTTTCCCTACACGACGCTCTTCCGATCTNNNATCAAGTGCACTGTCTGACCTT

GAGCACACGTCTGAACTCTTTCCCTACACGACGCTCTTCCGATCTNNNAGAGAGAGAGAGAGAGAGA

AGCACACGTCTGAACTCTTTCCCTACACGACGCTCTTCCGATCTNNNCCCAGCCCAAGGATGAGTATAT

AGCACACGTCTGAACTCTTTCCCTACACGACGCTCTTCCGATCTNNNTTTTCTACACACACACACACAC

GCACACGTCTGAACTCTTTCCCTACACGACGCTCTTCCGATCTNNNTGCTTGTTCACACACACAAAAAA

GAGCACACGTCTGAACTCTTTCCCTACACGACGCTCTTCCGATCTNNNTCCTTCCTTCCTTCTCTTTCTCC

AGCACACGTCTGAACTCTTTCCCTACACGACGCTCTTCCGATCTNNNTTGCTTCCTTCCTTTCCTTCCTTC

AGCACACGTCTGAACTCTTTCCCTACACGACGCTCTTCCGATCTNNNGACAGTGACTTCTGGAGTTCT

CACACGTCTGAACTCTTTCCCTACACGACGCTCTTCCGATCTNNNGCATTACTTGTCTATTGGCCCACC

AGCACACGTCTGAACTCTTTCCCTACACGACGCTCTTCCGATCTNNNCCTGCAGAACTGTGAGTCAATG

GCACACGTCTGAACTCTTTCCCTACACGACGCTCTTCCGATCTNNNTCCTACTGAGACTCATGAGCTGA

GAGCACACGTCTGAACTCTTTCCCTACACGACGCTCTTCCGATCTNNNCCAAGAGAAAGGACGAGAG

AGCACACGTCTGAACTCTTTCCCTACACGACGCTCTTCCGATCTNNNCAGTGAGCTGAGATCATGTCA

AGAGCACACGTCTGAACTCTTTCCCTACACGACGCTCTTCCGATCTNNNTGCAGTGAGCCAAAATATC

AGCACACGTCTGAACTCTTTCCCTACACGACGCTCTTCCGATCTNNNCTTGCATCACCATACCCCTAGTT

GCACACGTCTGAACTCTTTCCCTACACGACGCTCTTCCGATCTNNNGTGTGTGTGTGTGTGTGTAT

GAGCACACGTCTGAACTCTTTCCCTACACGACGCTCTTCCGATCTNNNGAGACTGGATGGAAAAGGA

GAGCACACGTCTGAACTCTTTCCCTACACGACGCTCTTCCGATCTNNNCCTGAGTTGGGTGAACAGA

AGCACACGTCTGAACTCTTTCCCTACACGACGCTCTTCCGATCTNNNTGGGGTCTCTCTCTCTCTTTT

AGCACACGTCTGAACTCTTTCCCTACACGACGCTCTTCCGATCTNNNAGCTGGCTAATAACTAGTGTG

AGCACACGTCTGAACTCTTTCCCTACACGACGCTCTTCCGATCTNNNCTCCTTCCTTCCTTCCTTCCTTCC

AGCACACGTCTGAACTCTTTCCCTACACGACGCTCTTCCGATCTNNNAGAACAGTGTCTACCTGAACTC

GAGCACACGTCTGAACTCTTTCCCTACACGACGCTCTTCCGATCTNNNCGGAAAGGCTCTAGGAAACA
AGCACACGTCTGAACTCTTTCCCTACACGACGCTCTTCCGATCTNNNCTTCCTTCCTTCTTTCCTCCCTCC
GCACACGTCTGAACTCTTTCCCTACACGACGCTCTTCCGATCTNNNAGTTTCCTCCATGCCATTCTCATC
AGCACACGTCTGAACTCTTTCCCTACACGACGCTCTTCCGATCTNNNTTGAAGTTTCCTTCCTCCCTTC
GCACACGTCTGAACTCTTTCCCTACACGACGCTCTTCCGATCTNNNGTCTGTATGTGTGTGTGTGT
AGCACACGTCTGAACTCTTTCCCTACACGACGCTCTTCCGATCTNNNGAGAACAAAGTGAGAAACAG
GAGCACACGTCTGAACTCTTTCCCTACACGACGCTCTTCCGATCTNNNATATTGCACACACACACACAC
GAGCACACGTCTGAACTCTTTCCCTACACGACGCTCTTCCGATCTNNNCTGTACAACCTGGAGGATTA
AGCACACGTCTGAACTCTTTCCCTACACGACGCTCTTCCGATCTNNNGAAGGAGGAAGGAGAGAGA
AGCACACGTCTGAACTCTTTCCCTACACGACGCTCTTCCGATCTNNNGTGTGTGTGAGAGAGAG
AAGAGCACACGTCTGAACTCTTTCCCTACACGACGCTCTTCCGATCTNNNTCCGTAGCACAGCAGATTT
GAGCACACGTCTGAACTCTTTCCCTACACGACGCTCTTCCGATCTNNNGCCTAACCATGGCCTATCGTA
GCACACGTCTGAACTCTTTCCCTACACGACGCTCTTCCGATCTNNNAGAGAGGGAGATAGAGGGAG
GCACACGTCTGAACTCTTTCCCTACACGACGCTCTTCCGATCTNNNGCTCTTTTCCTTCCCTTTTCCACCA
GCACACGTCTGAACTCTTTCCCTACACGACGCTCTTCCGATCTNNNTGCATATCAAAGCACATTCTTGT
AGCACACGTCTGAACTCTTTCCCTACACGACGCTCTTCCGATCTNNNACACAGCCTATCTACCTACCTC
AGCACACGTCTGAACTCTTTCCCTACACGACGCTCTTCCGATCTNNNTGTCACATATCTGGCACCTCAA
AGCACACGTCTGAACTCTTTCCCTACACGACGCTCTTCCGATCTNNNCTTTCCTTCCTTCCCTTCCCTTC
AGAGCACACGTCTGAACTCTTTCCCTACACGACGCTCTTCCGATCTNNNGACAGAGTGAGAGCTTAC
AGCACACGTCTGAACTCTTTCCCTACACGACGCTCTTCCGATCTNNNCAGAAGGACTAGTGGATGCA
GAGCACACGTCTGAACTCTTTCCCTACACGACGCTCTTCCGATCTNNNTTCTATGATAGAGTGGAGG
AGCACACGTCTGAACTCTTTCCCTACACGACGCTCTTCCGATCTNNNAGTGAGCTGAGATTGCACAA
AGCACACGTCTGAACTCTTTCCCTACACGACGCTCTTCCGATCTNNNTTGAGGCAGAGGTTGTAGTA
CACACGTCTGAACTCTTTCCCTACACGACGCTCTTCCGATCTNNNGCCACATTCTGAATTGAGGACATC
AGCACACGTCTGAACTCTTTCCCTACACGACGCTCTTCCGATCTNNNCTCTCCTCTCTCTCTCTCTCA
GCACACGTCTGAACTCTTTCCCTACACGACGCTCTTCCGATCTNNNTGATTGTGAGTGAGCTGTCATG
GCACACGTCTGAACTCTTTCCCTACACGACGCTCTTCCGATCTNNNTAAGCTGGGAGAATTGCTTGA
AGCACACGTCTGAACTCTTTCCCTACACGACGCTCTTCCGATCTNNNAGGCTGGAAACCTCATCTGAA
AGAGCACACGTCTGAACTCTTTCCCTACACGACGCTCTTCCGATCTNNNTGGACACCTGTAATTCCAG
AGCACACGTCTGAACTCTTTCCCTACACGACGCTCTTCCGATCTNNNATTCGGTAATCGTAGATGCTG
GCACACGTCTGAACTCTTTCCCTACACGACGCTCTTCCGATCTNNNGTGTTCATGTTCACTGTACAGG
GAGCACACGTCTGAACTCTTTCCCTACACGACGCTCTTCCGATCTNNNTTTAGTGGAGGATGTGGG
AGCACACGTCTGAACTCTTTCCCTACACGACGCTCTTCCGATCTNNNACAGCAAAACTCTGCCTCAAA
AGAGCACACGTCTGAACTCTTTCCCTACACGACGCTCTTCCGATCTNNNCTAGACAACTTTGCATGGC
AGCACACGTCTGAACTCTTTCCCTACACGACGCTCTTCCGATCTNNNTCAGTGGGCTGAGTAAGAAA
GCACACGTCTGAACTCTTTCCCTACACGACGCTCTTCCGATCTNNNTTCTCTAGCTTTGGAAAGTCCC
AGCACACGTCTGAACTCTTTCCCTACACGACGCTCTTCCGATCTNNNAGAGAATTCCTCAAACCTGGG
AGCACACGTCTGAACTCTTTCCCTACACGACGCTCTTCCGATCTNNNTAGTGCCTTTGTAGCTGCCTTT
CACACGTCTGAACTCTTTCCCTACACGACGCTCTTCCGATCTNNNAAAGAGGCTGAGAACCCTATGTG
AGCACACGTCTGAACTCTTTCCCTACACGACGCTCTTCCGATCTNNNTCACGTCTGAAGAACTTTTCCC
GCACACGTCTGAACTCTTTCCCTACACGACGCTCTTCCGATCTNNNCACAAACCATGCCCACATAAGAG
AGCACACGTCTGAACTCTTTCCCTACACGACGCTCTTCCGATCTNNNAGGTACCATACTCTTGGACAG
AGCACACGTCTGAACTCTTTCCCTACACGACGCTCTTCCGATCTNNNAGGCACAGAGAAATTAGGTTG

AGCACACGTCTGAACTCTTTCCCTACACGACGCTCTTCCGATCTNNNCATGGAGAGAGACAGACACA
AGCACACGTCTGAACTCTTTCCCTACACGACGCTCTTCCGATCTNNNAATCCATGCTAATTCCCCTCCT
AGAGCACACGTCTGAACTCTTTCCCTACACGACGCTCTTCCGATCTNNNCTGACTGCAGAAAATGGC
AGCACACGTCTGAACTCTTTCCCTACACGACGCTCTTCCGATCTNNNCTCTTTCTCCCTTCCTTCCTTCC
GAGCACACGTCTGAACTCTTTCCCTACACGACGCTCTTCCGATCTNNNAGGGATTGCTTGATCCAAGA
GAGCACACGTCTGAACTCTTTCCCTACACGACGCTCTTCCGATCTNNNAGCGAAATATTGTGGTGTTT
AGAGCACACGTCTGAACTCTTTCCCTACACGACGCTCTTCCGATCTNNNAAGCCCCAATTTTTCAATGT
GAGCACACGTCTGAACTCTTTCCCTACACGACGCTCTTCCGATCTNNNGAAGGCCAAATCAAAGTGT
GAGCACACGTCTGAACTCTTTCCCTACACGACGCTCTTCCGATCTNNNAACCAGTACTGTGCTAGCTT
GAGCACACGTCTGAACTCTTTCCCTACACGACGCTCTTCCGATCTNNNCTGTAATCCCAGCTACTCAGG
AAGAGCACACGTCTGAACTCTTTCCCTACACGACGCTCTTCCGATCTNNNGAGAAGTCCAGGAAATTC
GCACACGTCTGAACTCTTTCCCTACACGACGCTCTTCCGATCTNNNCCTCCAACTTCCATCCTCTTCTCC
AGCACACGTCTGAACTCTTTCCCTACACGACGCTCTTCCGATCTNNNAGTTTGGGTGAAAAAGGGAG
AGCACACGTCTGAACTCTTTCCCTACACGACGCTCTTCCGATCTNNNGTTTCGACAGAGCCAAACTTG
GAGCACACGTCTGAACTCTTTCCCTACACGACGCTCTTCCGATCTNNNTTACTACCCCTAGGCCTGAAG
GCACACGTCTGAACTCTTTCCCTACACGACGCTCTTCCGATCTNNNAATCATCTGGAGGGTTGACAT
AGCACACGTCTGAACTCTTTCCCTACACGACGCTCTTCCGATCTNNNTGGCCCGCATCTATTTTTTTGTT
AGCACACGTCTGAACTCTTTCCCTACACGACGCTCTTCCGATCTNNNCTCTCTCTCTCTCTCTCTCCCC
GCACACGTCTGAACTCTTTCCCTACACGACGCTCTTCCGATCTNNNTTTTCGCATAGAATCTTCGCAG
GCACACGTCTGAACTCTTTCCCTACACGACGCTCTTCCGATCTNNNGTCTCTGTCTTCCTCTGTCTCTGC
CACACGTCTGAACTCTTTCCCTACACGACGCTCTTCCGATCTNNNAACACTCAATGGAACCACTGACTC
GCACACGTCTGAACTCTTTCCCTACACGACGCTCTTCCGATCTNNNCCTTGAGTTTCTTCACCAACTGC
GCACACGTCTGAACTCTTTCCCTACACGACGCTCTTCCGATCTNNNTTCATATCTCCTGTCTCACCCCAC
AGCACACGTCTGAACTCTTTCCCTACACGACGCTCTTCCGATCTNNNACTCCACACTGATTCCTCTCCTA
GCACACGTCTGAACTCTTTCCCTACACGACGCTCTTCCGATCTNNNTATTCCTGAAAAGAAGGAGGGG
GCACACGTCTGAACTCTTTCCCTACACGACGCTCTTCCGATCTNNNTCCCTCTTCCTTTCTTCTTTTCCTC
AGCACACGTCTGAACTCTTTCCCTACACGACGCTCTTCCGATCTNNNGTACAGTGGCACAGTAGCTTA
AGCACACGTCTGAACTCTTTCCCTACACGACGCTCTTCCGATCTNNNAGATCAGTGCCATTCAGACAT
AGCACACGTCTGAACTCTTTCCCTACACGACGCTCTTCCGATCTNNNCTGGGCAACAAGAGCAAAACT
AGCACACGTCTGAACTCTTTCCCTACACGACGCTCTTCCGATCTNNNGTGTGTGTTTGTGTGTGTGTG
AGAGCACACGTCTGAACTCTTTCCCTACACGACGCTCTTCCGATCTNNNCCCAGCCCAATATGCATTTC
GCACACGTCTGAACTCTTTCCCTACACGACGCTCTTCCGATCTNNNATGAGACCAAGAACCCACCAATT
CACACGTCTGAACTCTTTCCCTACACGACGCTCTTCCGATCTNNNGTTCTCCTCTAAAGCACGGTGTA
GCACACGTCTGAACTCTTTCCCTACACGACGCTCTTCCGATCTNNNAGATTTCACCCTTGCTGTTCTCA
GAGCACACGTCTGAACTCTTTCCCTACACGACGCTCTTCCGATCTNNNCATGGTGGAGAGAGAGAGA
GAGCACACGTCTGAACTCTTTCCCTACACGACGCTCTTCCGATCTNNNTCTCTCTCTATGTGTGTGTGT
AGCACACGTCTGAACTCTTTCCCTACACGACGCTCTTCCGATCTNNNGGCTGAGGCAGGATAATTGTT
CACACGTCTGAACTCTTTCCCTACACGACGCTCTTCCGATCTNNNACTTCCTTTGTGAGAGGGAATGG
AGCACACGTCTGAACTCTTTCCCTACACGACGCTCTTCCGATCTNNNTTACTGAAAACGTTTCCCTCCC
CACACGTCTGAACTCTTTCCCTACACGACGCTCTTCCGATCTNNNTGAGCCACTATATGCCTCTCTCTCA
GCACACGTCTGAACTCTTTCCCTACACGACGCTCTTCCGATCTNNNGCAGGACTTGGCTCTGAATTTT
AGCACACGTCTGAACTCTTTCCCTACACGACGCTCTTCCGATCTNNNGAAGAAGGAAGGAAGGAAG
AGCACACGTCTGAACTCTTTCCCTACACGACGCTCTTCCGATCTNNNGATCGTGTCACTGTACTCTAGC

GAGCACACGTCTGAACTCTTTCCCTACACGACGCTCTTCCGATCTNNNATACCATGGGAGAGGATCAA
AGAGCACACGTCTGAACTCTTTCCCTACACGACGCTCTTCCGATCTNNNCCGTAATTGTGCACTCCAGA
AGCACACGTCTGAACTCTTTCCCTACACGACGCTCTTCCGATCTNNNTGGGGACTTTTGCTGAAGTAA
AGCACACGTCTGAACTCTTTCCCTACACGACGCTCTTCCGATCTNNNAAACTTTGTGTGTGTGTGTGTG
AGCACACGTCTGAACTCTTTCCCTACACGACGCTCTTCCGATCTNNNGGTTGCAGTGAGCCTAAATCAT
GAGCACACGTCTGAACTCTTTCCCTACACGACGCTCTTCCGATCTNNNGGTGTAGAAGTCGTCATTGA
AGCACACGTCTGAACTCTTTCCCTACACGACGCTCTTCCGATCTNNNGAGATGGAGAATAAGGCTGA
GAGCACACGTCTGAACTCTTTCCCTACACGACGCTCTTCCGATCTNNNTTTGCAAACCTAGGACCAGAA
AGAGCACACGTCTGAACTCTTTCCCTACACGACGCTCTTCCGATCTNNNTCCCAAAGTGTTAGGGTTACA
AGCACACGTCTGAACTCTTTCCCTACACGACGCTCTTCCGATCTNNNTCCCTATATTGCCTGTTGCCTTT
GAGCACACGTCTGAACTCTTTCCCTACACGACGCTCTTCCGATCTNNNAAGGAAAACACACACACACAC
AGCACACGTCTGAACTCTTTCCCTACACGACGCTCTTCCGATCTNNNCTCTCTCTCTCTCTCTCCTCGCA
AGCACACGTCTGAACTCTTTCCCTACACGACGCTCTTCCGATCTNNNCTGCATGTAGACAGCAACTTCA
GAGCACACGTCTGAACTCTTTCCCTACACGACGCTCTTCCGATCTNNNTATGTTGTGTGTGCCTGTATC
GAGCACACGTCTGAACTCTTTCCCTACACGACGCTCTTCCGATCTNNNCCATGTCCTGCAGATTCACATT
AGAGCACACGTCTGAACTCTTTCCCTACACGACGCTCTTCCGATCTNNNCATAGCGAGTGCCTATAAT
GAGCACACGTCTGAACTCTTTCCCTACACGACGCTCTTCCGATCTNNNGATAGTGCTACTGTACTCCAG
GAGCACACGTCTGAACTCTTTCCCTACACGACGCTCTTCCGATCTNNNTCTCTCTCTGTCTCTCTCTCTCT
AGCACACGTCTGAACTCTTTCCCTACACGACGCTCTTCCGATCTNNNTGATTGGATCATGGGGATGGTT
AGCACACGTCTGAACTCTTTCCCTACACGACGCTCTTCCGATCTNNNTTTGGTTCAGGGAGTGGTTTAC
AGCACACGTCTGAACTCTTTCCCTACACGACGCTCTTCCGATCTNNNCTTCCTTCCTTCCTTCCTTCCTCCA
AGAGCACACGTCTGAACTCTTTCCCTACACGACGCTCTTCCGATCTNNNACTGGTATCCCCTGTTCCGTA
AGAGCACACGTCTGAACTCTTTCCCTACACGACGCTCTTCCGATCTNNNAGGCTAGAAGTCCAACATCA
GAGCACACGTCTGAACTCTTTCCCTACACGACGCTCTTCCGATCTNNNCATTCGACCCAGCAATCTCAAT
AGCACACGTCTGAACTCTTTCCCTACACGACGCTCTTCCGATCTNNNAAAAGTTTACAGCTCCCTCACCT
GAGCACACGTCTGAACTCTTTCCCTACACGACGCTCTTCCGATCTNNNCCGAAGCTTAGAAAGAGGGC
AGAGCACACGTCTGAACTCTTTCCCTACACGACGCTCTTCCGATCTNNNATTGTTTGAACCCACCTAGGA
AGAGCACACGTCTGAACTCTTTCCCTACACGACGCTCTTCCGATCTNNNTGTCTCTGTCTCTGTCTCTGT
GCACACGTCTGAACTCTTTCCCTACACGACGCTCTTCCGATCTNNNCTGTACGTGTGTGTTCGTTATGG
GCACACGTCTGAACTCTTTCCCTACACGACGCTCTTCCGATCTNNNCTGTACGTGTGTGTTCGTTATGG
AGAGCACACGTCTGAACTCTTTCCCTACACGACGCTCTTCCGATCTNNNTAAACTATGATCGCTCCACTG
AGCACACGTCTGAACTCTTTCCCTACACGACGCTCTTCCGATCTNNNGTCTCTTGAGAAGACAGGTCTG
AGAGCACACGTCTGAACTCTTTCCCTACACGACGCTCTTCCGATCTNNNTGGGGCCCTGTCAAAAAAAA
GAGCACACGTCTGAACTCTTTCCCTACACGACGCTCTTCCGATCTNNNATTTTCTACCCCAGTCCTTGCTT
AGCACACGTCTGAACTCTTTCCCTACACGACGCTCTTCCGATCTNNNAAATCCCTTGTGGCTTAGTGCTA
AGAGCACACGTCTGAACTCTTTCCCTACACGACGCTCTTCCGATCTNNNGGAAAATTCTCCTGCCCCTTT
GAGCACACGTCTGAACTCTTTCCCTACACGACGCTCTTCCGATCTNNNTATGACAGGGAGAGAGAGG
AGAGCACACGTCTGAACTCTTTCCCTACACGACGCTCTTCCGATCTNNNAGGTACACAAGTTGAGGGT
GAGCACACGTCTGAACTCTTTCCCTACACGACGCTCTTCCGATCTNNNCTGAACCTGCAGCAGAAAGA
AGAGCACACGTCTGAACTCTTTCCCTACACGACGCTCTTCCGATCTNNNTAGAACTTAGTTTCCCCACCA
AGAGCACACGTCTGAACTCTTTCCCTACACGACGCTCTTCCGATCTNNNTTGAAGATGGGTACAAAGG
GAGCACACGTCTGAACTCTTTCCCTACACGACGCTCTTCCGATCTNNNCTTGACTGGAAGGGGTTTGA
AGCACACGTCTGAACTCTTTCCCTACACGACGCTCTTCCGATCTNNNAATATGCACAGACACACACACA

AGCACACGTCTGAACTCTTTCCCTACACGACGCTCTTCCGATCTNNNCCGGGACTTCTCAACCTCAATA
GCACACGTCTGAACTCTTTCCCTACACGACGCTCTTCCGATCTNNNAAATAAATGTGTGTGGCAGCT
AGCACACGTCTGAACTCTTTCCCTACACGACGCTCTTCCGATCTNNNCTGAGGCAGGAGAATCACTTG
GCACACGTCTGAACTCTTTCCCTACACGACGCTCTTCCGATCTNNNACAGAGAGAGAGAGAGAGAGA
GAGCACACGTCTGAACTCTTTCCCTACACGACGCTCTTCCGATCTNNNTCTCTGCCTTTATTTTTCCCC
AGCACACGTCTGAACTCTTTCCCTACACGACGCTCTTCCGATCTNNNTCACTTTAGTTAGCCCAGACCA
GAGCACACGTCTGAACTCTTTCCCTACACGACGCTCTTCCGATCTNNNGCCAAGCATCTCAGTGAGG
GAGCACACGTCTGAACTCTTTCCCTACACGACGCTCTTCCGATCTNNNACAGAGAGAGAGAGAAGAG
GCACACGTCTGAACTCTTTCCCTACACGACGCTCTTCCGATCTNNNAGAGTTATCGAGAGACTGACCC
GCACACGTCTGAACTCTTTCCCTACACGACGCTCTTCCGATCTNNNCTGTCATCTCTTGCCTGTCTCTGC
AGCACACGTCTGAACTCTTTCCCTACACGACGCTCTTCCGATCTNNNCCATTAGTGACCTGTAACAGC
AGCACACGTCTGAACTCTTTCCCTACACGACGCTCTTCCGATCTNNNAGATGATCAACATGCCTGCAA
AGCACACGTCTGAACTCTTTCCCTACACGACGCTCTTCCGATCTNNNAGTCTTCTAACACGGAAATGG
AGAGCACACGTCTGAACTCTTTCCCTACACGACGCTCTTCCGATCTNNNAGGTCCGGTAGGTGTAACT
GCACACGTCTGAACTCTTTCCCTACACGACGCTCTTCCGATCTNNNTCATGAAGAATGAGAAAGGGG
GCACACGTCTGAACTCTTTCCCTACACGACGCTCTTCCGATCTNNNACACACACAGAGAGAGAGAGA
AGCACACGTCTGAACTCTTTCCCTACACGACGCTCTTCCGATCTNNNGATAGGTTTCTTAGGGGTGG
AGCACACGTCTGAACTCTTTCCCTACACGACGCTCTTCCGATCTNNNCACACACACACACACACATAT
AGCACACGTCTGAACTCTTTCCCTACACGACGCTCTTCCGATCTNNNCATGGAGAGCAGTGTTTTTCC
GAGCACACGTCTGAACTCTTTCCCTACACGACGCTCTTCCGATCTNNNATGGGCAACGAAACAAGATT
AGAGCACACGTCTGAACTCTTTCCCTACACGACGCTCTTCCGATCTNNNGCTTGGATCCAATGAGAGG
GAGCACACGTCTGAACTCTTTCCCTACACGACGCTCTTCCGATCTNNNATGAGCCAGGTGAAGAATGA
CACACGTCTGAACTCTTTCCCTACACGACGCTCTTCCGATCTNNNACTTTCTGCTATCCACATCCCTGC
AGCACACGTCTGAACTCTTTCCCTACACGACGCTCTTCCGATCTNNNTGTGAACCCAGAAATCTCCACA
GAGCACACGTCTGAACTCTTTCCCTACACGACGCTCTTCCGATCTNNNTCCATAAATCTGCCACACACT
AGCACACGTCTGAACTCTTTCCCTACACGACGCTCTTCCGATCTNNNGTCGTAGATAAGGGTGCTGGT
GAGCACACGTCTGAACTCTTTCCCTACACGACGCTCTTCCGATCTNNNGACAGCTTTTGTTGCTACCTC
AGCACACGTCTGAACTCTTTCCCTACACGACGCTCTTCCGATCTNNNCTTGTAGGGCCAATTCTCCTCT
GCACACGTCTGAACTCTTTCCCTACACGACGCTCTTCCGATCTNNNTGTATGCCTGTAATCCCAGCTA
AGCACACGTCTGAACTCTTTCCCTACACGACGCTCTTCCGATCTNNNGAGCAAGTCTCTCTGTCTCTCA
AGCACACGTCTGAACTCTTTCCCTACACGACGCTCTTCCGATCTNNNCAGGGAAATGAGAATGCCCT
AGCACACGTCTGAACTCTTTCCCTACACGACGCTCTTCCGATCTNNNTGACCATCCATTACACTTGCTG
GCACACGTCTGAACTCTTTCCCTACACGACGCTCTTCCGATCTNNNATGTGGCTGAACTTCAAGAGG
AGCACACGTCTGAACTCTTTCCCTACACGACGCTCTTCCGATCTNNNTTCTCTGGTTTACCTCATCTGC
AGCACACGTCTGAACTCTTTCCCTACACGACGCTCTTCCGATCTNNNTATTGAGCAATGATGGTGGAG
AGCACACGTCTGAACTCTTTCCCTACACGACGCTCTTCCGATCTNNNGCTGAGGTATACAGAGAATG
GCACACGTCTGAACTCTTTCCCTACACGACGCTCTTCCGATCTNNNAGACCAGAGAGAGATAGCCAA
GAGCACACGTCTGAACTCTTTCCCTACACGACGCTCTTCCGATCTNNNTCTCTCTCTCTCTCTCTCCCT
AGCACACGTCTGAACTCTTTCCCTACACGACGCTCTTCCGATCTNNNACTACATGCTAGTGTGTGTGTG
GCACACGTCTGAACTCTTTCCCTACACGACGCTCTTCCGATCTNNNATTGCCACCATAATTGCCTTTCC
GCACACGTCTGAACTCTTTCCCTACACGACGCTCTTCCGATCTNNNAGATTGGCATGAGTCAGTGAA
CACACGTCTGAACTCTTTCCCTACACGACGCTCTTCCGATCTNNNTCTTTGGACCAGGGGAATACAG
GCACACGTCTGAACTCTTTCCCTACACGACGCTCTTCCGATCTNNNTTTCTTCTAAGGGGTGTGTGTG

GAGCACACGTCTGAACTCTTTCCCTACACGACGCTCTTCCGATCTNNNGACGCTGTCTTTTCAAGAAGC
AGCACACGTCTGAACTCTTTCCCTACACGACGCTCTTCCGATCTNNNAGGGCGCCAATAGTTTAGGTA
GAGCACACGTCTGAACTCTTTCCCTACACGACGCTCTTCCGATCTNNNAGGAGAGGGAAAGAGAGAG
GAGCACACGTCTGAACTCTTTCCCTACACGACGCTCTTCCGATCTNNNTCAGGCATCCCCATAGAAACA
GCACACGTCTGAACTCTTTCCCTACACGACGCTCTTCCGATCTNNNCTCAACTCCAAGAGAGAGGCATT
GCACACGTCTGAACTCTTTCCCTACACGACGCTCTTCCGATCTNNNGCAGACTTCTACCCTGATTGACAC
AGAGCACACGTCTGAACTCTTTCCCTACACGACGCTCTTCCGATCTNNNGGCGGGGGAAGATTCAAGT
GAGCACACGTCTGAACTCTTTCCCTACACGACGCTCTTCCGATCTNNNGTGTGTGTGAGAGAGAGA
AGCACACGTCTGAACTCTTTCCCTACACGACGCTCTTCCGATCTNNNATGGGGTAAGGTCTACAGAAT
GAGCACACGTCTGAACTCTTTCCCTACACGACGCTCTTCCGATCTNNNCTCATCCAGATCACTGTGGTAC
AGCACACGTCTGAACTCTTTCCCTACACGACGCTCTTCCGATCTNNNAACAGAATCACAGAACCAATGC
AGAGCACACGTCTGAACTCTTTCCCTACACGACGCTCTTCCGATCTNNNCTCAACAGCTTGGAAATCCCA
GAGCACACGTCTGAACTCTTTCCCTACACGACGCTCTTCCGATCTNNNAGAGGTGAGGAGAGATGGAC
AGAGCACACGTCTGAACTCTTTCCCTACACGACGCTCTTCCGATCTNNNGATGTCTGACTCAAGCTGGC
GCACACGTCTGAACTCTTTCCCTACACGACGCTCTTCCGATCTNNNACAAAGGCTTGACCACTCCTAATC
AGAGCACACGTCTGAACTCTTTCCCTACACGACGCTCTTCCGATCTNNNTTAGGTTGTTGGGTTGGAAC
GAGCACACGTCTGAACTCTTTCCCTACACGACGCTCTTCCGATCTNNNTCCTGGGAAACATAACCTGTG
GAGCACACGTCTGAACTCTTTCCCTACACGACGCTCTTCCGATCTNNNGTCCTAGCTACATGAGACATC
GAGCACACGTCTGAACTCTTTCCCTACACGACGCTCTTCCGATCTNNNGGGGGTTGGATGAATAAGAA
AGCACACGTCTGAACTCTTTCCCTACACGACGCTCTTCCGATCTNNNCTCTTTTCTCTTCTGGGGACTCCC
AGAGCACACGTCTGAACTCTTTCCCTACACGACGCTCTTCCGATCTNNNGAAAGTGAATCCCAGGAGA
AGCACACGTCTGAACTCTTTCCCTACACGACGCTCTTCCGATCTNNNTACTCATGTGTCTGGTGGTTGA
GAGCACACGTCTGAACTCTTTCCCTACACGACGCTCTTCCGATCTNNNGTTAGCCTGGTAATATTCCCCC
GAGCACACGTCTGAACTCTTTCCCTACACGACGCTCTTCCGATCTNNNTCTCTCATCCTCCAGTAGGCTA
AGCACACGTCTGAACTCTTTCCCTACACGACGCTCTTCCGATCTNNNTCTAGTATGGTGGCTTCAGGGT
GAGCACACGTCTGAACTCTTTCCCTACACGACGCTCTTCCGATCTNNNCCCTCAAAGCCGTAGAGTAGA
GAGCACACGTCTGAACTCTTTCCCTACACGACGCTCTTCCGATCTNNNTAAAGTGCTGGGATTACAGGC
GAGCACACGTCTGAACTCTTTCCCTACACGACGCTCTTCCGATCTNNNAATCACTTGAACCCCAGAGTCA
AGCACACGTCTGAACTCTTTCCCTACACGACGCTCTTCCGATCTNNNTTCTCTCTGCAAGGACTGTGAA
GAGCACACGTCTGAACTCTTTCCCTACACGACGCTCTTCCGATCTNNNTGGATTCCAGGCATCATTCAGA
AGAGCACACGTCTGAACTCTTTCCCTACACGACGCTCTTCCGATCTNNNCGAGTTTGGGAAGCAGTCGT
GAGCACACGTCTGAACTCTTTCCCTACACGACGCTCTTCCGATCTNNNTTTGCCAAGCATCATTTTGACC
AGAGCACACGTCTGAACTCTTTCCCTACACGACGCTCTTCCGATCTNNNAGGGAAGAGGTGATAAGAC
AGAGCACACGTCTGAACTCTTTCCCTACACGACGCTCTTCCGATCTNNNTGTTTGACACAGTGAGACTC
AGCACACGTCTGAACTCTTTCCCTACACGACGCTCTTCCGATCTNNNACCTGAGCTGTGAGTTTCTGTTT
AGCACACGTCTGAACTCTTTCCCTACACGACGCTCTTCCGATCTNNNGGTTTCCCCAATAAATCCCCTCTC
AGCACACGTCTGAACTCTTTCCCTACACGACGCTCTTCCGATCTNNNTAGACTTGAGACCCTCACCAGAT
GAGCACACGTCTGAACTCTTTCCCTACACGACGCTCTTCCGATCTNNNAATCATGATGGAAGGTGAAG
AGCACACGTCTGAACTCTTTCCCTACACGACGCTCTTCCGATCTNNNAATTTCCACAGCCCCAAAATTGT
AGCACACGTCTGAACTCTTTCCCTACACGACGCTCTTCCGATCTNNNGTTGGGATATCTGACATCCGAC
AGCACACGTCTGAACTCTTTCCCTACACGACGCTCTTCCGATCTNNNGATGTCACGAAGGGAAGATCAC
GAGCACACGTCTGAACTCTTTCCCTACACGACGCTCTTCCGATCTNNNAGGGTTTTGTGTGTGTGTTTC
AGCACACGTCTGAACTCTTTCCCTACACGACGCTCTTCCGATCTNNNCTGTGTGTATCCTGTGCTGAAT

GAGCACACGTCTGAACTCTTTCCCTACACGACGCTCTTCCGATCTNNNAGAGAGAGAGAGAGAGAGA
GAGCACACGTCTGAACTCTTTCCCTACACGACGCTCTTCCGATCTNNNGGGAAGGATGATTGGTGTG
AGCACACGTCTGAACTCTTTCCCTACACGACGCTCTTCCGATCTNNNCCTCAAAAATCAGCACTGGTGA
AGCACACGTCTGAACTCTTTCCCTACACGACGCTCTTCCGATCTNNNGATTTCTGTTGGGATCCTTGG
AGCACACGTCTGAACTCTTTCCCTACACGACGCTCTTCCGATCTNNNAAATGGCTACAGTGTCCAGAA
AGCACACGTCTGAACTCTTTCCCTACACGACGCTCTTCCGATCTNNNAACTGAGTCCCACTCTAAGCAA
AGAGCACACGTCTGAACTCTTTCCCTACACGACGCTCTTCCGATCTNNNGTGAGGCCCCATCTCAAAAA
AGCACACGTCTGAACTCTTTCCCTACACGACGCTCTTCCGATCTNNNATCAGGTTCAGGGAGGTCATCT
GAGCACACGTCTGAACTCTTTCCCTACACGACGCTCTTCCGATCTNNNCTGGGGGGGAGAGGATGATA
AGAGCACACGTCTGAACTCTTTCCCTACACGACGCTCTTCCGATCTNNNAGGGACATATCAGAGTCAC
AGCACACGTCTGAACTCTTTCCCTACACGACGCTCTTCCGATCTNNNAAGTCAAAGATCCCACCTACACG
GAGCACACGTCTGAACTCTTTCCCTACACGACGCTCTTCCGATCTNNNGTTCAAGCCATTCTCATGCCTT
AGCACACGTCTGAACTCTTTCCCTACACGACGCTCTTCCGATCTNNNCCATCATGCCCAGCTAATTTTGT
AGCACACGTCTGAACTCTTTCCCTACACGACGCTCTTCCGATCTNNNTTCCAGGTTCAAGCAATTCTCCT
AGAGCACACGTCTGAACTCTTTCCCTACACGACGCTCTTCCGATCTNNNTGTTCTGAAGTAGGCCACAA
AGAGCACACGTCTGAACTCTTTCCCTACACGACGCTCTTCCGATCTNNNAGTTTGGGCTACAAGTCCTA
AGCACACGTCTGAACTCTTTCCCTACACGACGCTCTTCCGATCTNNNTTTCATCCTGTTTCTTGTGCCC
GAGCACACGTCTGAACTCTTTCCCTACACGACGCTCTTCCGATCTNNNTTGAAATTGTCCCGTTTAGGC
AGCACACGTCTGAACTCTTTCCCTACACGACGCTCTTCCGATCTNNNTATTTTATTGCTTGGAGCCAGC
GAGCACACGTCTGAACTCTTTCCCTACACGACGCTCTTCCGATCTNNNAGGGCATAGCACACAACAAA
GAGCACACGTCTGAACTCTTTCCCTACACGACGCTCTTCCGATCTNNNTCAGTTCTCATGGCCATTTGGA
GAGCACACGTCTGAACTCTTTCCCTACACGACGCTCTTCCGATCTNNNCTGGGTGAATGCTGTGCAAAT
GAGCACACGTCTGAACTCTTTCCCTACACGACGCTCTTCCGATCTNNNCTGAGATTGTGCCACTGAACT
GAGCACACGTCTGAACTCTTTCCCTACACGACGCTCTTCCGATCTNNNGCTGCAGATCCAAAATAGTGT
GAGCACACGTCTGAACTCTTTCCCTACACGACGCTCTTCCGATCTNNNTTCAGGGAGTTGGTCTCTAGC
AGCACACGTCTGAACTCTTTCCCTACACGACGCTCTTCCGATCTNNNATCTAGGAGGTTGAGGCTGTA
AGCACACGTCTGAACTCTTTCCCTACACGACGCTCTTCCGATCTNNNACCCAGTAACAACAATTCCCCAA
GAGCACACGTCTGAACTCTTTCCCTACACGACGCTCTTCCGATCTNNNTTTCAATCCACTTCACATGCCAC
AGCACACGTCTGAACTCTTTCCCTACACGACGCTCTTCCGATCTNNNCAGTCTCCACACAACAAGCAAA
AGCACACGTCTGAACTCTTTCCCTACACGACGCTCTTCCGATCTNNNAAAATCAAGGTTCATGGCTGTG
AGCACACGTCTGAACTCTTTCCCTACACGACGCTCTTCCGATCTNNNGGTAGAGGTCTTCCCATAGAG
AGCACACGTCTGAACTCTTTCCCTACACGACGCTCTTCCGATCTNNNGACAAGGTGTCATTGGTTTCA
GAGCACACGTCTGAACTCTTTCCCTACACGACGCTCTTCCGATCTNNNGTCCATCTGAAGTTTCCTTTG
AGCACACGTCTGAACTCTTTCCCTACACGACGCTCTTCCGATCTNNNATACTTAGCAGACCAGGAAAGG
GAGCACACGTCTGAACTCTTTCCCTACACGACGCTCTTCCGATCTNNNGGGTTTGGGGCAAATAACACA
AGCACACGTCTGAACTCTTTCCCTACACGACGCTCTTCCGATCTNNNGGGTAAGGGGAGGTTCATCAT
GAGCACACGTCTGAACTCTTTCCCTACACGACGCTCTTCCGATCTNNNCCCTTGTAGGAAAGCCTCAGT
AGCACACGTCTGAACTCTTTCCCTACACGACGCTCTTCCGATCTNNNACAGGGTTTCACTGTAGATGG
AGCACACGTCTGAACTCTTTCCCTACACGACGCTCTTCCGATCTNNNATTGTTCCTAACATCCAGGGGTG
AGCACACGTCTGAACTCTTTCCCTACACGACGCTCTTCCGATCTNNNCCTTCCTTCCCTCCTTTTTCCTCA
GAGCACACGTCTGAACTCTTTCCCTACACGACGCTCTTCCGATCTNNNCTGGGCCTCGATTTTTAGAA
GAGCACACGTCTGAACTCTTTCCCTACACGACGCTCTTCCGATCTNNNTGCCAACATGTTCTTGTATGC
GCACACGTCTGAACTCTTTCCCTACACGACGCTCTTCCGATCTNNNGGCCACATCAAGAAGTTGACAAA

AGCACACGTCTGAACTCTTTCCCTACACGACGCTCTTCCGATCTNNNTGAGATTGTGCCATTACACTCCA
AGCACACGTCTGAACTCTTTCCCTACACGACGCTCTTCCGATCTNNNAGAAAGGAAAGAGGTTCACGT
GAGCACACGTCTGAACTCTTTCCCTACACGACGCTCTTCCGATCTNNNGAGAATGGCATGAACCCGAG
GAGCACACGTCTGAACTCTTTCCCTACACGACGCTCTTCCGATCTNNNTTCCTAAGCGTAATGACTGGG
AGCACACGTCTGAACTCTTTCCCTACACGACGCTCTTCCGATCTNNNGTGAAGCCCCGTCTCTACTAAA
GAGCACACGTCTGAACTCTTTCCCTACACGACGCTCTTCCGATCTNNNCTTCCTAAAAAGCTCCAAGCCA
AGCACACGTCTGAACTCTTTCCCTACACGACGCTCTTCCGATCTNNNGATGAGAGTTGAAGGTGGCCT
GAGCACACGTCTGAACTCTTTCCCTACACGACGCTCTTCCGATCTNNNCCCTCTTGTGTGTGATCTCTCT
AAGAGCACACGTCTGAACTCTTTCCCTACACGACGCTCTTCCGATCTNNNCAATGTGCTGTGTCTCACTT
AGAGCACACGTCTGAACTCTTTCCCTACACGACGCTCTTCCGATCTNNNTACTTTCTACAGCAGGATGT
AGAGCACACGTCTGAACTCTTTCCCTACACGACGCTCTTCCGATCTNNNTGGATGATAATGCTGCCAT
GAGCACACGTCTGAACTCTTTCCCTACACGACGCTCTTCCGATCTNNNCCCAAAAGAAAAGTAGTGCA
AGAGCACACGTCTGAACTCTTTCCCTACACGACGCTCTTCCGATCTNNNGGGAAAGGGAAGACAGTG
GAGCACACGTCTGAACTCTTTCCCTACACGACGCTCTTCCGATCTNNNGCCAGCCATATACTGTACTCCT
AGCACACGTCTGAACTCTTTCCCTACACGACGCTCTTCCGATCTNNNCACCTGTAATCCCAGCTACTCAG
AGCACACGTCTGAACTCTTTCCCTACACGACGCTCTTCCGATCTNNNTCTCTCTCTCTCTCACACACAC
AGCACACGTCTGAACTCTTTCCCTACACGACGCTCTTCCGATCTNNNGCCTCAACTTGGCTTTTTTGATA
AGCACACGTCTGAACTCTTTCCCTACACGACGCTCTTCCGATCTNNNATGTGCCACATTAGGGTAGGAA
AGCACACGTCTGAACTCTTTCCCTACACGACGCTCTTCCGATCTNNNTTTGCTTGAATTCCTGCCAGAA
AGCACACGTCTGAACTCTTTCCCTACACGACGCTCTTCCGATCTNNNCCCAGAACTTTGAAAGGCTGAA
AGCACACGTCTGAACTCTTTCCCTACACGACGCTCTTCCGATCTNNNTCCCCACCAACCTCAATTATCAG
AGAGCACACGTCTGAACTCTTTCCCTACACGACGCTCTTCCGATCTNNNGGTGGAAGAGCCTCTTGTTC
AGCACACGTCTGAACTCTTTCCCTACACGACGCTCTTCCGATCTNNNCTGAGGCATGAAAATCGCTTGA
GAGCACACGTCTGAACTCTTTCCCTACACGACGCTCTTCCGATCTNNNCTGGGTTCAAGCATTTCTCCAT
GAGCACACGTCTGAACTCTTTCCCTACACGACGCTCTTCCGATCTNNNAAGATCTCTCCCTCTCCTTCTCC
GAGCACACGTCTGAACTCTTTCCCTACACGACGCTCTTCCGATCTNNNCACTTGGTTTCTGTGTGTTCCT
AGCACACGTCTGAACTCTTTCCCTACACGACGCTCTTCCGATCTNNNAGATGTGGTCTTGCTCTGTCTT
AGCACACGTCTGAACTCTTTCCCTACACGACGCTCTTCCGATCTNNNAGCCTAACAACAAGAAAAAGC
GAGCACACGTCTGAACTCTTTCCCTACACGACGCTCTTCCGATCTNNNGTCCAGCTTCTTGGTTAGCTTT
AGCACACGTCTGAACTCTTTCCCTACACGACGCTCTTCCGATCTNNNTCTTCTCGCCTTTTTTCCCCTTCCA
AGCACACGTCTGAACTCTTTCCCTACACGACGCTCTTCCGATCTNNNGGCACACAAAAAGGCAAAACA
AGCACACGTCTGAACTCTTTCCCTACACGACGCTCTTCCGATCTNNNGCCCTGAATCTGCTTTTTAGCAA
AGCACACGTCTGAACTCTTTCCCTACACGACGCTCTTCCGATCTNNNACCAAACGTTCATTAGTGCCAA
AGCACACGTCTGAACTCTTTCCCTACACGACGCTCTTCCGATCTNNNGAATGTGCTCATGGAGGCATTT
GAGCACACGTCTGAACTCTTTCCCTACACGACGCTCTTCCGATCTNNNAATTCCAAATGCTAGAGGTG
AGCACACGTCTGAACTCTTTCCCTACACGACGCTCTTCCGATCTNNNGTTGAGACACTGGCCAAGATA
AGCACACGTCTGAACTCTTTCCCTACACGACGCTCTTCCGATCTNNNGCACTGCCACAAATACCTTGTA
AGCACACGTCTGAACTCTTTCCCTACACGACGCTCTTCCGATCTNNNTGTACACCCACTGCAATATCGA
GAGCACACGTCTGAACTCTTTCCCTACACGACGCTCTTCCGATCTNNNAGAAACACACAACCTCCCAAG
GAGCACACGTCTGAACTCTTTCCCTACACGACGCTCTTCCGATCTNNNTGAGGCAGGAGAATCACTTGA
GAGCACACGTCTGAACTCTTTCCCTACACGACGCTCTTCCGATCTNNNTGGAGACCTTGTGAGAAATG
AGCACACGTCTGAACTCTTTCCCTACACGACGCTCTTCCGATCTNNNAAAACGGAAGAGGCAGGAAAA
AGCACACGTCTGAACTCTTTCCCTACACGACGCTCTTCCGATCTNNNTGCCTGTAATCCCAACTACTTGG

GAGCACACGTCTGAACTCTTTCCCTACACGACGCTCTTCCGATCTNNNACAAAAATTAGCTGGGCATG
AGCACACGTCTGAACTCTTTCCCTACACGACGCTCTTCCGATCTNNNTTTCAAGCGATTCTCCTGTCTCA
GAGCACACGTCTGAACTCTTTCCCTACACGACGCTCTTCCGATCTNNNGCTTTCCTCAGGTTTTCTTCCA
AGAGCACACGTCTGAACTCTTTCCCTACACGACGCTCTTCCGATCTNNNTTCGAAGAGCTTGCAACTG
GAGCACACGTCTGAACTCTTTCCCTACACGACGCTCTTCCGATCTNNNAAAAATAAATACACAGGCCG
GAGCACACGTCTGAACTCTTTCCCTACACGACGCTCTTCCGATCTNNNTCCTACCCAACAAAAGTCACT
AGCACACGTCTGAACTCTTTCCCTACACGACGCTCTTCCGATCTNNNTTCTCTCTCTGCAACAGTGTCTC
AGCACACGTCTGAACTCTTTCCCTACACGACGCTCTTCCGATCTNNNGGTGGTGGCTAGAGGATGTT
GAGCACACGTCTGAACTCTTTCCCTACACGACGCTCTTCCGATCTNNNCCTCCCCACCATTCATATGTTG
GCACACGTCTGAACTCTTTCCCTACACGACGCTCTTCCGATCTNNNATTGCATGAAGTCAGAGACACC
AGCACACGTCTGAACTCTTTCCCTACACGACGCTCTTCCGATCTNNNTGGATCAGTGGTCTCCAAAACA
AGAGCACACGTCTGAACTCTTTCCCTACACGACGCTCTTCCGATCTNNNGTGTACAACCCCCTGTGATTT
GAGCACACGTCTGAACTCTTTCCCTACACGACGCTCTTCCGATCTNNNCTCCATCCTGTTCTTCCAAATG
GAGCACACGTCTGAACTCTTTCCCTACACGACGCTCTTCCGATCTNNNCTTCTGCCCCTTTTTTGTCTCT
AGAGCACACGTCTGAACTCTTTCCCTACACGACGCTCTTCCGATCTNNNTTTTTAGTAGAGGCGGGGT
GAGCACACGTCTGAACTCTTTCCCTACACGACGCTCTTCCGATCTNNNGGGGCCTCAGTGTTAGAAGA
AGCACACGTCTGAACTCTTTCCCTACACGACGCTCTTCCGATCTNNNTAGAGGTTGCAGTGAGCCAAA
GAGCACACGTCTGAACTCTTTCCCTACACGACGCTCTTCCGATCTNNNTTCATCCTGACTTGTCTCACCT
AGCACACGTCTGAACTCTTTCCCTACACGACGCTCTTCCGATCTNNNTCTGGAAGTTTTGTCTCAGAGG
AGAGCACACGTCTGAACTCTTTCCCTACACGACGCTCTTCCGATCTNNNGGATTACAGGTGTGAGCCT
AGAGCACACGTCTGAACTCTTTCCCTACACGACGCTCTTCCGATCTNNNAGGTATGCCTGACACATAAG
GAGCACACGTCTGAACTCTTTCCCTACACGACGCTCTTCCGATCTNNNCAGCCTGCTCACACTCATTTAT
AGCACACGTCTGAACTCTTTCCCTACACGACGCTCTTCCGATCTNNNGATAGGTGATGTTCCCGTACTC
GAGCACACGTCTGAACTCTTTCCCTACACGACGCTCTTCCGATCTNNNTCTCTCTCTCTCTCTCTCCCT
GAGCACACGTCTGAACTCTTTCCCTACACGACGCTCTTCCGATCTNNNCATCCTTTTCCTCCCTGGGTAT
GCACACGTCTGAACTCTTTCCCTACACGACGCTCTTCCGATCTNNNGCTACCATGCCCAGCTAATTTTC
GAGCACACGTCTGAACTCTTTCCCTACACGACGCTCTTCCGATCTNNNTCTCTGTGTATGTGAGACATG
GAGCACACGTCTGAACTCTTTCCCTACACGACGCTCTTCCGATCTNNNTCGTCTAGTGACAGAGTGAG
AGCACACGTCTGAACTCTTTCCCTACACGACGCTCTTCCGATCTNNNAGTTCATTCATCAGGTCTCCTC
AGCACACGTCTGAACTCTTTCCCTACACGACGCTCTTCCGATCTNNNAATTCACTTTCAGGAAGGCAT
AGAGCACACGTCTGAACTCTTTCCCTACACGACGCTCTTCCGATCTNNNTCCCATTGGGTTTCATCAACT
AGCACACGTCTGAACTCTTTCCCTACACGACGCTCTTCCGATCTNNNCCTTGTTTGGCCCAACTTACAA
AGCACACGTCTGAACTCTTTCCCTACACGACGCTCTTCCGATCTNNNTTTGATTCTCAGCCAAGGGAC
GAGCACACGTCTGAACTCTTTCCCTACACGACGCTCTTCCGATCTNNNCAGGAAAGGAGAATGGACCT
GAGCACACGTCTGAACTCTTTCCCTACACGACGCTCTTCCGATCTNNNGCATATTCGAATCACCTGGTG
AGCACACGTCTGAACTCTTTCCCTACACGACGCTCTTCCGATCTNNNAGAGAAAGCTAACCCCTGACA
GAGCACACGTCTGAACTCTTTCCCTACACGACGCTCTTCCGATCTNNNTGGCTTTGGAAATAGAGGG
GAGCACACGTCTGAACTCTTTCCCTACACGACGCTCTTCCGATCTNNNTGACTATGACACATGAGACGG
AGCACACGTCTGAACTCTTTCCCTACACGACGCTCTTCCGATCTNNNGTTTTAGTTGTCACAGCTGGA
AGCACACGTCTGAACTCTTTCCCTACACGACGCTCTTCCGATCTNNNTGAACTCCTGAGCTCAAGTGA
GAGCACACGTCTGAACTCTTTCCCTACACGACGCTCTTCCGATCTNNNCTGTAATCCCAGCTACTCGTG
GAGCACACGTCTGAACTCTTTCCCTACACGACGCTCTTCCGATCTNNNGAACAAAACCTTCCAGGGAAT
AGAGCACACGTCTGAACTCTTTCCCTACACGACGCTCTTCCGATCTNNNTATGAATGTCACCACTCAAG

AGCACACGTCTGAACTCTTTCCCTACACGACGCTCTTCCGATCTNNNTTTGTGTAGTCTGCAGATGTG
AGCACACGTCTGAACTCTTTCCCTACACGACGCTCTTCCGATCTNNNGATCTCTTGTGACCTTCCTGG
GAGCACACGTCTGAACTCTTTCCCTACACGACGCTCTTCCGATCTNNNTCACTGTATTGAGACCAGGT
AGCACACGTCTGAACTCTTTCCCTACACGACGCTCTTCCGATCTNNNACCCTGCAGTCTCTGTCTTTTT
AGCACACGTCTGAACTCTTTCCCTACACGACGCTCTTCCGATCTNNNTATTTTAGCAGAGACGGGG
GAGCACACGTCTGAACTCTTTCCCTACACGACGCTCTTCCGATCTNNNTTCGAGACCAGACTGACCAA
GCACACGTCTGAACTCTTTCCCTACACGACGCTCTTCCGATCTNNNCTCCAGAGTCATAGAACCGTCTG
GCACACGTCTGAACTCTTTCCCTACACGACGCTCTTCCGATCTNNNCCACGCCCAGCTAATTTTTGTAT
GAGCACACGTCTGAACTCTTTCCCTACACGACGCTCTTCCGATCTNNNTAATAAAACACCAGTGGGGA
AGCACACGTCTGAACTCTTTCCCTACACGACGCTCTTCCGATCTNNNAGGGTCCTGTTCTGTCAAGA
GAGCACACGTCTGAACTCTTTCCCTACACGACGCTCTTCCGATCTNNNGTAAAGGCCAACTCATGAG
GAGCACACGTCTGAACTCTTTCCCTACACGACGCTCTTCCGATCTNNNCGCATTGTTAGTCTTGCTCC
GCACACGTCTGAACTCTTTCCCTACACGACGCTCTTCCGATCTNNNGCGCCTAGATAAGGGTTTAGGA
AGCACACGTCTGAACTCTTTCCCTACACGACGCTCTTCCGATCTNNNGCTTCACAATCTCCTGAGACT
AGCACACGTCTGAACTCTTTCCCTACACGACGCTCTTCCGATCTNNNAGAGAACAATCCGACTTCCAC
AGCACACGTCTGAACTCTTTCCCTACACGACGCTCTTCCGATCTNNNGAATTGGGAGGTACCTGCTT
AGCACACGTCTGAACTCTTTCCCTACACGACGCTCTTCCGATCTNNNCTCACATCTGTAATCCCAGCACT
AGAGCACACGTCTGAACTCTTTCCCTACACGACGCTCTTCCGATCTNNNTGGACTGGGGAGTTGGTA
AGCACACGTCTGAACTCTTTCCCTACACGACGCTCTTCCGATCTNNNAGAGACCTGGACCTAGACTTG
AGCACACGTCTGAACTCTTTCCCTACACGACGCTCTTCCGATCTNNNGATCTTCCGCGGATAGGAGTG
AGCACACGTCTGAACTCTTTCCCTACACGACGCTCTTCCGATCTNNNCGGACTGGGGAACAAGAGTA
AGCACACGTCTGAACTCTTTCCCTACACGACGCTCTTCCGATCTNNNTTTGTTTTCTCGCAGTTCTGGA
GAGCACACGTCTGAACTCTTTCCCTACACGACGCTCTTCCGATCTNNNTTTGTGAAAAGACTGTCCCTC
GCACACGTCTGAACTCTTTCCCTACACGACGCTCTTCCGATCTNNNTTATCTCCTCTTGTCTGCCCACC
CACACGTCTGAACTCTTTCCCTACACGACGCTCTTCCGATCTNNNGCCTATCTCTTTCACATGCCTCTC
AGCACACGTCTGAACTCTTTCCCTACACGACGCTCTTCCGATCTNNNATTCTACCAGCTCAGATTGCA
GCACACGTCTGAACTCTTTCCCTACACGACGCTCTTCCGATCTNNNAAGGGACTCGATGGGAACAAA
AGCACACGTCTGAACTCTTTCCCTACACGACGCTCTTCCGATCTNNNCTCGGGAAAGGATTACTCAGC
GAGCACACGTCTGAACTCTTTCCCTACACGACGCTCTTCCGATCTNNNTAAAGGGTCATCCATGCTCA
AGCACACGTCTGAACTCTTTCCCTACACGACGCTCTTCCGATCTNNNGCTCCAAATCCAACCTCTTTCC
GAGCACACGTCTGAACTCTTTCCCTACACGACGCTCTTCCGATCTNNNCACTTGAGGTCAGGAGTTC
AGCACACGTCTGAACTCTTTCCCTACACGACGCTCTTCCGATCTNNNCTTGAACTCCTGACCTCAAGTG
GCACACGTCTGAACTCTTTCCCTACACGACGCTCTTCCGATCTNNNGTCCTGTTACTCCCTCATCAGTG
AGCACACGTCTGAACTCTTTCCCTACACGACGCTCTTCCGATCTNNNTGTCTCTCTCTCTGTCTCTCTC
GAGCACACGTCTGAACTCTTTCCCTACACGACGCTCTTCCGATCTNNNAGATGGGTCAATCGGAACT
AGCACACGTCTGAACTCTTTCCCTACACGACGCTCTTCCGATCTNNNCTTGCTCTCTGCTGATTTGACA
CACACGTCTGAACTCTTTCCCTACACGACGCTCTTCCGATCTNNNAAGCCAGGCTCTTGGAATATTCC
GAGCACACGTCTGAACTCTTTCCCTACACGACGCTCTTCCGATCTNNNTCTTTGGCTTTTACCCACTGA
AGCACACGTCTGAACTCTTTCCCTACACGACGCTCTTCCGATCTNNNTCATGTCAATCCCACTTTGCCT
GCACACGTCTGAACTCTTTCCCTACACGACGCTCTTCCGATCTNNNAGCCACTTCATCTAGCAACAAG
CACACGTCTGAACTCTTTCCCTACACGACGCTCTTCCGATCTNNNAAGAGCCACTTCATCTAGCAACAC
AGCACACGTCTGAACTCTTTCCCTACACGACGCTCTTCCGATCTNNNGGCTGGAGGAAGAACAAACT
GAGCACACGTCTGAACTCTTTCCCTACACGACGCTCTTCCGATCTNNNGGTCCCTCTTTGCTCTCTAGA

AGAGCACACGTCTGAACTCTTTCCCTACACGACGCTCTTCCGATCTNNNCACATGGGATTTGGAAAA
AGCACACGTCTGAACTCTTTCCCTACACGACGCTCTTCCGATCTNNNGAGCCAGTGATTGAGACAGA
GCACACGTCTGAACTCTTTCCCTACACGACGCTCTTCCGATCTNNNGTGATGGCTTTCTTACTGGCTTG
AGCACACGTCTGAACTCTTTCCCTACACGACGCTCTTCCGATCTNNNACCATTCATTCAGCCAATTTGC
GCACACGTCTGAACTCTTTCCCTACACGACGCTCTTCCGATCTNNNGTTTCAGTGAGCCAAGTTCAAG
GCACACGTCTGAACTCTTTCCCTACACGACGCTCTTCCGATCTNNNTTTTGGTTTACATCATGTCGGGC
AGCACACGTCTGAACTCTTTCCCTACACGACGCTCTTCCGATCTNNNAAGAGGCAGAGTGGGAGATT
AGCACACGTCTGAACTCTTTCCCTACACGACGCTCTTCCGATCTNNNATTATGTTTGCGTGTGTGTGT
AGCACACGTCTGAACTCTTTCCCTACACGACGCTCTTCCGATCTNNNAAGCACACATGAGACAGAGA
AGAGCACACGTCTGAACTCTTTCCCTACACGACGCTCTTCCGATCTNNNGAAGGGTAGCAGAGAGAG
AGCACACGTCTGAACTCTTTCCCTACACGACGCTCTTCCGATCTNNNCCGAGAGAGAGAGAGAGAG
GCACACGTCTGAACTCTTTCCCTACACGACGCTCTTCCGATCTNNNTCTCTCTCTCTCTGTGTGTGT
AGCACACGTCTGAACTCTTTCCCTACACGACGCTCTTCCGATCTNNNGCATAACTCCAGCATCTGCAT
GAGCACACGTCTGAACTCTTTCCCTACACGACGCTCTTCCGATCTNNNCAGCTCACATCCTGAAGTTAG
AGCACACGTCTGAACTCTTTCCCTACACGACGCTCTTCCGATCTNNNCAGCCCCCATATACATCAGGTT
GCACACGTCTGAACTCTTTCCCTACACGACGCTCTTCCGATCTNNNGAGCAAGTCTGGTGTTCTTTCA
AGCACACGTCTGAACTCTTTCCCTACACGACGCTCTTCCGATCTNNNACTCATGACCTATTCCTGGGG
AGCACACGTCTGAACTCTTTCCCTACACGACGCTCTTCCGATCTNNNCAGCAAGACCGTGACTCAAAA
GCACACGTCTGAACTCTTTCCCTACACGACGCTCTTCCGATCTNNNTGCCATGTATGATAGGGTAAG
GCACACGTCTGAACTCTTTCCCTACACGACGCTCTTCCGATCTNNNTCCACAGAGCTTTTCTTATGGGT
GCACACGTCTGAACTCTTTCCCTACACGACGCTCTTCCGATCTNNNCAGCTCTTCCTGATTCTGCAGTA
GCACACGTCTGAACTCTTTCCCTACACGACGCTCTTCCGATCTNNNGCCTTGATGGTCCTGGATAAGA
AGCACACGTCTGAACTCTTTCCCTACACGACGCTCTTCCGATCTNNNAGGCATCTCAGAAGGTTTTTG
GCACACGTCTGAACTCTTTCCCTACACGACGCTCTTCCGATCTNNNAGGAGAGAGAGAGAGAGGAG
GCACACGTCTGAACTCTTTCCCTACACGACGCTCTTCCGATCTNNNACGATTTGGCAAGACACAAGAT
AGCACACGTCTGAACTCTTTCCCTACACGACGCTCTTCCGATCTNNNAATTGTCAGGGGGTTGATGAT
AGAGCACACGTCTGAACTCTTTCCCTACACGACGCTCTTCCGATCTNNNGTCTGGAGTAGTTTGGGG
AGAGCACACGTCTGAACTCTTTCCCTACACGACGCTCTTCCGATCTNNNTTTTCCCAGACGAGGCTTCA
AGCACACGTCTGAACTCTTTCCCTACACGACGCTCTTCCGATCTNNNCAGTTATGGTGGAAGGCAAA
GAGCACACGTCTGAACTCTTTCCCTACACGACGCTCTTCCGATCTNNNTCTCTCTCTCTCTCTCTCCCT
AGCACACGTCTGAACTCTTTCCCTACACGACGCTCTTCCGATCTNNNCTCTCTCTCTGTCTCTCTCCCTCC
CACACGTCTGAACTCTTTCCCTACACGACGCTCTTCCGATCTNNNCACTCCCTCCCTCCTTTTTTTCTCAT
AGCACACGTCTGAACTCTTTCCCTACACGACGCTCTTCCGATCTNNNTCACCTCTCTGTTTCTCTGTGTC
GCACACGTCTGAACTCTTTCCCTACACGACGCTCTTCCGATCTNNNATTGAATCTTGGGGGCAAACTTC
GCACACGTCTGAACTCTTTCCCTACACGACGCTCTTCCGATCTNNNTGACTGCTCAACTGATAAGCTG
GCACACGTCTGAACTCTTTCCCTACACGACGCTCTTCCGATCTNNNTGACCTGGCCAAATACTGATGA
GCACACGTCTGAACTCTTTCCCTACACGACGCTCTTCCGATCTNNNGTTCCTTCCTTCCTTCCTTCCTTC
GCACACGTCTGAACTCTTTCCCTACACGACGCTCTTCCGATCTNNNCTTTCTCTCTGGCCTCTCTTCTCCA
AGCACACGTCTGAACTCTTTCCCTACACGACGCTCTTCCGATCTNNNGAGGAAGAGGAGGGTTTTG
GCACACGTCTGAACTCTTTCCCTACACGACGCTCTTCCGATCTNNNCCATATGTGTGTGTGTGTGTGT
AGAGCACACGTCTGAACTCTTTCCCTACACGACGCTCTTCCGATCTNNNCTGAGGCAGGAGAATTGC
AGCACACGTCTGAACTCTTTCCCTACACGACGCTCTTCCGATCTNNNAGGTACAGAAAGGCAATGTG
GCACACGTCTGAACTCTTTCCCTACACGACGCTCTTCCGATCTNNNACTTCCTCAAGTAAGTCAGCCTG

AGAGCACACGTCTGAACTCTTTCCCTACACGACGCTCTTCCGATCTNNNTGGGAAATGGCCAGCAATA
GAGCACACGTCTGAACTCTTTCCCTACACGACGCTCTTCCGATCTNNNATATGTGTGTGTGTGTGTGTG
AAGAGCACACGTCTGAACTCTTTCCCTACACGACGCTCTTCCGATCTNNNGAGGAAACATAGCAGCATC
AGAGCACACGTCTGAACTCTTTCCCTACACGACGCTCTTCCGATCTNNNAAAATGTCTTCACATGGTGG
GAGCACACGTCTGAACTCTTTCCCTACACGACGCTCTTCCGATCTNNNGGGCAGCGCTTGTATTATCTA
GAGCACACGTCTGAACTCTTTCCCTACACGACGCTCTTCCGATCTNNNGCACTTCCACTTCCAGAAATGT
AGAGCACACGTCTGAACTCTTTCCCTACACGACGCTCTTCCGATCTNNNTGATGCTGGAAAGATGGGA
GAGCACACGTCTGAACTCTTTCCCTACACGACGCTCTTCCGATCTNNNAGGCATCTCTCTCTTTCCTCT
CACACGTCTGAACTCTTTCCCTACACGACGCTCTTCCGATCTNNNGAAACATCCAATCCTGCTTTGCTCA
GAGCACACGTCTGAACTCTTTCCCTACACGACGCTCTTCCGATCTNNNCTTCTCTCTTCTCCCTCTCATGCC
GAGCACACGTCTGAACTCTTTCCCTACACGACGCTCTTCCGATCTNNNCAATTGCTCCAGGTCTCTCTTCT
AGAGCACACGTCTGAACTCTTTCCCTACACGACGCTCTTCCGATCTNNNCAAGAGCAAAACCGTCTCAG
AGAGCACACGTCTGAACTCTTTCCCTACACGACGCTCTTCCGATCTNNNAGACAGGAGAATCGCTTGAA
AGCACACGTCTGAACTCTTTCCCTACACGACGCTCTTCCGATCTNNNTGCAATTACAGTGGAAGGCAA
GAGCACACGTCTGAACTCTTTCCCTACACGACGCTCTTCCGATCTNNNGAGAGAGAGAGAGAGAGAG
GAGCACACGTCTGAACTCTTTCCCTACACGACGCTCTTCCGATCTNNNATGTGTGTGAGGAGAGAGAG
AGAGCACACGTCTGAACTCTTTCCCTACACGACGCTCTTCCGATCTNNNAAGCCGCTGGAAAATGTAGT
GAGCACACGTCTGAACTCTTTCCCTACACGACGCTCTTCCGATCTNNNAAAGGGAACATAGAAGGGAG
GAGCACACGTCTGAACTCTTTCCCTACACGACGCTCTTCCGATCTNNNGGGTTGTTCAGAGAAACAGAA
GAGCACACGTCTGAACTCTTTCCCTACACGACGCTCTTCCGATCTNNNAGTAGAAGCACCACAGCTAAG
GAGCACACGTCTGAACTCTTTCCCTACACGACGCTCTTCCGATCTNNNTCCATATGTGTCTCAGTGGCAA
GAGCACACGTCTGAACTCTTTCCCTACACGACGCTCTTCCGATCTNNNTTCATACACCTCTCTCCATCCCT
GAGCACACGTCTGAACTCTTTCCCTACACGACGCTCTTCCGATCTNNNGGAGAACTTTAGGAGGCAAA
GAGCACACGTCTGAACTCTTTCCCTACACGACGCTCTTCCGATCTNNNCCTCTCCATCTCTCTCTCCATCT
AGCACACGTCTGAACTCTTTCCCTACACGACGCTCTTCCGATCTNNNATAAAAGGAAGCCTGCCTGCAT
CACACGTCTGAACTCTTTCCCTACACGACGCTCTTCCGATCTNNNCTCCCTCCTTCCTTCCTTCTTTCCATC
GAGCACACGTCTGAACTCTTTCCCTACACGACGCTCTTCCGATCTNNNACCTCTCTGTCTGTCTCTCTCTC
AGCACACGTCTGAACTCTTTCCCTACACGACGCTCTTCCGATCTNNNCAAGCCATCCTTCTGTTTCAGTG
AGCACACGTCTGAACTCTTTCCCTACACGACGCTCTTCCGATCTNNNTGGGCACATGTTCTTAGGATCTC
GAGCACACGTCTGAACTCTTTCCCTACACGACGCTCTTCCGATCTNNNGAAGGGGAAGCAAGCACATA
AGCACACGTCTGAACTCTTTCCCTACACGACGCTCTTCCGATCTNNNATTATGCTGTGTGTGTGTGTGT
GAGCACACGTCTGAACTCTTTCCCTACACGACGCTCTTCCGATCTNNNAGTGAACTGAGATCATGCCAC
AGCACACGTCTGAACTCTTTCCCTACACGACGCTCTTCCGATCTNNNGGAGGAGAGGAGAGAGAGAG
GCACACGTCTGAACTCTTTCCCTACACGACGCTCTTCCGATCTNNNAGAAAGAGAGAGACAGAGAGG
AAGAGCACACGTCTGAACTCTTTCCCTACACGACGCTCTTCCGATCTNNNACTCAGCAGAGATAATGTG
GAGCACACGTCTGAACTCTTTCCCTACACGACGCTCTTCCGATCTNNNTGAGGCAGGAGAATCTCTTG
AGAGCACACGTCTGAACTCTTTCCCTACACGACGCTCTTCCGATCTNNNCACCTCCAGCCTTTCTTACTTC
GCACACGTCTGAACTCTTTCCCTACACGACGCTCTTCCGATCTNNNAGGAGTTTCAGGCTGTCATGAATG
AGCACACGTCTGAACTCTTTCCCTACACGACGCTCTTCCGATCTNNNTGTCTACTTCAGTCTGGTCTTGC
AGCACACGTCTGAACTCTTTCCCTACACGACGCTCTTCCGATCTNNNCTCTTTCCTTCCTTCCTTCCTCCCA
AGCACACGTCTGAACTCTTTCCCTACACGACGCTCTTCCGATCTNNNCCCTTTCTCTCTTCCTTCCTTCCCA
GAGCACACGTCTGAACTCTTTCCCTACACGACGCTCTTCCGATCTNNNAACATGGGGTAATGGAATGT
AGCACACGTCTGAACTCTTTCCCTACACGACGCTCTTCCGATCTNNNTCCGTCTCTTTCTGTCTCTCTCTC

AGCACACGTCTGAACTCTTTCCCTACACGACGCTCTTCCGATCTNNNAGATTGCAATGAGCCGAGATA
AGAGCACACGTCTGAACTCTTTCCCTACACGACGCTCTTCCGATCTNNNGGCATGAGAAATTAGGATG
AGCACACGTCTGAACTCTTTCCCTACACGACGCTCTTCCGATCTNNNAGACTCCATCTCAAGGACAAAAA
AGAGCACACGTCTGAACTCTTTCCCTACACGACGCTCTTCCGATCTNNNCAGCACAACTATGGCAAAGT
AGCACACGTCTGAACTCTTTCCCTACACGACGCTCTTCCGATCTNNNTTGAAAGTGTGTGTGTGTG
GAGCACACGTCTGAACTCTTTCCCTACACGACGCTCTTCCGATCTNNNCAAAGAGAGAGGGAGAGAG
AGAGCACACGTCTGAACTCTTTCCCTACACGACGCTCTTCCGATCTNNNGAGCTAGACTCTGTCTCAAAA
AGCACACGTCTGAACTCTTTCCCTACACGACGCTCTTCCGATCTNNNTGATCTCTAAGGCATCTTCCAG
GAGCACACGTCTGAACTCTTTCCCTACACGACGCTCTTCCGATCTNNNAGCAGGTTGGGTATAAACAG
GCACACGTCTGAACTCTTTCCCTACACGACGCTCTTCCGATCTNNNGACAGAGAGAGAGAGGGAGA
GAGCACACGTCTGAACTCTTTCCCTACACGACGCTCTTCCGATCTNNNTGGGACTATCAATAAGGCTG
GAGCACACGTCTGAACTCTTTCCCTACACGACGCTCTTCCGATCTNNNCCTTCTCCCTTTCTCTGTTCCT
GCACACGTCTGAACTCTTTCCCTACACGACGCTCTTCCGATCTNNNCAGGGGGAAAAATAGTTGAAG
GCACACGTCTGAACTCTTTCCCTACACGACGCTCTTCCGATCTNNNGAACTCCTGACCTCAAGTGATCCCA
GCACACGTCTGAACTCTTTCCCTACACGACGCTCTTCCGATCTNNNGTCTGTGGGATCTTGTGTCATCT
AGCACACGTCTGAACTCTTTCCCTACACGACGCTCTTCCGATCTNNNTGCAGCAATACATGTAACTCTC
GAGCACACGTCTGAACTCTTTCCCTACACGACGCTCTTCCGATCTNNNCCTAGGCATTTATCCCAGGGA
GAGCACACGTCTGAACTCTTTCCCTACACGACGCTCTTCCGATCTNNNAGCTTGCAGAGAGTAGATCAT
GAGCACACGTCTGAACTCTTTCCCTACACGACGCTCTTCCGATCTNNNCCCTTGTAGTTCCAGCTACTCT
AGCACACGTCTGAACTCTTTCCCTACACGACGCTCTTCCGATCTNNNTTTCATGCAATGCTTTTGCCAAA
GAGCACACGTCTGAACTCTTTCCCTACACGACGCTCTTCCGATCTNNNTTGGGGGAACAAGAAGTTACA
GAGCACACGTCTGAACTCTTTCCCTACACGACGCTCTTCCGATCTNNNCTGAGGCAGGAGAATTGCTT
AGCACACGTCTGAACTCTTTCCCTACACGACGCTCTTCCGATCTNNNGGGAGCTACAATTCAAGGTGA
AAGAGCACACGTCTGAACTCTTTCCCTACACGACGCTCTTCCGATCTNNNCACCTGAGGTCATGAGTTT
GAGCACACGTCTGAACTCTTTCCCTACACGACGCTCTTCCGATCTNNNGGAGAGACAGCAAGGAAGAA
AGAGCACACGTCTGAACTCTTTCCCTACACGACGCTCTTCCGATCTNNNCACACACACACACACACACAT
GAGCACACGTCTGAACTCTTTCCCTACACGACGCTCTTCCGATCTNNNGGCAAAAGACTGAGACCCTG
AGAGCACACGTCTGAACTCTTTCCCTACACGACGCTCTTCCGATCTNNNAGTGAGCTGTGATTATGCCA
AGAGCACACGTCTGAACTCTTTCCCTACACGACGCTCTTCCGATCTNNNTGAGGCGTTTTGTTGAACTT
GAGCACACGTCTGAACTCTTTCCCTACACGACGCTCTTCCGATCTNNNTCACCAAAAAATCACTGGAAA
AGCACACGTCTGAACTCTTTCCCTACACGACGCTCTTCCGATCTNNNGATCCAAAATGTTCCCCCAATG
GCACACGTCTGAACTCTTTCCCTACACGACGCTCTTCCGATCTNNNCCTTTTCCTTCACTACACCCAGACA
AGCACACGTCTGAACTCTTTCCCTACACGACGCTCTTCCGATCTNNNTCAAAAACAAATGGGCTCCTCA
GCACACGTCTGAACTCTTTCCCTACACGACGCTCTTCCGATCTNNNGCTCTCTGATCTCCACTTACACCC
GCACACGTCTGAACTCTTTCCCTACACGACGCTCTTCCGATCTNNNACAGCTGTATATGCAAGTGTCCA
AGCACACGTCTGAACTCTTTCCCTACACGACGCTCTTCCGATCTNNNGGTCCTTTGGGAGGTAATTA
AGCACACGTCTGAACTCTTTCCCTACACGACGCTCTTCCGATCTNNNACACTGTATTGGAGATGCTAG
AGAGCACACGTCTGAACTCTTTCCCTACACGACGCTCTTCCGATCTNNNCCCAGGTTGGTGATAAAAT
AAGAGCACACGTCTGAACTCTTTCCCTACACGACGCTCTTCCGATCTNNNGGAGGCTGAGGTAGAAGA
AGAGCACACGTCTGAACTCTTTCCCTACACGACGCTCTTCCGATCTNNNGTCTAACAGACAGGCAGAA
GAGCACACGTCTGAACTCTTTCCCTACACGACGCTCTTCCGATCTNNNGAAGTTTCCCAGCATGACTGT
GAGCACACGTCTGAACTCTTTCCCTACACGACGCTCTTCCGATCTNNNAGGGAGGAAAAGAGAGAGA
GAGCACACGTCTGAACTCTTTCCCTACACGACGCTCTTCCGATCTNNNAGAGAGAAAGGAAGAAGGA

GAGCACACGTCTGAACTCTTTCCCTACACGACGCTCTTCCGATCTNNNAGAAATGCCGCTGATTTTGT
AGCACACGTCTGAACTCTTTCCCTACACGACGCTCTTCCGATCTNNNTTGTCAAAGTGTGGCTGTCTTT
AGCACACGTCTGAACTCTTTCCCTACACGACGCTCTTCCGATCTNNNCAGACACTCAGCATGGCTATTT
CACACGTCTGAACTCTTTCCCTACACGACGCTCTTCCGATCTNNNTCTCTCTCTCTCCCTCTCTCTCATC
AGCACACGTCTGAACTCTTTCCCTACACGACGCTCTTCCGATCTNNNAGACAGAAGAATCGCTTGAAC
AGAGCACACGTCTGAACTCTTTCCCTACACGACGCTCTTCCGATCTNNNACACGGCCTATTTCCAATACA
GAGCACACGTCTGAACTCTTTCCCTACACGACGCTCTTCCGATCTNNNGTACACACTGGTATCCCCTGT
GAGCACACGTCTGAACTCTTTCCCTACACGACGCTCTTCCGATCTNNNAAAATGTGTGTGTGTGTGTGT
AGCACACGTCTGAACTCTTTCCCTACACGACGCTCTTCCGATCTNNNGTGCTGAGTCTCACTGTTCTAG
GAGCACACGTCTGAACTCTTTCCCTACACGACGCTCTTCCGATCTNNNGCCACGTAAGAAGTCTAGCCA
AGAGCACACGTCTGAACTCTTTCCCTACACGACGCTCTTCCGATCTNNNTGCTGGCTACTGGTGATAAA
AGCACACGTCTGAACTCTTTCCCTACACGACGCTCTTCCGATCTNNNGTGGGAAAAGACAGAGGAGA
GAGCACACGTCTGAACTCTTTCCCTACACGACGCTCTTCCGATCTNNNGAGTGCAGTGGTACCTTGAA
GCACACGTCTGAACTCTTTCCCTACACGACGCTCTTCCGATCTNNNTCTCTCTCTCTCTCTCCCTCTCAT
GAGCACACGTCTGAACTCTTTCCCTACACGACGCTCTTCCGATCTNNNTTTTGTTGAAGAAAGCCTGG
AGAGCACACGTCTGAACTCTTTCCCTACACGACGCTCTTCCGATCTNNNAGAGCAATCCCTTCCCAGAAA
AGCACACGTCTGAACTCTTTCCCTACACGACGCTCTTCCGATCTNNNCTGTTACGTTCCGTTTCAAACCTC
AAGAGCACACGTCTGAACTCTTTCCCTACACGACGCTCTTCCGATCTNNNGTGCAGTGAGCCCTTATCA
AGCACACGTCTGAACTCTTTCCCTACACGACGCTCTTCCGATCTNNNAAAAGATACACGCACACACACA
AGCACACGTCTGAACTCTTTCCCTACACGACGCTCTTCCGATCTNNNAGGGAGAGAGAGAGAGAGAG
AGCACACGTCTGAACTCTTTCCCTACACGACGCTCTTCCGATCTNNNAGGAGAATCACCTGAACCTAG
AGCACACGTCTGAACTCTTTCCCTACACGACGCTCTTCCGATCTNNNAGCCTGTTCTTATTCACCAGTGT
AGCACACGTCTGAACTCTTTCCCTACACGACGCTCTTCCGATCTNNNCCCTCCCTTTCTCTTTCTTCCTCCA
GAGCACACGTCTGAACTCTTTCCCTACACGACGCTCTTCCGATCTNNNAAGGTTGCAGTGAGCTAAGA
AGCACACGTCTGAACTCTTTCCCTACACGACGCTCTTCCGATCTNNNACTCTGACTCTCTCTCTCTCTCA
GCACACGTCTGAACTCTTTCCCTACACGACGCTCTTCCGATCTNNNTCTCCCTTGCTCTTCATTCTTCCCA
AGCACACGTCTGAACTCTTTCCCTACACGACGCTCTTCCGATCTNNNTGGTCTCAAACTCCTGACTTCAG
GCACACGTCTGAACTCTTTCCCTACACGACGCTCTTCCGATCTNNNATGGGCTCCAAAGGGAAAAAAAG
GAGCACACGTCTGAACTCTTTCCCTACACGACGCTCTTCCGATCTNNNGCAACAGAGCAAGACTGTCA
GCACACGTCTGAACTCTTTCCCTACACGACGCTCTTCCGATCTNNNGACTTCCAGCGTGATGGTATTTG
AGCACACGTCTGAACTCTTTCCCTACACGACGCTCTTCCGATCTNNNAGAGAGGGACCCAGATAGAAC
GAGCACACGTCTGAACTCTTTCCCTACACGACGCTCTTCCGATCTNNNTTCATGCTGTGGTTCTCCTACT
GCACACGTCTGAACTCTTTCCCTACACGACGCTCTTCCGATCTNNNTCCCTGTCTCTCTCTCTGTTTCTCA
GAGCACACGTCTGAACTCTTTCCCTACACGACGCTCTTCCGATCTNNNTTTCTTTGGTGCAAGCTTGAGA
GAGCACACGTCTGAACTCTTTCCCTACACGACGCTCTTCCGATCTNNNCTAAGGCAGGAGGATCACTTC
GAGCACACGTCTGAACTCTTTCCCTACACGACGCTCTTCCGATCTNNNCACTCCAACCTGGGTGATAAG
AGCACACGTCTGAACTCTTTCCCTACACGACGCTCTTCCGATCTNNNGATGGGAGAGACAGAGAGAGT
GCACACGTCTGAACTCTTTCCCTACACGACGCTCTTCCGATCTNNNGCCAATGCTCTTGAGTTGAGAACC
GAGCACACGTCTGAACTCTTTCCCTACACGACGCTCTTCCGATCTNNNAGGATCATCAAAAGCCACTCT
GCACACGTCTGAACTCTTTCCCTACACGACGCTCTTCCGATCTNNNTCTCTCCCTCTTTCTCCTCTCCAT
AGCACACGTCTGAACTCTTTCCCTACACGACGCTCTTCCGATCTNNNCAGAGTGAGACCCTTCCTTTCTC
GAGCACACGTCTGAACTCTTTCCCTACACGACGCTCTTCCGATCTNNNGCACACACTCTCTCTTTCTCTCTC
AGCACACGTCTGAACTCTTTCCCTACACGACGCTCTTCCGATCTNNNCCAAGTCTGTCTGTCTCTGTCTCC

GAGCACACGTCTGAACTCTTTCCCTACACGACGCTCTTCCGATCTNNNACAGCATTACCCTGGTACCAA
GAGCACACGTCTGAACTCTTTCCCTACACGACGCTCTTCCGATCTNNNTGTAATCGCACCACTGTACTT
AGCACACGTCTGAACTCTTTCCCTACACGACGCTCTTCCGATCTNNNCACACGAGAGAGAGAGAGAG
GCACACGTCTGAACTCTTTCCCTACACGACGCTCTTCCGATCTNNNTTCAGAGACCAAGATGATGCCA
AGAGCACACGTCTGAACTCTTTCCCTACACGACGCTCTTCCGATCTNNNGAAATTGAAAGCCCAGTG
AGCACACGTCTGAACTCTTTCCCTACACGACGCTCTTCCGATCTNNNTAATTGAAGTAGCCCACCTGA
AGAGCACACGTCTGAACTCTTTCCCTACACGACGCTCTTCCGATCTNNNGCAGGAGAATCATTTGAA
GAGCACACGTCTGAACTCTTTCCCTACACGACGCTCTTCCGATCTNNNGAAAGACCAGCAGAGAGG
GAGCACACGTCTGAACTCTTTCCCTACACGACGCTCTTCCGATCTNNNTTAGGGAGCAGTGAGAGAG
GCACACGTCTGAACTCTTTCCCTACACGACGCTCTTCCGATCTNNNCTTCCTACAAAAGTCATGGCGT
CACACGTCTGAACTCTTTCCCTACACGACGCTCTTCCGATCTNNNGGTCCCTGAGCGGTAATTAGAAT
AGCACACGTCTGAACTCTTTCCCTACACGACGCTCTTCCGATCTNNNCTTTTCTTCCCTTTCCCTCCCTCC
GAGCACACGTCTGAACTCTTTCCCTACACGACGCTCTTCCGATCTNNNTAAGAAGAGAAACCAGAGC
AGCACACGTCTGAACTCTTTCCCTACACGACGCTCTTCCGATCTNNNTTCAAGGCTGCAGTAAGCTAA
GCACACGTCTGAACTCTTTCCCTACACGACGCTCTTCCGATCTNNNATTGCTTGAGTCCAGGAGGTCT
AGAGCACACGTCTGAACTCTTTCCCTACACGACGCTCTTCCGATCTNNNGAGGCAGGCAGATCATTT
AGCACACGTCTGAACTCTTTCCCTACACGACGCTCTTCCGATCTNNNCATATATTTGGGGAGTGGAG
GAGCACACGTCTGAACTCTTTCCCTACACGACGCTCTTCCGATCTNNNTGTGTTTGTGTGTGTGTGTG
AGCACACGTCTGAACTCTTTCCCTACACGACGCTCTTCCGATCTNNNATTGAATGTTGCTAGGACTGT
GAGCACACGTCTGAACTCTTTCCCTACACGACGCTCTTCCGATCTNNNATATGTGTGTGTGTGTGTGT
AGCACACGTCTGAACTCTTTCCCTACACGACGCTCTTCCGATCTNNNGAATGTGTGTGGGTGGAAA
GCACACGTCTGAACTCTTTCCCTACACGACGCTCTTCCGATCTNNNTAGAAGGTTACGCAGTTTTGG
AGCACACGTCTGAACTCTTTCCCTACACGACGCTCTTCCGATCTNNNCCACTTACACACACACACATAC
GCACACGTCTGAACTCTTTCCCTACACGACGCTCTTCCGATCTNNNACAGACCAAGGCTTCATCTCAA
GAGCACACGTCTGAACTCTTTCCCTACACGACGCTCTTCCGATCTNNNTGTCTCTGTCTGTCTCTCTCT
GAGCACACGTCTGAACTCTTTCCCTACACGACGCTCTTCCGATCTNNNACGACTTTTTATAGAGGCTCT
GAGCACACGTCTGAACTCTTTCCCTACACGACGCTCTTCCGATCTNNNGGTGAGAAGAAAGTTTCCAG
GAGCACACGTCTGAACTCTTTCCCTACACGACGCTCTTCCGATCTNNNTATAGGGGGCAGTAAGACAT
GCACACGTCTGAACTCTTTCCCTACACGACGCTCTTCCGATCTNNNCCGTCTTTCTTTCTCTCCTCCTCC
AGCACACGTCTGAACTCTTTCCCTACACGACGCTCTTCCGATCTNNNCTTCCTCCATCTTCAAAGCCAA
AGCACACGTCTGAACTCTTTCCCTACACGACGCTCTTCCGATCTNNNTAGGGTGCCTTACTGTGACTTT
GCACACGTCTGAACTCTTTCCCTACACGACGCTCTTCCGATCTNNNTGATCTTGGTCACCTCAACTCAG
CACACGTCTGAACTCTTTCCCTACACGACGCTCTTCCGATCTNNNTAGGTCATGGAGCGGATTGTTA
CACACGTCTGAACTCTTTCCCTACACGACGCTCTTCCGATCTNNNCACTCTGGTCTAACACAAGGGAA
CACACGTCTGAACTCTTTCCCTACACGACGCTCTTCCGATCTNNNGTGGGGCCCTCATTATAGATCTG
AGCACACGTCTGAACTCTTTCCCTACACGACGCTCTTCCGATCTNNNGAGACAGAGAGACAGAGAGA
AGCACACGTCTGAACTCTTTCCCTACACGACGCTCTTCCGATCTNNNACACACAGACACAGAGTGAG
AAGAGCACACGTCTGAACTCTTTCCCTACACGACGCTCTTCCGATCTNNNTGGATTTTATGACTGTGG
AGAGCACACGTCTGAACTCTTTCCCTACACGACGCTCTTCCGATCTNNNAAGAGAAGAGAAAAGGCC
AAGAGCACACGTCTGAACTCTTTCCCTACACGACGCTCTTCCGATCTNNNAGAGAGCGAGACTGTCA
GCACACGTCTGAACTCTTTCCCTACACGACGCTCTTCCGATCTNNNCCATGCCCAGCTAATTTGTTTGT
AGCACACGTCTGAACTCTTTCCCTACACGACGCTCTTCCGATCTNNNGAGATGCTTGTGTCACTCCTCT
AGCACACGTCTGAACTCTTTCCCTACACGACGCTCTTCCGATCTNNNACACACATTCTCTCTCACACAC

GAGCACACGTCTGAACTCTTTCCCTACACGACGCTCTTCCGATCTNNNGTGTGTGTGAGAGAGAGA
GCACACGTCTGAACTCTTTCCCTACACGACGCTCTTCCGATCTNNNCACATGAAAGGAGAGAGAGGGA
GAGCACACGTCTGAACTCTTTCCCTACACGACGCTCTTCCGATCTNNNCCGCTTTATGCCCACTATGAAA
GAGCACACGTCTGAACTCTTTCCCTACACGACGCTCTTCCGATCTNNNCCTCTTAGGCTAAGGGTTTCT
AGAGCACACGTCTGAACTCTTTCCCTACACGACGCTCTTCCGATCTNNNGAGGTTGCAGTGAATGGAG
AGCACACGTCTGAACTCTTTCCCTACACGACGCTCTTCCGATCTNNNAGTGTCAGTCAATTTGAGCCAA
GAGCACACGTCTGAACTCTTTCCCTACACGACGCTCTTCCGATCTNNNCTCTAGTTCCAGGCCATTGACT
GCACACGTCTGAACTCTTTCCCTACACGACGCTCTTCCGATCTNNNTATCTACCAGGCTCTGCTCTAGGC
AGCACACGTCTGAACTCTTTCCCTACACGACGCTCTTCCGATCTNNNTGATTCCCTGCCTCCCTATTCTAC
AGCACACGTCTGAACTCTTTCCCTACACGACGCTCTTCCGATCTNNNTGCCTTCTACAGTTTGAGTACCC
GAGCACACGTCTGAACTCTTTCCCTACACGACGCTCTTCCGATCTNNNACCCGAGTTGCAAAGGAAAAA
GAGCACACGTCTGAACTCTTTCCCTACACGACGCTCTTCCGATCTNNNTAGTGAGAAGGGGAGGAAC
CACACGTCTGAACTCTTTCCCTACACGACGCTCTTCCGATCTNNNACAAGTCTTAGGCCTACCATTGGC
GAGCACACGTCTGAACTCTTTCCCTACACGACGCTCTTCCGATCTNNNCACACACACACACACACACATA
AGCACACGTCTGAACTCTTTCCCTACACGACGCTCTTCCGATCTNNNTACAATCATGGTGGAAGGGGA
GAGCACACGTCTGAACTCTTTCCCTACACGACGCTCTTCCGATCTNNNGTTACCAGAAACATGTTCACGC
AAGAGCACACGTCTGAACTCTTTCCCTACACGACGCTCTTCCGATCTNNNCAAGGCTGCAGAAAACCAT
GAGCACACGTCTGAACTCTTTCCCTACACGACGCTCTTCCGATCTNNNGGCTTGCAAACAGAGAGAGAA
AGCACACGTCTGAACTCTTTCCCTACACGACGCTCTTCCGATCTNNNCTCCCAAAGTGCTAGGTGATAGC
GAGCACACGTCTGAACTCTTTCCCTACACGACGCTCTTCCGATCTNNNTCCCCTGAAGACATTTGTGTTG
GAGCACACGTCTGAACTCTTTCCCTACACGACGCTCTTCCGATCTNNNTGAGAGAGAAAGAGGGAGA
AGCACACGTCTGAACTCTTTCCCTACACGACGCTCTTCCGATCTNNNCCAATGCTGATGAAGTCAAGCA
AAGAGCACACGTCTGAACTCTTTCCCTACACGACGCTCTTCCGATCTNNNTCTCCCCCTTAGGAAAAGA
GAGCACACGTCTGAACTCTTTCCCTACACGACGCTCTTCCGATCTNNNCCCCATCTCAGTCAATCAATCG
AAGAGCACACGTCTGAACTCTTTCCCTACACGACGCTCTTCCGATCTNNNTGAGGCAGGAGAATCACT
AGCACACGTCTGAACTCTTTCCCTACACGACGCTCTTCCGATCTNNNCCACATGGGGCATGTCCATACA
CACACGTCTGAACTCTTTCCCTACACGACGCTCTTCCGATCTNNNCAACTACCATTACACCCCAGTCCCAT
AGCACACGTCTGAACTCTTTCCCTACACGACGCTCTTCCGATCTNNNTGCCCCAAGAAGACATGGTAGT
AGCACACGTCTGAACTCTTTCCCTACACGACGCTCTTCCGATCTNNNAGAGAACCACTGCTGCCAGTAT
GCACACGTCTGAACTCTTTCCCTACACGACGCTCTTCCGATCTNNNAGGTCACATTCATACCCGAAGGA
AAGAGCACACGTCTGAACTCTTTCCCTACACGACGCTCTTCCGATCTNNNCTTCCTTTCCCCTCCCGCTTA
AGCACACGTCTGAACTCTTTCCCTACACGACGCTCTTCCGATCTNNNCGGGATTTCAAACACAGACAGC
GAGCACACGTCTGAACTCTTTCCCTACACGACGCTCTTCCGATCTNNNTCCTAGGAAGCCACCTTCTCTG
AGCACACGTCTGAACTCTTTCCCTACACGACGCTCTTCCGATCTNNNTTCCACCATCACCGAAAGTTCTC
GCACACGTCTGAACTCTTTCCCTACACGACGCTCTTCCGATCTNNNTAAAACCTGCCAGGGAACTCAAC
GCACACGTCTGAACTCTTTCCCTACACGACGCTCTTCCGATCTNNNTGAATGAAGCCATCTTGGACACC
GCACACGTCTGAACTCTTTCCCTACACGACGCTCTTCCGATCTNNNCGCCTTCTTCTCTGCTCTGTAAGCA
GAGCACACGTCTGAACTCTTTCCCTACACGACGCTCTTCCGATCTNNNCAGAAACCTGAGTGCTGACATC
GAGCACACGTCTGAACTCTTTCCCTACACGACGCTCTTCCGATCTNNNTCTTAGATCACTTGCTTGGGGC
GCACACGTCTGAACTCTTTCCCTACACGACGCTCTTCCGATCTNNNGTTGAGTGCAACTCTGGCTTCTAC
AGCACACGTCTGAACTCTTTCCCTACACGACGCTCTTCCGATCTNNNCTGTACTGGCTGACCCCTTAGTC
AGCACACGTCTGAACTCTTTCCCTACACGACGCTCTTCCGATCTNNNGAACGATGTTATGCCCTCCAGA
GCACACGTCTGAACTCTTTCCCTACACGACGCTCTTCCGATCTNNNCCAGGAAGACTGAGGTGTGTGTA

AGCACACGTCTGAACTCTTTCCCTACACGACGCTCTTCCGATCTNNNGCATATGTCCTCTGGGGTGTAT
AGCACACGTCTGAACTCTTTCCCTACACGACGCTCTTCCGATCTNNNCGACTTTCTGGAAGAGGAAGCA
GAGCACACGTCTGAACTCTTTCCCTACACGACGCTCTTCCGATCTNNNTTGCGACGAGTTCTGGAGTAC
GAGCACACGTCTGAACTCTTTCCCTACACGACGCTCTTCCGATCTNNNCACACGCTGACATACCGACATG
AGCACACGTCTGAACTCTTTCCCTACACGACGCTCTTCCGATCTNNNCACACCTTGCTGGTTCCTTCAGCA
GCACACGTCTGAACTCTTTCCCTACACGACGCTCTTCCGATCTNNNAATTCCTCATTCTCTGGGCCCTTCA
AGCACACGTCTGAACTCTTTCCCTACACGACGCTCTTCCGATCTNNNCTCCATGTTACTTCTGCTTCGCTC
AGAGCACACGTCTGAACTCTTTCCCTACACGACGCTCTTCCGATCTNNNCTGCAGCTTCCCAGAAGCAA
GCACACGTCTGAACTCTTTCCCTACACGACGCTCTTCCGATCTNNNGCAGGCAATCAGAGATTGTGGA
AGCACACGTCTGAACTCTTTCCCTACACGACGCTCTTCCGATCTNNNACAAAACAGAAGCTCTTTGGGG
GCACACGTCTGAACTCTTTCCCTACACGACGCTCTTCCGATCTNNNTTCAAGAAACTTCCCCTGCCTTCCA
GCACACGTCTGAACTCTTTCCCTACACGACGCTCTTCCGATCTNNNTGGAATCCACCTACTTCCAGGAGC
AGCACACGTCTGAACTCTTTCCCTACACGACGCTCTTCCGATCTNNNAGGTTTGCCAGTTTTCTTCCCAA
CACACGTCTGAACTCTTTCCCTACACGACGCTCTTCCGATCTNNNTCTCTGGGTCTCTGACGGTTTCCAT
GCACACGTCTGAACTCTTTCCCTACACGACGCTCTTCCGATCTNNNGGTGCACCTGGCCTTTATTCTTAC
AGCACACGTCTGAACTCTTTCCCTACACGACGCTCTTCCGATCTNNNAAAGTCAGAGACAGGAAGCGG
AGCACACGTCTGAACTCTTTCCCTACACGACGCTCTTCCGATCTNNNCTGGACCTGCAATTGCTCTACTT
GAGCACACGTCTGAACTCTTTCCCTACACGACGCTCTTCCGATCTNNNCCAGATCCAAGCCAGACAGAT
AGCACACGTCTGAACTCTTTCCCTACACGACGCTCTTCCGATCTNNNTTTGGCTCCTGCTTCATCTTCTCC
GAGCACACGTCTGAACTCTTTCCCTACACGACGCTCTTCCGATCTNNNTTGCTCTACCCTTCTGTCTCTCC
GAGCACACGTCTGAACTCTTTCCCTACACGACGCTCTTCCGATCTNNNTTAGGGGATTCTGGGGAGAA
GCACACGTCTGAACTCTTTCCCTACACGACGCTCTTCCGATCTNNNGATTGGTGCCCTCTGGAAACATC
AGCACACGTCTGAACTCTTTCCCTACACGACGCTCTTCCGATCTNNNCTTCTACCCAATGTGCCAAGGTTC
AGCACACGTCTGAACTCTTTCCCTACACGACGCTCTTCCGATCTNNNCATTGACGCTTGGAAGGGAAAA
AGCACACGTCTGAACTCTTTCCCTACACGACGCTCTTCCGATCTNNNCAACCTCTTGTCCAGTGTTGGA
AGCACACGTCTGAACTCTTTCCCTACACGACGCTCTTCCGATCTNNNTGTCACACCACTATAGACAGACC
GCACACGTCTGAACTCTTTCCCTACACGACGCTCTTCCGATCTNNNTGCTTGAGCTTTCCTTCTTCCTCCA
AGAGCACACGTCTGAACTCTTTCCCTACACGACGCTCTTCCGATCTNNNCCTTCCGTCCTTTTCTCTCCCC
GCACACGTCTGAACTCTTTCCCTACACGACGCTCTTCCGATCTNNNTGTTACGCAAATTGAGCGAGAG
AGCACACGTCTGAACTCTTTCCCTACACGACGCTCTTCCGATCTNNNTTAGGGGGTTAGTTCTTTCCTG
GCACACGTCTGAACTCTTTCCCTACACGACGCTCTTCCGATCTNNNTCCTTTCCCACATGAGATTCCCTCA
AGCACACGTCTGAACTCTTTCCCTACACGACGCTCTTCCGATCTNNNTTTCCATAAGGCTAGTGTGCAG
AGCACACGTCTGAACTCTTTCCCTACACGACGCTCTTCCGATCTNNNATCCAAATTCACTGGGTGCATC
GAGCACACGTCTGAACTCTTTCCCTACACGACGCTCTTCCGATCTNNNTATTTCCACCGTCCCTGTCATCT
GCACACGTCTGAACTCTTTCCCTACACGACGCTCTTCCGATCTNNNAGACCATGTTCTCAACAGGACACC
AGCACACGTCTGAACTCTTTCCCTACACGACGCTCTTCCGATCTNNNACAGAGCTATATGCCTCCCAGAG
GAGCACACGTCTGAACTCTTTCCCTACACGACGCTCTTCCGATCTNNNTGCCCCATGCAATTTGACAGAT
GAGCACACGTCTGAACTCTTTCCCTACACGACGCTCTTCCGATCTNNNGATCAAGCTTCTGCTCTGCACT
AGCACACGTCTGAACTCTTTCCCTACACGACGCTCTTCCGATCTNNNGAGTTCCATGGACACATGCACAG
GCACACGTCTGAACTCTTTCCCTACACGACGCTCTTCCGATCTNNNATAGATGGGGGCCAAAATGTTC
AGCACACGTCTGAACTCTTTCCCTACACGACGCTCTTCCGATCTNNNGGGAGAAATCATGGGAACCCC
GAGCACACGTCTGAACTCTTTCCCTACACGACGCTCTTCCGATCTNNNTTAGTGGAGGTCAGTAAGTGC
GAGCACACGTCTGAACTCTTTCCCTACACGACGCTCTTCCGATCTNNNCCCCATCAGAGCTCCCTGTAGC

GCACACGTCTGAACTCTTTCCCTACACGACGCTCTTCCGATCTNNNTTTGGCAAGTTGACCAGGTTGAT
AGCACACGTCTGAACTCTTTCCCTACACGACGCTCTTCCGATCTNNNCCGATCTCACCTCACGGCAAGCA
AGCACACGTCTGAACTCTTTCCCTACACGACGCTCTTCCGATCTNNNTTTGAGATGTCAGCAGTCTCTG
GCACACGTCTGAACTCTTTCCCTACACGACGCTCTTCCGATCTNNNGAGGTTGAGGGTACAGCGAGTA
CACACGTCTGAACTCTTTCCCTACACGACGCTCTTCCGATCTNNNTCAAACTCCACTCTGCTTCTGACCAT
GCACACGTCTGAACTCTTTCCCTACACGACGCTCTTCCGATCTNNNTCATCAGGTTTGGAAGTTGTGGG
GAGCACACGTCTGAACTCTTTCCCTACACGACGCTCTTCCGATCTNNNAAAAATTTAGCCAGGTGGGT
GAGCACACGTCTGAACTCTTTCCCTACACGACGCTCTTCCGATCTNNNAAGGGACTTCTGTTCCTCCTA
GCACACGTCTGAACTCTTTCCCTACACGACGCTCTTCCGATCTNNNAGCCAGGAACTGTGATGAAAAC
GCACACGTCTGAACTCTTTCCCTACACGACGCTCTTCCGATCTNNNACATTTGACTGAAAAGCTCGGGT
AGCACACGTCTGAACTCTTTCCCTACACGACGCTCTTCCGATCTNNNTGAGTAGCTAGTACCATGCCCT
GCACACGTCTGAACTCTTTCCCTACACGACGCTCTTCCGATCTNNNAAGAAAGAACAGCTAAGATGGCA
GCACACGTCTGAACTCTTTCCCTACACGACGCTCTTCCGATCTNNNCAAATTTTTCCAGCTTTGGCCTCT
AGCACACGTCTGAACTCTTTCCCTACACGACGCTCTTCCGATCTNNNGACTAGACTTGTGTCAGGGCTT
AGCACACGTCTGAACTCTTTCCCTACACGACGCTCTTCCGATCTNNNAAACACTGTGGATGCTTACCAG
AGCACACGTCTGAACTCTTTCCCTACACGACGCTCTTCCGATCTNNNTCAATCGTGACGGTTTTATGCCT
AGCACACGTCTGAACTCTTTCCCTACACGACGCTCTTCCGATCTNNNAGTTCAGTTGGCACTGTTGTTGA
GAGCACACGTCTGAACTCTTTCCCTACACGACGCTCTTCCGATCTNNNGAAAGGAAGTTTGAGCTCCC
AGCACACGTCTGAACTCTTTCCCTACACGACGCTCTTCCGATCTNNNTGCTCGGACCCTGATTACTATGT
GCACACGTCTGAACTCTTTCCCTACACGACGCTCTTCCGATCTNNNTTCTTACCTGGCCTTTGCTTCAGCA
AGCACACGTCTGAACTCTTTCCCTACACGACGCTCTTCCGATCTNNNGTGAGTGCACACGAGTATGTG
AGCACACGTCTGAACTCTTTCCCTACACGACGCTCTTCCGATCTNNNGGCTAACCTGTGTATGCCACTG
GCACACGTCTGAACTCTTTCCCTACACGACGCTCTTCCGATCTNNNGGACCACAGCCCATAGCTAATTT
AGCACACGTCTGAACTCTTTCCCTACACGACGCTCTTCCGATCTNNNTGGGTAAAGAGAACCCTCACCA
GAGCACACGTCTGAACTCTTTCCCTACACGACGCTCTTCCGATCTNNNGCACATTTCAGTTCCTTTCAG
AGCACACGTCTGAACTCTTTCCCTACACGACGCTCTTCCGATCTNNNGTGTATTGCCTGTGTGTATGT
GCACACGTCTGAACTCTTTCCCTACACGACGCTCTTCCGATCTNNNAGCCAAAGTGTCCTGATATGCTG
CACACGTCTGAACTCTTTCCCTACACGACGCTCTTCCGATCTNNNCCTTTCCCCCTATATGTGTGTGTGCA
GCACACGTCTGAACTCTTTCCCTACACGACGCTCTTCCGATCTNNNCAACCTTTGGTGCTCAAGTGATTC
AGCACACGTCTGAACTCTTTCCCTACACGACGCTCTTCCGATCTNNNACCTAAACCTGATGTCTCCACCA
GCACACGTCTGAACTCTTTCCCTACACGACGCTCTTCCGATCTNNNTCACTCTCTATTGCTCCCCCAAACA
GCACACGTCTGAACTCTTTCCCTACACGACGCTCTTCCGATCTNNNTTTTTCTTTGGCAGCTGATCCTGCA
AGCACACGTCTGAACTCTTTCCCTACACGACGCTCTTCCGATCTNNNTGTTTGTGTGCATGCATTGTGT
AGAGCACACGTCTGAACTCTTTCCCTACACGACGCTCTTCCGATCTNNNCAGATCTCTGGGATTCCTGT
AGCACACGTCTGAACTCTTTCCCTACACGACGCTCTTCCGATCTNNNTTTTGTCTGCCTTGTGCTCTCAT
GAGCACACGTCTGAACTCTTTCCCTACACGACGCTCTTCCGATCTNNNATGTTTCAGGCAAAGGCCATA
GAGCACACGTCTGAACTCTTTCCCTACACGACGCTCTTCCGATCTNNNTAGCTTGTTCCTGTGTCCTCTGA
AGCACACGTCTGAACTCTTTCCCTACACGACGCTCTTCCGATCTNNNTGAAACCCACTCCTCTCTCTCTCC
AGCACACGTCTGAACTCTTTCCCTACACGACGCTCTTCCGATCTNNNGCTGAAGACACTTTGGTAGTGC
AGCACACGTCTGAACTCTTTCCCTACACGACGCTCTTCCGATCTNNNTTGGCATGGCATTCTCTCAGTTT
GAGCACACGTCTGAACTCTTTCCCTACACGACGCTCTTCCGATCTNNNGCCAGCAGTGTACTGTATCACA
CACACGTCTGAACTCTTTCCCTACACGACGCTCTTCCGATCTNNNAGGTCTCTGTATGCTTTGTGCATCA
AGCACACGTCTGAACTCTTTCCCTACACGACGCTCTTCCGATCTNNNAAGTGAAACTCACTGGGGCAAA

AGCACACGTCTGAACTCTTTCCCTACACGACGCTCTTCCGATCTNNNGTTTTGTGTGTGGCTGGAGA
AGCACACGTCTGAACTCTTTCCCTACACGACGCTCTTCCGATCTNNNTGAAATCCAGCAACCCCATTTCT
GCACACGTCTGAACTCTTTCCCTACACGACGCTCTTCCGATCTNNNCCTACTTGTTTCTTTCCGTCCCC
GCACACGTCTGAACTCTTTCCCTACACGACGCTCTTCCGATCTNNNTGCCTTTGACAACAGCTTTCATTC
AGCACACGTCTGAACTCTTTCCCTACACGACGCTCTTCCGATCTNNNAAACTTGACAGTGGCTTTCCTG
AGCACACGTCTGAACTCTTTCCCTACACGACGCTCTTCCGATCTNNNGACAGCGATTGACGTGTAGTT
AGCACACGTCTGAACTCTTTCCCTACACGACGCTCTTCCGATCTNNNAACTCTCCAGGTAAGTTGGCTG
GAGCACACGTCTGAACTCTTTCCCTACACGACGCTCTTCCGATCTNNNATGAAATCAGTGTTGCCTTG
AGCACACGTCTGAACTCTTTCCCTACACGACGCTCTTCCGATCTNNNGGCATCTATTCCAAGAGGTAG
GCACACGTCTGAACTCTTTCCCTACACGACGCTCTTCCGATCTNNNATAGACGTGTGTCTGTGTGTGT
GCACACGTCTGAACTCTTTCCCTACACGACGCTCTTCCGATCTNNNCCTGAGTTTGAGTTCCATCCTG
AGCACACGTCTGAACTCTTTCCCTACACGACGCTCTTCCGATCTNNNGCACTGGCTCACTGAATGGAA
AGCACACGTCTGAACTCTTTCCCTACACGACGCTCTTCCGATCTNNNAGCAAATTGCAGAGCCGTATG
GAGCACACGTCTGAACTCTTTCCCTACACGACGCTCTTCCGATCTNNNGTCTGATCCAGCACAGACAC
GCACACGTCTGAACTCTTTCCCTACACGACGCTCTTCCGATCTNNNATGTGAGTTTCATACCCAGGTGC
AGCACACGTCTGAACTCTTTCCCTACACGACGCTCTTCCGATCTNNNTGTACACTGTGCAGAGGTAAG
GCACACGTCTGAACTCTTTCCCTACACGACGCTCTTCCGATCTNNNCTAGAAGGCCTTGTTGCCTGTAG
CACACGTCTGAACTCTTTCCCTACACGACGCTCTTCCGATCTNNNTGCAATGTTCATTCCTGGTTGAGA
GCACACGTCTGAACTCTTTCCCTACACGACGCTCTTCCGATCTNNNATTGGGTACACTGAATGCTTGC
GAGCACACGTCTGAACTCTTTCCCTACACGACGCTCTTCCGATCTNNNTTGTAAAGTGGCAATGCAAG
GAGCACACGTCTGAACTCTTTCCCTACACGACGCTCTTCCGATCTNNNGCTTGGAAGAAAGTTCCGTG
GCACACGTCTGAACTCTTTCCCTACACGACGCTCTTCCGATCTNNNTGCAAGCTGGTCAGAAACTCAA
GCACACGTCTGAACTCTTTCCCTACACGACGCTCTTCCGATCTNNNTAAGTCTCAGCCAGACTTGGACA
GAGCACACGTCTGAACTCTTTCCCTACACGACGCTCTTCCGATCTNNNGCTTCAGTTTTGGGGAAGGG
GCACACGTCTGAACTCTTTCCCTACACGACGCTCTTCCGATCTNNNGGTGAAACACCTGTGATGGGA
GCACACGTCTGAACTCTTTCCCTACACGACGCTCTTCCGATCTNNNTCATTAGACTTCTCGTGGTTGCG
GCACACGTCTGAACTCTTTCCCTACACGACGCTCTTCCGATCTNNNGAGTGTCACTTGAGCTGGGAA
GCACACGTCTGAACTCTTTCCCTACACGACGCTCTTCCGATCTNNNATTCTTTGGATTCTGGGGTCCCT
GCACACGTCTGAACTCTTTCCCTACACGACGCTCTTCCGATCTNNNCTTGTGTTGTGTATGTGTGCGT
GCACACGTCTGAACTCTTTCCCTACACGACGCTCTTCCGATCTNNNCTGTGAACTGCACACGTGAGTG
AGCACACGTCTGAACTCTTTCCCTACACGACGCTCTTCCGATCTNNNTATTGCAATGAGTGGGGGAG
GCACACGTCTGAACTCTTTCCCTACACGACGCTCTTCCGATCTNNNTGGGCACTTCCAGTATCATCTCA
GAGCACACGTCTGAACTCTTTCCCTACACGACGCTCTTCCGATCTNNNCTGACCTGGCCAACAGAGAA
GCACACGTCTGAACTCTTTCCCTACACGACGCTCTTCCGATCTNNNTCCCAGCCTAACCTAGTTCAACA
GCACACGTCTGAACTCTTTCCCTACACGACGCTCTTCCGATCTNNNTGTCCATGGTCTGACTTGTCTCT
AGCACACGTCTGAACTCTTTCCCTACACGACGCTCTTCCGATCTNNNCCAGCTGGAGTTCAGAATAAC
CACACGTCTGAACTCTTTCCCTACACGACGCTCTTCCGATCTNNNCCCAGGATTCCAGATTCTGAGTTC
GCACACGTCTGAACTCTTTCCCTACACGACGCTCTTCCGATCTNNNTTCCAGACTTTCTCCTGTGCCTAC
GCACACGTCTGAACTCTTTCCCTACACGACGCTCTTCCGATCTNNNGCTGGTTGTTCAAAGCAAACTC
AGCACACGTCTGAACTCTTTCCCTACACGACGCTCTTCCGATCTNNNGGGAAGGTTGTTGCCATAGA
AGCACACGTCTGAACTCTTTCCCTACACGACGCTCTTCCGATCTNNNCATTGCTGCCATAGCAAGTGT
GCACACGTCTGAACTCTTTCCCTACACGACGCTCTTCCGATCTNNNCCCCAAGGTACAATCTACAGGG
AGCACACGTCTGAACTCTTTCCCTACACGACGCTCTTCCGATCTNNNGTGCTGTTTTGTGTGTTGTGT

GCACACGTCTGAACTCTTTCCCTACACGACGCTCTTCCGATCTNNNTCTTGTGGTTCTGGATTACTGCC
AGCACACGTCTGAACTCTTTCCCTACACGACGCTCTTCCGATCTNNNTAATACACATTCCTGGGCCATG
GAGCACACGTCTGAACTCTTTCCCTACACGACGCTCTTCCGATCTNNNAGCAGTGTGAATCATGTGTGT
AGAGCACACGTCTGAACTCTTTCCCTACACGACGCTCTTCCGATCTNNNAACTTTGTGACTGTGTGCTT
AGCACACGTCTGAACTCTTTCCCTACACGACGCTCTTCCGATCTNNNTATTGACATGTGTGTGTGTGG
GAGCACACGTCTGAACTCTTTCCCTACACGACGCTCTTCCGATCTNNNGAGTATGCCTGGGAACCTTTG
GAGCACACGTCTGAACTCTTTCCCTACACGACGCTCTTCCGATCTNNNTGGATGTGGTAGGATGAAGG
AGAGCACACGTCTGAACTCTTTCCCTACACGACGCTCTTCCGATCTNNNGAGTGTGCAGTGAATCGAG
GAGCACACGTCTGAACTCTTTCCCTACACGACGCTCTTCCGATCTNNNAGAGTGGAAAGGGGTATGTG
GCACACGTCTGAACTCTTTCCCTACACGACGCTCTTCCGATCTNNNCAAGAGGATTGGTGGATGGATG
GAGCACACGTCTGAACTCTTTCCCTACACGACGCTCTTCCGATCTNNNGGGCCATATTCACATGCACAA
GCACACGTCTGAACTCTTTCCCTACACGACGCTCTTCCGATCTNNNGGGAAGAGTTTTTCTCCTCTGG
GAGCACACGTCTGAACTCTTTCCCTACACGACGCTCTTCCGATCTNNNATGTCAATGAGGGGGTTGAA
GAGCACACGTCTGAACTCTTTCCCTACACGACGCTCTTCCGATCTNNNTTTGCTGCCAGAATTTGAGAC
AGCACACGTCTGAACTCTTTCCCTACACGACGCTCTTCCGATCTNNNCGTGCACAGACACGCAGAAAAT
GCACACGTCTGAACTCTTTCCCTACACGACGCTCTTCCGATCTNNNACGTATTTTGTGTGTGGCTGTT
GCACACGTCTGAACTCTTTCCCTACACGACGCTCTTCCGATCTNNNATGGCTGTTTGAACCAGTGACA
AGCACACGTCTGAACTCTTTCCCTACACGACGCTCTTCCGATCTNNNTCCTCCCTAGGGTCAGATAAAG
AGCACACGTCTGAACTCTTTCCCTACACGACGCTCTTCCGATCTNNNGTTTTGAAGGTGGTGAGGACG
AGCACACGTCTGAACTCTTTCCCTACACGACGCTCTTCCGATCTNNNAGGATGCACAGGGCTAAGTAG
GAGCACACGTCTGAACTCTTTCCCTACACGACGCTCTTCCGATCTNNNGACTAGCTGCTCTGGGTGTTT
AGCACACGTCTGAACTCTTTCCCTACACGACGCTCTTCCGATCTNNNAATGCCTCCCATCAAAGCATCA
GCACACGTCTGAACTCTTTCCCTACACGACGCTCTTCCGATCTNNNTTATTGCATCCCCATCAGTGTCCC
GCACACGTCTGAACTCTTTCCCTACACGACGCTCTTCCGATCTNNNTACCCTCAACCTACAGAAGCACA
GCACACGTCTGAACTCTTTCCCTACACGACGCTCTTCCGATCTNNNTATAAATCAGATCCCCCGACCCCC
AGCACACGTCTGAACTCTTTCCCTACACGACGCTCTTCCGATCTNNNTTAGGTTACCAAGGGATGGGA
AGCACACGTCTGAACTCTTTCCCTACACGACGCTCTTCCGATCTNNNAAGTGGGTCATTCCCACAGACT
GAGCACACGTCTGAACTCTTTCCCTACACGACGCTCTTCCGATCTNNNGCAGTTTTCATGTGCCTTTCA
GCACACGTCTGAACTCTTTCCCTACACGACGCTCTTCCGATCTNNNTACAGTTGTCTCTTGGTGTGTGC
AGCACACGTCTGAACTCTTTCCCTACACGACGCTCTTCCGATCTNNNTCTCACAAGTGCCTTGGCTATT
GCACACGTCTGAACTCTTTCCCTACACGACGCTCTTCCGATCTNNNGTTTCACTCACCCCATTTCCTGTC
AGCACACGTCTGAACTCTTTCCCTACACGACGCTCTTCCGATCTNNNACATTCATTTCAGGCACACAGG
GAGCACACGTCTGAACTCTTTCCCTACACGACGCTCTTCCGATCTNNNTTGGCCTTGAAGTGTGTATGG
GCACACGTCTGAACTCTTTCCCTACACGACGCTCTTCCGATCTNNNCTGGACTTCCAACCTGCTTTGTA
AGCACACGTCTGAACTCTTTCCCTACACGACGCTCTTCCGATCTNNNGTTGTGACCACAAGGTCTGGA
GAGCACACGTCTGAACTCTTTCCCTACACGACGCTCTTCCGATCTNNNATTCGGGACCCCAGTGAATTT
GCACACGTCTGAACTCTTTCCCTACACGACGCTCTTCCGATCTNNNTTCTATCTATCCCTGGAGGCTGC
CACACGTCTGAACTCTTTCCCTACACGACGCTCTTCCGATCTNNNACCAAAGAAACTGTGGTGTGTGT
GCACACGTCTGAACTCTTTCCCTACACGACGCTCTTCCGATCTNNNGAGATCAATCAAAGCGGTCCTC
AGCACACGTCTGAACTCTTTCCCTACACGACGCTCTTCCGATCTNNNTCTGGAAGTGATGGAAGTGAA
GAGCACACGTCTGAACTCTTTCCCTACACGACGCTCTTCCGATCTNNNCTCCCTCCTGGGGGAAAAGA
GAGCACACGTCTGAACTCTTTCCCTACACGACGCTCTTCCGATCTNNNCATCGCTGTTTCTGAGTGAGT
GAGCACACGTCTGAACTCTTTCCCTACACGACGCTCTTCCGATCTNNNGGTATCCATGAGTGTGTACCT

AGCACACGTCTGAACTCTTTCCCTACACGACGCTCTTCCGATCTNNNAAGGCAAGTAGCTGGGTTAG

CACACGTCTGAACTCTTTCCCTACACGACGCTCTTCCGATCTNNNAATCAAAGGTACTCGGAGGCTG

AGCACACGTCTGAACTCTTTCCCTACACGACGCTCTTCCGATCTNNNCGGACTTTTATCCAGATGCCT

GCACACGTCTGAACTCTTTCCCTACACGACGCTCTTCCGATCTNNNTCTGTCCCACCCAAACATCTACTC

CACACGTCTGAACTCTTTCCCTACACGACGCTCTTCCGATCTNNNTTTCTGAGGCGGGTTCTCACTTT

GAGCACACGTCTGAACTCTTTCCCTACACGACGCTCTTCCGATCTNNNTTACAGTTGCAGTTATCCTGC

CACACGTCTGAACTCTTTCCCTACACGACGCTCTTCCGATCTNNNCCCTTCCCTTGGATTGGATTGAGC

AGCACACGTCTGAACTCTTTCCCTACACGACGCTCTTCCGATCTNNNGACAGTGCCTTTTCCCCTTCAG

GCACACGTCTGAACTCTTTCCCTACACGACGCTCTTCCGATCTNNNAGGGAAGAGAAGGTCCAAGG

CACACGTCTGAACTCTTTCCCTACACGACGCTCTTCCGATCTNNNGACAGGAAGGAAGAAAGAGGGT

GCACACGTCTGAACTCTTTCCCTACACGACGCTCTTCCGATCTNNNTCTGCCTGGGGGAAAAATAGTT

AGCACACGTCTGAACTCTTTCCCTACACGACGCTCTTCCGATCTNNNGTCAGATCCACAAGACCCCTTT

GAGCACACGTCTGAACTCTTTCCCTACACGACGCTCTTCCGATCTNNNTGTCTCCTGTCACTGAGGCT

GCACACGTCTGAACTCTTTCCCTACACGACGCTCTTCCGATCTNNNTCGTCCCTCTCCTCCATACCTAAC

GCACACGTCTGAACTCTTTCCCTACACGACGCTCTTCCGATCTNNNCGTACTCATGTGGGGAAACAA

GCACACGTCTGAACTCTTTCCCTACACGACGCTCTTCCGATCTNNNATCATCCAGCAAAGCACTAGAC

GCACACGTCTGAACTCTTTCCCTACACGACGCTCTTCCGATCTNNNTGTAGATCCCCCTACACTTCCTC

AGAGCACACGTCTGAACTCTTTCCCTACACGACGCTCTTCCGATCTNNNTGAATTCATAGATGGCTCC

AGCACACGTCTGAACTCTTTCCCTACACGACGCTCTTCCGATCTNNNGGATGCTGAAGTGTGAGTGG

GCACACGTCTGAACTCTTTCCCTACACGACGCTCTTCCGATCTNNNTGACGTTGTATGTTTCGCTGTCT

GCACACGTCTGAACTCTTTCCCTACACGACGCTCTTCCGATCTNNNCGCTTTCCAACACTGAAACCAAT

CACACGTCTGAACTCTTTCCCTACACGACGCTCTTCCGATCTNNNAGCAAGTAAAGACATCTGAGCC

AGCACACGTCTGAACTCTTTCCCTACACGACGCTCTTCCGATCTNNNACTGTTCCAGGGCATAGTCTC

GCACACGTCTGAACTCTTTCCCTACACGACGCTCTTCCGATCTNNNCAATAGGATGAGGGGTGTGTG

CACACGTCTGAACTCTTTCCCTACACGACGCTCTTCCGATCTNNNAGCTGCTGTGTGGTGTATACTTG

CACACGTCTGAACTCTTTCCCTACACGACGCTCTTCCGATCTNNNCTCCTCTTACCCCCTCCTTCCTCAT

AGCACACGTCTGAACTCTTTCCCTACACGACGCTCTTCCGATCTNNNCTGTGGACCAAAACTGACAGG

AGCACACGTCTGAACTCTTTCCCTACACGACGCTCTTCCGATCTNNNCACACTAGGCCCAGGTCTTTTT

AGCACACGTCTGAACTCTTTCCCTACACGACGCTCTTCCGATCTNNNATCTGCCCACAGAGCCTACAAT

AGCACACGTCTGAACTCTTTCCCTACACGACGCTCTTCCGATCTNNNGGTCATCAGTTAGGCTTGCAG

CACACGTCTGAACTCTTTCCCTACACGACGCTCTTCCGATCTNNNTTTGGTGCTAATGCTGATGGTCT

AGCACACGTCTGAACTCTTTCCCTACACGACGCTCTTCCGATCTNNNGGCATCTGATCTGGGAAACAG

GCACACGTCTGAACTCTTTCCCTACACGACGCTCTTCCGATCTNNNGACACACCTCATCAAGATGGG

CACACGTCTGAACTCTTTCCCTACACGACGCTCTTCCGATCTNNNAGCCCTCTTCACACCCATCTAATC

CACACGTCTGAACTCTTTCCCTACACGACGCTCTTCCGATCTNNNTCAGAATCCTTTCAGGGGTGTGT

AGCACACGTCTGAACTCTTTCCCTACACGACGCTCTTCCGATCTNNNATTCCACATTCATGCCCACAGT

CACACGTCTGAACTCTTTCCCTACACGACGCTCTTCCGATCTNNNGGGTTCCTATTTATCAGGCAGGC

CACACGTCTGAACTCTTTCCCTACACGACGCTCTTCCGATCTNNNCTGTGATGTGGAATGAAAGCAG

GCACACGTCTGAACTCTTTCCCTACACGACGCTCTTCCGATCTNNNAGAAAAGTAGGCCAGTGTGTG

CACACGTCTGAACTCTTTCCCTACACGACGCTCTTCCGATCTNNNCCAGGTCTGTTTCTTCTTCTCCCC

AGCACACGTCTGAACTCTTTCCCTACACGACGCTCTTCCGATCTNNNGTTTGCCTTTTTATCCCTGTGG

GCACACGTCTGAACTCTTTCCCTACACGACGCTCTTCCGATCTNNNATGCTCTTGGTATGAGTGATGG

AGCACACGTCTGAACTCTTTCCCTACACGACGCTCTTCCGATCTNNNGAGGCTGAGGTGAACCGTAAT

GCACACGTCTGAACTCTTTCCCTACACGACGCTCTTCCGATCTNNNTGCCACCTATGAACCAAGGACTAC
AGCACACGTCTGAACTCTTTCCCTACACGACGCTCTTCCGATCTNNNTCCGGTTCCCGAAAGTTTCTTTT
AGCACACGTCTGAACTCTTTCCCTACACGACGCTCTTCCGATCTNNNGGCAAAACCACCTAATGATGCA
GCACACGTCTGAACTCTTTCCCTACACGACGCTCTTCCGATCTNNNAAATATTTGCTGACGGCCTCCCCA
GCACACGTCTGAACTCTTTCCCTACACGACGCTCTTCCGATCTNNNGGGGTGGGGATACAAACATCAT
CACACGTCTGAACTCTTTCCCTACACGACGCTCTTCCGATCTNNNTCCAAAAGCTAGTGTGTGTGTGTCA
GCACACGTCTGAACTCTTTCCCTACACGACGCTCTTCCGATCTNNNTTCCACCTAGCAGATTGGGTACAC
CACACGTCTGAACTCTTTCCCTACACGACGCTCTTCCGATCTNNNCCAAACTCTGTGTCTGGGTTTCACA
AGCACACGTCTGAACTCTTTCCCTACACGACGCTCTTCCGATCTNNNGCTTAGTGCATAGGACAGGCTTA
GCACACGTCTGAACTCTTTCCCTACACGACGCTCTTCCGATCTNNNTGTGTTTTGGCCAAGAGGTCAAT
GCACACGTCTGAACTCTTTCCCTACACGACGCTCTTCCGATCTNNNGGAGTGGCAACTTTCTCTTTCCAC
AGCACACGTCTGAACTCTTTCCCTACACGACGCTCTTCCGATCTNNNGCTCTTTCTTAGGGGTTTCTGG
GAGCACACGTCTGAACTCTTTCCCTACACGACGCTCTTCCGATCTNNNCCATGTGAATTGAAGAGGGG
GAGCACACGTCTGAACTCTTTCCCTACACGACGCTCTTCCGATCTNNNTGTGTGAGTGTACGAGAGAG
AGCACACGTCTGAACTCTTTCCCTACACGACGCTCTTCCGATCTNNNACCAGTTATGGGGTACTGGACA
AGCACACGTCTGAACTCTTTCCCTACACGACGCTCTTCCGATCTNNNACAGATTCATTCATGATGCCACA
AGCACACGTCTGAACTCTTTCCCTACACGACGCTCTTCCGATCTNNNCAGCTAATGCTGCCTCCTTAGAG
AGCACACGTCTGAACTCTTTCCCTACACGACGCTCTTCCGATCTNNNAACTCCTACCTCCTCTGGGAAAC
AGAGCACACGTCTGAACTCTTTCCCTACACGACGCTCTTCCGATCTNNNTAGAGCCCTTGAACAAAGCT
AGCACACGTCTGAACTCTTTCCCTACACGACGCTCTTCCGATCTNNNACGTTCCTCTGGTATATTCGGGT
GCACACGTCTGAACTCTTTCCCTACACGACGCTCTTCCGATCTNNNGATACGGCTTTGCACAATCTTGG
AGCACACGTCTGAACTCTTTCCCTACACGACGCTCTTCCGATCTNNNCCCACTTTCTCTTCTCTGTGTCG
GAGCACACGTCTGAACTCTTTCCCTACACGACGCTCTTCCGATCTNNNGCAGGCCCCTCATCTTGAATA
GAGCACACGTCTGAACTCTTTCCCTACACGACGCTCTTCCGATCTNNNCACTGGAGATGCATGAAGTTG
GCACACGTCTGAACTCTTTCCCTACACGACGCTCTTCCGATCTNNNTTCTTGCTGAATTCAAGCTGGGTC
AGCACACGTCTGAACTCTTTCCCTACACGACGCTCTTCCGATCTNNNCCCGGCCCTGGGTATACATATT
GCACACGTCTGAACTCTTTCCCTACACGACGCTCTTCCGATCTNNNAGGCTTTTACATTCCCTGTGCTTC
AGCACACGTCTGAACTCTTTCCCTACACGACGCTCTTCCGATCTNNNTCTGCAAGAAGTAAGTTGGCTC
AGCACACGTCTGAACTCTTTCCCTACACGACGCTCTTCCGATCTNNNGGGAGAAGGGATGTTCAGCTTC
AGCACACGTCTGAACTCTTTCCCTACACGACGCTCTTCCGATCTNNNAATTCTCCCAGCCTCGCCTCTTCA
CACACGTCTGAACTCTTTCCCTACACGACGCTCTTCCGATCTNNNCATACACATACCTGCCACAACCACA
AGCACACGTCTGAACTCTTTCCCTACACGACGCTCTTCCGATCTNNNGTAGGAAGAGGAGGGGTGGAT
AGCACACGTCTGAACTCTTTCCCTACACGACGCTCTTCCGATCTNNNTGCTTGCATGCTGTGATGTCTAT
AGCACACGTCTGAACTCTTTCCCTACACGACGCTCTTCCGATCTNNNGGAAGACTGGCTTTGCATTCTCT
AGCACACGTCTGAACTCTTTCCCTACACGACGCTCTTCCGATCTNNNACCAAGTCCCAATGTCTTCAGTC
GCACACGTCTGAACTCTTTCCCTACACGACGCTCTTCCGATCTNNNCTGGCCCTGGTTTTCTTATCCATCA
GAGCACACGTCTGAACTCTTTCCCTACACGACGCTCTTCCGATCTNNNGAGCTCATGGGTGTTTGAGGA
GCACACGTCTGAACTCTTTCCCTACACGACGCTCTTCCGATCTNNNCTCGCTCCCTCCATCTCTCTTATCAT
AGCACACGTCTGAACTCTTTCCCTACACGACGCTCTTCCGATCTNNNAGGCACACGAAGGGACATTTAT
GAGCACACGTCTGAACTCTTTCCCTACACGACGCTCTTCCGATCTNNNCAGCTGGAGGAATTGTGCTC
AGCACACGTCTGAACTCTTTCCCTACACGACGCTCTTCCGATCTNNNCTTCCAGCATCTCGGACAACAGC
GAGCACACGTCTGAACTCTTTCCCTACACGACGCTCTTCCGATCTNNNGGGGGTAAGGCAAATCAAAA
AGCACACGTCTGAACTCTTTCCCTACACGACGCTCTTCCGATCTNNNTTTGGGGTAAAAATGCGTGTGT

GAGCACACGTCTGAACTCTTTCCCTACACGACGCTCTTCCGATCTNNNGTGTAGAAAAACAAGCCACAG

GCACACGTCTGAACTCTTTCCCTACACGACGCTCTTCCGATCTNNNATATAGCCCTCTGTGTCTGGTCCCA

CACACGTCTGAACTCTTTCCCTACACGACGCTCTTCCGATCTNNNACTGTTTTCATCAGTTTTCATGCGAC

AGCACACGTCTGAACTCTTTCCCTACACGACGCTCTTCCGATCTNNNTGTTCTGGAGTTCCTCGACTTCT

GCACACGTCTGAACTCTTTCCCTACACGACGCTCTTCCGATCTNNNCTGTGATGCAGTTCATCTCCTCCC

AGCACACGTCTGAACTCTTTCCCTACACGACGCTCTTCCGATCTNNNAATATGCCCTGTTGTCCCTCTGA

CACACGTCTGAACTCTTTCCCTACACGACGCTCTTCCGATCTNNNGATGGATCCAAGGGCTCAAACAAC

AGCACACGTCTGAACTCTTTCCCTACACGACGCTCTTCCGATCTNNNTTGCACCTGTGAGTATGTCTGTG

GCACACGTCTGAACTCTTTCCCTACACGACGCTCTTCCGATCTNNNCACCCTAGTTGGCTGATGTTGAA

GCACACGTCTGAACTCTTTCCCTACACGACGCTCTTCCGATCTNNNAGTAAATAACGAAAGCCCCAGCC

GCACACGTCTGAACTCTTTCCCTACACGACGCTCTTCCGATCTNNNGGACTGGGAGGCAAATAACTCTC

GAGCACACGTCTGAACTCTTTCCCTACACGACGCTCTTCCGATCTNNNAGGGGCACCTCCCTGTCTATAA

CACACGTCTGAACTCTTTCCCTACACGACGCTCTTCCGATCTNNNCGCGTGGCCCACTATTTTCTTATCA

AGCACACGTCTGAACTCTTTCCCTACACGACGCTCTTCCGATCTNNNAGTGACCTTTCACTTTTTCCCCC

AGCACACGTCTGAACTCTTTCCCTACACGACGCTCTTCCGATCTNNNGCCCTATCTGAATCCCTTTTGCCC

GCACACGTCTGAACTCTTTCCCTACACGACGCTCTTCCGATCTNNNCCTCGACAGTTCAGTCCCCATAGCA

AGCACACGTCTGAACTCTTTCCCTACACGACGCTCTTCCGATCTNNNCTCCATACAGCTTTCCATGGGTC

AGCACACGTCTGAACTCTTTCCCTACACGACGCTCTTCCGATCTNNNAGAGAATGAGCACTGGCTCTGA

GAGCACACGTCTGAACTCTTTCCCTACACGACGCTCTTCCGATCTNNNATGTAACACCAACACACCAGCA

GAGCACACGTCTGAACTCTTTCCCTACACGACGCTCTTCCGATCTNNNCAGCTTGGGTAACTTGCTGAA

GCACACGTCTGAACTCTTTCCCTACACGACGCTCTTCCGATCTNNNGAAGTAGTGCAGGTGATCAAGCA

GCACACGTCTGAACTCTTTCCCTACACGACGCTCTTCCGATCTNNNTTACAGTGCTTCAGGCCTCAGATC

GCACACGTCTGAACTCTTTCCCTACACGACGCTCTTCCGATCTNNNGAGGAGAGCAGCAGTCTAGTTG

CACACGTCTGAACTCTTTCCCTACACGACGCTCTTCCGATCTNNNACCCAACTGAACTTGAGAACTGGCA

GCACACGTCTGAACTCTTTCCCTACACGACGCTCTTCCGATCTNNNCTGAAGCCTCAGACCTTGGAAAG

GCACACGTCTGAACTCTTTCCCTACACGACGCTCTTCCGATCTNNNTTGTGGTCACCTTGCTCTCTTTACA

AGCACACGTCTGAACTCTTTCCCTACACGACGCTCTTCCGATCTNNNAGTTCAGAAAAATGCACGCACTC

AGCACACGTCTGAACTCTTTCCCTACACGACGCTCTTCCGATCTNNNTCAGGCCCCTTCTGTCAATGTAC

AGCACACGTCTGAACTCTTTCCCTACACGACGCTCTTCCGATCTNNNTTACAGACAGCATATCGTGGTGC

GCACACGTCTGAACTCTTTCCCTACACGACGCTCTTCCGATCTNNNCCAGCTTAGCACAGAATGGGAAT

CACACGTCTGAACTCTTTCCCTACACGACGCTCTTCCGATCTNNNGGTTGTTTGAAAGTGTGTGACTGA

GCACACGTCTGAACTCTTTCCCTACACGACGCTCTTCCGATCTNNNCTTATGCTGTGTTTAGCATGCAC

GAGCACACGTCTGAACTCTTTCCCTACACGACGCTCTTCCGATCTNNNTGTGTACCCAGAGAGAAGAGA

GAGCACACGTCTGAACTCTTTCCCTACACGACGCTCTTCCGATCTNNNTGAAGCTGTGTTGTTCAGAGA

GAGCACACGTCTGAACTCTTTCCCTACACGACGCTCTTCCGATCTNNNGTCTGCAAATTTTGGACAGCT

AGCACACGTCTGAACTCTTTCCCTACACGACGCTCTTCCGATCTNNNGATTCTACCTGCCCATCTTAGCG

CACACGTCTGAACTCTTTCCCTACACGACGCTCTTCCGATCTNNNTTCTTGGGGTCAAGTTATGCACTC

CACACGTCTGAACTCTTTCCCTACACGACGCTCTTCCGATCTNNNGCACCTGGTTTTTCCTATGCAACCA

CACACGTCTGAACTCTTTCCCTACACGACGCTCTTCCGATCTNNNAAGTTGGCTTGTGGTTACAACTTC

AGCACACGTCTGAACTCTTTCCCTACACGACGCTCTTCCGATCTNNNATGAGGGGCCCTGAAGAAGTAA

GCACACGTCTGAACTCTTTCCCTACACGACGCTCTTCCGATCTNNNTTCAATGTCTCTAACACACGGCTCA

AGCACACGTCTGAACTCTTTCCCTACACGACGCTCTTCCGATCTNNNAACTGACAACAGAACAGTGGA

AGCACACGTCTGAACTCTTTCCCTACACGACGCTCTTCCGATCTNNNTTGCAATTGTGTGTGAATGTGC

AGCACACGTCTGAACTCTTTCCCTACACGACGCTCTTCCGATCTNNNGAGAAGGGGAGGAAGAAAG
AGCACACGTCTGAACTCTTTCCCTACACGACGCTCTTCCGATCTNNNAAAGGGGCTCTCTTTCTCTGTC
CACACGTCTGAACTCTTTCCCTACACGACGCTCTTCCGATCTNNNTTAGCCTAGCATACCTGAAGGGG
GCACACGTCTGAACTCTTTCCCTACACGACGCTCTTCCGATCTNNNTCCTTGCCTCTGACTTCCCATTAC
AGCACACGTCTGAACTCTTTCCCTACACGACGCTCTTCCGATCTNNNAAGAGGAAGCAGTGCATAGA
AGCACACGTCTGAACTCTTTCCCTACACGACGCTCTTCCGATCTNNNCTAAGGATGCTTGTGTTGCCT
CACACGTCTGAACTCTTTCCCTACACGACGCTCTTCCGATCTNNNCCGGTTTGGTGTTTGATTGACAG
GAGCACACGTCTGAACTCTTTCCCTACACGACGCTCTTCCGATCTNNNCGGCCTGCTTGAATCACAA
CACACGTCTGAACTCTTTCCCTACACGACGCTCTTCCGATCTNNNGTGGTTTCTGTAGCGTGTGTACT
GCACACGTCTGAACTCTTTCCCTACACGACGCTCTTCCGATCTNNNATGTTTTCTTCCAACCTCTGGCT
GCACACGTCTGAACTCTTTCCCTACACGACGCTCTTCCGATCTNNNTCCATTTCCCCTCCCTCATCAAT
GCACACGTCTGAACTCTTTCCCTACACGACGCTCTTCCGATCTNNNCCCATGCATTTGAACTAACTGG
CACACGTCTGAACTCTTTCCCTACACGACGCTCTTCCGATCTNNNTTGGACCAGTGTCCCATGAGTAT
AGCACACGTCTGAACTCTTTCCCTACACGACGCTCTTCCGATCTNNNTTTTTCTCCTGCAGCTTCCCTC
AGCACACGTCTGAACTCTTTCCCTACACGACGCTCTTCCGATCTNNNAAAAAGAGGGAGAGGTAGCA
GCACACGTCTGAACTCTTTCCCTACACGACGCTCTTCCGATCTNNNGTGGTGATGGAGCAAGACCCT
GCACACGTCTGAACTCTTTCCCTACACGACGCTCTTCCGATCTNNNAGCTACTTGTACCCCAAAAGCTG
CACACGTCTGAACTCTTTCCCTACACGACGCTCTTCCGATCTNNNAGAGAGAGAGATCAGAGTGTG
GCACACGTCTGAACTCTTTCCCTACACGACGCTCTTCCGATCTNNNAACAAAGGCCATGTGTCCTCAC
GCACACGTCTGAACTCTTTCCCTACACGACGCTCTTCCGATCTNNNAGGACAGGGCATATGAGAGAG
CACACGTCTGAACTCTTTCCCTACACGACGCTCTTCCGATCTNNNAGAAGGGCTGGACACCTTACTTA
GCACACGTCTGAACTCTTTCCCTACACGACGCTCTTCCGATCTNNNAACTCTCCTGTCTTCCTTTCCCAC
ACACGTCTGAACTCTTTCCCTACACGACGCTCTTCCGATCTNNNGTTGTTTGCTAGGAGATGGCAGA
CACACGTCTGAACTCTTTCCCTACACGACGCTCTTCCGATCTNNNAGGATTTGGCTAGCAGACTGAG
AGCACACGTCTGAACTCTTTCCCTACACGACGCTCTTCCGATCTNNNGGGGTCATGTCATGTGTCAA
AGCACACGTCTGAACTCTTTCCCTACACGACGCTCTTCCGATCTNNNGTTGTAAGCTGGTCCCCCTAAT
AGCACACGTCTGAACTCTTTCCCTACACGACGCTCTTCCGATCTNNNGCCCAGATTCATTTGTTTCTGC
GCACACGTCTGAACTCTTTCCCTACACGACGCTCTTCCGATCTNNNCTGCAGGCAGTGATGCTATCTA
AGCACACGTCTGAACTCTTTCCCTACACGACGCTCTTCCGATCTNNNGGGCATGCTATATGCAAGTGA
AGCACACGTCTGAACTCTTTCCCTACACGACGCTCTTCCGATCTNNNAGAGAGAAAGAAGACACCA
CACACGTCTGAACTCTTTCCCTACACGACGCTCTTCCGATCTNNNCCTTCTCAAGGAAGGAGGTGAG
AGCACACGTCTGAACTCTTTCCCTACACGACGCTCTTCCGATCTNNNCATCTCAGAGGGTGGGTGGA
GAGCACACGTCTGAACTCTTTCCCTACACGACGCTCTTCCGATCTNNNTGGATCTAAGCTACCTCCAGT
AGCACACGTCTGAACTCTTTCCCTACACGACGCTCTTCCGATCTNNNGAGAACTTCTTCCACTCCCCCAC
GCACACGTCTGAACTCTTTCCCTACACGACGCTCTTCCGATCTNNNCTCTACAGTGTGATGCTGGCTTT
GCACACGTCTGAACTCTTTCCCTACACGACGCTCTTCCGATCTNNNGCTTACACAGAAACAGCTGCAA
GCACACGTCTGAACTCTTTCCCTACACGACGCTCTTCCGATCTNNNTCCCAGTTTGTCATGCAATTAGA
GCACACGTCTGAACTCTTTCCCTACACGACGCTCTTCCGATCTNNNTAAAGTCACCCCAGTCCTCAGTT
GAGCACACGTCTGAACTCTTTCCCTACACGACGCTCTTCCGATCTNNNCACATTGGTTGCCCAGGAGA
AGCACACGTCTGAACTCTTTCCCTACACGACGCTCTTCCGATCTNNNGAGAATGGGGAGGTGGTGGT
GAGCACACGTCTGAACTCTTTCCCTACACGACGCTCTTCCGATCTNNNTACAGCTCTGACTGTGCTGA
CACACGTCTGAACTCTTTCCCTACACGACGCTCTTCCGATCTNNNGGTCCATGTGCAGGTTGTATGTA
GAGCACACGTCTGAACTCTTTCCCTACACGACGCTCTTCCGATCTNNNCTTGGCCATGTGTAGCCTTT

GCACACGTCTGAACTCTTTCCCTACACGACGCTCTTCCGATCTNNNTTCCAGACTGTTGCTGATTTCTTC
GAGCACACGTCTGAACTCTTTCCCTACACGACGCTCTTCCGATCTNNNCTCGGCTGTGTTTATGATGGCA
GAGCACACGTCTGAACTCTTTCCCTACACGACGCTCTTCCGATCTNNNTGCGTATGCATCCACATGAAAA
GAGCACACGTCTGAACTCTTTCCCTACACGACGCTCTTCCGATCTNNNGCCTCACCTTCAGGGGATAAAA
GAGCACACGTCTGAACTCTTTCCCTACACGACGCTCTTCCGATCTNNNGGACAAGTGTCAGCAGGATT
AGCACACGTCTGAACTCTTTCCCTACACGACGCTCTTCCGATCTNNNGGCTTGTGTGGAGGTAAGTCA
GCACACGTCTGAACTCTTTCCCTACACGACGCTCTTCCGATCTNNNCAGGAAACAGAACCAACAAGATC
AGCACACGTCTGAACTCTTTCCCTACACGACGCTCTTCCGATCTNNNGCATTTCAACCTCAACTTGCTGG
AGCACACGTCTGAACTCTTTCCCTACACGACGCTCTTCCGATCTNNNGCATTTCAGGACCTGTCCCTATG
GCACACGTCTGAACTCTTTCCCTACACGACGCTCTTCCGATCTNNNACTCAACCTCAAGAGAGAGGAGC
GAGCACACGTCTGAACTCTTTCCCTACACGACGCTCTTCCGATCTNNNCTGTACTGACTGCAGACACGA
GAGCACACGTCTGAACTCTTTCCCTACACGACGCTCTTCCGATCTNNNTCCACTGGAAGAGACATAGTT
AGCACACGTCTGAACTCTTTCCCTACACGACGCTCTTCCGATCTNNNTAATAGCACAACAGCAGCAGCA
GCACACGTCTGAACTCTTTCCCTACACGACGCTCTTCCGATCTNNNAGCACTGGCTGACATTTAGTAGC
GCACACGTCTGAACTCTTTCCCTACACGACGCTCTTCCGATCTNNNGAGATTAGGGCTCCCACCTATGA
GCACACGTCTGAACTCTTTCCCTACACGACGCTCTTCCGATCTNNNGCTGTTCCCCTTACCTAGAATGCCA
AGCACACGTCTGAACTCTTTCCCTACACGACGCTCTTCCGATCTNNNCTTCTGGGGAGAAAGAAGGCTC
GAGCACACGTCTGAACTCTTTCCCTACACGACGCTCTTCCGATCTNNNGTACTACCAGCTTGGAAACCAC
GAGCACACGTCTGAACTCTTTCCCTACACGACGCTCTTCCGATCTNNNGCATGTATGATTGACAGCAGC
GCACACGTCTGAACTCTTTCCCTACACGACGCTCTTCCGATCTNNNGCCTCTATCTCTTTCTGCACTGGC
AGCACACGTCTGAACTCTTTCCCTACACGACGCTCTTCCGATCTNNNTCTACAACTGCCAGCATGCAAAT
GCACACGTCTGAACTCTTTCCCTACACGACGCTCTTCCGATCTNNNGTGCCAGGGTATATGTGACTGTC
GCACACGTCTGAACTCTTTCCCTACACGACGCTCTTCCGATCTNNNCTGCCCATTTACCTGTCTGATGGC
AGCACACGTCTGAACTCTTTCCCTACACGACGCTCTTCCGATCTNNNTTGTCTTGAGCTCACACAGTGG
GAGCACACGTCTGAACTCTTTCCCTACACGACGCTCTTCCGATCTNNNGAAATTGGCATGGCAAAGGC
GCACACGTCTGAACTCTTTCCCTACACGACGCTCTTCCGATCTNNNGACAAGGATTCAACCATTGCCGC
AGCACACGTCTGAACTCTTTCCCTACACGACGCTCTTCCGATCTNNNCTAGACATGCGTAGAGGCACAC
GCACACGTCTGAACTCTTTCCCTACACGACGCTCTTCCGATCTNNNTTTACAGATCAGGACCCCCTACGC
GAGCACACGTCTGAACTCTTTCCCTACACGACGCTCTTCCGATCTNNNATGCCAGATTCACAGTCCTCCTC
GCACACGTCTGAACTCTTTCCCTACACGACGCTCTTCCGATCTNNNGTGCTTCCTGTAGACAGAGTTGC
AGCACACGTCTGAACTCTTTCCCTACACGACGCTCTTCCGATCTNNNAGTCTCATATCACACACACACATC
GCACACGTCTGAACTCTTTCCCTACACGACGCTCTTCCGATCTNNNGCTGTACCATACACCCATCTACCCA
AGCACACGTCTGAACTCTTTCCCTACACGACGCTCTTCCGATCTNNNGTTGTGGTGCTTAGGGAAGAGA
AGCACACGTCTGAACTCTTTCCCTACACGACGCTCTTCCGATCTNNNCTGGTTACACCCCCTTCCCTAACA
AGCACACGTCTGAACTCTTTCCCTACACGACGCTCTTCCGATCTNNNAGGGCCTCACTCAGCTTATAGAT
AGCACACGTCTGAACTCTTTCCCTACACGACGCTCTTCCGATCTNNNTACACTTCAAGTCCAGCCCTACAC
GAGCACACGTCTGAACTCTTTCCCTACACGACGCTCTTCCGATCTNNNGGATTCAGCATGCTCCTAGCA
AGCACACGTCTGAACTCTTTCCCTACACGACGCTCTTCCGATCTNNNCAGCACGAGATTTCTAAGTCCCC
AGAGCACACGTCTGAACTCTTTCCCTACACGACGCTCTTCCGATCTNNNGATGATATATGGGCGGGCAC
AGCACACGTCTGAACTCTTTCCCTACACGACGCTCTTCCGATCTNNNCTGAGATTTCACCAGGCTGCAC
GCACACGTCTGAACTCTTTCCCTACACGACGCTCTTCCGATCTNNNACACATTAGCTGTCACCAAAGCAC
AGCACACGTCTGAACTCTTTCCCTACACGACGCTCTTCCGATCTNNNGCTGAGAATCAGAGGCAAAATC
AGCACACGTCTGAACTCTTTCCCTACACGACGCTCTTCCGATCTNNNCCTAGTGCCTGGCATATGTGTAC

AGCACACGTCTGAACTCTTTCCCTACACGACGCTCTTCCGATCTNNNCAGCCTGCAAAACCTTTCAGTGC
AGCACACGTCTGAACTCTTTCCCTACACGACGCTCTTCCGATCTNNNGACTTGCTCAAGGTCTCTGCAT
GAGCACACGTCTGAACTCTTTCCCTACACGACGCTCTTCCGATCTNNNTCTAAGCCGTACGCTCCAAAAC
AGCACACGTCTGAACTCTTTCCCTACACGACGCTCTTCCGATCTNNNTCAAACCCGGGTGTATTTCAAG
AGCACACGTCTGAACTCTTTCCCTACACGACGCTCTTCCGATCTNNNGACTGGAGTGTAGTGGTACCGT
GCACACGTCTGAACTCTTTCCCTACACGACGCTCTTCCGATCTNNNAGTCAAAAGTTGAGGTCGGTGTC
CACACGTCTGAACTCTTTCCCTACACGACGCTCTTCCGATCTNNNTGCTTCTTCCCTCTTCTTCTTCACATC
GCACACGTCTGAACTCTTTCCCTACACGACGCTCTTCCGATCTNNNCCTCCTTTGGTTTCTAGGAGCTGC
GAGCACACGTCTGAACTCTTTCCCTACACGACGCTCTTCCGATCTNNNTATTTGATGCCAAACAGAGGG
AGCACACGTCTGAACTCTTTCCCTACACGACGCTCTTCCGATCTNNNGAAGGGTCACTTGAGCCTGAAA
GCACACGTCTGAACTCTTTCCCTACACGACGCTCTTCCGATCTNNNTCCAATTAGATCTGACCAGGCCCC
AGCACACGTCTGAACTCTTTCCCTACACGACGCTCTTCCGATCTNNNCTCACTGTCCTTCCCTGGCTACCA
AGCACACGTCTGAACTCTTTCCCTACACGACGCTCTTCCGATCTNNNAAATAGCTTAGGGCAGAAGCAC
GCACACGTCTGAACTCTTTCCCTACACGACGCTCTTCCGATCTNNNCTGGGGAACTCCTAACCTCTTCACA
AGCACACGTCTGAACTCTTTCCCTACACGACGCTCTTCCGATCTNNNGAGCAATAAGGGCTGGGTTTGT
AGCACACGTCTGAACTCTTTCCCTACACGACGCTCTTCCGATCTNNNCATCACTGGGAAGAGAGAGGGA
AGCACACGTCTGAACTCTTTCCCTACACGACGCTCTTCCGATCTNNNCTGAAACTCTGCCCCGTGAAATA
GAGCACACGTCTGAACTCTTTCCCTACACGACGCTCTTCCGATCTNNNACACAACTTCTTCCACCACCTTC
AGCACACGTCTGAACTCTTTCCCTACACGACGCTCTTCCGATCTNNNCTCATTCAGGTCCCCAAGGGAG
AGCACACGTCTGAACTCTTTCCCTACACGACGCTCTTCCGATCTNNNAAAAATCTCCCCAGACCTGCCCC
CACACGTCTGAACTCTTTCCCTACACGACGCTCTTCCGATCTNNNAAGAAAGACAGCTTCCACACTGGC
AGCACACGTCTGAACTCTTTCCCTACACGACGCTCTTCCGATCTNNNCTCCCACTCCATAAAGCACATCC
CACACGTCTGAACTCTTTCCCTACACGACGCTCTTCCGATCTNNNCTCGCATTAGAGGAGAAGAAGGC
AGCACACGTCTGAACTCTTTCCCTACACGACGCTCTTCCGATCTNNNGAGGCATCCAAACCCACTTCAAC
GCACACGTCTGAACTCTTTCCCTACACGACGCTCTTCCGATCTNNNCAAAGTTTAGCTTTCCCATGCCCCA
GCACACGTCTGAACTCTTTCCCTACACGACGCTCTTCCGATCTNNNTGATGCAGGAGCACAACTAAGA
GCACACGTCTGAACTCTTTCCCTACACGACGCTCTTCCGATCTNNNAATTGTAAAGCCTCGGCTAGGTGC
AGCACACGTCTGAACTCTTTCCCTACACGACGCTCTTCCGATCTNNNTCCCATTCAAAGAGTACTACCCA
AGCACACGTCTGAACTCTTTCCCTACACGACGCTCTTCCGATCTNNNGACAGGGTCTCTCTCTGGTTTCT
AGCACACGTCTGAACTCTTTCCCTACACGACGCTCTTCCGATCTNNNCGTGACTATTTGTGCTTGCATCC
GCACACGTCTGAACTCTTTCCCTACACGACGCTCTTCCGATCTNNNGCCGAGGCATCTGTCTCAAAATAC
CACACGTCTGAACTCTTTCCCTACACGACGCTCTTCCGATCTNNNATGACAGGTTATGGTAGCCCCTCCA
GCACACGTCTGAACTCTTTCCCTACACGACGCTCTTCCGATCTNNNCTGTGGTTTAGCCCAGGAATCTCC
AGAGCACACGTCTGAACTCTTTCCCTACACGACGCTCTTCCGATCTNNNCATCTTTGAGCTTTCTGCAGC
AGCACACGTCTGAACTCTTTCCCTACACGACGCTCTTCCGATCTNNNCTCTTCCTCTAGTTTTTCCCCGCC
CACACGTCTGAACTCTTTCCCTACACGACGCTCTTCCGATCTNNNGATGACAGGATGTCCTCTTCAGGC
GAGCACACGTCTGAACTCTTTCCCTACACGACGCTCTTCCGATCTNNNCACCACCTTCTGTCATGCTGATC
GAGCACACGTCTGAACTCTTTCCCTACACGACGCTCTTCCGATCTNNNCAGCCCAGCACTGTATCTGTTT
AGCACACGTCTGAACTCTTTCCCTACACGACGCTCTTCCGATCTNNNGGACAGTGACTTGGGTCCATTTT
GCACACGTCTGAACTCTTTCCCTACACGACGCTCTTCCGATCTNNNCTTTCCTATGGGGAGAGGGAGA
AGCACACGTCTGAACTCTTTCCCTACACGACGCTCTTCCGATCTNNNTTTCAGGGGGAAGGTTGGTTTT
AGAGCACACGTCTGAACTCTTTCCCTACACGACGCTCTTCCGATCTNNNCAAAAGCGAGACCCTGTCCTC
GCACACGTCTGAACTCTTTCCCTACACGACGCTCTTCCGATCTNNNTTTTGCCTTGTTTTCAAGCCTCACA

GAGCACACGTCTGAACTCTTTCCCTACACGACGCTCTTCCGATCTNNNCCACCGCACCAGCTGTAAAATA
GCACACGTCTGAACTCTTTCCCTACACGACGCTCTTCCGATCTNNNCGACTGTCTGTAAAGTGGACCAA
AGCACACGTCTGAACTCTTTCCCTACACGACGCTCTTCCGATCTNNNAGCTCTGTGCTGGAAATTCTGA
AGCACACGTCTGAACTCTTTCCCTACACGACGCTCTTCCGATCTNNNCTCATTGCAGACTACCCTATGCCC
AGCACACGTCTGAACTCTTTCCCTACACGACGCTCTTCCGATCTNNNTGACGGCAGAAGGGAAAAGAA
GCACACGTCTGAACTCTTTCCCTACACGACGCTCTTCCGATCTNNNTGTGGGGATTAGGAGTTCCAGT
GCACACGTCTGAACTCTTTCCCTACACGACGCTCTTCCGATCTNNNGGGTTGTAAGGGTGTGTTGATG
AGCACACGTCTGAACTCTTTCCCTACACGACGCTCTTCCGATCTNNNGTCTGAGGATGAGAAACCTTGG
AGCACACGTCTGAACTCTTTCCCTACACGACGCTCTTCCGATCTNNNGGAAAAATGGCCAATGGAACGA
CACACGTCTGAACTCTTTCCCTACACGACGCTCTTCCGATCTNNNATTCCTCAACCATTGACACACCCCAT
AGCACACGTCTGAACTCTTTCCCTACACGACGCTCTTCCGATCTNNNAAAAAAAAAAAAGCAGCAGCCTC
CACACGTCTGAACTCTTTCCCTACACGACGCTCTTCCGATCTNNNGCCCCGGAGTTTTCTTATCTGTACA
AGAGCACACGTCTGAACTCTTTCCCTACACGACGCTCTTCCGATCTNNNTCAGTCAGAGAAACACGAGA
GCACACGTCTGAACTCTTTCCCTACACGACGCTCTTCCGATCTNNNTTCTTGGAAGGCACCAATTTTCCCA
AGCACACGTCTGAACTCTTTCCCTACACGACGCTCTTCCGATCTNNNGAGTCACCCATACCAAGGAAGC
GCACACGTCTGAACTCTTTCCCTACACGACGCTCTTCCGATCTNNNATCGATACCATCCTGGCTAACACC
CACACGTCTGAACTCTTTCCCTACACGACGCTCTTCCGATCTNNNCTCTCTCTCTCTCTCTCTCCCCATCC
AGCACACGTCTGAACTCTTTCCCTACACGACGCTCTTCCGATCTNNNGAACTTCCTAGCCTCCAGAAGAC
GCACACGTCTGAACTCTTTCCCTACACGACGCTCTTCCGATCTNNNCTCAAAGAAACTCAGGCAGCTTG
GAGCACACGTCTGAACTCTTTCCCTACACGACGCTCTTCCGATCTNNNCAAGTGAGCCTCTGTAGTCCA
AGAGCACACGTCTGAACTCTTTCCCTACACGACGCTCTTCCGATCTNNNAGTGAGCCGAGATTGTGACA
GCACACGTCTGAACTCTTTCCCTACACGACGCTCTTCCGATCTNNNCATGCCACTACACCCAGATGATTC
AGCACACGTCTGAACTCTTTCCCTACACGACGCTCTTCCGATCTNNNCTGGATTACAGGGGCATGATCTT
AGCACACGTCTGAACTCTTTCCCTACACGACGCTCTTCCGATCTNNNGTGACAAAGTGAGACCCTGTCT
AGCACACGTCTGAACTCTTTCCCTACACGACGCTCTTCCGATCTNNNACAACCTCTCTGTCTCTCTCTCTC
GCACACGTCTGAACTCTTTCCCTACACGACGCTCTTCCGATCTNNNAGAGAGACAGAGAGACAGAGAG
AGCACACGTCTGAACTCTTTCCCTACACGACGCTCTTCCGATCTNNNACTAGATCATTTCCCCTCCCTCAC
GCACACGTCTGAACTCTTTCCCTACACGACGCTCTTCCGATCTNNNTCCATGTGCTATAGACCACCAATCA
GCACACGTCTGAACTCTTTCCCTACACGACGCTCTTCCGATCTNNNGGTTCAAGTGACTCTCCTACCTCCA
AGCACACGTCTGAACTCTTTCCCTACACGACGCTCTTCCGATCTNNNCTTTCACTTACATAAGCAGGGG
AGCACACGTCTGAACTCTTTCCCTACACGACGCTCTTCCGATCTNNNCTGAATGCAAAACAGTACAGCC
AGAGCACACGTCTGAACTCTTTCCCTACACGACGCTCTTCCGATCTNNNGAGATTGCAGTGAGCCGAG
GAGCACACGTCTGAACTCTTTCCCTACACGACGCTCTTCCGATCTNNNCTTGCAGTAAGCCAAGATCGT
AGAGCACACGTCTGAACTCTTTCCCTACACGACGCTCTTCCGATCTNNNGAGGCAGAAGAATCGCTTGA
GAGCACACGTCTGAACTCTTTCCCTACACGACGCTCTTCCGATCTNNNGAGGCAGGAGAATAGCGTGA
AGCACACGTCTGAACTCTTTCCCTACACGACGCTCTTCCGATCTNNNAAAGATATGAGAGAGGCCTGAC
AGCACACGTCTGAACTCTTTCCCTACACGACGCTCTTCCGATCTNNNTATTTCTCCACATCCTCTCCAGCC
GCACACGTCTGAACTCTTTCCCTACACGACGCTCTTCCGATCTNNNTCTGCCTCCATCTTTACATGGTGCA
GCACACGTCTGAACTCTTTCCCTACACGACGCTCTTCCGATCTNNNGCGAATTTAGGCTCAGAGTTTGC
GCACACGTCTGAACTCTTTCCCTACACGACGCTCTTCCGATCTNNNGATTGGTGATGATGCTGTGTGAC
GCACACGTCTGAACTCTTTCCCTACACGACGCTCTTCCGATCTNNNGTTCTGTTTCAGCTGTTCCTGACCA
GAGCACACGTCTGAACTCTTTCCCTACACGACGCTCTTCCGATCTNNNGAACTCCTGACCTCAGTGATCC
GAGCACACGTCTGAACTCTTTCCCTACACGACGCTCTTCCGATCTNNNGGTGAGCTGAGATCGTGCTAT

GCACACGTCTGAACTCTTTCCCTACACGACGCTCTTCCGATCTNNNGTGCCTGTAATCCCAGCTACTTC
AGCACACGTCTGAACTCTTTCCCTACACGACGCTCTTCCGATCTNNNGAGATCACGAGATGCACTCCA
GCACACGTCTGAACTCTTTCCCTACACGACGCTCTTCCGATCTNNNGAGAATGGTGTGAACCTGGGA
AGCACACGTCTGAACTCTTTCCCTACACGACGCTCTTCCGATCTNNNGCTGAGGCAGAATCACTTGA
GCACACGTCTGAACTCTTTCCCTACACGACGCTCTTCCGATCTNNNGTGCCTGTAATCCCAGCTACTTC
AGCACACGTCTGAACTCTTTCCCTACACGACGCTCTTCCGATCTNNNATCAAATTGGACCAGGAGAGT
CACACGTCTGAACTCTTTCCCTACACGACGCTCTTCCGATCTNNNCTGTCTCCCTTATGTCCAACCTTCA
GCACACGTCTGAACTCTTTCCCTACACGACGCTCTTCCGATCTNNNCCTCTCCCTGCCCAATAAGTAAAC
AGCACACGTCTGAACTCTTTCCCTACACGACGCTCTTCCGATCTNNNGAGAATCGCTTGAACCCAGGA
GAGCACACGTCTGAACTCTTTCCCTACACGACGCTCTTCCGATCTNNNCTTGAGGTCAGGAGTTCGAG
GCACACGTCTGAACTCTTTCCCTACACGACGCTCTTCCGATCTNNNCACCTGTAATCCCAGCTACTCAG
CACACGTCTGAACTCTTTCCCTACACGACGCTCTTCCGATCTNNNTTGCCTGAAAGTATGCAACCTTG
AGCACACGTCTGAACTCTTTCCCTACACGACGCTCTTCCGATCTNNNCTCCATTGCTTCCTCTCTCCTGC
AGCACACGTCTGAACTCTTTCCCTACACGACGCTCTTCCGATCTNNNAGAACTGCTTGAAACTAGGAG
AGCACACGTCTGAACTCTTTCCCTACACGACGCTCTTCCGATCTNNNCACCTGTAGTCCCAGCTACTTG
AGCACACGTCTGAACTCTTTCCCTACACGACGCTCTTCCGATCTNNNTGTAATCCTAGCACTTTGGGA
AGCACACGTCTGAACTCTTTCCCTACACGACGCTCTTCCGATCTNNNTGGGTGACAGAACAAGACTCT
GAGCACACGTCTGAACTCTTTCCCTACACGACGCTCTTCCGATCTNNNGAGAGTCACTTGAACTGGG
GAGCACACGTCTGAACTCTTTCCCTACACGACGCTCTTCCGATCTNNNAGAATCACTTGAACCCGGGA
AGAGCACACGTCTGAACTCTTTCCCTACACGACGCTCTTCCGATCTNNNCACACTACTGTACTCCAGCC
GCACACGTCTGAACTCTTTCCCTACACGACGCTCTTCCGATCTNNNGATTTGTCTGGGAGGGGTACA
GCACACGTCTGAACTCTTTCCCTACACGACGCTCTTCCGATCTNNNGTGCCTGTAGTCCAAGCTACTC
GCACACGTCTGAACTCTTTCCCTACACGACGCTCTTCCGATCTNNNAGGCAACGACAGATGTACATCTT
CACACGTCTGAACTCTTTCCCTACACGACGCTCTTCCGATCTNNNGCCCGGCTAATTTTTGCATTTTTC
GCACACGTCTGAACTCTTTCCCTACACGACGCTCTTCCGATCTNNNCTCCCGAGTAGCTGGTATTACAG
GAGCACACGTCTGAACTCTTTCCCTACACGACGCTCTTCCGATCTNNNGAGGCATGAGAATCGCTTGA
AGCACACGTCTGAACTCTTTCCCTACACGACGCTCTTCCGATCTNNNGAGGTTGCAGTGAGCTGAGA
GCACACGTCTGAACTCTTTCCCTACACGACGCTCTTCCGATCTNNNCTACTTGGGAGATTGAGGCAG
AGCACACGTCTGAACTCTTTCCCTACACGACGCTCTTCCGATCTNNNCAAGATCGTGCCACTACACTCC
GCACACGTCTGAACTCTTTCCCTACACGACGCTCTTCCGATCTNNNACTCAAGAGAATCGCTTGAACC
GCACACGTCTGAACTCTTTCCCTACACGACGCTCTTCCGATCTNNNTGCATCATTCCAGTCTTCAAGGA
GCACACGTCTGAACTCTTTCCCTACACGACGCTCTTCCGATCTNNNCACACACACACACACACATATC
GCACACGTCTGAACTCTTTCCCTACACGACGCTCTTCCGATCTNNNGGCACCCAGACACCATTATGAT
GAGCACACGTCTGAACTCTTTCCCTACACGACGCTCTTCCGATCTNNNAAAGTGCTGGGATTACAGGC
GCACACGTCTGAACTCTTTCCCTACACGACGCTCTTCCGATCTNNNCTGCAGGAGGCTATGAGGTATT
CACACGTCTGAACTCTTTCCCTACACGACGCTCTTCCGATCTNNNTAGAGCTCGGTGCCTATTGAAAA
GCACACGTCTGAACTCTTTCCCTACACGACGCTCTTCCGATCTNNNGCCTGTAATCCCAGCTACTCTG
AGCACACGTCTGAACTCTTTCCCTACACGACGCTCTTCCGATCTNNNATGTGCCTGTAATCCCAGCTAC
AGCACACGTCTGAACTCTTTCCCTACACGACGCTCTTCCGATCTNNNGAGATCACACCATTGCACTCCA
GAGCACACGTCTGAACTCTTTCCCTACACGACGCTCTTCCGATCTNNNGAGAACTGCTTGAACCCAGG
AGCACACGTCTGAACTCTTTCCCTACACGACGCTCTTCCGATCTNNNCTGGGTGACAAGAGCAAGAC
AGCACACGTCTGAACTCTTTCCCTACACGACGCTCTTCCGATCTNNNGGGCAACAAGAGCAAAACTCA
AGCACACGTCTGAACTCTTTCCCTACACGACGCTCTTCCGATCTNNNGTGTGCCTGTAATCCCAGCTAC

GCACACGTCTGAACTCTTTCCCTACACGACGCTCTTCCGATCTNNNCTTACAGTGAGCCGAGACTTCG
GCACACGTCTGAACTCTTTCCCTACACGACGCTCTTCCGATCTNNNAAAGAATACAGACGGTCCCTGA
GCACACGTCTGAACTCTTTCCCTACACGACGCTCTTCCGATCTNNNTTGAGCCCAGGAATTTGAGGTT
GCACACGTCTGAACTCTTTCCCTACACGACGCTCTTCCGATCTNNNACACACATGTATTAGAAGTTGG
GCACACGTCTGAACTCTTTCCCTACACGACGCTCTTCCGATCTNNNTGTATTTCTCTTGGTCCCTAGGG
AGCACACGTCTGAACTCTTTCCCTACACGACGCTCTTCCGATCTNNNATCTCCCCACTTCATGCCATTCC
CACACGTCTGAACTCTTTCCCTACACGACGCTCTTCCGATCTNNNGTGTGTGTGTGTGTGTGTATA
CACACGTCTGAACTCTTTCCCTACACGACGCTCTTCCGATCTNNNAAATTACCCAGTCTCAGGTAGCC
GCACACGTCTGAACTCTTTCCCTACACGACGCTCTTCCGATCTNNNCACGCCCAGCTAATTCCTTGTAC
AGCACACGTCTGAACTCTTTCCCTACACGACGCTCTTCCGATCTNNNACCTGTAGTCCCAGCTACTAGG
GCACACGTCTGAACTCTTTCCCTACACGACGCTCTTCCGATCTNNNGGTCTCAAACTCCTGACCTCAGC
AGCACACGTCTGAACTCTTTCCCTACACGACGCTCTTCCGATCTNNNGAGCTTGCAGTGAGCCAAGA
GCACACGTCTGAACTCTTTCCCTACACGACGCTCTTCCGATCTNNNGAGAATGGCATGAACCCAGGA
CACACGTCTGAACTCTTTCCCTACACGACGCTCTTCCGATCTNNNGAGAATCTCTTGAACCTGGGAG
GAGCACACGTCTGAACTCTTTCCCTACACGACGCTCTTCCGATCTNNNGAAGGTGGGAGGATTGCTT
GCACACGTCTGAACTCTTTCCCTACACGACGCTCTTCCGATCTNNNGGGAGATGGAAGCACGAGAA
GAGCACACGTCTGAACTCTTTCCCTACACGACGCTCTTCCGATCTNNNGAGCTTGCAGTGAGTCGAG
AGCACACGTCTGAACTCTTTCCCTACACGACGCTCTTCCGATCTNNNCTCCCTTGCTCAAGAGATCCTC
AGCACACGTCTGAACTCTTTCCCTACACGACGCTCTTCCGATCTNNNATTCCGGATCCAACCCTGTTTG
GCACACGTCTGAACTCTTTCCCTACACGACGCTCTTCCGATCTNNNTGCCTGCAAATACCTGTATGAGT
GCACACGTCTGAACTCTTTCCCTACACGACGCTCTTCCGATCTNNNCCAGTCTGAAGGAGAGATTGAG
CACACGTCTGAACTCTTTCCCTACACGACGCTCTTCCGATCTNNNCTGAACTCCTAACCTCGTGATCCC
GCACACGTCTGAACTCTTTCCCTACACGACGCTCTTCCGATCTNNNGGTCTTGAACTCCTGACCTCAGC
CACACGTCTGAACTCTTTCCCTACACGACGCTCTTCCGATCTNNNCGCCGGCTGATCATGGAATATAT
GCACACGTCTGAACTCTTTCCCTACACGACGCTCTTCCGATCTNNNGGAGATTGCAGTGAGCTGAGA
GAGCACACGTCTGAACTCTTTCCCTACACGACGCTCTTCCGATCTNNNGTGCCTGTAATCCCAGCTAC
AGCACACGTCTGAACTCTTTCCCTACACGACGCTCTTCCGATCTNNNGAGATCACATCACTGCACTCCA
GCACACGTCTGAACTCTTTCCCTACACGACGCTCTTCCGATCTNNNGTGATGGTGAACGCCTGTAATC
AGCACACGTCTGAACTCTTTCCCTACACGACGCTCTTCCGATCTNNNGGAGAATCGCTAGAATCTGGG
AGCACACGTCTGAACTCTTTCCCTACACGACGCTCTTCCGATCTNNNGGTGACAGAGTGAGACTCCAT
AGCACACGTCTGAACTCTTTCCCTACACGACGCTCTTCCGATCTNNNGAGCTTGCAGTGAGCCAAGAT
AGCACACGTCTGAACTCTTTCCCTACACGACGCTCTTCCGATCTNNNCACCTGTAATCCCAGCTACTCG
AGCACACGTCTGAACTCTTTCCCTACACGACGCTCTTCCGATCTNNNTAGGAGAATTGCTTGAACCCA
AGCACACGTCTGAACTCTTTCCCTACACGACGCTCTTCCGATCTNNNCACCTGTAGTCCCAGCTACTTG
GAGCACACGTCTGAACTCTTTCCCTACACGACGCTCTTCCGATCTNNNGAGAATTGCTTGAAGCCGAG
AGAGCACACGTCTGAACTCTTTCCCTACACGACGCTCTTCCGATCTNNNGAGCTTGCAATGAGCCAAG
AGCACACGTCTGAACTCTTTCCCTACACGACGCTCTTCCGATCTNNNGAGGTTGCAGTGTGTGGAG
AGCACACGTCTGAACTCTTTCCCTACACGACGCTCTTCCGATCTNNNCCCAGCAGGTGATTCTGATACA
CACACGTCTGAACTCTTTCCCTACACGACGCTCTTCCGATCTNNNTGCTGTTGAGAAGTCAGAAGTC
GCACACGTCTGAACTCTTTCCCTACACGACGCTCTTCCGATCTNNNATCCCACCTGCTAATGCTTTACAC
AGCACACGTCTGAACTCTTTCCCTACACGACGCTCTTCCGATCTNNNAGAATGGGGAAGGAAATCTG
AGCACACGTCTGAACTCTTTCCCTACACGACGCTCTTCCGATCTNNNGGGTTAGTCAGGGCTCTGTT
AGCACACGTCTGAACTCTTTCCCTACACGACGCTCTTCCGATCTNNNTAAGTAGCTGGGATTACAGG

GCACACGTCTGAACTCTTTCCCTACACGACGCTCTTCCGATCTNNNTCGGCTGGAATCATTTTGCTTTT
GAGCACACGTCTGAACTCTTTCCCTACACGACGCTCTTCCGATCTNNNCACGAGAATTGCTTGAACCTG
AGCACACGTCTGAACTCTTTCCCTACACGACGCTCTTCCGATCTNNNCTGTGTTTGTCTCACTGCACTC
GAGCACACGTCTGAACTCTTTCCCTACACGACGCTCTTCCGATCTNNNGTTGCAGTGAGCTGAGATTG
AGCACACGTCTGAACTCTTTCCCTACACGACGCTCTTCCGATCTNNNATTGAGACCATCCTGGCTGATA
AGCACACGTCTGAACTCTTTCCCTACACGACGCTCTTCCGATCTNNNGAGAATGGCATGAACCTGGGA
GAGCACACGTCTGAACTCTTTCCCTACACGACGCTCTTCCGATCTNNNGCAGGAGAATTGCTTGAACCT
AGAGCACACGTCTGAACTCTTTCCCTACACGACGCTCTTCCGATCTNNNGAGGCAGAAGAATCGCTTG
AGCACACGTCTGAACTCTTTCCCTACACGACGCTCTTCCGATCTNNNCGACAGAACAAGACTCCCTCTC
GCACACGTCTGAACTCTTTCCCTACACGACGCTCTTCCGATCTNNNTCATGGGGTATTTTGGGTGAGA
AGCACACGTCTGAACTCTTTCCCTACACGACGCTCTTCCGATCTNNNAAAGTGCTGGGATTACAGGCA
GCACACGTCTGAACTCTTTCCCTACACGACGCTCTTCCGATCTNNNTGCAACACTTACTATGCCCTCATC
GCACACGTCTGAACTCTTTCCCTACACGACGCTCTTCCGATCTNNNCCGCACCCTACATTCACTATCTTC
AGCACACGTCTGAACTCTTTCCCTACACGACGCTCTTCCGATCTNNNGAACTCCAGACCTCAGGTGATC
GCACACGTCTGAACTCTTTCCCTACACGACGCTCTTCCGATCTNNNTGCCTCTATTGATCTCTGCAGAA
AGCACACGTCTGAACTCTTTCCCTACACGACGCTCTTCCGATCTNNNCCGGCTTCCTGAATTTCCTTATC
GCACACGTCTGAACTCTTTCCCTACACGACGCTCTTCCGATCTNNNTTTCTATCCATTCTGGCTTTCCAC
GCACACGTCTGAACTCTTTCCCTACACGACGCTCTTCCGATCTNNNAGATGATCAAAGAAGCCTGCCAT
AGCACACGTCTGAACTCTTTCCCTACACGACGCTCTTCCGATCTNNNGGATTACAGGCGTGAACCACTA
AGCACACGTCTGAACTCTTTCCCTACACGACGCTCTTCCGATCTNNNGGTTCTCCATTCCTGAGTCACTT
GCACACGTCTGAACTCTTTCCCTACACGACGCTCTTCCGATCTNNNAACAGACAACCACTAAGAGTCCC
GCACACGTCTGAACTCTTTCCCTACACGACGCTCTTCCGATCTNNNATGCAGAATGTCCACCTGTTTTG
GCACACGTCTGAACTCTTTCCCTACACGACGCTCTTCCGATCTNNNTACCCCTCTGACAGTATACCACA
AGCACACGTCTGAACTCTTTCCCTACACGACGCTCTTCCGATCTNNNTGGAACATTGACCCCTTTCTCAT
GCACACGTCTGAACTCTTTCCCTACACGACGCTCTTCCGATCTNNNTTCAATAGTGGTGCAAGGATCCA
GCACACGTCTGAACTCTTTCCCTACACGACGCTCTTCCGATCTNNNCTCAAGTGGCCGTAACAAAAACA
GAGCACACGTCTGAACTCTTTCCCTACACGACGCTCTTCCGATCTNNNGAGGTTTTGCTATGTTGGCCA
GAGCACACGTCTGAACTCTTTCCCTACACGACGCTCTTCCGATCTNNNCACGAGAATTGCTTGAACCCA
GAGCACACGTCTGAACTCTTTCCCTACACGACGCTCTTCCGATCTNNNGACCCCCAATCACTGCAAAAA
AGCACACGTCTGAACTCTTTCCCTACACGACGCTCTTCCGATCTNNNCGACAGAGTAAGGCTCCATCTC
AGCACACGTCTGAACTCTTTCCCTACACGACGCTCTTCCGATCTNNNGAGCTTGCAGTGAGTCGAGAT
GAGCACACGTCTGAACTCTTTCCCTACACGACGCTCTTCCGATCTNNNCGGAGCGAGACTTCCAAAAA
AGCACACGTCTGAACTCTTTCCCTACACGACGCTCTTCCGATCTNNNGAGAATGGTGTGAACCCAGGA
AGAGCACACGTCTGAACTCTTTCCCTACACGACGCTCTTCCGATCTNNNCAGGAGAATGGCGTGAAC
GAGCACACGTCTGAACTCTTTCCCTACACGACGCTCTTCCGATCTNNNACCTGTAGTCCCAGCTACTAG
GCACACGTCTGAACTCTTTCCCTACACGACGCTCTTCCGATCTNNNGATGGGTTGGATCATCTGAGGT
AGCACACGTCTGAACTCTTTCCCTACACGACGCTCTTCCGATCTNNNGCAGGAGAATCGCTTGAACTT
AGCACACGTCTGAACTCTTTCCCTACACGACGCTCTTCCGATCTNNNGCAGGAGAATTGCTTGAACCTG
GCACACGTCTGAACTCTTTCCCTACACGACGCTCTTCCGATCTNNNGGATCACTTGAGGCTAGGAGTT
GAGCACACGTCTGAACTCTTTCCCTACACGACGCTCTTCCGATCTNNNGAGACGGAGCAAGACTCTTGT
AGCACACGTCTGAACTCTTTCCCTACACGACGCTCTTCCGATCTNNNGAGAATGGCATGAACCCAGGA
GAGCACACGTCTGAACTCTTTCCCTACACGACGCTCTTCCGATCTNNNGAGAATCGCTTGAACCCAGG
AGCACACGTCTGAACTCTTTCCCTACACGACGCTCTTCCGATCTNNNAAGATAATGGCTTCCAGCTCCA

GCACACGTCTGAACTCTTTCCCTACACGACGCTCTTCCGATCTNNNTCTCCCAGGTTCAAGTGATTCTC
CACACGTCTGAACTCTTTCCCTACACGACGCTCTTCCGATCTNNNAGTGTGCATACTCCAAACTCACTC
GCACACGTCTGAACTCTTTCCCTACACGACGCTCTTCCGATCTNNNTGTGGCCTTTGCATCTTTCAATT
GCACACGTCTGAACTCTTTCCCTACACGACGCTCTTCCGATCTNNNACTGTAAATTCCATGAGGGCAG
GCACACGTCTGAACTCTTTCCCTACACGACGCTCTTCCGATCTNNNCTCGGTCTCAGTACGTCAATCAA
AGCACACGTCTGAACTCTTTCCCTACACGACGCTCTTCCGATCTNNNGCGAAACCACGTCTCTACTAAA
AGCACACGTCTGAACTCTTTCCCTACACGACGCTCTTCCGATCTNNNGCAACAGAATGAGACCCTGTCT
GAGCACACGTCTGAACTCTTTCCCTACACGACGCTCTTCCGATCTNNNTACACGCCTGTAATCCCAGCT
AGCACACGTCTGAACTCTTTCCCTACACGACGCTCTTCCGATCTNNNCTACTCAGGAGGCTAAGGCAA
AGCACACGTCTGAACTCTTTCCCTACACGACGCTCTTCCGATCTNNNGCCTGGCAATAGAGAGAGAC
AGCACACGTCTGAACTCTTTCCCTACACGACGCTCTTCCGATCTNNNGAGACAGGAGAGCCACTTGA
CACACGTCTGAACTCTTTCCCTACACGACGCTCTTCCGATCTNNNCTCAAACTATTCTCCCACCCCAACA
GCACACGTCTGAACTCTTTCCCTACACGACGCTCTTCCGATCTNNNGCTCAAGCGATTCTCCTACTTTG
GCACACGTCTGAACTCTTTCCCTACACGACGCTCTTCCGATCTNNNTTTGGCTCCACTGGTGAATTCTA
GAGCACACGTCTGAACTCTTTCCCTACACGACGCTCTTCCGATCTNNNGAGGCAGGAGAACCACTAGA
AGCACACGTCTGAACTCTTTCCCTACACGACGCTCTTCCGATCTNNNCCTATAGTCCCAGCTACTCGG
AGCACACGTCTGAACTCTTTCCCTACACGACGCTCTTCCGATCTNNNGAGAATCGCTTGAACCCAGGA
AGCACACGTCTGAACTCTTTCCCTACACGACGCTCTTCCGATCTNNNCGTAGCTGGGATTACAGGTCTG
GCACACGTCTGAACTCTTTCCCTACACGACGCTCTTCCGATCTNNNTGTGTGTCCTCTCTTGACAGAAG
CACACGTCTGAACTCTTTCCCTACACGACGCTCTTCCGATCTNNNCCCTTTGCTCTCCAAGAAGTTTGC
AGCACACGTCTGAACTCTTTCCCTACACGACGCTCTTCCGATCTNNNAGTCCCTTGTGTACTTCAGCTT
AGCACACGTCTGAACTCTTTCCCTACACGACGCTCTTCCGATCTNNNGATTACAGGTGTGAGCCACTG
AGCACACGTCTGAACTCTTTCCCTACACGACGCTCTTCCGATCTNNNGAGGTTGCAGTGAACTGAGAC
AGCACACGTCTGAACTCTTTCCCTACACGACGCTCTTCCGATCTNNNGCCTGTAGTCCCAGCTACTTGC
AGCACACGTCTGAACTCTTTCCCTACACGACGCTCTTCCGATCTNNNCAGGAGAATCGCTTGAACCCAA
GAGCACACGTCTGAACTCTTTCCCTACACGACGCTCTTCCGATCTNNNCACCTGTAATCCCAGCTACTCG
AGCACACGTCTGAACTCTTTCCCTACACGACGCTCTTCCGATCTNNNGAGAATGGTGTGAACCCAGGA
CACACGTCTGAACTCTTTCCCTACACGACGCTCTTCCGATCTNNNGCTTGTGTTTCCTTTTCCTCCTCCA
GCACACGTCTGAACTCTTTCCCTACACGACGCTCTTCCGATCTNNNGTCCTTCAGGTTAGGAACTGTG
GCACACGTCTGAACTCTTTCCCTACACGACGCTCTTCCGATCTNNNGGTTGACAATTTTCCTCTACAGC
AGCACACGTCTGAACTCTTTCCCTACACGACGCTCTTCCGATCTNNNTTACATGAAAAGTACCTCCCCG
CACACGTCTGAACTCTTTCCCTACACGACGCTCTTCCGATCTNNNCTCCTCTGCACCCTCAAAGTATTCA
GCACACGTCTGAACTCTTTCCCTACACGACGCTCTTCCGATCTNNNAGCACTAAGGTTTGAAGTCCCTC
AGCACACGTCTGAACTCTTTCCCTACACGACGCTCTTCCGATCTNNNCTTGCAGTGAGCTGATATCGTG
GAGCACACGTCTGAACTCTTTCCCTACACGACGCTCTTCCGATCTNNNGCCTGTAATCCCAGCTACTCG
AGCACACGTCTGAACTCTTTCCCTACACGACGCTCTTCCGATCTNNNGTTGCAGTGAGCCAAGAAGA
AGCACACGTCTGAACTCTTTCCCTACACGACGCTCTTCCGATCTNNNGGCGACAGAGAGATACTCCAT
AGCACACGTCTGAACTCTTTCCCTACACGACGCTCTTCCGATCTNNNGCAAGAGAATGGCTTGAACCT
GAGCACACGTCTGAACTCTTTCCCTACACGACGCTCTTCCGATCTNNNGCGACAGAGTGAGATTCCAT
AGAGCACACGTCTGAACTCTTTCCCTACACGACGCTCTTCCGATCTNNNAGAGGCAGGAGAATTGCTT
AGCACACGTCTGAACTCTTTCCCTACACGACGCTCTTCCGATCTNNNCTCCTGGGTTCAAGCGATTCTC
GCACACGTCTGAACTCTTTCCCTACACGACGCTCTTCCGATCTNNNGGGGTTTCTATTTCTGCTACCGA
GCACACGTCTGAACTCTTTCCCTACACGACGCTCTTCCGATCTNNNAGGCTTTCTACTAGAAGCAGCAA

GCACACGTCTGAACTCTTTCCCTACACGACGCTCTTCCGATCTNNNGGAGTTGTTTTTGCCTCAGGATC
GCACACGTCTGAACTCTTTCCCTACACGACGCTCTTCCGATCTNNNTTTTTAGTGGAGACGGGGTTTCA
GCACACGTCTGAACTCTTTCCCTACACGACGCTCTTCCGATCTNNNCGCACCACATACTGTTGTTTTCTCA
AGCACACGTCTGAACTCTTTCCCTACACGACGCTCTTCCGATCTNNNTGTATTTGCAATCAATCCCCAAG
AGCACACGTCTGAACTCTTTCCCTACACGACGCTCTTCCGATCTNNNGAGCTTGCAGTGAGCCTAGATC
AGAGCACACGTCTGAACTCTTTCCCTACACGACGCTCTTCCGATCTNNNAAGAGGTTGAGGTTGCAGC
GAGCACACGTCTGAACTCTTTCCCTACACGACGCTCTTCCGATCTNNNGGTGGATCACAAGGTCAAGA
GAGCACACGTCTGAACTCTTTCCCTACACGACGCTCTTCCGATCTNNNTGAGCCAAGATCACACCACTG
AGCACACGTCTGAACTCTTTCCCTACACGACGCTCTTCCGATCTNNNGCAGGAGAATTGCTTGAACCT
AGCACACGTCTGAACTCTTTCCCTACACGACGCTCTTCCGATCTNNNAGAGATCACACCACTGCATTCC
AGAGCACACGTCTGAACTCTTTCCCTACACGACGCTCTTCCGATCTNNNTAGTCCCAGCTACTCAGGAG
AGAGCACACGTCTGAACTCTTTCCCTACACGACGCTCTTCCGATCTNNNCCTATAGTCCCAGCTACTCG
AGCACACGTCTGAACTCTTTCCCTACACGACGCTCTTCCGATCTNNNGAGGCAGAAGAATCGCTTGAAC
GAGCACACGTCTGAACTCTTTCCCTACACGACGCTCTTCCGATCTNNNATGCCTGCAATCCCAACTACTC
AGCACACGTCTGAACTCTTTCCCTACACGACGCTCTTCCGATCTNNNAATGAAGCAAGTGGGAGAACA
AGCACACGTCTGAACTCTTTCCCTACACGACGCTCTTCCGATCTNNNGATCTCCTGACCTCGTGATCTG
GCACACGTCTGAACTCTTTCCCTACACGACGCTCTTCCGATCTNNNCCTGGGCAATATAGCAAGACACT
AGCACACGTCTGAACTCTTTCCCTACACGACGCTCTTCCGATCTNNNAAACAGCAGAGATGAGTGGCT
GCACACGTCTGAACTCTTTCCCTACACGACGCTCTTCCGATCTNNNAAATCCATTCAGCTACACTGCACCA
GCACACGTCTGAACTCTTTCCCTACACGACGCTCTTCCGATCTNNNTTGTAGCAGTTTGGCATTTCCTGC
AGCACACGTCTGAACTCTTTCCCTACACGACGCTCTTCCGATCTNNNCTACTTGGTAAGCTGAGGCAG
AGCACACGTCTGAACTCTTTCCCTACACGACGCTCTTCCGATCTNNNGCGAAAGAGCGAGATTCTGTC
GAGCACACGTCTGAACTCTTTCCCTACACGACGCTCTTCCGATCTNNNGAGGCAGAAGAATCGCTTGA
GAGCACACGTCTGAACTCTTTCCCTACACGACGCTCTTCCGATCTNNNGAGATCATGTCACTGCACTCCA
GCACACGTCTGAACTCTTTCCCTACACGACGCTCTTCCGATCTNNNGTGCCTATAGTCCCAGCTACTCCA
GAGCACACGTCTGAACTCTTTCCCTACACGACGCTCTTCCGATCTNNNGAAGCAGGAGAATGGCATGA
GAGCACACGTCTGAACTCTTTCCCTACACGACGCTCTTCCGATCTNNNGGTGACAGAACGAGACTCCAT
GCACACGTCTGAACTCTTTCCCTACACGACGCTCTTCCGATCTNNNCCACTGTGCCTGGCCATATATATCA
AGCACACGTCTGAACTCTTTCCCTACACGACGCTCTTCCGATCTNNNCTGTCATGGTGGCACTAATTCAC
GCACACGTCTGAACTCTTTCCCTACACGACGCTCTTCCGATCTNNNTTCCCCTGTGTGATAGTCCATTCCA
AGCACACGTCTGAACTCTTTCCCTACACGACGCTCTTCCGATCTNNNGGTCTTGAACTCCTGACCTCAG
AGCACACGTCTGAACTCTTTCCCTACACGACGCTCTTCCGATCTNNNTGACCCAGTGAGTCTCTGTCTC
GAGCACACGTCTGAACTCTTTCCCTACACGACGCTCTTCCGATCTNNNGAGAATCGCTTGAACCTGGGA
GAGCACACGTCTGAACTCTTTCCCTACACGACGCTCTTCCGATCTNNNCACAGTGTGCACAACTACAGT
GAGCACACGTCTGAACTCTTTCCCTACACGACGCTCTTCCGATCTNNNGAGAATGGCATGAACCCAGG
AGCACACGTCTGAACTCTTTCCCTACACGACGCTCTTCCGATCTNNNGTTGCAGTGAGCTGAGATTGT
AGCACACGTCTGAACTCTTTCCCTACACGACGCTCTTCCGATCTNNNGAGAATCGCTTGAACCTGGGA
AGCACACGTCTGAACTCTTTCCCTACACGACGCTCTTCCGATCTNNNCTTGCAGTGAGCCAAGATTGTG
GAGCACACGTCTGAACTCTTTCCCTACACGACGCTCTTCCGATCTNNNGAGGCAGGAGAATTGCTTCAA
AGCACACGTCTGAACTCTTTCCCTACACGACGCTCTTCCGATCTNNNGGAGGGAAATCACCAACCTACT
AAGAGCACACGTCTGAACTCTTTCCCTACACGACGCTCTTCCGATCTNNNAGGAGAATGGCGTGAACC
GAGCACACGTCTGAACTCTTTCCCTACACGACGCTCTTCCGATCTNNNGCACCTGTAGTACCAGCTACTC
AGAGCACACGTCTGAACTCTTTCCCTACACGACGCTCTTCCGATCTNNNGAGAATAGCTTGAACCCGG

GAGCACACGTCTGAACTCTTTCCCTACACGACGCTCTTCCGATCTNNNGAGACTGCAGTGAGCTGAGA
AGCACACGTCTGAACTCTTTCCCTACACGACGCTCTTCCGATCTNNNGAGCTTGCAGTGAGTCGAGAT
AGCACACGTCTGAACTCTTTCCCTACACGACGCTCTTCCGATCTNNNCATGAACCAGAGACAGAGACA
GAGCACACGTCTGAACTCTTTCCCTACACGACGCTCTTCCGATCTNNNCTGTCTCTTCTGGGTAAGGTG
AGCACACGTCTGAACTCTTTCCCTACACGACGCTCTTCCGATCTNNNTGCCAAAGTTTACGCAATTCTG
GCACACGTCTGAACTCTTTCCCTACACGACGCTCTTCCGATCTNNNGAAGTCTGGACAAGGGGCTTA
AGCACACGTCTGAACTCTTTCCCTACACGACGCTCTTCCGATCTNNNGAGACAGCAAGACTAACCTCT
AGCACACGTCTGAACTCTTTCCCTACACGACGCTCTTCCGATCTNNNTTTGAACATGCCCTATGCCAAA
GCACACGTCTGAACTCTTTCCCTACACGACGCTCTTCCGATCTNNNTCCTGAGTAGCTGAGACTACAGG
AGCACACGTCTGAACTCTTTCCCTACACGACGCTCTTCCGATCTNNNCTTGCAGTGAGGCGAGATTAC
AGCACACGTCTGAACTCTTTCCCTACACGACGCTCTTCCGATCTNNNGTTGCAGTGAGCTGAGATCAT
AGCACACGTCTGAACTCTTTCCCTACACGACGCTCTTCCGATCTNNNCTTGCAGTAAGCCAAGATCAC
GAGCACACGTCTGAACTCTTTCCCTACACGACGCTCTTCCGATCTNNNAGGCAAGAGAATGGTGTGA
AGAGCACACGTCTGAACTCTTTCCCTACACGACGCTCTTCCGATCTNNNCTGGGAGTTGGATTTTGCA
GAGCACACGTCTGAACTCTTTCCCTACACGACGCTCTTCCGATCTNNNGATAGCATTCTATCCAGCCTG
GAGCACACGTCTGAACTCTTTCCCTACACGACGCTCTTCCGATCTNNNGAGGTTGCAGTGAGCTGAGA
AGCACACGTCTGAACTCTTTCCCTACACGACGCTCTTCCGATCTNNNGTTGCAGTGAGCTGAGATTGT
GAGCACACGTCTGAACTCTTTCCCTACACGACGCTCTTCCGATCTNNNCACCTGTGATCCCAGCTACTT
CACACGTCTGAACTCTTTCCCTACACGACGCTCTTCCGATCTNNNCCCTTTGTAATTGTTTCCATAGGGC
GCACACGTCTGAACTCTTTCCCTACACGACGCTCTTCCGATCTNNNAGTGCTGGGATTACAGATGTGA
GCACACGTCTGAACTCTTTCCCTACACGACGCTCTTCCGATCTNNNGCAAGACAGCTTCACCTAACTCA
GCACACGTCTGAACTCTTTCCCTACACGACGCTCTTCCGATCTNNNACACCTATTACTATGCTGGCTGG
GCACACGTCTGAACTCTTTCCCTACACGACGCTCTTCCGATCTNNNAGTTTTCAGAGCACAGGTCTTCA
AGCACACGTCTGAACTCTTTCCCTACACGACGCTCTTCCGATCTNNNCTCCTGGGTTCAAGCAGTTCTC
AGCACACGTCTGAACTCTTTCCCTACACGACGCTCTTCCGATCTNNNCAAAACCCCAGTGGATGAAATG
GCACACGTCTGAACTCTTTCCCTACACGACGCTCTTCCGATCTNNNAGAGACTGGGTTTCTCCATGTT
AGCACACGTCTGAACTCTTTCCCTACACGACGCTCTTCCGATCTNNNGAGGTTGCAGTGAGCTTAGA
GAGCACACGTCTGAACTCTTTCCCTACACGACGCTCTTCCGATCTNNNGACAGAATGAGACCCCTGAC
GAGCACACGTCTGAACTCTTTCCCTACACGACGCTCTTCCGATCTNNNGAGCTTGCAGTTAGCCGAGA
GCACACGTCTGAACTCTTTCCCTACACGACGCTCTTCCGATCTNNNGAGGCAGAAGAATCGCTTGAAC
AGAGCACACGTCTGAACTCTTTCCCTACACGACGCTCTTCCGATCTNNNCACCTGTAGTCCCAGCTACTT
GCACACGTCTGAACTCTTTCCCTACACGACGCTCTTCCGATCTNNNCTGAGGTGGGAAAATTGCTTG
GAGCACACGTCTGAACTCTTTCCCTACACGACGCTCTTCCGATCTNNNATTGCTTGAGCTCAGGAGTTC
GAGCACACGTCTGAACTCTTTCCCTACACGACGCTCTTCCGATCTNNNCATGGAGAAACCCTGTCCCTA
GCACACGTCTGAACTCTTTCCCTACACGACGCTCTTCCGATCTNNNGCGACAGAGCAAGACATCATCT
AGCACACGTCTGAACTCTTTCCCTACACGACGCTCTTCCGATCTNNNGACGCAAGAGAATGGTGTGA
GCACACGTCTGAACTCTTTCCCTACACGACGCTCTTCCGATCTNNNTGACAGAGCAAGACACCATCTC
GCACACGTCTGAACTCTTTCCCTACACGACGCTCTTCCGATCTNNNCTGGGCAACAAGAGTGAAACTT
GCACACGTCTGAACTCTTTCCCTACACGACGCTCTTCCGATCTNNNGGTAAGACTTCGTTCAGGACCAT
GCACACGTCTGAACTCTTTCCCTACACGACGCTCTTCCGATCTNNNAGTGCATTTTCTCTCTCTCTCTCC
AGCACACGTCTGAACTCTTTCCCTACACGACGCTCTTCCGATCTNNNTTGTGGGGTTTTTTCTTTCTCT
GCACACGTCTGAACTCTTTCCCTACACGACGCTCTTCCGATCTNNNCCCTGGAAATATTGGCAATGGTG
GCACACGTCTGAACTCTTTCCCTACACGACGCTCTTCCGATCTNNNCCTTCGAGATACCAACAGGCATT

AGCACACGTCTGAACTCTTTCCCTACACGACGCTCTTCCGATCTNNNGAAACTGGACCTGGTGACTA
GCACACGTCTGAACTCTTTCCCTACACGACGCTCTTCCGATCTNNNATTTGGATTTCTTCTGCACACAG
CACACGTCTGAACTCTTTCCCTACACGACGCTCTTCCGATCTNNNGTTTGGGGACTGCCATTAGTTTG
AGCACACGTCTGAACTCTTTCCCTACACGACGCTCTTCCGATCTNNNCTGAGATCACACCACTGCACT
AGCACACGTCTGAACTCTTTCCCTACACGACGCTCTTCCGATCTNNNGAGAATCGCTTGAACCTGGGA
AGCACACGTCTGAACTCTTTCCCTACACGACGCTCTTCCGATCTNNNAGAATCGCTTGAACCAGGGA
AGCACACGTCTGAACTCTTTCCCTACACGACGCTCTTCCGATCTNNNAGGAGGTGGAGATTGCAATG
AGCACACGTCTGAACTCTTTCCCTACACGACGCTCTTCCGATCTNNNTGTAGTCCCAGCTACTCGGAA
GCACACGTCTGAACTCTTTCCCTACACGACGCTCTTCCGATCTNNNAATGAGCCGAGATCACACCATT
GCACACGTCTGAACTCTTTCCCTACACGACGCTCTTCCGATCTNNNGAGAATGACGTGAACCCAGGA
AGCACACGTCTGAACTCTTTCCCTACACGACGCTCTTCCGATCTNNNACCTGTAGTCCCTGCTACTCA
AGCACACGTCTGAACTCTTTCCCTACACGACGCTCTTCCGATCTNNNCACGAGAATTGCTTGAACCCA
GAGCACACGTCTGAACTCTTTCCCTACACGACGCTCTTCCGATCTNNNCACCTGTAGTCCCAGCTACTT
GCACACGTCTGAACTCTTTCCCTACACGACGCTCTTCCGATCTNNNTAGTCCCAGCTACTCGGTACTCC
AGCACACGTCTGAACTCTTTCCCTACACGACGCTCTTCCGATCTNNNGTGAACTGGGAGATGGAGCT
AGCACACGTCTGAACTCTTTCCCTACACGACGCTCTTCCGATCTNNNCCAAAACGCTCTTTTTCAAAGCC
GCACACGTCTGAACTCTTTCCCTACACGACGCTCTTCCGATCTNNNCAGCCTGATCTCCTCCTAAACATC
GCACACGTCTGAACTCTTTCCCTACACGACGCTCTTCCGATCTNNNGCCCTATATTGCATTTTCCCAGG
CACACGTCTGAACTCTTTCCCTACACGACGCTCTTCCGATCTNNNCCAAAACAGACAGTGATGTTGCT
GCACACGTCTGAACTCTTTCCCTACACGACGCTCTTCCGATCTNNNTGCTCTGACTCACAGTTTGGAAT
AGCACACGTCTGAACTCTTTCCCTACACGACGCTCTTCCGATCTNNNGTGCCTGTAATCCCAGCTACTT
AGCACACGTCTGAACTCTTTCCCTACACGACGCTCTTCCGATCTNNNCATGCCTGTAATCCCAGCTACT
AGCACACGTCTGAACTCTTTCCCTACACGACGCTCTTCCGATCTNNNGCAGGAGAATTGCTTGAACC
AGAGCACACGTCTGAACTCTTTCCCTACACGACGCTCTTCCGATCTNNNCGCCTATAGTCCCAGCTACT
AGCACACGTCTGAACTCTTTCCCTACACGACGCTCTTCCGATCTNNNCGCCTATAGTCCCAGCTACTT
AGCACACGTCTGAACTCTTTCCCTACACGACGCTCTTCCGATCTNNNGAGCTTGCAATGAGCCGAGAT
AGAGCACACGTCTGAACTCTTTCCCTACACGACGCTCTTCCGATCTNNNGCTCCTGTAATCCCAGCTACT
AGCACACGTCTGAACTCTTTCCCTACACGACGCTCTTCCGATCTNNNGTGAGCTGAGATTGTGCCATTG
GCACACGTCTGAACTCTTTCCCTACACGACGCTCTTCCGATCTNNNCTCCTGAGCTCAAGGGATCTTCC
AGCACACGTCTGAACTCTTTCCCTACACGACGCTCTTCCGATCTNNNGAGTTTGCAGTGAGCCGAGA
AGCACACGTCTGAACTCTTTCCCTACACGACGCTCTTCCGATCTNNNGGGACCCTTGTCATACATGGT
AGCACACGTCTGAACTCTTTCCCTACACGACGCTCTTCCGATCTNNNTAGGCTAGCCTCATTCACATCTG
AGCACACGTCTGAACTCTTTCCCTACACGACGCTCTTCCGATCTNNNCAGAAGAGAGAGAACAATGC
GAGCACACGTCTGAACTCTTTCCCTACACGACGCTCTTCCGATCTNNNGAGAGAGGGAGAGAGAGA
AGCACACGTCTGAACTCTTTCCCTACACGACGCTCTTCCGATCTNNNATTGGTGAGACATTTGCATG
AGCACACGTCTGAACTCTTTCCCTACACGACGCTCTTCCGATCTNNNGAGGTAGGAGAATGGCGTGA
GAGCACACGTCTGAACTCTTTCCCTACACGACGCTCTTCCGATCTNNNGAGTTTGCAGTGAGCCAAGA
AGCACACGTCTGAACTCTTTCCCTACACGACGCTCTTCCGATCTNNNTGGCTAACAAACCCTGTCTCTA
AGCACACGTCTGAACTCTTTCCCTACACGACGCTCTTCCGATCTNNNGAATGGCATGAACCCAGGAG
AGCACACGTCTGAACTCTTTCCCTACACGACGCTCTTCCGATCTNNNGAGAATGGTGTGAACCTGGG
GCACACGTCTGAACTCTTTCCCTACACGACGCTCTTCCGATCTNNNAATAAGCAATGCTGAATTCCGC
AGCACACGTCTGAACTCTTTCCCTACACGACGCTCTTCCGATCTNNNAGGGAACTTTCTAGGCCATGA
GCACACGTCTGAACTCTTTCCCTACACGACGCTCTTCCGATCTNNNACCACTGCACCAAAATGAGAAA

GCACACGTCTGAACTCTTTCCCTACACGACGCTCTTCCGATCTNNNCACTGCACTCGACCTAGATAGAT
GCACACGTCTGAACTCTTTCCCTACACGACGCTCTTCCGATCTNNNCACCTTCCTCACAGACTATGGTCC
GCACACGTCTGAACTCTTTCCCTACACGACGCTCTTCCGATCTNNNCTCAGGCTCAAATGATCCTTCCA
GAGCACACGTCTGAACTCTTTCCCTACACGACGCTCTTCCGATCTNNNAAGATCACACAACTGCACTCCA
AGCACACGTCTGAACTCTTTCCCTACACGACGCTCTTCCGATCTNNNCTTGCAGTAAGCCAAGATCACG
GAGCACACGTCTGAACTCTTTCCCTACACGACGCTCTTCCGATCTNNNTGTAGTCTCAGCTACTCAGGA
AGCACACGTCTGAACTCTTTCCCTACACGACGCTCTTCCGATCTNNNCACCTGTAGTCCCAGCTACTTGC
GAGCACACGTCTGAACTCTTTCCCTACACGACGCTCTTCCGATCTNNNCACCTGTAATCCCAGCTACTTG
GAGCACACGTCTGAACTCTTTCCCTACACGACGCTCTTCCGATCTNNNCTCATTGCAACCTTCGTCTCCT
AGCACACGTCTGAACTCTTTCCCTACACGACGCTCTTCCGATCTNNNCCAACGTGAAACCTAGGTCTCTA
AGCACACGTCTGAACTCTTTCCCTACACGACGCTCTTCCGATCTNNNGAGCTTGCAGTGAGTCGAGAT
GAGCACACGTCTGAACTCTTTCCCTACACGACGCTCTTCCGATCTNNNGGAGTTTGCATGAGCCAAGA
AGAGCACACGTCTGAACTCTTTCCCTACACGACGCTCTTCCGATCTNNNGAGCTTGCAGTGAGTGGAC
GAGCACACGTCTGAACTCTTTCCCTACACGACGCTCTTCCGATCTNNNGTTGCAGTGAGCTGAGATTG
AGCACACGTCTGAACTCTTTCCCTACACGACGCTCTTCCGATCTNNNCTCCTCAATGGACTCTATCAGG
AGCACACGTCTGAACTCTTTCCCTACACGACGCTCTTCCGATCTNNNAAAGTGCTGGGATTACAGGCA
GAGCACACGTCTGAACTCTTTCCCTACACGACGCTCTTCCGATCTNNNGCCTGTAATCCCAGCTACTTG
GAGCACACGTCTGAACTCTTTCCCTACACGACGCTCTTCCGATCTNNNTAGCTTGCAGTGAGCCGAGA
AGCACACGTCTGAACTCTTTCCCTACACGACGCTCTTCCGATCTNNNTGTAGTCCCAGCTACTCAGGAG
GAGCACACGTCTGAACTCTTTCCCTACACGACGCTCTTCCGATCTNNNGAGGCAGGAGAATTGCTTGA
AGCACACGTCTGAACTCTTTCCCTACACGACGCTCTTCCGATCTNNNGAGAATGGTGTGAACCTGGG
GAGCACACGTCTGAACTCTTTCCCTACACGACGCTCTTCCGATCTNNNGTTGCAGTCAGCTGAGATTGT
GAGCACACGTCTGAACTCTTTCCCTACACGACGCTCTTCCGATCTNNNGCCTTTTCTTACCTGGTGTGT
AGCACACGTCTGAACTCTTTCCCTACACGACGCTCTTCCGATCTNNNCTGGAGCTGAAGATGGAGGA
GCACACGTCTGAACTCTTTCCCTACACGACGCTCTTCCGATCTNNNACATGTTGTTTTGAAATCGGAAC
GCACACGTCTGAACTCTTTCCCTACACGACGCTCTTCCGATCTNNNAGTCATGGAGTTGACAGGGATT
AGCACACGTCTGAACTCTTTCCCTACACGACGCTCTTCCGATCTNNNTAGCCTGCTGAGGTATAAGGA
GCACACGTCTGAACTCTTTCCCTACACGACGCTCTTCCGATCTNNNAACCTTGATCTCCACATCCACTTC
AGCACACGTCTGAACTCTTTCCCTACACGACGCTCTTCCGATCTNNNAAAAACTCTGGTGGTTTTGGG
GAGCACACGTCTGAACTCTTTCCCTACACGACGCTCTTCCGATCTNNNTGAGCTATGATCACACCACTG
GAGCACACGTCTGAACTCTTTCCCTACACGACGCTCTTCCGATCTNNNGAGAATGGTGTGAACCCAGG
GAGCACACGTCTGAACTCTTTCCCTACACGACGCTCTTCCGATCTNNNCACCTGTAGTTCCAGCTACTCG
AGCACACGTCTGAACTCTTTCCCTACACGACGCTCTTCCGATCTNNNGAGGTTGCAGTGAGCTGAGA
GAGCACACGTCTGAACTCTTTCCCTACACGACGCTCTTCCGATCTNNNGAGATCGTATCACTGCACTCC
AGCACACGTCTGAACTCTTTCCCTACACGACGCTCTTCCGATCTNNNTCCACTACAGTTCTAGGAGGCA
AGCACACGTCTGAACTCTTTCCCTACACGACGCTCTTCCGATCTNNNGCCCCTAATCCAGTTCTCTAGTG
GCACACGTCTGAACTCTTTCCCTACACGACGCTCTTCCGATCTNNNGATGAGAAGGTCTGGAACTAG
AGCACACGTCTGAACTCTTTCCCTACACGACGCTCTTCCGATCTNNNTTCAAGTGATTCTCCTGCCTCAG
AGCACACGTCTGAACTCTTTCCCTACACGACGCTCTTCCGATCTNNNCTCTACAGTAGCTGGGACTCCA
AGCACACGTCTGAACTCTTTCCCTACACGACGCTCTTCCGATCTNNNAATTCTTCTGCCTCAGTCTCCTG
AGCACACGTCTGAACTCTTTCCCTACACGACGCTCTTCCGATCTNNNTAGGAGAATTGCTTGAACCTG
GCACACGTCTGAACTCTTTCCCTACACGACGCTCTTCCGATCTNNNGCAGCTGCCATAGTCCTATCTACC
GCACACGTCTGAACTCTTTCCCTACACGACGCTCTTCCGATCTNNNCACTCCAATGGACTATGTTCCCAC

GAGCACACGTCTGAACTCTTTCCCTACACGACGCTCTTCCGATCTNNNTGCTAGGATTACAGGCATGAG
AGCACACGTCTGAACTCTTTCCCTACACGACGCTCTTCCGATCTNNNCTTGGAGCAATCTCAATTCCAGC
CACACGTCTGAACTCTTTCCCTACACGACGCTCTTCCGATCTNNNCTTGAGGTCAGGAGTTCGAAACTC
GCACACGTCTGAACTCTTTCCCTACACGACGCTCTTCCGATCTNNNTGGTCTTTGTCCCCAGTTTCTAGCA
CACACGTCTGAACTCTTTCCCTACACGACGCTCTTCCGATCTNNNTGTTGTTTCACTGCAACCGATTTCA
GCACACGTCTGAACTCTTTCCCTACACGACGCTCTTCCGATCTNNNGCTTGGGAGGAACAAGAAAAGAG
GAGCACACGTCTGAACTCTTTCCCTACACGACGCTCTTCCGATCTNNNGAGAATGGTGTGAACCTGGC
AGCACACGTCTGAACTCTTTCCCTACACGACGCTCTTCCGATCTNNNGAGGTTGCAGTGAGCTGAGATT
AGCACACGTCTGAACTCTTTCCCTACACGACGCTCTTCCGATCTNNNCGGATGCCTATAATCCCAGCTAC
GCACACGTCTGAACTCTTTCCCTACACGACGCTCTTCCGATCTNNNGGAGAATCACTTGAACCTGGGAA
GAGCACACGTCTGAACTCTTTCCCTACACGACGCTCTTCCGATCTNNNGGAGGCAGGAGAATCACTTG
AGCACACGTCTGAACTCTTTCCCTACACGACGCTCTTCCGATCTNNNGAGAATCGCTTGAACCCAGGAG
AGCACACGTCTGAACTCTTTCCCTACACGACGCTCTTCCGATCTNNNGAAGCAGGAGAATCGCTTGAA
GCACACGTCTGAACTCTTTCCCTACACGACGCTCTTCCGATCTNNNGCTCCCTAACCTCAGCACATTATC
GCACACGTCTGAACTCTTTCCCTACACGACGCTCTTCCGATCTNNNGAGGTAGGAGAATGGCGTGAA
AGCACACGTCTGAACTCTTTCCCTACACGACGCTCTTCCGATCTNNNCTTGCAGTGAGCTGAGATTGTG
GAGCACACGTCTGAACTCTTTCCCTACACGACGCTCTTCCGATCTNNNAGGTCTCACAGAAGTGGTATT
CACACGTCTGAACTCTTTCCCTACACGACGCTCTTCCGATCTNNNACTTTGTCTCCTTTTTGTAAGCAACA
GCACACGTCTGAACTCTTTCCCTACACGACGCTCTTCCGATCTNNNTAGCACTGAAATTTTCGCAACCCC
AGCACACGTCTGAACTCTTTCCCTACACGACGCTCTTCCGATCTNNNTCTCTATCTCCTGACCACGTGATC
AGCACACGTCTGAACTCTTTCCCTACACGACGCTCTTCCGATCTNNNTGGGATTACAAGCATAAGCCAC
GAGCACACGTCTGAACTCTTTCCCTACACGACGCTCTTCCGATCTNNNATCTGTCCTCCTCTACCCAGACC
GCACACGTCTGAACTCTTTCCCTACACGACGCTCTTCCGATCTNNNTATGGAGTTTGTAGGGTGCAGT
ACACGTCTGAACTCTTTCCCTACACGACGCTCTTCCGATCTNNNAGAAACACCACAGTTTGATTACAGT
GCACACGTCTGAACTCTTTCCCTACACGACGCTCTTCCGATCTNNNTGCACAAAGTAGCCCCTCTTATTC
GCACACGTCTGAACTCTTTCCCTACACGACGCTCTTCCGATCTNNNACAAGTTTCTTTGTGCTGAGTGA
AGCACACGTCTGAACTCTTTCCCTACACGACGCTCTTCCGATCTNNNTGCACAGTGGTAATGCATGAAT
GAGCACACGTCTGAACTCTTTCCCTACACGACGCTCTTCCGATCTNNNGATCTCGGCTCACTACAAGCTC
AGCACACGTCTGAACTCTTTCCCTACACGACGCTCTTCCGATCTNNNTCTGGGTTTCCTACAACTTTCCA
GCACACGTCTGAACTCTTTCCCTACACGACGCTCTTCCGATCTNNNTTGTAAATGTCGTGCACTGTTGG
GCACACGTCTGAACTCTTTCCCTACACGACGCTCTTCCGATCTNNNCAGGGACACAGATCACAAGAGG
AGCACACGTCTGAACTCTTTCCCTACACGACGCTCTTCCGATCTNNNGAGGTTGCAGTGAGCTGAGAT
AGCACACGTCTGAACTCTTTCCCTACACGACGCTCTTCCGATCTNNNGAGGCTGGAATGAGCTGAGAT
GAGCACACGTCTGAACTCTTTCCCTACACGACGCTCTTCCGATCTNNNGCAGGAGAATTGCTTGAACCT
GCACACGTCTGAACTCTTTCCCTACACGACGCTCTTCCGATCTNNNGTTGCAGTGAGCCAAGATCATGC
GCACACGTCTGAACTCTTTCCCTACACGACGCTCTTCCGATCTNNNGTAATTCCAGCTACTAGGGAGGC
AGCACACGTCTGAACTCTTTCCCTACACGACGCTCTTCCGATCTNNNGATTGAGACCATCCTGGCTAACA
GAGCACACGTCTGAACTCTTTCCCTACACGACGCTCTTCCGATCTNNNGTGCCTATAGTCCCAGCTACTC
AGCACACGTCTGAACTCTTTCCCTACACGACGCTCTTCCGATCTNNNCTTGCACTCTAGCCTGGGTTAC
GAGCACACGTCTGAACTCTTTCCCTACACGACGCTCTTCCGATCTNNNGAGCTTACAGTGAGCCGAGAT
AGCACACGTCTGAACTCTTTCCCTACACGACGCTCTTCCGATCTNNNGCAGGAAAATTTCTTGAACCCC
GAGCACACGTCTGAACTCTTTCCCTACACGACGCTCTTCCGATCTNNNACACCTGTGGTCCTAGCTACTC
GCACACGTCTGAACTCTTTCCCTACACGACGCTCTTCCGATCTNNNCAAGATCATGCCACTGAACACTCC

GAGCACACGTCTGAACTCTTTCCCTACACGACGCTCTTCCGATCTNNNCTACTCAGGAGGCTGAAGCA
AGCACACGTCTGAACTCTTTCCCTACACGACGCTCTTCCGATCTNNNGTTGCAGTGAGCCAAGATTGT
GAGCACACGTCTGAACTCTTTCCCTACACGACGCTCTTCCGATCTNNNCAAGATCACACCCACTGCATT
GCACACGTCTGAACTCTTTCCCTACACGACGCTCTTCCGATCTNNNAACTTGCATCTAAACACTGCACC
CACACGTCTGAACTCTTTCCCTACACGACGCTCTTCCGATCTNNNTCCACAACATATTTTCATGCCCCA
CACACGTCTGAACTCTTTCCCTACACGACGCTCTTCCGATCTNNNTTGAAGTGATTCTCCTGCCTCAGC
GCACACGTCTGAACTCTTTCCCTACACGACGCTCTTCCGATCTNNNTCTTCACTAGCAGACAAGGTTG
AGCACACGTCTGAACTCTTTCCCTACACGACGCTCTTCCGATCTNNNCCAAAATGCTGGGATTACAGG
GCACACGTCTGAACTCTTTCCCTACACGACGCTCTTCCGATCTNNNGTGAAAATCCGCAAGTGTCTCT
GCACACGTCTGAACTCTTTCCCTACACGACGCTCTTCCGATCTNNNTATCACCCTCACCCCATACTTCTC
AGCACACGTCTGAACTCTTTCCCTACACGACGCTCTTCCGATCTNNNCAGGAGGTAGAGGTTTCAGT
AGCACACGTCTGAACTCTTTCCCTACACGACGCTCTTCCGATCTNNNGATGCCTCTAATCCCAGCTACT
AGCACACGTCTGAACTCTTTCCCTACACGACGCTCTTCCGATCTNNNGCCTGTAATCCCAGCTACTTGG
GCACACGTCTGAACTCTTTCCCTACACGACGCTCTTCCGATCTNNNTAATCCCAGCTACTCAGGAGGT
AGCACACGTCTGAACTCTTTCCCTACACGACGCTCTTCCGATCTNNNGAGAATCGTGTGAACCTGGGA
AGCACACGTCTGAACTCTTTCCCTACACGACGCTCTTCCGATCTNNNCTGAGGCAGGATAATCGCTT
GAGCACACGTCTGAACTCTTTCCCTACACGACGCTCTTCCGATCTNNNAATTAGCCGGGCATGGTTGA
AGCACACGTCTGAACTCTTTCCCTACACGACGCTCTTCCGATCTNNNCAACAGGGCAAGACTCTGTCT
AGCACACGTCTGAACTCTTTCCCTACACGACGCTCTTCCGATCTNNNAACAAGAGCGAGACTGTGAC
GCACACGTCTGAACTCTTTCCCTACACGACGCTCTTCCGATCTNNNCATAGTGTAAAACCGAGCAGCT
GCACACGTCTGAACTCTTTCCCTACACGACGCTCTTCCGATCTNNNCTCCGAGTAGCTGGGACTACAG
GCACACGTCTGAACTCTTTCCCTACACGACGCTCTTCCGATCTNNNCAAAGGGTCAGGACTTGTTCAA
GCACACGTCTGAACTCTTTCCCTACACGACGCTCTTCCGATCTNNNAATGCTTTAGGGGGAACTGAAG
AGCACACGTCTGAACTCTTTCCCTACACGACGCTCTTCCGATCTNNNCTTCCCAAAGATGACCTGCTGA
AGCACACGTCTGAACTCTTTCCCTACACGACGCTCTTCCGATCTNNNGACTCCCGAGTAGCTGTGACTA
AGCACACGTCTGAACTCTTTCCCTACACGACGCTCTTCCGATCTNNNCAAACAAAAGTCAGGGAAGG
GCACACGTCTGAACTCTTTCCCTACACGACGCTCTTCCGATCTNNNTCTCATTCCTTCCTTCTGGTGTG
CACACGTCTGAACTCTTTCCCTACACGACGCTCTTCCGATCTNNNCTTTCAGGTACCCCAATCAGTCAC
GCACACGTCTGAACTCTTTCCCTACACGACGCTCTTCCGATCTNNNTCTAGTGGAATGGGTCACTTTGG
CACACGTCTGAACTCTTTCCCTACACGACGCTCTTCCGATCTNNNTTCCTTGCCTAATATCCTGAAAGAA
AGCACACGTCTGAACTCTTTCCCTACACGACGCTCTTCCGATCTNNNATTCTCCAGCCTCAGCCTGTAG
GCACACGTCTGAACTCTTTCCCTACACGACGCTCTTCCGATCTNNNATACTACCCAGCAACCAAACAGT
CACACGTCTGAACTCTTTCCCTACACGACGCTCTTCCGATCTNNNAGTTCTGGGATTACAGGCATCAG
GCACACGTCTGAACTCTTTCCCTACACGACGCTCTTCCGATCTNNNGTATTGGCCTTGTGAAAACAGCT
GCACACGTCTGAACTCTTTCCCTACACGACGCTCTTCCGATCTNNNCTTCTCCTTCTTGGTTTCCAGGA
CACACGTCTGAACTCTTTCCCTACACGACGCTCTTCCGATCTNNNCCATCTTTTGGTTCAGATTGGCAC
AGCACACGTCTGAACTCTTTCCCTACACGACGCTCTTCCGATCTNNNGCAACAGAATGAGACTCCGT
ACACGTCTGAACTCTTTCCCTACACGACGCTCTTCCGATCTNNNTTCTTCTTTTTACCAGCTAGATGCC
AGCACACGTCTGAACTCTTTCCCTACACGACGCTCTTCCGATCTNNNCGACAGAGCAAGATTCCGTCT
GCACACGTCTGAACTCTTTCCCTACACGACGCTCTTCCGATCTNNNGTAGTGGCTCACTCCTGTAATCC
GCACACGTCTGAACTCTTTCCCTACACGACGCTCTTCCGATCTNNNGATGCAGGAGAATTGCTTGAAC
AGCACACGTCTGAACTCTTTCCCTACACGACGCTCTTCCGATCTNNNGAGGTTGCAGTGAGCTGAGA
AGCACACGTCTGAACTCTTTCCCTACACGACGCTCTTCCGATCTNNNCGAGATCACACCACTGTACTCC

GAGCACACGTCTGAACTCTTTCCCTACACGACGCTCTTCCGATCTNNNTAGTCCCAACTACTCAGGAGG
AGCACACGTCTGAACTCTTTCCCTACACGACGCTCTTCCGATCTNNNGAGCTTGCAATGACCCGAGATC
GAGCACACGTCTGAACTCTTTCCCTACACGACGCTCTTCCGATCTNNNGAGAATGGCATGAACTCGGGA
AGCACACGTCTGAACTCTTTCCCTACACGACGCTCTTCCGATCTNNNAAGAAGATCACTTGAGCCTGG
AGCACACGTCTGAACTCTTTCCCTACACGACGCTCTTCCGATCTNNNCAGAGCTTGCAGTGAGAGATC
GAGCACACGTCTGAACTCTTTCCCTACACGACGCTCTTCCGATCTNNNAAAAAAAAGCTTGCATTGAGC
GAGCACACGTCTGAACTCTTTCCCTACACGACGCTCTTCCGATCTNNNCAAGAAAATGCCTGTCGGTGA
GAGCACACGTCTGAACTCTTTCCCTACACGACGCTCTTCCGATCTNNNTAACTAGGCAGTTGGAGCTG
AGCACACGTCTGAACTCTTTCCCTACACGACGCTCTTCCGATCTNNNTAGTAGAGACGGGGTTTCTCCA
GCACACGTCTGAACTCTTTCCCTACACGACGCTCTTCCGATCTNNNTACAAATCTGCCCCAAGGACTAGC
GCACACGTCTGAACTCTTTCCCTACACGACGCTCTTCCGATCTNNNTGACCGTTTCCTTTTGTTTTGCTCA
AGCACACGTCTGAACTCTTTCCCTACACGACGCTCTTCCGATCTNNNGGCTCACTACAACCTCTACTTCCC
GCACACGTCTGAACTCTTTCCCTACACGACGCTCTTCCGATCTNNNACAGAGTCTTACAATGTTGCCCAC
AGCACACGTCTGAACTCTTTCCCTACACGACGCTCTTCCGATCTNNNAGACAGGTTTTCTCCATGTTGGT
GCACACGTCTGAACTCTTTCCCTACACGACGCTCTTCCGATCTNNNAATAGCTGGAACTACAGGAGCAC
GCACACGTCTGAACTCTTTCCCTACACGACGCTCTTCCGATCTNNNCCGTAAACTGGTTCATCTGTTCAC
GCACACGTCTGAACTCTTTCCCTACACGACGCTCTTCCGATCTNNNGAAGGACAAGCAAAAGCACATC
CACACGTCTGAACTCTTTCCCTACACGACGCTCTTCCGATCTNNNTCAATCCTTTCAAATCCTGCTGCCA
AGCACACGTCTGAACTCTTTCCCTACACGACGCTCTTCCGATCTNNNTTCAAGTGATTTCCCCTGCCTCCA
GCACACGTCTGAACTCTTTCCCTACACGACGCTCTTCCGATCTNNNCCTACTGAAACCTTTTGCCCATTCA
GCACACGTCTGAACTCTTTCCCTACACGACGCTCTTCCGATCTNNNTGACAAAGTGAGGCTCCATCTCCA
AGCACACGTCTGAACTCTTTCCCTACACGACGCTCTTCCGATCTNNNTGAACCTGGGAAGTGGAAAGT
AGAGCACACGTCTGAACTCTTTCCCTACACGACGCTCTTCCGATCTNNNGTGCCTGTAATCCCAGCTACT
AGCACACGTCTGAACTCTTTCCCTACACGACGCTCTTCCGATCTNNNGTTCCAGTGAGCCAAGATTGTC
GAGCACACGTCTGAACTCTTTCCCTACACGACGCTCTTCCGATCTNNNGAGGCACAAGAATCGGTTGAA
GAGCACACGTCTGAACTCTTTCCCTACACGACGCTCTTCCGATCTNNNGTTGCAGTGAGCTGAGATTGT
GAGCACACGTCTGAACTCTTTCCCTACACGACGCTCTTCCGATCTNNNCGACAGAGTGAGACTGCATCT
GAGCACACGTCTGAACTCTTTCCCTACACGACGCTCTTCCGATCTNNNGCGACAGAGTGAGATTTCGT
AGCACACGTCTGAACTCTTTCCCTACACGACGCTCTTCCGATCTNNNCTCCTGTAGTCCCAGCTACTCGC
GAGCACACGTCTGAACTCTTTCCCTACACGACGCTCTTCCGATCTNNNGAGCTTGCAGTGAGTCGAGA
GAGCACACGTCTGAACTCTTTCCCTACACGACGCTCTTCCGATCTNNNGTTGCAGTGAGCCAAGATTGA
GAGCACACGTCTGAACTCTTTCCCTACACGACGCTCTTCCGATCTNNNGCGATACAGCGAGATTCCATC
AGAGCACACGTCTGAACTCTTTCCCTACACGACGCTCTTCCGATCTNNNGCAAAAGAGTGAGACTCCG
GAGCACACGTCTGAACTCTTTCCCTACACGACGCTCTTCCGATCTNNNGAGAATGGTGTGAACCTGGC
AGCACACGTCTGAACTCTTTCCCTACACGACGCTCTTCCGATCTNNNCGCCTATAGTCCCAGCTACTTGC
AGAGCACACGTCTGAACTCTTTCCCTACACGACGCTCTTCCGATCTNNNCTTTGGGAGATGGAGGTGG
GAGCACACGTCTGAACTCTTTCCCTACACGACGCTCTTCCGATCTNNNGAGAATGGTGTGAACCTGGG
AGCACACGTCTGAACTCTTTCCCTACACGACGCTCTTCCGATCTNNNCACCTGTAGTCCCAGCTACATGC
AGAGCACACGTCTGAACTCTTTCCCTACACGACGCTCTTCCGATCTNNNGAGCTTTCAGTGAGCCGAG
GCACACGTCTGAACTCTTTCCCTACACGACGCTCTTCCGATCTNNNCACCTGTAATCCCAGCTACTTGGC
AGCACACGTCTGAACTCTTTCCCTACACGACGCTCTTCCGATCTNNNCTTGTAGTGAGCCGAGATCGAG
AGCACACGTCTGAACTCTTTCCCTACACGACGCTCTTCCGATCTNNNGAGTAGCTGAGATTACAGGCCC
GCACACGTCTGAACTCTTTCCCTACACGACGCTCTTCCGATCTNNNAGCTTTGAGCCAGTAAGAACAGA

GCACACGTCTGAACTCTTTCCCTACACGACGCTCTTCCGATCTNNNTGTGCCTCCACCTTTTTCTTTTC

CACACGTCTGAACTCTTTCCCTACACGACGCTCTTCCGATCTNNNAGTGCTGGGATTATAGGCATGA

CACACGTCTGAACTCTTTCCCTACACGACGCTCTTCCGATCTNNNACTAGGACTACAGGTGTGAGAC

CACACGTCTGAACTCTTTCCCTACACGACGCTCTTCCGATCTNNNTTGAGGCTGCCAGGAGTTTATAG

GCACACGTCTGAACTCTTTCCCTACACGACGCTCTTCCGATCTNNNTCCCAAAGTGCCGGGATTATAG

GCACACGTCTGAACTCTTTCCCTACACGACGCTCTTCCGATCTNNNAGAAGTACACCACTCCAAAAGCT

CACACGTCTGAACTCTTTCCCTACACGACGCTCTTCCGATCTNNNATACCTGCCAAAATGCCAATCTGC

CACACGTCTGAACTCTTTCCCTACACGACGCTCTTCCGATCTNNNGTTACCTTGTCCAAGATGACCCTC

GCACACGTCTGAACTCTTTCCCTACACGACGCTCTTCCGATCTNNNAATGTACAGTTTTTGCCTCAGCC

GCACACGTCTGAACTCTTTCCCTACACGACGCTCTTCCGATCTNNNTTCAAGCAATTCTCCTGCCTCAG

CACACGTCTGAACTCTTTCCCTACACGACGCTCTTCCGATCTNNNCCAATTTGCCCTTTGCCAATTTCC

AGCACACGTCTGAACTCTTTCCCTACACGACGCTCTTCCGATCTNNNGAGGTTGCAGTGAGCTGAGA

AGCACACGTCTGAACTCTTTCCCTACACGACGCTCTTCCGATCTNNNGGTGACAGAGCCAGACTTCA

CACACGTCTGAACTCTTTCCCTACACGACGCTCTTCCGATCTNNNGGATCACTTGAGGTCAGGAGTT

AGCACACGTCTGAACTCTTTCCCTACACGACGCTCTTCCGATCTNNNGGTTGCAGTGACCTGAGATCA

GCACACGTCTGAACTCTTTCCCTACACGACGCTCTTCCGATCTNNNCACCTGTAGACCCAGCTACTTG

GCACACGTCTGAACTCTTTCCCTACACGACGCTCTTCCGATCTNNNATGCCTGTTGTCTCAGCTACTCC

GAGCACACGTCTGAACTCTTTCCCTACACGACGCTCTTCCGATCTNNNTGGGTGACAGAGTGAGACT

AGCACACGTCTGAACTCTTTCCCTACACGACGCTCTTCCGATCTNNNGAGCTTACAGTGAGCGGAGA

AGCACACGTCTGAACTCTTTCCCTACACGACGCTCTTCCGATCTNNNGAGAATGGTGTGAACCTGGG

GCACACGTCTGAACTCTTTCCCTACACGACGCTCTTCCGATCTNNNGAGCTTGCAGTGAGTCGAGAT

CACACGTCTGAACTCTTTCCCTACACGACGCTCTTCCGATCTNNNTCTGGCATGACTTACTTCCTTCAC

GCACACGTCTGAACTCTTTCCCTACACGACGCTCTTCCGATCTNNNTGTGCACATGTCTCCTACACAG

AGCACACGTCTGAACTCTTTCCCTACACGACGCTCTTCCGATCTNNNCTCCCAGGTTCAAGCGATTCTC

AGCACACGTCTGAACTCTTTCCCTACACGACGCTCTTCCGATCTNNNCTCCCAGGTTCAAGCAATTCTG

AGCACACGTCTGAACTCTTTCCCTACACGACGCTCTTCCGATCTNNNCTCTGCCAGGAATATCTGACC

AGCACACGTCTGAACTCTTTCCCTACACGACGCTCTTCCGATCTNNNAATCTAATGCCGAAGCTCAGG

CACACGTCTGAACTCTTTCCCTACACGACGCTCTTCCGATCTNNNAACAGCATATGGTCAGCACAGT

GCACACGTCTGAACTCTTTCCCTACACGACGCTCTTCCGATCTNNNCCACAAGACTGACTCTAGGGTTT

AGCACACGTCTGAACTCTTTCCCTACACGACGCTCTTCCGATCTNNNCTCCAGAGTTCACGCCATTCTC

GCACACGTCTGAACTCTTTCCCTACACGACGCTCTTCCGATCTNNNAGAATTGCTTGAACCTGGGAG

AGCACACGTCTGAACTCTTTCCCTACACGACGCTCTTCCGATCTNNNGAGGTTGCAGTGAGCTGAG

AGCACACGTCTGAACTCTTTCCCTACACGACGCTCTTCCGATCTNNNGAAGGCTGAGGTAGGAGGA

GCACACGTCTGAACTCTTTCCCTACACGACGCTCTTCCGATCTNNNCCTATCCTGGCTAACACGGTAAA

GCACACGTCTGAACTCTTTCCCTACACGACGCTCTTCCGATCTNNNGGGTCCTGTGGTCTACAAGATT

AGCACACGTCTGAACTCTTTCCCTACACGACGCTCTTCCGATCTNNNCAGGAGTTTGAGACCAGCCTA

AGCACACGTCTGAACTCTTTCCCTACACGACGCTCTTCCGATCTNNNTGACAGAGTGAGACTCCCTCT

CACACGTCTGAACTCTTTCCCTACACGACGCTCTTCCGATCTNNNCAAGGCCTTGTATGAGGAGAAGT

GCACACGTCTGAACTCTTTCCCTACACGACGCTCTTCCGATCTNNNCCGTCTTTACTGCAGTCTCTGAA

ACACGTCTGAACTCTTTCCCTACACGACGCTCTTCCGATCTNNNACTCCCATTTTCCTGATATCCTGGCA

GCACACGTCTGAACTCTTTCCCTACACGACGCTCTTCCGATCTNNNGCACAGGGGAAATTCACATAAG

CACGTCTGAACTCTTTCCCTACACGACGCTCTTCCGATCTNNNACAACAACAAAAAGACACAAAAACA

CACACGTCTGAACTCTTTCCCTACACGACGCTCTTCCGATCTNNNTATCTTGGTTGTATGGCTGCAGT

GAGCACACGTCTGAACTCTTTCCCTACACGACGCTCTTCCGATCTNNNTGAGCTGAGATTGTGCCACTG
AGCACACGTCTGAACTCTTTCCCTACACGACGCTCTTCCGATCTNNNGAGAATCGCTTGAACCTGGGA
GAGCACACGTCTGAACTCTTTCCCTACACGACGCTCTTCCGATCTNNNCACCTGTAGTCCCAGCTATTCG
GAGCACACGTCTGAACTCTTTCCCTACACGACGCTCTTCCGATCTNNNGAGGCACGAGAATTGCTTGA
AGCACACGTCTGAACTCTTTCCCTACACGACGCTCTTCCGATCTNNNGAGCTTGCATTGAGCCAAGATC
AGCACACGTCTGAACTCTTTCCCTACACGACGCTCTTCCGATCTNNNGTTGCAGTGAGCCAAGATCAT
GAGCACACGTCTGAACTCTTTCCCTACACGACGCTCTTCCGATCTNNNGAGAATGGCATAAACCCGGG
AGCACACGTCTGAACTCTTTCCCTACACGACGCTCTTCCGATCTNNNGTGAGCCAAGATTGCATCACTG
GAGCACACGTCTGAACTCTTTCCCTACACGACGCTCTTCCGATCTNNNTTCTGGGTTCAAGCCATTCTCC
GCACACGTCTGAACTCTTTCCCTACACGACGCTCTTCCGATCTNNNTGGGGCTTCATTTTCATGACCTAC
GCACACGTCTGAACTCTTTCCCTACACGACGCTCTTCCGATCTNNNTGGGCAAAGGATATGAACAGACA
AGCACACGTCTGAACTCTTTCCCTACACGACGCTCTTCCGATCTNNNTGCATCCAGTTCATGAACATTGT
GCACACGTCTGAACTCTTTCCCTACACGACGCTCTTCCGATCTNNNCTCATTCCATCTTTCACCCACCTCA
AGCACACGTCTGAACTCTTTCCCTACACGACGCTCTTCCGATCTNNNTCACTGCAATTTCCACCTCCTAGC
GCACACGTCTGAACTCTTTCCCTACACGACGCTCTTCCGATCTNNNACCTCAGCATGTCTCAAAAGTCAC
GCACACGTCTGAACTCTTTCCCTACACGACGCTCTTCCGATCTNNNGTTTGCAATGTCTTTCGGCTTTGC
GCACACGTCTGAACTCTTTCCCTACACGACGCTCTTCCGATCTNNNCCTAGATATCCTCTCTCACGGGTC
GAGCACACGTCTGAACTCTTTCCCTACACGACGCTCTTCCGATCTNNNCAAGATCGAGCCACTGTACAC
GAGCACACGTCTGAACTCTTTCCCTACACGACGCTCTTCCGATCTNNNGAGAATGGTGTGAACCTGGG
GAGCACACGTCTGAACTCTTTCCCTACACGACGCTCTTCCGATCTNNNGCGACAAAACGAGACTCCATCT
GCACACGTCTGAACTCTTTCCCTACACGACGCTCTTCCGATCTNNNTGTGTAATCCCAGCACTTTGAGA
GAGCACACGTCTGAACTCTTTCCCTACACGACGCTCTTCCGATCTNNNGAGCTTACAGTGAGCCGAGA
AGCACACGTCTGAACTCTTTCCCTACACGACGCTCTTCCGATCTNNNGAGAATGGCATGAACCTGGGAC
AGCACACGTCTGAACTCTTTCCCTACACGACGCTCTTCCGATCTNNNGAGAATGGTGTGAACCTGGGA
AGCACACGTCTGAACTCTTTCCCTACACGACGCTCTTCCGATCTNNNGAGCTTACAGTGAGCCGAGATT
GCACACGTCTGAACTCTTTCCCTACACGACGCTCTTCCGATCTNNNCATGCCTGTAATCCCAGCTACTCCA
AGCACACGTCTGAACTCTTTCCCTACACGACGCTCTTCCGATCTNNNAGGTTGTGATGAGCTGAGATC
GCACACGTCTGAACTCTTTCCCTACACGACGCTCTTCCGATCTNNNGACACAGATGATGGGTTGAGTC
AGCACACGTCTGAACTCTTTCCCTACACGACGCTCTTCCGATCTNNNCATTGGTCCTACAGTCTCCCATT
GCACACGTCTGAACTCTTTCCCTACACGACGCTCTTCCGATCTNNNGTAGTCACATCTTCAGGCTCCATC
AGCACACGTCTGAACTCTTTCCCTACACGACGCTCTTCCGATCTNNNTGAAATTGTGAGGGAGAAGAC
GCACACGTCTGAACTCTTTCCCTACACGACGCTCTTCCGATCTNNNCCCACCAACAGTGTAAAAGCATTC
GCACACGTCTGAACTCTTTCCCTACACGACGCTCTTCCGATCTNNNGCTGCTTCCTGATCATCTACCTTC
GAGCACACGTCTGAACTCTTTCCCTACACGACGCTCTTCCGATCTNNNCATGATCTCAGCTCACTGCAAG
GCACACGTCTGAACTCTTTCCCTACACGACGCTCTTCCGATCTNNNGCTCAAGCAATCCCAAAGATCTCC
AGCACACGTCTGAACTCTTTCCCTACACGACGCTCTTCCGATCTNNNCTAATACTCTGGGAGGCTGAGC
GAGCACACGTCTGAACTCTTTCCCTACACGACGCTCTTCCGATCTNNNGAATGGCATGAACCCAGGAGT
GAGCACACGTCTGAACTCTTTCCCTACACGACGCTCTTCCGATCTNNNGAGGTTGCAGTGAGTCGAGA
GAGCACACGTCTGAACTCTTTCCCTACACGACGCTCTTCCGATCTNNNCACCTGTAGTCCCAGCTACTTG
GAGCACACGTCTGAACTCTTTCCCTACACGACGCTCTTCCGATCTNNNCTGCCTGTAGTCCCAGCTACTC
GAGCACACGTCTGAACTCTTTCCCTACACGACGCTCTTCCGATCTNNNGGTCAAGGCTCCATGTGATCA
AGAGCACACGTCTGAACTCTTTCCCTACACGACGCTCTTCCGATCTNNNATAGTGATGGGCCCCTGTAC
AGCACACGTCTGAACTCTTTCCCTACACGACGCTCTTCCGATCTNNNAAGAGAATGGCGTGAACCTGA

GAGCACACGTCTGAACTCTTTCCCTACACGACGCTCTTCCGATCTNNNCAGGAGTTCAAGACCAGCCTA
AGCACACGTCTGAACTCTTTCCCTACACGACGCTCTTCCGATCTNNNCAGGAGATGGAGGTTTCAGTG
GCACACGTCTGAACTCTTTCCCTACACGACGCTCTTCCGATCTNNNGGAGGAGAACCCATCTGTACAA
GAGCACACGTCTGAACTCTTTCCCTACACGACGCTCTTCCGATCTNNNCATGCCTGTATCCCAGCTACTC
GAGCACACGTCTGAACTCTTTCCCTACACGACGCTCTTCCGATCTNNNCATCAGATGTTGCTTCCAGCTG
AGCACACGTCTGAACTCTTTCCCTACACGACGCTCTTCCGATCTNNNCTGAGATCACACCACTGCACTCC
GCACACGTCTGAACTCTTTCCCTACACGACGCTCTTCCGATCTNNNCAGCAAACCCCATCTCTACAGAAC
GCACACGTCTGAACTCTTTCCCTACACGACGCTCTTCCGATCTNNNGAAGTTGCAGTAAGCCGAGATTC
AGCACACGTCTGAACTCTTTCCCTACACGACGCTCTTCCGATCTNNNGGTTTCACCAGGATTCAGAATG
GCACACGTCTGAACTCTTTCCCTACACGACGCTCTTCCGATCTNNNGTCCCAGAAAACATGGTACCTCTC
GCACACGTCTGAACTCTTTCCCTACACGACGCTCTTCCGATCTNNNATTCTCATAACAATCTGGCCCCAC
GCACACGTCTGAACTCTTTCCCTACACGACGCTCTTCCGATCTNNNCCCCATGTCTTTGTTCATGTTGTCA
GCACACGTCTGAACTCTTTCCCTACACGACGCTCTTCCGATCTNNNTAAAGTGCTGGGATTACAGGCAT
AGCACACGTCTGAACTCTTTCCCTACACGACGCTCTTCCGATCTNNNTGTGGATAAAATGAGATCCCGCA
AGCACACGTCTGAACTCTTTCCCTACACGACGCTCTTCCGATCTNNNGTTTACCTGAGGTCCGGAGTTC
AGCACACGTCTGAACTCTTTCCCTACACGACGCTCTTCCGATCTNNNCCAGGCAGGAAGATCTGTTGAA
AAGAGCACACGTCTGAACTCTTTCCCTACACGACGCTCTTCCGATCTNNNGAGGATCACTTGAACCCAG
GAGCACACGTCTGAACTCTTTCCCTACACGACGCTCTTCCGATCTNNNGTTGTGGTGAGCTGAAATCGT
AGCACACGTCTGAACTCTTTCCCTACACGACGCTCTTCCGATCTNNNAGGAGAATCACTAGAACCTGGC
GAGCACACGTCTGAACTCTTTCCCTACACGACGCTCTTCCGATCTNNNGAGCTTACAGTGAGCCGAGAT
AGCACACGTCTGAACTCTTTCCCTACACGACGCTCTTCCGATCTNNNCCAGAGGTCAGGAGTTTCAGAC
GAGCACACGTCTGAACTCTTTCCCTACACGACGCTCTTCCGATCTNNNGAGCTTGCAGTGAGTGGAGAT
GCACACGTCTGAACTCTTTCCCTACACGACGCTCTTCCGATCTNNNCTTGAACTCCTGGACTCAAGTGA
GCACACGTCTGAACTCTTTCCCTACACGACGCTCTTCCGATCTNNNTCAGGCTAAATCTTCGGACACAAC
AGCACACGTCTGAACTCTTTCCCTACACGACGCTCTTCCGATCTNNNTTGGGCAACAAGAGCAAAACTC
GCACACGTCTGAACTCTTTCCCTACACGACGCTCTTCCGATCTNNNCTTCCACCCCAGACATACTTCAACA
AGCACACGTCTGAACTCTTTCCCTACACGACGCTCTTCCGATCTNNNTCACAGGTTTCCAGGATTAGGA
GCACACGTCTGAACTCTTTCCCTACACGACGCTCTTCCGATCTNNNTGCCTAGTAATAGGATGCCAGGA
GCACACGTCTGAACTCTTTCCCTACACGACGCTCTTCCGATCTNNNCCAGCTGTAGGCATAAGATCTGTC
AGCACACGTCTGAACTCTTTCCCTACACGACGCTCTTCCGATCTNNNACTAGGGCAAGAGAACTTCACA
AGCACACGTCTGAACTCTTTCCCTACACGACGCTCTTCCGATCTNNNTGACTTGGTGTCCTCCATTTTCA
GCACACGTCTGAACTCTTTCCCTACACGACGCTCTTCCGATCTNNNAGCGTGTCAATTTCTGCAATGAA
GCACACGTCTGAACTCTTTCCCTACACGACGCTCTTCCGATCTNNNTCTATAGCACCAACAATCCCACCC
AGAGCACACGTCTGAACTCTTTCCCTACACGACGCTCTTCCGATCTNNNCTGCATTTAGTGGTGATGGT
GAGCACACGTCTGAACTCTTTCCCTACACGACGCTCTTCCGATCTNNNCTTGCAGTAAGCCGAGATTCCC
AGCACACGTCTGAACTCTTTCCCTACACGACGCTCTTCCGATCTNNNGAGAATGGTGTGAACCTGGGA
GAGCACACGTCTGAACTCTTTCCCTACACGACGCTCTTCCGATCTNNNCTGCTGTGAGCCATGATTACAC
GAGCACACGTCTGAACTCTTTCCCTACACGACGCTCTTCCGATCTNNNGAGTTTGCAGTGAACCGAGAT
GAGCACACGTCTGAACTCTTTCCCTACACGACGCTCTTCCGATCTNNNGTGCCTGTAATCCCAGCTACTC
GAGCACACGTCTGAACTCTTTCCCTACACGACGCTCTTCCGATCTNNNGAGAATGGCATGAACCCAGG
AGCACACGTCTGAACTCTTTCCCTACACGACGCTCTTCCGATCTNNNTAGTCCCAGCTACTCAGGAGAC
GAGCACACGTCTGAACTCTTTCCCTACACGACGCTCTTCCGATCTNNNTGTGCAACAGAGAGACACTCA
GAGCACACGTCTGAACTCTTTCCCTACACGACGCTCTTCCGATCTNNNGAGGTTGCAGTGAGCTGAGA

AGCACACGTCTGAACTCTTTCCCTACACGACGCTCTTCCGATCTNNNGAGAATGGCATGAACCCAGGA

GCACACGTCTGAACTCTTTCCCTACACGACGCTCTTCCGATCTNNNAGGTTGAGGCAGGAGAATCATT

GCACACGTCTGAACTCTTTCCCTACACGACGCTCTTCCGATCTNNNTCTCCAGTATGTTTCAGAGGCAC

GAGCACACGTCTGAACTCTTTCCCTACACGACGCTCTTCCGATCTNNNCTTGAGAAAGTAAGCATGGG

GAGCACACGTCTGAACTCTTTCCCTACACGACGCTCTTCCGATCTNNNCATTTCTCAAGCACAGCCCTTG

GCACACGTCTGAACTCTTTCCCTACACGACGCTCTTCCGATCTNNNCTTTCTCTATCTACTGCCCCCACC

GCACACGTCTGAACTCTTTCCCTACACGACGCTCTTCCGATCTNNNTTTTTGTGTGTGGTGGGAACAT

GCACACGTCTGAACTCTTTCCCTACACGACGCTCTTCCGATCTNNNATCATCCATGCCCCTATCTTTCCCA

AGCACACGTCTGAACTCTTTCCCTACACGACGCTCTTCCGATCTNNNAGTTTTACTCCCTTGTGATTGG

GCACACGTCTGAACTCTTTCCCTACACGACGCTCTTCCGATCTNNNCTGAGCAAAGCTTGAGCAAAGA

GCACACGTCTGAACTCTTTCCCTACACGACGCTCTTCCGATCTNNNCCCAACTCCTACTTTTACACATGC

GCACACGTCTGAACTCTTTCCCTACACGACGCTCTTCCGATCTNNNATCCCCACCAACCAAGGTATTTC

GCACACGTCTGAACTCTTTCCCTACACGACGCTCTTCCGATCTNNNCATGAACTCCTGGATTCAAGGGA

GCACACGTCTGAACTCTTTCCCTACACGACGCTCTTCCGATCTNNNGAGAATCGCTTGAACCCAGGAG

GAGCACACGTCTGAACTCTTTCCCTACACGACGCTCTTCCGATCTNNNGCAGGAGAATTGCTTGAACCT

GCACACGTCTGAACTCTTTCCCTACACGACGCTCTTCCGATCTNNNATGAGAATGGCATGAATCCAGG

GAGCACACGTCTGAACTCTTTCCCTACACGACGCTCTTCCGATCTNNNATCATGAGGTCAGGAGATCG

GAGCACACGTCTGAACTCTTTCCCTACACGACGCTCTTCCGATCTNNNGAGGGAGGAGAATGGCATG

AGCACACGTCTGAACTCTTTCCCTACACGACGCTCTTCCGATCTNNNCCTGGGTGACATAGTGAGACA

GAGCACACGTCTGAACTCTTTCCCTACACGACGCTCTTCCGATCTNNNCCAGGTAACAGTGTGAGACT

AGAGCACACGTCTGAACTCTTTCCCTACACGACGCTCTTCCGATCTNNNCTTGCAGTGAGCCGAGATT

GAGCACACGTCTGAACTCTTTCCCTACACGACGCTCTTCCGATCTNNNGCCTGTAGTCCCAGCTACTTG

GAGCACACGTCTGAACTCTTTCCCTACACGACGCTCTTCCGATCTNNNCGACAGAACAAGATTCCATCG

GCACACGTCTGAACTCTTTCCCTACACGACGCTCTTCCGATCTNNNGAGCTTGCAGTGAGTCGAGAT

GCACACGTCTGAACTCTTTCCCTACACGACGCTCTTCCGATCTNNNACCTTGCAGTGAGCTGAGATCCA

GAGCACACGTCTGAACTCTTTCCCTACACGACGCTCTTCCGATCTNNNGAGACCAGACTGACCAAGAA

AGCACACGTCTGAACTCTTTCCCTACACGACGCTCTTCCGATCTNNNCTGGGCAACAAGAGCAAAACT

GAGCACACGTCTGAACTCTTTCCCTACACGACGCTCTTCCGATCTNNNGAGGTTGCAGTGAGCTGAGA

GCACACGTCTGAACTCTTTCCCTACACGACGCTCTTCCGATCTNNNCTGAGATTGCACCATCGCATTCC

AGCACACGTCTGAACTCTTTCCCTACACGACGCTCTTCCGATCTNNNGCCTTTGATTCTAGCACTTTGG

AGCACACGTCTGAACTCTTTCCCTACACGACGCTCTTCCGATCTNNNGGAATTCCTTGAACCAGGGAG

AGCACACGTCTGAACTCTTTCCCTACACGACGCTCTTCCGATCTNNNGACTTCACAGAAAGGACCTGAC

AGCACACGTCTGAACTCTTTCCCTACACGACGCTCTTCCGATCTNNNCACATTTTTGCTTGAGTCCCTGT

GAGCACACGTCTGAACTCTTTCCCTACACGACGCTCTTCCGATCTNNNCATCTCAGCCTCCCAAATAGC

GAGCACACGTCTGAACTCTTTCCCTACACGACGCTCTTCCGATCTNNNGAAGCAGGAGAATCGCTTGA

GAGCACACGTCTGAACTCTTTCCCTACACGACGCTCTTCCGATCTNNNGTTGCAGTGAGCCAAGATCA

AGCACACGTCTGAACTCTTTCCCTACACGACGCTCTTCCGATCTNNNGAGATTGCAGTGAGCCAAGAT

GAGCACACGTCTGAACTCTTTCCCTACACGACGCTCTTCCGATCTNNNGAGAATCGCTTGAATTCGGG

GAGCACACGTCTGAACTCTTTCCCTACACGACGCTCTTCCGATCTNNNCCAGTCAGTCTAGGCAACAG

GAGCACACGTCTGAACTCTTTCCCTACACGACGCTCTTCCGATCTNNNGAGGTTTCAGTGAGCCGAGA

GAGCACACGTCTGAACTCTTTCCCTACACGACGCTCTTCCGATCTNNNGCGACAAAGTGAGACTCCAT

GCACACGTCTGAACTCTTTCCCTACACGACGCTCTTCCGATCTNNNGATGAGACGGGCTATAAGAAGC

AGCACACGTCTGAACTCTTTCCCTACACGACGCTCTTCCGATCTNNNGGAAGTACTGAGACTGAGTCA

GCACACGTCTGAACTCTTTCCCTACACGACGCTCTTCCGATCTNNNTAGAGCTGGAAAGAAGCCAAGA
GCACACGTCTGAACTCTTTCCCTACACGACGCTCTTCCGATCTNNNCACGTTTGCTTTGGTCTCCTTAT
AGCACACGTCTGAACTCTTTCCCTACACGACGCTCTTCCGATCTNNNGTCCTCCTTCTGACACTGTTAGG
GCACACGTCTGAACTCTTTCCCTACACGACGCTCTTCCGATCTNNNAAAGTGCTGGGATTACAGGCAT
AGCACACGTCTGAACTCTTTCCCTACACGACGCTCTTCCGATCTNNNCAAAGTGTTGGGATTACAGGC
GCACACGTCTGAACTCTTTCCCTACACGACGCTCTTCCGATCTNNNCCACACTTGGCCAACCTACTTATC
GCACACGTCTGAACTCTTTCCCTACACGACGCTCTTCCGATCTNNNCTGAATTATTGCATGAGCCCCAG
CACACGTCTGAACTCTTTCCCTACACGACGCTCTTCCGATCTNNNCACCCTCAAGCCTTTACTCAAACCA
GCACACGTCTGAACTCTTTCCCTACACGACGCTCTTCCGATCTNNNGACAGAGTATGACTGGGATAGG
GCACACGTCTGAACTCTTTCCCTACACGACGCTCTTCCGATCTNNNGGAAAATGACCAGTTCCTCCAAC
AGCACACGTCTGAACTCTTTCCCTACACGACGCTCTTCCGATCTNNNTTTAGTAGAGACAGGGTTTCGC
GCACACGTCTGAACTCTTTCCCTACACGACGCTCTTCCGATCTNNNCAAGGTCACACCACTGTACTCCC
GAGCACACGTCTGAACTCTTTCCCTACACGACGCTCTTCCGATCTNNNCACCTGTAGTCCCAGCTACTT
GCACACGTCTGAACTCTTTCCCTACACGACGCTCTTCCGATCTNNNTGAGAATCACTTGAACCTGGGA
GCACACGTCTGAACTCTTTCCCTACACGACGCTCTTCCGATCTNNNCTGAGGCAGGAGAATCACTTGA
AGCACACGTCTGAACTCTTTCCCTACACGACGCTCTTCCGATCTNNNGAGGATCACTTGAGCTTGGGA
GAGCACACGTCTGAACTCTTTCCCTACACGACGCTCTTCCGATCTNNNGTGACAGAGAGAGACTCCG
AGCACACGTCTGAACTCTTTCCCTACACGACGCTCTTCCGATCTNNNGAGAATGGCGTGAAACCAGA
AGCACACGTCTGAACTCTTTCCCTACACGACGCTCTTCCGATCTNNNGAGGCAGGAAAATGACGTGA
GAGCACACGTCTGAACTCTTTCCCTACACGACGCTCTTCCGATCTNNNATGCCTGTAGTCCCAGCTACT
AGAGCACACGTCTGAACTCTTTCCCTACACGACGCTCTTCCGATCTNNNCACCTGTAGTCCCAGCTACT
AGCACACGTCTGAACTCTTTCCCTACACGACGCTCTTCCGATCTNNNGAGAATGGTGTGAACCTGGGA
AGCACACGTCTGAACTCTTTCCCTACACGACGCTCTTCCGATCTNNNGACGCAGGAGAATTGCTTGAA
GAGCACACGTCTGAACTCTTTCCCTACACGACGCTCTTCCGATCTNNNCACCTGTAGTCCCAGCTACTT
GAGCACACGTCTGAACTCTTTCCCTACACGACGCTCTTCCGATCTNNNGAGGCTGAGGCAGGAGAAT
AGCACACGTCTGAACTCTTTCCCTACACGACGCTCTTCCGATCTNNNCGTAGTCCCAGCTACTTTGGAG
AGCACACGTCTGAACTCTTTCCCTACACGACGCTCTTCCGATCTNNNGCAGGAGAATTGCTTGAACCTG
AGAGCACACGTCTGAACTCTTTCCCTACACGACGCTCTTCCGATCTNNNGTTGCAGTAAGCCGAGATC
AGAGCACACGTCTGAACTCTTTCCCTACACGACGCTCTTCCGATCTNNNCTTGCAGTGAGCCGAGATA
GAGCACACGTCTGAACTCTTTCCCTACACGACGCTCTTCCGATCTNNNGAGATTGCACCACTGTACTCC
GAGCACACGTCTGAACTCTTTCCCTACACGACGCTCTTCCGATCTNNNGAGAATGGCATGAACCCAG
GAGCACACGTCTGAACTCTTTCCCTACACGACGCTCTTCCGATCTNNNGTGACACAGCAAGACTCTGT
GAGCACACGTCTGAACTCTTTCCCTACACGACGCTCTTCCGATCTNNNCAAGTGCCTGGGATTCTAGG
CACACGTCTGAACTCTTTCCCTACACGACGCTCTTCCGATCTNNNGAGATTAGGACTGGGGCTGTAA
AGCACACGTCTGAACTCTTTCCCTACACGACGCTCTTCCGATCTNNNATTCCTTGCAGAGCATTGTGA
GCACACGTCTGAACTCTTTCCCTACACGACGCTCTTCCGATCTNNNTGTTGTTGTTGTTTTGGTGGGT
AGCACACGTCTGAACTCTTTCCCTACACGACGCTCTTCCGATCTNNNCTCCTAGGCTCAAGCAATCCTC
AGCACACGTCTGAACTCTTTCCCTACACGACGCTCTTCCGATCTNNNATAGGATGTGTGTCAAGGACA
GCACACGTCTGAACTCTTTCCCTACACGACGCTCTTCCGATCTNNNCTCAAGTGATCCTCCCATCTCAG
AGCACACGTCTGAACTCTTTCCCTACACGACGCTCTTCCGATCTNNNGTAACATAGCGAGACCCCGTA
ACACGTCTGAACTCTTTCCCTACACGACGCTCTTCCGATCTNNNACAAGTATTTGTGACAGAGACAAC
AGCACACGTCTGAACTCTTTCCCTACACGACGCTCTTCCGATCTNNNGAGATTGCATCACTGGACTCC
AGCACACGTCTGAACTCTTTCCCTACACGACGCTCTTCCGATCTNNNCAACAGAGTGAGGCTCCATCT

AGCACACGTCTGAACTCTTTCCCTACACGACGCTCTTCCGATCTNNNCACTTGAGGTCAGGAGTTCAAG
AGCACACGTCTGAACTCTTTCCCTACACGACGCTCTTCCGATCTNNNAAGATCACACCACTGCACTTCAC
GCACACGTCTGAACTCTTTCCCTACACGACGCTCTTCCGATCTNNNTGTCTTTTGGACCCTTAGATCGG
AGCACACGTCTGAACTCTTTCCCTACACGACGCTCTTCCGATCTNNNCTCCCAGGTTCAAGTCATCCTC
GCACACGTCTGAACTCTTTCCCTACACGACGCTCTTCCGATCTNNNTTGGATCCAAAGGAAACATGCA
AGCACACGTCTGAACTCTTTCCCTACACGACGCTCTTCCGATCTNNNGTCTGTTGTCCCAGCTACTCAG
AGCACACGTCTGAACTCTTTCCCTACACGACGCTCTTCCGATCTNNNCAACAGAGTGAGACTCCAGCT
GAGCACACGTCTGAACTCTTTCCCTACACGACGCTCTTCCGATCTNNNAAGACCATCCTGGCTAACATT
GAGCACACGTCTGAACTCTTTCCCTACACGACGCTCTTCCGATCTNNNCAGGAGTTCAAGACCAGCCTA
AGCACACGTCTGAACTCTTTCCCTACACGACGCTCTTCCGATCTNNNGAGCAAGCTGCCAAAATTTGA
GCACACGTCTGAACTCTTTCCCTACACGACGCTCTTCCGATCTNNNTGCCCACATCTGAAAGGTTTTTCC
GCACACGTCTGAACTCTTTCCCTACACGACGCTCTTCCGATCTNNNTTGAGGTGTATTTCTCTCTGCCC
GCACACGTCTGAACTCTTTCCCTACACGACGCTCTTCCGATCTNNNATGTTTTCCATTTGCATAGCGCT
GCACACGTCTGAACTCTTTCCCTACACGACGCTCTTCCGATCTNNNAGATGTTGGGTCCTCAATGTGA
AGAGCACACGTCTGAACTCTTTCCCTACACGACGCTCTTCCGATCTNNNGTAGTCCAGCTACTCTTGCA
AGCACACGTCTGAACTCTTTCCCTACACGACGCTCTTCCGATCTNNNGAGCTTTCGGTGAGCCAAGAT
AGCACACGTCTGAACTCTTTCCCTACACGACGCTCTTCCGATCTNNNACACCCATAATCCCAGCACTTTC
AGCACACGTCTGAACTCTTTCCCTACACGACGCTCTTCCGATCTNNNGATCAGCTGAGGTCAGGACTT
AGCACACGTCTGAACTCTTTCCCTACACGACGCTCTTCCGATCTNNNGACCAGATCTTGAAAGTCCCCT
GCACACGTCTGAACTCTTTCCCTACACGACGCTCTTCCGATCTNNNAAGAGCAGCTTCTACCAGCAAA
AGCACACGTCTGAACTCTTTCCCTACACGACGCTCTTCCGATCTNNNATCTGGTCCGTTCTTCACACTC
AGAGCACACGTCTGAACTCTTTCCCTACACGACGCTCTTCCGATCTNNNGTGACAAAGTGAGACCCTG
AGCACACGTCTGAACTCTTTCCCTACACGACGCTCTTCCGATCTNNNAGATTGCCCCATTGCACTCTAG
GAGCACACGTCTGAACTCTTTCCCTACACGACGCTCTTCCGATCTNNNAGTGAGACGAGATTGCACCA
GAGCACACGTCTGAACTCTTTCCCTACACGACGCTCTTCCGATCTNNNTTGGAAAGGGAAGAGACTTG
GCACACGTCTGAACTCTTTCCCTACACGACGCTCTTCCGATCTNNNGGTTCAAACAATTCTCCTGCCTC
GCACACGTCTGAACTCTTTCCCTACACGACGCTCTTCCGATCTNNNGTACCTCCATAGACTGCTTCCACC
CACACGTCTGAACTCTTTCCCTACACGACGCTCTTCCGATCTNNNTGTTGTGCTCTATTTGTTCTGCAC
CACACGTCTGAACTCTTTCCCTACACGACGCTCTTCCGATCTNNNAGTTTACTAGCACAGCTCACCAAC
GCACACGTCTGAACTCTTTCCCTACACGACGCTCTTCCGATCTNNNCAGTATCCAAAGGTAACAGGCCT
AGCACACGTCTGAACTCTTTCCCTACACGACGCTCTTCCGATCTNNNTCAAGCAATTTTCCTGCCTTAG
GAGCACACGTCTGAACTCTTTCCCTACACGACGCTCTTCCGATCTNNNGAGAATGGCATGAACCTGGG
GAGCACACGTCTGAACTCTTTCCCTACACGACGCTCTTCCGATCTNNNCACCTGTAATCCCAGCTACTC
GCACACGTCTGAACTCTTTCCCTACACGACGCTCTTCCGATCTNNNGCACCTGTAGTCTCAGCTACTCC
GAGCACACGTCTGAACTCTTTCCCTACACGACGCTCTTCCGATCTNNNCTTGCAGTGAGCTGAGATTG
AGCACACGTCTGAACTCTTTCCCTACACGACGCTCTTCCGATCTNNNACTTTGGAAGGACAAGGTGA
GAGCACACGTCTGAACTCTTTCCCTACACGACGCTCTTCCGATCTNNNTCACTTGAGCTCAGGAGTTTG
GCACACGTCTGAACTCTTTCCCTACACGACGCTCTTCCGATCTNNNAGTAGCTGGAATTAGCAGACTG
GCACACGTCTGAACTCTTTCCCTACACGACGCTCTTCCGATCTNNNTGCTTTGGGTTTTTTTGTTGTTG
GCACACGTCTGAACTCTTTCCCTACACGACGCTCTTCCGATCTNNNAGGGCCAAAGTCTTAGTTATCT
AGCACACGTCTGAACTCTTTCCCTACACGACGCTCTTCCGATCTNNNTGGCCAGAGTAAAAAGAAGCA
GCACACGTCTGAACTCTTTCCCTACACGACGCTCTTCCGATCTNNNTTAGAACAGCCACGTCTAAGGA
GCACACGTCTGAACTCTTTCCCTACACGACGCTCTTCCGATCTNNNGAGTAGCTGGGACTACAGACAT

GAGCACACGTCTGAACTCTTTCCCTACACGACGCTCTTCCGATCTNNNCTTGCAGTGAGCCGAGATAG
GCACACGTCTGAACTCTTTCCCTACACGACGCTCTTCCGATCTNNNCTGGGCAACAAGAGCAAAACTC
GCACACGTCTGAACTCTTTCCCTACACGACGCTCTTCCGATCTNNNGGAGCTTGTAGTGAGCGAGAT
GCACACGTCTGAACTCTTTCCCTACACGACGCTCTTCCGATCTNNNGAAGCAGGAGAATGGCATGAA
AGCACACGTCTGAACTCTTTCCCTACACGACGCTCTTCCGATCTNNNGAGCTTGCAGTGAGTCGAGA
GCACACGTCTGAACTCTTTCCCTACACGACGCTCTTCCGATCTNNNCCATCTACTAGGGAGGCTGAG
GCACACGTCTGAACTCTTTCCCTACACGACGCTCTTCCGATCTNNNGTTGCAGTGAGCTGAGATCATG
AGCACACGTCTGAACTCTTTCCCTACACGACGCTCTTCCGATCTNNNGAGAATCCCTTGAACCCAGGA
AGCACACGTCTGAACTCTTTCCCTACACGACGCTCTTCCGATCTNNNCTGAGGCAGAATGGTGTGAA
AGCACACGTCTGAACTCTTTCCCTACACGACGCTCTTCCGATCTNNNCATGATGGCAGGCATCTGTAA
AGCACACGTCTGAACTCTTTCCCTACACGACGCTCTTCCGATCTNNNGCCTGTAGTCCCAGCTACTTG
CACACGTCTGAACTCTTTCCCTACACGACGCTCTTCCGATCTNNNGATCACGAGGTCAGGAGATTGA
GCACACGTCTGAACTCTTTCCCTACACGACGCTCTTCCGATCTNNNCACCTGTAGTCCCAGCTACTTG
GAGCACACGTCTGAACTCTTTCCCTACACGACGCTCTTCCGATCTNNNGAGGATAGCTTGAACCTGG
GCACACGTCTGAACTCTTTCCCTACACGACGCTCTTCCGATCTNNNGAGCTCACTCCTGTAATCCTAG
GCACACGTCTGAACTCTTTCCCTACACGACGCTCTTCCGATCTNNNCCTCCTGGATTGAAGTGATT
CACACGTCTGAACTCTTTCCCTACACGACGCTCTTCCGATCTNNNGCCTGGTTCAAATCTAGCTAGGT
AGCACACGTCTGAACTCTTTCCCTACACGACGCTCTTCCGATCTNNNCCCACACCCTTTTCCGAAAATTT
AGCACACGTCTGAACTCTTTCCCTACACGACGCTCTTCCGATCTNNNATTCCTGTATGTATTCCCCTCG
AGCACACGTCTGAACTCTTTCCCTACACGACGCTCTTCCGATCTNNNGATTACTTGAGCCCAGGAGTT
AGCACACGTCTGAACTCTTTCCCTACACGACGCTCTTCCGATCTNNNGAGGCAGGAAAATCACTTGAA
AGCACACGTCTGAACTCTTTCCCTACACGACGCTCTTCCGATCTNNNGGAGGTTGCAGTAAGCTGAG
AGCACACGTCTGAACTCTTTCCCTACACGACGCTCTTCCGATCTNNNCTGAGGCAGGAGAATCACTTG
GCACACGTCTGAACTCTTTCCCTACACGACGCTCTTCCGATCTNNNGAGTAGCTGGGACTACAGGCT
GAGCACACGTCTGAACTCTTTCCCTACACGACGCTCTTCCGATCTNNNAAGTAGCTGGGATTACAGG
AGAGCACACGTCTGAACTCTTTCCCTACACGACGCTCTTCCGATCTNNNAGAAGGTCAAGGATGCAG
GAGCACACGTCTGAACTCTTTCCCTACACGACGCTCTTCCGATCTNNNGGTGACACAGTGAGACTCCA
AGCACACGTCTGAACTCTTTCCCTACACGACGCTCTTCCGATCTNNNGAGCTTGCAGTGAGCCAAGA
GCACACGTCTGAACTCTTTCCCTACACGACGCTCTTCCGATCTNNNAGTGAGCACCGATAACACCATT
GCACACGTCTGAACTCTTTCCCTACACGACGCTCTTCCGATCTNNNGTTGCAGTGAGCCAAGATTGT
GCACACGTCTGAACTCTTTCCCTACACGACGCTCTTCCGATCTNNNAAGACTGCAGTGAGCTGAGAT
GCACACGTCTGAACTCTTTCCCTACACGACGCTCTTCCGATCTNNNGAGCTTGCAGTGAGCCTAGATT
GCACACGTCTGAACTCTTTCCCTACACGACGCTCTTCCGATCTNNNGGAGAGAGAGCAAGACTTTGT
AGCACACGTCTGAACTCTTTCCCTACACGACGCTCTTCCGATCTNNNGGAGGTTGCAGTAAGCTGAG
GAGCACACGTCTGAACTCTTTCCCTACACGACGCTCTTCCGATCTNNNAGGAGACTGAGGTAGGAGG
AGCACACGTCTGAACTCTTTCCCTACACGACGCTCTTCCGATCTNNNTAGTCCTAGCTACTCAGGAGG
CACACGTCTGAACTCTTTCCCTACACGACGCTCTTCCGATCTNNNTCCTGTTCTATCCACCGCCTATAC
CACACGTCTGAACTCTTTCCCTACACGACGCTCTTCCGATCTNNNTCTGAAAGTACTGGGGTTACAG
CACACGTCTGAACTCTTTCCCTACACGACGCTCTTCCGATCTNNNTAACTGGATGACTGGCTTTCTCC
CACACGTCTGAACTCTTTCCCTACACGACGCTCTTCCGATCTNNNGATTATTGCAGTAGCCAGGCTTG
GCACACGTCTGAACTCTTTCCCTACACGACGCTCTTCCGATCTNNNAACCCAGGCCACCATTATCAAAA
GCACACGTCTGAACTCTTTCCCTACACGACGCTCTTCCGATCTNNNATCCTACTTTGTTGCCCCAGATC
CACACGTCTGAACTCTTTCCCTACACGACGCTCTTCCGATCTNNNGCAATGGGATGCTTACACACTAC

GCACACGTCTGAACTCTTTCCCTACACGACGCTCTTCCGATCTNNNCACTCAGTTTGAACAGCACTTCA
AGCACACGTCTGAACTCTTTCCCTACACGACGCTCTTCCGATCTNNNATTGTTGGCCATCTTTTCATGG
GAGCACACGTCTGAACTCTTTCCCTACACGACGCTCTTCCGATCTNNNGGTCACCTGAAATCCCTCTGT
GCACACGTCTGAACTCTTTCCCTACACGACGCTCTTCCGATCTNNNTCTGTGTAATCTGTGTGCCTCAA
AGCACACGTCTGAACTCTTTCCCTACACGACGCTCTTCCGATCTNNNTTCTTGCTGTTTCATGTACCACT
AGCACACGTCTGAACTCTTTCCCTACACGACGCTCTTCCGATCTNNNGTCAAAGGTCAAACTGTGTGCT
AGCACACGTCTGAACTCTTTCCCTACACGACGCTCTTCCGATCTNNNCAGTACGCCCAGCTCATTTTCC
AGCACACGTCTGAACTCTTTCCCTACACGACGCTCTTCCGATCTNNNTCAACAAGGAGGGCAAGACATA
AGCACACGTCTGAACTCTTTCCCTACACGACGCTCTTCCGATCTNNNTAGTTGATGGGCTCTTGAGTT
GCACACGTCTGAACTCTTTCCCTACACGACGCTCTTCCGATCTNNNATGTTGAGCAAGTTTTCACATGC
AGCACACGTCTGAACTCTTTCCCTACACGACGCTCTTCCGATCTNNNGAGAATGGCATGAACCTGAGA
AGCACACGTCTGAACTCTTTCCCTACACGACGCTCTTCCGATCTNNNGAGAATGGCGAGAACACAGG
GAGCACACGTCTGAACTCTTTCCCTACACGACGCTCTTCCGATCTNNNGAGTATCGCTTGAACCCAGGA
GAGCACACGTCTGAACTCTTTCCCTACACGACGCTCTTCCGATCTNNNCACCTGTAGTCCCAGCTACTT
AGAGCACACGTCTGAACTCTTTCCCTACACGACGCTCTTCCGATCTNNNGAGAATCGCTTGAACCTGGG
AGCACACGTCTGAACTCTTTCCCTACACGACGCTCTTCCGATCTNNNCAGAGGCTGAGGCAGAAGAAT
GAGCACACGTCTGAACTCTTTCCCTACACGACGCTCTTCCGATCTNNNGCAGGAGAATTGCTTGAACCT
AGCACACGTCTGAACTCTTTCCCTACACGACGCTCTTCCGATCTNNNGACCTGTAGTCCCAGCTACTCG
AGCACACGTCTGAACTCTTTCCCTACACGACGCTCTTCCGATCTNNNGAGGTTGCAGTGAGCTGAGAT
GAGCACACGTCTGAACTCTTTCCCTACACGACGCTCTTCCGATCTNNNGAGGCAGGAGACTCTCTTGA
GAGCACACGTCTGAACTCTTTCCCTACACGACGCTCTTCCGATCTNNNGAGATTGTACCACTGCACTCCA
GCACACGTCTGAACTCTTTCCCTACACGACGCTCTTCCGATCTNNNTCTCAGTATGTTGCCAAGTAGCC
AGCACACGTCTGAACTCTTTCCCTACACGACGCTCTTCCGATCTNNNGGTTTCTAGGTCAAAGAGGCG
AGCACACGTCTGAACTCTTTCCCTACACGACGCTCTTCCGATCTNNNCCTGTTCCTTCAGAGACAGCTA
GCACACGTCTGAACTCTTTCCCTACACGACGCTCTTCCGATCTNNNAAACTCCTGACCTCAAGTGATCCC
GCACACGTCTGAACTCTTTCCCTACACGACGCTCTTCCGATCTNNNCGTCCCAAAGTTACATTACAGGC
AGCACACGTCTGAACTCTTTCCCTACACGACGCTCTTCCGATCTNNNATCAATGAGAAGTGGGTGAGT
GCACACGTCTGAACTCTTTCCCTACACGACGCTCTTCCGATCTNNNTGATGTCATTCACCCACAAAATG
AGCACACGTCTGAACTCTTTCCCTACACGACGCTCTTCCGATCTNNNCTATGTTGCCCAGTCTGGTCTG
GCACACGTCTGAACTCTTTCCCTACACGACGCTCTTCCGATCTNNNTCCTGAAAGTGATCATGGATGG
GCACACGTCTGAACTCTTTCCCTACACGACGCTCTTCCGATCTNNNTCAACCATGTTGTCACAAATGGC
AGCACACGTCTGAACTCTTTCCCTACACGACGCTCTTCCGATCTNNNTTAGTAAGTGTGTGTGTGTGT
AGAGCACACGTCTGAACTCTTTCCCTACACGACGCTCTTCCGATCTNNNTGCCTGTAGTCCCAGCTACT
GAGCACACGTCTGAACTCTTTCCCTACACGACGCTCTTCCGATCTNNNGAGAATCGCTTGAACCCAGGA
GAGCACACGTCTGAACTCTTTCCCTACACGACGCTCTTCCGATCTNNNAACATGGCAAAACCCGTCTCTA
AGCACACGTCTGAACTCTTTCCCTACACGACGCTCTTCCGATCTNNNCTGAGATCGTGCCACTGTACTC
AGAGCACACGTCTGAACTCTTTCCCTACACGACGCTCTTCCGATCTNNNCGTGGACAACAGAGCAAGA
GAGCACACGTCTGAACTCTTTCCCTACACGACGCTCTTCCGATCTNNNGTAAGCCAAGATGGTGCCATT
GAGCACACGTCTGAACTCTTTCCCTACACGACGCTCTTCCGATCTNNNGAGGTTGCATTGAGCCAAGAT
GAGCACACGTCTGAACTCTTTCCCTACACGACGCTCTTCCGATCTNNNGTTGCAGTGAGCTGAGATTG
GCACACGTCTGAACTCTTTCCCTACACGACGCTCTTCCGATCTNNNCTGAGGCAGGAGAATCGTTTGA
AGCACACGTCTGAACTCTTTCCCTACACGACGCTCTTCCGATCTNNNGCAAAGCCCCGTCTCTACTAAA
AGCACACGTCTGAACTCTTTCCCTACACGACGCTCTTCCGATCTNNNGAGGTTGTGGTGAGCTGAGAT

GAGCACACGTCTGAACTCTTTCCCTACACGACGCTCTTCCGATCTNNNGAGGCAGAAGAATCGCTTGA
AGCACACGTCTGAACTCTTTCCCTACACGACGCTCTTCCGATCTNNNAGATTGTGCCACTGTACTCTAG
GCACACGTCTGAACTCTTTCCCTACACGACGCTCTTCCGATCTNNNGGGTGACAGAGTAAGACTCCAT
GCACACGTCTGAACTCTTTCCCTACACGACGCTCTTCCGATCTNNNTGTAAACTGGGAACAAAAGCAG
CACACGTCTGAACTCTTTCCCTACACGACGCTCTTCCGATCTNNNAGATGGCCAACATTGTATATGGAG
AGCACACGTCTGAACTCTTTCCCTACACGACGCTCTTCCGATCTNNNCTCCCAGGTTCAAGCGATTCTCC
CACACGTCTGAACTCTTTCCCTACACGACGCTCTTCCGATCTNNNAGTGCTGGCTCATCTTCATTCTTCA
AGCACACGTCTGAACTCTTTCCCTACACGACGCTCTTCCGATCTNNNTTTTCAGCCTCTGAGTATCTGG
AGCACACGTCTGAACTCTTTCCCTACACGACGCTCTTCCGATCTNNNGCACCTGTAGTCTCAGCTACTG
GAGCACACGTCTGAACTCTTTCCCTACACGACGCTCTTCCGATCTNNNCGAGATCACACCACTGTACTC
GAGCACACGTCTGAACTCTTTCCCTACACGACGCTCTTCCGATCTNNNGAGAATGGCATGAACCTGG
AGAGCACACGTCTGAACTCTTTCCCTACACGACGCTCTTCCGATCTNNNGAGACTGAGACGGGAGAA
AGCACACGTCTGAACTCTTTCCCTACACGACGCTCTTCCGATCTNNNGAGAATGACGTGAACCCAGGA
AGCACACGTCTGAACTCTTTCCCTACACGACGCTCTTCCGATCTNNNATGCCATGGAAAACCTTTCTGT
AGCACACGTCTGAACTCTTTCCCTACACGACGCTCTTCCGATCTNNNAGATCCTGTCCCTTACTCATGT
GAGCACACGTCTGAACTCTTTCCCTACACGACGCTCTTCCGATCTNNNCAGGTTCAAGCAATTCTCCTG
AGCACACGTCTGAACTCTTTCCCTACACGACGCTCTTCCGATCTNNNGTGTGTGTGTGTGTGTGTGTA
AGCACACGTCTGAACTCTTTCCCTACACGACGCTCTTCCGATCTNNNTCTACTTAGAGCAGCACCTAGG
GCACACGTCTGAACTCTTTCCCTACACGACGCTCTTCCGATCTNNNCCCAACCCAGTTCCATTCTCTTACA
AGCACACGTCTGAACTCTTTCCCTACACGACGCTCTTCCGATCTNNNACTAGTAATAGTCTGCTGGCA
CACACGTCTGAACTCTTTCCCTACACGACGCTCTTCCGATCTNNNTCAAGATGGATAAAGCAGGCTAC
GCACACGTCTGAACTCTTTCCCTACACGACGCTCTTCCGATCTNNNAATTGCAACTTACTTCAGCCTCG
CACACGTCTGAACTCTTTCCCTACACGACGCTCTTCCGATCTNNNTCCTTCAAGAAGTGCATGAGAAGA
GCACACGTCTGAACTCTTTCCCTACACGACGCTCTTCCGATCTNNNAAGCTTTTCTTCCAGGAAGTTGC
AGCACACGTCTGAACTCTTTCCCTACACGACGCTCTTCCGATCTNNNTACATTTCCAGCTCCGTGAAGA
GCACACGTCTGAACTCTTTCCCTACACGACGCTCTTCCGATCTNNNAAAGAAGTAGCTCACTCACTGGG
GCACACGTCTGAACTCTTTCCCTACACGACGCTCTTCCGATCTNNNCACTCAGCAGTACAGAAGCAATG
GCACACGTCTGAACTCTTTCCCTACACGACGCTCTTCCGATCTNNNCCAGTGAAAGAGAATAGCAAGC
GAGCACACGTCTGAACTCTTTCCCTACACGACGCTCTTCCGATCTNNNGAGGTTGCAGTGAGCTGAG
GCACACGTCTGAACTCTTTCCCTACACGACGCTCTTCCGATCTNNNGAGAATGGCATGAACCCAGGAG
AGCACACGTCTGAACTCTTTCCCTACACGACGCTCTTCCGATCTNNNCTGTAGTCCCAGCTACTCGAAA
AGCACACGTCTGAACTCTTTCCCTACACGACGCTCTTCCGATCTNNNGAGGTTGCAGTGAGCTGAGAT
GAGCACACGTCTGAACTCTTTCCCTACACGACGCTCTTCCGATCTNNNGGACCACAGGAGCAAAAAAC
AGCACACGTCTGAACTCTTTCCCTACACGACGCTCTTCCGATCTNNNAAGGCTGAAGAATCGCTTGAAT
AGAGCACACGTCTGAACTCTTTCCCTACACGACGCTCTTCCGATCTNNNGAGAATCGCTTGAACCCAGG
GCACACGTCTGAACTCTTTCCCTACACGACGCTCTTCCGATCTNNNCTGGGTTGACAGAGTGAGACTC
AGCACACGTCTGAACTCTTTCCCTACACGACGCTCTTCCGATCTNNNGGAATTGCTTGAACCTGGGAG
AGCACACGTCTGAACTCTTTCCCTACACGACGCTCTTCCGATCTNNNGAGAATGGTGTGAACCTGGGA
AGAGCACACGTCTGAACTCTTTCCCTACACGACGCTCTTCCGATCTNNNGAGGCAGGAGAATGGAGT
GAGCACACGTCTGAACTCTTTCCCTACACGACGCTCTTCCGATCTNNNCACCTGTAGTCCCAGCTACTTG
AGCACACGTCTGAACTCTTTCCCTACACGACGCTCTTCCGATCTNNNGAGAATTGCTGGAACCTGGGA
GAGCACACGTCTGAACTCTTTCCCTACACGACGCTCTTCCGATCTNNNCCTGTAATCCCAGCTAATCGG
AGCACACGTCTGAACTCTTTCCCTACACGACGCTCTTCCGATCTNNNGAGCTTGCAGTGAGCCAAGAT

GAGCACACGTCTGAACTCTTTCCCTACACGACGCTCTTCCGATCTNNNGAGAATGACGTGAACCTGGG
GAGCACACGTCTGAACTCTTTCCCTACACGACGCTCTTCCGATCTNNNCTGGGCAACAAGAGCAAAACT
AGCACACGTCTGAACTCTTTCCCTACACGACGCTCTTCCGATCTNNNGAGAATGGCATGAACTCGGGA
AGAGCACACGTCTGAACTCTTTCCCTACACGACGCTCTTCCGATCTNNNGAGCTTGCAGTGAGTCGAG
AGCACACGTCTGAACTCTTTCCCTACACGACGCTCTTCCGATCTNNNCCTTTGTCTTTTAGCTTGTGGCA
AGCACACGTCTGAACTCTTTCCCTACACGACGCTCTTCCGATCTNNNCAAGCATTTCCTGAGCACTCACC
GCACACGTCTGAACTCTTTCCCTACACGACGCTCTTCCGATCTNNNGAGGAGGTTGAGTGACTTCTTG
GCACACGTCTGAACTCTTTCCCTACACGACGCTCTTCCGATCTNNNATGCTGTGTACATGTGTGTTCAT
AGCACACGTCTGAACTCTTTCCCTACACGACGCTCTTCCGATCTNNNACCACACCCGGCTAATTTTGTA
GAGCACACGTCTGAACTCTTTCCCTACACGACGCTCTTCCGATCTNNNCTCCTGGGTTCATGCCATTCTC
GCACACGTCTGAACTCTTTCCCTACACGACGCTCTTCCGATCTNNNGTGTGTGTGTGTGTGTGTGTATT
GCACACGTCTGAACTCTTTCCCTACACGACGCTCTTCCGATCTNNNTGGCCCGAAAAGAAGCTAAGTAT
GCACACGTCTGAACTCTTTCCCTACACGACGCTCTTCCGATCTNNNGAAGCTGAATGATGCCTTGGTT
GCACACGTCTGAACTCTTTCCCTACACGACGCTCTTCCGATCTNNNTGGGGTTTCACCATACTGATCAG
GCACACGTCTGAACTCTTTCCCTACACGACGCTCTTCCGATCTNNNAATCTCCTGACCTTGTGATCCACC
GAGCACACGTCTGAACTCTTTCCCTACACGACGCTCTTCCGATCTNNNGAGAATGGCATGAACCCAGG
AGCACACGTCTGAACTCTTTCCCTACACGACGCTCTTCCGATCTNNNCTTGCAGTGAGCCAAGATTGT
GAGCACACGTCTGAACTCTTTCCCTACACGACGCTCTTCCGATCTNNNCTCCCAGGCTAAAGTGATCCT
AGCACACGTCTGAACTCTTTCCCTACACGACGCTCTTCCGATCTNNNCACAGTGAAACCCCATCTCTACT
AGCACACGTCTGAACTCTTTCCCTACACGACGCTCTTCCGATCTNNNGAGAAACGCTTGAACACAGGA
CACACGTCTGAACTCTTTCCCTACACGACGCTCTTCCGATCTNNNGGGTAACAGAGTGAGACTCCATC
GAGCACACGTCTGAACTCTTTCCCTACACGACGCTCTTCCGATCTNNNAGGAGGCTGAGGTAGGAGA
AGAGCACACGTCTGAACTCTTTCCCTACACGACGCTCTTCCGATCTNNNGAGGCAGAAGAATCGCTTG
AGCACACGTCTGAACTCTTTCCCTACACGACGCTCTTCCGATCTNNNCGACAGAGGTAGACTCCGTCT
AGCACACGTCTGAACTCTTTCCCTACACGACGCTCTTCCGATCTNNNGTGACAGAGCAAGACCCTACTC
AGCACACGTCTGAACTCTTTCCCTACACGACGCTCTTCCGATCTNNNGAGAATCGCTTGAACCAGGGA
AGCACACGTCTGAACTCTTTCCCTACACGACGCTCTTCCGATCTNNNCTGGGCAACAAGAGTGAAACTG
AGCACACGTCTGAACTCTTTCCCTACACGACGCTCTTCCGATCTNNNAGGAGAATCGCTTGAATCTGG
AGAGCACACGTCTGAACTCTTTCCCTACACGACGCTCTTCCGATCTNNNGAGAATCGCTTGAACCTGG
GAGCACACGTCTGAACTCTTTCCCTACACGACGCTCTTCCGATCTNNNGATGACCGAGCGAGACTCCA
GAGCACACGTCTGAACTCTTTCCCTACACGACGCTCTTCCGATCTNNNGAATTGTTTGAACCCAGGAG
GAGCACACGTCTGAACTCTTTCCCTACACGACGCTCTTCCGATCTNNNCACCTGTAATCCCAGCTACTCG
GAGCACACGTCTGAACTCTTTCCCTACACGACGCTCTTCCGATCTNNNGAGAATGGCATGAACCTGGG
AGAGCACACGTCTGAACTCTTTCCCTACACGACGCTCTTCCGATCTNNNCGCCTGTAATCCCAGCTACT
AGCACACGTCTGAACTCTTTCCCTACACGACGCTCTTCCGATCTNNNATGCCTGTAATCCCAGCTACTTG
AGCACACGTCTGAACTCTTTCCCTACACGACGCTCTTCCGATCTNNNTCCATTGGTGATATTGAGCCTC
GCACACGTCTGAACTCTTTCCCTACACGACGCTCTTCCGATCTNNNTGTGGTAGAGGCAGTAATGTTG
GCACACGTCTGAACTCTTTCCCTACACGACGCTCTTCCGATCTNNNTCGTCAGTGAAGTTTGCTTACCT
AGCACACGTCTGAACTCTTTCCCTACACGACGCTCTTCCGATCTNNNCAAACCGGAGTCCTTGGGATA
GAGCACACGTCTGAACTCTTTCCCTACACGACGCTCTTCCGATCTNNNGAGTCTCACTCTGTTGCTCAG
AGCACACGTCTGAACTCTTTCCCTACACGACGCTCTTCCGATCTNNNACTACGCCTGGCAAATTTTTTGT
GCACACGTCTGAACTCTTTCCCTACACGACGCTCTTCCGATCTNNNTATTTTTTTGTAGAGACGGCGGC
AGCACACGTCTGAACTCTTTCCCTACACGACGCTCTTCCGATCTNNNAAATTCTATTAGGTGCAGGCTG

GCACACGTCTGAACTCTTTCCCTACACGACGCTCTTCCGATCTNNNGGTTCAGTGTGTCTATAGCCCTTC
AGCACACGTCTGAACTCTTTCCCTACACGACGCTCTTCCGATCTNNNCTACACGGACACTACTCCCTTGC
GCACACGTCTGAACTCTTTCCCTACACGACGCTCTTCCGATCTNNNTGGATTTCCTGTGCCTGTTCATAC
GCACACGTCTGAACTCTTTCCCTACACGACGCTCTTCCGATCTNNNCTGCAGCCCATTCTCAATATTGG
GCACACGTCTGAACTCTTTCCCTACACGACGCTCTTCCGATCTNNNCACCACACCTGGCTAATTTGTAC
GCACACGTCTGAACTCTTTCCCTACACGACGCTCTTCCGATCTNNNCAGGCTAGAAAACTCTGGTGAC
GCACACGTCTGAACTCTTTCCCTACACGACGCTCTTCCGATCTNNNATGGGGCAGAGTAGAAAATCTG
GCACACGTCTGAACTCTTTCCCTACACGACGCTCTTCCGATCTNNNCTCTCTCTCTCTCTCCACCTCCAT
GAGCACACGTCTGAACTCTTTCCCTACACGACGCTCTTCCGATCTNNNAAACTCTTGCCCTCAGGTGAT
GCACACGTCTGAACTCTTTCCCTACACGACGCTCTTCCGATCTNNNGCTAAGTTGAACCCTCCAGAGAA
GCACACGTCTGAACTCTTTCCCTACACGACGCTCTTCCGATCTNNNGAGAATCGCTTGAACCTGGGAG
AGCACACGTCTGAACTCTTTCCCTACACGACGCTCTTCCGATCTNNNGAGAATGGCGTGAATCTGGG
AGCACACGTCTGAACTCTTTCCCTACACGACGCTCTTCCGATCTNNNACCCTCGTAATCCCAGCTACTC
GAGCACACGTCTGAACTCTTTCCCTACACGACGCTCTTCCGATCTNNNCGAGGTAGGTGGATCACTTG
GCACACGTCTGAACTCTTTCCCTACACGACGCTCTTCCGATCTNNNCTACAGTGAACCATGATTGCACC
GAGCACACGTCTGAACTCTTTCCCTACACGACGCTCTTCCGATCTNNNGAGGCAGAAGAATCGCTTGA
GAGCACACGTCTGAACTCTTTCCCTACACGACGCTCTTCCGATCTNNNGCAACAGAGTGAGACTCCAT
GAGCACACGTCTGAACTCTTTCCCTACACGACGCTCTTCCGATCTNNNGAAATTGCAGTGAGCCGAGA
GAGCACACGTCTGAACTCTTTCCCTACACGACGCTCTTCCGATCTNNNGAAGCAGGAGAATCGCATG
AGAGCACACGTCTGAACTCTTTCCCTACACGACGCTCTTCCGATCTNNNAGAAGAATGGCGTGAACC
GCACACGTCTGAACTCTTTCCCTACACGACGCTCTTCCGATCTNNNGATGGTCTTGATCTCCTGAGCT
GAGCACACGTCTGAACTCTTTCCCTACACGACGCTCTTCCGATCTNNNAAAGTGCTGGGATTACAGGC
AGCACACGTCTGAACTCTTTCCCTACACGACGCTCTTCCGATCTNNNTAGGCTGCTTGGGAGGTATTTA
GCACACGTCTGAACTCTTTCCCTACACGACGCTCTTCCGATCTNNNTCTAAGACTGGGGGCTTTCTAGA
GCACACGTCTGAACTCTTTCCCTACACGACGCTCTTCCGATCTNNNGTTGTGTTGATGTGGGACATAG
GCACACGTCTGAACTCTTTCCCTACACGACGCTCTTCCGATCTNNNAGCAAGCATTAGGAAACTCTGG
GCACACGTCTGAACTCTTTCCCTACACGACGCTCTTCCGATCTNNNTGCTAGCAGTGAACTCTTTCAGA
AGCACACGTCTGAACTCTTTCCCTACACGACGCTCTTCCGATCTNNNTTTGAGACAGTCTCGCTCCAT
AGCACACGTCTGAACTCTTTCCCTACACGACGCTCTTCCGATCTNNNCCACAGCTTTTTGAGATGCAG
AGCACACGTCTGAACTCTTTCCCTACACGACGCTCTTCCGATCTNNNGCTGTGTGACTTTGAAGGCTA
GCACACGTCTGAACTCTTTCCCTACACGACGCTCTTCCGATCTNNNTTGGTGGCATCTTATCACAGGAA
AGCACACGTCTGAACTCTTTCCCTACACGACGCTCTTCCGATCTNNNTACCTGTAGTCCCAGCTACTTGA
GCACACGTCTGAACTCTTTCCCTACACGACGCTCTTCCGATCTNNNAGAATCGCTTGAATCTGGGAGT
GAGCACACGTCTGAACTCTTTCCCTACACGACGCTCTTCCGATCTNNNGCAGGAGAATTGCTTGAACC
AGCACACGTCTGAACTCTTTCCCTACACGACGCTCTTCCGATCTNNNGGGACAGAGTGAGACTTTT
AGAGCACACGTCTGAACTCTTTCCCTACACGACGCTCTTCCGATCTNNNCGACAGAGTAAGACTCCGTC
GAGCACACGTCTGAACTCTTTCCCTACACGACGCTCTTCCGATCTNNNGATTGCTTGAGCCCAGGAGT
GAGCACACGTCTGAACTCTTTCCCTACACGACGCTCTTCCGATCTNNNGAGGTAGGAGAATCGCTTGA
GAGCACACGTCTGAACTCTTTCCCTACACGACGCTCTTCCGATCTNNNGAGGTTGTGGTGAGCAGAG
GCACACGTCTGAACTCTTTCCCTACACGACGCTCTTCCGATCTNNNGCAGGAGAATTGCTTGAACCTG
AGCACACGTCTGAACTCTTTCCCTACACGACGCTCTTCCGATCTNNNGAGCTTGCAGTGAGCCAAGAT
AGCACACGTCTGAACTCTTTCCCTACACGACGCTCTTCCGATCTNNNTAGTCCCAGCTACTCAGGAGA
GAGCACACGTCTGAACTCTTTCCCTACACGACGCTCTTCCGATCTNNNAGAATCACTTGAACCCGGGA

AGCACACGTCTGAACTCTTTCCCTACACGACGCTCTTCCGATCTNNNCGCCTATAGTCCCAGCTACTTG
AGCACACGTCTGAACTCTTTCCCTACACGACGCTCTTCCGATCTNNNGAGGCAGGAAAATGGTGTG
GAGCACACGTCTGAACTCTTTCCCTACACGACGCTCTTCCGATCTNNNCTGGGCAACAAGAGCAAAAC
AGCACACGTCTGAACTCTTTCCCTACACGACGCTCTTCCGATCTNNNGAGAATGGCATGAACCTGGG
GAGCACACGTCTGAACTCTTTCCCTACACGACGCTCTTCCGATCTNNNGAGAATGGCGTGAATCCGA
AGCACACGTCTGAACTCTTTCCCTACACGACGCTCTTCCGATCTNNNACCTGTAGTCCCAGCTACTCTG
GAGCACACGTCTGAACTCTTTCCCTACACGACGCTCTTCCGATCTNNNGTGCCTGTAGTCTCAGCTAC
GAGCACACGTCTGAACTCTTTCCCTACACGACGCTCTTCCGATCTNNNGTGGCTGAGTGAGAATCTG
GCACACGTCTGAACTCTTTCCCTACACGACGCTCTTCCGATCTNNNCGATAGAGCGAGACTTCCTCTCC
AGCACACGTCTGAACTCTTTCCCTACACGACGCTCTTCCGATCTNNNGTTGCAGTGAGCTGAGATCAT
GAGCACACGTCTGAACTCTTTCCCTACACGACGCTCTTCCGATCTNNNGCTACAGAGTGAGACTCCG
AGCACACGTCTGAACTCTTTCCCTACACGACGCTCTTCCGATCTNNNCAACAGCGCAAGATTCTGTCT
AGCACACGTCTGAACTCTTTCCCTACACGACGCTCTTCCGATCTNNNGGAGAATTGTTTGAATCCGGG
GCACACGTCTGAACTCTTTCCCTACACGACGCTCTTCCGATCTNNNGCACATCAGTCTGCTCCATCTAA
CACACGTCTGAACTCTTTCCCTACACGACGCTCTTCCGATCTNNNAGAAGGAGGAGGAAGGAAGAA
AGCACACGTCTGAACTCTTTCCCTACACGACGCTCTTCCGATCTNNNAAAGCGTGATCCACATCCTCT
AGCACACGTCTGAACTCTTTCCCTACACGACGCTCTTCCGATCTNNNTCTGCAGATTCTCTGGCTTCTTT
GCACACGTCTGAACTCTTTCCCTACACGACGCTCTTCCGATCTNNNGAACACTTGGCTCTCAATGCAGC
GAGCACACGTCTGAACTCTTTCCCTACACGACGCTCTTCCGATCTNNNGTTCAAGCGATTCTCATGCCT
AGCACACGTCTGAACTCTTTCCCTACACGACGCTCTTCCGATCTNNNCCAGCCCATTTGCTTATTCTTT
CACACGTCTGAACTCTTTCCCTACACGACGCTCTTCCGATCTNNNACAGTGCCCAGCCACATTTTATAC
CACACGTCTGAACTCTTTCCCTACACGACGCTCTTCCGATCTNNNCCATGCCCGGCTAATTTTTGTATC
AGCACACGTCTGAACTCTTTCCCTACACGACGCTCTTCCGATCTNNNCTTTGGATGTCCTTTCCTGCCT
AGCACACGTCTGAACTCTTTCCCTACACGACGCTCTTCCGATCTNNNGGTGACATCCTGAGTCTCACA
GAGCACACGTCTGAACTCTTTCCCTACACGACGCTCTTCCGATCTNNNAGGGAGAGATGAGGACAG
GCACACGTCTGAACTCTTTCCCTACACGACGCTCTTCCGATCTNNNAACCACTTTGGGGAGGAAGAA
AGCACACGTCTGAACTCTTTCCCTACACGACGCTCTTCCGATCTNNNGTAGCTGCTTGAAGAGGCAAA
AGCACACGTCTGAACTCTTTCCCTACACGACGCTCTTCCGATCTNNNGAGGTCTCTCAGCTTCCAACTC
GCACACGTCTGAACTCTTTCCCTACACGACGCTCTTCCGATCTNNNCCAGTTGGAGGTTGTAGTGAGT
AGCACACGTCTGAACTCTTTCCCTACACGACGCTCTTCCGATCTNNNCTCACTGAAACCTCTGTCTCCCC
GCACACGTCTGAACTCTTTCCCTACACGACGCTCTTCCGATCTNNNCAAAGTGCTGGAATTACAGGC
CACACGTCTGAACTCTTTCCCTACACGACGCTCTTCCGATCTNNNTGCAGGTACTTCAACCAATTCTTC
AGCACACGTCTGAACTCTTTCCCTACACGACGCTCTTCCGATCTNNNCTCTCCCTTTTAGCTTCTTCCCTC
CACACGTCTGAACTCTTTCCCTACACGACGCTCTTCCGATCTNNNCCACTGTGTAGCCTGTTTCCTAAC
AGCACACGTCTGAACTCTTTCCCTACACGACGCTCTTCCGATCTNNNCTTGAGGTGAGGAGTTCGAG
CACACGTCTGAACTCTTTCCCTACACGACGCTCTTCCGATCTNNNGAAGTTTTGTCCGTGCCATGAAT
AGCACACGTCTGAACTCTTTCCCTACACGACGCTCTTCCGATCTNNNGAGATCACACCACTGCAATCC
CACACGTCTGAACTCTTTCCCTACACGACGCTCTTCCGATCTNNNGTTGTGTTGGTGCTATGCTGACC
GAGCACACGTCTGAACTCTTTCCCTACACGACGCTCTTCCGATCTNNNGAGTCCAGGAGTTTGAGAC
AGCACACGTCTGAACTCTTTCCCTACACGACGCTCTTCCGATCTNNNCTTGCAGTGAGCTGAGATTG
GCACACGTCTGAACTCTTTCCCTACACGACGCTCTTCCGATCTNNNCTCGAATTCCTGACCTCGTGATA
AGCACACGTCTGAACTCTTTCCCTACACGACGCTCTTCCGATCTNNNGCTGGTGCAATTCAGAATCAT
AGCACACGTCTGAACTCTTTCCCTACACGACGCTCTTCCGATCTNNNTGATTCCAGCCTAGACTGAAAC

GAGCACACGTCTGAACTCTTTCCCTACACGACGCTCTTCCGATCTNNNCCAAGCAGCCACTAGAGAAGA

GCACACGTCTGAACTCTTTCCCTACACGACGCTCTTCCGATCTNNNCTTCTAAGAGAAGCGCCTACCTTC

GCACACGTCTGAACTCTTTCCCTACACGACGCTCTTCCGATCTNNNATGTGAGATTTTCTTTGCCTGGCC

GAGCACACGTCTGAACTCTTTCCCTACACGACGCTCTTCCGATCTNNNGATGCTGAGCCTGTGGAAGTA

GCACACGTCTGAACTCTTTCCCTACACGACGCTCTTCCGATCTNNNAGATCAAGGCACCAGCAGATTTA

AGCACACGTCTGAACTCTTTCCCTACACGACGCTCTTCCGATCTNNNCGACAAGAACGAAACTCCATCTC

GAGCACACGTCTGAACTCTTTCCCTACACGACGCTCTTCCGATCTNNNGCCTGAGCAACAAGAGTGAA

AGCACACGTCTGAACTCTTTCCCTACACGACGCTCTTCCGATCTNNNGTGCCTGTAATCCCAGCTACTTC

GCACACGTCTGAACTCTTTCCCTACACGACGCTCTTCCGATCTNNNGAGATCATGCCACTGGACTCTAG

GAGCACACGTCTGAACTCTTTCCCTACACGACGCTCTTCCGATCTNNNCATCATTGCCTGATCGGGGAA

GAGCACACGTCTGAACTCTTTCCCTACACGACGCTCTTCCGATCTNNNGAGGCAAGAGAATCGCTTGA

GCACACGTCTGAACTCTTTCCCTACACGACGCTCTTCCGATCTNNNCTAAGTCCGCTTTTCCTTCTCCTCAT

AGCACACGTCTGAACTCTTTCCCTACACGACGCTCTTCCGATCTNNNGAGATTGCAGTGAGCCGAGAT

GCACACGTCTGAACTCTTTCCCTACACGACGCTCTTCCGATCTNNNCTTTTCCCTTAGGGCAAAGCTTTCA

AGCACACGTCTGAACTCTTTCCCTACACGACGCTCTTCCGATCTNNNAAAGTGCTGGGATTACAGGCA

GAGCACACGTCTGAACTCTTTCCCTACACGACGCTCTTCCGATCTNNNAGCCTGGACAGAGTGAGACTA

GCACACGTCTGAACTCTTTCCCTACACGACGCTCTTCCGATCTNNNTGAGGCAGAAGAATTGCTTGAAC

GAGCACACGTCTGAACTCTTTCCCTACACGACGCTCTTCCGATCTNNNCCGGTTTCAAGCAATTCTCCTC

GAGCACACGTCTGAACTCTTTCCCTACACGACGCTCTTCCGATCTNNNACTGAGCCAAGATCACGTCAT

AGCACACGTCTGAACTCTTTCCCTACACGACGCTCTTCCGATCTNNNCCCACTTCAGCCTTCAGAGTAGC

AGCACACGTCTGAACTCTTTCCCTACACGACGCTCTTCCGATCTNNNATCACTTGATCCCAGGAGTTTGC

AGCACACGTCTGAACTCTTTCCCTACACGACGCTCTTCCGATCTNNNGGCTGAAGCAGGAGAATCACT

CACACGTCTGAACTCTTTCCCTACACGACGCTCTTCCGATCTNNNTTCTACCGCTCCCCTTCCAGATCATC

GCACACGTCTGAACTCTTTCCCTACACGACGCTCTTCCGATCTNNNGGATAGGGAAGTGGGTGTATG

AGCACACGTCTGAACTCTTTCCCTACACGACGCTCTTCCGATCTNNNCGAGAGAGAGAGAGAGAGAG

GCACACGTCTGAACTCTTTCCCTACACGACGCTCTTCCGATCTNNNAGGAGTTTGAAGTTGCAATGAGC

GCACACGTCTGAACTCTTTCCCTACACGACGCTCTTCCGATCTNNNGAACTTCCAACCTCAGGTGATCTC

AGCACACGTCTGAACTCTTTCCCTACACGACGCTCTTCCGATCTNNNCATAGTGCTTGAGTCCAGGAGT

GAGCACACGTCTGAACTCTTTCCCTACACGACGCTCTTCCGATCTNNNGCAGGAGAATTGCTTGAACCT

AGCACACGTCTGAACTCTTTCCCTACACGACGCTCTTCCGATCTNNNCATTCTCTTGCCTCAACCTCCCCA

GCACACGTCTGAACTCTTTCCCTACACGACGCTCTTCCGATCTNNNGTCTTGAACTCCTGACCTTGTGAC

AGCACACGTCTGAACTCTTTCCCTACACGACGCTCTTCCGATCTNNNTAGGGAGAGCTATCACAGGGAA

AGCACACGTCTGAACTCTTTCCCTACACGACGCTCTTCCGATCTNNNTATGTGGAATGGAAGTGGCAA

AGCACACGTCTGAACTCTTTCCCTACACGACGCTCTTCCGATCTNNNCCACGCCCAGCTAATTTTTGTATC

GCACACGTCTGAACTCTTTCCCTACACGACGCTCTTCCGATCTNNNTGAGAGAGGGAATGTGTGTATC

GAGCACACGTCTGAACTCTTTCCCTACACGACGCTCTTCCGATCTNNNGCAATCGTAACACAATGGGAA

GCACACGTCTGAACTCTTTCCCTACACGACGCTCTTCCGATCTNNNCCCTTAGCCACATATTCTGTCCTCA

AGCACACGTCTGAACTCTTTCCCTACACGACGCTCTTCCGATCTNNNTCATTCCTCTATCTCCCACATGTC

GCACACGTCTGAACTCTTTCCCTACACGACGCTCTTCCGATCTNNNATGATTCTCTTACCTCAGCCTCCCA

CACACGTCTGAACTCTTTCCCTACACGACGCTCTTCCGATCTNNNTACAATGACATCCTGGTTCCCTCCA

AGCACACGTCTGAACTCTTTCCCTACACGACGCTCTTCCGATCTNNNGTTCAAGCAATTCTCCTGCCTCC

GCACACGTCTGAACTCTTTCCCTACACGACGCTCTTCCGATCTNNNTGCTGATTTAGAGGGAACAGAG

CACACGTCTGAACTCTTTCCCTACACGACGCTCTTCCGATCTNNNACACCATAACTCCTTCTAGGCTCCA

GCACACGTCTGAACTCTTTCCCTACACGACGCTCTTCCGATCTNNNTCTTGCTCCAAGTTTCTCCCTTTCA
AGCACACGTCTGAACTCTTTCCCTACACGACGCTCTTCCGATCTNNNGATCAAGACCATCCTGGCTAA
GAGCACACGTCTGAACTCTTTCCCTACACGACGCTCTTCCGATCTNNNGCAGGAGAATTGCTTGAACT
AGCACACGTCTGAACTCTTTCCCTACACGACGCTCTTCCGATCTNNNGAGAATGGTGTGAACCTGGGA
AGCACACGTCTGAACTCTTTCCCTACACGACGCTCTTCCGATCTNNNTGAGCTCAGGAGTTCAAGATCA
AGCACACGTCTGAACTCTTTCCCTACACGACGCTCTTCCGATCTNNNAAAGAGGAGGAGGGAATAGA
GCACACGTCTGAACTCTTTCCCTACACGACGCTCTTCCGATCTNNNGTTCTCCCTCTCACTTGGACTTTCA
GAGCACACGTCTGAACTCTTTCCCTACACGACGCTCTTCCGATCTNNNGAGAATCGCTTGAACCCAGGA
GCACACGTCTGAACTCTTTCCCTACACGACGCTCTTCCGATCTNNNGCTCATTCGTCCTTTGCCAAATACA
AGCACACGTCTGAACTCTTTCCCTACACGACGCTCTTCCGATCTNNNTCAAATAGAATGGGGTGTGAG
GAAGAGCACACGTCTGAACTCTTTCCCTACACGACGCTCTTCCGATCTNNNGAGGCTGAGGCAGGAGA
AGCACACGTCTGAACTCTTTCCCTACACGACGCTCTTCCGATCTNNNGAGGCAGGAGAATAGCGTGAA
AGCACACGTCTGAACTCTTTCCCTACACGACGCTCTTCCGATCTNNNGCCTGTAGTCCCAGATACTTG
AGCACACGTCTGAACTCTTTCCCTACACGACGCTCTTCCGATCTNNNGAGCTTGCAGTGAGTCGAGAT
AGCACACGTCTGAACTCTTTCCCTACACGACGCTCTTCCGATCTNNNGTGTGTAGTCCCAGCTACTCAG
GAGCACACGTCTGAACTCTTTCCCTACACGACGCTCTTCCGATCTNNNGTGAGCCAAGATCGCATCAT
GAGCACACGTCTGAACTCTTTCCCTACACGACGCTCTTCCGATCTNNNGCAGGAGAATTGCTTGAACCA
GAGCACACGTCTGAACTCTTTCCCTACACGACGCTCTTCCGATCTNNNCAACATGGCAAAACCCCAACT
AGAGCACACGTCTGAACTCTTTCCCTACACGACGCTCTTCCGATCTNNNGAGAATGGTGTGAACCTGG
AGCACACGTCTGAACTCTTTCCCTACACGACGCTCTTCCGATCTNNNCACCTGTAATCCCAGCTACTCG
CACACGTCTGAACTCTTTCCCTACACGACGCTCTTCCGATCTNNNCTGTAGTACCAGCTACTTGGGAGC
AGCACACGTCTGAACTCTTTCCCTACACGACGCTCTTCCGATCTNNNGAGAGGCTGAGACAGGAGAAT
GAGCACACGTCTGAACTCTTTCCCTACACGACGCTCTTCCGATCTNNNGAGCTTGTAGTGAGCCGAGA
AGCACACGTCTGAACTCTTTCCCTACACGACGCTCTTCCGATCTNNNGGAATCACTTGAACCCAGGAG
AGCACACGTCTGAACTCTTTCCCTACACGACGCTCTTCCGATCTNNNGGTTGCAGTGAGTCCAAATCA
AGCACACGTCTGAACTCTTTCCCTACACGACGCTCTTCCGATCTNNNCTTGCAGTGAGCTGAGATTGT
GAGCACACGTCTGAACTCTTTCCCTACACGACGCTCTTCCGATCTNNNGTTGCAGTGAGCCAAGATCA
GCACACGTCTGAACTCTTTCCCTACACGACGCTCTTCCGATCTNNNGGGTTACAGTGCTAGACAAAAG
AGCACACGTCTGAACTCTTTCCCTACACGACGCTCTTCCGATCTNNNTGTCTTCACATGGTTCAAGAGG
GCACACGTCTGAACTCTTTCCCTACACGACGCTCTTCCGATCTNNNCTCAGTCTAGTAGGAATGCCTGC
AGCACACGTCTGAACTCTTTCCCTACACGACGCTCTTCCGATCTNNNGGCAATTTCTTCCTCTTTCCGTTC
AGCACACGTCTGAACTCTTTCCCTACACGACGCTCTTCCGATCTNNNCCTTTGCAATGAGCCATTCAGA
GCACACGTCTGAACTCTTTCCCTACACGACGCTCTTCCGATCTNNNTCTTCTTATACATGCACCTGGCCCA
GAGCACACGTCTGAACTCTTTCCCTACACGACGCTCTTCCGATCTNNNGGGTTTGAGTTCATCACCTAG
AGCACACGTCTGAACTCTTTCCCTACACGACGCTCTTCCGATCTNNNCTATGCAAGCTGGACACAAAGT
GCACACGTCTGAACTCTTTCCCTACACGACGCTCTTCCGATCTNNNAACTATACGGAACACCTGTGACC
AGCACACGTCTGAACTCTTTCCCTACACGACGCTCTTCCGATCTNNNTGCTACAGAAAGCTTGCTCCTAA
GCACACGTCTGAACTCTTTCCCTACACGACGCTCTTCCGATCTNNNCTGTGTTCCTATTGGGCTTTTGG
AGCACACGTCTGAACTCTTTCCCTACACGACGCTCTTCCGATCTNNNCTATGTTGCCCAGACTGGTCTT
GCACACGTCTGAACTCTTTCCCTACACGACGCTCTTCCGATCTNNNAACCTAGCCTCATCTTACTCCCTCA
AGCACACGTCTGAACTCTTTCCCTACACGACGCTCTTCCGATCTNNNGGAGTAGCTGGGACTACAGGT
GCACACGTCTGAACTCTTTCCCTACACGACGCTCTTCCGATCTNNNCCAGGCCTTTTATAACTGGTTCC
GCACACGTCTGAACTCTTTCCCTACACGACGCTCTTCCGATCTNNNATTTGTTTGTTTTTGTGGGGGA

CACACGTCTGAACTCTTTCCCTACACGACGCTCTTCCGATCTNNNTTTGGAGCTTCTTCTCAGTCCAGC
CACACGTCTGAACTCTTTCCCTACACGACGCTCTTCCGATCTNNNCGGATATGGGGTGTCACATGATT
GCACACGTCTGAACTCTTTCCCTACACGACGCTCTTCCGATCTNNNTCCCAAAGTGCTGAGATTATCGG
GCACACGTCTGAACTCTTTCCCTACACGACGCTCTTCCGATCTNNNAGGTTCAAGCAATTCTTTTGCCT
AGCACACGTCTGAACTCTTTCCCTACACGACGCTCTTCCGATCTNNNGAGAATGCTGTGAACCTGGG
CACACGTCTGAACTCTTTCCCTACACGACGCTCTTCCGATCTNNNCGCCTATAATTCCAGCACTTTGG
GCACACGTCTGAACTCTTTCCCTACACGACGCTCTTCCGATCTNNNGTCGCGATTTGGTTTATTCTGCT
AGCACACGTCTGAACTCTTTCCCTACACGACGCTCTTCCGATCTNNNCTGTAGTCCCAGCTACTCAGG
GCACACGTCTGAACTCTTTCCCTACACGACGCTCTTCCGATCTNNNGAGAATCGCTTGAACCTGGGAG
AGCACACGTCTGAACTCTTTCCCTACACGACGCTCTTCCGATCTNNNGAGGTTGCAGTGAGCTGAGA
GCACACGTCTGAACTCTTTCCCTACACGACGCTCTTCCGATCTNNNGATCTTGCAGTGAGCCGAGATT
CACACGTCTGAACTCTTTCCCTACACGACGCTCTTCCGATCTNNNAAAAAAATTAGCCGGGTGTGTG
GAGCACACGTCTGAACTCTTTCCCTACACGACGCTCTTCCGATCTNNNAGGTTGCAGTGAGCCAAGAT
AGCACACGTCTGAACTCTTTCCCTACACGACGCTCTTCCGATCTNNNGAGCTTGTAGTGAGCCGAGA
GAGCACACGTCTGAACTCTTTCCCTACACGACGCTCTTCCGATCTNNNGAATGGCATGAACCTGGGA
GAGCACACGTCTGAACTCTTTCCCTACACGACGCTCTTCCGATCTNNNGAGAATGGCATGAACCTGG
AGCACACGTCTGAACTCTTTCCCTACACGACGCTCTTCCGATCTNNNCTTCAGTGAGCCAAGATCGTA
AGCACACGTCTGAACTCTTTCCCTACACGACGCTCTTCCGATCTNNNGATTGAGACCATCCTGGCTA
GCACACGTCTGAACTCTTTCCCTACACGACGCTCTTCCGATCTNNNGAGAGTCACTTGAACCTGGGA
GCACACGTCTGAACTCTTTCCCTACACGACGCTCTTCCGATCTNNNGAAGCAGGAGAATGGCATGA
AGCACACGTCTGAACTCTTTCCCTACACGACGCTCTTCCGATCTNNNCTGAGATTGTGCCGCTGTACT
GAGCACACGTCTGAACTCTTTCCCTACACGACGCTCTTCCGATCTNNNCTTGCAGTGAGCCAAGATTG
GCACACGTCTGAACTCTTTCCCTACACGACGCTCTTCCGATCTNNNGAGAATCGCTTGAACCAGGAG
AGCACACGTCTGAACTCTTTCCCTACACGACGCTCTTCCGATCTNNNGAGCTTGTAGTGAGCCGAGA
AGCACACGTCTGAACTCTTTCCCTACACGACGCTCTTCCGATCTNNNGAAGCAGGAGAATGGCATGA
GAGCACACGTCTGAACTCTTTCCCTACACGACGCTCTTCCGATCTNNNGGTGAGCCAAGATTGAGCTA
GAGCACACGTCTGAACTCTTTCCCTACACGACGCTCTTCCGATCTNNNGCAGGAGAATTGCTTGAACC
GAGCACACGTCTGAACTCTTTCCCTACACGACGCTCTTCCGATCTNNNGAAGGCTGAGTCAGGAGAA
CACACGTCTGAACTCTTTCCCTACACGACGCTCTTCCGATCTNNNGCTTCTGTAATCCCAGCTACTGTC
CACACGTCTGAACTCTTTCCCTACACGACGCTCTTCCGATCTNNNGTGGGTGGATAGCTCAACTCTAG
GCACACGTCTGAACTCTTTCCCTACACGACGCTCTTCCGATCTNNNGTGCCTGTAATCCCAGCTACTT
AGCACACGTCTGAACTCTTTCCCTACACGACGCTCTTCCGATCTNNNCTTGCAGTAAGCTGAGATCGT
GCACACGTCTGAACTCTTTCCCTACACGACGCTCTTCCGATCTNNNCTCTCACTGCAGTGTTAGTGT
GCACACGTCTGAACTCTTTCCCTACACGACGCTCTTCCGATCTNNNGGTGTTAGCTGAGATGGTCTTG
GCACACGTCTGAACTCTTTCCCTACACGACGCTCTTCCGATCTNNNTTGCAGTGAGCCAAGATAGCG
AGCACACGTCTGAACTCTTTCCCTACACGACGCTCTTCCGATCTNNNAGGTAACAGAACTGAGACACA
CACACGTCTGAACTCTTTCCCTACACGACGCTCTTCCGATCTNNNCAAACCTTTCTTGGGGATGCATT
GCACACGTCTGAACTCTTTCCCTACACGACGCTCTTCCGATCTNNNGAGATCGCTTCACTGTACTCCAC
AGCACACGTCTGAACTCTTTCCCTACACGACGCTCTTCCGATCTNNNTAGTCCCAGCTACTCAGGAGA
AGCACACGTCTGAACTCTTTCCCTACACGACGCTCTTCCGATCTNNNAAGAATTGCTTGAGCCTGGGA
GCACACGTCTGAACTCTTTCCCTACACGACGCTCTTCCGATCTNNNACTTGTGGTCCCAGCTTCTTAG
AGCACACGTCTGAACTCTTTCCCTACACGACGCTCTTCCGATCTNNNTGAGAACCATGGCAGAAGAG
AGCACACGTCTGAACTCTTTCCCTACACGACGCTCTTCCGATCTNNNACTTACCTCCATACAGTAGTGC

AGCACACGTCTGAACTCTTTCCCTACACGACGCTCTTCCGATCTNNNGTGCTTGTAGTCCCAGCTACTT
GCACACGTCTGAACTCTTTCCCTACACGACGCTCTTCCGATCTNNNAAAGGAATTCGAGACCATCCTG
AGCACACGTCTGAACTCTTTCCCTACACGACGCTCTTCCGATCTNNNAAGGACATAGACATCACCACC
GCACACGTCTGAACTCTTTCCCTACACGACGCTCTTCCGATCTNNNGAGGCACCAGAATTGCTTGAAC
AGCACACGTCTGAACTCTTTCCCTACACGACGCTCTTCCGATCTNNNGTAGTCCCAGCTACTCAGGAG
GCACACGTCTGAACTCTTTCCCTACACGACGCTCTTCCGATCTNNNTGTAGTCCCAGCTACTCAGGAG
AGCACACGTCTGAACTCTTTCCCTACACGACGCTCTTCCGATCTNNNGAGAATGGCATGAACCTGGGA
CACACGTCTGAACTCTTTCCCTACACGACGCTCTTCCGATCTNNNAGAGAAAAGCTTCTAGTGGAGCC
AGCACACGTCTGAACTCTTTCCCTACACGACGCTCTTCCGATCTNNNCTGAGCAACAAGAGCAAGACT
GAGCACACGTCTGAACTCTTTCCCTACACGACGCTCTTCCGATCTNNNGAGAATGGCATGAACCCAGG
GAGCACACGTCTGAACTCTTTCCCTACACGACGCTCTTCCGATCTNNNGAGAATGGTGTGAACCCAG
GCACACGTCTGAACTCTTTCCCTACACGACGCTCTTCCGATCTNNNCTCAATGCAACCTCAAACTCCTGC
AGCACACGTCTGAACTCTTTCCCTACACGACGCTCTTCCGATCTNNNATGCTGAGTCCAGTTCCCGATA
AGCACACGTCTGAACTCTTTCCCTACACGACGCTCTTCCGATCTNNNGAGCATCACTTGAACCTGGGAG
AGCACACGTCTGAACTCTTTCCCTACACGACGCTCTTCCGATCTNNNCAGGCAGGAAGAGGTGGTAT
AGCACACGTCTGAACTCTTTCCCTACACGACGCTCTTCCGATCTNNNCACCTGTAATCCCAGCTACTCG
GCACACGTCTGAACTCTTTCCCTACACGACGCTCTTCCGATCTNNNTGGGTGACAGAGAGAGACTGT
AGAGCACACGTCTGAACTCTTTCCCTACACGACGCTCTTCCGATCTNNNCTCAGTAATCCCAGCGACTC
GCACACGTCTGAACTCTTTCCCTACACGACGCTCTTCCGATCTNNNGGATTATAGGTGTGAGCCACTGT
GCACACGTCTGAACTCTTTCCCTACACGACGCTCTTCCGATCTNNNAAACTGCATGATCACAGGAAAC
AGCACACGTCTGAACTCTTTCCCTACACGACGCTCTTCCGATCTNNNGACAGCAGAGAGTCACTTGAG
CACACGTCTGAACTCTTTCCCTACACGACGCTCTTCCGATCTNNNGAGCCAACTAGGAAACTGAAGG
GCACACGTCTGAACTCTTTCCCTACACGACGCTCTTCCGATCTNNNGATCCCTAGGCAGCCAAATTAGA
CACACGTCTGAACTCTTTCCCTACACGACGCTCTTCCGATCTNNNTTGTCTCACTGGAAGGTCTTCAGC
AGCACACGTCTGAACTCTTTCCCTACACGACGCTCTTCCGATCTNNNCAAAACATTAGCTGAGCCAGGT
AGCACACGTCTGAACTCTTTCCCTACACGACGCTCTTCCGATCTNNNCCAGGCCTGGCTAATTTTGTTTT
GAGCACACGTCTGAACTCTTTCCCTACACGACGCTCTTCCGATCTNNNAAAGTGCTGGGATTACAGGC
GCACACGTCTGAACTCTTTCCCTACACGACGCTCTTCCGATCTNNNTGTCAACTTAGTGCCCTTGGAAG
GCACACGTCTGAACTCTTTCCCTACACGACGCTCTTCCGATCTNNNCTCCCTCTCCTTCTCAGCATTTTCA
AGCACACGTCTGAACTCTTTCCCTACACGACGCTCTTCCGATCTNNNATAAGCCATGCCTGAAAGTAGC
AGCACACGTCTGAACTCTTTCCCTACACGACGCTCTTCCGATCTNNNCAAGCAAAGTTCTCCCAGATCA
GCACACGTCTGAACTCTTTCCCTACACGACGCTCTTCCGATCTNNNCACACTGTCTTCCACAATGGTTG
GAGCACACGTCTGAACTCTTTCCCTACACGACGCTCTTCCGATCTNNNGTAAGCCAAGATTGCACCACT
GCACACGTCTGAACTCTTTCCCTACACGACGCTCTTCCGATCTNNNGAGAATCGCTTGAACTGGGGAG
AGCACACGTCTGAACTCTTTCCCTACACGACGCTCTTCCGATCTNNNAGTGAGCTGTGATTGTACCACT
AGCACACGTCTGAACTCTTTCCCTACACGACGCTCTTCCGATCTNNNTAACAGAGTGAGACCCCATCCT
CACACGTCTGAACTCTTTCCCTACACGACGCTCTTCCGATCTNNNACTGATGACTAGGAAGGGTGGAT
GAGCACACGTCTGAACTCTTTCCCTACACGACGCTCTTCCGATCTNNNCAAGATCACACCACTGCATTC
AGCACACGTCTGAACTCTTTCCCTACACGACGCTCTTCCGATCTNNNGAGAATGGTGTGAACCCAGG
GAGCACACGTCTGAACTCTTTCCCTACACGACGCTCTTCCGATCTNNNGAGGTTGAAGTGAGCCGAG
AGCACACGTCTGAACTCTTTCCCTACACGACGCTCTTCCGATCTNNNACCCGAGCAGAAAAAGCAAAA
GCACACGTCTGAACTCTTTCCCTACACGACGCTCTTCCGATCTNNNTTGATACTGGGAGGTGAAGGTT
GAGCACACGTCTGAACTCTTTCCCTACACGACGCTCTTCCGATCTNNNACACCTGTAATCCCAGCACTTC

GAGCACACGTCTGAACTCTTTCCCTACACGACGCTCTTCCGATCTNNNGCAACAGAGTGAGACTCCAT
GCACACGTCTGAACTCTTTCCCTACACGACGCTCTTCCGATCTNNNTTTCTCTCAGCTCTTTCTAGGCCC
AGCACACGTCTGAACTCTTTCCCTACACGACGCTCTTCCGATCTNNNGAGGTTGCAGTGAGTCGAGA
AGCACACGTCTGAACTCTTTCCCTACACGACGCTCTTCCGATCTNNNGAGGTTGCAGTGAGCTGAGAT
GCACACGTCTGAACTCTTTCCCTACACGACGCTCTTCCGATCTNNNATCATTCCGTAGTGTTCTGAGT
CACACGTCTGAACTCTTTCCCTACACGACGCTCTTCCGATCTNNNTGCAGTAGCTAAGCATCAGTGAA
AGCACACGTCTGAACTCTTTCCCTACACGACGCTCTTCCGATCTNNNCTGGCCAACAAGAGCAAAACT
GCACACGTCTGAACTCTTTCCCTACACGACGCTCTTCCGATCTNNNGATCACTTGAGCCTGAGAGGTC
GCACACGTCTGAACTCTTTCCCTACACGACGCTCTTCCGATCTNNNCCCTACATCAACAGTCGTAGCTTC
CACACGTCTGAACTCTTTCCCTACACGACGCTCTTCCGATCTNNNCCCGAGGACTTGTTCTACTTTCTC
CACACGTCTGAACTCTTTCCCTACACGACGCTCTTCCGATCTNNNCCCATGTCTCAAACCATCCTGTAC
AGCACACGTCTGAACTCTTTCCCTACACGACGCTCTTCCGATCTNNNATCTGGTCCGTTCTTCACACTCC
CACACGTCTGAACTCTTTCCCTACACGACGCTCTTCCGATCTNNNTCATACCTGTAATGCCAGCACTGC
CACACGTCTGAACTCTTTCCCTACACGACGCTCTTCCGATCTNNNGGTGCTGAGTGTCTGTTTCTTTC
AGCACACGTCTGAACTCTTTCCCTACACGACGCTCTTCCGATCTNNNTGAGGCAGGAGAATAGTGTG
AGCACACGTCTGAACTCTTTCCCTACACGACGCTCTTCCGATCTNNNTTAGCCTGGCTAACATGGTGA
AGCACACGTCTGAACTCTTTCCCTACACGACGCTCTTCCGATCTNNNCAGTGAAACCCCGTCTGTACTA
AGCACACGTCTGAACTCTTTCCCTACACGACGCTCTTCCGATCTNNNTGAGGCAGAGAATGGAGTG
AGCACACGTCTGAACTCTTTCCCTACACGACGCTCTTCCGATCTNNNATTGTGTGTGTGTGTGTGTGT
CACACGTCTGAACTCTTTCCCTACACGACGCTCTTCCGATCTNNNCTCAAGCAATCATTCCACCTTGGC
GAGCACACGTCTGAACTCTTTCCCTACACGACGCTCTTCCGATCTNNNGAGGCAGGACAATTGCTTGA
GAGCACACGTCTGAACTCTTTCCCTACACGACGCTCTTCCGATCTNNNCTGTAGTCCCAGCTACTCAGG
CACACGTCTGAACTCTTTCCCTACACGACGCTCTTCCGATCTNNNACCACCACCACTATCACCAAATTC
CACACGTCTGAACTCTTTCCCTACACGACGCTCTTCCGATCTNNNAAAAAAATTTAGCCGGGCATGG
GCACACGTCTGAACTCTTTCCCTACACGACGCTCTTCCGATCTNNNGAGAATAGCGTGAACCCAGGAG
GAGCACACGTCTGAACTCTTTCCCTACACGACGCTCTTCCGATCTNNNGTGCCTGTAATCCCACCTTCT
AGCACACGTCTGAACTCTTTCCCTACACGACGCTCTTCCGATCTNNNGTGATCATGCCACTGTACTCCA
AGCACACGTCTGAACTCTTTCCCTACACGACGCTCTTCCGATCTNNNCACGATCTCAGCTTACTGCAAG
AGCACACGTCTGAACTCTTTCCCTACACGACGCTCTTCCGATCTNNNGCCTGTAGTCCCAGCTACTTG
GCACACGTCTGAACTCTTTCCCTACACGACGCTCTTCCGATCTNNNCATGCCTAGGAAAGAGACCATG
AGCACACGTCTGAACTCTTTCCCTACACGACGCTCTTCCGATCTNNNGGAGCCATGTGTGTTCTTTTAC
AGCACACGTCTGAACTCTTTCCCTACACGACGCTCTTCCGATCTNNNCAGAGTGCAGTGATCCCATA
CACACGTCTGAACTCTTTCCCTACACGACGCTCTTCCGATCTNNNTCTCACTGTGGAGTTGTTATGCA
GAGCACACGTCTGAACTCTTTCCCTACACGACGCTCTTCCGATCTNNNCAGGTTCAAGCAATTCTCCTG
AGCACACGTCTGAACTCTTTCCCTACACGACGCTCTTCCGATCTNNNTCCCTCACTTTGCTCTCCTTTCC
AGAGCACACGTCTGAACTCTTTCCCTACACGACGCTCTTCCGATCTNNNGTAGCAGAACCCAGTCTCT
AGCACACGTCTGAACTCTTTCCCTACACGACGCTCTTCCGATCTNNNAGTGGGTCTTGCATATGTACG
GCACACGTCTGAACTCTTTCCCTACACGACGCTCTTCCGATCTNNNAACCTCCATCCTTCCATCCTCTAC
GCACACGTCTGAACTCTTTCCCTACACGACGCTCTTCCGATCTNNNAGAGCAAGGAGACAGACAGAA
GAGCACACGTCTGAACTCTTTCCCTACACGACGCTCTTCCGATCTNNNGATGACCTCTCTTTCTCCTTC
AGCACACGTCTGAACTCTTTCCCTACACGACGCTCTTCCGATCTNNNAGAGGTAGAGACAGAGAAG
GCACACGTCTGAACTCTTTCCCTACACGACGCTCTTCCGATCTNNNGACACTGCCTTGCATGTGAATAG
GCACACGTCTGAACTCTTTCCCTACACGACGCTCTTCCGATCTNNNGGTTGCAGTGAAGCACAATCAT

AGCACACGTCTGAACTCTTTCCCTACACGACGCTCTTCCGATCTNNNGCAACAGAGTGAGATCCTGTCT
AGCACACGTCTGAACTCTTTCCCTACACGACGCTCTTCCGATCTNNNCCATTTGCTGCTCATAAGGCTACC
GAGCACACGTCTGAACTCTTTCCCTACACGACGCTCTTCCGATCTNNNGCGATGTAGTGAGACACTGTC
GCACACGTCTGAACTCTTTCCCTACACGACGCTCTTCCGATCTNNNCATAACCAAGGCGACATGCTTAG
GAGCACACGTCTGAACTCTTTCCCTACACGACGCTCTTCCGATCTNNNGGATTACCAGCATGAGTCACCA
AGAGCACACGTCTGAACTCTTTCCCTACACGACGCTCTTCCGATCTNNNGAGGCAGGAGAATTGCATG
GAGCACACGTCTGAACTCTTTCCCTACACGACGCTCTTCCGATCTNNNCACTGCAACCTCTACCTCTCAGC
AGAGCACACGTCTGAACTCTTTCCCTACACGACGCTCTTCCGATCTNNNCTTGGGTGAGAACTAGTGCT
GAGCACACGTCTGAACTCTTTCCCTACACGACGCTCTTCCGATCTNNNGCAGGAGAATTGCTTGAACCT
GAGCACACGTCTGAACTCTTTCCCTACACGACGCTCTTCCGATCTNNNGAGGTTGCAGTTAGCCGAGAT
CACACGTCTGAACTCTTTCCCTACACGACGCTCTTCCGATCTNNNGAAGTTTGCAGTGAGCCAAGATCC
GCACACGTCTGAACTCTTTCCCTACACGACGCTCTTCCGATCTNNNTGTTTCTACCCTACTCTCCCCCA
AGCACACGTCTGAACTCTTTCCCTACACGACGCTCTTCCGATCTNNNTTGTTGTTTGTAGCACAGGGAA
GCACACGTCTGAACTCTTTCCCTACACGACGCTCTTCCGATCTNNNCTGGGCAACAAGAGCAAAACTCC
GAGCACACGTCTGAACTCTTTCCCTACACGACGCTCTTCCGATCTNNNGTTGCAGTGAGCTGAGATCA
GCACACGTCTGAACTCTTTCCCTACACGACGCTCTTCCGATCTNNNATTGTATTTGGAGATGGGGGAG
AGCACACGTCTGAACTCTTTCCCTACACGACGCTCTTCCGATCTNNNAGCAGCAGCCAAAAGAGTATG
GAGCACACGTCTGAACTCTTTCCCTACACGACGCTCTTCCGATCTNNNCTGCAGTGAACCGAGATCACC
AGCACACGTCTGAACTCTTTCCCTACACGACGCTCTTCCGATCTNNNACAGTGACTCTCTCTCTCTCTC
GAGCACACGTCTGAACTCTTTCCCTACACGACGCTCTTCCGATCTNNNATGAGCTGATAGCCTGAACTCA
GCACACGTCTGAACTCTTTCCCTACACGACGCTCTTCCGATCTNNNAAGCACTGGGATTACAGGCATAA
GAGCACACGTCTGAACTCTTTCCCTACACGACGCTCTTCCGATCTNNNCTGGACGACAAGAGCAAGAC
GCACACGTCTGAACTCTTTCCCTACACGACGCTCTTCCGATCTNNNTAGAAATAGTGGTTGCCTCTGGG
GAGCACACGTCTGAACTCTTTCCCTACACGACGCTCTTCCGATCTNNNGTTGCAGTGAGCCAAGATCAT
GAGCACACGTCTGAACTCTTTCCCTACACGACGCTCTTCCGATCTNNNACAAAAAATTCACAGGCTGGT
AGCACACGTCTGAACTCTTTCCCTACACGACGCTCTTCCGATCTNNNGACAACAGAGTGAGACTCCGTC
GAGCACACGTCTGAACTCTTTCCCTACACGACGCTCTTCCGATCTNNNGAGACAGACCAAGACACACAC
AGAGCACACGTCTGAACTCTTTCCCTACACGACGCTCTTCCGATCTNNNAGAAGCAGAAGCAGAAGCA
GCACACGTCTGAACTCTTTCCCTACACGACGCTCTTCCGATCTNNNCCCTGATTGGTATGGTCATGGAA
GAGCACACGTCTGAACTCTTTCCCTACACGACGCTCTTCCGATCTNNNCTACTTGCTACTTGGGAGGGTC
AGCACACGTCTGAACTCTTTCCCTACACGACGCTCTTCCGATCTNNNTGCTGGGCAATACTACTTGGTAC
GCACACGTCTGAACTCTTTCCCTACACGACGCTCTTCCGATCTNNNGAGCCGTACTTCTTTCTCAGTCACA
AGCACACGTCTGAACTCTTTCCCTACACGACGCTCTTCCGATCTNNNGAGAATGGCATGAACCTGGGA
GCACACGTCTGAACTCTTTCCCTACACGACGCTCTTCCGATCTNNNGTAGGAAGTGGAAGCAGAGCTAC
AGCACACGTCTGAACTCTTTCCCTACACGACGCTCTTCCGATCTNNNTAGCTGATGAAGTTGTAGGAGC
AGCACACGTCTGAACTCTTTCCCTACACGACGCTCTTCCGATCTNNNGAGTGCAATTGTTCAGCCACAT
AGCACACGTCTGAACTCTTTCCCTACACGACGCTCTTCCGATCTNNNCACTGGTACCCGAGAACTTTCCC
GCACACGTCTGAACTCTTTCCCTACACGACGCTCTTCCGATCTNNNTACCTCTCTCAATTCCGTGTTCCCA
AGCACACGTCTGAACTCTTTCCCTACACGACGCTCTTCCGATCTNNNAGCTACCACACCTGGCTAATTTT
AGAGCACACGTCTGAACTCTTTCCCTACACGACGCTCTTCCGATCTNNNGAGGTCACAGTGAGCCAAGA
AGCACACGTCTGAACTCTTTCCCTACACGACGCTCTTCCGATCTNNNCGAGATCATGTCACTGAACTCCA
AGCACACGTCTGAACTCTTTCCCTACACGACGCTCTTCCGATCTNNNGCAACAGAGTGAGACTCCATCT
GAGCACACGTCTGAACTCTTTCCCTACACGACGCTCTTCCGATCTNNNACTGAGCCAAGATCACGTCAT

GCACACGTCTGAACTCTTTCCCTACACGACGCTCTTCCGATCTNNNGAGCGAAACTCCGTCTCAAAAAA
AGAGCACACGTCTGAACTCTTTCCCTACACGACGCTCTTCCGATCTNNNCAATCCAGCCTAGGTGACAGA
GAGCACACGTCTGAACTCTTTCCCTACACGACGCTCTTCCGATCTNNNGTCTTGGCTAACACAGTGAAA
GAGCACACGTCTGAACTCTTTCCCTACACGACGCTCTTCCGATCTNNNGTGTGTGTGTCTGGAAGGCTA
GCACACGTCTGAACTCTTTCCCTACACGACGCTCTTCCGATCTNNNCCTATGTCCAAGGAATCCTACCG
GCACACGTCTGAACTCTTTCCCTACACGACGCTCTTCCGATCTNNNCTGAGGTTGAAGAATGGCTTGA
AGCACACGTCTGAACTCTTTCCCTACACGACGCTCTTCCGATCTNNNGCAACACAGTGAGACTCTGTCT
AGCACACGTCTGAACTCTTTCCCTACACGACGCTCTTCCGATCTNNNAGAGGATATGAGACTCCCAGT
GAGCACACGTCTGAACTCTTTCCCTACACGACGCTCTTCCGATCTNNNATGAGCTGATAGCCTGAACTCA
GCACACGTCTGAACTCTTTCCCTACACGACGCTCTTCCGATCTNNNGTGTTGGGATTATAGGCATGAG
GCACACGTCTGAACTCTTTCCCTACACGACGCTCTTCCGATCTNNNTTTCAGAGACCTCCATCTGCATCC
AGCACACGTCTGAACTCTTTCCCTACACGACGCTCTTCCGATCTNNNAAAGACTGTCTATGGGAGATG
GCACACGTCTGAACTCTTTCCCTACACGACGCTCTTCCGATCTNNNCTCAAGCAATCTTCCTGCCTTAGC
GAGCACACGTCTGAACTCTTTCCCTACACGACGCTCTTCCGATCTNNNCTCAACCTCCTGGACTCAAGTG
AGCACACGTCTGAACTCTTTCCCTACACGACGCTCTTCCGATCTNNNGATAGAGCCTTCTGTGGGGGT
GCACACGTCTGAACTCTTTCCCTACACGACGCTCTTCCGATCTNNNTATCATCGGCTCTCTTGGAAACA
GCACACGTCTGAACTCTTTCCCTACACGACGCTCTTCCGATCTNNNTTCTCAGAGATTACTGCAGCCTC
GAGCACACGTCTGAACTCTTTCCCTACACGACGCTCTTCCGATCTNNNGAGGTTGTGGTGAGTCGAGA
AGCACACGTCTGAACTCTTTCCCTACACGACGCTCTTCCGATCTNNNACAGGTAATCCCAGCACTTCAG
AGCACACGTCTGAACTCTTTCCCTACACGACGCTCTTCCGATCTNNNCCTTCAGCTCCAAAATGTCCATG
GAGCACACGTCTGAACTCTTTCCCTACACGACGCTCTTCCGATCTNNNAGGCTCTCTTTCTCTCTCTCTCT
AGCACACGTCTGAACTCTTTCCCTACACGACGCTCTTCCGATCTNNNGAGACTGCAGTGACCCAAGAT
AGCACACGTCTGAACTCTTTCCCTACACGACGCTCTTCCGATCTNNNCTCAAAGAGATCCACCTGCCTT
GAGCACACGTCTGAACTCTTTCCCTACACGACGCTCTTCCGATCTNNNCTGAGTGAGTCAAGACCACT
GCACACGTCTGAACTCTTTCCCTACACGACGCTCTTCCGATCTNNNCGTTATGGAAACCAATTCTGCCA
AGCACACGTCTGAACTCTTTCCCTACACGACGCTCTTCCGATCTNNNGGACCACAGGTACACTCAATCA
GAGCACACGTCTGAACTCTTTCCCTACACGACGCTCTTCCGATCTNNNGACTTTGTGCCATTGACTCCA
AGCACACGTCTGAACTCTTTCCCTACACGACGCTCTTCCGATCTNNNTTTTCAGCAAGCATACCAAGACT
AGCACACGTCTGAACTCTTTCCCTACACGACGCTCTTCCGATCTNNNCACAGCCCTGAATTGAGCAATA
GAGCACACGTCTGAACTCTTTCCCTACACGACGCTCTTCCGATCTNNNCACCAAAAGCTAGCTCCTCTCC
GCACACGTCTGAACTCTTTCCCTACACGACGCTCTTCCGATCTNNNTCTCTAATTGTTTCCCCTGGAGC
AGCACACGTCTGAACTCTTTCCCTACACGACGCTCTTCCGATCTNNNCATTTGGTCAGTTGTGTGGTCC
GAGCACACGTCTGAACTCTTTCCCTACACGACGCTCTTCCGATCTNNNCTCATGTGGTCGCTGTGTTCTA
GAGCACACGTCTGAACTCTTTCCCTACACGACGCTCTTCCGATCTNNNCCAGGTCCAGGTGACTTAGAA
GAGCACACGTCTGAACTCTTTCCCTACACGACGCTCTTCCGATCTNNNCAGACTGGACACCTTTCCATCA
AGCACACGTCTGAACTCTTTCCCTACACGACGCTCTTCCGATCTNNNGCGACTTCTATGTAGATGAGG
GCACACGTCTGAACTCTTTCCCTACACGACGCTCTTCCGATCTNNNAGGGATCTTCTCACCAGTGTTTC
GCACACGTCTGAACTCTTTCCCTACACGACGCTCTTCCGATCTNNNGCCTCAATTTCCCCCTCTAGAAAC
AGCACACGTCTGAACTCTTTCCCTACACGACGCTCTTCCGATCTNNNCCACTGGCTCAATAAAGCAGAG
GAGCACACGTCTGAACTCTTTCCCTACACGACGCTCTTCCGATCTNNNCAGGAGAGTGGCTTGAACCT
AGCACACGTCTGAACTCTTTCCCTACACGACGCTCTTCCGATCTNNNGGCGAAACCTCGTCTCTACTAAA
GAGCACACGTCTGAACTCTTTCCCTACACGACGCTCTTCCGATCTNNNGCTTGGAAAAGCAGCAGATGA
GAGCACACGTCTGAACTCTTTCCCTACACGACGCTCTTCCGATCTNNNCTAGTGGTTCACCTGAGGTCA

AGCACACGTCTGAACTCTTTCCCTACACGACGCTCTTCCGATCTNNNTGCCCAGCTAATTTTTGTGTG
AGCACACGTCTGAACTCTTTCCCTACACGACGCTCTTCCGATCTNNNCTGATGGTGAACACACACACA
GAGCACACGTCTGAACTCTTTCCCTACACGACGCTCTTCCGATCTNNNGAGGATTGCTTGAACCTGGG
CACACGTCTGAACTCTTTCCCTACACGACGCTCTTCCGATCTNNNCCGGAAGTCTGGATTTCTGATGT
GAGCACACGTCTGAACTCTTTCCCTACACGACGCTCTTCCGATCTNNNGAGAATCGCTTGAACCCAGG
AGCACACGTCTGAACTCTTTCCCTACACGACGCTCTTCCGATCTNNNCTGTCCAACTGCAATAACCTGC
AGAGCACACGTCTGAACTCTTTCCCTACACGACGCTCTTCCGATCTNNNAGGAGAATCGCTTGAATCT
GCACACGTCTGAACTCTTTCCCTACACGACGCTCTTCCGATCTNNNGTATGGTTTCTGCTGGGTTGTT
GAGCACACGTCTGAACTCTTTCCCTACACGACGCTCTTCCGATCTNNNTGGGTGACAGAGCAAGACT
GCACACGTCTGAACTCTTTCCCTACACGACGCTCTTCCGATCTNNNTATGCTGAGGATGGGGAGGA
GCACACGTCTGAACTCTTTCCCTACACGACGCTCTTCCGATCTNNNGATTGCTTGAGCTTGGGAGATG
CACACGTCTGAACTCTTTCCCTACACGACGCTCTTCCGATCTNNNAGAGAATGTCAGGATTTTGCCCT
AGCACACGTCTGAACTCTTTCCCTACACGACGCTCTTCCGATCTNNNCATGATCTCAGCTCCCTGTGAG
GCACACGTCTGAACTCTTTCCCTACACGACGCTCTTCCGATCTNNNCTGGGCAATAAGAGCGAAACT
GAGCACACGTCTGAACTCTTTCCCTACACGACGCTCTTCCGATCTNNNGAGGCAGGAAAATCGGTTG
GCACACGTCTGAACTCTTTCCCTACACGACGCTCTTCCGATCTNNNACAGAGGATATGAGACTCCCA
CACACGTCTGAACTCTTTCCCTACACGACGCTCTTCCGATCTNNNCAAAACTCCATCACACACACACA
GCACACGTCTGAACTCTTTCCCTACACGACGCTCTTCCGATCTNNNCGTGTGTGTGTGTATATGTGTG
GCACACGTCTGAACTCTTTCCCTACACGACGCTCTTCCGATCTNNNAGTTGTCCCACCTTTCCAGAAG
AGCACACGTCTGAACTCTTTCCCTACACGACGCTCTTCCGATCTNNNATCTGGTCCGTTCTTCACACTC
CACACGTCTGAACTCTTTCCCTACACGACGCTCTTCCGATCTNNNGTAGCTAGTCAGACATGAGCAG
AGCACACGTCTGAACTCTTTCCCTACACGACGCTCTTCCGATCTNNNAAAGTGCTGGGATTACAGGA
GCACACGTCTGAACTCTTTCCCTACACGACGCTCTTCCGATCTNNNCGAGATCACACCACTGTACTCC
CACACGTCTGAACTCTTTCCCTACACGACGCTCTTCCGATCTNNNCTCCTGAGTAGCTGGGATTACAG
AGCACACGTCTGAACTCTTTCCCTACACGACGCTCTTCCGATCTNNNCTCCTGAGTAGCTGGGATTAC
GAGCACACGTCTGAACTCTTTCCCTACACGACGCTCTTCCGATCTNNNAGTGAACTGAGATCGTGCC
CACACGTCTGAACTCTTTCCCTACACGACGCTCTTCCGATCTNNNGAAGGGCTAAGTAAAGAAGGGC
AGCACACGTCTGAACTCTTTCCCTACACGACGCTCTTCCGATCTNNNATGTGAAGATGGAGGGAACA
GAGCACACGTCTGAACTCTTTCCCTACACGACGCTCTTCCGATCTNNNCTGGGTGTCAAGAGTGAGA
CACACGTCTGAACTCTTTCCCTACACGACGCTCTTCCGATCTNNNTAAGTCTGGTCCTCCTGTGAGAT
GCACACGTCTGAACTCTTTCCCTACACGACGCTCTTCCGATCTNNNGAGGTTGCAGTGAGCTGAGAT
GCACACGTCTGAACTCTTTCCCTACACGACGCTCTTCCGATCTNNNCACCTGGGTGTTCCGAAAAGGA
CACACGTCTGAACTCTTTCCCTACACGACGCTCTTCCGATCTNNNTCTCTCTCCCTCTCTCTCTCTCATC
AGCACACGTCTGAACTCTTTCCCTACACGACGCTCTTCCGATCTNNNGCCATCTTGAGTGGTACCCAA
GCACACGTCTGAACTCTTTCCCTACACGACGCTCTTCCGATCTNNNGAACTTGCAGTGAGCCGAGATC
AGCACACGTCTGAACTCTTTCCCTACACGACGCTCTTCCGATCTNNNGACGGACAGAGACAGAAAG
AGCACACGTCTGAACTCTTTCCCTACACGACGCTCTTCCGATCTNNNCTTCAACAGAGGACAGCTGTC
AGCACACGTCTGAACTCTTTCCCTACACGACGCTCTTCCGATCTNNNGAAAAATTGGTGCCCTGTGTC
AGCACACGTCTGAACTCTTTCCCTACACGACGCTCTTCCGATCTNNNGGACACGAGAAAGGTCATTT
AGCACACGTCTGAACTCTTTCCCTACACGACGCTCTTCCGATCTNNNGCCTAGATGTGCTTGTCATGA
CACACGTCTGAACTCTTTCCCTACACGACGCTCTTCCGATCTNNNGGTTTGATCCCTCCGTAACTCTCC
AGCACACGTCTGAACTCTTTCCCTACACGACGCTCTTCCGATCTNNNGGACCCATCAGCTCTTTCATA
CACACGTCTGAACTCTTTCCCTACACGACGCTCTTCCGATCTNNNACTCTGAAGAGCTAGAATCCCCT

GAGCACACGTCTGAACTCTTTCCCTACACGACGCTCTTCCGATCTNNNGCTGAGTCTGTAGTGTGTGCA
AGCACACGTCTGAACTCTTTCCCTACACGACGCTCTTCCGATCTNNNATGTAACAGCCCTGAGGAACAT
GCACACGTCTGAACTCTTTCCCTACACGACGCTCTTCCGATCTNNNGTTGCAGTGAACCGAGATCATG
CACACGTCTGAACTCTTTCCCTACACGACGCTCTTCCGATCTNNNGAGGTTGCAGTGAGCTGAGATCC
AGCACACGTCTGAACTCTTTCCCTACACGACGCTCTTCCGATCTNNNCAATCTTGGCTCACTGCAATCTC
GCACACGTCTGAACTCTTTCCCTACACGACGCTCTTCCGATCTNNNAGTGAGCTGAGATCATGCCATTG
GCACACGTCTGAACTCTTTCCCTACACGACGCTCTTCCGATCTNNNGGCTCACTCCTGTAATATCAGCA
GAGCACACGTCTGAACTCTTTCCCTACACGACGCTCTTCCGATCTNNNGAGAATGGCATGAACCTGGG
AGAGCACACGTCTGAACTCTTTCCCTACACGACGCTCTTCCGATCTNNNGTGAGCTGAGATTTTGCCAC
AGCACACGTCTGAACTCTTTCCCTACACGACGCTCTTCCGATCTNNNACAGCAATGGGAAACTGATACA
AGAGCACACGTCTGAACTCTTTCCCTACACGACGCTCTTCCGATCTNNNTGACAACAGCGAGACTCATC
GAGCACACGTCTGAACTCTTTCCCTACACGACGCTCTTCCGATCTNNNGAGAATCGCTTGAACTCGGG
GAGCACACGTCTGAACTCTTTCCCTACACGACGCTCTTCCGATCTNNNCATCTGTGGACCCAGCTACTT
AGCACACGTCTGAACTCTTTCCCTACACGACGCTCTTCCGATCTNNNGCAGGAGAATTGCTTGAACCT
GCACACGTCTGAACTCTTTCCCTACACGACGCTCTTCCGATCTNNNTCTTTCCTCTGACTTTCTCAGGCC
GCACACGTCTGAACTCTTTCCCTACACGACGCTCTTCCGATCTNNNGGGTTCTCATTTCTCCACATCCTC
AGCACACGTCTGAACTCTTTCCCTACACGACGCTCTTCCGATCTNNNATATGTGTGTGTGTGTGTGTG
AGCACACGTCTGAACTCTTTCCCTACACGACGCTCTTCCGATCTNNNGGGCAACAAAGCAAGACTTCAT
GAGCACACGTCTGAACTCTTTCCCTACACGACGCTCTTCCGATCTNNNCGTGGTGGACCTCATATTGAA
GAGCACACGTCTGAACTCTTTCCCTACACGACGCTCTTCCGATCTNNNCAGTTAGCCATCATCGCAACA
GAGCACACGTCTGAACTCTTTCCCTACACGACGCTCTTCCGATCTNNNACACACACACACACACTCTCAG
GAGCACACGTCTGAACTCTTTCCCTACACGACGCTCTTCCGATCTNNNGAGTAGCTGGGATTACAGGC
GCACACGTCTGAACTCTTTCCCTACACGACGCTCTTCCGATCTNNNCATGTTGAAGGTTGTGAAGCCT
AGCACACGTCTGAACTCTTTCCCTACACGACGCTCTTCCGATCTNNNCATGCCTCAACCTCCCAAGTAGC
GAGCACACGTCTGAACTCTTTCCCTACACGACGCTCTTCCGATCTNNNGAGAATCGCTTGAACCTGGGA
GAGCACACGTCTGAACTCTTTCCCTACACGACGCTCTTCCGATCTNNNGGCGAAACCCTGTCTCTACTAA
GAGCACACGTCTGAACTCTTTCCCTACACGACGCTCTTCCGATCTNNNGAGGTTGCAGTGAGCCAAGA
AGCACACGTCTGAACTCTTTCCCTACACGACGCTCTTCCGATCTNNNATCATGAGCTCACTGAGGGTTG
GAGCACACGTCTGAACTCTTTCCCTACACGACGCTCTTCCGATCTNNNATACTGCACACACATTGCACA
AGCACACGTCTGAACTCTTTCCCTACACGACGCTCTTCCGATCTNNNGTTTCACCCCTATCTGCCTAGGC
GCACACGTCTGAACTCTTTCCCTACACGACGCTCTTCCGATCTNNNTTCAGGAGTTCGAAACCAGTCTG
AGCACACGTCTGAACTCTTTCCCTACACGACGCTCTTCCGATCTNNNCACACACGGAGGATATGAGACT
GAGCACACGTCTGAACTCTTTCCCTACACGACGCTCTTCCGATCTNNNACTATCTCAGCTCACTGCAAGC
AGCACACGTCTGAACTCTTTCCCTACACGACGCTCTTCCGATCTNNNGCTTGCAGTGAGCTGAAATTGT
GAGCACACGTCTGAACTCTTTCCCTACACGACGCTCTTCCGATCTNNNATGAGCTGATAGCCTGAACTCA
GCACACGTCTGAACTCTTTCCCTACACGACGCTCTTCCGATCTNNNATTCTCCCCACTCTCATCTATCTCA
GCACACGTCTGAACTCTTTCCCTACACGACGCTCTTCCGATCTNNNTACATTCATTTCTGGACGCATCCC
GAGCACACGTCTGAACTCTTTCCCTACACGACGCTCTTCCGATCTNNNATCATGAGGTCAGGAGTTTGA
GCACACGTCTGAACTCTTTCCCTACACGACGCTCTTCCGATCTNNNGTTCATTGCTAACAGGGTGTCAC
AGCACACGTCTGAACTCTTTCCCTACACGACGCTCTTCCGATCTNNNCCCTGCCTCAATACACTGTACTT
GAGCACACGTCTGAACTCTTTCCCTACACGACGCTCTTCCGATCTNNNGCCTAGGTACAATCCACAGGA
GCACACGTCTGAACTCTTTCCCTACACGACGCTCTTCCGATCTNNNCCTCCCCAAGTGCTAGAATTACACA
AGCACACGTCTGAACTCTTTCCCTACACGACGCTCTTCCGATCTNNNCTGAGCAACATAGTGAGACTCCA

AGCACACGTCTGAACTCTTTCCCTACACGACGCTCTTCCGATCTNNNCTGAGCAACATAGTGAGACTCC
GAGCACACGTCTGAACTCTTTCCCTACACGACGCTCTTCCGATCTNNNTGGGACACAGAATGAGACTC
GCACACGTCTGAACTCTTTCCCTACACGACGCTCTTCCGATCTNNNTACTGCTGCGTACGTAACAAGTA
GAGCACACGTCTGAACTCTTTCCCTACACGACGCTCTTCCGATCTNNNGTTGCACTGAGCCAAGATCAA
AGCACACGTCTGAACTCTTTCCCTACACGACGCTCTTCCGATCTNNNTAATCCCAGCTACTCAAGAGGCT
GAGCACACGTCTGAACTCTTTCCCTACACGACGCTCTTCCGATCTNNNCTAGGCGACAGAGTGAGACTC
AGCACACGTCTGAACTCTTTCCCTACACGACGCTCTTCCGATCTNNNGAATCGCTTGAACCTGGGAGAT
GCACACGTCTGAACTCTTTCCCTACACGACGCTCTTCCGATCTNNNCACCTGTAATCCCAGCTACTCAGC
GCACACGTCTGAACTCTTTCCCTACACGACGCTCTTCCGATCTNNNCAAAATGATGCCAGTGGGATG
GCACACGTCTGAACTCTTTCCCTACACGACGCTCTTCCGATCTNNNTCATGGGTAACACAGAAAGCAGA
GCACACGTCTGAACTCTTTCCCTACACGACGCTCTTCCGATCTNNNGAGTCTCTCTCTCTCCCTCTCTCCA
GCACACGTCTGAACTCTTTCCCTACACGACGCTCTTCCGATCTNNNGGCCTAGCACAGTATCACAAGAA
AGCACACGTCTGAACTCTTTCCCTACACGACGCTCTTCCGATCTNNNCTGTACTTCAGCCTGGAGACAG
GCACACGTCTGAACTCTTTCCCTACACGACGCTCTTCCGATCTNNNGACGGACAGAGACAGAAAGAG
GCACACGTCTGAACTCTTTCCCTACACGACGCTCTTCCGATCTNNNTCACTCCCTCATTTCCATTCTTGACA
GAGCACACGTCTGAACTCTTTCCCTACACGACGCTCTTCCGATCTNNNATTTTGGAGCCAGGTGGTTCT
AGAGCACACGTCTGAACTCTTTCCCTACACGACGCTCTTCCGATCTNNNAGACATTGGACAGTGCTCTC
GCACACGTCTGAACTCTTTCCCTACACGACGCTCTTCCGATCTNNNGGTGTGCCATTTATTGACTCTGG
AGCACACGTCTGAACTCTTTCCCTACACGACGCTCTTCCGATCTNNNGAGCTTGTCTCAAAGAGGAGG
GAGCACACGTCTGAACTCTTTCCCTACACGACGCTCTTCCGATCTNNNCAGTCTTTTCTCCTCCCTCACCC
GAGCACACGTCTGAACTCTTTCCCTACACGACGCTCTTCCGATCTNNNGAGAAGCATTGGAGCAGACT
GCACACGTCTGAACTCTTTCCCTACACGACGCTCTTCCGATCTNNNCTGCAGGATTCTGGAAGCATTTC
AGCACACGTCTGAACTCTTTCCCTACACGACGCTCTTCCGATCTNNNCCATCTGAGGCCTCCATATTCTC
GAGCACACGTCTGAACTCTTTCCCTACACGACGCTCTTCCGATCTNNNACCCCACCAAAACACATATGT
GCACACGTCTGAACTCTTTCCCTACACGACGCTCTTCCGATCTNNNGCCAGAACTTAGTGGGAGAGAG
AGCACACGTCTGAACTCTTTCCCTACACGACGCTCTTCCGATCTNNNGAGTAGCTGGGACTACATGCA
GCACACGTCTGAACTCTTTCCCTACACGACGCTCTTCCGATCTNNNGAGTAGCTGGGATTACAGGCATG
AGAGCACACGTCTGAACTCTTTCCCTACACGACGCTCTTCCGATCTNNNCTGTACTCCCAGCTACTCAGG
GCACACGTCTGAACTCTTTCCCTACACGACGCTCTTCCGATCTNNNGGGCAACAAGAGTAAGACTCCAT
GCACACGTCTGAACTCTTTCCCTACACGACGCTCTTCCGATCTNNNAGGAGACCGGAATAAGACTCTCT
AGCACACGTCTGAACTCTTTCCCTACACGACGCTCTTCCGATCTNNNGTAGCTGGGATTACAGGCACTT
AGCACACGTCTGAACTCTTTCCCTACACGACGCTCTTCCGATCTNNNTGTCTTATCTGCAAGTCTCGACT
GCACACGTCTGAACTCTTTCCCTACACGACGCTCTTCCGATCTNNNAGTCCACCTCTGGTTTTTGGTAAC
GCACACGTCTGAACTCTTTCCCTACACGACGCTCTTCCGATCTNNNAACGAGGACATCCAGTTAGATCG
AGCACACGTCTGAACTCTTTCCCTACACGACGCTCTTCCGATCTNNNCTGGGCAACAAGAGCAAAACTC
AGCACACGTCTGAACTCTTTCCCTACACGACGCTCTTCCGATCTNNNAACTGTGGTTGTAGCTACTGG
GCACACGTCTGAACTCTTTCCCTACACGACGCTCTTCCGATCTNNNAGCAGGGAGTTTTTGGGTAAGA
GCACACGTCTGAACTCTTTCCCTACACGACGCTCTTCCGATCTNNNACTGCTGGTGGGAGTATAAACTC
GCACACGTCTGAACTCTTTCCCTACACGACGCTCTTCCGATCTNNNCTTGCTACAGATGCCTTTTTGCTC
GAGCACACGTCTGAACTCTTTCCCTACACGACGCTCTTCCGATCTNNNCTTGAGCCCAGGAGTTCAAGA
AGCACACGTCTGAACTCTTTCCCTACACGACGCTCTTCCGATCTNNNAAAGTGCTGGGATTACAGGCA
AGCACACGTCTGAACTCTTTCCCTACACGACGCTCTTCCGATCTNNNGTGAGCTGAGATTGTGCCATTG
GCACACGTCTGAACTCTTTCCCTACACGACGCTCTTCCGATCTNNNATTCAAACTAGGTAGCCTGGCTCC

AGCACACGTCTGAACTCTTTCCCTACACGACGCTCTTCCGATCTNNNAATGGCAAAACCCCATCTCTAC
GCACACGTCTGAACTCTTTCCCTACACGACGCTCTTCCGATCTNNNCTGGGCAACAAGAGCAAAACT
AGCACACGTCTGAACTCTTTCCCTACACGACGCTCTTCCGATCTNNNATGCCTGTAATCCCAGCACTTT
GAGCACACGTCTGAACTCTTTCCCTACACGACGCTCTTCCGATCTNNNGAGGTTGCAGTGAGCCAAG
GCACACGTCTGAACTCTTTCCCTACACGACGCTCTTCCGATCTNNNATTTCAATTCCGAATGGTGGTG
GCACACGTCTGAACTCTTTCCCTACACGACGCTCTTCCGATCTNNNGTTTGAGACCAGTCTGTGCAAC
GCACACGTCTGAACTCTTTCCCTACACGACGCTCTTCCGATCTNNNAGTAAGCCGAGATCACACCATG
AGCACACGTCTGAACTCTTTCCCTACACGACGCTCTTCCGATCTNNNGACAGGGATTCATGGTGACCA
GCACACGTCTGAACTCTTTCCCTACACGACGCTCTTCCGATCTNNNATAATCTCAGCTCACTGCAACCT
AGCACACGTCTGAACTCTTTCCCTACACGACGCTCTTCCGATCTNNNCTCAACCTCCCAGACTCAAGTC
AGCACACGTCTGAACTCTTTCCCTACACGACGCTCTTCCGATCTNNNGTAGTCCTAGCTGCTTGGGAA
CACACGTCTGAACTCTTTCCCTACACGACGCTCTTCCGATCTNNNAAAGAGTCACCCATGTTGTTGTG
GCACACGTCTGAACTCTTTCCCTACACGACGCTCTTCCGATCTNNNGGTGTTCGCCTAGCCAAGATAG
CACACGTCTGAACTCTTTCCCTACACGACGCTCTTCCGATCTNNNATGTGAGTTGGGAGGTGATGA
AGCACACGTCTGAACTCTTTCCCTACACGACGCTCTTCCGATCTNNNGAGAATCCCTTGAACCCAGGA
AGCACACGTCTGAACTCTTTCCCTACACGACGCTCTTCCGATCTNNNGCCTCCTGAGTAGTTGGGAC
GCACACGTCTGAACTCTTTCCCTACACGACGCTCTTCCGATCTNNNGTAATTCTGGCACTTTGGAAGG
GCACACGTCTGAACTCTTTCCCTACACGACGCTCTTCCGATCTNNNAAAATTTTTCTCGTGCCCTGTGT
AGCACACGTCTGAACTCTTTCCCTACACGACGCTCTTCCGATCTNNNGACAGCAGGAGACTCCATCAA
GCACACGTCTGAACTCTTTCCCTACACGACGCTCTTCCGATCTNNNTCTCACTTTGATGTCACCGTTCT
AGCACACGTCTGAACTCTTTCCCTACACGACGCTCTTCCGATCTNNNCAGTAAAGTTGCGTGGGACA
CACACGTCTGAACTCTTTCCCTACACGACGCTCTTCCGATCTNNNGGCCTTCTGGTGGAAAGTTTCTA
AGCACACGTCTGAACTCTTTCCCTACACGACGCTCTTCCGATCTNNNCTTGGGTGACAGAGTGAGAT
GCACACGTCTGAACTCTTTCCCTACACGACGCTCTTCCGATCTNNNCCAAAGTGCTGGGATTACAAGC
GCACACGTCTGAACTCTTTCCCTACACGACGCTCTTCCGATCTNNNGTGCTGTCTTGTTTGTCCAGAAA
AGCACACGTCTGAACTCTTTCCCTACACGACGCTCTTCCGATCTNNNCTGATCACGAGGTCAGCAGAT
AGCACACGTCTGAACTCTTTCCCTACACGACGCTCTTCCGATCTNNNTTCTACAAAAAAATTGGCCGG
GAGCACACGTCTGAACTCTTTCCCTACACGACGCTCTTCCGATCTNNNCAGGTTCAGGCAATTCTTCT
GCACACGTCTGAACTCTTTCCCTACACGACGCTCTTCCGATCTNNNGAGGCTTTAGGGATAAGGAGA
CACACGTCTGAACTCTTTCCCTACACGACGCTCTTCCGATCTNNNCCTGCTCTCTGAGAAAAGCTACA
GCACACGTCTGAACTCTTTCCCTACACGACGCTCTTCCGATCTNNNCGCCACCACACTTAGCTATTTTT
AGCACACGTCTGAACTCTTTCCCTACACGACGCTCTTCCGATCTNNNGTGAGCCAAGATCATGCCATT
GAGCACACGTCTGAACTCTTTCCCTACACGACGCTCTTCCGATCTNNNGGTTACAGTGAGCCAAGAT
GAGCACACGTCTGAACTCTTTCCCTACACGACGCTCTTCCGATCTNNNTGAGGCTGTGTCTGAGTGA
AGAGCACACGTCTGAACTCTTTCCCTACACGACGCTCTTCCGATCTNNNCTTGAGGAGAGGGCTGT
CACACGTCTGAACTCTTTCCCTACACGACGCTCTTCCGATCTNNNTCTCTCTCTCTCTCTCTCCCTCAT
GCACACGTCTGAACTCTTTCCCTACACGACGCTCTTCCGATCTNNNGAGAAACAGAACCAATATGCG
AGCACACGTCTGAACTCTTTCCCTACACGACGCTCTTCCGATCTNNNACCTGTAGTCCCAGCTACTCTG
AGCACACGTCTGAACTCTTTCCCTACACGACGCTCTTCCGATCTNNNGAGATCACACCACTCACTCCA
AGCACACGTCTGAACTCTTTCCCTACACGACGCTCTTCCGATCTNNNGAGATCACGCCACTGTACTAC
GCACACGTCTGAACTCTTTCCCTACACGACGCTCTTCCGATCTNNNTCTCTCCCTTCCTTCCTTTCTCA
GAGCACACGTCTGAACTCTTTCCCTACACGACGCTCTTCCGATCTNNNCATGAGAGTGAGGACCAGC
GCACACGTCTGAACTCTTTCCCTACACGACGCTCTTCCGATCTNNNGACGGACAGAGACAGAAAGAG

AGCACACGTCTGAACTCTTTCCCTACACGACGCTCTTCCGATCTNNNGACGGACAGAGACAGAAAGAG
AGCACACGTCTGAACTCTTTCCCTACACGACGCTCTTCCGATCTNNNGTGTGTGTGTGAGAGAGAGAG
AGCACACGTCTGAACTCTTTCCCTACACGACGCTCTTCCGATCTNNNCAGAACTGGTCTCAGGGAAGT
AGCACACGTCTGAACTCTTTCCCTACACGACGCTCTTCCGATCTNNNGTTGCAGTGAGCCAAGATCAT
AGCACACGTCTGAACTCTTTCCCTACACGACGCTCTTCCGATCTNNNGACTCGGTTCACATCCCAAGAT
GCACACGTCTGAACTCTTTCCCTACACGACGCTCTTCCGATCTNNNGAGAGAGAGATGTGTGTGTG
AAGAGCACACGTCTGAACTCTTTCCCTACACGACGCTCTTCCGATCTNNNAAGAATCGCTTGAACCTGG
AGCACACGTCTGAACTCTTTCCCTACACGACGCTCTTCCGATCTNNNAGCTACTCAGGAGGCTAAGGT
AGCACACGTCTGAACTCTTTCCCTACACGACGCTCTTCCGATCTNNNGACGGACAGAGACAGAAAGAG
AGCACACGTCTGAACTCTTTCCCTACACGACGCTCTTCCGATCTNNNGACGGACAGAGACAGAAAGAG
AGCACACGTCTGAACTCTTTCCCTACACGACGCTCTTCCGATCTNNNGAGGTTGCAGTGAGTGGAGAT
GAGCACACGTCTGAACTCTTTCCCTACACGACGCTCTTCCGATCTNNNGTGGTTGTATCACTCCACCCTC
GCACACGTCTGAACTCTTTCCCTACACGACGCTCTTCCGATCTNNNCATCTTATAGAACACGGGGCGG
GCACACGTCTGAACTCTTTCCCTACACGACGCTCTTCCGATCTNNNAGTTTAGGCTACCACACCATGAG
AGCACACGTCTGAACTCTTTCCCTACACGACGCTCTTCCGATCTNNNTCTCTCTCTCTCAGGTATGCA
AGCACACGTCTGAACTCTTTCCCTACACGACGCTCTTCCGATCTNNNCTTGAGGCCAGGAGTTTGAGA
AGAGCACACGTCTGAACTCTTTCCCTACACGACGCTCTTCCGATCTNNNGTTGCAGTGAGTCGAGATTG
AGCACACGTCTGAACTCTTTCCCTACACGACGCTCTTCCGATCTNNNCTTCCCTTCCTTTTCTCCCTCCCAT
CACACGTCTGAACTCTTTCCCTACACGACGCTCTTCCGATCTNNNAGATGATTTGGGGAAACTCTGCTCA
GAGCACACGTCTGAACTCTTTCCCTACACGACGCTCTTCCGATCTNNNCTGGGCAACAAGAGCAAAACT
AGCACACGTCTGAACTCTTTCCCTACACGACGCTCTTCCGATCTNNNCATTCCAGCCTAGGTGACAGAG
GAGCACACGTCTGAACTCTTTCCCTACACGACGCTCTTCCGATCTNNNGAGGAGGAGAGAGAAGGAG
AGCACACGTCTGAACTCTTTCCCTACACGACGCTCTTCCGATCTNNNGACGGACAGAGACAGAAAGAG
AGCACACGTCTGAACTCTTTCCCTACACGACGCTCTTCCGATCTNNNGACGGACAGAGACAGAAAGAG
GAGCACACGTCTGAACTCTTTCCCTACACGACGCTCTTCCGATCTNNNGGTCTCTGTGAAGTCCCTTCTG
GAGCACACGTCTGAACTCTTTCCCTACACGACGCTCTTCCGATCTNNNTCACTCCCTTTCTCTAGCGAGA
GCACACGTCTGAACTCTTTCCCTACACGACGCTCTTCCGATCTNNNGAAAAAGAAAGGTCACTCCCTGC
AGCACACGTCTGAACTCTTTCCCTACACGACGCTCTTCCGATCTNNNGGGAGGCTGAGGAACAAGAAT
GAGCACACGTCTGAACTCTTTCCCTACACGACGCTCTTCCGATCTNNNGGCTTGAACTCAGAGAGCGA
CACACGTCTGAACTCTTTCCCTACACGACGCTCTTCCGATCTNNNGCTTTGGGTCTGTTGAACTCTTCCA
AGCACACGTCTGAACTCTTTCCCTACACGACGCTCTTCCGATCTNNNGGCTCTTTTCTCTCTCTCCCAAA
AGCACACGTCTGAACTCTTTCCCTACACGACGCTCTTCCGATCTNNNGGAGAGACCGTGCTAAAGCTAA
GCACACGTCTGAACTCTTTCCCTACACGACGCTCTTCCGATCTNNNTGCAATCTTCCTTTTGTCATCATGT
GCACACGTCTGAACTCTTTCCCTACACGACGCTCTTCCGATCTNNNTCCAAAGTGCTCGGATTACAGG
GAGCACACGTCTGAACTCTTTCCCTACACGACGCTCTTCCGATCTNNNCTGGGTTTCACTGTACAGGGA
GAGCACACGTCTGAACTCTTTCCCTACACGACGCTCTTCCGATCTNNNGAGGCAGAAGAATCGCTTGA
AGCACACGTCTGAACTCTTTCCCTACACGACGCTCTTCCGATCTNNNCTCAATGCTTGTGGGCACTATC
CACACGTCTGAACTCTTTCCCTACACGACGCTCTTCCGATCTNNNTCACAACTAGACAGTGCTCCAAACA
AGAGCACACGTCTGAACTCTTTCCCTACACGACGCTCTTCCGATCTNNNGAGAATCACTCGAACCTGGG
AGCACACGTCTGAACTCTTTCCCTACACGACGCTCTTCCGATCTNNNGACGGACAGAGACAGAAAGAG
AGCACACGTCTGAACTCTTTCCCTACACGACGCTCTTCCGATCTNNNGACGGACAGAGACAGAAAGAG
AGCACACGTCTGAACTCTTTCCCTACACGACGCTCTTCCGATCTNNNGACGGACAGAGACAGAAAGAG
GAGCACACGTCTGAACTCTTTCCCTACACGACGCTCTTCCGATCTNNNCTTTTCTGTGTCTCCAGCTTGC

AGCACACGTCTGAACTCTTTCCCTACACGACGCTCTTCCGATCTNNNCAGCTCCAGGAACCTTAGAACA
AGCACACGTCTGAACTCTTTCCCTACACGACGCTCTTCCGATCTNNNGAAGCAGGAGAATCGCTTGA
GCACACGTCTGAACTCTTTCCCTACACGACGCTCTTCCGATCTNNNCTTTTCCCTGTGGCTTTGGAAA
GCACACGTCTGAACTCTTTCCCTACACGACGCTCTTCCGATCTNNNCCCAGGTTCAAGCAATTCTCCTA
AGCACACGTCTGAACTCTTTCCCTACACGACGCTCTTCCGATCTNNNTAGAACATTCCCACTGCAGACA
GAGCACACGTCTGAACTCTTTCCCTACACGACGCTCTTCCGATCTNNNCTACGGAAGGAAGTGCAGA
GCACACGTCTGAACTCTTTCCCTACACGACGCTCTTCCGATCTNNNGATGCAAATGGACCACTCAAG
GCACACGTCTGAACTCTTTCCCTACACGACGCTCTTCCGATCTNNNAACGTGACGCTCTTCCTTACTGC
AGCACACGTCTGAACTCTTTCCCTACACGACGCTCTTCCGATCTNNNCATCTTTCAAACCCACCGAGTC
GCACACGTCTGAACTCTTTCCCTACACGACGCTCTTCCGATCTNNNCAACAGCAACCAACAGTCACTA
GCACACGTCTGAACTCTTTCCCTACACGACGCTCTTCCGATCTNNNAGGTCACAGATGTGGAAACTGA
AGCACACGTCTGAACTCTTTCCCTACACGACGCTCTTCCGATCTNNNCCCTGATTGTGCTAGGACAAG
GCACACGTCTGAACTCTTTCCCTACACGACGCTCTTCCGATCTNNNAGTTACGTGTGTTTACTGGAGC
GCACACGTCTGAACTCTTTCCCTACACGACGCTCTTCCGATCTNNNCATGGAGGTATTACAAGTGCCC
AGCACACGTCTGAACTCTTTCCCTACACGACGCTCTTCCGATCTNNNTGATGTGCTGGCTTCAGATCT
ACACGTCTGAACTCTTTCCCTACACGACGCTCTTCCGATCTNNNTTCTTCCACATCTCTCTCTCCCTCATC
GCACACGTCTGAACTCTTTCCCTACACGACGCTCTTCCGATCTNNNCCTCCTTCCTTCCTTCCTTCTTTCAT
GCACACGTCTGAACTCTTTCCCTACACGACGCTCTTCCGATCTNNNGCTTGCAGTGAGCTGAGATAGT
GCACACGTCTGAACTCTTTCCCTACACGACGCTCTTCCGATCTNNNCATACACACACACACACACACACC
AAGAGCACACGTCTGAACTCTTTCCCTACACGACGCTCTTCCGATCTNNNCACGCCCACCTGATAGTAT
AGCACACGTCTGAACTCTTTCCCTACACGACGCTCTTCCGATCTNNNAAGTCTATGCTAGCACAGTGG
GCACACGTCTGAACTCTTTCCCTACACGACGCTCTTCCGATCTNNNGACGGACAGAGACAGAAAGAG
AGAGCACACGTCTGAACTCTTTCCCTACACGACGCTCTTCCGATCTNNNAGCAGGTTGAGGTTACAG
GAGCACACGTCTGAACTCTTTCCCTACACGACGCTCTTCCGATCTNNNTCTGTAATCCCAGGTACTCAG
AGCACACGTCTGAACTCTTTCCCTACACGACGCTCTTCCGATCTNNNTGAAGAATGTCAACAAACAGG
GAGCACACGTCTGAACTCTTTCCCTACACGACGCTCTTCCGATCTNNNCTATGACAGTGTGTGTGTGT
GAGCACACGTCTGAACTCTTTCCCTACACGACGCTCTTCCGATCTNNNGAATGCTTCTTTCGGCACAAG
AGCACACGTCTGAACTCTTTCCCTACACGACGCTCTTCCGATCTNNNTTCCCCCACATGTTCAGAACTTC
GCACACGTCTGAACTCTTTCCCTACACGACGCTCTTCCGATCTNNNGTAGTGGAGAGCAAGGTCAGT
CACACGTCTGAACTCTTTCCCTACACGACGCTCTTCCGATCTNNNCCCATATACTAGCCAGAGGTCTGC
GCACACGTCTGAACTCTTTCCCTACACGACGCTCTTCCGATCTNNNCTCAAGAGATCCACCTACCTCAGC
CACACGTCTGAACTCTTTCCCTACACGACGCTCTTCCGATCTNNNCAGATGTTGGGAAGGTCTGAGA
AGCACACGTCTGAACTCTTTCCCTACACGACGCTCTTCCGATCTNNNGATGGGAAAGGCAAGAACAC
GAGCACACGTCTGAACTCTTTCCCTACACGACGCTCTTCCGATCTNNNGATGCAGGAAACTCACTGAG
AGCACACGTCTGAACTCTTTCCCTACACGACGCTCTTCCGATCTNNNTGGGTGACAGACTGAGATTCT
GAGCACACGTCTGAACTCTTTCCCTACACGACGCTCTTCCGATCTNNNGGTAGCGAGGAGTTGATACA
GCACACGTCTGAACTCTTTCCCTACACGACGCTCTTCCGATCTNNNGCTCTCAGTGCTCACTATCCAGC
AGCACACGTCTGAACTCTTTCCCTACACGACGCTCTTCCGATCTNNNCTTGAGGTGGACGTCAGTTTCT
GCACACGTCTGAACTCTTTCCCTACACGACGCTCTTCCGATCTNNNCGATAATGGGGGAGAGGGAAA
GCACACGTCTGAACTCTTTCCCTACACGACGCTCTTCCGATCTNNNCCTCTCTCCCTCTCTCTCTCTCTCA
AGCACACGTCTGAACTCTTTCCCTACACGACGCTCTTCCGATCTNNNGATCACTTGAGCCCAGGAAGT
GCACACGTCTGAACTCTTTCCCTACACGACGCTCTTCCGATCTNNNCTGAGATCACACCACTGCACTCC
GCACACGTCTGAACTCTTTCCCTACACGACGCTCTTCCGATCTNNNATCCTGGAGGTATATGTGTGTG

GCACACGTCTGAACTCTTTCCCTACACGACGCTCTTCCGATCTNNNTCTTTTGTGTGTGTGTGTGTGTG
AGCACACGTCTGAACTCTTTCCCTACACGACGCTCTTCCGATCTNNNGACATTCACTGCAATGCACACTT
AGCACACGTCTGAACTCTTTCCCTACACGACGCTCTTCCGATCTNNNGACGGACAGAGACAGAAAGAG
AGCACACGTCTGAACTCTTTCCCTACACGACGCTCTTCCGATCTNNNGAAACAGAGGCAGAGAGAGA
AGCACACGTCTGAACTCTTTCCCTACACGACGCTCTTCCGATCTNNNGTACTGGTTGCTCTCTAGGTCTG
AGCACACGTCTGAACTCTTTCCCTACACGACGCTCTTCCGATCTNNNGACCTTGTCTGATACCTGTCCTG
GCACACGTCTGAACTCTTTCCCTACACGACGCTCTTCCGATCTNNNTTACAGGCACAAACCACTGCATCA
GCACACGTCTGAACTCTTTCCCTACACGACGCTCTTCCGATCTNNNGAAATGAGGTTCCACGTTGAAG
GCACACGTCTGAACTCTTTCCCTACACGACGCTCTTCCGATCTNNNGGAACATGAAGGTGGACTAGGA
AGCACACGTCTGAACTCTTTCCCTACACGACGCTCTTCCGATCTNNNCACTGGTATCCCATGTACCCTGCA
CACACGTCTGAACTCTTTCCCTACACGACGCTCTTCCGATCTNNNGGGGAAAGCAGACAAAAAGAA
AGCACACGTCTGAACTCTTTCCCTACACGACGCTCTTCCGATCTNNNGCAACATAAGGAGACCCCATCT
GAGCACACGTCTGAACTCTTTCCCTACACGACGCTCTTCCGATCTNNNACTGTGGTGAGGAAATGGAA
GAGCACACGTCTGAACTCTTTCCCTACACGACGCTCTTCCGATCTNNNGCTCCCTCCAGTCTCTGTTATG
GAGCACACGTCTGAACTCTTTCCCTACACGACGCTCTTCCGATCTNNNCACCTGTAGTTCCAGCTACTCG
AGCACACGTCTGAACTCTTTCCCTACACGACGCTCTTCCGATCTNNNGAAAGAGAGGGAGAGGGGAG
GCACACGTCTGAACTCTTTCCCTACACGACGCTCTTCCGATCTNNNGAGGATCACTTGAGCTCAGAAGT
AGAGCACACGTCTGAACTCTTTCCCTACACGACGCTCTTCCGATCTNNNCTACAGTGAGCCGAGATAGT
GCACACGTCTGAACTCTTTCCCTACACGACGCTCTTCCGATCTNNNTCTTTCTTTCTCTCTCCCTCCCTCAT
GCACACGTCTGAACTCTTTCCCTACACGACGCTCTTCCGATCTNNNTCCCTCTCTCTCTCTCTCTCTCAT
GCACACGTCTGAACTCTTTCCCTACACGACGCTCTTCCGATCTNNNTTTGTAAGTGGCATAGTCTCGGT
GAGCACACGTCTGAACTCTTTCCCTACACGACGCTCTTCCGATCTNNNTGGAGGAAGTCAGTGTTGCT
AGCACACGTCTGAACTCTTTCCCTACACGACGCTCTTCCGATCTNNNTTCGTAATGCTGTTCTTGCTGTG
GAGCACACGTCTGAACTCTTTCCCTACACGACGCTCTTCCGATCTNNNCATTGCACTCCAGTCTCAGTGA
GCACACGTCTGAACTCTTTCCCTACACGACGCTCTTCCGATCTNNNGGTTCAAAATGGCTGCCAAGATCA
GCACACGTCTGAACTCTTTCCCTACACGACGCTCTTCCGATCTNNNGAAGATGTGCAGTCCAAGGCTAA
GAGCACACGTCTGAACTCTTTCCCTACACGACGCTCTTCCGATCTNNNGCCCTACTTCTTGGCTCCTCAC
GCACACGTCTGAACTCTTTCCCTACACGACGCTCTTCCGATCTNNNTTATCCCAGAATCAGCTGGCAAC
GCACACGTCTGAACTCTTTCCCTACACGACGCTCTTCCGATCTNNNCATAGAGGGAACCCCTACCACTG
CACACGTCTGAACTCTTTCCCTACACGACGCTCTTCCGATCTNNNACAGTTTACCAGCAGTCTAAGCGCA
CACACGTCTGAACTCTTTCCCTACACGACGCTCTTCCGATCTNNNACTTATACTTCCACCTGGGGCTTCA
GCACACGTCTGAACTCTTTCCCTACACGACGCTCTTCCGATCTNNNGTAACACAAAGCAGGGCTGAAA
CACACGTCTGAACTCTTTCCCTACACGACGCTCTTCCGATCTNNNCTTGCCCAAGGTCATAAAAGGGTCA
AGCACACGTCTGAACTCTTTCCCTACACGACGCTCTTCCGATCTNNNAATCAGCAAACACAGATGACCCA
GCACACGTCTGAACTCTTTCCCTACACGACGCTCTTCCGATCTNNNCAGCACTTCATACCCATGGTCTCCA
CACACGTCTGAACTCTTTCCCTACACGACGCTCTTCCGATCTNNNCTGGGTACCCTGTCTTCCTCCATCA
AGCACACGTCTGAACTCTTTCCCTACACGACGCTCTTCCGATCTNNNCCAGGTGGGCCAAATAAACAGT
CACACGTCTGAACTCTTTCCCTACACGACGCTCTTCCGATCTNNNATATTTCTCCTAGTGCCACAGCCCAT
GCACACGTCTGAACTCTTTCCCTACACGACGCTCTTCCGATCTNNNGTCTGTGTCTCTCAGTTGTGCATC
CACACGTCTGAACTCTTTCCCTACACGACGCTCTTCCGATCTNNNCCGTGCCTAGCCAGAAATTGTATCA
AGAGCACACGTCTGAACTCTTTCCCTACACGACGCTCTTCCGATCTNNNCTTCCTTTCCCCTCCCGCTTAC
GCACACGTCTGAACTCTTTCCCTACACGACGCTCTTCCGATCTNNNTGGTCTCGTTGGCTTAGTAGTGA
GCACACGTCTGAACTCTTTCCCTACACGACGCTCTTCCGATCTNNNTCTCCCCGTTTTCTCTTTTGGTGCA

ACACGTCTGAACTCTTTCCCTACACGACGCTCTTCCGATCTNNNGTTTTCCTCCTGGCAGGTAGATGC
CACACGTCTGAACTCTTTCCCTACACGACGCTCTTCCGATCTNNNAATGTCCATTTCCTCTTCCCCACCA
GCACACGTCTGAACTCTTTCCCTACACGACGCTCTTCCGATCTNNNAATAACAAACGGGAGGAGGAA
CACACGTCTGAACTCTTTCCCTACACGACGCTCTTCCGATCTNNNTTATTATAGCTCCCCCTCCCCCTCAT
GCACACGTCTGAACTCTTTCCCTACACGACGCTCTTCCGATCTNNNGCTGCTTCAGCACATCATGGAG
GCACACGTCTGAACTCTTTCCCTACACGACGCTCTTCCGATCTNNNAGGAGAGGGAGAAAGGGAAG
AGCACACGTCTGAACTCTTTCCCTACACGACGCTCTTCCGATCTNNNGATAGCTAGTCCCGGCGTGCTA
GCACACGTCTGAACTCTTTCCCTACACGACGCTCTTCCGATCTNNNCAGCCCCACATCAGTGACATCCA
CACACGTCTGAACTCTTTCCCTACACGACGCTCTTCCGATCTNNNTTCTTCTCCCTAGTGGATTTGCGC
CACACGTCTGAACTCTTTCCCTACACGACGCTCTTCCGATCTNNNCATCAGTGTGAACCAGATGGACC
CACACGTCTGAACTCTTTCCCTACACGACGCTCTTCCGATCTNNNCCAGATTGAGGGAAGCTTCTTGG
GCACACGTCTGAACTCTTTCCCTACACGACGCTCTTCCGATCTNNNCAGAGACATTCCAGGGGTCCAG
CACACGTCTGAACTCTTTCCCTACACGACGCTCTTCCGATCTNNNTTAGCAAAAGCCCCAGAGATTCC
GCACACGTCTGAACTCTTTCCCTACACGACGCTCTTCCGATCTNNNATAAGCAAGGGCCCCAATTTCT
GCACACGTCTGAACTCTTTCCCTACACGACGCTCTTCCGATCTNNNGTGCTTATGGCAGCAGAGAGG
CACACGTCTGAACTCTTTCCCTACACGACGCTCTTCCGATCTNNNTCAAGAGGTGGTTTTGTGTTGCT
GCACACGTCTGAACTCTTTCCCTACACGACGCTCTTCCGATCTNNNCCCAGCAGCTTATGTGAATGTC
GCACACGTCTGAACTCTTTCCCTACACGACGCTCTTCCGATCTNNNATCAACCCCCAGCATTCTGATTCC
CACACGTCTGAACTCTTTCCCTACACGACGCTCTTCCGATCTNNNATGTCATAGCTCAAGTGGCAAGG
CACACGTCTGAACTCTTTCCCTACACGACGCTCTTCCGATCTNNNGGAGGAAGGGAGAGAAGATCTG
ACACGTCTGAACTCTTTCCCTACACGACGCTCTTCCGATCTNNNAGAAAACAGTGAAGCCATGGTAGA
GCACACGTCTGAACTCTTTCCCTACACGACGCTCTTCCGATCTNNNGAACATGTGTGGCCAGAACTAC
ACACGTCTGAACTCTTTCCCTACACGACGCTCTTCCGATCTNNNAGCTTTGAACTCCTAGGCTCTCTCA
GCACACGTCTGAACTCTTTCCCTACACGACGCTCTTCCGATCTNNNCCAGCCAGGTAAGGGGTGTAT
CACACGTCTGAACTCTTTCCCTACACGACGCTCTTCCGATCTNNNCGCTGTTGCTCTCTCTTTCTGTTC
CACACGTCTGAACTCTTTCCCTACACGACGCTCTTCCGATCTNNNAAGCTAGAACTTGAACTTGCCCC
GCACACGTCTGAACTCTTTCCCTACACGACGCTCTTCCGATCTNNNCTCCCCGCTCTAAGTGATTCTCC
CACACGTCTGAACTCTTTCCCTACACGACGCTCTTCCGATCTNNNAAACCATGAAACCTCCATGTCCC
AGCACACGTCTGAACTCTTTCCCTACACGACGCTCTTCCGATCTNNNAGGATTTAGGGATGGAGGTT
AGCACACGTCTGAACTCTTTCCCTACACGACGCTCTTCCGATCTNNNAGGCAAGTAGCCTGTTTTTGC
ACACGTCTGAACTCTTTCCCTACACGACGCTCTTCCGATCTNNNAATACCTCATCACAGCCTCAACCCAT
CACACGTCTGAACTCTTTCCCTACACGACGCTCTTCCGATCTNNNAGACCATGGATAAATGGCAACCG
CACACGTCTGAACTCTTTCCCTACACGACGCTCTTCCGATCTNNNGTTCATGCGATTCTCCTGGTTCTC
CACACGTCTGAACTCTTTCCCTACACGACGCTCTTCCGATCTNNNCAAAGCAGCTCTGTTTCCCTACTC
GCACACGTCTGAACTCTTTCCCTACACGACGCTCTTCCGATCTNNNAGCCACACGCAAAATGAAACAA
CACACGTCTGAACTCTTTCCCTACACGACGCTCTTCCGATCTNNNAAATCACGTGTCAGTCTCCTTCCC
GCACACGTCTGAACTCTTTCCCTACACGACGCTCTTCCGATCTNNNTTTGTTACCCCAACTCAAGAGG
ACACGTCTGAACTCTTTCCCTACACGACGCTCTTCCGATCTNNNTGGCTTTGGAAGTCAGAAGGAAG
CACACGTCTGAACTCTTTCCCTACACGACGCTCTTCCGATCTNNNGAAATCTGGCATAGGAGCCTGA
CACACGTCTGAACTCTTTCCCTACACGACGCTCTTCCGATCTNNNTCCATTGCTACATGTGCCTTTTTC
CACACGTCTGAACTCTTTCCCTACACGACGCTCTTCCGATCTNNNACCATATGAGCTTGTGTGTGCATC
CACACGTCTGAACTCTTTCCCTACACGACGCTCTTCCGATCTNNNGTAGCTTCAGGTTTGACCGCTAA
CACACGTCTGAACTCTTTCCCTACACGACGCTCTTCCGATCTNNNGCCACCGCACATAAATGTTCAAGC

CACACGTCTGAACTCTTTCCCTACACGACGCTCTTCCGATCTNNNGTCTTCCCAACTGCTCTTCTCTGC
GCACACGTCTGAACTCTTTCCCTACACGACGCTCTTCCGATCTNNNCCTGGTATCCATCTGGGTGACA
AGCACACGTCTGAACTCTTTCCCTACACGACGCTCTTCCGATCTNNNATGTTGGGTGAAAGCTCCTTG
CACACGTCTGAACTCTTTCCCTACACGACGCTCTTCCGATCTNNNGGATGACGAACTTGGCATTCTCT
GCACACGTCTGAACTCTTTCCCTACACGACGCTCTTCCGATCTNNNTAGTCAAGTGGTGTTGAGAGA
CACACGTCTGAACTCTTTCCCTACACGACGCTCTTCCGATCTNNNGTCAAGCACCAAATGGAAACCAAC
AGCACACGTCTGAACTCTTTCCCTACACGACGCTCTTCCGATCTNNNACTCCTTCATTGCACACATGGTT
CACACGTCTGAACTCTTTCCCTACACGACGCTCTTCCGATCTNNNTTTAGGGCCACTTACTGGCTTCC
AGCACACGTCTGAACTCTTTCCCTACACGACGCTCTTCCGATCTNNNAAGAGAGAAGGGGATGTGTG
CACACGTCTGAACTCTTTCCCTACACGACGCTCTTCCGATCTNNNCACTGTATCCCAACTGCACATCAC
CACACGTCTGAACTCTTTCCCTACACGACGCTCTTCCGATCTNNNTGGATGGGGCAAATAATCACAT
CACACGTCTGAACTCTTTCCCTACACGACGCTCTTCCGATCTNNNCACTTGTCAGTTCCTGCACCATAC
AGCACACGTCTGAACTCTTTCCCTACACGACGCTCTTCCGATCTNNNGAATGAGCAAAAACAGCCAGA
CACACGTCTGAACTCTTTCCCTACACGACGCTCTTCCGATCTNNNGGAGGCTCAAGTCTGTGTATGTG
GCACACGTCTGAACTCTTTCCCTACACGACGCTCTTCCGATCTNNNCCCAGCTGAAAGATAACTGGTGA
CACACGTCTGAACTCTTTCCCTACACGACGCTCTTCCGATCTNNNTACACACACACATACAGAGCCAG
CACACGTCTGAACTCTTTCCCTACACGACGCTCTTCCGATCTNNNCGTGTGTCTATCGACTGGGTTTT
ACACGTCTGAACTCTTTCCCTACACGACGCTCTTCCGATCTNNNTACCATACAGGCCTTCTGCACATCA
GCACACGTCTGAACTCTTTCCCTACACGACGCTCTTCCGATCTNNNTAGAGTGCGATTTGGAAATGG
GCACACGTCTGAACTCTTTCCCTACACGACGCTCTTCCGATCTNNNGATAGCTAGTCCCGGCGTGCTA
ACACGTCTGAACTCTTTCCCTACACGACGCTCTTCCGATCTNNNTGGCCAATCATAGATCCTTCCTGCA
GCACACGTCTGAACTCTTTCCCTACACGACGCTCTTCCGATCTNNNTCAGGCAATTACAGGAGGATTG
GCACACGTCTGAACTCTTTCCCTACACGACGCTCTTCCGATCTNNNGTAAGCAGATGATGCCAGAATG
GCACACGTCTGAACTCTTTCCCTACACGACGCTCTTCCGATCTNNNCAGCTCTAGAATCCTGGCTTTCC
ACACGTCTGAACTCTTTCCCTACACGACGCTCTTCCGATCTNNNTCTTCCGTGCTTGGCATCTAAAACA
AGCACACGTCTGAACTCTTTCCCTACACGACGCTCTTCCGATCTNNNTTCATGAACCAATCTCAGGAGC
GCACACGTCTGAACTCTTTCCCTACACGACGCTCTTCCGATCTNNNAAAACAGTGCTGGAACACCTAG
CACACGTCTGAACTCTTTCCCTACACGACGCTCTTCCGATCTNNNCAGCTGGTGCTTTTCTCGAACTTC
CACACGTCTGAACTCTTTCCCTACACGACGCTCTTCCGATCTNNNCCATTTGCGTTATCAGGCTCTCTCA
CACACGTCTGAACTCTTTCCCTACACGACGCTCTTCCGATCTNNNCACACATTTTACTCCGCTCTCCACAT
CACACGTCTGAACTCTTTCCCTACACGACGCTCTTCCGATCTNNNATCCACCACACAGGTGTGAAAAA
AGCACACGTCTGAACTCTTTCCCTACACGACGCTCTTCCGATCTNNNGGCAAAGCTTGCCTGCTAAAG
CACACGTCTGAACTCTTTCCCTACACGACGCTCTTCCGATCTNNNCTGGGAAAGAGAAATGGTAGGG
GCACACGTCTGAACTCTTTCCCTACACGACGCTCTTCCGATCTNNNGCCACCTCTTTGGCATTTGTATG
CACACGTCTGAACTCTTTCCCTACACGACGCTCTTCCGATCTNNNTCCTCATTTCTACCTGCAACGTGC
GCACACGTCTGAACTCTTTCCCTACACGACGCTCTTCCGATCTNNNGGCGACAGAGCTACTCTCAAAA
CACACGTCTGAACTCTTTCCCTACACGACGCTCTTCCGATCTNNNCTCAAGCTACCAAAGACCACTGTC
GCACACGTCTGAACTCTTTCCCTACACGACGCTCTTCCGATCTNNNTGAATGTGAGGAGGTCTACAG
GCACACGTCTGAACTCTTTCCCTACACGACGCTCTTCCGATCTNNNGTCAACTCAAAGGGCTCCTCATT
AGCACACGTCTGAACTCTTTCCCTACACGACGCTCTTCCGATCTNNNAGGGAGAACGGAAAGGTGTG
CACACGTCTGAACTCTTTCCCTACACGACGCTCTTCCGATCTNNNTGCAGAGACACAGACGTAGAGAG
GCACACGTCTGAACTCTTTCCCTACACGACGCTCTTCCGATCTNNNATTTTGTCTGCCATACATGCTGCC
CACACGTCTGAACTCTTTCCCTACACGACGCTCTTCCGATCTNNNCTGTGTTCTGCCTCCTCAAATCTC

CACACGTCTGAACTCTTTCCCTACACGACGCTCTTCCGATCTNNNTAGTTGAAAGGACACTTGGGGG

GCACACGTCTGAACTCTTTCCCTACACGACGCTCTTCCGATCTNNNGTGAGGGACTGTGCTATCTTTCC

ACACGTCTGAACTCTTTCCCTACACGACGCTCTTCCGATCTNNNGATCTCTTTCTGGAAAGGGGGTGC

GCACACGTCTGAACTCTTTCCCTACACGACGCTCTTCCGATCTNNNAGCTGTCTCGTACACACCCATCAT

AGCACACGTCTGAACTCTTTCCCTACACGACGCTCTTCCGATCTNNNGGAAATTCCCTCTCCTGCCACC

CACACGTCTGAACTCTTTCCCTACACGACGCTCTTCCGATCTNNNTTCAGTGTCGAATAAGGGCACAG

CACACGTCTGAACTCTTTCCCTACACGACGCTCTTCCGATCTNNNAGGGAGTAGAATGAGTTCTTGGC

AGCACACGTCTGAACTCTTTCCCTACACGACGCTCTTCCGATCTNNNGGGAAGGGTATGGTGACCAG

GCACACGTCTGAACTCTTTCCCTACACGACGCTCTTCCGATCTNNNCTCTGGCTTCTGTCACTGCAACCA

GCACACGTCTGAACTCTTTCCCTACACGACGCTCTTCCGATCTNNNCTCGCAACGCTAAACAGTGAAAA

CACACGTCTGAACTCTTTCCCTACACGACGCTCTTCCGATCTNNNAAGACACAGTTTTCCGAAGTCCCCA

GCACACGTCTGAACTCTTTCCCTACACGACGCTCTTCCGATCTNNNCAGCACACAAGAACCAGGTCATC

GCACACGTCTGAACTCTTTCCCTACACGACGCTCTTCCGATCTNNNACTCATTTCACATGGTTTCCCGTC

CACACGTCTGAACTCTTTCCCTACACGACGCTCTTCCGATCTNNNAGTGCAGCTGGATACAAGTTCAA

CACACGTCTGAACTCTTTCCCTACACGACGCTCTTCCGATCTNNNATCAACAGTGAGAAATGCCCTCCC

AGCACACGTCTGAACTCTTTCCCTACACGACGCTCTTCCGATCTNNNCAGGAGAGGCCAGAAGTTTG

GCACACGTCTGAACTCTTTCCCTACACGACGCTCTTCCGATCTNNNGGACTGCATGTCTTCCATCCATTC

CACACGTCTGAACTCTTTCCCTACACGACGCTCTTCCGATCTNNNGATCCATCTGCAGAGCTGTTCTTC

AGCACACGTCTGAACTCTTTCCCTACACGACGCTCTTCCGATCTNNNTAACTCTTCCTTCCCTCCCTTCCC

GCACACGTCTGAACTCTTTCCCTACACGACGCTCTTCCGATCTNNNCTGGAGGGCAACTCCTCCTTTCA

CACACGTCTGAACTCTTTCCCTACACGACGCTCTTCCGATCTNNNAGCCATCTTCATGTTTGAGGGTCC

GCACACGTCTGAACTCTTTCCCTACACGACGCTCTTCCGATCTNNNCATCAGTGTCTGGAGGCTTTTCAC

GCACACGTCTGAACTCTTTCCCTACACGACGCTCTTCCGATCTNNNCCTCCTGGAATGACACGTTTCTCC

CACACGTCTGAACTCTTTCCCTACACGACGCTCTTCCGATCTNNNAAACATACAGATTTTGGGGGCGAC

GCACACGTCTGAACTCTTTCCCTACACGACGCTCTTCCGATCTNNNAGGCTTCATGTTGCTCCATTTCTC

CACACGTCTGAACTCTTTCCCTACACGACGCTCTTCCGATCTNNNCCACACTTGGTTATGTCCTTCGTCA

GCACACGTCTGAACTCTTTCCCTACACGACGCTCTTCCGATCTNNNGAACACTGTTACCCATCAACTGCC

CACACGTCTGAACTCTTTCCCTACACGACGCTCTTCCGATCTNNNCAGGCTTTGTAAAAGCAAGGACCC

CACACGTCTGAACTCTTTCCCTACACGACGCTCTTCCGATCTNNNGGACACTAGGGTTCCTGTGAGAA

CACACGTCTGAACTCTTTCCCTACACGACGCTCTTCCGATCTNNNTCCTCTATCCCCACAACCAACTTCAT

AGCACACGTCTGAACTCTTTCCCTACACGACGCTCTTCCGATCTNNNCGCTCTGCCTCCTGCTCTATCA

GCACACGTCTGAACTCTTTCCCTACACGACGCTCTTCCGATCTNNNCTGGATCTCTCTGTTAGAGGGGG

CACACGTCTGAACTCTTTCCCTACACGACGCTCTTCCGATCTNNNCCCTTCCTTATGAGGTTGGTGTGC

CACACGTCTGAACTCTTTCCCTACACGACGCTCTTCCGATCTNNNAGCTCGAACTGCCTGGATAAAAG

GCACACGTCTGAACTCTTTCCCTACACGACGCTCTTCCGATCTNNNACAATAGGTGAGGGCCCCATTC

GCACACGTCTGAACTCTTTCCCTACACGACGCTCTTCCGATCTNNNCCCAGCAGTCATGAAAGAAGGA

AGCACACGTCTGAACTCTTTCCCTACACGACGCTCTTCCGATCTNNNGACCCCAAGTCAGCAAGTCAG

GCACACGTCTGAACTCTTTCCCTACACGACGCTCTTCCGATCTNNNGGGTGGATTTGGGACACTTATG

CACACGTCTGAACTCTTTCCCTACACGACGCTCTTCCGATCTNNNTATTCCTCCTAGGCTGTGTGTGTC

GCACACGTCTGAACTCTTTCCCTACACGACGCTCTTCCGATCTNNNCAAACCTAGTGCAGACCCAAAGT

GCACACGTCTGAACTCTTTCCCTACACGACGCTCTTCCGATCTNNNATTTCCGCAAGTACAGACTGAGG

GCACACGTCTGAACTCTTTCCCTACACGACGCTCTTCCGATCTNNNTCAAGGGTATTTGAAGTTTGGG

GCACACGTCTGAACTCTTTCCCTACACGACGCTCTTCCGATCTNNNTTGCTGGACATGGATGGATTCTG

CACACGTCTGAACTCTTTCCCTACACGACGCTCTTCCGATCTNNNCTCTCTCTCGTGGACACCTCACCA
AGCACACGTCTGAACTCTTTCCCTACACGACGCTCTTCCGATCTNNNGTCTTGGTGGATCTGGCTTGG
GCACACGTCTGAACTCTTTCCCTACACGACGCTCTTCCGATCTNNNCTTCCATCCTCCCTCCTCTTCTCCA
GCACACGTCTGAACTCTTTCCCTACACGACGCTCTTCCGATCTNNNCCCAGTCTTAGTCCCCACACTCAT
CACACGTCTGAACTCTTTCCCTACACGACGCTCTTCCGATCTNNNTCCCTGGCTTAGAAACAGATGAC
CACACGTCTGAACTCTTTCCCTACACGACGCTCTTCCGATCTNNNCATAAGCTGTGCATAGCAGGCAT
CACACGTCTGAACTCTTTCCCTACACGACGCTCTTCCGATCTNNNTTCCTTTTCTCTTTCCAGCCACCCA
CACACGTCTGAACTCTTTCCCTACACGACGCTCTTCCGATCTNNNCCATTCTTTATTTGGGGAGCGGA
CACACGTCTGAACTCTTTCCCTACACGACGCTCTTCCGATCTNNNAGGCTCAAAGTCTAACGCCAATC
GCACACGTCTGAACTCTTTCCCTACACGACGCTCTTCCGATCTNNNCCTGTGAAAGGAATGTTTGTGG
GCACACGTCTGAACTCTTTCCCTACACGACGCTCTTCCGATCTNNNTACCTGCAGTTCCCAGGAAAAAC
GCACACGTCTGAACTCTTTCCCTACACGACGCTCTTCCGATCTNNNTCCCTCTCCTCTTTCCAATCCTGCA
GCACACGTCTGAACTCTTTCCCTACACGACGCTCTTCCGATCTNNNGTGACCACGCCCCTTAGTCTCCA
CACACGTCTGAACTCTTTCCCTACACGACGCTCTTCCGATCTNNNAGAATTCCAGGGTGCAAGATGT
CACACGTCTGAACTCTTTCCCTACACGACGCTCTTCCGATCTNNNCGAGATCCCATCCGAACATACAC
CACACGTCTGAACTCTTTCCCTACACGACGCTCTTCCGATCTNNNATGCCCTGTACTGAGGAAAGAGA
CACACGTCTGAACTCTTTCCCTACACGACGCTCTTCCGATCTNNNTAGCGGAGAGCTAAGGTACTCAC
GCACACGTCTGAACTCTTTCCCTACACGACGCTCTTCCGATCTNNNGTGACAATCCTCAACCCTTTCCA
CACACGTCTGAACTCTTTCCCTACACGACGCTCTTCCGATCTNNNGAAGCAGAGCAAAAATGGCAGA
GAGCACACGTCTGAACTCTTTCCCTACACGACGCTCTTCCGATCTNNNCAGACAGCGTCTGGTCAGCT
AGCACACGTCTGAACTCTTTCCCTACACGACGCTCTTCCGATCTNNNAAATTGCTGGGAACAAGGACT
CACACGTCTGAACTCTTTCCCTACACGACGCTCTTCCGATCTNNNCACAGACAGACCTGGACTTGAACC
ACACGTCTGAACTCTTTCCCTACACGACGCTCTTCCGATCTNNNTAACTGTACCGGGAGCACCTAAAC
AGCACACGTCTGAACTCTTTCCCTACACGACGCTCTTCCGATCTNNNCGGCACTCCCTAAATGATCTCTC
GCACACGTCTGAACTCTTTCCCTACACGACGCTCTTCCGATCTNNNGGGCAAACTGTAGGTATTCAGC
ACACGTCTGAACTCTTTCCCTACACGACGCTCTTCCGATCTNNNGTGGATGGCTAGAGAAGGGAAG
CACACGTCTGAACTCTTTCCCTACACGACGCTCTTCCGATCTNNNCTGCTCCCAACTTCTTGAAACCTCA
CACACGTCTGAACTCTTTCCCTACACGACGCTCTTCCGATCTNNNTAGGCTGAGGTTGGTAAAGGTC
GCACACGTCTGAACTCTTTCCCTACACGACGCTCTTCCGATCTNNNAGTGCTGTGACTGGGATAGTT
ACACGTCTGAACTCTTTCCCTACACGACGCTCTTCCGATCTNNNGGGTGTATGCACTCCACAATACAC
GCACACGTCTGAACTCTTTCCCTACACGACGCTCTTCCGATCTNNNGTTTCAGCCATGTCAGCCAGGC
GCACACGTCTGAACTCTTTCCCTACACGACGCTCTTCCGATCTNNNCTCATCCATCTTGTTCCCTGCTCC
CACACGTCTGAACTCTTTCCCTACACGACGCTCTTCCGATCTNNNGAACTAAGCTACATGTGTGGGCA
ACACGTCTGAACTCTTTCCCTACACGACGCTCTTCCGATCTNNNTCTCTAGCCTGAAAGAGTGCTGTC
CACACGTCTGAACTCTTTCCCTACACGACGCTCTTCCGATCTNNNAAGAAAACTCACCCTGTTGTGCTC
CACACGTCTGAACTCTTTCCCTACACGACGCTCTTCCGATCTNNNGCACAAGGAGATAGGGTTCCTCT
CACACGTCTGAACTCTTTCCCTACACGACGCTCTTCCGATCTNNNTGTCTGTAGGCTTTGTACTGAAA
GCACACGTCTGAACTCTTTCCCTACACGACGCTCTTCCGATCTNNNAATGGTGATACCATGTGTGCCT
AGCACACGTCTGAACTCTTTCCCTACACGACGCTCTTCCGATCTNNNGAGGTGCTGGGATTCAGGGT
CACACGTCTGAACTCTTTCCCTACACGACGCTCTTCCGATCTNNNCATTGGAAAACGTTAGGTGGTGC
AGCACACGTCTGAACTCTTTCCCTACACGACGCTCTTCCGATCTNNNCAACATGGCGAGATCCTGTCT
CACACGTCTGAACTCTTTCCCTACACGACGCTCTTCCGATCTNNNACCAGGCAGAGACTCTTTTTCTTC
AGCACACGTCTGAACTCTTTCCCTACACGACGCTCTTCCGATCTNNNCAGACACACACACACACACACC

GAGCACACGTCTGAACTCTTTCCCTACACGACGCTCTTCCGATCTNNNCCTGGGGCAACAGAAAGAGA
GCACACGTCTGAACTCTTTCCCTACACGACGCTCTTCCGATCTNNNCCAGGCAGAAATTCTTGTTCTCC
CACACGTCTGAACTCTTTCCCTACACGACGCTCTTCCGATCTNNNCTCTCCAGAACAGACAGACACAAC
AGCACACGTCTGAACTCTTTCCCTACACGACGCTCTTCCGATCTNNNGAGATCACACCACTGCACTCCC
AGCACACGTCTGAACTCTTTCCCTACACGACGCTCTTCCGATCTNNNCACACACAGACACACACACACC
GCACACGTCTGAACTCTTTCCCTACACGACGCTCTTCCGATCTNNNTGAGCTAAGATTGTGGCACTAC
AGCACACGTCTGAACTCTTTCCCTACACGACGCTCTTCCGATCTNNNGAGTACAGTGGCTCGATCTCG
GCACACGTCTGAACTCTTTCCCTACACGACGCTCTTCCGATCTNNNATAGGAAGAGGGTCAAGGCTTT
ACACGTCTGAACTCTTTCCCTACACGACGCTCTTCCGATCTNNNTTTCCCATGCTGTTCTCGTGATACAT
GCACACGTCTGAACTCTTTCCCTACACGACGCTCTTCCGATCTNNNTGCCTGTAGTCCCAGCTACTCCA
AGCACACGTCTGAACTCTTTCCCTACACGACGCTCTTCCGATCTNNNCTTGCAGTGAGCTGAAATCGCC
AGCACACGTCTGAACTCTTTCCCTACACGACGCTCTTCCGATCTNNNGGTTGCAGTGAGCCAAGATT
GCACACGTCTGAACTCTTTCCCTACACGACGCTCTTCCGATCTNNNATCAGGAGGTTTTAGGACAGCT
GCACACGTCTGAACTCTTTCCCTACACGACGCTCTTCCGATCTNNNGCCATGCCCAGCTAATTTTTGTA
AGCACACGTCTGAACTCTTTCCCTACACGACGCTCTTCCGATCTNNNGTAGTCCCAGCTACTCCGGAG
GAGCACACGTCTGAACTCTTTCCCTACACGACGCTCTTCCGATCTNNNGCCTGTAGTCCCAGCTACTTG
AGAGCACACGTCTGAACTCTTTCCCTACACGACGCTCTTCCGATCTNNNGACTGAGAGACTGTCCCTG
GCACACGTCTGAACTCTTTCCCTACACGACGCTCTTCCGATCTNNNTGGGAGACAGAGTGAGATGAG
AGCACACGTCTGAACTCTTTCCCTACACGACGCTCTTCCGATCTNNNCACCTGTAGTGCCAGCTACTCC
CACACGTCTGAACTCTTTCCCTACACGACGCTCTTCCGATCTNNNGATGGTCTCGATCTCTTGACCTCC
AGCACACGTCTGAACTCTTTCCCTACACGACGCTCTTCCGATCTNNNTTCATGCCATTCTCCTGCCTCCA
ACACGTCTGAACTCTTTCCCTACACGACGCTCTTCCGATCTNNNACCCTCATATCAACAGCGATTTGCA
GCACACGTCTGAACTCTTTCCCTACACGACGCTCTTCCGATCTNNNTCCTGCTCCAAAGAACCTAGTTC
CACACGTCTGAACTCTTTCCCTACACGACGCTCTTCCGATCTNNNAATAATGGGATGACTGGGTCACG
GAGCACACGTCTGAACTCTTTCCCTACACGACGCTCTTCCGATCTNNNCTTGCAGTGAGCTGAGATTG
GCACACGTCTGAACTCTTTCCCTACACGACGCTCTTCCGATCTNNNGAGGCAGGAGAATGGTGTGA
CACACGTCTGAACTCTTTCCCTACACGACGCTCTTCCGATCTNNNTGCATTAGAGTTTCCTCCAGGTC
GCACACGTCTGAACTCTTTCCCTACACGACGCTCTTCCGATCTNNNAGGTGGCAGGAAAAAAGAAAT
GCACACGTCTGAACTCTTTCCCTACACGACGCTCTTCCGATCTNNNCAAAGTGCTGGGATTACAGGC
CACACGTCTGAACTCTTTCCCTACACGACGCTCTTCCGATCTNNNGAGGCTGAGGCAAGAGAATGG
GCACACGTCTGAACTCTTTCCCTACACGACGCTCTTCCGATCTNNNGATCACCAGGTCAGGAGATCGA
AGCACACGTCTGAACTCTTTCCCTACACGACGCTCTTCCGATCTNNNCTTGCAGTGAGCTGAGATCGC
ACACGTCTGAACTCTTTCCCTACACGACGCTCTTCCGATCTNNNCAAGAGGCCTAACTTTTTCAGCCCA
GCACACGTCTGAACTCTTTCCCTACACGACGCTCTTCCGATCTNNNCCTGTGTGTGATGTTCCCTTTCT
GCACACGTCTGAACTCTTTCCCTACACGACGCTCTTCCGATCTNNNGAGGCAGGAGAATGGTGTGAA
AGCACACGTCTGAACTCTTTCCCTACACGACGCTCTTCCGATCTNNNGAGGTTGCAGTGAGCTGAGA
AGAGCACACGTCTGAACTCTTTCCCTACACGACGCTCTTCCGATCTNNNCTCCTGTAGTTCCAGCCACT
GCACACGTCTGAACTCTTTCCCTACACGACGCTCTTCCGATCTNNNCCGCACCTGGCCTTACTATTATTC
GCACACGTCTGAACTCTTTCCCTACACGACGCTCTTCCGATCTNNNGTATGTGTGTGTGTGTGTGTGT
AGCACACGTCTGAACTCTTTCCCTACACGACGCTCTTCCGATCTNNNGAAGGTGGAGCTTGCAGTGA
AGCACACGTCTGAACTCTTTCCCTACACGACGCTCTTCCGATCTNNNGAGATCGCACCACTACACTCC
CACACGTCTGAACTCTTTCCCTACACGACGCTCTTCCGATCTNNNGAACCACATGTGAGCCAATCAAA
GCACACGTCTGAACTCTTTCCCTACACGACGCTCTTCCGATCTNNNAATGGGAATTTCCTTGCACAAG

GCACACGTCTGAACTCTTTCCCTACACGACGCTCTTCCGATCTNNNGGTCCCCTTGTAGCATCTCAAAT
GCACACGTCTGAACTCTTTCCCTACACGACGCTCTTCCGATCTNNNGCCACCACATCCAACTGATTTTC
AGAGCACACGTCTGAACTCTTTCCCTACACGACGCTCTTCCGATCTNNNGAGGTTGCAATGAGCCTG
GCACACGTCTGAACTCTTTCCCTACACGACGCTCTTCCGATCTNNNCTGTAGTCCCAGCTACTCGGAG
AGCACACGTCTGAACTCTTTCCCTACACGACGCTCTTCCGATCTNNNCGACAGAGTGAGACTCCGTCTC
GAGCACACGTCTGAACTCTTTCCCTACACGACGCTCTTCCGATCTNNNTGGGTGACAGAGTGAGACTC
GAGCACACGTCTGAACTCTTTCCCTACACGACGCTCTTCCGATCTNNNGCGACAGAGTGAGACTCCAT
AGCACACGTCTGAACTCTTTCCCTACACGACGCTCTTCCGATCTNNNGACACAGCAAGACTCCGTCTC
GCACACGTCTGAACTCTTTCCCTACACGACGCTCTTCCGATCTNNNTCTGCTTTGACCTGTTCCTATCC
GCACACGTCTGAACTCTTTCCCTACACGACGCTCTTCCGATCTNNNCTTTTTTTTTTGGGGCCCACAG
AGCACACGTCTGAACTCTTTCCCTACACGACGCTCTTCCGATCTNNNGCTGGGACTACAGGTGTACAC
CACACGTCTGAACTCTTTCCCTACACGACGCTCTTCCGATCTNNNAAATAGCTGGGACTACTCGTGTG
GAGCACACGTCTGAACTCTTTCCCTACACGACGCTCTTCCGATCTNNNGAGGTTGCAGTGAGCTGAGA
AGAGCACACGTCTGAACTCTTTCCCTACACGACGCTCTTCCGATCTNNNGCCTGTAGTCCCAGCTACTT
AGCACACGTCTGAACTCTTTCCCTACACGACGCTCTTCCGATCTNNNGATCGAGACCATCCTGCCTAAC
GCACACGTCTGAACTCTTTCCCTACACGACGCTCTTCCGATCTNNNTGTAGTACCAGTTACTCGGGAG
GCACACGTCTGAACTCTTTCCCTACACGACGCTCTTCCGATCTNNNAAAAATTAGGCATGGTGGCAG
AGCACACGTCTGAACTCTTTCCCTACACGACGCTCTTCCGATCTNNNGAGGCTGAGGCAAGAGAATG
GAGCACACGTCTGAACTCTTTCCCTACACGACGCTCTTCCGATCTNNNTGCCTGTAGTCCCAGCTACTC
AGCACACGTCTGAACTCTTTCCCTACACGACGCTCTTCCGATCTNNNGAGGCAGGAGAATGGTGTGA
AGCACACGTCTGAACTCTTTCCCTACACGACGCTCTTCCGATCTNNNCTAGGCGACAGAGCAAGACTC
CACACGTCTGAACTCTTTCCCTACACGACGCTCTTCCGATCTNNNACAAAGTGAGACCCTGTCTTTGTC
CACACGTCTGAACTCTTTCCCTACACGACGCTCTTCCGATCTNNNTTGAGTGATTCTCCTGCCTCATCCA
CACACGTCTGAACTCTTTCCCTACACGACGCTCTTCCGATCTNNNGTTCAAGCAATTCTCTGCCTCAGC
GCACACGTCTGAACTCTTTCCCTACACGACGCTCTTCCGATCTNNNCAATAGACACTTTGGTCGCTTCC
AGCACACGTCTGAACTCTTTCCCTACACGACGCTCTTCCGATCTNNNGTTTGCAGTGAGCCGAGATTG
AGCACACGTCTGAACTCTTTCCCTACACGACGCTCTTCCGATCTNNNGCTTGCAGTGAGCTGAGATTG
AGCACACGTCTGAACTCTTTCCCTACACGACGCTCTTCCGATCTNNNCTTGCAGTGAGCCGAGATTGT
AGCACACGTCTGAACTCTTTCCCTACACGACGCTCTTCCGATCTNNNGAGACAGGAGAATGGCGTGA
CACACGTCTGAACTCTTTCCCTACACGACGCTCTTCCGATCTNNNGGCGACAACAATGAAACTCCTCCA
ACACGTCTGAACTCTTTCCCTACACGACGCTCTTCCGATCTNNNGGGGAAAATCTTCATGACCTTGTC
CACACGTCTGAACTCTTTCCCTACACGACGCTCTTCCGATCTNNNGTCCCTCACTGAACAATGAAGGA
CACACGTCTGAACTCTTTCCCTACACGACGCTCTTCCGATCTNNNGTAGTCAAAGGACCTGAAACCCT
GCACACGTCTGAACTCTTTCCCTACACGACGCTCTTCCGATCTNNNGGTTTCCCAGACTCAAGCAATTC
CACACGTCTGAACTCTTTCCCTACACGACGCTCTTCCGATCTNNNACAAAAATTAGCAGAGCGTGGTG
GAGCACACGTCTGAACTCTTTCCCTACACGACGCTCTTCCGATCTNNNGAGGCAGGAGAATGACGTG
GCACACGTCTGAACTCTTTCCCTACACGACGCTCTTCCGATCTNNNTTTCCTCGGCTTACAGACACATCC
GCACACGTCTGAACTCTTTCCCTACACGACGCTCTTCCGATCTNNNCCCGCGTCTATCACCTTGTAATA
AGCACACGTCTGAACTCTTTCCCTACACGACGCTCTTCCGATCTNNNCTTGCAGTGAGCCGAGATCGC
GCACACGTCTGAACTCTTTCCCTACACGACGCTCTTCCGATCTNNNGCCTGTAGTCCCAGCTACTTGCA
AGCACACGTCTGAACTCTTTCCCTACACGACGCTCTTCCGATCTNNNGTAGTCCCAGCTACTCGGAGA
CACACGTCTGAACTCTTTCCCTACACGACGCTCTTCCGATCTNNNCCAATTGCTAAGTCCAGTCTCCTCA
GCACACGTCTGAACTCTTTCCCTACACGACGCTCTTCCGATCTNNNCCAGAAACATGGCTAGAGAGTG

GCACACGTCTGAACTCTTTCCCTACACGACGCTCTTCCGATCTNNNTGGATACAGCAAAGCAAGGATCT
GCACACGTCTGAACTCTTTCCCTACACGACGCTCTTCCGATCTNNNGCTCTATCTAACTGGGGGCATTT
GCACACGTCTGAACTCTTTCCCTACACGACGCTCTTCCGATCTNNNGATTGCTCTAGCCAGGACTTCTA
GAGCACACGTCTGAACTCTTTCCCTACACGACGCTCTTCCGATCTNNNACCTGTAGTCCCAGCTACTCG
GCACACGTCTGAACTCTTTCCCTACACGACGCTCTTCCGATCTNNNCATCCTTGCTAACACGGTGAAAC
AGCACACGTCTGAACTCTTTCCCTACACGACGCTCTTCCGATCTNNNCAAGAGAGCCGATTGAACCCCA
GAGCACACGTCTGAACTCTTTCCCTACACGACGCTCTTCCGATCTNNNGCCTGTAGTCCCAGCTACTTG
CACACGTCTGAACTCTTTCCCTACACGACGCTCTTCCGATCTNNNCTCTCGAGTAGCTGGGATTACAG
GAGCACACGTCTGAACTCTTTCCCTACACGACGCTCTTCCGATCTNNNCCGAGTAGCTGGGACTACAG
GCACACGTCTGAACTCTTTCCCTACACGACGCTCTTCCGATCTNNNTTTGGATGCCATTTCCCATATGCCA
GCACACGTCTGAACTCTTTCCCTACACGACGCTCTTCCGATCTNNNCCTTGTGCAAGAAAAGGCATTCT
GCACACGTCTGAACTCTTTCCCTACACGACGCTCTTCCGATCTNNNGAACAAGAGAGAGTGGCAGAAG
AGCACACGTCTGAACTCTTTCCCTACACGACGCTCTTCCGATCTNNNTACGCACCACCATACCTTGTCCAT
CACGTCTGAACTCTTTCCCTACACGACGCTCTTCCGATCTNNNGGTACAGTCAACAATTTCTAGCCCCAT
GCACACGTCTGAACTCTTTCCCTACACGACGCTCTTCCGATCTNNNGATCAAGACCATCCTGGCTAACAC
AGCACACGTCTGAACTCTTTCCCTACACGACGCTCTTCCGATCTNNNGGTTGCAGTGAGCCAAGATTG
AGCACACGTCTGAACTCTTTCCCTACACGACGCTCTTCCGATCTNNNGCCTGTAGTCCCAGCTACTTGCA
AGCACACGTCTGAACTCTTTCCCTACACGACGCTCTTCCGATCTNNNGAGAATCACGAGCTCAGGAGA
GAGCACACGTCTGAACTCTTTCCCTACACGACGCTCTTCCGATCTNNNCTTGCAGTGAGCCGAGATTG
GAGCACACGTCTGAACTCTTTCCCTACACGACGCTCTTCCGATCTNNNCCTGTAGTCCCAGCTACTTGG
AAGAGCACACGTCTGAACTCTTTCCCTACACGACGCTCTTCCGATCTNNNACCTGTAGTCCCAGCTACT
CACACGTCTGAACTCTTTCCCTACACGACGCTCTTCCGATCTNNNAGACAGAATCAGAAACAGAGCCT
AGCACACGTCTGAACTCTTTCCCTACACGACGCTCTTCCGATCTNNNGAGGCAGGAGAATGGTGTGAA
GAGCACACGTCTGAACTCTTTCCCTACACGACGCTCTTCCGATCTNNNGACAGAGCAAGACTCCGTCTC
GCACACGTCTGAACTCTTTCCCTACACGACGCTCTTCCGATCTNNNAAATTTTGAGCACCTACCAATGC
GCACACGTCTGAACTCTTTCCCTACACGACGCTCTTCCGATCTNNNATTCTGAGGCTCAGGAATCTCTG
GCACACGTCTGAACTCTTTCCCTACACGACGCTCTTCCGATCTNNNTGGTTTCTACAGGGGTAGGTACT
GCACACGTCTGAACTCTTTCCCTACACGACGCTCTTCCGATCTNNNCTGTATGTTTTCCATGCCCTCACC
GCACACGTCTGAACTCTTTCCCTACACGACGCTCTTCCGATCTNNNCCTGAACATGTTCCCCCTAAGAACA
CACACGTCTGAACTCTTTCCCTACACGACGCTCTTCCGATCTNNNCTCTTACACTGGTGGCCCTATTCCAT
GCACACGTCTGAACTCTTTCCCTACACGACGCTCTTCCGATCTNNNTGTCTCCAAGCAGCATTTTTTTGCA
AGAGCACACGTCTGAACTCTTTCCCTACACGACGCTCTTCCGATCTNNNGAGGCTGAGGCAGGAGAAT
AGCACACGTCTGAACTCTTTCCCTACACGACGCTCTTCCGATCTNNNGAAGTCAGGAGATCGAGACCA
GAGCACACGTCTGAACTCTTTCCCTACACGACGCTCTTCCGATCTNNNGCCTGTAGTCCCAGCTACTTG
GAGCACACGTCTGAACTCTTTCCCTACACGACGCTCTTCCGATCTNNNCTTGCAGTGAGCTGAGATTG
GAGCACACGTCTGAACTCTTTCCCTACACGACGCTCTTCCGATCTNNNGAGGCTAAGGCAGGAGAATG
GAGCACACGTCTGAACTCTTTCCCTACACGACGCTCTTCCGATCTNNNGCCTGTAGTCCCAGCTACTTG
AGCACACGTCTGAACTCTTTCCCTACACGACGCTCTTCCGATCTNNNTAATCCCAGCTACTCAGGAGTCT
GAGCACACGTCTGAACTCTTTCCCTACACGACGCTCTTCCGATCTNNNGTTTGCAGTGAGCCGAGATT
GCACACGTCTGAACTCTTTCCCTACACGACGCTCTTCCGATCTNNNACGCCCGGCTAATTTTTGTATTTC
CACACGTCTGAACTCTTTCCCTACACGACGCTCTTCCGATCTNNNGCTAGTCACCTATCACAGCTCATCA
CACACGTCTGAACTCTTTCCCTACACGACGCTCTTCCGATCTNNNTCTCCACATCTCAACACAGTTGTCAT
CACACGTCTGAACTCTTTCCCTACACGACGCTCTTCCGATCTNNNTCACTGCCTAAACAATCTGGTCACA

GCACACGTCTGAACTCTTTCCCTACACGACGCTCTTCCGATCTNNNAGGCTGTATTTAGATTTCCCAGT
GCACACGTCTGAACTCTTTCCCTACACGACGCTCTTCCGATCTNNNGATTTCTCTGGCTTCAGTTGTGT
GCACACGTCTGAACTCTTTCCCTACACGACGCTCTTCCGATCTNNNTCAAAAGTGAATACTGGCACACG
GCACACGTCTGAACTCTTTCCCTACACGACGCTCTTCCGATCTNNNATATGTTGTCCGTCGTTGACTGA
CACACGTCTGAACTCTTTCCCTACACGACGCTCTTCCGATCTNNNAGGAAAAACTTCAGTAACAGGGCT
GAGCACACGTCTGAACTCTTTCCCTACACGACGCTCTTCCGATCTNNNGAGGCAGGGAAATCGCTTG
AGCACACGTCTGAACTCTTTCCCTACACGACGCTCTTCCGATCTNNNGCCTGTAGTCCCAGCTACTTGC
GCACACGTCTGAACTCTTTCCCTACACGACGCTCTTCCGATCTNNNTGTAGTCCCAGCTACTCTAGAGG
AGCACACGTCTGAACTCTTTCCCTACACGACGCTCTTCCGATCTNNNGAGGCTGAAGCAGGAGAATG
GAGCACACGTCTGAACTCTTTCCCTACACGACGCTCTTCCGATCTNNNGCCTGTAGTCCCAGCTACTTG
GAGCACACGTCTGAACTCTTTCCCTACACGACGCTCTTCCGATCTNNNGACAGAGCGAAACTCCGTCT
GAGCACACGTCTGAACTCTTTCCCTACACGACGCTCTTCCGATCTNNNGACAGAGCAAGACTCCGTCTC
GCACACGTCTGAACTCTTTCCCTACACGACGCTCTTCCGATCTNNNTGTAGTCCCAGATACTCAGGAGG
AGCACACGTCTGAACTCTTTCCCTACACGACGCTCTTCCGATCTNNNCAAGATCGCGTCACTGCATTCC
GCACACGTCTGAACTCTTTCCCTACACGACGCTCTTCCGATCTNNNTTTGGTTAGGTTAGGCTGGTCTC
CACACGTCTGAACTCTTTCCCTACACGACGCTCTTCCGATCTNNNGCACCATGCCAGGTAATCTTTATC
GCACACGTCTGAACTCTTTCCCTACACGACGCTCTTCCGATCTNNNGCACTGAAAAGCTATGGGGATT
CACACGTCTGAACTCTTTCCCTACACGACGCTCTTCCGATCTNNNAGAGAGCTAAAGACCAAACTCCAC
GAGCACACGTCTGAACTCTTTCCCTACACGACGCTCTTCCGATCTNNNTTGCAGTGAGCCGAGATCAAA
GCACACGTCTGAACTCTTTCCCTACACGACGCTCTTCCGATCTNNNTGCATCTTGAATCTACTGGCCTTC
GCACACGTCTGAACTCTTTCCCTACACGACGCTCTTCCGATCTNNNCCAAGTATCTAAGACCCACCTCA
GCACACGTCTGAACTCTTTCCCTACACGACGCTCTTCCGATCTNNNAATAACTTCTGGGTAGGTCTGG
CACACGTCTGAACTCTTTCCCTACACGACGCTCTTCCGATCTNNNGATAAGTGAGGCCTGTCAGTCAG
CACACGTCTGAACTCTTTCCCTACACGACGCTCTTCCGATCTNNNTGTGGTCTCATTCATGTCAGTGTCA
GCACACGTCTGAACTCTTTCCCTACACGACGCTCTTCCGATCTNNNAGAAGCAGAAATTCACCTGGTCT
GCACACGTCTGAACTCTTTCCCTACACGACGCTCTTCCGATCTNNNTTTCACTAAAAGCTGGGTCTGG
AGCACACGTCTGAACTCTTTCCCTACACGACGCTCTTCCGATCTNNNTACAGCTCAGTTGTTGGGTGA
ACACGTCTGAACTCTTTCCCTACACGACGCTCTTCCGATCTNNNCGAATACCTGAGAATCCCTCCAGCAT
AGCACACGTCTGAACTCTTTCCCTACACGACGCTCTTCCGATCTNNNGCCTGTAGTCCCAGCTACTTGC
AGCACACGTCTGAACTCTTTCCCTACACGACGCTCTTCCGATCTNNNGGTGACAGACTGAGACCCTGT
AGCACACGTCTGAACTCTTTCCCTACACGACGCTCTTCCGATCTNNNGCCTGTAATCCCAGCTACTCGC
AGCACACGTCTGAACTCTTTCCCTACACGACGCTCTTCCGATCTNNNGTGACAGAACGAGACTCCGTC
GAGCACACGTCTGAACTCTTTCCCTACACGACGCTCTTCCGATCTNNNCTGTAGTCCCAGCTACTCGG
AGCACACGTCTGAACTCTTTCCCTACACGACGCTCTTCCGATCTNNNCTTGCAGTGAGCCGAGATTGT
AGCACACGTCTGAACTCTTTCCCTACACGACGCTCTTCCGATCTNNNCAATGAACCAAGGTCATGCCAC
AGCACACGTCTGAACTCTTTCCCTACACGACGCTCTTCCGATCTNNNGTGGATCACAAGGTCAGGAG
AGCACACGTCTGAACTCTTTCCCTACACGACGCTCTTCCGATCTNNNGCCTGTAGTCCCAGCTACTTGCA
AGCACACGTCTGAACTCTTTCCCTACACGACGCTCTTCCGATCTNNNTGGGAGACAGAGTGAGACTCC
AGCACACGTCTGAACTCTTTCCCTACACGACGCTCTTCCGATCTNNNGAGATCACACCACTGCACTCC
AGCACACGTCTGAACTCTTTCCCTACACGACGCTCTTCCGATCTNNNCTGGGCGAAAGAGGAAGACT
AGCACACGTCTGAACTCTTTCCCTACACGACGCTCTTCCGATCTNNNGAGGCAGGAGAATGGTGTGA
CACACGTCTGAACTCTTTCCCTACACGACGCTCTTCCGATCTNNNACTATTGGCCTGCAAAGTCACTAC
CACACGTCTGAACTCTTTCCCTACACGACGCTCTTCCGATCTNNNATTTCCGAGTTTCCGTCGACACAT

GCACACGTCTGAACTCTTTCCCTACACGACGCTCTTCCGATCTNNNACCATTTGCATCTTTTCTCCTCTC
ACACGTCTGAACTCTTTCCCTACACGACGCTCTTCCGATCTNNNTCAATTTGATCTGCTACCCACACA
ACACGTCTGAACTCTTTCCCTACACGACGCTCTTCCGATCTNNNTGCTTGGTCTGTGGTTAGTTAGGC
GAGCACACGTCTGAACTCTTTCCCTACACGACGCTCTTCCGATCTNNNGCCTGTAGTCCCAGCTACTTG
AGCACACGTCTGAACTCTTTCCCTACACGACGCTCTTCCGATCTNNNCTTGCAGTGAGCTGAGATCGT
AGCACACGTCTGAACTCTTTCCCTACACGACGCTCTTCCGATCTNNNGAGACAGGAGAATGGCGTGA
GCACACGTCTGAACTCTTTCCCTACACGACGCTCTTCCGATCTNNNGCTCTGTCCTCCTTTTTTCCCTC
GCACACGTCTGAACTCTTTCCCTACACGACGCTCTTCCGATCTNNNCCCCATGGTTTTCTAGTTGTGTC
AGCACACGTCTGAACTCTTTCCCTACACGACGCTCTTCCGATCTNNNTTCATGCCATTCTCCTGCCTCC
GCACACGTCTGAACTCTTTCCCTACACGACGCTCTTCCGATCTNNNCAACATAGCGAGGCAGCTTTGC
CACACGTCTGAACTCTTTCCCTACACGACGCTCTTCCGATCTNNNGTAGAGACGAGGTTTCACCATGT
CACACGTCTGAACTCTTTCCCTACACGACGCTCTTCCGATCTNNNGCTAGTCTCGAATTCCTGACCTCC
ACACGTCTGAACTCTTTCCCTACACGACGCTCTTCCGATCTNNNGGACATTCCCTGAAAATTTGGCTC
GCACACGTCTGAACTCTTTCCCTACACGACGCTCTTCCGATCTNNNGTAGTGAGAGCTGGGCTTGTC
GAGCACACGTCTGAACTCTTTCCCTACACGACGCTCTTCCGATCTNNNGTTGCAGTGAGCCGAGATTG
AGCACACGTCTGAACTCTTTCCCTACACGACGCTCTTCCGATCTNNNCTTGCAGTGAGCTGAGATCGA
ACACGTCTGAACTCTTTCCCTACACGACGCTCTTCCGATCTNNNCCTCCTTTTACATTCCCCTTCCACAT
GCACACGTCTGAACTCTTTCCCTACACGACGCTCTTCCGATCTNNNCCATACAACTTGGGTCAGGAAG
AGCACACGTCTGAACTCTTTCCCTACACGACGCTCTTCCGATCTNNNGCCTGTAGTCCCAGCTACTTGC
GCACACGTCTGAACTCTTTCCCTACACGACGCTCTTCCGATCTNNNGCCTGTAGTCCCAGCTACTTGC
AGCACACGTCTGAACTCTTTCCCTACACGACGCTCTTCCGATCTNNNACCTGTAGTCCCAGCTACTCGC
AGCACACGTCTGAACTCTTTCCCTACACGACGCTCTTCCGATCTNNNGAGCTTGCATTGAGCCGAAAT
GCACACGTCTGAACTCTTTCCCTACACGACGCTCTTCCGATCTNNNGCCTGGGCAATACAGTAAGATC
CACACGTCTGAACTCTTTCCCTACACGACGCTCTTCCGATCTNNNCTTTACCAAACACGGAAGACCTC
CACACGTCTGAACTCTTTCCCTACACGACGCTCTTCCGATCTNNNCCTGCTCAGGGATGCAATAGATA
ACACGTCTGAACTCTTTCCCTACACGACGCTCTTCCGATCTNNNAGCCTTTATAGCGGTTACTCACTCAT
CACACGTCTGAACTCTTTCCCTACACGACGCTCTTCCGATCTNNNTTTAGTACAGACGGGGTTTCACC
GCACACGTCTGAACTCTTTCCCTACACGACGCTCTTCCGATCTNNNTTCCTCTGGTGGTAGGAAGATC
AGCACACGTCTGAACTCTTTCCCTACACGACGCTCTTCCGATCTNNNCTTGCAGTGAGCTGAGATCG
AGCACACGTCTGAACTCTTTCCCTACACGACGCTCTTCCGATCTNNNCGACAGAGCAAGATCCGTCTC
GCACACGTCTGAACTCTTTCCCTACACGACGCTCTTCCGATCTNNNGAGAATGGCGTGAACCTGGAA
CACACGTCTGAACTCTTTCCCTACACGACGCTCTTCCGATCTNNNCAATCTCAGCTCACAGCAACTTCC
CACACGTCTGAACTCTTTCCCTACACGACGCTCTTCCGATCTNNNTCAGTCTGTCTTTGAGATGCGAA
GCACACGTCTGAACTCTTTCCCTACACGACGCTCTTCCGATCTNNNCTGAGTAGCTGGATTTACAGGC
ACACGTCTGAACTCTTTCCCTACACGACGCTCTTCCGATCTNNNAGTGTGAAGTGATATCTTGTGCTC
GAGCACACGTCTGAACTCTTTCCCTACACGACGCTCTTCCGATCTNNNGTGAACCTGGGAGACAGCTT
AGCACACGTCTGAACTCTTTCCCTACACGACGCTCTTCCGATCTNNNGCTTGCAGTGAGCTGAGATC
AGCACACGTCTGAACTCTTTCCCTACACGACGCTCTTCCGATCTNNNGACAGAACAAGCCTCCGTCTCC
GAGCACACGTCTGAACTCTTTCCCTACACGACGCTCTTCCGATCTNNNCCTGTAGTCCCAGCTACTTGG
GAGCACACGTCTGAACTCTTTCCCTACACGACGCTCTTCCGATCTNNNGCCTGTAGTCCCAGCTACTTG
GAGCACACGTCTGAACTCTTTCCCTACACGACGCTCTTCCGATCTNNNGCTTGCAGTGAGCTGAGAT
AGCACACGTCTGAACTCTTTCCCTACACGACGCTCTTCCGATCTNNNGATGGAGTGCGACTCTGTCT
GAGCACACGTCTGAACTCTTTCCCTACACGACGCTCTTCCGATCTNNNGCCTGTAGTCCCAGCTACTTG

AGCACACGTCTGAACTCTTTCCCTACACGACGCTCTTCCGATCTNNNGCCTGTAATCCCAGCTACTCGCA

GCACACGTCTGAACTCTTTCCCTACACGACGCTCTTCCGATCTNNNTGTCTCGTGATTCACAGAAGTGTC

GCACACGTCTGAACTCTTTCCCTACACGACGCTCTTCCGATCTNNNCTCTGACACCTGACACATATCCAC

AGCACACGTCTGAACTCTTTCCCTACACGACGCTCTTCCGATCTNNNTGGCCTATGGACATTTTCTTCCAC

GAGCACACGTCTGAACTCTTTCCCTACACGACGCTCTTCCGATCTNNNGCCTGTAGTCCCAGCTACTTGC

GAGCACACGTCTGAACTCTTTCCCTACACGACGCTCTTCCGATCTNNNGATCGCGCCACTACACTCTAG

GCACACGTCTGAACTCTTTCCCTACACGACGCTCTTCCGATCTNNNCGAGACCATCCTAGCTAACATGG

GCACACGTCTGAACTCTTTCCCTACACGACGCTCTTCCGATCTNNNATAGCCTAACCTGAAACAGCACG

CACACGTCTGAACTCTTTCCCTACACGACGCTCTTCCGATCTNNNCCACGCCCAGCTAATTTTGTATTCA

CACACGTCTGAACTCTTTCCCTACACGACGCTCTTCCGATCTNNNCTCCAGGGTTCAAGTGATTCTCCCA

GCACACGTCTGAACTCTTTCCCTACACGACGCTCTTCCGATCTNNNCAATCAGCCGCATTTTGTTGATTCA

CACACGTCTGAACTCTTTCCCTACACGACGCTCTTCCGATCTNNNGAGTATCTCCGAGGGTCTTTTCCCA

CACACGTCTGAACTCTTTCCCTACACGACGCTCTTCCGATCTNNNTTTCTTCCCTCCCTCCTTCTTTCCATCC

GCACACGTCTGAACTCTTTCCCTACACGACGCTCTTCCGATCTNNNAAAGTGCTGGGATTACAGGCATG

CACACGTCTGAACTCTTTCCCTACACGACGCTCTTCCGATCTNNNATCCTCTCCAGCATCTGTTGTTTCA

AGCACACGTCTGAACTCTTTCCCTACACGACGCTCTTCCGATCTNNNGCCTGTAGTCCCAGCTACTTGCA

GAGCACACGTCTGAACTCTTTCCCTACACGACGCTCTTCCGATCTNNNAGAATGGCATGAACCTGGGAC

AGCACACGTCTGAACTCTTTCCCTACACGACGCTCTTCCGATCTNNNGACAGAGCAAGACTCCGTCTCCA

GCACACGTCTGAACTCTTTCCCTACACGACGCTCTTCCGATCTNNNGCATGAAAGCAGCCATAGACAGC

GCACACGTCTGAACTCTTTCCCTACACGACGCTCTTCCGATCTNNNTCTCCTCCATTCACAAACCACTTCAT

GCACACGTCTGAACTCTTTCCCTACACGACGCTCTTCCGATCTNNNAAAGTGCTGGGATTGCAGGTATC

GCACACGTCTGAACTCTTTCCCTACACGACGCTCTTCCGATCTNNNGCACAGAAAGACCAATACCACACC

ACACGTCTGAACTCTTTCCCTACACGACGCTCTTCCGATCTNNNGCATAAGACCAATAAACGTGGCCCA

CACACGTCTGAACTCTTTCCCTACACGACGCTCTTCCGATCTNNNAATCACAACAGATGTCACTAGGACA

GCACACGTCTGAACTCTTTCCCTACACGACGCTCTTCCGATCTNNNTCACATTGACTTTTCCCCTCAGTCA

CACACGTCTGAACTCTTTCCCTACACGACGCTCTTCCGATCTNNNTTCAACAATCCCAGTTACCAACGCAT

CACACGTCTGAACTCTTTCCCTACACGACGCTCTTCCGATCTNNNACAGTGCTATGTTCAAGTAACCCTC

GCACACGTCTGAACTCTTTCCCTACACGACGCTCTTCCGATCTNNNGACATAACAGCCCAGTACTTGCT

CACACGTCTGAACTCTTTCCCTACACGACGCTCTTCCGATCTNNNAAATCACTGTACCAACCCTGTCACA

GCACACGTCTGAACTCTTTCCCTACACGACGCTCTTCCGATCTNNNTCTGTAGCTTGTCTTTTCGTCCTC

CACACGTCTGAACTCTTTCCCTACACGACGCTCTTCCGATCTNNNTGGTTGTCTTGAATCAGGTCCATCA

AGCACACGTCTGAACTCTTTCCCTACACGACGCTCTTCCGATCTNNNACCTGTAGTCCCAGCTACTCGCA

GAGCACACGTCTGAACTCTTTCCCTACACGACGCTCTTCCGATCTNNNCACCACTGCACTGCACTCTAGC

AGCACACGTCTGAACTCTTTCCCTACACGACGCTCTTCCGATCTNNNTTGCAGTGAGCTGAGATCGTG

AGAGCACACGTCTGAACTCTTTCCCTACACGACGCTCTTCCGATCTNNNGCCTGTAGTCCCAGCTACTTG

GCACACGTCTGAACTCTTTCCCTACACGACGCTCTTCCGATCTNNNGCCTGTAGTCCCAGCTACTTGCAT

AGCACACGTCTGAACTCTTTCCCTACACGACGCTCTTCCGATCTNNNGACAGAGCGAGCCTACATCTCC

AGAGCACACGTCTGAACTCTTTCCCTACACGACGCTCTTCCGATCTNNNCTTGCAGTGAGCCGAGATTG

AGCACACGTCTGAACTCTTTCCCTACACGACGCTCTTCCGATCTNNNGAGATCGGAGCTTGCAGTGAGC

AGAGCACACGTCTGAACTCTTTCCCTACACGACGCTCTTCCGATCTNNNTGGGTGACAGAGTGAGACTC

GAGCACACGTCTGAACTCTTTCCCTACACGACGCTCTTCCGATCTNNNGAGGCGAGAGAATGGTGTGA

GCACACGTCTGAACTCTTTCCCTACACGACGCTCTTCCGATCTNNNCAACAGCCAACATGAACTCTTGAC

GCACACGTCTGAACTCTTTCCCTACACGACGCTCTTCCGATCTNNNACCCTGTCTCGGAAAAAGGAAA

ACACGTCTGAACTCTTTCCCTACACGACGCTCTTCCGATCTNNNTGATATTGCAAATGCGTTCCCAACAT

GCACACGTCTGAACTCTTTCCCTACACGACGCTCTTCCGATCTNNNTAGCCAGTTCTCCTCTACTTCCACA

ACACGTCTGAACTCTTTCCCTACACGACGCTCTTCCGATCTNNNGCTATTTGGCAGGACATCTCCTACAT

AGCACACGTCTGAACTCTTTCCCTACACGACGCTCTTCCGATCTNNNGCAGTCTCAGCTACTTGGGAGC

GAGCACACGTCTGAACTCTTTCCCTACACGACGCTCTTCCGATCTNNNGCCTGTAGTCCCAGCTACTTGC

AGCACACGTCTGAACTCTTTCCCTACACGACGCTCTTCCGATCTNNNGCCTGTAGTCCCAGCTACTTGCA

GAGCACACGTCTGAACTCTTTCCCTACACGACGCTCTTCCGATCTNNNCTGAGGTCAGGAGTCCAAGA

AGCACACGTCTGAACTCTTTCCCTACACGACGCTCTTCCGATCTNNNGCCTGTAGTCCCAGCTACTTGCA

GCACACGTCTGAACTCTTTCCCTACACGACGCTCTTCCGATCTNNNATATCTCTCCTCGGCTCTACCAACA

GCACACGTCTGAACTCTTTCCCTACACGACGCTCTTCCGATCTNNNATTCAGTTTCATGTTCCTGCTGGC

GCACACGTCTGAACTCTTTCCCTACACGACGCTCTTCCGATCTNNNTGAACCAATTAGGCCTCCTTTCTC

GCACACGTCTGAACTCTTTCCCTACACGACGCTCTTCCGATCTNNNAGTGCTACAATGAACACATGCAC

GAGCACACGTCTGAACTCTTTCCCTACACGACGCTCTTCCGATCTNNNAGAAGAATGGCGTGAACCCA

CACACGTCTGAACTCTTTCCCTACACGACGCTCTTCCGATCTNNNTGATAGAGCCAGACCTCATCTCACA

AGAGCACACGTCTGAACTCTTTCCCTACACGACGCTCTTCCGATCTNNNGAGATCGTACCACTGCACTC

GAGCACACGTCTGAACTCTTTCCCTACACGACGCTCTTCCGATCTNNNGAGGCTGAGGCAGGAGAAT

ACACGTCTGAACTCTTTCCCTACACGACGCTCTTCCGATCTNNNGAACCACCTTTGCATTCCTCAAACAT

GAGCACACGTCTGAACTCTTTCCCTACACGACGCTCTTCCGATCTNNNCCTCCCAGGTTCAAGCGATTCC

GCACACGTCTGAACTCTTTCCCTACACGACGCTCTTCCGATCTNNNTCATTGCTGGAATTCTACTGCCTC

GCACACGTCTGAACTCTTTCCCTACACGACGCTCTTCCGATCTNNNAGTAAGTACGTAGGACCCTTGGA

CACACGTCTGAACTCTTTCCCTACACGACGCTCTTCCGATCTNNNTTCTCTCATGCTGTTTGCTGTTGCA

GAGCACACGTCTGAACTCTTTCCCTACACGACGCTCTTCCGATCTNNNCTTGCAGTGAGCCAAGAACAC

AGCACACGTCTGAACTCTTTCCCTACACGACGCTCTTCCGATCTNNNGCCTGTAGTCCCAGCTACTTGCA

AGAGCACACGTCTGAACTCTTTCCCTACACGACGCTCTTCCGATCTNNNGGTTGCAGTGAGCCAAGATT

AGCACACGTCTGAACTCTTTCCCTACACGACGCTCTTCCGATCTNNNCTTGCAGTGAGCCGAGATTGTC

GAGCACACGTCTGAACTCTTTCCCTACACGACGCTCTTCCGATCTNNNGCTTGCAGTGAGCTGAGATCA

GAGCACACGTCTGAACTCTTTCCCTACACGACGCTCTTCCGATCTNNNGCCTGTAGTCCCAGCTACTTG

AGAGCACACGTCTGAACTCTTTCCCTACACGACGCTCTTCCGATCTNNNGAGATCCAGCCACTGCACTC

GAGCACACGTCTGAACTCTTTCCCTACACGACGCTCTTCCGATCTNNNGCCTGTAGTCCCAGCTACTTGC

GAGCACACGTCTGAACTCTTTCCCTACACGACGCTCTTCCGATCTNNNGCCTGTAGTCCCAGTTACTCGC

AGCACACGTCTGAACTCTTTCCCTACACGACGCTCTTCCGATCTNNNGTAGTCCCAGCTACTCCGGAGC

GAGCACACGTCTGAACTCTTTCCCTACACGACGCTCTTCCGATCTNNNGAGTCAGGAGAATGGCGTGA

GAGCACACGTCTGAACTCTTTCCCTACACGACGCTCTTCCGATCTNNNGCCTGTAGTCCCAGCTACTTGC

CACACGTCTGAACTCTTTCCCTACACGACGCTCTTCCGATCTNNNACGCCCGGCTAATTTTTGTATTTCA

GCACACGTCTGAACTCTTTCCCTACACGACGCTCTTCCGATCTNNNAGATACCTCCCCGCATTAGTAGAC

GCACACGTCTGAACTCTTTCCCTACACGACGCTCTTCCGATCTNNNAACACATACACACACACACACACC

CACACGTCTGAACTCTTTCCCTACACGACGCTCTTCCGATCTNNNTGGGGTGAAACAAAGTCAAAACA

AGCACACGTCTGAACTCTTTCCCTACACGACGCTCTTCCGATCTNNNAAAGTGCTGGGATTACAGGCAT

GAGCACACGTCTGAACTCTTTCCCTACACGACGCTCTTCCGATCTNNNACCTGTAGTCCCAGCTACTCGC

GAGCACACGTCTGAACTCTTTCCCTACACGACGCTCTTCCGATCTNNNGAGGCTGAGGCAGGAGAATG

GAGCACACGTCTGAACTCTTTCCCTACACGACGCTCTTCCGATCTNNNTGAGCCAAGATCGTGTCACTG

AGAGCACACGTCTGAACTCTTTCCCTACACGACGCTCTTCCGATCTNNNGATCACGCCACTGCACTCTAC

AGCACACGTCTGAACTCTTTCCCTACACGACGCTCTTCCGATCTNNNGCCTGTAGTCCCAGCTACTTGCA

GCACACGTCTGAACTCTTTCCCTACACGACGCTCTTCCGATCTNNNGAGGCAGGAGAATGGTGTGAA
AGAGCACACGTCTGAACTCTTTCCCTACACGACGCTCTTCCGATCTNNNGCCTGTAGTCCCAGCTACTT
CACACGTCTGAACTCTTTCCCTACACGACGCTCTTCCGATCTNNNCGTCTCTGTCTTTGTCTCTGTCTCA
GCACACGTCTGAACTCTTTCCCTACACGACGCTCTTCCGATCTNNNTGAGTTCATATCATGTCCCAGGT
GCACACGTCTGAACTCTTTCCCTACACGACGCTCTTCCGATCTNNNTGAGTCTCTGGTAATAGCCCCTA
GCACACGTCTGAACTCTTTCCCTACACGACGCTCTTCCGATCTNNNGCTGGTCTTGAACTCCTGATCTT
CACACGTCTGAACTCTTTCCCTACACGACGCTCTTCCGATCTNNNGAGTGTAGTGGCTCGATCTCAGC
CACACGTCTGAACTCTTTCCCTACACGACGCTCTTCCGATCTNNNTCCACTATTTGCTAACCTGAGGCC
GCACACGTCTGAACTCTTTCCCTACACGACGCTCTTCCGATCTNNNCACAACACCATGTCCAGCTAATT
CACACGTCTGAACTCTTTCCCTACACGACGCTCTTCCGATCTNNNGGCTTTGGAAGTACTCCCTCTAGC
GCACACGTCTGAACTCTTTCCCTACACGACGCTCTTCCGATCTNNNGAGGGAGGCATGGTAGAAACC
AGCACACGTCTGAACTCTTTCCCTACACGACGCTCTTCCGATCTNNNGAGGTTGCAGTGAGCTGAGA
GAGCACACGTCTGAACTCTTTCCCTACACGACGCTCTTCCGATCTNNNGAGGCAGGAGAATGGTGTG
AGCACACGTCTGAACTCTTTCCCTACACGACGCTCTTCCGATCTNNNCATCCTGGCTAACACAGTGAAA
GCACACGTCTGAACTCTTTCCCTACACGACGCTCTTCCGATCTNNNAGGATCTCCTCTCTGATTCCCCCA
GCACACGTCTGAACTCTTTCCCTACACGACGCTCTTCCGATCTNNNACTGGTGAAGATGTGAACACTT
CACACGTCTGAACTCTTTCCCTACACGACGCTCTTCCGATCTNNNCTGACACTACAGCTTGATGACCTC
GCACACGTCTGAACTCTTTCCCTACACGACGCTCTTCCGATCTNNNTGAGTAGCTGAGATTACAGGC
ACACGTCTGAACTCTTTCCCTACACGACGCTCTTCCGATCTNNNTCCAGAAGCTGTGACTGTTTTTCA
CACACGTCTGAACTCTTTCCCTACACGACGCTCTTCCGATCTNNNCCTGCTTACGACATTCTCAACAACA
AGCACACGTCTGAACTCTTTCCCTACACGACGCTCTTCCGATCTNNNGCCTGTAGTCCCAGCTACTTGC
AAGAGCACACGTCTGAACTCTTTCCCTACACGACGCTCTTCCGATCTNNNGATGGTGTGAACCCAGGA
GCACACGTCTGAACTCTTTCCCTACACGACGCTCTTCCGATCTNNNGGCTAATACGATGAAACCCTGTC
GCACACGTCTGAACTCTTTCCCTACACGACGCTCTTCCGATCTNNNGTGACACAAGCGAAACTCCATCC
GAGCACACGTCTGAACTCTTTCCCTACACGACGCTCTTCCGATCTNNNGTGACAGAGCAAGAGTCCCT
AGCACACGTCTGAACTCTTTCCCTACACGACGCTCTTCCGATCTNNNTGAGCCATGATCGTGTCACTG
AGCACACGTCTGAACTCTTTCCCTACACGACGCTCTTCCGATCTNNNCAGGAGAATGGCGTGAACTT
AGCACACGTCTGAACTCTTTCCCTACACGACGCTCTTCCGATCTNNNGCCTGTAGTCCCAGCTACTTG
GAGCACACGTCTGAACTCTTTCCCTACACGACGCTCTTCCGATCTNNNGCAACAGAGTGAGACTCCGT
AGCACACGTCTGAACTCTTTCCCTACACGACGCTCTTCCGATCTNNNGACAGAGCAAGACTCCGTCTC
GAGCACACGTCTGAACTCTTTCCCTACACGACGCTCTTCCGATCTNNNGCCTGTAGTCCCAGCTACATG
AGCACACGTCTGAACTCTTTCCCTACACGACGCTCTTCCGATCTNNNGCCTGTAGTCCCAGCTACTTG
CACACGTCTGAACTCTTTCCCTACACGACGCTCTTCCGATCTNNNTCAAACTCCTGACCTCATGATCCC
GCACACGTCTGAACTCTTTCCCTACACGACGCTCTTCCGATCTNNNGGAAACCTCTACTCGCTTGCTAT
AGCACACGTCTGAACTCTTTCCCTACACGACGCTCTTCCGATCTNNNACAGCTTCAGTTTGTTTTGTCA
GCACACGTCTGAACTCTTTCCCTACACGACGCTCTTCCGATCTNNNTTTGAATCTGGCTGACCTTGGTA
GCACACGTCTGAACTCTTTCCCTACACGACGCTCTTCCGATCTNNNCCACCACGTTCCTGAATTTGATTC
CACACGTCTGAACTCTTTCCCTACACGACGCTCTTCCGATCTNNNGCTGCGTAACTTTCTAAGGTTGGC
CACACGTCTGAACTCTTTCCCTACACGACGCTCTTCCGATCTNNNCCCAGAGAACACCTGTGAGATACC
AGCACACGTCTGAACTCTTTCCCTACACGACGCTCTTCCGATCTNNNCTTGCAGTGAGCTGAGATTGC
AGCACACGTCTGAACTCTTTCCCTACACGACGCTCTTCCGATCTNNNCTGTGATCCCAGGACTTTGAGA
GAGCACACGTCTGAACTCTTTCCCTACACGACGCTCTTCCGATCTNNNGCCTGTAGTCCCAGCTACTTG
AGCACACGTCTGAACTCTTTCCCTACACGACGCTCTTCCGATCTNNNGACAGAGCGAAACTCCGTCTCC

GCACACGTCTGAACTCTTTCCCTACACGACGCTCTTCCGATCTNNNGAGATCACACCACTGCACTCCCA
CACACGTCTGAACTCTTTCCCTACACGACGCTCTTCCGATCTNNNAGTTGGGATCTCACTATGTTGCC
GCACACGTCTGAACTCTTTCCCTACACGACGCTCTTCCGATCTNNNGAGAGTCAAGCGTTTACCCTACT
GCACACGTCTGAACTCTTTCCCTACACGACGCTCTTCCGATCTNNNTGTGTGGGGACTTACAATCA
GCACACGTCTGAACTCTTTCCCTACACGACGCTCTTCCGATCTNNNCCTTACCATCAAAGTCCACTGGA
GCACACGTCTGAACTCTTTCCCTACACGACGCTCTTCCGATCTNNNTTTTTCTCCATGTTGTCCAGGCC
AGCACACGTCTGAACTCTTTCCCTACACGACGCTCTTCCGATCTNNNCACCTGTAGTCCCAGCTACTTG
GCACACGTCTGAACTCTTTCCCTACACGACGCTCTTCCGATCTNNNGCCTGTAGTCCCAGCTACTTGCA
GAGCACACGTCTGAACTCTTTCCCTACACGACGCTCTTCCGATCTNNNCTTGCAGTGAGCCGAGATCA
GAGCACACGTCTGAACTCTTTCCCTACACGACGCTCTTCCGATCTNNNCTTGCAGTGAGCCAGATCAT
GAGCACACGTCTGAACTCTTTCCCTACACGACGCTCTTCCGATCTNNNGAGGCTGAGGCAGGAGAAT
GAGCACACGTCTGAACTCTTTCCCTACACGACGCTCTTCCGATCTNNNGCAACAGAGTGAGACTCCGT
AGCACACGTCTGAACTCTTTCCCTACACGACGCTCTTCCGATCTNNNGGAACTTGCTTGAACCTGGG
GCACACGTCTGAACTCTTTCCCTACACGACGCTCTTCCGATCTNNNCGCTTTTCAGTTTCAGGTATA
CACACGTCTGAACTCTTTCCCTACACGACGCTCTTCCGATCTNNNAGCCTAAGCAACCTAAGAAGTGC
GCACACGTCTGAACTCTTTCCCTACACGACGCTCTTCCGATCTNNNACCCATCAGCCATAGTGTTCATT
CACACGTCTGAACTCTTTCCCTACACGACGCTCTTCCGATCTNNNCCATTTTCCCCTACATCCTGGATCA
GCACACGTCTGAACTCTTTCCCTACACGACGCTCTTCCGATCTNNNTGATTGGTGAGAATAGGCATCC
GAGCACACGTCTGAACTCTTTCCCTACACGACGCTCTTCCGATCTNNNGCTTCTCTGGTGGGGTTTTTC
ACACGTCTGAACTCTTTCCCTACACGACGCTCTTCCGATCTNNNCCATCCATGTTCCCACAAAAGACCAT
GCACACGTCTGAACTCTTTCCCTACACGACGCTCTTCCGATCTNNNCAGGTTCAAGCAATTCTCCTGCC
CACACGTCTGAACTCTTTCCCTACACGACGCTCTTCCGATCTNNNGGACAGTGCAGGCTTAGACTAAT
GCACACGTCTGAACTCTTTCCCTACACGACGCTCTTCCGATCTNNNATGGAAAAGAGATGTAGGCTG
CACACGTCTGAACTCTTTCCCTACACGACGCTCTTCCGATCTNNNTGCTGAGAAAAGTGACTTGAAGT
GAGCACACGTCTGAACTCTTTCCCTACACGACGCTCTTCCGATCTNNNGCCTGTAGTCCCAGCTACTTG
AGCACACGTCTGAACTCTTTCCCTACACGACGCTCTTCCGATCTNNNGAGGCAGGAGAATGGTGTGA
AGCACACGTCTGAACTCTTTCCCTACACGACGCTCTTCCGATCTNNNGCCTGTAGTCCCAGCTACTTG
AGCACACGTCTGAACTCTTTCCCTACACGACGCTCTTCCGATCTNNNGCCTGTAGTCCCAGCTACTTGC
AGCACACGTCTGAACTCTTTCCCTACACGACGCTCTTCCGATCTNNNGATGCCTGTAGTCCCAGCTAC
AGCACACGTCTGAACTCTTTCCCTACACGACGCTCTTCCGATCTNNNGCCTGTAGTCCCAGCTACTTG
GCACACGTCTGAACTCTTTCCCTACACGACGCTCTTCCGATCTNNNGCCTGTAGTCCCAGCTACTTGCA
GCACACGTCTGAACTCTTTCCCTACACGACGCTCTTCCGATCTNNNGCCTGTAGTCCCAGCTACTTGC
AGAGCACACGTCTGAACTCTTTCCCTACACGACGCTCTTCCGATCTNNNGAGGCAGGAGAATGGTGT
CACACGTCTGAACTCTTTCCCTACACGACGCTCTTCCGATCTNNNAGAAAATAGTGGAGCCCTATGCA
CACGTCTGAACTCTTTCCCTACACGACGCTCTTCCGATCTNNNGGATACCTTCCGATTTCTCTAGTTCAT
CACACGTCTGAACTCTTTCCCTACACGACGCTCTTCCGATCTNNNGGCTTTTCATTTGTTTGCAAGCAC
GCACACGTCTGAACTCTTTCCCTACACGACGCTCTTCCGATCTNNNGAGTAGCTGGGACTACAGGCA
GCACACGTCTGAACTCTTTCCCTACACGACGCTCTTCCGATCTNNNTTACTGGGCTTTTAGTTCCACCA
GCACACGTCTGAACTCTTTCCCTACACGACGCTCTTCCGATCTNNNAACCTTGACTGGACAAAGGAGA
CACACGTCTGAACTCTTTCCCTACACGACGCTCTTCCGATCTNNNGCCTTCAGCACATTGTGATCTTTC
AGCACACGTCTGAACTCTTTCCCTACACGACGCTCTTCCGATCTNNNCTTGCAGTGAGCCGAGATTGT
AGCACACGTCTGAACTCTTTCCCTACACGACGCTCTTCCGATCTNNNGCCTGTAGTCCCAGCTACTTGC
AGCACACGTCTGAACTCTTTCCCTACACGACGCTCTTCCGATCTNNNCTGTAAGTCCCAGCTACTCGG

GCACACGTCTGAACTCTTTCCCTACACGACGCTCTTCCGATCTNNNGATGGTCTGTGAGTATGCCCTTG
GCACACGTCTGAACTCTTTCCCTACACGACGCTCTTCCGATCTNNNCTTGGGTTGAACTGCATTGGTAC
ACACGTCTGAACTCTTTCCCTACACGACGCTCTTCCGATCTNNNCTAGAGAGAGAGTGAGGGGGTAG
GAGCACACGTCTGAACTCTTTCCCTACACGACGCTCTTCCGATCTNNNGCCTGTAGTCCCAGCTACTTG
ACACGTCTGAACTCTTTCCCTACACGACGCTCTTCCGATCTNNNCAAGACATTTATGTGCCTCAGCCCA
GAGCACACGTCTGAACTCTTTCCCTACACGACGCTCTTCCGATCTNNNCTGGGTGACAGCAAGACTCT
GAGCACACGTCTGAACTCTTTCCCTACACGACGCTCTTCCGATCTNNNCTGGGTGAGAGAGCAAGAC
AGCACACGTCTGAACTCTTTCCCTACACGACGCTCTTCCGATCTNNNCTGCAGTGAGCTGTGATTGTGC
CACACGTCTGAACTCTTTCCCTACACGACGCTCTTCCGATCTNNNGAGACTTGTGAAGAGCGAGAGA
AGCACACGTCTGAACTCTTTCCCTACACGACGCTCTTCCGATCTNNNCTGGGCAATAGAGCGAGACTC
AGCACACGTCTGAACTCTTTCCCTACACGACGCTCTTCCGATCTNNNAGAATGGCATGAACCTGGGA
AGCACACGTCTGAACTCTTTCCCTACACGACGCTCTTCCGATCTNNNCTATCACACCCCTGGAACCTCC
CACACGTCTGAACTCTTTCCCTACACGACGCTCTTCCGATCTNNNATCACTTTTGTAGCCTGGATGCTC
ACACGTCTGAACTCTTTCCCTACACGACGCTCTTCCGATCTNNNGTTCAACTGGGAGCTAACACAATCA
CACACGTCTGAACTCTTTCCCTACACGACGCTCTTCCGATCTNNNCTTCTATAGAAGGGTGCGCTCTC
AGCACACGTCTGAACTCTTTCCCTACACGACGCTCTTCCGATCTNNNGAGGTGGGAGAATGGCATGA
GCACACGTCTGAACTCTTTCCCTACACGACGCTCTTCCGATCTNNNTTCACATCATTCTCCTGCCTCAGC
AGCACACGTCTGAACTCTTTCCCTACACGACGCTCTTCCGATCTNNNGAGGTTGCAGTGAGCTGAGA
AGCACACGTCTGAACTCTTTCCCTACACGACGCTCTTCCGATCTNNNTTCAAGCGATTCTCCTGCCTCC
GCACACGTCTGAACTCTTTCCCTACACGACGCTCTTCCGATCTNNNGAGGCAGAAGAGTCGCTTGAA
CACACGTCTGAACTCTTTCCCTACACGACGCTCTTCCGATCTNNNAGGCAGGAGAATCACTTGAACC
GAGCACACGTCTGAACTCTTTCCCTACACGACGCTCTTCCGATCTNNNGAGATCGAGATCGCACCACT
CACACGTCTGAACTCTTTCCCTACACGACGCTCTTCCGATCTNNNTTTTTCTAATGCCTGACTTGCCCCA
CACGTCTGAACTCTTTCCCTACACGACGCTCTTCCGATCTNNNAGTGAGCTTTGGTAATTTTTGCCTCA
CACACGTCTGAACTCTTTCCCTACACGACGCTCTTCCGATCTNNNAACTCCTGACATCATGATCCACCC
CACACGTCTGAACTCTTTCCCTACACGACGCTCTTCCGATCTNNNAGACAATCATGTTCCCTGGGAAT
CACACGTCTGAACTCTTTCCCTACACGACGCTCTTCCGATCTNNNTGCCCAAGTGAGACTTTTATCCTC
CACACGTCTGAACTCTTTCCCTACACGACGCTCTTCCGATCTNNNCCAGTTTGCTCGTTTCCTTTCATCA
AGAGCACACGTCTGAACTCTTTCCCTACACGACGCTCTTCCGATCTNNNGACAGAGCAAGACTCCGTC
AGCACACGTCTGAACTCTTTCCCTACACGACGCTCTTCCGATCTNNNAGAATGGCGTGAATCCAGGA
AGCACACGTCTGAACTCTTTCCCTACACGACGCTCTTCCGATCTNNNGAGATCGCACCACTGTACTCCC
ACACGTCTGAACTCTTTCCCTACACGACGCTCTTCCGATCTNNNTCCCCATTTTTCTTCCCTTGACACAT
AGCACACGTCTGAACTCTTTCCCTACACGACGCTCTTCCGATCTNNNCTTGCAGTGAGCTGAGATCCC
ACACGTCTGAACTCTTTCCCTACACGACGCTCTTCCGATCTNNNAAACCCAATTCCAAGACCCTTTCCAT
CACACGTCTGAACTCTTTCCCTACACGACGCTCTTCCGATCTNNNACTTTGTTGGATGTTGGATCTGAC
ACACGTCTGAACTCTTTCCCTACACGACGCTCTTCCGATCTNNNAGCTACAAATCCAGCTGACTAAGC
CACACGTCTGAACTCTTTCCCTACACGACGCTCTTCCGATCTNNNCTTGAACTCCTGACCTCAAGTGA
AGCACACGTCTGAACTCTTTCCCTACACGACGCTCTTCCGATCTNNNGCGACAGAGTGAGACTCCAT
GAGCACACGTCTGAACTCTTTCCCTACACGACGCTCTTCCGATCTNNNACCTGTAGTCCCAGCTACTCG
AGCACACGTCTGAACTCTTTCCCTACACGACGCTCTTCCGATCTNNNGCAGTCCCAGCTACTCAGAAG
AGCACACGTCTGAACTCTTTCCCTACACGACGCTCTTCCGATCTNNNCTGAGGCAGGAGAATGGTGT
GAGCACACGTCTGAACTCTTTCCCTACACGACGCTCTTCCGATCTNNNGAGACTGAGGCAGGAGAAT
GAGCACACGTCTGAACTCTTTCCCTACACGACGCTCTTCCGATCTNNNGAGACTGAGGCAGGAGAAT

GCACACGTCTGAACTCTTTCCCTACACGACGCTCTTCCGATCTNNNGTGGATCATGAGGTCAGGAGATC
AGCACACGTCTGAACTCTTTCCCTACACGACGCTCTTCCGATCTNNNGCCTGTAGTCCCAGCTACTCTCA
GAGCACACGTCTGAACTCTTTCCCTACACGACGCTCTTCCGATCTNNNGAGGCAGGAGAATGGTGTG
AGCACACGTCTGAACTCTTTCCCTACACGACGCTCTTCCGATCTNNNCTGAGGCAGGAGAATCACTTGA
GCACACGTCTGAACTCTTTCCCTACACGACGCTCTTCCGATCTNNNACCTGTAATCCCAGCACTTTGAG
ACACGTCTGAACTCTTTCCCTACACGACGCTCTTCCGATCTNNNTACCTAGCCTGTTCACTGATTGGCAT
GCACACGTCTGAACTCTTTCCCTACACGACGCTCTTCCGATCTNNNGGAGGGAGAAGTGCCTAAAGAA
AGCACACGTCTGAACTCTTTCCCTACACGACGCTCTTCCGATCTNNNGGGAAGCTGAGGCAGAAGAAT
GAGCACACGTCTGAACTCTTTCCCTACACGACGCTCTTCCGATCTNNNCTTGCAGTGAGCCAAGATTGC
GAGCACACGTCTGAACTCTTTCCCTACACGACGCTCTTCCGATCTNNNTGAGCCAAGATCACACCACTG
GAGCACACGTCTGAACTCTTTCCCTACACGACGCTCTTCCGATCTNNNGCCTGTAGTCCCAGCTACTTGC
AGCACACGTCTGAACTCTTTCCCTACACGACGCTCTTCCGATCTNNNGCCTGTAGTCCCAGCTACTTGCA
CACACGTCTGAACTCTTTCCCTACACGACGCTCTTCCGATCTNNNTGAGATGGAAGTGTTTTAGGCAG
CACACGTCTGAACTCTTTCCCTACACGACGCTCTTCCGATCTNNNCTCAGGACTTTGCCTATGATCCACA
GCACACGTCTGAACTCTTTCCCTACACGACGCTCTTCCGATCTNNNATTTCTGCAGCCAGCTTGAATTTC
GCACACGTCTGAACTCTTTCCCTACACGACGCTCTTCCGATCTNNNCAAAACTCACAGTCAAGCAGTGA
ACACGTCTGAACTCTTTCCCTACACGACGCTCTTCCGATCTNNNTTGTGGCAGCATAACTTCACATGCAT
GAGCACACGTCTGAACTCTTTCCCTACACGACGCTCTTCCGATCTNNNGAGTCTGAGGCAGGAGAATG
CACACGTCTGAACTCTTTCCCTACACGACGCTCTTCCGATCTNNNCACCACTCCTGGCTAATTTTGTTCAT
GCACACGTCTGAACTCTTTCCCTACACGACGCTCTTCCGATCTNNNTGAAAAGTTTCTGTTCGCATCCTC
GCACACGTCTGAACTCTTTCCCTACACGACGCTCTTCCGATCTNNNCAGGAGACTCACGTCCTTGAAAT
AGCACACGTCTGAACTCTTTCCCTACACGACGCTCTTCCGATCTNNNCCTCAGCCTCCCAAGTAGTTGCA
CACACGTCTGAACTCTTTCCCTACACGACGCTCTTCCGATCTNNNACATGTTCAATACAGACCCAGCTCA
AGAGCACACGTCTGAACTCTTTCCCTACACGACGCTCTTCCGATCTNNNAGGAGAATGGCTTGAACCTC
AGCACACGTCTGAACTCTTTCCCTACACGACGCTCTTCCGATCTNNNGAGGCAGGAGAATCGCTTGAAC
GAGCACACGTCTGAACTCTTTCCCTACACGACGCTCTTCCGATCTNNNCGACAGAACGAGACTCCATCTC
GAGCACACGTCTGAACTCTTTCCCTACACGACGCTCTTCCGATCTNNNGTAGCAGTGAGGCGTGATCAC
GAGCACACGTCTGAACTCTTTCCCTACACGACGCTCTTCCGATCTNNNGAGGCAGGAGAATGGTGTGA
CACGTCTGAACTCTTTCCCTACACGACGCTCTTCCGATCTNNNGTTTCAAACTACTTTTCCCTCGCACATG
GCACACGTCTGAACTCTTTCCCTACACGACGCTCTTCCGATCTNNNCTGGGATTACAGGTGTGAGTCA
GCACACGTCTGAACTCTTTCCCTACACGACGCTCTTCCGATCTNNNTGAAGCTAAGGGAGAGTCTGAG
GCACACGTCTGAACTCTTTCCCTACACGACGCTCTTCCGATCTNNNAAATAATGTGTCCGGTAGCCCTCC
ACACGTCTGAACTCTTTCCCTACACGACGCTCTTCCGATCTNNNTTCAAGCAATTTTCCTGCCTCAGCAT
GCACACGTCTGAACTCTTTCCCTACACGACGCTCTTCCGATCTNNNATCTCACCTTGAAAACGTGCTTCC
GCACACGTCTGAACTCTTTCCCTACACGACGCTCTTCCGATCTNNNGACCCTGTCTCAAAAGCAAAACAC
GCACACGTCTGAACTCTTTCCCTACACGACGCTCTTCCGATCTNNNCCCCATGGCCATTTTCTTTTGTACA
AGCACACGTCTGAACTCTTTCCCTACACGACGCTCTTCCGATCTNNNCTTGCAGTGAGCTGAGATTGCC
AGAGCACACGTCTGAACTCTTTCCCTACACGACGCTCTTCCGATCTNNNCTTGCAGTGAGCTGAGATTG
AGCACACGTCTGAACTCTTTCCCTACACGACGCTCTTCCGATCTNNNCTGAGGCAGAGAATTGCTTGAA
GCACACGTCTGAACTCTTTCCCTACACGACGCTCTTCCGATCTNNNGAGGCTAAGGCAGGAGAATGG
AGCACACGTCTGAACTCTTTCCCTACACGACGCTCTTCCGATCTNNNTGTAATCCTAGCACTTTGGGAG
GAGCACACGTCTGAACTCTTTCCCTACACGACGCTCTTCCGATCTNNNGCCTGTAGTCCCACCTACTTGC
GAGCACACGTCTGAACTCTTTCCCTACACGACGCTCTTCCGATCTNNNGTCAGGAGTTTGAGACCAGC

CACACGTCTGAACTCTTTCCCTACACGACGCTCTTCCGATCTNNNCAGAGGCTGAAGCATGAGAATTG
CACACGTCTGAACTCTTTCCCTACACGACGCTCTTCCGATCTNNNGGCTCTGCTCCCGTTAGTTTATACA
GCACACGTCTGAACTCTTTCCCTACACGACGCTCTTCCGATCTNNNCAACCTTAGCCTCCTGAGTAGCC
GCACACGTCTGAACTCTTTCCCTACACGACGCTCTTCCGATCTNNNACACTTCAGAAAACTGTTTGGCA
GCACACGTCTGAACTCTTTCCCTACACGACGCTCTTCCGATCTNNNTCCAGCCATCATCTAGTCCAGTA
CACGTCTGAACTCTTTCCCTACACGACGCTCTTCCGATCTNNNTCATGTCATACAAGGCAAACAAGTCA
CACACGTCTGAACTCTTTCCCTACACGACGCTCTTCCGATCTNNNACTTTTGCTGGCTGGCATTTTATCA
AGCACACGTCTGAACTCTTTCCCTACACGACGCTCTTCCGATCTNNNTGAGCCAAGATCACACCACTGC
GAGCACACGTCTGAACTCTTTCCCTACACGACGCTCTTCCGATCTNNNGCCTGTAGTCCCAGCTACTTG
AGCACACGTCTGAACTCTTTCCCTACACGACGCTCTTCCGATCTNNNGAGATAGCACGACTGCACTCCC
AGCACACGTCTGAACTCTTTCCCTACACGACGCTCTTCCGATCTNNNGAGACAGGAGAATGGCGTGA
AGCACACGTCTGAACTCTTTCCCTACACGACGCTCTTCCGATCTNNNCTTGCAGTGAGCCGAGATCTC
GAGCACACGTCTGAACTCTTTCCCTACACGACGCTCTTCCGATCTNNNCGACAGTGCAAGACTCCTCTC
GAGCACACGTCTGAACTCTTTCCCTACACGACGCTCTTCCGATCTNNNGAGACTGAGGCAGGAGAAT
AGAGCACACGTCTGAACTCTTTCCCTACACGACGCTCTTCCGATCTNNNGCCTGTAGTCCCAGCTACTTG
AGCACACGTCTGAACTCTTTCCCTACACGACGCTCTTCCGATCTNNNGCTCCTGTAGTCCCAGCTACTC
AGCACACGTCTGAACTCTTTCCCTACACGACGCTCTTCCGATCTNNNGAGGTTGCAGTGAGCCAAGA
GAGCACACGTCTGAACTCTTTCCCTACACGACGCTCTTCCGATCTNNNCCTGTTGTCCCAGCTACTTGG
GCACACGTCTGAACTCTTTCCCTACACGACGCTCTTCCGATCTNNNGCCTGTAGTCCCAGCTACTTGCA
GAGCACACGTCTGAACTCTTTCCCTACACGACGCTCTTCCGATCTNNNTGCCTGTAGTCCCAGCTACTC
GCACACGTCTGAACTCTTTCCCTACACGACGCTCTTCCGATCTNNNTATGCATGGATGGTCCTTTCCTT
GCACACGTCTGAACTCTTTCCCTACACGACGCTCTTCCGATCTNNNCTCCCGAGTTCAAGCAATTCTGC
ACACGTCTGAACTCTTTCCCTACACGACGCTCTTCCGATCTNNNTCCTTGCCCCCATTTTACAGATTCAT
GCACACGTCTGAACTCTTTCCCTACACGACGCTCTTCCGATCTNNNGTACTGGTAAGCTGCTCTGGTGC
CACACGTCTGAACTCTTTCCCTACACGACGCTCTTCCGATCTNNNGTTCAAGCTATTCTCCTGCCTCTCA
CACACGTCTGAACTCTTTCCCTACACGACGCTCTTCCGATCTNNNCACGACTTCTCCCTCTTCTGAAACA
GCACACGTCTGAACTCTTTCCCTACACGACGCTCTTCCGATCTNNNACAGTTTTCAGGGTGGTGTCTA
ACACGTCTGAACTCTTTCCCTACACGACGCTCTTCCGATCTNNNGGTGGCAGTGGGGAGAAATATTA
AGCACACGTCTGAACTCTTTCCCTACACGACGCTCTTCCGATCTNNNGAGATCACACCACTGCACTCCC
AGCACACGTCTGAACTCTTTCCCTACACGACGCTCTTCCGATCTNNNTGAGCTGTGATTGAGCCACTG
CACACGTCTGAACTCTTTCCCTACACGACGCTCTTCCGATCTNNNGATCACGAGGTCAGGAGATCGA
GAGCACACGTCTGAACTCTTTCCCTACACGACGCTCTTCCGATCTNNNGCCTGTAATCCCAGCTACTCG
AGCACACGTCTGAACTCTTTCCCTACACGACGCTCTTCCGATCTNNNCGACAGAGTGAGACTCCGTCTC
GCACACGTCTGAACTCTTTCCCTACACGACGCTCTTCCGATCTNNNCCTGGACAACAAGAGCCAAATTG
AGCACACGTCTGAACTCTTTCCCTACACGACGCTCTTCCGATCTNNNACCTGTAGTCCCAGCTACTCGC
CACACGTCTGAACTCTTTCCCTACACGACGCTCTTCCGATCTNNNCTGCGTCTGGCCAGTATAATTTGC
GCACACGTCTGAACTCTTTCCCTACACGACGCTCTTCCGATCTNNNCTCCTGGGTTGATGCCATTCTCC
CACACGTCTGAACTCTTTCCCTACACGACGCTCTTCCGATCTNNNCCCTCTCCCATCCTTTTTGCTATCAT
GCACACGTCTGAACTCTTTCCCTACACGACGCTCTTCCGATCTNNNGGCTCATGACTGTAATCCTAGCA
ACACGTCTGAACTCTTTCCCTACACGACGCTCTTCCGATCTNNNCAGTGCTTTTCAGATGAGCAACACA
CACACGTCTGAACTCTTTCCCTACACGACGCTCTTCCGATCTNNNTTGAATCACTTTGGCCTAGCTCTC
AGCACACGTCTGAACTCTTTCCCTACACGACGCTCTTCCGATCTNNNGCCTGTAGTCCCAGCTACTTGC
GAGCACACGTCTGAACTCTTTCCCTACACGACGCTCTTCCGATCTNNNGCCTGTAGTCCCAGCTACTTG

AGCACACGTCTGAACTCTTTCCCTACACGACGCTCTTCCGATCTNNNGCCTGTAGTCCCAGCTACTTG
AGCACACGTCTGAACTCTTTCCCTACACGACGCTCTTCCGATCTNNNGCCTGTAGTCCCAGCTACTTGC
GCACACGTCTGAACTCTTTCCCTACACGACGCTCTTCCGATCTNNNAGAATGGCATGAACCAGGAGG
AGCACACGTCTGAACTCTTTCCCTACACGACGCTCTTCCGATCTNNNGCCTGTAGTCCCAGCTACTTGC
CACACGTCTGAACTCTTTCCCTACACGACGCTCTTCCGATCTNNNCTTGCAGTGAGCCAAGATTGTGC
AGCACACGTCTGAACTCTTTCCCTACACGACGCTCTTCCGATCTNNNTCTGTAGTCCCAGCTACTCGG
AGCACACGTCTGAACTCTTTCCCTACACGACGCTCTTCCGATCTNNNGCCTGTAGTCCCAGCTACTTG
CACACGTCTGAACTCTTTCCCTACACGACGCTCTTCCGATCTNNNGCAACATTCTGAGACCCCTGTCCA
GAGCACACGTCTGAACTCTTTCCCTACACGACGCTCTTCCGATCTNNNCCTGGGCAACAAGAGTGAA
AGCACACGTCTGAACTCTTTCCCTACACGACGCTCTTCCGATCTNNNGCCTGTAGTCCCAGCTACTTGC
GAGCACACGTCTGAACTCTTTCCCTACACGACGCTCTTCCGATCTNNNCTTGAGCCCAGGAGTTTGAG
AGCACACGTCTGAACTCTTTCCCTACACGACGCTCTTCCGATCTNNNGCGACAGAGTGAGACTCCATC
CACACGTCTGAACTCTTTCCCTACACGACGCTCTTCCGATCTNNNCCAGTCTGTTCTCAAACTCCTGACA
CACACGTCTGAACTCTTTCCCTACACGACGCTCTTCCGATCTNNNGAAGACAAACCCAGGTAGGAAGA
CACACGTCTGAACTCTTTCCCTACACGACGCTCTTCCGATCTNNNGAGTTTTGCTCTTGTTGCCTAGG
GCACACGTCTGAACTCTTTCCCTACACGACGCTCTTCCGATCTNNNTTTACATGGTGTCAGGCAAGAG
GCACACGTCTGAACTCTTTCCCTACACGACGCTCTTCCGATCTNNNAATTGGAGGGAAATGGGGAAT
GCACACGTCTGAACTCTTTCCCTACACGACGCTCTTCCGATCTNNNAATTGGAGGGAAATGGGGAAT
GCACACGTCTGAACTCTTTCCCTACACGACGCTCTTCCGATCTNNNACTATCGGCCATTATCCCTGATG
ACACGTCTGAACTCTTTCCCTACACGACGCTCTTCCGATCTNNNTCTAACACAAGGCGGCTTATTCTCA
GCACACGTCTGAACTCTTTCCCTACACGACGCTCTTCCGATCTNNNCCAGGACTGTATGCATGATCTCA
CACACGTCTGAACTCTTTCCCTACACGACGCTCTTCCGATCTNNNTCCAACCTTTGTCCCTTACTCAGC
CACACGTCTGAACTCTTTCCCTACACGACGCTCTTCCGATCTNNNAACGTCATCTGACAAGTTCCACTC
CACACGTCTGAACTCTTTCCCTACACGACGCTCTTCCGATCTNNNCTTTCCCCCATAGCACTCTGATC
CACACGTCTGAACTCTTTCCCTACACGACGCTCTTCCGATCTNNNTCCAGAAGTGTTCCCAAGAGAAC
AGAGCACACGTCTGAACTCTTTCCCTACACGACGCTCTTCCGATCTNNNCTTGCAGTGAGCCAAGATC
GAGCACACGTCTGAACTCTTTCCCTACACGACGCTCTTCCGATCTNNNTGGTGACAGAGTGAGACTCC
AGCACACGTCTGAACTCTTTCCCTACACGACGCTCTTCCGATCTNNNTGTAGTCCCAGCTACTCAGGA
GAGCACACGTCTGAACTCTTTCCCTACACGACGCTCTTCCGATCTNNNCTGGGTGAAAGAGCGAGAC
GCACACGTCTGAACTCTTTCCCTACACGACGCTCTTCCGATCTNNNGCCTGTAGTCCCAGCTACTTGCA
AGCACACGTCTGAACTCTTTCCCTACACGACGCTCTTCCGATCTNNNCTTGCAGTGAGCTGAGATTG
GCACACGTCTGAACTCTTTCCCTACACGACGCTCTTCCGATCTNNNGACAGAGCAAGACTCCGTCTCCA
GAGCACACGTCTGAACTCTTTCCCTACACGACGCTCTTCCGATCTNNNCTAGGCGACAGAGCAAGAC
AGCACACGTCTGAACTCTTTCCCTACACGACGCTCTTCCGATCTNNNGAGCTTGCAGTGAGTGGAGA
ACACGTCTGAACTCTTTCCCTACACGACGCTCTTCCGATCTNNNTGCAATGTTACTAACTGGAGTTGTC
GCACACGTCTGAACTCTTTCCCTACACGACGCTCTTCCGATCTNNNGGTGACAGAGTGAGACTCCATC
AGCACACGTCTGAACTCTTTCCCTACACGACGCTCTTCCGATCTNNNCGACAGAGAGAGACTCCGTCT
AGCACACGTCTGAACTCTTTCCCTACACGACGCTCTTCCGATCTNNNCTGCTCAGCTATCCCCAGGTC
GCACACGTCTGAACTCTTTCCCTACACGACGCTCTTCCGATCTNNNGAAGAGCTGGAGTCTGGAGGA
GCACACGTCTGAACTCTTTCCCTACACGACGCTCTTCCGATCTNNNATTTCAATGTTTCAGCAGGCAAC
AGCACACGTCTGAACTCTTTCCCTACACGACGCTCTTCCGATCTNNNCCACCTTGACCTCCCAAAGTGC
GAGCACACGTCTGAACTCTTTCCCTACACGACGCTCTTCCGATCTNNNGAGATCACGCTACTGCACTCC
CACACGTCTGAACTCTTTCCCTACACGACGCTCTTCCGATCTNNNATACAAACAGGAAAACACAGCCG

AGCACACGTCTGAACTCTTTCCCTACACGACGCTCTTCCGATCTNNNGCCTGTAGTCCCAGCTACTTG
CACACGTCTGAACTCTTTCCCTACACGACGCTCTTCCGATCTNNNCCAGGGTGCCACACTACATTTAG
AGCACACGTCTGAACTCTTTCCCTACACGACGCTCTTCCGATCTNNNTGAGCCGAGATCACACCAATG
AGCACACGTCTGAACTCTTTCCCTACACGACGCTCTTCCGATCTNNNCCACAGGGGTACTTGCATCTT
GCACACGTCTGAACTCTTTCCCTACACGACGCTCTTCCGATCTNNNAGCTTTTGTCCCTTTGAGAAG
AGCACACGTCTGAACTCTTTCCCTACACGACGCTCTTCCGATCTNNNCCTGCCAACACACCTAGCTAA
GCACACGTCTGAACTCTTTCCCTACACGACGCTCTTCCGATCTNNNGATGGGGTTCCCTTTGTTAGTG
CACACGTCTGAACTCTTTCCCTACACGACGCTCTTCCGATCTNNNCGGTTTGAGCTTGCAGATTCTTT
AGCACACGTCTGAACTCTTTCCCTACACGACGCTCTTCCGATCTNNNGCCTGTAGTCCCAGCTACTTGC
CACACGTCTGAACTCTTTCCCTACACGACGCTCTTCCGATCTNNNAAAGTGGTGGGATTACAGGTGT
ACACGTCTGAACTCTTTCCCTACACGACGCTCTTCCGATCTNNNTGTTTCTTAGGAGACAAGGGGTGC
GCACACGTCTGAACTCTTTCCCTACACGACGCTCTTCCGATCTNNNGGGTACTTGAGATTAGGGAGT
GCACACGTCTGAACTCTTTCCCTACACGACGCTCTTCCGATCTNNNAATGCAGTGGGATGATCTCAAC
CACACGTCTGAACTCTTTCCCTACACGACGCTCTTCCGATCTNNNACTCCCGAACTGATTTTCTCCAAC
CACACGTCTGAACTCTTTCCCTACACGACGCTCTTCCGATCTNNNAGTGAGGTAGATACAACCCTGGA
GAGCACACGTCTGAACTCTTTCCCTACACGACGCTCTTCCGATCTNNNCACAGAGCCAGATTCCGTCT
AGCACACGTCTGAACTCTTTCCCTACACGACGCTCTTCCGATCTNNNCTTGCAGTGAGCCGAGATTG
AGCACACGTCTGAACTCTTTCCCTACACGACGCTCTTCCGATCTNNNGCCTGTAGTCCCAGCTACTTG
GAGCACACGTCTGAACTCTTTCCCTACACGACGCTCTTCCGATCTNNNGTAGTGCCAGCTACACGAGA
GCACACGTCTGAACTCTTTCCCTACACGACGCTCTTCCGATCTNNNAGGTTGCAGTGAGCTGAGATC
GCACACGTCTGAACTCTTTCCCTACACGACGCTCTTCCGATCTNNNGAGGCACAGAATCGCTTGAACC
AGCACACGTCTGAACTCTTTCCCTACACGACGCTCTTCCGATCTNNNCTGTAATCCCAGCCACTCAGG
GCACACGTCTGAACTCTTTCCCTACACGACGCTCTTCCGATCTNNNCTCTAGTCCCAGCTACTCAGGAG
GCACACGTCTGAACTCTTTCCCTACACGACGCTCTTCCGATCTNNNTTTGATGTGCTGCTGGATTCAG
GCACACGTCTGAACTCTTTCCCTACACGACGCTCTTCCGATCTNNNGGAGACGGAGCTTGCAGTAAG
GCACACGTCTGAACTCTTTCCCTACACGACGCTCTTCCGATCTNNNCTCTCTCAAGGTAGCCACTTCTG
GCACACGTCTGAACTCTTTCCCTACACGACGCTCTTCCGATCTNNNGCTGGATAGGAGAGACAAGAA
GCACACGTCTGAACTCTTTCCCTACACGACGCTCTTCCGATCTNNNCCACCATGCCCAGCTAATTTTTC
GCACACGTCTGAACTCTTTCCCTACACGACGCTCTTCCGATCTNNNCTTGCAGTGAGCCAAGATTGCC
GCACACGTCTGAACTCTTTCCCTACACGACGCTCTTCCGATCTNNNTTGTTGTTCGAACTCCTTACAGA
AGAGCACACGTCTGAACTCTTTCCCTACACGACGCTCTTCCGATCTNNNGCCTGTAGTCCCAGCTACTT
AGAGCACACGTCTGAACTCTTTCCCTACACGACGCTCTTCCGATCTNNNCTTGCAGTGAGCCGAGATTG
AGCACACGTCTGAACTCTTTCCCTACACGACGCTCTTCCGATCTNNNGCCTGTAGTCCCAGCTACTTG
AGCACACGTCTGAACTCTTTCCCTACACGACGCTCTTCCGATCTNNNCTGGGTGACAGTGTGAGACTC
AGCACACGTCTGAACTCTTTCCCTACACGACGCTCTTCCGATCTNNNGAGGTTGCAGTGAGCCAAGA
AGCACACGTCTGAACTCTTTCCCTACACGACGCTCTTCCGATCTNNNGCCTGTAGTCCCAGCTACTTG
AGCACACGTCTGAACTCTTTCCCTACACGACGCTCTTCCGATCTNNNGAGGCTAAGGCAGGAGAATG
AGCACACGTCTGAACTCTTTCCCTACACGACGCTCTTCCGATCTNNNACCTGTAGTCCCAGCTACTCGC
CACACGTCTGAACTCTTTCCCTACACGACGCTCTTCCGATCTNNNTGCGTTTCTCTACCTACAACTGTCA
CACACGTCTGAACTCTTTCCCTACACGACGCTCTTCCGATCTNNNCCTGCATGACTCGAAGATGTAGT
AGCACACGTCTGAACTCTTTCCCTACACGACGCTCTTCCGATCTNNNTTCACCATTCTCCTGCCTCCA
GCACACGTCTGAACTCTTTCCCTACACGACGCTCTTCCGATCTNNNTCAAGAAGAAAACCCTCACCAG
GCACACGTCTGAACTCTTTCCCTACACGACGCTCTTCCGATCTNNNCCCCTGTATTTTATGTGGGGTG

AGCACACGTCTGAACTCTTTCCCTACACGACGCTCTTCCGATCTNNNCTATTTGGGAGGCTGAGGCA
GCACACGTCTGAACTCTTTCCCTACACGACGCTCTTCCGATCTNNNCAACAAGAGCGAGAGTCCATCT
GAGCACACGTCTGAACTCTTTCCCTACACGACGCTCTTCCGATCTNNNCTGTGGTCAGGAGTTCGAGA
AGCACACGTCTGAACTCTTTCCCTACACGACGCTCTTCCGATCTNNNAGGAGAATCGCTTGAACCCAG
GAGCACACGTCTGAACTCTTTCCCTACACGACGCTCTTCCGATCTNNNGAATGGCGTGAACCCAGGA
AGAGCACACGTCTGAACTCTTTCCCTACACGACGCTCTTCCGATCTNNNAGGAGAATGGCGTGAACCT
GCACACGTCTGAACTCTTTCCCTACACGACGCTCTTCCGATCTNNNAACAGAGTGAGACCCCGTCTCAT
GAGCACACGTCTGAACTCTTTCCCTACACGACGCTCTTCCGATCTNNNGAGGCAGGAGAATGGTGTG
AGCACACGTCTGAACTCTTTCCCTACACGACGCTCTTCCGATCTNNNGCCTGTAGTCCCAGCTACTTG
AGCACACGTCTGAACTCTTTCCCTACACGACGCTCTTCCGATCTNNNGAGGCTGAGGCAGGAGAAT
GAGCACACGTCTGAACTCTTTCCCTACACGACGCTCTTCCGATCTNNNGCCTGTAGTCCCAGCTACTTG
AGCACACGTCTGAACTCTTTCCCTACACGACGCTCTTCCGATCTNNNGCCTGTAATCCCAGCTACTCGC
AGCACACGTCTGAACTCTTTCCCTACACGACGCTCTTCCGATCTNNNCAGATCACGAGGTCAGGAGAT
AGCACACGTCTGAACTCTTTCCCTACACGACGCTCTTCCGATCTNNNATCATGAGGTCAGGAGATCGA
AGCACACGTCTGAACTCTTTCCCTACACGACGCTCTTCCGATCTNNNGAGGTTGCAGTGAGCTGAGAT
AGCACACGTCTGAACTCTTTCCCTACACGACGCTCTTCCGATCTNNNGCCTGTAGTCCCAGCTACTTG
GCACACGTCTGAACTCTTTCCCTACACGACGCTCTTCCGATCTNNNAGGAACCTCTGAAAGCCATCTTC
AGAGCACACGTCTGAACTCTTTCCCTACACGACGCTCTTCCGATCTNNNACCTGTAGTCCCAGCTACTC
AGCACACGTCTGAACTCTTTCCCTACACGACGCTCTTCCGATCTNNNAAGATCACACCACTGCACTCCC
AGCACACGTCTGAACTCTTTCCCTACACGACGCTCTTCCGATCTNNNGAGAATGGCATGAACTGGGG
GCACACGTCTGAACTCTTTCCCTACACGACGCTCTTCCGATCTNNNGTGAGATTTTTGAGGGGATGG
AGCACACGTCTGAACTCTTTCCCTACACGACGCTCTTCCGATCTNNNTGCCTGTAGTCCCAGCTACTCC
GCACACGTCTGAACTCTTTCCCTACACGACGCTCTTCCGATCTNNNCCCGGCCTTGTTTTTTTTTGTTTC
GCACACGTCTGAACTCTTTCCCTACACGACGCTCTTCCGATCTNNNAGTGCTGGGATTATAGGCATGA
CACACGTCTGAACTCTTTCCCTACACGACGCTCTTCCGATCTNNNGGGAGTCTAAGCCTCTTTGTAGG
CACACGTCTGAACTCTTTCCCTACACGACGCTCTTCCGATCTNNNCTCCTGAGTAGCTGGAACTACAG
CACACGTCTGAACTCTTTCCCTACACGACGCTCTTCCGATCTNNNGCTGCTGTTTATTGAGCACCTACC
AGCACACGTCTGAACTCTTTCCCTACACGACGCTCTTCCGATCTNNNGCCTGTAGTCCCAGCTACTTGC
AGCACACGTCTGAACTCTTTCCCTACACGACGCTCTTCCGATCTNNNGGTGAAGAAAGTGGAGGTGC
AGCACACGTCTGAACTCTTTCCCTACACGACGCTCTTCCGATCTNNNCTCTAGCTCTGTTGGGCTGAG
GCACACGTCTGAACTCTTTCCCTACACGACGCTCTTCCGATCTNNNCTTCCGAGTAGCTGGGATGACC
CACACGTCTGAACTCTTTCCCTACACGACGCTCTTCCGATCTNNNGTCTTGAACTCCTGACCTTGTGA
AGCACACGTCTGAACTCTTTCCCTACACGACGCTCTTCCGATCTNNNGCCTGTAGTCCCAGCTACTTG
ACACGTCTGAACTCTTTCCCTACACGACGCTCTTCCGATCTNNNGGCTCCTTGTCTGTTTCATTCTCCAT
CACACGTCTGAACTCTTTCCCTACACGACGCTCTTCCGATCTNNNCATCCCAAAGCCAGGTTTTCTGCA
GCACACGTCTGAACTCTTTCCCTACACGACGCTCTTCCGATCTNNNCCAAGTAGCTGGGACTACAACA
AGCACACGTCTGAACTCTTTCCCTACACGACGCTCTTCCGATCTNNNACTCCAAGTTGGGCAACAGAG
CACACGTCTGAACTCTTTCCCTACACGACGCTCTTCCGATCTNNNTGGCCATATGAGTATGCATTATGT
GCACACGTCTGAACTCTTTCCCTACACGACGCTCTTCCGATCTNNNCAGATCACGAGGTCAGGAGAT
GAGCACACGTCTGAACTCTTTCCCTACACGACGCTCTTCCGATCTNNNCTTGTAGTGAGCCAAGGTCG
AGCACACGTCTGAACTCTTTCCCTACACGACGCTCTTCCGATCTNNNCTGTCGCCAACAACAACAACAC
AGCACACGTCTGAACTCTTTCCCTACACGACGCTCTTCCGATCTNNNCTTGCAGTGAGCTGAGATTGC
GCACACGTCTGAACTCTTTCCCTACACGACGCTCTTCCGATCTNNNTCTCTCCCTCTCTCTCTCTCTCA

AGCACACGTCTGAACTCTTTCCCTACACGACGCTCTTCCGATCTNNNCTGCCTCAGCTTCCCAAGTAGC
GCACACGTCTGAACTCTTTCCCTACACGACGCTCTTCCGATCTNNNGCAGAGGTTATAGTGAGCCAAG
AGCACACGTCTGAACTCTTTCCCTACACGACGCTCTTCCGATCTNNNGCCTGTAGTCCCAGCTATTCGCA
CACACGTCTGAACTCTTTCCCTACACGACGCTCTTCCGATCTNNNGCTAACACAGTGAAACCCTGTTCC
CACACGTCTGAACTCTTTCCCTACACGACGCTCTTCCGATCTNNNAGAAGCAAAAGTGGCTCTGAGTA
GCACACGTCTGAACTCTTTCCCTACACGACGCTCTTCCGATCTNNNCTCTCTTGACCCAGCACCTTTCCA
GCACACGTCTGAACTCTTTCCCTACACGACGCTCTTCCGATCTNNNAAAGTGTTGGGGATTACAGGTG
GCACACGTCTGAACTCTTTCCCTACACGACGCTCTTCCGATCTNNNGCCTGTAGTCCCAGCTACTTGCA
AGCACACGTCTGAACTCTTTCCCTACACGACGCTCTTCCGATCTNNNGCCTGTAGTCCCAGCTACTTGCA
AGAGCACACGTCTGAACTCTTTCCCTACACGACGCTCTTCCGATCTNNNGACAGAGCAAGACTCCGTC
CACACGTCTGAACTCTTTCCCTACACGACGCTCTTCCGATCTNNNGGTTACAGAGCAAGACCTTGTCTC
GAGCACACGTCTGAACTCTTTCCCTACACGACGCTCTTCCGATCTNNNTGAACCGAGATCATGCCACTG
AGCACACGTCTGAACTCTTTCCCTACACGACGCTCTTCCGATCTNNNCAACTTCACACAGGTGGTTGCC
AGCACACGTCTGAACTCTTTCCCTACACGACGCTCTTCCGATCTNNNGCCTGTAGTCCCAGCTACTTGC
CACACGTCTGAACTCTTTCCCTACACGACGCTCTTCCGATCTNNNATGCTTGTAATCCCAGCACTTTGCA
AGCACACGTCTGAACTCTTTCCCTACACGACGCTCTTCCGATCTNNNCTTGCAGTGAGCCGAGATTGC
CACACGTCTGAACTCTTTCCCTACACGACGCTCTTCCGATCTNNNTGGGACAGACTTGGTGAATATCG
AGCACACGTCTGAACTCTTTCCCTACACGACGCTCTTCCGATCTNNNGCCTGTAGTCCCAGCTACTTGC
CACACGTCTGAACTCTTTCCCTACACGACGCTCTTCCGATCTNNNACACTTAGATTGAGCTGGTCTTGA
CACACGTCTGAACTCTTTCCCTACACGACGCTCTTCCGATCTNNNGGTAACAGAGAACCCACAGACAG
GCACACGTCTGAACTCTTTCCCTACACGACGCTCTTCCGATCTNNNCCTGATGCATTTTCTCAGCCTTTC
AGCACACGTCTGAACTCTTTCCCTACACGACGCTCTTCCGATCTNNNGTCCTGAGACTCTGCATCTTCCC
GCACACGTCTGAACTCTTTCCCTACACGACGCTCTTCCGATCTNNNGCCACTAGGAATTCGTGCCTTATC
GAGCACACGTCTGAACTCTTTCCCTACACGACGCTCTTCCGATCTNNNCTCTCCACGCTTCCTTGCATCCA
GCACACGTCTGAACTCTTTCCCTACACGACGCTCTTCCGATCTNNNTTCATGCCATTCTCCTGCCTCCAT
GCACACGTCTGAACTCTTTCCCTACACGACGCTCTTCCGATCTNNNTGAGCCAAGATCACACCACTGCA
CACACGTCTGAACTCTTTCCCTACACGACGCTCTTCCGATCTNNNAACTAGGTTAGGGTTGCCAGATCC
GAGCACACGTCTGAACTCTTTCCCTACACGACGCTCTTCCGATCTNNNGTTTGGATGCCCGTTTCAGA
GCACACGTCTGAACTCTTTCCCTACACGACGCTCTTCCGATCTNNNCATGATCTTGGCTCACTGCAAGC
CACACGTCTGAACTCTTTCCCTACACGACGCTCTTCCGATCTNNNAGGATCAACCAACTATATGCAGCTC
AGCACACGTCTGAACTCTTTCCCTACACGACGCTCTTCCGATCTNNNTGAGCCAAGATCATGCCACTGC
GCACACGTCTGAACTCTTTCCCTACACGACGCTCTTCCGATCTNNNAAGAGCCATGCTTCAGAACTCAA
AGCACACGTCTGAACTCTTTCCCTACACGACGCTCTTCCGATCTNNNAAAGAGGAGCCAGACTGAAG
ACACGTCTGAACTCTTTCCCTACACGACGCTCTTCCGATCTNNNACCGACTACAGTTCCATCACATTCAT
GCACACGTCTGAACTCTTTCCCTACACGACGCTCTTCCGATCTNNNGCCTAAGACTCTCCTCTCTCTCTCA
GAGCACACGTCTGAACTCTTTCCCTACACGACGCTCTTCCGATCTNNNCTACTCCCAGAAGTCCCTGCAC
GAGCACACGTCTGAACTCTTTCCCTACACGACGCTCTTCCGATCTNNNCTGCTGAGAAGCTGGCACTAC
CACACGTCTGAACTCTTTCCCTACACGACGCTCTTCCGATCTNNNCGTTGAGTATACCCACTGTAGGTC
GCACACGTCTGAACTCTTTCCCTACACGACGCTCTTCCGATCTNNNTTTCACACCATTCTCCTGCCTCCA
CACACGTCTGAACTCTTTCCCTACACGACGCTCTTCCGATCTNNNTCCAAAGAAGCAGAACAGACAGAC
AGCACACGTCTGAACTCTTTCCCTACACGACGCTCTTCCGATCTNNNCTTACAGTGAGCCGAGACTGGC
CACACGTCTGAACTCTTTCCCTACACGACGCTCTTCCGATCTNNNCCCATTGCTAACAACTGTTACCACA
CACACGTCTGAACTCTTTCCCTACACGACGCTCTTCCGATCTNNNTGTGTGTGTGTGAGAGACTACAG

GAGCACACGTCTGAACTCTTTCCCTACACGACGCTCTTCCGATCTNNNCCACCGAGTGAGACTCCATCT
CACACGTCTGAACTCTTTCCCTACACGACGCTCTTCCGATCTNNNCGACAGAGTGAGACTCCGTCTCAT
CACACGTCTGAACTCTTTCCCTACACGACGCTCTTCCGATCTNNNACAGAGCTGCACATTTCCATCTTCA
AGCACACGTCTGAACTCTTTCCCTACACGACGCTCTTCCGATCTNNNGACTTCTCCTCCCAGCTTCCCAT
AAGAGCACACGTCTGAACTCTTTCCCTACACGACGCTCTTCCGATCTNNNTTTGTGAGGGCCACTCTCT
AGCACACGTCTGAACTCTTTCCCTACACGACGCTCTTCCGATCTNNNCTAGGCGACAGAACGAGACTC
CACACGTCTGAACTCTTTCCCTACACGACGCTCTTCCGATCTNNNCTCCCTATAGCTCACTGGGATAC
AGCACACGTCTGAACTCTTTCCCTACACGACGCTCTTCCGATCTNNNGACTGACAGACTCGCTAGTGA
GCACACGTCTGAACTCTTTCCCTACACGACGCTCTTCCGATCTNNNATTCTCGTTTCCCCACAAAAAGG
GCACACGTCTGAACTCTTTCCCTACACGACGCTCTTCCGATCTNNNAGTGCCTTTTCTCAGCCATAAGA
CACACGTCTGAACTCTTTCCCTACACGACGCTCTTCCGATCTNNNCTCAATCTCAAGTGAACCTCCCAC
AGCACACGTCTGAACTCTTTCCCTACACGACGCTCTTCCGATCTNNNCACACACACACACACACACACCA
AGCACACGTCTGAACTCTTTCCCTACACGACGCTCTTCCGATCTNNNCAGGGTGACAGAGCAAGACT
GCACACGTCTGAACTCTTTCCCTACACGACGCTCTTCCGATCTNNNCCTGGCAAAAGAGCAAGACTCC
GCACACGTCTGAACTCTTTCCCTACACGACGCTCTTCCGATCTNNNGGTGAAACGCCGTCTCTACTAAA
GAGCACACGTCTGAACTCTTTCCCTACACGACGCTCTTCCGATCTNNNGAGATCGCGCCACTGAAATT
AGCACACGTCTGAACTCTTTCCCTACACGACGCTCTTCCGATCTNNNGCAAGAGAATGGCATGAACCC
CACACGTCTGAACTCTTTCCCTACACGACGCTCTTCCGATCTNNNTTCTCACCTCCAATCCCATGATCCAT
GCACACGTCTGAACTCTTTCCCTACACGACGCTCTTCCGATCTNNNCTTGCAGTGAGCTGAGATCGCC
AGCACACGTCTGAACTCTTTCCCTACACGACGCTCTTCCGATCTNNNGTCTAAATGTGCGCATCCTCGC
CACACGTCTGAACTCTTTCCCTACACGACGCTCTTCCGATCTNNNAATGACAACCCAGGCACTATGAA
AGCACACGTCTGAACTCTTTCCCTACACGACGCTCTTCCGATCTNNNCTCCCGGGTTCAAGGATTCTCC
CACACGTCTGAACTCTTTCCCTACACGACGCTCTTCCGATCTNNNAGGTCCTCGTGACTTTGAAAAGA
AGCACACGTCTGAACTCTTTCCCTACACGACGCTCTTCCGATCTNNNGGTGACAGCAAGACTCAGTCT
AGCACACGTCTGAACTCTTTCCCTACACGACGCTCTTCCGATCTNNNCAAGATCAGGCCACTGCATTC
CACACGTCTGAACTCTTTCCCTACACGACGCTCTTCCGATCTNNNTGTGTCCTAAATCCTGAAGGTGG
AGCACACGTCTGAACTCTTTCCCTACACGACGCTCTTCCGATCTNNNCTTCTCAACCCCCTCCAGAACCA
CACACGTCTGAACTCTTTCCCTACACGACGCTCTTCCGATCTNNNCATGCACTCTCTCTCTGTCTCTGCAT
AGAGCACACGTCTGAACTCTTTCCCTACACGACGCTCTTCCGATCTNNNCAGACCCCACATCACACTCT
AGCACACGTCTGAACTCTTTCCCTACACGACGCTCTTCCGATCTNNNCTCCCATCTGGCCTTTCTCTCCA
CACACGTCTGAACTCTTTCCCTACACGACGCTCTTCCGATCTNNNAATACAGTGACAGTCTGAGGTGC
AGCACACGTCTGAACTCTTTCCCTACACGACGCTCTTCCGATCTNNNCTTGCTGCCCCTTCAGTCTAGCA
CACACGTCTGAACTCTTTCCCTACACGACGCTCTTCCGATCTNNNTGAGCATTGCAAGTGGAGAGAAT
AGCACACGTCTGAACTCTTTCCCTACACGACGCTCTTCCGATCTNNNAAGGTGGGAGGATTGCTTGAG
GAGCACACGTCTGAACTCTTTCCCTACACGACGCTCTTCCGATCTNNNGAGGACCTCAGGACACGTCA
GCACACGTCTGAACTCTTTCCCTACACGACGCTCTTCCGATCTNNNGTCAGGAGTTCAAGACCAGCTTC
AGCACACGTCTGAACTCTTTCCCTACACGACGCTCTTCCGATCTNNNGGAAGTGGAGCTTGCAGTGA
CACACGTCTGAACTCTTTCCCTACACGACGCTCTTCCGATCTNNNACTAGGTCTCAACAACAGCAACAC
CACACGTCTGAACTCTTTCCCTACACGACGCTCTTCCGATCTNNNTAATCCTAACACTTTGGGAGGCCC
CACACGTCTGAACTCTTTCCCTACACGACGCTCTTCCGATCTNNNTGGAATAATGTCCTCTGGGTTGT
AGCACACGTCTGAACTCTTTCCCTACACGACGCTCTTCCGATCTNNNTGGGTGACAGAGATTCCATCT
CACACGTCTGAACTCTTTCCCTACACGACGCTCTTCCGATCTNNNTGAAAGGTCAACGAAAAACCCAA
CACACGTCTGAACTCTTTCCCTACACGACGCTCTTCCGATCTNNNTCCAATCTCTGCCTGTCTTCAAACA

CACACGTCTGAACTCTTTCCCTACACGACGCTCTTCCGATCTNNNTCAAGTGATTCTACTGCCTCAGCC
CACACGTCTGAACTCTTTCCCTACACGACGCTCTTCCGATCTNNNTCCCACAAATAGCTCCCACAATTCA
AGCACACGTCTGAACTCTTTCCCTACACGACGCTCTTCCGATCTNNNGAGATTGCGCCACTGTAGTCC
GAGCACACGTCTGAACTCTTTCCCTACACGACGCTCTTCCGATCTNNNCTGAGCTCAGGAGTTGGAG
CACACGTCTGAACTCTTTCCCTACACGACGCTCTTCCGATCTNNNGGCTTATGAATCATTTGGGGCTT
GAGCACACGTCTGAACTCTTTCCCTACACGACGCTCTTCCGATCTNNNGCGACAGGGTGACATTTGG
GCACACGTCTGAACTCTTTCCCTACACGACGCTCTTCCGATCTNNNATTTGCAAATCACATGGCTGAC
AGCACACGTCTGAACTCTTTCCCTACACGACGCTCTTCCGATCTNNNCTGTCACCCAGTGAGACTGTG
GCACACGTCTGAACTCTTTCCCTACACGACGCTCTTCCGATCTNNNGCAACAGAGTGAGACTCCATCTC
AGCACACGTCTGAACTCTTTCCCTACACGACGCTCTTCCGATCTNNNGAGCTTGAAGTGAGCCGAGAT
AGCACACGTCTGAACTCTTTCCCTACACGACGCTCTTCCGATCTNNNGTGTGTGTGTGTGTGTGT
AGCACACGTCTGAACTCTTTCCCTACACGACGCTCTTCCGATCTNNNAGTTATGTACCCAGGTTGTGC
CACACGTCTGAACTCTTTCCCTACACGACGCTCTTCCGATCTNNNCGCTTCGAGATCCCCAAATACTAC
GCACACGTCTGAACTCTTTCCCTACACGACGCTCTTCCGATCTNNNGTCCATGGTACTAACACACACCA
AGCACACGTCTGAACTCTTTCCCTACACGACGCTCTTCCGATCTNNNCTGCGCTGGTCTTCAAATGACC
CACACGTCTGAACTCTTTCCCTACACGACGCTCTTCCGATCTNNNCTCTGTGTGACAACCTTGAAAGCC
AAGAGCACACGTCTGAACTCTTTCCCTACACGACGCTCTTCCGATCTNNNGAGGCCAGGAGTTCAAGA
GCACACGTCTGAACTCTTTCCCTACACGACGCTCTTCCGATCTNNNGGGCAACATAGATCCCATCTCTA
AGCACACGTCTGAACTCTTTCCCTACACGACGCTCTTCCGATCTNNNAAGCCCAGGTTTTCTCGTTCA
AGCACACGTCTGAACTCTTTCCCTACACGACGCTCTTCCGATCTNNNCTGGCCCACTCTTCAGATGTGC
GCACACGTCTGAACTCTTTCCCTACACGACGCTCTTCCGATCTNNNCAAATTCATATGCAGACCCACCC
GCACACGTCTGAACTCTTTCCCTACACGACGCTCTTCCGATCTNNNTTGAATCTGATGCATTCCGTCTCC
AGCACACGTCTGAACTCTTTCCCTACACGACGCTCTTCCGATCTNNNCATGCCAAGTTGTCCAATCCGC
GAGCACACGTCTGAACTCTTTCCCTACACGACGCTCTTCCGATCTNNNGAGAATGAGGCAGGGAGG
CACACGTCTGAACTCTTTCCCTACACGACGCTCTTCCGATCTNNNTGTTATGACAGCCATTTAGCAGCC
GCACACGTCTGAACTCTTTCCCTACACGACGCTCTTCCGATCTNNNGTGTGTGTGTGTGTGTGTGC
GCACACGTCTGAACTCTTTCCCTACACGACGCTCTTCCGATCTNNNAAGAAAGGAGTAGGAGGAGGA
CACACGTCTGAACTCTTTCCCTACACGACGCTCTTCCGATCTNNNACTTCTAGTAGTTGTGTGTGTGA
CACACGTCTGAACTCTTTCCCTACACGACGCTCTTCCGATCTNNNCAAAACTGGGTTGTGAGCTCAAG
GCACACGTCTGAACTCTTTCCCTACACGACGCTCTTCCGATCTNNNTAGCTAGGATTACAGGTGTGTG
GAGCACACGTCTGAACTCTTTCCCTACACGACGCTCTTCCGATCTNNNGAGGTAGGAGAATGGCGTG
GCACACGTCTGAACTCTTTCCCTACACGACGCTCTTCCGATCTNNNGAGGCAGGAGAATCGCTTGAA
CACACGTCTGAACTCTTTCCCTACACGACGCTCTTCCGATCTNNNCCTCCGCAGACATCCATAACTGCA
GAGCACACGTCTGAACTCTTTCCCTACACGACGCTCTTCCGATCTNNNAGTGCAGTCCCAGTATTGGT
GAGCACACGTCTGAACTCTTTCCCTACACGACGCTCTTCCGATCTNNNGCTGAGTAGCTGGGACTACA
GAGCACACGTCTGAACTCTTTCCCTACACGACGCTCTTCCGATCTNNNAGAGGTTGCAGTGAGTCAA
GCACACGTCTGAACTCTTTCCCTACACGACGCTCTTCCGATCTNNNCTTGCAGTGAGCTGAGATCGCC
CACACGTCTGAACTCTTTCCCTACACGACGCTCTTCCGATCTNNNTGGGTGACAGAATGAGACCTTG
AGCACACGTCTGAACTCTTTCCCTACACGACGCTCTTCCGATCTNNNAGCTGGGATTACAGGTGTGTG
AGCACACGTCTGAACTCTTTCCCTACACGACGCTCTTCCGATCTNNNCCTGGGCAACAAGAGTGAAAC
GCACACGTCTGAACTCTTTCCCTACACGACGCTCTTCCGATCTNNNGAGCGAAACTCCATCTCAAAACA
GCACACGTCTGAACTCTTTCCCTACACGACGCTCTTCCGATCTNNNGGAGGATCACTTGAGCTCACAA
GCACACGTCTGAACTCTTTCCCTACACGACGCTCTTCCGATCTNNNCCACTGCACCTGGTCATAGATC

AGCACACGTCTGAACTCTTTCCCTACACGACGCTCTTCCGATCTNNNCTGAGGCCAGAGGACTGATT
AGCACACGTCTGAACTCTTTCCCTACACGACGCTCTTCCGATCTNNNGAGCTTGCAGTGAGTGGAGA
AGCACACGTCTGAACTCTTTCCCTACACGACGCTCTTCCGATCTNNNTGAGCTGAGATCGCACCATTG
CACACGTCTGAACTCTTTCCCTACACGACGCTCTTCCGATCTNNNAAGCATTCACTTGAGGAGAGAG
AGCACACGTCTGAACTCTTTCCCTACACGACGCTCTTCCGATCTNNNTGAGCTGAGATTGCACCACTG
CACACGTCTGAACTCTTTCCCTACACGACGCTCTTCCGATCTNNNTCTCTCTCTCTCTCTCTCCCTCAT
AGCACACGTCTGAACTCTTTCCCTACACGACGCTCTTCCGATCTNNNGAGGTTGCAGTGAGCCAAGA
AGCACACGTCTGAACTCTTTCCCTACACGACGCTCTTCCGATCTNNNCCTGGTGTTCAGTGCTTAGCAC
GCACACGTCTGAACTCTTTCCCTACACGACGCTCTTCCGATCTNNNAGTGAGCCAATATCATGCCACT
CACACGTCTGAACTCTTTCCCTACACGACGCTCTTCCGATCTNNNTCCCACAATTCTCTATCTCACACAC
GAGCACACGTCTGAACTCTTTCCCTACACGACGCTCTTCCGATCTNNNCTTGCAGTGAGCCAAGATTG
CACACGTCTGAACTCTTTCCCTACACGACGCTCTTCCGATCTNNNGGGATTACAGGTGTGAGCCACC
CACACGTCTGAACTCTTTCCCTACACGACGCTCTTCCGATCTNNNAGAAGGAAAGAAAGAAGGAAGG
AGCACACGTCTGAACTCTTTCCCTACACGACGCTCTTCCGATCTNNNCTCTCAACAGCCTCACCACAGC
GCACACGTCTGAACTCTTTCCCTACACGACGCTCTTCCGATCTNNNCTCATGCCTATAATCCCAGCACT
CACACGTCTGAACTCTTTCCCTACACGACGCTCTTCCGATCTNNNCCTGAGGTCTCAAGGCCTTTGCA
AAGAGCACACGTCTGAACTCTTTCCCTACACGACGCTCTTCCGATCTNNNCTGCTCTCCAGTCTGGGT
GCACACGTCTGAACTCTTTCCCTACACGACGCTCTTCCGATCTNNNTCACGGATAGGGCGATTTCTAT
CACACGTCTGAACTCTTTCCCTACACGACGCTCTTCCGATCTNNNGACACCACTATGAGGGACTTGAG
AGCACACGTCTGAACTCTTTCCCTACACGACGCTCTTCCGATCTNNNCATTGTAAAGGGTGGGAGGG
GAGCACACGTCTGAACTCTTTCCCTACACGACGCTCTTCCGATCTNNNGAAAACACCAAATGGCGGAT
CACACGTCTGAACTCTTTCCCTACACGACGCTCTTCCGATCTNNNTGCTACACGATAGCTGTTCTCAGC
GCACACGTCTGAACTCTTTCCCTACACGACGCTCTTCCGATCTNNNAGGTTGCAGTGAGCCAAGATC
CACACGTCTGAACTCTTTCCCTACACGACGCTCTTCCGATCTNNNTCCTTCCTTCCTTCTTTCTTTCTCA
GAGCACACGTCTGAACTCTTTCCCTACACGACGCTCTTCCGATCTNNNGATTGGAAAGAGGGGTGG
GCACACGTCTGAACTCTTTCCCTACACGACGCTCTTCCGATCTNNNCAGGTAGAAGAACTGCGTCAG
AGCACACGTCTGAACTCTTTCCCTACACGACGCTCTTCCGATCTNNNGGAGAAGGGGGCACATGTT
CACACGTCTGAACTCTTTCCCTACACGACGCTCTTCCGATCTNNNATCCACACAGAAGACTTCTCGACC
CACACGTCTGAACTCTTTCCCTACACGACGCTCTTCCGATCTNNNCATTGGGTAAGTGCAAGTGACT
AGAGCACACGTCTGAACTCTTTCCCTACACGACGCTCTTCCGATCTNNNGAGTCCTGGTTTCTGGCAG
CACACGTCTGAACTCTTTCCCTACACGACGCTCTTCCGATCTNNNTCTCTCCTTCCTTCCTTTCCTCAT
AGAGCACACGTCTGAACTCTTTCCCTACACGACGCTCTTCCGATCTNNNCTGCAGTGAGCCATGTTTG
AGCACACGTCTGAACTCTTTCCCTACACGACGCTCTTCCGATCTNNNGTGGCAGAGTGAGAGTCCAT
CACACGTCTGAACTCTTTCCCTACACGACGCTCTTCCGATCTNNNCCTCTCTCAAACTGCATCTGTCTC
CACACGTCTGAACTCTTTCCCTACACGACGCTCTTCCGATCTNNNCAATGCTACCTGGAACCTGTACCC
GCACACGTCTGAACTCTTTCCCTACACGACGCTCTTCCGATCTNNNTGCTGATGAAGTTACGGTAGCA
CACACGTCTGAACTCTTTCCCTACACGACGCTCTTCCGATCTNNNAAGGCATTTTTCCTCTCCTGTGAC
AGCACACGTCTGAACTCTTTCCCTACACGACGCTCTTCCGATCTNNNTTTTGGCTGTGCTCATTACCT
AGCACACGTCTGAACTCTTTCCCTACACGACGCTCTTCCGATCTNNNGTGTGTAGGGATGCCTGAGT
CACACGTCTGAACTCTTTCCCTACACGACGCTCTTCCGATCTNNNCTCCCTTTAGAGTGAGATGCTG
AGAGCACACGTCTGAACTCTTTCCCTACACGACGCTCTTCCGATCTNNNGAGGCTGAGGCAGGAGA
GCACACGTCTGAACTCTTTCCCTACACGACGCTCTTCCGATCTNNNCACCCAGACTCCTCAAAGGTGC
CACACGTCTGAACTCTTTCCCTACACGACGCTCTTCCGATCTNNNAGCAAATACCCAGAGGAACCAGC

AGCACACGTCTGAACTCTTTCCCTACACGACGCTCTTCCGATCTNNNCTCGAACCTCCTTCCACCAAGCA
GCACACGTCTGAACTCTTTCCCTACACGACGCTCTTCCGATCTNNNTATTCCGGGCGTCACTACTTTAGC
GCACACGTCTGAACTCTTTCCCTACACGACGCTCTTCCGATCTNNNGTAACTGCAATACTGGACGATGC
CACACGTCTGAACTCTTTCCCTACACGACGCTCTTCCGATCTNNNTAGAAAGAAGTGCTGAAGGAGG
GAGCACACGTCTGAACTCTTTCCCTACACGACGCTCTTCCGATCTNNNAGAATCGCTTGAACCTGGGA
GAGCACACGTCTGAACTCTTTCCCTACACGACGCTCTTCCGATCTNNNGGTTGCAGTGAGCCAAGATT
AGCACACGTCTGAACTCTTTCCCTACACGACGCTCTTCCGATCTNNNACTCCTTATCCTTGGCCTTGCCAT
AGAGCACACGTCTGAACTCTTTCCCTACACGACGCTCTTCCGATCTNNNACATCTGCCCAGTGTGTTCT
AGCACACGTCTGAACTCTTTCCCTACACGACGCTCTTCCGATCTNNNCAAGGCTACCAGGTGATCCACC
AGCACACGTCTGAACTCTTTCCCTACACGACGCTCTTCCGATCTNNNCGCACAGCAGGAAAACATCTC
GCACACGTCTGAACTCTTTCCCTACACGACGCTCTTCCGATCTNNNGCGTTGTACCAATGACTGACAAA
GCACACGTCTGAACTCTTTCCCTACACGACGCTCTTCCGATCTNNNTTCTCTGGTTTTTCTGCTCCAGTCA
GAGCACACGTCTGAACTCTTTCCCTACACGACGCTCTTCCGATCTNNNGAGAATGGTGTGAACCTGG
CACACGTCTGAACTCTTTCCCTACACGACGCTCTTCCGATCTNNNATGTCATTCACAAACGTCTGCAGC
GAGCACACGTCTGAACTCTTTCCCTACACGACGCTCTTCCGATCTNNNGATAGCAGGGGTGCTCAGA
GCACACGTCTGAACTCTTTCCCTACACGACGCTCTTCCGATCTNNNCTGACGAACACAGGTACCAAAG
AGCACACGTCTGAACTCTTTCCCTACACGACGCTCTTCCGATCTNNNGAGAACATCCCAGCTCCACACC
AGCACACGTCTGAACTCTTTCCCTACACGACGCTCTTCCGATCTNNNCCCATTCTGCACATCTTAGCTCT
GCACACGTCTGAACTCTTTCCCTACACGACGCTCTTCCGATCTNNNTGATTCTCTCCAGAGTCCCAGATC
GCACACGTCTGAACTCTTTCCCTACACGACGCTCTTCCGATCTNNNGAGCGAGACTCCTTTCAAAAAG
ACACGTCTGAACTCTTTCCCTACACGACGCTCTTCCGATCTNNNTCTTTTCTTCTCTCTCCCTCCCTCATCG
GAGCACACGTCTGAACTCTTTCCCTACACGACGCTCTTCCGATCTNNNGAGGTGAGAGGATCGCTTCA
ACACGTCTGAACTCTTTCCCTACACGACGCTCTTCCGATCTNNNTTTCTTGTGTGTGTGTGTGTGTGC
AGCACACGTCTGAACTCTTTCCCTACACGACGCTCTTCCGATCTNNNACAGAGTGAGGGAAAGAGAG
GCACACGTCTGAACTCTTTCCCTACACGACGCTCTTCCGATCTNNNTCTCCTTTCCTTCCTTCCTTCTTCA
CACACGTCTGAACTCTTTCCCTACACGACGCTCTTCCGATCTNNNGTTTCACCATGTTAGCCAGGATGC
GCACACGTCTGAACTCTTTCCCTACACGACGCTCTTCCGATCTNNNGTTGCAGTGAGCTGAGATTGTG
ACACGTCTGAACTCTTTCCCTACACGACGCTCTTCCGATCTNNNATCTGGGAGGACTTCAACTCCAACA
CACACGTCTGAACTCTTTCCCTACACGACGCTCTTCCGATCTNNNCAATCTCTTGGGCTTGGGTGATTC
CACGTCTGAACTCTTTCCCTACACGACGCTCTTCCGATCTNNNGTTGTGTTATGTCTCCCTCCACCCATC
GCACACGTCTGAACTCTTTCCCTACACGACGCTCTTCCGATCTNNNAGGGAGAATCCAGCCTGTAAGT
ACACGTCTGAACTCTTTCCCTACACGACGCTCTTCCGATCTNNNGCATAAAATCTGGAAAGCCACCGC
CACACGTCTGAACTCTTTCCCTACACGACGCTCTTCCGATCTNNNACACAAAGGGCAAACTCCTGATTC
CACGTCTGAACTCTTTCCCTACACGACGCTCTTCCGATCTNNNCTAGGCAATTCTGCCTCCATAGCCAT
ACACGTCTGAACTCTTTCCCTACACGACGCTCTTCCGATCTNNNTTTAGCTGCCCCCAACTTCATTTCATC
ACACGTCTGAACTCTTTCCCTACACGACGCTCTTCCGATCTNNNTTTGGTGCTAATGCTGATGGTCTCA
ACACGTCTGAACTCTTTCCCTACACGACGCTCTTCCGATCTNNNTCCTCACCTGTCACGAGGATAAACA
CACACGTCTGAACTCTTTCCCTACACGACGCTCTTCCGATCTNNNATAGTGAAAGTGTGTGTGTGTGC
GCACACGTCTGAACTCTTTCCCTACACGACGCTCTTCCGATCTNNNACATGTTTACTTGCTGCCTGAGA
CACACGTCTGAACTCTTTCCCTACACGACGCTCTTCCGATCTNNNCAATGTGTGTAAGCGTGTGTTGTC
CACGTCTGAACTCTTTCCCTACACGACGCTCTTCCGATCTNNNTAATGGGTTGTTCCTGCTTGGAGCA
CACACGTCTGAACTCTTTCCCTACACGACGCTCTTCCGATCTNNNCACTGGCACCAGACCATCAAAGCA
CACACGTCTGAACTCTTTCCCTACACGACGCTCTTCCGATCTNNNTTAGGATCTGGCTCACTTGGGTTC

AGCACACGTCTGAACTCTTTCCCTACACGACGCTCTTCCGATCTNNNCGCAGGCTCCAATTCAATCCAC
ACACGTCTGAACTCTTTCCCTACACGACGCTCTTCCGATCTNNNAGGTGATTCTCCACCTCTCAGTCCAT
CACACGTCTGAACTCTTTCCCTACACGACGCTCTTCCGATCTNNNTAGGCCTCTCTTCTCTAGACCATCA
CACACGTCTGAACTCTTTCCCTACACGACGCTCTTCCGATCTNNNAGGAGATAAAGACACAGGCCATT
CACACGTCTGAACTCTTTCCCTACACGACGCTCTTCCGATCTNNNCTTTTGCCAACATGATGCCTCGCA
CACACGTCTGAACTCTTTCCCTACACGACGCTCTTCCGATCTNNNTTGAGGAGGAAAGGCATGTTAG
GCACACGTCTGAACTCTTTCCCTACACGACGCTCTTCCGATCTNNNCGCTTCATGCTGACGAATGTTTT
GAGCACACGTCTGAACTCTTTCCCTACACGACGCTCTTCCGATCTNNNCTCTTGGCTGCTGCAGTCTCC
CACACGTCTGAACTCTTTCCCTACACGACGCTCTTCCGATCTNNNAGGTTAGATCAGAAGAGCCCCAG
CACACGTCTGAACTCTTTCCCTACACGACGCTCTTCCGATCTNNNAGGTATCTTGGCGTGACAACTTC
GCACACGTCTGAACTCTTTCCCTACACGACGCTCTTCCGATCTNNNGGCTCCCCTTCTGGATCCTTCATC
CACACGTCTGAACTCTTTCCCTACACGACGCTCTTCCGATCTNNNCTCCCCACGTTCTGCTTGAATAACA
GCACACGTCTGAACTCTTTCCCTACACGACGCTCTTCCGATCTNNNACAAACCCCAGCTCCACTCTCATC
AGCACACGTCTGAACTCTTTCCCTACACGACGCTCTTCCGATCTNNNGAAGCCCCAGACCCATCTGTC
ACACGTCTGAACTCTTTCCCTACACGACGCTCTTCCGATCTNNNGTACATGCAGAGTGAGTAGCTGG
CACACGTCTGAACTCTTTCCCTACACGACGCTCTTCCGATCTNNNCTTCCCCGCTGTCCATTCATTCATC
ACACGTCTGAACTCTTTCCCTACACGACGCTCTTCCGATCTNNNACAAAGACACCTAACTTCTCAGGCAT
ACACGTCTGAACTCTTTCCCTACACGACGCTCTTCCGATCTNNNCTACCACTGCTCAGAGGAACAGAC
CACACGTCTGAACTCTTTCCCTACACGACGCTCTTCCGATCTNNNCCGCCTGCATCTCCTAATAAACAC
CACACGTCTGAACTCTTTCCCTACACGACGCTCTTCCGATCTNNNGCGTGTGGGGTGTGTGTGTAGA
CACACGTCTGAACTCTTTCCCTACACGACGCTCTTCCGATCTNNNACATTCTCACACAGCTTCTGTCCCA
GCACACGTCTGAACTCTTTCCCTACACGACGCTCTTCCGATCTNNNCGCACATATTGCAACAGAACCCC
CACACGTCTGAACTCTTTCCCTACACGACGCTCTTCCGATCTNNNCCTGCTGGTCTCCCAATTCTTTGCA
ACACGTCTGAACTCTTTCCCTACACGACGCTCTTCCGATCTNNNTAGAAGGTGTTTGTGAAGTCGGCC
CACACGTCTGAACTCTTTCCCTACACGACGCTCTTCCGATCTNNNGTAACGCATCATCTACCGTCCTCC
AGCACACGTCTGAACTCTTTCCCTACACGACGCTCTTCCGATCTNNNCACACACAGGCTCACATGCCTC
ACACGTCTGAACTCTTTCCCTACACGACGCTCTTCCGATCTNNNCAGGACAAGCACGAGTACTACAGC
ACACGTCTGAACTCTTTCCCTACACGACGCTCTTCCGATCTNNNCGTGGTGGCGAAGAGAGTAAAAT
AGAGCACACGTCTGAACTCTTTCCCTACACGACGCTCTTCCGATCTNNNGAAGGGGTTTTCGGGAGC
CACACGTCTGAACTCTTTCCCTACACGACGCTCTTCCGATCTNNNGGAGCAGAAACTTTGCGTCTCTAC
GAGCACACGTCTGAACTCTTTCCCTACACGACGCTCTTCCGATCTNNNGGTTGCCGTGGGTTTGTTTG
AGCACACGTCTGAACTCTTTCCCTACACGACGCTCTTCCGATCTNNNTCACGCGTGGTCCTAAATGTCC
GAGCACACGTCTGAACTCTTTCCCTACACGACGCTCTTCCGATCTNNNGCGAATACACACGCCCTGACC
CACACGTCTGAACTCTTTCCCTACACGACGCTCTTCCGATCTNNNGTACCTCTGCTTCCACACATACCCA
ACACGTCTGAACTCTTTCCCTACACGACGCTCTTCCGATCTNNNATACACACTCTCTTGAGCCAAATCAT
GCACACGTCTGAACTCTTTCCCTACACGACGCTCTTCCGATCTNNNGGATGGTTGGGTTTTTCCGTACT
CACGTCTGAACTCTTTCCCTACACGACGCTCTTCCGATCTNNNCGACGCTTTCCTTTCATTTTTGCCATC
CACGTCTGAACTCTTTCCCTACACGACGCTCTTCCGATCTNNNCCCTTGGAGCATTTTGTCACTGGCAT
ACACGTCTGAACTCTTTCCCTACACGACGCTCTTCCGATCTNNNAAGAGAAATTACAGGCCACAGCCC
ACACGTCTGAACTCTTTCCCTACACGACGCTCTTCCGATCTNNNGCCCATCAGAAATAACCAACAGCCAT
CACACGTCTGAACTCTTTCCCTACACGACGCTCTTCCGATCTNNNAAGCAGACATAGAGGAGGGGAA
AGCACACGTCTGAACTCTTTCCCTACACGACGCTCTTCCGATCTNNNCAACCCCCTCCCTGACTGCATCAT
CACACGTCTGAACTCTTTCCCTACACGACGCTCTTCCGATCTNNNCTACTAGACTGATACCCAGGGGCC

ACACGTCTGAACTCTTTCCCTACACGACGCTCTTCCGATCTNNNCTAAAAAGGGAGTTGGACCCTCGCA
ACACGTCTGAACTCTTTCCCTACACGACGCTCTTCCGATCTNNNCGCACAAGAACAAATGACAGTCCCAT
CACACGTCTGAACTCTTTCCCTACACGACGCTCTTCCGATCTNNNCTGGGAGTTCACAGCTGTATGTGC
CACACGTCTGAACTCTTTCCCTACACGACGCTCTTCCGATCTNNNTAGCAGATCTTGATGTTGGCAGGC
AGCACACGTCTGAACTCTTTCCCTACACGACGCTCTTCCGATCTNNNCCGAGTAGCTGGGACTACAGG
GAGCACACGTCTGAACTCTTTCCCTACACGACGCTCTTCCGATCTNNNCGGTAATAGGGGCTAAGGCT
GCACACGTCTGAACTCTTTCCCTACACGACGCTCTTCCGATCTNNNCCTACGGGAAGCCCAAACTCCAT
CACACGTCTGAACTCTTTCCCTACACGACGCTCTTCCGATCTNNNGCTACCTCTTGATGGTTCTGAGTCA
GCACACGTCTGAACTCTTTCCCTACACGACGCTCTTCCGATCTNNNGAGGATCACAAGGTCAGGAGAT
CACGTCTGAACTCTTTCCCTACACGACGCTCTTCCGATCTNNNTCCAATCTGTCTGCGCTTTTTTTCATC
CACGTCTGAACTCTTTCCCTACACGACGCTCTTCCGATCTNNNAGTCCTGTAGGGAGGAATCATGGCA
CACACGTCTGAACTCTTTCCCTACACGACGCTCTTCCGATCTNNNGCCACTGTACATGATCACCCTAGCAT
CACGTCTGAACTCTTTCCCTACACGACGCTCTTCCGATCTNNNGTTTCAAATGGTTTCCCCCGTTTCATC
AGCACACGTCTGAACTCTTTCCCTACACGACGCTCTTCCGATCTNNNGAGCTGAGATCGCCTCACTGCA
GCACACGTCTGAACTCTTTCCCTACACGACGCTCTTCCGATCTNNNGAGGCTGAGGCAGGAGAATGCA
GAGCACACGTCTGAACTCTTTCCCTACACGACGCTCTTCCGATCTNNNAGGAGAATGGCGTGAACCTG
AGCACACGTCTGAACTCTTTCCCTACACGACGCTCTTCCGATCTNNNGAGGCTGAGGCAGGAGAATG
AGCACACGTCTGAACTCTTTCCCTACACGACGCTCTTCCGATCTNNNGAGGCTGAGGCAGGAGAATG
GAGCACACGTCTGAACTCTTTCCCTACACGACGCTCTTCCGATCTNNNGAGGTGGAGCTTGCAGTGAG
GCACACGTCTGAACTCTTTCCCTACACGACGCTCTTCCGATCTNNNGCAGAACCGCCTATGAACTTTCCA
GCACACGTCTGAACTCTTTCCCTACACGACGCTCTTCCGATCTNNNGAGGCTGAGGCAGGAGAATGCA
CACACGTCTGAACTCTTTCCCTACACGACGCTCTTCCGATCTNNNCACCAGAACATTCAGCACAATGTCA
CACGTCTGAACTCTTTCCCTACACGACGCTCTTCCGATCTNNNAAGTCCACAATTTGTCAAGGCTGCAT
GAGCACACGTCTGAACTCTTTCCCTACACGACGCTCTTCCGATCTNNNGAGGCTGAGGCAGGAGAAT
CACACGTCTGAACTCTTTCCCTACACGACGCTCTTCCGATCTNNNTTTACAAGTCTCGCTTCGGATGTCAT
GAGCACACGTCTGAACTCTTTCCCTACACGACGCTCTTCCGATCTNNNGAGGCTGAGGCAGGAGAATG
GAGCACACGTCTGAACTCTTTCCCTACACGACGCTCTTCCGATCTNNNCTTGCAGTGAGCCGAGATCAC
AGCACACGTCTGAACTCTTTCCCTACACGACGCTCTTCCGATCTNNNGAGGCTGAGGCAGGAGAATG
AGCACACGTCTGAACTCTTTCCCTACACGACGCTCTTCCGATCTNNNGAGGCTGAGGCAGGAGAATG
CACGTCTGAACTCTTTCCCTACACGACGCTCTTCCGATCTNNNATCCTTTGATTCTGGCTTACTGCCATC
AGCACACGTCTGAACTCTTTCCCTACACGACGCTCTTCCGATCTNNNGAGGCTGAGGCAGGAGAATG
GAGCACACGTCTGAACTCTTTCCCTACACGACGCTCTTCCGATCTNNNGAGGCTGAGGCAGGAGAATG
GCACACGTCTGAACTCTTTCCCTACACGACGCTCTTCCGATCTNNNGAGGCTGAGGCAGGAGAATGC
CACGTCTGAACTCTTTCCCTACACGACGCTCTTCCGATCTNNNATCCCTGGCTGTCCTAAGATACTCAT
GAGCACACGTCTGAACTCTTTCCCTACACGACGCTCTTCCGATCTNNNGAGGCTGAGGCAGGAGAAT
CACGTCTGAACTCTTTCCCTACACGACGCTCTTCCGATCTNNNGACTAAAGCTGCAGACCTTCAGACAT
CACACGTCTGAACTCTTTCCCTACACGACGCTCTTCCGATCTNNNTCTTATCTCATCTCCCTGCCTCTCATC
ACACGTCTGAACTCTTTCCCTACACGACGCTCTTCCGATCTNNNACCAAAGACAAAAACCAGCCAAACAT
ACACGTCTGAACTCTTTCCCTACACGACGCTCTTCCGATCTNNNTCATGTGAAGAAGAGGCAATACACA
CACGTCTGAACTCTTTCCCTACACGACGCTCTTCCGATCTNNNGGAGCACCAAAAATGTGAAATGCCA
GAGCACACGTCTGAACTCTTTCCCTACACGACGCTCTTCCGATCTNNNGAGGCTGAGGCAGGAGAAT
GCACACGTCTGAACTCTTTCCCTACACGACGCTCTTCCGATCTNNNGAGGCTGAGGCAGGAGAATGC
GAGCACACGTCTGAACTCTTTCCCTACACGACGCTCTTCCGATCTNNNGAGGCTGAGGCAGGAGAATG

AGCACACGTCTGAACTCTTTCCCTACACGACGCTCTTCCGATCTNNNCAGGAGAATGGCGTGAACCCC
AGCACACGTCTGAACTCTTTCCCTACACGACGCTCTTCCGATCTNNNCTTGCAGTGAGCCGAGATTGC
AGCACACGTCTGAACTCTTTCCCTACACGACGCTCTTCCGATCTNNNGAGGCTGAGGCAGGAGAATG
CACACGTCTGAACTCTTTCCCTACACGACGCTCTTCCGATCTNNNGTCTCAAACTCCTGACCTCATGACAT
CACGTCTGAACTCTTTCCCTACACGACGCTCTTCCGATCTNNNTCCTGGAAAAGAAAAGCTGGAGACA
GAGCACACGTCTGAACTCTTTCCCTACACGACGCTCTTCCGATCTNNNAGGAGAATGGCGTGAACCTG
CACGTCTGAACTCTTTCCCTACACGACGCTCTTCCGATCTNNNCTCTGTCAGAGTACAACATGCTCCAT
CACGTCTGAACTCTTTCCCTACACGACGCTCTTCCGATCTNNNTGTGAGATGATCCCTGTGTTGTTCAT
ACACGTCTGAACTCTTTCCCTACACGACGCTCTTCCGATCTNNNACTGAAAGGCTACCTCACAGAAACA
GCACACGTCTGAACTCTTTCCCTACACGACGCTCTTCCGATCTNNNGTCAGGCTCTACTGCTGAACCCAT
CACACGTCTGAACTCTTTCCCTACACGACGCTCTTCCGATCTNNNCCAAGATCAAACCACTGCACTACCA
ACACGTCTGAACTCTTTCCCTACACGACGCTCTTCCGATCTNNNTCTCTCTCTCTCTCTCTTCCTCATCG
CACGTCTGAACTCTTTCCCTACACGACGCTCTTCCGATCTNNNTTCCATTCCTTTGCTCAGTTCCTCATCG
ACACGTCTGAACTCTTTCCCTACACGACGCTCTTCCGATCTNNNTTCAGGAAGTCGTCTTGGAGAAGCA
ACACGTCTGAACTCTTTCCCTACACGACGCTCTTCCGATCTNNNCCCATATTTCTTGGAGGCTCTGTCAT
CACGTCTGAACTCTTTCCCTACACGACGCTCTTCCGATCTNNNCCTGTGGATTGACATATTCCGGACAT
AGCACACGTCTGAACTCTTTCCCTACACGACGCTCTTCCGATCTNNNCTTGCAGTGAGCCGAGATCCCA
AGAGCACACGTCTGAACTCTTTCCCTACACGACGCTCTTCCGATCTNNNCACCATTGCATTCCAGCCTGC
GAGCACACGTCTGAACTCTTTCCCTACACGACGCTCTTCCGATCTNNNGAGGCTGAGGCAGGAGAATG
AGCACACGTCTGAACTCTTTCCCTACACGACGCTCTTCCGATCTNNNGAGGCTGAGGCAGGAGAATG
AGCACACGTCTGAACTCTTTCCCTACACGACGCTCTTCCGATCTNNNGAGGCTGAGGCAGGAGAATG
GAGCACACGTCTGAACTCTTTCCCTACACGACGCTCTTCCGATCTNNNAGGAGAATGGCGTGAACCTG
CACGTCTGAACTCTTTCCCTACACGACGCTCTTCCGATCTNNNTTTCCTTCCCAGAGCTGAAAACTCAT
GAGCACACGTCTGAACTCTTTCCCTACACGACGCTCTTCCGATCTNNNGAGGCTGAGGCAGGAGAATG
GAGCACACGTCTGAACTCTTTCCCTACACGACGCTCTTCCGATCTNNNGAGGCTGAGGCAGGAGAAT
GAGCACACGTCTGAACTCTTTCCCTACACGACGCTCTTCCGATCTNNNAGGAGAATGGCGTGAACCT
AGCACACGTCTGAACTCTTTCCCTACACGACGCTCTTCCGATCTNNNGAGGCTGAGGCAGGAGAATG
AGCACACGTCTGAACTCTTTCCCTACACGACGCTCTTCCGATCTNNNGAGGCTGAGGCAGGAGAATGC
AGCACACGTCTGAACTCTTTCCCTACACGACGCTCTTCCGATCTNNNGAGGCTGAGGCAGGAGAATG
CACGTCTGAACTCTTTCCCTACACGACGCTCTTCCGATCTNNNTCCAGATGCTCTTCCTCCAGATACATC
CACGTCTGAACTCTTTCCCTACACGACGCTCTTCCGATCTNNNGTCAGACTTGCACCCTAGTAACACAT
CACACGTCTGAACTCTTTCCCTACACGACGCTCTTCCGATCTNNNTCTGCCAGATGTATTAGCACTCCCAT
CACGTCTGAACTCTTTCCCTACACGACGCTCTTCCGATCTNNNTTCTAAGGGCAGGAACCATATCACAT
ACACGTCTGAACTCTTTCCCTACACGACGCTCTTCCGATCTNNNCGACCGCAAATCCTGGTAATAAACAT
GAGCACACGTCTGAACTCTTTCCCTACACGACGCTCTTCCGATCTNNNGAATGGCGTGAACCCAGGAG
AGCACACGTCTGAACTCTTTCCCTACACGACGCTCTTCCGATCTNNNGAGGCTGAGGCAGGAGAATGC
AGCACACGTCTGAACTCTTTCCCTACACGACGCTCTTCCGATCTNNNGACAGAGCGAGACTCCGTCTCAT
GAGCACACGTCTGAACTCTTTCCCTACACGACGCTCTTCCGATCTNNNCTTGCAGTGAGCCGAGATCCCA
AGAGCACACGTCTGAACTCTTTCCCTACACGACGCTCTTCCGATCTNNNGAGGCTGAGGCAGGAGAAT
GAGCACACGTCTGAACTCTTTCCCTACACGACGCTCTTCCGATCTNNNTGGGTGACAGAGCAAGACTC
GAGCACACGTCTGAACTCTTTCCCTACACGACGCTCTTCCGATCTNNNAGGAGAATGGCGTGAACCTGC
GAGCACACGTCTGAACTCTTTCCCTACACGACGCTCTTCCGATCTNNNGAGGCTGAGGCAGGAGAAT
AGCACACGTCTGAACTCTTTCCCTACACGACGCTCTTCCGATCTNNNGAGGCTGAGGCAGGAGAATGC

AGCACACGTCTGAACTCTTTCCCTACACGACGCTCTTCCGATCTNNNCGACAGAACGAGACTCCGTCCA
ACACGTCTGAACTCTTTCCCTACACGACGCTCTTCCGATCTNNNAGAAGGCGTCTGAGATAGGTTTCC
CACACGTCTGAACTCTTTCCCTACACGACGCTCTTCCGATCTNNNGCTGGCTTTCTGCTTTCTTTCTTCAT
CACACGTCTGAACTCTTTCCCTACACGACGCTCTTCCGATCTNNNACCAACCAACGATAGAAAGATCAC
CACGTCTGAACTCTTTCCCTACACGACGCTCTTCCGATCTNNNAAAGCTCAGGACCAGAAGAGTTTCA
CACGTCTGAACTCTTTCCCTACACGACGCTCTTCCGATCTNNNATTCTTCCACCCCCATCTTTGTTCATC
GAGCACACGTCTGAACTCTTTCCCTACACGACGCTCTTCCGATCTNNNCTTGCAGTGAGCCGAGATCA
GAGCACACGTCTGAACTCTTTCCCTACACGACGCTCTTCCGATCTNNNCTTGCAGTGAGCCGAGATTG
AGCACACGTCTGAACTCTTTCCCTACACGACGCTCTTCCGATCTNNNGAGGCTGAGGCAGGAGAATG
GAGCACACGTCTGAACTCTTTCCCTACACGACGCTCTTCCGATCTNNNGAAGAGCGAGACCCTGTCTG
GAGCACACGTCTGAACTCTTTCCCTACACGACGCTCTTCCGATCTNNNGAGGCTGAGGCAGGAGAATG
CACGTCTGAACTCTTTCCCTACACGACGCTCTTCCGATCTNNNGAGTAGCTGGGATTATAGGCACCCA
CACGTCTGAACTCTTTCCCTACACGACGCTCTTCCGATCTNNNACCAATCCTCTTTCCACTCTTCCCATCG
GAGCACACGTCTGAACTCTTTCCCTACACGACGCTCTTCCGATCTNNNGTTCTCTTGCCTCAGCCTCCCA
CACACGTCTGAACTCTTTCCCTACACGACGCTCTTCCGATCTNNNAGGGGCTGTAGTTTGATAAAGCA
GCACACGTCTGAACTCTTTCCCTACACGACGCTCTTCCGATCTNNNCCGAGTAACCGGGATTACAGG
ACACGTCTGAACTCTTTCCCTACACGACGCTCTTCCGATCTNNNGATAACCTGAGGTCGGGAGTTTGC
GAGCACACGTCTGAACTCTTTCCCTACACGACGCTCTTCCGATCTNNNGAGGCTGAGGCAGGAGAAT
AGCACACGTCTGAACTCTTTCCCTACACGACGCTCTTCCGATCTNNNGAGGCTGAGGCAGGAGAATG
AGCACACGTCTGAACTCTTTCCCTACACGACGCTCTTCCGATCTNNNGAGGCTGAGGCAGGAGAATG
AGCACACGTCTGAACTCTTTCCCTACACGACGCTCTTCCGATCTNNNGAGCTTGCAGTGAGCCAAGA
AGCACACGTCTGAACTCTTTCCCTACACGACGCTCTTCCGATCTNNNGAGGCTGAGGCAGGAGAATG
CACACGTCTGAACTCTTTCCCTACACGACGCTCTTCCGATCTNNNTGACTTCCTCTTTCCCAATACGGC
ACACGTCTGAACTCTTTCCCTACACGACGCTCTTCCGATCTNNNACTAACATAAACAGCAGCATGGACA
CACACGTCTGAACTCTTTCCCTACACGACGCTCTTCCGATCTNNNTTTCAATCTTTGAGGCTGCTGACC
CACACGTCTGAACTCTTTCCCTACACGACGCTCTTCCGATCTNNNGTGCTTCCTGGAGGACCATATTTC
ACACGTCTGAACTCTTTCCCTACACGACGCTCTTCCGATCTNNNACCCAAAAAACGTGCTAATAGCCCA
AGCACACGTCTGAACTCTTTCCCTACACGACGCTCTTCCGATCTNNNGAGCTGAGATCCTGCCACTGCA
GCACACGTCTGAACTCTTTCCCTACACGACGCTCTTCCGATCTNNNGACAGAGCAAGACTCCGTCTCCA
AGCACACGTCTGAACTCTTTCCCTACACGACGCTCTTCCGATCTNNNGAGGCTGAGGCAGGAGAATG
CACGTCTGAACTCTTTCCCTACACGACGCTCTTCCGATCTNNNCAGGGGTATATCAACTCTCCAGCCAT
CACGTCTGAACTCTTTCCCTACACGACGCTCTTCCGATCTNNNTAGGCATTCCTCATCATTCTCCCCATC
CACACGTCTGAACTCTTTCCCTACACGACGCTCTTCCGATCTNNNATGATCACGTCACTACACTCCAGCA
CACGTCTGAACTCTTTCCCTACACGACGCTCTTCCGATCTNNNTAAATGTCTTCTTGGGCTCCAGTCAT
AGCACACGTCTGAACTCTTTCCCTACACGACGCTCTTCCGATCTNNNGAGGCTGAGGCAGGAGAATG
GAGCACACGTCTGAACTCTTTCCCTACACGACGCTCTTCCGATCTNNNGAGGCTGAGGCAGGAGAAT
AGCACACGTCTGAACTCTTTCCCTACACGACGCTCTTCCGATCTNNNCTTGCAGTGAGCCGAGATCCCA
GAGCACACGTCTGAACTCTTTCCCTACACGACGCTCTTCCGATCTNNNAGGAGAATGGCGTGAACCTG
GAGCACACGTCTGAACTCTTTCCCTACACGACGCTCTTCCGATCTNNNAGGAGAATGGCGTGAACCTG
CACACGTCTGAACTCTTTCCCTACACGACGCTCTTCCGATCTNNNTCTCTAACTCCTGACCTCGTGATCA
GAGCACACGTCTGAACTCTTTCCCTACACGACGCTCTTCCGATCTNNNGCCTGTAGTCCCAGCTACTG
CACGTCTGAACTCTTTCCCTACACGACGCTCTTCCGATCTNNNTCAATCGCCAGAACTGCTAAGAACAT
CACGTCTGAACTCTTTCCCTACACGACGCTCTTCCGATCTNNNTCCTTCAGTCATTTCTTCGCTCTCATC

CACGTCTGAACTCTTTCCCTACACGACGCTCTTCCGATCTNNNTGTGATTCTAGGGAACACGTGAGCA
AGCACACGTCTGAACTCTTTCCCTACACGACGCTCTTCCGATCTNNNGAGGCTGAGGCAGGAGAATG
AGCACACGTCTGAACTCTTTCCCTACACGACGCTCTTCCGATCTNNNGAGGCTGAGGCAGGAGAATG
AGAGCACACGTCTGAACTCTTTCCCTACACGACGCTCTTCCGATCTNNNGAGGCTGAGGCAGGAGAA
GAGCACACGTCTGAACTCTTTCCCTACACGACGCTCTTCCGATCTNNNCTTGCAGTGAGCCGAGATTG
AGCACACGTCTGAACTCTTTCCCTACACGACGCTCTTCCGATCTNNNGAGGCTGAGGCAGGAGAATG
GAGCACACGTCTGAACTCTTTCCCTACACGACGCTCTTCCGATCTNNNGAGGCTGAGGCAGGAGAAT
AGCACACGTCTGAACTCTTTCCCTACACGACGCTCTTCCGATCTNNNAGGAGAATGGCGTGAACCTG
CACGTCTGAACTCTTTCCCTACACGACGCTCTTCCGATCTNNNGAGAAATGTCTCTCTGCCCACTCCAT
AGAGCACACGTCTGAACTCTTTCCCTACACGACGCTCTTCCGATCTNNNCTTGCAGTGAGCCGAGATT
GCACACGTCTGAACTCTTTCCCTACACGACGCTCTTCCGATCTNNNGTTTGAACCTGGGAGGCAGAG
GCACACGTCTGAACTCTTTCCCTACACGACGCTCTTCCGATCTNNNCGTTCATCAATCCACGACATGAA
CACACGTCTGAACTCTTTCCCTACACGACGCTCTTCCGATCTNNNCAAAGGCTTCCTCAGACCTCTTTCA
ACACGTCTGAACTCTTTCCCTACACGACGCTCTTCCGATCTNNNTGGCTTTCTGCAACGAAAAATTGC
CACGTCTGAACTCTTTCCCTACACGACGCTCTTCCGATCTNNNTCACACTCTACCCTGAATTGCAACAT
GAGCACACGTCTGAACTCTTTCCCTACACGACGCTCTTCCGATCTNNNCTGGGCAACAAGAGCGAAA
GAGCACACGTCTGAACTCTTTCCCTACACGACGCTCTTCCGATCTNNNGAGGCTGAGGCAGGAGAAT
AGCACACGTCTGAACTCTTTCCCTACACGACGCTCTTCCGATCTNNNGAGGCTGAGGCAGGAGAATG
CACGTCTGAACTCTTTCCCTACACGACGCTCTTCCGATCTNNNAATGCTGGGATTACAGGTCTGAGCA
ACACGTCTGAACTCTTTCCCTACACGACGCTCTTCCGATCTNNNTTTCAGGAATGCCAATGAGTCATCA
AGCACACGTCTGAACTCTTTCCCTACACGACGCTCTTCCGATCTNNNTGATCCTGGCTCACTGCAAGC
CACGTCTGAACTCTTTCCCTACACGACGCTCTTCCGATCTNNNGTGGCACTCAAATAAAAAGCCCTCAT
ACACGTCTGAACTCTTTCCCTACACGACGCTCTTCCGATCTNNNGGACCCAGACTTCTCTCCTTTTTCAT
GAGCACACGTCTGAACTCTTTCCCTACACGACGCTCTTCCGATCTNNNGAGGCTGAGGCAGGAGAAT
AGCACACGTCTGAACTCTTTCCCTACACGACGCTCTTCCGATCTNNNAGGAGAATGGCGTGAACTCG
AGCACACGTCTGAACTCTTTCCCTACACGACGCTCTTCCGATCTNNNGAGGCTGAGGCAGGAGAATG
AGAGCACACGTCTGAACTCTTTCCCTACACGACGCTCTTCCGATCTNNNCTTGCAGTGAGCCGAGATC
AGCACACGTCTGAACTCTTTCCCTACACGACGCTCTTCCGATCTNNNGAGGCTGAGGCAGGAGAATG
CACGTCTGAACTCTTTCCCTACACGACGCTCTTCCGATCTNNNCTCCTGAGTAGCTGGGATAACAGCA
CACGTCTGAACTCTTTCCCTACACGACGCTCTTCCGATCTNNNTTGATTTGAGGTAAACCGTTGGGCA
ACACGTCTGAACTCTTTCCCTACACGACGCTCTTCCGATCTNNNTTCTGGATTCTCTTCCTTTGGGGCA
AGCACACGTCTGAACTCTTTCCCTACACGACGCTCTTCCGATCTNNNGAGGCTGAGGCAGGAGAATG
AGCACACGTCTGAACTCTTTCCCTACACGACGCTCTTCCGATCTNNNCTGGGCAACAAGAGCGAAAC
GAGCACACGTCTGAACTCTTTCCCTACACGACGCTCTTCCGATCTNNNGAGGCTGAGGCAGGAGAAT
GAGCACACGTCTGAACTCTTTCCCTACACGACGCTCTTCCGATCTNNNGAGGCTGAGGCAGGAGAAT
GAGCACACGTCTGAACTCTTTCCCTACACGACGCTCTTCCGATCTNNNGAGGCTGAGGCAGGAGAAT
GAGCACACGTCTGAACTCTTTCCCTACACGACGCTCTTCCGATCTNNNGAGGCTGAGGCAGGAGAAT
AGCACACGTCTGAACTCTTTCCCTACACGACGCTCTTCCGATCTNNNAGAGCTGTCCTGGTCCTGAGCA
CACGTCTGAACTCTTTCCCTACACGACGCTCTTCCGATCTNNNTTGGAAAAAGTGTTCACAGGATCCA
ACACGTCTGAACTCTTTCCCTACACGACGCTCTTCCGATCTNNNCAGTGGATAAAAGGAGAGAGGCA
ACACGTCTGAACTCTTTCCCTACACGACGCTCTTCCGATCTNNNTGGGTGACAGAATGAGACTCTGC
CACACGTCTGAACTCTTTCCCTACACGACGCTCTTCCGATCTNNNCTGCACACAAAGACCATCAGATTC
AGCACACGTCTGAACTCTTTCCCTACACGACGCTCTTCCGATCTNNNCTTGCAGTGAGCCGAGATCGC

AGCACACGTCTGAACTCTTTCCCTACACGACGCTCTTCCGATCTNNNGAGGCTGAGGCAGGAGAATG
AGCACACGTCTGAACTCTTTCCCTACACGACGCTCTTCCGATCTNNNCTTGCAGTGAGCCGAGATCCC
GAGCACACGTCTGAACTCTTTCCCTACACGACGCTCTTCCGATCTNNNGAGGCTGAGGCAGGAGAAT
AGCACACGTCTGAACTCTTTCCCTACACGACGCTCTTCCGATCTNNNCTTGCAGTGAGCCGAGATTG
AGCACACGTCTGAACTCTTTCCCTACACGACGCTCTTCCGATCTNNNAGGAGAATGGCGTGAACCTG
GAGCACACGTCTGAACTCTTTCCCTACACGACGCTCTTCCGATCTNNNAGGAGAATGGCGTGAACCT
AGCACACGTCTGAACTCTTTCCCTACACGACGCTCTTCCGATCTNNNGAGGCTGAGGCAGGAGAATG
ACACGTCTGAACTCTTTCCCTACACGACGCTCTTCCGATCTNNNAATACAAAAAATTAGCCGGGCGTC
AGCACACGTCTGAACTCTTTCCCTACACGACGCTCTTCCGATCTNNNGAGGCTGAGGCAGGAGAATG
ACACGTCTGAACTCTTTCCCTACACGACGCTCTTCCGATCTNNNTTCCCTCTTGGTTCTATTCTGGCCAT
ACACGTCTGAACTCTTTCCCTACACGACGCTCTTCCGATCTNNNACCGACAAAAGTGGATCTCATGAC
CACGTCTGAACTCTTTCCCTACACGACGCTCTTCCGATCTNNNTTCAGTTTTGCGTCTGTATGAAACAT
ACACGTCTGAACTCTTTCCCTACACGACGCTCTTCCGATCTNNNAGGTTCAAGCAATTCTCCTGTCTCA
ACACGTCTGAACTCTTTCCCTACACGACGCTCTTCCGATCTNNNGCCCAGTGTAGTTCTCTGTAGTTCA
AGCACACGTCTGAACTCTTTCCCTACACGACGCTCTTCCGATCTNNNTGGGTGACAGAGCGAGACTC
ACACGTCTGAACTCTTTCCCTACACGACGCTCTTCCGATCTNNNCAGAGATCACACCACTGTACTCCCA
AGCACACGTCTGAACTCTTTCCCTACACGACGCTCTTCCGATCTNNNGACAGAGCGAGACTCCGTCTCA
GAGCACACGTCTGAACTCTTTCCCTACACGACGCTCTTCCGATCTNNNGCCTGTAGTCCCAGCTCTTGC
AGCACACGTCTGAACTCTTTCCCTACACGACGCTCTTCCGATCTNNNGAGGCTGAGGCAGGAGAATG
AGCACACGTCTGAACTCTTTCCCTACACGACGCTCTTCCGATCTNNNCTTTCTTCTGCCTCAGCCTCCCAT
ACACGTCTGAACTCTTTCCCTACACGACGCTCTTCCGATCTNNNGGGAACTCTCATTACTGCAAGGAC
AGCACACGTCTGAACTCTTTCCCTACACGACGCTCTTCCGATCTNNNAGCAGAGCGGGATTTCATCTCC
AGCACACGTCTGAACTCTTTCCCTACACGACGCTCTTCCGATCTNNNGAGGCTGAGGCAGGAGAATG
CACGTCTGAACTCTTTCCCTACACGACGCTCTTCCGATCTNNNTTTGGTAGAGATGGGGTTTCACCCA
AGCACACGTCTGAACTCTTTCCCTACACGACGCTCTTCCGATCTNNNAGGTTCACGCCATTCTCCTGCA
CACGTCTGAACTCTTTCCCTACACGACGCTCTTCCGATCTNNNAGGCCTCAGGAAACTTATCATGGCA
ACACGTCTGAACTCTTTCCCTACACGACGCTCTTCCGATCTNNNTGCTCTGAAAACCAAACCATTGTCA
AGCACACGTCTGAACTCTTTCCCTACACGACGCTCTTCCGATCTNNNGTTGCAGTGAGCCGAGATGGC
ACACGTCTGAACTCTTTCCCTACACGACGCTCTTCCGATCTNNNTAAGAATAAATGGGCTGGGTGGG
GAGCACACGTCTGAACTCTTTCCCTACACGACGCTCTTCCGATCTNNNGAGGCTGAGGCAGGAGAAT
AGCACACGTCTGAACTCTTTCCCTACACGACGCTCTTCCGATCTNNNCAGGAAAATGGCGTGAACCCC
CACGTCTGAACTCTTTCCCTACACGACGCTCTTCCGATCTNNNTCACACTGTGAGAACATGGTCTTCAT
CACGTCTGAACTCTTTCCCTACACGACGCTCTTCCGATCTNNNTGTGCTTGGTGGTAGAATACAGACA
CACGTCTGAACTCTTTCCCTACACGACGCTCTTCCGATCTNNNAAAACACGTAAAGCACCGATGACCA
CACGTCTGAACTCTTTCCCTACACGACGCTCTTCCGATCTNNNGCCTCCAAAACTGTTGGGATTACCAT
ACACGTCTGAACTCTTTCCCTACACGACGCTCTTCCGATCTNNNTGAGGTAGGAGAATCGCTTGAAC
CACGTCTGAACTCTTTCCCTACACGACGCTCTTCCGATCTNNNCTTTGGGATTCAGCTTCGACATCCAT
CACACGTCTGAACTCTTTCCCTACACGACGCTCTTCCGATCTNNNCATTCTGACAGGTATGGGATGGT
CACACGTCTGAACTCTTTCCCTACACGACGCTCTTCCGATCTNNNCTGAGTTCTGCCACACACAAAAAC
ACACGTCTGAACTCTTTCCCTACACGACGCTCTTCCGATCTNNNGTGTTAGCCAGGATGATCTCGATC
AGCACACGTCTGAACTCTTTCCCTACACGACGCTCTTCCGATCTNNNCTCACTGCAACCTCCACCTCCATC
ACACGTCTGAACTCTTTCCCTACACGACGCTCTTCCGATCTNNNCCTGTCCTCAATCCTGACTCATTCAT
AGCACACGTCTGAACTCTTTCCCTACACGACGCTCTTCCGATCTNNNGAGGCTGAGGCAGGAGAATG

AGCACACGTCTGAACTCTTTCCCTACACGACGCTCTTCCGATCTNNNGAGGCTGAGGCAGGAGAAT
AGCACACGTCTGAACTCTTTCCCTACACGACGCTCTTCCGATCTNNNGAGGTGGAGCTTGCAGTGAG
AGCACACGTCTGAACTCTTTCCCTACACGACGCTCTTCCGATCTNNNGAGGCTGAGGCAGGAGAATG
AGCACACGTCTGAACTCTTTCCCTACACGACGCTCTTCCGATCTNNNGAGGCTGAGGCAGGAGAATG
AGCACACGTCTGAACTCTTTCCCTACACGACGCTCTTCCGATCTNNNGAGGCTGAGGCAGGAGAAT
AGCACACGTCTGAACTCTTTCCCTACACGACGCTCTTCCGATCTNNNGAGGCTGAGGCAGGAGAATG
AGCACACGTCTGAACTCTTTCCCTACACGACGCTCTTCCGATCTNNNCTTGCAGTGAGCCGAGATCCC
AGCACACGTCTGAACTCTTTCCCTACACGACGCTCTTCCGATCTNNNGCCTGTAGTCCCAGCTACACC
GAGCACACGTCTGAACTCTTTCCCTACACGACGCTCTTCCGATCTNNNCTTGCAGTGAGCCGAGATCC
GAGCACACGTCTGAACTCTTTCCCTACACGACGCTCTTCCGATCTNNNGAGGCTGAGGCAGGAGAAT
CACGTCTGAACTCTTTCCCTACACGACGCTCTTCCGATCTNNNATGAGTTTAGTCAGAATCCCCGGCAT
CACGTCTGAACTCTTTCCCTACACGACGCTCTTCCGATCTNNNTTGCGTTCTGCTAGCTTTTGAATCAT
CACGTCTGAACTCTTTCCCTACACGACGCTCTTCCGATCTNNNTCAGGGTTATTGTCTTCGTGGTTCAT
CACGTCTGAACTCTTTCCCTACACGACGCTCTTCCGATCTNNNTAGACTCCCATTGGCTGAATCTGCAT
AGCACACGTCTGAACTCTTTCCCTACACGACGCTCTTCCGATCTNNNGAGGCTGAGGCAGGAGAAT
AGCACACGTCTGAACTCTTTCCCTACACGACGCTCTTCCGATCTNNNGAGGCTGAGGCAGGAGAAT
GCACACGTCTGAACTCTTTCCCTACACGACGCTCTTCCGATCTNNNGAGGCTGAGGCAGGAGAATG
AGCACACGTCTGAACTCTTTCCCTACACGACGCTCTTCCGATCTNNNGAGGCTGAGGCAGGAGAATG
CACGTCTGAACTCTTTCCCTACACGACGCTCTTCCGATCTNNNGGTGGAGGAATAGGATCACAGTCC
ACACGTCTGAACTCTTTCCCTACACGACGCTCTTCCGATCTNNNGCTTCTCAAAGTGCTGGGATTACC
CACGTCTGAACTCTTTCCCTACACGACGCTCTTCCGATCTNNNGCTGTGTGTTCCATTTTTGGGAACAT
ACACGTCTGAACTCTTTCCCTACACGACGCTCTTCCGATCTNNNATACTCACAGCATTACCCAGTCGCAT
AGCACACGTCTGAACTCTTTCCCTACACGACGCTCTTCCGATCTNNNGAGGCTGAGGCAGGAGAATG
GAGCACACGTCTGAACTCTTTCCCTACACGACGCTCTTCCGATCTNNNGAGGCTGAGGCAGGAGAAT
CACGTCTGAACTCTTTCCCTACACGACGCTCTTCCGATCTNNNGTCCCTGGGCAACTACTAATCTGCAT
ACACGTCTGAACTCTTTCCCTACACGACGCTCTTCCGATCTNNNTTTAGTAGAGACGGGGTTTCACCCA
ACACGTCTGAACTCTTTCCCTACACGACGCTCTTCCGATCTNNNTTTTATACCTCCTGGGAAGTTCCCA
ACACGTCTGAACTCTTTCCCTACACGACGCTCTTCCGATCTNNNAGAACCCCAGATTTCTAGAACAAC
CACGTCTGAACTCTTTCCCTACACGACGCTCTTCCGATCTNNNAGATGGCCTCAGCTCTTGTTTTACAT
AGCACACGTCTGAACTCTTTCCCTACACGACGCTCTTCCGATCTNNNAGGAGAATGGCGTGAACCTGC
GCACACGTCTGAACTCTTTCCCTACACGACGCTCTTCCGATCTNNNGAGGCTGAGGCAGGAGAATG
GAGCACACGTCTGAACTCTTTCCCTACACGACGCTCTTCCGATCTNNNTGGACAACAGAGCGAGACT
GAGCACACGTCTGAACTCTTTCCCTACACGACGCTCTTCCGATCTNNNGAGGCTGAGGCAGGAGAAT
AGCACACGTCTGAACTCTTTCCCTACACGACGCTCTTCCGATCTNNNGAGGCTGAGGCAGGAGAATG
ACACGTCTGAACTCTTTCCCTACACGACGCTCTTCCGATCTNNNCAATTTCCTGAGTAGCTGGGACTCA
CACGTCTGAACTCTTTCCCTACACGACGCTCTTCCGATCTNNNACCAAAAGAGACAGAGTGGTGAAC
ACACGTCTGAACTCTTTCCCTACACGACGCTCTTCCGATCTNNNGGCATACATGTTGATTGCACACAC
CACGTCTGAACTCTTTCCCTACACGACGCTCTTCCGATCTNNNCCCAAAGTGCTGAGATTACAAGCCAT
ACACGTCTGAACTCTTTCCCTACACGACGCTCTTCCGATCTNNNAGCATGACATTTTTGTGAAGCCTCA
CACGTCTGAACTCTTTCCCTACACGACGCTCTTCCGATCTNNNCACACCCAACCTGATTGTTTTCTCAT
GCACACGTCTGAACTCTTTCCCTACACGACGCTCTTCCGATCTNNNGAGATCCTGCCACTGCACTCCAT
CACGTCTGAACTCTTTCCCTACACGACGCTCTTCCGATCTNNNGGCTACTCTAGAATTGCCCTAGCCAT
ACACGTCTGAACTCTTTCCCTACACGACGCTCTTCCGATCTNNNTCCCAAAGTGCTAGGATTACAGGCA

AGCACACGTCTGAACTCTTTCCCTACACGACGCTCTTCCGATCTNNNGAGGCTGAGGCAGGAGAATG
AGCACACGTCTGAACTCTTTCCCTACACGACGCTCTTCCGATCTNNNCTTGCAGTGAGCCGAGATCGCA
AGCACACGTCTGAACTCTTTCCCTACACGACGCTCTTCCGATCTNNNGAGGCTGAGGCAGGAGAATG
AGCACACGTCTGAACTCTTTCCCTACACGACGCTCTTCCGATCTNNNCTTGCAGTGAGCCGAGATTGCA
GAGCACACGTCTGAACTCTTTCCCTACACGACGCTCTTCCGATCTNNNGAGGCTGAGGCAGGAGAATG
CACACGTCTGAACTCTTTCCCTACACGACGCTCTTCCGATCTNNNTATCCATGTGTTCCTGTGGCTTTCAT
ACACGTCTGAACTCTTTCCCTACACGACGCTCTTCCGATCTNNNTACCAAGGCCTTGACTAGAACAACAT
ACACGTCTGAACTCTTTCCCTACACGACGCTCTTCCGATCTNNNTCCCCACATCCCAATTTCATCTTCATCG
GAGCACACGTCTGAACTCTTTCCCTACACGACGCTCTTCCGATCTNNNGAGGCTGAGGCAGGAGAATG
AGAGCACACGTCTGAACTCTTTCCCTACACGACGCTCTTCCGATCTNNNGAGCCAAGATCGTGCCACTG
GAGCACACGTCTGAACTCTTTCCCTACACGACGCTCTTCCGATCTNNNGAGGCTGAGGCAGGAGAATG
GAGCACACGTCTGAACTCTTTCCCTACACGACGCTCTTCCGATCTNNNGAGGCTGAGGCAGGAGAATG
CACACGTCTGAACTCTTTCCCTACACGACGCTCTTCCGATCTNNNTCATTCACTCAGGTCTTCCCTTGCAT
AGCACACGTCTGAACTCTTTCCCTACACGACGCTCTTCCGATCTNNNGAGGCTGAGGCAGGAGAATG
ACACGTCTGAACTCTTTCCCTACACGACGCTCTTCCGATCTNNNTCAAGTGATCTTCCTGTCTCAGCCAT
ACACGTCTGAACTCTTTCCCTACACGACGCTCTTCCGATCTNNNTCCTGGAATCCTGTGCAAAATCTCAT
ACACGTCTGAACTCTTTCCCTACACGACGCTCTTCCGATCTNNNTACTCCAATTTTGCTCCTCTGCTCATCC
CACACGTCTGAACTCTTTCCCTACACGACGCTCTTCCGATCTNNNACCCGGCCTCCTTTTTTTTTTTTCATC
GAGCACACGTCTGAACTCTTTCCCTACACGACGCTCTTCCGATCTNNNCTGTTGTCCCAGCTACTCGGC
AGCACACGTCTGAACTCTTTCCCTACACGACGCTCTTCCGATCTNNNGAGGCTGAGGCAGGAGAATG
GCACACGTCTGAACTCTTTCCCTACACGACGCTCTTCCGATCTNNNGCAACAAAAGCGAGACTCTGTCCA
GAGCACACGTCTGAACTCTTTCCCTACACGACGCTCTTCCGATCTNNNGAGGCTGAGGCAGGAGAAT
GAGCACACGTCTGAACTCTTTCCCTACACGACGCTCTTCCGATCTNNNGAGGCTGAGGCAGGAGAATG
AGAGCACACGTCTGAACTCTTTCCCTACACGACGCTCTTCCGATCTNNNCTTGCAGTGAGCCGAGATTG
ACACGTCTGAACTCTTTCCCTACACGACGCTCTTCCGATCTNNNTTAGCAGTGTGAGAACAGACTAACAT
CACACGTCTGAACTCTTTCCCTACACGACGCTCTTCCGATCTNNNGGGCATATGGGAACTCTGTACTACA
ACACGTCTGAACTCTTTCCCTACACGACGCTCTTCCGATCTNNNATGGCAAAGTGACCCATACAGATCAT
ACACGTCTGAACTCTTTCCCTACACGACGCTCTTCCGATCTNNNCATATTTCATCCCTGCAGCAAGGCATC
ACACGTCTGAACTCTTTCCCTACACGACGCTCTTCCGATCTNNNTGCCACAGTTTCATCTAAGACCTCATC
ACACGTCTGAACTCTTTCCCTACACGACGCTCTTCCGATCTNNNCTAAGTGCTGGGATTACACACCTCAT
ACACGTCTGAACTCTTTCCCTACACGACGCTCTTCCGATCTNNNTCTTCCAAAGTGCTGGGATTACACAT
ACACGTCTGAACTCTTTCCCTACACGACGCTCTTCCGATCTNNNGCTCATTCATGTAAGCTGCATGTCATC
AGAGCACACGTCTGAACTCTTTCCCTACACGACGCTCTTCCGATCTNNNCTGCAGTGAGCCGAGATCAT
AGCACACGTCTGAACTCTTTCCCTACACGACGCTCTTCCGATCTNNNCTGTGGTCCCAGCTACTTGGCAT
AGAGCACACGTCTGAACTCTTTCCCTACACGACGCTCTTCCGATCTNNNAGGAGAATGGCATGAACCC
GAGCACACGTCTGAACTCTTTCCCTACACGACGCTCTTCCGATCTNNNGAGGCTGAGGCAGGAGAATG
GCACACGTCTGAACTCTTTCCCTACACGACGCTCTTCCGATCTNNNGAGATCGCACCACTGCTCTCCATC
GAGCACACGTCTGAACTCTTTCCCTACACGACGCTCTTCCGATCTNNNGAGGCTGAGGCAGGAGAATG
GAGCACACGTCTGAACTCTTTCCCTACACGACGCTCTTCCGATCTNNNCTTGCAGTGAGCCGAGATTG
AGCACACGTCTGAACTCTTTCCCTACACGACGCTCTTCCGATCTNNNGAGGCTGAGGCAGGAGAATGC
ACACGTCTGAACTCTTTCCCTACACGACGCTCTTCCGATCTNNNTCCCATCGCTTTCCTCTTACAAACATCC
ACACGTCTGAACTCTTTCCCTACACGACGCTCTTCCGATCTNNNATCACACTCACTCTTGTAGCACCCATC
ACACGTCTGAACTCTTTCCCTACACGACGCTCTTCCGATCTNNNTGGCCTATGAAGAGTTGCTTTCTCAT

AGCACACGTCTGAACTCTTTCCCTACACGACGCTCTTCCGATCTNNNGAGGCTGAGGCAGGAGAAT
GCACACGTCTGAACTCTTTCCCTACACGACGCTCTTCCGATCTNNNGAGGCTGAGGCAGGAGAATG
GAGCACACGTCTGAACTCTTTCCCTACACGACGCTCTTCCGATCTNNNGAGGCTGAGGCAGGAGAAT
GAGCACACGTCTGAACTCTTTCCCTACACGACGCTCTTCCGATCTNNNGAGATAGCGCCACTGCACTC
CACGTCTGAACTCTTTCCCTACACGACGCTCTTCCGATCTNNNGCTACGGTTTTAGCACATCTTCACAT
ACACGTCTGAACTCTTTCCCTACACGACGCTCTTCCGATCTNNNATTACACACGTGCCACAATTTGTCAT
ACACGTCTGAACTCTTTCCCTACACGACGCTCTTCCGATCTNNNAGTTCTGGAGTGGAAAGAATGCTC
CACACGTCTGAACTCTTTCCCTACACGACGCTCTTCCGATCTNNNAACCTGCATATGTACCCCTGAACC
ACACGTCTGAACTCTTTCCCTACACGACGCTCTTCCGATCTNNNCCAGCCGTTGGTAATTCTTTCTGCA
ACACGTCTGAACTCTTTCCCTACACGACGCTCTTCCGATCTNNNTCAGGTGGCAGATATGTTCAACAC
CACACGTCTGAACTCTTTCCCTACACGACGCTCTTCCGATCTNNNCTATGTTGGCCAGACTGGTTTTG
GAGCACACGTCTGAACTCTTTCCCTACACGACGCTCTTCCGATCTNNNCTGCAGTCCCAGCTACTTGG
GCACACGTCTGAACTCTTTCCCTACACGACGCTCTTCCGATCTNNNGAGGCTGAGGCAGGAGAATGC
CACACGTCTGAACTCTTTCCCTACACGACGCTCTTCCGATCTNNNCACCTGTAGTCCCAGCTACTTGCA
AGCACACGTCTGAACTCTTTCCCTACACGACGCTCTTCCGATCTNNNGAGGCTGAGGCAGGAGAAT
AGCACACGTCTGAACTCTTTCCCTACACGACGCTCTTCCGATCTNNNGAGGCTGAGGCAGGAGAAT
AGCACACGTCTGAACTCTTTCCCTACACGACGCTCTTCCGATCTNNNGAGGCTGAGGCAGGAGAAT
AGCACACGTCTGAACTCTTTCCCTACACGACGCTCTTCCGATCTNNNGAGGCTGAGGCAGGAGAAT
GAGCACACGTCTGAACTCTTTCCCTACACGACGCTCTTCCGATCTNNNCAGGAGAATGGCGTGAACCC
CACGTCTGAACTCTTTCCCTACACGACGCTCTTCCGATCTNNNTGAGTCCTTATCATGAGGCAAGTCAT
CACGTCTGAACTCTTTCCCTACACGACGCTCTTCCGATCTNNNACGCTGTCATCTTGATCTCAGATCAT
ACACGTCTGAACTCTTTCCCTACACGACGCTCTTCCGATCTNNNGGCAACAGAGTGAGATTCCATCTCA
CACGTCTGAACTCTTTCCCTACACGACGCTCTTCCGATCTNNNCCCATACATCTATGGCCAGTTGACAT
CACGTCTGAACTCTTTCCCTACACGACGCTCTTCCGATCTNNNGCAATAGAAGAGAGCTTGTGCAGCA
AGCACACGTCTGAACTCTTTCCCTACACGACGCTCTTCCGATCTNNNAGGAGAATGGCGTGAACCTG
AGCACACGTCTGAACTCTTTCCCTACACGACGCTCTTCCGATCTNNNGAGGCTGAGGCAGGAGAATG
GAGCACACGTCTGAACTCTTTCCCTACACGACGCTCTTCCGATCTNNNCCTGCAGTGAGCCGAGATTG
GAGCACACGTCTGAACTCTTTCCCTACACGACGCTCTTCCGATCTNNNGAGGCTGAGGCAGGAGAA
ACACGTCTGAACTCTTTCCCTACACGACGCTCTTCCGATCTNNNATTCCACCACGGATGTTACAGAACA
CACACGTCTGAACTCTTTCCCTACACGACGCTCTTCCGATCTNNNCAGGAGATAGAGACCATTCTGGC
AGCACACGTCTGAACTCTTTCCCTACACGACGCTCTTCCGATCTNNNGAGGCTGAGGCAGGAGAAT
AGCACACGTCTGAACTCTTTCCCTACACGACGCTCTTCCGATCTNNNGCAGAAGAATGGCGTGAACC
AGCACACGTCTGAACTCTTTCCCTACACGACGCTCTTCCGATCTNNNGAGGCTGAGGCAGGAGAAT
ACACGTCTGAACTCTTTCCCTACACGACGCTCTTCCGATCTNNNGGTTCCGATTTTTCCACATCCTCCAT
CACGTCTGAACTCTTTCCCTACACGACGCTCTTCCGATCTNNNAGAATGACTTTGCTCTCCTGAAACAT
CACGTCTGAACTCTTTCCCTACACGACGCTCTTCCGATCTNNNCTTGTCTTCTTCCCTCCAACCTTCATCG
AGCACACGTCTGAACTCTTTCCCTACACGACGCTCTTCCGATCTNNNGAGGCTGAGGCAGGAGAATG
AGCACACGTCTGAACTCTTTCCCTACACGACGCTCTTCCGATCTNNNTGCTATTGCACTCCAGCCTGC
GAGCACACGTCTGAACTCTTTCCCTACACGACGCTCTTCCGATCTNNNCGACAGAACGAGACTCCGTC
ACACGTCTGAACTCTTTCCCTACACGACGCTCTTCCGATCTNNNCTCTGTGTTGTATGCCTGCATTTCAT
CACGTCTGAACTCTTTCCCTACACGACGCTCTTCCGATCTNNNACTACTCAACACTCCCTTCCTCACATC
CACACGTCTGAACTCTTTCCCTACACGACGCTCTTCCGATCTNNNGGGGCTCACAGAAAAAAAAATGC
AGCACACGTCTGAACTCTTTCCCTACACGACGCTCTTCCGATCTNNNGAGGTGGAGCTTGCAGTGAG

AGCACACGTCTGAACTCTTTCCCTACACGACGCTCTTCCGATCTNNNGCCTGGTCAACATGGTGAAAC
CACGTCTGAACTCTTTCCCTACACGACGCTCTTCCGATCTNNNGGGGTATCCATCACATCAAGCATCAT
CACACGTCTGAACTCTTTCCCTACACGACGCTCTTCCGATCTNNNAGGGAAACTGGGATTCTAGGAG
CACGTCTGAACTCTTTCCCTACACGACGCTCTTCCGATCTNNNGACCTGCAAAAATATGCCCAACTCAT
GAGCACACGTCTGAACTCTTTCCCTACACGACGCTCTTCCGATCTNNNAGGAGAATGGCGTGAACCTG
AGCACACGTCTGAACTCTTTCCCTACACGACGCTCTTCCGATCTNNNCGACAGAACGAGACTCCGTCCA
CACGTCTGAACTCTTTCCCTACACGACGCTCTTCCGATCTNNNTTTTCTTTCATGCTGCAAGGTGTCAT
GAGCACACGTCTGAACTCTTTCCCTACACGACGCTCTTCCGATCTNNNGAGGCTGAGGCAGGAGAAT
AGCACACGTCTGAACTCTTTCCCTACACGACGCTCTTCCGATCTNNNAGGAGAATGGCGTGAACCTG
CACGTCTGAACTCTTTCCCTACACGACGCTCTTCCGATCTNNNTCCCGTGAAGCATGATTTAGGAACAT
CACACGTCTGAACTCTTTCCCTACACGACGCTCTTCCGATCTNNNGAACTCATGGGCTCAAACAATCCCA
GCACACGTCTGAACTCTTTCCCTACACGACGCTCTTCCGATCTNNNCTCACCATAGCTTCGACCTCCCAT
CACGTCTGAACTCTTTCCCTACACGACGCTCTTCCGATCTNNNTGTGTGTGCTTTTTTTGGTTTGGCAT
CACGTCTGAACTCTTTCCCTACACGACGCTCTTCCGATCTNNNAACAGAACAGAGCCCTCAGAAATCAT
CACACGTCTGAACTCTTTCCCTACACGACGCTCTTCCGATCTNNNCCAAGAAATGGGCAGGTAACTTCC
CACGTCTGAACTCTTTCCCTACACGACGCTCTTCCGATCTNNNGTGCTGCAAGGTGTTTCTATCACCAT
CACACGTCTGAACTCTTTCCCTACACGACGCTCTTCCGATCTNNNCCGGGATTTGTTACATGTCACACCA
AGAGCACACGTCTGAACTCTTTCCCTACACGACGCTCTTCCGATCTNNNGCACTCTAGCCTGGACAACA
AGCACACGTCTGAACTCTTTCCCTACACGACGCTCTTCCGATCTNNNGAGGCTGAGGCAGGAGAATG
GAGCACACGTCTGAACTCTTTCCCTACACGACGCTCTTCCGATCTNNNCTTGCAGTGAGCCGAGATCCC
CACGTCTGAACTCTTTCCCTACACGACGCTCTTCCGATCTNNNTGGGTTTGTTTATGAACACGCTTCAT
ACACGTCTGAACTCTTTCCCTACACGACGCTCTTCCGATCTNNNCGCCCGGCTTGTACTTTTTTATCAT
ACACGTCTGAACTCTTTCCCTACACGACGCTCTTCCGATCTNNNATATAATTCCAGTCGGGGGAAGCC
GCACACGTCTGAACTCTTTCCCTACACGACGCTCTTCCGATCTNNNCAGAGGCAGAGGGAGTTTTCTC
AGAGCACACGTCTGAACTCTTTCCCTACACGACGCTCTTCCGATCTNNNGAGGCTGAGGCAGGAGAAT
GAGCACACGTCTGAACTCTTTCCCTACACGACGCTCTTCCGATCTNNNGAGGCTGAGGCAGGAGAATG
GAGCACACGTCTGAACTCTTTCCCTACACGACGCTCTTCCGATCTNNNGAGGCTGAGGCAGGAGAATG
GAGCACACGTCTGAACTCTTTCCCTACACGACGCTCTTCCGATCTNNNGAGGCTGAGGCAGGAGAAT
CACACGTCTGAACTCTTTCCCTACACGACGCTCTTCCGATCTNNNCTGGGTGACAGTGCAAGATTCTC
GAGCACACGTCTGAACTCTTTCCCTACACGACGCTCTTCCGATCTNNNTGGGTGACAGAGTGAGACTC
GAGCACACGTCTGAACTCTTTCCCTACACGACGCTCTTCCGATCTNNNGAGGCTGAGGCAGGAGAAT
GAGCACACGTCTGAACTCTTTCCCTACACGACGCTCTTCCGATCTNNNGAGATGGAGCCACTGCACTCC
GAGCACACGTCTGAACTCTTTCCCTACACGACGCTCTTCCGATCTNNNGAGGCTGAGGCAGGAGAAT
CACACGTCTGAACTCTTTCCCTACACGACGCTCTTCCGATCTNNNTCATCAGTCAAGACCATACTGGCC
GAGCACACGTCTGAACTCTTTCCCTACACGACGCTCTTCCGATCTNNNGAGGCTGAGGCAGGAGAAT
GAGCACACGTCTGAACTCTTTCCCTACACGACGCTCTTCCGATCTNNNGAGGCTGAGGCAGGAGAATG
AGCACACGTCTGAACTCTTTCCCTACACGACGCTCTTCCGATCTNNNGAGGCTGAGGCAGGAGAATG
AGCACACGTCTGAACTCTTTCCCTACACGACGCTCTTCCGATCTNNNCTTGCAGTGAGCCGAGATTGC
ACACGTCTGAACTCTTTCCCTACACGACGCTCTTCCGATCTNNNGTGTCCTTCAGCTCCTTTCACTACATC
ACACGTCTGAACTCTTTCCCTACACGACGCTCTTCCGATCTNNNATACAGCATTGTGGAGAAAAGCTCA
CACGTCTGAACTCTTTCCCTACACGACGCTCTTCCGATCTNNNGGACAGGTGATCAAACCATAGGTCA
GAGCACACGTCTGAACTCTTTCCCTACACGACGCTCTTCCGATCTNNNGAGGCTGAGGCAGGAGAATC
AGCACACGTCTGAACTCTTTCCCTACACGACGCTCTTCCGATCTNNNCCTGCAGTGAGCCGAGATTGC

GCACACGTCTGAACTCTTTCCCTACACGACGCTCTTCCGATCTNNNGAGCCGAGATCATGCCACTGCA
GAGCACACGTCTGAACTCTTTCCCTACACGACGCTCTTCCGATCTNNNGAGGCTGAGGCAGGAGAAT
AGCACACGTCTGAACTCTTTCCCTACACGACGCTCTTCCGATCTNNNCTTGCAGTGAGGGGAGATCG
GAGCACACGTCTGAACTCTTTCCCTACACGACGCTCTTCCGATCTNNNGAGGCTGAGGCAGGAGAA
GCACACGTCTGAACTCTTTCCCTACACGACGCTCTTCCGATCTNNNGATTGAGACCATCCTGGCTAAC
GAGCACACGTCTGAACTCTTTCCCTACACGACGCTCTTCCGATCTNNNCTTGCAGTGAGCCGAGATCA
CACGTCTGAACTCTTTCCCTACACGACGCTCTTCCGATCTNNNGCCTTTCCCACCAATCTCTTTTCCATC
GCACACGTCTGAACTCTTTCCCTACACGACGCTCTTCCGATCTNNNGGCAATACCCACCAAGGAAGA
CACACGTCTGAACTCTTTCCCTACACGACGCTCTTCCGATCTNNNCTGGCCTAGAAGGGTGTTTTTCC
CACGTCTGAACTCTTTCCCTACACGACGCTCTTCCGATCTNNNTGACCTCTGATCTCTACGCACTACAT
ACACGTCTGAACTCTTTCCCTACACGACGCTCTTCCGATCTNNNTGCTGACTTGACAAAATGGCAAAC
CACGTCTGAACTCTTTCCCTACACGACGCTCTTCCGATCTNNNGAGTAGCTGGGATTACAGGCATGC
ACACGTCTGAACTCTTTCCCTACACGACGCTCTTCCGATCTNNNTGTTAGCAAACTAAGTCCAGCAAC
CACGTCTGAACTCTTTCCCTACACGACGCTCTTCCGATCTNNNATAAATCCAACAAACCCCCTTGCCAT
AGCACACGTCTGAACTCTTTCCCTACACGACGCTCTTCCGATCTNNNCTTGCAGTGAGCCGAGATTG
GAGCACACGTCTGAACTCTTTCCCTACACGACGCTCTTCCGATCTNNNGAGGCTGAGGCAGGAGAAT
AGCACACGTCTGAACTCTTTCCCTACACGACGCTCTTCCGATCTNNNGAGGCTGAGGCAGGAGAAT
GAGCACACGTCTGAACTCTTTCCCTACACGACGCTCTTCCGATCTNNNAGGAGAATGGCGTGAACCT
AGCACACGTCTGAACTCTTTCCCTACACGACGCTCTTCCGATCTNNNGAGGCTGAGGCAGGAGAATG
CACGTCTGAACTCTTTCCCTACACGACGCTCTTCCGATCTNNNAATTGCTAATGCGTAAGGCCAAACA
GCACACGTCTGAACTCTTTCCCTACACGACGCTCTTCCGATCTNNNGGAGAATGTGGTCCCAGCATT
CACGTCTGAACTCTTTCCCTACACGACGCTCTTCCGATCTNNNGAAAGGAAGCTCCCAACTACAGACA
ACACGTCTGAACTCTTTCCCTACACGACGCTCTTCCGATCTNNNAGACAAGGTTTCTCCATGTTGGTCA
ACACGTCTGAACTCTTTCCCTACACGACGCTCTTCCGATCTNNNCGTGCCTATGTCCTGAATGGTATCA
CACACGTCTGAACTCTTTCCCTACACGACGCTCTTCCGATCTNNNAGGGCAAGACTCCATCTCAAAT
ACACGTCTGAACTCTTTCCCTACACGACGCTCTTCCGATCTNNNAATGTGTCATGGAGGCTTTCAGAC
AGCACACGTCTGAACTCTTTCCCTACACGACGCTCTTCCGATCTNNNGAGATCGCACCACTGCACTCCA
GAGCACACGTCTGAACTCTTTCCCTACACGACGCTCTTCCGATCTNNNGAGATCGCACCACTGCACTCC
AGCACACGTCTGAACTCTTTCCCTACACGACGCTCTTCCGATCTNNNGAGGCTGAGGCAGGAGAATG
AGCACACGTCTGAACTCTTTCCCTACACGACGCTCTTCCGATCTNNNGACAGAGCGAGACTCCGTCTC
AGCACACGTCTGAACTCTTTCCCTACACGACGCTCTTCCGATCTNNNGAGGCTGAGGCAGGAGAATG
AGCACACGTCTGAACTCTTTCCCTACACGACGCTCTTCCGATCTNNNCTTGCAGTGAGCCGAGATCCC
GAGCACACGTCTGAACTCTTTCCCTACACGACGCTCTTCCGATCTNNNGAGGCTGAGGCAGGAGAAT
AGCACACGTCTGAACTCTTTCCCTACACGACGCTCTTCCGATCTNNNCTTGCAGTGAGCCGAGATCC
AGCACACGTCTGAACTCTTTCCCTACACGACGCTCTTCCGATCTNNNGAGGCTGAGGCAGGAGAAT
AGCACACGTCTGAACTCTTTCCCTACACGACGCTCTTCCGATCTNNNAACAGAGCGAGACACCCAAG
GCACACGTCTGAACTCTTTCCCTACACGACGCTCTTCCGATCTNNNGAGCTGAGATGGAGCCACTGC
GCACACGTCTGAACTCTTTCCCTACACGACGCTCTTCCGATCTNNNGAGGCTGAGGCAGGAGAATG
GAGCACACGTCTGAACTCTTTCCCTACACGACGCTCTTCCGATCTNNNAGGAGAATGGCGTGAACCT
CACGTCTGAACTCTTTCCCTACACGACGCTCTTCCGATCTNNNGCAGAAGCATTGACTAAACCTGACA
ACACGTCTGAACTCTTTCCCTACACGACGCTCTTCCGATCTNNNCGAACCACCTCCAATAACACCTACAT
GCACACGTCTGAACTCTTTCCCTACACGACGCTCTTCCGATCTNNNGAGAATCGCTTGATCCTGGGAG
GCACACGTCTGAACTCTTTCCCTACACGACGCTCTTCCGATCTNNNGAGGCTGAGGCAGGAGAATG

GCACACGTCTGAACTCTTTCCCTACACGACGCTCTTCCGATCTNNNAGGAGAATGGCGTGAACCTGCA
ACACGTCTGAACTCTTTCCCTACACGACGCTCTTCCGATCTNNNAGGCAGGAGAATCATTTGAACCTCA
AGCACACGTCTGAACTCTTTCCCTACACGACGCTCTTCCGATCTNNNGGAACCAAAGTGAGACCCTGCA
ACACGTCTGAACTCTTTCCCTACACGACGCTCTTCCGATCTNNNCAAAAGAGTGGATTGGTCACTGGCA
CACACGTCTGAACTCTTTCCCTACACGACGCTCTTCCGATCTNNNTCTTTTCTCGTAGGACCCTTAGTCAT
GAGCACACGTCTGAACTCTTTCCCTACACGACGCTCTTCCGATCTNNNTACTGTGTCACCCACTCCCACAT
GAGCACACGTCTGAACTCTTTCCCTACACGACGCTCTTCCGATCTNNNGAGGCTGAGGCAGGAGAATG
ACACGTCTGAACTCTTTCCCTACACGACGCTCTTCCGATCTNNNGCAACAGACCATCTCCAGACATACATG
GAGCACACGTCTGAACTCTTTCCCTACACGACGCTCTTCCGATCTNNNAGGAGAATGGCGTGAACCTG
GCACACGTCTGAACTCTTTCCCTACACGACGCTCTTCCGATCTNNNAAAGTGCTGGGATTACAGGCTTG
ACACGTCTGAACTCTTTCCCTACACGACGCTCTTCCGATCTNNNTTTCCTCAGGGCTCTTCTCTTTCCATCC
ACACGTCTGAACTCTTTCCCTACACGACGCTCTTCCGATCTNNNTGTTCCCTTTGTGTAGCTGAGAACAT
GCACACGTCTGAACTCTTTCCCTACACGACGCTCTTCCGATCTNNNAGAATGGCTTGAACCCAGGAGCA
GCACACGTCTGAACTCTTTCCCTACACGACGCTCTTCCGATCTNNNGAGGCTGAGGCAGGAGAATGCA
AGAGCACACGTCTGAACTCTTTCCCTACACGACGCTCTTCCGATCTNNNAGGAGAATGGCATGAACCC
ACACGTCTGAACTCTTTCCCTACACGACGCTCTTCCGATCTNNNTTGCACATTCCTTCCCTGAAGATCATG
GAGCACACGTCTGAACTCTTTCCCTACACGACGCTCTTCCGATCTNNNGAGGCTGAGGCAGGAGAATG
GAGCACACGTCTGAACTCTTTCCCTACACGACGCTCTTCCGATCTNNNGAGGCTGAGGCAGGAGAATG
ACACGTCTGAACTCTTTCCCTACACGACGCTCTTCCGATCTNNNCCCTTCTACCATGAGTGAAAGCTCATG
GCACACGTCTGAACTCTTTCCCTACACGACGCTCTTCCGATCTNNNCTGTCAAGCCCCTCTAGTGAATTCA
CACACGTCTGAACTCTTTCCCTACACGACGCTCTTCCGATCTNNNTACTGGGTTTCTCCATATTGGCCCA
ACACGTCTGAACTCTTTCCCTACACGACGCTCTTCCGATCTNNNCCCCACCATGTCAATCAACAAATCATC
GAGCACACGTCTGAACTCTTTCCCTACACGACGCTCTTCCGATCTNNNGAGGCTGAGGCAGGAGATTG
GAGCACACGTCTGAACTCTTTCCCTACACGACGCTCTTCCGATCTNNNGAGGCTGAGGCAGGAGAATG
AGAGCACACGTCTGAACTCTTTCCCTACACGACGCTCTTCCGATCTNNNGAGGCTGAGGCAGGAGAAT
GAGCACACGTCTGAACTCTTTCCCTACACGACGCTCTTCCGATCTNNNAGGAGAATGGCGTGAACCTGG
GAGCACACGTCTGAACTCTTTCCCTACACGACGCTCTTCCGATCTNNNGAACTTGCAGTGAGCCGAGA
ACACGTCTGAACTCTTTCCCTACACGACGCTCTTCCGATCTNNNAACAGCAGATCTCTCAGCAGAAACAT
ACACGTCTGAACTCTTTCCCTACACGACGCTCTTCCGATCTNNNTGTCTTCACTGCCTGCTTTAGATCATC
ACACGTCTGAACTCTTTCCCTACACGACGCTCTTCCGATCTNNNGGCTCAACACCTGAAATGAGATTCA
CACACGTCTGAACTCTTTCCCTACACGACGCTCTTCCGATCTNNNCCACACGAAATGTTTCCCTTAGGCA
CACACGTCTGAACTCTTTCCCTACACGACGCTCTTCCGATCTNNNGACAGCCTTTCCTAAACCTAGCTCA
ACACGTCTGAACTCTTTCCCTACACGACGCTCTTCCGATCTNNNGGAGCTTGCTCTCTCTCTTTCTTCATC
ACACGTCTGAACTCTTTCCCTACACGACGCTCTTCCGATCTNNNCCTCCAAATTTGTGGTTGCTACACATG
ACACGTCTGAACTCTTTCCCTACACGACGCTCTTCCGATCTNNNTCTAGTGATGCAGTCTGCTTTGACAT
AGCACACGTCTGAACTCTTTCCCTACACGACGCTCTTCCGATCTNNNGAGAACGCACCATGGCAGTCCA
GAGCACACGTCTGAACTCTTTCCCTACACGACGCTCTTCCGATCTNNNGTTGCAGTGAGCCGAGATTGC
AGAGCACACGTCTGAACTCTTTCCCTACACGACGCTCTTCCGATCTNNNGAGATCCCACCACTGCACTC
AGCACACGTCTGAACTCTTTCCCTACACGACGCTCTTCCGATCTNNNGAGGCTGAGGCAGGAGAATG
GCACACGTCTGAACTCTTTCCCTACACGACGCTCTTCCGATCTNNNCATGCCTGTAATCCCAGCTACTACA
CACACGTCTGAACTCTTTCCCTACACGACGCTCTTCCGATCTNNNTACAAAAAATTAGCCCGGCATGGC
AGCACACGTCTGAACTCTTTCCCTACACGACGCTCTTCCGATCTNNNGAGGCTGAGGCAGGAGAATG
GAGCACACGTCTGAACTCTTTCCCTACACGACGCTCTTCCGATCTNNNCTTGCAGTGAGCCGAGATTGC

CACACGTCTGAACTCTTTCCCTACACGACGCTCTTCCGATCTNNNAGGGGGCTTCTCTTTTGGAATTA
CACGTCTGAACTCTTTCCCTACACGACGCTCTTCCGATCTNNNCAGTGGTCTGATCATGTCTCCTTCAT
AGCACACGTCTGAACTCTTTCCCTACACGACGCTCTTCCGATCTNNNGACAGAGCGAGACTCCGTCTC
GCACACGTCTGAACTCTTTCCCTACACGACGCTCTTCCGATCTNNNCACTTTCGTCCCTGGGAGACCAT
AGCACACGTCTGAACTCTTTCCCTACACGACGCTCTTCCGATCTNNNAATGGCATGAACCCAGGAGG
AGCACACGTCTGAACTCTTTCCCTACACGACGCTCTTCCGATCTNNNTGGGCAACAGAGCAAGACTCC
AGCACACGTCTGAACTCTTTCCCTACACGACGCTCTTCCGATCTNNNCTTGCAGTGAGCCGAGATTGC
AGCACACGTCTGAACTCTTTCCCTACACGACGCTCTTCCGATCTNNNGAGACAGCGAGACTCCGTCTC
ACACGTCTGAACTCTTTCCCTACACGACGCTCTTCCGATCTNNNTGACTCTGACTCTAAGGACCACTCAT
CACGTCTGAACTCTTTCCCTACACGACGCTCTTCCGATCTNNNATGCAGACTGTGTGATTCTGACTCAT
AGCACACGTCTGAACTCTTTCCCTACACGACGCTCTTCCGATCTNNNGAGACTGAGGCGGGAGAATG
CACACGTCTGAACTCTTTCCCTACACGACGCTCTTCCGATCTNNNAAAGCTAGCAGTGGTGAAATTGG
GAGCACACGTCTGAACTCTTTCCCTACACGACGCTCTTCCGATCTNNNGAGGCTGAGGCAGGAGAAT
CACACGTCTGAACTCTTTCCCTACACGACGCTCTTCCGATCTNNNGACAGAGCGAGACTCCGTCTCAT
AGCACACGTCTGAACTCTTTCCCTACACGACGCTCTTCCGATCTNNNTGAGCCAAGATTGTGCCACTCA
CACGTCTGAACTCTTTCCCTACACGACGCTCTTCCGATCTNNNCCACAGATTGCTTGGCTAAAACACAT
GAGCACACGTCTGAACTCTTTCCCTACACGACGCTCTTCCGATCTNNNGGCGACAGTGAGAGACTCT
CACGTCTGAACTCTTTCCCTACACGACGCTCTTCCGATCTNNNTGGAAAGCAAAAATGGCAAATGCCA
AGCACACGTCTGAACTCTTTCCCTACACGACGCTCTTCCGATCTNNNGGTTGTGGTGAGCGGAGATC
AGCACACGTCTGAACTCTTTCCCTACACGACGCTCTTCCGATCTNNNCAGGAGAATGGCGTGAACCC
ACACGTCTGAACTCTTTCCCTACACGACGCTCTTCCGATCTNNNGTGGGTGTTTGTGTGTGTATACAC
AGCACACGTCTGAACTCTTTCCCTACACGACGCTCTTCCGATCTNNNAGGAGAATGGCATGAACCCG
ACACGTCTGAACTCTTTCCCTACACGACGCTCTTCCGATCTNNNACGGAGAGAGGGACATTTGTTTTC
ACACGTCTGAACTCTTTCCCTACACGACGCTCTTCCGATCTNNNACCTCGTCTCTACAAAACACACACAT
GAGCACACGTCTGAACTCTTTCCCTACACGACGCTCTTCCGATCTNNNAGGCCAACAGAGCAAGACT
GAGCACACGTCTGAACTCTTTCCCTACACGACGCTCTTCCGATCTNNNGAGGTGGAGGTTGTGGTGA
ACACGTCTGAACTCTTTCCCTACACGACGCTCTTCCGATCTNNNGTGAAAAGGGAATGTTGGTGCAT
AGCACACGTCTGAACTCTTTCCCTACACGACGCTCTTCCGATCTNNNCTTGCAGTGAGCCGACATCGC
CACACGTCTGAACTCTTTCCCTACACGACGCTCTTCCGATCTNNNACACGGTGAAACTCTGTCTCTACC
ACACGTCTGAACTCTTTCCCTACACGACGCTCTTCCGATCTNNNGTCAAGAGATCGAGACCATCCTGCA
AGCACACGTCTGAACTCTTTCCCTACACGACGCTCTTCCGATCTNNNGAGGCTGAGGCAGGAGAATG
AGCACACGTCTGAACTCTTTCCCTACACGACGCTCTTCCGATCTNNNCATAAACCTGGGAGGCGGAG
CACACGTCTGAACTCTTTCCCTACACGACGCTCTTCCGATCTNNNCGACAGAGCGAGATTCCGTCCAT
GAGCACACGTCTGAACTCTTTCCCTACACGACGCTCTTCCGATCTNNNCAGGAGAATGGCGTGAACCC
CACGTCTGAACTCTTTCCCTACACGACGCTCTTCCGATCTNNNTTGGGTGGCAGATGGTGTAATAACA
CACGTCTGAACTCTTTCCCTACACGACGCTCTTCCGATCTNNNCTATCATCCCCTGTGTCCTTCAGCATC
AGCACACGTCTGAACTCTTTCCCTACACGACGCTCTTCCGATCTNNNGAGACAGGTGTCCAGGTGTCC
AGAGCACACGTCTGAACTCTTTCCCTACACGACGCTCTTCCGATCTNNNCTGGAGGAAAGCGAGAGG
CACACGTCTGAACTCTTTCCCTACACGACGCTCTTCCGATCTNNNATCATTGACAAGTGGTTAGGGGG
CACACGTCTGAACTCTTTCCCTACACGACGCTCTTCCGATCTNNNGGCAGATCACAAGGTCAGGATAT
ACACGTCTGAACTCTTTCCCTACACGACGCTCTTCCGATCTNNNGAAACAGACATCTGATCAGCTGCCA
GCACACGTCTGAACTCTTTCCCTACACGACGCTCTTCCGATCTNNNTCTCTAACACACACACACACACA
GCACACGTCTGAACTCTTTCCCTACACGACGCTCTTCCGATCTNNNGATCGAGACCATCCTGGCTAAAA

CACACGTCTGAACTCTTTCCCTACACGACGCTCTTCCGATCTNNNCAGAGTAGCTGGGATTACAGGCC
GAGCACACGTCTGAACTCTTTCCCTACACGACGCTCTTCCGATCTNNNGAGGCTGAGGCAGGAGAAT
CACACGTCTGAACTCTTTCCCTACACGACGCTCTTCCGATCTNNNCTGGAGATTCGAGGGAAAACGA
GAGCACACGTCTGAACTCTTTCCCTACACGACGCTCTTCCGATCTNNNCAGTTGGCCTCAGTAGCTCCC
CACACGTCTGAACTCTTTCCCTACACGACGCTCTTCCGATCTNNNAGCCAGCATCTCATATCCTTGAGC
CACACGTCTGAACTCTTTCCCTACACGACGCTCTTCCGATCTNNNGATTGCATCAGGTGTGGAGCAG
AGCACACGTCTGAACTCTTTCCCTACACGACGCTCTTCCGATCTNNNGTTCCAGGCTGCAGGAAGAC
AGCACACGTCTGAACTCTTTCCCTACACGACGCTCTTCCGATCTNNNTTCACTGCAACCTCCACCTCCAT
AGCACACGTCTGAACTCTTTCCCTACACGACGCTCTTCCGATCTNNNCTGCCGAGTAGCTGGGACTA
AGCACACGTCTGAACTCTTTCCCTACACGACGCTCTTCCGATCTNNNCAAGTAGCTGGGACTACAGGC
GCACACGTCTGAACTCTTTCCCTACACGACGCTCTTCCGATCTNNNATGCTACTTCCAGAGGAATTGCA
AGCACACGTCTGAACTCTTTCCCTACACGACGCTCTTCCGATCTNNNCCAGGTCCACATGCATGGTG
CACACGTCTGAACTCTTTCCCTACACGACGCTCTTCCGATCTNNNCAAAATGCTGGGATTACAGGCAT
GCACACGTCTGAACTCTTTCCCTACACGACGCTCTTCCGATCTNNNTGTGTAGAGCTGTGTGTGTGTG
GCACACGTCTGAACTCTTTCCCTACACGACGCTCTTCCGATCTNNNCCATATCCTGGCTAACATGGTGA
ACACGTCTGAACTCTTTCCCTACACGACGCTCTTCCGATCTNNNGATGTAAGACTTTCCAGCCTCCACA
GCACACGTCTGAACTCTTTCCCTACACGACGCTCTTCCGATCTNNNGAGGCTGAGGCAGGAGAATG
GCACACGTCTGAACTCTTTCCCTACACGACGCTCTTCCGATCTNNNCAACACACACACCCACACACCAT
CACACGTCTGAACTCTTTCCCTACACGACGCTCTTCCGATCTNNNCCTGACGCTCAGCTAGGAAGCAT
GCACACGTCTGAACTCTTTCCCTACACGACGCTCTTCCGATCTNNNCCTGGGCAACAAGAGTGAAACC
GAGCACACGTCTGAACTCTTTCCCTACACGACGCTCTTCCGATCTNNNTGGGTGACAGAGCGAAACT
CACACGTCTGAACTCTTTCCCTACACGACGCTCTTCCGATCTNNNGTAAGCCATGTTTGCACCACTGCA
CACACGTCTGAACTCTTTCCCTACACGACGCTCTTCCGATCTNNNCCCAGCCTGGCTAGAGGAATATAC
AGCACACGTCTGAACTCTTTCCCTACACGACGCTCTTCCGATCTNNNGATGCAGAAAGGAGGGGCT
ACACGTCTGAACTCTTTCCCTACACGACGCTCTTCCGATCTNNNTAGATGGAGTCTCACTCTGTTGCCA
ACGTCTGAACTCTTTCCCTACACGACGCTCTTCCGATCTNNNTCTTTACTTTTTCCCACCCACCTCATCG
GCACACGTCTGAACTCTTTCCCTACACGACGCTCTTCCGATCTNNNCTACCCCCTCAGCTGACAGGCAT
GCACACGTCTGAACTCTTTCCCTACACGACGCTCTTCCGATCTNNNCACACATGTGCACACAGCTGCAT
CACACGTCTGAACTCTTTCCCTACACGACGCTCTTCCGATCTNNNCACGTACACGGAGTTTCACAACCA
GCACACGTCTGAACTCTTTCCCTACACGACGCTCTTCCGATCTNNNGAGCTTGCAGTGAACCGAGAC
AGCACACGTCTGAACTCTTTCCCTACACGACGCTCTTCCGATCTNNNTGGGCGAAAGAGCGAGACTC
ACACGTCTGAACTCTTTCCCTACACGACGCTCTTCCGATCTNNNTGGTCATCCACTTTGTAGCTAGTCA
AGCACACGTCTGAACTCTTTCCCTACACGACGCTCTTCCGATCTNNNCCTGTCTAATGCCTGGCTCAAC
ACACGTCTGAACTCTTTCCCTACACGACGCTCTTCCGATCTNNNACTCAAGGAAAAGGCTCTTCCAAC
GCACACGTCTGAACTCTTTCCCTACACGACGCTCTTCCGATCTNNNTAGTCCCTTTATCACCCCCTCATC
AGCACACGTCTGAACTCTTTCCCTACACGACGCTCTTCCGATCTNNNCTTGCAGTGAGCCGAGATCAC
AGCACACGTCTGAACTCTTTCCCTACACGACGCTCTTCCGATCTNNNGTTGCAGTGAGCCGAGATTG
AGCACACGTCTGAACTCTTTCCCTACACGACGCTCTTCCGATCTNNNGAGCGAAGATGTGGACGGA
ACACGTCTGAACTCTTTCCCTACACGACGCTCTTCCGATCTNNNGCCCATGAATTCTTCTATGCCACCAT
ACACGTCTGAACTCTTTCCCTACACGACGCTCTTCCGATCTNNNGGCAGTTTCATCATGTGTGAACACA
CACACGTCTGAACTCTTTCCCTACACGACGCTCTTCCGATCTNNNCATCCCGGCTAACATGGTGAAAC
AGCACACGTCTGAACTCTTTCCCTACACGACGCTCTTCCGATCTNNNGCACACACTCTCCCCAGTACCAT
ACACGTCTGAACTCTTTCCCTACACGACGCTCTTCCGATCTNNNCATCACAAAGCAAAGAATAGGCAC

AGAGCACACGTCTGAACTCTTTCCCTACACGACGCTCTTCCGATCTNNNGAGATGGCTCCAGTGCACT

CACACGTCTGAACTCTTTCCCTACACGACGCTCTTCCGATCTNNNGCTGTTATCAGTAGGCAGTTTGCC

AGCACACGTCTGAACTCTTTCCCTACACGACGCTCTTCCGATCTNNNCTGGGGAAGCTGTATGTCGGC

GCACACGTCTGAACTCTTTCCCTACACGACGCTCTTCCGATCTNNNCCGCTTGATAAATACGCAGGAT

AGCACACGTCTGAACTCTTTCCCTACACGACGCTCTTCCGATCTNNNCGAGAACATCCCAGCTCCACCA

ACACGTCTGAACTCTTTCCCTACACGACGCTCTTCCGATCTNNNCTGCTTCACCATCTTCCAGCAGCAT

GCACACGTCTGAACTCTTTCCCTACACGACGCTCTTCCGATCTNNNGACAAACAGCCCTCGCGTTGCAT

AGCACACGTCTGAACTCTTTCCCTACACGACGCTCTTCCGATCTNNNCCATAGAAGTCAGCGCTCAG

GCACACGTCTGAACTCTTTCCCTACACGACGCTCTTCCGATCTNNNGTGCAGAGACTGTACCGACCCA

GCACACGTCTGAACTCTTTCCCTACACGACGCTCTTCCGATCTNNNCACGGCTTTTTGACCCCCTGCAT

AGCACACGTCTGAACTCTTTCCCTACACGACGCTCTTCCGATCTNNNCAATCTCGGCTCACTGCAAGC

CACACGTCTGAACTCTTTCCCTACACGACGCTCTTCCGATCTNNNCTATCCTGAGCTACCCCTCACCCAT

AGCACACGTCTGAACTCTTTCCCTACACGACGCTCTTCCGATCTNNNCTGTGCGAGTGTCGGAGATG

GCACACGTCTGAACTCTTTCCCTACACGACGCTCTTCCGATCTNNNCAGACCCCATGAACCCCAGACA

ACACGTCTGAACTCTTTCCCTACACGACGCTCTTCCGATCTNNNAGTGTTGGGATTACAGGCGTAAG

CACACGTCTGAACTCTTTCCCTACACGACGCTCTTCCGATCTNNNCAACCCCAACCAAGGAAAACAGC

CACACGTCTGAACTCTTTCCCTACACGACGCTCTTCCGATCTNNNCTAGACGTCTCAGGCCACAAACC

AGCACACGTCTGAACTCTTTCCCTACACGACGCTCTTCCGATCTNNNCTCACCTGCTCCTGAGTGTGCA

GAGCACACGTCTGAACTCTTTCCCTACACGACGCTCTTCCGATCTNNNCTGGTAGTTCGGAGTCGGTG

ACACGTCTGAACTCTTTCCCTACACGACGCTCTTCCGATCTNNNGGCCACTGGGTTAGTAGAGAATCC

CACACGTCTGAACTCTTTCCCTACACGACGCTCTTCCGATCTNNNCCCCAAAACTGTGTAAGCCAATTC

AGAGCACACGTCTGAACTCTTTCCCTACACGACGCTCTTCCGATCTNNNGAGCTGTGATTGGGCCACT

AGCACACGTCTGAACTCTTTCCCTACACGACGCTCTTCCGATCTNNNGAGGCTGAGGCAGGAGAAT

CACACGTCTGAACTCTTTCCCTACACGACGCTCTTCCGATCTNNNAAGGAGAAAGGTACAGCTTGAG

AGCACACGTCTGAACTCTTTCCCTACACGACGCTCTTCCGATCTNNNCTTTCCTTGTCAGGCACAGCC

CACACGTCTGAACTCTTTCCCTACACGACGCTCTTCCGATCTNNNAGGAAGTTGAAGTGTTCAGGAC

ACACGTCTGAACTCTTTCCCTACACGACGCTCTTCCGATCTNNNGCCGCTTTATATAGAAACGCCTGC

ACACGTCTGAACTCTTTCCCTACACGACGCTCTTCCGATCTNNNCATCCTGGCTCCCTGTCTGAATACA

ACACGTCTGAACTCTTTCCCTACACGACGCTCTTCCGATCTNNNAGAGGTTGGTGTCAAATAGCTCAC

CACACGTCTGAACTCTTTCCCTACACGACGCTCTTCCGATCTNNNGAGTGAGACTCCACCTCTCTCTAC

GAGCACACGTCTGAACTCTTTCCCTACACGACGCTCTTCCGATCTNNNCATCGTCTGTGCCCCATGTCC

ACACGTCTGAACTCTTTCCCTACACGACGCTCTTCCGATCTNNNGGTGGCATATAGGGCAAAAGTCCC

ACACGTCTGAACTCTTTCCCTACACGACGCTCTTCCGATCTNNNCACCCGATAAAGTCAGTCAGTCTCA

ACACGTCTGAACTCTTTCCCTACACGACGCTCTTCCGATCTNNNCACAGAAAGAGACTGAGAGGACGC

GCACACGTCTGAACTCTTTCCCTACACGACGCTCTTCCGATCTNNNGTCACCTCCATGCGTCTCTCCATC

CACACGTCTGAACTCTTTCCCTACACGACGCTCTTCCGATCTNNNGTACCGTGTTCCATTCAAAGTGG

CACGTCTGAACTCTTTCCCTACACGACGCTCTTCCGATCTNNNCTCAACGCTAGAGCTTTTTGACCCAT

CACACGTCTGAACTCTTTCCCTACACGACGCTCTTCCGATCTNNNCTAGTCCTGACAACCGCAGATGCA

AGCACACGTCTGAACTCTTTCCCTACACGACGCTCTTCCGATCTNNNCAAAAAGCAAGCGCAGAACAA

CACACGTCTGAACTCTTTCCCTACACGACGCTCTTCCGATCTNNNCACAAATCAGGAACAGTGCCTTTC

ACACGTCTGAACTCTTTCCCTACACGACGCTCTTCCGATCTNNNAAGTTTCTGGGTCTTCCTCAGTCCAT

ACACGTCTGAACTCTTTCCCTACACGACGCTCTTCCGATCTNNNGACGAGGAGTATGAGCCCCACCAT

CACACGTCTGAACTCTTTCCCTACACGACGCTCTTCCGATCTNNNCCGGCAGCTATACACTGGTCCAT

GCACACGTCTGAACTCTTTCCCTACACGACGCTCTTCCGATCTNNNCAACACAGGTTGCCTGTTCCCCAT
GCACACGTCTGAACTCTTTCCCTACACGACGCTCTTCCGATCTNNNCCTGGGGCCAAGGAAATTGTGC
CACACGTCTGAACTCTTTCCCTACACGACGCTCTTCCGATCTNNNCGGCCAGGTCTCATGTGTTTTCAT
ACACGTCTGAACTCTTTCCCTACACGACGCTCTTCCGATCTNNNGTGGCTATGCCTTCGGAGTTTCCAT
CACACGTCTGAACTCTTTCCCTACACGACGCTCTTCCGATCTNNNCAGGTCTCAGGGAGAGGGAACC
AGCACACGTCTGAACTCTTTCCCTACACGACGCTCTTCCGATCTNNNACAACCAGGGCACAGGGAATC
GCACACGTCTGAACTCTTTCCCTACACGACGCTCTTCCGATCTNNNCTGAGAGTTGCGGTGGCTATGCA
ACACGTCTGAACTCTTTCCCTACACGACGCTCTTCCGATCTNNNGAGTTTCTGGACCCCCTCATGGCAT
GCACACGTCTGAACTCTTTCCCTACACGACGCTCTTCCGATCTNNNCTTAGCTGGCTACCAGGCTCTCA
CACACGTCTGAACTCTTTCCCTACACGACGCTCTTCCGATCTNNNGAGAGAGGGAACTGGGGAAGTG
GAGCACACGTCTGAACTCTTTCCCTACACGACGCTCTTCCGATCTNNNCTCTCTGTTCACCAGCAGCACA
CACGTCTGAACTCTTTCCCTACACGACGCTCTTCCGATCTNNNCTATCCCGAAGCTCCCCAAATCCATCG
CACGTCTGAACTCTTTCCCTACACGACGCTCTTCCGATCTNNNCACCCAGAAATTCTGCTTCAGCCATC
GAGCACACGTCTGAACTCTTTCCCTACACGACGCTCTTCCGATCTNNNGAGATTGCGCCATTGCACTCC
AGCACACGTCTGAACTCTTTCCCTACACGACGCTCTTCCGATCTNNNGCCTGTAGTCCCAGCTACTTGCA
GCACACGTCTGAACTCTTTCCCTACACGACGCTCTTCCGATCTNNNCAGATCACGAGGTCAGGAGATG
GAGCACACGTCTGAACTCTTTCCCTACACGACGCTCTTCCGATCTNNNCTTGCAGTGAGCCGAGATTGC
AGCACACGTCTGAACTCTTTCCCTACACGACGCTCTTCCGATCTNNNGCCTGTAGTCCCAGCTACTTGC
GCACACGTCTGAACTCTTTCCCTACACGACGCTCTTCCGATCTNNNCTTGCAGTGAGCCGAGATCCCAT
GCACACGTCTGAACTCTTTCCCTACACGACGCTCTTCCGATCTNNNCTGGCTGCTACAGTGAAAGGGC
GCACACGTCTGAACTCTTTCCCTACACGACGCTCTTCCGATCTNNNCACGTGCTGTCAGTGTCCTGCATC
GCACACGTCTGAACTCTTTCCCTACACGACGCTCTTCCGATCTNNNGGGACAGAGTGAGACTTCGTCTC
CACGTCTGAACTCTTTCCCTACACGACGCTCTTCCGATCTNNNGACTACAGGCACACACCATCATCATC
GCACACGTCTGAACTCTTTCCCTACACGACGCTCTTCCGATCTNNNGGCGACAGAGTGAGACACTGCA
AGCACACGTCTGAACTCTTTCCCTACACGACGCTCTTCCGATCTNNNGAGGAGGGGGATGAGTGAG
CACGTCTGAACTCTTTCCCTACACGACGCTCTTCCGATCTNNNCTGGGTTCAAGCGATTCTTCTGCATC
CACGTCTGAACTCTTTCCCTACACGACGCTCTTCCGATCTNNNCCATGCCCTGCAGCTAATTTTTCATCG
GCACACGTCTGAACTCTTTCCCTACACGACGCTCTTCCGATCTNNNCACACACACACACACACAACTGCA
AGCACACGTCTGAACTCTTTCCCTACACGACGCTCTTCCGATCTNNNCTGCCTCCACTTCCACTCTGCAT
GCACACGTCTGAACTCTTTCCCTACACGACGCTCTTCCGATCTNNNGATCGAGACCATCCTGGCTAACC
AGCACACGTCTGAACTCTTTCCCTACACGACGCTCTTCCGATCTNNNGTTTCACGCCATTCTCCTGCCAT
CACACGTCTGAACTCTTTCCCTACACGACGCTCTTCCGATCTNNNCCACCTCAGTCCCCCAAATAGCATC
GCACACGTCTGAACTCTTTCCCTACACGACGCTCTTCCGATCTNNNGTGATTGCTCCATTGCACTCCCAT
AGCACACGTCTGAACTCTTTCCCTACACGACGCTCTTCCGATCTNNNCTGCAGACGGCTACCCTTGGCAT
AGCACACGTCTGAACTCTTTCCCTACACGACGCTCTTCCGATCTNNNCACACACACACACACACACACCAT
GAGCACACGTCTGAACTCTTTCCCTACACGACGCTCTTCCGATCTNNNCTCCACCCCTTCATGAATGGCC
ACACGTCTGAACTCTTTCCCTACACGACGCTCTTCCGATCTNNNGTGGTTCTGGATATCCCAACCCCATC
GCACACGTCTGAACTCTTTCCCTACACGACGCTCTTCCGATCTNNNCGCTTACGAAAGAGCCAACACCA
GCACACGTCTGAACTCTTTCCCTACACGACGCTCTTCCGATCTNNNCTCTCCTTCCCCTCCCCTTCCATCG
CACACGTCTGAACTCTTTCCCTACACGACGCTCTTCCGATCTNNNGGGGTCCTCACTTCCCAGTAGCAT
ACACGTCTGAACTCTTTCCCTACACGACGCTCTTCCGATCTNNNGTGCCATGCTCAAATTCTCGACCATC
AGCACACGTCTGAACTCTTTCCCTACACGACGCTCTTCCGATCTNNNAACCCACATCACAGTGTCCCCAT
AGCACACGTCTGAACTCTTTCCCTACACGACGCTCTTCCGATCTNNNGGTGGGAAGGTCAAGGTCAG

GAGCACACGTCTGAACTCTTTCCCTACACGACGCTCTTCCGATCTNNNGTCACTCAGAACCACCAGGGC
CACGTCTGAACTCTTTCCCTACACGACGCTCTTCCGATCTNNNATCACCAGCCATACCAAGCAACATCG
AGCACACGTCTGAACTCTTTCCCTACACGACGCTCTTCCGATCTNNNCTGTCCGGTGAGAATGTAGGC
AGCACACGTCTGAACTCTTTCCCTACACGACGCTCTTCCGATCTNNNGCACGTGTAGGCAAAGAAGCC
GCACACGTCTGAACTCTTTCCCTACACGACGCTCTTCCGATCTNNNCCCTACTGGAAAGTGAGGAGCC
GCACACGTCTGAACTCTTTCCCTACACGACGCTCTTCCGATCTNNNTCCCAGTAAGGTCAGCACCCCAT
CACACGTCTGAACTCTTTCCCTACACGACGCTCTTCCGATCTNNNTCAGGGACACCAGGTTTGTTCCAT
GCACACGTCTGAACTCTTTCCCTACACGACGCTCTTCCGATCTNNNCAGTGTCTGGTCAGAAGTGCTCC
CACACGTCTGAACTCTTTCCCTACACGACGCTCTTCCGATCTNNNCTCTTTACTCCCCGCAGCAAGCATC
CACACGTCTGAACTCTTTCCCTACACGACGCTCTTCCGATCTNNNCTGCTCTCTAGACCTCCCGTGCATC
CACACGTCTGAACTCTTTCCCTACACGACGCTCTTCCGATCTNNNCACAGAGCGCCCTTTGAGGAGCAT
CACACGTCTGAACTCTTTCCCTACACGACGCTCTTCCGATCTNNNCTGTGTCCAGCTCTCCTTGCCATCG
CACACGTCTGAACTCTTTCCCTACACGACGCTCTTCCGATCTNNNCTCTGCCCTCTTCCTTCCCCCCATCG
ACACGTCTGAACTCTTTCCCTACACGACGCTCTTCCGATCTNNNCGAAAGCCCACCTGACCATTGCATC
GCACACGTCTGAACTCTTTCCCTACACGACGCTCTTCCGATCTNNNCTGTCGTCCCAGCTACTTGGCATC
AGCACACGTCTGAACTCTTTCCCTACACGACGCTCTTCCGATCTNNNGAGGTGGAGCTTGCAGTGAG
CACACGTCTGAACTCTTTCCCTACACGACGCTCTTCCGATCTNNNGAGGCTGAGGCAGGAGAATGCAT
GCACACGTCTGAACTCTTTCCCTACACGACGCTCTTCCGATCTNNNCTGGGACTACAGGTGTGTGCCAT
CACACGTCTGAACTCTTTCCCTACACGACGCTCTTCCGATCTNNNGATGATCTGCCAGGGGGTATGCAT
CACACGTCTGAACTCTTTCCCTACACGACGCTCTTCCGATCTNNNGAGGCTGAGGCAGGAGAATGCA
CACACGTCTGAACTCTTTCCCTACACGACGCTCTTCCGATCTNNNCTCCCCTCAAACAGACACAGCATC
CACACGTCTGAACTCTTTCCCTACACGACGCTCTTCCGATCTNNNCCTGACGCTCAGCTAGGAAGCATC
AGCACACGTCTGAACTCTTTCCCTACACGACGCTCTTCCGATCTNNNCTTCCTTTCAGGGAGGAGGCCA
ACACGTCTGAACTCTTTCCCTACACGACGCTCTTCCGATCTNNNGAGGAAGGGTGTGGAAGACAGCAT
CACACGTCTGAACTCTTTCCCTACACGACGCTCTTCCGATCTNNNCCCTCACCTCCTGGATAGGGCATC
GCACACGTCTGAACTCTTTCCCTACACGACGCTCTTCCGATCTNNNCTGCACTCCTCAGCCCTTGGCATC
CACACGTCTGAACTCTTTCCCTACACGACGCTCTTCCGATCTNNNTGGGAACTGACCTCCTCTCTGCATC
CACACGTCTGAACTCTTTCCCTACACGACGCTCTTCCGATCTNNNCTGATTGGCAGGAAGCTCGCCATC
ACACGTCTGAACTCTTTCCCTACACGACGCTCTTCCGATCTNNNCAGAGCAAGGTCCCTGAGAAGCATC
GCACACGTCTGAACTCTTTCCCTACACGACGCTCTTCCGATCTNNNCTCTGTTCCACTCACGCTTCTCATC
AGCACACGTCTGAACTCTTTCCCTACACGACGCTCTTCCGATCTNNNCTTTTGCTCCCGCGCTACCCAT
GCACACGTCTGAACTCTTTCCCTACACGACGCTCTTCCGATCTNNNAAGCACACTGGGCAAAGCTGCAT
CACACGTCTGAACTCTTTCCCTACACGACGCTCTTCCGATCTNNNGCCGAAGGGATGTGCTCCTGCATC
ACACGTCTGAACTCTTTCCCTACACGACGCTCTTCCGATCTNNNAGCCTTAGGGCCCACACATCCATCG
CACACGTCTGAACTCTTTCCCTACACGACGCTCTTCCGATCTNNNCCTCCCTGTATCACCCCCTCCATCGG
CACACGTCTGAACTCTTTCCCTACACGACGCTCTTCCGATCTNNNAGTCCCCAACTCCCTGGTCCATCG
ACACGTCTGAACTCTTTCCCTACACGACGCTCTTCCGATCTNNNCCAACCCCGTTGTCCCCCTGCATCGG
CACACGTCTGAACTCTTTCCCTACACGACGCTCTTCCGATCTNNNCTCCAGTTCCCGATCCACCCCATCG
ACACGTCTGAACTCTTTCCCTACACGACGCTCTTCCGATCTNNNCCGTGGCTTTTTACCCTCCACATCGG
CACACGTCTGAACTCTTTCCCTACACGACGCTCTTCCGATCTNNNGAGATGACCCTACAGCGGTGCATC
CACACGTCTGAACTCTTTCCCTACACGACGCTCTTCCGATCTNNNGGGTCACGTGATAGCCGAGGCAT
AGAGCACACGTCTGAACTCTTTCCCTACACGACGCTCTTCCGATCTNNNCAGAATGGATGAATATGTG
GAGCACACGTCTGAACTCTTTCCCTACACGACGCTCTTCCGATCTNNNACTCCTTTTTTTTGTTTTGTTT

GAGCACACGTCTGAACTCTTTCCCTACACGACGCTCTTCCGATCTNNNTGTCAAAAGTAAAAAGAAAA
AGCACACGTCTGAACTCTTTCCCTACACGACGCTCTTCCGATCTNNNCATTCCTTTTTATGGCTGAGT
AGAGCACACGTCTGAACTCTTTCCCTACACGACGCTCTTCCGATCTNNNTCCTTTCTTCTTTTTTTCCT
GAGCACACGTCTGAACTCTTTCCCTACACGACGCTCTTCCGATCTNNNACTTTGTTAGTGGGCATGTA
AGAGCACACGTCTGAACTCTTTCCCTACACGACGCTCTTCCGATCTNNNTGTGTGTGTATGTAAATGT
AAGAGCACACGTCTGAACTCTTTCCCTACACGACGCTCTTCCGATCTNNNAGCAGTGTGAAAATGGA
AGAGCACACGTCTGAACTCTTTCCCTACACGACGCTCTTCCGATCTNNNAGTCGCTATAAGTATCACA
AGAGCACACGTCTGAACTCTTTCCCTACACGACGCTCTTCCGATCTNNNCAACCCTTTACCTAGACTAA
AGAGCACACGTCTGAACTCTTTCCCTACACGACGCTCTTCCGATCTNNNTCGGACCACTATCTGACTA
AAGAGCACACGTCTGAACTCTTTCCCTACACGACGCTCTTCCGATCTNNNATATCACCTCATGTCCATT
GGAAGAGCACACGTCTGAACTCTTTCCCTACACGACGCTCTTCCGATCTNNNAGAACTAGATAACCAA
AGAGCACACGTCTGAACTCTTTCCCTACACGACGCTCTTCCGATCTNNNTGGACACAGAAAGACAAAA
GCACACGTCTGAACTCTTTCCCTACACGACGCTCTTCCGATCTNNNACACACACACAATGTATTTATTC
AGAGCACACGTCTGAACTCTTTCCCTACACGACGCTCTTCCGATCTNNNTGTATTAGAGTTCTCCAGA
GAGCACACGTCTGAACTCTTTCCCTACACGACGCTCTTCCGATCTNNNGCTCATACTCACTAAGGGAAG
AGAGCACACGTCTGAACTCTTTCCCTACACGACGCTCTTCCGATCTNNNCCAGAATCAAATGTTTCTCT
GAGCACACGTCTGAACTCTTTCCCTACACGACGCTCTTCCGATCTNNNTGAAGACATGACATAGTTAG
AGAGCACACGTCTGAACTCTTTCCCTACACGACGCTCTTCCGATCTNNNTCATCAGAGTTTTGTAGTTT
AGAGCACACGTCTGAACTCTTTCCCTACACGACGCTCTTCCGATCTNNNGGGAAAGCTAAGTTGATTC
GAGCACACGTCTGAACTCTTTCCCTACACGACGCTCTTCCGATCTNNNAGCTGTTCTCATCATTTTTTCC
AGCACACGTCTGAACTCTTTCCCTACACGACGCTCTTCCGATCTNNNTGTGTTCAAATTACTTTGTGGT
AAGAGCACACGTCTGAACTCTTTCCCTACACGACGCTCTTCCGATCTNNNGCCCCTGCAATCATCAAT
AAGAGCACACGTCTGAACTCTTTCCCTACACGACGCTCTTCCGATCTNNNTGAGATGGTCACAAGAA
AAGAGCACACGTCTGAACTCTTTCCCTACACGACGCTCTTCCGATCTNNNACATTTGCACCGACCTAAG
GAAGAGCACACGTCTGAACTCTTTCCCTACACGACGCTCTTCCGATCTNNNAAACAATCATGGCAAGG
AGAGCACACGTCTGAACTCTTTCCCTACACGACGCTCTTCCGATCTNNNTGACACACAGTAATTGTAG
AAGAGCACACGTCTGAACTCTTTCCCTACACGACGCTCTTCCGATCTNNNCAGGATTTCATTCCATTGT
GAGCACACGTCTGAACTCTTTCCCTACACGACGCTCTTCCGATCTNNNAGTAAGTTTTAGTGTGTTTCT
AAGAGCACACGTCTGAACTCTTTCCCTACACGACGCTCTTCCGATCTNNNGTTGGGTCAATCAGTCAGG
AAGAGCACACGTCTGAACTCTTTCCCTACACGACGCTCTTCCGATCTNNNGCATCTTCTGGTTGATGCC
GCACACGTCTGAACTCTTTCCCTACACGACGCTCTTCCGATCTNNNACATCAGTCAATAGAAGAAAGA
GAGCACACGTCTGAACTCTTTCCCTACACGACGCTCTTCCGATCTNNNACTCTCATGATGGTACATTTT
AAGAGCACACGTCTGAACTCTTTCCCTACACGACGCTCTTCCGATCTNNNACAATCAGCCCTCTATAA
GAAGAGCACACGTCTGAACTCTTTCCCTACACGACGCTCTTCCGATCTNNNATCCCCTCCTGTACCTTGT
AGCACACGTCTGAACTCTTTCCCTACACGACGCTCTTCCGATCTNNNGCCAAATAAAATAAAAAGCAG
GAGCACACGTCTGAACTCTTTCCCTACACGACGCTCTTCCGATCTNNNACAGATAAGAAGACTGAGT
GCACACGTCTGAACTCTTTCCCTACACGACGCTCTTCCGATCTNNNGCTGATATTCTGGGTATCTTTTC
GAAGAGCACACGTCTGAACTCTTTCCCTACACGACGCTCTTCCGATCTNNNGCTGTGTTGGTCGTTGT
AGCACACGTCTGAACTCTTTCCCTACACGACGCTCTTCCGATCTNNNTGCTTAGCTTTTGAATATGAAC
GCACACGTCTGAACTCTTTCCCTACACGACGCTCTTCCGATCTNNNGCTGAGTATTATATTTCCATATT
AAGAGCACACGTCTGAACTCTTTCCCTACACGACGCTCTTCCGATCTNNNGAACTCTGTGGAACTTTG
GAAGAGCACACGTCTGAACTCTTTCCCTACACGACGCTCTTCCGATCTNNNCTATGTGGTGAGTTTCA
GAAGAGCACACGTCTGAACTCTTTCCCTACACGACGCTCTTCCGATCTNNNTCTTGCTTGGACCTCTC

AAGAGCACACGTCTGAACTCTTTCCCTACACGACGCTCTTCCGATCTNNNTTAGAGCCAGGAGATAGA
GAAGAGCACACGTCTGAACTCTTTCCCTACACGACGCTCTTCCGATCTNNNCAGATACCCTGTTCCCCT
GAGCACACGTCTGAACTCTTTCCCTACACGACGCTCTTCCGATCTNNNAGTGAAATTCCATCTCTCTCTC
GAAGAGCACACGTCTGAACTCTTTCCCTACACGACGCTCTTCCGATCTNNNTGGGGCAAAAACCCTAG
GAGCACACGTCTGAACTCTTTCCCTACACGACGCTCTTCCGATCTNNNTTTTACCTAAGATAATGGCCT
AGCACACGTCTGAACTCTTTCCCTACACGACGCTCTTCCGATCTNNNCCCATTTATTACAACGAACTTAT
GCACACGTCTGAACTCTTTCCCTACACGACGCTCTTCCGATCTNNNTGAGTTCATTGTAATCAAACAAA
AGAGCACACGTCTGAACTCTTTCCCTACACGACGCTCTTCCGATCTNNNGCAGACAATGGTTTCTTTTT
AGCACACGTCTGAACTCTTTCCCTACACGACGCTCTTCCGATCTNNNACTAGATGATCTCTTCCTCTAGA
GAAGAGCACACGTCTGAACTCTTTCCCTACACGACGCTCTTCCGATCTNNNAGCTTTTGCCTATGCTTC
AAGAGCACACGTCTGAACTCTTTCCCTACACGACGCTCTTCCGATCTNNNGCTGATCCAGAAAGAGGTA
GAAGAGCACACGTCTGAACTCTTTCCCTACACGACGCTCTTCCGATCTNNNTTCCAGGGTTGGATATGT
GAGCACACGTCTGAACTCTTTCCCTACACGACGCTCTTCCGATCTNNNGTCCCCAGTTACTTTGGAGAC
GGAAGAGCACACGTCTGAACTCTTTCCCTACACGACGCTCTTCCGATCTNNNTCACTAGACAAGCTCAA
GAGCACACGTCTGAACTCTTTCCCTACACGACGCTCTTCCGATCTNNNAATTTTCTCCCATTCTGTTGTCT
AGAGCACACGTCTGAACTCTTTCCCTACACGACGCTCTTCCGATCTNNNCGCAAATAGGCTCAAAACA
GAAGAGCACACGTCTGAACTCTTTCCCTACACGACGCTCTTCCGATCTNNNTTCAAGCCACAGCATTTC
GAAGAGCACACGTCTGAACTCTTTCCCTACACGACGCTCTTCCGATCTNNNATGGTAGCACTAGAATC
AAGAGCACACGTCTGAACTCTTTCCCTACACGACGCTCTTCCGATCTNNNACATGATGTTTGGCTGAA
GAAGAGCACACGTCTGAACTCTTTCCCTACACGACGCTCTTCCGATCTNNNACGCATCCTTATTGCTTTT
AGAGCACACGTCTGAACTCTTTCCCTACACGACGCTCTTCCGATCTNNNAGTCATGCTTCTGTTATAG
GAGCACACGTCTGAACTCTTTCCCTACACGACGCTCTTCCGATCTNNNGTTTTGAATTATTTCTAGTACT
AGCACACGTCTGAACTCTTTCCCTACACGACGCTCTTCCGATCTNNNCCAGGATTTCTGTTATTTGAATT
GAGCACACGTCTGAACTCTTTCCCTACACGACGCTCTTCCGATCTNNNAGATTCATATTTCCCAATAAGA
GAGCACACGTCTGAACTCTTTCCCTACACGACGCTCTTCCGATCTNNNTCTTTCTGTATTTCCAATTTTCT
GAGCACACGTCTGAACTCTTTCCCTACACGACGCTCTTCCGATCTNNNGCCAACTAGACATAACAGTTTA
GAGCACACGTCTGAACTCTTTCCCTACACGACGCTCTTCCGATCTNNNCTCAGAATGCTACCATAATTCA
GAAGAGCACACGTCTGAACTCTTTCCCTACACGACGCTCTTCCGATCTNNNCCTCAGGCCCAACATAGA
AGCACACGTCTGAACTCTTTCCCTACACGACGCTCTTCCGATCTNNNTGTTCCCTTCTAATAAGACAAAA
AGCACACGTCTGAACTCTTTCCCTACACGACGCTCTTCCGATCTNNNAGCAAACATAAGATTGATGCAA
GAGCACACGTCTGAACTCTTTCCCTACACGACGCTCTTCCGATCTNNNCCTAGTTTTAGAAGTTTCAATA
AGCACACGTCTGAACTCTTTCCCTACACGACGCTCTTCCGATCTNNNATAGTACTTTTTCTCTCTCTCTCT
AAGAGCACACGTCTGAACTCTTTCCCTACACGACGCTCTTCCGATCTNNNATGGGAGAGAAAACACTA
GAAGAGCACACGTCTGAACTCTTTCCCTACACGACGCTCTTCCGATCTNNNAAGCTGTGGGATCTCTC
AAGAGCACACGTCTGAACTCTTTCCCTACACGACGCTCTTCCGATCTNNNTGTGACCTTGGGCTTTCAC
AAGAGCACACGTCTGAACTCTTTCCCTACACGACGCTCTTCCGATCTNNNATACATTGAAAACTCCCCT
AAGAGCACACGTCTGAACTCTTTCCCTACACGACGCTCTTCCGATCTNNNTCTCAAGAATTTCGGATAT
GAGCACACGTCTGAACTCTTTCCCTACACGACGCTCTTCCGATCTNNNACAATTGTATACCTAAAACTG
GAGCACACGTCTGAACTCTTTCCCTACACGACGCTCTTCCGATCTNNNACCCATGATATACAATACACAT
GAAGAGCACACGTCTGAACTCTTTCCCTACACGACGCTCTTCCGATCTNNNGCTACGAGTGAATACTT
GGAAGAGCACACGTCTGAACTCTTTCCCTACACGACGCTCTTCCGATCTNNNTTCCTTATGTCCCATAGG
GAAGAGCACACGTCTGAACTCTTTCCCTACACGACGCTCTTCCGATCTNNNCATATTCCTGGGTTTGG
GAGCACACGTCTGAACTCTTTCCCTACACGACGCTCTTCCGATCTNNNTGAAATGAAGAAGTGTAGGA

AGAGCACACGTCTGAACTCTTTCCCTACACGACGCTCTTCCGATCTNNNTCCTGTGTCTTTTACTTTTGC
GAAGAGCACACGTCTGAACTCTTTCCCTACACGACGCTCTTCCGATCTNNNCAGTGGTTCCCAACCTTA
GAAGAGCACACGTCTGAACTCTTTCCCTACACGACGCTCTTCCGATCTNNNCACGTGCTTTTCTTTGAG
AAGAGCACACGTCTGAACTCTTTCCCTACACGACGCTCTTCCGATCTNNNGGAACAATATAGGGGAAA
GAAGAGCACACGTCTGAACTCTTTCCCTACACGACGCTCTTCCGATCTNNNAGCTTGTAGTGAGCTGA
GAAGAGCACACGTCTGAACTCTTTCCCTACACGACGCTCTTCCGATCTNNNACTGAAGCATTTAGGTCG
AGCACACGTCTGAACTCTTTCCCTACACGACGCTCTTCCGATCTNNNGCCAGGATATATATGTATGTAT
AAGAGCACACGTCTGAACTCTTTCCCTACACGACGCTCTTCCGATCTNNNAATAAGTGGCAGAACTGG
AAGAGCACACGTCTGAACTCTTTCCCTACACGACGCTCTTCCGATCTNNNCCGAATGGTGCTGCAATA
GAAGAGCACACGTCTGAACTCTTTCCCTACACGACGCTCTTCCGATCTNNNCTACTGAGTGTCTACCCA
GAAGAGCACACGTCTGAACTCTTTCCCTACACGACGCTCTTCCGATCTNNNTTGCGCACGTATGTTTAT
GAAGAGCACACGTCTGAACTCTTTCCCTACACGACGCTCTTCCGATCTNNNCCAGAAAAGGGTCACAC
AGAGCACACGTCTGAACTCTTTCCCTACACGACGCTCTTCCGATCTNNNTGACCATAAAAGTCCCTCCA
AGAGCACACGTCTGAACTCTTTCCCTACACGACGCTCTTCCGATCTNNNTTCTGTAGCTATCATCTGTCTA
GAAGAGCACACGTCTGAACTCTTTCCCTACACGACGCTCTTCCGATCTNNNGGGACATAACTTGGGCC
AAGAGCACACGTCTGAACTCTTTCCCTACACGACGCTCTTCCGATCTNNNACCGTGTAAGGTGTATGTA
CGGAAGAGCACACGTCTGAACTCTTTCCCTACACGACGCTCTTCCGATCTNNNGGATGAATGGATGGA
AAGAGCACACGTCTGAACTCTTTCCCTACACGACGCTCTTCCGATCTNNNAGCAGACTCTCCCTGTATCT
GAGCACACGTCTGAACTCTTTCCCTACACGACGCTCTTCCGATCTNNNACCTTGAAAACATTATGCTAAA
AAGAGCACACGTCTGAACTCTTTCCCTACACGACGCTCTTCCGATCTNNNGGGTTCTTTCCCTTTAGCTC
GAAGAGCACACGTCTGAACTCTTTCCCTACACGACGCTCTTCCGATCTNNNTGGTGGGTAGGTATGTG
GAGCACACGTCTGAACTCTTTCCCTACACGACGCTCTTCCGATCTNNNGCAAGCAAAAATATGAAATTC
GAAGAGCACACGTCTGAACTCTTTCCCTACACGACGCTCTTCCGATCTNNNTAAGCACAAGGTGGTGA
GAGCACACGTCTGAACTCTTTCCCTACACGACGCTCTTCCGATCTNNNTGTGTGTATATAGTGTTTGTA
TCGGAAGAGCACACGTCTGAACTCTTTCCCTACACGACGCTCTTCCGATCTNNNTACCTATAGTCCCAGC
AGCACACGTCTGAACTCTTTCCCTACACGACGCTCTTCCGATCTNNNTGTGTGCGTGTATATACATAAAT
GAAGAGCACACGTCTGAACTCTTTCCCTACACGACGCTCTTCCGATCTNNNTCAGAGGCCTGACATAAC
AGAGCACACGTCTGAACTCTTTCCCTACACGACGCTCTTCCGATCTNNNAGCCTGTATAGTATACACAC
GAGCACACGTCTGAACTCTTTCCCTACACGACGCTCTTCCGATCTNNNACATGATACAGAAATAAGAGA
GAAGAGCACACGTCTGAACTCTTTCCCTACACGACGCTCTTCCGATCTNNNAAGTCCTAGCCAGAGCAA
AGAGCACACGTCTGAACTCTTTCCCTACACGACGCTCTTCCGATCTNNNCTCTCTCTCTCTCTGTCCAC
GAGCACACGTCTGAACTCTTTCCCTACACGACGCTCTTCCGATCTNNNAAAACTATTTTTATGTTGCAC
AGCACACGTCTGAACTCTTTCCCTACACGACGCTCTTCCGATCTNNNAGACTGCTATCTAGACTAACAA
GGAAGAGCACACGTCTGAACTCTTTCCCTACACGACGCTCTTCCGATCTNNNTGCTCCCTTGAAATCCGT
GAAGAGCACACGTCTGAACTCTTTCCCTACACGACGCTCTTCCGATCTNNNGCAGGAGAATCACTTGAA
AGCACACGTCTGAACTCTTTCCCTACACGACGCTCTTCCGATCTNNNCACGTGTGTGTATATGTATATAT
GAAGAGCACACGTCTGAACTCTTTCCCTACACGACGCTCTTCCGATCTNNNTCCCAACTGACAAATGAG
GGAAGAGCACACGTCTGAACTCTTTCCCTACACGACGCTCTTCCGATCTNNNGATATGACCTCAAAAGC
AGAGCACACGTCTGAACTCTTTCCCTACACGACGCTCTTCCGATCTNNNAGAAGTTGGTTCTTCAAAAA
AGCACACGTCTGAACTCTTTCCCTACACGACGCTCTTCCGATCTNNNTCAAGAATAGTTCAAGAGAGGA
GAGCACACGTCTGAACTCTTTCCCTACACGACGCTCTTCCGATCTNNNACAGTAATCCTAGTGATTTAT
AAGAGCACACGTCTGAACTCTTTCCCTACACGACGCTCTTCCGATCTNNNCTGAGGCAGGAGAATAGC
AGCACACGTCTGAACTCTTTCCCTACACGACGCTCTTCCGATCTNNNTCTCTTGCTTCTCTAAGTTTTTTT

AAGAGCACACGTCTGAACTCTTTCCCTACACGACGCTCTTCCGATCTNNNACCTAATGCTAGATGACAC
GAGCACACGTCTGAACTCTTTCCCTACACGACGCTCTTCCGATCTNNNACTCCAAAACATTATGTTGTAC
AGCACACGTCTGAACTCTTTCCCTACACGACGCTCTTCCGATCTNNNTGCTAATATTTCTCCCATTTTTT
AGCACACGTCTGAACTCTTTCCCTACACGACGCTCTTCCGATCTNNNTCTTCTTTCCTCTTTCTTTCTTTCT
AGCACACGTCTGAACTCTTTCCCTACACGACGCTCTTCCGATCTNNNTCTTCTTCCTTTCTCTCTCTTCTT
GCACACGTCTGAACTCTTTCCCTACACGACGCTCTTCCGATCTNNNTGCATATTTCTTCACTTTTACTTA
GAAGAGCACACGTCTGAACTCTTTCCCTACACGACGCTCTTCCGATCTNNNATTCAAGACCCAAGAAGC
AGCACACGTCTGAACTCTTTCCCTACACGACGCTCTTCCGATCTNNNTGAGAATTCTCCTGATATTCTG
AGCACACGTCTGAACTCTTTCCCTACACGACGCTCTTCCGATCTNNNTTCTAGTACACATAGTCATTTA
AGCACACGTCTGAACTCTTTCCCTACACGACGCTCTTCCGATCTNNNTTCTAGTACACATAGTCATTTA
AGCACACGTCTGAACTCTTTCCCTACACGACGCTCTTCCGATCTNNNGTTTCTCAAATTCTGGATTTTT
GCACACGTCTGAACTCTTTCCCTACACGACGCTCTTCCGATCTNNNTGATTTTGTAACCTGAAACTTTA
GAAGAGCACACGTCTGAACTCTTTCCCTACACGACGCTCTTCCGATCTNNNAGAATCACTTGAACCCAG
AAGAGCACACGTCTGAACTCTTTCCCTACACGACGCTCTTCCGATCTNNNACAATCATGGAGATTCCCT
GAAGAGCACACGTCTGAACTCTTTCCCTACACGACGCTCTTCCGATCTNNNCAGAATTGCTTGAAACTG
AGAGCACACGTCTGAACTCTTTCCCTACACGACGCTCTTCCGATCTNNNTGGACAAAGGGTAATATAGT
GAAGAGCACACGTCTGAACTCTTTCCCTACACGACGCTCTTCCGATCTNNNGGAGGCTAAGACAGGA
AGAGCACACGTCTGAACTCTTTCCCTACACGACGCTCTTCCGATCTNNNCAGAAGCTAACCATTTGTTC
GAAGAGCACACGTCTGAACTCTTTCCCTACACGACGCTCTTCCGATCTNNNCACGCCTGTAATCCCAAC
GGAAGAGCACACGTCTGAACTCTTTCCCTACACGACGCTCTTCCGATCTNNNCTGAGACAGGAGAATC
AGAGCACACGTCTGAACTCTTTCCCTACACGACGCTCTTCCGATCTNNNACTGTCACTAAAAGGCAAAT
AAGAGCACACGTCTGAACTCTTTCCCTACACGACGCTCTTCCGATCTNNNGGTGCCTGTAATCTCAGCT
GAAGAGCACACGTCTGAACTCTTTCCCTACACGACGCTCTTCCGATCTNNNCTGAGGTGGGTGGATAG
GAAGAGCACACGTCTGAACTCTTTCCCTACACGACGCTCTTCCGATCTNNNGAGAATTGCTTGAACCTG
GAAGAGCACACGTCTGAACTCTTTCCCTACACGACGCTCTTCCGATCTNNNTGCCTCTTTTTGAATGCTT
AGAGCACACGTCTGAACTCTTTCCCTACACGACGCTCTTCCGATCTNNNACAGATTCGAAACACTCTCT
GGAAGAGCACACGTCTGAACTCTTTCCCTACACGACGCTCTTCCGATCTNNNGAGGCAGGAGAATTGT
GAGCACACGTCTGAACTCTTTCCCTACACGACGCTCTTCCGATCTNNNATAAAGGGAGAGGAAAAAG
GGAAGAGCACACGTCTGAACTCTTTCCCTACACGACGCTCTTCCGATCTNNNAGGCAAGAGAATCACT
AGCACACGTCTGAACTCTTTCCCTACACGACGCTCTTCCGATCTNNNAGACTGCTATCTAGACTAACAA
GGAAGAGCACACGTCTGAACTCTTTCCCTACACGACGCTCTTCCGATCTNNNTAATCCCAGCTACTTGG
GAAGAGCACACGTCTGAACTCTTTCCCTACACGACGCTCTTCCGATCTNNNCATCTGTGATCCCAGCTA
GGAAGAGCACACGTCTGAACTCTTTCCCTACACGACGCTCTTCCGATCTNNNAACAAAAATAGCCAGGT
GAGCACACGTCTGAACTCTTTCCCTACACGACGCTCTTCCGATCTNNNAGACAAATTTTTTGAAACACA
AGCACACGTCTGAACTCTTTCCCTACACGACGCTCTTCCGATCTNNNTGAGGTAAAAGACTGATATAAC
GGAAGAGCACACGTCTGAACTCTTTCCCTACACGACGCTCTTCCGATCTNNNTGGAAAAACCTAAGGAC
GAGCACACGTCTGAACTCTTTCCCTACACGACGCTCTTCCGATCTNNNCCATCAAATTGGATGAATCTCA
GAAGAGCACACGTCTGAACTCTTTCCCTACACGACGCTCTTCCGATCTNNNCACAATCTCCTCCAGAACA
GGAAGAGCACACGTCTGAACTCTTTCCCTACACGACGCTCTTCCGATCTNNNAAATGCTGCAACACTGG
AGCACACGTCTGAACTCTTTCCCTACACGACGCTCTTCCGATCTNNNGGGAAATCCTTATAAACTATTG
GGAAGAGCACACGTCTGAACTCTTTCCCTACACGACGCTCTTCCGATCTNNNTCTGTAGTCCCAGCTATT
GAAGAGCACACGTCTGAACTCTTTCCCTACACGACGCTCTTCCGATCTNNNTGAATACTGCCGCTATGA
AGCACACGTCTGAACTCTTTCCCTACACGACGCTCTTCCGATCTNNNTCTTCTTTACAAAAGCTATCAGT

GCACACGTCTGAACTCTTTCCCTACACGACGCTCTTCCGATCTNNNCCATACCTTTATTTCAGGCTAT
GAAGAGCACACGTCTGAACTCTTTCCCTACACGACGCTCTTCCGATCTNNNCACGCATGTCTATTGCA
AAGAGCACACGTCTGAACTCTTTCCCTACACGACGCTCTTCCGATCTNNNTGAATGGGCTTAGGAGA
AAGAGCACACGTCTGAACTCTTTCCCTACACGACGCTCTTCCGATCTNNNTGAATGGGCTTAGGAGA
AGCACACGTCTGAACTCTTTCCCTACACGACGCTCTTCCGATCTNNNGCTACATACTCTTGATGTATTT
AGCACACGTCTGAACTCTTTCCCTACACGACGCTCTTCCGATCTNNNAGAGCCTATTATTTGAAAATT
AGCACACGTCTGAACTCTTTCCCTACACGACGCTCTTCCGATCTNNNGTATATATATACACACACACAC
GCACACGTCTGAACTCTTTCCCTACACGACGCTCTTCCGATCTNNNCACACGTGTGTATATGTATATAT
AGCACACGTCTGAACTCTTTCCCTACACGACGCTCTTCCGATCTNNNACACACCATATATATATACGCA
GAGCACACGTCTGAACTCTTTCCCTACACGACGCTCTTCCGATCTNNNTGTGAGTGTATATATATG
AGCACACGTCTGAACTCTTTCCCTACACGACGCTCTTCCGATCTNNNAGAGCCTATTATTTGAAAATT
AAGAGCACACGTCTGAACTCTTTCCCTACACGACGCTCTTCCGATCTNNNACATTTGAAAAAGCCCTCT
GGAAGAGCACACGTCTGAACTCTTTCCCTACACGACGCTCTTCCGATCTNNNAGAATTGCTTGAAATT
AAGAGCACACGTCTGAACTCTTTCCCTACACGACGCTCTTCCGATCTNNNTGCTTCATGGGTAGTGT
AGCACACGTCTGAACTCTTTCCCTACACGACGCTCTTCCGATCTNNNTGGAATAGTAATACAGTTATT
AAGAGCACACGTCTGAACTCTTTCCCTACACGACGCTCTTCCGATCTNNNAGAGATGGGGTTTCTCC
AGCACACGTCTGAACTCTTTCCCTACACGACGCTCTTCCGATCTNNNCGTCGATATATTTATGTATGT
GAAGAGCACACGTCTGAACTCTTTCCCTACACGACGCTCTTCCGATCTNNNAGACAACCAAAAGCTGA
AGCACACGTCTGAACTCTTTCCCTACACGACGCTCTTCCGATCTNNNACATATATAAGTTGTGTGTGT
GAAGAGCACACGTCTGAACTCTTTCCCTACACGACGCTCTTCCGATCTNNNAGGGTTCTCTAGAGGG
AAGAGCACACGTCTGAACTCTTTCCCTACACGACGCTCTTCCGATCTNNNTCTATGAGGGGAGATCAT
AGAGCACACGTCTGAACTCTTTCCCTACACGACGCTCTTCCGATCTNNNTGGTCTTGGGAGGGTTTAT
AAGAGCACACGTCTGAACTCTTTCCCTACACGACGCTCTTCCGATCTNNNGACCTCATTATTGGCTTGC
GAAGAGCACACGTCTGAACTCTTTCCCTACACGACGCTCTTCCGATCTNNNAGTAGCCATAGGGAAAG
GAAGAGCACACGTCTGAACTCTTTCCCTACACGACGCTCTTCCGATCTNNNTAGCTGGGAGGTAGGA
AGCACACGTCTGAACTCTTTCCCTACACGACGCTCTTCCGATCTNNNTGTTTCTTAGGTAATAGAATGT
AAGAGCACACGTCTGAACTCTTTCCCTACACGACGCTCTTCCGATCTNNNCTGAGGCAGGAGAATAG
GGAAGAGCACACGTCTGAACTCTTTCCCTACACGACGCTCTTCCGATCTNNNGTGCAAATGTAAAATG
GAGCACACGTCTGAACTCTTTCCCTACACGACGCTCTTCCGATCTNNNACATAGATTGTCCTGAGATAC
GAAGAGCACACGTCTGAACTCTTTCCCTACACGACGCTCTTCCGATCTNNNGACCCACTACAGCAAAA
AGCACACGTCTGAACTCTTTCCCTACACGACGCTCTTCCGATCTNNNTGGATTTCTCTGATAATCACTG
AGAGCACACGTCTGAACTCTTTCCCTACACGACGCTCTTCCGATCTNNNTCTGTCACTTGATTTTTCTTT
AGCACACGTCTGAACTCTTTCCCTACACGACGCTCTTCCGATCTNNNTCTTGGTCTAGCTAAAGTTTTAT
AAGAGCACACGTCTGAACTCTTTCCCTACACGACGCTCTTCCGATCTNNNACTGCTCAATACCATGGT
GAGCACACGTCTGAACTCTTTCCCTACACGACGCTCTTCCGATCTNNNTCTCTTTGTATTAGTCTAGAC
AGCACACGTCTGAACTCTTTCCCTACACGACGCTCTTCCGATCTNNNTGTGCTTAGAATCACACTTTCTT
AGCACACGTCTGAACTCTTTCCCTACACGACGCTCTTCCGATCTNNNTCTTCTTCCTTTCTCTCTCTTCTT
GAGCACACGTCTGAACTCTTTCCCTACACGACGCTCTTCCGATCTNNNCTTTCTCTTTCTTCCTTCTTTCT
AGCACACGTCTGAACTCTTTCCCTACACGACGCTCTTCCGATCTNNNGTTCCTTTTGAAAAGAATGTAC
AGCACACGTCTGAACTCTTTCCCTACACGACGCTCTTCCGATCTNNNACATTAGGGTTATTACATCTGC
AGCACACGTCTGAACTCTTTCCCTACACGACGCTCTTCCGATCTNNNACTGTTGGTGTAATGTAAATTA
AGCACACGTCTGAACTCTTTCCCTACACGACGCTCTTCCGATCTNNNTCTCTCTTTCTTTCTTTCTTTCGT
GCACACGTCTGAACTCTTTCCCTACACGACGCTCTTCCGATCTNNNTGCCCAGTTTTTAGTAGAACTAT

GAGCACACGTCTGAACTCTTTCCCTACACGACGCTCTTCCGATCTNNNTGGAAGAAACTATAAAACTTG
AGCACACGTCTGAACTCTTTCCCTACACGACGCTCTTCCGATCTNNNTCTTCTTTCCTCTTTCTTTCTTTCT
GAGCACACGTCTGAACTCTTTCCCTACACGACGCTCTTCCGATCTNNNTCAGTTTTATTTAGTCCAGGA
AGCACACGTCTGAACTCTTTCCCTACACGACGCTCTTCCGATCTNNNACCACAATACATGTTTGTTATTT
GAGCACACGTCTGAACTCTTTCCCTACACGACGCTCTTCCGATCTNNNGTTCACAAAACTTTAGTATTCT
GGAAGAGCACACGTCTGAACTCTTTCCCTACACGACGCTCTTCCGATCTNNNTCCCAAGGACAAAAATG
GAGCACACGTCTGAACTCTTTCCCTACACGACGCTCTTCCGATCTNNNACACTAAAAACTATAAACGTAC
GAAGAGCACACGTCTGAACTCTTTCCCTACACGACGCTCTTCCGATCTNNNGCAAAACCCTGCCTCATGA
GAGCACACGTCTGAACTCTTTCCCTACACGACGCTCTTCCGATCTNNNGCTATGATTATGTCACTTTTCA
AGCACACGTCTGAACTCTTTCCCTACACGACGCTCTTCCGATCTNNNTCTCTTTCTTTCTTTCTTTCTTTTG
GAGCACACGTCTGAACTCTTTCCCTACACGACGCTCTTCCGATCTNNNTCCTGCAACTTTACAGAATTTA
AGCACACGTCTGAACTCTTTCCCTACACGACGCTCTTCCGATCTNNNTGATTGTAGTTTTTGCAGGATA
GAGCACACGTCTGAACTCTTTCCCTACACGACGCTCTTCCGATCTNNNTCACTGAATATCTACTCTTCCA
GAGCACACGTCTGAACTCTTTCCCTACACGACGCTCTTCCGATCTNNNTCATATTCTTCCATCATCAGTCT
GGAAGAGCACACGTCTGAACTCTTTCCCTACACGACGCTCTTCCGATCTNNNGCAAGAAAAGTTCCTGT
GAAGAGCACACGTCTGAACTCTTTCCCTACACGACGCTCTTCCGATCTNNNTAACAGCAGAATTCGCAA
GGAAGAGCACACGTCTGAACTCTTTCCCTACACGACGCTCTTCCGATCTNNNTGCTCCCTTGAAATCCGT
GCACACGTCTGAACTCTTTCCCTACACGACGCTCTTCCGATCTNNNGGGTACTACAGATATATGTTTAG
CGGAAGAGCACACGTCTGAACTCTTTCCCTACACGACGCTCTTCCGATCTNNNACACAAAACACCCAGA
GAAGAGCACACGTCTGAACTCTTTCCCTACACGACGCTCTTCCGATCTNNNCTTAGCACCAAGAGGGA
GAGCACACGTCTGAACTCTTTCCCTACACGACGCTCTTCCGATCTNNNTGCATCTATGTATCTATCTGTC
GAAGAGCACACGTCTGAACTCTTTCCCTACACGACGCTCTTCCGATCTNNNAAGCAGATCACTGGTTGA
AAGAGCACACGTCTGAACTCTTTCCCTACACGACGCTCTTCCGATCTNNNTAAATTTGACCTGCCCTCCT
AAGAGCACACGTCTGAACTCTTTCCCTACACGACGCTCTTCCGATCTNNNCCAGGAATGTGTAAGGAAA
AGAGCACACGTCTGAACTCTTTCCCTACACGACGCTCTTCCGATCTNNNGAGAAGGCTGATGTCTTTGC
AGAGCACACGTCTGAACTCTTTCCCTACACGACGCTCTTCCGATCTNNNAAGACGCATAAGTAAAGAAA
AAGAGCACACGTCTGAACTCTTTCCCTACACGACGCTCTTCCGATCTNNNTGGGGGAAAATTCCCAACC
GAAGAGCACACGTCTGAACTCTTTCCCTACACGACGCTCTTCCGATCTNNNAGATAACTTGAAGGGGA
AGAGCACACGTCTGAACTCTTTCCCTACACGACGCTCTTCCGATCTNNNTGAAGTCAGGACTGTTAGG
AGAGCACACGTCTGAACTCTTTCCCTACACGACGCTCTTCCGATCTNNNTGAAAAGACAAGATGCATAC
AAGAGCACACGTCTGAACTCTTTCCCTACACGACGCTCTTCCGATCTNNNGAGATAGGAGGTACCCAG
AGCACACGTCTGAACTCTTTCCCTACACGACGCTCTTCCGATCTNNNTGGAGAAAACATAGCCCTATTAC
AAGAGCACACGTCTGAACTCTTTCCCTACACGACGCTCTTCCGATCTNNNTATGCTGCTGGACTTAGTTT
AAGAGCACACGTCTGAACTCTTTCCCTACACGACGCTCTTCCGATCTNNNCCGGTTGCTGCATCCATTAT
AGAGCACACGTCTGAACTCTTTCCCTACACGACGCTCTTCCGATCTNNNAGGATGAGTAAGAGTTTTCT
AGAGCACACGTCTGAACTCTTTCCCTACACGACGCTCTTCCGATCTNNNACGTGCCAAAGACATATTTC
AGAGCACACGTCTGAACTCTTTCCCTACACGACGCTCTTCCGATCTNNNAGTGAGAGAAAAGAGAACA
AAGAGCACACGTCTGAACTCTTTCCCTACACGACGCTCTTCCGATCTNNNTCCCACTTCAGCCTCCTAAT
AGCACACGTCTGAACTCTTTCCCTACACGACGCTCTTCCGATCTNNNGCCACAGATCTTATAGAGATGT
AGAGCACACGTCTGAACTCTTTCCCTACACGACGCTCTTCCGATCTNNNTGCTAACGTGGAGAGAGAC
GAAGAGCACACGTCTGAACTCTTTCCCTACACGACGCTCTTCCGATCTNNNCTATTTGGGAGGATGA
GAAGAGCACACGTCTGAACTCTTTCCCTACACGACGCTCTTCCGATCTNNNAAATTGAGCAGGGTGAG
AAGAGCACACGTCTGAACTCTTTCCCTACACGACGCTCTTCCGATCTNNNCTAGTCCCTGCCAGGTGAA

AGAGCACACGTCTGAACTCTTTCCCTACACGACGCTCTTCCGATCTNNNTTGCTAAAGGCAAATTCTAG

GCACACGTCTGAACTCTTTCCCTACACGACGCTCTTCCGATCTNNNAGACTTTCTCCACTTCAATTTTCCCT

GAAGAGCACACGTCTGAACTCTTTCCCTACACGACGCTCTTCCGATCTNNNTGGCCTTTGTATCTTGAC

AAGAGCACACGTCTGAACTCTTTCCCTACACGACGCTCTTCCGATCTNNNAGAGTAGCACCCAGGAAT

AGAGCACACGTCTGAACTCTTTCCCTACACGACGCTCTTCCGATCTNNNACAAGCTCTTCAGTTGCAAA

GAGCACACGTCTGAACTCTTTCCCTACACGACGCTCTTCCGATCTNNNGGCCTAGTTTCAAAGAAGATA

GAGCACACGTCTGAACTCTTTCCCTACACGACGCTCTTCCGATCTNNNAGAGAATAAAGAGAGTATTTC

AGAGCACACGTCTGAACTCTTTCCCTACACGACGCTCTTCCGATCTNNNCGTGCCTTTTGATAATGCAG

AAGAGCACACGTCTGAACTCTTTCCCTACACGACGCTCTTCCGATCTNNNAGTGAGACACTTCAGGGAA

AAGAGCACACGTCTGAACTCTTTCCCTACACGACGCTCTTCCGATCTNNNGGGGTTTCTCTCAGGTCTA

AAGAGCACACGTCTGAACTCTTTCCCTACACGACGCTCTTCCGATCTNNNCCCCAATGGTAACATCCTC

AGCACACGTCTGAACTCTTTCCCTACACGACGCTCTTCCGATCTNNNAGCCTTATAGGGATAACATTGA

AGCACACGTCTGAACTCTTTCCCTACACGACGCTCTTCCGATCTNNNACAAACTATTTGAGAGTGAGTTC

AGCACACGTCTGAACTCTTTCCCTACACGACGCTCTTCCGATCTNNNTGCAAGCATAGTATAAGAAACTC

GAGCACACGTCTGAACTCTTTCCCTACACGACGCTCTTCCGATCTNNNGTGAAAAATGTCAAAGTGCA

GAAGAGCACACGTCTGAACTCTTTCCCTACACGACGCTCTTCCGATCTNNNAATTTGCTAGCAGGGTAC

GAAGAGCACACGTCTGAACTCTTTCCCTACACGACGCTCTTCCGATCTNNNACCCGTAACAAGCAGCTAA

GAAGAGCACACGTCTGAACTCTTTCCCTACACGACGCTCTTCCGATCTNNNAAGCTCAAGCTGTCAACCT

GAAGAGCACACGTCTGAACTCTTTCCCTACACGACGCTCTTCCGATCTNNNCTTGAGCACTTTTTACTTG

AGAGCACACGTCTGAACTCTTTCCCTACACGACGCTCTTCCGATCTNNNTCTCTCTCTAGACAAATGAT

AAGAGCACACGTCTGAACTCTTTCCCTACACGACGCTCTTCCGATCTNNNATCTTGGGGAGGAGTACAC

AAGAGCACACGTCTGAACTCTTTCCCTACACGACGCTCTTCCGATCTNNNTCAAGTCAAGAACTGAAAC

AGAGCACACGTCTGAACTCTTTCCCTACACGACGCTCTTCCGATCTNNNCCATGCAGTAGCATAGATGA

GAGCACACGTCTGAACTCTTTCCCTACACGACGCTCTTCCGATCTNNNCCCCATGAGAAGAACTCAGTA

AGAGCACACGTCTGAACTCTTTCCCTACACGACGCTCTTCCGATCTNNNTTCACTCACTCAACCAGCTAA

AAGAGCACACGTCTGAACTCTTTCCCTACACGACGCTCTTCCGATCTNNNATGATACCACTTTTCCCCTC

AAGAGCACACGTCTGAACTCTTTCCCTACACGACGCTCTTCCGATCTNNNGCCAAATCCCATCTCTCTCG

AGAGCACACGTCTGAACTCTTTCCCTACACGACGCTCTTCCGATCTNNNTAGCCCGGTCAACAAGAGCA

GAAGAGCACACGTCTGAACTCTTTCCCTACACGACGCTCTTCCGATCTNNNTGCTGGGTCTTGATTTCC

GAAGAGCACACGTCTGAACTCTTTCCCTACACGACGCTCTTCCGATCTNNNTTGGACATAAGACAACCCT

AGAGCACACGTCTGAACTCTTTCCCTACACGACGCTCTTCCGATCTNNNTTCCTGTCTACCTTTGCTGAT

AAGAGCACACGTCTGAACTCTTTCCCTACACGACGCTCTTCCGATCTNNNAGTGTTGGGAAATCCCCTG

AGAGCACACGTCTGAACTCTTTCCCTACACGACGCTCTTCCGATCTNNNTTGAATTTCTGCTCCACCAC

AAGAGCACACGTCTGAACTCTTTCCCTACACGACGCTCTTCCGATCTNNNTGTTTTCCAAAGCTACTGT

AAGAGCACACGTCTGAACTCTTTCCCTACACGACGCTCTTCCGATCTNNNAATGTGTCAGACGTTGTGC

AGAGCACACGTCTGAACTCTTTCCCTACACGACGCTCTTCCGATCTNNNAGAAACACACCCAAATCAGTA

AAGAGCACACGTCTGAACTCTTTCCCTACACGACGCTCTTCCGATCTNNNCTGAAAAATGAACAGAGCT

AAGAGCACACGTCTGAACTCTTTCCCTACACGACGCTCTTCCGATCTNNNTCCAAGTTATCTTCCCAAAC

AAGAGCACACGTCTGAACTCTTTCCCTACACGACGCTCTTCCGATCTNNNGAATACATTGCTCTGGGG

AGCACACGTCTGAACTCTTTCCCTACACGACGCTCTTCCGATCTNNNAAAGAAGAAAGAACCAGAAAA

AAGAGCACACGTCTGAACTCTTTCCCTACACGACGCTCTTCCGATCTNNNCTGCTGGTTTTCAGTGAACC

GAAGAGCACACGTCTGAACTCTTTCCCTACACGACGCTCTTCCGATCTNNNCACCTCACTTCTGTCTTCAC

AGCACACGTCTGAACTCTTTCCCTACACGACGCTCTTCCGATCTNNNACAGTCTTTCTGAAACATTCAAC

AGCACACGTCTGAACTCTTTCCCTACACGACGCTCTTCCGATCTNNNTGCACACATATATAGGCATCTAT
GAGCACACGTCTGAACTCTTTCCCTACACGACGCTCTTCCGATCTNNNTGTGTCAGGAAAGAAAATACA
GAAGAGCACACGTCTGAACTCTTTCCCTACACGACGCTCTTCCGATCTNNNAAGTGGCTTCTGTAGAG
GAGCACACGTCTGAACTCTTTCCCTACACGACGCTCTTCCGATCTNNNTTGCTTTCTGAGATGGGACAC
AGCACACGTCTGAACTCTTTCCCTACACGACGCTCTTCCGATCTNNNACCTATAAGACGGTGTGCTAAT
GAGCACACGTCTGAACTCTTTCCCTACACGACGCTCTTCCGATCTNNNACCTGCTAAAAAGTTCTAGGA
GAAGAGCACACGTCTGAACTCTTTCCCTACACGACGCTCTTCCGATCTNNNTGTGTGTATCCATCCATC
AGCACACGTCTGAACTCTTTCCCTACACGACGCTCTTCCGATCTNNNTGTTGTCTGTGTACTCATAATTG
AGAGCACACGTCTGAACTCTTTCCCTACACGACGCTCTTCCGATCTNNNACCCAGTTTTATTTGCTTAGC
GAGCACACGTCTGAACTCTTTCCCTACACGACGCTCTTCCGATCTNNNACTGGGAAAGAAAACTCTCTA
AGAGCACACGTCTGAACTCTTTCCCTACACGACGCTCTTCCGATCTNNNAGCTTTTGCAATTCTGATTCT
GAAGAGCACACGTCTGAACTCTTTCCCTACACGACGCTCTTCCGATCTNNNGGCCCATCATTTCTTGTG
GAAGAGCACACGTCTGAACTCTTTCCCTACACGACGCTCTTCCGATCTNNNGTCGTTTCATATCTCCACA
AAGAGCACACGTCTGAACTCTTTCCCTACACGACGCTCTTCCGATCTNNNTTTGGCAGAGCTTTAGCT
GGAAGAGCACACGTCTGAACTCTTTCCCTACACGACGCTCTTCCGATCTNNNCCTGAAGCCTTGCTGAA
GAAGAGCACACGTCTGAACTCTTTCCCTACACGACGCTCTTCCGATCTNNNTGCATCAGAAAAGCATTC
AGAGCACACGTCTGAACTCTTTCCCTACACGACGCTCTTCCGATCTNNNAATTTCCAGAATGAATGAAT
AGAGCACACGTCTGAACTCTTTCCCTACACGACGCTCTTCCGATCTNNNCCCCAAGTAACCTCATAGGTT
AGAGCACACGTCTGAACTCTTTCCCTACACGACGCTCTTCCGATCTNNNGGCTGTTTAGTGTTCAACCCA
GAGCACACGTCTGAACTCTTTCCCTACACGACGCTCTTCCGATCTNNNACAATTCATATGGGGTCTTATC
AGCACACGTCTGAACTCTTTCCCTACACGACGCTCTTCCGATCTNNNACGATGACTGGATAAAAACAAT
AAGAGCACACGTCTGAACTCTTTCCCTACACGACGCTCTTCCGATCTNNNACTCACTTTGCTTTCCATTTC
AGAGCACACGTCTGAACTCTTTCCCTACACGACGCTCTTCCGATCTNNNTCAAATTAGCTTTTTGCTGGC
AGCACACGTCTGAACTCTTTCCCTACACGACGCTCTTCCGATCTNNNTGCCAAAAATATTCTTTTCTTCTC
AAGAGCACACGTCTGAACTCTTTCCCTACACGACGCTCTTCCGATCTNNNTCACTCCCATGAGTATTGAA
AGAGCACACGTCTGAACTCTTTCCCTACACGACGCTCTTCCGATCTNNNGGGATCTTCCTCTTGGGATC
AAGAGCACACGTCTGAACTCTTTCCCTACACGACGCTCTTCCGATCTNNNTCTTGAGACAGCTCAAAAT
AAGAGCACACGTCTGAACTCTTTCCCTACACGACGCTCTTCCGATCTNNNATGATCAGCCCACTACACTC
GAAGAGCACACGTCTGAACTCTTTCCCTACACGACGCTCTTCCGATCTNNNACAGTGCCTAGCCTAGAA
GAAGAGCACACGTCTGAACTCTTTCCCTACACGACGCTCTTCCGATCTNNNTTGTGCAAAACTGGGTAA
AAGAGCACACGTCTGAACTCTTTCCCTACACGACGCTCTTCCGATCTNNNGGCCAGAAGAGGCAACTT
AGAGCACACGTCTGAACTCTTTCCCTACACGACGCTCTTCCGATCTNNNTCCTATTGTTATCCCTCCCCA
GAGCACACGTCTGAACTCTTTCCCTACACGACGCTCTTCCGATCTNNNACCATGCCCAGCCACTTATTAT
AAGAGCACACGTCTGAACTCTTTCCCTACACGACGCTCTTCCGATCTNNNGAGAGGGTTGCTCAGTAG
GAGCACACGTCTGAACTCTTTCCCTACACGACGCTCTTCCGATCTNNNTGTCATTAGTTACACTGAATG
AAGAGCACACGTCTGAACTCTTTCCCTACACGACGCTCTTCCGATCTNNNTAACGAGAGATTGGGGCA
AGAGCACACGTCTGAACTCTTTCCCTACACGACGCTCTTCCGATCTNNNGTGTCAGTGGCAATACTCC
AAGAGCACACGTCTGAACTCTTTCCCTACACGACGCTCTTCCGATCTNNNTTCAAATCCCACTCACTCGT
AGAGCACACGTCTGAACTCTTTCCCTACACGACGCTCTTCCGATCTNNNACTTGCCTTTGGATAGCAAG
AGAGCACACGTCTGAACTCTTTCCCTACACGACGCTCTTCCGATCTNNNTGCATGTCATCACTTCCTTTC
AGAGCACACGTCTGAACTCTTTCCCTACACGACGCTCTTCCGATCTNNNGAAGGCACCGGATAATTGA
AAGAGCACACGTCTGAACTCTTTCCCTACACGACGCTCTTCCGATCTNNNGTTTGGCCCATTATAGGCA
GGAAGAGCACACGTCTGAACTCTTTCCCTACACGACGCTCTTCCGATCTNNNCTACAGCCTGGGAGTTC

AAGAGCACACGTCTGAACTCTTTCCCTACACGACGCTCTTCCGATCTNNNGTGAGCCTGTAATGTCAG
GAGCACACGTCTGAACTCTTTCCCTACACGACGCTCTTCCGATCTNNNCGAGTCACATATAAGGGAAT
AAGAGCACACGTCTGAACTCTTTCCCTACACGACGCTCTTCCGATCTNNNAATAGCTTGAATCTGGGA
GAGCACACGTCTGAACTCTTTCCCTACACGACGCTCTTCCGATCTNNNTCTAGGAATCACATTACCTAA
AAGAGCACACGTCTGAACTCTTTCCCTACACGACGCTCTTCCGATCTNNNAGACAATCTATGGTCACA
AGCACACGTCTGAACTCTTTCCCTACACGACGCTCTTCCGATCTNNNCCTGACTTCAAATTGTACTACA
AAGAGCACACGTCTGAACTCTTTCCCTACACGACGCTCTTCCGATCTNNNTGTAATCCCAGCTACTTTG
AAGAGCACACGTCTGAACTCTTTCCCTACACGACGCTCTTCCGATCTNNNCCCAATTAGAAATGGCAA
GAGCACACGTCTGAACTCTTTCCCTACACGACGCTCTTCCGATCTNNNTACTCTGGAGGCTAAGGAA
AAGAGCACACGTCTGAACTCTTTCCCTACACGACGCTCTTCCGATCTNNNTCAGGATAATCGCTTGAA
AAGAGCACACGTCTGAACTCTTTCCCTACACGACGCTCTTCCGATCTNNNGAGAATGGTGTGAACCCA
AAGAGCACACGTCTGAACTCTTTCCCTACACGACGCTCTTCCGATCTNNNTTGGAGGTATCCCCCCTA
AGAGCACACGTCTGAACTCTTTCCCTACACGACGCTCTTCCGATCTNNNCTGTAATCCCAGCTACTCAG
AAGAGCACACGTCTGAACTCTTTCCCTACACGACGCTCTTCCGATCTNNNATACAAAAATTAGCTGGG
AGCACACGTCTGAACTCTTTCCCTACACGACGCTCTTCCGATCTNNNTCAAATGTACCCCAGAAACATA
AAGAGCACACGTCTGAACTCTTTCCCTACACGACGCTCTTCCGATCTNNNCTGTAGTCCAAGCTATTC
GAAGAGCACACGTCTGAACTCTTTCCCTACACGACGCTCTTCCGATCTNNNTCTGTAATCCCAGCTACT
GCACACGTCTGAACTCTTTCCCTACACGACGCTCTTCCGATCTNNNGGTCCTAACATCATTGTATAGG
AAGAGCACACGTCTGAACTCTTTCCCTACACGACGCTCTTCCGATCTNNNGAGAATGGCATGAACTC
GAAGAGCACACGTCTGAACTCTTTCCCTACACGACGCTCTTCCGATCTNNNGAAAATGGCATGAACCC
AAGAGCACACGTCTGAACTCTTTCCCTACACGACGCTCTTCCGATCTNNNTGTCTTTTGAAATAAGCC
AGAGCACACGTCTGAACTCTTTCCCTACACGACGCTCTTCCGATCTNNNGAGGCAGAAGAATCACTTG
AAGAGCACACGTCTGAACTCTTTCCCTACACGACGCTCTTCCGATCTNNNAATCACTTGAAGCCAGG
GAAGAGCACACGTCTGAACTCTTTCCCTACACGACGCTCTTCCGATCTNNNAAAAAAAAAATTAGCCG
AAGAGCACACGTCTGAACTCTTTCCCTACACGACGCTCTTCCGATCTNNNACTTGGTAGGTTGAGGTA
AAGAGCACACGTCTGAACTCTTTCCCTACACGACGCTCTTCCGATCTNNNTTATCCTTCCCTCCTCCAAC
AAGAGCACACGTCTGAACTCTTTCCCTACACGACGCTCTTCCGATCTNNNAGCTTTCAACATTGGCAC
AGAGCACACGTCTGAACTCTTTCCCTACACGACGCTCTTCCGATCTNNNATAGAACGCAATAGAGAG
AAGAGCACACGTCTGAACTCTTTCCCTACACGACGCTCTTCCGATCTNNNTGAGGCAGAAGAATTGCT
AGCACACGTCTGAACTCTTTCCCTACACGACGCTCTTCCGATCTNNNACTCTGTTAGATGTTGAGGAT
AAGAGCACACGTCTGAACTCTTTCCCTACACGACGCTCTTCCGATCTNNNTGCATTGGCGGCTATATTT
AGCACACGTCTGAACTCTTTCCCTACACGACGCTCTTCCGATCTNNNTGGAATATCCATTACCTCAAA
AGCACACGTCTGAACTCTTTCCCTACACGACGCTCTTCCGATCTNNNACCACTACACACCTATTAGAAT
AAGAGCACACGTCTGAACTCTTTCCCTACACGACGCTCTTCCGATCTNNNATAATCCCAGCTATTCAGG
AAGAGCACACGTCTGAACTCTTTCCCTACACGACGCTCTTCCGATCTNNNACAACAAACCCTTATGACA
AAGAGCACACGTCTGAACTCTTTCCCTACACGACGCTCTTCCGATCTNNNTGAAGCATGAGAATCGCT
GAGCACACGTCTGAACTCTTTCCCTACACGACGCTCTTCCGATCTNNNACAAAATTCATGAAAGGGTG
AAGAGCACACGTCTGAACTCTTTCCCTACACGACGCTCTTCCGATCTNNNATCAGTTCAGCTTGGCAGT
GAAGAGCACACGTCTGAACTCTTTCCCTACACGACGCTCTTCCGATCTNNNGGGAAGTGAAGTTTGT
AGCACACGTCTGAACTCTTTCCCTACACGACGCTCTTCCGATCTNNNAGTTGTTACACTGTTTTGTTTA
GCACACGTCTGAACTCTTTCCCTACACGACGCTCTTCCGATCTNNNTGTTTATCAGGAATGTGTTTTGA
GCACACGTCTGAACTCTTTCCCTACACGACGCTCTTCCGATCTNNNTGAACTTTACATTCTCCTTTTGAC
AAGAGCACACGTCTGAACTCTTTCCCTACACGACGCTCTTCCGATCTNNNGTAATTCACATCCCTCCCT

GAGCACACGTCTGAACTCTTTCCCTACACGACGCTCTTCCGATCTNNNTGAAGGCTTTGTCTTTTAGTA
AGAGCACACGTCTGAACTCTTTCCCTACACGACGCTCTTCCGATCTNNNCACCCGGCCTGAATTCATTT
AGCACACGTCTGAACTCTTTCCCTACACGACGCTCTTCCGATCTNNNTCTCTACCTTTGTATTTTCATCT
AGAGCACACGTCTGAACTCTTTCCCTACACGACGCTCTTCCGATCTNNNTCAGTAGGGGAACAAATAA
GCACACGTCTGAACTCTTTCCCTACACGACGCTCTTCCGATCTNNNATGCATACATATGTATACATGCA
AGCACACGTCTGAACTCTTTCCCTACACGACGCTCTTCCGATCTNNNTCTTTCTTCTCTCTTTCTCCCTTT
GGAAGAGCACACGTCTGAACTCTTTCCCTACACGACGCTCTTCCGATCTNNNTGCAGACTTCAGTTAG
GCACACGTCTGAACTCTTTCCCTACACGACGCTCTTCCGATCTNNNAGCATGCCTCTATATAGCATATAT
GAGCACACGTCTGAACTCTTTCCCTACACGACGCTCTTCCGATCTNNNAAACAAAAAATATACACACA
AGCACACGTCTGAACTCTTTCCCTACACGACGCTCTTCCGATCTNNNGTCCAACAGGAAAATATCACAA
AGAGCACACGTCTGAACTCTTTCCCTACACGACGCTCTTCCGATCTNNNCACCCGGCCTGAATTCATTT
AGCACACGTCTGAACTCTTTCCCTACACGACGCTCTTCCGATCTNNNATGTATAGACACAAACATGCAT
AGAGCACACGTCTGAACTCTTTCCCTACACGACGCTCTTCCGATCTNNNTCTTGCTTTTTATGGCTGCAT
GAGCACACGTCTGAACTCTTTCCCTACACGACGCTCTTCCGATCTNNNCCATGGTGTGTGTATATGTAT
AAGAGCACACGTCTGAACTCTTTCCCTACACGACGCTCTTCCGATCTNNNAAGGGGTGATAGGAGTG
AGAGCACACGTCTGAACTCTTTCCCTACACGACGCTCTTCCGATCTNNNAGAATTGCTTGAATCTGGG
AGAGCACACGTCTGAACTCTTTCCCTACACGACGCTCTTCCGATCTNNNGGCTCCTAGCTTGGAAAATA
AAGAGCACACGTCTGAACTCTTTCCCTACACGACGCTCTTCCGATCTNNNCACTCATGAAGGGTCCTGA
GAGCACACGTCTGAACTCTTTCCCTACACGACGCTCTTCCGATCTNNNTCACAGTTGCAAAGATAAGGA
AAGAGCACACGTCTGAACTCTTTCCCTACACGACGCTCTTCCGATCTNNNTGAGGCAGGAGAATAGTG
GAGCACACGTCTGAACTCTTTCCCTACACGACGCTCTTCCGATCTNNNTGTATATATGTGTGTGTGTGT
AGCACACGTCTGAACTCTTTCCCTACACGACGCTCTTCCGATCTNNNTGTGTGTAAGTTGCTAAAACAA
AGCACACGTCTGAACTCTTTCCCTACACGACGCTCTTCCGATCTNNNACAATTTTGCCTTTTTTTTCTTTC
AAGAGCACACGTCTGAACTCTTTCCCTACACGACGCTCTTCCGATCTNNNGAGGTTGCAGTGAACTGA
AGAGCACACGTCTGAACTCTTTCCCTACACGACGCTCTTCCGATCTNNNACTAGAAAGCAGCAAGGAA
GAAGAGCACACGTCTGAACTCTTTCCCTACACGACGCTCTTCCGATCTNNNTTCACACACACACATACAC
AAGAGCACACGTCTGAACTCTTTCCCTACACGACGCTCTTCCGATCTNNNTCTACACTCCCATGTTCATT
AAGAGCACACGTCTGAACTCTTTCCCTACACGACGCTCTTCCGATCTNNNCCAAGTAGTCACAATCCTG
AGAGCACACGTCTGAACTCTTTCCCTACACGACGCTCTTCCGATCTNNNTGTAAATTTCATGCAATTTCC
GGAAGAGCACACGTCTGAACTCTTTCCCTACACGACGCTCTTCCGATCTNNNGCTGCTCCTCCTTACTTC
AGCACACGTCTGAACTCTTTCCCTACACGACGCTCTTCCGATCTNNNGCCATAAAAAGGAATGAATCA
AAGAGCACACGTCTGAACTCTTTCCCTACACGACGCTCTTCCGATCTNNNAATATGGAACCAGCCTAAC
AGAGCACACGTCTGAACTCTTTCCCTACACGACGCTCTTCCGATCTNNNGAACCAACCCAAATGTCCAT
AGCACACGTCTGAACTCTTTCCCTACACGACGCTCTTCCGATCTNNNCACACACACATATATACACACACA
AAGAGCACACGTCTGAACTCTTTCCCTACACGACGCTCTTCCGATCTNNNTCCCACCCAGCATACATAAC
GAGCACACGTCTGAACTCTTTCCCTACACGACGCTCTTCCGATCTNNNTCTTTGTTTCAATTCATCCTTTT
GAGCACACGTCTGAACTCTTTCCCTACACGACGCTCTTCCGATCTNNNACCATACTGTTTTGATCATCAT
AGAGCACACGTCTGAACTCTTTCCCTACACGACGCTCTTCCGATCTNNNTCAAATCATTGGAAAGCATT
AGCACACGTCTGAACTCTTTCCCTACACGACGCTCTTCCGATCTNNNTCCAGTTTTATTTGAAGTGAGGA
AAGAGCACACGTCTGAACTCTTTCCCTACACGACGCTCTTCCGATCTNNNTTGTCTCTTTGTCTTTACCC
AGCACACGTCTGAACTCTTTCCCTACACGACGCTCTTCCGATCTNNNTGAAAGAATGTTTTGTGGCATT
AGCACACGTCTGAACTCTTTCCCTACACGACGCTCTTCCGATCTNNNTGGTTGGTTACAATTATACAGT
GAAGAGCACACGTCTGAACTCTTTCCCTACACGACGCTCTTCCGATCTNNNTTGCATTTCCACCAGCAAT

AGCACACGTCTGAACTCTTTCCCTACACGACGCTCTTCCGATCTNNNCAGTCATTTCATAGTGTTGATG
AGCACACGTCTGAACTCTTTCCCTACACGACGCTCTTCCGATCTNNNACATTTTGCTTTATTTTCTTTGGC
AGCACACGTCTGAACTCTTTCCCTACACGACGCTCTTCCGATCTNNNTGAATGAGCTTTCATAATTTGT
GAGCACACGTCTGAACTCTTTCCCTACACGACGCTCTTCCGATCTNNNCCTTTTGCAAAGTTCAAGTTTA
GCACACGTCTGAACTCTTTCCCTACACGACGCTCTTCCGATCTNNNCAGTGTCTCATTACTTGTATATCC
GAGCACACGTCTGAACTCTTTCCCTACACGACGCTCTTCCGATCTNNNACTGTTTACTGTTGAATAAGGA
AGCACACGTCTGAACTCTTTCCCTACACGACGCTCTTCCGATCTNNNTGCCATATTGAAAAAAAATGAC
AGCACACGTCTGAACTCTTTCCCTACACGACGCTCTTCCGATCTNNNTGAAAGTATTATACCCAGGAGT
AGCACACGTCTGAACTCTTTCCCTACACGACGCTCTTCCGATCTNNNAGCTTCATTTGTTATTGTCCATTC
AGAGCACACGTCTGAACTCTTTCCCTACACGACGCTCTTCCGATCTNNNTTCAGCAACCAAGTCCTGTAC
AAGAGCACACGTCTGAACTCTTTCCCTACACGACGCTCTTCCGATCTNNNTGTGTTTTGCAGTTCTGCTT
AGCACACGTCTGAACTCTTTCCCTACACGACGCTCTTCCGATCTNNNTGAGACAGACTTTTTTTGAGAC
GAGCACACGTCTGAACTCTTTCCCTACACGACGCTCTTCCGATCTNNNGCAGTAATAGCTTTTTGCATTT
AGCACACGTCTGAACTCTTTCCCTACACGACGCTCTTCCGATCTNNNGGTTTCAGAAACAATTTGTTATC
AGCACACGTCTGAACTCTTTCCCTACACGACGCTCTTCCGATCTNNNATCCGTATAACAGAAAATTTGC
GAGCACACGTCTGAACTCTTTCCCTACACGACGCTCTTCCGATCTNNNTCTCTGAGATGTAAATATTGC
AGCACACGTCTGAACTCTTTCCCTACACGACGCTCTTCCGATCTNNNTGTCTGATATTAGGATTGCAACT
GAGCACACGTCTGAACTCTTTCCCTACACGACGCTCTTCCGATCTNNNAGAATGATAGCAAAATCCATA
AGAGCACACGTCTGAACTCTTTCCCTACACGACGCTCTTCCGATCTNNNTGCCTAAAAGCTTACTTACTT
AAGAGCACACGTCTGAACTCTTTCCCTACACGACGCTCTTCCGATCTNNNGCAAAAAATATGGAACCAC
GAGCACACGTCTGAACTCTTTCCCTACACGACGCTCTTCCGATCTNNNTGAATCCTTCAACTTCACTGAA
AAGAGCACACGTCTGAACTCTTTCCCTACACGACGCTCTTCCGATCTNNNAAAAAATTAGCTGGACGTG
AGCACACGTCTGAACTCTTTCCCTACACGACGCTCTTCCGATCTNNNACTTGTTTGTTGTCCATTTGTAT
GAAGAGCACACGTCTGAACTCTTTCCCTACACGACGCTCTTCCGATCTNNNGCTAAGGTAGGAGGATT
AGCACACGTCTGAACTCTTTCCCTACACGACGCTCTTCCGATCTNNNTCTATGTATAAGAGGAATGGAC
AGAGCACACGTCTGAACTCTTTCCCTACACGACGCTCTTCCGATCTNNNACAAGTAGAGTTCTTAGCCT
GAAGAGCACACGTCTGAACTCTTTCCCTACACGACGCTCTTCCGATCTNNNGAAGAAATTGCACTGACT
AAGAGCACACGTCTGAACTCTTTCCCTACACGACGCTCTTCCGATCTNNNCCATGTAAGTGGATCACTA
GAAGAGCACACGTCTGAACTCTTTCCCTACACGACGCTCTTCCGATCTNNNGGTGCCTGTAATCTCAGC
AGCACACGTCTGAACTCTTTCCCTACACGACGCTCTTCCGATCTNNNTCTTCAGTTGGTTGATTGTCTTT
AAGAGCACACGTCTGAACTCTTTCCCTACACGACGCTCTTCCGATCTNNNTTTTCTCTCTCTCTCCCCC
GGAAGAGCACACGTCTGAACTCTTTCCCTACACGACGCTCTTCCGATCTNNNAAGATGGAAGGAAGG
AAGAGCACACGTCTGAACTCTTTCCCTACACGACGCTCTTCCGATCTNNNCCATTCCTCTGTATTGGTGC
AGAGCACACGTCTGAACTCTTTCCCTACACGACGCTCTTCCGATCTNNNAAAGGTTATGAATGCCAGGC
AGAGCACACGTCTGAACTCTTTCCCTACACGACGCTCTTCCGATCTNNNAAGTTGGGATGCAGGGTAC
GCACACGTCTGAACTCTTTCCCTACACGACGCTCTTCCGATCTNNNGCCAGTATAGACAATAATTGGCC
AGAGCACACGTCTGAACTCTTTCCCTACACGACGCTCTTCCGATCTNNNCAAAACTTCGTCTCTCTCCCC
AGAGCACACGTCTGAACTCTTTCCCTACACGACGCTCTTCCGATCTNNNAGTATTTTGGCATGCTGAGG
AAGAGCACACGTCTGAACTCTTTCCCTACACGACGCTCTTCCGATCTNNNAAGTAATGAGCTTCAGCTG
AGAGCACACGTCTGAACTCTTTCCCTACACGACGCTCTTCCGATCTNNNGTTGACAGGCTGTAGTTTGC
AGCACACGTCTGAACTCTTTCCCTACACGACGCTCTTCCGATCTNNNCGCCAACATATCTGTGTATTTG
AGAGCACACGTCTGAACTCTTTCCCTACACGACGCTCTTCCGATCTNNNAGCAAAAATGAACAAACTG
AAGAGCACACGTCTGAACTCTTTCCCTACACGACGCTCTTCCGATCTNNNACTCAGGCATCATTCAGAT

AGAGCACACGTCTGAACTCTTTCCCTACACGACGCTCTTCCGATCTNNNCTACAGTTCTGGGTTCTTGGC
AGCACACGTCTGAACTCTTTCCCTACACGACGCTCTTCCGATCTNNNACCTGGCTATTAGTGAGTTACT
AAGAGCACACGTCTGAACTCTTTCCCTACACGACGCTCTTCCGATCTNNNACCAAACTCATCCAAAGACA
AGAGCACACGTCTGAACTCTTTCCCTACACGACGCTCTTCCGATCTNNNTAGGTACTCATTGCTTCTGGC
GAGCACACGTCTGAACTCTTTCCCTACACGACGCTCTTCCGATCTNNNGCCTGGCCAAGTATCACTTTT
GAGCACACGTCTGAACTCTTTCCCTACACGACGCTCTTCCGATCTNNNTGGCAAATGACAGGACTTCTTA
GAGCACACGTCTGAACTCTTTCCCTACACGACGCTCTTCCGATCTNNNAGATGTTTTCTCTGATGGTCT
GCACACGTCTGAACTCTTTCCCTACACGACGCTCTTCCGATCTNNNGGGAATTTCTAGGGTGTCTTTTT
AGAGCACACGTCTGAACTCTTTCCCTACACGACGCTCTTCCGATCTNNNGGCTACAAGCATCGTATAGTC
AAGAGCACACGTCTGAACTCTTTCCCTACACGACGCTCTTCCGATCTNNNTAAGTGCTCATCATCCCACA
AGAGCACACGTCTGAACTCTTTCCCTACACGACGCTCTTCCGATCTNNNGAATCTGGCTCAGGGCAACT
AGCACACGTCTGAACTCTTTCCCTACACGACGCTCTTCCGATCTNNNACCCAATTAGGGTGTATTAGTC
GAGCACACGTCTGAACTCTTTCCCTACACGACGCTCTTCCGATCTNNNAAGAGATGTGGTCTTGCCGTC
GAGCACACGTCTGAACTCTTTCCCTACACGACGCTCTTCCGATCTNNNACATCACCTACATCACACATACC
AGCACACGTCTGAACTCTTTCCCTACACGACGCTCTTCCGATCTNNNACTCACCTAGGAATATGGCTTCT
GAGCACACGTCTGAACTCTTTCCCTACACGACGCTCTTCCGATCTNNNTGTCGCATATTTCAGAATTCC
GAGCACACGTCTGAACTCTTTCCCTACACGACGCTCTTCCGATCTNNNTGCTTACTACATCCATCACTTC
GAGCACACGTCTGAACTCTTTCCCTACACGACGCTCTTCCGATCTNNNTTTGTGCCTACGGTACAGTTC
AGAGCACACGTCTGAACTCTTTCCCTACACGACGCTCTTCCGATCTNNNTCACTAACTGGCAGAAAATG
GCACACGTCTGAACTCTTTCCCTACACGACGCTCTTCCGATCTNNNAAGGCTTCATAATACATTTGTGGA
GAGCACACGTCTGAACTCTTTCCCTACACGACGCTCTTCCGATCTNNNTGGAGTGAGACTATGTCTCGT
AGCACACGTCTGAACTCTTTCCCTACACGACGCTCTTCCGATCTNNNACATGTGTGTACATATACTTCTG
GCACACGTCTGAACTCTTTCCCTACACGACGCTCTTCCGATCTNNNTCCTGAAGCCAAAAACCAAATTTC
AGAGCACACGTCTGAACTCTTTCCCTACACGACGCTCTTCCGATCTNNNTAGCTTACTTCCCCTGTAGCA
GAGCACACGTCTGAACTCTTTCCCTACACGACGCTCTTCCGATCTNNNTGCCATATAACTGGAAATGCA
AGAGCACACGTCTGAACTCTTTCCCTACACGACGCTCTTCCGATCTNNNTGTATGAGTACTTGCACACA
AGAGCACACGTCTGAACTCTTTCCCTACACGACGCTCTTCCGATCTNNNACTTGAGAGAAAGAGAGGG
AAGAGCACACGTCTGAACTCTTTCCCTACACGACGCTCTTCCGATCTNNNGAGACTGGCTTTCTGCTTCT
GAGCACACGTCTGAACTCTTTCCCTACACGACGCTCTTCCGATCTNNNGCCCACTTTCTGATGGAGATGA
GAAGAGCACACGTCTGAACTCTTTCCCTACACGACGCTCTTCCGATCTNNNTTTGTCACTGGTTGCTGTC
GAGCACACGTCTGAACTCTTTCCCTACACGACGCTCTTCCGATCTNNNAAACAAAAGCAGTTCCTGGAC
AAGAGCACACGTCTGAACTCTTTCCCTACACGACGCTCTTCCGATCTNNNCGAAGGCATAGAGGGCAA
AGAGCACACGTCTGAACTCTTTCCCTACACGACGCTCTTCCGATCTNNNGTCTTGAGCCTGTCTCGAAC
GAGCACACGTCTGAACTCTTTCCCTACACGACGCTCTTCCGATCTNNNAGATCAGAGGCATACATGTAA
GAGCACACGTCTGAACTCTTTCCCTACACGACGCTCTTCCGATCTNNNACTGATTACCACTCTGTGCTTG
GAGCACACGTCTGAACTCTTTCCCTACACGACGCTCTTCCGATCTNNNTCTTTCCTGTAGATTGTGTTCT
AGCACACGTCTGAACTCTTTCCCTACACGACGCTCTTCCGATCTNNNGCAGATCGAATATGGCAAAAG
AGAGCACACGTCTGAACTCTTTCCCTACACGACGCTCTTCCGATCTNNNCCCCACCCCAGTTTAGGAGAT
AGAGCACACGTCTGAACTCTTTCCCTACACGACGCTCTTCCGATCTNNNTTCTACCCACTAGGCCATGTC
AGAGCACACGTCTGAACTCTTTCCCTACACGACGCTCTTCCGATCTNNNAGGGGATATATGTGATGGA
AGAGCACACGTCTGAACTCTTTCCCTACACGACGCTCTTCCGATCTNNNATGCTTGTTTGCTGAGAAAG
AGCACACGTCTGAACTCTTTCCCTACACGACGCTCTTCCGATCTNNNTCTTCTCACATGGTAACATTTTCC
AGCACACGTCTGAACTCTTTCCCTACACGACGCTCTTCCGATCTNNNACTTGGAAAGAAACCAAACATC

AGAGCACACGTCTGAACTCTTTCCCTACACGACGCTCTTCCGATCTNNNGAGTAGCTAGGACCGTGGG
AGAGCACACGTCTGAACTCTTTCCCTACACGACGCTCTTCCGATCTNNNCTGCAAAGAGGCATTTTTCC
GAAGAGCACACGTCTGAACTCTTTCCCTACACGACGCTCTTCCGATCTNNNATGGGGGAACCAGTTACA
GAGCACACGTCTGAACTCTTTCCCTACACGACGCTCTTCCGATCTNNNAAATGATAGATAACGCCCCAC
AAGAGCACACGTCTGAACTCTTTCCCTACACGACGCTCTTCCGATCTNNNTGGGCTGGTTACAACCTCT
AGCACACGTCTGAACTCTTTCCCTACACGACGCTCTTCCGATCTNNNCCAACTTTTTGTCTTGTTGGTCC
GAGCACACGTCTGAACTCTTTCCCTACACGACGCTCTTCCGATCTNNNAGTCTGTTGCCAAACTTTTCAT
GAGCACACGTCTGAACTCTTTCCCTACACGACGCTCTTCCGATCTNNNGCAGACTTGTAGATACCACCTT
AGAGCACACGTCTGAACTCTTTCCCTACACGACGCTCTTCCGATCTNNNGACCCTATTCTCCTGCTGCAC
AGAGCACACGTCTGAACTCTTTCCCTACACGACGCTCTTCCGATCTNNNGAGGGATAATTTCACACAG
AAGAGCACACGTCTGAACTCTTTCCCTACACGACGCTCTTCCGATCTNNNGCGTAAATCCACGATACCA
AGCACACGTCTGAACTCTTTCCCTACACGACGCTCTTCCGATCTNNNTTGAGGGTATATATGTGTGTGT
AGAGCACACGTCTGAACTCTTTCCCTACACGACGCTCTTCCGATCTNNNTGAGAACGGACTAATCCACC
GAGCACACGTCTGAACTCTTTCCCTACACGACGCTCTTCCGATCTNNNCCTGTACCTATTGGCCTTTCCA
AGAGCACACGTCTGAACTCTTTCCCTACACGACGCTCTTCCGATCTNNNTGCACTTGGGTTCATCAGGC
AAGAGCACACGTCTGAACTCTTTCCCTACACGACGCTCTTCCGATCTNNNAATCACTCTTTGCCTAACCC
AAGAGCACACGTCTGAACTCTTTCCCTACACGACGCTCTTCCGATCTNNNCGGCTGTGAATCCATCATAT
AAGAGCACACGTCTGAACTCTTTCCCTACACGACGCTCTTCCGATCTNNNTGCAACTACACCTCAGAGA
AGAGCACACGTCTGAACTCTTTCCCTACACGACGCTCTTCCGATCTNNNCCCCCTCTACAATGCCTGTTTT
AAGAGCACACGTCTGAACTCTTTCCCTACACGACGCTCTTCCGATCTNNNCCTTTGTATCTGGTCTGGG
AAGAGCACACGTCTGAACTCTTTCCCTACACGACGCTCTTCCGATCTNNNACTGGCTAACAGTGAACTA
AGAGCACACGTCTGAACTCTTTCCCTACACGACGCTCTTCCGATCTNNNCCAGCAATATCCAGGACCAG
GAGCACACGTCTGAACTCTTTCCCTACACGACGCTCTTCCGATCTNNNAAGAAGTTTCCTCCACCAAATA
AGAGCACACGTCTGAACTCTTTCCCTACACGACGCTCTTCCGATCTNNNGGCTATCTTCTTGGGCCTTC
GAAGAGCACACGTCTGAACTCTTTCCCTACACGACGCTCTTCCGATCTNNNGCACAGACTAACCAGCTCT
AAGAGCACACGTCTGAACTCTTTCCCTACACGACGCTCTTCCGATCTNNNCTGAGGTGTGTACCATTCA
AGAGCACACGTCTGAACTCTTTCCCTACACGACGCTCTTCCGATCTNNNGCCAGTGCTCCAACTTTTTCT
AGAGCACACGTCTGAACTCTTTCCCTACACGACGCTCTTCCGATCTNNNCAAATACCTGTGAGCGTTGC
AGCACACGTCTGAACTCTTTCCCTACACGACGCTCTTCCGATCTNNNCCTCTTTTATCCAAGACTAGGAC
AGAGCACACGTCTGAACTCTTTCCCTACACGACGCTCTTCCGATCTNNNTGCATAAAGAATAGGGAGG
AGAGCACACGTCTGAACTCTTTCCCTACACGACGCTCTTCCGATCTNNNAAAGCACCGCTTCTAAGAGG
AAGAGCACACGTCTGAACTCTTTCCCTACACGACGCTCTTCCGATCTNNNTATTGCACCTTGCCAGGTA
GAGCACACGTCTGAACTCTTTCCCTACACGACGCTCTTCCGATCTNNNCAGTTTGCTCAGATATGCCCT
AGAGCACACGTCTGAACTCTTTCCCTACACGACGCTCTTCCGATCTNNNCACATACCAGGAGATTGCCT
AGCACACGTCTGAACTCTTTCCCTACACGACGCTCTTCCGATCTNNNGACATCATGATCTACCTGCTTCC
CACACGTCTGAACTCTTTCCCTACACGACGCTCTTCCGATCTNNNTTTATTTTCCTCCTCCTCCTTCTTCTT
AAGAGCACACGTCTGAACTCTTTCCCTACACGACGCTCTTCCGATCTNNNACAAATTCAAAGCCAGCAC
AGAGCACACGTCTGAACTCTTTCCCTACACGACGCTCTTCCGATCTNNNAGAAGCTCACCTTACCACAG
GAGCACACGTCTGAACTCTTTCCCTACACGACGCTCTTCCGATCTNNNTGAGGAGCAAAAGGAAGTGA
GAGCACACGTCTGAACTCTTTCCCTACACGACGCTCTTCCGATCTNNNTTCCATAAGGTCTGAGTGTGT
GAGCACACGTCTGAACTCTTTCCCTACACGACGCTCTTCCGATCTNNNACTTATACCCTGATGAACTGG
GAGCACACGTCTGAACTCTTTCCCTACACGACGCTCTTCCGATCTNNNAATTTGTGAGCACTAAGATTGA
GAGCACACGTCTGAACTCTTTCCCTACACGACGCTCTTCCGATCTNNNTCAAGGTTTTCGAGTTGCCCAT

AAGAGCACACGTCTGAACTCTTTCCCTACACGACGCTCTTCCGATCTNNNGCCCACAAAATTGACAAA
AGAGCACACGTCTGAACTCTTTCCCTACACGACGCTCTTCCGATCTNNNTTATCTCTGTCCACTGGTCC
AGCACACGTCTGAACTCTTTCCCTACACGACGCTCTTCCGATCTNNNCAGGAATAATGGAGTTTACCC
AAGAGCACACGTCTGAACTCTTTCCCTACACGACGCTCTTCCGATCTNNNTCTGTTGTCAAGGCTGAAG
AGAGCACACGTCTGAACTCTTTCCCTACACGACGCTCTTCCGATCTNNNCCGAGTTCTTCCAAGAAACA
AGCACACGTCTGAACTCTTTCCCTACACGACGCTCTTCCGATCTNNNACATGTATCTCCTGTTATTGTGT
GAGCACACGTCTGAACTCTTTCCCTACACGACGCTCTTCCGATCTNNNAGTGCAGTCTTTGATTACTCC
AAGAGCACACGTCTGAACTCTTTCCCTACACGACGCTCTTCCGATCTNNNAGAAAGAGACTGAGGGA
AGAGCACACGTCTGAACTCTTTCCCTACACGACGCTCTTCCGATCTNNNTTATTCAATGCACATGGCCA
GAGCACACGTCTGAACTCTTTCCCTACACGACGCTCTTCCGATCTNNNAGTACTGTCTGTTTCGTTTCAC
GCACACGTCTGAACTCTTTCCCTACACGACGCTCTTCCGATCTNNNATGATTATTTCATCTGAGCACTTG
AGAGCACACGTCTGAACTCTTTCCCTACACGACGCTCTTCCGATCTNNNGCTAACAATGCTGTTTGCGA
AGAGCACACGTCTGAACTCTTTCCCTACACGACGCTCTTCCGATCTNNNGCACACAAACACACAAGACA
GAGCACACGTCTGAACTCTTTCCCTACACGACGCTCTTCCGATCTNNNCGGGAAACAGCAAGGACTAA
AGAGCACACGTCTGAACTCTTTCCCTACACGACGCTCTTCCGATCTNNNTTTGCCCACACCCTCACATAT
GAGCACACGTCTGAACTCTTTCCCTACACGACGCTCTTCCGATCTNNNGCTTTACAAACCAGAAATGAG
AGAGCACACGTCTGAACTCTTTCCCTACACGACGCTCTTCCGATCTNNNTATCACCTCCTCCAAACAGACA
AAGAGCACACGTCTGAACTCTTTCCCTACACGACGCTCTTCCGATCTNNNAGTGGCTATTCAACTGGTC
AGAGCACACGTCTGAACTCTTTCCCTACACGACGCTCTTCCGATCTNNNCAGTGTGTAAGTCTTTACCT
GAGCACACGTCTGAACTCTTTCCCTACACGACGCTCTTCCGATCTNNNAAACTGCTCTAGCCTCTTTCAC
GAGCACACGTCTGAACTCTTTCCCTACACGACGCTCTTCCGATCTNNNGAAGAAAAGTAAGGATGCCA
AAGAGCACACGTCTGAACTCTTTCCCTACACGACGCTCTTCCGATCTNNNGAAATACTGTGCTCATGTG
GAAGAGCACACGTCTGAACTCTTTCCCTACACGACGCTCTTCCGATCTNNNAGCAGACATTTCGGTGA
AGAGCACACGTCTGAACTCTTTCCCTACACGACGCTCTTCCGATCTNNNAGATTGAGAGGCAATAACT
AGAGCACACGTCTGAACTCTTTCCCTACACGACGCTCTTCCGATCTNNNCAGTGTTCAATGTCCTCACA
GAGCACACGTCTGAACTCTTTCCCTACACGACGCTCTTCCGATCTNNNTCATGTCCATGTGAACAGTAG
GAGCACACGTCTGAACTCTTTCCCTACACGACGCTCTTCCGATCTNNNCAGCTGAAGGGTATTTCTCTC
AGAGCACACGTCTGAACTCTTTCCCTACACGACGCTCTTCCGATCTNNNTCCCTTCAGGTCCAAACAGA
AGAGCACACGTCTGAACTCTTTCCCTACACGACGCTCTTCCGATCTNNNACAGCAAACCAAATCCAAAG
AGCACACGTCTGAACTCTTTCCCTACACGACGCTCTTCCGATCTNNNAGACATACTGTTGAGAAAACTG
AGAGCACACGTCTGAACTCTTTCCCTACACGACGCTCTTCCGATCTNNNTGATGACATCCCTCTCAACG
AGAGCACACGTCTGAACTCTTTCCCTACACGACGCTCTTCCGATCTNNNGCTCCTTTGGAAGAAAGTT
GAAGAGCACACGTCTGAACTCTTTCCCTACACGACGCTCTTCCGATCTNNNCTCCAGCAGCCAAAGTAC
AGAGCACACGTCTGAACTCTTTCCCTACACGACGCTCTTCCGATCTNNNCAAGAGGGAGGGACAGAG
AGCACACGTCTGAACTCTTTCCCTACACGACGCTCTTCCGATCTNNNACATACCTACACACATTCATACA
AGCACACGTCTGAACTCTTTCCCTACACGACGCTCTTCCGATCTNNNTGGATGTTTGATAGGATTGCA
AGAGCACACGTCTGAACTCTTTCCCTACACGACGCTCTTCCGATCTNNNAGATCCTCTGCAGAAATGTA
AGCACACGTCTGAACTCTTTCCCTACACGACGCTCTTCCGATCTNNNACTACCATTTGCAAAAACATTCC
AGCACACGTCTGAACTCTTTCCCTACACGACGCTCTTCCGATCTNNNTCAGCTTAGAGTTTTGAATGAT
AGAGCACACGTCTGAACTCTTTCCCTACACGACGCTCTTCCGATCTNNNTGCAGGAGAACGTTTTAGG
AAGAGCACACGTCTGAACTCTTTCCCTACACGACGCTCTTCCGATCTNNNGAGGAAGGTCTCCCAACAA
AGCACACGTCTGAACTCTTTCCCTACACGACGCTCTTCCGATCTNNNACTCTTTGGTGAAACATTTCTCT
AGCACACGTCTGAACTCTTTCCCTACACGACGCTCTTCCGATCTNNNCCTAGAACCATGTATGGAACAA

GAGCACACGTCTGAACTCTTTCCCTACACGACGCTCTTCCGATCTNNNCCCCATCAAAATTTATCCATC
AGCACACGTCTGAACTCTTTCCCTACACGACGCTCTTCCGATCTNNNTCTAATAGGATTTGGAACACA
GAGCACACGTCTGAACTCTTTCCCTACACGACGCTCTTCCGATCTNNNTGGTATCTGATCCAAGTACCA
AGAGCACACGTCTGAACTCTTTCCCTACACGACGCTCTTCCGATCTNNNCTAGCACAGAGCTTAGAAC
AGCACACGTCTGAACTCTTTCCCTACACGACGCTCTTCCGATCTNNNACATTTAGGTTTGTCTGCTTTG
AGAGCACACGTCTGAACTCTTTCCCTACACGACGCTCTTCCGATCTNNNGATCCCAAACCGTTGAAGG
GAGCACACGTCTGAACTCTTTCCCTACACGACGCTCTTCCGATCTNNNAGGTAAAAGCTCAAGAAAGC
GCACACGTCTGAACTCTTTCCCTACACGACGCTCTTCCGATCTNNNAGACTTGTTAGTGTTTGCAAAG
AAGAGCACACGTCTGAACTCTTTCCCTACACGACGCTCTTCCGATCTNNNGAGAGTGGTCCAGAGGT
AGAGCACACGTCTGAACTCTTTCCCTACACGACGCTCTTCCGATCTNNNCCCAAACTGTGGAAATTGC
GAGCACACGTCTGAACTCTTTCCCTACACGACGCTCTTCCGATCTNNNTTATTGCCTGCTCTCTCTCCTG
AGAGCACACGTCTGAACTCTTTCCCTACACGACGCTCTTCCGATCTNNNGACCTGTGTGGAAACTGGC
AGAGCACACGTCTGAACTCTTTCCCTACACGACGCTCTTCCGATCTNNNTGTATATCTCCTAATGCTGG
AGCACACGTCTGAACTCTTTCCCTACACGACGCTCTTCCGATCTNNNGTGTCTGTTCACATCCTTTGGC
AGAGCACACGTCTGAACTCTTTCCCTACACGACGCTCTTCCGATCTNNNGGAGTGTAGTGAACTCCTG
AGAGCACACGTCTGAACTCTTTCCCTACACGACGCTCTTCCGATCTNNNTCGAGCGTGCTGTGTATTAC
AGCACACGTCTGAACTCTTTCCCTACACGACGCTCTTCCGATCTNNNGAGTAGGTGGGATTTCAGGC
GAGCACACGTCTGAACTCTTTCCCTACACGACGCTCTTCCGATCTNNNTGGAGGTACCAACCTATCCA
AGAGCACACGTCTGAACTCTTTCCCTACACGACGCTCTTCCGATCTNNNACTCATCGTCATGACAACCA
AGAGCACACGTCTGAACTCTTTCCCTACACGACGCTCTTCCGATCTNNNCACTGCAATCATGGGATTC
GAGCACACGTCTGAACTCTTTCCCTACACGACGCTCTTCCGATCTNNNTGTGGCAATGCTTTCTTGGA
AGAGCACACGTCTGAACTCTTTCCCTACACGACGCTCTTCCGATCTNNNATGAGGAATGAGGACAAG
AAGAGCACACGTCTGAACTCTTTCCCTACACGACGCTCTTCCGATCTNNNCTCCTGCAGTTCTCCTACAA
AGAGCACACGTCTGAACTCTTTCCCTACACGACGCTCTTCCGATCTNNNTTACAGGTGGATGACACAA
AGCACACGTCTGAACTCTTTCCCTACACGACGCTCTTCCGATCTNNNAGTAACAGTCTTACACTGCCAA
AAGAGCACACGTCTGAACTCTTTCCCTACACGACGCTCTTCCGATCTNNNTCTGAACACACTGAGAGC
AAGAGCACACGTCTGAACTCTTTCCCTACACGACGCTCTTCCGATCTNNNATGGTGTTTTGTGATGTT
AGAGCACACGTCTGAACTCTTTCCCTACACGACGCTCTTCCGATCTNNNTGAGGACCCAAGGGCTAAA
AGAGCACACGTCTGAACTCTTTCCCTACACGACGCTCTTCCGATCTNNNATTGGAGATAGTGCTTTGT
GAGCACACGTCTGAACTCTTTCCCTACACGACGCTCTTCCGATCTNNNAGCACTATTGCTGGATTGATG
GAAGAGCACACGTCTGAACTCTTTCCCTACACGACGCTCTTCCGATCTNNNGAGACAACAGCCTTACA
AGAGCACACGTCTGAACTCTTTCCCTACACGACGCTCTTCCGATCTNNNCCACTTGGTAAAAAGGATG
AAGAGCACACGTCTGAACTCTTTCCCTACACGACGCTCTTCCGATCTNNNAGTGAGCCAATGTAGTGC
AGAGCACACGTCTGAACTCTTTCCCTACACGACGCTCTTCCGATCTNNNCAGTGCATGCTCCATCAAG
AGAGCACACGTCTGAACTCTTTCCCTACACGACGCTCTTCCGATCTNNNTTGCTTCCTTAGCATCTGGTT
AAGAGCACACGTCTGAACTCTTTCCCTACACGACGCTCTTCCGATCTNNNTAGGAGCAAACTTCTCAG
AGAGCACACGTCTGAACTCTTTCCCTACACGACGCTCTTCCGATCTNNNATCTCCTAGTGTGCAGGTCT
AGAGCACACGTCTGAACTCTTTCCCTACACGACGCTCTTCCGATCTNNNACTATCCCCAGTCTCATCCA
AGAGCACACGTCTGAACTCTTTCCCTACACGACGCTCTTCCGATCTNNNCACCTTACTGATGGGCTATC
AGAGCACACGTCTGAACTCTTTCCCTACACGACGCTCTTCCGATCTNNNCCACCACCATACCACTTTTG
AGCACACGTCTGAACTCTTTCCCTACACGACGCTCTTCCGATCTNNNCTATGTATTTTCCGGGGTTGG
AGAGCACACGTCTGAACTCTTTCCCTACACGACGCTCTTCCGATCTNNNACAGGTCCTCTTGGTACAAT
AGAGCACACGTCTGAACTCTTTCCCTACACGACGCTCTTCCGATCTNNNTGTAATTCCAGCTAATCGGG

AGAGCACACGTCTGAACTCTTTCCCTACACGACGCTCTTCCGATCTNNNGCAATTGCAAAAATGTGGA
AGAGCACACGTCTGAACTCTTTCCCTACACGACGCTCTTCCGATCTNNNGTGGGCTAGCATGCAGTAA
AAGAGCACACGTCTGAACTCTTTCCCTACACGACGCTCTTCCGATCTNNNGGAACAACAGGAAGAGTA
GAGCACACGTCTGAACTCTTTCCCTACACGACGCTCTTCCGATCTNNNGGTATCTCACCTATTTGTCATC
GAGCACACGTCTGAACTCTTTCCCTACACGACGCTCTTCCGATCTNNNTCTATATCTCCCTGAAATGATG
GAAGAGCACACGTCTGAACTCTTTCCCTACACGACGCTCTTCCGATCTNNNAGTGAGCTACGATCATGT
AGAGCACACGTCTGAACTCTTTCCCTACACGACGCTCTTCCGATCTNNNTCCTCCCATCTCATCCTCTTAC
AGCACACGTCTGAACTCTTTCCCTACACGACGCTCTTCCGATCTNNNTGAAGCACTTCCATACTATTCTC
GAGCACACGTCTGAACTCTTTCCCTACACGACGCTCTTCCGATCTNNNTCCCTTTGTGATTGATATCTCT
GAGCACACGTCTGAACTCTTTCCCTACACGACGCTCTTCCGATCTNNNACTGAGTCACAAACAACATCA
AGAGCACACGTCTGAACTCTTTCCCTACACGACGCTCTTCCGATCTNNNACCTTTACTCATTGATGGAGA
AGAGCACACGTCTGAACTCTTTCCCTACACGACGCTCTTCCGATCTNNNCAACTGAGTACACATACACA
GAAGAGCACACGTCTGAACTCTTTCCCTACACGACGCTCTTCCGATCTNNNGTGAGAGAGTGCGTGTC
AGCACACGTCTGAACTCTTTCCCTACACGACGCTCTTCCGATCTNNNGCAATTGCAAAAATATGAGACC
AGCACACGTCTGAACTCTTTCCCTACACGACGCTCTTCCGATCTNNNTGTAGATACAGACCAAATCAAC
AGAGCACACGTCTGAACTCTTTCCCTACACGACGCTCTTCCGATCTNNNAAGAATTGCCTGAACCTGG
AGCACACGTCTGAACTCTTTCCCTACACGACGCTCTTCCGATCTNNNACCTAAATGTAAGGTATGCTTGA
GAGCACACGTCTGAACTCTTTCCCTACACGACGCTCTTCCGATCTNNNAGTGTGGGATAATTATGGCTA
GAGCACACGTCTGAACTCTTTCCCTACACGACGCTCTTCCGATCTNNNTGAAACAATGGAAGATGTAGC
AGAGCACACGTCTGAACTCTTTCCCTACACGACGCTCTTCCGATCTNNNTTATAACAGAGGTGGGTGA
GAGCACACGTCTGAACTCTTTCCCTACACGACGCTCTTCCGATCTNNNAGCCTTGTAAAATGAACTGGA
GAGCACACGTCTGAACTCTTTCCCTACACGACGCTCTTCCGATCTNNNTGTGCTTTGTATCCTGTATCCT
AGAGCACACGTCTGAACTCTTTCCCTACACGACGCTCTTCCGATCTNNNAGCATAGTTATTTATGGCCT
AGAGCACACGTCTGAACTCTTTCCCTACACGACGCTCTTCCGATCTNNNACTGTGGTTATGTAAGAGGT
AGAGCACACGTCTGAACTCTTTCCCTACACGACGCTCTTCCGATCTNNNTGTTTGTTTCCTTGCATCTGCT
AGCACACGTCTGAACTCTTTCCCTACACGACGCTCTTCCGATCTNNNTCCCTCTCTCTCTCTCTCTCTCA
AGCACACGTCTGAACTCTTTCCCTACACGACGCTCTTCCGATCTNNNAGAAAGAGAGAGACAGAGAG
GCACACGTCTGAACTCTTTCCCTACACGACGCTCTTCCGATCTNNNAGTGTTCTTTTCCTCCAAAGTCTC
AGAGCACACGTCTGAACTCTTTCCCTACACGACGCTCTTCCGATCTNNNGATACACACACACACACACAC
AGAGCACACGTCTGAACTCTTTCCCTACACGACGCTCTTCCGATCTNNNGTGAGACCTCTCTCTCTCTCT
GCACACGTCTGAACTCTTTCCCTACACGACGCTCTTCCGATCTNNNTGGTGGAGAAATTGTATTTTTGC
AGCACACGTCTGAACTCTTTCCCTACACGACGCTCTTCCGATCTNNNACAGTGATTATTACAGTGACTG
AGAGCACACGTCTGAACTCTTTCCCTACACGACGCTCTTCCGATCTNNNCAGTCAATTGGCCTTGTAAA
GAGCACACGTCTGAACTCTTTCCCTACACGACGCTCTTCCGATCTNNNAGGCATATATATGTGTGTGTC
AGCACACGTCTGAACTCTTTCCCTACACGACGCTCTTCCGATCTNNNTGTGAATTCCTATACCATCCACAC
AAGAGCACACGTCTGAACTCTTTCCCTACACGACGCTCTTCCGATCTNNNAATGCAGAGTAAACGTGTC
AGAGCACACGTCTGAACTCTTTCCCTACACGACGCTCTTCCGATCTNNNTTTCAGCTTGCAAATAGCCTC
AGAGCACACGTCTGAACTCTTTCCCTACACGACGCTCTTCCGATCTNNNTAGTTCTCTAGCTTGCAGACA
GAGCACACGTCTGAACTCTTTCCCTACACGACGCTCTTCCGATCTNNNACTTTCTGTGTGAAGCAATCA
GAGCACACGTCTGAACTCTTTCCCTACACGACGCTCTTCCGATCTNNNACTCCATAATAAGTCAAGGACC
AGCACACGTCTGAACTCTTTCCCTACACGACGCTCTTCCGATCTNNNTTGGATCCCAACTATTTCTTTCTC
AGAGCACACGTCTGAACTCTTTCCCTACACGACGCTCTTCCGATCTNNNTAGAGTGGAGATGACAGA
AAGAGCACACGTCTGAACTCTTTCCCTACACGACGCTCTTCCGATCTNNNTCCTGTATTGTGCATCTCTC

GAGCACACGTCTGAACTCTTTCCCTACACGACGCTCTTCCGATCTNNNTTTACCTTTACTACAGCCATAT
GAGCACACGTCTGAACTCTTTCCCTACACGACGCTCTTCCGATCTNNNTGTCTTTCTGTATCTCTCTCTCT
GGAAGAGCACACGTCTGAACTCTTTCCCTACACGACGCTCTTCCGATCTNNNTTTCCCTTTAGGCTCATG
GGAAGAGCACACGTCTGAACTCTTTCCCTACACGACGCTCTTCCGATCTNNNAAGGGAGGAAGAAAG
AAGAGCACACGTCTGAACTCTTTCCCTACACGACGCTCTTCCGATCTNNNCTGTAGGGCTGCAAGTCAT
AAGAGCACACGTCTGAACTCTTTCCCTACACGACGCTCTTCCGATCTNNNCTGTAATTCCAGCTACTCG
AGAGCACACGTCTGAACTCTTTCCCTACACGACGCTCTTCCGATCTNNNCAGGAGCATGAAAATAGCA
AAGAGCACACGTCTGAACTCTTTCCCTACACGACGCTCTTCCGATCTNNNGAGGCAGTAGAATTGCTT
GAGCACACGTCTGAACTCTTTCCCTACACGACGCTCTTCCGATCTNNNTGGCATAGAGTTATTTCTCCC
AGAGCACACGTCTGAACTCTTTCCCTACACGACGCTCTTCCGATCTNNNATACAAAAAATTAGCCGGG
AGAGCACACGTCTGAACTCTTTCCCTACACGACGCTCTTCCGATCTNNNTGGCCTGGCAGATATAAAC
AGAGCACACGTCTGAACTCTTTCCCTACACGACGCTCTTCCGATCTNNNAGCAAATGAGTGACTTTGT
GAGCACACGTCTGAACTCTTTCCCTACACGACGCTCTTCCGATCTNNNTCATCAAAGTTACAGAGTTCTC
AGCACACGTCTGAACTCTTTCCCTACACGACGCTCTTCCGATCTNNNTGTAGAGATTTTTCACCTCTCC
AGCACACGTCTGAACTCTTTCCCTACACGACGCTCTTCCGATCTNNNGCTAATCTATATCCTGTGGACA
AGCACACGTCTGAACTCTTTCCCTACACGACGCTCTTCCGATCTNNNTTTGCATCTCATTTAGTAAGCA
AGAGCACACGTCTGAACTCTTTCCCTACACGACGCTCTTCCGATCTNNNTGTAGTCTCAGCTACTCAG
AGAGCACACGTCTGAACTCTTTCCCTACACGACGCTCTTCCGATCTNNNCTGTAATCCCAGCTACTCCA
AGCACACGTCTGAACTCTTTCCCTACACGACGCTCTTCCGATCTNNNGGAGAATTTTCCTTTTGCTTGA
GAGCACACGTCTGAACTCTTTCCCTACACGACGCTCTTCCGATCTNNNTCAGAGACTAAGATTGTAACC
GAGCACACGTCTGAACTCTTTCCCTACACGACGCTCTTCCGATCTNNNTGAGGTAGGAGAATTGCTTG
GCACACGTCTGAACTCTTTCCCTACACGACGCTCTTCCGATCTNNNGCAAACAGTTCTTCTAAAGCTTT
GAGCACACGTCTGAACTCTTTCCCTACACGACGCTCTTCCGATCTNNNTGGCTTTGAATTTATCTGTTC
AGCACACGTCTGAACTCTTTCCCTACACGACGCTCTTCCGATCTNNNTCCTGCCTTTTTTTTTGTATTG
AGAGCACACGTCTGAACTCTTTCCCTACACGACGCTCTTCCGATCTNNNAGTACAACATAAGGCTGAG
GAAGAGCACACGTCTGAACTCTTTCCCTACACGACGCTCTTCCGATCTNNNGAGAATCGCTTGAACCCA
AGCACACGTCTGAACTCTTTCCCTACACGACGCTCTTCCGATCTNNNAGACTGCTATCAAAACCAAACT
AGAGCACACGTCTGAACTCTTTCCCTACACGACGCTCTTCCGATCTNNNAGCCAGACTAACAAAAGAGG
AGAGCACACGTCTGAACTCTTTCCCTACACGACGCTCTTCCGATCTNNNTGTAATCTCAGCACTTTGGG
AAGAGCACACGTCTGAACTCTTTCCCTACACGACGCTCTTCCGATCTNNNCACCTGTAGTCCCAGCTAA
AAGAGCACACGTCTGAACTCTTTCCCTACACGACGCTCTTCCGATCTNNNGCCTCTTGTGTCTGGCTTAT
GAGCACACGTCTGAACTCTTTCCCTACACGACGCTCTTCCGATCTNNNTGCGTATTCTGCATAAAATTG
GGAAGAGCACACGTCTGAACTCTTTCCCTACACGACGCTCTTCCGATCTNNNGTTCTCCTAGTGGTGAC
AAGAGCACACGTCTGAACTCTTTCCCTACACGACGCTCTTCCGATCTNNNCTGTAGTCACAGCTACTCA
AAGAGCACACGTCTGAACTCTTTCCCTACACGACGCTCTTCCGATCTNNNAGAAAACACCAGAGATCC
AAGAGCACACGTCTGAACTCTTTCCCTACACGACGCTCTTCCGATCTNNNGTAATCCCAGCTACTCAAG
GAGCACACGTCTGAACTCTTTCCCTACACGACGCTCTTCCGATCTNNNTGGTTTTCTGGTATTTTGTTG
AGAGCACACGTCTGAACTCTTTCCCTACACGACGCTCTTCCGATCTNNNGTAATCCCAGCTACTCAGAA
AGAGCACACGTCTGAACTCTTTCCCTACACGACGCTCTTCCGATCTNNNAAGCTGAGACAGGAGAAT
AGCACACGTCTGAACTCTTTCCCTACACGACGCTCTTCCGATCTNNNACATTTCAACCTCTAAATCATGC
GCACACGTCTGAACTCTTTCCCTACACGACGCTCTTCCGATCTNNNACAAGTCATCAAGATTCAACAAA
AAGAGCACACGTCTGAACTCTTTCCCTACACGACGCTCTTCCGATCTNNNTGTAATTCCAGCTACTTGG
AAGAGCACACGTCTGAACTCTTTCCCTACACGACGCTCTTCCGATCTNNNCAAGGCAGGAGAATCGTT

AGAGCACACGTCTGAACTCTTTCCCTACACGACGCTCTTCCGATCTNNNAAAAATAAAGTGGGTGTG
GAGCACACGTCTGAACTCTTTCCCTACACGACGCTCTTCCGATCTNNNTGTGGACTAGAACATACATG
GGAAGAGCACACGTCTGAACTCTTTCCCTACACGACGCTCTTCCGATCTNNNGAAGCAGGAGAATCG
AGAGCACACGTCTGAACTCTTTCCCTACACGACGCTCTTCCGATCTNNNTGTAATTCCAGCTACTTGGG
AAGAGCACACGTCTGAACTCTTTCCCTACACGACGCTCTTCCGATCTNNNGTGCCTGTAATCCTAGCTA
GAAGAGCACACGTCTGAACTCTTTCCCTACACGACGCTCTTCCGATCTNNNGAGAATGGTGTGAACCC
GGAAGAGCACACGTCTGAACTCTTTCCCTACACGACGCTCTTCCGATCTNNNAAGAGGCTGAGACAGG
AAGAGCACACGTCTGAACTCTTTCCCTACACGACGCTCTTCCGATCTNNNAGGAGAATTGCCTGAAC
AGAGCACACGTCTGAACTCTTTCCCTACACGACGCTCTTCCGATCTNNNACATTCATAAAGGCCAAAC
AGAGCACACGTCTGAACTCTTTCCCTACACGACGCTCTTCCGATCTNNNCATGAGAATTGCTTGAACC
GAGCACACGTCTGAACTCTTTCCCTACACGACGCTCTTCCGATCTNNNAAAAAATTAGCCAGGTGTG
AAGAGCACACGTCTGAACTCTTTCCCTACACGACGCTCTTCCGATCTNNNGCCCAATGGTATGGTGTT
GAGCACACGTCTGAACTCTTTCCCTACACGACGCTCTTCCGATCTNNNTGTGCTTATAGTCCCAGCAA
AGAGCACACGTCTGAACTCTTTCCCTACACGACGCTCTTCCGATCTNNNCTGTAATCCCAGCTACTTCG
GAGCACACGTCTGAACTCTTTCCCTACACGACGCTCTTCCGATCTNNNTACAAAAACATTAGCTGGGC
AGAGCACACGTCTGAACTCTTTCCCTACACGACGCTCTTCCGATCTNNNAAGATGGTGCCATTGTACT
AGAGCACACGTCTGAACTCTTTCCCTACACGACGCTCTTCCGATCTNNNAGAAGAATCGTTTGAACCC
AAGAGCACACGTCTGAACTCTTTCCCTACACGACGCTCTTCCGATCTNNNAATGAGCTATGATCGCGTC
AGAGCACACGTCTGAACTCTTTCCCTACACGACGCTCTTCCGATCTNNNGGAGGCTGAGGCAGAATA
GAGCACACGTCTGAACTCTTTCCCTACACGACGCTCTTCCGATCTNNNTTTAGAAAACACTAAAGACTC
AAGAGCACACGTCTGAACTCTTTCCCTACACGACGCTCTTCCGATCTNNNACAGAGTCAAGCTGATAG
AGAGCACACGTCTGAACTCTTTCCCTACACGACGCTCTTCCGATCTNNNAGGAAGGAGAATTGCTTGA
AAGAGCACACGTCTGAACTCTTTCCCTACACGACGCTCTTCCGATCTNNNAAAAAAATTAGCCGGGCG
AAGAGCACACGTCTGAACTCTTTCCCTACACGACGCTCTTCCGATCTNNNGATGCCTGTAGTCTCAGCT
AGAGCACACGTCTGAACTCTTTCCCTACACGACGCTCTTCCGATCTNNNTCTCAAGGAATCTAGAGGA
AAGAGCACACGTCTGAACTCTTTCCCTACACGACGCTCTTCCGATCTNNNGAGTTTGCAGTGAGCTGA
AAGAGCACACGTCTGAACTCTTTCCCTACACGACGCTCTTCCGATCTNNNAGGCTGAGGTAGAATTG
AAGAGCACACGTCTGAACTCTTTCCCTACACGACGCTCTTCCGATCTNNNCTGTAATTCCAGCTACTCG
GAGCACACGTCTGAACTCTTTCCCTACACGACGCTCTTCCGATCTNNNACTTTATGTCCACCATACTGC
GAGCACACGTCTGAACTCTTTCCCTACACGACGCTCTTCCGATCTNNNACTTTATGTCCACCATACTGC
GAGCACACGTCTGAACTCTTTCCCTACACGACGCTCTTCCGATCTNNNACTTTATGTCCACCATACTGC
GAGCACACGTCTGAACTCTTTCCCTACACGACGCTCTTCCGATCTNNNACTTTATGTCCACCATACTGC
GAGCACACGTCTGAACTCTTTCCCTACACGACGCTCTTCCGATCTNNNACTTTATGTCCACCATACTGC
GAGCACACGTCTGAACTCTTTCCCTACACGACGCTCTTCCGATCTNNNACTTTATGTCCACCATACTGC
GAGCACACGTCTGAACTCTTTCCCTACACGACGCTCTTCCGATCTNNNACTTTATGTCCACCATACTGC
GAGCACACGTCTGAACTCTTTCCCTACACGACGCTCTTCCGATCTNNNACTTTATGTCCACCATACTGC
GCACACGTCTGAACTCTTTCCCTACACGACGCTCTTCCGATCTNNNTGGACCTCTGTTTCTGTTCTTCTC
GCACACGTCTGAACTCTTTCCCTACACGACGCTCTTCCGATCTNNNTGGACCTCTGTTTCTGTTCTTCTC
GCACACGTCTGAACTCTTTCCCTACACGACGCTCTTCCGATCTNNNTGGACCTCTGTTTCTGTTCTTCTC
GCACACGTCTGAACTCTTTCCCTACACGACGCTCTTCCGATCTNNNTGGACCTCTGTTTCTGTTCTTCTC
GCACACGTCTGAACTCTTTCCCTACACGACGCTCTTCCGATCTNNNTGGACCTCTGTTTCTGTTCTTCTC
GCACACGTCTGAACTCTTTCCCTACACGACGCTCTTCCGATCTNNNTGGACCTCTGTTTCTGTTCTTCTC
GCACACGTCTGAACTCTTTCCCTACACGACGCTCTTCCGATCTNNNTGGACCTCTGTTTCTGTTCTTCTC

GCACACGTCTGAACTCTTTCCCTACACGACGCTCTTCCGATCTNNNTGGACCTCTGTTTCTGTTCTTCT
AGCACACGTCTGAACTCTTTCCCTACACGACGCTCTTCCGATCTNNNCAAGATCTCACCACTGCACTCT
AGCACACGTCTGAACTCTTTCCCTACACGACGCTCTTCCGATCTNNNCAAGATCTCACCACTGCACTCT
AGCACACGTCTGAACTCTTTCCCTACACGACGCTCTTCCGATCTNNNCAAGATCTCACCACTGCACTCT
AGCACACGTCTGAACTCTTTCCCTACACGACGCTCTTCCGATCTNNNCAAGATCTCACCACTGCACTCT
AGCACACGTCTGAACTCTTTCCCTACACGACGCTCTTCCGATCTNNNCAAGATCTCACCACTGCACTCT
AGCACACGTCTGAACTCTTTCCCTACACGACGCTCTTCCGATCTNNNCAAGATCTCACCACTGCACTCT
AGCACACGTCTGAACTCTTTCCCTACACGACGCTCTTCCGATCTNNNCAAGATCTCACCACTGCACTCT
AGCACACGTCTGAACTCTTTCCCTACACGACGCTCTTCCGATCTNNNCAAGATCTCACCACTGCACTCT
AGCACACGTCTGAACTCTTTCCCTACACGACGCTCTTCCGATCTNNNCTGAGGCAGAAGAATCACCTG
AGCACACGTCTGAACTCTTTCCCTACACGACGCTCTTCCGATCTNNNCTGAGGCAGAAGAATCACCTG
AGCACACGTCTGAACTCTTTCCCTACACGACGCTCTTCCGATCTNNNCTGAGGCAGAAGAATCACCTG
AGCACACGTCTGAACTCTTTCCCTACACGACGCTCTTCCGATCTNNNCTGAGGCAGAAGAATCACCTG
AGCACACGTCTGAACTCTTTCCCTACACGACGCTCTTCCGATCTNNNCTGAGGCAGAAGAATCACCTG
AGCACACGTCTGAACTCTTTCCCTACACGACGCTCTTCCGATCTNNNCTGAGGCAGAAGAATCACCTG
AGCACACGTCTGAACTCTTTCCCTACACGACGCTCTTCCGATCTNNNCTGAGGCAGAAGAATCACCTG
AGCACACGTCTGAACTCTTTCCCTACACGACGCTCTTCCGATCTNNNCTGAGGCAGAAGAATCACCTG
GCACACGTCTGAACTCTTTCCCTACACGACGCTCTTCCGATCTNNNTCACCTGTAATCCCAGCTACTTG
GCACACGTCTGAACTCTTTCCCTACACGACGCTCTTCCGATCTNNNTCACCTGTAATCCCAGCTACTTG
GCACACGTCTGAACTCTTTCCCTACACGACGCTCTTCCGATCTNNNTCACCTGTAATCCCAGCTACTTG
GCACACGTCTGAACTCTTTCCCTACACGACGCTCTTCCGATCTNNNTCACCTGTAATCCCAGCTACTTG
GCACACGTCTGAACTCTTTCCCTACACGACGCTCTTCCGATCTNNNTCACCTGTAATCCCAGCTACTTG
GCACACGTCTGAACTCTTTCCCTACACGACGCTCTTCCGATCTNNNTCACCTGTAATCCCAGCTACTTG
GCACACGTCTGAACTCTTTCCCTACACGACGCTCTTCCGATCTNNNTCACCTGTAATCCCAGCTACTTG
GCACACGTCTGAACTCTTTCCCTACACGACGCTCTTCCGATCTNNNTCACCTGTAATCCCAGCTACTTG
GAGCACACGTCTGAACTCTTTCCCTACACGACGCTCTTCCGATCTNNNCTAGGTTTACAGCTGTGAGC
GAGCACACGTCTGAACTCTTTCCCTACACGACGCTCTTCCGATCTNNNCTAGGTTTACAGCTGTGAGC
GAGCACACGTCTGAACTCTTTCCCTACACGACGCTCTTCCGATCTNNNCTAGGTTTACAGCTGTGAGC
GAGCACACGTCTGAACTCTTTCCCTACACGACGCTCTTCCGATCTNNNCTAGGTTTACAGCTGTGAGC
GAGCACACGTCTGAACTCTTTCCCTACACGACGCTCTTCCGATCTNNNCTAGGTTTACAGCTGTGAGC
GAGCACACGTCTGAACTCTTTCCCTACACGACGCTCTTCCGATCTNNNCTAGGTTTACAGCTGTGAGC
GAGCACACGTCTGAACTCTTTCCCTACACGACGCTCTTCCGATCTNNNCTAGGTTTACAGCTGTGAGC
GAGCACACGTCTGAACTCTTTCCCTACACGACGCTCTTCCGATCTNNNCTAGGTTTACAGCTGTGAGC
AGCACACGTCTGAACTCTTTCCCTACACGACGCTCTTCCGATCTNNNATACATACTTGCACACACACCCT
AGCACACGTCTGAACTCTTTCCCTACACGACGCTCTTCCGATCTNNNATACATACTTGCACACACACCCT
AGCACACGTCTGAACTCTTTCCCTACACGACGCTCTTCCGATCTNNNATACATACTTGCACACACACCCT
AGCACACGTCTGAACTCTTTCCCTACACGACGCTCTTCCGATCTNNNATACATACTTGCACACACACCCT
AGCACACGTCTGAACTCTTTCCCTACACGACGCTCTTCCGATCTNNNATACATACTTGCACACACACCCT
AGCACACGTCTGAACTCTTTCCCTACACGACGCTCTTCCGATCTNNNATACATACTTGCACACACACCCT
AGCACACGTCTGAACTCTTTCCCTACACGACGCTCTTCCGATCTNNNATACATACTTGCACACACACCCT
AGCACACGTCTGAACTCTTTCCCTACACGACGCTCTTCCGATCTNNNATACATACTTGCACACACACCCT
AGCACACGTCTGAACTCTTTCCCTACACGACGCTCTTCCGATCTNNNATAGAGCAAGCATCTGCATAC
AGCACACGTCTGAACTCTTTCCCTACACGACGCTCTTCCGATCTNNNATAGAGCAAGCATCTGCATAC

AGCACACGTCTGAACTCTTTCCCTACACGACGCTCTTCCGATCTNNNATAGAGCAAGCATCTGCATAC
AGCACACGTCTGAACTCTTTCCCTACACGACGCTCTTCCGATCTNNNATAGAGCAAGCATCTGCATAC
AGCACACGTCTGAACTCTTTCCCTACACGACGCTCTTCCGATCTNNNATAGAGCAAGCATCTGCATAC
AGCACACGTCTGAACTCTTTCCCTACACGACGCTCTTCCGATCTNNNATAGAGCAAGCATCTGCATAC
AGCACACGTCTGAACTCTTTCCCTACACGACGCTCTTCCGATCTNNNATAGAGCAAGCATCTGCATAC
AGCACACGTCTGAACTCTTTCCCTACACGACGCTCTTCCGATCTNNNATAGAGCAAGCATCTGCATAC
AGCACACGTCTGAACTCTTTCCCTACACGACGCTCTTCCGATCTNNNCCACCATACTTCTCTGCCTCAAA
AGCACACGTCTGAACTCTTTCCCTACACGACGCTCTTCCGATCTNNNCCACCATACTTCTCTGCCTCAAA
AGCACACGTCTGAACTCTTTCCCTACACGACGCTCTTCCGATCTNNNCCACCATACTTCTCTGCCTCAAA
AGCACACGTCTGAACTCTTTCCCTACACGACGCTCTTCCGATCTNNNCCACCATACTTCTCTGCCTCAAA
AGCACACGTCTGAACTCTTTCCCTACACGACGCTCTTCCGATCTNNNCCACCATACTTCTCTGCCTCAAA
AGCACACGTCTGAACTCTTTCCCTACACGACGCTCTTCCGATCTNNNCCACCATACTTCTCTGCCTCAAA
AGCACACGTCTGAACTCTTTCCCTACACGACGCTCTTCCGATCTNNNCCACCATACTTCTCTGCCTCAAA
AGCACACGTCTGAACTCTTTCCCTACACGACGCTCTTCCGATCTNNNCCACCATACTTCTCTGCCTCAAA
AGCACACGTCTGAACTCTTTCCCTACACGACGCTCTTCCGATCTNNNGAGAATCACTTGAACCCGGGA
AGCACACGTCTGAACTCTTTCCCTACACGACGCTCTTCCGATCTNNNGAGAATCACTTGAACCCGGGA
AGCACACGTCTGAACTCTTTCCCTACACGACGCTCTTCCGATCTNNNGAGAATCACTTGAACCCGGGA
AGCACACGTCTGAACTCTTTCCCTACACGACGCTCTTCCGATCTNNNGAGAATCACTTGAACCCGGGA
AGCACACGTCTGAACTCTTTCCCTACACGACGCTCTTCCGATCTNNNGAGAATCACTTGAACCCGGGA
AGCACACGTCTGAACTCTTTCCCTACACGACGCTCTTCCGATCTNNNGAGAATCACTTGAACCCGGGA
AGCACACGTCTGAACTCTTTCCCTACACGACGCTCTTCCGATCTNNNGAGAATCACTTGAACCCGGGA
AGCACACGTCTGAACTCTTTCCCTACACGACGCTCTTCCGATCTNNNGAGAATCACTTGAACCCGGGA
AGCACACGTCTGAACTCTTTCCCTACACGACGCTCTTCCGATCTNNNTCCTGGAGAACAAGAGCAAAA
AGCACACGTCTGAACTCTTTCCCTACACGACGCTCTTCCGATCTNNNTCCTGGAGAACAAGAGCAAAA
AGCACACGTCTGAACTCTTTCCCTACACGACGCTCTTCCGATCTNNNTCCTGGAGAACAAGAGCAAAA
AGCACACGTCTGAACTCTTTCCCTACACGACGCTCTTCCGATCTNNNTCCTGGAGAACAAGAGCAAAA
AGCACACGTCTGAACTCTTTCCCTACACGACGCTCTTCCGATCTNNNTCCTGGAGAACAAGAGCAAAA
AGCACACGTCTGAACTCTTTCCCTACACGACGCTCTTCCGATCTNNNTCCTGGAGAACAAGAGCAAAA
AGCACACGTCTGAACTCTTTCCCTACACGACGCTCTTCCGATCTNNNTCCTGGAGAACAAGAGCAAAA
AGCACACGTCTGAACTCTTTCCCTACACGACGCTCTTCCGATCTNNNTCCTGGAGAACAAGAGCAAAA
AGCACACGTCTGAACTCTTTCCCTACACGACGCTCTTCCGATCTNNNATCACAGAGGTAGTGGGAGG
AGCACACGTCTGAACTCTTTCCCTACACGACGCTCTTCCGATCTNNNATCACAGAGGTAGTGGGAGG
AGCACACGTCTGAACTCTTTCCCTACACGACGCTCTTCCGATCTNNNATCACAGAGGTAGTGGGAGG
AGCACACGTCTGAACTCTTTCCCTACACGACGCTCTTCCGATCTNNNATCACAGAGGTAGTGGGAGG
AGCACACGTCTGAACTCTTTCCCTACACGACGCTCTTCCGATCTNNNATCACAGAGGTAGTGGGAGG
AGCACACGTCTGAACTCTTTCCCTACACGACGCTCTTCCGATCTNNNATCACAGAGGTAGTGGGAGG
AGCACACGTCTGAACTCTTTCCCTACACGACGCTCTTCCGATCTNNNATCACAGAGGTAGTGGGAGG
AGCACACGTCTGAACTCTTTCCCTACACGACGCTCTTCCGATCTNNNATCACAGAGGTAGTGGGAGG
GAGCACACGTCTGAACTCTTTCCCTACACGACGCTCTTCCGATCTNNNTGAGGTAGGAGAATGGCAT
GAGCACACGTCTGAACTCTTTCCCTACACGACGCTCTTCCGATCTNNNTGAGGTAGGAGAATGGCAT
GAGCACACGTCTGAACTCTTTCCCTACACGACGCTCTTCCGATCTNNNTGAGGTAGGAGAATGGCAT
GAGCACACGTCTGAACTCTTTCCCTACACGACGCTCTTCCGATCTNNNTGAGGTAGGAGAATGGCAT
GAGCACACGTCTGAACTCTTTCCCTACACGACGCTCTTCCGATCTNNNTGAGGTAGGAGAATGGCAT

AGCACACGTCTGAACTCTTTCCCTACACGACGCTCTTCCGATCTNNNTGAGGTAGGAGAATGGCAT
AGCACACGTCTGAACTCTTTCCCTACACGACGCTCTTCCGATCTNNNTGAGGTAGGAGAATGGCAT
AGCACACGTCTGAACTCTTTCCCTACACGACGCTCTTCCGATCTNNNTGAGGTAGGAGAATGGCAT
GAGCACACGTCTGAACTCTTTCCCTACACGACGCTCTTCCGATCTNNNTATACTGGAGGCTGAGGTA
GAGCACACGTCTGAACTCTTTCCCTACACGACGCTCTTCCGATCTNNNTATACTGGAGGCTGAGGTA
GAGCACACGTCTGAACTCTTTCCCTACACGACGCTCTTCCGATCTNNNTATACTGGAGGCTGAGGTA
GAGCACACGTCTGAACTCTTTCCCTACACGACGCTCTTCCGATCTNNNTATACTGGAGGCTGAGGTA
GAGCACACGTCTGAACTCTTTCCCTACACGACGCTCTTCCGATCTNNNTATACTGGAGGCTGAGGTA
GAGCACACGTCTGAACTCTTTCCCTACACGACGCTCTTCCGATCTNNNTATACTGGAGGCTGAGGTA
GAGCACACGTCTGAACTCTTTCCCTACACGACGCTCTTCCGATCTNNNTATACTGGAGGCTGAGGTA
GAGCACACGTCTGAACTCTTTCCCTACACGACGCTCTTCCGATCTNNNTATACTGGAGGCTGAGGTA
GAGCACACGTCTGAACTCTTTCCCTACACGACGCTCTTCCGATCTNNNTAAAAATTCAACCATATCTGG
GAGCACACGTCTGAACTCTTTCCCTACACGACGCTCTTCCGATCTNNNTAAAAATTCAACCATATCTGG
GAGCACACGTCTGAACTCTTTCCCTACACGACGCTCTTCCGATCTNNNTAAAAATTCAACCATATCTGG
GAGCACACGTCTGAACTCTTTCCCTACACGACGCTCTTCCGATCTNNNTAAAAATTCAACCATATCTGG
GAGCACACGTCTGAACTCTTTCCCTACACGACGCTCTTCCGATCTNNNTAAAAATTCAACCATATCTGG
GAGCACACGTCTGAACTCTTTCCCTACACGACGCTCTTCCGATCTNNNTAAAAATTCAACCATATCTGG
GAGCACACGTCTGAACTCTTTCCCTACACGACGCTCTTCCGATCTNNNTAAAAATTCAACCATATCTGG
GAGCACACGTCTGAACTCTTTCCCTACACGACGCTCTTCCGATCTNNNTAAAAATTCAACCATATCTGG
AGCACACGTCTGAACTCTTTCCCTACACGACGCTCTTCCGATCTNNNCAGTATTCTCAGGCAGTGTGG
AGCACACGTCTGAACTCTTTCCCTACACGACGCTCTTCCGATCTNNNCAGTATTCTCAGGCAGTGTGG
AGCACACGTCTGAACTCTTTCCCTACACGACGCTCTTCCGATCTNNNCAGTATTCTCAGGCAGTGTGG
AGCACACGTCTGAACTCTTTCCCTACACGACGCTCTTCCGATCTNNNCAGTATTCTCAGGCAGTGTGG
AGCACACGTCTGAACTCTTTCCCTACACGACGCTCTTCCGATCTNNNCAGTATTCTCAGGCAGTGTGG
AGCACACGTCTGAACTCTTTCCCTACACGACGCTCTTCCGATCTNNNCAGTATTCTCAGGCAGTGTGG
AGCACACGTCTGAACTCTTTCCCTACACGACGCTCTTCCGATCTNNNCAGTATTCTCAGGCAGTGTGG
AGCACACGTCTGAACTCTTTCCCTACACGACGCTCTTCCGATCTNNNCAGTATTCTCAGGCAGTGTGG
AGCACACGTCTGAACTCTTTCCCTACACGACGCTCTTCCGATCTNNNACAAACAACGGAAAGATGTG
AGCACACGTCTGAACTCTTTCCCTACACGACGCTCTTCCGATCTNNNACAAACAACGGAAAGATGTG
AGCACACGTCTGAACTCTTTCCCTACACGACGCTCTTCCGATCTNNNACAAACAACGGAAAGATGTG
AGCACACGTCTGAACTCTTTCCCTACACGACGCTCTTCCGATCTNNNACAAACAACGGAAAGATGTG
AGCACACGTCTGAACTCTTTCCCTACACGACGCTCTTCCGATCTNNNACAAACAACGGAAAGATGTG
AGCACACGTCTGAACTCTTTCCCTACACGACGCTCTTCCGATCTNNNACAAACAACGGAAAGATGTG
AGCACACGTCTGAACTCTTTCCCTACACGACGCTCTTCCGATCTNNNACAAACAACGGAAAGATGTG
AGCACACGTCTGAACTCTTTCCCTACACGACGCTCTTCCGATCTNNNACAAACAACGGAAAGATGTG
ACACGTCTGAACTCTTTCCCTACACGACGCTCTTCCGATCTNNNAGCTGTGAAATTACTCAGATGTTG
ACACGTCTGAACTCTTTCCCTACACGACGCTCTTCCGATCTNNNAGCTGTGAAATTACTCAGATGTTG
ACACGTCTGAACTCTTTCCCTACACGACGCTCTTCCGATCTNNNAGCTGTGAAATTACTCAGATGTTG
ACACGTCTGAACTCTTTCCCTACACGACGCTCTTCCGATCTNNNAGCTGTGAAATTACTCAGATGTTG
ACACGTCTGAACTCTTTCCCTACACGACGCTCTTCCGATCTNNNAGCTGTGAAATTACTCAGATGTTG
ACACGTCTGAACTCTTTCCCTACACGACGCTCTTCCGATCTNNNAGCTGTGAAATTACTCAGATGTTG
ACACGTCTGAACTCTTTCCCTACACGACGCTCTTCCGATCTNNNAGCTGTGAAATTACTCAGATGTTG
ACACGTCTGAACTCTTTCCCTACACGACGCTCTTCCGATCTNNNAGCTGTGAAATTACTCAGATGTTG

AGCACACGTCTGAACTCTTTCCCTACACGACGCTCTTCCGATCTNNNTGCAGAGAGCCAGATAAAGACA
AGCACACGTCTGAACTCTTTCCCTACACGACGCTCTTCCGATCTNNNTGCAGAGAGCCAGATAAAGACA
AGCACACGTCTGAACTCTTTCCCTACACGACGCTCTTCCGATCTNNNTGCAGAGAGCCAGATAAAGACA
AGCACACGTCTGAACTCTTTCCCTACACGACGCTCTTCCGATCTNNNTGCAGAGAGCCAGATAAAGACA
AGCACACGTCTGAACTCTTTCCCTACACGACGCTCTTCCGATCTNNNTGCAGAGAGCCAGATAAAGACA
AGCACACGTCTGAACTCTTTCCCTACACGACGCTCTTCCGATCTNNNTGCAGAGAGCCAGATAAAGACA
AGCACACGTCTGAACTCTTTCCCTACACGACGCTCTTCCGATCTNNNTGCAGAGAGCCAGATAAAGACA
AGCACACGTCTGAACTCTTTCCCTACACGACGCTCTTCCGATCTNNNTGCAGAGAGCCAGATAAAGACA
CACGTCTGAACTCTTTCCCTACACGACGCTCTTCCGATCTNNNTTGCCACGTTCGTGCTGATGCATCGGA
CACGTCTGAACTCTTTCCCTACACGACGCTCTTCCGATCTNNNTTGCCACGTTCGTGCTGATGCATCGGA
CACGTCTGAACTCTTTCCCTACACGACGCTCTTCCGATCTNNNTTGCCACGTTCGTGCTGATGCATCGGA
CACGTCTGAACTCTTTCCCTACACGACGCTCTTCCGATCTNNNTTGCCACGTTCGTGCTGATGCATCGGA
CACGTCTGAACTCTTTCCCTACACGACGCTCTTCCGATCTNNNTTGCCACGTTCGTGCTGATGCATCGGA
CACGTCTGAACTCTTTCCCTACACGACGCTCTTCCGATCTNNNTTGCCACGTTCGTGCTGATGCATCGGA
CACGTCTGAACTCTTTCCCTACACGACGCTCTTCCGATCTNNNTTGCCACGTTCGTGCTGATGCATCGGA
CACGTCTGAACTCTTTCCCTACACGACGCTCTTCCGATCTNNNTTGCCACGTTCGTGCTGATGCATCGGA
AGAGCACACGTCTGAACTCTTTCCCTACACGACGCTCTTCCGATCTNNNCCTTGTACCCATTAGCACTCA
AGAGCACACGTCTGAACTCTTTCCCTACACGACGCTCTTCCGATCTNNNCCTTGTACCCATTAGCACTCA
AGAGCACACGTCTGAACTCTTTCCCTACACGACGCTCTTCCGATCTNNNCCTTGTACCCATTAGCACTCA
AGAGCACACGTCTGAACTCTTTCCCTACACGACGCTCTTCCGATCTNNNCCTTGTACCCATTAGCACTCA
AGAGCACACGTCTGAACTCTTTCCCTACACGACGCTCTTCCGATCTNNNCCTTGTACCCATTAGCACTCA
AGAGCACACGTCTGAACTCTTTCCCTACACGACGCTCTTCCGATCTNNNCCTTGTACCCATTAGCACTCA
AGAGCACACGTCTGAACTCTTTCCCTACACGACGCTCTTCCGATCTNNNCCTTGTACCCATTAGCACTCA
AGAGCACACGTCTGAACTCTTTCCCTACACGACGCTCTTCCGATCTNNNCCTTGTACCCATTAGCACTCA
GAGCACACGTCTGAACTCTTTCCCTACACGACGCTCTTCCGATCTNNNAACAGCACTGCAAAAGCATGA
GAGCACACGTCTGAACTCTTTCCCTACACGACGCTCTTCCGATCTNNNAACAGCACTGCAAAAGCATGA
GAGCACACGTCTGAACTCTTTCCCTACACGACGCTCTTCCGATCTNNNAACAGCACTGCAAAAGCATGA
GAGCACACGTCTGAACTCTTTCCCTACACGACGCTCTTCCGATCTNNNAACAGCACTGCAAAAGCATGA
GAGCACACGTCTGAACTCTTTCCCTACACGACGCTCTTCCGATCTNNNAACAGCACTGCAAAAGCATGA
GAGCACACGTCTGAACTCTTTCCCTACACGACGCTCTTCCGATCTNNNAACAGCACTGCAAAAGCATGA
GAGCACACGTCTGAACTCTTTCCCTACACGACGCTCTTCCGATCTNNNAACAGCACTGCAAAAGCATGA
GAGCACACGTCTGAACTCTTTCCCTACACGACGCTCTTCCGATCTNNNAACAGCACTGCAAAAGCATGA
AGCACACGTCTGAACTCTTTCCCTACACGACGCTCTTCCGATCTNNNTTTCCTTCTGCTTCTTTCTCTCCCA
AGCACACGTCTGAACTCTTTCCCTACACGACGCTCTTCCGATCTNNNTTTCCTTCTGCTTCTTTCTCTCCCA
AGCACACGTCTGAACTCTTTCCCTACACGACGCTCTTCCGATCTNNNTTTCCTTCTGCTTCTTTCTCTCCCA
AGCACACGTCTGAACTCTTTCCCTACACGACGCTCTTCCGATCTNNNTTTCCTTCTGCTTCTTTCTCTCCCA
AGCACACGTCTGAACTCTTTCCCTACACGACGCTCTTCCGATCTNNNTTTCCTTCTGCTTCTTTCTCTCCCA
AGCACACGTCTGAACTCTTTCCCTACACGACGCTCTTCCGATCTNNNTTTCCTTCTGCTTCTTTCTCTCCCA
AGCACACGTCTGAACTCTTTCCCTACACGACGCTCTTCCGATCTNNNTTTCCTTCTGCTTCTTTCTCTCCCA
AGCACACGTCTGAACTCTTTCCCTACACGACGCTCTTCCGATCTNNNTTTCCTTCTGCTTCTTTCTCTCCCA
GCACACGTCTGAACTCTTTCCCTACACGACGCTCTTCCGATCTNNNGCCCAGAAACAAATCCATGCATAC
GCACACGTCTGAACTCTTTCCCTACACGACGCTCTTCCGATCTNNNGCCCAGAAACAAATCCATGCATAC
GCACACGTCTGAACTCTTTCCCTACACGACGCTCTTCCGATCTNNNGCCCAGAAACAAATCCATGCATAC

CACACGTCTGAACTCTTTCCCTACACGACGCTCTTCCGATCTNNNGCCCAGAAACAAATCCATGCATAC
CACACGTCTGAACTCTTTCCCTACACGACGCTCTTCCGATCTNNNGCCCAGAAACAAATCCATGCATAC
CACACGTCTGAACTCTTTCCCTACACGACGCTCTTCCGATCTNNNGCCCAGAAACAAATCCATGCATAC
CACACGTCTGAACTCTTTCCCTACACGACGCTCTTCCGATCTNNNGCCCAGAAACAAATCCATGCATAC
CACACGTCTGAACTCTTTCCCTACACGACGCTCTTCCGATCTNNNGCCCAGAAACAAATCCATGCATAC
GAGCACACGTCTGAACTCTTTCCCTACACGACGCTCTTCCGATCTNNNGAGGGAAGGACAGATGAAG
GAGCACACGTCTGAACTCTTTCCCTACACGACGCTCTTCCGATCTNNNGAGGGAAGGACAGATGAAG
GAGCACACGTCTGAACTCTTTCCCTACACGACGCTCTTCCGATCTNNNGAGGGAAGGACAGATGAAG
GAGCACACGTCTGAACTCTTTCCCTACACGACGCTCTTCCGATCTNNNGAGGGAAGGACAGATGAAG
GAGCACACGTCTGAACTCTTTCCCTACACGACGCTCTTCCGATCTNNNGAGGGAAGGACAGATGAAG
GAGCACACGTCTGAACTCTTTCCCTACACGACGCTCTTCCGATCTNNNGAGGGAAGGACAGATGAAG
GAGCACACGTCTGAACTCTTTCCCTACACGACGCTCTTCCGATCTNNNGAGGGAAGGACAGATGAAG
GAGCACACGTCTGAACTCTTTCCCTACACGACGCTCTTCCGATCTNNNGAGGGAAGGACAGATGAAG
AGAGCACACGTCTGAACTCTTTCCCTACACGACGCTCTTCCGATCTNNNAGCTCAGAATTTGATGAAA
AGAGCACACGTCTGAACTCTTTCCCTACACGACGCTCTTCCGATCTNNNAGCTCAGAATTTGATGAAA
AGAGCACACGTCTGAACTCTTTCCCTACACGACGCTCTTCCGATCTNNNAGCTCAGAATTTGATGAAA
AGAGCACACGTCTGAACTCTTTCCCTACACGACGCTCTTCCGATCTNNNAGCTCAGAATTTGATGAAA
AGAGCACACGTCTGAACTCTTTCCCTACACGACGCTCTTCCGATCTNNNAGCTCAGAATTTGATGAAA
AGAGCACACGTCTGAACTCTTTCCCTACACGACGCTCTTCCGATCTNNNAGCTCAGAATTTGATGAAA
AGAGCACACGTCTGAACTCTTTCCCTACACGACGCTCTTCCGATCTNNNAGCTCAGAATTTGATGAAA
AGAGCACACGTCTGAACTCTTTCCCTACACGACGCTCTTCCGATCTNNNAGCTCAGAATTTGATGAAA
GAGCACACGTCTGAACTCTTTCCCTACACGACGCTCTTCCGATCTNNNAATCCATGTTGAGAGAACAG
GAGCACACGTCTGAACTCTTTCCCTACACGACGCTCTTCCGATCTNNNAATCCATGTTGAGAGAACAG
GAGCACACGTCTGAACTCTTTCCCTACACGACGCTCTTCCGATCTNNNAATCCATGTTGAGAGAACAG
GAGCACACGTCTGAACTCTTTCCCTACACGACGCTCTTCCGATCTNNNAATCCATGTTGAGAGAACAG
GAGCACACGTCTGAACTCTTTCCCTACACGACGCTCTTCCGATCTNNNAATCCATGTTGAGAGAACAG
GAGCACACGTCTGAACTCTTTCCCTACACGACGCTCTTCCGATCTNNNAATCCATGTTGAGAGAACAG
GAGCACACGTCTGAACTCTTTCCCTACACGACGCTCTTCCGATCTNNNAATCCATGTTGAGAGAACAG
GAGCACACGTCTGAACTCTTTCCCTACACGACGCTCTTCCGATCTNNNAATCCATGTTGAGAGAACAG
AGCACACGTCTGAACTCTTTCCCTACACGACGCTCTTCCGATCTNNNCGCCTGGCTTTCCTTCATTTATT
AGCACACGTCTGAACTCTTTCCCTACACGACGCTCTTCCGATCTNNNCGCCTGGCTTTCCTTCATTTATT
AGCACACGTCTGAACTCTTTCCCTACACGACGCTCTTCCGATCTNNNCGCCTGGCTTTCCTTCATTTATT
AGCACACGTCTGAACTCTTTCCCTACACGACGCTCTTCCGATCTNNNCGCCTGGCTTTCCTTCATTTATT
AGCACACGTCTGAACTCTTTCCCTACACGACGCTCTTCCGATCTNNNCGCCTGGCTTTCCTTCATTTATT
AGCACACGTCTGAACTCTTTCCCTACACGACGCTCTTCCGATCTNNNCGCCTGGCTTTCCTTCATTTATT
AGCACACGTCTGAACTCTTTCCCTACACGACGCTCTTCCGATCTNNNCGCCTGGCTTTCCTTCATTTATT
AGCACACGTCTGAACTCTTTCCCTACACGACGCTCTTCCGATCTNNNCGCCTGGCTTTCCTTCATTTATT
AGCACACGTCTGAACTCTTTCCCTACACGACGCTCTTCCGATCTNNNTCTCTATTTCTCTCCCTCCCTCCC
AGCACACGTCTGAACTCTTTCCCTACACGACGCTCTTCCGATCTNNNTCTCTATTTCTCTCCCTCCCTCCC
AGCACACGTCTGAACTCTTTCCCTACACGACGCTCTTCCGATCTNNNTCTCTATTTCTCTCCCTCCCTCCC
AGCACACGTCTGAACTCTTTCCCTACACGACGCTCTTCCGATCTNNNTCTCTATTTCTCTCCCTCCCTCCC
AGCACACGTCTGAACTCTTTCCCTACACGACGCTCTTCCGATCTNNNTCTCTATTTCTCTCCCTCCCTCCC
AGCACACGTCTGAACTCTTTCCCTACACGACGCTCTTCCGATCTNNNTCTCTATTTCTCTCCCTCCCTCCC

AGCACACGTCTGAACTCTTTCCCTACACGACGCTCTTCCGATCTNNNTCTCTATTTCTCTCCCTCCCTCCCA
AGCACACGTCTGAACTCTTTCCCTACACGACGCTCTTCCGATCTNNNTCTCTATTTCTCTCCCTCCCTCCCA
AGCACACGTCTGAACTCTTTCCCTACACGACGCTCTTCCGATCTNNNGATTCCAGTGCTTCCCTGTTTTG
AGCACACGTCTGAACTCTTTCCCTACACGACGCTCTTCCGATCTNNNGATTCCAGTGCTTCCCTGTTTTG
AGCACACGTCTGAACTCTTTCCCTACACGACGCTCTTCCGATCTNNNGATTCCAGTGCTTCCCTGTTTTG
AGCACACGTCTGAACTCTTTCCCTACACGACGCTCTTCCGATCTNNNGATTCCAGTGCTTCCCTGTTTTG
AGCACACGTCTGAACTCTTTCCCTACACGACGCTCTTCCGATCTNNNGATTCCAGTGCTTCCCTGTTTTG
AGCACACGTCTGAACTCTTTCCCTACACGACGCTCTTCCGATCTNNNGATTCCAGTGCTTCCCTGTTTTG
AGCACACGTCTGAACTCTTTCCCTACACGACGCTCTTCCGATCTNNNGATTCCAGTGCTTCCCTGTTTTG
AGCACACGTCTGAACTCTTTCCCTACACGACGCTCTTCCGATCTNNNGATTCCAGTGCTTCCCTGTTTTG
GAGCACACGTCTGAACTCTTTCCCTACACGACGCTCTTCCGATCTNNNAGAGAACTGCTTGAATCTGGG
GAGCACACGTCTGAACTCTTTCCCTACACGACGCTCTTCCGATCTNNNAGAGAACTGCTTGAATCTGGG
GAGCACACGTCTGAACTCTTTCCCTACACGACGCTCTTCCGATCTNNNAGAGAACTGCTTGAATCTGGG
GAGCACACGTCTGAACTCTTTCCCTACACGACGCTCTTCCGATCTNNNAGAGAACTGCTTGAATCTGGG
GAGCACACGTCTGAACTCTTTCCCTACACGACGCTCTTCCGATCTNNNAGAGAACTGCTTGAATCTGGG
GAGCACACGTCTGAACTCTTTCCCTACACGACGCTCTTCCGATCTNNNAGAGAACTGCTTGAATCTGGG
GAGCACACGTCTGAACTCTTTCCCTACACGACGCTCTTCCGATCTNNNAGAGAACTGCTTGAATCTGGG
GAGCACACGTCTGAACTCTTTCCCTACACGACGCTCTTCCGATCTNNNAGAGAACTGCTTGAATCTGGG
AGAGCACACGTCTGAACTCTTTCCCTACACGACGCTCTTCCGATCTNNNCGACAGAGTGAAAGTCCATG
AGAGCACACGTCTGAACTCTTTCCCTACACGACGCTCTTCCGATCTNNNCGACAGAGTGAAAGTCCATG
AGAGCACACGTCTGAACTCTTTCCCTACACGACGCTCTTCCGATCTNNNCGACAGAGTGAAAGTCCATG
AGAGCACACGTCTGAACTCTTTCCCTACACGACGCTCTTCCGATCTNNNCGACAGAGTGAAAGTCCATG
AGAGCACACGTCTGAACTCTTTCCCTACACGACGCTCTTCCGATCTNNNCGACAGAGTGAAAGTCCATG
AGAGCACACGTCTGAACTCTTTCCCTACACGACGCTCTTCCGATCTNNNCGACAGAGTGAAAGTCCATG
AGAGCACACGTCTGAACTCTTTCCCTACACGACGCTCTTCCGATCTNNNCGACAGAGTGAAAGTCCATG
AGAGCACACGTCTGAACTCTTTCCCTACACGACGCTCTTCCGATCTNNNCGACAGAGTGAAAGTCCATG
GAGCACACGTCTGAACTCTTTCCCTACACGACGCTCTTCCGATCTNNNTGTTCTCCACCTATCGGAAACT
GAGCACACGTCTGAACTCTTTCCCTACACGACGCTCTTCCGATCTNNNTGTTCTCCACCTATCGGAAACT
GAGCACACGTCTGAACTCTTTCCCTACACGACGCTCTTCCGATCTNNNTGTTCTCCACCTATCGGAAACT
GAGCACACGTCTGAACTCTTTCCCTACACGACGCTCTTCCGATCTNNNTGTTCTCCACCTATCGGAAACT
GAGCACACGTCTGAACTCTTTCCCTACACGACGCTCTTCCGATCTNNNTGTTCTCCACCTATCGGAAACT
GAGCACACGTCTGAACTCTTTCCCTACACGACGCTCTTCCGATCTNNNTGTTCTCCACCTATCGGAAACT
GAGCACACGTCTGAACTCTTTCCCTACACGACGCTCTTCCGATCTNNNTGTTCTCCACCTATCGGAAACT
GAGCACACGTCTGAACTCTTTCCCTACACGACGCTCTTCCGATCTNNNTGTTCTCCACCTATCGGAAACT
GAGCACACGTCTGAACTCTTTCCCTACACGACGCTCTTCCGATCTNNNGAGAATGGAGTGAACCAGGG
GAGCACACGTCTGAACTCTTTCCCTACACGACGCTCTTCCGATCTNNNGAGAATGGAGTGAACCAGGG
GAGCACACGTCTGAACTCTTTCCCTACACGACGCTCTTCCGATCTNNNGAGAATGGAGTGAACCAGGG
GAGCACACGTCTGAACTCTTTCCCTACACGACGCTCTTCCGATCTNNNGAGAATGGAGTGAACCAGGG
GAGCACACGTCTGAACTCTTTCCCTACACGACGCTCTTCCGATCTNNNGAGAATGGAGTGAACCAGGG
GAGCACACGTCTGAACTCTTTCCCTACACGACGCTCTTCCGATCTNNNGAGAATGGAGTGAACCAGGG
GAGCACACGTCTGAACTCTTTCCCTACACGACGCTCTTCCGATCTNNNGAGAATGGAGTGAACCAGGG
GAGCACACGTCTGAACTCTTTCCCTACACGACGCTCTTCCGATCTNNNGAGAATGGAGTGAACCAGGG
AGAGCACACGTCTGAACTCTTTCCCTACACGACGCTCTTCCGATCTNNNGAGGCATTAGAATCGCTTGA

AGAGCACACGTCTGAACTCTTTCCCTACACGACGCTCTTCCGATCTNNNGAGGCATTAGAATCGCTTG
AGAGCACACGTCTGAACTCTTTCCCTACACGACGCTCTTCCGATCTNNNGAGGCATTAGAATCGCTTG
AGAGCACACGTCTGAACTCTTTCCCTACACGACGCTCTTCCGATCTNNNGAGGCATTAGAATCGCTTG
AGAGCACACGTCTGAACTCTTTCCCTACACGACGCTCTTCCGATCTNNNGAGGCATTAGAATCGCTTG
AGAGCACACGTCTGAACTCTTTCCCTACACGACGCTCTTCCGATCTNNNGAGGCATTAGAATCGCTTG
AGAGCACACGTCTGAACTCTTTCCCTACACGACGCTCTTCCGATCTNNNGAGGCATTAGAATCGCTTG
AGAGCACACGTCTGAACTCTTTCCCTACACGACGCTCTTCCGATCTNNNGAGGCATTAGAATCGCTTG
GAGCACACGTCTGAACTCTTTCCCTACACGACGCTCTTCCGATCTNNNAGGAGAATCGCTTGAACCCG
GAGCACACGTCTGAACTCTTTCCCTACACGACGCTCTTCCGATCTNNNAGGAGAATCGCTTGAACCCG
GAGCACACGTCTGAACTCTTTCCCTACACGACGCTCTTCCGATCTNNNAGGAGAATCGCTTGAACCCG
GAGCACACGTCTGAACTCTTTCCCTACACGACGCTCTTCCGATCTNNNAGGAGAATCGCTTGAACCCG
GAGCACACGTCTGAACTCTTTCCCTACACGACGCTCTTCCGATCTNNNAGGAGAATCGCTTGAACCCG
GAGCACACGTCTGAACTCTTTCCCTACACGACGCTCTTCCGATCTNNNAGGAGAATCGCTTGAACCCG
GAGCACACGTCTGAACTCTTTCCCTACACGACGCTCTTCCGATCTNNNAGGAGAATCGCTTGAACCCG
GAGCACACGTCTGAACTCTTTCCCTACACGACGCTCTTCCGATCTNNNAGGAGAATCGCTTGAACCCG
GAGCACACGTCTGAACTCTTTCCCTACACGACGCTCTTCCGATCTNNNGCAGGTCCTAACAGAAGAA
GAGCACACGTCTGAACTCTTTCCCTACACGACGCTCTTCCGATCTNNNGCAGGTCCTAACAGAAGAA
GAGCACACGTCTGAACTCTTTCCCTACACGACGCTCTTCCGATCTNNNGCAGGTCCTAACAGAAGAA
GAGCACACGTCTGAACTCTTTCCCTACACGACGCTCTTCCGATCTNNNGCAGGTCCTAACAGAAGAA
GAGCACACGTCTGAACTCTTTCCCTACACGACGCTCTTCCGATCTNNNGCAGGTCCTAACAGAAGAA
GAGCACACGTCTGAACTCTTTCCCTACACGACGCTCTTCCGATCTNNNGCAGGTCCTAACAGAAGAA
GAGCACACGTCTGAACTCTTTCCCTACACGACGCTCTTCCGATCTNNNGCAGGTCCTAACAGAAGAA
GAGCACACGTCTGAACTCTTTCCCTACACGACGCTCTTCCGATCTNNNGCAGGTCCTAACAGAAGAA
AGCACACGTCTGAACTCTTTCCCTACACGACGCTCTTCCGATCTNNNTTCAAGCTATTCTCCTGCCTCAG
AGCACACGTCTGAACTCTTTCCCTACACGACGCTCTTCCGATCTNNNTTCAAGCTATTCTCCTGCCTCAG
AGCACACGTCTGAACTCTTTCCCTACACGACGCTCTTCCGATCTNNNTTCAAGCTATTCTCCTGCCTCAG
AGCACACGTCTGAACTCTTTCCCTACACGACGCTCTTCCGATCTNNNTTCAAGCTATTCTCCTGCCTCAG
AGCACACGTCTGAACTCTTTCCCTACACGACGCTCTTCCGATCTNNNTTCAAGCTATTCTCCTGCCTCAG
AGCACACGTCTGAACTCTTTCCCTACACGACGCTCTTCCGATCTNNNTTCAAGCTATTCTCCTGCCTCAG
AGCACACGTCTGAACTCTTTCCCTACACGACGCTCTTCCGATCTNNNTTCAAGCTATTCTCCTGCCTCAG
AGCACACGTCTGAACTCTTTCCCTACACGACGCTCTTCCGATCTNNNTTCAAGCTATTCTCCTGCCTCAG
AGCACACGTCTGAACTCTTTCCCTACACGACGCTCTTCCGATCTNNNCCTACTTCTCTGAGGTGGTCCT
AGCACACGTCTGAACTCTTTCCCTACACGACGCTCTTCCGATCTNNNCCTACTTCTCTGAGGTGGTCCT
AGCACACGTCTGAACTCTTTCCCTACACGACGCTCTTCCGATCTNNNCCTACTTCTCTGAGGTGGTCCT
AGCACACGTCTGAACTCTTTCCCTACACGACGCTCTTCCGATCTNNNCCTACTTCTCTGAGGTGGTCCT
AGCACACGTCTGAACTCTTTCCCTACACGACGCTCTTCCGATCTNNNCCTACTTCTCTGAGGTGGTCCT
AGCACACGTCTGAACTCTTTCCCTACACGACGCTCTTCCGATCTNNNCCTACTTCTCTGAGGTGGTCCT
AGCACACGTCTGAACTCTTTCCCTACACGACGCTCTTCCGATCTNNNCCTACTTCTCTGAGGTGGTCCT
AGCACACGTCTGAACTCTTTCCCTACACGACGCTCTTCCGATCTNNNCCTACTTCTCTGAGGTGGTCCT
GCACACGTCTGAACTCTTTCCCTACACGACGCTCTTCCGATCTNNNACCTAGTTGTCACCTTTGTATAA
GCACACGTCTGAACTCTTTCCCTACACGACGCTCTTCCGATCTNNNACCTAGTTGTCACCTTTGTATAA
GCACACGTCTGAACTCTTTCCCTACACGACGCTCTTCCGATCTNNNACCTAGTTGTCACCTTTGTATAA
GCACACGTCTGAACTCTTTCCCTACACGACGCTCTTCCGATCTNNNACCTAGTTGTCACCTTTGTATAA

GCACACGTCTGAACTCTTTCCCTACACGACGCTCTTCCGATCTNNNACCTAGTTGTCACCTTTGTATAAC
GCACACGTCTGAACTCTTTCCCTACACGACGCTCTTCCGATCTNNNACCTAGTTGTCACCTTTGTATAAC
GCACACGTCTGAACTCTTTCCCTACACGACGCTCTTCCGATCTNNNACCTAGTTGTCACCTTTGTATAAC
GCACACGTCTGAACTCTTTCCCTACACGACGCTCTTCCGATCTNNNACCTAGTTGTCACCTTTGTATAAC
AGCACACGTCTGAACTCTTTCCCTACACGACGCTCTTCCGATCTNNNGAAGATGGTTTGGTTGTCAGGT
AGCACACGTCTGAACTCTTTCCCTACACGACGCTCTTCCGATCTNNNGAAGATGGTTTGGTTGTCAGGT
AGCACACGTCTGAACTCTTTCCCTACACGACGCTCTTCCGATCTNNNGAAGATGGTTTGGTTGTCAGGT
AGCACACGTCTGAACTCTTTCCCTACACGACGCTCTTCCGATCTNNNGAAGATGGTTTGGTTGTCAGGT
AGCACACGTCTGAACTCTTTCCCTACACGACGCTCTTCCGATCTNNNGAAGATGGTTTGGTTGTCAGGT
AGCACACGTCTGAACTCTTTCCCTACACGACGCTCTTCCGATCTNNNGAAGATGGTTTGGTTGTCAGGT
AGCACACGTCTGAACTCTTTCCCTACACGACGCTCTTCCGATCTNNNGAAGATGGTTTGGTTGTCAGGT
AGCACACGTCTGAACTCTTTCCCTACACGACGCTCTTCCGATCTNNNGAAGATGGTTTGGTTGTCAGGT
GAGCACACGTCTGAACTCTTTCCCTACACGACGCTCTTCCGATCTNNNACAATCCTCTTCATGTACTTGCA
GAGCACACGTCTGAACTCTTTCCCTACACGACGCTCTTCCGATCTNNNACAATCCTCTTCATGTACTTGCA
GAGCACACGTCTGAACTCTTTCCCTACACGACGCTCTTCCGATCTNNNACAATCCTCTTCATGTACTTGCA
GAGCACACGTCTGAACTCTTTCCCTACACGACGCTCTTCCGATCTNNNACAATCCTCTTCATGTACTTGCA
GAGCACACGTCTGAACTCTTTCCCTACACGACGCTCTTCCGATCTNNNACAATCCTCTTCATGTACTTGCA
GAGCACACGTCTGAACTCTTTCCCTACACGACGCTCTTCCGATCTNNNACAATCCTCTTCATGTACTTGCA
GAGCACACGTCTGAACTCTTTCCCTACACGACGCTCTTCCGATCTNNNACAATCCTCTTCATGTACTTGCA
GAGCACACGTCTGAACTCTTTCCCTACACGACGCTCTTCCGATCTNNNACAATCCTCTTCATGTACTTGCA
GAGCACACGTCTGAACTCTTTCCCTACACGACGCTCTTCCGATCTNNNCAAGACCCCCATCTCTACCAAA
GAGCACACGTCTGAACTCTTTCCCTACACGACGCTCTTCCGATCTNNNCAAGACCCCCATCTCTACCAAA
GAGCACACGTCTGAACTCTTTCCCTACACGACGCTCTTCCGATCTNNNCAAGACCCCCATCTCTACCAAA
GAGCACACGTCTGAACTCTTTCCCTACACGACGCTCTTCCGATCTNNNCAAGACCCCCATCTCTACCAAA
GAGCACACGTCTGAACTCTTTCCCTACACGACGCTCTTCCGATCTNNNCAAGACCCCCATCTCTACCAAA
GAGCACACGTCTGAACTCTTTCCCTACACGACGCTCTTCCGATCTNNNCAAGACCCCCATCTCTACCAAA
GAGCACACGTCTGAACTCTTTCCCTACACGACGCTCTTCCGATCTNNNCAAGACCCCCATCTCTACCAAA
GAGCACACGTCTGAACTCTTTCCCTACACGACGCTCTTCCGATCTNNNCAAGACCCCCATCTCTACCAAA
GAGCACACGTCTGAACTCTTTCCCTACACGACGCTCTTCCGATCTNNNTACGCCTGTAATCCCAGCTATTG
GAGCACACGTCTGAACTCTTTCCCTACACGACGCTCTTCCGATCTNNNTACGCCTGTAATCCCAGCTATTG
GAGCACACGTCTGAACTCTTTCCCTACACGACGCTCTTCCGATCTNNNTACGCCTGTAATCCCAGCTATTG
GAGCACACGTCTGAACTCTTTCCCTACACGACGCTCTTCCGATCTNNNTACGCCTGTAATCCCAGCTATTG
GAGCACACGTCTGAACTCTTTCCCTACACGACGCTCTTCCGATCTNNNTACGCCTGTAATCCCAGCTATTG
GAGCACACGTCTGAACTCTTTCCCTACACGACGCTCTTCCGATCTNNNTACGCCTGTAATCCCAGCTATTG
GAGCACACGTCTGAACTCTTTCCCTACACGACGCTCTTCCGATCTNNNTACGCCTGTAATCCCAGCTATTG
GAGCACACGTCTGAACTCTTTCCCTACACGACGCTCTTCCGATCTNNNTACGCCTGTAATCCCAGCTATTG
AGCACACGTCTGAACTCTTTCCCTACACGACGCTCTTCCGATCTNNNATTTGAACTCCAGTCCATTTGG
AGCACACGTCTGAACTCTTTCCCTACACGACGCTCTTCCGATCTNNNATTTGAACTCCAGTCCATTTGG
AGCACACGTCTGAACTCTTTCCCTACACGACGCTCTTCCGATCTNNNATTTGAACTCCAGTCCATTTGG
AGCACACGTCTGAACTCTTTCCCTACACGACGCTCTTCCGATCTNNNATTTGAACTCCAGTCCATTTGG
AGCACACGTCTGAACTCTTTCCCTACACGACGCTCTTCCGATCTNNNATTTGAACTCCAGTCCATTTGG
AGCACACGTCTGAACTCTTTCCCTACACGACGCTCTTCCGATCTNNNATTTGAACTCCAGTCCATTTGG
AGCACACGTCTGAACTCTTTCCCTACACGACGCTCTTCCGATCTNNNATTTGAACTCCAGTCCATTTGG

AGCACACGTCTGAACTCTTTCCCTACACGACGCTCTTCCGATCTNNNATTTGAACTCCAGTCCATTTGG
GAGCACACGTCTGAACTCTTTCCCTACACGACGCTCTTCCGATCTNNNGCTTACCAGGTAGCTGGGAGT
GAGCACACGTCTGAACTCTTTCCCTACACGACGCTCTTCCGATCTNNNGCTTACCAGGTAGCTGGGAGT
GAGCACACGTCTGAACTCTTTCCCTACACGACGCTCTTCCGATCTNNNGCTTACCAGGTAGCTGGGAGT
GAGCACACGTCTGAACTCTTTCCCTACACGACGCTCTTCCGATCTNNNGCTTACCAGGTAGCTGGGAGT
GAGCACACGTCTGAACTCTTTCCCTACACGACGCTCTTCCGATCTNNNGCTTACCAGGTAGCTGGGAGT
GAGCACACGTCTGAACTCTTTCCCTACACGACGCTCTTCCGATCTNNNGCTTACCAGGTAGCTGGGAGT
GAGCACACGTCTGAACTCTTTCCCTACACGACGCTCTTCCGATCTNNNGCTTACCAGGTAGCTGGGAGT
GAGCACACGTCTGAACTCTTTCCCTACACGACGCTCTTCCGATCTNNNGCTTACCAGGTAGCTGGGAGT
GCACACGTCTGAACTCTTTCCCTACACGACGCTCTTCCGATCTNNNTCACTTGAGTCCAGGACTTCAAGC
GCACACGTCTGAACTCTTTCCCTACACGACGCTCTTCCGATCTNNNTCACTTGAGTCCAGGACTTCAAGC
GCACACGTCTGAACTCTTTCCCTACACGACGCTCTTCCGATCTNNNTCACTTGAGTCCAGGACTTCAAGC
GCACACGTCTGAACTCTTTCCCTACACGACGCTCTTCCGATCTNNNTCACTTGAGTCCAGGACTTCAAGC
GCACACGTCTGAACTCTTTCCCTACACGACGCTCTTCCGATCTNNNTCACTTGAGTCCAGGACTTCAAGC
GCACACGTCTGAACTCTTTCCCTACACGACGCTCTTCCGATCTNNNTCACTTGAGTCCAGGACTTCAAGC
GCACACGTCTGAACTCTTTCCCTACACGACGCTCTTCCGATCTNNNTCACTTGAGTCCAGGACTTCAAGC
GCACACGTCTGAACTCTTTCCCTACACGACGCTCTTCCGATCTNNNTCACTTGAGTCCAGGACTTCAAGC
GAGCACACGTCTGAACTCTTTCCCTACACGACGCTCTTCCGATCTNNNAGATCCTGTTTCTGAATTGATC
GAGCACACGTCTGAACTCTTTCCCTACACGACGCTCTTCCGATCTNNNAGATCCTGTTTCTGAATTGATC
GAGCACACGTCTGAACTCTTTCCCTACACGACGCTCTTCCGATCTNNNAGATCCTGTTTCTGAATTGATC
GAGCACACGTCTGAACTCTTTCCCTACACGACGCTCTTCCGATCTNNNAGATCCTGTTTCTGAATTGATC
GAGCACACGTCTGAACTCTTTCCCTACACGACGCTCTTCCGATCTNNNAGATCCTGTTTCTGAATTGATC
GAGCACACGTCTGAACTCTTTCCCTACACGACGCTCTTCCGATCTNNNAGATCCTGTTTCTGAATTGATC
GAGCACACGTCTGAACTCTTTCCCTACACGACGCTCTTCCGATCTNNNAGATCCTGTTTCTGAATTGATC
GAGCACACGTCTGAACTCTTTCCCTACACGACGCTCTTCCGATCTNNNAGATCCTGTTTCTGAATTGATC
CACACGTCTGAACTCTTTCCCTACACGACGCTCTTCCGATCTNNNTATGTGTGAGAGCGTCTGTGAGAC
CACACGTCTGAACTCTTTCCCTACACGACGCTCTTCCGATCTNNNTATGTGTGAGAGCGTCTGTGAGAC
CACACGTCTGAACTCTTTCCCTACACGACGCTCTTCCGATCTNNNTATGTGTGAGAGCGTCTGTGAGAC
CACACGTCTGAACTCTTTCCCTACACGACGCTCTTCCGATCTNNNTATGTGTGAGAGCGTCTGTGAGAC
CACACGTCTGAACTCTTTCCCTACACGACGCTCTTCCGATCTNNNTATGTGTGAGAGCGTCTGTGAGAC
CACACGTCTGAACTCTTTCCCTACACGACGCTCTTCCGATCTNNNTATGTGTGAGAGCGTCTGTGAGAC
CACACGTCTGAACTCTTTCCCTACACGACGCTCTTCCGATCTNNNTATGTGTGAGAGCGTCTGTGAGAC
CACACGTCTGAACTCTTTCCCTACACGACGCTCTTCCGATCTNNNTATGTGTGAGAGCGTCTGTGAGAC
AGCACACGTCTGAACTCTTTCCCTACACGACGCTCTTCCGATCTNNNATATTGCCCATGCTGGTGTCAA
AGCACACGTCTGAACTCTTTCCCTACACGACGCTCTTCCGATCTNNNATATTGCCCATGCTGGTGTCAA
AGCACACGTCTGAACTCTTTCCCTACACGACGCTCTTCCGATCTNNNATATTGCCCATGCTGGTGTCAA
AGCACACGTCTGAACTCTTTCCCTACACGACGCTCTTCCGATCTNNNATATTGCCCATGCTGGTGTCAA
AGCACACGTCTGAACTCTTTCCCTACACGACGCTCTTCCGATCTNNNATATTGCCCATGCTGGTGTCAA
AGCACACGTCTGAACTCTTTCCCTACACGACGCTCTTCCGATCTNNNATATTGCCCATGCTGGTGTCAA
AGCACACGTCTGAACTCTTTCCCTACACGACGCTCTTCCGATCTNNNATATTGCCCATGCTGGTGTCAA
AGCACACGTCTGAACTCTTTCCCTACACGACGCTCTTCCGATCTNNNATATTGCCCATGCTGGTGTCAA
AGCACACGTCTGAACTCTTTCCCTACACGACGCTCTTCCGATCTNNNTTTGTTCTGGGGAGGATCATCT
AGCACACGTCTGAACTCTTTCCCTACACGACGCTCTTCCGATCTNNNTTTGTTCTGGGGAGGATCATCT

AGCACACGTCTGAACTCTTTCCCTACACGACGCTCTTCCGATCTNNNTTTGTTCTGGGGAGGATCATCT
AGCACACGTCTGAACTCTTTCCCTACACGACGCTCTTCCGATCTNNNTTTGTTCTGGGGAGGATCATCT
AGCACACGTCTGAACTCTTTCCCTACACGACGCTCTTCCGATCTNNNTTTGTTCTGGGGAGGATCATCT
AGCACACGTCTGAACTCTTTCCCTACACGACGCTCTTCCGATCTNNNTTTGTTCTGGGGAGGATCATCT
AGCACACGTCTGAACTCTTTCCCTACACGACGCTCTTCCGATCTNNNTTTGTTCTGGGGAGGATCATCT
AGCACACGTCTGAACTCTTTCCCTACACGACGCTCTTCCGATCTNNNTTTGTTCTGGGGAGGATCATCT
AGAGCACACGTCTGAACTCTTTCCCTACACGACGCTCTTCCGATCTNNNAGACCTTTAGATGTAGCTG
AGAGCACACGTCTGAACTCTTTCCCTACACGACGCTCTTCCGATCTNNNAGACCTTTAGATGTAGCTG
AGAGCACACGTCTGAACTCTTTCCCTACACGACGCTCTTCCGATCTNNNAGACCTTTAGATGTAGCTG
AGAGCACACGTCTGAACTCTTTCCCTACACGACGCTCTTCCGATCTNNNAGACCTTTAGATGTAGCTG
AGAGCACACGTCTGAACTCTTTCCCTACACGACGCTCTTCCGATCTNNNAGACCTTTAGATGTAGCTG
AGAGCACACGTCTGAACTCTTTCCCTACACGACGCTCTTCCGATCTNNNAGACCTTTAGATGTAGCTG
AGAGCACACGTCTGAACTCTTTCCCTACACGACGCTCTTCCGATCTNNNAGACCTTTAGATGTAGCTG
AGAGCACACGTCTGAACTCTTTCCCTACACGACGCTCTTCCGATCTNNNAGACCTTTAGATGTAGCTG
GAGCACACGTCTGAACTCTTTCCCTACACGACGCTCTTCCGATCTNNNCTGCCAATGCCCAATATGATCT
GAGCACACGTCTGAACTCTTTCCCTACACGACGCTCTTCCGATCTNNNCTGCCAATGCCCAATATGATCT
GAGCACACGTCTGAACTCTTTCCCTACACGACGCTCTTCCGATCTNNNCTGCCAATGCCCAATATGATCT
GAGCACACGTCTGAACTCTTTCCCTACACGACGCTCTTCCGATCTNNNCTGCCAATGCCCAATATGATCT
GAGCACACGTCTGAACTCTTTCCCTACACGACGCTCTTCCGATCTNNNCTGCCAATGCCCAATATGATCT
GAGCACACGTCTGAACTCTTTCCCTACACGACGCTCTTCCGATCTNNNCTGCCAATGCCCAATATGATCT
GAGCACACGTCTGAACTCTTTCCCTACACGACGCTCTTCCGATCTNNNCTGCCAATGCCCAATATGATCT
GAGCACACGTCTGAACTCTTTCCCTACACGACGCTCTTCCGATCTNNNCTGCCAATGCCCAATATGATCT
AGCACACGTCTGAACTCTTTCCCTACACGACGCTCTTCCGATCTNNNCCTTTGTCAGGTGCGTAGTTTT
AGCACACGTCTGAACTCTTTCCCTACACGACGCTCTTCCGATCTNNNCCTTTGTCAGGTGCGTAGTTTT
AGCACACGTCTGAACTCTTTCCCTACACGACGCTCTTCCGATCTNNNCCTTTGTCAGGTGCGTAGTTTT
AGCACACGTCTGAACTCTTTCCCTACACGACGCTCTTCCGATCTNNNCCTTTGTCAGGTGCGTAGTTTT
AGCACACGTCTGAACTCTTTCCCTACACGACGCTCTTCCGATCTNNNCCTTTGTCAGGTGCGTAGTTTT
AGCACACGTCTGAACTCTTTCCCTACACGACGCTCTTCCGATCTNNNCCTTTGTCAGGTGCGTAGTTTT
AGCACACGTCTGAACTCTTTCCCTACACGACGCTCTTCCGATCTNNNCCTTTGTCAGGTGCGTAGTTTT
AGCACACGTCTGAACTCTTTCCCTACACGACGCTCTTCCGATCTNNNCCTTTGTCAGGTGCGTAGTTTT
AGCACACGTCTGAACTCTTTCCCTACACGACGCTCTTCCGATCTNNNTAGAACAATTAGCGGGGAATG
AGCACACGTCTGAACTCTTTCCCTACACGACGCTCTTCCGATCTNNNTAGAACAATTAGCGGGGAATG
AGCACACGTCTGAACTCTTTCCCTACACGACGCTCTTCCGATCTNNNTAGAACAATTAGCGGGGAATG
AGCACACGTCTGAACTCTTTCCCTACACGACGCTCTTCCGATCTNNNTAGAACAATTAGCGGGGAATG
AGCACACGTCTGAACTCTTTCCCTACACGACGCTCTTCCGATCTNNNTAGAACAATTAGCGGGGAATG
AGCACACGTCTGAACTCTTTCCCTACACGACGCTCTTCCGATCTNNNTAGAACAATTAGCGGGGAATG
AGCACACGTCTGAACTCTTTCCCTACACGACGCTCTTCCGATCTNNNTAGAACAATTAGCGGGGAATG
AGCACACGTCTGAACTCTTTCCCTACACGACGCTCTTCCGATCTNNNTAGAACAATTAGCGGGGAATG
GCACACGTCTGAACTCTTTCCCTACACGACGCTCTTCCGATCTNNNTTACCTTCTCGGGAACTCCAATG
GCACACGTCTGAACTCTTTCCCTACACGACGCTCTTCCGATCTNNNTTACCTTCTCGGGAACTCCAATG
GCACACGTCTGAACTCTTTCCCTACACGACGCTCTTCCGATCTNNNTTACCTTCTCGGGAACTCCAATG
GCACACGTCTGAACTCTTTCCCTACACGACGCTCTTCCGATCTNNNTTACCTTCTCGGGAACTCCAATG
GCACACGTCTGAACTCTTTCCCTACACGACGCTCTTCCGATCTNNNTTACCTTCTCGGGAACTCCAATG

GCACACGTCTGAACTCTTTCCCTACACGACGCTCTTCCGATCTNNNTTACCTTCTCGGGAACTCCAATG
GCACACGTCTGAACTCTTTCCCTACACGACGCTCTTCCGATCTNNNTTACCTTCTCGGGAACTCCAATG
GCACACGTCTGAACTCTTTCCCTACACGACGCTCTTCCGATCTNNNTTACCTTCTCGGGAACTCCAATG
AGCACACGTCTGAACTCTTTCCCTACACGACGCTCTTCCGATCTNNNCGTGTATGATGAAATGAGTAA
AGCACACGTCTGAACTCTTTCCCTACACGACGCTCTTCCGATCTNNNCGTGTATGATGAAATGAGTAA
AGCACACGTCTGAACTCTTTCCCTACACGACGCTCTTCCGATCTNNNCGTGTATGATGAAATGAGTAA
AGCACACGTCTGAACTCTTTCCCTACACGACGCTCTTCCGATCTNNNCGTGTATGATGAAATGAGTAA
AGCACACGTCTGAACTCTTTCCCTACACGACGCTCTTCCGATCTNNNCGTGTATGATGAAATGAGTAA
AGCACACGTCTGAACTCTTTCCCTACACGACGCTCTTCCGATCTNNNCGTGTATGATGAAATGAGTAA
AGCACACGTCTGAACTCTTTCCCTACACGACGCTCTTCCGATCTNNNCGTGTATGATGAAATGAGTAA
AGCACACGTCTGAACTCTTTCCCTACACGACGCTCTTCCGATCTNNNCGTGTATGATGAAATGAGTAA
GCACACGTCTGAACTCTTTCCCTACACGACGCTCTTCCGATCTNNNTGGAAGTTGAATTCTCCCACTT
GCACACGTCTGAACTCTTTCCCTACACGACGCTCTTCCGATCTNNNTGGAAGTTGAATTCTCCCACTT
GCACACGTCTGAACTCTTTCCCTACACGACGCTCTTCCGATCTNNNTGGAAGTTGAATTCTCCCACTT
GCACACGTCTGAACTCTTTCCCTACACGACGCTCTTCCGATCTNNNTGGAAGTTGAATTCTCCCACTT
GCACACGTCTGAACTCTTTCCCTACACGACGCTCTTCCGATCTNNNTGGAAGTTGAATTCTCCCACTT
GCACACGTCTGAACTCTTTCCCTACACGACGCTCTTCCGATCTNNNTGGAAGTTGAATTCTCCCACTT
GCACACGTCTGAACTCTTTCCCTACACGACGCTCTTCCGATCTNNNTGGAAGTTGAATTCTCCCACTT
GCACACGTCTGAACTCTTTCCCTACACGACGCTCTTCCGATCTNNNTGGAAGTTGAATTCTCCCACTT
GCACACGTCTGAACTCTTTCCCTACACGACGCTCTTCCGATCTNNNTTCCAAGTAGCTGGGATTACAG
GCACACGTCTGAACTCTTTCCCTACACGACGCTCTTCCGATCTNNNTTCCAAGTAGCTGGGATTACAG
GCACACGTCTGAACTCTTTCCCTACACGACGCTCTTCCGATCTNNNTTCCAAGTAGCTGGGATTACAG
GCACACGTCTGAACTCTTTCCCTACACGACGCTCTTCCGATCTNNNTTCCAAGTAGCTGGGATTACAG
GCACACGTCTGAACTCTTTCCCTACACGACGCTCTTCCGATCTNNNTTCCAAGTAGCTGGGATTACAG
GCACACGTCTGAACTCTTTCCCTACACGACGCTCTTCCGATCTNNNTTCCAAGTAGCTGGGATTACAG
GCACACGTCTGAACTCTTTCCCTACACGACGCTCTTCCGATCTNNNTTCCAAGTAGCTGGGATTACAG
GCACACGTCTGAACTCTTTCCCTACACGACGCTCTTCCGATCTNNNTTCCAAGTAGCTGGGATTACAG
AGCACACGTCTGAACTCTTTCCCTACACGACGCTCTTCCGATCTNNNACCATCTTCCACACATCAGTCTC
AGCACACGTCTGAACTCTTTCCCTACACGACGCTCTTCCGATCTNNNACCATCTTCCACACATCAGTCTC
AGCACACGTCTGAACTCTTTCCCTACACGACGCTCTTCCGATCTNNNACCATCTTCCACACATCAGTCTC
AGCACACGTCTGAACTCTTTCCCTACACGACGCTCTTCCGATCTNNNACCATCTTCCACACATCAGTCTC
AGCACACGTCTGAACTCTTTCCCTACACGACGCTCTTCCGATCTNNNACCATCTTCCACACATCAGTCTC
AGCACACGTCTGAACTCTTTCCCTACACGACGCTCTTCCGATCTNNNACCATCTTCCACACATCAGTCTC
AGCACACGTCTGAACTCTTTCCCTACACGACGCTCTTCCGATCTNNNACCATCTTCCACACATCAGTCTC
AGCACACGTCTGAACTCTTTCCCTACACGACGCTCTTCCGATCTNNNACCATCTTCCACACATCAGTCTC
GAGCACACGTCTGAACTCTTTCCCTACACGACGCTCTTCCGATCTNNNAAAAAATTAGCCAGGCGTG
GAGCACACGTCTGAACTCTTTCCCTACACGACGCTCTTCCGATCTNNNAAAAAATTAGCCAGGCGTG
GAGCACACGTCTGAACTCTTTCCCTACACGACGCTCTTCCGATCTNNNAAAAAATTAGCCAGGCGTG
GAGCACACGTCTGAACTCTTTCCCTACACGACGCTCTTCCGATCTNNNAAAAAATTAGCCAGGCGTG
GAGCACACGTCTGAACTCTTTCCCTACACGACGCTCTTCCGATCTNNNAAAAAATTAGCCAGGCGTG
GAGCACACGTCTGAACTCTTTCCCTACACGACGCTCTTCCGATCTNNNAAAAAATTAGCCAGGCGTG
GAGCACACGTCTGAACTCTTTCCCTACACGACGCTCTTCCGATCTNNNAAAAAATTAGCCAGGCGTG
GAGCACACGTCTGAACTCTTTCCCTACACGACGCTCTTCCGATCTNNNAAAAAATTAGCCAGGCGTG

GAGCACACGTCTGAACTCTTTCCCTACACGACGCTCTTCCGATCTNNNTCACTAATCATTAGGGCAGTG
GAGCACACGTCTGAACTCTTTCCCTACACGACGCTCTTCCGATCTNNNTCACTAATCATTAGGGCAGTG
GAGCACACGTCTGAACTCTTTCCCTACACGACGCTCTTCCGATCTNNNTCACTAATCATTAGGGCAGTG
GAGCACACGTCTGAACTCTTTCCCTACACGACGCTCTTCCGATCTNNNTCACTAATCATTAGGGCAGTG
GAGCACACGTCTGAACTCTTTCCCTACACGACGCTCTTCCGATCTNNNTCACTAATCATTAGGGCAGTG
GAGCACACGTCTGAACTCTTTCCCTACACGACGCTCTTCCGATCTNNNTCACTAATCATTAGGGCAGTG
GAGCACACGTCTGAACTCTTTCCCTACACGACGCTCTTCCGATCTNNNTCACTAATCATTAGGGCAGTG
GAGCACACGTCTGAACTCTTTCCCTACACGACGCTCTTCCGATCTNNNTCACTAATCATTAGGGCAGTG
AGCACACGTCTGAACTCTTTCCCTACACGACGCTCTTCCGATCTNNNCTCCATCTCAGATCATCAGGCAT
AGCACACGTCTGAACTCTTTCCCTACACGACGCTCTTCCGATCTNNNCTCCATCTCAGATCATCAGGCAT
AGCACACGTCTGAACTCTTTCCCTACACGACGCTCTTCCGATCTNNNCTCCATCTCAGATCATCAGGCAT
AGCACACGTCTGAACTCTTTCCCTACACGACGCTCTTCCGATCTNNNCTCCATCTCAGATCATCAGGCAT
AGCACACGTCTGAACTCTTTCCCTACACGACGCTCTTCCGATCTNNNCTCCATCTCAGATCATCAGGCAT
AGCACACGTCTGAACTCTTTCCCTACACGACGCTCTTCCGATCTNNNCTCCATCTCAGATCATCAGGCAT
AGCACACGTCTGAACTCTTTCCCTACACGACGCTCTTCCGATCTNNNCTCCATCTCAGATCATCAGGCAT
AGCACACGTCTGAACTCTTTCCCTACACGACGCTCTTCCGATCTNNNCTCCATCTCAGATCATCAGGCAT
AGCACACGTCTGAACTCTTTCCCTACACGACGCTCTTCCGATCTNNNTCTTCAGCTGTGGTAAGTCAGT
AGCACACGTCTGAACTCTTTCCCTACACGACGCTCTTCCGATCTNNNTCTTCAGCTGTGGTAAGTCAGT
AGCACACGTCTGAACTCTTTCCCTACACGACGCTCTTCCGATCTNNNTCTTCAGCTGTGGTAAGTCAGT
AGCACACGTCTGAACTCTTTCCCTACACGACGCTCTTCCGATCTNNNTCTTCAGCTGTGGTAAGTCAGT
AGCACACGTCTGAACTCTTTCCCTACACGACGCTCTTCCGATCTNNNTCTTCAGCTGTGGTAAGTCAGT
AGCACACGTCTGAACTCTTTCCCTACACGACGCTCTTCCGATCTNNNTCTTCAGCTGTGGTAAGTCAGT
AGCACACGTCTGAACTCTTTCCCTACACGACGCTCTTCCGATCTNNNTCTTCAGCTGTGGTAAGTCAGT
AGCACACGTCTGAACTCTTTCCCTACACGACGCTCTTCCGATCTNNNTCTTCAGCTGTGGTAAGTCAGT
GAGCACACGTCTGAACTCTTTCCCTACACGACGCTCTTCCGATCTNNNGGGACCATGTCTTATTCACCCT
GAGCACACGTCTGAACTCTTTCCCTACACGACGCTCTTCCGATCTNNNGGGACCATGTCTTATTCACCCT
GAGCACACGTCTGAACTCTTTCCCTACACGACGCTCTTCCGATCTNNNGGGACCATGTCTTATTCACCCT
GAGCACACGTCTGAACTCTTTCCCTACACGACGCTCTTCCGATCTNNNGGGACCATGTCTTATTCACCCT
GAGCACACGTCTGAACTCTTTCCCTACACGACGCTCTTCCGATCTNNNGGGACCATGTCTTATTCACCCT
GAGCACACGTCTGAACTCTTTCCCTACACGACGCTCTTCCGATCTNNNGGGACCATGTCTTATTCACCCT
GAGCACACGTCTGAACTCTTTCCCTACACGACGCTCTTCCGATCTNNNGGGACCATGTCTTATTCACCCT
GAGCACACGTCTGAACTCTTTCCCTACACGACGCTCTTCCGATCTNNNGGGACCATGTCTTATTCACCCT
AGCACACGTCTGAACTCTTTCCCTACACGACGCTCTTCCGATCTNNNTTTAGACTGTGGTCCCAATCAC
AGCACACGTCTGAACTCTTTCCCTACACGACGCTCTTCCGATCTNNNTTTAGACTGTGGTCCCAATCAC
AGCACACGTCTGAACTCTTTCCCTACACGACGCTCTTCCGATCTNNNTTTAGACTGTGGTCCCAATCAC
AGCACACGTCTGAACTCTTTCCCTACACGACGCTCTTCCGATCTNNNTTTAGACTGTGGTCCCAATCAC
AGCACACGTCTGAACTCTTTCCCTACACGACGCTCTTCCGATCTNNNTTTAGACTGTGGTCCCAATCAC
AGCACACGTCTGAACTCTTTCCCTACACGACGCTCTTCCGATCTNNNTTTAGACTGTGGTCCCAATCAC
AGCACACGTCTGAACTCTTTCCCTACACGACGCTCTTCCGATCTNNNTTTAGACTGTGGTCCCAATCAC
AGCACACGTCTGAACTCTTTCCCTACACGACGCTCTTCCGATCTNNNTTTAGACTGTGGTCCCAATCAC
AGCACACGTCTGAACTCTTTCCCTACACGACGCTCTTCCGATCTNNNCATCCTCAGCTCCACTCTACACCA
AGCACACGTCTGAACTCTTTCCCTACACGACGCTCTTCCGATCTNNNCATCCTCAGCTCCACTCTACACCA
AGCACACGTCTGAACTCTTTCCCTACACGACGCTCTTCCGATCTNNNCATCCTCAGCTCCACTCTACACCA

AGCACACGTCTGAACTCTTTCCCTACACGACGCTCTTCCGATCTNNNCATCCTCAGCTCCACTCTACACCA
AGCACACGTCTGAACTCTTTCCCTACACGACGCTCTTCCGATCTNNNCATCCTCAGCTCCACTCTACACCA
AGCACACGTCTGAACTCTTTCCCTACACGACGCTCTTCCGATCTNNNCATCCTCAGCTCCACTCTACACCA
AGCACACGTCTGAACTCTTTCCCTACACGACGCTCTTCCGATCTNNNCATCCTCAGCTCCACTCTACACCA
AGCACACGTCTGAACTCTTTCCCTACACGACGCTCTTCCGATCTNNNCATCCTCAGCTCCACTCTACACCA
AGAGCACACGTCTGAACTCTTTCCCTACACGACGCTCTTCCGATCTNNNGAGGTGGAGGTTGCAGTG
AGAGCACACGTCTGAACTCTTTCCCTACACGACGCTCTTCCGATCTNNNGAGGTGGAGGTTGCAGTG
AGAGCACACGTCTGAACTCTTTCCCTACACGACGCTCTTCCGATCTNNNGAGGTGGAGGTTGCAGTG
AGAGCACACGTCTGAACTCTTTCCCTACACGACGCTCTTCCGATCTNNNGAGGTGGAGGTTGCAGTG
AGAGCACACGTCTGAACTCTTTCCCTACACGACGCTCTTCCGATCTNNNGAGGTGGAGGTTGCAGTG
AGAGCACACGTCTGAACTCTTTCCCTACACGACGCTCTTCCGATCTNNNGAGGTGGAGGTTGCAGTG
AGAGCACACGTCTGAACTCTTTCCCTACACGACGCTCTTCCGATCTNNNGAGGTGGAGGTTGCAGTG
AGAGCACACGTCTGAACTCTTTCCCTACACGACGCTCTTCCGATCTNNNGAGGTGGAGGTTGCAGTG
AGCACACGTCTGAACTCTTTCCCTACACGACGCTCTTCCGATCTNNNCAGCTGTTTTGGAGACTGAGTT
AGCACACGTCTGAACTCTTTCCCTACACGACGCTCTTCCGATCTNNNCAGCTGTTTTGGAGACTGAGTT
AGCACACGTCTGAACTCTTTCCCTACACGACGCTCTTCCGATCTNNNCAGCTGTTTTGGAGACTGAGTT
AGCACACGTCTGAACTCTTTCCCTACACGACGCTCTTCCGATCTNNNCAGCTGTTTTGGAGACTGAGTT
AGCACACGTCTGAACTCTTTCCCTACACGACGCTCTTCCGATCTNNNCAGCTGTTTTGGAGACTGAGTT
AGCACACGTCTGAACTCTTTCCCTACACGACGCTCTTCCGATCTNNNCAGCTGTTTTGGAGACTGAGTT
AGCACACGTCTGAACTCTTTCCCTACACGACGCTCTTCCGATCTNNNCAGCTGTTTTGGAGACTGAGTT
AGCACACGTCTGAACTCTTTCCCTACACGACGCTCTTCCGATCTNNNCAGCTGTTTTGGAGACTGAGTT
GAGCACACGTCTGAACTCTTTCCCTACACGACGCTCTTCCGATCTNNNGGTGACAGAGTGAGACTCCA
GAGCACACGTCTGAACTCTTTCCCTACACGACGCTCTTCCGATCTNNNGGTGACAGAGTGAGACTCCA
GAGCACACGTCTGAACTCTTTCCCTACACGACGCTCTTCCGATCTNNNGGTGACAGAGTGAGACTCCA
GAGCACACGTCTGAACTCTTTCCCTACACGACGCTCTTCCGATCTNNNGGTGACAGAGTGAGACTCCA
GAGCACACGTCTGAACTCTTTCCCTACACGACGCTCTTCCGATCTNNNGGTGACAGAGTGAGACTCCA
GAGCACACGTCTGAACTCTTTCCCTACACGACGCTCTTCCGATCTNNNGGTGACAGAGTGAGACTCCA
GAGCACACGTCTGAACTCTTTCCCTACACGACGCTCTTCCGATCTNNNGGTGACAGAGTGAGACTCCA
GAGCACACGTCTGAACTCTTTCCCTACACGACGCTCTTCCGATCTNNNGGTGACAGAGTGAGACTCCA
GAGCACACGTCTGAACTCTTTCCCTACACGACGCTCTTCCGATCTNNNCTGAGGCAGGAGAATCACTTG
GAGCACACGTCTGAACTCTTTCCCTACACGACGCTCTTCCGATCTNNNCTGAGGCAGGAGAATCACTTG
GAGCACACGTCTGAACTCTTTCCCTACACGACGCTCTTCCGATCTNNNCTGAGGCAGGAGAATCACTTG
GAGCACACGTCTGAACTCTTTCCCTACACGACGCTCTTCCGATCTNNNCTGAGGCAGGAGAATCACTTG
GAGCACACGTCTGAACTCTTTCCCTACACGACGCTCTTCCGATCTNNNCTGAGGCAGGAGAATCACTTG
GAGCACACGTCTGAACTCTTTCCCTACACGACGCTCTTCCGATCTNNNCTGAGGCAGGAGAATCACTTG
GAGCACACGTCTGAACTCTTTCCCTACACGACGCTCTTCCGATCTNNNCTGAGGCAGGAGAATCACTTG
GAGCACACGTCTGAACTCTTTCCCTACACGACGCTCTTCCGATCTNNNCTGAGGCAGGAGAATCACTTG
AGCACACGTCTGAACTCTTTCCCTACACGACGCTCTTCCGATCTNNNTGTGGAGTCGTTAGTGTTTTCCA
AGCACACGTCTGAACTCTTTCCCTACACGACGCTCTTCCGATCTNNNTGTGGAGTCGTTAGTGTTTTCCA
AGCACACGTCTGAACTCTTTCCCTACACGACGCTCTTCCGATCTNNNTGTGGAGTCGTTAGTGTTTTCCA
AGCACACGTCTGAACTCTTTCCCTACACGACGCTCTTCCGATCTNNNTGTGGAGTCGTTAGTGTTTTCCA
AGCACACGTCTGAACTCTTTCCCTACACGACGCTCTTCCGATCTNNNTGTGGAGTCGTTAGTGTTTTCCA
AGCACACGTCTGAACTCTTTCCCTACACGACGCTCTTCCGATCTNNNTGTGGAGTCGTTAGTGTTTTCCA

GCACACGTCTGAACTCTTTCCCTACACGACGCTCTTCCGATCTNNNTGTGGAGTCGTTAGTGTTTTCC

GCACACGTCTGAACTCTTTCCCTACACGACGCTCTTCCGATCTNNNTGTGGAGTCGTTAGTGTTTTCC

GCACACGTCTGAACTCTTTCCCTACACGACGCTCTTCCGATCTNNNGGTAACTGTGGTCAAGAGGTA

GCACACGTCTGAACTCTTTCCCTACACGACGCTCTTCCGATCTNNNGGTAACTGTGGTCAAGAGGTA

GCACACGTCTGAACTCTTTCCCTACACGACGCTCTTCCGATCTNNNGGTAACTGTGGTCAAGAGGTA

GCACACGTCTGAACTCTTTCCCTACACGACGCTCTTCCGATCTNNNGGTAACTGTGGTCAAGAGGTA

GCACACGTCTGAACTCTTTCCCTACACGACGCTCTTCCGATCTNNNGGTAACTGTGGTCAAGAGGTA

GCACACGTCTGAACTCTTTCCCTACACGACGCTCTTCCGATCTNNNGGTAACTGTGGTCAAGAGGTA

GCACACGTCTGAACTCTTTCCCTACACGACGCTCTTCCGATCTNNNGGTAACTGTGGTCAAGAGGTA

GCACACGTCTGAACTCTTTCCCTACACGACGCTCTTCCGATCTNNNGGTAACTGTGGTCAAGAGGTA

GCACACGTCTGAACTCTTTCCCTACACGACGCTCTTCCGATCTNNNGCAGCTTATAGTGATTTAGGATA

GCACACGTCTGAACTCTTTCCCTACACGACGCTCTTCCGATCTNNNGCAGCTTATAGTGATTTAGGATA

GCACACGTCTGAACTCTTTCCCTACACGACGCTCTTCCGATCTNNNGCAGCTTATAGTGATTTAGGATA

GCACACGTCTGAACTCTTTCCCTACACGACGCTCTTCCGATCTNNNGCAGCTTATAGTGATTTAGGATA

GCACACGTCTGAACTCTTTCCCTACACGACGCTCTTCCGATCTNNNGCAGCTTATAGTGATTTAGGATA

GCACACGTCTGAACTCTTTCCCTACACGACGCTCTTCCGATCTNNNGCAGCTTATAGTGATTTAGGATA

GCACACGTCTGAACTCTTTCCCTACACGACGCTCTTCCGATCTNNNGCAGCTTATAGTGATTTAGGATA

GCACACGTCTGAACTCTTTCCCTACACGACGCTCTTCCGATCTNNNGCAGCTTATAGTGATTTAGGATA

AGCACACGTCTGAACTCTTTCCCTACACGACGCTCTTCCGATCTNNNCTGTAATCCAGCTACTCAGG

AGCACACGTCTGAACTCTTTCCCTACACGACGCTCTTCCGATCTNNNCTGTAATCCAGCTACTCAGG

AGCACACGTCTGAACTCTTTCCCTACACGACGCTCTTCCGATCTNNNCTGTAATCCAGCTACTCAGG

AGCACACGTCTGAACTCTTTCCCTACACGACGCTCTTCCGATCTNNNCTGTAATCCAGCTACTCAGG

AGCACACGTCTGAACTCTTTCCCTACACGACGCTCTTCCGATCTNNNCTGTAATCCAGCTACTCAGG

AGCACACGTCTGAACTCTTTCCCTACACGACGCTCTTCCGATCTNNNCTGTAATCCAGCTACTCAGG

AGCACACGTCTGAACTCTTTCCCTACACGACGCTCTTCCGATCTNNNCTGTAATCCAGCTACTCAGG

AGCACACGTCTGAACTCTTTCCCTACACGACGCTCTTCCGATCTNNNCTGTAATCCAGCTACTCAGG

AGCACACGTCTGAACTCTTTCCCTACACGACGCTCTTCCGATCTNNNGAGGCAGGAGAATCGCTTGA

AGCACACGTCTGAACTCTTTCCCTACACGACGCTCTTCCGATCTNNNGAGGCAGGAGAATCGCTTGA

AGCACACGTCTGAACTCTTTCCCTACACGACGCTCTTCCGATCTNNNGAGGCAGGAGAATCGCTTGA

AGCACACGTCTGAACTCTTTCCCTACACGACGCTCTTCCGATCTNNNGAGGCAGGAGAATCGCTTGA

AGCACACGTCTGAACTCTTTCCCTACACGACGCTCTTCCGATCTNNNGAGGCAGGAGAATCGCTTGA

AGCACACGTCTGAACTCTTTCCCTACACGACGCTCTTCCGATCTNNNGAGGCAGGAGAATCGCTTGA

AGCACACGTCTGAACTCTTTCCCTACACGACGCTCTTCCGATCTNNNGAGGCAGGAGAATCGCTTGA

AGCACACGTCTGAACTCTTTCCCTACACGACGCTCTTCCGATCTNNNGAGGCAGGAGAATCGCTTGA

AGAGCACACGTCTGAACTCTTTCCCTACACGACGCTCTTCCGATCTNNNCCTCAGATCAATCCTCCCAG

AGAGCACACGTCTGAACTCTTTCCCTACACGACGCTCTTCCGATCTNNNCCTCAGATCAATCCTCCCAG

AGAGCACACGTCTGAACTCTTTCCCTACACGACGCTCTTCCGATCTNNNCCTCAGATCAATCCTCCCAG

AGAGCACACGTCTGAACTCTTTCCCTACACGACGCTCTTCCGATCTNNNCCTCAGATCAATCCTCCCAG

AGAGCACACGTCTGAACTCTTTCCCTACACGACGCTCTTCCGATCTNNNCCTCAGATCAATCCTCCCAG

AGAGCACACGTCTGAACTCTTTCCCTACACGACGCTCTTCCGATCTNNNCCTCAGATCAATCCTCCCAG

AGAGCACACGTCTGAACTCTTTCCCTACACGACGCTCTTCCGATCTNNNCCTCAGATCAATCCTCCCAG

AGAGCACACGTCTGAACTCTTTCCCTACACGACGCTCTTCCGATCTNNNCCTCAGATCAATCCTCCCAG

CACACGTCTGAACTCTTTCCCTACACGACGCTCTTCCGATCTNNNGTGTTGGAATTACAGACGTGAG

GCACACGTCTGAACTCTTTCCCTACACGACGCTCTTCCGATCTNNNGTGTTGGAATTACAGACGTGAGC
GCACACGTCTGAACTCTTTCCCTACACGACGCTCTTCCGATCTNNNGTGTTGGAATTACAGACGTGAGC
GCACACGTCTGAACTCTTTCCCTACACGACGCTCTTCCGATCTNNNGTGTTGGAATTACAGACGTGAGC
GCACACGTCTGAACTCTTTCCCTACACGACGCTCTTCCGATCTNNNGTGTTGGAATTACAGACGTGAGC
GCACACGTCTGAACTCTTTCCCTACACGACGCTCTTCCGATCTNNNGTGTTGGAATTACAGACGTGAGC
GCACACGTCTGAACTCTTTCCCTACACGACGCTCTTCCGATCTNNNGTGTTGGAATTACAGACGTGAGC
GCACACGTCTGAACTCTTTCCCTACACGACGCTCTTCCGATCTNNNGTGTTGGAATTACAGACGTGAGC
AGCACACGTCTGAACTCTTTCCCTACACGACGCTCTTCCGATCTNNNCCCAAAGTCCTTTGTGTCATTCC
AGCACACGTCTGAACTCTTTCCCTACACGACGCTCTTCCGATCTNNNCCCAAAGTCCTTTGTGTCATTCC
AGCACACGTCTGAACTCTTTCCCTACACGACGCTCTTCCGATCTNNNCCCAAAGTCCTTTGTGTCATTCC
AGCACACGTCTGAACTCTTTCCCTACACGACGCTCTTCCGATCTNNNCCCAAAGTCCTTTGTGTCATTCC
AGCACACGTCTGAACTCTTTCCCTACACGACGCTCTTCCGATCTNNNCCCAAAGTCCTTTGTGTCATTCC
AGCACACGTCTGAACTCTTTCCCTACACGACGCTCTTCCGATCTNNNCCCAAAGTCCTTTGTGTCATTCC
AGCACACGTCTGAACTCTTTCCCTACACGACGCTCTTCCGATCTNNNCCCAAAGTCCTTTGTGTCATTCC
AGCACACGTCTGAACTCTTTCCCTACACGACGCTCTTCCGATCTNNNCCCAAAGTCCTTTGTGTCATTCC
AGCACACGTCTGAACTCTTTCCCTACACGACGCTCTTCCGATCTNNNACTAAAACTCAGCGATACTGCCA
AGCACACGTCTGAACTCTTTCCCTACACGACGCTCTTCCGATCTNNNACTAAAACTCAGCGATACTGCCA
AGCACACGTCTGAACTCTTTCCCTACACGACGCTCTTCCGATCTNNNACTAAAACTCAGCGATACTGCCA
AGCACACGTCTGAACTCTTTCCCTACACGACGCTCTTCCGATCTNNNACTAAAACTCAGCGATACTGCCA
AGCACACGTCTGAACTCTTTCCCTACACGACGCTCTTCCGATCTNNNACTAAAACTCAGCGATACTGCCA
AGCACACGTCTGAACTCTTTCCCTACACGACGCTCTTCCGATCTNNNACTAAAACTCAGCGATACTGCCA
AGCACACGTCTGAACTCTTTCCCTACACGACGCTCTTCCGATCTNNNACTAAAACTCAGCGATACTGCCA
AGCACACGTCTGAACTCTTTCCCTACACGACGCTCTTCCGATCTNNNACTAAAACTCAGCGATACTGCCA
GAGCACACGTCTGAACTCTTTCCCTACACGACGCTCTTCCGATCTNNNGAGGTTGCAGTGAGTCGAGA
GAGCACACGTCTGAACTCTTTCCCTACACGACGCTCTTCCGATCTNNNGAGGTTGCAGTGAGTCGAGA
GAGCACACGTCTGAACTCTTTCCCTACACGACGCTCTTCCGATCTNNNGAGGTTGCAGTGAGTCGAGA
GAGCACACGTCTGAACTCTTTCCCTACACGACGCTCTTCCGATCTNNNGAGGTTGCAGTGAGTCGAGA
GAGCACACGTCTGAACTCTTTCCCTACACGACGCTCTTCCGATCTNNNGAGGTTGCAGTGAGTCGAGA
GAGCACACGTCTGAACTCTTTCCCTACACGACGCTCTTCCGATCTNNNGAGGTTGCAGTGAGTCGAGA
GAGCACACGTCTGAACTCTTTCCCTACACGACGCTCTTCCGATCTNNNGAGGTTGCAGTGAGTCGAGA
GAGCACACGTCTGAACTCTTTCCCTACACGACGCTCTTCCGATCTNNNGAGGTTGCAGTGAGTCGAGA
AGCACACGTCTGAACTCTTTCCCTACACGACGCTCTTCCGATCTNNNCTAATTTGGAAATAGCTGGGGC
AGCACACGTCTGAACTCTTTCCCTACACGACGCTCTTCCGATCTNNNCTAATTTGGAAATAGCTGGGGC
AGCACACGTCTGAACTCTTTCCCTACACGACGCTCTTCCGATCTNNNCTAATTTGGAAATAGCTGGGGC
AGCACACGTCTGAACTCTTTCCCTACACGACGCTCTTCCGATCTNNNCTAATTTGGAAATAGCTGGGGC
AGCACACGTCTGAACTCTTTCCCTACACGACGCTCTTCCGATCTNNNCTAATTTGGAAATAGCTGGGGC
AGCACACGTCTGAACTCTTTCCCTACACGACGCTCTTCCGATCTNNNCTAATTTGGAAATAGCTGGGGC
AGCACACGTCTGAACTCTTTCCCTACACGACGCTCTTCCGATCTNNNCTAATTTGGAAATAGCTGGGGC
AGCACACGTCTGAACTCTTTCCCTACACGACGCTCTTCCGATCTNNNCTAATTTGGAAATAGCTGGGGC

AGCATCGGACTCATCAGGAAGC
AGCATCGGACTCATCAGGAAGC
AGCATCGGACTCATCAGGAAGC
CTTGCATCGGACTCATCAGGAAGC
CTTGCATCGGACTCATCAGGAAGC
CTTGCATCGGACTCATCAGGAAGC
CTTGCATCGGACTCATCAGGAAGC
CTTGCATCGGACTCATCAGGAAGC
CTTGCATCGGACTCATCAGGAAGC
CTTGCATCGGACTCATCAGGAAGC
CATCGGACTCATCAGGAAGC
CATCGGACTCATCAGGAAGC
CATCGGACTCATCAGGAAGC
CATCGGACTCATCAGGAAGC
CATCGGACTCATCAGGAAGC
CATCGGACTCATCAGGAAGC
CATCGGACTCATCAGGAAGC
CATCATCGGACTCATCAGGAAGC
CATCATCGGACTCATCAGGAAGC
CATCATCGGACTCATCAGGAAGC
CATCATCGGACTCATCAGGAAGC
CATCATCGGACTCATCAGGAAGC
CATCATCGGACTCATCAGGAAGC
CATCATCGGACTCATCAGGAAGC
GAACATCGGACTCATCAGGAAGC
GAACATCGGACTCATCAGGAAGC
GAACATCGGACTCATCAGGAAGC
GAACATCGGACTCATCAGGAAGC
GAACATCGGACTCATCAGGAAGC
GAACATCGGACTCATCAGGAAGC
GAACATCGGACTCATCAGGAAGC
CAGGCACATCGGACTCATCAGGAAG
CAGGCACATCGGACTCATCAGGAAG
CAGGCACATCGGACTCATCAGGAAG
CAGGCACATCGGACTCATCAGGAAG
CAGGCACATCGGACTCATCAGGAAG
CAGGCACATCGGACTCATCAGGAAG
CAGGCACATCGGACTCATCAGGAAG
CATCGGACTCATCAGGAAGC
CATCGGACTCATCAGGAAGC
CATCGGACTCATCAGGAAGC
CATCGGACTCATCAGGAAGC
CATCGGACTCATCAGGAAGC

CATCGGACTCATCAGGAAGC
CATCGGACTCATCAGGAAGC
ATCGGACTCATCAGGAAGC
ATCGGACTCATCAGGAAGC
ATCGGACTCATCAGGAAGC
ATCGGACTCATCAGGAAGC
ATCGGACTCATCAGGAAGC
ATCGGACTCATCAGGAAGC
ATCGGACTCATCAGGAAGC
GAGGACATCGGACTCATCAGGAAGC
GAGGACATCGGACTCATCAGGAAGC
GAGGACATCGGACTCATCAGGAAGC
GAGGACATCGGACTCATCAGGAAGC
GAGGACATCGGACTCATCAGGAAGC
GAGGACATCGGACTCATCAGGAAGC
GAGGACATCGGACTCATCAGGAAGC
TCATCGGACTCATCAGGAAGC
TCATCGGACTCATCAGGAAGC
TCATCGGACTCATCAGGAAGC
TCATCGGACTCATCAGGAAGC
TCATCGGACTCATCAGGAAGC
TCATCGGACTCATCAGGAAGC
TCATCGGACTCATCAGGAAGC
AGGCATCGGACTCATCAGGAAGC
AGGCATCGGACTCATCAGGAAGC
AGGCATCGGACTCATCAGGAAGC
AGGCATCGGACTCATCAGGAAGC
AGGCATCGGACTCATCAGGAAGC
AGGCATCGGACTCATCAGGAAGC
AGGCATCGGACTCATCAGGAAGC
GGTACATCGGACTCATCAGGAAGC
GGTACATCGGACTCATCAGGAAGC
GGTACATCGGACTCATCAGGAAGC
GGTACATCGGACTCATCAGGAAGC
GGTACATCGGACTCATCAGGAAGC
GGTACATCGGACTCATCAGGAAGC
GGTACATCGGACTCATCAGGAAGC
CATCGGACTCATCAGGAAGC
CATCGGACTCATCAGGAAGC
CATCGGACTCATCAGGAAGC
CATCGGACTCATCAGGAAGC
CATCGGACTCATCAGGAAGC
CATCGGACTCATCAGGAAGC

CACATCGGACTCATCAGGAAGC
TCGGACTCATCAGGAAGC
TCGGACTCATCAGGAAGC
TCGGACTCATCAGGAAGC
GGCACATCGGACTCATCAGGAAGC
GGCACATCGGACTCATCAGGAAGC
GGCACATCGGACTCATCAGGAAGC
ATCATCGGACTCATCAGGAAGC
ATCATCGGACTCATCAGGAAGC
ATCATCGGACTCATCAGGAAGC
CATCGGACTCATCAGGAAGC
CATCGGACTCATCAGGAAGC
CATCGGACTCATCAGGAAGC
CCATCGGACTCATCAGGAAGC
CCATCGGACTCATCAGGAAGC
CCATCGGACTCATCAGGAAGC
GGGGCATCGGACTCATCAGGAAGC
GGGGCATCGGACTCATCAGGAAGC
GGGGCATCGGACTCATCAGGAAGC
AGTCATCGGACTCATCAGGAAGC
AGTCATCGGACTCATCAGGAAGC
AGTCATCGGACTCATCAGGAAGC
TCCATCGGACTCATCAGGAAGC
TCCATCGGACTCATCAGGAAGC
TCCATCGGACTCATCAGGAAGC
GGGCATCGGACTCATCAGGAAGC
GGGCATCGGACTCATCAGGAAGC
GGGCATCGGACTCATCAGGAAGC
ACTGGACATCGGACTCATCAGGAAG
ACTGGACATCGGACTCATCAGGAAG
ACTGGACATCGGACTCATCAGGAAG
CACATCGGACTCATCAGGAAGC
CACATCGGACTCATCAGGAAGC
CACATCGGACTCATCAGGAAGC
GAACATCGGACTCATCAGGAAGC
GAACATCGGACTCATCAGGAAGC
GAACATCGGACTCATCAGGAAGC
GCCATCGGACTCATCAGGAAGC
GCCATCGGACTCATCAGGAAGC
GCCATCGGACTCATCAGGAAGC
CCATCGGACTCATCAGGAAGC
CCATCGGACTCATCAGGAAGC
CCATCGGACTCATCAGGAAGC

CTCATCGGACTCATCAGGAAGC
CTCATCGGACTCATCAGGAAGC
CTCATCGGACTCATCAGGAAGC
AGCATCGGACTCATCAGGAAGC
AGCATCGGACTCATCAGGAAGC
AGCATCGGACTCATCAGGAAGC
GGCATCGGACTCATCAGGAAGC
GGCATCGGACTCATCAGGAAGC
GGCATCGGACTCATCAGGAAGC
GATCAGACCATCGGACTCATCAGGAAG
GATCAGACCATCGGACTCATCAGGAAG
GATCAGACCATCGGACTCATCAGGAAG
ATCGGACTCATCAGGAAGC
ATCGGACTCATCAGGAAGC
ATCGGACTCATCAGGAAGC
TTCATCGGACTCATCAGGAAGC
TTCATCGGACTCATCAGGAAGC
TTCATCGGACTCATCAGGAAGC
GAAGACATCGGACTCATCAGGAAGC
GAAGACATCGGACTCATCAGGAAGC
GAAGACATCGGACTCATCAGGAAGC
AGCCATCGGACTCATCAGGAAGC
AGCCATCGGACTCATCAGGAAGC
AGCCATCGGACTCATCAGGAAGC
ATCGGACTCATCAGGAAGC
ATCGGACTCATCAGGAAGC
ATCGGACTCATCAGGAAGC
TCGGACTCATCAGGAAGC
TCGGACTCATCAGGAAGC
TCGGACTCATCAGGAAGC
ATCGGACTCATCAGGAAGC
ATCGGACTCATCAGGAAGC
ATCGGACTCATCAGGAAGC
AAAGTCATCGGACTCATCAGGAAGC
AAAGTCATCGGACTCATCAGGAAGC
AAAGTCATCGGACTCATCAGGAAGC
GACATCGGACTCATCAGGAAGC
GACATCGGACTCATCAGGAAGC
GACATCGGACTCATCAGGAAGC
AGCATCGGACTCATCAGGAAGC
AGCATCGGACTCATCAGGAAGC
AGCATCGGACTCATCAGGAAGC
ACCTCATCGGACTCATCAGGAAGC

CATCGGACTCATCAGGAAGC
CATCGGACTCATCAGGAAGC
TCCATCGGACTCATCAGGAAGC
TCCATCGGACTCATCAGGAAGC
TCCATCGGACTCATCAGGAAGC
TGGGCCATCGGACTCATCAGGAAGC
TGGGCCATCGGACTCATCAGGAAGC
TGGGCCATCGGACTCATCAGGAAGC
GAATCATCGGACTCATCAGGAAGC
GAATCATCGGACTCATCAGGAAGC
GAATCATCGGACTCATCAGGAAGC
GGAGCATCGGACTCATCAGGAAGC
GGAGCATCGGACTCATCAGGAAGC
GGAGCATCGGACTCATCAGGAAGC
AACATCGGACTCATCAGGAAGC
AACATCGGACTCATCAGGAAGC
AACATCGGACTCATCAGGAAGC
GTCCATCGGACTCATCAGGAAGC
GTCCATCGGACTCATCAGGAAGC
GTCCATCGGACTCATCAGGAAGC
CATCGGACTCATCAGGAAGC
CATCGGACTCATCAGGAAGC
CATCGGACTCATCAGGAAGC
CATCGGACTCATCAGGAAGC
CATCGGACTCATCAGGAAGC
CATCGGACTCATCAGGAAGC
TGGCCATCGGACTCATCAGGAAGC
TGGCCATCGGACTCATCAGGAAGC
TGGCCATCGGACTCATCAGGAAGC
CATCGGACTCATCAGGAAGC
CATCGGACTCATCAGGAAGC
CATCGGACTCATCAGGAAGC
GTAGCATCGGACTCATCAGGAAGC
GTAGCATCGGACTCATCAGGAAGC
GTAGCATCGGACTCATCAGGAAGC
ACATCGGACTCATCAGGAAGC
ACATCGGACTCATCAGGAAGC
ACATCGGACTCATCAGGAAGC
AGTCATCGGACTCATCAGGAAGC
AGTCATCGGACTCATCAGGAAGC
AGTCATCGGACTCATCAGGAAGC
CCCATCGGACTCATCAGGAAGC
CCCATCGGACTCATCAGGAAGC

TGCATCGGACTCATCAGGAAGC
TGCATCGGACTCATCAGGAAGC
TGCATCGGACTCATCAGGAAGC
TGCATCGGACTCATCAGGAAGC
GCATCGGACTCATCAGGAAGC
GCATCGGACTCATCAGGAAGC
GCATCGGACTCATCAGGAAGC
GCATCGGACTCATCAGGAAGC
GCATCGGACTCATCAGGAAGC
GCATCGGACTCATCAGGAAGC
GCATCGGACTCATCAGGAAGC
GCATCGGACTCATCAGGAAGC
GCATCGGACTCATCAGGAAGC
CATCGGACTCATCAGGAAGC
CATCGGACTCATCAGGAAGC
CATCGGACTCATCAGGAAGC
CATCGGACTCATCAGGAAGC
CATCGGACTCATCAGGAAGC
CATCGGACTCATCAGGAAGC
CATCGGACTCATCAGGAAGC
CATCGGACTCATCAGGAAGC
CATCGGACTCATCAGGAAGC
TTCATCGGACTCATCAGGAAGC
TTCATCGGACTCATCAGGAAGC
TTCATCGGACTCATCAGGAAGC
TTCATCGGACTCATCAGGAAGC
TTCATCGGACTCATCAGGAAGC
TTCATCGGACTCATCAGGAAGC
TTCATCGGACTCATCAGGAAGC
TTCATCGGACTCATCAGGAAGC
TTCATCGGACTCATCAGGAAGC
ACTCATCGGACTCATCAGGAAGC
ACTCATCGGACTCATCAGGAAGC
ACTCATCGGACTCATCAGGAAGC
ACTCATCGGACTCATCAGGAAGC
ACTCATCGGACTCATCAGGAAGC
ACTCATCGGACTCATCAGGAAGC
ACTCATCGGACTCATCAGGAAGC
ACTCATCGGACTCATCAGGAAGC
ACTCATCGGACTCATCAGGAAGC
GAGCTCATCGGACTCATCAGGAAG
GAGCTCATCGGACTCATCAGGAAG
GAGCTCATCGGACTCATCAGGAAG

GAGCTCATCGGACTCATCAGGAAG
GAGCTCATCGGACTCATCAGGAAG
GAGCTCATCGGACTCATCAGGAAG
GAGCTCATCGGACTCATCAGGAAG
GAGCTCATCGGACTCATCAGGAAG
GAGCTCATCGGACTCATCAGGAAG
ATGCATCGGACTCATCAGGAAGC
ATGCATCGGACTCATCAGGAAGC
ATGCATCGGACTCATCAGGAAGC
ATGCATCGGACTCATCAGGAAGC
ATGCATCGGACTCATCAGGAAGC
ATGCATCGGACTCATCAGGAAGC
ATGCATCGGACTCATCAGGAAGC
ATGCATCGGACTCATCAGGAAGC
ATGCATCGGACTCATCAGGAAGC
CATCGGACTCATCAGGAAGC
CATCGGACTCATCAGGAAGC
CATCGGACTCATCAGGAAGC
CATCGGACTCATCAGGAAGC
CATCGGACTCATCAGGAAGC
CATCGGACTCATCAGGAAGC
CATCGGACTCATCAGGAAGC
CATCGGACTCATCAGGAAGC
CATCGGACTCATCAGGAAGC
AACATCGGACTCATCAGGAAGC
AACATCGGACTCATCAGGAAGC
AACATCGGACTCATCAGGAAGC
AACATCGGACTCATCAGGAAGC
AACATCGGACTCATCAGGAAGC
AACATCGGACTCATCAGGAAGC
AACATCGGACTCATCAGGAAGC
AACATCGGACTCATCAGGAAGC
AACATCGGACTCATCAGGAAGC
CATCGGACTCATCAGGAAGC
CATCGGACTCATCAGGAAGC
CATCGGACTCATCAGGAAGC
CATCGGACTCATCAGGAAGC
CATCGGACTCATCAGGAAGC
CATCGGACTCATCAGGAAGC
CATCGGACTCATCAGGAAGC
CATCGGACTCATCAGGAAGC
CATCGGACTCATCAGGAAGC
AGCATCGGACTCATCAGGAAGC

AGCATCGGACTCATCAGGAAGC
AGCATCGGACTCATCAGGAAGC
AGCATCGGACTCATCAGGAAGC
AGCATCGGACTCATCAGGAAGC
AGCATCGGACTCATCAGGAAGC
AGCATCGGACTCATCAGGAAGC
AGCATCGGACTCATCAGGAAGC
AGCATCGGACTCATCAGGAAGC
CCATCGGACTCATCAGGAAGC
CCATCGGACTCATCAGGAAGC
CCATCGGACTCATCAGGAAGC
CCATCGGACTCATCAGGAAGC
CCATCGGACTCATCAGGAAGC
CCATCGGACTCATCAGGAAGC
CCATCGGACTCATCAGGAAGC
CCATCGGACTCATCAGGAAGC
CCATCGGACTCATCAGGAAGC
TCATCGGACTCATCAGGAAGC
TCATCGGACTCATCAGGAAGC
TCATCGGACTCATCAGGAAGC
TCATCGGACTCATCAGGAAGC
TCATCGGACTCATCAGGAAGC
TCATCGGACTCATCAGGAAGC
TCATCGGACTCATCAGGAAGC
TCATCGGACTCATCAGGAAGC
TCATCGGACTCATCAGGAAGC
TCTCATCGGACTCATCAGGAAGC
TCTCATCGGACTCATCAGGAAGC
TCTCATCGGACTCATCAGGAAGC
TCTCATCGGACTCATCAGGAAGC
TCTCATCGGACTCATCAGGAAGC
TCTCATCGGACTCATCAGGAAGC
TCTCATCGGACTCATCAGGAAGC
TCTCATCGGACTCATCAGGAAGC
TCTCATCGGACTCATCAGGAAGC
GTAGTCCATCGGACTCATCAGGAAG
GTAGTCCATCGGACTCATCAGGAAG
GTAGTCCATCGGACTCATCAGGAAG
GTAGTCCATCGGACTCATCAGGAAG
GTAGTCCATCGGACTCATCAGGAAG
GTAGTCCATCGGACTCATCAGGAAG
GTAGTCCATCGGACTCATCAGGAAG
GTAGTCCATCGGACTCATCAGGAAG

ATAACATCGGACTCATCAGGAAGC
ATAACATCGGACTCATCAGGAAGC
ATAACATCGGACTCATCAGGAAGC
GACTCATCGGACTCATCAGGAAGC
GACTCATCGGACTCATCAGGAAGC
GACTCATCGGACTCATCAGGAAGC
GACTCATCGGACTCATCAGGAAGC
CATCGGACTCATCAGGAAGC
CATCGGACTCATCAGGAAGC
CATCGGACTCATCAGGAAGC
CATCGGACTCATCAGGAAGC
CCATCGGACTCATCAGGAAGC
CCATCGGACTCATCAGGAAGC
CCATCGGACTCATCAGGAAGC
CCATCGGACTCATCAGGAAGC
ATCCATCGGACTCATCAGGAAGC
ATCCATCGGACTCATCAGGAAGC
ATCCATCGGACTCATCAGGAAGC
ATCCATCGGACTCATCAGGAAGC
CGTCATCGGACTCATCAGGAAGC
CGTCATCGGACTCATCAGGAAGC
CGTCATCGGACTCATCAGGAAGC
CGTCATCGGACTCATCAGGAAGC
GCATCGGACTCATCAGGAAGC
GCATCGGACTCATCAGGAAGC
GCATCGGACTCATCAGGAAGC
GCATCGGACTCATCAGGAAGC
ACGCATCGGACTCATCAGGAAGC
ACGCATCGGACTCATCAGGAAGC
ACGCATCGGACTCATCAGGAAGC
ACGCATCGGACTCATCAGGAAGC
AACATCGGACTCATCAGGAAGC
AACATCGGACTCATCAGGAAGC
AACATCGGACTCATCAGGAAGC
AACATCGGACTCATCAGGAAGC
ACCATCGGACTCATCAGGAAGC
ACCATCGGACTCATCAGGAAGC
ACCATCGGACTCATCAGGAAGC
ACCATCGGACTCATCAGGAAGC
TACATCGGACTCATCAGGAAGC
TACATCGGACTCATCAGGAAGC
TACATCGGACTCATCAGGAAGC
TACATCGGACTCATCAGGAAGC

CATCGGACTCATCAGGAAGC
CATCGGACTCATCAGGAAGC
TCCTCATCGGACTCATCAGGAAGC
TCCTCATCGGACTCATCAGGAAGC
CTGCATCGGACTCATCAGGAAGC
CTGCATCGGACTCATCAGGAAGC
CACTCATCGGACTCATCAGGAAGC
CACTCATCGGACTCATCAGGAAGC
CATCGGACTCATCAGGAAGC
CATCGGACTCATCAGGAAGC
TTCATCGGACTCATCAGGAAGC
TTCATCGGACTCATCAGGAAGC
ACATCGGACTCATCAGGAAGC
ACATCGGACTCATCAGGAAGC
ATCGGACTCATCAGGAAGC
ATCGGACTCATCAGGAAGC
GACTCATCGGACTCATCAGGAAGC
GACTCATCGGACTCATCAGGAAGC
TGACATCGGACTCATCAGGAAGC
TGACATCGGACTCATCAGGAAGC
AGCATCGGACTCATCAGGAAGC
AGCATCGGACTCATCAGGAAGC
ATCGGACTCATCAGGAAGC
ATCGGACTCATCAGGAAGC
CGCACATCGGACTCATCAGGAAGC
CGCACATCGGACTCATCAGGAAGC
AGAGCATCGGACTCATCAGGAAGC
AGAGCATCGGACTCATCAGGAAGC
TACATCGGACTCATCAGGAAGC
TACATCGGACTCATCAGGAAGC
CAACATCGGACTCATCAGGAAGC
CAACATCGGACTCATCAGGAAGC
TGGCATCGGACTCATCAGGAAGC
TGGCATCGGACTCATCAGGAAGC
CATCGGACTCATCAGGAAGC
CATCGGACTCATCAGGAAGC
ATCGGACTCATCAGGAAGC
ATCGGACTCATCAGGAAGC
GGAACATCGGACTCATCAGGAAGC
GGAACATCGGACTCATCAGGAAGC
ACATCGGACTCATCAGGAAGC
ACATCGGACTCATCAGGAAGC
ACATCGGACTCATCAGGAAGC

GCATCGGACTCATCAGGAAGC
CATCGGACTCATCAGGAAGC
CATCGGACTCATCAGGAAGC
TCATCGGACTCATCAGGAAGC
TCATCGGACTCATCAGGAAGC
GATCATCGGACTCATCAGGAAGC
GATCATCGGACTCATCAGGAAGC
CCATCGGACTCATCAGGAAGC
CCATCGGACTCATCAGGAAGC
AGCATCGGACTCATCAGGAAGC
AGCATCGGACTCATCAGGAAGC
CATCGGACTCATCAGGAAGC
CATCGGACTCATCAGGAAGC
CATCCATCGGACTCATCAGGAAGC
CATCCATCGGACTCATCAGGAAGC
CCATCGGACTCATCAGGAAGC
CCATCGGACTCATCAGGAAGC
ATGGCATCGGACTCATCAGGAAGC
ATGGCATCGGACTCATCAGGAAGC
CATCGGACTCATCAGGAAGC
CATCGGACTCATCAGGAAGC
AATCATCGGACTCATCAGGAAGC
AATCATCGGACTCATCAGGAAGC
GCCATCGGACTCATCAGGAAGC
GCCATCGGACTCATCAGGAAGC
CCCATCGGACTCATCAGGAAGC
CCCATCGGACTCATCAGGAAGC
GACATCGGACTCATCAGGAAGC
GACATCGGACTCATCAGGAAGC
CATCGGACTCATCAGGAAGC
CATCGGACTCATCAGGAAGC
TTCATCGGACTCATCAGGAAGC
TTCATCGGACTCATCAGGAAGC
TCATCGGACTCATCAGGAAGC
TCATCGGACTCATCAGGAAGC
GCATCGGACTCATCAGGAAGC
GCATCGGACTCATCAGGAAGC
CCCATCGGACTCATCAGGAAGC
CCCATCGGACTCATCAGGAAGC
CTCCGTCATCGGACTCATCAGGAAG
CTCCGTCATCGGACTCATCAGGAAG
CATCGGACTCATCAGGAAGC
CATCGGACTCATCAGGAAGC

CCATCGGACTCATCAGGAAGC

CCATCGGACTCATCAGGAAGC

GCATCGGACTCATCAGGAAGC

AGCATCGGACTCATCAGGAAGC
TGCATCGGACTCATCAGGAAGC
TGCATCGGACTCATCAGGAAGC
ATGTCCATCGGACTCATCAGGAAGC
ATGTCCATCGGACTCATCAGGAAGC
AATGCACATCGGACTCATCAGGAAG
AATGCACATCGGACTCATCAGGAAG
CATCGGACTCATCAGGAAGC
CATCGGACTCATCAGGAAGC
ACATCGGACTCATCAGGAAGC
ACATCGGACTCATCAGGAAGC
CATCGGACTCATCAGGAAGC
CATCGGACTCATCAGGAAGC
ACATCGGACTCATCAGGAAGC
ACATCGGACTCATCAGGAAGC
CATCGGACTCATCAGGAAGC
CATCGGACTCATCAGGAAGC
ATCATCGGACTCATCAGGAAGC
ATCATCGGACTCATCAGGAAGC
ATCGGACTCATCAGGAAGC
ATCGGACTCATCAGGAAGC
TCATCGGACTCATCAGGAAGC
TCATCGGACTCATCAGGAAGC
TCATCGGACTCATCAGGAAGC
TCATCGGACTCATCAGGAAGC
GGCATCGGACTCATCAGGAAGC
GGCATCGGACTCATCAGGAAGC
TCCGCATCGGACTCATCAGGAAGC
TCCGCATCGGACTCATCAGGAAGC
GCATCGGACTCATCAGGAAGC
GCATCGGACTCATCAGGAAGC
CCATCGGACTCATCAGGAAGC
CCATCGGACTCATCAGGAAGC
TCATCGGACTCATCAGGAAGC
TCATCGGACTCATCAGGAAGC
TCGGACTCATCAGGAAGC
TCGGACTCATCAGGAAGC
CCATCGGACTCATCAGGAAGC
CCATCGGACTCATCAGGAAGC
ATCGGACTCATCAGGAAGC
ATCGGACTCATCAGGAAGC
CATCGGACTCATCAGGAAGC
CATCGGACTCATCAGGAAGC

CATCGGACTCATCAGGAAGC
CTGCATCGGACTCATCAGGAAGC
CTGCATCGGACTCATCAGGAAGC
AACATCGGACTCATCAGGAAGC
AACATCGGACTCATCAGGAAGC
TGCATCGGACTCATCAGGAAGC
TGCATCGGACTCATCAGGAAGC
AGTCATCGGACTCATCAGGAAGC
AGTCATCGGACTCATCAGGAAGC
CATCGGACTCATCAGGAAGC
CATCGGACTCATCAGGAAGC
CATCGGACTCATCAGGAAGC
CATCGGACTCATCAGGAAGC
TCATCGGACTCATCAGGAAGC
TCATCGGACTCATCAGGAAGC
GTCATCGGACTCATCAGGAAGC
GTCATCGGACTCATCAGGAAGC
GCATCGGACTCATCAGGAAGC
GCATCGGACTCATCAGGAAGC
AGCATCGGACTCATCAGGAAGC
AGCATCGGACTCATCAGGAAGC
GACATCGGACTCATCAGGAAGC
GACATCGGACTCATCAGGAAGC
TCGGACTCATCAGGAAGC
TCGGACTCATCAGGAAGC
GCATCGGACTCATCAGGAAGC
GCATCGGACTCATCAGGAAGC
CTCCCATCGGACTCATCAGGAAGC
CTCCCATCGGACTCATCAGGAAGC
CCATCGGACTCATCAGGAAGC
CCATCGGACTCATCAGGAAGC
CCATCGGACTCATCAGGAAGC
CCATCGGACTCATCAGGAAGC
TCATCGGACTCATCAGGAAGC
TCATCGGACTCATCAGGAAGC
TCACCCATCGGACTCATCAGGAAGC
TCACCCATCGGACTCATCAGGAAGC
GAAGCATCGGACTCATCAGGAAGC
GAAGCATCGGACTCATCAGGAAGC
GCTGTCCATCGGACTCATCAGGAAG
GCTGTCCATCGGACTCATCAGGAAG
TACATCGGACTCATCAGGAAGC
TACATCGGACTCATCAGGAAGC

AGCATCGGACTCATCAGGAAGC

AGCATCGGACTCATCAGGAAGC

GACTGCATCGGACTCATCAGGAAGC

GACTGCATCGGACTCATCAGGAAGC

AGCATCGGACTCATCAGGAAGC

AGCATCGGACTCATCAGGAAGC

GGACATCGGACTCATCAGGAAGC

GGACATCGGACTCATCAGGAAGC

TACATCGGACTCATCAGGAAGC

TACATCGGACTCATCAGGAAGC

GTGATCATCGGACTCATCAGGAAGC

GTGATCATCGGACTCATCAGGAAGC

TATCATCGGACTCATCAGGAAGC

TATCATCGGACTCATCAGGAAGC

CAGCATCGGACTCATCAGGAAGC

CAGCATCGGACTCATCAGGAAGC

TCTCATCGGACTCATCAGGAAGC

TCTCATCGGACTCATCAGGAAGC

CATCGGACTCATCAGGAAGC
CATCGGACTCATCAGGAAGC
CATCGGACTCATCAGGAAGC
CTCCATCGGACTCATCAGGAAGC
CTCCATCGGACTCATCAGGAAGC
TTCATCGGACTCATCAGGAAGC
TTCATCGGACTCATCAGGAAGC
CATCGGACTCATCAGGAAGC
CATCGGACTCATCAGGAAGC
CATCGGACTCATCAGGAAGC
CATCGGACTCATCAGGAAGC
CATCGGACTCATCAGGAAGC
CATCGGACTCATCAGGAAGC
GCATCGGACTCATCAGGAAGC
GCATCGGACTCATCAGGAAGC
CCTGTGCATCGGACTCATCAGGAAG
CCTGTGCATCGGACTCATCAGGAAG
TCATCGGACTCATCAGGAAGC
TCATCGGACTCATCAGGAAGC
TTCATCGGACTCATCAGGAAGC
TTCATCGGACTCATCAGGAAGC
CTCATCGGACTCATCAGGAAGC
CTCATCGGACTCATCAGGAAGC
GGCATCGGACTCATCAGGAAGC
GGCATCGGACTCATCAGGAAGC
GCATCGGACTCATCAGGAAGC
GCATCGGACTCATCAGGAAGC
TCATCGGACTCATCAGGAAGC
TCATCGGACTCATCAGGAAGC
TCATCGGACTCATCAGGAAGC
TCATCGGACTCATCAGGAAGC
TTCACATCGGACTCATCAGGAAGC
TTCACATCGGACTCATCAGGAAGC
GAACATCGGACTCATCAGGAAGC
GAACATCGGACTCATCAGGAAGC
CTCATCGGACTCATCAGGAAGC
CTCATCGGACTCATCAGGAAGC
ACATCGGACTCATCAGGAAGC
ACATCGGACTCATCAGGAAGC
GTGACATCGGACTCATCAGGAAGC
GTGACATCGGACTCATCAGGAAGC
CCATCGGACTCATCAGGAAGC
CCATCGGACTCATCAGGAAGC

TGGTGCATCGGACTCATCAGGAAGC
TGGTGCATCGGACTCATCAGGAAGC
CATCGGACTCATCAGGAAGC
CATCGGACTCATCAGGAAGC
GCATCGGACTCATCAGGAAGC
GCATCGGACTCATCAGGAAGC
TCATCGGACTCATCAGGAAGC
TCATCGGACTCATCAGGAAGC
AGCATCGGACTCATCAGGAAGC
AGCATCGGACTCATCAGGAAGC
TGCATCGGACTCATCAGGAAGC
TGCATCGGACTCATCAGGAAGC
CCATCGGACTCATCAGGAAGC
CCATCGGACTCATCAGGAAGC
GCATCGGACTCATCAGGAAGC
GCATCGGACTCATCAGGAAGC
GCATCGGACTCATCAGGAAGC
GCATCGGACTCATCAGGAAGC
GAACATCGGACTCATCAGGAAGC
GAACATCGGACTCATCAGGAAGC
TACATCGGACTCATCAGGAAGC
TACATCGGACTCATCAGGAAGC
CATCGGACTCATCAGGAAGC
CATCGGACTCATCAGGAAGC
ATCGGACTCATCAGGAAGC
ATCGGACTCATCAGGAAGC
CTCATCGGACTCATCAGGAAGC
CTCATCGGACTCATCAGGAAGC
TCTCCATCGGACTCATCAGGAAGC
TCTCCATCGGACTCATCAGGAAGC
TCCATCGGACTCATCAGGAAGC
TCCATCGGACTCATCAGGAAGC
AAACATCGGACTCATCAGGAAGC
AAACATCGGACTCATCAGGAAGC
GGCCATCGGACTCATCAGGAAGC
GGCCATCGGACTCATCAGGAAGC
CATCGGACTCATCAGGAAGC
CATCGGACTCATCAGGAAGC
CATCGGACTCATCAGGAAGC
CATCGGACTCATCAGGAAGC
GGAGACATCGGACTCATCAGGAAGC
GGAGACATCGGACTCATCAGGAAGC
ACATCGGACTCATCAGGAAGC

ACATCGGACTCATCAGGAAGC

CATCGGACTCATCAGGAAGC

CATCGGACTCATCAGGAAGC

ATGCTCATCGGACTCATCAGGAAG

ATGCTCATCGGACTCATCAGGAAG

ACCATCGGACTCATCAGGAAGC

ACCATCGGACTCATCAGGAAGC

CTGCATCGGACTCATCAGGAAGC

CTGCATCGGACTCATCAGGAAGC

ACATCGGACTCATCAGGAAGC

TGATGCATCGGACTCATCAGGAAGC

TGATGCATCGGACTCATCAGGAAGC

CCATCGGACTCATCAGGAAGC

CCATCGGACTCATCAGGAAGC

CCATCGGACTCATCAGGAAGC

CCATCGGACTCATCAGGAAGC

CATCGGACTCATCAGGAAGC

CATCGGACTCATCAGGAAGC

ACATCGGACTCATCAGGAAGC

ACATCGGACTCATCAGGAAGC

TCGGACTCATCAGGAAGC

TCGGACTCATCAGGAAGC

TACATCGGACTCATCAGGAAGC

TACATCGGACTCATCAGGAAGC

ATTGAACATCGGACTCATCAGGAAGC

ATTGAACATCGGACTCATCAGGAAGC

ATCGGACTCATCAGGAAGC

ATCGGACTCATCAGGAAGC

ACATCGGACTCATCAGGAAGC

ACATCGGACTCATCAGGAAGC

ACTAGACATCGGACTCATCAGGAAG

ACTAGACATCGGACTCATCAGGAAG

AAAGCATCGGACTCATCAGGAAGC

AAAGCATCGGACTCATCAGGAAGC

CCATCGGACTCATCAGGAAGC

CCATCGGACTCATCAGGAAGC

AGCATCGGACTCATCAGGAAGC

AGCATCGGACTCATCAGGAAGC

CATCGGACTCATCAGGAAGC

CATCGGACTCATCAGGAAGC
CATCGGACTCATCAGGAAGC
CATCGGACTCATCAGGAAGC
CATCGGACTCATCAGGAAGC
CATCGGACTCATCAGGAAGC
CATCGGACTCATCAGGAAGC
CATCGGACTCATCAGGAAGC
CGGTAGCATCGGACTCATCAGGAAG
CGGTAGCATCGGACTCATCAGGAAG
CATCGGACTCATCAGGAAGC
CATCGGACTCATCAGGAAGC
GCATCGGACTCATCAGGAAGC
GCATCGGACTCATCAGGAAGC
CATCGGACTCATCAGGAAGC
CATCGGACTCATCAGGAAGC
ATCGGACTCATCAGGAAGC
ATCGGACTCATCAGGAAGC
GCATCGGACTCATCAGGAAGC
GCATCGGACTCATCAGGAAGC
GCATCGGACTCATCAGGAAGC
GCATCGGACTCATCAGGAAGC
GCATCGGACTCATCAGGAAGC
GCATCGGACTCATCAGGAAGC
GTCATCGGACTCATCAGGAAGC
GTCATCGGACTCATCAGGAAGC
TAACATCGGACTCATCAGGAAGC
TAACATCGGACTCATCAGGAAGC
GCATCGGACTCATCAGGAAGC
GCATCGGACTCATCAGGAAGC
GCCATCGGACTCATCAGGAAGC
GCCATCGGACTCATCAGGAAGC
CCTCATCGGACTCATCAGGAAGC
CCTCATCGGACTCATCAGGAAGC
CATCGGACTCATCAGGAAGC
CATCGGACTCATCAGGAAGC
CATCGGACTCATCAGGAAGC
CATCGGACTCATCAGGAAGC
CATACATCGGACTCATCAGGAAGC
CATACATCGGACTCATCAGGAAGC
CATCGGACTCATCAGGAAGC
CATCGGACTCATCAGGAAGC
AACATCGGACTCATCAGGAAGC
AACATCGGACTCATCAGGAAGC

GCATCGGACTCATCAGGAAGC

GCATCGGACTCATCAGGAAGC

GAACATCGGACTCATCAGGAAGC

GAACATCGGACTCATCAGGAAGC

GATGACATCGGACTCATCAGGAAGC

GATGACATCGGACTCATCAGGAAGC

AACATCGGACTCATCAGGAAGC

AACATCGGACTCATCAGGAAGC

AGGCATCGGACTCATCAGGAAGC

TTGCATCGGACTCATCAGGAAGC
TCATCGGACTCATCAGGAAGC
TCATCGGACTCATCAGGAAGC
CCATCGGACTCATCAGGAAGC
CCATCGGACTCATCAGGAAGC
CATCGGACTCATCAGGAAGC
CATCGGACTCATCAGGAAGC
GTCCATCGGACTCATCAGGAAGC
GTCCATCGGACTCATCAGGAAGC
ACATCGGACTCATCAGGAAGC
ACATCGGACTCATCAGGAAGC
CCATCATCGGACTCATCAGGAAGC
CCATCATCGGACTCATCAGGAAGC
ATTTGCCATCGGACTCATCAGGAAG
ATTTGCCATCGGACTCATCAGGAAG
CATCGGACTCATCAGGAAGC
CATCGGACTCATCAGGAAGC
TGAACATCGGACTCATCAGGAAGC
TGAACATCGGACTCATCAGGAAGC
GCGTCATCGGACTCATCAGGAAGC
GCGTCATCGGACTCATCAGGAAGC
CATCGGACTCATCAGGAAGC
CATCGGACTCATCAGGAAGC
GGCATCGGACTCATCAGGAAGC
GGCATCGGACTCATCAGGAAGC
TCATCGGACTCATCAGGAAGC
TCATCGGACTCATCAGGAAGC
TTTCATCGGACTCATCAGGAAGC
TTTCATCGGACTCATCAGGAAGC
TCTCATCGGACTCATCAGGAAGC
TCTCATCGGACTCATCAGGAAGC
ATCGGACTCATCAGGAAGC
ATCGGACTCATCAGGAAGC
AACCCATCGGACTCATCAGGAAGC
AACCCATCGGACTCATCAGGAAGC
AGCATCGGACTCATCAGGAAGC
AGCATCGGACTCATCAGGAAGC
CCCATCGGACTCATCAGGAAGC
CCCATCGGACTCATCAGGAAGC
AACATCGGACTCATCAGGAAGC
AACATCGGACTCATCAGGAAGC
CATCGGACTCATCAGGAAGC
CATCGGACTCATCAGGAAGC

TCATCGGACTCATCAGGAAGC

TCATCGGACTCATCAGGAAGC

TCATCGGACTCATCAGGAAGC

TCATCGGACTCATCAGGAAGC

TCATCGGACTCATCAGGAAGC

TCGGACTCATCAGGAAGC

TCGGACTCATCAGGAAGC

TATCTCATCGGACTCATCAGGAAGC

TATCTCATCGGACTCATCAGGAAGC

GAGACATCGGACTCATCAGGAAGC

ATCGGACTCATCAGGAAGC
ATCGGACTCATCAGGAAGC
AGTTCATCGGACTCATCAGGAAGC
AGTTCATCGGACTCATCAGGAAGC
GTGCATCGGACTCATCAGGAAGC
GTGCATCGGACTCATCAGGAAGC
AGATGCATCGGACTCATCAGGAAGC
AGATGCATCGGACTCATCAGGAAGC
ACATCGGACTCATCAGGAAGC
ACATCGGACTCATCAGGAAGC
TAACATCGGACTCATCAGGAAGC
TAACATCGGACTCATCAGGAAGC
AACATCGGACTCATCAGGAAGC
AACATCGGACTCATCAGGAAGC
AACATCGGACTCATCAGGAAGC
AACATCGGACTCATCAGGAAGC
GCCATCGGACTCATCAGGAAGC
GCCATCGGACTCATCAGGAAGC
CATCGGACTCATCAGGAAGC
CATCGGACTCATCAGGAAGC
CCATCGGACTCATCAGGAAGC
CCATCGGACTCATCAGGAAGC
TCATCGGACTCATCAGGAAGC
TCATCGGACTCATCAGGAAGC
CCATCGGACTCATCAGGAAGC
CCATCGGACTCATCAGGAAGC
ATTGCATCGGACTCATCAGGAAGC
ATTGCATCGGACTCATCAGGAAGC
GATACATCGGACTCATCAGGAAGC
GATACATCGGACTCATCAGGAAGC
ATTCATCGGACTCATCAGGAAGC
ATTCATCGGACTCATCAGGAAGC
GAGGACATCGGACTCATCAGGAAGC
GAGGACATCGGACTCATCAGGAAGC
GGACATCGGACTCATCAGGAAGC
GGACATCGGACTCATCAGGAAGC
CTGCCATCGGACTCATCAGGAAGC
CTGCCATCGGACTCATCAGGAAGC
CATCGGACTCATCAGGAAGC
CATCGGACTCATCAGGAAGC
CTCATCGGACTCATCAGGAAGC
CTCATCGGACTCATCAGGAAGC
ACACATCGGACTCATCAGGAAGC

AGAACACATCGGACTCATCAGGAAG
AGAACACATCGGACTCATCAGGAAG
AGAACACATCGGACTCATCAGGAAG
AGAACACATCGGACTCATCAGGAAG
AGAACACATCGGACTCATCAGGAAG
TGCCATCGGACTCATCAGGAAGC
TGCCATCGGACTCATCAGGAAGC
TGCCATCGGACTCATCAGGAAGC
TGCCATCGGACTCATCAGGAAGC
TGCCATCGGACTCATCAGGAAGC
AGCATCGGACTCATCAGGAAGC
AGCATCGGACTCATCAGGAAGC
AGCATCGGACTCATCAGGAAGC
AGCATCGGACTCATCAGGAAGC
AGCATCGGACTCATCAGGAAGC
GAGCATCGGACTCATCAGGAAGC
GAGCATCGGACTCATCAGGAAGC
GAGCATCGGACTCATCAGGAAGC
GAGCATCGGACTCATCAGGAAGC
GAGCATCGGACTCATCAGGAAGC
TTTCCATCGGACTCATCAGGAAGC
TTTCCATCGGACTCATCAGGAAGC
TTTCCATCGGACTCATCAGGAAGC
TTTCCATCGGACTCATCAGGAAGC
TTTCCATCGGACTCATCAGGAAGC
CCATCGGACTCATCAGGAAGC
CCATCGGACTCATCAGGAAGC
CCATCGGACTCATCAGGAAGC
CCATCGGACTCATCAGGAAGC
CCATCGGACTCATCAGGAAGC
CATTCATCGGACTCATCAGGAAGC
CATTCATCGGACTCATCAGGAAGC
CATTCATCGGACTCATCAGGAAGC
CATTCATCGGACTCATCAGGAAGC
CATTCATCGGACTCATCAGGAAGC
ACTTCATCGGACTCATCAGGAAGC
ACTTCATCGGACTCATCAGGAAGC
ACTTCATCGGACTCATCAGGAAGC
ACTTCATCGGACTCATCAGGAAGC
ACTTCATCGGACTCATCAGGAAGC
AGTTCATCGGACTCATCAGGAAGC
AGTTCATCGGACTCATCAGGAAGC
AGTTCATCGGACTCATCAGGAAGC

TGGCATCGGACTCATCAGGAAGC
TGGCATCGGACTCATCAGGAAGC
TGGCATCGGACTCATCAGGAAGC
TGGCATCGGACTCATCAGGAAGC
TGGCATCGGACTCATCAGGAAGC
GACATCGGACTCATCAGGAAGC
GACATCGGACTCATCAGGAAGC
GACATCGGACTCATCAGGAAGC
GACATCGGACTCATCAGGAAGC
GACATCGGACTCATCAGGAAGC
ACATCGGACTCATCAGGAAGC
ACATCGGACTCATCAGGAAGC
ACATCGGACTCATCAGGAAGC
ACATCGGACTCATCAGGAAGC
ACATCGGACTCATCAGGAAGC
CATCGGACTCATCAGGAAGC
CATCGGACTCATCAGGAAGC
CATCGGACTCATCAGGAAGC
CATCGGACTCATCAGGAAGC
CATCGGACTCATCAGGAAGC
ACATCGGACTCATCAGGAAGC
ACATCGGACTCATCAGGAAGC
ACATCGGACTCATCAGGAAGC
ACATCGGACTCATCAGGAAGC
ACATCGGACTCATCAGGAAGC
GCTGTCCATCGGACTCATCAGGAAG
GCTGTCCATCGGACTCATCAGGAAG
GCTGTCCATCGGACTCATCAGGAAG
GCTGTCCATCGGACTCATCAGGAAG
GCTGTCCATCGGACTCATCAGGAAG
CATCGGACTCATCAGGAAGC
CATCGGACTCATCAGGAAGC
CATCGGACTCATCAGGAAGC
CATCGGACTCATCAGGAAGC
CATCGGACTCATCAGGAAGC
CATCGGACTCATCAGGAAGC
CATCGGACTCATCAGGAAGC
CATCGGACTCATCAGGAAGC
CATCGGACTCATCAGGAAGC
CATCGGACTCATCAGGAAGC
TGGTCATCGGACTCATCAGGAAGC
TGGTCATCGGACTCATCAGGAAGC
TGGTCATCGGACTCATCAGGAAGC

TCATCGGACTCATCAGGAAGC
TCATCGGACTCATCAGGAAGC
TCATCGGACTCATCAGGAAGC
CCATCGGACTCATCAGGAAGC
CCATCGGACTCATCAGGAAGC
CCATCGGACTCATCAGGAAGC
CCATCGGACTCATCAGGAAGC
CCATCGGACTCATCAGGAAGC
GTATAGGTCATCGGACTCATCAGGAA
GTTTCATCGGACTCATCAGGAAG
TTTATGCCATCGGACTCATCAGGAAG
TGAGGCATCGGACTCATCAGGAAG
GAGCACATCGGACTCATCAGGAAG
TCATGTCATCGGACTCATCAGGAAG
TGTTGTCATCGGACTCATCAGGAAG
GTTTACACATCGGACTCATCAGGAAG
CCATGCCATCGGACTCATCAGGAAG
ACACTCACATCGGACTCATCAGGAAG
TAAGACCATCGGACTCATCAGGAAG
AAGACATCGGACTCATCAGGAAG
AAAGATCATCGGACTCATCAGGAAGC
ATGGCACATCGGACTCATCAGGAAGC
CCATGTCATCGGACTCATCAGGAAGC
ATTCCATCGGACTCATCAGGAAGC
CCTTCACATCGGACTCATCAGGAAGC
GTCCATCGGACTCATCAGGAAGC
TGCCACATCGGACTCATCAGGAAGC
TTGAAACATCGGACTCATCAGGAAGC
ATGACACATCGGACTCATCAGGAAGC
AAGAAGCATCGGACTCATCAGGAAGC
ATGCAGCATCGGACTCATCAGGAAGC
AGGGCACATCGGACTCATCAGGAAGC
ACAATCATCGGACTCATCAGGAAGC
GACCACATCGGACTCATCAGGAAGC
GTACTGACATCGGACTCATCAGGAAGC
TAGCGTCATCGGACTCATCAGGAAGC
TATTGCTCATCGGACTCATCAGGAAGC
AAATGCATCGGACTCATCAGGAAGC
ATACGACATCGGACTCATCAGGAAGC
CCGTGCATCGGACTCATCAGGAAGC
TCCTCATCGGACTCATCAGGAAGC
AGGGCCATCGGACTCATCAGGAAGC
CTCCATCGGACTCATCAGGAAGC

GCTGGACATCGGACTCATCAGGAAGC
ACAGGACATCGGACTCATCAGGAAGC
GTATCTCATCGGACTCATCAGGAAGC
TCCTTGACATCGGACTCATCAGGAAGC
AGTATGTCATCGGACTCATCAGGAAGC
CACCTCATCGGACTCATCAGGAAGC
CGCAACATCGGACTCATCAGGAAGC
ATTGGTCATCGGACTCATCAGGAAGC
GGCACATCGGACTCATCAGGAAGC
AGCCATCGGACTCATCAGGAAGC
TCCCCATCGGACTCATCAGGAAGC
TTCTGACATCGGACTCATCAGGAAGC
TGTGTCATCGGACTCATCAGGAAGC
GAACCCATCGGACTCATCAGGAAGC
GGCCATCGGACTCATCAGGAAGC
CAACACATCGGACTCATCAGGAAGC
ATACACATCGGACTCATCAGGAAGC
TTGCTCATCGGACTCATCAGGAAGC
TGTGCCATCGGACTCATCAGGAAGC
AGAGGCATCGGACTCATCAGGAAGC
CTACTGCATCGGACTCATCAGGAAGC
TCCATCATCGGACTCATCAGGAAGC
CTACCTCATCGGACTCATCAGGAAGC
GACCACATCGGACTCATCAGGAAGC
TAGCTGCATCGGACTCATCAGGAAGC
AGGATCATCGGACTCATCAGGAAGC
ATACGCATCGGACTCATCAGGAAGC
TTCTCATCGGACTCATCAGGAAGC
ACCATGACATCGGACTCATCAGGAAGC
AAGCCATCGGACTCATCAGGAAGC
TATGAGCCATCGGACTCATCAGGAAGC
TGTGTCATCGGACTCATCAGGAAGC
CGAACTCACATCGGACTCATCAGGAAGC
AGTTACATCGGACTCATCAGGAAGC
CCAGCCATCGGACTCATCAGGAAGC
CACTAACATCGGACTCATCAGGAAGC
GGCTTCATCGGACTCATCAGGAAGC
TGCTGTGCATCGGACTCATCAGGAAGC
ACACATCATCGGACTCATCAGGAAGC
ACATACACATCGGACTCATCAGGAAGC
TTCAGCCATCGGACTCATCAGGAAGC
TTGTCATCGGACTCATCAGGAAGC
TTTGTCATCGGACTCATCAGGAAGC

TTGTTTCATCGGACTCATCAGGAAGC
AAAGGCTCATCGGACTCATCAGGAAGC
CGCACGCATCGGACTCATCAGGAAGC
AATGCCCATCGGACTCATCAGGAAGC
TGGAATGCATCGGACTCATCAGGAAGC
AGCAGACATCGGACTCATCAGGAAGC
ATATGACATCGGACTCATCAGGAAGC
TCACATCGGACTCATCAGGAAGC
CGTTCATCGGACTCATCAGGAAGC
GCAACATCGGACTCATCAGGAAGC
GCTAGGCATCGGACTCATCAGGAAGC
TCCACCATCGGACTCATCAGGAAGC
GGGGGCATCGGACTCATCAGGAAGC
CTCCATCGGACTCATCAGGAAGC
TATAAGTCATCGGACTCATCAGGAAGC
CTCCACCATCGGACTCATCAGGAAGC
TGCAGACATCGGACTCATCAGGAAGC
AATCACATCGGACTCATCAGGAAGC
GCCCCTCATCGGACTCATCAGGAAGC
GTGCCCATCGGACTCATCAGGAAGC
TTCTCATCGGACTCATCAGGAAGC
CAAATACATCGGACTCATCAGGAAGC
TGAGTGCATCGGACTCATCAGGAAGC
CTGCTCATCGGACTCATCAGGAAGC
ACAAAACATCGGACTCATCAGGAAGC
GCTCTTCATCGGACTCATCAGGAAGC
CTTATCATCGGACTCATCAGGAAGC
CAGGCATCATCGGACTCATCAGGAAGC
GCTCATCGGACTCATCAGGAAGC
GTGTCATCGGACTCATCAGGAAGC
TGACTACATCGGACTCATCAGGAAGC
GTCAGTTCATCGGACTCATCAGGAAGC
TCCATCATCGGACTCATCAGGAAGC
AAAAAGCATCGGACTCATCAGGAAGC
ACCACGCATCGGACTCATCAGGAAGC
GCTCGCATCGGACTCATCAGGAAGC
GTGTGTCATCGGACTCATCAGGAAGC
TCTGCATCGGACTCATCAGGAAGC
ACACACATCGGACTCATCAGGAAGC
GCAAACATCGGACTCATCAGGAAGC
TAATGACATCGGACTCATCAGGAAGC
GTGTGCATCGGACTCATCAGGAAGC
GGATGTCATCGGACTCATCAGGAAGC

ATGTGTGCATCGGACTCATCAGGAAGC
CAATCATCGGACTCATCAGGAAGC
GAGCGGCATCGGACTCATCAGGAAGC
TCACACATCGGACTCATCAGGAAGC
AGCAGACATCGGACTCATCAGGAAGC
ATATGCGCATCGGACTCATCAGGAAGC
GGAAACATCGGACTCATCAGGAAGC
GACACATCGGACTCATCAGGAAGC
GTTTTCGCATCGGACTCATCAGGAAGC
GTGTCATCGGACTCATCAGGAAGC
CTTCCATCGGACTCATCAGGAAGC
GGAGATCATCGGACTCATCAGGAAGC
CACCGCCATCGGACTCATCAGGAAGC
GCTTATCATCGGACTCATCAGGAAGC
CAATCGCATCGGACTCATCAGGAAGC
ATTCCCATCGGACTCATCAGGAAGC
CATAGTCATCGGACTCATCAGGAAGC
TATGCCATCGGACTCATCAGGAAGC
TCACTGCATCGGACTCATCAGGAAGC
GAACACATCGGACTCATCAGGAAGC
AAGCGCATCGGACTCATCAGGAAGC
GATAGGCATCGGACTCATCAGGAAGC
GGTGTCTCATCGGACTCATCAGGAAGC
GAATTTGCATCGGACTCATCAGGAAGC
CTCTCATCGGACTCATCAGGAAGC
TCAACCTCATCGGACTCATCAGGAAGC
AACCCCACATCGGACTCATCAGGAAGC
ACACACATCGGACTCATCAGGAAGC
AGCCATCATCGGACTCATCAGGAAGC
ATGGTCATCGGACTCATCAGGAAGC
CTCAAGCATCGGACTCATCAGGAAGC
TGCAACATCGGACTCATCAGGAAGC
CAATCATCGGACTCATCAGGAAGC
ATGGAACATCGGACTCATCAGGAAGC
CTCTCTCATCGGACTCATCAGGAAGC
GCAGGTCATCGGACTCATCAGGAAGC
AGCCCCATCGGACTCATCAGGAAGC
CTGTCATCGGACTCATCAGGAAGC
TTACTCACATCGGACTCATCAGGAAGC
ATGCTCCATCGGACTCATCAGGAAGC
GCCATACATCGGACTCATCAGGAAGC
AGAGGCCATCGGACTCATCAGGAAGC
TGTCATCGGACTCATCAGGAAGC

GATAGACATCGGACTCATCAGGAAGC
TGTGTCATCGGACTCATCAGGAAGC
TAACACATCGGACTCATCAGGAAGC
CCCCACCATCGGACTCATCAGGAAGC
TGGGTTCATCGGACTCATCAGGAAGC
GTTCACCATCGGACTCATCAGGAAGC
GTGTCATCGGACTCATCAGGAAGC
AAACACATCGGACTCATCAGGAAGC
AACCACATCGGACTCATCAGGAAGC
TGACATCGGACTCATCAGGAAGC
CAACCACATCGGACTCATCAGGAAGC
TAAGACATCGGACTCATCAGGAAGC
ACCTGCATCGGACTCATCAGGAAGC
GATTACACATCGGACTCATCAGGAAGC
CTGTCATCGGACTCATCAGGAAGC
TCTCAACATCGGACTCATCAGGAAGC
ACACAGCATCGGACTCATCAGGAAGC
TAGGGTCATCGGACTCATCAGGAAGC
CATTTCATCGGACTCATCAGGAAGC
CCATCTCATCGGACTCATCAGGAAGC
AGTGGCCATCGGACTCATCAGGAAGC
CAGTCATCGGACTCATCAGGAAGC
TACTGACATCGGACTCATCAGGAAGC
TTGCTCATCGGACTCATCAGGAAGC
GCTGCATCGGACTCATCAGGAAGC
GCTCCATCGGACTCATCAGGAAGC
GGACACATCGGACTCATCAGGAAGC
TCTCCACATCGGACTCATCAGGAAGC
TACACATCGGACTCATCAGGAAGC
AAGGCACATCGGACTCATCAGGAAGC
ATGCCATCGGACTCATCAGGAAGC
TTGGCCATCGGACTCATCAGGAAGC
GTGACCATCGGACTCATCAGGAAGC
GTAGTCATCGGACTCATCAGGAAGC
ACTACATCGGACTCATCAGGAAGC
TATAACATCGGACTCATCAGGAAGC
TTTGGTCATCGGACTCATCAGGAAGC
TGAGGTTCATCGGACTCATCAGGAAGC
CTGGCCATCGGACTCATCAGGAAGC
CTCAGCATCGGACTCATCAGGAAGC
TCCCCATCGGACTCATCAGGAAGC
CCTAACATCGGACTCATCAGGAAGC
CTCTCATCGGACTCATCAGGAAGC

CCTACATCGGACTCATCAGGAAGC
GCTCTACATCGGACTCATCAGGAAGC
TCAGCATCGGACTCATCAGGAAGC
CTCATCGGACTCATCAGGAAGC
TGGGTCATCGGACTCATCAGGAAGC
GTCCACATCGGACTCATCAGGAAGC
TCACCACCCATCGGACTCATCAGGAAGC
ACACCATCGGACTCATCAGGAAGC
ATTGCCTCATCGGACTCATCAGGAAGC
GAACATCGGACTCATCAGGAAGC
ATGCCCATCGGACTCATCAGGAAGC
GATCCCACATCGGACTCATCAGGAAGC
ATAGCATCGGACTCATCAGGAAGC
TCTCTCATCGGACTCATCAGGAAGC
GATTGACCATCGGACTCATCAGGAAGC
TGAATCCCATCGGACTCATCAGGAAGC
AACAGACATCGGACTCATCAGGAAGC
ACAGAGCATCGGACTCATCAGGAAGC
GTGCATCATCGGACTCATCAGGAAGC
ACCCCCCATCGGACTCATCAGGAAGC
ACTGCCTCATCGGACTCATCAGGAAGC
TAGACCACATCGGACTCATCAGGAAGC
ATTCCACATCGGACTCATCAGGAAGC
TCTCTCATCGGACTCATCAGGAAGC
TAAGAACATCGGACTCATCAGGAAGC
CTACGCATCGGACTCATCAGGAAGC
TCTGGCATCGGACTCATCAGGAAGC
AGCATCATCGGACTCATCAGGAAGC
TAATACATCGGACTCATCAGGAAGC
CCCCATCGGACTCATCAGGAAGC
GTCTCATCGGACTCATCAGGAAGC
AAGGCCATCGGACTCATCAGGAAGC
AACTCACATCGGACTCATCAGGAAGC
GTCCACATCGGACTCATCAGGAAGC
TCAACATCGGACTCATCAGGAAGC
GCACACATCGGACTCATCAGGAAGC
GATAGACATCGGACTCATCAGGAAGC
GCTGACATCGGACTCATCAGGAAGC
ATGCCCATCGGACTCATCAGGAAGC
CAATTACCATCGGACTCATCAGGAAGC
GACAACATCGGACTCATCAGGAAGC
TCGCACATCGGACTCATCAGGAAGC
AAACAGACATCGGACTCATCAGGAAGC

CCTTCACATCGGACTCATCAGGAAGC
AGTCACATCGGACTCATCAGGAAGC
TTTTGTCATCGGACTCATCAGGAAGC
TGGAACATCGGACTCATCAGGAAGC
ACATGCCATCGGACTCATCAGGAAGC
TCGCTCATCGGACTCATCAGGAAGC
ACAATACATCGGACTCATCAGGAAGC
TCATCCATCGGACTCATCAGGAAGC
GTGTGTTCATCGGACTCATCAGGAAGC
GTGTCATCGGACTCATCAGGAAGC
GCACACATCGGACTCATCAGGAAGC
AAGTCCATCGGACTCATCAGGAAGC
CATTGACATCGGACTCATCAGGAAGC
CACCCCATCGGACTCATCAGGAAGC
TCTCTCCATCGGACTCATCAGGAAGC
CCATCCATCGGACTCATCAGGAAGC
GTGGAACATCGGACTCATCAGGAAGC
TGCAGTCTCATCGGACTCATCAGGAAGC
AAATGCCATCGGACTCATCAGGAAGC
AAGCACATCGGACTCATCAGGAAGC
TTTCCCATCGGACTCATCAGGAAGC
TTCCACATCGGACTCATCAGGAAGC
AGTCACATCGGACTCATCAGGAAGC
AATAACATCGGACTCATCAGGAAGC
TGTGTGCATCGGACTCATCAGGAAGC
TATGCCATCGGACTCATCAGGAAGC
TACTCATCGGACTCATCAGGAAGC
AACTCATCGGACTCATCAGGAAGC
GAGACAGCATCGGACTCATCAGGAAGC
GCTGGCATCGGACTCATCAGGAAGC
TTAGTCATCGGACTCATCAGGAAGC
TCACACATCGGACTCATCAGGAAGC
TCCCTCATCGGACTCATCAGGAAGC
TATGTCATCGGACTCATCAGGAAGC
TGGACATCGGACTCATCAGGAAGC
GTCCACATCGGACTCATCAGGAAGC
GGGTCATCGGACTCATCAGGAAGC
GGTCAGCATCGGACTCATCAGGAAGC
TATACCATCGGACTCATCAGGAAGC
CAGCATCATCGGACTCATCAGGAAGC
ATGCTTCATCGGACTCATCAGGAAGC
GAAAGCCATCGGACTCATCAGGAAGC
AGAGGCATCGGACTCATCAGGAAGC

CTGTGCGCATCGGACTCATCAGGAAGC
CAGAAACATCGGACTCATCAGGAAGC
ACCCACATCGGACTCATCAGGAAGC
CTTCACATCGGACTCATCAGGAAGC
ACTATCCATCGGACTCATCAGGAAGC
AAGCTACATCGGACTCATCAGGAAGC
GTCCACATCGGACTCATCAGGAAGC
GTGTGGCATCGGACTCATCAGGAAGC
ACCAGCCATCGGACTCATCAGGAAGC
GGTCCTTCATCGGACTCATCAGGAAGC
GGTGCCCCATCGGACTCATCAGGAAGC
CTTGGCATCGGACTCATCAGGAAGC
GATTCAACATCGGACTCATCAGGAAGC
TGCATCATCGGACTCATCAGGAAGC
GTCTCATCGGACTCATCAGGAAGC
TCTCTCATCGGACTCATCAGGAAGC
GATGGCCATCGGACTCATCAGGAAGC
TTGTTTCATCGGACTCATCAGGAAGC
CTGCATCGGACTCATCAGGAAGC
GTGTGTCATCGGACTCATCAGGAAGC
TGCTCATCGGACTCATCAGGAAGC
AATGCTCATCGGACTCATCAGGAAGC
GTTGACCATCGGACTCATCAGGAAGC
CACACCCATCGGACTCATCAGGAAGC
TAGGCACATCGGACTCATCAGGAAGC
TCAGTCATCGGACTCATCAGGAAGC
ACTTGAACATCGGACTCATCAGGAAGC
GAGAACCCGCATCGGACTCATCAGGAAGC
TATTTCATCGGACTCATCAGGAAGC
TGGGGACATCGGACTCATCAGGAAGC
AACTTGCATCGGACTCATCAGGAAGC
GGGAAGCATCGGACTCATCAGGAAGC
ATCCGGCATCGGACTCATCAGGAAGC
ACTTTTGCATCGGACTCATCAGGAAGC
TCTCTCATCGGACTCATCAGGAAGC
ACCTGGACATCGGACTCATCAGGAAGC
TCCTGCCATCGGACTCATCAGGAAGC
GATCATCATCGGACTCATCAGGAAGC
GAGGCCATCGGACTCATCAGGAAGC
GATGGACATCGGACTCATCAGGAAGC
ACGTCATCGGACTCATCAGGAAGC
CATGTCATCGGACTCATCAGGAAGC
GGGAGGCATCGGACTCATCAGGAAGC

ACTGTACCATCGGACTCATCAGGAAGC
TAAGACATCGGACTCATCAGGAAGC
GGCCACATCGGACTCATCAGGAAGC
ACAGTCATCGGACTCATCAGGAAGC
AGAAGCATCGGACTCATCAGGAAGC
ACATCTCATCGGACTCATCAGGAAGC
TGGAATCATCGGACTCATCAGGAAGC
ACACACATCGGACTCATCAGGAAGC
GTTGGCCATCGGACTCATCAGGAAGC
AGATTGCATCGGACTCATCAGGAAGC
TCATGATGCATCGGACTCATCAGGAAGC
TCTGAACCATCGGACTCATCAGGAAGC
TGCCATGCCATCGGACTCATCAGGAAGC
TATGAACCATCGGACTCATCAGGAAGC
TCTCATCGGACTCATCAGGAAGC
AGTGAGCATCGGACTCATCAGGAAGC
CTCTCTCATCGGACTCATCAGGAAGC
AGGAGGCATCGGACTCATCAGGAAGC
TGAGTCATCGGACTCATCAGGAAGC
GTATGTCATCGGACTCATCAGGAAGC
AAGATTGCATCGGACTCATCAGGAAGC
TTTACATCGGACTCATCAGGAAGC
TACATCTCATCGGACTCATCAGGAAGC
ACACACATCGGACTCATCAGGAAGC
TATCCACATCGGACTCATCAGGAAGC
GTGTGGCATCGGACTCATCAGGAAGC
TTTCTGCATCGGACTCATCAGGAAGC
GTTTCACATCGGACTCATCAGGAAGC
ATCTCCATCGGACTCATCAGGAAGC
GTGTGTCATCGGACTCATCAGGAAGC
AAACCACATCGGACTCATCAGGAAGC
AGACACATCGGACTCATCAGGAAGC
CTGACATCGGACTCATCAGGAAGC
AAACACATCGGACTCATCAGGAAGC
TGTGTCATCGGACTCATCAGGAAGC
AGACTCATCGGACTCATCAGGAAGC
GTGTGTCATCGGACTCATCAGGAAGC
CTGTGCACATCGGACTCATCAGGAAGC
GCAGGACATCGGACTCATCAGGAAGC
TCTAGGACATCGGACTCATCAGGAAGC
TGACACATCGGACTCATCAGGAAGC
ATCTGACATCGGACTCATCAGGAAGC
ATTTGTCATCGGACTCATCAGGAAGC

TCTCCATCGGACTCATCAGGAAGC
TGTCCATCGGACTCATCAGGAAGC
TGTTCATCGGACTCATCAGGAAGC
TTCCTCACATCGGACTCATCAGGAAGC
GTCACTGCATCGGACTCATCAGGAAGC
ACGAGGCATCGGACTCATCAGGAAGC
ACCTGCATCGGACTCATCAGGAAGC
CTCCCATCGGACTCATCAGGAAGC
AAGAAAGACATCGGACTCATCAGGAAGC
AGGACATCGGACTCATCAGGAAGC
AGAGAGATTCATCGGACTCATCAGGAAGC
ACCTATGATCATCGGACTCATCAGGAAGC
TTCTCATCGGACTCATCAGGAAGC
GCAGGCATCGGACTCATCAGGAAGC
AAAAAAACCACATCGGACTCATCAGGAAGC
TGGTCATCGGACTCATCAGGAAGC
TAGTCATCGGACTCATCAGGAAGC
TCATTCATCGGACTCATCAGGAAGC
GTCTTCATCGGACTCATCAGGAAGC
TTCATCGGACTCATCAGGAAGC
CACATCGGACTCATCAGGAAGC
CATGATCATCGGACTCATCAGGAAGC
AGAAAGACATCGGACTCATCAGGAAGC
TCTTCATCGGACTCATCAGGAAGC
CTCATCGGACTCATCAGGAAGC
AGTCCATCGGACTCATCAGGAAGC
CTAGCATCGGACTCATCAGGAAGC
TCTCATCGGACTCATCAGGAAGC
CAGTAAGCCCATCGGACTCATCAGGAAGC
GTAGGCATCGGACTCATCAGGAAGC
CTAGTCATCGGACTCATCAGGAAGC
AGTTCCATCGGACTCATCAGGAAGC
TTGGCACATCGGACTCATCAGGAAGC
GGGGGCATCGGACTCATCAGGAAGC
CTCAACATCGGACTCATCAGGAAGC
GAGAGAGCATCGGACTCATCAGGAAGC
TACACATCGGACTCATCAGGAAGC
CAAAACATCGGACTCATCAGGAAGC
AGGCACATCGGACTCATCAGGAAGC
TATGCATCGGACTCATCAGGAAGC
TGCTCATCGGACTCATCAGGAAGC
TAAAACATCGGACTCATCAGGAAGC
TTGGAGCATCGGACTCATCAGGAAGC

TGAAACATCGGACTCATCAGGAAGC
AGGGACATCGGACTCATCAGGAAGC
TAGCCATCGGACTCATCAGGAAGC
GCAGACATCGGACTCATCAGGAAGC
CCTGACATCGGACTCATCAGGAAGC
GTTGACATCGGACTCATCAGGAAGC
ACCAACATCGGACTCATCAGGAAGC
TGGACATCGGACTCATCAGGAAGC
TCAGACATCGGACTCATCAGGAAGC
ACCTGTCATCGGACTCATCAGGAAGC
TCAGCATCGGACTCATCAGGAAGC
TCAGGGCATCGGACTCATCAGGAAGC
CTCCCACATCGGACTCATCAGGAAGC
CACCCACATCGGACTCATCAGGAAGC
TGTTTCATCGGACTCATCAGGAAGC
GAGCGACATCGGACTCATCAGGAAGC
GAGTGTCATCGGACTCATCAGGAAGC
AGGCATCGGACTCATCAGGAAGC
ACACATCGGACTCATCAGGAAGC
TGTCATCGGACTCATCAGGAAGC
CGTGACATCGGACTCATCAGGAAGC
ACACTCATCGGACTCATCAGGAAGC
GTAGTGCATCGGACTCATCAGGAAGC
GTACATCGGACTCATCAGGAAGC
ATAACACATCGGACTCATCAGGAAGC
TGGGCATCGGACTCATCAGGAAGC
GCTGATCATCGGACTCATCAGGAAGC
GAGACATCGGACTCATCAGGAAGC
AGAGTGACATCGGACTCATCAGGAAGC
AGCATCATCGGACTCATCAGGAAGC
ATACACATCGGACTCATCAGGAAGC
CACACATCGGACTCATCAGGAAGC
CCCCATCGGACTCATCAGGAAGC
CTCATCATCGGACTCATCAGGAAGC
TATGTGCATCGGACTCATCAGGAAGC
GCGCTCCATCGGACTCATCAGGAAGC
CTGTCATCGGACTCATCAGGAAGC
TGTTCATCGGACTCATCAGGAAGC
GTGACATCGGACTCATCAGGAAGC
TTTGCATCGGACTCATCAGGAAGC
GTATCCATCGGACTCATCAGGAAGC
GCCTGTCATCGGACTCATCAGGAAGC
GAGCTACATCGGACTCATCAGGAAGC

ATGAAACATCGGACTCATCAGGAAGC
AATGCTCATCGGACTCATCAGGAAGC
TTGACATCGGACTCATCAGGAAGC
ATCTGACATCGGACTCATCAGGAAGC
GTGTGTCATCGGACTCATCAGGAAGC
TTCACTCATCGGACTCATCAGGAAGC
TCATGTCATCGGACTCATCAGGAAGC
TAGACATCGGACTCATCAGGAAGC
TGTCATCGGACTCATCAGGAAGC
AAGACATCGGACTCATCAGGAAGC
CTCCTCATCGGACTCATCAGGAAGC
ATTGACATCGGACTCATCAGGAAGC
TCTCCATCGGACTCATCAGGAAGC
GGCAAACATCGGACTCATCAGGAAGC
TGTCATCGGACTCATCAGGAAGC
AGATCATCGGACTCATCAGGAAGC
GTTCATCGGACTCATCAGGAAGC
GCTACATCGGACTCATCAGGAAGC
TTTTCATCGGACTCATCAGGAAGC
CACCTGCATCGGACTCATCAGGAAGC
GACACACATCGGACTCATCAGGAAGC
AAAAACATCGGACTCATCAGGAAGC
CCCCATCGGACTCATCAGGAAGC
GTACATCGGACTCATCAGGAAGC
TTTGATCATCGGACTCATCAGGAAGC
AGGCACATCGGACTCATCAGGAAGC
CCCATCGGACTCATCAGGAAGC
TGCCTCCATCGGACTCATCAGGAAGC
ACCCCTCATCGGACTCATCAGGAAGC
AAACATCGGACTCATCAGGAAGC
TGCCCCCATCGGACTCATCAGGAAGC
AAATCATCGGACTCATCAGGAAGC
TGAAGACATCGGACTCATCAGGAAGC
AGAGCATCGGACTCATCAGGAAGC
TCGATACATCGGACTCATCAGGAAGC
ATTGTCCATCGGACTCATCAGGAAGC
CCAACATCGGACTCATCAGGAAGC
ACACATCGGACTCATCAGGAAGC
GTTGCCATCGGACTCATCAGGAAGC
GTCATCATCGGACTCATCAGGAAGC
CCTACATCGGACTCATCAGGAAGC
CTTTCCATCGGACTCATCAGGAAGC
TGGCATCGGACTCATCAGGAAGC

ATGCAACATCGGACTCATCAGGAAGC
CAGGCCATCGGACTCATCAGGAAGC
AGCCACCATCGGACTCATCAGGAAGC
TCTCATCGGACTCATCAGGAAGC
GCACATCGGACTCATCAGGAAGC
CCAATCATCGGACTCATCAGGAAGC
TGCATCGGACTCATCAGGAAGC
CAGCACATCGGACTCATCAGGAAGC
AGCCCATCGGACTCATCAGGAAGC
ATACCACATCGGACTCATCAGGAAGC
TCTGGCATCGGACTCATCAGGAAGC
ACCTCATCGGACTCATCAGGAAGC
GTGTGCCATCGGACTCATCAGGAAGC
AATGTTCATCGGACTCATCAGGAAGC
TATGCCATCGGACTCATCAGGAAGC
TGAGAACCATCGGACTCATCAGGAAGC
TGTGCATCGGACTCATCAGGAAGC
GCACACATCGGACTCATCAGGAAGC
CCCCTACATCGGACTCATCAGGAAGC
GTGCATCGGACTCATCAGGAAGC
TCCCCATCGGACTCATCAGGAAGC
TTCCGTTCATCGGACTCATCAGGAAGC
CAGCATCGGACTCATCAGGAAGC
AGTCAGCATCGGACTCATCAGGAAGC
ATCACATCGGACTCATCAGGAAGC
CATTCATCGGACTCATCAGGAAGC
CCTTCATCGGACTCATCAGGAAGC
TCAAACATCGGACTCATCAGGAAGC
GTGGCTCATCGGACTCATCAGGAAGC
ATTCCACATCGGACTCATCAGGAAGC
AACATCGGACTCATCAGGAAGC
ATCCCACATCGGACTCATCAGGAAGC
GTGGCATCGGACTCATCAGGAAGC
GGAAACATCGGACTCATCAGGAAGC
AACCCCATCGGACTCATCAGGAAGC
CAATACATCGGACTCATCAGGAAGC
GCTCCATCGGACTCATCAGGAAGC
TGACTACATCGGACTCATCAGGAAGC
CAAATCATCGGACTCATCAGGAAGC
AGCAGAGCATCGGACTCATCAGGAAGC
CCCCATCGGACTCATCAGGAAGC
TCCTGACATCGGACTCATCAGGAAGC
TGCCATCGGACTCATCAGGAAGC

CTGTCATCGGACTCATCAGGAAGC
CTATCCATCGGACTCATCAGGAAGC
AGTGACCATCGGACTCATCAGGAAGC
AATACATCGGACTCATCAGGAAGC
AAAACATCGGACTCATCAGGAAGC
GGAGCCATCGGACTCATCAGGAAGC
GAGACCATCGGACTCATCAGGAAGC
GAATCATCGGACTCATCAGGAAGC
TCTCATCGGACTCATCAGGAAGC
TCAGTCATCGGACTCATCAGGAAGC
TTCCATCGGACTCATCAGGAAGC
CTCTCATCGGACTCATCAGGAAGC
TTCTACATCGGACTCATCAGGAAGC
CTGCAACATCGGACTCATCAGGAAGC
CAAACATCGGACTCATCAGGAAGC
TGTCATCGGACTCATCAGGAAGC
AATGCATCGGACTCATCAGGAAGC
GCAAACATCGGACTCATCAGGAAGC
GTAGTTCATCGGACTCATCAGGAAGC
ATCCATCGGACTCATCAGGAAGC
GAACATCGGACTCATCAGGAAGC
GCCACATCGGACTCATCAGGAAGC
ACTGTGCATCGGACTCATCAGGAAGC
GTGTTTCATCGGACTCATCAGGAAGC
ATTTCACATCGGACTCATCAGGAAGC
TATGTGCATCGGACTCATCAGGAAGC
TGTCATCGGACTCATCAGGAAGC
ATATGCATCGGACTCATCAGGAAGC
GCCCATCGGACTCATCAGGAAGC
GCTATCCATCGGACTCATCAGGAAGC
ATCTCTCATCGGACTCATCAGGAAGC
CTCCATCGGACTCATCAGGAAGC
TATGCCATCGGACTCATCAGGAAGC
GTATCATCGGACTCATCAGGAAGC
TAAACATCGGACTCATCAGGAAGC
TTCTCCATCGGACTCATCAGGAAGC
ACATCGGACTCATCAGGAAGC
ATGGCTCATCGGACTCATCAGGAAGC
ATGTTTACATCGGACTCATCAGGAAGC
TACATCGGACTCATCAGGAAGC
ACTCCCATCGGACTCATCAGGAAGC
AGAGAGCATCGGACTCATCAGGAAGC
CCATCGGACTCATCAGGAAGC

GGTCTCATCGGACTCATCAGGAAGC
TGGTCATCGGACTCATCAGGAAGC
TCTCCCATCGGACTCATCAGGAAGC
GCATCATCGGACTCATCAGGAAGC
CAACACATCGGACTCATCAGGAAGC
CTTGCCATCGGACTCATCAGGAAGC
CCACATCGGACTCATCAGGAAGC
GAAGACATCGGACTCATCAGGAAGC
AACACATCGGACTCATCAGGAAGC
ATGTTCATCGGACTCATCAGGAAGC
CCCACATCGGACTCATCAGGAAGC
CTCATCGGACTCATCAGGAAGC
CTCATCGGACTCATCAGGAAGC
TTATTCATCGGACTCATCAGGAAGC
TTGTGCATCGGACTCATCAGGAAGC
TGCAAACATCGGACTCATCAGGAAGC
GCAACATCGGACTCATCAGGAAGC
GTTTCATCGGACTCATCAGGAAGC
TAGGCACATCGGACTCATCAGGAAGC
CGAGCATCGGACTCATCAGGAAGC
GAGAACCATCGGACTCATCAGGAAGC
CACACATCGGACTCATCAGGAAGC
GAGCCATCGGACTCATCAGGAAGC
GGATCATCGGACTCATCAGGAAGC
GCCATCGGACTCATCAGGAAGC
GATGCATCGGACTCATCAGGAAGC
GACACATCGGACTCATCAGGAAGC
GTTACATCGGACTCATCAGGAAGC
AGGTTCATCGGACTCATCAGGAAGC
GACATCGGACTCATCAGGAAGC
AGGGCATCGGACTCATCAGGAAGC
TTCACATCGGACTCATCAGGAAGC
AGCCATCGGACTCATCAGGAAGC
CTCTCATCGGACTCATCAGGAAGC
TTCGTTCATCGGACTCATCAGGAAGC
TACCCATCGGACTCATCAGGAAGC
CTGACATCGGACTCATCAGGAAGC
ACACATCGGACTCATCAGGAAGC
GAGAGCATCGGACTCATCAGGAAGC
TGCCATCGGACTCATCAGGAAGC
TGACACATCGGACTCATCAGGAAGC
AAGACATCGGACTCATCAGGAAGC
TGTCATCGGACTCATCAGGAAGC

GAAGAAGCCATCGGACTCATCAGGAAGC
ACAGACATCGGACTCATCAGGAAGC
AGCACATCGGACTCATCAGGAAGC
ATGCCATCGGACTCATCAGGAAGC
TGTGAGACATCGGACTCATCAGGAAGC
TGGGTCATCGGACTCATCAGGAAGC
ACCCCATCGGACTCATCAGGAAGC
TAGCCATCGGACTCATCAGGAAGC
GCCAAACATCGGACTCATCAGGAAGC
TCCTCATCGGACTCATCAGGAAGC
GAGGAACATCGGACTCATCAGGAAGC
GGACATCGGACTCATCAGGAAGC
ATCCCATCGGACTCATCAGGAAGC
CTTCATCGGACTCATCAGGAAGC
CTCAACATCGGACTCATCAGGAAGC
CCCATCGGACTCATCAGGAAGC
GCTCATCGGACTCATCAGGAAGC
AGTGCATCGGACTCATCAGGAAGC
CTTCCACATCGGACTCATCAGGAAGC
AAGCCATCGGACTCATCAGGAAGC
ACAGGCATCGGACTCATCAGGAAGC
TTCATCGGACTCATCAGGAAGC
TCTATAACATCGGACTCATCAGGAAGC
TGTCCCATCGGACTCATCAGGAAGC
CACGCCATCGGACTCATCAGGAAGC
CTGTCATCGGACTCATCAGGAAGC
CTCCACATCGGACTCATCAGGAAGC
CACTCATCGGACTCATCAGGAAGC
TGAACATCGGACTCATCAGGAAGC
GCCCACATCGGACTCATCAGGAAGC
TAGTCTCATCGGACTCATCAGGAAGC
CGCTGCATCGGACTCATCAGGAAGC
GATTTCATCGGACTCATCAGGAAGC
CCTGCATCGGACTCATCAGGAAGC
TTCCTCATCGGACTCATCAGGAAGC
CTCCATCGGACTCATCAGGAAGC
ATGCATCGGACTCATCAGGAAGC
ATATGACATCGGACTCATCAGGAAGC
TTGTCATCGGACTCATCAGGAAGC
TCACATCGGACTCATCAGGAAGC
GGAGACATCGGACTCATCAGGAAGC
AGAGACATCGGACTCATCAGGAAGC
TCTCTCATCGGACTCATCAGGAAGC

AAACACATCGGACTCATCAGGAAGC
AGAAGACATCGGACTCATCAGGAAGC
ACATTGGCATCGGACTCATCAGGAAGC
TCGCACATCGGACTCATCAGGAAGC
AAACACATCGGACTCATCAGGAAGC
GTCCATCGGACTCATCAGGAAGC
AGCACATCGGACTCATCAGGAAGC
AGTCACATCGGACTCATCAGGAAGC
ACTGCCATCGGACTCATCAGGAAGC
GCTAAACATCGGACTCATCAGGAAGC
AATGGTCATCGGACTCATCAGGAAGC
ACACATCGGACTCATCAGGAAGC
AGTGTCATCGGACTCATCAGGAAGC
TGACCATCGGACTCATCAGGAAGC
TAGCCATCGGACTCATCAGGAAGC
AGGCATCATCGGACTCATCAGGAAGC
TAGGTCATCGGACTCATCAGGAAGC
AACCTCATCGGACTCATCAGGAAGC
AACGCATCGGACTCATCAGGAAGC
TACTGCATCGGACTCATCAGGAAGC
ACCACATCGGACTCATCAGGAAGC
GTCCATCGGACTCATCAGGAAGC
ATGGACATCGGACTCATCAGGAAGC
ATACTCATCGGACTCATCAGGAAGC
ATTTCATCGGACTCATCAGGAAGC
AGTCATCGGACTCATCAGGAAGC
CGCACATCGGACTCATCAGGAAGC
AGCTGACATCGGACTCATCAGGAAGC
TACATCGGACTCATCAGGAAGC
TTCTCATCGGACTCATCAGGAAGC
AAATGCATCGGACTCATCAGGAAGC
CATTCATCGGACTCATCAGGAAGC
AGCACATCGGACTCATCAGGAAGC
TGTACATCGGACTCATCAGGAAGC
GGATGCATCGGACTCATCAGGAAGC
TTTCCCCATCGGACTCATCAGGAAGC
AAGGGGGCATCGGACTCATCAGGAAGC
ACACATCGGACTCATCAGGAAGC
TGTAAACATCGGACTCATCAGGAAGC
AGTGCATCGGACTCATCAGGAAGC
GTTGATCATCGGACTCATCAGGAAGC
CCCATCGGACTCATCAGGAAGC
TATCACATCGGACTCATCAGGAAGC

AGGTCATCGGACTCATCAGGAAGC
ACTGCCATCGGACTCATCAGGAAGC
TGCCATCGGACTCATCAGGAAGC
AGACATCGGACTCATCAGGAAGC
CCACATCGGACTCATCAGGAAGC
CTATCATCGGACTCATCAGGAAGC
AGATGGACATCGGACTCATCAGGAAGC
AGTCAGCATCGGACTCATCAGGAAGC
CCCAACATCGGACTCATCAGGAAGC
CTGCCATCGGACTCATCAGGAAGC
TCCCCATCGGACTCATCAGGAAGC
CTTTCATCGGACTCATCAGGAAGC
AGAACATCGGACTCATCAGGAAGC
TGTCATCGGACTCATCAGGAAGC
TACGGACATCGGACTCATCAGGAAGC
CACATCGGACTCATCAGGAAGC
ATTTACATCGGACTCATCAGGAAGC
GGCATCGGACTCATCAGGAAGC
TCTCATCGGACTCATCAGGAAGC
TGTGCATCGGACTCATCAGGAAGC
AACCCCATCGGACTCATCAGGAAGC
ACCCACATCGGACTCATCAGGAAGC
AGGCTCATCGGACTCATCAGGAAGC
TTCAGCATCGGACTCATCAGGAAGC
ACCACTCATCGGACTCATCAGGAAGC
GAGAATCATCGGACTCATCAGGAAGC
GCCATCGGACTCATCAGGAAGC
GGACATCGGACTCATCAGGAAGC
TCCAGCATCGGACTCATCAGGAAGC
CTTTCTCATCGGACTCATCAGGAAGC
CTGTCATCGGACTCATCAGGAAGC
TAAGCACATCGGACTCATCAGGAAGC
CTGATCATCGGACTCATCAGGAAGC
TGATCATCGGACTCATCAGGAAGC
CAGCATCGGACTCATCAGGAAGC
AATCATCGGACTCATCAGGAAGC
ACAGTCATCGGACTCATCAGGAAGC
TTGACATCGGACTCATCAGGAAGC
TCCTCATCGGACTCATCAGGAAGC
CAGTCCATCGGACTCATCAGGAAGC
TTGCCCATCGGACTCATCAGGAAGC
ACAGACATCGGACTCATCAGGAAGC
GGGCATCGGACTCATCAGGAAGC

GCCTCCATCGGACTCATCAGGAAGC
CTTTGACATCGGACTCATCAGGAAGC
CAGAGGACATCGGACTCATCAGGAAGC
TCAAACATCGGACTCATCAGGAAGC
GTCATCATCGGACTCATCAGGAAGC
TGGCATCGGACTCATCAGGAAGC
CACACCATCGGACTCATCAGGAAGC
CTGACCATCGGACTCATCAGGAAGC
AAAAGGCATCGGACTCATCAGGAAGC
CAGTCATCGGACTCATCAGGAAGC
AGGGGACATCGGACTCATCAGGAAGC
TGTACATCGGACTCATCAGGAAGC
GTCTACATCGGACTCATCAGGAAGC
CCAACATCGGACTCATCAGGAAGC
GGACCATCGGACTCATCAGGAAGC
AGCCCCATCGGACTCATCAGGAAGC
AGGGCATCGGACTCATCAGGAAGC
GCAGGCATCGGACTCATCAGGAAGC
ATCATCATCGGACTCATCAGGAAGC
TCTCATCGGACTCATCAGGAAGC
ACCCCATCGGACTCATCAGGAAGC
GTCTCATCGGACTCATCAGGAAGC
GAAAGAACATCGGACTCATCAGGAAGC
AAGACATCGGACTCATCAGGAAGC
CTTGAACATCGGACTCATCAGGAAGC
AACACATCGGACTCATCAGGAAGC
CTTGCATCGGACTCATCAGGAAGC
GAAGGCATCGGACTCATCAGGAAGC
AGTTACATCGGACTCATCAGGAAGC
GTGTGAACATCGGACTCATCAGGAAGC
TGTCATCGGACTCATCAGGAAGC
TTCGCCATCGGACTCATCAGGAAGC
GTGTGCATCGGACTCATCAGGAAGC
TAACACATCGGACTCATCAGGAAGC
GATTGCATCGGACTCATCAGGAAGC
TCCCATCGGACTCATCAGGAAGC
TTCACATCGGACTCATCAGGAAGC
GGTGACATCGGACTCATCAGGAAGC
TAGGGCATCGGACTCATCAGGAAGC
TCGGCATCGGACTCATCAGGAAGC
GAGGCATCGGACTCATCAGGAAGC
TGGGGACATCGGACTCATCAGGAAGC
TACTTGCATCGGACTCATCAGGAAGC

GAGATACATCGGACTCATCAGGAAGC
TCAACATCGGACTCATCAGGAAGC
TGTCACATCGGACTCATCAGGAAGC
GCTGCATCGGACTCATCAGGAAGC
GGCTCATCGGACTCATCAGGAAGC
GAGTATTCATCGGACTCATCAGGAAGC
ACACCATCGGACTCATCAGGAAGC
ATTAGTCATCGGACTCATCAGGAAGC
CATATCATCGGACTCATCAGGAAGC
ATACCATCGGACTCATCAGGAAGC
TGTCATCGGACTCATCAGGAAGC
ATCACATCGGACTCATCAGGAAGC
CTCCATCGGACTCATCAGGAAGC
CTACATCGGACTCATCAGGAAGC
TTGTGCATCGGACTCATCAGGAAGC
ACCCCAACATCGGACTCATCAGGAAGC
CCAGACATCGGACTCATCAGGAAGC
CCACATCGGACTCATCAGGAAGC
CCGGACATCGGACTCATCAGGAAGC
ACTGGGCATCGGACTCATCAGGAAGC
GTGCATCGGACTCATCAGGAAGC
TTGGCATCGGACTCATCAGGAAGC
TGCCATCGGACTCATCAGGAAGC
AAACATCGGACTCATCAGGAAGC
CTCTCATCGGACTCATCAGGAAGC
ACTCATCGGACTCATCAGGAAGC
TCATGGCATCGGACTCATCAGGAAGC
AACCCCATCGGACTCATCAGGAAGC
TTGAACATCGGACTCATCAGGAAGC
CCTGGGCATCGGACTCATCAGGAAGC
ATACCTCATCGGACTCATCAGGAAGC
GTGTCATCGGACTCATCAGGAAGC
TCCCACATCGGACTCATCAGGAAGC
TAACATCGGACTCATCAGGAAGC
TGCCATCGGACTCATCAGGAAGC
AATCATCGGACTCATCAGGAAGC
TGCCATCGGACTCATCAGGAAGC
CTCATCGGACTCATCAGGAAGC
GACTCATCGGACTCATCAGGAAGC
CACTCATCGGACTCATCAGGAAGC
AAAACATCGGACTCATCAGGAAGC
AGACTCATCGGACTCATCAGGAAGC
AGCACATCGGACTCATCAGGAAGC

TGTCATCGGACTCATCAGGAAGC
ACACATCGGACTCATCAGGAAGC
CTTCACATCGGACTCATCAGGAAGC
GGCTCACATCGGACTCATCAGGAAGC
GAGCCATCGGACTCATCAGGAAGC
ACAGCATCGGACTCATCAGGAAGC
TGTGCATCGGACTCATCAGGAAGC
CTTCATCGGACTCATCAGGAAGC
CTCCATCGGACTCATCAGGAAGC
GAACACATCGGACTCATCAGGAAGC
ATTCACATCGGACTCATCAGGAAGC
TGTTCATCGGACTCATCAGGAAGC
GCCTCATCGGACTCATCAGGAAGC
GGCTCATCGGACTCATCAGGAAGC
GGACATCGGACTCATCAGGAAGC
GTGTCATCGGACTCATCAGGAAGC
CATGTCATCGGACTCATCAGGAAGC
TAGTCATCGGACTCATCAGGAAGC
GTCCATCGGACTCATCAGGAAGC
GCAACATCGGACTCATCAGGAAGC
CTAACATCGGACTCATCAGGAAGC
ACCCCATCGGACTCATCAGGAAGC
AAGCATCGGACTCATCAGGAAGC
GAGCATCGGACTCATCAGGAAGC
TCTTCATCGGACTCATCAGGAAGC
GAAACATCGGACTCATCAGGAAGC
AAGGCATCGGACTCATCAGGAAGC
TACTCATCGGACTCATCAGGAAGC
CACTACATCGGACTCATCAGGAAGC
CTCCCATCGGACTCATCAGGAAGC
TGGCTCATCGGACTCATCAGGAAGC
CACATCGGACTCATCAGGAAGC
CACTATCATCGGACTCATCAGGAAGC
TGAACATCGGACTCATCAGGAAGC
CACCATCGGACTCATCAGGAAGC
ATGCCATCGGACTCATCAGGAAGC
ATAGACATCGGACTCATCAGGAAGC
TGTTCATCGGACTCATCAGGAAGC
AACAGCATCGGACTCATCAGGAAGC
GATGCCATCGGACTCATCAGGAAGC
GCACCATCGGACTCATCAGGAAGC
GGAGAGCATCGGACTCATCAGGAAGC
AGGGACATCGGACTCATCAGGAAGC

CATGACATCGGACTCATCAGGAAGC
CAAACATCGGACTCATCAGGAAGC
CTCATCGGACTCATCAGGAAGC
GTCATCGGACTCATCAGGAAGC
GCATCGGACTCATCAGGAAGC
CTCATCGGACTCATCAGGAAGC
TGACCATCGGACTCATCAGGAAGC
AGACATCGGACTCATCAGGAAGC
AAGGCATCGGACTCATCAGGAAGC
CCATCGGACTCATCAGGAAGC
TACATCGGACTCATCAGGAAGC
AAGGCATCGGACTCATCAGGAAGC
ACATCGGACTCATCAGGAAGC
ACATCATCGGACTCATCAGGAAGC
TCATCGGACTCATCAGGAAGC
CTCATCGGACTCATCAGGAAGC
TTTACATCGGACTCATCAGGAAGC
ATAAAGCATCGGACTCATCAGGAAGC
CACCATCGGACTCATCAGGAAGC
ATCATCGGACTCATCAGGAAGC
ACCTCATCGGACTCATCAGGAAGC
CTCATCGGACTCATCAGGAAGC
CACATCGGACTCATCAGGAAGC
ATACATCGGACTCATCAGGAAGC
ATTACATCGGACTCATCAGGAAGC
TGAAGCATCGGACTCATCAGGAAGC
CAGCATCGGACTCATCAGGAAGC
ATTGCATCGGACTCATCAGGAAGC
AGGGCATCGGACTCATCAGGAAGC
CCCCATCGGACTCATCAGGAAGC
TGCCATCGGACTCATCAGGAAGC
TTCCATCGGACTCATCAGGAAGC
CCCATCGGACTCATCAGGAAGC
TACAACATCGGACTCATCAGGAAGC
AGACATCGGACTCATCAGGAAGC
CACATCGGACTCATCAGGAAGC
CATCATCGGACTCATCAGGAAGC
CACCCATCGGACTCATCAGGAAGC
GCATCGGACTCATCAGGAAGC
AGTCATCGGACTCATCAGGAAGC
ATACATCGGACTCATCAGGAAGC
GGTGGCATCGGACTCATCAGGAAGC
AGACATCGGACTCATCAGGAAGC

TGAGACATCGGACTCATCAGGAAGC
CACATCGGACTCATCAGGAAGC
AGATGGTCATCGGACTCATCAGGAAGC
GGCACATCGGACTCATCAGGAAGC
TGCCCCATCGGACTCATCAGGAAGC
ATTTCATCGGACTCATCAGGAAGC
AGGACATCGGACTCATCAGGAAGC
GGATCATCGGACTCATCAGGAAGC
GCTCATCGGACTCATCAGGAAGC
ACACATCGGACTCATCAGGAAGC
GAGACATCGGACTCATCAGGAAGC
CTCCATCGGACTCATCAGGAAGC
GCACATCGGACTCATCAGGAAGC
GGCATCGGACTCATCAGGAAGC
CTGGACATCGGACTCATCAGGAAGC
CTCCCATCGGACTCATCAGGAAGC
GTAACATCGGACTCATCAGGAAGC
TGTCATCGGACTCATCAGGAAGC
ATTCATCGGACTCATCAGGAAGC
GTCGCATCGGACTCATCAGGAAGC
AGGACATCGGACTCATCAGGAAGC
TACTCATCGGACTCATCAGGAAGC
ACAACATCGGACTCATCAGGAAGC
GTCCCATCGGACTCATCAGGAAGC
GTTTCATCGGACTCATCAGGAAGC
AGGACATCGGACTCATCAGGAAGC
GGGGCATCGGACTCATCAGGAAGC
TTCCATCGGACTCATCAGGAAGC
ATGACATCGGACTCATCAGGAAGC
GGTCATCGGACTCATCAGGAAGC
TGGCATCGGACTCATCAGGAAGC
GGAATCATCGGACTCATCAGGAAGC
TCCCATCGGACTCATCAGGAAGC
GCATACATCGGACTCATCAGGAAGC
GTGTCCATCGGACTCATCAGGAAGC
TTGCATCGGACTCATCAGGAAGC
TGGGGTCATCGGACTCATCAGGAAGC
CGAGCATCGGACTCATCAGGAAGC
TTGACATCGGACTCATCAGGAAGC
TGATCATCGGACTCATCAGGAAGC
GGTTCATCGGACTCATCAGGAAGC
TAAACATCGGACTCATCAGGAAGC
ACTACATCGGACTCATCAGGAAGC

CCGCATCGGACTCATCAGGAAGC
CTCCATCGGACTCATCAGGAAGC
CAGGCATCGGACTCATCAGGAAGC
TCTCCATCGGACTCATCAGGAAGC
TTTACATCGGACTCATCAGGAAGC
TGCCATCGGACTCATCAGGAAGC
CTACATCGGACTCATCAGGAAGC
CCACATCGGACTCATCAGGAAGC
AGGTCATCGGACTCATCAGGAAGC
GTCCATCGGACTCATCAGGAAGC
TGTCATCGGACTCATCAGGAAGC
AAGACATCGGACTCATCAGGAAGC
AGACATCGGACTCATCAGGAAGC
TGTCATCGGACTCATCAGGAAGC
GGTACATCGGACTCATCAGGAAGC
CTCCATCGGACTCATCAGGAAGC
CAGCATCGGACTCATCAGGAAGC
CTGGTCATCGGACTCATCAGGAAGC
GAACATCGGACTCATCAGGAAGC
GGCATCGGACTCATCAGGAAGC
TGGCATCGGACTCATCAGGAAGC
CCAGGCATCGGACTCATCAGGAAGC
AATCCATCGGACTCATCAGGAAGC
AGCCATCGGACTCATCAGGAAGC
GGTACATCGGACTCATCAGGAAGC
CACATCGGACTCATCAGGAAGC
GCATCGGACTCATCAGGAAGC
CAACATCGGACTCATCAGGAAGC
TCCATCGGACTCATCAGGAAGC
TATCACATCGGACTCATCAGGAAGC
GGCATCGGACTCATCAGGAAGC
CACATCGGACTCATCAGGAAGC
GCACATCGGACTCATCAGGAAGC
CCTGTCATCGGACTCATCAGGAAGC
TAGTCCATCGGACTCATCAGGAAGC
TCATCGGACTCATCAGGAAGC
TGGTCATCGGACTCATCAGGAAGC
TACACATCGGACTCATCAGGAAGC
AAGCCATCGGACTCATCAGGAAGC
GTCTCATCGGACTCATCAGGAAGC
CAGCATCGGACTCATCAGGAAGC
CTACACATCGGACTCATCAGGAAGC
TACACATCGGACTCATCAGGAAGC

CTTTCATCGGACTCATCAGGAAGC
CCCATCGGACTCATCAGGAAGC
GATGGCATCGGACTCATCAGGAAGC
TGACATCGGACTCATCAGGAAGC
ATGGGGCATCGGACTCATCAGGAAGC
AGTGCATCGGACTCATCAGGAAGC
CCTCATCGGACTCATCAGGAAGC
AGTGTCATCGGACTCATCAGGAAGC
CAAGGCATCGGACTCATCAGGAAGC
GTGTCATCGGACTCATCAGGAAGC
ATGAACATCGGACTCATCAGGAAGC
CAAAACATCGGACTCATCAGGAAGC
GGTACTCATCGGACTCATCAGGAAGC
AAAGACATCGGACTCATCAGGAAGC
GTTTTCATCGGACTCATCAGGAAGC
ATGTCATCGGACTCATCAGGAAGC
CAGAACATCGGACTCATCAGGAAGC
GCTCATCGGACTCATCAGGAAGC
CCTGCATCGGACTCATCAGGAAGC
TGGACATCGGACTCATCAGGAAGC
CAACCATCGGACTCATCAGGAAGC
GTGCCATCGGACTCATCAGGAAGC
TGATATGCATCGGACTCATCAGGAAGC
CCCCACATCGGACTCATCAGGAAGC
GACATCGGACTCATCAGGAAGC
AACGCATCGGACTCATCAGGAAGC
TGTGTCATCGGACTCATCAGGAAGC
AGCACATCGGACTCATCAGGAAGC
TTGCATCGGACTCATCAGGAAGC
GGCACATCGGACTCATCAGGAAGC
TTCATCATCGGACTCATCAGGAAGC
GAACATCGGACTCATCAGGAAGC
AAGACATCGGACTCATCAGGAAGC
GGTCATCGGACTCATCAGGAAGC
CTCCATCGGACTCATCAGGAAGC
GTGTGCATCGGACTCATCAGGAAGC
ATAACTCATCGGACTCATCAGGAAGC
CTCATCGGACTCATCAGGAAGC
GGGTCATCGGACTCATCAGGAAGC
AGGTCATCGGACTCATCAGGAAGC
CACATCGGACTCATCAGGAAGC
AACATCGGACTCATCAGGAAGC
CACATCGGACTCATCAGGAAGC

GAGGGCATCGGACTCATCAGGAAGC
AAACATCGGACTCATCAGGAAGC
ACACATCGGACTCATCAGGAAGC
TGGACATCGGACTCATCAGGAAGC
GGCATCGGACTCATCAGGAAGC
CATCATCGGACTCATCAGGAAGC
CTCATCGGACTCATCAGGAAGC
AAAGTCATCGGACTCATCAGGAAGC
AACACATCGGACTCATCAGGAAGC
TGAGCATCGGACTCATCAGGAAGC
TCATCATCGGACTCATCAGGAAGC
CATCATCGGACTCATCAGGAAGC
TCATCGGACTCATCAGGAAGC
CACATCGGACTCATCAGGAAGC
GACGCATCGGACTCATCAGGAAGC
AAGACATCGGACTCATCAGGAAGC
AGCCATCGGACTCATCAGGAAGC
CTCCATCGGACTCATCAGGAAGC
TTACATCGGACTCATCAGGAAGC
ATGCCATCGGACTCATCAGGAAGC
CCATCGGACTCATCAGGAAGC
TGCATCGGACTCATCAGGAAGC
ACATCGGACTCATCAGGAAGC
CACATCGGACTCATCAGGAAGC
GACCATCGGACTCATCAGGAAGC
CACATCGGACTCATCAGGAAGC
TGGACATCGGACTCATCAGGAAGC
GCCATCGGACTCATCAGGAAGC
GGGACATCGGACTCATCAGGAAGC
GAACATCGGACTCATCAGGAAGC
GCACATCGGACTCATCAGGAAGC
GACATCGGACTCATCAGGAAGC
ACATCGGACTCATCAGGAAGC
CTGACACATCGGACTCATCAGGAAGC
TGCCATCGGACTCATCAGGAAGC
ACCACATCGGACTCATCAGGAAGC
GAGAGCATCGGACTCATCAGGAAGC
ACCACATCGGACTCATCAGGAAGC
AAAACATCGGACTCATCAGGAAGC
TACATCGGACTCATCAGGAAGC
ATCCCATCGGACTCATCAGGAAGC
CTCATCGGACTCATCAGGAAGC
AGACATCGGACTCATCAGGAAGC

ACACATCGGACTCATCAGGAAGC
TCGCATCGGACTCATCAGGAAGC
TCCATCGGACTCATCAGGAAGC
GTCATCGGACTCATCAGGAAGC
CCATCGGACTCATCAGGAAGC
GCCAGCCATCGGACTCATCAGGAAGC
GTCATCGGACTCATCAGGAAGC
AACATCGGACTCATCAGGAAGC
CTACCATCGGACTCATCAGGAAGC
TCCATCGGACTCATCAGGAAGC
GGTCATCGGACTCATCAGGAAGC
CACCATCGGACTCATCAGGAAGC
CCACATCGGACTCATCAGGAAGC
GGTGGCATCGGACTCATCAGGAAGC
TGGGCATCGGACTCATCAGGAAGC
TACATCGGACTCATCAGGAAGC
GCCTGCATCGGACTCATCAGGAAGC
TCCCATCGGACTCATCAGGAAGC
CTCTCATCGGACTCATCAGGAAGC
CAGCATCGGACTCATCAGGAAGC
CTAAATCATCGGACTCATCAGGAAGC
CAACATCGGACTCATCAGGAAGC
CCCATCGGACTCATCAGGAAGC
CTCATCGGACTCATCAGGAAGC
CACATCGGACTCATCAGGAAGC
CCACATCGGACTCATCAGGAAGC
GTCACATCGGACTCATCAGGAAGC
CCCATCGGACTCATCAGGAAGC
TGCATCGGACTCATCAGGAAGC
TTGCATCGGACTCATCAGGAAGC
TCCCATCGGACTCATCAGGAAGC
CACACATCGGACTCATCAGGAAGC
TCAGCATCGGACTCATCAGGAAGC
TTTCATCGGACTCATCAGGAAGC
TCTCATCGGACTCATCAGGAAGC
AACACATCGGACTCATCAGGAAGC
CCCATCGGACTCATCAGGAAGC
TAGTCGCATCGGACTCATCAGGAAGC
AGCACATCGGACTCATCAGGAAGC
CACATCGGACTCATCAGGAAGC
AACTCCATCGGACTCATCAGGAAGC
ATGTACATCGGACTCATCAGGAAGC
CGGACATCGGACTCATCAGGAAGC

AACTCATCGGACTCATCAGGAAGC
GTCATCGGACTCATCAGGAAGC
AAGCCATCGGACTCATCAGGAAGC
ACACATCGGACTCATCAGGAAGC
ATGCCATCGGACTCATCAGGAAGC
ACATCGGACTCATCAGGAAGC
AAGACATCGGACTCATCAGGAAGC
AGAACCATCGGACTCATCAGGAAGC
TGAACCATCGGACTCATCAGGAAGC
ACCTCATCGGACTCATCAGGAAGC
CCAACATCGGACTCATCAGGAAGC
CTCATCGGACTCATCAGGAAGC
TCACATCGGACTCATCAGGAAGC
CCCATCGGACTCATCAGGAAGC
CAAACATCGGACTCATCAGGAAGC
GTTCTCCATCGGACTCATCAGGAAGC
TGTCATCGGACTCATCAGGAAGC
ATTCATCGGACTCATCAGGAAGC
CCCATCGGACTCATCAGGAAGC
TTCCTCATCGGACTCATCAGGAAGC
CAGTCATCGGACTCATCAGGAAGC
GCCATCGGACTCATCAGGAAGC
GAAACATCGGACTCATCAGGAAGC
CCTCATCGGACTCATCAGGAAGC
GCTCATCGGACTCATCAGGAAGC
CAGCCATCGGACTCATCAGGAAGC
CCACAACATCGGACTCATCAGGAAGC
CTCTCATCGGACTCATCAGGAAGC
GGACATCGGACTCATCAGGAAGC
GTGTCATCGGACTCATCAGGAAGC
GGCCATCGGACTCATCAGGAAGC
CTCCATCGGACTCATCAGGAAGC
CATCATCGGACTCATCAGGAAGC
GGTCATCGGACTCATCAGGAAGC
ACACACATCGGACTCATCAGGAAGC
CTCATCGGACTCATCAGGAAGC
AGCCATCGGACTCATCAGGAAGC
GAAACATCGGACTCATCAGGAAGC
ACTCTCATCGGACTCATCAGGAAGC
AGTACGCATCGGACTCATCAGGAAGC
CAGGCCATCGGACTCATCAGGAAGC
GGTCATCGGACTCATCAGGAAGC
AAGCATCGGACTCATCAGGAAGC

CATCCATCGGACTCATCAGGAAGC
AGATACATCGGACTCATCAGGAAGC
GTCTCATCGGACTCATCAGGAAGC
GCCATCGGACTCATCAGGAAGC
GGGACATCGGACTCATCAGGAAGC
AGACATCGGACTCATCAGGAAGC
GGCTACATCGGACTCATCAGGAAGC
TCTGACATCGGACTCATCAGGAAGC
TCCCATCGGACTCATCAGGAAGC
TGTCATCGGACTCATCAGGAAGC
GTGCGCATCGGACTCATCAGGAAGC
TAAGGCATCGGACTCATCAGGAAGC
AGGGCATCGGACTCATCAGGAAGC
GCCATCGGACTCATCAGGAAGC
ACAGCATCGGACTCATCAGGAAGC
GCACATCGGACTCATCAGGAAGC
CCATCGGACTCATCAGGAAGC
CTTTCATCGGACTCATCAGGAAGC
AAAGCCATCGGACTCATCAGGAAGC
ACACCATCGGACTCATCAGGAAGC
AGTGCATCGGACTCATCAGGAAGC
GAGCATCGGACTCATCAGGAAGC
GCACATCGGACTCATCAGGAAGC
TGGGACATCGGACTCATCAGGAAGC
TTTCATCGGACTCATCAGGAAGC
AGGACATCGGACTCATCAGGAAGC
TCCATCGGACTCATCAGGAAGC
AGGGCATCGGACTCATCAGGAAGC
TGCAGTCATCGGACTCATCAGGAAGC
GTATGCATCGGACTCATCAGGAAGC
TGTTTCATCGGACTCATCAGGAAGC
AAAAGCCATCGGACTCATCAGGAAGC
CTCATCGGACTCATCAGGAAGC
AGACATCGGACTCATCAGGAAGC
ATCAGCATCGGACTCATCAGGAAGC
AAGAACATCGGACTCATCAGGAAGC
CTCCATCGGACTCATCAGGAAGC
ATGTCATCGGACTCATCAGGAAGC
ATGCCACATCGGACTCATCAGGAAGC
CATCATCGGACTCATCAGGAAGC
TCCATCGGACTCATCAGGAAGC
TGCAACATCGGACTCATCAGGAAGC
AGGCATCGGACTCATCAGGAAGC

AGGCTGCATCGGACTCATCAGGAAGC
TCTCACATCGGACTCATCAGGAAGC
TCCCATCGGACTCATCAGGAAGC
TAACATCATCGGACTCATCAGGAAGC
TCCAGTCATCGGACTCATCAGGAAGC
TGAGCCATCGGACTCATCAGGAAGC
ACAACATCGGACTCATCAGGAAGC
TCTGGACATCGGACTCATCAGGAAGC
ACTGTCATCGGACTCATCAGGAAGC
CCATGACATCGGACTCATCAGGAAGC
AAGCACATCGGACTCATCAGGAAGC
TGTGTGCATCGGACTCATCAGGAAGC
TGAGCATCGGACTCATCAGGAAGC
CAGCATCATCGGACTCATCAGGAAGC
TTTTCATCGGACTCATCAGGAAGC
AATGGCATCGGACTCATCAGGAAGC
AGTCTCATCGGACTCATCAGGAAGC
AGTGAACATCGGACTCATCAGGAAGC
CTGGTCATCGGACTCATCAGGAAGC
TGGGACATCGGACTCATCAGGAAGC
GTCCCCATCGGACTCATCAGGAAGC
TGCTGGCATCGGACTCATCAGGAAGC
TTTGGCATCGGACTCATCAGGAAGC
ACAGCACATCGGACTCATCAGGAAGC
CCCTCATCGGACTCATCAGGAAGC
ATACATCATCGGACTCATCAGGAAGC
ATTTCACATCGGACTCATCAGGAAGC
ACATTCCATCGGACTCATCAGGAAGC
AGAAACATCGGACTCATCAGGAAGC
AATAGCCATCGGACTCATCAGGAAGC
AGCAGTCATCGGACTCATCAGGAAGC
CTCCATCGGACTCATCAGGAAGC
TAGCACATCGGACTCATCAGGAAGC
TCTCTCCATCGGACTCATCAGGAAGC
CTTACCATCGGACTCATCAGGAAGC
TCCCCATCGGACTCATCAGGAAGC
TGGGGCATCGGACTCATCAGGAAGC
GTGTAACATCGGACTCATCAGGAAGC
TAACCCCATCGGACTCATCAGGAAGC
GGGTGACATCGGACTCATCAGGAAGC
GCAACCCATCGGACTCATCAGGAAGC
CCACTTCATCGGACTCATCAGGAAGC
AGTGACACATCGGACTCATCAGGAAGC

TCCACATCGGACTCATCAGGAAGC
AGGACACATCGGACTCATCAGGAAGC
GGCTTTCCATCGGACTCATCAGGAAGC
CAACATCGGACTCATCAGGAAGC
TTGTCATCGGACTCATCAGGAAGC
AAGAAACATCGGACTCATCAGGAAGC
CTCCTCATCGGACTCATCAGGAAGC
GCCACATCGGACTCATCAGGAAGC
GTAGCCATCGGACTCATCAGGAAGC
ATGTGGCATCGGACTCATCAGGAAGC
TTGGTCATCGGACTCATCAGGAAGC
TAACTCATCGGACTCATCAGGAAGC
AACCTCATCGGACTCATCAGGAAGC
AGGACATCGGACTCATCAGGAAGC
ATTTGCCATCGGACTCATCAGGAAGC
GTGTGTCATCGGACTCATCAGGAAGC
TGGTCACATCGGACTCATCAGGAAGC
TTGTAACATCGGACTCATCAGGAAGC
AAATGCATCGGACTCATCAGGAAGC
GCTCACATCGGACTCATCAGGAAGC
TGTGCCATCGGACTCATCAGGAAGC
GCACCATCGGACTCATCAGGAAGC
GTTGACATCGGACTCATCAGGAAGC
GTCCATCGGACTCATCAGGAAGC
AGCACCATCGGACTCATCAGGAAGC
CCATCATCGGACTCATCAGGAAGC
TCCCTCATCGGACTCATCAGGAAGC
CTAGCCATCGGACTCATCAGGAAGC
ATGCTTCATCGGACTCATCAGGAAGC
AGTCCACATCGGACTCATCAGGAAGC
TATGACATCGGACTCATCAGGAAGC
AGTCATCGGACTCATCAGGAAGC
TTTATCATCGGACTCATCAGGAAGC
GCTAGACATCGGACTCATCAGGAAGC
GCTCTCATCGGACTCATCAGGAAGC
AAACCCATCGGACTCATCAGGAAGC
CTTTCATCGGACTCATCAGGAAGC
TGTCCATCGGACTCATCAGGAAGC
ACCCATCGGACTCATCAGGAAGC
CTCATCGGACTCATCAGGAAGC
GTGTCATCGGACTCATCAGGAAGC
ATCCACATCGGACTCATCAGGAAGC
CCGTTCCATCGGACTCATCAGGAAGC

TAGAAGGCATCGGACTCATCAGGAAGC
TGTAACATCGGACTCATCAGGAAGC
CTTAGACATCGGACTCATCAGGAAGC
GCCCAACATCGGACTCATCAGGAAGC
TTCATCGGACTCATCAGGAAGC
CACACATCGGACTCATCAGGAAGC
GTGTGCATCGGACTCATCAGGAAGC
CTTGAACATCGGACTCATCAGGAAGC
TGTACATCGGACTCATCAGGAAGC
GGGCATCGGACTCATCAGGAAGC
TGGCCATCGGACTCATCAGGAAGC
CACTCCATCGGACTCATCAGGAAGC
ATACATCGGACTCATCAGGAAGC
AGAACATCGGACTCATCAGGAAGC
ACCACATCGGACTCATCAGGAAGC
ATTCATCGGACTCATCAGGAAGC
ACTGCACATCGGACTCATCAGGAAGC
GGGGCATCGGACTCATCAGGAAGC
AGTGGACATCGGACTCATCAGGAAGC
CAGTTCATCGGACTCATCAGGAAGC
ACGTCATCGGACTCATCAGGAAGC
TCAGGCATCGGACTCATCAGGAAGC
CTCATCGGACTCATCAGGAAGC
TGTGTCATCGGACTCATCAGGAAGC
TATAGCCATCGGACTCATCAGGAAGC
GTCATCGGACTCATCAGGAAGC
GCGCTCATCGGACTCATCAGGAAGC
GACTGCATCGGACTCATCAGGAAGC
GCCAGCATCGGACTCATCAGGAAGC
AATGACATCGGACTCATCAGGAAGC
TCAGTCATCGGACTCATCAGGAAGC
ATGCACATCGGACTCATCAGGAAGC
TGTATACATCGGACTCATCAGGAAGC
CACACATCGGACTCATCAGGAAGC
AAGGCATCGGACTCATCAGGAAGC
CTTGAGCATCGGACTCATCAGGAAGC
TTTCACATCGGACTCATCAGGAAGC
GAGCATCGGACTCATCAGGAAGC
CACCATCGGACTCATCAGGAAGC
TTATGCATCGGACTCATCAGGAAGC
GTGCCATCGGACTCATCAGGAAGC
TGAACCCATCGGACTCATCAGGAAGC
TGACATCGGACTCATCAGGAAGC

ACGCTCATCGGACTCATCAGGAAGC
CACCATCGGACTCATCAGGAAGC
CAGGCATCGGACTCATCAGGAAGC
GTACGCATCGGACTCATCAGGAAGC
GAACCATCGGACTCATCAGGAAGC
GCCATCGGACTCATCAGGAAGC
TTGACATCGGACTCATCAGGAAGC
CCAGAGCATCGGACTCATCAGGAAGC
CACCATCGGACTCATCAGGAAGC
CTGTCATCGGACTCATCAGGAAGC
TCATCGGACTCATCAGGAAGC
CATCATCGGACTCATCAGGAAGC
AGACCATCGGACTCATCAGGAAGC
ATAGACATCGGACTCATCAGGAAGC
TACCATCGGACTCATCAGGAAGC
GCGACATCGGACTCATCAGGAAGC
ACCATCGGACTCATCAGGAAGC
CTTCGCATCGGACTCATCAGGAAGC
TGTGCCATCGGACTCATCAGGAAGC
GCCATCGGACTCATCAGGAAGC
ATTCCACATCGGACTCATCAGGAAGC
GTTGGCATCGGACTCATCAGGAAGC
AAAACCTCATCGGACTCATCAGGAAGC
GGCACATCGGACTCATCAGGAAGC
GGTCATCGGACTCATCAGGAAGC
GTGACATCGGACTCATCAGGAAGC
GGGCATCGGACTCATCAGGAAGC
AGAACATCGGACTCATCAGGAAGC
GTAAGCCATCGGACTCATCAGGAAGC
GCAACATCGGACTCATCAGGAAGC
TCACATCGGACTCATCAGGAAGC
TGGCCATCGGACTCATCAGGAAGC
TGAATCATCGGACTCATCAGGAAGC
GAGAATCATCGGACTCATCAGGAAGC
CACATCGGACTCATCAGGAAGC
TGACCATCGGACTCATCAGGAAGC
AAAAGTCATCGGACTCATCAGGAAGC
CGTCATCGGACTCATCAGGAAGC
TGGTCATCGGACTCATCAGGAAGC
GAAATGCATCGGACTCATCAGGAAGC
AACTCCATCGGACTCATCAGGAAGC
TCTCATCGGACTCATCAGGAAGC
CCACATCGGACTCATCAGGAAGC

CCTCATCGGACTCATCAGGAAGC
CACACATCGGACTCATCAGGAAGC
ACAACATCGGACTCATCAGGAAGC
CCAGGCATCGGACTCATCAGGAAGC
GACCATCGGACTCATCAGGAAGC
AACTCATCGGACTCATCAGGAAGC
CGACACATCGGACTCATCAGGAAGC
TGACATCGGACTCATCAGGAAGC
AAATGCCATCGGACTCATCAGGAAGC
TTCCCATCGGACTCATCAGGAAGC
GTCATCGGACTCATCAGGAAGC
AATACATCGGACTCATCAGGAAGC
CCCTCATCGGACTCATCAGGAAGC
TATCATCGGACTCATCAGGAAGC
GACCTCATCGGACTCATCAGGAAGC
CTTTCATCGGACTCATCAGGAAGC
ATTTGCATCGGACTCATCAGGAAGC
GTGCATCGGACTCATCAGGAAGC
CTGCATCGGACTCATCAGGAAGC
TGGACATCGGACTCATCAGGAAGC
TACCATCGGACTCATCAGGAAGC
CACACCATCGGACTCATCAGGAAGC
AACCCATCGGACTCATCAGGAAGC
TGCCCATCGGACTCATCAGGAAGC
AAGCCATCGGACTCATCAGGAAGC
CACTACATCGGACTCATCAGGAAGC
TTCCGTCATCGGACTCATCAGGAAGC
GTAACATCGGACTCATCAGGAAGC
GGCTCATCGGACTCATCAGGAAGC
GGCCATCGGACTCATCAGGAAGC
GGCACCATCGGACTCATCAGGAAGC
GCCTCATCGGACTCATCAGGAAGC
AACTGCATCGGACTCATCAGGAAGC
TACACATCGGACTCATCAGGAAGC
TCTCATCGGACTCATCAGGAAGC
CAGGACATCGGACTCATCAGGAAGC
CTCCATCGGACTCATCAGGAAGC
CTGCATCGGACTCATCAGGAAGC
CACCCATCGGACTCATCAGGAAGC
TGTCATCGGACTCATCAGGAAGC
GGCATCGGACTCATCAGGAAGC
GTAATCATCGGACTCATCAGGAAGC
GAGGCATCGGACTCATCAGGAAGC

GTGGCCATCGGACTCATCAGGAAGC
TAGCTCATCGGACTCATCAGGAAGC
CTTGAACCATCGGACTCATCAGGAAGC
GAGGCATCGGACTCATCAGGAAGC
ACTCGCATCGGACTCATCAGGAAGC
AGGAGCATCGGACTCATCAGGAAGC
GAGAATTGCATCGGACTCATCAGGAAGC
CCAGCATCGGACTCATCAGGAAGC
ACAGGCATCGGACTCATCAGGAAGC
CAGGCATCGGACTCATCAGGAAGC
GTGGCATCGGACTCATCAGGAAGC
TTCAACATCGGACTCATCAGGAAGC
CTTCATCGGACTCATCAGGAAGC
GAGCATCGGACTCATCAGGAAGC
ACGCATCGGACTCATCAGGAAGC
CCAGCATCGGACTCATCAGGAAGC
AGGACATCGGACTCATCAGGAAGC
CACTCATCGGACTCATCAGGAAGC
ATCATCATCGGACTCATCAGGAAGC
CTGCCCATCGGACTCATCAGGAAGC
CCAAACATCGGACTCATCAGGAAGC
AACCTCATCGGACTCATCAGGAAGC
GTAGTGCATCGGACTCATCAGGAAGC
ACTCCATCGGACTCATCAGGAAGC
CACCTCATCGGACTCATCAGGAAGC
GATTGCATCGGACTCATCAGGAAGC
CTTCAACATCGGACTCATCAGGAAGC
GGAGCATCGGACTCATCAGGAAGC
GTCATCGGACTCATCAGGAAGC
GTCATCGGACTCATCAGGAAGC
GTCATCGGACTCATCAGGAAGC
GTCATCGGACTCATCAGGAAGC
GTCATCGGACTCATCAGGAAGC
GTCATCGGACTCATCAGGAAGC
GTCATCGGACTCATCAGGAAGC
GTCATCGGACTCATCAGGAAGC
CATCGGACTCATCAGGAAGC
CATCGGACTCATCAGGAAGC
CATCGGACTCATCAGGAAGC
CATCGGACTCATCAGGAAGC
CATCGGACTCATCAGGAAGC
CATCGGACTCATCAGGAAGC
CATCGGACTCATCAGGAAGC

CATCGGACTCATCAGGAAGC
ACATCGGACTCATCAGGAAGC
ACATCGGACTCATCAGGAAGC
ACATCGGACTCATCAGGAAGC
ACATCGGACTCATCAGGAAGC
ACATCGGACTCATCAGGAAGC
ACATCGGACTCATCAGGAAGC
ACATCGGACTCATCAGGAAGC
ACATCGGACTCATCAGGAAGC
AACATCGGACTCATCAGGAAGC
AACATCGGACTCATCAGGAAGC
AACATCGGACTCATCAGGAAGC
AACATCGGACTCATCAGGAAGC
AACATCGGACTCATCAGGAAGC
AACATCGGACTCATCAGGAAGC
AACATCGGACTCATCAGGAAGC
AACATCGGACTCATCAGGAAGC
CATCGGACTCATCAGGAAGC
CATCGGACTCATCAGGAAGC
CATCGGACTCATCAGGAAGC
CATCGGACTCATCAGGAAGC
CATCGGACTCATCAGGAAGC
CATCGGACTCATCAGGAAGC
CATCGGACTCATCAGGAAGC
CATCGGACTCATCAGGAAGC
CACATCGGACTCATCAGGAAGC
CACATCGGACTCATCAGGAAGC
CACATCGGACTCATCAGGAAGC
CACATCGGACTCATCAGGAAGC
CACATCGGACTCATCAGGAAGC
CACATCGGACTCATCAGGAAGC
CACATCGGACTCATCAGGAAGC
CACATCGGACTCATCAGGAAGC
TCATCGGACTCATCAGGAAGC
TCATCGGACTCATCAGGAAGC
TCATCGGACTCATCAGGAAGC
TCATCGGACTCATCAGGAAGC
TCATCGGACTCATCAGGAAGC
TCATCGGACTCATCAGGAAGC
TCATCGGACTCATCAGGAAGC
TCATCGGACTCATCAGGAAGC
CACATCGGACTCATCAGGAAGC
CACATCGGACTCATCAGGAAGC

CACATCGGACTCATCAGGAAGC
CACATCGGACTCATCAGGAAGC
CACATCGGACTCATCAGGAAGC
CACATCGGACTCATCAGGAAGC
CACATCGGACTCATCAGGAAGC
CACATCGGACTCATCAGGAAGC
CATCGGACTCATCAGGAAGC
CATCGGACTCATCAGGAAGC
CATCGGACTCATCAGGAAGC
CATCGGACTCATCAGGAAGC
CATCGGACTCATCAGGAAGC
CATCGGACTCATCAGGAAGC
CATCGGACTCATCAGGAAGC
CATCGGACTCATCAGGAAGC
AAGCATCGGACTCATCAGGAAGC
AAGCATCGGACTCATCAGGAAGC
AAGCATCGGACTCATCAGGAAGC
AAGCATCGGACTCATCAGGAAGC
AAGCATCGGACTCATCAGGAAGC
AAGCATCGGACTCATCAGGAAGC
AAGCATCGGACTCATCAGGAAGC
AAGCATCGGACTCATCAGGAAGC
ACTCATCGGACTCATCAGGAAGC
ACTCATCGGACTCATCAGGAAGC
ACTCATCGGACTCATCAGGAAGC
ACTCATCGGACTCATCAGGAAGC
ACTCATCGGACTCATCAGGAAGC
ACTCATCGGACTCATCAGGAAGC
ACTCATCGGACTCATCAGGAAGC
ACTCATCGGACTCATCAGGAAGC
TATCATCGGACTCATCAGGAAGC
TATCATCGGACTCATCAGGAAGC
TATCATCGGACTCATCAGGAAGC
TATCATCGGACTCATCAGGAAGC
TATCATCGGACTCATCAGGAAGC
TATCATCGGACTCATCAGGAAGC
TATCATCGGACTCATCAGGAAGC
TATCATCGGACTCATCAGGAAGC
GAACCATCGGACTCATCAGGAAGC
GAACCATCGGACTCATCAGGAAGC
GAACCATCGGACTCATCAGGAAGC
GAACCATCGGACTCATCAGGAAGC
GAACCATCGGACTCATCAGGAAGC

GAACCATCGGACTCATCAGGAAGC
GAACCATCGGACTCATCAGGAAGC
GAACCATCGGACTCATCAGGAAGC
GGAGCATCGGACTCATCAGGAAGC
GGAGCATCGGACTCATCAGGAAGC
GGAGCATCGGACTCATCAGGAAGC
GGAGCATCGGACTCATCAGGAAGC
GGAGCATCGGACTCATCAGGAAGC
GGAGCATCGGACTCATCAGGAAGC
GGAGCATCGGACTCATCAGGAAGC
GGAGCATCGGACTCATCAGGAAGC
ACCATCGGACTCATCAGGAAGC
ACCATCGGACTCATCAGGAAGC
ACCATCGGACTCATCAGGAAGC
ACCATCGGACTCATCAGGAAGC
ACCATCGGACTCATCAGGAAGC
ACCATCGGACTCATCAGGAAGC
ACCATCGGACTCATCAGGAAGC
ACCATCGGACTCATCAGGAAGC
ACCATCGGACTCATCAGGAAGC
ACCATCGGACTCATCAGGAAGC
ACCATCGGACTCATCAGGAAGC
ACCATCGGACTCATCAGGAAGC
ACCATCGGACTCATCAGGAAGC
ACCATCGGACTCATCAGGAAGC
ACCATCGGACTCATCAGGAAGC
ACCATCGGACTCATCAGGAAGC
GTCCATCGGACTCATCAGGAAGC
GTCCATCGGACTCATCAGGAAGC
GTCCATCGGACTCATCAGGAAGC
GTCCATCGGACTCATCAGGAAGC
GTCCATCGGACTCATCAGGAAGC
GTCCATCGGACTCATCAGGAAGC
GTCCATCGGACTCATCAGGAAGC
GTCCATCGGACTCATCAGGAAGC
ACATCGGACTCATCAGGAAGC
ACATCGGACTCATCAGGAAGC
ACATCGGACTCATCAGGAAGC
ACATCGGACTCATCAGGAAGC
ACATCGGACTCATCAGGAAGC
ACATCGGACTCATCAGGAAGC
ACATCGGACTCATCAGGAAGC
ACATCGGACTCATCAGGAAGC

AACATCGGACTCATCAGGAAGC
AACATCGGACTCATCAGGAAGC
AACATCGGACTCATCAGGAAGC
AACATCGGACTCATCAGGAAGC
AACATCGGACTCATCAGGAAGC
AACATCGGACTCATCAGGAAGC
AACATCGGACTCATCAGGAAGC
AACATCGGACTCATCAGGAAGC
ACTCATCAGGAAGC
ACTCATCAGGAAGC
ACTCATCAGGAAGC
ACTCATCAGGAAGC
ACTCATCAGGAAGC
ACTCATCAGGAAGC
ACTCATCAGGAAGC
ACTCATCAGGAAGC
CTCATCGGACTCATCAGGAAGC
CTCATCGGACTCATCAGGAAGC
CTCATCGGACTCATCAGGAAGC
CTCATCGGACTCATCAGGAAGC
CTCATCGGACTCATCAGGAAGC
CTCATCGGACTCATCAGGAAGC
CTCATCGGACTCATCAGGAAGC
CTCATCGGACTCATCAGGAAGC
AACATCGGACTCATCAGGAAGC
AACATCGGACTCATCAGGAAGC
AACATCGGACTCATCAGGAAGC
AACATCGGACTCATCAGGAAGC
AACATCGGACTCATCAGGAAGC
AACATCGGACTCATCAGGAAGC
AACATCGGACTCATCAGGAAGC
AACATCGGACTCATCAGGAAGC
ATCGGACTCATCAGGAAGC
ATCGGACTCATCAGGAAGC
ATCGGACTCATCAGGAAGC
ATCGGACTCATCAGGAAGC
ATCGGACTCATCAGGAAGC
ATCGGACTCATCAGGAAGC
ATCGGACTCATCAGGAAGC
ATCGGACTCATCAGGAAGC
CATCGGACTCATCAGGAAGC
CATCGGACTCATCAGGAAGC
CATCGGACTCATCAGGAAGC

CATCGGACTCATCAGGAAGC
CATCGGACTCATCAGGAAGC
CATCGGACTCATCAGGAAGC
CATCGGACTCATCAGGAAGC
CATCGGACTCATCAGGAAGC
GGAAGCATCGGACTCATCAGGAAGC
GGAAGCATCGGACTCATCAGGAAGC
GGAAGCATCGGACTCATCAGGAAGC
GGAAGCATCGGACTCATCAGGAAGC
GGAAGCATCGGACTCATCAGGAAGC
GGAAGCATCGGACTCATCAGGAAGC
GGAAGCATCGGACTCATCAGGAAGC
GGAAGCATCGGACTCATCAGGAAGC
CGCACATCGGACTCATCAGGAAGC
CGCACATCGGACTCATCAGGAAGC
CGCACATCGGACTCATCAGGAAGC
CGCACATCGGACTCATCAGGAAGC
CGCACATCGGACTCATCAGGAAGC
CGCACATCGGACTCATCAGGAAGC
CGCACATCGGACTCATCAGGAAGC
CGCACATCGGACTCATCAGGAAGC
GCACATCGGACTCATCAGGAAGC
GCACATCGGACTCATCAGGAAGC
GCACATCGGACTCATCAGGAAGC
GCACATCGGACTCATCAGGAAGC
GCACATCGGACTCATCAGGAAGC
GCACATCGGACTCATCAGGAAGC
GCACATCGGACTCATCAGGAAGC
GCACATCGGACTCATCAGGAAGC
TCATCGGACTCATCAGGAAGC
TCATCGGACTCATCAGGAAGC
TCATCGGACTCATCAGGAAGC
TCATCGGACTCATCAGGAAGC
TCATCGGACTCATCAGGAAGC
TCATCGGACTCATCAGGAAGC
TCATCGGACTCATCAGGAAGC
TCATCGGACTCATCAGGAAGC
ATCGGACTCATCAGGAAGC
ATCGGACTCATCAGGAAGC
ATCGGACTCATCAGGAAGC
ATCGGACTCATCAGGAAGC
ATCGGACTCATCAGGAAGC
ATCGGACTCATCAGGAAGC
